# A genome-wide association study of mass spectrometry proteomics using the Seer Proteograph platform

**DOI:** 10.1101/2024.05.27.596028

**Authors:** Karsten Suhre, Qingwen Chen, Anna Halama, Kevin Mendez, Amber Dahlin, Nisha Stephan, Gaurav Thareja, Hina Sarwath, Harendra Guturu, Varun B. Dwaraka, Serafim Batzoglou, Frank Schmidt, Jessica A. Lasky-Su

## Abstract

Genome-wide association studies (GWAS) with proteomics are essential tools for drug discovery. To date, most studies have used affinity proteomics platforms, which have limited discovery to protein panels covered by the available affinity binders. Furthermore, it is not clear to which extent protein epitope changing variants interfere with the detection of protein quantitative trait loci (pQTLs). Mass spectrometry-based (MS) proteomics can overcome some of these limitations. Here we report a GWAS using the MS-based Seer Proteograph^TM^ platform with blood samples from a discovery cohort of 1,260 American participants and a replication in 325 individuals from Asia, with diverse ethnic backgrounds. We analysed 1,980 proteins quantified in at least 80% of the samples, out of 5,753 proteins quantified across the discovery cohort. We identified 252 and replicated 90 pQTLs, where 30 of the replicated pQTLs have not been reported before. We further investigated 200 of the strongest associated cis-pQTLs previously identified using the SOMAscan and the Olink platforms and found that up to one third of the affinity proteomics pQTLs may be affected by epitope effects, while another third were confirmed by MS proteomics to be consistent with the hypothesis that genetic variants induce changes in protein expression. The present study demonstrates the complementarity of the different proteomics approaches and reports pQTLs not accessible to affinity proteomics, suggesting that many more pQTLs remain to be discovered using MS-based platforms.

**Graphical Abstract:** Summarizing the approach taken to identify potential epitope effects.

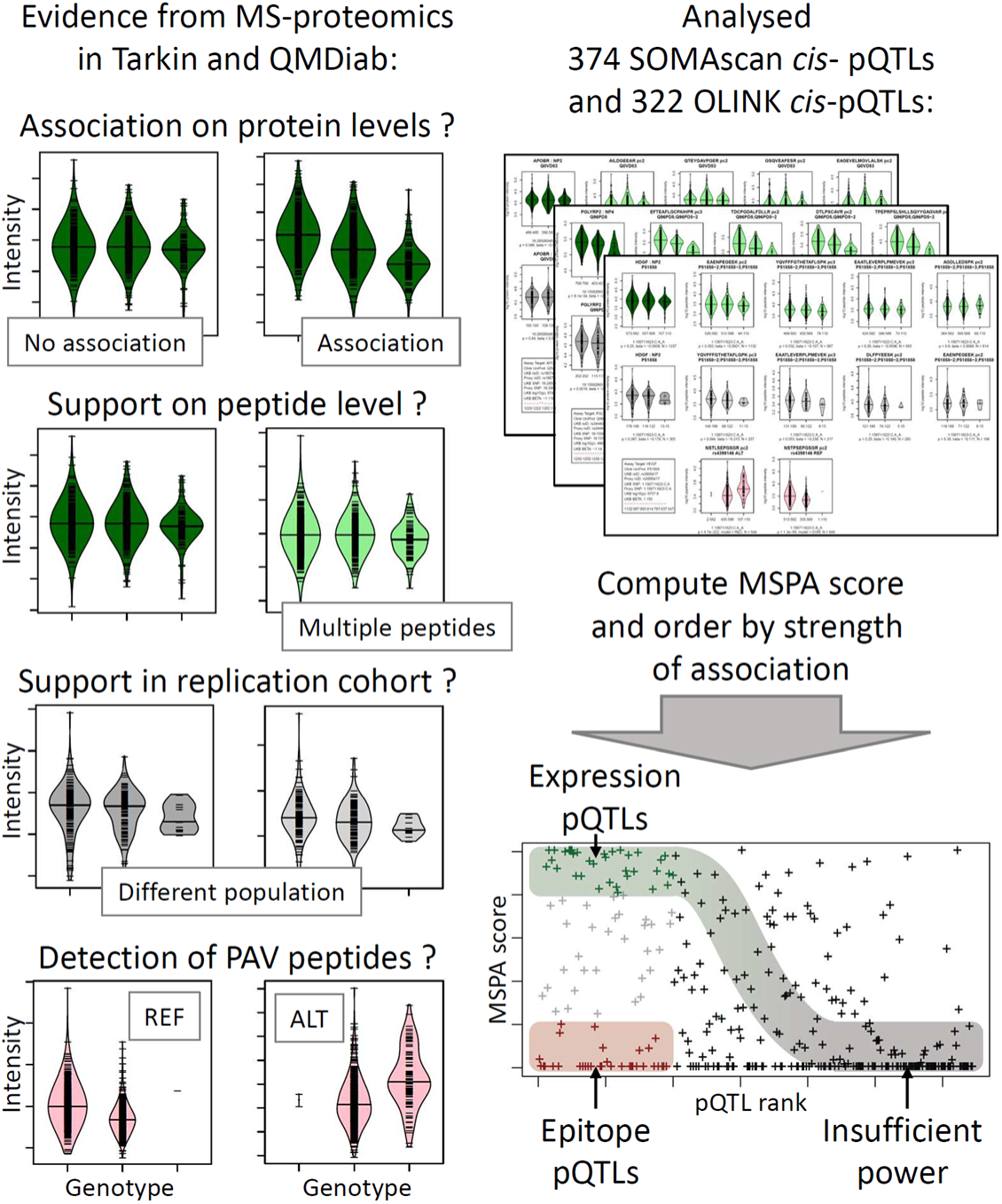

## INTRODUCTION

Genome-wide association studies (GWAS) with proteomics leverage the naturally occurring genetic variation in human populations and link differences between individual genomes to their effects on the proteome and beyond ^1^. Protein quantitative trait loci (pQTLs) are central in the drug discovery process as they provide supporting evidence for drug target identification and hypothesis generation on their modes of action ^2^. Most large-scale GWAS so far have been conducted using affinity proteomics platforms ^3-12^. The largest studies to date are from deCODE using the SOMAscan platform with 4,907 aptamers in 35,559 Icelandic samples from Icelanders ^13^ and from the UKB-PPP consortium using the Olink platform with dual antibodies targeting 2,923 proteins in 54,219 samples from participants of the UK Biobank ^14^. These studies reported thousands of pQTLs that are now available for further exploitation, such as Mendelian randomization (MR) experiments, to identify new drug targets and to further their development.

However, it should be noted that SOMAscan and Olink pQTLs represent genetic associations with protein binding affinity, rather than protein expression. This is because protein altering variants (PAVs) can modify the affinity binding epitopes of the proteins, thereby leading to genotype-dependent read-outs that do not correspond to real changes in protein levels ^15^. Such epitope effects can invalidate conclusions drawn from MR experiments, as their basic hypothesis requires that changes in the exposure, i.e. the protein expression levels, are causal for changes in the disease outcome. Epitope effects can also skew the prediction of protein levels using polygenic scores and confound correlations with other omics modalities. Therefore, it is important to validate key pQTLs on an independent platform that is immune against epitope effects, such as MS-based proteomics.

A few smaller-scale MS-based GWAS have been previously reported. Johansson et al. ^16^ identified and quantified the abundance of 1,056 tryptic-digested peptides, representing 163 proteins in the plasma of 1,060 individuals from two population-based cohorts. Xu et al. ^17^ conducted a GWAS for 304 proteins measured by SWATH-MS in blood serum from 2,958 Han Chinese individuals. Niu et al. ^18^ performed a GWAS for 420 proteins using MS-based proteomics in blood plasma from 1,914 children and adolescents with a replication in 558 adults. However, these studies were all limited to a small number of proteins.

Here we conduct a GWAS using the MS-based Proteograph™ platform (*Seer, Inc*.) for deep, unbiased proteomics ^15,19,20^. The Seer technology increases proteome coverage by using nanoparticle enrichment, followed by a data-independent acquisition protocol implemented on a Bruker timsTOF Pro 2 mass spectrometer (*Bruker Daltonics*). We have previously shown that the protein readouts of the Proteograph platform can reliably distinguish between epitope- and protein expression QTLs when a specific data analysis protocol is applied that eliminates PAV-containing peptides. We extended this approach here to a larger study cohort and a full GWAS ^15^. We analysed 1,980 proteins that were quantified in at least 80% of the samples, out of 5,753 proteins quantified across a discovery cohort of 1,260 participants of a diverse American ethnic background and a replication phase using 325 samples from participants of mainly Arab, Indian, and Filipino ethnic backgrounds. We then validated the protein associations with age and sex against those reported by the two largest affinity proteomics GWAS. Finally, we investigated the lead cis-pQTLs reported by the SOMAscan and Olink platforms for potential epitope effects.

## RESULTS

### A GWAS with the Seer Proteograph platform identified 252 and replicated 90 pQTLs

We conducted a GWAS with 1,980 proteins detected in >80% of the 1,260 samples from the Tarkin study (**Table S1**, see Methods). A total of 252 independent protein associations reached a Bonferroni level of significance (p < 2.5x10^-11^), involving 224 genetic loci for 152 different proteins (**Figure 1** and **Table S2**). Replication was attempted using 325 samples from the QMDiab study with matching genotype and proteomics data. A pQTL was considered replicated if its association reached a significance level of p < 0.05 / 252 and had concordant effect direction. A total of 90 pQTLs satisfied these criteria. Of the 94 pQTLs calculated to have 80% replication power, determined by sampling, a total of 60 pQTLs (63.8%) replicated, and most of the non-replicated pQTLs also had concordant directionality (**Figure 2A**). Differences in allele frequencies of the pQTLs between Tarkin and QMDiab were generally in the range of +/-10% allele frequency (**Figure 2B**).

**Figure 1:**
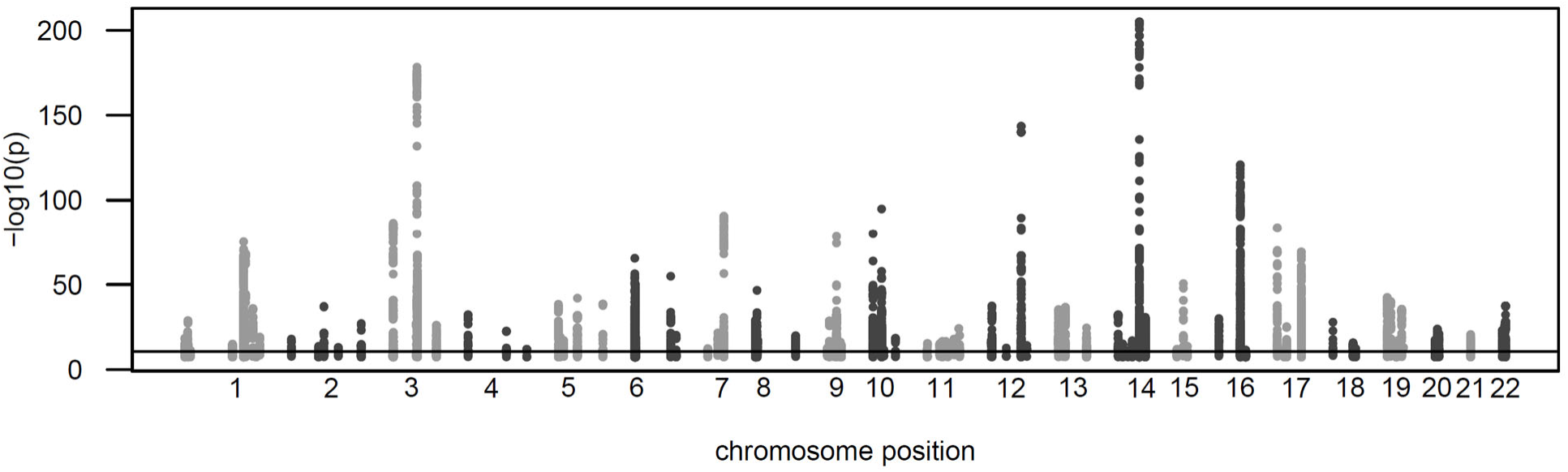
Manhattan plot. Shown are all protein associations that reached a significance level p < 5x10^-8^ in the discovery study.

**Figure 2:**
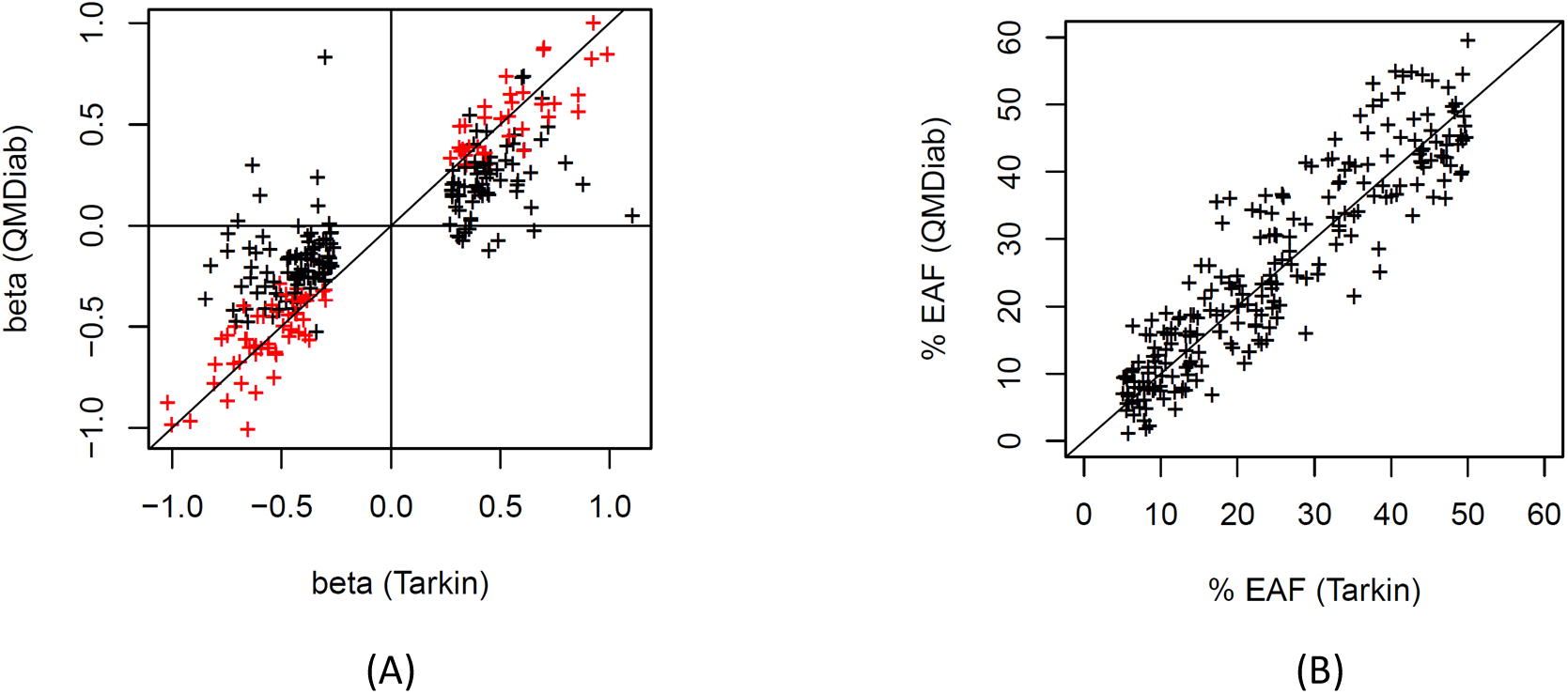
Effect size and effect allele frequencies. Scatterplot of the pQTL effect sizes from the discovery (Tarkin) and the replication (QMDiab) study, replicated loci are shown in red (A); Scatterplot of the effect allele frequencies (EAF), Tarkin versus QMDiab (B).

MS-based proteomics platforms generate a rich set of readouts, including quantifications of multiple peptides issued from a same protein, often repeatedly measured at multiple precursor charges. We generated visualizations of this data stratified by genotype for each of the 252 pQTLs (**Figure S1**). An example is given in **Figure 3**, where the associations with the four most frequently quantified peptides per protein are shown together with the derived protein level associations, both for Tarkin and QMDiab. Additionally, we present plots of PAV containing peptides, in cases they were detected for the protein in question. These plots provide individual verifiable experimental evidence that further supports the validity of these pQTLs.

**Figure 3:**
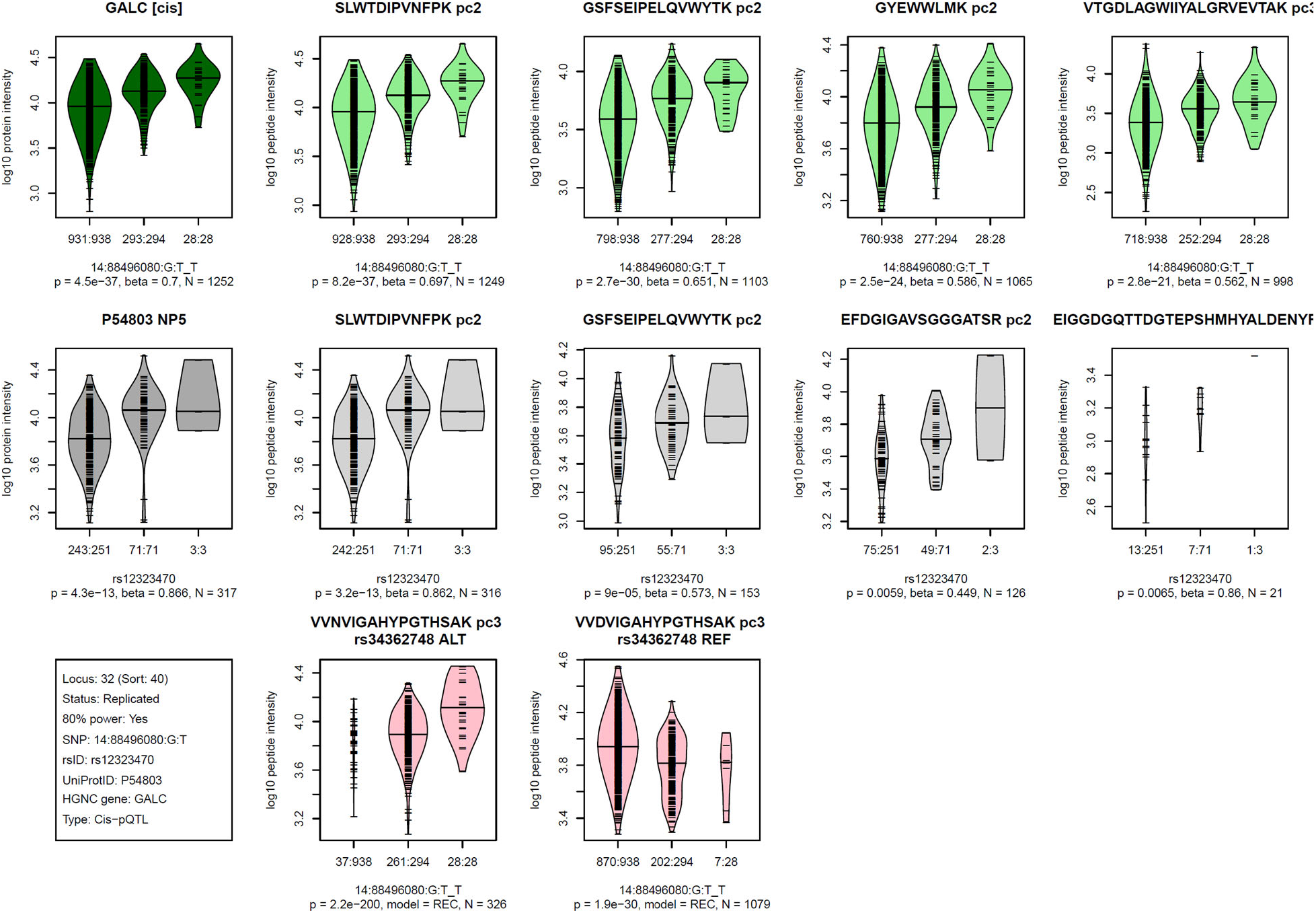
**Visualization of the Proteograph MS-proteomics data for a prototypical pQTL**. Violin plots of log-scaled engine-normalized protein and peptide intensities by genotype for the indicated protein and genetic variant (bottom left box, details are in **Table S2**); Top row (green): Tarkin data, using *PAV-exclusive* library, Middle row (grey): QMDiab data, using *PAV-exclusive* library, Bottom row (red): Tarkin data, using *PAV-inclusive* library, limited to PAV containing peptides; Plot titles indicate the protein UniProt identifier and the pQTL type (*cis*/*trans*) above the Tarkin protein plot, and HGNC gene name and nanoparticle number are above the QMDiab protein plot; Titles of the peptide plots indicate the respective peptide sequences and precursor charges (pc), and additionally the PAV variant rsID and allele type (REF/ALT) for the PAV peptide plots; Summary statistics are reported below the plots based on linear models with residualized and inverse-normal scaled data for associations with the *PAV-exclusive* library data (top two rows) and Fisher’s exact test for *PAV-inclusive* library data (bottom row); Associations with peptide data are sorted by increasing p-values from left to right and limited to a maximum of four plots; Whenever peptides were detected at multiple precursor charge values, only the strongest association was plotted; Numbers at the x-axis tick marks indicate the number of detected peptides by genotype followed by the number a samples with the corresponding genotype (e.g. 662:665); Genotypes are ordered as (1) other allele, (2) heterozygote, (3) effect allele, where the effect allele is indicated following the SNP name (chr:pos:ref:alt_eff, e.g. 3:49721532:G:A_A). Similar plots for all 252 pQTLs are provided as **Figure S2**.

We then asked which MS-based pQTLs had been identified before, and which were novel. Summary statistics from the deCODE SOMAscan study ^13^ and the UKB-PPP Olink study ^14^ were used to identify overlapping associations with previously reported pQTLs. Of the 252 pQTLs identified in this study, 65 were found by both platforms, 43 were reported by SOMAscan alone and 33 were seen by Olink alone. Four pQTLs were not reported by either of these studies, but were identified using PhenoScanner ^21^ as a pQTL in some other study. This leaves 107 pQTLs that have not been reported before, including 30 of the 90 replicated pQTLs (Table 1), suggesting an expected novel discovery rate when using the Proteograph platform of about one in three compared to existing affinity pQTLs.

**Table 1.**
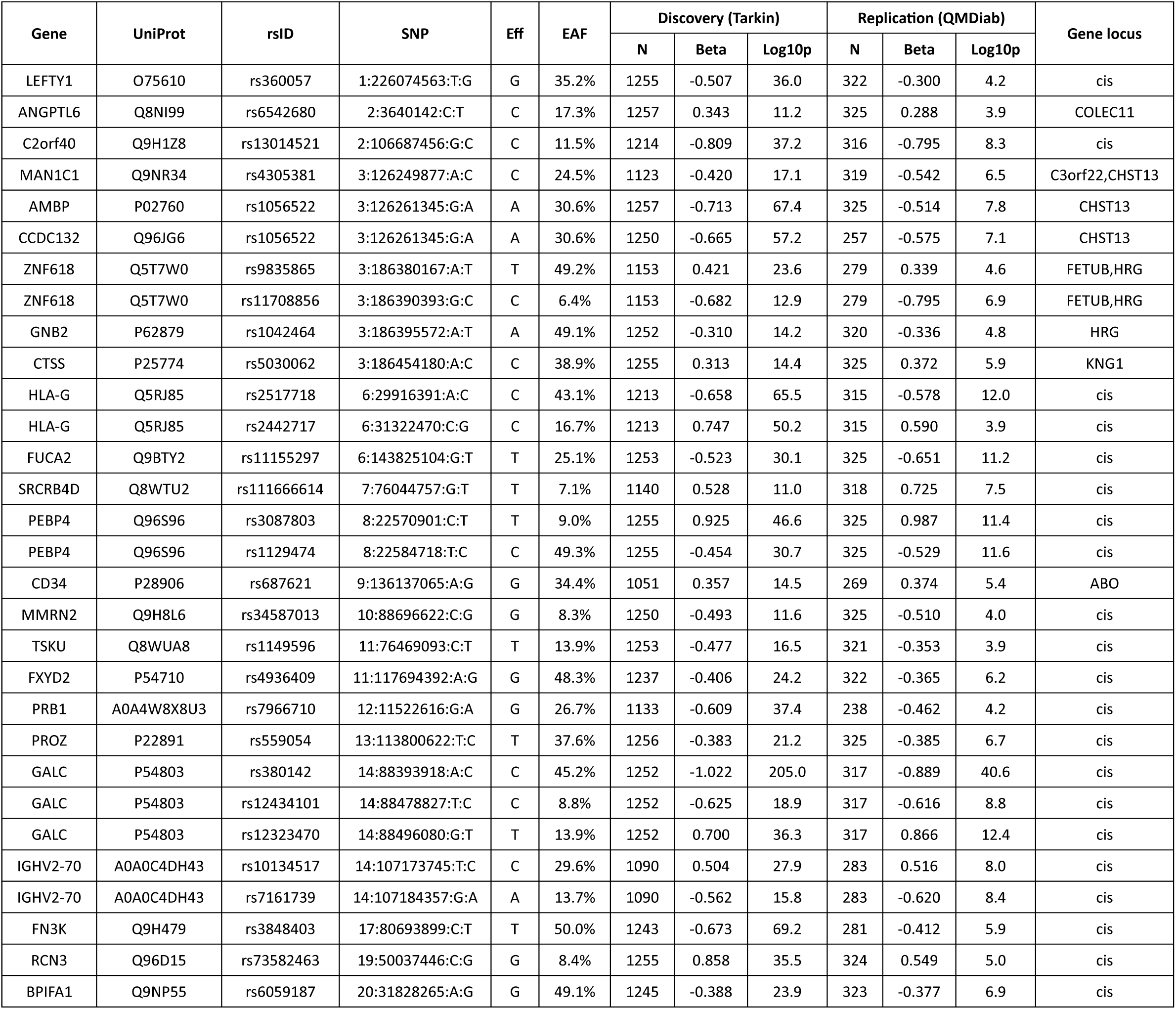
List of 30 previously unreported replicated pQTLs discovered using the Seer Proteograph platform.

### Age and sex associations were concordant between the affinity and MS based proteomics platforms

We computed the associations between all 1,980 protein readouts with age, sex, and the ten genotype principal components (**Table S3**). We then asked whether the associations with age and sex were concordant between the two affinity platforms and between the affinity and the MS proteomics platforms. A total of 507 proteins were quantified on all three platforms (**Figure S1A and Table S4**). The overall effect sizes were concordant for most of the proteins between all three platforms (**Figure 4**), although with some exceptions for the associations with age. Overall, 62 proteins shared significant associations with sex across all three platforms (**Figure S1B**), while 110 proteins exhibited significant associations with age (**Figure S1C**).

**Figure 4:**
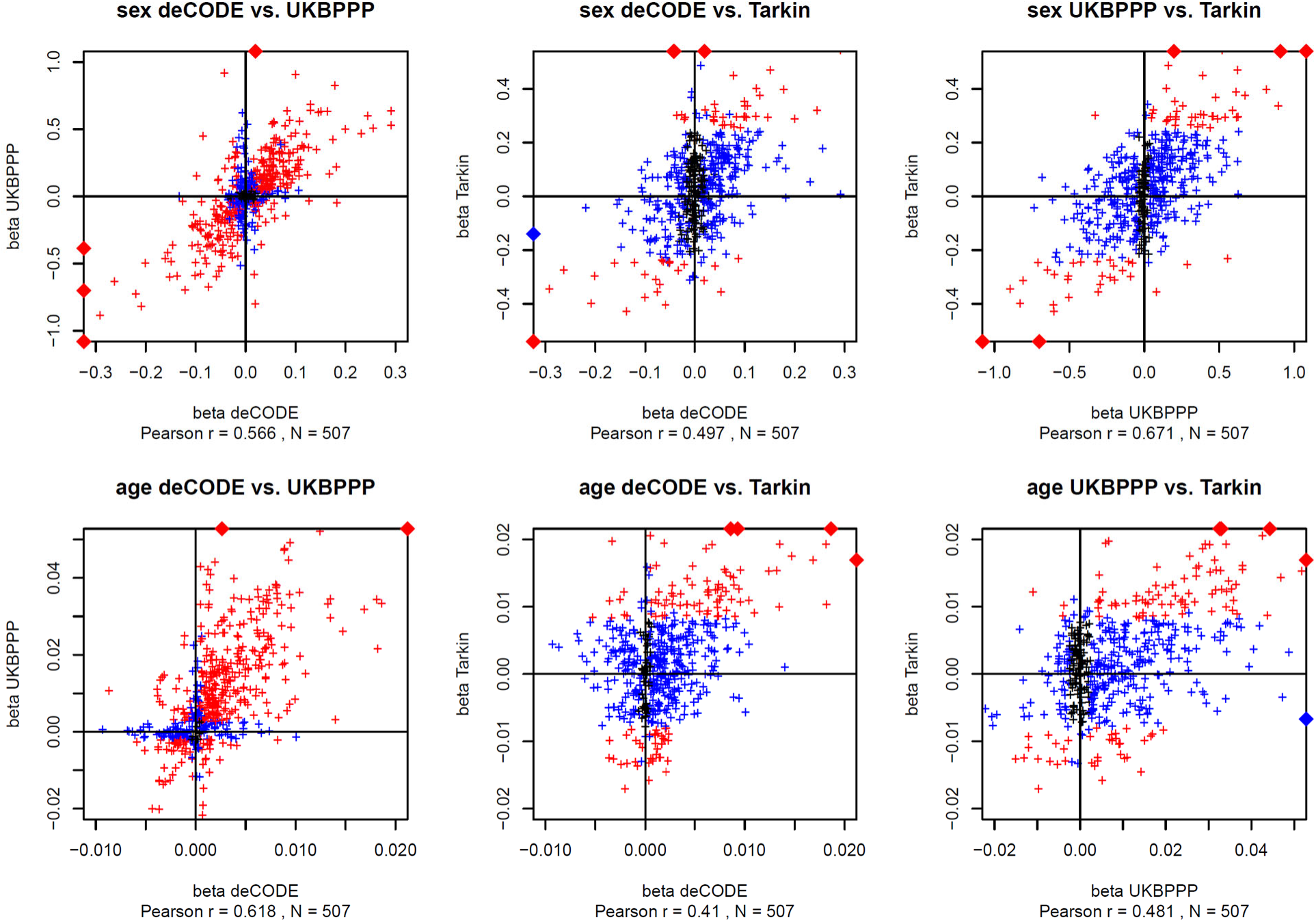
Scatterplot of the effect sizes for the protein associations with sex and age. Summary statistics for the associations with affinity proteomics were from the respective GWAS studies. Associations that reached Bonferroni significance in both respective studies are in red and in one study are in blue (p < 0.05 / number of reported associations). The effect sizes (beta) are reported in units of standard deviations (s.d.). Data points outside the plotting window are indicated by diamonds on the plot frames. Plot data are available in **Table ST4**. Scatterplots are limited to 507 unique proteins that were reported by all three studies. In cases where data for multiple affinity binders was reported, the most significant association was retained.

### One third of affinity proteomics pQTLs are potentially affected by epitope effects

The deCODE SOMAscan study ^13^ and the UKB-PPP Olink study ^14^ reported 1,881 and 1,917 cis-pQTLs, respectively (**Table S5 & S6**). For 1,041 and 1,415 of these pQTLs matching genetic variants were available in Tarkin and QMDiab. Out of these, 322 and 374 pQTLs also had the matching protein quantified in Tarkin with a missing rate of less than 20%. For these pQTLs the genetic associations with the matching protein and all corresponding peptide levels were computed using the Proteograph data, both for the Tarkin and the QMDiab study (**Table S7 & S8**). Note that most pQTLs without a matching variant in Tarkin had minor allele frequencies below 5% and were excluded from our analysis.

Quantification of protein levels from peptide intensities is sometimes a limiting factor in MS proteomics and prone to large uncertainties. We therefore integrated information from individual peptide measurements. For each pQTL the association data was integrated into what we refer to here as the MS peptide association (MSPA) score. This score is designed as a proxy for the likelihood of a protein expression signal being observed in the MS proteomics data and combines association signals from all peptides observed for a given protein. Note that PAV containing peptides had been removed from our MS peptide library. We defined the MSPA score as follows: All peptides were attributed a weight using the number of individual peptide detections for that protein, divided by the sum of all peptide detections for that protein. Repeated peptide detections with different precursor charges were excluded, retaining the peptide with the highest number of detections. The weights of all peptide associations where the 99% confidence interval of the beta estimate did not contain the zero were then summed up. An MSPA score of one therefore represents a case where all peptides support a genetic association with the protein expression level, and an MSPA score of zero indicates cases where the data provides no statistical support for such an association, suggesting either a lack of statistical power to detect an association or the presence of an epitope effect. As association directionality was not included into the score, and to identify possible artifacts, the associations for all analysed pQTLs were visualized for additional manual inspections; no inconsistencies were found (**Figure S2&S3**).

The MSPA score of each pQTL was then plotted against the rank of the p-value of the original pQTL from the affinity proteomics study. As shown in **Figure 5**, a sigmoid distribution running from the upper left to the lower right corner of the plot may be discerned for both studies. This curve approximately represents the MSPA score expected for a protein expression QTL as a function of pQTL rank as statistical power decreases with rank. Roughly, the first 100 pQTLs of each study appear to have sufficient power to replicate in Tarkin. Within these 100 strongest *cis-*pQTLs, both studies have almost the same proportions of protein expression QTLs (35% SOMA, 37% Olink) and the same number of likely epitope effect driven pQTLs (33% for SOMA and Olink). Hence, these plots suggest that about one third of the affinity proteomics pQTLs are possibly due to epitope effects, one third are reproducible using MS proteomics in a study of our current size, and one third fall into a “grey zone” where power may be an issue, but hybrids of epitope- and expression-pQTLs are also possible, that is, cases where a genetic variant interferes with the affinity binding, but at the same time affects protein expression via some biological feedback mechanism.

**Figure 5:**
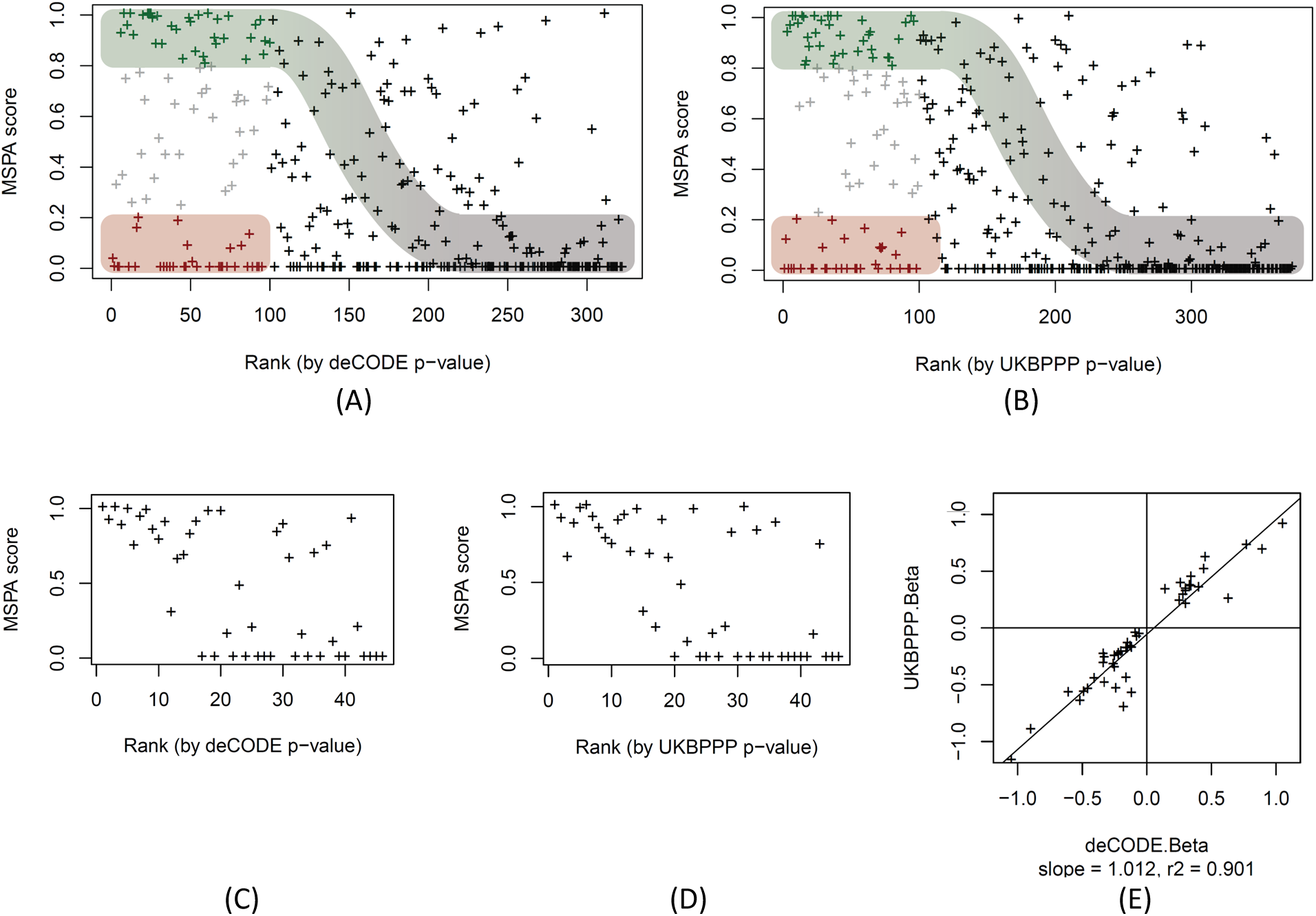
MS-peptide association score plotted by pQTL rank. Scatterplot of the MSPA scores against the rank of the affinity proteomics pQTLs of the deCODE SOMAscan (panel A, data in **Table S7**) and the UKB-PPP OLINK (panel B, data in **Table S8**) studies, ranked starting with the lowest p-value. The first 100 pQTLs (out of 322 pQTLs for SOMAscan and 374 pQTLs for Olink) are coloured to indicate likely protein expression QTLs (MSPA score > 0.8; green) and likely epitope effect driven pQTLs (MSPA score < 0.2; red), the sigmoid curve indicates the assumed dependence of power to detect a pQTL as a function of the strength of the association, approximated by the rank of the pQTL in the respective study; MSPA scores limited to 46 pQTLs that were reported on the same variant in deCODE (panel C) and UKBPPP (panel D); scatterplot of the effect size (beta) of the 46 pQTLs reported deCODE and UKBPPP (panel E, data in **Table S9**).

To further support the validity of the MSPA score as a proxy for the detection/non-detection of a genetic association and its potential to identify true positive protein expression pQTLs, we selected all pQTLs that were reported on the same genetic variant by deCODE and UKBPPP, and for which also matching protein and genetic data were also available in Tarkin and QMDiab. We used this set of 46 pQTLs as a gold standard for comparing effect sizes and directionality without the caveat of having to use proxy SNPs (**Table S9**). A few things were notable when looking at this set of pQTLs. Firstly, in contrast to the overall distribution of the MSPA scores in **Figures 5A and 5B**, where there were many high ranking pQTLs with low and zero MSPA scores present, the top ranking pQTLs in this set all had high MSPA scores (**Figures 5C and 5D)**, suggesting that none of these pQTLs were affected by epitope effects. This inference is further supported by the near perfect correlation of the effect sizes between the Olink and the SOMAscan platforms (**Figure 5E**). Considering that these pQTLs had been pre-selected based on the requirement that they were detected on both affinity platforms, it appears that a pQTL being detected by both affinity platforms is also a strong indicator for a true protein expression QTL and the absence of an epitope effect. This is a reasonable assumption because there is low likelihood that two different affinity binders target the same surface area of a protein and lead to a similar epitope readout.

If these 46 pQTLs are mostly devoid of epitope effects, the dependence of the MSPA values on the association strength (rank) should follow the power of each pQTL to be detected by the Tarkin study, supporting our earlier assumption that the high-ranking pQTLs with low MSPA scores in **Figure 5A** and **5B** are epitope-effect driven pQTLs. We therefore asked whether a candidate epitope-changing variant was identifiable in all these cases (the pQTLs in the red zone of **Figure 5A** and **5B**). We manually queried the Ensembl database ^22^ for coding variants that were potentially epitope-changing (**Table S10**) and found that for 23 out of 33 SOMA and 25 out of 33 Olink pQTLs such a variant had been reported (requiring LD r^2^>0.8). The lead pQTL SNP or a SNP in perfect LD of r^2^=1 was apparently epitope-changing in all but four of the 48 cases. Additionally, in 11 out of these 48 cases, a PAV-containing peptide was also detected on the Proteograph platform, with heterozygotes having about half the protein level, showing that the absence of a protein expression pQTL is not due to limitations in the quantification of the peptides, and confirming the presence of the protein variant in blood. In five cases (CPN2, SERPINA1, ENO3, HDGF, APOBR) both alleles, reference (REF) and alternate (ALT), were detected and showed significant associations with the coding variant, while all other peptides did not associate with the variant (for an example see **Figure 7**).

**Figure 6:**
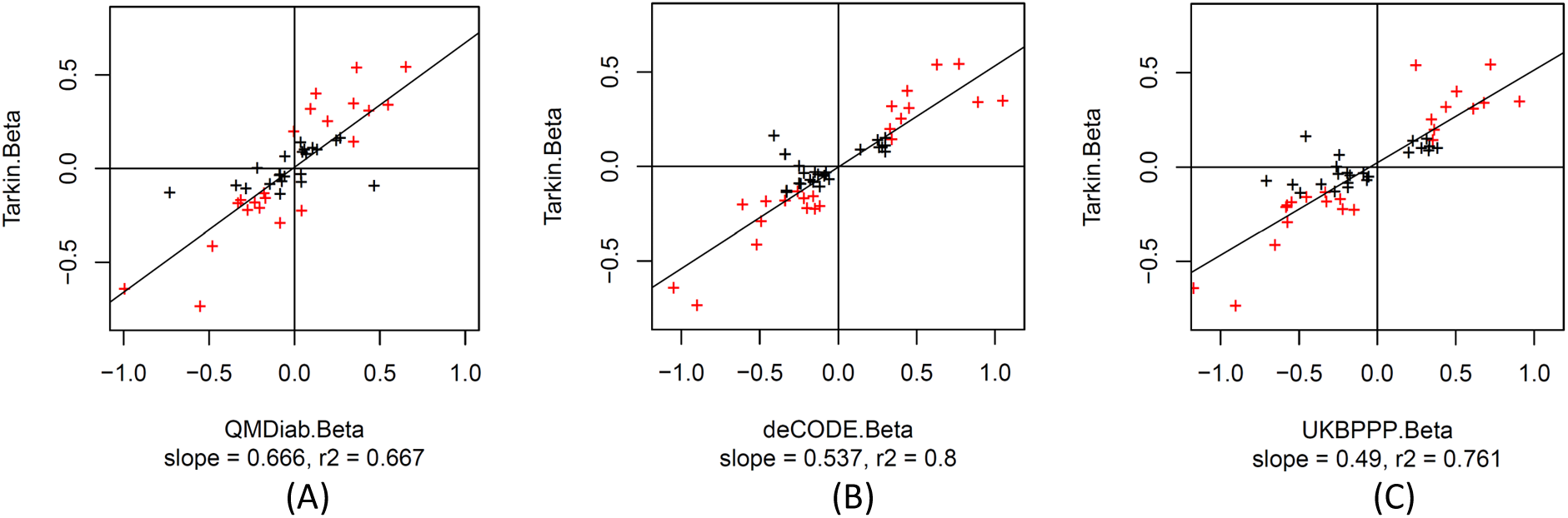
Scatterplot of the effect sizes (beta) of 46 pQTLs that were reported by deCODE and UKBPPP on the same SNP and for which association data was also available for Tarkin and QMDiab; Tarkin vs. QMDiab (A), Tarkin vs. deCODE (B), Tarkin vs. UKBPPP (C); associations that reach a significance level of p < 0.05 / 46 in Tarkin are in red (**Table S9**).

**Figure 7:**
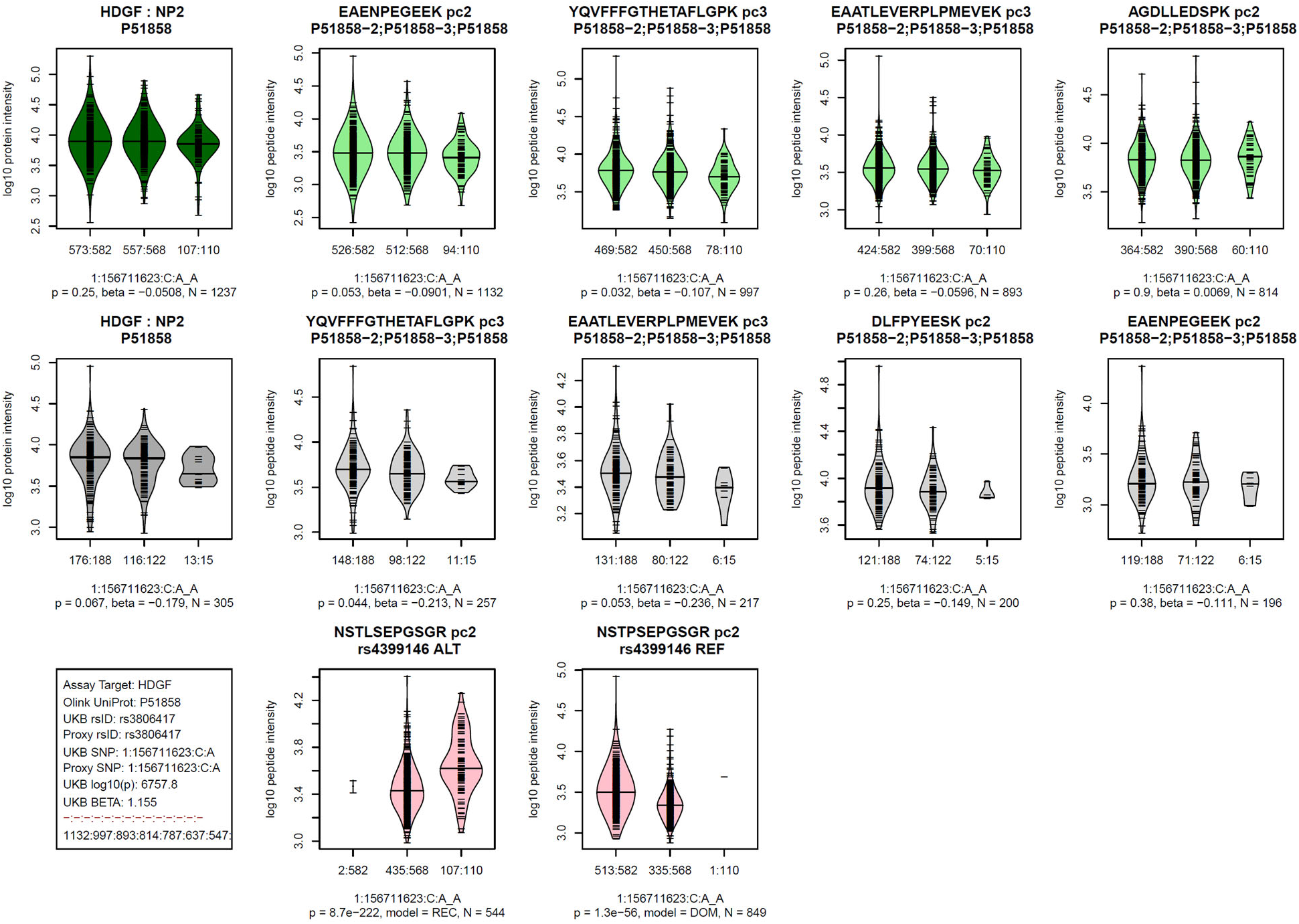
Example of a pQTL that is likely affected by an epitope effect. See legend of Figure 3 for legend; similar plots for all 374 pQTLs with OLINK data and for all 322 pQTLs with SOMAscan data are provided as **Figures S3** and **S4**; data is in **Tables S6** and **S7**.

## DISCUSSION

This study presents the first comprehensive GWAS using the MS-based Seer Proteograph platform, accompanied by a full replication, and employing a proteomics data analysis protocol that accounts for genetic variants within the analysed peptide ^15^. Our methodology not only identified new pQTLs on proteins previously unassessed by affinity proteomics platforms, but also examined previously reported affinity pQTLs for confounding due to epitope-altering variants. We estimate that about one-third of the 200 pQTLs evaluated were influenced by such effects. However, due to limitations in statistical power that restricted evaluation to only the strongest affinity pQTLs, which are the ones most likely to be enriched for epitope effects due to ascertainment bias, this estimate should be considered as an upper bound.

We reported a total of 252 pQTLs, with 90 successfully replicated in an independent population. While the replication rate of 63.8% for sufficiently powered pQTLs is high, it falls below the theoretically expected 80%. We attribute this difference to genetic and probably also lifestyle differences between the discovery and replication cohorts, which are ethnically very distinct. Nonetheless, this diversity also enhances the robustness and translatability of the replicated pQTLs across populations.

The use of a proteome library that accounts for PAVs was central to our study. Without using this approach, a very high number of false positive pQTLs would have been detected, as we previously discussed ^15^. Traditional proteomics data analysis methods often rely on a limited peptide library for protein quantification, and the presence of a single peptide with a large effect can skew the quantification. A PAV containing peptide would not be detected in homozygotes of the alternate allele and heterozygotes would have half the peptide level. The inclusion of PAV containing peptides into the protein quantification would hence lead to the equivalent of an epitope effect, that is, a pQTL signal where in reality, there is no genotype dependence of the protein expression level. Also, PAV peptides corresponding to the alternate allele are not present in an in-silico digest library of the standard UniProt database. Indeed, using data from a standard proteomics data processing run, we observed cases where fragment spectra of these peptides miss-matched to peptides from other proteins and in extreme cases led to false protein identifications.

We used the associations with sex and age to externally validate the comparability of the affinity and MS proteomics readouts to predict non-genetic outcomes. Although our MS-based study was less powered compared to the two much larger affinity proteomics studies, the scatterplots of the effect sizes between the two large studies and one of the large studies and our study are largely comparable. Key signals, such as the association of sex hormone binding globulin (SHBG) with sex and leptin (LEP) with age, were significant on all platforms. There are more age-related associations with conflicting effect sizes. These may be population specific and lifestyle related effects. Noteworthy is also the higher number of positive associations with age that is present across all three platforms, which we can only speculate about.

Our study also has some caveats. The interpretation of absence of pQTL signal in MS proteomics as indicative of epitope effects necessitates formal power analysis. However, such an approach is challenging because it requires precisely estimating the effect sizes for MS-based data from associations using affinity proteomics, whereas effect sizes are measured in relative units of standard deviations within the study population that do not translate between platforms and studies. Thus, we limited our analysis to the 100 strongest pQTLs in each study, under the assumption that these are sufficiently powered to replicate using MS proteomics, if the signal is truly based on protein expression rather than affinity binding. We support this argument using the distribution of the MSPA scores as discussed around **Figure 5**.

As we interpret the absence of a pQTL signal using an MS platform as an indicator of a potential epitope effect, we applied a stringent missingness criterion (<20%). The total number of 1,980 proteins analysed in this GWAS are therefore lower than the 5,753 proteins that were quantified in any sample across the study.

Additionally, the matching of proteins between platforms using UniProt identifiers can be challenging, as affinity proteomics platforms sometimes report multiple protein identifiers when they are targeting protein complexes or use the UniProt identifier specific to the proteoform employed to generate the affinity binder while alternate proteoforms may be included in the MS library, occasionally leading to annotations by protein groups rather than specific proteins. Mapping of UniProt ids to gene names can also be challenging, especially when the versions of the underlying databases do not match between studies.

While we reported many novel pQTLs, about one third of the identified pQTLs, we opted not to highlight any new biologically relevant findings. We believe that the biomedical relevance of pQTLs has already been amply shown in many of the previous pQTL studies, and that cherry-picking one or two new highlights would distract from the central message of our paper, which is to demonstrate the complementarity of MS proteomics to affinity approaches in (1) validating associations that may be driven by epitope effects and (2) substantially extending the panel of proteins accessible to pQTL studies.

## METHODS

### Ethics

The original Tarkin study was reviewed and approved by the IRCM IRB and the Mass General Brigham IRB. Participants provided written informed consent to take part in the study. The WCMQ IRB determined that use of the Tarkin data for the present project does not meet the definition of human research for this study (IRB document HRP-532). The QMDiab study was approved by the institutional research boards of Weill Cornell Medicine – Qatar under protocol #2011-0012 and of Hamad Medical Corporation under protocol #11131/11 and complies with all relevant ethical regulations. For forthgoing work with the study a non-human subject research determination was obtained. The study design and conduct complied with all relevant regulations regarding the use of human study participants and was conducted in accordance with the criteria set by the Declaration of Helsinki.

### Cohorts

The samples used in the Tarkin study were obtained from the Massachusetts General Brigham (MGB) Biobank. Joint phenotype and genotype data were available for 1,260 samples, comprised of 662 females and 598 males with an age range of 23-99 years (median 70 years, mean 67.2 years). 1,057 of the participants were white. This subset of MGB samples together with its deep omics characterization is referred to here as the “Tarkin study” in this paper ^23^. For replication, a total of 345 previously unthawed citrate blood plasma samples from participants of the Qatar Metabolomics study of Diabetes (QMDiab), including adult female and male participants of predominantly Arab, Indian, and Filipino ethnic background, with and without diabetes in an age range from 18-80 were assayed using the Proteograph platform (*Seer Inc*.) ^15,24,25^.

### Genotyping

Imputed genotype data for 1,980 samples of the Tarkin study was received on a per-chromosome basis in vcf format (build 37, imputed using Minimac3, no indels). The genotype data was filtered for biallelic variants and variant names were standardized using bcftools (version 1.16), converted to plink format and filtered using plink2 (version v2.00a5LM) with the options --geno 0.1 --mac 10 –maf 0.05 --hwe 1E-15. For 1,260 samples proteomics data was available. These samples were merged into a single genotype file and further filtered using plink2 with the options --maf 0.05 --hwe 1E-6 --geno 0.02, leaving data for 5,461,287 genetic variants. The first ten genetic principal components were then computed using plink2 with the --pca option. QMDiab samples were genotyped using the Illumina Omni 2.5 array (version 8) and imputed using the SHAPEIT software with 1000 Genomes (phase3) haplotypes. Genotyping data was available for 325 of the 345 samples with proteomics data.

### Proteomics

The workflow used for the proteomics analyses of the Tarkin and the QMDiab samples were essentially identical and have been described in detail before ^15^. Briefly, plasma samples were prepared using the Proteograph workflow ^19,20^ (*Seer, Inc.*) to generate purified peptides that were then analyzed using a dia-PASEF method ^26^ on a timsTOF Pro 2 mass spectrometer *(Bruker Daltonics)*. Each study was conducted at independent times using two mass spectrometers. One mass spectrometer coincidentally was reused in both studies. DIA-NN (1.8.1) ^27^ was used to derive peptide and protein intensities. A library-free search based on UniProt version UP000005640_9606 was used, processing the data a second time using the match between runs (MBR) option. Two additional libraries were created, one excluding common (MAF > 10%) protein-altering variant (PAV) containing peptides, and one injecting the alternate alleles into the reference protein sequences. These libraries were referred to as the *PAV-exclusive* library and the *PAV-inclusive* library, respectively. Details of this library generation process have been described elsewhere ^15^. DIA-NN’s normalized intensities (PG.Normalised) were used as protein readout. 5,753 unique protein groups were quantified, 4,109 were detected in at least 20% of the samples, and 1,980 had less than 20% of missing values. Wherever a protein was detected at this level in more than one nanoparticle run, the one with the largest sum of protein intensities was retained. Protein levels were then log-scaled, residualized using age, sex and the ten first genotype principal components, and finally inverse-normal scaled.

### Genome-wide association

Genetic associations of 1,980 residualized and inverse-normal scaled protein levels with 5,461,287 genetic variants in 1,260 samples were evaluated using linear models (plink v1.90b7.1, option --linear). Missing data points were excluded. The Bonferroni significance level for protein associations was p_Bonf_ = 5x10^-8^/1,980 = 2.5x10^-11^. For six proteins, the residualization did not entirely remove association with these confounders. Three of these proteins presented with inflated GWAS statistics and the corresponding pQTLs were removed (UniProt V9GYE7, B1AKG0, and O15230). Protein associations on correlated variants were clumped on a per-trait basis using an LD cut-off of r^2^=0.1 For each cluster the trait sentinel association was identified. Sentinel associations were then clumped using an LD cut-off of r^2^=0.9 into loci.

### Replication

Replication was attempted using data from 325 samples of the QMDiab study that had joint genotype and proteomics information. Power to replicate 80% of the pQTLs was determined as the 80% quantile of the p-values obtained from 1,000 random samples from Tarkin data set using the number of samples available in the QMDiab for the respective genotype-protein pair.

### Overlap with previous OLINK and SOMAscan pQTLs

Summary statistics from UKB-PPP Olink study ^14^ and deCODE SOMAscan study ^13^ were downloaded from the respective sites. Associations were reported for 4,660 proteins by deCODE and for 2,908 proteins by UKB-PPP. All protein associations on variants matching one of the Proteograph pQTLs were retrieved. An association reported by deCODE and Olink was considered as significant at levels of p < 0.05/139/4660 and p < 0.05/117/2908, respectively, accounting for the number of evaluated variants and proteins on the panel.

### Evaluation of age and sex associations

Summary statistics for age and sex associations were retrieved from the from the supplementary Excel files of Ferkingstad *et al*. ^13^ for deCODE SOMAscan (sheet ST01) and of Sun *et al*. ^14^ for UKB-PPP Olink (sheet ST5). Associations with age and sex for the Tarkin study were computed with PLINK using identical datasets and models as for the genome-wide association (option -- linear no-snp). Proteins were matched using UniProt identifiers. In rare cases when there were matches to multiple protein groups, the strongest association was retained.

### Evaluation of OLINK and SOMAscan cis-pQTLs

Cis-pQTLs were obtained from the supplementary tables of the respective studies (ST02 from Ferkingstad *et al*. ^13^ for deCODE SOMAscan and ST10 from Sun *et al*. ^14^ for UKB-PPP OLINK). pQTLs were limited to variants that were located on autologous chromosomes, had a suitable replication SNP available in both, Tarkin and QMDiab (LD r^2^ > 0.8), and a minor allele frequency greater than 5% in the Tarkin study. Matching of proteins between platforms was done using Uniprot IDs, allowing for matches to protein groups that contained the Uniprot ID. Ambiguous cases where more than one matching protein group was found were omitted. In cases where protein readouts for multiple nanoparticle runs where available, the one with the highest single number of peptide detections was used. Cases where the number of quantified proteins in Tarkin was below 80% were excluded.

### Annotation of pQTLs

snipa.org ^28^, phenoscanner.medschl.cam.ac.uk ^21^ and omicsciences.org ^29^ were used to annotate pQTLs with overlapping information on disease GWAS, gene expression and metabolomics QTLs. Protein epitope changing variants were identified using Ensembl ^22^.

## DATA AVAILABILITY STATEMENT

The MS-proteomics data for the Tarkin study has been deposited with ProteomeXchange under project accession code PXD048709 and will be made public at time of publication. The MS-proteomics data of QMDiab is already available on ProteomeXchange with identifier PXD042852. Summary statistics of the GWAS will be deposited in the GWAS catalog. Consent obtained from the QMDiab study participants does not allow deposition of genetic information in public databases. Researcher affiliated with a research institution may request access to genetic data on an individual basis from the corresponding author (Karsten Suhre, Weill Cornell Medicine - Qatar, Doha, Qatar). Access is subject to approval by the institutional research board of Weill Cornell Medicine - Qatar.

## ACKNOWLEDGEMENTS

We are grateful to all study participants for providing their time and blood. K.S. and F.S. are supported by the Biomedical Research Program at Weill Cornell Medicine in Qatar, a program funded by the Qatar Foundation. K.S. is also supported by Qatar National Research Fund (QNRF) grants NPRP11C-0115-180010 and ARG01-0420-230007. J.L-S. is supported by the National Institute of Health grants R01HL123915, R01HL155742, and U19AI168643. The statements made herein are solely the responsibility of the authors.

We thank Guhan Venkataraman, Amir Alavi and Jian Wang for developing software workflows that were used in the proteogenomic analyses in this work.

## AUTHOR CONTRIBUTION STATEMENT

Financial Support: K.S., J.L-S

Study design: K.S., J.L-S.

Data analysis: K.S., H.G., S.B.

Provided Materials and Conducted Experiments: Q.C., A.H., K.M., A.D., N.S., G.T., H.S., V.B.D., F.S.

Manuscript writing: K.S., S.B., J.L-S.

All authors contributed to the interpretation of the results and critically reviewed the manuscript.

## Supplementary Material

**Supplementary Tables** [provided as a multi-sheet MS Excel file].

|  | A | B |
| --- | --- | --- |
| 1 | **Supplementary Tables** |  |
| 2 |  |  |
| 3 | **Table** | **Content** |
| 4 | ST1 | Annotated list of 1,980 proteins assayed in this GWAS |
| 5 | ST2 | Annotated list of 252 pQTLs identified in this study |
| 6 | ST3 | Summary statistics for association of the 1,980 with age, sex, and ten genotype principle components (PCs) |
| 7 | ST4 | 6498 proteins with age & sex associations in at least one of deCODE, UKBPPP, and Tarkin |
| 8 | ST5 | 1,881 pQTLs from deCODE, annotated with association data with proteins and peptides in Tarkin and QMDiab |
| 9 | ST6 | 1,917 pQTLs from UKB-PPP, annotated with association data with proteins and peptides in Tarkin and QMDiab |
| 10 | ST7 | 322 pQTLs from deCODE, scored with MSPA |
| 11 | ST8 | 374 pQTLs from UKB-PPP, scored with MSPA |
| 12 | ST9 | 46 cis-pQTLs reported both by deCODE and UKBPPP, evaluated on same SNPs in Tarkin and QMDiab |
| 13 | ST10 | 66 cis-pQTLs with potential epitope-effect generating variants |

**Figure S1:**
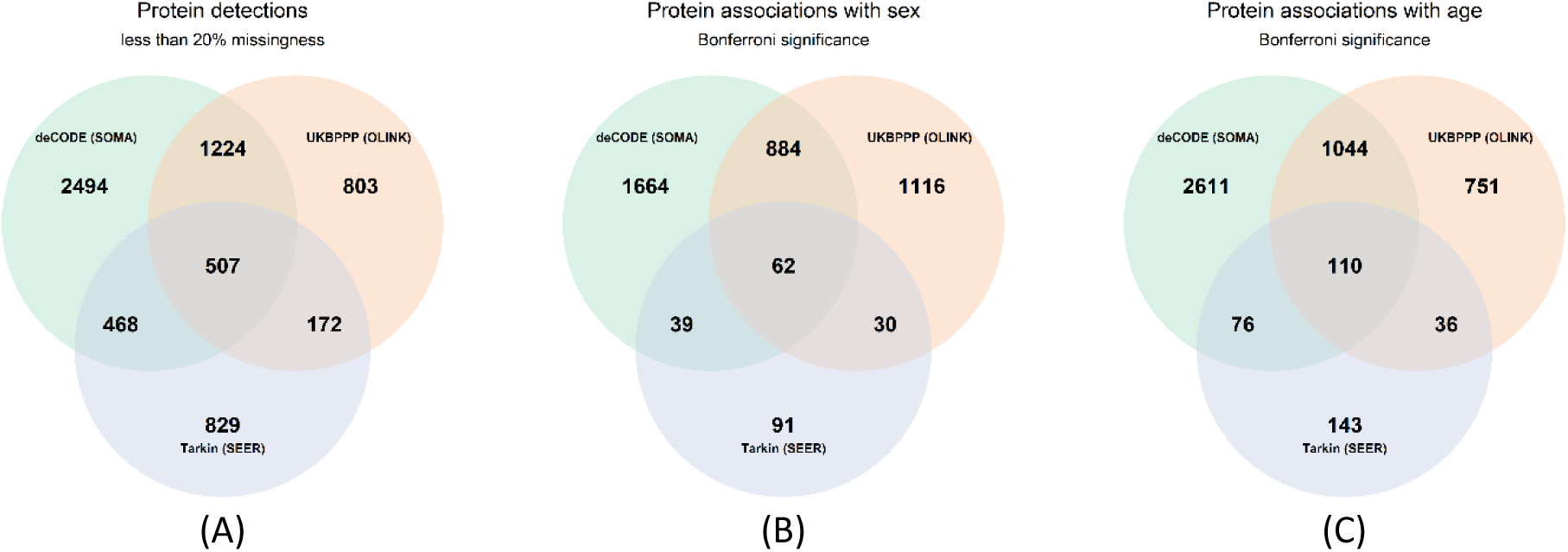
Venn diagrams. Proteins reported by the three studies, 1,980 proteins with <20% missingness for Tarkin, out of 5,753 proteins quantified across the discovery cohort (A); Proteins associated with sex (B) and age (C) at a significance level of 0.05 / number of proteins reported.

The following Figures are provided as separate PDF files:

**Figure S2: Violin plots of protein and peptide levels by genotype for 252 pQTLs identified in this study.** See Figure 3 for legend. File: Figure_S2_PGWAS_replicate_check.20240523.pdf

**Figure S3: Violin plots of protein and peptide levels by genotype for 322 pQTLs identified by the deCODE OLINK study.** See Figure 7 for legend. File: Figure_S3_PGWAS_test_epitope_SOMA.20240523.pdf

**Figure S4: Violin plots of protein and peptide levels by genotype for 374 pQTLs identified by the UKB-PPP SOMAscan study.** See Figure 7 for legend. File: Figure_S4_PGWAS_test_epitope_OLINK.20240523.pdf

