## Supplementary material for "A genome-wide association study of mass spectrometry proteomics using the Seer Proteograph platform": Figure S3

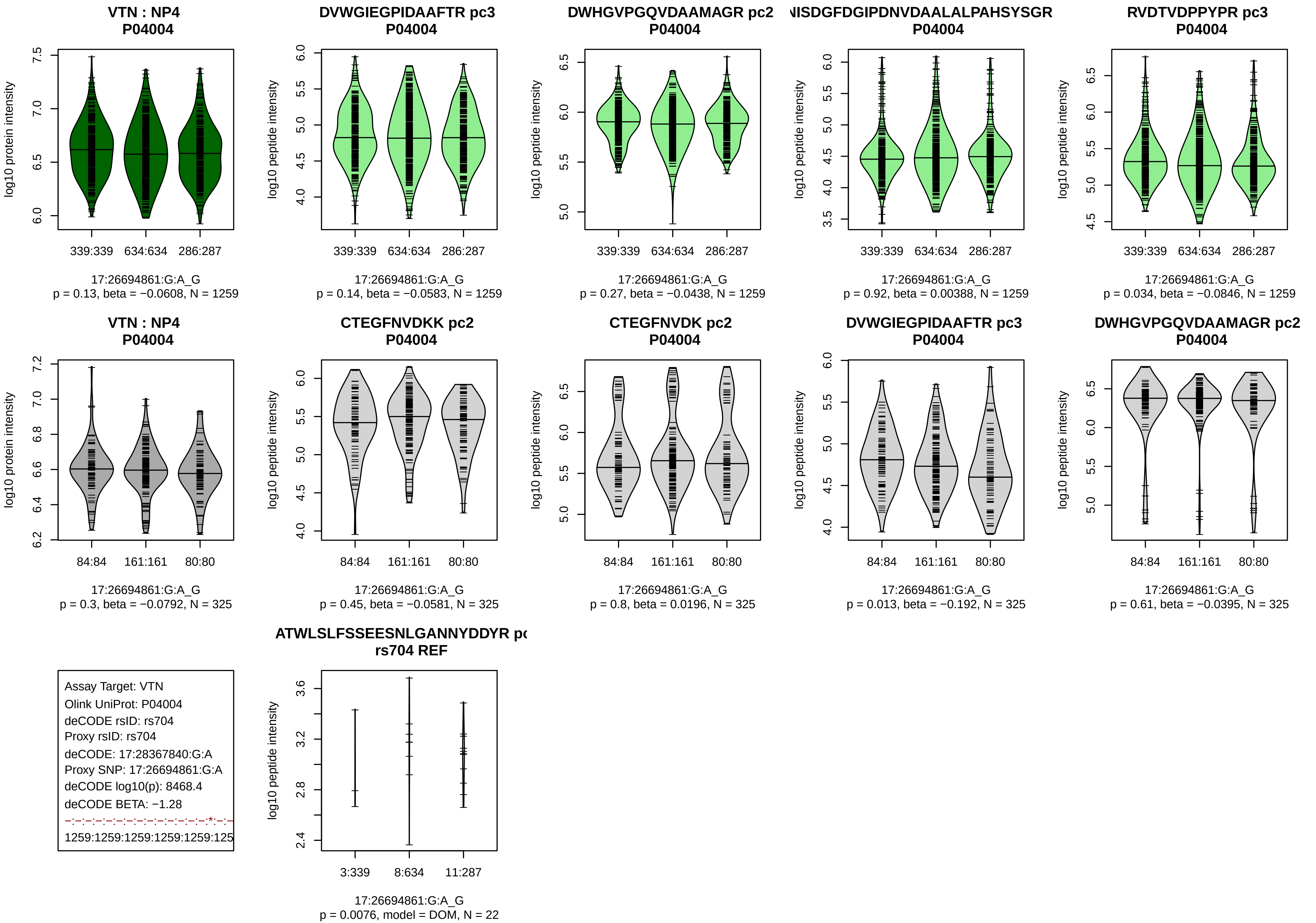
VTN : NP4
P04004
log10 protein intensity
339:339 634:634 286:287
17:26694861:G:A_G
p = 0.13, beta = −0.0608, N = 1259
DVWGIEGPIDAAFTR pc3
P04004
log10 peptide intensity
339:339 634:634 286:287
17:26694861:G:A_G
p = 0.14, beta = −0.0583, N = 1259
DWHGVPGQVDAAMAGR pc2
P04004
log10 peptide intensity
339:339 634:634 286:287
17:26694861:G:A_G
p = 0.27, beta = −0.0438, N = 1259
NISDGFDGIPDNVDAALALPAHSYSGR
P04004
log10 peptide intensity
339:339 634:634 286:287
17:26694861:G:A_G
p = 0.92, beta = 0.00388, N = 1259
RVDTVDPPYPR pc3
P04004
log10 peptide intensity
339:339 634:634 286:287
17:26694861:G:A_G
p = 0.034, beta = −0.0846, N = 1259
VTN : NP4
P04004
log10 protein intensity
84:84 161:161 80:80
17:26694861:G:A_G
p = 0.3, beta = −0.0792, N = 325
CTEGFNVDKK pc2
P04004
log10 peptide intensity
84:84 161:161 80:80
17:26694861:G:A_G
p = 0.45, beta = −0.0581, N = 325
CTEGFNVDK pc2
P04004
log10 peptide intensity
84:84 161:161 80:80
17:26694861:G:A_G
p = 0.8, beta = 0.0196, N = 325
DVWGIEGPIDAAFTR pc3
P04004
log10 peptide intensity
84:84 161:161 80:80
17:26694861:G:A_G
p = 0.013, beta = −0.192, N = 325
DWHGVPGQVDAAMAGR pc2
P04004
log10 peptide intensity
84:84 161:161 80:80
17:26694861:G:A_G
p = 0.61, beta = −0.0395, N = 325
Assay Target: VTN
Olink UniProt: P04004
deCODE rsID: rs704
Proxy rsID: rs704
deCODE: 17:28367840:G:A
Proxy SNP: 17:26694861:G:A
deCODE log10(p): 8468.4
deCODE BETA: −1.28
1259:1259:1259:1259:1259:125
ATWLSLFSSEESNLGANNYDDYR pc
rs704 REF
log10 peptide intensity
3:339 8:634 11:287
17:26694861:G:A_G
p = 0.0076, model = DOM, N = 22

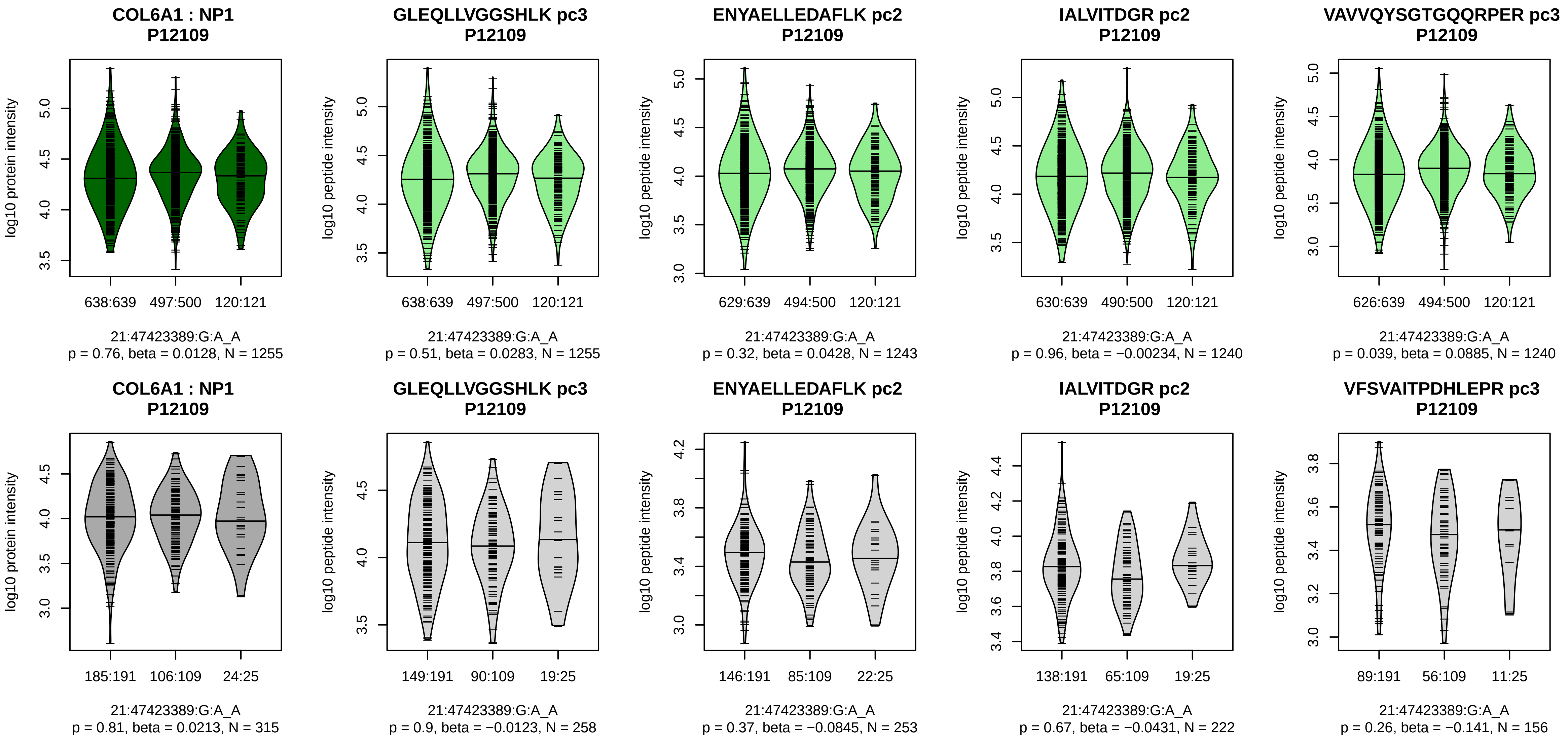


Assay Target: COL6A1
Olink UniProt: P12109
deCODE rsID: rs1053312
Proxy rsID: rs1053312
deCODE: 21:46003475:A:G
Proxy SNP: 21:47423389:G:A
deCODE log10(p): 7739.5
deCODE BETA: −1.23
–:–:–:–:–:–:–:–:–:–:–:–:–:–:–:–:
1255:1251:1243:1240:1240:124

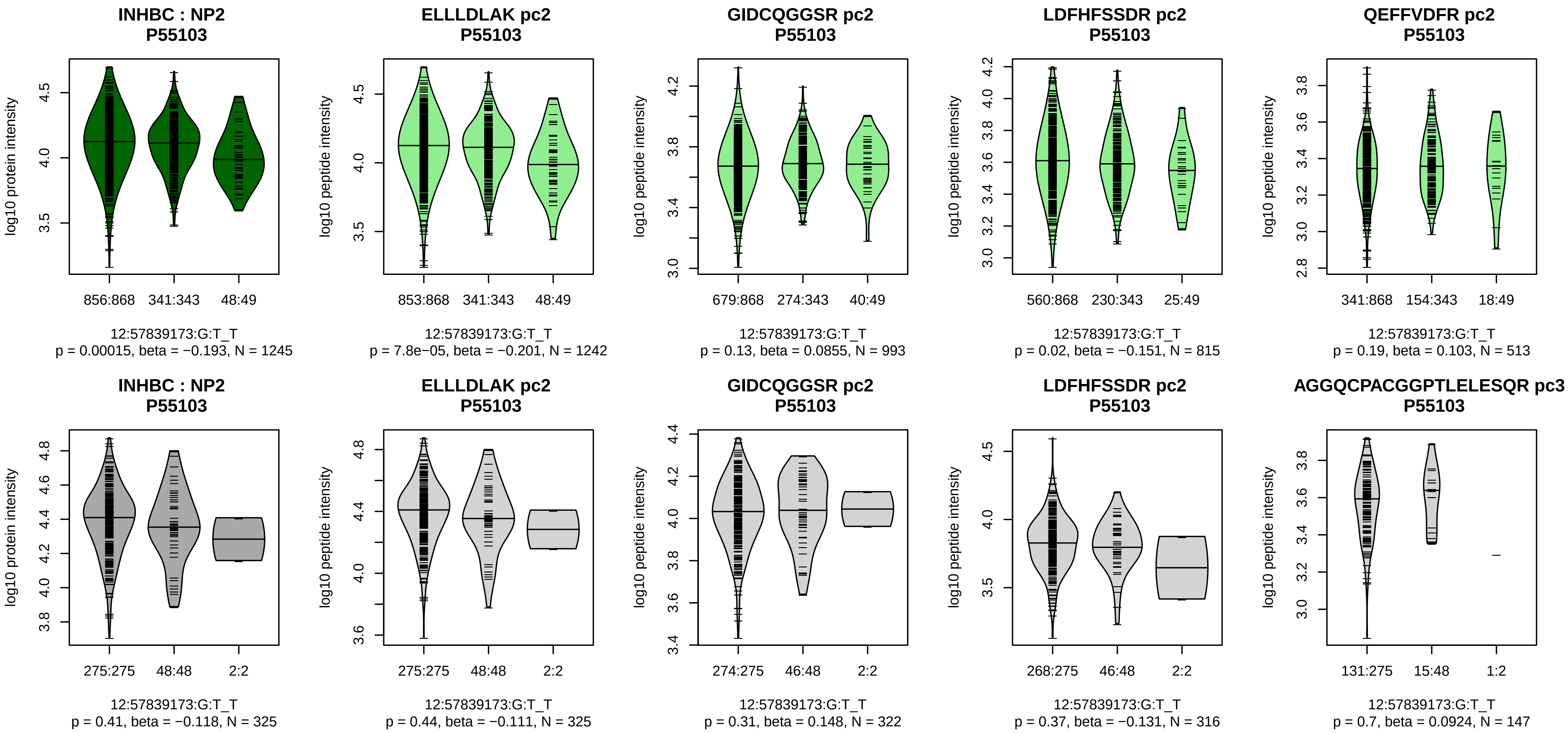


Assay Target: INHBC
Olink UniProt: P55103
deCODE rsID: rs61352607
Proxy rsID: rs61352607
deCODE: 12:57445381:GAAAA
Proxy SNP: 12:57839173:G:T
deCODE log10(p): 7136.9
deCODE BETA: −1.25
*:−:−:−:−
1242:993:815:513:260

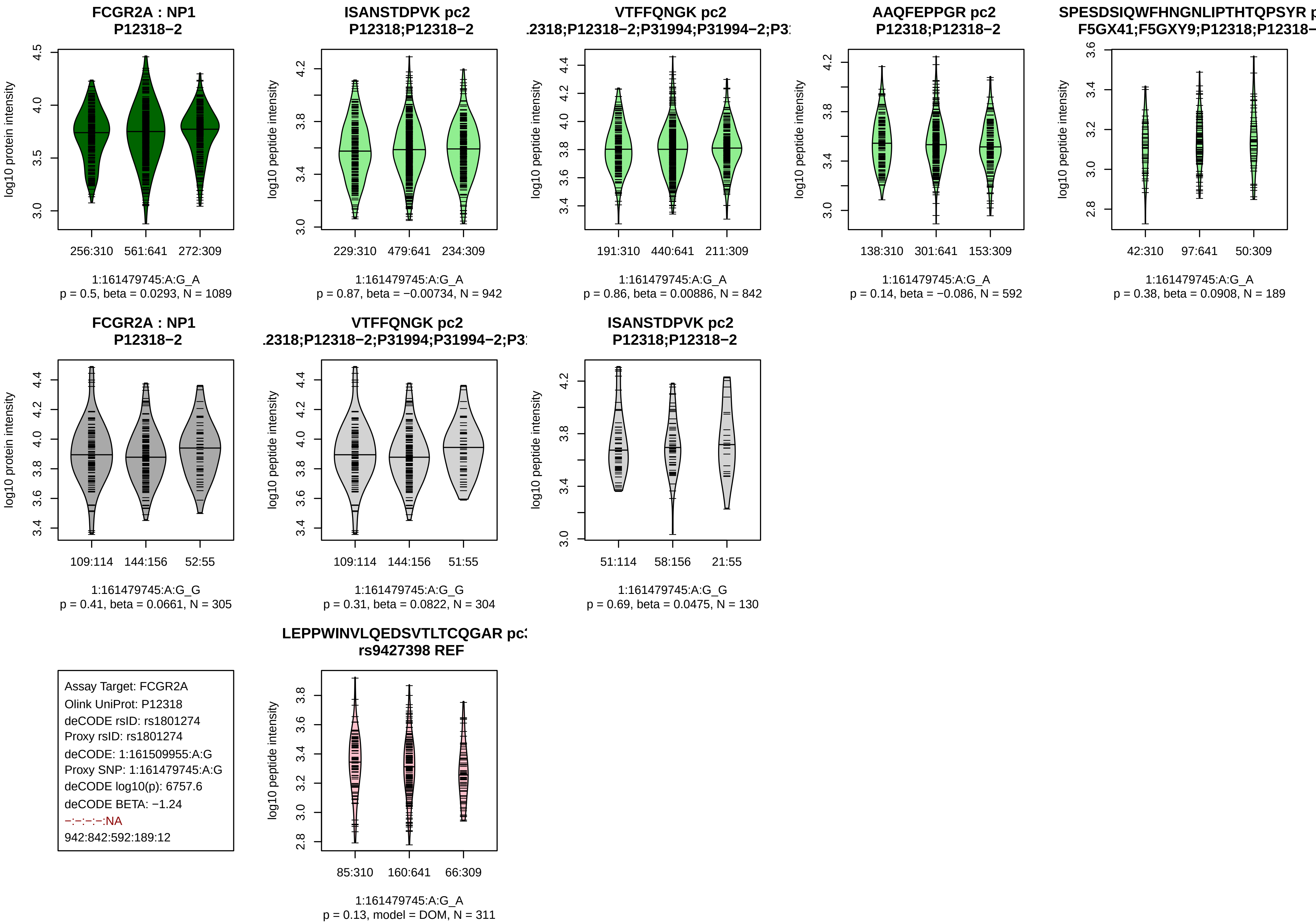
FCGR2A : NP1
P12318−2
log10 protein intensity
256:310 561:641 272:309
1:161479745:A:G_A
p = 0.5, beta = 0.0293, N = 1089
ISANSTDPVK pc2
P12318;P12318−2
log10 peptide intensity
229:310 479:641 234:309
1:161479745:A:G_A
p = 0.87, beta = −0.00734, N = 942
VTFFQNGK pc2
.2318;P12318−2;P31994;P31994−2;P3
log10 peptide intensity
191:310 440:641 211:309
1:161479745:A:G_A
p = 0.86, beta = 0.00886, N = 842
AAQFEPPGR pc2
P12318;P12318−2
log10 peptide intensity
138:310 301:641 153:309
1:161479745:A:G_A
p = 0.14, beta = −0.086, N = 592
SPESDSIQWFHNGNLIPTHTQPSYR p
F5GX41;F5GXY9;P12318;P12318−
log10 peptide intensity
42:310 97:641 50:309
1:161479745:A:G_A
p = 0.38, beta = 0.0908, N = 189
FCGR2A : NP1
P12318−2
log10 protein intensity
109:114 144:156 52:55
1:161479745:A:G_G
p = 0.41, beta = 0.0661, N = 305
VTFFQNGK pc2
.2318;P12318−2;P31994;P31994−2;P3
log10 peptide intensity
109:114 144:156 51:55
1:161479745:A:G_G
p = 0.31, beta = 0.0822, N = 304
ISANSTDPVK pc2
P12318;P12318−2
log10 peptide intensity
51:114 58:156 21:55
1:161479745:A:G_G
p = 0.69, beta = 0.0475, N = 130
Assay Target: FCGR2A
Olink UniProt: P12318
deCODE rsID: rs1801274
Proxy rsID: rs1801274
deCODE: 1:161509955:A:G
Proxy SNP: 1:161479745:A:G
deCODE log10(p): 6757.6
deCODE BETA: −1.24
−:−:−:−:NA
942:842:592:189:12
LEPPWINVLQEDSVTLTCQGAR pc
rs9427398 REF
log10 peptide intensity
85:310 160:641 66:309
1:161479745:A:G_A
p = 0.13, model = DOM, N = 311

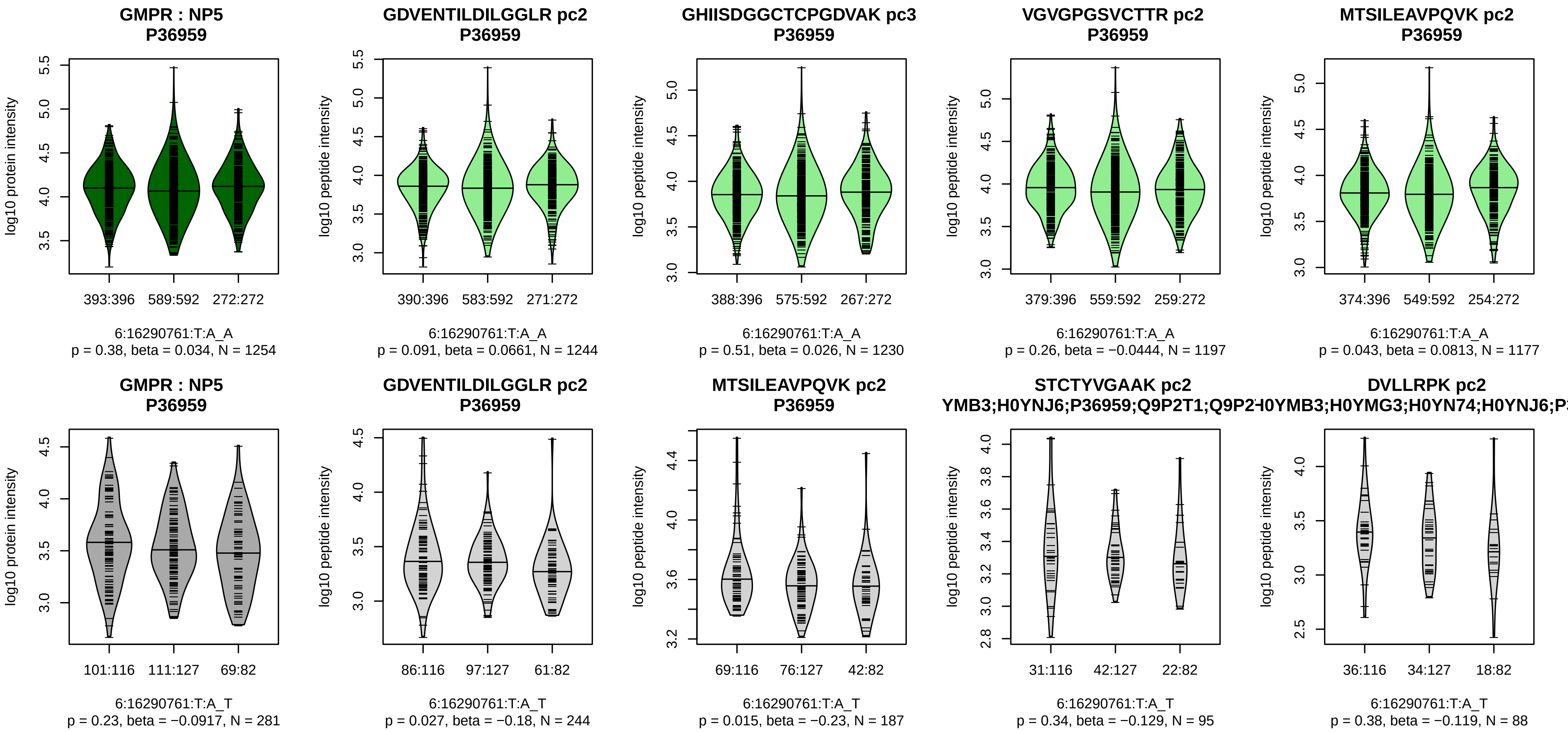


Assay Target: GMPR
Olink UniProt: P36959
deCODE rsID: rs71535075
Proxy rsID: rs1042391
deCODE: 6:16289677:GTC:G
Proxy SNP: 6:16290761:T:A
deCODE log10(p): 6478
deCODE BETA: 1.16
−:−:−:−:−:−:−:−:−:−:−:−:−:−:−:NA
1244:1230:1197:1177:1174:109

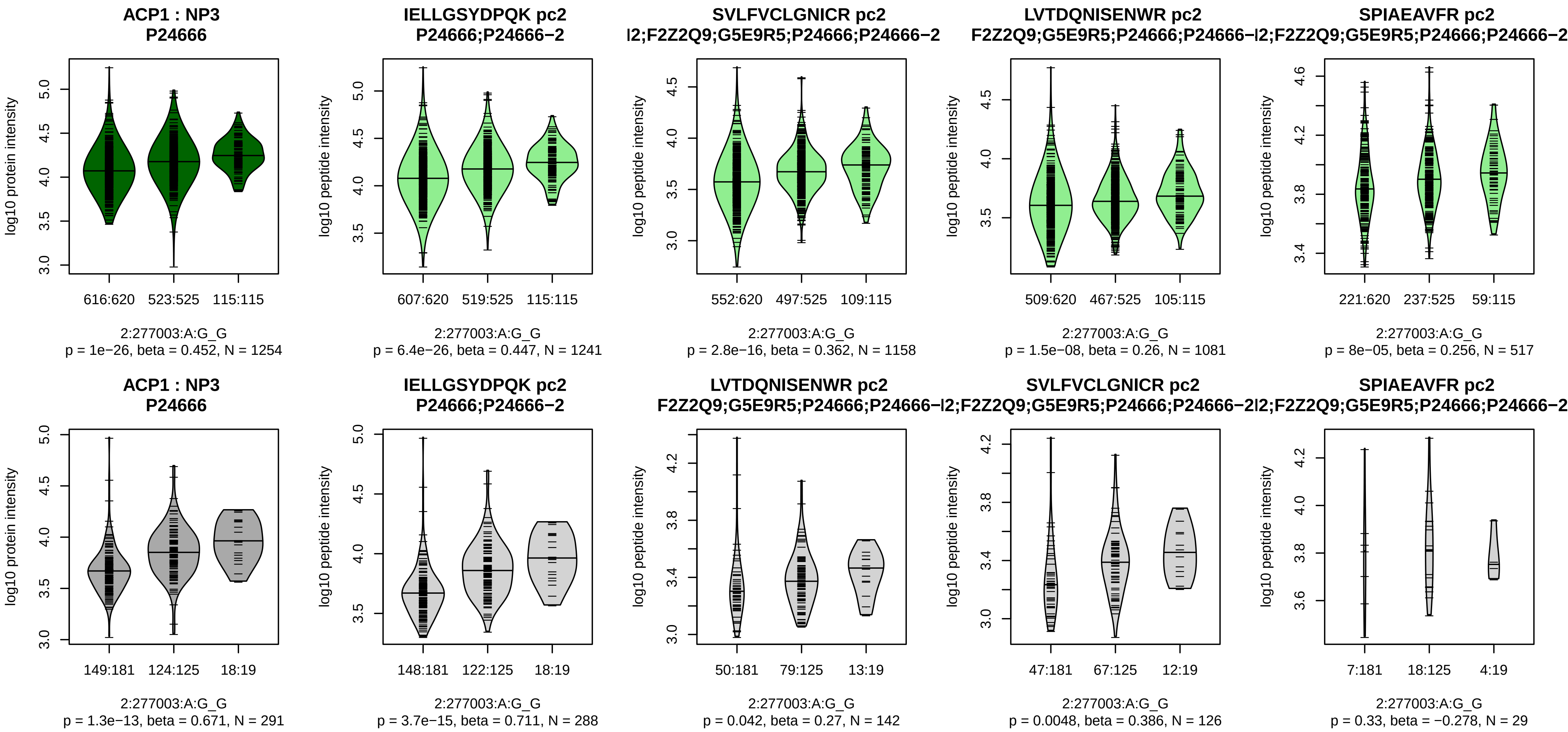


Assay Target: ACP1
Olink UniProt: P24666
deCODE rsID: rs79716074
Proxy rsID: rs79716074
deCODE: 2:277003:G:A
Proxy SNP: 2:277003:A:G
deCODE log10(p): 5886.7
deCODE BETA: 1.09
*:*:*:*:*:−:−:−:−:NA:NA:NA
1241:1158:1081:517:226:165:11

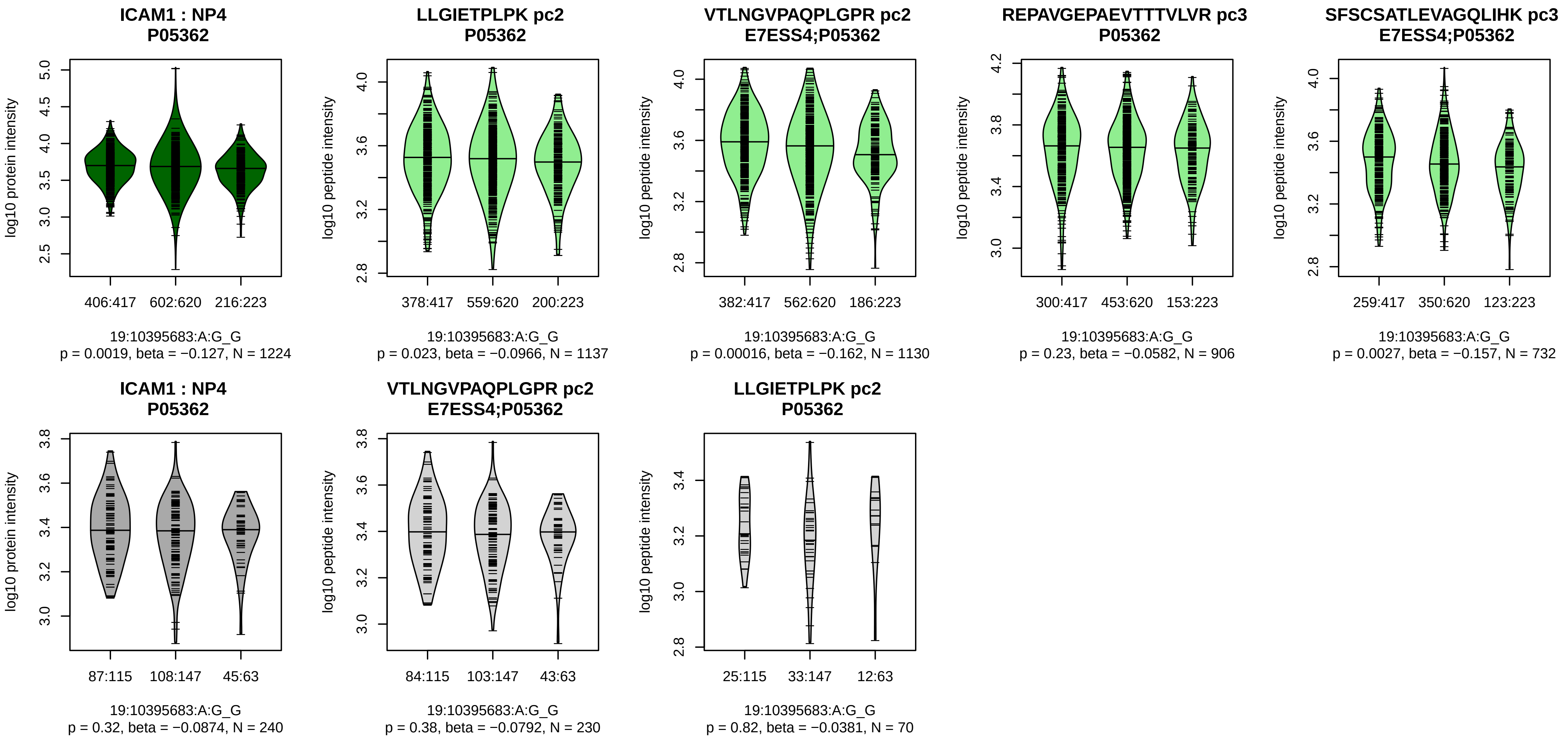


Assay Target: ICAM1
Olink UniProt: P05362
deCODE rsID: rs5498
Proxy rsID: rs5498
deCODE: 19:10285007:G:A
Proxy SNP: 19:10395683:A:G
deCODE log10(p): 5538.2
deCODE BETA: −1.14
−:*:−:*:−:−
1137:1130:906:732:664:564

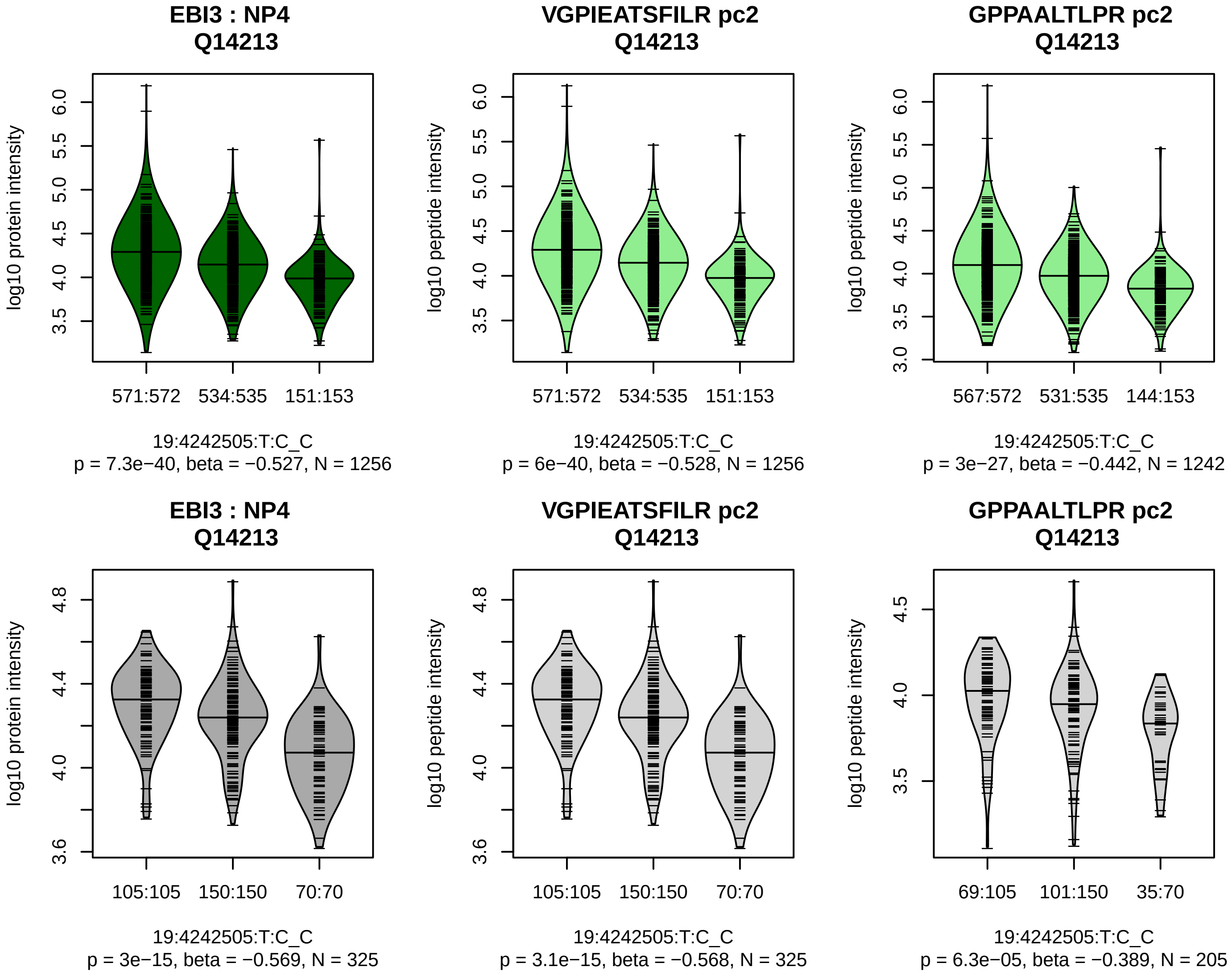


Assay Target: EBI3
Olink UniProt: Q14213
deCODE rsID: rs353696
Proxy rsID: rs353696
deCODE: 19:4242508:C:T
Proxy SNP: 19:4242505:T:C
deCODE log10(p): 4454.4
deCODE BETA: −1.12
*:*
1256:1242

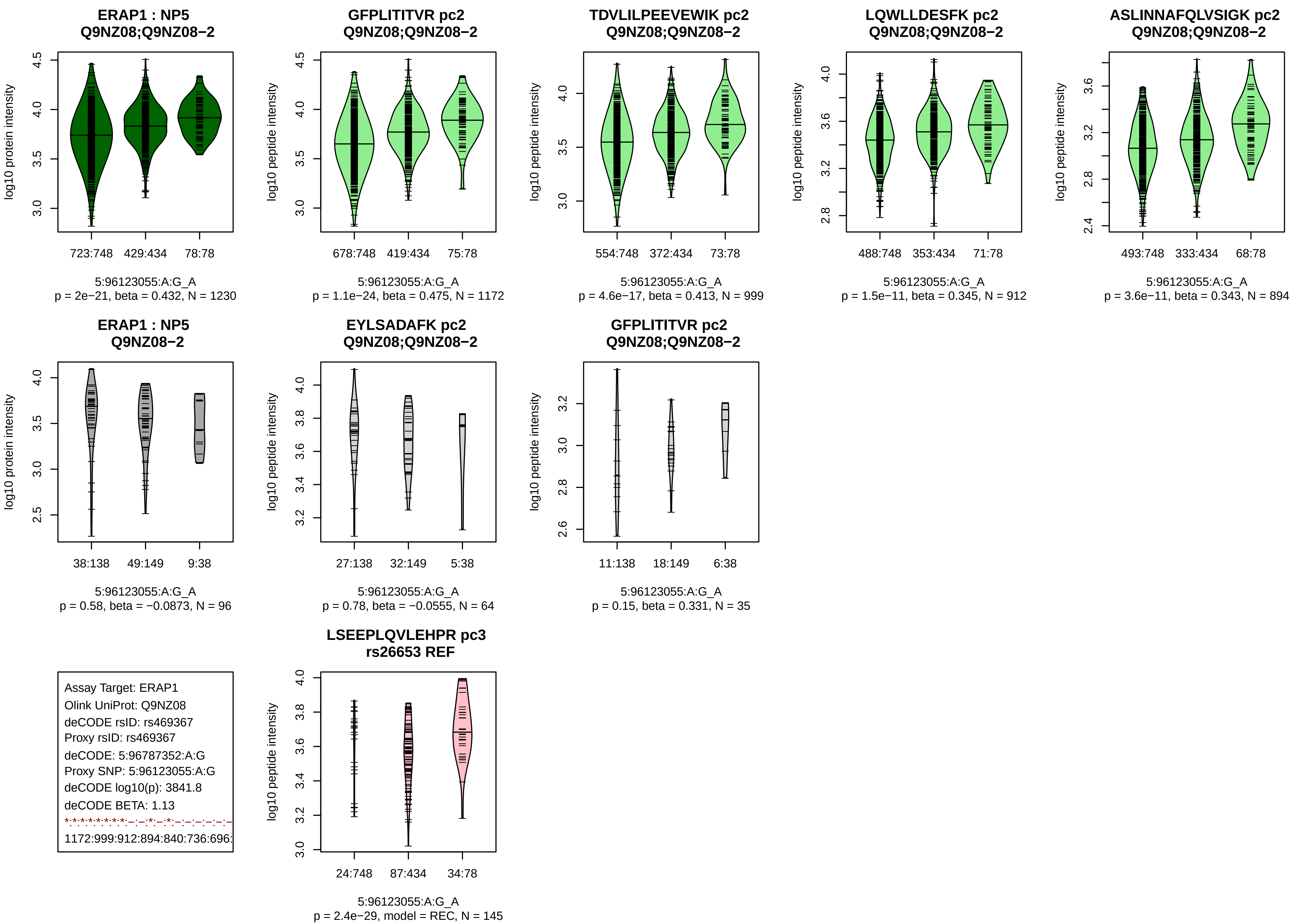
ERAP1 : NP5
Q9NZ08;Q9NZ08−2
log10 protein intensity
723:748 429:434 78:78
5:96123055:A:G_A
p = 2e−21, beta = 0.432, N = 1230
GFPLITITVR pc2
Q9NZ08;Q9NZ08−2
log10 peptide intensity
678:748 419:434 75:78
5:96123055:A:G_A
p = 1.1e−24, beta = 0.475, N = 1172
TDVLILPEEVEWIK pc2
Q9NZ08;Q9NZ08−2
log10 peptide intensity
554:748 372:434 73:78
5:96123055:A:G_A
p = 4.6e−17, beta = 0.413, N = 999
LQWLLDESFK pc2
Q9NZ08;Q9NZ08−2
log10 peptide intensity
488:748 353:434 71:78
5:96123055:A:G_A
p = 1.5e−11, beta = 0.345, N = 912
ASLINNAFQLVSIGK pc2
Q9NZ08;Q9NZ08−2
log10 peptide intensity
493:748 333:434 68:78
5:96123055:A:G_A
p = 3.6e−11, beta = 0.343, N = 894
ERAP1 : NP5
Q9NZ08−2
log10 protein intensity
38:138 49:149 9:38
5:96123055:A:G_A
p = 0.58, beta = −0.0873, N = 96
EYLSADAFK pc2
Q9NZ08;Q9NZ08−2
log10 peptide intensity
27:138 32:149 5:38
5:96123055:A:G_A
p = 0.78, beta = −0.0555, N = 64
GFPLITITVR pc2
Q9NZ08;Q9NZ08−2
log10 peptide intensity
11:138 18:149 6:38
5:96123055:A:G_A
p = 0.15, beta = 0.331, N = 35
Assay Target: ERAP1
Olink UniProt: Q9NZ08
deCODE rsID: rs469367
Proxy rsID: rs469367
deCODE: 5:96787352:A:G
Proxy SNP: 5:96123055:A:G
deCODE log10(p): 3841.8
deCODE BETA: 1.13
1172:999:912:894:840:736:696:
LSEEPLQVLEHPR pc3
rs26653 REF
log10 peptide intensity
24:748 87:434 34:78
5:96123055:A:G_A
p = 2.4e−29, model = REC, N = 145

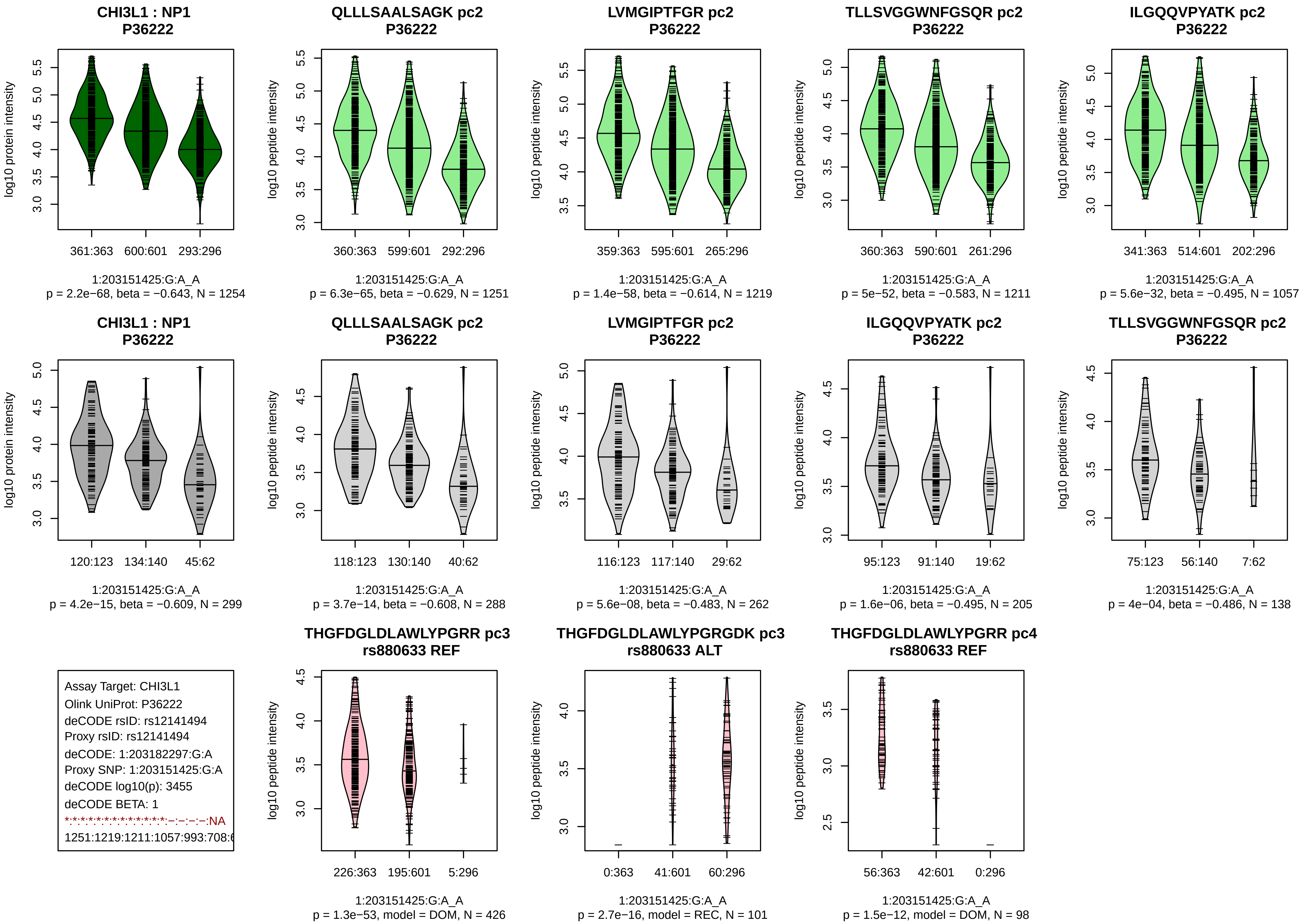
CHI3L1 : NP1
P36222
log10 protein intensity
361:363
600:601
293:296
1:203151425:G:A_A
p = 2.2e−68, beta = −0.643, N = 1254
QLLLSAALSAGK pc2
P36222
log10 peptide intensity
360:363
599:601
292:296
1:203151425:G:A_A
p = 6.3e−65, beta = −0.629, N = 1251
LVMGIPTFGR pc2
P36222
log10 peptide intensity
359:363
595:601
265:296
1:203151425:G:A_A
p = 1.4e−58, beta = −0.614, N = 1219
TLLSVGGWNFGSQR pc2
P36222
log10 peptide intensity
360:363
590:601
261:296
1:203151425:G:A_A
p = 5e−52, beta = −0.583, N = 1211
ILGQQVPYATK pc2
P36222
log10 peptide intensity
341:363
514:601
202:296
1:203151425:G:A_A
p = 5.6e−32, beta = −0.495, N = 1057
CHI3L1 : NP1
P36222
log10 protein intensity
120:123
134:140
45:62
1:203151425:G:A_A
p = 4.2e−15, beta = −0.609, N = 299
QLLLSAALSAGK pc2
P36222
log10 peptide intensity
118:123
130:140
40:62
1:203151425:G:A_A
p = 3.7e−14, beta = −0.608, N = 288
LVMGIPTFGR pc2
P36222
log10 peptide intensity
116:123
117:140
29:62
1:203151425:G:A_A
p = 5.6e−08, beta = −0.483, N = 262
ILGQQVPYATK pc2
P36222
log10 peptide intensity
95:123
91:140
19:62
1:203151425:G:A_A
p = 1.6e−06, beta = −0.495, N = 205
TLLSVGGWNFGSQR pc2
P36222
log10 peptide intensity
75:123
56:140
7:62
1:203151425:G:A_A
p = 4e−04, beta = −0.486, N = 138
Assay Target: CHI3L1
Olink UniProt: P36222
deCODE rsID: rs12141494
Proxy rsID: rs12141494
deCODE: 1:203182297:G:A
Proxy SNP: 1:203151425:G:A
deCODE log10(p): 3455
deCODE BETA: 1
*:*:*:*:*:*:*:*:*:*:–:–:–:–:NA
1251:1219:1211:1057:993:708:6
THGFDGLDLAWLYPGRR pc3
rs880633 REF
log10 peptide intensity
226:363
195:601
5:296
1:203151425:G:A_A
p = 1.3e−53, model = DOM, N = 426
THGFDGLDLAWLYPGRGDK pc3
rs880633 ALT
log10 peptide intensity
0:363
41:601
60:296
1:203151425:G:A_A
p = 2.7e−16, model = REC, N = 101
THGFDGLDLAWLYPGRR pc4
rs880633 REF
log10 peptide intensity
56:363
42:601
0:296
1:203151425:G:A_A
p = 1.5e−12, model = DOM, N = 98

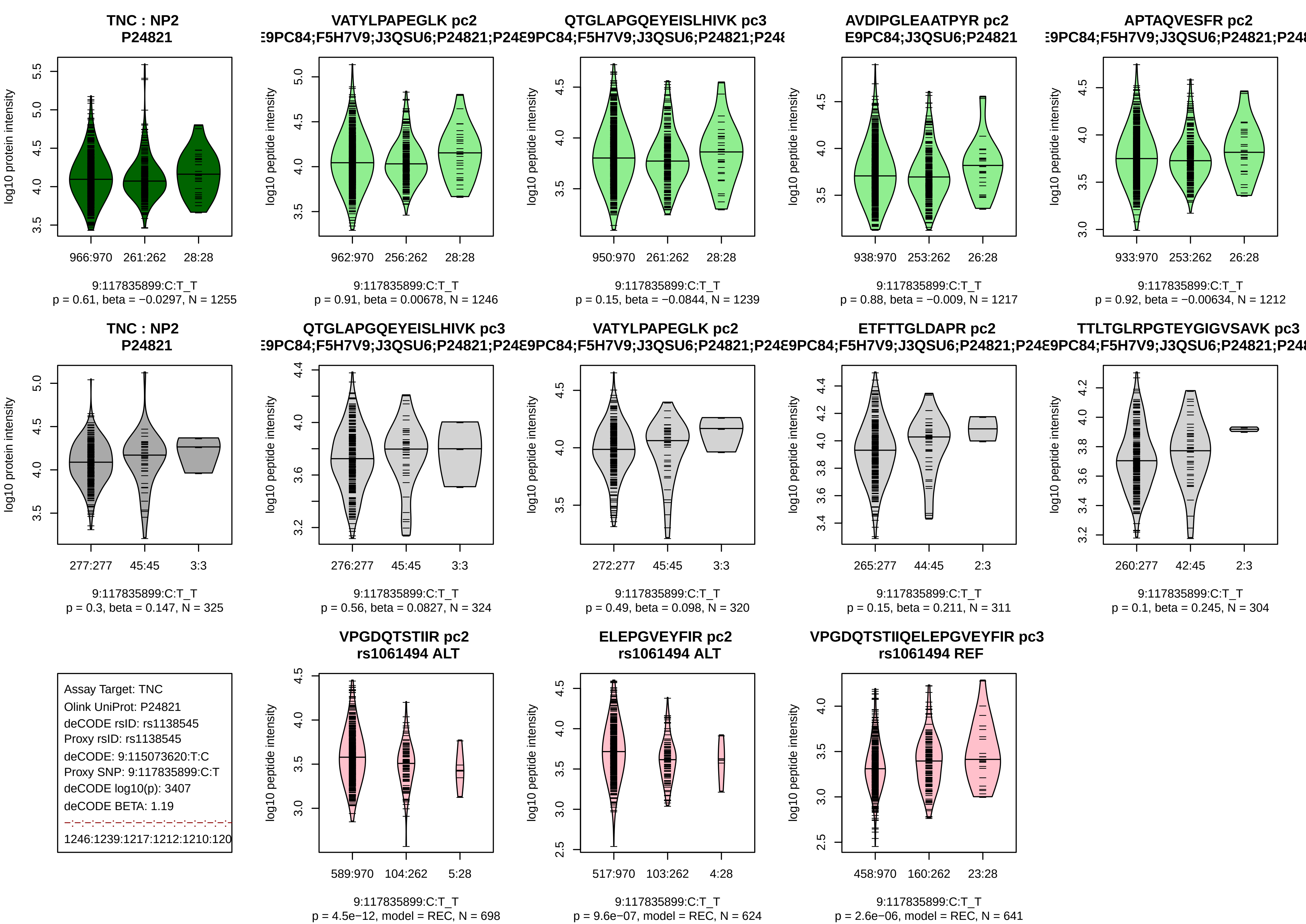

TNC : NP2
P24821
966:970
261:262
28:28
9:117835899:C:T_T
p = 0.61, beta = −0.0297, N = 1255
VATYLPAPEGLK pc2
9:117835899:C:T_T
p = 0.91, beta = 0.00678, N = 1246
QTGLAPGQEYEISLHIVK pc3
9:117835899:C:T_T
p = 0.15, beta = −0.0844, N = 1239
AVDIPGLEAATPYR pc2
E9PC84;J3QSU6;P24821
9:117835899:C:T_T
p = 0.88, beta = −0.009, N = 1217
APTAQVESFR pc2
9:117835899:C:T_T
p = 0.92, beta = −0.00634, N = 1212
TNC : NP2
P24821
9:117835899:C:T_T
p = 0.3, beta = 0.147, N = 325
QTGLAPGQEYEISLHIVK pc3
9:117835899:C:T_T
p = 0.56, beta = 0.0827, N = 324
VATYLPAPEGLK pc2
9:117835899:C:T_T
p = 0.49, beta = 0.098, N = 320
ETFTTGLDAPR pc2
9:117835899:C:T_T
p = 0.15, beta = 0.211, N = 311
TTLTGLRPGTEYGIGVSAVK pc3
9:117835899:C:T_T
p = 0.1, beta = 0.245, N = 304
Assay Target: TNC
Olink UniProt: P24821
deCODE rsID: rs1138545
Proxy rsID: rs1138545
deCODE: 9:115073620:T:C
Proxy SNP: 9:117835899:C:T
deCODE log10(p): 3407
deCODE BETA: 1.19
VPGDQTSTIIR pc2
rs1061494 ALT
9:117835899:C:T_T
p = 4.5e−12, model = REC, N = 698
ELEPGVEYFIR pc2
rs1061494 ALT
9:117835899:C:T_T
p = 9.6e−07, model = REC, N = 624
VPGDQTSTIIQELEPGVEYFIR pc3
rs1061494 REF
9:117835899:C:T_T
p = 2.6e−06, model = REC, N = 641
log10 protein intensity
log10 peptide intensity

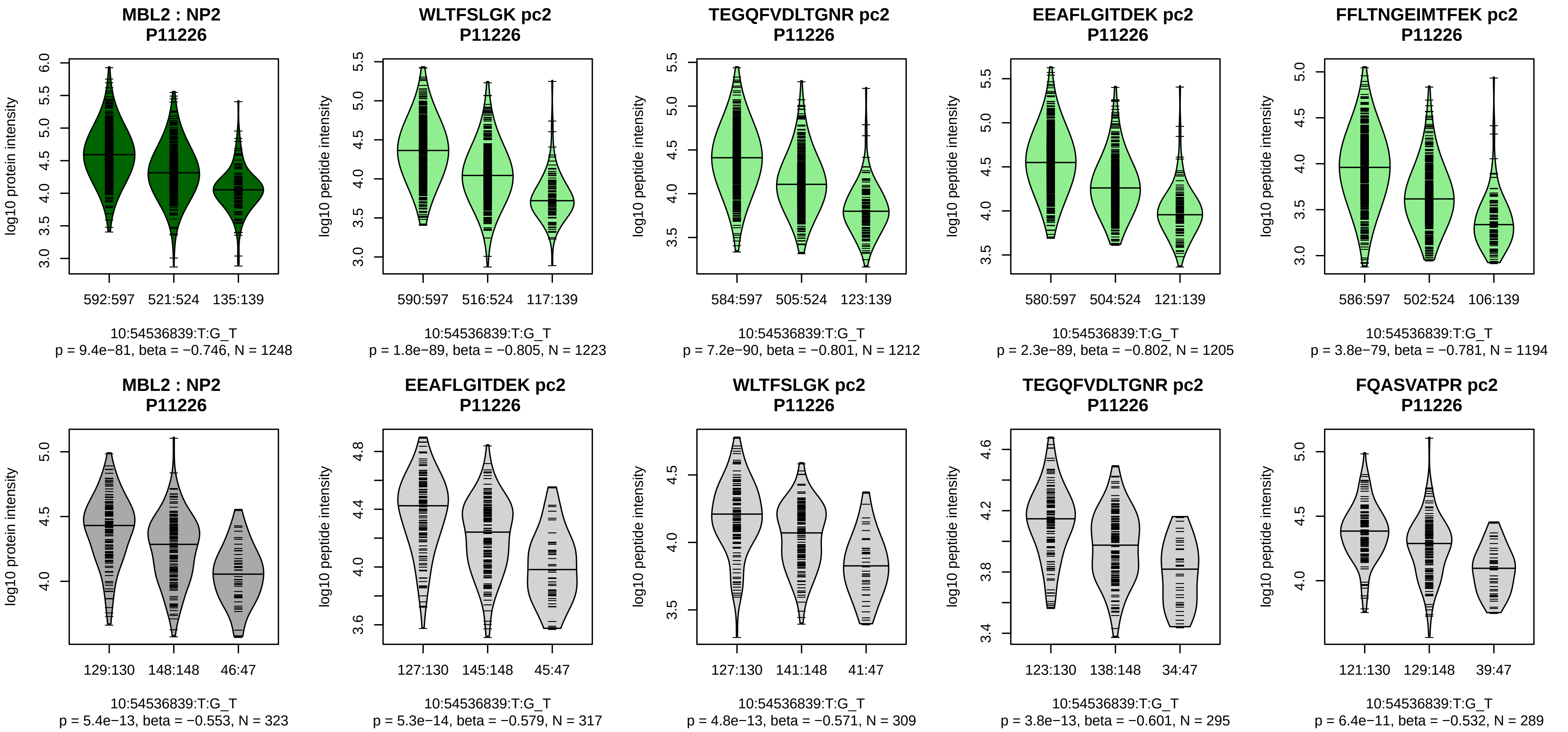


Assay Target: MBL2
Olink UniProt: P11226
deCODE rsID: rs7899547
Proxy rsID: rs7899547
deCODE: 10:52777079:T:G
Proxy SNP: 10:54536839:T:G
deCODE log10(p): 3017.5
deCODE BETA: −0.9
*:*:*:*:*:*:*:*:NA
1223:1212:1205:1194:1164:106

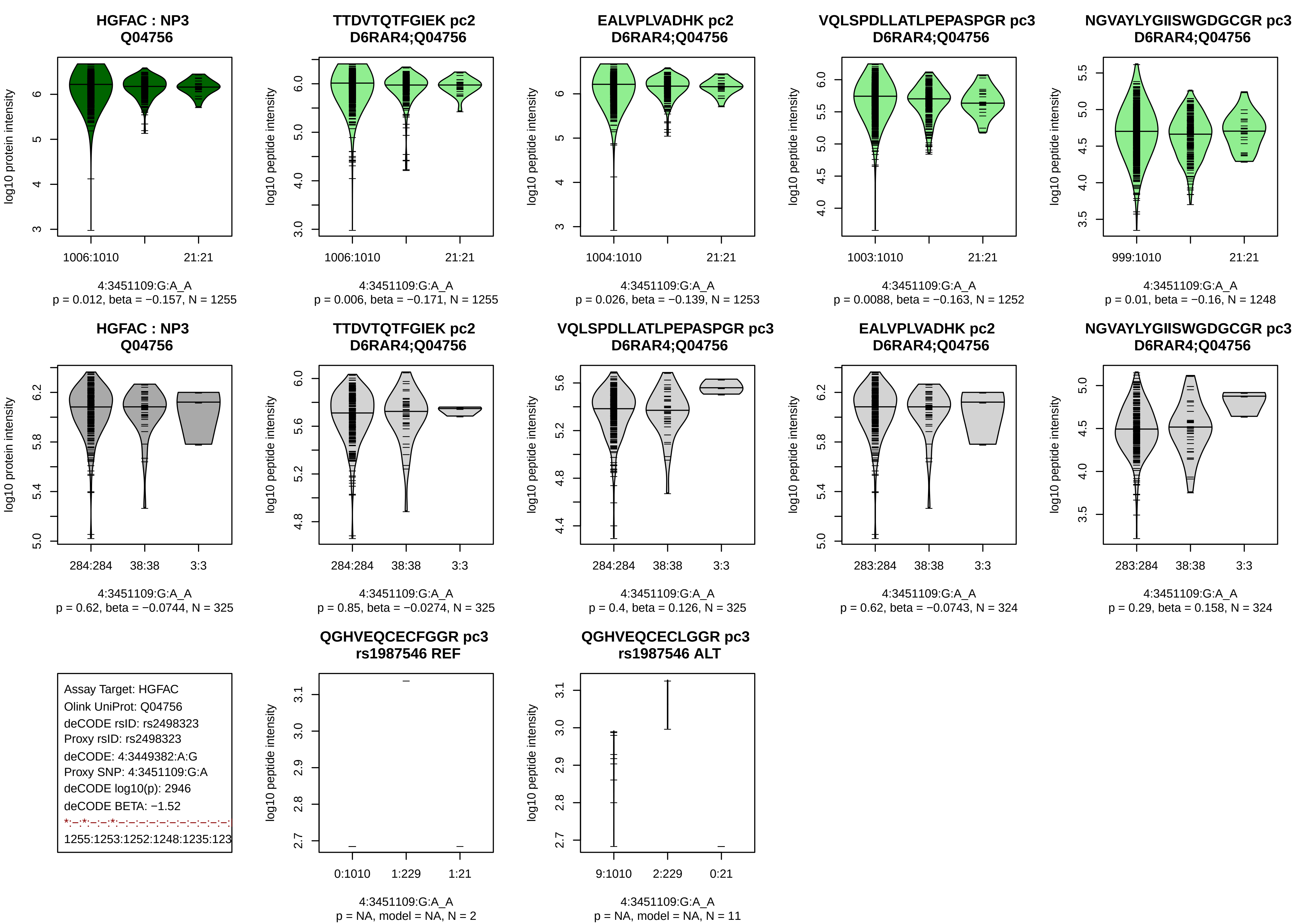
HGFAC : NP3
Q04756
log10 protein intensity
1006:1010
21:21
4:3451109:G:A_A
p = 0.012, beta = −0.157, N = 1255
TTDVTQTFGIEK pc2
D6RAR4;Q04756
log10 peptide intensity
1006:1010
21:21
4:3451109:G:A_A
p = 0.006, beta = −0.171, N = 1255
EALVPLVADHK pc2
D6RAR4;Q04756
log10 peptide intensity
1004:1010
21:21
4:3451109:G:A_A
p = 0.026, beta = −0.139, N = 1253
VQLSPDLLATLPEPASPGR pc3
D6RAR4;Q04756
log10 peptide intensity
1003:1010
21:21
4:3451109:G:A_A
p = 0.0088, beta = −0.163, N = 1252
NGVAYLYGIISWGDGCGR pc3
D6RAR4;Q04756
log10 peptide intensity
999:1010
21:21
4:3451109:G:A_A
p = 0.01, beta = −0.16, N = 1248
HGFAC : NP3
Q04756
log10 protein intensity
284:284
38:38
3:3
4:3451109:G:A_A
p = 0.62, beta = −0.0744, N = 325
TTDVTQTFGIEK pc2
D6RAR4;Q04756
log10 peptide intensity
284:284
38:38
3:3
4:3451109:G:A_A
p = 0.85, beta = −0.0274, N = 325
VQLSPDLLATLPEPASPGR pc3
D6RAR4;Q04756
log10 peptide intensity
284:284
38:38
3:3
4:3451109:G:A_A
p = 0.4, beta = 0.126, N = 325
EALVPLVADHK pc2
D6RAR4;Q04756
log10 peptide intensity
283:284
38:38
3:3
4:3451109:G:A_A
p = 0.62, beta = −0.0743, N = 324
NGVAYLYGIISWGDGCGR pc3
D6RAR4;Q04756
log10 peptide intensity
283:284
38:38
3:3
4:3451109:G:A_A
p = 0.29, beta = 0.158, N = 324
Assay Target: HGFAC
Olink UniProt: Q04756
deCODE rsID: rs2498323
Proxy rsID: rs2498323
deCODE: 4:3449382:A:G
Proxy SNP: 4:3451109:G:A
deCODE log10(p): 2946
deCODE BETA: −1.52
*:–:*:–:–:*:–:–:–:–:–:–:–:–:–:–:–:
1255:1253:1252:1248:1235:123
QGHVEQCECFGGR pc3
rs1987546 REF
log10 peptide intensity
0:1010
1:229
1:21
4:3451109:G:A_A
p = NA, model = NA, N = 2
QGHVEQCECLGGR pc3
rs1987546 ALT
log10 peptide intensity
9:1010
2:229
0:21
4:3451109:G:A_A
p = NA, model = NA, N = 11

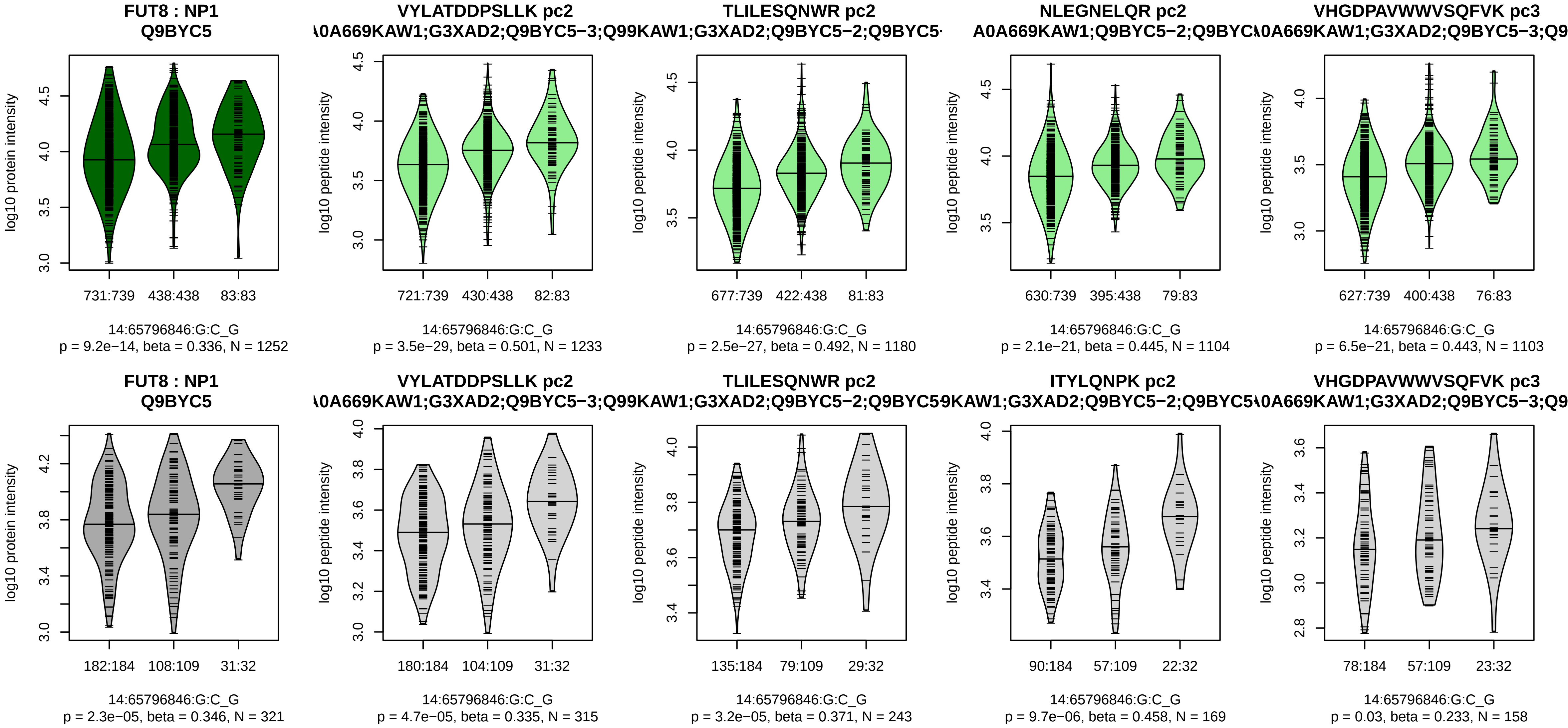


Assay Target: FUT8
Olink UniProt: Q9BYC5
deCODE rsID: rs2127870
Proxy rsID: rs2127870
deCODE: 14:65330128:G:C
Proxy SNP: 14:65796846:G:C
deCODE log10(p): 2568.3
deCODE BETA: 1.05
1233:1180:1104:1103:1050:979

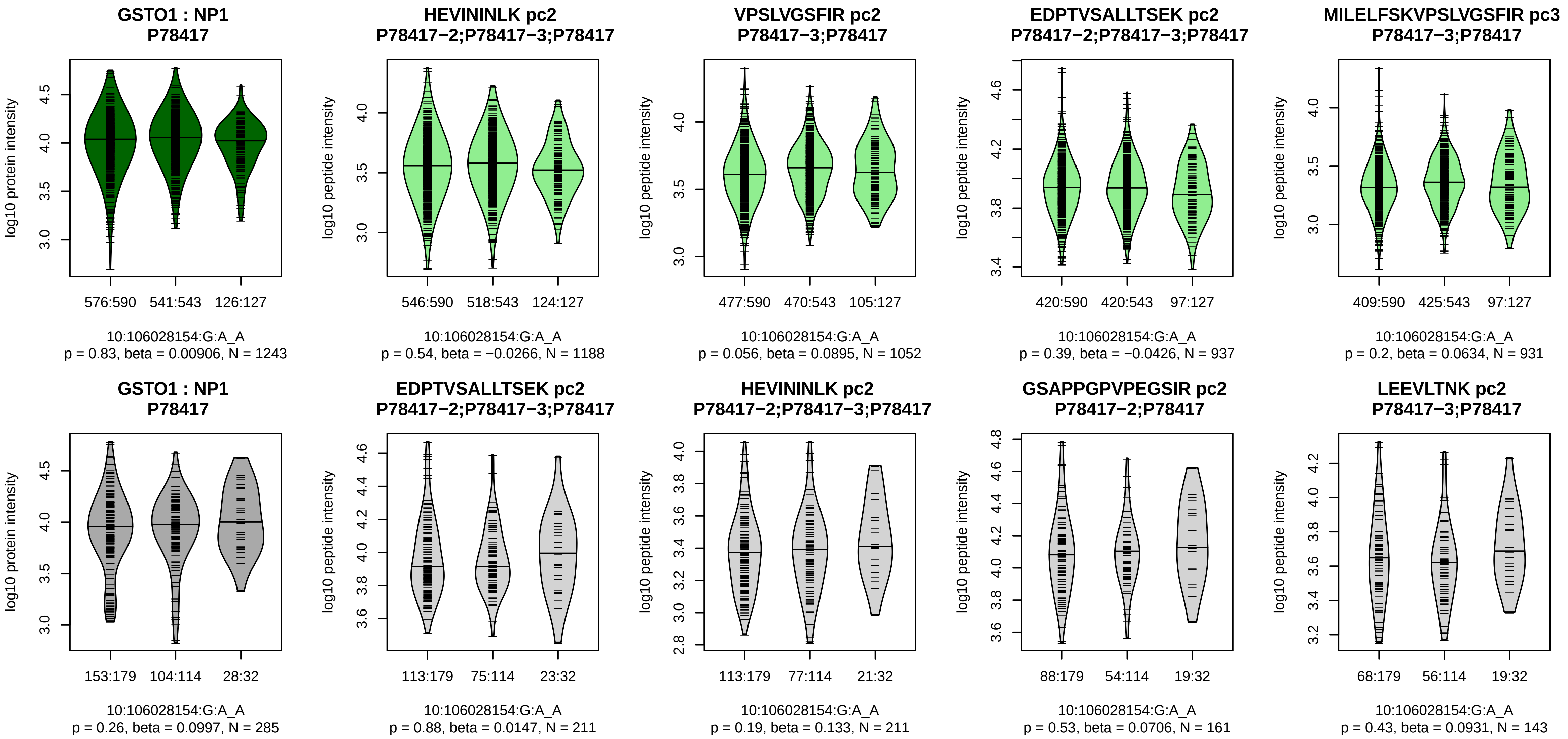


Assay Target: GSTO1
Olink UniProt: P78417
deCODE rsID: rs45596840
Proxy rsID: rs45596840
deCODE: 10:104268396:A:G
Proxy SNP: 10:106028154:G:A
deCODE log10(p): 2443.9
deCODE BETA: −0.82
−:−:−:−:−:−:−:−:−:−:−:−:−:−:NA
1188:1052:937:931:893:831:77[illegible]

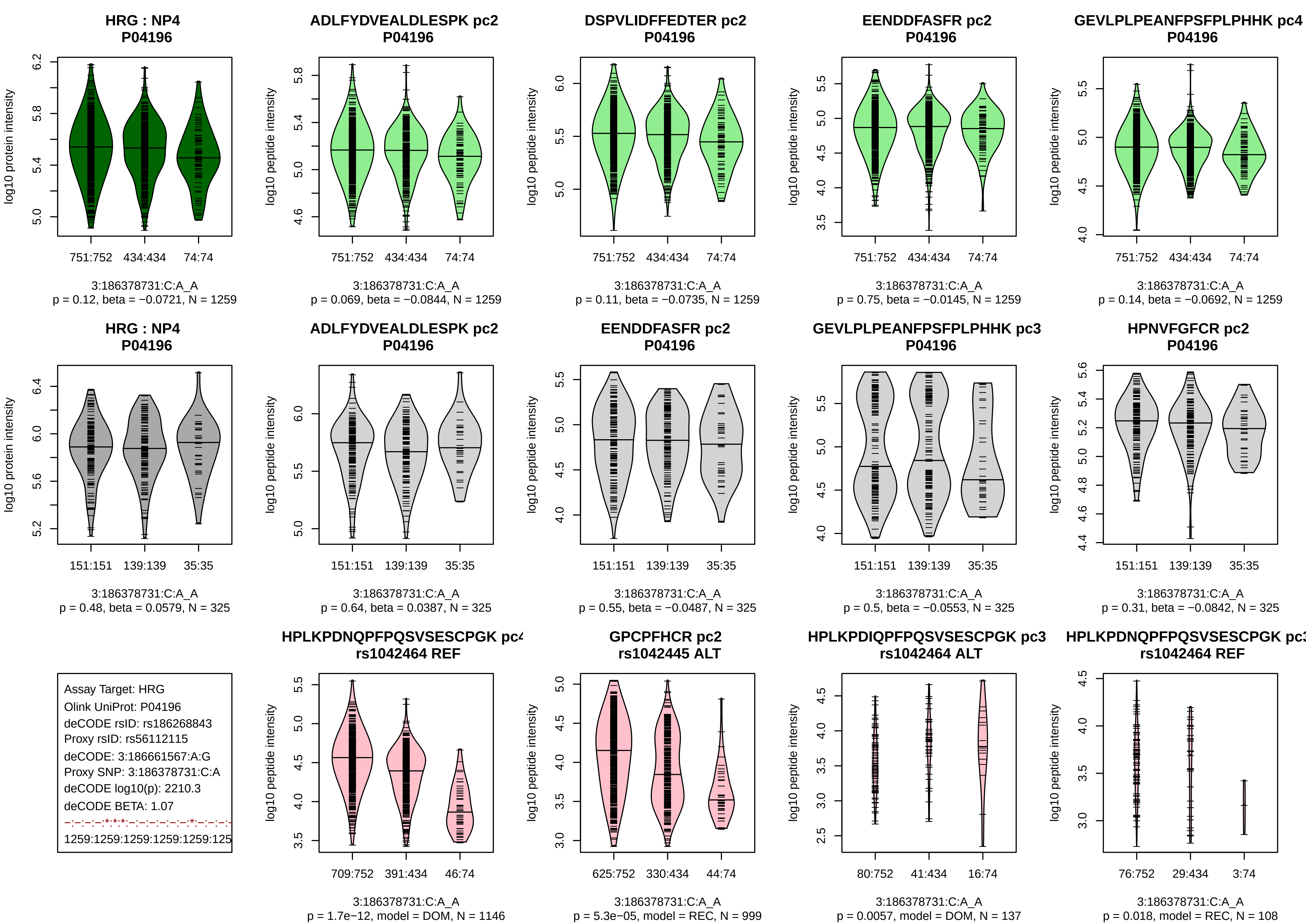
HRG : NP4
P04196
log10 protein intensity
751:752 434:434 74:74
3:186378731:C:A_A
p = 0.12, beta = −0.0721, N = 1259
ADLFYDVEALDLESPK pc2
P04196
log10 peptide intensity
751:752 434:434 74:74
3:186378731:C:A_A
p = 0.069, beta = −0.0844, N = 1259
DSPVLIDFFEDTER pc2
P04196
log10 peptide intensity
751:752 434:434 74:74
3:186378731:C:A_A
p = 0.11, beta = −0.0735, N = 1259
EENDDFASFR pc2
P04196
log10 peptide intensity
751:752 434:434 74:74
3:186378731:C:A_A
p = 0.75, beta = −0.0145, N = 1259
GEVLPLPEANFPSFPLPHHK pc4
P04196
log10 peptide intensity
751:752 434:434 74:74
3:186378731:C:A_A
p = 0.14, beta = −0.0692, N = 1259
HRG : NP4
P04196
log10 protein intensity
151:151 139:139 35:35
3:186378731:C:A_A
p = 0.48, beta = 0.0579, N = 325
ADLFYDVEALDLESPK pc2
P04196
log10 peptide intensity
151:151 139:139 35:35
3:186378731:C:A_A
p = 0.64, beta = 0.0387, N = 325
EENDDFASFR pc2
P04196
log10 peptide intensity
151:151 139:139 35:35
3:186378731:C:A_A
p = 0.55, beta = −0.0487, N = 325
GEVLPLPEANFPSFPLPHHK pc3
P04196
log10 peptide intensity
151:151 139:139 35:35
3:186378731:C:A_A
p = 0.5, beta = −0.0553, N = 325
HPNVFGFCR pc2
P04196
log10 peptide intensity
151:151 139:139 35:35
3:186378731:C:A_A
p = 0.31, beta = −0.0842, N = 325
Assay Target: HRG
Olink UniProt: P04196
deCODE rsID: rs186268843
Proxy rsID: rs56112115
deCODE: 3:186661567:A:G
Proxy SNP: 3:186378731:C:A
deCODE log10(p): 2210.3
deCODE BETA: 1.07
−:−:−:−:*:*:*:−:−:−:−:−:−:*:−:−:−:
1259:1259:1259:1259:1259:125
HPLKPDNQPFPQSVSESCPGK pc4
rs1042464 REF
log10 peptide intensity
709:752 391:434 46:74
3:186378731:C:A_A
p = 1.7e−12, model = DOM, N = 1146
GPCPFHCR pc2
rs1042445 ALT
log10 peptide intensity
625:752 330:434 44:74
3:186378731:C:A_A
p = 5.3e−05, model = REC, N = 999
HPLKPDIQPFPQSVSESCPGK pc3
rs1042464 ALT
log10 peptide intensity
80:752 41:434 16:74
3:186378731:C:A_A
p = 0.0057, model = DOM, N = 137
HPLKPDNQPFPQSVSESCPGK pc3
rs1042464 REF
log10 peptide intensity
76:752 29:434 3:74
3:186378731:C:A_A
p = 0.018, model = REC, N = 108

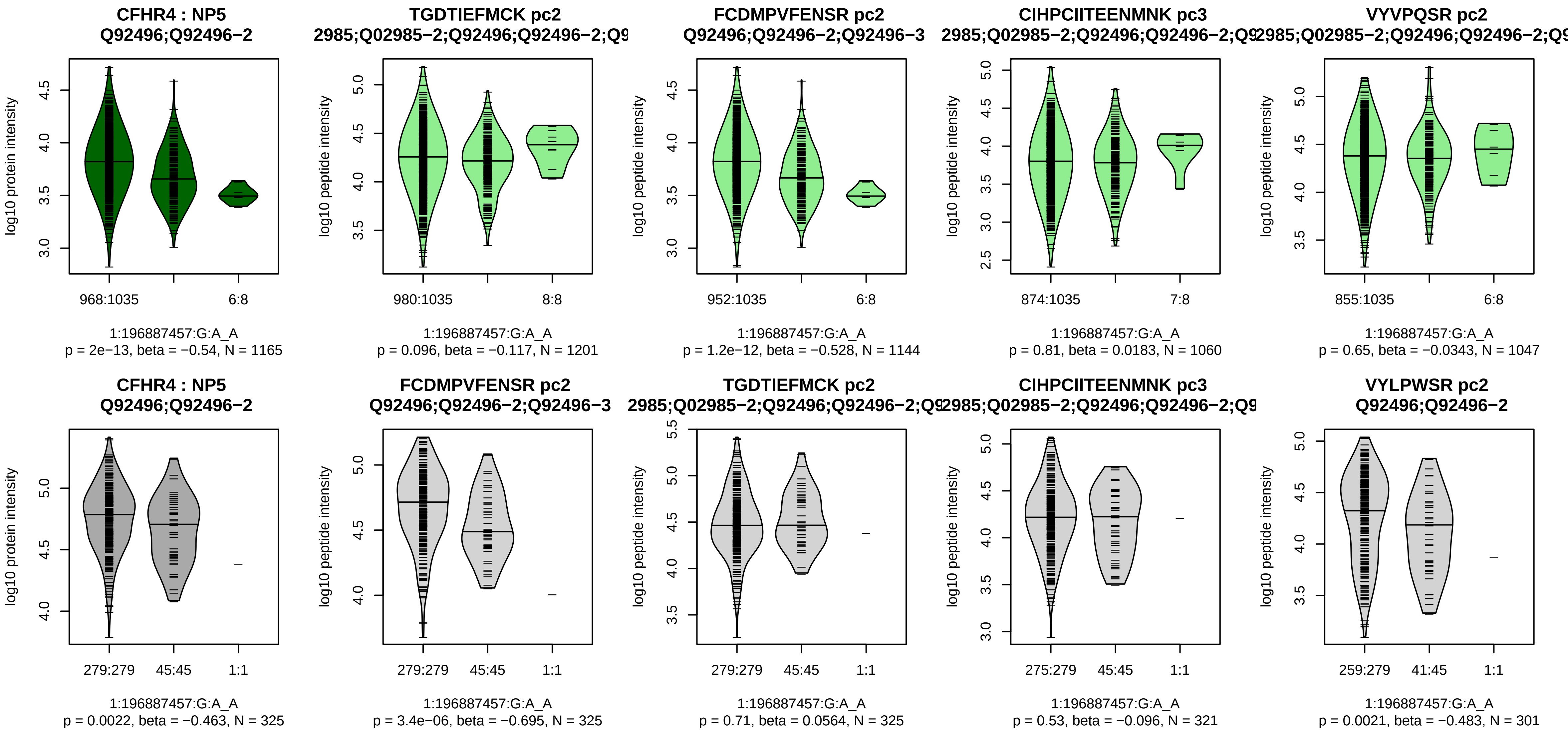


Assay Target: CFHR4
Olink UniProt: Q92496
deCODE rsID: rs10494745
Proxy rsID: rs10494745
deCODE: 1:196918327:A:G
Proxy SNP: 1:196887457:G:A
deCODE log10(p): 2098.5
deCODE BETA: −0.97
−:*:−:−:−:−:−:−
1201:1144:1060:1047:690:585:1

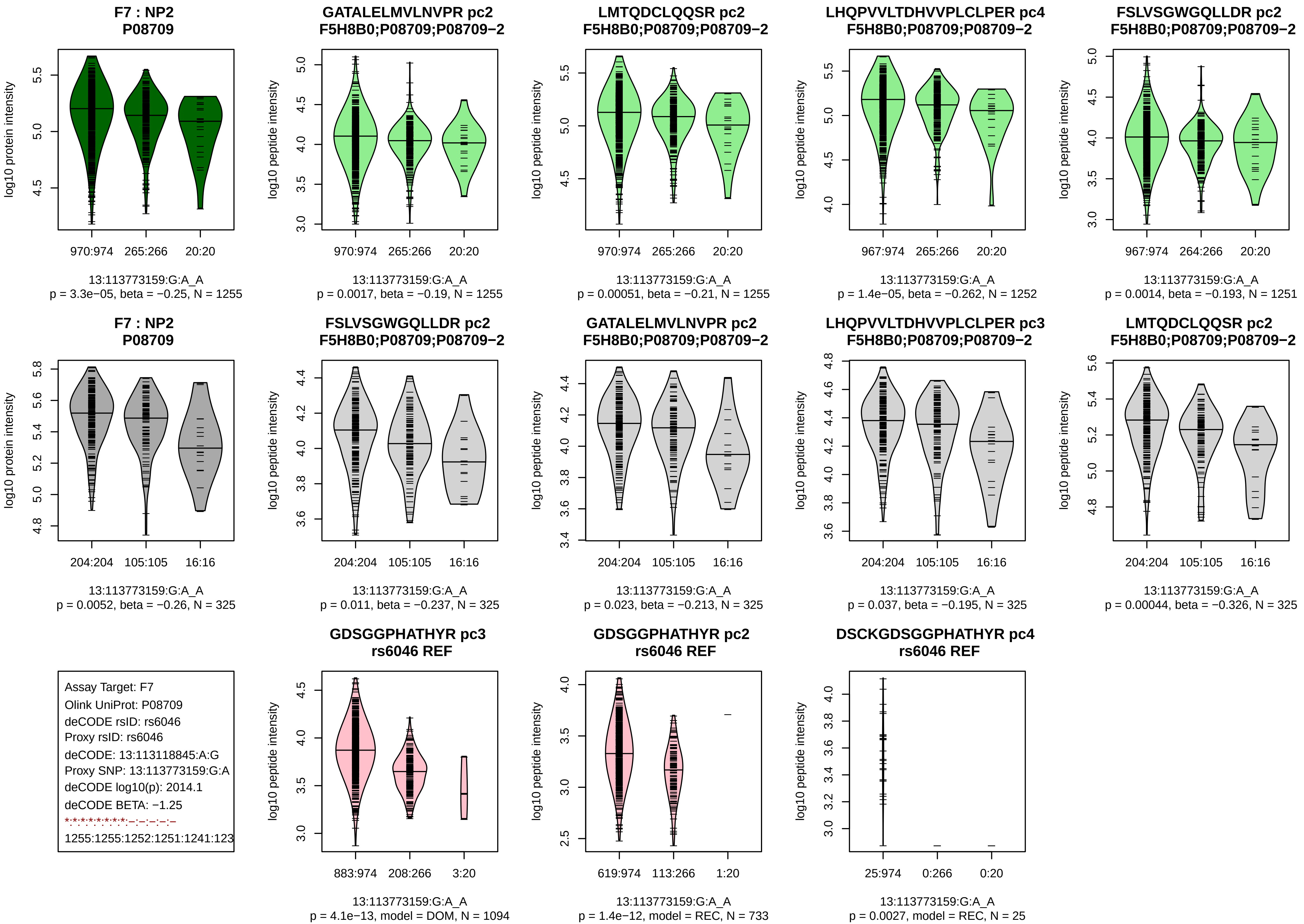
F7 : NP2
P08709
log10 protein intensity
970:974 265:266 20:20
13:113773159:G:A_A
p = 3.3e−05, beta = −0.25, N = 1255
GATALELMVLNVPR pc2
F5H8B0;P08709;P08709−2
log10 peptide intensity
970:974 265:266 20:20
13:113773159:G:A_A
p = 0.0017, beta = −0.19, N = 1255
LMTQDCLQQSR pc2
F5H8B0;P08709;P08709−2
log10 peptide intensity
970:974 265:266 20:20
13:113773159:G:A_A
p = 0.00051, beta = −0.21, N = 1255
LHQPVVLTDHVVPLCLPER pc4
F5H8B0;P08709;P08709−2
log10 peptide intensity
967:974 265:266 20:20
13:113773159:G:A_A
p = 1.4e−05, beta = −0.262, N = 1252
FSLVSGWGQLLDR pc2
F5H8B0;P08709;P08709−2
log10 peptide intensity
967:974 264:266 20:20
13:113773159:G:A_A
p = 0.0014, beta = −0.193, N = 1251
F7 : NP2
P08709
log10 protein intensity
204:204 105:105 16:16
13:113773159:G:A_A
p = 0.0052, beta = −0.26, N = 325
FSLVSGWGQLLDR pc2
F5H8B0;P08709;P08709−2
log10 peptide intensity
204:204 105:105 16:16
13:113773159:G:A_A
p = 0.011, beta = −0.237, N = 325
GATALELMVLNVPR pc2
F5H8B0;P08709;P08709−2
log10 peptide intensity
204:204 105:105 16:16
13:113773159:G:A_A
p = 0.023, beta = −0.213, N = 325
LHQPVVLTDHVVPLCLPER pc3
F5H8B0;P08709;P08709−2
log10 peptide intensity
204:204 105:105 16:16
13:113773159:G:A_A
p = 0.037, beta = −0.195, N = 325
LMTQDCLQQSR pc2
F5H8B0;P08709;P08709−2
log10 peptide intensity
204:204 105:105 16:16
13:113773159:G:A_A
p = 0.00044, beta = −0.326, N = 325
Assay Target: F7
Olink UniProt: P08709
deCODE rsID: rs6046
Proxy rsID: rs6046
deCODE: 13:113118845:A:G
Proxy SNP: 13:113773159:G:A
deCODE log10(p): 2014.1
deCODE BETA: −1.25
*:*:*:*:*:*:*:*:–:–:–:–:–:–
1255:1255:1252:1251:1241:123
GDSGGPHATHYR pc3
rs6046 REF
log10 peptide intensity
883:974 208:266 3:20
13:113773159:G:A_A
p = 4.1e−13, model = DOM, N = 1094
GDSGGPHATHYR pc2
rs6046 REF
log10 peptide intensity
619:974 113:266 1:20
13:113773159:G:A_A
p = 1.4e−12, model = REC, N = 733
DSCKGDSGGPHATHYR pc4
rs6046 REF
log10 peptide intensity
25:974 0:266 0:20
13:113773159:G:A_A
p = 0.0027, model = REC, N = 25

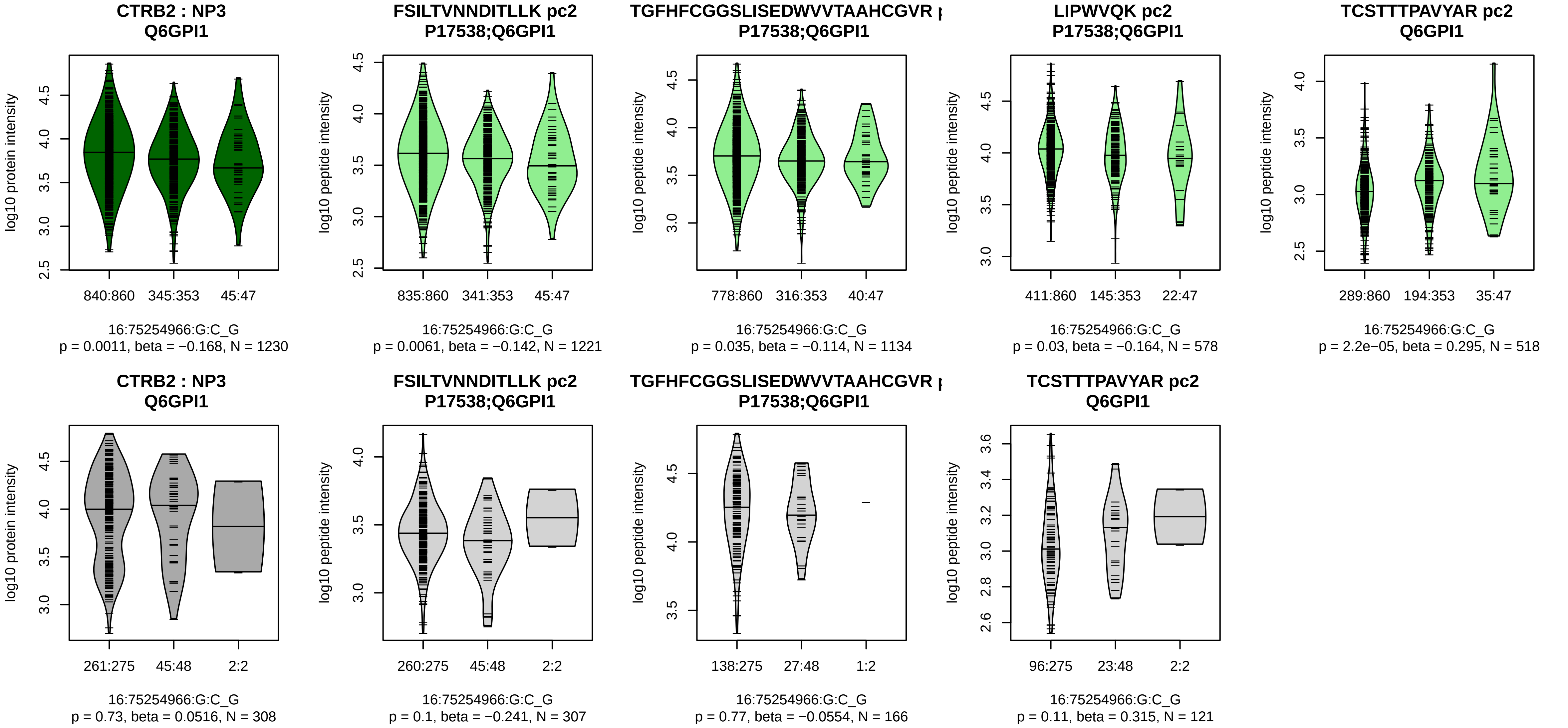


Assay Target: CTRB2
Olink UniProt: Q6GPI1
deCODE rsID: rs8048956
Proxy rsID: rs8048956
deCODE: 16:75221068:G:C
Proxy SNP: 16:75254966:G:C
deCODE log10(p): 1800.8
deCODE BETA: 0.86
*:−:−:*:−:−
1221:1134:578:518:370:84

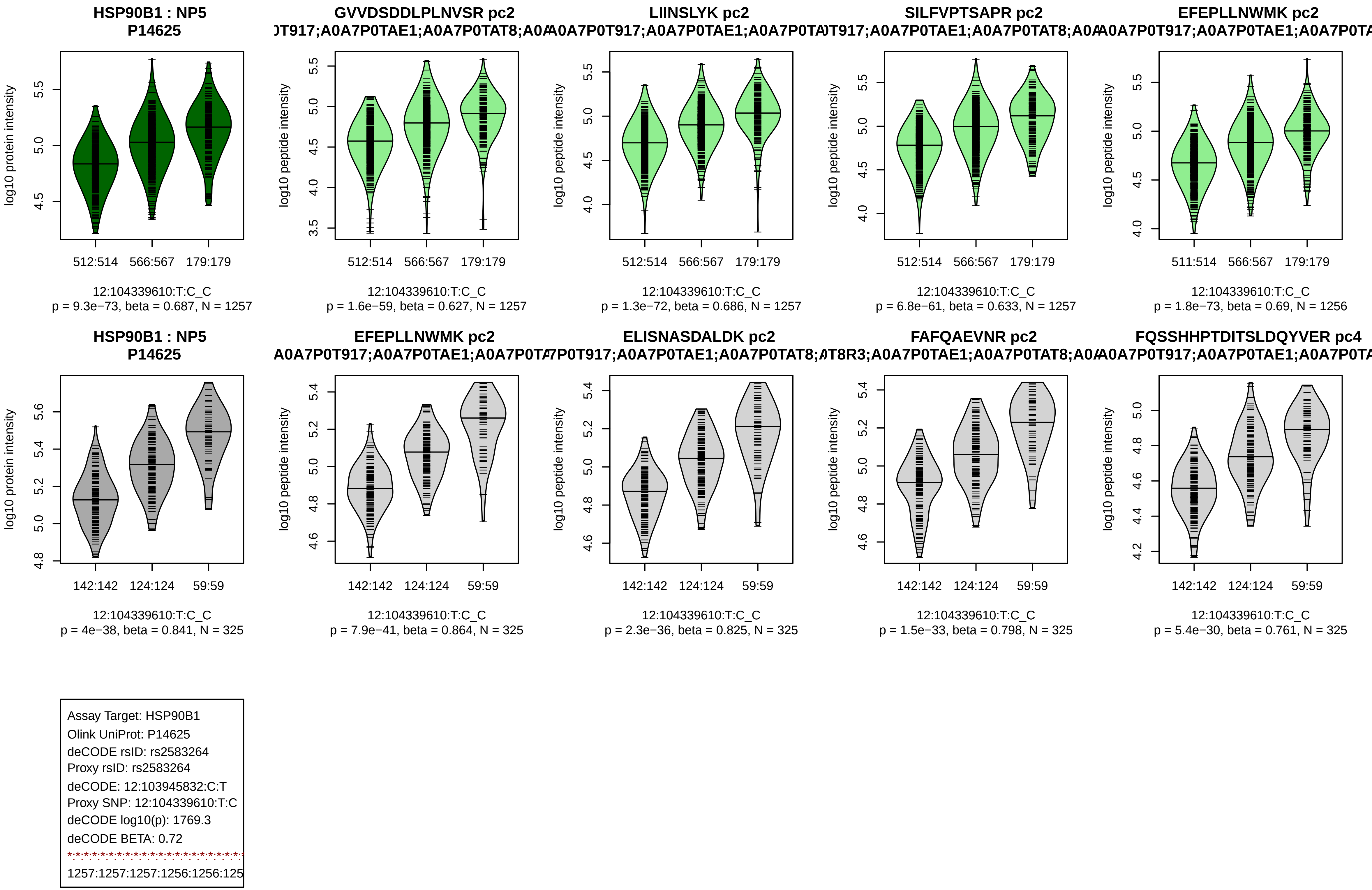
HSP90B1 : NP5
P14625
log10 protein intensity
512:514
566:567
179:179
12:104339610:T:C_C
p = 9.3e−73, beta = 0.687, N = 1257
GVVDSDDLPLNVSR pc2
log10 peptide intensity
12:104339610:T:C_C
p = 1.6e−59, beta = 0.627, N = 1257
LIINSLYK pc2
12:104339610:T:C_C
p = 1.3e−72, beta = 0.686, N = 1257
SILFVPTSAPR pc2
12:104339610:T:C_C
p = 6.8e−61, beta = 0.633, N = 1257
EFEPLLNWMK pc2
511:514
12:104339610:T:C_C
p = 1.8e−73, beta = 0.69, N = 1256
HSP90B1 : NP5
P14625
142:142
124:124
59:59
12:104339610:T:C_C
p = 4e−38, beta = 0.841, N = 325
EFEPLLNWMK pc2
12:104339610:T:C_C
p = 7.9e−41, beta = 0.864, N = 325
ELISNASDALDK pc2
12:104339610:T:C_C
p = 2.3e−36, beta = 0.825, N = 325
FAFQAEVNR pc2
12:104339610:T:C_C
p = 1.5e−33, beta = 0.798, N = 325
FQSSHHPTDITSLDQYVER pc4
12:104339610:T:C_C
p = 5.4e−30, beta = 0.761, N = 325
Assay Target: HSP90B1
Olink UniProt: P14625
deCODE rsID: rs2583264
Proxy rsID: rs2583264
deCODE: 12:103945832:C:T
Proxy SNP: 12:104339610:T:C
deCODE log10(p): 1769.3
deCODE BETA: 0.72
1257:1257:1257:1256:1256:125

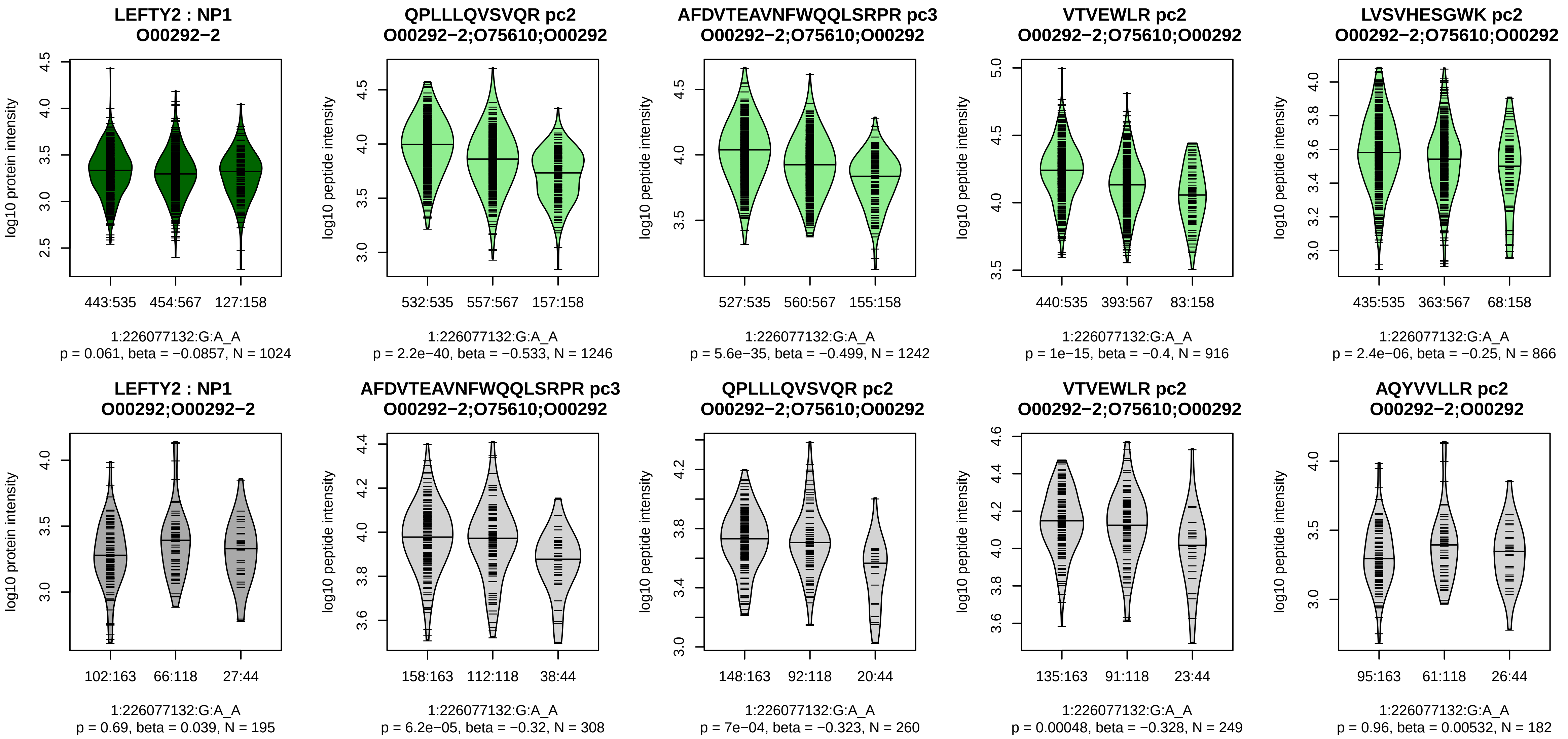


Assay Target: LEFTY2
Olink UniProt: O00292
deCODE rsID: rs360059
Proxy rsID: rs360059
deCODE: 1:225889432:A:G
Proxy SNP: 1:226077132:G:A
deCODE log10(p): 1700
deCODE BETA: −0.71
*:*:*:*:–:–:–:–:–
1246:1242:916:866:856:628:33[illegible]

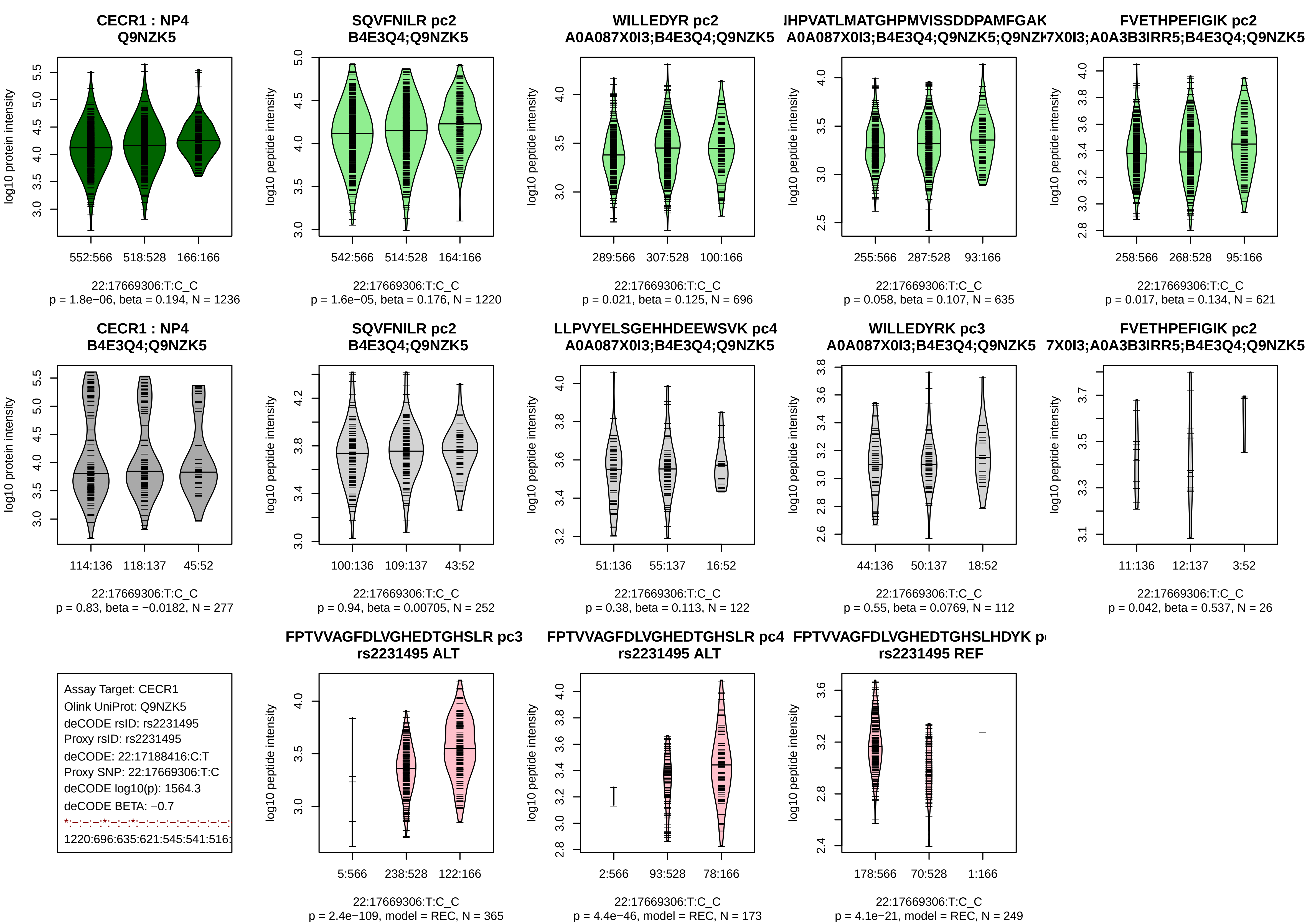
CECR1 : NP4
Q9NZK5
log10 protein intensity
552:566
518:528
166:166
22:17669306:T:C_C
p = 1.8e−06, beta = 0.194, N = 1236
SQVFNILR pc2
B4E3Q4;Q9NZK5
log10 peptide intensity
542:566
514:528
164:166
22:17669306:T:C_C
p = 1.6e−05, beta = 0.176, N = 1220
WILLEDYR pc2
A0A087X0I3;B4E3Q4;Q9NZK5
log10 peptide intensity
289:566
307:528
100:166
22:17669306:T:C_C
p = 0.021, beta = 0.125, N = 696
IHPVATLMATGHPMVISSDDPAMFGAK
A0A087X0I3;B4E3Q4;Q9NZK5;Q9NZI
log10 peptide intensity
255:566
287:528
93:166
22:17669306:T:C_C
p = 0.058, beta = 0.107, N = 635
FVETHPEFIGIK pc2
7X0I3;A0A3B3IRR5;B4E3Q4;Q9NZK5
log10 peptide intensity
258:566
268:528
95:166
22:17669306:T:C_C
p = 0.017, beta = 0.134, N = 621
CECR1 : NP4
B4E3Q4;Q9NZK5
log10 protein intensity
114:136
118:137
45:52
22:17669306:T:C_C
p = 0.83, beta = −0.0182, N = 277
SQVFNILR pc2
B4E3Q4;Q9NZK5
log10 peptide intensity
100:136
109:137
43:52
22:17669306:T:C_C
p = 0.94, beta = 0.00705, N = 252
LLPVYELSGEHHDEEWSVK pc4
A0A087X0I3;B4E3Q4;Q9NZK5
log10 peptide intensity
51:136
55:137
16:52
22:17669306:T:C_C
p = 0.38, beta = 0.113, N = 122
WILLEDYRK pc3
A0A087X0I3;B4E3Q4;Q9NZK5
log10 peptide intensity
44:136
50:137
18:52
22:17669306:T:C_C
p = 0.55, beta = 0.0769, N = 112
FVETHPEFIGIK pc2
7X0I3;A0A3B3IRR5;B4E3Q4;Q9NZK5
log10 peptide intensity
11:136
12:137
3:52
22:17669306:T:C_C
p = 0.042, beta = 0.537, N = 26
Assay Target: CECR1
Olink UniProt: Q9NZK5
deCODE rsID: rs2231495
Proxy rsID: rs2231495
deCODE: 22:17188416:C:T
Proxy SNP: 22:17669306:T:C
deCODE log10(p): 1564.3
deCODE BETA: −0.7
*:−:−:−:*:−:−:*:−:−:−:−:−:−:−:−:
1220:696:635:621:545:541:516:
FPTVVAGFDLVGHEDTGHSLR pc3
rs2231495 ALT
log10 peptide intensity
5:566
238:528
122:166
22:17669306:T:C_C
p = 2.4e−109, model = REC, N = 365
FPTVVAGFDLVGHEDTGHSLR pc4
rs2231495 ALT
log10 peptide intensity
2:566
93:528
78:166
22:17669306:T:C_C
p = 4.4e−46, model = REC, N = 173
FPTVVAGFDLVGHEDTGHSLHDYK p
rs2231495 REF
log10 peptide intensity
178:566
70:528
1:166
22:17669306:T:C_C
p = 4.1e−21, model = REC, N = 249

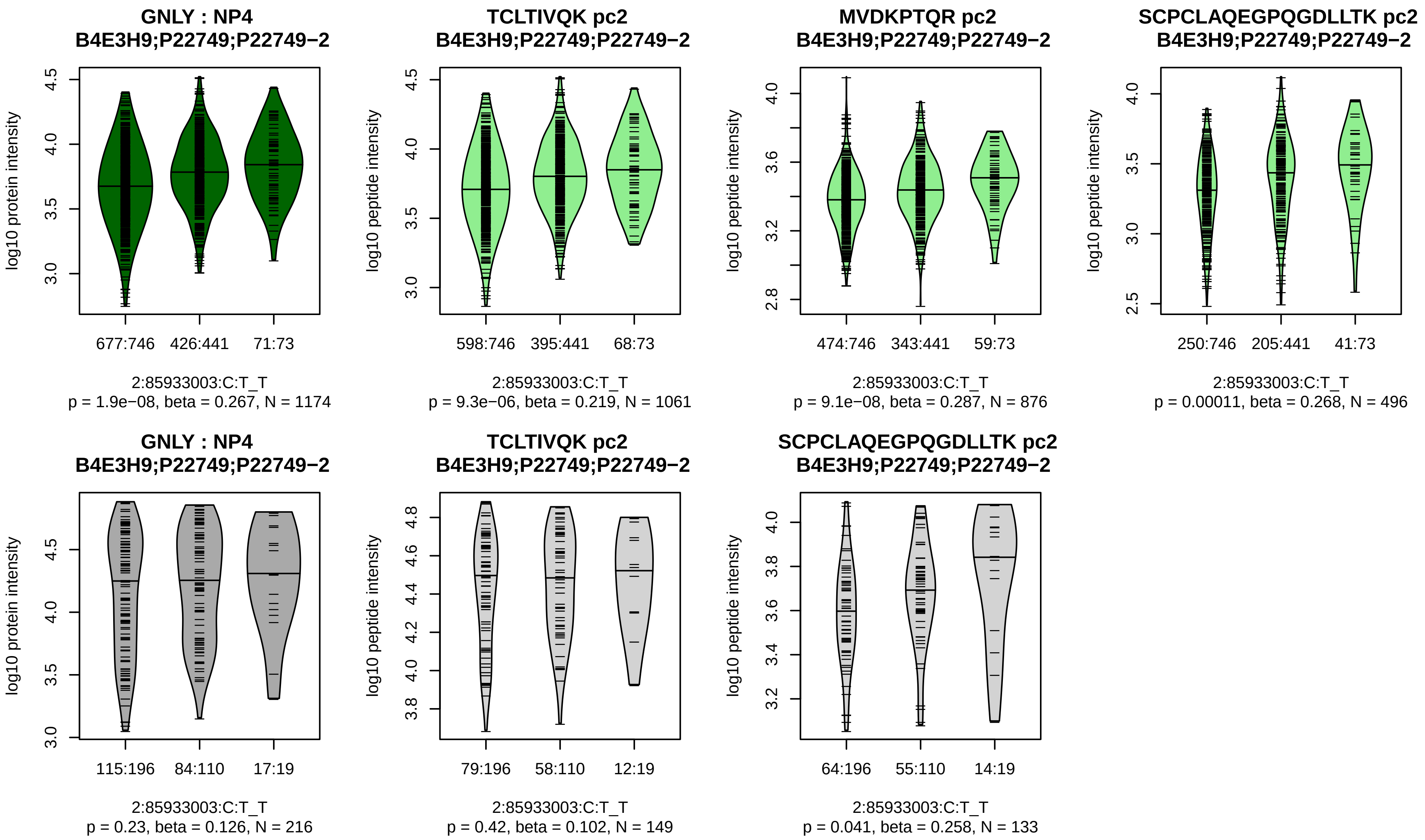


Assay Target: GNLY
Olink UniProt: P22749
deCODE rsID: rs12151742
Proxy rsID: rs12151742
deCODE: 2:85705880:T:C
Proxy SNP: 2:85933003:C:T
deCODE log10(p): 1527.6
deCODE BETA: 0.81
*:*:*
1061:876:496

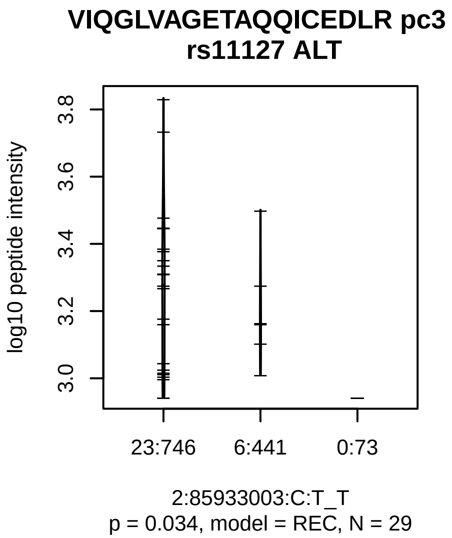

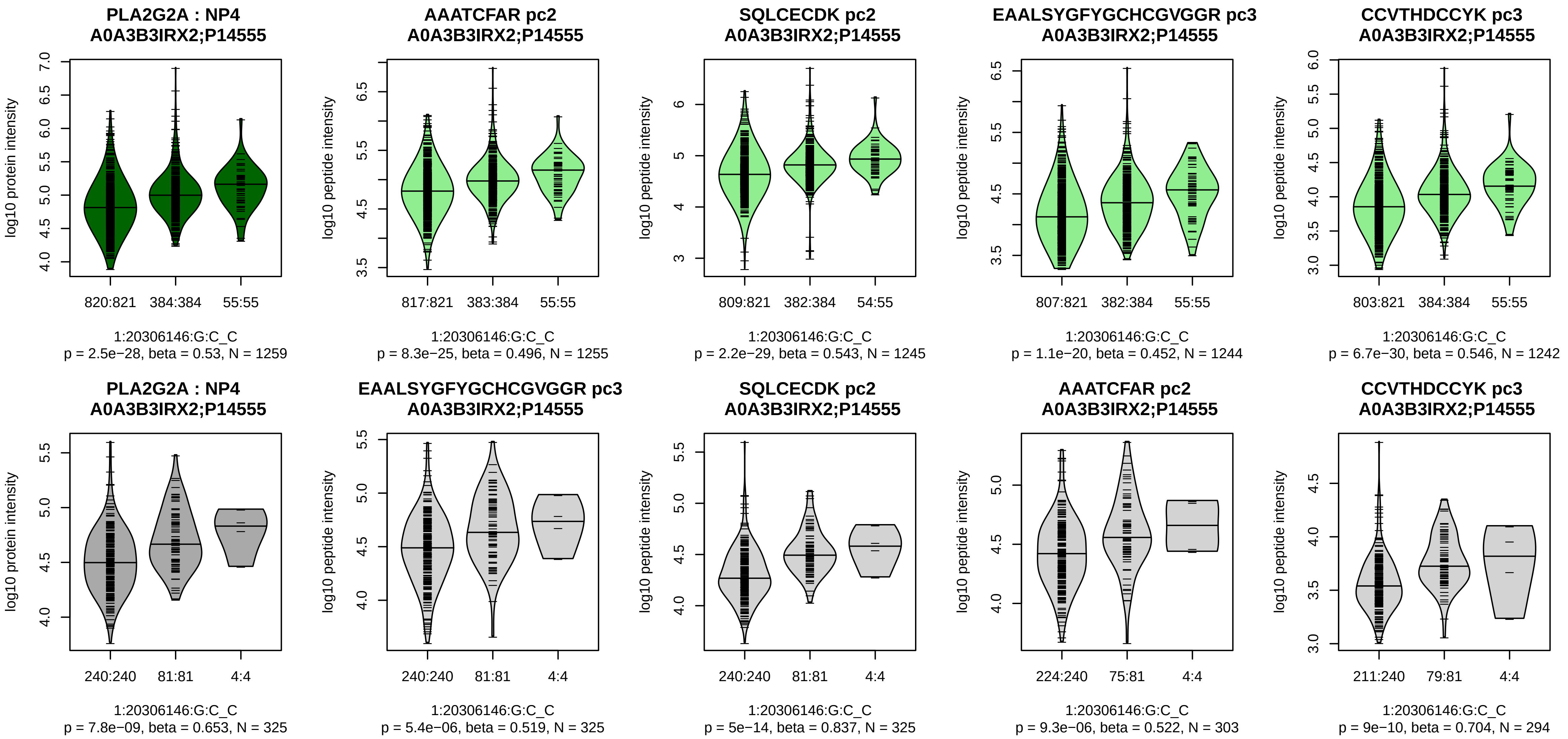


Assay Target: PLA2G2A
Olink UniProt: P14555
deCODE rsID: rs11573156
Proxy rsID: rs11573156
deCODE: 1:19979653:C:G
Proxy SNP: 1:20306146:G:C
deCODE log10(p): 1497.1
deCODE BETA: 0.77
*:*:*:*:*:*
1255:1245:1244:1242:1180:114

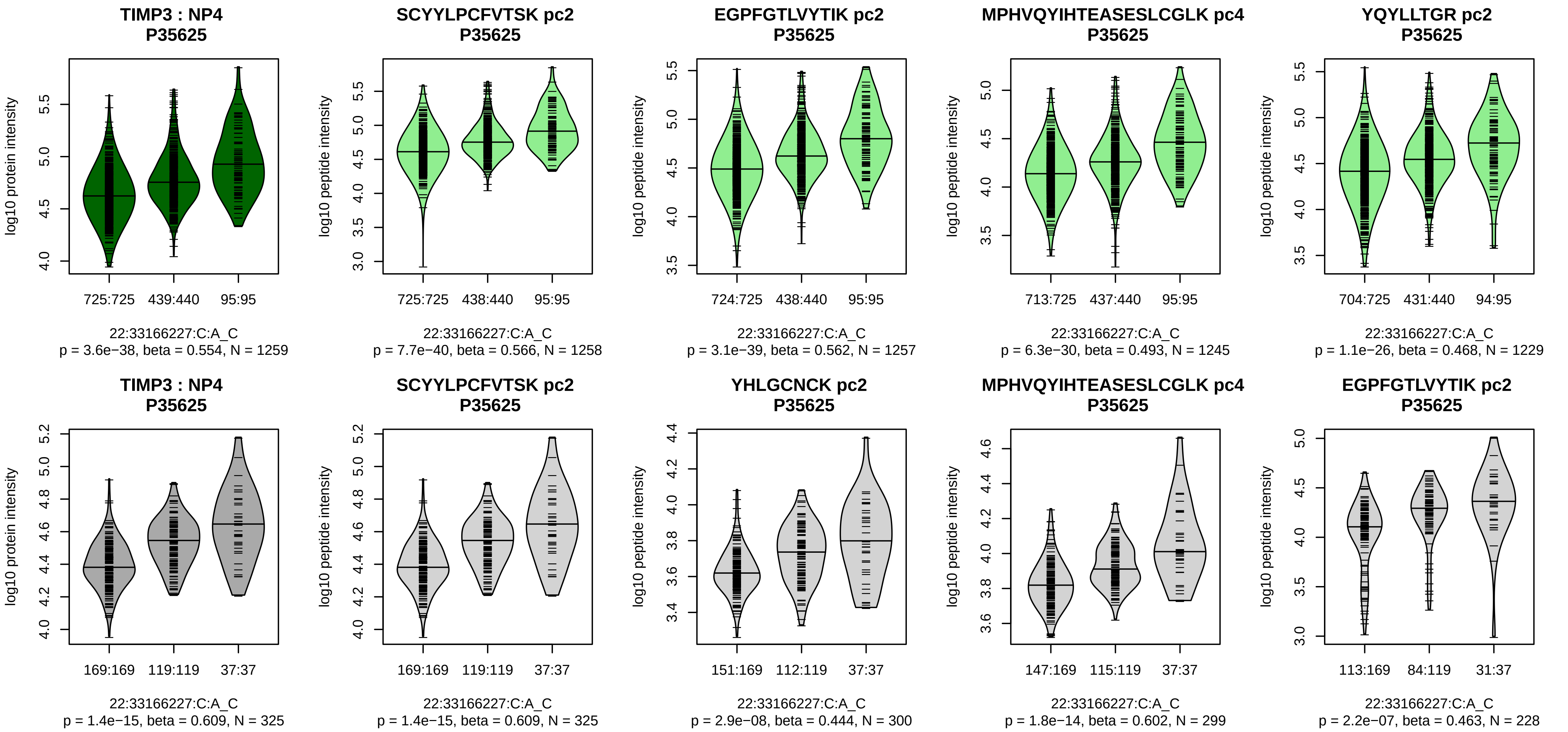


Assay Target: TIMP3
Olink UniProt: P35625
deCODE rsID: rs4821102
Proxy rsID: rs4821102
deCODE: 22:32770241:C:A
Proxy SNP: 22:33166227:C:A
deCODE log10(p): 1471
deCODE BETA: 0.78
*:*:*:*:*:*:*:*
1258:1257:1245:1229:1218:121

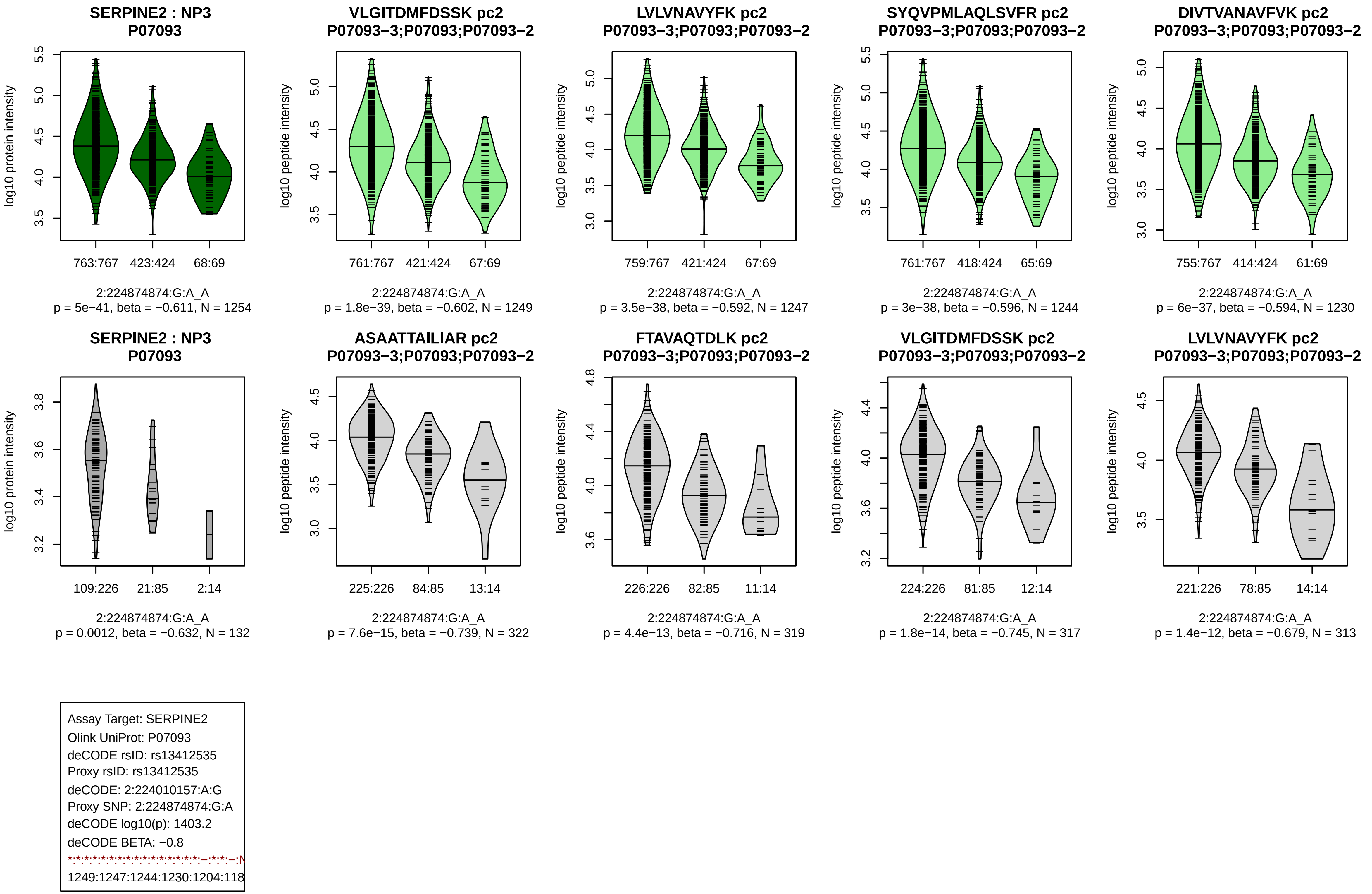
SERPINE2 : NP3
P07093
log10 protein intensity
763:767
423:424
68:69
2:224874874:G:A_A
p = 5e−41, beta = −0.611, N = 1254
VLGITDMFDSSK pc2
P07093−3;P07093;P07093−2
log10 peptide intensity
761:767
421:424
67:69
2:224874874:G:A_A
p = 1.8e−39, beta = −0.602, N = 1249
LVLVNAVYFK pc2
P07093−3;P07093;P07093−2
log10 peptide intensity
759:767
421:424
67:69
2:224874874:G:A_A
p = 3.5e−38, beta = −0.592, N = 1247
SYQVPMLAQLSVFR pc2
P07093−3;P07093;P07093−2
log10 peptide intensity
761:767
418:424
65:69
2:224874874:G:A_A
p = 3e−38, beta = −0.596, N = 1244
DIVTVANAVFVK pc2
P07093−3;P07093;P07093−2
log10 peptide intensity
755:767
414:424
61:69
2:224874874:G:A_A
p = 6e−37, beta = −0.594, N = 1230
SERPINE2 : NP3
P07093
log10 protein intensity
109:226
21:85
2:14
2:224874874:G:A_A
p = 0.0012, beta = −0.632, N = 132
ASAATTAILIAR pc2
P07093−3;P07093;P07093−2
log10 peptide intensity
225:226
84:85
13:14
2:224874874:G:A_A
p = 7.6e−15, beta = −0.739, N = 322
FTAVAQTDLK pc2
P07093−3;P07093;P07093−2
log10 peptide intensity
226:226
82:85
11:14
2:224874874:G:A_A
p = 4.4e−13, beta = −0.716, N = 319
VLGITDMFDSSK pc2
P07093−3;P07093;P07093−2
log10 peptide intensity
224:226
81:85
12:14
2:224874874:G:A_A
p = 1.8e−14, beta = −0.745, N = 317
LVLVNAVYFK pc2
P07093−3;P07093;P07093−2
log10 peptide intensity
221:226
78:85
14:14
2:224874874:G:A_A
p = 1.4e−12, beta = −0.679, N = 313
Assay Target: SERPINE2
Olink UniProt: P07093
deCODE rsID: rs13412535
Proxy rsID: rs13412535
deCODE: 2:224010157:A:G
Proxy SNP: 2:224874874:G:A
deCODE log10(p): 1403.2
deCODE BETA: −0.8
1249:1247:1244:1230:1204:118

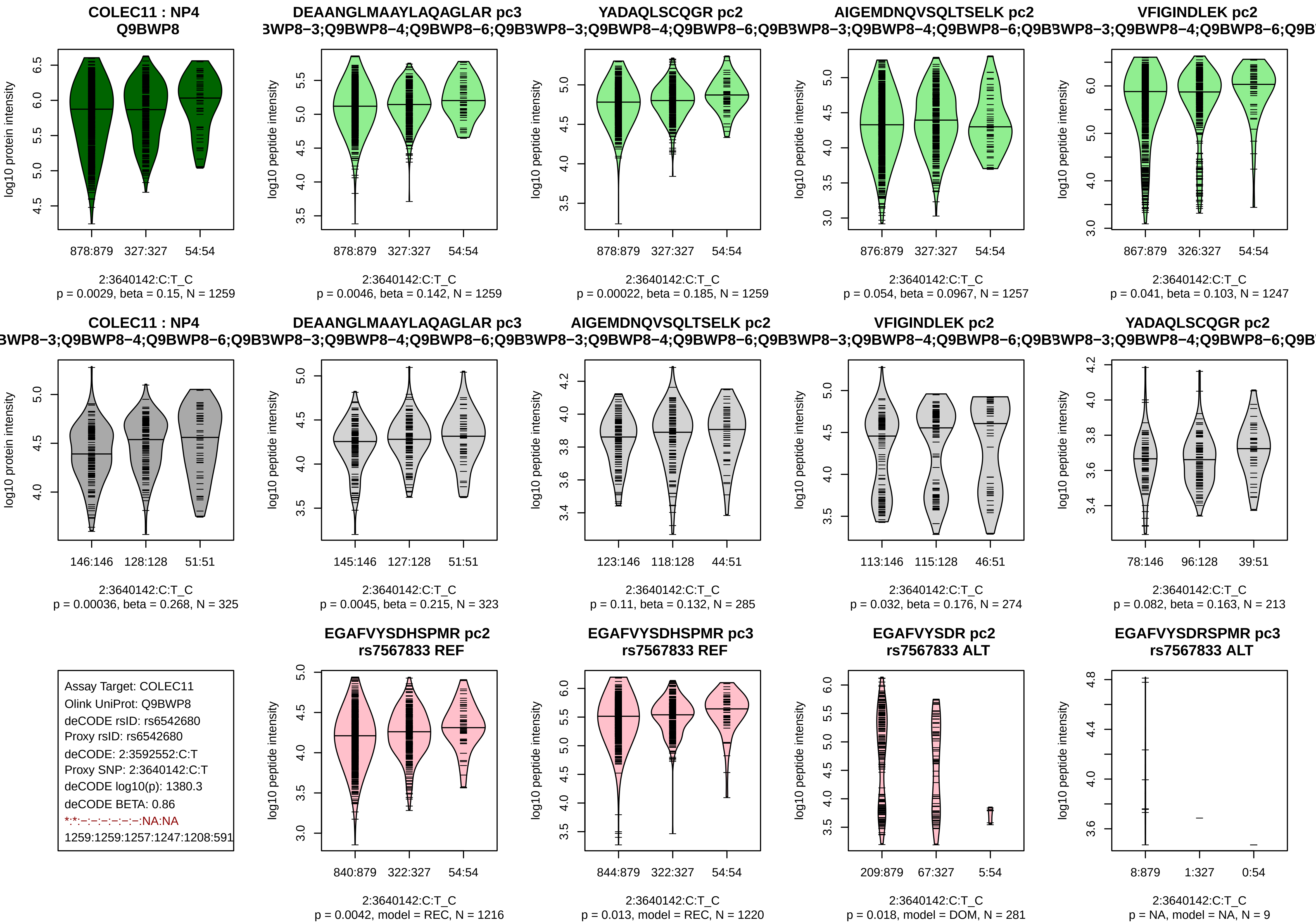
COLEC11 : NP4
Q9BWP8
log10 protein intensity
878:879 327:327 54:54
2:3640142:C:T_C
p = 0.0029, beta = 0.15, N = 1259
DEAANGLMAAYLAQAGLAR pc3
3WP8−3;Q9BWP8−4;Q9BWP8−6;Q9B
log10 peptide intensity
878:879 327:327 54:54
2:3640142:C:T_C
p = 0.0046, beta = 0.142, N = 1259
YADAQLSCQGR pc2
3WP8−3;Q9BWP8−4;Q9BWP8−6;Q9B
log10 peptide intensity
878:879 327:327 54:54
2:3640142:C:T_C
p = 0.00022, beta = 0.185, N = 1259
AIGEMDNQVSQLTSELK pc2
3WP8−3;Q9BWP8−4;Q9BWP8−6;Q9B
log10 peptide intensity
876:879 327:327 54:54
2:3640142:C:T_C
p = 0.054, beta = 0.0967, N = 1257
VFIGINDLEK pc2
3WP8−3;Q9BWP8−4;Q9BWP8−6;Q9B
log10 peptide intensity
867:879 326:327 54:54
2:3640142:C:T_C
p = 0.041, beta = 0.103, N = 1247
COLEC11 : NP4
3WP8−3;Q9BWP8−4;Q9BWP8−6;Q9B
log10 protein intensity
146:146 128:128 51:51
2:3640142:C:T_C
p = 0.00036, beta = 0.268, N = 325
DEAANGLMAAYLAQAGLAR pc3
3WP8−3;Q9BWP8−4;Q9BWP8−6;Q9B
log10 peptide intensity
145:146 127:128 51:51
2:3640142:C:T_C
p = 0.0045, beta = 0.215, N = 323
AIGEMDNQVSQLTSELK pc2
3WP8−3;Q9BWP8−4;Q9BWP8−6;Q9B
log10 peptide intensity
123:146 118:128 44:51
2:3640142:C:T_C
p = 0.11, beta = 0.132, N = 285
VFIGINDLEK pc2
3WP8−3;Q9BWP8−4;Q9BWP8−6;Q9B
log10 peptide intensity
113:146 115:128 46:51
2:3640142:C:T_C
p = 0.032, beta = 0.176, N = 274
YADAQLSCQGR pc2
3WP8−3;Q9BWP8−4;Q9BWP8−6;Q9B
log10 peptide intensity
78:146 96:128 39:51
2:3640142:C:T_C
p = 0.082, beta = 0.163, N = 213
Assay Target: COLEC11
Olink UniProt: Q9BWP8
deCODE rsID: rs6542680
Proxy rsID: rs6542680
deCODE: 2:3592552:C:T
Proxy SNP: 2:3640142:C:T
deCODE log10(p): 1380.3
deCODE BETA: 0.86
*:*:−:−:−:−:−:−:NA:NA
1259:1259:1257:1247:1208:591
EGAFVYSDHSPMR pc2
rs7567833 REF
log10 peptide intensity
840:879 322:327 54:54
2:3640142:C:T_C
p = 0.0042, model = REC, N = 1216
EGAFVYSDHSPMR pc3
rs7567833 REF
log10 peptide intensity
844:879 322:327 54:54
2:3640142:C:T_C
p = 0.013, model = REC, N = 1220
EGAFVYSDR pc2
rs7567833 ALT
log10 peptide intensity
209:879 67:327 5:54
2:3640142:C:T_C
p = 0.018, model = DOM, N = 281
EGAFVYSDRSPMR pc3
rs7567833 ALT
log10 peptide intensity
8:879 1:327 0:54
2:3640142:C:T_C
p = NA, model = NA, N = 9

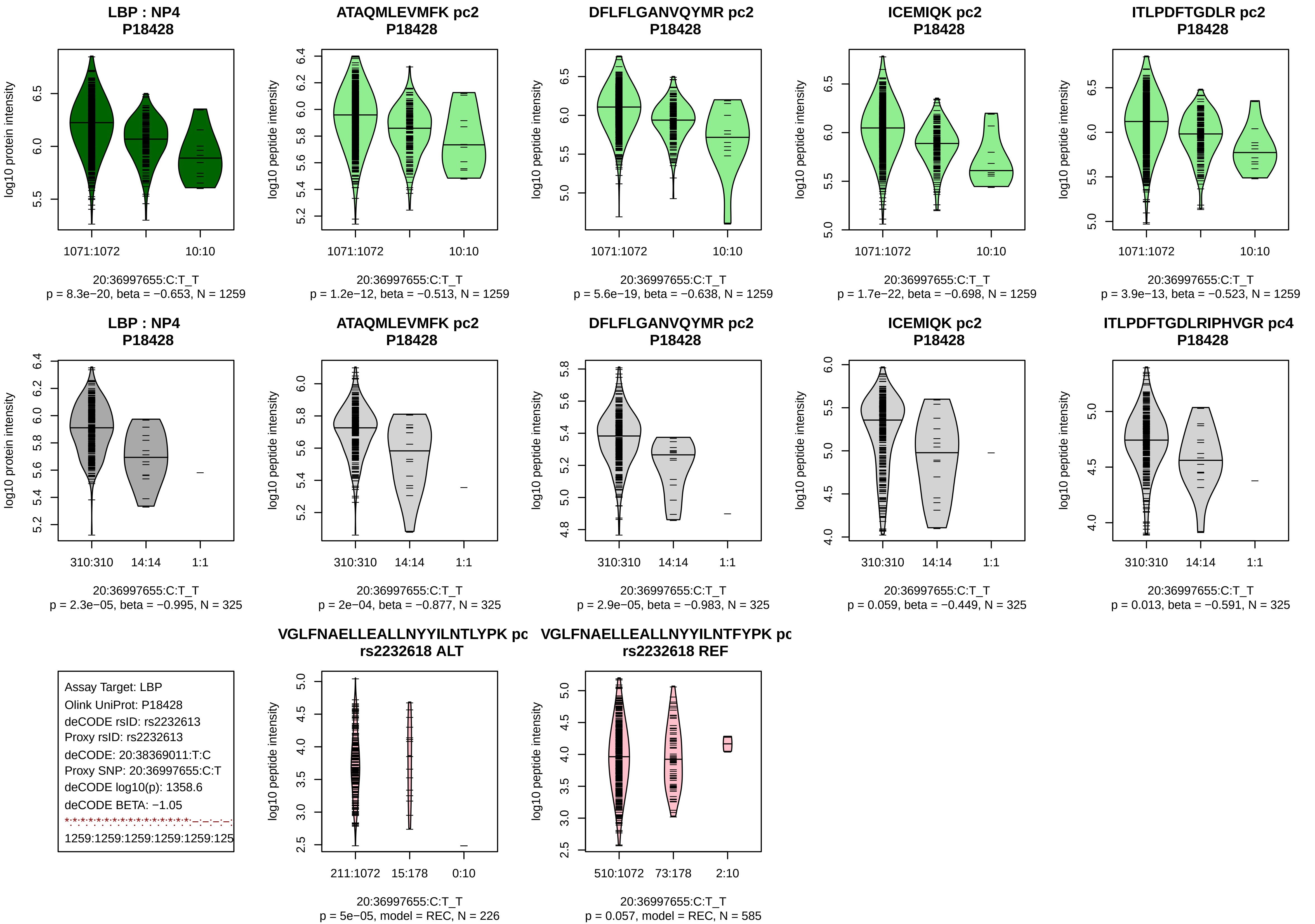

LBP : NP4
P18428
ATAQMLEVMFK pc2
P18428
DFLFLGANVQYMR pc2
P18428
ICEMIQK pc2
P18428
ITLPDFTGDLR pc2
P18428
log10 protein intensity
log10 peptide intensity
1071:1072
10:10
20:36997655:C:T_T
p = 8.3e−20, beta = −0.653, N = 1259
p = 1.2e−12, beta = −0.513, N = 1259
p = 5.6e−19, beta = −0.638, N = 1259
p = 1.7e−22, beta = −0.698, N = 1259
p = 3.9e−13, beta = −0.523, N = 1259
ITLPDFTGDLRIPHVGR pc4
310:310
14:14
1:1
p = 2.3e−05, beta = −0.995, N = 325
p = 2e−04, beta = −0.877, N = 325
p = 2.9e−05, beta = −0.983, N = 325
p = 0.059, beta = −0.449, N = 325
p = 0.013, beta = −0.591, N = 325
Assay Target: LBP
Olink UniProt: P18428
deCODE rsID: rs2232613
Proxy rsID: rs2232613
deCODE: 20:38369011:T:C
Proxy SNP: 20:36997655:C:T
deCODE log10(p): 1358.6
deCODE BETA: −1.05
1259:1259:1259:1259:1259:125
VGLFNAELLEALLNYYILNTLYPK pc
rs2232618 ALT
VGLFNAELLEALLNYYILNTFYPK pc
rs2232618 REF
211:1072
15:178
0:10
510:1072
73:178
2:10
p = 5e−05, model = REC, N = 226
p = 0.057, model = REC, N = 585

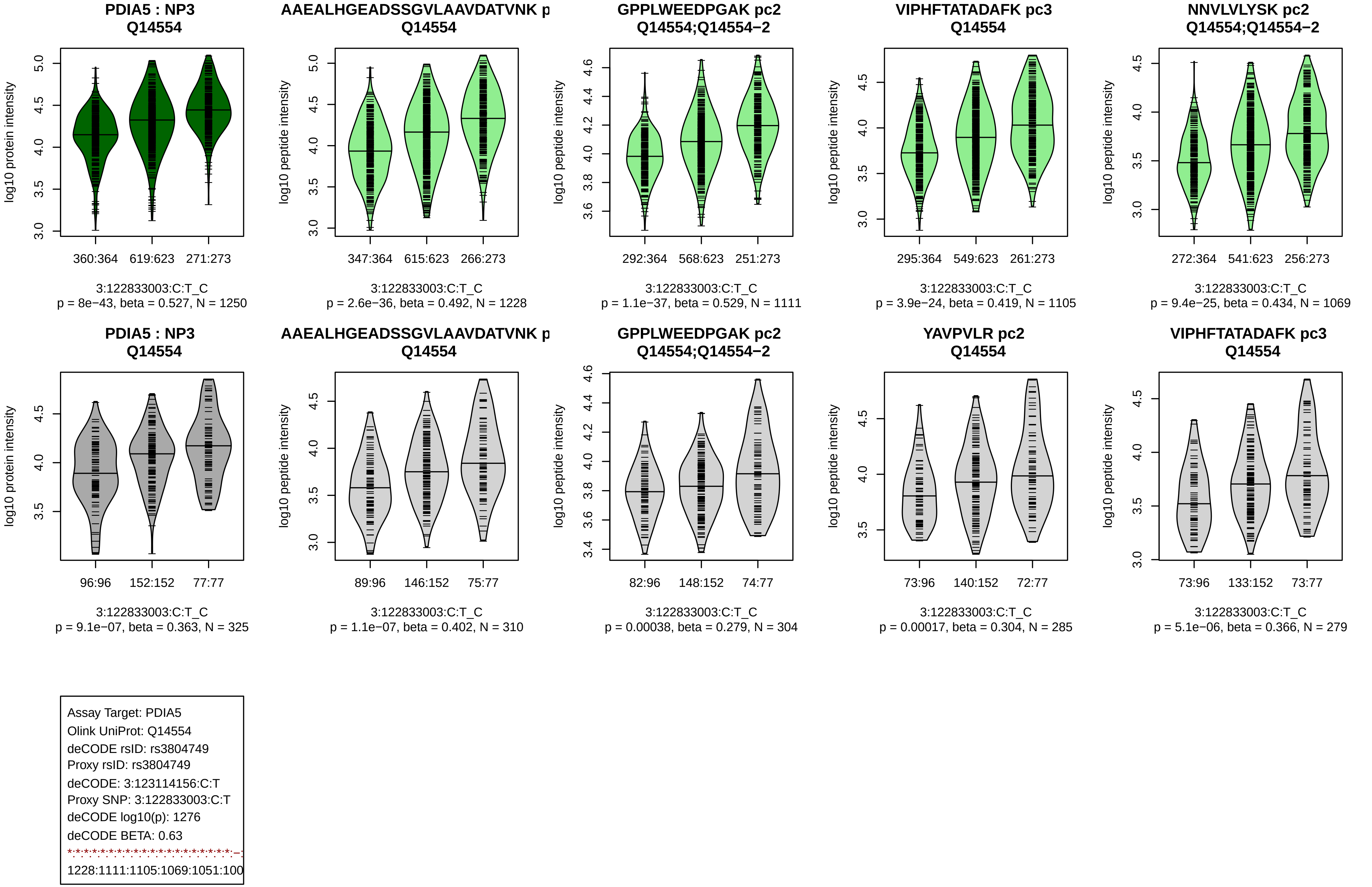
PDIA5 : NP3
Q14554
log10 protein intensity
360:364
619:623
271:273
3:122833003:C:T_C
p = 8e−43, beta = 0.527, N = 1250
AAEALHGEADSSGVLAAVDATVNK p
Q14554
log10 peptide intensity
347:364
615:623
266:273
3:122833003:C:T_C
p = 2.6e−36, beta = 0.492, N = 1228
GPPLWEEDPGAK pc2
Q14554;Q14554−2
log10 peptide intensity
292:364
568:623
251:273
3:122833003:C:T_C
p = 1.1e−37, beta = 0.529, N = 1111
VIPHFTATADAFK pc3
Q14554
log10 peptide intensity
295:364
549:623
261:273
3:122833003:C:T_C
p = 3.9e−24, beta = 0.419, N = 1105
NNVLVLYSK pc2
Q14554;Q14554−2
log10 peptide intensity
272:364
541:623
256:273
3:122833003:C:T_C
p = 9.4e−25, beta = 0.434, N = 1069
PDIA5 : NP3
Q14554
log10 protein intensity
96:96
152:152
77:77
3:122833003:C:T_C
p = 9.1e−07, beta = 0.363, N = 325
AAEALHGEADSSGVLAAVDATVNK p
Q14554
log10 peptide intensity
89:96
146:152
75:77
3:122833003:C:T_C
p = 1.1e−07, beta = 0.402, N = 310
GPPLWEEDPGAK pc2
Q14554;Q14554−2
log10 peptide intensity
82:96
148:152
74:77
3:122833003:C:T_C
p = 0.00038, beta = 0.279, N = 304
YAVPVLR pc2
Q14554
log10 peptide intensity
73:96
140:152
72:77
3:122833003:C:T_C
p = 0.00017, beta = 0.304, N = 285
VIPHFTATADAFK pc3
Q14554
log10 peptide intensity
73:96
133:152
73:77
3:122833003:C:T_C
p = 5.1e−06, beta = 0.366, N = 279
Assay Target: PDIA5
Olink UniProt: Q14554
deCODE rsID: rs3804749
Proxy rsID: rs3804749
deCODE: 3:123114156:C:T
Proxy SNP: 3:122833003:C:T
deCODE log10(p): 1276
deCODE BETA: 0.63
1228:1111:1105:1069:1051:100

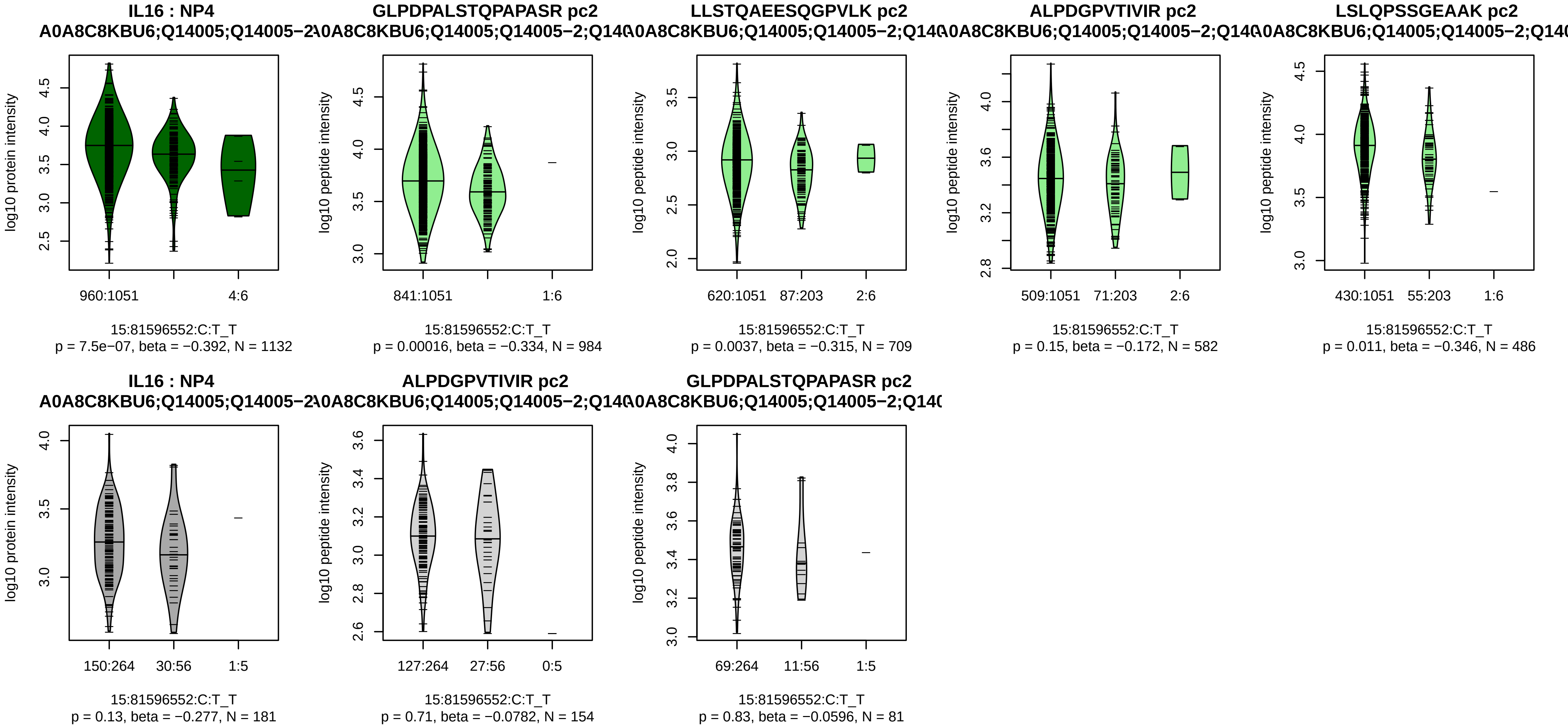


Assay Target: IL16
Olink UniProt: Q14005
deCODE rsID: rs17875523
Proxy rsID: rs17875523
deCODE: 15:81304211:T:C
Proxy SNP: 15:81596552:C:T
deCODE log10(p): 1235
deCODE BETA: −1.06
*:*:−:−:−:−:−:−
984:709:582:486:235:210:91:41

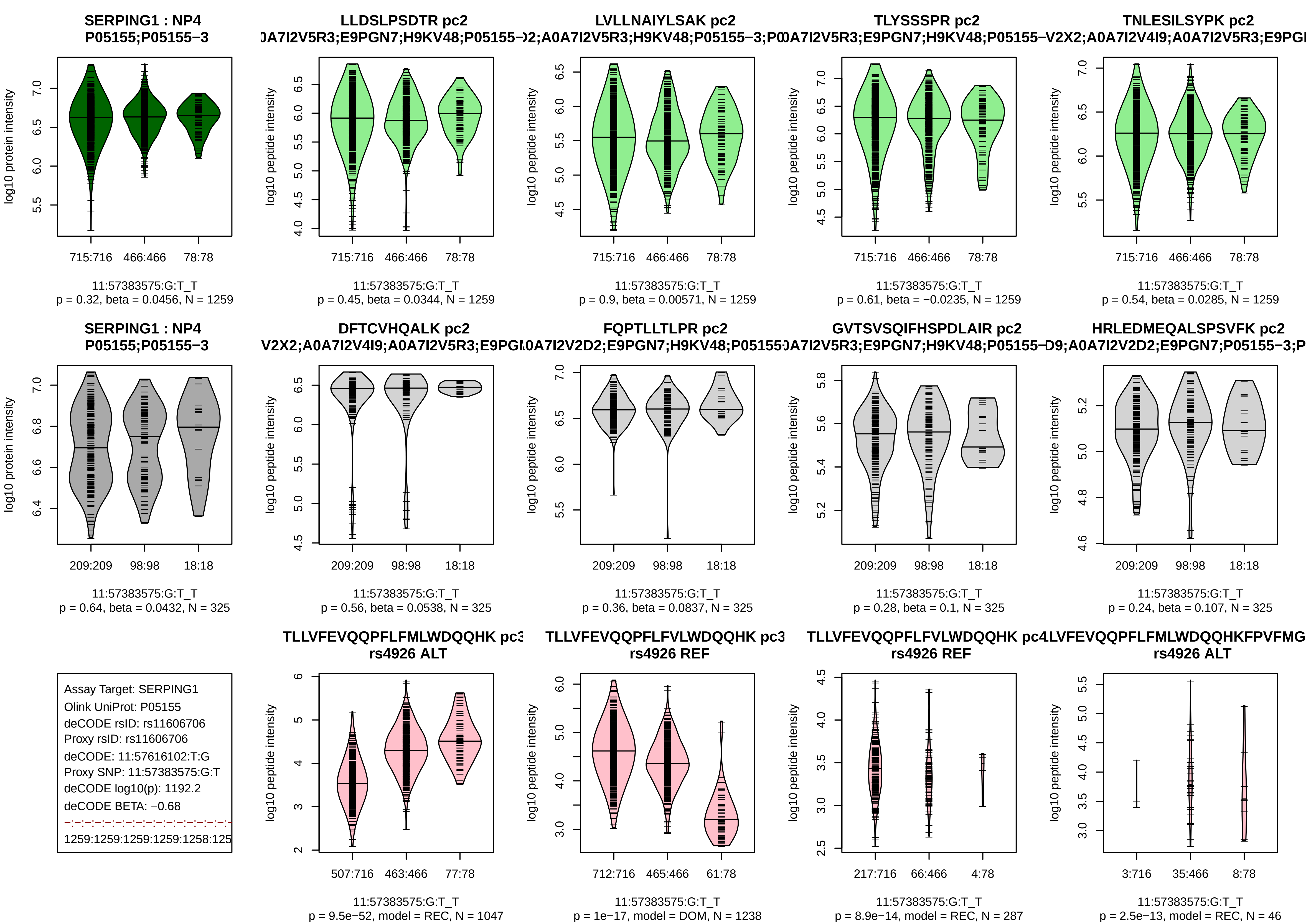
SERPING1 : NP4
P05155;P05155−3
log10 protein intensity
715:716 466:466 78:78
11:57383575:G:T_T
p = 0.32, beta = 0.0456, N = 1259
LLDSLPSDTR pc2
log10 peptide intensity
p = 0.45, beta = 0.0344, N = 1259
LVLLNAIYLSAK pc2
p = 0.9, beta = 0.00571, N = 1259
TLYSSSPR pc2
p = 0.61, beta = −0.0235, N = 1259
TNLESILSYPK pc2
p = 0.54, beta = 0.0285, N = 1259
209:209 98:98 18:18
p = 0.64, beta = 0.0432, N = 325
DFTCVHQALK pc2
p = 0.56, beta = 0.0538, N = 325
FQPTLLTLPR pc2
p = 0.36, beta = 0.0837, N = 325
GVTSVSQIFHSPDLAIR pc2
p = 0.28, beta = 0.1, N = 325
HRLEDMEQALSPSVFK pc2
p = 0.24, beta = 0.107, N = 325
Assay Target: SERPING1
Olink UniProt: P05155
deCODE rsID: rs11606706
Proxy rsID: rs11606706
deCODE: 11:57616102:T:G
Proxy SNP: 11:57383575:G:T
deCODE log10(p): 1192.2
deCODE BETA: −0.68
1259:1259:1259:1259:1258:125
rs4926 ALT
507:716 463:466 77:78
p = 9.5e−52, model = REC, N = 1047
rs4926 REF
712:716 465:466 61:78
p = 1e−17, model = DOM, N = 1238
TLLVFEVQQPFLFVLWDQQHK pc4
rs4926 REF
217:716 66:466 4:78
p = 8.9e−14, model = REC, N = 287
rs4926 ALT
3:716 35:466 8:78
p = 2.5e−13, model = REC, N = 46

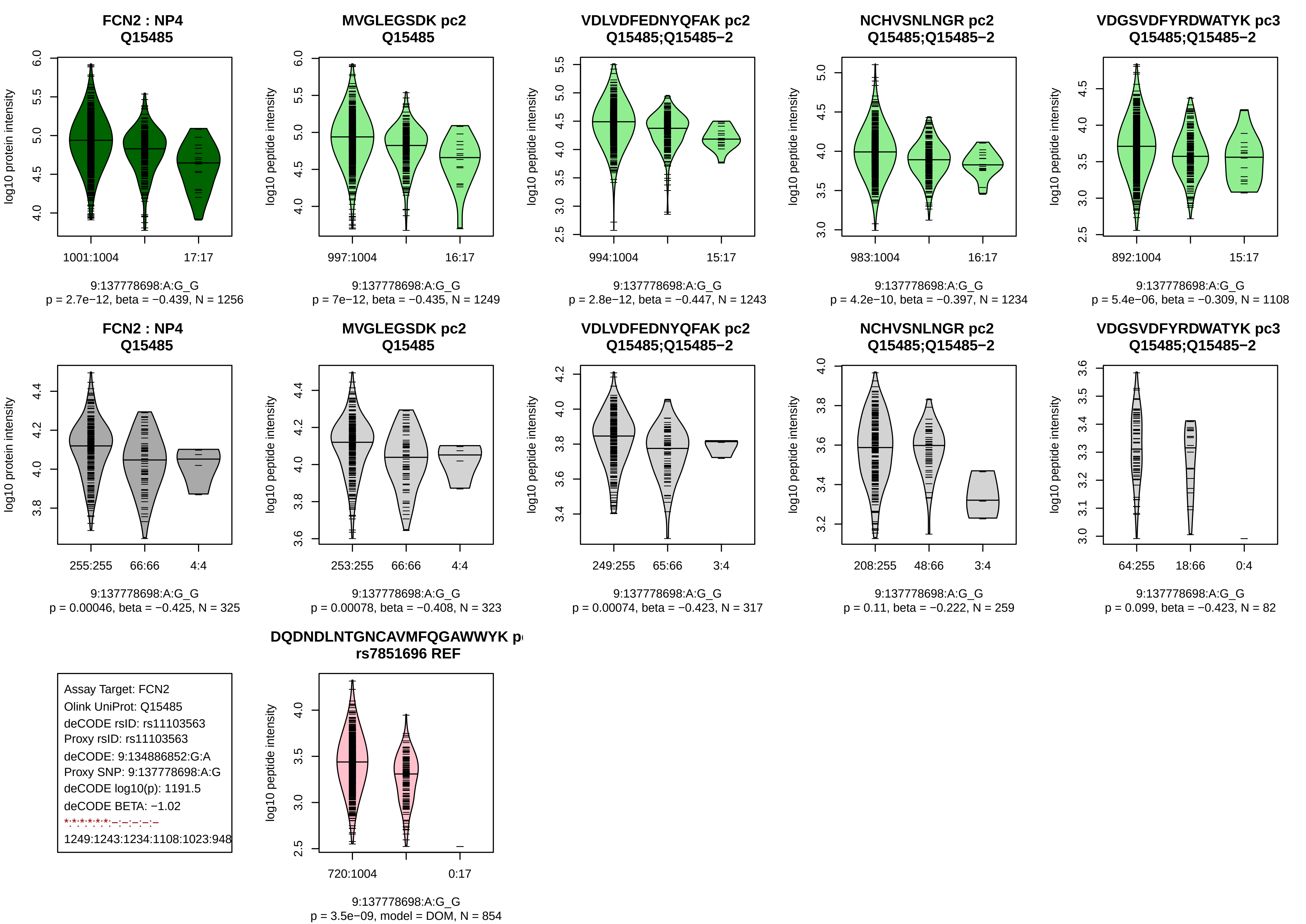
FCN2 : NP4
Q15485
log10 protein intensity
1001:1004
17:17
9:137778698:A:G_G
p = 2.7e−12, beta = −0.439, N = 1256
MVGLEGSDK pc2
Q15485
log10 peptide intensity
997:1004
16:17
9:137778698:A:G_G
p = 7e−12, beta = −0.435, N = 1249
VDLVDFEDNYQFAK pc2
Q15485;Q15485−2
log10 peptide intensity
994:1004
15:17
9:137778698:A:G_G
p = 2.8e−12, beta = −0.447, N = 1243
NCHVSNLNGR pc2
Q15485;Q15485−2
log10 peptide intensity
983:1004
16:17
9:137778698:A:G_G
p = 4.2e−10, beta = −0.397, N = 1234
VDGSVDFYRDWATYK pc3
Q15485;Q15485−2
log10 peptide intensity
892:1004
15:17
9:137778698:A:G_G
p = 5.4e−06, beta = −0.309, N = 1108
FCN2 : NP4
Q15485
log10 protein intensity
255:255
66:66
4:4
9:137778698:A:G_G
p = 0.00046, beta = −0.425, N = 325
MVGLEGSDK pc2
Q15485
log10 peptide intensity
253:255
66:66
4:4
9:137778698:A:G_G
p = 0.00078, beta = −0.408, N = 323
VDLVDFEDNYQFAK pc2
Q15485;Q15485−2
log10 peptide intensity
249:255
65:66
3:4
9:137778698:A:G_G
p = 0.00074, beta = −0.423, N = 317
NCHVSNLNGR pc2
Q15485;Q15485−2
log10 peptide intensity
208:255
48:66
3:4
9:137778698:A:G_G
p = 0.11, beta = −0.222, N = 259
VDGSVDFYRDWATYK pc3
Q15485;Q15485−2
log10 peptide intensity
64:255
18:66
0:4
9:137778698:A:G_G
p = 0.099, beta = −0.423, N = 82
Assay Target: FCN2
Olink UniProt: Q15485
deCODE rsID: rs11103563
Proxy rsID: rs11103563
deCODE: 9:134886852:G:A
Proxy SNP: 9:137778698:A:G
deCODE log10(p): 1191.5
deCODE BETA: −1.02
*:*:*:*:*:*:–:–:–:–:–
1249:1243:1234:1108:1023:948
DQDNDLNTGNCAVMFQGAWWYK p
rs7851696 REF
log10 peptide intensity
720:1004
0:17
9:137778698:A:G_G
p = 3.5e−09, model = DOM, N = 854

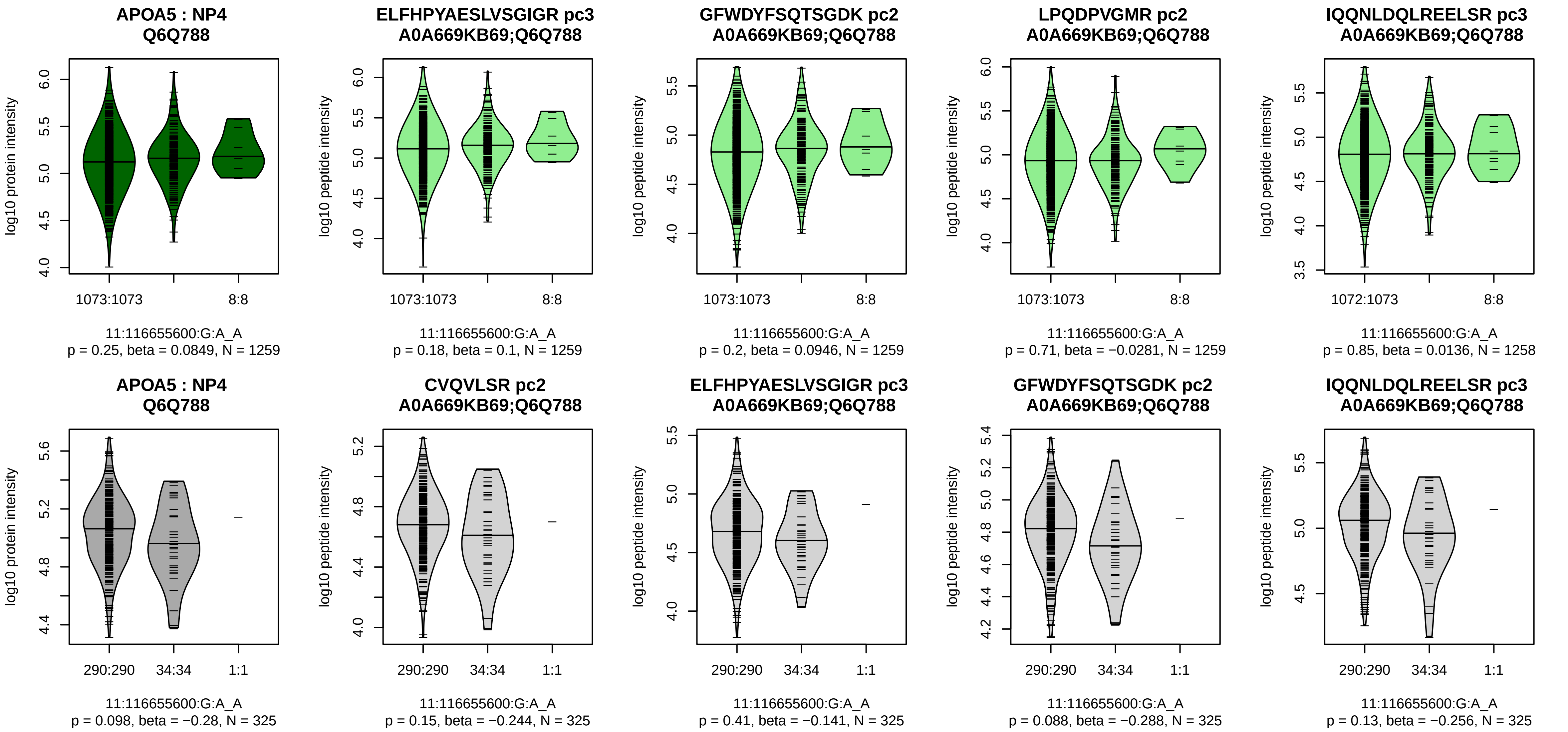


Assay Target: APOA5
Olink UniProt: Q6Q788
deCODE rsID: rs35120633
Proxy rsID: rs35120633
deCODE: 11:116784884:A:G
Proxy SNP: 11:116655600:G:A
deCODE log10(p): 1180.2
deCODE BETA: 1.26
–:–:–:–:–:–:–:–:–:–:–:–:–:–:–:–:
1259:1259:1259:1258:1258:125

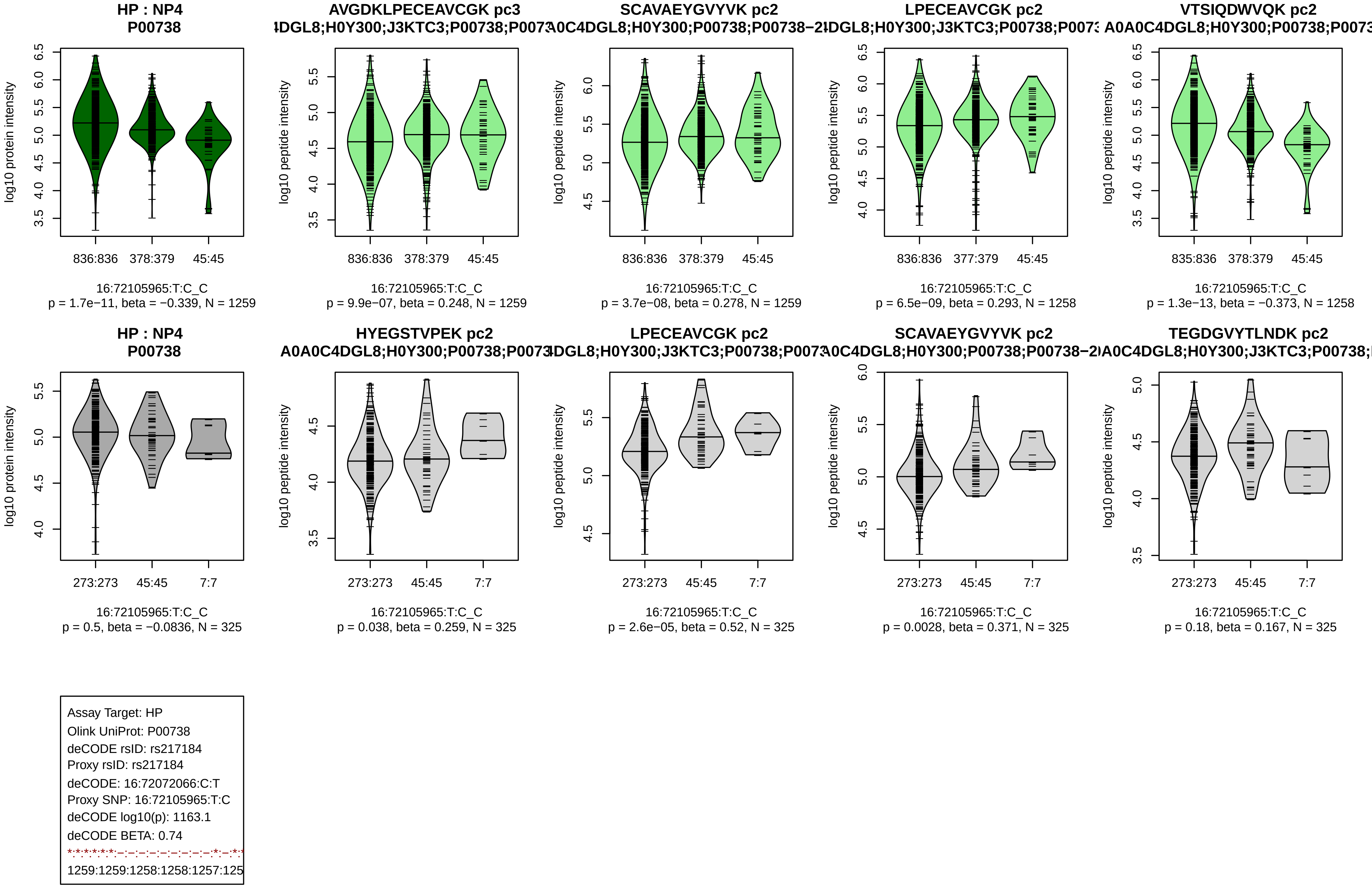

HP : NP4
P00738
log10 protein intensity
836:836
378:379
45:45
16:72105965:T:C_C
p = 1.7e−11, beta = −0.339, N = 1259
AVGDKLPECEAVCGK pc3
4DGL8;H0Y300;J3KTC3;P00738;P0073
log10 peptide intensity
836:836
378:379
45:45
16:72105965:T:C_C
p = 9.9e−07, beta = 0.248, N = 1259
SCAVAEYGVYVK pc2
A0C4DGL8;H0Y300;P00738;P00738−2
log10 peptide intensity
836:836
378:379
45:45
16:72105965:T:C_C
p = 3.7e−08, beta = 0.278, N = 1259
LPECEAVCGK pc2
4DGL8;H0Y300;J3KTC3;P00738;P0073
log10 peptide intensity
836:836
377:379
45:45
16:72105965:T:C_C
p = 6.5e−09, beta = 0.293, N = 1258
VTSIQDWVQK pc2
A0A0C4DGL8;H0Y300;P00738;P0073
log10 peptide intensity
835:836
378:379
45:45
16:72105965:T:C_C
p = 1.3e−13, beta = −0.373, N = 1258
HP : NP4
P00738
log10 protein intensity
273:273
45:45
7:7
16:72105965:T:C_C
p = 0.5, beta = −0.0836, N = 325
HYEGSTVPEK pc2
A0A0C4DGL8;H0Y300;P00738;P0073
log10 peptide intensity
273:273
45:45
7:7
16:72105965:T:C_C
p = 0.038, beta = 0.259, N = 325
LPECEAVCGK pc2
4DGL8;H0Y300;J3KTC3;P00738;P0073
log10 peptide intensity
273:273
45:45
7:7
16:72105965:T:C_C
p = 2.6e−05, beta = 0.52, N = 325
SCAVAEYGVYVK pc2
A0C4DGL8;H0Y300;P00738;P00738−2
log10 peptide intensity
273:273
45:45
7:7
16:72105965:T:C_C
p = 0.0028, beta = 0.371, N = 325
TEGDGVYTLNDK pc2
A0A0C4DGL8;H0Y300;J3KTC3;P00738;
log10 peptide intensity
273:273
45:45
7:7
16:72105965:T:C_C
p = 0.18, beta = 0.167, N = 325
Assay Target: HP
Olink UniProt: P00738
deCODE rsID: rs217184
Proxy rsID: rs217184
deCODE: 16:72072066:C:T
Proxy SNP: 16:72105965:T:C
deCODE log10(p): 1163.1
deCODE BETA: 0.74
*:*:*:*:*:*:–:–:–:–:–:–:–:–:–:*:–:*:*
1259:1259:1258:1258:1257:125

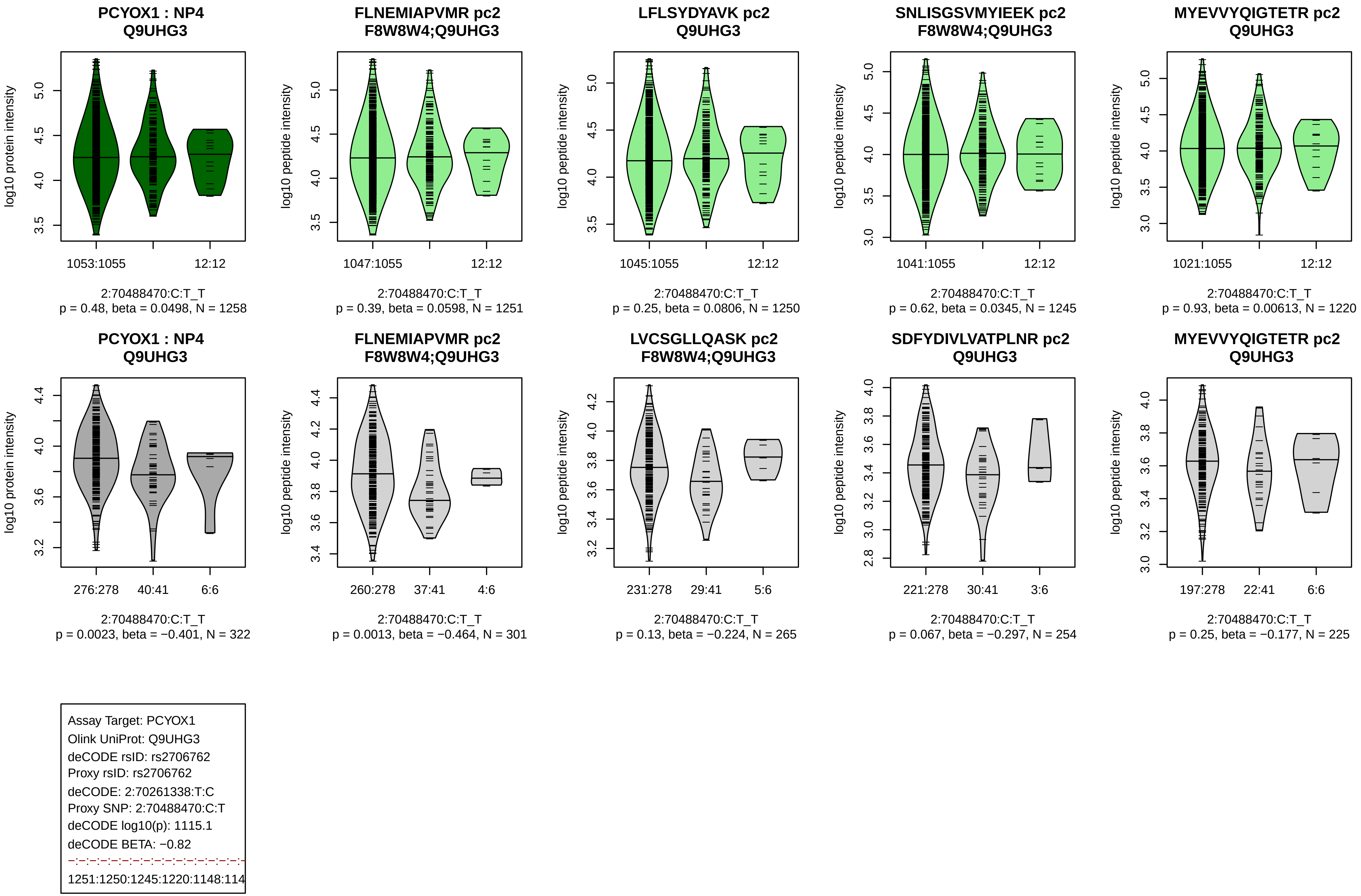
PCYOX1 : NP4
Q9UHG3
log10 protein intensity
1053:1055
12:12
2:70488470:C:T_T
p = 0.48, beta = 0.0498, N = 1258
FLNEMIAPVMR pc2
F8W8W4;Q9UHG3
log10 peptide intensity
1047:1055
12:12
2:70488470:C:T_T
p = 0.39, beta = 0.0598, N = 1251
LFLSYDYAVK pc2
Q9UHG3
log10 peptide intensity
1045:1055
12:12
2:70488470:C:T_T
p = 0.25, beta = 0.0806, N = 1250
SNLISGSVMYIEEK pc2
F8W8W4;Q9UHG3
log10 peptide intensity
1041:1055
12:12
2:70488470:C:T_T
p = 0.62, beta = 0.0345, N = 1245
MYEVVYQIGTETR pc2
Q9UHG3
log10 peptide intensity
1021:1055
12:12
2:70488470:C:T_T
p = 0.93, beta = 0.00613, N = 1220
PCYOX1 : NP4
Q9UHG3
log10 protein intensity
276:278
40:41
6:6
2:70488470:C:T_T
p = 0.0023, beta = −0.401, N = 322
FLNEMIAPVMR pc2
F8W8W4;Q9UHG3
log10 peptide intensity
260:278
37:41
4:6
2:70488470:C:T_T
p = 0.0013, beta = −0.464, N = 301
LVCSGLLQASK pc2
F8W8W4;Q9UHG3
log10 peptide intensity
231:278
29:41
5:6
2:70488470:C:T_T
p = 0.13, beta = −0.224, N = 265
SDFYDIVLVATPLNR pc2
Q9UHG3
log10 peptide intensity
221:278
30:41
3:6
2:70488470:C:T_T
p = 0.067, beta = −0.297, N = 254
MYEVVYQIGTETR pc2
Q9UHG3
log10 peptide intensity
197:278
22:41
6:6
2:70488470:C:T_T
p = 0.25, beta = −0.177, N = 225
Assay Target: PCYOX1
Olink UniProt: Q9UHG3
deCODE rsID: rs2706762
Proxy rsID: rs2706762
deCODE: 2:70261338:T:C
Proxy SNP: 2:70488470:C:T
deCODE log10(p): 1115.1
deCODE BETA: −0.82
1251:1250:1245:1220:1148:114

**KLK10 : NP3**
**O43240**

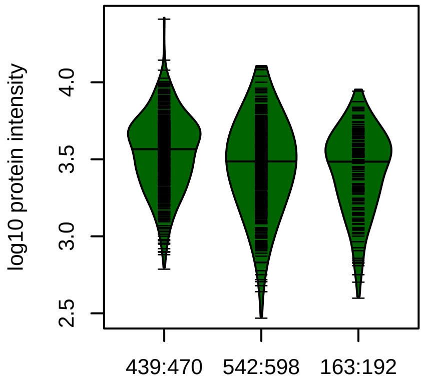


19:51523203:T:C_T
p = 6.1e−07, beta = −0.213, N = 1144

**GLTCSSITILSPK pc2**
**O43240**

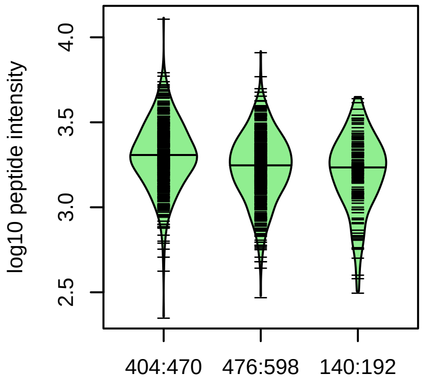


19:51523203:T:C_T
p = 2.3e−07, beta = −0.235, N = 1020

**ALQLPYR pc2**
**O43240**

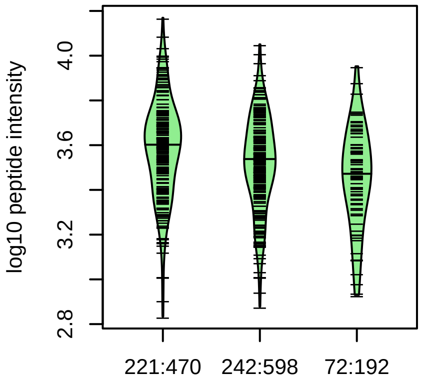


19:51523203:T:C_T
p = 1.2e−05, beta = −0.272, N = 535

**RTDEHDLMLLK pc3**
**O43240**

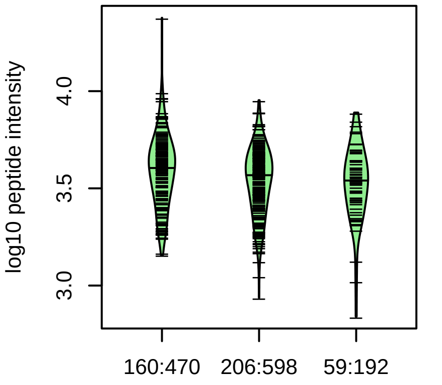


19:51523203:T:C_T
p = 0.005, beta = −0.198, N = 425

**YHQGSGPILPR pc2**
**O43240**

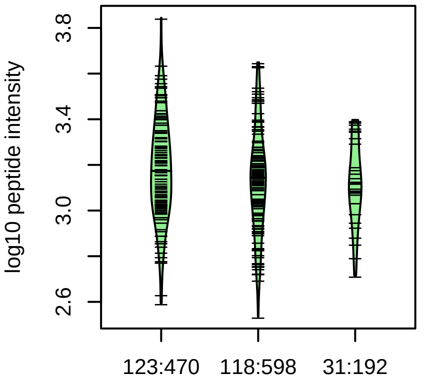


19:51523203:T:C_T
p = 0.059, beta = −0.168, N = 272

Assay Target: KLK10
Olink UniProt: O43240
deCODE rsID: rs2569454
Proxy rsID: rs2569454
deCODE: 19:51019947:T:C
Proxy SNP: 19:51523203:T:C
deCODE log10(p): 1063.6
deCODE BETA: −0.61
*:*:*:−:−:−:−:NA:NA
1020:535:425:272:248:86:76:5:4

**LDPEAYGAPCAR pc2**
**rs3745535 ALT**

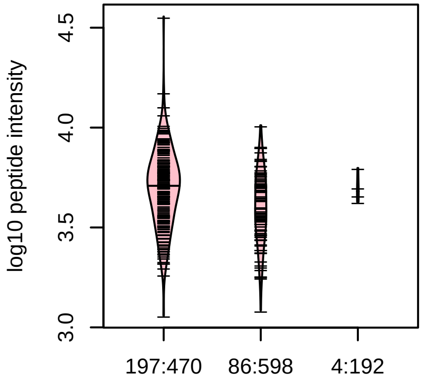


19:51523203:T:C_T
p = 6.1e−35, model = REC, N = 287

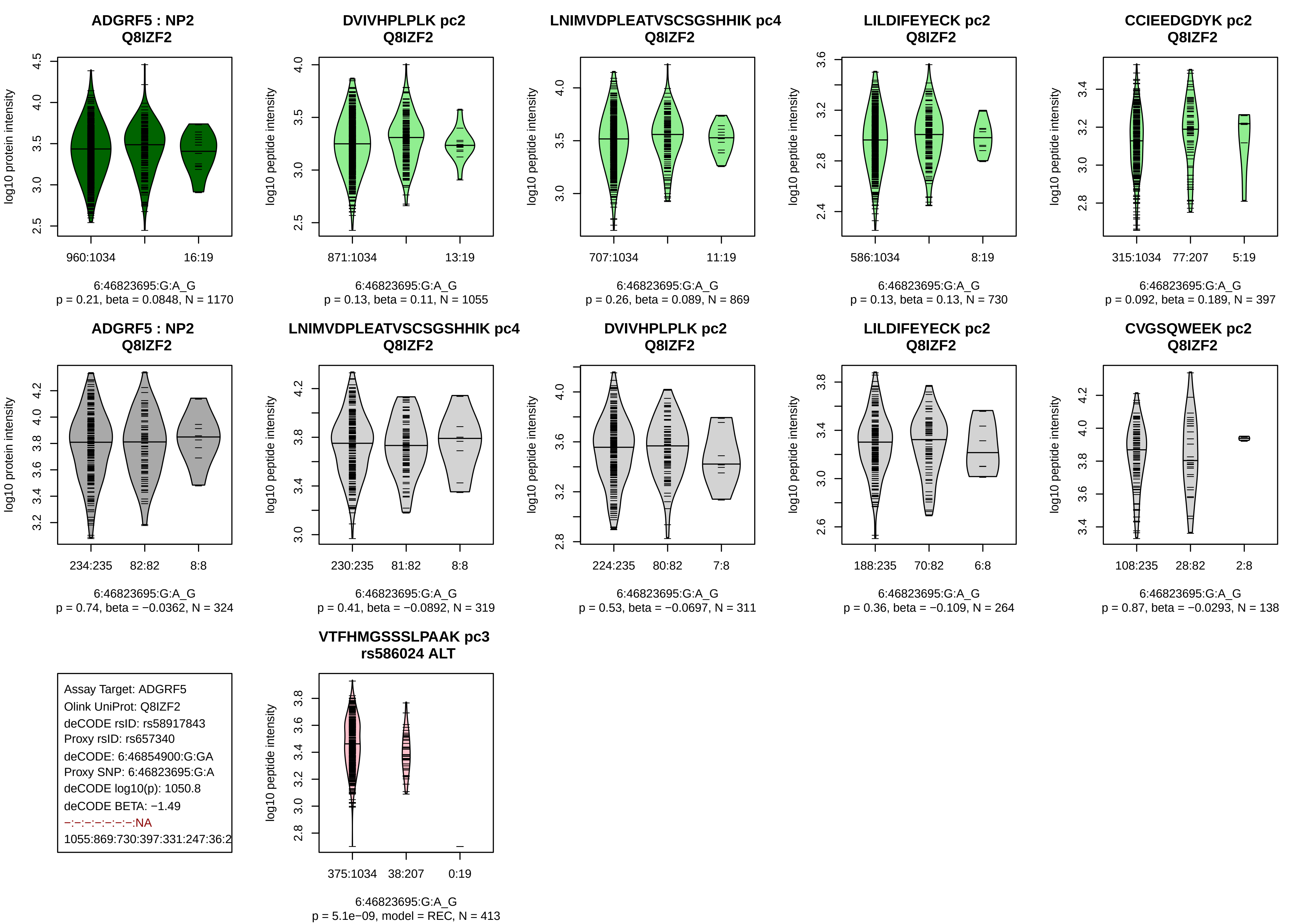
ADGRF5 : NP2
Q8IZF2
log10 protein intensity
960:1034
16:19
6:46823695:G:A_G
p = 0.21, beta = 0.0848, N = 1170
DVIVHPLPLK pc2
Q8IZF2
log10 peptide intensity
871:1034
13:19
6:46823695:G:A_G
p = 0.13, beta = 0.11, N = 1055
LNIMVDPLEATVSCSGSHHIK pc4
Q8IZF2
707:1034
11:19
6:46823695:G:A_G
p = 0.26, beta = 0.089, N = 869
LILDIFEYECK pc2
Q8IZF2
586:1034
8:19
6:46823695:G:A_G
p = 0.13, beta = 0.13, N = 730
CCIEEDGDYK pc2
Q8IZF2
315:1034
77:207
5:19
6:46823695:G:A_G
p = 0.092, beta = 0.189, N = 397
ADGRF5 : NP2
Q8IZF2
234:235
82:82
8:8
6:46823695:G:A_G
p = 0.74, beta = −0.0362, N = 324
LNIMVDPLEATVSCSGSHHIK pc4
Q8IZF2
230:235
81:82
8:8
6:46823695:G:A_G
p = 0.41, beta = −0.0892, N = 319
DVIVHPLPLK pc2
Q8IZF2
224:235
80:82
7:8
6:46823695:G:A_G
p = 0.53, beta = −0.0697, N = 311
LILDIFEYECK pc2
Q8IZF2
188:235
70:82
6:8
6:46823695:G:A_G
p = 0.36, beta = −0.109, N = 264
CVGSQWEEK pc2
Q8IZF2
108:235
28:82
2:8
6:46823695:G:A_G
p = 0.87, beta = −0.0293, N = 138
Assay Target: ADGRF5
Olink UniProt: Q8IZF2
deCODE rsID: rs58917843
Proxy rsID: rs657340
deCODE: 6:46854900:G:GA
Proxy SNP: 6:46823695:G:A
deCODE log10(p): 1050.8
deCODE BETA: −1.49
−:−:−:−:−:−:−:NA
1055:869:730:397:331:247:36:2
VTFHMGSSSLPAAK pc3
rs586024 ALT
log10 peptide intensity
375:1034
38:207
0:19
6:46823695:G:A_G
p = 5.1e−09, model = REC, N = 413

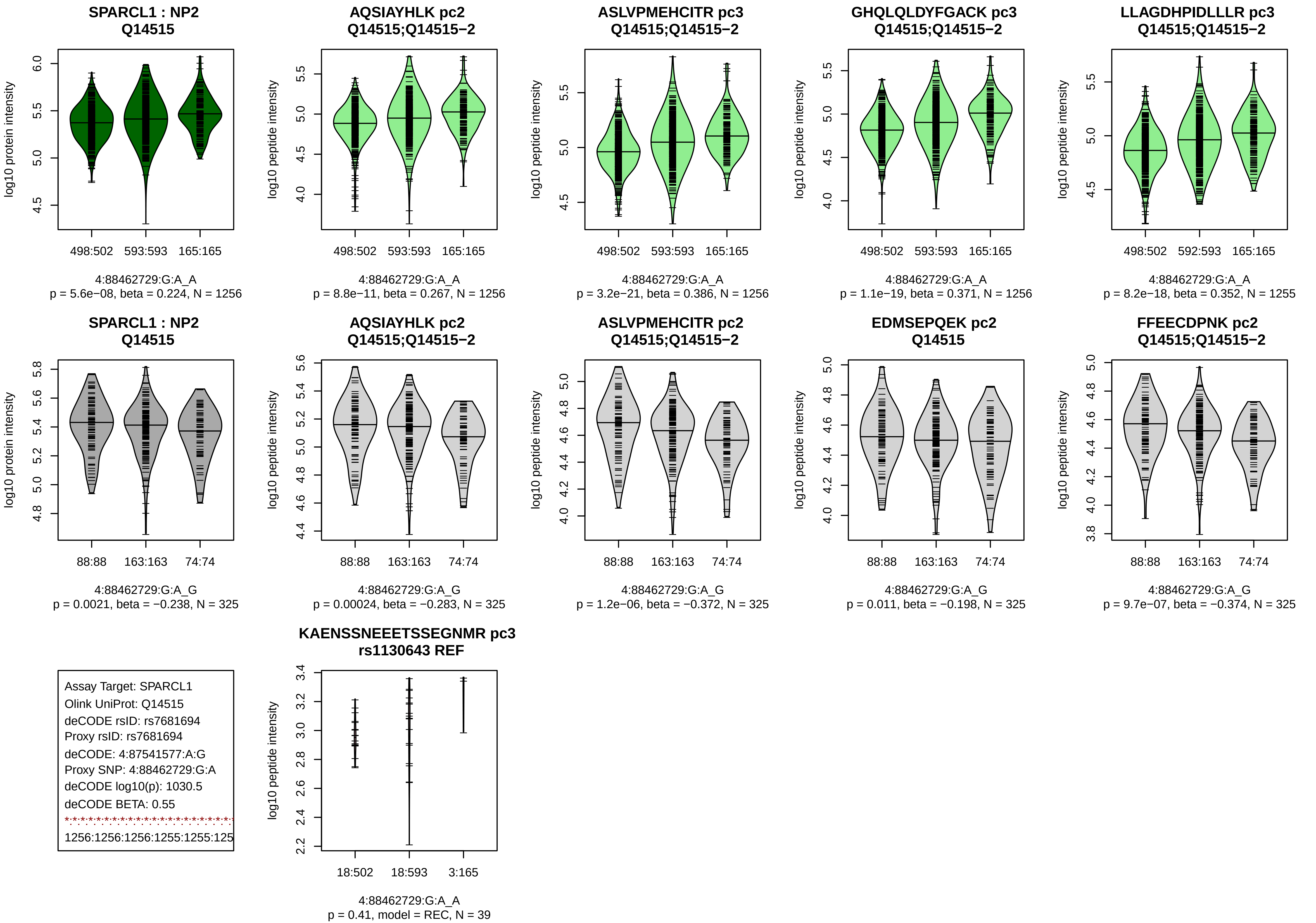

SPARCL1 : NP2
Q14515
log10 protein intensity
498:502
593:593
165:165
4:88462729:G:A_A
p = 5.6e−08, beta = 0.224, N = 1256
AQSIAYHLK pc2
Q14515;Q14515−2
log10 peptide intensity
498:502
593:593
165:165
4:88462729:G:A_A
p = 8.8e−11, beta = 0.267, N = 1256
ASLVPMEHCITR pc3
Q14515;Q14515−2
log10 peptide intensity
498:502
593:593
165:165
4:88462729:G:A_A
p = 3.2e−21, beta = 0.386, N = 1256
GHQLQLDYFGACK pc3
Q14515;Q14515−2
log10 peptide intensity
498:502
593:593
165:165
4:88462729:G:A_A
p = 1.1e−19, beta = 0.371, N = 1256
LLAGDHPIDLLLR pc3
Q14515;Q14515−2
log10 peptide intensity
498:502
592:593
165:165
4:88462729:G:A_A
p = 8.2e−18, beta = 0.352, N = 1255
SPARCL1 : NP2
Q14515
log10 protein intensity
88:88
163:163
74:74
4:88462729:G:A_G
p = 0.0021, beta = −0.238, N = 325
AQSIAYHLK pc2
Q14515;Q14515−2
log10 peptide intensity
88:88
163:163
74:74
4:88462729:G:A_G
p = 0.00024, beta = −0.283, N = 325
ASLVPMEHCITR pc2
Q14515;Q14515−2
log10 peptide intensity
88:88
163:163
74:74
4:88462729:G:A_G
p = 1.2e−06, beta = −0.372, N = 325
EDMSEPQEK pc2
Q14515
log10 peptide intensity
88:88
163:163
74:74
4:88462729:G:A_G
p = 0.011, beta = −0.198, N = 325
FFEECDPNK pc2
Q14515;Q14515−2
log10 peptide intensity
88:88
163:163
74:74
4:88462729:G:A_G
p = 9.7e−07, beta = −0.374, N = 325
Assay Target: SPARCL1
Olink UniProt: Q14515
deCODE rsID: rs7681694
Proxy rsID: rs7681694
deCODE: 4:87541577:A:G
Proxy SNP: 4:88462729:G:A
deCODE log10(p): 1030.5
deCODE BETA: 0.55
1256:1256:1256:1255:1255:125
KAENSSNEEETSSEGNMR pc3
rs1130643 REF
log10 peptide intensity
18:502
18:593
3:165
4:88462729:G:A_A
p = 0.41, model = REC, N = 39

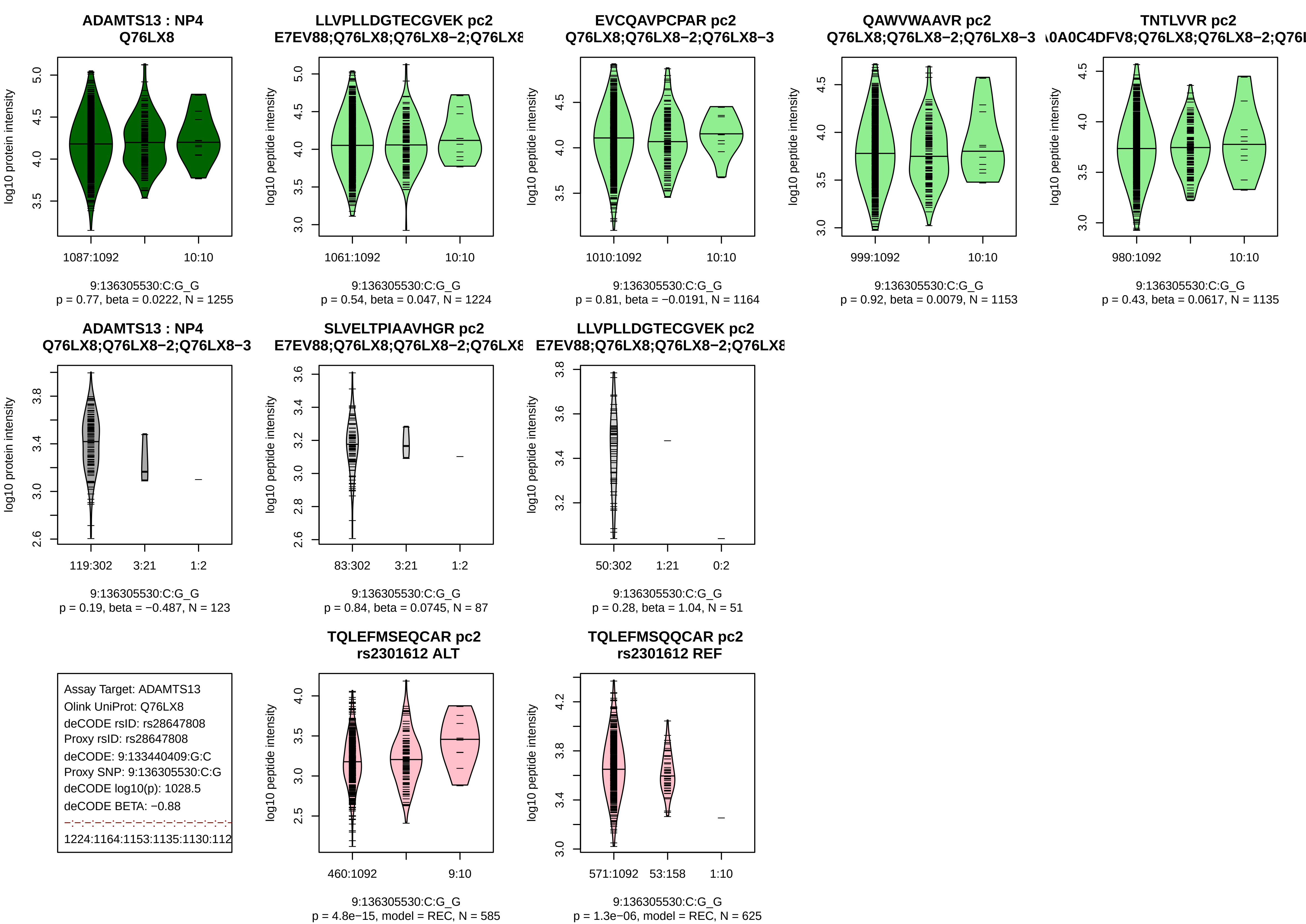

ADAMTS13 : NP4
Q76LX8
log10 protein intensity
1087:1092
10:10
9:136305530:C:G_G
p = 0.77, beta = 0.0222, N = 1255
LLVPLLDGTECGVEK pc2
E7EV88;Q76LX8;Q76LX8−2;Q76LX8
log10 peptide intensity
1061:1092
10:10
9:136305530:C:G_G
p = 0.54, beta = 0.047, N = 1224
EVCQAVPCPAR pc2
Q76LX8;Q76LX8−2;Q76LX8−3
1010:1092
10:10
9:136305530:C:G_G
p = 0.81, beta = −0.0191, N = 1164
QAWVWAAVR pc2
Q76LX8;Q76LX8−2;Q76LX8−3
999:1092
10:10
9:136305530:C:G_G
p = 0.92, beta = 0.0079, N = 1153
TNTLVVR pc2
A0A0C4DFV8;Q76LX8;Q76LX8−2;Q76L
980:1092
10:10
9:136305530:C:G_G
p = 0.43, beta = 0.0617, N = 1135
ADAMTS13 : NP4
Q76LX8;Q76LX8−2;Q76LX8−3
119:302
3:21
1:2
9:136305530:C:G_G
p = 0.19, beta = −0.487, N = 123
SLVELTPIAAVHGR pc2
E7EV88;Q76LX8;Q76LX8−2;Q76LX8
83:302
3:21
1:2
9:136305530:C:G_G
p = 0.84, beta = 0.0745, N = 87
LLVPLLDGTECGVEK pc2
E7EV88;Q76LX8;Q76LX8−2;Q76LX8
50:302
1:21
0:2
9:136305530:C:G_G
p = 0.28, beta = 1.04, N = 51
Assay Target: ADAMTS13
Olink UniProt: Q76LX8
deCODE rsID: rs28647808
Proxy rsID: rs28647808
deCODE: 9:133440409:G:C
Proxy SNP: 9:136305530:C:G
deCODE log10(p): 1028.5
deCODE BETA: −0.88
1224:1164:1153:1135:1130:112
TQLEFMSEQCAR pc2
rs2301612 ALT
460:1092
9:10
9:136305530:C:G_G
p = 4.8e−15, model = REC, N = 585
TQLEFMSQQCAR pc2
rs2301612 REF
571:1092
53:158
1:10
9:136305530:C:G_G
p = 1.3e−06, model = REC, N = 625

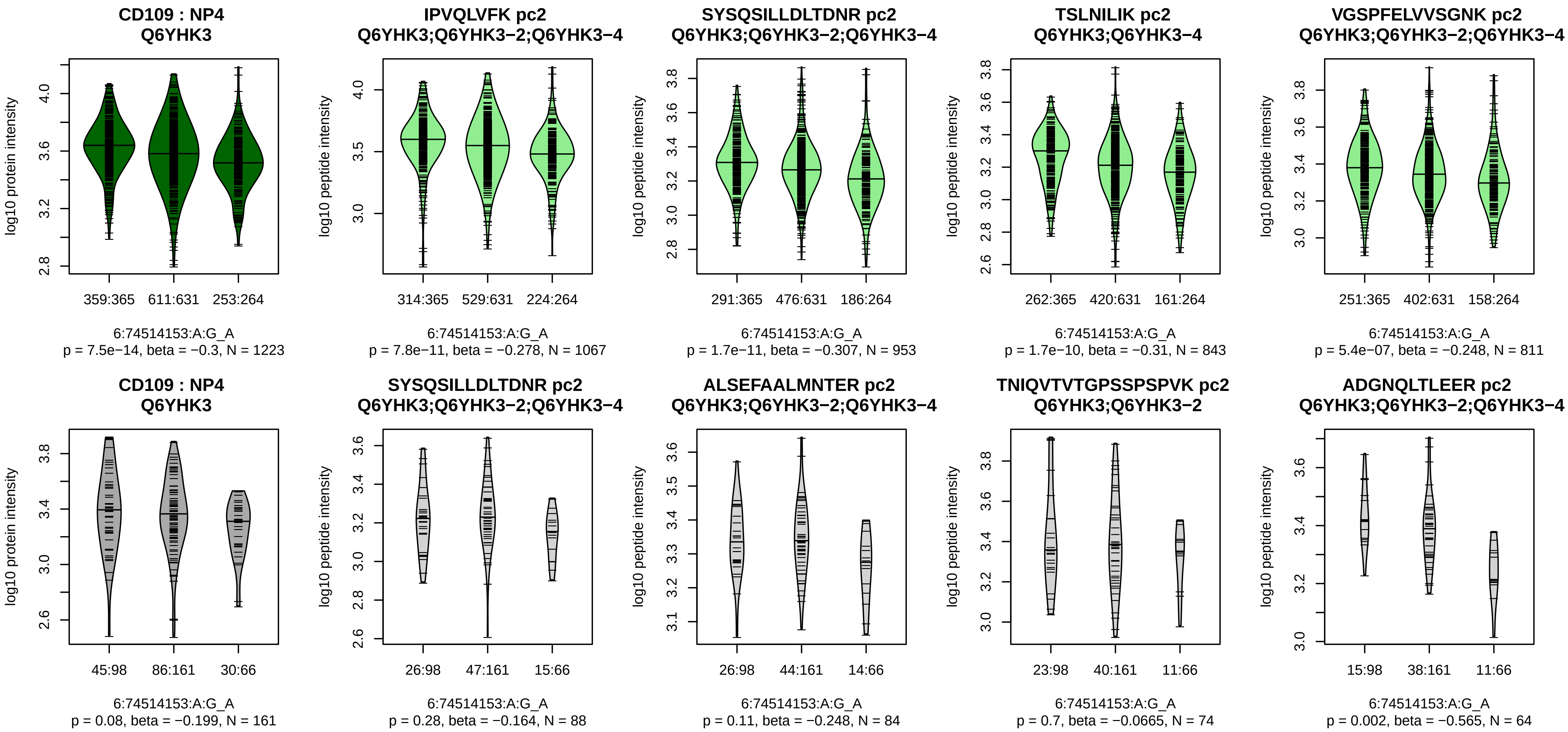


Assay Target: CD109
Olink UniProt: Q6YHK3
deCODE rsID: rs4708089
Proxy rsID: rs4708089
deCODE: 6:73804430:G:A
Proxy SNP: 6:74514153:A:G
deCODE log10(p): 1021.6
deCODE BETA: 0.52
*:*:*:*:*:–:–:–:–:*:*:*:–:–:–:–:–:–:–:
1067:953:843:811:609:533:491:

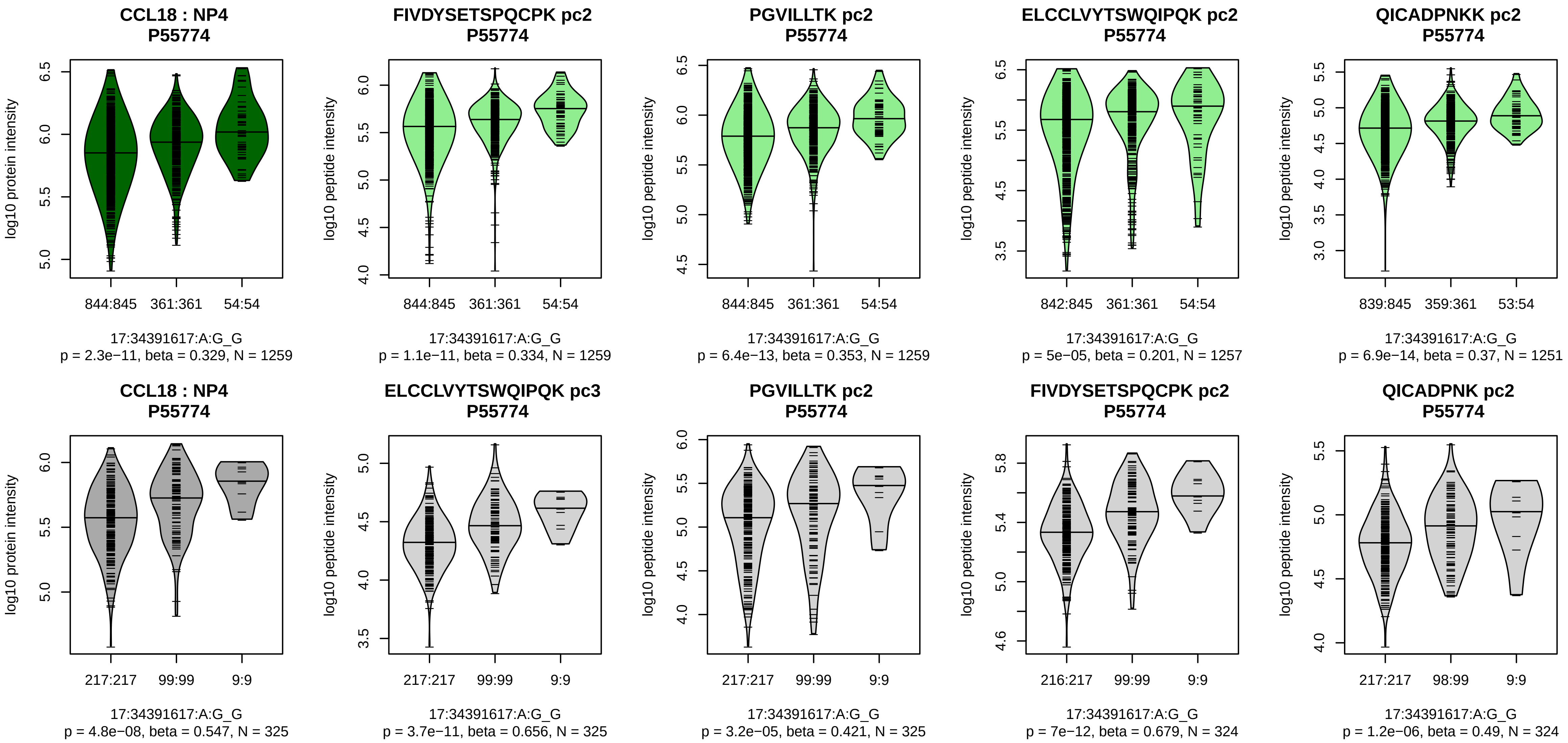


Assay Target: CCL18
Olink UniProt: P55774
deCODE rsID: rs2015086
Proxy rsID: rs2015086
deCODE: 17:36064257:G:A
Proxy SNP: 17:34391617:A:G
deCODE log10(p): 1016.6
deCODE BETA: 0.89
*:*:*:*:*:*:–
1259:1259:1257:1257:1251:124

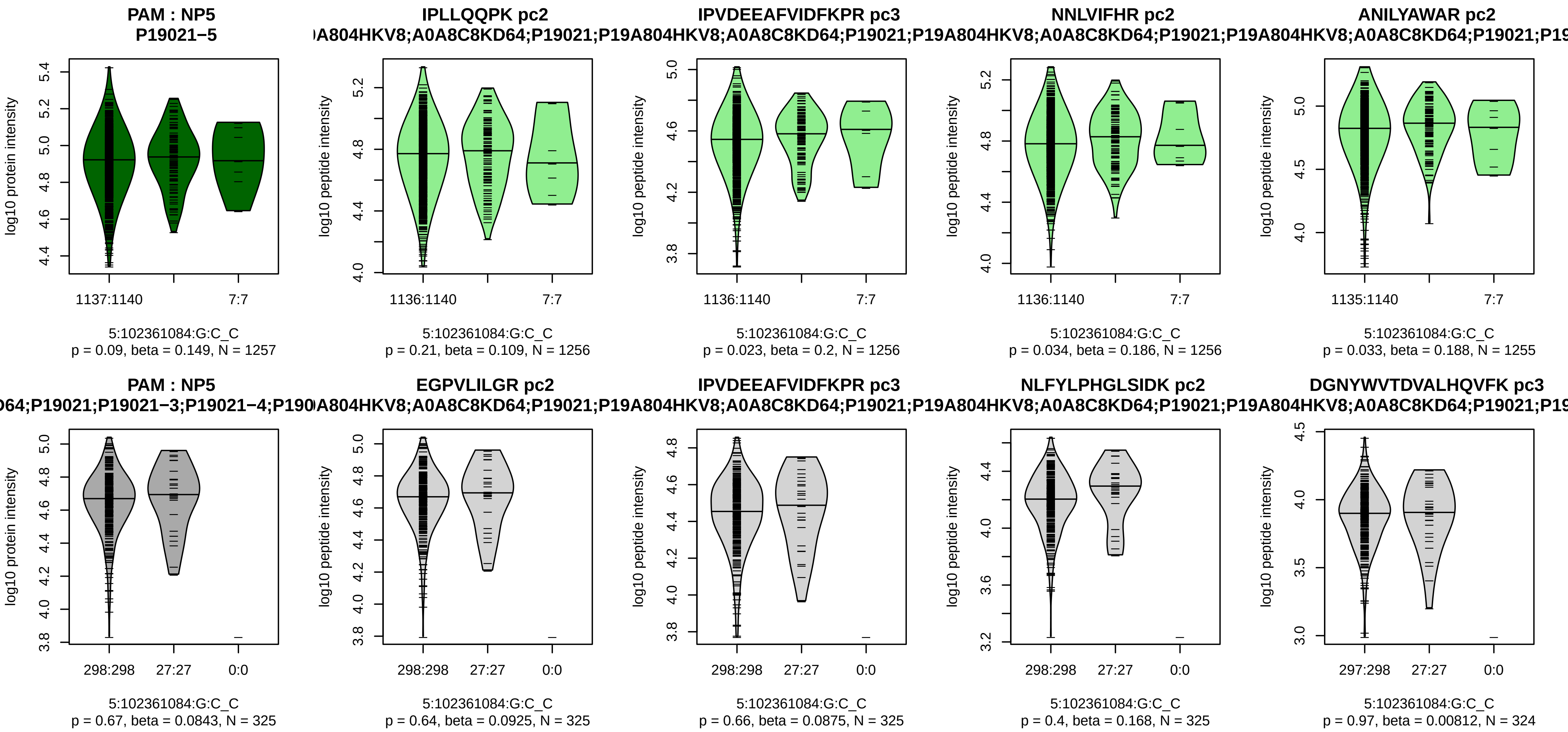


Assay Target: PAM
Olink UniProt: P19021
deCODE rsID: rs41300805
Proxy rsID: rs41300805
deCODE: 5:103025380:C:G
Proxy SNP: 5:102361084:G:C
deCODE log10(p): 1011.9
deCODE BETA: 0.7
–:–:–:–:–:*:–:–:–:*:–:–:–:*:*:–:*:–
1256:1256:1256:1255:1255:125

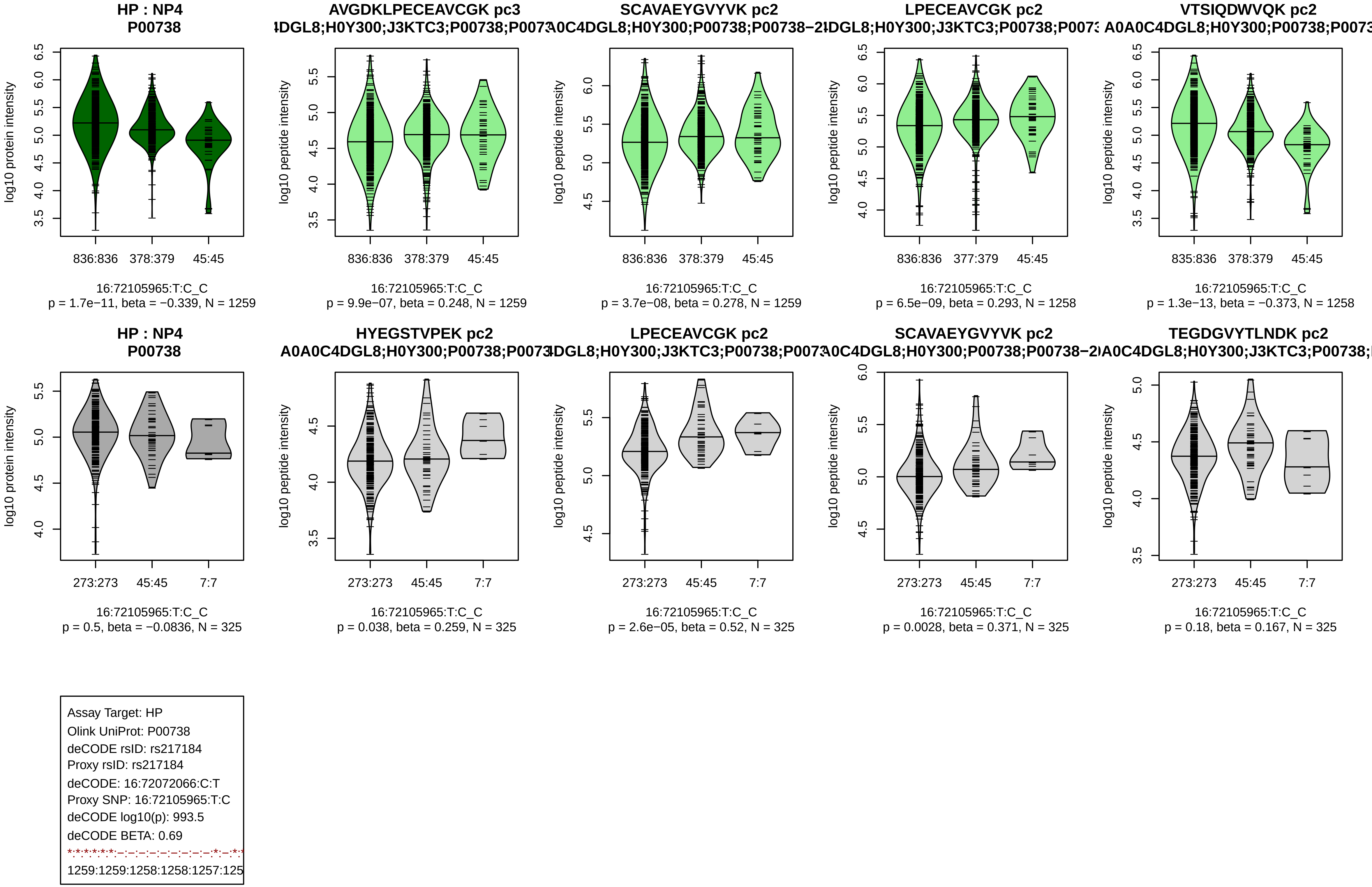
HP : NP4
P00738
AVGDKLPECEAVCGK pc3
4DGL8;H0Y300;J3KTC3;P00738;P0073
SCAVAEYGVYVK pc2
A0C4DGL8;H0Y300;P00738;P00738−2
LPECEAVCGK pc2
4DGL8;H0Y300;J3KTC3;P00738;P007
VTSIQDWVQK pc2
A0A0C4DGL8;H0Y300;P00738;P0073
log10 protein intensity
log10 peptide intensity
836:836
378:379
45:45
377:379
835:836
16:72105965:T:C_C
p = 1.7e−11, beta = −0.339, N = 1259
p = 9.9e−07, beta = 0.248, N = 1259
p = 3.7e−08, beta = 0.278, N = 1259
p = 6.5e−09, beta = 0.293, N = 1258
p = 1.3e−13, beta = −0.373, N = 1258
HP : NP4
P00738
HYEGSTVPEK pc2
A0A0C4DGL8;H0Y300;P00738;P0073
LPECEAVCGK pc2
4DGL8;H0Y300;J3KTC3;P00738;P0073
SCAVAEYGVYVK pc2
A0C4DGL8;H0Y300;P00738;P00738−2
TEGDGVYTLNDK pc2
A0A0C4DGL8;H0Y300;J3KTC3;P00738;
273:273
45:45
7:7
p = 0.5, beta = −0.0836, N = 325
p = 0.038, beta = 0.259, N = 325
p = 2.6e−05, beta = 0.52, N = 325
p = 0.0028, beta = 0.371, N = 325
p = 0.18, beta = 0.167, N = 325
Assay Target: HP
Olink UniProt: P00738
deCODE rsID: rs217184
Proxy rsID: rs217184
deCODE: 16:72072066:C:T
Proxy SNP: 16:72105965:T:C
deCODE log10(p): 993.5
deCODE BETA: 0.69
*:*:*:*:*:*:–:–:–:–:–:–:–:–:–:*:–:*:*
1259:1259:1258:1258:1257:125

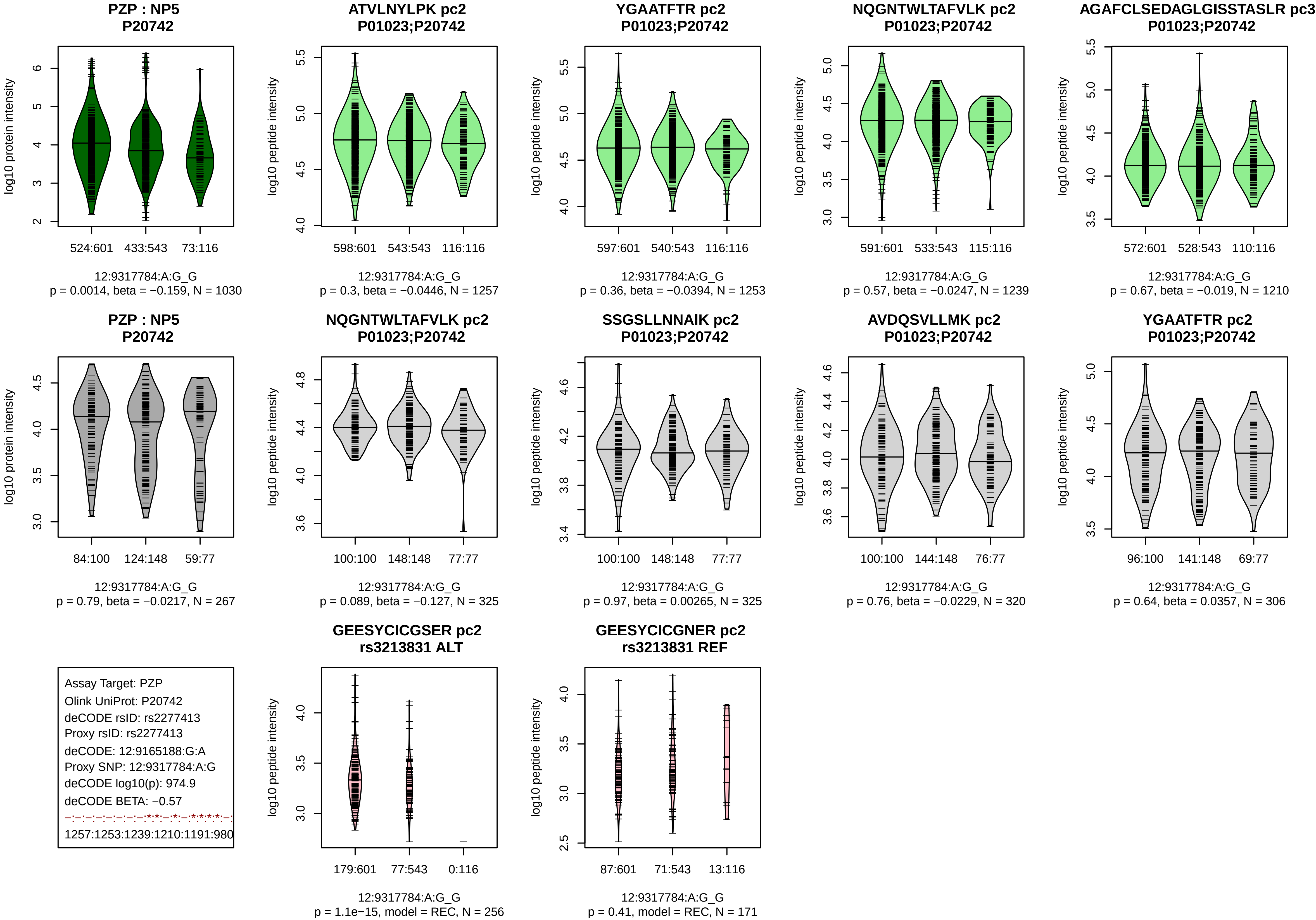
PZP : NP5
P20742
log10 protein intensity
524:601
433:543
73:116
12:9317784:A:G_G
p = 0.0014, beta = −0.159, N = 1030
ATVLNYLPK pc2
P01023;P20742
log10 peptide intensity
598:601
543:543
116:116
12:9317784:A:G_G
p = 0.3, beta = −0.0446, N = 1257
YGAATFTR pc2
P01023;P20742
log10 peptide intensity
597:601
540:543
116:116
12:9317784:A:G_G
p = 0.36, beta = −0.0394, N = 1253
NQGNTWLTAFVLK pc2
P01023;P20742
log10 peptide intensity
591:601
533:543
115:116
12:9317784:A:G_G
p = 0.57, beta = −0.0247, N = 1239
AGAFCLSEDAGLGISSTASLR pc3
P01023;P20742
log10 peptide intensity
572:601
528:543
110:116
12:9317784:A:G_G
p = 0.67, beta = −0.019, N = 1210
PZP : NP5
P20742
log10 protein intensity
84:100
124:148
59:77
12:9317784:A:G_G
p = 0.79, beta = −0.0217, N = 267
NQGNTWLTAFVLK pc2
P01023;P20742
log10 peptide intensity
100:100
148:148
77:77
12:9317784:A:G_G
p = 0.089, beta = −0.127, N = 325
SSGSLLNNAIK pc2
P01023;P20742
log10 peptide intensity
100:100
148:148
77:77
12:9317784:A:G_G
p = 0.97, beta = 0.00265, N = 325
AVDQSVLLMK pc2
P01023;P20742
log10 peptide intensity
100:100
144:148
76:77
12:9317784:A:G_G
p = 0.76, beta = −0.0229, N = 320
YGAATFTR pc2
P01023;P20742
log10 peptide intensity
96:100
141:148
69:77
12:9317784:A:G_G
p = 0.64, beta = 0.0357, N = 306
Assay Target: PZP
Olink UniProt: P20742
deCODE rsID: rs2277413
Proxy rsID: rs2277413
deCODE: 12:9165188:G:A
Proxy SNP: 12:9317784:A:G
deCODE log10(p): 974.9
deCODE BETA: −0.57
−:−:−:−:−:−:−:*:*:−:*:−:*:*:*:*:−:
1257:1253:1239:1210:1191:980
GEESYCICGSER pc2
rs3213831 ALT
log10 peptide intensity
179:601
77:543
0:116
12:9317784:A:G_G
p = 1.1e−15, model = REC, N = 256
GEESYCICGNER pc2
rs3213831 REF
log10 peptide intensity
87:601
71:543
13:116
12:9317784:A:G_G
p = 0.41, model = REC, N = 171

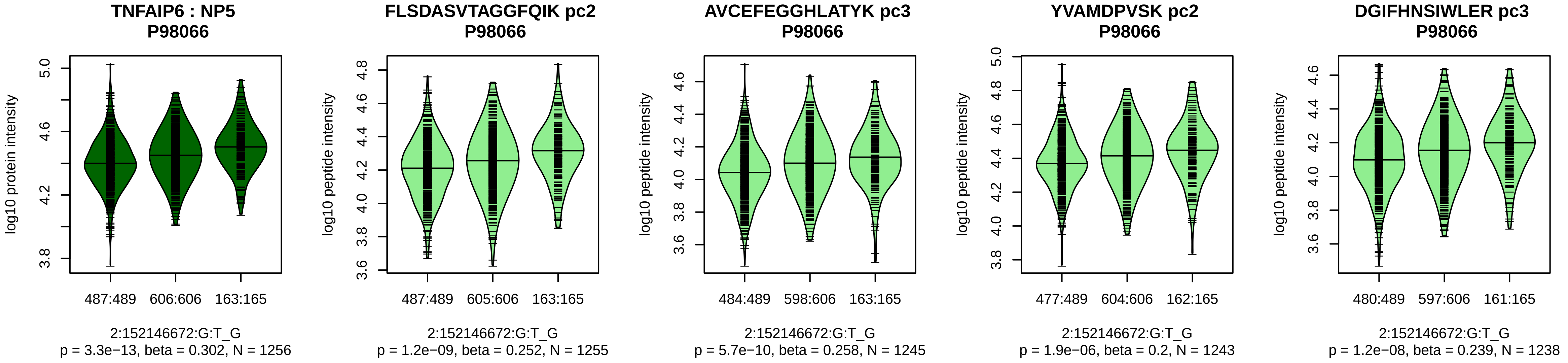


Assay Target: TNFAIP6
Olink UniProt: P98066
deCODE rsID: rs2278089
Proxy rsID: rs2278089
deCODE: 2:151290158:G:T
Proxy SNP: 2:152146672:G:T
deCODE log10(p): 959.2
deCODE BETA: 0.52
*:*:*:*:*:*:*:*:–:*:–:–:–:–:–
1255:1245:1243:1238:1158:110

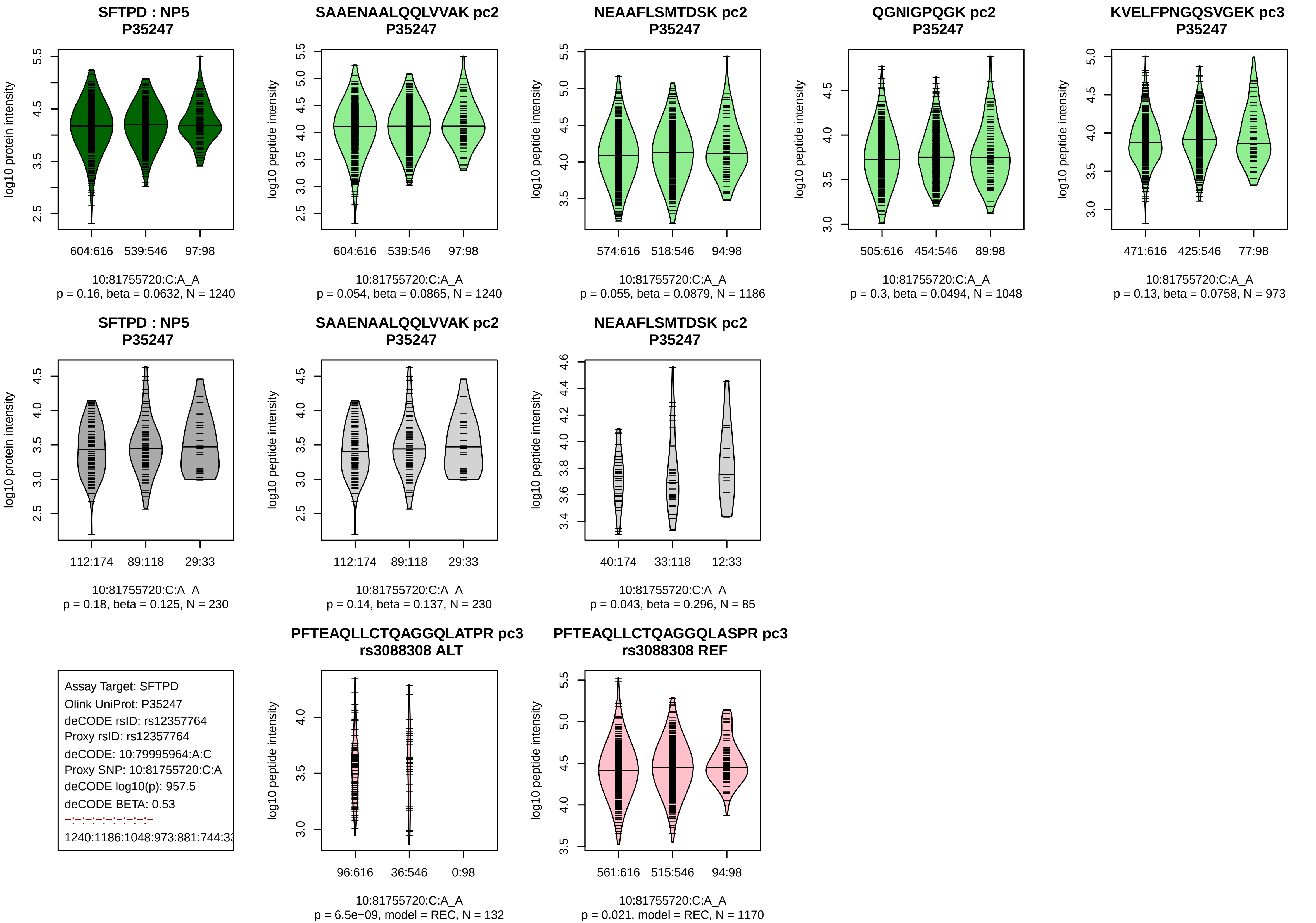

SFTPD : NP5
P35247
log10 protein intensity
604:616
539:546
97:98
10:81755720:C:A_A
p = 0.16, beta = 0.0632, N = 1240
SAAENAALQQLVVAK pc2
P35247
log10 peptide intensity
604:616
539:546
97:98
10:81755720:C:A_A
p = 0.054, beta = 0.0865, N = 1240
NEAAFLSMTDSK pc2
P35247
log10 peptide intensity
574:616
518:546
94:98
10:81755720:C:A_A
p = 0.055, beta = 0.0879, N = 1186
QGNIGPQGK pc2
P35247
log10 peptide intensity
505:616
454:546
89:98
10:81755720:C:A_A
p = 0.3, beta = 0.0494, N = 1048
KVELFPNGQSVGEK pc3
P35247
log10 peptide intensity
471:616
425:546
77:98
10:81755720:C:A_A
p = 0.13, beta = 0.0758, N = 973
SFTPD : NP5
P35247
log10 protein intensity
112:174
89:118
29:33
10:81755720:C:A_A
p = 0.18, beta = 0.125, N = 230
SAAENAALQQLVVAK pc2
P35247
log10 peptide intensity
112:174
89:118
29:33
10:81755720:C:A_A
p = 0.14, beta = 0.137, N = 230
NEAAFLSMTDSK pc2
P35247
log10 peptide intensity
40:174
33:118
12:33
10:81755720:C:A_A
p = 0.043, beta = 0.296, N = 85
Assay Target: SFTPD
Olink UniProt: P35247
deCODE rsID: rs12357764
Proxy rsID: rs12357764
deCODE: 10:79995964:A:C
Proxy SNP: 10:81755720:C:A
deCODE log10(p): 957.5
deCODE BETA: 0.53
–:–:–:–:–:–:–:–:–
1240:1186:1048:973:881:744:33
PFTEAQLLCTQAGGQLATPR pc3
rs3088308 ALT
log10 peptide intensity
96:616
36:546
0:98
10:81755720:C:A_A
p = 6.5e−09, model = REC, N = 132
PFTEAQLLCTQAGGQLASPR pc3
rs3088308 REF
log10 peptide intensity
561:616
515:546
94:98
10:81755720:C:A_A
p = 0.021, model = REC, N = 1170

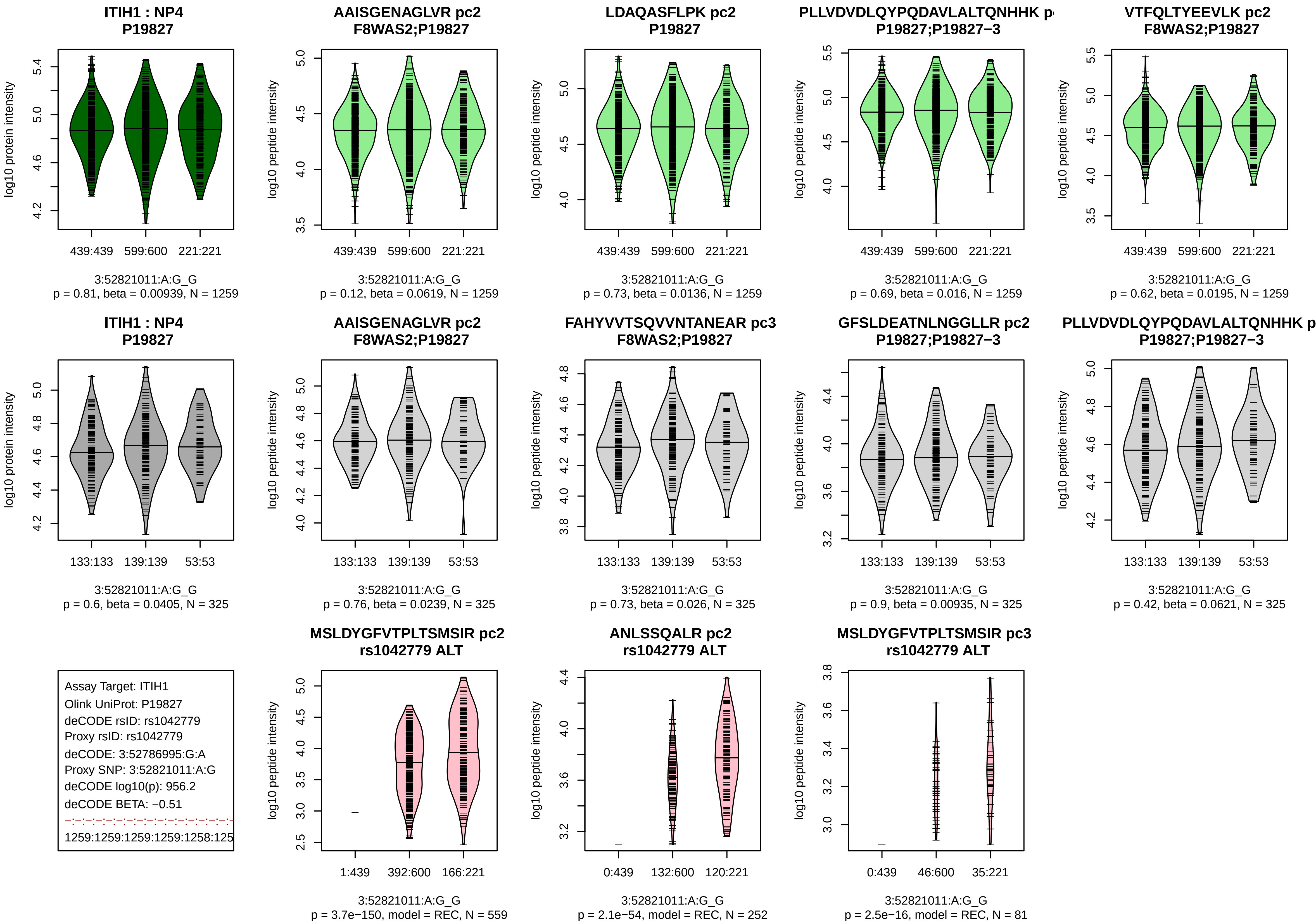
ITIH1 : NP4
P19827
log10 protein intensity
439:439 599:600 221:221
3:52821011:A:G_G
p = 0.81, beta = 0.00939, N = 1259
AAISGENAGLVR pc2
F8WAS2;P19827
log10 peptide intensity
439:439 599:600 221:221
3:52821011:A:G_G
p = 0.12, beta = 0.0619, N = 1259
LDAQASFLPK pc2
P19827
log10 peptide intensity
439:439 599:600 221:221
3:52821011:A:G_G
p = 0.73, beta = 0.0136, N = 1259
PLLVDVDLQYPQDAVLALTQNHHK p
P19827;P19827−3
log10 peptide intensity
439:439 599:600 221:221
3:52821011:A:G_G
p = 0.69, beta = 0.016, N = 1259
VTFQLTYEEVLK pc2
F8WAS2;P19827
log10 peptide intensity
439:439 599:600 221:221
3:52821011:A:G_G
p = 0.62, beta = 0.0195, N = 1259
ITIH1 : NP4
P19827
log10 protein intensity
133:133 139:139 53:53
3:52821011:A:G_G
p = 0.6, beta = 0.0405, N = 325
AAISGENAGLVR pc2
F8WAS2;P19827
log10 peptide intensity
133:133 139:139 53:53
3:52821011:A:G_G
p = 0.76, beta = 0.0239, N = 325
FAHYVVTSQVVNTANEAR pc3
F8WAS2;P19827
log10 peptide intensity
133:133 139:139 53:53
3:52821011:A:G_G
p = 0.73, beta = 0.026, N = 325
GFSLDEATNLNGGLLR pc2
P19827;P19827−3
log10 peptide intensity
133:133 139:139 53:53
3:52821011:A:G_G
p = 0.9, beta = 0.00935, N = 325
PLLVDVDLQYPQDAVLALTQNHHK p
P19827;P19827−3
log10 peptide intensity
133:133 139:139 53:53
3:52821011:A:G_G
p = 0.42, beta = 0.0621, N = 325
Assay Target: ITIH1
Olink UniProt: P19827
deCODE rsID: rs1042779
Proxy rsID: rs1042779
deCODE: 3:52786995:G:A
Proxy SNP: 3:52821011:A:G
deCODE log10(p): 956.2
deCODE BETA: −0.51
1259:1259:1259:1259:1258:125
MSLDYGFVTPLTSMSIR pc2
rs1042779 ALT
log10 peptide intensity
1:439 392:600 166:221
3:52821011:A:G_G
p = 3.7e−150, model = REC, N = 559
ANLSSQALR pc2
rs1042779 ALT
log10 peptide intensity
0:439 132:600 120:221
3:52821011:A:G_G
p = 2.1e−54, model = REC, N = 252
MSLDYGFVTPLTSMSIR pc3
rs1042779 ALT
log10 peptide intensity
0:439 46:600 35:221
3:52821011:A:G_G
p = 2.5e−16, model = REC, N = 81

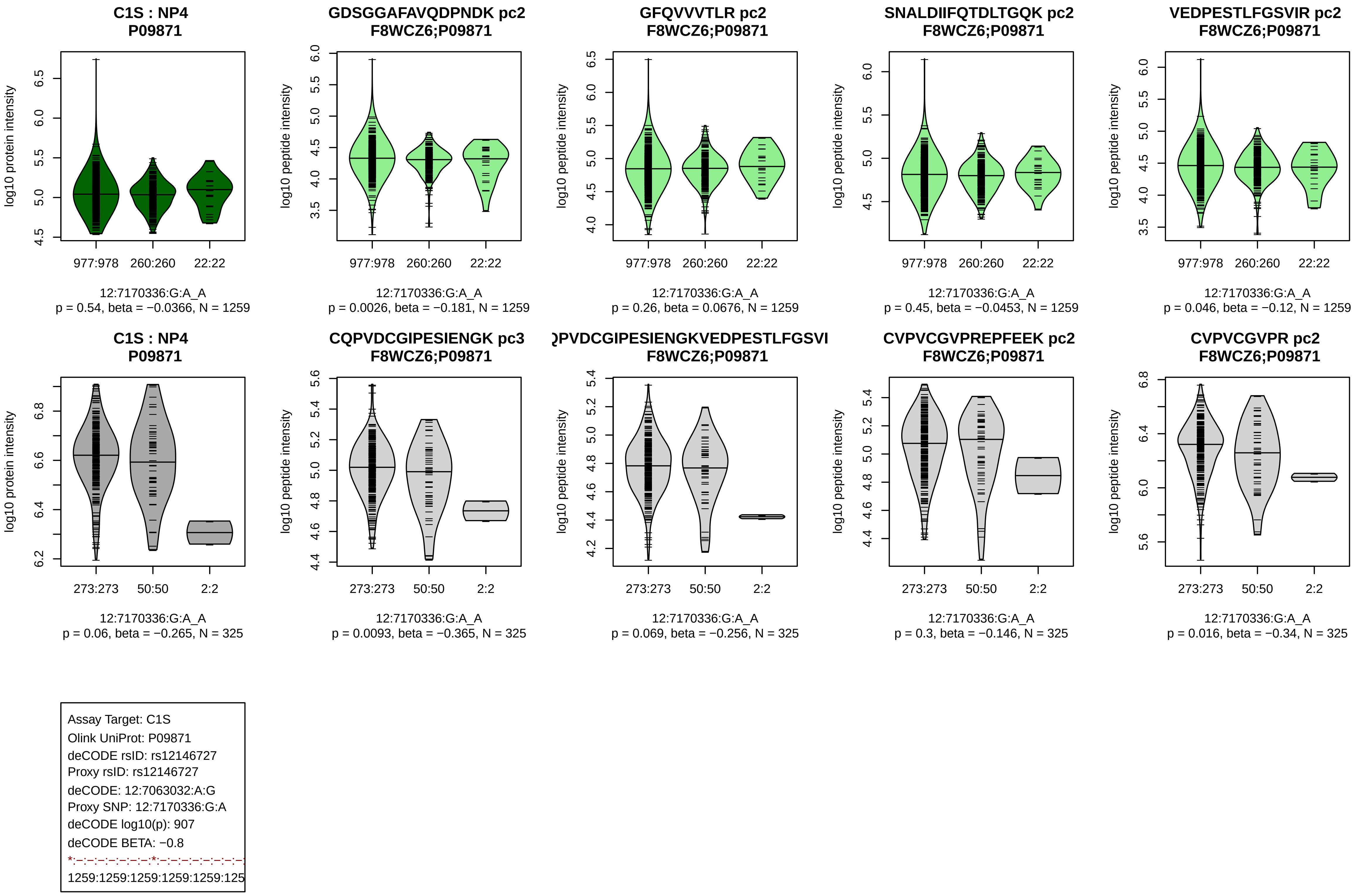

C1S : NP4
P09871
log10 protein intensity
977:978
260:260
22:22
12:7170336:G:A_A
p = 0.54, beta = −0.0366, N = 1259
GDSGGAFAVQDPNDK pc2
F8WCZ6;P09871
log10 peptide intensity
12:7170336:G:A_A
p = 0.0026, beta = −0.181, N = 1259
GFQVVVTLR pc2
F8WCZ6;P09871
12:7170336:G:A_A
p = 0.26, beta = 0.0676, N = 1259
SNALDIIFQTDLTGQK pc2
F8WCZ6;P09871
12:7170336:G:A_A
p = 0.45, beta = −0.0453, N = 1259
VEDPESTLFGSVIR pc2
F8WCZ6;P09871
12:7170336:G:A_A
p = 0.046, beta = −0.12, N = 1259
C1S : NP4
P09871
273:273
50:50
2:2
12:7170336:G:A_A
p = 0.06, beta = −0.265, N = 325
CQPVDCGIPESIENGK pc3
F8WCZ6;P09871
12:7170336:G:A_A
p = 0.0093, beta = −0.365, N = 325
QPVDCGIPESIENGKVEDPESTLFGSVI
F8WCZ6;P09871
12:7170336:G:A_A
p = 0.069, beta = −0.256, N = 325
CVPVCGVPREPFEEK pc2
F8WCZ6;P09871
12:7170336:G:A_A
p = 0.3, beta = −0.146, N = 325
CVPVCGVPR pc2
F8WCZ6;P09871
12:7170336:G:A_A
p = 0.016, beta = −0.34, N = 325
Assay Target: C1S
Olink UniProt: P09871
deCODE rsID: rs12146727
Proxy rsID: rs12146727
deCODE: 12:7063032:A:G
Proxy SNP: 12:7170336:G:A
deCODE log10(p): 907
deCODE BETA: −0.8
*:–:–:–:–:–:–:*:–:–:–:–:–:–:–:–:
1259:1259:1259:1259:1259:125

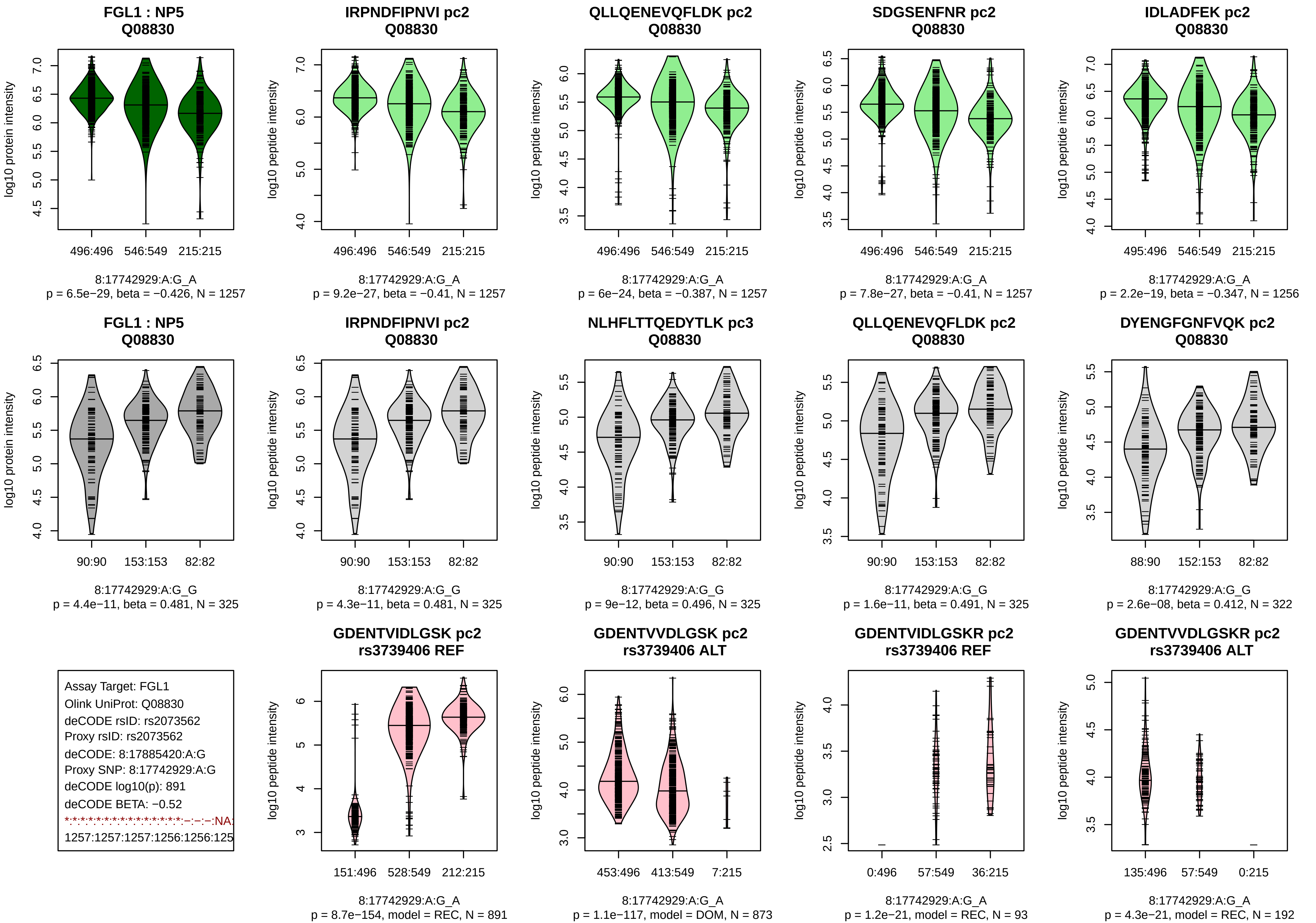

FGL1 : NP5
Q08830
496:496 546:549 215:215
8:17742929:A:G_A
p = 6.5e−29, beta = −0.426, N = 1257
IRPNDFIPNVI pc2
Q08830
496:496 546:549 215:215
8:17742929:A:G_A
p = 9.2e−27, beta = −0.41, N = 1257
QLLQENEVQFLDK pc2
Q08830
496:496 546:549 215:215
8:17742929:A:G_A
p = 6e−24, beta = −0.387, N = 1257
SDGSENFNR pc2
Q08830
496:496 546:549 215:215
8:17742929:A:G_A
p = 7.8e−27, beta = −0.41, N = 1257
IDLADFEK pc2
Q08830
495:496 546:549 215:215
8:17742929:A:G_A
p = 2.2e−19, beta = −0.347, N = 1256
FGL1 : NP5
Q08830
90:90 153:153 82:82
8:17742929:A:G_G
p = 4.4e−11, beta = 0.481, N = 325
IRPNDFIPNVI pc2
Q08830
90:90 153:153 82:82
8:17742929:A:G_G
p = 4.3e−11, beta = 0.481, N = 325
NLHFLTTQEDYTLK pc3
Q08830
90:90 153:153 82:82
8:17742929:A:G_G
p = 9e−12, beta = 0.496, N = 325
QLLQENEVQFLDK pc2
Q08830
90:90 153:153 82:82
8:17742929:A:G_G
p = 1.6e−11, beta = 0.491, N = 325
DYENGFGNFVQK pc2
Q08830
88:90 152:153 82:82
8:17742929:A:G_G
p = 2.6e−08, beta = 0.412, N = 322
Assay Target: FGL1
Olink UniProt: Q08830
deCODE rsID: rs2073562
Proxy rsID: rs2073562
deCODE: 8:17885420:A:G
Proxy SNP: 8:17742929:A:G
deCODE log10(p): 891
deCODE BETA: −0.52
*:*:*:*:*:*:*:*:*:*:*:*:*:*:*:*:*:*:–:–:–:NA:
1257:1257:1257:1256:1256:125
GDENTVIDLGSK pc2
rs3739406 REF
151:496 528:549 212:215
8:17742929:A:G_A
p = 8.7e−154, model = REC, N = 891
GDENTVVDLGSK pc2
rs3739406 ALT
453:496 413:549 7:215
8:17742929:A:G_A
p = 1.1e−117, model = DOM, N = 873
GDENTVIDLGSKR pc2
rs3739406 REF
0:496 57:549 36:215
8:17742929:A:G_A
p = 1.2e−21, model = REC, N = 93
GDENTVVDLGSKR pc2
rs3739406 ALT
135:496 57:549 0:215
8:17742929:A:G_A
p = 4.3e−21, model = REC, N = 192
log10 protein intensity
log10 peptide intensity

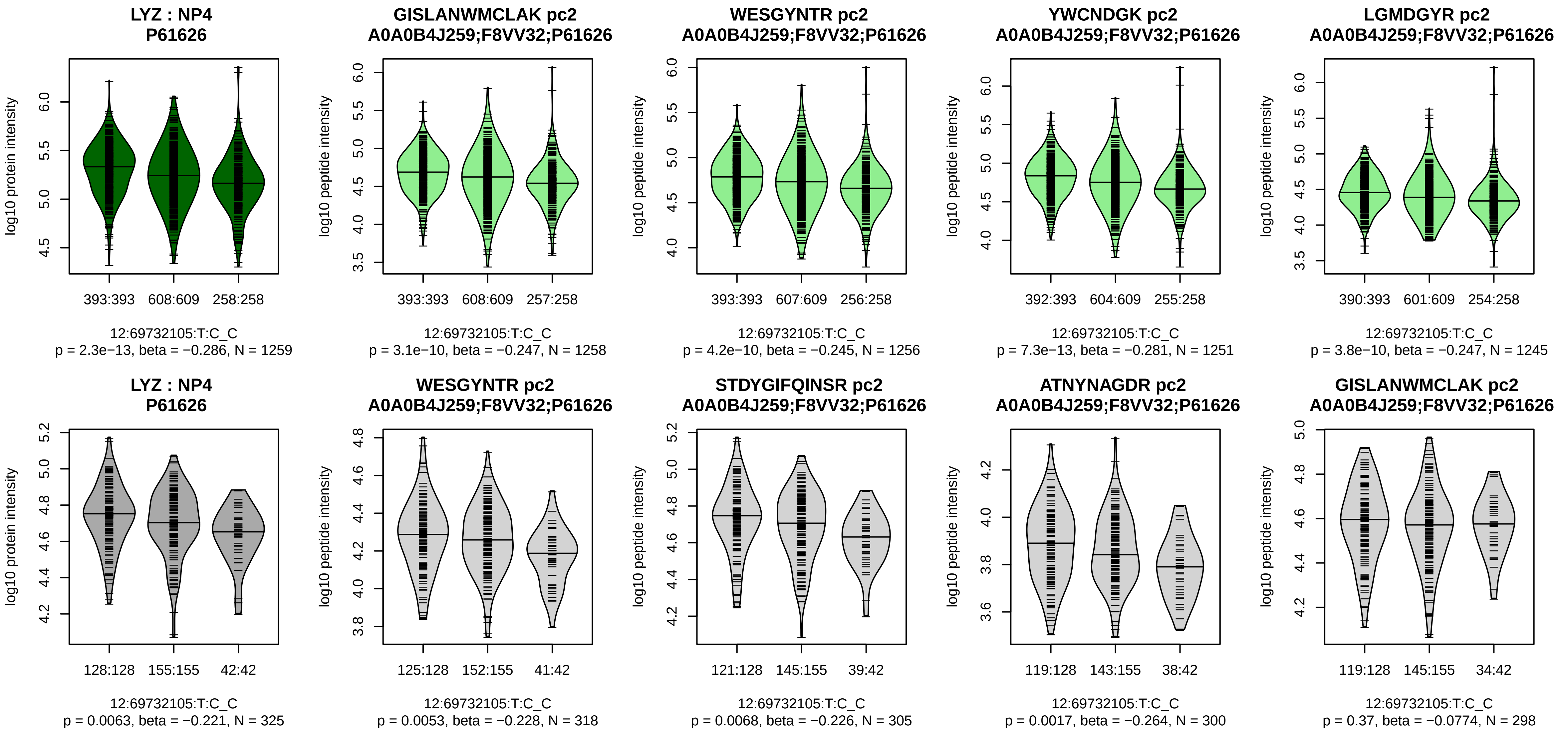


Assay Target: LYZ
Olink UniProt: P61626
deCODE rsID: rs4761234
Proxy rsID: rs4761234
deCODE: 12:69338325:C:T
Proxy SNP: 12:69732105:T:C
deCODE log10(p): 861.2
deCODE BETA: −0.49
*:*:*:*:*:*:−:−:−:NA
1258:1256:1251:1245:1244:122

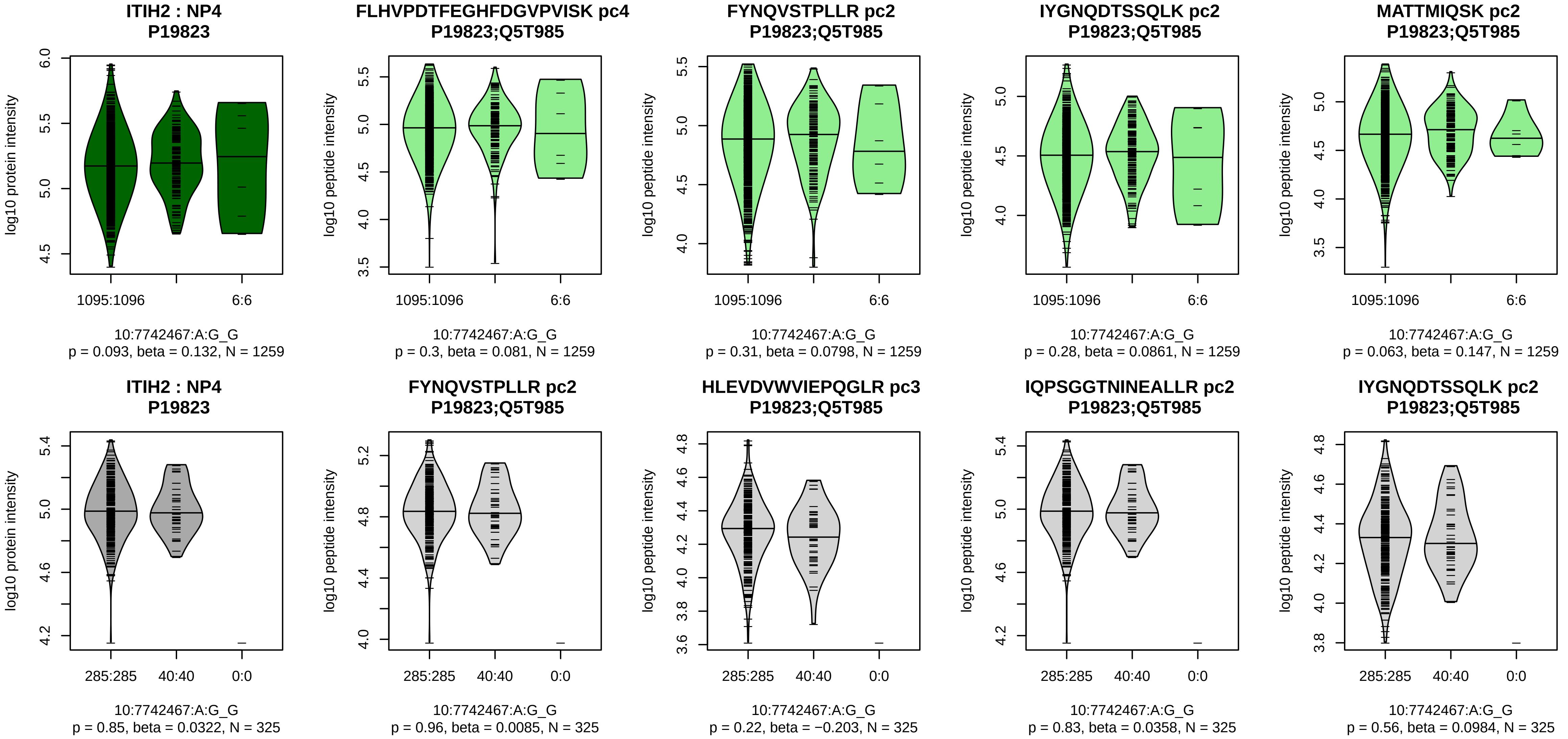


Assay Target: ITIH2
Olink UniProt: P19823
deCODE rsID: rs77938199
Proxy rsID: rs77938199
deCODE: 10:7700504:G:A
Proxy SNP: 10:7742467:A:G
deCODE log10(p): 855.8
deCODE BETA: 1.01
–·–·–·–·–·–·–·–·–·–·–·–·–·–·–·–·–·
1259:1259:1259:1259:1258:125

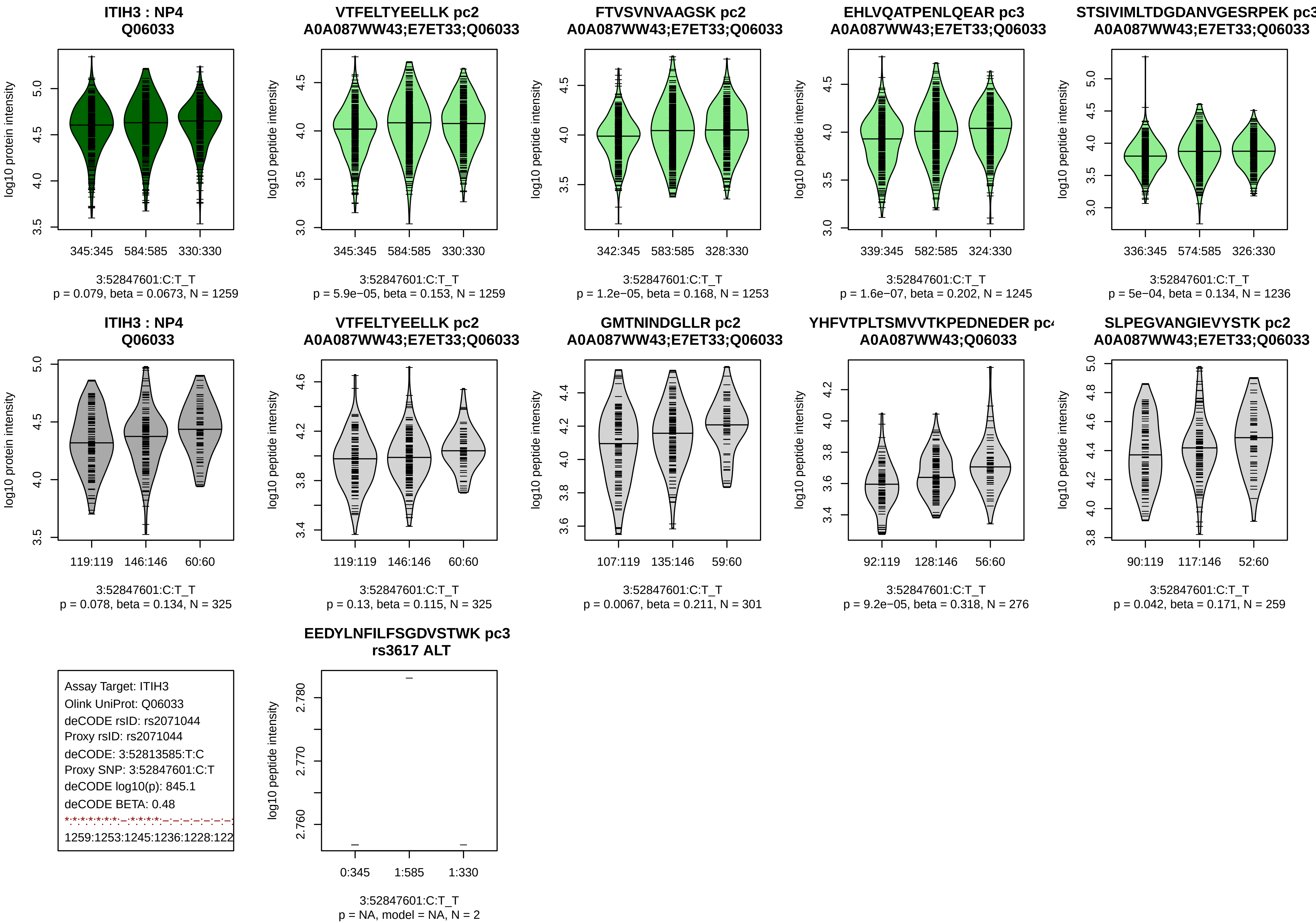
ITIH3 : NP4
Q06033
log10 protein intensity
345:345
584:585
330:330
3:52847601:C:T_T
p = 0.079, beta = 0.0673, N = 1259
VTFELTYEELLK pc2
A0A087WW43;E7ET33;Q06033
log10 peptide intensity
345:345
584:585
330:330
3:52847601:C:T_T
p = 5.9e−05, beta = 0.153, N = 1259
FTVSVNVAAGSK pc2
A0A087WW43;E7ET33;Q06033
log10 peptide intensity
342:345
583:585
328:330
3:52847601:C:T_T
p = 1.2e−05, beta = 0.168, N = 1253
EHLVQATPENLQEAR pc3
A0A087WW43;E7ET33;Q06033
log10 peptide intensity
339:345
582:585
324:330
3:52847601:C:T_T
p = 1.6e−07, beta = 0.202, N = 1245
STSIVIMLTDGDANVGESRPEK pc3
A0A087WW43;E7ET33;Q06033
log10 peptide intensity
336:345
574:585
326:330
3:52847601:C:T_T
p = 5e−04, beta = 0.134, N = 1236
ITIH3 : NP4
Q06033
log10 protein intensity
119:119
146:146
60:60
3:52847601:C:T_T
p = 0.078, beta = 0.134, N = 325
VTFELTYEELLK pc2
A0A087WW43;E7ET33;Q06033
log10 peptide intensity
119:119
146:146
60:60
3:52847601:C:T_T
p = 0.13, beta = 0.115, N = 325
GMTNINDGLLR pc2
A0A087WW43;E7ET33;Q06033
log10 peptide intensity
107:119
135:146
59:60
3:52847601:C:T_T
p = 0.0067, beta = 0.211, N = 301
YHFVTPLTSMVVTKPEDNEDER pc
A0A087WW43;Q06033
log10 peptide intensity
92:119
128:146
56:60
3:52847601:C:T_T
p = 9.2e−05, beta = 0.318, N = 276
SLPEGVANGIEVYSTK pc2
A0A087WW43;E7ET33;Q06033
log10 peptide intensity
90:119
117:146
52:60
3:52847601:C:T_T
p = 0.042, beta = 0.171, N = 259
Assay Target: ITIH3
Olink UniProt: Q06033
deCODE rsID: rs2071044
Proxy rsID: rs2071044
deCODE: 3:52813585:T:C
Proxy SNP: 3:52847601:C:T
deCODE log10(p): 845.1
deCODE BETA: 0.48
1259:1253:1245:1236:1228:122
EEDYLNFILFSGDVSTWK pc3
rs3617 ALT
log10 peptide intensity
0:345
1:585
1:330
3:52847601:C:T_T
p = NA, model = NA, N = 2

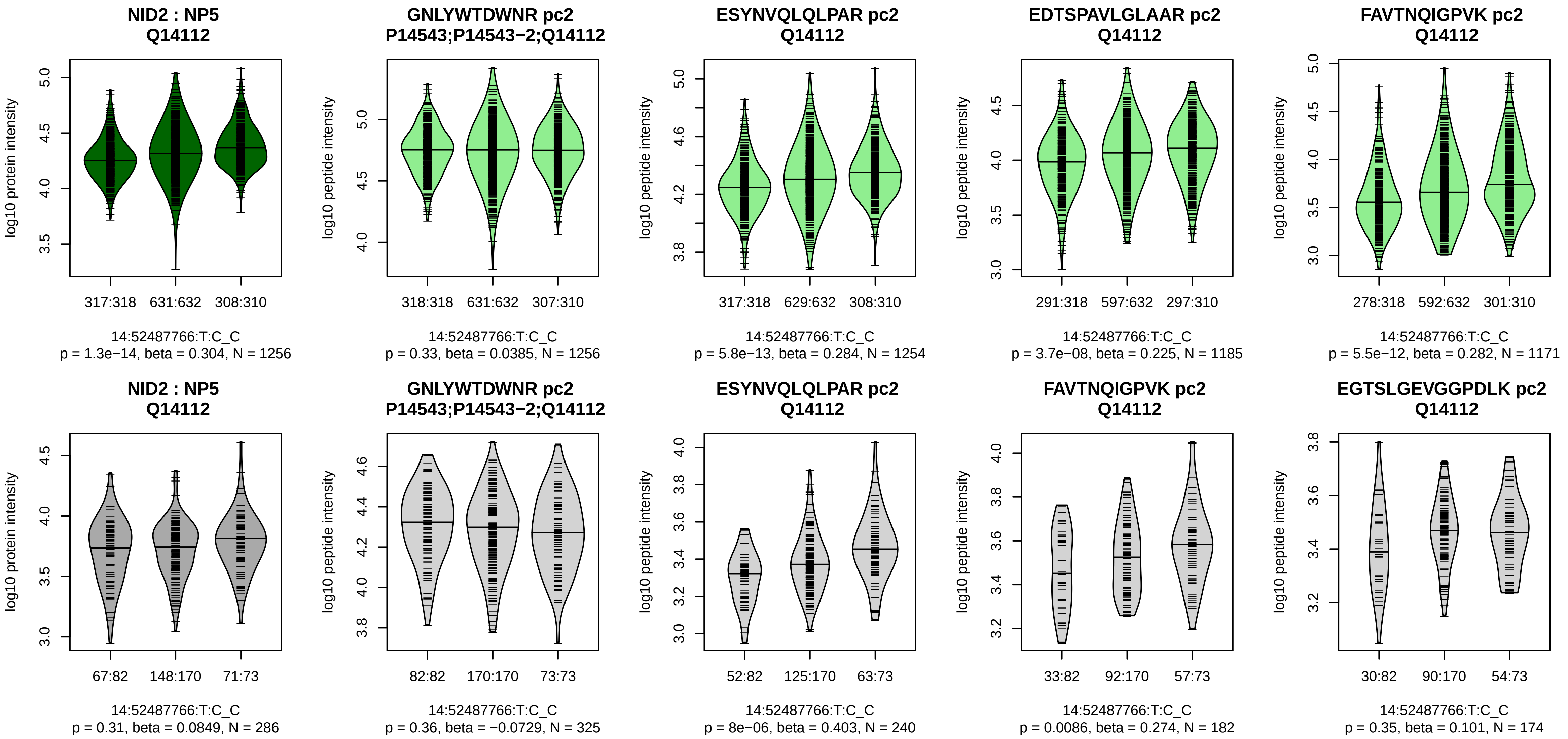


Assay Target: NID2
Olink UniProt: Q14112
deCODE rsID: rs2516600
Proxy rsID: rs2516600
deCODE: 14:52021048:T:C
Proxy SNP: 14:52487766:T:C
deCODE log10(p): 829.6
deCODE BETA: −0.49

1256:1254:1185:1171:1164:107

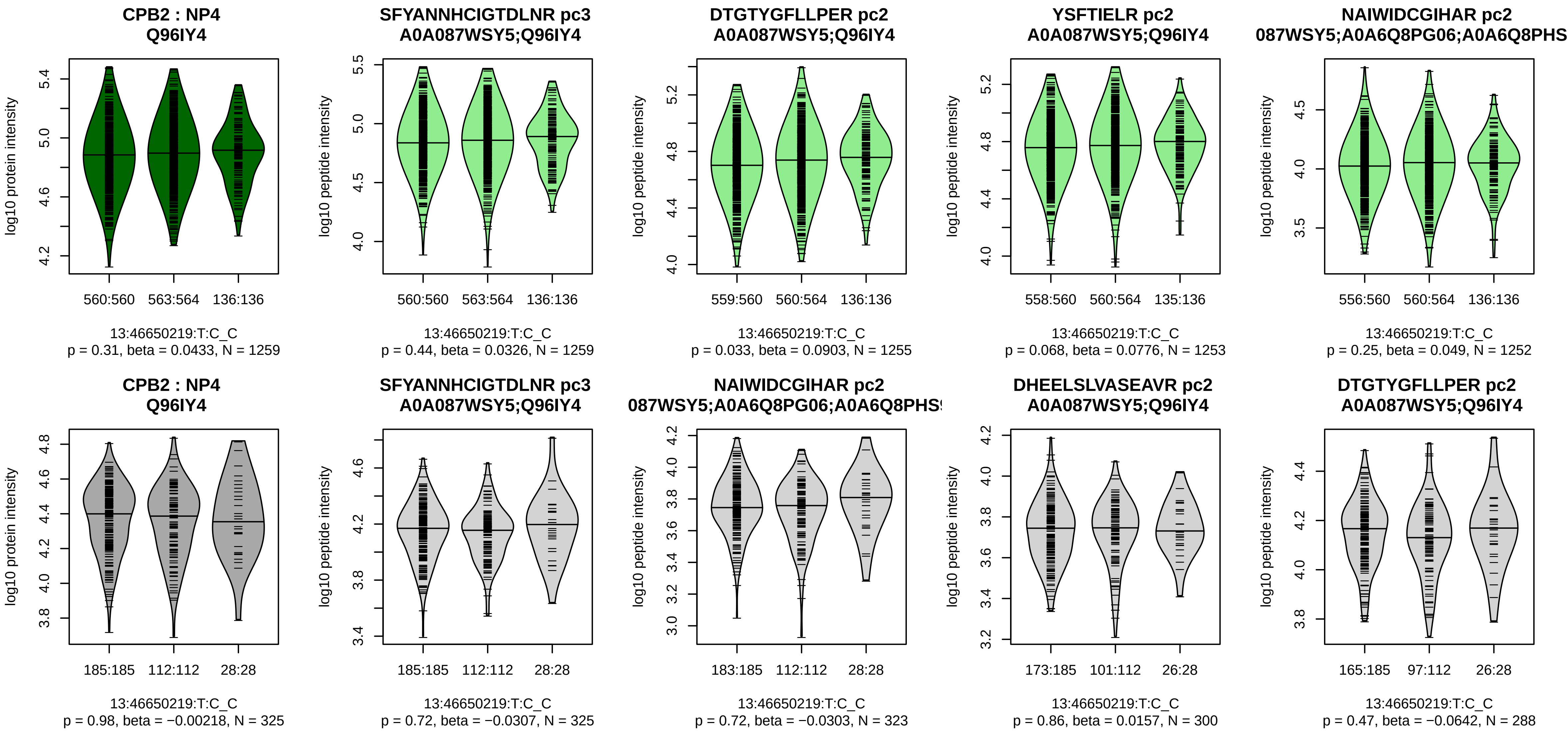


Assay Target: CPB2
Olink UniProt: Q96IY4
deCODE rsID: rs9534312
Proxy rsID: rs9534312
deCODE: 13:46076084:C:T
Proxy SNP: 13:46650219:T:C
deCODE log10(p): 764.1
deCODE BETA: 0.5
–:–:–:–:–:–:–:–:–:–:–:–:–:–:–:–:
1259:1255:1253:1252:1247:124

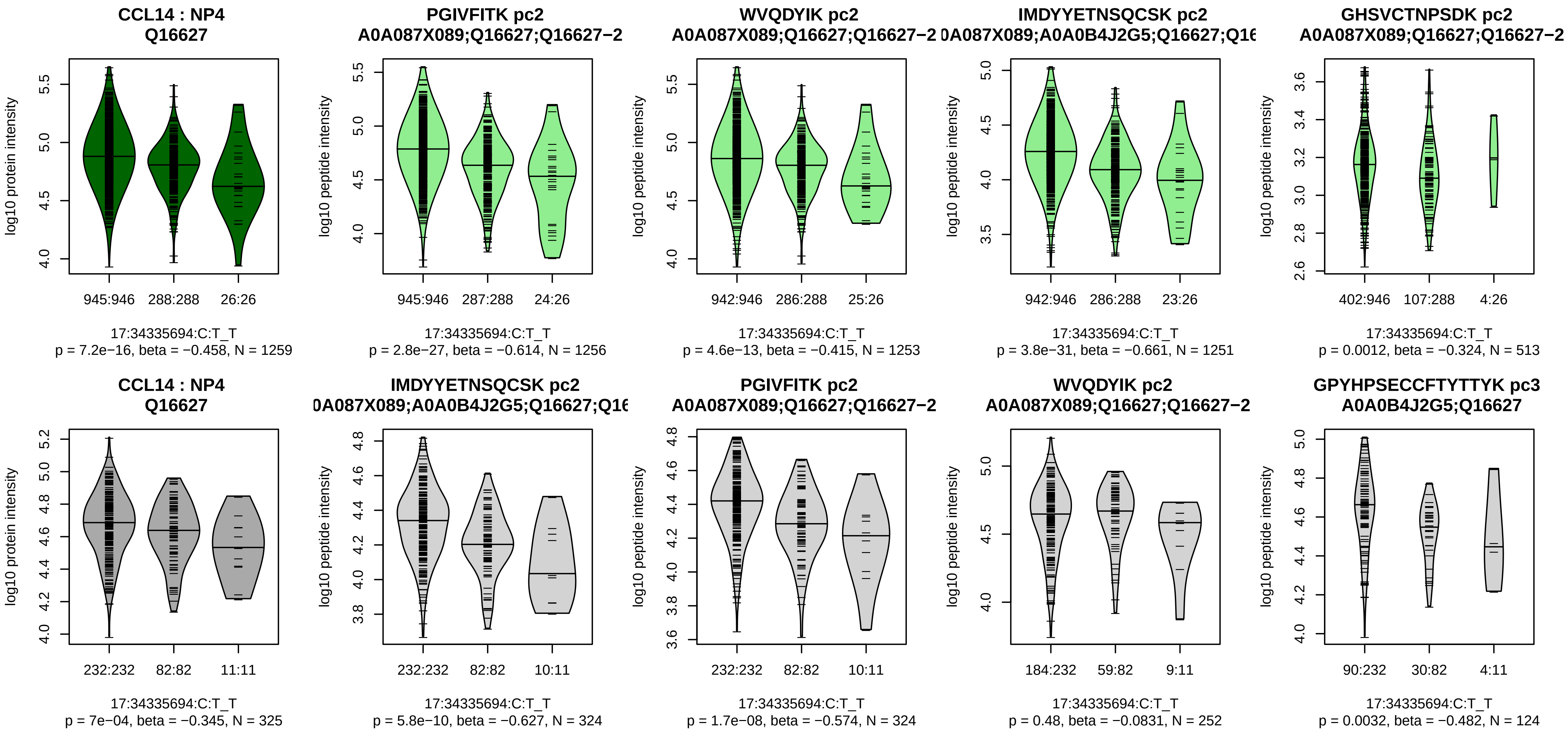


Assay Target: CCL14
Olink UniProt: Q16627
deCODE rsID: rs7222922
Proxy rsID: rs7222922
deCODE: 17:36008654:T:C
Proxy SNP: 17:34335694:C:T
deCODE log10(p): 737.6
deCODE BETA: −0.86
*:*:*:*:*:–
1256:1253:1251:513:239:32

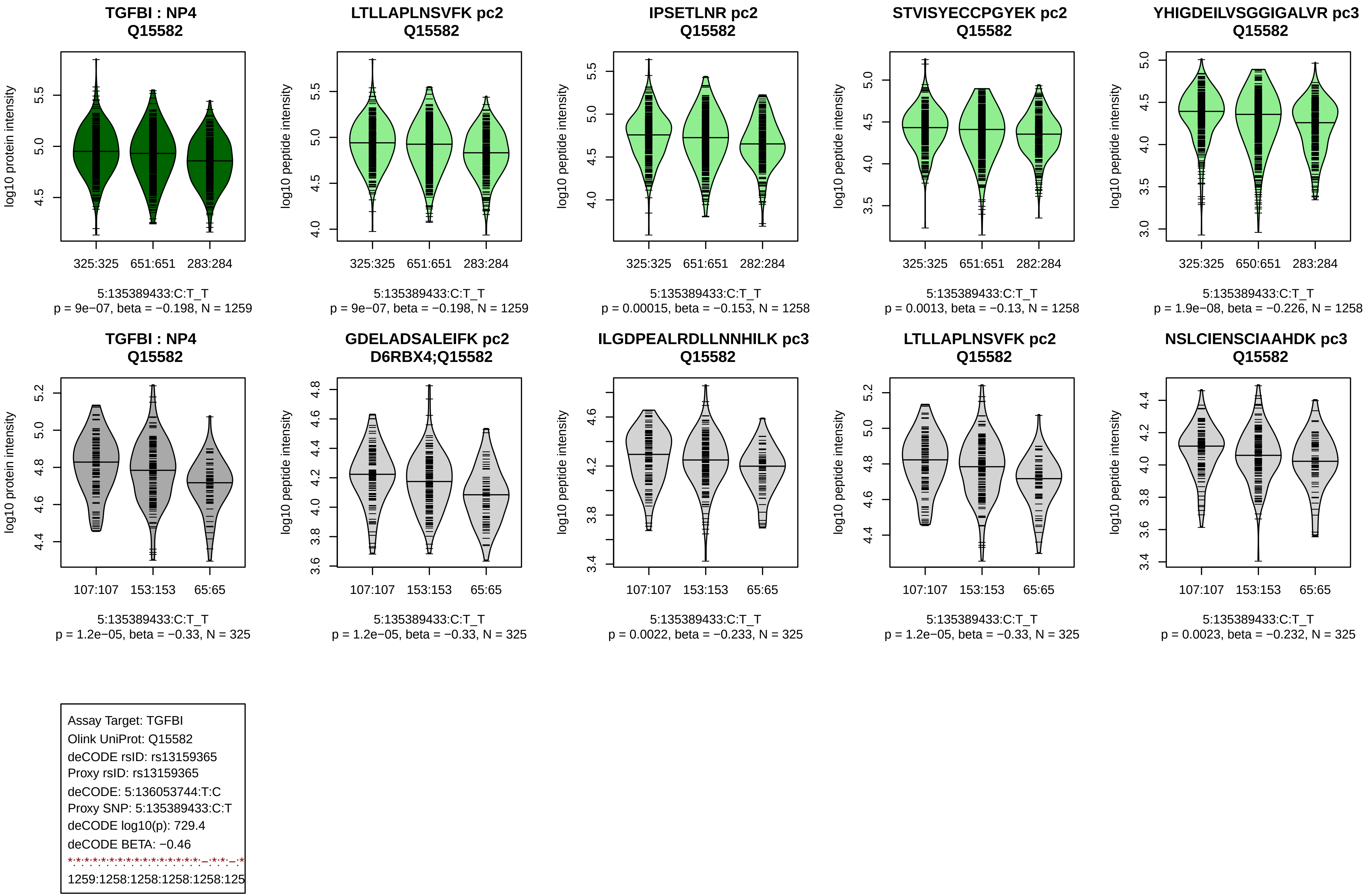
TGFBI : NP4
Q15582
log10 protein intensity
325:325 651:651 283:284
5:135389433:C:T_T
p = 9e−07, beta = −0.198, N = 1259
LTLLAPLNSVFK pc2
Q15582
log10 peptide intensity
325:325 651:651 283:284
5:135389433:C:T_T
p = 9e−07, beta = −0.198, N = 1259
IPSETLNR pc2
Q15582
log10 peptide intensity
325:325 651:651 282:284
5:135389433:C:T_T
p = 0.00015, beta = −0.153, N = 1258
STVISYECCPGYEK pc2
Q15582
log10 peptide intensity
325:325 651:651 282:284
5:135389433:C:T_T
p = 0.0013, beta = −0.13, N = 1258
YHIGDEILVSGGIGALVR pc3
Q15582
log10 peptide intensity
325:325 650:651 283:284
5:135389433:C:T_T
p = 1.9e−08, beta = −0.226, N = 1258
TGFBI : NP4
Q15582
log10 protein intensity
107:107 153:153 65:65
5:135389433:C:T_T
p = 1.2e−05, beta = −0.33, N = 325
GDELADSALEIFK pc2
D6RBX4;Q15582
log10 peptide intensity
107:107 153:153 65:65
5:135389433:C:T_T
p = 1.2e−05, beta = −0.33, N = 325
ILGDPEALRDLLNNHILK pc3
Q15582
log10 peptide intensity
107:107 153:153 65:65
5:135389433:C:T_T
p = 0.0022, beta = −0.233, N = 325
LTLLAPLNSVFK pc2
Q15582
log10 peptide intensity
107:107 153:153 65:65
5:135389433:C:T_T
p = 1.2e−05, beta = −0.33, N = 325
NSLCIENSCIAAHDK pc3
Q15582
log10 peptide intensity
107:107 153:153 65:65
5:135389433:C:T_T
p = 0.0023, beta = −0.232, N = 325
Assay Target: TGFBI
Olink UniProt: Q15582
deCODE rsID: rs13159365
Proxy rsID: rs13159365
deCODE: 5:136053744:T:C
Proxy SNP: 5:135389433:C:T
deCODE log10(p): 729.4
deCODE BETA: −0.46
1259:1258:1258:1258:1258:125

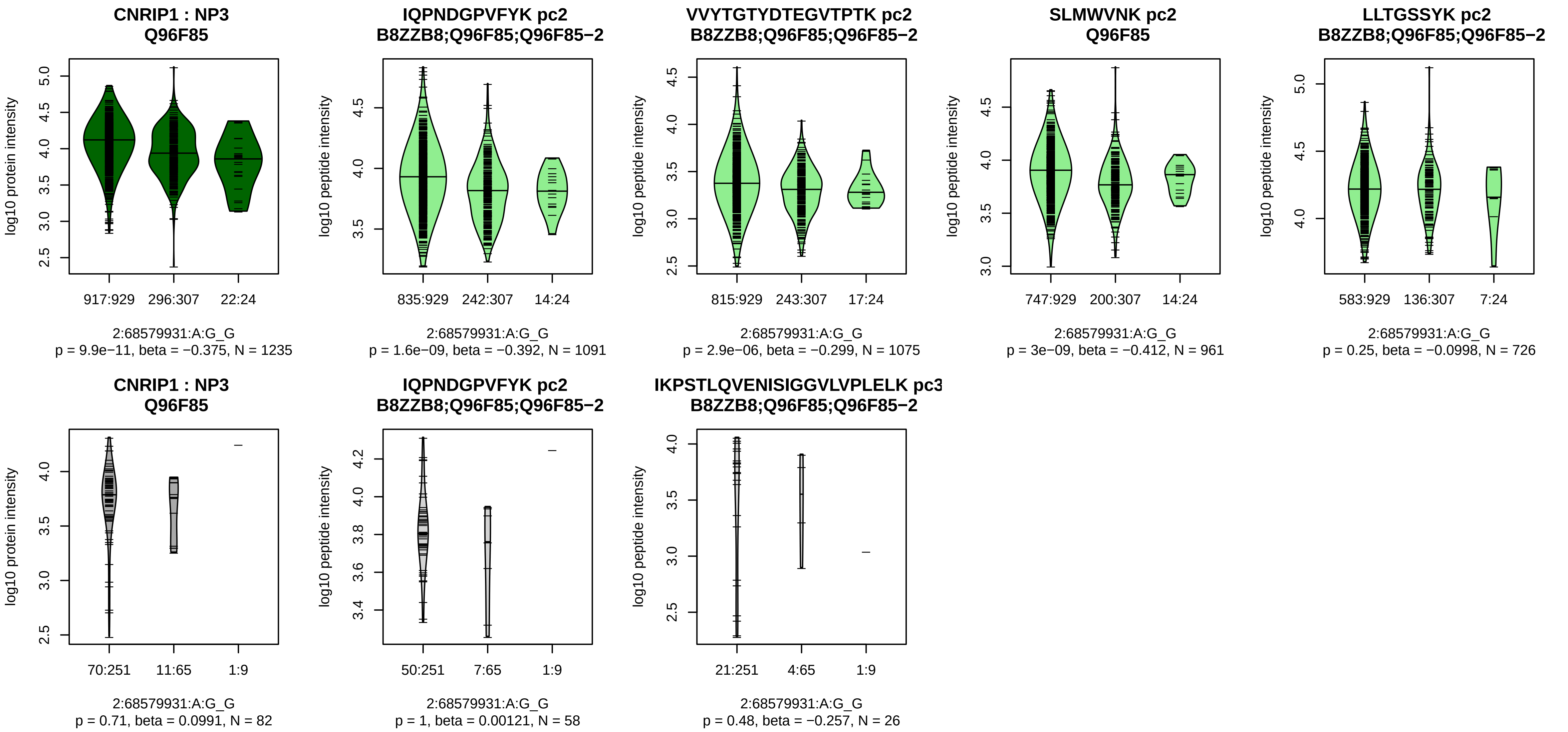


Assay Target: CNRIP1
Olink UniProt: Q96F85
deCODE rsID: rs7578047
Proxy rsID: rs7578047
deCODE: 2:68352799:G:A
Proxy SNP: 2:68579931:A:G
deCODE log10(p): 707.8
deCODE BETA: −0.66
*:*:*:−:−:−:−:NA
1091:1075:961:726:424:225:64:

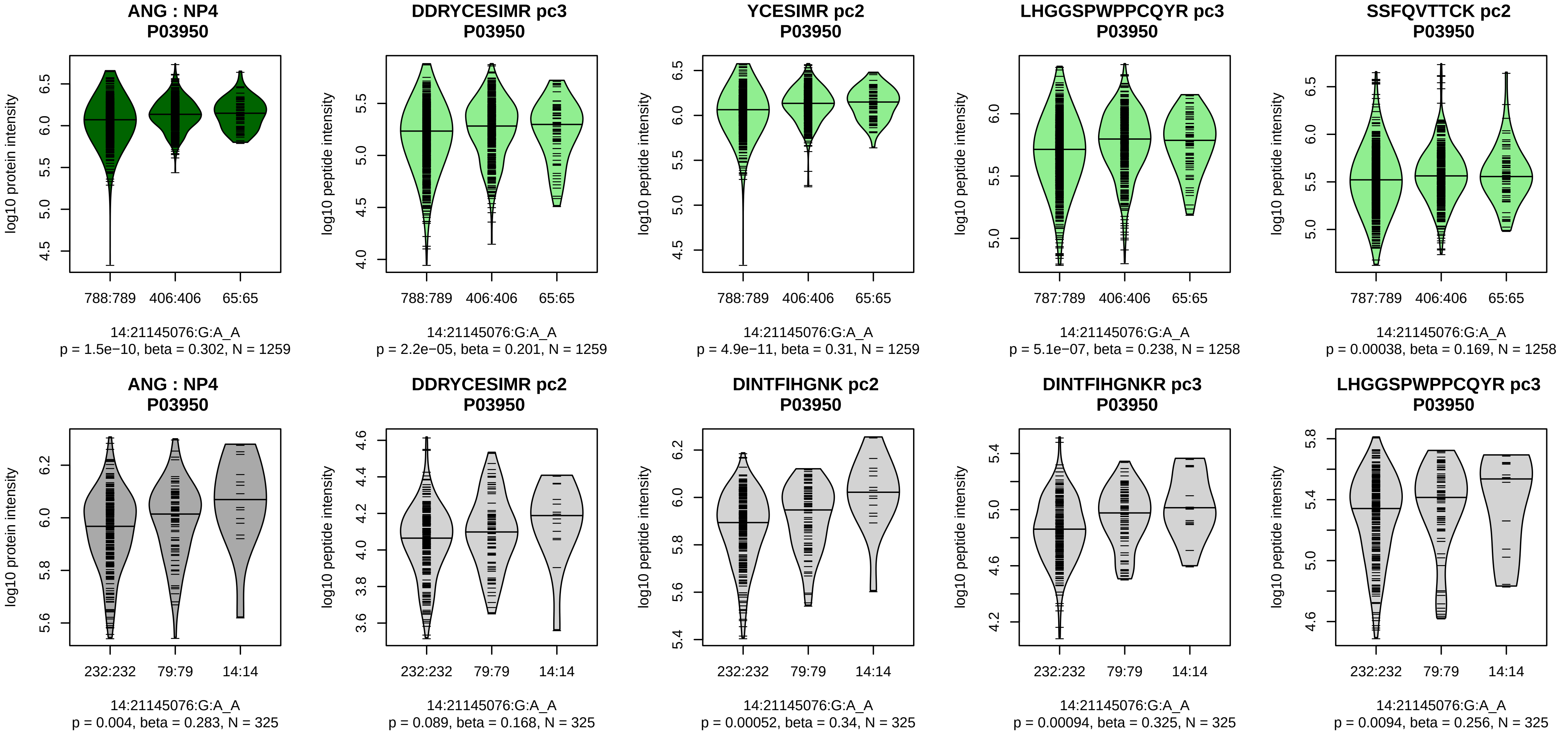


Assay Target: ANG
Olink UniProt: P03950
deCODE rsID: rs28535078
Proxy rsID: rs28535078
deCODE: 14:20676917:A:G
Proxy SNP: 14:21145076:G:A
deCODE log10(p): 686
deCODE BETA: 0.56
*:*:*:*:*:*:*:*:*:*:–:*:–:*:–:*:–
1259:1259:1258:1258:1253:124

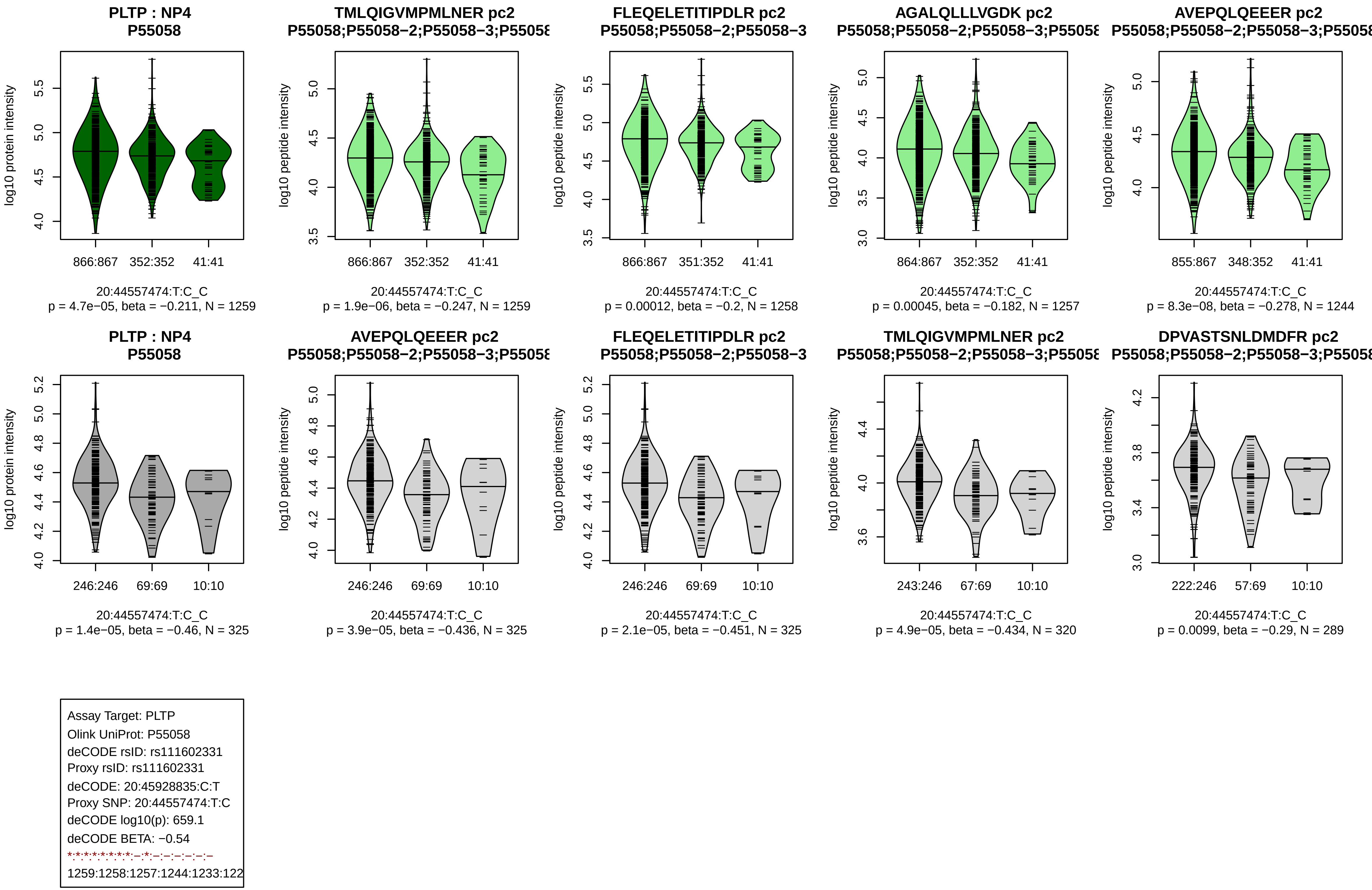
PLTP : NP4
P55058
log10 protein intensity
866:867
352:352
41:41
20:44557474:T:C_C
p = 4.7e−05, beta = −0.211, N = 1259
TMLQIGVMPMLNER pc2
P55058;P55058−2;P55058−3;P5505
log10 peptide intensity
866:867
352:352
41:41
20:44557474:T:C_C
p = 1.9e−06, beta = −0.247, N = 1259
FLEQELETITIPDLR pc2
P55058;P55058−2;P55058−3
log10 peptide intensity
866:867
351:352
41:41
20:44557474:T:C_C
p = 0.00012, beta = −0.2, N = 1258
AGALQLLLVGDK pc2
P55058;P55058−2;P55058−3;P5505
log10 peptide intensity
864:867
352:352
41:41
20:44557474:T:C_C
p = 0.00045, beta = −0.182, N = 1257
AVEPQLQEEER pc2
P55058;P55058−2;P55058−3;P5505
log10 peptide intensity
855:867
348:352
41:41
20:44557474:T:C_C
p = 8.3e−08, beta = −0.278, N = 1244
PLTP : NP4
P55058
log10 protein intensity
246:246
69:69
10:10
20:44557474:T:C_C
p = 1.4e−05, beta = −0.46, N = 325
AVEPQLQEEER pc2
P55058;P55058−2;P55058−3;P5505
log10 peptide intensity
246:246
69:69
10:10
20:44557474:T:C_C
p = 3.9e−05, beta = −0.436, N = 325
FLEQELETITIPDLR pc2
P55058;P55058−2;P55058−3
log10 peptide intensity
246:246
69:69
10:10
20:44557474:T:C_C
p = 2.1e−05, beta = −0.451, N = 325
TMLQIGVMPMLNER pc2
P55058;P55058−2;P55058−3;P5505
log10 peptide intensity
243:246
67:69
10:10
20:44557474:T:C_C
p = 4.9e−05, beta = −0.434, N = 320
DPVASTSNLDMDFR pc2
P55058;P55058−2;P55058−3;P5505
log10 peptide intensity
222:246
57:69
10:10
20:44557474:T:C_C
p = 0.0099, beta = −0.29, N = 289
Assay Target: PLTP
Olink UniProt: P55058
deCODE rsID: rs111602331
Proxy rsID: rs111602331
deCODE: 20:45928835:C:T
Proxy SNP: 20:44557474:T:C
deCODE log10(p): 659.1
deCODE BETA: −0.54
*:*:*:*:*:*:*:*:–:*:–:–:–:–:–
1259:1258:1257:1244:1233:122

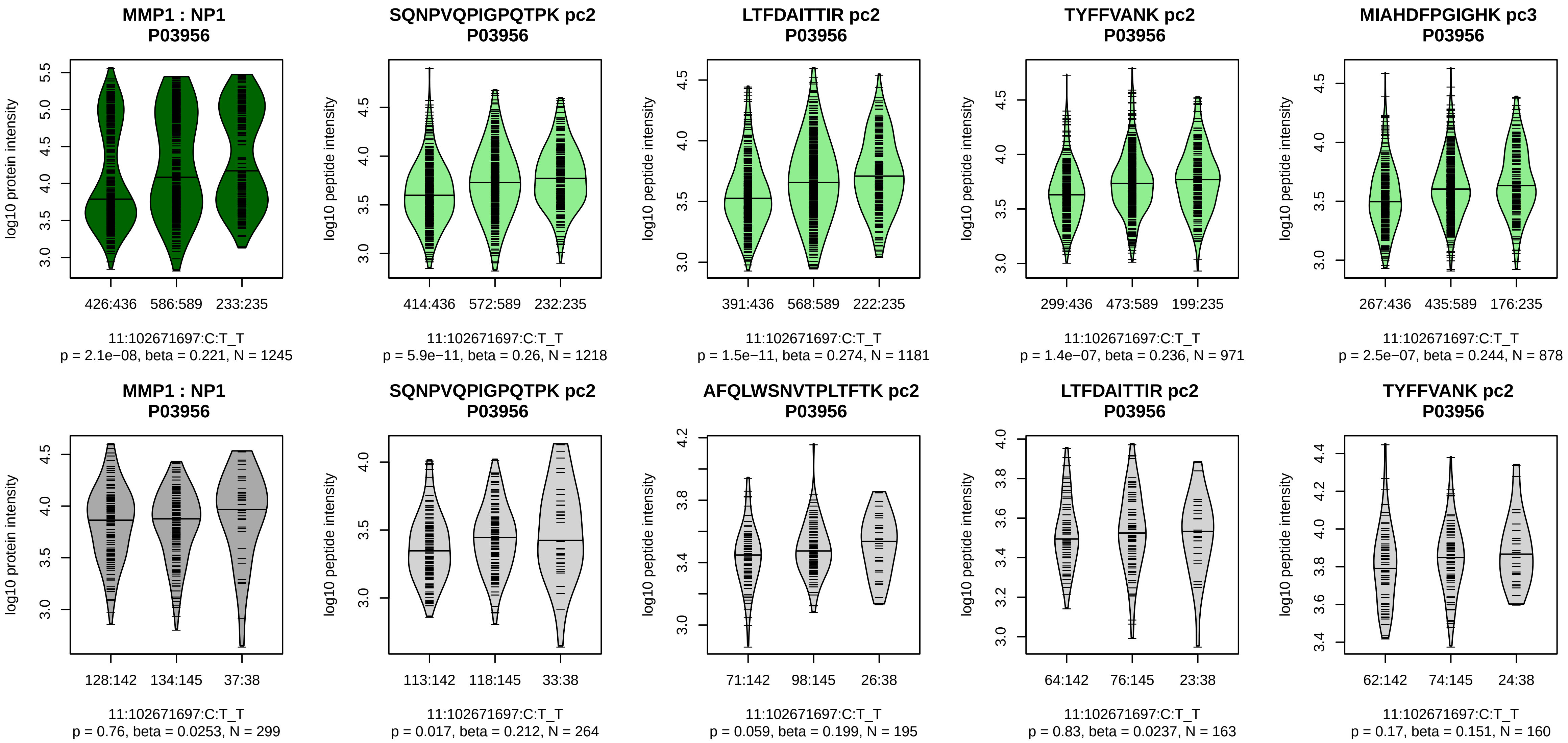


Assay Target: MMP1
Olink UniProt: P03956
deCODE rsID: rs534191
Proxy rsID: rs534191
deCODE: 11:102800966:T:C
Proxy SNP: 11:102671697:C:T
deCODE log10(p): 589.7
deCODE BETA: 0.43
*:*:*:*:*:*:_:*:*:_:*:_:_:_:_:_:_:*:*:
1218:1181:971:878:685:652:639

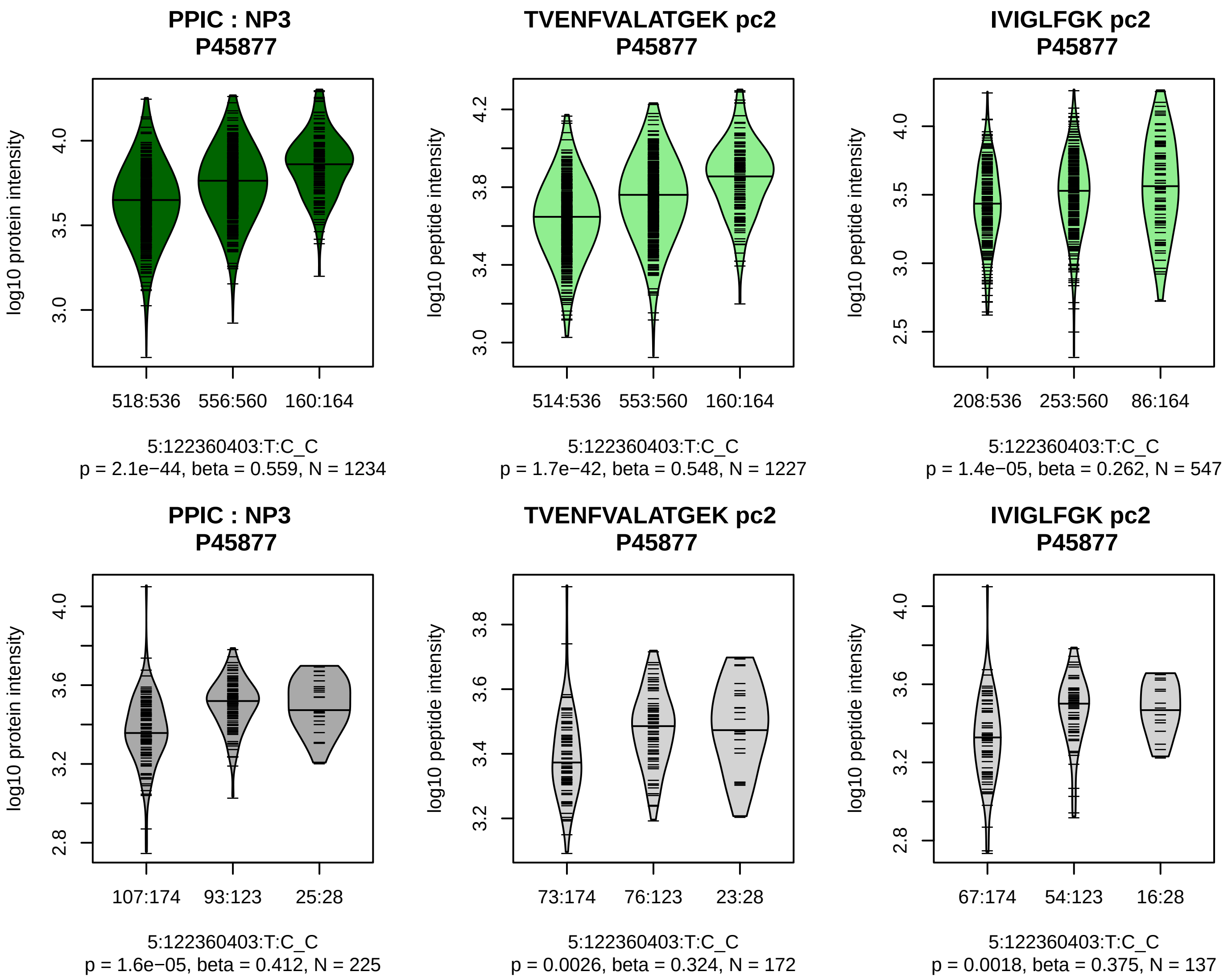


Assay Target: PPIC
Olink UniProt: P45877
deCODE rsID: rs17388251
Proxy rsID: rs17388251
deCODE: 5:123024708:C:T
Proxy SNP: 5:122360403:T:C
deCODE log10(p): 585.3
deCODE BETA: −0.42
*:*
1227:547

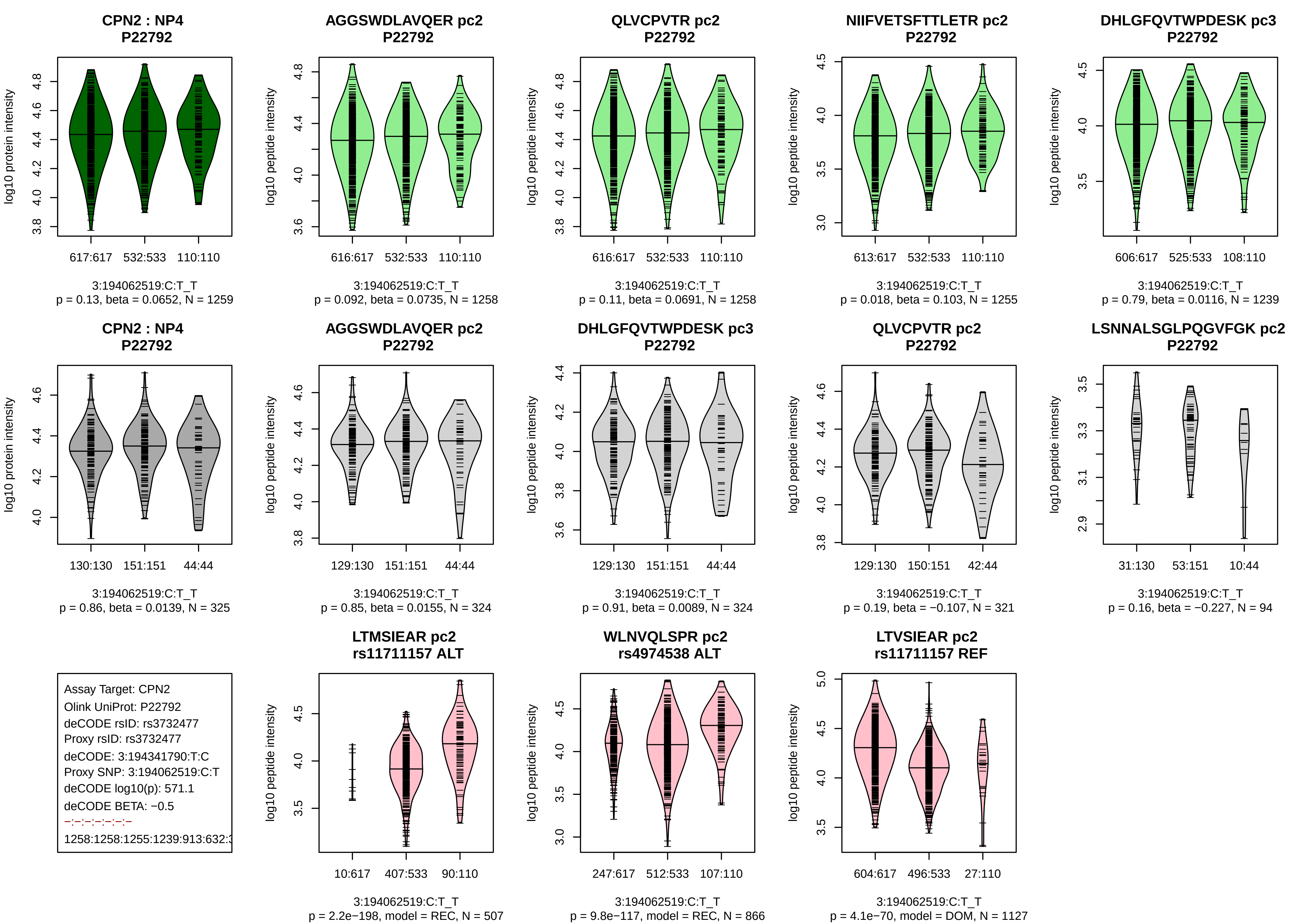
CPN2 : NP4
P22792
log10 protein intensity
617:617
532:533
110:110
3:194062519:C:T_T
p = 0.13, beta = 0.0652, N = 1259
AGGSWDLAVQER pc2
P22792
log10 peptide intensity
616:617
532:533
110:110
3:194062519:C:T_T
p = 0.092, beta = 0.0735, N = 1258
QLVCPVTR pc2
P22792
log10 peptide intensity
616:617
532:533
110:110
3:194062519:C:T_T
p = 0.11, beta = 0.0691, N = 1258
NIIFVETSFTTLETR pc2
P22792
log10 peptide intensity
613:617
532:533
110:110
3:194062519:C:T_T
p = 0.018, beta = 0.103, N = 1255
DHLGFQVTWPDESK pc3
P22792
log10 peptide intensity
606:617
525:533
108:110
3:194062519:C:T_T
p = 0.79, beta = 0.0116, N = 1239
CPN2 : NP4
P22792
log10 protein intensity
130:130
151:151
44:44
3:194062519:C:T_T
p = 0.86, beta = 0.0139, N = 325
AGGSWDLAVQER pc2
P22792
log10 peptide intensity
129:130
151:151
44:44
3:194062519:C:T_T
p = 0.85, beta = 0.0155, N = 324
DHLGFQVTWPDESK pc3
P22792
log10 peptide intensity
129:130
151:151
44:44
3:194062519:C:T_T
p = 0.91, beta = 0.0089, N = 324
QLVCPVTR pc2
P22792
log10 peptide intensity
129:130
150:151
42:44
3:194062519:C:T_T
p = 0.19, beta = −0.107, N = 321
LSNNALSGLPQGVFGK pc2
P22792
log10 peptide intensity
31:130
53:151
10:44
3:194062519:C:T_T
p = 0.16, beta = −0.227, N = 94
Assay Target: CPN2
Olink UniProt: P22792
deCODE rsID: rs3732477
Proxy rsID: rs3732477
deCODE: 3:194341790:T:C
Proxy SNP: 3:194062519:C:T
deCODE log10(p): 571.1
deCODE BETA: −0.5
−:−:−:−:−:−:−
1258:1258:1255:1239:913:632:3
LTMSIEAR pc2
rs11711157 ALT
log10 peptide intensity
10:617
407:533
90:110
3:194062519:C:T_T
p = 2.2e−198, model = REC, N = 507
WLNVQLSPR pc2
rs4974538 ALT
log10 peptide intensity
247:617
512:533
107:110
3:194062519:C:T_T
p = 9.8e−117, model = REC, N = 866
LTVSIEAR pc2
rs11711157 REF
log10 peptide intensity
604:617
496:533
27:110
3:194062519:C:T_T
p = 4.1e−70, model = DOM, N = 1127

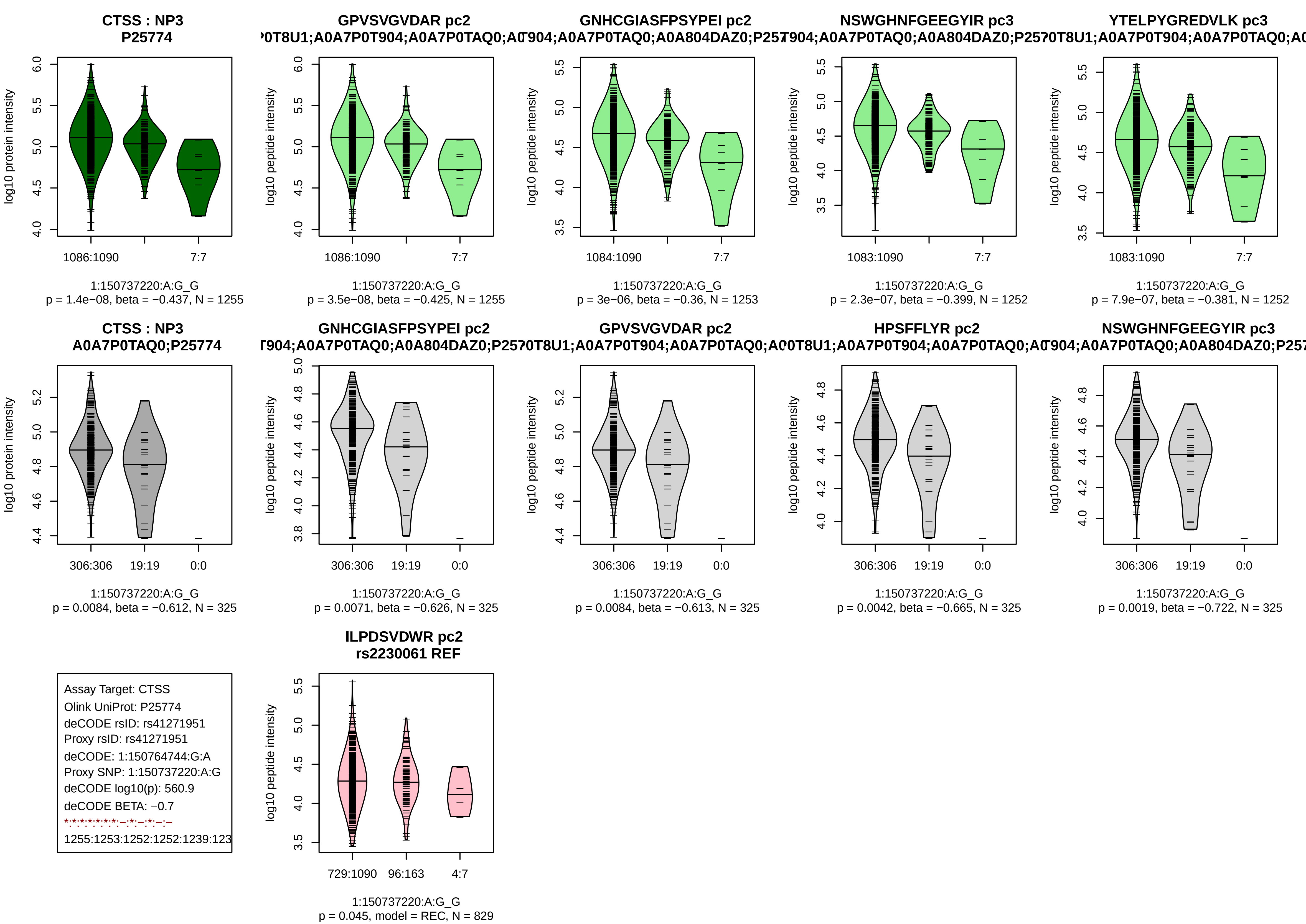
CTSS : NP3
P25774
log10 protein intensity
1086:1090
7:7
1:150737220:A:G_G
p = 1.4e−08, beta = −0.437, N = 1255
GPVSVGVDAR pc2
log10 peptide intensity
1086:1090
7:7
1:150737220:A:G_G
p = 3.5e−08, beta = −0.425, N = 1255
GNHCGIASFPSYPEI pc2
log10 peptide intensity
1084:1090
7:7
1:150737220:A:G_G
p = 3e−06, beta = −0.36, N = 1253
NSWGHNFGEEGYIR pc3
log10 peptide intensity
1083:1090
7:7
1:150737220:A:G_G
p = 2.3e−07, beta = −0.399, N = 1252
YTELPYGREDVLK pc3
log10 peptide intensity
1083:1090
7:7
1:150737220:A:G_G
p = 7.9e−07, beta = −0.381, N = 1252
CTSS : NP3
A0A7P0TAQ0;P25774
log10 protein intensity
306:306
19:19
0:0
1:150737220:A:G_G
p = 0.0084, beta = −0.612, N = 325
GNHCGIASFPSYPEI pc2
log10 peptide intensity
306:306
19:19
0:0
1:150737220:A:G_G
p = 0.0071, beta = −0.626, N = 325
GPVSVGVDAR pc2
log10 peptide intensity
306:306
19:19
0:0
1:150737220:A:G_G
p = 0.0084, beta = −0.613, N = 325
HPSFFLYR pc2
log10 peptide intensity
306:306
19:19
0:0
1:150737220:A:G_G
p = 0.0042, beta = −0.665, N = 325
NSWGHNFGEEGYIR pc3
log10 peptide intensity
306:306
19:19
0:0
1:150737220:A:G_G
p = 0.0019, beta = −0.722, N = 325
Assay Target: CTSS
Olink UniProt: P25774
deCODE rsID: rs41271951
Proxy rsID: rs41271951
deCODE: 1:150764744:G:A
Proxy SNP: 1:150737220:A:G
deCODE log10(p): 560.9
deCODE BETA: −0.7
1255:1253:1252:1252:1239:123
ILPDSVDWR pc2
rs2230061 REF
log10 peptide intensity
729:1090
96:163
4:7
1:150737220:A:G_G
p = 0.045, model = REC, N = 829

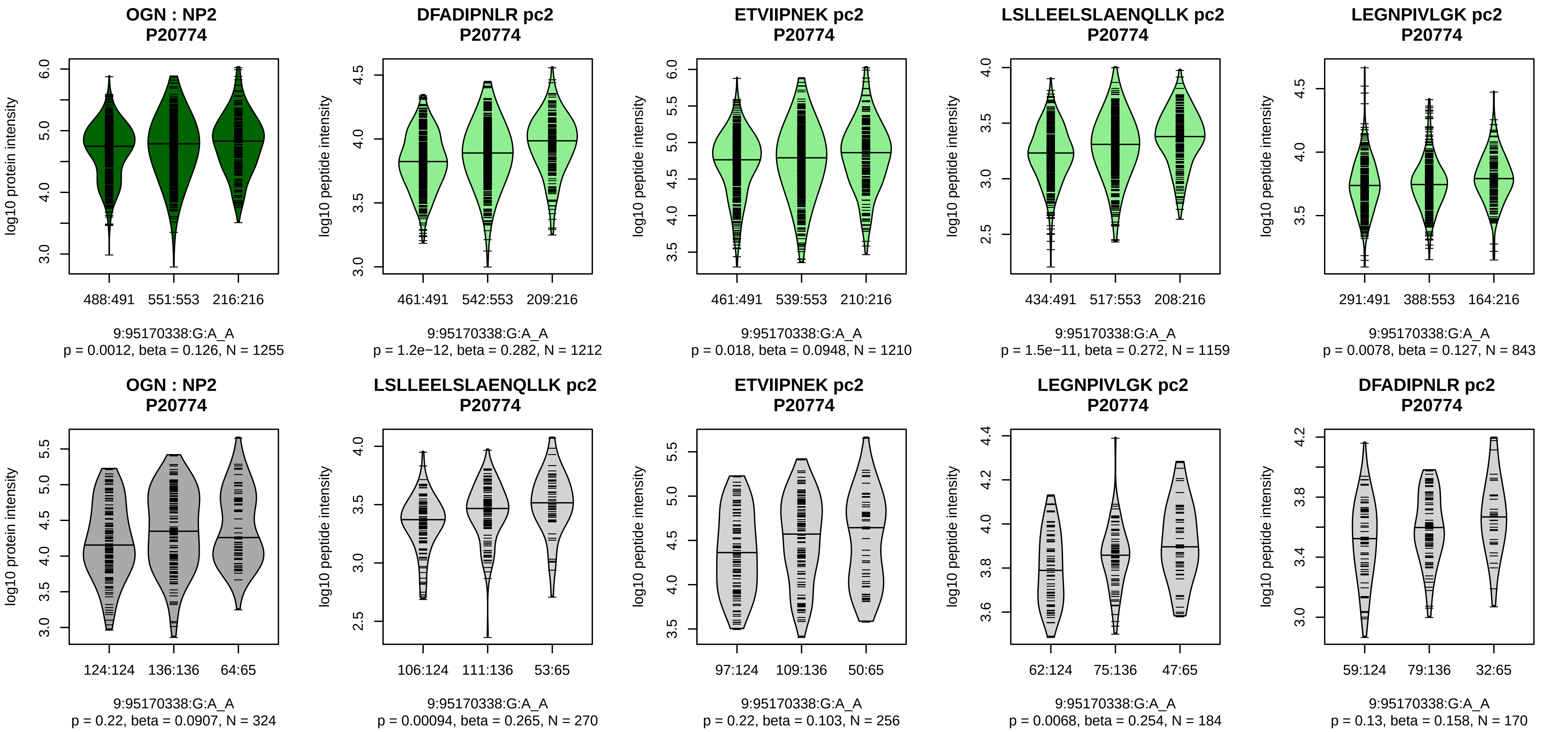


Assay Target: OGN
Olink UniProt: P20774
deCODE rsID: rs9299404
Proxy rsID: rs9299404
deCODE: 9:92408056:A:G
Proxy SNP: 9:95170338:G:A
deCODE log10(p): 550.6
deCODE BETA: 0.39
*:–:*:*:–:–:–:–:–
1212:1210:1159:843:383:235:22

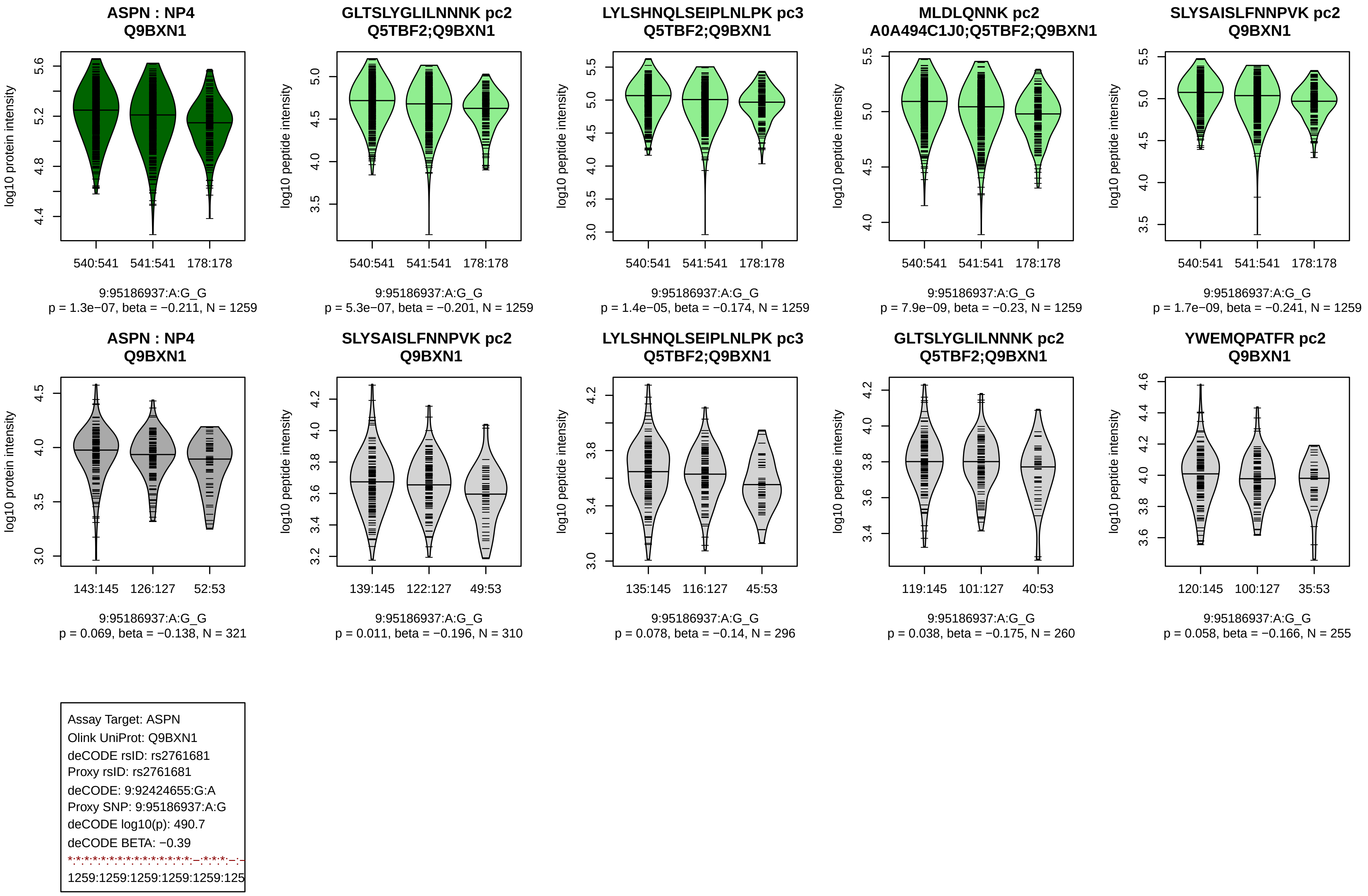
ASPN : NP4
Q9BXN1
log10 protein intensity
540:541
541:541
178:178
9:95186937:A:G_G
p = 1.3e−07, beta = −0.211, N = 1259
GLTSLYGLILNNNK pc2
Q5TBF2;Q9BXN1
log10 peptide intensity
540:541
541:541
178:178
9:95186937:A:G_G
p = 5.3e−07, beta = −0.201, N = 1259
LYLSHNQLSEIPLNLPK pc3
Q5TBF2;Q9BXN1
log10 peptide intensity
540:541
541:541
178:178
9:95186937:A:G_G
p = 1.4e−05, beta = −0.174, N = 1259
MLDLQNNK pc2
A0A494C1J0;Q5TBF2;Q9BXN1
log10 peptide intensity
540:541
541:541
178:178
9:95186937:A:G_G
p = 7.9e−09, beta = −0.23, N = 1259
SLYSAISLFNNPVK pc2
Q9BXN1
log10 peptide intensity
540:541
541:541
178:178
9:95186937:A:G_G
p = 1.7e−09, beta = −0.241, N = 1259
ASPN : NP4
Q9BXN1
log10 protein intensity
143:145
126:127
52:53
9:95186937:A:G_G
p = 0.069, beta = −0.138, N = 321
SLYSAISLFNNPVK pc2
Q9BXN1
log10 peptide intensity
139:145
122:127
49:53
9:95186937:A:G_G
p = 0.011, beta = −0.196, N = 310
LYLSHNQLSEIPLNLPK pc3
Q5TBF2;Q9BXN1
log10 peptide intensity
135:145
116:127
45:53
9:95186937:A:G_G
p = 0.078, beta = −0.14, N = 296
GLTSLYGLILNNNK pc2
Q5TBF2;Q9BXN1
log10 peptide intensity
119:145
101:127
40:53
9:95186937:A:G_G
p = 0.038, beta = −0.175, N = 260
YWEMQPATFR pc2
Q9BXN1
log10 peptide intensity
120:145
100:127
35:53
9:95186937:A:G_G
p = 0.058, beta = −0.166, N = 255
Assay Target: ASPN
Olink UniProt: Q9BXN1
deCODE rsID: rs2761681
Proxy rsID: rs2761681
deCODE: 9:92424655:G:A
Proxy SNP: 9:95186937:A:G
deCODE log10(p): 490.7
deCODE BETA: −0.39
1259:1259:1259:1259:1259:125

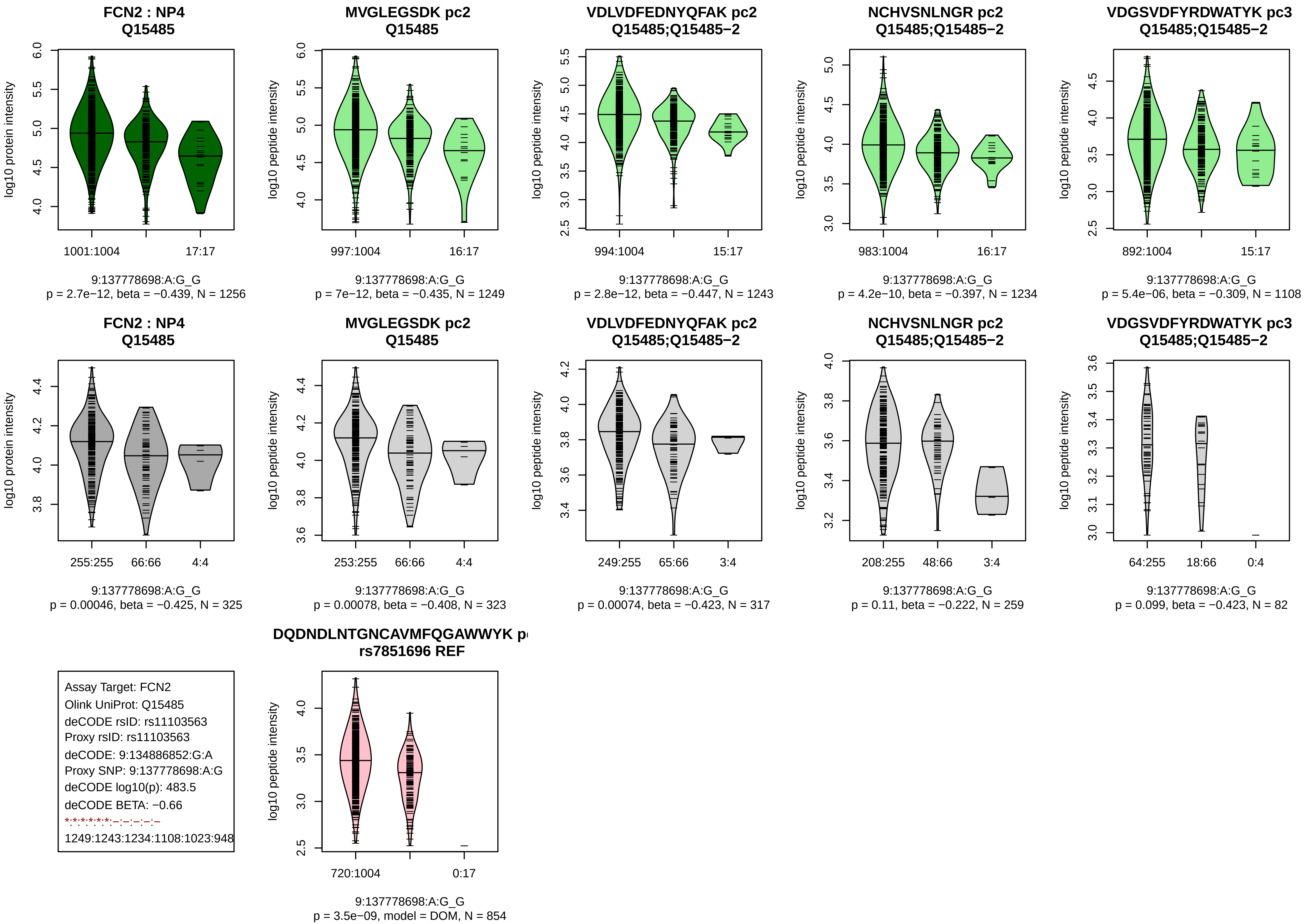
FCN2 : NP4
Q15485
MVGLEGSDK pc2
Q15485
VDLVDFEDNYQFAK pc2
Q15485;Q15485−2
NCHVSNLNGR pc2
Q15485;Q15485−2
VDGSVDFYRDWATYK pc3
Q15485;Q15485−2
log10 protein intensity
log10 peptide intensity
1001:1004
17:17
997:1004
16:17
994:1004
15:17
983:1004
16:17
892:1004
15:17
9:137778698:A:G_G
p = 2.7e−12, beta = −0.439, N = 1256
p = 7e−12, beta = −0.435, N = 1249
p = 2.8e−12, beta = −0.447, N = 1243
p = 4.2e−10, beta = −0.397, N = 1234
p = 5.4e−06, beta = −0.309, N = 1108
255:255
66:66
4:4
253:255
66:66
4:4
249:255
65:66
3:4
208:255
48:66
3:4
64:255
18:66
0:4
p = 0.00046, beta = −0.425, N = 325
p = 0.00078, beta = −0.408, N = 323
p = 0.00074, beta = −0.423, N = 317
p = 0.11, beta = −0.222, N = 259
p = 0.099, beta = −0.423, N = 82
Assay Target: FCN2
Olink UniProt: Q15485
deCODE rsID: rs11103563
Proxy rsID: rs11103563
deCODE: 9:134886852:G:A
Proxy SNP: 9:137778698:A:G
deCODE log10(p): 483.5
deCODE BETA: −0.66
*:*:*:*:*:*:–:–:–:–:–
1249:1243:1234:1108:1023:948
DQDNDLNTGNCAVMFQGAWWYK p
rs7851696 REF
720:1004
0:17
9:137778698:A:G_G
p = 3.5e−09, model = DOM, N = 854

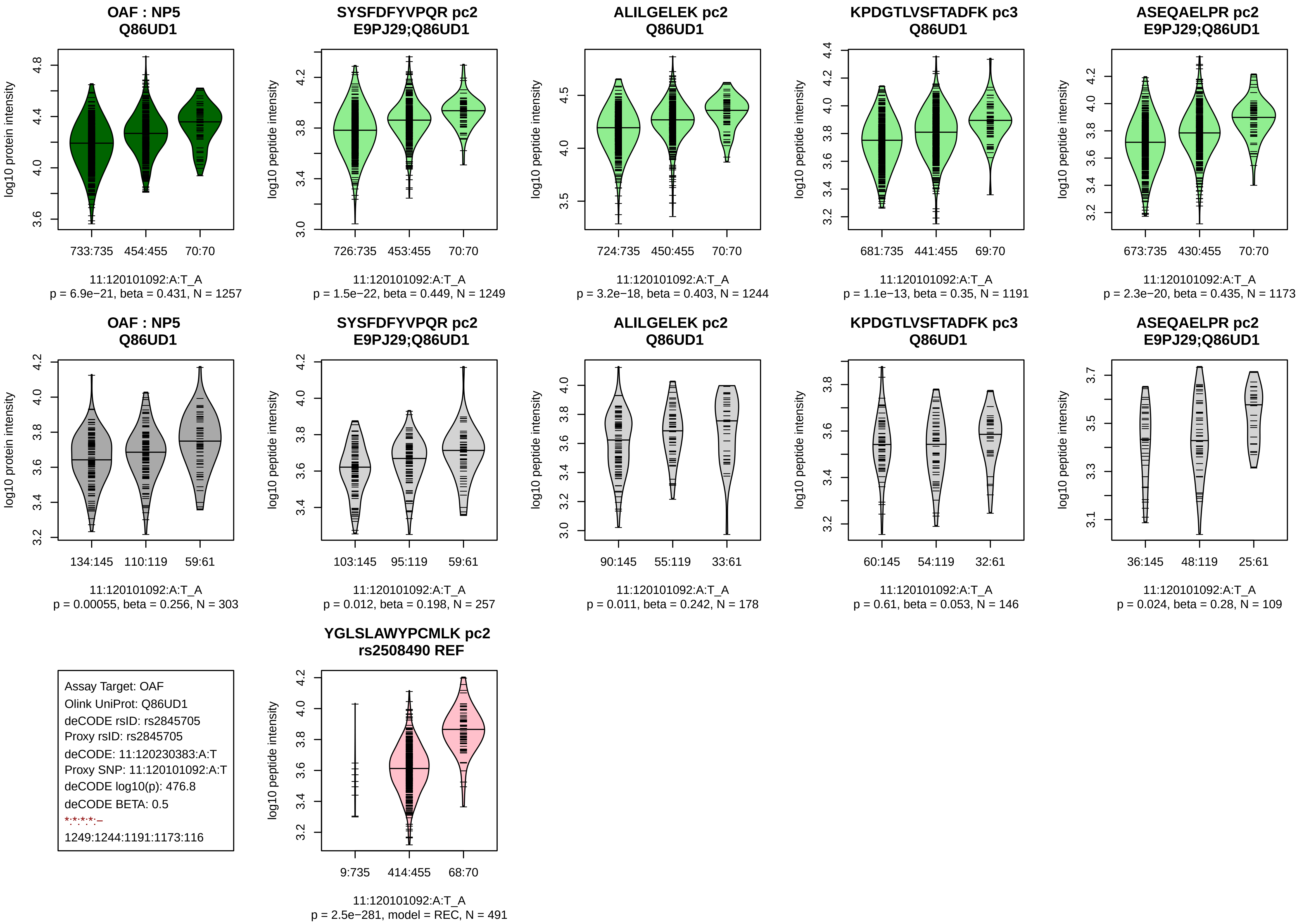

OAF : NP5
Q86UD1
log10 protein intensity
733:735
454:455
70:70
11:120101092:A:T_A
p = 6.9e−21, beta = 0.431, N = 1257
SYSFDFYVPQR pc2
E9PJ29;Q86UD1
log10 peptide intensity
726:735
453:455
70:70
11:120101092:A:T_A
p = 1.5e−22, beta = 0.449, N = 1249
ALILGELEK pc2
Q86UD1
log10 peptide intensity
724:735
450:455
70:70
11:120101092:A:T_A
p = 3.2e−18, beta = 0.403, N = 1244
KPDGTLVSFTADFK pc3
Q86UD1
log10 peptide intensity
681:735
441:455
69:70
11:120101092:A:T_A
p = 1.1e−13, beta = 0.35, N = 1191
ASEQAELPR pc2
E9PJ29;Q86UD1
log10 peptide intensity
673:735
430:455
70:70
11:120101092:A:T_A
p = 2.3e−20, beta = 0.435, N = 1173
OAF : NP5
Q86UD1
log10 protein intensity
134:145
110:119
59:61
11:120101092:A:T_A
p = 0.00055, beta = 0.256, N = 303
SYSFDFYVPQR pc2
E9PJ29;Q86UD1
log10 peptide intensity
103:145
95:119
59:61
11:120101092:A:T_A
p = 0.012, beta = 0.198, N = 257
ALILGELEK pc2
Q86UD1
log10 peptide intensity
90:145
55:119
33:61
11:120101092:A:T_A
p = 0.011, beta = 0.242, N = 178
KPDGTLVSFTADFK pc3
Q86UD1
log10 peptide intensity
60:145
54:119
32:61
11:120101092:A:T_A
p = 0.61, beta = 0.053, N = 146
ASEQAELPR pc2
E9PJ29;Q86UD1
log10 peptide intensity
36:145
48:119
25:61
11:120101092:A:T_A
p = 0.024, beta = 0.28, N = 109
Assay Target: OAF
Olink UniProt: Q86UD1
deCODE rsID: rs2845705
Proxy rsID: rs2845705
deCODE: 11:120230383:A:T
Proxy SNP: 11:120101092:A:T
deCODE log10(p): 476.8
deCODE BETA: 0.5
*:*:*:*:–
1249:1244:1191:1173:116
YGLSLAWYPCMLK pc2
rs2508490 REF
log10 peptide intensity
9:735
414:455
68:70
11:120101092:A:T_A
p = 2.5e−281, model = REC, N = 491

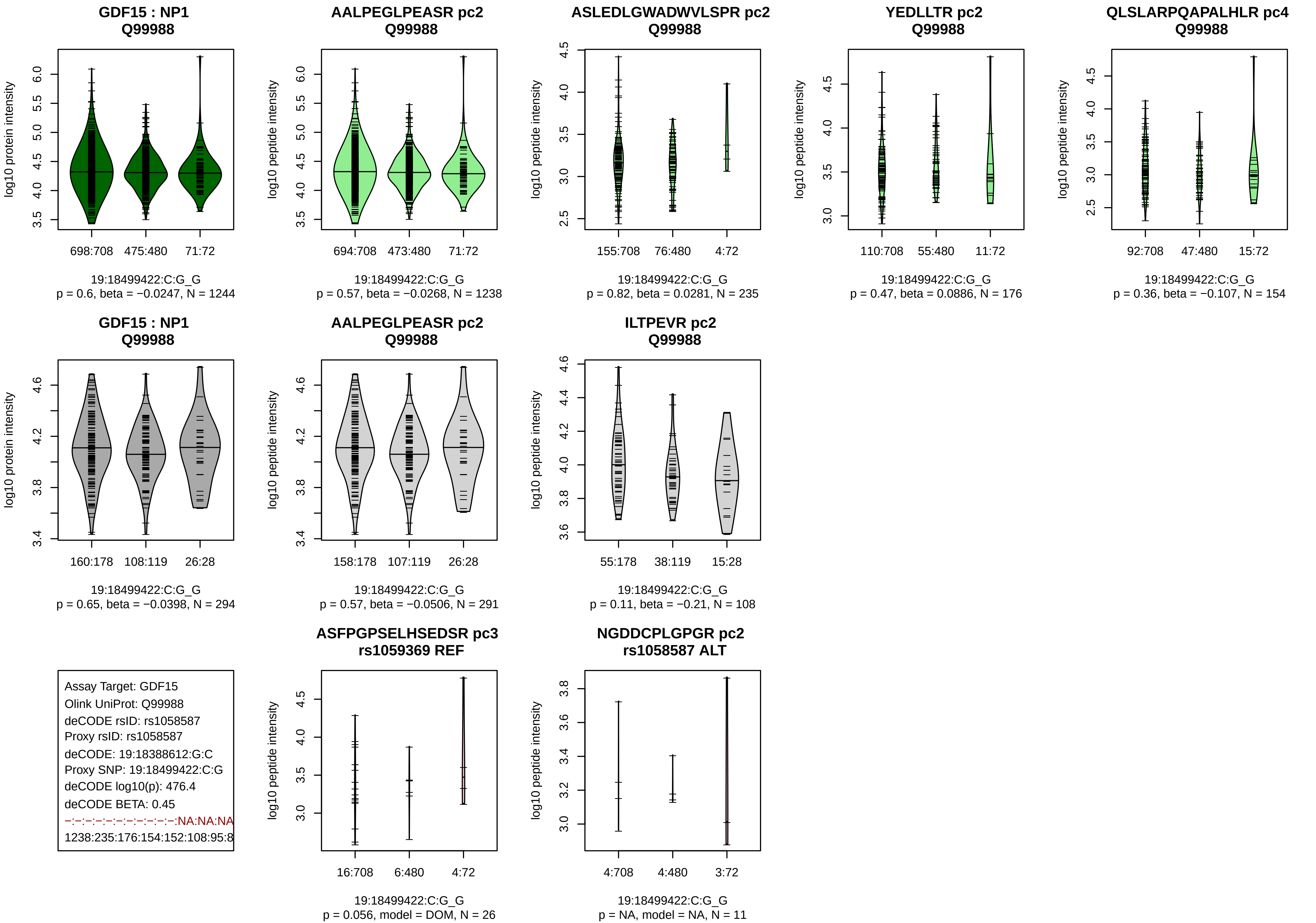

GDF15 : NP1
Q99988
log10 protein intensity
698:708 475:480 71:72
19:18499422:C:G_G
p = 0.6, beta = −0.0247, N = 1244
AALPEGLPEASR pc2
Q99988
log10 peptide intensity
694:708 473:480 71:72
19:18499422:C:G_G
p = 0.57, beta = −0.0268, N = 1238
ASLEDLGWADWVLSPR pc2
Q99988
155:708 76:480 4:72
19:18499422:C:G_G
p = 0.82, beta = 0.0281, N = 235
YEDLLTR pc2
Q99988
110:708 55:480 11:72
19:18499422:C:G_G
p = 0.47, beta = 0.0886, N = 176
QLSLARPQAPALHLR pc4
Q99988
92:708 47:480 15:72
19:18499422:C:G_G
p = 0.36, beta = −0.107, N = 154
GDF15 : NP1
Q99988
160:178 108:119 26:28
19:18499422:C:G_G
p = 0.65, beta = −0.0398, N = 294
AALPEGLPEASR pc2
Q99988
158:178 107:119 26:28
19:18499422:C:G_G
p = 0.57, beta = −0.0506, N = 291
ILTPEVR pc2
Q99988
55:178 38:119 15:28
19:18499422:C:G_G
p = 0.11, beta = −0.21, N = 108
Assay Target: GDF15
Olink UniProt: Q99988
deCODE rsID: rs1058587
Proxy rsID: rs1058587
deCODE: 19:18388612:G:C
Proxy SNP: 19:18499422:C:G
deCODE log10(p): 476.4
deCODE BETA: 0.45
−:−:−:−:−:−:−:−:−:−:−:NA:NA:NA
1238:235:176:154:152:108:95:8
ASFPGPSELHSEDSR pc3
rs1059369 REF
16:708 6:480 4:72
19:18499422:C:G_G
p = 0.056, model = DOM, N = 26
NGDDCPLGPGR pc2
rs1058587 ALT
4:708 4:480 3:72
19:18499422:C:G_G
p = NA, model = NA, N = 11

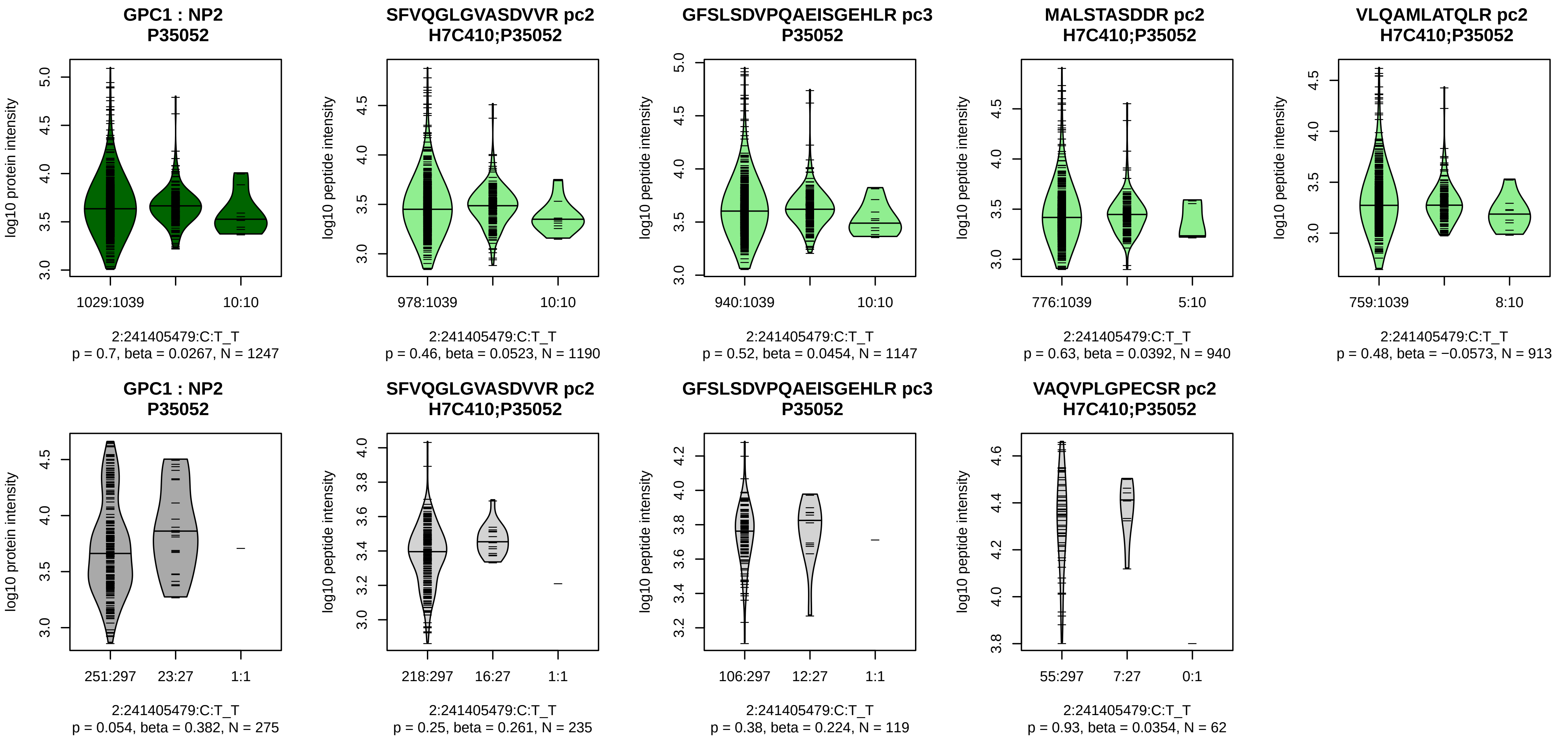


Assay Target: GPC1
Olink UniProt: P35052
deCODE rsID: rs1126920
Proxy rsID: rs1126920
deCODE: 2:240466062:T:C
Proxy SNP: 2:241405479:C:T
deCODE log10(p): 475.1
deCODE BETA: 0.63
−:−:−:−:−:−:−:−:−:−:−:−:−:−:NA:NA
1190:1147:940:913:582:472:319

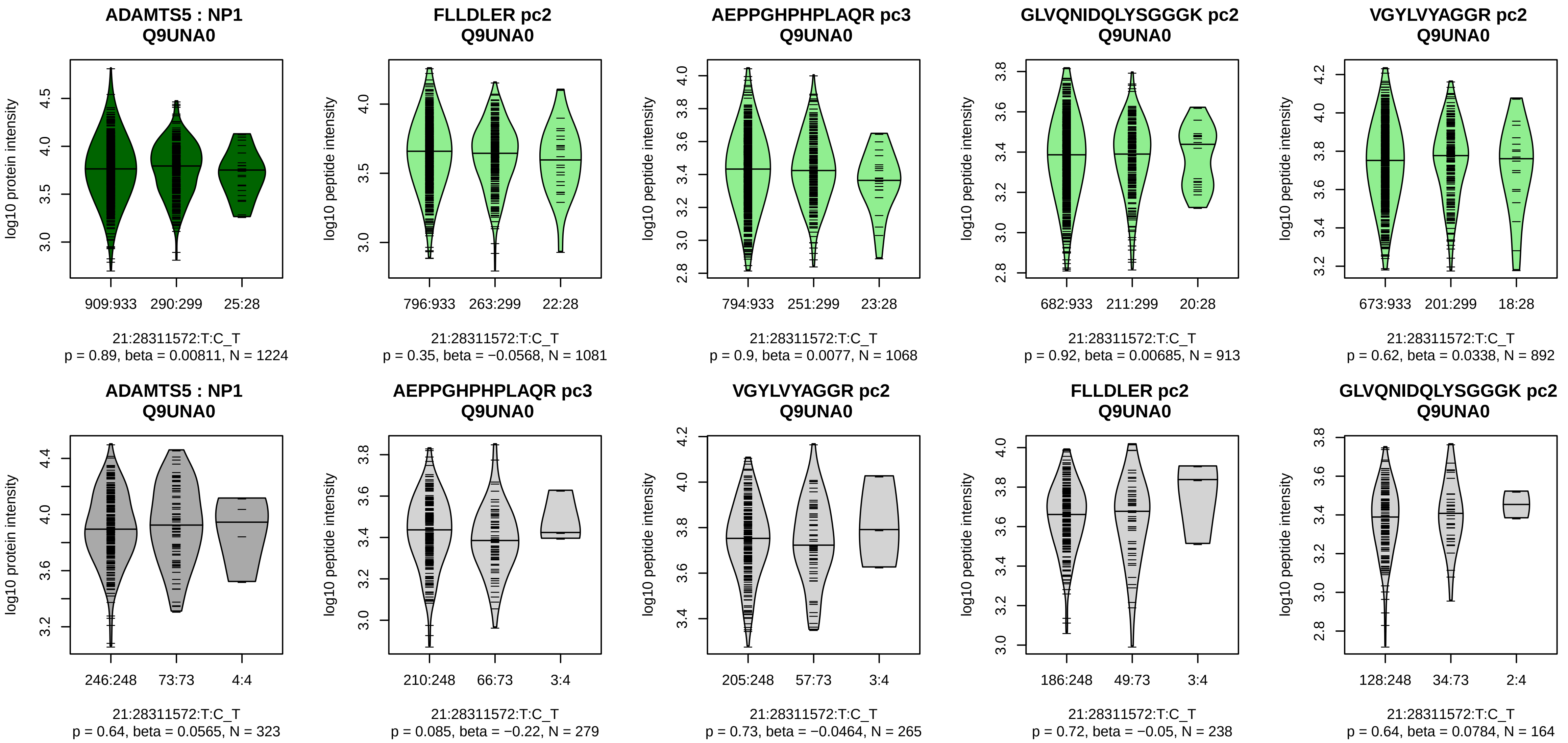


Assay Target: ADAMTS5
Olink UniProt: Q9UNA0
deCODE rsID: rs151058
Proxy rsID: rs151058
deCODE: 21:26939253:T:C
Proxy SNP: 21:28311572:T:C
deCODE log10(p): 469
deCODE BETA: −0.56
−:−:−:−:−:−
1081:1068:913:892:364:140

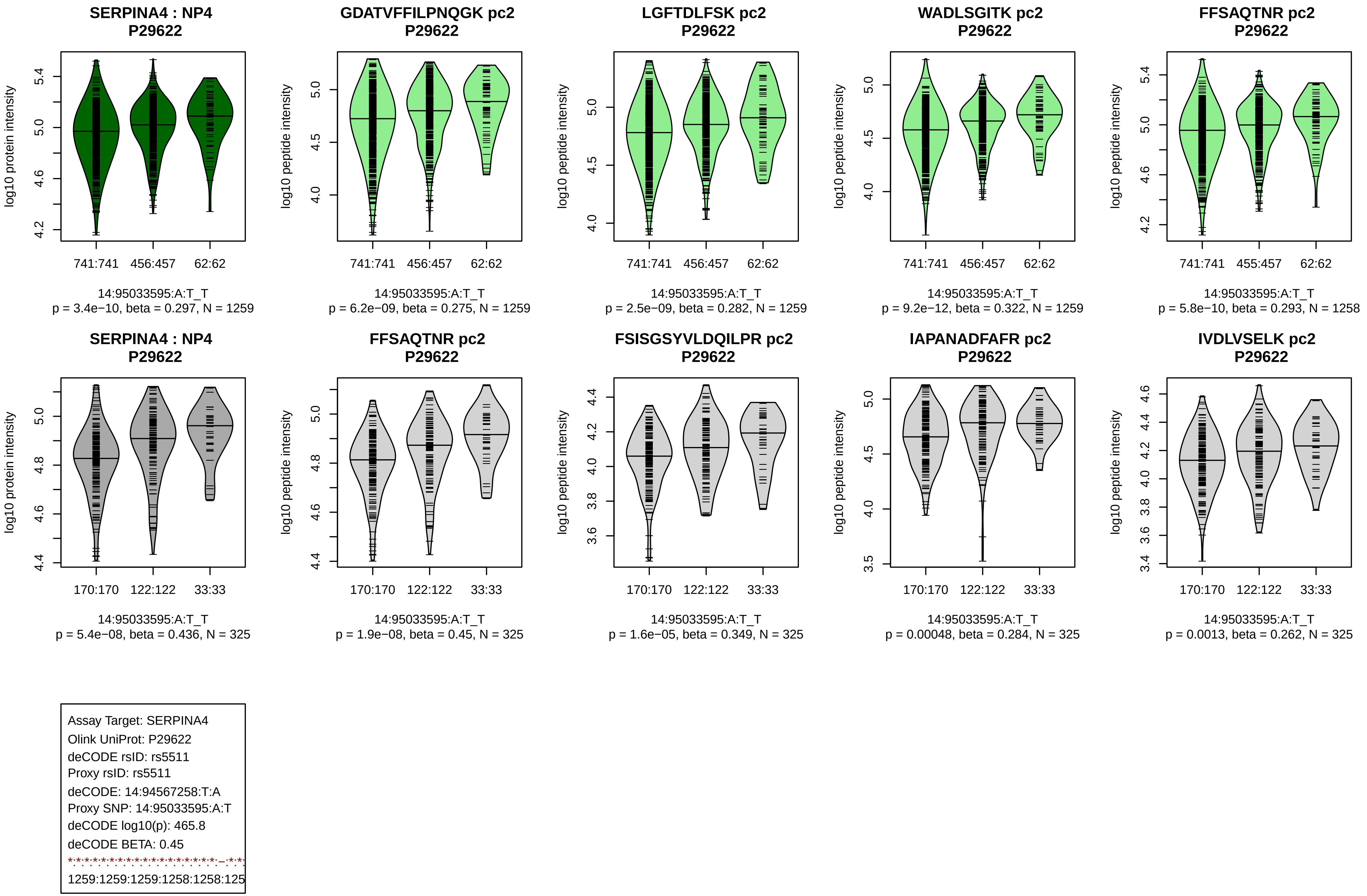

SERPINA4 : NP4
P29622
log10 protein intensity
741:741
456:457
62:62
14:95033595:A:T_T
p = 3.4e−10, beta = 0.297, N = 1259
GDATVFFILPNQGK pc2
P29622
log10 peptide intensity
741:741
456:457
62:62
14:95033595:A:T_T
p = 6.2e−09, beta = 0.275, N = 1259
LGFTDLFSK pc2
P29622
log10 peptide intensity
741:741
456:457
62:62
14:95033595:A:T_T
p = 2.5e−09, beta = 0.282, N = 1259
WADLSGITK pc2
P29622
log10 peptide intensity
741:741
456:457
62:62
14:95033595:A:T_T
p = 9.2e−12, beta = 0.322, N = 1259
FFSAQTNR pc2
P29622
log10 peptide intensity
741:741
455:457
62:62
14:95033595:A:T_T
p = 5.8e−10, beta = 0.293, N = 1258
SERPINA4 : NP4
P29622
log10 protein intensity
170:170
122:122
33:33
14:95033595:A:T_T
p = 5.4e−08, beta = 0.436, N = 325
FFSAQTNR pc2
P29622
log10 peptide intensity
170:170
122:122
33:33
14:95033595:A:T_T
p = 1.9e−08, beta = 0.45, N = 325
FSISGSYVLDQILPR pc2
P29622
log10 peptide intensity
170:170
122:122
33:33
14:95033595:A:T_T
p = 1.6e−05, beta = 0.349, N = 325
IAPANADFAFR pc2
P29622
log10 peptide intensity
170:170
122:122
33:33
14:95033595:A:T_T
p = 0.00048, beta = 0.284, N = 325
IVDLVSELK pc2
P29622
log10 peptide intensity
170:170
122:122
33:33
14:95033595:A:T_T
p = 0.0013, beta = 0.262, N = 325
Assay Target: SERPINA4
Olink UniProt: P29622
deCODE rsID: rs5511
Proxy rsID: rs5511
deCODE: 14:94567258:T:A
Proxy SNP: 14:95033595:A:T
deCODE log10(p): 465.8
deCODE BETA: 0.45
1259:1259:1259:1258:1258:125

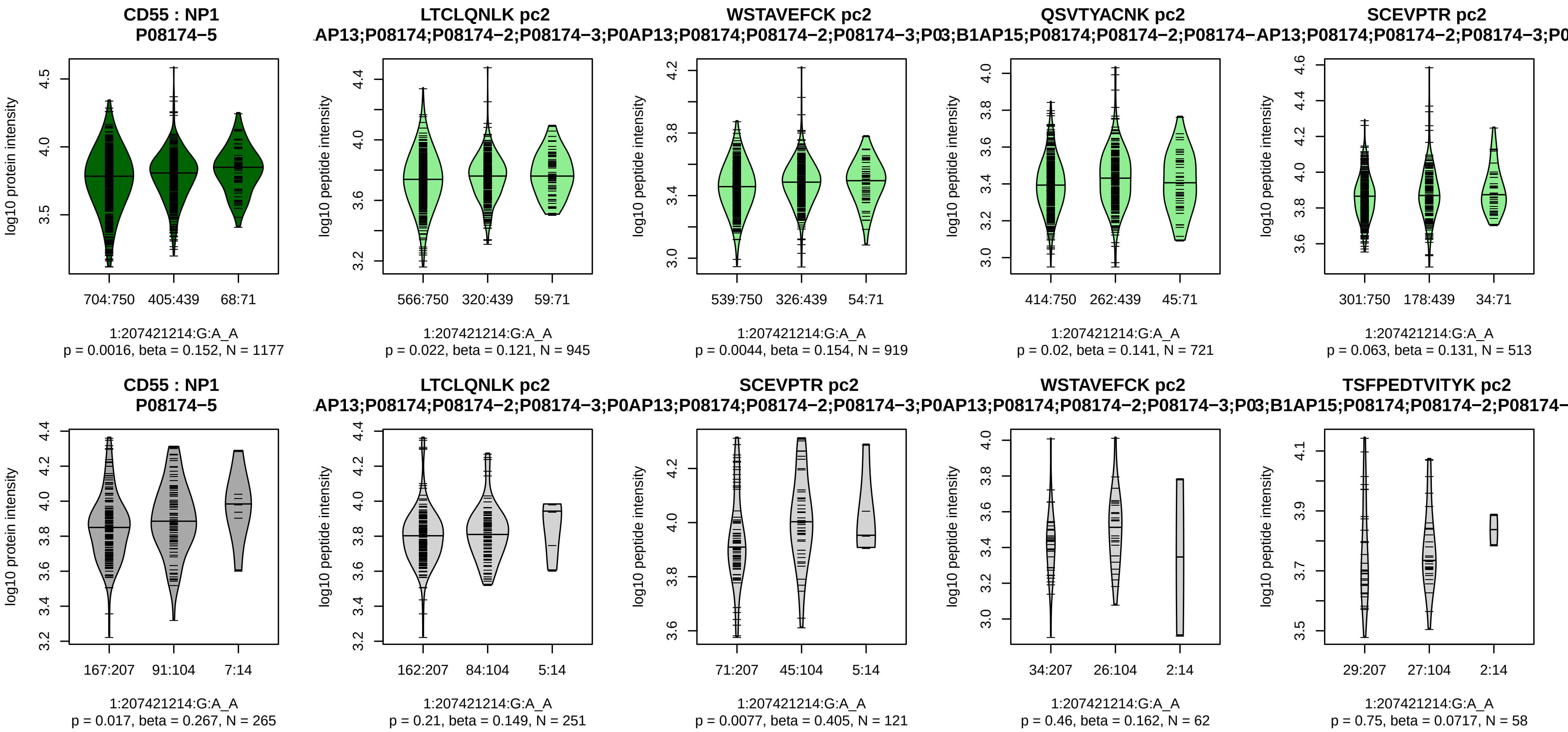


Assay Target: CD55
Olink UniProt: P08174
deCODE rsID: rs61822669
Proxy rsID: rs61822669
deCODE: 1:207247869:A:G
Proxy SNP: 1:207421214:G:A
deCODE log10(p): 433.9
deCODE BETA: −0.41
−:*:−:−:−:−:*:−
945:919:721:513:406:389:366:5

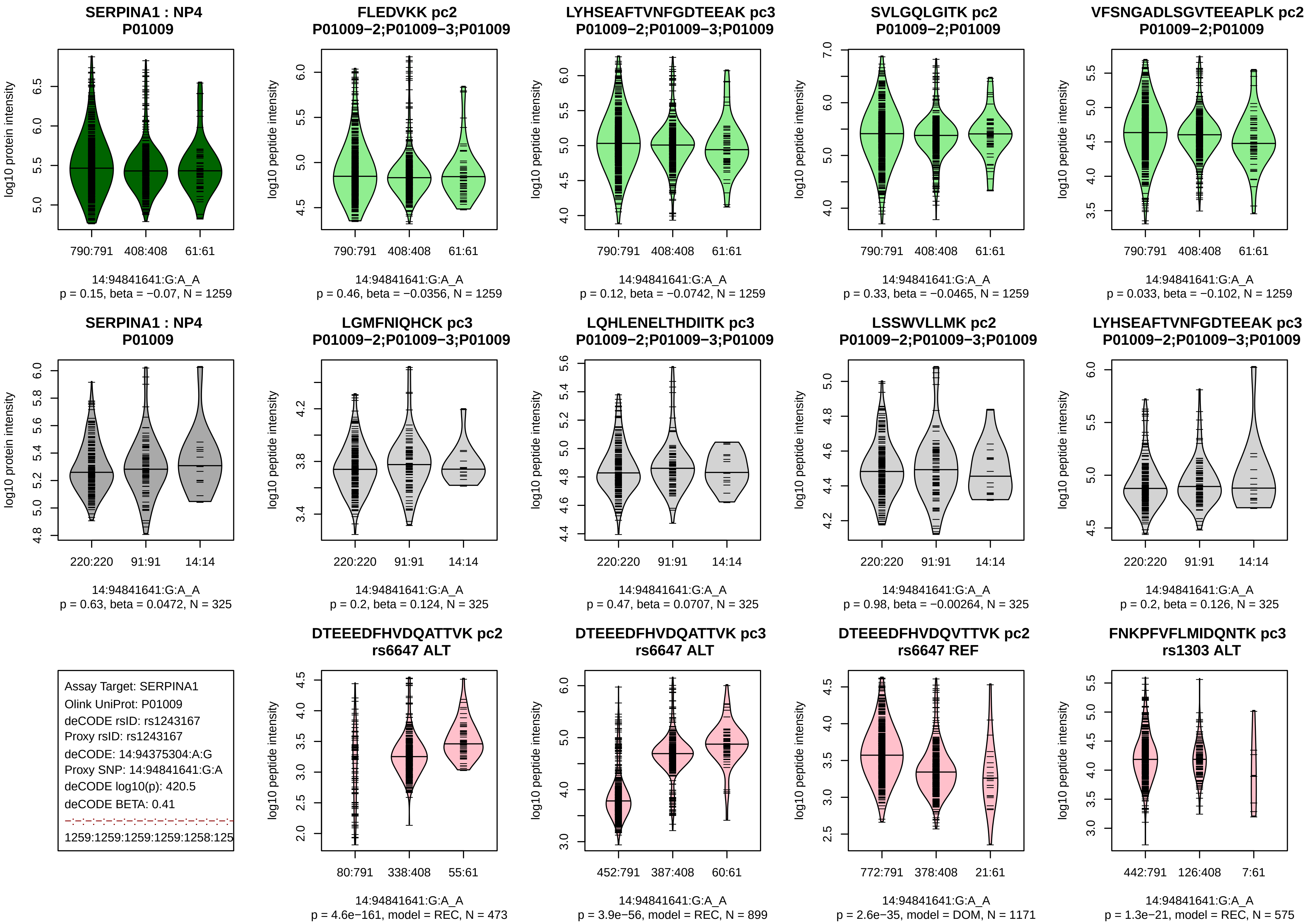
SERPINA1 : NP4
P01009
log10 protein intensity
790:791 408:408 61:61
14:94841641:G:A_A
p = 0.15, beta = −0.07, N = 1259
FLEDVKK pc2
P01009−2;P01009−3;P01009
log10 peptide intensity
790:791 408:408 61:61
14:94841641:G:A_A
p = 0.46, beta = −0.0356, N = 1259
LYHSEAFTVNFGDTEEAK pc3
P01009−2;P01009−3;P01009
log10 peptide intensity
790:791 408:408 61:61
14:94841641:G:A_A
p = 0.12, beta = −0.0742, N = 1259
SVLGQLGITK pc2
P01009−2;P01009
log10 peptide intensity
790:791 408:408 61:61
14:94841641:G:A_A
p = 0.33, beta = −0.0465, N = 1259
VFSNGADLSGVTEEAPLK pc2
P01009−2;P01009
log10 peptide intensity
790:791 408:408 61:61
14:94841641:G:A_A
p = 0.033, beta = −0.102, N = 1259
SERPINA1 : NP4
P01009
log10 protein intensity
220:220 91:91 14:14
14:94841641:G:A_A
p = 0.63, beta = 0.0472, N = 325
LGMFNIQHCK pc3
P01009−2;P01009−3;P01009
log10 peptide intensity
220:220 91:91 14:14
14:94841641:G:A_A
p = 0.2, beta = 0.124, N = 325
LQHLENELTHDIITK pc3
P01009−2;P01009−3;P01009
log10 peptide intensity
220:220 91:91 14:14
14:94841641:G:A_A
p = 0.47, beta = 0.0707, N = 325
LSSWVLLMK pc2
P01009−2;P01009−3;P01009
log10 peptide intensity
220:220 91:91 14:14
14:94841641:G:A_A
p = 0.98, beta = −0.00264, N = 325
LYHSEAFTVNFGDTEEAK pc3
P01009−2;P01009−3;P01009
log10 peptide intensity
220:220 91:91 14:14
14:94841641:G:A_A
p = 0.2, beta = 0.126, N = 325
Assay Target: SERPINA1
Olink UniProt: P01009
deCODE rsID: rs1243167
Proxy rsID: rs1243167
deCODE: 14:94375304:A:G
Proxy SNP: 14:94841641:G:A
deCODE log10(p): 420.5
deCODE BETA: 0.41
1259:1259:1259:1259:1258:125
DTEEEDFHVDQATTVK pc2
rs6647 ALT
log10 peptide intensity
80:791 338:408 55:61
14:94841641:G:A_A
p = 4.6e−161, model = REC, N = 473
DTEEEDFHVDQATTVK pc3
rs6647 ALT
log10 peptide intensity
452:791 387:408 60:61
14:94841641:G:A_A
p = 3.9e−56, model = REC, N = 899
DTEEEDFHVDQVTTVK pc2
rs6647 REF
log10 peptide intensity
772:791 378:408 21:61
14:94841641:G:A_A
p = 2.6e−35, model = DOM, N = 1171
FNKPFVFLMIDQNTK pc3
rs1303 ALT
log10 peptide intensity
442:791 126:408 7:61
14:94841641:G:A_A
p = 1.3e−21, model = REC, N = 575

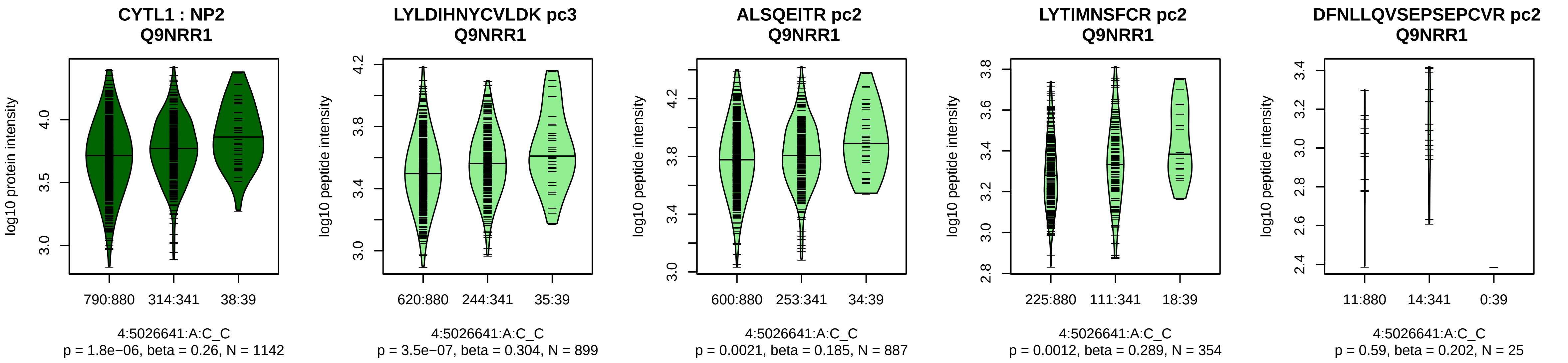


Assay Target: CYTL1
Olink UniProt: Q9NRR1
deCODE rsID: rs9998211
Proxy rsID: rs9998211
deCODE: 4:5024914:C:!
Proxy SNP: 4:5026641:A:C
deCODE log10(p): 411.7
deCODE BETA: 0.45
*:*:*:–
899:887:354:25

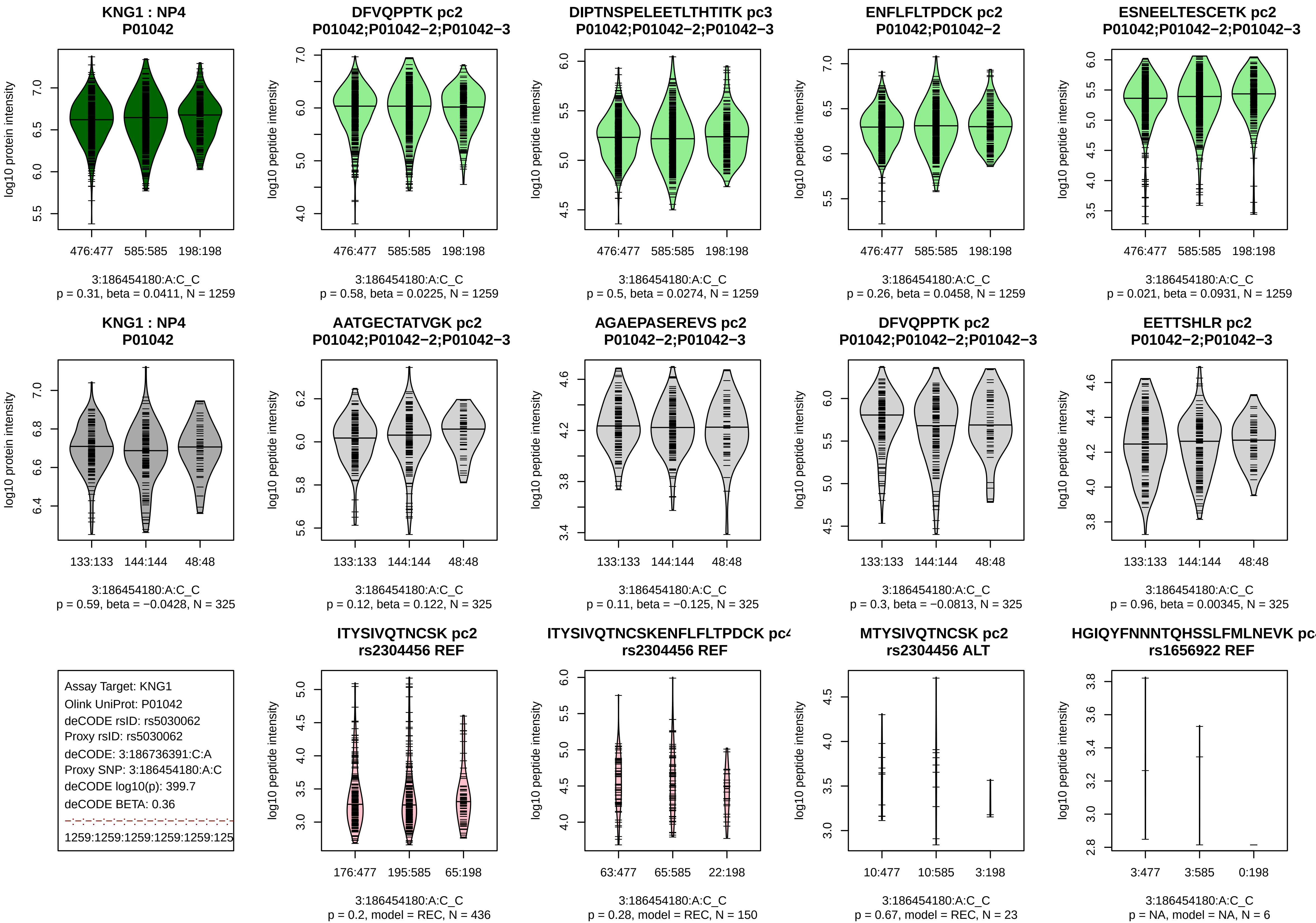
KNG1 : NP4
P01042
log10 protein intensity
476:477
585:585
198:198
3:186454180:A:C_C
p = 0.31, beta = 0.0411, N = 1259
DFVQPPTK pc2
P01042;P01042−2;P01042−3
log10 peptide intensity
476:477
585:585
198:198
3:186454180:A:C_C
p = 0.58, beta = 0.0225, N = 1259
DIPTNSPELEETLTHTITK pc3
P01042;P01042−2;P01042−3
log10 peptide intensity
476:477
585:585
198:198
3:186454180:A:C_C
p = 0.5, beta = 0.0274, N = 1259
ENFLFLTPDCK pc2
P01042;P01042−2
log10 peptide intensity
476:477
585:585
198:198
3:186454180:A:C_C
p = 0.26, beta = 0.0458, N = 1259
ESNEELTESCETK pc2
P01042;P01042−2;P01042−3
log10 peptide intensity
476:477
585:585
198:198
3:186454180:A:C_C
p = 0.021, beta = 0.0931, N = 1259
KNG1 : NP4
P01042
log10 protein intensity
133:133
144:144
48:48
3:186454180:A:C_C
p = 0.59, beta = −0.0428, N = 325
AATGECTATVGK pc2
P01042;P01042−2;P01042−3
log10 peptide intensity
133:133
144:144
48:48
3:186454180:A:C_C
p = 0.12, beta = 0.122, N = 325
AGAEPASEREVS pc2
P01042−2;P01042−3
log10 peptide intensity
133:133
144:144
48:48
3:186454180:A:C_C
p = 0.11, beta = −0.125, N = 325
DFVQPPTK pc2
P01042;P01042−2;P01042−3
log10 peptide intensity
133:133
144:144
48:48
3:186454180:A:C_C
p = 0.3, beta = −0.0813, N = 325
EETTSHLR pc2
P01042−2;P01042−3
log10 peptide intensity
133:133
144:144
48:48
3:186454180:A:C_C
p = 0.96, beta = 0.00345, N = 325
Assay Target: KNG1
Olink UniProt: P01042
deCODE rsID: rs5030062
Proxy rsID: rs5030062
deCODE: 3:186736391:C:A
Proxy SNP: 3:186454180:A:C
deCODE log10(p): 399.7
deCODE BETA: 0.36
1259:1259:1259:1259:1259:125
ITYSIVQTNCSK pc2
rs2304456 REF
log10 peptide intensity
176:477
195:585
65:198
3:186454180:A:C_C
p = 0.2, model = REC, N = 436
ITYSIVQTNCSKENFLFLTPDCK pc4
rs2304456 REF
log10 peptide intensity
63:477
65:585
22:198
3:186454180:A:C_C
p = 0.28, model = REC, N = 150
MTYSIVQTNCSK pc2
rs2304456 ALT
log10 peptide intensity
10:477
10:585
3:198
3:186454180:A:C_C
p = 0.67, model = REC, N = 23
HGIQYFNNNTQHSSLFMLNEVK pc
rs1656922 REF
log10 peptide intensity
3:477
3:585
0:198
3:186454180:A:C_C
p = NA, model = NA, N = 6

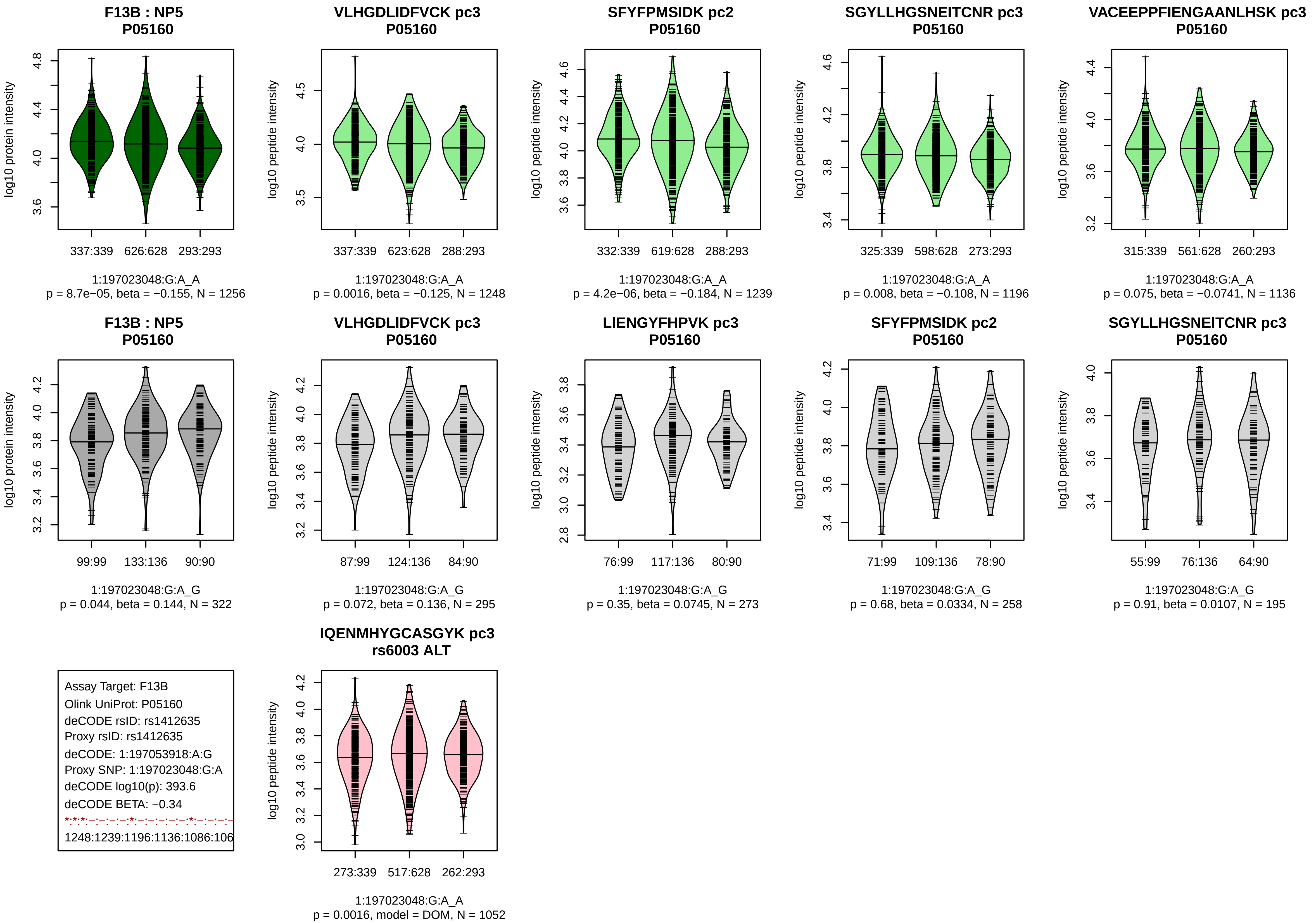
F13B : NP5
P05160
log10 protein intensity
337:339 626:628 293:293
1:197023048:G:A_A
p = 8.7e−05, beta = −0.155, N = 1256
VLHGDLIDFVCK pc3
P05160
log10 peptide intensity
337:339 623:628 288:293
1:197023048:G:A_A
p = 0.0016, beta = −0.125, N = 1248
SFYFPMSIDK pc2
P05160
332:339 619:628 288:293
1:197023048:G:A_A
p = 4.2e−06, beta = −0.184, N = 1239
SGYLLHGSNEITCNR pc3
P05160
325:339 598:628 273:293
1:197023048:G:A_A
p = 0.008, beta = −0.108, N = 1196
VACEEPPFIENGAANLHSK pc3
P05160
315:339 561:628 260:293
1:197023048:G:A_A
p = 0.075, beta = −0.0741, N = 1136
F13B : NP5
P05160
99:99 133:136 90:90
1:197023048:G:A_G
p = 0.044, beta = 0.144, N = 322
VLHGDLIDFVCK pc3
P05160
87:99 124:136 84:90
1:197023048:G:A_G
p = 0.072, beta = 0.136, N = 295
LIENGYFHPVK pc3
P05160
76:99 117:136 80:90
1:197023048:G:A_G
p = 0.35, beta = 0.0745, N = 273
SFYFPMSIDK pc2
P05160
71:99 109:136 78:90
1:197023048:G:A_G
p = 0.68, beta = 0.0334, N = 258
SGYLLHGSNEITCNR pc3
P05160
55:99 76:136 64:90
1:197023048:G:A_G
p = 0.91, beta = 0.0107, N = 195
Assay Target: F13B
Olink UniProt: P05160
deCODE rsID: rs1412635
Proxy rsID: rs1412635
deCODE: 1:197053918:A:G
Proxy SNP: 1:197023048:G:A
deCODE log10(p): 393.6
deCODE BETA: −0.34
*:*:*:–:–:–:–:*:–:–:–:–:–:*:–:–:–:–
1248:1239:1196:1136:1086:106
IQENMHYGCASGYK pc3
rs6003 ALT
273:339 517:628 262:293
1:197023048:G:A_A
p = 0.0016, model = DOM, N = 1052

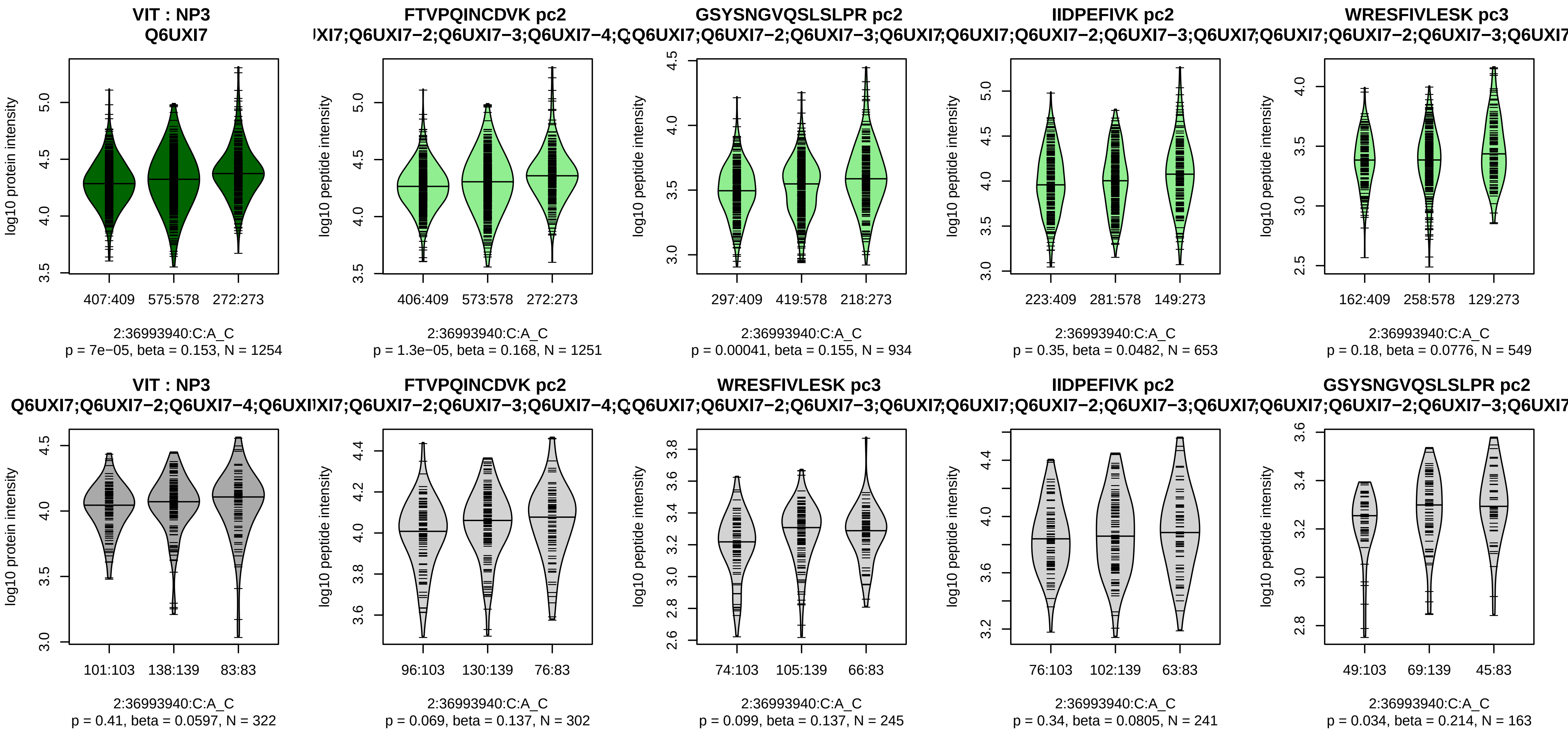


Assay Target: VIT
Olink UniProt: Q6UXI7
deCODE rsID: rs1468810
Proxy rsID: rs1468810
deCODE: 2:36766797:C:A
Proxy SNP: 2:36993940:C:A
deCODE log10(p): 389.4
deCODE BETA: 0.36
*:*:−:−:−:−:−:−:−:−:−:NA:NA:NA
1251:934:653:549:480:465:415:

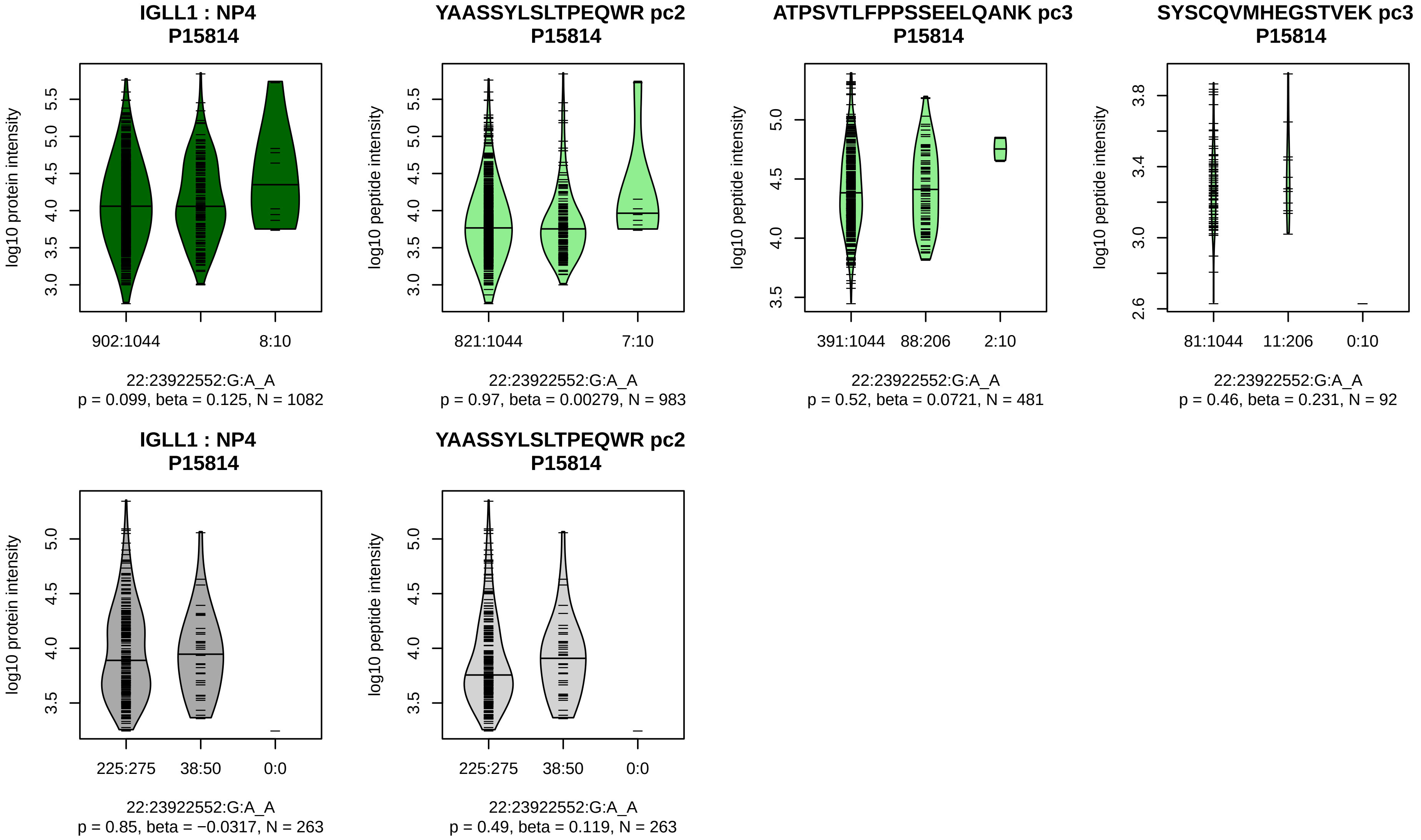


Assay Target: IGLL1
Olink UniProt: P15814
deCODE rsID: rs9624216
Proxy rsID: rs9624216
deCODE: 22:23580365:A:G
Proxy SNP: 22:23922552:G:A
deCODE log10(p): 374.6
deCODE BETA: −0.6
−:−:−:−
983:481:92:35

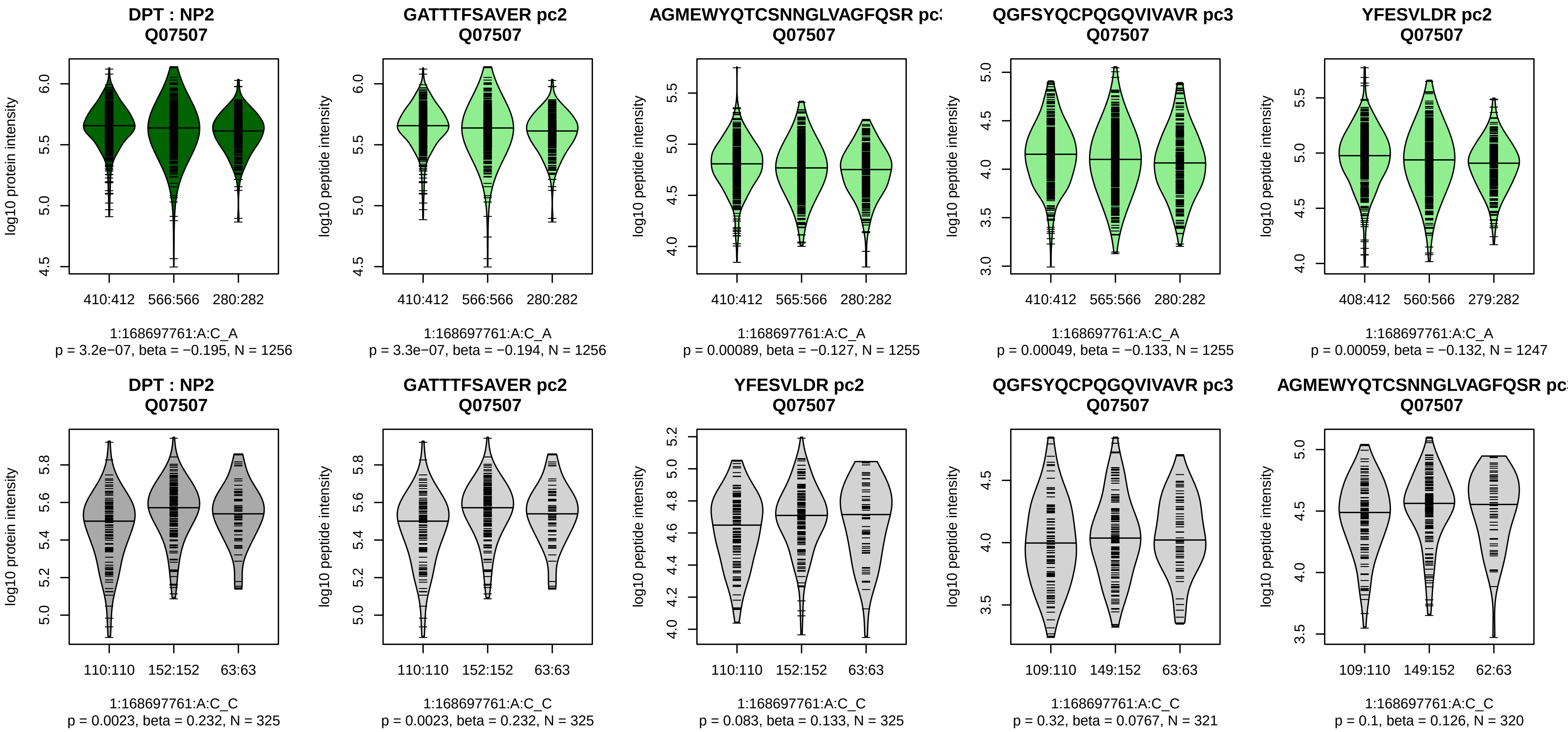


Assay Target: DPT
Olink UniProt: Q07507
deCODE rsID: rs1018454
Proxy rsID: rs1018454
deCODE: 1:168728523:A:C
Proxy SNP: 1:168697761:A:C
deCODE log10(p): 370
deCODE BETA: −0.34
*:*:*:*:–:–:–:–
1256:1255:1255:1247:1169:858

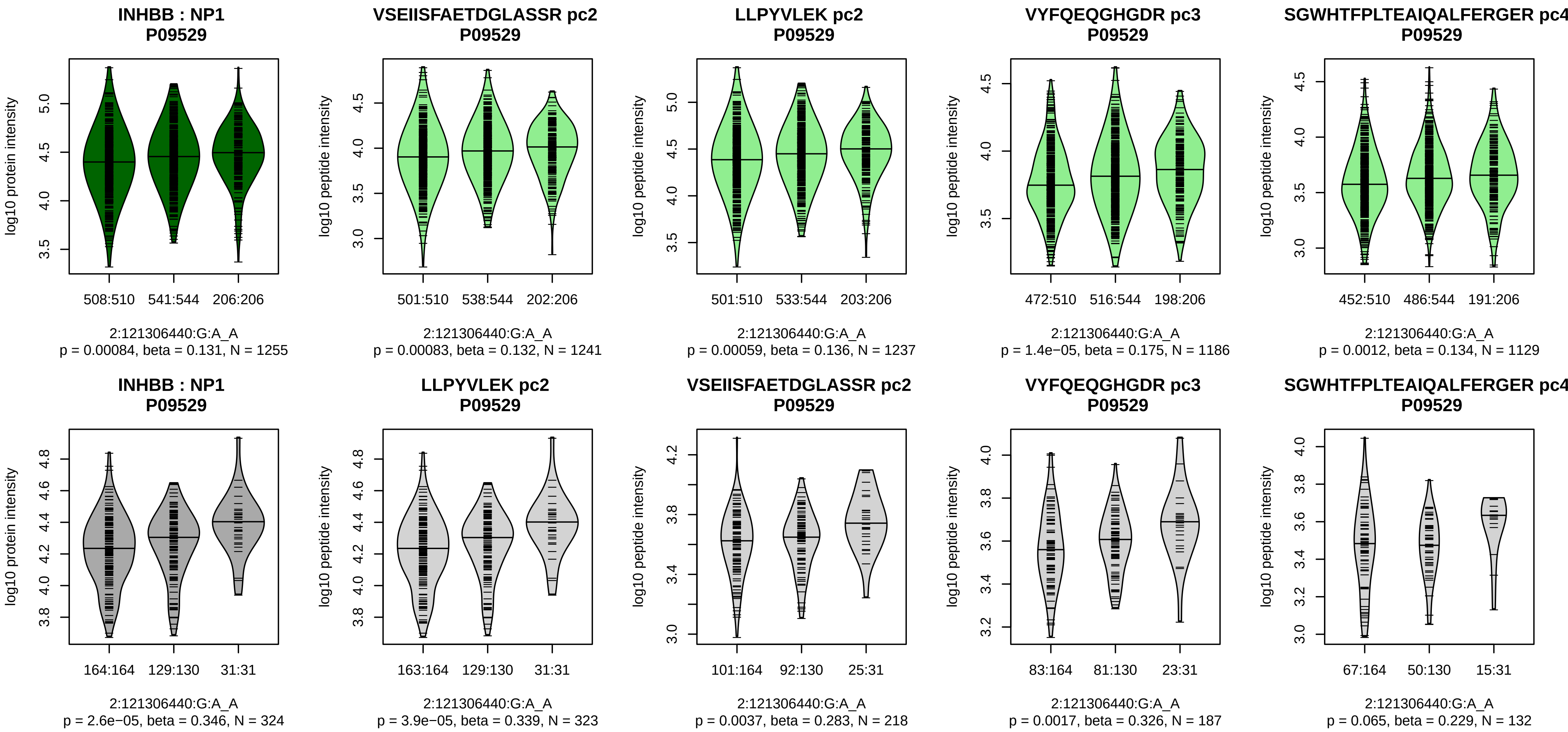


Assay Target: INHBB
Olink UniProt: P09529
deCODE rsID: rs17050272
Proxy rsID: rs17050272
deCODE: 2:120548864:A:G
Proxy SNP: 2:121306440:G:A
deCODE log10(p): 367.3
deCODE BETA: 0.34
*:*:*:*:−:−:−:−:−:−:−:−:−:NA
1241:1237:1186:1129:751:500:3

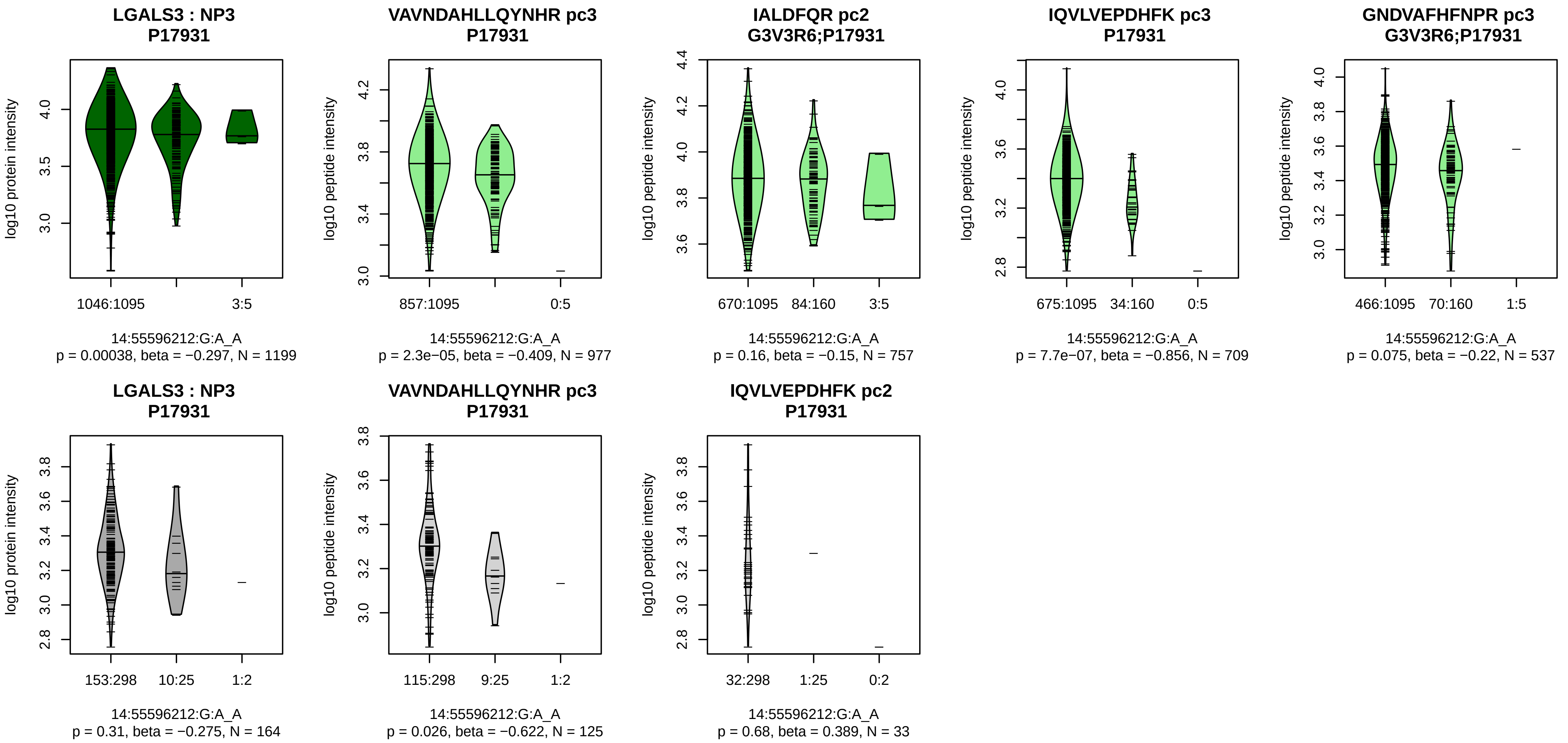


Assay Target: LGALS3
Olink UniProt: P17931
deCODE rsID: rs118083722
Proxy rsID: rs118083722
deCODE: 14:55129494:A:G
Proxy SNP: 14:55596212:G:A
deCODE log10(p): 357.8
deCODE BETA: −0.57
*:–:*:*:–:–:–:–:–:–
977:757:709:658:537:280:231:1

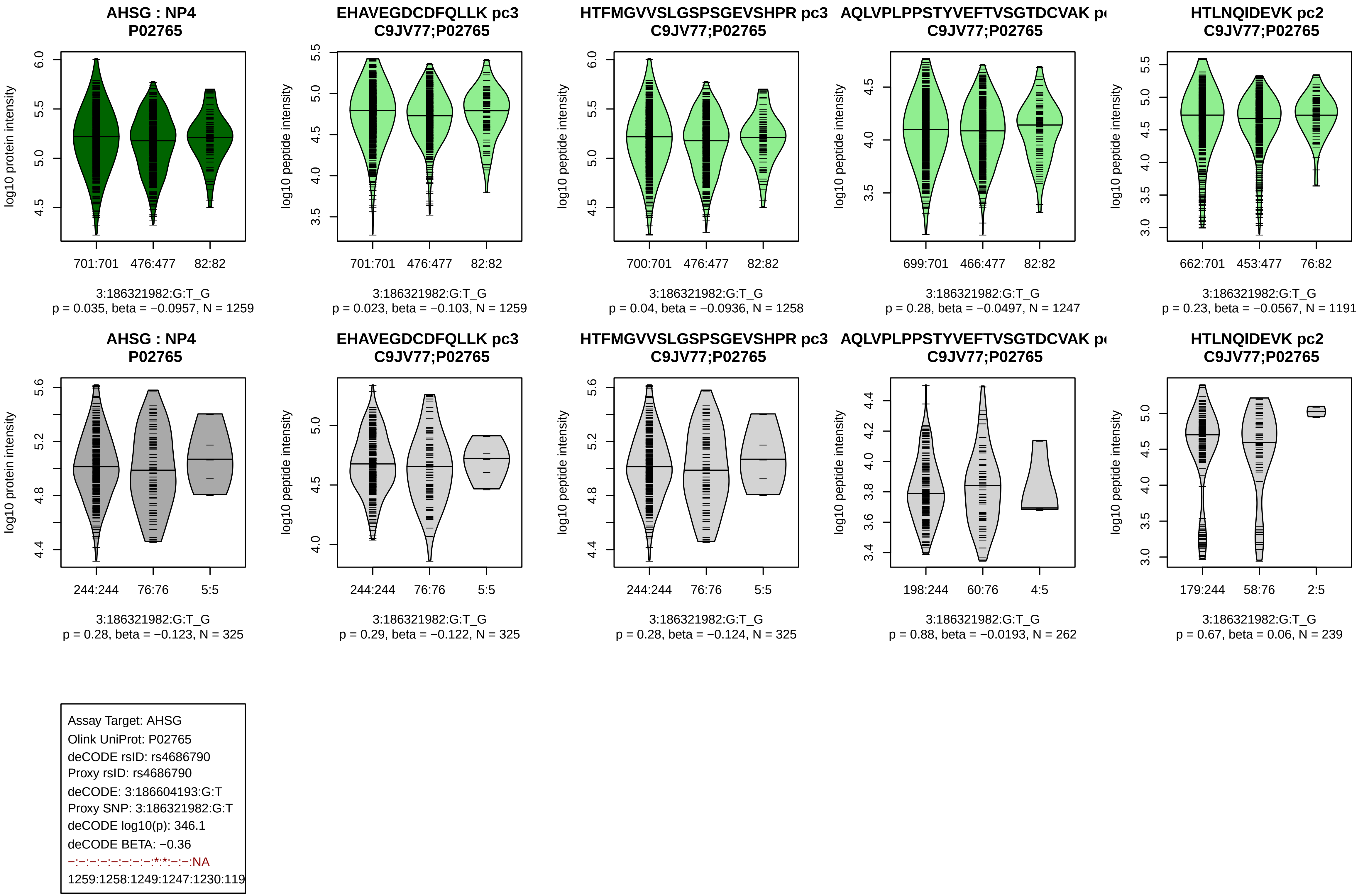
AHSG : NP4
P02765
log10 protein intensity
701:701
476:477
82:82
3:186321982:G:T_G
p = 0.035, beta = −0.0957, N = 1259
EHAVEGDCDFQLLK pc3
C9JV77;P02765
log10 peptide intensity
701:701
476:477
82:82
3:186321982:G:T_G
p = 0.023, beta = −0.103, N = 1259
HTFMGVVSLGSPSGEVSHPR pc3
C9JV77;P02765
log10 peptide intensity
700:701
476:477
82:82
3:186321982:G:T_G
p = 0.04, beta = −0.0936, N = 1258
AQLVPLPPSTYVEFTVSGTDCVAK p
C9JV77;P02765
log10 peptide intensity
699:701
466:477
82:82
3:186321982:G:T_G
p = 0.28, beta = −0.0497, N = 1247
HTLNQIDEVK pc2
C9JV77;P02765
log10 peptide intensity
662:701
453:477
76:82
3:186321982:G:T_G
p = 0.23, beta = −0.0567, N = 1191
AHSG : NP4
P02765
log10 protein intensity
244:244
76:76
5:5
3:186321982:G:T_G
p = 0.28, beta = −0.123, N = 325
EHAVEGDCDFQLLK pc3
C9JV77;P02765
log10 peptide intensity
244:244
76:76
5:5
3:186321982:G:T_G
p = 0.29, beta = −0.122, N = 325
HTFMGVVSLGSPSGEVSHPR pc3
C9JV77;P02765
log10 peptide intensity
244:244
76:76
5:5
3:186321982:G:T_G
p = 0.28, beta = −0.124, N = 325
AQLVPLPPSTYVEFTVSGTDCVAK p
C9JV77;P02765
log10 peptide intensity
198:244
60:76
4:5
3:186321982:G:T_G
p = 0.88, beta = −0.0193, N = 262
HTLNQIDEVK pc2
C9JV77;P02765
log10 peptide intensity
179:244
58:76
2:5
3:186321982:G:T_G
p = 0.67, beta = 0.06, N = 239
Assay Target: AHSG
Olink UniProt: P02765
deCODE rsID: rs4686790
Proxy rsID: rs4686790
deCODE: 3:186604193:G:T
Proxy SNP: 3:186321982:G:T
deCODE log10(p): 346.1
deCODE BETA: −0.36
−:−:−:−:−:−:−:−:*:*:−:−:NA
1259:1258:1249:1247:1230:119

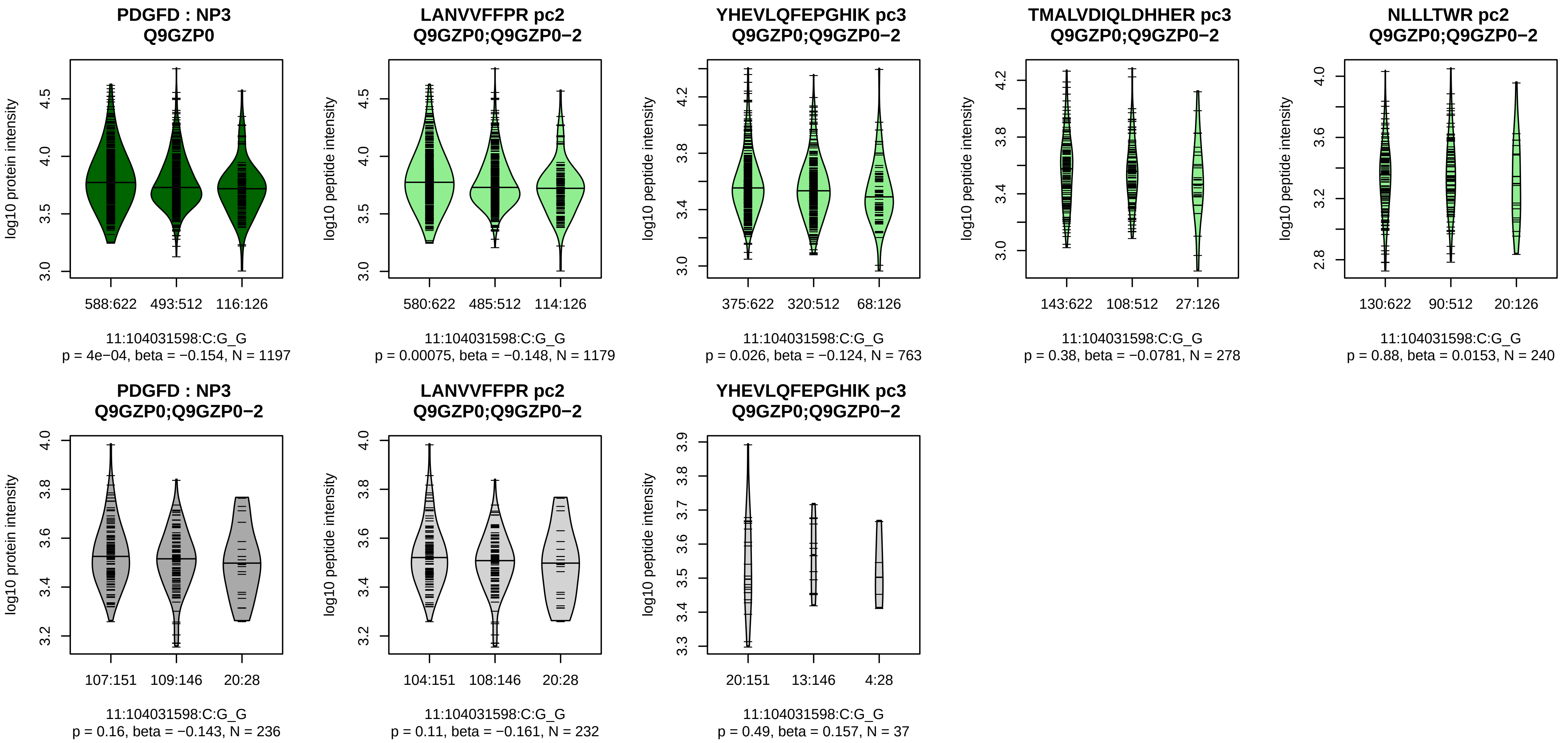


Assay Target: PDGFD
Olink UniProt: Q9GZP0
deCODE rsID: rs7950273
Proxy rsID: rs7950273
deCODE: 11:104160870:G:C
Proxy SNP: 11:104031598:C:G
deCODE log10(p): 342.9
deCODE BETA: −0.35
*:−:−:−:−:NA:NA:NA:NA
1179:763:278:240:198:8:7:13:3

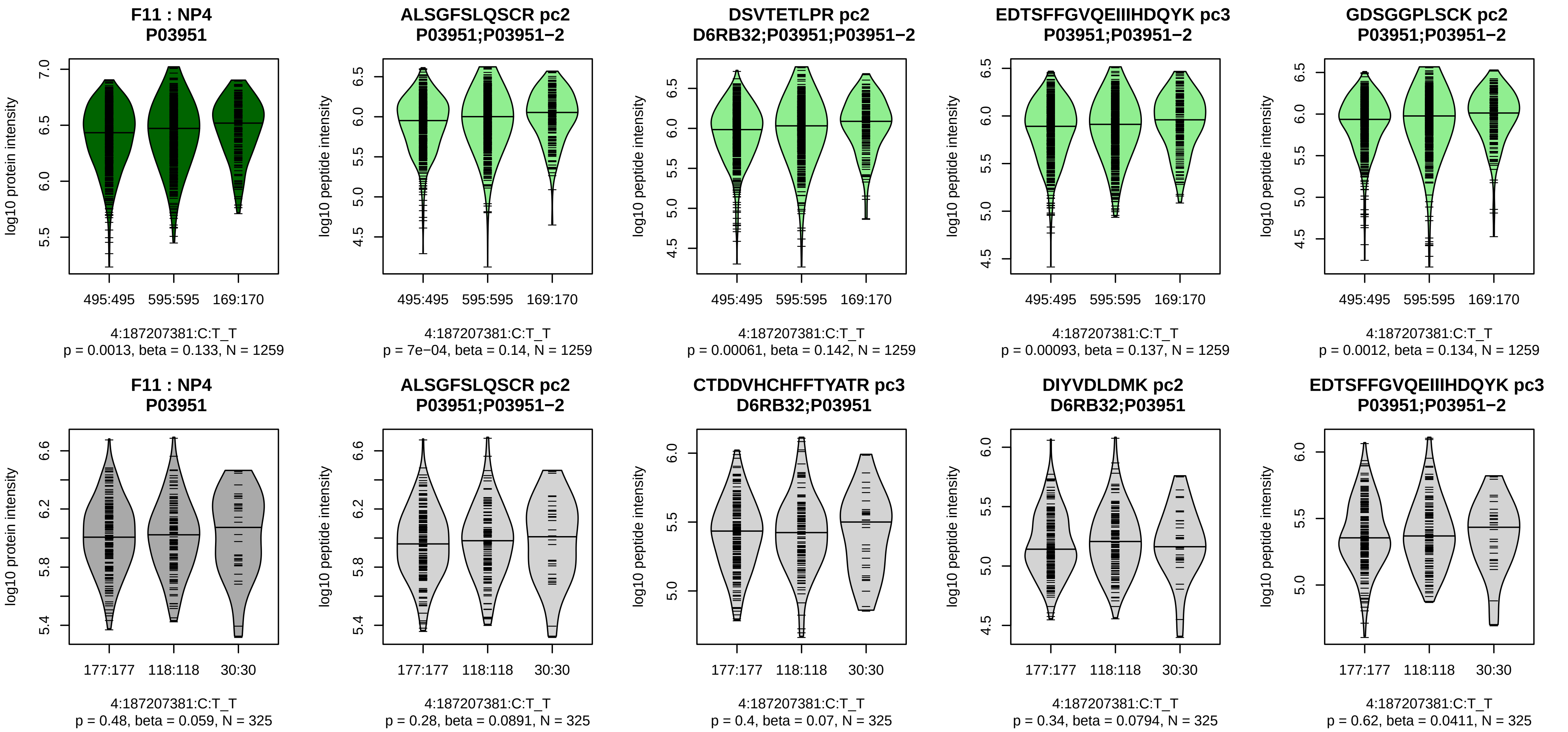


Assay Target: F11
Olink UniProt: P03951
deCODE rsID: rs2289252
Proxy rsID: rs2289252
deCODE: 4:186286227:T:C
Proxy SNP: 4:187207381:C:T
deCODE log10(p): 331.7
deCODE BETA: 0.33
*:*:*:*:*:*:*:*:*:*:*:*:*:*:_:*:*:*:_:*:*
1259:1259:1259:1259:1259:125

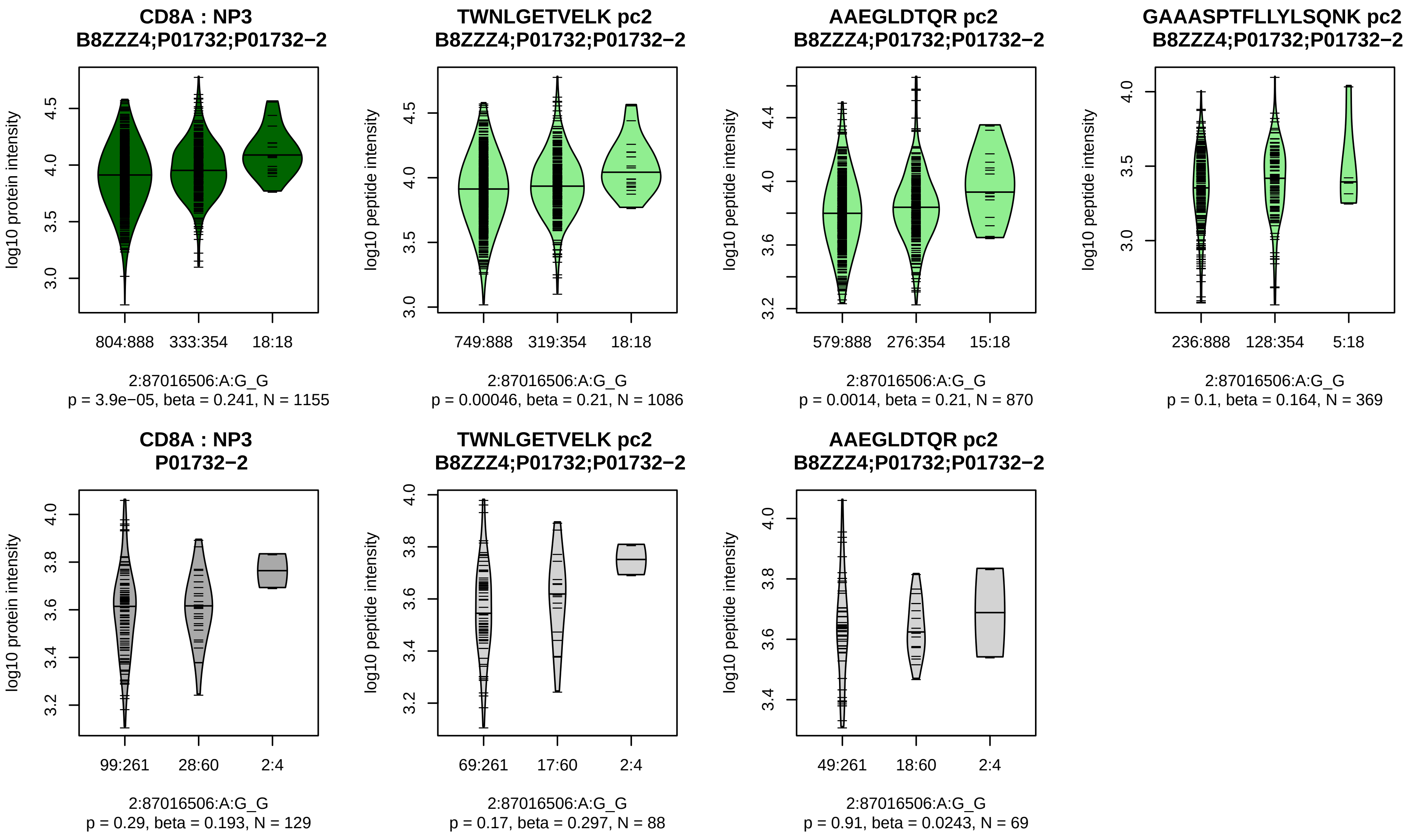


Assay Target: CD8A
Olink UniProt: P01732
deCODE rsID: rs3020726
Proxy rsID: rs3020726
deCODE: 2:86789383:G:A
Proxy SNP: 2:87016506:A:G
deCODE log10(p): 329.6
deCODE BETA: 0.4
*:*:−:−
1086:870:369:65

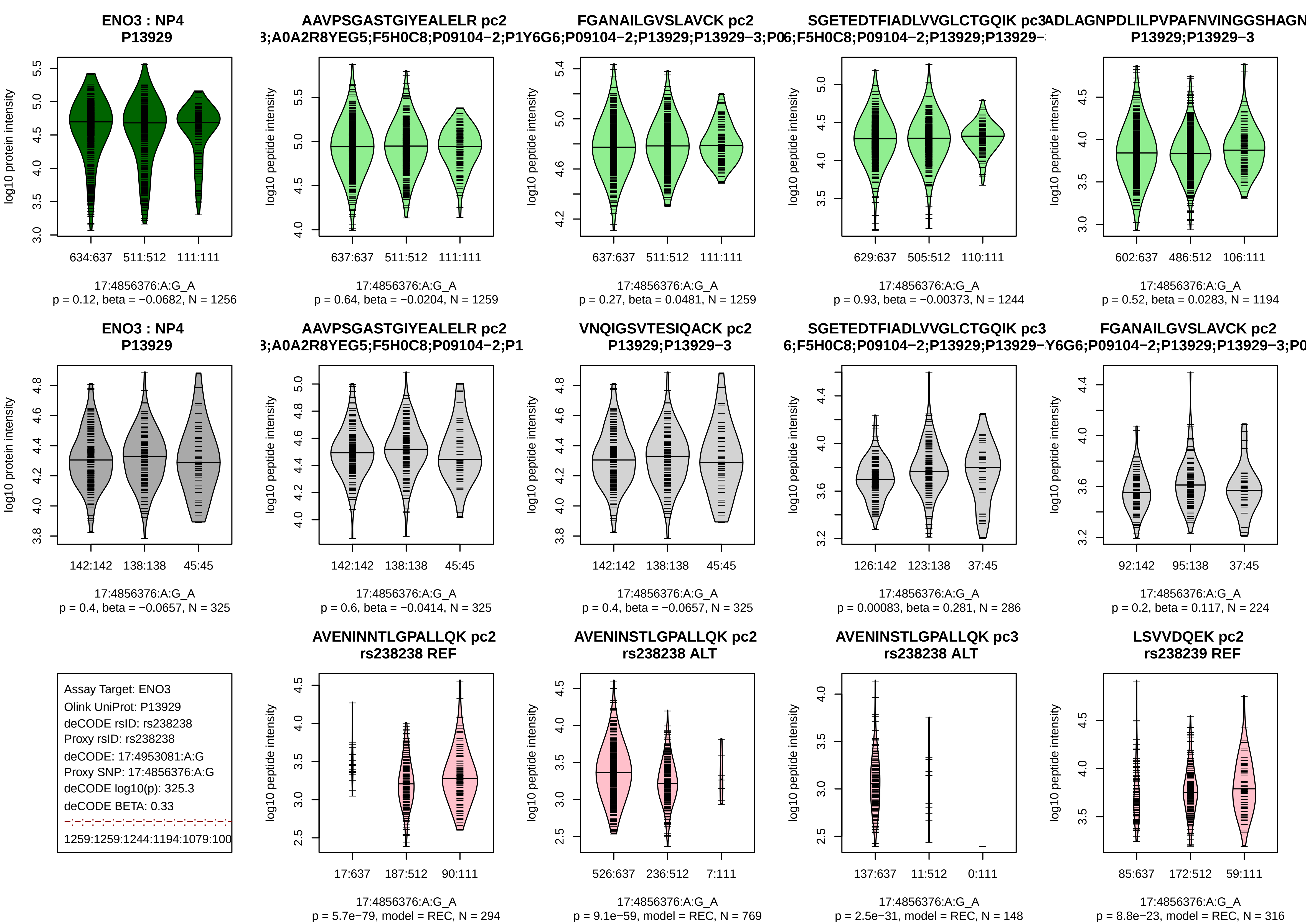
ENO3 : NP4
P13929
log10 protein intensity
634:637 511:512 111:111
17:4856376:A:G_A
p = 0.12, beta = −0.0682, N = 1256
AAVPSGASTGIYEALELR pc2
3;A0A2R8YEG5;F5H0C8;P09104−2;P1
log10 peptide intensity
637:637 511:512 111:111
17:4856376:A:G_A
p = 0.64, beta = −0.0204, N = 1259
FGANAILGVSLAVCK pc2
Y6G6;P09104−2;P13929;P13929−3;P0
log10 peptide intensity
637:637 511:512 111:111
17:4856376:A:G_A
p = 0.27, beta = 0.0481, N = 1259
SGETEDTFIADLVVGLCTGQIK pc3
6;F5H0C8;P09104−2;P13929;P13929−
log10 peptide intensity
629:637 505:512 110:111
17:4856376:A:G_A
p = 0.93, beta = −0.00373, N = 1244
ADLAGNPDLILPVPAFNVINGGSHAGN
P13929;P13929−3
log10 peptide intensity
602:637 486:512 106:111
17:4856376:A:G_A
p = 0.52, beta = 0.0283, N = 1194
ENO3 : NP4
P13929
log10 protein intensity
142:142 138:138 45:45
17:4856376:A:G_A
p = 0.4, beta = −0.0657, N = 325
AAVPSGASTGIYEALELR pc2
3;A0A2R8YEG5;F5H0C8;P09104−2;P1
log10 peptide intensity
142:142 138:138 45:45
17:4856376:A:G_A
p = 0.6, beta = −0.0414, N = 325
VNQIGSVTESIQACK pc2
P13929;P13929−3
log10 peptide intensity
142:142 138:138 45:45
17:4856376:A:G_A
p = 0.4, beta = −0.0657, N = 325
SGETEDTFIADLVVGLCTGQIK pc3
6;F5H0C8;P09104−2;P13929;P13929−
log10 peptide intensity
126:142 123:138 37:45
17:4856376:A:G_A
p = 0.00083, beta = 0.281, N = 286
FGANAILGVSLAVCK pc2
Y6G6;P09104−2;P13929;P13929−3;P0
log10 peptide intensity
92:142 95:138 37:45
17:4856376:A:G_A
p = 0.2, beta = 0.117, N = 224
Assay Target: ENO3
Olink UniProt: P13929
deCODE rsID: rs238238
Proxy rsID: rs238238
deCODE: 17:4953081:A:G
Proxy SNP: 17:4856376:A:G
deCODE log10(p): 325.3
deCODE BETA: 0.33
1259:1259:1244:1194:1079:100
AVENINNTLGPALLQK pc2
rs238238 REF
log10 peptide intensity
17:637 187:512 90:111
17:4856376:A:G_A
p = 5.7e−79, model = REC, N = 294
AVENINSTLGPALLQK pc2
rs238238 ALT
log10 peptide intensity
526:637 236:512 7:111
17:4856376:A:G_A
p = 9.1e−59, model = REC, N = 769
AVENINSTLGPALLQK pc3
rs238238 ALT
log10 peptide intensity
137:637 11:512 0:111
17:4856376:A:G_A
p = 2.5e−31, model = REC, N = 148
LSVVDQEK pc2
rs238239 REF
log10 peptide intensity
85:637 172:512 59:111
17:4856376:A:G_A
p = 8.8e−23, model = REC, N = 316

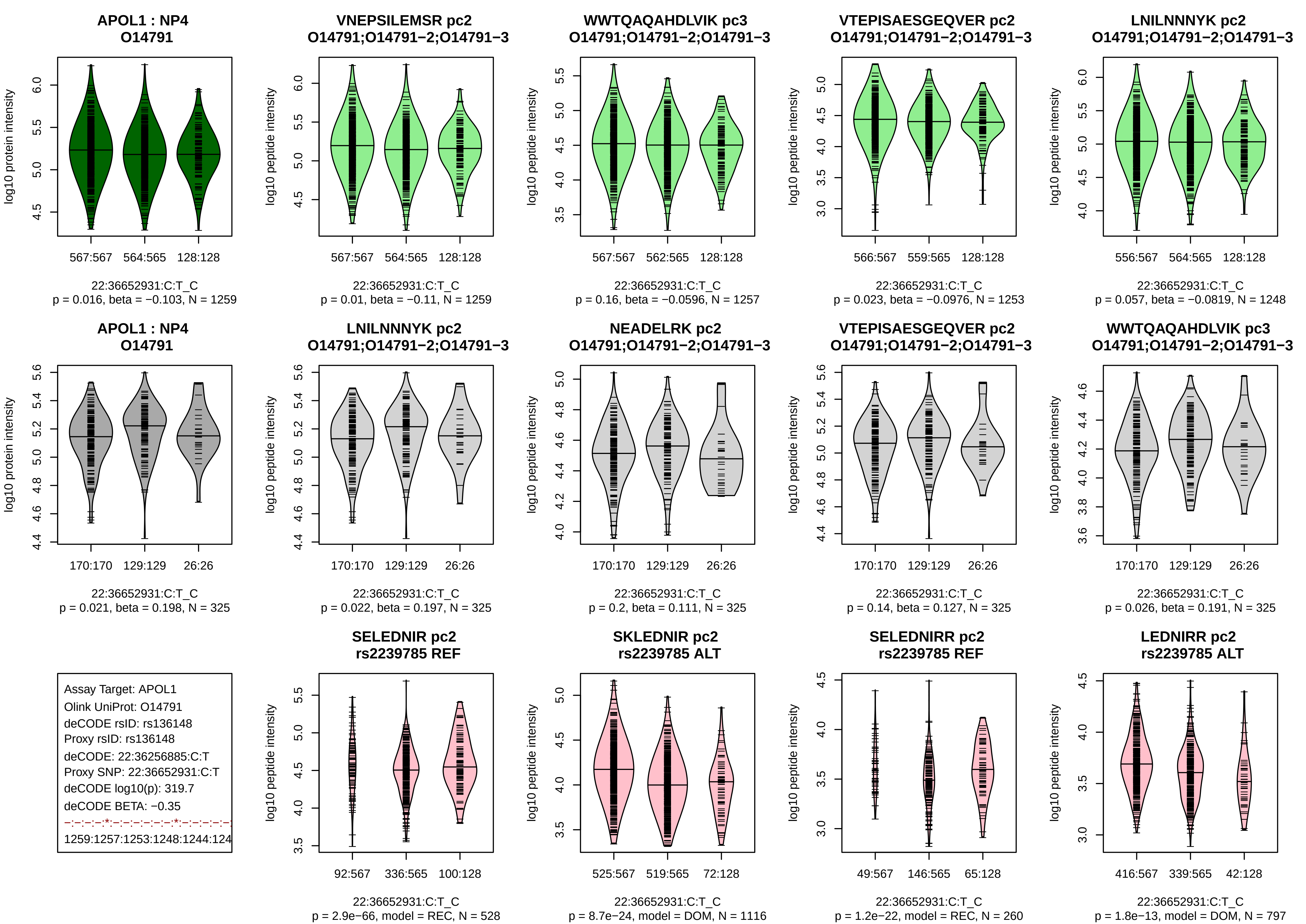
APOL1 : NP4
O14791
log10 protein intensity
567:567
564:565
128:128
22:36652931:C:T_C
p = 0.016, beta = −0.103, N = 1259
VNEPSILEMSR pc2
O14791;O14791−2;O14791−3
log10 peptide intensity
567:567
564:565
128:128
22:36652931:C:T_C
p = 0.01, beta = −0.11, N = 1259
WWTQAQAHDLVIK pc3
O14791;O14791−2;O14791−3
log10 peptide intensity
567:567
562:565
128:128
22:36652931:C:T_C
p = 0.16, beta = −0.0596, N = 1257
VTEPISAESGEQVER pc2
O14791;O14791−2;O14791−3
log10 peptide intensity
566:567
559:565
128:128
22:36652931:C:T_C
p = 0.023, beta = −0.0976, N = 1253
LNILNNNYK pc2
O14791;O14791−2;O14791−3
log10 peptide intensity
556:567
564:565
128:128
22:36652931:C:T_C
p = 0.057, beta = −0.0819, N = 1248
APOL1 : NP4
O14791
log10 protein intensity
170:170
129:129
26:26
22:36652931:C:T_C
p = 0.021, beta = 0.198, N = 325
LNILNNNYK pc2
O14791;O14791−2;O14791−3
log10 peptide intensity
170:170
129:129
26:26
22:36652931:C:T_C
p = 0.022, beta = 0.197, N = 325
NEADELRK pc2
O14791;O14791−2;O14791−3
log10 peptide intensity
170:170
129:129
26:26
22:36652931:C:T_C
p = 0.2, beta = 0.111, N = 325
VTEPISAESGEQVER pc2
O14791;O14791−2;O14791−3
log10 peptide intensity
170:170
129:129
26:26
22:36652931:C:T_C
p = 0.14, beta = 0.127, N = 325
WWTQAQAHDLVIK pc3
O14791;O14791−2;O14791−3
log10 peptide intensity
170:170
129:129
26:26
22:36652931:C:T_C
p = 0.026, beta = 0.191, N = 325
Assay Target: APOL1
Olink UniProt: O14791
deCODE rsID: rs136148
Proxy rsID: rs136148
deCODE: 22:36256885:C:T
Proxy SNP: 22:36652931:C:T
deCODE log10(p): 319.7
deCODE BETA: −0.35
−:−:−:−:*:−:−:−:−:−:−:*:−:−:−:−:−:−
1259:1257:1253:1248:1244:124
SELEDNIR pc2
rs2239785 REF
log10 peptide intensity
92:567
336:565
100:128
22:36652931:C:T_C
p = 2.9e−66, model = REC, N = 528
SKLEDNIR pc2
rs2239785 ALT
log10 peptide intensity
525:567
519:565
72:128
22:36652931:C:T_C
p = 8.7e−24, model = DOM, N = 1116
SELEDNIRR pc2
rs2239785 REF
log10 peptide intensity
49:567
146:565
65:128
22:36652931:C:T_C
p = 1.2e−22, model = REC, N = 260
LEDNIRR pc2
rs2239785 ALT
log10 peptide intensity
416:567
339:565
42:128
22:36652931:C:T_C
p = 1.8e−13, model = DOM, N = 797

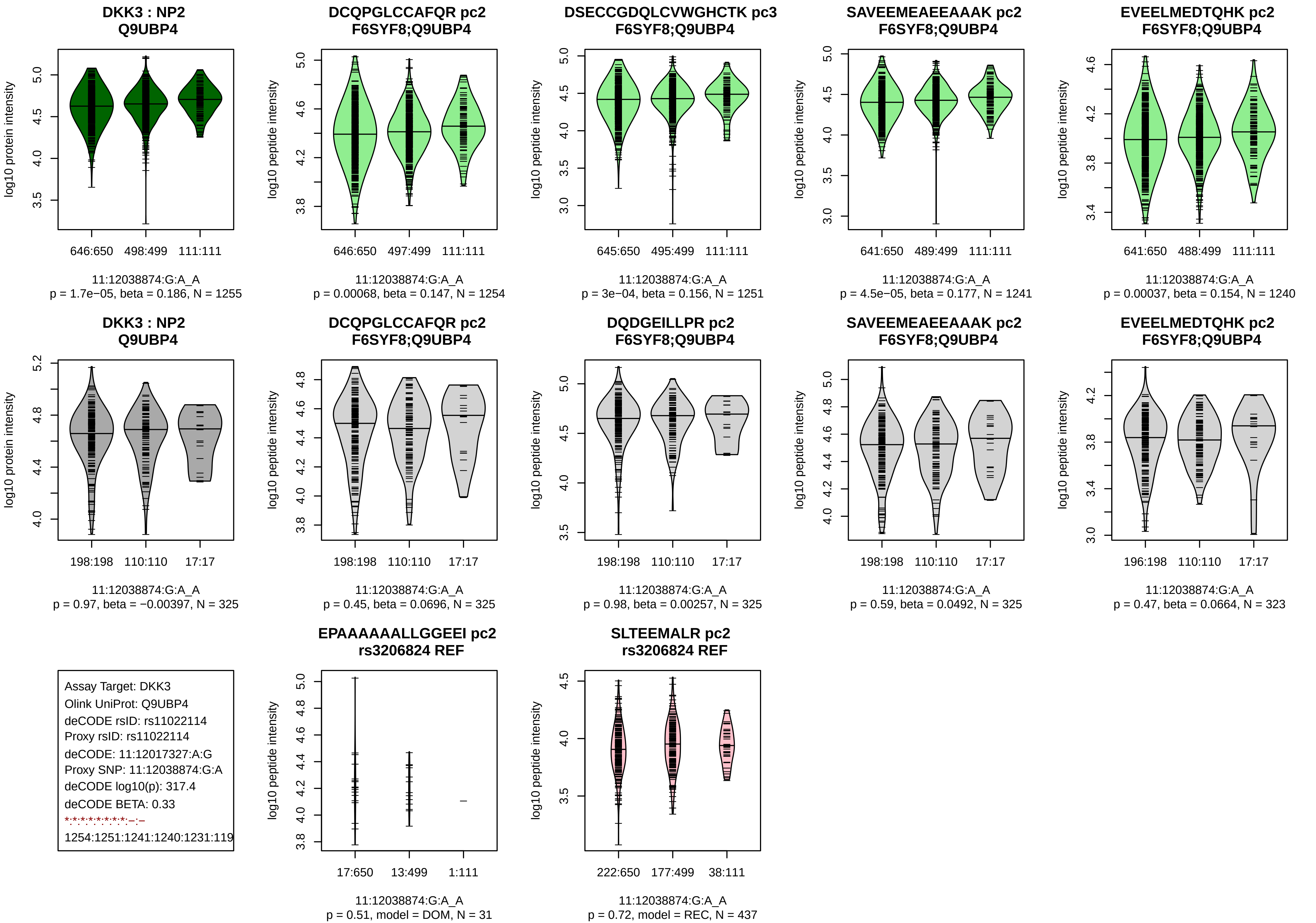
DKK3 : NP2
Q9UBP4
log10 protein intensity
646:650 498:499 111:111
11:12038874:G:A_A
p = 1.7e−05, beta = 0.186, N = 1255
DCQPGLCCAFQR pc2
F6SYF8;Q9UBP4
log10 peptide intensity
646:650 497:499 111:111
11:12038874:G:A_A
p = 0.00068, beta = 0.147, N = 1254
DSECCGDQLCVWGHCTK pc3
F6SYF8;Q9UBP4
log10 peptide intensity
645:650 495:499 111:111
11:12038874:G:A_A
p = 3e−04, beta = 0.156, N = 1251
SAVEEMEAEEAAAK pc2
F6SYF8;Q9UBP4
log10 peptide intensity
641:650 489:499 111:111
11:12038874:G:A_A
p = 4.5e−05, beta = 0.177, N = 1241
EVEELMEDTQHK pc2
F6SYF8;Q9UBP4
log10 peptide intensity
641:650 488:499 111:111
11:12038874:G:A_A
p = 0.00037, beta = 0.154, N = 1240
DKK3 : NP2
Q9UBP4
log10 protein intensity
198:198 110:110 17:17
11:12038874:G:A_A
p = 0.97, beta = −0.00397, N = 325
DCQPGLCCAFQR pc2
F6SYF8;Q9UBP4
log10 peptide intensity
198:198 110:110 17:17
11:12038874:G:A_A
p = 0.45, beta = 0.0696, N = 325
DQDGEILLPR pc2
F6SYF8;Q9UBP4
log10 peptide intensity
198:198 110:110 17:17
11:12038874:G:A_A
p = 0.98, beta = 0.00257, N = 325
SAVEEMEAEEAAAK pc2
F6SYF8;Q9UBP4
log10 peptide intensity
198:198 110:110 17:17
11:12038874:G:A_A
p = 0.59, beta = 0.0492, N = 325
EVEELMEDTQHK pc2
F6SYF8;Q9UBP4
log10 peptide intensity
196:198 110:110 17:17
11:12038874:G:A_A
p = 0.47, beta = 0.0664, N = 323
Assay Target: DKK3
Olink UniProt: Q9UBP4
deCODE rsID: rs11022114
Proxy rsID: rs11022114
deCODE: 11:12017327:A:G
Proxy SNP: 11:12038874:G:A
deCODE log10(p): 317.4
deCODE BETA: 0.33
1254:1251:1241:1240:1231:119
EPAAAAAALLGGEEI pc2
rs3206824 REF
log10 peptide intensity
17:650 13:499 1:111
11:12038874:G:A_A
p = 0.51, model = DOM, N = 31
SLTEEMALR pc2
rs3206824 REF
log10 peptide intensity
222:650 177:499 38:111
11:12038874:G:A_A
p = 0.72, model = REC, N = 437

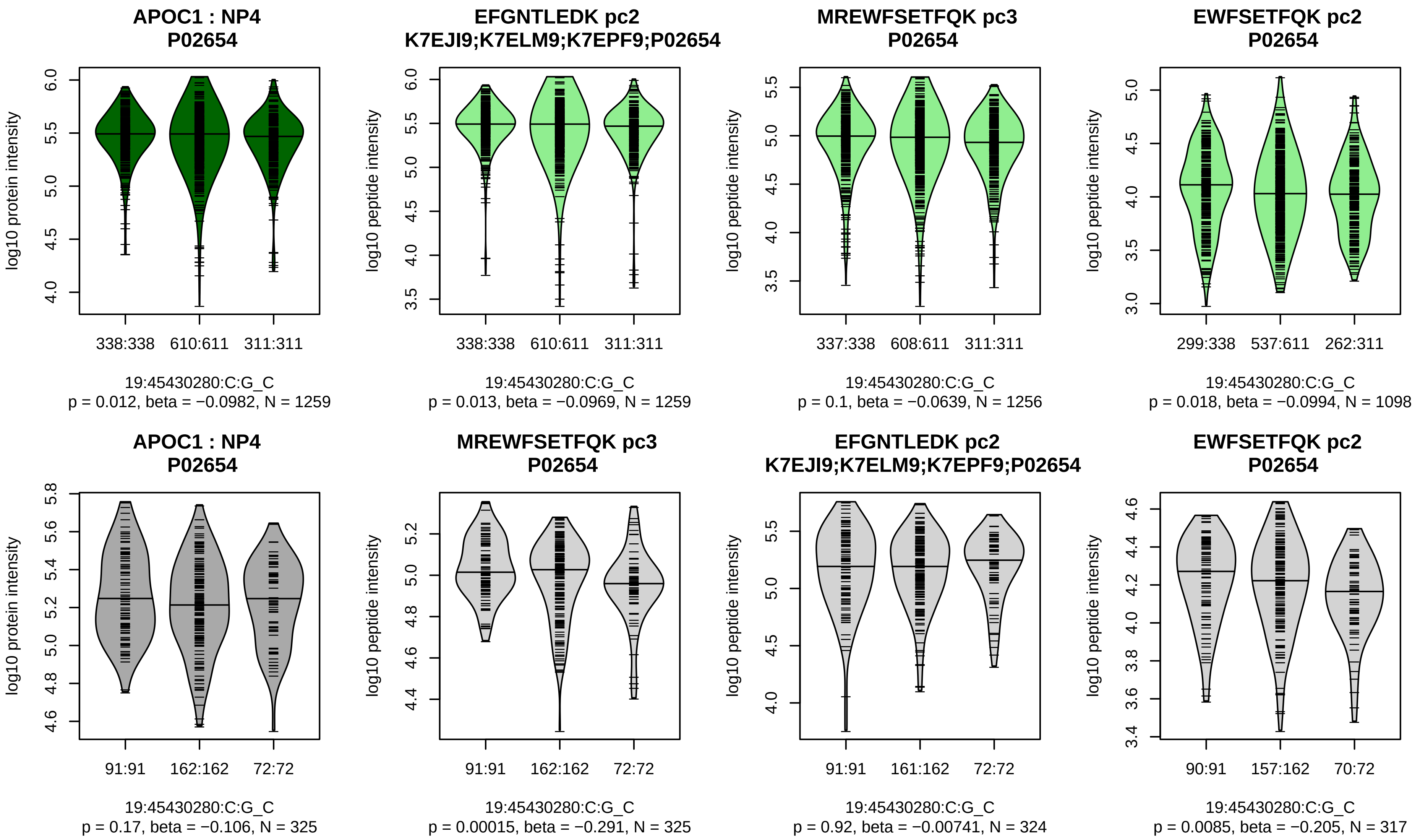


Assay Target: APOC1
Olink UniProt: P02654
deCODE rsID: rs5112
Proxy rsID: rs5112
deCODE: 19:44927023:G:C
Proxy SNP: 19:45430280:C:G
deCODE log10(p): 298.6
deCODE BETA: 0.3
−:−:−:−:NA:NA:NA:NA
1259:1256:1143:1098:6:7:4:10

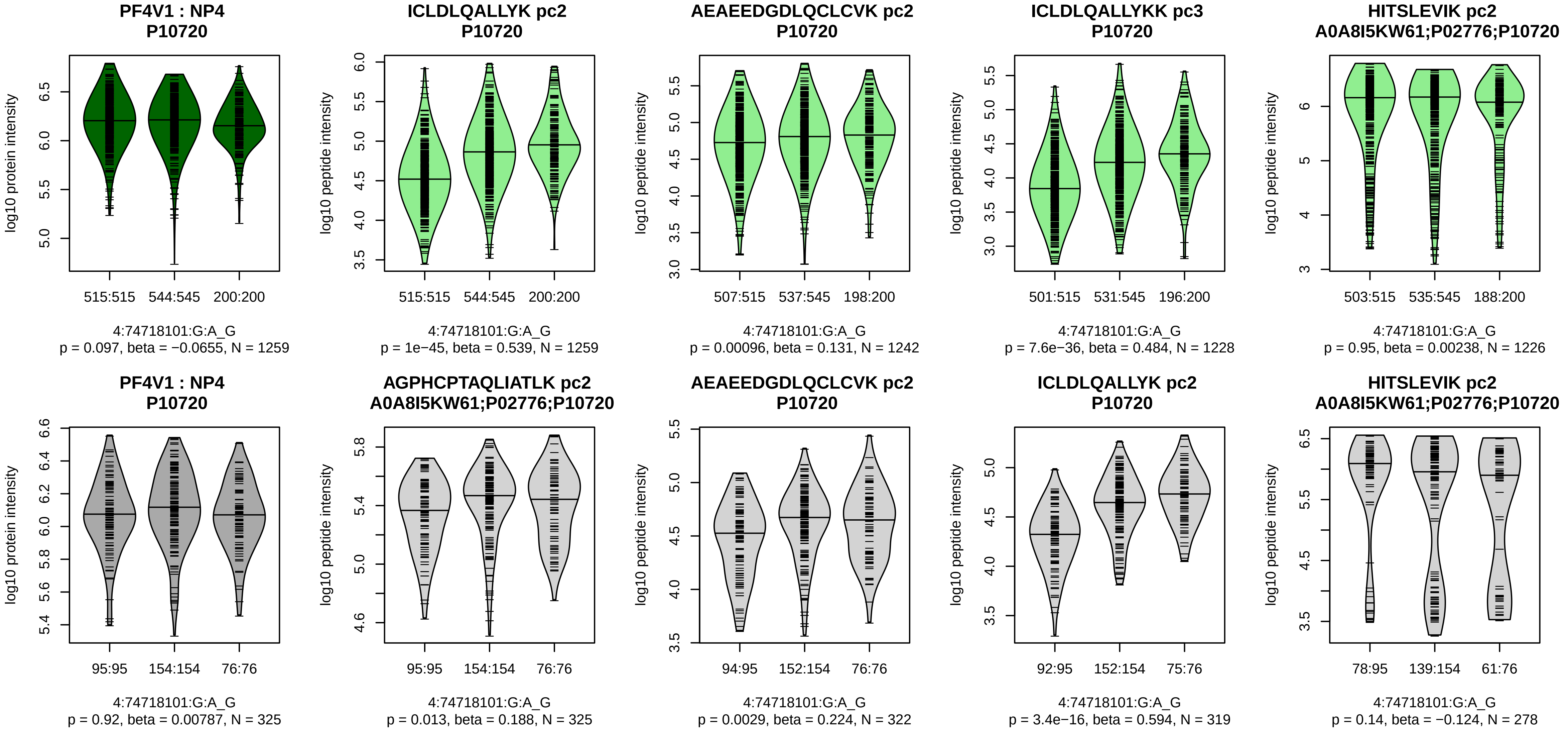


Assay Target: PF4V1
Olink UniProt: P10720
deCODE rsID: rs872914
Proxy rsID: rs872914
deCODE: 4:73852384:G:A
Proxy SNP: 4:74718101:G:A
deCODE log10(p): 297.5
deCODE BETA: 0.32
*:*:*:–:–:–:*:–:*:–
1259:1242:1228:1226:1209:120

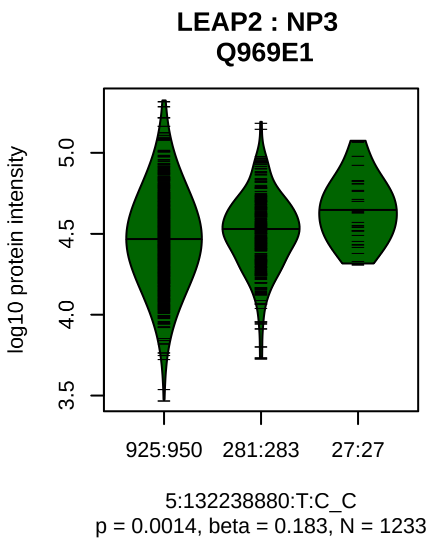


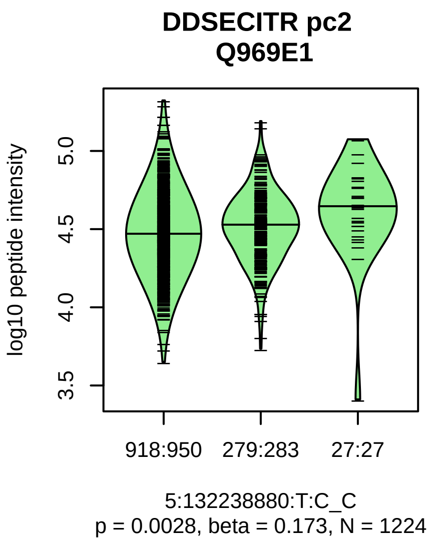


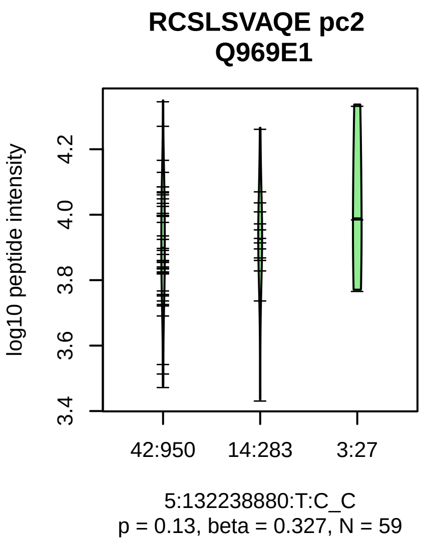


Assay Target: LEAP2
Olink UniProt: Q969E1
deCODE rsID: rs12515756
Proxy rsID: rs12515756
deCODE: 5:132903188:C:T
Proxy SNP: 5:132238880:T:C
deCODE log10(p): 296.6
deCODE BETA: 0.4
*:–
1224:59

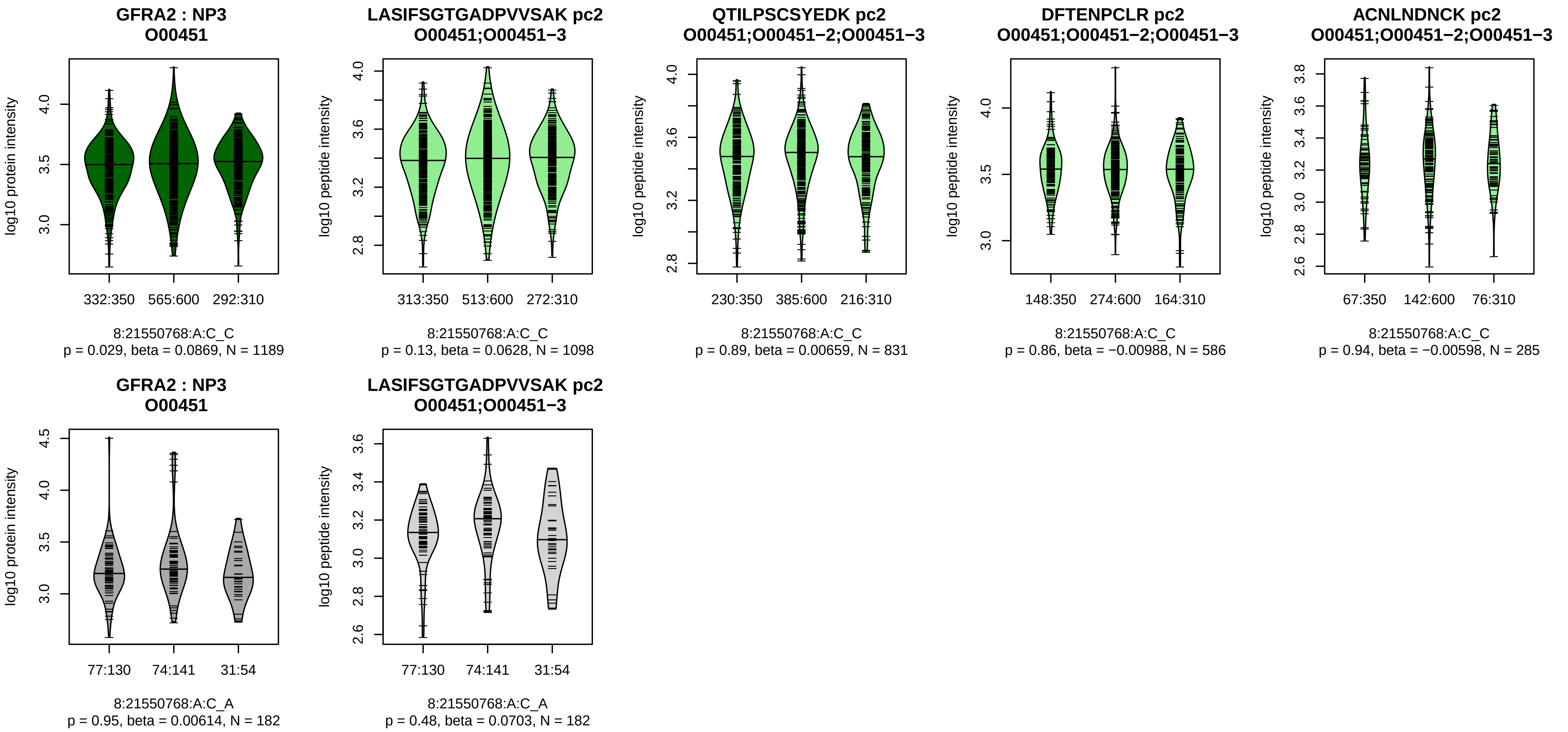


Assay Target: GFRA2
Olink UniProt: O00451
deCODE rsID: rs15881
Proxy rsID: rs15881
deCODE: 8:21693256:A:C
Proxy SNP: 8:21550768:A:C
deCODE log10(p): 292.7
deCODE BETA: −0.31
−:−:−:−:−:−:−
1098:831:586:285:211:169:139

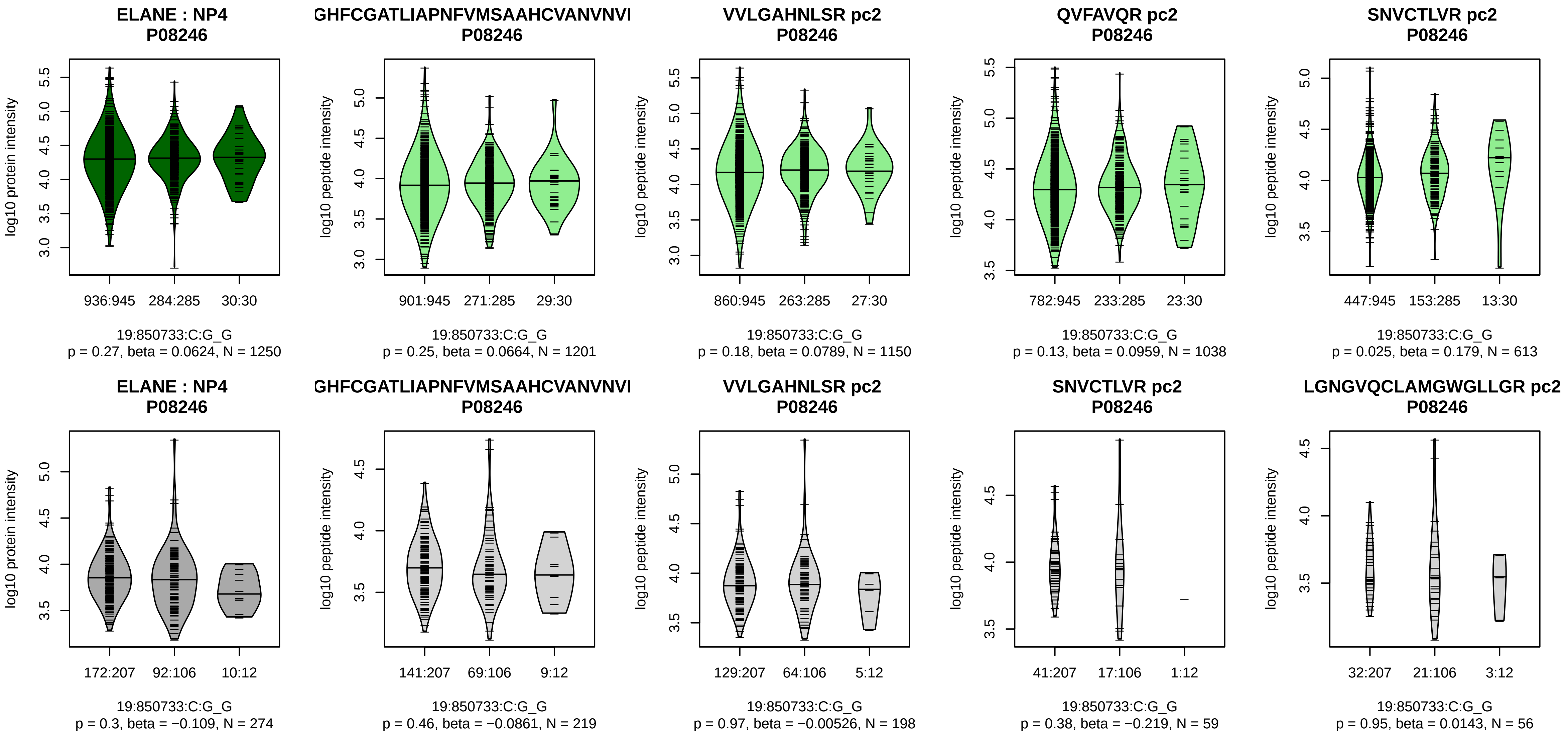


Assay Target: ELANE
Olink UniProt: P08246
deCODE rsID: rs10409474
Proxy rsID: rs10409474
deCODE: 19:850733:G:C
Proxy SNP: 19:850733:C:G
deCODE log10(p): 290.7
deCODE BETA: 0.45
–:–:–:–:–:–
1201:1150:1038:613:426:417

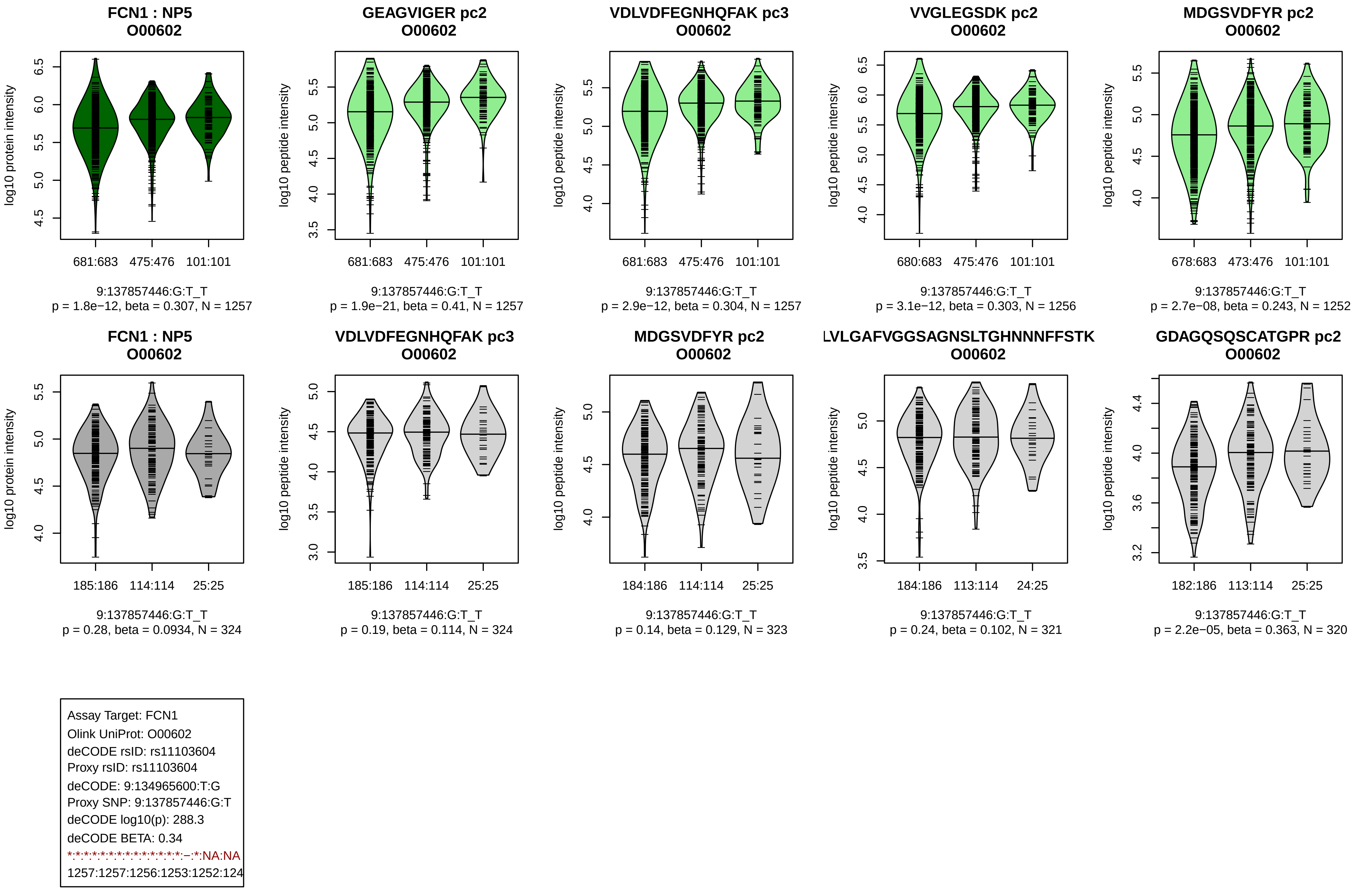
FCN1 : NP5
O00602
log10 protein intensity
681:683 475:476 101:101
9:137857446:G:T_T
p = 1.8e−12, beta = 0.307, N = 1257
GEAGVIGER pc2
O00602
log10 peptide intensity
681:683 475:476 101:101
9:137857446:G:T_T
p = 1.9e−21, beta = 0.41, N = 1257
VDLVDFEGNHQFAK pc3
O00602
log10 peptide intensity
681:683 475:476 101:101
9:137857446:G:T_T
p = 2.9e−12, beta = 0.304, N = 1257
VVGLEGSDK pc2
O00602
log10 peptide intensity
680:683 475:476 101:101
9:137857446:G:T_T
p = 3.1e−12, beta = 0.303, N = 1256
MDGSVDFYR pc2
O00602
log10 peptide intensity
678:683 473:476 101:101
9:137857446:G:T_T
p = 2.7e−08, beta = 0.243, N = 1252
FCN1 : NP5
O00602
log10 protein intensity
185:186 114:114 25:25
9:137857446:G:T_T
p = 0.28, beta = 0.0934, N = 324
VDLVDFEGNHQFAK pc3
O00602
log10 peptide intensity
185:186 114:114 25:25
9:137857446:G:T_T
p = 0.19, beta = 0.114, N = 324
MDGSVDFYR pc2
O00602
log10 peptide intensity
184:186 114:114 25:25
9:137857446:G:T_T
p = 0.14, beta = 0.129, N = 323
LVLGAFVGGSAGNSLTGHNNNFFSTK
O00602
log10 peptide intensity
184:186 113:114 24:25
9:137857446:G:T_T
p = 0.24, beta = 0.102, N = 321
GDAGQSQSCATGPR pc2
O00602
log10 peptide intensity
182:186 113:114 25:25
9:137857446:G:T_T
p = 2.2e−05, beta = 0.363, N = 320
Assay Target: FCN1
Olink UniProt: O00602
deCODE rsID: rs11103604
Proxy rsID: rs11103604
deCODE: 9:134965600:T:G
Proxy SNP: 9:137857446:G:T
deCODE log10(p): 288.3
deCODE BETA: 0.34
*:*:*:*:*:*:*:*:*:*:*:*:*:*:−:*:NA:NA
1257:1257:1256:1253:1252:124

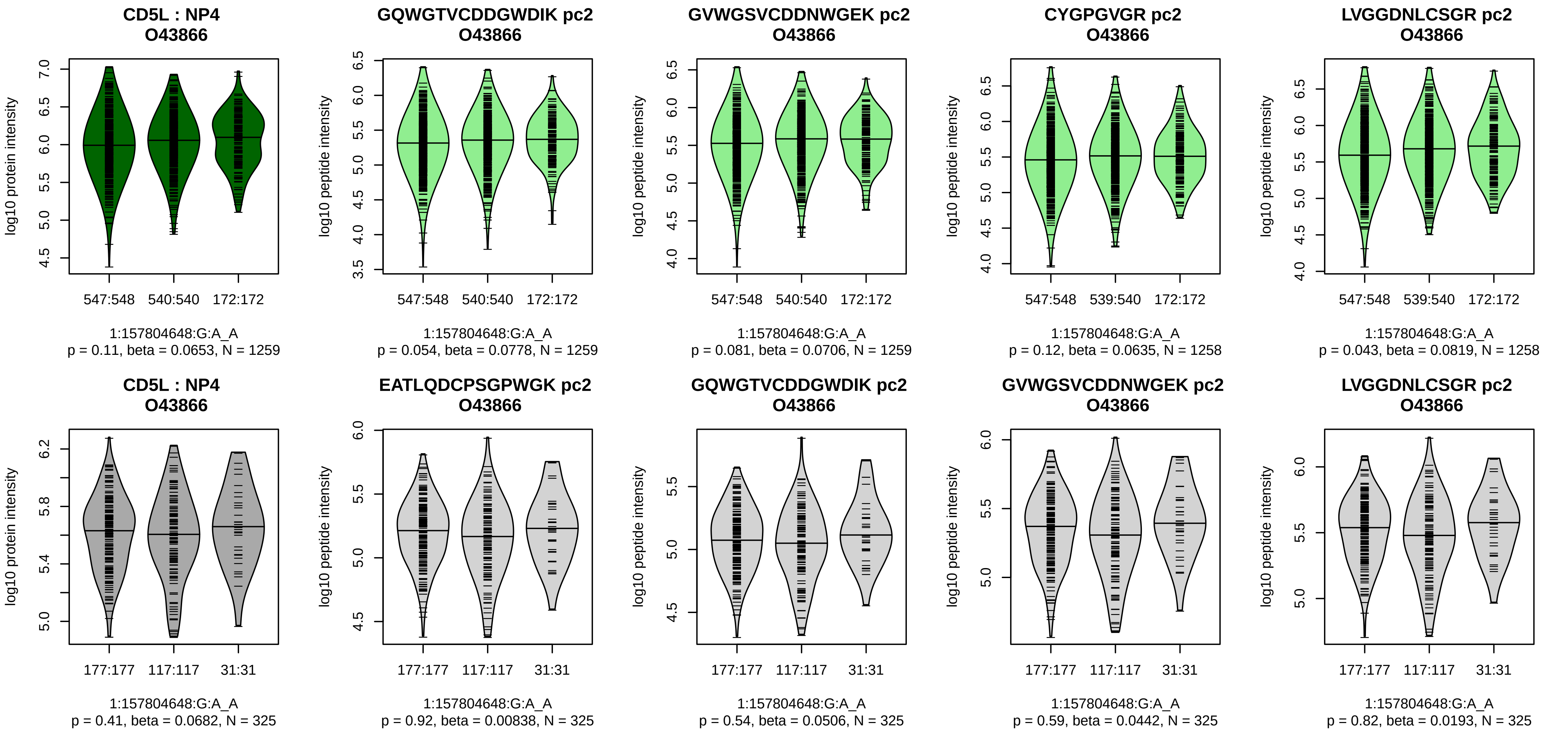


Assay Target: CD5L
Olink UniProt: O43866
deCODE rsID: rs2765501
Proxy rsID: rs2765501
deCODE: 1:157834858:A:G
Proxy SNP: 1:157804648:G:A
deCODE log10(p): 284
deCODE BETA: 0.3
–:–:–:–:–:–:–:–:–:–:–:–:–:–:–:–:
1259:1259:1258:1258:1254:124

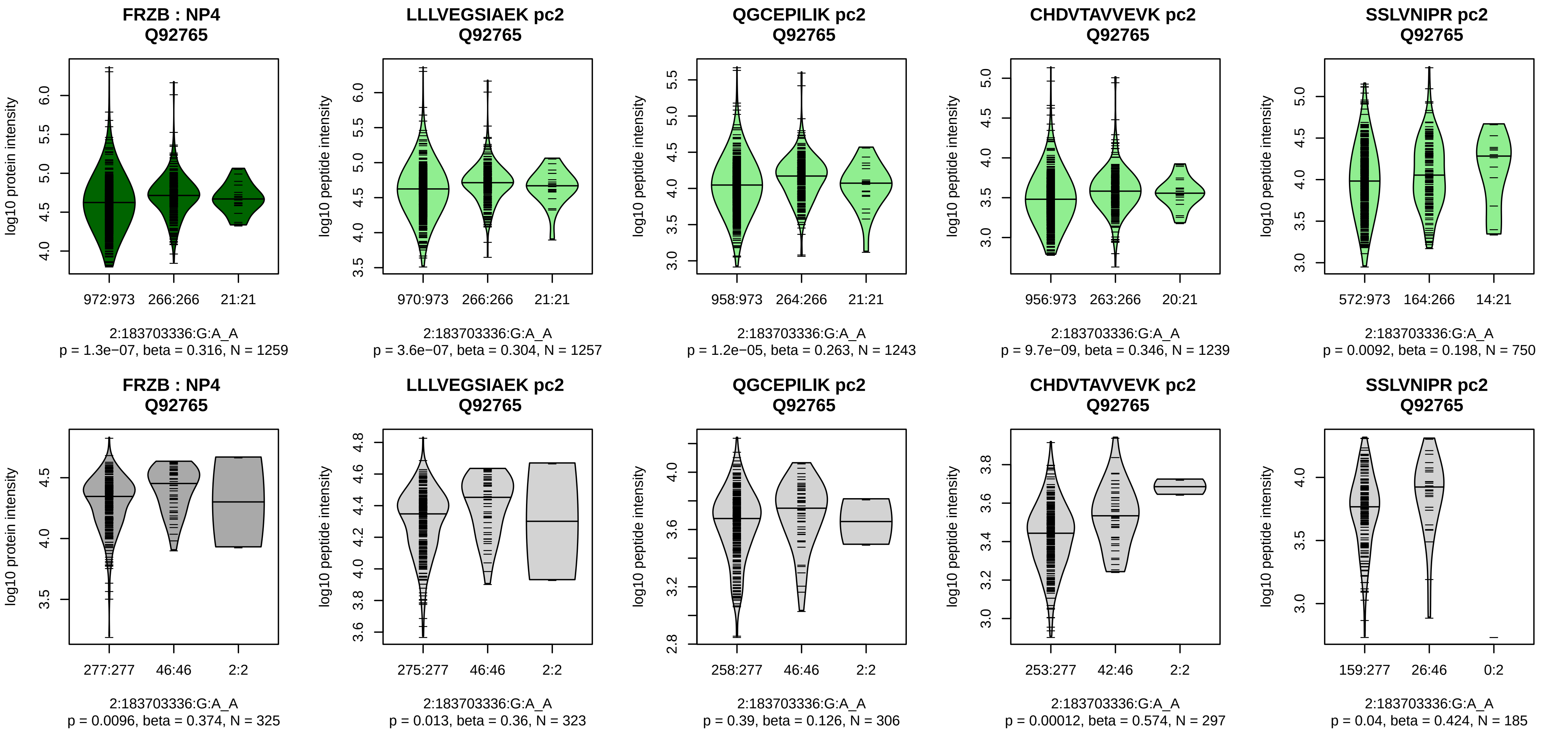


Assay Target: FRZB
Olink UniProt: Q92765
deCODE rsID: rs288326
Proxy rsID: rs288326
deCODE: 2:182838608:A:G
Proxy SNP: 2:183703336:G:A
deCODE log10(p): 280.4
deCODE BETA: 0.54
*:*:*:*:−:−:−:NA
1257:1243:1239:750:522:294:53

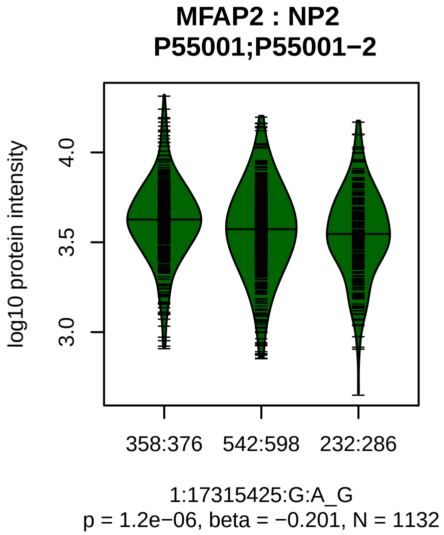


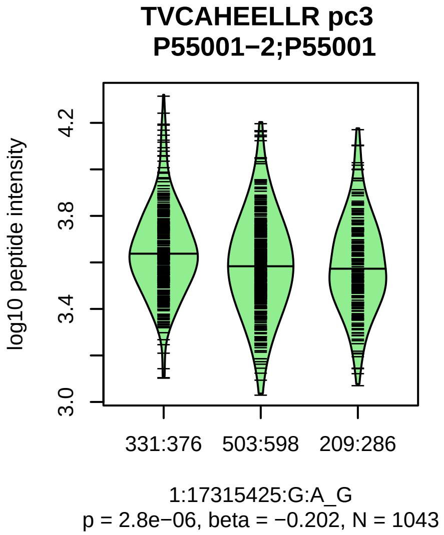


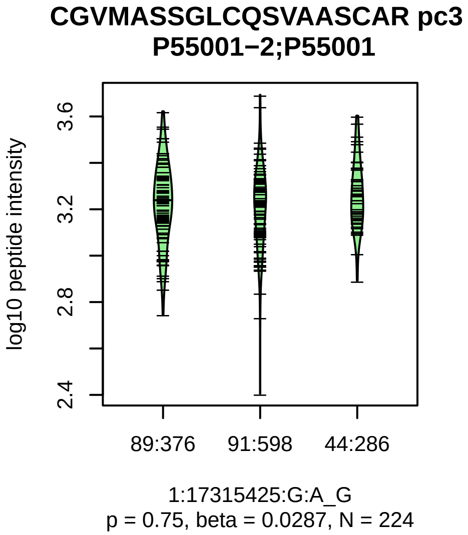


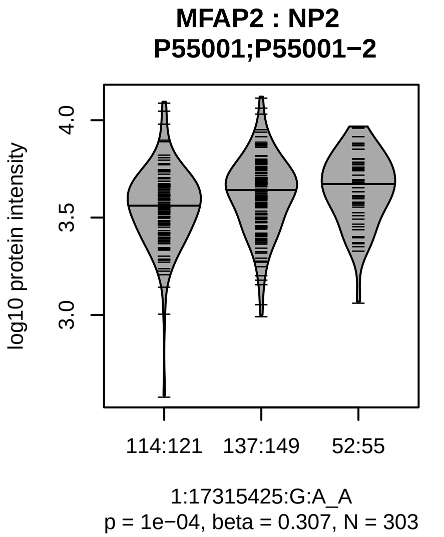


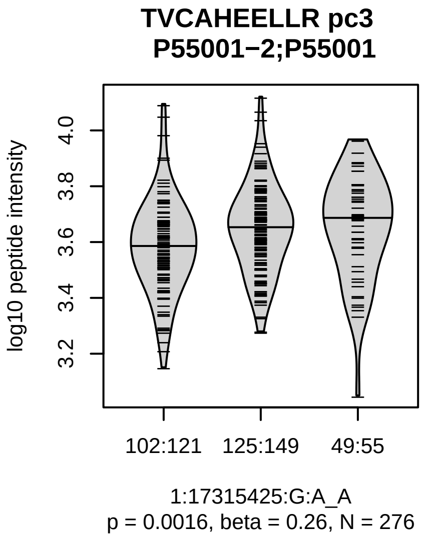


Assay Target: MFAP2
Olink UniProt: P55001
deCODE rsID: rs4920605
Proxy rsID: rs4920605
deCODE: 1:16988925:!:TGCCC
Proxy SNP: 1:17315425:G:A
deCODE log10(p): 275.9
deCODE BETA: −0.29
*:*:−
1043:779:224

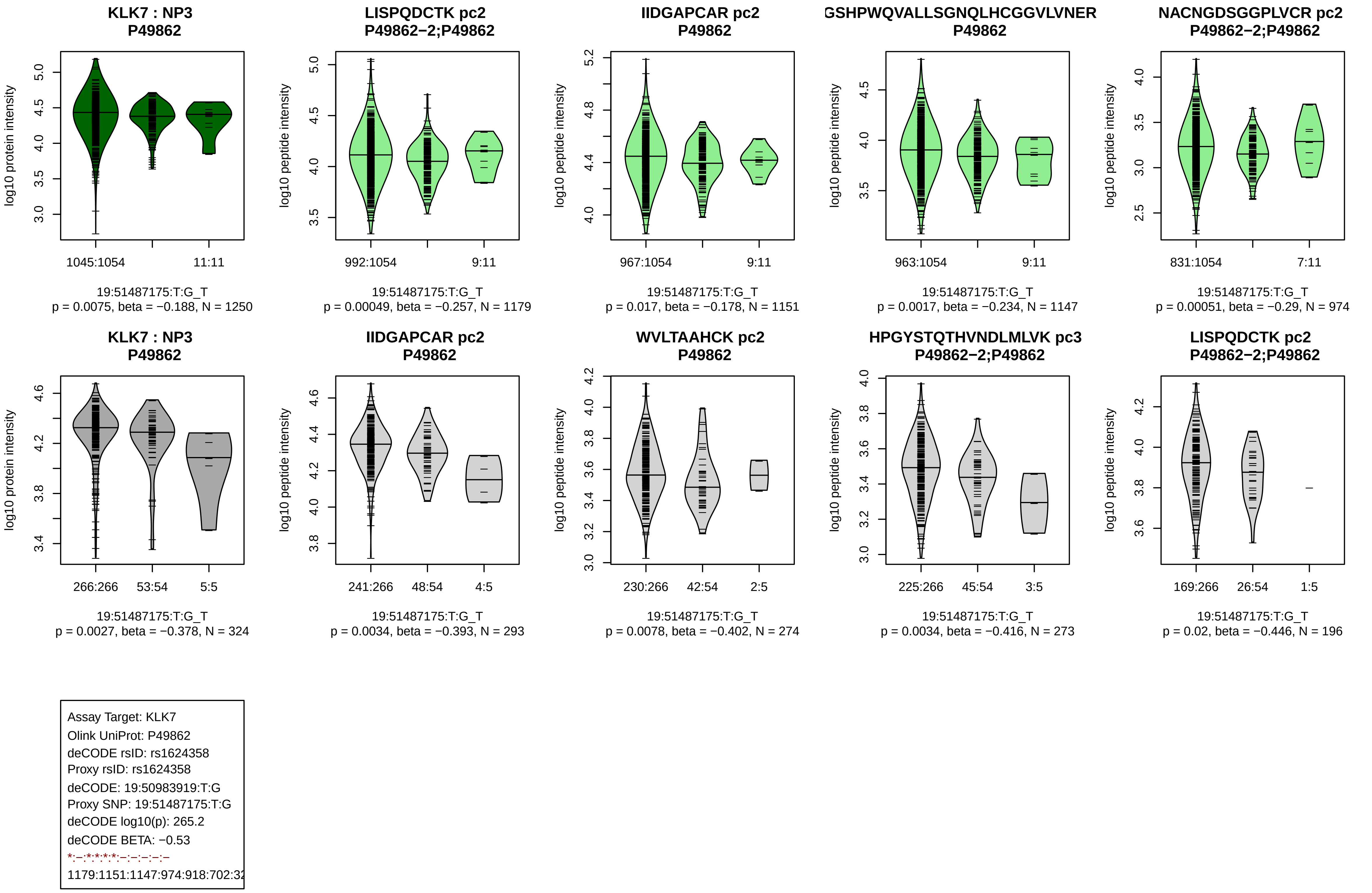

KLK7 : NP3
P49862
log10 protein intensity
1045:1054
11:11
19:51487175:T:G_T
p = 0.0075, beta = −0.188, N = 1250
LISPQDCTK pc2
P49862−2;P49862
log10 peptide intensity
992:1054
9:11
19:51487175:T:G_T
p = 0.00049, beta = −0.257, N = 1179
IIDGAPCAR pc2
P49862
log10 peptide intensity
967:1054
9:11
19:51487175:T:G_T
p = 0.017, beta = −0.178, N = 1151
GSHPWQVALLSGNQLHCGGVLVNER
P49862
log10 peptide intensity
963:1054
9:11
19:51487175:T:G_T
p = 0.0017, beta = −0.234, N = 1147
NACNGDSGGPLVCR pc2
P49862−2;P49862
log10 peptide intensity
831:1054
7:11
19:51487175:T:G_T
p = 0.00051, beta = −0.29, N = 974
KLK7 : NP3
P49862
log10 protein intensity
266:266
53:54
5:5
19:51487175:T:G_T
p = 0.0027, beta = −0.378, N = 324
IIDGAPCAR pc2
P49862
log10 peptide intensity
241:266
48:54
4:5
19:51487175:T:G_T
p = 0.0034, beta = −0.393, N = 293
WVLTAAHCK pc2
P49862
log10 peptide intensity
230:266
42:54
2:5
19:51487175:T:G_T
p = 0.0078, beta = −0.402, N = 274
HPGYSTQTHVNDLMLVK pc3
P49862−2;P49862
log10 peptide intensity
225:266
45:54
3:5
19:51487175:T:G_T
p = 0.0034, beta = −0.416, N = 273
LISPQDCTK pc2
P49862−2;P49862
log10 peptide intensity
169:266
26:54
1:5
19:51487175:T:G_T
p = 0.02, beta = −0.446, N = 196
Assay Target: KLK7
Olink UniProt: P49862
deCODE rsID: rs1624358
Proxy rsID: rs1624358
deCODE: 19:50983919:T:G
Proxy SNP: 19:51487175:T:G
deCODE log10(p): 265.2
deCODE BETA: −0.53
*:–:*:*:*:*:–:–:–:–:–
1179:1151:1147:974:918:702:32

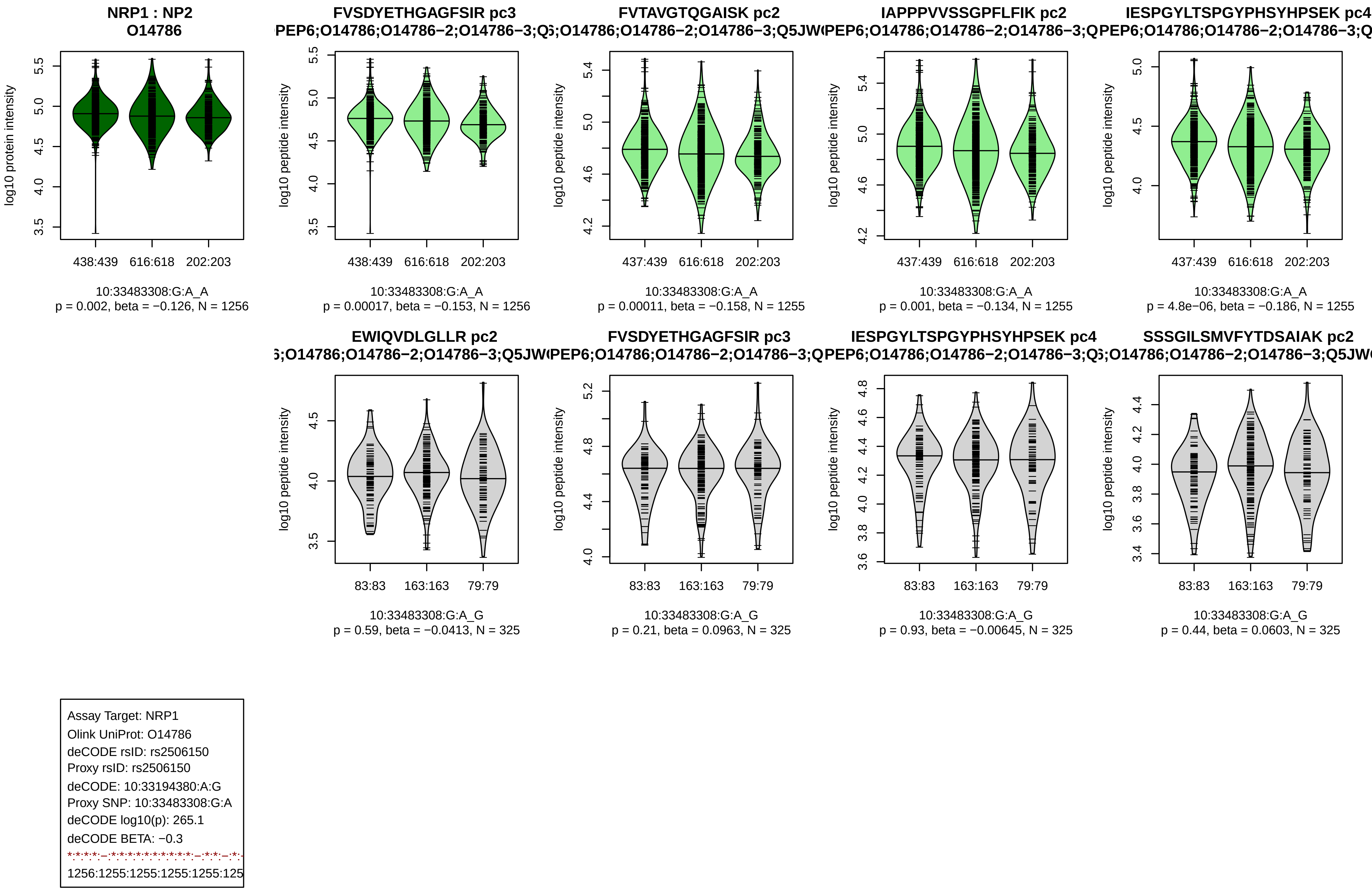
NRP1 : NP2
O14786
log10 protein intensity
438:439
616:618
202:203
10:33483308:G:A_A
p = 0.002, beta = −0.126, N = 1256
FVSDYETHGAGFSIR pc3
log10 peptide intensity
p = 0.00017, beta = −0.153, N = 1256
FVTAVGTQGAISK pc2
437:439
p = 0.00011, beta = −0.158, N = 1255
IAPPPVVSSGPFLFIK pc2
p = 0.001, beta = −0.134, N = 1255
IESPGYLTSPGYPHSYHPSEK pc4
p = 4.8e−06, beta = −0.186, N = 1255
EWIQVDLGLLR pc2
83:83
163:163
79:79
10:33483308:G:A_G
p = 0.59, beta = −0.0413, N = 325
FVSDYETHGAGFSIR pc3
p = 0.21, beta = 0.0963, N = 325
IESPGYLTSPGYPHSYHPSEK pc4
p = 0.93, beta = −0.00645, N = 325
SSSGILSMVFYTDSAIAK pc2
p = 0.44, beta = 0.0603, N = 325
Assay Target: NRP1
Olink UniProt: O14786
deCODE rsID: rs2506150
Proxy rsID: rs2506150
deCODE: 10:33194380:A:G
Proxy SNP: 10:33483308:G:A
deCODE log10(p): 265.1
deCODE BETA: −0.3
1256:1255:1255:1255:1255:125

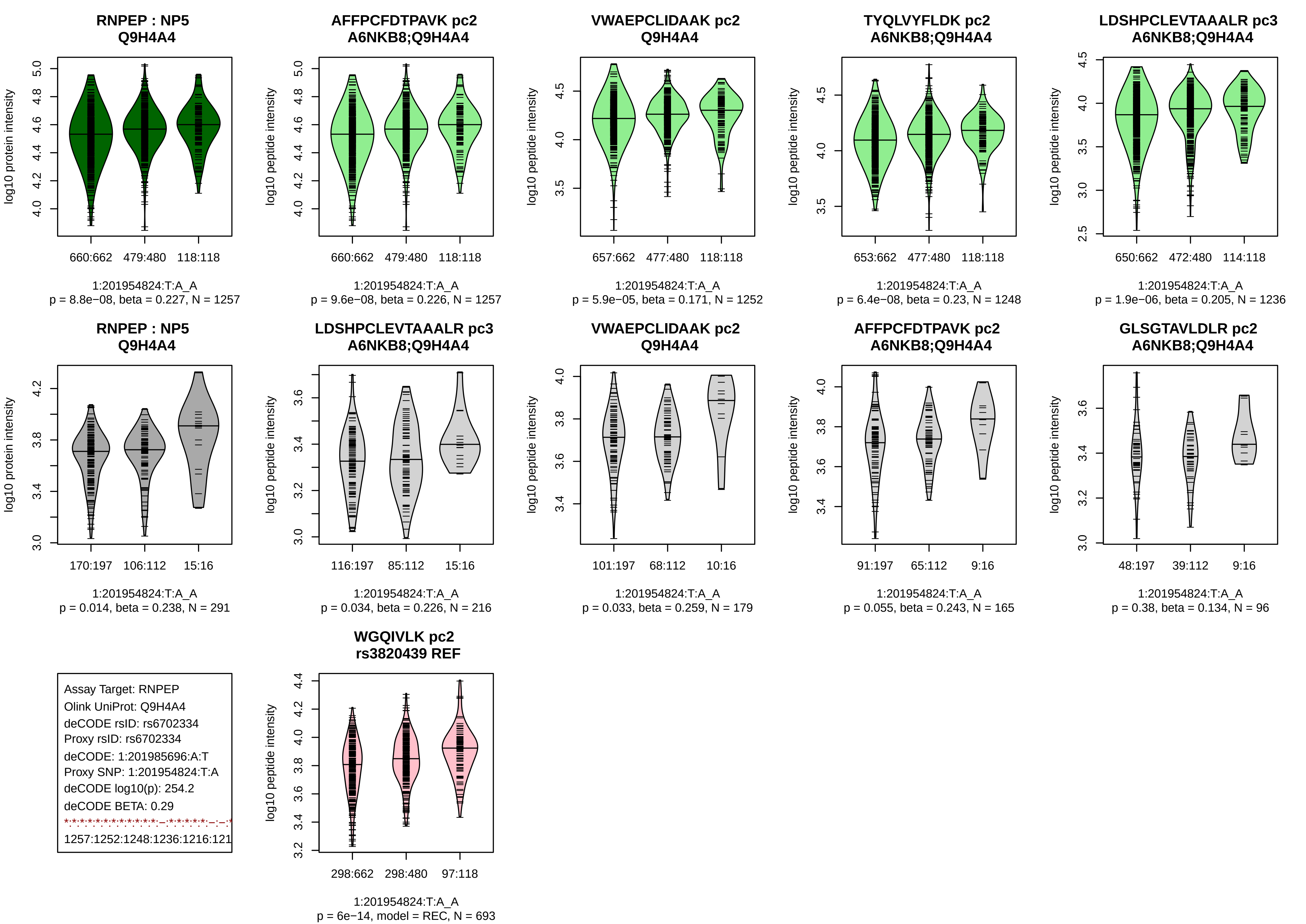
RNPEP : NP5
Q9H4A4
log10 protein intensity
660:662 479:480 118:118
1:201954824:T:A_A
p = 8.8e−08, beta = 0.227, N = 1257
AFFPCFDTPAVK pc2
A6NKB8;Q9H4A4
log10 peptide intensity
660:662 479:480 118:118
1:201954824:T:A_A
p = 9.6e−08, beta = 0.226, N = 1257
VWAEPCLIDAAK pc2
Q9H4A4
log10 peptide intensity
657:662 477:480 118:118
1:201954824:T:A_A
p = 5.9e−05, beta = 0.171, N = 1252
TYQLVYFLDK pc2
A6NKB8;Q9H4A4
log10 peptide intensity
653:662 477:480 118:118
1:201954824:T:A_A
p = 6.4e−08, beta = 0.23, N = 1248
LDSHPCLEVTAAALR pc3
A6NKB8;Q9H4A4
log10 peptide intensity
650:662 472:480 114:118
1:201954824:T:A_A
p = 1.9e−06, beta = 0.205, N = 1236
RNPEP : NP5
Q9H4A4
log10 protein intensity
170:197 106:112 15:16
1:201954824:T:A_A
p = 0.014, beta = 0.238, N = 291
LDSHPCLEVTAAALR pc3
A6NKB8;Q9H4A4
log10 peptide intensity
116:197 85:112 15:16
1:201954824:T:A_A
p = 0.034, beta = 0.226, N = 216
VWAEPCLIDAAK pc2
Q9H4A4
log10 peptide intensity
101:197 68:112 10:16
1:201954824:T:A_A
p = 0.033, beta = 0.259, N = 179
AFFPCFDTPAVK pc2
A6NKB8;Q9H4A4
log10 peptide intensity
91:197 65:112 9:16
1:201954824:T:A_A
p = 0.055, beta = 0.243, N = 165
GLSGTAVLDLR pc2
A6NKB8;Q9H4A4
log10 peptide intensity
48:197 39:112 9:16
1:201954824:T:A_A
p = 0.38, beta = 0.134, N = 96
Assay Target: RNPEP
Olink UniProt: Q9H4A4
deCODE rsID: rs6702334
Proxy rsID: rs6702334
deCODE: 1:201985696:A:T
Proxy SNP: 1:201954824:T:A
deCODE log10(p): 254.2
deCODE BETA: 0.29
1257:1252:1248:1236:1216:121
WGQIVLK pc2
rs3820439 REF
log10 peptide intensity
298:662 298:480 97:118
1:201954824:T:A_A
p = 6e−14, model = REC, N = 693

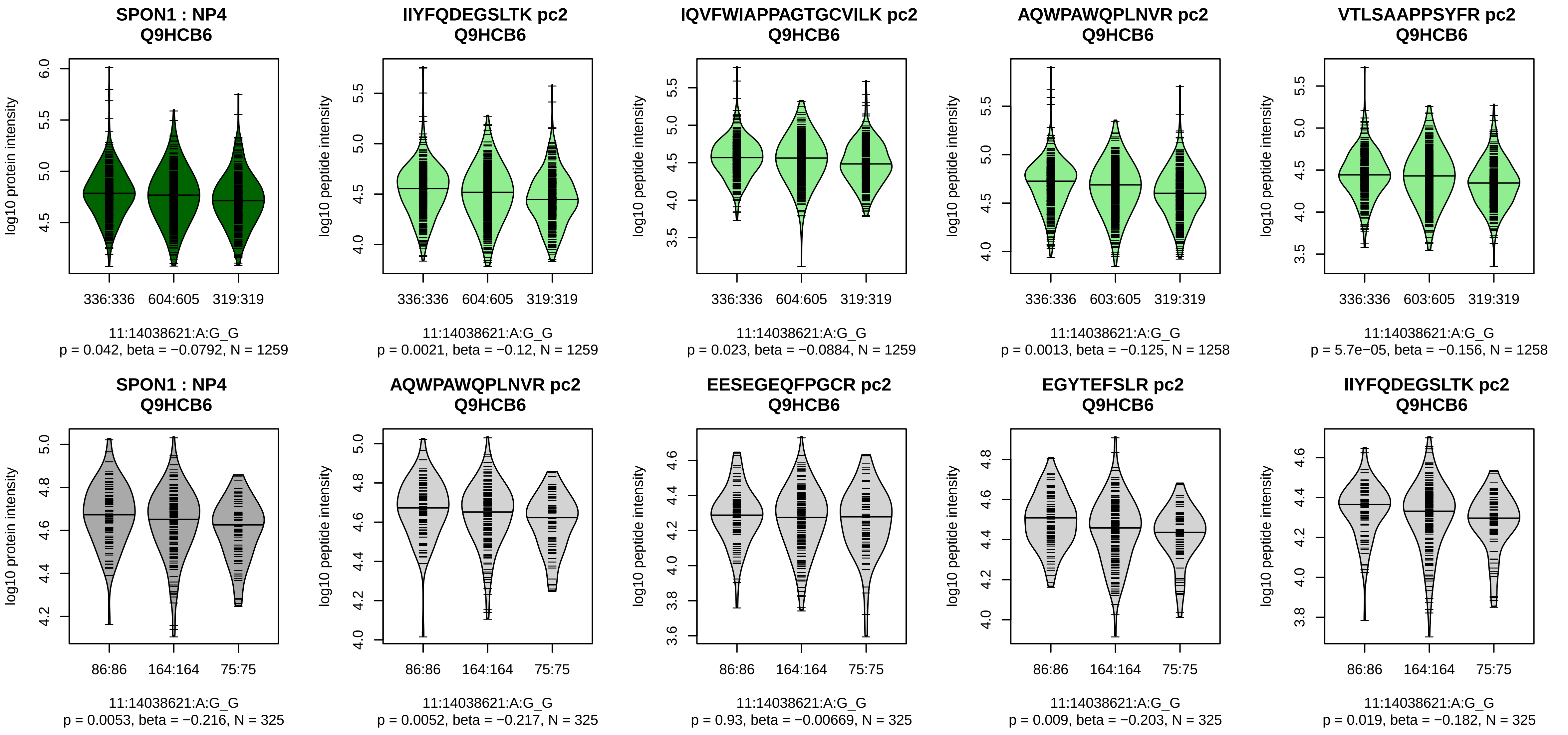


Assay Target: SPON1
Olink UniProt: Q9HCB6
deCODE rsID: rs1969539
Proxy rsID: rs1969539
deCODE: 11:14017074:A:G
Proxy SNP: 11:14038621:A:G
deCODE log10(p): 253.1
deCODE BETA: 0.28
*:–:*:*:*:–:–:*:*:–:*:–:*:–:–:–:–:–:
1259:1259:1258:1258:1246:124

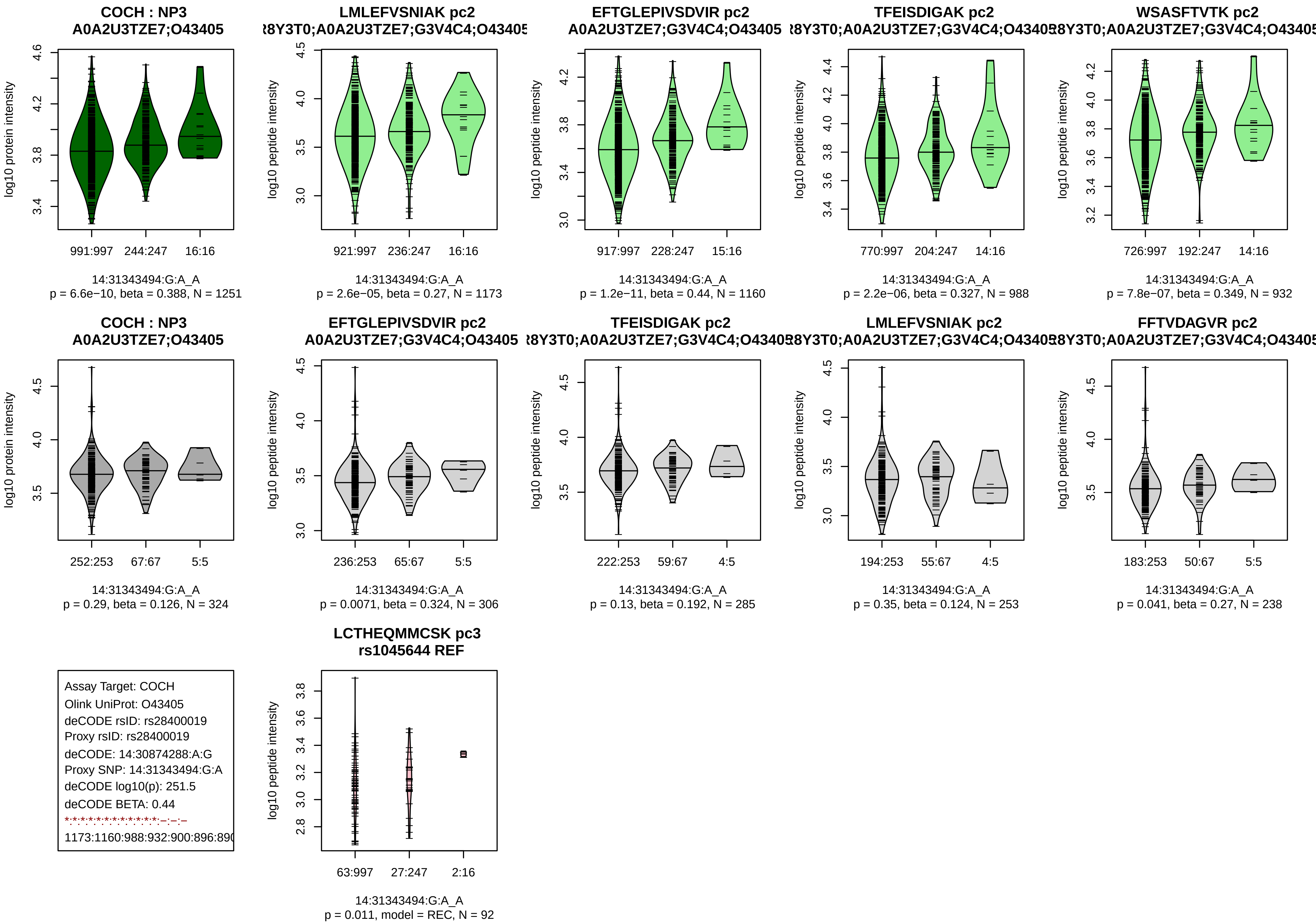
COCH : NP3
A0A2U3TZE7;O43405
log10 protein intensity
991:997
244:247
16:16
14:31343494:G:A_A
p = 6.6e−10, beta = 0.388, N = 1251
LMLEFVSNIAK pc2
R8Y3T0;A0A2U3TZE7;G3V4C4;O43405
log10 peptide intensity
921:997
236:247
16:16
14:31343494:G:A_A
p = 2.6e−05, beta = 0.27, N = 1173
EFTGLEPIVSDVIR pc2
A0A2U3TZE7;G3V4C4;O43405
log10 peptide intensity
917:997
228:247
15:16
14:31343494:G:A_A
p = 1.2e−11, beta = 0.44, N = 1160
TFEISDIGAK pc2
R8Y3T0;A0A2U3TZE7;G3V4C4;O43405
log10 peptide intensity
770:997
204:247
14:16
14:31343494:G:A_A
p = 2.2e−06, beta = 0.327, N = 988
WSASFTVTK pc2
R8Y3T0;A0A2U3TZE7;G3V4C4;O43405
log10 peptide intensity
726:997
192:247
14:16
14:31343494:G:A_A
p = 7.8e−07, beta = 0.349, N = 932
COCH : NP3
A0A2U3TZE7;O43405
log10 protein intensity
252:253
67:67
5:5
14:31343494:G:A_A
p = 0.29, beta = 0.126, N = 324
EFTGLEPIVSDVIR pc2
A0A2U3TZE7;G3V4C4;O43405
log10 peptide intensity
236:253
65:67
5:5
14:31343494:G:A_A
p = 0.0071, beta = 0.324, N = 306
TFEISDIGAK pc2
R8Y3T0;A0A2U3TZE7;G3V4C4;O43405
log10 peptide intensity
222:253
59:67
4:5
14:31343494:G:A_A
p = 0.13, beta = 0.192, N = 285
LMLEFVSNIAK pc2
R8Y3T0;A0A2U3TZE7;G3V4C4;O43405
log10 peptide intensity
194:253
55:67
4:5
14:31343494:G:A_A
p = 0.35, beta = 0.124, N = 253
FFTVDAGVR pc2
R8Y3T0;A0A2U3TZE7;G3V4C4;O43405
log10 peptide intensity
183:253
50:67
5:5
14:31343494:G:A_A
p = 0.041, beta = 0.27, N = 238
Assay Target: COCH
Olink UniProt: O43405
deCODE rsID: rs28400019
Proxy rsID: rs28400019
deCODE: 14:30874288:A:G
Proxy SNP: 14:31343494:G:A
deCODE log10(p): 251.5
deCODE BETA: 0.44
1173:1160:988:932:900:896:89
LCTHEQMMCSK pc3
rs1045644 REF
log10 peptide intensity
63:997
27:247
2:16
14:31343494:G:A_A
p = 0.011, model = REC, N = 92

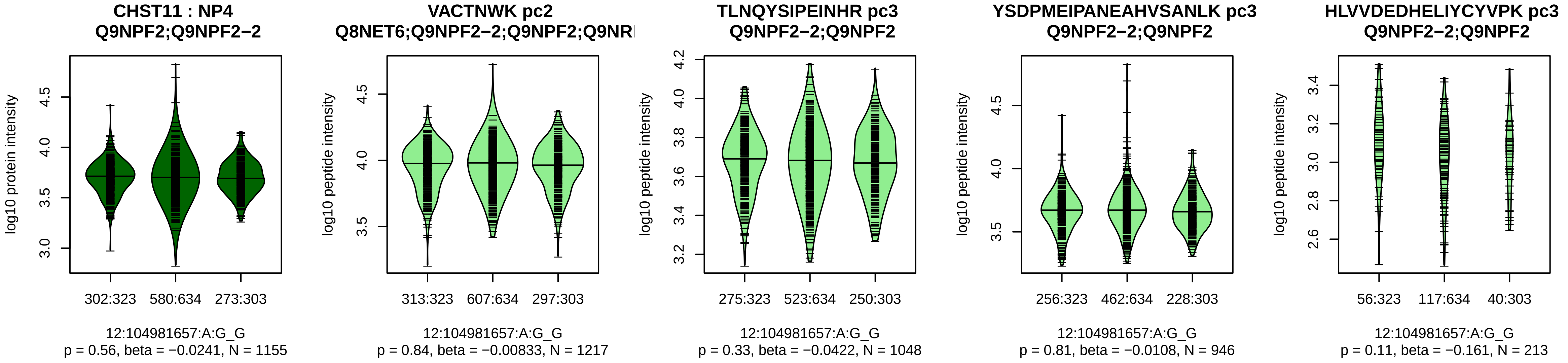


Assay Target: CHST11
Olink UniProt: Q9NPF2
deCODE rsID: rs1650132
Proxy rsID: rs1650132
deCODE: 12:104587879:G:A
Proxy SNP: 12:104981657:A:G
deCODE log10(p): 250.7
deCODE BETA: −0.28
−:−:−:−
1217:1048:946:213

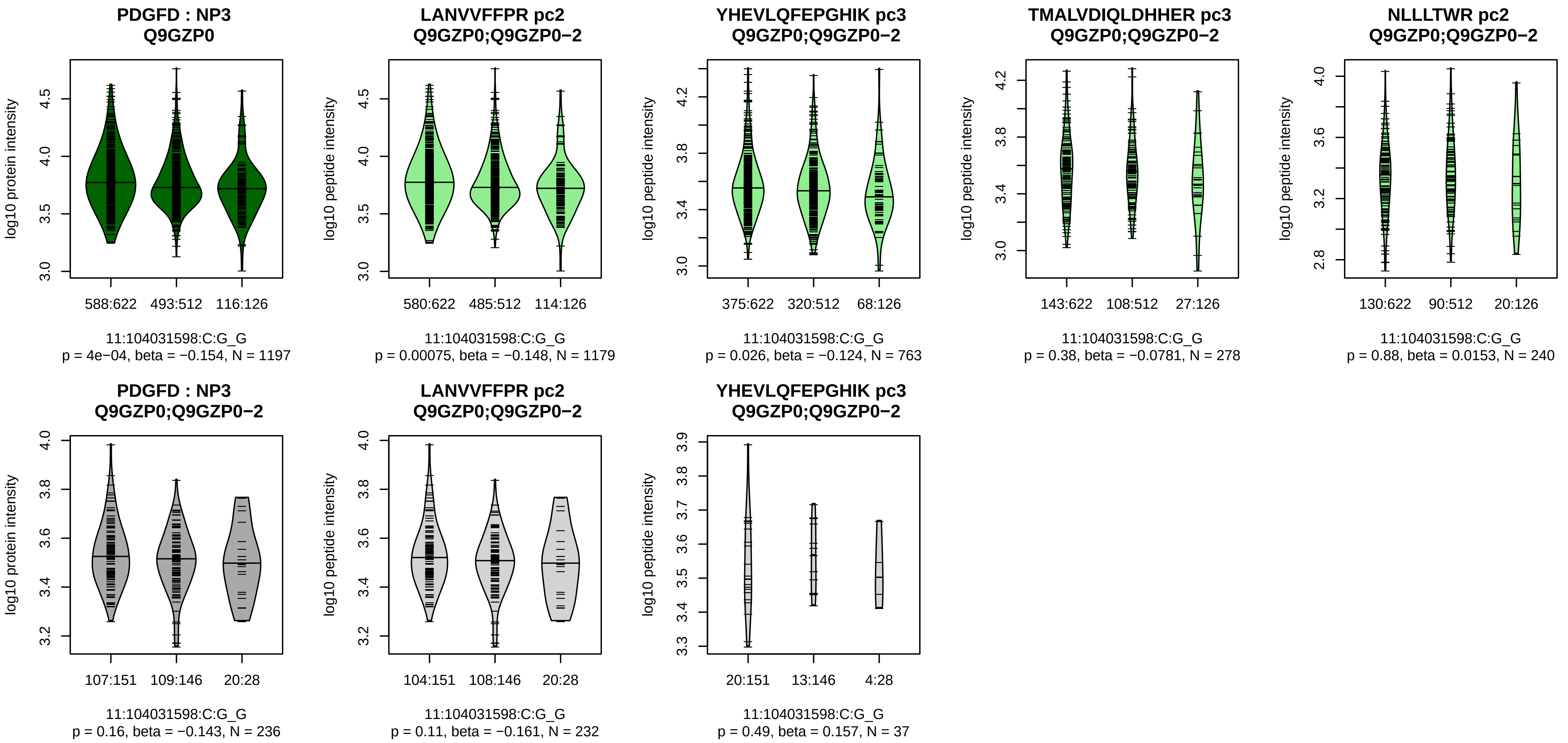


Assay Target: PDGFD
Olink UniProt: Q9GZP0
deCODE rsID: rs7950273
Proxy rsID: rs7950273
deCODE: 11:104160870:G:C
Proxy SNP: 11:104031598:C:G
deCODE log10(p): 250.6
deCODE BETA: −0.3
*:−:−:−:−:NA:NA:NA:NA
1179:763:278:240:198:8:7:13:3

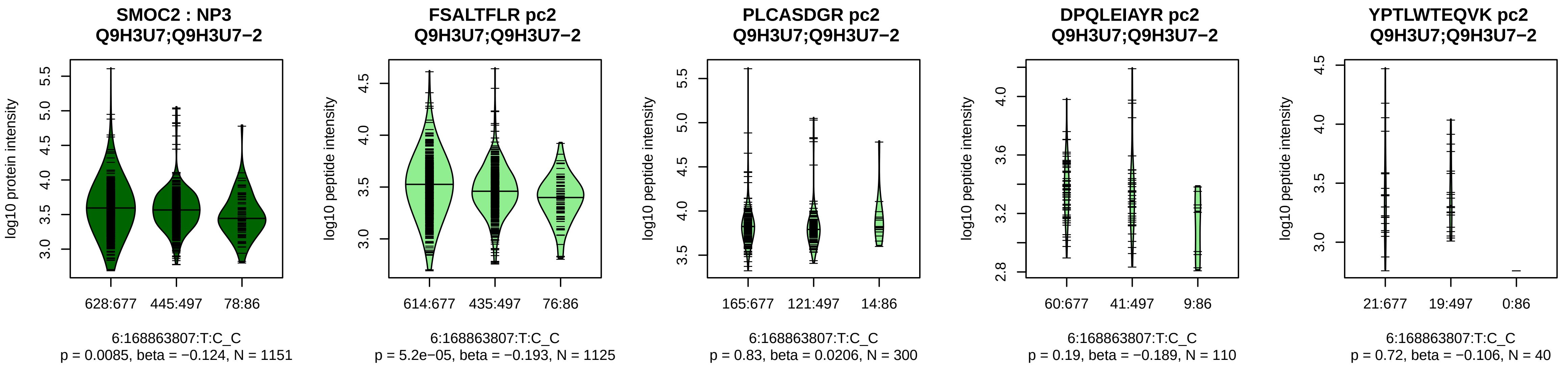


Assay Target: SMOC2
Olink UniProt: Q9H3U7
deCODE rsID: rs56296467
Proxy rsID: rs56296467
deCODE: 6:168463127:C:T
Proxy SNP: 6:168863807:T:C
deCODE log10(p): 243.7
deCODE BETA: −0.31
*:−:−:−:−:−:NA:NA:NA:NA:NA:NA
1125:300:110:40:38:21:15:11:3:

**PI3 : NP3**
**P19957**

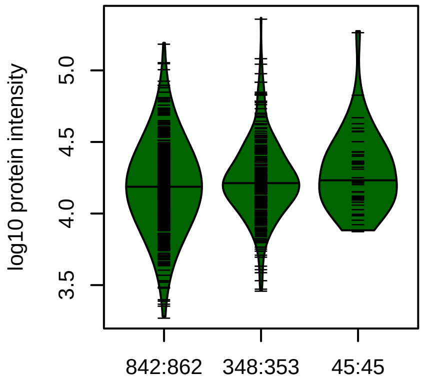


20:43777506:C:T_T
p = 7.2e−05, beta = 0.204, N = 1235

**PGSCPIILIR pc2**
**P19957**

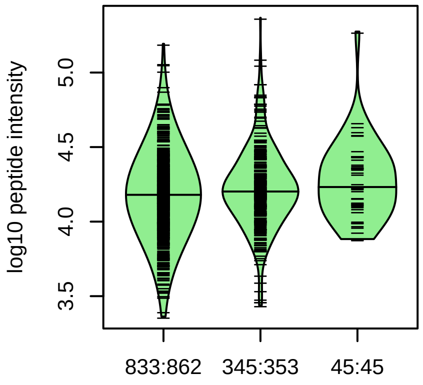


20:43777506:C:T_T
p = 0.00022, beta = 0.19, N = 1223

**CAMLNPPNR pc2**
**P19957**

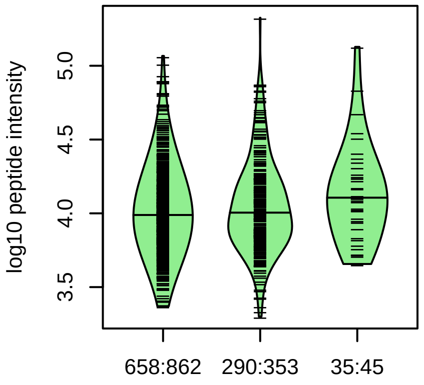


20:43777506:C:T_T
p = 0.0037, beta = 0.167, N = 983

**VPFNGQDPVK pc2**
**P19957**

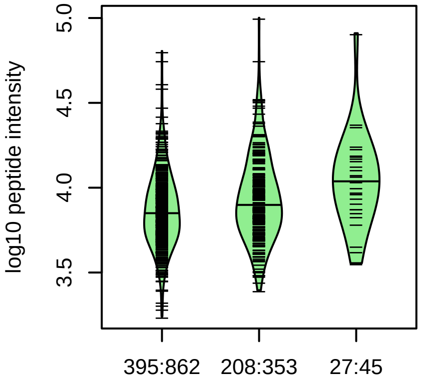


20:43777506:C:T_T
p = 1.3e−07, beta = 0.36, N = 630

**DTDCPGIK pc2**
**P19957**

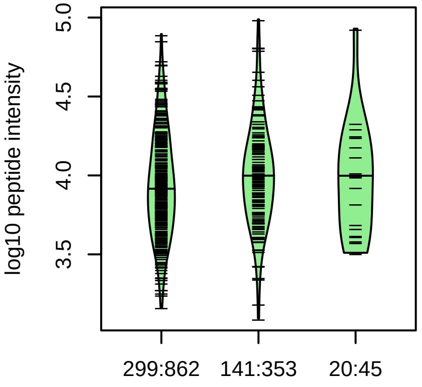


20:43777506:C:T_T
p = 0.063, beta = 0.15, N = 460

**PI3 : NP3**
**P19957**

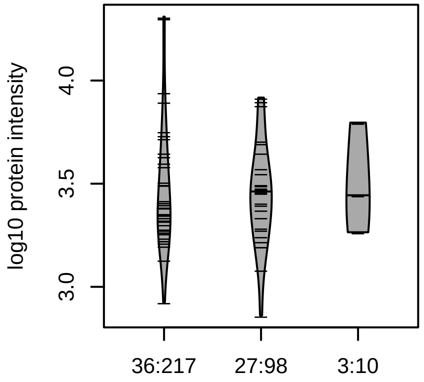


20:43777506:C:T_T
p = 0.49, beta = −0.14, N = 66

**PGSCPIILIR pc2**
**P19957**

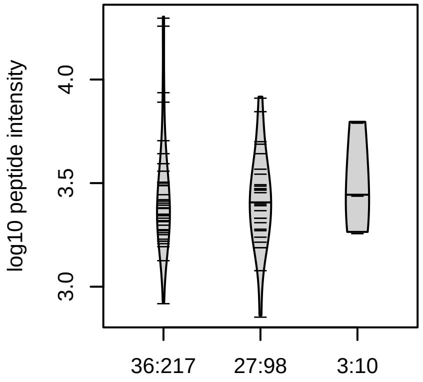


20:43777506:C:T_T
p = 0.46, beta = −0.15, N = 66

Assay Target: PI3
Olink UniProt: P19957
deCODE rsID: rs17332620
Proxy rsID: rs17332620
deCODE: 20:45148865:T:C
Proxy SNP: 20:43777506:C:T
deCODE log10(p): 241.6
deCODE BETA: 0.34
*:*:*:–:–
1223:983:630:460:31

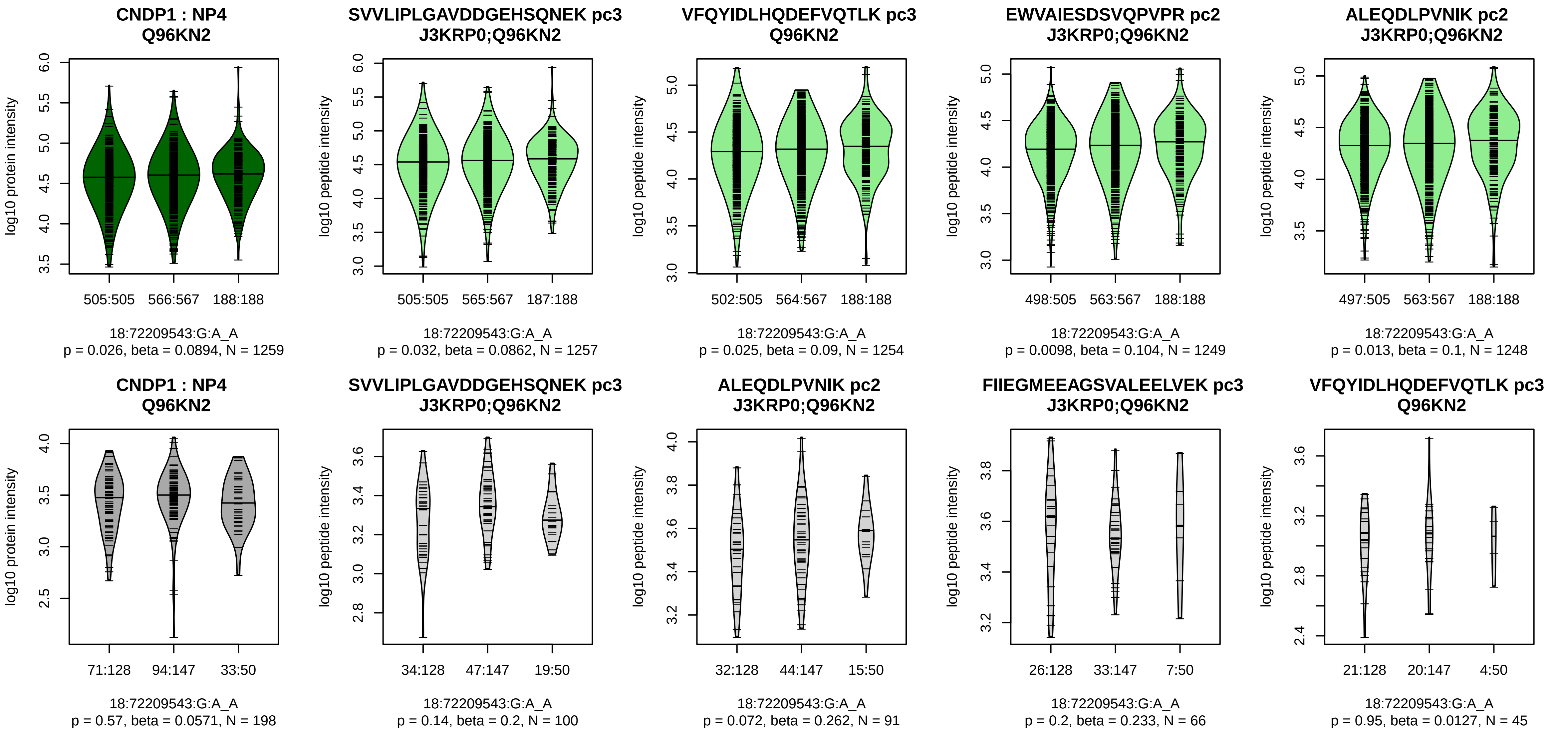


Assay Target: CNDP1
Olink UniProt: Q96KN2
deCODE rsID: rs17817077
Proxy rsID: rs17817077
deCODE: 18:74542308:A:G
Proxy SNP: 18:72209543:G:A
deCODE log10(p): 240.1
deCODE BETA: 0.28
–:–:*:–:–:–:*:–:–:–:–:–:–:–:*:–:–:
1257:1254:1249:1248:1235:123

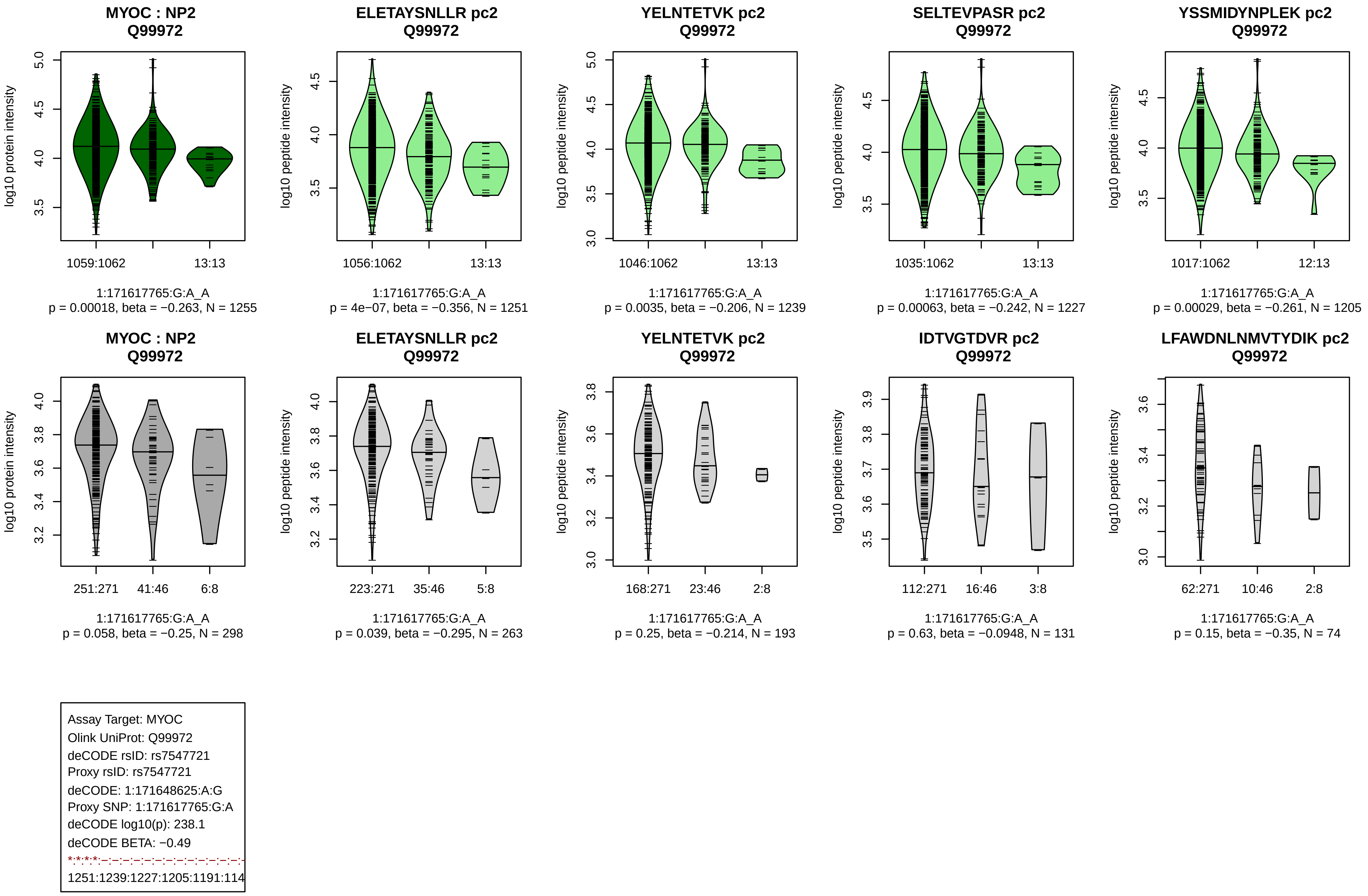
MYOC : NP2
Q99972
log10 protein intensity
1059:1062
13:13
1:171617765:G:A_A
p = 0.00018, beta = −0.263, N = 1255
ELETAYSNLLR pc2
Q99972
log10 peptide intensity
1056:1062
13:13
1:171617765:G:A_A
p = 4e−07, beta = −0.356, N = 1251
YELNTETVK pc2
Q99972
log10 peptide intensity
1046:1062
13:13
1:171617765:G:A_A
p = 0.0035, beta = −0.206, N = 1239
SELTEVPASR pc2
Q99972
log10 peptide intensity
1035:1062
13:13
1:171617765:G:A_A
p = 0.00063, beta = −0.242, N = 1227
YSSMIDYNPLEK pc2
Q99972
log10 peptide intensity
1017:1062
12:13
1:171617765:G:A_A
p = 0.00029, beta = −0.261, N = 1205
MYOC : NP2
Q99972
log10 protein intensity
251:271
41:46
6:8
1:171617765:G:A_A
p = 0.058, beta = −0.25, N = 298
ELETAYSNLLR pc2
Q99972
log10 peptide intensity
223:271
35:46
5:8
1:171617765:G:A_A
p = 0.039, beta = −0.295, N = 263
YELNTETVK pc2
Q99972
log10 peptide intensity
168:271
23:46
2:8
1:171617765:G:A_A
p = 0.25, beta = −0.214, N = 193
IDTVGTDVR pc2
Q99972
log10 peptide intensity
112:271
16:46
3:8
1:171617765:G:A_A
p = 0.63, beta = −0.0948, N = 131
LFAWDNLNMVTYDIK pc2
Q99972
log10 peptide intensity
62:271
10:46
2:8
1:171617765:G:A_A
p = 0.15, beta = −0.35, N = 74
Assay Target: MYOC
Olink UniProt: Q99972
deCODE rsID: rs7547721
Proxy rsID: rs7547721
deCODE: 1:171648625:A:G
Proxy SNP: 1:171617765:G:A
deCODE log10(p): 238.1
deCODE BETA: −0.49
1251:1239:1227:1205:1191:114

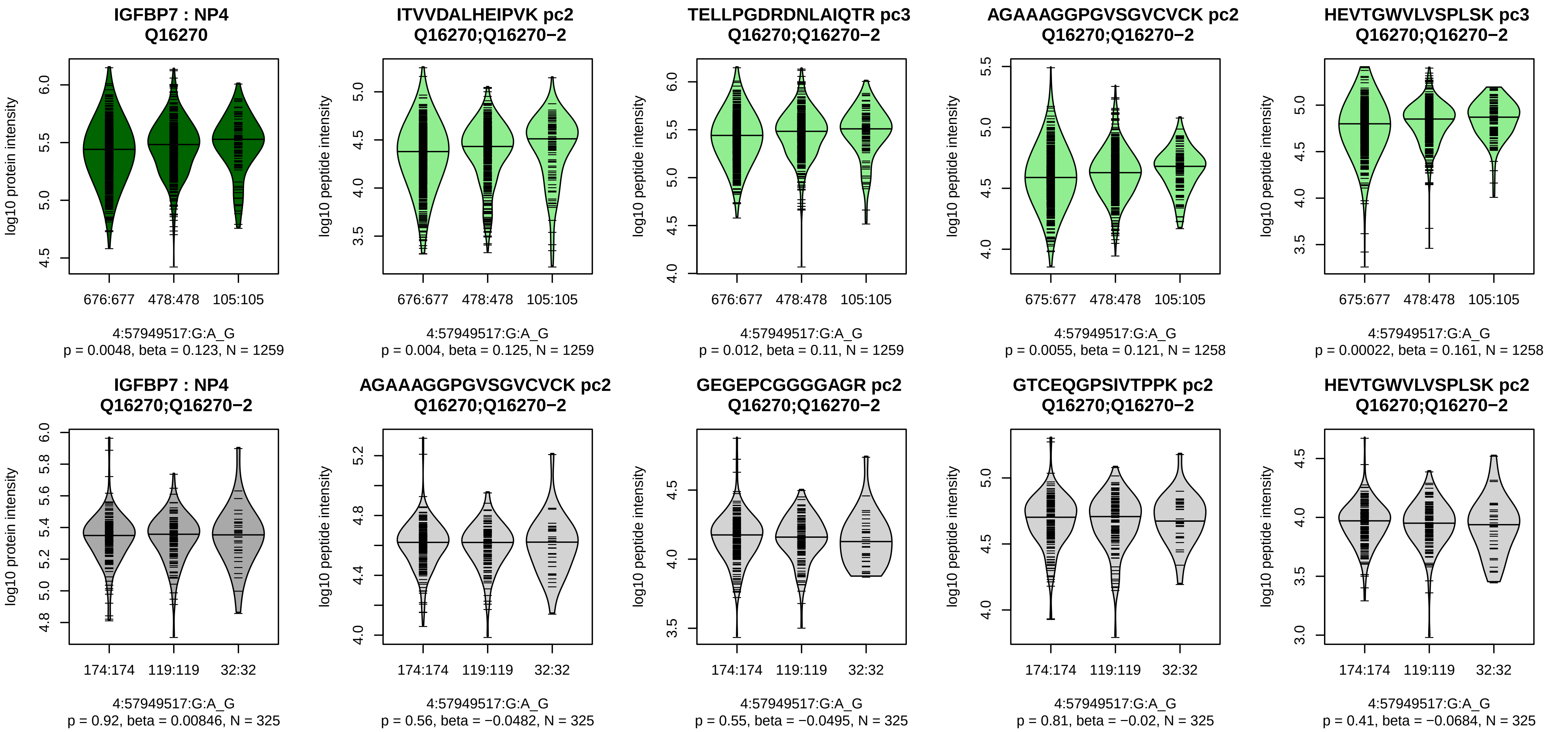


Assay Target: IGFBP7
Olink UniProt: Q16270
deCODE rsID: rs1718860
Proxy rsID: rs1718860
deCODE: 4:57083351:G:A
Proxy SNP: 4:57949517:G:A
deCODE log10(p): 235.7
deCODE BETA: 0.3
*:−:*:*:*:−:*:*:*:*:−:−:NA
1259:1259:1258:1258:1258:125

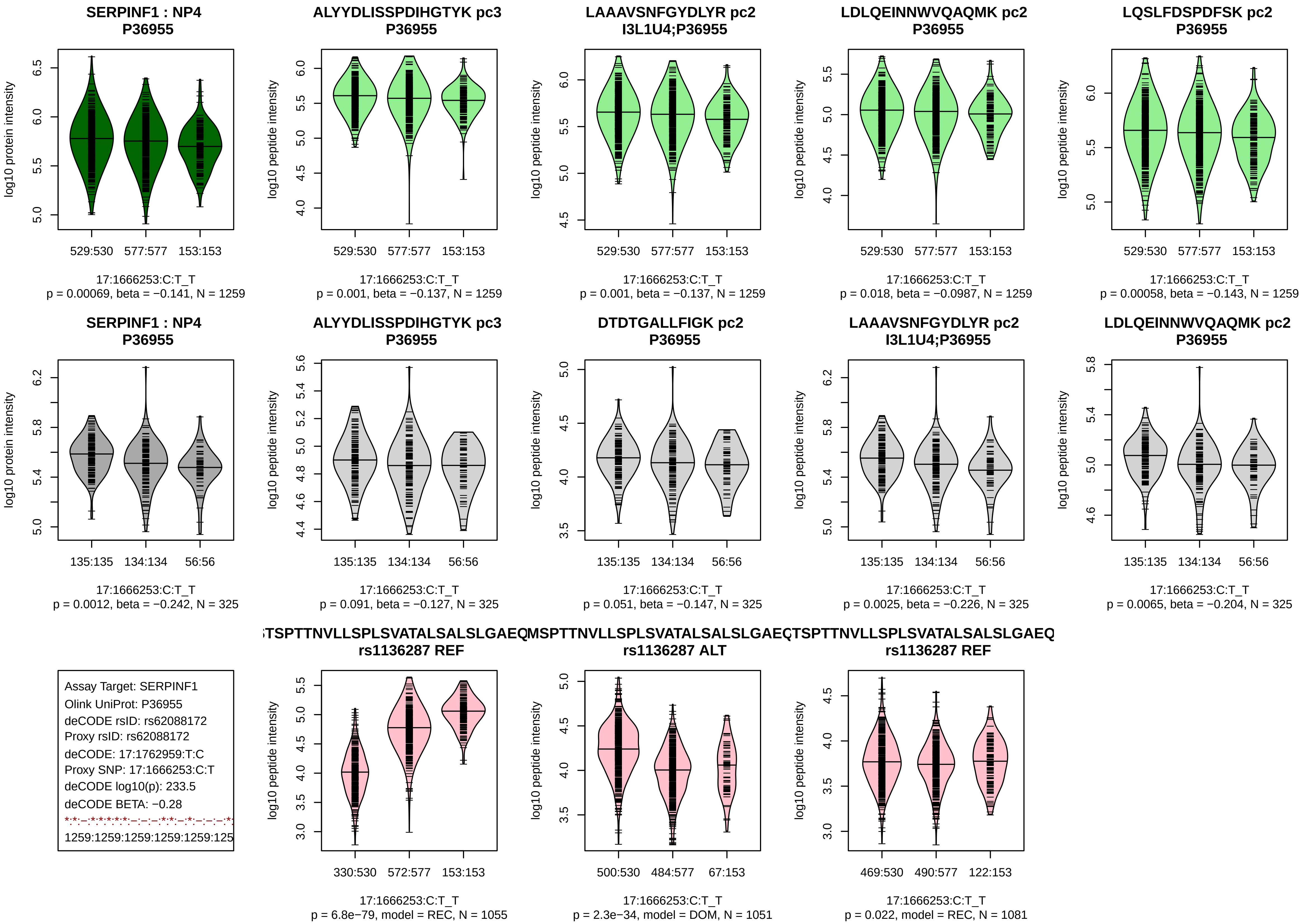

SERPINF1 : NP4
P36955
log10 protein intensity
529:530
577:577
153:153
17:1666253:C:T_T
p = 0.00069, beta = −0.141, N = 1259
ALYYDLISSPDIHGTYK pc3
P36955
log10 peptide intensity
529:530
577:577
153:153
17:1666253:C:T_T
p = 0.001, beta = −0.137, N = 1259
LAAAVSNFGYDLYR pc2
I3L1U4;P36955
log10 peptide intensity
529:530
577:577
153:153
17:1666253:C:T_T
p = 0.001, beta = −0.137, N = 1259
LDLQEINNWVQAQMK pc2
P36955
log10 peptide intensity
529:530
577:577
153:153
17:1666253:C:T_T
p = 0.018, beta = −0.0987, N = 1259
LQSLFDSPDFSK pc2
P36955
log10 peptide intensity
529:530
577:577
153:153
17:1666253:C:T_T
p = 0.00058, beta = −0.143, N = 1259
SERPINF1 : NP4
P36955
log10 protein intensity
135:135
134:134
56:56
17:1666253:C:T_T
p = 0.0012, beta = −0.242, N = 325
ALYYDLISSPDIHGTYK pc3
P36955
log10 peptide intensity
135:135
134:134
56:56
17:1666253:C:T_T
p = 0.091, beta = −0.127, N = 325
DTDTGALLFIGK pc2
P36955
log10 peptide intensity
135:135
134:134
56:56
17:1666253:C:T_T
p = 0.051, beta = −0.147, N = 325
LAAAVSNFGYDLYR pc2
I3L1U4;P36955
log10 peptide intensity
135:135
134:134
56:56
17:1666253:C:T_T
p = 0.0025, beta = −0.226, N = 325
LDLQEINNWVQAQMK pc2
P36955
log10 peptide intensity
135:135
134:134
56:56
17:1666253:C:T_T
p = 0.0065, beta = −0.204, N = 325
Assay Target: SERPINF1
Olink UniProt: P36955
deCODE rsID: rs62088172
Proxy rsID: rs62088172
deCODE: 17:1762959:T:C
Proxy SNP: 17:1666253:C:T
deCODE log10(p): 233.5
deCODE BETA: −0.28
1259:1259:1259:1259:1259:125
TSPTTNVLLSPLSVATALSALSLGAEQ
rs1136287 REF
log10 peptide intensity
330:530
572:577
153:153
17:1666253:C:T_T
p = 6.8e−79, model = REC, N = 1055
MSPTTNVLLSPLSVATALSALSLGAEQ
rs1136287 ALT
log10 peptide intensity
500:530
484:577
67:153
17:1666253:C:T_T
p = 2.3e−34, model = DOM, N = 1051
TSPTTNVLLSPLSVATALSALSLGAEQ
rs1136287 REF
log10 peptide intensity
469:530
490:577
122:153
17:1666253:C:T_T
p = 0.022, model = REC, N = 1081

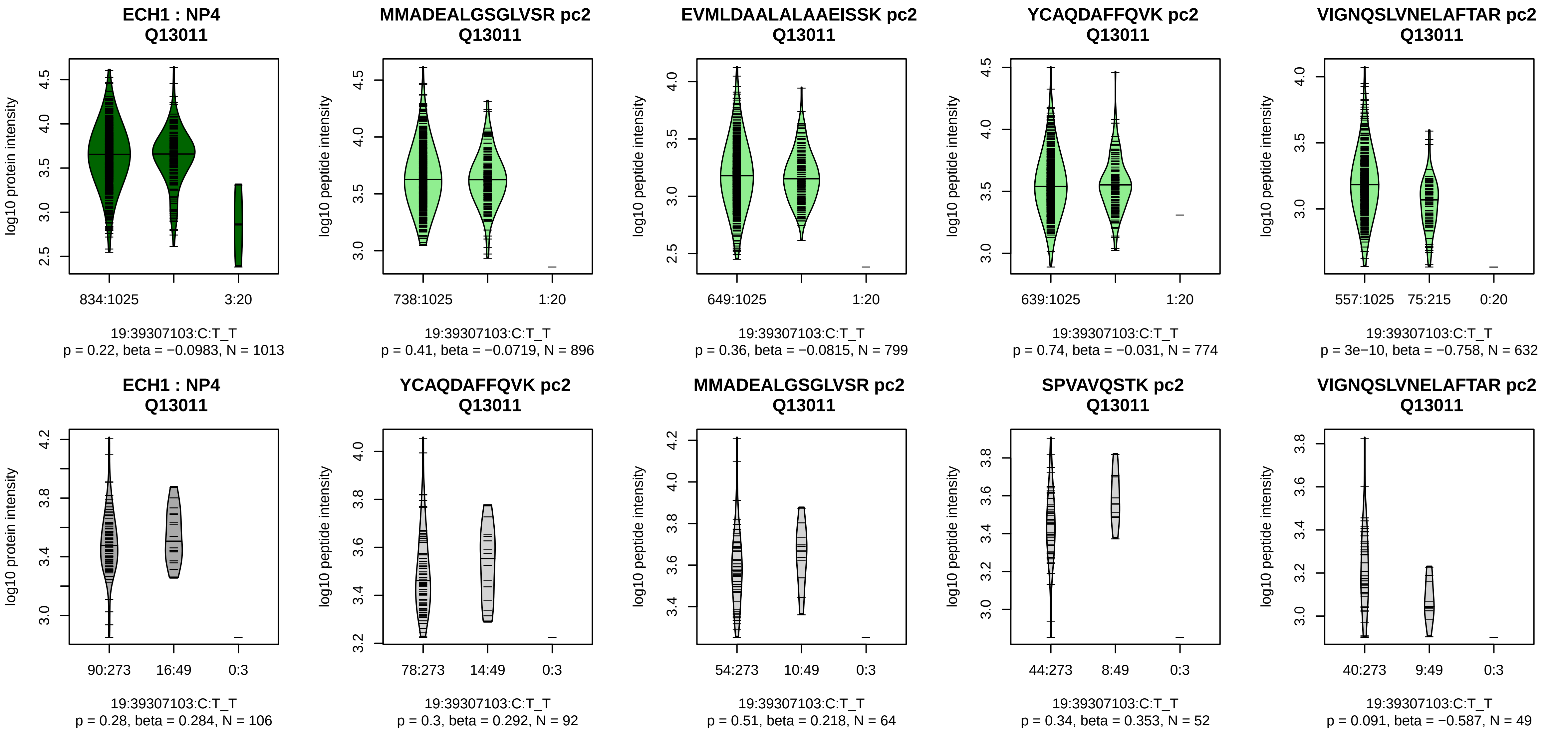


Assay Target: ECH1
Olink UniProt: Q13011
deCODE rsID: rs2229259
Proxy rsID: rs2229259
deCODE: 19:38816463:T:C
Proxy SNP: 19:39307103:C:T
deCODE log10(p): 230.5
deCODE BETA: 0.45
−:−:−:*:−:−:−:−:−:−:NA
896:799:774:632:570:552:527:3

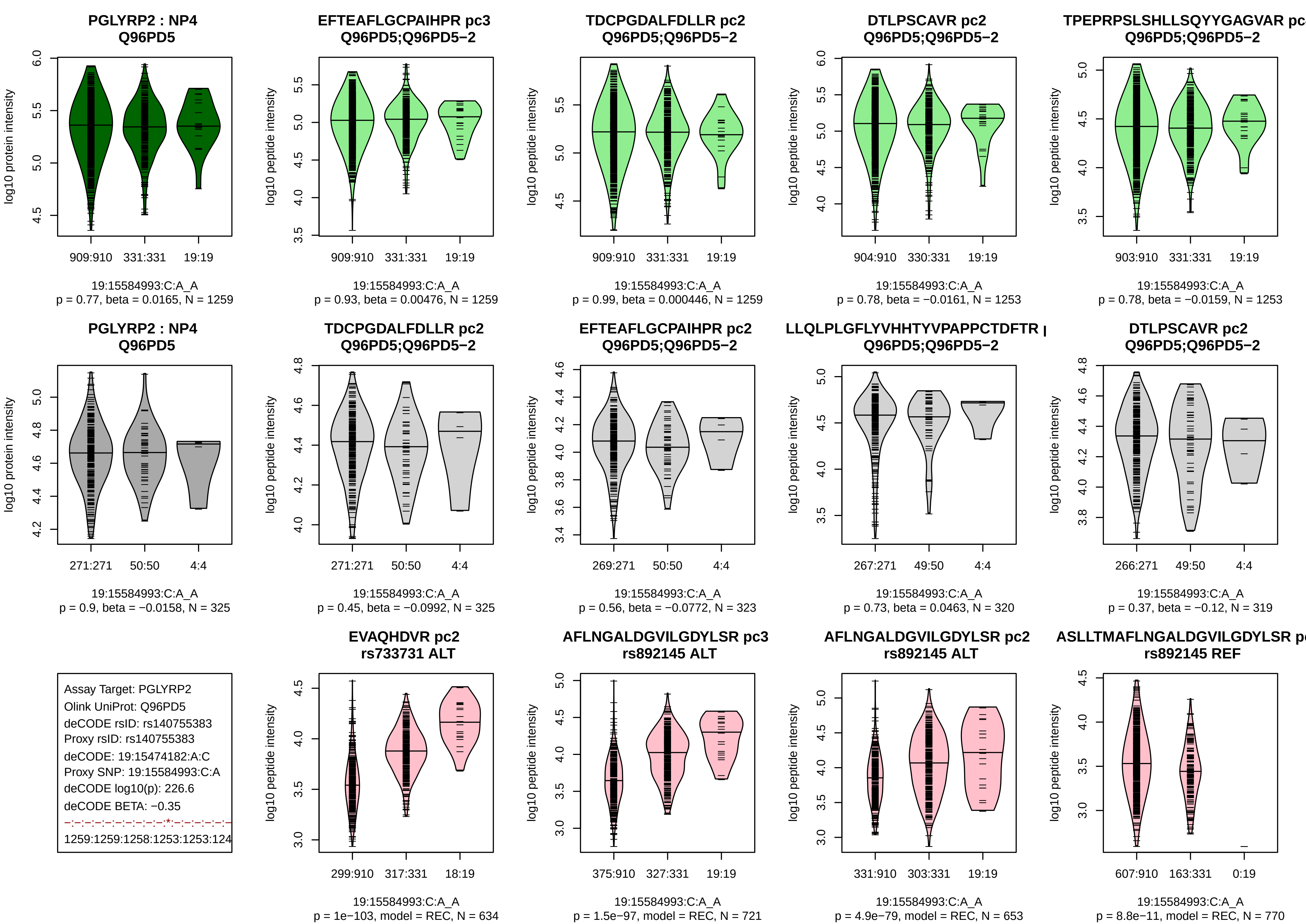
PGLYRP2 : NP4
Q96PD5
log10 protein intensity
909:910 331:331 19:19
19:15584993:C:A_A
p = 0.77, beta = 0.0165, N = 1259
EFTEAFLGCPAIHPR pc3
Q96PD5;Q96PD5−2
log10 peptide intensity
909:910 331:331 19:19
19:15584993:C:A_A
p = 0.93, beta = 0.00476, N = 1259
TDCPGDALFDLLR pc2
Q96PD5;Q96PD5−2
log10 peptide intensity
909:910 331:331 19:19
19:15584993:C:A_A
p = 0.99, beta = 0.000446, N = 1259
DTLPSCAVR pc2
Q96PD5;Q96PD5−2
log10 peptide intensity
904:910 330:331 19:19
19:15584993:C:A_A
p = 0.78, beta = −0.0161, N = 1253
TPEPRPSLSHLLSQYYGAGVAR pc
Q96PD5;Q96PD5−2
log10 peptide intensity
903:910 331:331 19:19
19:15584993:C:A_A
p = 0.78, beta = −0.0159, N = 1253
PGLYRP2 : NP4
Q96PD5
log10 protein intensity
271:271 50:50 4:4
19:15584993:C:A_A
p = 0.9, beta = −0.0158, N = 325
TDCPGDALFDLLR pc2
Q96PD5;Q96PD5−2
log10 peptide intensity
271:271 50:50 4:4
19:15584993:C:A_A
p = 0.45, beta = −0.0992, N = 325
EFTEAFLGCPAIHPR pc2
Q96PD5;Q96PD5−2
log10 peptide intensity
269:271 50:50 4:4
19:15584993:C:A_A
p = 0.56, beta = −0.0772, N = 323
LLQLPLGFLYVHHTYVPAPPCTDFTR
Q96PD5;Q96PD5−2
log10 peptide intensity
267:271 49:50 4:4
19:15584993:C:A_A
p = 0.73, beta = 0.0463, N = 320
DTLPSCAVR pc2
Q96PD5;Q96PD5−2
log10 peptide intensity
266:271 49:50 4:4
19:15584993:C:A_A
p = 0.37, beta = −0.12, N = 319
Assay Target: PGLYRP2
Olink UniProt: Q96PD5
deCODE rsID: rs140755383
Proxy rsID: rs140755383
deCODE: 19:15474182:A:C
Proxy SNP: 19:15584993:C:A
deCODE log10(p): 226.6
deCODE BETA: −0.35
−:−:−:−:−:−:−:−:−:*:−:−:−:−:−:−
1259:1259:1258:1253:1253:124
EVAQHDVR pc2
rs733731 ALT
log10 peptide intensity
299:910 317:331 18:19
19:15584993:C:A_A
p = 1e−103, model = REC, N = 634
AFLNGALDGVILGDYLSR pc3
rs892145 ALT
log10 peptide intensity
375:910 327:331 19:19
19:15584993:C:A_A
p = 1.5e−97, model = REC, N = 721
AFLNGALDGVILGDYLSR pc2
rs892145 ALT
log10 peptide intensity
331:910 303:331 19:19
19:15584993:C:A_A
p = 4.9e−79, model = REC, N = 653
ASLLTMAFLNGALDGVILGDYLSR p
rs892145 REF
log10 peptide intensity
607:910 163:331 0:19
19:15584993:C:A_A
p = 8.8e−11, model = REC, N = 770

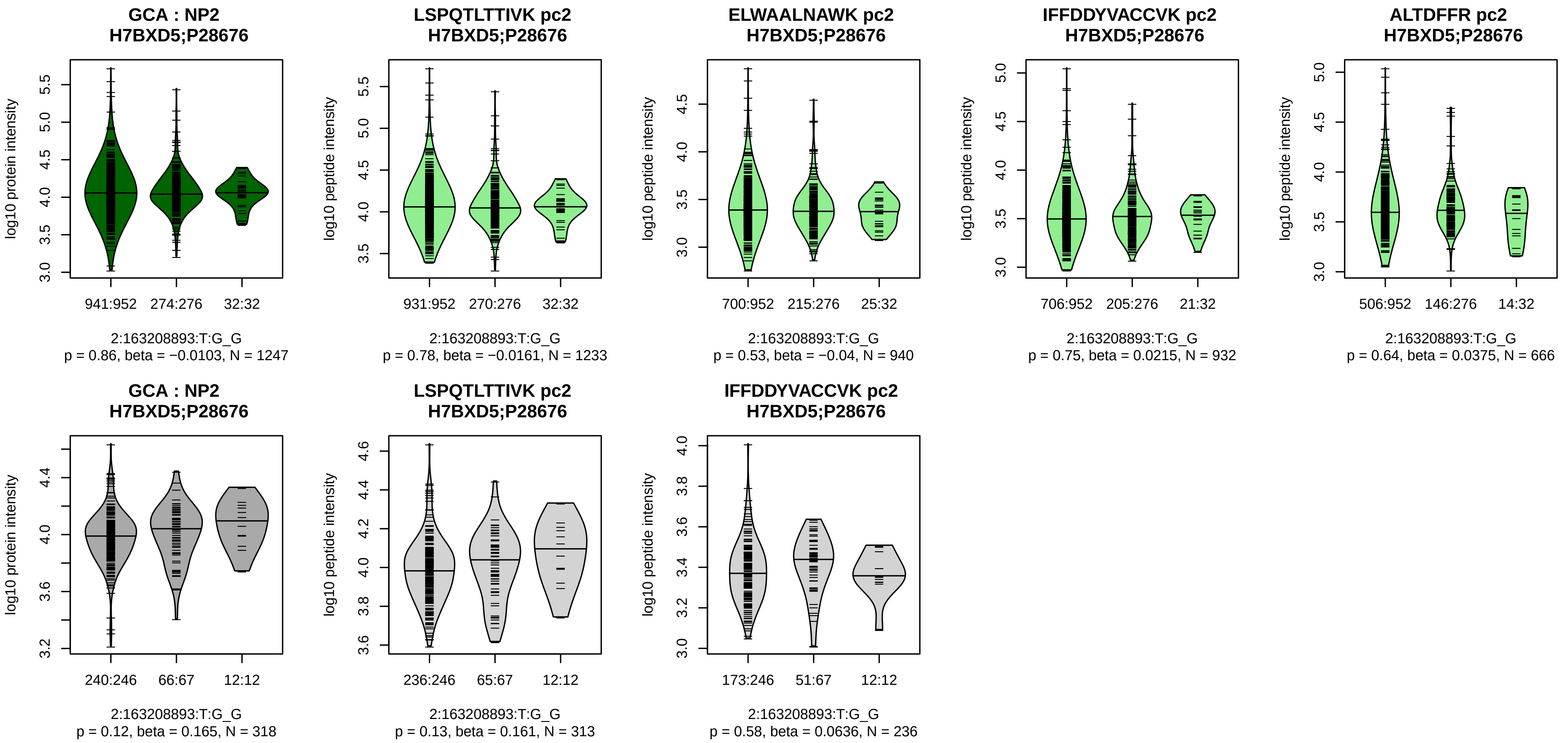


Assay Target: GCA
Olink UniProt: P28676
deCODE rsID: rs17783344
Proxy rsID: rs17783344
deCODE: 2:162352383:G:T
Proxy SNP: 2:163208893:T:G
deCODE log10(p): 222.1
deCODE BETA: −0.43
−:−:−:−:−:−:−:−
1233:940:932:666:349:185:128:

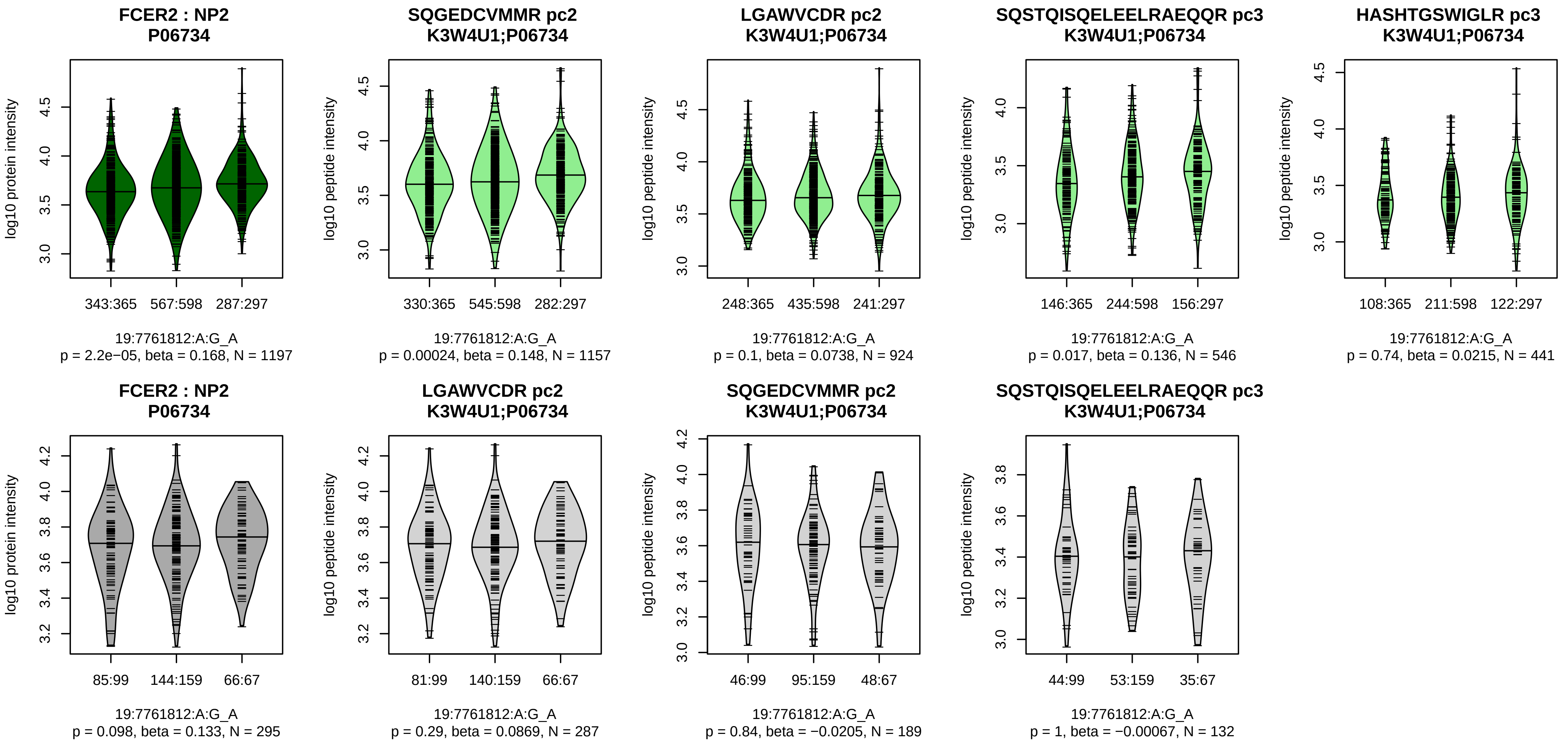


Assay Target: FCER2
Olink UniProt: P06734
deCODE rsID: rs2277989
Proxy rsID: rs2277989
deCODE: 19:7696926:G:A
Proxy SNP: 19:7761812:A:G
deCODE log10(p): 217.9
deCODE BETA: −0.27
*:−:−:−:−
1157:924:546:441:205

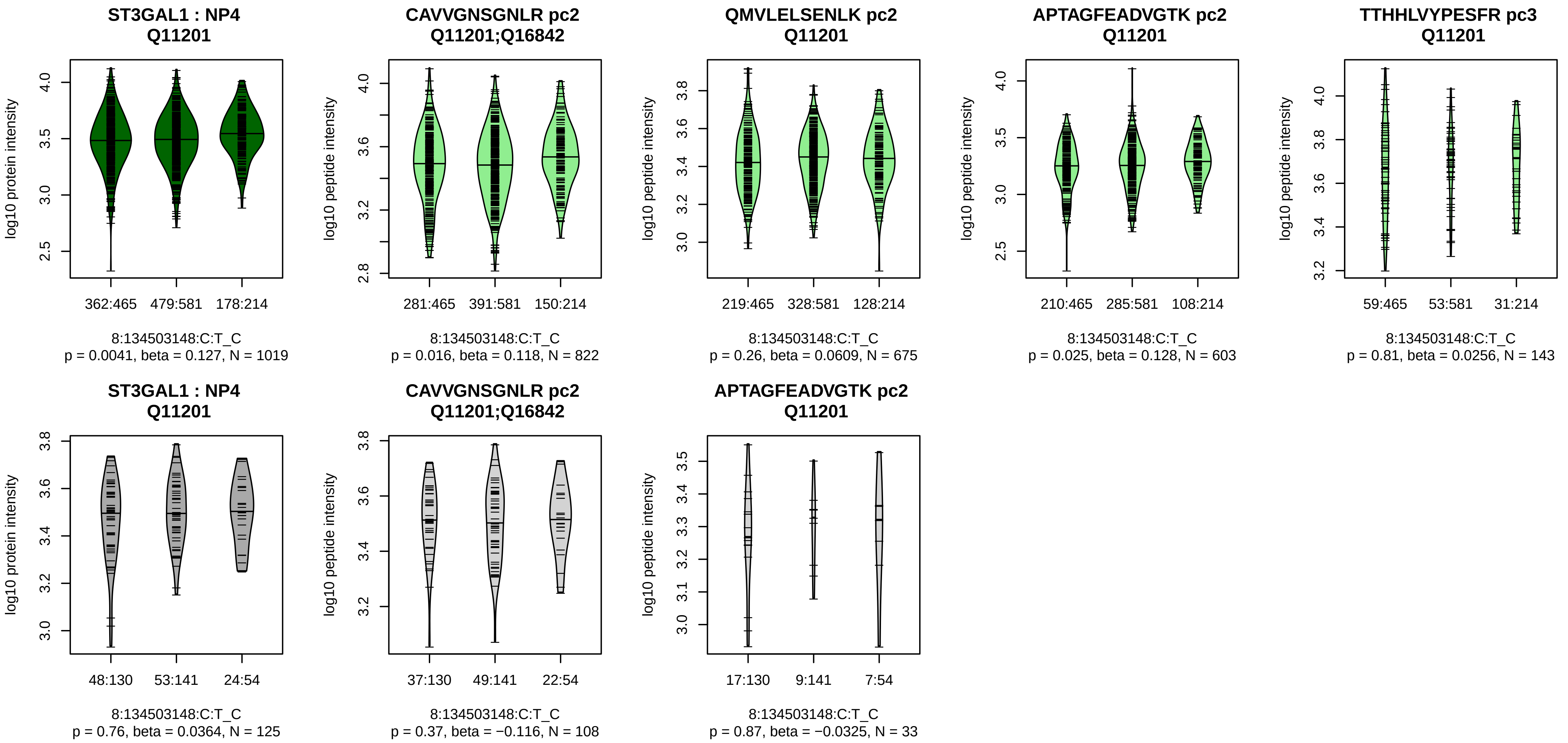


Assay Target: ST3GAL1
Olink UniProt: Q11201
deCODE rsID: rs9643300
Proxy rsID: rs9643300
deCODE: 8:133490905:C:T
Proxy SNP: 8:134503148:C:T
deCODE log10(p): 207.5
deCODE BETA: 0.25
–:–:–:–
822:675:603:143

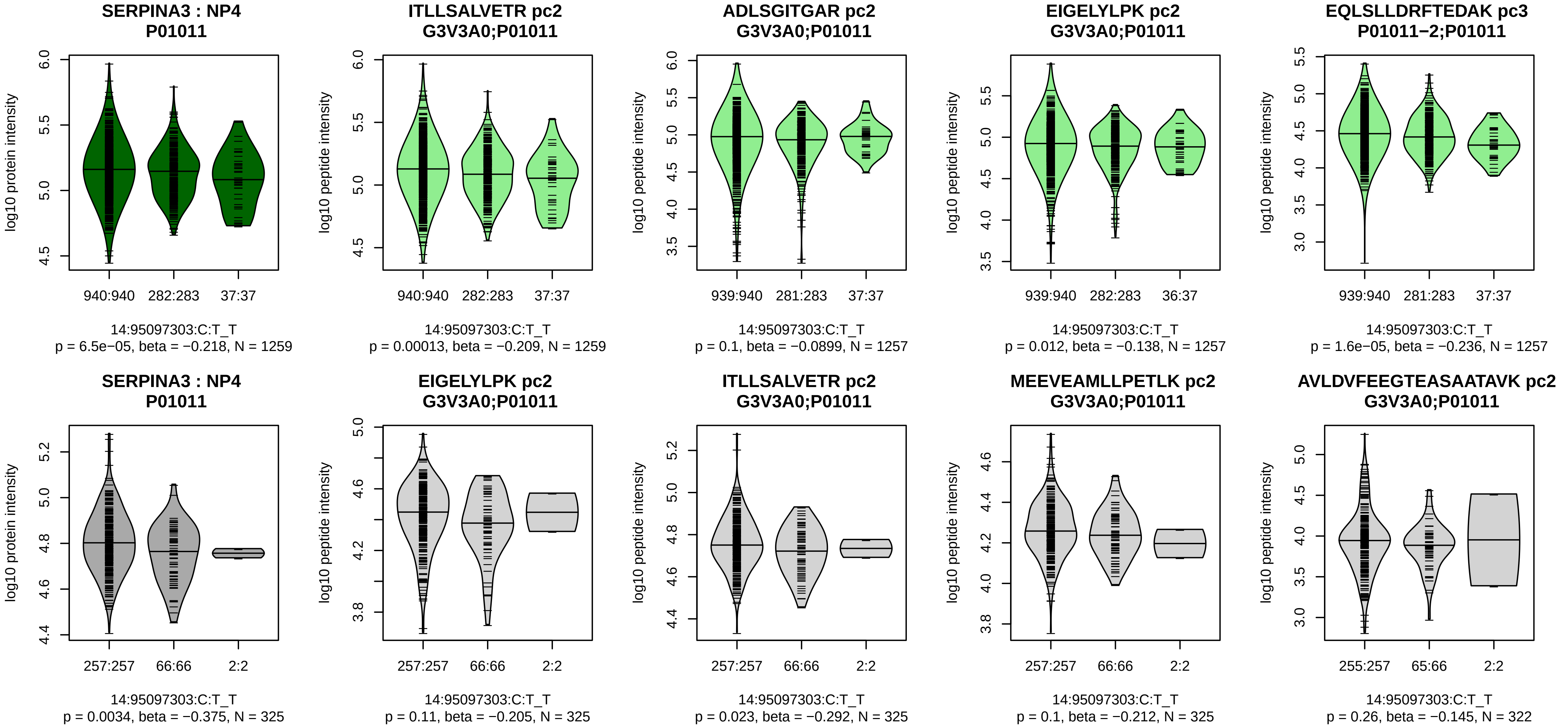


Assay Target: SERPINA3
Olink UniProt: P01011
deCODE rsID: rs140735783
Proxy rsID: rs6575449
deCODE: 14:94634064:TC:T
Proxy SNP: 14:95097303:C:T
deCODE log10(p): 206.5
deCODE BETA: −0.32
*:–:–:*:*:–:*:*:–:*:–:–:–:–:–:–:–:–
1259:1257:1257:1257:1248:124

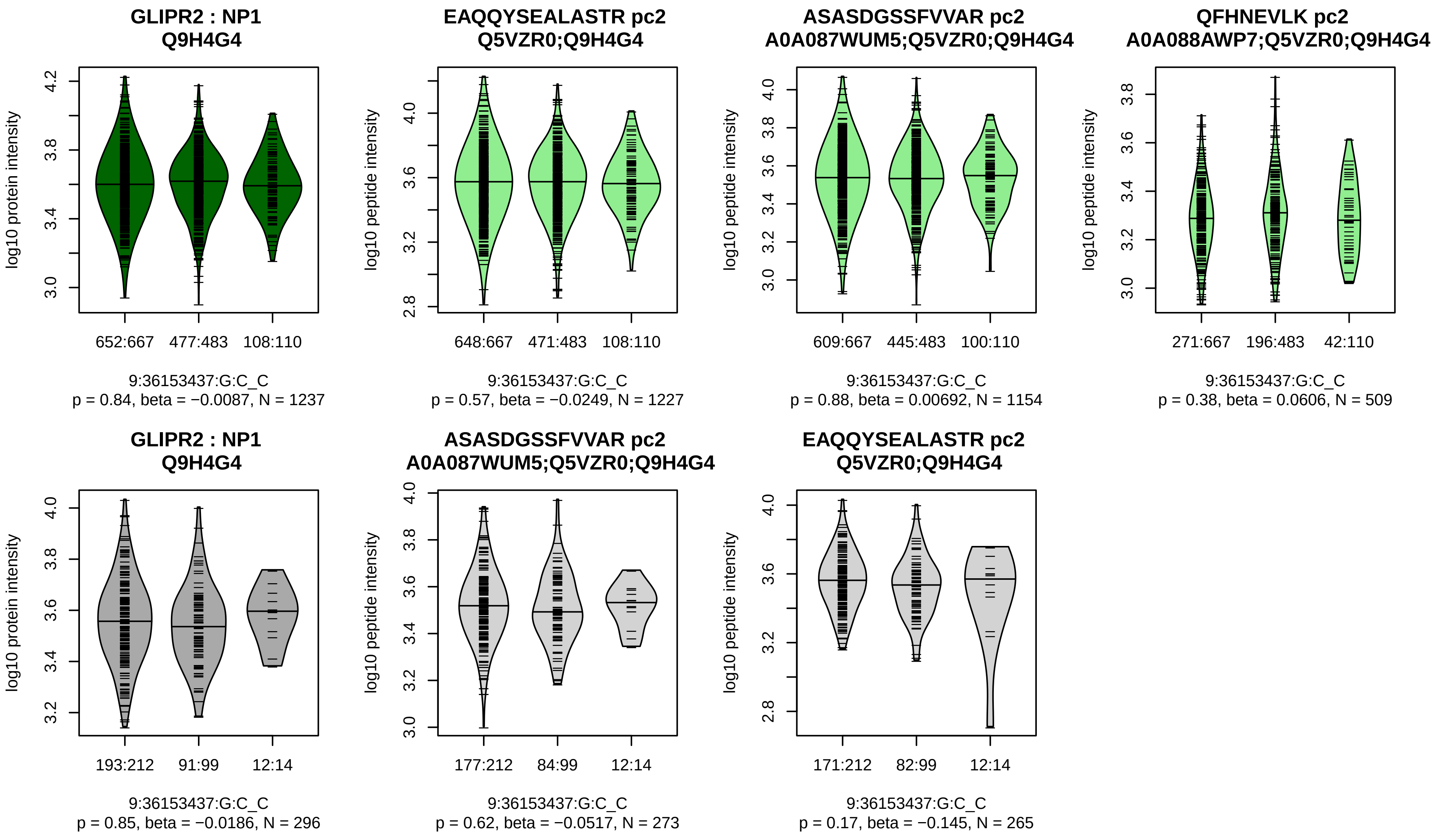


Assay Target: GLIPR2
Olink UniProt: Q9H4G4
deCODE rsID: rs4878643
Proxy rsID: rs4878643
deCODE: 9:36153440:C:G
Proxy SNP: 9:36153437:G:C
deCODE log10(p): 204.4
deCODE BETA: −0.28
−:−:−:NA:NA
1227:1154:509:6:7

**MFAP2 : NP2**
**P55001;P55001−2**

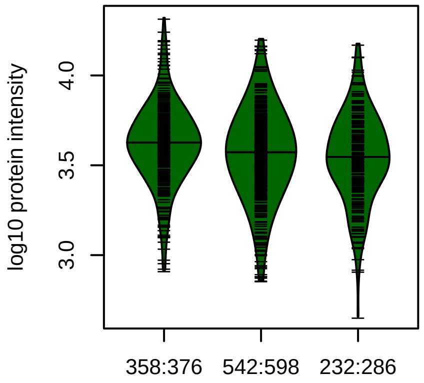


1:17315425:G:A_G
p = 1.2e−06, beta = −0.201, N = 1132

**TVCAHEELLR pc3**
**P55001−2;P55001**

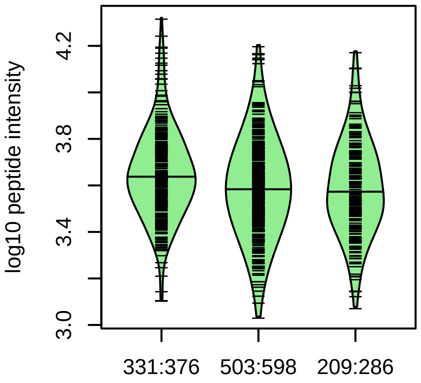


1:17315425:G:A_G
p = 2.8e−06, beta = −0.202, N = 1043

**CGVMASSGLCQSVAASCAR pc3**
**P55001−2;P55001**

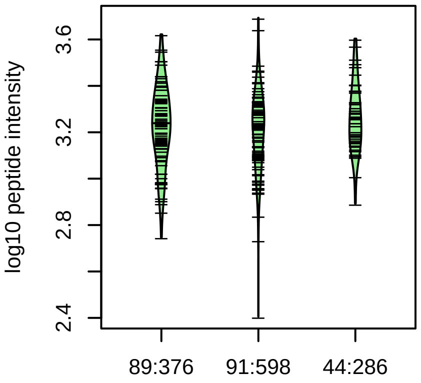


1:17315425:G:A_G
p = 0.75, beta = 0.0287, N = 224

**MFAP2 : NP2**
**P55001;P55001−2**

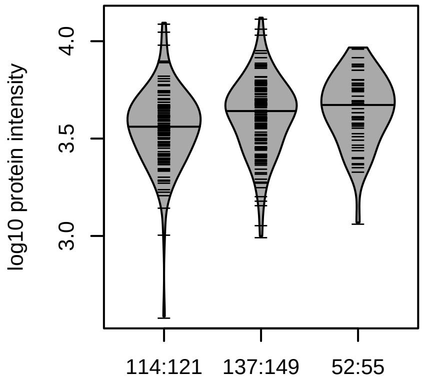


1:17315425:G:A_A
p = 1e−04, beta = 0.307, N = 303

**TVCAHEELLR pc3**
**P55001−2;P55001**

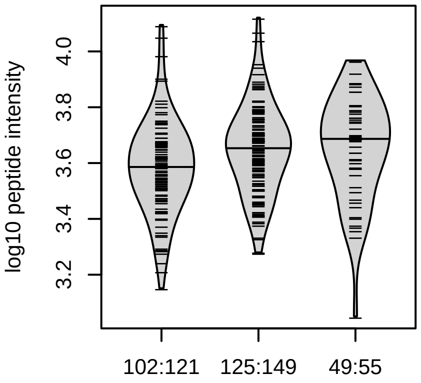


1:17315425:G:A_A
p = 0.0016, beta = 0.26, N = 276

Assay Target: MFAP2
Olink UniProt: P55001
deCODE rsID: rs4920605
Proxy rsID: rs4920605
deCODE: 1:16988925:!:TGCCC
Proxy SNP: 1:17315425:G:A
deCODE log10(p): 199.1
deCODE BETA: −0.25
*:*:−
1043:779:224

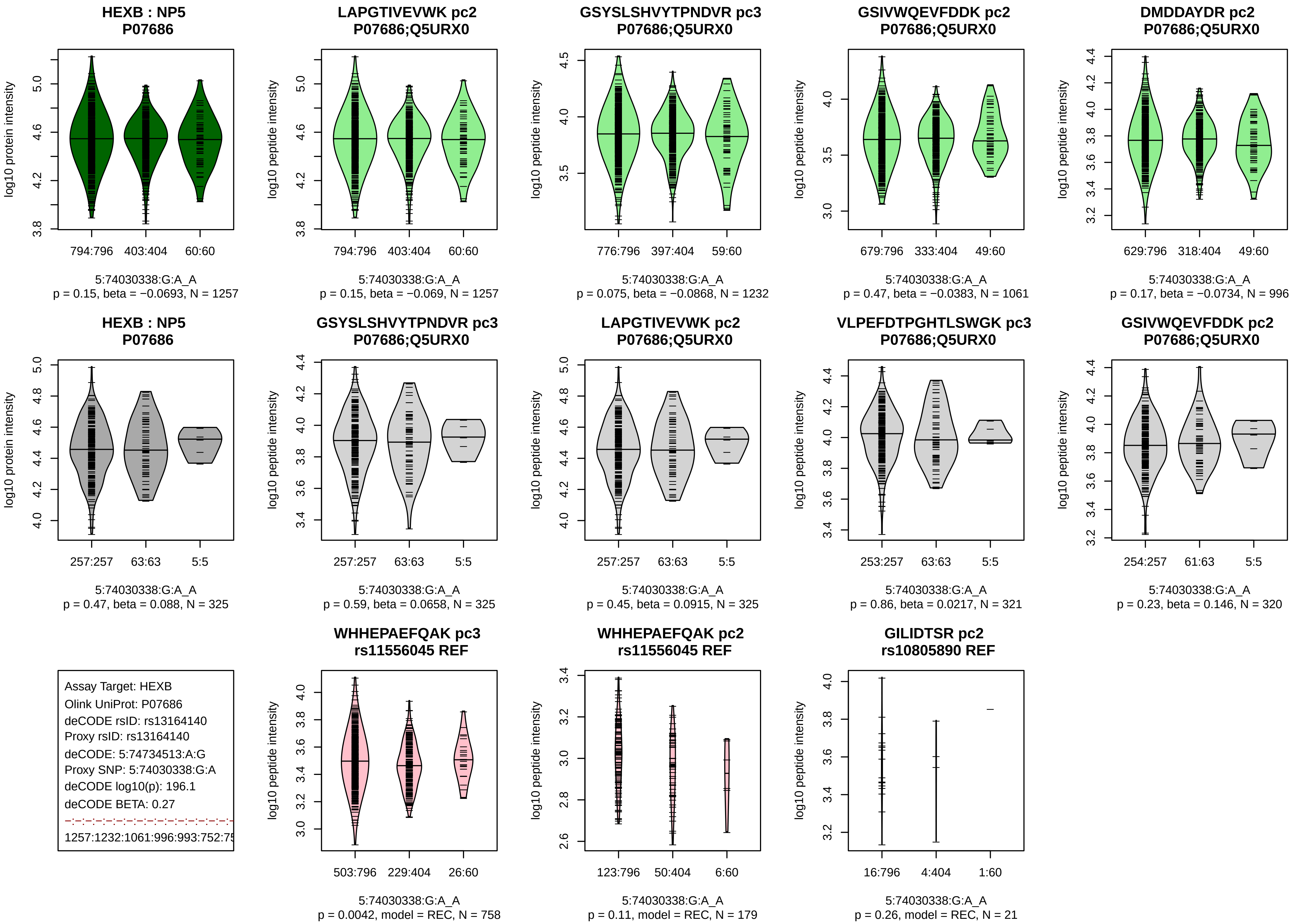
HEXB : NP5
P07686
log10 protein intensity
794:796
403:404
60:60
5:74030338:G:A_A
p = 0.15, beta = −0.0693, N = 1257
LAPGTIVEVWK pc2
P07686;Q5URX0
log10 peptide intensity
794:796
403:404
60:60
5:74030338:G:A_A
p = 0.15, beta = −0.069, N = 1257
GSYSLSHVYTPNDVR pc3
P07686;Q5URX0
log10 peptide intensity
776:796
397:404
59:60
5:74030338:G:A_A
p = 0.075, beta = −0.0868, N = 1232
GSIVWQEVFDDK pc2
P07686;Q5URX0
log10 peptide intensity
679:796
333:404
49:60
5:74030338:G:A_A
p = 0.47, beta = −0.0383, N = 1061
DMDDAYDR pc2
P07686;Q5URX0
log10 peptide intensity
629:796
318:404
49:60
5:74030338:G:A_A
p = 0.17, beta = −0.0734, N = 996
HEXB : NP5
P07686
log10 protein intensity
257:257
63:63
5:5
5:74030338:G:A_A
p = 0.47, beta = 0.088, N = 325
GSYSLSHVYTPNDVR pc3
P07686;Q5URX0
log10 peptide intensity
257:257
63:63
5:5
5:74030338:G:A_A
p = 0.59, beta = 0.0658, N = 325
LAPGTIVEVWK pc2
P07686;Q5URX0
log10 peptide intensity
257:257
63:63
5:5
5:74030338:G:A_A
p = 0.45, beta = 0.0915, N = 325
VLPEFDTPGHTLSWGK pc3
P07686;Q5URX0
log10 peptide intensity
253:257
63:63
5:5
5:74030338:G:A_A
p = 0.86, beta = 0.0217, N = 321
GSIVWQEVFDDK pc2
P07686;Q5URX0
log10 peptide intensity
254:257
61:63
5:5
5:74030338:G:A_A
p = 0.23, beta = 0.146, N = 320
Assay Target: HEXB
Olink UniProt: P07686
deCODE rsID: rs13164140
Proxy rsID: rs13164140
deCODE: 5:74734513:A:G
Proxy SNP: 5:74030338:G:A
deCODE log10(p): 196.1
deCODE BETA: 0.27
1257:1232:1061:996:993:752:75
WHHEPAEFQAK pc3
rs11556045 REF
log10 peptide intensity
503:796
229:404
26:60
5:74030338:G:A_A
p = 0.0042, model = REC, N = 758
WHHEPAEFQAK pc2
rs11556045 REF
log10 peptide intensity
123:796
50:404
6:60
5:74030338:G:A_A
p = 0.11, model = REC, N = 179
GILIDTSR pc2
rs10805890 REF
log10 peptide intensity
16:796
4:404
1:60
5:74030338:G:A_A
p = 0.26, model = REC, N = 21

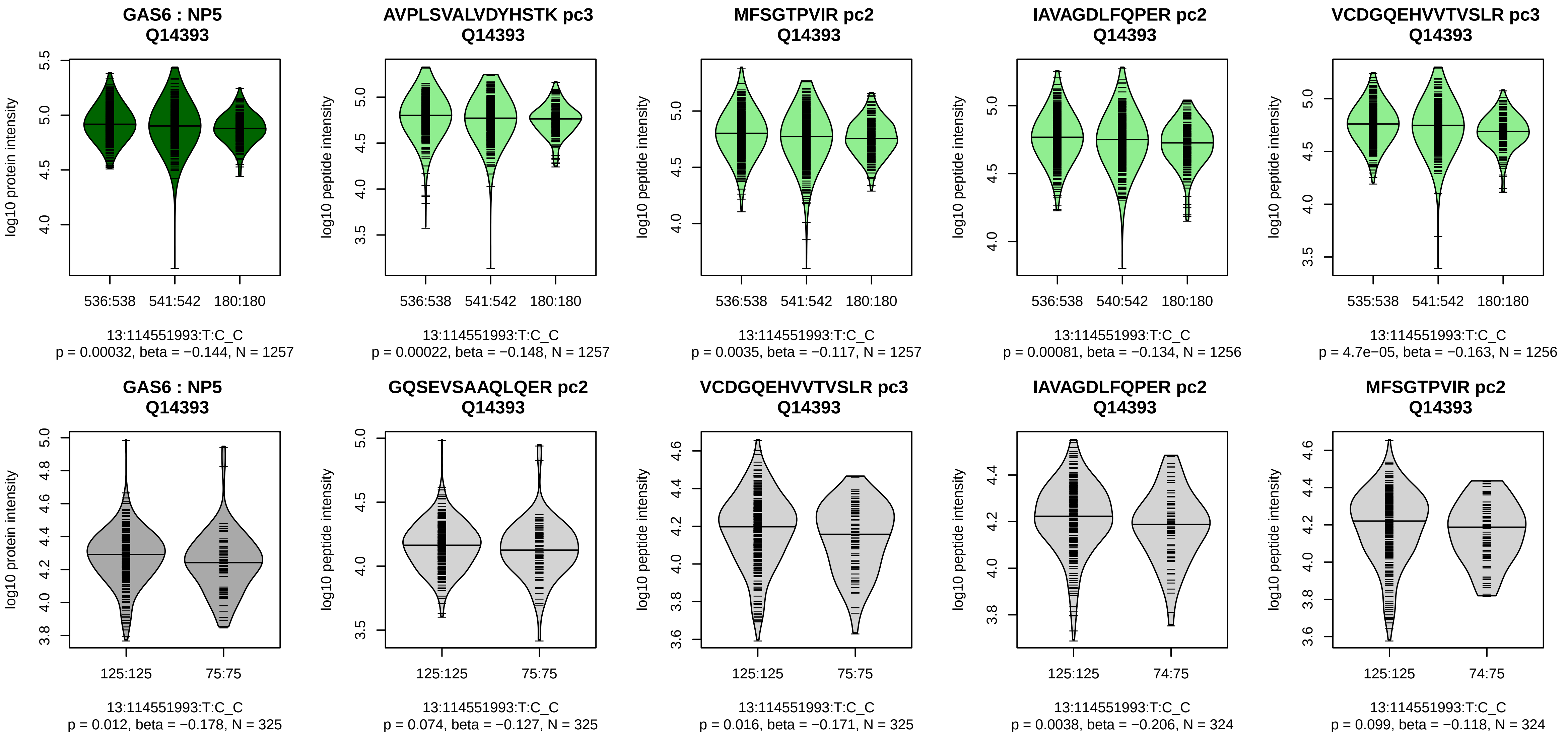


Assay Target: GAS6
Olink UniProt: Q14393
deCODE rsID: rs6602909
Proxy rsID: rs6602909
deCODE: 13:113849020:C:T
Proxy SNP: 13:114551993:T:C
deCODE log10(p): 194.7
deCODE BETA: −0.26
*:*:*:*:*:–:*:*:*:*:*:–:–:–:–:–:–:–:*:–:–
1257:1257:1256:1256:1256:125

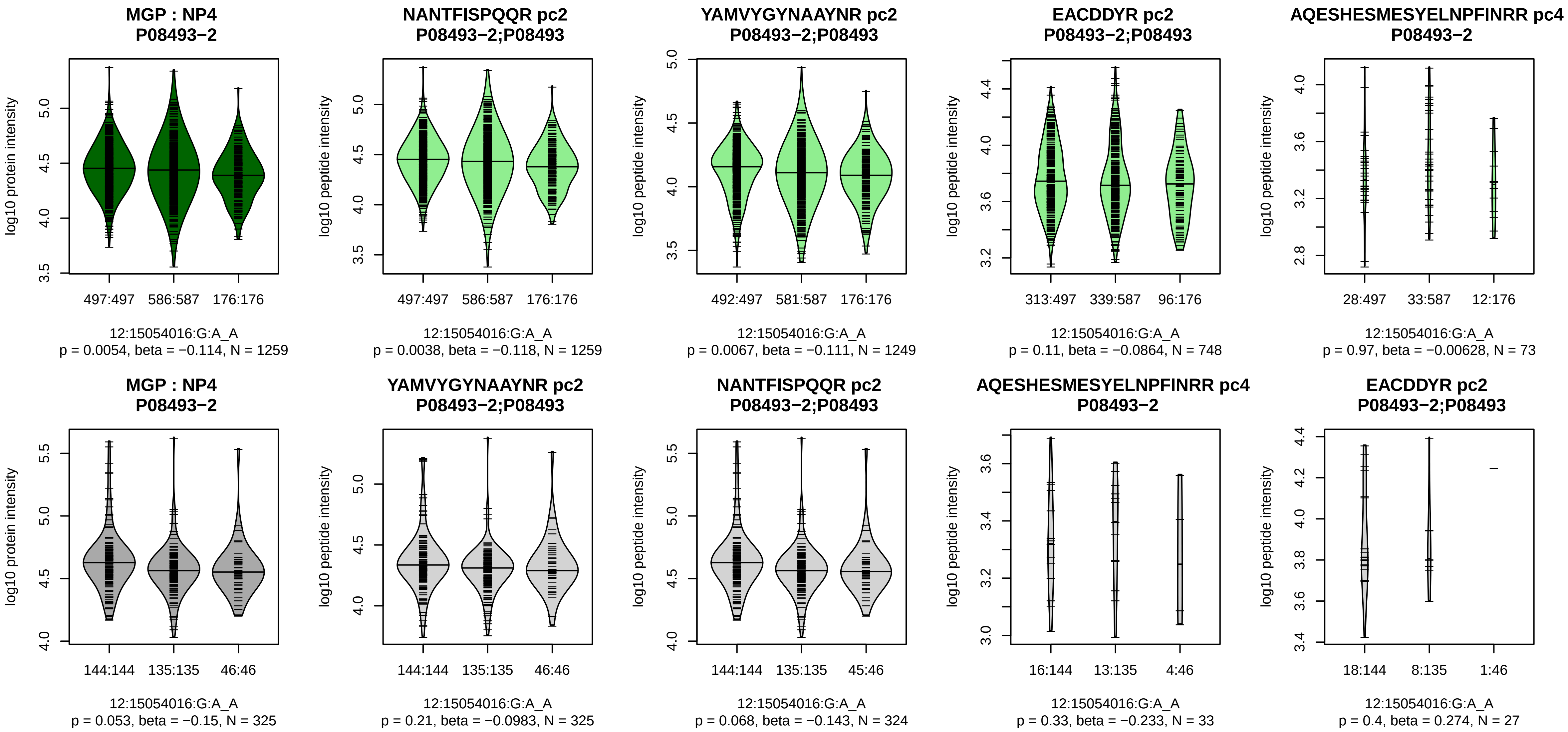


Assay Target: MGP
Olink UniProt: P08493
deCODE rsID: rs7294636
Proxy rsID: rs7294636
deCODE: 12:14901082:A:G
Proxy SNP: 12:15054016:G:A
deCODE log10(p): 192.8
deCODE BETA: −0.25
*:*:−:−:NA:NA
1259:1249:748:73:3:2

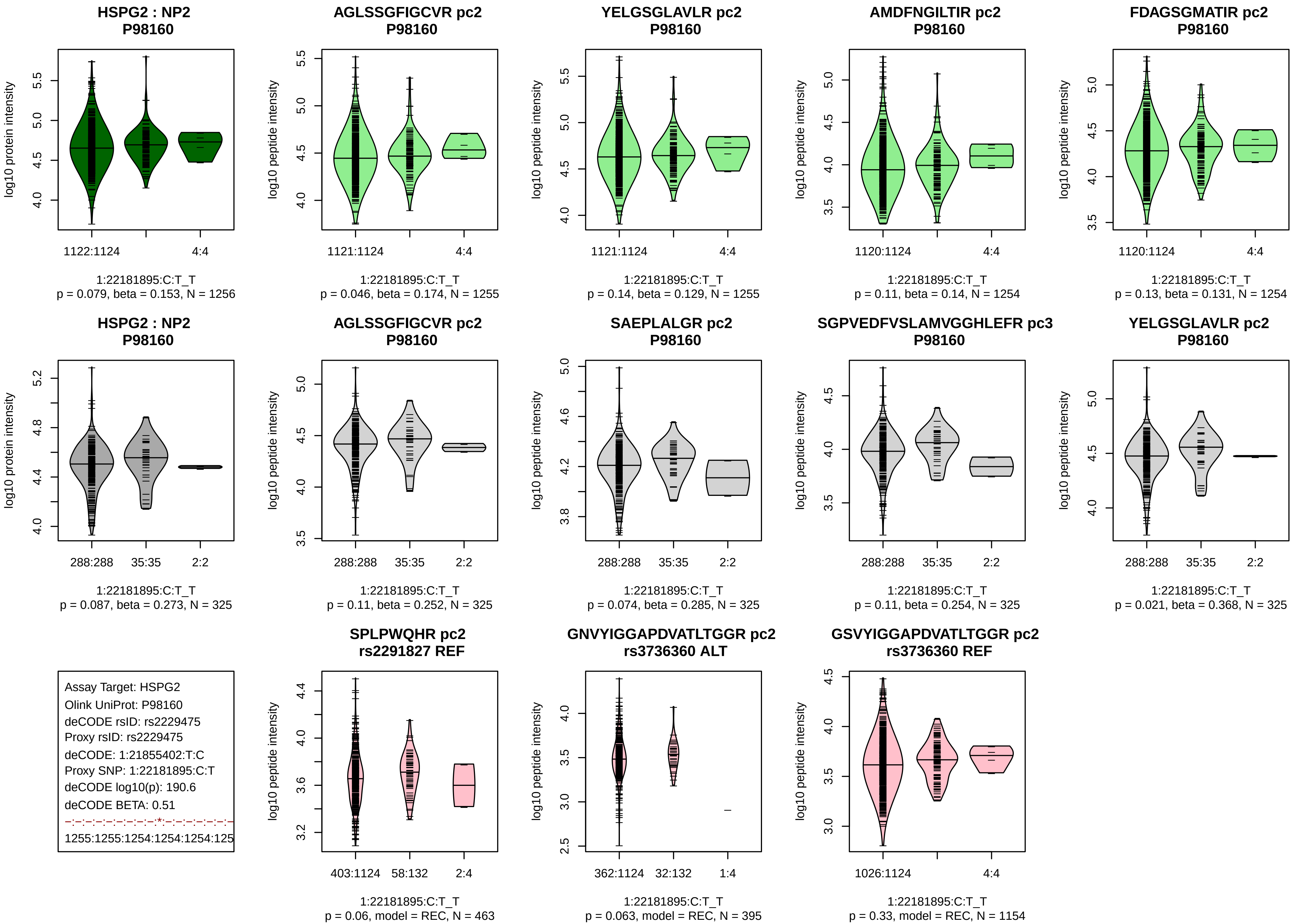

HSPG2 : NP2
P98160
log10 protein intensity
1122:1124
4:4
1:22181895:C:T_T
p = 0.079, beta = 0.153, N = 1256
AGLSSGFIGCVR pc2
P98160
log10 peptide intensity
1121:1124
4:4
1:22181895:C:T_T
p = 0.046, beta = 0.174, N = 1255
YELGSGLAVLR pc2
P98160
log10 peptide intensity
1121:1124
4:4
1:22181895:C:T_T
p = 0.14, beta = 0.129, N = 1255
AMDFNGILTIR pc2
P98160
log10 peptide intensity
1120:1124
4:4
1:22181895:C:T_T
p = 0.11, beta = 0.14, N = 1254
FDAGSGMATIR pc2
P98160
log10 peptide intensity
1120:1124
4:4
1:22181895:C:T_T
p = 0.13, beta = 0.131, N = 1254
HSPG2 : NP2
P98160
log10 protein intensity
288:288
35:35
2:2
1:22181895:C:T_T
p = 0.087, beta = 0.273, N = 325
AGLSSGFIGCVR pc2
P98160
log10 peptide intensity
288:288
35:35
2:2
1:22181895:C:T_T
p = 0.11, beta = 0.252, N = 325
SAEPLALGR pc2
P98160
log10 peptide intensity
288:288
35:35
2:2
1:22181895:C:T_T
p = 0.074, beta = 0.285, N = 325
SGPVEDFVSLAMVGGHLEFR pc3
P98160
log10 peptide intensity
288:288
35:35
2:2
1:22181895:C:T_T
p = 0.11, beta = 0.254, N = 325
YELGSGLAVLR pc2
P98160
log10 peptide intensity
288:288
35:35
2:2
1:22181895:C:T_T
p = 0.021, beta = 0.368, N = 325
Assay Target: HSPG2
Olink UniProt: P98160
deCODE rsID: rs2229475
Proxy rsID: rs2229475
deCODE: 1:21855402:T:C
Proxy SNP: 1:22181895:C:T
deCODE log10(p): 190.6
deCODE BETA: 0.51
-:-:-:-:-:-:-:-:-*:-:-:-:-:-:-:-
1255:1255:1254:1254:1254:125
SPLPWQHR pc2
rs2291827 REF
log10 peptide intensity
403:1124
58:132
2:4
1:22181895:C:T_T
p = 0.06, model = REC, N = 463
GNVYIGGAPDVATLTGGR pc2
rs3736360 ALT
log10 peptide intensity
362:1124
32:132
1:4
1:22181895:C:T_T
p = 0.063, model = REC, N = 395
GSVYIGGAPDVATLTGGR pc2
rs3736360 REF
log10 peptide intensity
1026:1124
4:4
1:22181895:C:T_T
p = 0.33, model = REC, N = 1154

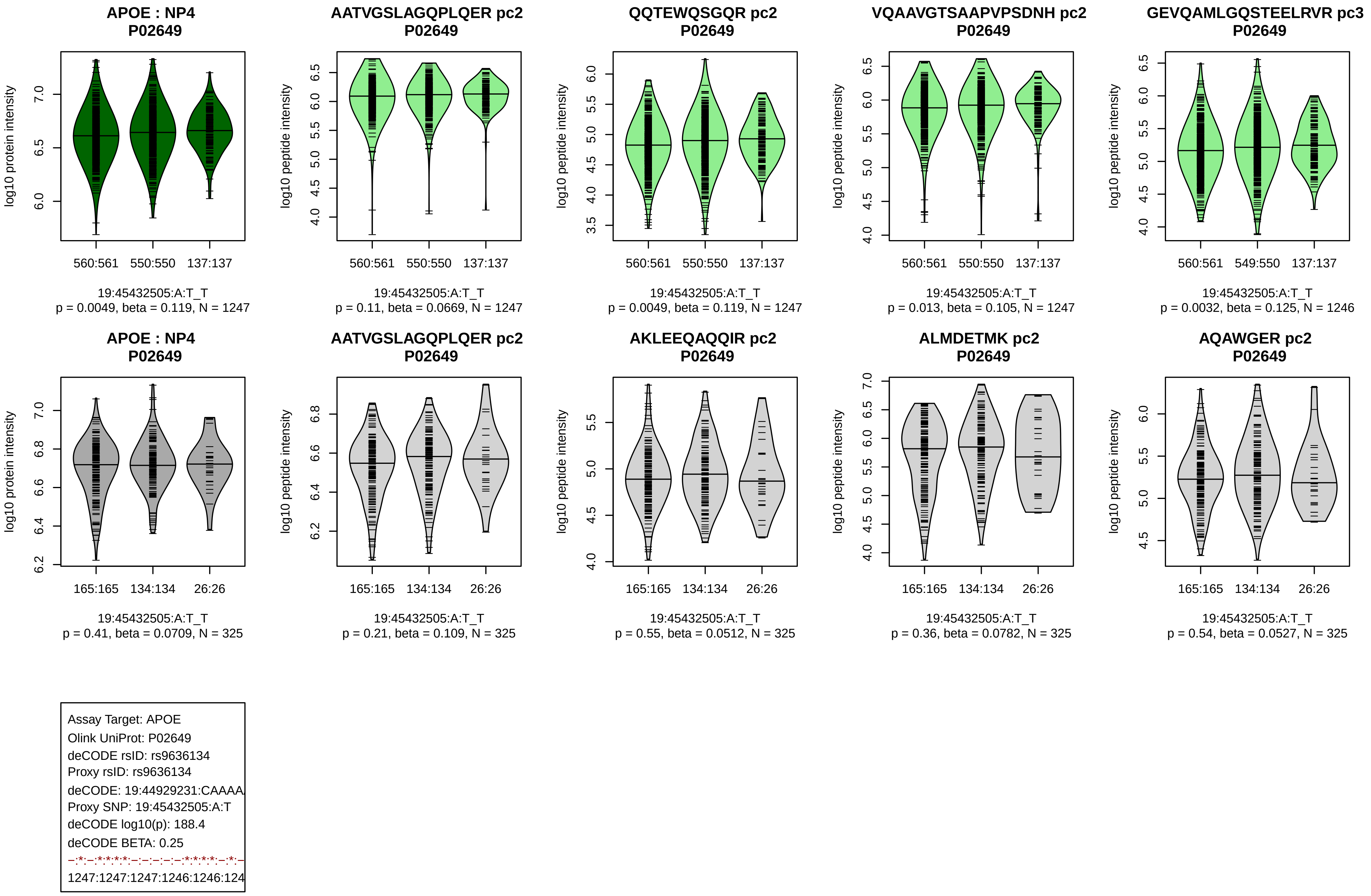
APOE : NP4
P02649
log10 protein intensity
560:561 550:550 137:137
19:45432505:A:T_T
p = 0.0049, beta = 0.119, N = 1247
AATVGSLAGQPLQER pc2
P02649
log10 peptide intensity
560:561 550:550 137:137
19:45432505:A:T_T
p = 0.11, beta = 0.0669, N = 1247
QQTEWQSGQR pc2
P02649
log10 peptide intensity
560:561 550:550 137:137
19:45432505:A:T_T
p = 0.0049, beta = 0.119, N = 1247
VQAAVGTSAAPVPSDNH pc2
P02649
log10 peptide intensity
560:561 550:550 137:137
19:45432505:A:T_T
p = 0.013, beta = 0.105, N = 1247
GEVQAMLGQSTEELRVR pc3
P02649
log10 peptide intensity
560:561 549:550 137:137
19:45432505:A:T_T
p = 0.0032, beta = 0.125, N = 1246
APOE : NP4
P02649
log10 protein intensity
165:165 134:134 26:26
19:45432505:A:T_T
p = 0.41, beta = 0.0709, N = 325
AATVGSLAGQPLQER pc2
P02649
log10 peptide intensity
165:165 134:134 26:26
19:45432505:A:T_T
p = 0.21, beta = 0.109, N = 325
AKLEEQAQQIR pc2
P02649
log10 peptide intensity
165:165 134:134 26:26
19:45432505:A:T_T
p = 0.55, beta = 0.0512, N = 325
ALMDETMK pc2
P02649
log10 peptide intensity
165:165 134:134 26:26
19:45432505:A:T_T
p = 0.36, beta = 0.0782, N = 325
AQAWGER pc2
P02649
log10 peptide intensity
165:165 134:134 26:26
19:45432505:A:T_T
p = 0.54, beta = 0.0527, N = 325
Assay Target: APOE
Olink UniProt: P02649
deCODE rsID: rs9636134
Proxy rsID: rs9636134
deCODE: 19:44929231:CAAAA
Proxy SNP: 19:45432505:A:T
deCODE log10(p): 188.4
deCODE BETA: 0.25
–:*:–:*:*:*:*:–:–:–:–:–:*:*:*:*:–:*:–
1247:1247:1247:1246:1246:124

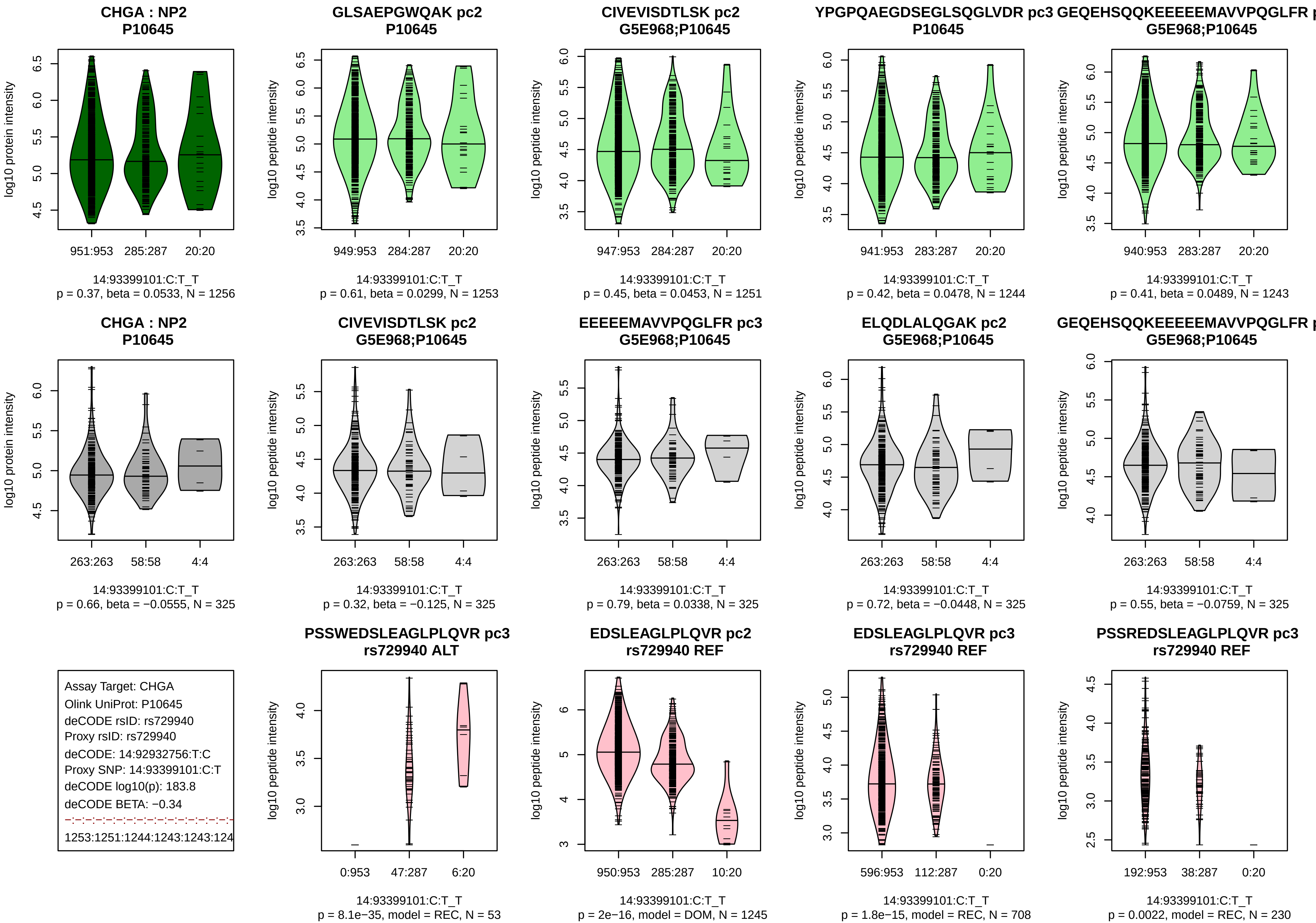
CHGA : NP2
P10645
log10 protein intensity
951:953 285:287 20:20
14:93399101:C:T_T
p = 0.37, beta = 0.0533, N = 1256
GLSAEPGWQAK pc2
P10645
log10 peptide intensity
949:953 284:287 20:20
14:93399101:C:T_T
p = 0.61, beta = 0.0299, N = 1253
CIVEVISDTLSK pc2
G5E968;P10645
log10 peptide intensity
947:953 284:287 20:20
14:93399101:C:T_T
p = 0.45, beta = 0.0453, N = 1251
YPGPQAEGDSEGLSQGLVDR pc3
P10645
log10 peptide intensity
941:953 283:287 20:20
14:93399101:C:T_T
p = 0.42, beta = 0.0478, N = 1244
GEQEHSQQKEEEEEMAVVPQGLFR p
G5E968;P10645
log10 peptide intensity
940:953 283:287 20:20
14:93399101:C:T_T
p = 0.41, beta = 0.0489, N = 1243
CHGA : NP2
P10645
log10 protein intensity
263:263 58:58 4:4
14:93399101:C:T_T
p = 0.66, beta = −0.0555, N = 325
CIVEVISDTLSK pc2
G5E968;P10645
log10 peptide intensity
263:263 58:58 4:4
14:93399101:C:T_T
p = 0.32, beta = −0.125, N = 325
EEEEEMAVVPQGLFR pc3
G5E968;P10645
log10 peptide intensity
263:263 58:58 4:4
14:93399101:C:T_T
p = 0.79, beta = 0.0338, N = 325
ELQDLALQGAK pc2
G5E968;P10645
log10 peptide intensity
263:263 58:58 4:4
14:93399101:C:T_T
p = 0.72, beta = −0.0448, N = 325
GEQEHSQQKEEEEEMAVVPQGLFR p
G5E968;P10645
log10 peptide intensity
263:263 58:58 4:4
14:93399101:C:T_T
p = 0.55, beta = −0.0759, N = 325
Assay Target: CHGA
Olink UniProt: P10645
deCODE rsID: rs729940
Proxy rsID: rs729940
deCODE: 14:92932756:T:C
Proxy SNP: 14:93399101:C:T
deCODE log10(p): 183.8
deCODE BETA: −0.34
1253:1251:1244:1243:1243:124
PSSWEDSLEAGLPLQVR pc3
rs729940 ALT
log10 peptide intensity
0:953 47:287 6:20
14:93399101:C:T_T
p = 8.1e−35, model = REC, N = 53
EDSLEAGLPLQVR pc2
rs729940 REF
log10 peptide intensity
950:953 285:287 10:20
14:93399101:C:T_T
p = 2e−16, model = DOM, N = 1245
EDSLEAGLPLQVR pc3
rs729940 REF
log10 peptide intensity
596:953 112:287 0:20
14:93399101:C:T_T
p = 1.8e−15, model = REC, N = 708
PSSREDSLEAGLPLQVR pc3
rs729940 REF
log10 peptide intensity
192:953 38:287 0:20
14:93399101:C:T_T
p = 0.0022, model = REC, N = 230

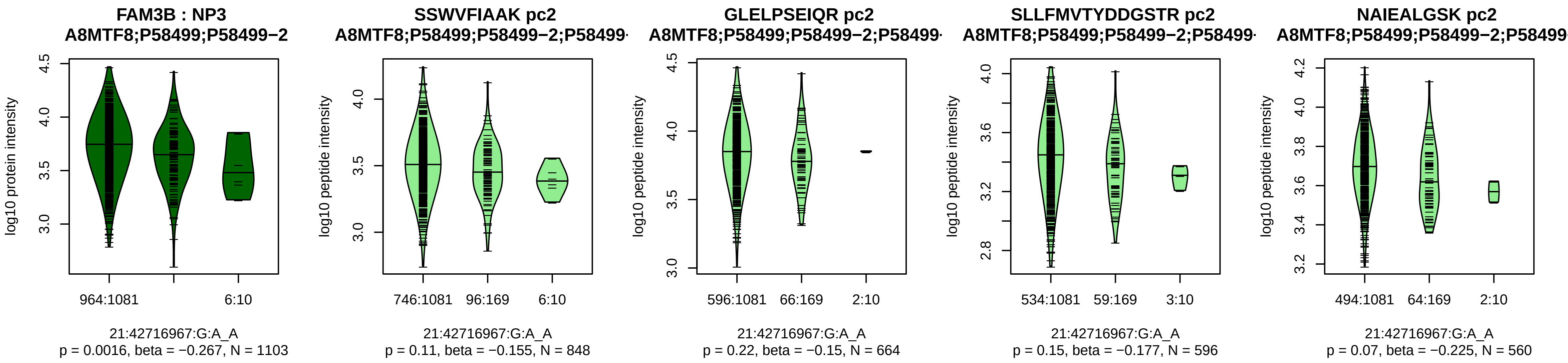


Assay Target: FAM3B
Olink UniProt: P58499
deCODE rsID: rs60265870
Proxy rsID: rs60265870
deCODE: 21:41345040:A:G
Proxy SNP: 21:42716967:G:A
deCODE log10(p): 181.2
deCODE BETA: −0.55
−:−:−:−:*:−
848:664:596:560:520:92

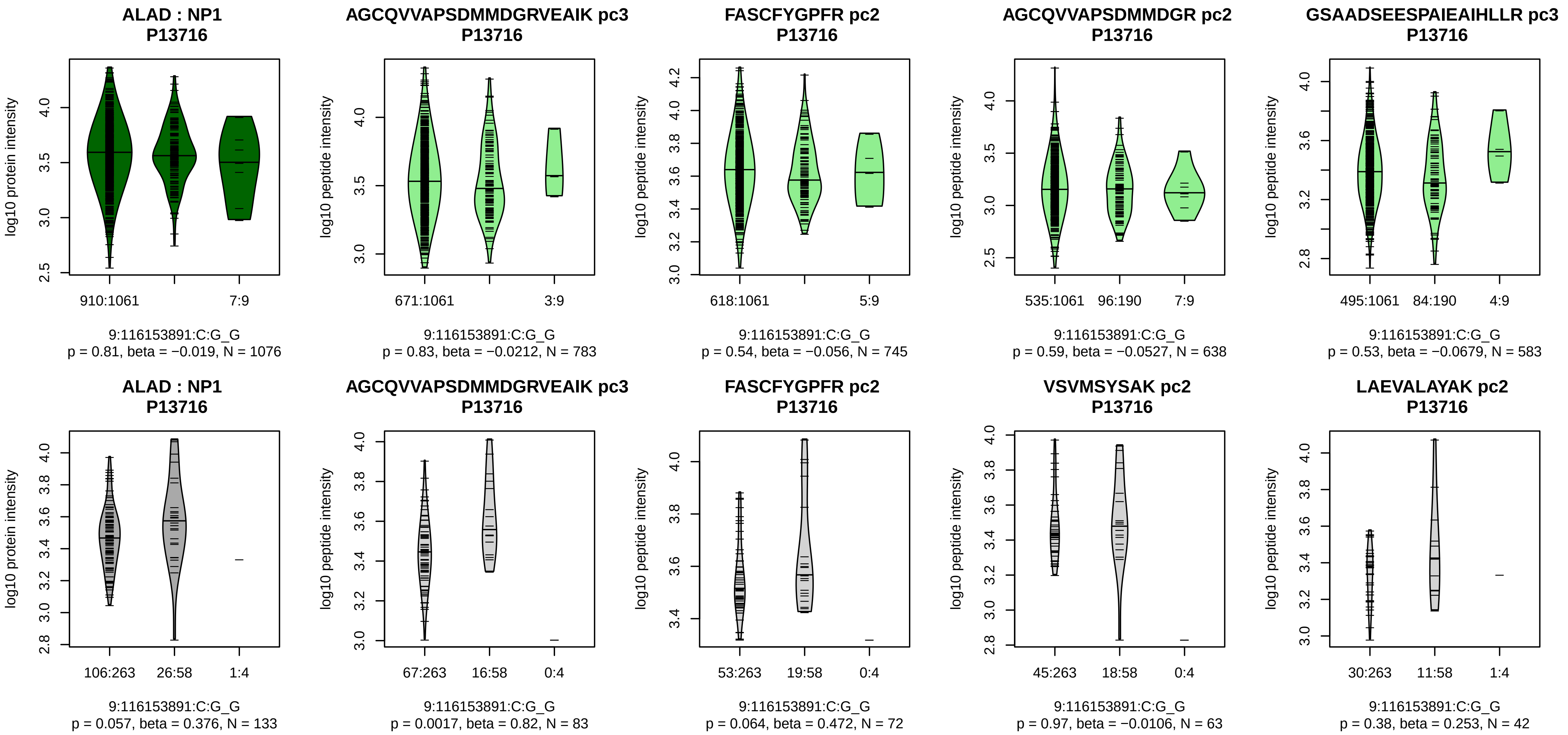


Assay Target: ALAD
Olink UniProt: P13716
deCODE rsID: rs1800435
Proxy rsID: rs1800435
deCODE: 9:113391611:G:C
Proxy SNP: 9:116153891:C:G
deCODE log10(p): 178.9
deCODE BETA: −0.42
−:−:−:−:−:−:−:−:−:−:−
783:745:638:583:535:512:430:4

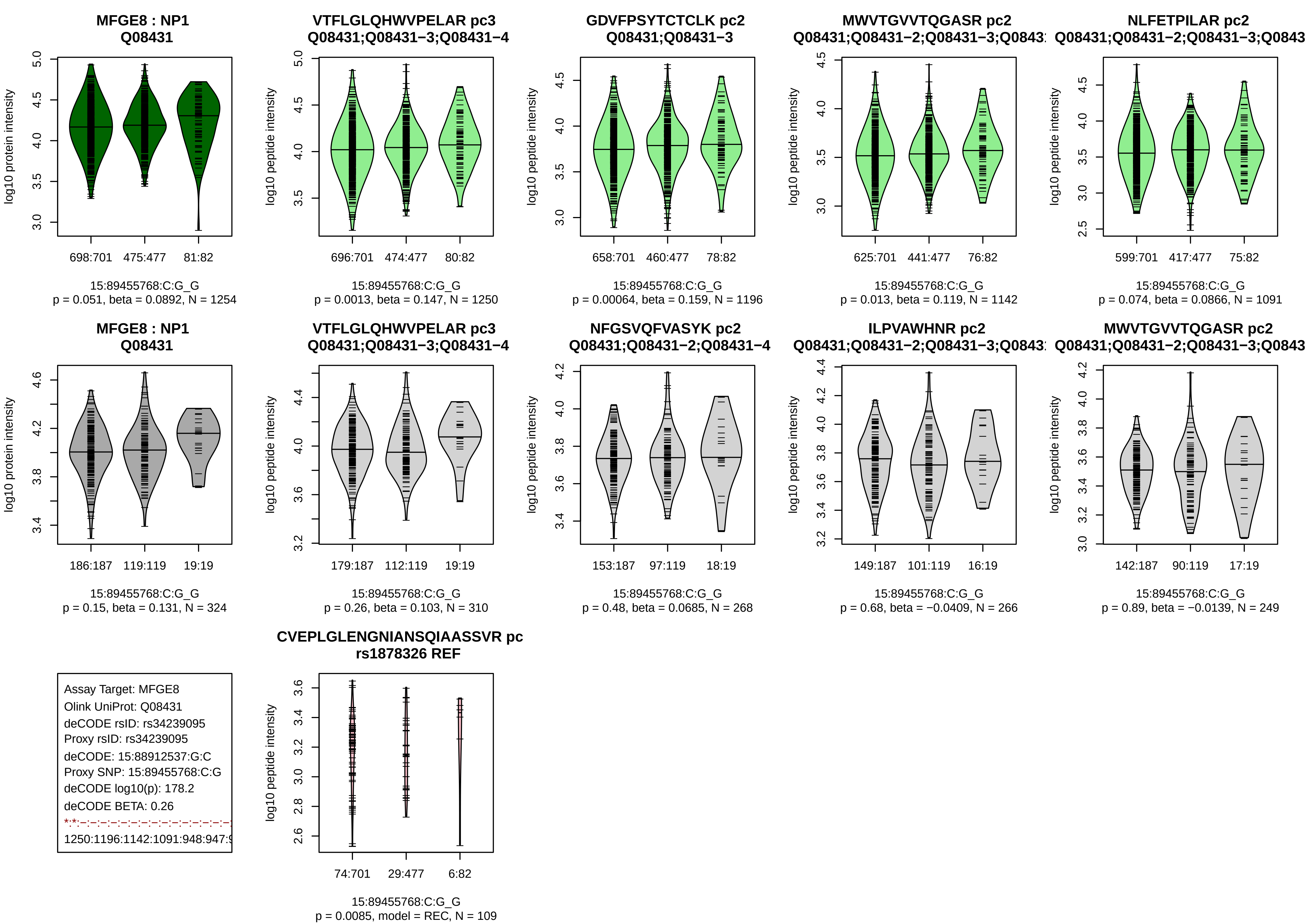
MFGE8 : NP1
Q08431
log10 protein intensity
698:701
475:477
81:82
15:89455768:C:G_G
p = 0.051, beta = 0.0892, N = 1254
VTFLGLQHWVPELAR pc3
Q08431;Q08431−3;Q08431−4
log10 peptide intensity
696:701
474:477
80:82
15:89455768:C:G_G
p = 0.0013, beta = 0.147, N = 1250
GDVFPSYTCTCLK pc2
Q08431;Q08431−3
658:701
460:477
78:82
15:89455768:C:G_G
p = 0.00064, beta = 0.159, N = 1196
MWVTGVVTQGASR pc2
Q08431;Q08431−2;Q08431−3;Q0843
625:701
441:477
76:82
15:89455768:C:G_G
p = 0.013, beta = 0.119, N = 1142
NLFETPILAR pc2
Q08431;Q08431−2;Q08431−3;Q0843
599:701
417:477
75:82
15:89455768:C:G_G
p = 0.074, beta = 0.0866, N = 1091
MFGE8 : NP1
Q08431
186:187
119:119
19:19
15:89455768:C:G_G
p = 0.15, beta = 0.131, N = 324
VTFLGLQHWVPELAR pc3
Q08431;Q08431−3;Q08431−4
179:187
112:119
19:19
15:89455768:C:G_G
p = 0.26, beta = 0.103, N = 310
NFGSVQFVASYK pc2
Q08431;Q08431−2;Q08431−4
153:187
97:119
18:19
15:89455768:C:G_G
p = 0.48, beta = 0.0685, N = 268
ILPVAWHNR pc2
Q08431;Q08431−2;Q08431−3;Q0843
149:187
101:119
16:19
15:89455768:C:G_G
p = 0.68, beta = −0.0409, N = 266
MWVTGVVTQGASR pc2
Q08431;Q08431−2;Q08431−3;Q0843
142:187
90:119
17:19
15:89455768:C:G_G
p = 0.89, beta = −0.0139, N = 249
Assay Target: MFGE8
Olink UniProt: Q08431
deCODE rsID: rs34239095
Proxy rsID: rs34239095
deCODE: 15:88912537:G:C
Proxy SNP: 15:89455768:C:G
deCODE log10(p): 178.2
deCODE BETA: 0.26
1250:1196:1142:1091:948:947:
CVEPLGLENGNIANSQIAASSVR pc
rs1878326 REF
74:701
29:477
6:82
15:89455768:C:G_G
p = 0.0085, model = REC, N = 109

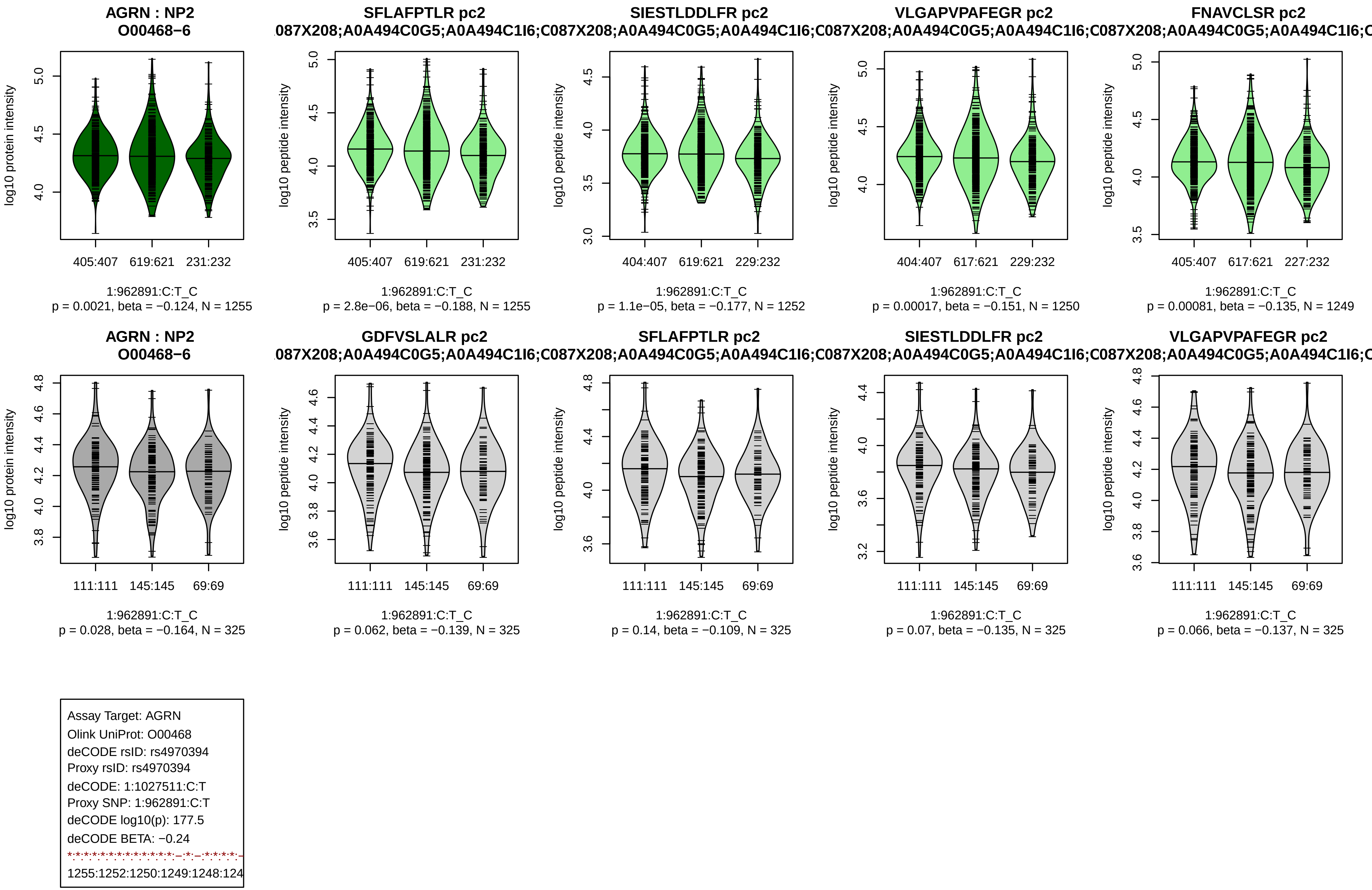
AGRN : NP2
O00468−6
log10 protein intensity
405:407 619:621 231:232
1:962891:C:T_C
p = 0.0021, beta = −0.124, N = 1255
SFLAFPTLR pc2
log10 peptide intensity
405:407 619:621 231:232
1:962891:C:T_C
p = 2.8e−06, beta = −0.188, N = 1255
SIESTLDDLFR pc2
404:407 619:621 229:232
1:962891:C:T_C
p = 1.1e−05, beta = −0.177, N = 1252
VLGAPVPAFEGR pc2
404:407 617:621 229:232
1:962891:C:T_C
p = 0.00017, beta = −0.151, N = 1250
FNAVCLSR pc2
405:407 617:621 227:232
1:962891:C:T_C
p = 0.00081, beta = −0.135, N = 1249
.087X208;A0A494C0G5;A0A494C1I6;C087X208;A0A494C0G5;A0A494C1I6;C087X208;A0A494C0G5;A0A494C1I6;C087X208;A0A494C0G5;A0A494C1I6;C
AGRN : NP2
O00468−6
111:111 145:145 69:69
1:962891:C:T_C
p = 0.028, beta = −0.164, N = 325
GDFVSLALR pc2
1:962891:C:T_C
p = 0.062, beta = −0.139, N = 325
SFLAFPTLR pc2
1:962891:C:T_C
p = 0.14, beta = −0.109, N = 325
SIESTLDDLFR pc2
1:962891:C:T_C
p = 0.07, beta = −0.135, N = 325
VLGAPVPAFEGR pc2
1:962891:C:T_C
p = 0.066, beta = −0.137, N = 325
Assay Target: AGRN
Olink UniProt: O00468
deCODE rsID: rs4970394
Proxy rsID: rs4970394
deCODE: 1:1027511:C:T
Proxy SNP: 1:962891:C:T
deCODE log10(p): 177.5
deCODE BETA: −0.24
1255:1252:1250:1249:1248:124

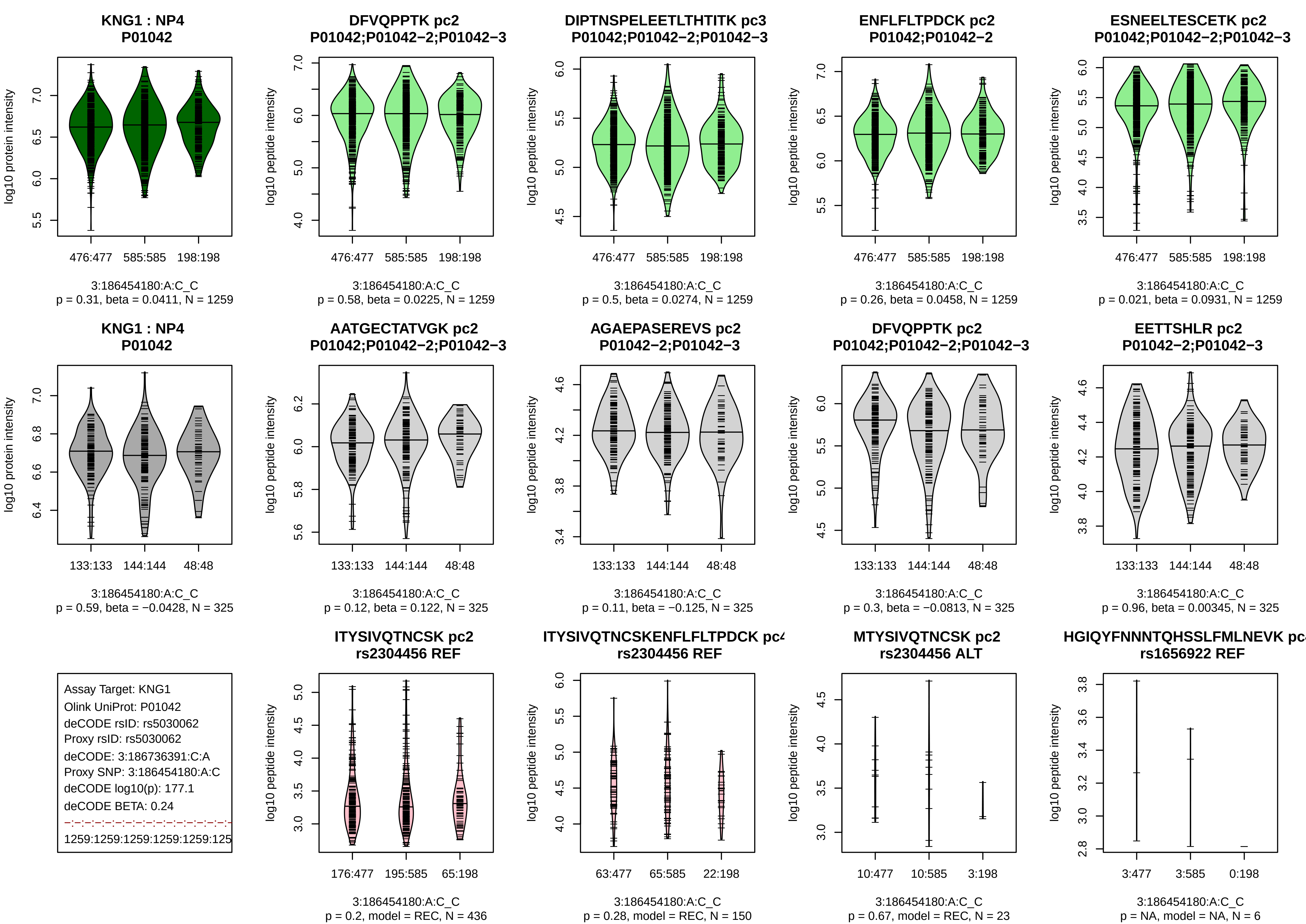
KNG1 : NP4
P01042
log10 protein intensity
476:477
585:585
198:198
3:186454180:A:C_C
p = 0.31, beta = 0.0411, N = 1259
DFVQPPTK pc2
P01042;P01042−2;P01042−3
log10 peptide intensity
476:477
585:585
198:198
3:186454180:A:C_C
p = 0.58, beta = 0.0225, N = 1259
DIPTNSPELEETLTHTITK pc3
P01042;P01042−2;P01042−3
log10 peptide intensity
476:477
585:585
198:198
3:186454180:A:C_C
p = 0.5, beta = 0.0274, N = 1259
ENFLFLTPDCK pc2
P01042;P01042−2
log10 peptide intensity
476:477
585:585
198:198
3:186454180:A:C_C
p = 0.26, beta = 0.0458, N = 1259
ESNEELTESCETK pc2
P01042;P01042−2;P01042−3
log10 peptide intensity
476:477
585:585
198:198
3:186454180:A:C_C
p = 0.021, beta = 0.0931, N = 1259
KNG1 : NP4
P01042
log10 protein intensity
133:133
144:144
48:48
3:186454180:A:C_C
p = 0.59, beta = −0.0428, N = 325
AATGECTATVGK pc2
P01042;P01042−2;P01042−3
log10 peptide intensity
133:133
144:144
48:48
3:186454180:A:C_C
p = 0.12, beta = 0.122, N = 325
AGAEPASEREVS pc2
P01042−2;P01042−3
log10 peptide intensity
133:133
144:144
48:48
3:186454180:A:C_C
p = 0.11, beta = −0.125, N = 325
DFVQPPTK pc2
P01042;P01042−2;P01042−3
log10 peptide intensity
133:133
144:144
48:48
3:186454180:A:C_C
p = 0.3, beta = −0.0813, N = 325
EETTSHLR pc2
P01042−2;P01042−3
log10 peptide intensity
133:133
144:144
48:48
3:186454180:A:C_C
p = 0.96, beta = 0.00345, N = 325
Assay Target: KNG1
Olink UniProt: P01042
deCODE rsID: rs5030062
Proxy rsID: rs5030062
deCODE: 3:186736391:C:A
Proxy SNP: 3:186454180:A:C
deCODE log10(p): 177.1
deCODE BETA: 0.24
1259:1259:1259:1259:1259:125
ITYSIVQTNCSK pc2
rs2304456 REF
log10 peptide intensity
176:477
195:585
65:198
3:186454180:A:C_C
p = 0.2, model = REC, N = 436
ITYSIVQTNCSKENFLFLTPDCK pc4
rs2304456 REF
log10 peptide intensity
63:477
65:585
22:198
3:186454180:A:C_C
p = 0.28, model = REC, N = 150
MTYSIVQTNCSK pc2
rs2304456 ALT
log10 peptide intensity
10:477
10:585
3:198
3:186454180:A:C_C
p = 0.67, model = REC, N = 23
HGIQYFNNNTQHSSLFMLNEVK pc
rs1656922 REF
log10 peptide intensity
3:477
3:585
0:198
3:186454180:A:C_C
p = NA, model = NA, N = 6

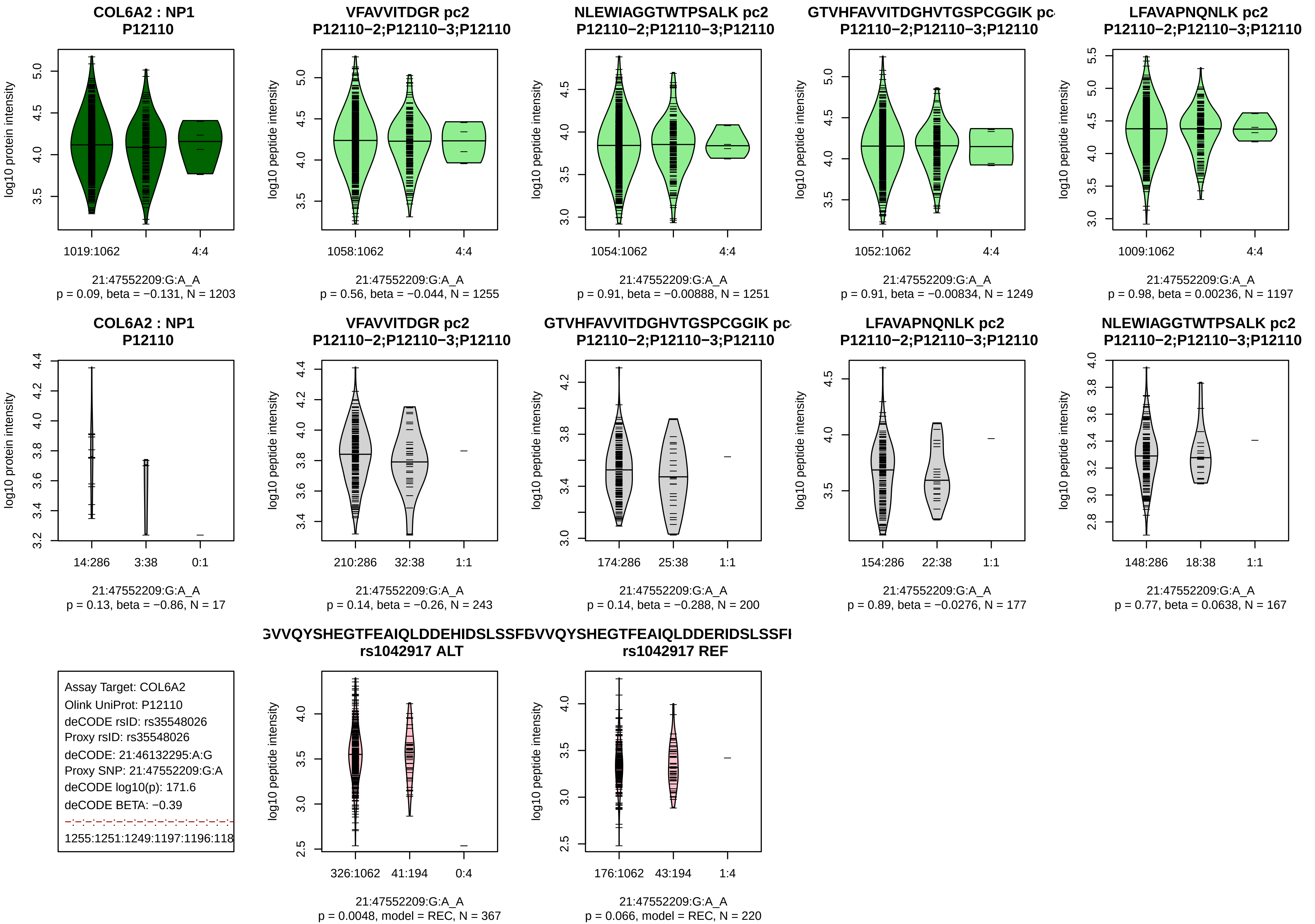
COL6A2 : NP1
P12110
log10 protein intensity
1019:1062
4:4
21:47552209:G:A_A
p = 0.09, beta = −0.131, N = 1203
VFAVVITDGR pc2
P12110−2;P12110−3;P12110
log10 peptide intensity
1058:1062
4:4
21:47552209:G:A_A
p = 0.56, beta = −0.044, N = 1255
NLEWIAGGTWTPSALK pc2
P12110−2;P12110−3;P12110
log10 peptide intensity
1054:1062
4:4
21:47552209:G:A_A
p = 0.91, beta = −0.00888, N = 1251
GTVHFAVVITDGHVTGSPCGGIK pc
P12110−2;P12110−3;P12110
log10 peptide intensity
1052:1062
4:4
21:47552209:G:A_A
p = 0.91, beta = −0.00834, N = 1249
LFAVAPNQNLK pc2
P12110−2;P12110−3;P12110
log10 peptide intensity
1009:1062
4:4
21:47552209:G:A_A
p = 0.98, beta = 0.00236, N = 1197
COL6A2 : NP1
P12110
log10 protein intensity
14:286
3:38
0:1
21:47552209:G:A_A
p = 0.13, beta = −0.86, N = 17
VFAVVITDGR pc2
P12110−2;P12110−3;P12110
log10 peptide intensity
210:286
32:38
1:1
21:47552209:G:A_A
p = 0.14, beta = −0.26, N = 243
GTVHFAVVITDGHVTGSPCGGIK pc
P12110−2;P12110−3;P12110
log10 peptide intensity
174:286
25:38
1:1
21:47552209:G:A_A
p = 0.14, beta = −0.288, N = 200
LFAVAPNQNLK pc2
P12110−2;P12110−3;P12110
log10 peptide intensity
154:286
22:38
1:1
21:47552209:G:A_A
p = 0.89, beta = −0.0276, N = 177
NLEWIAGGTWTPSALK pc2
P12110−2;P12110−3;P12110
log10 peptide intensity
148:286
18:38
1:1
21:47552209:G:A_A
p = 0.77, beta = 0.0638, N = 167
Assay Target: COL6A2
Olink UniProt: P12110
deCODE rsID: rs35548026
Proxy rsID: rs35548026
deCODE: 21:46132295:A:G
Proxy SNP: 21:47552209:G:A
deCODE log10(p): 171.6
deCODE BETA: −0.39
1255:1251:1249:1197:1196:118
GVVQYSHEGTFEAIQLDDEHIDSLSSF
rs1042917 ALT
log10 peptide intensity
326:1062
41:194
0:4
21:47552209:G:A_A
p = 0.0048, model = REC, N = 367
GVVQYSHEGTFEAIQLDDERIDSLSSFI
rs1042917 REF
log10 peptide intensity
176:1062
43:194
1:4
21:47552209:G:A_A
p = 0.066, model = REC, N = 220

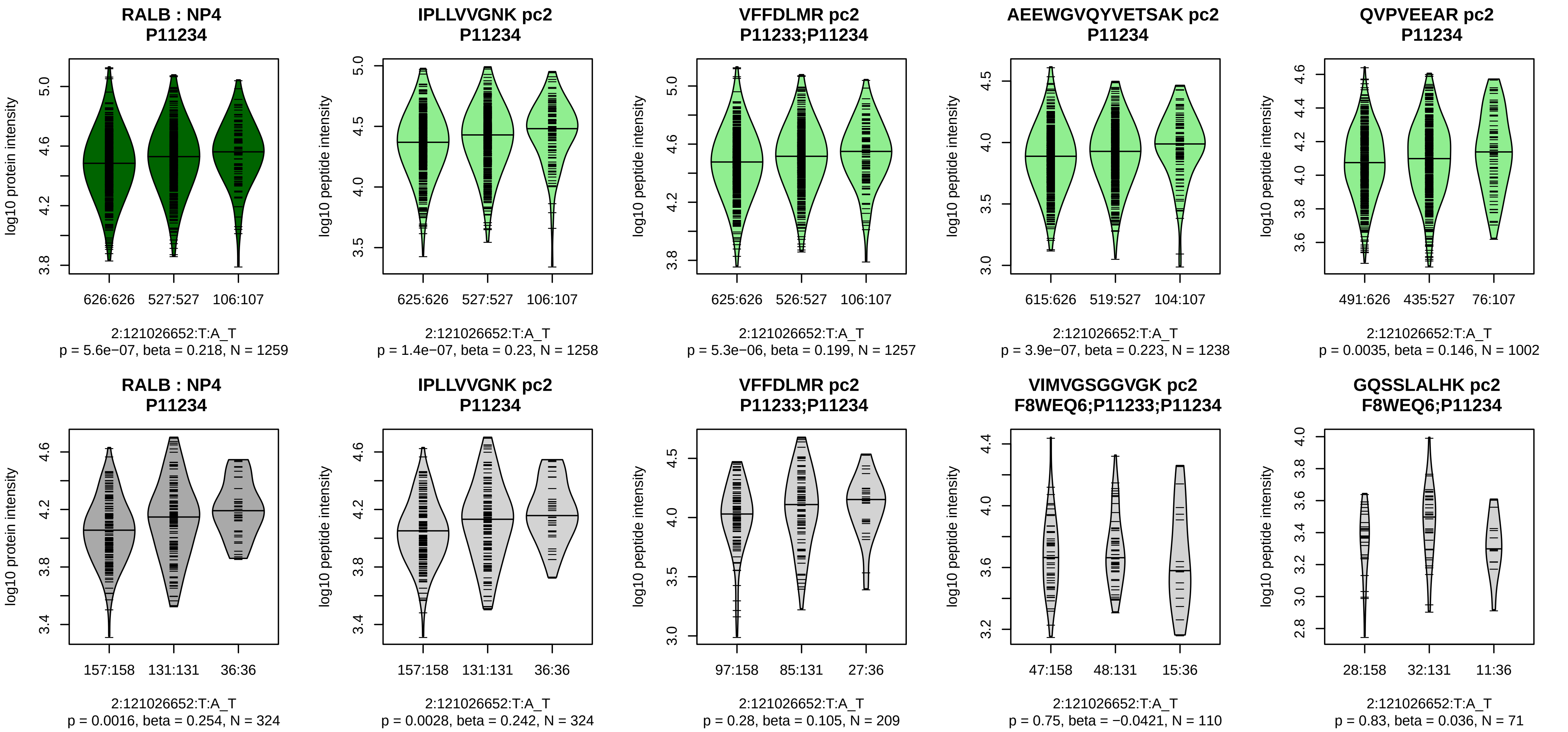


Assay Target: RALB
Olink UniProt: P11234
deCODE rsID: rs11448973
Proxy rsID: rs7593660
deCODE: 2:120268935:!:CAAT
Proxy SNP: 2:121026652:T:A
deCODE log10(p): 168.9
deCODE BETA: 0.24
*:*:*:*:*:–:–:–:–:–
1258:1257:1238:1002:831:464:1

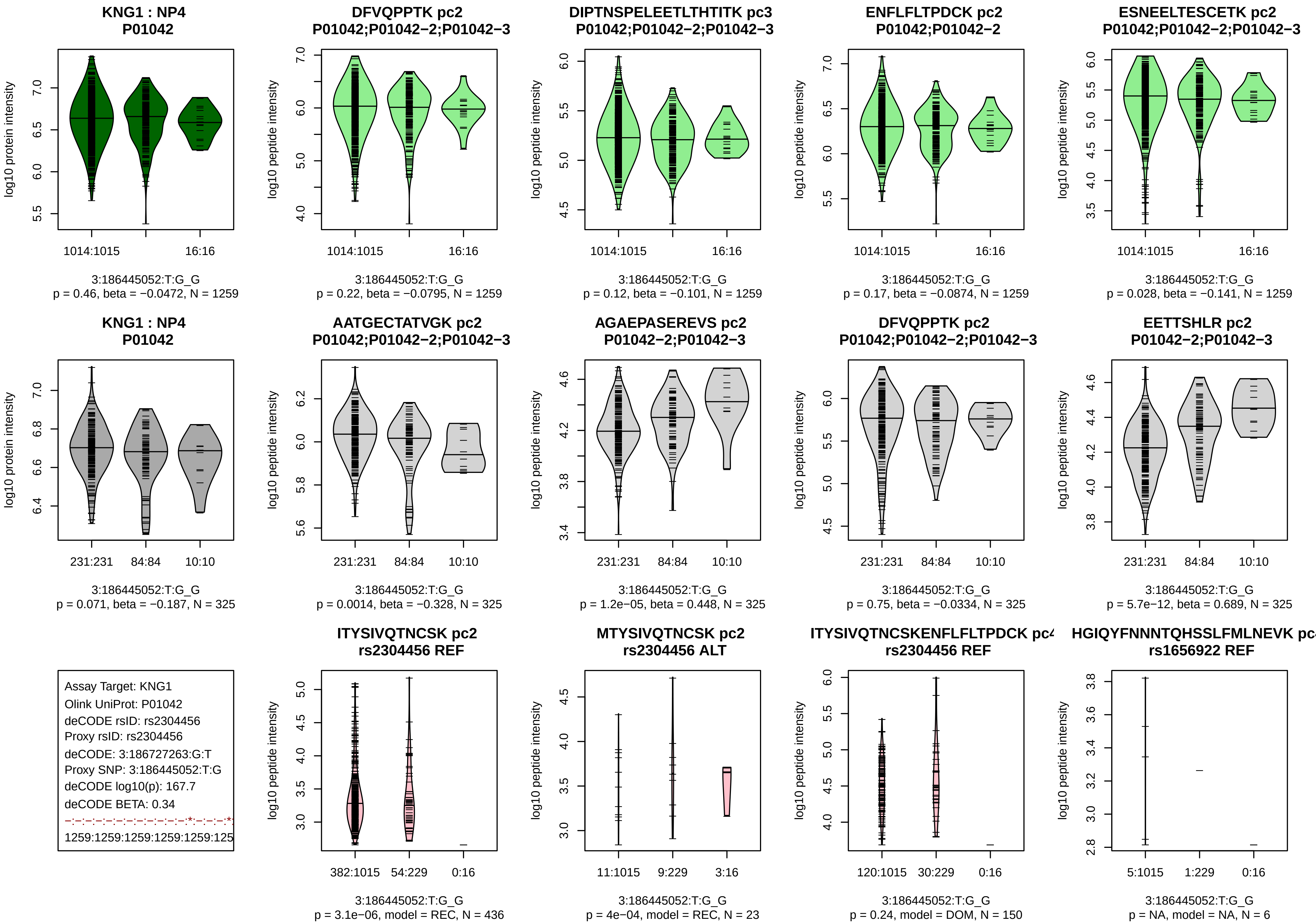

KNG1 : NP4
P01042
log10 protein intensity
1014:1015
16:16
3:186445052:T:G_G
p = 0.46, beta = −0.0472, N = 1259
DFVQPPTK pc2
P01042;P01042−2;P01042−3
log10 peptide intensity
1014:1015
16:16
3:186445052:T:G_G
p = 0.22, beta = −0.0795, N = 1259
DIPTNSPELEETLTHTITK pc3
P01042;P01042−2;P01042−3
log10 peptide intensity
1014:1015
16:16
3:186445052:T:G_G
p = 0.12, beta = −0.101, N = 1259
ENFLFLTPDCK pc2
P01042;P01042−2
log10 peptide intensity
1014:1015
16:16
3:186445052:T:G_G
p = 0.17, beta = −0.0874, N = 1259
ESNEELTESCETK pc2
P01042;P01042−2;P01042−3
log10 peptide intensity
1014:1015
16:16
3:186445052:T:G_G
p = 0.028, beta = −0.141, N = 1259
KNG1 : NP4
P01042
log10 protein intensity
231:231
84:84
10:10
3:186445052:T:G_G
p = 0.071, beta = −0.187, N = 325
AATGECTATVGK pc2
P01042;P01042−2;P01042−3
log10 peptide intensity
231:231
84:84
10:10
3:186445052:T:G_G
p = 0.0014, beta = −0.328, N = 325
AGAEPASEREVS pc2
P01042−2;P01042−3
log10 peptide intensity
231:231
84:84
10:10
3:186445052:T:G_G
p = 1.2e−05, beta = 0.448, N = 325
DFVQPPTK pc2
P01042;P01042−2;P01042−3
log10 peptide intensity
231:231
84:84
10:10
3:186445052:T:G_G
p = 0.75, beta = −0.0334, N = 325
EETTSHLR pc2
P01042−2;P01042−3
log10 peptide intensity
231:231
84:84
10:10
3:186445052:T:G_G
p = 5.7e−12, beta = 0.689, N = 325
Assay Target: KNG1
Olink UniProt: P01042
deCODE rsID: rs2304456
Proxy rsID: rs2304456
deCODE: 3:186727263:G:T
Proxy SNP: 3:186445052:T:G
deCODE log10(p): 167.7
deCODE BETA: 0.34
−:−:−:−:−:−:−:−:−:−:−:*:−:−:−:*
1259:1259:1259:1259:1259:125
ITYSIVQTNCSK pc2
rs2304456 REF
log10 peptide intensity
382:1015
54:229
0:16
3:186445052:T:G_G
p = 3.1e−06, model = REC, N = 436
MTYSIVQTNCSK pc2
rs2304456 ALT
log10 peptide intensity
11:1015
9:229
3:16
3:186445052:T:G_G
p = 4e−04, model = REC, N = 23
ITYSIVQTNCSKENFLFLTPDCK pc
rs2304456 REF
log10 peptide intensity
120:1015
30:229
0:16
3:186445052:T:G_G
p = 0.24, model = DOM, N = 150
HGIQYFNNNTQHSSLFMLNEVK pc
rs1656922 REF
log10 peptide intensity
5:1015
1:229
0:16
3:186445052:T:G_G
p = NA, model = NA, N = 6

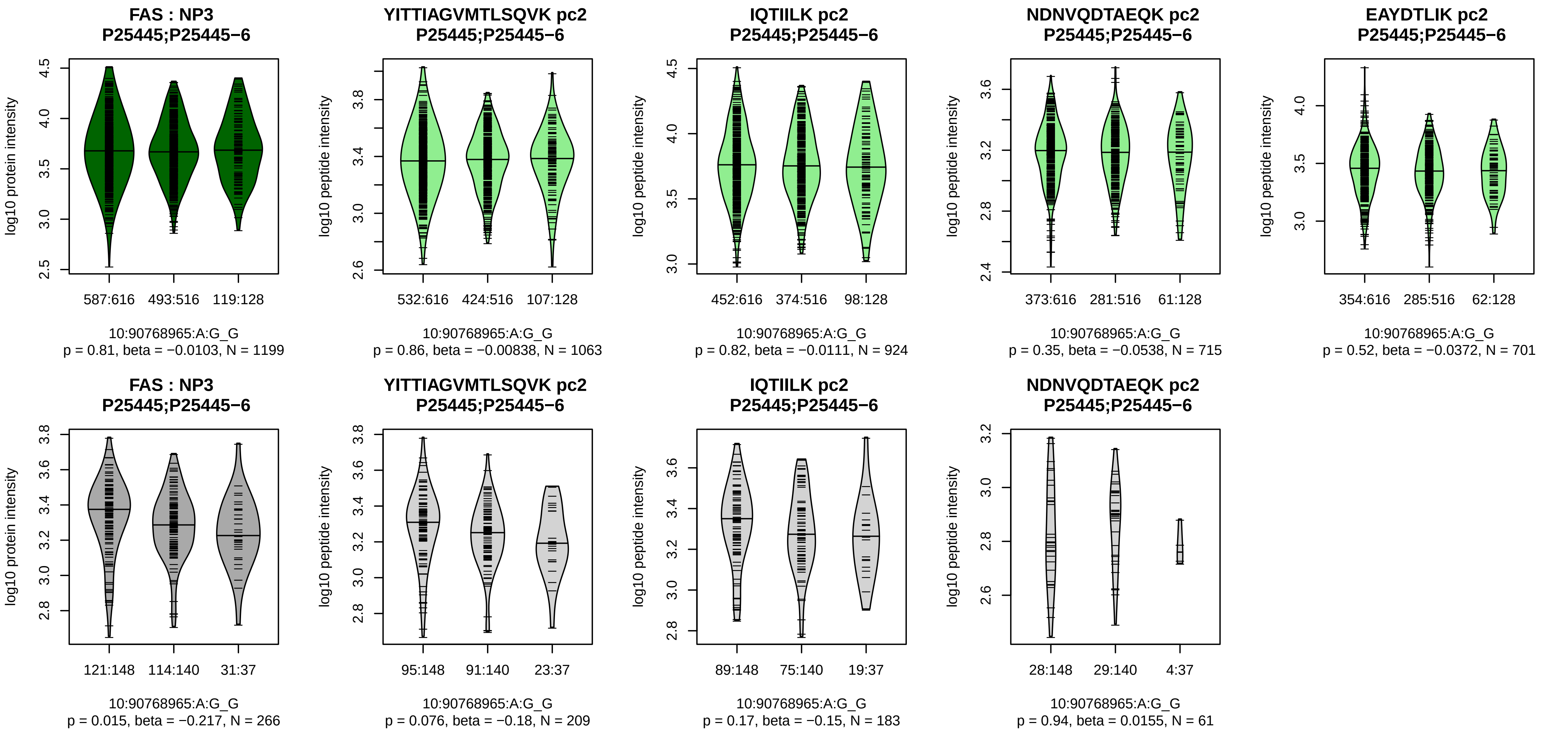


Assay Target: FAS
Olink UniProt: P25445
deCODE rsID: rs7911226
Proxy rsID: rs7911226
deCODE: 10:89009208:G:A
Proxy SNP: 10:90768965:A:G
deCODE log10(p): 166.6
deCODE BETA: −0.25
−:−:−:−
1063:924:715:701

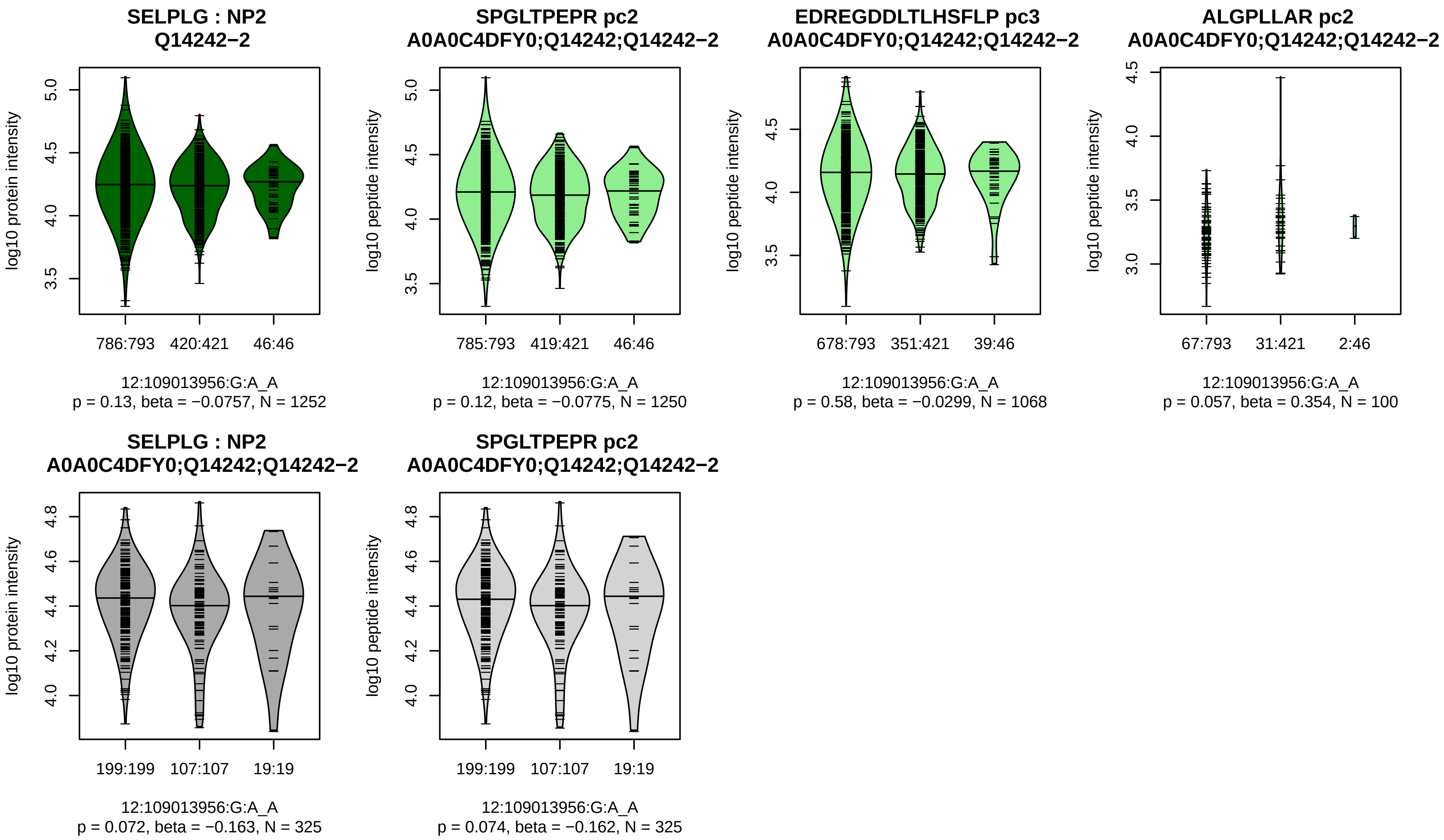


Assay Target: SELPLG
Olink UniProt: Q14242
deCODE rsID: rs73191242
Proxy rsID: rs73191242
deCODE: 12:108620180:A:G
Proxy SNP: 12:109013956:G:A
deCODE log10(p): 166.5
deCODE BETA: −0.28
−:−:−
1250:1068:100

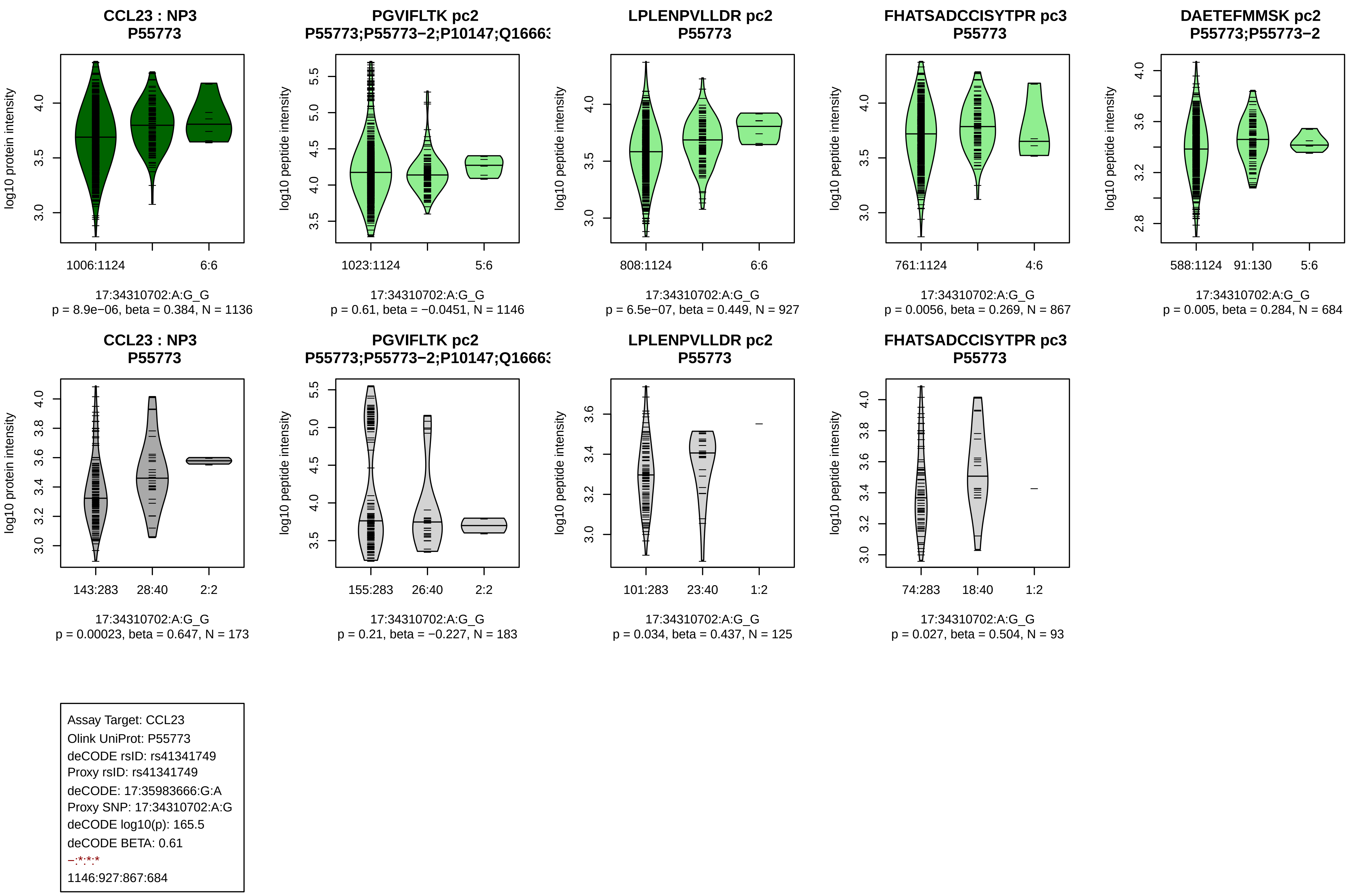
CCL23 : NP3
P55773
log10 protein intensity
1006:1124
6:6
17:34310702:A:G_G
p = 8.9e−06, beta = 0.384, N = 1136
PGVIFLTK pc2
P55773;P55773−2;P10147;Q16663
log10 peptide intensity
1023:1124
5:6
17:34310702:A:G_G
p = 0.61, beta = −0.0451, N = 1146
LPLENPVLLDR pc2
P55773
log10 peptide intensity
808:1124
6:6
17:34310702:A:G_G
p = 6.5e−07, beta = 0.449, N = 927
FHATSADCCISYTPR pc3
P55773
log10 peptide intensity
761:1124
4:6
17:34310702:A:G_G
p = 0.0056, beta = 0.269, N = 867
DAETEFMMSK pc2
P55773;P55773−2
log10 peptide intensity
588:1124
91:130
5:6
17:34310702:A:G_G
p = 0.005, beta = 0.284, N = 684
CCL23 : NP3
P55773
log10 protein intensity
143:283
28:40
2:2
17:34310702:A:G_G
p = 0.00023, beta = 0.647, N = 173
PGVIFLTK pc2
P55773;P55773−2;P10147;Q16663
log10 peptide intensity
155:283
26:40
2:2
17:34310702:A:G_G
p = 0.21, beta = −0.227, N = 183
LPLENPVLLDR pc2
P55773
log10 peptide intensity
101:283
23:40
1:2
17:34310702:A:G_G
p = 0.034, beta = 0.437, N = 125
FHATSADCCISYTPR pc3
P55773
log10 peptide intensity
74:283
18:40
1:2
17:34310702:A:G_G
p = 0.027, beta = 0.504, N = 93
Assay Target: CCL23
Olink UniProt: P55773
deCODE rsID: rs41341749
Proxy rsID: rs41341749
deCODE: 17:35983666:G:A
Proxy SNP: 17:34310702:A:G
deCODE log10(p): 165.5
deCODE BETA: 0.61
−:*:*:*
1146:927:867:684

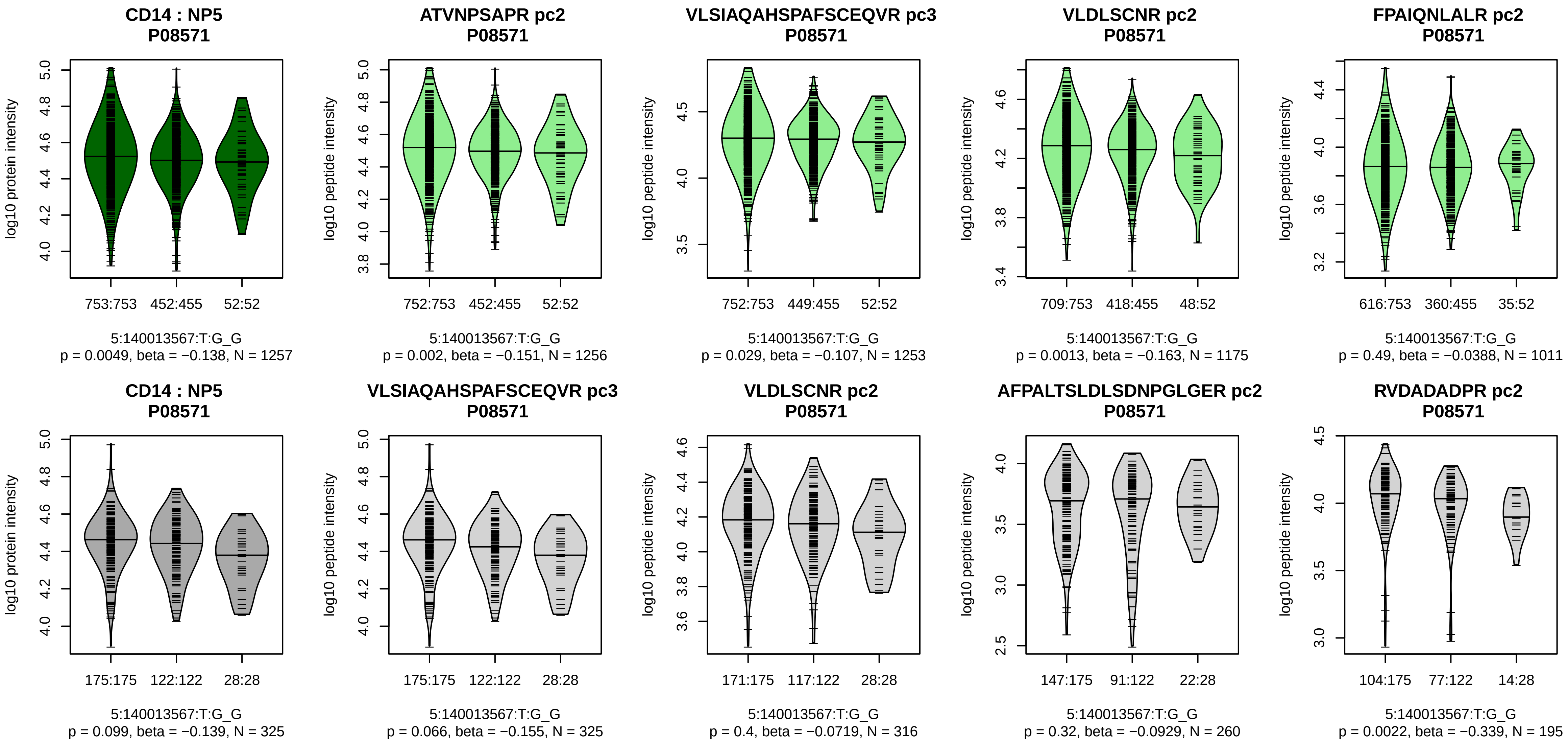


Assay Target: CD14
Olink UniProt: P08571
deCODE rsID: rs5744454
Proxy rsID: rs5744454
deCODE: 5:140633982:G:T
Proxy SNP: 5:140013567:T:G
deCODE log10(p): 160.4
deCODE BETA: −0.24
*:−:*:−:−:−:−:−:−:−:−:−:−
1256:1253:1175:1011:982:929:8

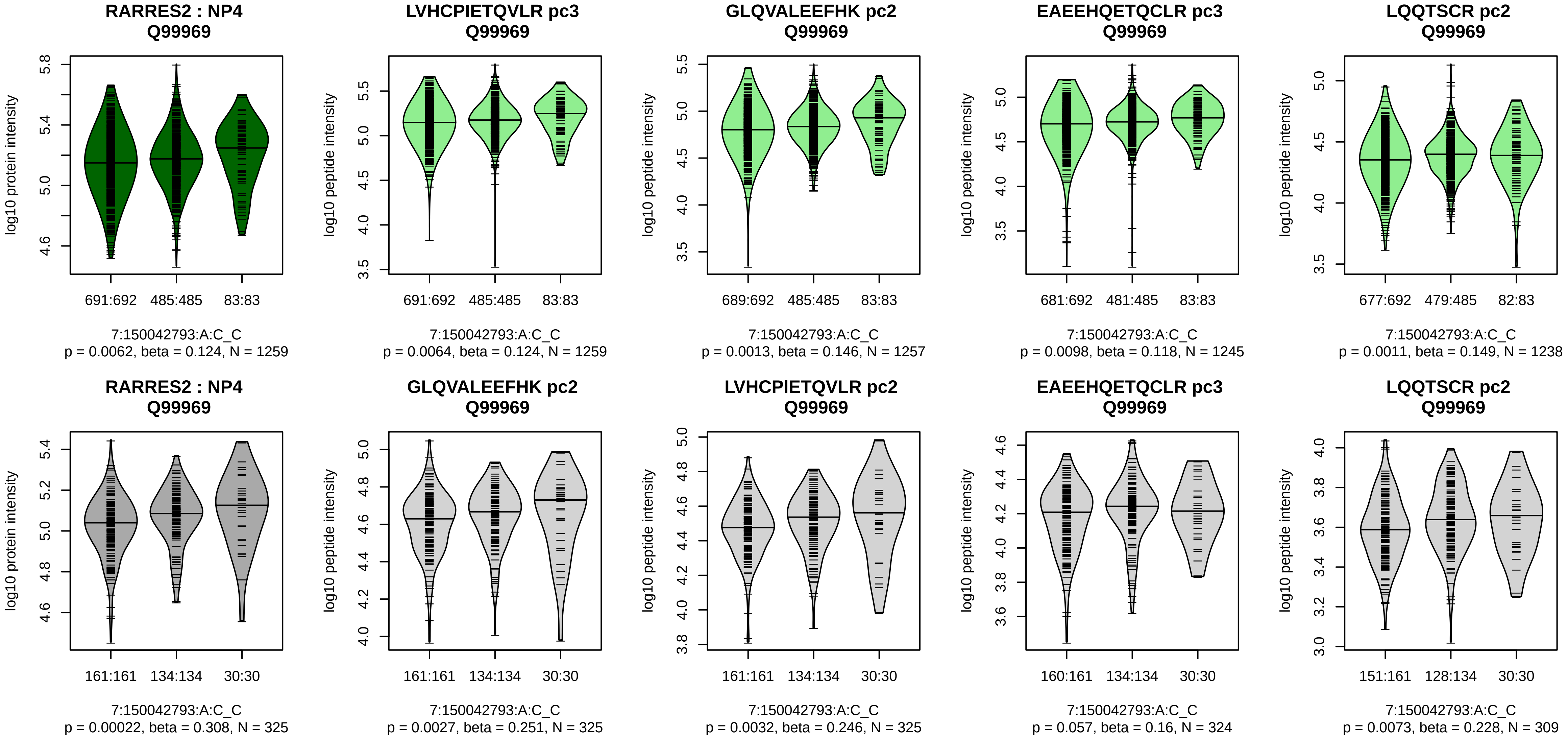


Assay Target: RARRES2
Olink UniProt: Q99969
deCODE rsID: rs883138
Proxy rsID: rs883138
deCODE: 7:150345704:C:A
Proxy SNP: 7:150042793:A:C
deCODE log10(p): 158.7
deCODE BETA: 0.25
*:*:*:*:*:*:–:–:–:–
1259:1257:1257:1250:1245:123

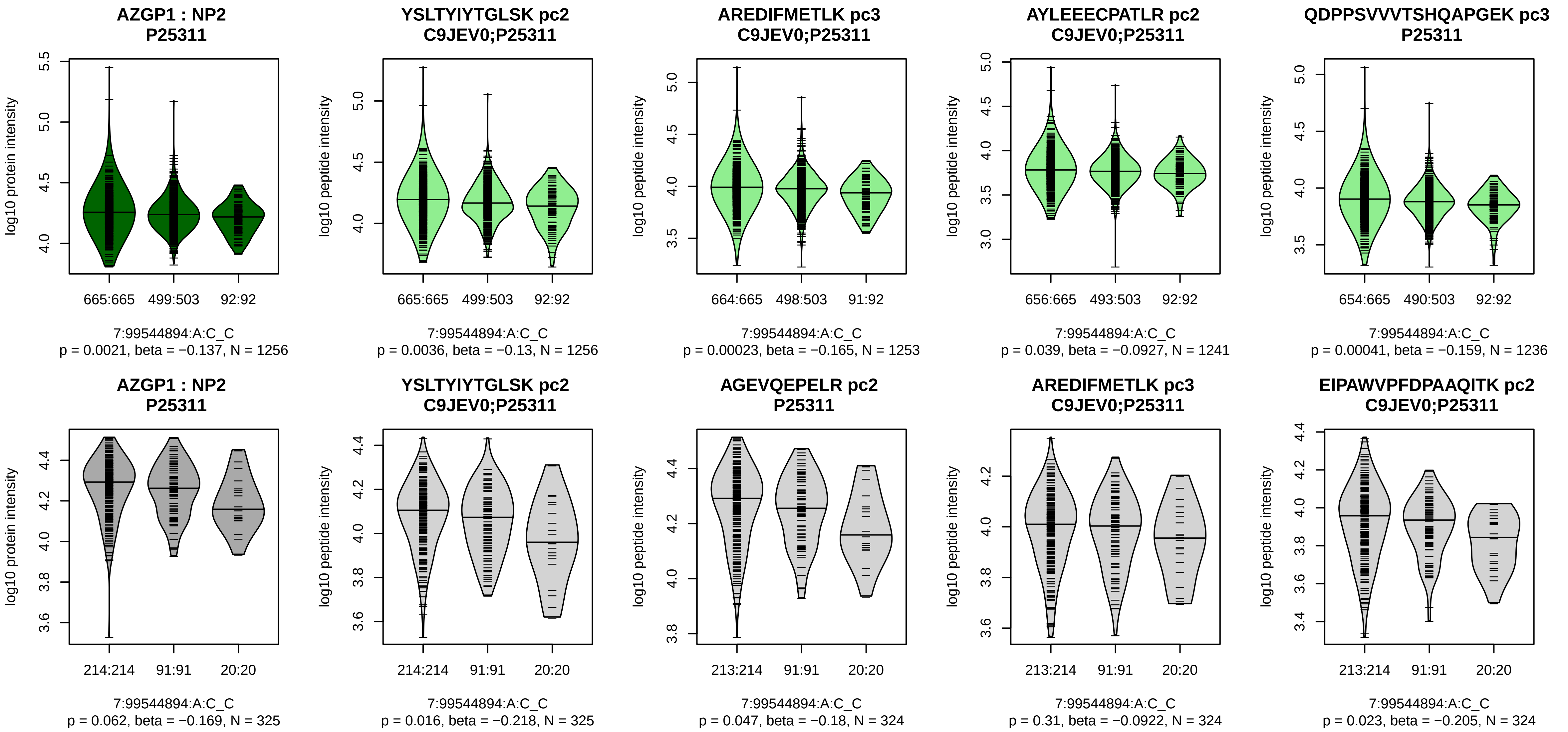


Assay Target: AZGP1
Olink UniProt: P25311
deCODE rsID: rs2525546
Proxy rsID: rs2040787
deCODE: 7:99954321:T:C
Proxy SNP: 7:99544894:A:C
deCODE log10(p): 157.9
deCODE BETA: −0.24
*:*:−:*:*:−:*:−:−:−:−:−:*:−:−:−:NA
1256:1253:1241:1236:1231:118

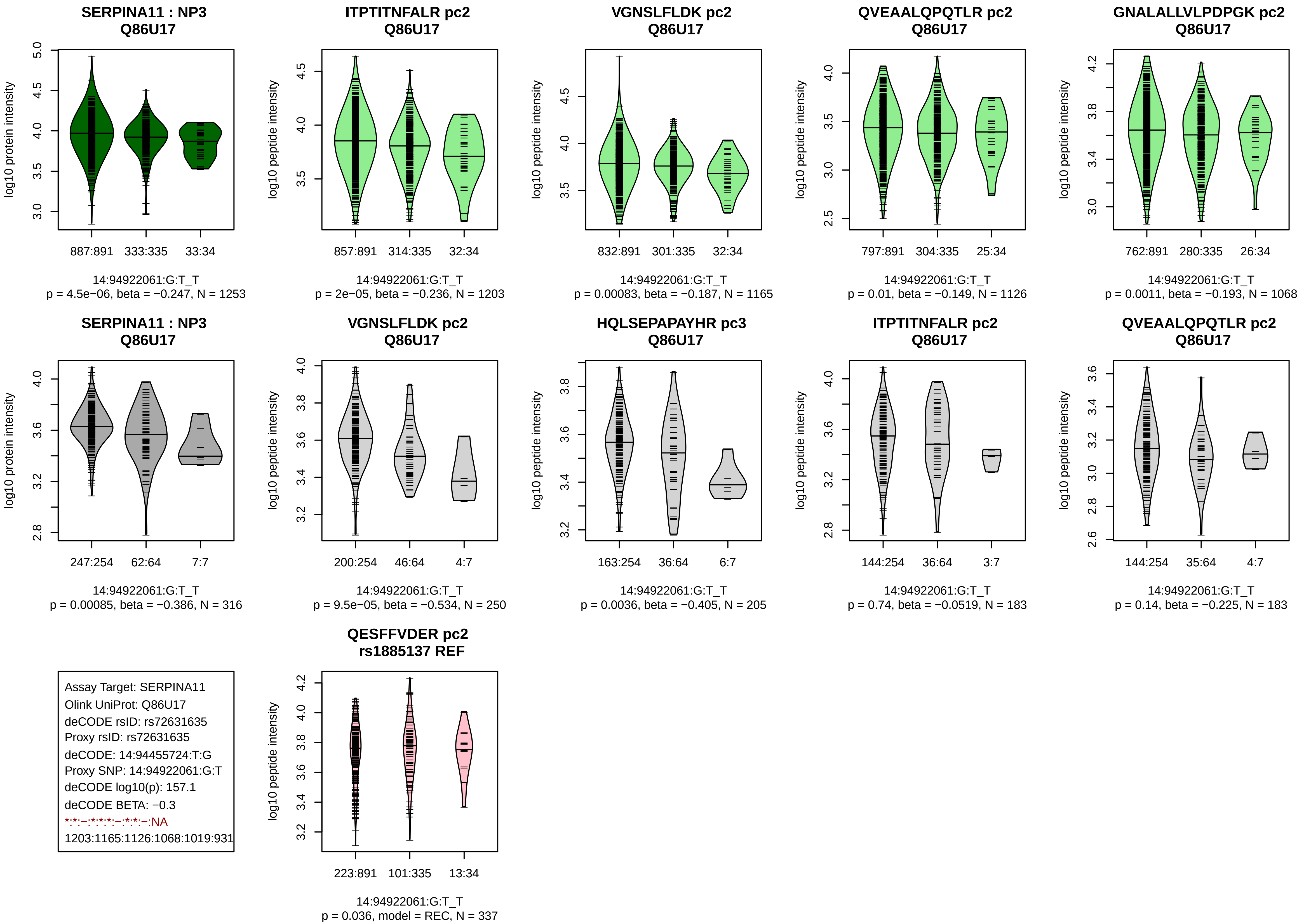

SERPINA11 : NP3
Q86U17
log10 protein intensity
887:891 333:335 33:34
14:94922061:G:T_T
p = 4.5e−06, beta = −0.247, N = 1253
ITPTITNFALR pc2
Q86U17
log10 peptide intensity
857:891 314:335 32:34
14:94922061:G:T_T
p = 2e−05, beta = −0.236, N = 1203
VGNSLFLDK pc2
Q86U17
log10 peptide intensity
832:891 301:335 32:34
14:94922061:G:T_T
p = 0.00083, beta = −0.187, N = 1165
QVEAALQPQTLR pc2
Q86U17
log10 peptide intensity
797:891 304:335 25:34
14:94922061:G:T_T
p = 0.01, beta = −0.149, N = 1126
GNALALLVLPDPGK pc2
Q86U17
log10 peptide intensity
762:891 280:335 26:34
14:94922061:G:T_T
p = 0.0011, beta = −0.193, N = 1068
SERPINA11 : NP3
Q86U17
log10 protein intensity
247:254 62:64 7:7
14:94922061:G:T_T
p = 0.00085, beta = −0.386, N = 316
VGNSLFLDK pc2
Q86U17
log10 peptide intensity
200:254 46:64 4:7
14:94922061:G:T_T
p = 9.5e−05, beta = −0.534, N = 250
HQLSEPAPAYHR pc3
Q86U17
log10 peptide intensity
163:254 36:64 6:7
14:94922061:G:T_T
p = 0.0036, beta = −0.405, N = 205
ITPTITNFALR pc2
Q86U17
log10 peptide intensity
144:254 36:64 3:7
14:94922061:G:T_T
p = 0.74, beta = −0.0519, N = 183
QVEAALQPQTLR pc2
Q86U17
log10 peptide intensity
144:254 35:64 4:7
14:94922061:G:T_T
p = 0.14, beta = −0.225, N = 183
Assay Target: SERPINA11
Olink UniProt: Q86U17
deCODE rsID: rs72631635
Proxy rsID: rs72631635
deCODE: 14:94455724:T:G
Proxy SNP: 14:94922061:G:T
deCODE log10(p): 157.1
deCODE BETA: −0.3
*:*:−:*:*:*:−:*:*:−:NA
1203:1165:1126:1068:1019:931
QESFFVDER pc2
rs1885137 REF
log10 peptide intensity
223:891 101:335 13:34
14:94922061:G:T_T
p = 0.036, model = REC, N = 337

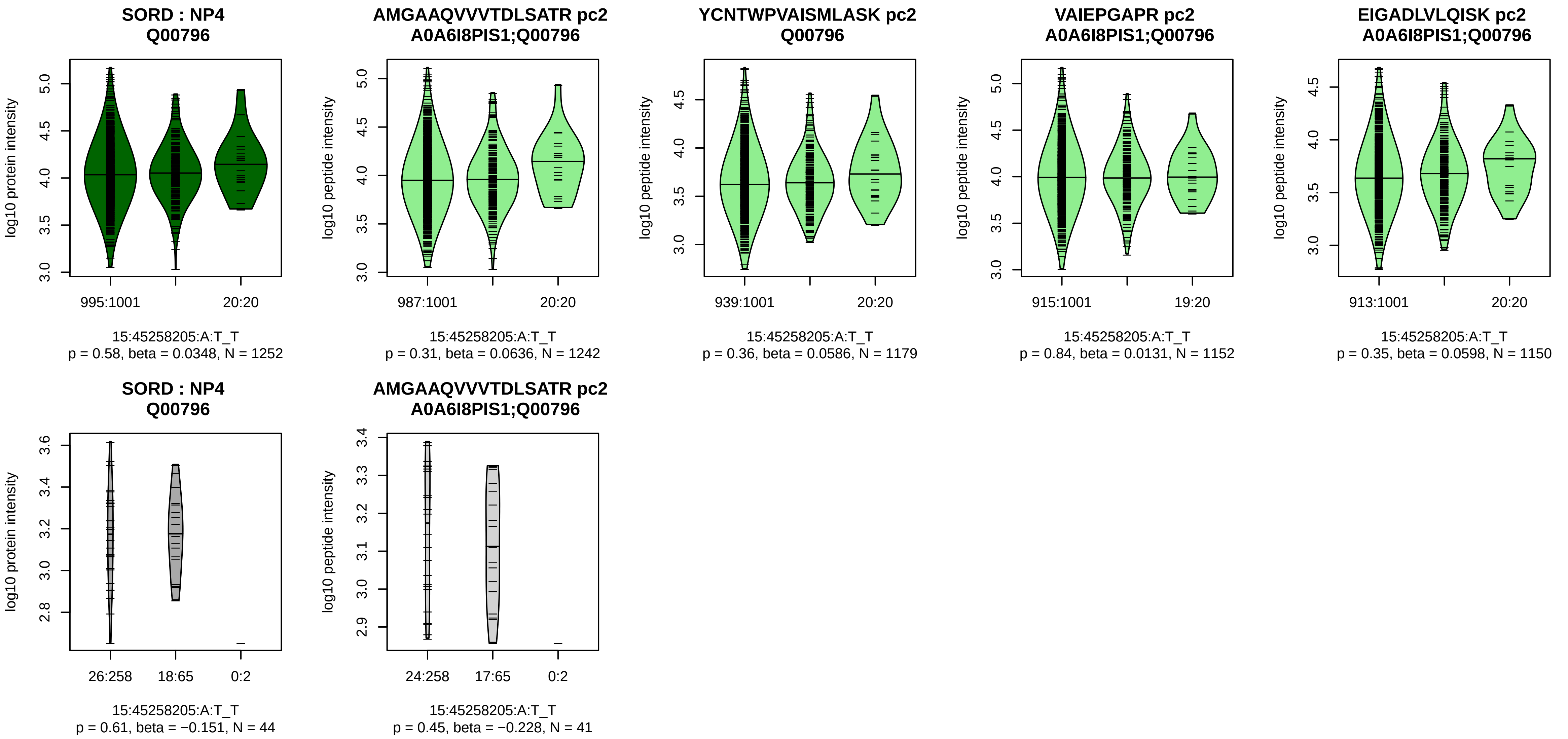


Assay Target: SORD
Olink UniProt: Q00796
deCODE rsID: rs72722045
Proxy rsID: rs72722045
deCODE: 15:44966007:T:A
Proxy SNP: 15:45258205:A:T
deCODE log10(p): 152.1
deCODE BETA: 0.3
–:–:–:–:–:–:–:–:–:–:–:–:–:–:–:–:–:
1242:1179:1152:1150:995:979:[illegible]

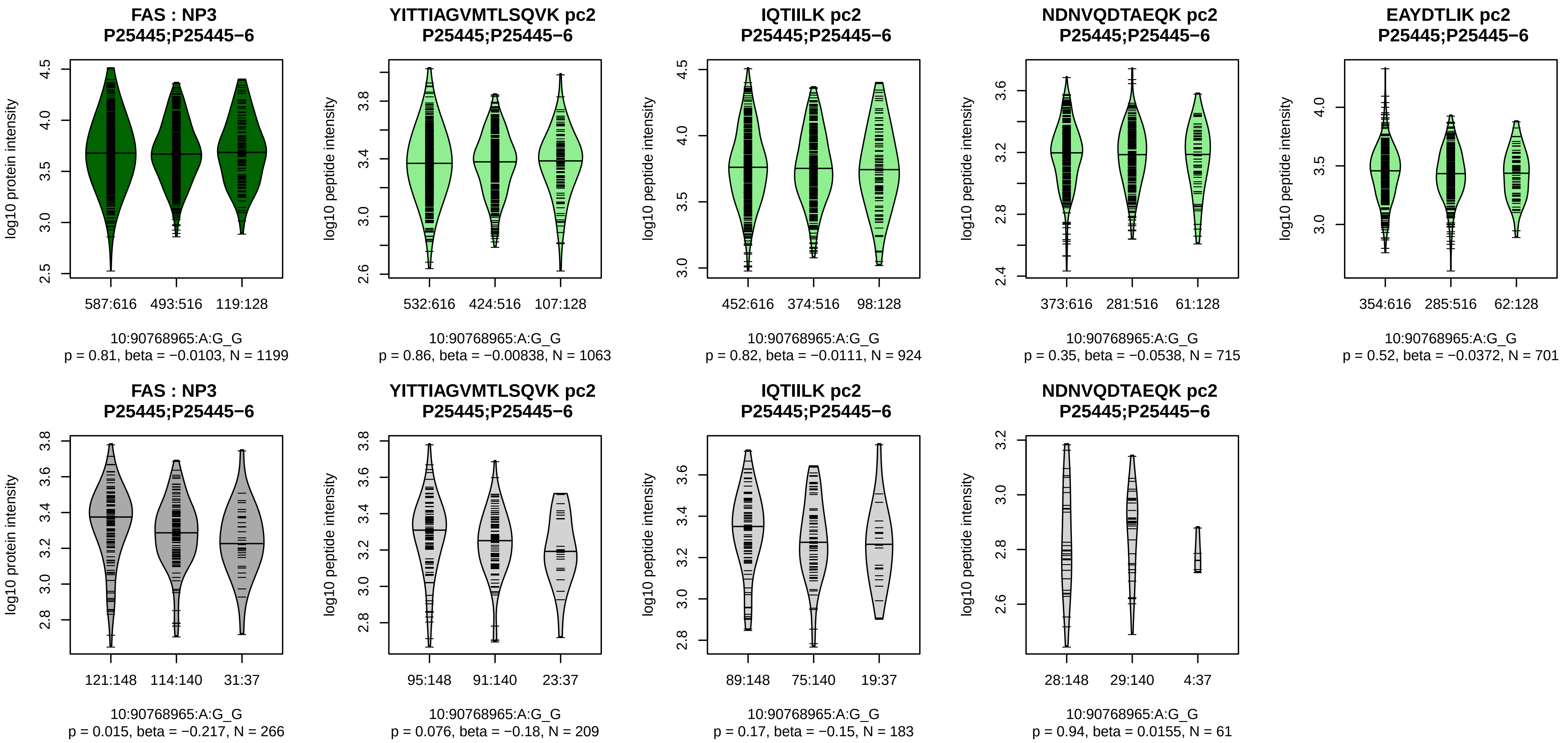


Assay Target: FAS
Olink UniProt: P25445
deCODE rsID: rs7911226
Proxy rsID: rs7911226
deCODE: 10:89009208:G:A
Proxy SNP: 10:90768965:A:G
deCODE log10(p): 152
deCODE BETA: −0.24
−:−:−:−
1063:924:715:701

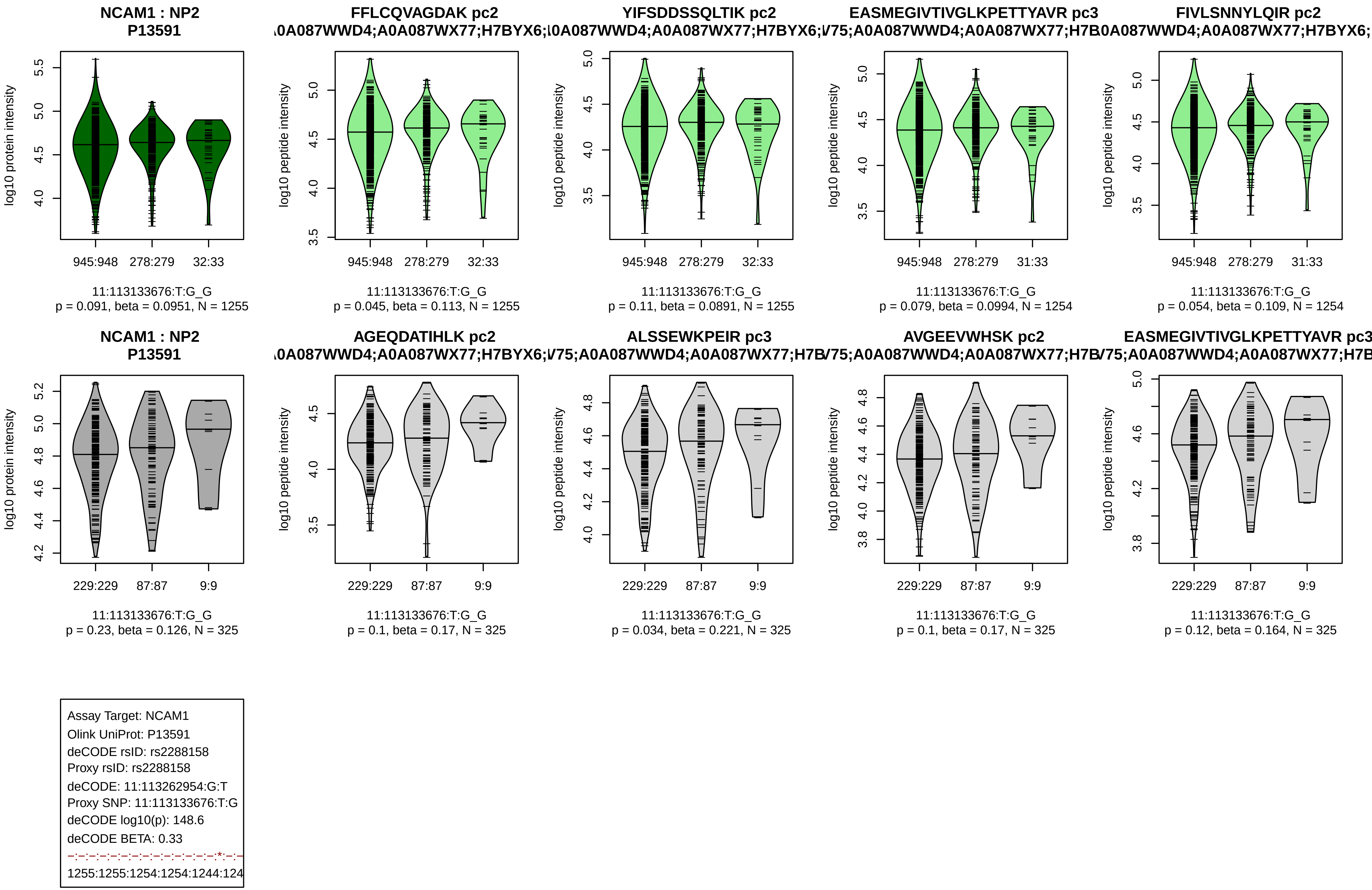
NCAM1 : NP2
P13591
log10 protein intensity
945:948
278:279
32:33
11:113133676:T:G_G
p = 0.091, beta = 0.0951, N = 1255
FFLCQVAGDAK pc2
A0A087WWD4;A0A087WX77;H7BYX6;
log10 peptide intensity
945:948
278:279
32:33
11:113133676:T:G_G
p = 0.045, beta = 0.113, N = 1255
YIFSDDSSQLTIK pc2
A0A087WWD4;A0A087WX77;H7BYX6;
log10 peptide intensity
945:948
278:279
32:33
11:113133676:T:G_G
p = 0.11, beta = 0.0891, N = 1255
EASMEGIVTIVGLKPETTYAVR pc3
V75;A0A087WWD4;A0A087WX77;H7B
log10 peptide intensity
945:948
278:279
31:33
11:113133676:T:G_G
p = 0.079, beta = 0.0994, N = 1254
FIVLSNNYLQIR pc2
A0A087WWD4;A0A087WX77;H7BYX6;
log10 peptide intensity
945:948
278:279
31:33
11:113133676:T:G_G
p = 0.054, beta = 0.109, N = 1254
NCAM1 : NP2
P13591
log10 protein intensity
229:229
87:87
9:9
11:113133676:T:G_G
p = 0.23, beta = 0.126, N = 325
AGEQDATIHLK pc2
A0A087WWD4;A0A087WX77;H7BYX6;
log10 peptide intensity
229:229
87:87
9:9
11:113133676:T:G_G
p = 0.1, beta = 0.17, N = 325
ALSSEWKPEIR pc3
V75;A0A087WWD4;A0A087WX77;H7B
log10 peptide intensity
229:229
87:87
9:9
11:113133676:T:G_G
p = 0.034, beta = 0.221, N = 325
AVGEEVWHSK pc2
V75;A0A087WWD4;A0A087WX77;H7B
log10 peptide intensity
229:229
87:87
9:9
11:113133676:T:G_G
p = 0.1, beta = 0.17, N = 325
EASMEGIVTIVGLKPETTYAVR pc3
V75;A0A087WWD4;A0A087WX77;H7B
log10 peptide intensity
229:229
87:87
9:9
11:113133676:T:G_G
p = 0.12, beta = 0.164, N = 325
Assay Target: NCAM1
Olink UniProt: P13591
deCODE rsID: rs2288158
Proxy rsID: rs2288158
deCODE: 11:113262954:G:T
Proxy SNP: 11:113133676:T:G
deCODE log10(p): 148.6
deCODE BETA: 0.33
-:-:-:-:-:-:-:-:-:-:-:-:-:-:*:-:-
1255:1255:1254:1254:1244:124

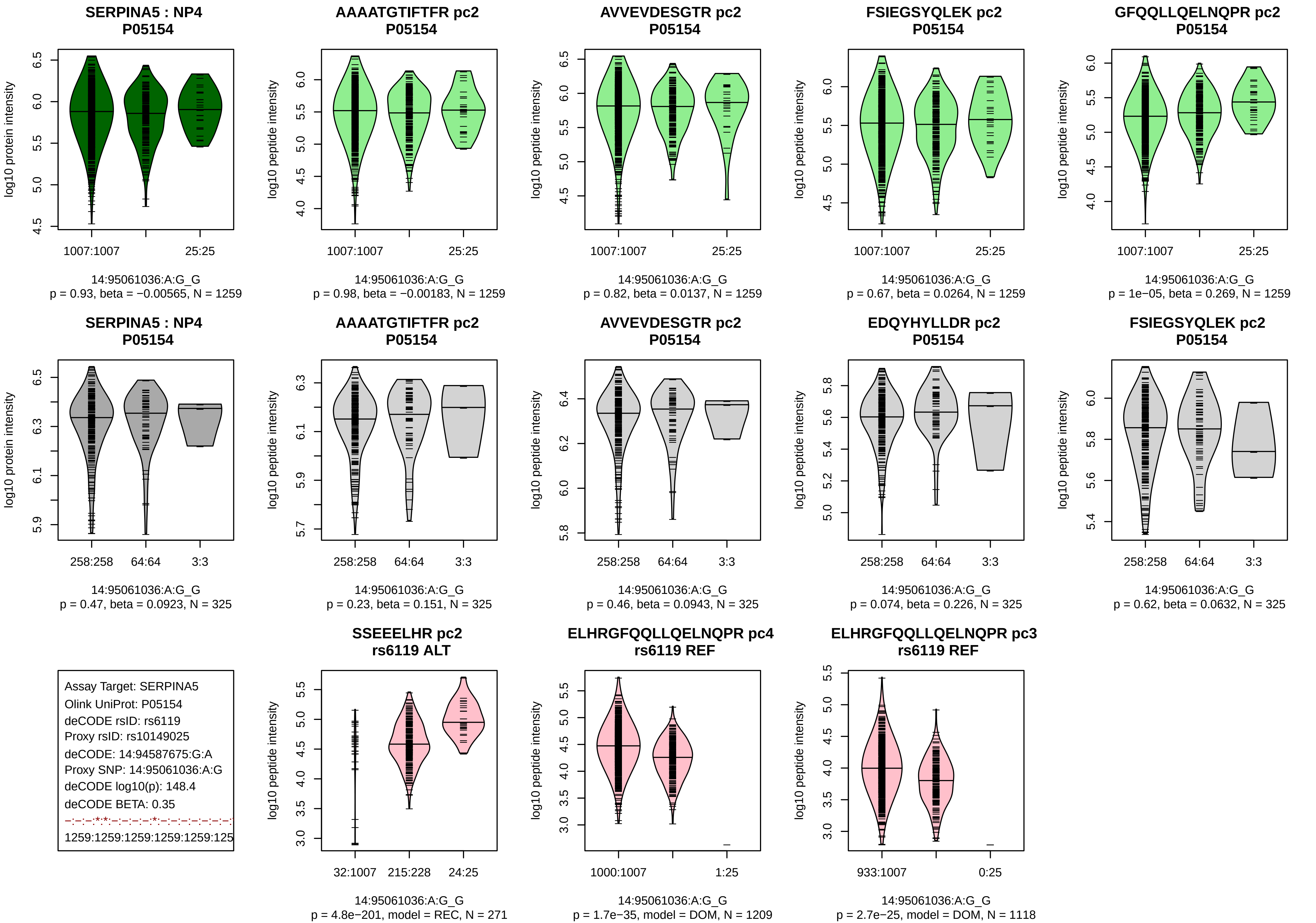
SERPINA5 : NP4
P05154
log10 protein intensity
1007:1007
25:25
14:95061036:A:G_G
p = 0.93, beta = −0.00565, N = 1259
AAAATGTIFTFR pc2
P05154
log10 peptide intensity
14:95061036:A:G_G
p = 0.98, beta = −0.00183, N = 1259
AVVEVDESGTR pc2
P05154
14:95061036:A:G_G
p = 0.82, beta = 0.0137, N = 1259
FSIEGSYQLEK pc2
P05154
14:95061036:A:G_G
p = 0.67, beta = 0.0264, N = 1259
GFQQLLQELNQPR pc2
P05154
14:95061036:A:G_G
p = 1e−05, beta = 0.269, N = 1259
SERPINA5 : NP4
P05154
258:258
64:64
3:3
14:95061036:A:G_G
p = 0.47, beta = 0.0923, N = 325
AAAATGTIFTFR pc2
P05154
14:95061036:A:G_G
p = 0.23, beta = 0.151, N = 325
AVVEVDESGTR pc2
P05154
14:95061036:A:G_G
p = 0.46, beta = 0.0943, N = 325
EDQYHYLLDR pc2
P05154
14:95061036:A:G_G
p = 0.074, beta = 0.226, N = 325
FSIEGSYQLEK pc2
P05154
14:95061036:A:G_G
p = 0.62, beta = 0.0632, N = 325
Assay Target: SERPINA5
Olink UniProt: P05154
deCODE rsID: rs6119
Proxy rsID: rs10149025
deCODE: 14:94587675:G:A
Proxy SNP: 14:95061036:A:G
deCODE log10(p): 148.4
deCODE BETA: 0.35
−:−:−:*:*:−:−:−:−:*:−:−:−:−:−:−:−:−:
1259:1259:1259:1259:1259:125
SSEEELHR pc2
rs6119 ALT
32:1007
215:228
24:25
14:95061036:A:G_G
p = 4.8e−201, model = REC, N = 271
ELHRGFQQLLQELNQPR pc4
rs6119 REF
1000:1007
1:25
14:95061036:A:G_G
p = 1.7e−35, model = DOM, N = 1209
ELHRGFQQLLQELNQPR pc3
rs6119 REF
933:1007
0:25
14:95061036:A:G_G
p = 2.7e−25, model = DOM, N = 1118

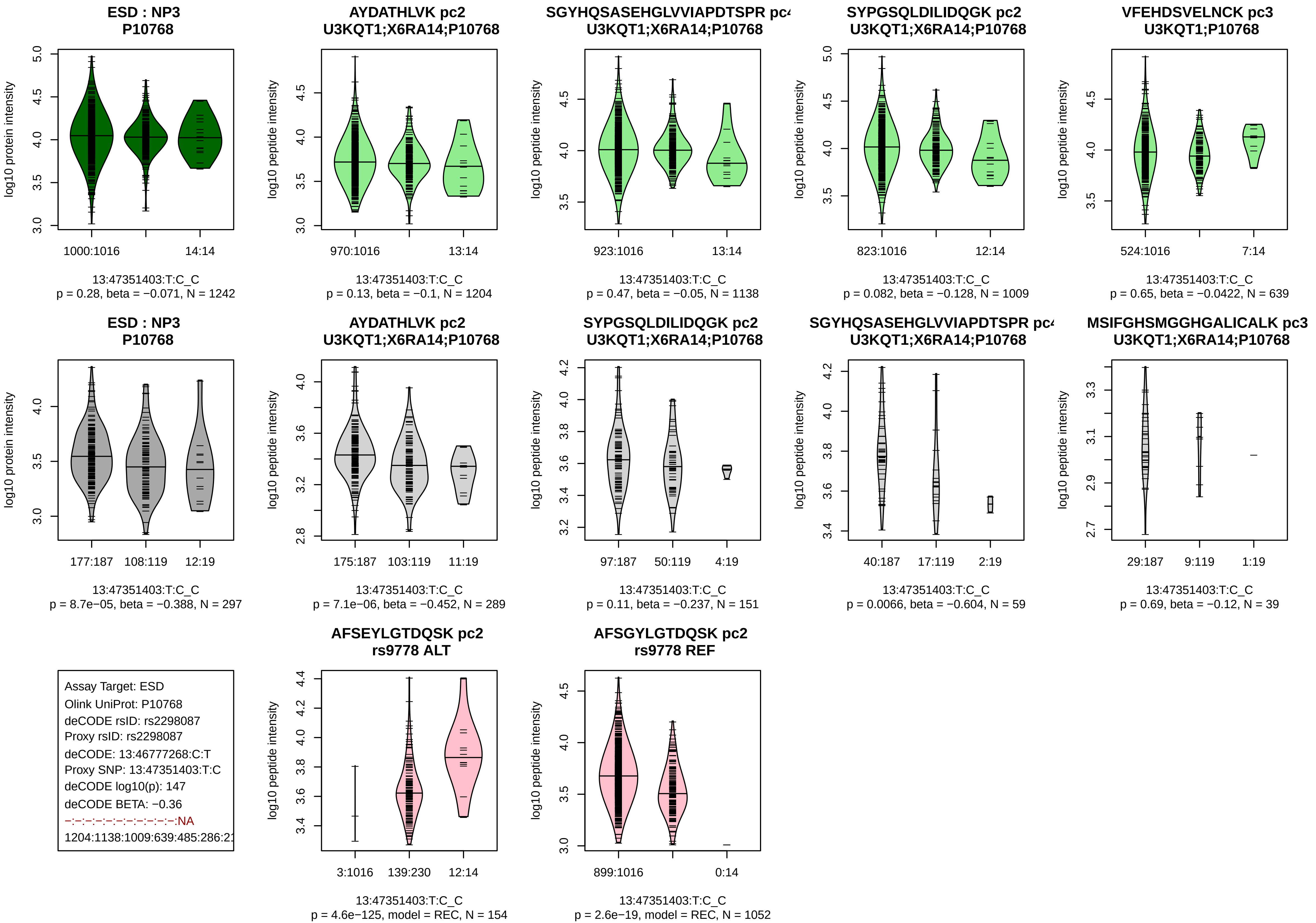
ESD : NP3
P10768
log10 protein intensity
1000:1016
14:14
13:47351403:T:C_C
p = 0.28, beta = −0.071, N = 1242
AYDATHLVK pc2
U3KQT1;X6RA14;P10768
log10 peptide intensity
970:1016
13:14
13:47351403:T:C_C
p = 0.13, beta = −0.1, N = 1204
SGYHQSASEHGLVVIAPDTSPR pc
U3KQT1;X6RA14;P10768
log10 peptide intensity
923:1016
13:14
13:47351403:T:C_C
p = 0.47, beta = −0.05, N = 1138
SYPGSQLDILIDQGK pc2
U3KQT1;X6RA14;P10768
log10 peptide intensity
823:1016
12:14
13:47351403:T:C_C
p = 0.082, beta = −0.128, N = 1009
VFEHDSVELNCK pc3
U3KQT1;P10768
log10 peptide intensity
524:1016
7:14
13:47351403:T:C_C
p = 0.65, beta = −0.0422, N = 639
ESD : NP3
P10768
log10 protein intensity
177:187
108:119
12:19
13:47351403:T:C_C
p = 8.7e−05, beta = −0.388, N = 297
AYDATHLVK pc2
U3KQT1;X6RA14;P10768
log10 peptide intensity
175:187
103:119
11:19
13:47351403:T:C_C
p = 7.1e−06, beta = −0.452, N = 289
SYPGSQLDILIDQGK pc2
U3KQT1;X6RA14;P10768
log10 peptide intensity
97:187
50:119
4:19
13:47351403:T:C_C
p = 0.11, beta = −0.237, N = 151
SGYHQSASEHGLVVIAPDTSPR pc
U3KQT1;X6RA14;P10768
log10 peptide intensity
40:187
17:119
2:19
13:47351403:T:C_C
p = 0.0066, beta = −0.604, N = 59
MSIFGHSMGGHGALICALK pc3
U3KQT1;X6RA14;P10768
log10 peptide intensity
29:187
9:119
1:19
13:47351403:T:C_C
p = 0.69, beta = −0.12, N = 39
Assay Target: ESD
Olink UniProt: P10768
deCODE rsID: rs2298087
Proxy rsID: rs2298087
deCODE: 13:46777268:C:T
Proxy SNP: 13:47351403:T:C
deCODE log10(p): 147
deCODE BETA: −0.36
−:−:−:−:−:−:−:−:−:−:−:NA
1204:1138:1009:639:485:286:2
AFSEYLGTDQSK pc2
rs9778 ALT
log10 peptide intensity
3:1016
139:230
12:14
13:47351403:T:C_C
p = 4.6e−125, model = REC, N = 154
AFSGYLGTDQSK pc2
rs9778 REF
log10 peptide intensity
899:1016
0:14
13:47351403:T:C_C
p = 2.6e−19, model = REC, N = 1052

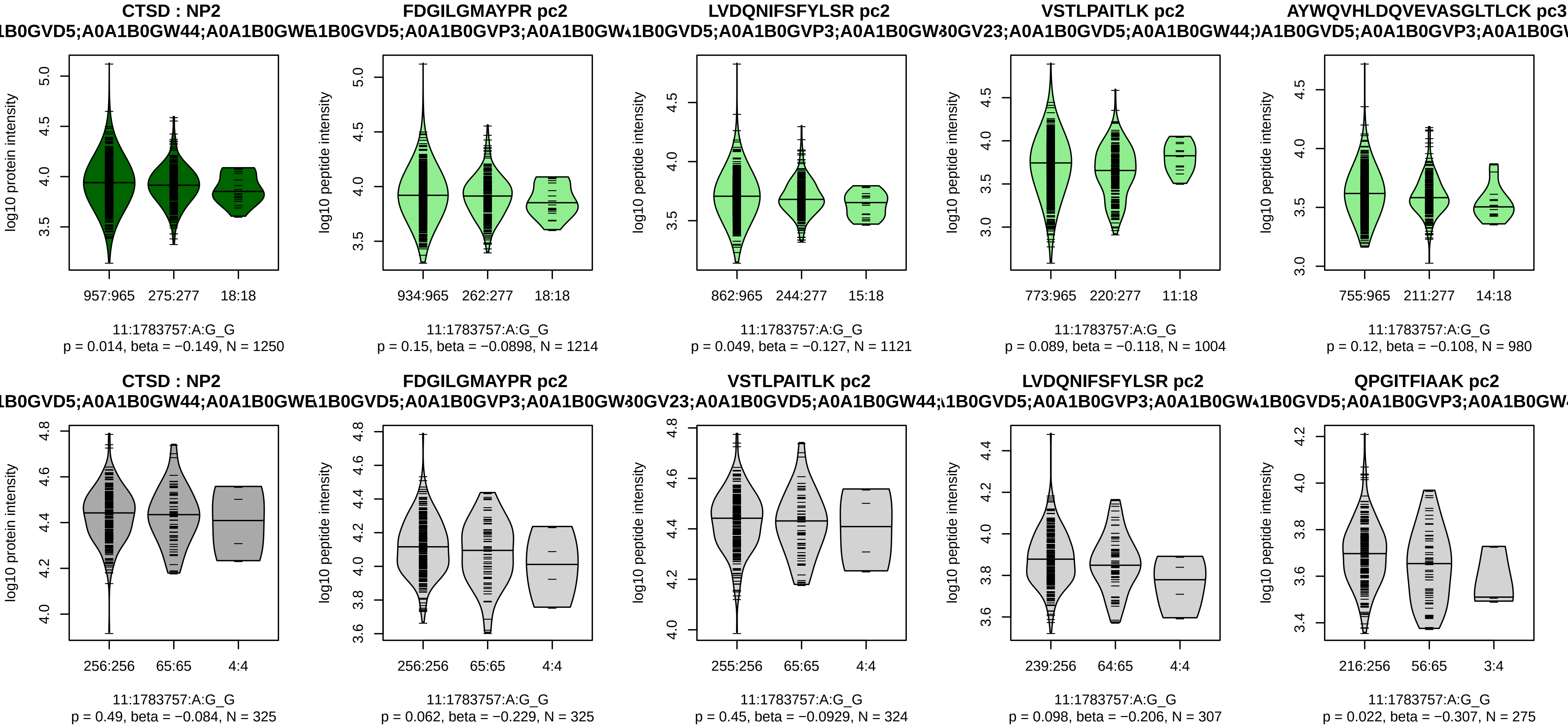


Assay Target: CTSD
Olink UniProt: P07339
deCODE rsID: rs55861089
Proxy rsID: rs55861089
deCODE: 11:1762527:G:A
Proxy SNP: 11:1783757:A:G
deCODE log10(p): 141.9
deCODE BETA: −0.33
−:−:−:−:−:−:−:−:−:−:−:−:−:−:NA
1214:1121:1004:980:964:486:40

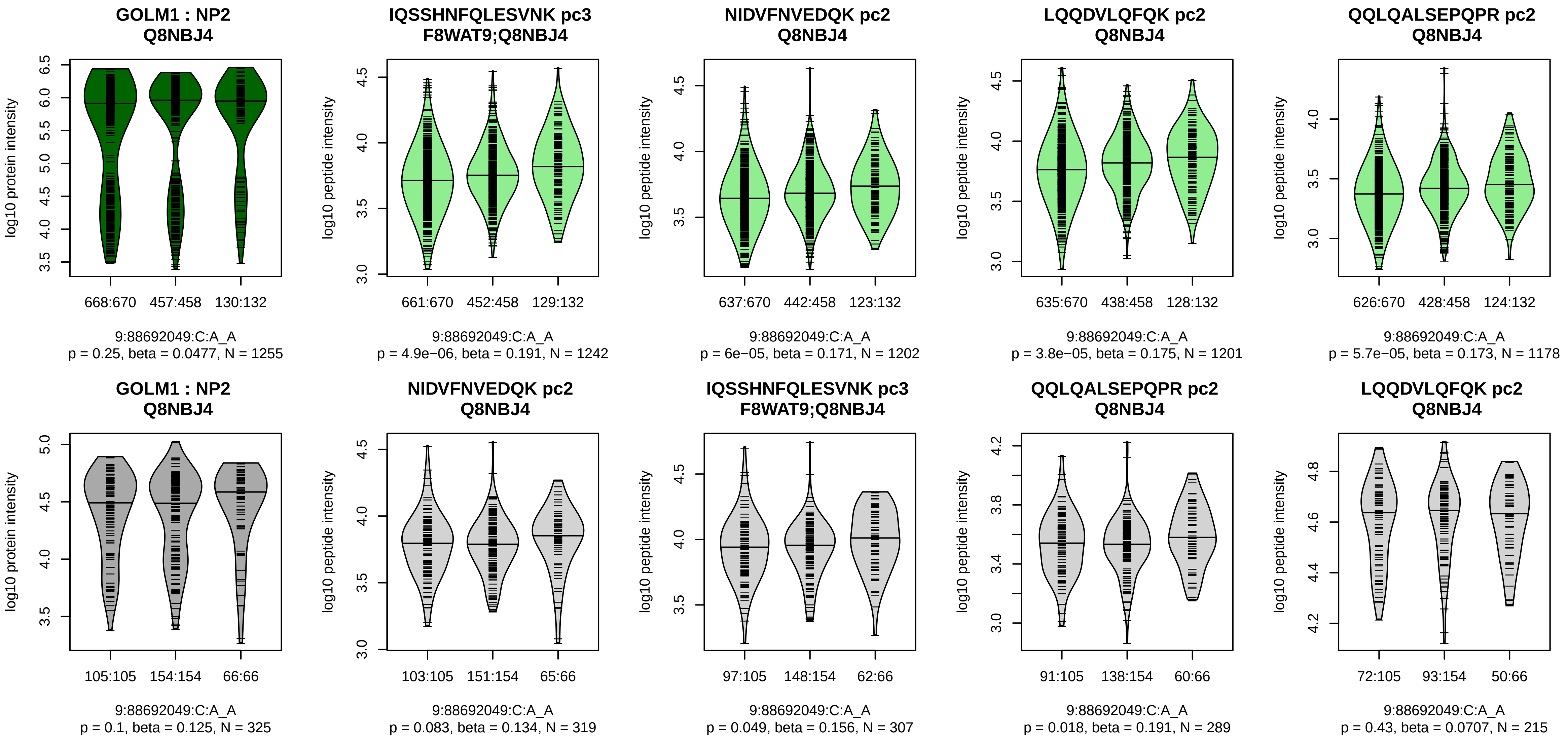


Assay Target: GOLM1
Olink UniProt: Q8NBJ4
deCODE rsID: rs4333693
Proxy rsID: rs4333693
deCODE: 9:86077134:A:C
Proxy SNP: 9:88692049:C:A
deCODE log10(p): 140.8
deCODE BETA: 0.26
*:*:*:*:*:−:−:*:−:−:NA
1242:1202:1201:1178:1081:950

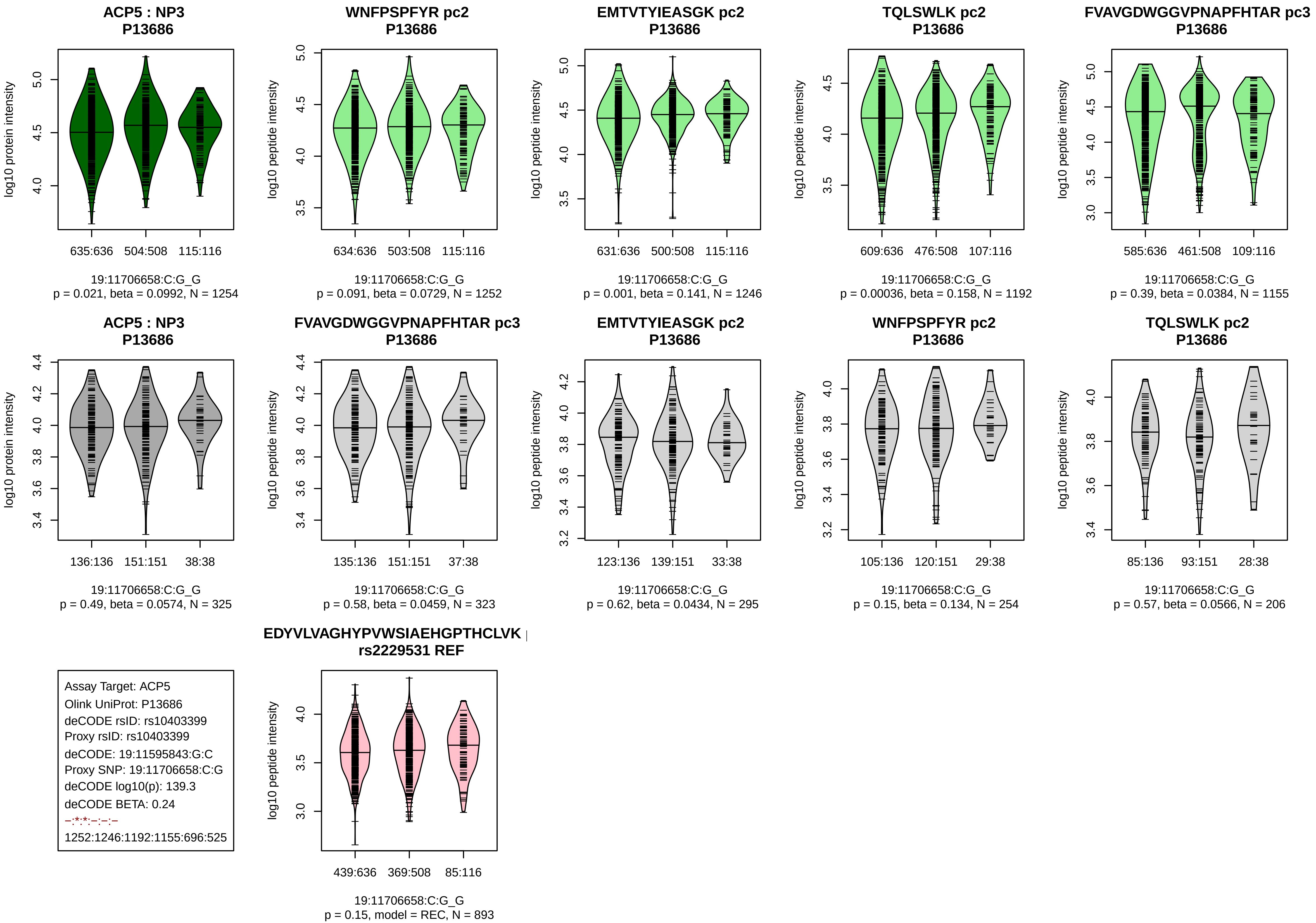
ACP5 : NP3
P13686
log10 protein intensity
635:636
504:508
115:116
19:11706658:C:G_G
p = 0.021, beta = 0.0992, N = 1254
WNFPSPFYR pc2
P13686
log10 peptide intensity
634:636
503:508
115:116
19:11706658:C:G_G
p = 0.091, beta = 0.0729, N = 1252
EMTVTYIEASGK pc2
P13686
log10 peptide intensity
631:636
500:508
115:116
19:11706658:C:G_G
p = 0.001, beta = 0.141, N = 1246
TQLSWLK pc2
P13686
log10 peptide intensity
609:636
476:508
107:116
19:11706658:C:G_G
p = 0.00036, beta = 0.158, N = 1192
FVAVGDWGGVPNAPFHTAR pc3
P13686
log10 peptide intensity
585:636
461:508
109:116
19:11706658:C:G_G
p = 0.39, beta = 0.0384, N = 1155
ACP5 : NP3
P13686
log10 protein intensity
136:136
151:151
38:38
19:11706658:C:G_G
p = 0.49, beta = 0.0574, N = 325
FVAVGDWGGVPNAPFHTAR pc3
P13686
log10 peptide intensity
135:136
151:151
37:38
19:11706658:C:G_G
p = 0.58, beta = 0.0459, N = 323
EMTVTYIEASGK pc2
P13686
log10 peptide intensity
123:136
139:151
33:38
19:11706658:C:G_G
p = 0.62, beta = 0.0434, N = 295
WNFPSPFYR pc2
P13686
log10 peptide intensity
105:136
120:151
29:38
19:11706658:C:G_G
p = 0.15, beta = 0.134, N = 254
TQLSWLK pc2
P13686
log10 peptide intensity
85:136
93:151
28:38
19:11706658:C:G_G
p = 0.57, beta = 0.0566, N = 206
Assay Target: ACP5
Olink UniProt: P13686
deCODE rsID: rs10403399
Proxy rsID: rs10403399
deCODE: 19:11595843:G:C
Proxy SNP: 19:11706658:C:G
deCODE log10(p): 139.3
deCODE BETA: 0.24
–:*:*:–:–:–
1252:1246:1192:1155:696:525
EDYVLVAGHYPVWSIAEHGPTHCLVK |
rs2229531 REF
log10 peptide intensity
439:636
369:508
85:116
19:11706658:C:G_G
p = 0.15, model = REC, N = 893

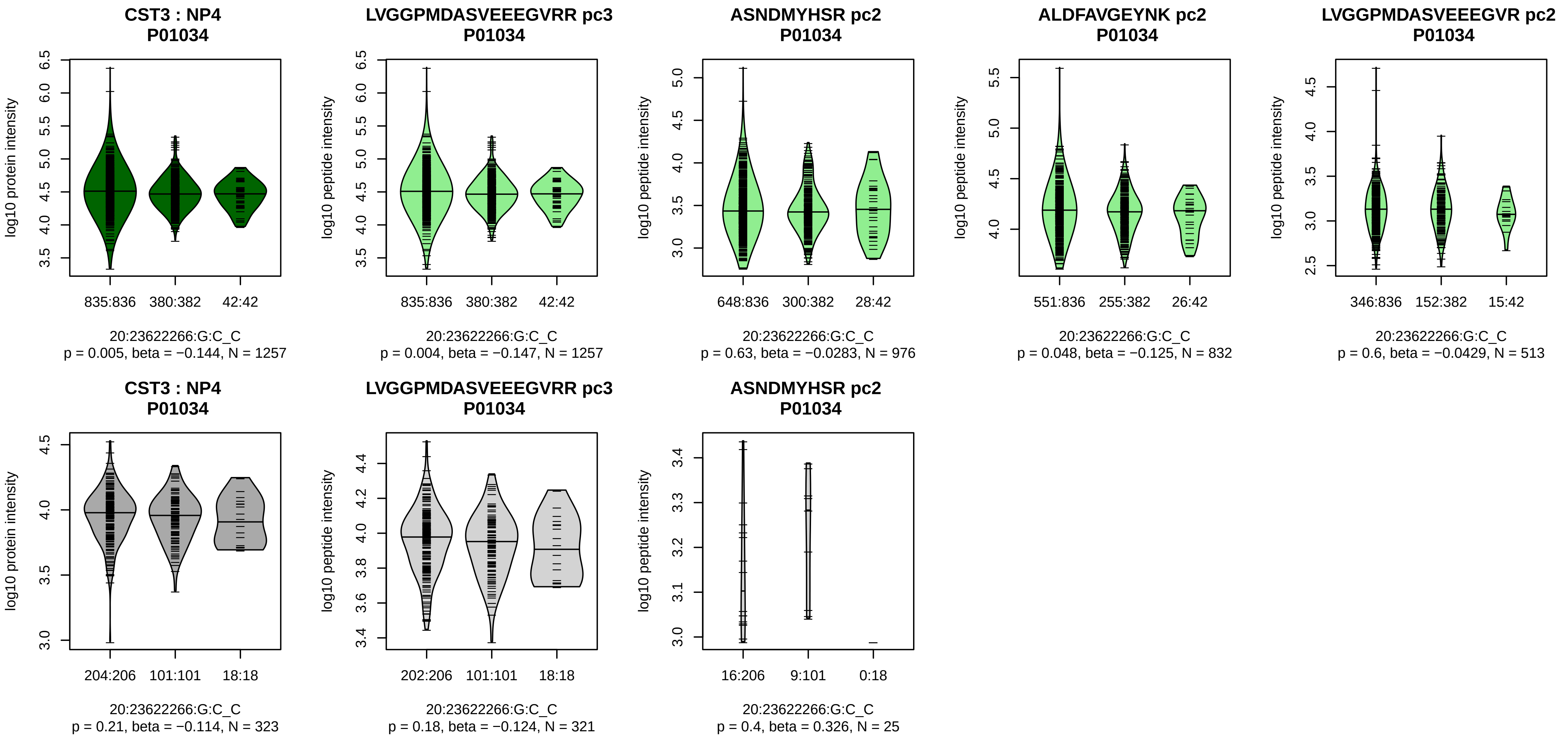


Assay Target: CST3
Olink UniProt: P01034
deCODE rsID: rs6114209
Proxy rsID: rs6114209
deCODE: 20:23641629:C:G
Proxy SNP: 20:23622266:G:C
deCODE log10(p): 137.9
deCODE BETA: −0.25
*:−:−:−:−:−:−:−:−:−:NA:NA:NA
1257:976:832:513:510:184:160:

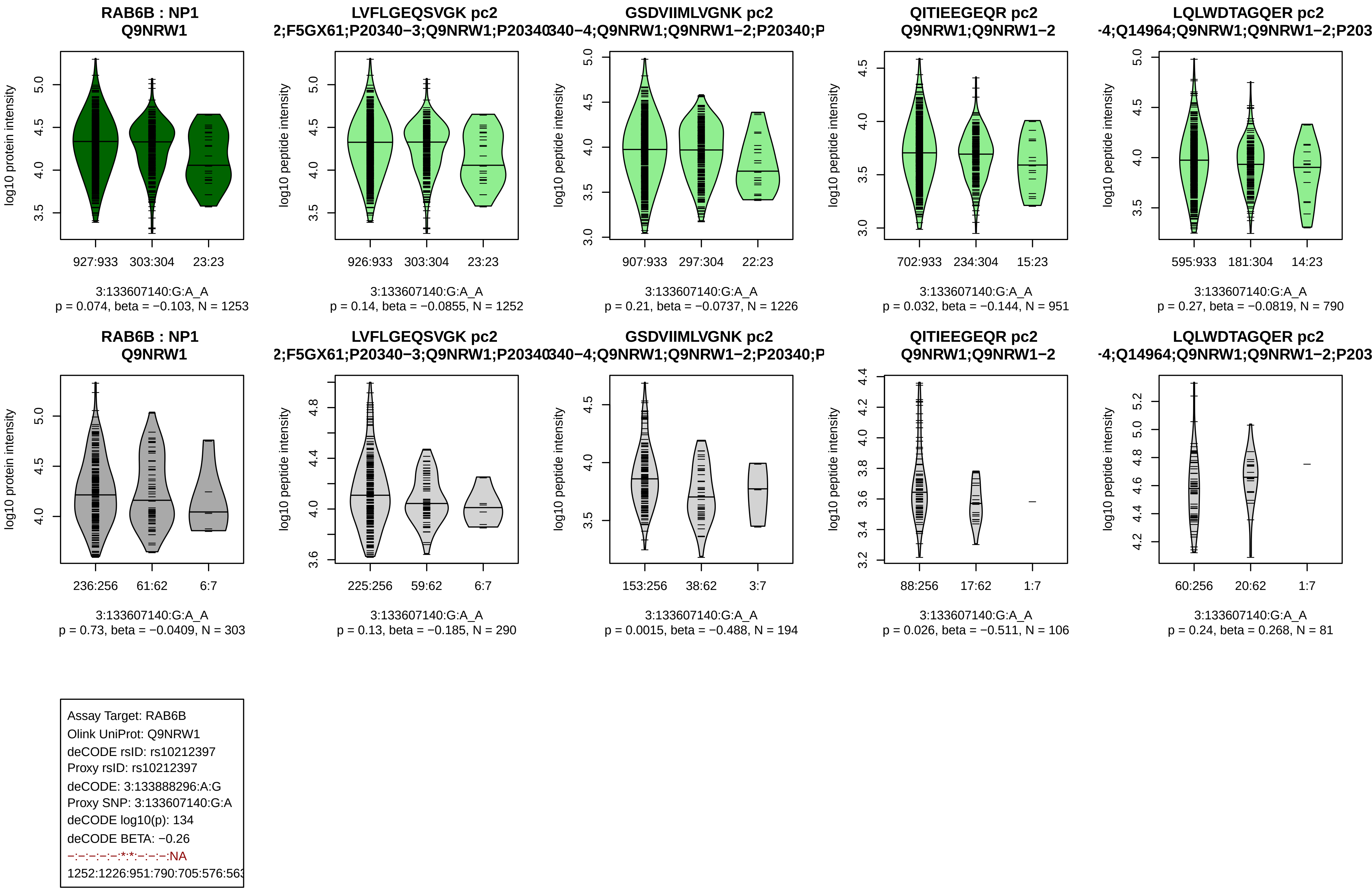
RAB6B : NP1
Q9NRW1
log10 protein intensity
927:933 303:304 23:23
3:133607140:G:A_A
p = 0.074, beta = −0.103, N = 1253
LVFLGEQSVGK pc2
2;F5GX61;P20340−3;Q9NRW1;P20340
log10 peptide intensity
926:933 303:304 23:23
3:133607140:G:A_A
p = 0.14, beta = −0.0855, N = 1252
GSDVIIMLVGNK pc2
340−4;Q9NRW1;Q9NRW1−2;P20340;P
log10 peptide intensity
907:933 297:304 22:23
3:133607140:G:A_A
p = 0.21, beta = −0.0737, N = 1226
QITIEEGEQR pc2
Q9NRW1;Q9NRW1−2
log10 peptide intensity
702:933 234:304 15:23
3:133607140:G:A_A
p = 0.032, beta = −0.144, N = 951
LQLWDTAGQER pc2
-4;Q14964;Q9NRW1;Q9NRW1−2;P203
log10 peptide intensity
595:933 181:304 14:23
3:133607140:G:A_A
p = 0.27, beta = −0.0819, N = 790
RAB6B : NP1
Q9NRW1
log10 protein intensity
236:256 61:62 6:7
3:133607140:G:A_A
p = 0.73, beta = −0.0409, N = 303
LVFLGEQSVGK pc2
2;F5GX61;P20340−3;Q9NRW1;P20340
log10 peptide intensity
225:256 59:62 6:7
3:133607140:G:A_A
p = 0.13, beta = −0.185, N = 290
GSDVIIMLVGNK pc2
340−4;Q9NRW1;Q9NRW1−2;P20340;P
log10 peptide intensity
153:256 38:62 3:7
3:133607140:G:A_A
p = 0.0015, beta = −0.488, N = 194
QITIEEGEQR pc2
Q9NRW1;Q9NRW1−2
log10 peptide intensity
88:256 17:62 1:7
3:133607140:G:A_A
p = 0.026, beta = −0.511, N = 106
LQLWDTAGQER pc2
-4;Q14964;Q9NRW1;Q9NRW1−2;P203
log10 peptide intensity
60:256 20:62 1:7
3:133607140:G:A_A
p = 0.24, beta = 0.268, N = 81
Assay Target: RAB6B
Olink UniProt: Q9NRW1
deCODE rsID: rs10212397
Proxy rsID: rs10212397
deCODE: 3:133888296:A:G
Proxy SNP: 3:133607140:G:A
deCODE log10(p): 134
deCODE BETA: −0.26
−:−:−:−:−:*:*:−:−:−:NA
1252:1226:951:790:705:576:563

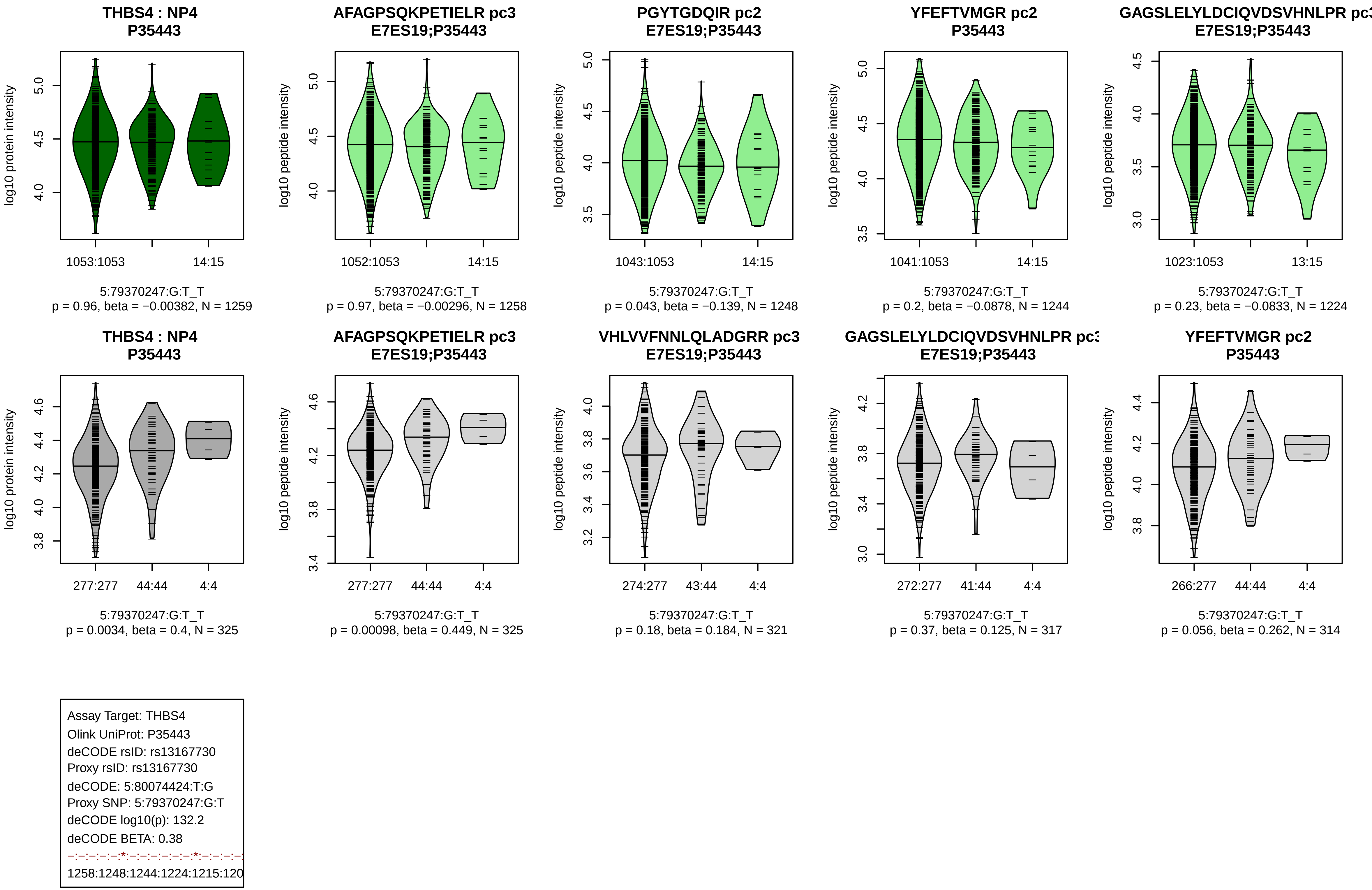
THBS4 : NP4
P35443
log10 protein intensity
1053:1053
14:15
5:79370247:G:T_T
p = 0.96, beta = −0.00382, N = 1259
AFAGPSQKPETIELR pc3
E7ES19;P35443
log10 peptide intensity
1052:1053
14:15
5:79370247:G:T_T
p = 0.97, beta = −0.00296, N = 1258
PGYTGDQIR pc2
E7ES19;P35443
log10 peptide intensity
1043:1053
14:15
5:79370247:G:T_T
p = 0.043, beta = −0.139, N = 1248
YFEFTVMGR pc2
P35443
log10 peptide intensity
1041:1053
14:15
5:79370247:G:T_T
p = 0.2, beta = −0.0878, N = 1244
GAGSLELYLDCIQVDSVHNLPR pc3
E7ES19;P35443
log10 peptide intensity
1023:1053
13:15
5:79370247:G:T_T
p = 0.23, beta = −0.0833, N = 1224
THBS4 : NP4
P35443
log10 protein intensity
277:277
44:44
4:4
5:79370247:G:T_T
p = 0.0034, beta = 0.4, N = 325
AFAGPSQKPETIELR pc3
E7ES19;P35443
log10 peptide intensity
277:277
44:44
4:4
5:79370247:G:T_T
p = 0.00098, beta = 0.449, N = 325
VHLVVFNNLQLADGRR pc3
E7ES19;P35443
log10 peptide intensity
274:277
43:44
4:4
5:79370247:G:T_T
p = 0.18, beta = 0.184, N = 321
GAGSLELYLDCIQVDSVHNLPR pc3
E7ES19;P35443
log10 peptide intensity
272:277
41:44
4:4
5:79370247:G:T_T
p = 0.37, beta = 0.125, N = 317
YFEFTVMGR pc2
P35443
log10 peptide intensity
266:277
44:44
4:4
5:79370247:G:T_T
p = 0.056, beta = 0.262, N = 314
Assay Target: THBS4
Olink UniProt: P35443
deCODE rsID: rs13167730
Proxy rsID: rs13167730
deCODE: 5:80074424:T:G
Proxy SNP: 5:79370247:G:T
deCODE log10(p): 132.2
deCODE BETA: 0.38
1258:1248:1244:1224:1215:120

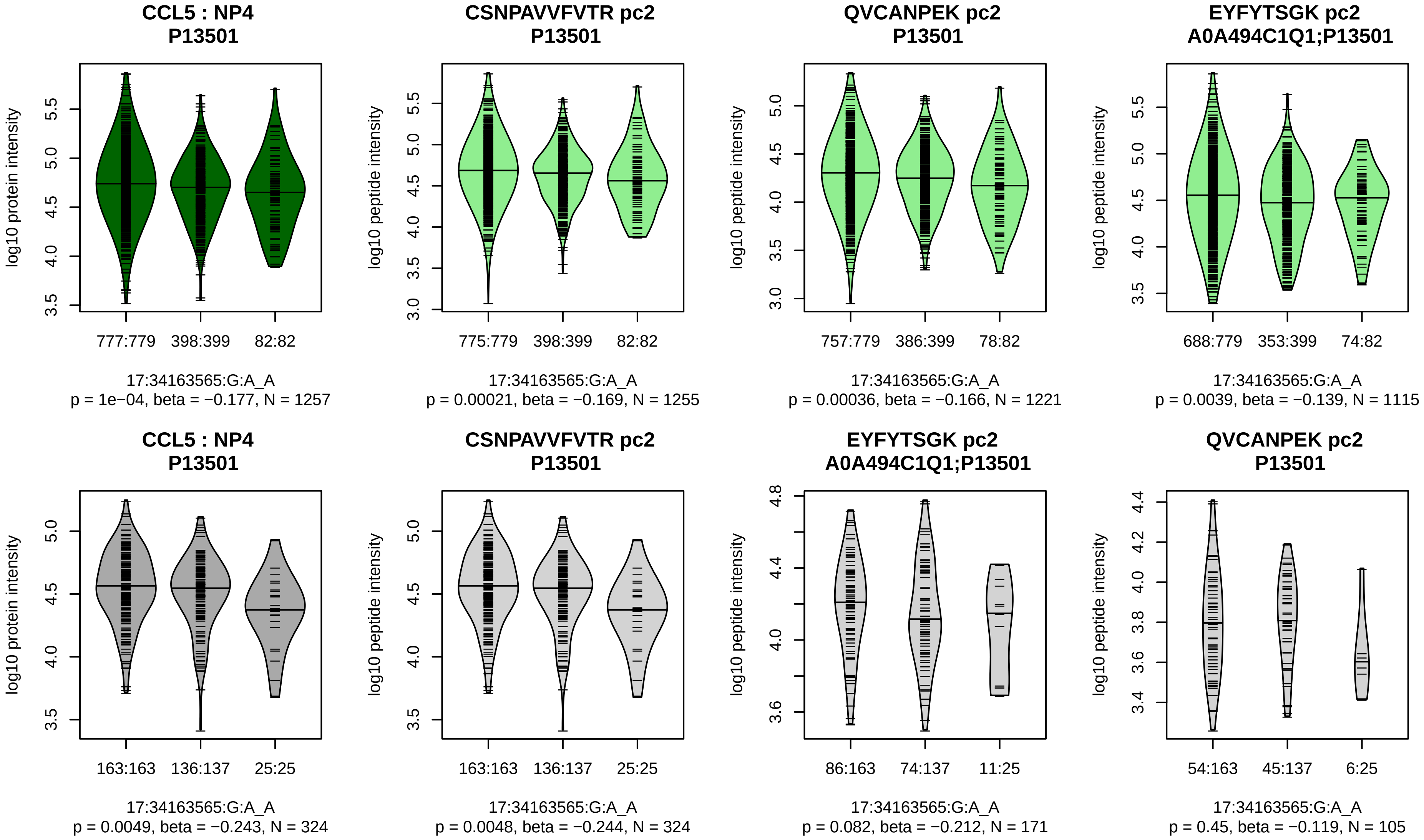


Assay Target: CCL5
Olink UniProt: P13501
deCODE rsID: rs4239252
Proxy rsID: rs4239252
deCODE: 17:35836561:A:G
Proxy SNP: 17:34163565:G:A
deCODE log10(p): 132
deCODE BETA: −0.26
*:*:*
1255:1221:1115

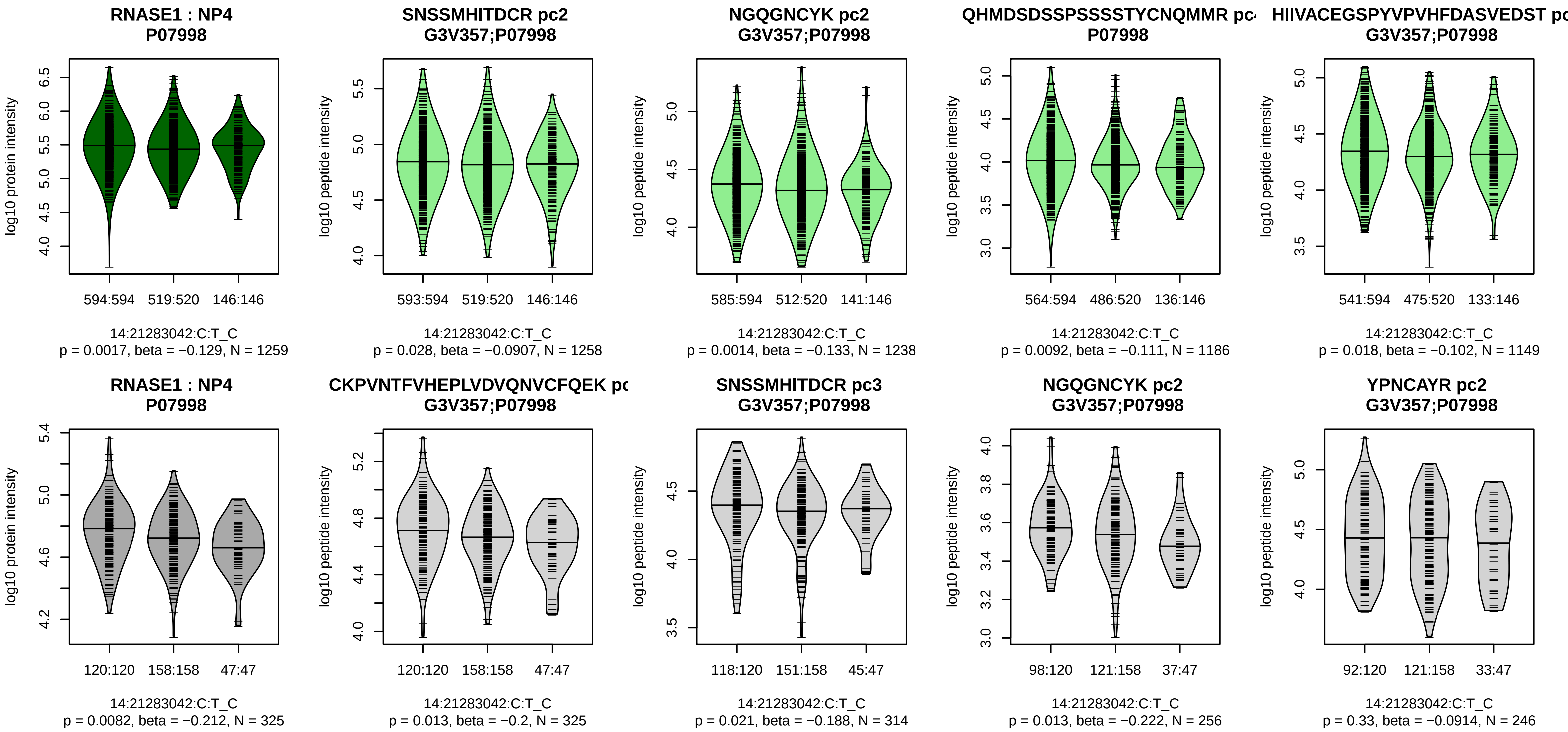


Assay Target: RNASE1
Olink UniProt: P07998
deCODE rsID: rs12897030
Proxy rsID: rs12897030
deCODE: 14:20814883:C:T
Proxy SNP: 14:21283042:C:T
deCODE log10(p): 130.1
deCODE BETA: −0.22
–:–:*:*:–:–:–:–
1258:1256:1238:1186:1149:114

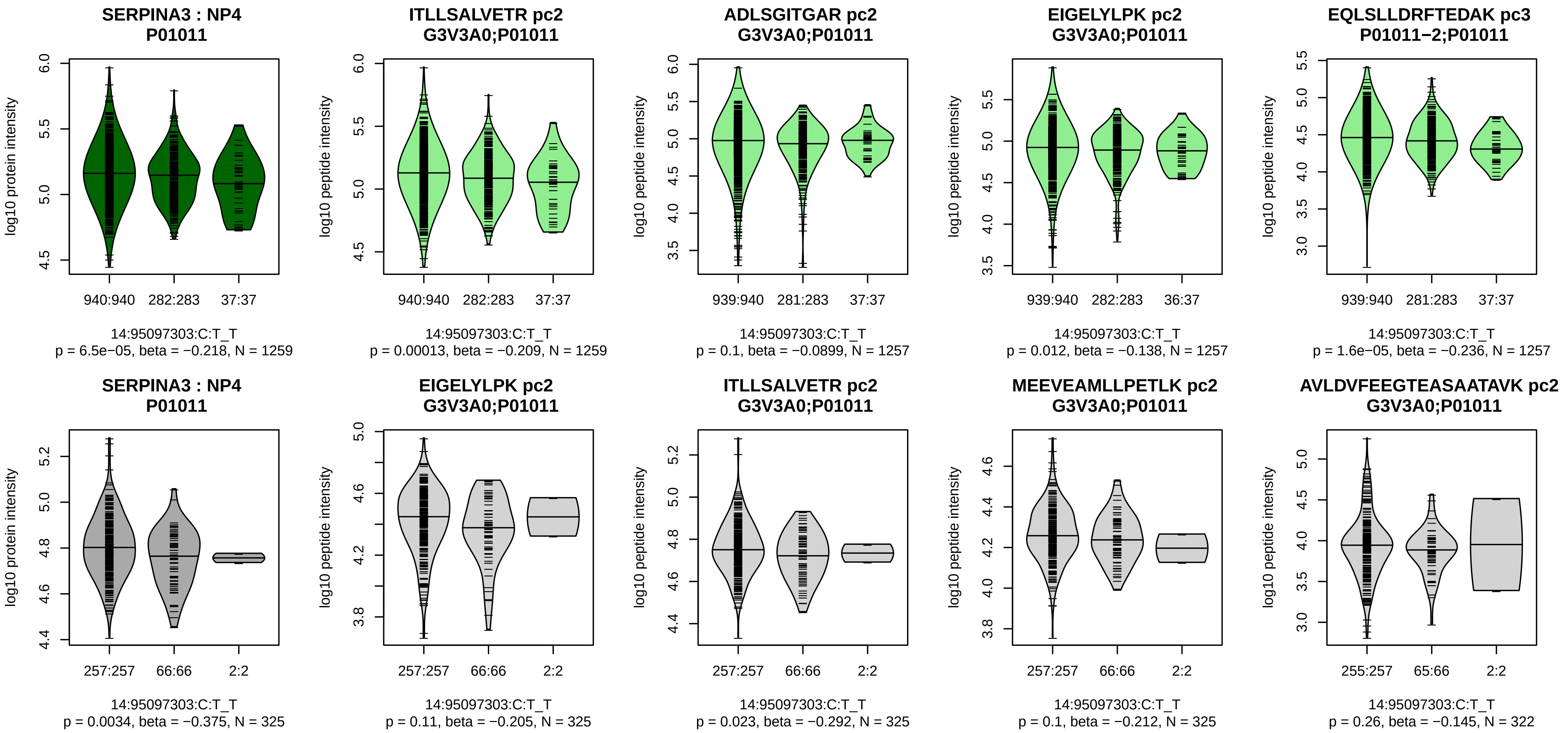


Assay Target: SERPINA3
Olink UniProt: P01011
deCODE rsID: rs140735783
Proxy rsID: rs6575449
deCODE: 14:94634064:TC:T
Proxy SNP: 14:95097303:C:T
deCODE log10(p): 128.2
deCODE BETA: −0.24
*:–:–:*:*:–:*:*:–:*:–:–:–:–:–:–:–:–
1259:1257:1257:1257:1248:124

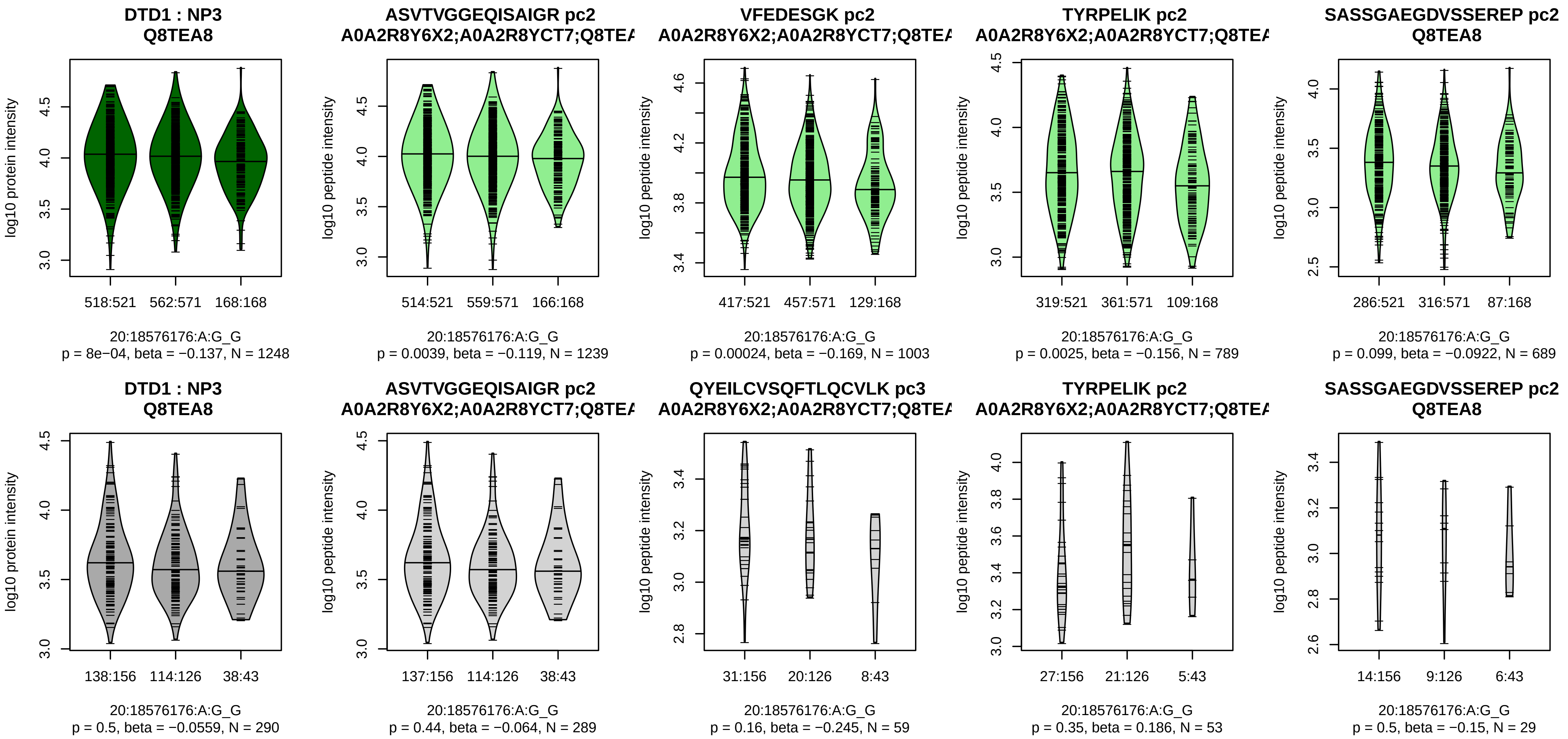


Assay Target: DTD1
Olink UniProt: Q8TEA8
deCODE rsID: rs6081231
Proxy rsID: rs6081235
deCODE: 20:18590320:A:G
Proxy SNP: 20:18576176:A:G
deCODE log10(p): 127.8
deCODE BETA: −0.2
*:*:*:–:*:–:–
1239:1003:789:689:604:456:254

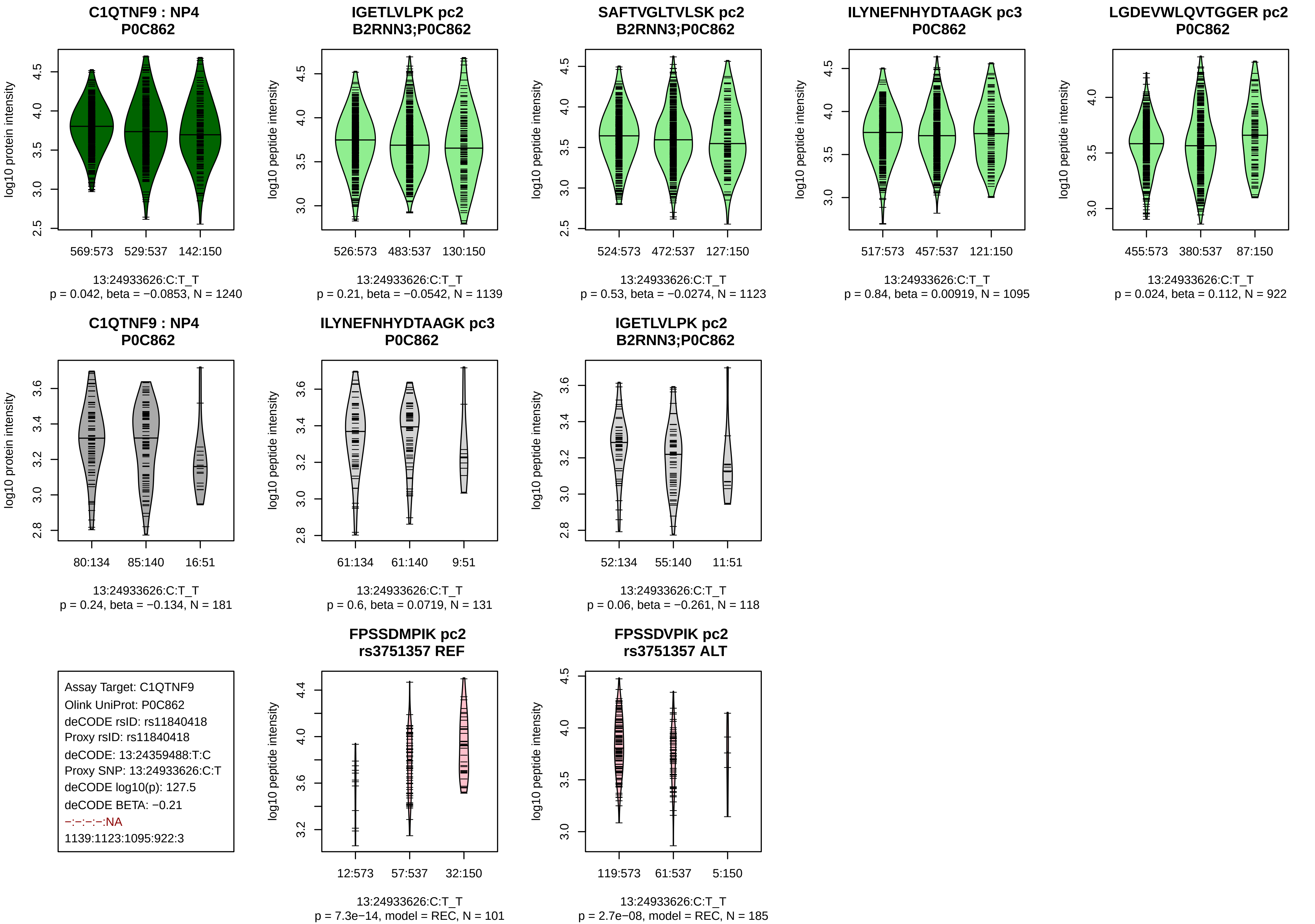
C1QTNF9 : NP4
P0C862
log10 protein intensity
569:573
529:537
142:150
13:24933626:C:T_T
p = 0.042, beta = −0.0853, N = 1240
IGETLVLPK pc2
B2RNN3;P0C862
log10 peptide intensity
526:573
483:537
130:150
13:24933626:C:T_T
p = 0.21, beta = −0.0542, N = 1139
SAFTVGLTVLSK pc2
B2RNN3;P0C862
log10 peptide intensity
524:573
472:537
127:150
13:24933626:C:T_T
p = 0.53, beta = −0.0274, N = 1123
ILYNEFNHYDTAAGK pc3
P0C862
log10 peptide intensity
517:573
457:537
121:150
13:24933626:C:T_T
p = 0.84, beta = 0.00919, N = 1095
LGDEVWLQVTGGER pc2
P0C862
log10 peptide intensity
455:573
380:537
87:150
13:24933626:C:T_T
p = 0.024, beta = 0.112, N = 922
C1QTNF9 : NP4
P0C862
log10 protein intensity
80:134
85:140
16:51
13:24933626:C:T_T
p = 0.24, beta = −0.134, N = 181
ILYNEFNHYDTAAGK pc3
P0C862
log10 peptide intensity
61:134
61:140
9:51
13:24933626:C:T_T
p = 0.6, beta = 0.0719, N = 131
IGETLVLPK pc2
B2RNN3;P0C862
log10 peptide intensity
52:134
55:140
11:51
13:24933626:C:T_T
p = 0.06, beta = −0.261, N = 118
Assay Target: C1QTNF9
Olink UniProt: P0C862
deCODE rsID: rs11840418
Proxy rsID: rs11840418
deCODE: 13:24359488:T:C
Proxy SNP: 13:24933626:C:T
deCODE log10(p): 127.5
deCODE BETA: −0.21
−:−:−:−:NA
1139:1123:1095:922:3
FPSSDMPIK pc2
rs3751357 REF
log10 peptide intensity
12:573
57:537
32:150
13:24933626:C:T_T
p = 7.3e−14, model = REC, N = 101
FPSSDVPIK pc2
rs3751357 ALT
log10 peptide intensity
119:573
61:537
5:150
13:24933626:C:T_T
p = 2.7e−08, model = REC, N = 185

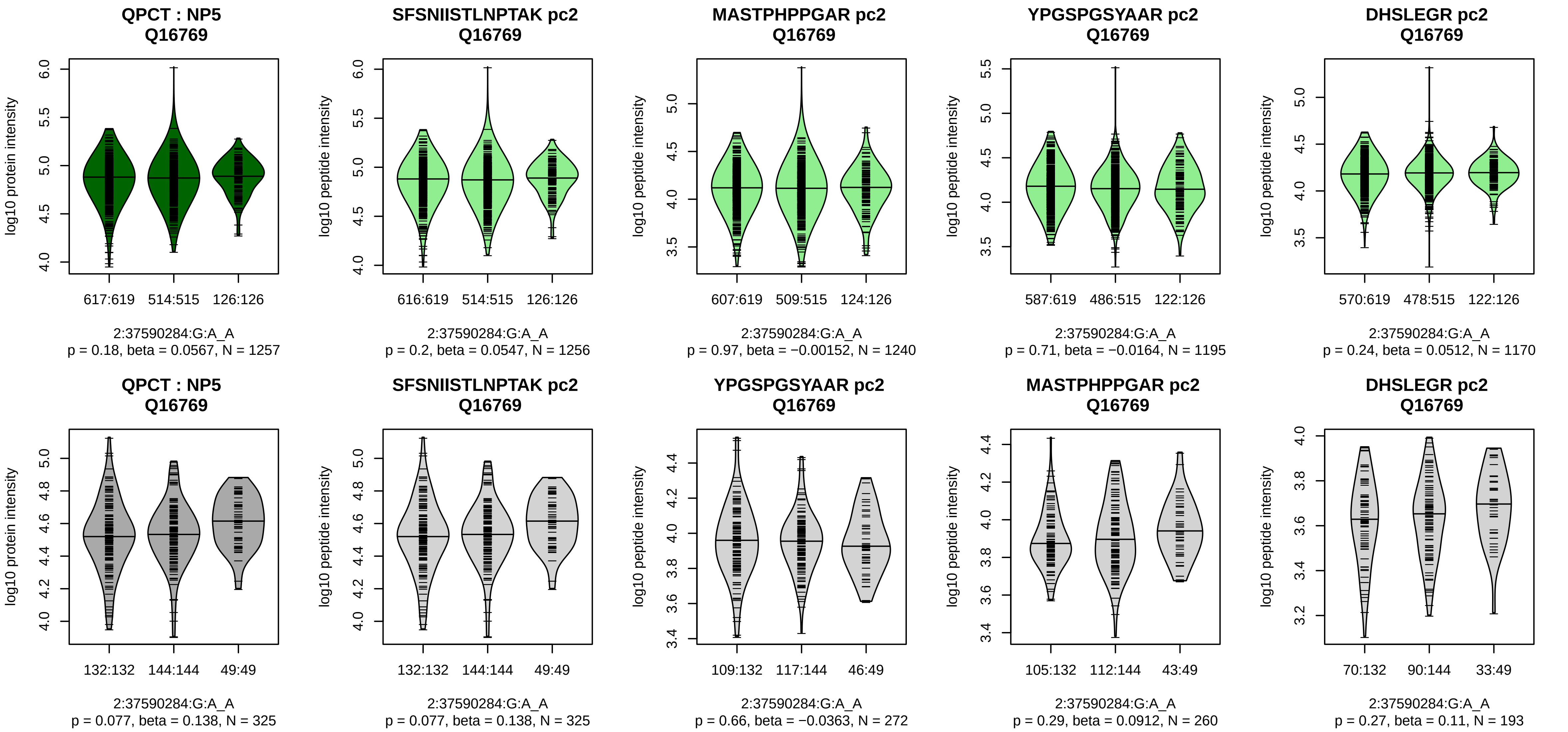


Assay Target: QPCT
Olink UniProt: Q16769
deCODE rsID: rs4384764
Proxy rsID: rs4384764
deCODE: 2:37363141:A:G
Proxy SNP: 2:37590284:G:A
deCODE log10(p): 126.5
deCODE BETA: 0.22
–:–:–:–:–:–:–:–:–:–
1256:1240:1195:1170:1112:973

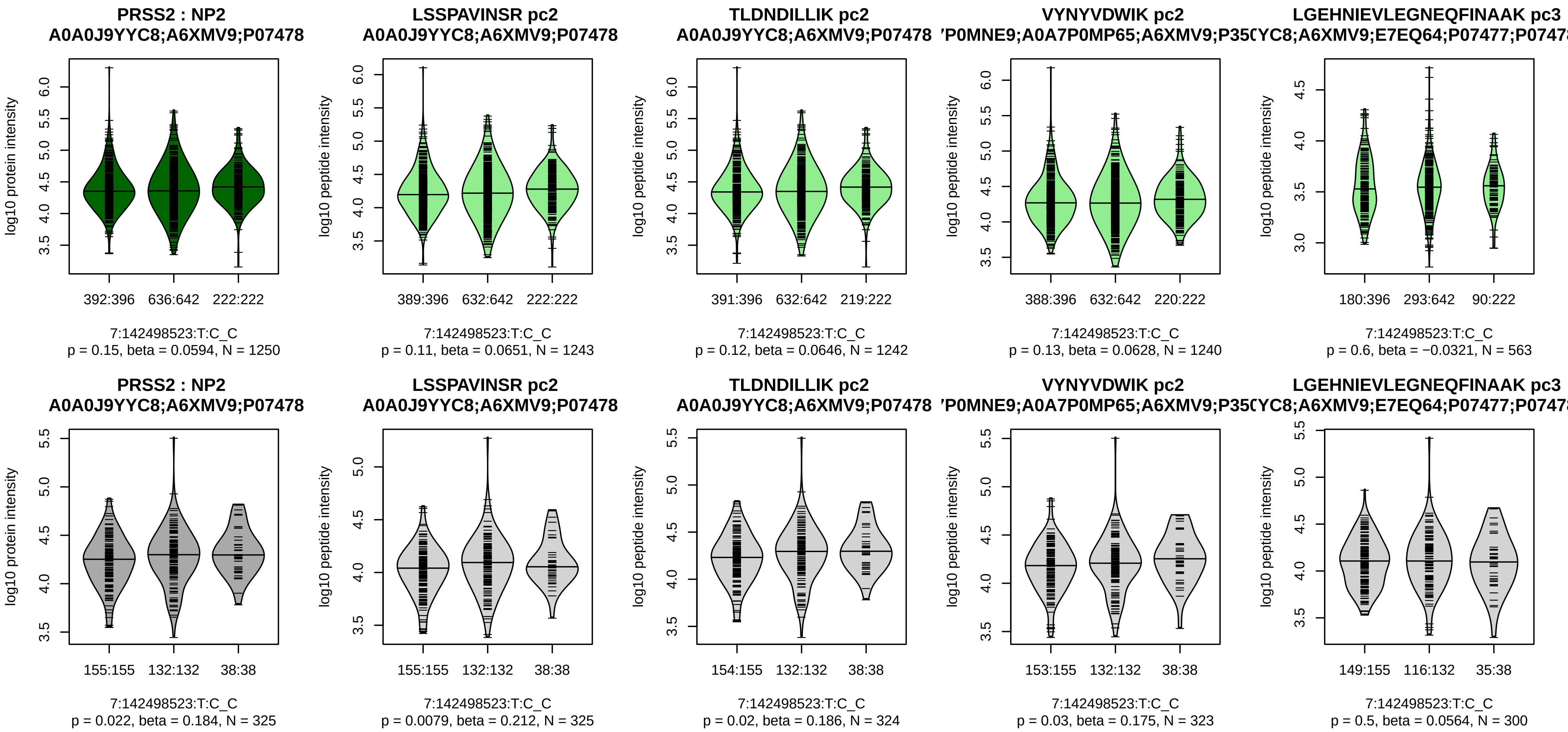


Assay Target: PRSS2
Olink UniProt: P07478
deCODE rsID: rs1799886
Proxy rsID: rs1799886
deCODE: 7:142800839:C:T
Proxy SNP: 7:142498523:T:C
deCODE log10(p): 123.9
deCODE BETA: 0.2
–:–:–:–:–
1243:1242:1240:563:425

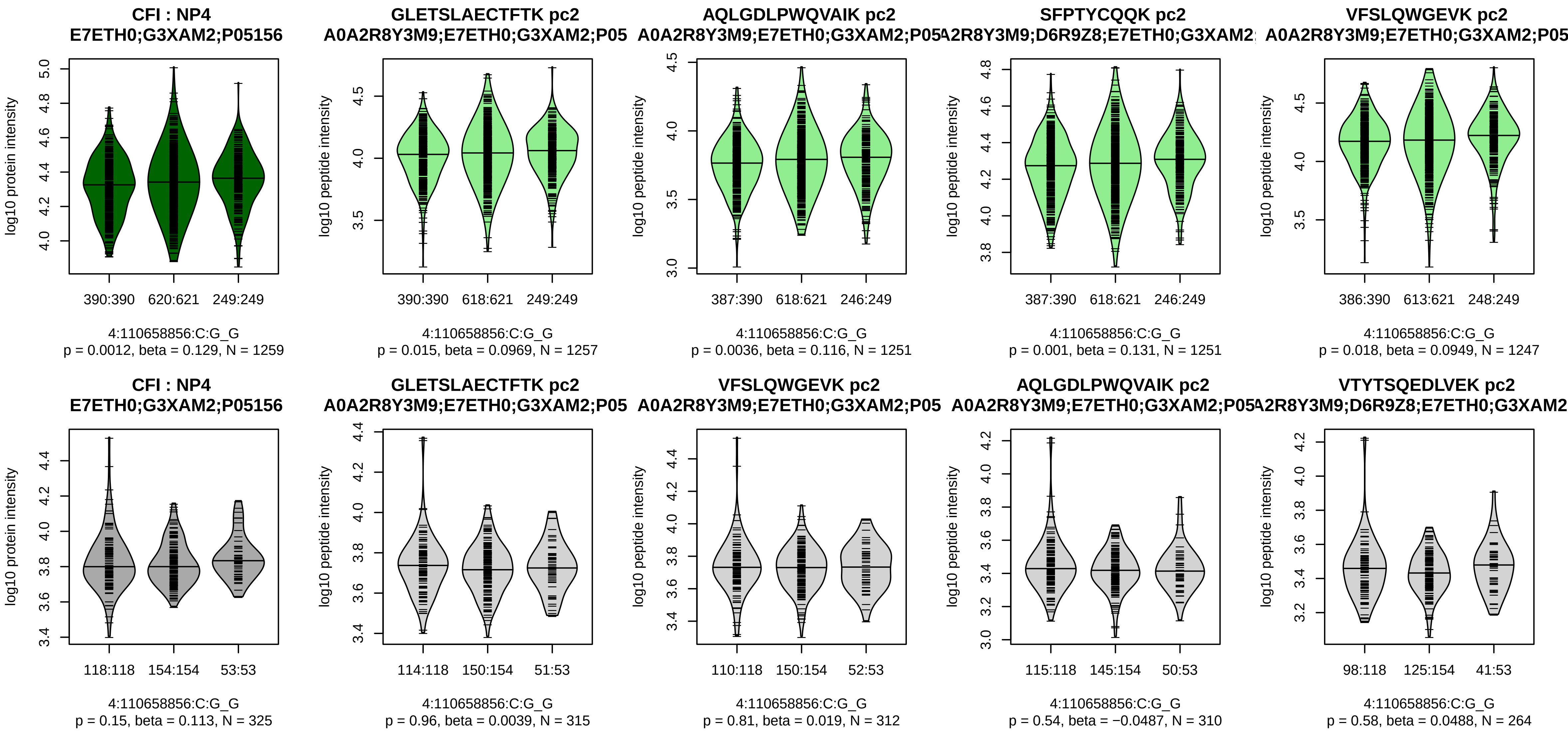


Assay Target: CFI
Olink UniProt: P05156
deCODE rsID: rs13117504
Proxy rsID: rs13117504
deCODE: 4:109737700:G:C
Proxy SNP: 4:110658856:C:G
deCODE log10(p): 123.5
deCODE BETA: 0.2
–:*:*:–:–:*:*:–:–:–:–:–:–:–:–:–:–:
1257:1251:1251:1247:1234:122

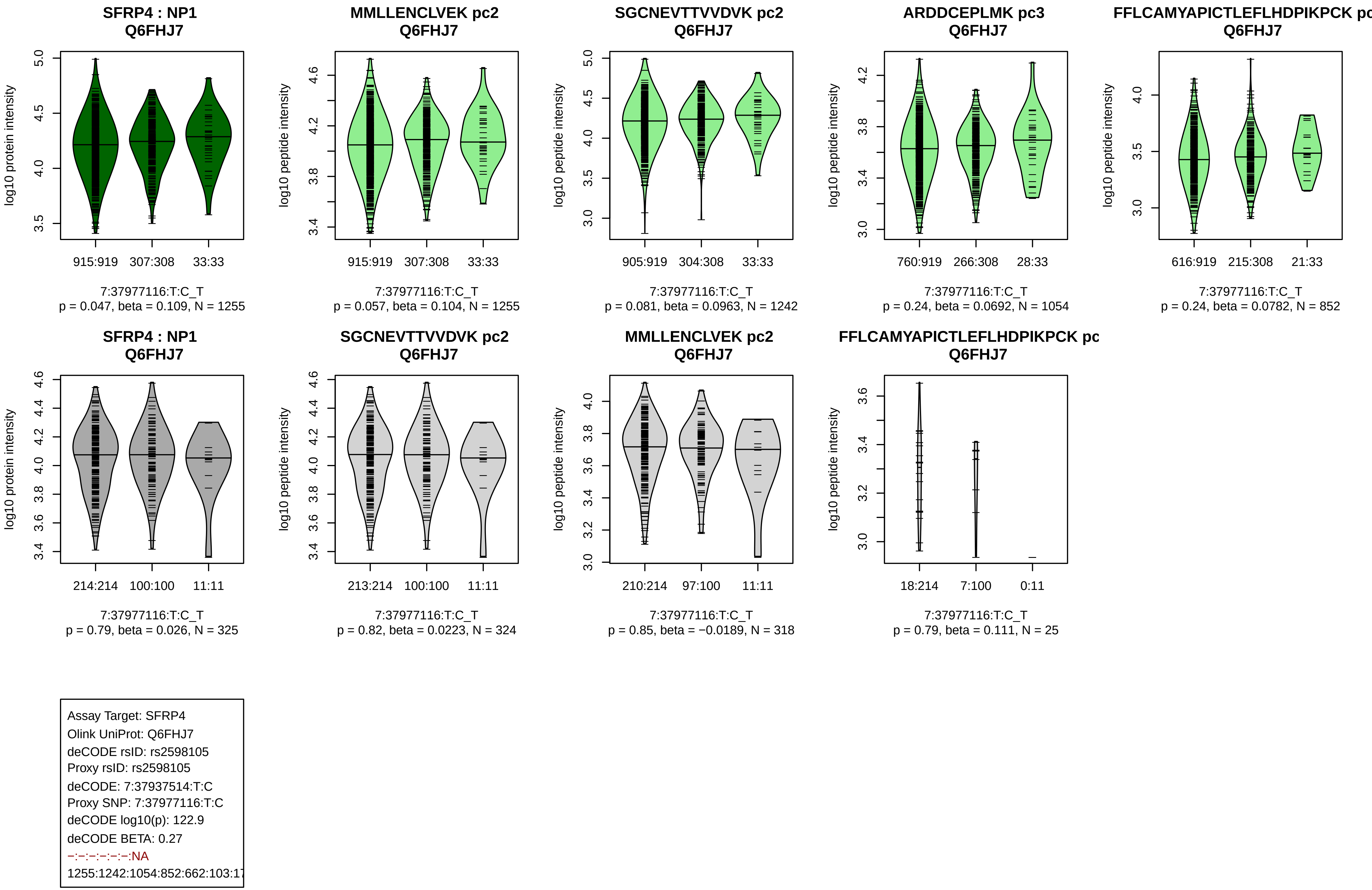
SFRP4 : NP1
Q6FHJ7
log10 protein intensity
915:919 307:308 33:33
7:37977116:T:C_T
p = 0.047, beta = 0.109, N = 1255
MMLLENCLVEK pc2
Q6FHJ7
log10 peptide intensity
915:919 307:308 33:33
7:37977116:T:C_T
p = 0.057, beta = 0.104, N = 1255
SGCNEVTTVVDVK pc2
Q6FHJ7
log10 peptide intensity
905:919 304:308 33:33
7:37977116:T:C_T
p = 0.081, beta = 0.0963, N = 1242
ARDDCEPLMK pc3
Q6FHJ7
log10 peptide intensity
760:919 266:308 28:33
7:37977116:T:C_T
p = 0.24, beta = 0.0692, N = 1054
FFLCAMYAPICTLEFLHDPIKPCK pc
Q6FHJ7
log10 peptide intensity
616:919 215:308 21:33
7:37977116:T:C_T
p = 0.24, beta = 0.0782, N = 852
SFRP4 : NP1
Q6FHJ7
log10 protein intensity
214:214 100:100 11:11
7:37977116:T:C_T
p = 0.79, beta = 0.026, N = 325
SGCNEVTTVVDVK pc2
Q6FHJ7
log10 peptide intensity
213:214 100:100 11:11
7:37977116:T:C_T
p = 0.82, beta = 0.0223, N = 324
MMLLENCLVEK pc2
Q6FHJ7
log10 peptide intensity
210:214 97:100 11:11
7:37977116:T:C_T
p = 0.85, beta = −0.0189, N = 318
FFLCAMYAPICTLEFLHDPIKPCK pc
Q6FHJ7
log10 peptide intensity
18:214 7:100 0:11
7:37977116:T:C_T
p = 0.79, beta = 0.111, N = 25
Assay Target: SFRP4
Olink UniProt: Q6FHJ7
deCODE rsID: rs2598105
Proxy rsID: rs2598105
deCODE: 7:37937514:T:C
Proxy SNP: 7:37977116:T:C
deCODE log10(p): 122.9
deCODE BETA: 0.27
−:−:−:−:−:−:NA
1255:1242:1054:852:662:103:17

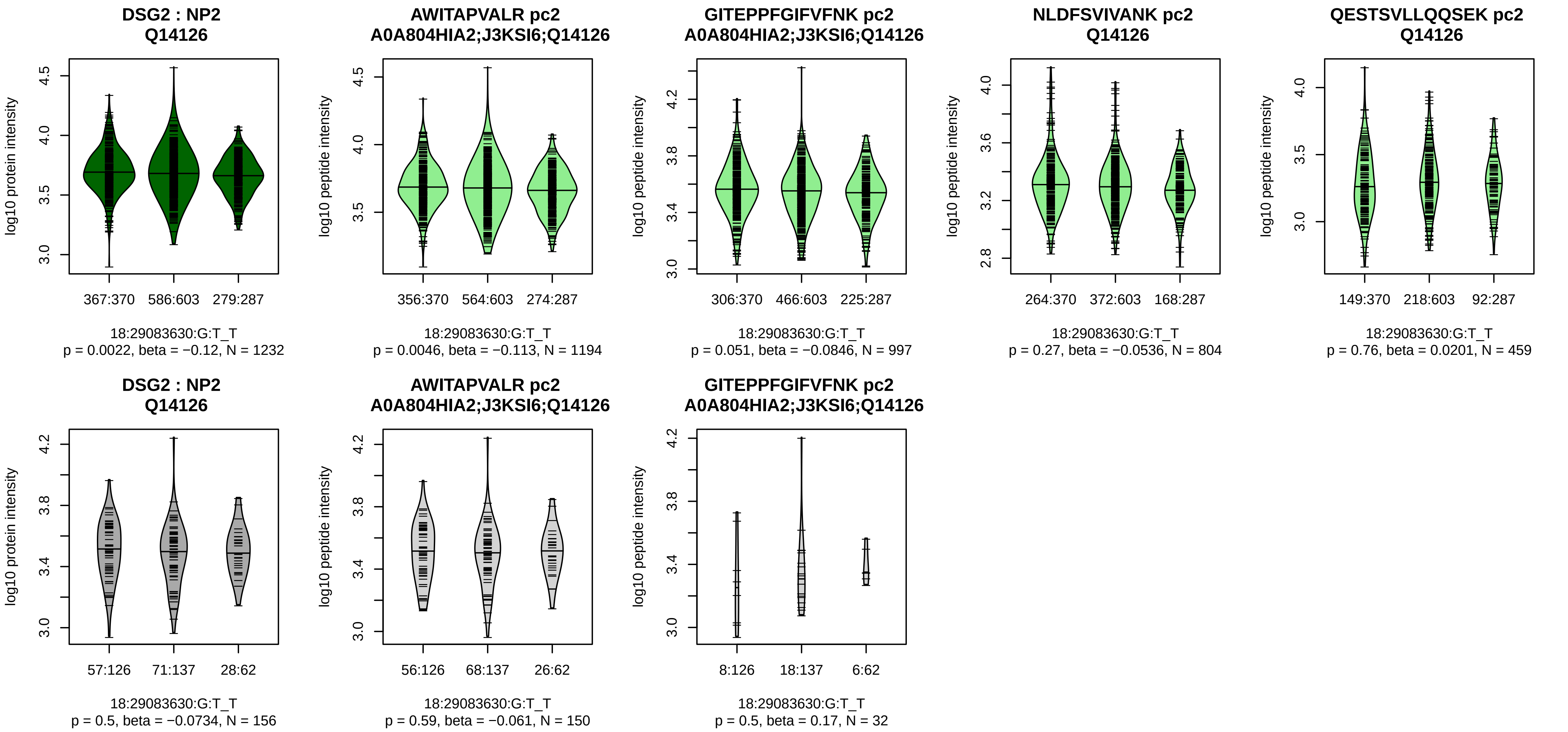


Assay Target: DSG2
Olink UniProt: Q14126
deCODE rsID: rs9304098
Proxy rsID: rs9304098
deCODE: 18:31503667:T:G
Proxy SNP: 18:29083630:G:T
deCODE log10(p): 118.8
deCODE BETA: −0.19
*:−:−:−:−:−:−:−
1194:997:804:459:313:275:252:

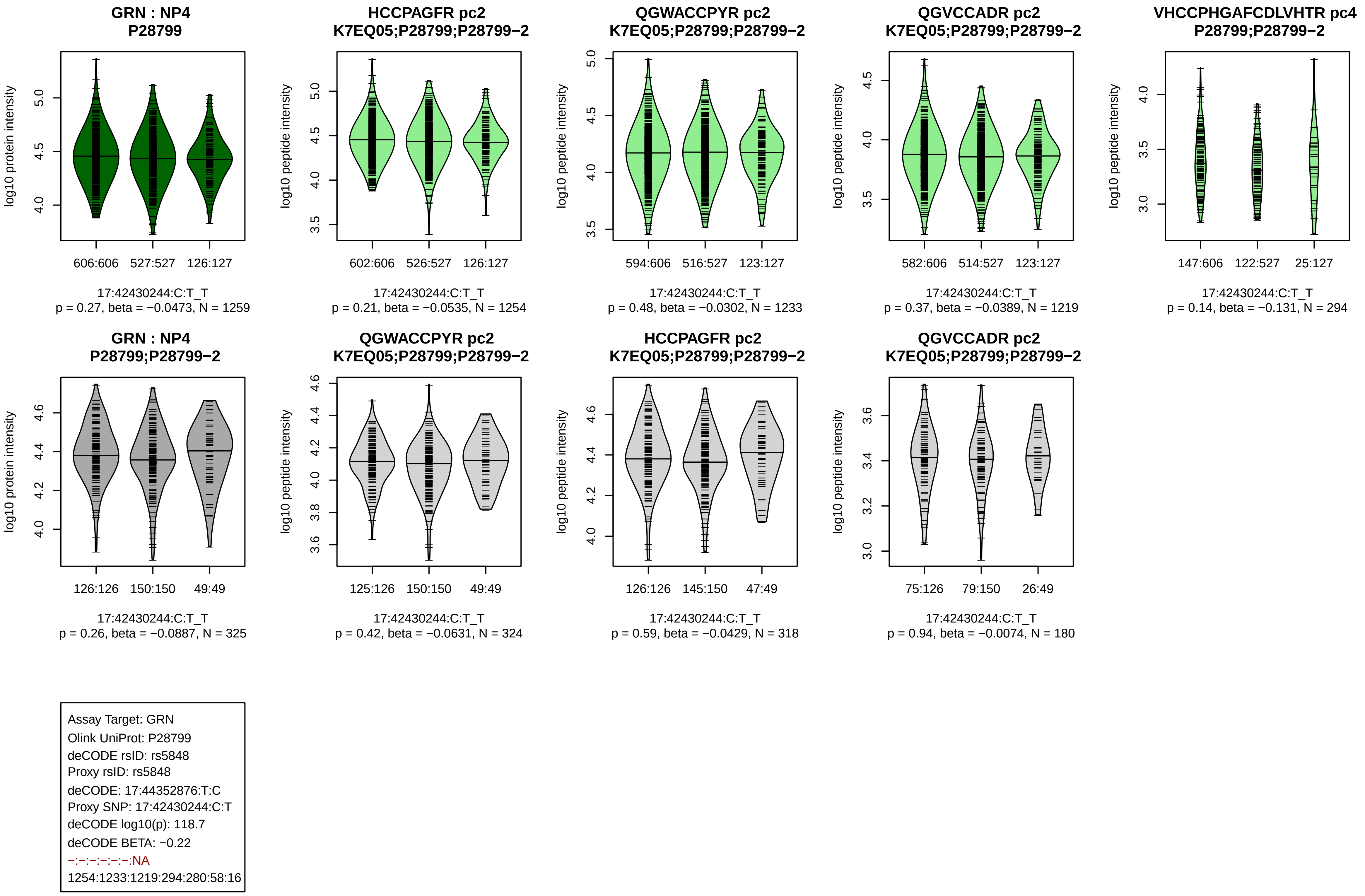

GRN : NP4
P28799
log10 protein intensity
606:606
527:527
126:127
17:42430244:C:T_T
p = 0.27, beta = −0.0473, N = 1259
HCCPAGFR pc2
K7EQ05;P28799;P28799−2
log10 peptide intensity
602:606
526:527
126:127
17:42430244:C:T_T
p = 0.21, beta = −0.0535, N = 1254
QGWACCPYR pc2
K7EQ05;P28799;P28799−2
log10 peptide intensity
594:606
516:527
123:127
17:42430244:C:T_T
p = 0.48, beta = −0.0302, N = 1233
QGVCCADR pc2
K7EQ05;P28799;P28799−2
log10 peptide intensity
582:606
514:527
123:127
17:42430244:C:T_T
p = 0.37, beta = −0.0389, N = 1219
VHCCPHGAFCDLVHTR pc4
P28799;P28799−2
log10 peptide intensity
147:606
122:527
25:127
17:42430244:C:T_T
p = 0.14, beta = −0.131, N = 294
GRN : NP4
P28799;P28799−2
log10 protein intensity
126:126
150:150
49:49
17:42430244:C:T_T
p = 0.26, beta = −0.0887, N = 325
QGWACCPYR pc2
K7EQ05;P28799;P28799−2
log10 peptide intensity
125:126
150:150
49:49
17:42430244:C:T_T
p = 0.42, beta = −0.0631, N = 324
HCCPAGFR pc2
K7EQ05;P28799;P28799−2
log10 peptide intensity
126:126
145:150
47:49
17:42430244:C:T_T
p = 0.59, beta = −0.0429, N = 318
QGVCCADR pc2
K7EQ05;P28799;P28799−2
log10 peptide intensity
75:126
79:150
26:49
17:42430244:C:T_T
p = 0.94, beta = −0.0074, N = 180
Assay Target: GRN
Olink UniProt: P28799
deCODE rsID: rs5848
Proxy rsID: rs5848
deCODE: 17:44352876:T:C
Proxy SNP: 17:42430244:C:T
deCODE log10(p): 118.7
deCODE BETA: −0.22
−:−:−:−:−:−:NA
1254:1233:1219:294:280:58:16

**MAN1C1 : NP2**
**Q9NR34**

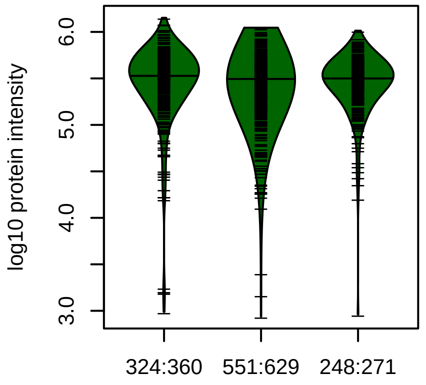


1:26008659:T:C_C
p = 0.11, beta = −0.0663, N = 1123

**ELAAQITK pc2**
**A0A087WZ31;A6NGN6;B1AJZ5;Q9NI**

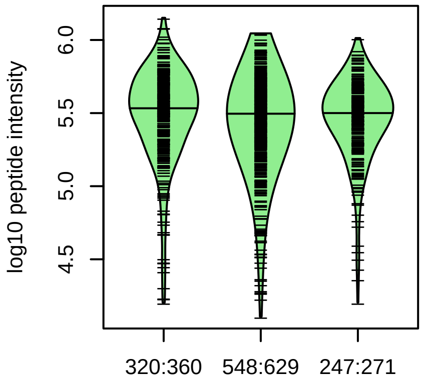


1:26008659:T:C_C
p = 0.073, beta = −0.0754, N = 1115

**TQQPGLEVVAEIAGHAPAR pc3**
**Q9NR34**

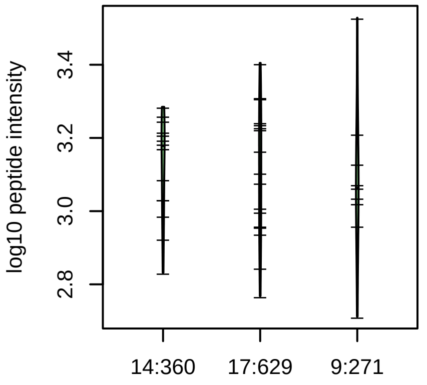


1:26008659:T:C_C
p = 0.63, beta = −0.0978, N = 40

**LLPAFNTPTGIPK pc2**
**A0A087WZ31;A6NGN6;B1AJZ5;Q9NI**

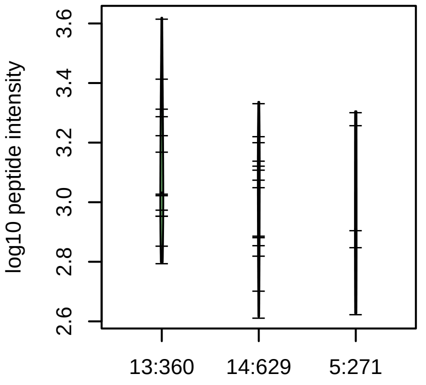


1:26008659:T:C_C
p = 0.28, beta = −0.25, N = 32

**MAN1C1 : NP2**
**A6NGN6;B1AJZ5;Q9NR34**

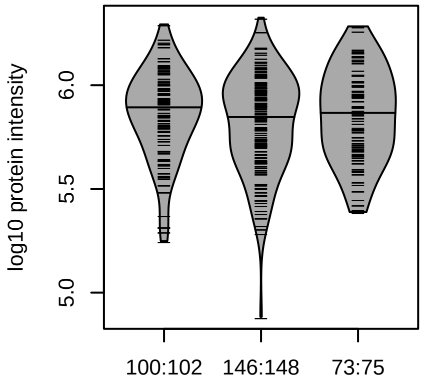


1:26008659:T:C_C
p = 0.43, beta = 0.0602, N = 319

**ELAAQITK pc2**
**A0A087WZ31;A6NGN6;B1AJZ5;Q9NI**

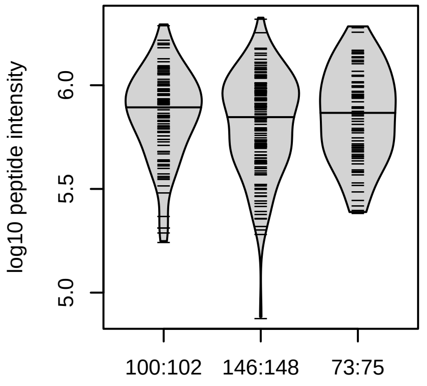


1:26008659:T:C_C
p = 0.43, beta = 0.0602, N = 319

Assay Target: MAN1C1
Olink UniProt: Q9NR34
deCODE rsID: rs11247595
Proxy rsID: rs11247595
deCODE: 1:25682168:T:C
Proxy SNP: 1:26008659:T:C
deCODE log10(p): 116.6
deCODE BETA: 0.19
−:−:−
1115:40:32

**SPOCK2 : NP2**
**Q92563**

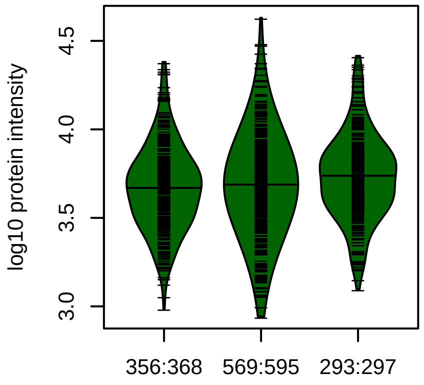


10:73845884:G:A_A
p = 0.028, beta = 0.0858, N = 1218

**LEQQACLSSK pc2**
**Q92563**

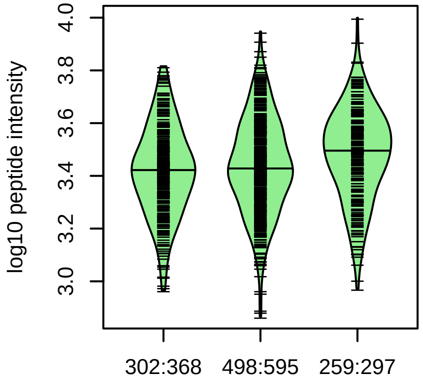


10:73845884:G:A_A
p = 0.0032, beta = 0.124, N = 1059

**DSIGWMFSK pc2**
**Q92563**

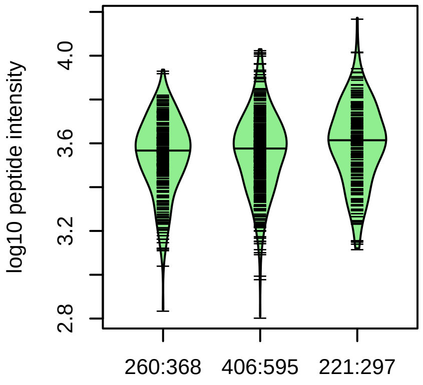


10:73845884:G:A_A
p = 0.01, beta = 0.116, N = 887

**VCIAQGYQR pc2**
**Q92563**

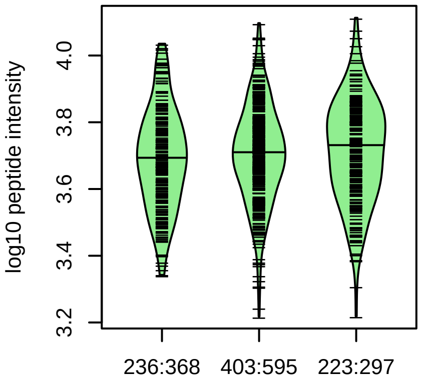


10:73845884:G:A_A
p = 0.46, beta = 0.0345, N = 862

**PCHMAQLASVCGSDGHTYSSVCK p**
**Q92563**

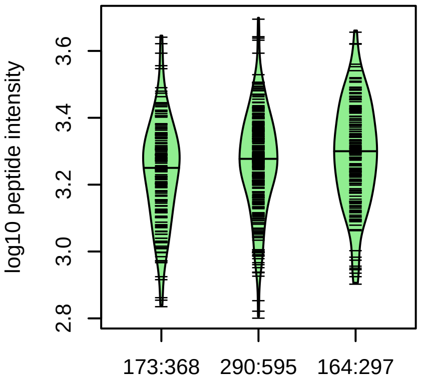


10:73845884:G:A_A
p = 0.0055, beta = 0.15, N = 627

**SPOCK2 : NP2**
**Q92563**

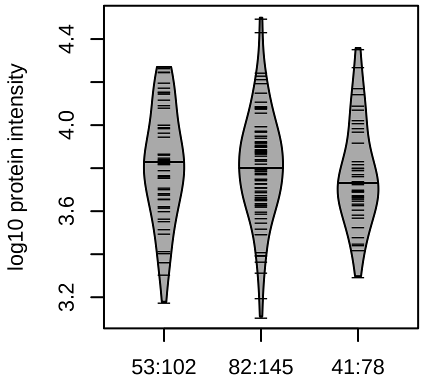


10:73845884:G:A_G
p = 0.42, beta = −0.0823, N = 176

**DSIGWMFSK pc2**
**Q92563**

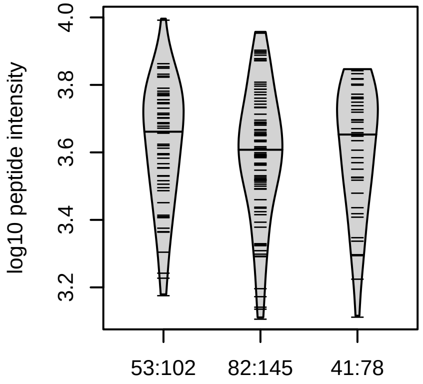


10:73845884:G:A_G
p = 0.7, beta = −0.0398, N = 176

Assay Target: SPOCK2
Olink UniProt: Q92563
deCODE rsID: rs1245548
Proxy rsID: rs1245548
deCODE: 10:72086126:A:G
Proxy SNP: 10:73845884:G:A
deCODE log10(p): 116
deCODE BETA: 0.19
*:–:–:*:*:–
1059:887:862:627:541:243

**S100A12 : NP5**
**P80511**

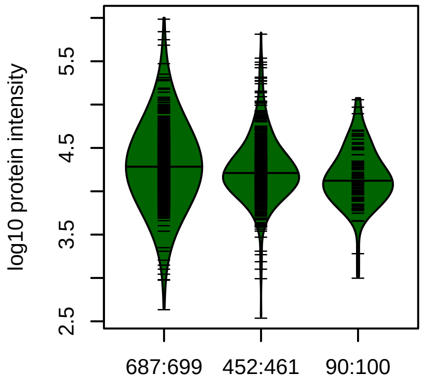


1:153337943:G:A_A
p = 6.2e−05, beta = −0.18, N = 1229

**GHFDTLSK pc2**
**P80511**

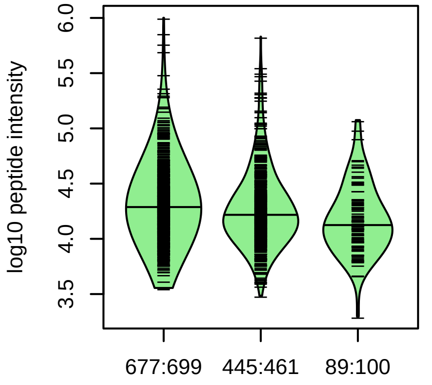


1:153337943:G:A_A
p = 2e−05, beta = −0.193, N = 1211

**TKLEEHLEGIVNIFHQYSVR pc4**
**P80511**

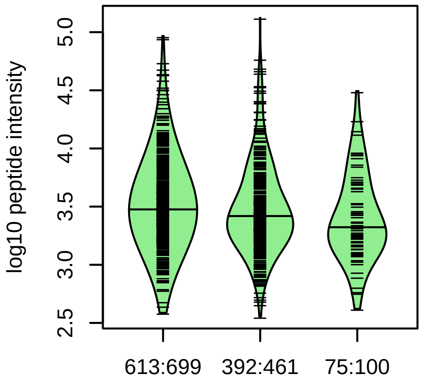


1:153337943:G:A_A
p = 0.00067, beta = −0.165, N = 1080

**LEEHLEGIVNIFHQYSVRK pc4**
**P80511**

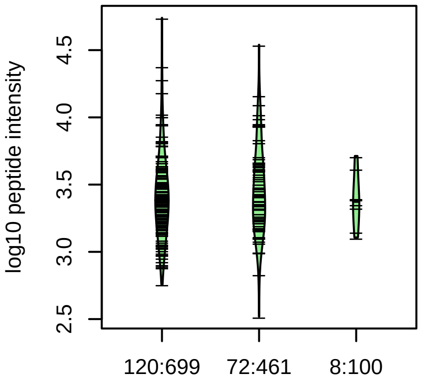


1:153337943:G:A_A
p = 0.81, beta = 0.03, N = 200

**LEEHLEGIVNIFHQYSVR pc4**
**P80511**

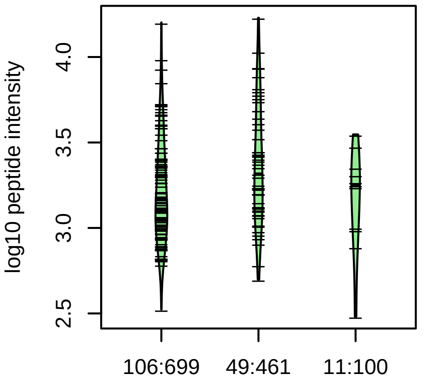


1:153337943:G:A_A
p = 0.29, beta = 0.13, N = 166

**S100A12 : NP5**
**P80511**

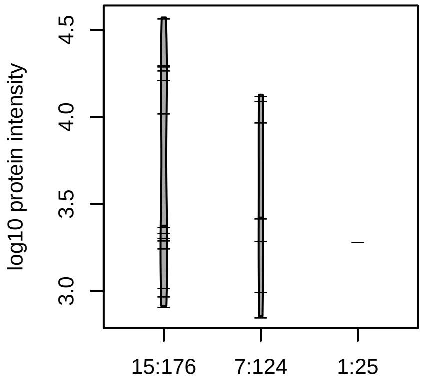


1:153337943:G:A_A
p = 0.35, beta = −0.314, N = 23

Assay Target: S100A12
Olink UniProt: P80511
deCODE rsID: rs3014874
Proxy rsID: rs3014874
deCODE: 1:153365467:A:G
Proxy SNP: 1:153337943:G:A
deCODE log10(p): 114.8
deCODE BETA: −0.22
*:*:−:−:−:−
1211:1080:200:166:55:39

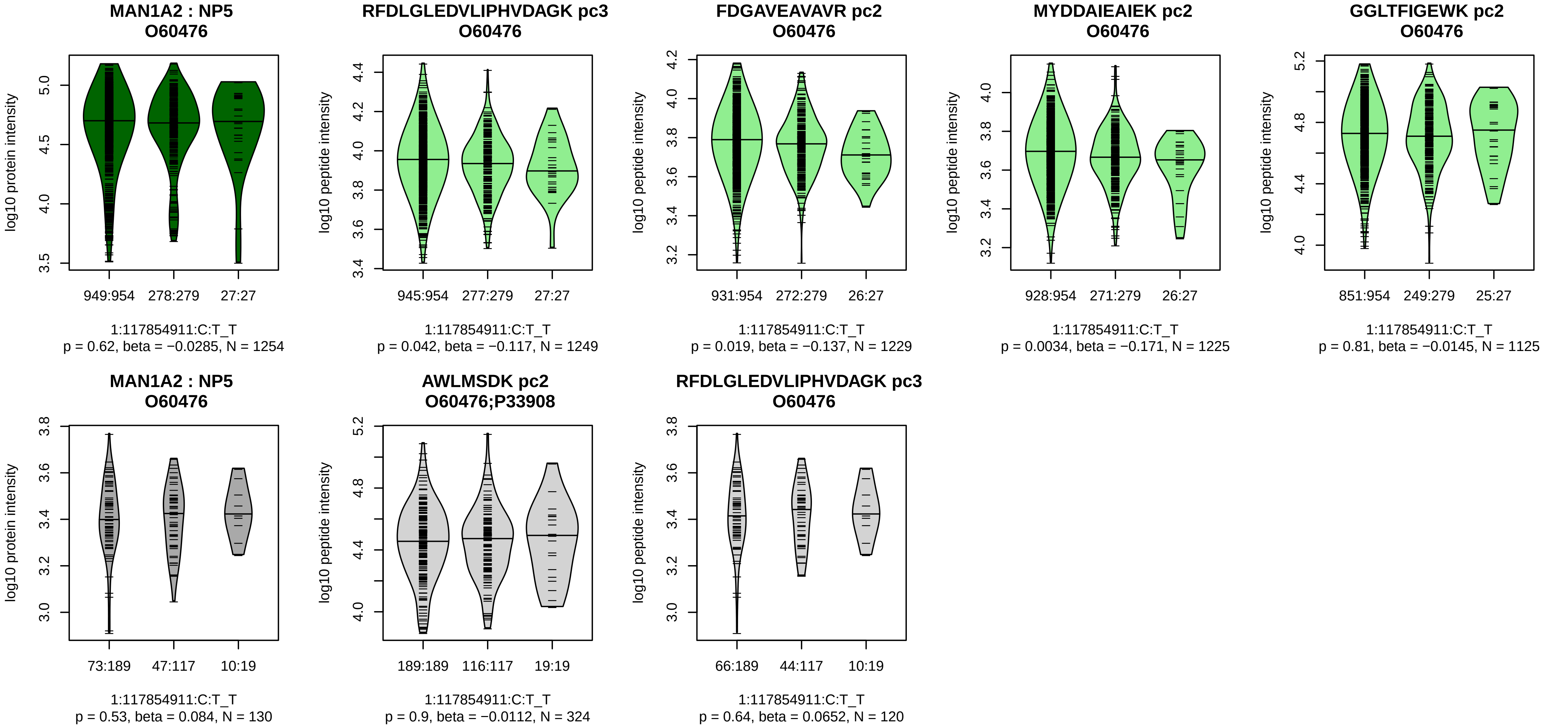


Assay Target: MAN1A2
Olink UniProt: O60476
deCODE rsID: rs73013841
Proxy rsID: rs73013841
deCODE: 1:117312289:T:C
Proxy SNP: 1:117854911:C:T
deCODE log10(p): 109.1
deCODE BETA: −0.3
−:−:*:−:−:−:−:−:−:−:−:−
1249:1229:1225:1125:1124:998

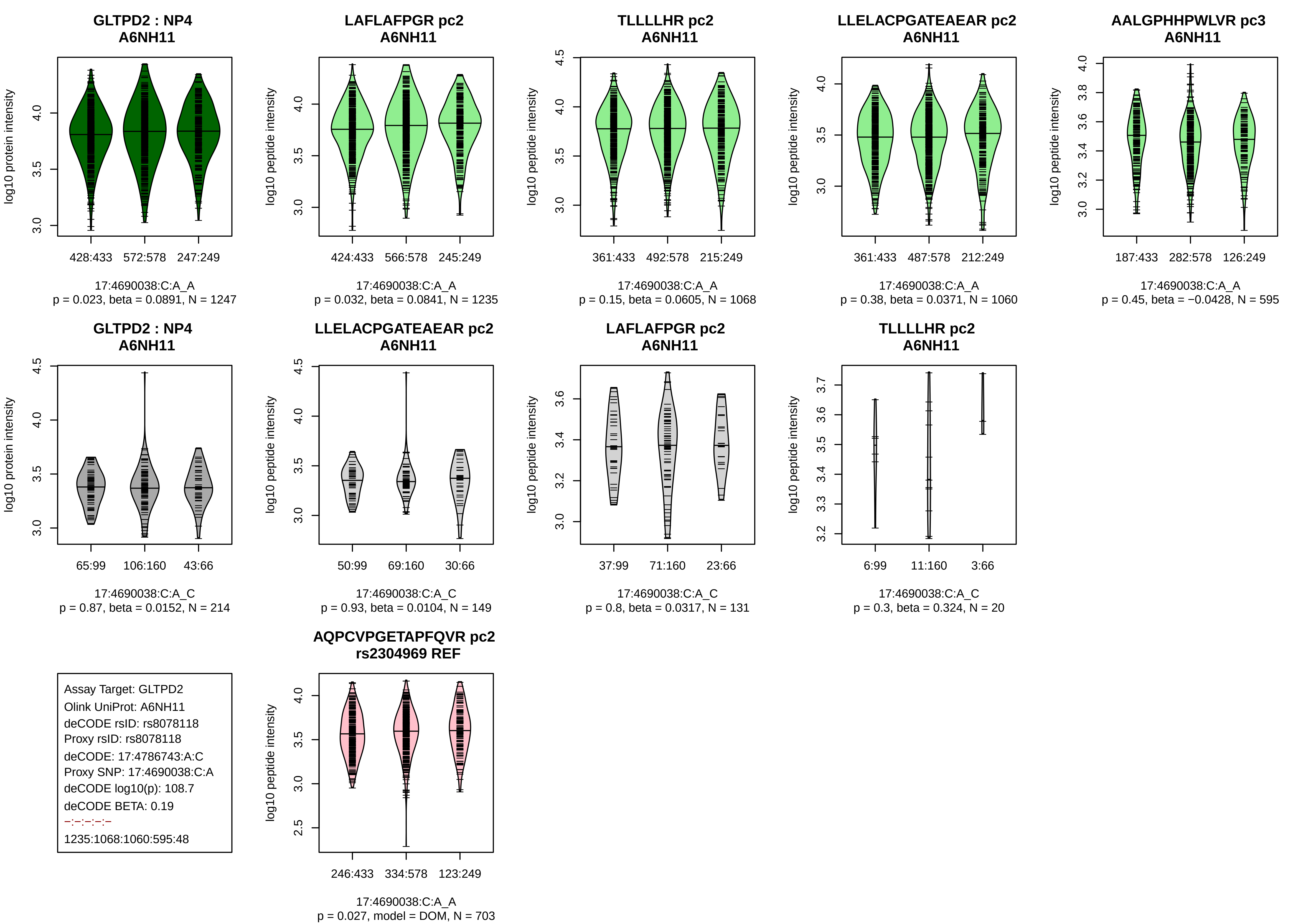
GLTPD2 : NP4
A6NH11
log10 protein intensity
428:433
572:578
247:249
17:4690038:C:A_A
p = 0.023, beta = 0.0891, N = 1247
LAFLAFPGR pc2
A6NH11
log10 peptide intensity
424:433
566:578
245:249
17:4690038:C:A_A
p = 0.032, beta = 0.0841, N = 1235
TLLLLHR pc2
A6NH11
log10 peptide intensity
361:433
492:578
215:249
17:4690038:C:A_A
p = 0.15, beta = 0.0605, N = 1068
LLELACPGATEAEAR pc2
A6NH11
log10 peptide intensity
361:433
487:578
212:249
17:4690038:C:A_A
p = 0.38, beta = 0.0371, N = 1060
AALGPHHPWLVR pc3
A6NH11
log10 peptide intensity
187:433
282:578
126:249
17:4690038:C:A_A
p = 0.45, beta = −0.0428, N = 595
GLTPD2 : NP4
A6NH11
log10 protein intensity
65:99
106:160
43:66
17:4690038:C:A_C
p = 0.87, beta = 0.0152, N = 214
LLELACPGATEAEAR pc2
A6NH11
log10 peptide intensity
50:99
69:160
30:66
17:4690038:C:A_C
p = 0.93, beta = 0.0104, N = 149
LAFLAFPGR pc2
A6NH11
log10 peptide intensity
37:99
71:160
23:66
17:4690038:C:A_C
p = 0.8, beta = 0.0317, N = 131
TLLLLHR pc2
A6NH11
log10 peptide intensity
6:99
11:160
3:66
17:4690038:C:A_C
p = 0.3, beta = 0.324, N = 20
Assay Target: GLTPD2
Olink UniProt: A6NH11
deCODE rsID: rs8078118
Proxy rsID: rs8078118
deCODE: 17:4786743:A:C
Proxy SNP: 17:4690038:C:A
deCODE log10(p): 108.7
deCODE BETA: 0.19
–:–:–:–:–
1235:1068:1060:595:48
AQPCVPGETAPFQVR pc2
rs2304969 REF
log10 peptide intensity
246:433
334:578
123:249
17:4690038:C:A_A
p = 0.027, model = DOM, N = 703

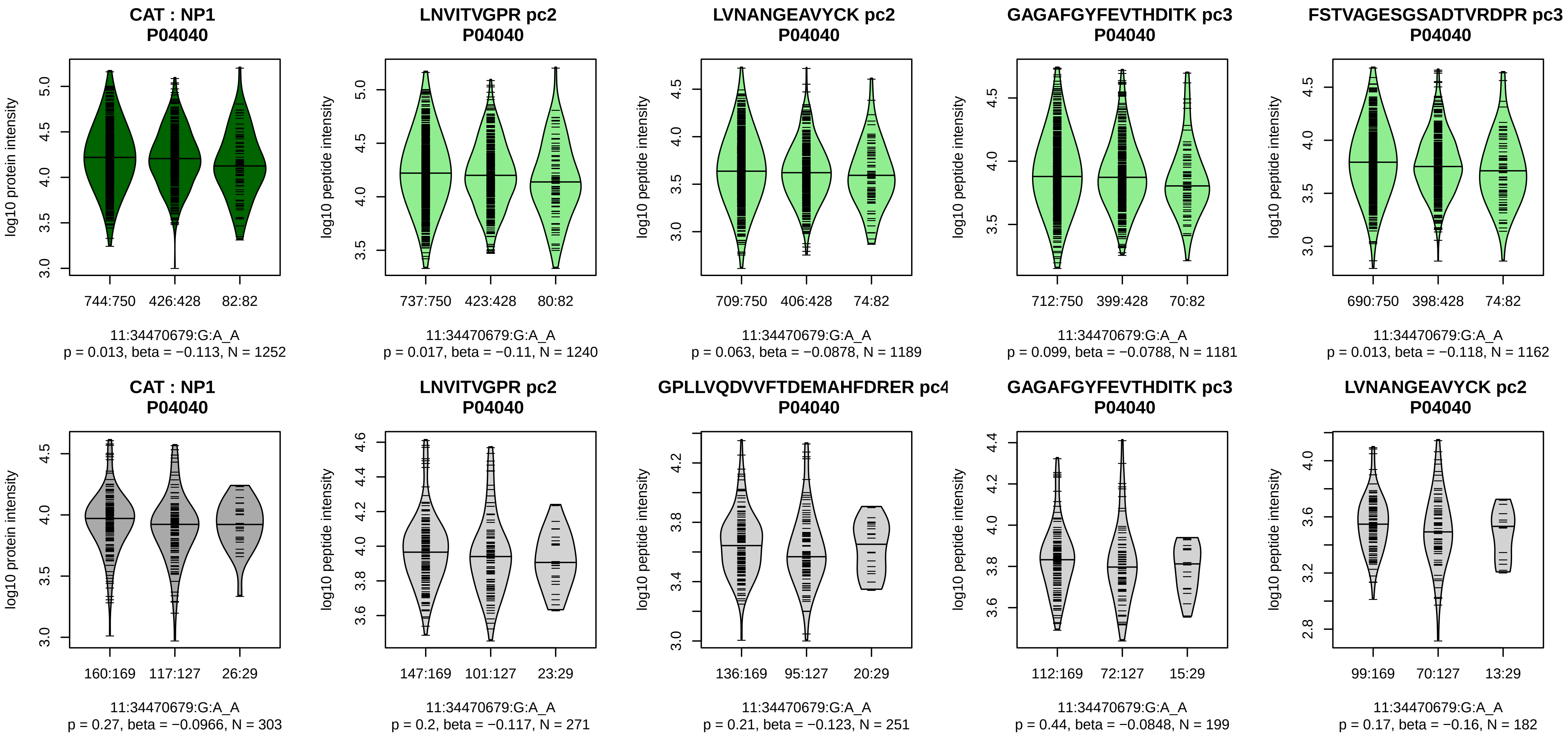


Assay Target: CAT
Olink UniProt: P04040
deCODE rsID: rs769218
Proxy rsID: rs769218
deCODE: 11:34449132:A:G
Proxy SNP: 11:34470679:G:A
deCODE log10(p): 106.5
deCODE BETA: −0.21
-:-:-:-:-:-:-:-:-:-:-:-:-:-:-:-:-:-
1240:1189:1181:1162:1087:105

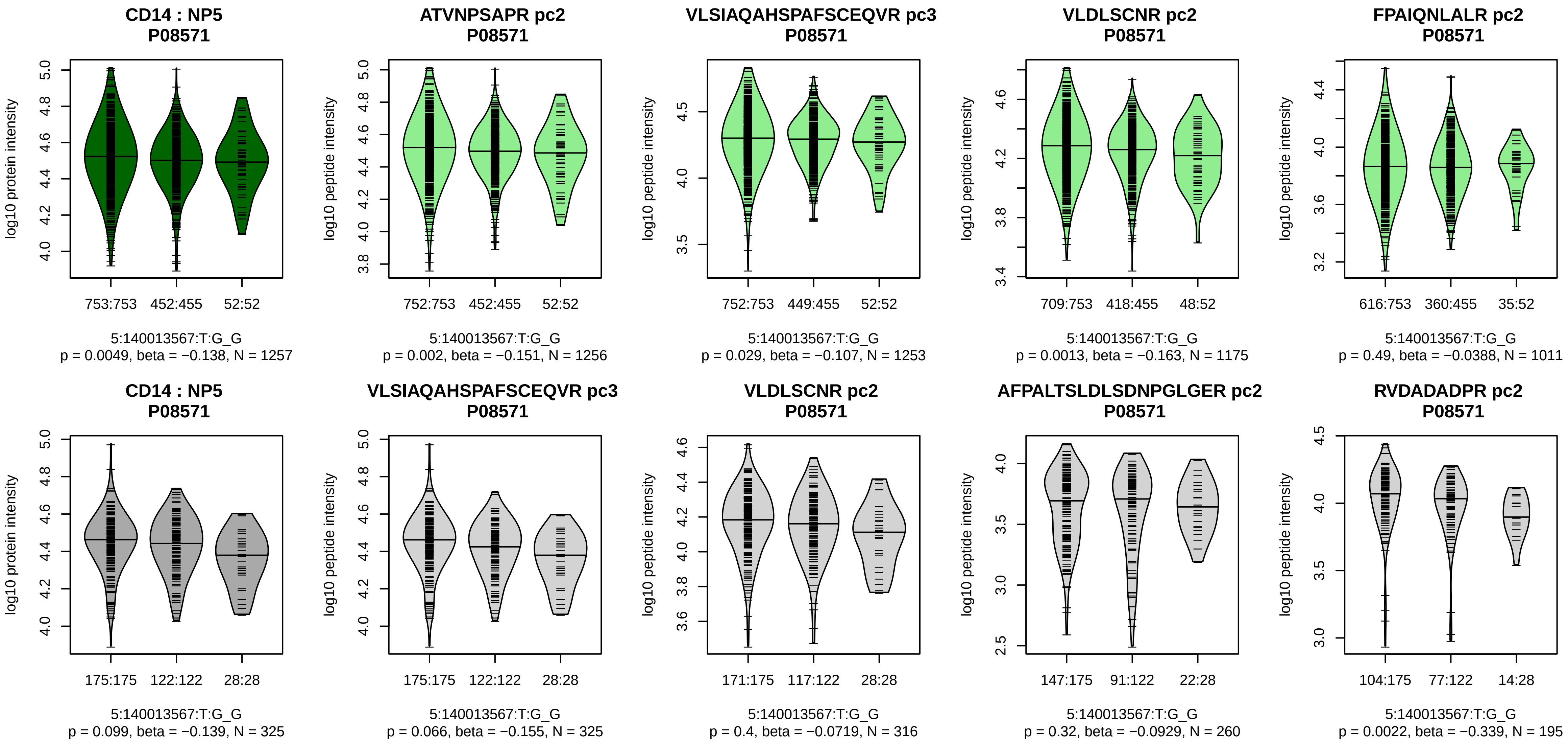


Assay Target: CD14
Olink UniProt: P08571
deCODE rsID: rs5744454
Proxy rsID: rs5744454
deCODE: 5:140633982:G:T
Proxy SNP: 5:140013567:T:G
deCODE log10(p): 104
deCODE BETA: −0.19
*:–:*:–:–:–:–:–:–:–:–:–:–:–
1256:1253:1175:1011:982:929:8

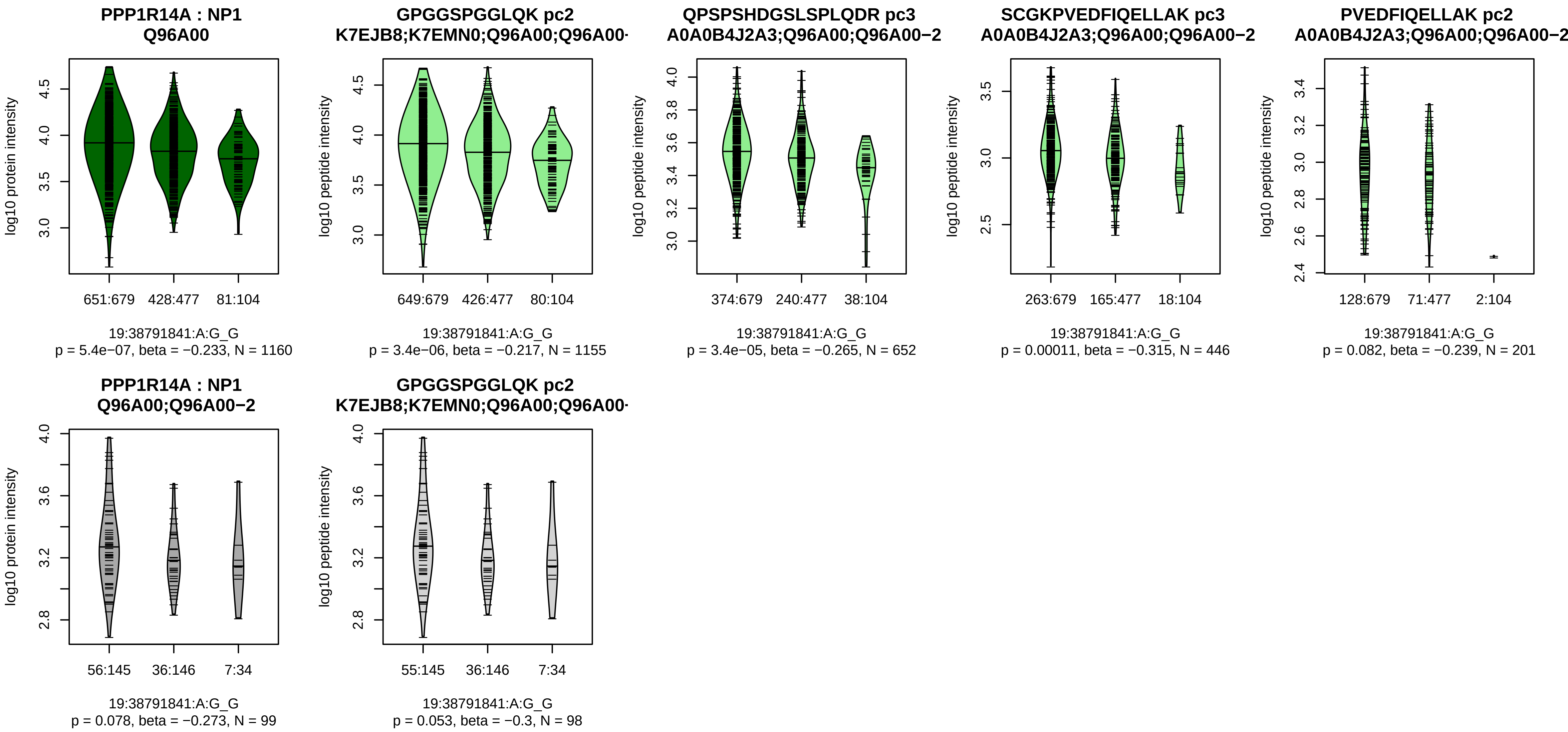


Assay Target: PPP1R14A
Olink UniProt: Q96A00
deCODE rsID: rs71354995
Proxy rsID: rs71354995
deCODE: 19:38301201:G:A
Proxy SNP: 19:38791841:A:G
deCODE log10(p): 103.5
deCODE BETA: −0.2
*:*:*:–:–
1155:652:446:201:90

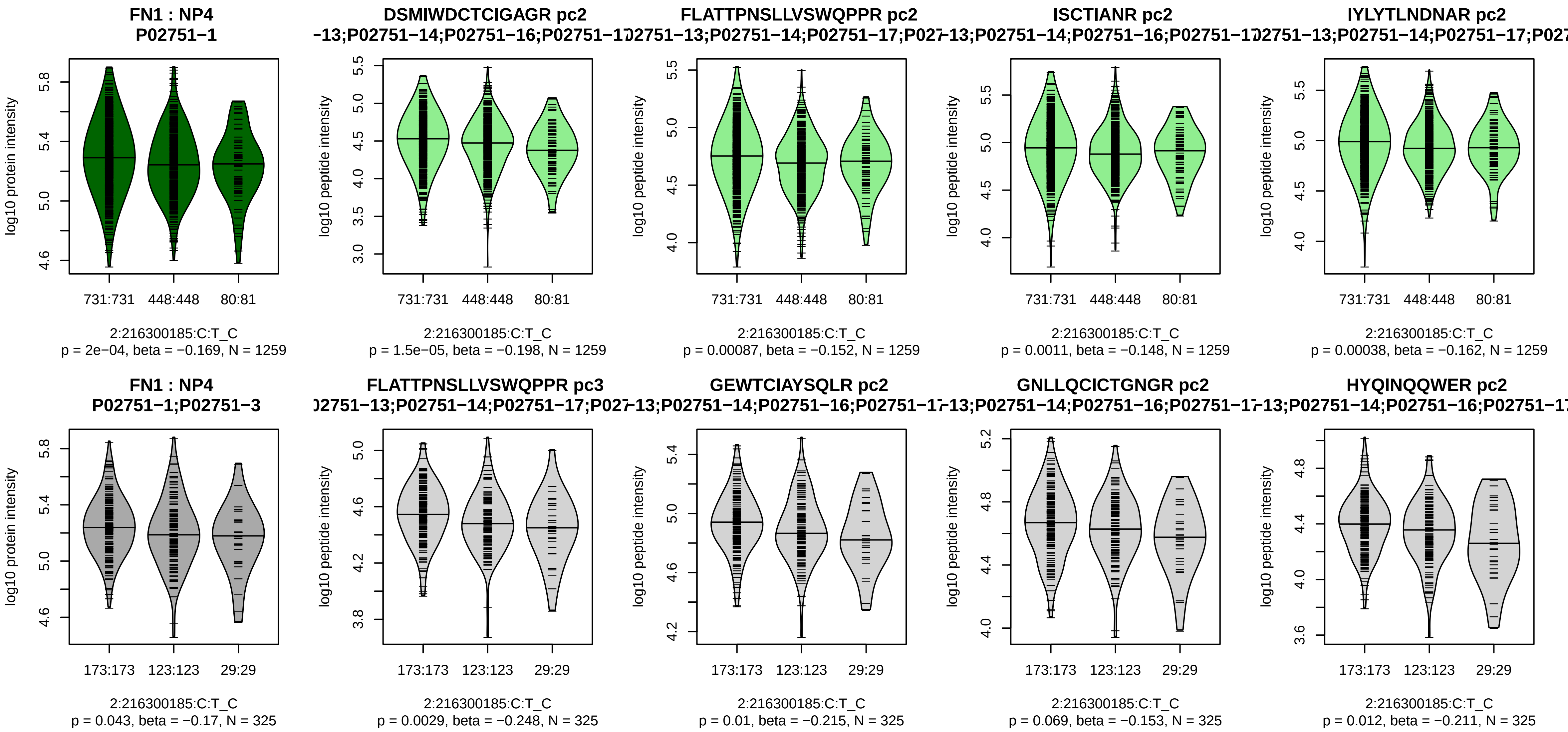


Assay Target: FN1
Olink UniProt: P02751
deCODE rsID: rs1250258
Proxy rsID: rs1250258
deCODE: 2:215435462:C:T
Proxy SNP: 2:216300185:C:T
deCODE log10(p): 101.7
deCODE BETA: −0.18
1259:1259:1259:1259:1259:125

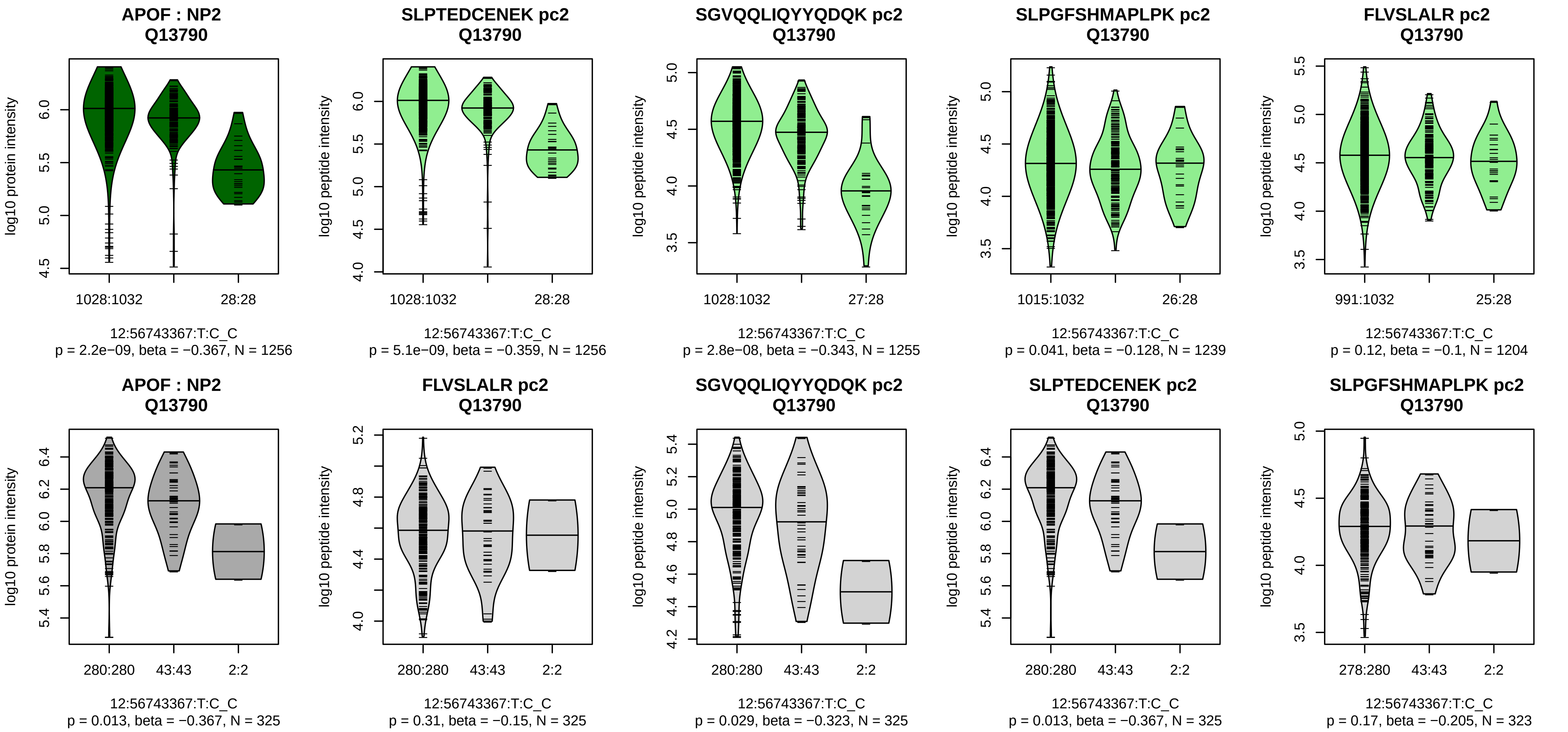


Assay Target: APOF
Olink UniProt: Q13790
deCODE rsID: rs2020854
Proxy rsID: rs2020854
deCODE: 12:56349583:C:T
Proxy SNP: 12:56743367:T:C
deCODE log10(p): 101.5
deCODE BETA: −0.31
*:*:*:–:–:–:–:–:–
1256:1255:1254:1239:1204:625

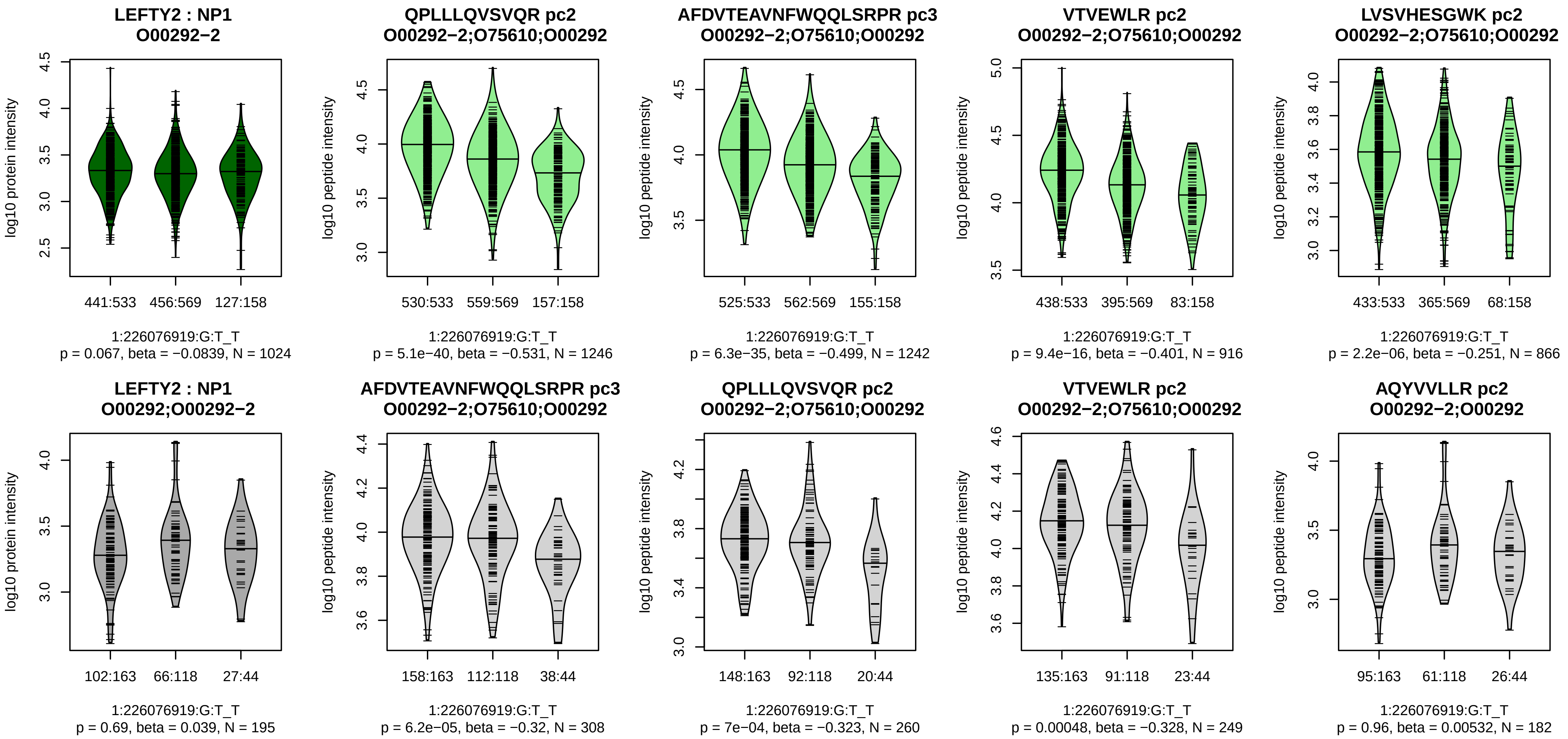


Assay Target: LEFTY2
Olink UniProt: O00292
deCODE rsID: rs360058
Proxy rsID: rs360058
deCODE: 1:225889219:T:G
Proxy SNP: 1:226076919:G:T
deCODE log10(p): 101
deCODE BETA: −0.18
*:*:*:*:–:–:–:–:–
1246:1242:916:866:856:628:337

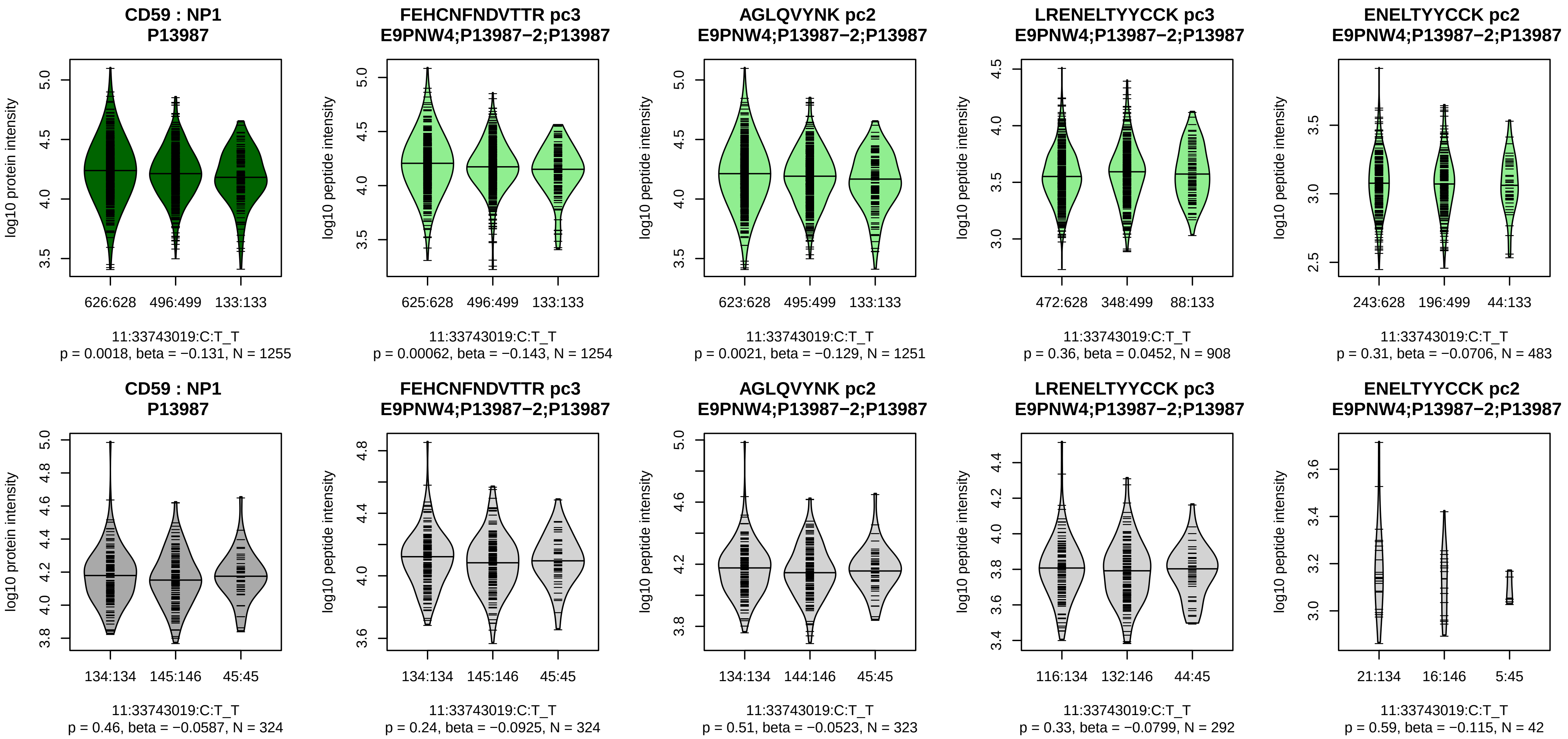


Assay Target: CD59
Olink UniProt: P13987
deCODE rsID: rs831630
Proxy rsID: rs831630
deCODE: 11:33721473:T:C
Proxy SNP: 11:33743019:C:T
deCODE log10(p): 100.3
deCODE BETA: −0.18
*:*:−:−:−:−
1254:1251:908:589:483:60

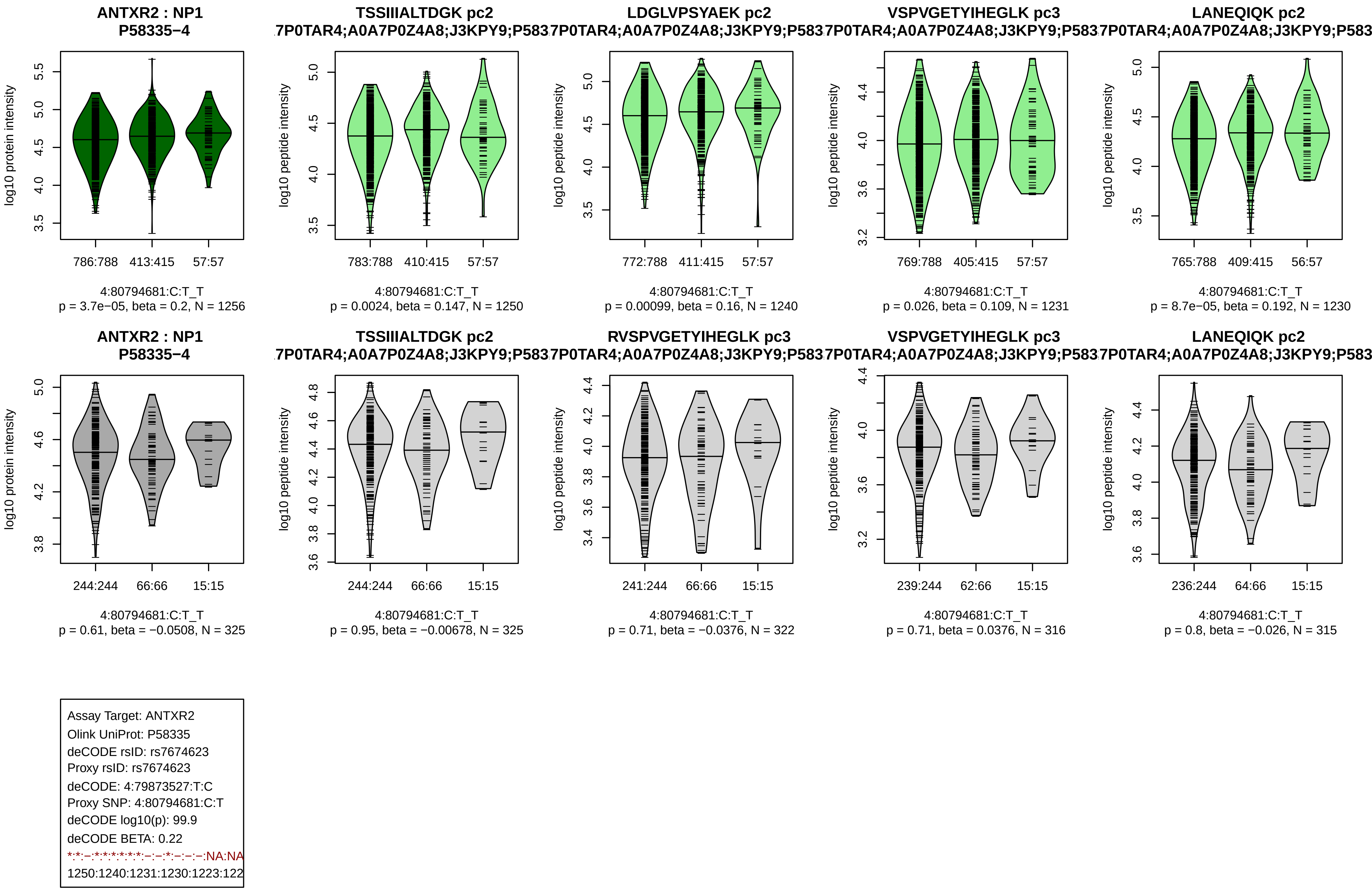
ANTXR2 : NP1
P58335−4
log10 protein intensity
786:788
413:415
57:57
4:80794681:C:T_T
p = 3.7e−05, beta = 0.2, N = 1256
TSSIIIALTDGK pc2
.7P0TAR4;A0A7P0Z4A8;J3KPY9;P583
log10 peptide intensity
783:788
410:415
57:57
4:80794681:C:T_T
p = 0.0024, beta = 0.147, N = 1250
LDGLVPSYAEK pc2
7P0TAR4;A0A7P0Z4A8;J3KPY9;P583
log10 peptide intensity
772:788
411:415
57:57
4:80794681:C:T_T
p = 0.00099, beta = 0.16, N = 1240
VSPVGETYIHEGLK pc3
7P0TAR4;A0A7P0Z4A8;J3KPY9;P583
log10 peptide intensity
769:788
405:415
57:57
4:80794681:C:T_T
p = 0.026, beta = 0.109, N = 1231
LANEQIQK pc2
7P0TAR4;A0A7P0Z4A8;J3KPY9;P583
log10 peptide intensity
765:788
409:415
56:57
4:80794681:C:T_T
p = 8.7e−05, beta = 0.192, N = 1230
ANTXR2 : NP1
P58335−4
log10 protein intensity
244:244
66:66
15:15
4:80794681:C:T_T
p = 0.61, beta = −0.0508, N = 325
TSSIIIALTDGK pc2
.7P0TAR4;A0A7P0Z4A8;J3KPY9;P583
log10 peptide intensity
244:244
66:66
15:15
4:80794681:C:T_T
p = 0.95, beta = −0.00678, N = 325
RVSPVGETYIHEGLK pc3
7P0TAR4;A0A7P0Z4A8;J3KPY9;P583
log10 peptide intensity
241:244
66:66
15:15
4:80794681:C:T_T
p = 0.71, beta = −0.0376, N = 322
VSPVGETYIHEGLK pc3
7P0TAR4;A0A7P0Z4A8;J3KPY9;P583
log10 peptide intensity
239:244
62:66
15:15
4:80794681:C:T_T
p = 0.71, beta = 0.0376, N = 316
LANEQIQK pc2
7P0TAR4;A0A7P0Z4A8;J3KPY9;P583
log10 peptide intensity
236:244
64:66
15:15
4:80794681:C:T_T
p = 0.8, beta = −0.026, N = 315
Assay Target: ANTXR2
Olink UniProt: P58335
deCODE rsID: rs7674623
Proxy rsID: rs7674623
deCODE: 4:79873527:T:C
Proxy SNP: 4:80794681:C:T
deCODE log10(p): 99.9
deCODE BETA: 0.22
*:*:−:*:*:*:*:*:*:−:−:*:−:−:−:NA:NA
1250:1240:1231:1230:1223:122

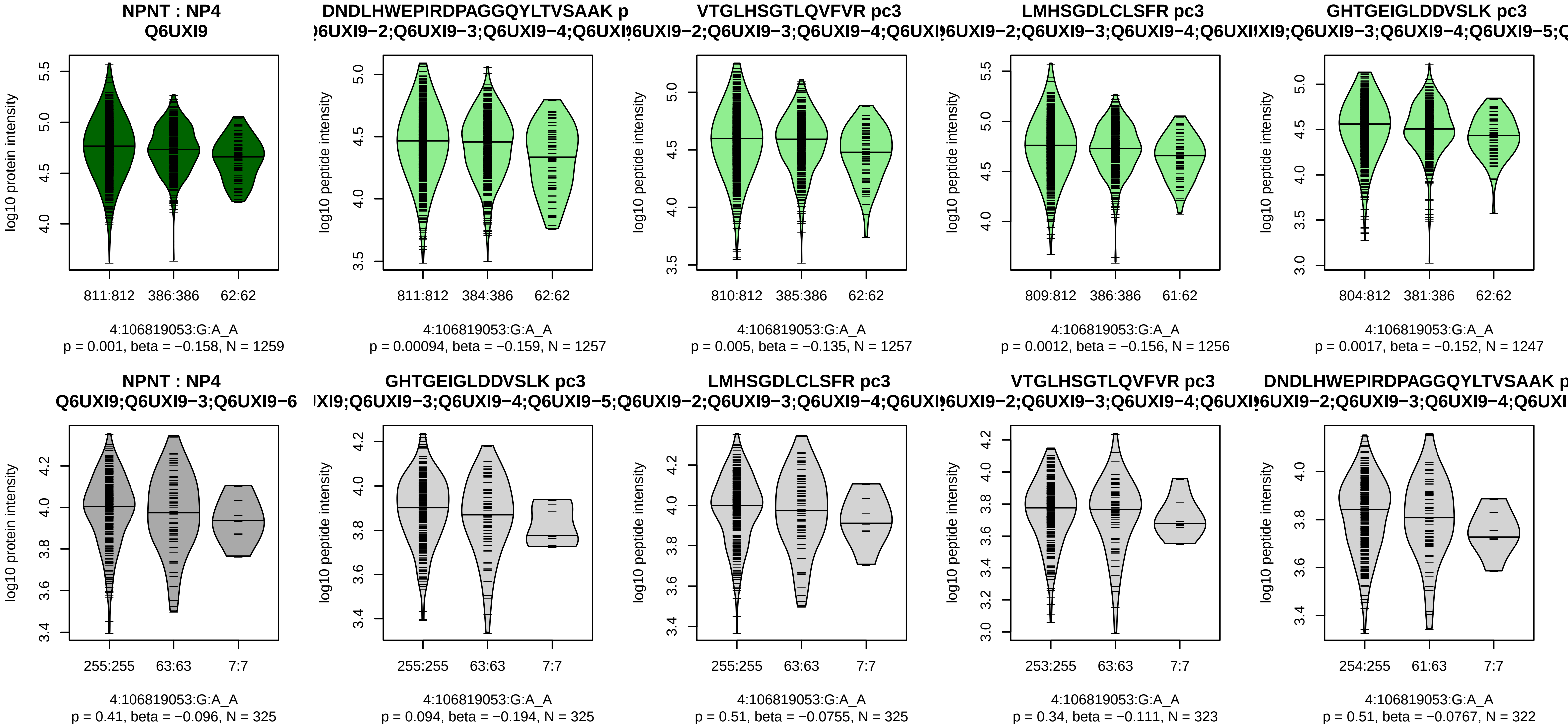


Assay Target: NPNT
Olink UniProt: Q6UXI9
deCODE rsID: rs34712979
Proxy rsID: rs34712979
deCODE: 4:105897896:A:G
Proxy SNP: 4:106819053:G:A
deCODE log10(p): 99.7
deCODE BETA: −0.2
*:*:*:*:−:*:*:−:−:−:−:*:−:−:−:−:NA:
1257:1257:1256:1247:1170:116

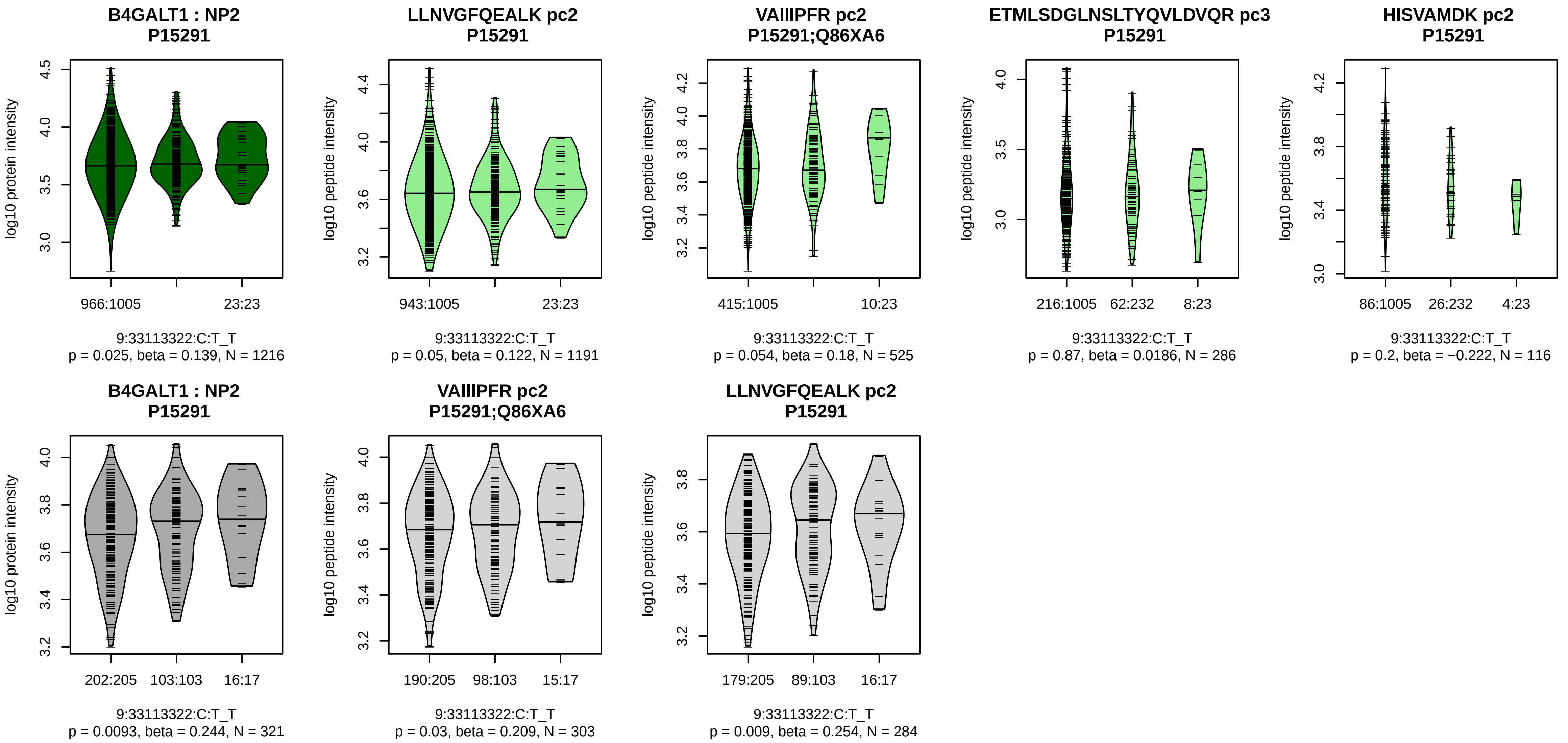


Assay Target: B4GALT1
Olink UniProt: P15291
deCODE rsID: rs7019909
Proxy rsID: rs7019909
deCODE: 9:33113324:T:C
Proxy SNP: 9:33113322:C:T
deCODE log10(p): 98.3
deCODE BETA: 0.3
−:−:−:−:−:−:NA
1191:525:286:116:89:74:9

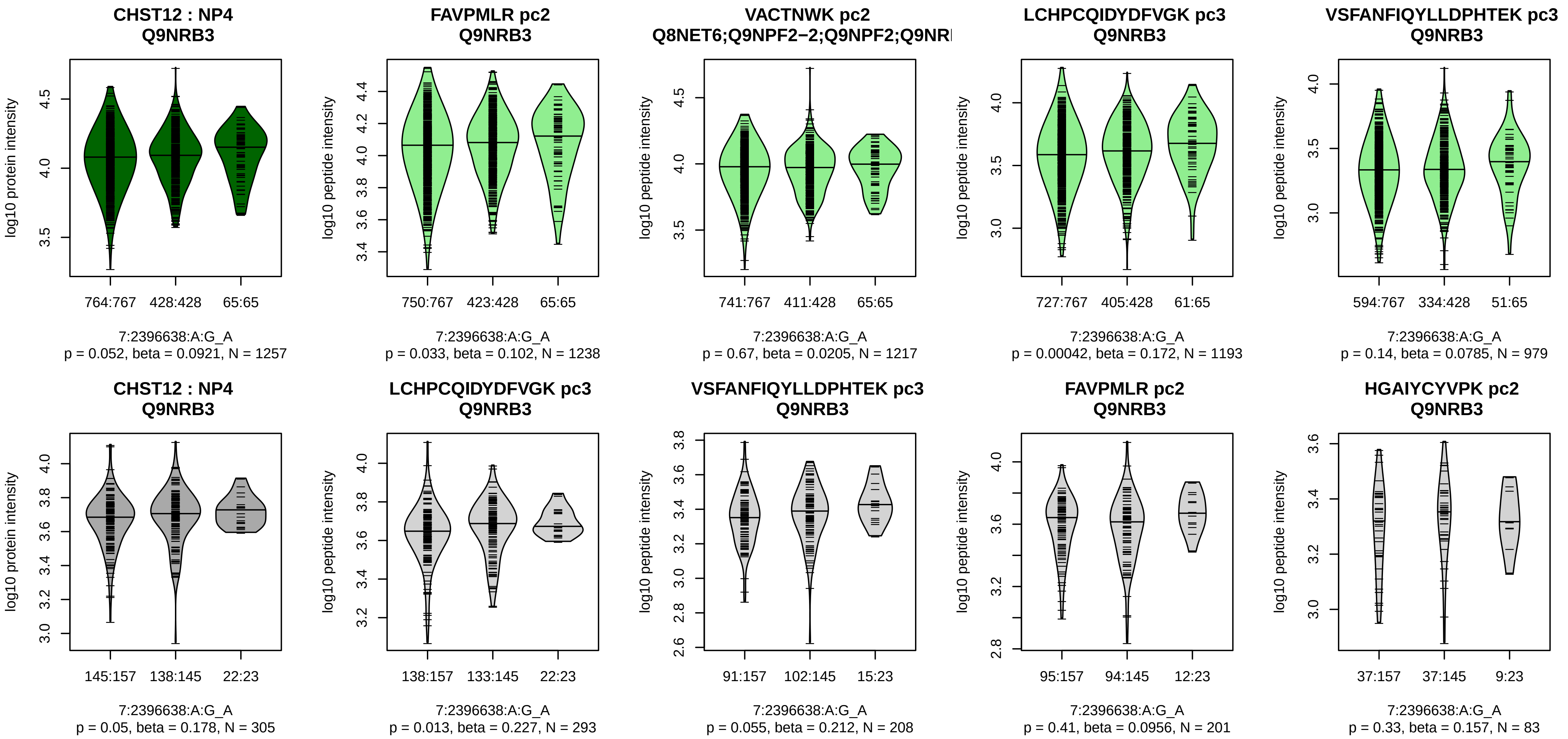


Assay Target: CHST12
Olink UniProt: Q9NRB3
deCODE rsID: rs6952433
Proxy rsID: rs6952433
deCODE: 7:2357003:A:G
Proxy SNP: 7:2396638:A:G
deCODE log10(p): 95
deCODE BETA: 0.2
−:−:*:−:−:−:−:−:−:−:NA
1238:1217:1193:979:807:685:67

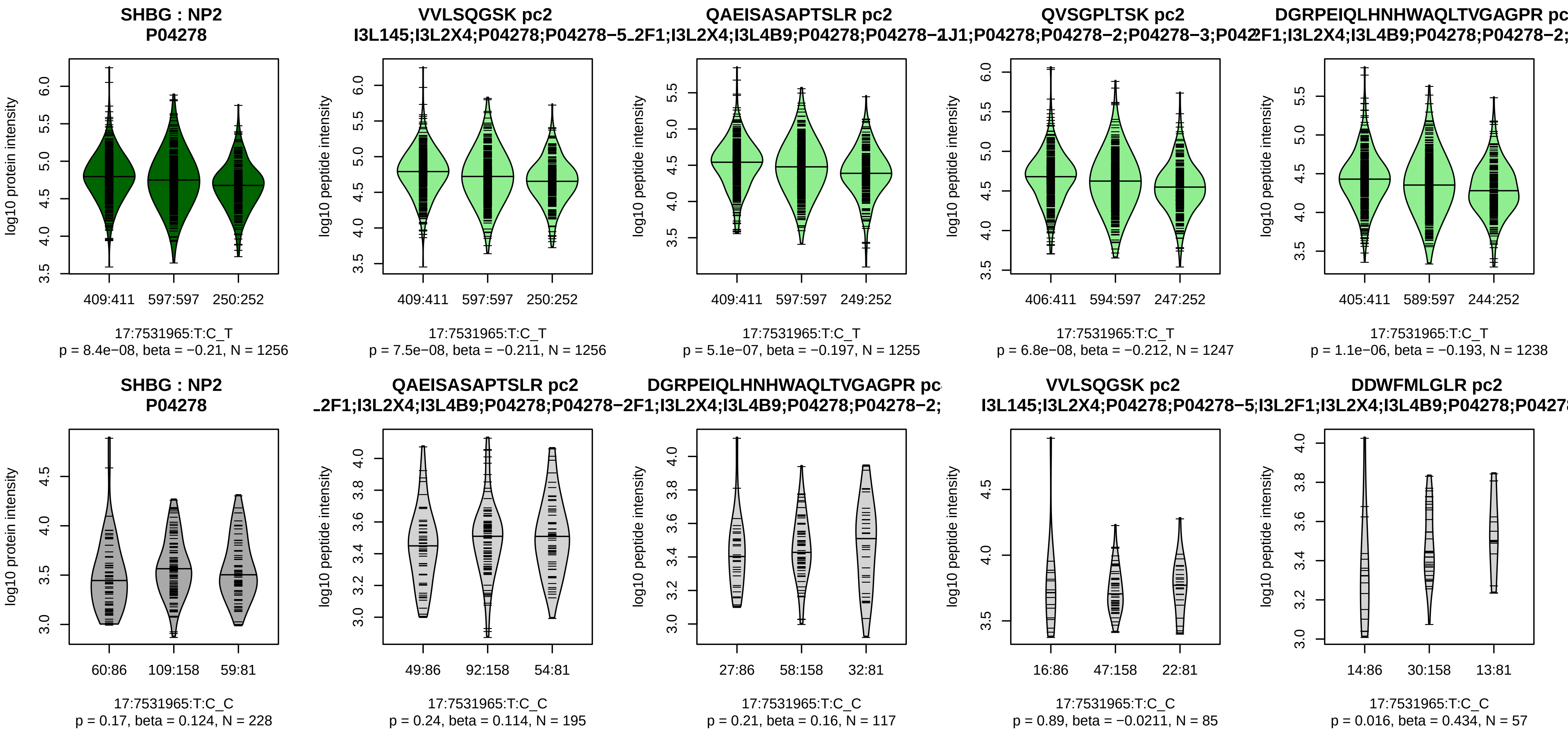


Assay Target: SHBG
Olink UniProt: P04278
deCODE rsID: rs858519
Proxy rsID: rs858519
deCODE: 17:7628647:T:C
Proxy SNP: 17:7531965:T:C
deCODE log10(p): 94
deCODE BETA: −0.17
*:*:*:*:*:*:*:*:*:*:*:*:*:*:*:*:*:*:–:–:–:–:–:–
1256:1255:1247:1238:1238:123

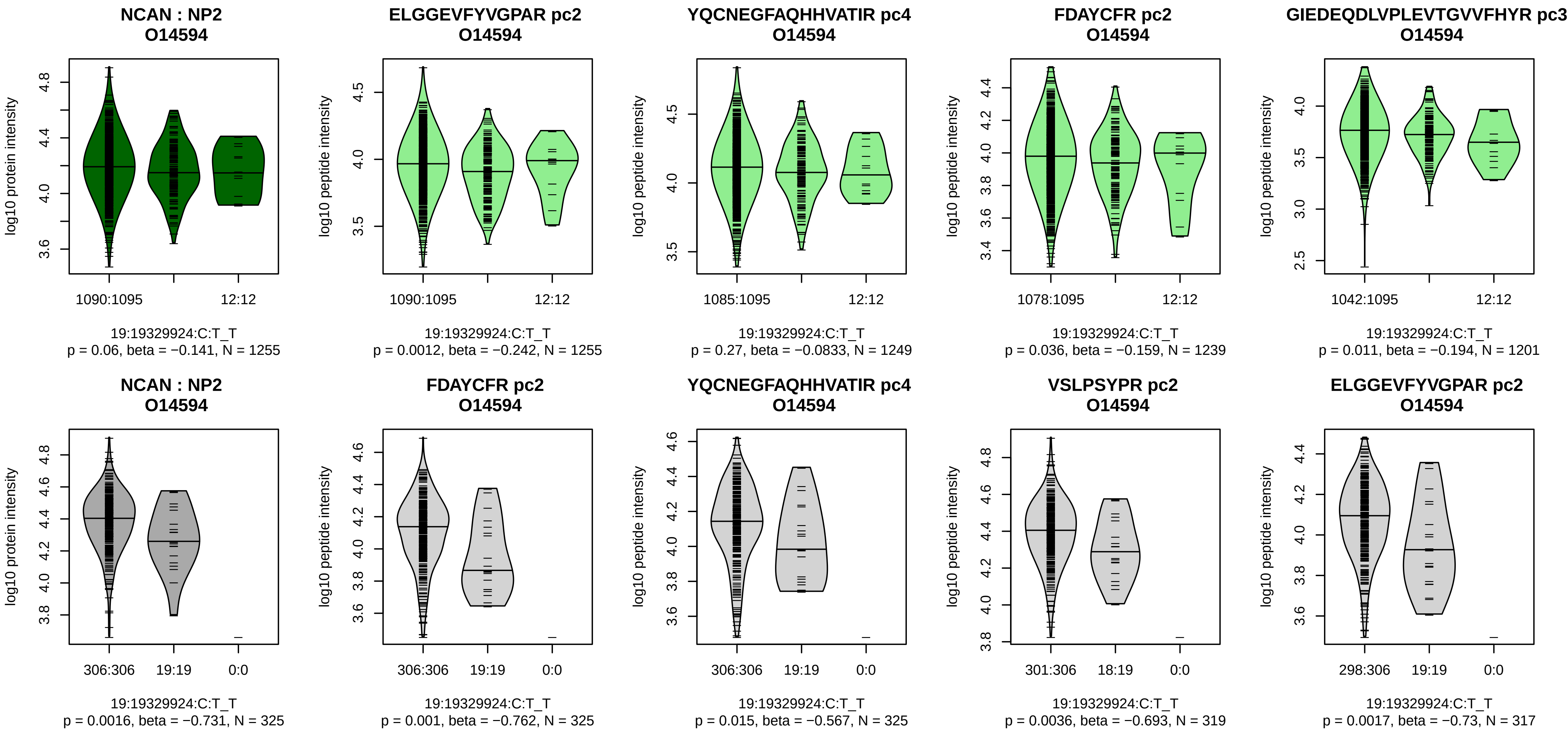


Assay Target: NCAN
Olink UniProt: O14594
deCODE rsID: rs2228603
Proxy rsID: rs2228603
deCODE: 19:19219115:T:C
Proxy SNP: 19:19329924:C:T
deCODE log10(p): 93.2
deCODE BETA: −0.33
*:–:–:–:–:–:–:–:–:*:–:–:–:–:*:–:–:
1255:1249:1239:1201:1189:117

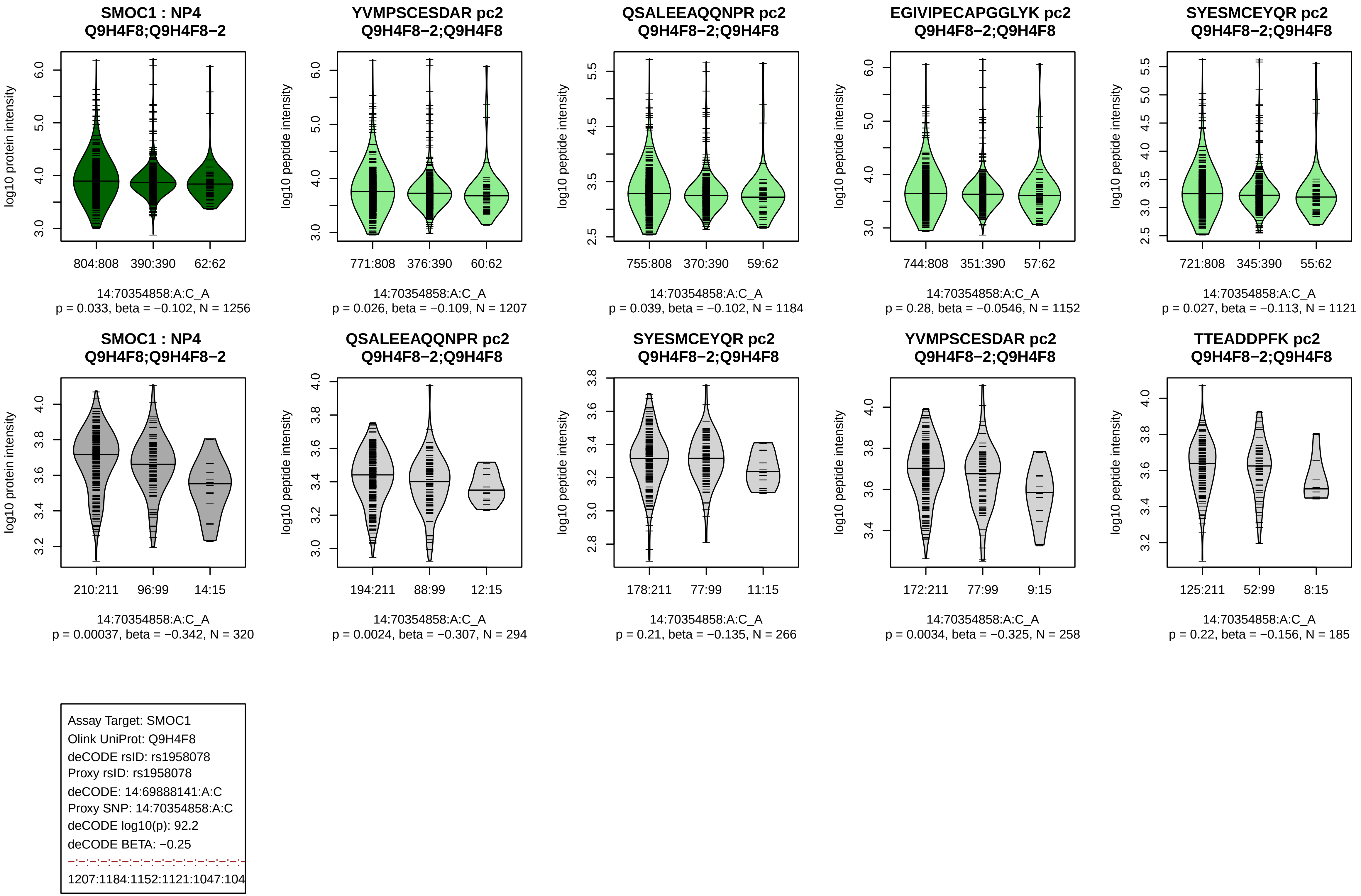
SMOC1 : NP4
Q9H4F8;Q9H4F8−2
log10 protein intensity
804:808 390:390 62:62
14:70354858:A:C_A
p = 0.033, beta = −0.102, N = 1256
YVMPSCESDAR pc2
Q9H4F8−2;Q9H4F8
log10 peptide intensity
771:808 376:390 60:62
14:70354858:A:C_A
p = 0.026, beta = −0.109, N = 1207
QSALEEAQQNPR pc2
Q9H4F8−2;Q9H4F8
log10 peptide intensity
755:808 370:390 59:62
14:70354858:A:C_A
p = 0.039, beta = −0.102, N = 1184
EGIVIPECAPGGLYK pc2
Q9H4F8−2;Q9H4F8
log10 peptide intensity
744:808 351:390 57:62
14:70354858:A:C_A
p = 0.28, beta = −0.0546, N = 1152
SYESMCEYQR pc2
Q9H4F8−2;Q9H4F8
log10 peptide intensity
721:808 345:390 55:62
14:70354858:A:C_A
p = 0.027, beta = −0.113, N = 1121
SMOC1 : NP4
Q9H4F8;Q9H4F8−2
log10 protein intensity
210:211 96:99 14:15
14:70354858:A:C_A
p = 0.00037, beta = −0.342, N = 320
QSALEEAQQNPR pc2
Q9H4F8−2;Q9H4F8
log10 peptide intensity
194:211 88:99 12:15
14:70354858:A:C_A
p = 0.0024, beta = −0.307, N = 294
SYESMCEYQR pc2
Q9H4F8−2;Q9H4F8
log10 peptide intensity
178:211 77:99 11:15
14:70354858:A:C_A
p = 0.21, beta = −0.135, N = 266
YVMPSCESDAR pc2
Q9H4F8−2;Q9H4F8
log10 peptide intensity
172:211 77:99 9:15
14:70354858:A:C_A
p = 0.0034, beta = −0.325, N = 258
TTEADDPFK pc2
Q9H4F8−2;Q9H4F8
log10 peptide intensity
125:211 52:99 8:15
14:70354858:A:C_A
p = 0.22, beta = −0.156, N = 185
Assay Target: SMOC1
Olink UniProt: Q9H4F8
deCODE rsID: rs1958078
Proxy rsID: rs1958078
deCODE: 14:69888141:A:C
Proxy SNP: 14:70354858:A:C
deCODE log10(p): 92.2
deCODE BETA: −0.25
1207:1184:1152:1121:1047:104

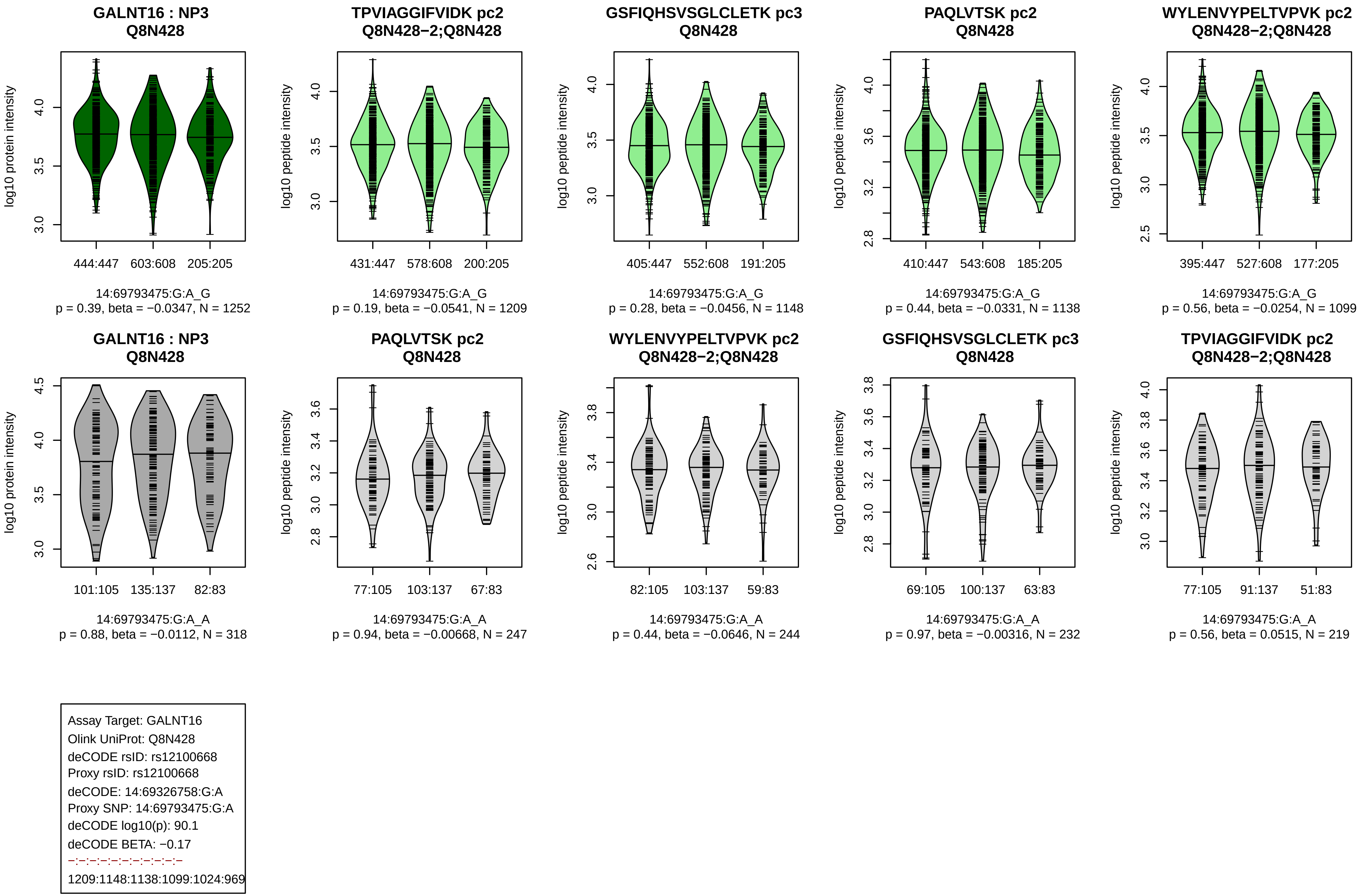
GALNT16 : NP3
Q8N428
log10 protein intensity
444:447 603:608 205:205
14:69793475:G:A_G
p = 0.39, beta = −0.0347, N = 1252
TPVIAGGIFVIDK pc2
Q8N428−2;Q8N428
log10 peptide intensity
431:447 578:608 200:205
14:69793475:G:A_G
p = 0.19, beta = −0.0541, N = 1209
GSFIQHSVSGLCLETK pc3
Q8N428
log10 peptide intensity
405:447 552:608 191:205
14:69793475:G:A_G
p = 0.28, beta = −0.0456, N = 1148
PAQLVTSK pc2
Q8N428
log10 peptide intensity
410:447 543:608 185:205
14:69793475:G:A_G
p = 0.44, beta = −0.0331, N = 1138
WYLENVYPELTVPVK pc2
Q8N428−2;Q8N428
log10 peptide intensity
395:447 527:608 177:205
14:69793475:G:A_G
p = 0.56, beta = −0.0254, N = 1099
GALNT16 : NP3
Q8N428
log10 protein intensity
101:105 135:137 82:83
14:69793475:G:A_A
p = 0.88, beta = −0.0112, N = 318
PAQLVTSK pc2
Q8N428
log10 peptide intensity
77:105 103:137 67:83
14:69793475:G:A_A
p = 0.94, beta = −0.00668, N = 247
WYLENVYPELTVPVK pc2
Q8N428−2;Q8N428
log10 peptide intensity
82:105 103:137 59:83
14:69793475:G:A_A
p = 0.44, beta = −0.0646, N = 244
GSFIQHSVSGLCLETK pc3
Q8N428
log10 peptide intensity
69:105 100:137 63:83
14:69793475:G:A_A
p = 0.97, beta = −0.00316, N = 232
TPVIAGGIFVIDK pc2
Q8N428−2;Q8N428
log10 peptide intensity
77:105 91:137 51:83
14:69793475:G:A_A
p = 0.56, beta = 0.0515, N = 219
Assay Target: GALNT16
Olink UniProt: Q8N428
deCODE rsID: rs12100668
Proxy rsID: rs12100668
deCODE: 14:69326758:G:A
Proxy SNP: 14:69793475:G:A
deCODE log10(p): 90.1
deCODE BETA: −0.17
−:−:−:−:−:−:−:−:−:−:−
1209:1148:1138:1099:1024:969

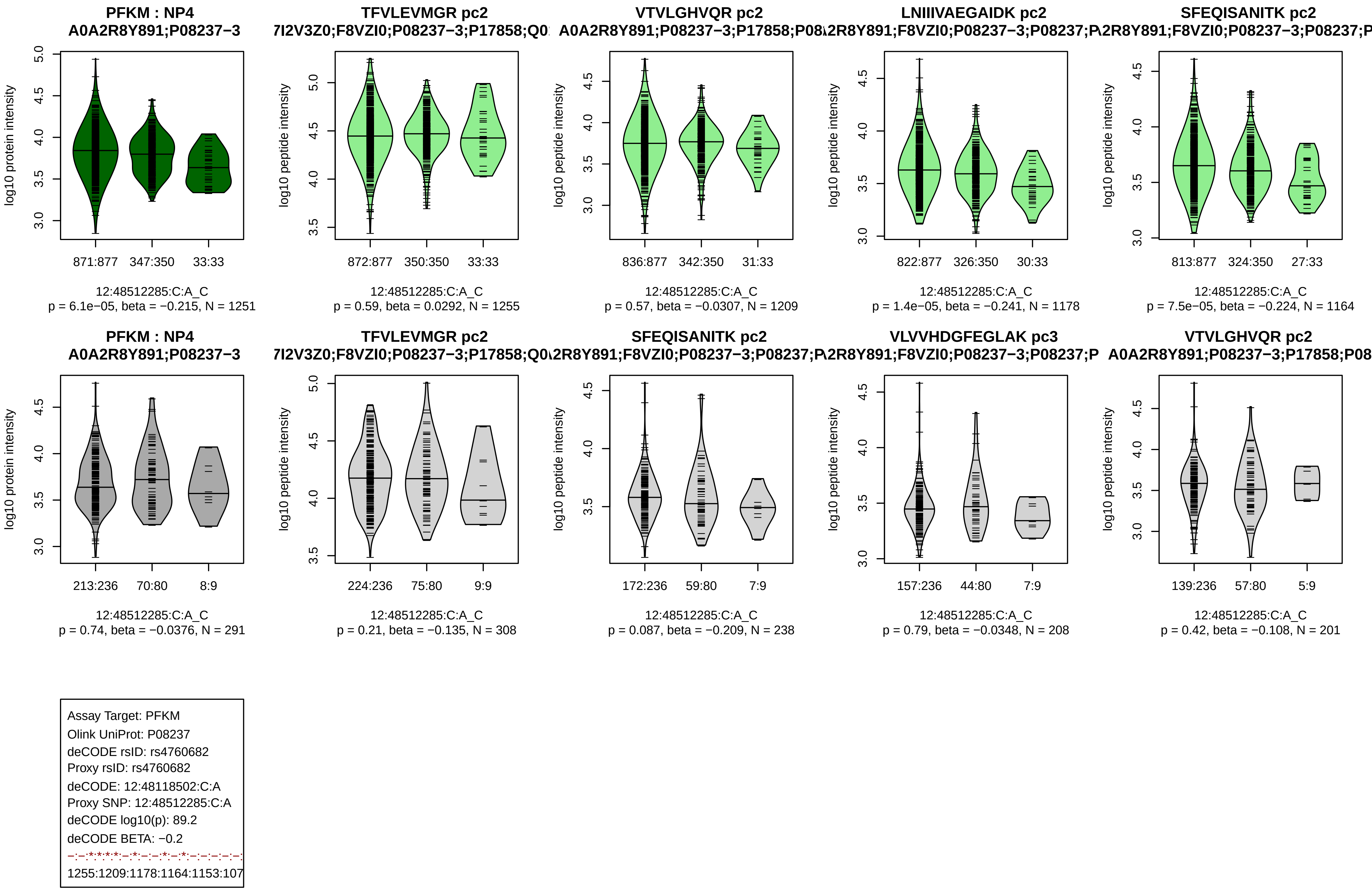
PFKM : NP4
A0A2R8Y891;P08237−3
log10 protein intensity
871:877 347:350 33:33
12:48512285:C:A_C
p = 6.1e−05, beta = −0.215, N = 1251
TFVLEVMGR pc2
7I2V3Z0;F8VZI0;P08237−3;P17858;Q0
log10 peptide intensity
872:877 350:350 33:33
12:48512285:C:A_C
p = 0.59, beta = 0.0292, N = 1255
VTVLGHVQR pc2
A0A2R8Y891;P08237−3;P17858;P08
log10 peptide intensity
836:877 342:350 31:33
12:48512285:C:A_C
p = 0.57, beta = −0.0307, N = 1209
LNIIIVAEGAIDK pc2
\2R8Y891;F8VZI0;P08237−3;P08237;P
log10 peptide intensity
822:877 326:350 30:33
12:48512285:C:A_C
p = 1.4e−05, beta = −0.241, N = 1178
SFEQISANITK pc2
\2R8Y891;F8VZI0;P08237−3;P08237;P
log10 peptide intensity
813:877 324:350 27:33
12:48512285:C:A_C
p = 7.5e−05, beta = −0.224, N = 1164
PFKM : NP4
A0A2R8Y891;P08237−3
log10 protein intensity
213:236 70:80 8:9
12:48512285:C:A_C
p = 0.74, beta = −0.0376, N = 291
TFVLEVMGR pc2
7I2V3Z0;F8VZI0;P08237−3;P17858;Q0
log10 peptide intensity
224:236 75:80 9:9
12:48512285:C:A_C
p = 0.21, beta = −0.135, N = 308
SFEQISANITK pc2
\2R8Y891;F8VZI0;P08237−3;P08237;P
log10 peptide intensity
172:236 59:80 7:9
12:48512285:C:A_C
p = 0.087, beta = −0.209, N = 238
VLVVHDGFEGLAK pc3
\2R8Y891;F8VZI0;P08237−3;P08237;P
log10 peptide intensity
157:236 44:80 7:9
12:48512285:C:A_C
p = 0.79, beta = −0.0348, N = 208
VTVLGHVQR pc2
A0A2R8Y891;P08237−3;P17858;P08
log10 peptide intensity
139:236 57:80 5:9
12:48512285:C:A_C
p = 0.42, beta = −0.108, N = 201
Assay Target: PFKM
Olink UniProt: P08237
deCODE rsID: rs4760682
Proxy rsID: rs4760682
deCODE: 12:48118502:C:A
Proxy SNP: 12:48512285:C:A
deCODE log10(p): 89.2
deCODE BETA: −0.2
−:−:*:*:*:*:−:*:−:−:*:−:*:−:−:−:−:−:
1255:1209:1178:1164:1153:107

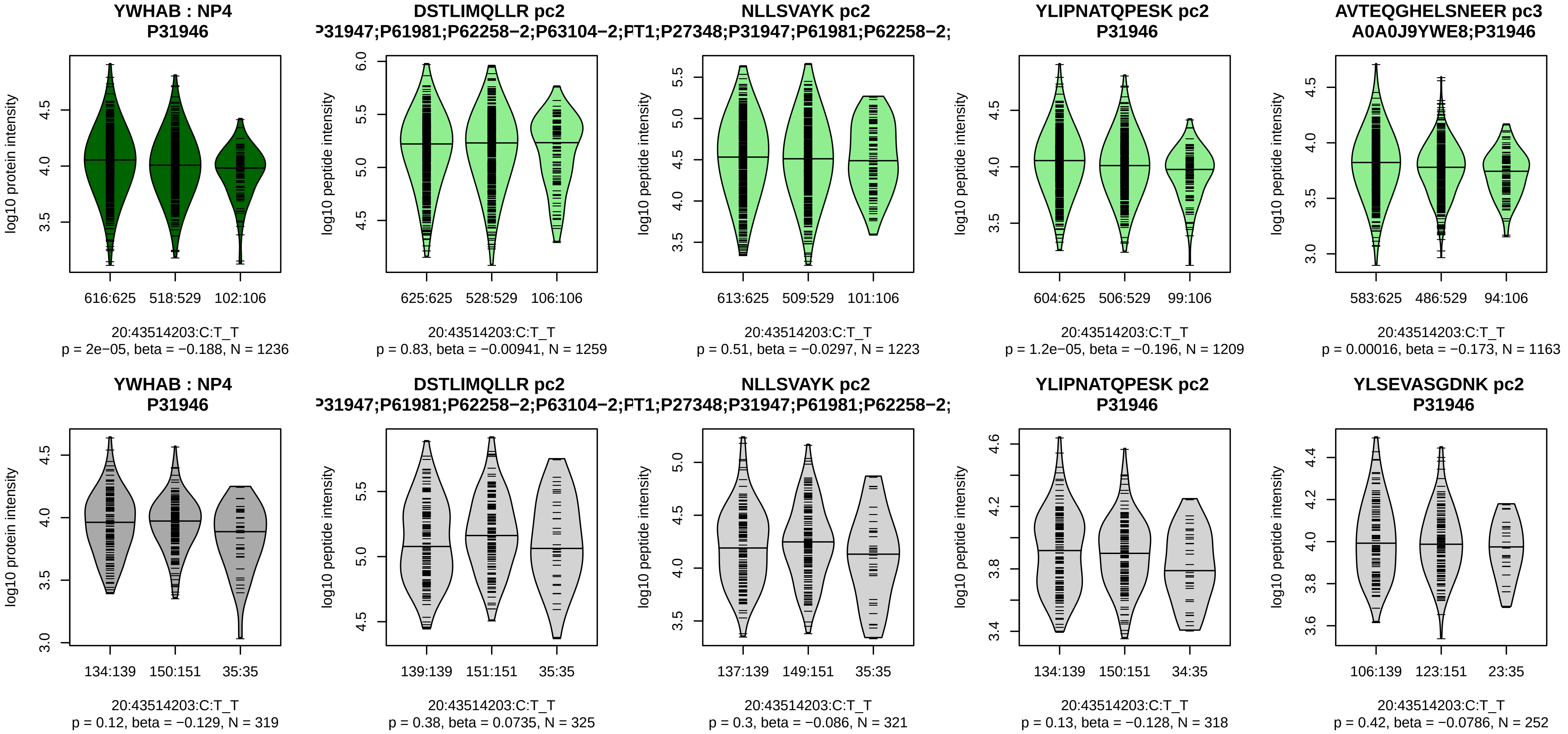


Assay Target: YWHAB
Olink UniProt: P31946
deCODE rsID: rs6031847
Proxy rsID: rs6031847
deCODE: 20:44885562:T:C
Proxy SNP: 20:43514203:C:T
deCODE log10(p): 86.5
deCODE BETA: −0.18
−:−:*:*:−:−:*:−:−:−:−:−:−
1259:1223:1209:1163:1074:101

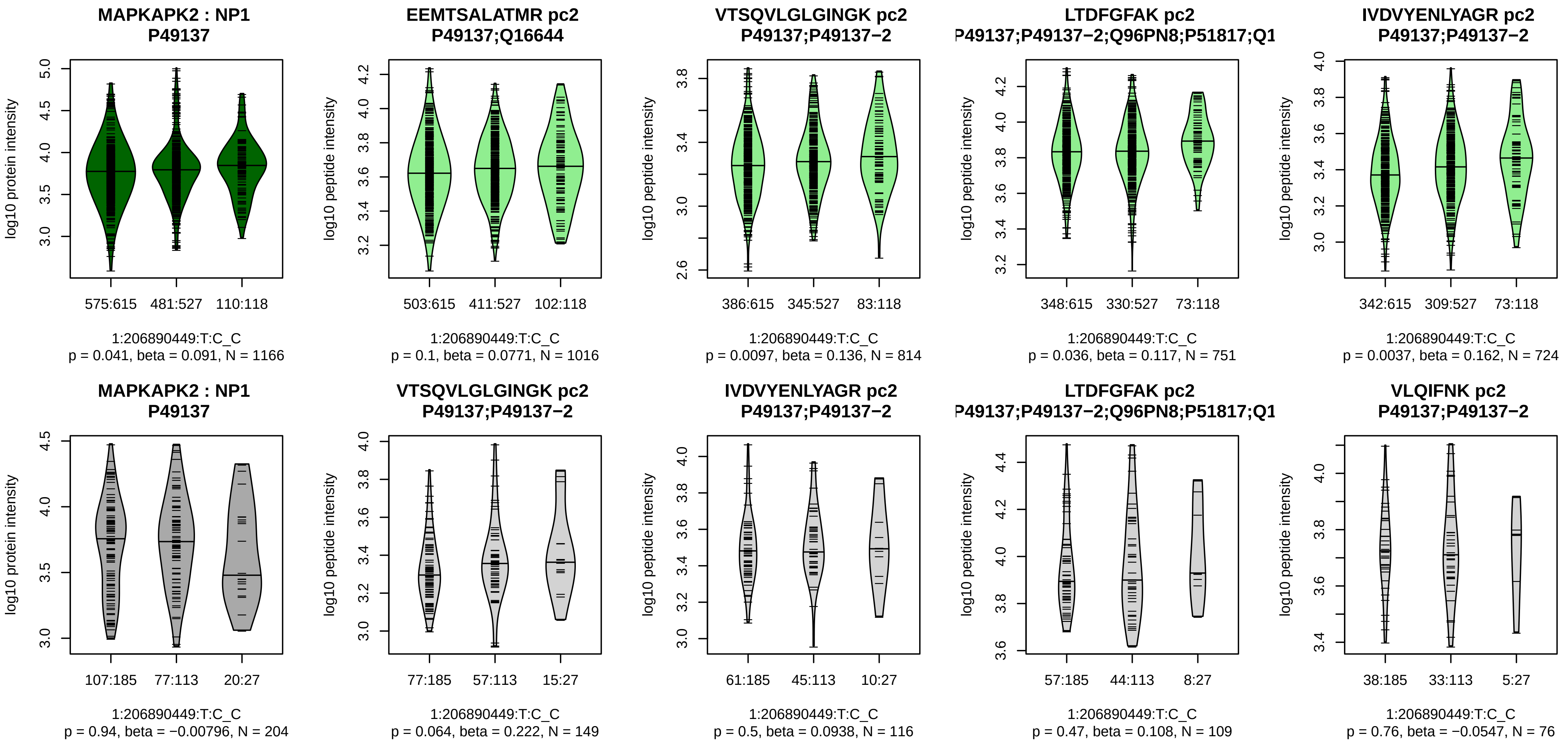


Assay Target: MAPKAPK2
Olink UniProt: P49137
deCODE rsID: rs4256810
Proxy rsID: rs4256810
deCODE: 1:206717104:C:T
Proxy SNP: 1:206890449:T:C
deCODE log10(p): 83.3
deCODE BETA: 0.18
–·*·–·*·*·–·–·–·–·–·–·–·*·–·–·–·–·
1016:814:751:724:696:666:437:

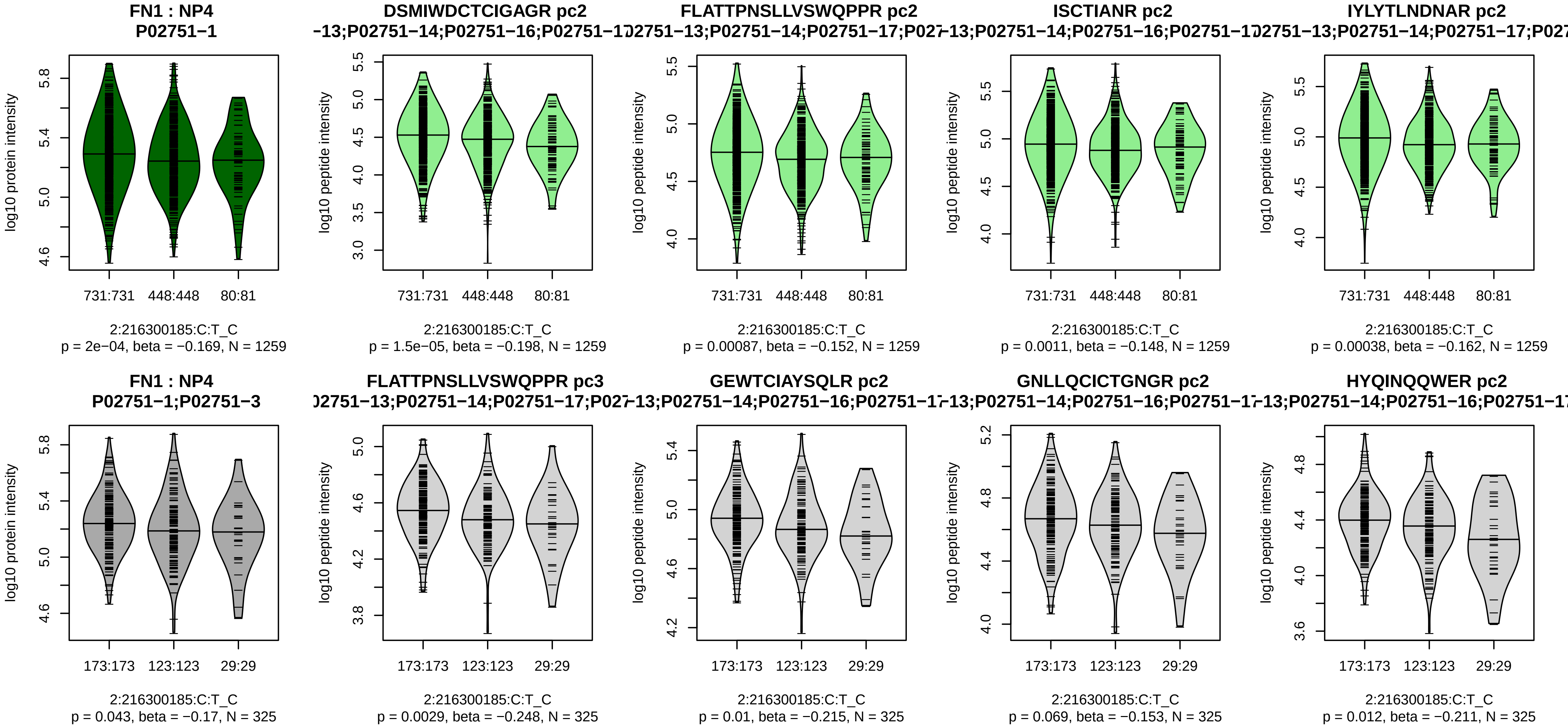


Assay Target: FN1
Olink UniProt: P02751
deCODE rsID: rs1250258
Proxy rsID: rs1250258
deCODE: 2:215435462:C:T
Proxy SNP: 2:216300185:C:T
deCODE log10(p): 82.3
deCODE BETA: −0.16
**************************
1259:1259:1259:1259:1259:125

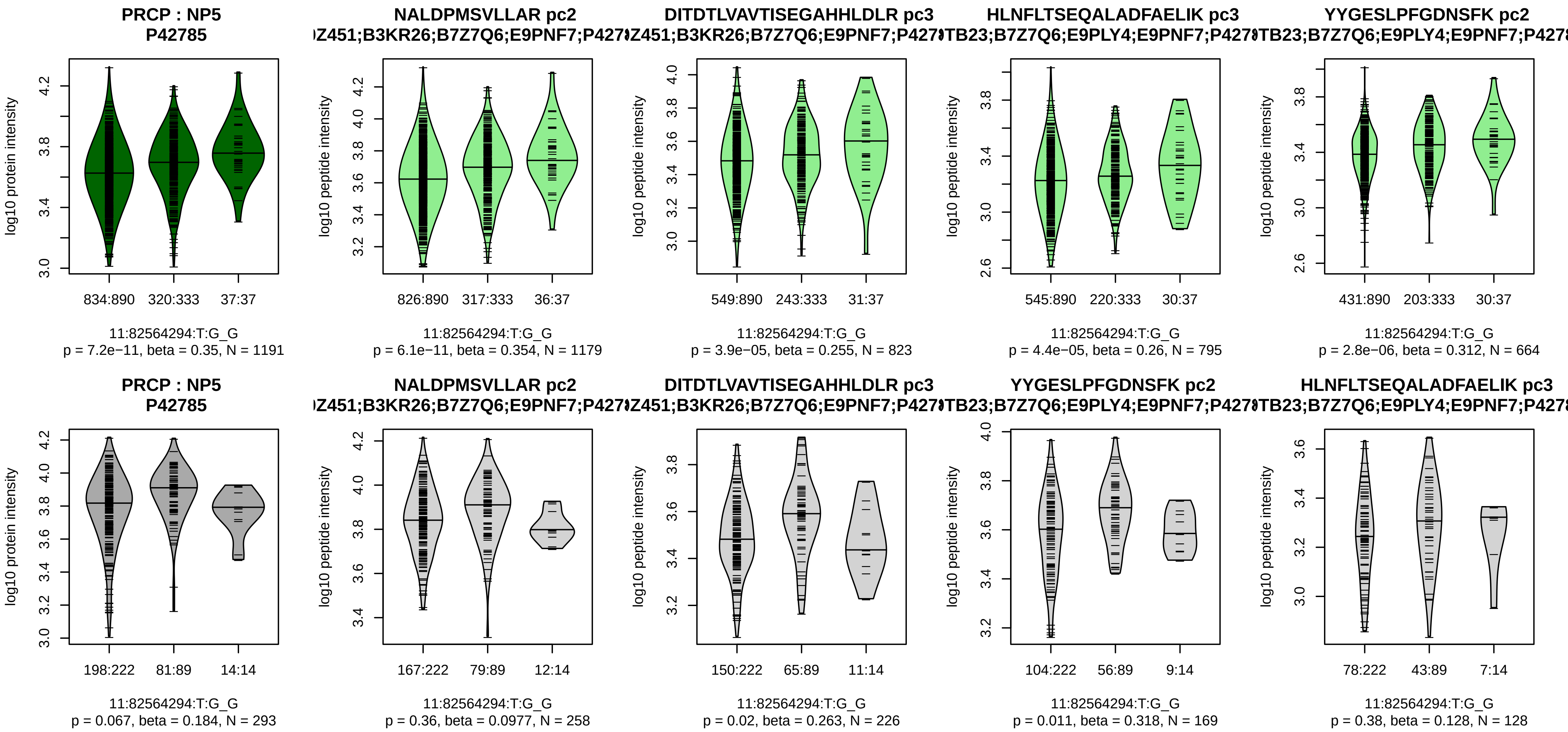


Assay Target: PRCP
Olink UniProt: P42785
deCODE rsID: rs2229437
Proxy rsID: rs2229437
deCODE: 11:82853252:G:T
Proxy SNP: 11:82564294:T:G
deCODE log10(p): 82.2
deCODE BETA: 0.22
*:*:*:*:–:–:–:–
1179:823:795:664:216:87:63:39

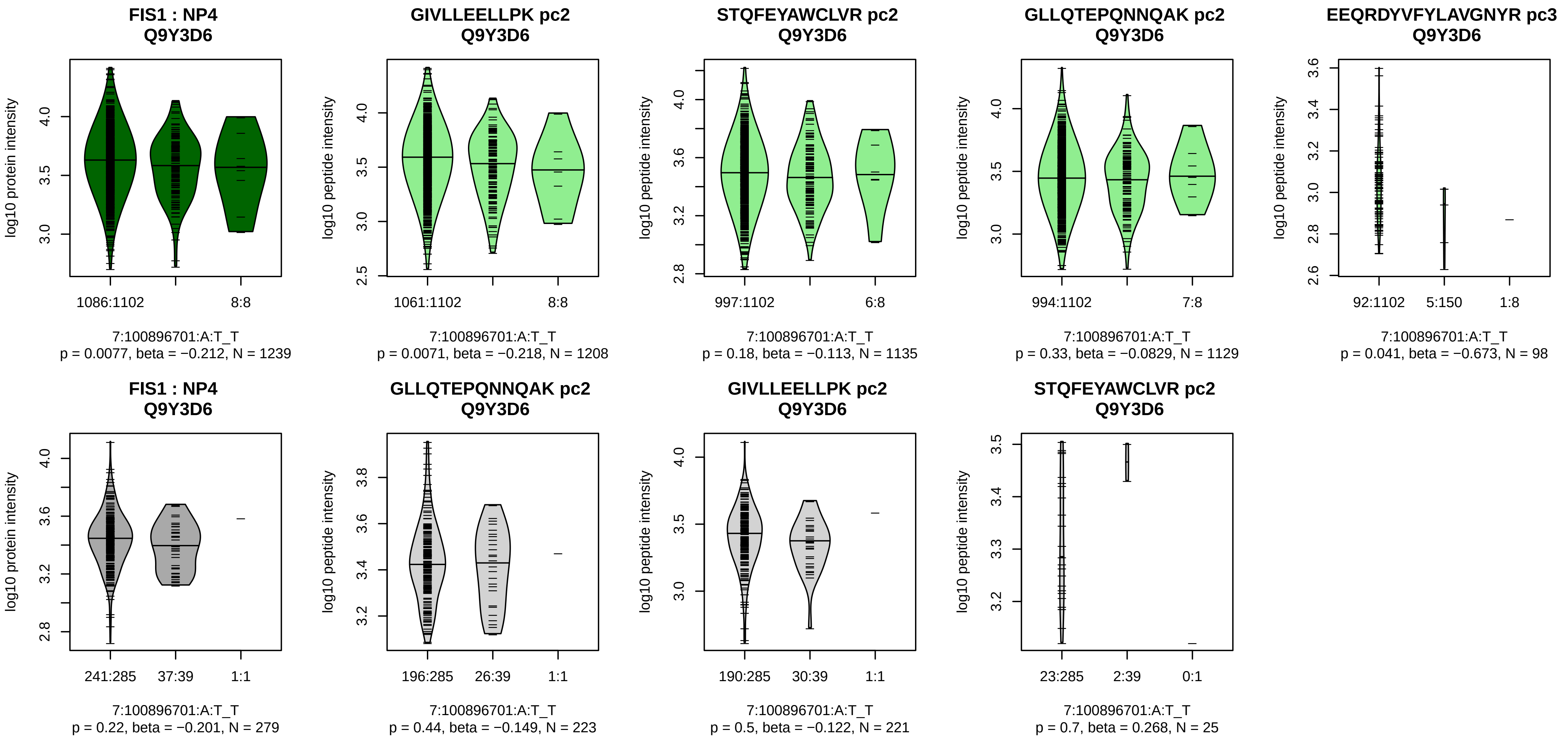


Assay Target: FIS1
Olink UniProt: Q9Y3D6
deCODE rsID: rs75487681
Proxy rsID: rs75487681
deCODE: 7:101253420:T:A
Proxy SNP: 7:100896701:A:T
deCODE log10(p): 80.3
deCODE BETA: −0.32
*:−:−:−
1208:1135:1129:98

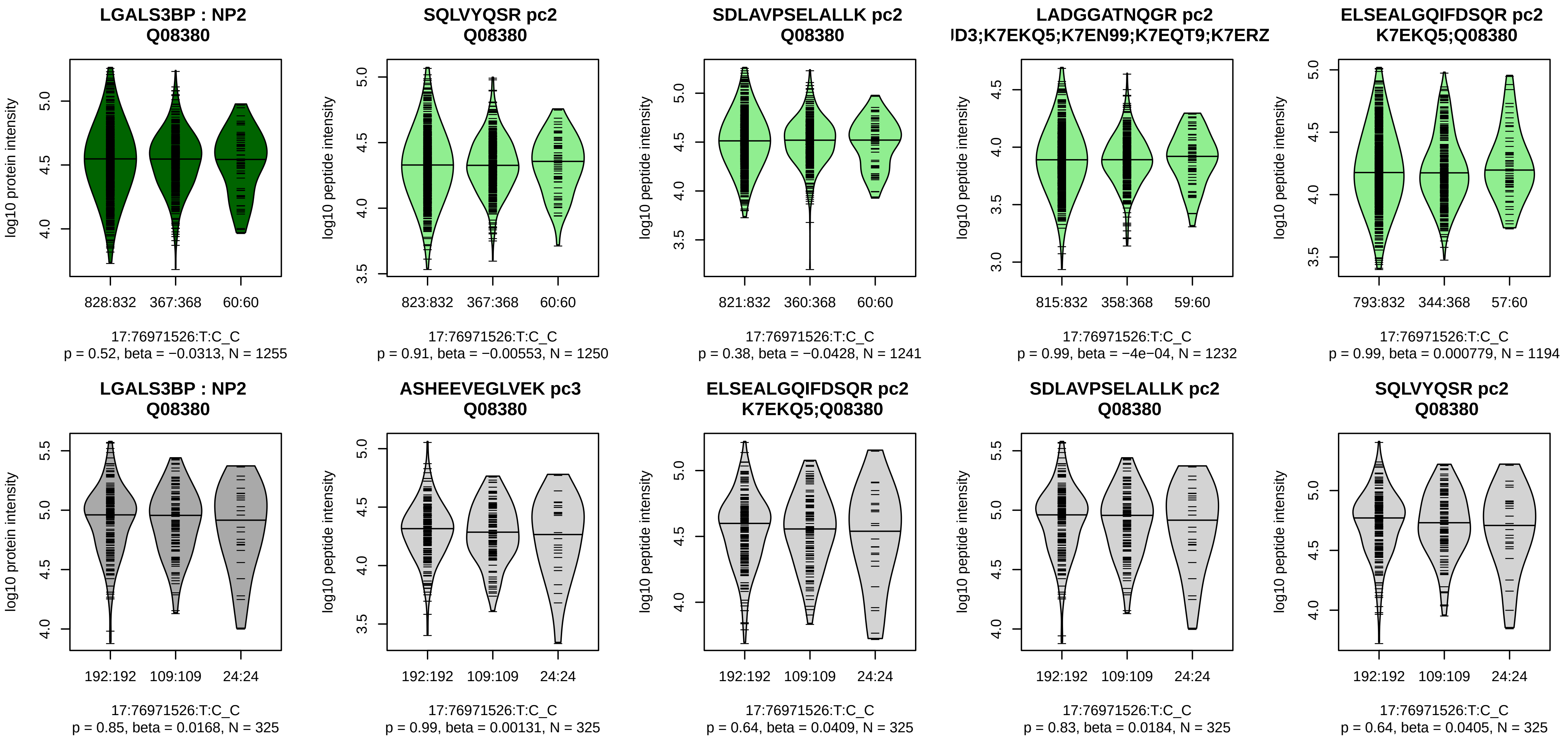


Assay Target: LGALS3BP
Olink UniProt: Q08380
deCODE rsID: rs3826311
Proxy rsID: rs3826311
deCODE: 17:78975444:C:T
Proxy SNP: 17:76971526:T:C
deCODE log10(p): 80.3
deCODE BETA: 0.2
–:–:–:–:–:–:–:–:–:–:–:–:–:–:–:–:
1250:1241:1232:1194:1169:113

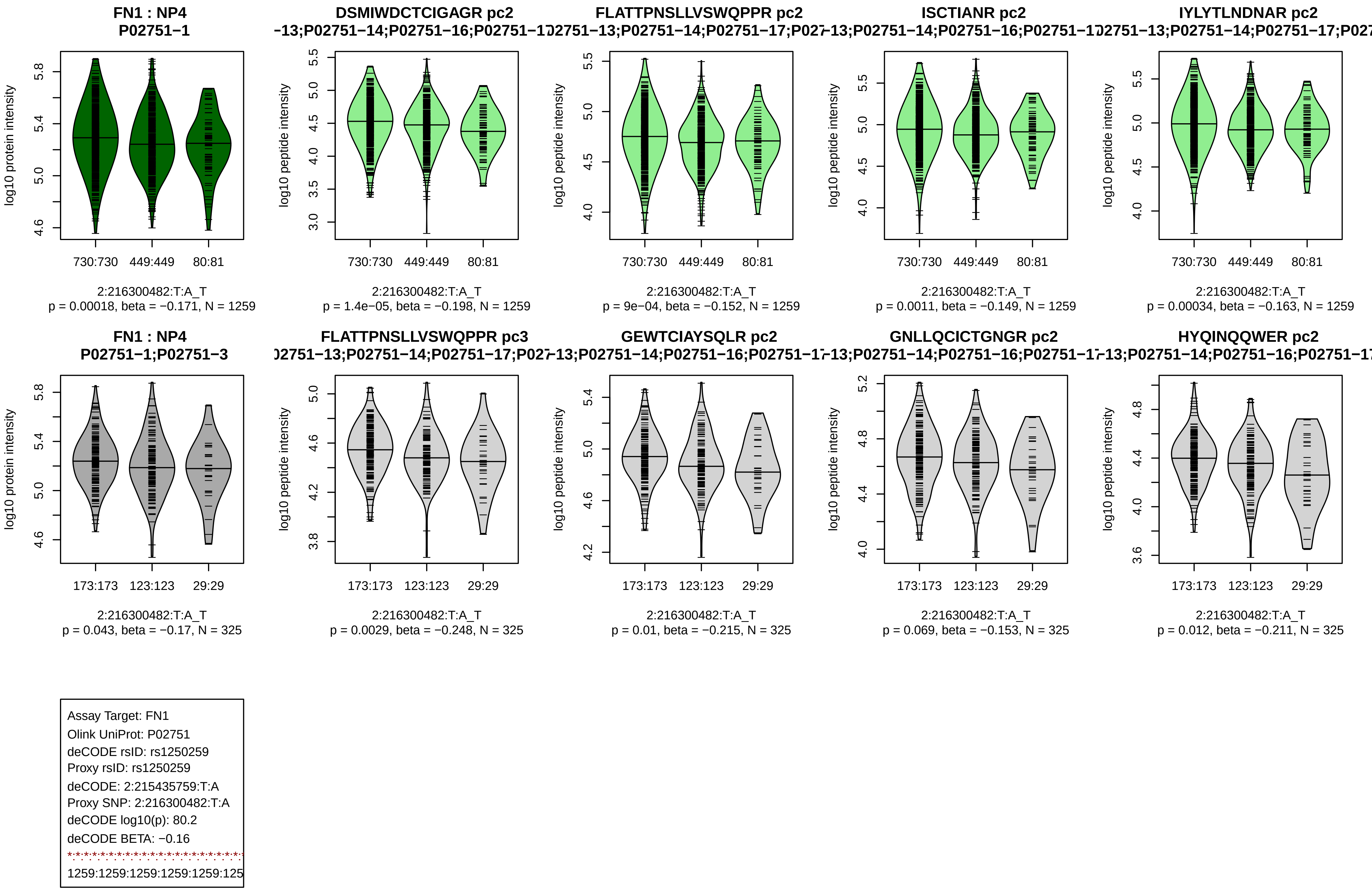
FN1 : NP4
P02751−1
log10 protein intensity
730:730 449:449 80:81
2:216300482:T:A_T
p = 0.00018, beta = −0.171, N = 1259
DSMIWDCTCIGAGR pc2
−13;P02751−14;P02751−16;P02751−1
log10 peptide intensity
730:730 449:449 80:81
2:216300482:T:A_T
p = 1.4e−05, beta = −0.198, N = 1259
FLATTPNSLLVSWQPPR pc2
)2751−13;P02751−14;P02751−17;P027
log10 peptide intensity
730:730 449:449 80:81
2:216300482:T:A_T
p = 9e−04, beta = −0.152, N = 1259
ISCTIANR pc2
−13;P02751−14;P02751−16;P02751−1
log10 peptide intensity
730:730 449:449 80:81
2:216300482:T:A_T
p = 0.0011, beta = −0.149, N = 1259
IYLYTLNDNAR pc2
)2751−13;P02751−14;P02751−17;P027
log10 peptide intensity
730:730 449:449 80:81
2:216300482:T:A_T
p = 0.00034, beta = −0.163, N = 1259
FN1 : NP4
P02751−1;P02751−3
log10 protein intensity
173:173 123:123 29:29
2:216300482:T:A_T
p = 0.043, beta = −0.17, N = 325
FLATTPNSLLVSWQPPR pc3
)2751−13;P02751−14;P02751−17;P027
log10 peptide intensity
173:173 123:123 29:29
2:216300482:T:A_T
p = 0.0029, beta = −0.248, N = 325
GEWTCIAYSQLR pc2
−13;P02751−14;P02751−16;P02751−1
log10 peptide intensity
173:173 123:123 29:29
2:216300482:T:A_T
p = 0.01, beta = −0.215, N = 325
GNLLQCICTGNGR pc2
−13;P02751−14;P02751−16;P02751−1
log10 peptide intensity
173:173 123:123 29:29
2:216300482:T:A_T
p = 0.069, beta = −0.153, N = 325
HYQINQQWER pc2
−13;P02751−14;P02751−16;P02751−1
log10 peptide intensity
173:173 123:123 29:29
2:216300482:T:A_T
p = 0.012, beta = −0.211, N = 325
Assay Target: FN1
Olink UniProt: P02751
deCODE rsID: rs1250259
Proxy rsID: rs1250259
deCODE: 2:215435759:T:A
Proxy SNP: 2:216300482:T:A
deCODE log10(p): 80.2
deCODE BETA: −0.16
1259:1259:1259:1259:1259:125

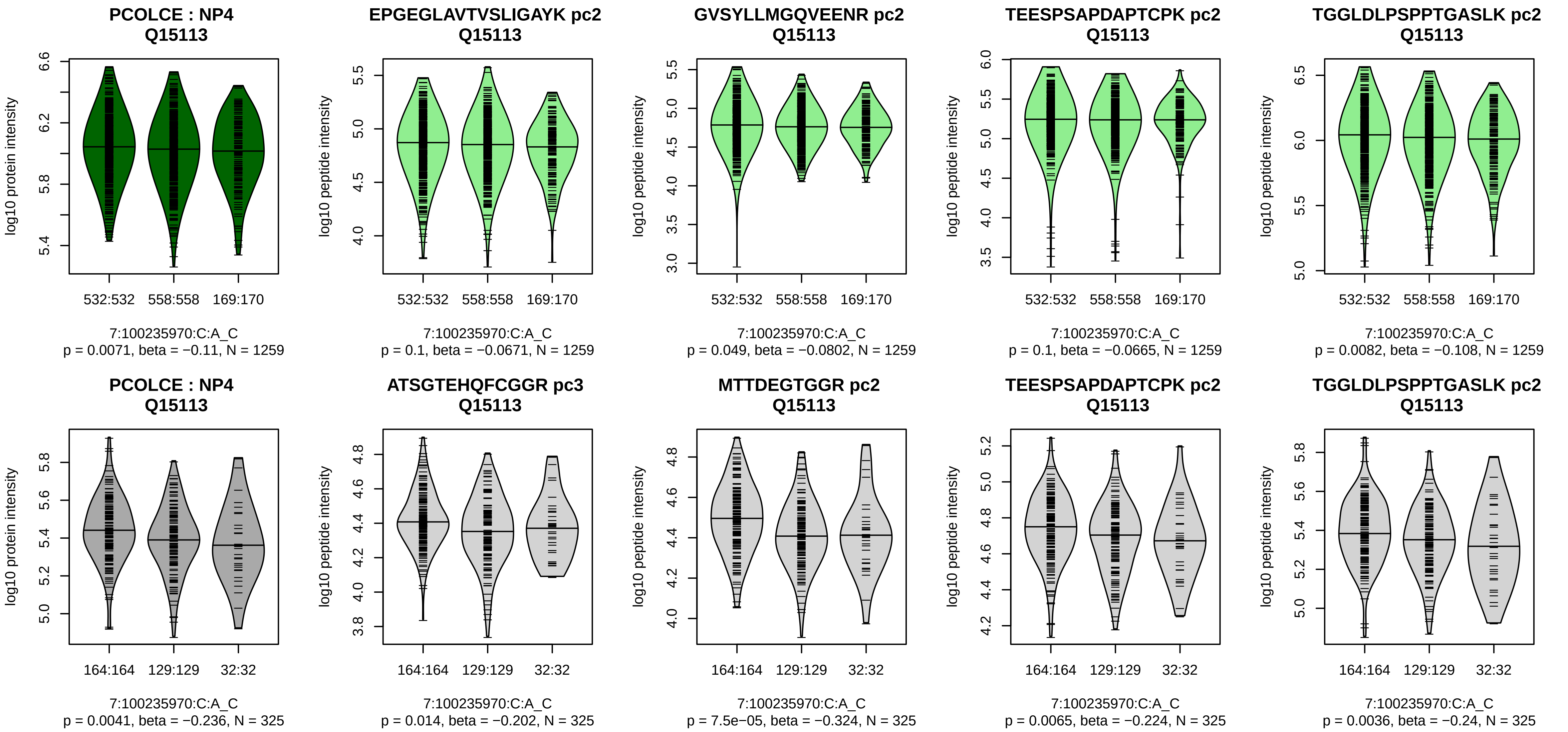


Assay Target: PCOLCE
Olink UniProt: Q15113
deCODE rsID: rs7385804
Proxy rsID: rs7385804
deCODE: 7:100638347:C:A
Proxy SNP: 7:100235970:C:A
deCODE log10(p): 77.9
deCODE BETA: −0.16
−:−:−:*:*:−:−:−:−:−:−:−:−:−:−:−:−:−
1259:1259:1259:1259:1259:125

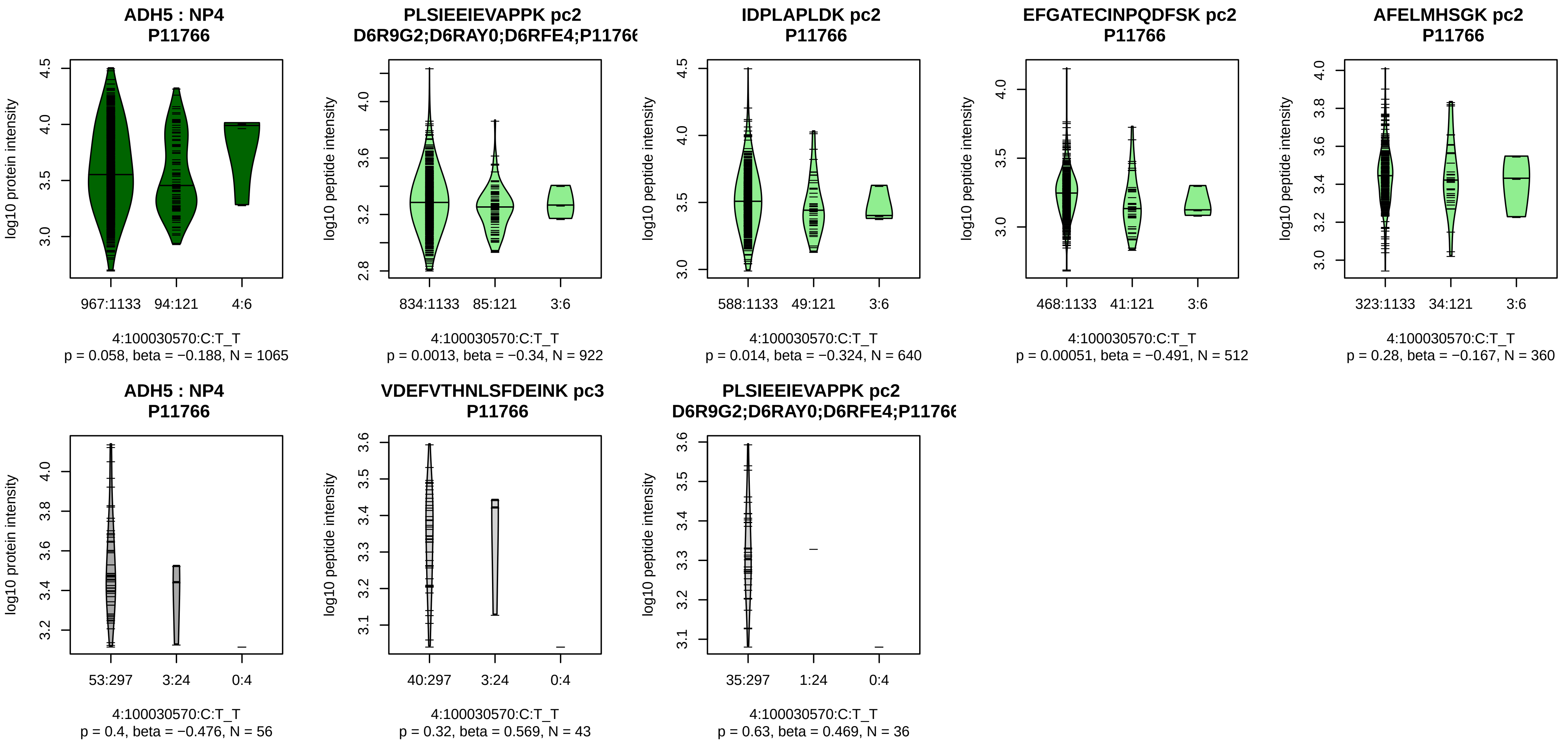


Assay Target: ADH5
Olink UniProt: P11766
deCODE rsID: rs28894371
Proxy rsID: rs75238346
deCODE: 4:99083469:A:G
Proxy SNP: 4:100030570:C:T
deCODE log10(p): 76.8
deCODE BETA: −0.36
*:−:*:−:−:−:−:−:−:−:−:NA
922:640:512:360:309:256:230:2

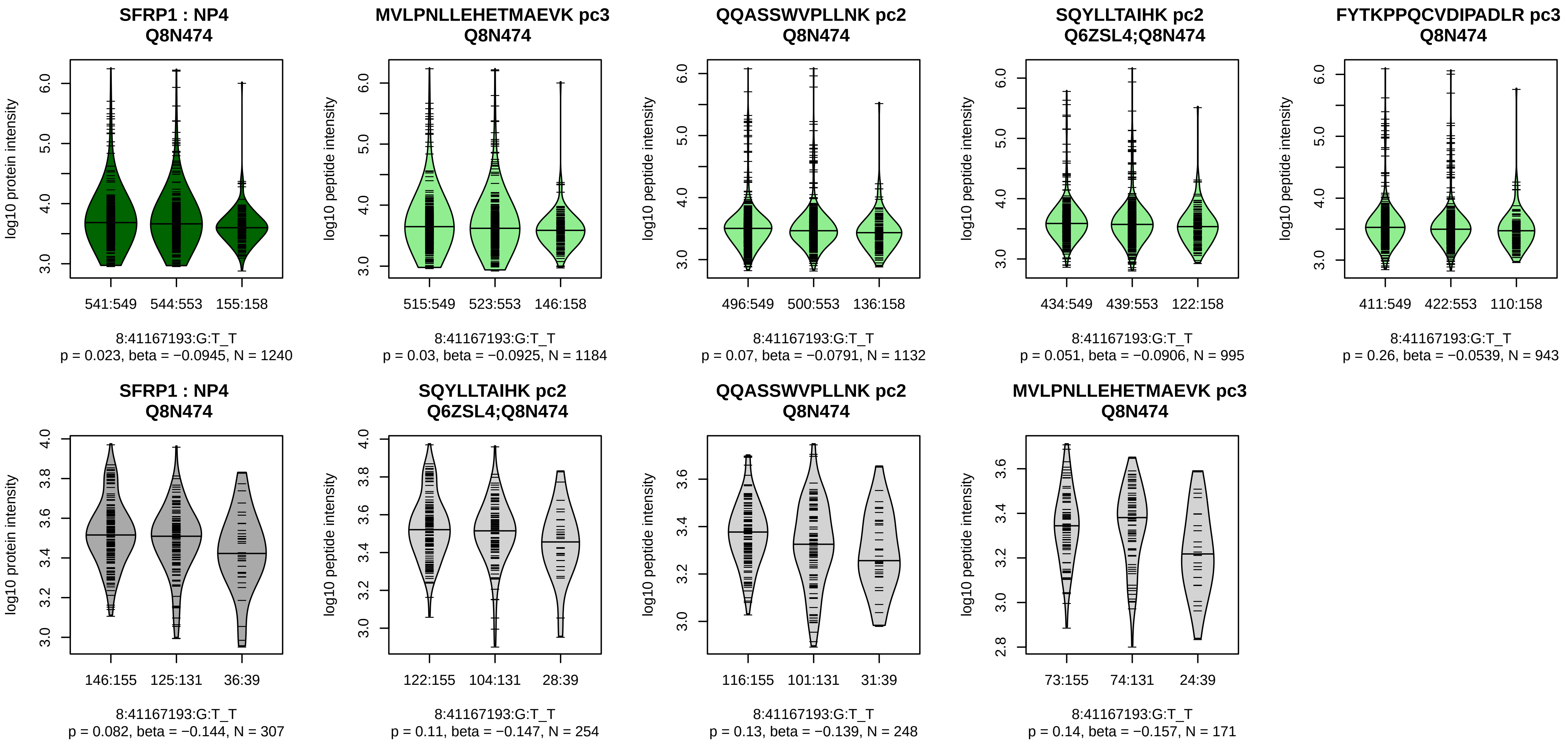


Assay Target: SFRP1
Olink UniProt: Q8N474
deCODE rsID: rs72643819
Proxy rsID: rs72643819
deCODE: 8:41309674:T:G
Proxy SNP: 8:41167193:G:T
deCODE log10(p): 76.2
deCODE BETA: −0.16
−:−:−:−:−:−:−:−:−:−:−:−:−:NA:NA
1184:1132:995:943:637:260:82:

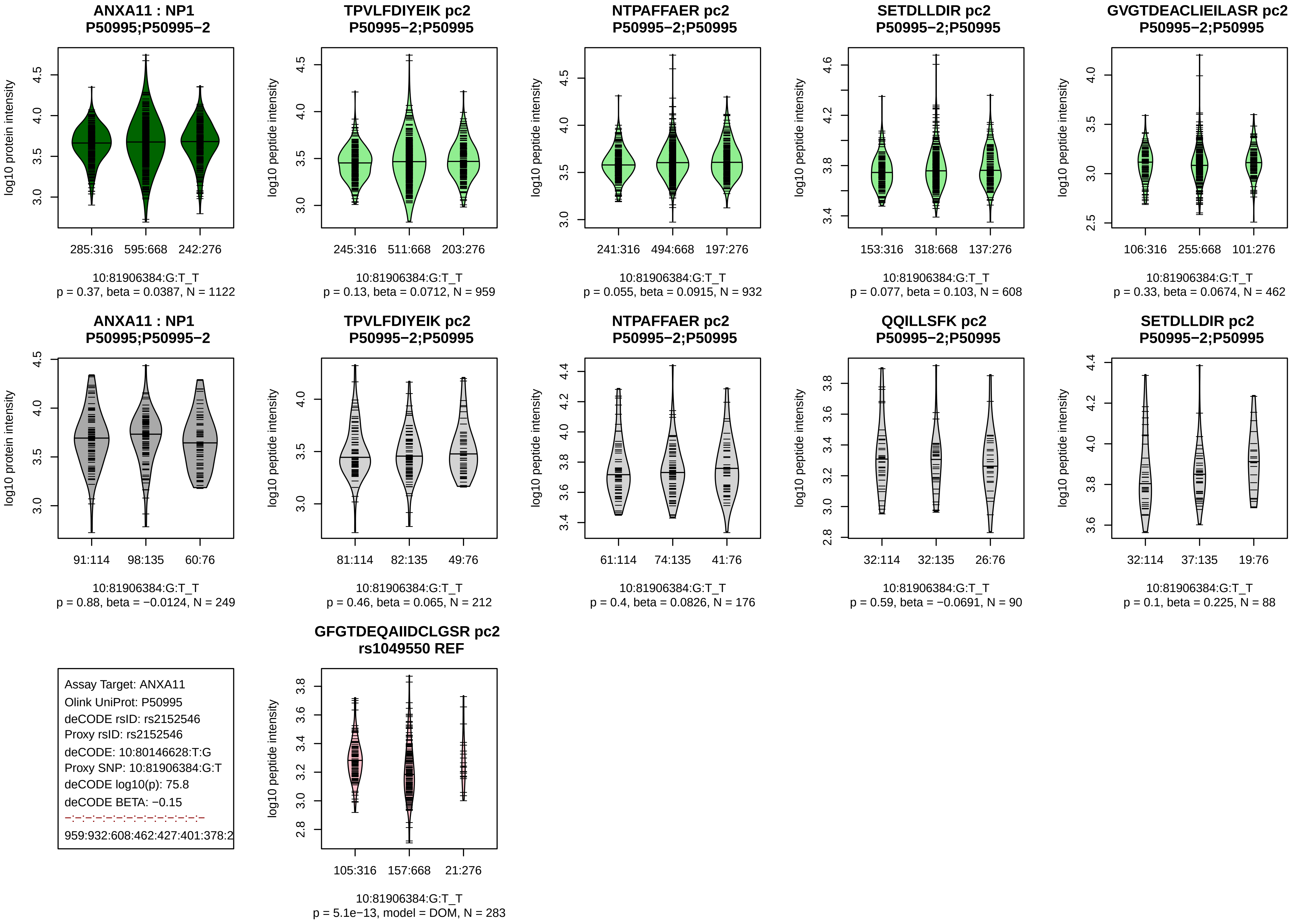
ANXA11 : NP1
P50995;P50995−2
log10 protein intensity
285:316
595:668
242:276
10:81906384:G:T_T
p = 0.37, beta = 0.0387, N = 1122
TPVLFDIYEIK pc2
P50995−2;P50995
log10 peptide intensity
245:316
511:668
203:276
10:81906384:G:T_T
p = 0.13, beta = 0.0712, N = 959
NTPAFFAER pc2
P50995−2;P50995
241:316
494:668
197:276
10:81906384:G:T_T
p = 0.055, beta = 0.0915, N = 932
SETDLLDIR pc2
P50995−2;P50995
153:316
318:668
137:276
10:81906384:G:T_T
p = 0.077, beta = 0.103, N = 608
GVGTDEACLIEILASR pc2
P50995−2;P50995
106:316
255:668
101:276
10:81906384:G:T_T
p = 0.33, beta = 0.0674, N = 462
ANXA11 : NP1
P50995;P50995−2
91:114
98:135
60:76
10:81906384:G:T_T
p = 0.88, beta = −0.0124, N = 249
TPVLFDIYEIK pc2
P50995−2;P50995
81:114
82:135
49:76
10:81906384:G:T_T
p = 0.46, beta = 0.065, N = 212
NTPAFFAER pc2
P50995−2;P50995
61:114
74:135
41:76
10:81906384:G:T_T
p = 0.4, beta = 0.0826, N = 176
QQILLSFK pc2
P50995−2;P50995
32:114
32:135
26:76
10:81906384:G:T_T
p = 0.59, beta = −0.0691, N = 90
SETDLLDIR pc2
P50995−2;P50995
32:114
37:135
19:76
10:81906384:G:T_T
p = 0.1, beta = 0.225, N = 88
Assay Target: ANXA11
Olink UniProt: P50995
deCODE rsID: rs2152546
Proxy rsID: rs2152546
deCODE: 10:80146628:T:G
Proxy SNP: 10:81906384:G:T
deCODE log10(p): 75.8
deCODE BETA: −0.15
959:932:608:462:427:401:378:2
GFGTDEQAIIDCLGSR pc2
rs1049550 REF
105:316
157:668
21:276
10:81906384:G:T_T
p = 5.1e−13, model = DOM, N = 283

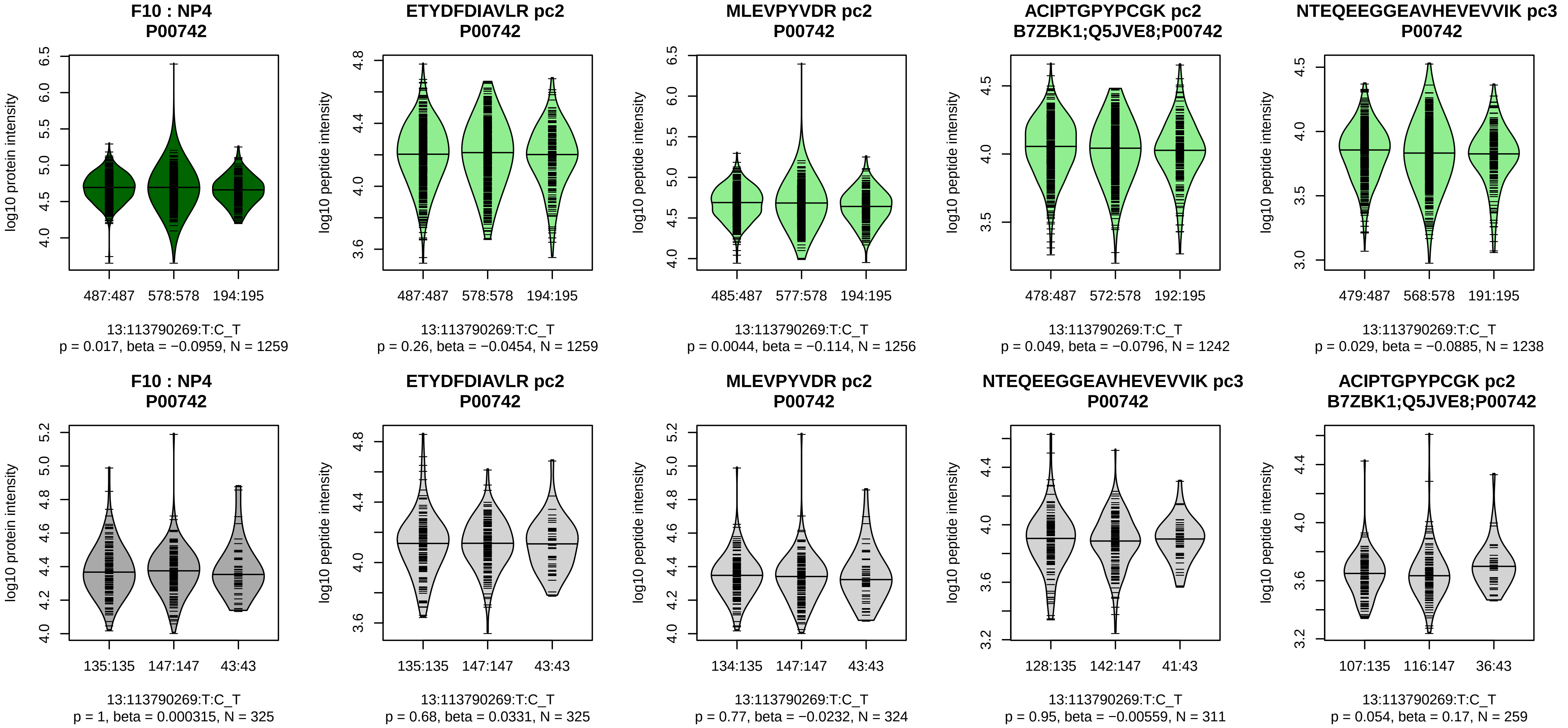


Assay Target: F10
Olink UniProt: P00742
deCODE rsID: rs3211757
Proxy rsID: rs3211757
deCODE: 13:113135954:!:AC
Proxy SNP: 13:113790269:T:C
deCODE log10(p): 75
deCODE BETA: −0.16
−:*:−:−:−:−:−:−:−:−:−:−:−:−:−:−:NA
1259:1256:1242:1238:1216:106

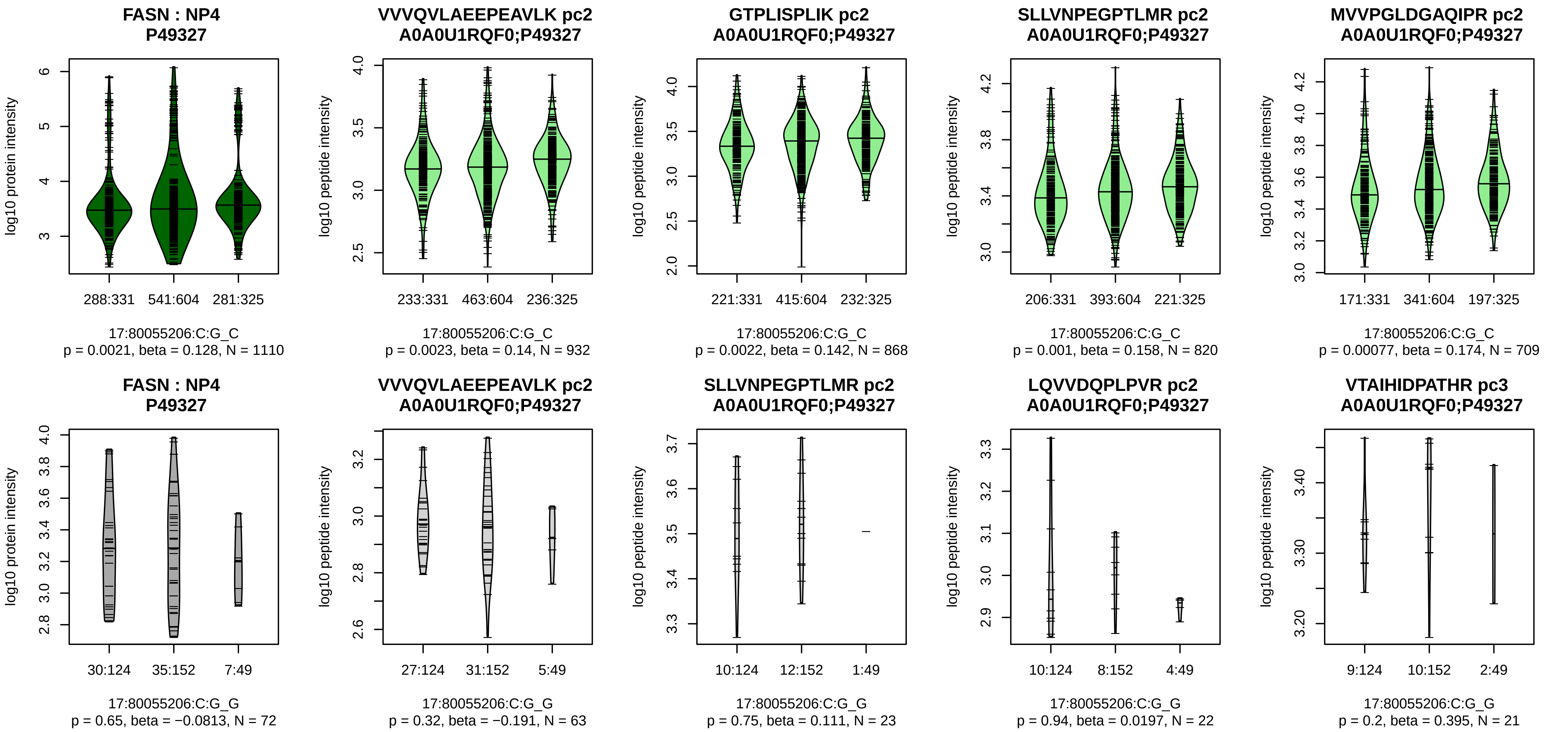


Assay Target: FASN
Olink UniProt: P49327
deCODE rsID: rs62078747
Proxy rsID: rs62078747
deCODE: 17:82097330:C:G
Proxy SNP: 17:80055206:C:G
deCODE log10(p): 74.2
deCODE BETA: 0.15
*:*:*:*:*:–:*:–:–:–:–:–:–:–:–:–:–:–:–
932:868:820:709:641:621:615:6

**RNASE2 : NP1**
**P10153**

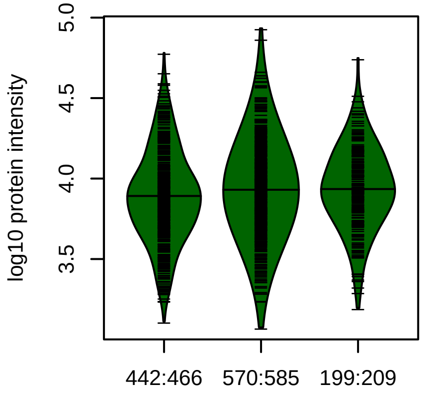


14:21359808:C:A_A
p = 0.013, beta = 0.102, N = 1211

**RDPPQYPVVPVHLDR pc3**
**P10153**

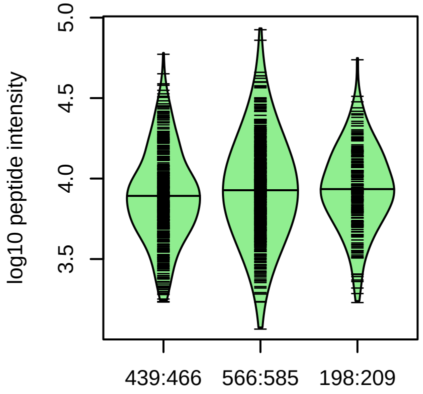


14:21359808:C:A_A
p = 0.017, beta = 0.0976, N = 1203

**YAQTPANMFYIVACDNR pc3**
**P10153**

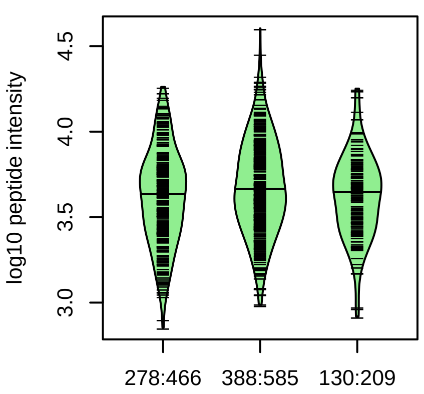


14:21359808:C:A_A
p = 0.49, beta = 0.0348, N = 796

**RNASE2 : NP1**
**P10153**

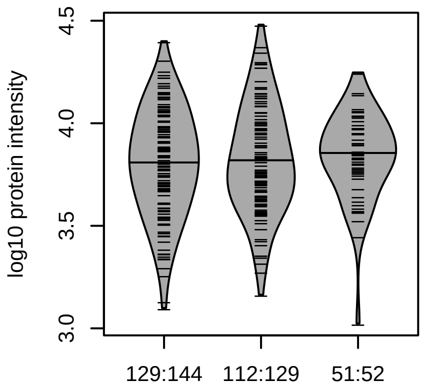


14:21359808:C:A_A
p = 0.5, beta = 0.0526, N = 292

**RDPPQYPVVPVHLDR pc3**
**P10153**

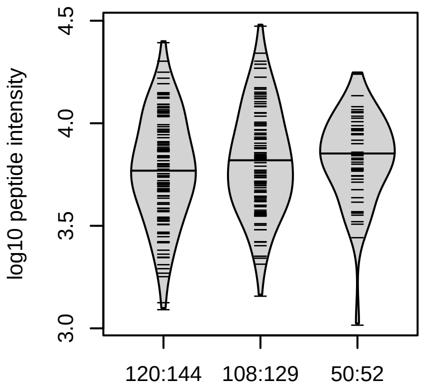


14:21359808:C:A_A
p = 0.19, beta = 0.106, N = 278

Assay Target: RNASE2
Olink UniProt: P10153
deCODE rsID: rs2233859
Proxy rsID: rs2233859
deCODE: 14:20891649:A:C
Proxy SNP: 14:21359808:C:A
deCODE log10(p): 73.8
deCODE BETA: 0.14
–:–:–
1203:862:796

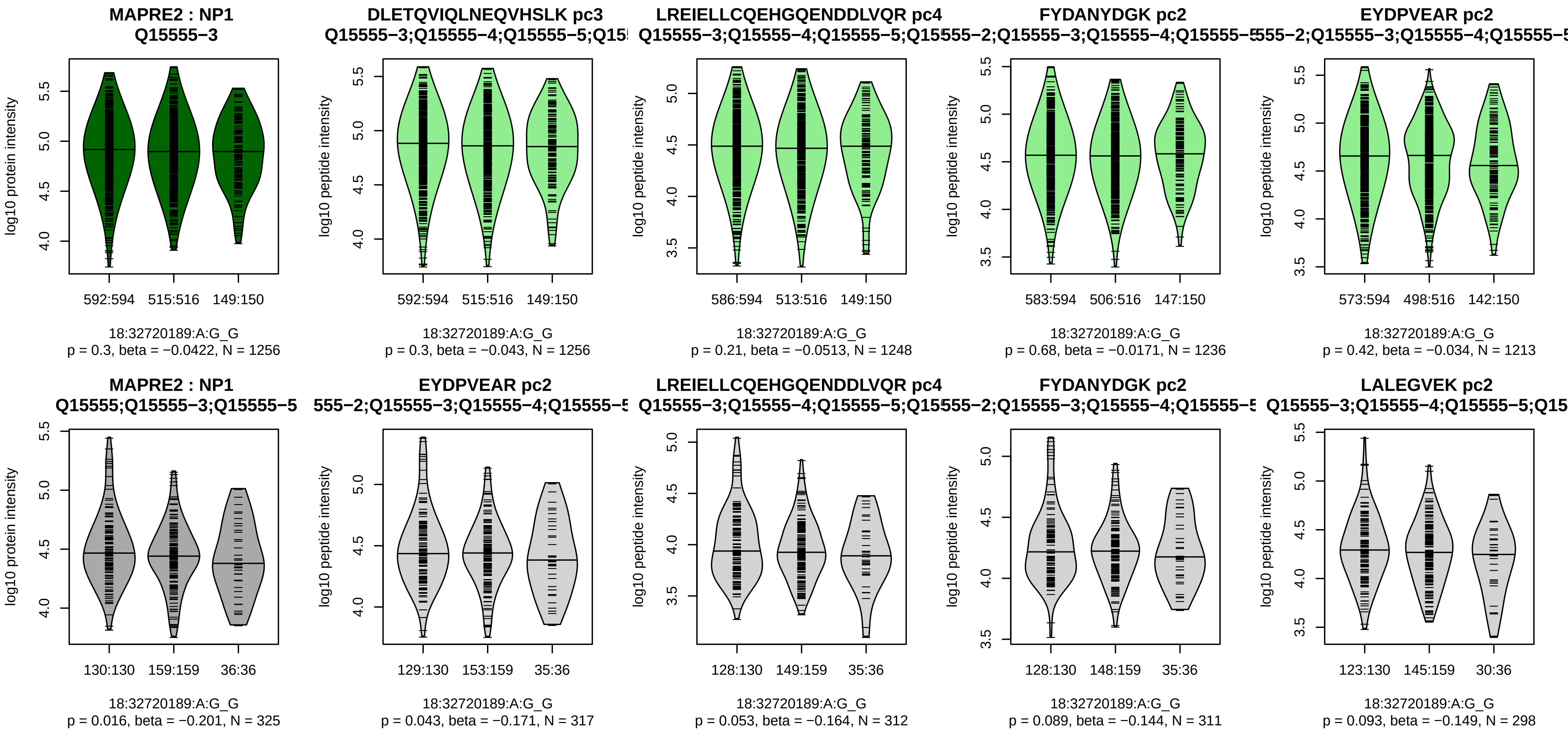


Assay Target: MAPRE2
Olink UniProt: Q15555
deCODE rsID: rs3786314
Proxy rsID: rs3786314
deCODE: 18:35140225:G:A
Proxy SNP: 18:32720189:A:G
deCODE log10(p): 72.9
deCODE BETA: −0.15
−:−:−:−:−:−:−:−:−:−:−:−:−:−:−
1256:1248:1236:1213:1209:120

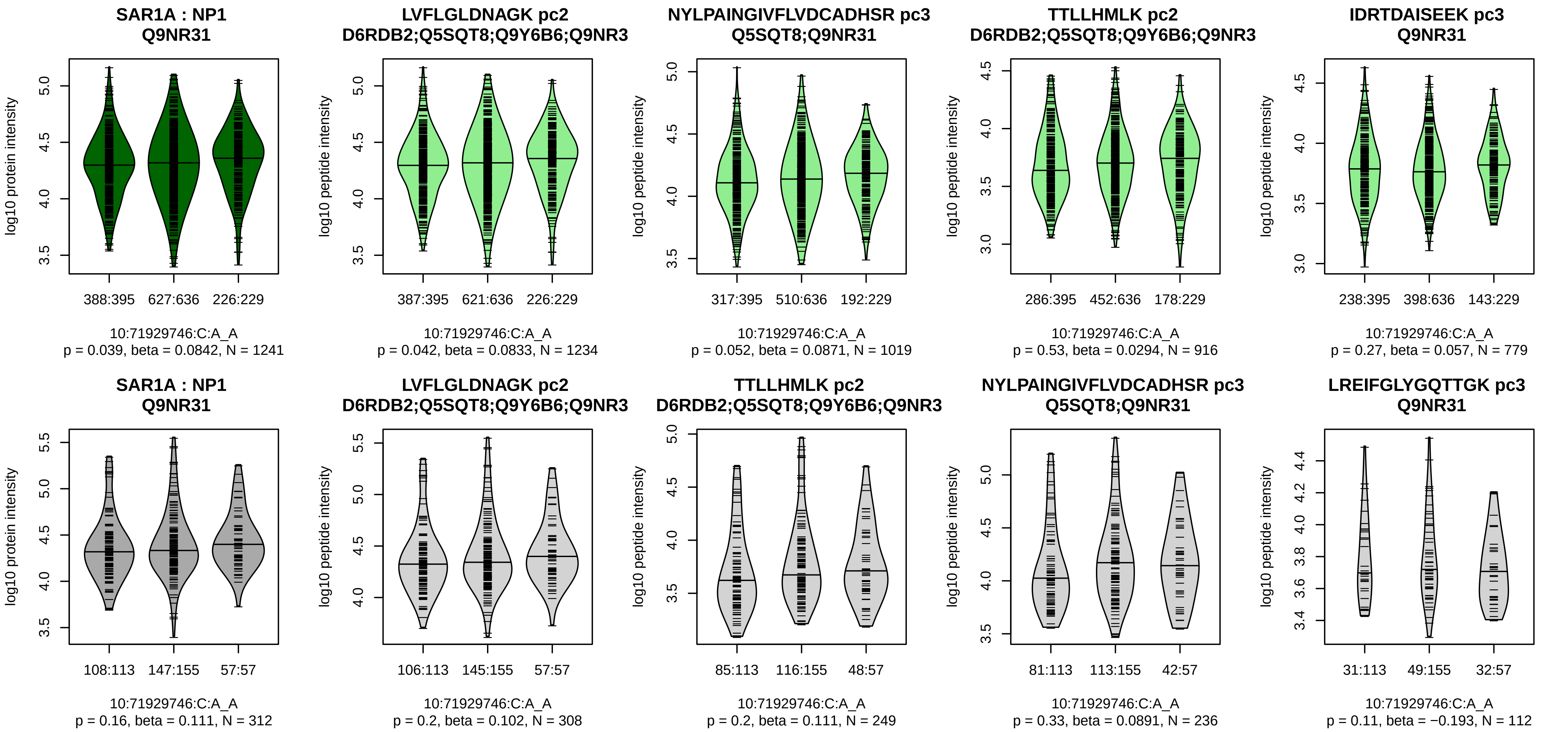


Assay Target: SAR1A
Olink UniProt: Q9NR31
deCODE rsID: rs870801
Proxy rsID: rs870801
deCODE: 10:70169990:A:C
Proxy SNP: 10:71929746:C:A
deCODE log10(p): 72.2
deCODE BETA: 0.15
–:–:–:–:*:–:–:–:–:–
1234:1019:916:779:502:470:38[illegible]

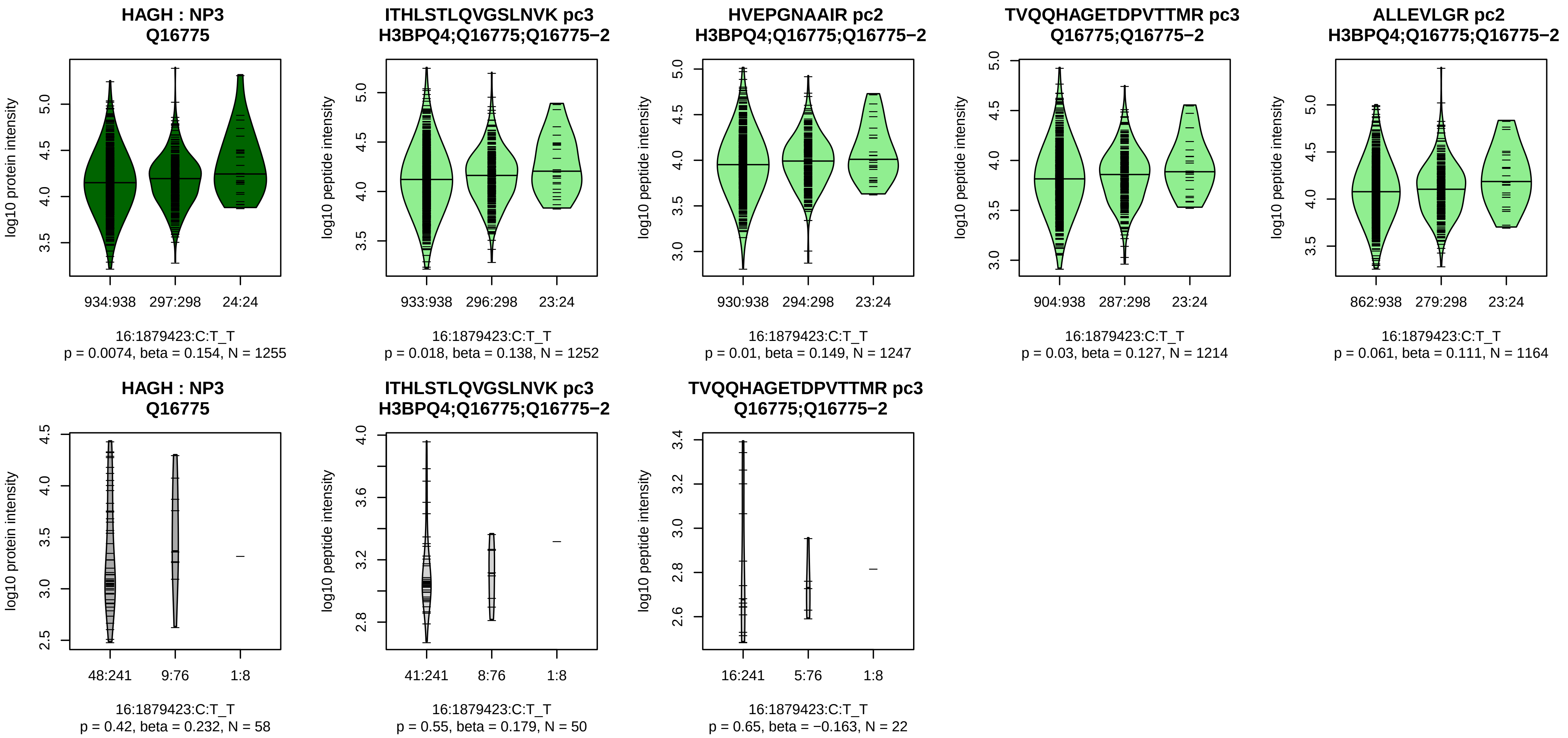


Assay Target: HAGH
Olink UniProt: Q16775
deCODE rsID: rs116869551
Proxy rsID: rs7185299
deCODE: 16:1826618:T:C
Proxy SNP: 16:1879423:C:T
deCODE log10(p): 72.2
deCODE BETA: 0.21
−:−:−:−:−:−:−:−:−:−:−:NA:NA:NA
1252:1247:1214:1164:988:798:3

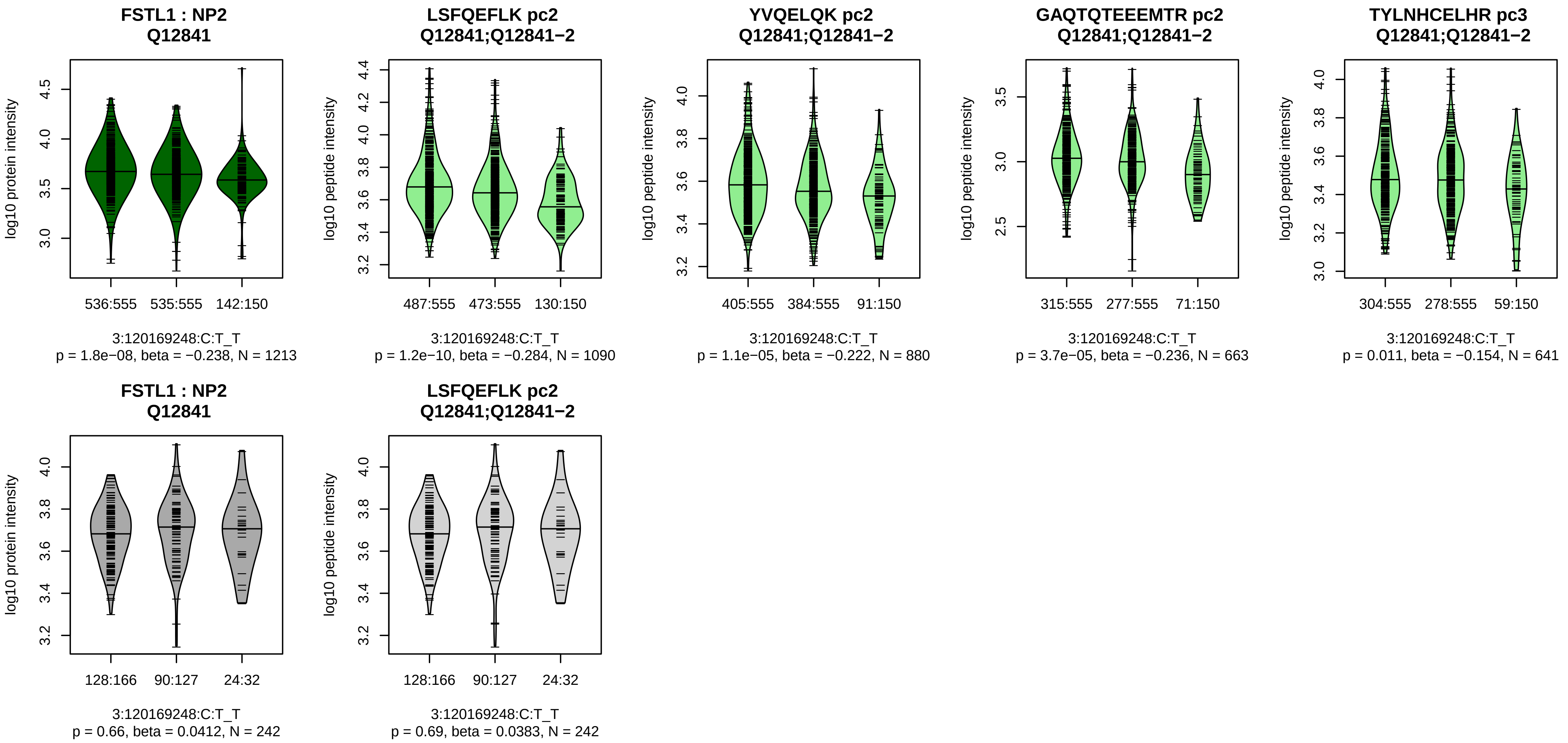


Assay Target: FSTL1
Olink UniProt: Q12841
deCODE rsID: rs1147707
Proxy rsID: rs1147707
deCODE: 3:120450401:T:C
Proxy SNP: 3:120169248:C:T
deCODE log10(p): 69.8
deCODE BETA: −0.15
*:*:*:–:*:*:–:–:–
1090:880:663:641:545:538:361:

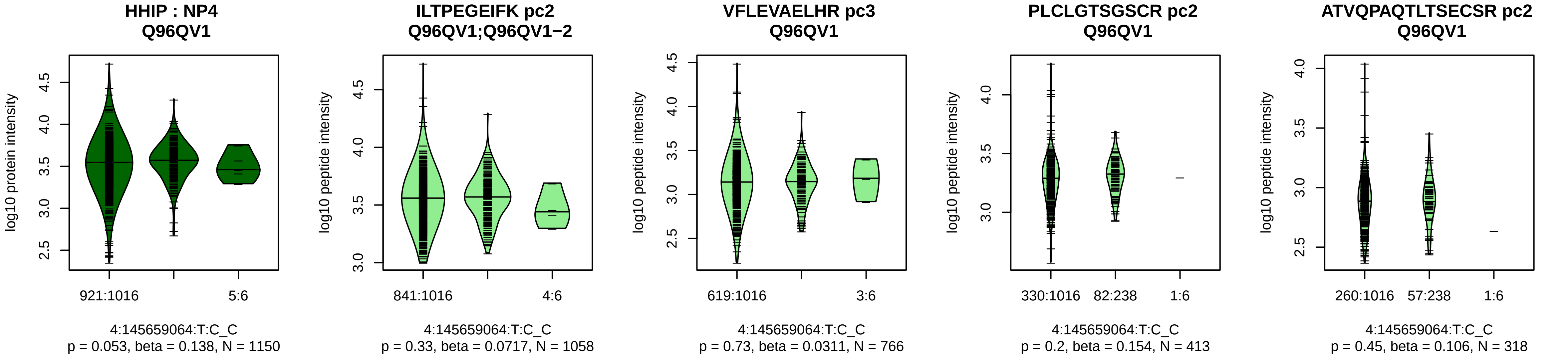


Assay Target: HHIP
Olink UniProt: Q96QV1
deCODE rsID: rs11727676
Proxy rsID: rs11727676
deCODE: 4:144737912:C:T
Proxy SNP: 4:145659064:T:C
deCODE log10(p): 69.3
deCODE BETA: 0.24
−:-:−:-:−:-:−:-:−:-:−:-:−:-:−:-:−:N
1058:766:413:318:268:236:222:

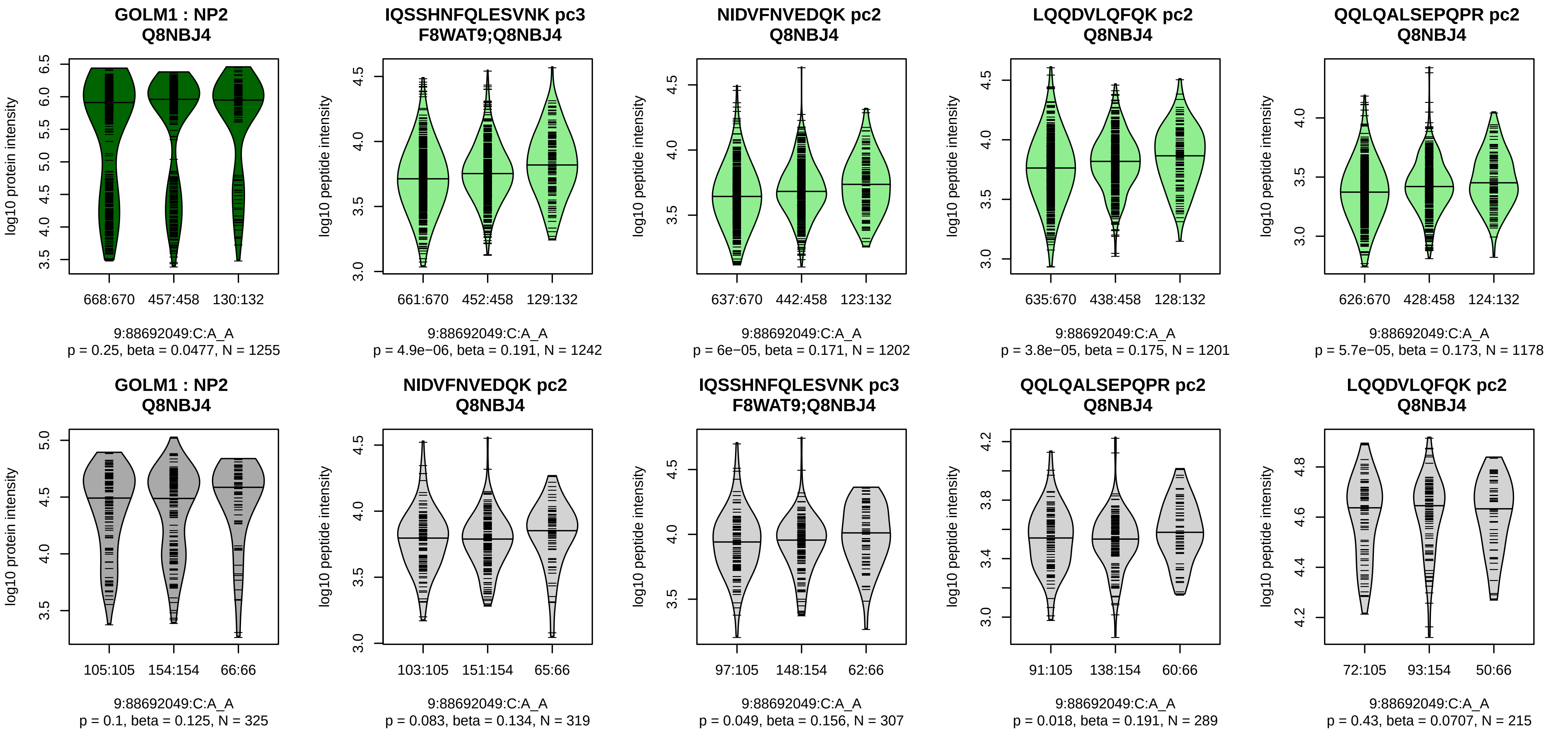


Assay Target: GOLM1
Olink UniProt: Q8NBJ4
deCODE rsID: rs4333693
Proxy rsID: rs4333693
deCODE: 9:86077134:A:C
Proxy SNP: 9:88692049:C:A
deCODE log10(p): 68.7
deCODE BETA: 0.18
*:*:*:*:*:−:−:*:−:−:NA
1242:1202:1201:1178:1081:950

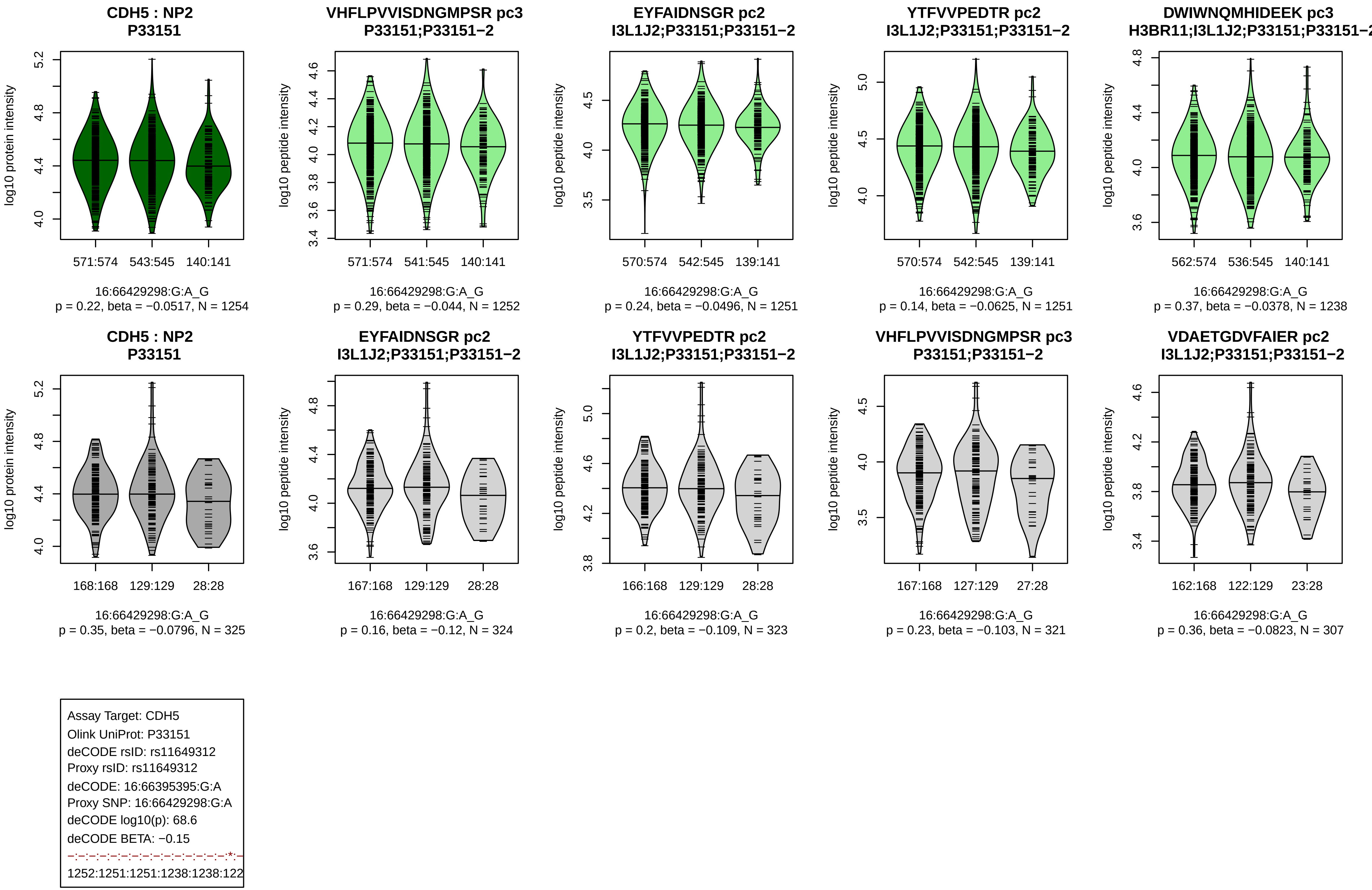
CDH5 : NP2
P33151
log10 protein intensity
571:574 543:545 140:141
16:66429298:G:A_G
p = 0.22, beta = −0.0517, N = 1254
VHFLPVVISDNGMPSR pc3
P33151;P33151−2
log10 peptide intensity
571:574 541:545 140:141
16:66429298:G:A_G
p = 0.29, beta = −0.044, N = 1252
EYFAIDNSGR pc2
I3L1J2;P33151;P33151−2
570:574 542:545 139:141
16:66429298:G:A_G
p = 0.24, beta = −0.0496, N = 1251
YTFVVPEDTR pc2
I3L1J2;P33151;P33151−2
570:574 542:545 139:141
16:66429298:G:A_G
p = 0.14, beta = −0.0625, N = 1251
DWIWNQMHIDEEK pc3
H3BR11;I3L1J2;P33151;P33151−2
562:574 536:545 140:141
16:66429298:G:A_G
p = 0.37, beta = −0.0378, N = 1238
CDH5 : NP2
P33151
168:168 129:129 28:28
16:66429298:G:A_G
p = 0.35, beta = −0.0796, N = 325
EYFAIDNSGR pc2
I3L1J2;P33151;P33151−2
167:168 129:129 28:28
16:66429298:G:A_G
p = 0.16, beta = −0.12, N = 324
YTFVVPEDTR pc2
I3L1J2;P33151;P33151−2
166:168 129:129 28:28
16:66429298:G:A_G
p = 0.2, beta = −0.109, N = 323
VHFLPVVISDNGMPSR pc3
P33151;P33151−2
167:168 127:129 27:28
16:66429298:G:A_G
p = 0.23, beta = −0.103, N = 321
VDAETGDVFAIER pc2
I3L1J2;P33151;P33151−2
162:168 122:129 23:28
16:66429298:G:A_G
p = 0.36, beta = −0.0823, N = 307
Assay Target: CDH5
Olink UniProt: P33151
deCODE rsID: rs11649312
Proxy rsID: rs11649312
deCODE: 16:66395395:G:A
Proxy SNP: 16:66429298:G:A
deCODE log10(p): 68.6
deCODE BETA: −0.15
−:−:−:−:−:−:−:−:−:−:−:−:−:−:−:*:−
1252:1251:1251:1238:1238:122

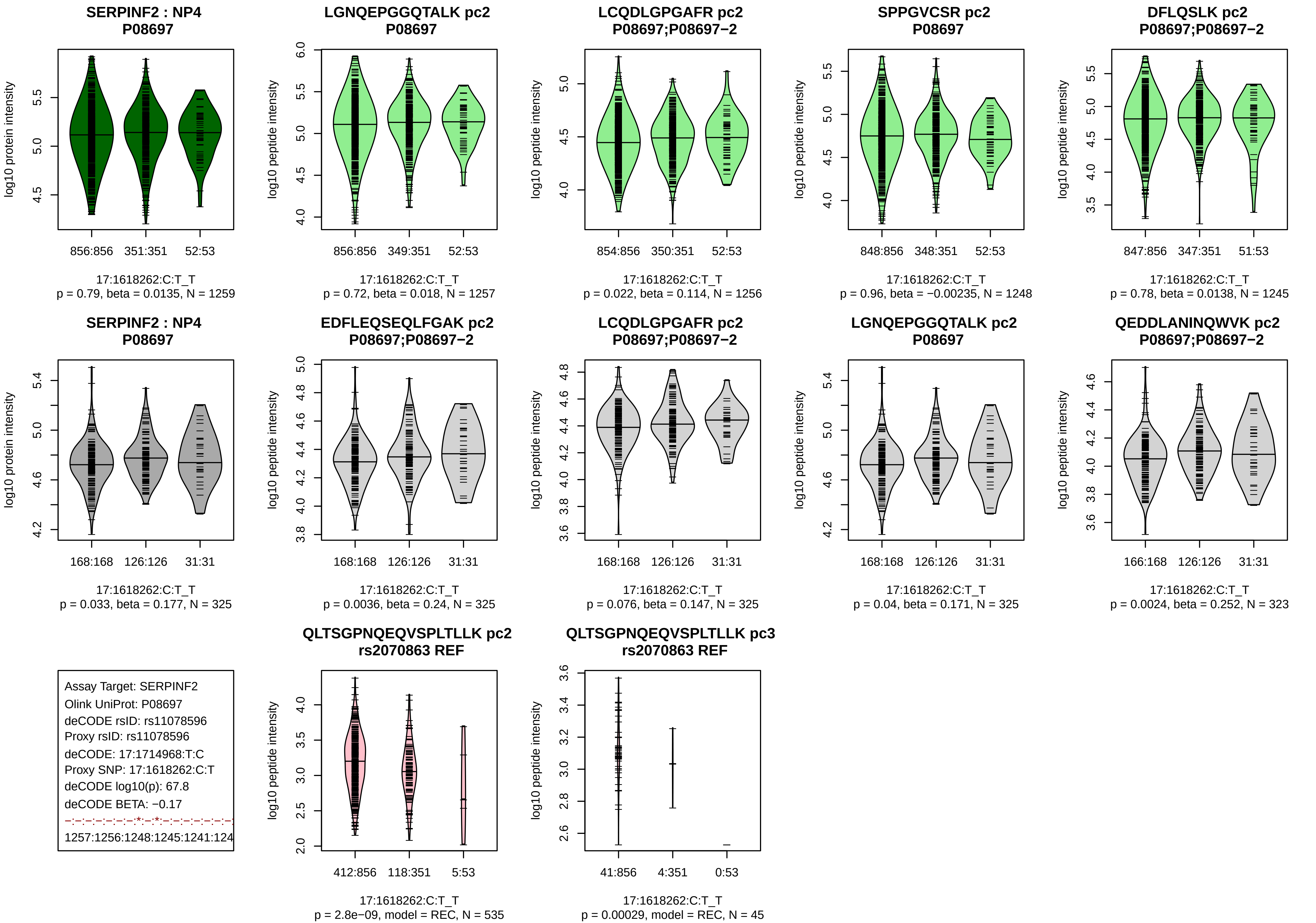

SERPINF2 : NP4
P08697
log10 protein intensity
856:856
351:351
52:53
17:1618262:C:T_T
p = 0.79, beta = 0.0135, N = 1259
LGNQEPGGQTALK pc2
P08697
log10 peptide intensity
856:856
349:351
52:53
17:1618262:C:T_T
p = 0.72, beta = 0.018, N = 1257
LCQDLGPGAFR pc2
P08697;P08697−2
854:856
350:351
52:53
17:1618262:C:T_T
p = 0.022, beta = 0.114, N = 1256
SPPGVCSR pc2
P08697
848:856
348:351
52:53
17:1618262:C:T_T
p = 0.96, beta = −0.00235, N = 1248
DFLQSLK pc2
P08697;P08697−2
847:856
347:351
51:53
17:1618262:C:T_T
p = 0.78, beta = 0.0138, N = 1245
SERPINF2 : NP4
P08697
168:168
126:126
31:31
17:1618262:C:T_T
p = 0.033, beta = 0.177, N = 325
EDFLEQSEQLFGAK pc2
P08697;P08697−2
168:168
126:126
31:31
17:1618262:C:T_T
p = 0.0036, beta = 0.24, N = 325
LCQDLGPGAFR pc2
P08697;P08697−2
168:168
126:126
31:31
17:1618262:C:T_T
p = 0.076, beta = 0.147, N = 325
LGNQEPGGQTALK pc2
P08697
168:168
126:126
31:31
17:1618262:C:T_T
p = 0.04, beta = 0.171, N = 325
QEDDLANINQWVK pc2
P08697;P08697−2
166:168
126:126
31:31
17:1618262:C:T_T
p = 0.0024, beta = 0.252, N = 323
Assay Target: SERPINF2
Olink UniProt: P08697
deCODE rsID: rs11078596
Proxy rsID: rs11078596
deCODE: 17:1714968:T:C
Proxy SNP: 17:1618262:C:T
deCODE log10(p): 67.8
deCODE BETA: −0.17
−:−:−:−:−:−:−:*:−:*:−:−:−:−:−:−:−:−
1257:1256:1248:1245:1241:124
QLTSGPNQEQVSPLTLLK pc2
rs2070863 REF
412:856
118:351
5:53
17:1618262:C:T_T
p = 2.8e−09, model = REC, N = 535
QLTSGPNQEQVSPLTLLK pc3
rs2070863 REF
41:856
4:351
0:53
17:1618262:C:T_T
p = 0.00029, model = REC, N = 45

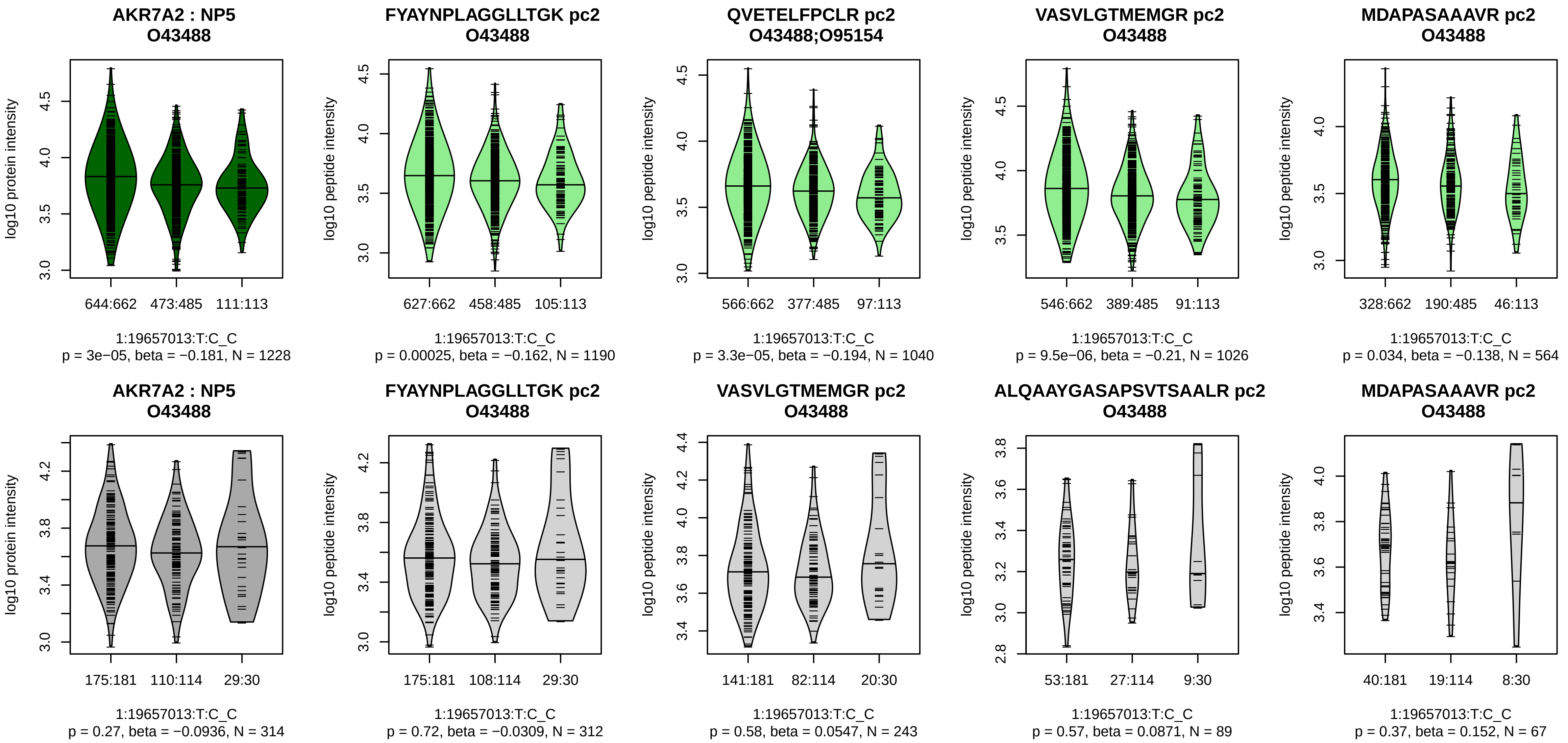


Assay Target: AKR7A2
Olink UniProt: O43488
deCODE rsID: rs61766662
Proxy rsID: rs61766662
deCODE: 1:19330519:C:T
Proxy SNP: 1:19657013:T:C
deCODE log10(p): 67.7
deCODE BETA: −0.15
*:*:*:–:–:–:–:–:–
1190:1040:1026:564:537:263:78

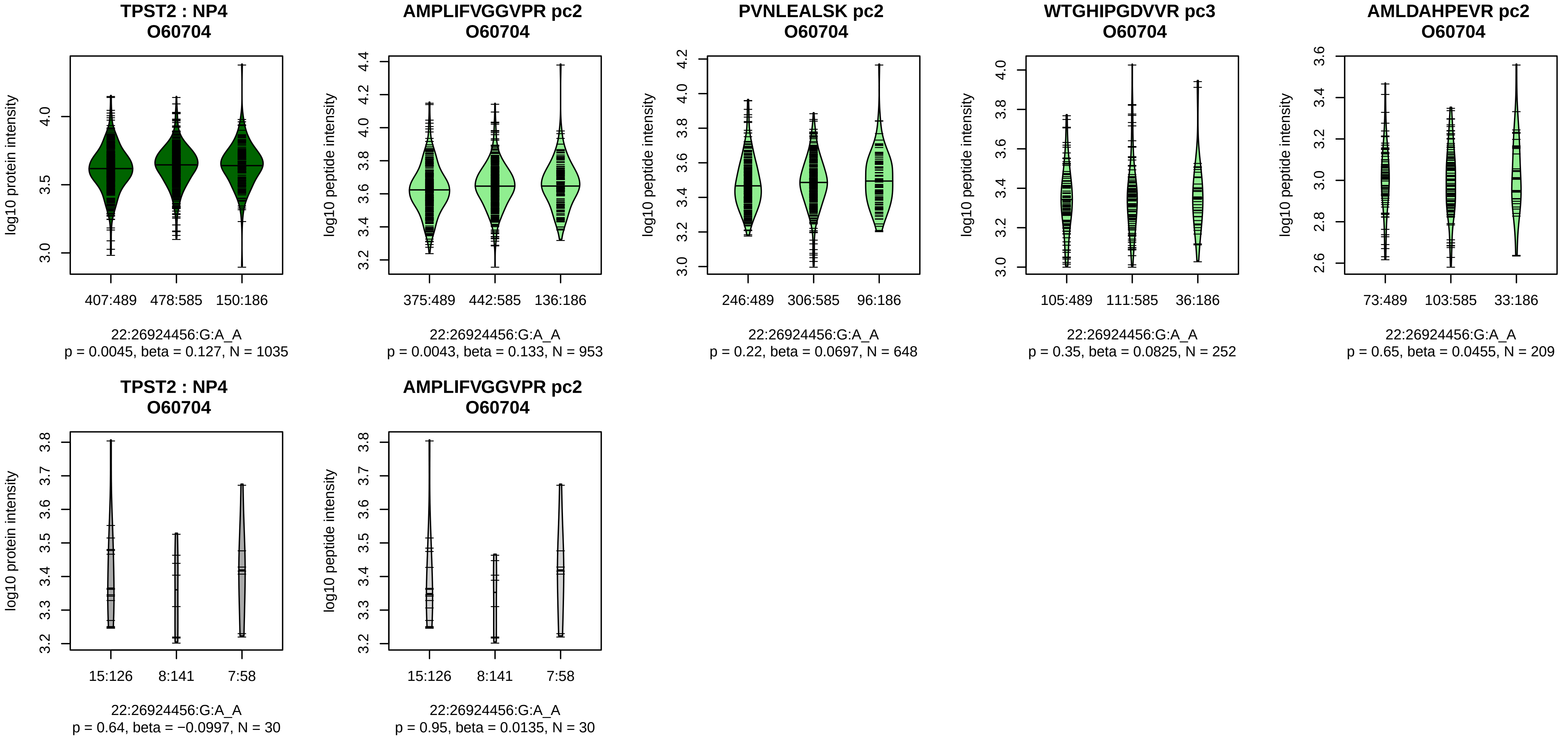


Assay Target: TPST2
Olink UniProt: O60704
deCODE rsID: rs2283824
Proxy rsID: rs2283824
deCODE: 22:26528490:A:G
Proxy SNP: 22:26924456:G:A
deCODE log10(p): 67
deCODE BETA: 0.15
*:−:−:−:−:−:−:−
953:648:252:209:129:118:116:5

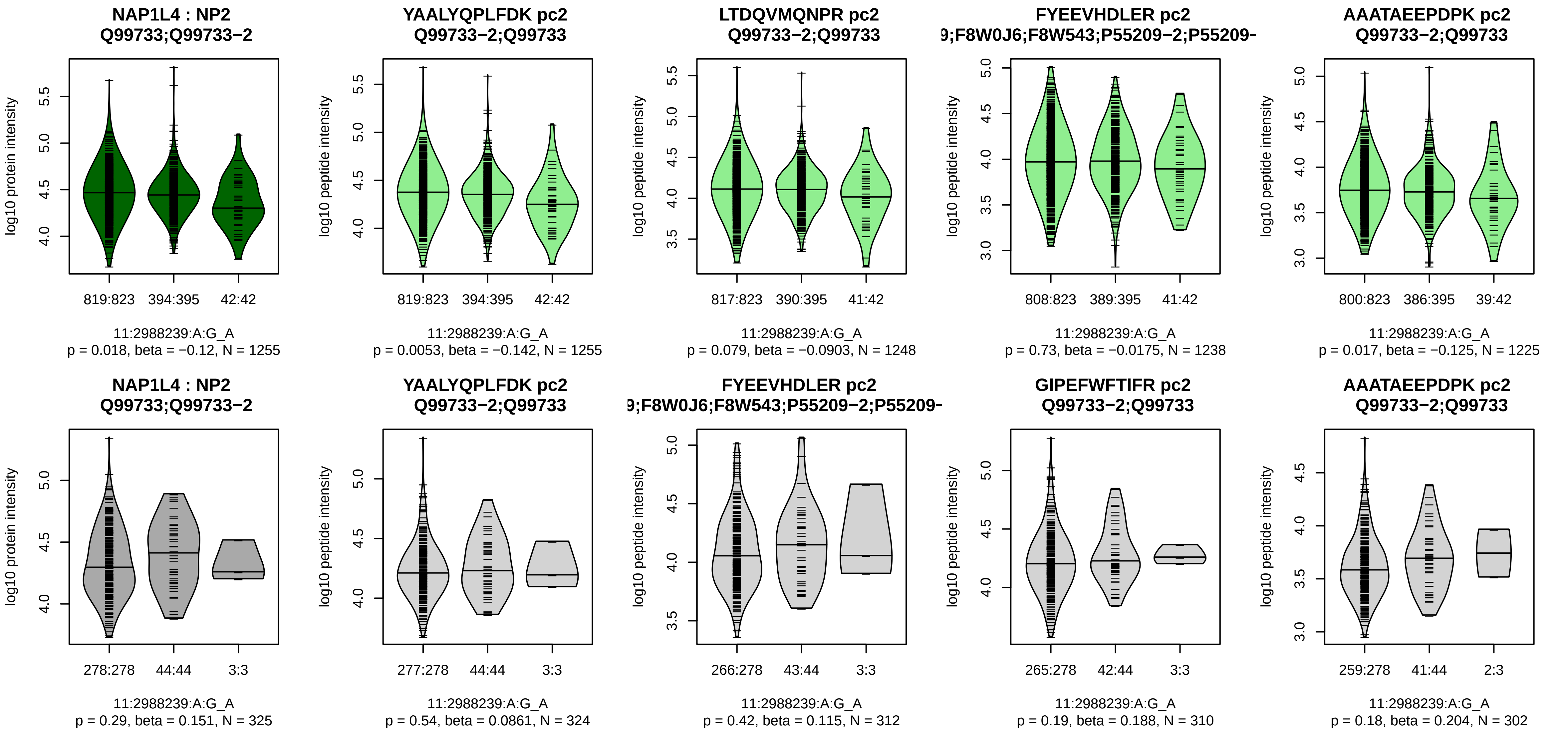


Assay Target: NAP1L4
Olink UniProt: Q99733
deCODE rsID: rs57928937
Proxy rsID: rs7109587
deCODE: 11:2969023:C:CT
Proxy SNP: 11:2988239:A:G
deCODE log10(p): 66.1
deCODE BETA: −0.18
*:−:−:−:−:−:*:−:−:−:−:−:−:NA
1255:1248:1238:1225:1225:118

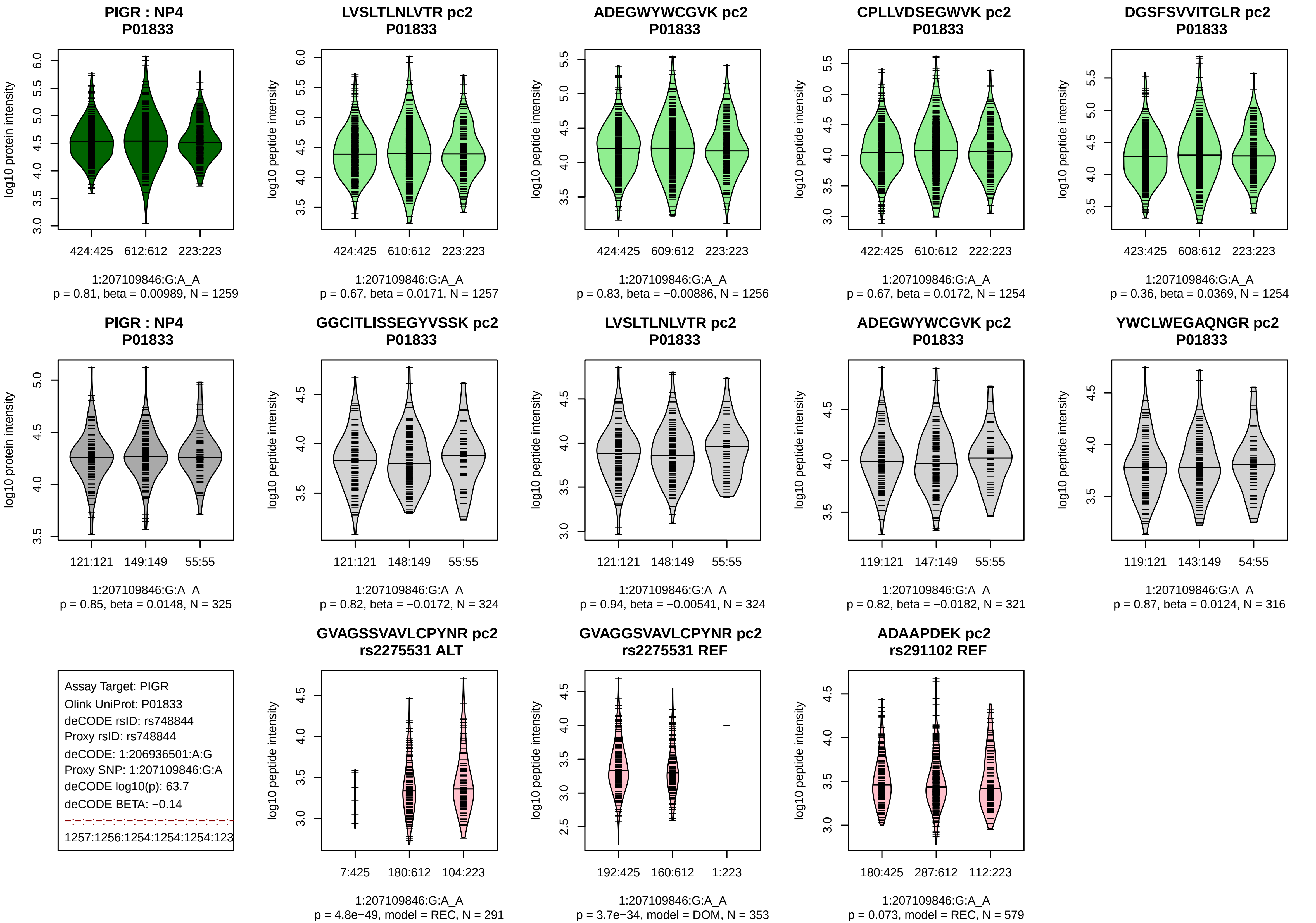
PIGR : NP4
P01833
log10 protein intensity
424:425
612:612
223:223
1:207109846:G:A_A
p = 0.81, beta = 0.00989, N = 1259
LVSLTLNLVTR pc2
P01833
log10 peptide intensity
424:425
610:612
223:223
1:207109846:G:A_A
p = 0.67, beta = 0.0171, N = 1257
ADEGWYWCGVK pc2
P01833
log10 peptide intensity
424:425
609:612
223:223
1:207109846:G:A_A
p = 0.83, beta = −0.00886, N = 1256
CPLLVDSEGWVK pc2
P01833
log10 peptide intensity
422:425
610:612
222:223
1:207109846:G:A_A
p = 0.67, beta = 0.0172, N = 1254
DGSFSVVITGLR pc2
P01833
log10 peptide intensity
423:425
608:612
223:223
1:207109846:G:A_A
p = 0.36, beta = 0.0369, N = 1254
PIGR : NP4
P01833
log10 protein intensity
121:121
149:149
55:55
1:207109846:G:A_A
p = 0.85, beta = 0.0148, N = 325
GGCITLISSEGYVSSK pc2
P01833
log10 peptide intensity
121:121
148:149
55:55
1:207109846:G:A_A
p = 0.82, beta = −0.0172, N = 324
LVSLTLNLVTR pc2
P01833
log10 peptide intensity
121:121
148:149
55:55
1:207109846:G:A_A
p = 0.94, beta = −0.00541, N = 324
ADEGWYWCGVK pc2
P01833
log10 peptide intensity
119:121
147:149
55:55
1:207109846:G:A_A
p = 0.82, beta = −0.0182, N = 321
YWCLWEGAQNGR pc2
P01833
log10 peptide intensity
119:121
143:149
54:55
1:207109846:G:A_A
p = 0.87, beta = 0.0124, N = 316
Assay Target: PIGR
Olink UniProt: P01833
deCODE rsID: rs748844
Proxy rsID: rs748844
deCODE: 1:206936501:A:G
Proxy SNP: 1:207109846:G:A
deCODE log10(p): 63.7
deCODE BETA: −0.14
1257:1256:1254:1254:1254:123
GVAGSSVAVLCPYNR pc2
rs2275531 ALT
log10 peptide intensity
7:425
180:612
104:223
1:207109846:G:A_A
p = 4.8e−49, model = REC, N = 291
GVAGGSVAVLCPYNR pc2
rs2275531 REF
log10 peptide intensity
192:425
160:612
1:223
1:207109846:G:A_A
p = 3.7e−34, model = DOM, N = 353
ADAAPDEK pc2
rs291102 REF
log10 peptide intensity
180:425
287:612
112:223
1:207109846:G:A_A
p = 0.073, model = REC, N = 579

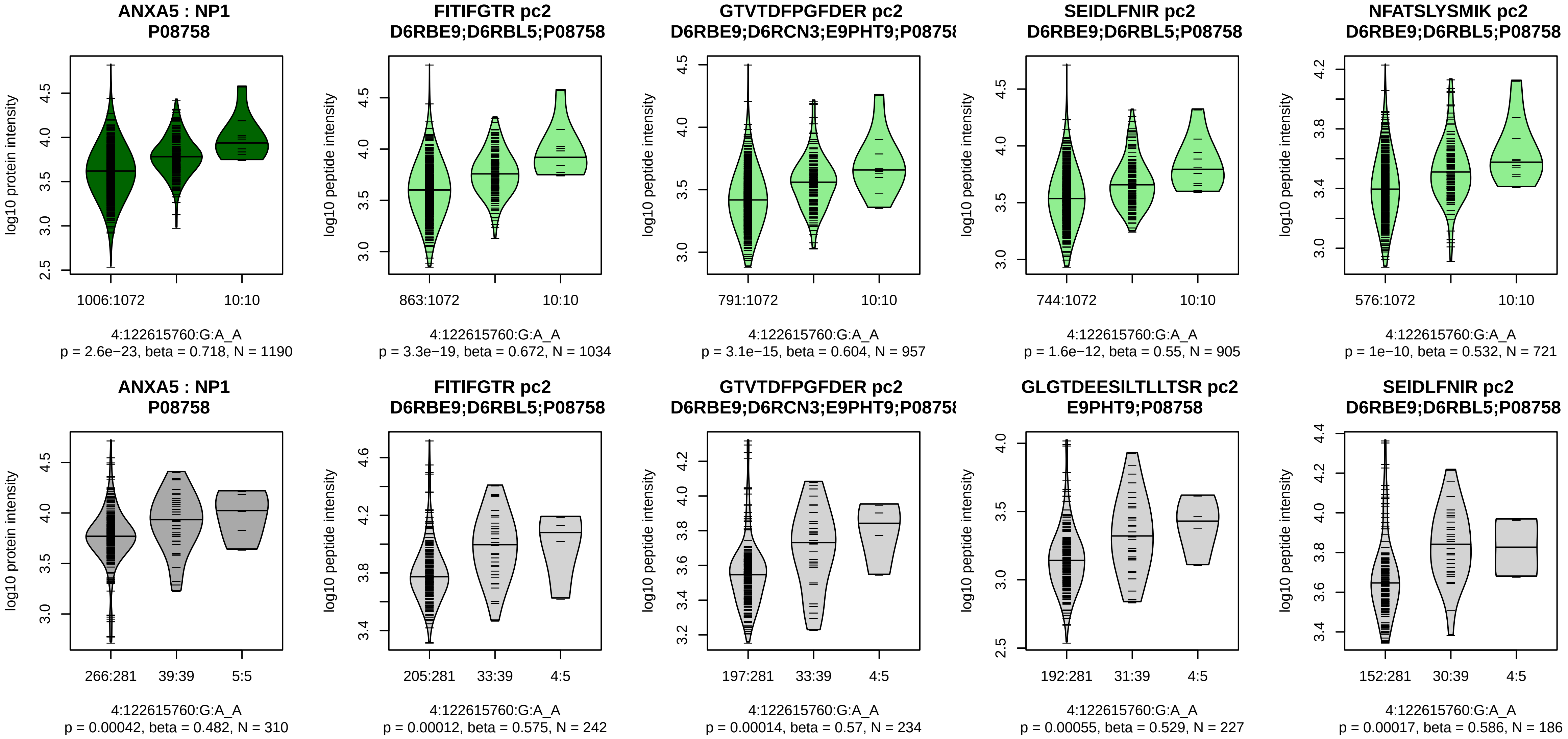


Assay Target: ANXA5
Olink UniProt: P08758
deCODE rsID: rs9995591
Proxy rsID: rs9995591
deCODE: 4:121694605:A:G
Proxy SNP: 4:122615760:G:A
deCODE log10(p): 63.4
deCODE BETA: 0.26
*:*:*:*:*:*:*:–:–:–:*:–:–:–:NA
1034:957:905:721:641:579:208:

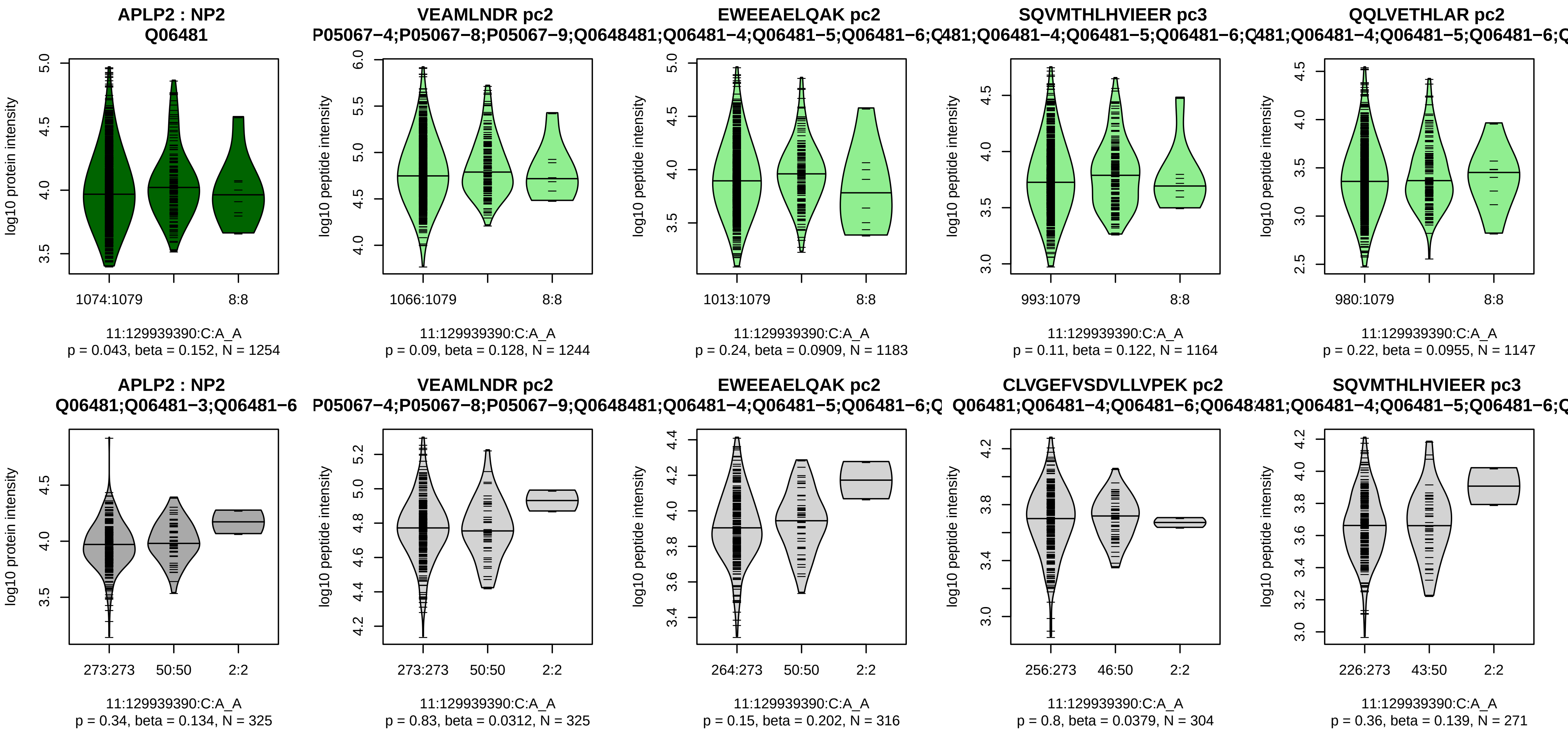


Assay Target: APLP2
Olink UniProt: Q06481
deCODE rsID: rs73583419
Proxy rsID: rs73583419
deCODE: 11:130069495:A:C
Proxy SNP: 11:129939390:C:A
deCODE log10(p): 62.6
deCODE BETA: 0.28
−:−:−:−:−:−:−:−:−:−:−:−:−:−:−:
1244:1183:1164:1147:1121:110

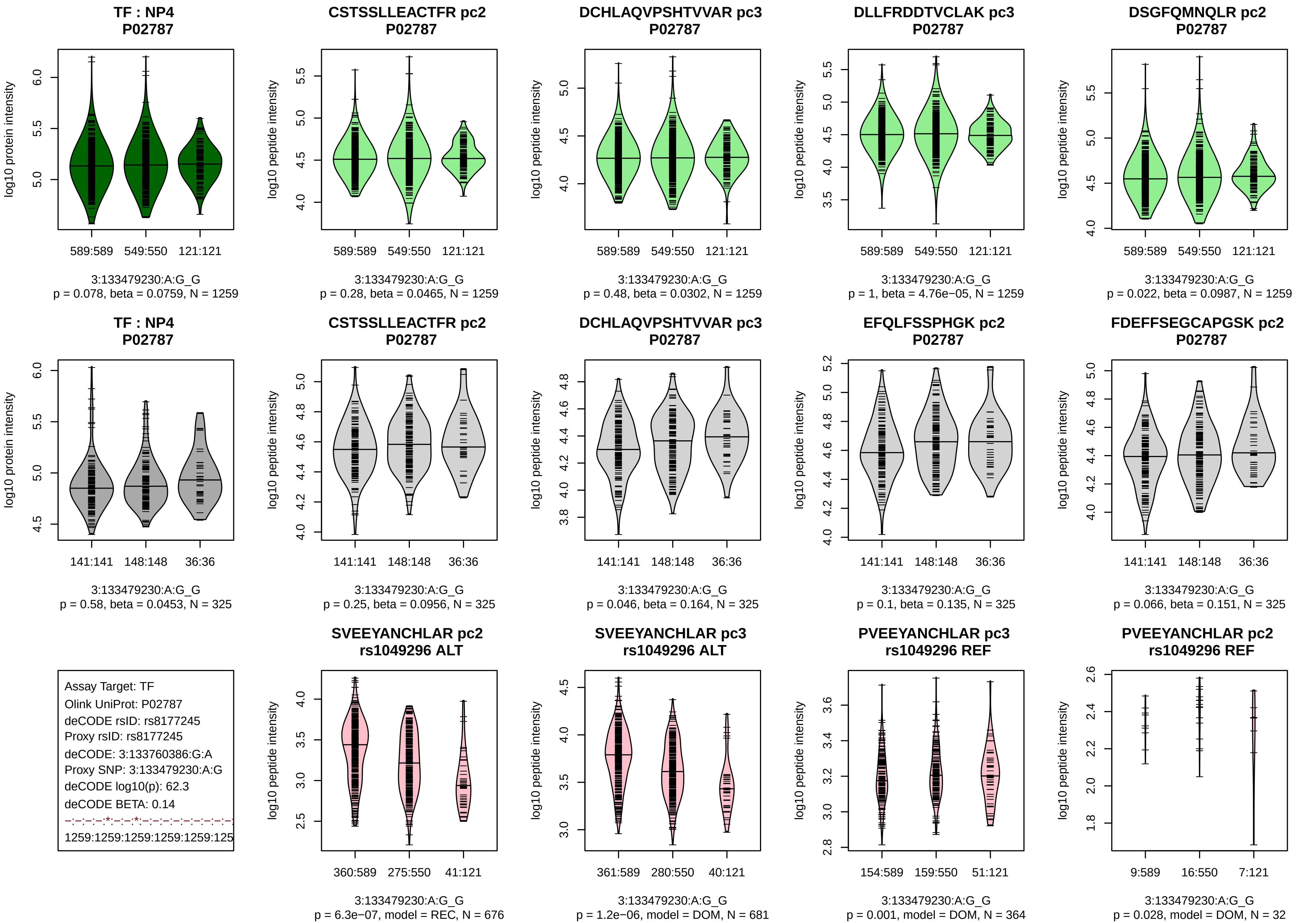

TF : NP4
P02787
log10 protein intensity
589:589
549:550
121:121
3:133479230:A:G_G
p = 0.078, beta = 0.0759, N = 1259
CSTSSLLEACTFR pc2
P02787
log10 peptide intensity
589:589
549:550
121:121
3:133479230:A:G_G
p = 0.28, beta = 0.0465, N = 1259
DCHLAQVPSHTVVAR pc3
P02787
log10 peptide intensity
589:589
549:550
121:121
3:133479230:A:G_G
p = 0.48, beta = 0.0302, N = 1259
DLLFRDDTVCLAK pc3
P02787
log10 peptide intensity
589:589
549:550
121:121
3:133479230:A:G_G
p = 1, beta = 4.76e−05, N = 1259
DSGFQMNQLR pc2
P02787
log10 peptide intensity
589:589
549:550
121:121
3:133479230:A:G_G
p = 0.022, beta = 0.0987, N = 1259
TF : NP4
P02787
log10 protein intensity
141:141
148:148
36:36
3:133479230:A:G_G
p = 0.58, beta = 0.0453, N = 325
CSTSSLLEACTFR pc2
P02787
log10 peptide intensity
141:141
148:148
36:36
3:133479230:A:G_G
p = 0.25, beta = 0.0956, N = 325
DCHLAQVPSHTVVAR pc3
P02787
log10 peptide intensity
141:141
148:148
36:36
3:133479230:A:G_G
p = 0.046, beta = 0.164, N = 325
EFQLFSSPHGK pc2
P02787
log10 peptide intensity
141:141
148:148
36:36
3:133479230:A:G_G
p = 0.1, beta = 0.135, N = 325
FDEFFSEGCAPGSK pc2
P02787
log10 peptide intensity
141:141
148:148
36:36
3:133479230:A:G_G
p = 0.066, beta = 0.151, N = 325
Assay Target: TF
Olink UniProt: P02787
deCODE rsID: rs8177245
Proxy rsID: rs8177245
deCODE: 3:133760386:G:A
Proxy SNP: 3:133479230:A:G
deCODE log10(p): 62.3
deCODE BETA: 0.14
−:−:−:*:−:−:*:−:−:−:−:−:−:−:−:−
1259:1259:1259:1259:1259:125
SVEEYANCHLAR pc2
rs1049296 ALT
log10 peptide intensity
360:589
275:550
41:121
3:133479230:A:G_G
p = 6.3e−07, model = REC, N = 676
SVEEYANCHLAR pc3
rs1049296 ALT
log10 peptide intensity
361:589
280:550
40:121
3:133479230:A:G_G
p = 1.2e−06, model = DOM, N = 681
PVEEYANCHLAR pc3
rs1049296 REF
log10 peptide intensity
154:589
159:550
51:121
3:133479230:A:G_G
p = 0.001, model = DOM, N = 364
PVEEYANCHLAR pc2
rs1049296 REF
log10 peptide intensity
9:589
16:550
7:121
3:133479230:A:G_G
p = 0.028, model = DOM, N = 32

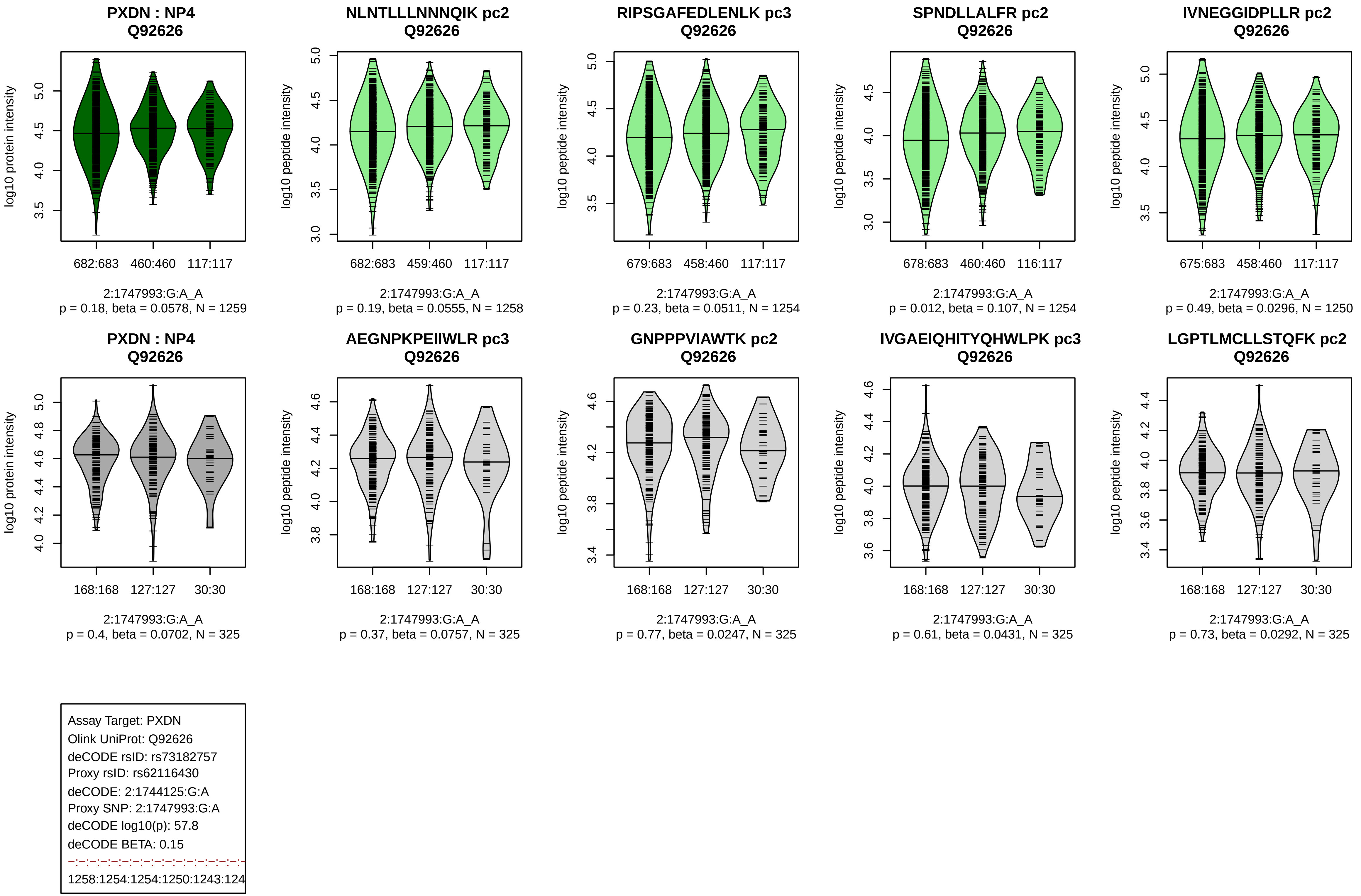

PXDN : NP4
Q92626
log10 protein intensity
682:683 460:460 117:117
2:1747993:G:A_A
p = 0.18, beta = 0.0578, N = 1259
NLNTLLLNNNQIK pc2
Q92626
log10 peptide intensity
682:683 459:460 117:117
2:1747993:G:A_A
p = 0.19, beta = 0.0555, N = 1258
RIPSGAFEDLENLK pc3
Q92626
log10 peptide intensity
679:683 458:460 117:117
2:1747993:G:A_A
p = 0.23, beta = 0.0511, N = 1254
SPNDLLALFR pc2
Q92626
log10 peptide intensity
678:683 460:460 116:117
2:1747993:G:A_A
p = 0.012, beta = 0.107, N = 1254
IVNEGGIDPLLR pc2
Q92626
log10 peptide intensity
675:683 458:460 117:117
2:1747993:G:A_A
p = 0.49, beta = 0.0296, N = 1250
PXDN : NP4
Q92626
log10 protein intensity
168:168 127:127 30:30
2:1747993:G:A_A
p = 0.4, beta = 0.0702, N = 325
AEGNPKPEIIWLR pc3
Q92626
log10 peptide intensity
168:168 127:127 30:30
2:1747993:G:A_A
p = 0.37, beta = 0.0757, N = 325
GNPPPVIAWTK pc2
Q92626
log10 peptide intensity
168:168 127:127 30:30
2:1747993:G:A_A
p = 0.77, beta = 0.0247, N = 325
IVGAEIQHITYQHWLPK pc3
Q92626
log10 peptide intensity
168:168 127:127 30:30
2:1747993:G:A_A
p = 0.61, beta = 0.0431, N = 325
LGPTLMCLLSTQFK pc2
Q92626
log10 peptide intensity
168:168 127:127 30:30
2:1747993:G:A_A
p = 0.73, beta = 0.0292, N = 325
Assay Target: PXDN
Olink UniProt: Q92626
deCODE rsID: rs73182757
Proxy rsID: rs62116430
deCODE: 2:1744125:G:A
Proxy SNP: 2:1747993:G:A
deCODE log10(p): 57.8
deCODE BETA: 0.15
1258:1254:1254:1250:1243:124

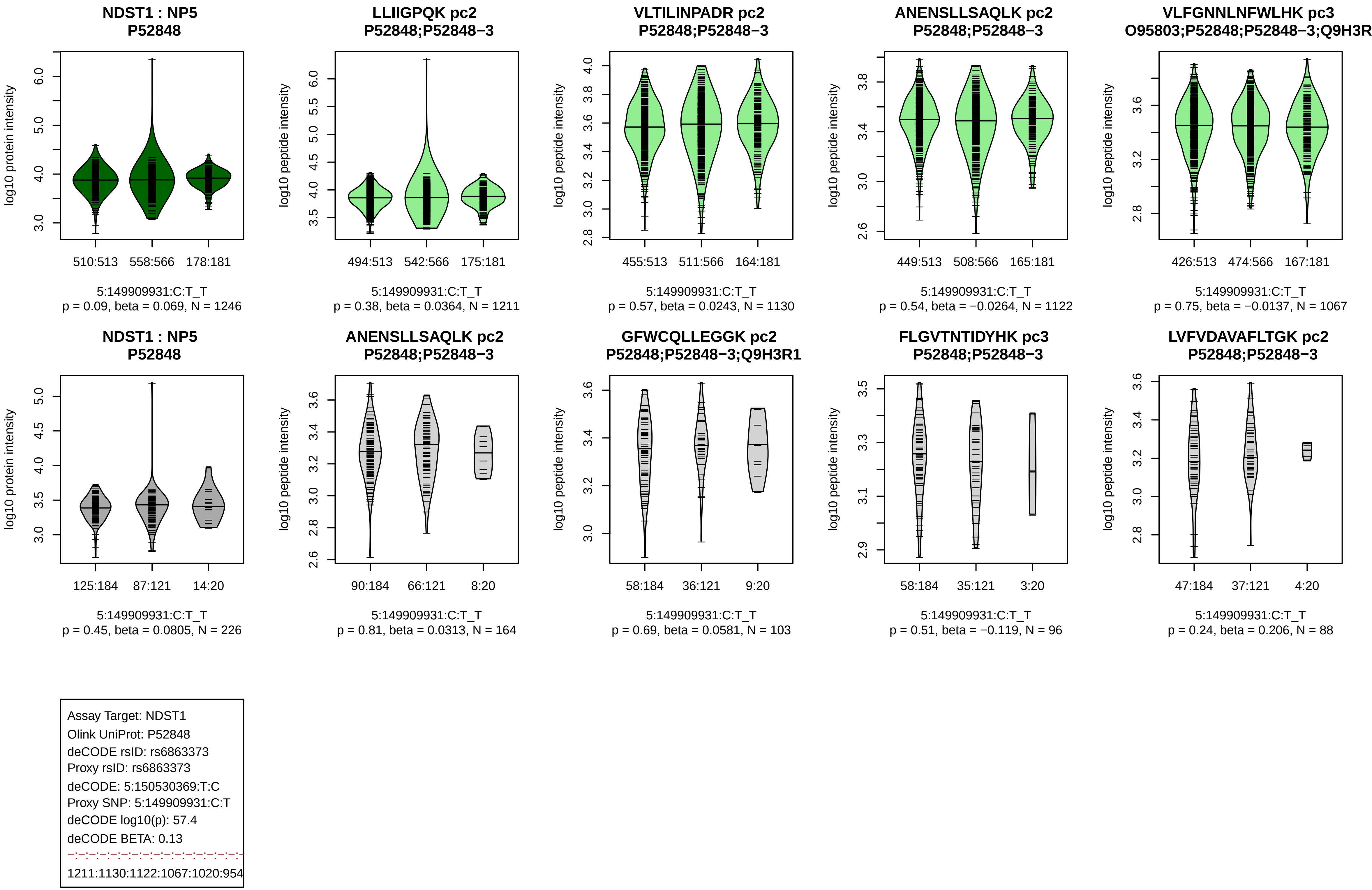
NDST1 : NP5
P52848
log10 protein intensity
510:513
558:566
178:181
5:149909931:C:T_T
p = 0.09, beta = 0.069, N = 1246
LLIIGPQK pc2
P52848;P52848−3
log10 peptide intensity
494:513
542:566
175:181
5:149909931:C:T_T
p = 0.38, beta = 0.0364, N = 1211
VLTILINPADR pc2
P52848;P52848−3
log10 peptide intensity
455:513
511:566
164:181
5:149909931:C:T_T
p = 0.57, beta = 0.0243, N = 1130
ANENSLLSAQLK pc2
P52848;P52848−3
log10 peptide intensity
449:513
508:566
165:181
5:149909931:C:T_T
p = 0.54, beta = −0.0264, N = 1122
VLFGNNLNFWLHK pc3
O95803;P52848;P52848−3;Q9H3R
log10 peptide intensity
426:513
474:566
167:181
5:149909931:C:T_T
p = 0.75, beta = −0.0137, N = 1067
NDST1 : NP5
P52848
log10 protein intensity
125:184
87:121
14:20
5:149909931:C:T_T
p = 0.45, beta = 0.0805, N = 226
ANENSLLSAQLK pc2
P52848;P52848−3
log10 peptide intensity
90:184
66:121
8:20
5:149909931:C:T_T
p = 0.81, beta = 0.0313, N = 164
GFWCQLLEGGK pc2
P52848;P52848−3;Q9H3R1
log10 peptide intensity
58:184
36:121
9:20
5:149909931:C:T_T
p = 0.69, beta = 0.0581, N = 103
FLGVTNTIDYHK pc3
P52848;P52848−3
log10 peptide intensity
58:184
35:121
3:20
5:149909931:C:T_T
p = 0.51, beta = −0.119, N = 96
LVFVDAVAFLTGK pc2
P52848;P52848−3
log10 peptide intensity
47:184
37:121
4:20
5:149909931:C:T_T
p = 0.24, beta = 0.206, N = 88
Assay Target: NDST1
Olink UniProt: P52848
deCODE rsID: rs6863373
Proxy rsID: rs6863373
deCODE: 5:150530369:T:C
Proxy SNP: 5:149909931:C:T
deCODE log10(p): 57.4
deCODE BETA: 0.13
1211:1130:1122:1067:1020:954

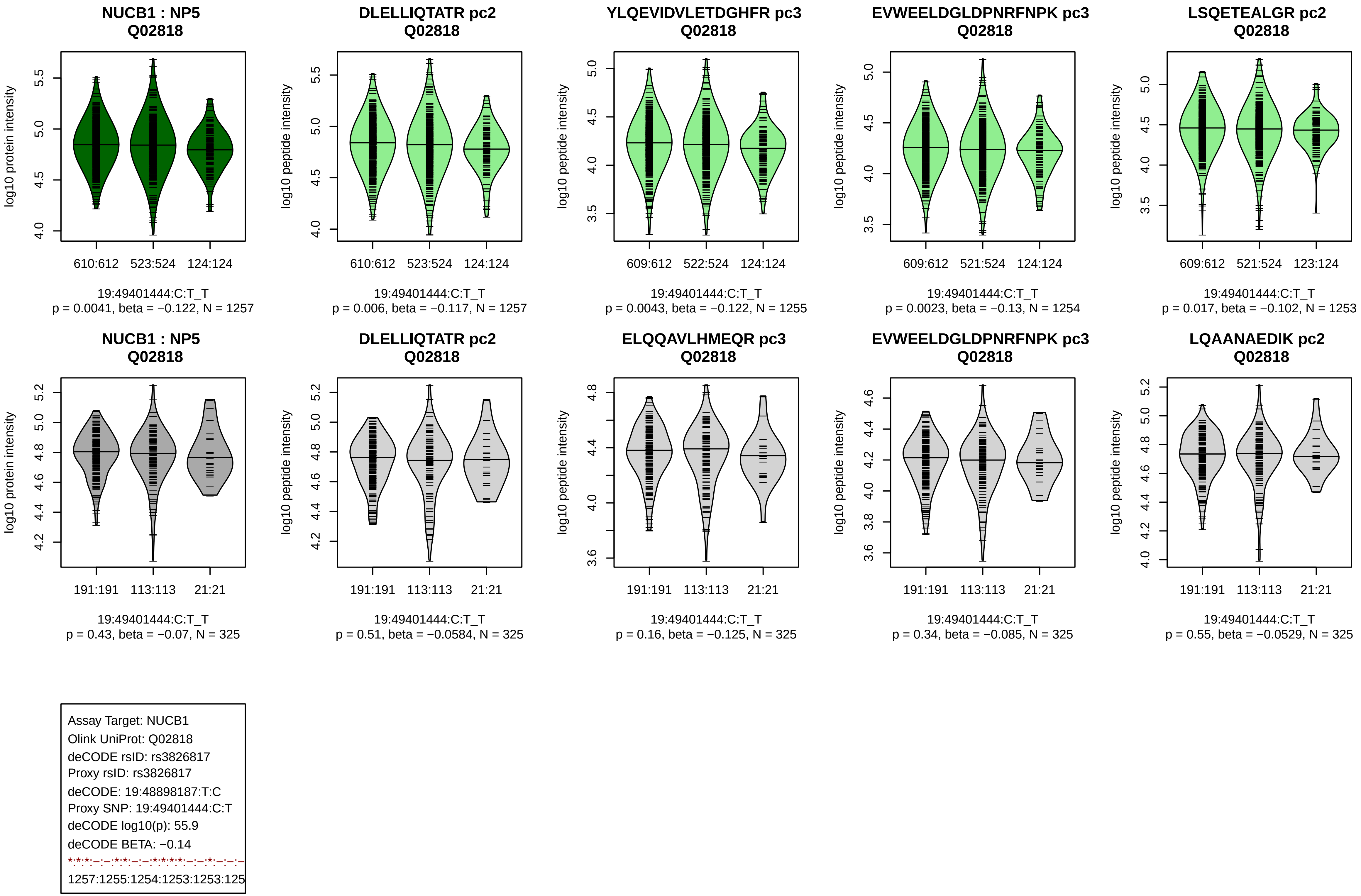
NUCB1 : NP5
Q02818
log10 protein intensity
610:612
523:524
124:124
19:49401444:C:T_T
p = 0.0041, beta = −0.122, N = 1257
DLELLIQTATR pc2
Q02818
log10 peptide intensity
610:612
523:524
124:124
19:49401444:C:T_T
p = 0.006, beta = −0.117, N = 1257
YLQEVIDVLETDGHFR pc3
Q02818
log10 peptide intensity
609:612
522:524
124:124
19:49401444:C:T_T
p = 0.0043, beta = −0.122, N = 1255
EVWEELDGLDPNRFNPK pc3
Q02818
log10 peptide intensity
609:612
521:524
124:124
19:49401444:C:T_T
p = 0.0023, beta = −0.13, N = 1254
LSQETEALGR pc2
Q02818
log10 peptide intensity
609:612
521:524
123:124
19:49401444:C:T_T
p = 0.017, beta = −0.102, N = 1253
NUCB1 : NP5
Q02818
log10 protein intensity
191:191
113:113
21:21
19:49401444:C:T_T
p = 0.43, beta = −0.07, N = 325
DLELLIQTATR pc2
Q02818
log10 peptide intensity
191:191
113:113
21:21
19:49401444:C:T_T
p = 0.51, beta = −0.0584, N = 325
ELQQAVLHMEQR pc3
Q02818
log10 peptide intensity
191:191
113:113
21:21
19:49401444:C:T_T
p = 0.16, beta = −0.125, N = 325
EVWEELDGLDPNRFNPK pc3
Q02818
log10 peptide intensity
191:191
113:113
21:21
19:49401444:C:T_T
p = 0.34, beta = −0.085, N = 325
LQAANAEDIK pc2
Q02818
log10 peptide intensity
191:191
113:113
21:21
19:49401444:C:T_T
p = 0.55, beta = −0.0529, N = 325
Assay Target: NUCB1
Olink UniProt: Q02818
deCODE rsID: rs3826817
Proxy rsID: rs3826817
deCODE: 19:48898187:T:C
Proxy SNP: 19:49401444:C:T
deCODE log10(p): 55.9
deCODE BETA: −0.14
1257:1255:1254:1253:1253:125

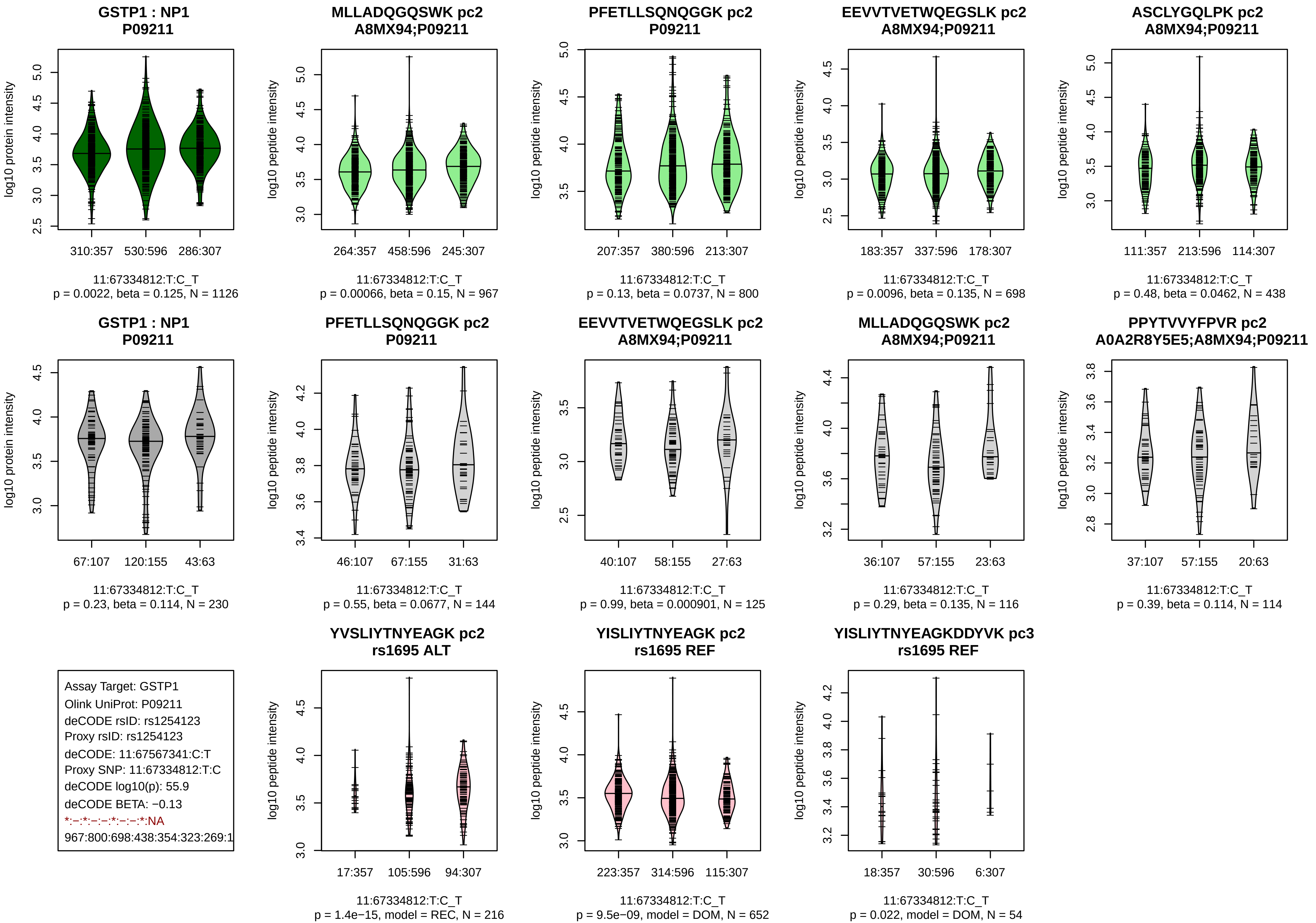
GSTP1 : NP1
P09211
log10 protein intensity
310:357
530:596
286:307
11:67334812:T:C_T
p = 0.0022, beta = 0.125, N = 1126
MLLADQGQSWK pc2
A8MX94;P09211
log10 peptide intensity
264:357
458:596
245:307
11:67334812:T:C_T
p = 0.00066, beta = 0.15, N = 967
PFETLLSQNQGGK pc2
P09211
log10 peptide intensity
207:357
380:596
213:307
11:67334812:T:C_T
p = 0.13, beta = 0.0737, N = 800
EEVVTVETWQEGSLK pc2
A8MX94;P09211
log10 peptide intensity
183:357
337:596
178:307
11:67334812:T:C_T
p = 0.0096, beta = 0.135, N = 698
ASCLYGQLPK pc2
A8MX94;P09211
log10 peptide intensity
111:357
213:596
114:307
11:67334812:T:C_T
p = 0.48, beta = 0.0462, N = 438
GSTP1 : NP1
P09211
log10 protein intensity
67:107
120:155
43:63
11:67334812:T:C_T
p = 0.23, beta = 0.114, N = 230
PFETLLSQNQGGK pc2
P09211
log10 peptide intensity
46:107
67:155
31:63
11:67334812:T:C_T
p = 0.55, beta = 0.0677, N = 144
EEVVTVETWQEGSLK pc2
A8MX94;P09211
log10 peptide intensity
40:107
58:155
27:63
11:67334812:T:C_T
p = 0.99, beta = 0.000901, N = 125
MLLADQGQSWK pc2
A8MX94;P09211
log10 peptide intensity
36:107
57:155
23:63
11:67334812:T:C_T
p = 0.29, beta = 0.135, N = 116
PPYTVVYFPVR pc2
A0A2R8Y5E5;A8MX94;P09211
log10 peptide intensity
37:107
57:155
20:63
11:67334812:T:C_T
p = 0.39, beta = 0.114, N = 114
Assay Target: GSTP1
Olink UniProt: P09211
deCODE rsID: rs1254123
Proxy rsID: rs1254123
deCODE: 11:67567341:C:T
Proxy SNP: 11:67334812:T:C
deCODE log10(p): 55.9
deCODE BETA: −0.13
*:−:*:−:−:*:−:−:*:NA
967:800:698:438:354:323:269:1
YVSLIYTNYEAGK pc2
rs1695 ALT
log10 peptide intensity
17:357
105:596
94:307
11:67334812:T:C_T
p = 1.4e−15, model = REC, N = 216
YISLIYTNYEAGK pc2
rs1695 REF
log10 peptide intensity
223:357
314:596
115:307
11:67334812:T:C_T
p = 9.5e−09, model = DOM, N = 652
YISLIYTNYEAGKDDYVK pc3
rs1695 REF
log10 peptide intensity
18:357
30:596
6:307
11:67334812:T:C_T
p = 0.022, model = DOM, N = 54

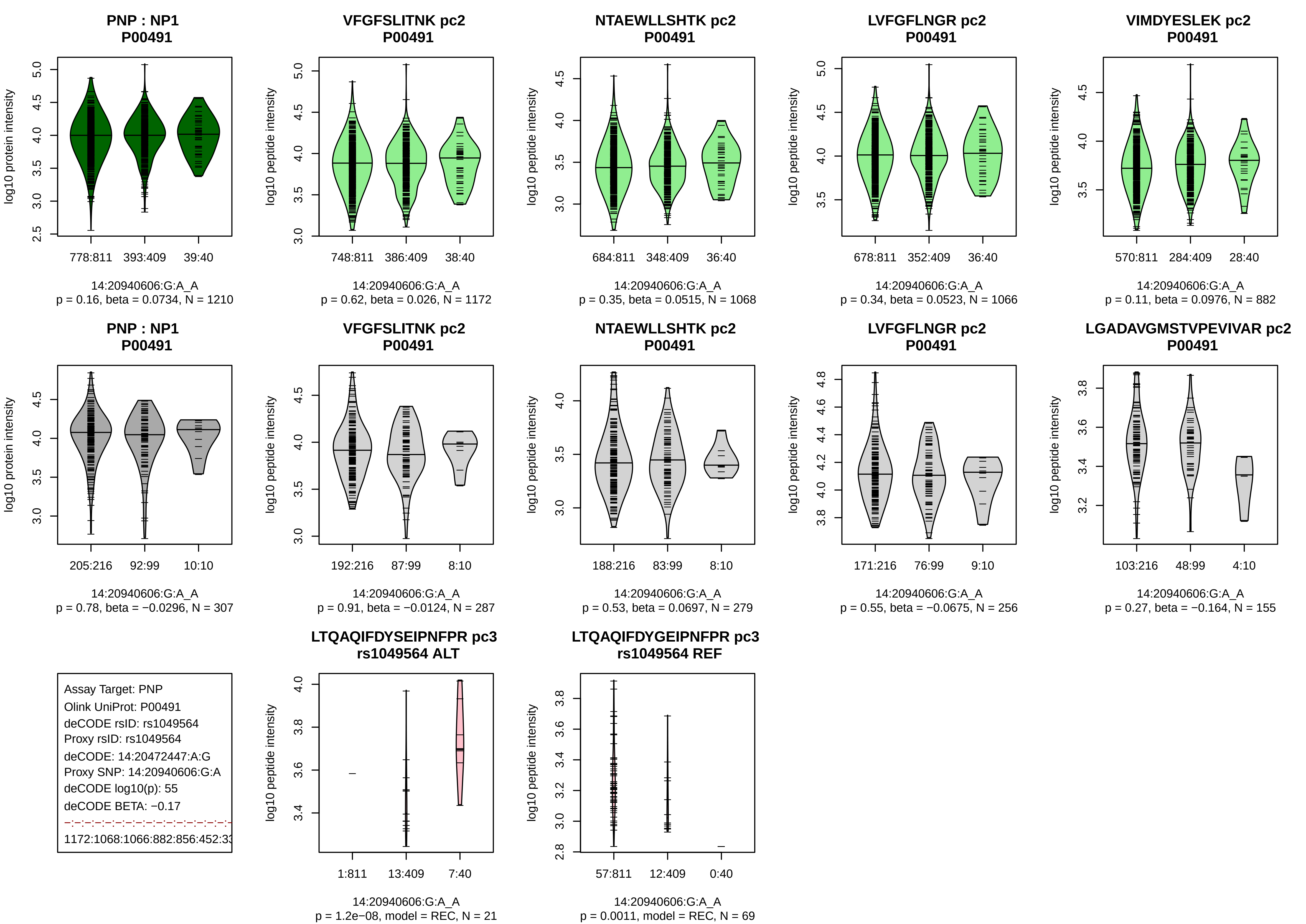

PNP : NP1
P00491
log10 protein intensity
778:811 393:409 39:40
14:20940606:G:A_A
p = 0.16, beta = 0.0734, N = 1210
VFGFSLITNK pc2
P00491
log10 peptide intensity
748:811 386:409 38:40
14:20940606:G:A_A
p = 0.62, beta = 0.026, N = 1172
NTAEWLLSHTK pc2
P00491
log10 peptide intensity
684:811 348:409 36:40
14:20940606:G:A_A
p = 0.35, beta = 0.0515, N = 1068
LVFGFLNGR pc2
P00491
log10 peptide intensity
678:811 352:409 36:40
14:20940606:G:A_A
p = 0.34, beta = 0.0523, N = 1066
VIMDYESLEK pc2
P00491
log10 peptide intensity
570:811 284:409 28:40
14:20940606:G:A_A
p = 0.11, beta = 0.0976, N = 882
PNP : NP1
P00491
log10 protein intensity
205:216 92:99 10:10
14:20940606:G:A_A
p = 0.78, beta = −0.0296, N = 307
VFGFSLITNK pc2
P00491
log10 peptide intensity
192:216 87:99 8:10
14:20940606:G:A_A
p = 0.91, beta = −0.0124, N = 287
NTAEWLLSHTK pc2
P00491
log10 peptide intensity
188:216 83:99 8:10
14:20940606:G:A_A
p = 0.53, beta = 0.0697, N = 279
LVFGFLNGR pc2
P00491
log10 peptide intensity
171:216 76:99 9:10
14:20940606:G:A_A
p = 0.55, beta = −0.0675, N = 256
LGADAVGMSTVPEVIVAR pc2
P00491
log10 peptide intensity
103:216 48:99 4:10
14:20940606:G:A_A
p = 0.27, beta = −0.164, N = 155
Assay Target: PNP
Olink UniProt: P00491
deCODE rsID: rs1049564
Proxy rsID: rs1049564
deCODE: 14:20472447:A:G
Proxy SNP: 14:20940606:G:A
deCODE log10(p): 55
deCODE BETA: −0.17
1172:1068:1066:882:856:452:33
LTQAQIFDYSEIPNFPR pc3
rs1049564 ALT
log10 peptide intensity
1:811 13:409 7:40
14:20940606:G:A_A
p = 1.2e−08, model = REC, N = 21
LTQAQIFDYGEIPNFPR pc3
rs1049564 REF
log10 peptide intensity
57:811 12:409 0:40
14:20940606:G:A_A
p = 0.0011, model = REC, N = 69

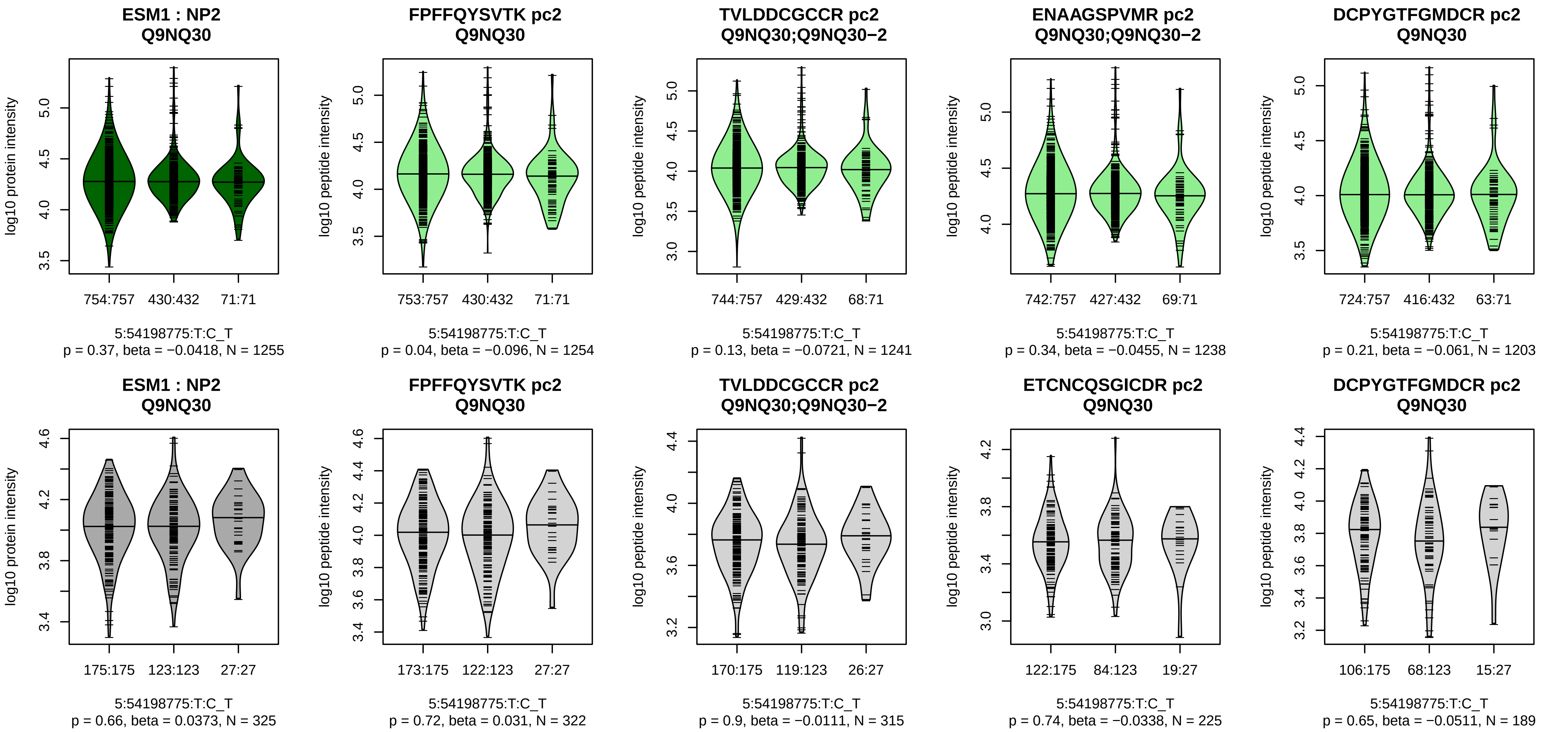


Assay Target: ESM1
Olink UniProt: Q9NQ30
deCODE rsID: rs4242051
Proxy rsID: rs4242051
deCODE: 5:54902947:T:C
Proxy SNP: 5:54198775:T:C
deCODE log10(p): 54.7
deCODE BETA: −0.15
−:−:−:−:−:−
1254:1241:1238:1203:1146:998

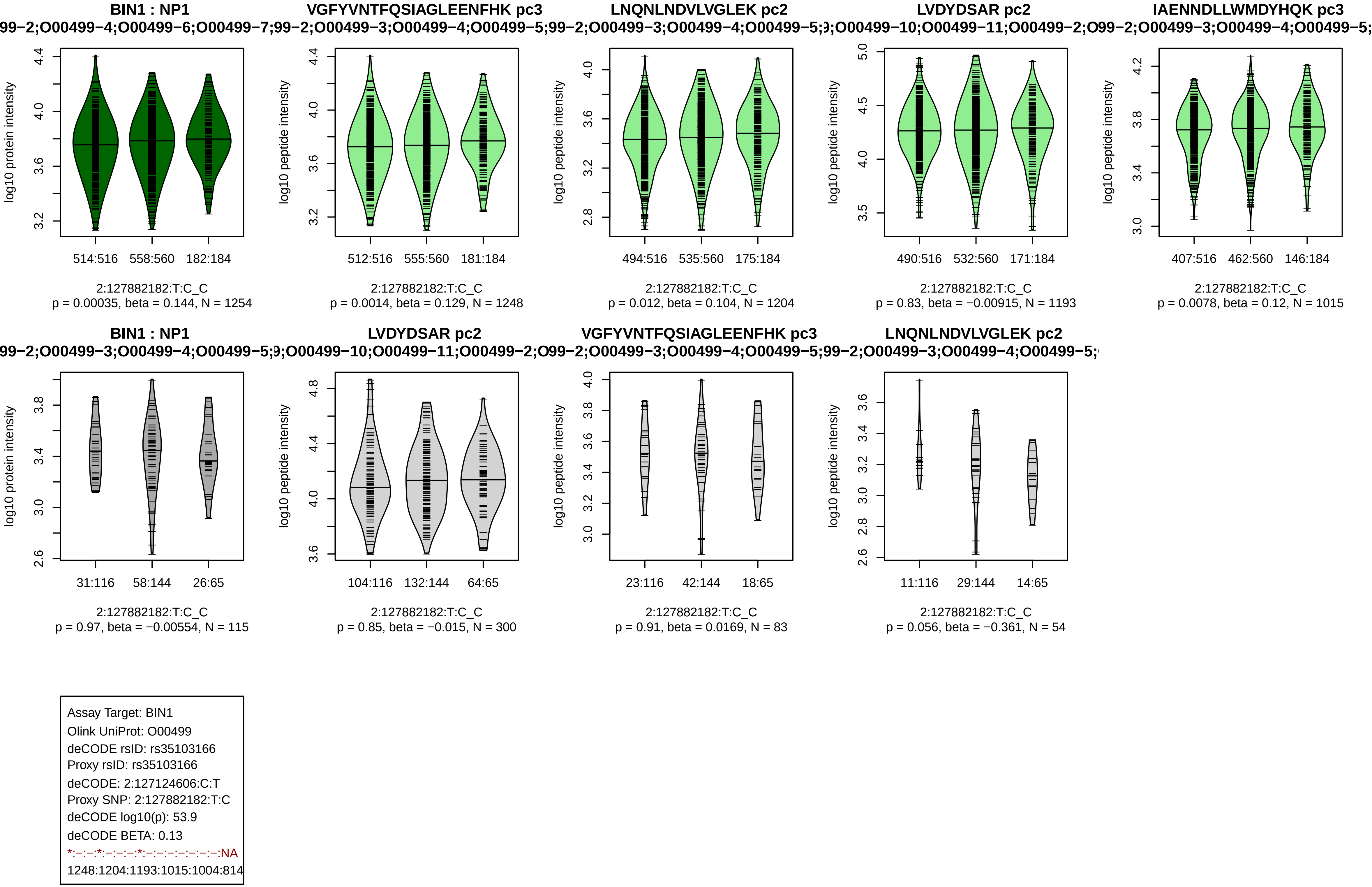

BIN1 : NP1
99−2;O00499−4;O00499−6;O00499−7;99−2;O00499−3;O00499−4;O00499−5;99−2;O00499−3;O00499−4;O00499−5;9;O00499−10;O00499−11;O00499−2;O99−2;O00499−3;O00499−4;O00499−5;
log10 protein intensity
514:516
558:560
182:184
2:127882182:T:C_C
p = 0.00035, beta = 0.144, N = 1254
VGFYVNTFQSIAGLEENFHK pc3
log10 peptide intensity
512:516
555:560
181:184
2:127882182:T:C_C
p = 0.0014, beta = 0.129, N = 1248
LNQNLNDVLVGLEK pc2
log10 peptide intensity
494:516
535:560
175:184
2:127882182:T:C_C
p = 0.012, beta = 0.104, N = 1204
LVDYDSAR pc2
log10 peptide intensity
490:516
532:560
171:184
2:127882182:T:C_C
p = 0.83, beta = −0.00915, N = 1193
IAENNDLLWMDYHQK pc3
log10 peptide intensity
407:516
462:560
146:184
2:127882182:T:C_C
p = 0.0078, beta = 0.12, N = 1015
BIN1 : NP1
99−2;O00499−3;O00499−4;O00499−5;9;O00499−10;O00499−11;O00499−2;O99−2;O00499−3;O00499−4;O00499−5;99−2;O00499−3;O00499−4;O00499−5;
log10 protein intensity
31:116
58:144
26:65
2:127882182:T:C_C
p = 0.97, beta = −0.00554, N = 115
LVDYDSAR pc2
log10 peptide intensity
104:116
132:144
64:65
2:127882182:T:C_C
p = 0.85, beta = −0.015, N = 300
VGFYVNTFQSIAGLEENFHK pc3
log10 peptide intensity
23:116
42:144
18:65
2:127882182:T:C_C
p = 0.91, beta = 0.0169, N = 83
LNQNLNDVLVGLEK pc2
log10 peptide intensity
11:116
29:144
14:65
2:127882182:T:C_C
p = 0.056, beta = −0.361, N = 54
Assay Target: BIN1
Olink UniProt: O00499
deCODE rsID: rs35103166
Proxy rsID: rs35103166
deCODE: 2:127124606:C:T
Proxy SNP: 2:127882182:T:C
deCODE log10(p): 53.9
deCODE BETA: 0.13
*:−:−:*:−:−:−:*:−:−:−:−:−:−:−:NA
1248:1204:1193:1015:1004:814

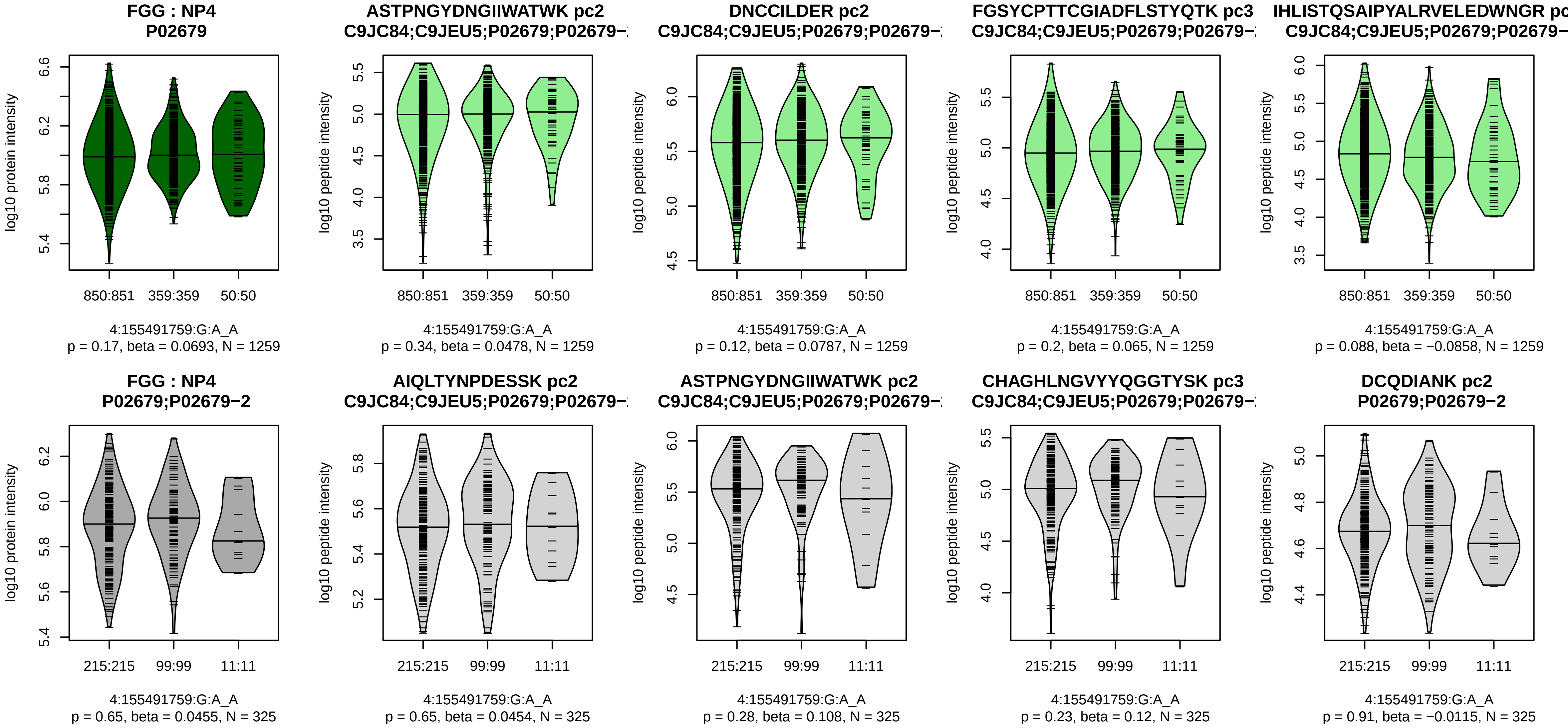


Assay Target: FGG
Olink UniProt: P02679
deCODE rsID: rs4220
Proxy rsID: rs4220
deCODE: 4:154570607:A:G
Proxy SNP: 4:155491759:G:A
deCODE log10(p): 53.5
deCODE BETA: 0.18
−:−:−:−:−:−:−:−:−:−:−:−:−:−:−:−:
1259:1259:1259:1259:1259:125

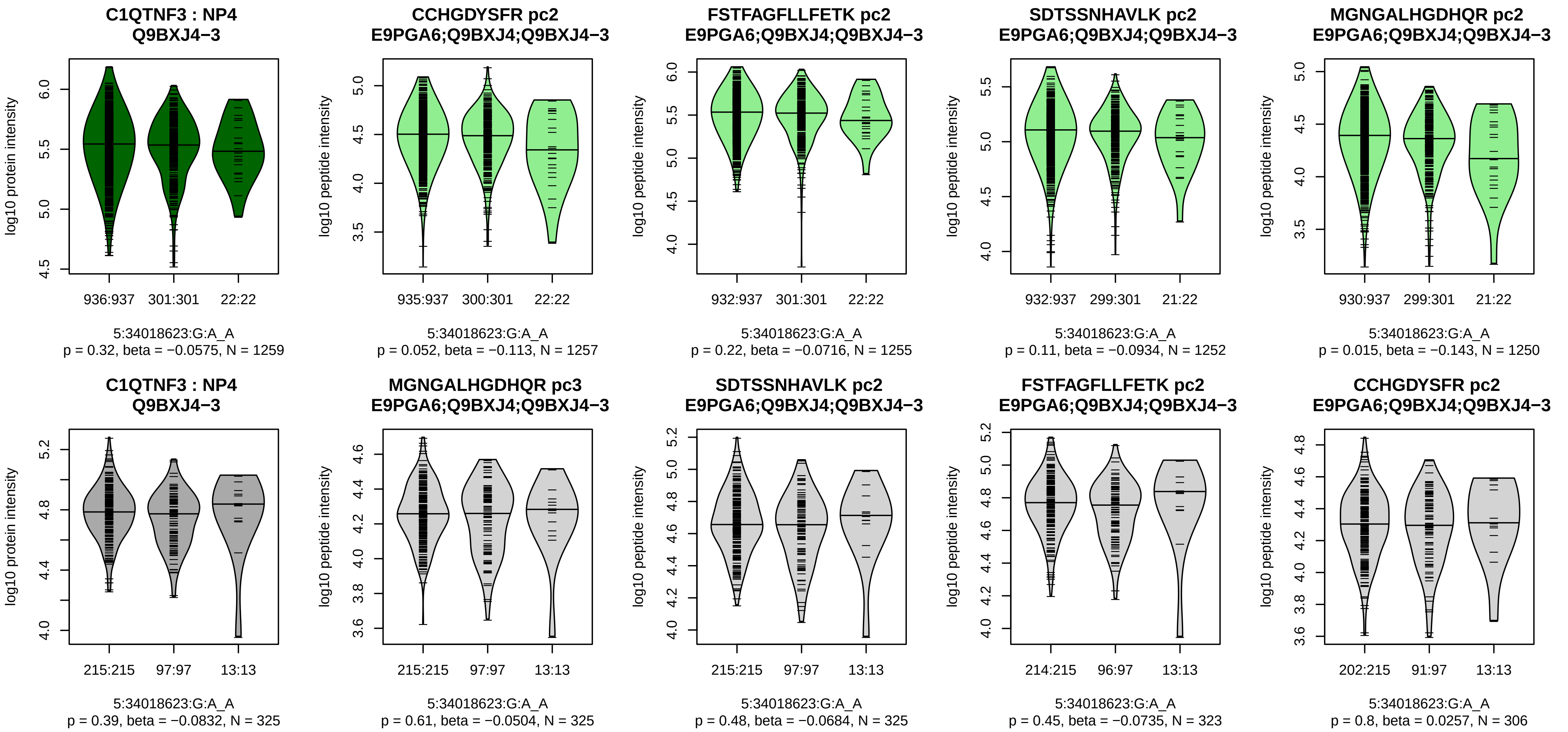


Assay Target: C1QTNF3
Olink UniProt: Q9BXJ4
deCODE rsID: rs840390
Proxy rsID: rs840390
deCODE: 5:34018518:A:G
Proxy SNP: 5:34018623:G:A
deCODE log10(p): 53.4
deCODE BETA: −0.18
−:−:−:−:−:−:−:−:−:−:−:−:−:−:−:NA
1257:1255:1255:1252:1250:125

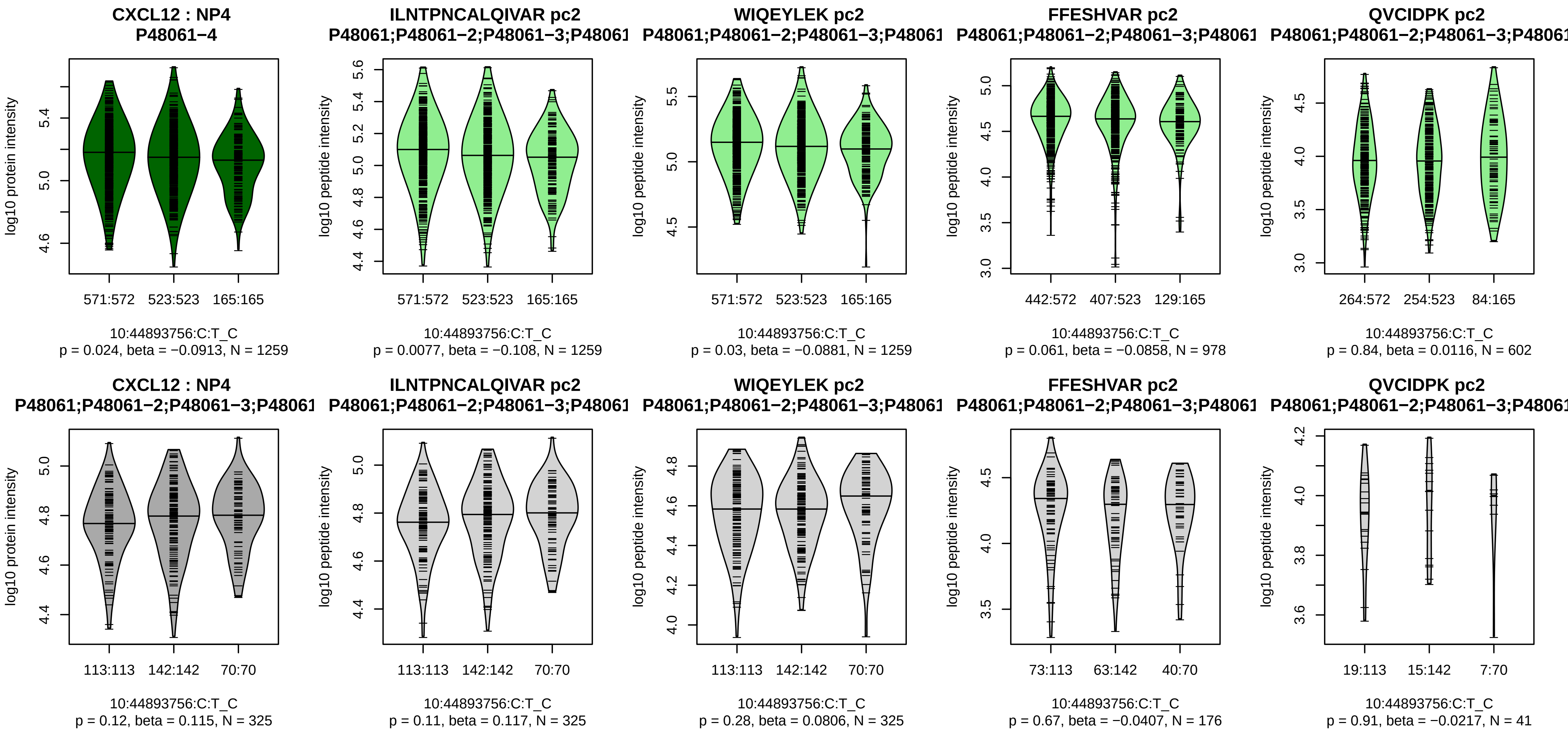


Assay Target: CXCL12
Olink UniProt: P48061
deCODE rsID: rs1023264
Proxy rsID: rs1023264
deCODE: 10:44398308:C:T
Proxy SNP: 10:44893756:C:T
deCODE log10(p): 53.1
deCODE BETA: 0.14
*:−:−:−
1259:1259:978:602

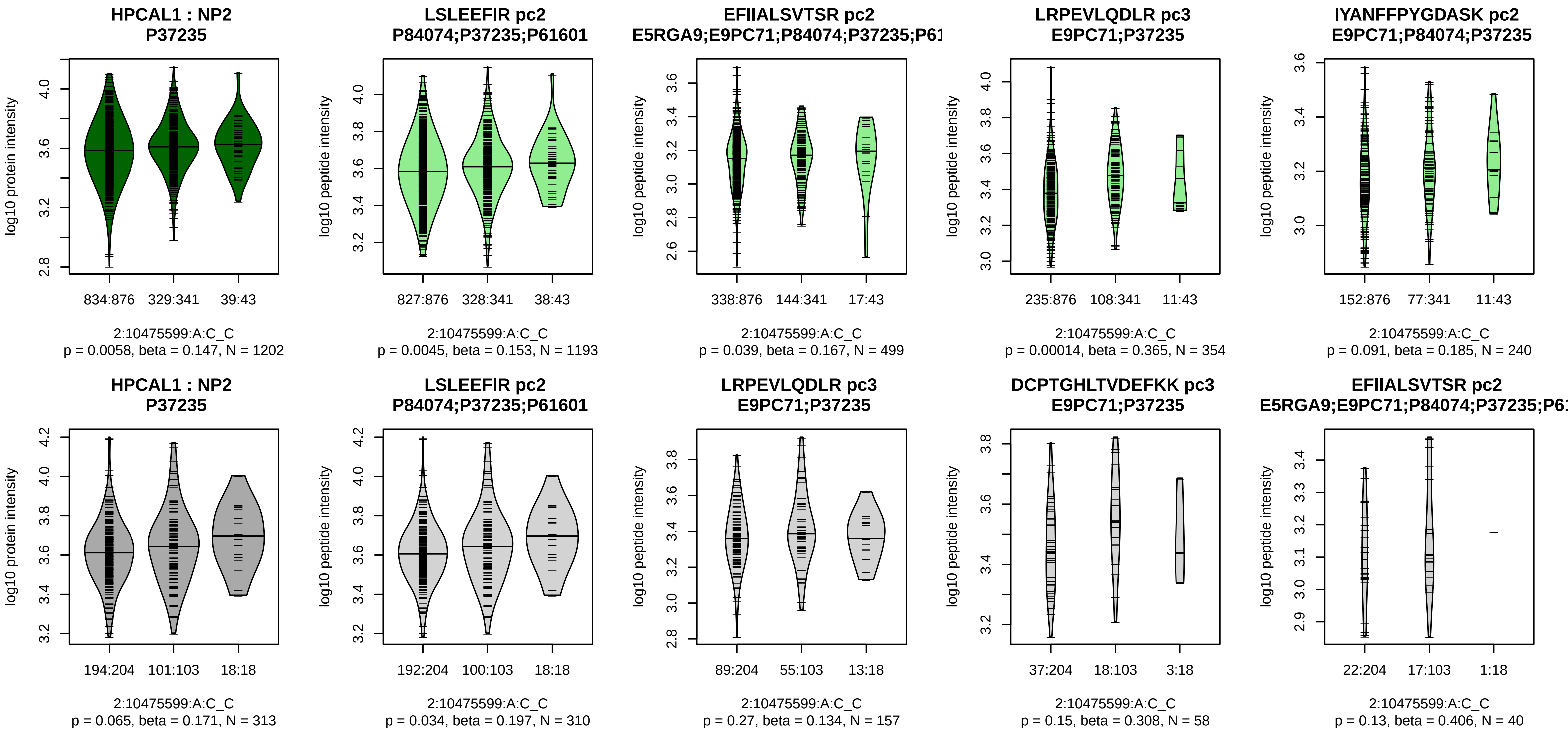


Assay Target: HPCAL1
Olink UniProt: P37235
deCODE rsID: rs6737972
Proxy rsID: rs11891604
deCODE: 2:10334724:C:T
Proxy SNP: 2:10475599:A:C
deCODE log10(p): 52.1
deCODE BETA: 0.16
*:−:*:−:−:−:NA
1193:499:354:240:130:95:2

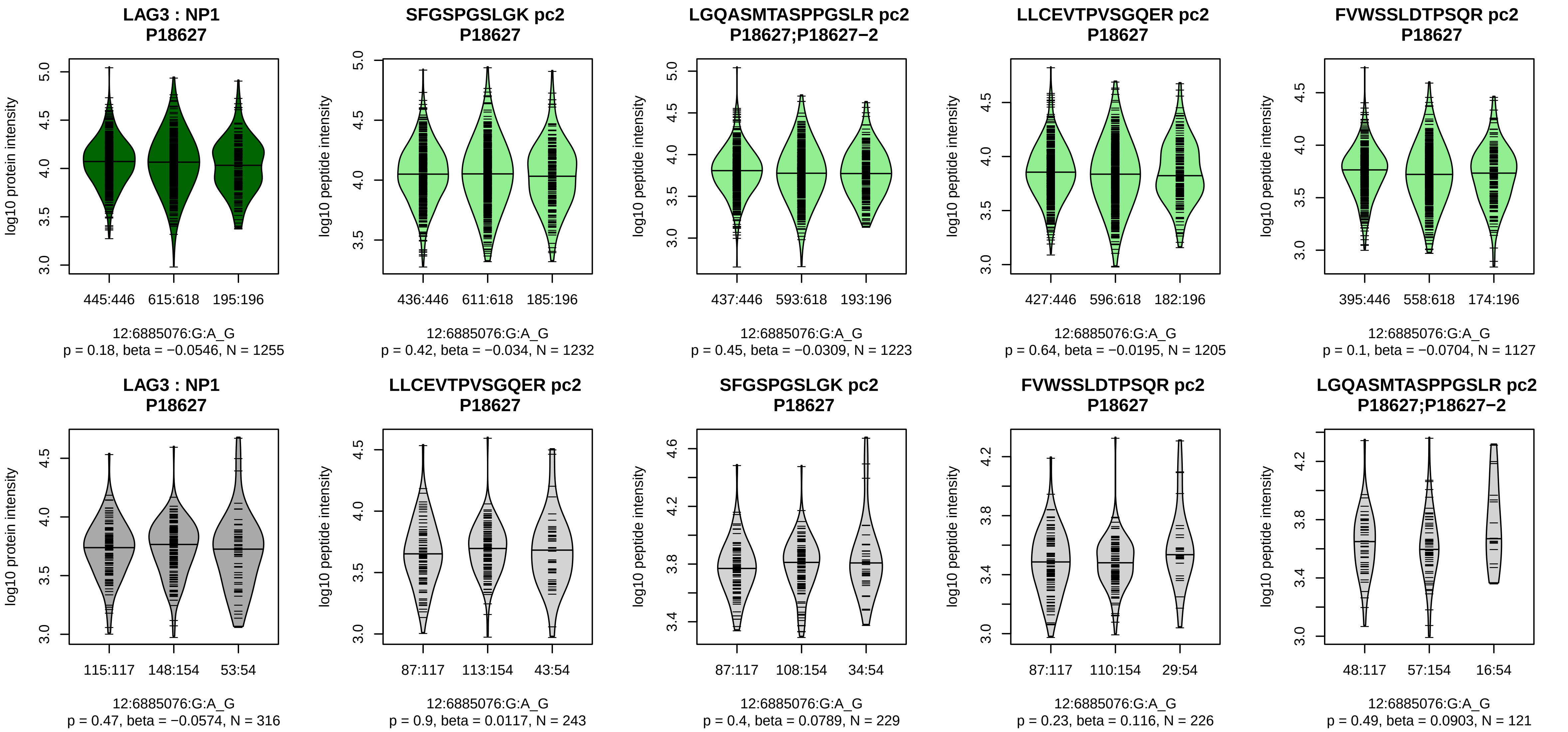


Assay Target: LAG3
Olink UniProt: P18627
deCODE rsID: rs3782735
Proxy rsID: rs3782735
deCODE: 12:6775910:G:A
Proxy SNP: 12:6885076:G:A
deCODE log10(p): 51.1
deCODE BETA: −0.13
−:−:−:−:−:−:−
1232:1223:1205:1127:651:92:5

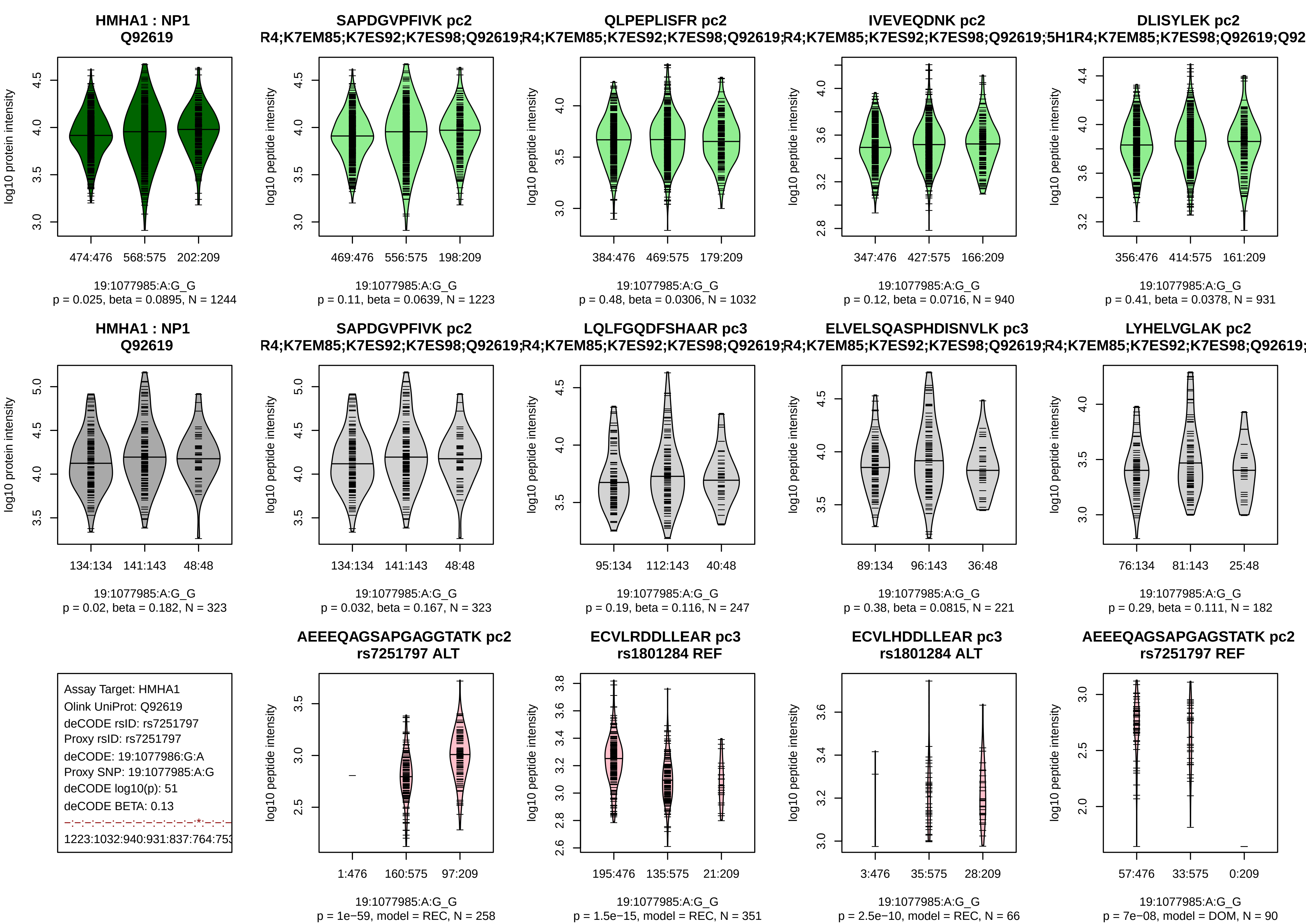

HMHA1 : NP1
Q92619
log10 protein intensity
474:476 568:575 202:209
19:1077985:A:G_G
p = 0.025, beta = 0.0895, N = 1244
SAPDGVPFIVK pc2
R4;K7EM85;K7ES92;K7ES98;Q92619;
log10 peptide intensity
469:476 556:575 198:209
19:1077985:A:G_G
p = 0.11, beta = 0.0639, N = 1223
QLPEPLISFR pc2
R4;K7EM85;K7ES92;K7ES98;Q92619;
384:476 469:575 179:209
19:1077985:A:G_G
p = 0.48, beta = 0.0306, N = 1032
IVEVEQDNK pc2
R4;K7EM85;K7ES92;K7ES98;Q92619;5H1
347:476 427:575 166:209
19:1077985:A:G_G
p = 0.12, beta = 0.0716, N = 940
DLISYLEK pc2
R4;K7EM85;K7ES98;Q92619;Q92
356:476 414:575 161:209
19:1077985:A:G_G
p = 0.41, beta = 0.0378, N = 931
HMHA1 : NP1
Q92619
134:134 141:143 48:48
19:1077985:A:G_G
p = 0.02, beta = 0.182, N = 323
SAPDGVPFIVK pc2
R4;K7EM85;K7ES92;K7ES98;Q92619;
134:134 141:143 48:48
19:1077985:A:G_G
p = 0.032, beta = 0.167, N = 323
LQLFGQDFSHAAR pc3
R4;K7EM85;K7ES92;K7ES98;Q92619;
95:134 112:143 40:48
19:1077985:A:G_G
p = 0.19, beta = 0.116, N = 247
ELVELSQASPHDISNVLK pc3
R4;K7EM85;K7ES92;K7ES98;Q92619;
89:134 96:143 36:48
19:1077985:A:G_G
p = 0.38, beta = 0.0815, N = 221
LYHELVGLAK pc2
R4;K7EM85;K7ES92;K7ES98;Q92619;
76:134 81:143 25:48
19:1077985:A:G_G
p = 0.29, beta = 0.111, N = 182
Assay Target: HMHA1
Olink UniProt: Q92619
deCODE rsID: rs7251797
Proxy rsID: rs7251797
deCODE: 19:1077986:G:A
Proxy SNP: 19:1077985:A:G
deCODE log10(p): 51
deCODE BETA: 0.13
1223:1032:940:931:837:764:753
AEEEQAGSAPGAGGTATK pc2
rs7251797 ALT
1:476 160:575 97:209
19:1077985:A:G_G
p = 1e−59, model = REC, N = 258
ECVLRDDLLEAR pc3
rs1801284 REF
195:476 135:575 21:209
19:1077985:A:G_G
p = 1.5e−15, model = REC, N = 351
ECVLHDDLLEAR pc3
rs1801284 ALT
3:476 35:575 28:209
19:1077985:A:G_G
p = 2.5e−10, model = REC, N = 66
AEEEQAGSAPGAGSTATK pc2
rs7251797 REF
57:476 33:575 0:209
19:1077985:A:G_G
p = 7e−08, model = DOM, N = 90

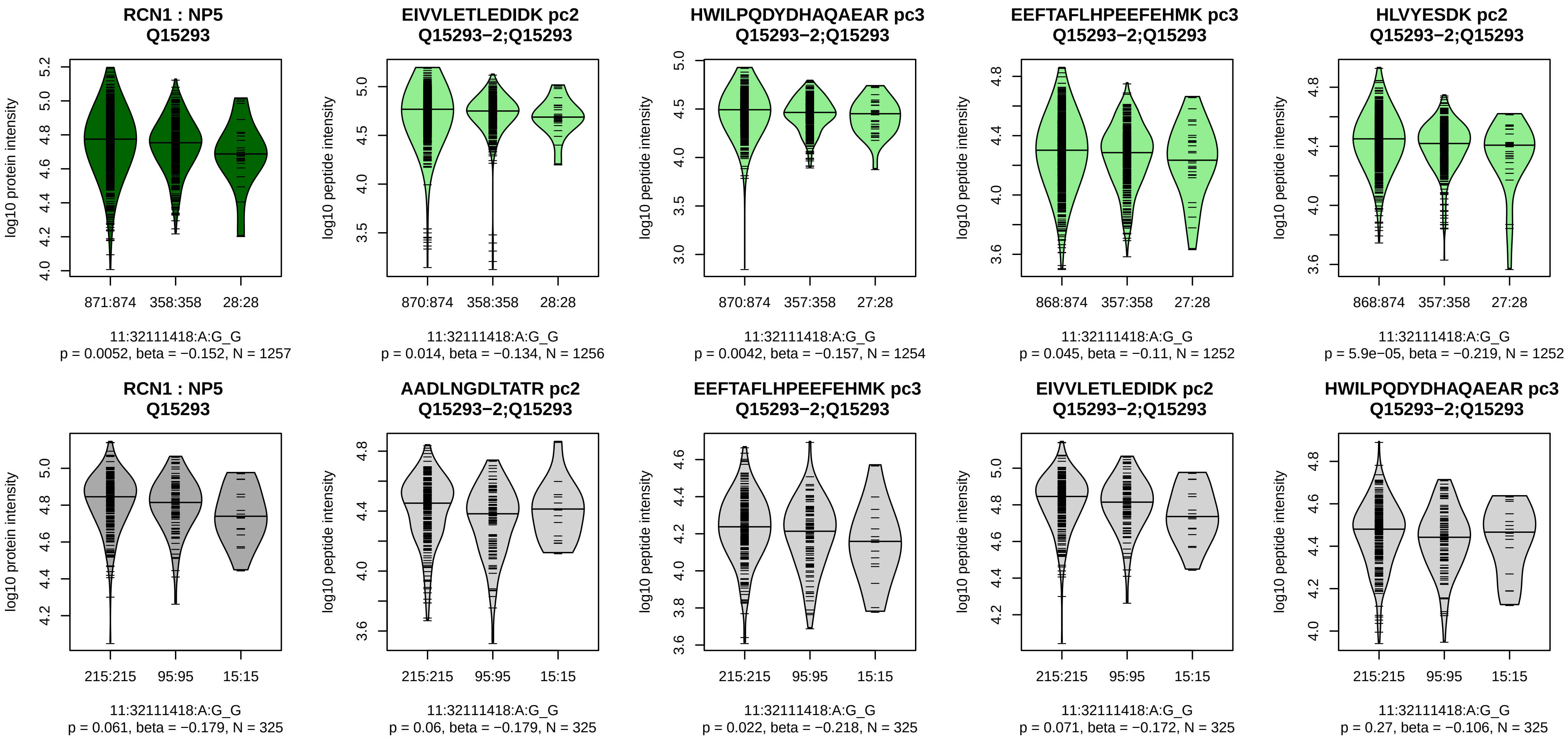


Assay Target: RCN1
Olink UniProt: Q15293
deCODE rsID: rs79314307
Proxy rsID: rs531603
deCODE: 11:32089698:CTT:C
Proxy SNP: 11:32111418:A:G
deCODE log10(p): 50.5
deCODE BETA: −0.17
−:*:−:*:*:−:−:−:−:−:−:−:−:−:−:−:−
1256:1254:1252:1252:1250:124

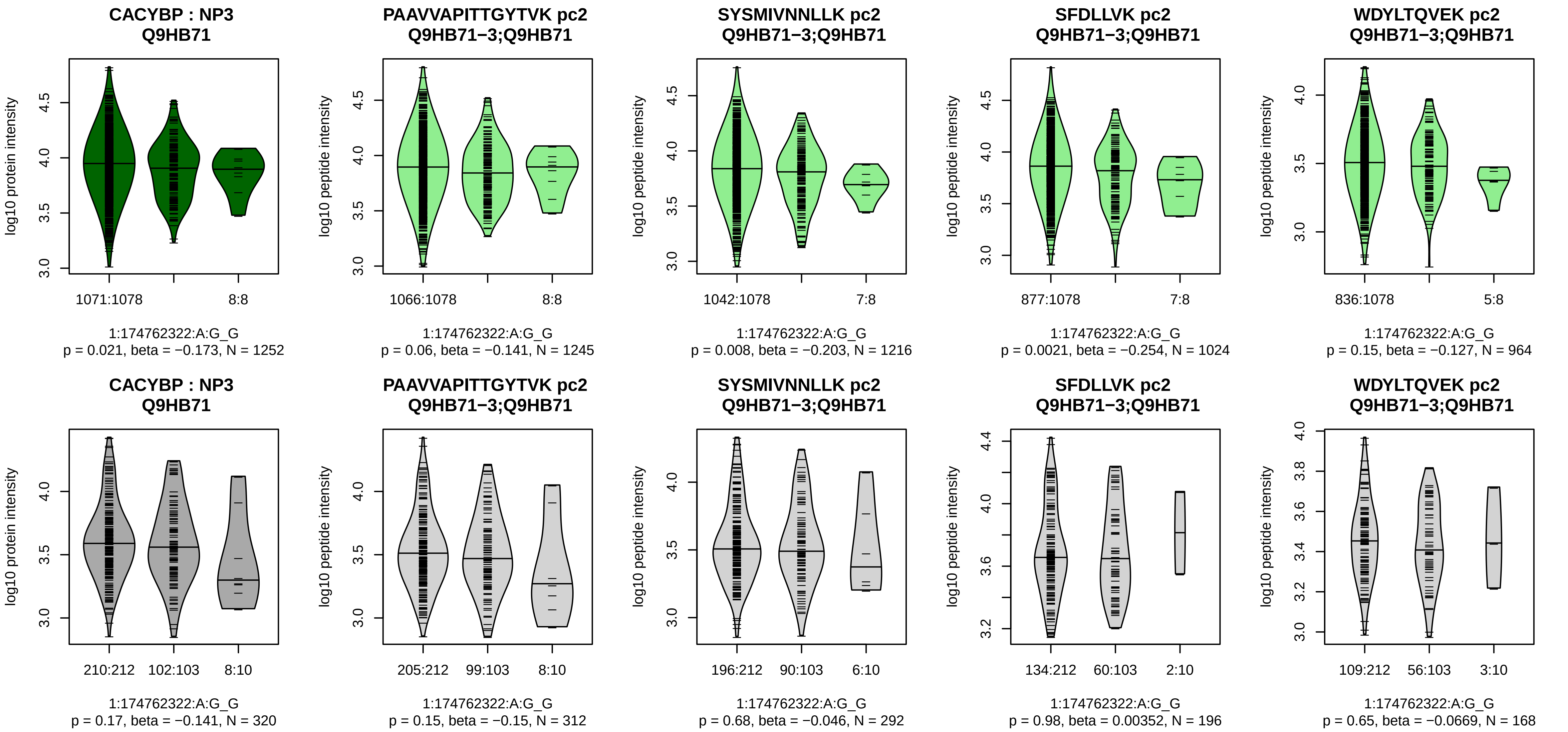


Assay Target: CACYBP
Olink UniProt: Q9HB71
deCODE rsID: rs139185768
Proxy rsID: rs16847450
deCODE: 1:174814998:GTATT[illegible]
Proxy SNP: 1:174762322:A:G
deCODE log10(p): 49.5
deCODE BETA: −0.2
−:*:*:−:−:−:−:−:−:−:−:−:NA
1245:1216:1024:964:914:691:4[illegible]

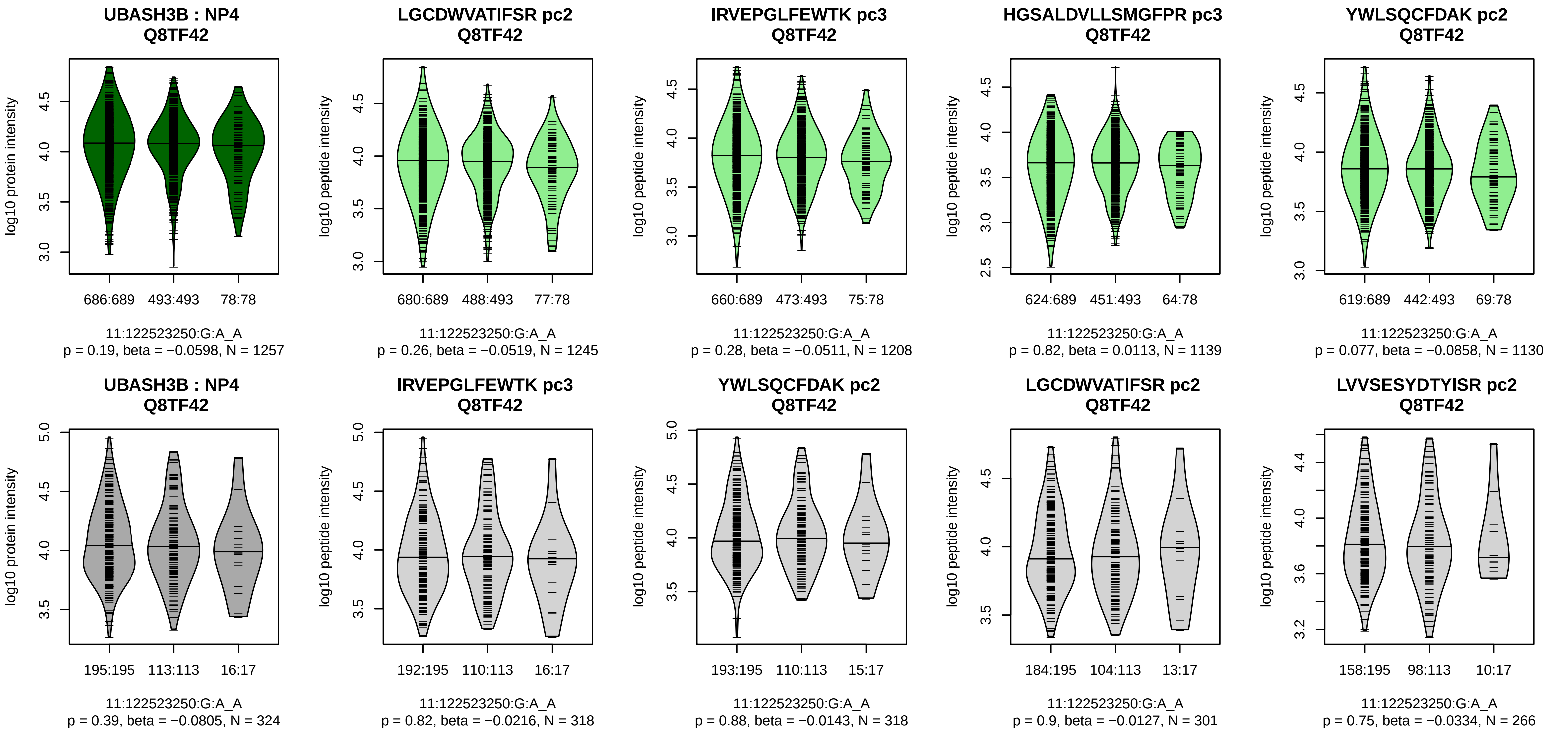


Assay Target: UBASH3B
Olink UniProt: Q8TF42
deCODE rsID: rs11218735
Proxy rsID: rs11218735
deCODE: 11:122652542:A:G
Proxy SNP: 11:122523250:G:A
deCODE log10(p): 48.5
deCODE BETA: −0.13
−:−:−:−:−:−:−:−:−:−:−:−:−:−:−:−:−
1245:1208:1139:1130:1119:105

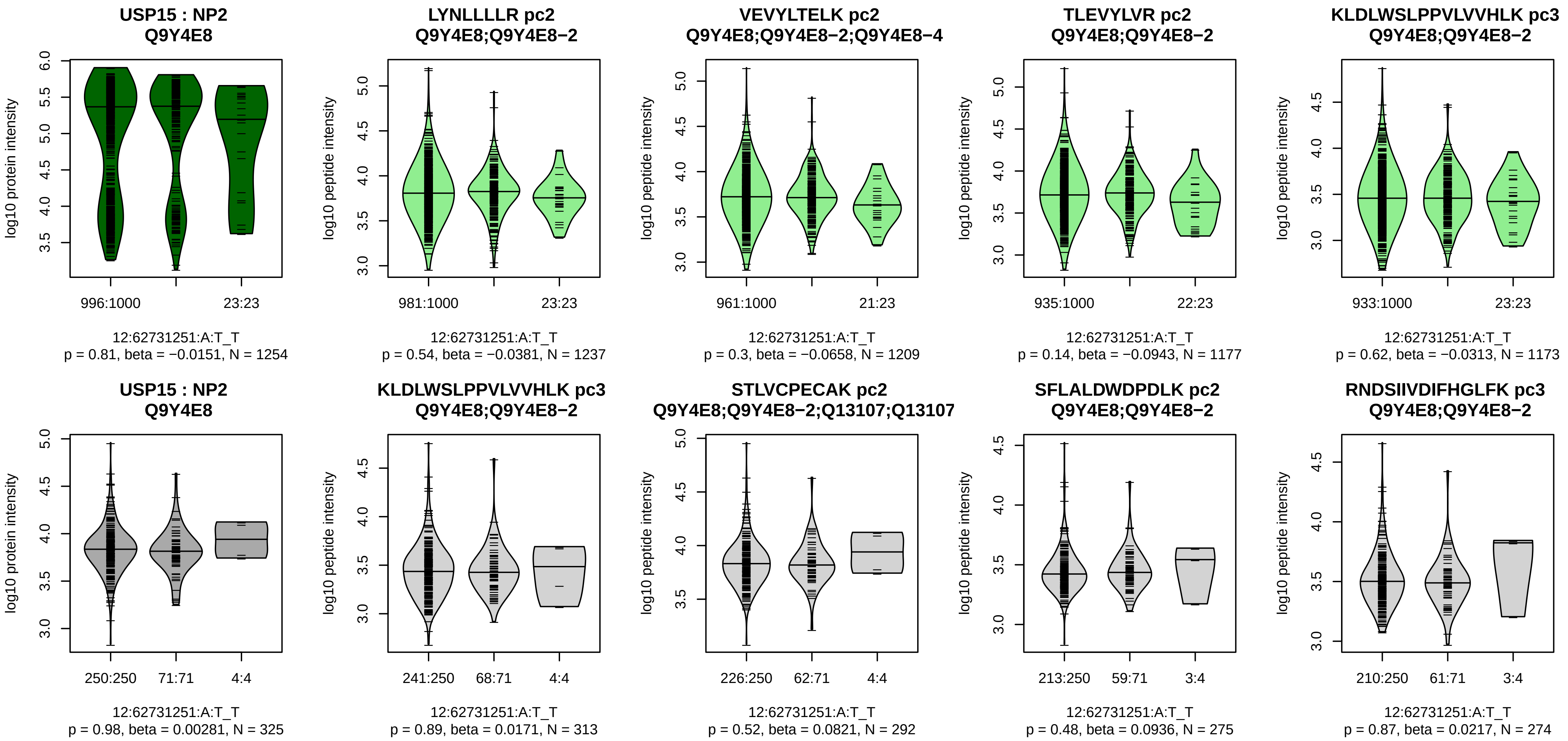


Assay Target: USP15
Olink UniProt: Q9Y4E8
deCODE rsID: rs73136831
Proxy rsID: rs73136831
deCODE: 12:62337470:T:A
Proxy SNP: 12:62731251:A:T
deCODE log10(p): 48
deCODE BETA: 0.2
−:−:−:−:−:−:−:−:−:−:−:−:−:−:−:−:
1237:1209:1177:1173:1167:115

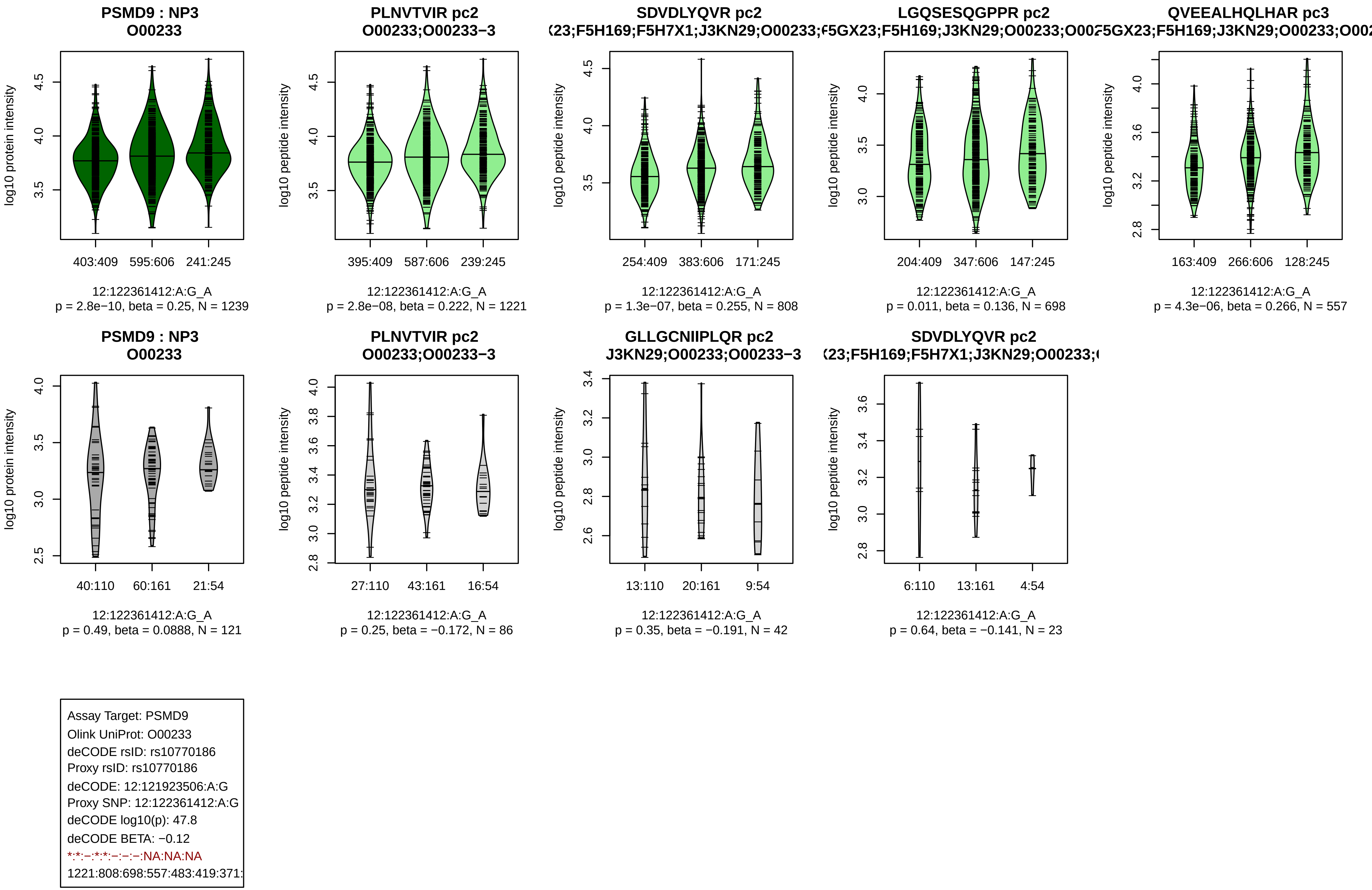
PSMD9 : NP3
O00233
log10 protein intensity
403:409 595:606 241:245
12:122361412:A:G_A
p = 2.8e−10, beta = 0.25, N = 1239
PLNVTVIR pc2
O00233;O00233−3
log10 peptide intensity
395:409 587:606 239:245
12:122361412:A:G_A
p = 2.8e−08, beta = 0.222, N = 1221
SDVDLYQVR pc2
(23;F5H169;F5H7X1;J3KN29;O00233;(
log10 peptide intensity
254:409 383:606 171:245
12:122361412:A:G_A
p = 1.3e−07, beta = 0.255, N = 808
LGQSESQGPPR pc2
F5GX23;F5H169;J3KN29;O00233;O002
log10 peptide intensity
204:409 347:606 147:245
12:122361412:A:G_A
p = 0.011, beta = 0.136, N = 698
QVEEALHQLHAR pc3
F5GX23;F5H169;J3KN29;O00233;O002
log10 peptide intensity
163:409 266:606 128:245
12:122361412:A:G_A
p = 4.3e−06, beta = 0.266, N = 557
PSMD9 : NP3
O00233
log10 protein intensity
40:110 60:161 21:54
12:122361412:A:G_A
p = 0.49, beta = 0.0888, N = 121
PLNVTVIR pc2
O00233;O00233−3
log10 peptide intensity
27:110 43:161 16:54
12:122361412:A:G_A
p = 0.25, beta = −0.172, N = 86
GLLGCNIIPLQR pc2
J3KN29;O00233;O00233−3
log10 peptide intensity
13:110 20:161 9:54
12:122361412:A:G_A
p = 0.35, beta = −0.191, N = 42
SDVDLYQVR pc2
(23;F5H169;F5H7X1;J3KN29;O00233;(
log10 peptide intensity
6:110 13:161 4:54
12:122361412:A:G_A
p = 0.64, beta = −0.141, N = 23
Assay Target: PSMD9
Olink UniProt: O00233
deCODE rsID: rs10770186
Proxy rsID: rs10770186
deCODE: 12:121923506:A:G
Proxy SNP: 12:122361412:A:G
deCODE log10(p): 47.8
deCODE BETA: −0.12
*:*:−:*:*:−:−:−:NA:NA:NA
1221:808:698:557:483:419:371:

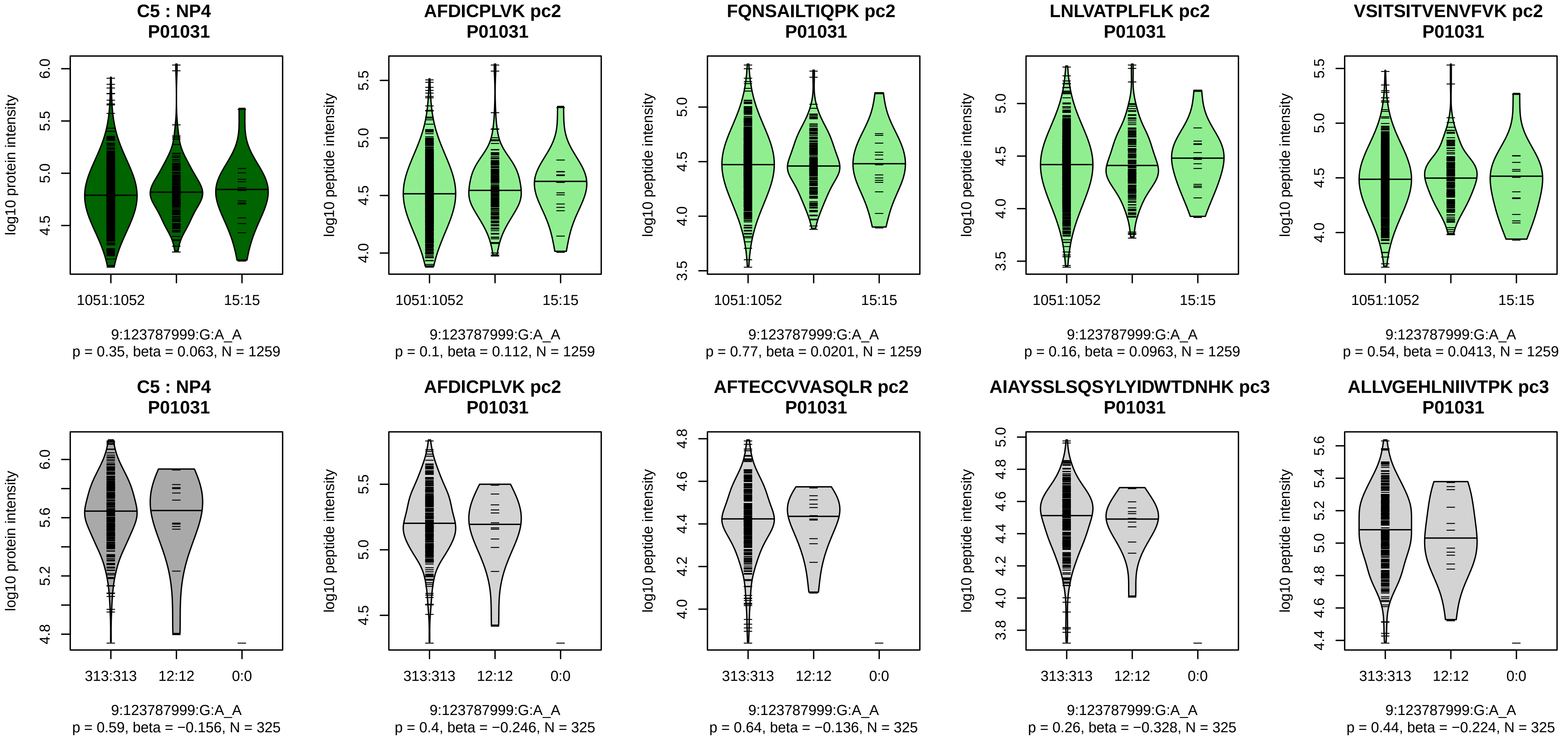


Assay Target: C5
Olink UniProt: P01031
deCODE rsID: rs17220750
Proxy rsID: rs17220750
deCODE: 9:121025721:A:G
Proxy SNP: 9:123787999:G:A
deCODE log10(p): 47.4
deCODE BETA: 0.19
−:−:−:−:−:−:−:−:−:−:−:−:−:−:−:−:
1259:1259:1259:1259:1258:125

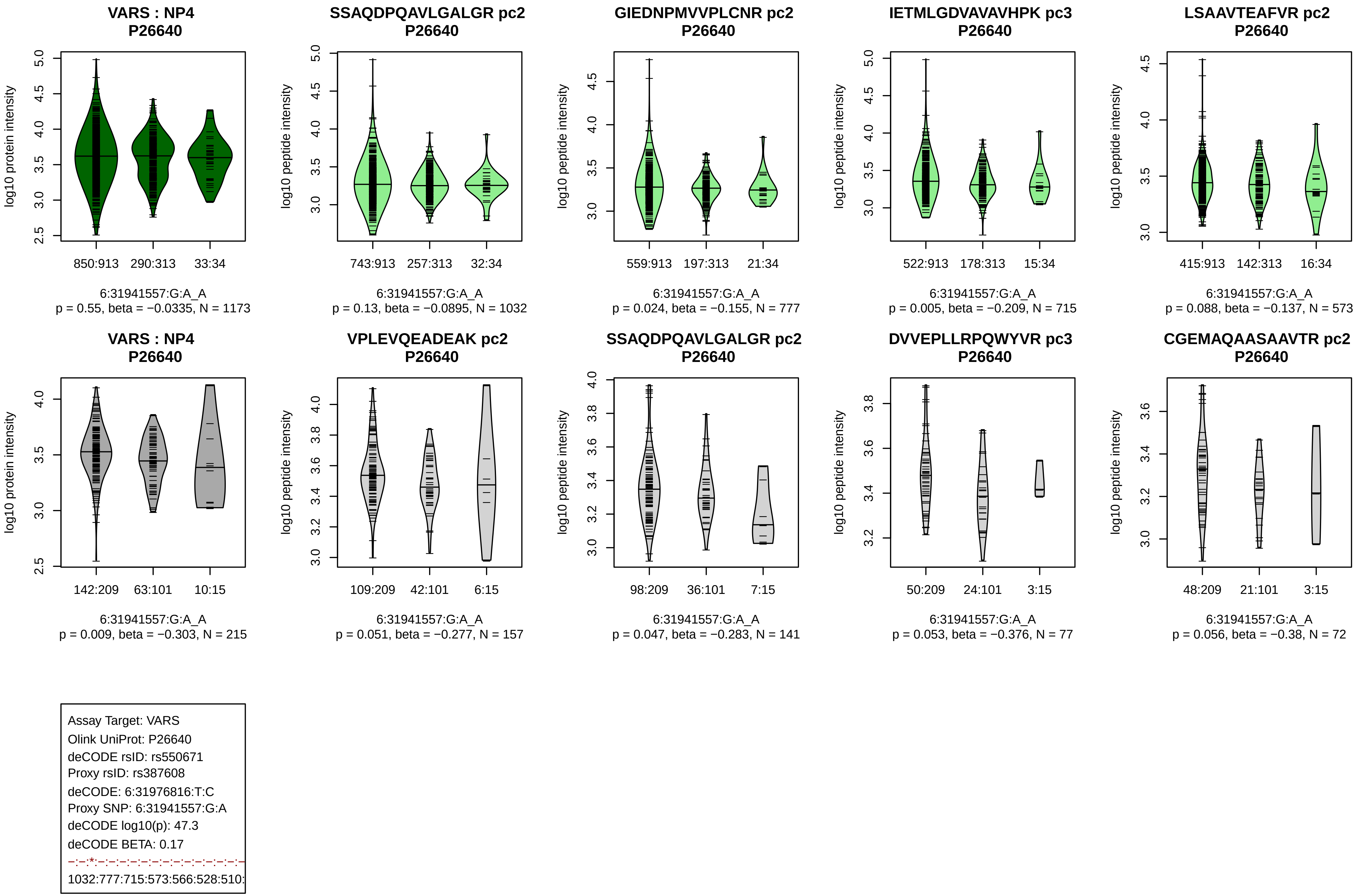
VARS : NP4
P26640
log10 protein intensity
850:913
290:313
33:34
6:31941557:G:A_A
p = 0.55, beta = −0.0335, N = 1173
SSAQDPQAVLGALGR pc2
P26640
log10 peptide intensity
743:913
257:313
32:34
6:31941557:G:A_A
p = 0.13, beta = −0.0895, N = 1032
GIEDNPMVVPLCNR pc2
P26640
log10 peptide intensity
559:913
197:313
21:34
6:31941557:G:A_A
p = 0.024, beta = −0.155, N = 777
IETMLGDVAVAVHPK pc3
P26640
log10 peptide intensity
522:913
178:313
15:34
6:31941557:G:A_A
p = 0.005, beta = −0.209, N = 715
LSAAVTEAFVR pc2
P26640
log10 peptide intensity
415:913
142:313
16:34
6:31941557:G:A_A
p = 0.088, beta = −0.137, N = 573
VARS : NP4
P26640
log10 protein intensity
142:209
63:101
10:15
6:31941557:G:A_A
p = 0.009, beta = −0.303, N = 215
VPLEVQEADEAK pc2
P26640
log10 peptide intensity
109:209
42:101
6:15
6:31941557:G:A_A
p = 0.051, beta = −0.277, N = 157
SSAQDPQAVLGALGR pc2
P26640
log10 peptide intensity
98:209
36:101
7:15
6:31941557:G:A_A
p = 0.047, beta = −0.283, N = 141
DVVEPLLRPQWYVR pc3
P26640
log10 peptide intensity
50:209
24:101
3:15
6:31941557:G:A_A
p = 0.053, beta = −0.376, N = 77
CGEMAQAASAAVTR pc2
P26640
log10 peptide intensity
48:209
21:101
3:15
6:31941557:G:A_A
p = 0.056, beta = −0.38, N = 72
Assay Target: VARS
Olink UniProt: P26640
deCODE rsID: rs550671
Proxy rsID: rs387608
deCODE: 6:31976816:T:C
Proxy SNP: 6:31941557:G:A
deCODE log10(p): 47.3
deCODE BETA: 0.17
1032:777:715:573:566:528:510:

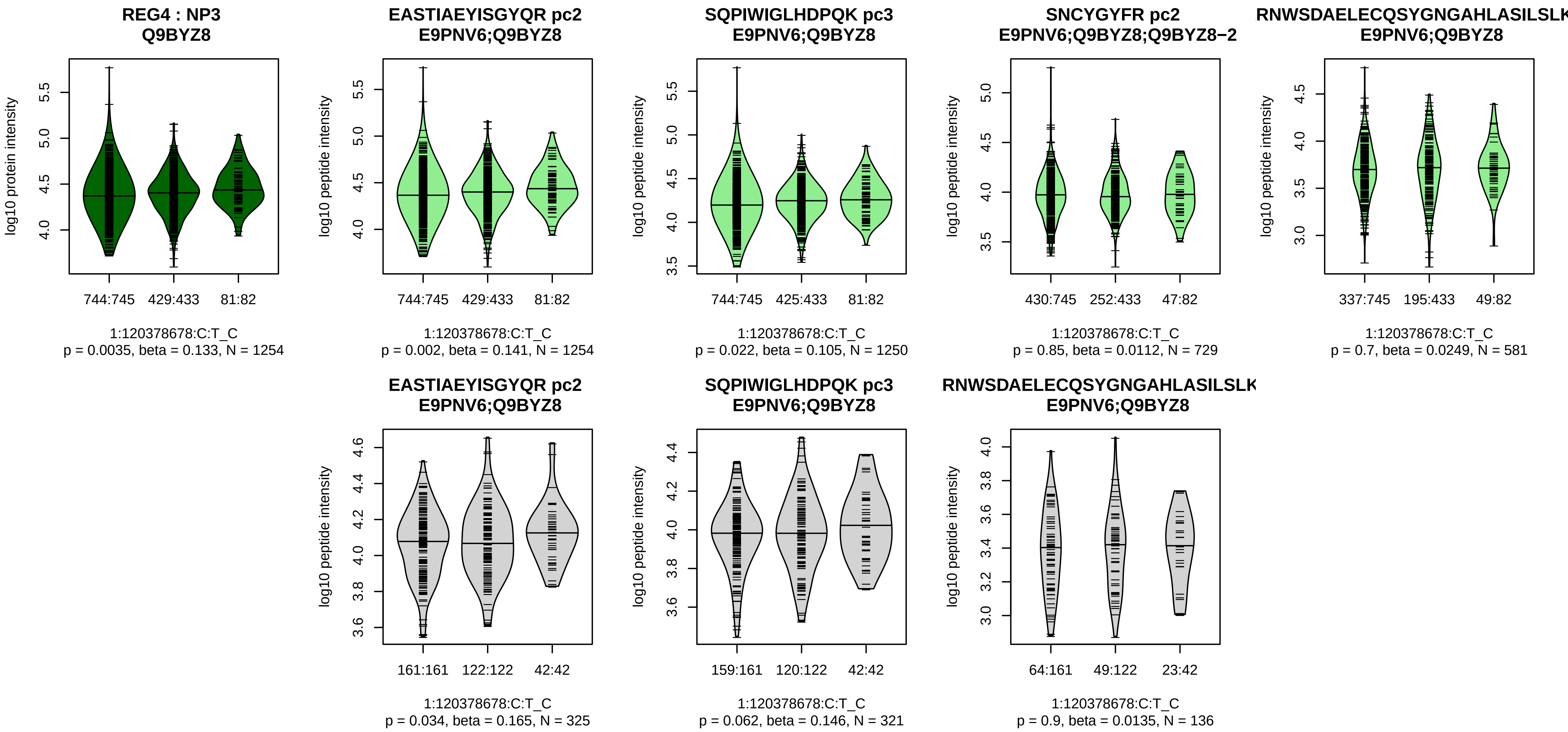


Assay Target: REG4
Olink UniProt: Q9BYZ8
deCODE rsID: rs3009182
Proxy rsID: rs3009182
deCODE: 1:119836055:C:T
Proxy SNP: 1:120378678:C:T
deCODE log10(p): 45.6
deCODE BETA: 0.14
*:−:*:−:−:−:−:−:−:−:NA
1254:1250:1175:1156:729:581:4

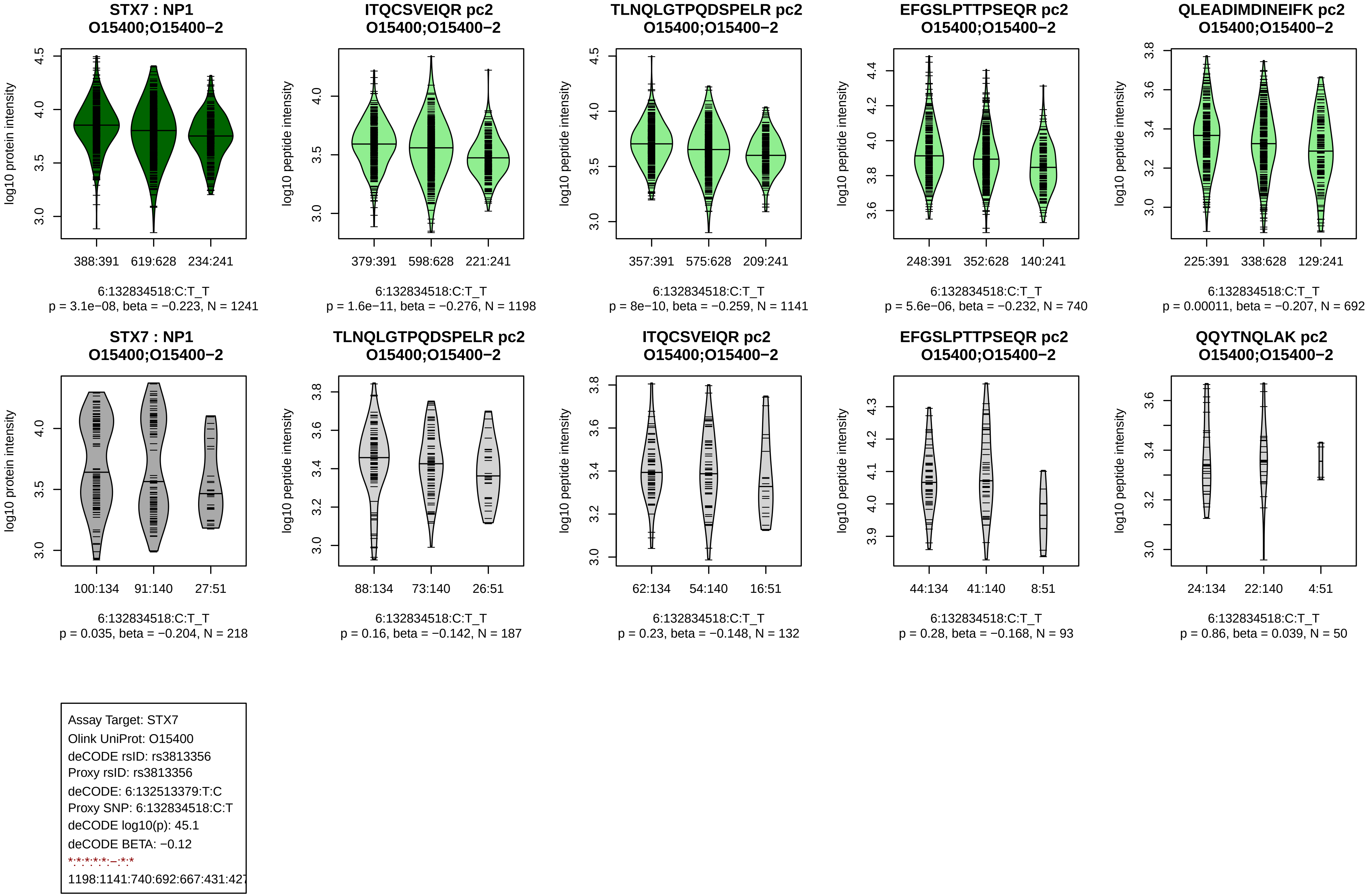
STX7 : NP1
O15400;O15400−2
log10 protein intensity
388:391
619:628
234:241
6:132834518:C:T_T
p = 3.1e−08, beta = −0.223, N = 1241
ITQCSVEIQR pc2
O15400;O15400−2
log10 peptide intensity
379:391
598:628
221:241
6:132834518:C:T_T
p = 1.6e−11, beta = −0.276, N = 1198
TLNQLGTPQDSPELR pc2
O15400;O15400−2
log10 peptide intensity
357:391
575:628
209:241
6:132834518:C:T_T
p = 8e−10, beta = −0.259, N = 1141
EFGSLPTTPSEQR pc2
O15400;O15400−2
log10 peptide intensity
248:391
352:628
140:241
6:132834518:C:T_T
p = 5.6e−06, beta = −0.232, N = 740
QLEADIMDINEIFK pc2
O15400;O15400−2
log10 peptide intensity
225:391
338:628
129:241
6:132834518:C:T_T
p = 0.00011, beta = −0.207, N = 692
STX7 : NP1
O15400;O15400−2
log10 protein intensity
100:134
91:140
27:51
6:132834518:C:T_T
p = 0.035, beta = −0.204, N = 218
TLNQLGTPQDSPELR pc2
O15400;O15400−2
log10 peptide intensity
88:134
73:140
26:51
6:132834518:C:T_T
p = 0.16, beta = −0.142, N = 187
ITQCSVEIQR pc2
O15400;O15400−2
log10 peptide intensity
62:134
54:140
16:51
6:132834518:C:T_T
p = 0.23, beta = −0.148, N = 132
EFGSLPTTPSEQR pc2
O15400;O15400−2
log10 peptide intensity
44:134
41:140
8:51
6:132834518:C:T_T
p = 0.28, beta = −0.168, N = 93
QQYTNQLAK pc2
O15400;O15400−2
log10 peptide intensity
24:134
22:140
4:51
6:132834518:C:T_T
p = 0.86, beta = 0.039, N = 50
Assay Target: STX7
Olink UniProt: O15400
deCODE rsID: rs3813356
Proxy rsID: rs3813356
deCODE: 6:132513379:T:C
Proxy SNP: 6:132834518:C:T
deCODE log10(p): 45.1
deCODE BETA: −0.12
*:*:*:*:*:–:*:*
1198:1141:740:692:667:431:427

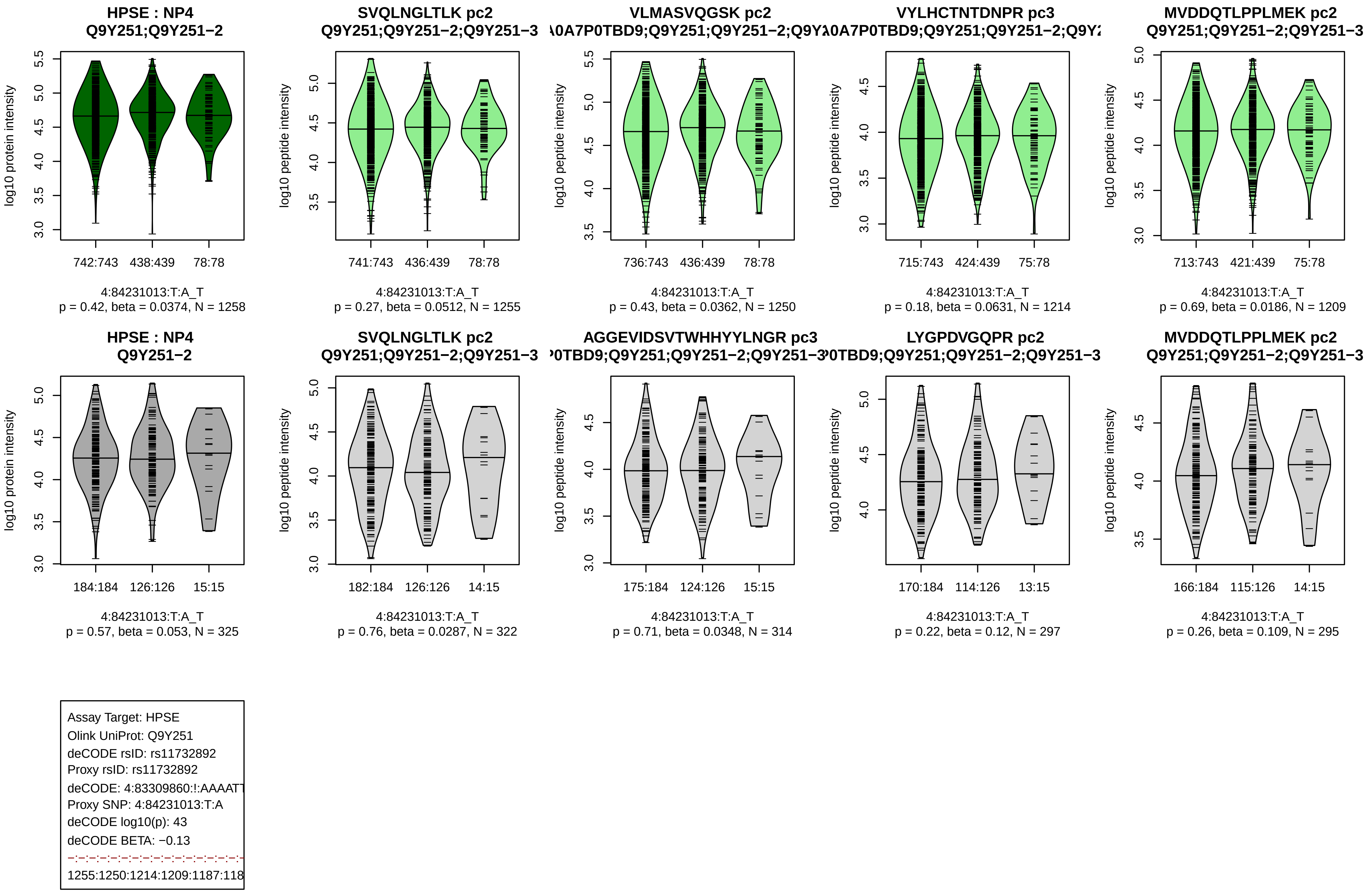

HPSE : NP4
Q9Y251;Q9Y251−2
log10 protein intensity
742:743 438:439 78:78
4:84231013:T:A_T
p = 0.42, beta = 0.0374, N = 1258
SVQLNGLTLK pc2
Q9Y251;Q9Y251−2;Q9Y251−3
log10 peptide intensity
741:743 436:439 78:78
4:84231013:T:A_T
p = 0.27, beta = 0.0512, N = 1255
VLMASVQGSK pc2
A0A7P0TBD9;Q9Y251;Q9Y251−2;Q9Y
log10 peptide intensity
736:743 436:439 78:78
4:84231013:T:A_T
p = 0.43, beta = 0.0362, N = 1250
VYLHCTNTDNPR pc3
A0A7P0TBD9;Q9Y251;Q9Y251−2;Q9Y
log10 peptide intensity
715:743 424:439 75:78
4:84231013:T:A_T
p = 0.18, beta = 0.0631, N = 1214
MVDDQTLPPLMEK pc2
Q9Y251;Q9Y251−2;Q9Y251−3
log10 peptide intensity
713:743 421:439 75:78
4:84231013:T:A_T
p = 0.69, beta = 0.0186, N = 1209
HPSE : NP4
Q9Y251−2
log10 protein intensity
184:184 126:126 15:15
4:84231013:T:A_T
p = 0.57, beta = 0.053, N = 325
SVQLNGLTLK pc2
Q9Y251;Q9Y251−2;Q9Y251−3
log10 peptide intensity
182:184 126:126 14:15
4:84231013:T:A_T
p = 0.76, beta = 0.0287, N = 322
AGGEVIDSVTWHHYYLNGR pc3
P0TBD9;Q9Y251;Q9Y251−2;Q9Y251−3
log10 peptide intensity
175:184 124:126 15:15
4:84231013:T:A_T
p = 0.71, beta = 0.0348, N = 314
LYGPDVGQPR pc2
P0TBD9;Q9Y251;Q9Y251−2;Q9Y251−3
log10 peptide intensity
170:184 114:126 13:15
4:84231013:T:A_T
p = 0.22, beta = 0.12, N = 297
MVDDQTLPPLMEK pc2
Q9Y251;Q9Y251−2;Q9Y251−3
log10 peptide intensity
166:184 115:126 14:15
4:84231013:T:A_T
p = 0.26, beta = 0.109, N = 295
Assay Target: HPSE
Olink UniProt: Q9Y251
deCODE rsID: rs11732892
Proxy rsID: rs11732892
deCODE: 4:83309860:!:AAAATT
Proxy SNP: 4:84231013:T:A
deCODE log10(p): 43
deCODE BETA: −0.13
1255:1250:1214:1209:1187:118

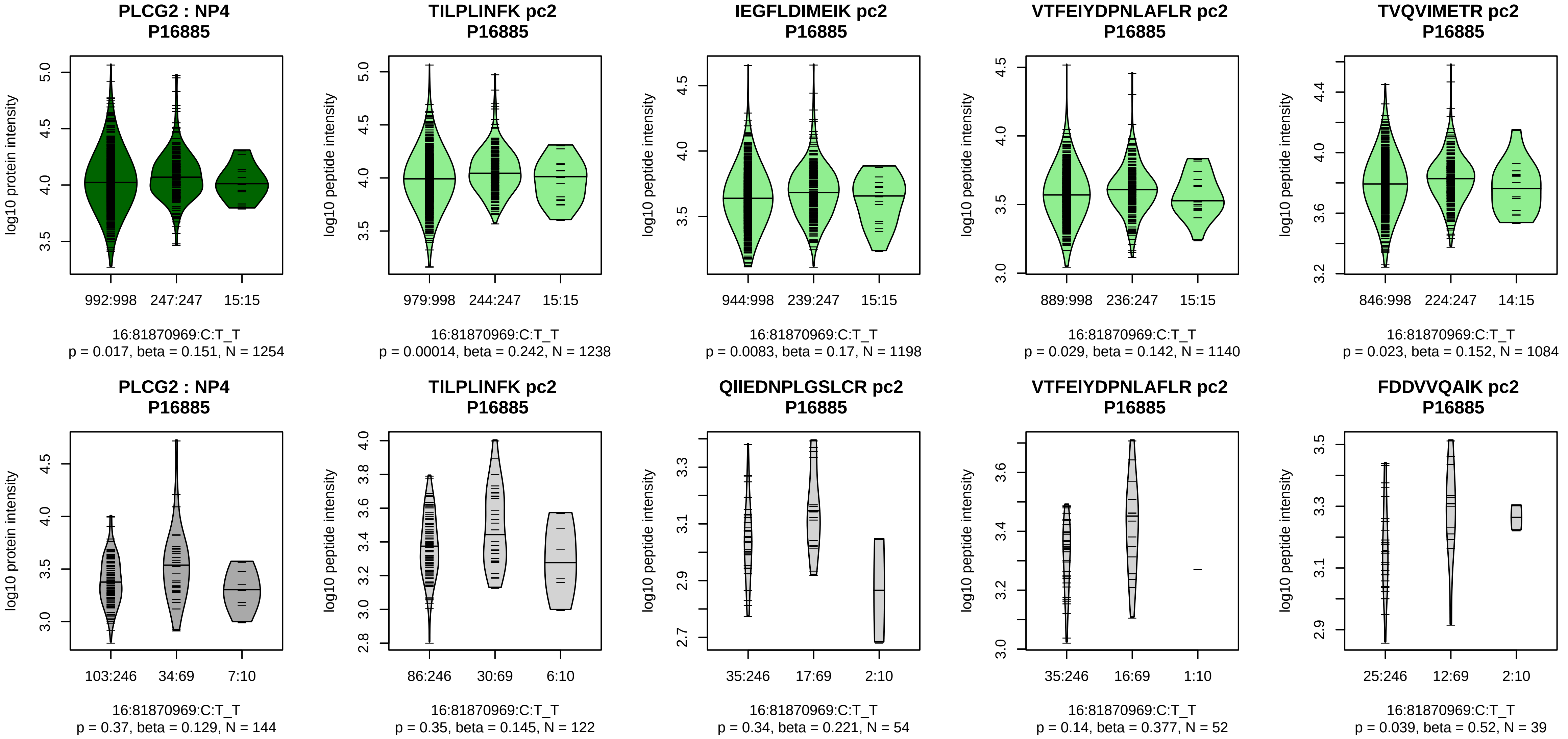


Assay Target: PLCG2
Olink UniProt: P16885
deCODE rsID: rs12445050
Proxy rsID: rs12445050
deCODE: 16:81837364:T:C
Proxy SNP: 16:81870969:C:T
deCODE log10(p): 42.9
deCODE BETA: 0.18
*:*:–:–:–:–:–:*:–:–:–:*:–:–:–:–:–:
1238:1198:1140:1084:1082:104

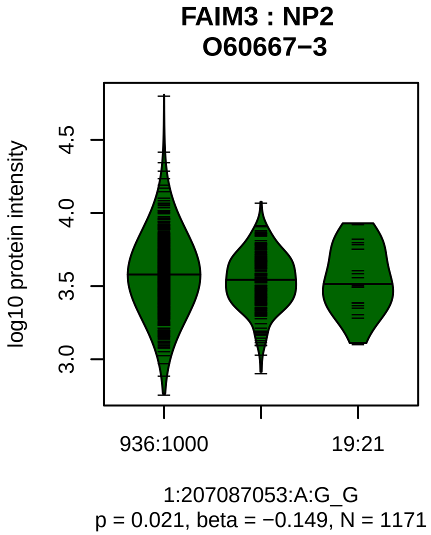


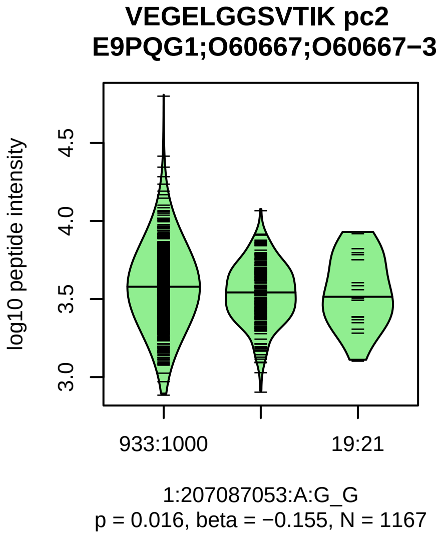


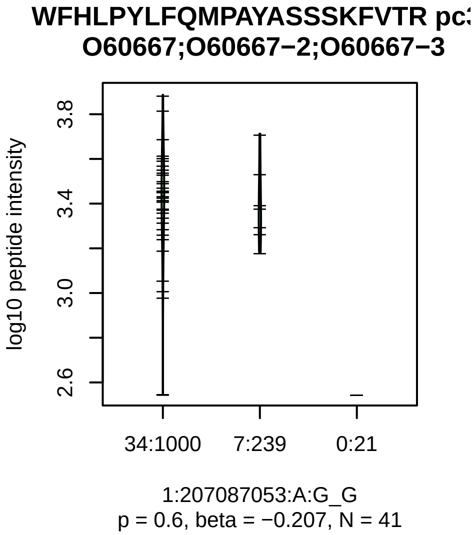


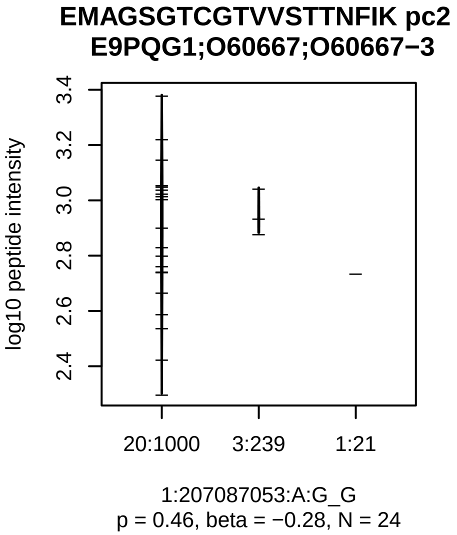


Assay Target: FAIM3
Olink UniProt: O60667
deCODE rsID: rs72758947
Proxy rsID: rs72758947
deCODE: 1:206913708:G:A
Proxy SNP: 1:207087053:A:G
deCODE log10(p): 42.2
deCODE BETA: −0.16
−:−:−
1167:41:24

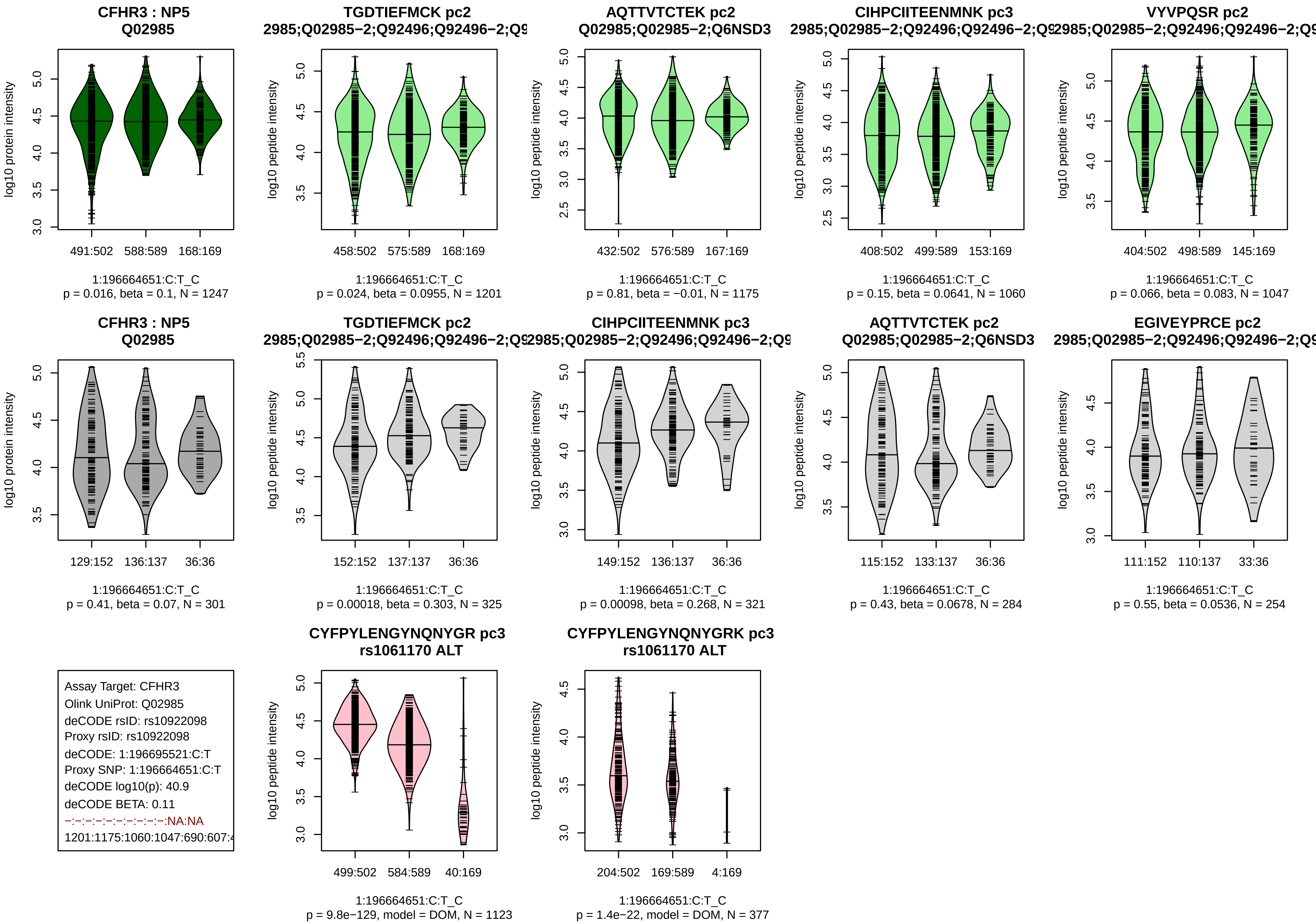
CFHR3 : NP5
Q02985
log10 protein intensity
491:502 588:589 168:169
1:196664651:C:T_C
p = 0.016, beta = 0.1, N = 1247
TGDTIEFMCK pc2
2985;Q02985−2;Q92496;Q92496−2;Q9
log10 peptide intensity
458:502 575:589 168:169
1:196664651:C:T_C
p = 0.024, beta = 0.0955, N = 1201
AQTTVTCTEK pc2
Q02985;Q02985−2;Q6NSD3
log10 peptide intensity
432:502 576:589 167:169
1:196664651:C:T_C
p = 0.81, beta = −0.01, N = 1175
CIHPCIITEENMNK pc3
2985;Q02985−2;Q92496;Q92496−2;Q9
log10 peptide intensity
408:502 499:589 153:169
1:196664651:C:T_C
p = 0.15, beta = 0.0641, N = 1060
VYVPQSR pc2
2985;Q02985−2;Q92496;Q92496−2;Q9
log10 peptide intensity
404:502 498:589 145:169
1:196664651:C:T_C
p = 0.066, beta = 0.083, N = 1047
CFHR3 : NP5
Q02985
log10 protein intensity
129:152 136:137 36:36
1:196664651:C:T_C
p = 0.41, beta = 0.07, N = 301
TGDTIEFMCK pc2
2985;Q02985−2;Q92496;Q92496−2;Q9
log10 peptide intensity
152:152 137:137 36:36
1:196664651:C:T_C
p = 0.00018, beta = 0.303, N = 325
CIHPCIITEENMNK pc3
2985;Q02985−2;Q92496;Q92496−2;Q9
log10 peptide intensity
149:152 136:137 36:36
1:196664651:C:T_C
p = 0.00098, beta = 0.268, N = 321
AQTTVTCTEK pc2
Q02985;Q02985−2;Q6NSD3
log10 peptide intensity
115:152 133:137 36:36
1:196664651:C:T_C
p = 0.43, beta = 0.0678, N = 284
EGIVEYPRCE pc2
2985;Q02985−2;Q92496;Q92496−2;Q9
log10 peptide intensity
111:152 110:137 33:36
1:196664651:C:T_C
p = 0.55, beta = 0.0536, N = 254
Assay Target: CFHR3
Olink UniProt: Q02985
deCODE rsID: rs10922098
Proxy rsID: rs10922098
deCODE: 1:196695521:C:T
Proxy SNP: 1:196664651:C:T
deCODE log10(p): 40.9
deCODE BETA: 0.11
−:−:−:−:−:−:−:−:−:−:NA:NA
1201:1175:1060:1047:690:607:4
CYFPYLENGYNQNYGR pc3
rs1061170 ALT
log10 peptide intensity
499:502 584:589 40:169
1:196664651:C:T_C
p = 9.8e−129, model = DOM, N = 1123
CYFPYLENGYNQNYGRK pc3
rs1061170 ALT
log10 peptide intensity
204:502 169:589 4:169
1:196664651:C:T_C
p = 1.4e−22, model = DOM, N = 377

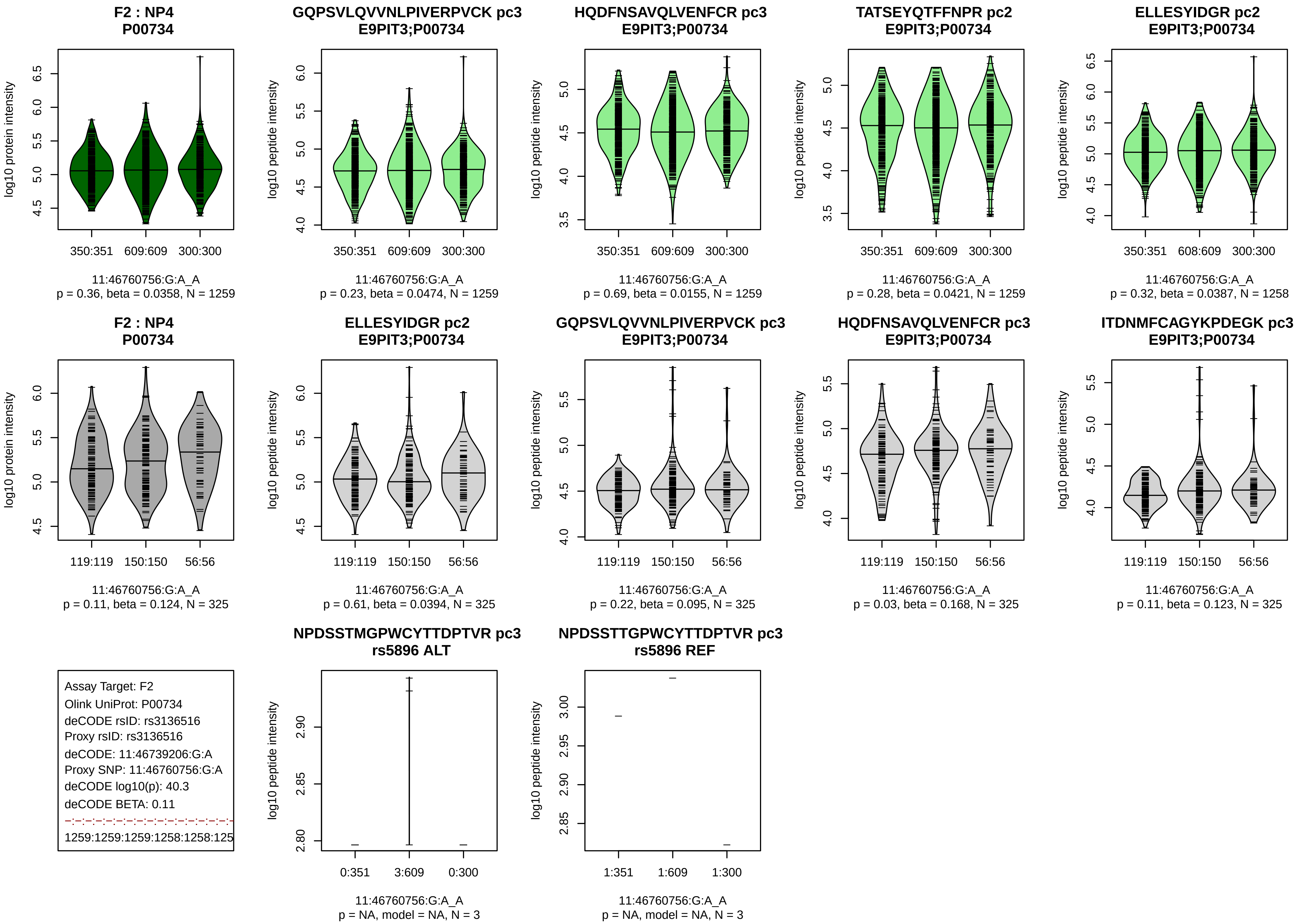

F2 : NP4
P00734
log10 protein intensity
350:351 609:609 300:300
11:46760756:G:A_A
p = 0.36, beta = 0.0358, N = 1259
GQPSVLQVVNLPIVERPVCK pc3
E9PIT3;P00734
log10 peptide intensity
350:351 609:609 300:300
11:46760756:G:A_A
p = 0.23, beta = 0.0474, N = 1259
HQDFNSAVQLVENFCR pc3
E9PIT3;P00734
log10 peptide intensity
350:351 609:609 300:300
11:46760756:G:A_A
p = 0.69, beta = 0.0155, N = 1259
TATSEYQTFFNPR pc2
E9PIT3;P00734
log10 peptide intensity
350:351 609:609 300:300
11:46760756:G:A_A
p = 0.28, beta = 0.0421, N = 1259
ELLESYIDGR pc2
E9PIT3;P00734
log10 peptide intensity
350:351 608:609 300:300
11:46760756:G:A_A
p = 0.32, beta = 0.0387, N = 1258
F2 : NP4
P00734
log10 protein intensity
119:119 150:150 56:56
11:46760756:G:A_A
p = 0.11, beta = 0.124, N = 325
ELLESYIDGR pc2
E9PIT3;P00734
log10 peptide intensity
119:119 150:150 56:56
11:46760756:G:A_A
p = 0.61, beta = 0.0394, N = 325
GQPSVLQVVNLPIVERPVCK pc3
E9PIT3;P00734
log10 peptide intensity
119:119 150:150 56:56
11:46760756:G:A_A
p = 0.22, beta = 0.095, N = 325
HQDFNSAVQLVENFCR pc3
E9PIT3;P00734
log10 peptide intensity
119:119 150:150 56:56
11:46760756:G:A_A
p = 0.03, beta = 0.168, N = 325
ITDNMFCAGYKPDEGK pc3
E9PIT3;P00734
log10 peptide intensity
119:119 150:150 56:56
11:46760756:G:A_A
p = 0.11, beta = 0.123, N = 325
Assay Target: F2
Olink UniProt: P00734
deCODE rsID: rs3136516
Proxy rsID: rs3136516
deCODE: 11:46739206:G:A
Proxy SNP: 11:46760756:G:A
deCODE log10(p): 40.3
deCODE BETA: 0.11
1259:1259:1259:1258:1258:125
NPDSSTMGPWCYTTDPTVR pc3
rs5896 ALT
log10 peptide intensity
0:351 3:609 0:300
11:46760756:G:A_A
p = NA, model = NA, N = 3
NPDSSTTGPWCYTTDPTVR pc3
rs5896 REF
log10 peptide intensity
1:351 1:609 1:300
11:46760756:G:A_A
p = NA, model = NA, N = 3

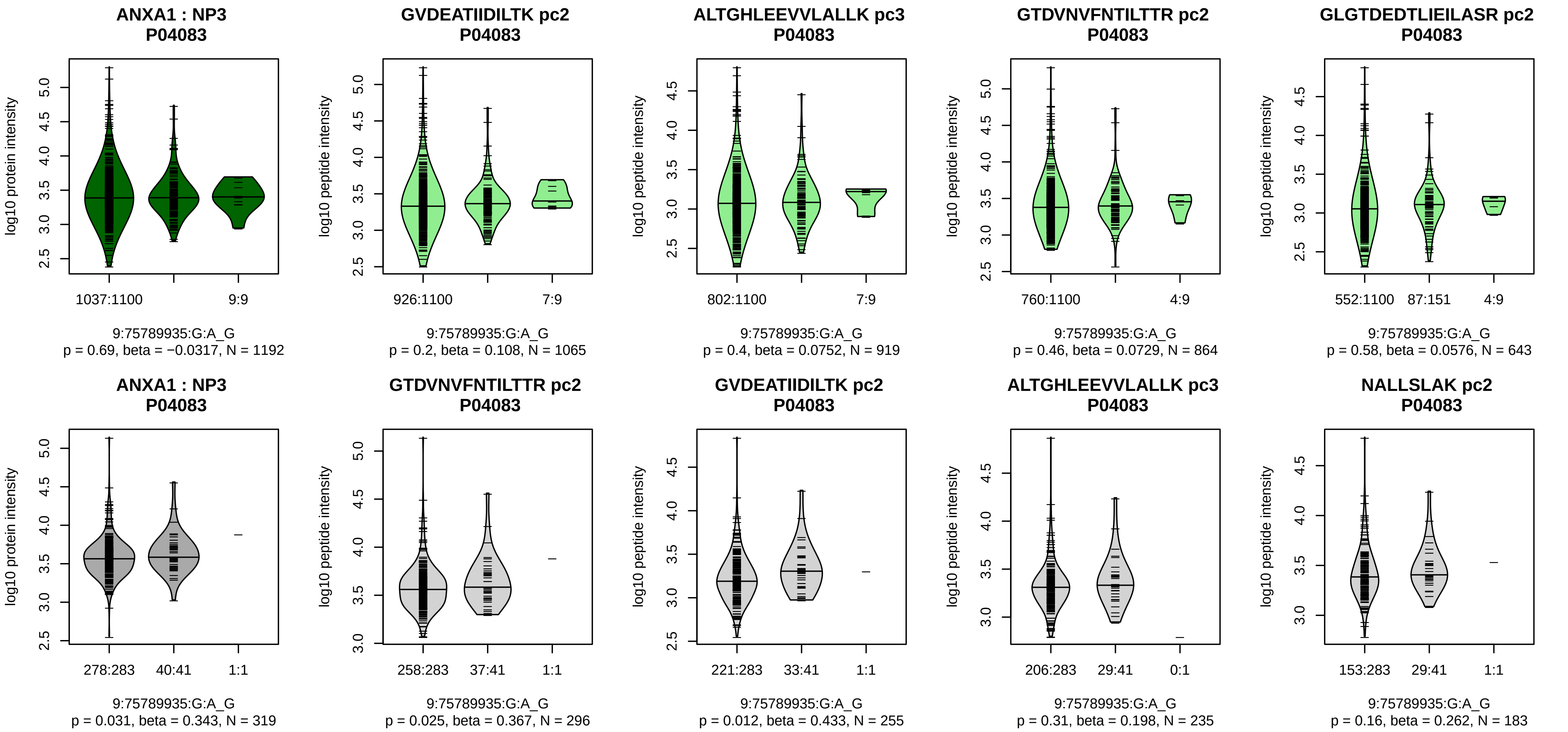


Assay Target: ANXA1
Olink UniProt: P04083
deCODE rsID: rs7357731
Proxy rsID: rs7357731
deCODE: 9:73175019:G:A
Proxy SNP: 9:75789935:G:A
deCODE log10(p): 39.6
deCODE BETA: 0.34
–:–:–:–:–:–:–:–:–:–:–:–:–:–:–:–:
1065:919:864:643:345:333:269:

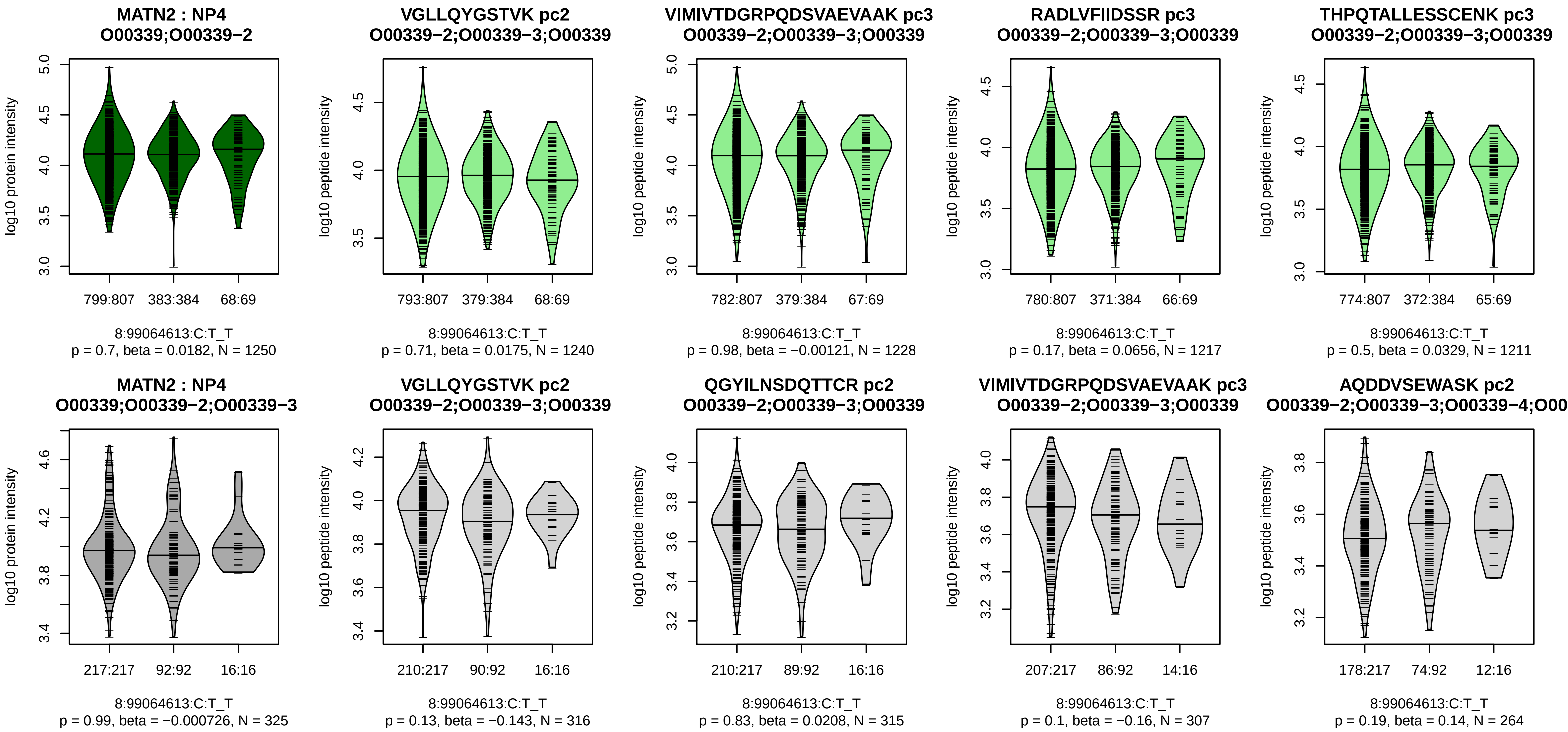


Assay Target: MATN2
Olink UniProt: O00339
deCODE rsID: rs78548919
Proxy rsID: rs28522929
deCODE: 8:98052647:T:C
Proxy SNP: 8:99064613:C:T
deCODE log10(p): 39.4
deCODE BETA: 0.14

1240:1228:1217:1211:1205:114

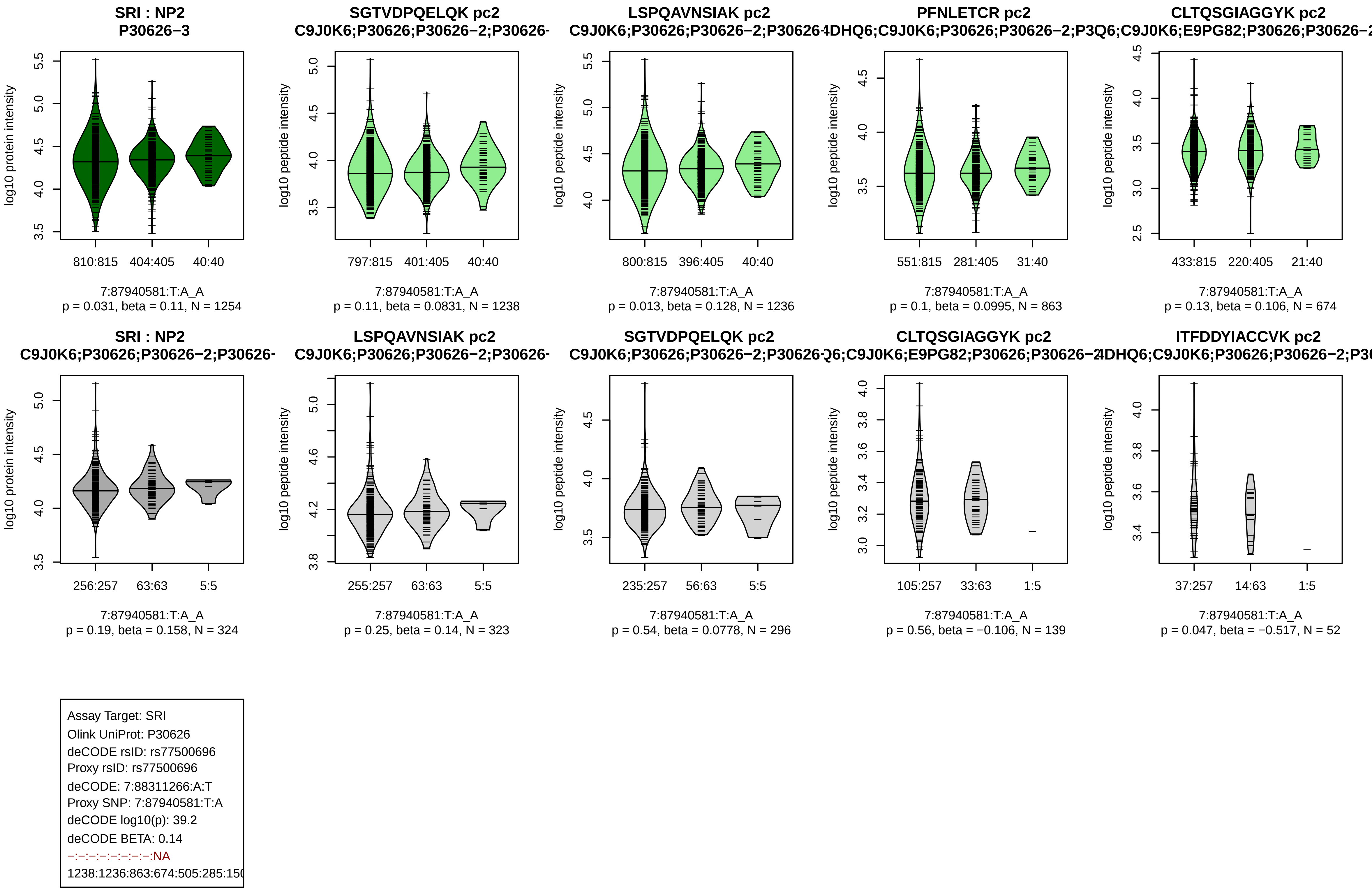
SRI : NP2
P30626−3
log10 protein intensity
810:815
404:405
40:40
7:87940581:T:A_A
p = 0.031, beta = 0.11, N = 1254
SGTVDPQELQK pc2
C9J0K6;P30626;P30626−2;P30626-
log10 peptide intensity
797:815
401:405
40:40
7:87940581:T:A_A
p = 0.11, beta = 0.0831, N = 1238
LSPQAVNSIAK pc2
C9J0K6;P30626;P30626−2;P30626
log10 peptide intensity
800:815
396:405
40:40
7:87940581:T:A_A
p = 0.013, beta = 0.128, N = 1236
PFNLETCR pc2
4DHQ6;C9J0K6;P30626;P30626−2;P3
log10 peptide intensity
551:815
281:405
31:40
7:87940581:T:A_A
p = 0.1, beta = 0.0995, N = 863
CLTQSGIAGGYK pc2
Q6;C9J0K6;E9PG82;P30626;P30626−2
log10 peptide intensity
433:815
220:405
21:40
7:87940581:T:A_A
p = 0.13, beta = 0.106, N = 674
SRI : NP2
C9J0K6;P30626;P30626−2;P30626-
log10 protein intensity
256:257
63:63
5:5
7:87940581:T:A_A
p = 0.19, beta = 0.158, N = 324
LSPQAVNSIAK pc2
C9J0K6;P30626;P30626−2;P30626-
log10 peptide intensity
255:257
63:63
5:5
7:87940581:T:A_A
p = 0.25, beta = 0.14, N = 323
SGTVDPQELQK pc2
C9J0K6;P30626;P30626−2;P30626
log10 peptide intensity
235:257
56:63
5:5
7:87940581:T:A_A
p = 0.54, beta = 0.0778, N = 296
CLTQSGIAGGYK pc2
Q6;C9J0K6;E9PG82;P30626;P30626−2
log10 peptide intensity
105:257
33:63
1:5
7:87940581:T:A_A
p = 0.56, beta = −0.106, N = 139
ITFDDYIACCVK pc2
4DHQ6;C9J0K6;P30626;P30626−2;P3
log10 peptide intensity
37:257
14:63
1:5
7:87940581:T:A_A
p = 0.047, beta = −0.517, N = 52
Assay Target: SRI
Olink UniProt: P30626
deCODE rsID: rs77500696
Proxy rsID: rs77500696
deCODE: 7:88311266:A:T
Proxy SNP: 7:87940581:T:A
deCODE log10(p): 39.2
deCODE BETA: 0.14
−:−:−:−:−:−:−:−:NA
1238:1236:863:674:505:285:150

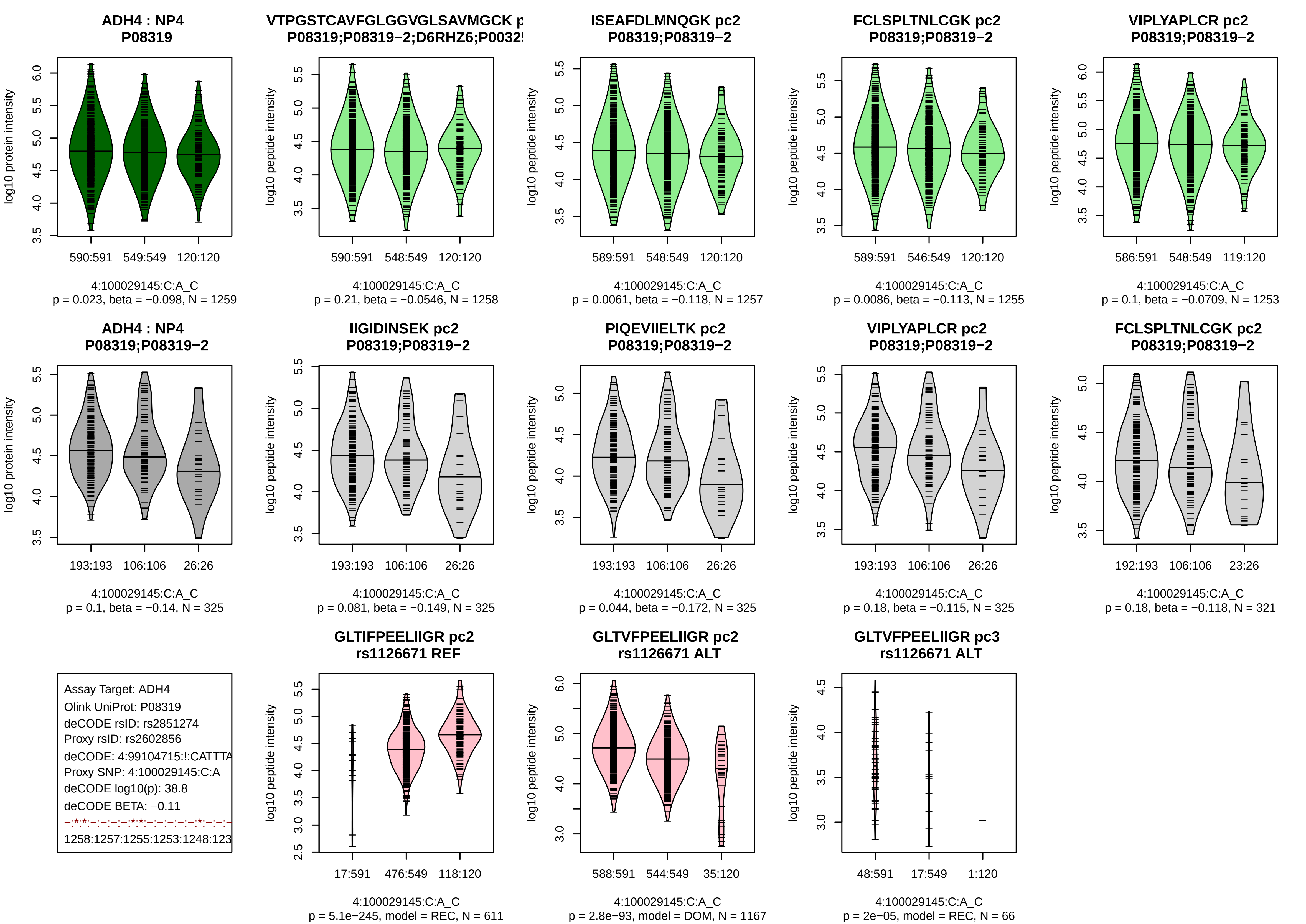
ADH4 : NP4
P08319
log10 protein intensity
590:591
549:549
120:120
4:100029145:C:A_C
p = 0.023, beta = −0.098, N = 1259
VTPGSTCAVFGLGGVGLSAVMGCK p
P08319;P08319−2;D6RHZ6;P0032
log10 peptide intensity
590:591
548:549
120:120
4:100029145:C:A_C
p = 0.21, beta = −0.0546, N = 1258
ISEAFDLMNQGK pc2
P08319;P08319−2
log10 peptide intensity
589:591
548:549
120:120
4:100029145:C:A_C
p = 0.0061, beta = −0.118, N = 1257
FCLSPLTNLCGK pc2
P08319;P08319−2
log10 peptide intensity
589:591
546:549
120:120
4:100029145:C:A_C
p = 0.0086, beta = −0.113, N = 1255
VIPLYAPLCR pc2
P08319;P08319−2
log10 peptide intensity
586:591
548:549
119:120
4:100029145:C:A_C
p = 0.1, beta = −0.0709, N = 1253
ADH4 : NP4
P08319;P08319−2
log10 protein intensity
193:193
106:106
26:26
4:100029145:C:A_C
p = 0.1, beta = −0.14, N = 325
IIGIDINSEK pc2
P08319;P08319−2
log10 peptide intensity
193:193
106:106
26:26
4:100029145:C:A_C
p = 0.081, beta = −0.149, N = 325
PIQEVIIELTK pc2
P08319;P08319−2
log10 peptide intensity
193:193
106:106
26:26
4:100029145:C:A_C
p = 0.044, beta = −0.172, N = 325
VIPLYAPLCR pc2
P08319;P08319−2
log10 peptide intensity
193:193
106:106
26:26
4:100029145:C:A_C
p = 0.18, beta = −0.115, N = 325
FCLSPLTNLCGK pc2
P08319;P08319−2
log10 peptide intensity
192:193
106:106
23:26
4:100029145:C:A_C
p = 0.18, beta = −0.118, N = 321
Assay Target: ADH4
Olink UniProt: P08319
deCODE rsID: rs2851274
Proxy rsID: rs2602856
deCODE: 4:99104715:!:CATTTA
Proxy SNP: 4:100029145:C:A
deCODE log10(p): 38.8
deCODE BETA: −0.11
−:*:*:−:−:−:−:*:*:−:−:−:−:−:*:−:−:−
1258:1257:1255:1253:1248:123
GLTIFPEELIIGR pc2
rs1126671 REF
log10 peptide intensity
17:591
476:549
118:120
4:100029145:C:A_C
p = 5.1e−245, model = REC, N = 611
GLTVFPEELIIGR pc2
rs1126671 ALT
log10 peptide intensity
588:591
544:549
35:120
4:100029145:C:A_C
p = 2.8e−93, model = DOM, N = 1167
GLTVFPEELIIGR pc3
rs1126671 ALT
log10 peptide intensity
48:591
17:549
1:120
4:100029145:C:A_C
p = 2e−05, model = REC, N = 66

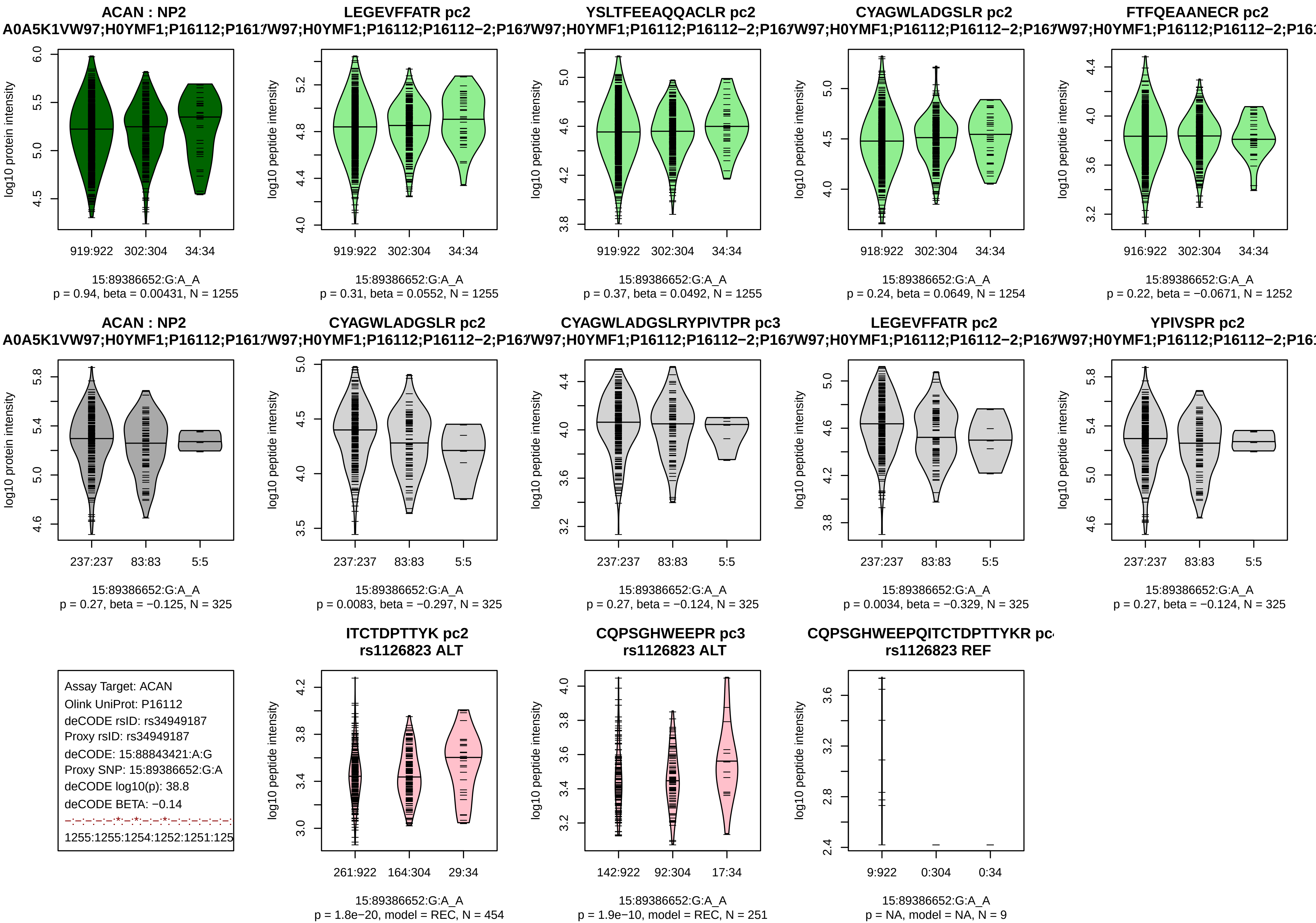
ACAN : NP2
A0A5K1VW97;H0YMF1;P16112;P161
log10 protein intensity
919:922 302:304 34:34
15:89386652:G:A_A
p = 0.94, beta = 0.00431, N = 1255
LEGEVFFATR pc2
W97;H0YMF1;P16112;P16112−2;P16
log10 peptide intensity
919:922 302:304 34:34
15:89386652:G:A_A
p = 0.31, beta = 0.0552, N = 1255
YSLTFEEAQQACLR pc2
W97;H0YMF1;P16112;P16112−2;P16
log10 peptide intensity
919:922 302:304 34:34
15:89386652:G:A_A
p = 0.37, beta = 0.0492, N = 1255
CYAGWLADGSLR pc2
W97;H0YMF1;P16112;P16112−2;P16
log10 peptide intensity
918:922 302:304 34:34
15:89386652:G:A_A
p = 0.24, beta = 0.0649, N = 1254
FTFQEAANECR pc2
W97;H0YMF1;P16112;P16112−2;P16
log10 peptide intensity
916:922 302:304 34:34
15:89386652:G:A_A
p = 0.22, beta = −0.0671, N = 1252
ACAN : NP2
A0A5K1VW97;H0YMF1;P16112;P161
log10 protein intensity
237:237 83:83 5:5
15:89386652:G:A_A
p = 0.27, beta = −0.125, N = 325
CYAGWLADGSLR pc2
W97;H0YMF1;P16112;P16112−2;P16
log10 peptide intensity
237:237 83:83 5:5
15:89386652:G:A_A
p = 0.0083, beta = −0.297, N = 325
CYAGWLADGSLRYPIVTPR pc3
W97;H0YMF1;P16112;P16112−2;P16
log10 peptide intensity
237:237 83:83 5:5
15:89386652:G:A_A
p = 0.27, beta = −0.124, N = 325
LEGEVFFATR pc2
W97;H0YMF1;P16112;P16112−2;P16
log10 peptide intensity
237:237 83:83 5:5
15:89386652:G:A_A
p = 0.0034, beta = −0.329, N = 325
YPIVSPR pc2
W97;H0YMF1;P16112;P16112−2;P16
log10 peptide intensity
237:237 83:83 5:5
15:89386652:G:A_A
p = 0.27, beta = −0.124, N = 325
Assay Target: ACAN
Olink UniProt: P16112
deCODE rsID: rs34949187
Proxy rsID: rs34949187
deCODE: 15:88843421:A:G
Proxy SNP: 15:89386652:G:A
deCODE log10(p): 38.8
deCODE BETA: −0.14
−:−:−:−:−:*:−:*:−:−:*:−:−:−:−:−:−:
1255:1255:1254:1252:1251:125
ITCTDPTTYK pc2
rs1126823 ALT
log10 peptide intensity
261:922 164:304 29:34
15:89386652:G:A_A
p = 1.8e−20, model = REC, N = 454
CQPSGHWEEPR pc3
rs1126823 ALT
log10 peptide intensity
142:922 92:304 17:34
15:89386652:G:A_A
p = 1.9e−10, model = REC, N = 251
CQPSGHWEEPQITCTDPTTYKR pc
rs1126823 REF
log10 peptide intensity
9:922 0:304 0:34
15:89386652:G:A_A
p = NA, model = NA, N = 9

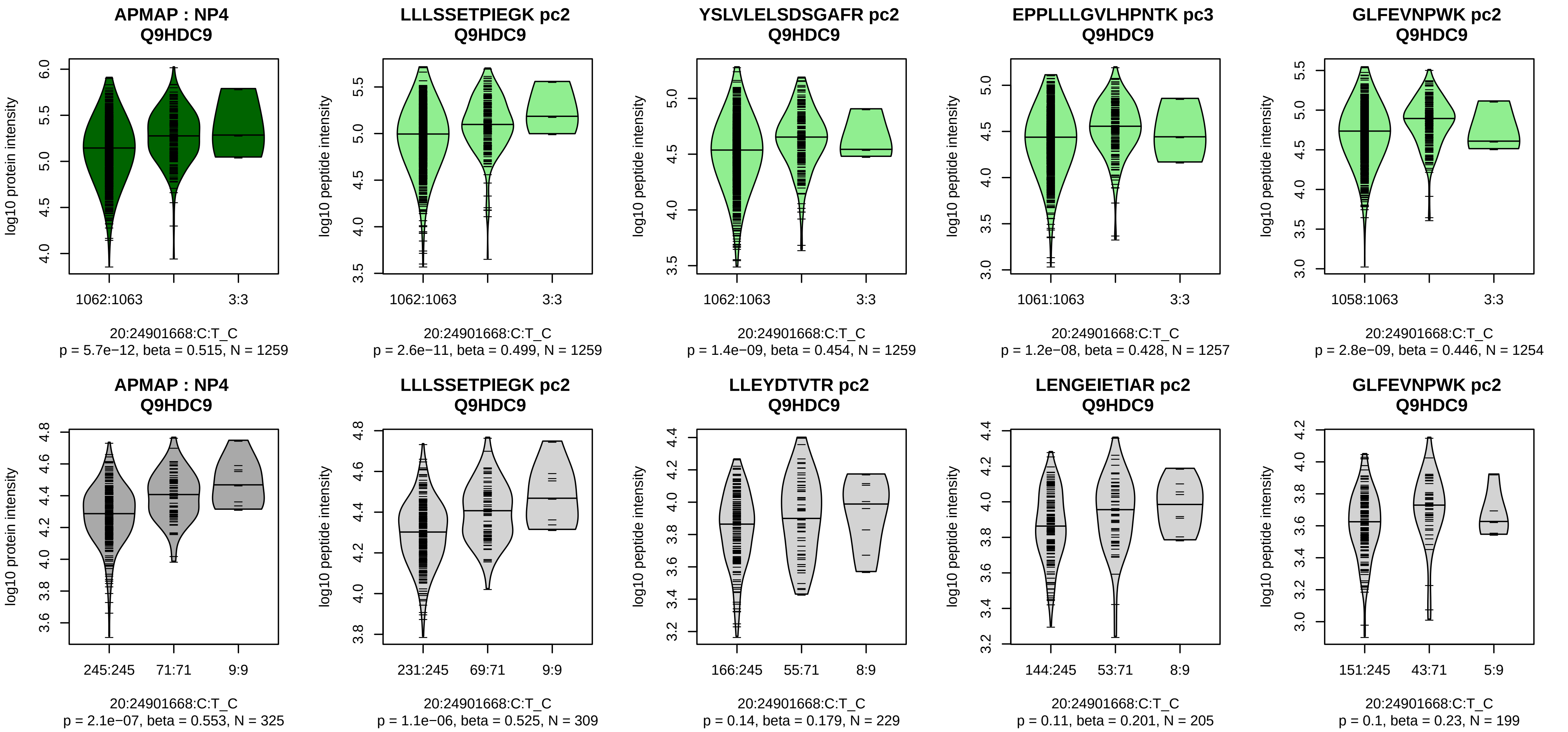


Assay Target: APMAP
Olink UniProt: Q9HDC9
deCODE rsID: rs6036977
Proxy rsID: rs6036977
deCODE: 20:24921032:C:T
Proxy SNP: 20:24901668:C:T
deCODE log10(p): 37.6
deCODE BETA: 0.16
*:*:*:*:*:*:*:*:*:*:*:*:*:*:*:*:–:*:–:*:–:
1259:1259:1257:1254:1250:124

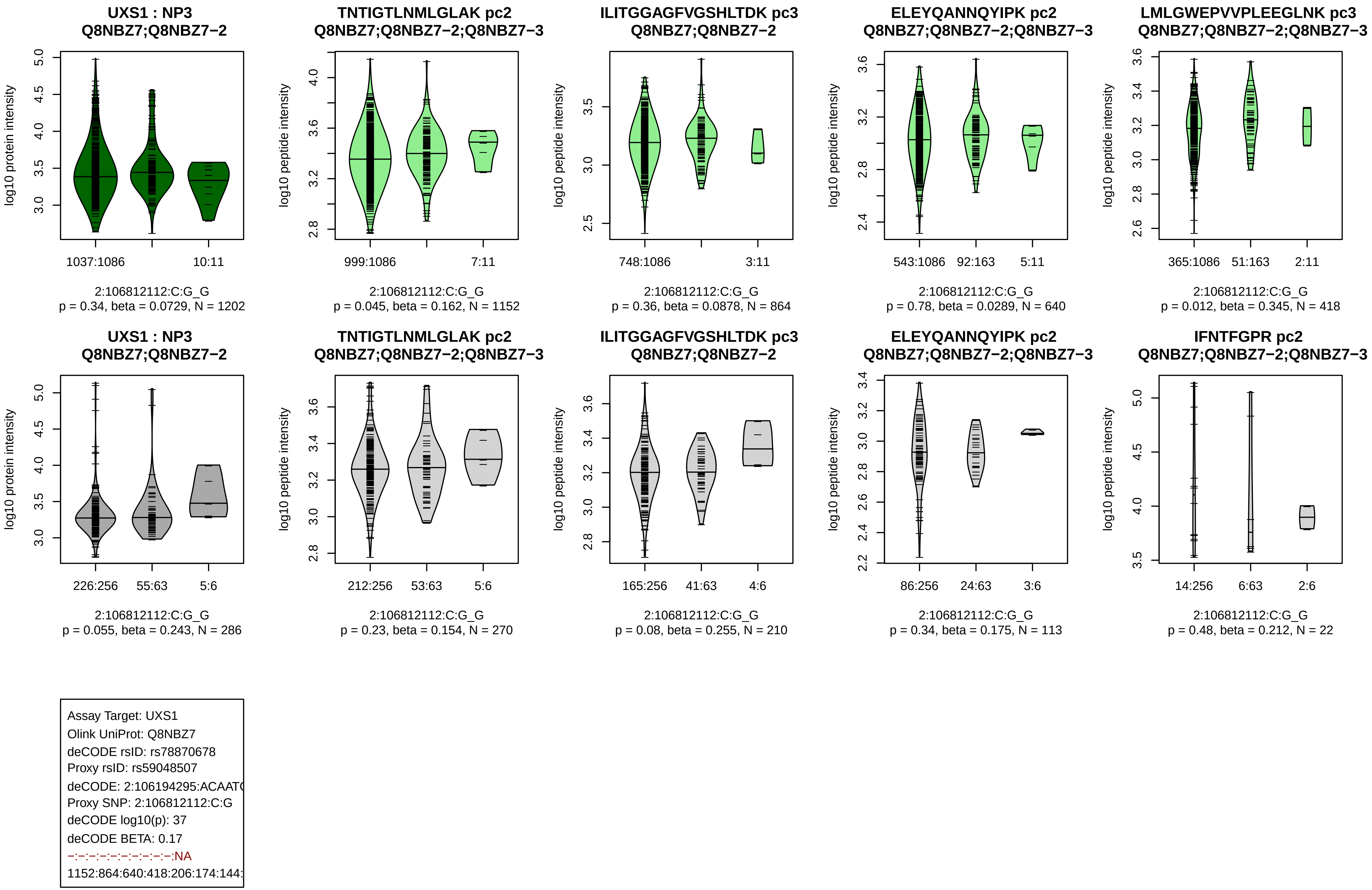
UXS1 : NP3
Q8NBZ7;Q8NBZ7−2
log10 protein intensity
1037:1086
10:11
2:106812112:C:G_G
p = 0.34, beta = 0.0729, N = 1202
TNTIGTLNMLGLAK pc2
Q8NBZ7;Q8NBZ7−2;Q8NBZ7−3
log10 peptide intensity
999:1086
7:11
2:106812112:C:G_G
p = 0.045, beta = 0.162, N = 1152
ILITGGAGFVGSHLTDK pc3
Q8NBZ7;Q8NBZ7−2
log10 peptide intensity
748:1086
3:11
2:106812112:C:G_G
p = 0.36, beta = 0.0878, N = 864
ELEYQANNQYIPK pc2
Q8NBZ7;Q8NBZ7−2;Q8NBZ7−3
log10 peptide intensity
543:1086
92:163
5:11
2:106812112:C:G_G
p = 0.78, beta = 0.0289, N = 640
LMLGWEPVVPLEEGLNK pc3
Q8NBZ7;Q8NBZ7−2;Q8NBZ7−3
log10 peptide intensity
365:1086
51:163
2:11
2:106812112:C:G_G
p = 0.012, beta = 0.345, N = 418
UXS1 : NP3
Q8NBZ7;Q8NBZ7−2
log10 protein intensity
226:256
55:63
5:6
2:106812112:C:G_G
p = 0.055, beta = 0.243, N = 286
TNTIGTLNMLGLAK pc2
Q8NBZ7;Q8NBZ7−2;Q8NBZ7−3
log10 peptide intensity
212:256
53:63
5:6
2:106812112:C:G_G
p = 0.23, beta = 0.154, N = 270
ILITGGAGFVGSHLTDK pc3
Q8NBZ7;Q8NBZ7−2
log10 peptide intensity
165:256
41:63
4:6
2:106812112:C:G_G
p = 0.08, beta = 0.255, N = 210
ELEYQANNQYIPK pc2
Q8NBZ7;Q8NBZ7−2;Q8NBZ7−3
log10 peptide intensity
86:256
24:63
3:6
2:106812112:C:G_G
p = 0.34, beta = 0.175, N = 113
IFNTFGPR pc2
Q8NBZ7;Q8NBZ7−2;Q8NBZ7−3
log10 peptide intensity
14:256
6:63
2:6
2:106812112:C:G_G
p = 0.48, beta = 0.212, N = 22
Assay Target: UXS1
Olink UniProt: Q8NBZ7
deCODE rsID: rs78870678
Proxy rsID: rs59048507
deCODE: 2:106194295:ACAAT
Proxy SNP: 2:106812112:C:G
deCODE log10(p): 37
deCODE BETA: 0.17
−:−:−:−:−:−:−:−:−:−:NA
1152:864:640:418:206:174:144:

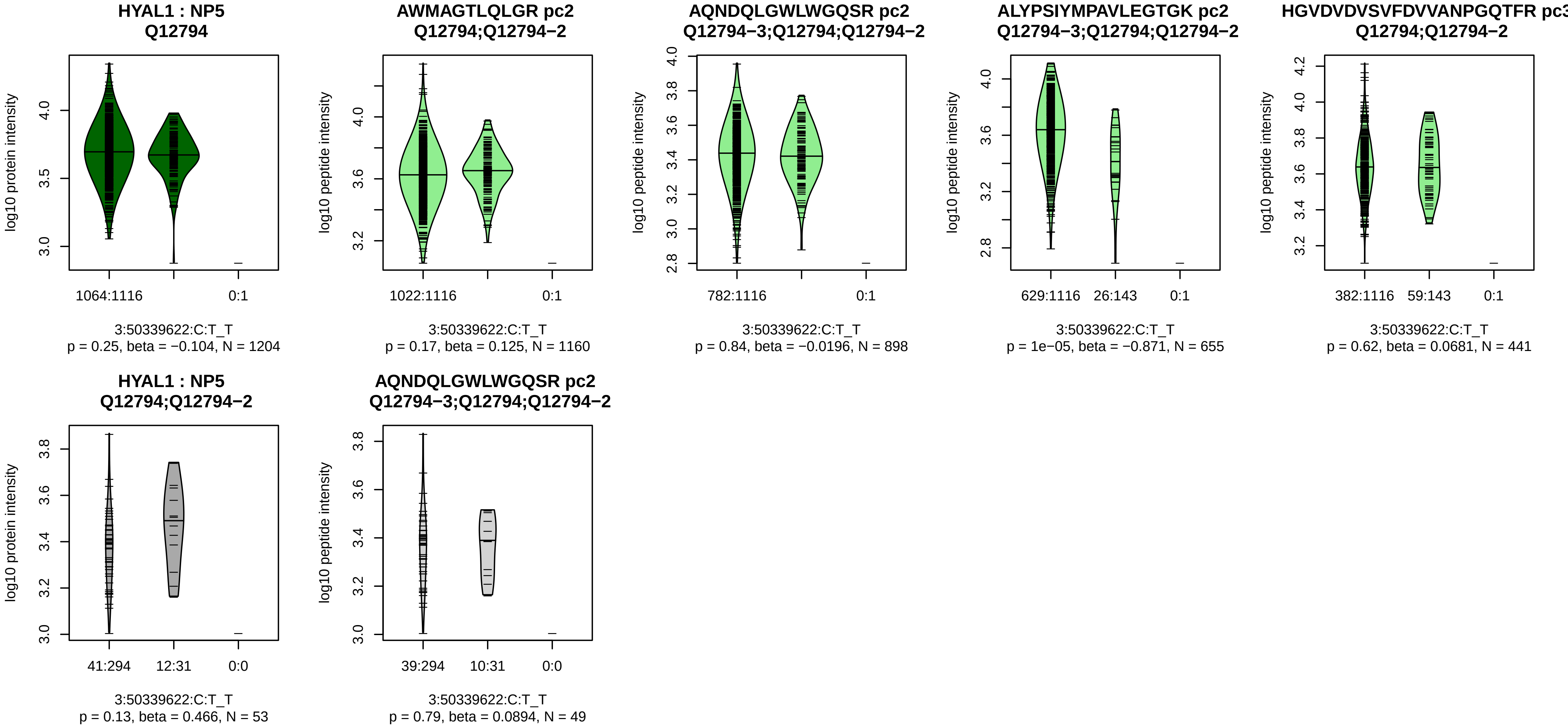


Assay Target: HYAL1
Olink UniProt: Q12794
deCODE rsID: rs116482870
Proxy rsID: rs116482870
deCODE: 3:50302191:T:C
Proxy SNP: 3:50339622:C:T
deCODE log10(p): 35.8
deCODE BETA: −0.24
−:−:*:−:−
1160:898:655:441:146

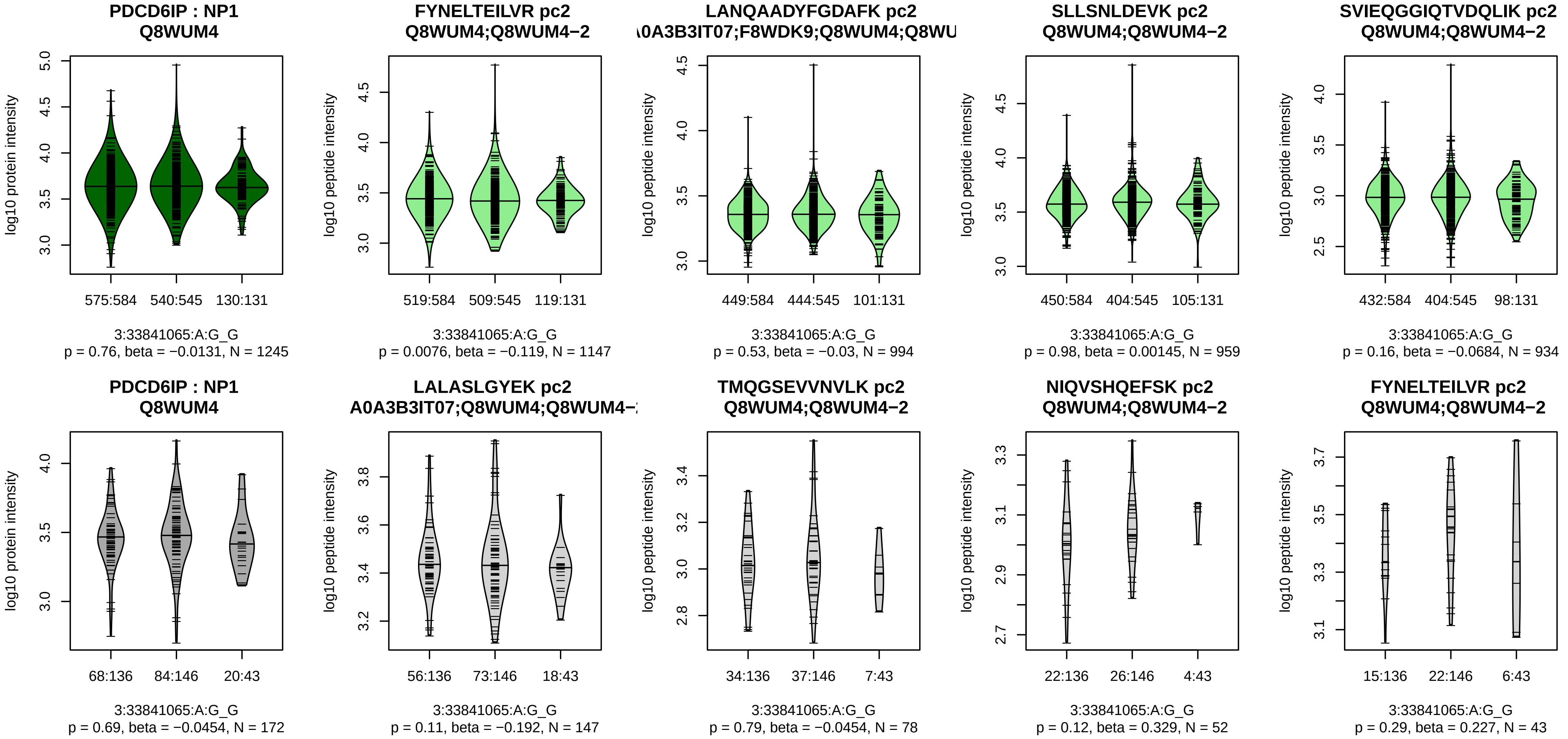


Assay Target: PDCD6IP
Olink UniProt: Q8WUM4
deCODE rsID: rs9876986
Proxy rsID: rs9876986
deCODE: 3:33799573:G:A
Proxy SNP: 3:33841065:A:G
deCODE log10(p): 34.2
deCODE BETA: −0.11
*:−:−:−:*:−:−:−:−:−:−:−:−:−:−:−:−:−:
1147:994:959:934:926:895:740:

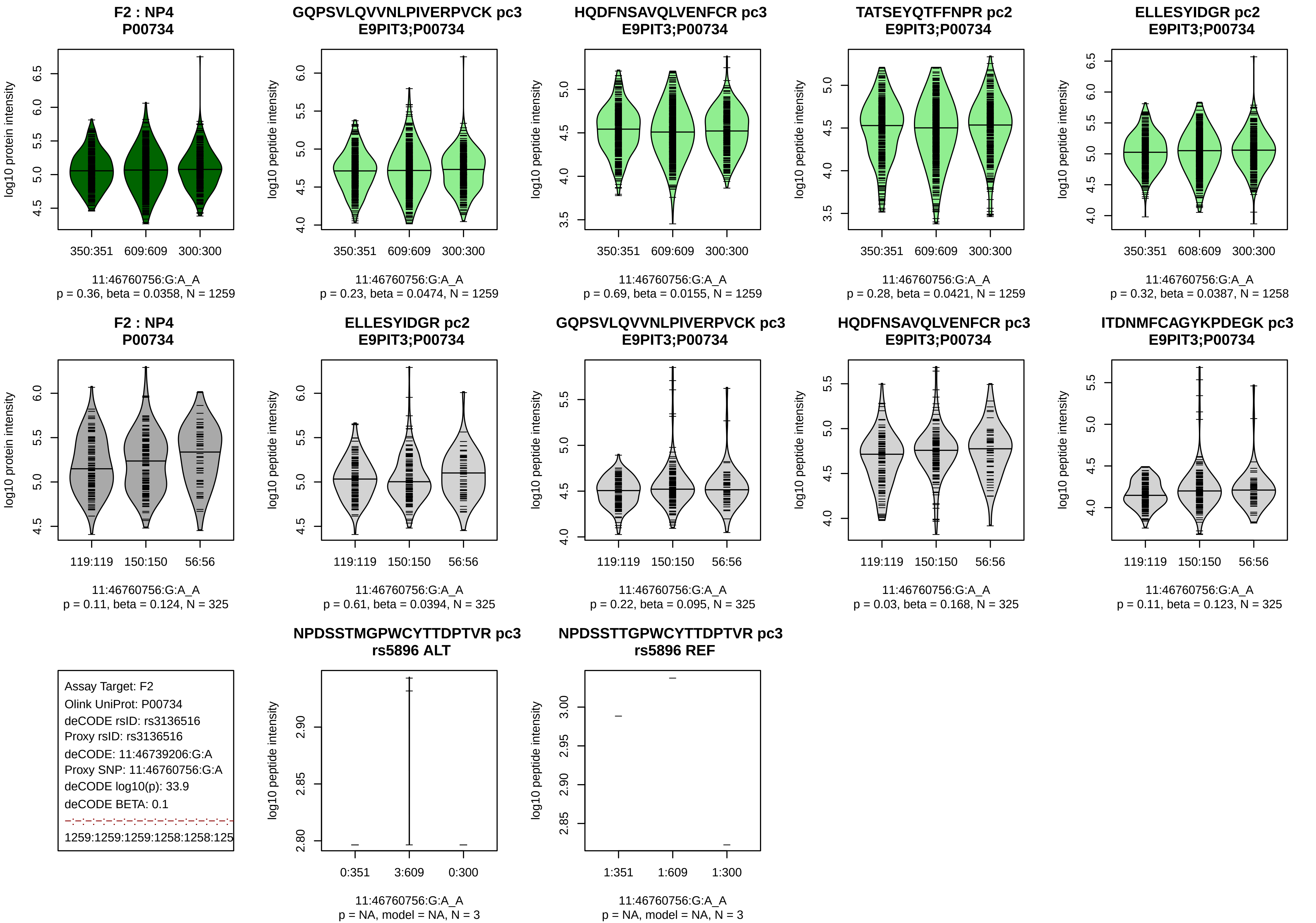

F2 : NP4
P00734
log10 protein intensity
350:351 609:609 300:300
11:46760756:G:A_A
p = 0.36, beta = 0.0358, N = 1259
GQPSVLQVVNLPIVERPVCK pc3
E9PIT3;P00734
log10 peptide intensity
350:351 609:609 300:300
11:46760756:G:A_A
p = 0.23, beta = 0.0474, N = 1259
HQDFNSAVQLVENFCR pc3
E9PIT3;P00734
log10 peptide intensity
350:351 609:609 300:300
11:46760756:G:A_A
p = 0.69, beta = 0.0155, N = 1259
TATSEYQTFFNPR pc2
E9PIT3;P00734
log10 peptide intensity
350:351 609:609 300:300
11:46760756:G:A_A
p = 0.28, beta = 0.0421, N = 1259
ELLESYIDGR pc2
E9PIT3;P00734
log10 peptide intensity
350:351 608:609 300:300
11:46760756:G:A_A
p = 0.32, beta = 0.0387, N = 1258
F2 : NP4
P00734
log10 protein intensity
119:119 150:150 56:56
11:46760756:G:A_A
p = 0.11, beta = 0.124, N = 325
ELLESYIDGR pc2
E9PIT3;P00734
log10 peptide intensity
119:119 150:150 56:56
11:46760756:G:A_A
p = 0.61, beta = 0.0394, N = 325
GQPSVLQVVNLPIVERPVCK pc3
E9PIT3;P00734
log10 peptide intensity
119:119 150:150 56:56
11:46760756:G:A_A
p = 0.22, beta = 0.095, N = 325
HQDFNSAVQLVENFCR pc3
E9PIT3;P00734
log10 peptide intensity
119:119 150:150 56:56
11:46760756:G:A_A
p = 0.03, beta = 0.168, N = 325
ITDNMFCAGYKPDEGK pc3
E9PIT3;P00734
log10 peptide intensity
119:119 150:150 56:56
11:46760756:G:A_A
p = 0.11, beta = 0.123, N = 325
Assay Target: F2
Olink UniProt: P00734
deCODE rsID: rs3136516
Proxy rsID: rs3136516
deCODE: 11:46739206:G:A
Proxy SNP: 11:46760756:G:A
deCODE log10(p): 33.9
deCODE BETA: 0.1
1259:1259:1259:1258:1258:125
NPDSSTMGPWCYTTDPTVR pc3
rs5896 ALT
log10 peptide intensity
0:351 3:609 0:300
11:46760756:G:A_A
p = NA, model = NA, N = 3
NPDSSTTGPWCYTTDPTVR pc3
rs5896 REF
log10 peptide intensity
1:351 1:609 1:300
11:46760756:G:A_A
p = NA, model = NA, N = 3

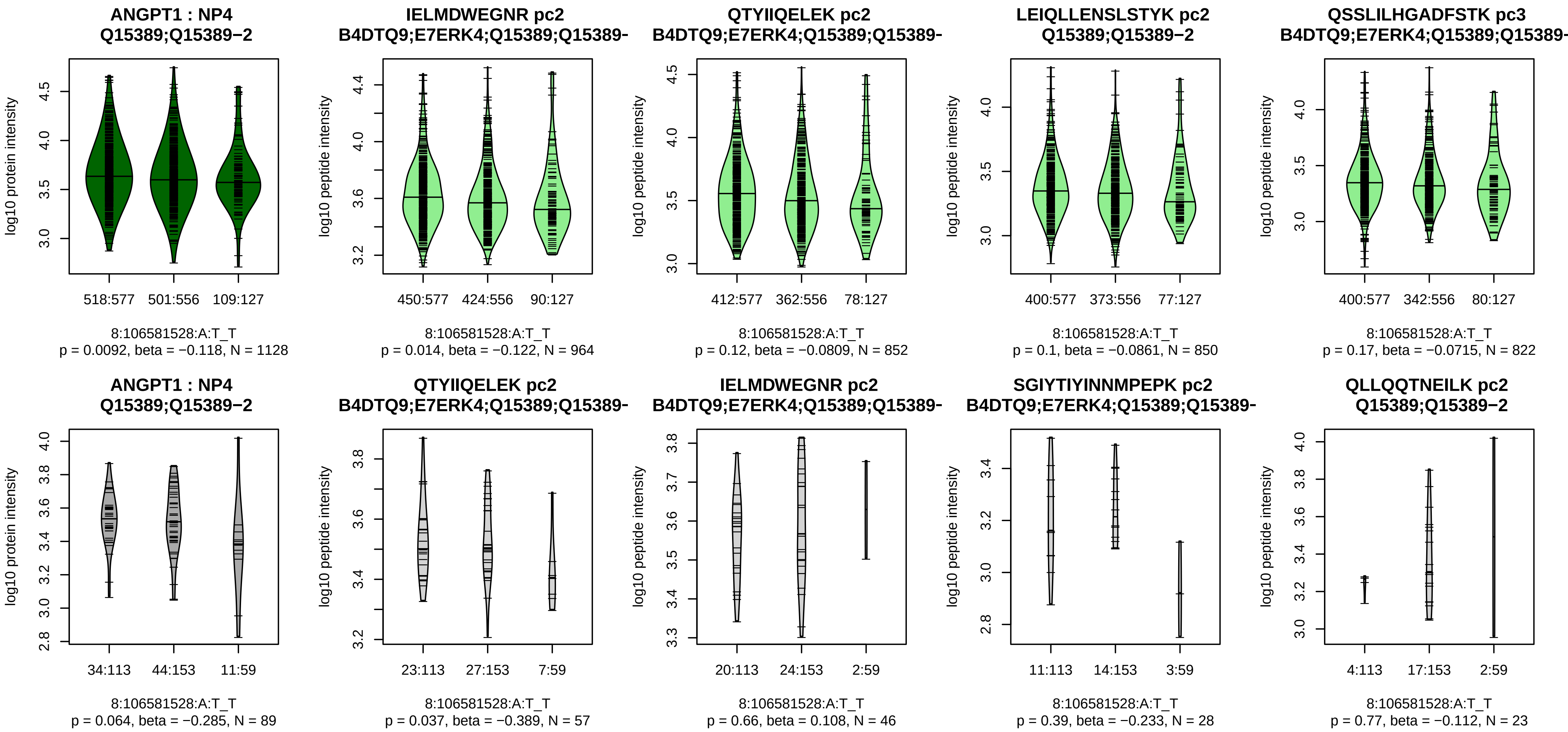


Assay Target: ANGPT1
Olink UniProt: Q15389
deCODE rsID: rs6993770
Proxy rsID: rs6993770
deCODE: 8:105569300:T:A
Proxy SNP: 8:106581528:A:T
deCODE log10(p): 31.9
deCODE BETA: −0.12
−:−:−:−:−:−:−:−:−:−:−:−:−:−:−:−:
964:852:850:822:732:624:507:4

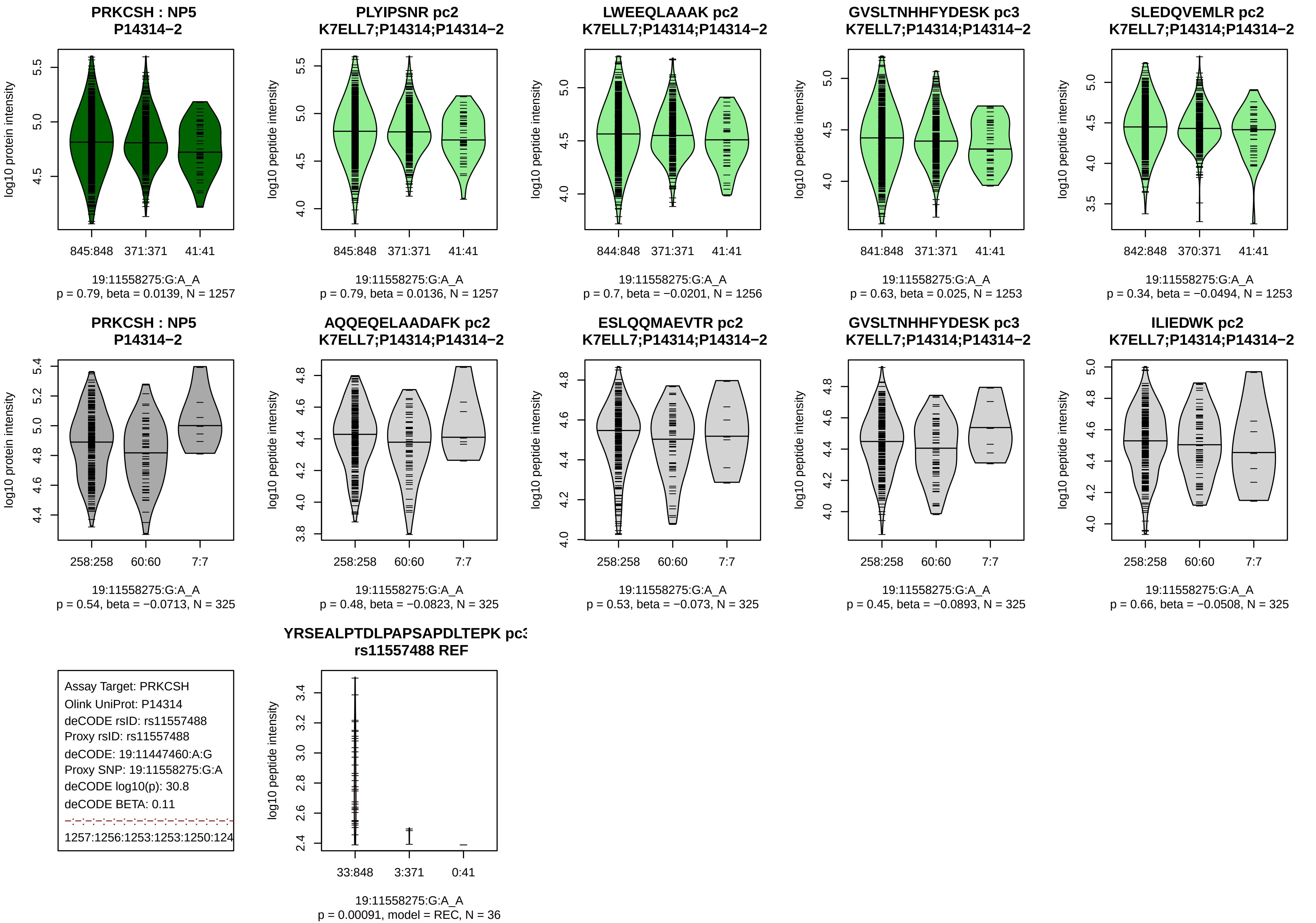
PRKCSH : NP5
P14314−2
log10 protein intensity
845:848
371:371
41:41
19:11558275:G:A_A
p = 0.79, beta = 0.0139, N = 1257
PLYIPSNR pc2
K7ELL7;P14314;P14314−2
log10 peptide intensity
845:848
371:371
41:41
19:11558275:G:A_A
p = 0.79, beta = 0.0136, N = 1257
LWEEQLAAAK pc2
K7ELL7;P14314;P14314−2
log10 peptide intensity
844:848
371:371
41:41
19:11558275:G:A_A
p = 0.7, beta = −0.0201, N = 1256
GVSLTNHHFYDESK pc3
K7ELL7;P14314;P14314−2
log10 peptide intensity
841:848
371:371
41:41
19:11558275:G:A_A
p = 0.63, beta = 0.025, N = 1253
SLEDQVEMLR pc2
K7ELL7;P14314;P14314−2
log10 peptide intensity
842:848
370:371
41:41
19:11558275:G:A_A
p = 0.34, beta = −0.0494, N = 1253
PRKCSH : NP5
P14314−2
log10 protein intensity
258:258
60:60
7:7
19:11558275:G:A_A
p = 0.54, beta = −0.0713, N = 325
AQQEQELAADAFK pc2
K7ELL7;P14314;P14314−2
log10 peptide intensity
258:258
60:60
7:7
19:11558275:G:A_A
p = 0.48, beta = −0.0823, N = 325
ESLQQMAEVTR pc2
K7ELL7;P14314;P14314−2
log10 peptide intensity
258:258
60:60
7:7
19:11558275:G:A_A
p = 0.53, beta = −0.073, N = 325
GVSLTNHHFYDESK pc3
K7ELL7;P14314;P14314−2
log10 peptide intensity
258:258
60:60
7:7
19:11558275:G:A_A
p = 0.45, beta = −0.0893, N = 325
ILIEDWK pc2
K7ELL7;P14314;P14314−2
log10 peptide intensity
258:258
60:60
7:7
19:11558275:G:A_A
p = 0.66, beta = −0.0508, N = 325
Assay Target: PRKCSH
Olink UniProt: P14314
deCODE rsID: rs11557488
Proxy rsID: rs11557488
deCODE: 19:11447460:A:G
Proxy SNP: 19:11558275:G:A
deCODE log10(p): 30.8
deCODE BETA: 0.11
1257:1256:1253:1253:1250:124
YRSEALPTDLPAPSAPDLTEPK pc3
rs11557488 REF
log10 peptide intensity
33:848
3:371
0:41
19:11558275:G:A_A
p = 0.00091, model = REC, N = 36

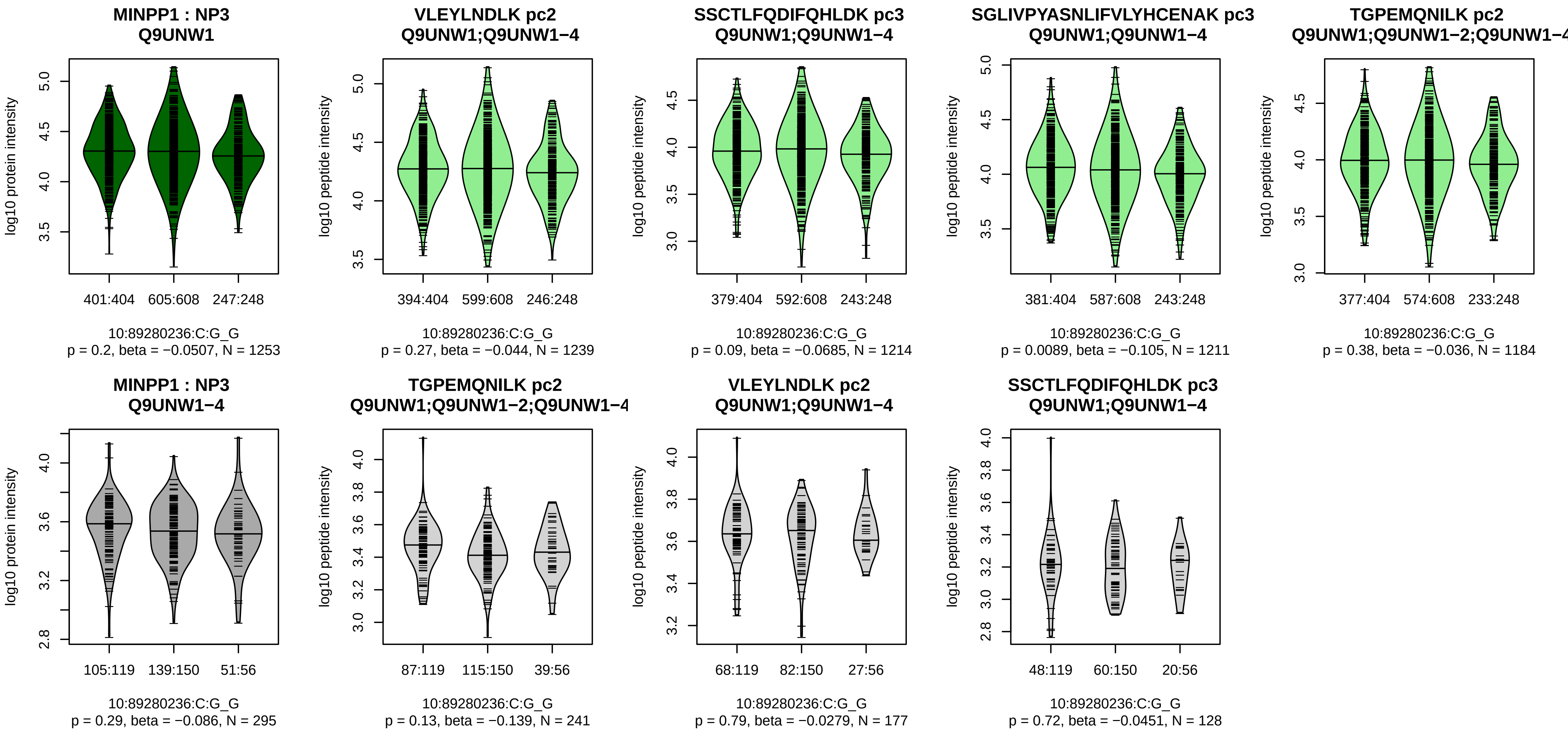


Assay Target: MINPP1
Olink UniProt: Q9UNW1
deCODE rsID: rs59980852
Proxy rsID: rs12356542
deCODE: 10:87515989:T:C
Proxy SNP: 10:89280236:C:G
deCODE log10(p): 30.2
deCODE BETA: −0.1
−:−:*:−:−:−:−:−:−:−:−:−:−:−
1239:1214:1211:1184:1174:987

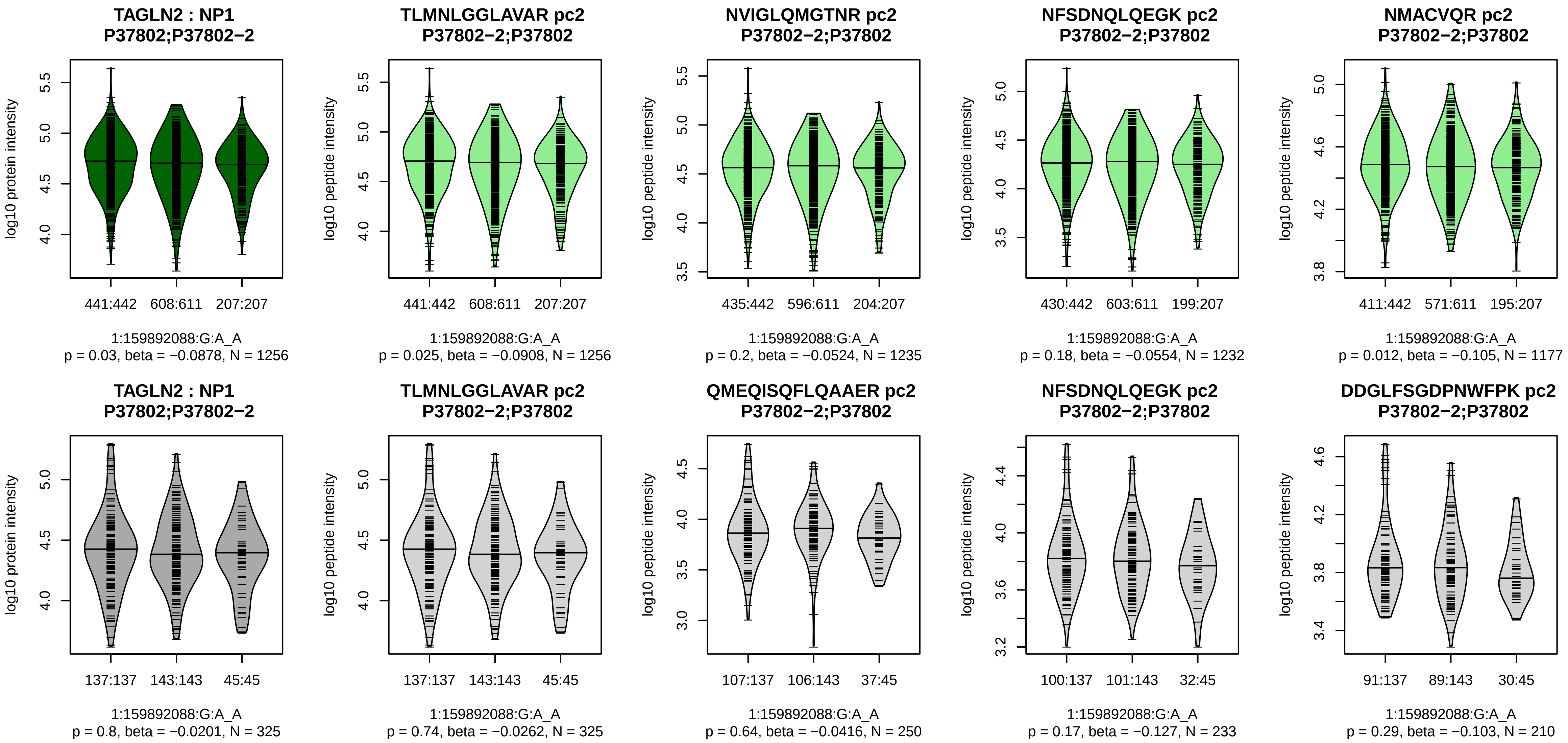


Assay Target: TAGLN2
Olink UniProt: P37802
deCODE rsID: rs2789422
Proxy rsID: rs2789422
deCODE: 1:159922298:A:G
Proxy SNP: 1:159892088:G:A
deCODE log10(p): 28.7
deCODE BETA: −0.09
−:−:−:−:−:−:*:−:−:−:−:−:*:−:−:−:N
1256:1235:1232:1177:1149:112

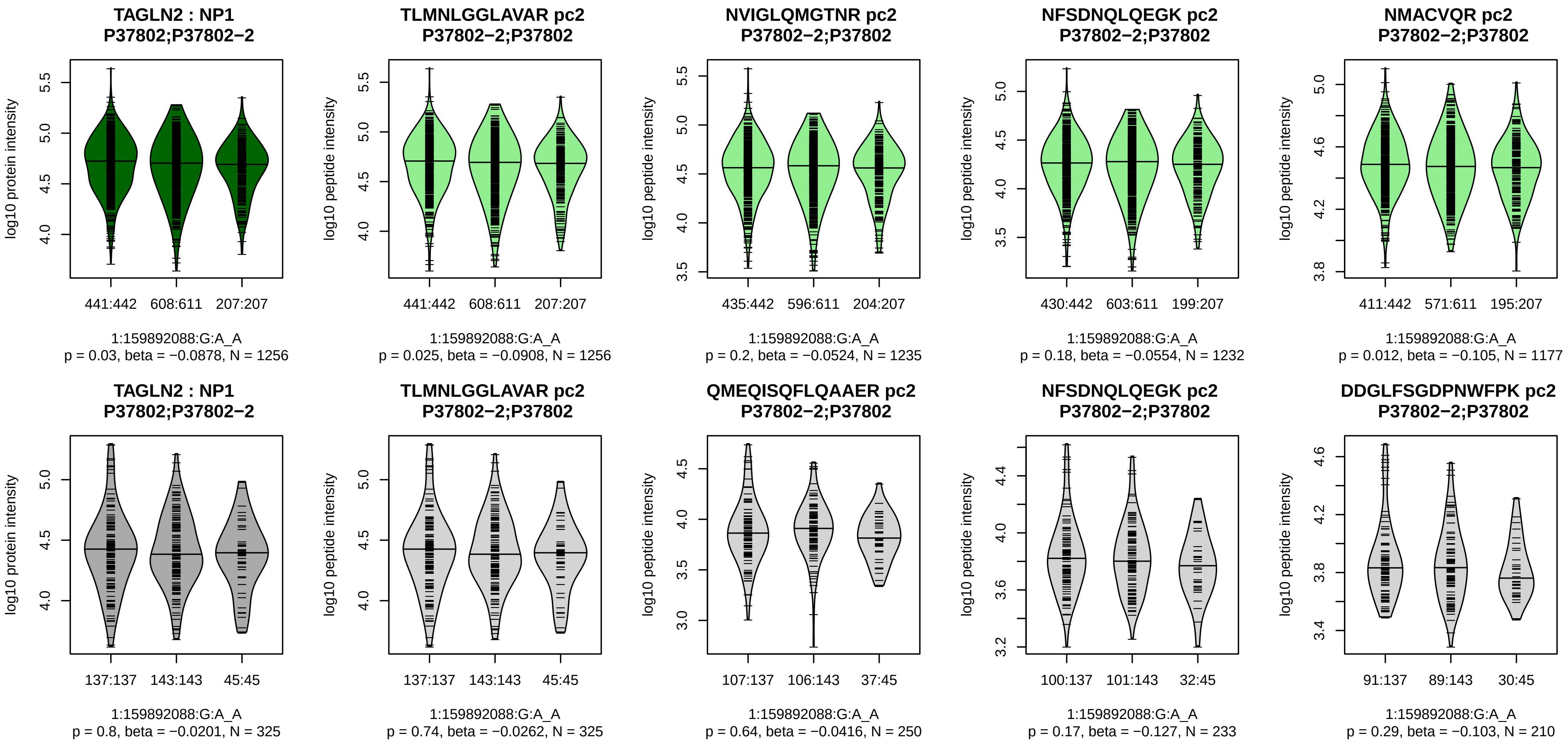


Assay Target: TAGLN2
Olink UniProt: P37802
deCODE rsID: rs2789422
Proxy rsID: rs2789422
deCODE: 1:159922298:A:G
Proxy SNP: 1:159892088:G:A
deCODE log10(p): 28.6
deCODE BETA: −0.09
−:−:−:−:−:−:*:−:−:−:−:−:*:−:−:−:N
1256:1235:1232:1177:1149:112

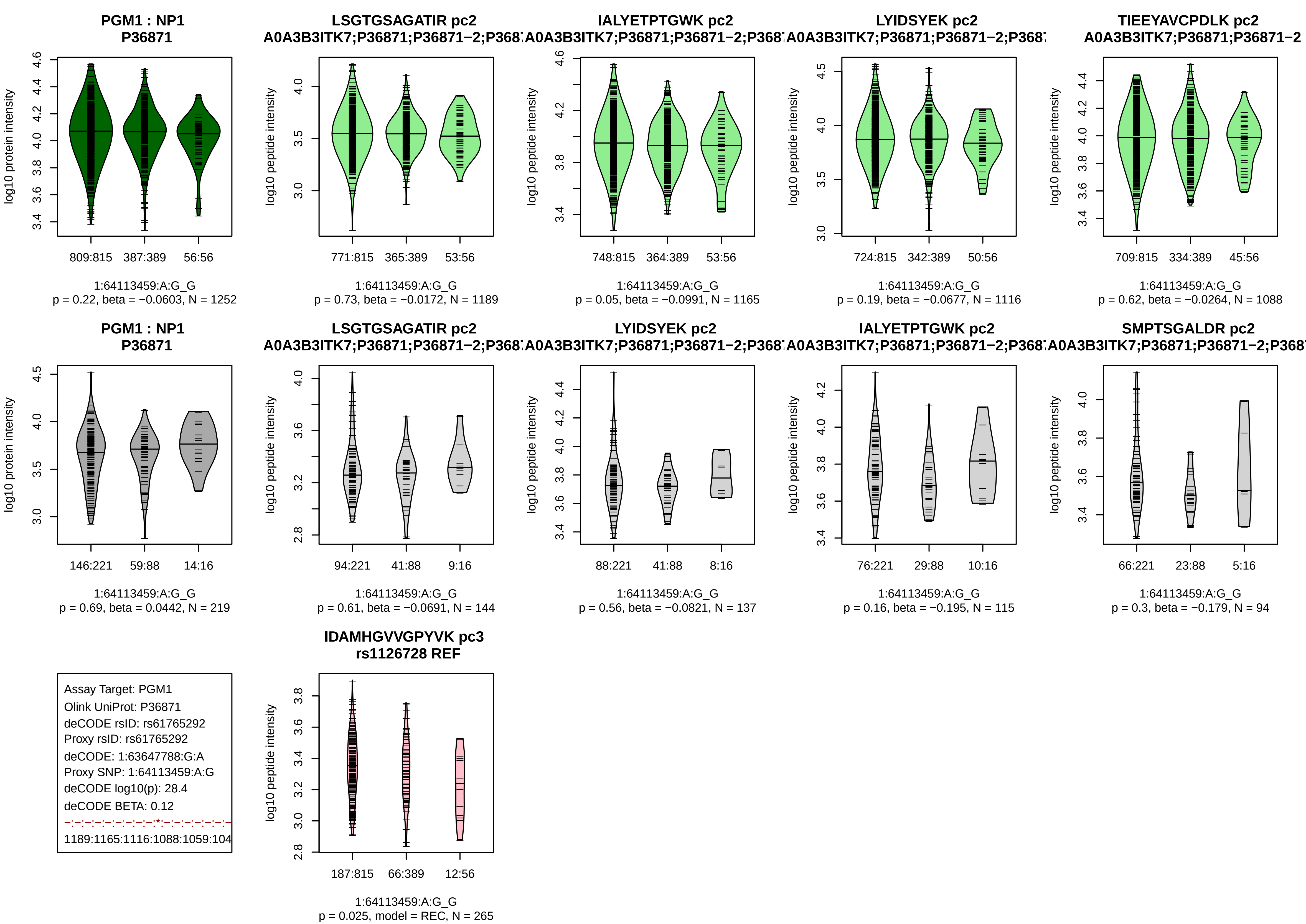

PGM1 : NP1
P36871
log10 protein intensity
809:815
387:389
56:56
1:64113459:A:G_G
p = 0.22, beta = −0.0603, N = 1252
LSGTGSAGATIR pc2
A0A3B3ITK7;P36871;P36871−2;P368
log10 peptide intensity
771:815
365:389
53:56
1:64113459:A:G_G
p = 0.73, beta = −0.0172, N = 1189
IALYETPTGWK pc2
A0A3B3ITK7;P36871;P36871−2;P368
log10 peptide intensity
748:815
364:389
53:56
1:64113459:A:G_G
p = 0.05, beta = −0.0991, N = 1165
LYIDSYEK pc2
A0A3B3ITK7;P36871;P36871−2;P368
log10 peptide intensity
724:815
342:389
50:56
1:64113459:A:G_G
p = 0.19, beta = −0.0677, N = 1116
TIEEYAVCPDLK pc2
A0A3B3ITK7;P36871;P36871−2
log10 peptide intensity
709:815
334:389
45:56
1:64113459:A:G_G
p = 0.62, beta = −0.0264, N = 1088
PGM1 : NP1
P36871
log10 protein intensity
146:221
59:88
14:16
1:64113459:A:G_G
p = 0.69, beta = 0.0442, N = 219
LSGTGSAGATIR pc2
A0A3B3ITK7;P36871;P36871−2;P368
log10 peptide intensity
94:221
41:88
9:16
1:64113459:A:G_G
p = 0.61, beta = −0.0691, N = 144
LYIDSYEK pc2
A0A3B3ITK7;P36871;P36871−2;P368
log10 peptide intensity
88:221
41:88
8:16
1:64113459:A:G_G
p = 0.56, beta = −0.0821, N = 137
IALYETPTGWK pc2
A0A3B3ITK7;P36871;P36871−2;P368
log10 peptide intensity
76:221
29:88
10:16
1:64113459:A:G_G
p = 0.16, beta = −0.195, N = 115
SMPTSGALDR pc2
A0A3B3ITK7;P36871;P36871−2;P368
log10 peptide intensity
66:221
23:88
5:16
1:64113459:A:G_G
p = 0.3, beta = −0.179, N = 94
Assay Target: PGM1
Olink UniProt: P36871
deCODE rsID: rs61765292
Proxy rsID: rs61765292
deCODE: 1:63647788:G:A
Proxy SNP: 1:64113459:A:G
deCODE log10(p): 28.4
deCODE BETA: 0.12
−:−:−:−:−:−:−:−:*:−:−:−:−:−:−:−
1189:1165:1116:1088:1059:104
IDAMHGVVGPYVK pc3
rs1126728 REF
log10 peptide intensity
187:815
66:389
12:56
1:64113459:A:G_G
p = 0.025, model = REC, N = 265

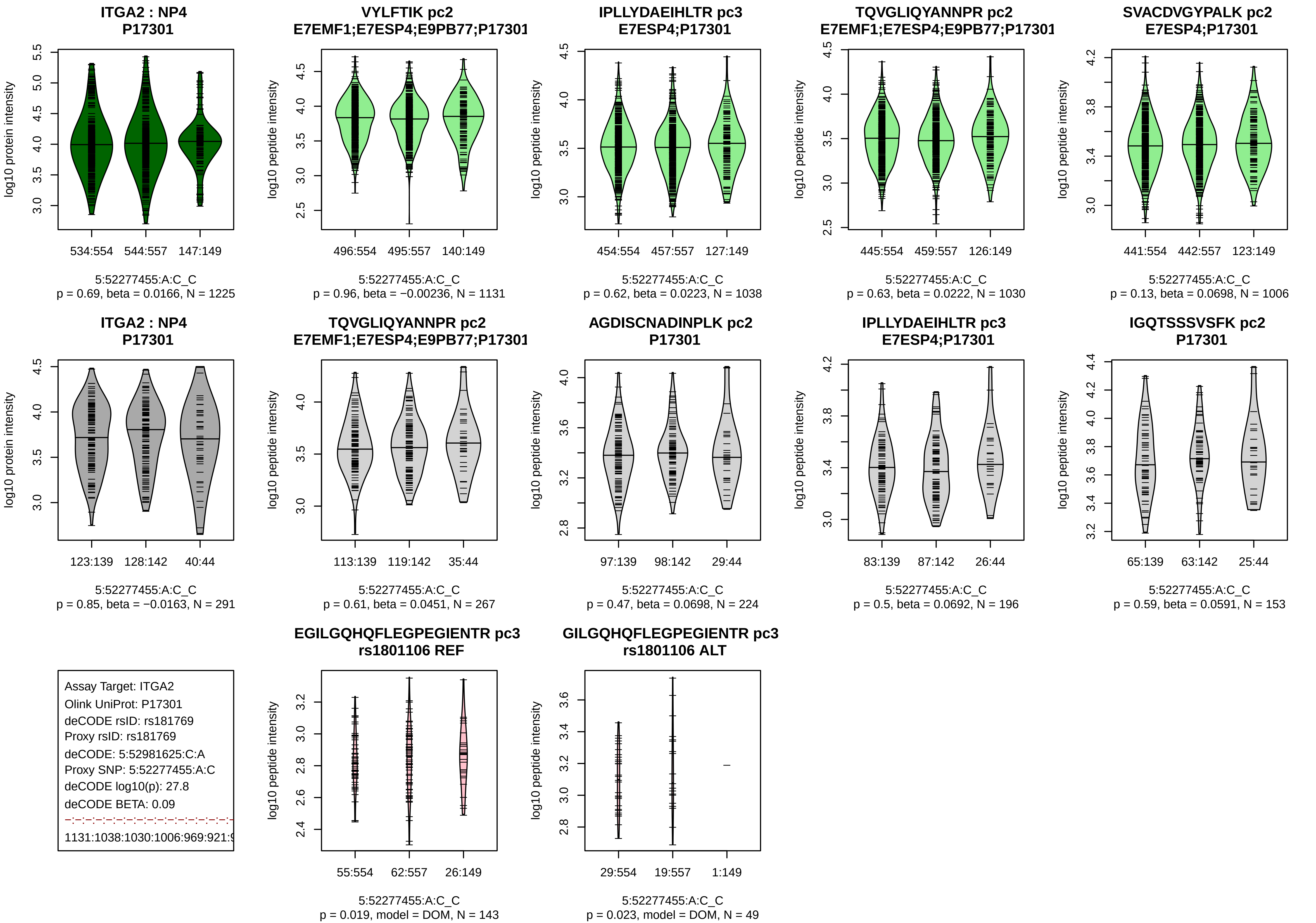

ITGA2 : NP4
P17301
log10 protein intensity
534:554
544:557
147:149
5:52277455:A:C_C
p = 0.69, beta = 0.0166, N = 1225
VYLFTIK pc2
E7EMF1;E7ESP4;E9PB77;P17301
log10 peptide intensity
496:554
495:557
140:149
5:52277455:A:C_C
p = 0.96, beta = −0.00236, N = 1131
IPLLYDAEIHLTR pc3
E7ESP4;P17301
454:554
457:557
127:149
5:52277455:A:C_C
p = 0.62, beta = 0.0223, N = 1038
TQVGLIQYANNPR pc2
E7EMF1;E7ESP4;E9PB77;P17301
445:554
459:557
126:149
5:52277455:A:C_C
p = 0.63, beta = 0.0222, N = 1030
SVACDVGYPALK pc2
E7ESP4;P17301
441:554
442:557
123:149
5:52277455:A:C_C
p = 0.13, beta = 0.0698, N = 1006
ITGA2 : NP4
P17301
123:139
128:142
40:44
5:52277455:A:C_C
p = 0.85, beta = −0.0163, N = 291
TQVGLIQYANNPR pc2
E7EMF1;E7ESP4;E9PB77;P17301
113:139
119:142
35:44
5:52277455:A:C_C
p = 0.61, beta = 0.0451, N = 267
AGDISCNADINPLK pc2
P17301
97:139
98:142
29:44
5:52277455:A:C_C
p = 0.47, beta = 0.0698, N = 224
IPLLYDAEIHLTR pc3
E7ESP4;P17301
83:139
87:142
26:44
5:52277455:A:C_C
p = 0.5, beta = 0.0692, N = 196
IGQTSSSVSFK pc2
P17301
65:139
63:142
25:44
5:52277455:A:C_C
p = 0.59, beta = 0.0591, N = 153
Assay Target: ITGA2
Olink UniProt: P17301
deCODE rsID: rs181769
Proxy rsID: rs181769
deCODE: 5:52981625:C:A
Proxy SNP: 5:52277455:A:C
deCODE log10(p): 27.8
deCODE BETA: 0.09
1131:1038:1030:1006:969:921:
EGILGQHQFLEGPEGIENTR pc3
rs1801106 REF
55:554
62:557
26:149
5:52277455:A:C_C
p = 0.019, model = DOM, N = 143
GILGQHQFLEGPEGIENTR pc3
rs1801106 ALT
29:554
19:557
1:149
5:52277455:A:C_C
p = 0.023, model = DOM, N = 49

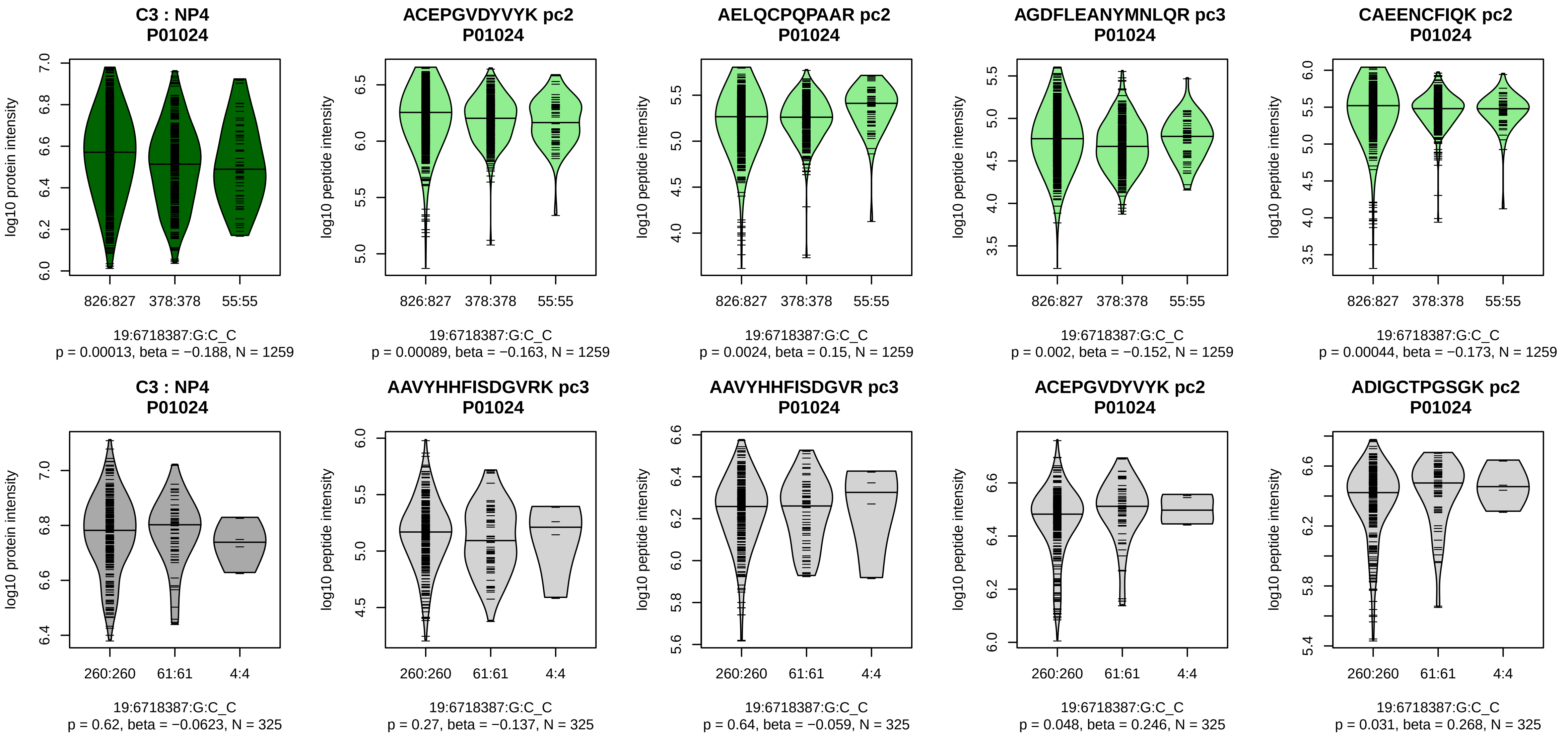


Assay Target: C3
Olink UniProt: P01024
deCODE rsID: rs2230199
Proxy rsID: rs2230199
deCODE: 19:6718376:C:G
Proxy SNP: 19:6718387:G:C
deCODE log10(p): 27.4
deCODE BETA: 0.1
*:*:*:*:*:*:*:*:*:*:*:_:*:*:*:*:*:*:_:*:*
1259:1259:1259:1259:1259:125

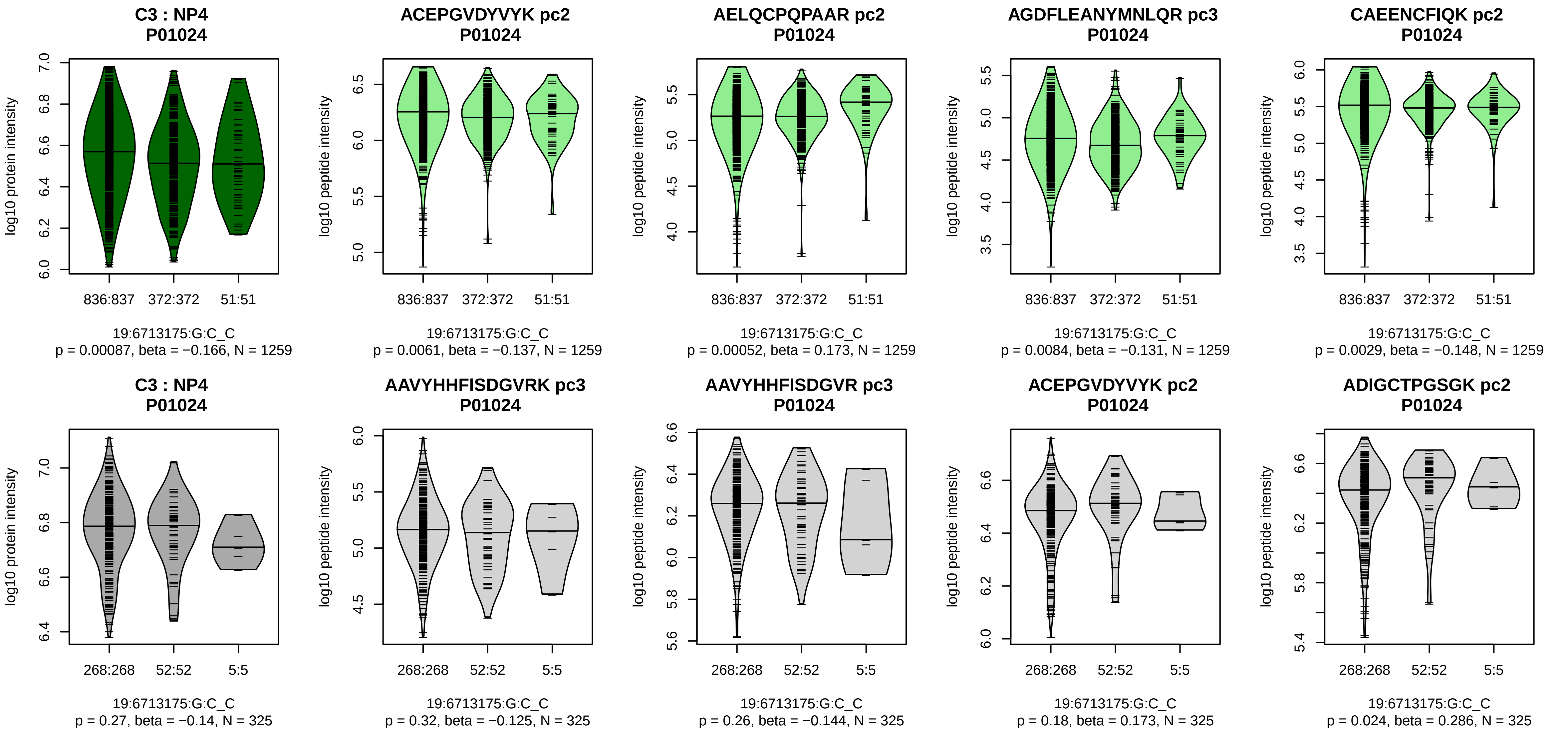


Assay Target: C3
Olink UniProt: P01024
deCODE rsID: rs11085197
Proxy rsID: rs11085197
deCODE: 19:6713164:C:G
Proxy SNP: 19:6713175:G:C
deCODE log10(p): 27.2
deCODE BETA: −0.1
*:*:*:*:*:*:–:*:–:–:*:–:*:*:*:–:*:*:–:
1259:1259:1259:1259:1259:125

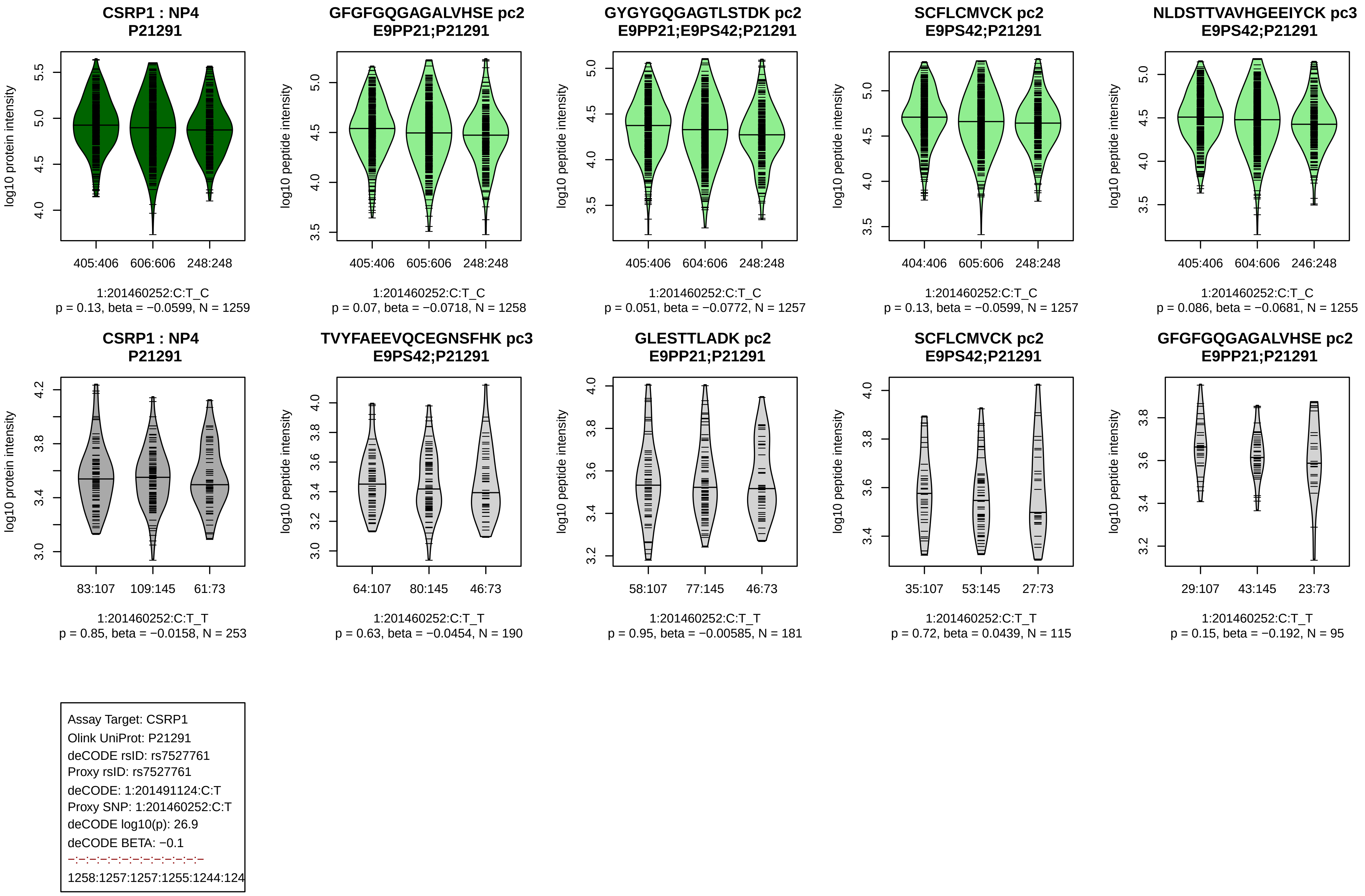
CSRP1 : NP4
P21291
log10 protein intensity
405:406
606:606
248:248
1:201460252:C:T_C
p = 0.13, beta = −0.0599, N = 1259
GFGFGQGAGALVHSE pc2
E9PP21;P21291
log10 peptide intensity
405:406
605:606
248:248
1:201460252:C:T_C
p = 0.07, beta = −0.0718, N = 1258
GYGYGQGAGTLSTDK pc2
E9PP21;E9PS42;P21291
log10 peptide intensity
405:406
604:606
248:248
1:201460252:C:T_C
p = 0.051, beta = −0.0772, N = 1257
SCFLCMVCK pc2
E9PS42;P21291
log10 peptide intensity
404:406
605:606
248:248
1:201460252:C:T_C
p = 0.13, beta = −0.0599, N = 1257
NLDSTTVAVHGEEIYCK pc3
E9PS42;P21291
log10 peptide intensity
405:406
604:606
246:248
1:201460252:C:T_C
p = 0.086, beta = −0.0681, N = 1255
CSRP1 : NP4
P21291
log10 protein intensity
83:107
109:145
61:73
1:201460252:C:T_T
p = 0.85, beta = −0.0158, N = 253
TVYFAEEVQCEGNSFHK pc3
E9PS42;P21291
log10 peptide intensity
64:107
80:145
46:73
1:201460252:C:T_T
p = 0.63, beta = −0.0454, N = 190
GLESTTLADK pc2
E9PP21;P21291
log10 peptide intensity
58:107
77:145
46:73
1:201460252:C:T_T
p = 0.95, beta = −0.00585, N = 181
SCFLCMVCK pc2
E9PS42;P21291
log10 peptide intensity
35:107
53:145
27:73
1:201460252:C:T_T
p = 0.72, beta = 0.0439, N = 115
GFGFGQGAGALVHSE pc2
E9PP21;P21291
log10 peptide intensity
29:107
43:145
23:73
1:201460252:C:T_T
p = 0.15, beta = −0.192, N = 95
Assay Target: CSRP1
Olink UniProt: P21291
deCODE rsID: rs7527761
Proxy rsID: rs7527761
deCODE: 1:201491124:C:T
Proxy SNP: 1:201460252:C:T
deCODE log10(p): 26.9
deCODE BETA: −0.1
−:−:−:−:−:−:−:−:−:−:−:−
1258:1257:1257:1255:1244:124

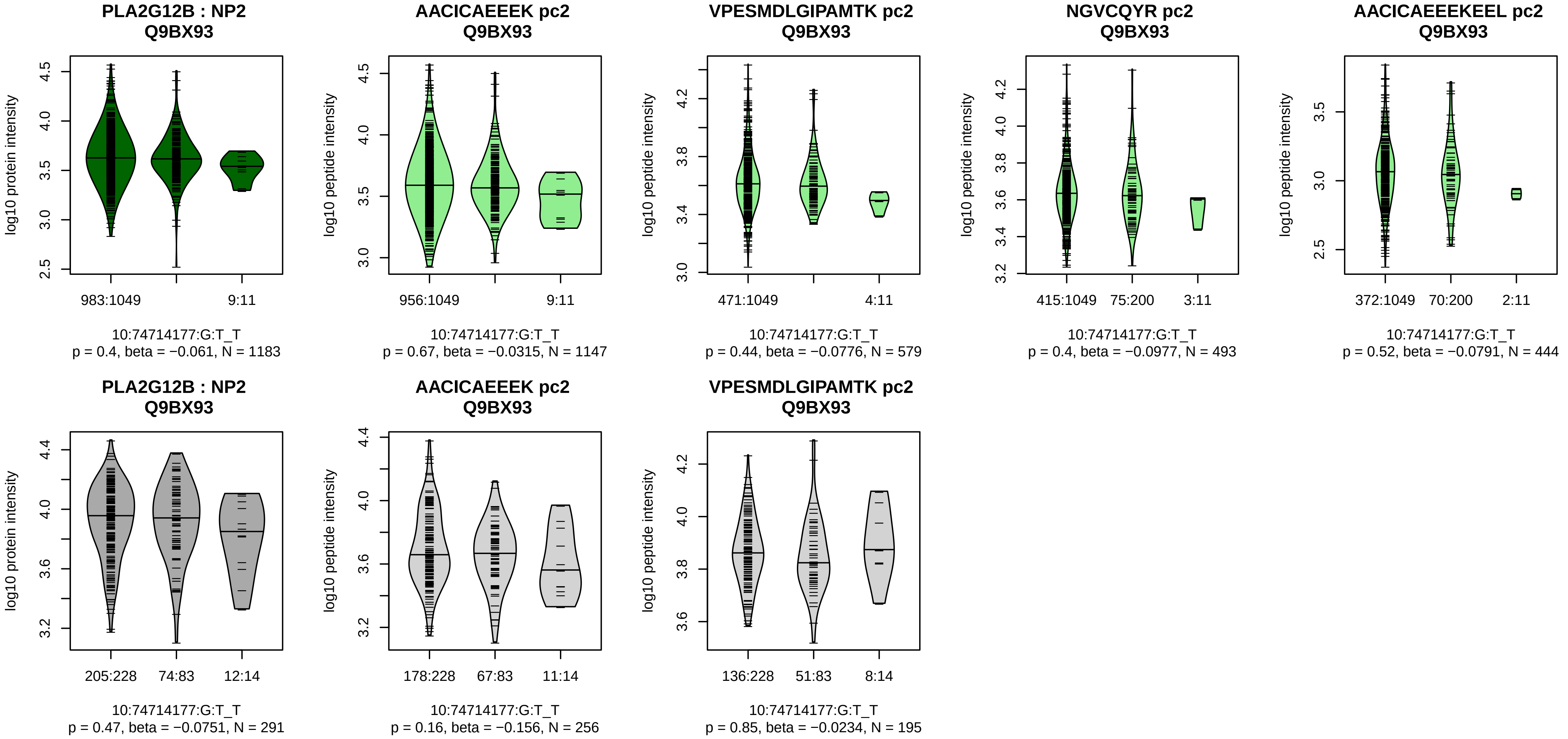


Assay Target: PLA2G12B
Olink UniProt: Q9BX93
deCODE rsID: rs3829126
Proxy rsID: rs3829126
deCODE: 10:72954419:T:G
Proxy SNP: 10:74714177:G:T
deCODE log10(p): 26.9
deCODE BETA: −0.15
−:−:−:−:−
1147:579:493:444:93

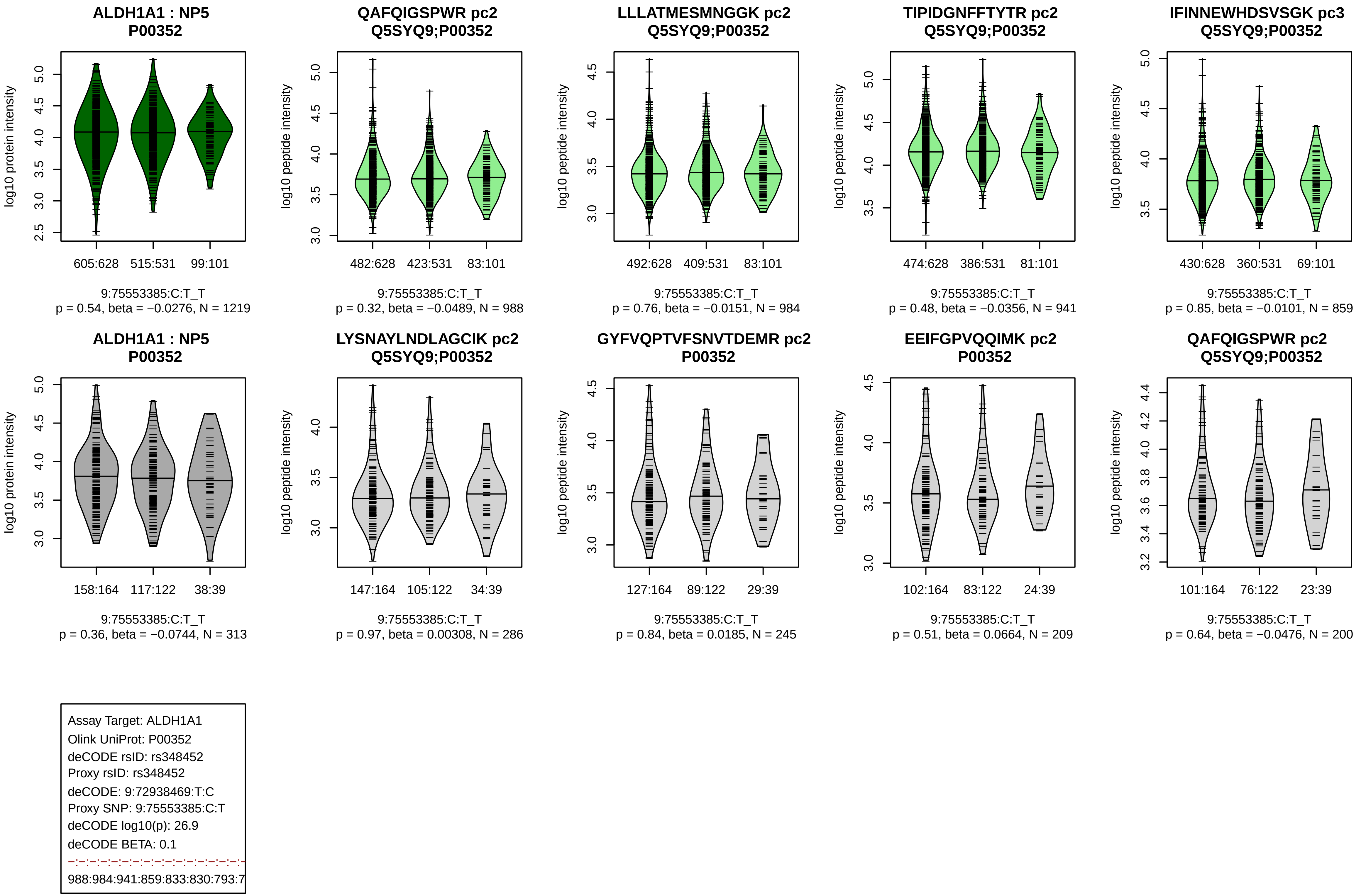
ALDH1A1 : NP5
P00352
log10 protein intensity
605:628 515:531 99:101
9:75553385:C:T_T
p = 0.54, beta = −0.0276, N = 1219
QAFQIGSPWR pc2
Q5SYQ9;P00352
log10 peptide intensity
482:628 423:531 83:101
9:75553385:C:T_T
p = 0.32, beta = −0.0489, N = 988
LLLATMESMNGGK pc2
Q5SYQ9;P00352
log10 peptide intensity
492:628 409:531 83:101
9:75553385:C:T_T
p = 0.76, beta = −0.0151, N = 984
TIPIDGNFFTYTR pc2
Q5SYQ9;P00352
log10 peptide intensity
474:628 386:531 81:101
9:75553385:C:T_T
p = 0.48, beta = −0.0356, N = 941
IFINNEWHDSVSGK pc3
Q5SYQ9;P00352
log10 peptide intensity
430:628 360:531 69:101
9:75553385:C:T_T
p = 0.85, beta = −0.0101, N = 859
ALDH1A1 : NP5
P00352
log10 protein intensity
158:164 117:122 38:39
9:75553385:C:T_T
p = 0.36, beta = −0.0744, N = 313
LYSNAYLNDLAGCIK pc2
Q5SYQ9;P00352
log10 peptide intensity
147:164 105:122 34:39
9:75553385:C:T_T
p = 0.97, beta = 0.00308, N = 286
GYFVQPTVFSNVTDEMR pc2
P00352
log10 peptide intensity
127:164 89:122 29:39
9:75553385:C:T_T
p = 0.84, beta = 0.0185, N = 245
EEIFGPVQQIMK pc2
P00352
log10 peptide intensity
102:164 83:122 24:39
9:75553385:C:T_T
p = 0.51, beta = 0.0664, N = 209
QAFQIGSPWR pc2
Q5SYQ9;P00352
log10 peptide intensity
101:164 76:122 23:39
9:75553385:C:T_T
p = 0.64, beta = −0.0476, N = 200
Assay Target: ALDH1A1
Olink UniProt: P00352
deCODE rsID: rs348452
Proxy rsID: rs348452
deCODE: 9:72938469:T:C
Proxy SNP: 9:75553385:C:T
deCODE log10(p): 26.9
deCODE BETA: 0.1
988:984:941:859:833:830:793:7

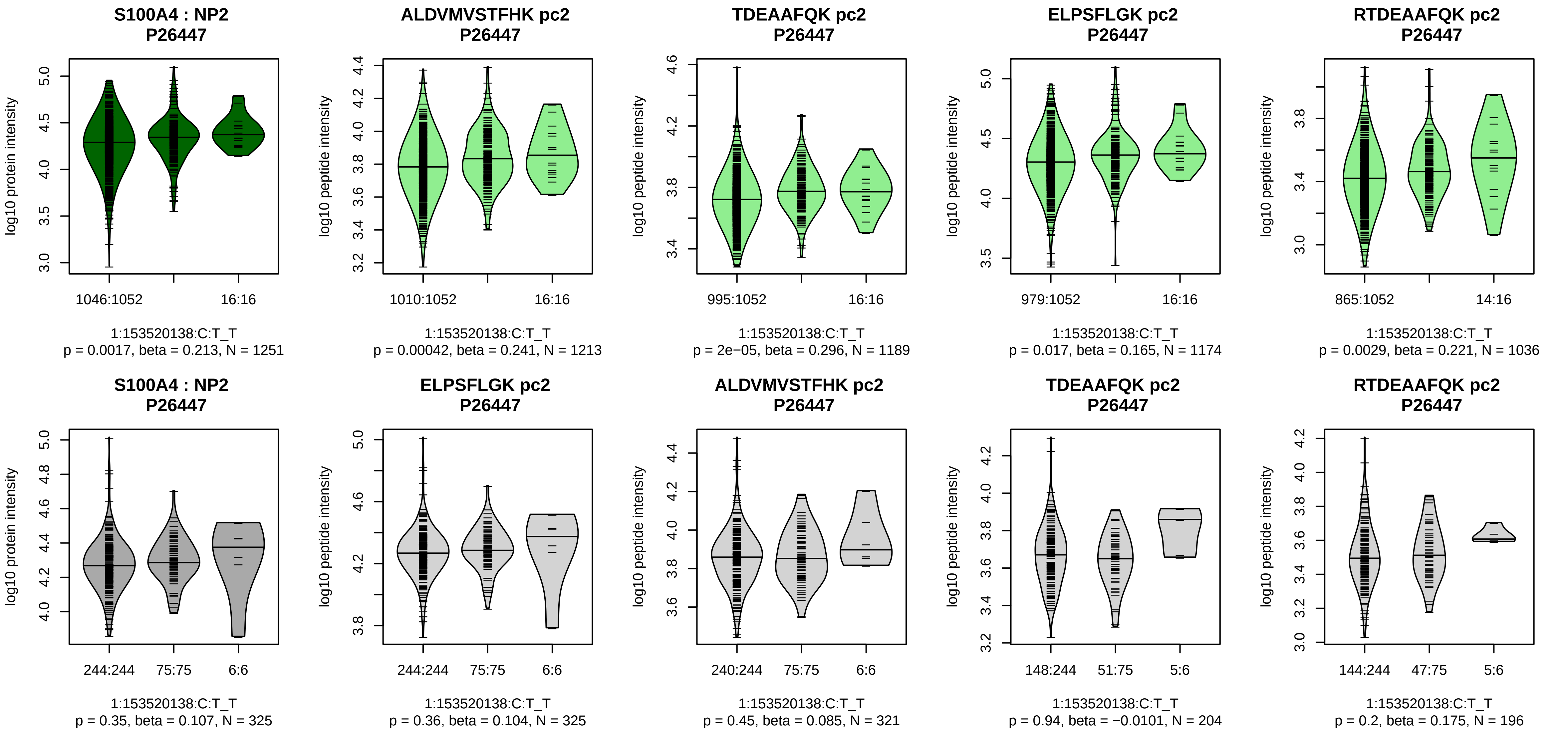


Assay Target: S100A4
Olink UniProt: P26447
deCODE rsID: rs41265162
Proxy rsID: rs41265162
deCODE: 1:153547662:T:C
Proxy SNP: 1:153520138:C:T
deCODE log10(p): 26.4
deCODE BETA: 0.21
*:*:−:*:NA
1213:1189:1174:1036:17

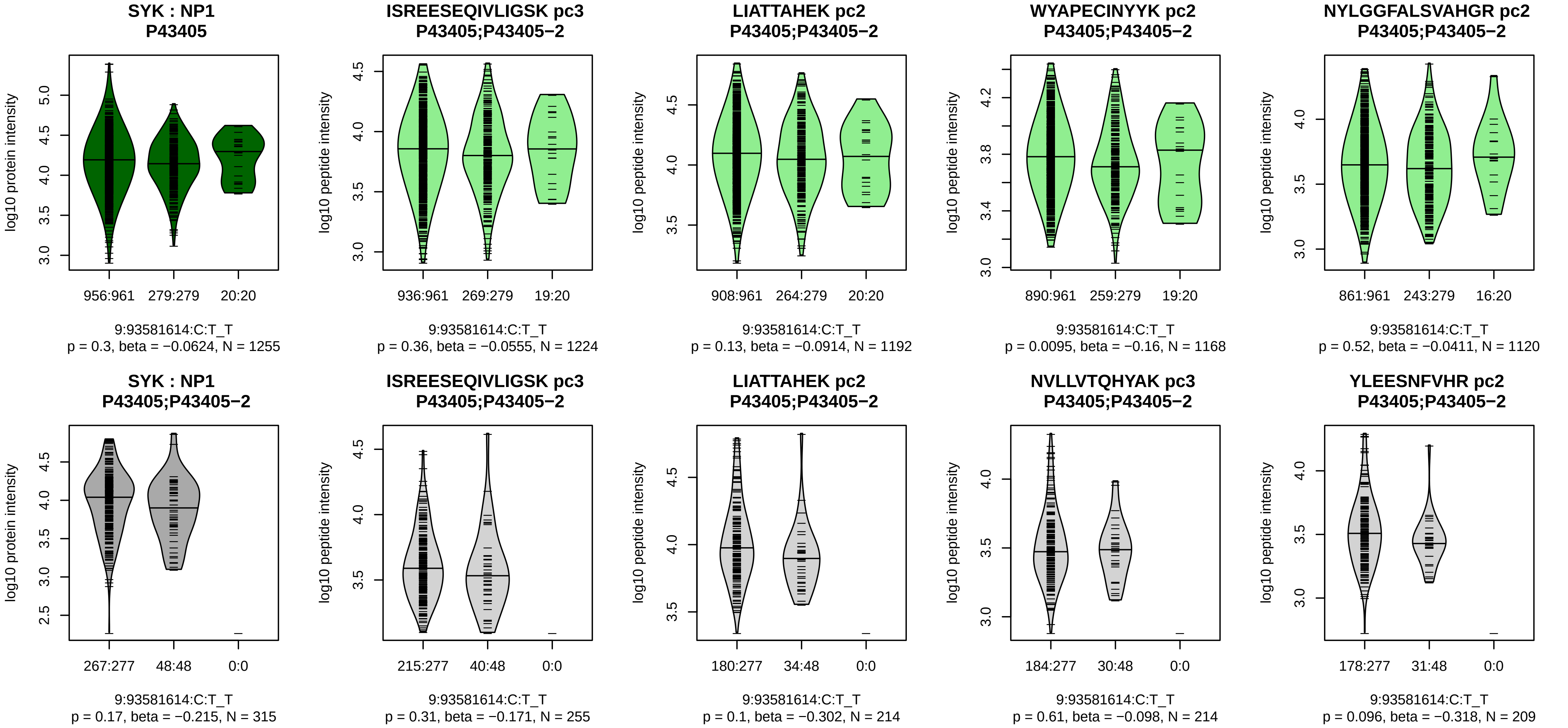


Assay Target: SYK
Olink UniProt: P43405
deCODE rsID: rs75505307
Proxy rsID: rs75505307
deCODE: 9:90819332:T:C
Proxy SNP: 9:93581614:C:T
deCODE log10(p): 26.1
deCODE BETA: −0.13
−:−:*:−:−:−:−:−:−:−:−:−:−:−:−:−:−
1224:1192:1168:1120:1101:109

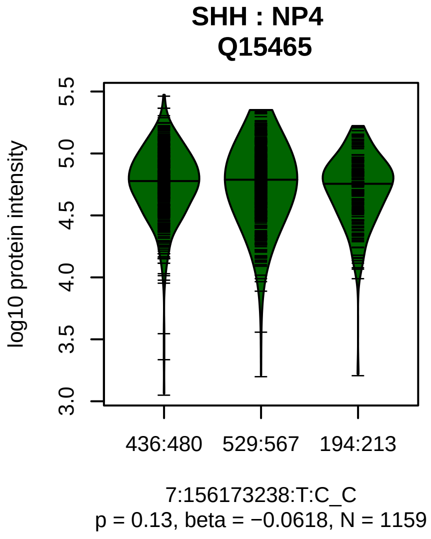


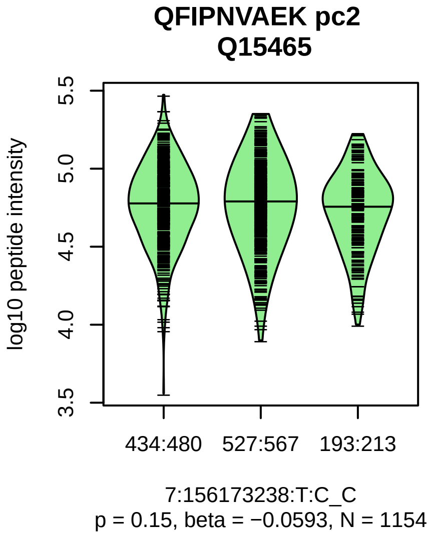


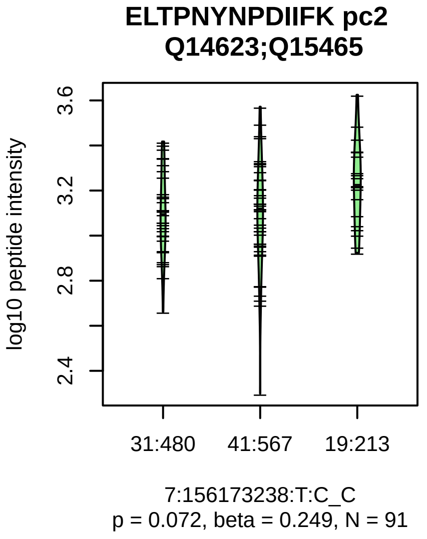


Assay Target: SHH
Olink UniProt: Q15465
deCODE rsID: rs10225292
Proxy rsID: rs10225292
deCODE: 7:156380544:C:T
Proxy SNP: 7:156173238:T:C
deCODE log10(p): 25.7
deCODE BETA: −0.09
−:−
1154:91

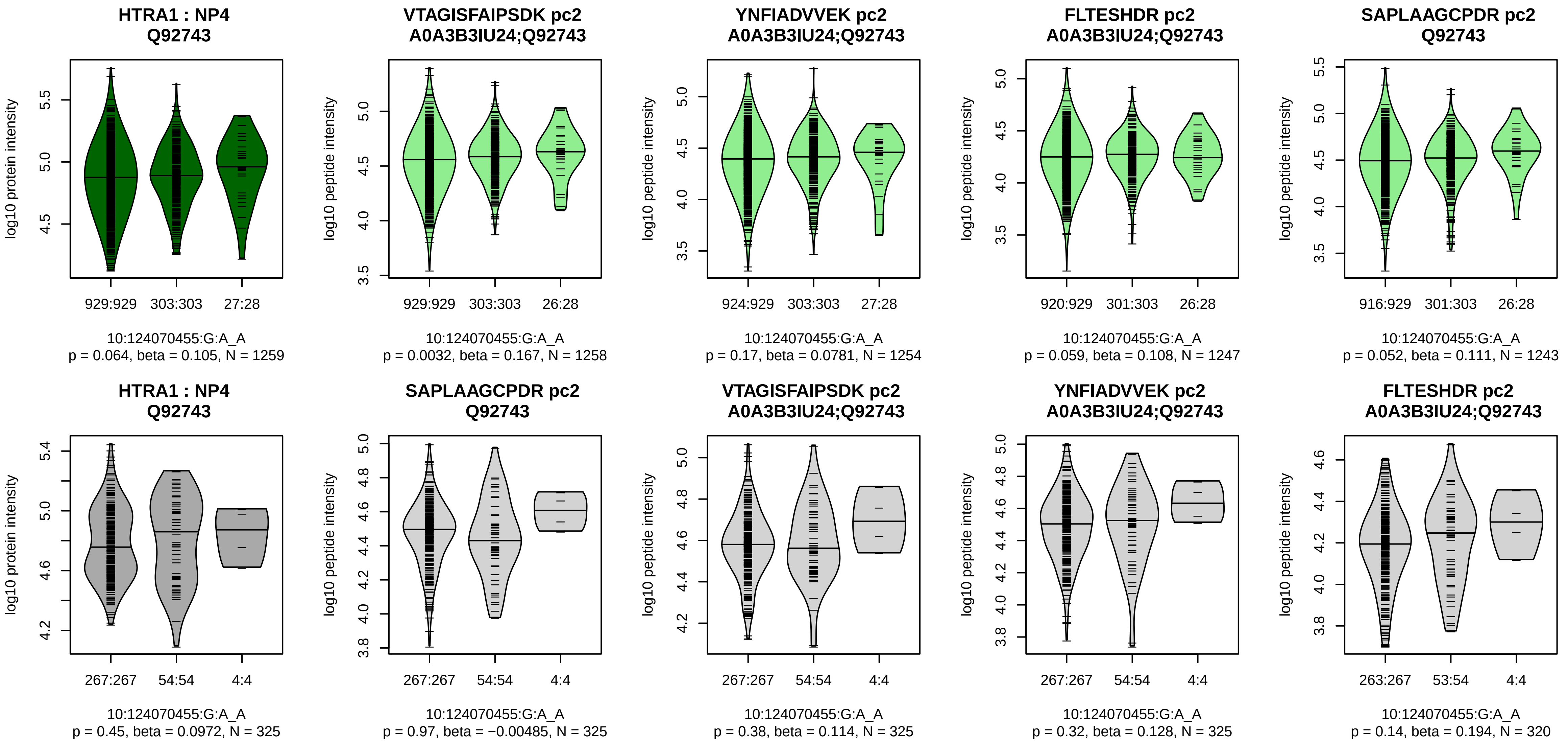


Assay Target: HTRA1
Olink UniProt: Q92743
deCODE rsID: rs61871680
Proxy rsID: rs61871680
deCODE: 10:122310940:A:G
Proxy SNP: 10:124070455:G:A
deCODE log10(p): 25.1
deCODE BETA: 0.11
*:–:–:–:–:–:–:–:–:–:–:–:–:–:–:–
1258:1254:1247:1243:1239:123

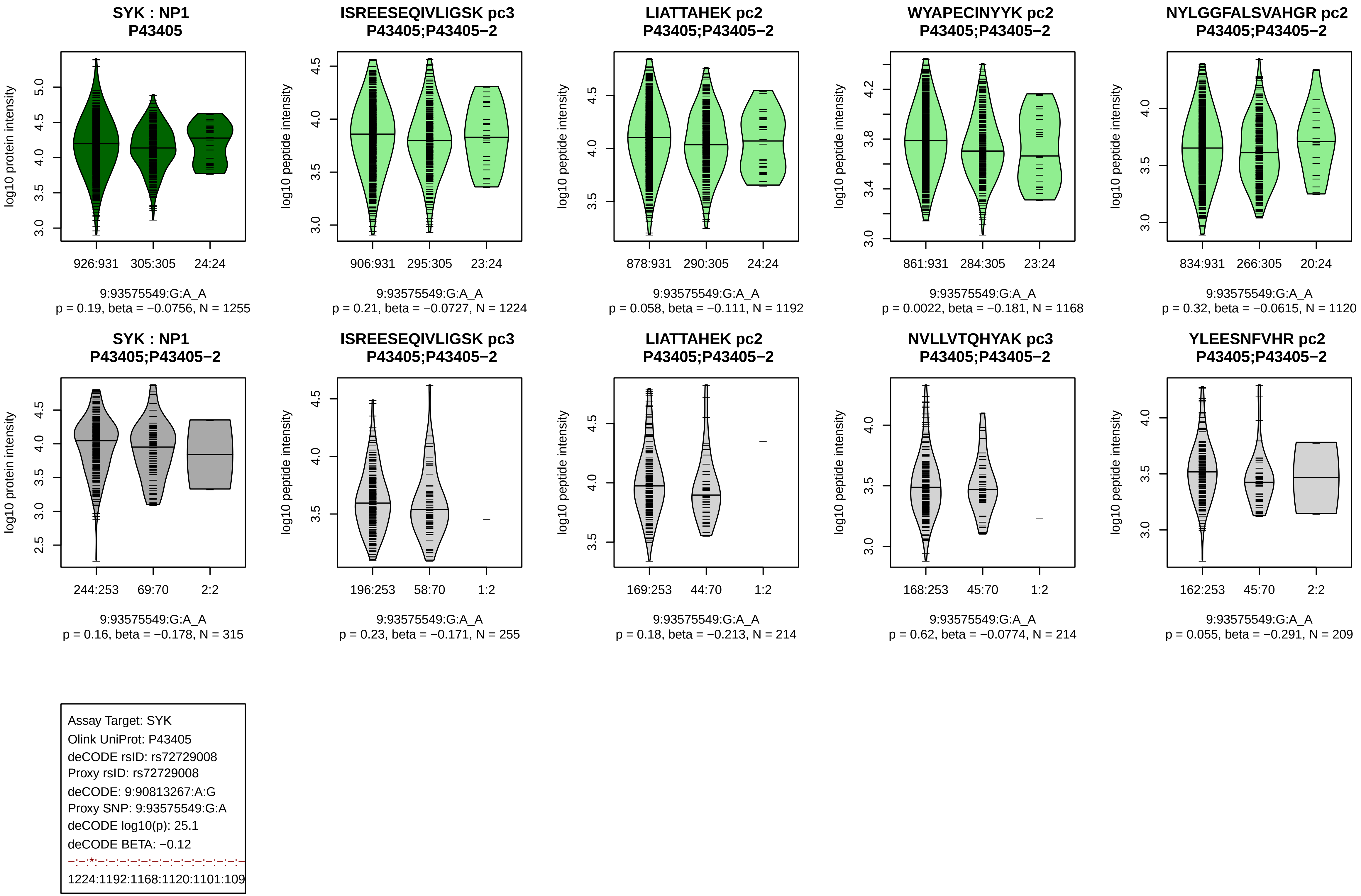
SYK : NP1
P43405
926:931 305:305 24:24
9:93575549:G:A_A
p = 0.19, beta = −0.0756, N = 1255
ISREESEQIVLIGSK pc3
P43405;P43405−2
906:931 295:305 23:24
9:93575549:G:A_A
p = 0.21, beta = −0.0727, N = 1224
LIATTAHEK pc2
P43405;P43405−2
878:931 290:305 24:24
9:93575549:G:A_A
p = 0.058, beta = −0.111, N = 1192
WYAPECINYYK pc2
P43405;P43405−2
861:931 284:305 23:24
9:93575549:G:A_A
p = 0.0022, beta = −0.181, N = 1168
NYLGGFALSVAHGR pc2
P43405;P43405−2
834:931 266:305 20:24
9:93575549:G:A_A
p = 0.32, beta = −0.0615, N = 1120
SYK : NP1
P43405;P43405−2
244:253 69:70 2:2
9:93575549:G:A_A
p = 0.16, beta = −0.178, N = 315
ISREESEQIVLIGSK pc3
P43405;P43405−2
196:253 58:70 1:2
9:93575549:G:A_A
p = 0.23, beta = −0.171, N = 255
LIATTAHEK pc2
P43405;P43405−2
169:253 44:70 1:2
9:93575549:G:A_A
p = 0.18, beta = −0.213, N = 214
NVLLVTQHYAK pc3
P43405;P43405−2
168:253 45:70 1:2
9:93575549:G:A_A
p = 0.62, beta = −0.0774, N = 214
YLEESNFVHR pc2
P43405;P43405−2
162:253 45:70 2:2
9:93575549:G:A_A
p = 0.055, beta = −0.291, N = 209
log10 protein intensity
log10 peptide intensity
Assay Target: SYK
Olink UniProt: P43405
deCODE rsID: rs72729008
Proxy rsID: rs72729008
deCODE: 9:90813267:A:G
Proxy SNP: 9:93575549:G:A
deCODE log10(p): 25.1
deCODE BETA: −0.12
1224:1192:1168:1120:1101:109

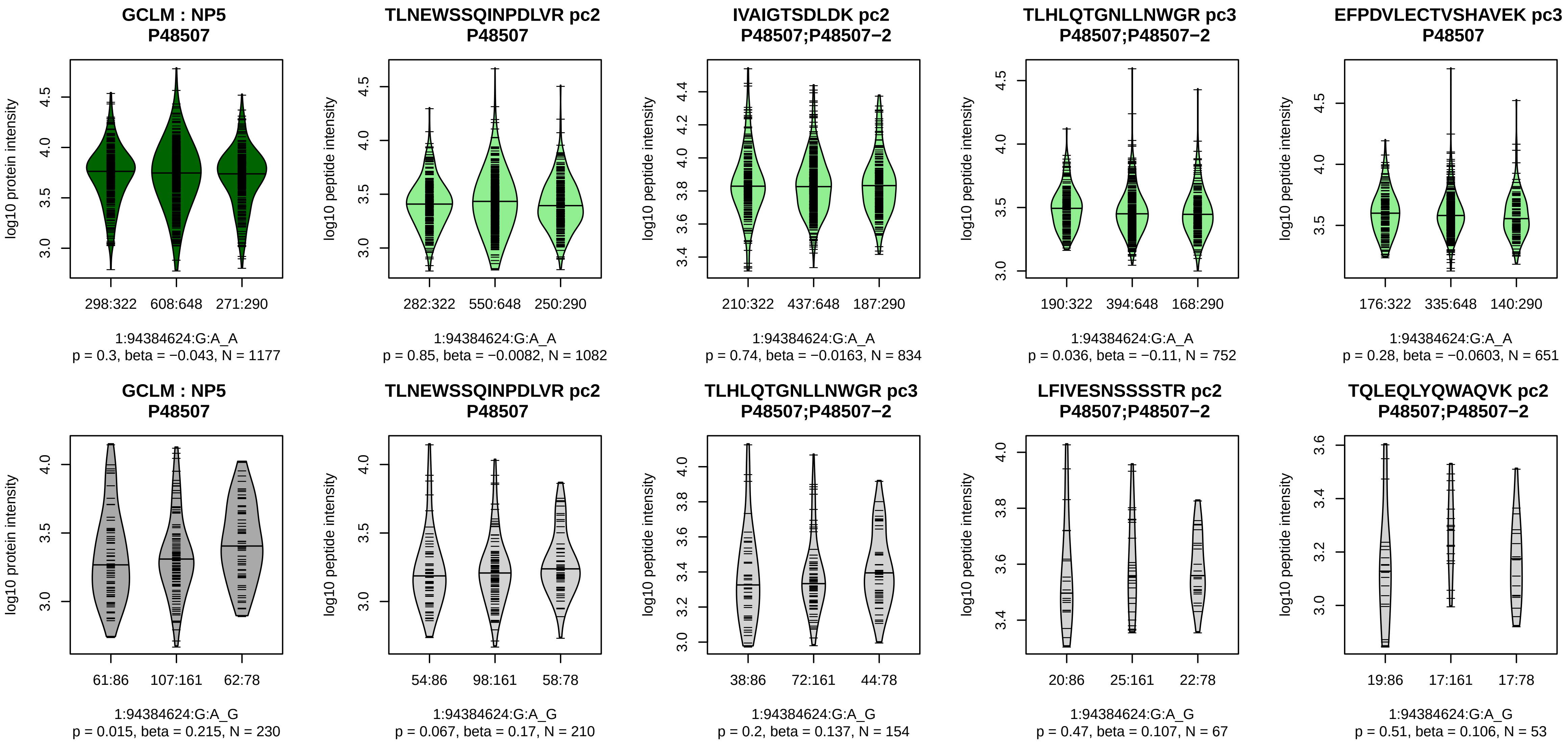


Assay Target: GCLM
Olink UniProt: P48507
deCODE rsID: rs12563495
Proxy rsID: rs12563495
deCODE: 1:93919068:A:G
Proxy SNP: 1:94384624:G:A
deCODE log10(p): 25
deCODE BETA: −0.08
−:−:−:−:−:−:−:−:−:−:−:−:NA:NA
1082:834:752:651:444:227:219:

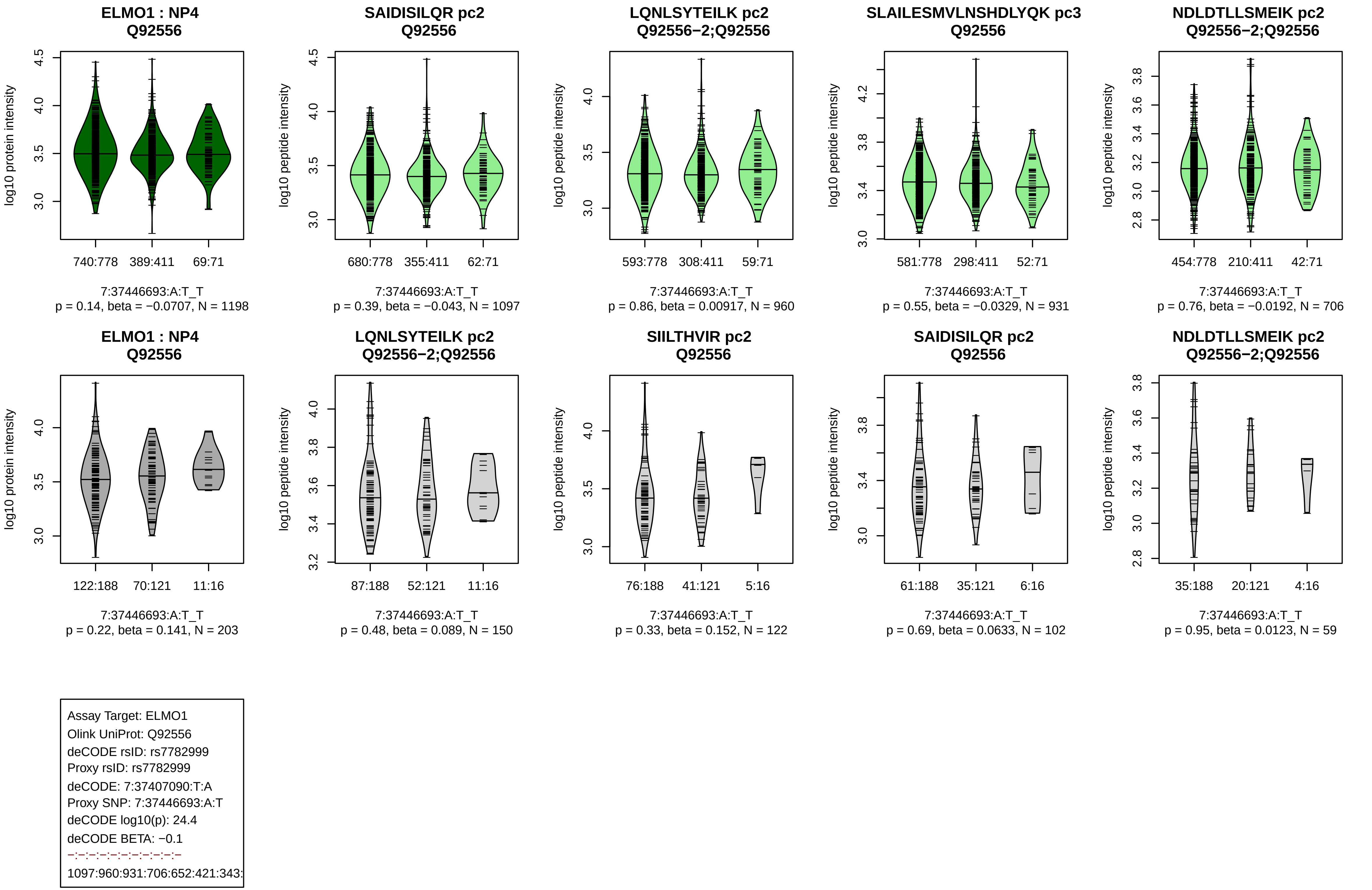
ELMO1 : NP4
Q92556
log10 protein intensity
740:778 389:411 69:71
7:37446693:A:T_T
p = 0.14, beta = −0.0707, N = 1198
SAIDISILQR pc2
Q92556
log10 peptide intensity
680:778 355:411 62:71
7:37446693:A:T_T
p = 0.39, beta = −0.043, N = 1097
LQNLSYTEILK pc2
Q92556−2;Q92556
log10 peptide intensity
593:778 308:411 59:71
7:37446693:A:T_T
p = 0.86, beta = 0.00917, N = 960
SLAILESMVLNSHDLYQK pc3
Q92556
log10 peptide intensity
581:778 298:411 52:71
7:37446693:A:T_T
p = 0.55, beta = −0.0329, N = 931
NDLDTLLSMEIK pc2
Q92556−2;Q92556
log10 peptide intensity
454:778 210:411 42:71
7:37446693:A:T_T
p = 0.76, beta = −0.0192, N = 706
ELMO1 : NP4
Q92556
log10 protein intensity
122:188 70:121 11:16
7:37446693:A:T_T
p = 0.22, beta = 0.141, N = 203
LQNLSYTEILK pc2
Q92556−2;Q92556
log10 peptide intensity
87:188 52:121 11:16
7:37446693:A:T_T
p = 0.48, beta = 0.089, N = 150
SIILTHVIR pc2
Q92556
log10 peptide intensity
76:188 41:121 5:16
7:37446693:A:T_T
p = 0.33, beta = 0.152, N = 122
SAIDISILQR pc2
Q92556
log10 peptide intensity
61:188 35:121 6:16
7:37446693:A:T_T
p = 0.69, beta = 0.0633, N = 102
NDLDTLLSMEIK pc2
Q92556−2;Q92556
log10 peptide intensity
35:188 20:121 4:16
7:37446693:A:T_T
p = 0.95, beta = 0.0123, N = 59
Assay Target: ELMO1
Olink UniProt: Q92556
deCODE rsID: rs7782999
Proxy rsID: rs7782999
deCODE: 7:37407090:T:A
Proxy SNP: 7:37446693:A:T
deCODE log10(p): 24.4
deCODE BETA: −0.1
−:−:−:−:−:−:−:−:−:−
1097:960:931:706:652:421:343:

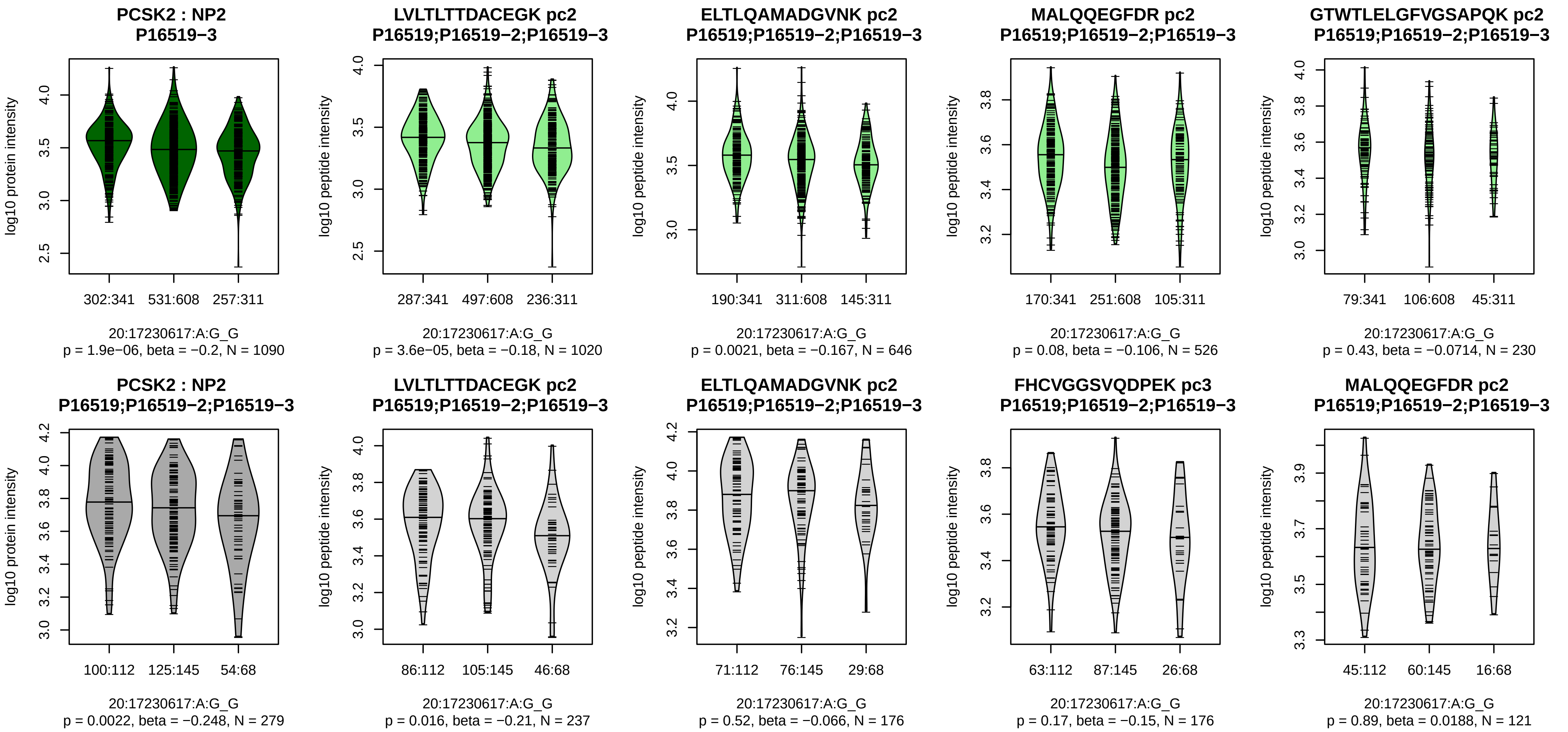


Assay Target: PCSK2
Olink UniProt: P16519
deCODE rsID: rs6044699
Proxy rsID: rs6044699
deCODE: 20:17249972:A:G
Proxy SNP: 20:17230617:A:G
deCODE log10(p): 23.9
deCODE BETA: 0.08
*:*:−:−:−:−
1020:646:526:230:226:199

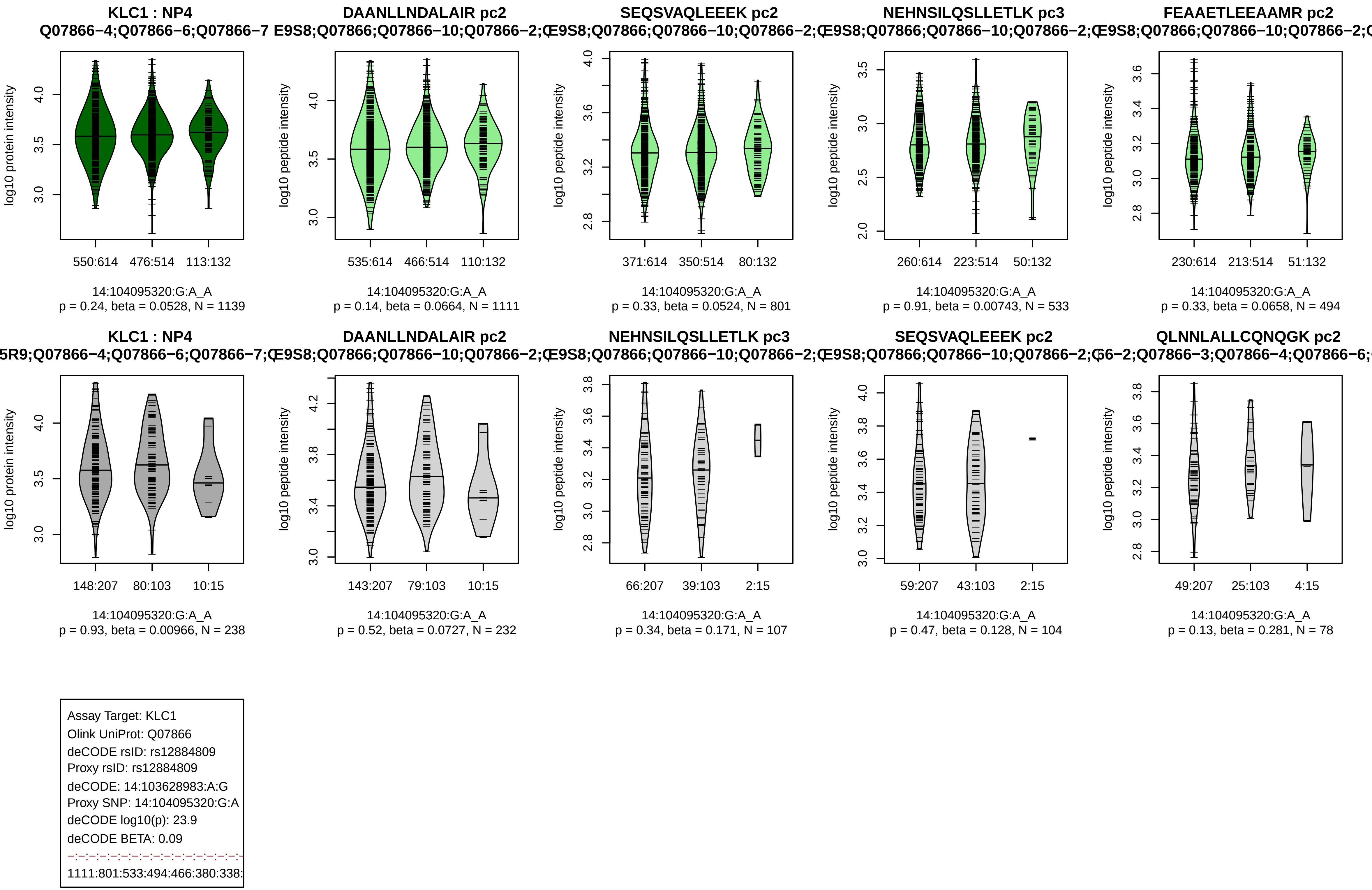
KLC1 : NP4
Q07866−4;Q07866−6;Q07866−7
log10 protein intensity
550:614 476:514 113:132
14:104095320:G:A_A
p = 0.24, beta = 0.0528, N = 1139
DAANLLNDALAIR pc2
E9S8;Q07866;Q07866−10;Q07866−2;
log10 peptide intensity
535:614 466:514 110:132
14:104095320:G:A_A
p = 0.14, beta = 0.0664, N = 1111
SEQSVAQLEEEK pc2
E9S8;Q07866;Q07866−10;Q07866−2;
371:614 350:514 80:132
14:104095320:G:A_A
p = 0.33, beta = 0.0524, N = 801
NEHNSILQSLLETLK pc3
E9S8;Q07866;Q07866−10;Q07866−2;
260:614 223:514 50:132
14:104095320:G:A_A
p = 0.91, beta = 0.00743, N = 533
FEAAETLEEAAMR pc2
E9S8;Q07866;Q07866−10;Q07866−2;
230:614 213:514 51:132
14:104095320:G:A_A
p = 0.33, beta = 0.0658, N = 494
KLC1 : NP4
5R9;Q07866−4;Q07866−6;Q07866−7;
148:207 80:103 10:15
14:104095320:G:A_A
p = 0.93, beta = 0.00966, N = 238
DAANLLNDALAIR pc2
E9S8;Q07866;Q07866−10;Q07866−2;
143:207 79:103 10:15
14:104095320:G:A_A
p = 0.52, beta = 0.0727, N = 232
NEHNSILQSLLETLK pc3
E9S8;Q07866;Q07866−10;Q07866−2;
66:207 39:103 2:15
14:104095320:G:A_A
p = 0.34, beta = 0.171, N = 107
SEQSVAQLEEEK pc2
E9S8;Q07866;Q07866−10;Q07866−2;
59:207 43:103 2:15
14:104095320:G:A_A
p = 0.47, beta = 0.128, N = 104
QLNNLALLCQNQGK pc2
66−2;Q07866−3;Q07866−4;Q07866−6;
49:207 25:103 4:15
14:104095320:G:A_A
p = 0.13, beta = 0.281, N = 78
Assay Target: KLC1
Olink UniProt: Q07866
deCODE rsID: rs12884809
Proxy rsID: rs12884809
deCODE: 14:103628983:A:G
Proxy SNP: 14:104095320:G:A
deCODE log10(p): 23.9
deCODE BETA: 0.09
1111:801:533:494:466:380:338:

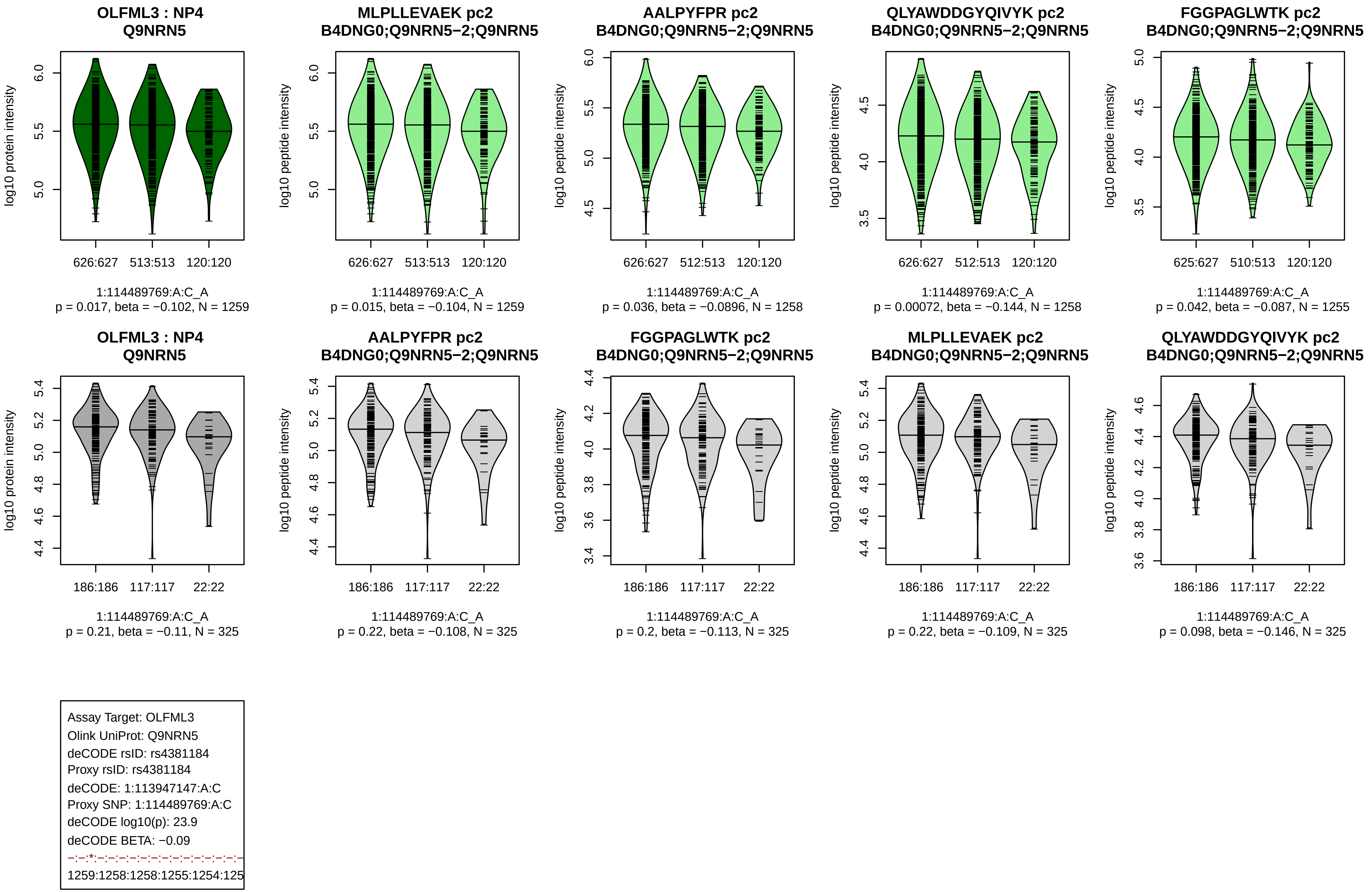
OLFML3 : NP4
Q9NRN5
log10 protein intensity
626:627 513:513 120:120
1:114489769:A:C_A
p = 0.017, beta = −0.102, N = 1259
MLPLLEVAEK pc2
B4DNG0;Q9NRN5−2;Q9NRN5
log10 peptide intensity
626:627 513:513 120:120
1:114489769:A:C_A
p = 0.015, beta = −0.104, N = 1259
AALPYFPR pc2
B4DNG0;Q9NRN5−2;Q9NRN5
log10 peptide intensity
626:627 512:513 120:120
1:114489769:A:C_A
p = 0.036, beta = −0.0896, N = 1258
QLYAWDDGYQIVYK pc2
B4DNG0;Q9NRN5−2;Q9NRN5
log10 peptide intensity
626:627 512:513 120:120
1:114489769:A:C_A
p = 0.00072, beta = −0.144, N = 1258
FGGPAGLWTK pc2
B4DNG0;Q9NRN5−2;Q9NRN5
log10 peptide intensity
625:627 510:513 120:120
1:114489769:A:C_A
p = 0.042, beta = −0.087, N = 1255
OLFML3 : NP4
Q9NRN5
log10 protein intensity
186:186 117:117 22:22
1:114489769:A:C_A
p = 0.21, beta = −0.11, N = 325
AALPYFPR pc2
B4DNG0;Q9NRN5−2;Q9NRN5
log10 peptide intensity
186:186 117:117 22:22
1:114489769:A:C_A
p = 0.22, beta = −0.108, N = 325
FGGPAGLWTK pc2
B4DNG0;Q9NRN5−2;Q9NRN5
log10 peptide intensity
186:186 117:117 22:22
1:114489769:A:C_A
p = 0.2, beta = −0.113, N = 325
MLPLLEVAEK pc2
B4DNG0;Q9NRN5−2;Q9NRN5
log10 peptide intensity
186:186 117:117 22:22
1:114489769:A:C_A
p = 0.22, beta = −0.109, N = 325
QLYAWDDGYQIVYK pc2
B4DNG0;Q9NRN5−2;Q9NRN5
log10 peptide intensity
186:186 117:117 22:22
1:114489769:A:C_A
p = 0.098, beta = −0.146, N = 325
Assay Target: OLFML3
Olink UniProt: Q9NRN5
deCODE rsID: rs4381184
Proxy rsID: rs4381184
deCODE: 1:113947147:A:C
Proxy SNP: 1:114489769:A:C
deCODE log10(p): 23.9
deCODE BETA: −0.09
−:−:*:−:−:−:−:−:−:−:−:−:−:−:−:−:−
1259:1258:1258:1255:1254:125

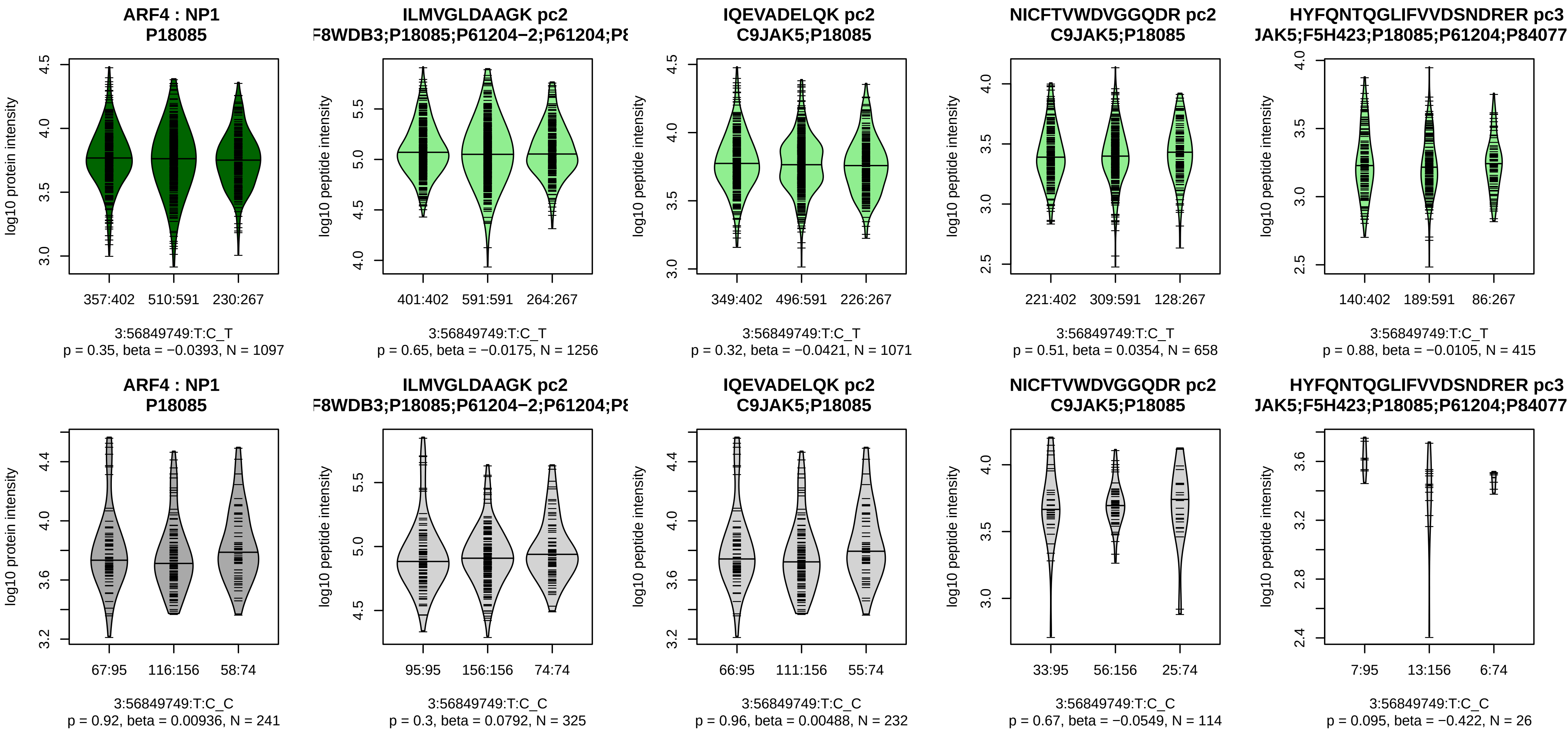


Assay Target: ARF4
Olink UniProt: P18085
deCODE rsID: rs1354034
Proxy rsID: rs1354034
deCODE: 3:56815721:T:C
Proxy SNP: 3:56849749:T:C
deCODE log10(p): 23.8
deCODE BETA: −0.08
−:−:−:−:−:−:−:−:NA
1256:1071:658:415:319:285:62:

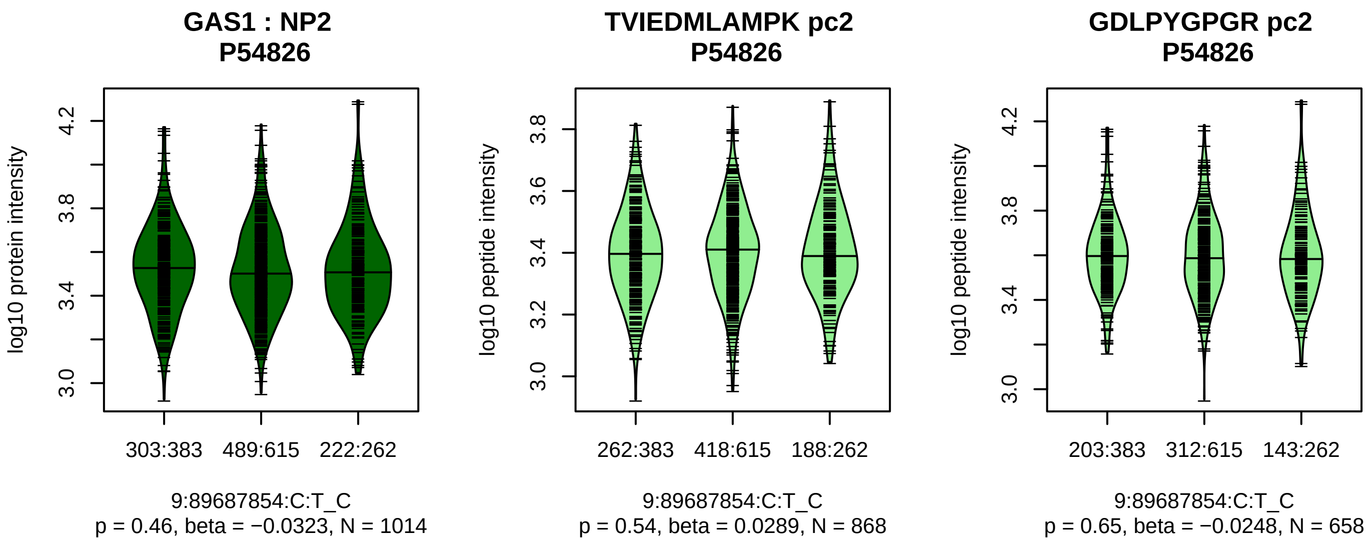


Assay Target: GAS1
Olink UniProt: P54826
deCODE rsID: rs4878043
Proxy rsID: rs4878043
deCODE: 9:87072939:C:T
Proxy SNP: 9:89687854:C:T
deCODE log10(p): 23.1
deCODE BETA: −0.08
−:−
868:658

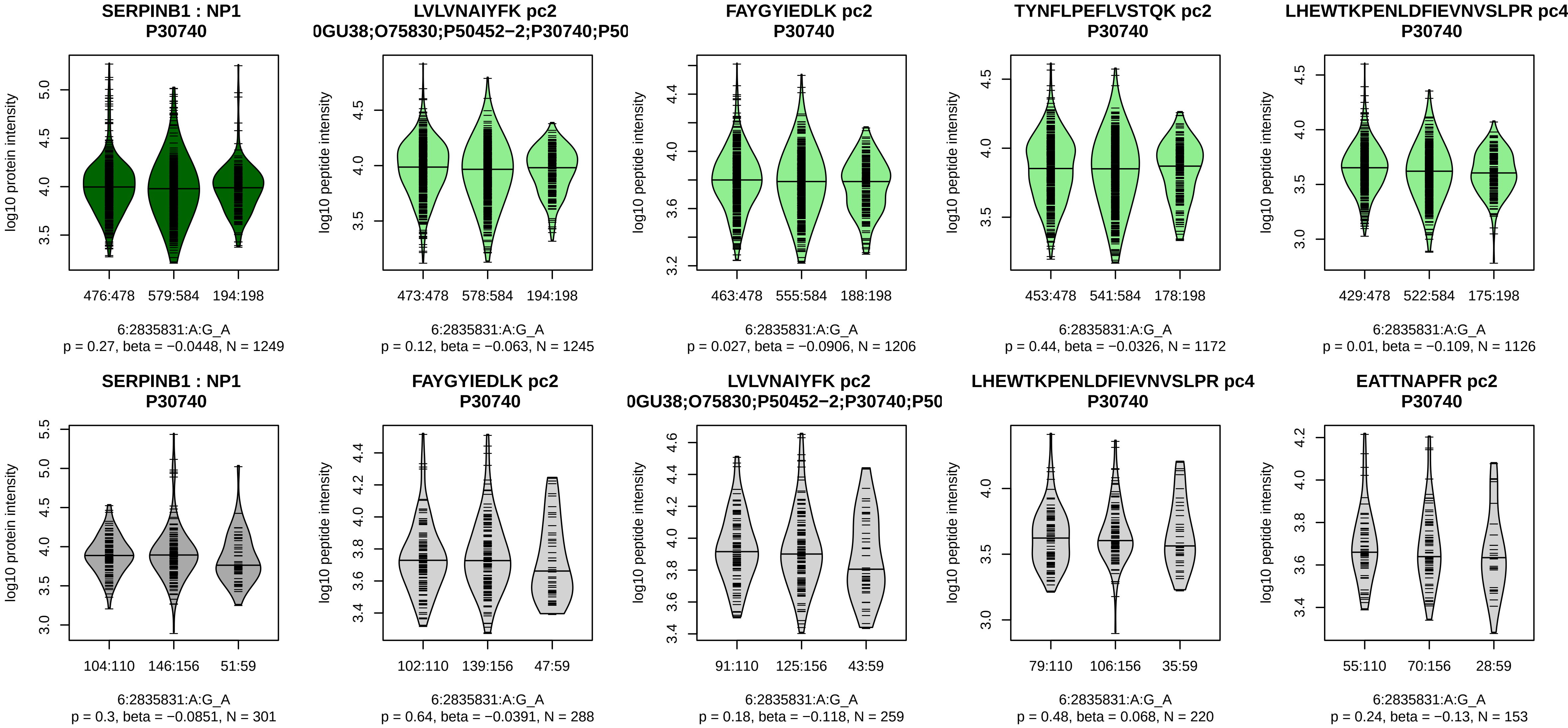


Assay Target: SERPINB1
Olink UniProt: P30740
deCODE rsID: rs2293772
Proxy rsID: rs2293772
deCODE: 6:2835597:A:G
Proxy SNP: 6:2835831:A:G
deCODE log10(p): 21.2
deCODE BETA: −0.08

1245:1206:1172:1126:892:822:7

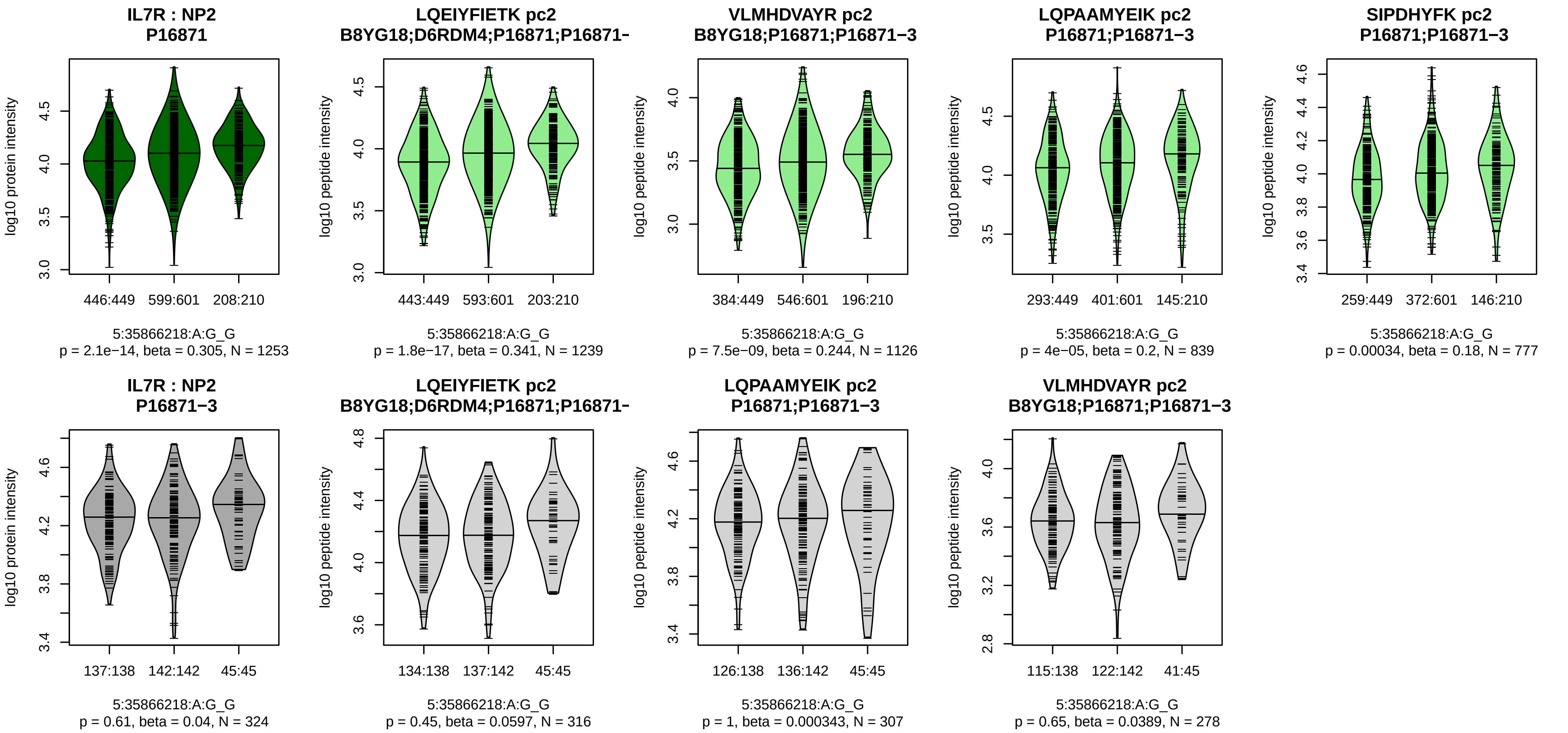


Assay Target: IL7R
Olink UniProt: P16871
deCODE rsID: rs10058572
Proxy rsID: rs6451229
deCODE: 5:35880754:!:TTTTTC
Proxy SNP: 5:35866218:A:G
deCODE log10(p): 20.9
deCODE BETA: 0.08
*:*:*:*:–:–
1239:1126:839:777:84:38

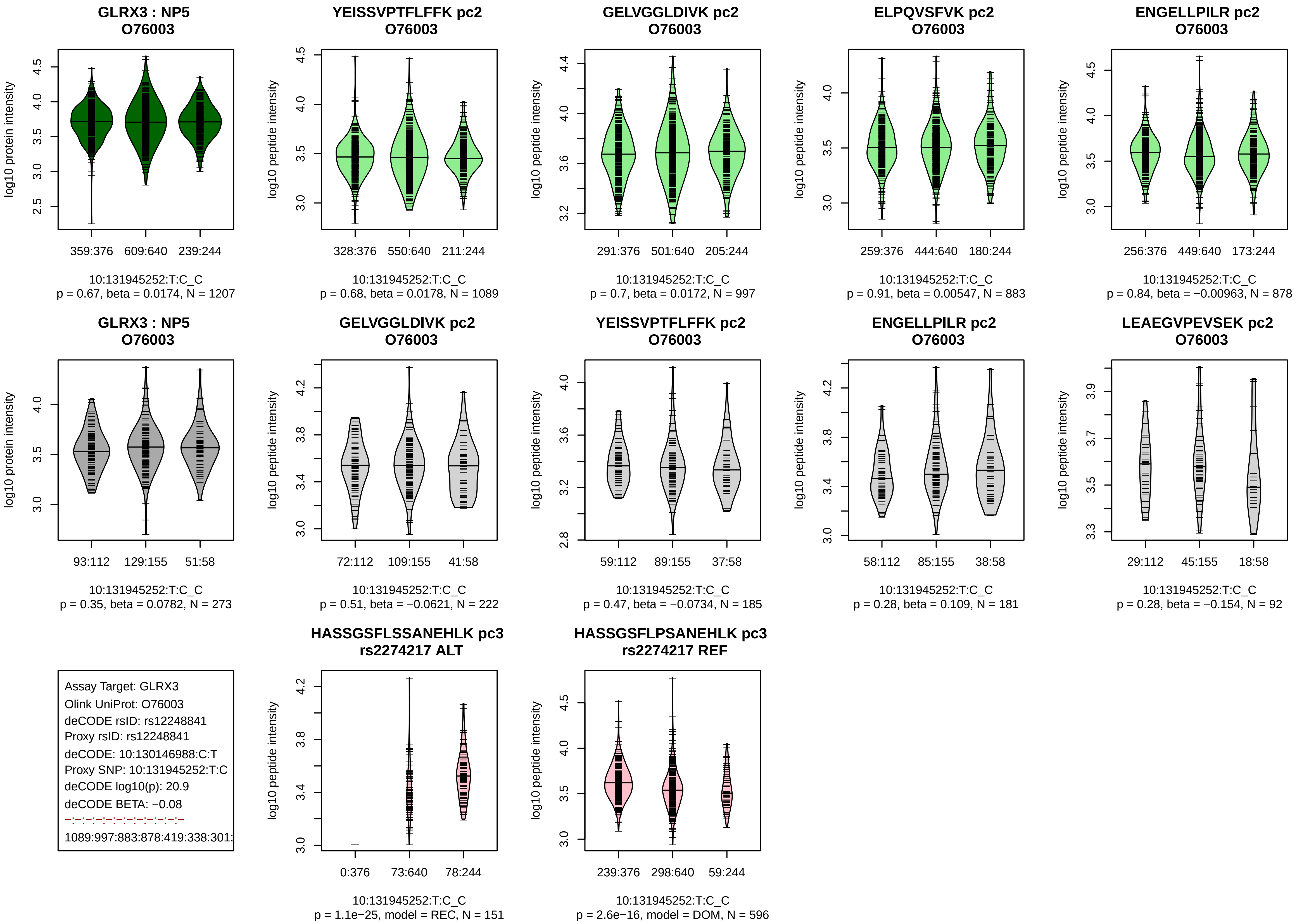
GLRX3 : NP5
O76003
log10 protein intensity
359:376
609:640
239:244
10:131945252:T:C_C
p = 0.67, beta = 0.0174, N = 1207
YEISSVPTFLFFK pc2
O76003
log10 peptide intensity
328:376
550:640
211:244
10:131945252:T:C_C
p = 0.68, beta = 0.0178, N = 1089
GELVGGLDIVK pc2
O76003
291:376
501:640
205:244
10:131945252:T:C_C
p = 0.7, beta = 0.0172, N = 997
ELPQVSFVK pc2
O76003
259:376
444:640
180:244
10:131945252:T:C_C
p = 0.91, beta = 0.00547, N = 883
ENGELLPILR pc2
O76003
256:376
449:640
173:244
10:131945252:T:C_C
p = 0.84, beta = −0.00963, N = 878
GLRX3 : NP5
O76003
93:112
129:155
51:58
10:131945252:T:C_C
p = 0.35, beta = 0.0782, N = 273
GELVGGLDIVK pc2
O76003
72:112
109:155
41:58
10:131945252:T:C_C
p = 0.51, beta = −0.0621, N = 222
YEISSVPTFLFFK pc2
O76003
59:112
89:155
37:58
10:131945252:T:C_C
p = 0.47, beta = −0.0734, N = 185
ENGELLPILR pc2
O76003
58:112
85:155
38:58
10:131945252:T:C_C
p = 0.28, beta = 0.109, N = 181
LEAEGVPEVSEK pc2
O76003
29:112
45:155
18:58
10:131945252:T:C_C
p = 0.28, beta = −0.154, N = 92
Assay Target: GLRX3
Olink UniProt: O76003
deCODE rsID: rs12248841
Proxy rsID: rs12248841
deCODE: 10:130146988:C:T
Proxy SNP: 10:131945252:T:C
deCODE log10(p): 20.9
deCODE BETA: −0.08
1089:997:883:878:419:338:301:
HASSGSFLSSANEHLK pc3
rs2274217 ALT
0:376
73:640
78:244
10:131945252:T:C_C
p = 1.1e−25, model = REC, N = 151
HASSGSFLPSANEHLK pc3
rs2274217 REF
239:376
298:640
59:244
10:131945252:T:C_C
p = 2.6e−16, model = DOM, N = 596

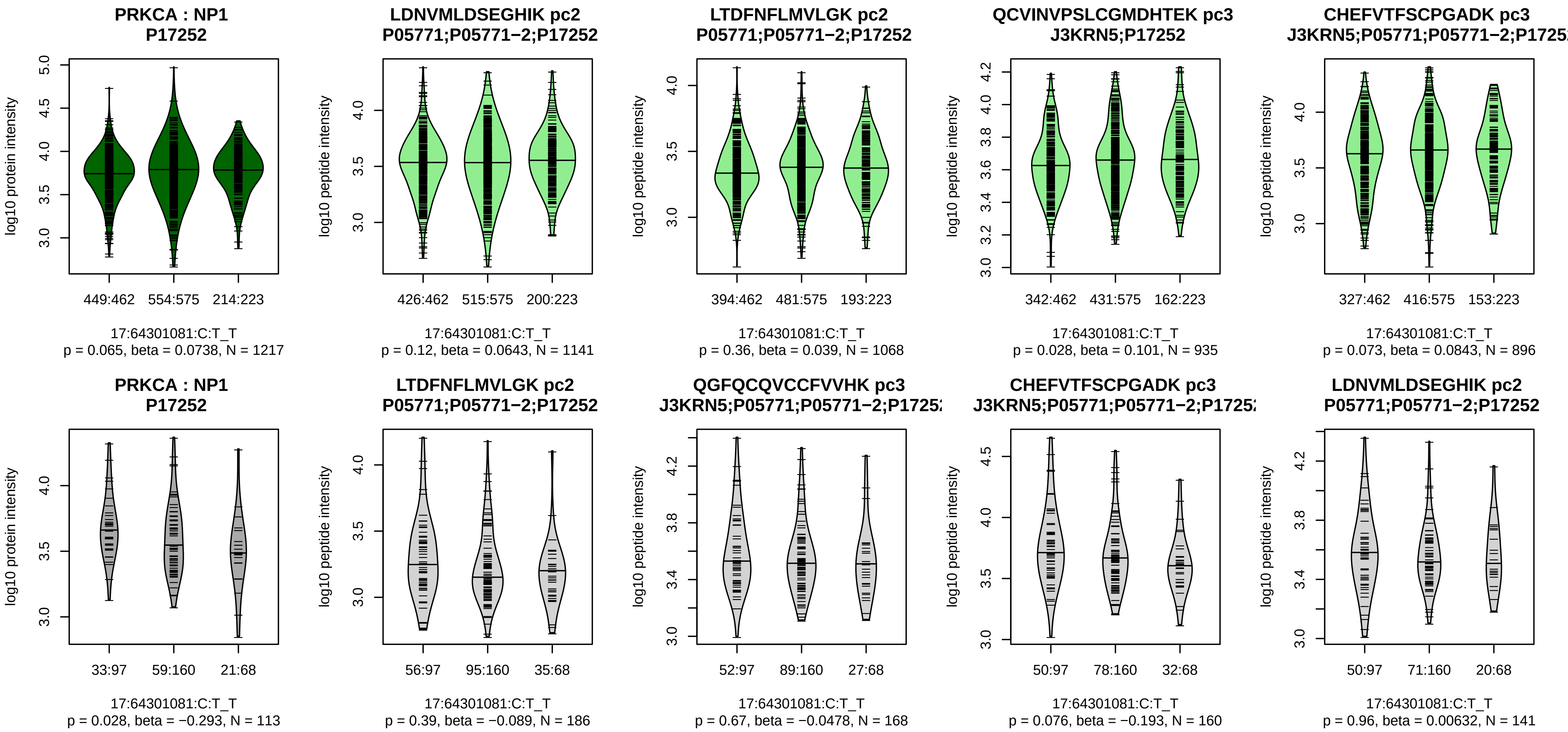


Assay Target: PRKCA
Olink UniProt: P17252
deCODE rsID: rs61762372
Proxy rsID: rs67700546
deCODE: 17:66302564:A:G
Proxy SNP: 17:64301081:C:T
deCODE log10(p): 20.8
deCODE BETA: 0.08
−:−:−:−:−:−:−:−:−:−:−:−:−:−:−:−:
1141:1068:935:896:812:791:766

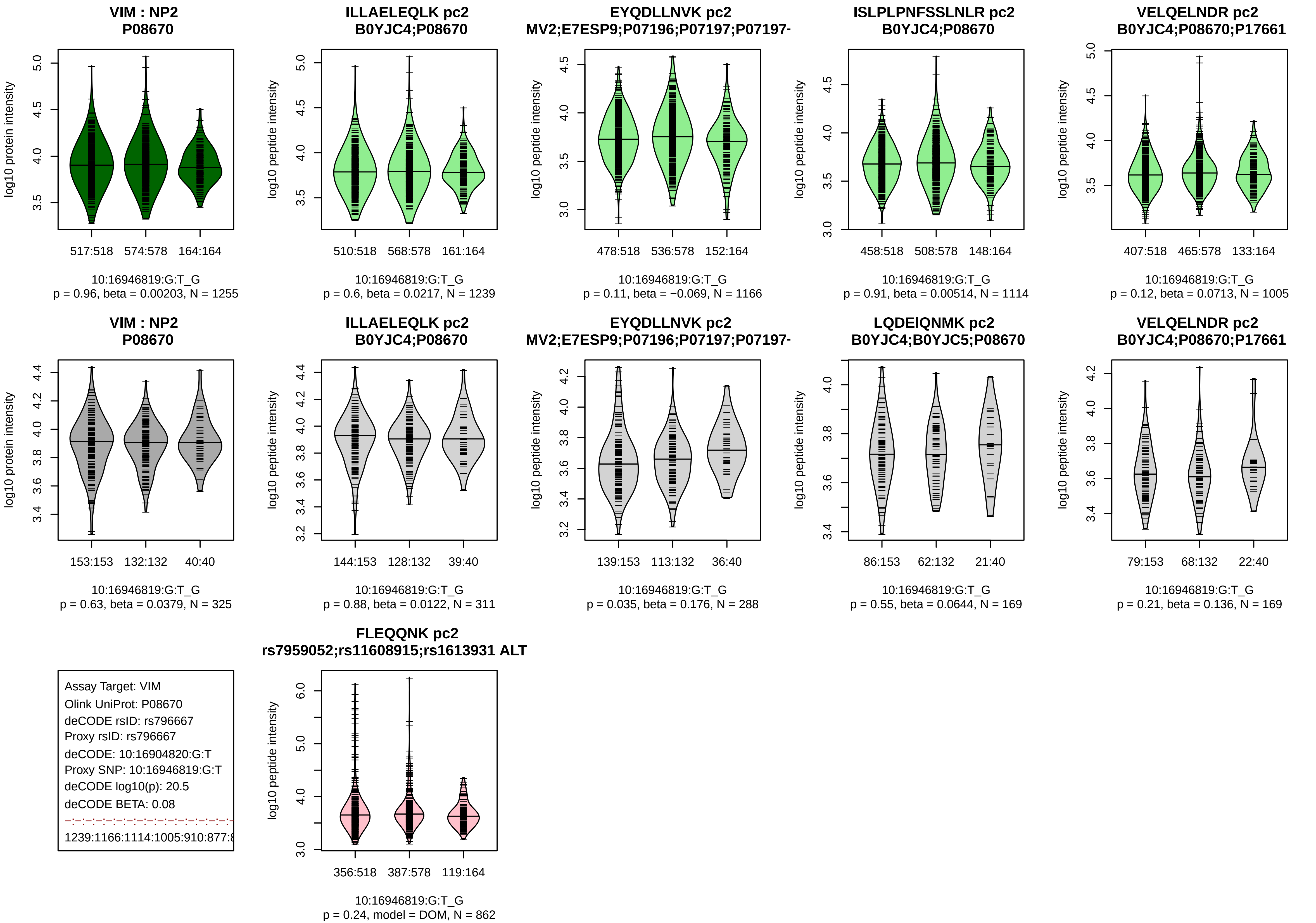

VIM : NP2
P08670
log10 protein intensity
517:518
574:578
164:164
10:16946819:G:T_G
p = 0.96, beta = 0.00203, N = 1255
ILLAELEQLK pc2
B0YJC4;P08670
log10 peptide intensity
510:518
568:578
161:164
10:16946819:G:T_G
p = 0.6, beta = 0.0217, N = 1239
EYQDLLNVK pc2
MV2;E7ESP9;P07196;P07197;P07197-
log10 peptide intensity
478:518
536:578
152:164
10:16946819:G:T_G
p = 0.11, beta = −0.069, N = 1166
ISLPLPNFSSLNLR pc2
B0YJC4;P08670
log10 peptide intensity
458:518
508:578
148:164
10:16946819:G:T_G
p = 0.91, beta = 0.00514, N = 1114
VELQELNDR pc2
B0YJC4;P08670;P17661
log10 peptide intensity
407:518
465:578
133:164
10:16946819:G:T_G
p = 0.12, beta = 0.0713, N = 1005
VIM : NP2
P08670
log10 protein intensity
153:153
132:132
40:40
10:16946819:G:T_G
p = 0.63, beta = 0.0379, N = 325
ILLAELEQLK pc2
B0YJC4;P08670
log10 peptide intensity
144:153
128:132
39:40
10:16946819:G:T_G
p = 0.88, beta = 0.0122, N = 311
EYQDLLNVK pc2
MV2;E7ESP9;P07196;P07197;P07197-
log10 peptide intensity
139:153
113:132
36:40
10:16946819:G:T_G
p = 0.035, beta = 0.176, N = 288
LQDEIQNMK pc2
B0YJC4;B0YJC5;P08670
log10 peptide intensity
86:153
62:132
21:40
10:16946819:G:T_G
p = 0.55, beta = 0.0644, N = 169
VELQELNDR pc2
B0YJC4;P08670;P17661
log10 peptide intensity
79:153
68:132
22:40
10:16946819:G:T_G
p = 0.21, beta = 0.136, N = 169
Assay Target: VIM
Olink UniProt: P08670
deCODE rsID: rs796667
Proxy rsID: rs796667
deCODE: 10:16904820:G:T
Proxy SNP: 10:16946819:G:T
deCODE log10(p): 20.5
deCODE BETA: 0.08
1239:1166:1114:1005:910:877:8
FLEQQNK pc2
rs7959052;rs11608915;rs1613931 ALT
log10 peptide intensity
356:518
387:578
119:164
10:16946819:G:T_G
p = 0.24, model = DOM, N = 862

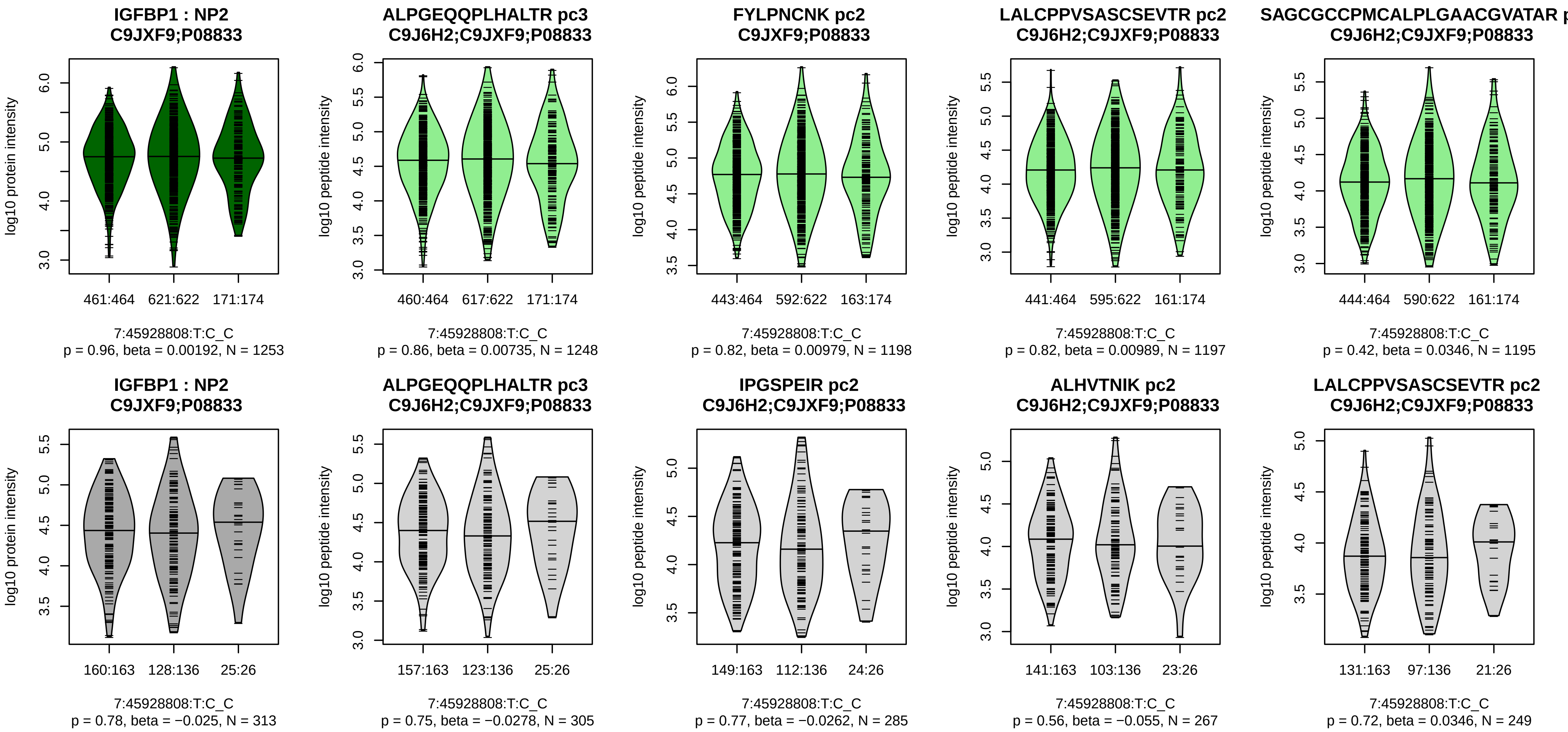


Assay Target: IGFBP1
Olink UniProt: P08833
deCODE rsID: rs3828998
Proxy rsID: rs3828998
deCODE: 7:45889209:C:T
Proxy SNP: 7:45928808:T:C
deCODE log10(p): 20.4
deCODE BETA: −0.08
–:–:–:–:–:–:–:–:–:–
1248:1206:1198:1197:1195:119

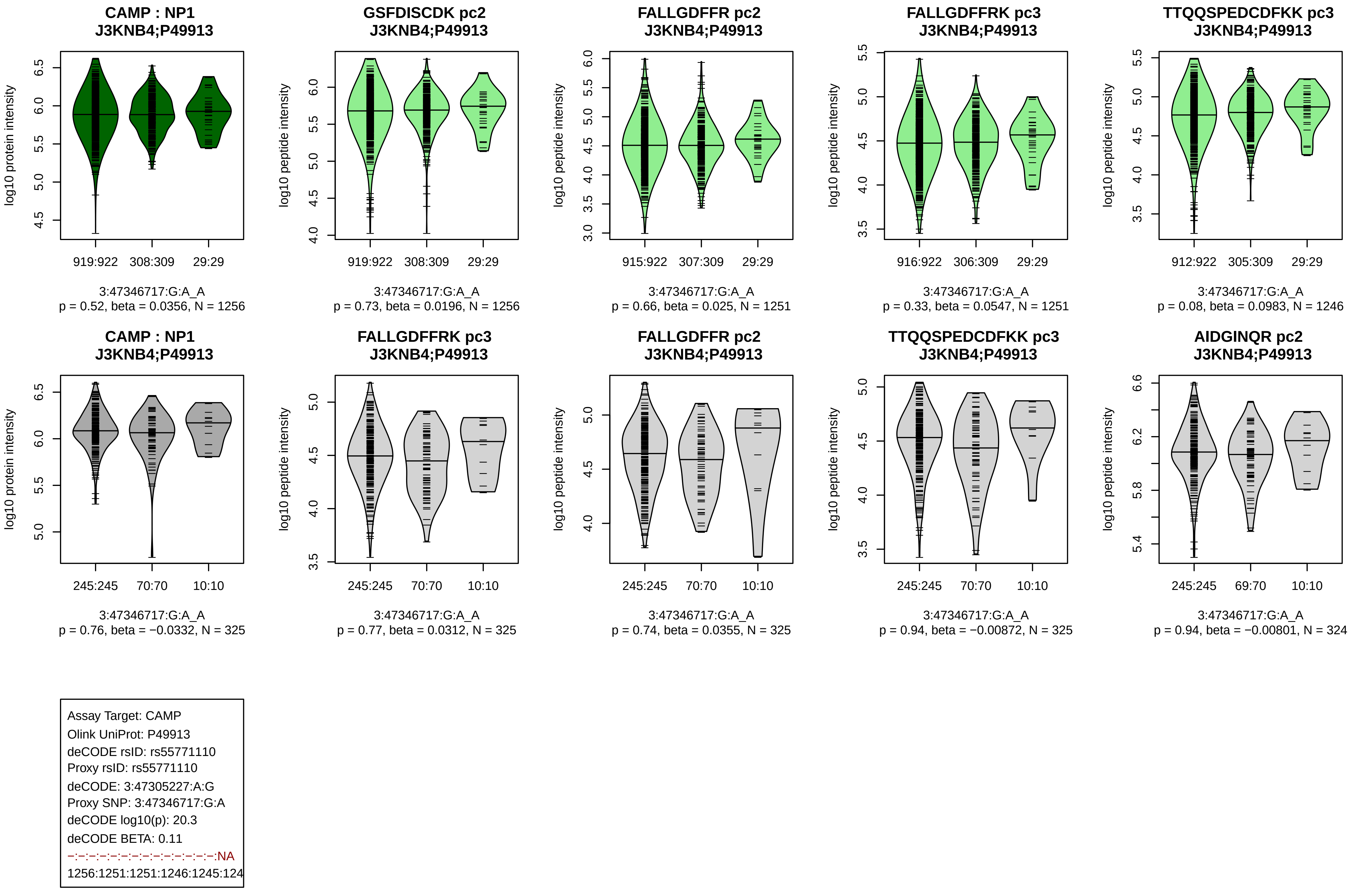
CAMP : NP1
J3KNB4;P49913
log10 protein intensity
919:922 308:309 29:29
3:47346717:G:A_A
p = 0.52, beta = 0.0356, N = 1256
GSFDISCDK pc2
J3KNB4;P49913
log10 peptide intensity
919:922 308:309 29:29
3:47346717:G:A_A
p = 0.73, beta = 0.0196, N = 1256
FALLGDFFR pc2
J3KNB4;P49913
log10 peptide intensity
915:922 307:309 29:29
3:47346717:G:A_A
p = 0.66, beta = 0.025, N = 1251
FALLGDFFRK pc3
J3KNB4;P49913
log10 peptide intensity
916:922 306:309 29:29
3:47346717:G:A_A
p = 0.33, beta = 0.0547, N = 1251
TTQQSPEDCDFKK pc3
J3KNB4;P49913
log10 peptide intensity
912:922 305:309 29:29
3:47346717:G:A_A
p = 0.08, beta = 0.0983, N = 1246
CAMP : NP1
J3KNB4;P49913
log10 protein intensity
245:245 70:70 10:10
3:47346717:G:A_A
p = 0.76, beta = −0.0332, N = 325
FALLGDFFRK pc3
J3KNB4;P49913
log10 peptide intensity
245:245 70:70 10:10
3:47346717:G:A_A
p = 0.77, beta = 0.0312, N = 325
FALLGDFFR pc2
J3KNB4;P49913
log10 peptide intensity
245:245 70:70 10:10
3:47346717:G:A_A
p = 0.74, beta = 0.0355, N = 325
TTQQSPEDCDFKK pc3
J3KNB4;P49913
log10 peptide intensity
245:245 70:70 10:10
3:47346717:G:A_A
p = 0.94, beta = −0.00872, N = 325
AIDGINQR pc2
J3KNB4;P49913
log10 peptide intensity
245:245 69:70 10:10
3:47346717:G:A_A
p = 0.94, beta = −0.00801, N = 324
Assay Target: CAMP
Olink UniProt: P49913
deCODE rsID: rs55771110
Proxy rsID: rs55771110
deCODE: 3:47305227:A:G
Proxy SNP: 3:47346717:G:A
deCODE log10(p): 20.3
deCODE BETA: 0.11
−:−:−:−:−:−:−:−:−:−:−:−:−:−:NA
1256:1251:1251:1246:1245:124

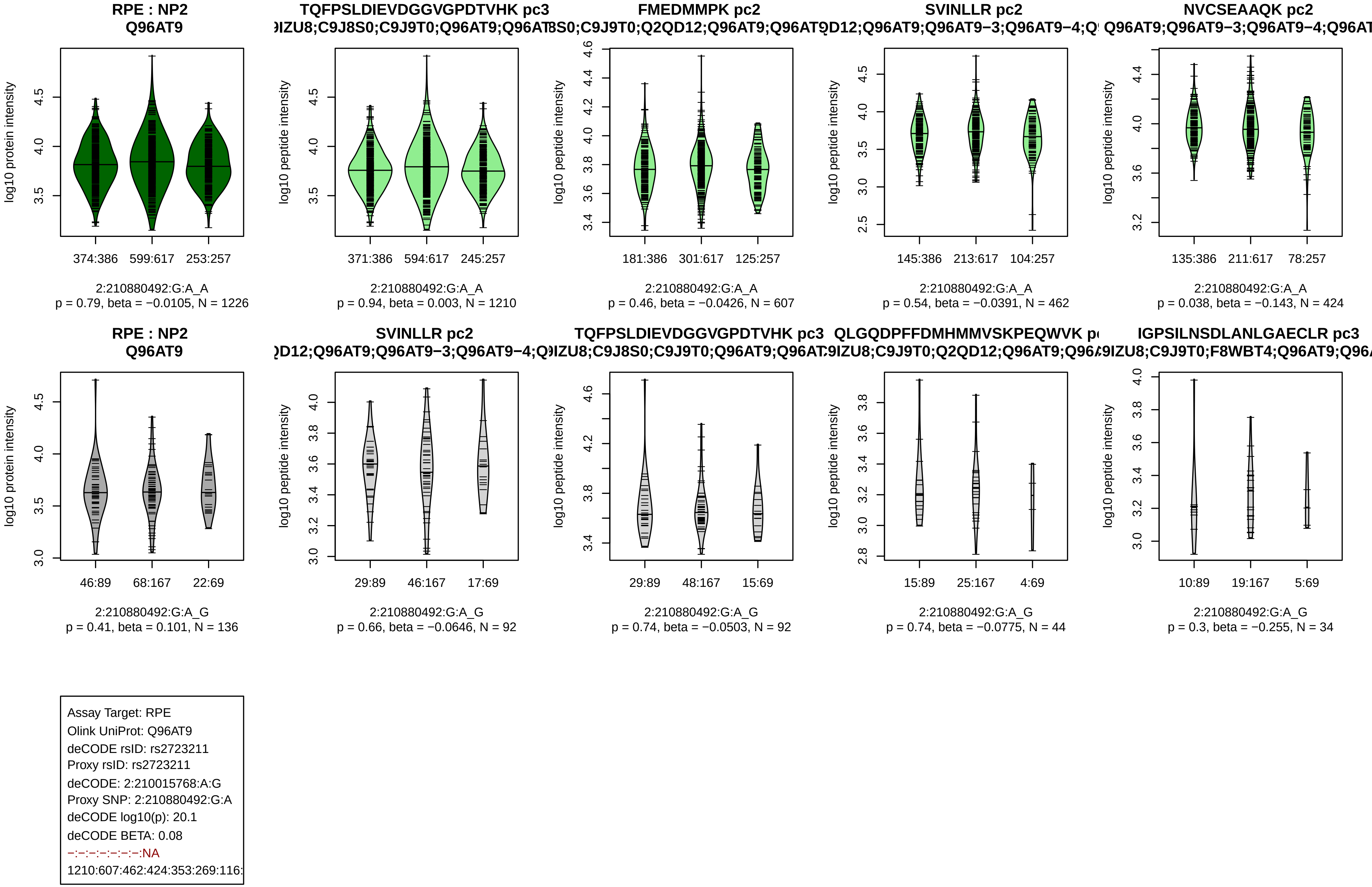

RPE : NP2
Q96AT9
log10 protein intensity
374:386 599:617 253:257
2:210880492:G:A_A
p = 0.79, beta = −0.0105, N = 1226
TQFPSLDIEVDGGVGPDTVHK pc3
IZU8;C9J8S0;C9J9T0;Q96AT9;Q96AT
log10 peptide intensity
371:386 594:617 245:257
2:210880492:G:A_A
p = 0.94, beta = 0.003, N = 1210
FMEDMMPK pc2
8S0;C9J9T0;Q2QD12;Q96AT9;Q96AT9
log10 peptide intensity
181:386 301:617 125:257
2:210880492:G:A_A
p = 0.46, beta = −0.0426, N = 607
SVINLLR pc2
D12;Q96AT9;Q96AT9−3;Q96AT9−4;Q
log10 peptide intensity
145:386 213:617 104:257
2:210880492:G:A_A
p = 0.54, beta = −0.0391, N = 462
NVCSEAAQK pc2
Q96AT9;Q96AT9−3;Q96AT9−4;Q96AT
log10 peptide intensity
135:386 211:617 78:257
2:210880492:G:A_A
p = 0.038, beta = −0.143, N = 424
RPE : NP2
Q96AT9
log10 protein intensity
46:89 68:167 22:69
2:210880492:G:A_G
p = 0.41, beta = 0.101, N = 136
SVINLLR pc2
D12;Q96AT9;Q96AT9−3;Q96AT9−4;Q
log10 peptide intensity
29:89 46:167 17:69
2:210880492:G:A_G
p = 0.66, beta = −0.0646, N = 92
TQFPSLDIEVDGGVGPDTVHK pc3
IZU8;C9J8S0;C9J9T0;Q96AT9;Q96AT
log10 peptide intensity
29:89 48:167 15:69
2:210880492:G:A_G
p = 0.74, beta = −0.0503, N = 92
QLGQDPFFDMHMMVSKPEQWVK p
9IZU8;C9J9T0;Q2QD12;Q96AT9;Q96A
log10 peptide intensity
15:89 25:167 4:69
2:210880492:G:A_G
p = 0.74, beta = −0.0775, N = 44
IGPSILNSDLANLGAECLR pc3
9IZU8;C9J9T0;F8WBT4;Q96AT9;Q96A
log10 peptide intensity
10:89 19:167 5:69
2:210880492:G:A_G
p = 0.3, beta = −0.255, N = 34
Assay Target: RPE
Olink UniProt: Q96AT9
deCODE rsID: rs2723211
Proxy rsID: rs2723211
deCODE: 2:210015768:A:G
Proxy SNP: 2:210880492:G:A
deCODE log10(p): 20.1
deCODE BETA: 0.08
−:−:−:−:−:−:−:NA
1210:607:462:424:353:269:116:

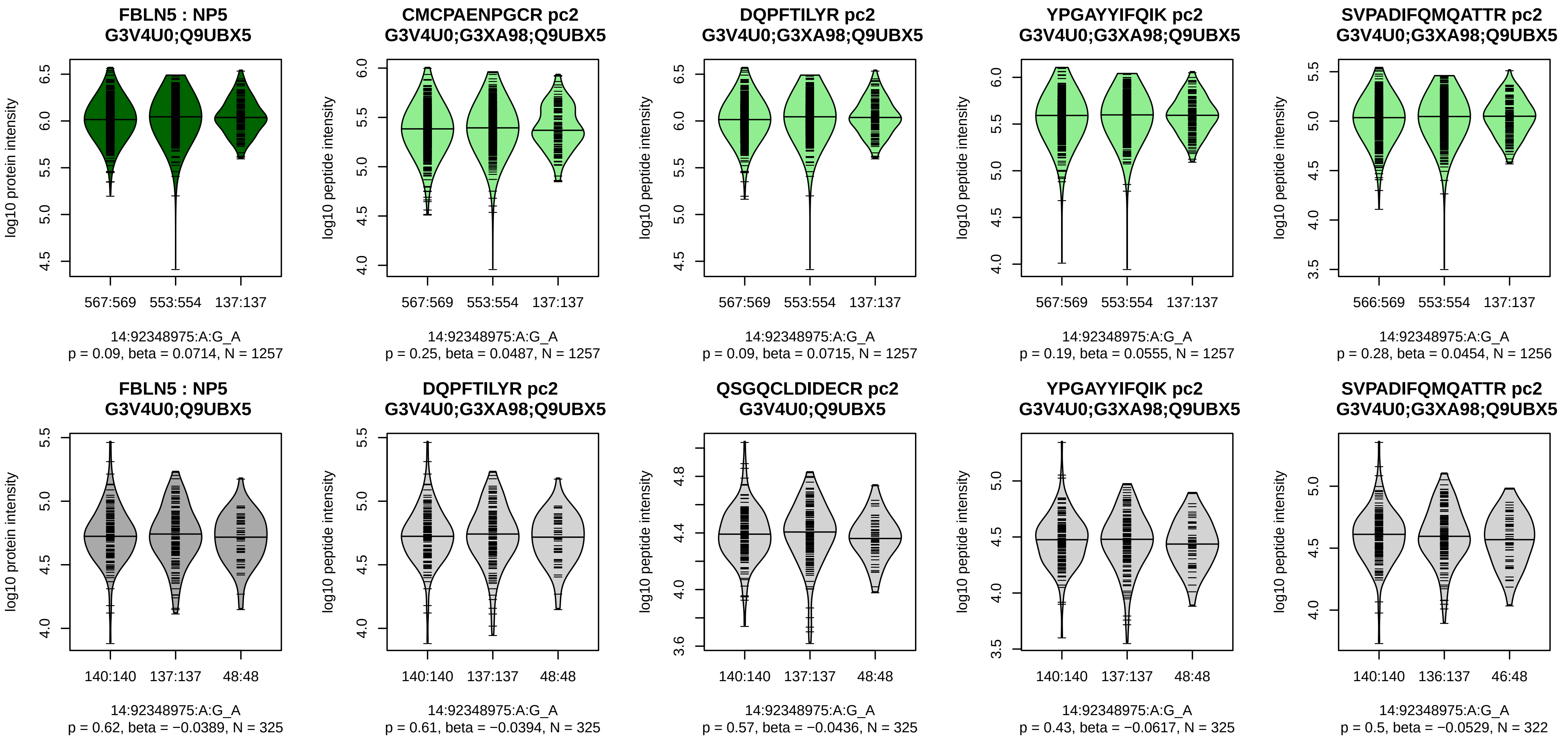


Assay Target: FBLN5
Olink UniProt: Q9UBX5
deCODE rsID: rs2430349
Proxy rsID: rs2430349
deCODE: 14:91882631:A:G
Proxy SNP: 14:92348975:A:G
deCODE log10(p): 19.1
deCODE BETA: 0.08
–:–:–:–:–:–:–:–:–:–:–:–:–:–:–:–:
1257:1257:1257:1256:1255:125

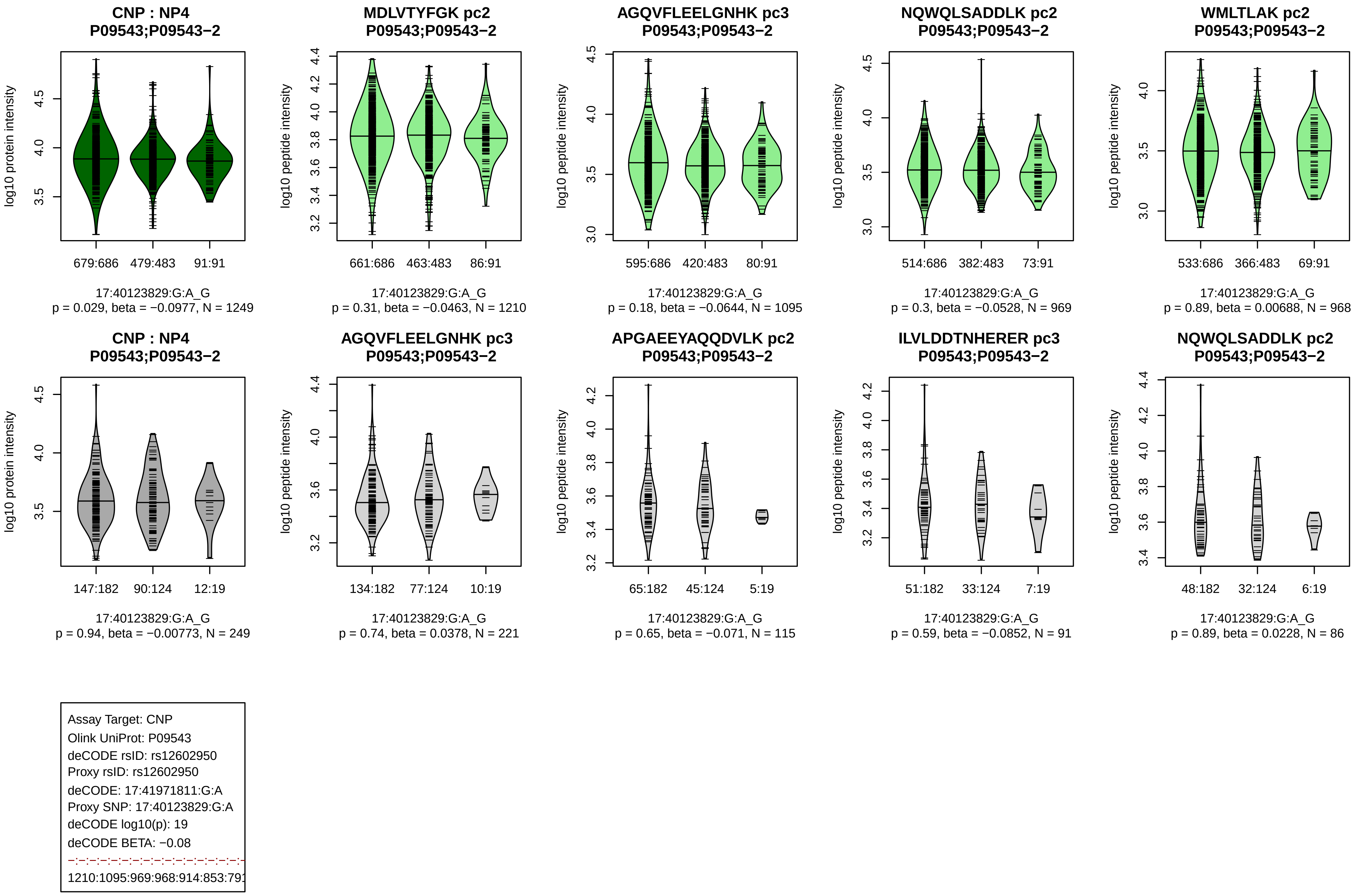

CNP : NP4
P09543;P09543−2
log10 protein intensity
679:686 479:483 91:91
17:40123829:G:A_G
p = 0.029, beta = −0.0977, N = 1249
MDLVTYFGK pc2
P09543;P09543−2
log10 peptide intensity
661:686 463:483 86:91
17:40123829:G:A_G
p = 0.31, beta = −0.0463, N = 1210
AGQVFLEELGNHK pc3
P09543;P09543−2
log10 peptide intensity
595:686 420:483 80:91
17:40123829:G:A_G
p = 0.18, beta = −0.0644, N = 1095
NQWQLSADDLK pc2
P09543;P09543−2
log10 peptide intensity
514:686 382:483 73:91
17:40123829:G:A_G
p = 0.3, beta = −0.0528, N = 969
WMLTLAK pc2
P09543;P09543−2
log10 peptide intensity
533:686 366:483 69:91
17:40123829:G:A_G
p = 0.89, beta = 0.00688, N = 968
CNP : NP4
P09543;P09543−2
log10 protein intensity
147:182 90:124 12:19
17:40123829:G:A_G
p = 0.94, beta = −0.00773, N = 249
AGQVFLEELGNHK pc3
P09543;P09543−2
log10 peptide intensity
134:182 77:124 10:19
17:40123829:G:A_G
p = 0.74, beta = 0.0378, N = 221
APGAEEYAQQDVLK pc2
P09543;P09543−2
log10 peptide intensity
65:182 45:124 5:19
17:40123829:G:A_G
p = 0.65, beta = −0.071, N = 115
ILVLDDTNHERER pc3
P09543;P09543−2
log10 peptide intensity
51:182 33:124 7:19
17:40123829:G:A_G
p = 0.59, beta = −0.0852, N = 91
NQWQLSADDLK pc2
P09543;P09543−2
log10 peptide intensity
48:182 32:124 6:19
17:40123829:G:A_G
p = 0.89, beta = 0.0228, N = 86
Assay Target: CNP
Olink UniProt: P09543
deCODE rsID: rs12602950
Proxy rsID: rs12602950
deCODE: 17:41971811:G:A
Proxy SNP: 17:40123829:G:A
deCODE log10(p): 19
deCODE BETA: −0.08
1210:1095:969:968:914:853:791

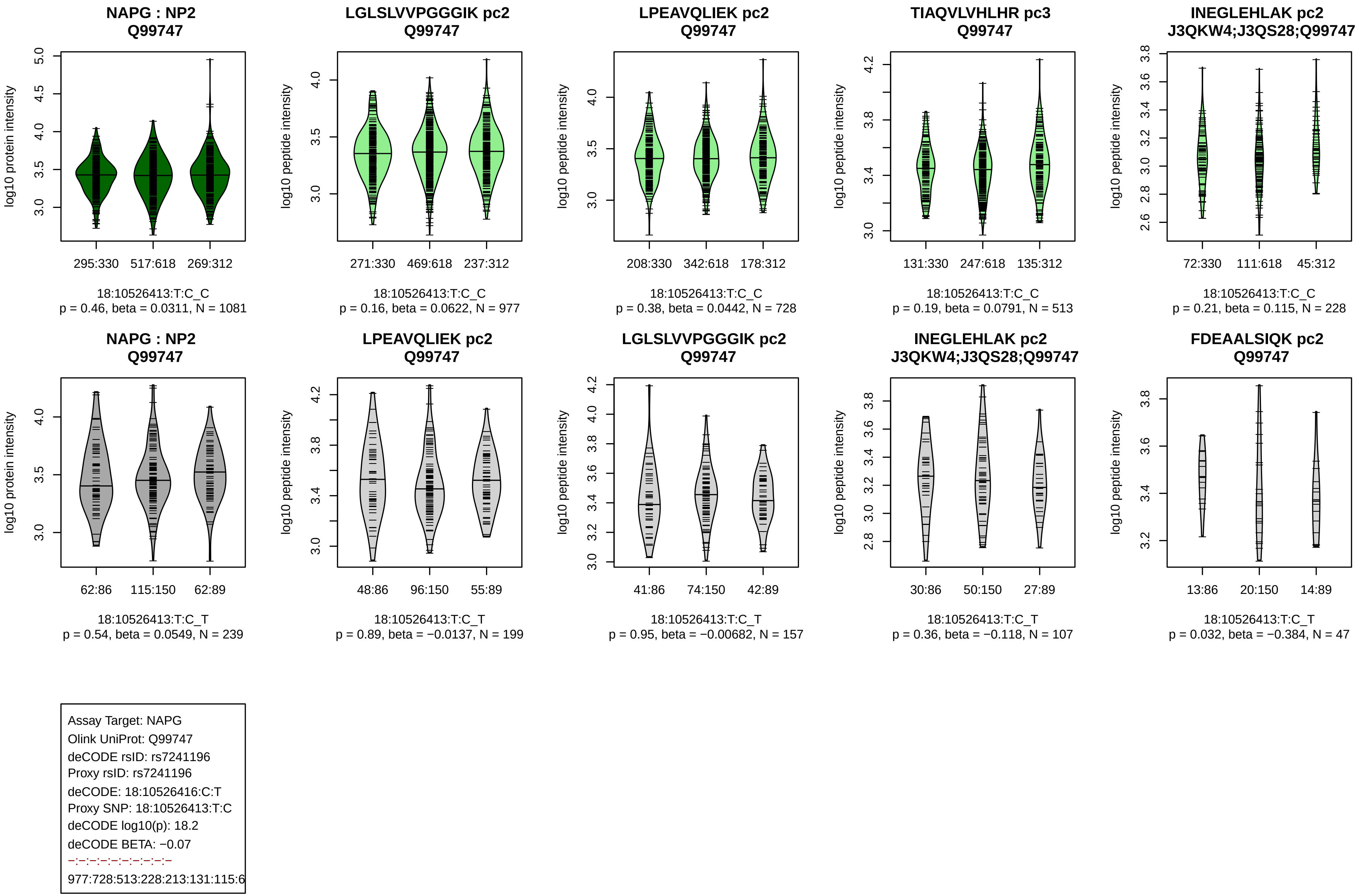
NAPG : NP2
Q99747
log10 protein intensity
295:330
517:618
269:312
18:10526413:T:C_C
p = 0.46, beta = 0.0311, N = 1081
LGLSLVVPGGGIK pc2
Q99747
log10 peptide intensity
271:330
469:618
237:312
18:10526413:T:C_C
p = 0.16, beta = 0.0622, N = 977
LPEAVQLIEK pc2
Q99747
log10 peptide intensity
208:330
342:618
178:312
18:10526413:T:C_C
p = 0.38, beta = 0.0442, N = 728
TIAQVLVHLHR pc3
Q99747
log10 peptide intensity
131:330
247:618
135:312
18:10526413:T:C_C
p = 0.19, beta = 0.0791, N = 513
INEGLEHLAK pc2
J3QKW4;J3QS28;Q99747
log10 peptide intensity
72:330
111:618
45:312
18:10526413:T:C_C
p = 0.21, beta = 0.115, N = 228
NAPG : NP2
Q99747
log10 protein intensity
62:86
115:150
62:89
18:10526413:T:C_T
p = 0.54, beta = 0.0549, N = 239
LPEAVQLIEK pc2
Q99747
log10 peptide intensity
48:86
96:150
55:89
18:10526413:T:C_T
p = 0.89, beta = −0.0137, N = 199
LGLSLVVPGGGIK pc2
Q99747
log10 peptide intensity
41:86
74:150
42:89
18:10526413:T:C_T
p = 0.95, beta = −0.00682, N = 157
INEGLEHLAK pc2
J3QKW4;J3QS28;Q99747
log10 peptide intensity
30:86
50:150
27:89
18:10526413:T:C_T
p = 0.36, beta = −0.118, N = 107
FDEAALSIQK pc2
Q99747
log10 peptide intensity
13:86
20:150
14:89
18:10526413:T:C_T
p = 0.032, beta = −0.384, N = 47
Assay Target: NAPG
Olink UniProt: Q99747
deCODE rsID: rs7241196
Proxy rsID: rs7241196
deCODE: 18:10526416:C:T
Proxy SNP: 18:10526413:T:C
deCODE log10(p): 18.2
deCODE BETA: −0.07
−:−:−:−:−:−:−:−:−
977:728:513:228:213:131:115:6

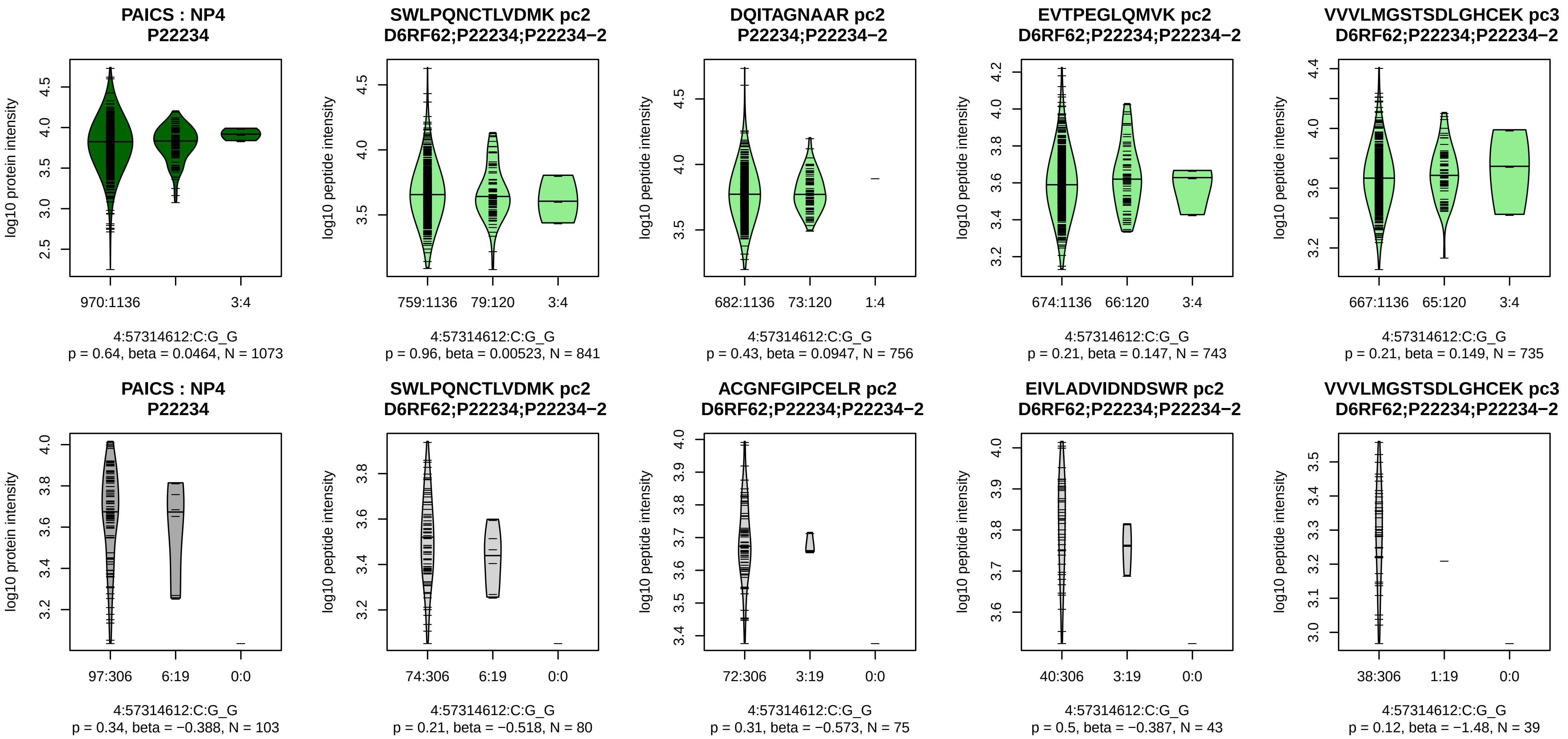


Assay Target: PAICS
Olink UniProt: P22234
deCODE rsID: rs138933625
Proxy rsID: rs138933625
deCODE: 4:56448446:G:C
Proxy SNP: 4:57314612:C:G
deCODE log10(p): 18.2
deCODE BETA: −0.19
−:−:−:−:−:−:−:−:−:−:−:−:*:−
841:756:743:735:661:568:566:4

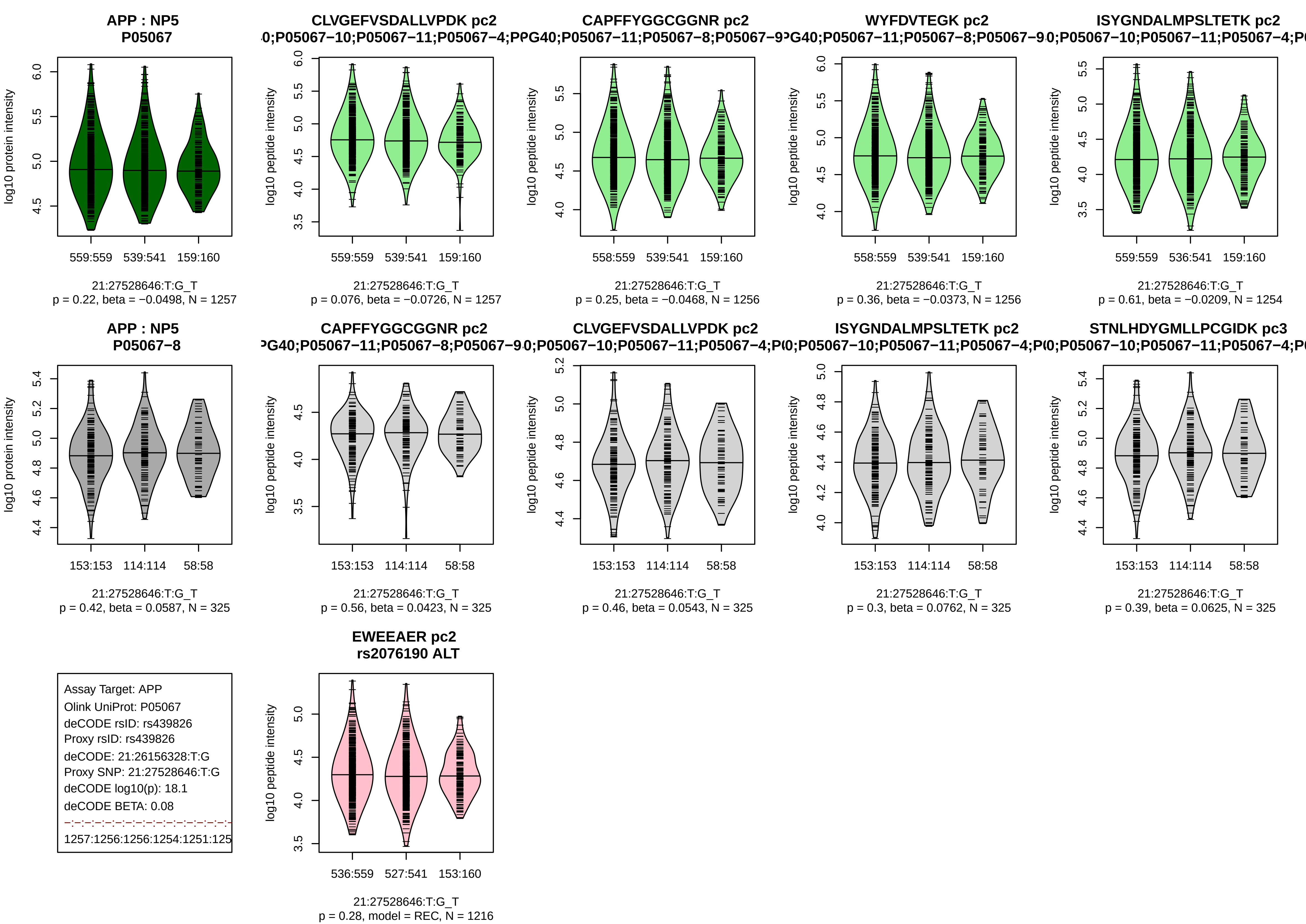

APP : NP5
P05067
log10 protein intensity
559:559 539:541 159:160
21:27528646:T:G_T
p = 0.22, beta = −0.0498, N = 1257
CLVGEFVSDALLVPDK pc2
0;P05067−10;P05067−11;P05067−4;P
log10 peptide intensity
559:559 539:541 159:160
21:27528646:T:G_T
p = 0.076, beta = −0.0726, N = 1257
CAPFFYGGCGGNR pc2
PG40;P05067−11;P05067−8;P05067−9
558:559 539:541 159:160
21:27528646:T:G_T
p = 0.25, beta = −0.0468, N = 1256
WYFDVTEGK pc2
PG40;P05067−11;P05067−8;P05067−9
558:559 539:541 159:160
21:27528646:T:G_T
p = 0.36, beta = −0.0373, N = 1256
ISYGNDALMPSLTETK pc2
0;P05067−10;P05067−11;P05067−4;P
559:559 536:541 159:160
21:27528646:T:G_T
p = 0.61, beta = −0.0209, N = 1254
APP : NP5
P05067−8
153:153 114:114 58:58
21:27528646:T:G_T
p = 0.42, beta = 0.0587, N = 325
CAPFFYGGCGGNR pc2
PG40;P05067−11;P05067−8;P05067−9
21:27528646:T:G_T
p = 0.56, beta = 0.0423, N = 325
CLVGEFVSDALLVPDK pc2
0;P05067−10;P05067−11;P05067−4;P
21:27528646:T:G_T
p = 0.46, beta = 0.0543, N = 325
ISYGNDALMPSLTETK pc2
0;P05067−10;P05067−11;P05067−4;P
21:27528646:T:G_T
p = 0.3, beta = 0.0762, N = 325
STNLHDYGMLLPCGIDK pc3
0;P05067−10;P05067−11;P05067−4;P
21:27528646:T:G_T
p = 0.39, beta = 0.0625, N = 325
Assay Target: APP
Olink UniProt: P05067
deCODE rsID: rs439826
Proxy rsID: rs439826
deCODE: 21:26156328:T:G
Proxy SNP: 21:27528646:T:G
deCODE log10(p): 18.1
deCODE BETA: 0.08
1257:1256:1256:1254:1251:125
EWEEAER pc2
rs2076190 ALT
536:559 527:541 153:160
21:27528646:T:G_T
p = 0.28, model = REC, N = 1216

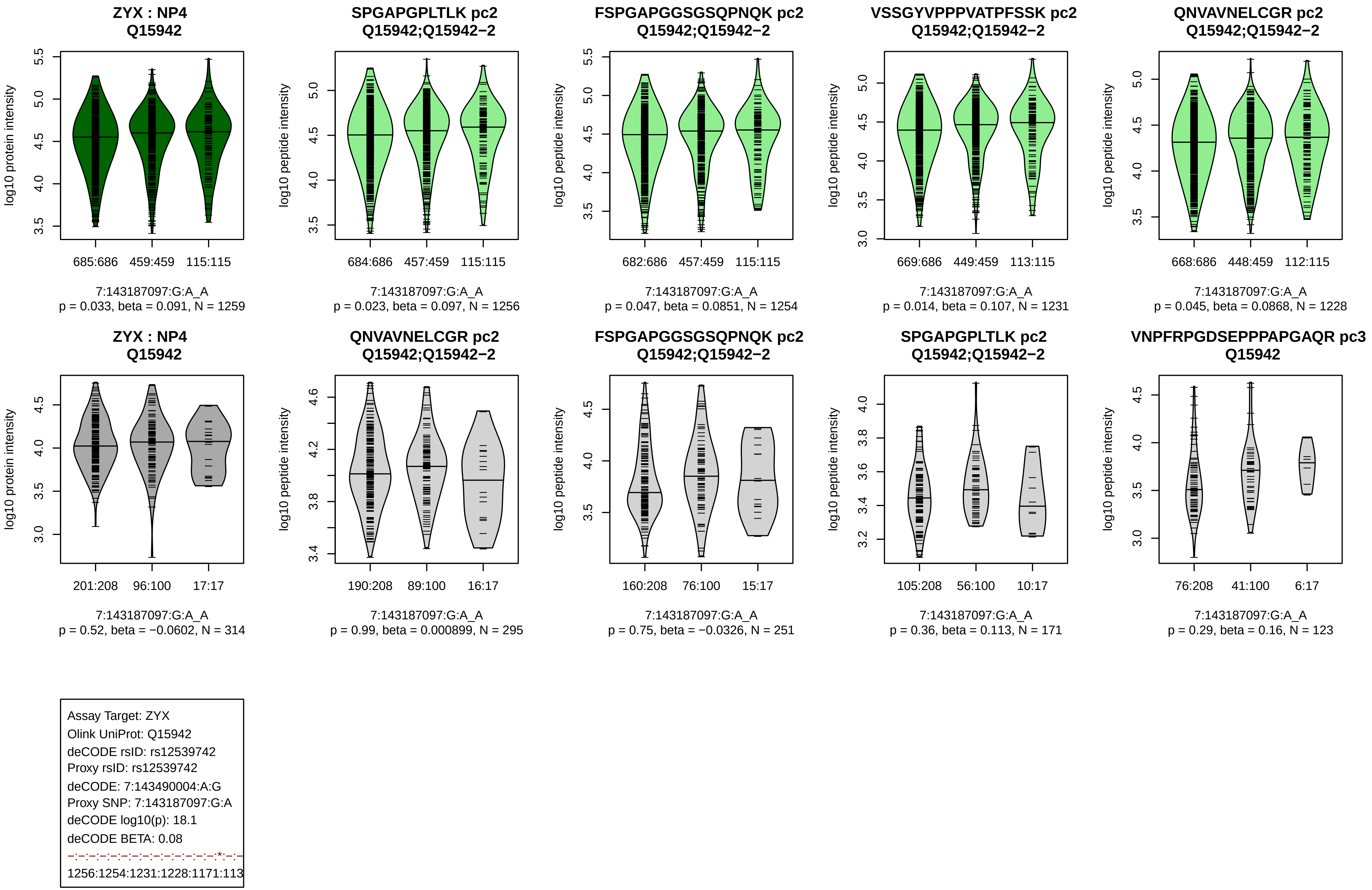

ZYX : NP4
Q15942
log10 protein intensity
685:686
459:459
115:115
7:143187097:G:A_A
p = 0.033, beta = 0.091, N = 1259
SPGAPGPLTLK pc2
Q15942;Q15942−2
log10 peptide intensity
684:686
457:459
115:115
7:143187097:G:A_A
p = 0.023, beta = 0.097, N = 1256
FSPGAPGGSGSQPNQK pc2
Q15942;Q15942−2
log10 peptide intensity
682:686
457:459
115:115
7:143187097:G:A_A
p = 0.047, beta = 0.0851, N = 1254
VSSGYVPPPVATPFSSK pc2
Q15942;Q15942−2
log10 peptide intensity
669:686
449:459
113:115
7:143187097:G:A_A
p = 0.014, beta = 0.107, N = 1231
QNVAVNELCGR pc2
Q15942;Q15942−2
log10 peptide intensity
668:686
448:459
112:115
7:143187097:G:A_A
p = 0.045, beta = 0.0868, N = 1228
ZYX : NP4
Q15942
log10 protein intensity
201:208
96:100
17:17
7:143187097:G:A_A
p = 0.52, beta = −0.0602, N = 314
QNVAVNELCGR pc2
Q15942;Q15942−2
log10 peptide intensity
190:208
89:100
16:17
7:143187097:G:A_A
p = 0.99, beta = 0.000899, N = 295
FSPGAPGGSGSQPNQK pc2
Q15942;Q15942−2
log10 peptide intensity
160:208
76:100
15:17
7:143187097:G:A_A
p = 0.75, beta = −0.0326, N = 251
SPGAPGPLTLK pc2
Q15942;Q15942−2
log10 peptide intensity
105:208
56:100
10:17
7:143187097:G:A_A
p = 0.36, beta = 0.113, N = 171
VNPFRPGDSEPPPAPGAQR pc3
Q15942
log10 peptide intensity
76:208
41:100
6:17
7:143187097:G:A_A
p = 0.29, beta = 0.16, N = 123
Assay Target: ZYX
Olink UniProt: Q15942
deCODE rsID: rs12539742
Proxy rsID: rs12539742
deCODE: 7:143490004:A:G
Proxy SNP: 7:143187097:G:A
deCODE log10(p): 18.1
deCODE BETA: 0.08
−:−:−:−:−:−:−:−:−:−:−:−:−:−:*:−:−
1256:1254:1231:1228:1171:113

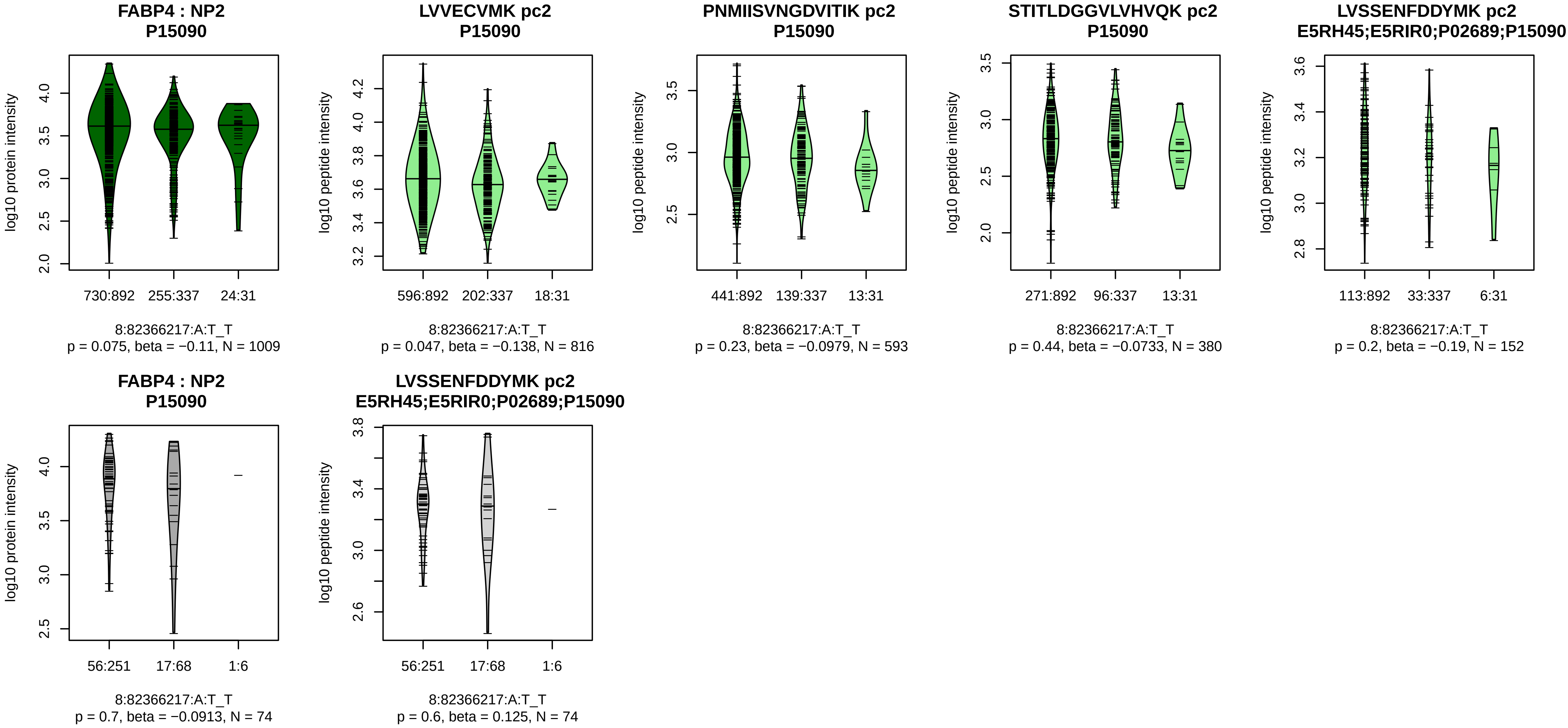


Assay Target: FABP4
Olink UniProt: P15090
deCODE rsID: rs16909138
Proxy rsID: rs16909138
deCODE: 8:81453981:TT:!
Proxy SNP: 8:82366217:A:T
deCODE log10(p): 17.4
deCODE BETA: −0.1
–:–:–:–:*
816:593:380:152:149

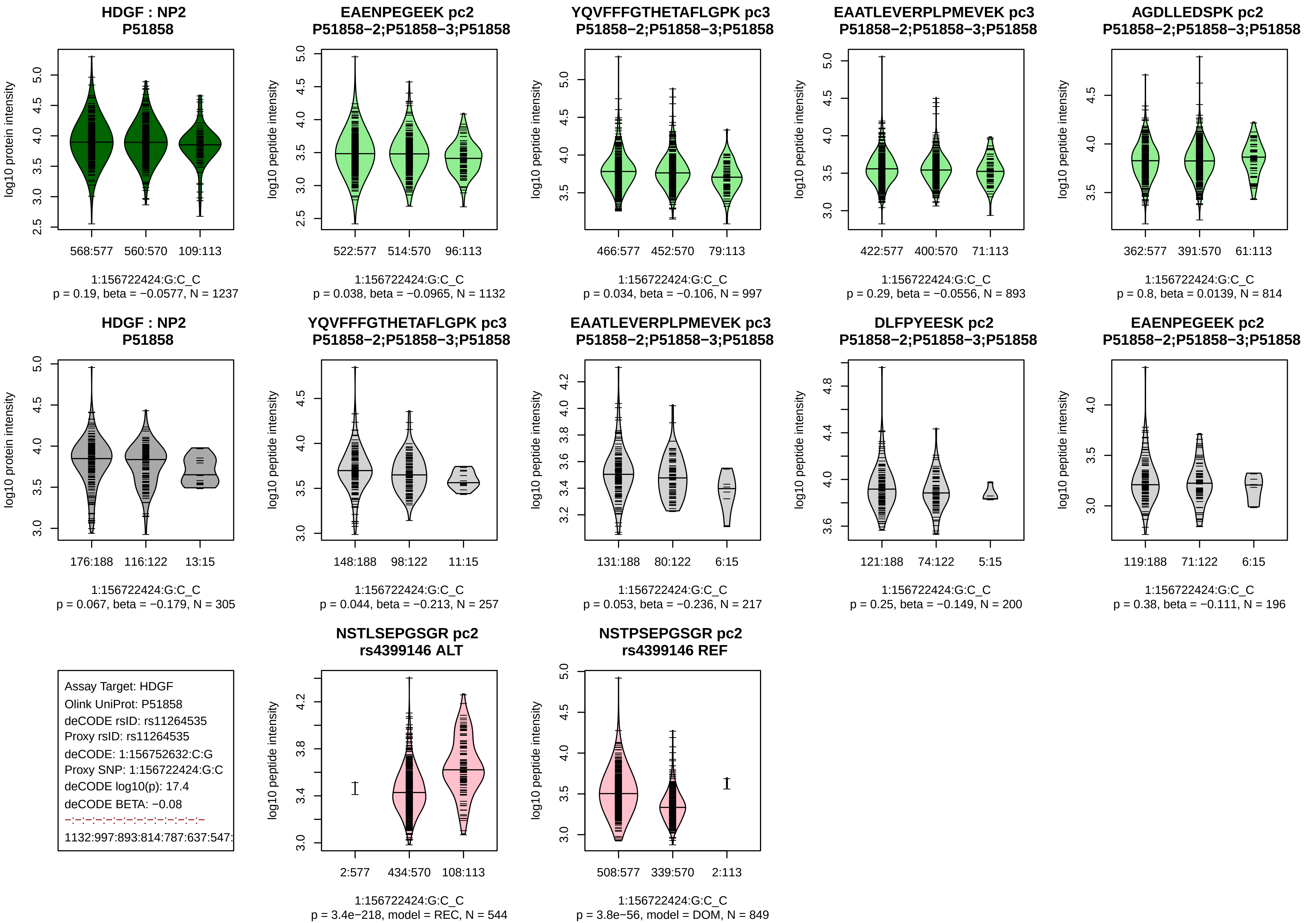
HDGF : NP2
P51858
log10 protein intensity
568:577
560:570
109:113
1:156722424:G:C_C
p = 0.19, beta = −0.0577, N = 1237
EAENPEGEEK pc2
P51858−2;P51858−3;P51858
log10 peptide intensity
522:577
514:570
96:113
1:156722424:G:C_C
p = 0.038, beta = −0.0965, N = 1132
YQVFFFGTHETAFLGPK pc3
P51858−2;P51858−3;P51858
log10 peptide intensity
466:577
452:570
79:113
1:156722424:G:C_C
p = 0.034, beta = −0.106, N = 997
EAATLEVERPLPMEVEK pc3
P51858−2;P51858−3;P51858
log10 peptide intensity
422:577
400:570
71:113
1:156722424:G:C_C
p = 0.29, beta = −0.0556, N = 893
AGDLLEDSPK pc2
P51858−2;P51858−3;P51858
log10 peptide intensity
362:577
391:570
61:113
1:156722424:G:C_C
p = 0.8, beta = 0.0139, N = 814
HDGF : NP2
P51858
log10 protein intensity
176:188
116:122
13:15
1:156722424:G:C_C
p = 0.067, beta = −0.179, N = 305
YQVFFFGTHETAFLGPK pc3
P51858−2;P51858−3;P51858
log10 peptide intensity
148:188
98:122
11:15
1:156722424:G:C_C
p = 0.044, beta = −0.213, N = 257
EAATLEVERPLPMEVEK pc3
P51858−2;P51858−3;P51858
log10 peptide intensity
131:188
80:122
6:15
1:156722424:G:C_C
p = 0.053, beta = −0.236, N = 217
DLFPYEESK pc2
P51858−2;P51858−3;P51858
log10 peptide intensity
121:188
74:122
5:15
1:156722424:G:C_C
p = 0.25, beta = −0.149, N = 200
EAENPEGEEK pc2
P51858−2;P51858−3;P51858
log10 peptide intensity
119:188
71:122
6:15
1:156722424:G:C_C
p = 0.38, beta = −0.111, N = 196
Assay Target: HDGF
Olink UniProt: P51858
deCODE rsID: rs11264535
Proxy rsID: rs11264535
deCODE: 1:156752632:C:G
Proxy SNP: 1:156722424:G:C
deCODE log10(p): 17.4
deCODE BETA: −0.08
1132:997:893:814:787:637:547:
NSTLSEPGSGR pc2
rs4399146 ALT
log10 peptide intensity
2:577
434:570
108:113
1:156722424:G:C_C
p = 3.4e−218, model = REC, N = 544
NSTPSEPGSGR pc2
rs4399146 REF
log10 peptide intensity
508:577
339:570
2:113
1:156722424:G:C_C
p = 3.8e−56, model = DOM, N = 849

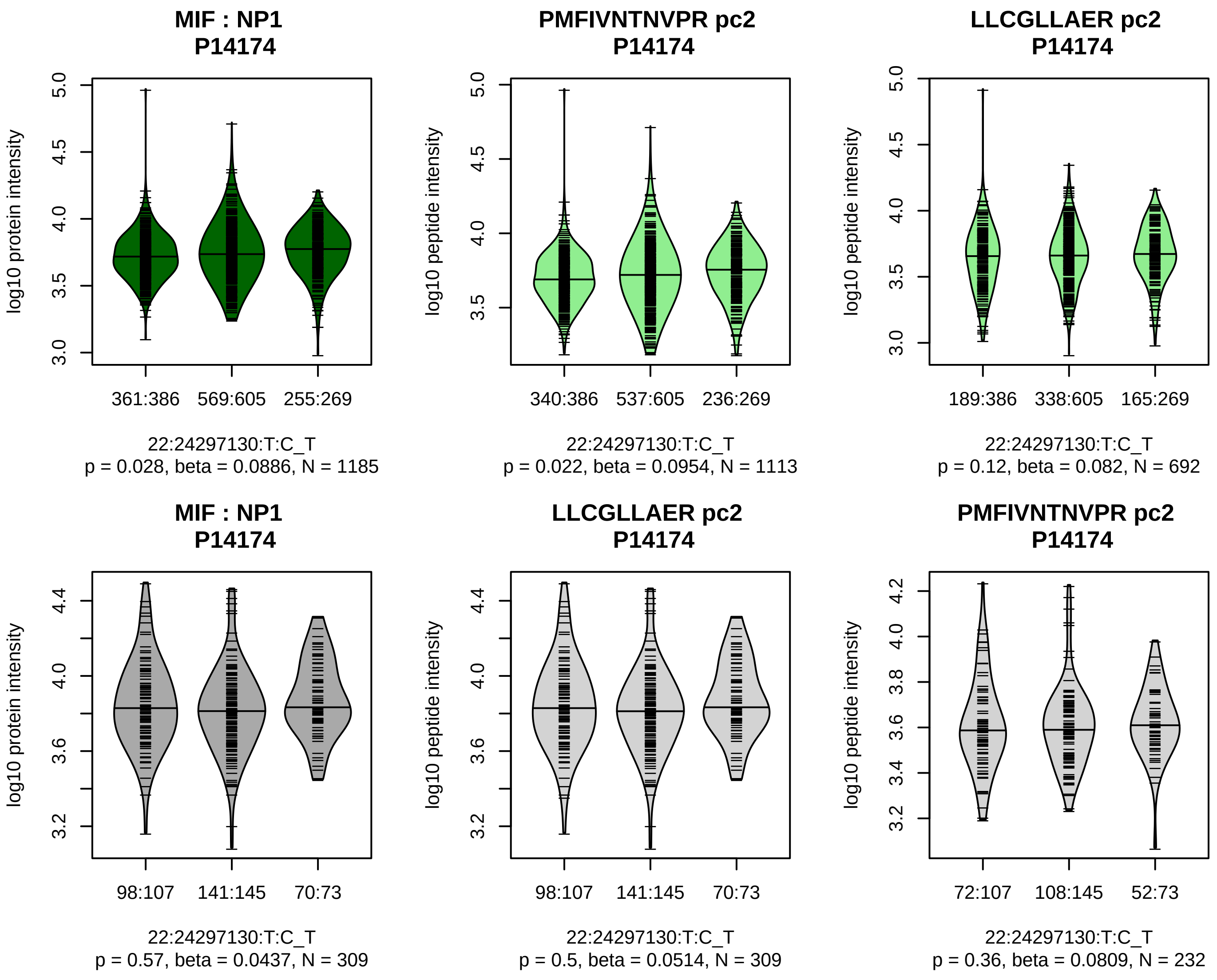


Assay Target: MIF
Olink UniProt: P14174
deCODE rsID: rs2739338
Proxy rsID: rs2739338
deCODE: 22:23954943:T:C
Proxy SNP: 22:24297130:T:C
deCODE log10(p): 17.3
deCODE BETA: 0.07
–:–
1113:692

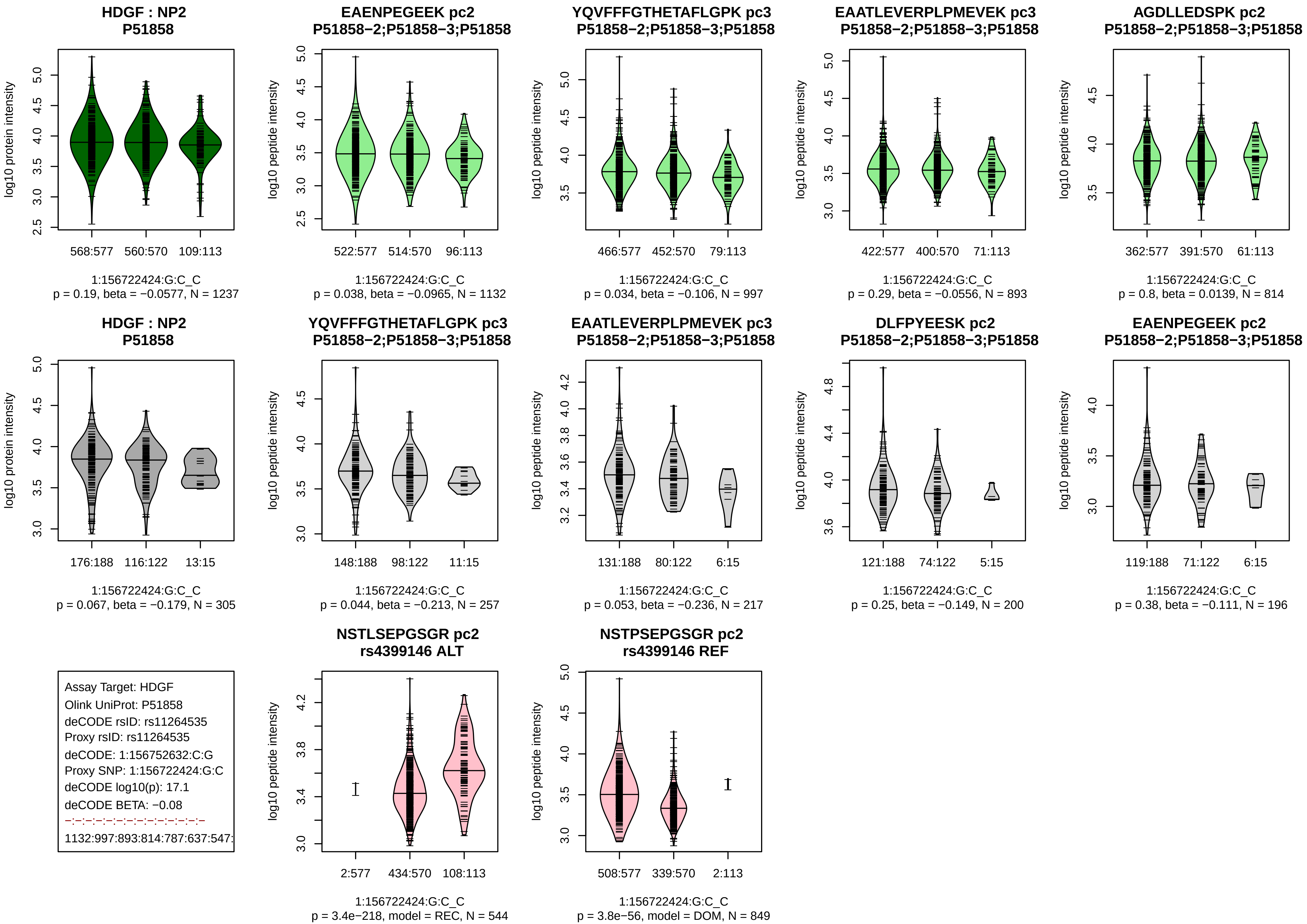
HDGF : NP2
P51858
log10 protein intensity
568:577 560:570 109:113
1:156722424:G:C_C
p = 0.19, beta = −0.0577, N = 1237
EAENPEGEEK pc2
P51858−2;P51858−3;P51858
log10 peptide intensity
522:577 514:570 96:113
1:156722424:G:C_C
p = 0.038, beta = −0.0965, N = 1132
YQVFFFGTHETAFLGPK pc3
P51858−2;P51858−3;P51858
466:577 452:570 79:113
1:156722424:G:C_C
p = 0.034, beta = −0.106, N = 997
EAATLEVERPLPMEVEK pc3
P51858−2;P51858−3;P51858
422:577 400:570 71:113
1:156722424:G:C_C
p = 0.29, beta = −0.0556, N = 893
AGDLLEDSPK pc2
P51858−2;P51858−3;P51858
362:577 391:570 61:113
1:156722424:G:C_C
p = 0.8, beta = 0.0139, N = 814
HDGF : NP2
P51858
176:188 116:122 13:15
1:156722424:G:C_C
p = 0.067, beta = −0.179, N = 305
YQVFFFGTHETAFLGPK pc3
P51858−2;P51858−3;P51858
148:188 98:122 11:15
1:156722424:G:C_C
p = 0.044, beta = −0.213, N = 257
EAATLEVERPLPMEVEK pc3
P51858−2;P51858−3;P51858
131:188 80:122 6:15
1:156722424:G:C_C
p = 0.053, beta = −0.236, N = 217
DLFPYEESK pc2
P51858−2;P51858−3;P51858
121:188 74:122 5:15
1:156722424:G:C_C
p = 0.25, beta = −0.149, N = 200
EAENPEGEEK pc2
P51858−2;P51858−3;P51858
119:188 71:122 6:15
1:156722424:G:C_C
p = 0.38, beta = −0.111, N = 196
Assay Target: HDGF
Olink UniProt: P51858
deCODE rsID: rs11264535
Proxy rsID: rs11264535
deCODE: 1:156752632:C:G
Proxy SNP: 1:156722424:G:C
deCODE log10(p): 17.1
deCODE BETA: −0.08
−:−:−:−:−:−:−:−:−:−:−:−:−
1132:997:893:814:787:637:547:
NSTLSEPGSGR pc2
rs4399146 ALT
2:577 434:570 108:113
1:156722424:G:C_C
p = 3.4e−218, model = REC, N = 544
NSTPSEPGSGR pc2
rs4399146 REF
508:577 339:570 2:113
1:156722424:G:C_C
p = 3.8e−56, model = DOM, N = 849

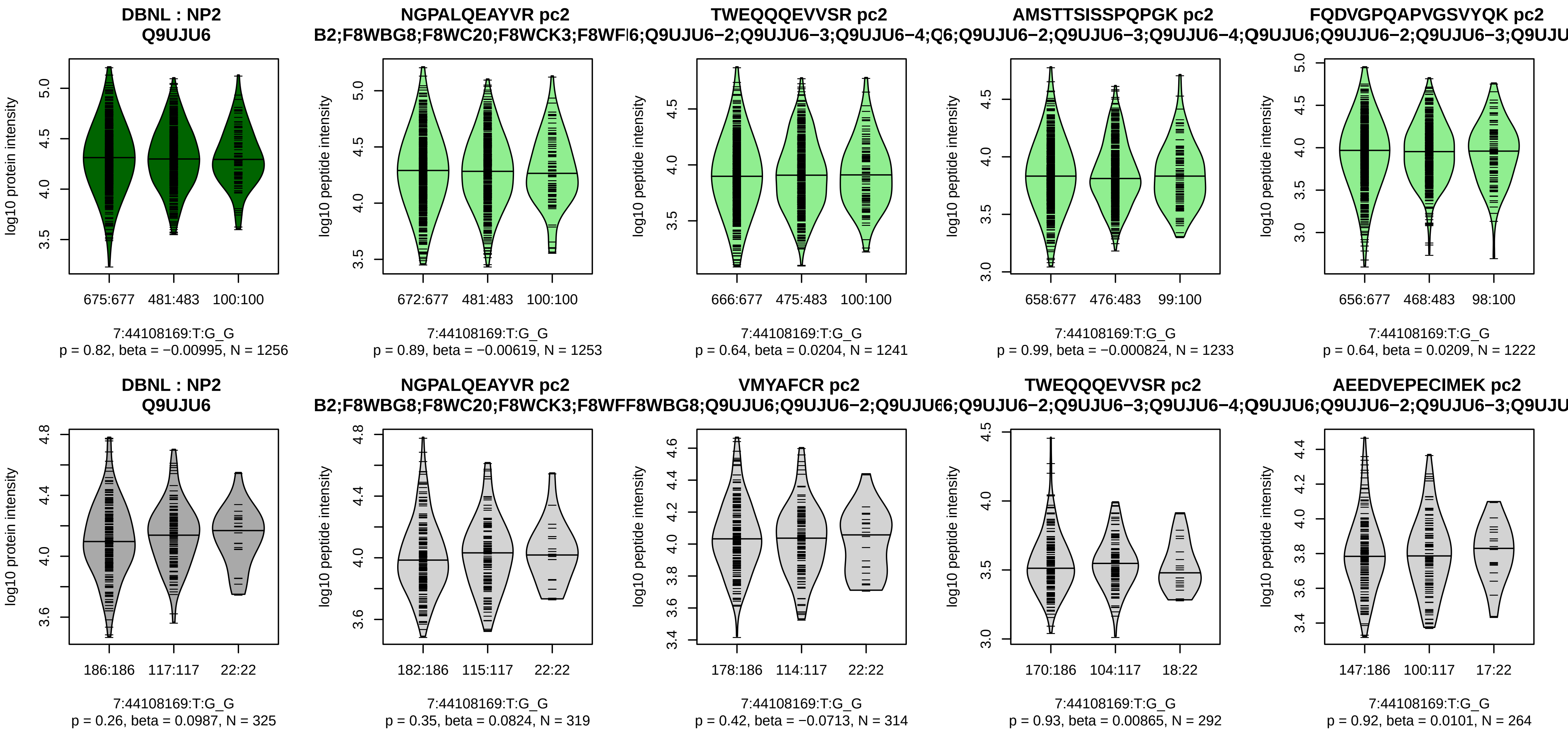


Assay Target: DBNL
Olink UniProt: Q9UJU6
deCODE rsID: rs56665454
Proxy rsID: rs10236567
deCODE: 7:44037871:T:C
Proxy SNP: 7:44108169:T:G
deCODE log10(p): 16.7
deCODE BETA: 0.07
−:−:−:−:−:−:−:−:−:−:−:−:−:−:−:−:
1253:1241:1233:1222:1182:112

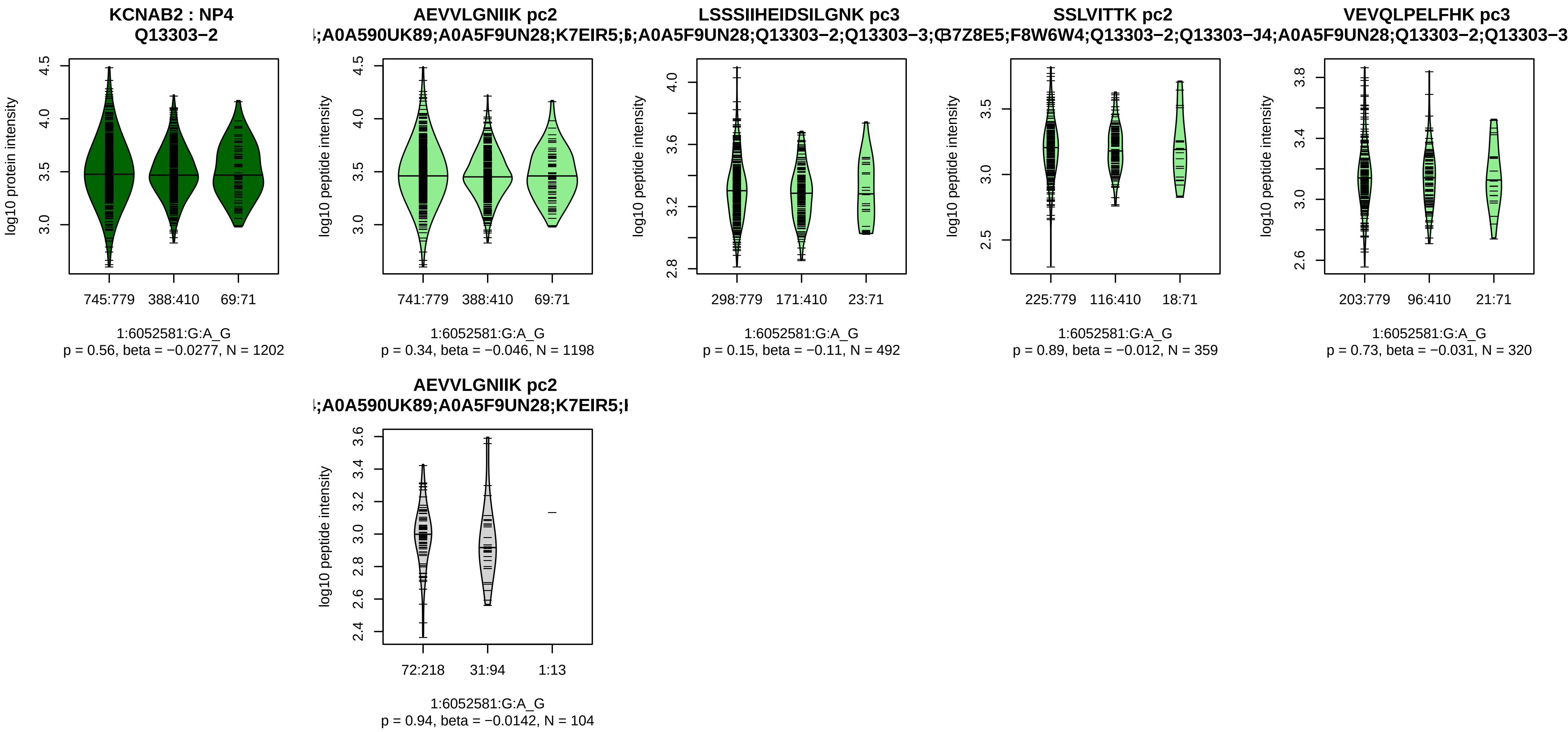


Assay Target: KCNAB2
Olink UniProt: Q13303
deCODE rsID: rs806109
Proxy rsID: rs806109
deCODE: 1:5992521:G:A
Proxy SNP: 1:6052581:G:A
deCODE log10(p): 16.2
deCODE BETA: −0.08
−:−:−:−:−:−:−:−:−
1198:492:359:320:311:274:228:

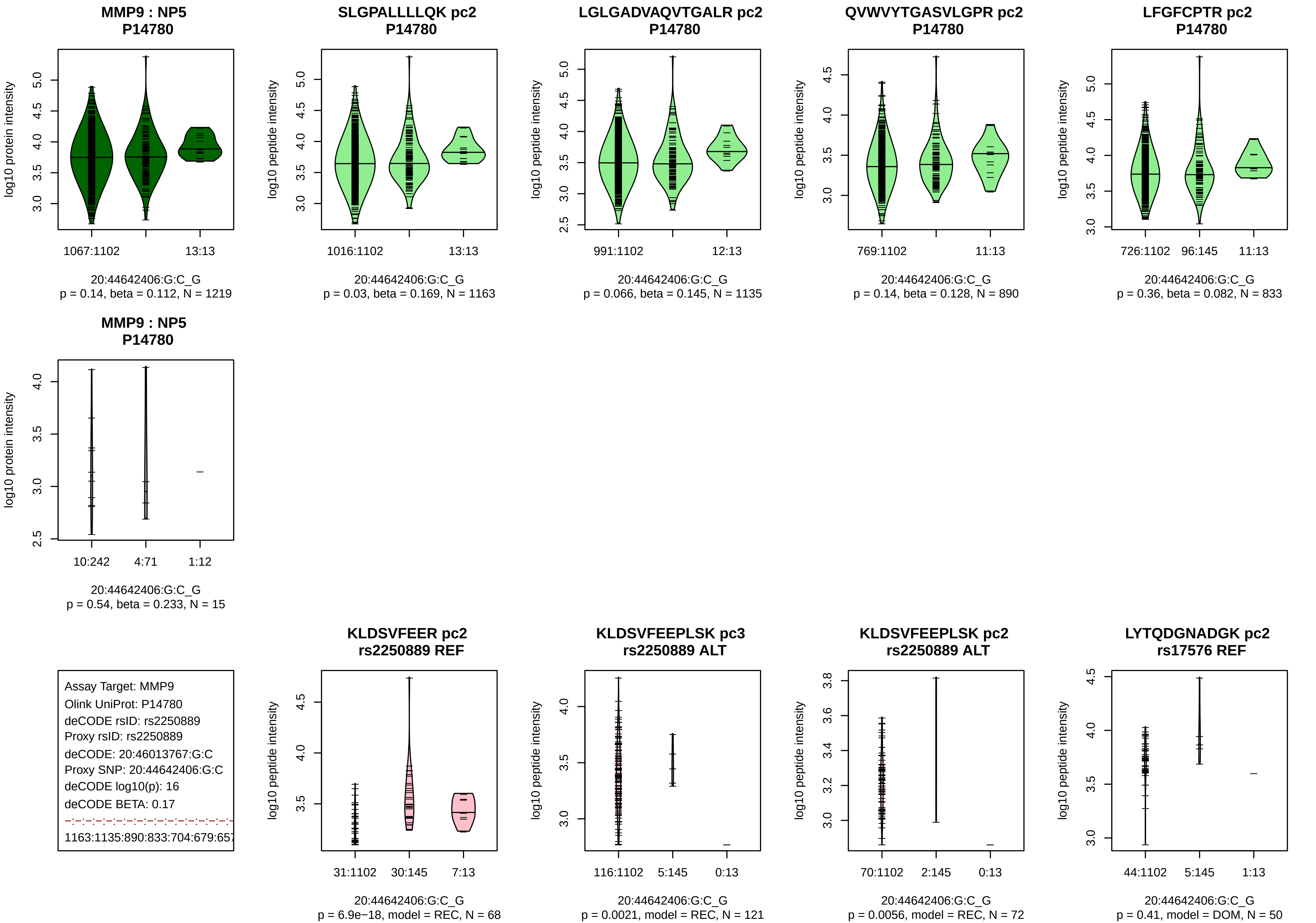

MMP9 : NP5
P14780
log10 protein intensity
1067:1102
13:13
20:44642406:G:C_G
p = 0.14, beta = 0.112, N = 1219
SLGPALLLLQK pc2
P14780
log10 peptide intensity
1016:1102
13:13
20:44642406:G:C_G
p = 0.03, beta = 0.169, N = 1163
LGLGADVAQVTGALR pc2
P14780
log10 peptide intensity
991:1102
12:13
20:44642406:G:C_G
p = 0.066, beta = 0.145, N = 1135
QVWVYTGASVLGPR pc2
P14780
log10 peptide intensity
769:1102
11:13
20:44642406:G:C_G
p = 0.14, beta = 0.128, N = 890
LFGFCPTR pc2
P14780
log10 peptide intensity
726:1102
96:145
11:13
20:44642406:G:C_G
p = 0.36, beta = 0.082, N = 833
MMP9 : NP5
P14780
log10 protein intensity
10:242
4:71
1:12
20:44642406:G:C_G
p = 0.54, beta = 0.233, N = 15
Assay Target: MMP9
Olink UniProt: P14780
deCODE rsID: rs2250889
Proxy rsID: rs2250889
deCODE: 20:46013767:G:C
Proxy SNP: 20:44642406:G:C
deCODE log10(p): 16
deCODE BETA: 0.17
1163:1135:890:833:704:679:657
KLDSVFEER pc2
rs2250889 REF
log10 peptide intensity
31:1102
30:145
7:13
20:44642406:G:C_G
p = 6.9e−18, model = REC, N = 68
KLDSVFEEPLSK pc3
rs2250889 ALT
log10 peptide intensity
116:1102
5:145
0:13
20:44642406:G:C_G
p = 0.0021, model = REC, N = 121
KLDSVFEEPLSK pc2
rs2250889 ALT
log10 peptide intensity
70:1102
2:145
0:13
20:44642406:G:C_G
p = 0.0056, model = REC, N = 72
LYTQDGNADGK pc2
rs17576 REF
log10 peptide intensity
44:1102
5:145
1:13
20:44642406:G:C_G
p = 0.41, model = DOM, N = 50

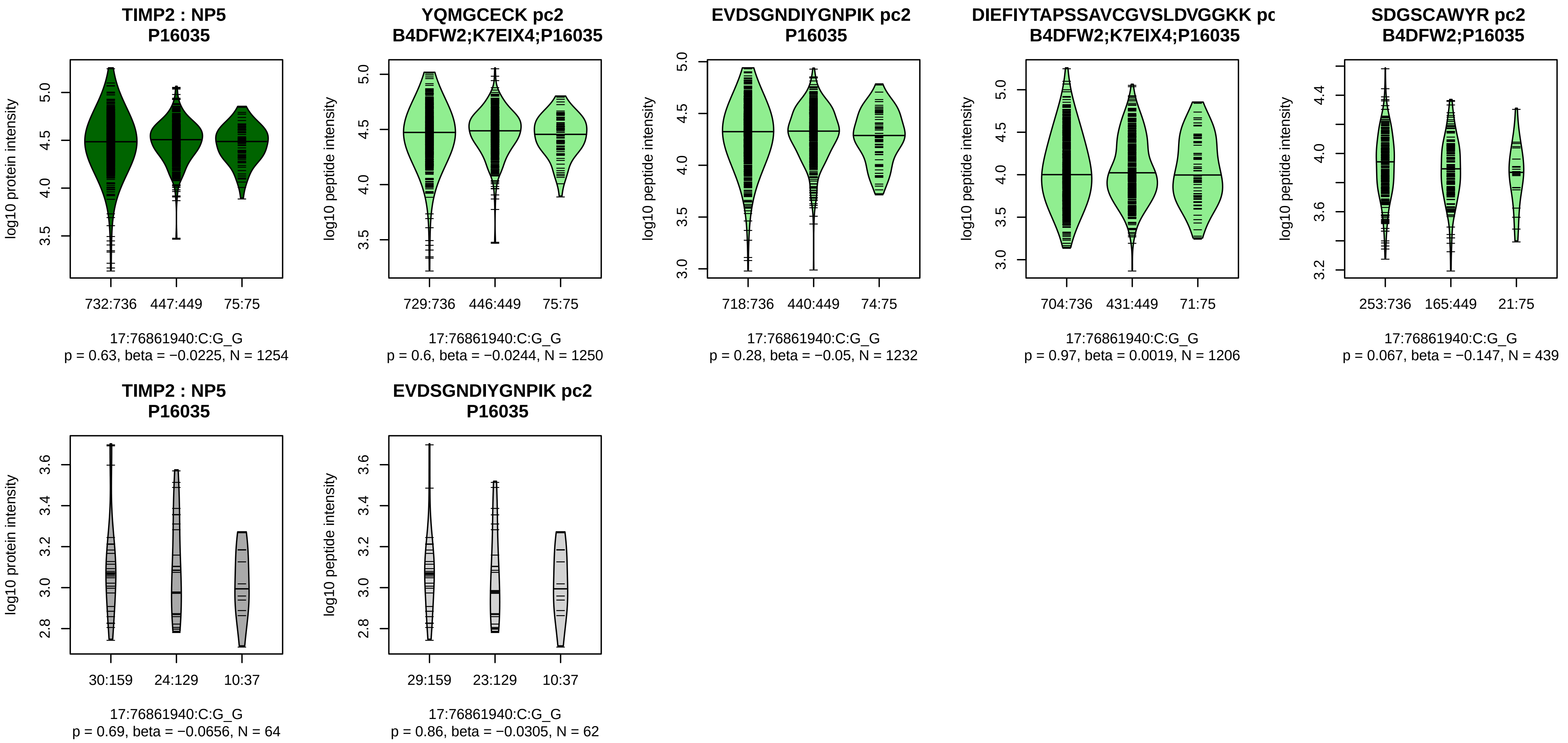


Assay Target: TIMP2
Olink UniProt: P16035
deCODE rsID: rs9894212
Proxy rsID: rs9894212
deCODE: 17:78865858:G:C
Proxy SNP: 17:76861940:C:G
deCODE log10(p): 15.8
deCODE BETA: −0.08
−:−:−:−:−:−:−:NA
1250:1232:1206:439:237:34:24:

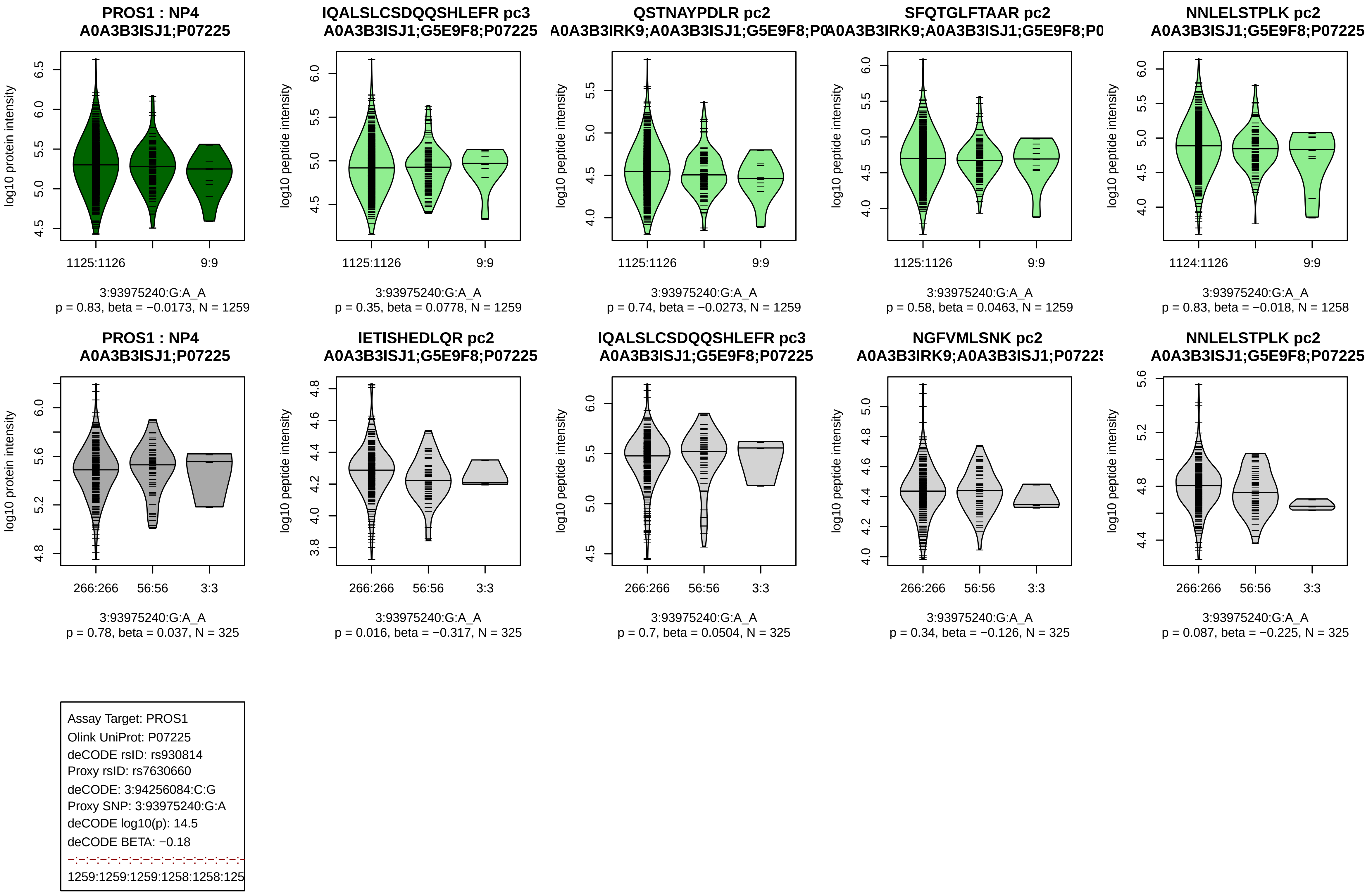
PROS1 : NP4
A0A3B3ISJ1;P07225
log10 protein intensity
1125:1126
9:9
3:93975240:G:A_A
p = 0.83, beta = −0.0173, N = 1259
IQALSLCSDQQSHLEFR pc3
A0A3B3ISJ1;G5E9F8;P07225
log10 peptide intensity
1125:1126
9:9
3:93975240:G:A_A
p = 0.35, beta = 0.0778, N = 1259
QSTNAYPDLR pc2
A0A3B3IRK9;A0A3B3ISJ1;G5E9F8;P0
log10 peptide intensity
1125:1126
9:9
3:93975240:G:A_A
p = 0.74, beta = −0.0273, N = 1259
SFQTGLFTAAR pc2
A0A3B3IRK9;A0A3B3ISJ1;G5E9F8;P0
log10 peptide intensity
1125:1126
9:9
3:93975240:G:A_A
p = 0.58, beta = 0.0463, N = 1259
NNLELSTPLK pc2
A0A3B3ISJ1;G5E9F8;P07225
log10 peptide intensity
1124:1126
9:9
3:93975240:G:A_A
p = 0.83, beta = −0.018, N = 1258
PROS1 : NP4
A0A3B3ISJ1;P07225
log10 protein intensity
266:266
56:56
3:3
3:93975240:G:A_A
p = 0.78, beta = 0.037, N = 325
IETISHEDLQR pc2
A0A3B3ISJ1;G5E9F8;P07225
log10 peptide intensity
266:266
56:56
3:3
3:93975240:G:A_A
p = 0.016, beta = −0.317, N = 325
IQALSLCSDQQSHLEFR pc3
A0A3B3ISJ1;G5E9F8;P07225
log10 peptide intensity
266:266
56:56
3:3
3:93975240:G:A_A
p = 0.7, beta = 0.0504, N = 325
NGFVMLSNK pc2
A0A3B3IRK9;A0A3B3ISJ1;P07225
log10 peptide intensity
266:266
56:56
3:3
3:93975240:G:A_A
p = 0.34, beta = −0.126, N = 325
NNLELSTPLK pc2
A0A3B3ISJ1;G5E9F8;P07225
log10 peptide intensity
266:266
56:56
3:3
3:93975240:G:A_A
p = 0.087, beta = −0.225, N = 325
Assay Target: PROS1
Olink UniProt: P07225
deCODE rsID: rs930814
Proxy rsID: rs7630660
deCODE: 3:94256084:C:G
Proxy SNP: 3:93975240:G:A
deCODE log10(p): 14.5
deCODE BETA: −0.18
1259:1259:1259:1258:1258:125

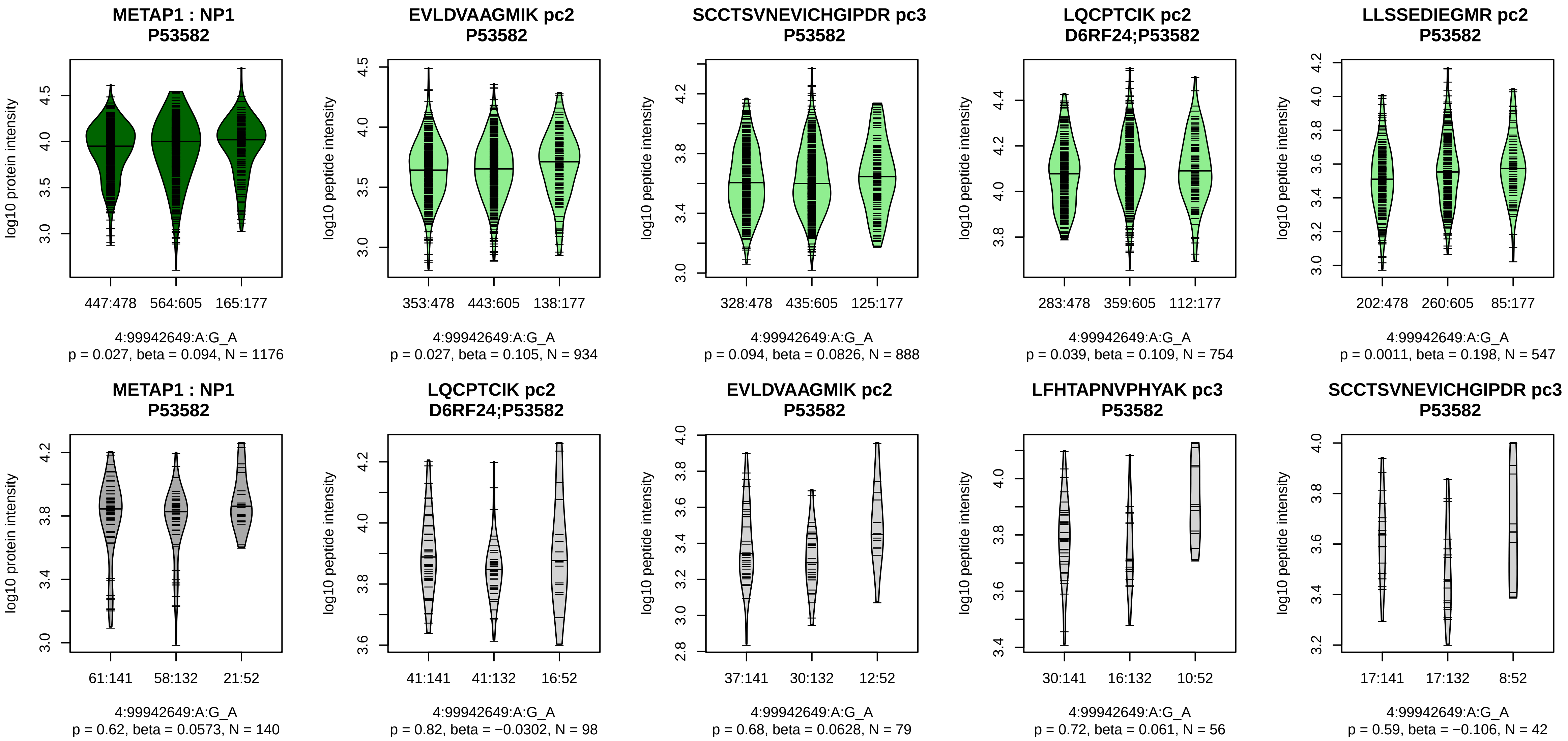


Assay Target: METAP1
Olink UniProt: P53582
deCODE rsID: rs7676259
Proxy rsID: rs7676259
deCODE: 4:99021498:A:G
Proxy SNP: 4:99942649:A:G
deCODE log10(p): 14.4
deCODE BETA: 0.06
–:–:–:*:–:–:–:–:–:–:–:–:–:–
934:888:754:547:531:477:454:4

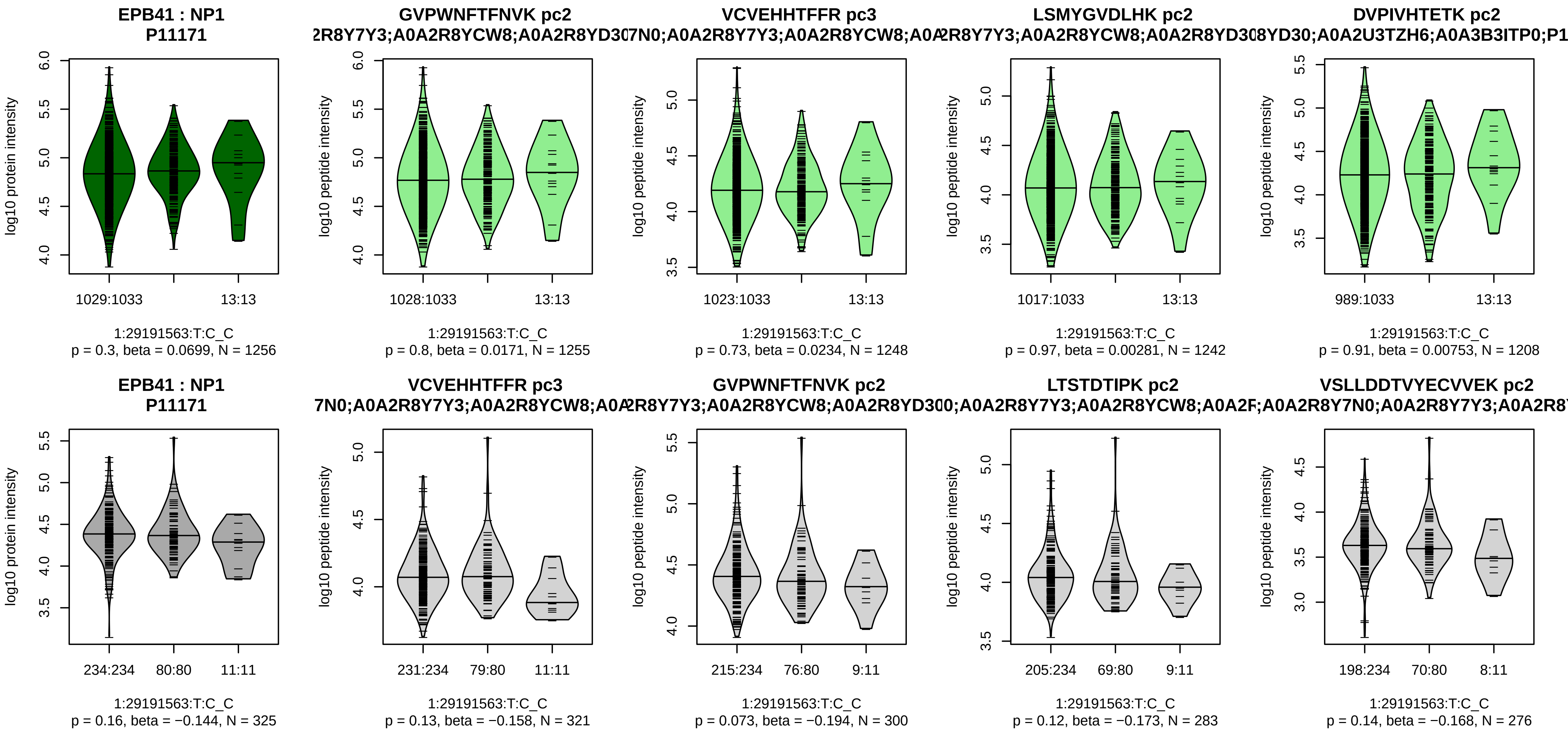


Assay Target: EPB41
Olink UniProt: P11171
deCODE rsID: rs204074
Proxy rsID: rs204074
deCODE: 1:28865051:C:T
Proxy SNP: 1:29191563:T:C
deCODE log10(p): 14.2
deCODE BETA: −0.1

1255:1248:1242:1208:1205:120

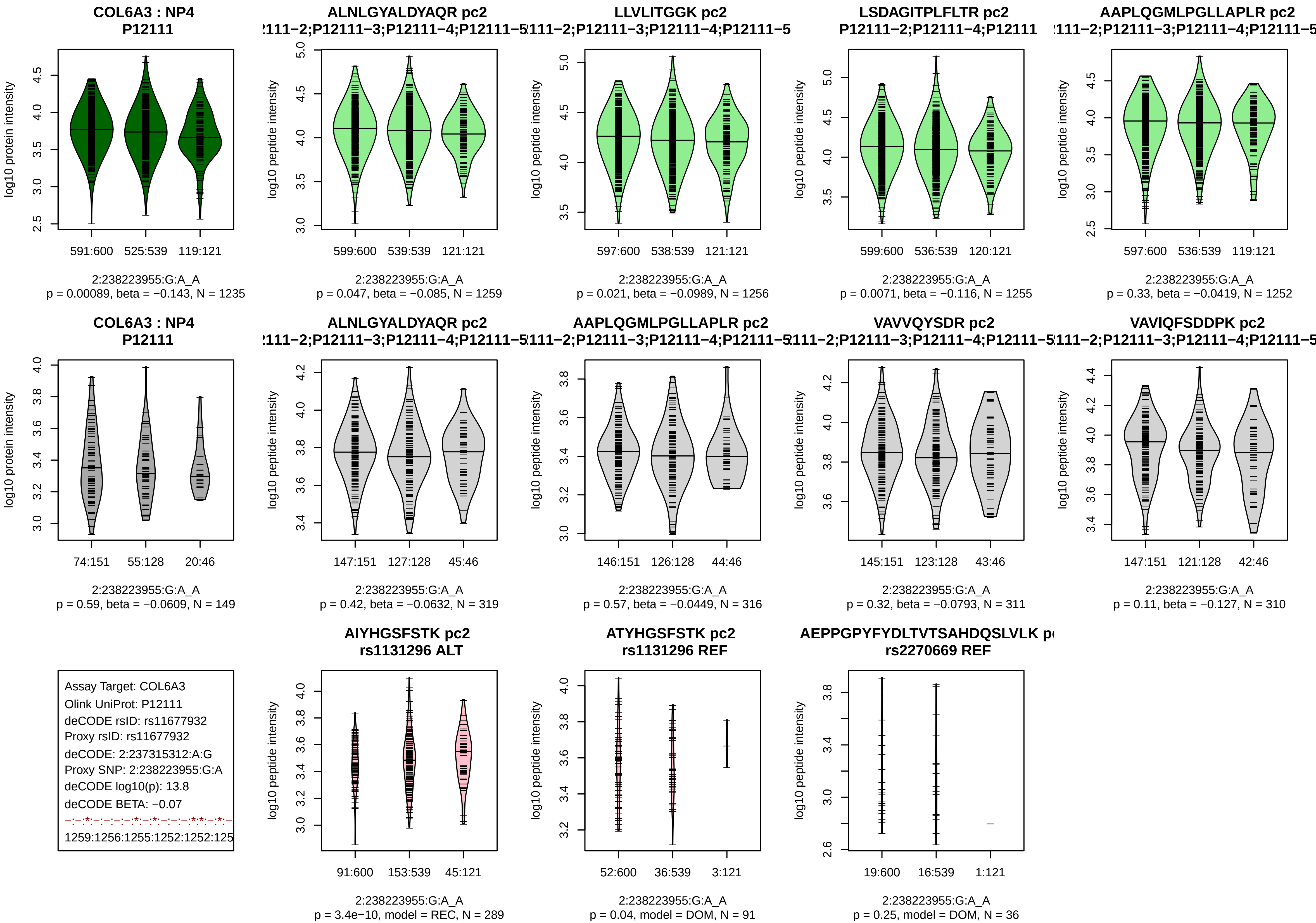
COL6A3 : NP4
P12111
log10 protein intensity
591:600 525:539 119:121
2:238223955:G:A_A
p = 0.00089, beta = −0.143, N = 1235
ALNLGYALDYAQR pc2
:111−2;P12111−3;P12111−4;P12111−5
log10 peptide intensity
599:600 539:539 121:121
2:238223955:G:A_A
p = 0.047, beta = −0.085, N = 1259
LLVLITGGK pc2
:111−2;P12111−3;P12111−4;P12111−5
log10 peptide intensity
597:600 538:539 121:121
2:238223955:G:A_A
p = 0.021, beta = −0.0989, N = 1256
LSDAGITPLFLTR pc2
P12111−2;P12111−4;P12111
log10 peptide intensity
599:600 536:539 120:121
2:238223955:G:A_A
p = 0.0071, beta = −0.116, N = 1255
AAPLQGMLPGLLAPLR pc2
:111−2;P12111−3;P12111−4;P12111−5
log10 peptide intensity
597:600 536:539 119:121
2:238223955:G:A_A
p = 0.33, beta = −0.0419, N = 1252
COL6A3 : NP4
P12111
log10 protein intensity
74:151 55:128 20:46
2:238223955:G:A_A
p = 0.59, beta = −0.0609, N = 149
ALNLGYALDYAQR pc2
:111−2;P12111−3;P12111−4;P12111−5
log10 peptide intensity
147:151 127:128 45:46
2:238223955:G:A_A
p = 0.42, beta = −0.0632, N = 319
AAPLQGMLPGLLAPLR pc2
:111−2;P12111−3;P12111−4;P12111−5
log10 peptide intensity
146:151 126:128 44:46
2:238223955:G:A_A
p = 0.57, beta = −0.0449, N = 316
VAVVQYSDR pc2
:111−2;P12111−3;P12111−4;P12111−5
log10 peptide intensity
145:151 123:128 43:46
2:238223955:G:A_A
p = 0.32, beta = −0.0793, N = 311
VAVIQFSDDPK pc2
:111−2;P12111−3;P12111−4;P12111−5
log10 peptide intensity
147:151 121:128 42:46
2:238223955:G:A_A
p = 0.11, beta = −0.127, N = 310
Assay Target: COL6A3
Olink UniProt: P12111
deCODE rsID: rs11677932
Proxy rsID: rs11677932
deCODE: 2:237315312:A:G
Proxy SNP: 2:238223955:G:A
deCODE log10(p): 13.8
deCODE BETA: −0.07
−:−:*:−:−:−:−:*:−:*:−:−:−:*:*:−:*:−
1259:1256:1255:1252:1252:125
AIYHGSFSTK pc2
rs1131296 ALT
log10 peptide intensity
91:600 153:539 45:121
2:238223955:G:A_A
p = 3.4e−10, model = REC, N = 289
ATYHGSFSTK pc2
rs1131296 REF
log10 peptide intensity
52:600 36:539 3:121
2:238223955:G:A_A
p = 0.04, model = DOM, N = 91
AEPPGPYFYDLTVTSAHDQSLVLK pc
rs2270669 REF
log10 peptide intensity
19:600 16:539 1:121
2:238223955:G:A_A
p = 0.25, model = DOM, N = 36

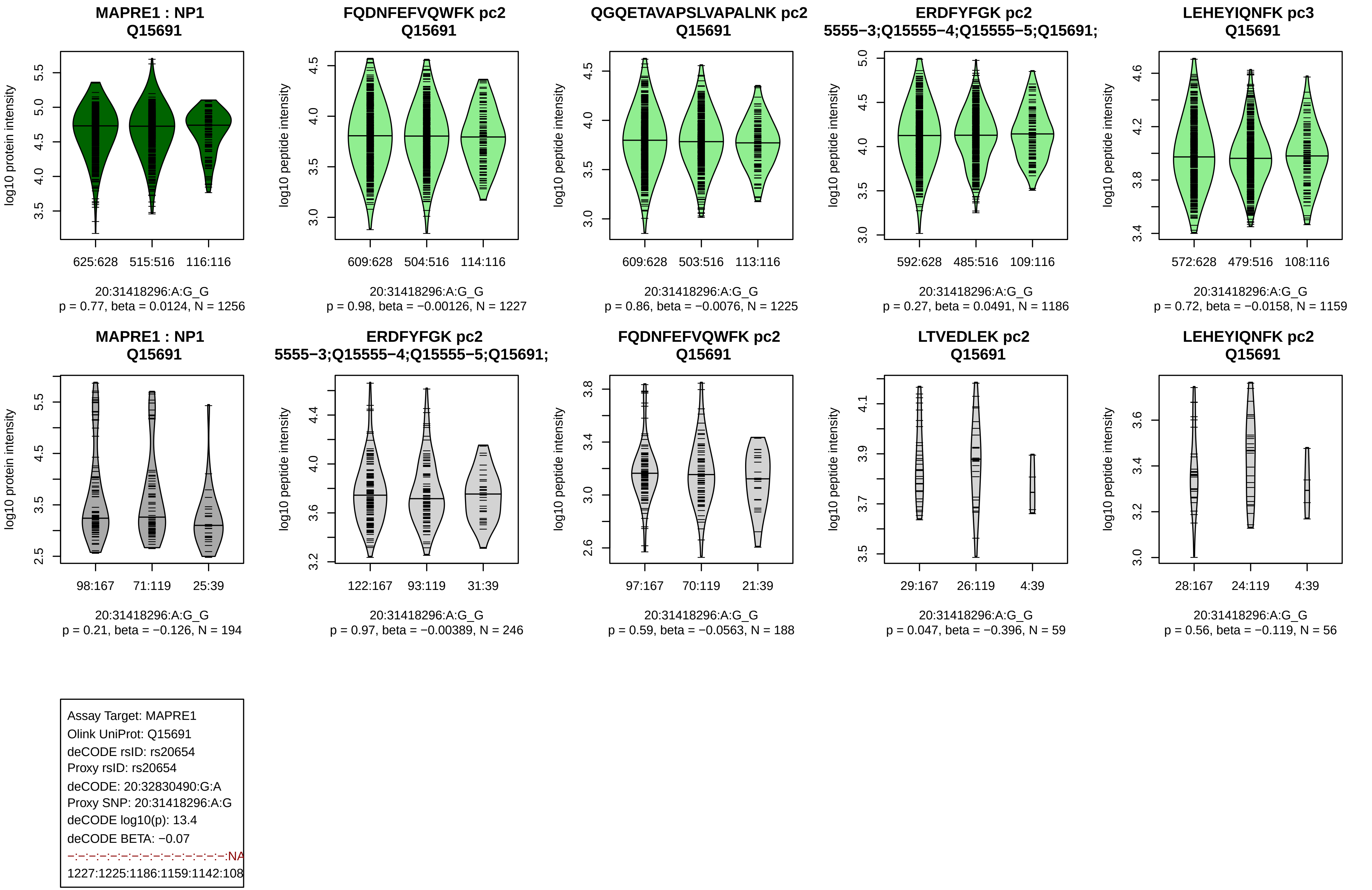

MAPRE1 : NP1
Q15691
log10 protein intensity
625:628
515:516
116:116
20:31418296:A:G_G
p = 0.77, beta = 0.0124, N = 1256
FQDNFEFVQWFK pc2
Q15691
log10 peptide intensity
609:628
504:516
114:116
20:31418296:A:G_G
p = 0.98, beta = −0.00126, N = 1227
QGQETAVAPSLVAPALNK pc2
Q15691
log10 peptide intensity
609:628
503:516
113:116
20:31418296:A:G_G
p = 0.86, beta = −0.0076, N = 1225
ERDFYFGK pc2
5555−3;Q15555−4;Q15555−5;Q15691;
log10 peptide intensity
592:628
485:516
109:116
20:31418296:A:G_G
p = 0.27, beta = 0.0491, N = 1186
LEHEYIQNFK pc3
Q15691
log10 peptide intensity
572:628
479:516
108:116
20:31418296:A:G_G
p = 0.72, beta = −0.0158, N = 1159
MAPRE1 : NP1
Q15691
log10 protein intensity
98:167
71:119
25:39
20:31418296:A:G_G
p = 0.21, beta = −0.126, N = 194
ERDFYFGK pc2
5555−3;Q15555−4;Q15555−5;Q15691;
log10 peptide intensity
122:167
93:119
31:39
20:31418296:A:G_G
p = 0.97, beta = −0.00389, N = 246
FQDNFEFVQWFK pc2
Q15691
log10 peptide intensity
97:167
70:119
21:39
20:31418296:A:G_G
p = 0.59, beta = −0.0563, N = 188
LTVEDLEK pc2
Q15691
log10 peptide intensity
29:167
26:119
4:39
20:31418296:A:G_G
p = 0.047, beta = −0.396, N = 59
LEHEYIQNFK pc2
Q15691
log10 peptide intensity
28:167
24:119
4:39
20:31418296:A:G_G
p = 0.56, beta = −0.119, N = 56
Assay Target: MAPRE1
Olink UniProt: Q15691
deCODE rsID: rs20654
Proxy rsID: rs20654
deCODE: 20:32830490:G:A
Proxy SNP: 20:31418296:A:G
deCODE log10(p): 13.4
deCODE BETA: −0.07
−:−:−:−:−:−:−:−:−:−:−:−:−:−:−:NA
1227:1225:1186:1159:1142:108

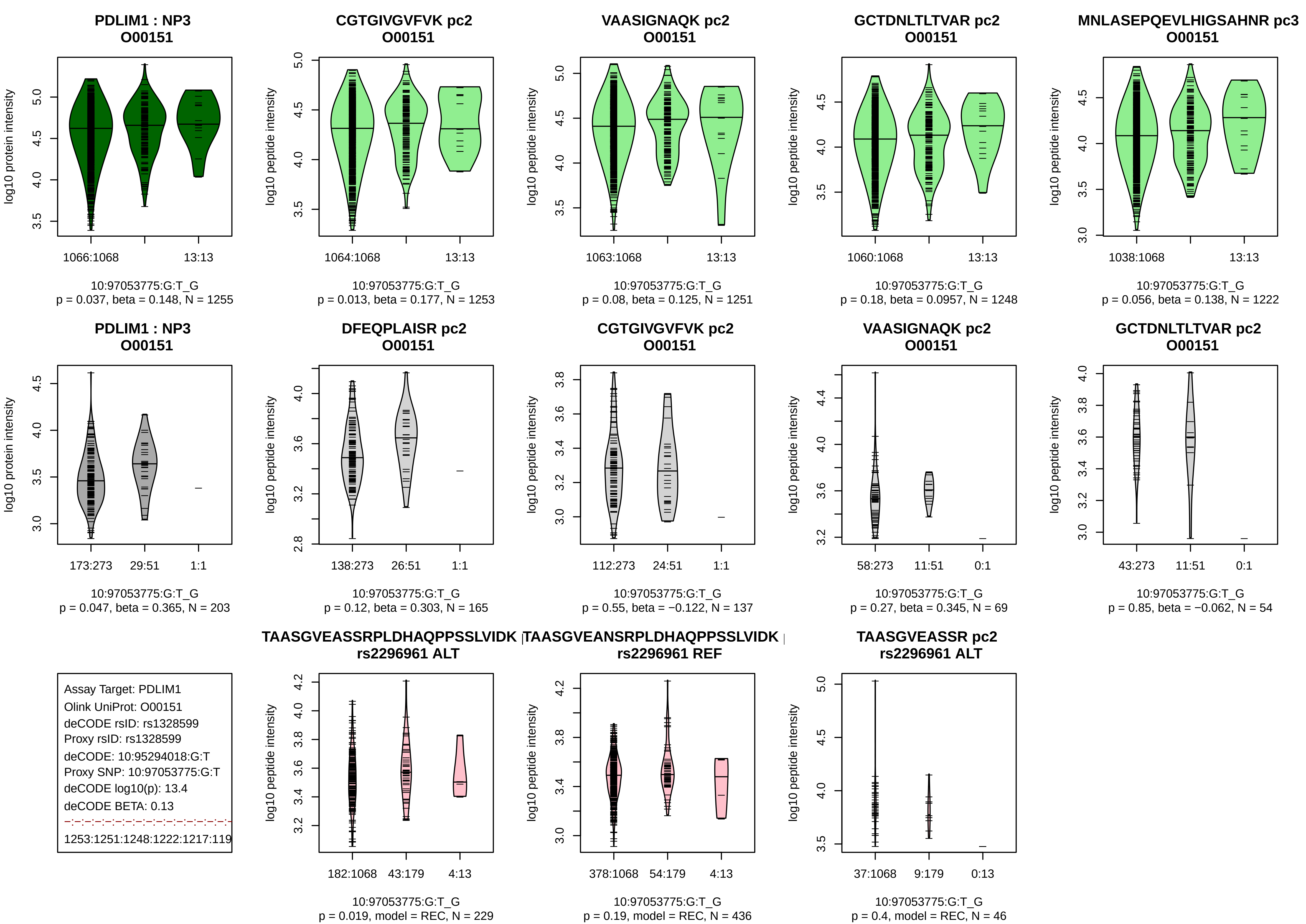
PDLIM1 : NP3
O00151
log10 protein intensity
1066:1068
13:13
10:97053775:G:T_G
p = 0.037, beta = 0.148, N = 1255
CGTGIVGVFVK pc2
O00151
log10 peptide intensity
1064:1068
13:13
10:97053775:G:T_G
p = 0.013, beta = 0.177, N = 1253
VAASIGNAQK pc2
O00151
log10 peptide intensity
1063:1068
13:13
10:97053775:G:T_G
p = 0.08, beta = 0.125, N = 1251
GCTDNLTLTVAR pc2
O00151
log10 peptide intensity
1060:1068
13:13
10:97053775:G:T_G
p = 0.18, beta = 0.0957, N = 1248
MNLASEPQEVLHIGSAHNR pc3
O00151
log10 peptide intensity
1038:1068
13:13
10:97053775:G:T_G
p = 0.056, beta = 0.138, N = 1222
PDLIM1 : NP3
O00151
log10 protein intensity
173:273
29:51
1:1
10:97053775:G:T_G
p = 0.047, beta = 0.365, N = 203
DFEQPLAISR pc2
O00151
log10 peptide intensity
138:273
26:51
1:1
10:97053775:G:T_G
p = 0.12, beta = 0.303, N = 165
CGTGIVGVFVK pc2
O00151
log10 peptide intensity
112:273
24:51
1:1
10:97053775:G:T_G
p = 0.55, beta = −0.122, N = 137
VAASIGNAQK pc2
O00151
log10 peptide intensity
58:273
11:51
0:1
10:97053775:G:T_G
p = 0.27, beta = 0.345, N = 69
GCTDNLTLTVAR pc2
O00151
log10 peptide intensity
43:273
11:51
0:1
10:97053775:G:T_G
p = 0.85, beta = −0.062, N = 54
Assay Target: PDLIM1
Olink UniProt: O00151
deCODE rsID: rs1328599
Proxy rsID: rs1328599
deCODE: 10:95294018:G:T
Proxy SNP: 10:97053775:G:T
deCODE log10(p): 13.4
deCODE BETA: 0.13
1253:1251:1248:1222:1217:119
TAASGVEASSRPLDHAQPPSSLVIDK
rs2296961 ALT
log10 peptide intensity
182:1068
43:179
4:13
10:97053775:G:T_G
p = 0.019, model = REC, N = 229
TAASGVEANSRPLDHAQPPSSLVIDK
rs2296961 REF
log10 peptide intensity
378:1068
54:179
4:13
10:97053775:G:T_G
p = 0.19, model = REC, N = 436
TAASGVEASSR pc2
rs2296961 ALT
log10 peptide intensity
37:1068
9:179
0:13
10:97053775:G:T_G
p = 0.4, model = REC, N = 46

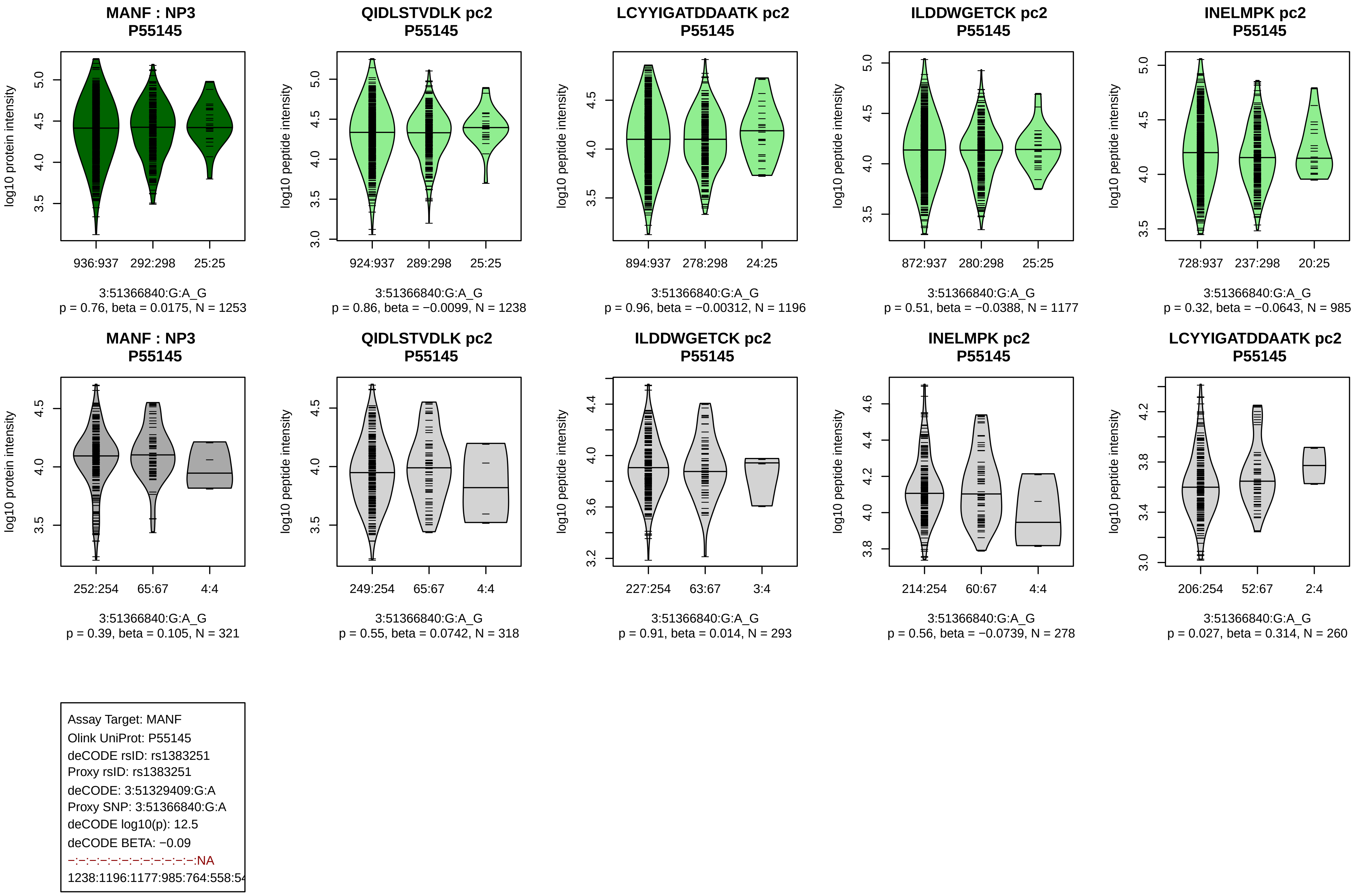
MANF : NP3
P55145
log10 protein intensity
936:937 292:298 25:25
3:51366840:G:A_G
p = 0.76, beta = 0.0175, N = 1253
QIDLSTVDLK pc2
P55145
log10 peptide intensity
924:937 289:298 25:25
3:51366840:G:A_G
p = 0.86, beta = −0.0099, N = 1238
LCYYIGATDDAATK pc2
P55145
log10 peptide intensity
894:937 278:298 24:25
3:51366840:G:A_G
p = 0.96, beta = −0.00312, N = 1196
ILDDWGETCK pc2
P55145
log10 peptide intensity
872:937 280:298 25:25
3:51366840:G:A_G
p = 0.51, beta = −0.0388, N = 1177
INELMPK pc2
P55145
log10 peptide intensity
728:937 237:298 20:25
3:51366840:G:A_G
p = 0.32, beta = −0.0643, N = 985
MANF : NP3
P55145
log10 protein intensity
252:254 65:67 4:4
3:51366840:G:A_G
p = 0.39, beta = 0.105, N = 321
QIDLSTVDLK pc2
P55145
log10 peptide intensity
249:254 65:67 4:4
3:51366840:G:A_G
p = 0.55, beta = 0.0742, N = 318
ILDDWGETCK pc2
P55145
log10 peptide intensity
227:254 63:67 3:4
3:51366840:G:A_G
p = 0.91, beta = 0.014, N = 293
INELMPK pc2
P55145
log10 peptide intensity
214:254 60:67 4:4
3:51366840:G:A_G
p = 0.56, beta = −0.0739, N = 278
LCYYIGATDDAATK pc2
P55145
log10 peptide intensity
206:254 52:67 2:4
3:51366840:G:A_G
p = 0.027, beta = 0.314, N = 260
Assay Target: MANF
Olink UniProt: P55145
deCODE rsID: rs1383251
Proxy rsID: rs1383251
deCODE: 3:51329409:G:A
Proxy SNP: 3:51366840:G:A
deCODE log10(p): 12.5
deCODE BETA: −0.09
−:−:−:−:−:−:−:−:−:−:−:−:−:NA
1238:1196:1177:985:764:558:54

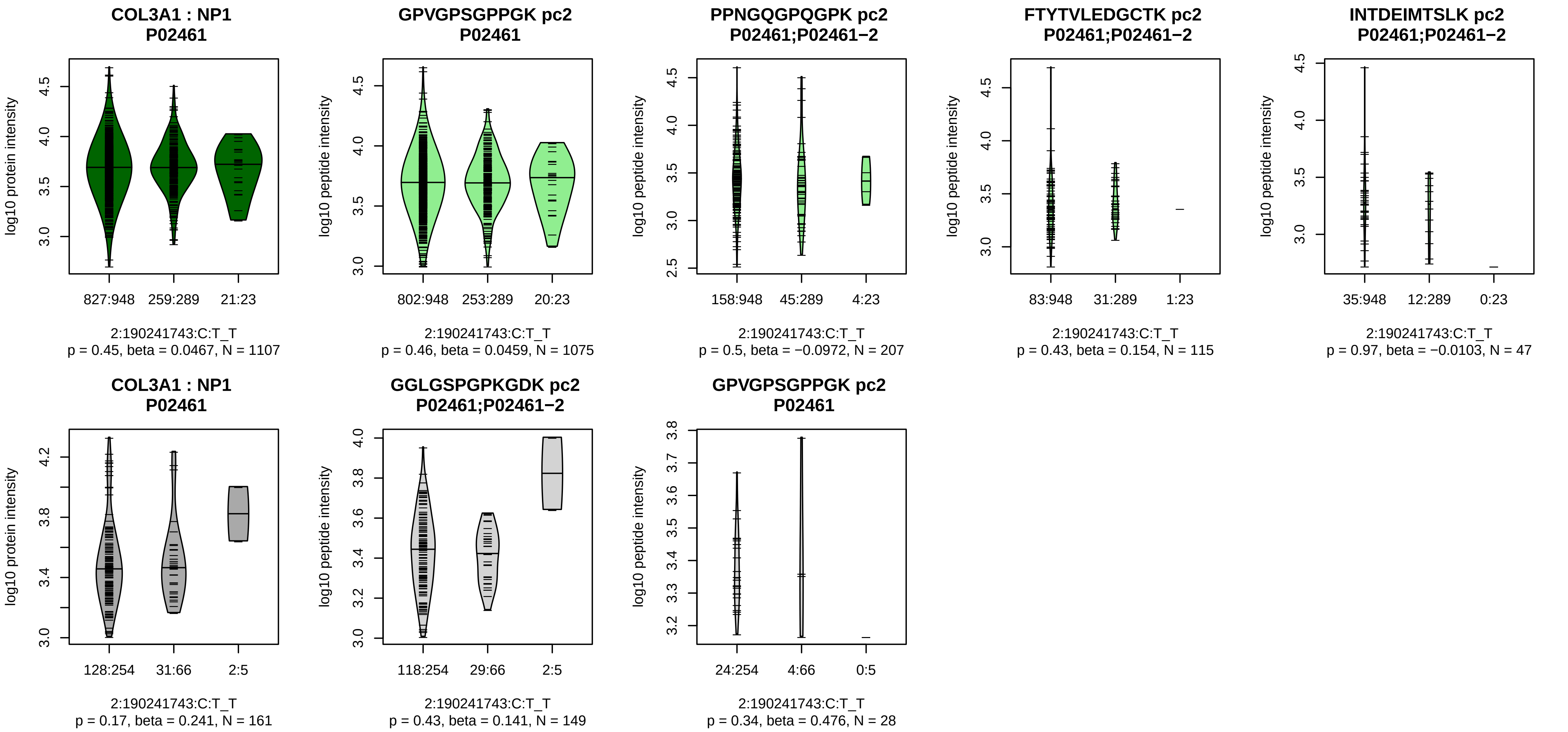


Assay Target: COL3A1
Olink UniProt: P02461
deCODE rsID: rs72914144
Proxy rsID: rs72914144
deCODE: 2:189377017:T:C
Proxy SNP: 2:190241743:C:T
deCODE log10(p): 11.9
deCODE BETA: 0.08
−:−:−:−:−:NA:NA:NA
1075:207:115:47:26:6:8:11

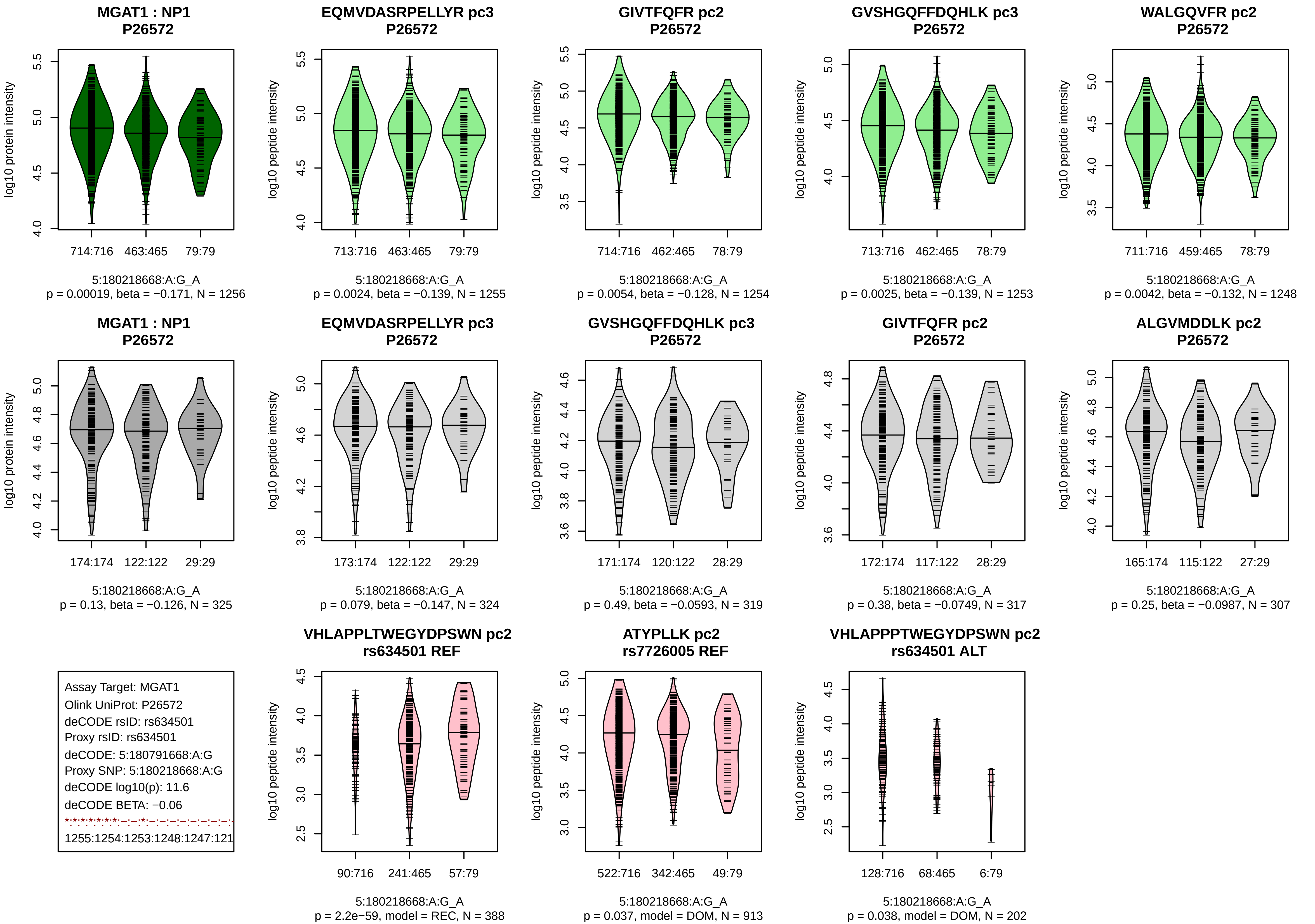
MGAT1 : NP1
P26572
log10 protein intensity
714:716
463:465
79:79
5:180218668:A:G_A
p = 0.00019, beta = −0.171, N = 1256
EQMVDASRPELLYR pc3
P26572
log10 peptide intensity
713:716
463:465
79:79
5:180218668:A:G_A
p = 0.0024, beta = −0.139, N = 1255
GIVTFQFR pc2
P26572
log10 peptide intensity
714:716
462:465
78:79
5:180218668:A:G_A
p = 0.0054, beta = −0.128, N = 1254
GVSHGQFFDQHLK pc3
P26572
log10 peptide intensity
713:716
462:465
78:79
5:180218668:A:G_A
p = 0.0025, beta = −0.139, N = 1253
WALGQVFR pc2
P26572
log10 peptide intensity
711:716
459:465
78:79
5:180218668:A:G_A
p = 0.0042, beta = −0.132, N = 1248
MGAT1 : NP1
P26572
log10 protein intensity
174:174
122:122
29:29
5:180218668:A:G_A
p = 0.13, beta = −0.126, N = 325
EQMVDASRPELLYR pc3
P26572
log10 peptide intensity
173:174
122:122
29:29
5:180218668:A:G_A
p = 0.079, beta = −0.147, N = 324
GVSHGQFFDQHLK pc3
P26572
log10 peptide intensity
171:174
120:122
28:29
5:180218668:A:G_A
p = 0.49, beta = −0.0593, N = 319
GIVTFQFR pc2
P26572
log10 peptide intensity
172:174
117:122
28:29
5:180218668:A:G_A
p = 0.38, beta = −0.0749, N = 317
ALGVMDDLK pc2
P26572
log10 peptide intensity
165:174
115:122
27:29
5:180218668:A:G_A
p = 0.25, beta = −0.0987, N = 307
Assay Target: MGAT1
Olink UniProt: P26572
deCODE rsID: rs634501
Proxy rsID: rs634501
deCODE: 5:180791668:A:G
Proxy SNP: 5:180218668:A:G
deCODE log10(p): 11.6
deCODE BETA: −0.06
1255:1254:1253:1248:1247:121
VHLAPPLTWEGYDPSWN pc2
rs634501 REF
log10 peptide intensity
90:716
241:465
57:79
5:180218668:A:G_A
p = 2.2e−59, model = REC, N = 388
ATYPLLK pc2
rs7726005 REF
log10 peptide intensity
522:716
342:465
49:79
5:180218668:A:G_A
p = 0.037, model = DOM, N = 913
VHLAPPPTWEGYDPSWN pc2
rs634501 ALT
log10 peptide intensity
128:716
68:465
6:79
5:180218668:A:G_A
p = 0.038, model = DOM, N = 202

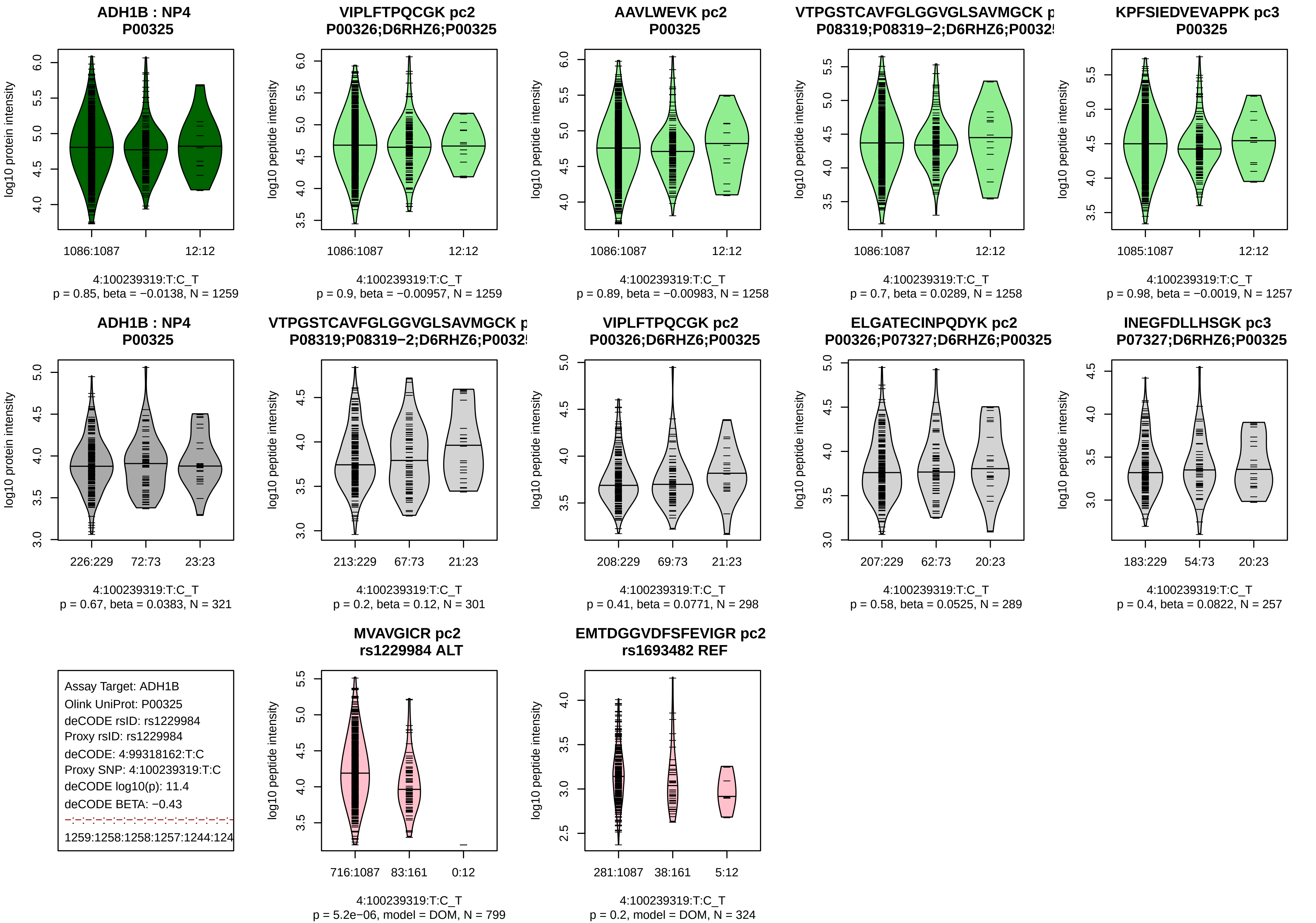
ADH1B : NP4
P00325
log10 protein intensity
1086:1087
12:12
4:100239319:T:C_T
p = 0.85, beta = −0.0138, N = 1259
VIPLFTPQCGK pc2
P00326;D6RHZ6;P00325
log10 peptide intensity
1086:1087
12:12
4:100239319:T:C_T
p = 0.9, beta = −0.00957, N = 1259
AAVLWEVK pc2
P00325
log10 peptide intensity
1086:1087
12:12
4:100239319:T:C_T
p = 0.89, beta = −0.00983, N = 1258
VTPGSTCAVFGLGGVGLSAVMGCK p
P08319;P08319−2;D6RHZ6;P0032
log10 peptide intensity
1086:1087
12:12
4:100239319:T:C_T
p = 0.7, beta = 0.0289, N = 1258
KPFSIEDVEVAPPK pc3
P00325
log10 peptide intensity
1085:1087
12:12
4:100239319:T:C_T
p = 0.98, beta = −0.0019, N = 1257
ADH1B : NP4
P00325
log10 protein intensity
226:229
72:73
23:23
4:100239319:T:C_T
p = 0.67, beta = 0.0383, N = 321
VTPGSTCAVFGLGGVGLSAVMGCK p
P08319;P08319−2;D6RHZ6;P0032
log10 peptide intensity
213:229
67:73
21:23
4:100239319:T:C_T
p = 0.2, beta = 0.12, N = 301
VIPLFTPQCGK pc2
P00326;D6RHZ6;P00325
log10 peptide intensity
208:229
69:73
21:23
4:100239319:T:C_T
p = 0.41, beta = 0.0771, N = 298
ELGATECINPQDYK pc2
P00326;P07327;D6RHZ6;P00325
log10 peptide intensity
207:229
62:73
20:23
4:100239319:T:C_T
p = 0.58, beta = 0.0525, N = 289
INEGFDLLHSGK pc3
P07327;D6RHZ6;P00325
log10 peptide intensity
183:229
54:73
20:23
4:100239319:T:C_T
p = 0.4, beta = 0.0822, N = 257
Assay Target: ADH1B
Olink UniProt: P00325
deCODE rsID: rs1229984
Proxy rsID: rs1229984
deCODE: 4:99318162:T:C
Proxy SNP: 4:100239319:T:C
deCODE log10(p): 11.4
deCODE BETA: −0.43
1259:1258:1258:1257:1244:124
MVAVGICR pc2
rs1229984 ALT
log10 peptide intensity
716:1087
83:161
0:12
4:100239319:T:C_T
p = 5.2e−06, model = DOM, N = 799
EMTDGGVDFSFEVIGR pc2
rs1693482 REF
log10 peptide intensity
281:1087
38:161
5:12
4:100239319:T:C_T
p = 0.2, model = DOM, N = 324

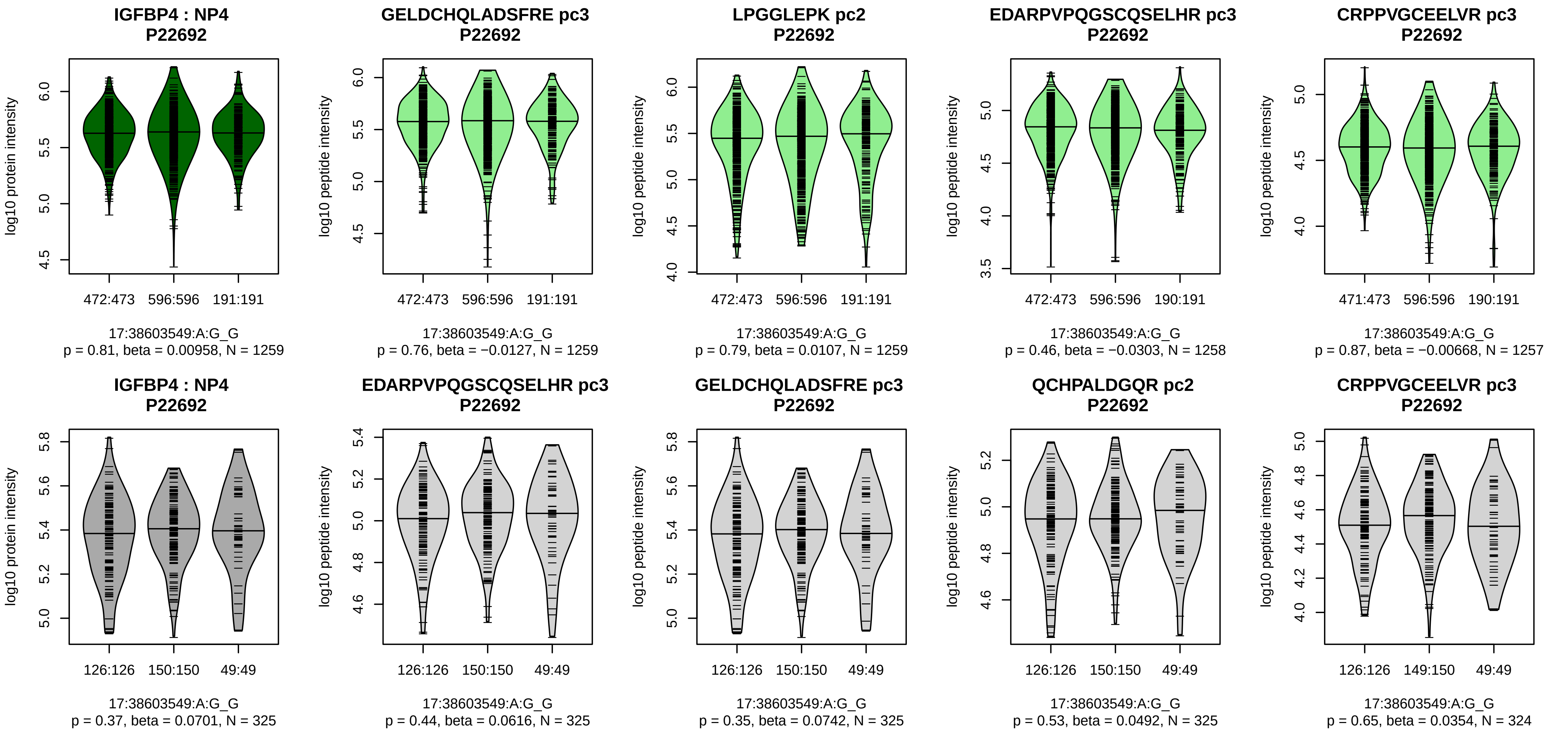


Assay Target: IGFBP4
Olink UniProt: P22692
deCODE rsID: rs4890114
Proxy rsID: rs4890114
deCODE: 17:40447297:G:A
Proxy SNP: 17:38603549:A:G
deCODE log10(p): 11.3
deCODE BETA: 0.06
-:-:-:-:-:-:-:-:-:-:-:-:-:-:-:-:-:
1259:1259:1258:1257:1256:125

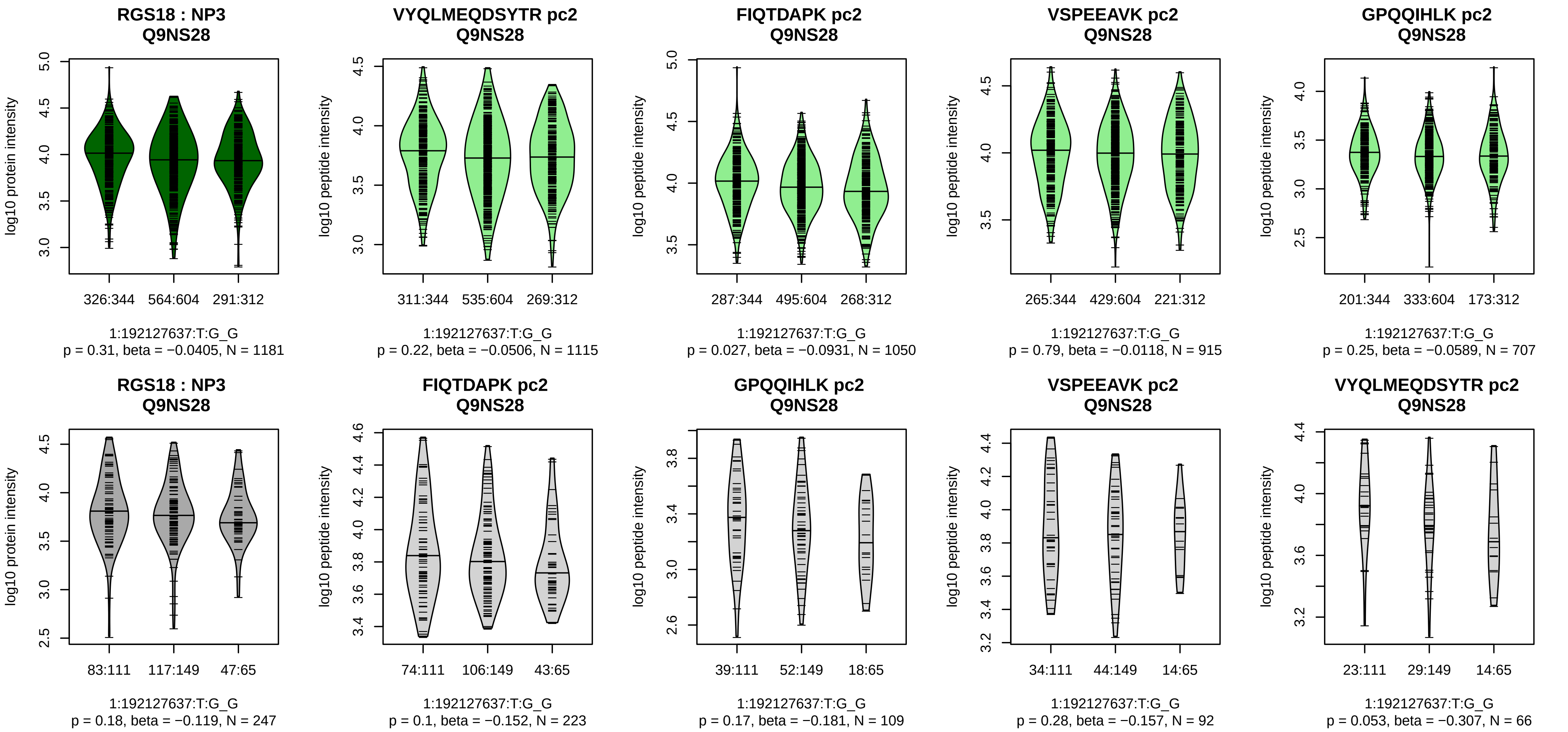


Assay Target: RGS18
Olink UniProt: Q9NS28
deCODE rsID: rs4495675
Proxy rsID: rs4495675
deCODE: 1:192158507:T:G
Proxy SNP: 1:192127637:T:G
deCODE log10(p): 11.1
deCODE BETA: 0.06
–:–:–:–:–:–:–:–:–:–:–
1115:1050:915:707:467:344:307

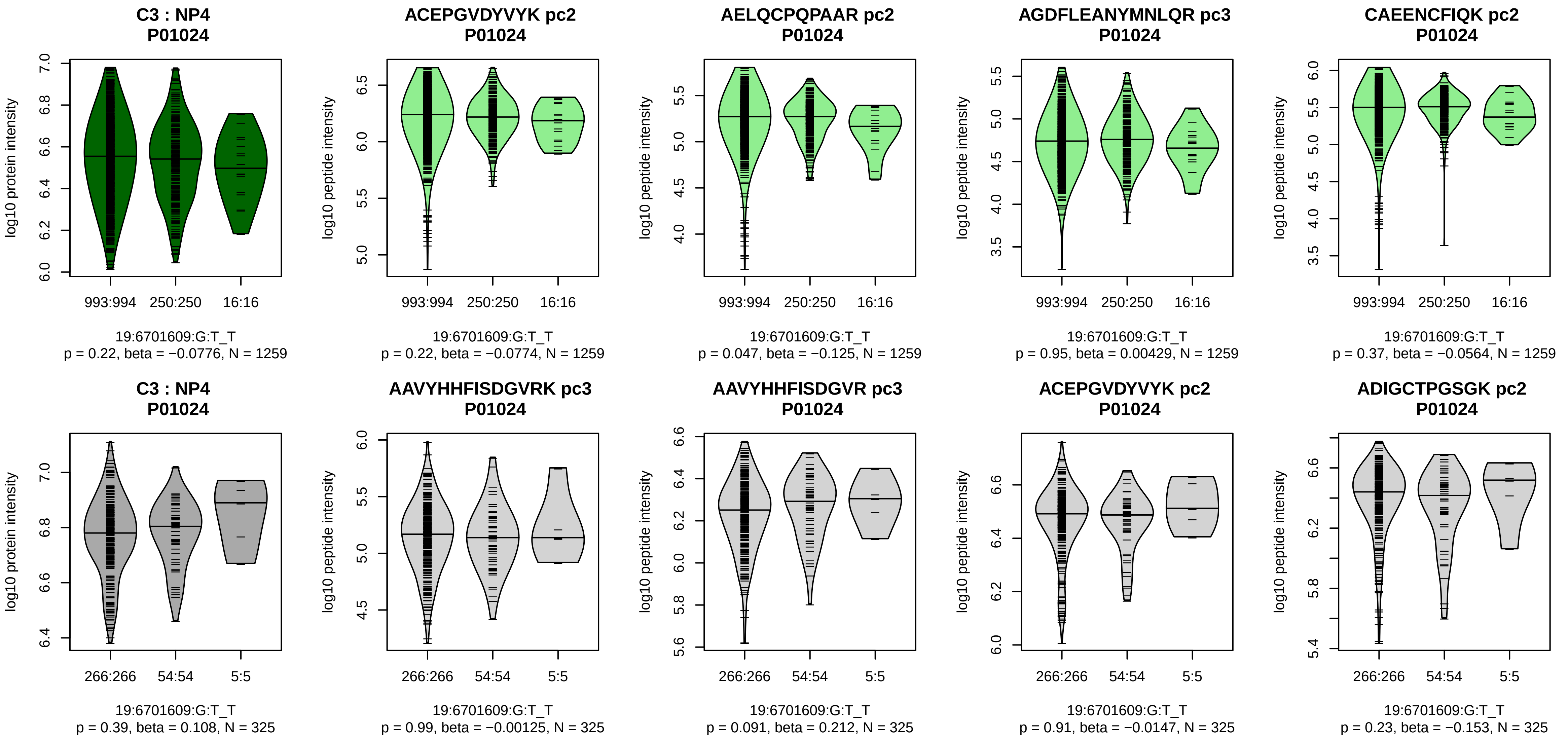


Assay Target: C3
Olink UniProt: P01024
deCODE rsID: rs11569466
Proxy rsID: rs11569466
deCODE: 19:6701598:T:!
Proxy SNP: 19:6701609:G:T
deCODE log10(p): 11.1
deCODE BETA: −0.08
–:–:–:–:–:–:–:–:–:–:–:–:–:–:*:–
1259:1259:1259:1259:1259:125

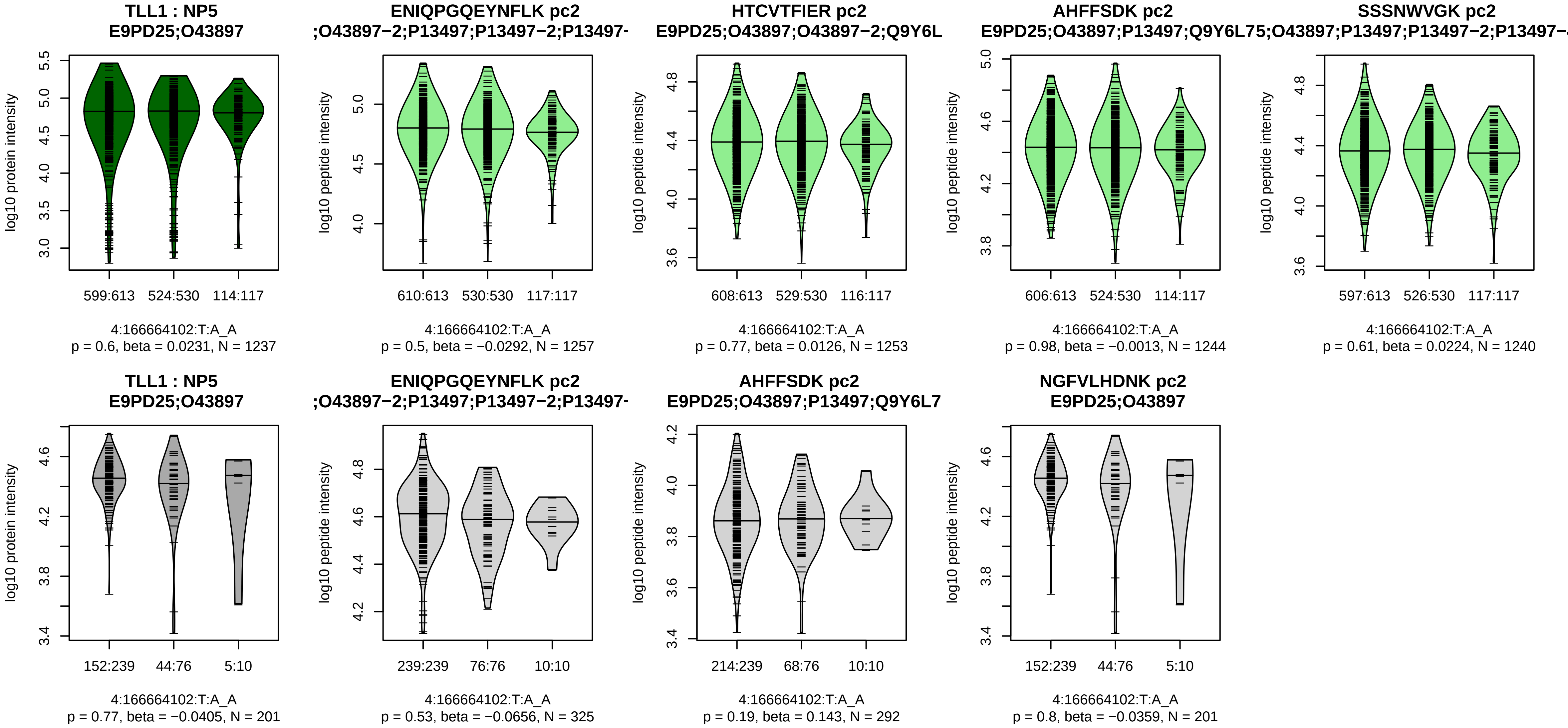


Assay Target: TLL1
Olink UniProt: O43897
deCODE rsID: rs1903176
Proxy rsID: rs1903176
deCODE: 4:165742950:A:T
Proxy SNP: 4:166664102:T:A
deCODE log10(p): 10.9
deCODE BETA: −0.06
−:−:−:−:−:−:−:−:−:−:−:−
1257:1253:1244:1240:1157:716

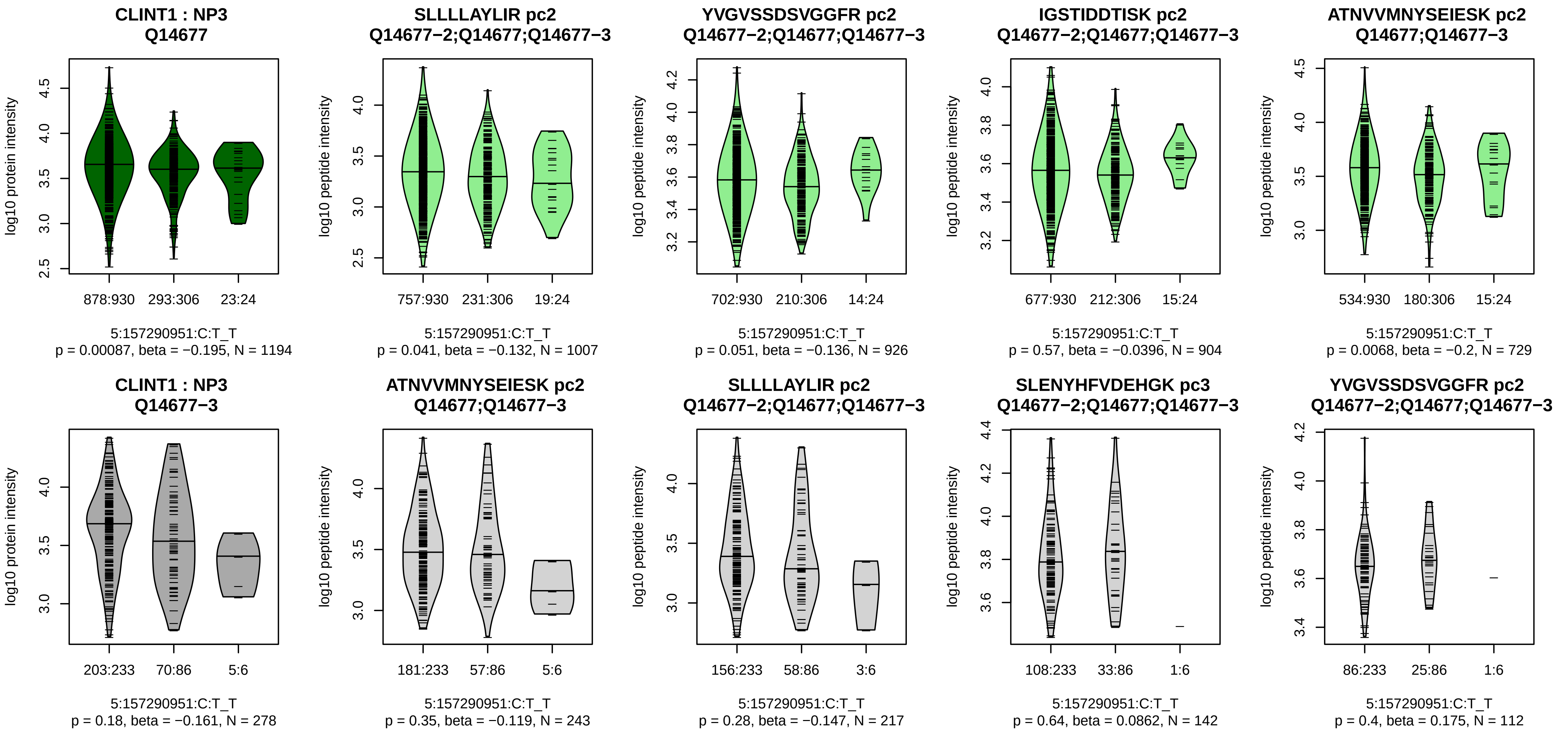


Assay Target: CLINT1
Olink UniProt: Q14677
deCODE rsID: rs62389053
Proxy rsID: rs62389053
deCODE: 5:157863943:T:C
Proxy SNP: 5:157290951:C:T
deCODE log10(p): 10.7
deCODE BETA: −0.08
−:−:−:*:−:−:−:−:*:−:−:−:−:−:−:−:−:−
1007:926:904:729:654:413:280:

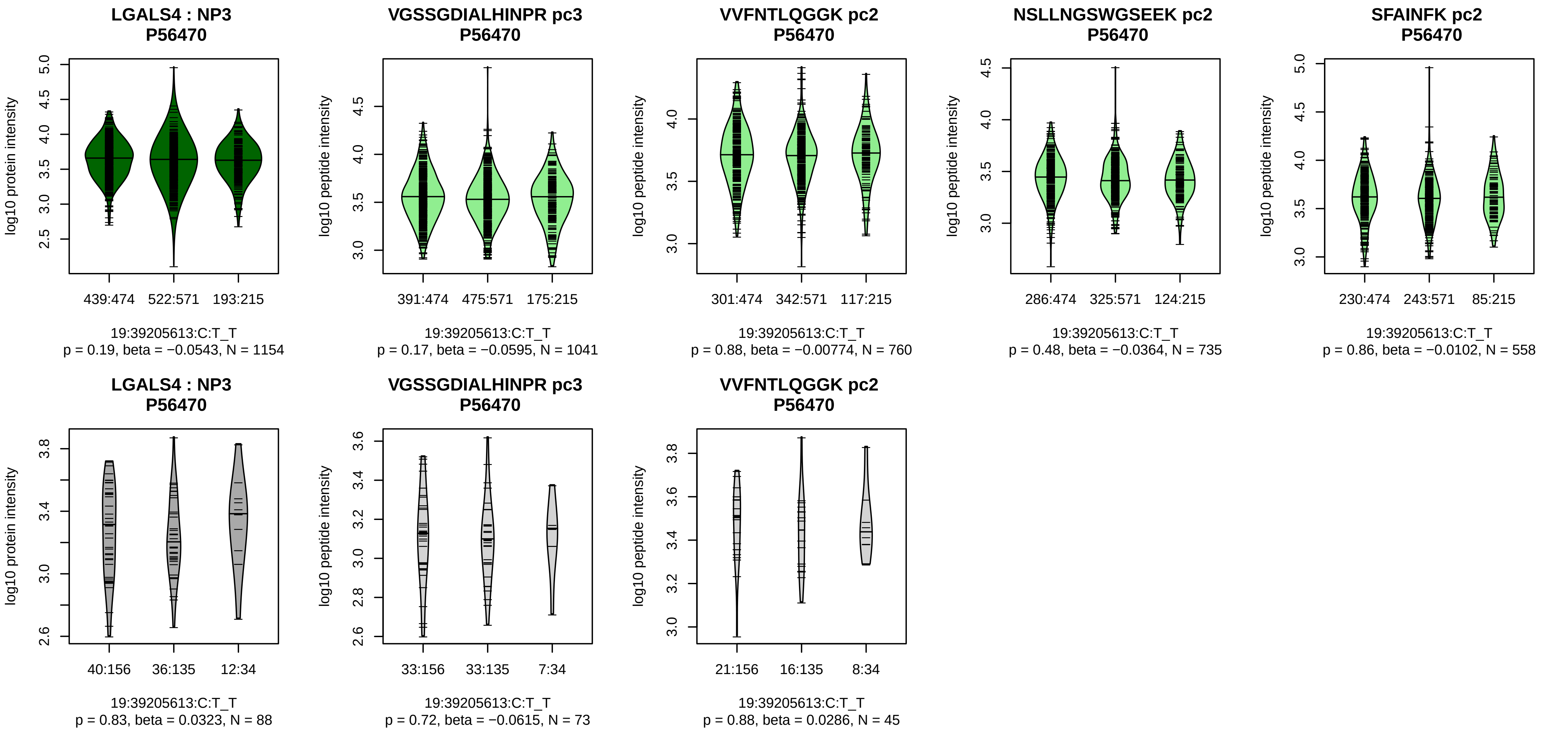


Assay Target: LGALS4
Olink UniProt: P56470
deCODE rsID: rs883394
Proxy rsID: rs883394
deCODE: 19:38714973:T:C
Proxy SNP: 19:39205613:C:T
deCODE log10(p): 10.4
deCODE BETA: −0.05
−:−:−:−:*:−:−
1041:760:735:558:342:153:29

**WFDC5 : NP1**
**Q8TCV5;Q8TCV5−2**

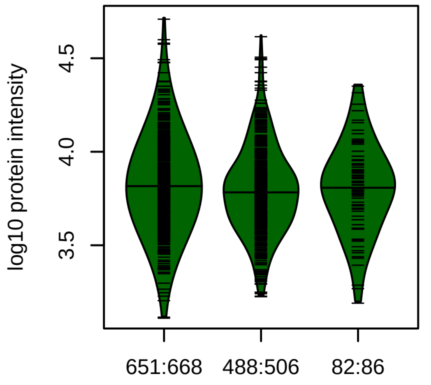


20:43727443:G:C_C
p = 0.0077, beta = −0.123, N = 1221

**LGSCPEDQLR pc2**
**Q8TCV5;Q8TCV5−2**

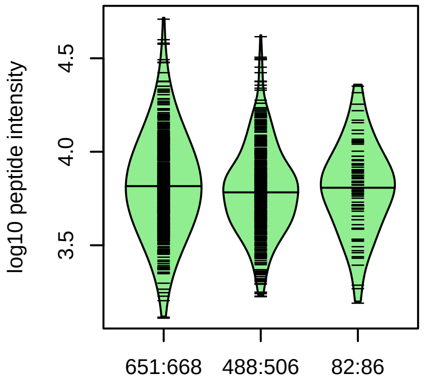


20:43727443:G:C_C
p = 0.0077, beta = −0.123, N = 1221

**WFDC5 : NP1**
**Q8TCV5;Q8TCV5−2**

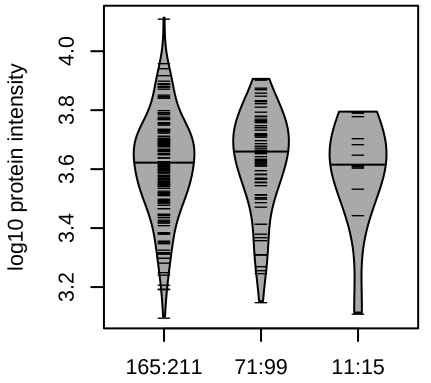


20:43727443:G:C_C
p = 0.88, beta = −0.0169, N = 247

**LGSCPEDQLR pc2**
**Q8TCV5;Q8TCV5−2**

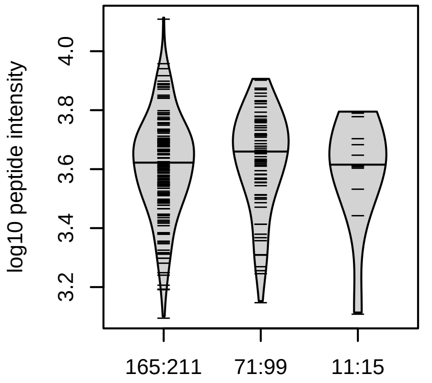


20:43727443:G:C_C
p = 0.88, beta = −0.0169, N = 247

Assay Target: WFDC5
Olink UniProt: Q8TCV5
deCODE rsID: rs35456522
Proxy rsID: rs35456522
deCODE: 20:45098802:C:G
Proxy SNP: 20:43727443:G:C
deCODE log10(p): 10.2
deCODE BETA: −0.06
*
1221

**CLSPMNHLCHK pc3**
**rs17422688 REF**

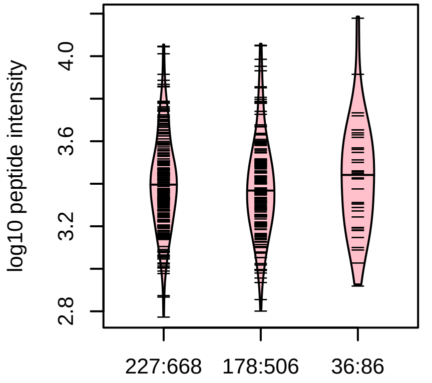


20:43727443:G:C_C
p = 0.2, model = DOM, N = 441

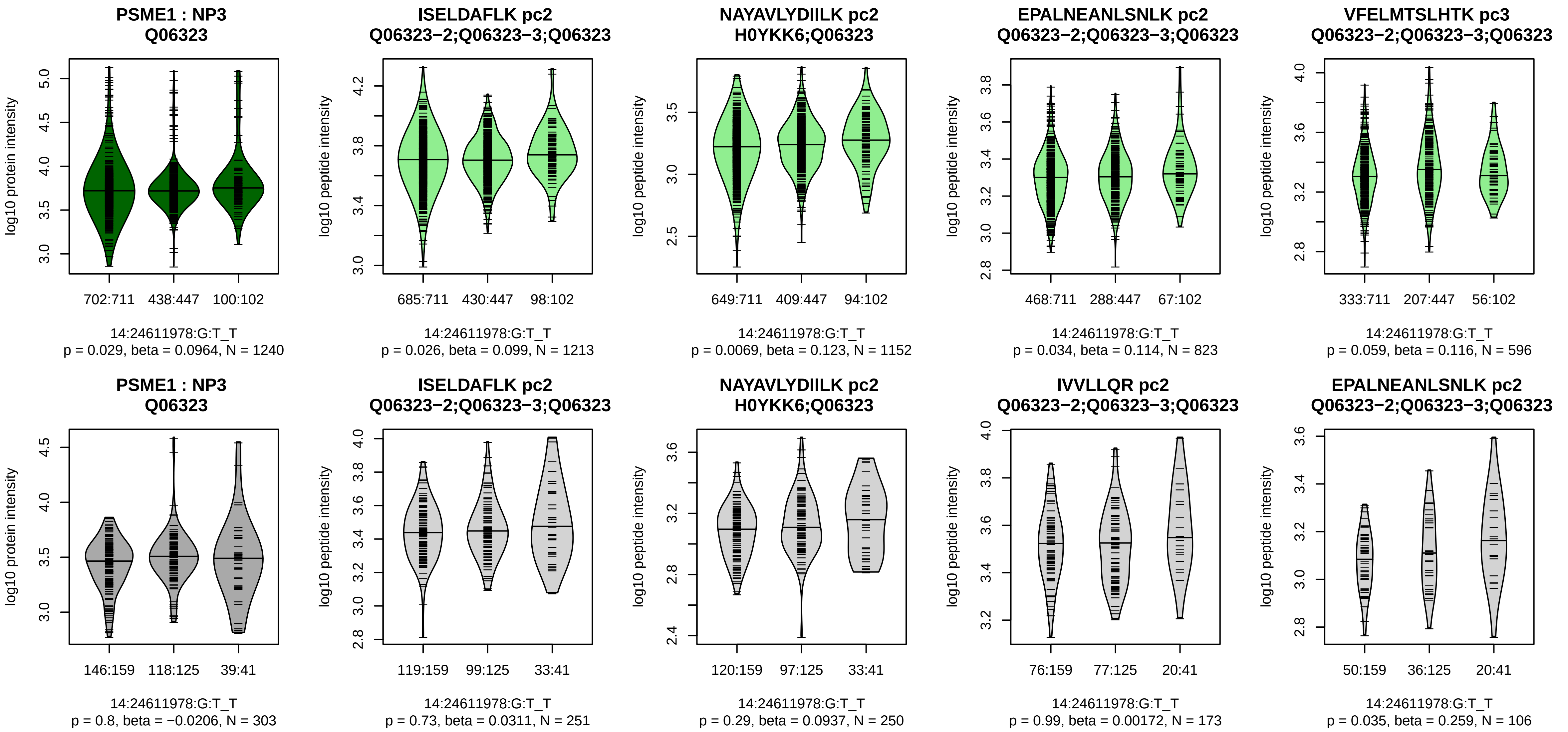


Assay Target: PSME1
Olink UniProt: Q06323
deCODE rsID: rs3742500
Proxy rsID: rs3742500
deCODE: 14:24142769:T:G
Proxy SNP: 14:24611978:G:T
deCODE log10(p): 9.9
deCODE BETA: 0.06
−:*:−:−:*:−:−:−:−:−:−:−:−:−:−:−:NA
1213:1152:823:596:552:510:502

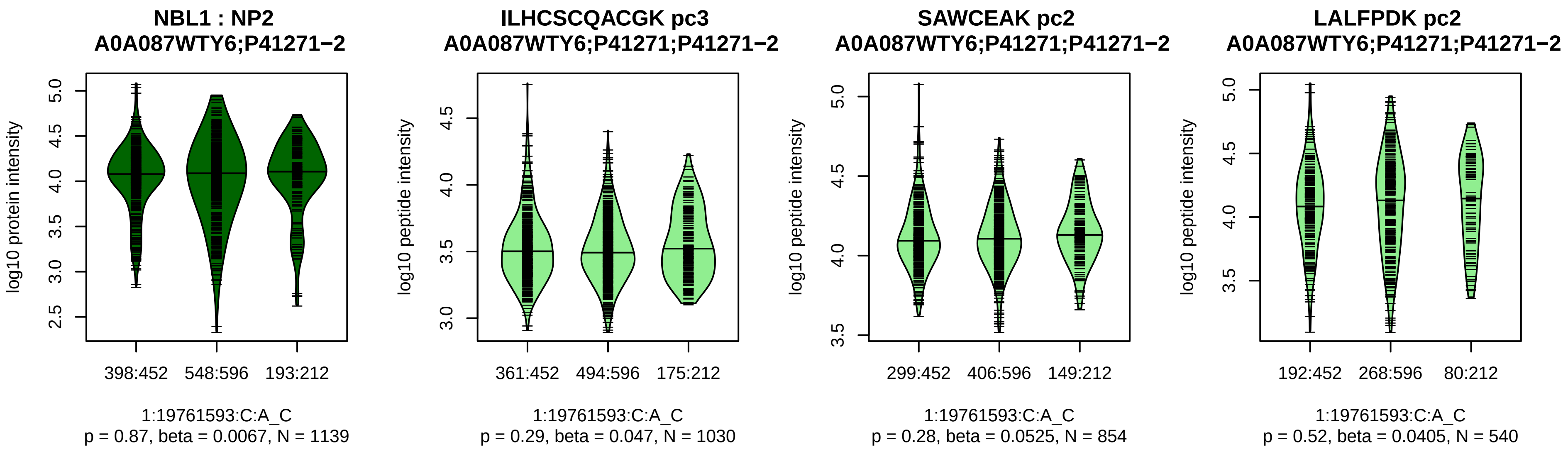


Assay Target: NBL1
Olink UniProt: P41271
deCODE rsID: rs4911994
Proxy rsID: rs4911994
deCODE: 1:19435099:C:A
Proxy SNP: 1:19761593:C:A
deCODE log10(p): 9.8
deCODE BETA: 0.05
−:−:−:−
1030:854:540:517

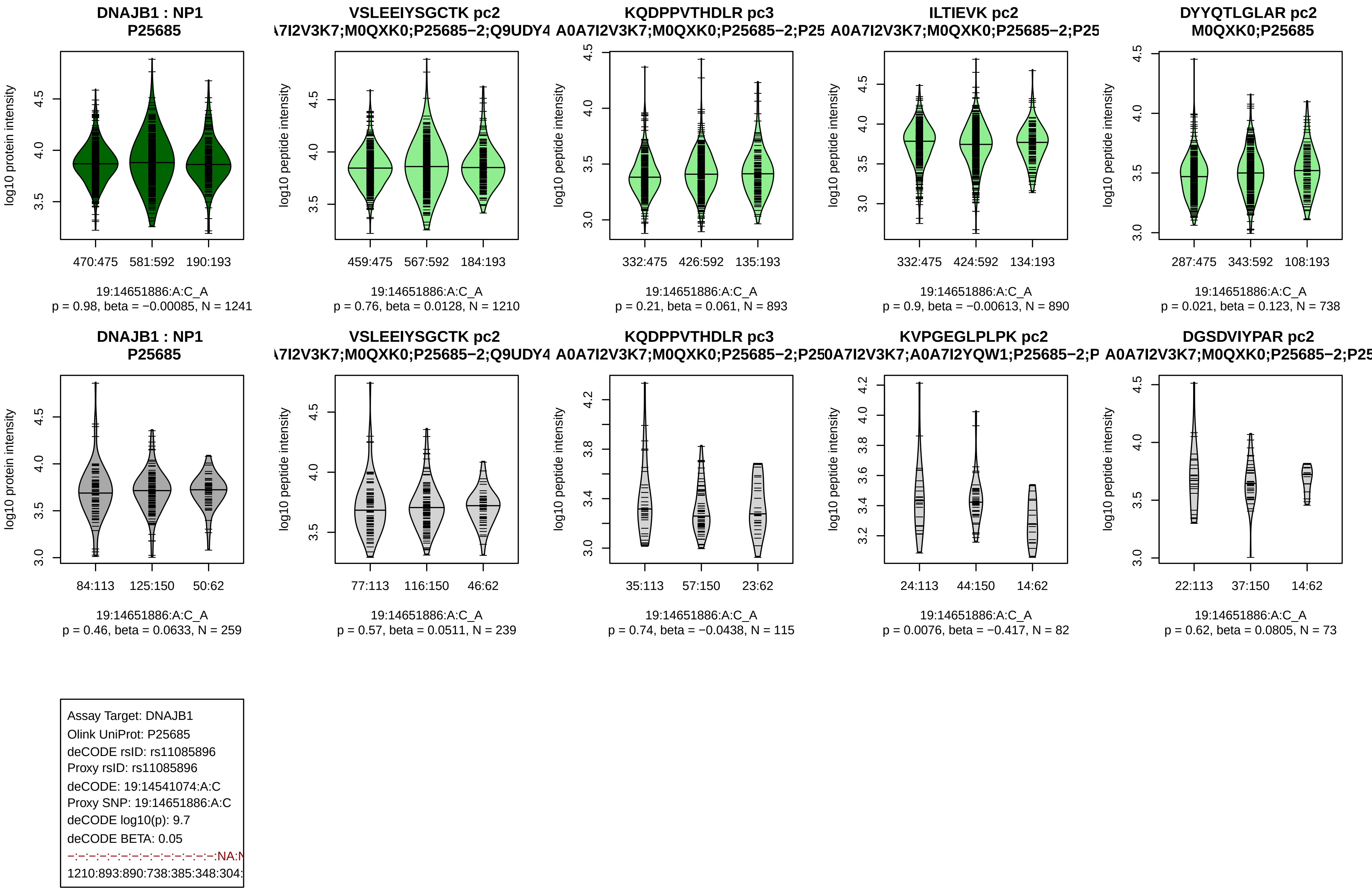
DNAJB1 : NP1
P25685
log10 protein intensity
470:475 581:592 190:193
19:14651886:A:C_A
p = 0.98, beta = −0.00085, N = 1241
VSLEEIYSGCTK pc2
A7I2V3K7;M0QXK0;P25685−2;Q9UDY4
log10 peptide intensity
459:475 567:592 184:193
19:14651886:A:C_A
p = 0.76, beta = 0.0128, N = 1210
KQDPPVTHDLR pc3
A0A7I2V3K7;M0QXK0;P25685−2;P25
log10 peptide intensity
332:475 426:592 135:193
19:14651886:A:C_A
p = 0.21, beta = 0.061, N = 893
ILTIEVK pc2
A0A7I2V3K7;M0QXK0;P25685−2;P25
log10 peptide intensity
332:475 424:592 134:193
19:14651886:A:C_A
p = 0.9, beta = −0.00613, N = 890
DYYQTLGLAR pc2
M0QXK0;P25685
log10 peptide intensity
287:475 343:592 108:193
19:14651886:A:C_A
p = 0.021, beta = 0.123, N = 738
DNAJB1 : NP1
P25685
log10 protein intensity
84:113 125:150 50:62
19:14651886:A:C_A
p = 0.46, beta = 0.0633, N = 259
VSLEEIYSGCTK pc2
A7I2V3K7;M0QXK0;P25685−2;Q9UDY4
log10 peptide intensity
77:113 116:150 46:62
19:14651886:A:C_A
p = 0.57, beta = 0.0511, N = 239
KQDPPVTHDLR pc3
A0A7I2V3K7;M0QXK0;P25685−2;P25
log10 peptide intensity
35:113 57:150 23:62
19:14651886:A:C_A
p = 0.74, beta = −0.0438, N = 115
KVPGEGLPLPK pc2
0A7I2V3K7;A0A7I2YQW1;P25685−2;P
log10 peptide intensity
24:113 44:150 14:62
19:14651886:A:C_A
p = 0.0076, beta = −0.417, N = 82
DGSDVIYPAR pc2
A0A7I2V3K7;M0QXK0;P25685−2;P25
log10 peptide intensity
22:113 37:150 14:62
19:14651886:A:C_A
p = 0.62, beta = 0.0805, N = 73
Assay Target: DNAJB1
Olink UniProt: P25685
deCODE rsID: rs11085896
Proxy rsID: rs11085896
deCODE: 19:14541074:A:C
Proxy SNP: 19:14651886:A:C
deCODE log10(p): 9.7
deCODE BETA: 0.05
−:−:−:−:−:−:−:−:−:−:−:−:−:−:NA:N
1210:893:890:738:385:348:304:

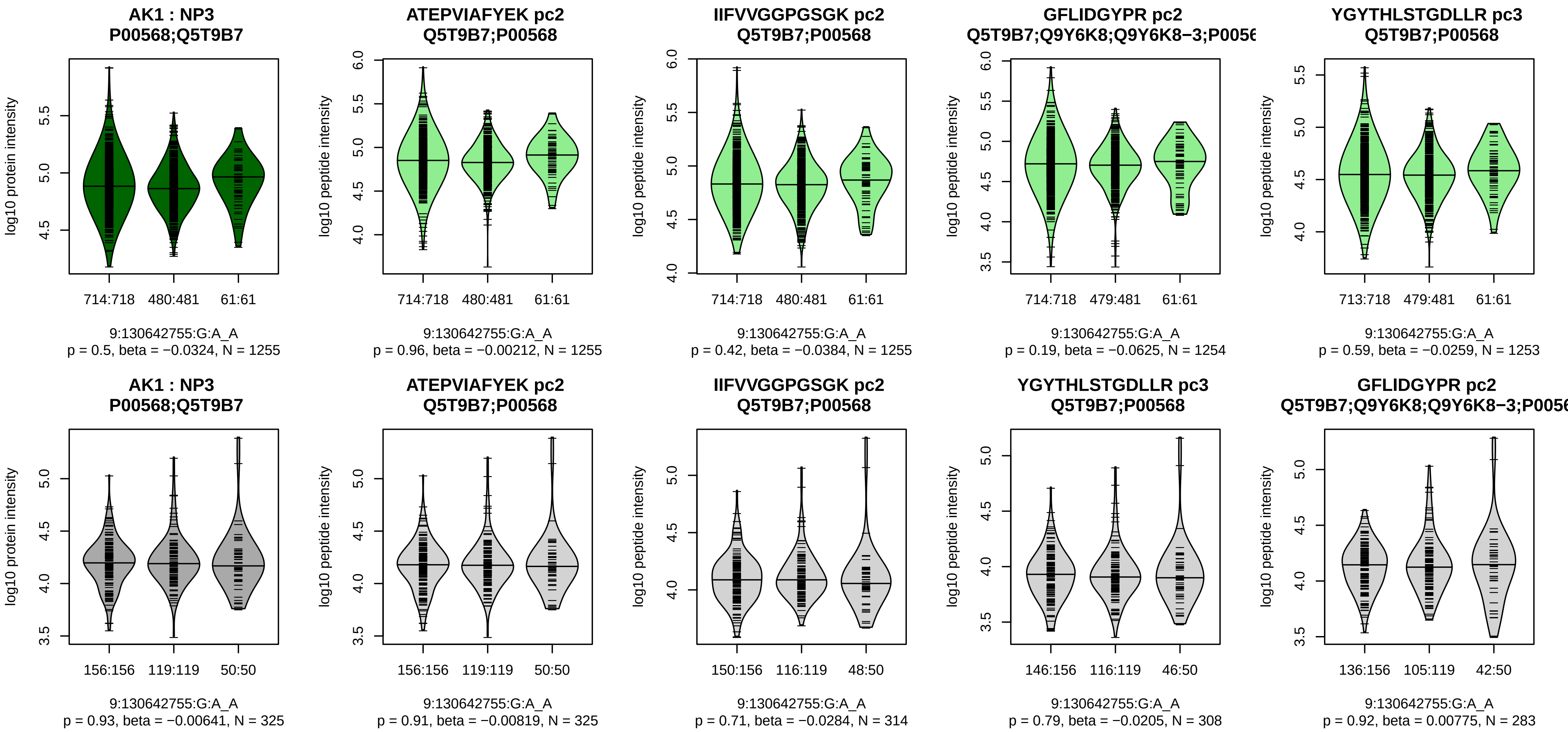


Assay Target: AK1
Olink UniProt: P00568
deCODE rsID: rs9919018
Proxy rsID: rs9919018
deCODE: 9:127880476:A:G
Proxy SNP: 9:130642755:G:A
deCODE log10(p): 9.7
deCODE BETA: −0.06
−:−:−:−:−:−:−:−:−:−:−:−:−:−:−:
1255:1255:1254:1253:1224:121

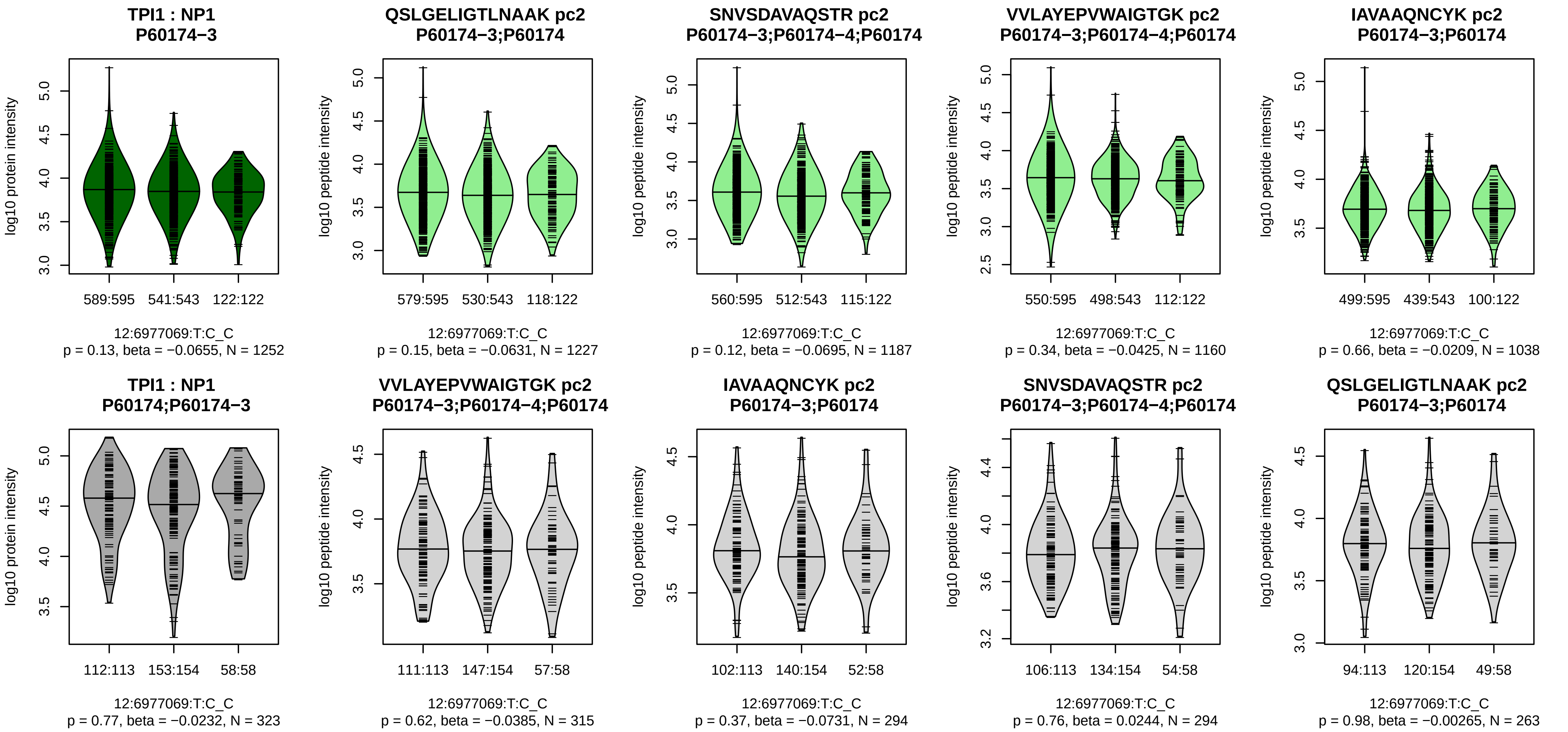


Assay Target: TPI1
Olink UniProt: P60174
deCODE rsID: rs2071065
Proxy rsID: rs2071065
deCODE: 12:6867905:C:T
Proxy SNP: 12:6977069:T:C
deCODE log10(p): 9.7
deCODE BETA: −0.06
−:−:−:−:−:−:−:−:−:−:−:−:−:−:−:
1227:1187:1160:1038:947:934:[illegible]

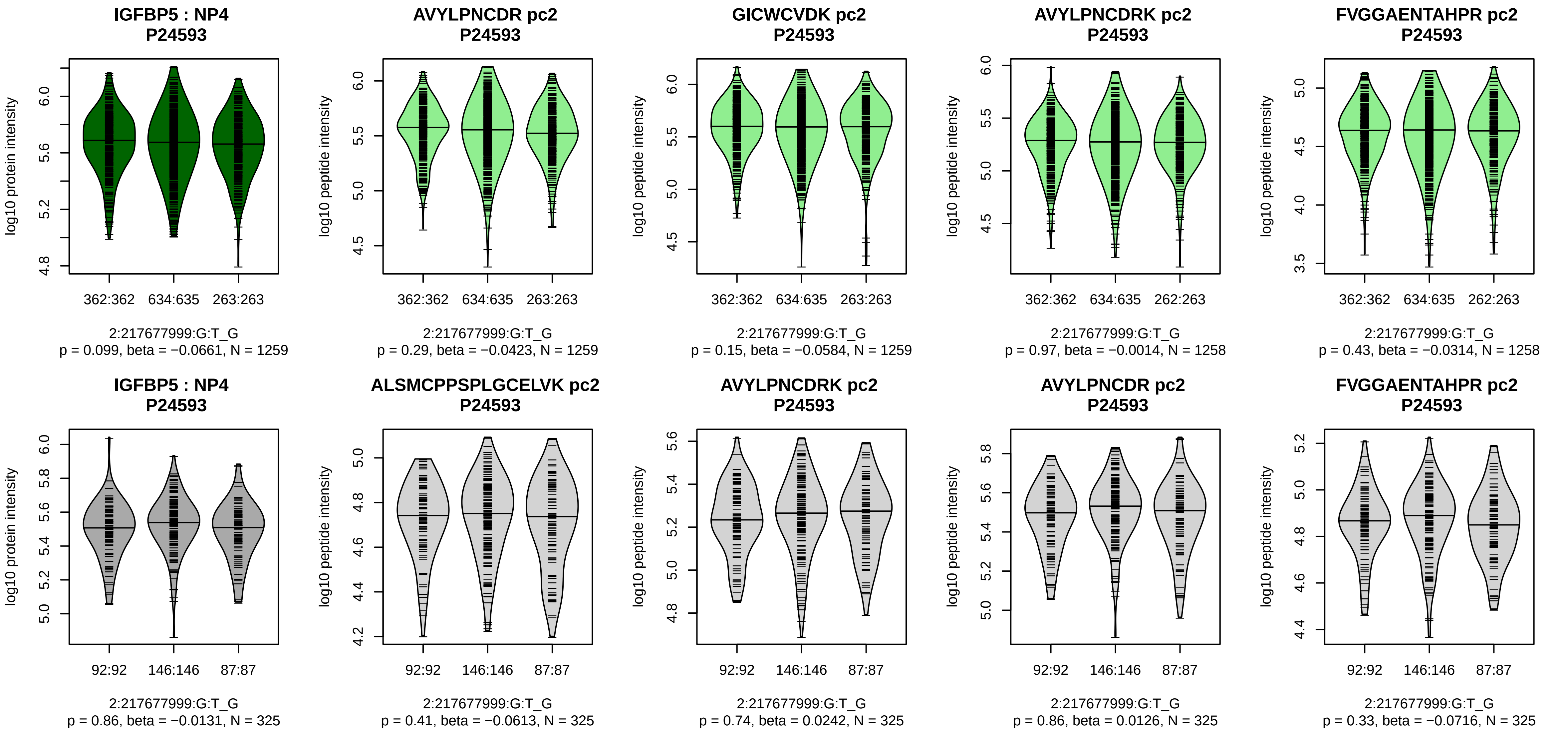


Assay Target: IGFBP5
Olink UniProt: P24593
deCODE rsID: rs7598172
Proxy rsID: rs7598172
deCODE: 2:216813276:G:T
Proxy SNP: 2:217677999:G:T
deCODE log10(p): 9.6
deCODE BETA: 0.05

1259:1259:1258:1258:1253:125

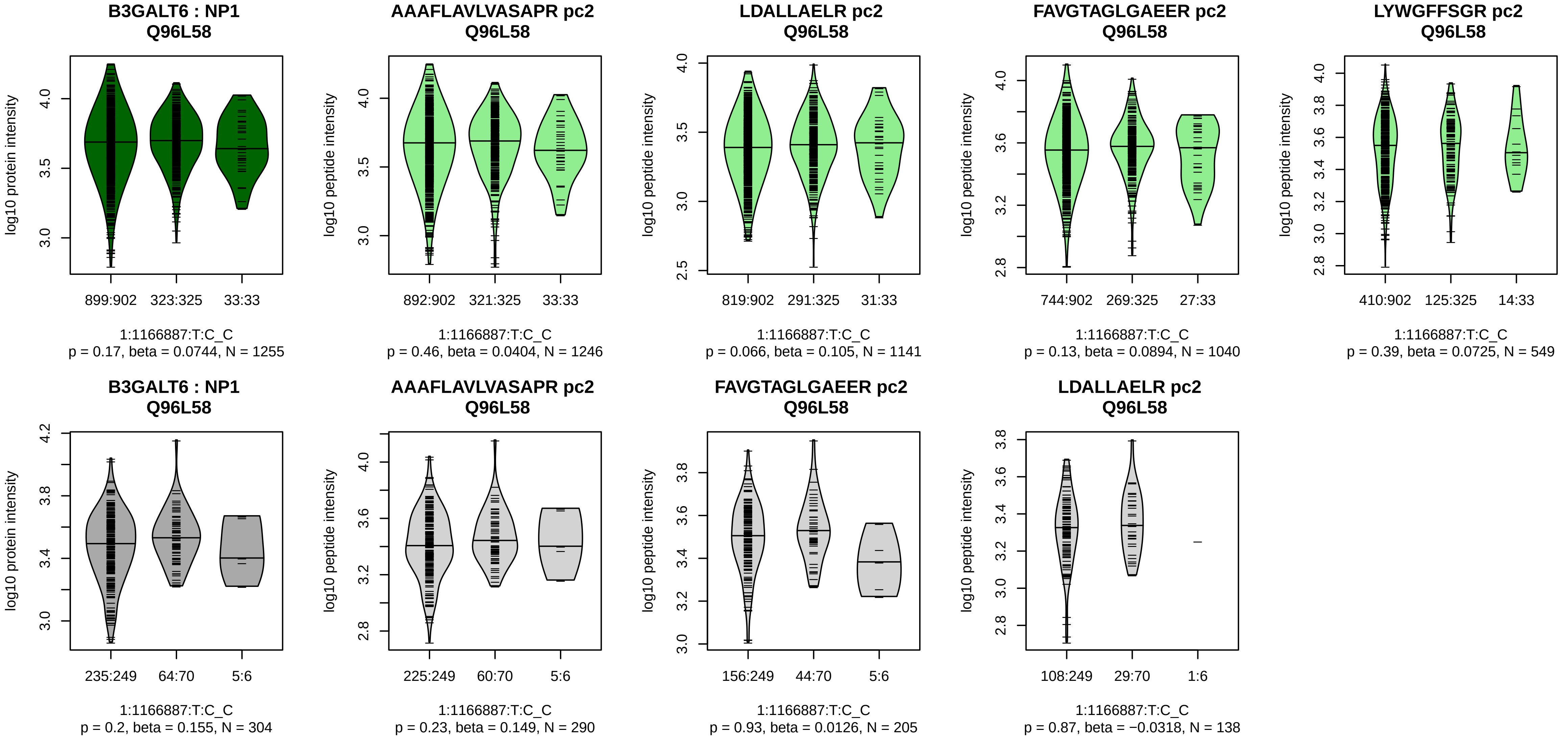


Assay Target: B3GALT6
Olink UniProt: Q96L58
deCODE rsID: rs60252802
Proxy rsID: rs60252802
deCODE: 1:1231507:C:T
Proxy SNP: 1:1166887:T:C
deCODE log10(p): 9.4
deCODE BETA: −0.07
−:−:−:−:−:−:−:−
1246:1141:1040:549:362:305:1[illegible]

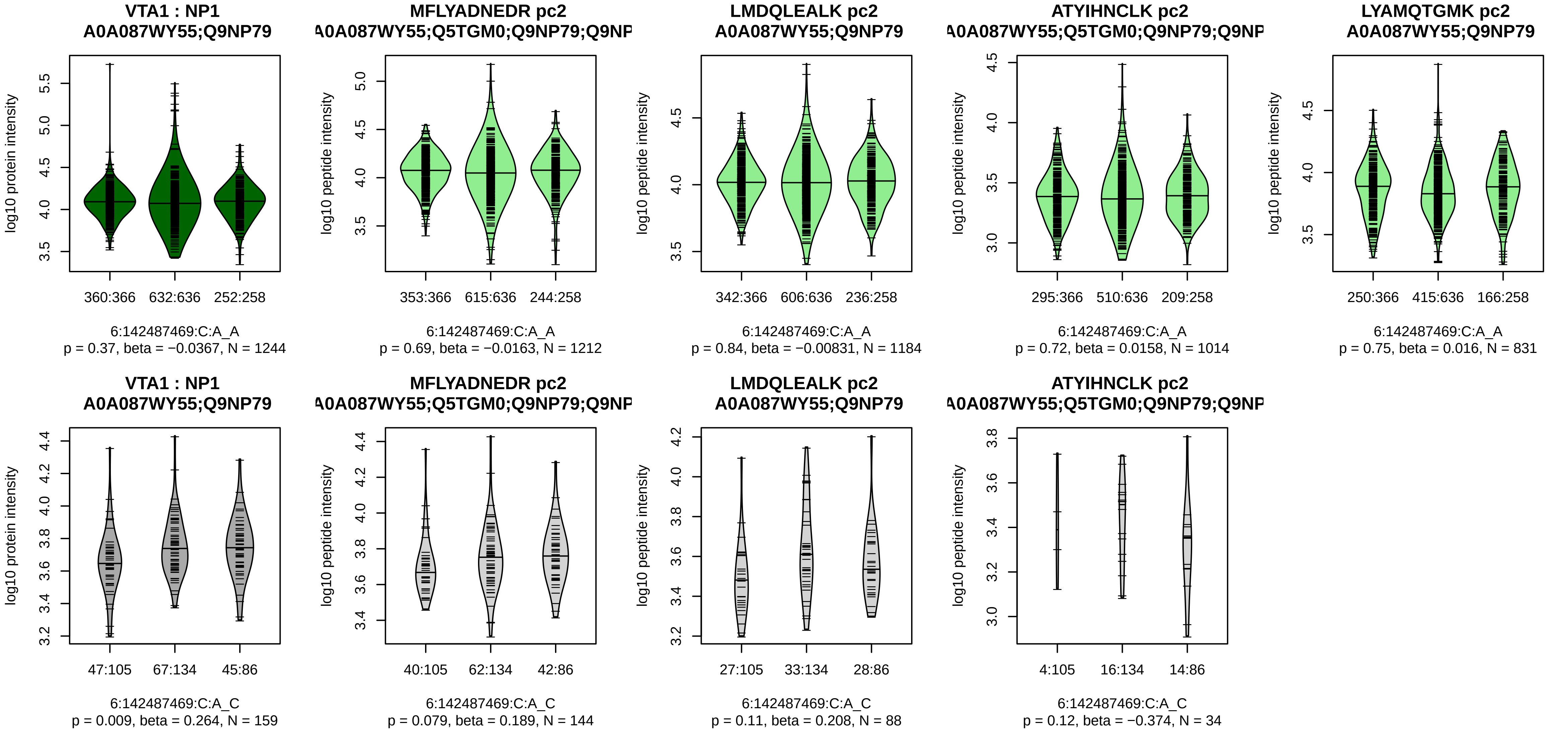


Assay Target: VTA1
Olink UniProt: Q9NP79
deCODE rsID: rs225656
Proxy rsID: rs225656
deCODE: 6:142166332:A:C
Proxy SNP: 6:142487469:C:A
deCODE log10(p): 9.1
deCODE BETA: −0.05
−:−:−:−:−:−:NA:NA
1212:1184:1014:831:146:89:12:

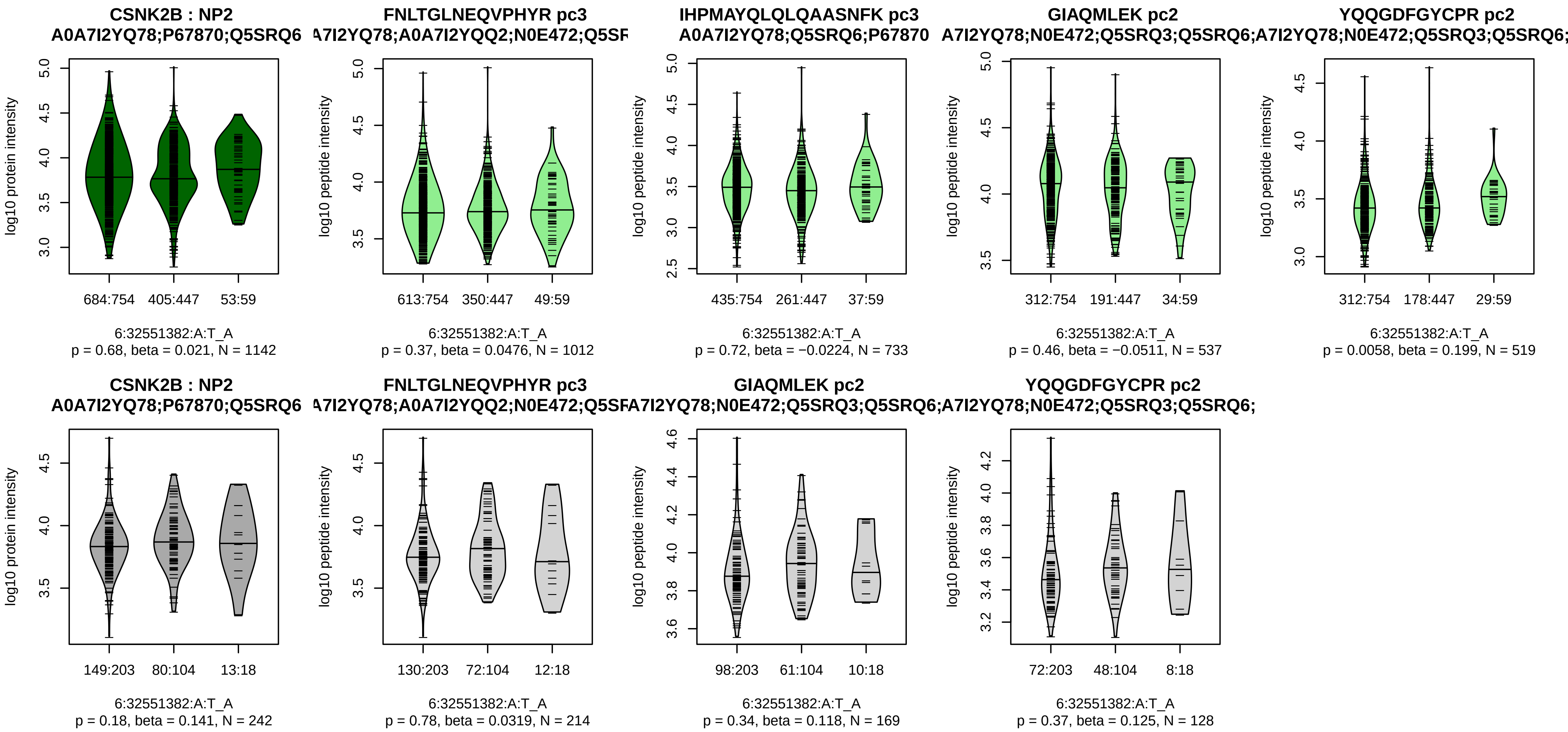


Assay Target: CSNK2B
Olink UniProt: P67870
deCODE rsID: rs78912960
Proxy rsID: rs9269911
deCODE: 6:32538131:G:A
Proxy SNP: 6:32551382:A:T
deCODE log10(p): 9
deCODE BETA: −0.05
−:−:−:*:NA
1012:733:537:519:9

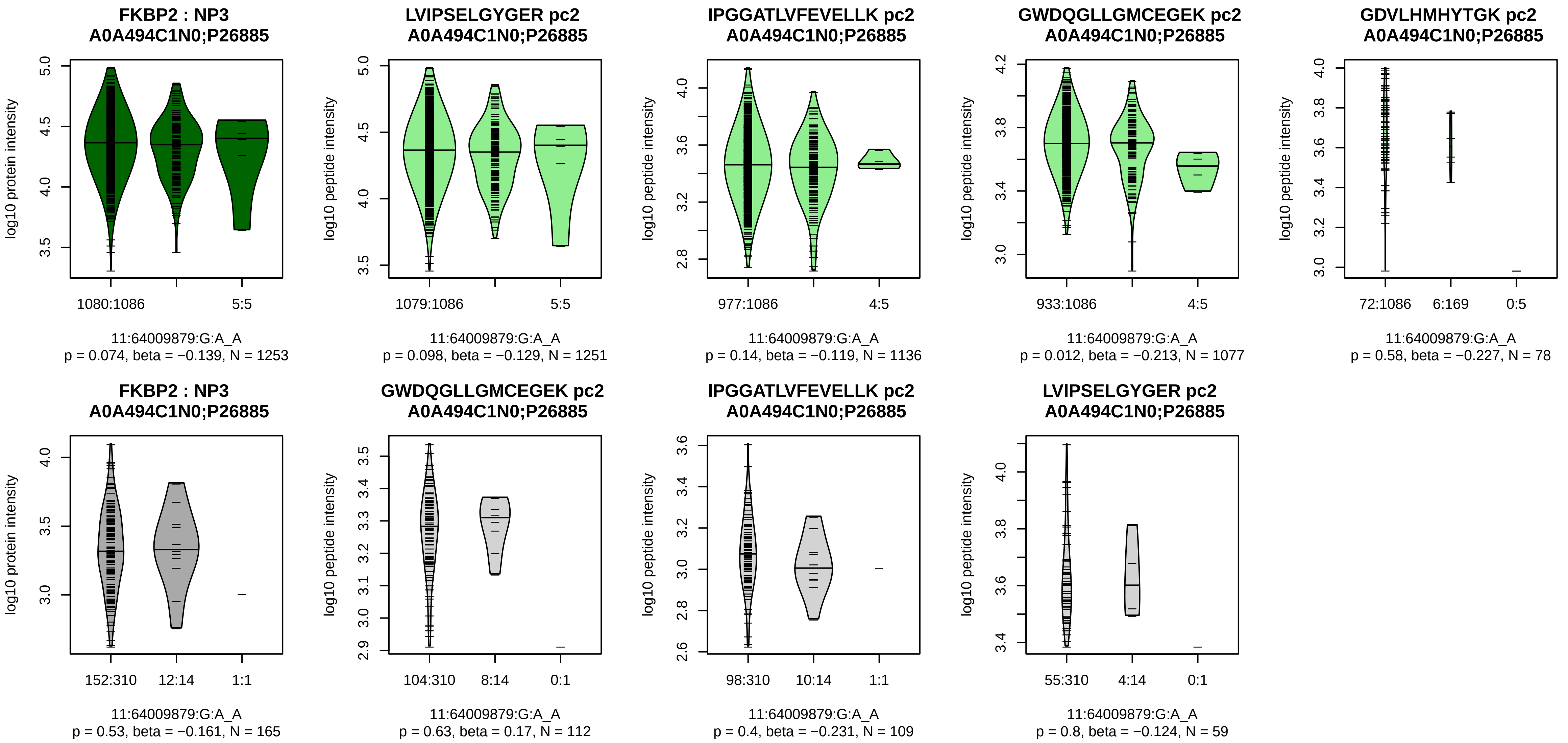


Assay Target: FKBP2
Olink UniProt: P26885
deCODE rsID: rs4672
Proxy rsID: rs4672
deCODE: 11:64242407:A:G
Proxy SNP: 11:64009879:G:A
deCODE log10(p): 8.9
deCODE BETA: −0.08
−:−:−:−:−
1251:1136:1077:78:57

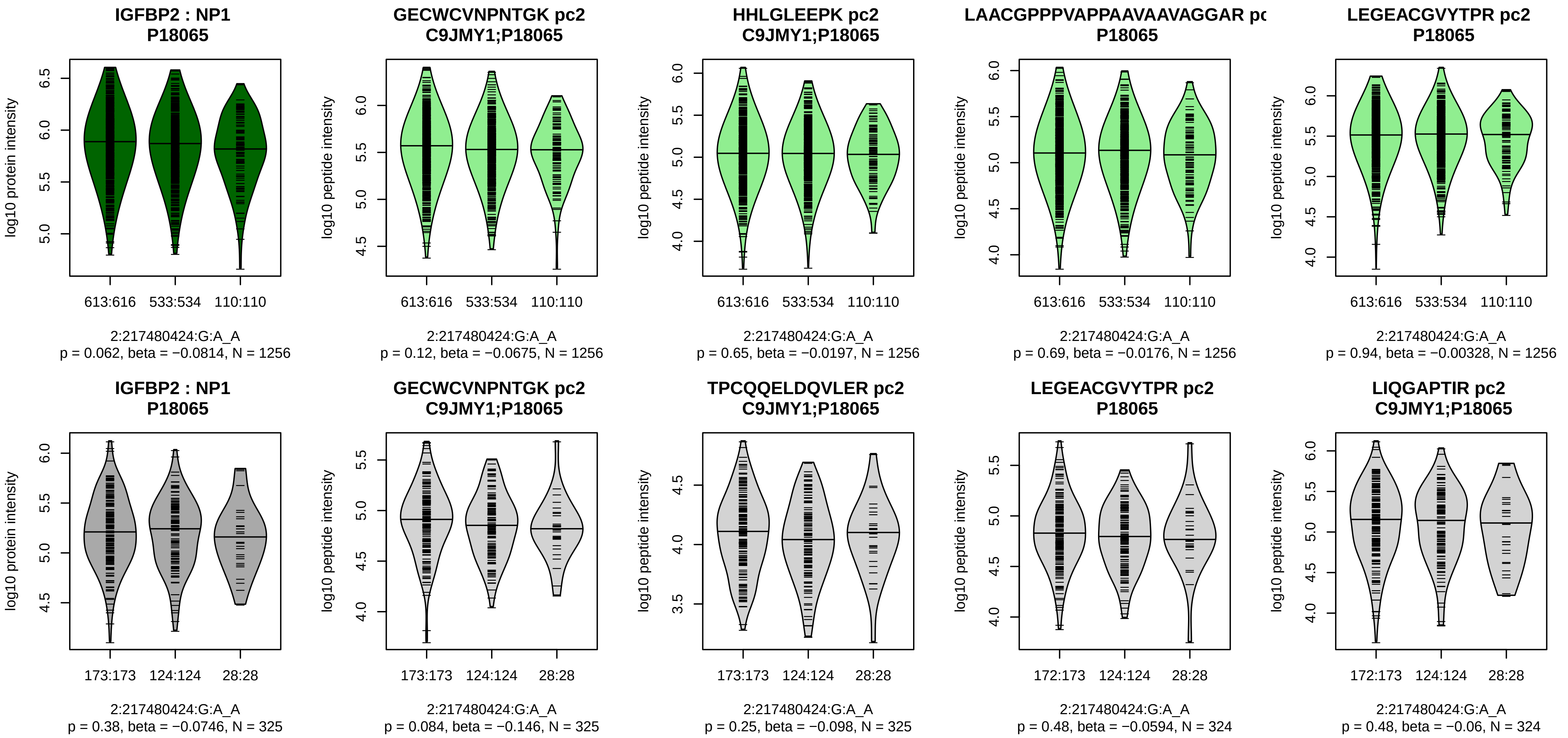


Assay Target: IGFBP2
Olink UniProt: P18065
deCODE rsID: rs4674100
Proxy rsID: rs4674100
deCODE: 2:216615701:A:G
Proxy SNP: 2:217480424:G:A
deCODE log10(p): 8.9
deCODE BETA: −0.06
–:–:–:–:–:–:–:–:–:–:–:–:–:–:–:–:
1256:1256:1256:1256:1255:125
