## Supplementary material for "A genome-wide association study of mass spectrometry proteomics using the Seer Proteograph platform": Figure S2

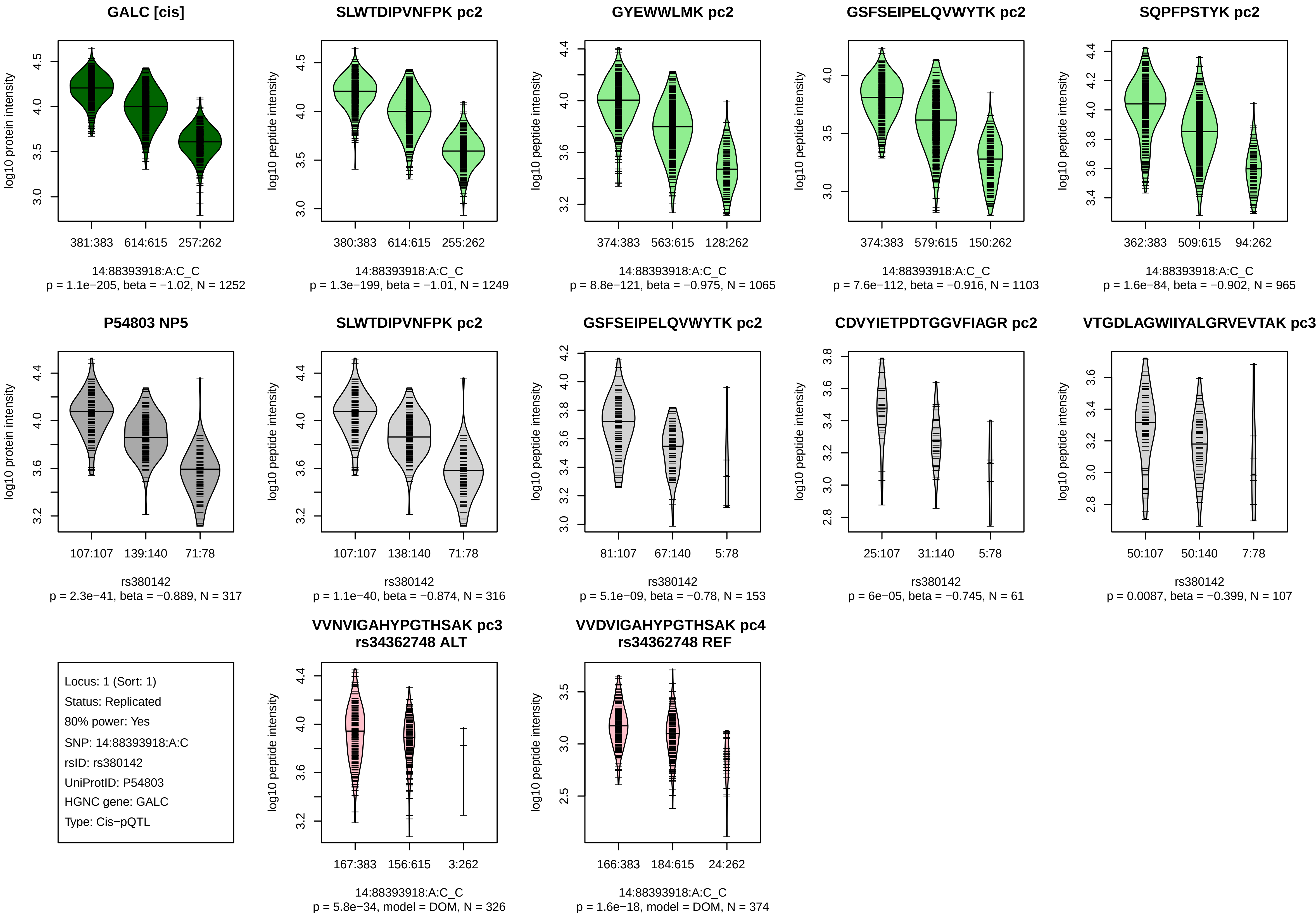
GALC [cis]
log10 protein intensity
381:383
614:615
257:262
14:88393918:A:C_C
p = 1.1e−205, beta = −1.02, N = 1252
SLWTDIPVNFPK pc2
log10 peptide intensity
380:383
614:615
255:262
14:88393918:A:C_C
p = 1.3e−199, beta = −1.01, N = 1249
GYEWWLMK pc2
log10 peptide intensity
374:383
563:615
128:262
14:88393918:A:C_C
p = 8.8e−121, beta = −0.975, N = 1065
GSFSEIPELQVWYTK pc2
log10 peptide intensity
374:383
579:615
150:262
14:88393918:A:C_C
p = 7.6e−112, beta = −0.916, N = 1103
SQPFPSTYK pc2
log10 peptide intensity
362:383
509:615
94:262
14:88393918:A:C_C
p = 1.6e−84, beta = −0.902, N = 965
P54803 NP5
log10 protein intensity
107:107
139:140
71:78
rs380142
p = 2.3e−41, beta = −0.889, N = 317
SLWTDIPVNFPK pc2
log10 peptide intensity
107:107
138:140
71:78
rs380142
p = 1.1e−40, beta = −0.874, N = 316
GSFSEIPELQVWYTK pc2
log10 peptide intensity
81:107
67:140
5:78
rs380142
p = 5.1e−09, beta = −0.78, N = 153
CDVYIETPDTGGVFIAGR pc2
log10 peptide intensity
25:107
31:140
5:78
rs380142
p = 6e−05, beta = −0.745, N = 61
VTGDLAGWIIYALGRVEVTAK pc3
log10 peptide intensity
50:107
50:140
7:78
rs380142
p = 0.0087, beta = −0.399, N = 107
Locus: 1 (Sort: 1)
Status: Replicated
80% power: Yes
SNP: 14:88393918:A:C
rsID: rs380142
UniProtID: P54803
HGNC gene: GALC
Type: Cis−pQTL
VVNVIGAHYPGTHSAK pc3
rs34362748 ALT
log10 peptide intensity
167:383
156:615
3:262
14:88393918:A:C_C
p = 5.8e−34, model = DOM, N = 326
VVDVIGAHYPGTHSAK pc4
rs34362748 REF
log10 peptide intensity
166:383
184:615
24:262
14:88393918:A:C_C
p = 1.6e−18, model = DOM, N = 374

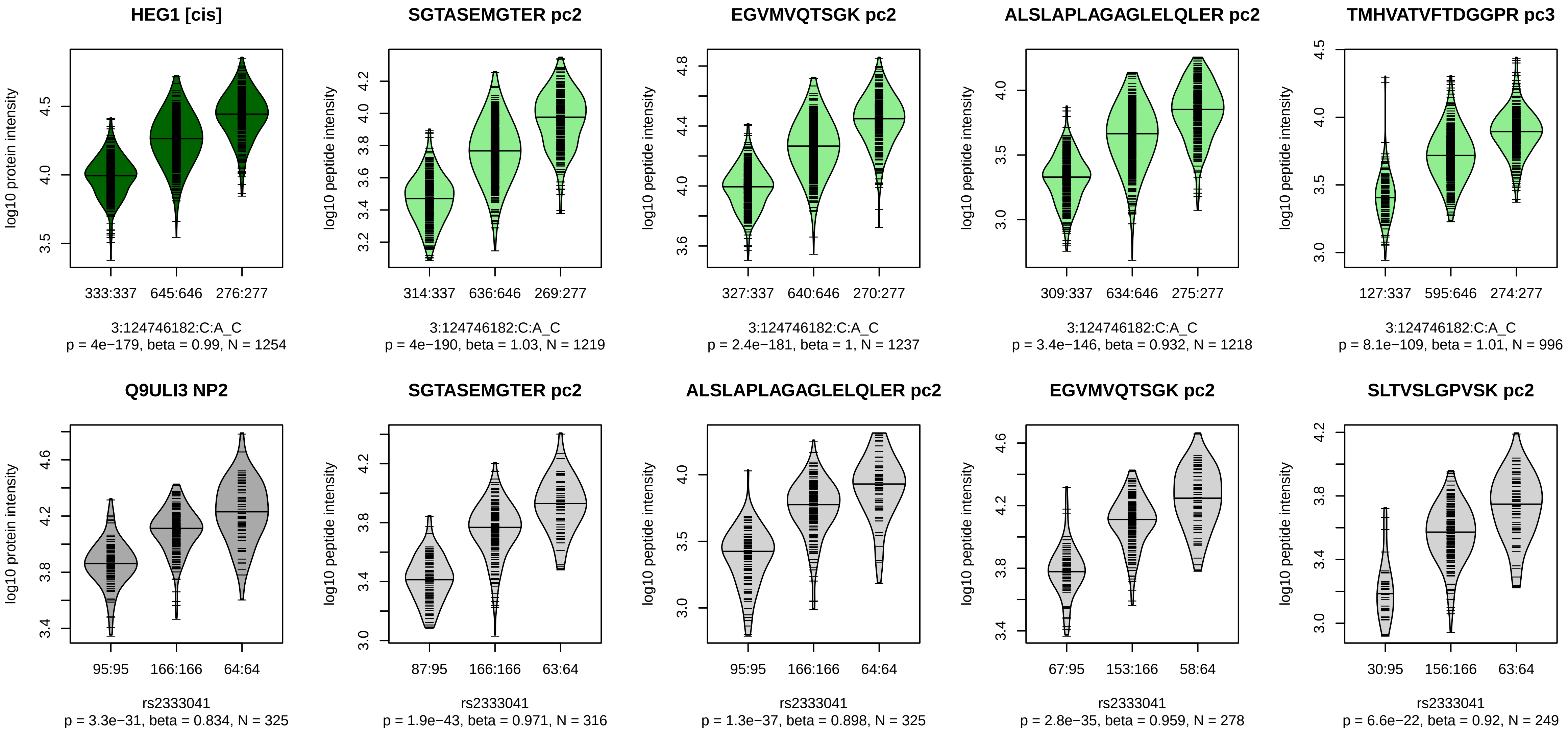


Locus: 2 (Sort: 2)
Status: Replicated
80% power: Yes
SNP: 3:124746182:C:A
rsID: rs2333041
UniProtID: Q9ULI3
HGNC gene: HEG1
Type: Cis−pQTL

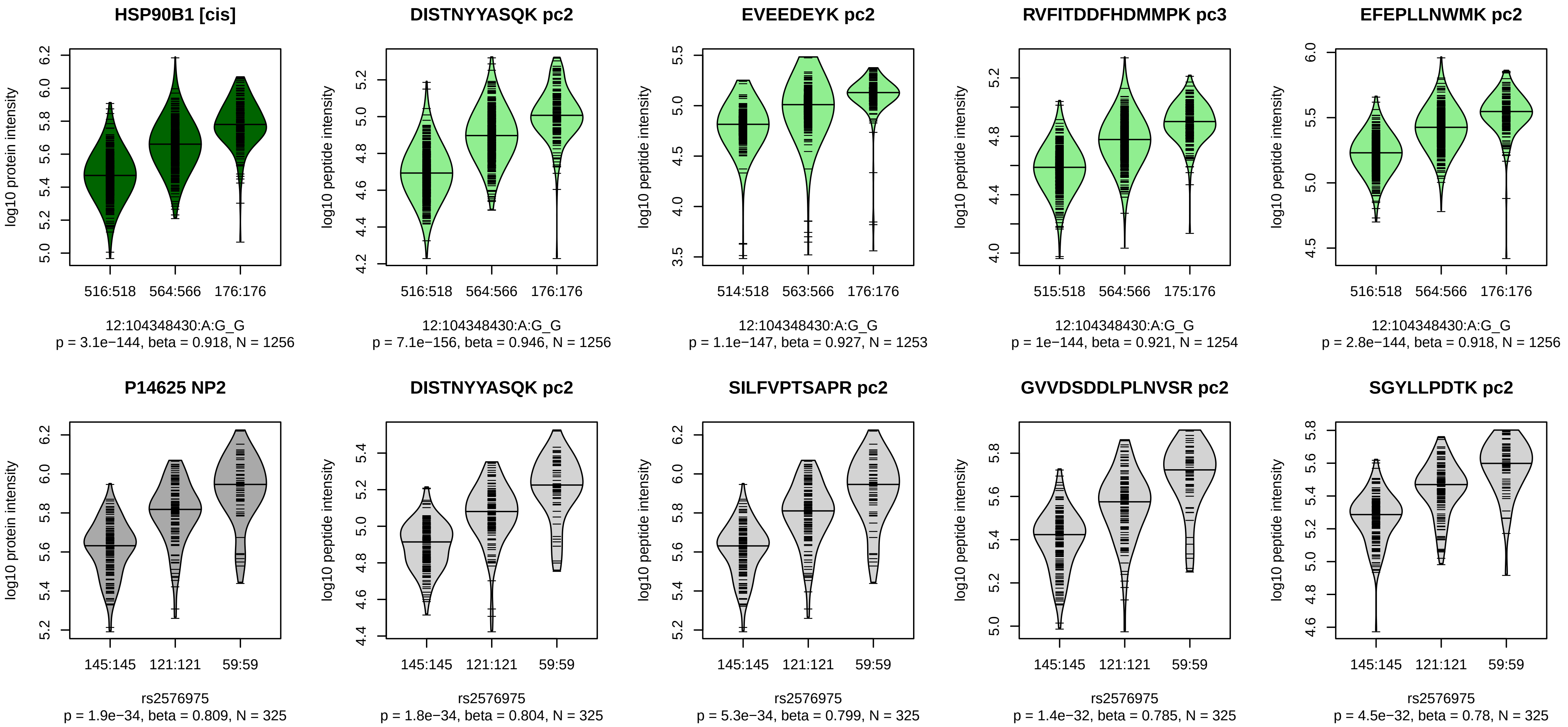


Locus: 3 (Sort: 3)

Status: Replicated

80% power: Yes

SNP: 12:104348430:A:G

rsID: rs2576975

UniProtID: P14625

HGNC gene: HSP90B1

Type: Cis−pQTL

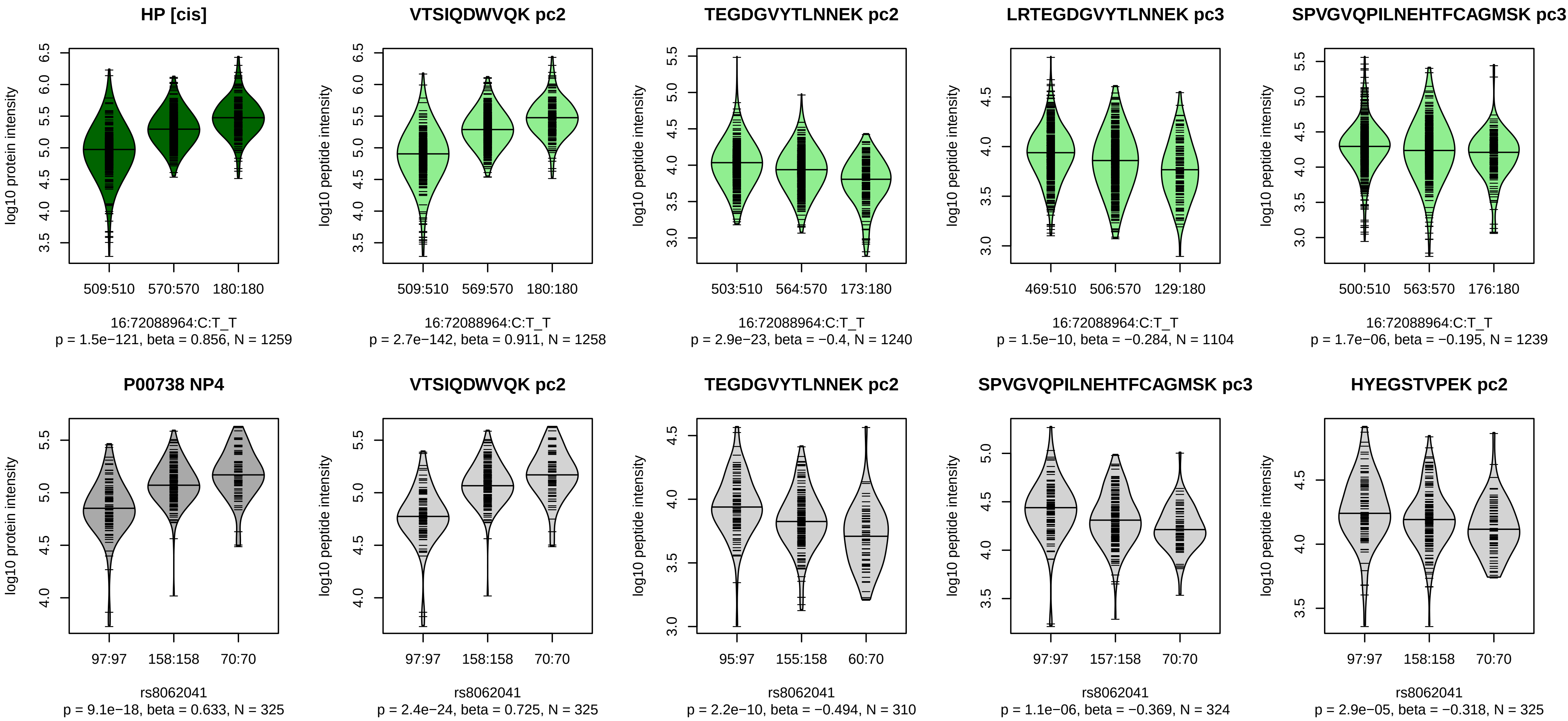


Locus: 4 (Sort: 4)
Status: Replicated
80% power: Yes
SNP: 16:72088964:C:T
rsID: rs8062041
UniProtID: P00738
HGNC gene: HP
Type: Cis−pQTL

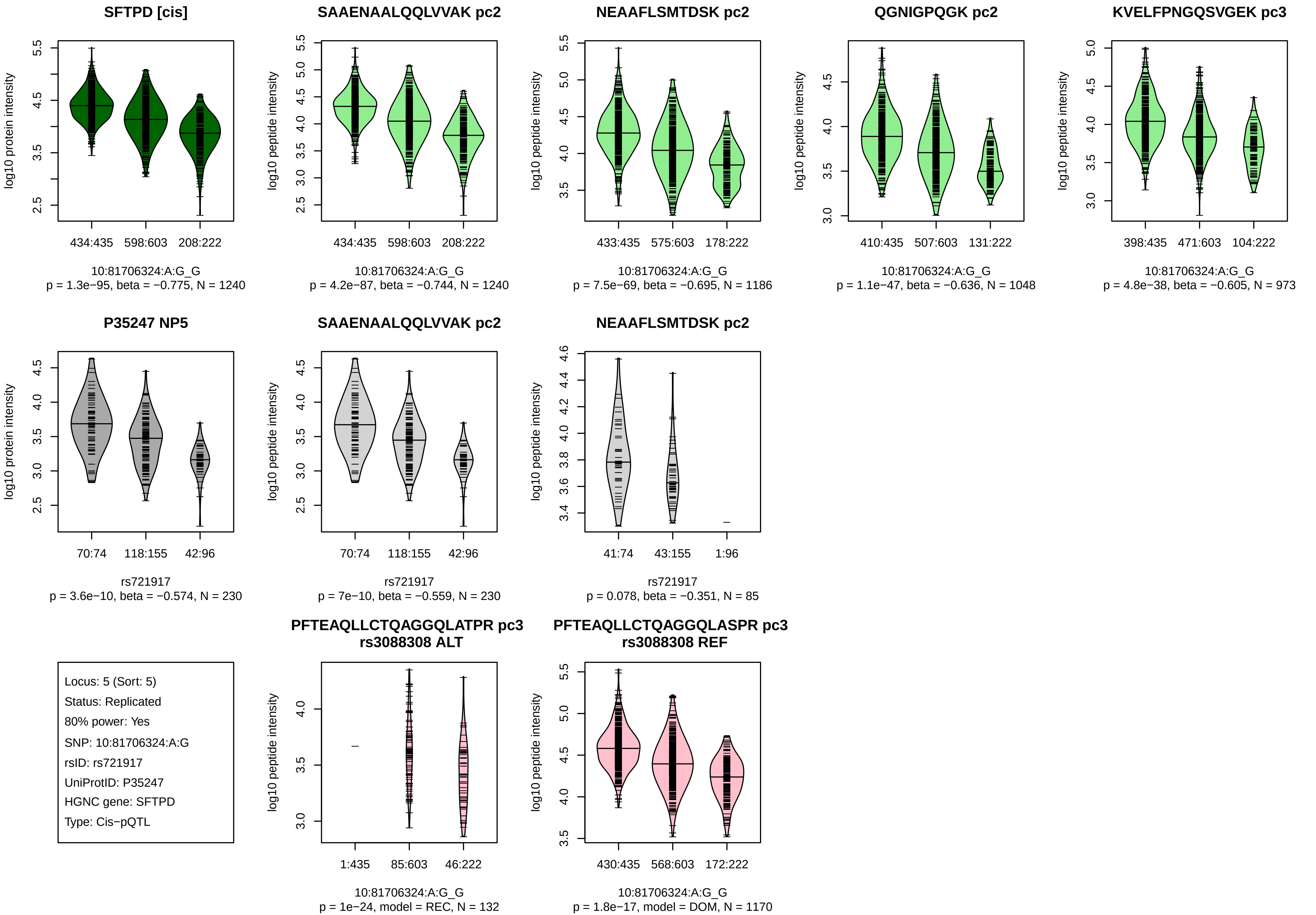
SFTPD [cis]
log10 protein intensity
434:435
598:603
208:222
10:81706324:A:G_G
p = 1.3e−95, beta = −0.775, N = 1240
SAAENAALQQLVVAK pc2
log10 peptide intensity
434:435
598:603
208:222
10:81706324:A:G_G
p = 4.2e−87, beta = −0.744, N = 1240
NEAAFLSMTDSK pc2
log10 peptide intensity
433:435
575:603
178:222
10:81706324:A:G_G
p = 7.5e−69, beta = −0.695, N = 1186
QGNIGPQGK pc2
log10 peptide intensity
410:435
507:603
131:222
10:81706324:A:G_G
p = 1.1e−47, beta = −0.636, N = 1048
KVELFPNGQSVGEK pc3
log10 peptide intensity
398:435
471:603
104:222
10:81706324:A:G_G
p = 4.8e−38, beta = −0.605, N = 973
P35247 NP5
log10 protein intensity
70:74
118:155
42:96
rs721917
p = 3.6e−10, beta = −0.574, N = 230
SAAENAALQQLVVAK pc2
log10 peptide intensity
70:74
118:155
42:96
rs721917
p = 7e−10, beta = −0.559, N = 230
NEAAFLSMTDSK pc2
log10 peptide intensity
41:74
43:155
1:96
rs721917
p = 0.078, beta = −0.351, N = 85
Locus: 5 (Sort: 5)
Status: Replicated
80% power: Yes
SNP: 10:81706324:A:G
rsID: rs721917
UniProtID: P35247
HGNC gene: SFTPD
Type: Cis−pQTL
PFTEAQLLCTQAGGQLATPR pc3
rs3088308 ALT
log10 peptide intensity
1:435
85:603
46:222
10:81706324:A:G_G
p = 1e−24, model = REC, N = 132
PFTEAQLLCTQAGGQLASPR pc3
rs3088308 REF
log10 peptide intensity
430:435
568:603
172:222
10:81706324:A:G_G
p = 1.8e−17, model = DOM, N = 1170

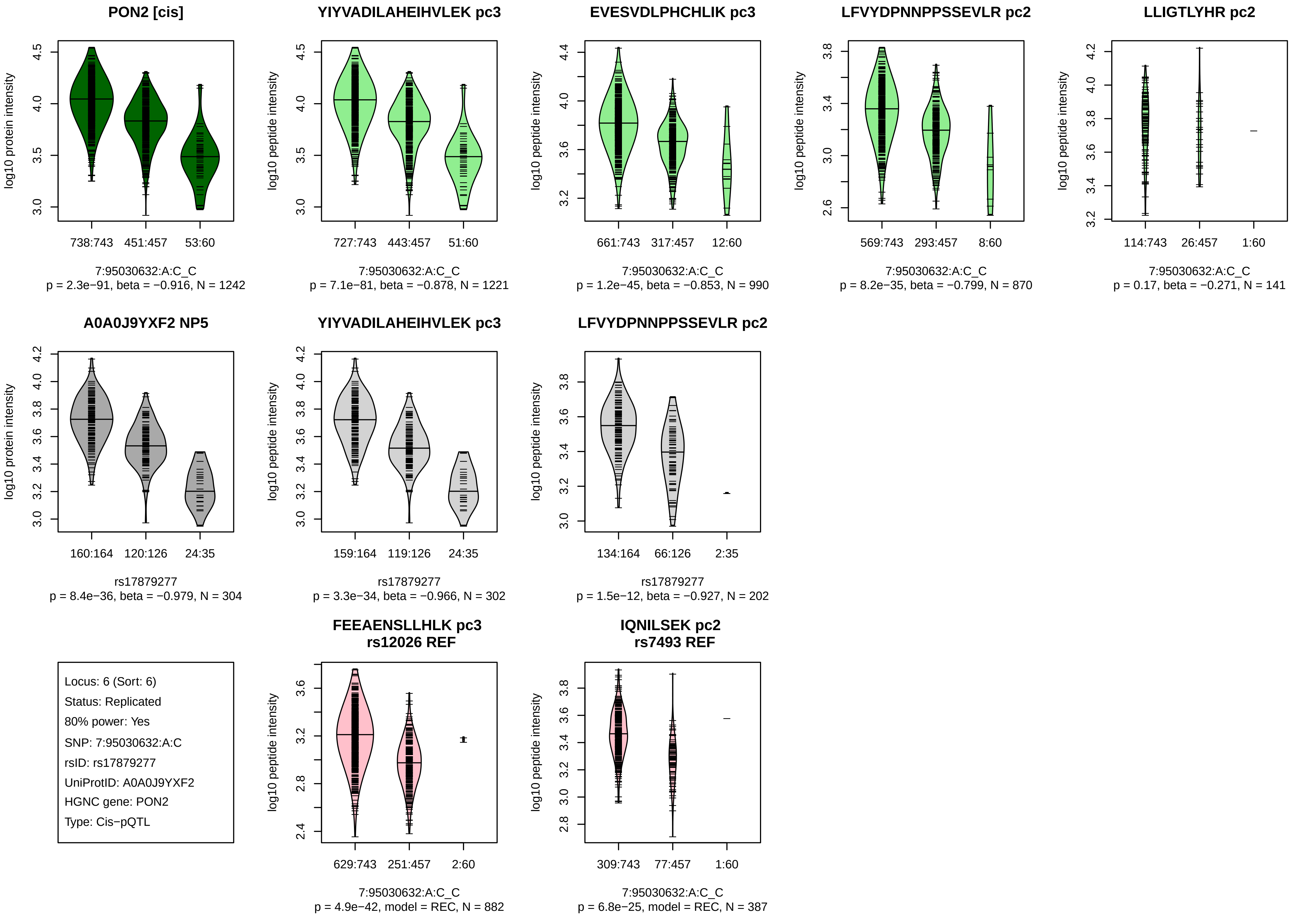

PON2 [cis]
log10 protein intensity
738:743
451:457
53:60
7:95030632:A:C_C
p = 2.3e−91, beta = −0.916, N = 1242
YIYVADILAHEIHVLEK pc3
log10 peptide intensity
727:743
443:457
51:60
7:95030632:A:C_C
p = 7.1e−81, beta = −0.878, N = 1221
EVESVDLPHCHLIK pc3
log10 peptide intensity
661:743
317:457
12:60
7:95030632:A:C_C
p = 1.2e−45, beta = −0.853, N = 990
LFVYDPNNPPSSEVLR pc2
log10 peptide intensity
569:743
293:457
8:60
7:95030632:A:C_C
p = 8.2e−35, beta = −0.799, N = 870
LLIGTLYHR pc2
log10 peptide intensity
114:743
26:457
1:60
7:95030632:A:C_C
p = 0.17, beta = −0.271, N = 141
A0A0J9YXF2 NP5
log10 protein intensity
160:164
120:126
24:35
rs17879277
p = 8.4e−36, beta = −0.979, N = 304
YIYVADILAHEIHVLEK pc3
log10 peptide intensity
159:164
119:126
24:35
rs17879277
p = 3.3e−34, beta = −0.966, N = 302
LFVYDPNNPPSSEVLR pc2
log10 peptide intensity
134:164
66:126
2:35
rs17879277
p = 1.5e−12, beta = −0.927, N = 202
Locus: 6 (Sort: 6)
Status: Replicated
80% power: Yes
SNP: 7:95030632:A:C
rsID: rs17879277
UniProtID: A0A0J9YXF2
HGNC gene: PON2
Type: Cis−pQTL
FEEAENSLLHLK pc3
rs12026 REF
log10 peptide intensity
629:743
251:457
2:60
7:95030632:A:C_C
p = 4.9e−42, model = REC, N = 882
IQNILSEK pc2
rs7493 REF
log10 peptide intensity
309:743
77:457
1:60
7:95030632:A:C_C
p = 6.8e−25, model = REC, N = 387

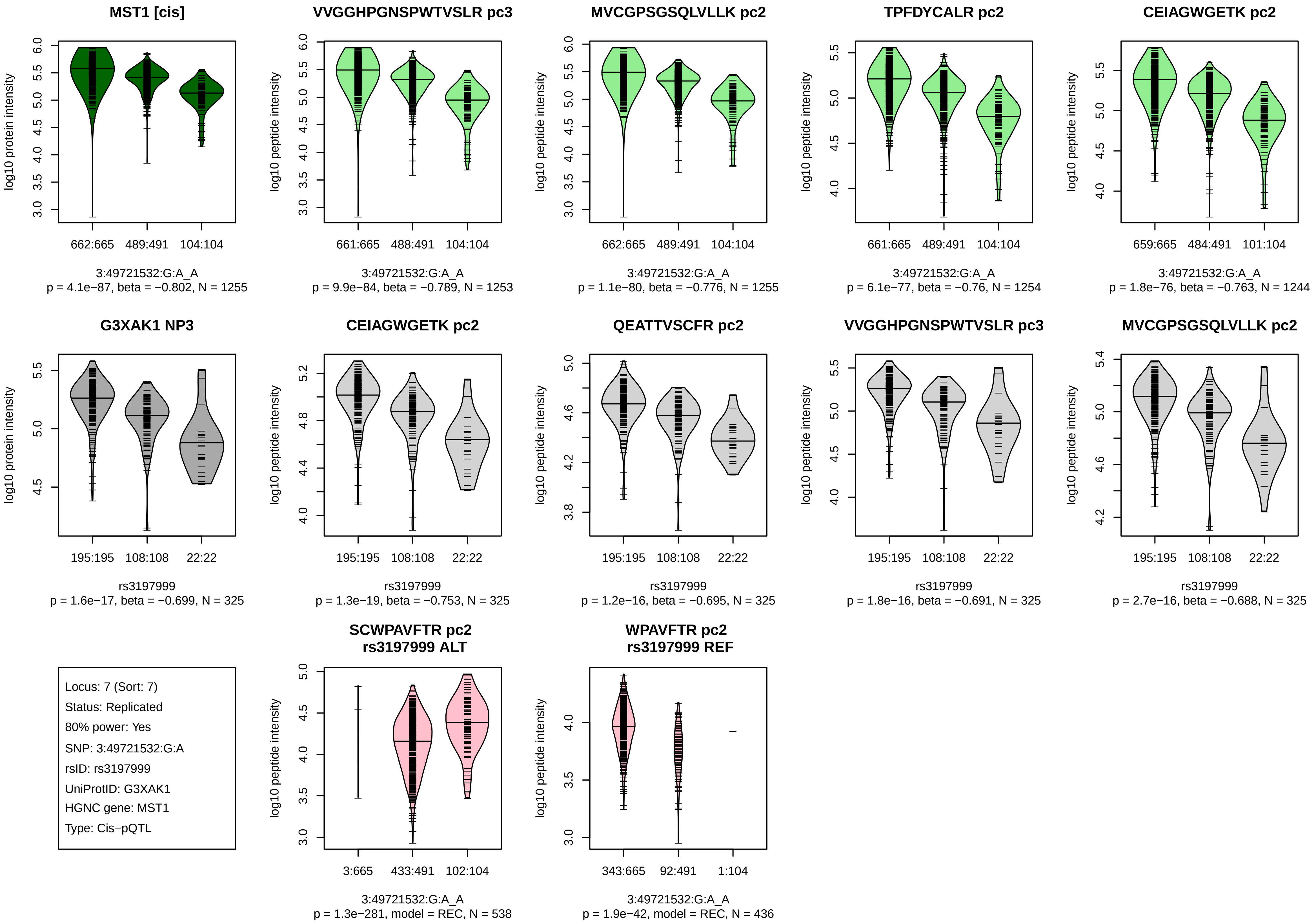
MST1 [cis]
log10 protein intensity
662:665 489:491 104:104
3:49721532:G:A_A
p = 4.1e−87, beta = −0.802, N = 1255
VVGGHPGNSPWTVSLR pc3
log10 peptide intensity
661:665 488:491 104:104
3:49721532:G:A_A
p = 9.9e−84, beta = −0.789, N = 1253
MVCGPSGSQLVLLK pc2
log10 peptide intensity
662:665 489:491 104:104
3:49721532:G:A_A
p = 1.1e−80, beta = −0.776, N = 1255
TPFDYCALR pc2
log10 peptide intensity
661:665 489:491 104:104
3:49721532:G:A_A
p = 6.1e−77, beta = −0.76, N = 1254
CEIAGWGETK pc2
log10 peptide intensity
659:665 484:491 101:104
3:49721532:G:A_A
p = 1.8e−76, beta = −0.763, N = 1244
G3XAK1 NP3
log10 protein intensity
195:195 108:108 22:22
rs3197999
p = 1.6e−17, beta = −0.699, N = 325
CEIAGWGETK pc2
log10 peptide intensity
195:195 108:108 22:22
rs3197999
p = 1.3e−19, beta = −0.753, N = 325
QEATTVSCFR pc2
log10 peptide intensity
195:195 108:108 22:22
rs3197999
p = 1.2e−16, beta = −0.695, N = 325
VVGGHPGNSPWTVSLR pc3
log10 peptide intensity
195:195 108:108 22:22
rs3197999
p = 1.8e−16, beta = −0.691, N = 325
MVCGPSGSQLVLLK pc2
log10 peptide intensity
195:195 108:108 22:22
rs3197999
p = 2.7e−16, beta = −0.688, N = 325
Locus: 7 (Sort: 7)
Status: Replicated
80% power: Yes
SNP: 3:49721532:G:A
rsID: rs3197999
UniProtID: G3XAK1
HGNC gene: MST1
Type: Cis−pQTL
SCWPAVFTR pc2
rs3197999 ALT
log10 peptide intensity
3:665 433:491 102:104
3:49721532:G:A_A
p = 1.3e−281, model = REC, N = 538
WPAVFTR pc2
rs3197999 REF
log10 peptide intensity
343:665 92:491 1:104
3:49721532:G:A_A
p = 1.9e−42, model = REC, N = 436

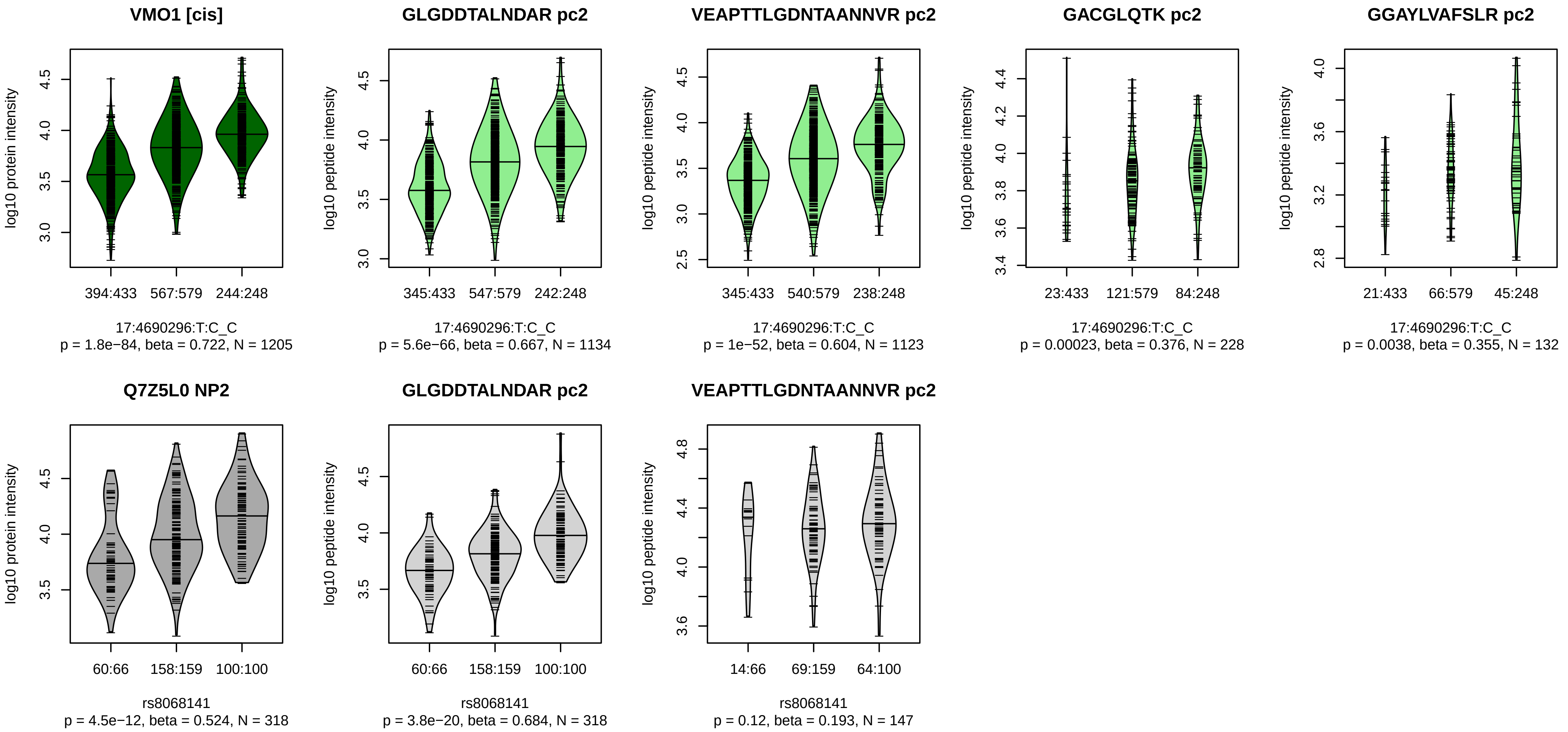


Locus: 8 (Sort: 8)
Status: Replicated
80% power: Yes
SNP: 17:4690296:T:C
rsID: rs8068141
UniProtID: Q7Z5L0
HGNC gene: VMO1
Type: Cis−pQTL

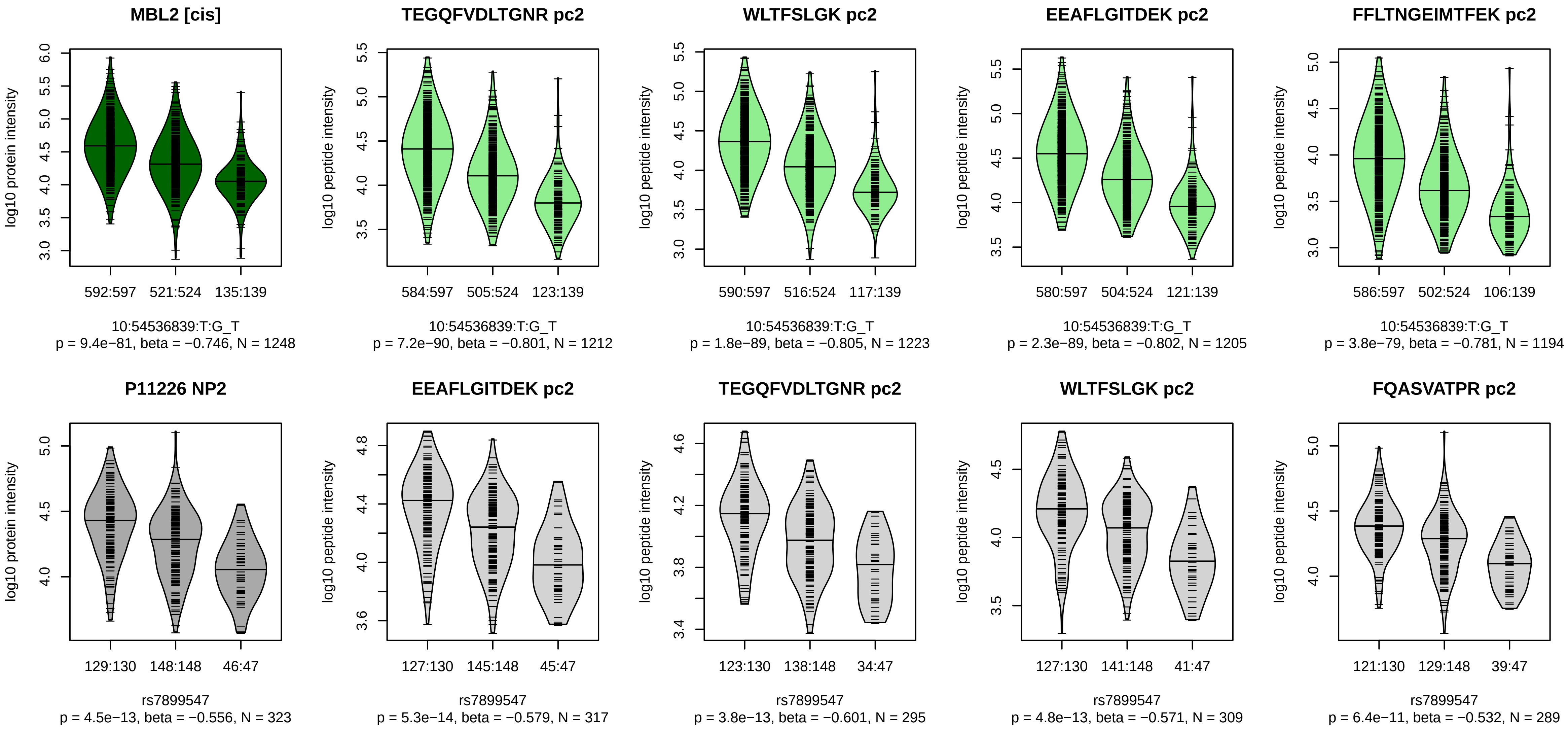


Locus: 9 (Sort: 9)

Status: Replicated

80% power: Yes

SNP: 10:54536839:T:G

rsID: rs7899547

UniProtID: P11226

HGNC gene: MBL2

Type: Cis−pQTL

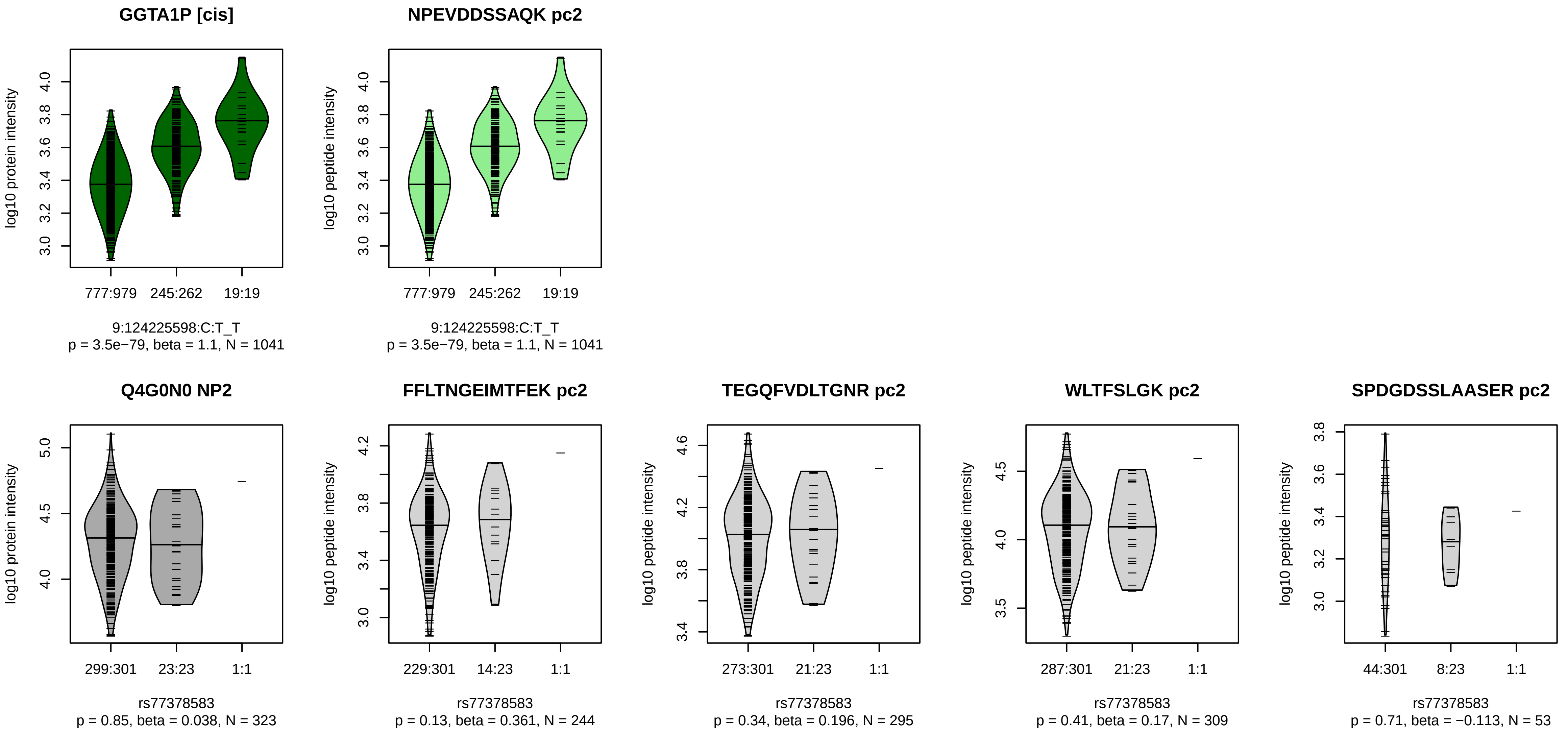


Locus: 10 (Sort: 10)
Status: Not replicated
80% power: Yes
SNP: 9:124225598:C:T
rsID: rs77378583
UniProtID: Q4G0N0
HGNC gene: GGTA1P
Type: Cis−pQTL

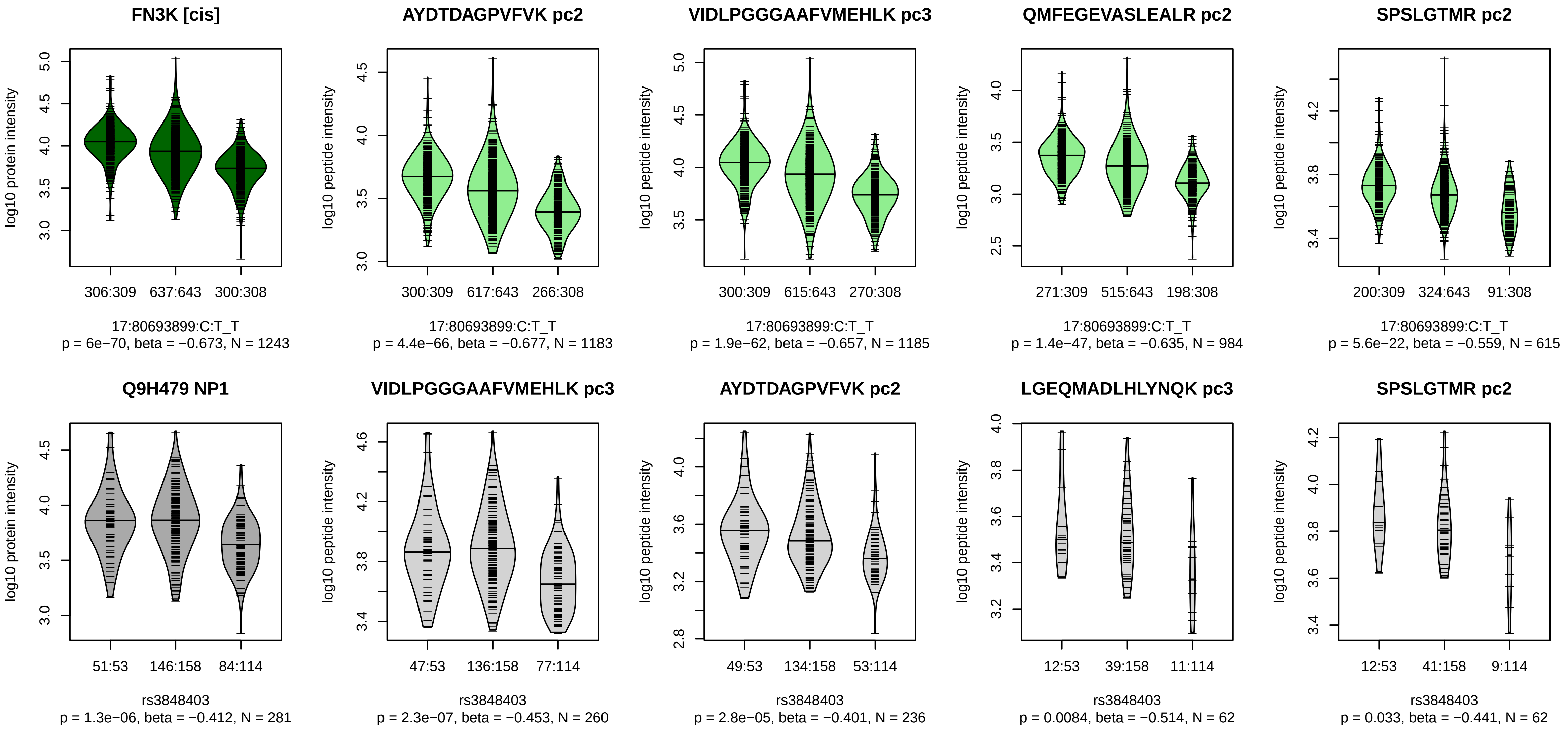


Locus: 11 (Sort: 11)
Status: Replicated
80% power: Yes
SNP: 17:80693899:C:T
rsID: rs3848403
UniProtID: Q9H479
HGNC gene: FN3K
Type: Cis−pQTL

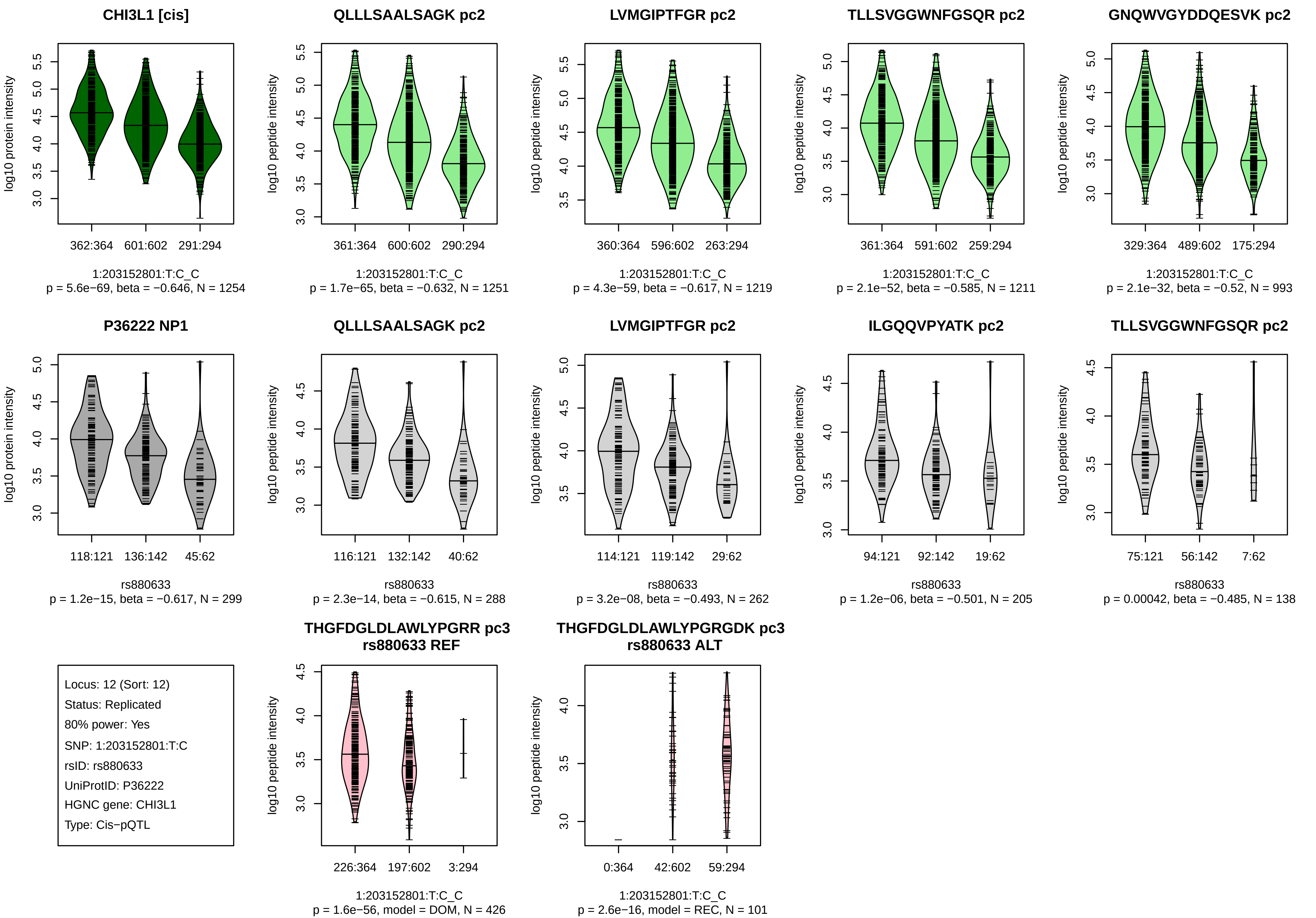
CHI3L1 [cis]
log10 protein intensity
362:364
601:602
291:294
1:203152801:T:C_C
p = 5.6e−69, beta = −0.646, N = 1254
QLLLSAALSAGK pc2
log10 peptide intensity
361:364
600:602
290:294
1:203152801:T:C_C
p = 1.7e−65, beta = −0.632, N = 1251
LVMGIPTFGR pc2
log10 peptide intensity
360:364
596:602
263:294
1:203152801:T:C_C
p = 4.3e−59, beta = −0.617, N = 1219
TLLSVGGWNFGSQR pc2
log10 peptide intensity
361:364
591:602
259:294
1:203152801:T:C_C
p = 2.1e−52, beta = −0.585, N = 1211
GNQWVGYDDQESVK pc2
log10 peptide intensity
329:364
489:602
175:294
1:203152801:T:C_C
p = 2.1e−32, beta = −0.52, N = 993
P36222 NP1
log10 protein intensity
118:121
136:142
45:62
rs880633
p = 1.2e−15, beta = −0.617, N = 299
QLLLSAALSAGK pc2
log10 peptide intensity
116:121
132:142
40:62
rs880633
p = 2.3e−14, beta = −0.615, N = 288
LVMGIPTFGR pc2
log10 peptide intensity
114:121
119:142
29:62
rs880633
p = 3.2e−08, beta = −0.493, N = 262
ILGQQVPYATK pc2
log10 peptide intensity
94:121
92:142
19:62
rs880633
p = 1.2e−06, beta = −0.501, N = 205
TLLSVGGWNFGSQR pc2
log10 peptide intensity
75:121
56:142
7:62
rs880633
p = 0.00042, beta = −0.485, N = 138
Locus: 12 (Sort: 12)
Status: Replicated
80% power: Yes
SNP: 1:203152801:T:C
rsID: rs880633
UniProtID: P36222
HGNC gene: CHI3L1
Type: Cis−pQTL
THGFDGLDLAWLYPGRR pc3
rs880633 REF
log10 peptide intensity
226:364
197:602
3:294
1:203152801:T:C_C
p = 1.6e−56, model = DOM, N = 426
THGFDGLDLAWLYPGRGDK pc3
rs880633 ALT
log10 peptide intensity
0:364
42:602
59:294
1:203152801:T:C_C
p = 2.6e−16, model = REC, N = 101

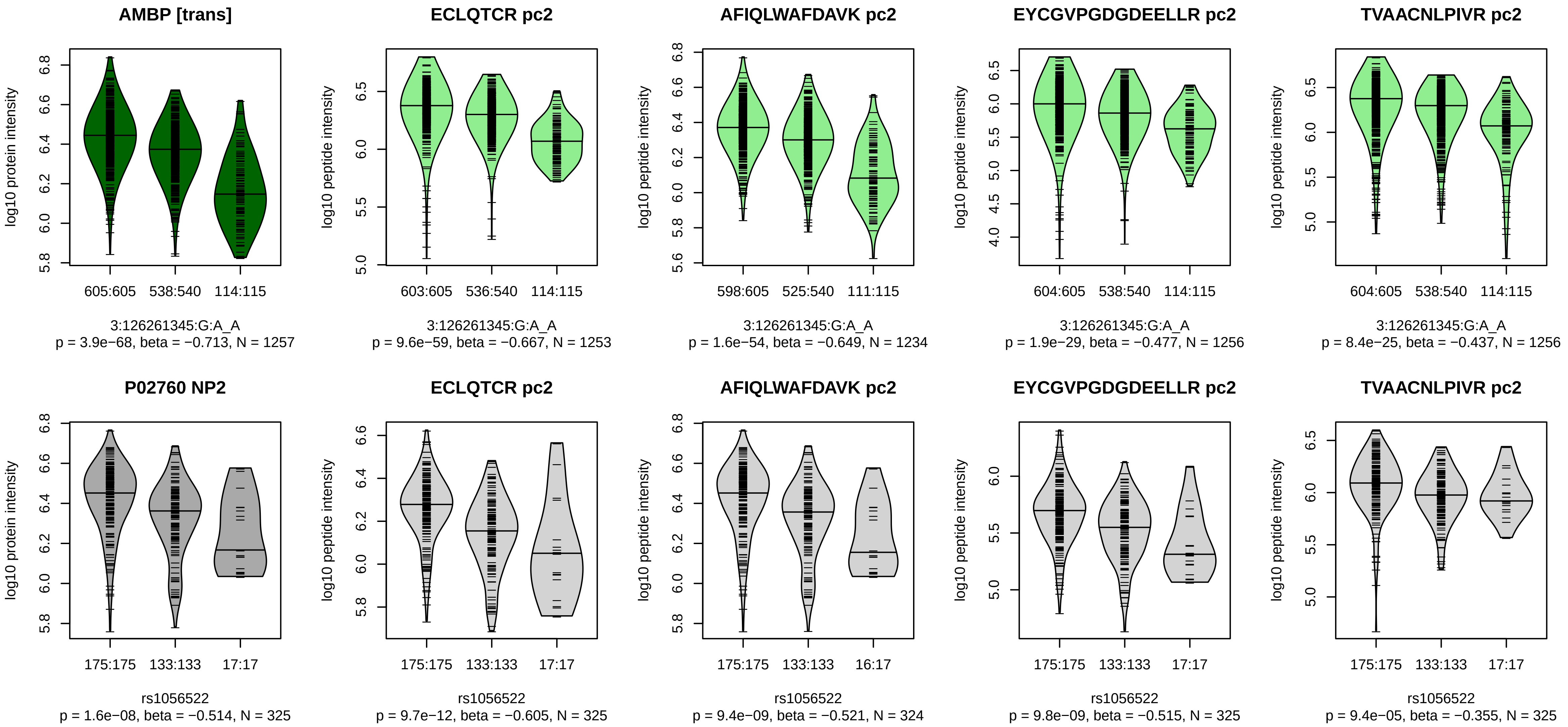


Locus: 13 (Sort: 13)
Status: Replicated
80% power: Yes
SNP: 3:126261345:G:A
rsID: rs1056522
UniProtID: P02760
HGNC gene: AMBP
Type: Trans−pQTL

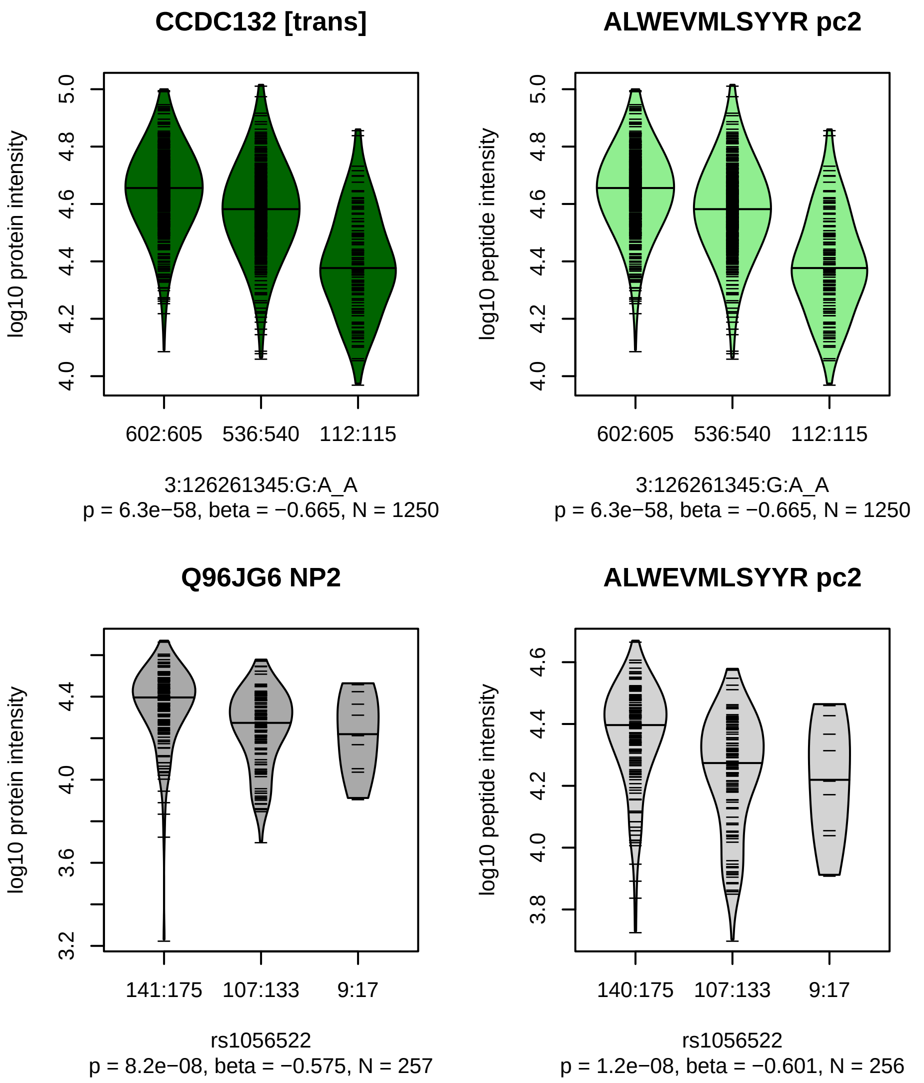


Locus: 13 (Sort: 14)

Status: Replicated

80% power: Yes

SNP: 3:126261345:G:A

rsID: rs1056522

UniProtID: Q96JG6

HGNC gene: CCDC132

Type: Trans−pQTL

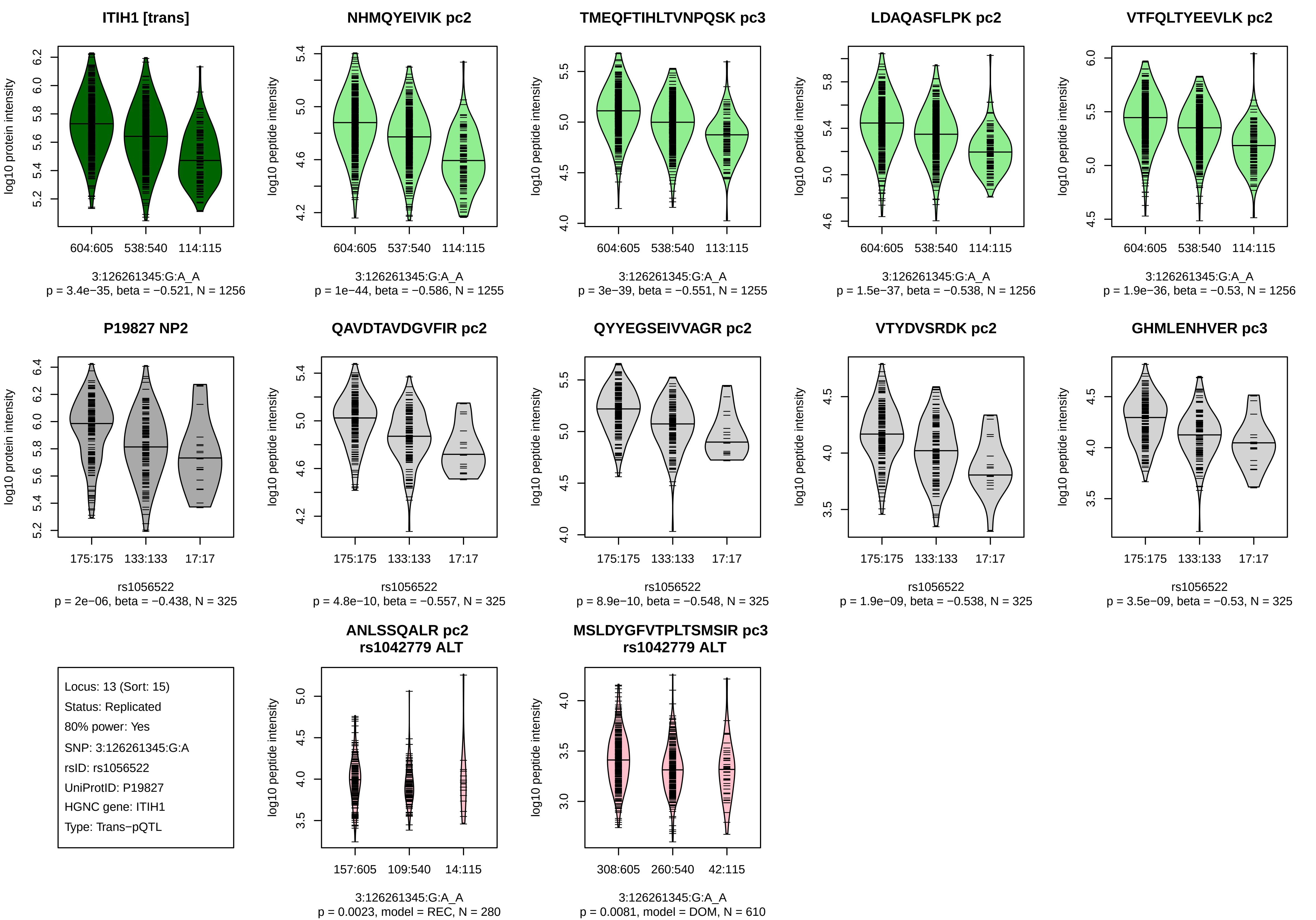
ITIH1 [trans]
log10 protein intensity
604:605 538:540 114:115
3:126261345:G:A_A
p = 3.4e−35, beta = −0.521, N = 1256
NHMQYEIVIK pc2
log10 peptide intensity
604:605 537:540 114:115
3:126261345:G:A_A
p = 1e−44, beta = −0.586, N = 1255
TMEQFTIHLTVNPQSK pc3
log10 peptide intensity
604:605 538:540 113:115
3:126261345:G:A_A
p = 3e−39, beta = −0.551, N = 1255
LDAQASFLPK pc2
log10 peptide intensity
604:605 538:540 114:115
3:126261345:G:A_A
p = 1.5e−37, beta = −0.538, N = 1256
VTFQLTYEEVLK pc2
log10 peptide intensity
604:605 538:540 114:115
3:126261345:G:A_A
p = 1.9e−36, beta = −0.53, N = 1256
P19827 NP2
log10 protein intensity
175:175 133:133 17:17
rs1056522
p = 2e−06, beta = −0.438, N = 325
QAVDTAVDGVFIR pc2
log10 peptide intensity
175:175 133:133 17:17
rs1056522
p = 4.8e−10, beta = −0.557, N = 325
QYYEGSEIVVAGR pc2
log10 peptide intensity
175:175 133:133 17:17
rs1056522
p = 8.9e−10, beta = −0.548, N = 325
VTYDVSRDK pc2
log10 peptide intensity
175:175 133:133 17:17
rs1056522
p = 1.9e−09, beta = −0.538, N = 325
GHMLENHVER pc3
log10 peptide intensity
175:175 133:133 17:17
rs1056522
p = 3.5e−09, beta = −0.53, N = 325
Locus: 13 (Sort: 15)
Status: Replicated
80% power: Yes
SNP: 3:126261345:G:A
rsID: rs1056522
UniProtID: P19827
HGNC gene: ITIH1
Type: Trans−pQTL
ANLSSQALR pc2
rs1042779 ALT
log10 peptide intensity
157:605 109:540 14:115
3:126261345:G:A_A
p = 0.0023, model = REC, N = 280
MSLDYGFVTPLTSMSIR pc3
rs1042779 ALT
log10 peptide intensity
308:605 260:540 42:115
3:126261345:G:A_A
p = 0.0081, model = DOM, N = 610

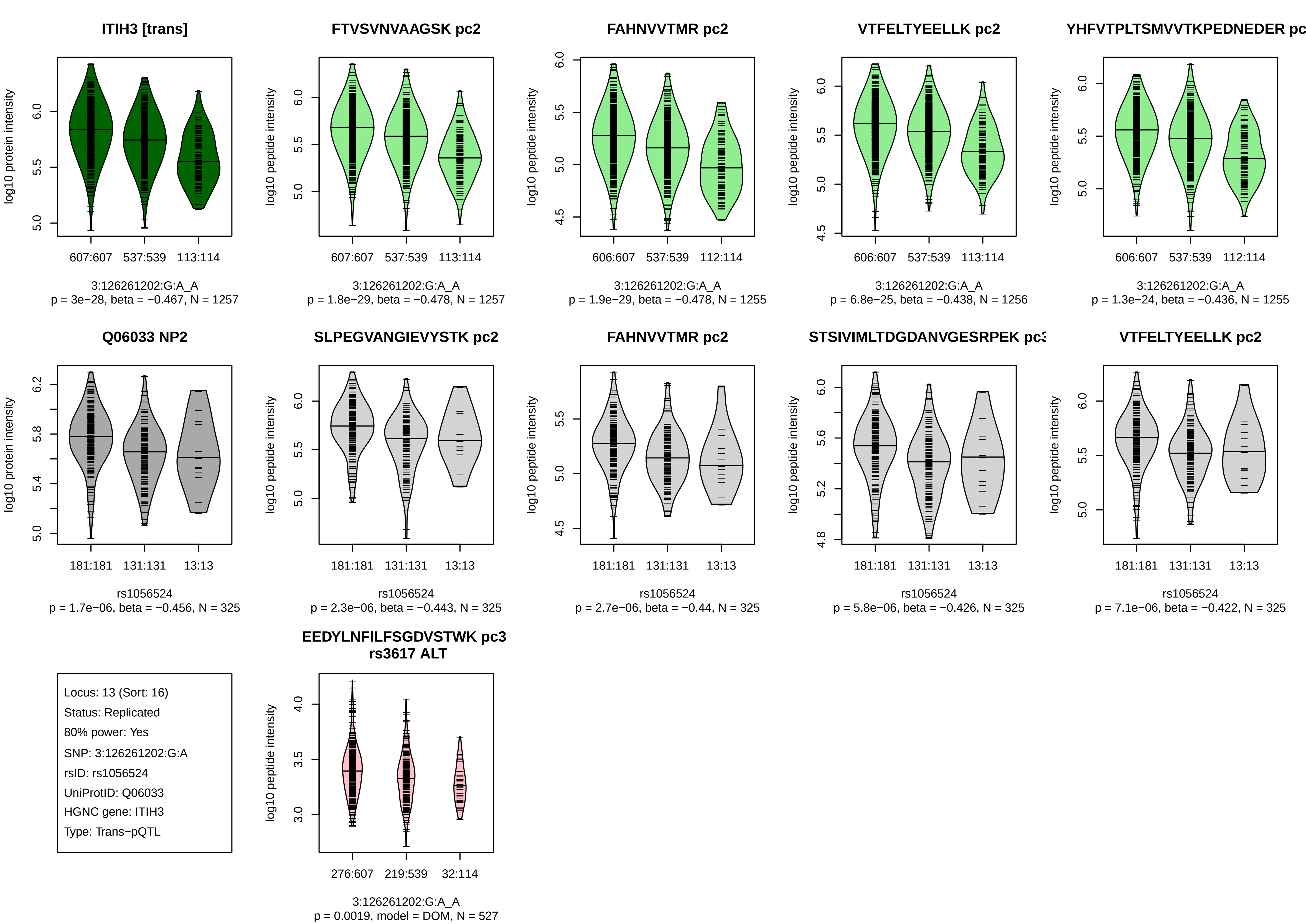
ITIH3 [trans]
log10 protein intensity
607:607 537:539 113:114
3:126261202:G:A_A
p = 3e−28, beta = −0.467, N = 1257
FTVSVNVAAGSK pc2
log10 peptide intensity
607:607 537:539 113:114
3:126261202:G:A_A
p = 1.8e−29, beta = −0.478, N = 1257
FAHNVVTMR pc2
log10 peptide intensity
606:607 537:539 112:114
3:126261202:G:A_A
p = 1.9e−29, beta = −0.478, N = 1255
VTFELTYEELLK pc2
log10 peptide intensity
606:607 537:539 113:114
3:126261202:G:A_A
p = 6.8e−25, beta = −0.438, N = 1256
YHFVTPLTSMVVTKPEDNEDER pc
log10 peptide intensity
606:607 537:539 112:114
3:126261202:G:A_A
p = 1.3e−24, beta = −0.436, N = 1255
Q06033 NP2
log10 protein intensity
181:181 131:131 13:13
rs1056524
p = 1.7e−06, beta = −0.456, N = 325
SLPEGVANGIEVYSTK pc2
log10 peptide intensity
181:181 131:131 13:13
rs1056524
p = 2.3e−06, beta = −0.443, N = 325
FAHNVVTMR pc2
log10 peptide intensity
181:181 131:131 13:13
rs1056524
p = 2.7e−06, beta = −0.44, N = 325
STSIVIMLTDGDANVGESRPEK pc3
log10 peptide intensity
181:181 131:131 13:13
rs1056524
p = 5.8e−06, beta = −0.426, N = 325
VTFELTYEELLK pc2
log10 peptide intensity
181:181 131:131 13:13
rs1056524
p = 7.1e−06, beta = −0.422, N = 325
Locus: 13 (Sort: 16)
Status: Replicated
80% power: Yes
SNP: 3:126261202:G:A
rsID: rs1056524
UniProtID: Q06033
HGNC gene: ITIH3
Type: Trans−pQTL
EEDYLNFILFSGDVSTWK pc3
rs3617 ALT
log10 peptide intensity
276:607 219:539 32:114
3:126261202:G:A_A
p = 0.0019, model = DOM, N = 527

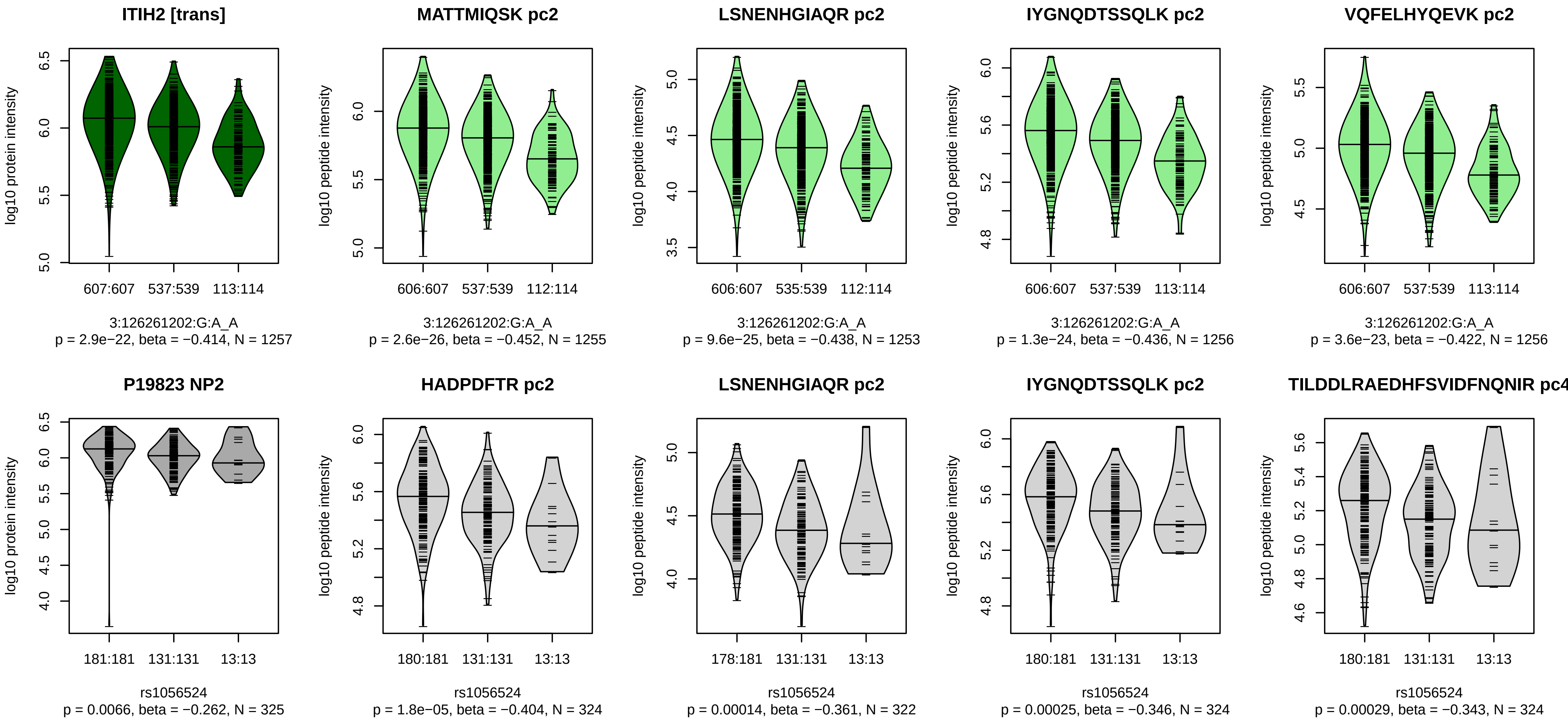


Locus: 13 (Sort: 17)

Status: Not replicated

80% power: Yes

SNP: 3:126261202:G:A

rsID: rs1056524

UniProtID: P19823

HGNC gene: ITIH2

Type: Trans−pQTL

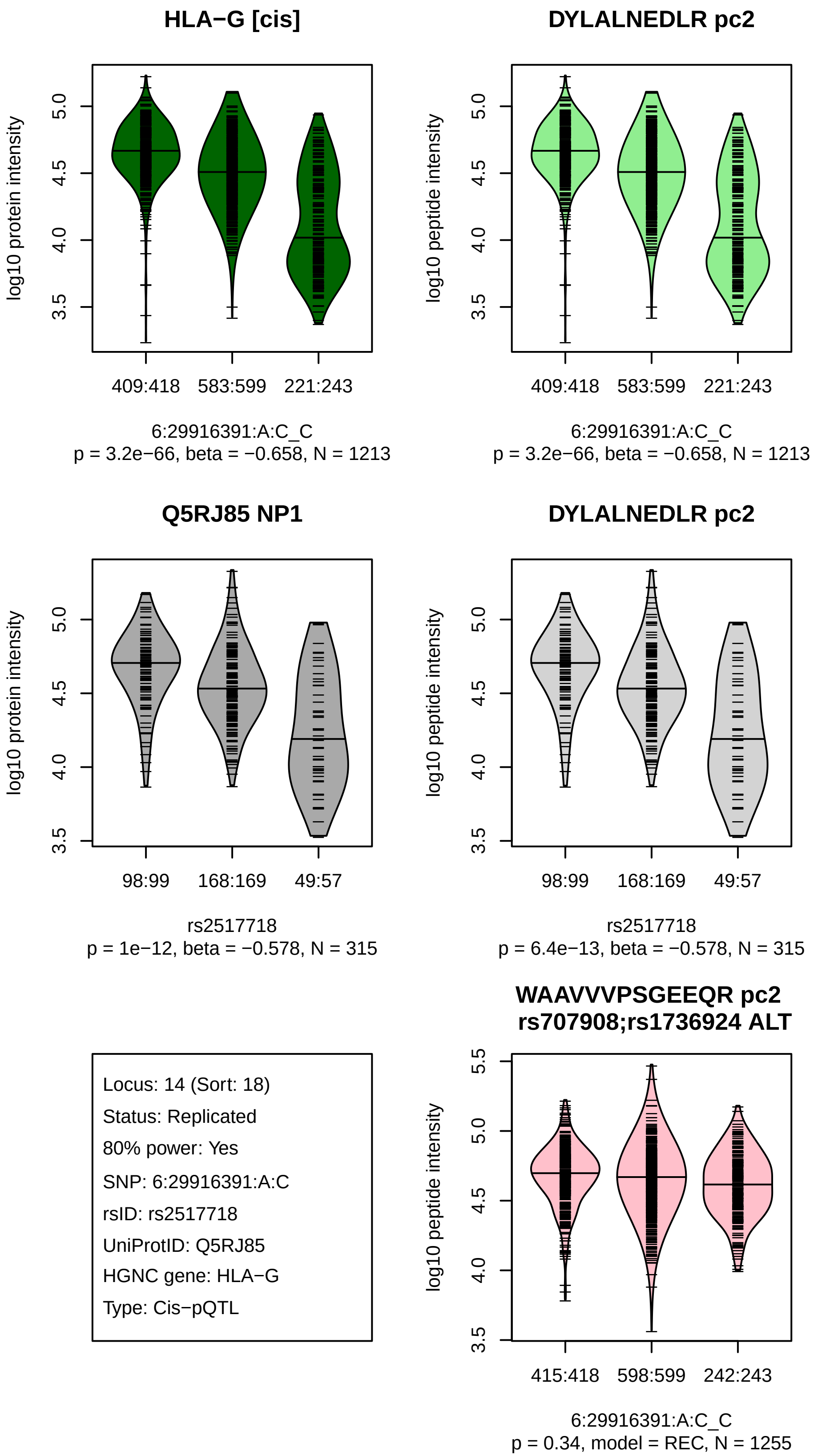
HLA−G [cis]
log10 protein intensity
3.5
4.0
4.5
5.0
409:418
583:599
221:243
6:29916391:A:C_C
p = 3.2e−66, beta = −0.658, N = 1213
DYLALNEDLR pc2
log10 peptide intensity
409:418
583:599
221:243
6:29916391:A:C_C
p = 3.2e−66, beta = −0.658, N = 1213
Q5RJ85 NP1
log10 protein intensity
98:99
168:169
49:57
rs2517718
p = 1e−12, beta = −0.578, N = 315
DYLALNEDLR pc2
log10 peptide intensity
98:99
168:169
49:57
rs2517718
p = 6.4e−13, beta = −0.578, N = 315
Locus: 14 (Sort: 18)
Status: Replicated
80% power: Yes
SNP: 6:29916391:A:C
rsID: rs2517718
UniProtID: Q5RJ85
HGNC gene: HLA−G
Type: Cis−pQTL
WAAVVVPSGEEQR pc2
rs707908;rs1736924 ALT
log10 peptide intensity
3.5
4.0
4.5
5.0
5.5
415:418
598:599
242:243
6:29916391:A:C_C
p = 0.34, model = REC, N = 1255

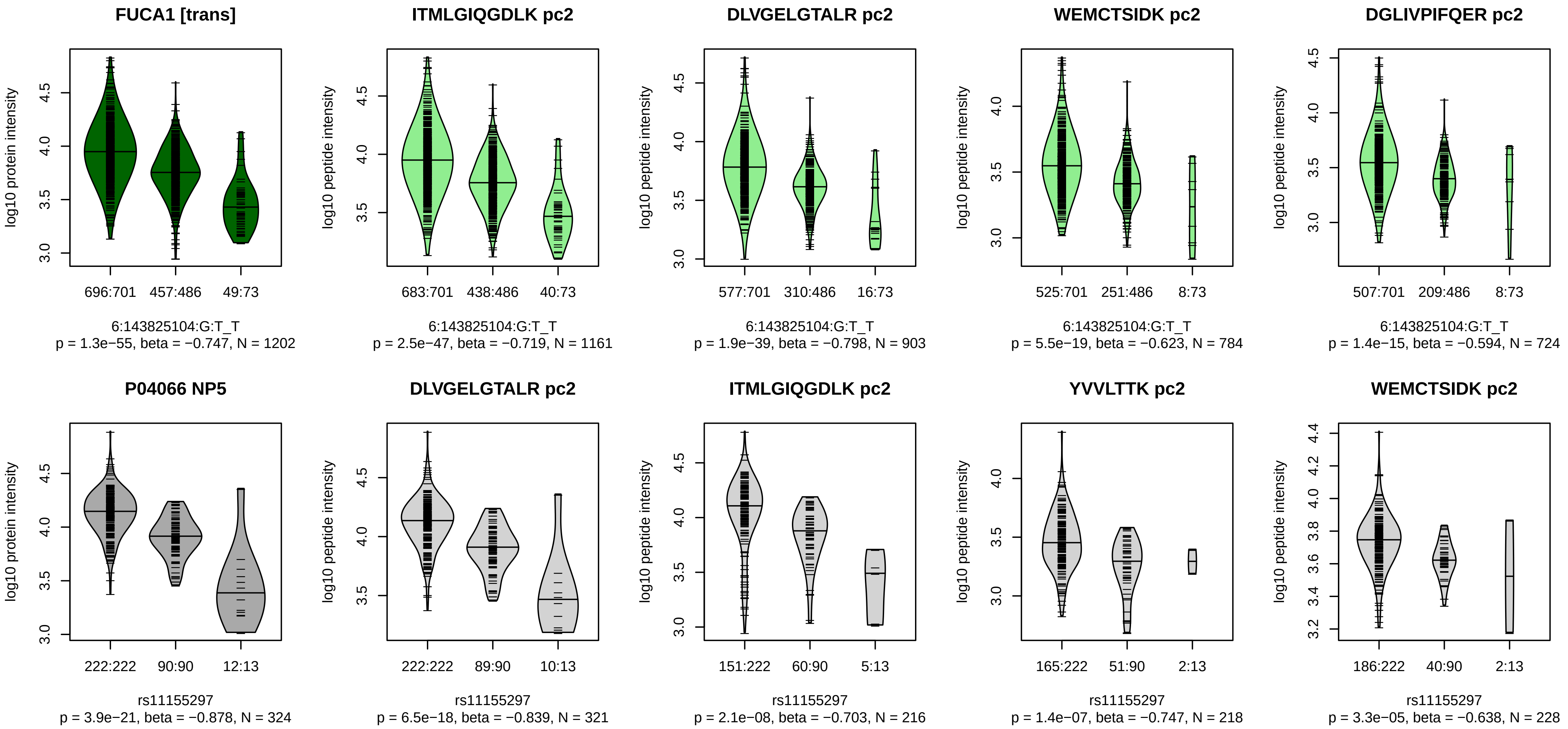


Locus: 15 (Sort: 19)

Status: Replicated

80% power: Yes

SNP: 6:143825104:G:T

rsID: rs11155297

UniProtID: P04066

HGNC gene: FUCA1

Type: Trans−pQTL

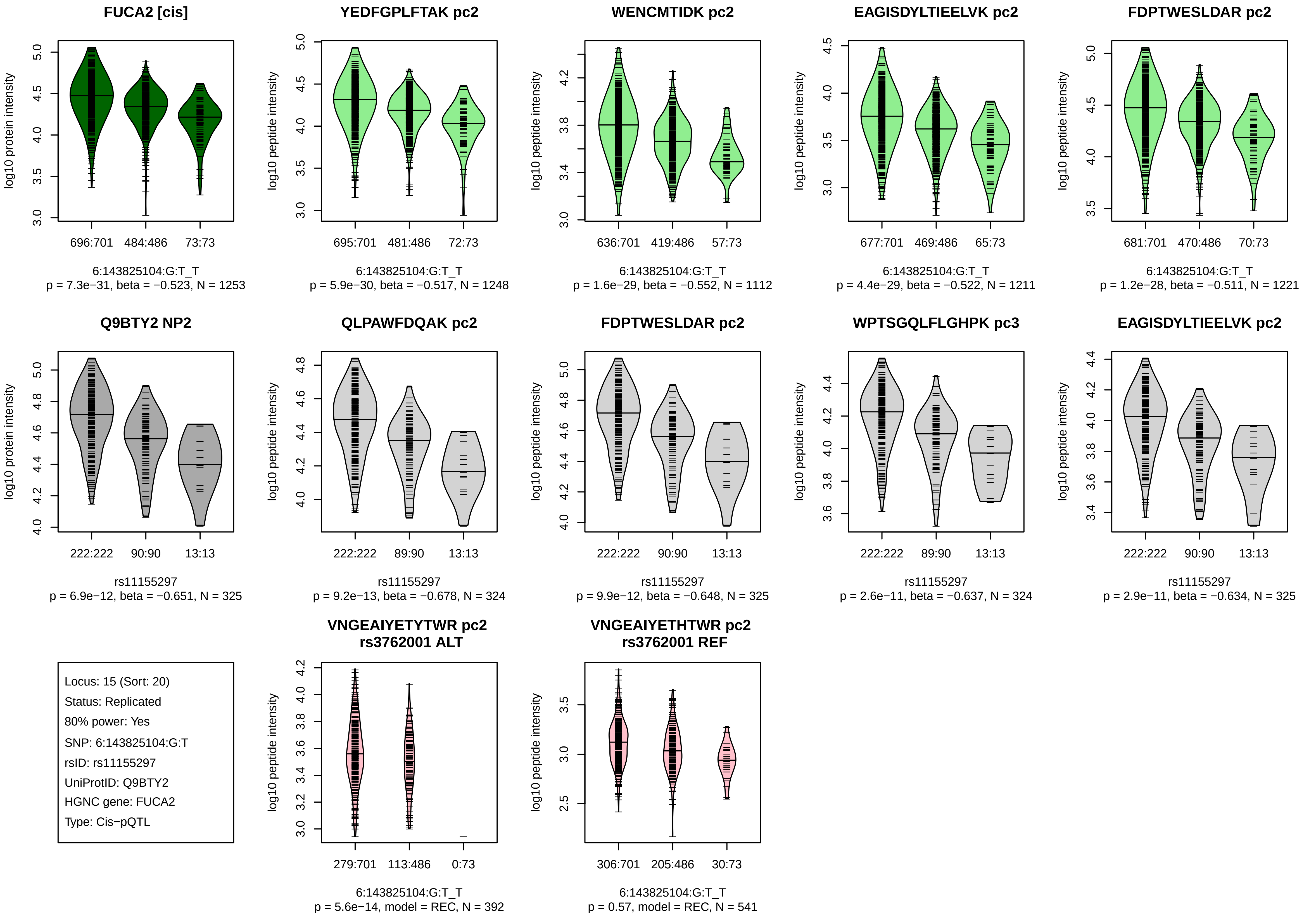
FUCA2 [cis]
YEDFGPLFTAK pc2
WENCMTIDK pc2
EAGISDYLTIEELVK pc2
FDPTWESLDAR pc2
log10 protein intensity
log10 peptide intensity
696:701
484:486
73:73
695:701
481:486
72:73
636:701
419:486
57:73
677:701
469:486
65:73
681:701
470:486
70:73
6:143825104:G:T_T
p = 7.3e−31, beta = −0.523, N = 1253
p = 5.9e−30, beta = −0.517, N = 1248
p = 1.6e−29, beta = −0.552, N = 1112
p = 4.4e−29, beta = −0.522, N = 1211
p = 1.2e−28, beta = −0.511, N = 1221
Q9BTY2 NP2
QLPAWFDQAK pc2
FDPTWESLDAR pc2
WPTSGQLFLGHPK pc3
EAGISDYLTIEELVK pc2
222:222
90:90
89:90
13:13
rs11155297
p = 6.9e−12, beta = −0.651, N = 325
p = 9.2e−13, beta = −0.678, N = 324
p = 9.9e−12, beta = −0.648, N = 325
p = 2.6e−11, beta = −0.637, N = 324
p = 2.9e−11, beta = −0.634, N = 325
Locus: 15 (Sort: 20)
Status: Replicated
80% power: Yes
SNP: 6:143825104:G:T
rsID: rs11155297
UniProtID: Q9BTY2
HGNC gene: FUCA2
Type: Cis−pQTL
VNGEAIYETYTWR pc2
rs3762001 ALT
VNGEAIYETHTWR pc2
rs3762001 REF
279:701
113:486
0:73
306:701
205:486
30:73
p = 5.6e−14, model = REC, N = 392
p = 0.57, model = REC, N = 541

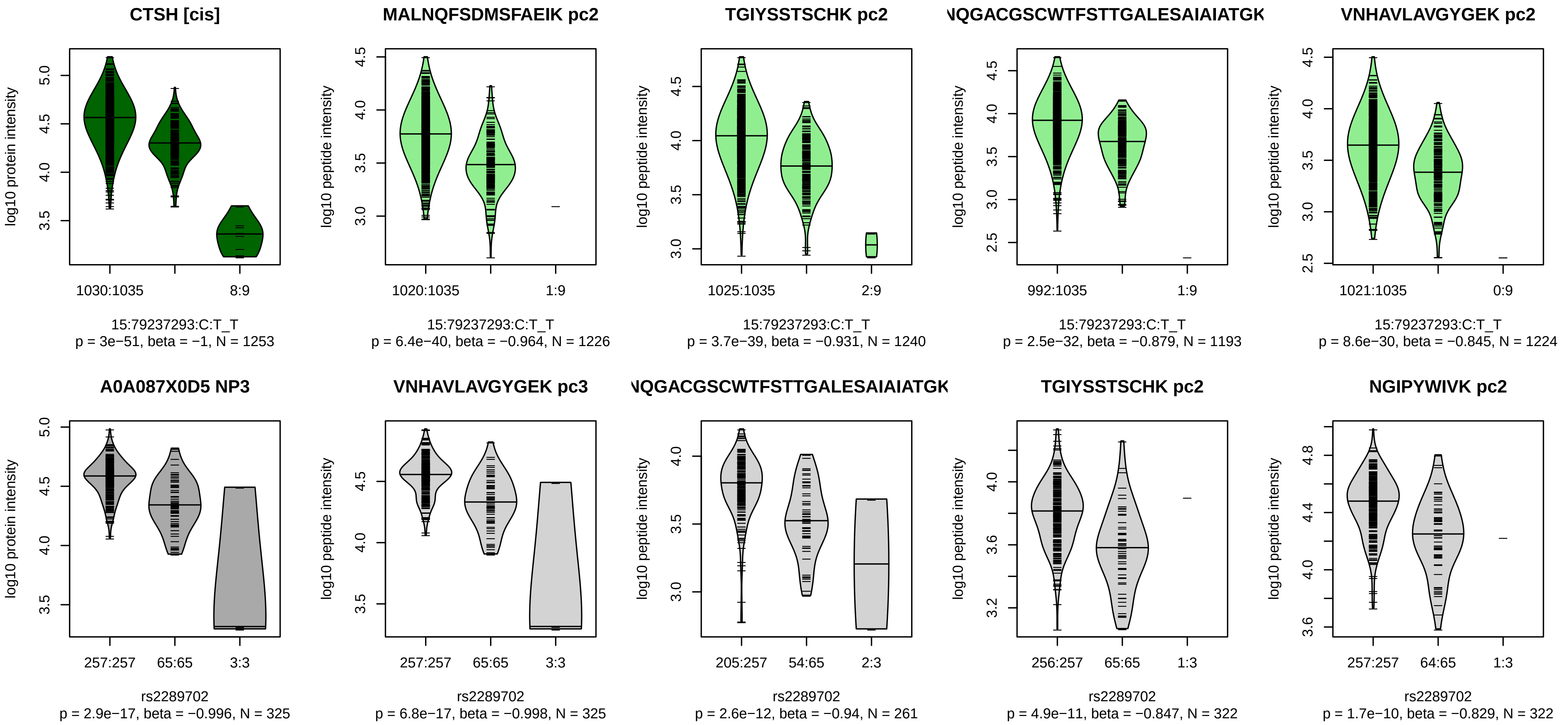


Locus: 16 (Sort: 21)
Status: Replicated
80% power: Yes
SNP: 15:79237293:C:T
rsID: rs2289702
UniProtID: A0A087X0D5
HGNC gene: CTSH
Type: Cis−pQTL

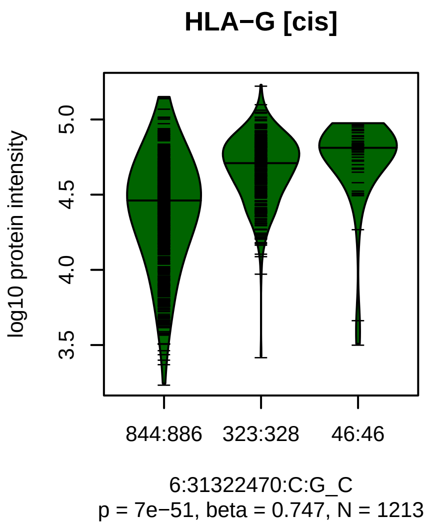


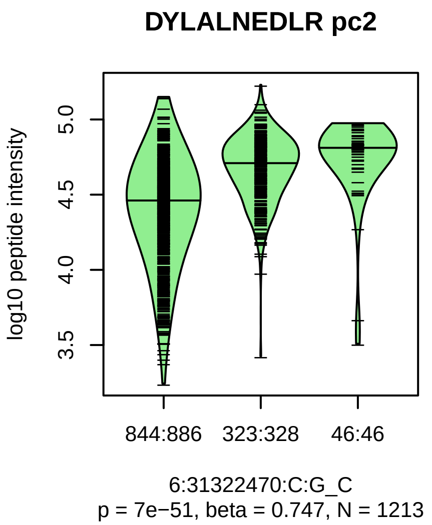


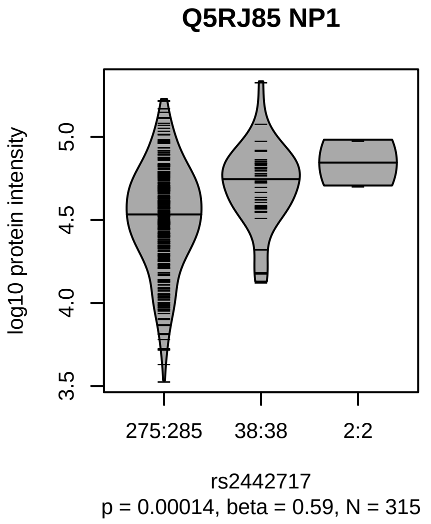


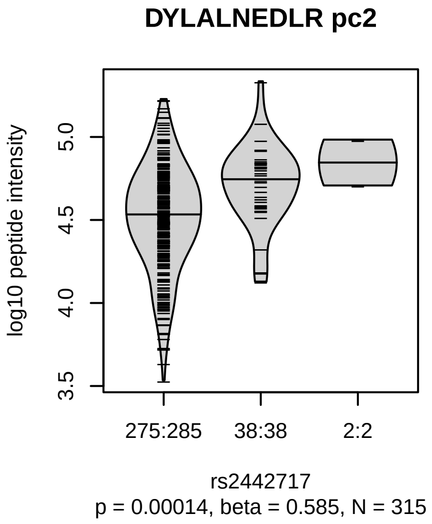


Locus: 17 (Sort: 22)

Status: Replicated

80% power: Yes

SNP: 6:31322470:C:G

rsID: rs2442717

UniProtID: Q5RJ85

HGNC gene: HLA−G

Type: Cis−pQTL

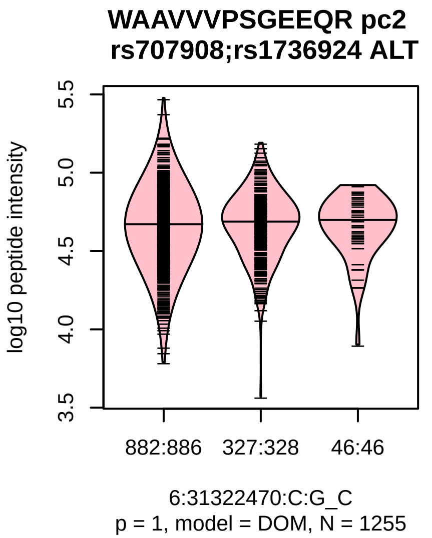

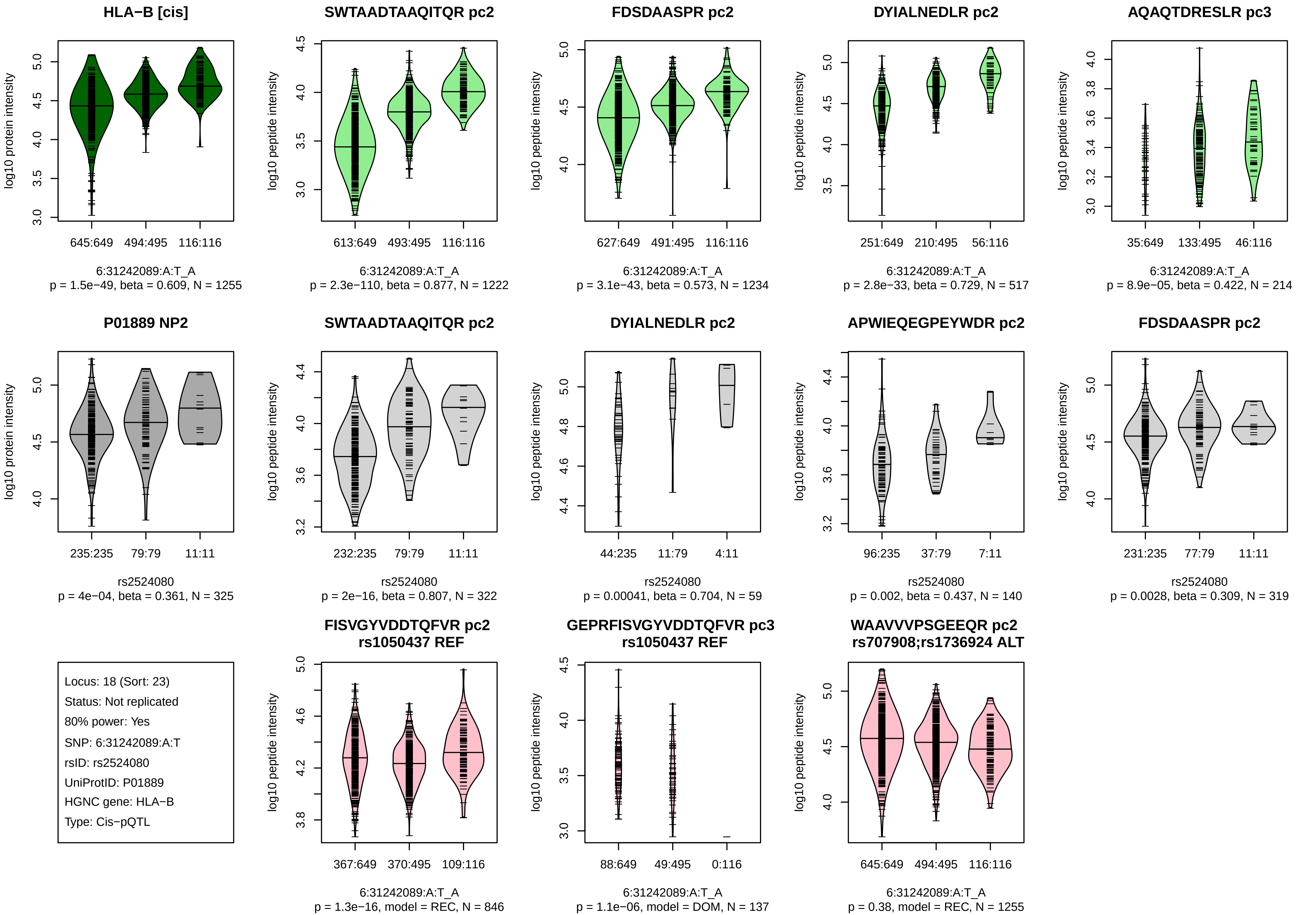
HLA−B [cis]
log10 protein intensity
645:649 494:495 116:116
6:31242089:A:T_A
p = 1.5e−49, beta = 0.609, N = 1255
SWTAADTAAQITQR pc2
log10 peptide intensity
613:649 493:495 116:116
6:31242089:A:T_A
p = 2.3e−110, beta = 0.877, N = 1222
FDSDAASPR pc2
log10 peptide intensity
627:649 491:495 116:116
6:31242089:A:T_A
p = 3.1e−43, beta = 0.573, N = 1234
DYIALNEDLR pc2
log10 peptide intensity
251:649 210:495 56:116
6:31242089:A:T_A
p = 2.8e−33, beta = 0.729, N = 517
AQAQTDRESLR pc3
log10 peptide intensity
35:649 133:495 46:116
6:31242089:A:T_A
p = 8.9e−05, beta = 0.422, N = 214
P01889 NP2
log10 protein intensity
235:235 79:79 11:11
rs2524080
p = 4e−04, beta = 0.361, N = 325
SWTAADTAAQITQR pc2
log10 peptide intensity
232:235 79:79 11:11
rs2524080
p = 2e−16, beta = 0.807, N = 322
DYIALNEDLR pc2
log10 peptide intensity
44:235 11:79 4:11
rs2524080
p = 0.00041, beta = 0.704, N = 59
APWIEQEGPEYWDR pc2
log10 peptide intensity
96:235 37:79 7:11
rs2524080
p = 0.002, beta = 0.437, N = 140
FDSDAASPR pc2
log10 peptide intensity
231:235 77:79 11:11
rs2524080
p = 0.0028, beta = 0.309, N = 319
Locus: 18 (Sort: 23)
Status: Not replicated
80% power: Yes
SNP: 6:31242089:A:T
rsID: rs2524080
UniProtID: P01889
HGNC gene: HLA−B
Type: Cis−pQTL
FISVGYVDDTQFVR pc2
rs1050437 REF
log10 peptide intensity
367:649 370:495 109:116
6:31242089:A:T_A
p = 1.3e−16, model = REC, N = 846
GEPRFISVGYVDDTQFVR pc3
rs1050437 REF
log10 peptide intensity
88:649 49:495 0:116
6:31242089:A:T_A
p = 1.1e−06, model = DOM, N = 137
WAAVVVPSGEEQR pc2
rs707908;rs1736924 ALT
log10 peptide intensity
645:649 494:495 116:116
6:31242089:A:T_A
p = 0.38, model = REC, N = 1255

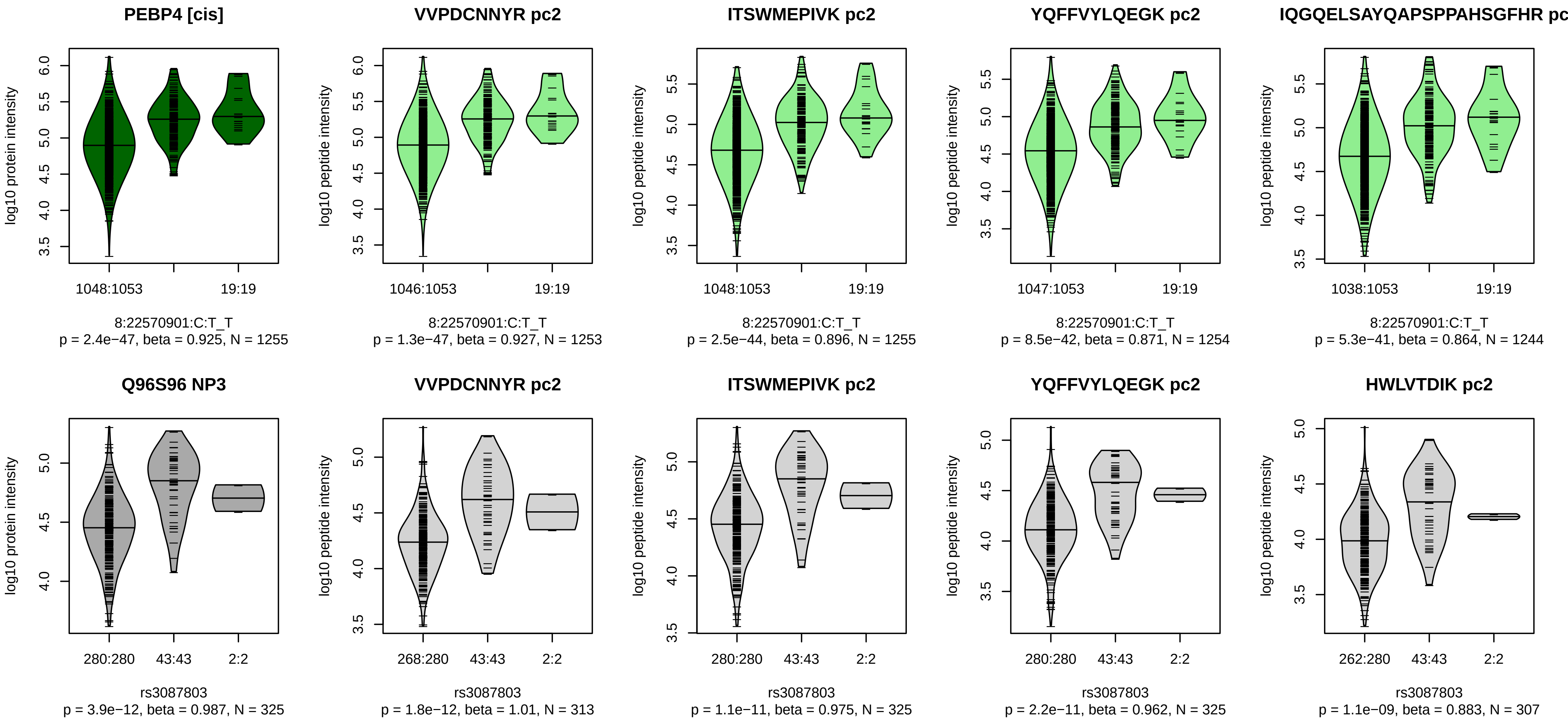


Locus: 19 (Sort: 24)

Status: Replicated

80% power: Yes

SNP: 8:22570901:C:T

rsID: rs3087803

UniProtID: Q96S96

HGNC gene: PEBP4

Type: Cis−pQTL

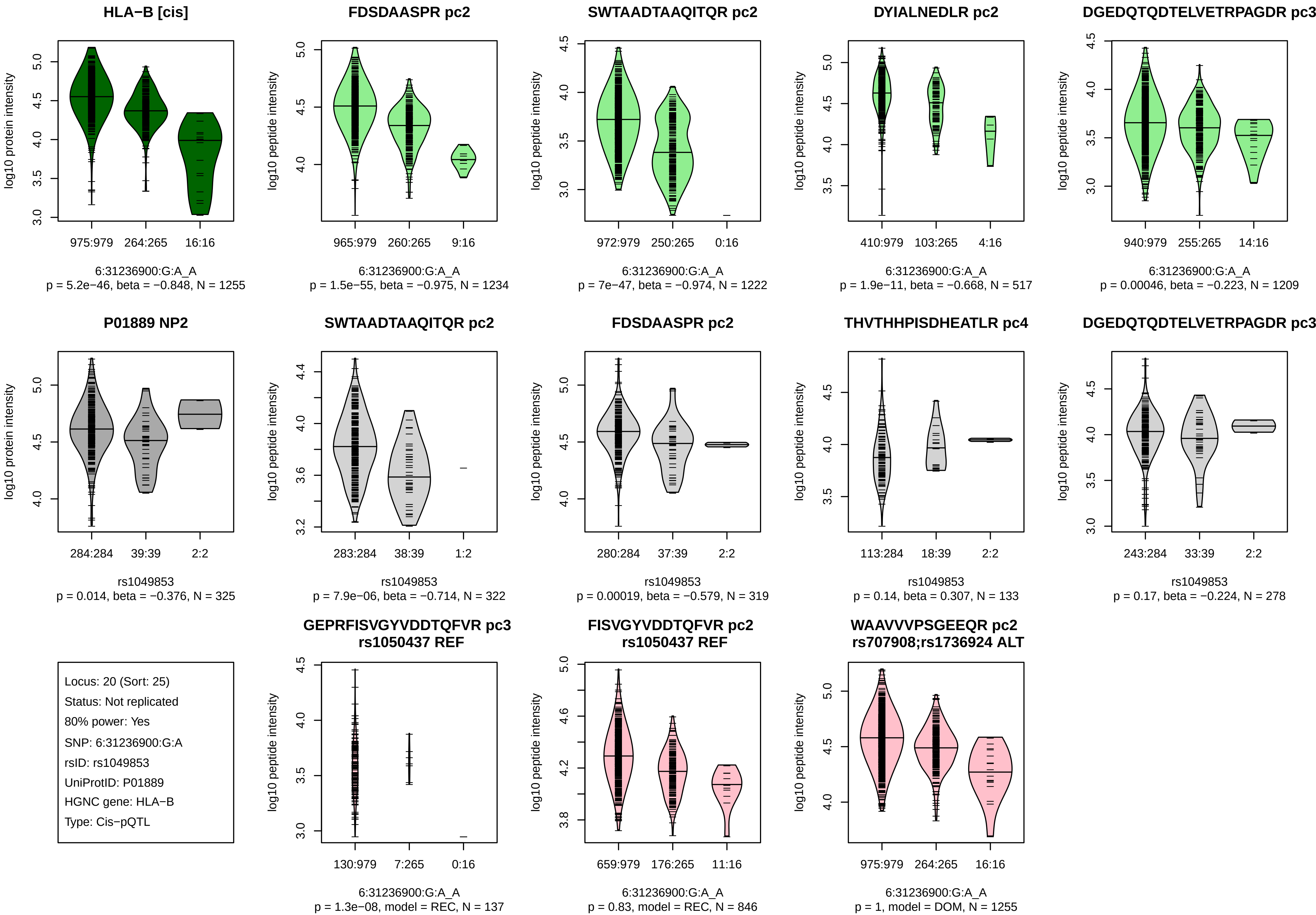
HLA−B [cis]
log10 protein intensity
975:979
264:265
16:16
6:31236900:G:A_A
p = 5.2e−46, beta = −0.848, N = 1255
FDSDAASPR pc2
log10 peptide intensity
965:979
260:265
9:16
6:31236900:G:A_A
p = 1.5e−55, beta = −0.975, N = 1234
SWTAADTAAQITQR pc2
972:979
250:265
0:16
6:31236900:G:A_A
p = 7e−47, beta = −0.974, N = 1222
DYIALNEDLR pc2
410:979
103:265
4:16
6:31236900:G:A_A
p = 1.9e−11, beta = −0.668, N = 517
DGEDQTQDTELVETRPAGDR pc3
940:979
255:265
14:16
6:31236900:G:A_A
p = 0.00046, beta = −0.223, N = 1209
P01889 NP2
284:284
39:39
2:2
rs1049853
p = 0.014, beta = −0.376, N = 325
SWTAADTAAQITQR pc2
283:284
38:39
1:2
rs1049853
p = 7.9e−06, beta = −0.714, N = 322
FDSDAASPR pc2
280:284
37:39
2:2
rs1049853
p = 0.00019, beta = −0.579, N = 319
THVTHHPISDHEATLR pc4
113:284
18:39
2:2
rs1049853
p = 0.14, beta = 0.307, N = 133
DGEDQTQDTELVETRPAGDR pc3
243:284
33:39
2:2
rs1049853
p = 0.17, beta = −0.224, N = 278
Locus: 20 (Sort: 25)
Status: Not replicated
80% power: Yes
SNP: 6:31236900:G:A
rsID: rs1049853
UniProtID: P01889
HGNC gene: HLA−B
Type: Cis−pQTL
GEPRFISVGYVDDTQFVR pc3
rs1050437 REF
130:979
7:265
0:16
6:31236900:G:A_A
p = 1.3e−08, model = REC, N = 137
FISVGYVDDTQFVR pc2
rs1050437 REF
659:979
176:265
11:16
6:31236900:G:A_A
p = 0.83, model = REC, N = 846
WAAVVVPSGEEQR pc2
rs707908;rs1736924 ALT
975:979
264:265
16:16
6:31236900:G:A_A
p = 1, model = DOM, N = 1255

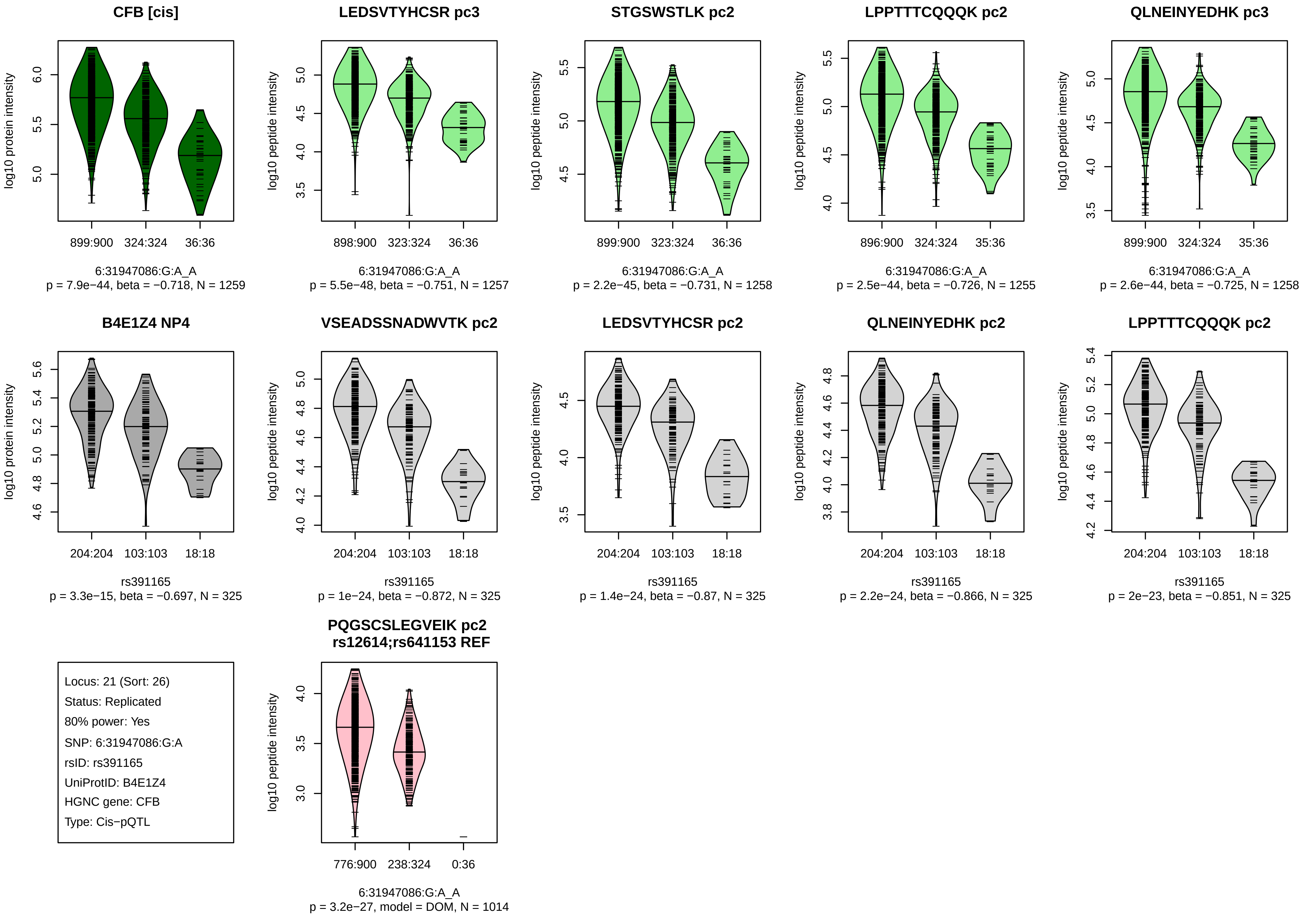
CFB [cis]
log10 protein intensity
899:900
324:324
36:36
6:31947086:G:A_A
p = 7.9e−44, beta = −0.718, N = 1259
LEDSVTYHCSR pc3
log10 peptide intensity
898:900
323:324
36:36
6:31947086:G:A_A
p = 5.5e−48, beta = −0.751, N = 1257
STGSWSTLK pc2
log10 peptide intensity
899:900
323:324
36:36
6:31947086:G:A_A
p = 2.2e−45, beta = −0.731, N = 1258
LPPTTTCQQQK pc2
log10 peptide intensity
896:900
324:324
35:36
6:31947086:G:A_A
p = 2.5e−44, beta = −0.726, N = 1255
QLNEINYEDHK pc3
log10 peptide intensity
899:900
324:324
35:36
6:31947086:G:A_A
p = 2.6e−44, beta = −0.725, N = 1258
B4E1Z4 NP4
log10 protein intensity
204:204
103:103
18:18
rs391165
p = 3.3e−15, beta = −0.697, N = 325
VSEADSSNADWVTK pc2
log10 peptide intensity
204:204
103:103
18:18
rs391165
p = 1e−24, beta = −0.872, N = 325
LEDSVTYHCSR pc2
log10 peptide intensity
204:204
103:103
18:18
rs391165
p = 1.4e−24, beta = −0.87, N = 325
QLNEINYEDHK pc2
log10 peptide intensity
204:204
103:103
18:18
rs391165
p = 2.2e−24, beta = −0.866, N = 325
LPPTTTCQQQK pc2
log10 peptide intensity
204:204
103:103
18:18
rs391165
p = 2e−23, beta = −0.851, N = 325
Locus: 21 (Sort: 26)
Status: Replicated
80% power: Yes
SNP: 6:31947086:G:A
rsID: rs391165
UniProtID: B4E1Z4
HGNC gene: CFB
Type: Cis−pQTL
PQGSCSLEGVEIK pc2
rs12614;rs641153 REF
log10 peptide intensity
776:900
238:324
0:36
6:31947086:G:A_A
p = 3.2e−27, model = DOM, N = 1014

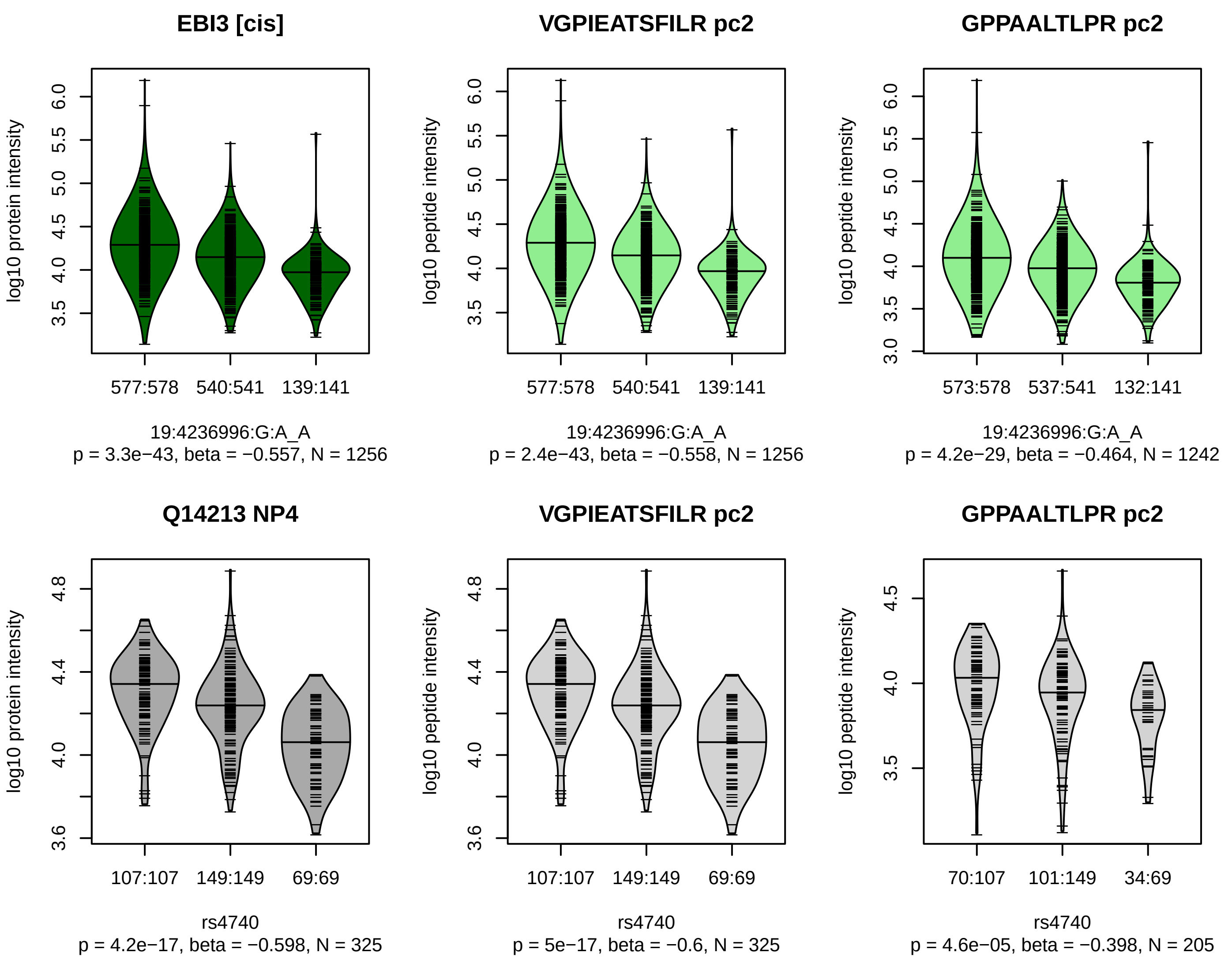


Locus: 22 (Sort: 27)

Status: Replicated

80% power: Yes

SNP: 19:4236996:G:A

rsID: rs4740

UniProtID: Q14213

HGNC gene: EBI3

Type: Cis−pQTL

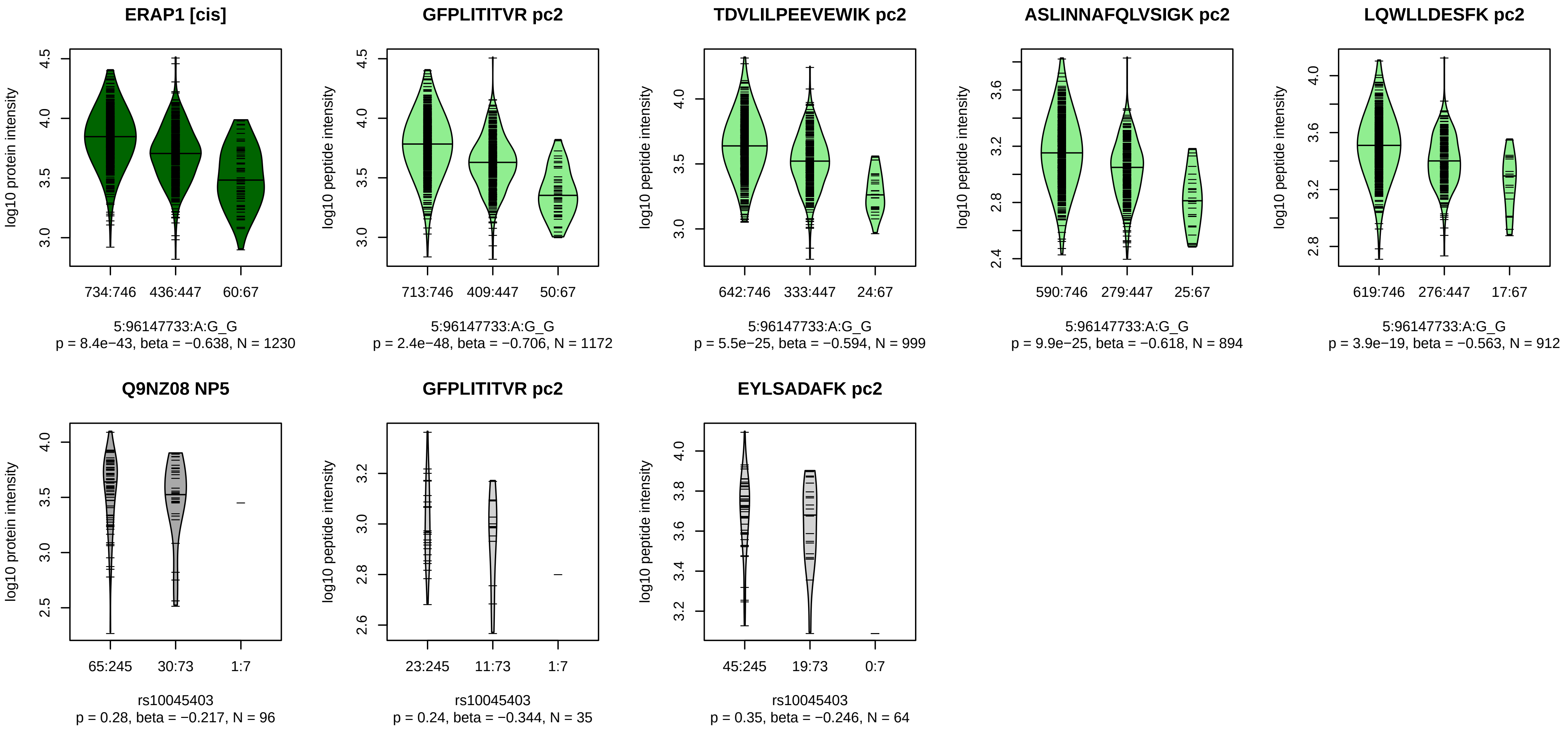


Locus: 23 (Sort: 28)
Status: Not replicated
80% power: No
SNP: 5:96147733:A:G
rsID: rs10045403
UniProtID: Q9NZ08
HGNC gene: ERAP1
Type: Cis−pQTL

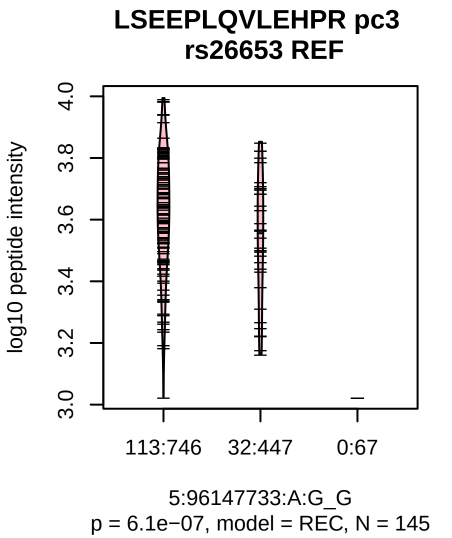

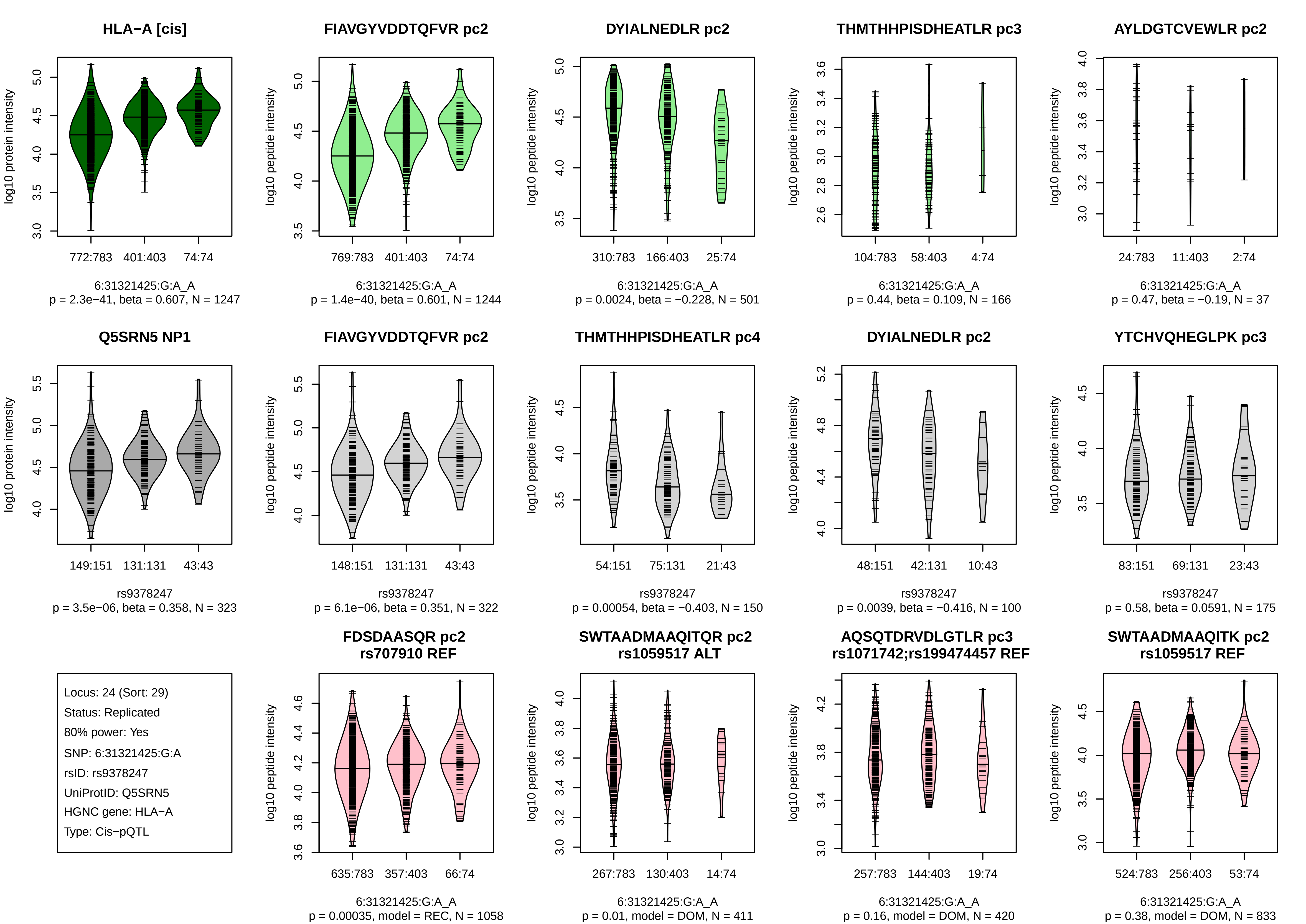
HLA−A [cis]
log10 protein intensity
772:783 401:403 74:74
6:31321425:G:A_A
p = 2.3e−41, beta = 0.607, N = 1247
FIAVGYVDDTQFVR pc2
log10 peptide intensity
769:783 401:403 74:74
6:31321425:G:A_A
p = 1.4e−40, beta = 0.601, N = 1244
DYIALNEDLR pc2
310:783 166:403 25:74
6:31321425:G:A_A
p = 0.0024, beta = −0.228, N = 501
THMTHHPISDHEATLR pc3
104:783 58:403 4:74
6:31321425:G:A_A
p = 0.44, beta = 0.109, N = 166
AYLDGTCVEWLR pc2
24:783 11:403 2:74
6:31321425:G:A_A
p = 0.47, beta = −0.19, N = 37
Q5SRN5 NP1
149:151 131:131 43:43
rs9378247
p = 3.5e−06, beta = 0.358, N = 323
FIAVGYVDDTQFVR pc2
148:151 131:131 43:43
rs9378247
p = 6.1e−06, beta = 0.351, N = 322
THMTHHPISDHEATLR pc4
54:151 75:131 21:43
rs9378247
p = 0.00054, beta = −0.403, N = 150
DYIALNEDLR pc2
48:151 42:131 10:43
rs9378247
p = 0.0039, beta = −0.416, N = 100
YTCHVQHEGLPK pc3
83:151 69:131 23:43
rs9378247
p = 0.58, beta = 0.0591, N = 175
Locus: 24 (Sort: 29)
Status: Replicated
80% power: Yes
SNP: 6:31321425:G:A
rsID: rs9378247
UniProtID: Q5SRN5
HGNC gene: HLA−A
Type: Cis−pQTL
FDSDAASQR pc2
rs707910 REF
635:783 357:403 66:74
6:31321425:G:A_A
p = 0.00035, model = REC, N = 1058
SWTAADMAAQITQR pc2
rs1059517 ALT
267:783 130:403 14:74
6:31321425:G:A_A
p = 0.01, model = DOM, N = 411
AQSQTDRVDLGTLR pc3
rs1071742;rs199474457 REF
257:783 144:403 19:74
6:31321425:G:A_A
p = 0.16, model = DOM, N = 420
SWTAADMAAQITK pc2
rs1059517 REF
524:783 256:403 53:74
6:31321425:G:A_A
p = 0.38, model = DOM, N = 833

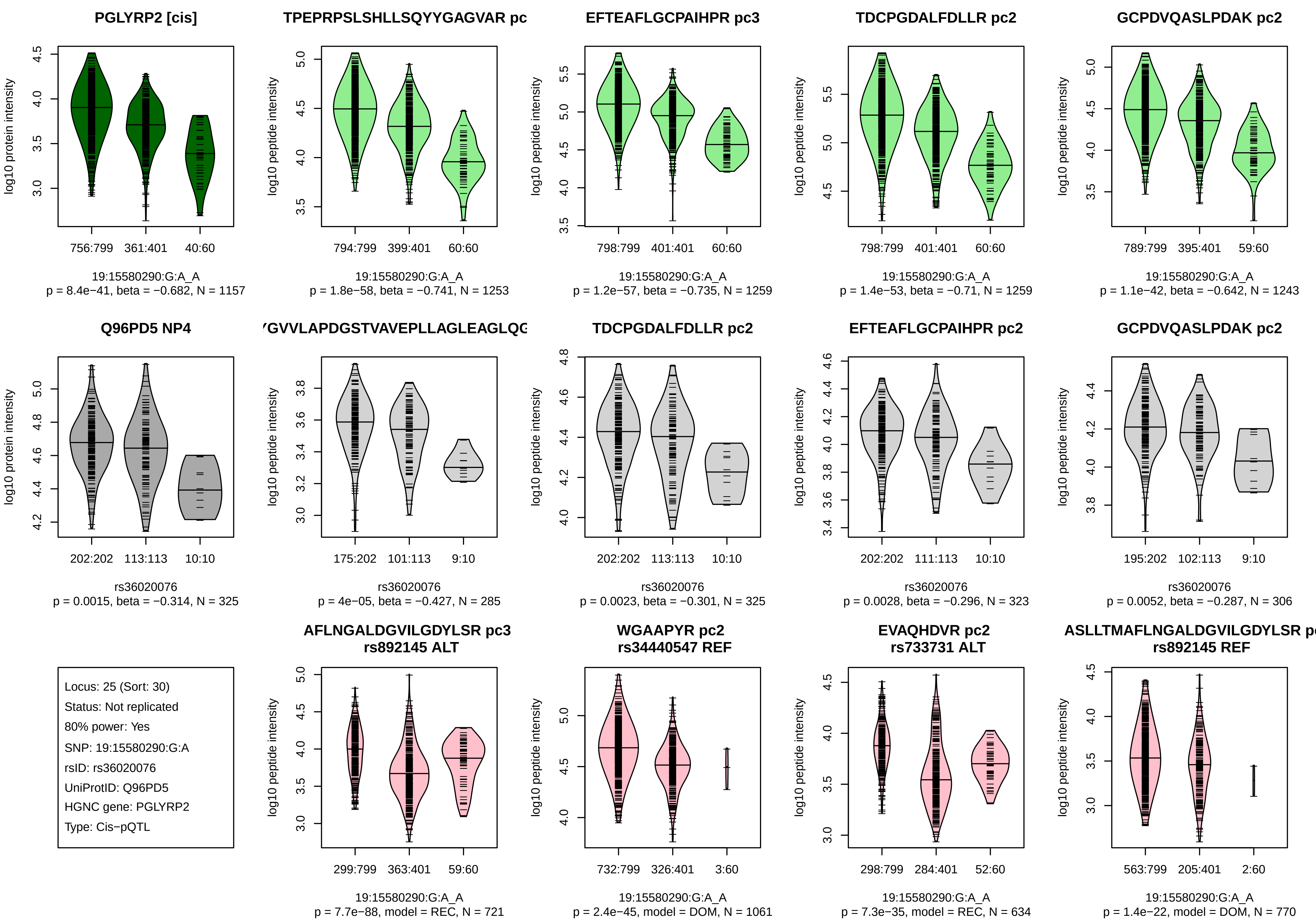
PGLYRP2 [cis]
log10 protein intensity
756:799 361:401 40:60
19:15580290:G:A_A
p = 8.4e−41, beta = −0.682, N = 1157
TPEPRPSLSHLLSQYYGAGVAR pc
log10 peptide intensity
794:799 399:401 60:60
19:15580290:G:A_A
p = 1.8e−58, beta = −0.741, N = 1253
EFTEAFLGCPAIHPR pc3
log10 peptide intensity
798:799 401:401 60:60
19:15580290:G:A_A
p = 1.2e−57, beta = −0.735, N = 1259
TDCPGDALFDLLR pc2
log10 peptide intensity
798:799 401:401 60:60
19:15580290:G:A_A
p = 1.4e−53, beta = −0.71, N = 1259
GCPDVQASLPDAK pc2
log10 peptide intensity
789:799 395:401 59:60
19:15580290:G:A_A
p = 1.1e−42, beta = −0.642, N = 1243
Q96PD5 NP4
log10 protein intensity
202:202 113:113 10:10
rs36020076
p = 0.0015, beta = −0.314, N = 325
GVVLAPDGSTVAVEPLLAGLEAGLQG
log10 peptide intensity
175:202 101:113 9:10
rs36020076
p = 4e−05, beta = −0.427, N = 285
TDCPGDALFDLLR pc2
log10 peptide intensity
202:202 113:113 10:10
rs36020076
p = 0.0023, beta = −0.301, N = 325
EFTEAFLGCPAIHPR pc2
log10 peptide intensity
202:202 111:113 10:10
rs36020076
p = 0.0028, beta = −0.296, N = 323
GCPDVQASLPDAK pc2
log10 peptide intensity
195:202 102:113 9:10
rs36020076
p = 0.0052, beta = −0.287, N = 306
Locus: 25 (Sort: 30)
Status: Not replicated
80% power: Yes
SNP: 19:15580290:G:A
rsID: rs36020076
UniProtID: Q96PD5
HGNC gene: PGLYRP2
Type: Cis−pQTL
AFLNGALDGVILGDYLSR pc3
rs892145 ALT
log10 peptide intensity
299:799 363:401 59:60
19:15580290:G:A_A
p = 7.7e−88, model = REC, N = 721
WGAAPYR pc2
rs34440547 REF
log10 peptide intensity
732:799 326:401 3:60
19:15580290:G:A_A
p = 2.4e−45, model = DOM, N = 1061
EVAQHDVR pc2
rs733731 ALT
log10 peptide intensity
298:799 284:401 52:60
19:15580290:G:A_A
p = 7.3e−35, model = REC, N = 634
ASLLTMAFLNGALDGVILGDYLSR p
rs892145 REF
log10 peptide intensity
563:799 205:401 2:60
19:15580290:G:A_A
p = 1.4e−22, model = DOM, N = 770

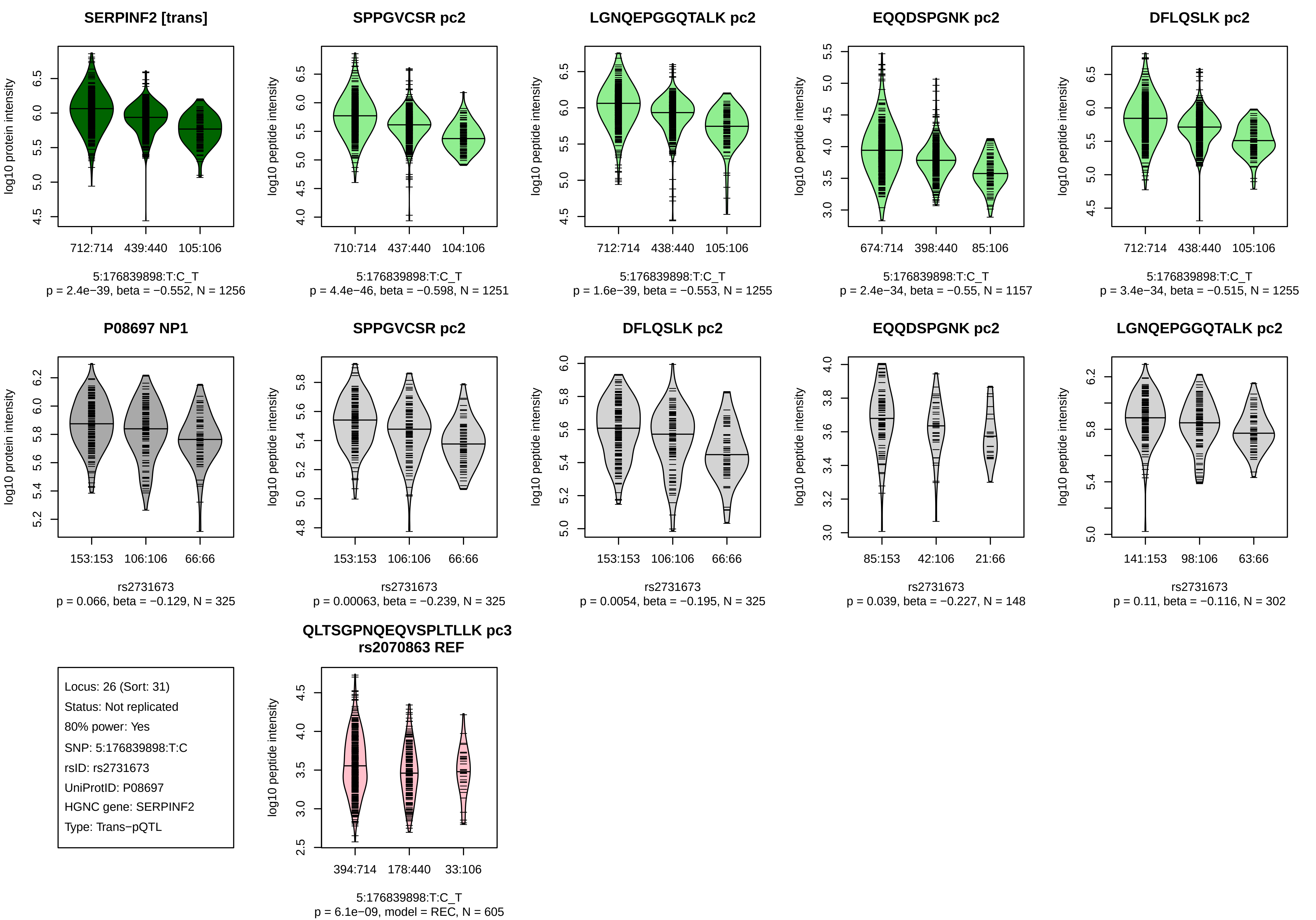
SERPINF2 [trans]
log10 protein intensity
712:714
439:440
105:106
5:176839898:T:C_T
p = 2.4e−39, beta = −0.552, N = 1256
SPPGVCSR pc2
log10 peptide intensity
710:714
437:440
104:106
5:176839898:T:C_T
p = 4.4e−46, beta = −0.598, N = 1251
LGNQEPGGQTALK pc2
log10 peptide intensity
712:714
438:440
105:106
5:176839898:T:C_T
p = 1.6e−39, beta = −0.553, N = 1255
EQQDSPGNK pc2
log10 peptide intensity
674:714
398:440
85:106
5:176839898:T:C_T
p = 2.4e−34, beta = −0.55, N = 1157
DFLQSLK pc2
log10 peptide intensity
712:714
438:440
105:106
5:176839898:T:C_T
p = 3.4e−34, beta = −0.515, N = 1255
P08697 NP1
log10 protein intensity
153:153
106:106
66:66
rs2731673
p = 0.066, beta = −0.129, N = 325
SPPGVCSR pc2
log10 peptide intensity
153:153
106:106
66:66
rs2731673
p = 0.00063, beta = −0.239, N = 325
DFLQSLK pc2
log10 peptide intensity
153:153
106:106
66:66
rs2731673
p = 0.0054, beta = −0.195, N = 325
EQQDSPGNK pc2
log10 peptide intensity
85:153
42:106
21:66
rs2731673
p = 0.039, beta = −0.227, N = 148
LGNQEPGGQTALK pc2
log10 peptide intensity
141:153
98:106
63:66
rs2731673
p = 0.11, beta = −0.116, N = 302
Locus: 26 (Sort: 31)
Status: Not replicated
80% power: Yes
SNP: 5:176839898:T:C
rsID: rs2731673
UniProtID: P08697
HGNC gene: SERPINF2
Type: Trans−pQTL
QLTSGPNQEQVSPLTLLK pc3
rs2070863 REF
log10 peptide intensity
394:714
178:440
33:106
5:176839898:T:C_T
p = 6.1e−09, model = REC, N = 605

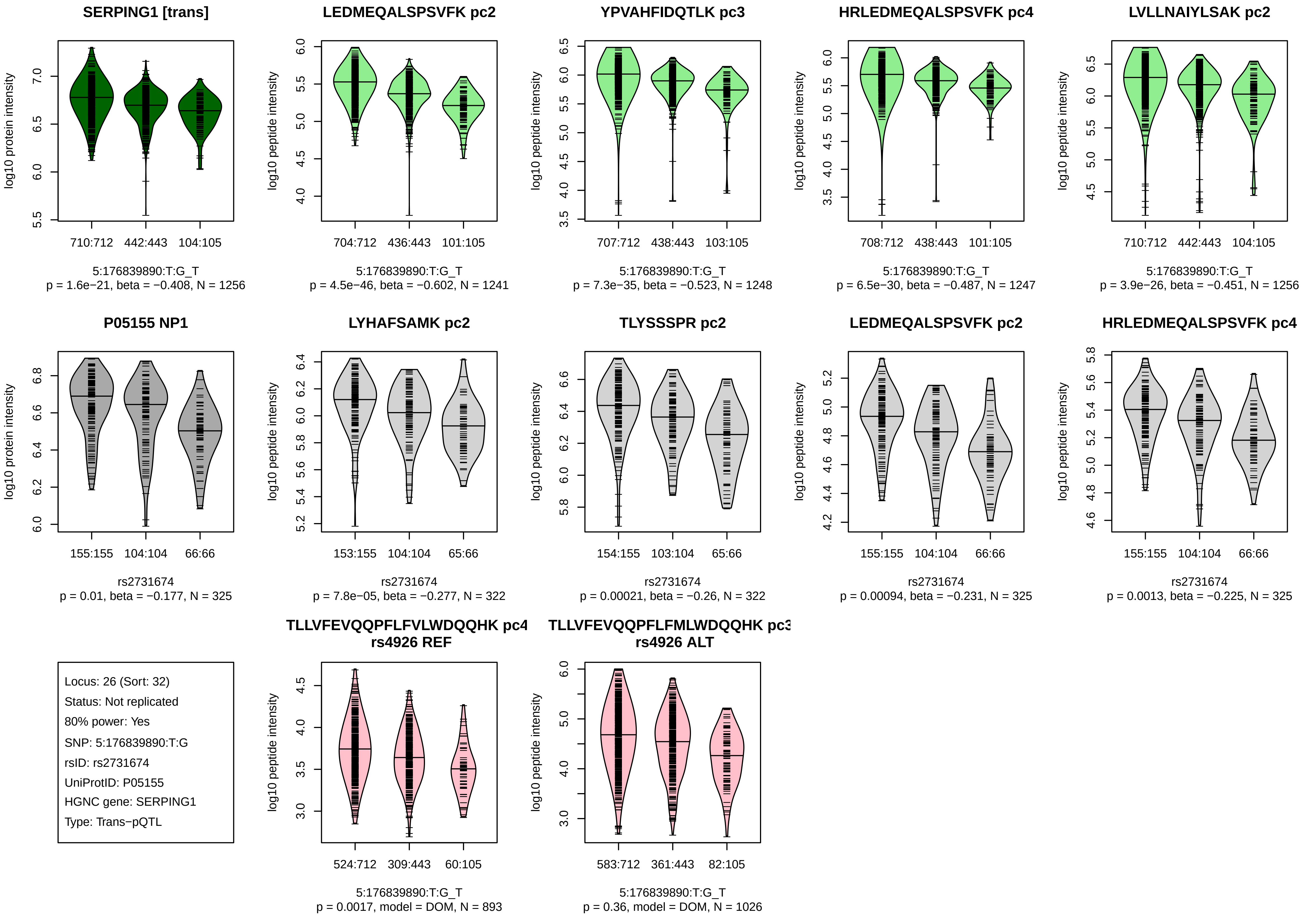
SERPING1 [trans]
log10 protein intensity
710:712 442:443 104:105
5:176839890:T:G_T
p = 1.6e−21, beta = −0.408, N = 1256
LEDMEQALSPSVFK pc2
log10 peptide intensity
704:712 436:443 101:105
5:176839890:T:G_T
p = 4.5e−46, beta = −0.602, N = 1241
YPVAHFIDQTLK pc3
log10 peptide intensity
707:712 438:443 103:105
5:176839890:T:G_T
p = 7.3e−35, beta = −0.523, N = 1248
HRLEDMEQALSPSVFK pc4
log10 peptide intensity
708:712 438:443 101:105
5:176839890:T:G_T
p = 6.5e−30, beta = −0.487, N = 1247
LVLLNAIYLSAK pc2
log10 peptide intensity
710:712 442:443 104:105
5:176839890:T:G_T
p = 3.9e−26, beta = −0.451, N = 1256
P05155 NP1
log10 protein intensity
155:155 104:104 66:66
rs2731674
p = 0.01, beta = −0.177, N = 325
LYHAFSAMK pc2
log10 peptide intensity
153:155 104:104 65:66
rs2731674
p = 7.8e−05, beta = −0.277, N = 322
TLYSSSPR pc2
log10 peptide intensity
154:155 103:104 65:66
rs2731674
p = 0.00021, beta = −0.26, N = 322
LEDMEQALSPSVFK pc2
log10 peptide intensity
155:155 104:104 66:66
rs2731674
p = 0.00094, beta = −0.231, N = 325
HRLEDMEQALSPSVFK pc4
log10 peptide intensity
155:155 104:104 66:66
rs2731674
p = 0.0013, beta = −0.225, N = 325
Locus: 26 (Sort: 32)
Status: Not replicated
80% power: Yes
SNP: 5:176839890:T:G
rsID: rs2731674
UniProtID: P05155
HGNC gene: SERPING1
Type: Trans−pQTL
TLLVFEVQQPFLFVLWDQQHK pc4
rs4926 REF
log10 peptide intensity
524:712 309:443 60:105
5:176839890:T:G_T
p = 0.0017, model = DOM, N = 893
TLLVFEVQQPFLFMLWDQQHK pc3
rs4926 ALT
log10 peptide intensity
583:712 361:443 82:105
5:176839890:T:G_T
p = 0.36, model = DOM, N = 1026

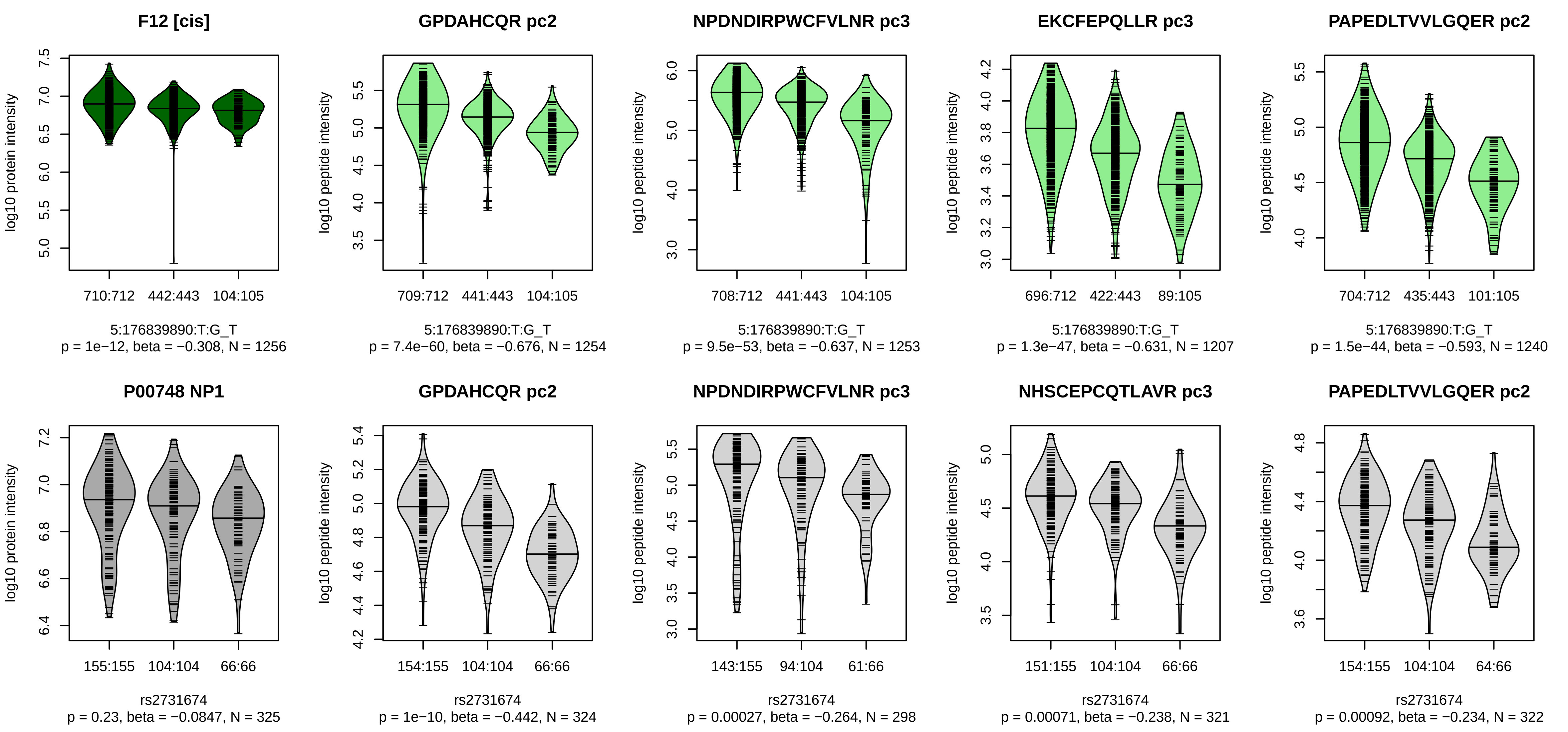


Locus: 26 (Sort: 33)

Status: Not replicated

80% power: No

SNP: 5:176839890:T:G

rsID: rs2731674

UniProtID: P00748

HGNC gene: F12

Type: Cis−pQTL

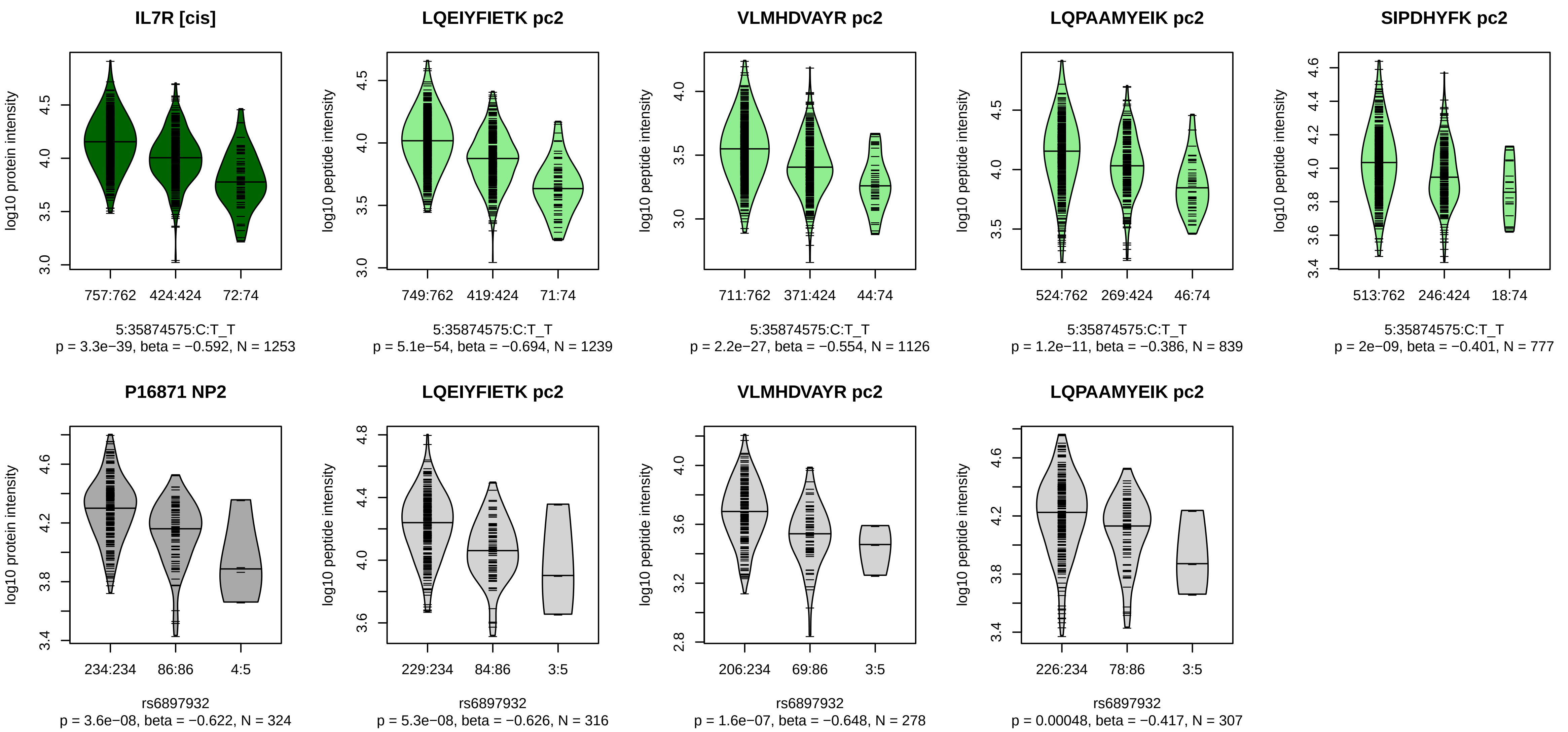


Locus: 27 (Sort: 34)
Status: Replicated
80% power: Yes
SNP: 5:35874575:C:T
rsID: rs6897932
UniProtID: P16871
HGNC gene: IL7R
Type: Cis−pQTL

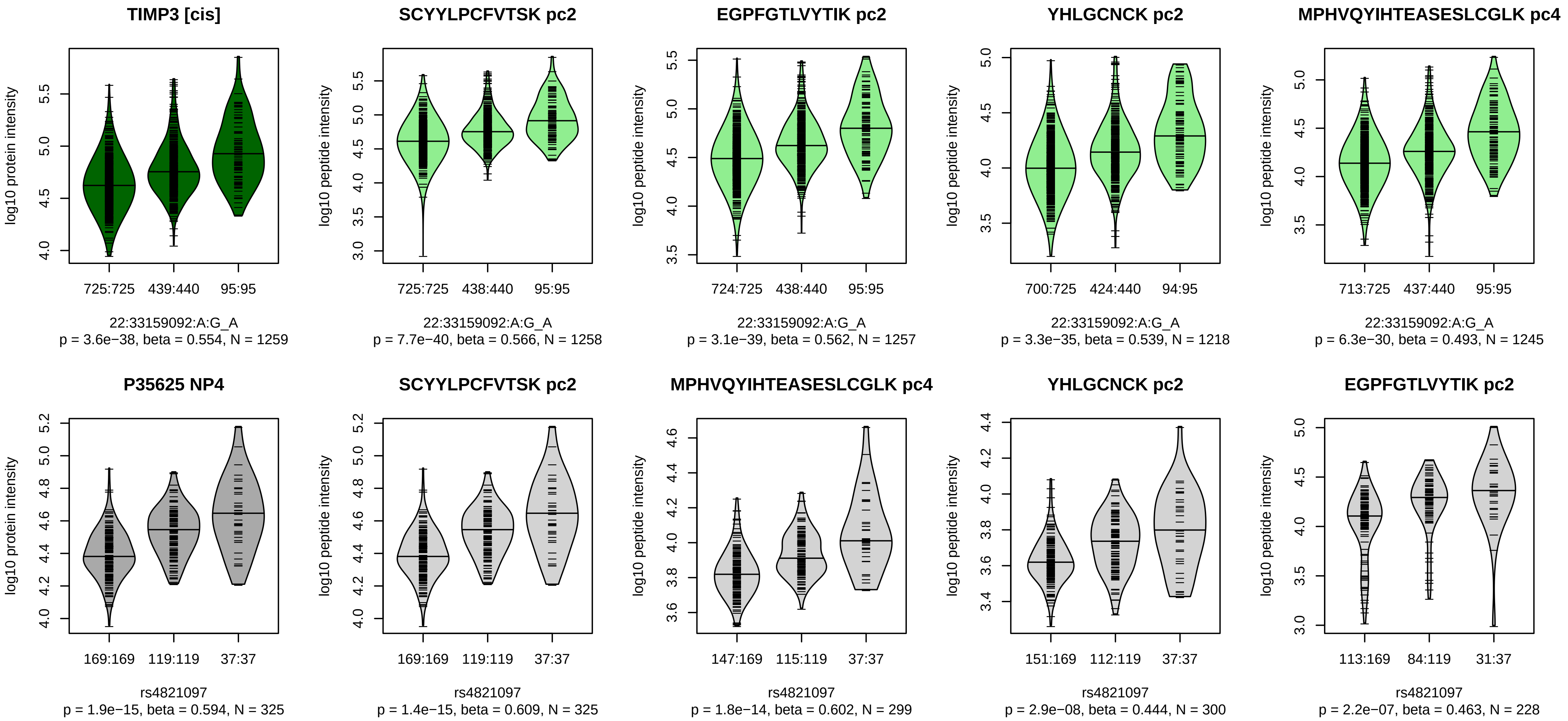


Locus: 28 (Sort: 35)
Status: Replicated
80% power: Yes
SNP: 22:33159092:A:G
rsID: rs4821097
UniProtID: P35625
HGNC gene: TIMP3
Type: Cis−pQTL

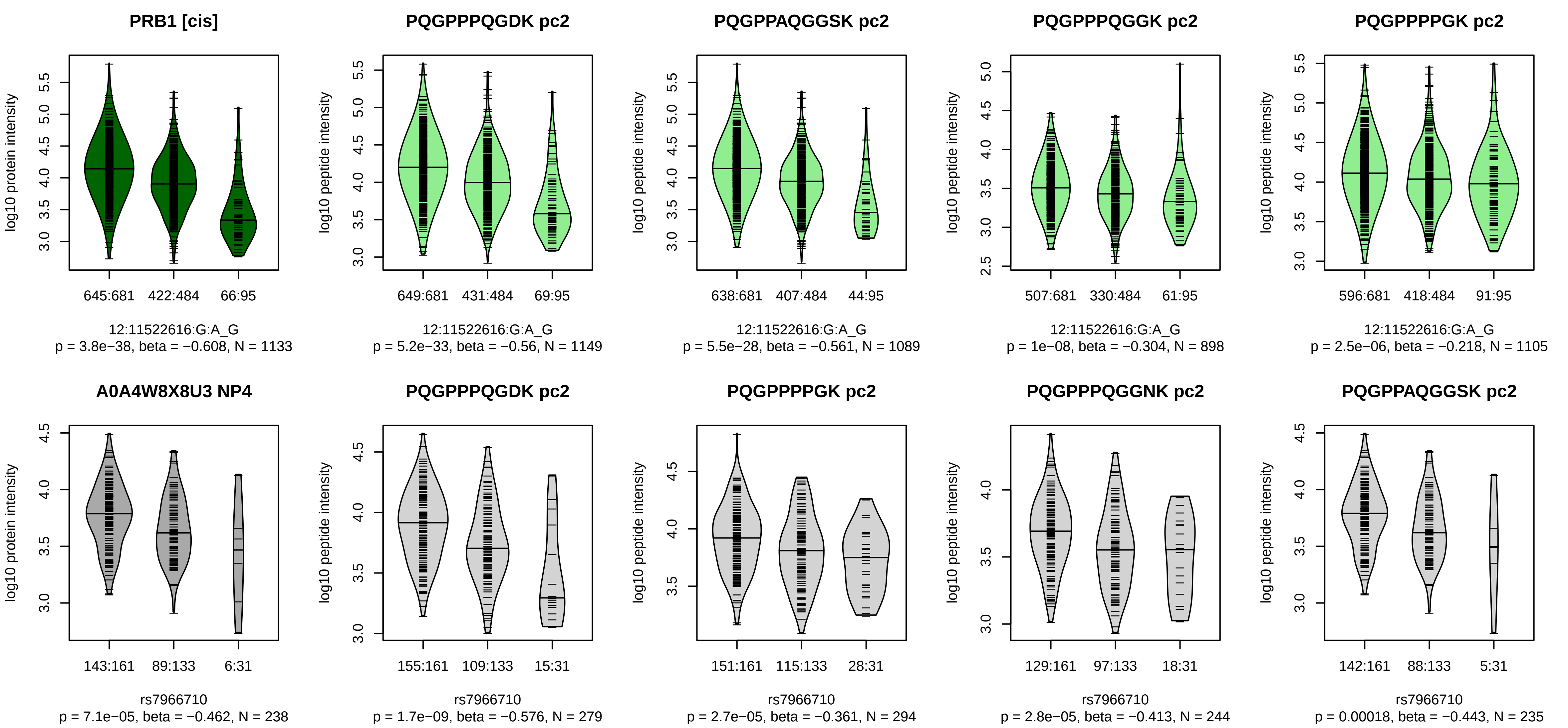


Locus: 29 (Sort: 36)
Status: Replicated
80% power: Yes
SNP: 12:11522616:G:A
rsID: rs7966710
UniProtID: A0A4W8X8U3
HGNC gene: PRB1
Type: Cis−pQTL

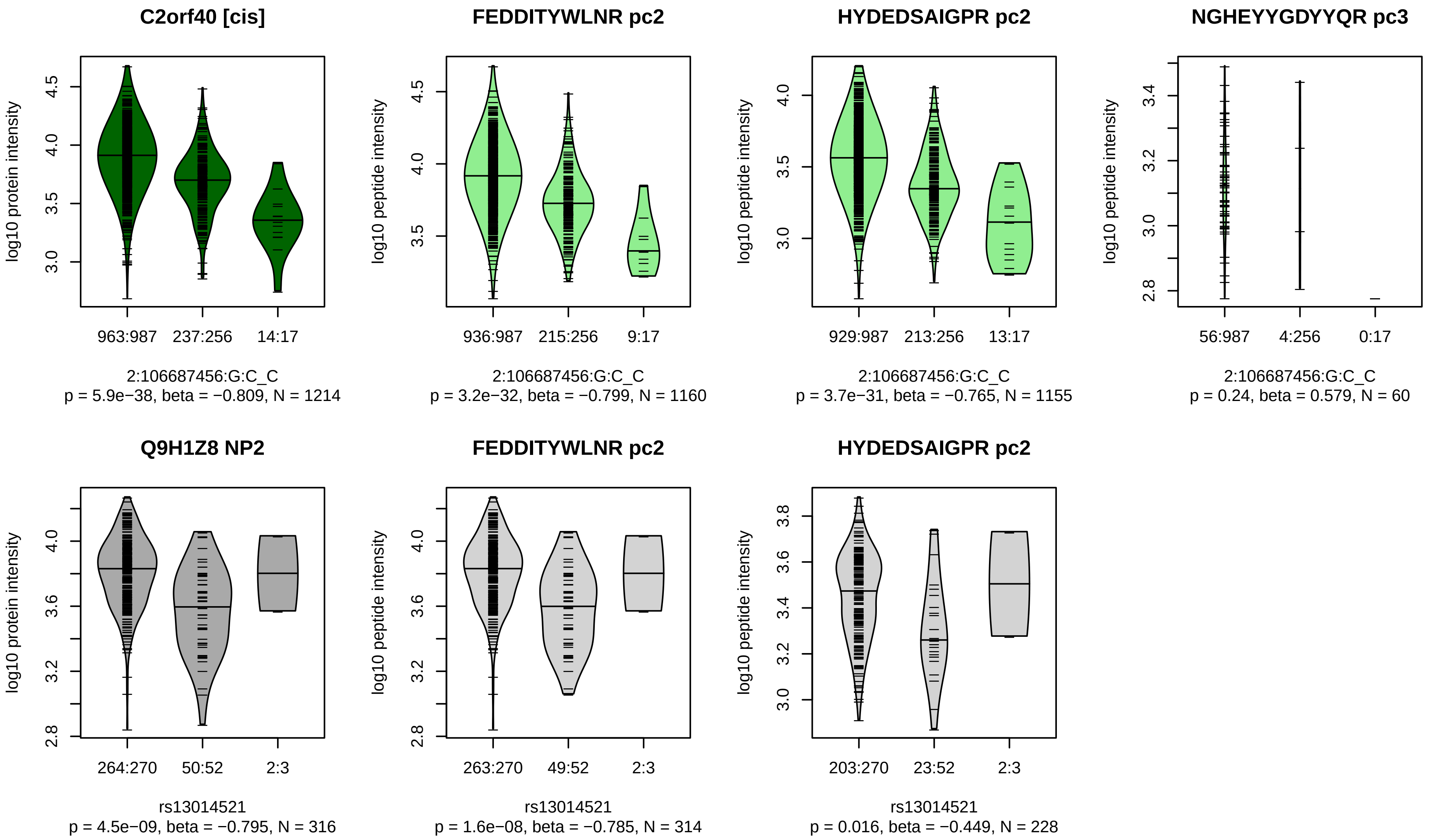


Locus: 30 (Sort: 37)

Status: Replicated

80% power: Yes

SNP: 2:106687456:G:C

rsID: rs13014521

UniProtID: Q9H1Z8

HGNC gene: C2orf40

Type: Cis−pQTL

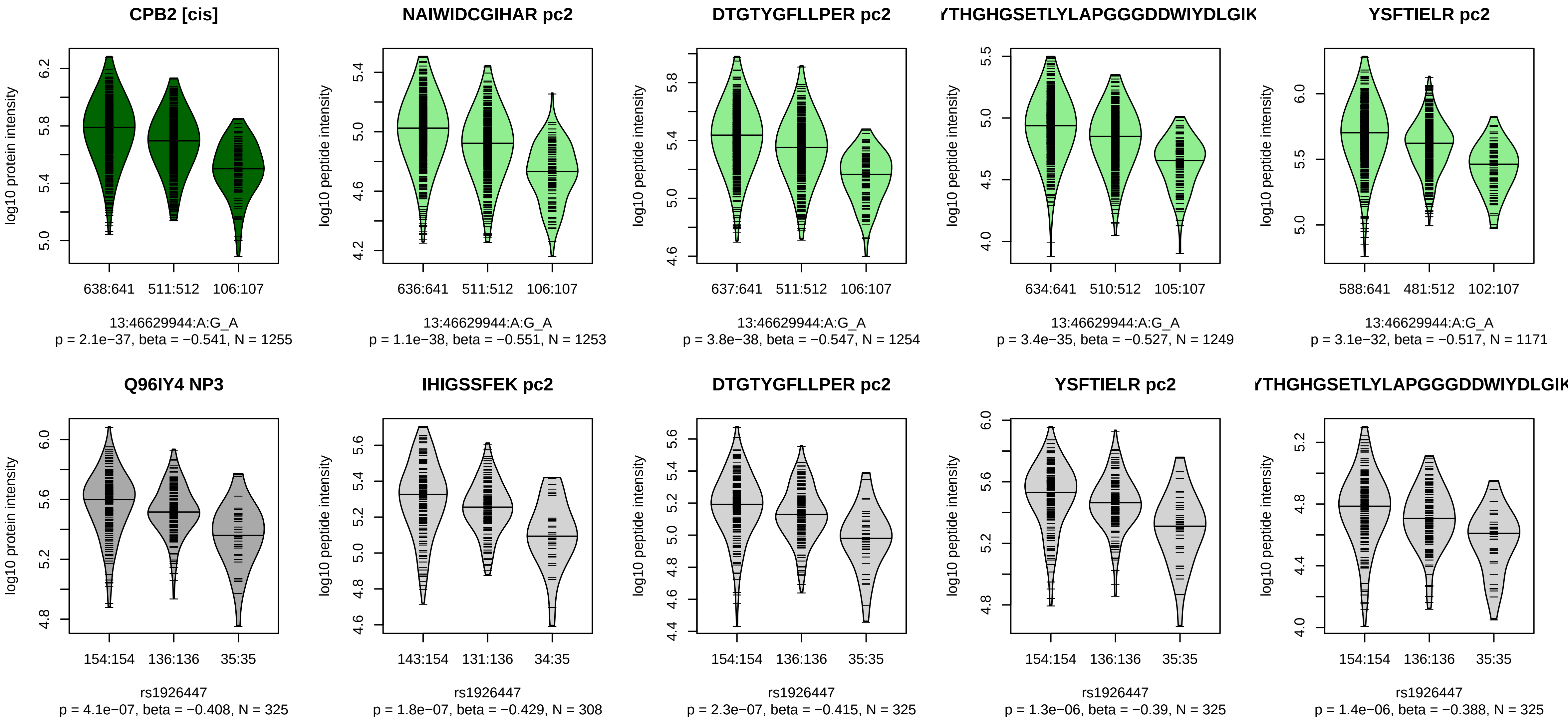


Locus: 31 (Sort: 38)
Status: Replicated
80% power: Yes
SNP: 13:46629944:A:G
rsID: rs1926447
UniProtID: Q96IY4
HGNC gene: CPB2
Type: Cis−pQTL

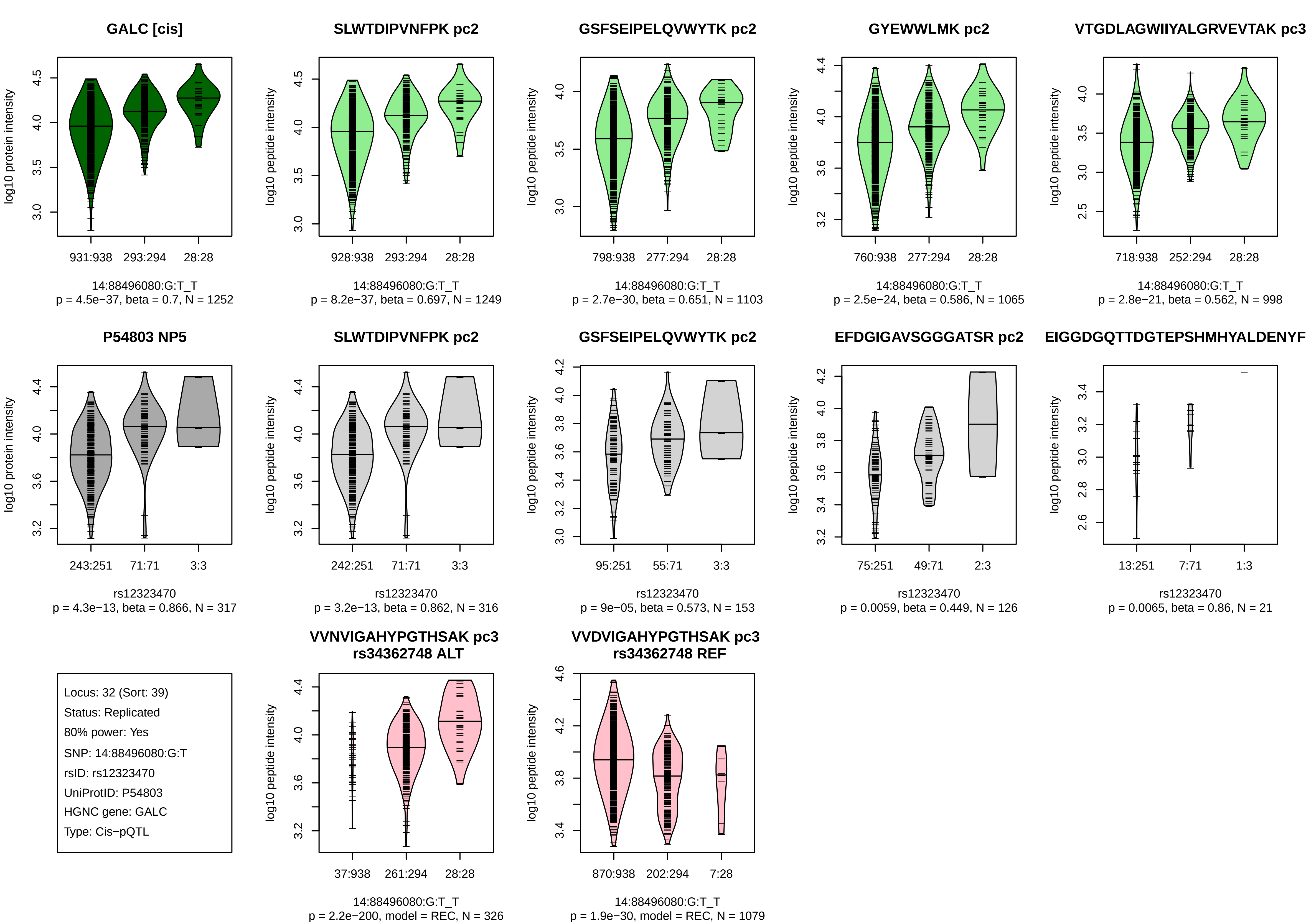
GALC [cis]
log10 protein intensity
931:938
293:294
28:28
14:88496080:G:T_T
p = 4.5e−37, beta = 0.7, N = 1252
SLWTDIPVNFPK pc2
log10 peptide intensity
928:938
293:294
28:28
14:88496080:G:T_T
p = 8.2e−37, beta = 0.697, N = 1249
GSFSEIPELQVWYTK pc2
log10 peptide intensity
798:938
277:294
28:28
14:88496080:G:T_T
p = 2.7e−30, beta = 0.651, N = 1103
GYEWWLMK pc2
log10 peptide intensity
760:938
277:294
28:28
14:88496080:G:T_T
p = 2.5e−24, beta = 0.586, N = 1065
VTGDLAGWIIYALGRVEVTAK pc3
log10 peptide intensity
718:938
252:294
28:28
14:88496080:G:T_T
p = 2.8e−21, beta = 0.562, N = 998
P54803 NP5
log10 protein intensity
243:251
71:71
3:3
rs12323470
p = 4.3e−13, beta = 0.866, N = 317
SLWTDIPVNFPK pc2
log10 peptide intensity
242:251
71:71
3:3
rs12323470
p = 3.2e−13, beta = 0.862, N = 316
GSFSEIPELQVWYTK pc2
log10 peptide intensity
95:251
55:71
3:3
rs12323470
p = 9e−05, beta = 0.573, N = 153
EFDGIGAVSGGGATSR pc2
log10 peptide intensity
75:251
49:71
2:3
rs12323470
p = 0.0059, beta = 0.449, N = 126
EIGGDGQTTDGTEPSHMHYALDENYF
log10 peptide intensity
13:251
7:71
1:3
rs12323470
p = 0.0065, beta = 0.86, N = 21
Locus: 32 (Sort: 39)
Status: Replicated
80% power: Yes
SNP: 14:88496080:G:T
rsID: rs12323470
UniProtID: P54803
HGNC gene: GALC
Type: Cis−pQTL
VVNVIGAHYPGTHSAK pc3
rs34362748 ALT
log10 peptide intensity
37:938
261:294
28:28
14:88496080:G:T_T
p = 2.2e−200, model = REC, N = 326
VVDVIGAHYPGTHSAK pc3
rs34362748 REF
log10 peptide intensity
870:938
202:294
7:28
14:88496080:G:T_T
p = 1.9e−30, model = REC, N = 1079

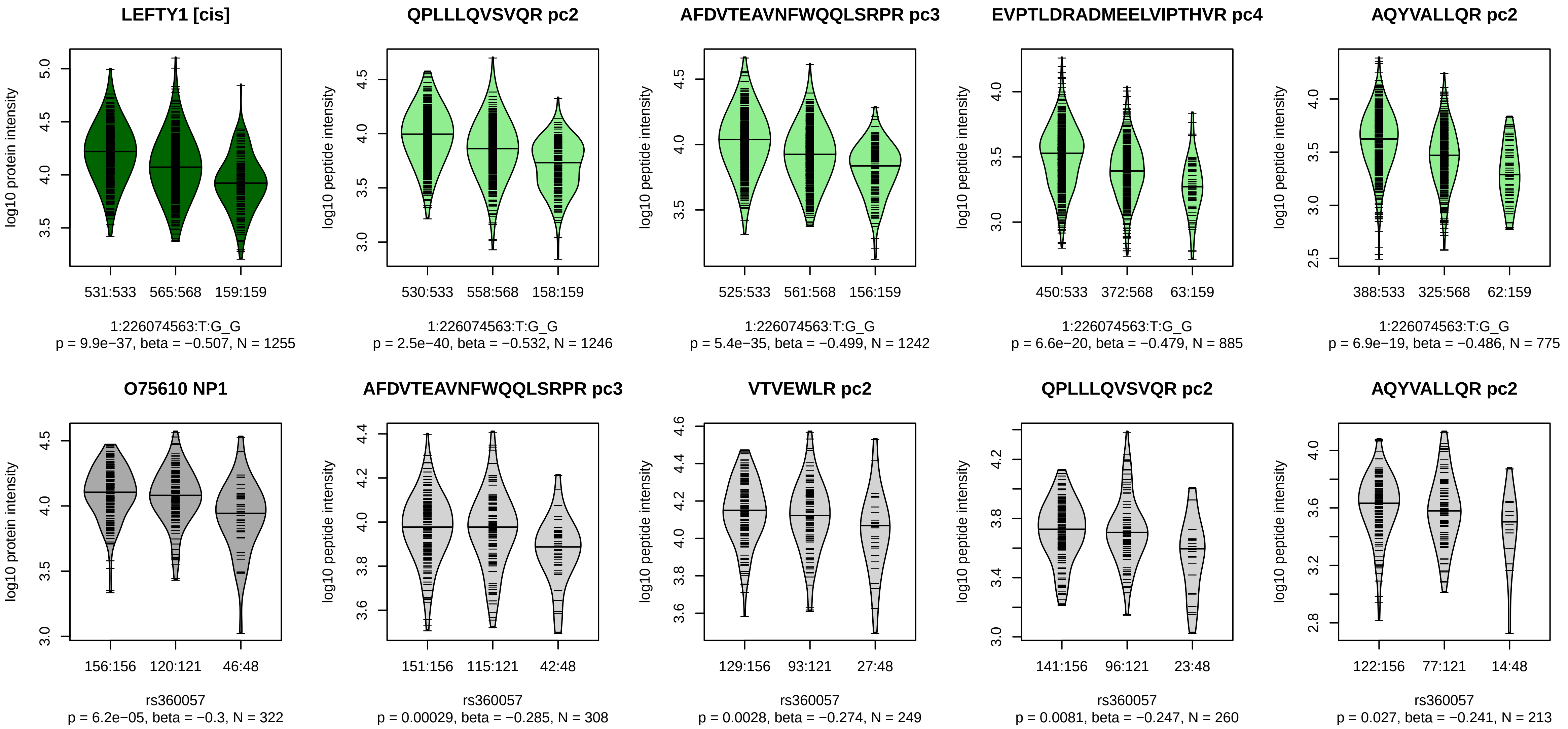


Locus: 33 (Sort: 40)
Status: Replicated
80% power: Yes
SNP: 1:226074563:T:G
rsID: rs360057
UniProtID: O75610
HGNC gene: LEFTY1
Type: Cis−pQTL

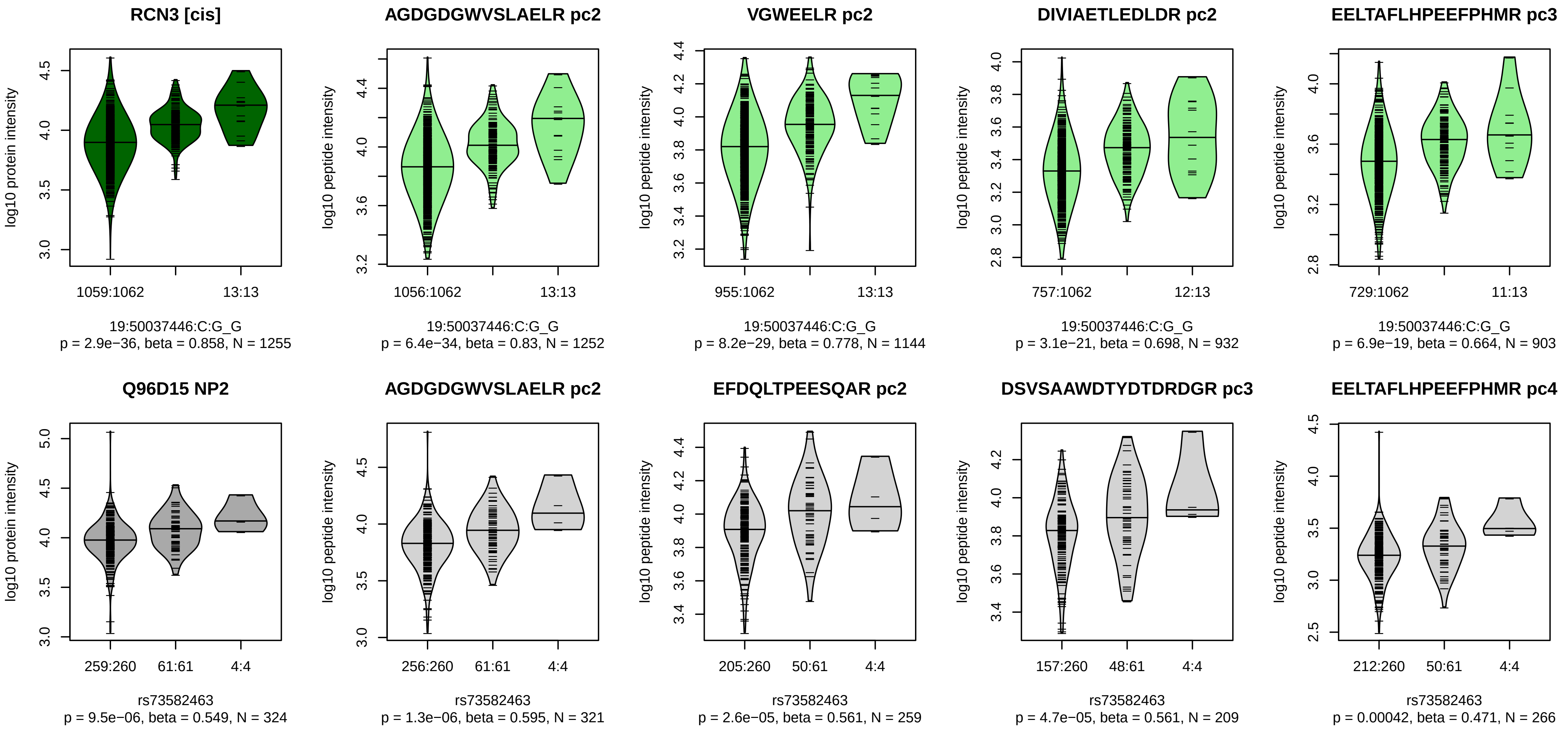


Locus: 34 (Sort: 41)
Status: Replicated
80% power: Yes
SNP: 19:50037446:C:G
rsID: rs73582463
UniProtID: Q96D15
HGNC gene: RCN3
Type: Cis−pQTL

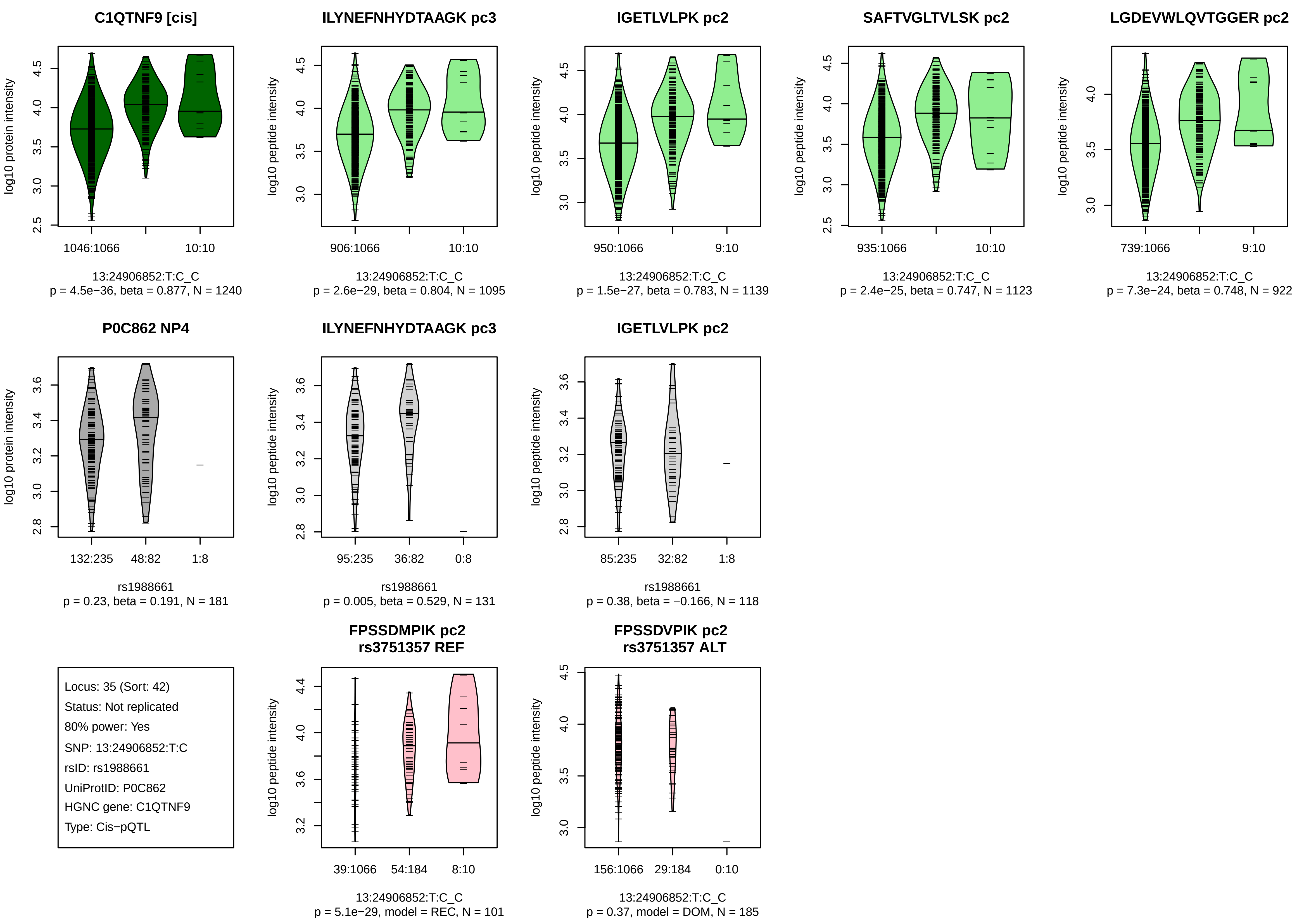

C1QTNF9 [cis]
log10 protein intensity
1046:1066
10:10
13:24906852:T:C_C
p = 4.5e−36, beta = 0.877, N = 1240
ILYNEFNHYDTAAGK pc3
log10 peptide intensity
906:1066
10:10
13:24906852:T:C_C
p = 2.6e−29, beta = 0.804, N = 1095
IGETLVLPK pc2
log10 peptide intensity
950:1066
9:10
13:24906852:T:C_C
p = 1.5e−27, beta = 0.783, N = 1139
SAFTVGLTVLSK pc2
log10 peptide intensity
935:1066
10:10
13:24906852:T:C_C
p = 2.4e−25, beta = 0.747, N = 1123
LGDEVWLQVTGGER pc2
log10 peptide intensity
739:1066
9:10
13:24906852:T:C_C
p = 7.3e−24, beta = 0.748, N = 922
P0C862 NP4
log10 protein intensity
132:235
48:82
1:8
rs1988661
p = 0.23, beta = 0.191, N = 181
ILYNEFNHYDTAAGK pc3
log10 peptide intensity
95:235
36:82
0:8
rs1988661
p = 0.005, beta = 0.529, N = 131
IGETLVLPK pc2
log10 peptide intensity
85:235
32:82
1:8
rs1988661
p = 0.38, beta = −0.166, N = 118
Locus: 35 (Sort: 42)
Status: Not replicated
80% power: Yes
SNP: 13:24906852:T:C
rsID: rs1988661
UniProtID: P0C862
HGNC gene: C1QTNF9
Type: Cis−pQTL
FPSSDMPIK pc2
rs3751357 REF
log10 peptide intensity
39:1066
54:184
8:10
13:24906852:T:C_C
p = 5.1e−29, model = REC, N = 101
FPSSDVPIK pc2
rs3751357 ALT
log10 peptide intensity
156:1066
29:184
0:10
13:24906852:T:C_C
p = 0.37, model = DOM, N = 185

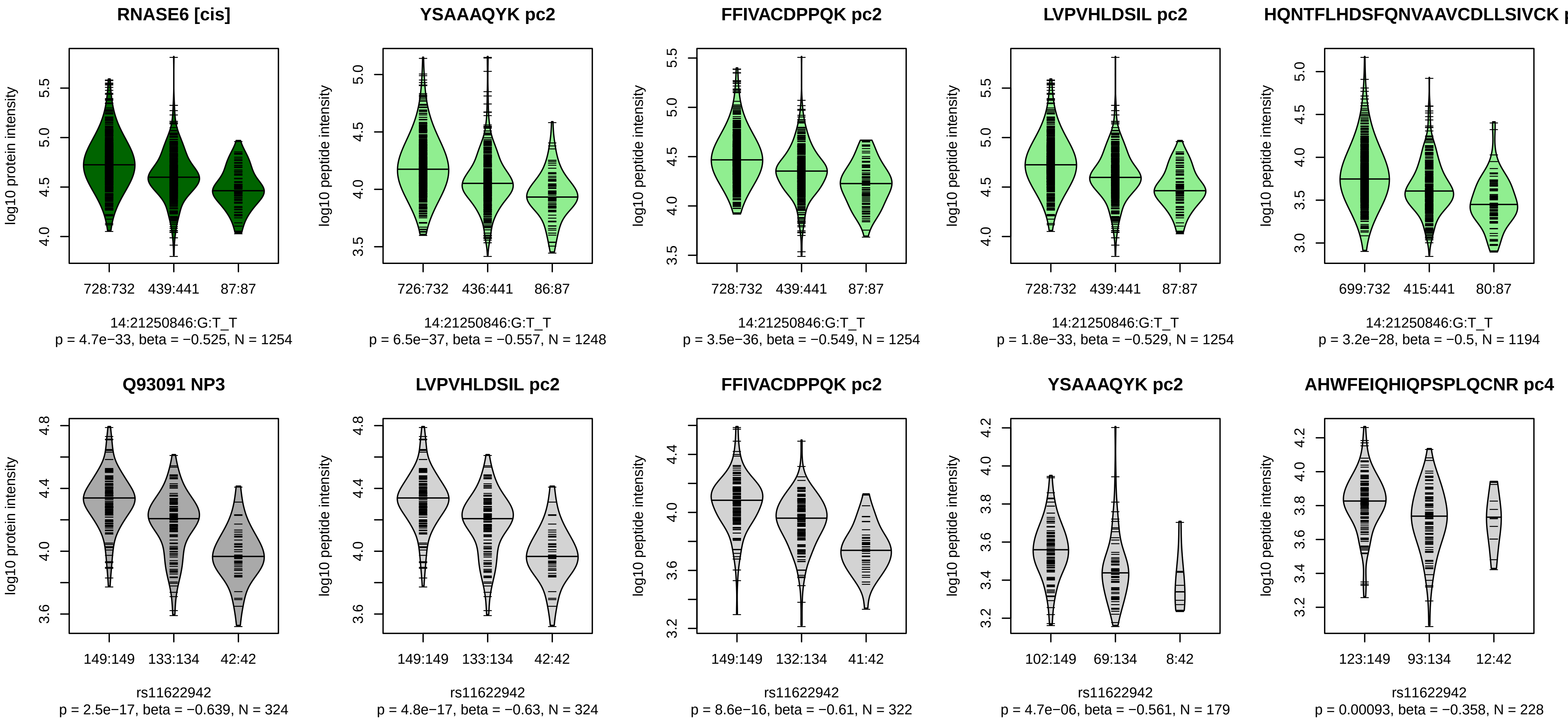


Locus: 36 (Sort: 43)

Status: Replicated

80% power: Yes

SNP: 14:21250846:G:T

rsID: rs11622942

UniProtID: Q93091

HGNC gene: RNASE6

Type: Cis−pQTL

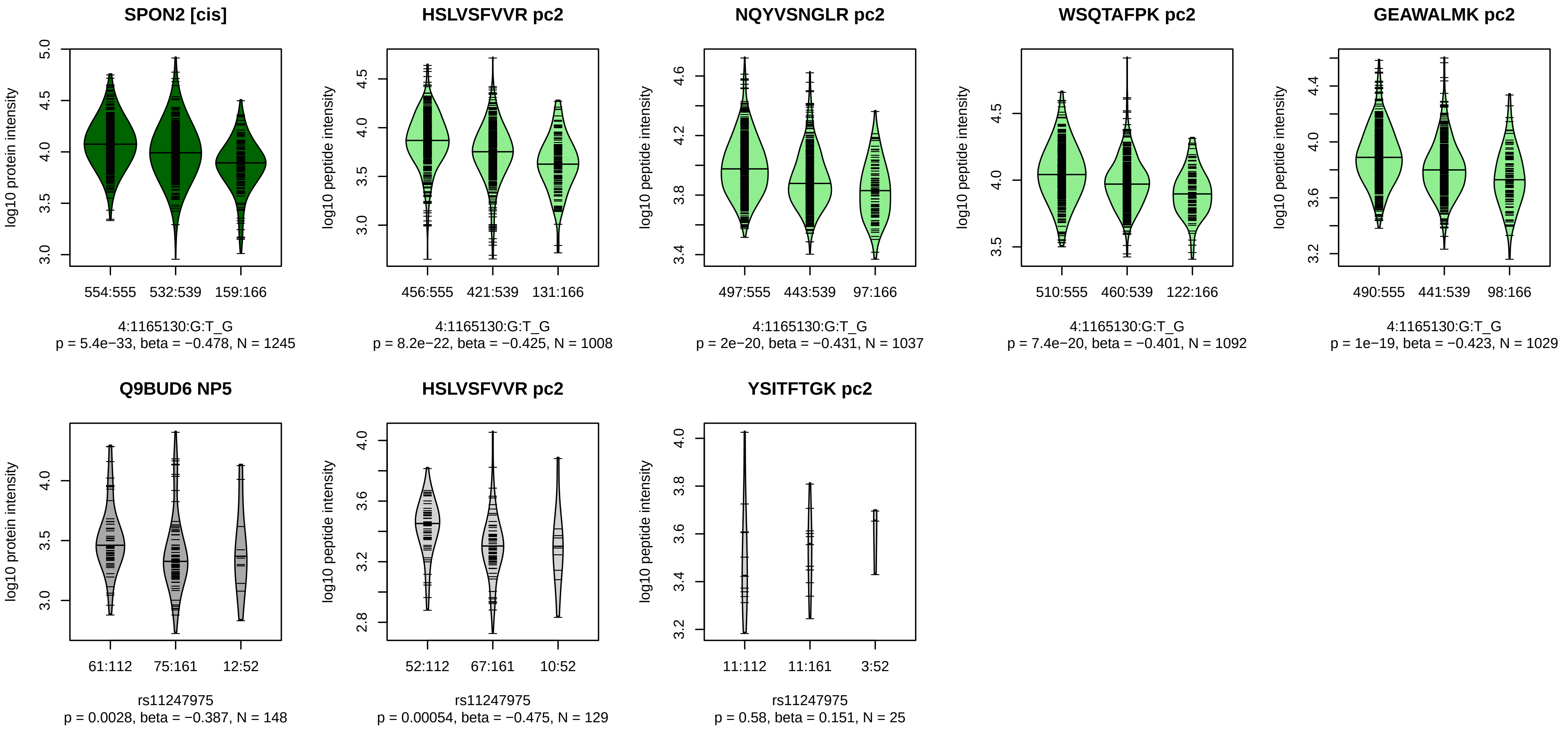


Locus: 37 (Sort: 44)
Status: Not replicated
80% power: No
SNP: 4:1165130:G:T
rsID: rs11247975
UniProtID: Q9BUD6
HGNC gene: SPON2
Type: Cis−pQTL

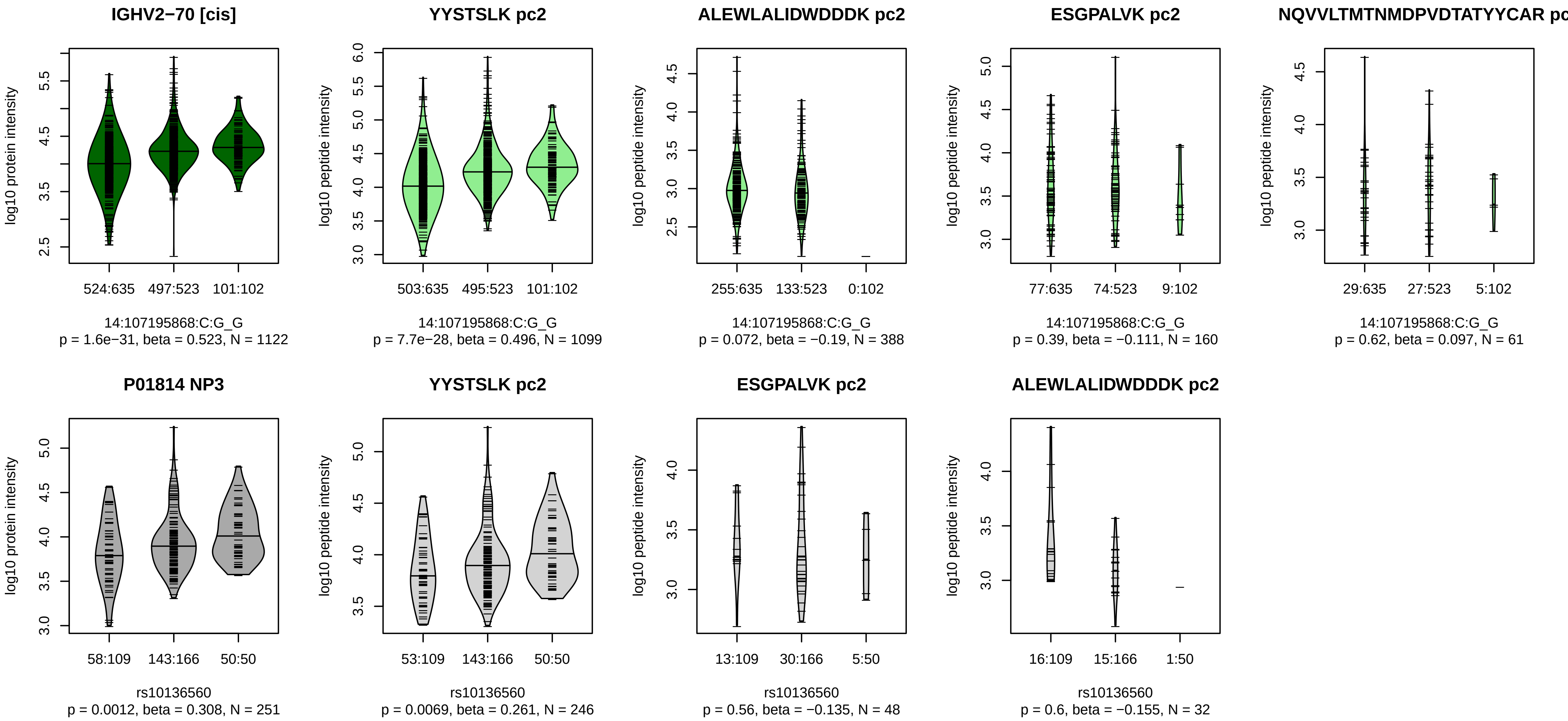


Locus: 38 (Sort: 45)

Status: Not replicated

80% power: Yes

SNP: 14:107195868:C:G

rsID: rs10136560

UniProtID: P01814

HGNC gene: IGHV2−70

Type: Cis−pQTL

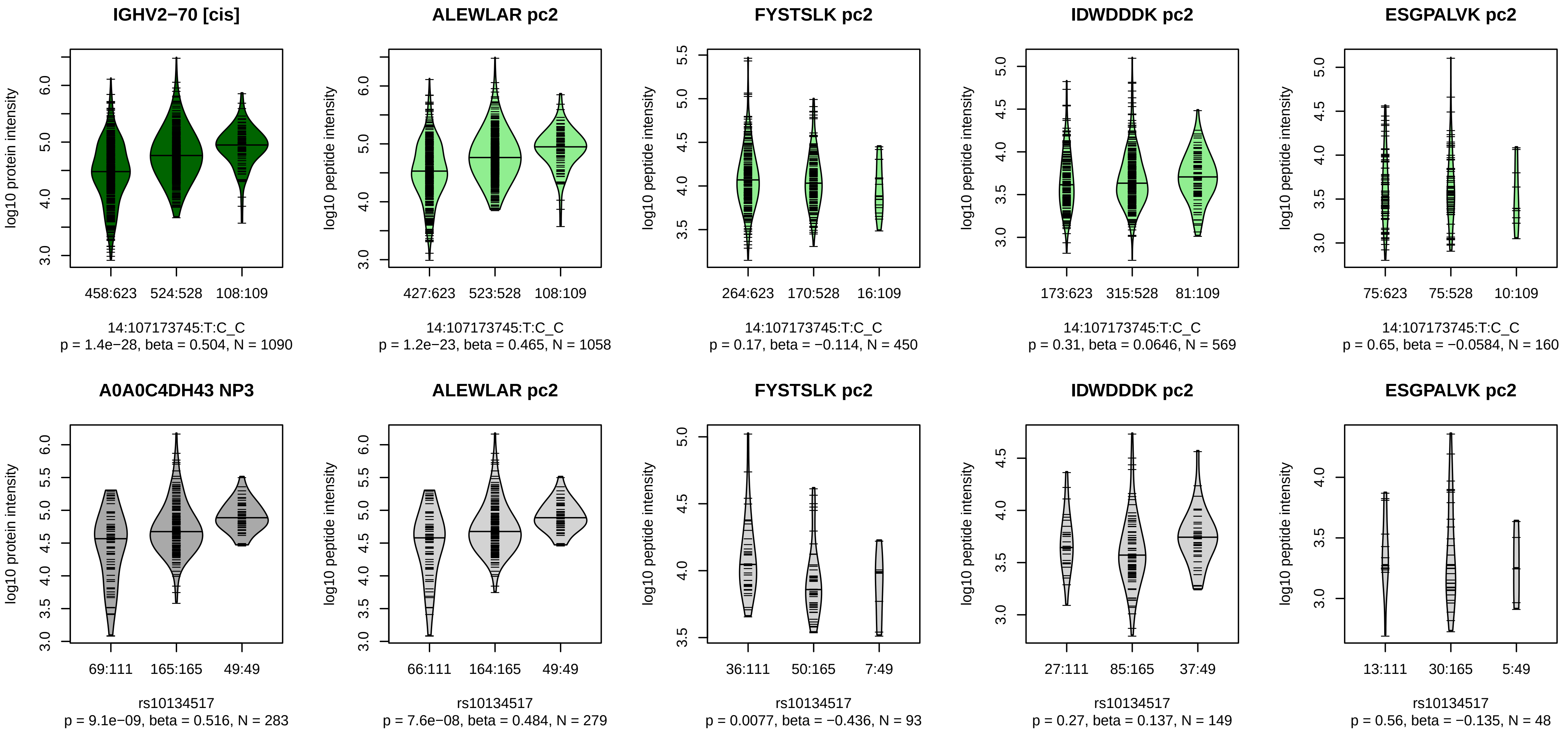


Locus: 38 (Sort: 46)
Status: Replicated
80% power: Yes
SNP: 14:107173745:T:C
rsID: rs10134517
UniProtID: A0A0C4DH43
HGNC gene: IGHV2−70
Type: Cis−pQTL

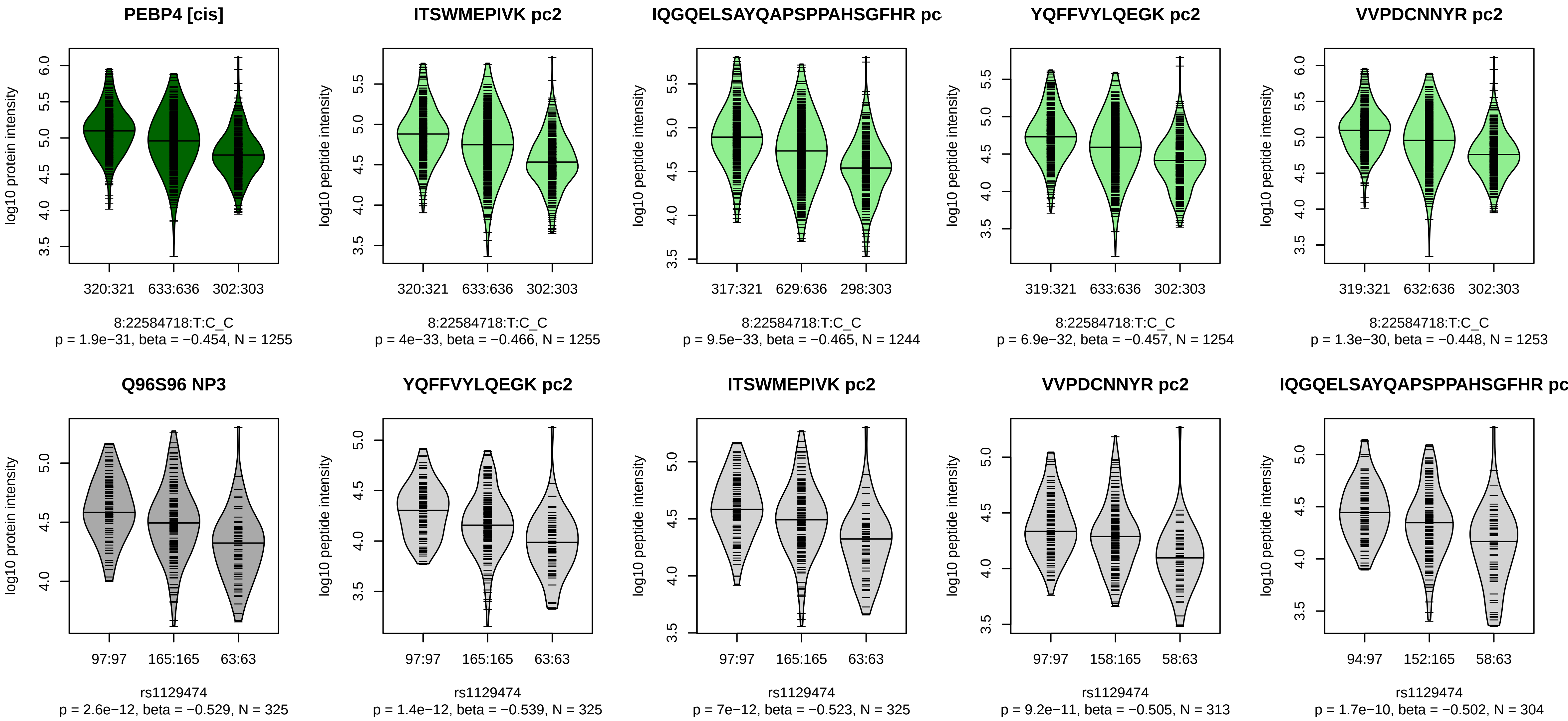


Locus: 39 (Sort: 47)
Status: Replicated
80% power: Yes
SNP: 8:22584718:T:C
rsID: rs1129474
UniProtID: Q96S96
HGNC gene: PEBP4
Type: Cis−pQTL

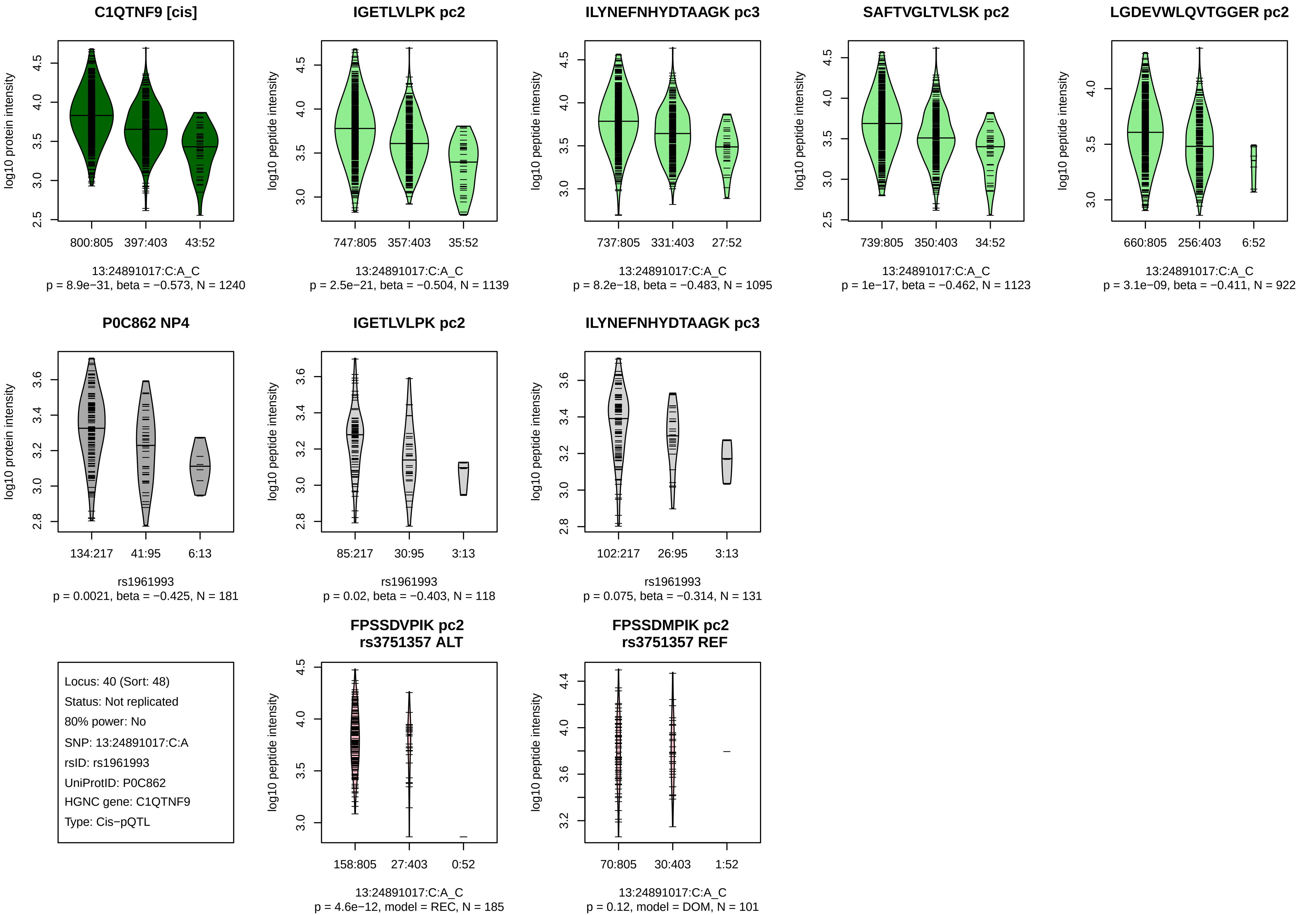

C1QTNF9 [cis]
log10 protein intensity
800:805
397:403
43:52
13:24891017:C:A_C
p = 8.9e−31, beta = −0.573, N = 1240
IGETLVLPK pc2
log10 peptide intensity
747:805
357:403
35:52
13:24891017:C:A_C
p = 2.5e−21, beta = −0.504, N = 1139
ILYNEFNHYDTAAGK pc3
log10 peptide intensity
737:805
331:403
27:52
13:24891017:C:A_C
p = 8.2e−18, beta = −0.483, N = 1095
SAFTVGLTVLSK pc2
log10 peptide intensity
739:805
350:403
34:52
13:24891017:C:A_C
p = 1e−17, beta = −0.462, N = 1123
LGDEVWLQVTGGER pc2
log10 peptide intensity
660:805
256:403
6:52
13:24891017:C:A_C
p = 3.1e−09, beta = −0.411, N = 922
P0C862 NP4
log10 protein intensity
134:217
41:95
6:13
rs1961993
p = 0.0021, beta = −0.425, N = 181
IGETLVLPK pc2
log10 peptide intensity
85:217
30:95
3:13
rs1961993
p = 0.02, beta = −0.403, N = 118
ILYNEFNHYDTAAGK pc3
log10 peptide intensity
102:217
26:95
3:13
rs1961993
p = 0.075, beta = −0.314, N = 131
Locus: 40 (Sort: 48)
Status: Not replicated
80% power: No
SNP: 13:24891017:C:A
rsID: rs1961993
UniProtID: P0C862
HGNC gene: C1QTNF9
Type: Cis−pQTL
FPSSDVPIK pc2
rs3751357 ALT
log10 peptide intensity
158:805
27:403
0:52
13:24891017:C:A_C
p = 4.6e−12, model = REC, N = 185
FPSSDMPIK pc2
rs3751357 REF
log10 peptide intensity
70:805
30:403
1:52
13:24891017:C:A_C
p = 0.12, model = DOM, N = 101

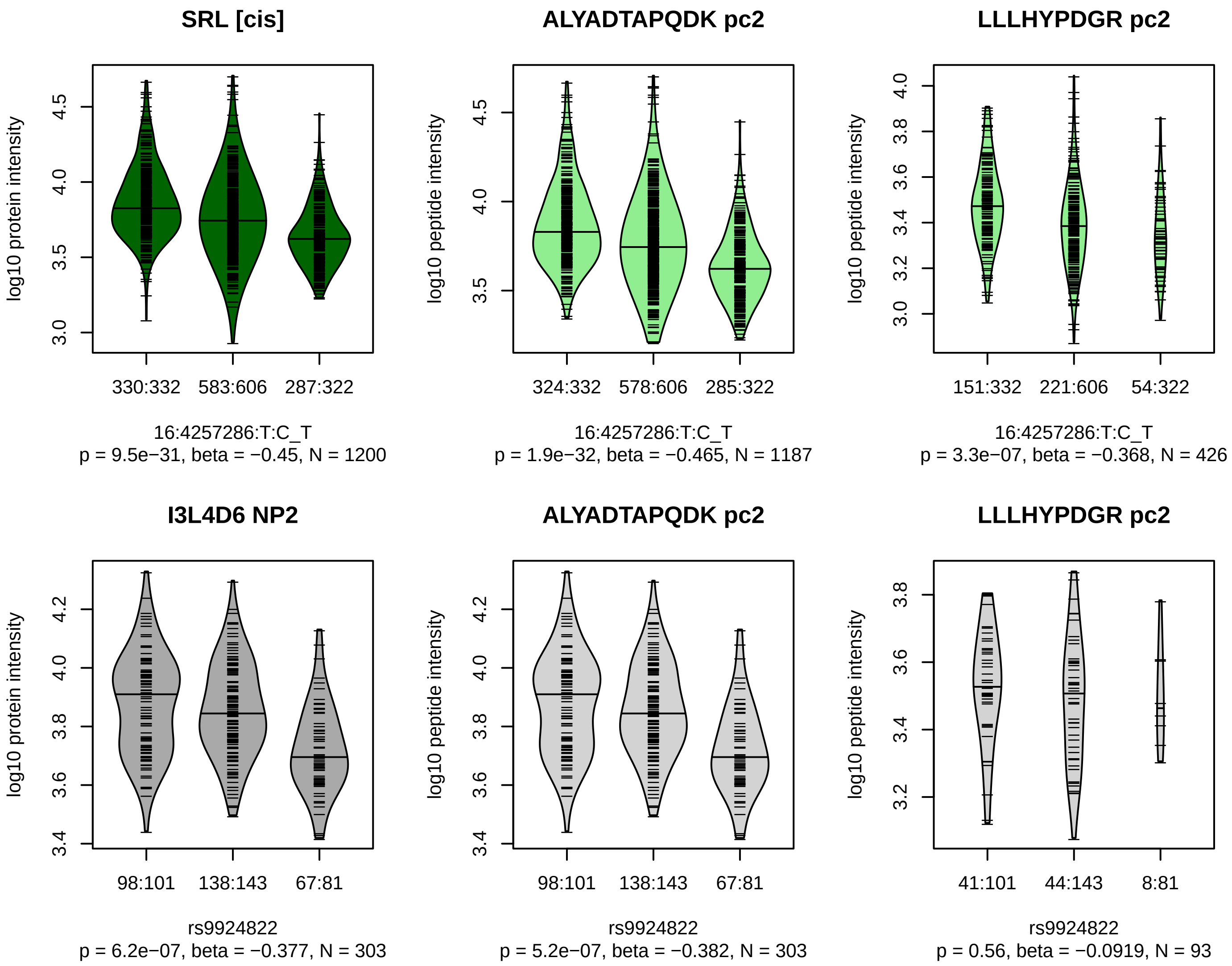


Locus: 41 (Sort: 49)

Status: Replicated

80% power: Yes

SNP: 16:4257286:T:C

rsID: rs9924822

UniProtID: I3L4D6

HGNC gene: SRL

Type: Cis−pQTL

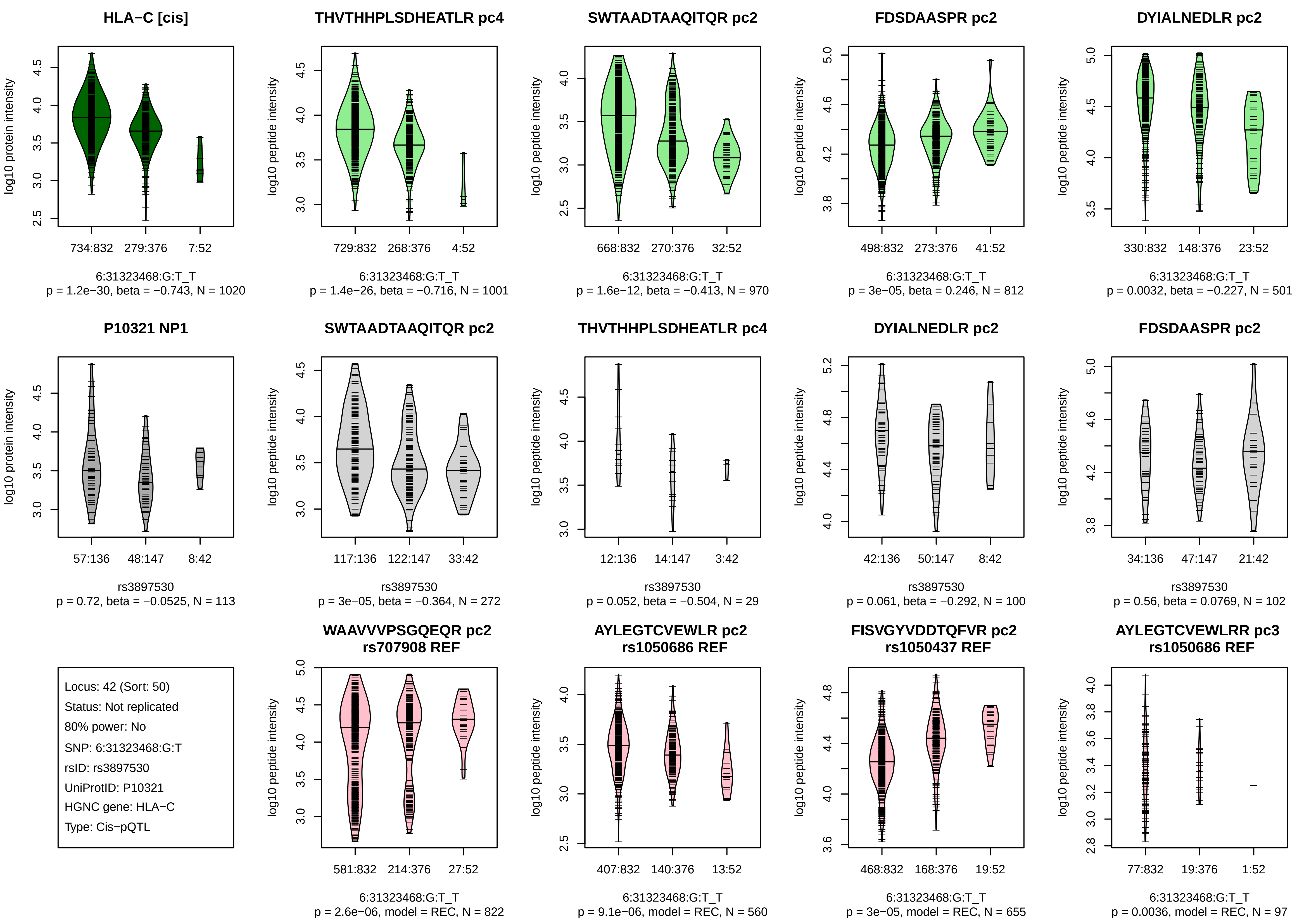
HLA−C [cis]
log10 protein intensity
734:832
279:376
7:52
6:31323468:G:T_T
p = 1.2e−30, beta = −0.743, N = 1020
THVTHHPLSDHEATLR pc4
log10 peptide intensity
729:832
268:376
4:52
6:31323468:G:T_T
p = 1.4e−26, beta = −0.716, N = 1001
SWTAADTAAQITQR pc2
log10 peptide intensity
668:832
270:376
32:52
6:31323468:G:T_T
p = 1.6e−12, beta = −0.413, N = 970
FDSDAASPR pc2
log10 peptide intensity
498:832
273:376
41:52
6:31323468:G:T_T
p = 3e−05, beta = 0.246, N = 812
DYIALNEDLR pc2
log10 peptide intensity
330:832
148:376
23:52
6:31323468:G:T_T
p = 0.0032, beta = −0.227, N = 501
P10321 NP1
log10 protein intensity
57:136
48:147
8:42
rs3897530
p = 0.72, beta = −0.0525, N = 113
SWTAADTAAQITQR pc2
log10 peptide intensity
117:136
122:147
33:42
rs3897530
p = 3e−05, beta = −0.364, N = 272
THVTHHPLSDHEATLR pc4
log10 peptide intensity
12:136
14:147
3:42
rs3897530
p = 0.052, beta = −0.504, N = 29
DYIALNEDLR pc2
log10 peptide intensity
42:136
50:147
8:42
rs3897530
p = 0.061, beta = −0.292, N = 100
FDSDAASPR pc2
log10 peptide intensity
34:136
47:147
21:42
rs3897530
p = 0.56, beta = 0.0769, N = 102
Locus: 42 (Sort: 50)
Status: Not replicated
80% power: No
SNP: 6:31323468:G:T
rsID: rs3897530
UniProtID: P10321
HGNC gene: HLA−C
Type: Cis−pQTL
WAAVVVPSGQEQR pc2
rs707908 REF
log10 peptide intensity
581:832
214:376
27:52
6:31323468:G:T_T
p = 2.6e−06, model = REC, N = 822
AYLEGTCVEWLR pc2
rs1050686 REF
log10 peptide intensity
407:832
140:376
13:52
6:31323468:G:T_T
p = 9.1e−06, model = REC, N = 560
FISVGYVDDTQFVR pc2
rs1050437 REF
log10 peptide intensity
468:832
168:376
19:52
6:31323468:G:T_T
p = 3e−05, model = REC, N = 655
AYLEGTCVEWLRR pc3
rs1050686 REF
log10 peptide intensity
77:832
19:376
1:52
6:31323468:G:T_T
p = 0.0036, model = REC, N = 97

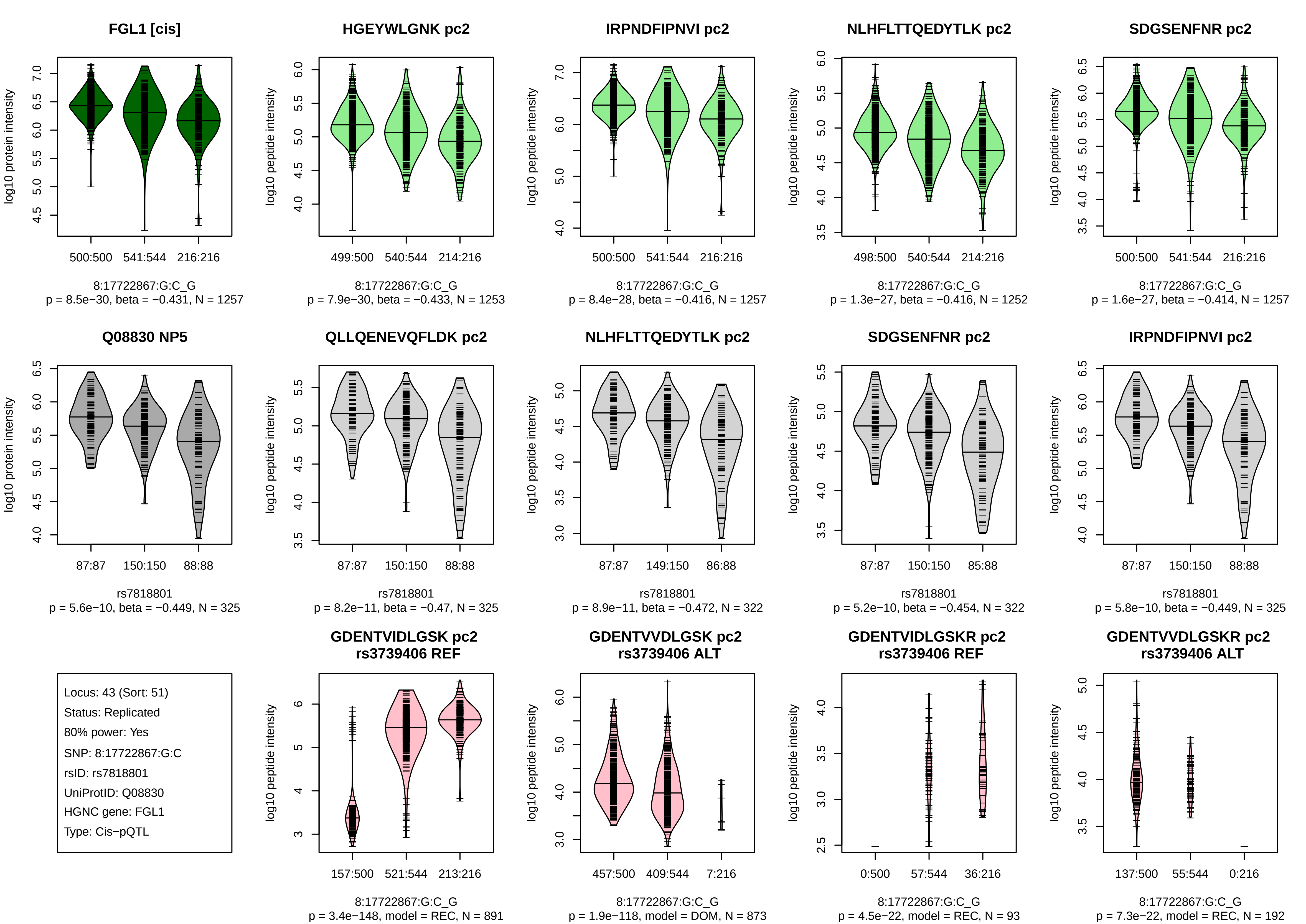

FGL1 [cis]
log10 protein intensity
500:500
541:544
216:216
8:17722867:G:C_G
p = 8.5e−30, beta = −0.431, N = 1257
HGEYWLGNK pc2
log10 peptide intensity
499:500
540:544
214:216
8:17722867:G:C_G
p = 7.9e−30, beta = −0.433, N = 1253
IRPNDFIPNVI pc2
log10 peptide intensity
500:500
541:544
216:216
8:17722867:G:C_G
p = 8.4e−28, beta = −0.416, N = 1257
NLHFLTTQEDYTLK pc2
log10 peptide intensity
498:500
540:544
214:216
8:17722867:G:C_G
p = 1.3e−27, beta = −0.416, N = 1252
SDGSENFNR pc2
log10 peptide intensity
500:500
541:544
216:216
8:17722867:G:C_G
p = 1.6e−27, beta = −0.414, N = 1257
Q08830 NP5
log10 protein intensity
87:87
150:150
88:88
rs7818801
p = 5.6e−10, beta = −0.449, N = 325
QLLQENEVQFLDK pc2
log10 peptide intensity
87:87
150:150
88:88
rs7818801
p = 8.2e−11, beta = −0.47, N = 325
NLHFLTTQEDYTLK pc2
log10 peptide intensity
87:87
149:150
86:88
rs7818801
p = 8.9e−11, beta = −0.472, N = 322
SDGSENFNR pc2
log10 peptide intensity
87:87
150:150
85:88
rs7818801
p = 5.2e−10, beta = −0.454, N = 322
IRPNDFIPNVI pc2
log10 peptide intensity
87:87
150:150
88:88
rs7818801
p = 5.8e−10, beta = −0.449, N = 325
Locus: 43 (Sort: 51)
Status: Replicated
80% power: Yes
SNP: 8:17722867:G:C
rsID: rs7818801
UniProtID: Q08830
HGNC gene: FGL1
Type: Cis−pQTL
GDENTVIDLGSK pc2
rs3739406 REF
log10 peptide intensity
157:500
521:544
213:216
8:17722867:G:C_G
p = 3.4e−148, model = REC, N = 891
GDENTVVDLGSK pc2
rs3739406 ALT
log10 peptide intensity
457:500
409:544
7:216
8:17722867:G:C_G
p = 1.9e−118, model = DOM, N = 873
GDENTVIDLGSKR pc2
rs3739406 REF
log10 peptide intensity
0:500
57:544
36:216
8:17722867:G:C_G
p = 4.5e−22, model = REC, N = 93
GDENTVVDLGSKR pc2
rs3739406 ALT
log10 peptide intensity
137:500
55:544
0:216
8:17722867:G:C_G
p = 7.3e−22, model = REC, N = 192

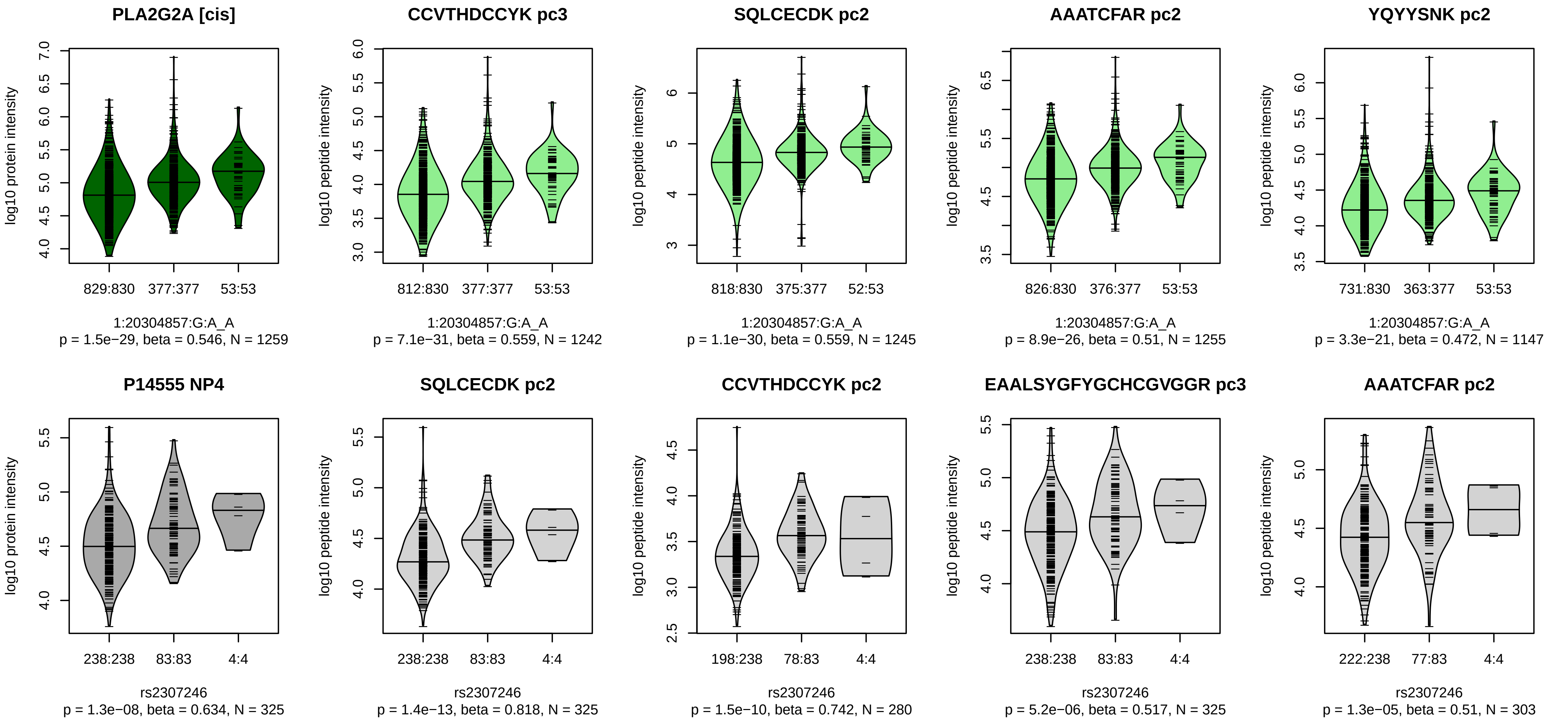


Locus: 44 (Sort: 52)
Status: Replicated
80% power: Yes
SNP: 1:20304857:G:A
rsID: rs2307246
UniProtID: P14555
HGNC gene: PLA2G2A
Type: Cis−pQTL

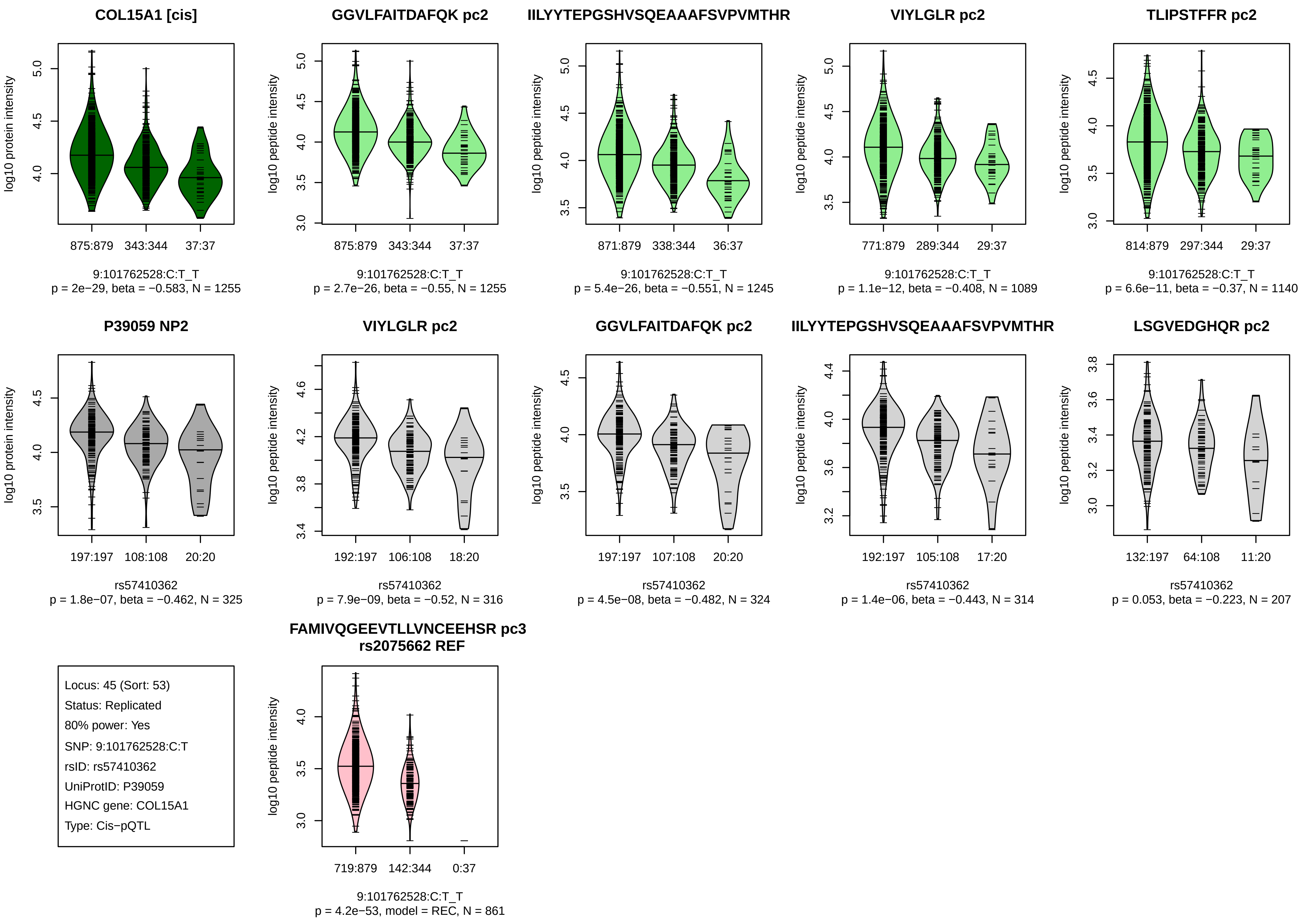
COL15A1 [cis]
log10 protein intensity
875:879
343:344
37:37
9:101762528:C:T_T
p = 2e−29, beta = −0.583, N = 1255
GGVLFAITDAFQK pc2
log10 peptide intensity
875:879
343:344
37:37
9:101762528:C:T_T
p = 2.7e−26, beta = −0.55, N = 1255
IILYYTEPGSHVSQEAAAFSVPVMTHR
log10 peptide intensity
871:879
338:344
36:37
9:101762528:C:T_T
p = 5.4e−26, beta = −0.551, N = 1245
VIYLGLR pc2
log10 peptide intensity
771:879
289:344
29:37
9:101762528:C:T_T
p = 1.1e−12, beta = −0.408, N = 1089
TLIPSTFFR pc2
log10 peptide intensity
814:879
297:344
29:37
9:101762528:C:T_T
p = 6.6e−11, beta = −0.37, N = 1140
P39059 NP2
log10 protein intensity
197:197
108:108
20:20
rs57410362
p = 1.8e−07, beta = −0.462, N = 325
VIYLGLR pc2
log10 peptide intensity
192:197
106:108
18:20
rs57410362
p = 7.9e−09, beta = −0.52, N = 316
GGVLFAITDAFQK pc2
log10 peptide intensity
197:197
107:108
20:20
rs57410362
p = 4.5e−08, beta = −0.482, N = 324
IILYYTEPGSHVSQEAAAFSVPVMTHR
log10 peptide intensity
192:197
105:108
17:20
rs57410362
p = 1.4e−06, beta = −0.443, N = 314
LSGVEDGHQR pc2
log10 peptide intensity
132:197
64:108
11:20
rs57410362
p = 0.053, beta = −0.223, N = 207
Locus: 45 (Sort: 53)
Status: Replicated
80% power: Yes
SNP: 9:101762528:C:T
rsID: rs57410362
UniProtID: P39059
HGNC gene: COL15A1
Type: Cis−pQTL
FAMIVQGEEVTLLVNCEEHSR pc3
rs2075662 REF
log10 peptide intensity
719:879
142:344
0:37
9:101762528:C:T_T
p = 4.2e−53, model = REC, N = 861

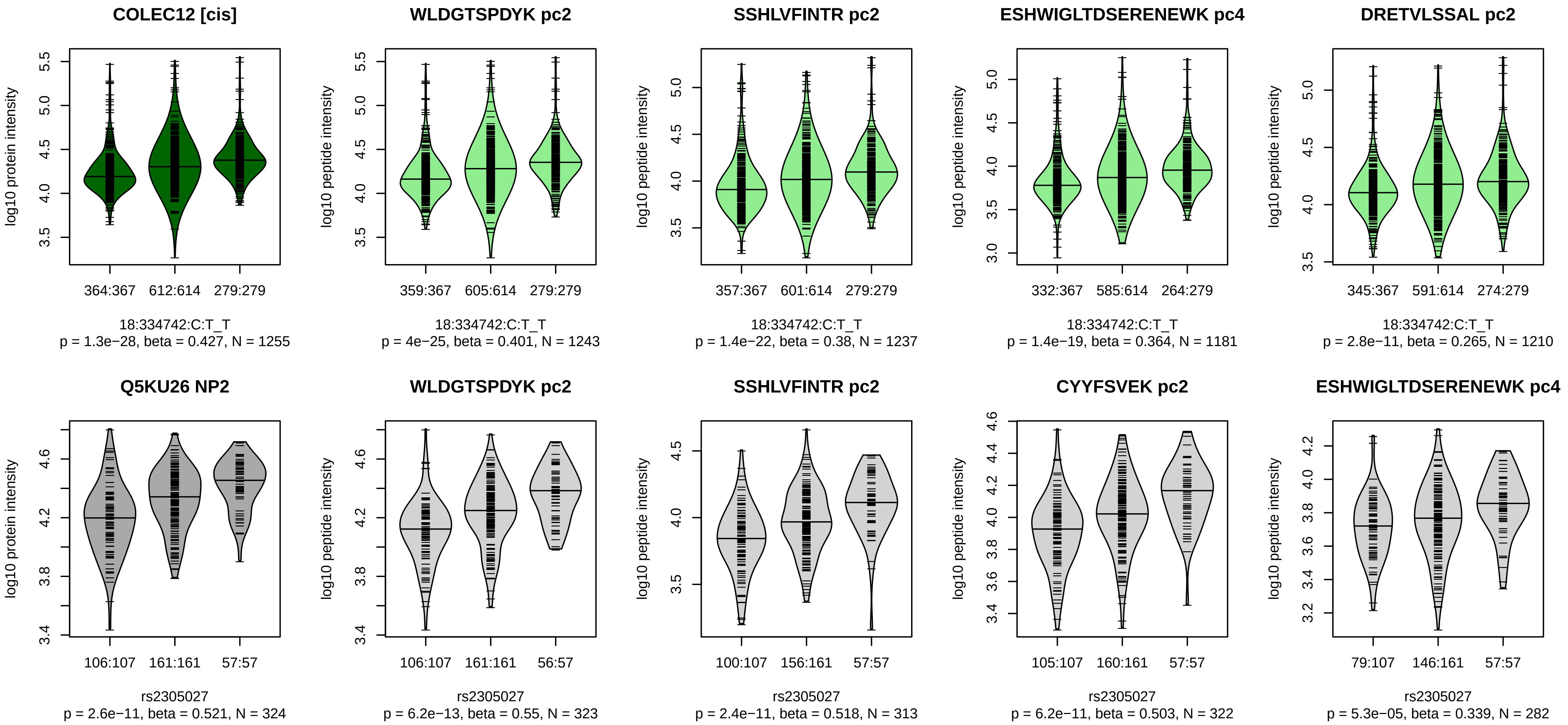


Locus: 46 (Sort: 54)
Status: Replicated
80% power: Yes
SNP: 18:334742:C:T
rsID: rs2305027
UniProtID: Q5KU26
HGNC gene: COLEC12
Type: Cis−pQTL

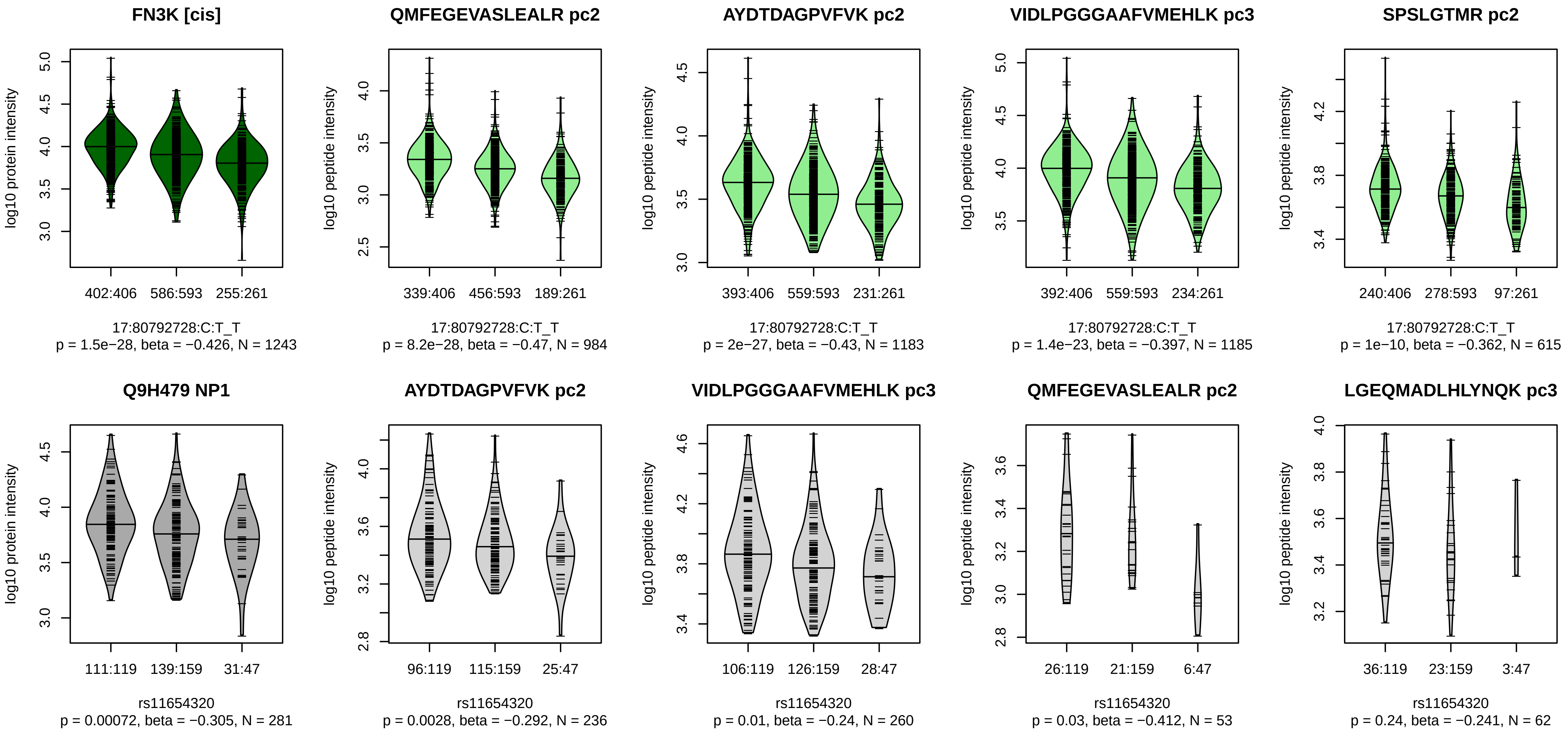


Locus: 47 (Sort: 55)
Status: Not replicated
80% power: Yes
SNP: 17:80792728:C:T
rsID: rs11654320
UniProtID: Q9H479
HGNC gene: FN3K
Type: Cis−pQTL

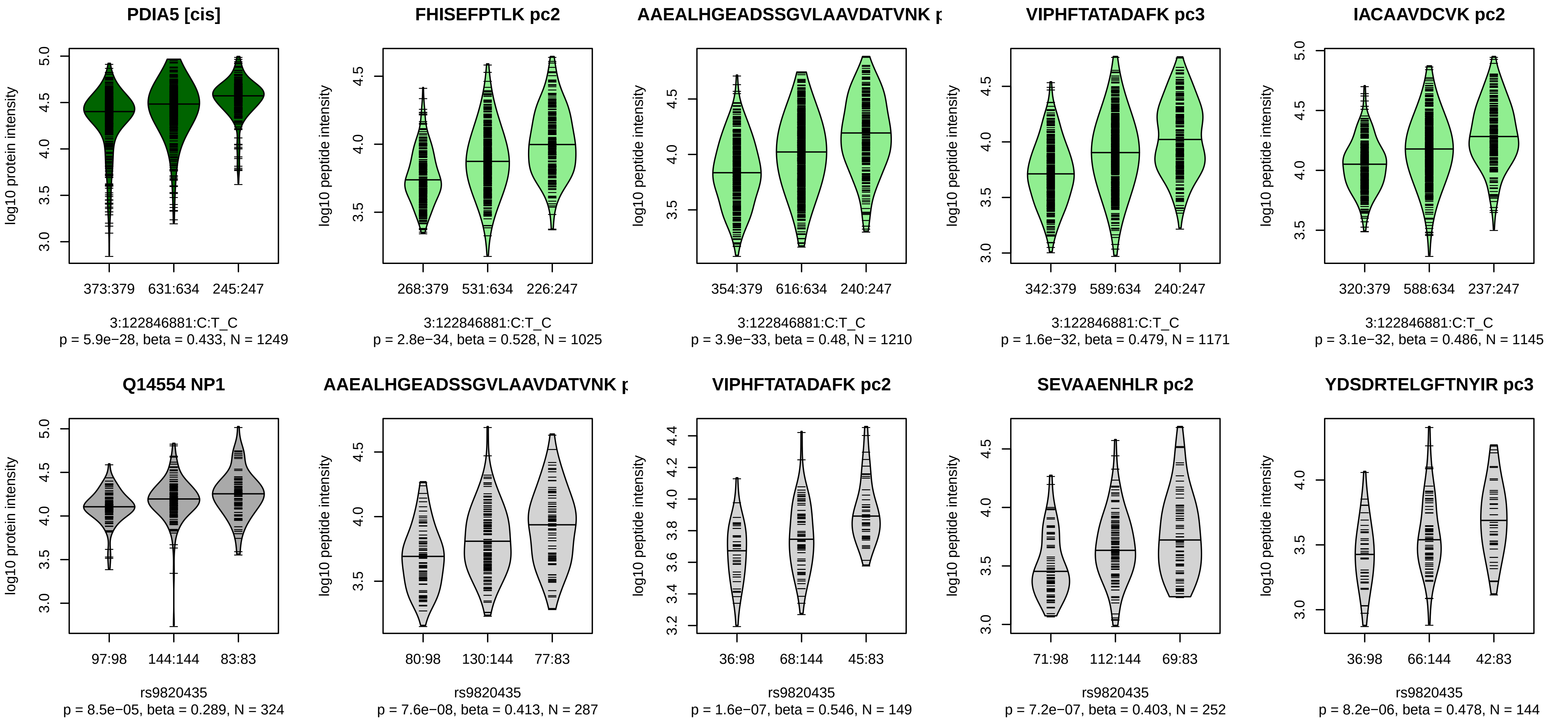


Locus: 48 (Sort: 56)

Status: Replicated

80% power: Yes

SNP: 3:122846881:C:T

rsID: rs9820435

UniProtID: Q14554

HGNC gene: PDIA5

Type: Cis−pQTL

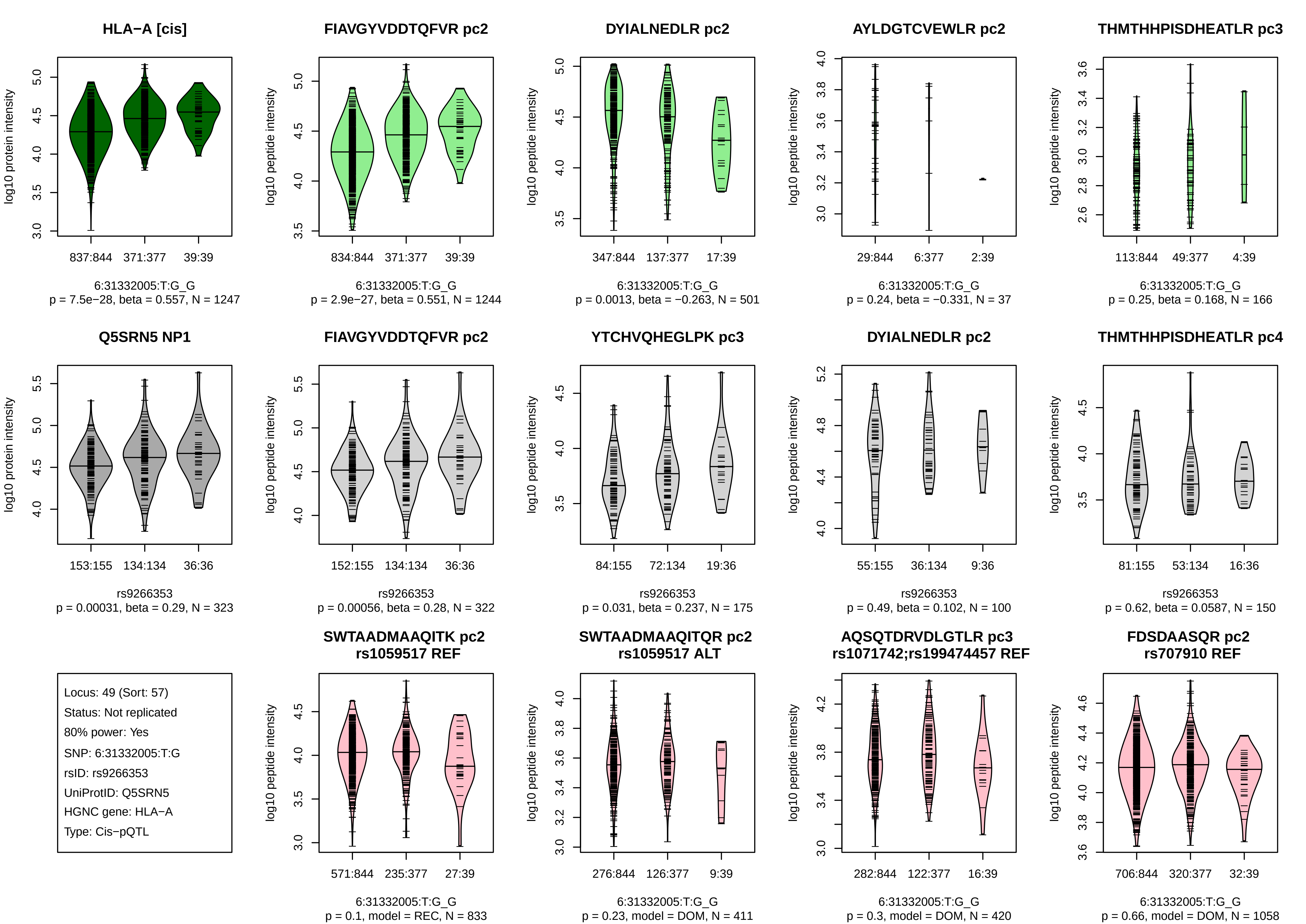
HLA−A [cis]
log10 protein intensity
837:844
371:377
39:39
6:31332005:T:G_G
p = 7.5e−28, beta = 0.557, N = 1247
FIAVGYVDDTQFVR pc2
log10 peptide intensity
834:844
371:377
39:39
6:31332005:T:G_G
p = 2.9e−27, beta = 0.551, N = 1244
DYIALNEDLR pc2
log10 peptide intensity
347:844
137:377
17:39
6:31332005:T:G_G
p = 0.0013, beta = −0.263, N = 501
AYLDGTCVEWLR pc2
log10 peptide intensity
29:844
6:377
2:39
6:31332005:T:G_G
p = 0.24, beta = −0.331, N = 37
THMTHHPISDHEATLR pc3
log10 peptide intensity
113:844
49:377
4:39
6:31332005:T:G_G
p = 0.25, beta = 0.168, N = 166
Q5SRN5 NP1
log10 protein intensity
153:155
134:134
36:36
rs9266353
p = 0.00031, beta = 0.29, N = 323
FIAVGYVDDTQFVR pc2
log10 peptide intensity
152:155
134:134
36:36
rs9266353
p = 0.00056, beta = 0.28, N = 322
YTCHVQHEGLPK pc3
log10 peptide intensity
84:155
72:134
19:36
rs9266353
p = 0.031, beta = 0.237, N = 175
DYIALNEDLR pc2
log10 peptide intensity
55:155
36:134
9:36
rs9266353
p = 0.49, beta = 0.102, N = 100
THMTHHPISDHEATLR pc4
log10 peptide intensity
81:155
53:134
16:36
rs9266353
p = 0.62, beta = 0.0587, N = 150
Locus: 49 (Sort: 57)
Status: Not replicated
80% power: Yes
SNP: 6:31332005:T:G
rsID: rs9266353
UniProtID: Q5SRN5
HGNC gene: HLA−A
Type: Cis−pQTL
SWTAADMAAQITK pc2
rs1059517 REF
log10 peptide intensity
571:844
235:377
27:39
6:31332005:T:G_G
p = 0.1, model = REC, N = 833
SWTAADMAAQITQR pc2
rs1059517 ALT
log10 peptide intensity
276:844
126:377
9:39
6:31332005:T:G_G
p = 0.23, model = DOM, N = 411
AQSQTDRVDLGTLR pc3
rs1071742;rs199474457 REF
log10 peptide intensity
282:844
122:377
16:39
6:31332005:T:G_G
p = 0.3, model = DOM, N = 420
FDSDAASQR pc2
rs707910 REF
log10 peptide intensity
706:844
320:377
32:39
6:31332005:T:G_G
p = 0.66, model = DOM, N = 1058

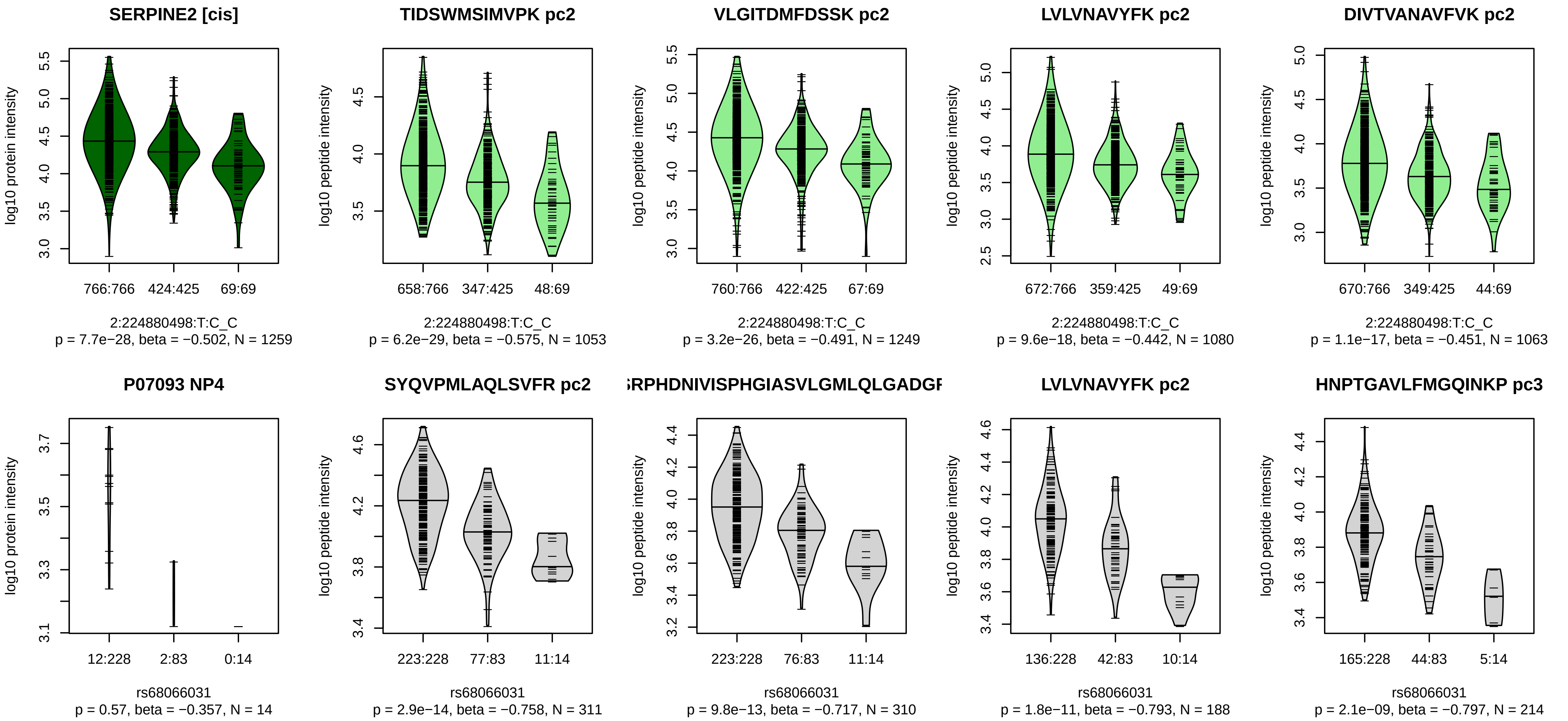


Locus: 50 (Sort: 58)
Status: Not replicated
80% power: No
SNP: 2:224880498:T:C
rsID: rs68066031
UniProtID: P07093
HGNC gene: SERPINE2
Type: Cis−pQTL

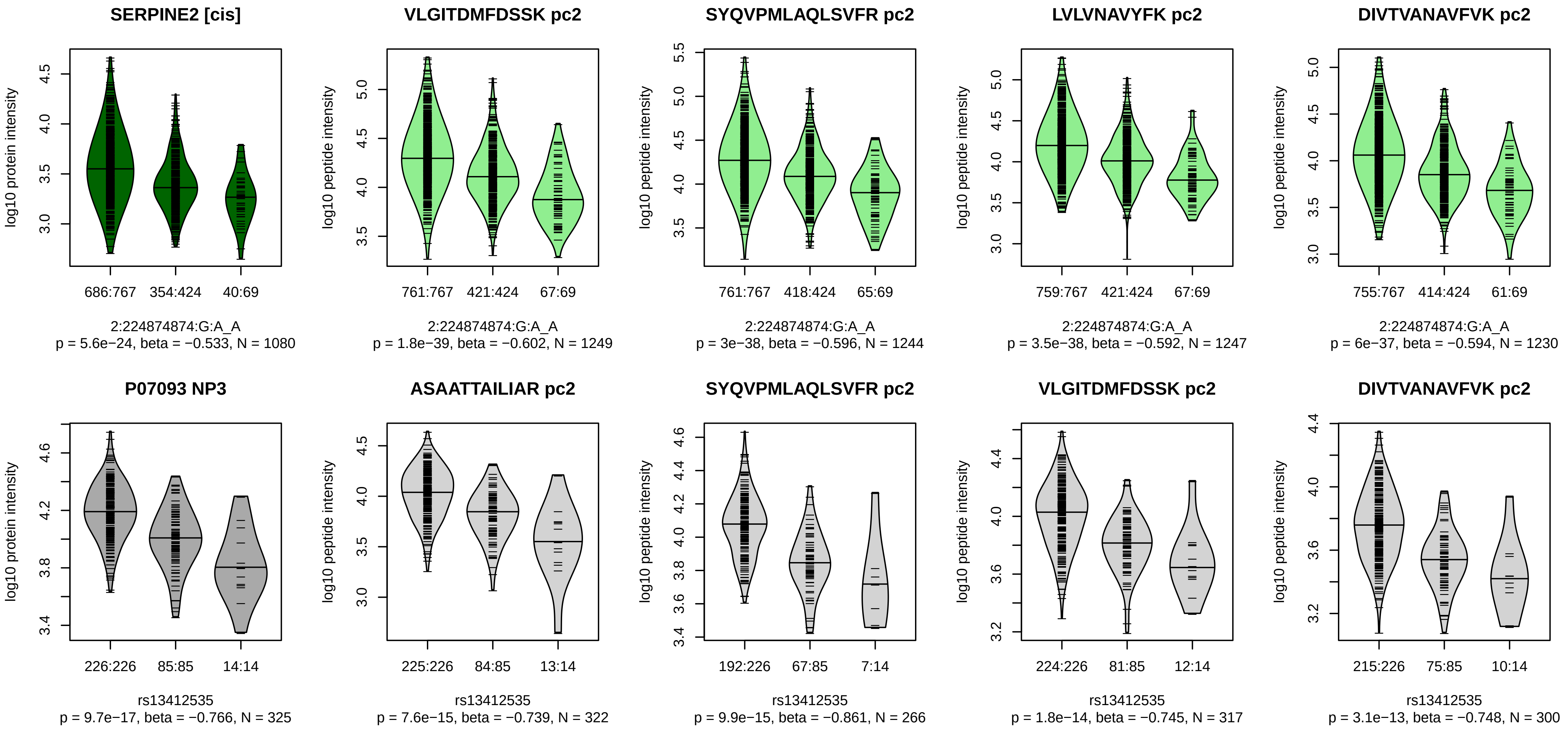


Locus: 50 (Sort: 59)

Status: Replicated

80% power: Yes

SNP: 2:224874874:G:A

rsID: rs13412535

UniProtID: P07093

HGNC gene: SERPINE2

Type: Cis−pQTL

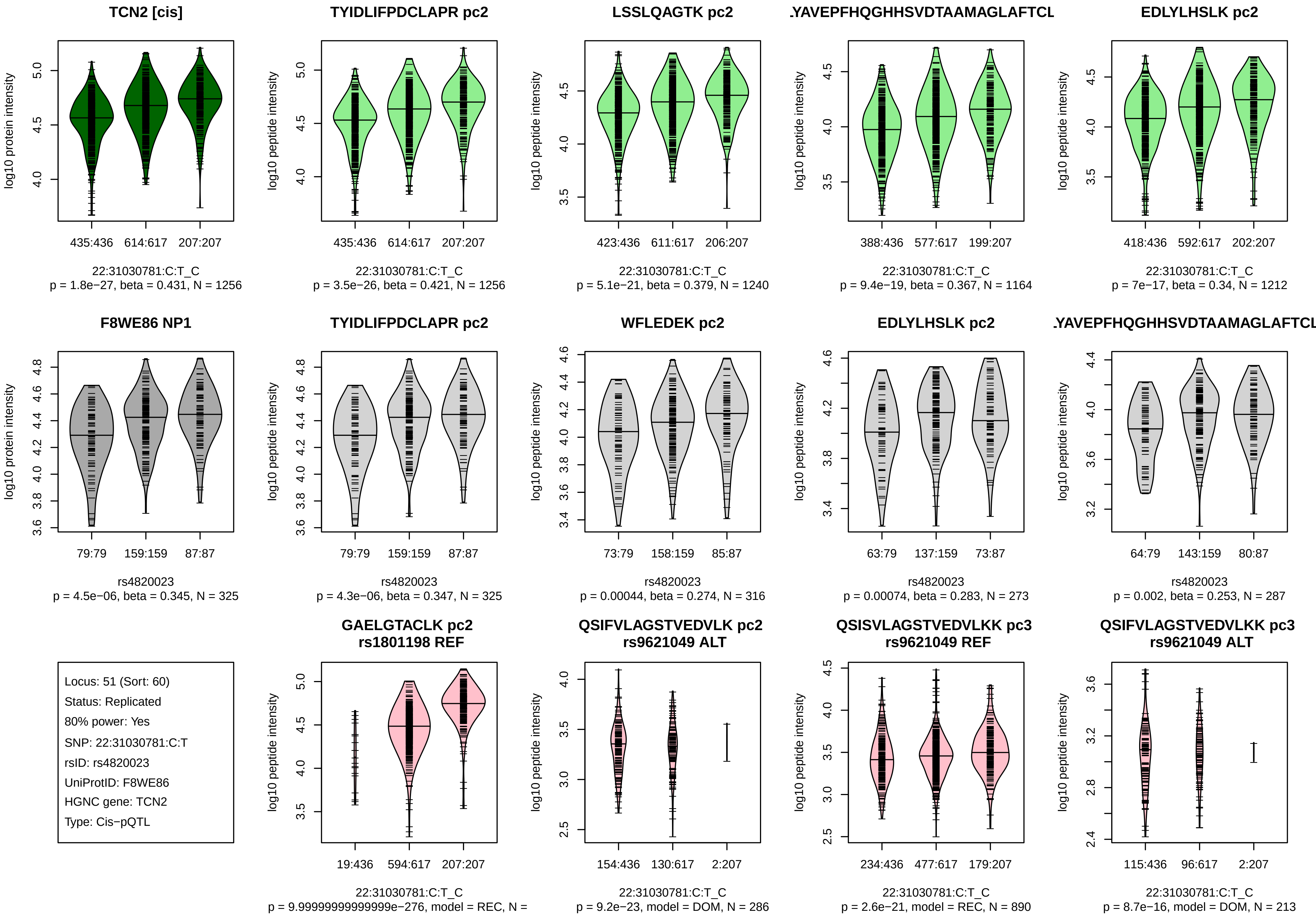
TCN2 [cis]
log10 protein intensity
435:436
614:617
207:207
22:31030781:C:T_C
p = 1.8e−27, beta = 0.431, N = 1256
TYIDLIFPDCLAPR pc2
log10 peptide intensity
435:436
614:617
207:207
22:31030781:C:T_C
p = 3.5e−26, beta = 0.421, N = 1256
LSSLQAGTK pc2
log10 peptide intensity
423:436
611:617
206:207
22:31030781:C:T_C
p = 5.1e−21, beta = 0.379, N = 1240
.YAVEPFHQGHHSVDTAAMAGLAFTCL
log10 peptide intensity
388:436
577:617
199:207
22:31030781:C:T_C
p = 9.4e−19, beta = 0.367, N = 1164
EDLYLHSLK pc2
log10 peptide intensity
418:436
592:617
202:207
22:31030781:C:T_C
p = 7e−17, beta = 0.34, N = 1212
F8WE86 NP1
log10 protein intensity
79:79
159:159
87:87
rs4820023
p = 4.5e−06, beta = 0.345, N = 325
TYIDLIFPDCLAPR pc2
log10 peptide intensity
79:79
159:159
87:87
rs4820023
p = 4.3e−06, beta = 0.347, N = 325
WFLEDEK pc2
log10 peptide intensity
73:79
158:159
85:87
rs4820023
p = 0.00044, beta = 0.274, N = 316
EDLYLHSLK pc2
log10 peptide intensity
63:79
137:159
73:87
rs4820023
p = 0.00074, beta = 0.283, N = 273
.YAVEPFHQGHHSVDTAAMAGLAFTCL
log10 peptide intensity
64:79
143:159
80:87
rs4820023
p = 0.002, beta = 0.253, N = 287
Locus: 51 (Sort: 60)
Status: Replicated
80% power: Yes
SNP: 22:31030781:C:T
rsID: rs4820023
UniProtID: F8WE86
HGNC gene: TCN2
Type: Cis−pQTL
GAELGTACLK pc2
rs1801198 REF
log10 peptide intensity
19:436
594:617
207:207
22:31030781:C:T_C
p = 9.99999999999999e−276, model = REC, N =
QSIFVLAGSTVEDVLK pc2
rs9621049 ALT
log10 peptide intensity
154:436
130:617
2:207
22:31030781:C:T_C
p = 9.2e−23, model = DOM, N = 286
QSISVLAGSTVEDVLKK pc3
rs9621049 REF
log10 peptide intensity
234:436
477:617
179:207
22:31030781:C:T_C
p = 2.6e−21, model = REC, N = 890
QSIFVLAGSTVEDVLKK pc3
rs9621049 ALT
log10 peptide intensity
115:436
96:617
2:207
22:31030781:C:T_C
p = 8.7e−16, model = DOM, N = 213

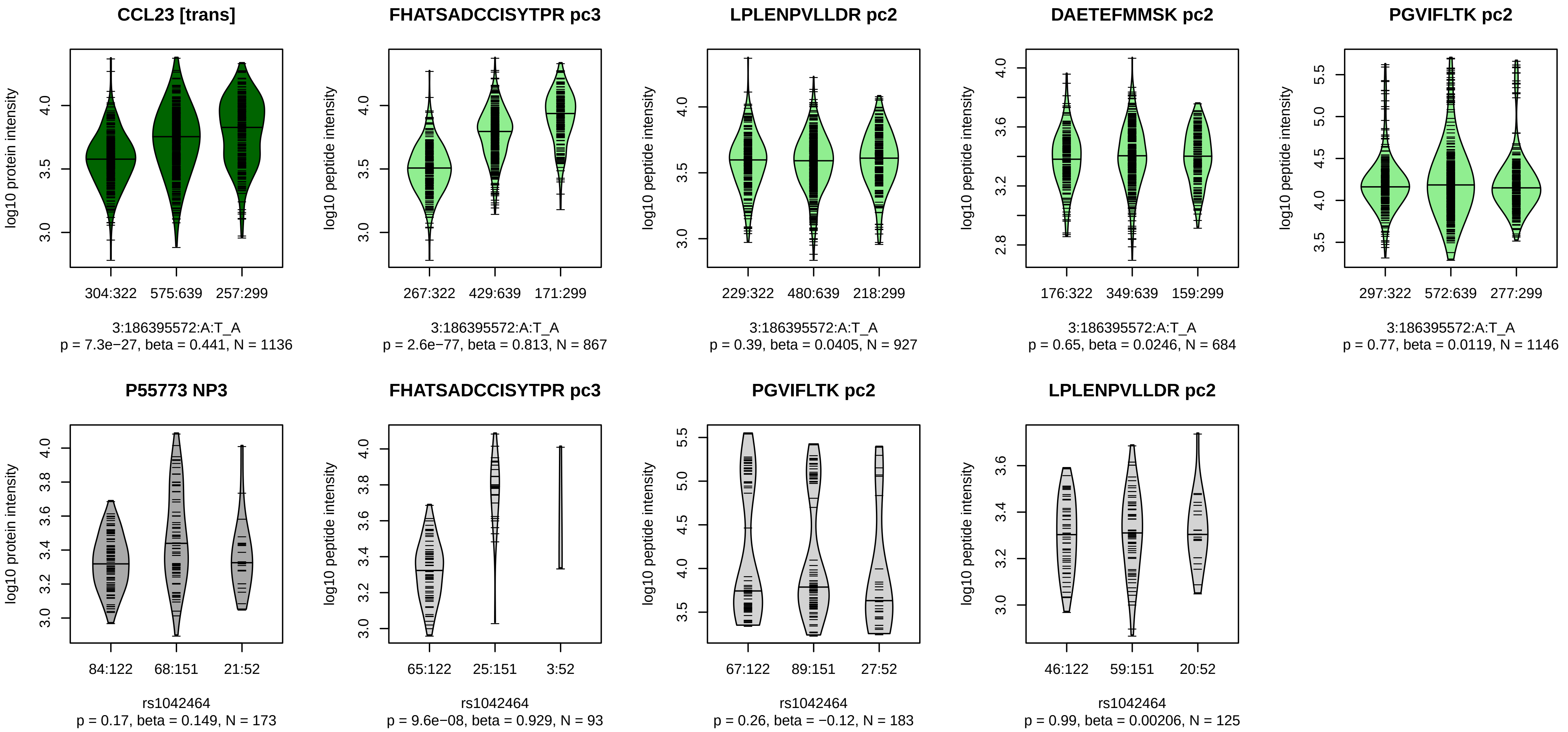


Locus: 52 (Sort: 61)

Status: Not replicated

80% power: No

SNP: 3:186395572:A:T

rsID: rs1042464

UniProtID: P55773

HGNC gene: CCL23

Type: Trans−pQTL

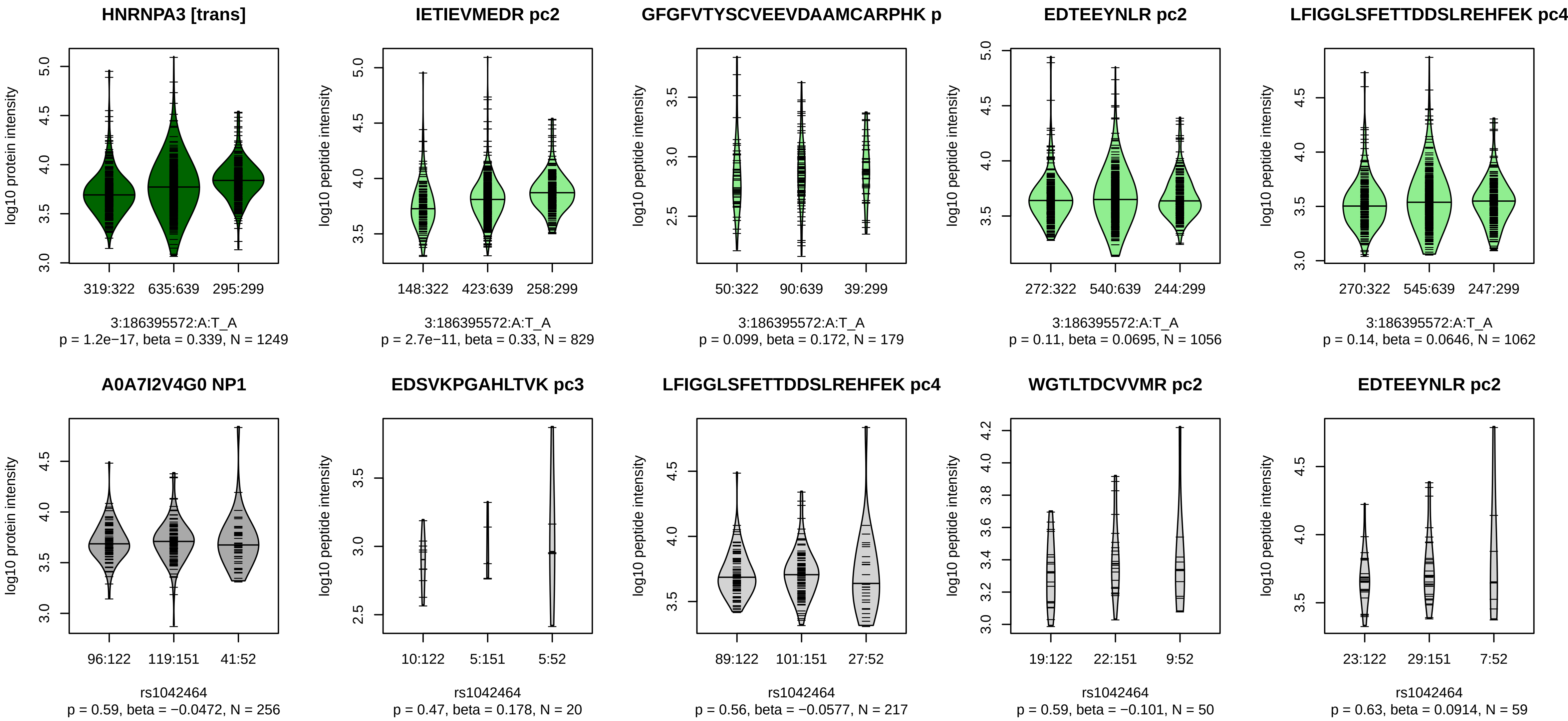


Locus: 52 (Sort: 62)

Status: Not replicated

80% power: No

SNP: 3:186395572:A:T

rsID: rs1042464

UniProtID: A0A7I2V4G0

HGNC gene: HNRNPA3

Type: Trans−pQTL

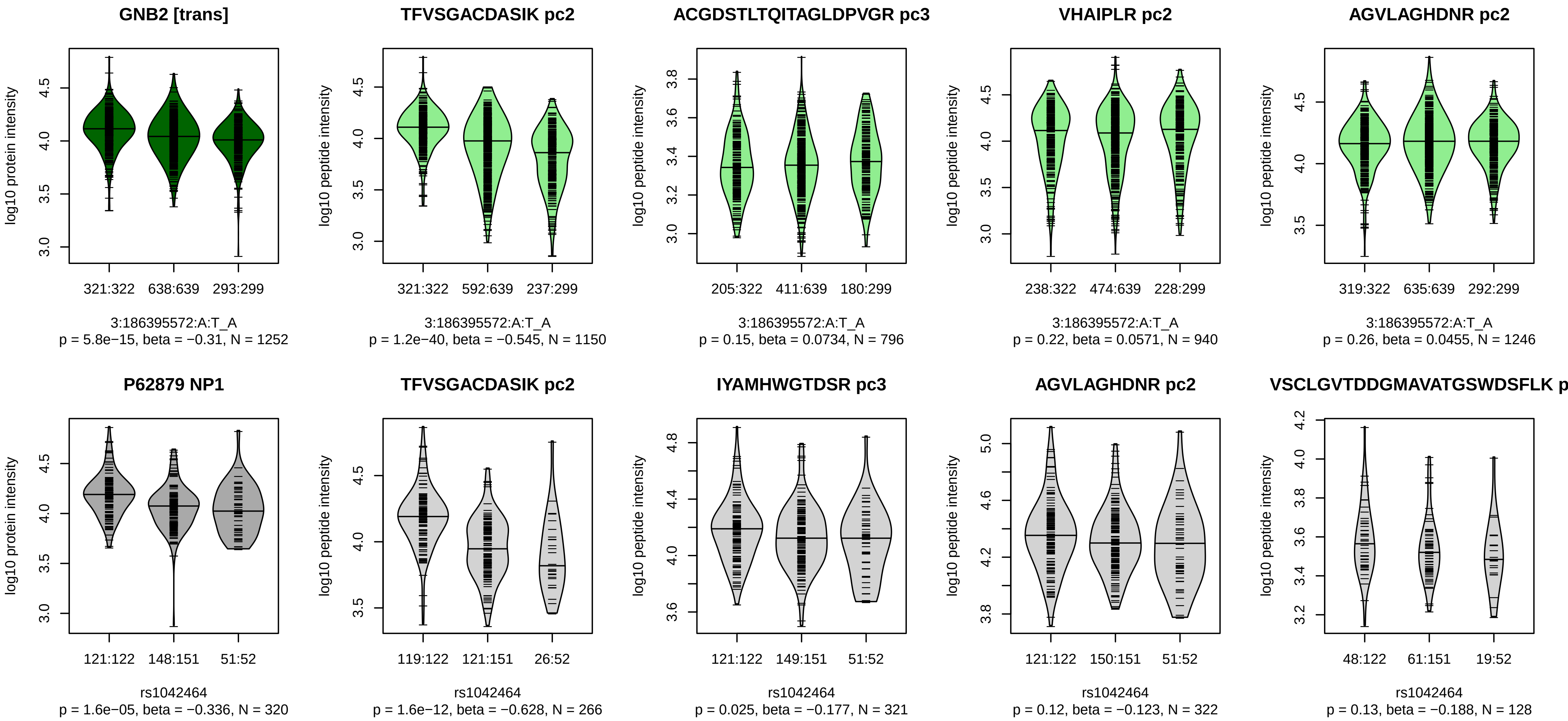


Locus: 52 (Sort: 63)

Status: Replicated

80% power: No

SNP: 3:186395572:A:T

rsID: rs1042464

UniProtID: P62879

HGNC gene: GNB2

Type: Trans−pQTL

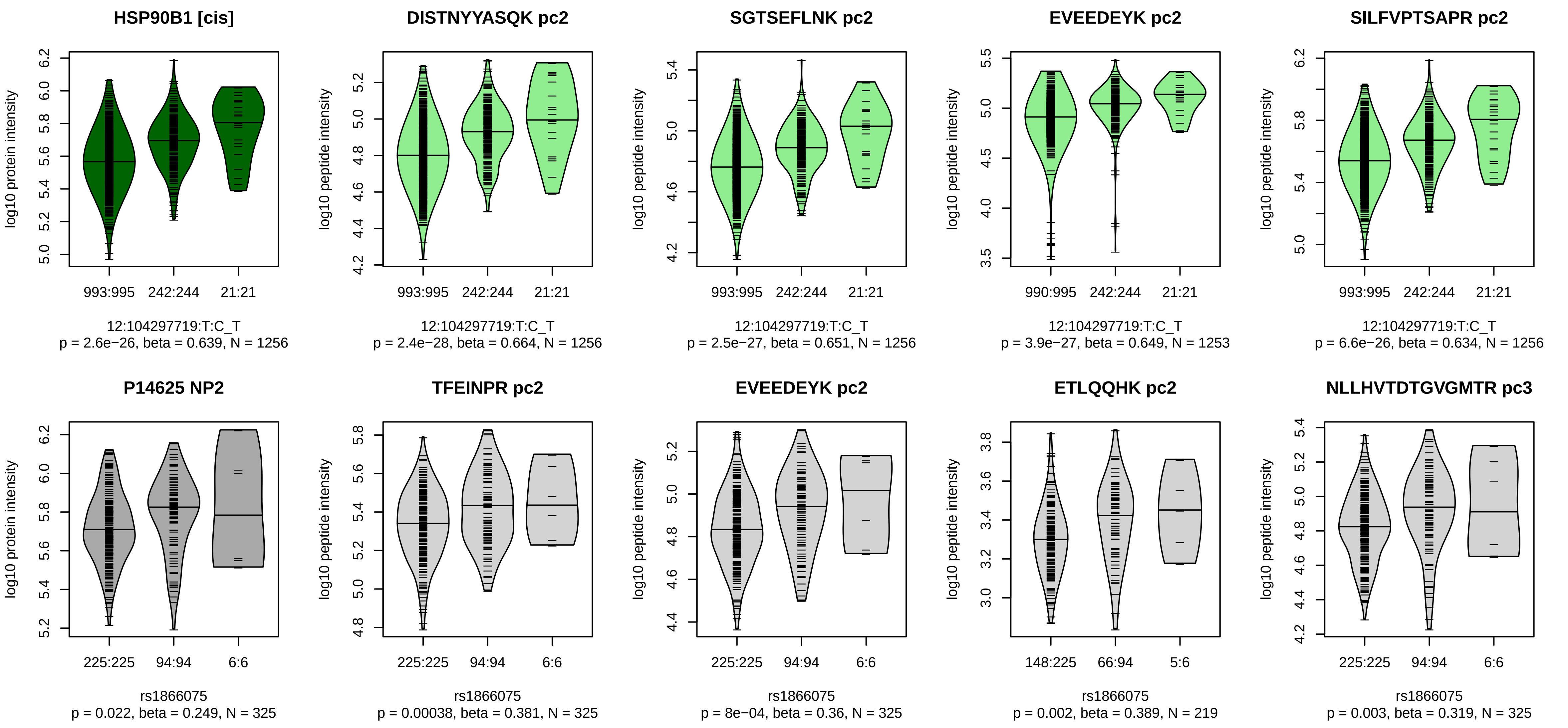


Locus: 53 (Sort: 64)
Status: Not replicated
80% power: Yes
SNP: 12:104297719:T:C
rsID: rs1866075
UniProtID: P14625
HGNC gene: HSP90B1
Type: Cis−pQTL

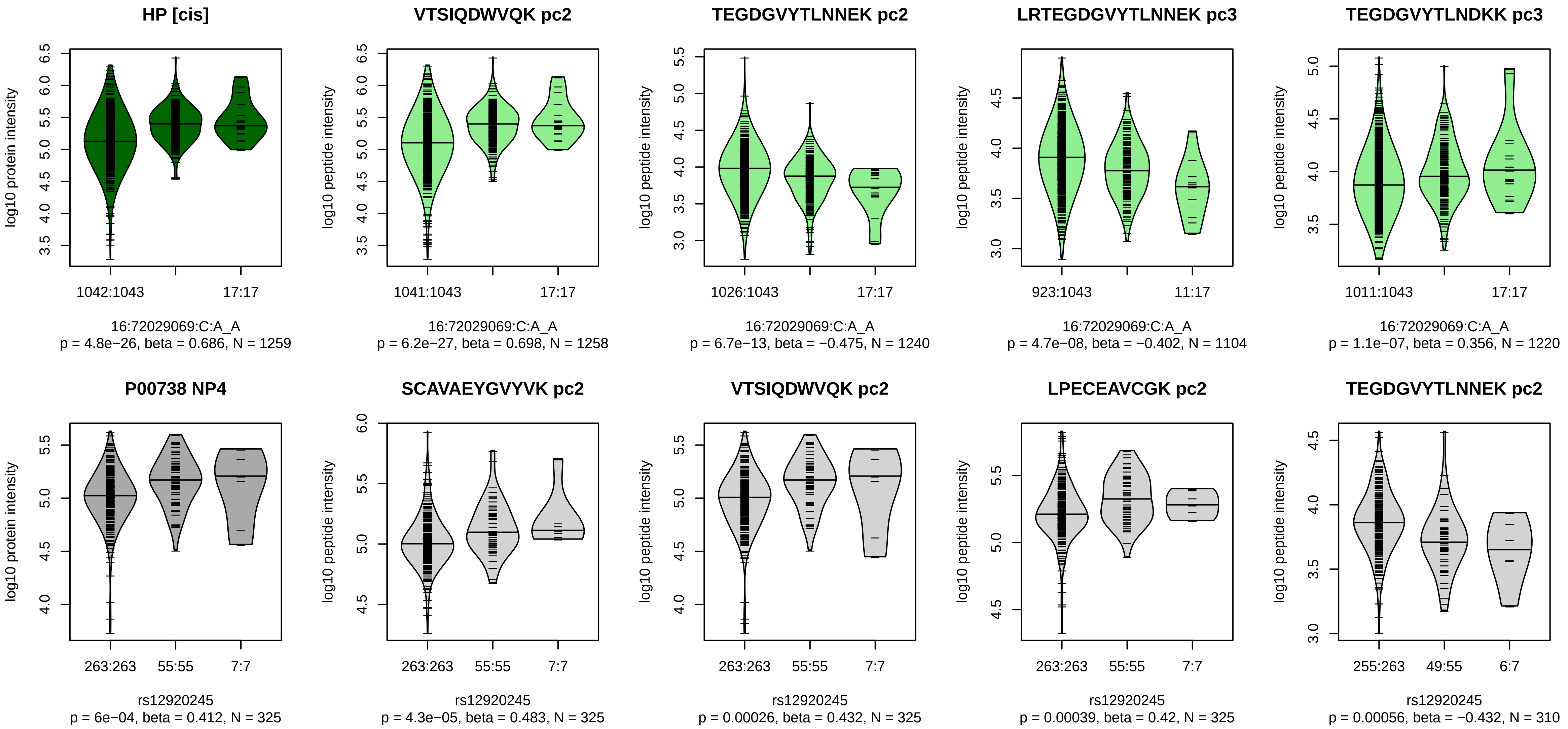


Locus: 54 (Sort: 65)
Status: Not replicated
80% power: Yes
SNP: 16:72029069:C:A
rsID: rs12920245
UniProtID: P00738
HGNC gene: HP
Type: Cis−pQTL

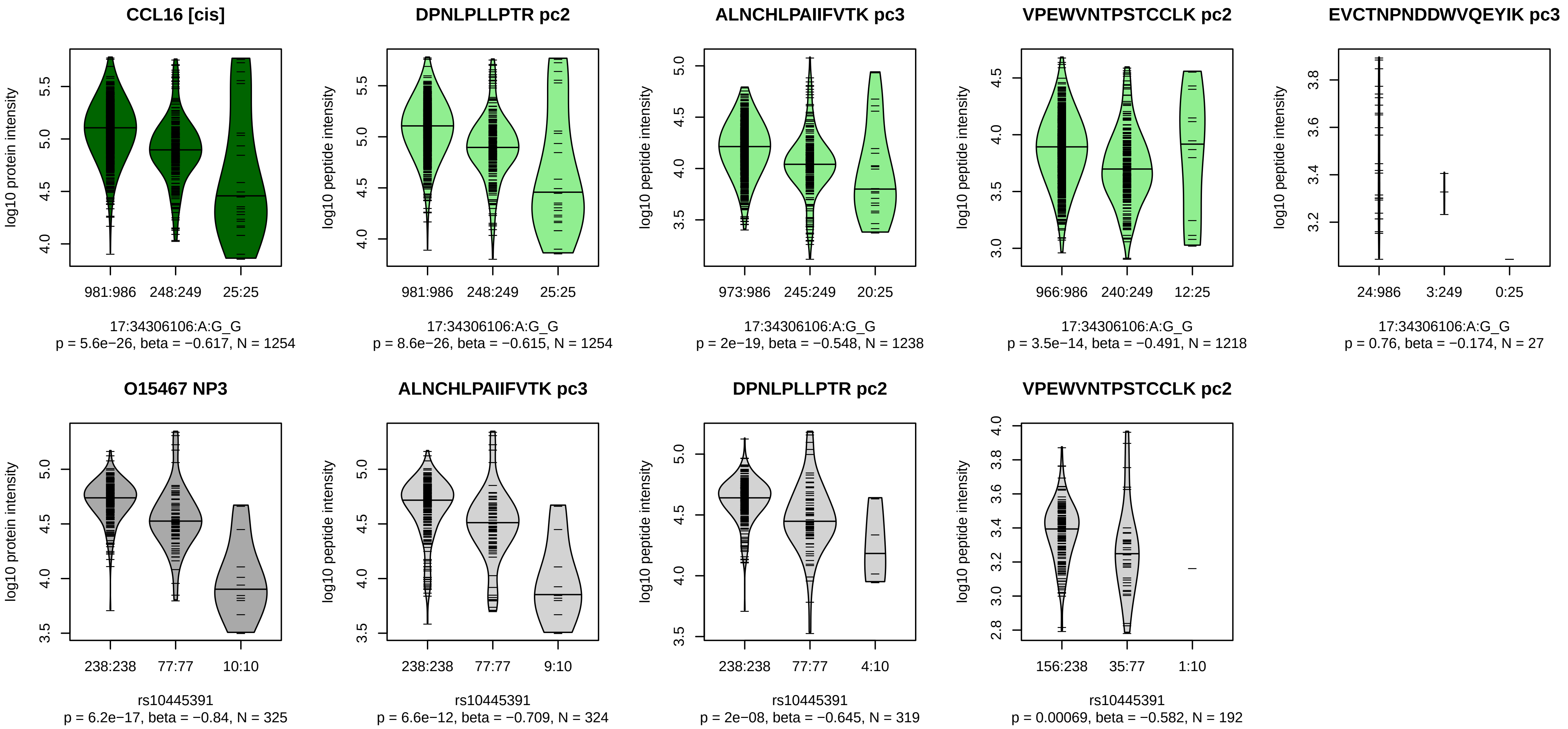


Locus: 55 (Sort: 66)

Status: Replicated

80% power: Yes

SNP: 17:34306106:A:G

rsID: rs10445391

UniProtID: O15467

HGNC gene: CCL16

Type: Cis−pQTL

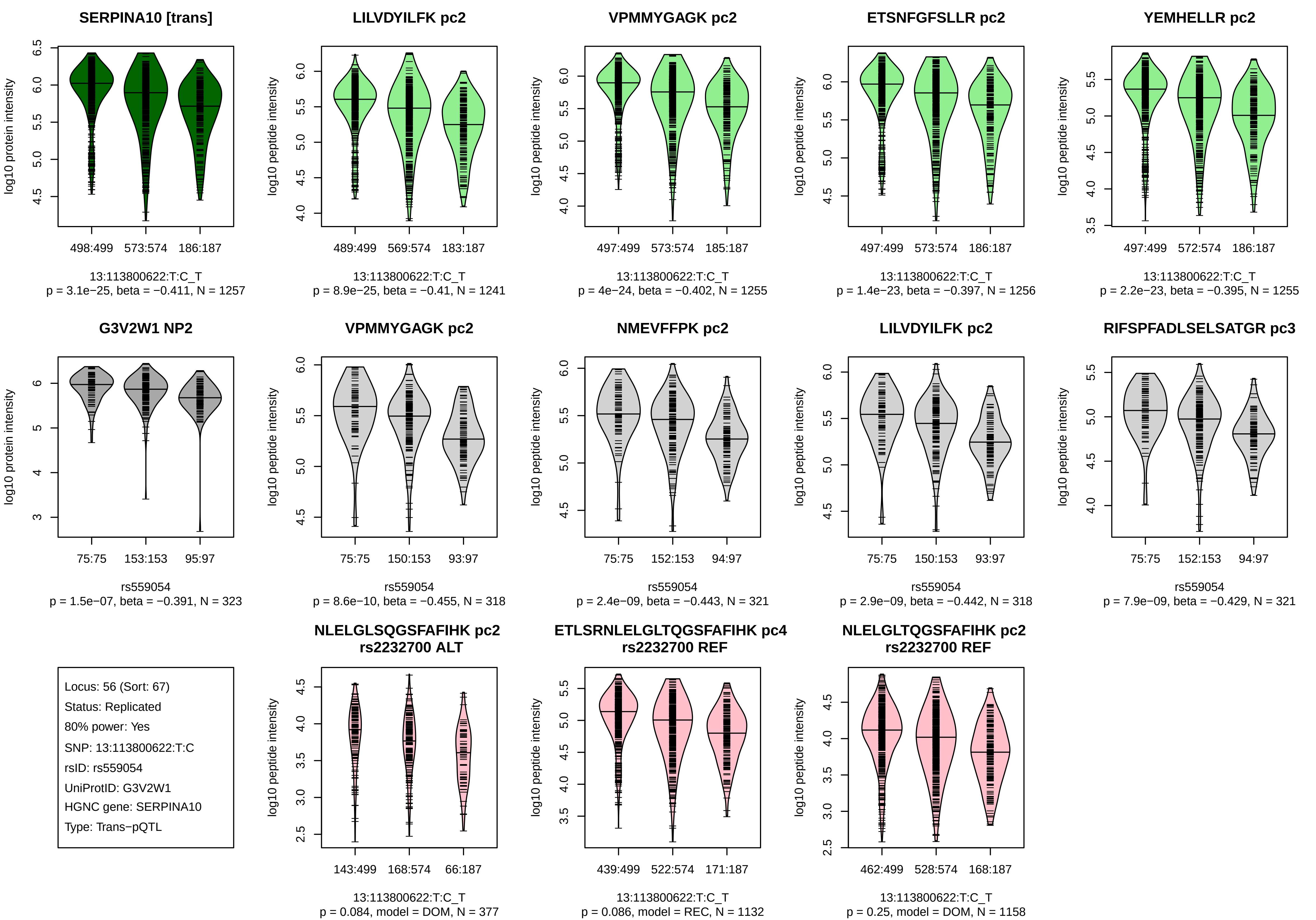
SERPINA10 [trans]
log10 protein intensity
498:499
573:574
186:187
13:113800622:T:C_T
p = 3.1e−25, beta = −0.411, N = 1257
LILVDYILFK pc2
log10 peptide intensity
489:499
569:574
183:187
13:113800622:T:C_T
p = 8.9e−25, beta = −0.41, N = 1241
VPMMYGAGK pc2
log10 peptide intensity
497:499
573:574
185:187
13:113800622:T:C_T
p = 4e−24, beta = −0.402, N = 1255
ETSNFGFSLLR pc2
log10 peptide intensity
497:499
573:574
186:187
13:113800622:T:C_T
p = 1.4e−23, beta = −0.397, N = 1256
YEMHELLR pc2
log10 peptide intensity
497:499
572:574
186:187
13:113800622:T:C_T
p = 2.2e−23, beta = −0.395, N = 1255
G3V2W1 NP2
log10 protein intensity
75:75
153:153
95:97
rs559054
p = 1.5e−07, beta = −0.391, N = 323
VPMMYGAGK pc2
log10 peptide intensity
75:75
150:153
93:97
rs559054
p = 8.6e−10, beta = −0.455, N = 318
NMEVFFPK pc2
log10 peptide intensity
75:75
152:153
94:97
rs559054
p = 2.4e−09, beta = −0.443, N = 321
LILVDYILFK pc2
log10 peptide intensity
75:75
150:153
93:97
rs559054
p = 2.9e−09, beta = −0.442, N = 318
RIFSPFADLSELSATGR pc3
log10 peptide intensity
75:75
152:153
94:97
rs559054
p = 7.9e−09, beta = −0.429, N = 321
Locus: 56 (Sort: 67)
Status: Replicated
80% power: Yes
SNP: 13:113800622:T:C
rsID: rs559054
UniProtID: G3V2W1
HGNC gene: SERPINA10
Type: Trans−pQTL
NLELGLSQGSFAFIHK pc2
rs2232700 ALT
log10 peptide intensity
143:499
168:574
66:187
13:113800622:T:C_T
p = 0.084, model = DOM, N = 377
ETLSRNLELGLTQGSFAFIHK pc4
rs2232700 REF
log10 peptide intensity
439:499
522:574
171:187
13:113800622:T:C_T
p = 0.086, model = REC, N = 1132
NLELGLTQGSFAFIHK pc2
rs2232700 REF
log10 peptide intensity
462:499
528:574
168:187
13:113800622:T:C_T
p = 0.25, model = DOM, N = 1158

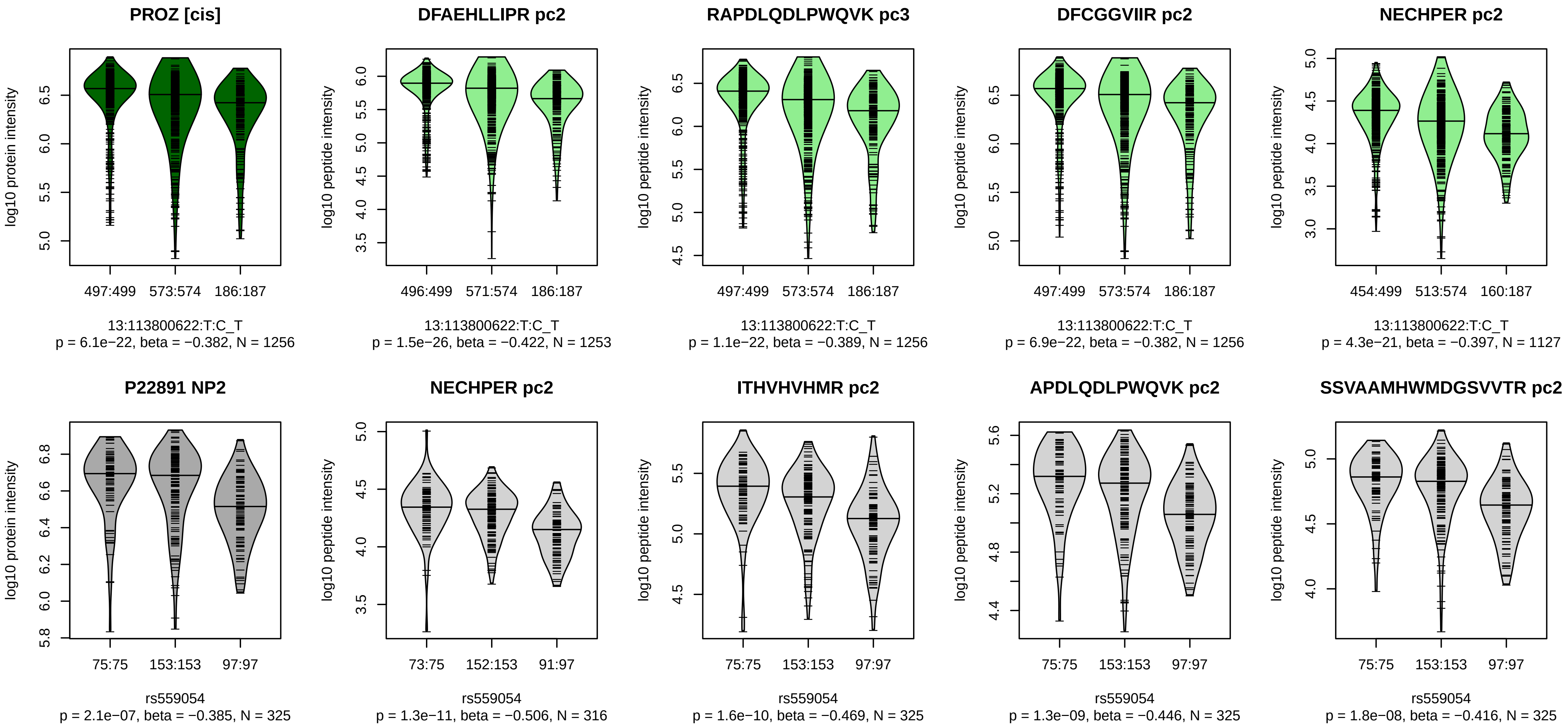


Locus: 56 (Sort: 68)
Status: Replicated
80% power: Yes
SNP: 13:113800622:T:C
rsID: rs559054
UniProtID: P22891
HGNC gene: PROZ
Type: Cis−pQTL

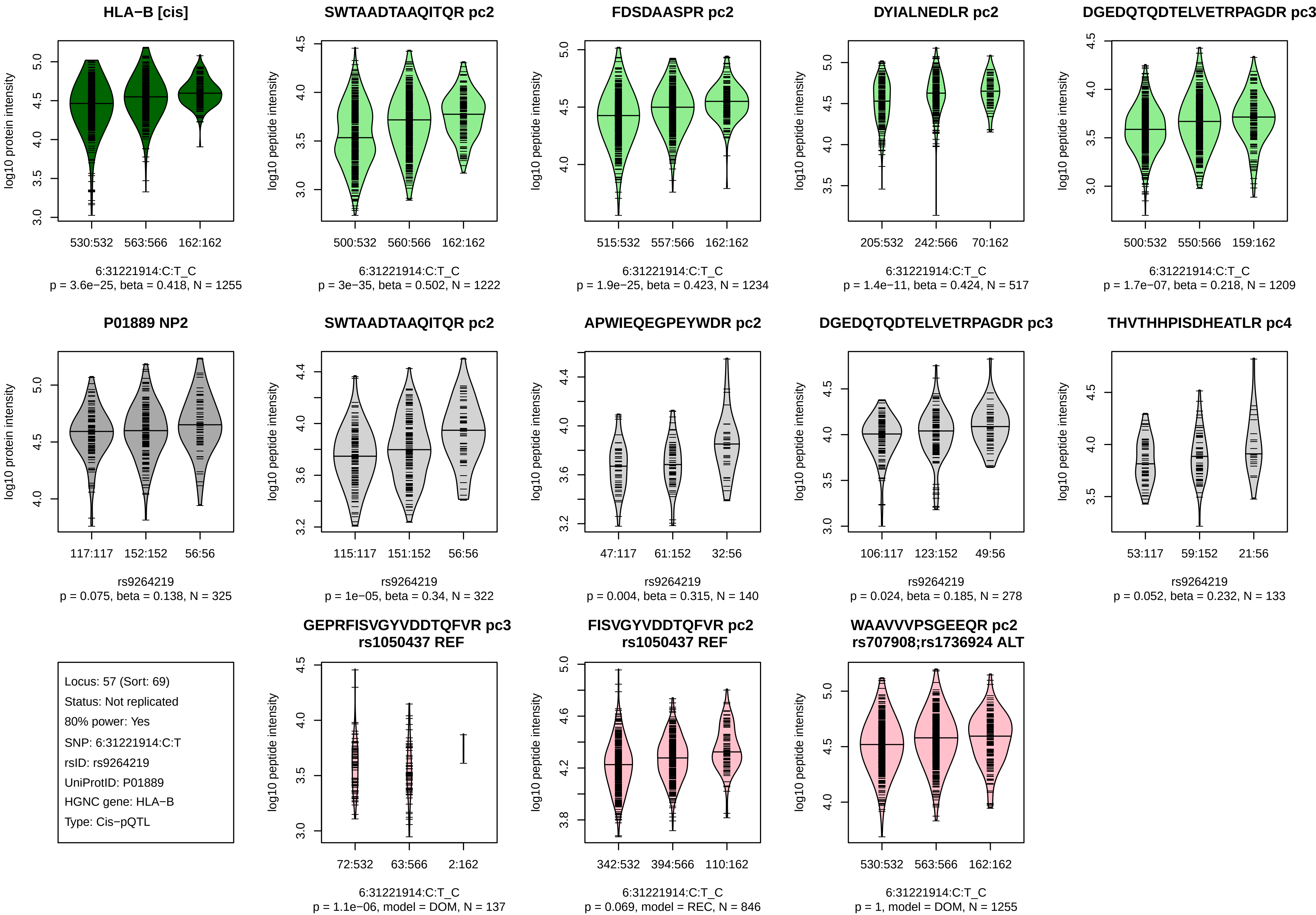
HLA−B [cis]
log10 protein intensity
530:532
563:566
162:162
6:31221914:C:T_C
p = 3.6e−25, beta = 0.418, N = 1255
SWTAADTAAQITQR pc2
log10 peptide intensity
500:532
560:566
162:162
6:31221914:C:T_C
p = 3e−35, beta = 0.502, N = 1222
FDSDAASPR pc2
log10 peptide intensity
515:532
557:566
162:162
6:31221914:C:T_C
p = 1.9e−25, beta = 0.423, N = 1234
DYIALNEDLR pc2
log10 peptide intensity
205:532
242:566
70:162
6:31221914:C:T_C
p = 1.4e−11, beta = 0.424, N = 517
DGEDQTQDTELVETRPAGDR pc3
log10 peptide intensity
500:532
550:566
159:162
6:31221914:C:T_C
p = 1.7e−07, beta = 0.218, N = 1209
P01889 NP2
log10 protein intensity
117:117
152:152
56:56
rs9264219
p = 0.075, beta = 0.138, N = 325
SWTAADTAAQITQR pc2
log10 peptide intensity
115:117
151:152
56:56
rs9264219
p = 1e−05, beta = 0.34, N = 322
APWIEQEGPEYWDR pc2
log10 peptide intensity
47:117
61:152
32:56
rs9264219
p = 0.004, beta = 0.315, N = 140
DGEDQTQDTELVETRPAGDR pc3
log10 peptide intensity
106:117
123:152
49:56
rs9264219
p = 0.024, beta = 0.185, N = 278
THVTHHPISDHEATLR pc4
log10 peptide intensity
53:117
59:152
21:56
rs9264219
p = 0.052, beta = 0.232, N = 133
Locus: 57 (Sort: 69)
Status: Not replicated
80% power: Yes
SNP: 6:31221914:C:T
rsID: rs9264219
UniProtID: P01889
HGNC gene: HLA−B
Type: Cis−pQTL
GEPRFISVGYVDDTQFVR pc3
rs1050437 REF
log10 peptide intensity
72:532
63:566
2:162
6:31221914:C:T_C
p = 1.1e−06, model = DOM, N = 137
FISVGYVDDTQFVR pc2
rs1050437 REF
log10 peptide intensity
342:532
394:566
110:162
6:31221914:C:T_C
p = 0.069, model = REC, N = 846
WAAVVVPSGEEQR pc2
rs707908;rs1736924 ALT
log10 peptide intensity
530:532
563:566
162:162
6:31221914:C:T_C
p = 1, model = DOM, N = 1255

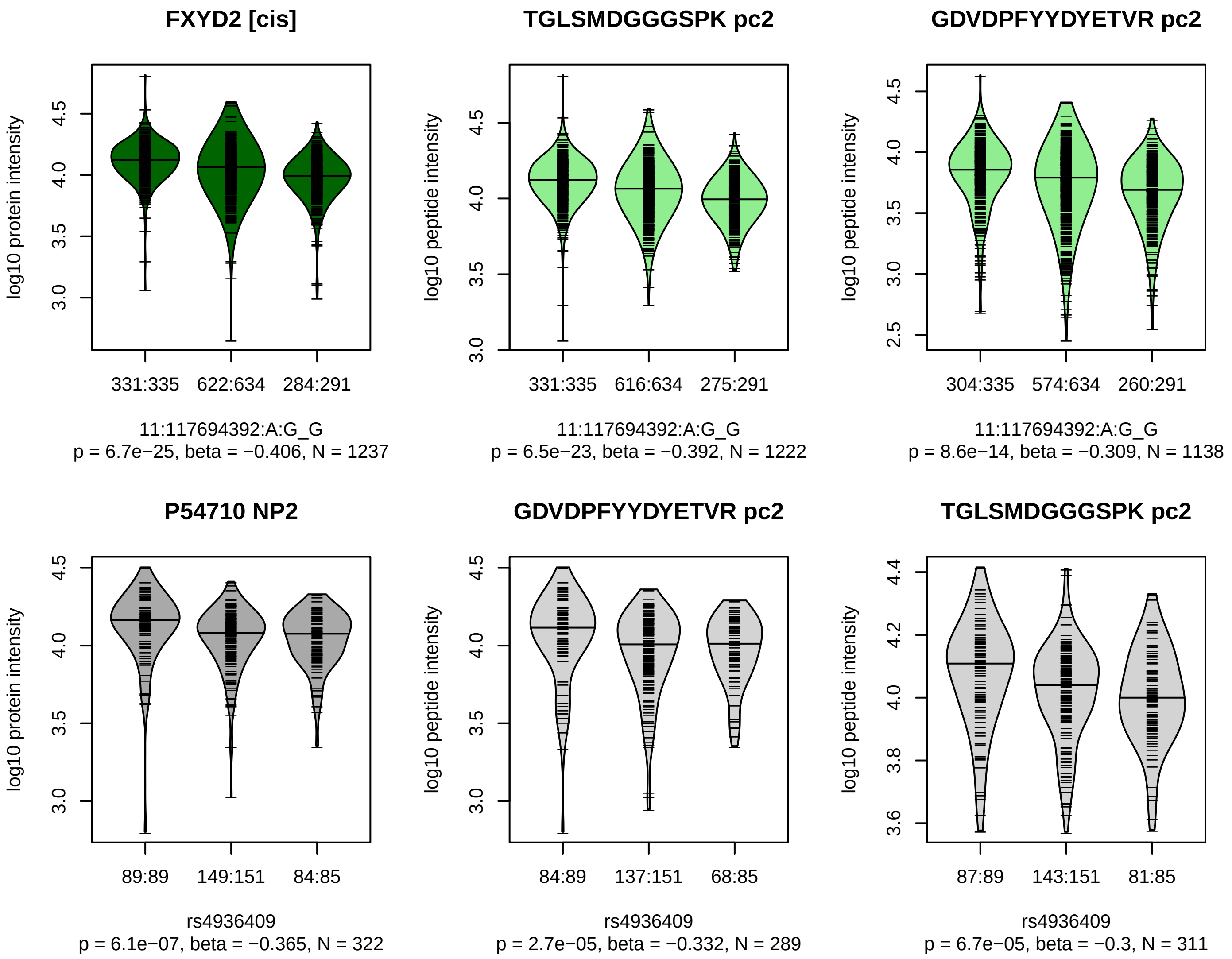


Locus: 58 (Sort: 70)

Status: Replicated

80% power: Yes

SNP: 11:117694392:A:G

rsID: rs4936409

UniProtID: P54710

HGNC gene: FXYD2

Type: Cis−pQTL

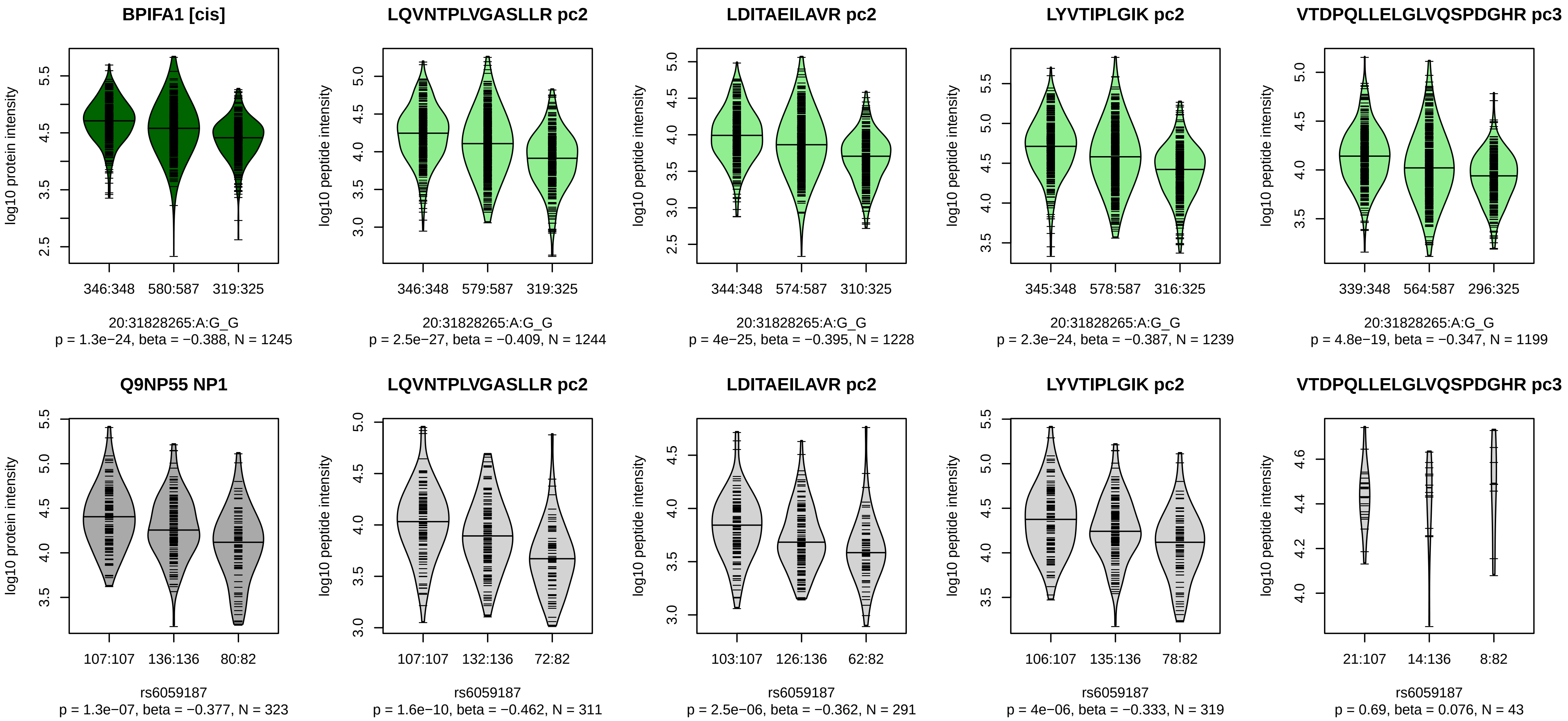


Locus: 59 (Sort: 71)

Status: Replicated

80% power: Yes

SNP: 20:31828265:A:G

rsID: rs6059187

UniProtID: Q9NP55

HGNC gene: BPIFA1

Type: Cis−pQTL

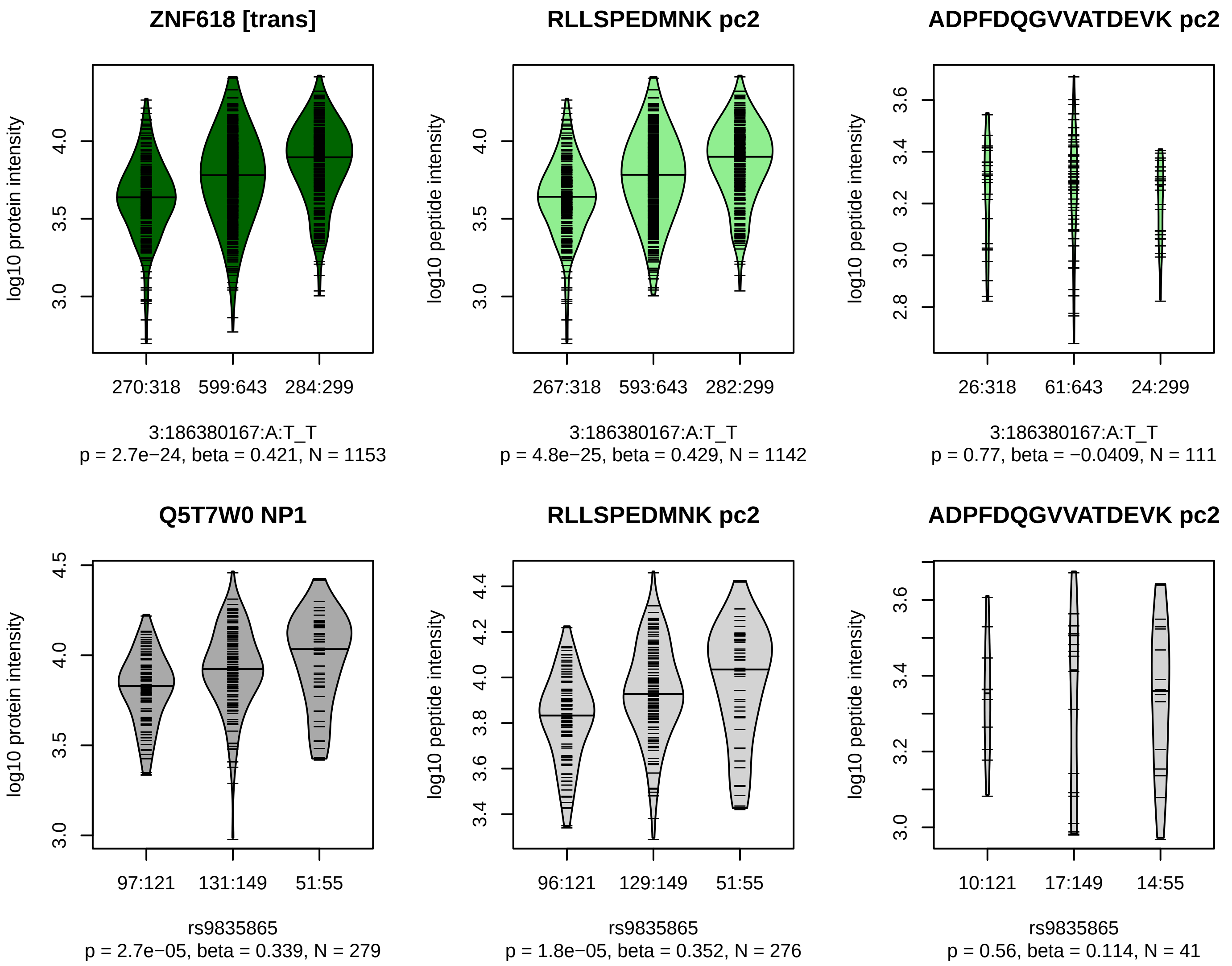


Locus: 60 (Sort: 72)

Status: Replicated

80% power: Yes

SNP: 3:186380167:A:T

rsID: rs9835865

UniProtID: Q5T7W0

HGNC gene: ZNF618

Type: Trans−pQTL

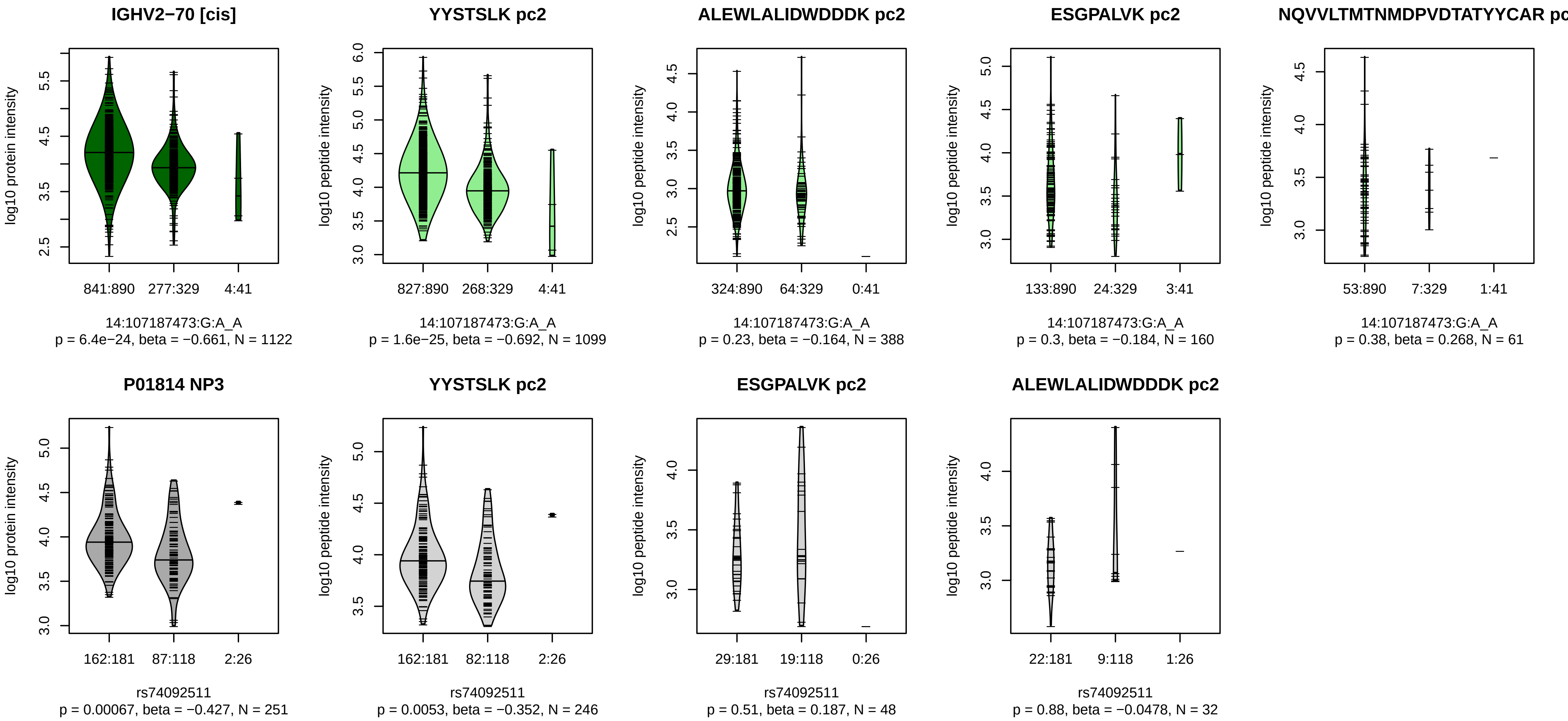


Locus: 61 (Sort: 73)
Status: Not replicated
80% power: Yes
SNP: 14:107187473:G:A
rsID: rs74092511
UniProtID: P01814
HGNC gene: IGHV2−70
Type: Cis−pQTL

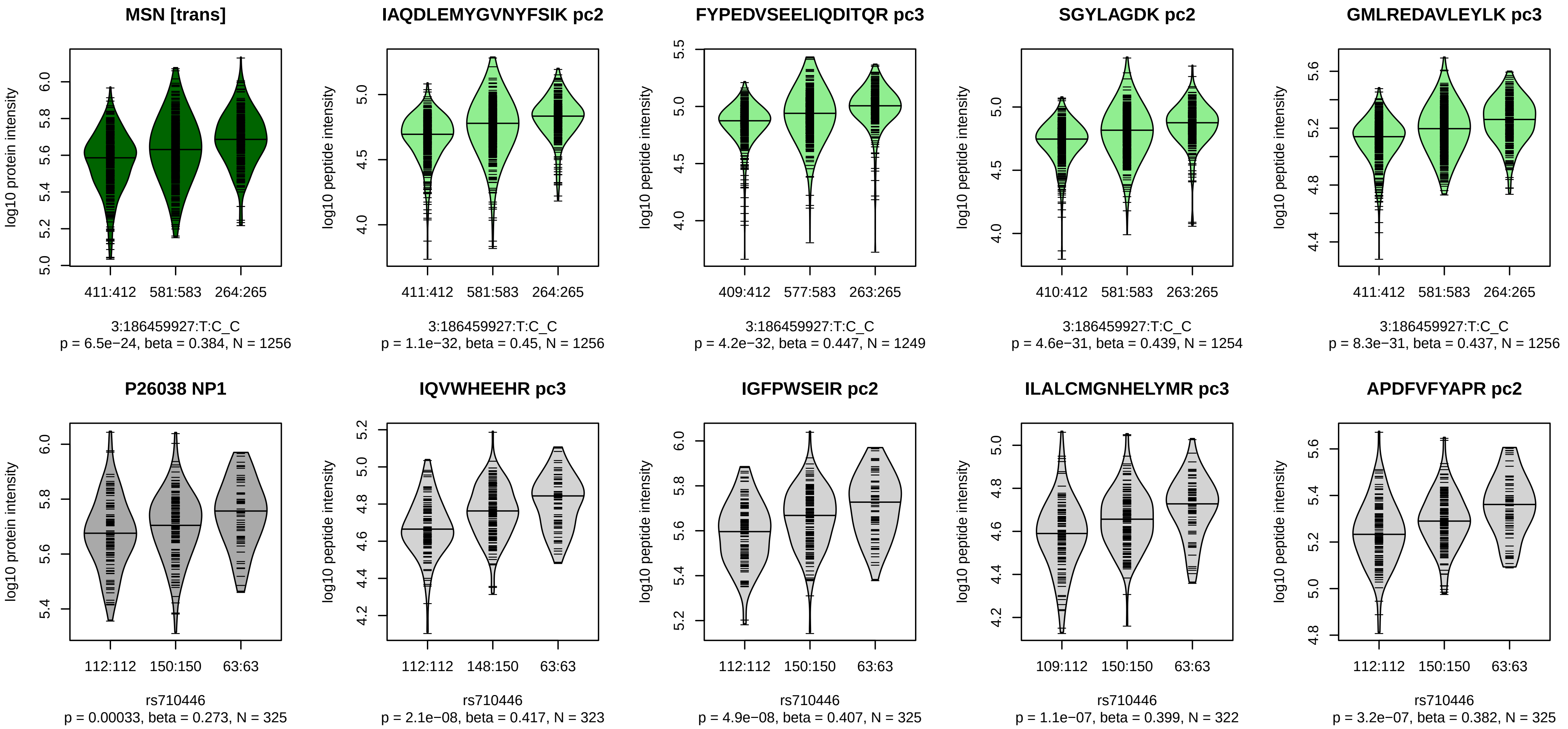


Locus: 62 (Sort: 74)

Status: Not replicated

80% power: Yes

SNP: 3:186459927:T:C

rsID: rs710446

UniProtID: P26038

HGNC gene: MSN

Type: Trans−pQTL

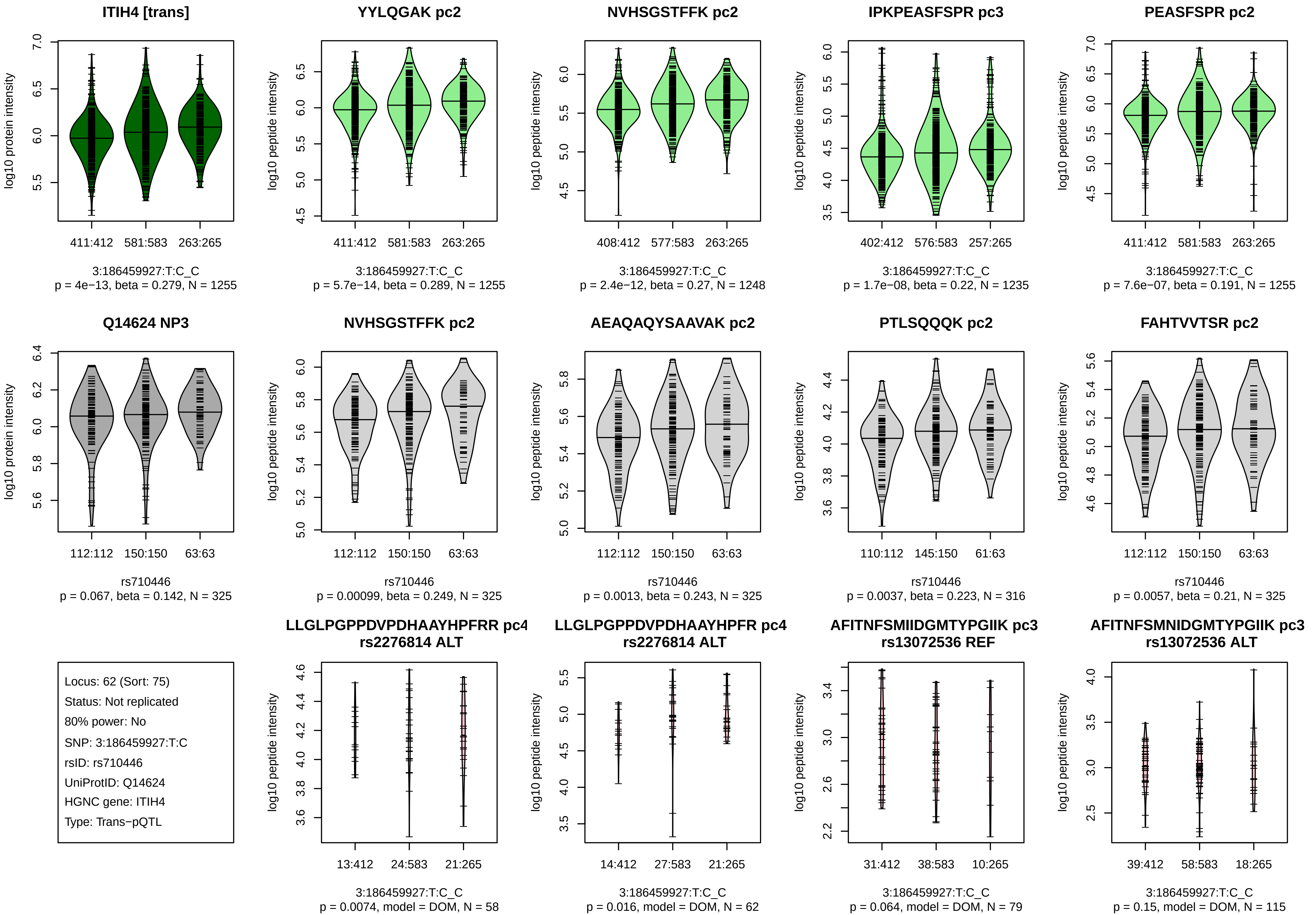
ITIH4 [trans]
log10 protein intensity
411:412
581:583
263:265
3:186459927:T:C_C
p = 4e−13, beta = 0.279, N = 1255
YYLQGAK pc2
log10 peptide intensity
411:412
581:583
263:265
3:186459927:T:C_C
p = 5.7e−14, beta = 0.289, N = 1255
NVHSGSTFFK pc2
log10 peptide intensity
408:412
577:583
263:265
3:186459927:T:C_C
p = 2.4e−12, beta = 0.27, N = 1248
IPKPEASFSPR pc3
log10 peptide intensity
402:412
576:583
257:265
3:186459927:T:C_C
p = 1.7e−08, beta = 0.22, N = 1235
PEASFSPR pc2
log10 peptide intensity
411:412
581:583
263:265
3:186459927:T:C_C
p = 7.6e−07, beta = 0.191, N = 1255
Q14624 NP3
log10 protein intensity
112:112
150:150
63:63
rs710446
p = 0.067, beta = 0.142, N = 325
NVHSGSTFFK pc2
log10 peptide intensity
112:112
150:150
63:63
rs710446
p = 0.00099, beta = 0.249, N = 325
AEAQAQYSAAVAK pc2
log10 peptide intensity
112:112
150:150
63:63
rs710446
p = 0.0013, beta = 0.243, N = 325
PTLSQQQK pc2
log10 peptide intensity
110:112
145:150
61:63
rs710446
p = 0.0037, beta = 0.223, N = 316
FAHTVVTSR pc2
log10 peptide intensity
112:112
150:150
63:63
rs710446
p = 0.0057, beta = 0.21, N = 325
Locus: 62 (Sort: 75)
Status: Not replicated
80% power: No
SNP: 3:186459927:T:C
rsID: rs710446
UniProtID: Q14624
HGNC gene: ITIH4
Type: Trans−pQTL
LLGLPGPPDVPDHAAYHPFRR pc4
rs2276814 ALT
log10 peptide intensity
13:412
24:583
21:265
3:186459927:T:C_C
p = 0.0074, model = DOM, N = 58
LLGLPGPPDVPDHAAYHPFR pc4
rs2276814 ALT
log10 peptide intensity
14:412
27:583
21:265
3:186459927:T:C_C
p = 0.016, model = DOM, N = 62
AFITNFSMIIDGMTYPGIIK pc3
rs13072536 REF
log10 peptide intensity
31:412
38:583
10:265
3:186459927:T:C_C
p = 0.064, model = DOM, N = 79
AFITNFSMNIDGMTYPGIIK pc3
rs13072536 ALT
log10 peptide intensity
39:412
58:583
18:265
3:186459927:T:C_C
p = 0.15, model = DOM, N = 115

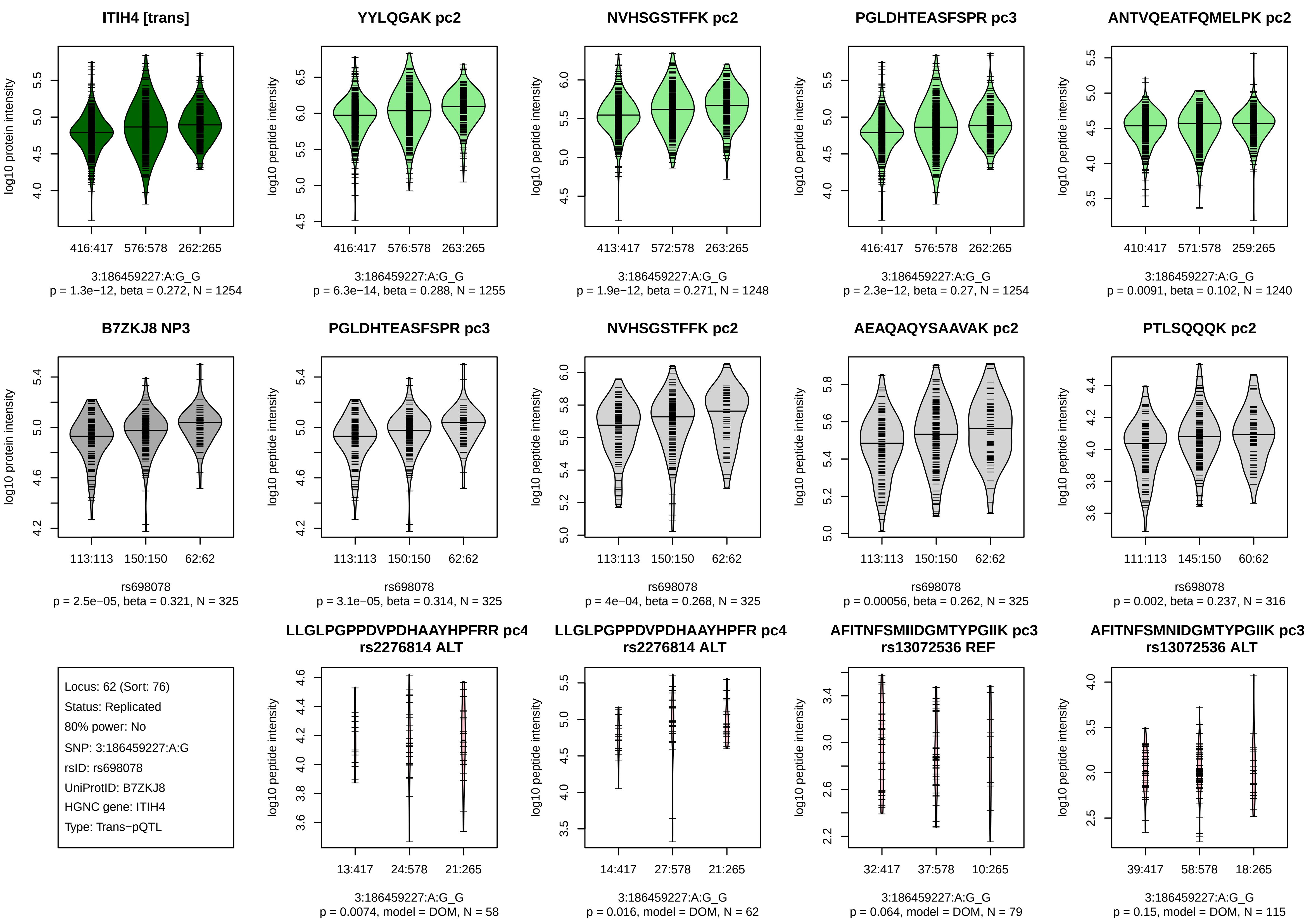
ITIH4 [trans]
log10 protein intensity
416:417 576:578 262:265
3:186459227:A:G_G
p = 1.3e−12, beta = 0.272, N = 1254
YYLQGAK pc2
log10 peptide intensity
416:417 576:578 263:265
3:186459227:A:G_G
p = 6.3e−14, beta = 0.288, N = 1255
NVHSGSTFFK pc2
log10 peptide intensity
413:417 572:578 263:265
3:186459227:A:G_G
p = 1.9e−12, beta = 0.271, N = 1248
PGLDHTEASFSPR pc3
log10 peptide intensity
416:417 576:578 262:265
3:186459227:A:G_G
p = 2.3e−12, beta = 0.27, N = 1254
ANTVQEATFQMELPK pc2
log10 peptide intensity
410:417 571:578 259:265
3:186459227:A:G_G
p = 0.0091, beta = 0.102, N = 1240
B7ZKJ8 NP3
log10 protein intensity
113:113 150:150 62:62
rs698078
p = 2.5e−05, beta = 0.321, N = 325
PGLDHTEASFSPR pc3
log10 peptide intensity
113:113 150:150 62:62
rs698078
p = 3.1e−05, beta = 0.314, N = 325
NVHSGSTFFK pc2
log10 peptide intensity
113:113 150:150 62:62
rs698078
p = 4e−04, beta = 0.268, N = 325
AEAQAQYSAAVAK pc2
log10 peptide intensity
113:113 150:150 62:62
rs698078
p = 0.00056, beta = 0.262, N = 325
PTLSQQQK pc2
log10 peptide intensity
111:113 145:150 60:62
rs698078
p = 0.002, beta = 0.237, N = 316
Locus: 62 (Sort: 76)
Status: Replicated
80% power: No
SNP: 3:186459227:A:G
rsID: rs698078
UniProtID: B7ZKJ8
HGNC gene: ITIH4
Type: Trans−pQTL
LLGLPGPPDVPDHAAYHPFRR pc4
rs2276814 ALT
log10 peptide intensity
13:417 24:578 21:265
3:186459227:A:G_G
p = 0.0074, model = DOM, N = 58
LLGLPGPPDVPDHAAYHPFR pc4
rs2276814 ALT
log10 peptide intensity
14:417 27:578 21:265
3:186459227:A:G_G
p = 0.016, model = DOM, N = 62
AFITNFSMIIDGMTYPGIIK pc3
rs13072536 REF
log10 peptide intensity
32:417 37:578 10:265
3:186459227:A:G_G
p = 0.064, model = DOM, N = 79
AFITNFSMNIDGMTYPGIIK pc3
rs13072536 ALT
log10 peptide intensity
39:417 58:578 18:265
3:186459227:A:G_G
p = 0.15, model = DOM, N = 115

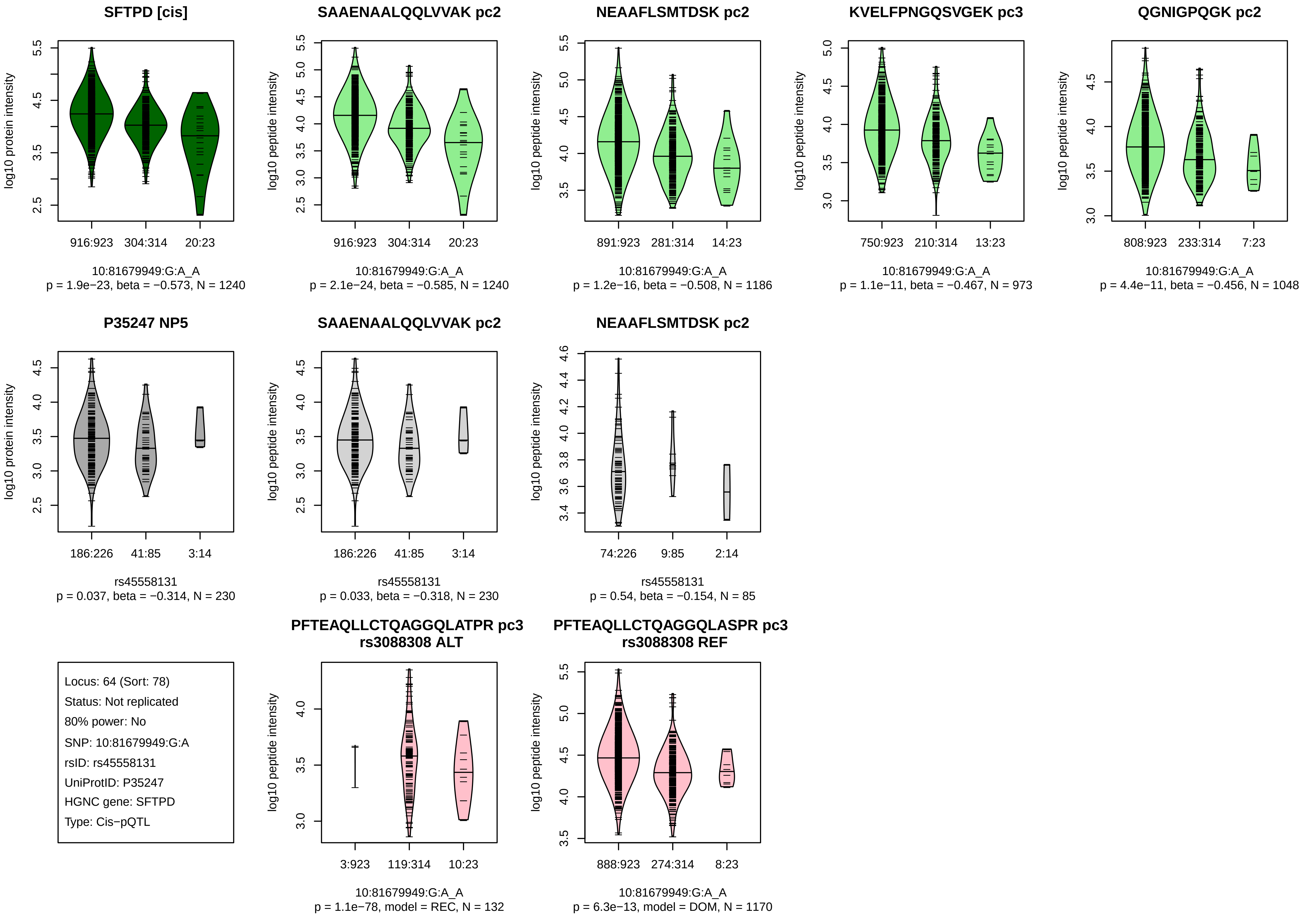
SFTPD [cis]
log10 protein intensity
916:923
304:314
20:23
10:81679949:G:A_A
p = 1.9e−23, beta = −0.573, N = 1240
SAAENAALQQLVVAK pc2
log10 peptide intensity
916:923
304:314
20:23
10:81679949:G:A_A
p = 2.1e−24, beta = −0.585, N = 1240
NEAAFLSMTDSK pc2
log10 peptide intensity
891:923
281:314
14:23
10:81679949:G:A_A
p = 1.2e−16, beta = −0.508, N = 1186
KVELFPNGQSVGEK pc3
log10 peptide intensity
750:923
210:314
13:23
10:81679949:G:A_A
p = 1.1e−11, beta = −0.467, N = 973
QGNIGPQGK pc2
log10 peptide intensity
808:923
233:314
7:23
10:81679949:G:A_A
p = 4.4e−11, beta = −0.456, N = 1048
P35247 NP5
log10 protein intensity
186:226
41:85
3:14
rs45558131
p = 0.037, beta = −0.314, N = 230
SAAENAALQQLVVAK pc2
log10 peptide intensity
186:226
41:85
3:14
rs45558131
p = 0.033, beta = −0.318, N = 230
NEAAFLSMTDSK pc2
log10 peptide intensity
74:226
9:85
2:14
rs45558131
p = 0.54, beta = −0.154, N = 85
Locus: 64 (Sort: 78)
Status: Not replicated
80% power: No
SNP: 10:81679949:G:A
rsID: rs45558131
UniProtID: P35247
HGNC gene: SFTPD
Type: Cis−pQTL
PFTEAQLLCTQAGGQLATPR pc3
rs3088308 ALT
log10 peptide intensity
3:923
119:314
10:23
10:81679949:G:A_A
p = 1.1e−78, model = REC, N = 132
PFTEAQLLCTQAGGQLASPR pc3
rs3088308 REF
log10 peptide intensity
888:923
274:314
8:23
10:81679949:G:A_A
p = 6.3e−13, model = DOM, N = 1170

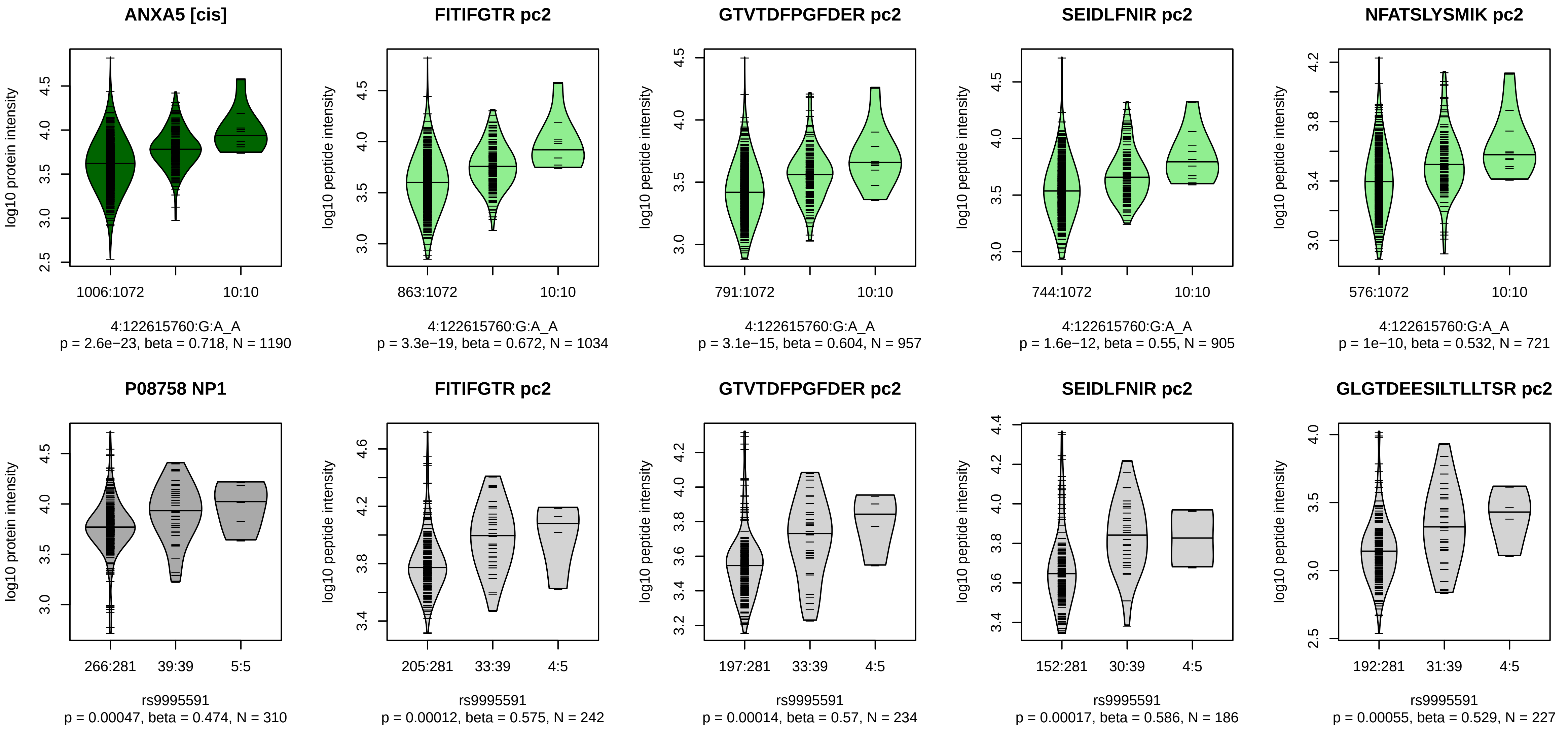


Locus: 65 (Sort: 79)
Status: Not replicated
80% power: Yes
SNP: 4:122615760:G:A
rsID: rs9995591
UniProtID: P08758
HGNC gene: ANXA5
Type: Cis−pQTL

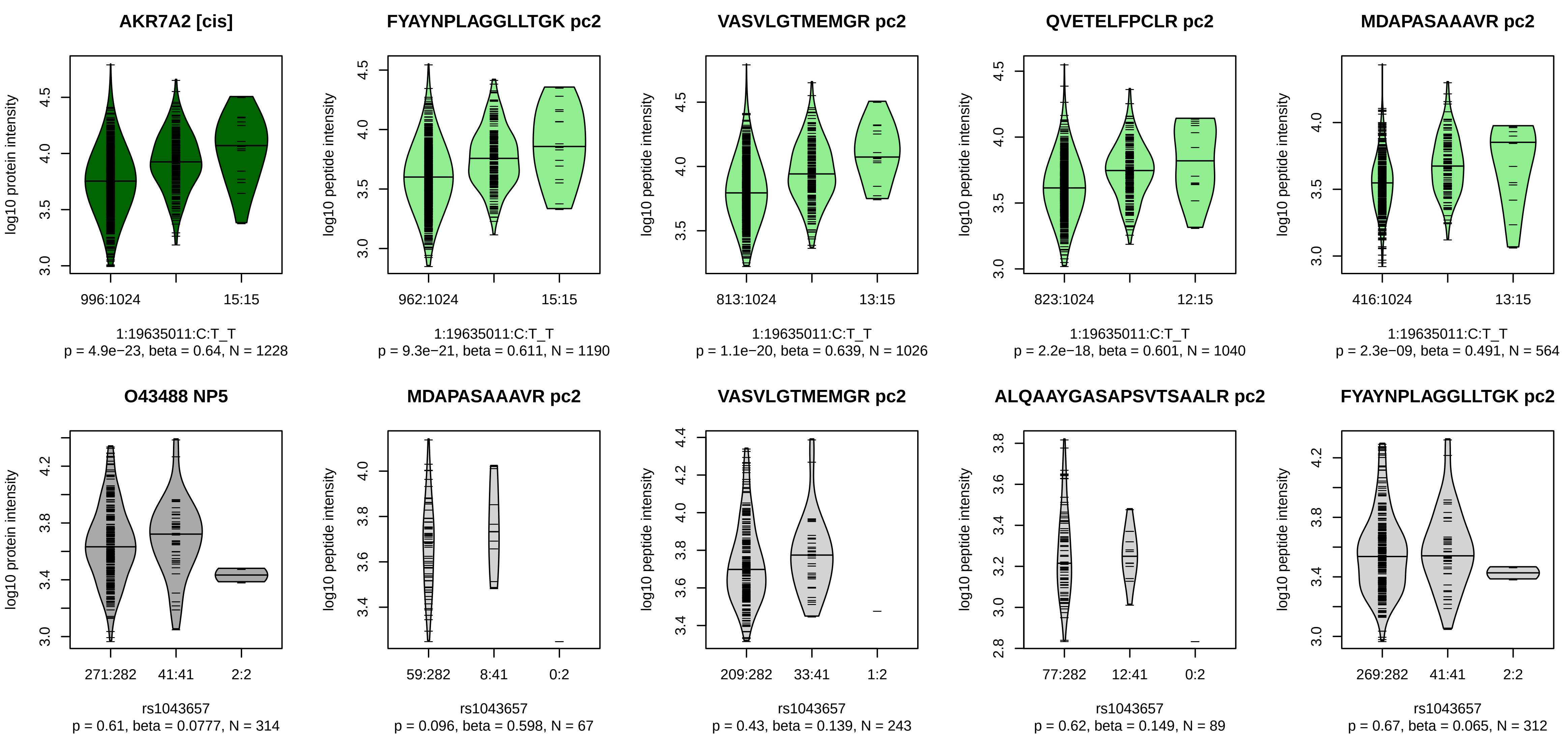


Locus: 66 (Sort: 80)
Status: Not replicated
80% power: Yes
SNP: 1:19635011:C:T
rsID: rs1043657
UniProtID: O43488
HGNC gene: AKR7A2
Type: Cis−pQTL

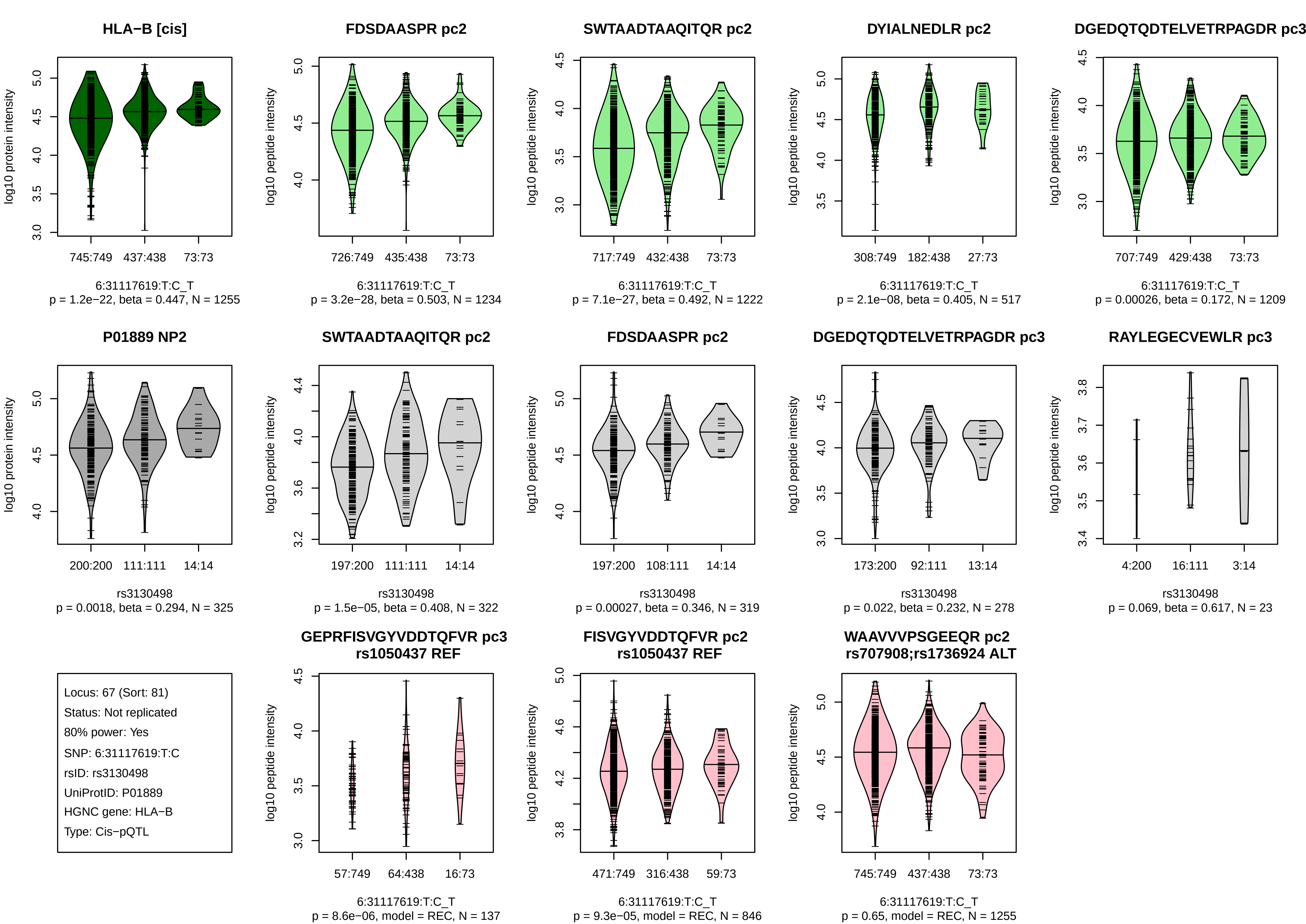
HLA−B [cis]
log10 protein intensity
745:749 437:438 73:73
6:31117619:T:C_T
p = 1.2e−22, beta = 0.447, N = 1255
FDSDAASPR pc2
log10 peptide intensity
726:749 435:438 73:73
6:31117619:T:C_T
p = 3.2e−28, beta = 0.503, N = 1234
SWTAADTAAQITQR pc2
log10 peptide intensity
717:749 432:438 73:73
6:31117619:T:C_T
p = 7.1e−27, beta = 0.492, N = 1222
DYIALNEDLR pc2
log10 peptide intensity
308:749 182:438 27:73
6:31117619:T:C_T
p = 2.1e−08, beta = 0.405, N = 517
DGEDQTQDTELVETRPAGDR pc3
log10 peptide intensity
707:749 429:438 73:73
6:31117619:T:C_T
p = 0.00026, beta = 0.172, N = 1209
P01889 NP2
log10 protein intensity
200:200 111:111 14:14
rs3130498
p = 0.0018, beta = 0.294, N = 325
SWTAADTAAQITQR pc2
log10 peptide intensity
197:200 111:111 14:14
rs3130498
p = 1.5e−05, beta = 0.408, N = 322
FDSDAASPR pc2
log10 peptide intensity
197:200 108:111 14:14
rs3130498
p = 0.00027, beta = 0.346, N = 319
DGEDQTQDTELVETRPAGDR pc3
log10 peptide intensity
173:200 92:111 13:14
rs3130498
p = 0.022, beta = 0.232, N = 278
RAYLEGECVEWLR pc3
log10 peptide intensity
4:200 16:111 3:14
rs3130498
p = 0.069, beta = 0.617, N = 23
Locus: 67 (Sort: 81)
Status: Not replicated
80% power: Yes
SNP: 6:31117619:T:C
rsID: rs3130498
UniProtID: P01889
HGNC gene: HLA−B
Type: Cis−pQTL
GEPRFISVGYVDDTQFVR pc3
rs1050437 REF
log10 peptide intensity
57:749 64:438 16:73
6:31117619:T:C_T
p = 8.6e−06, model = REC, N = 137
FISVGYVDDTQFVR pc2
rs1050437 REF
log10 peptide intensity
471:749 316:438 59:73
6:31117619:T:C_T
p = 9.3e−05, model = REC, N = 846
WAAVVVPSGEEQR pc2
rs707908;rs1736924 ALT
log10 peptide intensity
745:749 437:438 73:73
6:31117619:T:C_T
p = 0.65, model = REC, N = 1255

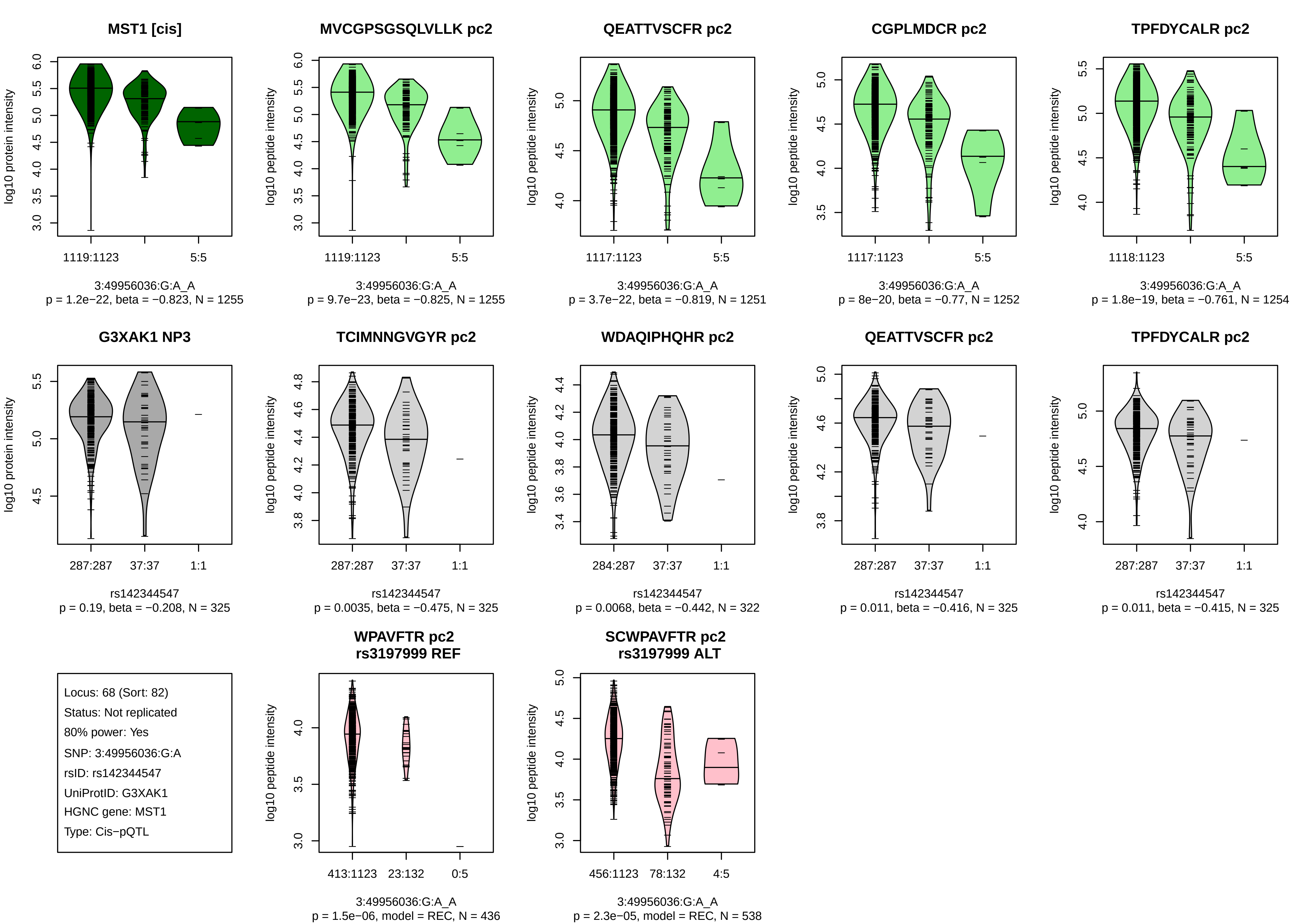
MST1 [cis]
log10 protein intensity
1119:1123
5:5
3:49956036:G:A_A
p = 1.2e−22, beta = −0.823, N = 1255
MVCGPSGSQLVLLK pc2
log10 peptide intensity
1119:1123
5:5
3:49956036:G:A_A
p = 9.7e−23, beta = −0.825, N = 1255
QEATTVSCFR pc2
log10 peptide intensity
1117:1123
5:5
3:49956036:G:A_A
p = 3.7e−22, beta = −0.819, N = 1251
CGPLMDCR pc2
log10 peptide intensity
1117:1123
5:5
3:49956036:G:A_A
p = 8e−20, beta = −0.77, N = 1252
TPFDYCALR pc2
log10 peptide intensity
1118:1123
5:5
3:49956036:G:A_A
p = 1.8e−19, beta = −0.761, N = 1254
G3XAK1 NP3
log10 protein intensity
287:287
37:37
1:1
rs142344547
p = 0.19, beta = −0.208, N = 325
TCIMNNGVGYR pc2
log10 peptide intensity
287:287
37:37
1:1
rs142344547
p = 0.0035, beta = −0.475, N = 325
WDAQIPHQHR pc2
log10 peptide intensity
284:287
37:37
1:1
rs142344547
p = 0.0068, beta = −0.442, N = 322
QEATTVSCFR pc2
log10 peptide intensity
287:287
37:37
1:1
rs142344547
p = 0.011, beta = −0.416, N = 325
TPFDYCALR pc2
log10 peptide intensity
287:287
37:37
1:1
rs142344547
p = 0.011, beta = −0.415, N = 325
Locus: 68 (Sort: 82)
Status: Not replicated
80% power: Yes
SNP: 3:49956036:G:A
rsID: rs142344547
UniProtID: G3XAK1
HGNC gene: MST1
Type: Cis−pQTL
WPAVFTR pc2
rs3197999 REF
log10 peptide intensity
413:1123
23:132
0:5
3:49956036:G:A_A
p = 1.5e−06, model = REC, N = 436
SCWPAVFTR pc2
rs3197999 ALT
log10 peptide intensity
456:1123
78:132
4:5
3:49956036:G:A_A
p = 2.3e−05, model = REC, N = 538

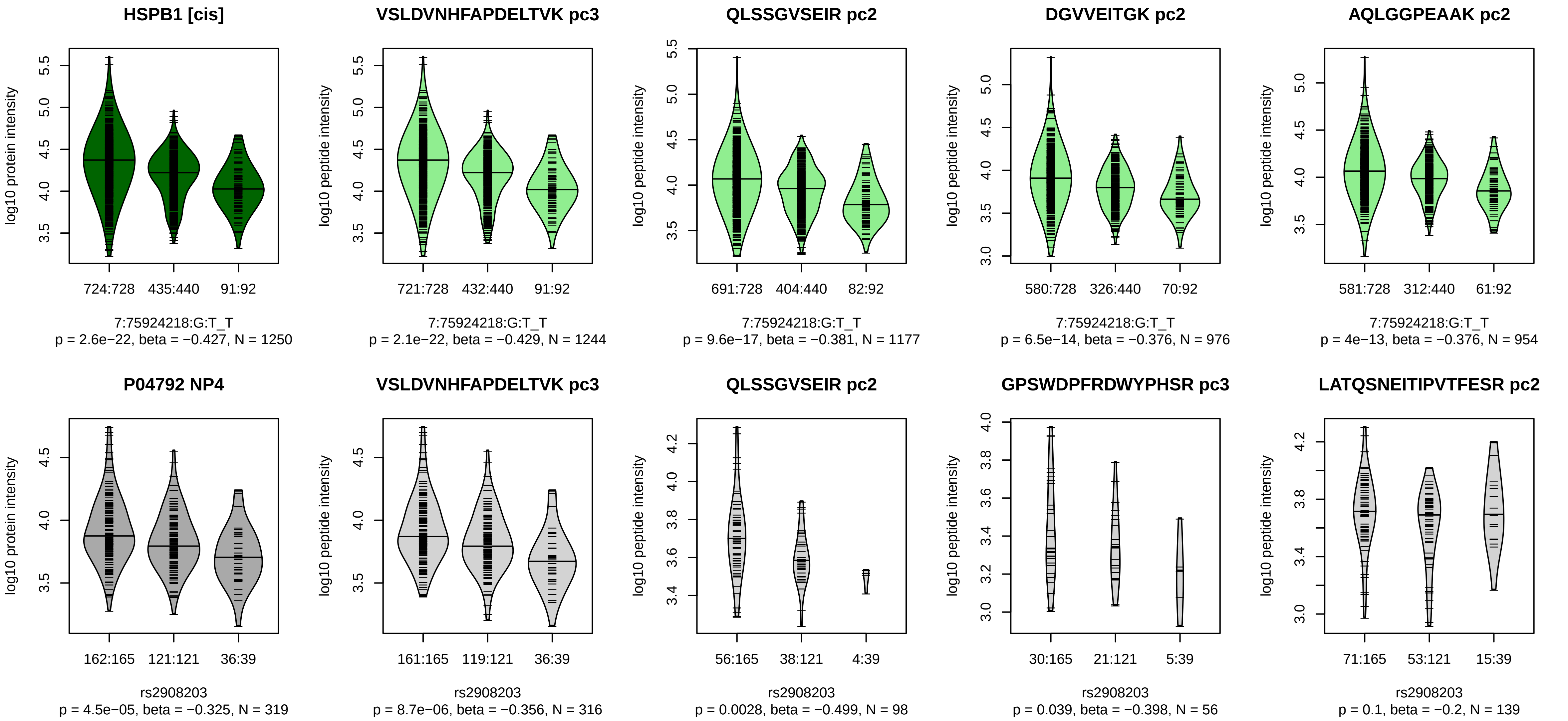


Locus: 69 (Sort: 83)
Status: Replicated
80% power: Yes
SNP: 7:75924218:G:T
rsID: rs2908203
UniProtID: P04792
HGNC gene: HSPB1
Type: Cis−pQTL

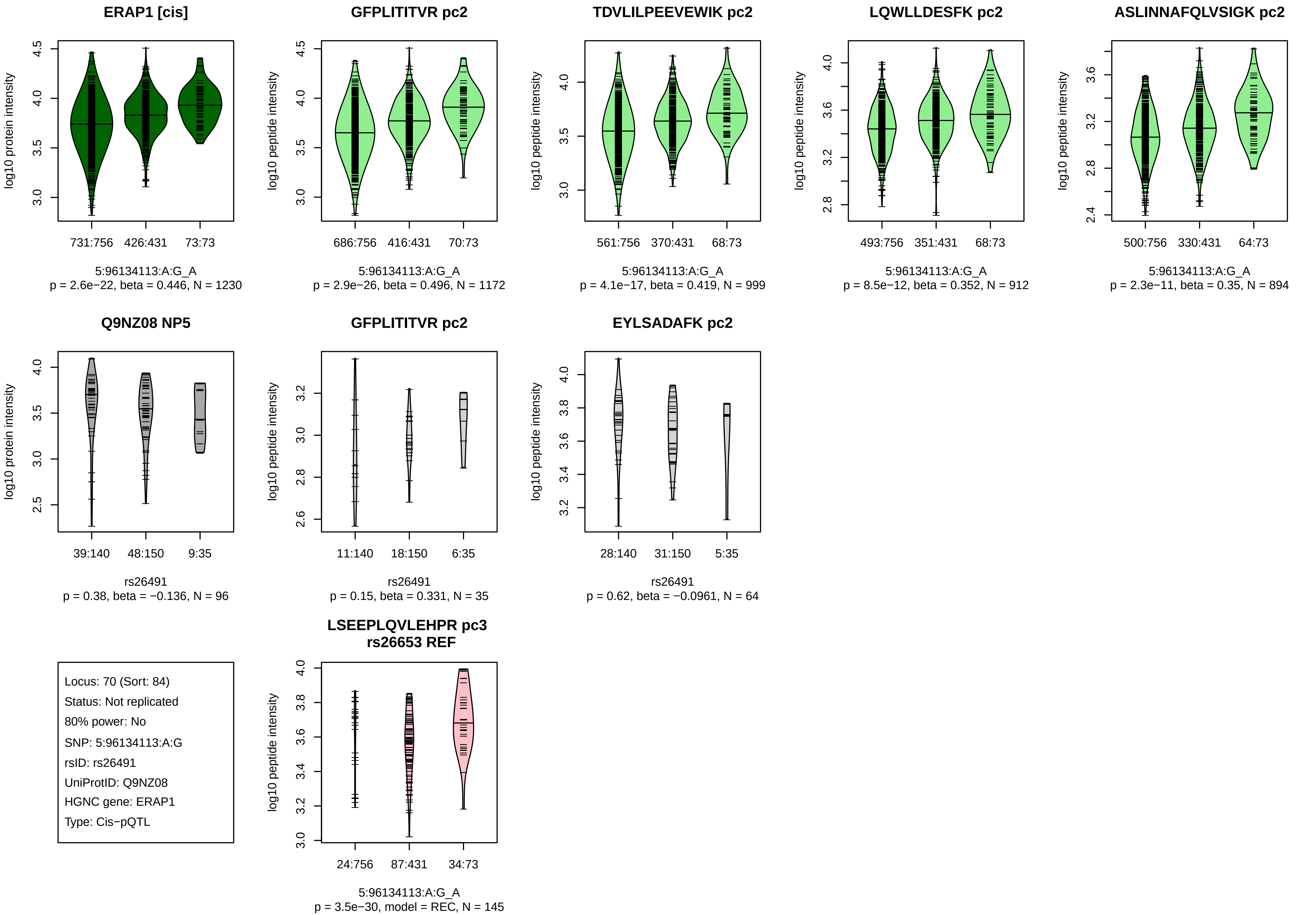
ERAP1 [cis]
log10 protein intensity
731:756
426:431
73:73
5:96134113:A:G_A
p = 2.6e−22, beta = 0.446, N = 1230
GFPLITITVR pc2
log10 peptide intensity
686:756
416:431
70:73
5:96134113:A:G_A
p = 2.9e−26, beta = 0.496, N = 1172
TDVLILPEEVEWIK pc2
log10 peptide intensity
561:756
370:431
68:73
5:96134113:A:G_A
p = 4.1e−17, beta = 0.419, N = 999
LQWLLDESFK pc2
log10 peptide intensity
493:756
351:431
68:73
5:96134113:A:G_A
p = 8.5e−12, beta = 0.352, N = 912
ASLINNAFQLVSIGK pc2
log10 peptide intensity
500:756
330:431
64:73
5:96134113:A:G_A
p = 2.3e−11, beta = 0.35, N = 894
Q9NZ08 NP5
log10 protein intensity
39:140
48:150
9:35
rs26491
p = 0.38, beta = −0.136, N = 96
GFPLITITVR pc2
log10 peptide intensity
11:140
18:150
6:35
rs26491
p = 0.15, beta = 0.331, N = 35
EYLSADAFK pc2
log10 peptide intensity
28:140
31:150
5:35
rs26491
p = 0.62, beta = −0.0961, N = 64
Locus: 70 (Sort: 84)
Status: Not replicated
80% power: No
SNP: 5:96134113:A:G
rsID: rs26491
UniProtID: Q9NZ08
HGNC gene: ERAP1
Type: Cis−pQTL
LSEEPLQVLEHPR pc3
rs26653 REF
log10 peptide intensity
24:756
87:431
34:73
5:96134113:A:G_A
p = 3.5e−30, model = REC, N = 145

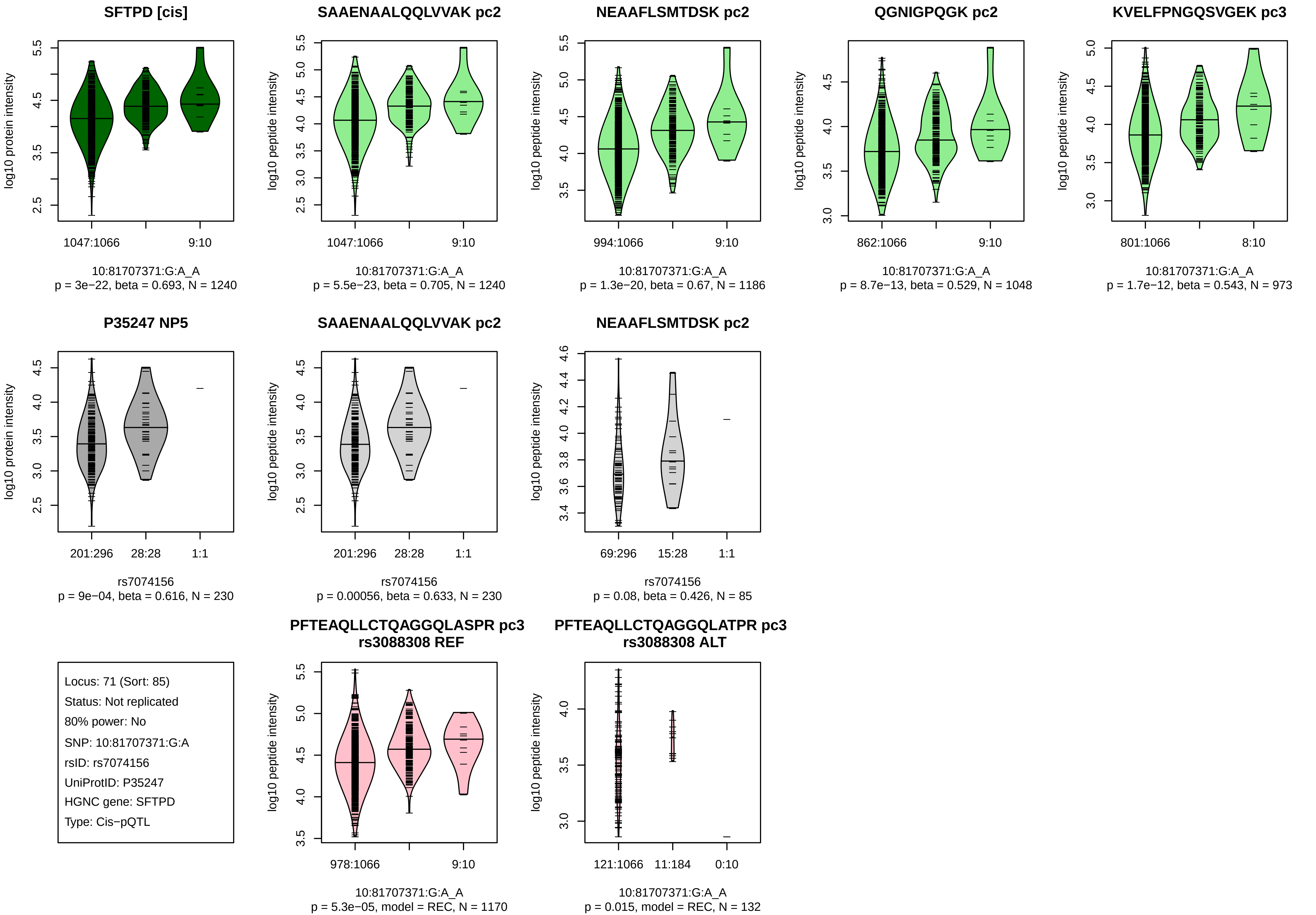
SFTPD [cis]
log10 protein intensity
1047:1066
9:10
10:81707371:G:A_A
p = 3e−22, beta = 0.693, N = 1240
SAAENAALQQLVVAK pc2
log10 peptide intensity
1047:1066
9:10
10:81707371:G:A_A
p = 5.5e−23, beta = 0.705, N = 1240
NEAAFLSMTDSK pc2
log10 peptide intensity
994:1066
9:10
10:81707371:G:A_A
p = 1.3e−20, beta = 0.67, N = 1186
QGNIGPQGK pc2
log10 peptide intensity
862:1066
9:10
10:81707371:G:A_A
p = 8.7e−13, beta = 0.529, N = 1048
KVELFPNGQSVGEK pc3
log10 peptide intensity
801:1066
8:10
10:81707371:G:A_A
p = 1.7e−12, beta = 0.543, N = 973
P35247 NP5
log10 protein intensity
201:296
28:28
1:1
rs7074156
p = 9e−04, beta = 0.616, N = 230
SAAENAALQQLVVAK pc2
log10 peptide intensity
201:296
28:28
1:1
rs7074156
p = 0.00056, beta = 0.633, N = 230
NEAAFLSMTDSK pc2
log10 peptide intensity
69:296
15:28
1:1
rs7074156
p = 0.08, beta = 0.426, N = 85
Locus: 71 (Sort: 85)
Status: Not replicated
80% power: No
SNP: 10:81707371:G:A
rsID: rs7074156
UniProtID: P35247
HGNC gene: SFTPD
Type: Cis−pQTL
PFTEAQLLCTQAGGQLASPR pc3
rs3088308 REF
log10 peptide intensity
978:1066
9:10
10:81707371:G:A_A
p = 5.3e−05, model = REC, N = 1170
PFTEAQLLCTQAGGQLATPR pc3
rs3088308 ALT
log10 peptide intensity
121:1066
11:184
0:10
10:81707371:G:A_A
p = 0.015, model = REC, N = 132

**HLA−G [cis]**

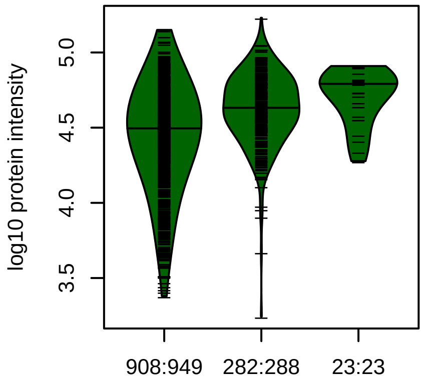


6:29815713:C:T_C
p = 4.6e−22, beta = 0.559, N = 1213

**DYLALNEDLR pc2**

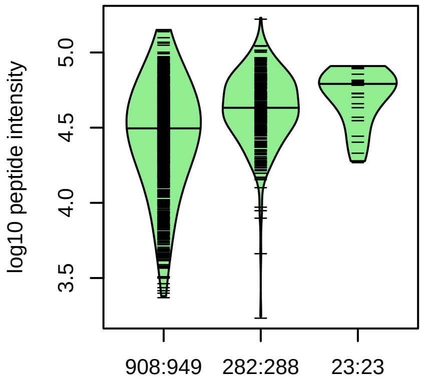


6:29815713:C:T_C
p = 4.6e−22, beta = 0.559, N = 1213

**Q5RJ85 NP1**

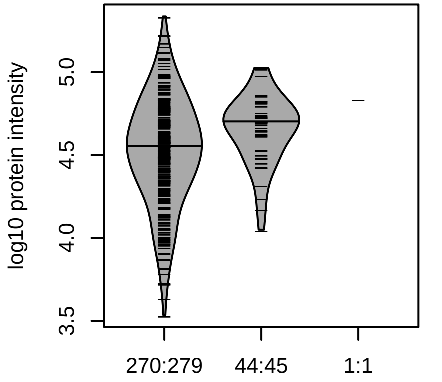


rs3115629
p = 0.011, beta = 0.391, N = 315

**DYLALNEDLR pc2**

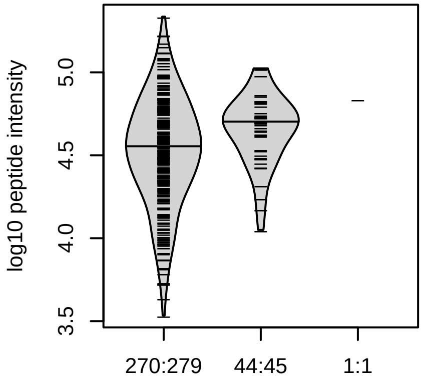


rs3115629
p = 0.011, beta = 0.391, N = 315

Locus: 72 (Sort: 86)
Status: Not replicated
80% power: Yes
SNP: 6:29815713:C:T
rsID: rs3115629
UniProtID: Q5RJ85
HGNC gene: HLA−G
Type: Cis−pQTL

**WAAVVVPSGEEQR pc2**
**rs707908;rs1736924 ALT**

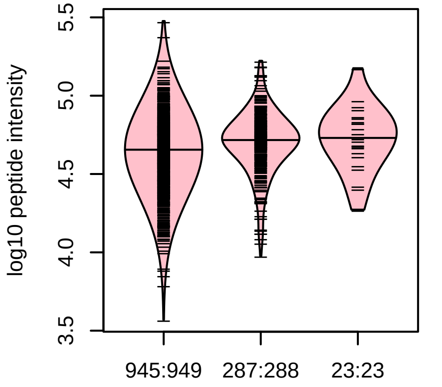


6:29815713:C:T_C
p = 1, model = DOM, N = 1255

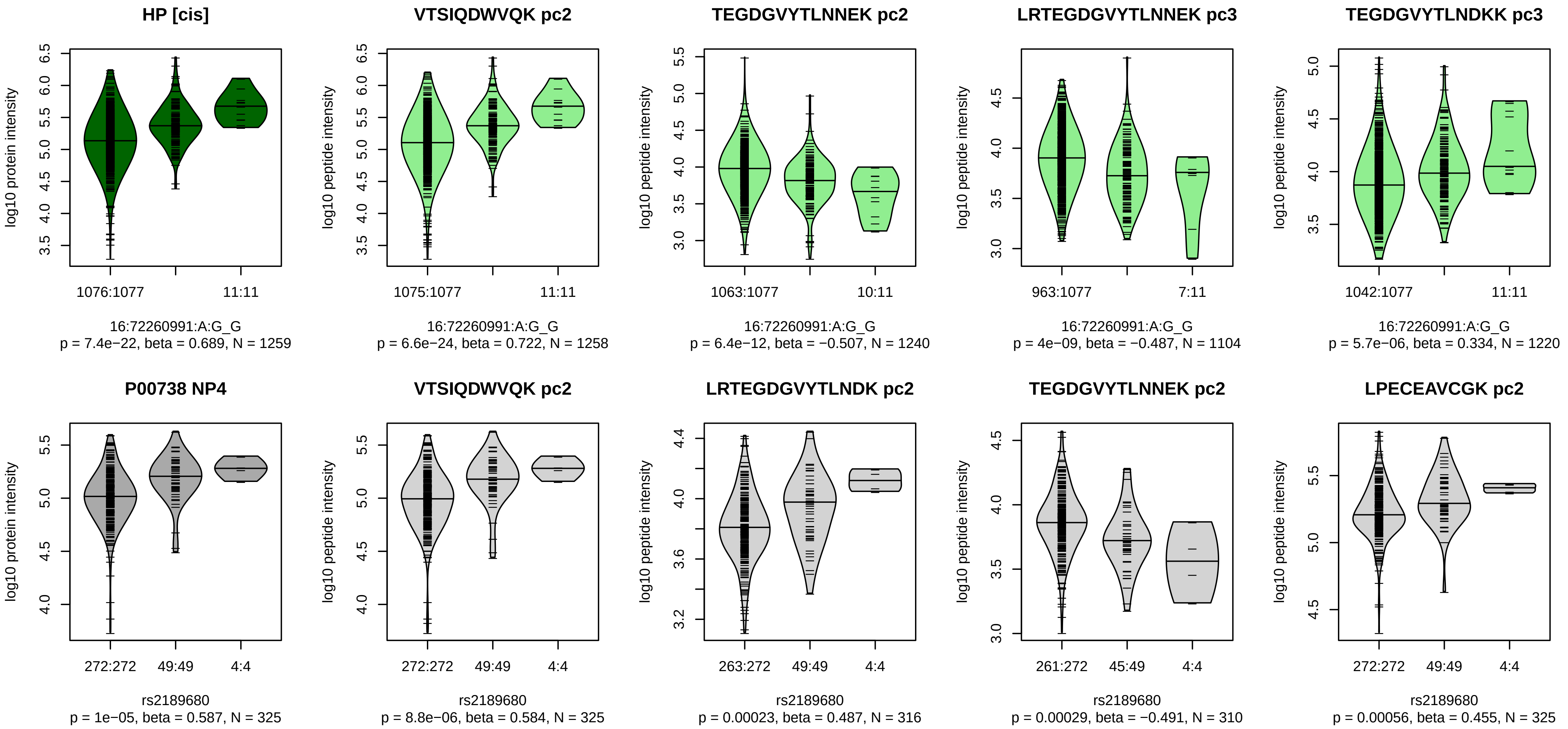


Locus: 73 (Sort: 87)

Status: Replicated

80% power: Yes

SNP: 16:72260991:A:G

rsID: rs2189680

UniProtID: P00738

HGNC gene: HP

Type: Cis−pQTL

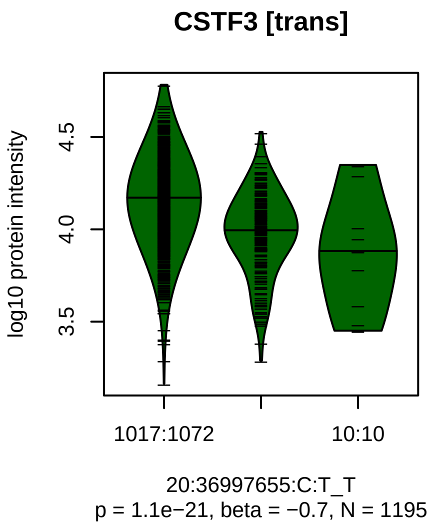


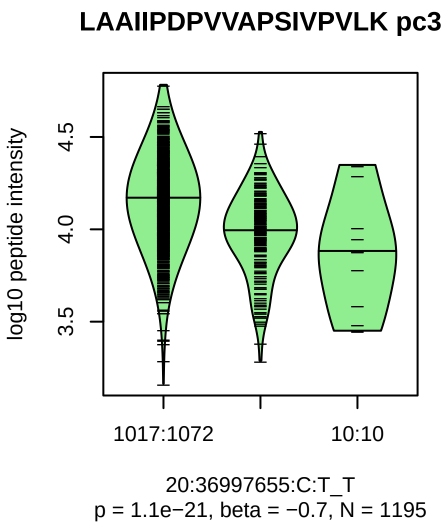


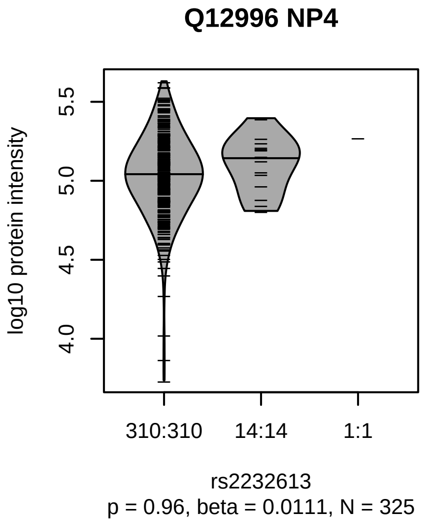


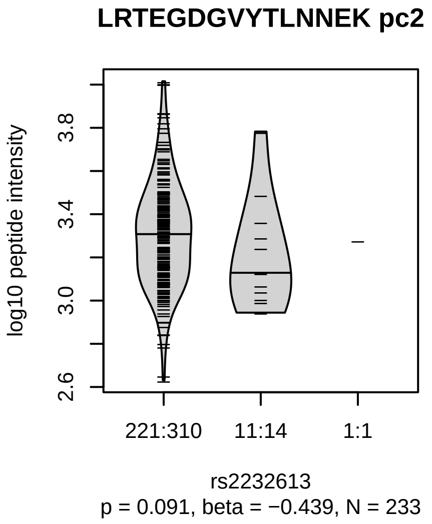


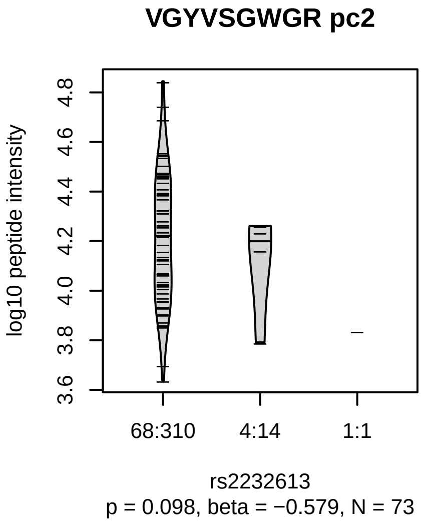


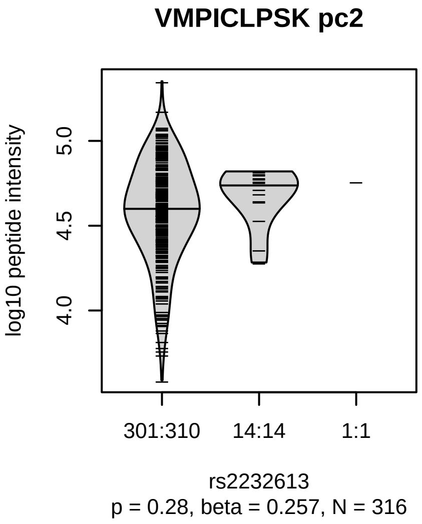


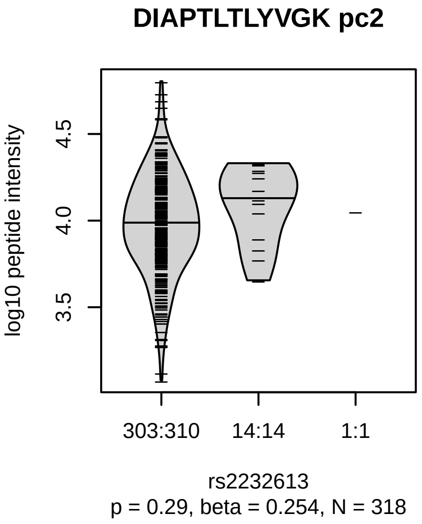


Locus: 74 (Sort: 88)
Status: Not replicated
80% power: Yes
SNP: 20:36997655:C:T
rsID: rs2232613
UniProtID: Q12996
HGNC gene: CSTF3
Type: Trans−pQTL

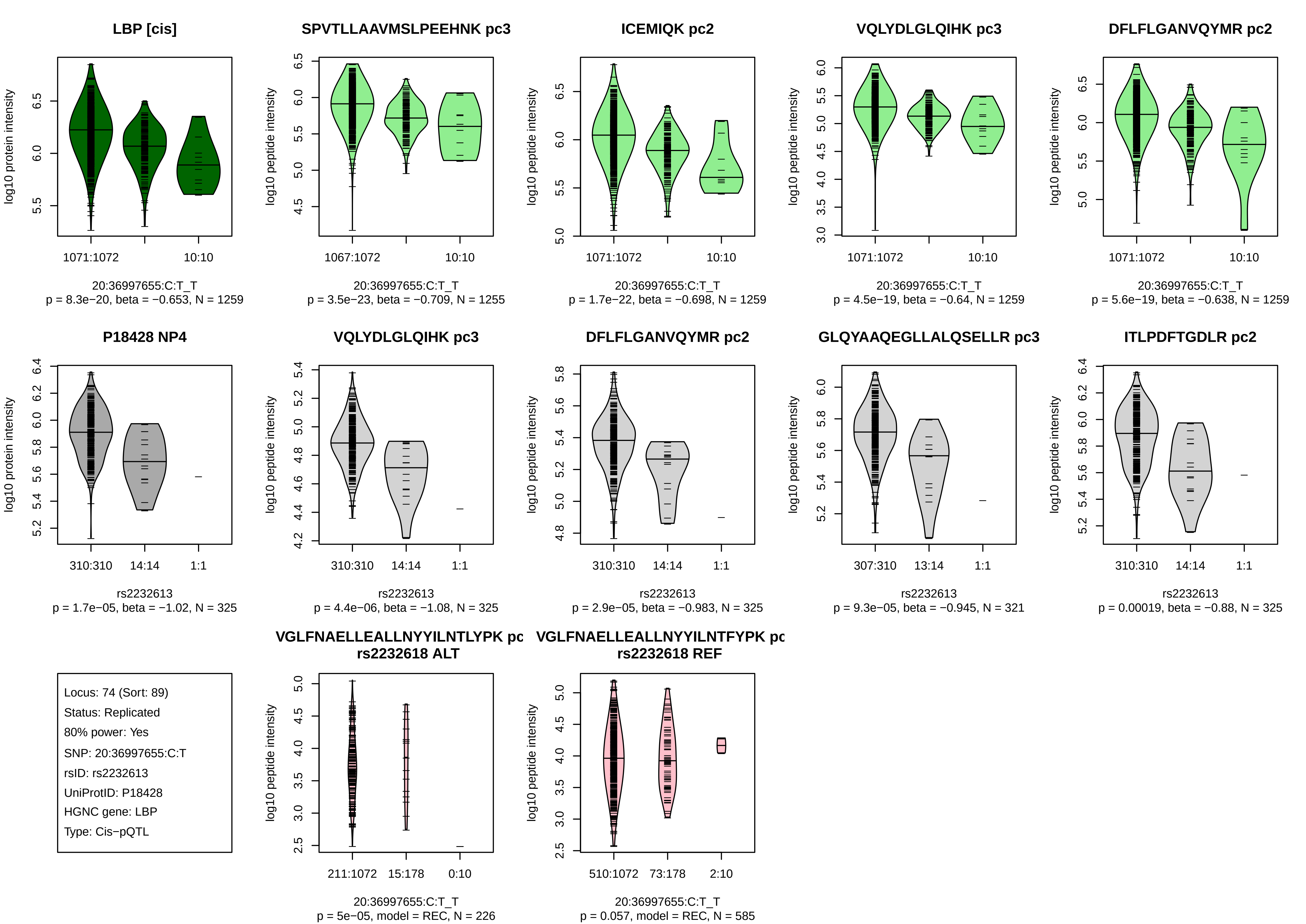

LBP [cis]
log10 protein intensity
1071:1072
10:10
20:36997655:C:T_T
p = 8.3e−20, beta = −0.653, N = 1259
SPVTLLAAVMSLPEEHNK pc3
log10 peptide intensity
1067:1072
10:10
20:36997655:C:T_T
p = 3.5e−23, beta = −0.709, N = 1255
ICEMIQK pc2
log10 peptide intensity
1071:1072
10:10
20:36997655:C:T_T
p = 1.7e−22, beta = −0.698, N = 1259
VQLYDLGLQIHK pc3
log10 peptide intensity
1071:1072
10:10
20:36997655:C:T_T
p = 4.5e−19, beta = −0.64, N = 1259
DFLFLGANVQYMR pc2
log10 peptide intensity
1071:1072
10:10
20:36997655:C:T_T
p = 5.6e−19, beta = −0.638, N = 1259
P18428 NP4
log10 protein intensity
310:310
14:14
1:1
rs2232613
p = 1.7e−05, beta = −1.02, N = 325
VQLYDLGLQIHK pc3
log10 peptide intensity
310:310
14:14
1:1
rs2232613
p = 4.4e−06, beta = −1.08, N = 325
DFLFLGANVQYMR pc2
log10 peptide intensity
310:310
14:14
1:1
rs2232613
p = 2.9e−05, beta = −0.983, N = 325
GLQYAAQEGLLALQSELLR pc3
log10 peptide intensity
307:310
13:14
1:1
rs2232613
p = 9.3e−05, beta = −0.945, N = 321
ITLPDFTGDLR pc2
log10 peptide intensity
310:310
14:14
1:1
rs2232613
p = 0.00019, beta = −0.88, N = 325
Locus: 74 (Sort: 89)
Status: Replicated
80% power: Yes
SNP: 20:36997655:C:T
rsID: rs2232613
UniProtID: P18428
HGNC gene: LBP
Type: Cis−pQTL
VGLFNAELLEALLNYYILNTLYPK pc
rs2232618 ALT
log10 peptide intensity
211:1072
15:178
0:10
20:36997655:C:T_T
p = 5e−05, model = REC, N = 226
VGLFNAELLEALLNYYILNTFYPK pc
rs2232618 REF
log10 peptide intensity
510:1072
73:178
2:10
20:36997655:C:T_T
p = 0.057, model = REC, N = 585

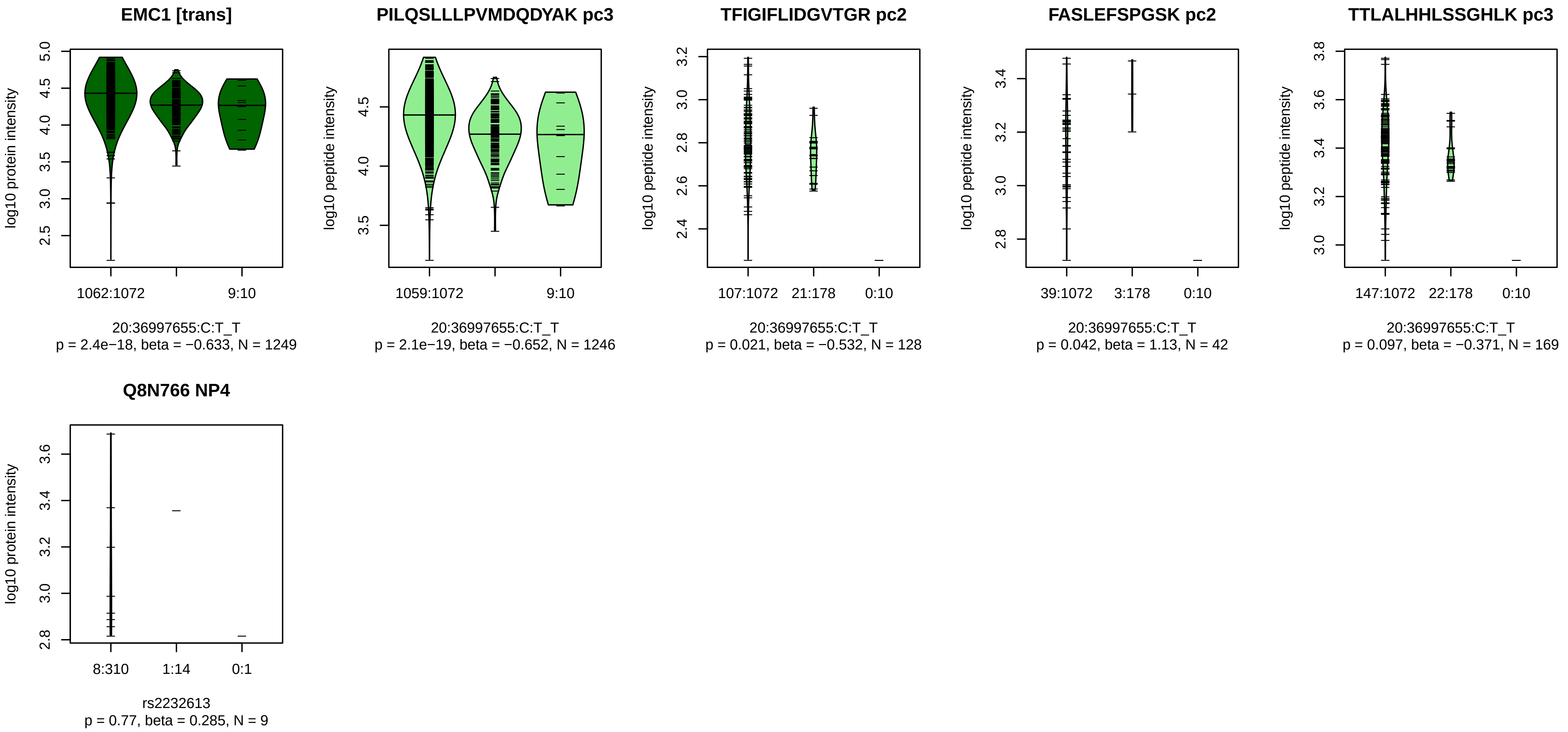


Locus: 74 (Sort: 90)

Status: Not replicated

80% power: No

SNP: 20:36997655:C:T

rsID: rs2232613

UniProtID: Q8N766

HGNC gene: EMC1

Type: Trans−pQTL

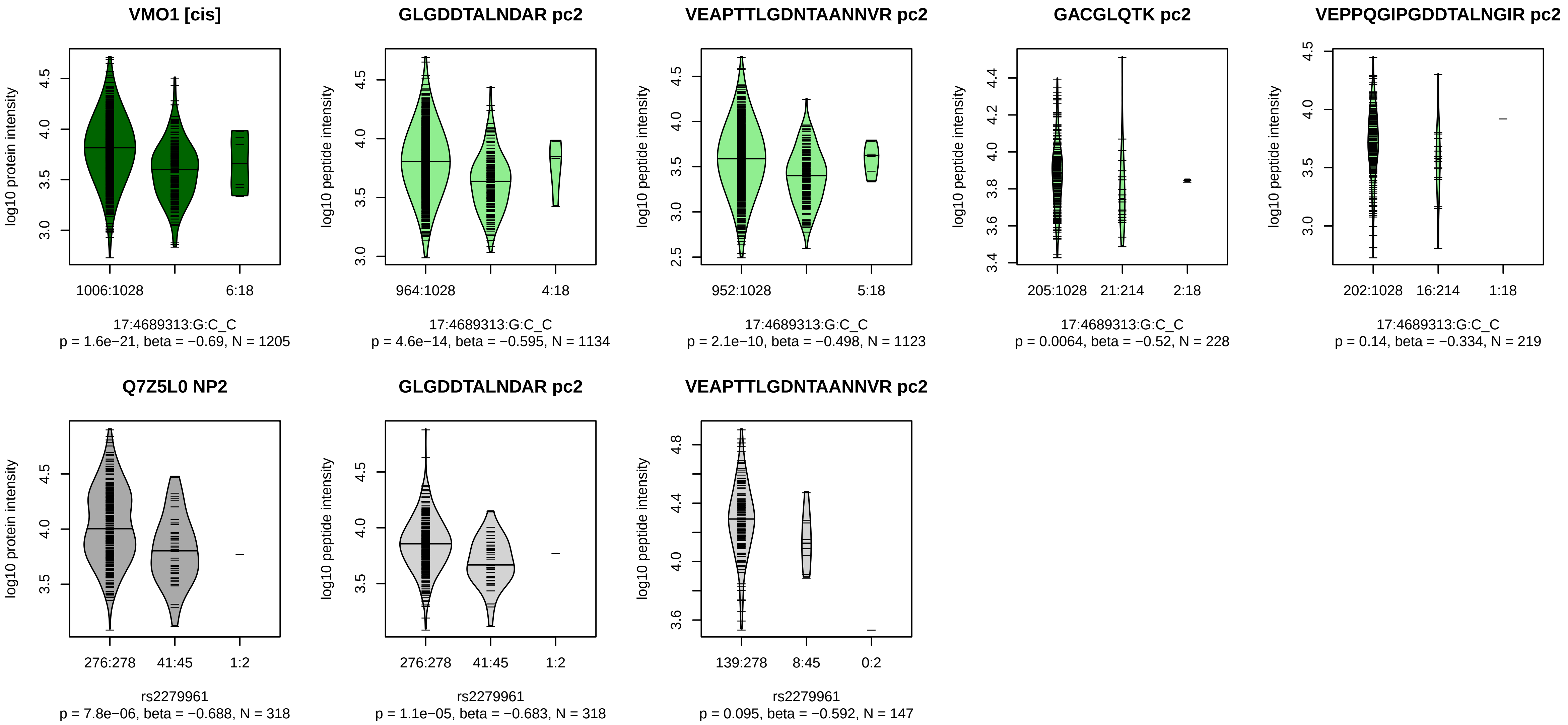


Locus: 75 (Sort: 91)
Status: Replicated
80% power: Yes
SNP: 17:4689313:G:C
rsID: rs2279961
UniProtID: Q7Z5L0
HGNC gene: VMO1
Type: Cis−pQTL

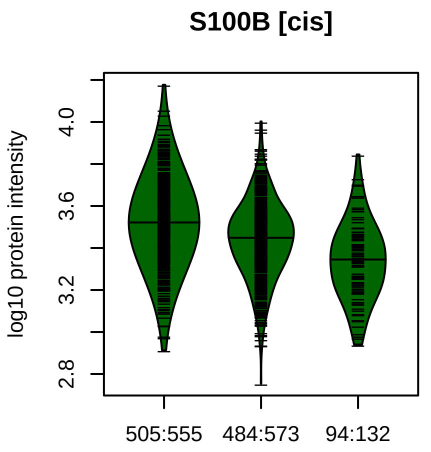


21:48025097:A:G_A
p = 2.3e−21, beta = −0.44, N = 1083

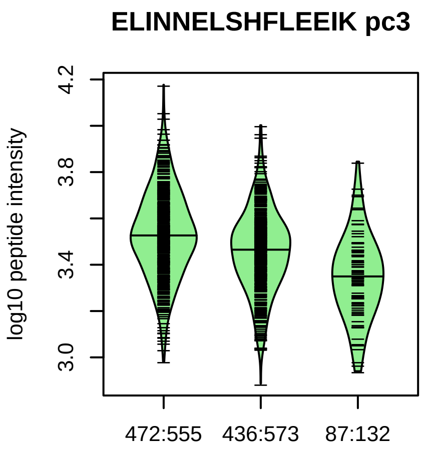


21:48025097:A:G_A
p = 3.3e−20, beta = −0.443, N = 995

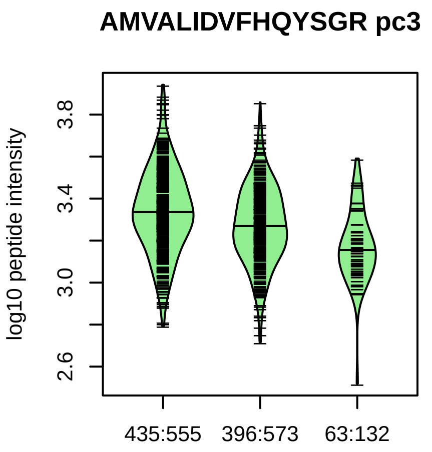


21:48025097:A:G_A
p = 6.7e−12, beta = −0.364, N = 894

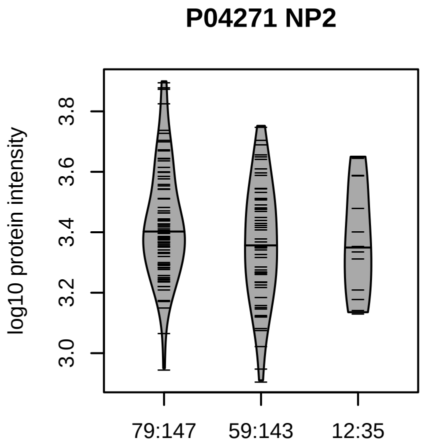


rs8128872
p = 0.025, beta = −0.276, N = 150

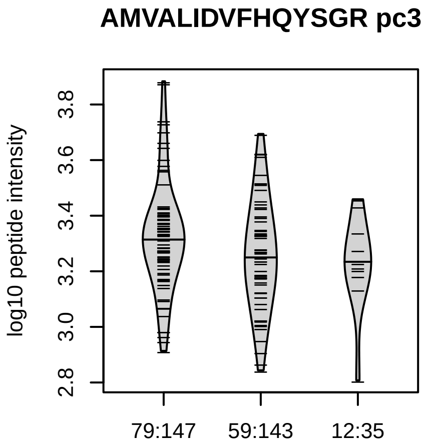


rs8128872
p = 0.019, beta = −0.29, N = 150

Locus: 76 (Sort: 92)
Status: Not replicated
80% power: No
SNP: 21:48025097:A:G
rsID: rs8128872
UniProtID: P04271
HGNC gene: S100B
Type: Cis−pQTL

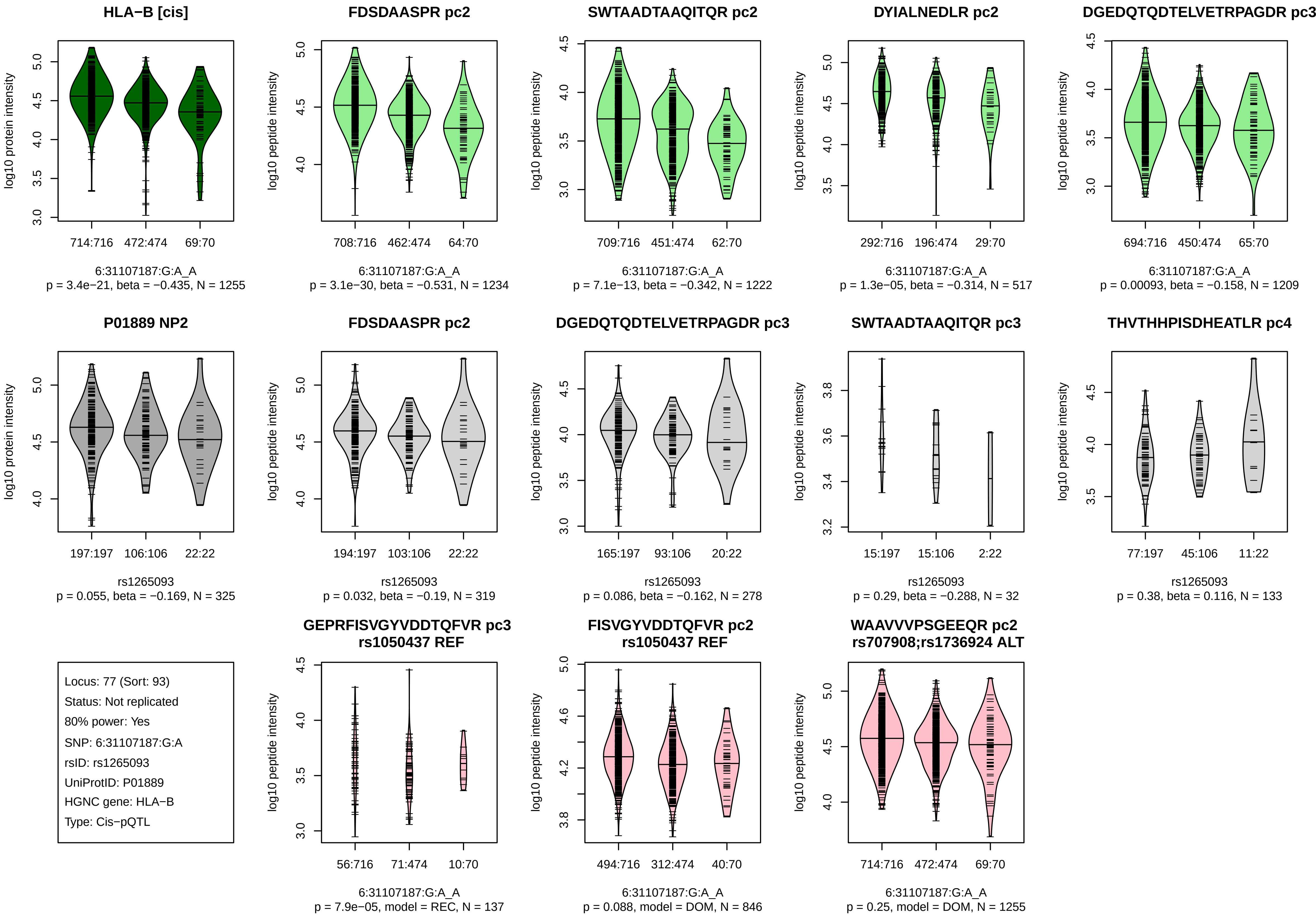
HLA−B [cis]
log10 protein intensity
714:716
472:474
69:70
6:31107187:G:A_A
p = 3.4e−21, beta = −0.435, N = 1255
FDSDAASPR pc2
log10 peptide intensity
708:716
462:474
64:70
6:31107187:G:A_A
p = 3.1e−30, beta = −0.531, N = 1234
SWTAADTAAQITQR pc2
log10 peptide intensity
709:716
451:474
62:70
6:31107187:G:A_A
p = 7.1e−13, beta = −0.342, N = 1222
DYIALNEDLR pc2
log10 peptide intensity
292:716
196:474
29:70
6:31107187:G:A_A
p = 1.3e−05, beta = −0.314, N = 517
DGEDQTQDTELVETRPAGDR pc3
log10 peptide intensity
694:716
450:474
65:70
6:31107187:G:A_A
p = 0.00093, beta = −0.158, N = 1209
P01889 NP2
log10 protein intensity
197:197
106:106
22:22
rs1265093
p = 0.055, beta = −0.169, N = 325
FDSDAASPR pc2
log10 peptide intensity
194:197
103:106
22:22
rs1265093
p = 0.032, beta = −0.19, N = 319
DGEDQTQDTELVETRPAGDR pc3
log10 peptide intensity
165:197
93:106
20:22
rs1265093
p = 0.086, beta = −0.162, N = 278
SWTAADTAAQITQR pc3
log10 peptide intensity
15:197
15:106
2:22
rs1265093
p = 0.29, beta = −0.288, N = 32
THVTHHPISDHEATLR pc4
log10 peptide intensity
77:197
45:106
11:22
rs1265093
p = 0.38, beta = 0.116, N = 133
Locus: 77 (Sort: 93)
Status: Not replicated
80% power: Yes
SNP: 6:31107187:G:A
rsID: rs1265093
UniProtID: P01889
HGNC gene: HLA−B
Type: Cis−pQTL
GEPRFISVGYVDDTQFVR pc3
rs1050437 REF
log10 peptide intensity
56:716
71:474
10:70
6:31107187:G:A_A
p = 7.9e−05, model = REC, N = 137
FISVGYVDDTQFVR pc2
rs1050437 REF
log10 peptide intensity
494:716
312:474
40:70
6:31107187:G:A_A
p = 0.088, model = DOM, N = 846
WAAVVVPSGEEQR pc2
rs707908;rs1736924 ALT
log10 peptide intensity
714:716
472:474
69:70
6:31107187:G:A_A
p = 0.25, model = DOM, N = 1255

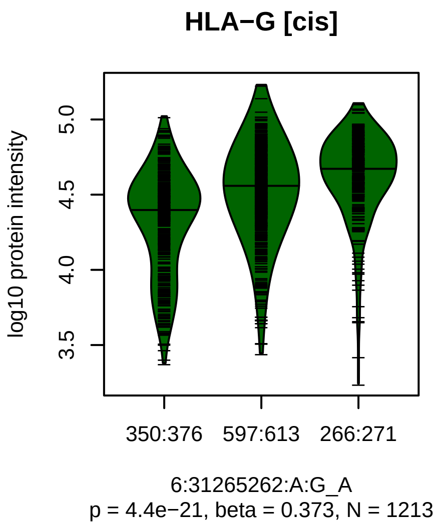


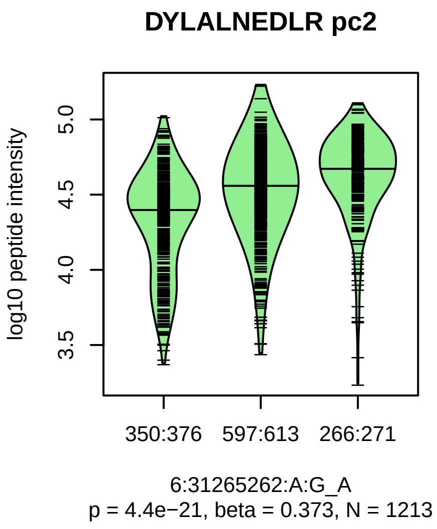


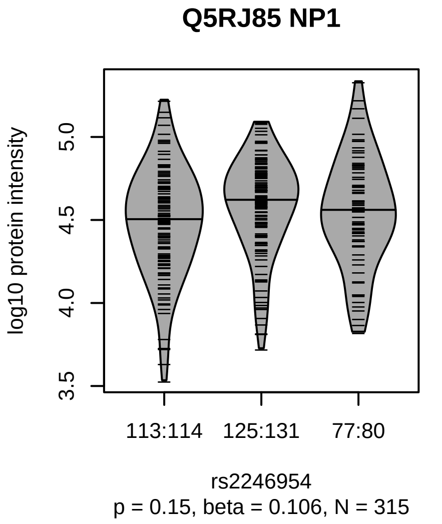


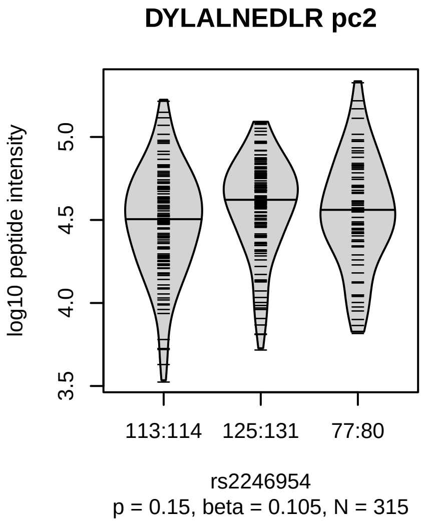


Locus: 78 (Sort: 94)
Status: Not replicated
80% power: Yes
SNP: 6:31265262:A:G
rsID: rs2246954
UniProtID: Q5RJ85
HGNC gene: HLA−G
Type: Cis−pQTL

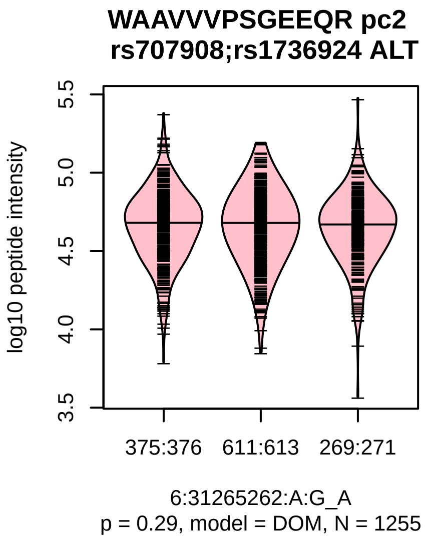

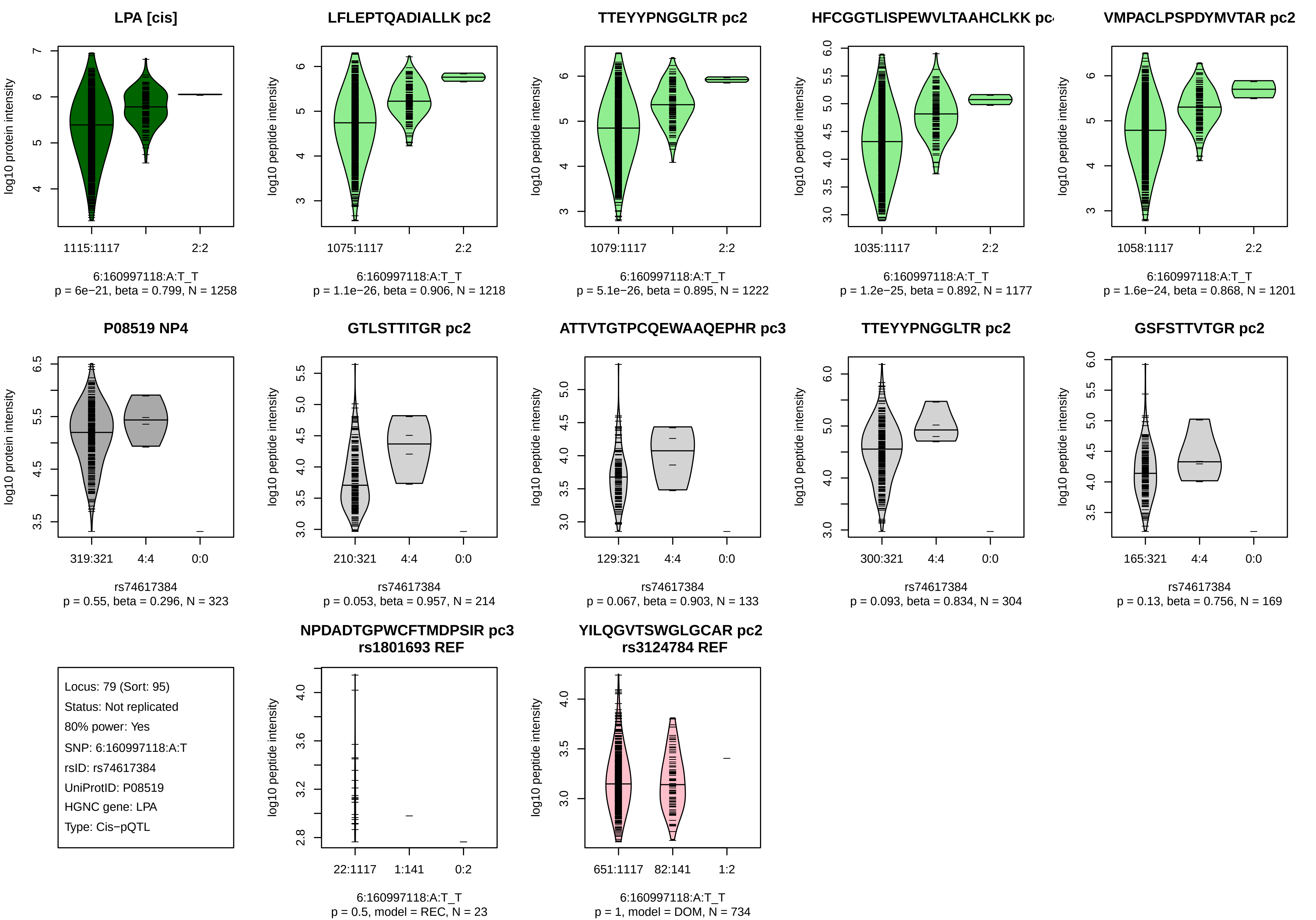

LPA [cis]
log10 protein intensity
1115:1117
2:2
6:160997118:A:T_T
p = 6e−21, beta = 0.799, N = 1258
LFLEPTQADIALLK pc2
log10 peptide intensity
1075:1117
2:2
6:160997118:A:T_T
p = 1.1e−26, beta = 0.906, N = 1218
TTEYYPNGGLTR pc2
1079:1117
2:2
6:160997118:A:T_T
p = 5.1e−26, beta = 0.895, N = 1222
HFCGGTLISPEWVLTAAHCLKK pc
1035:1117
2:2
6:160997118:A:T_T
p = 1.2e−25, beta = 0.892, N = 1177
VMPACLPSPDYMVTAR pc2
1058:1117
2:2
6:160997118:A:T_T
p = 1.6e−24, beta = 0.868, N = 1201
P08519 NP4
319:321
4:4
0:0
rs74617384
p = 0.55, beta = 0.296, N = 323
GTLSTTITGR pc2
210:321
4:4
0:0
rs74617384
p = 0.053, beta = 0.957, N = 214
ATTVTGTPCQEWAAQEPHR pc3
129:321
4:4
0:0
rs74617384
p = 0.067, beta = 0.903, N = 133
TTEYYPNGGLTR pc2
300:321
4:4
0:0
rs74617384
p = 0.093, beta = 0.834, N = 304
GSFSTTVTGR pc2
165:321
4:4
0:0
rs74617384
p = 0.13, beta = 0.756, N = 169
Locus: 79 (Sort: 95)
Status: Not replicated
80% power: Yes
SNP: 6:160997118:A:T
rsID: rs74617384
UniProtID: P08519
HGNC gene: LPA
Type: Cis−pQTL
NPDADTGPWCFTMDPSIR pc3
rs1801693 REF
22:1117
1:141
0:2
6:160997118:A:T_T
p = 0.5, model = REC, N = 23
YILQGVTSWGLGCAR pc2
rs3124784 REF
651:1117
82:141
1:2
6:160997118:A:T_T
p = 1, model = DOM, N = 734

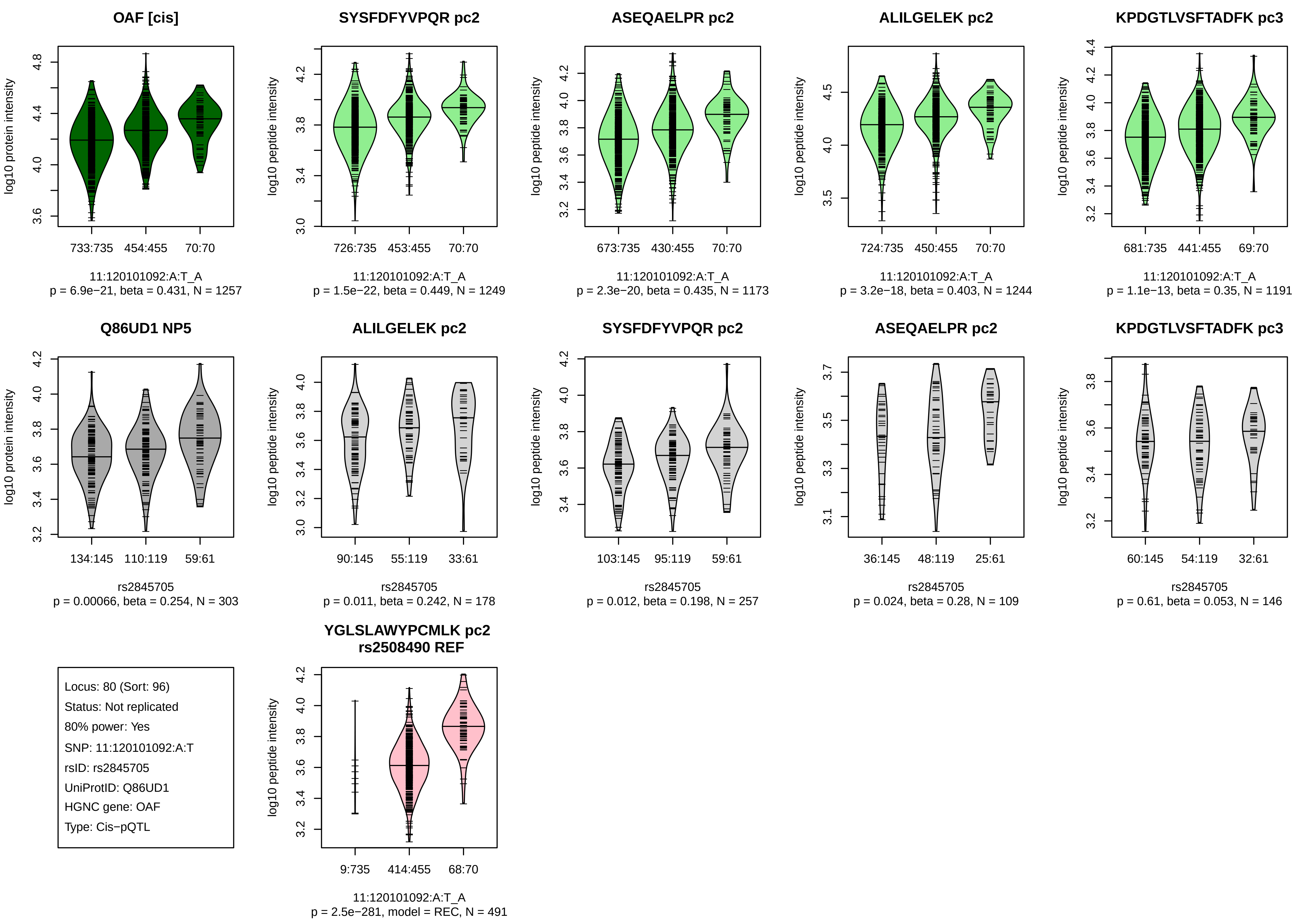
OAF [cis]
log10 protein intensity
733:735
454:455
70:70
11:120101092:A:T_A
p = 6.9e−21, beta = 0.431, N = 1257
SYSFDFYVPQR pc2
log10 peptide intensity
726:735
453:455
70:70
11:120101092:A:T_A
p = 1.5e−22, beta = 0.449, N = 1249
ASEQAELPR pc2
log10 peptide intensity
673:735
430:455
70:70
11:120101092:A:T_A
p = 2.3e−20, beta = 0.435, N = 1173
ALILGELEK pc2
log10 peptide intensity
724:735
450:455
70:70
11:120101092:A:T_A
p = 3.2e−18, beta = 0.403, N = 1244
KPDGTLVSFTADFK pc3
log10 peptide intensity
681:735
441:455
69:70
11:120101092:A:T_A
p = 1.1e−13, beta = 0.35, N = 1191
Q86UD1 NP5
log10 protein intensity
134:145
110:119
59:61
rs2845705
p = 0.00066, beta = 0.254, N = 303
ALILGELEK pc2
log10 peptide intensity
90:145
55:119
33:61
rs2845705
p = 0.011, beta = 0.242, N = 178
SYSFDFYVPQR pc2
log10 peptide intensity
103:145
95:119
59:61
rs2845705
p = 0.012, beta = 0.198, N = 257
ASEQAELPR pc2
log10 peptide intensity
36:145
48:119
25:61
rs2845705
p = 0.024, beta = 0.28, N = 109
KPDGTLVSFTADFK pc3
log10 peptide intensity
60:145
54:119
32:61
rs2845705
p = 0.61, beta = 0.053, N = 146
Locus: 80 (Sort: 96)
Status: Not replicated
80% power: Yes
SNP: 11:120101092:A:T
rsID: rs2845705
UniProtID: Q86UD1
HGNC gene: OAF
Type: Cis−pQTL
YGLSLAWYPCMLK pc2
rs2508490 REF
log10 peptide intensity
9:735
414:455
68:70
11:120101092:A:T_A
p = 2.5e−281, model = REC, N = 491

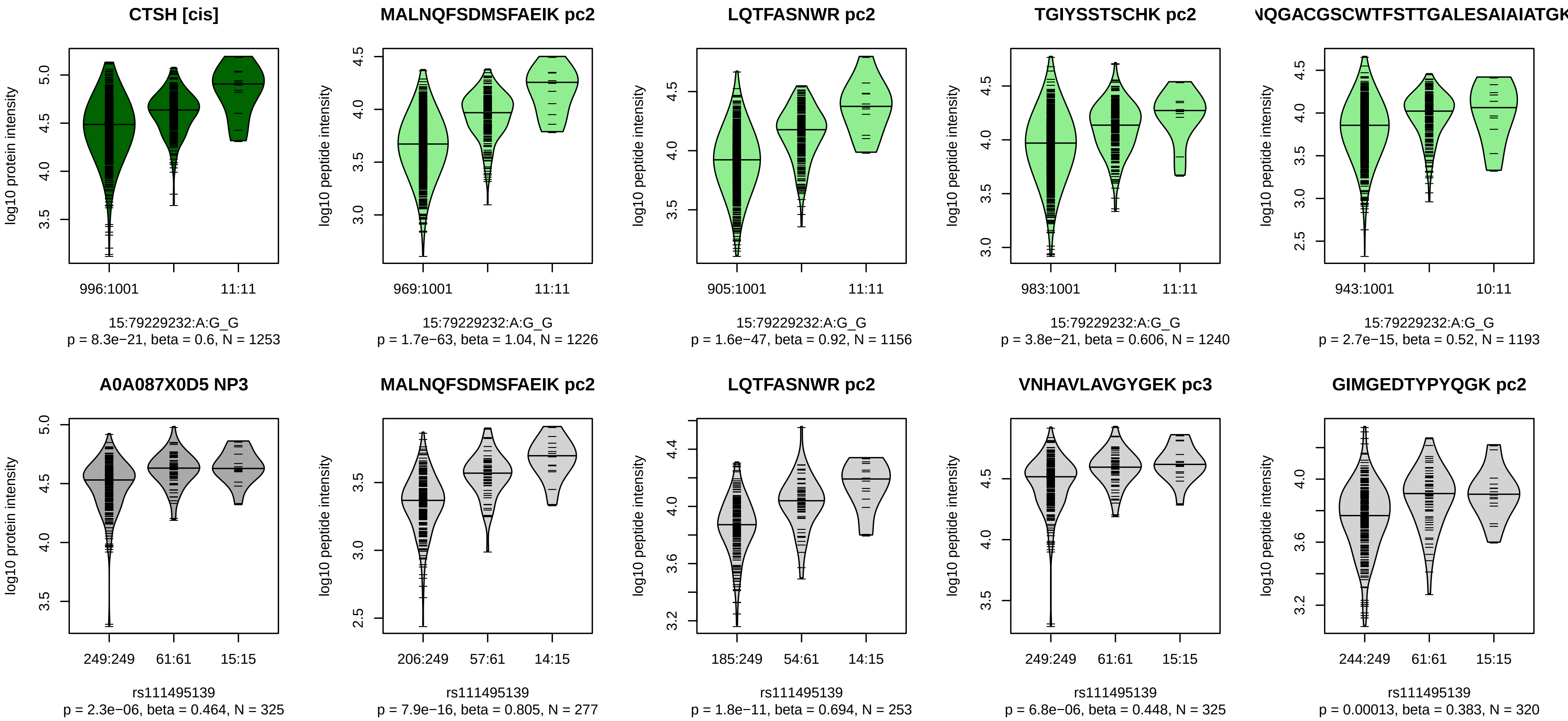


Locus: 81 (Sort: 97)
Status: Replicated
80% power: Yes
SNP: 15:79229232:A:G
rsID: rs111495139
UniProtID: A0A087X0D5
HGNC gene: CTSH
Type: Cis−pQTL

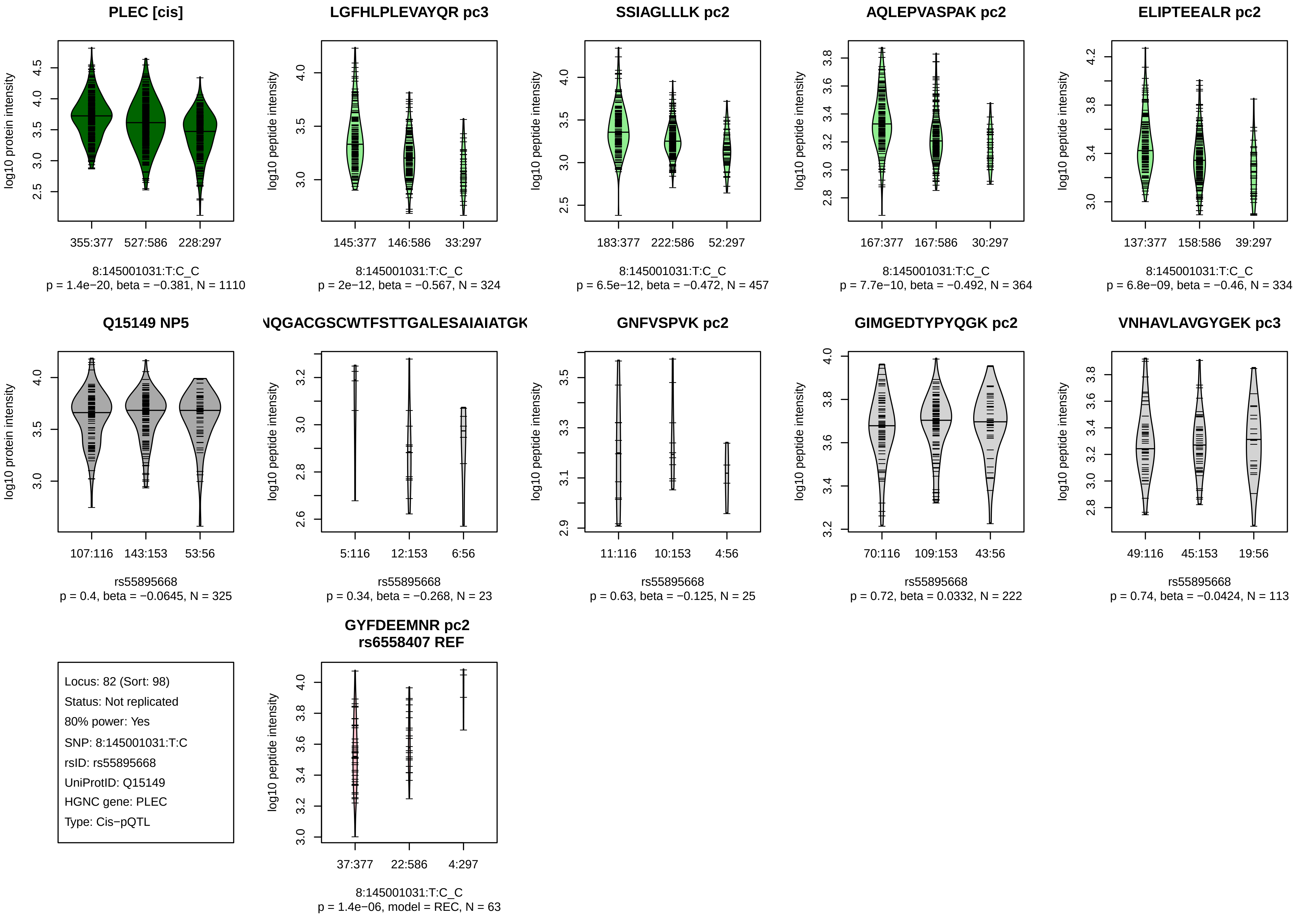
PLEC [cis]
log10 protein intensity
355:377
527:586
228:297
8:145001031:T:C_C
p = 1.4e−20, beta = −0.381, N = 1110
LGFHLPLEVAYQR pc3
log10 peptide intensity
145:377
146:586
33:297
8:145001031:T:C_C
p = 2e−12, beta = −0.567, N = 324
SSIAGLLLK pc2
log10 peptide intensity
183:377
222:586
52:297
8:145001031:T:C_C
p = 6.5e−12, beta = −0.472, N = 457
AQLEPVASPAK pc2
log10 peptide intensity
167:377
167:586
30:297
8:145001031:T:C_C
p = 7.7e−10, beta = −0.492, N = 364
ELIPTEEALR pc2
log10 peptide intensity
137:377
158:586
39:297
8:145001031:T:C_C
p = 6.8e−09, beta = −0.46, N = 334
Q15149 NP5
log10 protein intensity
107:116
143:153
53:56
rs55895668
p = 0.4, beta = −0.0645, N = 325
NQGACGSCWTFSTTGALESAIAIATGK
log10 peptide intensity
5:116
12:153
6:56
rs55895668
p = 0.34, beta = −0.268, N = 23
GNFVSPVK pc2
log10 peptide intensity
11:116
10:153
4:56
rs55895668
p = 0.63, beta = −0.125, N = 25
GIMGEDTYPYQGK pc2
log10 peptide intensity
70:116
109:153
43:56
rs55895668
p = 0.72, beta = 0.0332, N = 222
VNHAVLAVGYGEK pc3
log10 peptide intensity
49:116
45:153
19:56
rs55895668
p = 0.74, beta = −0.0424, N = 113
Locus: 82 (Sort: 98)
Status: Not replicated
80% power: Yes
SNP: 8:145001031:T:C
rsID: rs55895668
UniProtID: Q15149
HGNC gene: PLEC
Type: Cis−pQTL
GYFDEEMNR pc2
rs6558407 REF
log10 peptide intensity
37:377
22:586
4:297
8:145001031:T:C_C
p = 1.4e−06, model = REC, N = 63

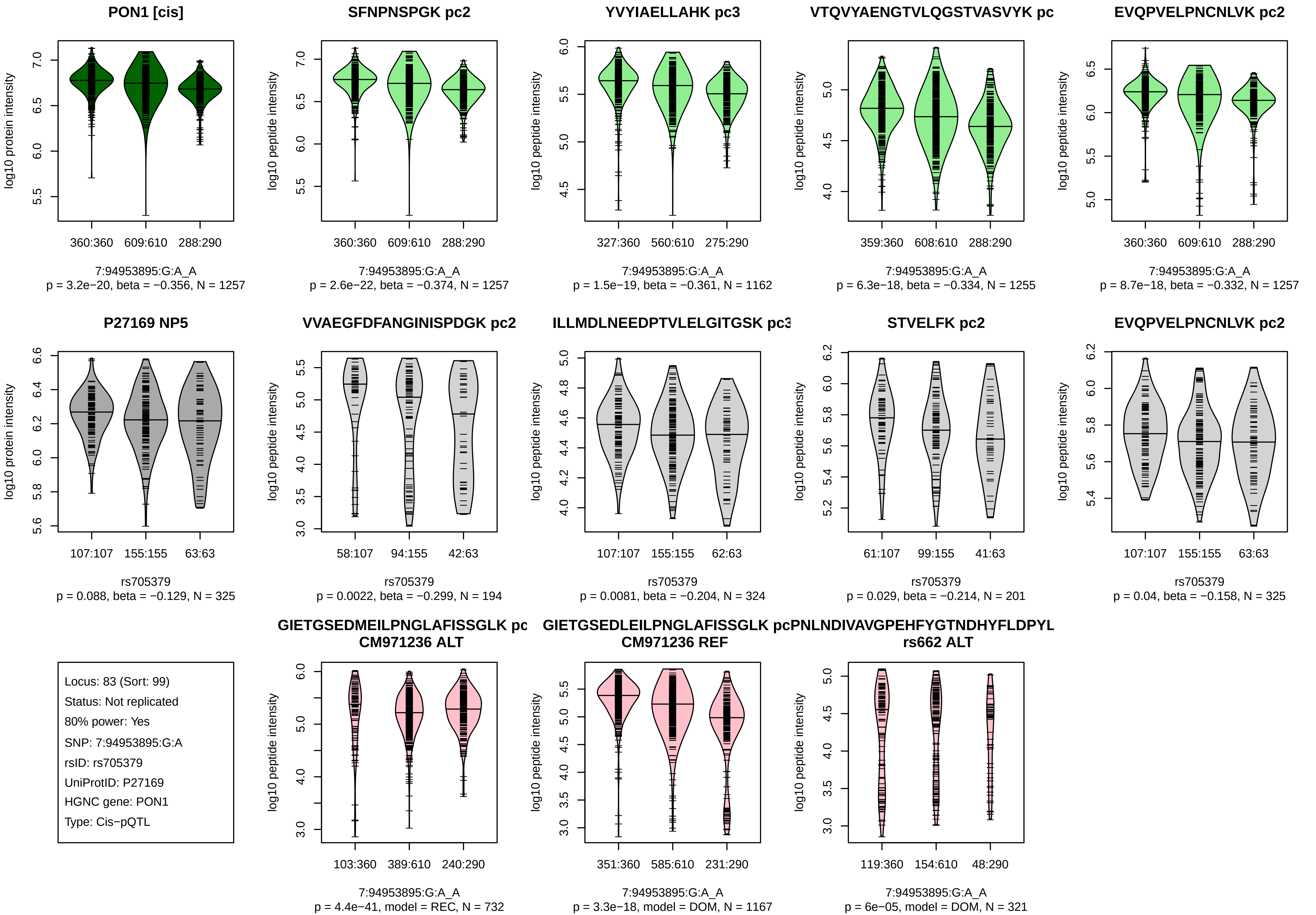
PON1 [cis]
log10 protein intensity
360:360 609:610 288:290
7:94953895:G:A_A
p = 3.2e−20, beta = −0.356, N = 1257
SFNPNSPGK pc2
log10 peptide intensity
360:360 609:610 288:290
7:94953895:G:A_A
p = 2.6e−22, beta = −0.374, N = 1257
YVYIAELLAHK pc3
log10 peptide intensity
327:360 560:610 275:290
7:94953895:G:A_A
p = 1.5e−19, beta = −0.361, N = 1162
VTQVYAENGTVLQGSTVASVYK pc
log10 peptide intensity
359:360 608:610 288:290
7:94953895:G:A_A
p = 6.3e−18, beta = −0.334, N = 1255
EVQPVELPNCNLVK pc2
log10 peptide intensity
360:360 609:610 288:290
7:94953895:G:A_A
p = 8.7e−18, beta = −0.332, N = 1257
P27169 NP5
log10 protein intensity
107:107 155:155 63:63
rs705379
p = 0.088, beta = −0.129, N = 325
VVAEGFDFANGINISPDGK pc2
log10 peptide intensity
58:107 94:155 42:63
rs705379
p = 0.0022, beta = −0.299, N = 194
ILLMDLNEEDPTVLELGITGSK pc3
log10 peptide intensity
107:107 155:155 62:63
rs705379
p = 0.0081, beta = −0.204, N = 324
STVELFK pc2
log10 peptide intensity
61:107 99:155 41:63
rs705379
p = 0.029, beta = −0.214, N = 201
EVQPVELPNCNLVK pc2
log10 peptide intensity
107:107 155:155 63:63
rs705379
p = 0.04, beta = −0.158, N = 325
Locus: 83 (Sort: 99)
Status: Not replicated
80% power: Yes
SNP: 7:94953895:G:A
rsID: rs705379
UniProtID: P27169
HGNC gene: PON1
Type: Cis−pQTL
GIETGSEDMEILPNGLAFISSGLK pc
CM971236 ALT
log10 peptide intensity
103:360 389:610 240:290
7:94953895:G:A_A
p = 4.4e−41, model = REC, N = 732
GIETGSEDLEILPNGLAFISSGLK pc
CM971236 REF
log10 peptide intensity
351:360 585:610 231:290
7:94953895:G:A_A
p = 3.3e−18, model = DOM, N = 1167
PNLNDIVAVGPEHFYGTNDHYFLDPYL
rs662 ALT
log10 peptide intensity
119:360 154:610 48:290
7:94953895:G:A_A
p = 6e−05, model = DOM, N = 321

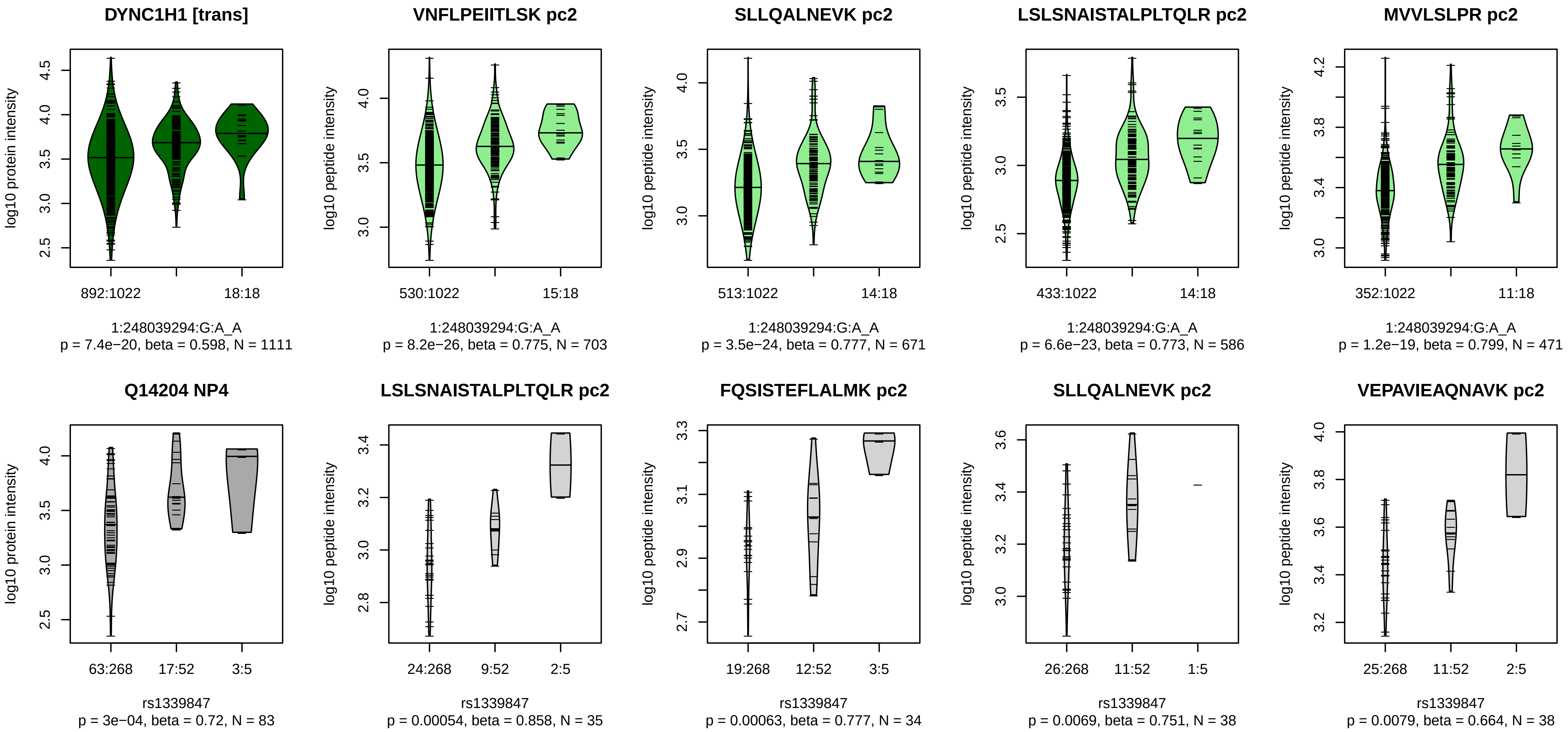


Locus: 84 (Sort: 100)

Status: Not replicated

80% power: No

SNP: 1:248039294:G:A

rsID: rs1339847

UniProtID: Q14204

HGNC gene: DYNC1H1

Type: Trans−pQTL

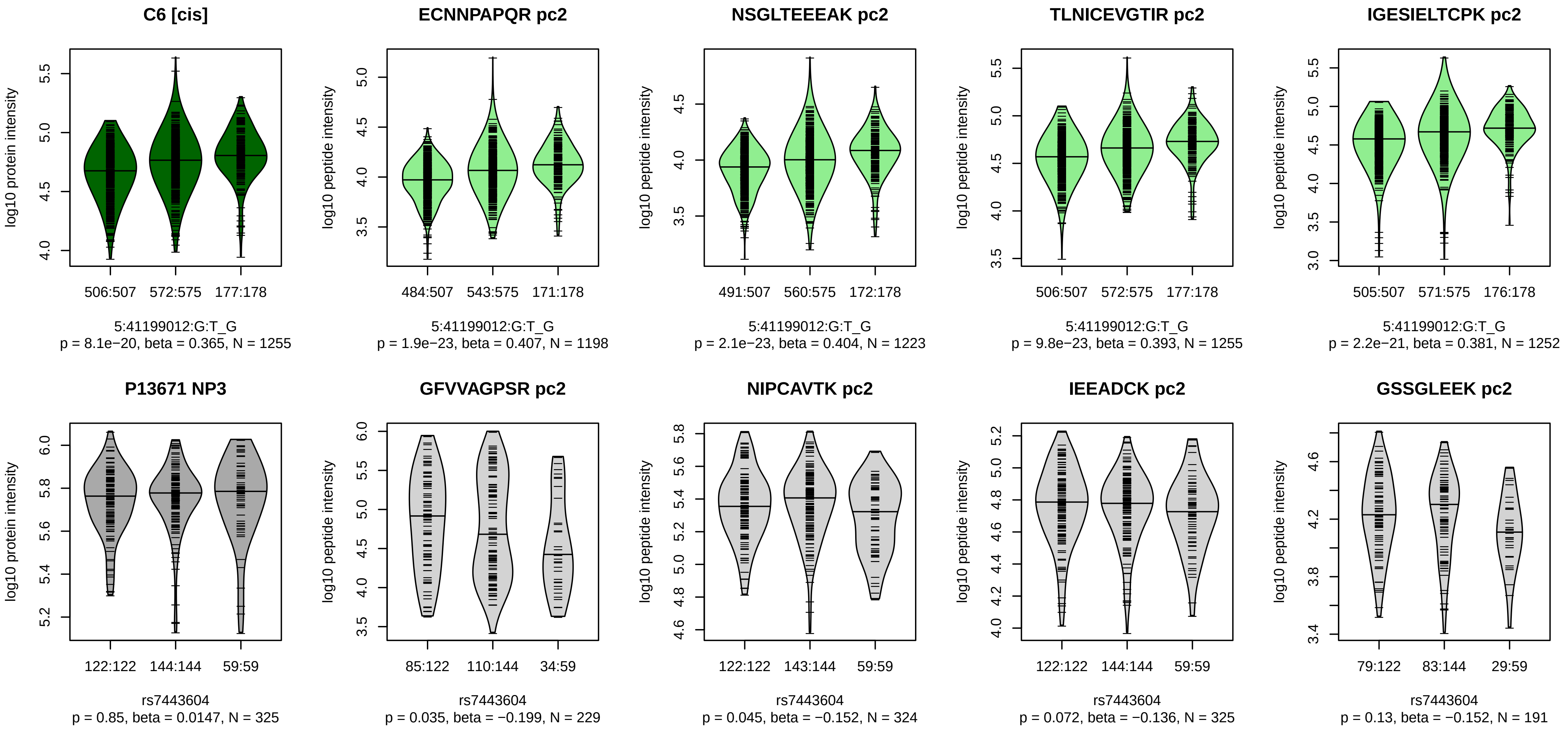


Locus: 85 (Sort: 101)
Status: Not replicated
80% power: Yes
SNP: 5:41199012:G:T
rsID: rs7443604
UniProtID: P13671
HGNC gene: C6
Type: Cis−pQTL

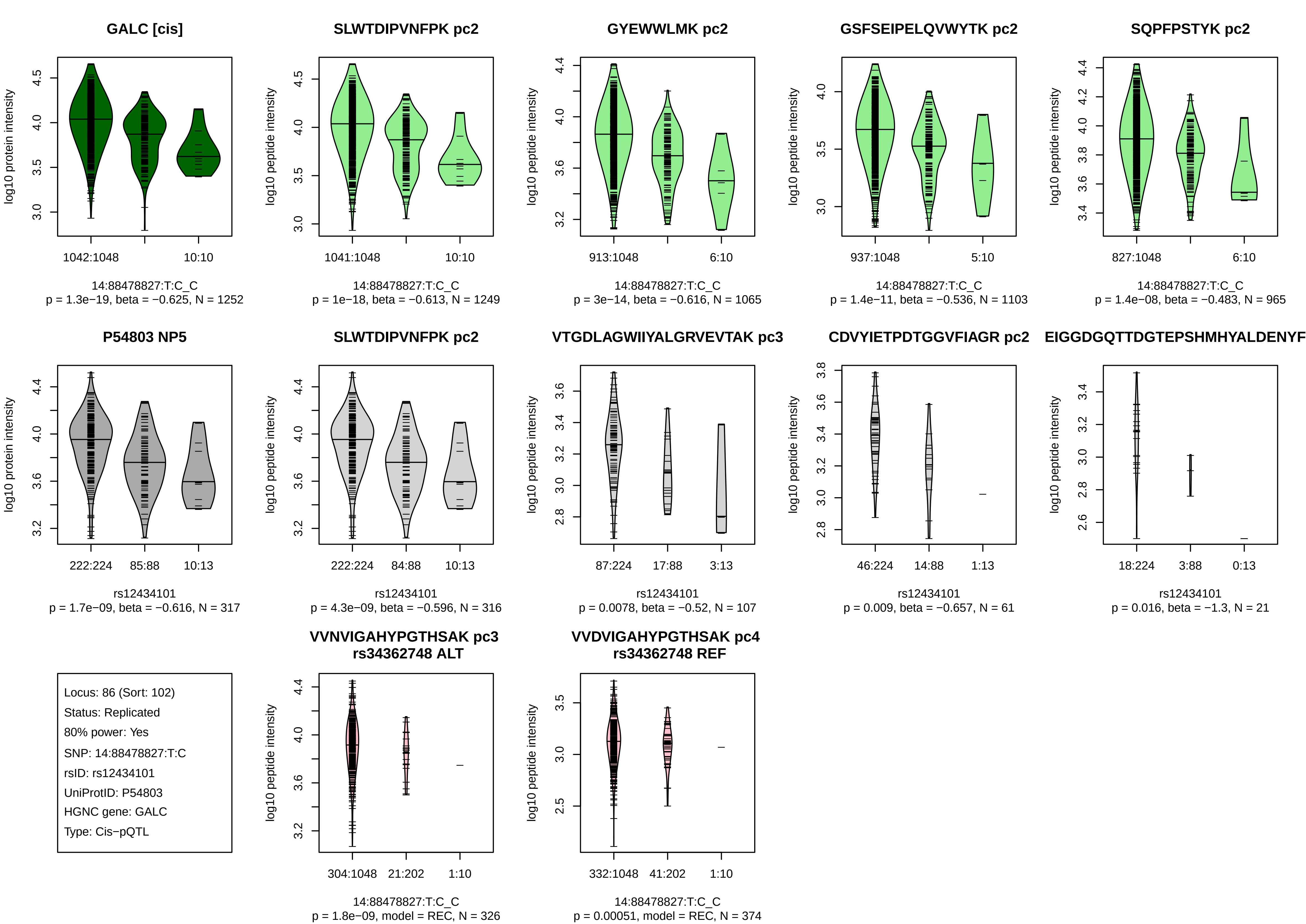

GALC [cis]
log10 protein intensity
1042:1048
10:10
14:88478827:T:C_C
p = 1.3e−19, beta = −0.625, N = 1252
SLWTDIPVNFPK pc2
log10 peptide intensity
1041:1048
10:10
14:88478827:T:C_C
p = 1e−18, beta = −0.613, N = 1249
GYEWWLMK pc2
log10 peptide intensity
913:1048
6:10
14:88478827:T:C_C
p = 3e−14, beta = −0.616, N = 1065
GSFSEIPELQVWYTK pc2
log10 peptide intensity
937:1048
5:10
14:88478827:T:C_C
p = 1.4e−11, beta = −0.536, N = 1103
SQPFPSTYK pc2
log10 peptide intensity
827:1048
6:10
14:88478827:T:C_C
p = 1.4e−08, beta = −0.483, N = 965
P54803 NP5
log10 protein intensity
222:224
85:88
10:13
rs12434101
p = 1.7e−09, beta = −0.616, N = 317
SLWTDIPVNFPK pc2
log10 peptide intensity
222:224
84:88
10:13
rs12434101
p = 4.3e−09, beta = −0.596, N = 316
VTGDLAGWIIYALGRVEVTAK pc3
log10 peptide intensity
87:224
17:88
3:13
rs12434101
p = 0.0078, beta = −0.52, N = 107
CDVYIETPDTGGVFIAGR pc2
log10 peptide intensity
46:224
14:88
1:13
rs12434101
p = 0.009, beta = −0.657, N = 61
EIGGDGQTTDGTEPSHMHYALDENYF
log10 peptide intensity
18:224
3:88
0:13
rs12434101
p = 0.016, beta = −1.3, N = 21
Locus: 86 (Sort: 102)
Status: Replicated
80% power: Yes
SNP: 14:88478827:T:C
rsID: rs12434101
UniProtID: P54803
HGNC gene: GALC
Type: Cis−pQTL
VVNVIGAHYPGTHSAK pc3
rs34362748 ALT
log10 peptide intensity
304:1048
21:202
1:10
14:88478827:T:C_C
p = 1.8e−09, model = REC, N = 326
VVDVIGAHYPGTHSAK pc4
rs34362748 REF
log10 peptide intensity
332:1048
41:202
1:10
14:88478827:T:C_C
p = 0.00051, model = REC, N = 374

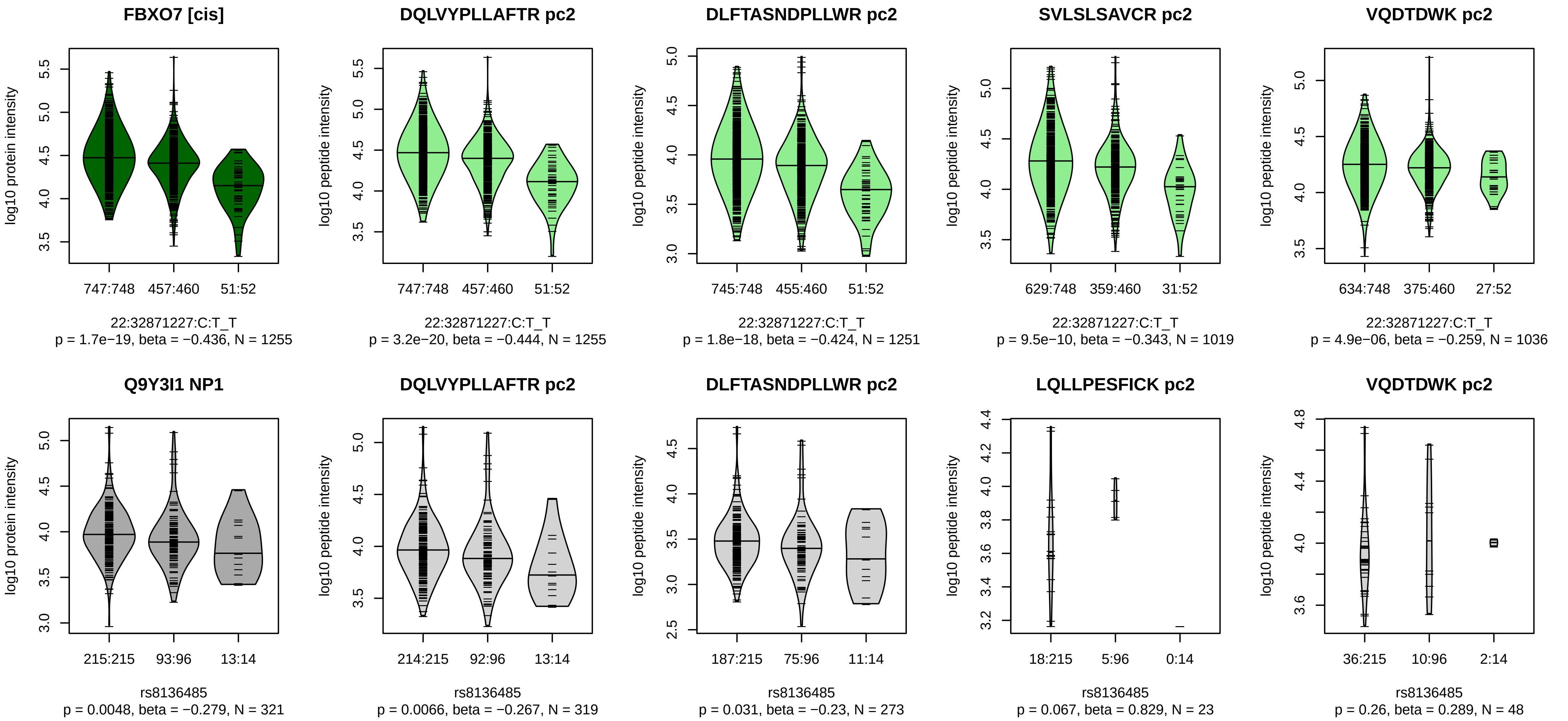


Locus: 87 (Sort: 103)
Status: Not replicated
80% power: Yes
SNP: 22:32871227:C:T
rsID: rs8136485
UniProtID: Q9Y3I1
HGNC gene: FBXO7
Type: Cis−pQTL

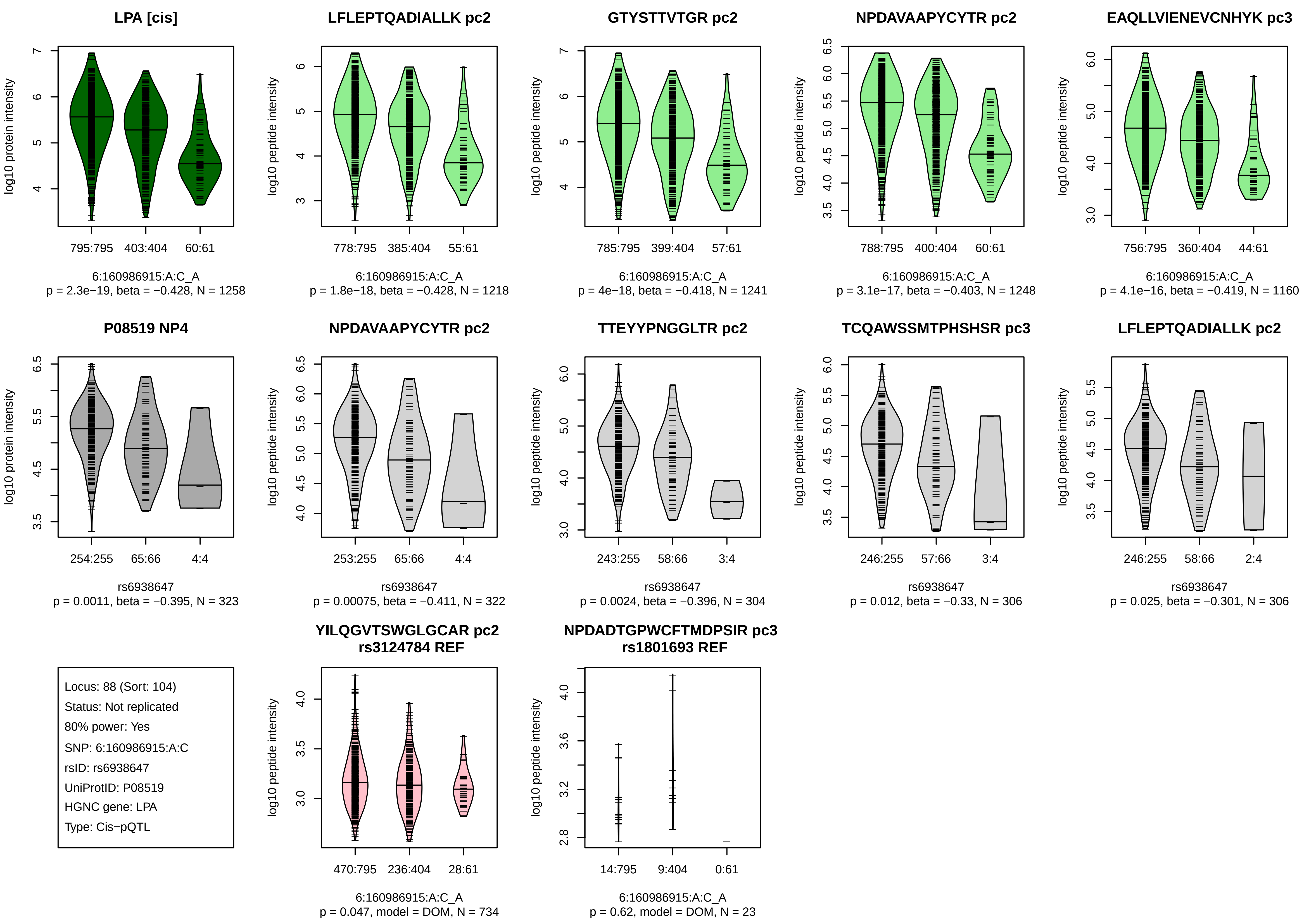
LPA [cis]
795:795 403:404 60:61
6:160986915:A:C_A
p = 2.3e−19, beta = −0.428, N = 1258
log10 protein intensity
LFLEPTQADIALLK pc2
778:795 385:404 55:61
6:160986915:A:C_A
p = 1.8e−18, beta = −0.428, N = 1218
log10 peptide intensity
GTYSTTVTGR pc2
785:795 399:404 57:61
6:160986915:A:C_A
p = 4e−18, beta = −0.418, N = 1241
NPDAVAAPYCYTR pc2
788:795 400:404 60:61
6:160986915:A:C_A
p = 3.1e−17, beta = −0.403, N = 1248
EAQLLVIENEVCNHYK pc3
756:795 360:404 44:61
6:160986915:A:C_A
p = 4.1e−16, beta = −0.419, N = 1160
P08519 NP4
254:255 65:66 4:4
rs6938647
p = 0.0011, beta = −0.395, N = 323
NPDAVAAPYCYTR pc2
253:255 65:66 4:4
rs6938647
p = 0.00075, beta = −0.411, N = 322
TTEYYPNGGLTR pc2
243:255 58:66 3:4
rs6938647
p = 0.0024, beta = −0.396, N = 304
TCQAWSSMTPHSHSR pc3
246:255 57:66 3:4
rs6938647
p = 0.012, beta = −0.33, N = 306
LFLEPTQADIALLK pc2
246:255 58:66 2:4
rs6938647
p = 0.025, beta = −0.301, N = 306
Locus: 88 (Sort: 104)
Status: Not replicated
80% power: Yes
SNP: 6:160986915:A:C
rsID: rs6938647
UniProtID: P08519
HGNC gene: LPA
Type: Cis−pQTL
YILQGVTSWGLGCAR pc2
rs3124784 REF
470:795 236:404 28:61
6:160986915:A:C_A
p = 0.047, model = DOM, N = 734
NPDADTGPWCFTMDPSIR pc3
rs1801693 REF
14:795 9:404 0:61
6:160986915:A:C_A
p = 0.62, model = DOM, N = 23

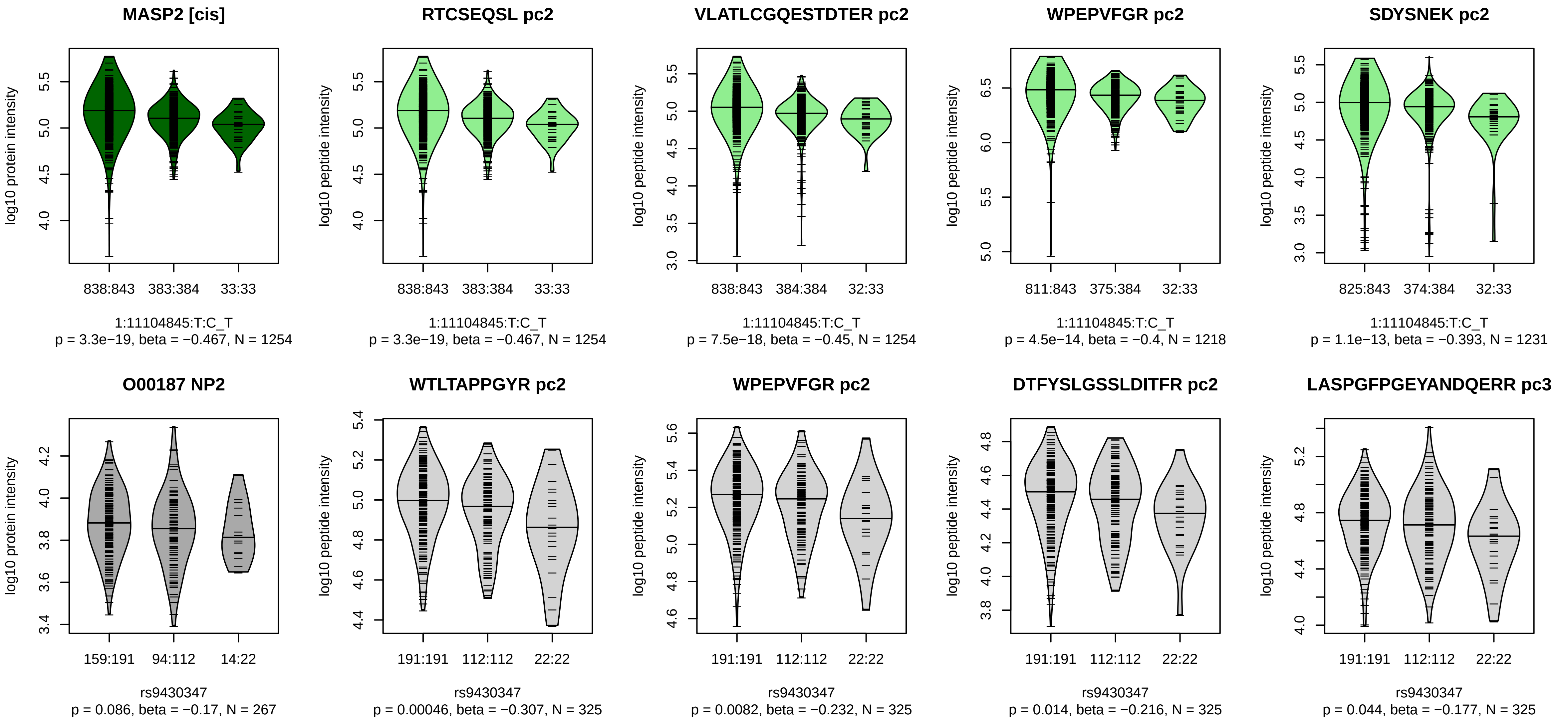


Locus: 89 (Sort: 105)

Status: Not replicated

80% power: No

SNP: 1:11104845:T:C

rsID: rs9430347

UniProtID: O00187

HGNC gene: MASP2

Type: Cis−pQTL

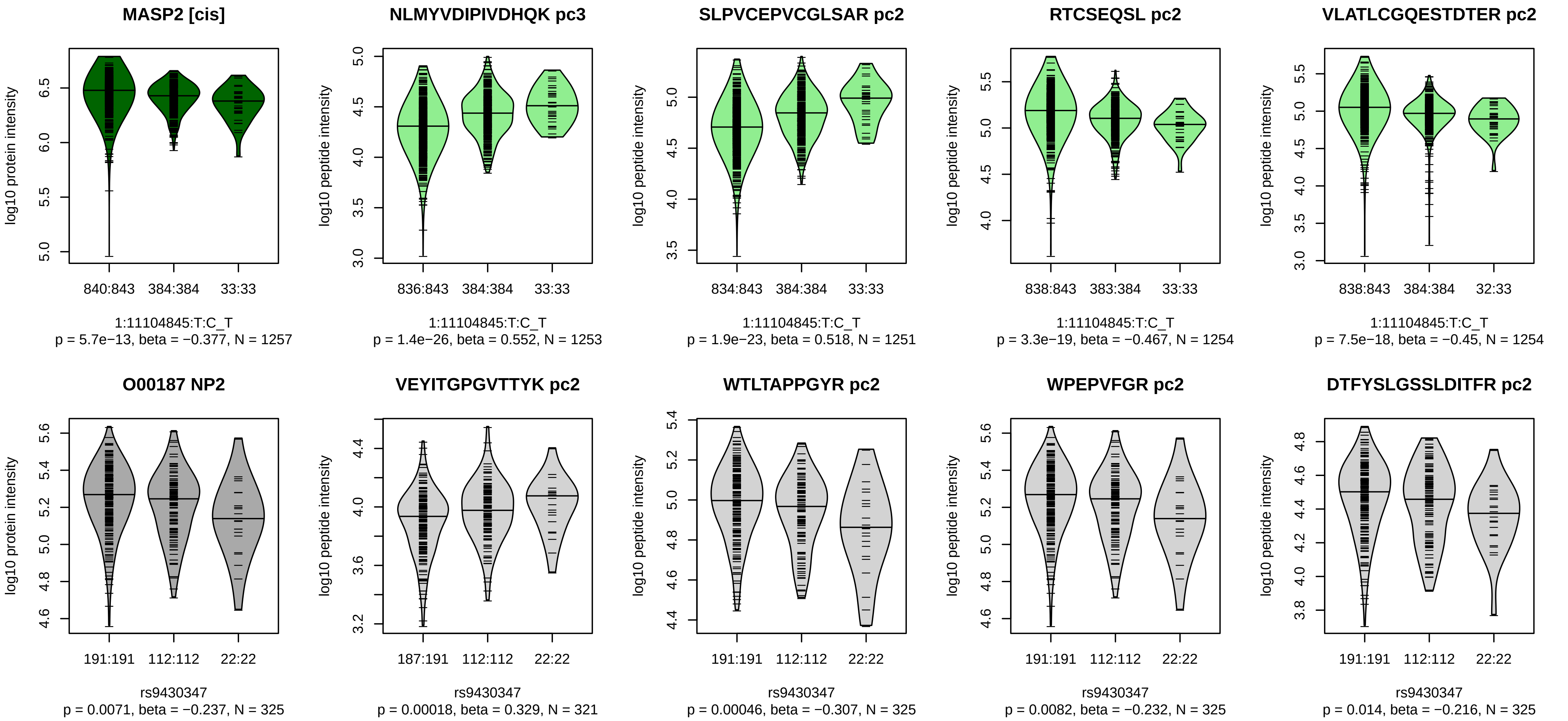


Locus: 89 (Sort: 106)
Status: Not replicated
80% power: No
SNP: 1:11104845:T:C
rsID: rs9430347
UniProtID: O00187
HGNC gene: MASP2
Type: Cis−pQTL

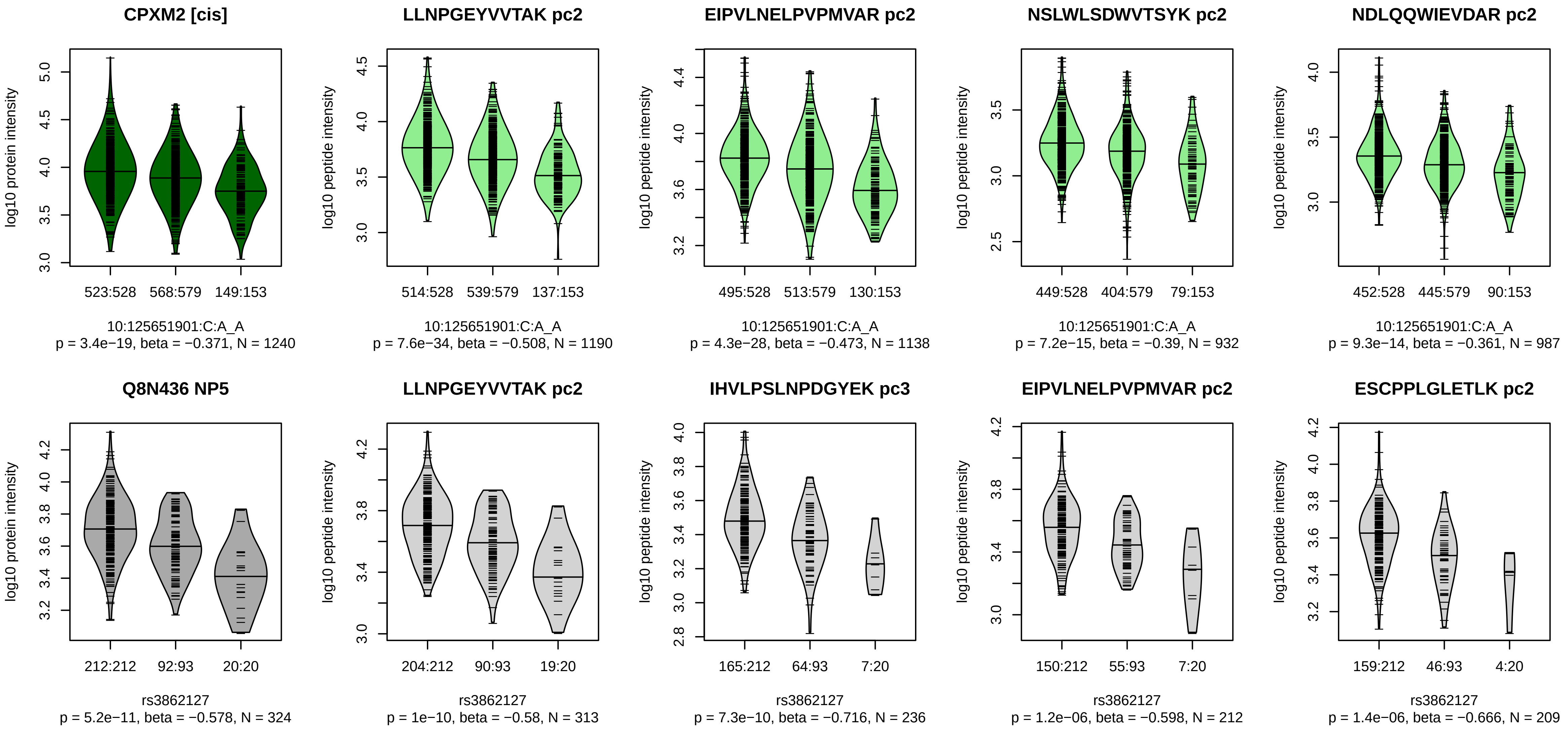


Locus: 90 (Sort: 107)
Status: Replicated
80% power: Yes
SNP: 10:125651901:C:A
rsID: rs3862127
UniProtID: Q8N436
HGNC gene: CPXM2
Type: Cis−pQTL

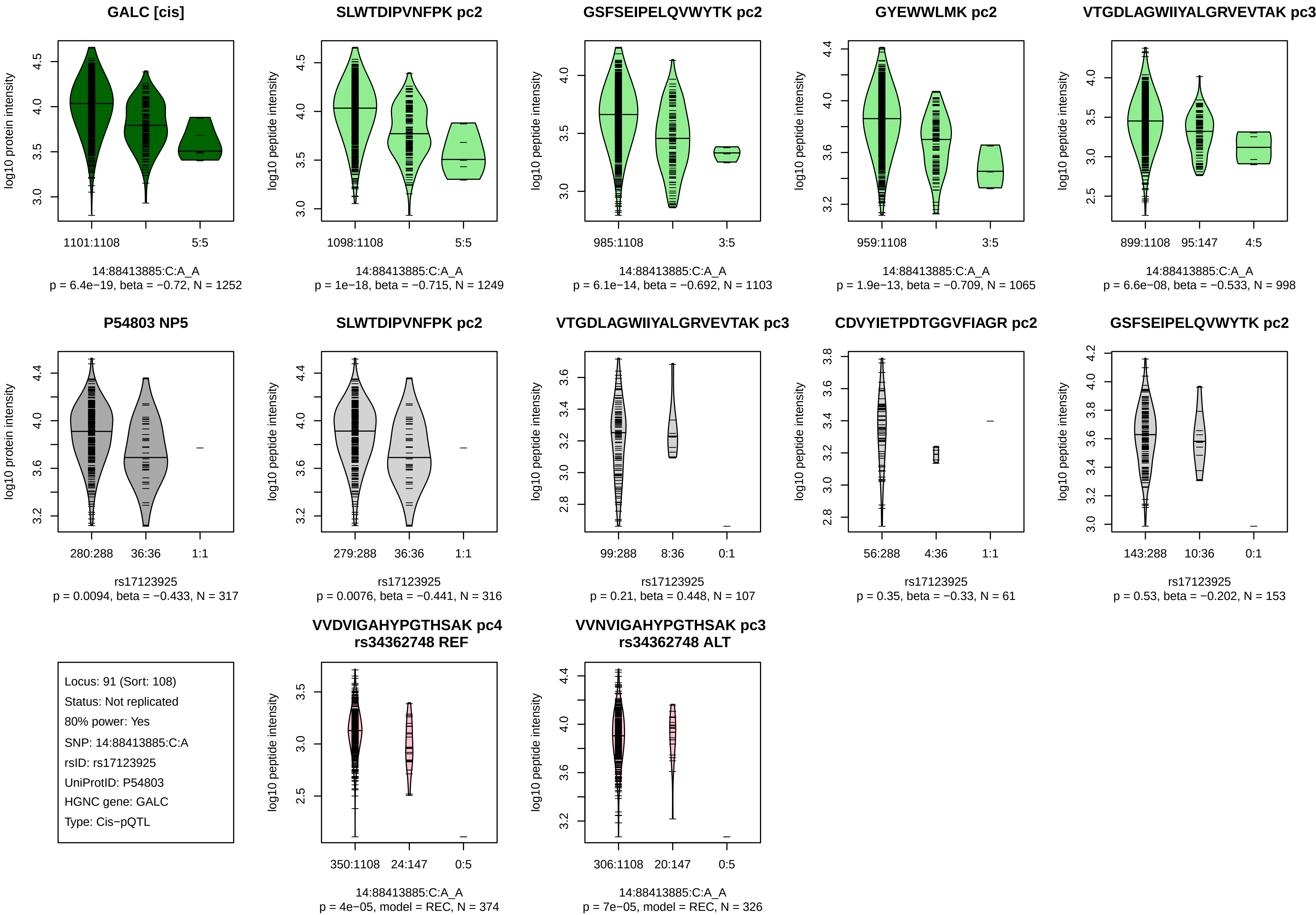
GALC [cis]
log10 protein intensity
1101:1108
5:5
14:88413885:C:A_A
p = 6.4e−19, beta = −0.72, N = 1252
SLWTDIPVNFPK pc2
log10 peptide intensity
1098:1108
5:5
14:88413885:C:A_A
p = 1e−18, beta = −0.715, N = 1249
GSFSEIPELQVWYTK pc2
log10 peptide intensity
985:1108
3:5
14:88413885:C:A_A
p = 6.1e−14, beta = −0.692, N = 1103
GYEWWLMK pc2
log10 peptide intensity
959:1108
3:5
14:88413885:C:A_A
p = 1.9e−13, beta = −0.709, N = 1065
VTGDLAGWIIYALGRVEVTAK pc3
log10 peptide intensity
899:1108
95:147
4:5
14:88413885:C:A_A
p = 6.6e−08, beta = −0.533, N = 998
P54803 NP5
log10 protein intensity
280:288
36:36
1:1
rs17123925
p = 0.0094, beta = −0.433, N = 317
SLWTDIPVNFPK pc2
log10 peptide intensity
279:288
36:36
1:1
rs17123925
p = 0.0076, beta = −0.441, N = 316
VTGDLAGWIIYALGRVEVTAK pc3
log10 peptide intensity
99:288
8:36
0:1
rs17123925
p = 0.21, beta = 0.448, N = 107
CDVYIETPDTGGVFIAGR pc2
log10 peptide intensity
56:288
4:36
1:1
rs17123925
p = 0.35, beta = −0.33, N = 61
GSFSEIPELQVWYTK pc2
log10 peptide intensity
143:288
10:36
0:1
rs17123925
p = 0.53, beta = −0.202, N = 153
Locus: 91 (Sort: 108)
Status: Not replicated
80% power: Yes
SNP: 14:88413885:C:A
rsID: rs17123925
UniProtID: P54803
HGNC gene: GALC
Type: Cis−pQTL
VVDVIGAHYPGTHSAK pc4
rs34362748 REF
log10 peptide intensity
350:1108
24:147
0:5
14:88413885:C:A_A
p = 4e−05, model = REC, N = 374
VVNVIGAHYPGTHSAK pc3
rs34362748 ALT
log10 peptide intensity
306:1108
20:147
0:5
14:88413885:C:A_A
p = 7e−05, model = REC, N = 326

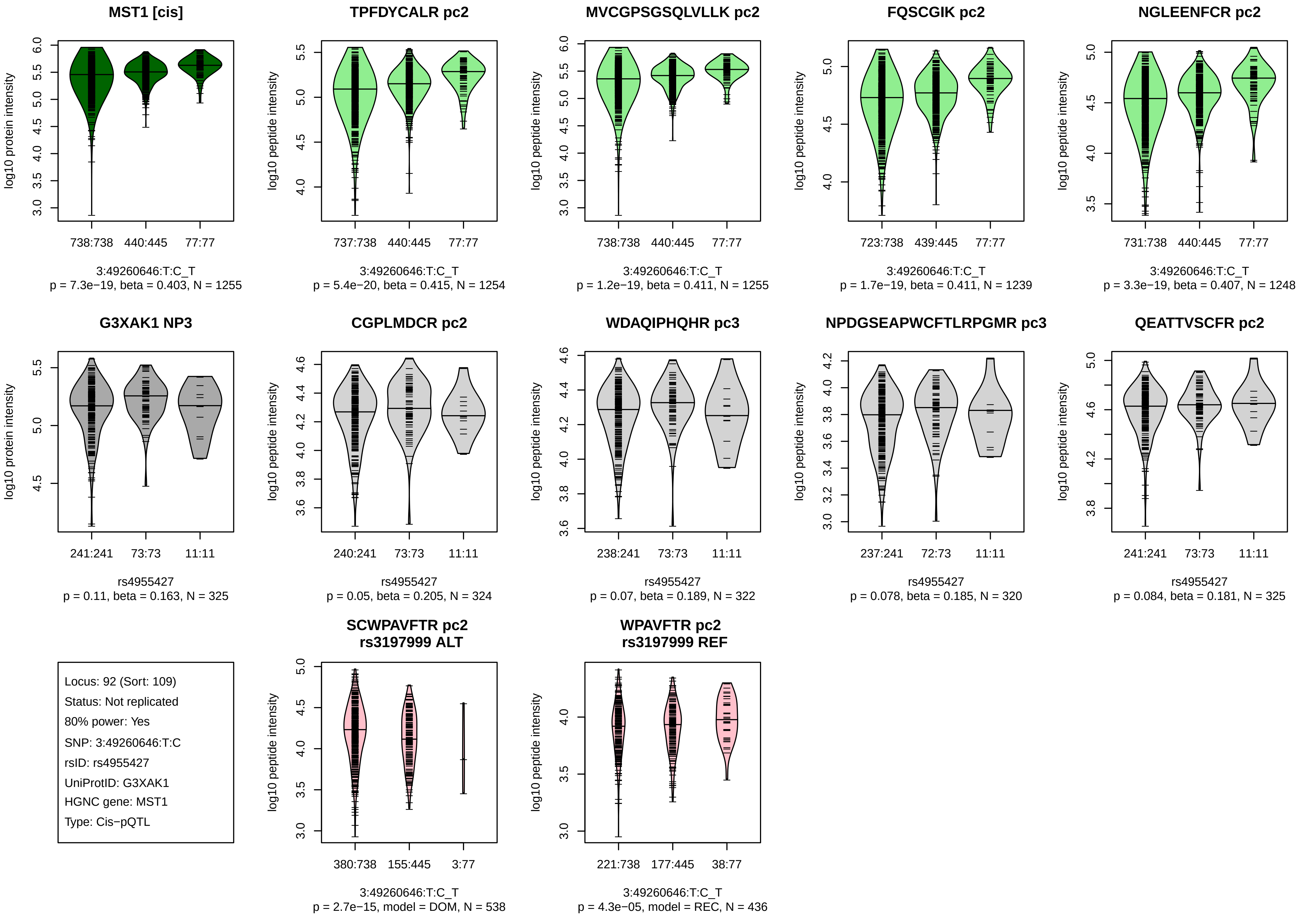
MST1 [cis]
log10 protein intensity
738:738 440:445 77:77
3:49260646:T:C_T
p = 7.3e−19, beta = 0.403, N = 1255
TPFDYCALR pc2
log10 peptide intensity
737:738 440:445 77:77
3:49260646:T:C_T
p = 5.4e−20, beta = 0.415, N = 1254
MVCGPSGSQLVLLK pc2
log10 peptide intensity
738:738 440:445 77:77
3:49260646:T:C_T
p = 1.2e−19, beta = 0.411, N = 1255
FQSCGIK pc2
log10 peptide intensity
723:738 439:445 77:77
3:49260646:T:C_T
p = 1.7e−19, beta = 0.411, N = 1239
NGLEENFCR pc2
log10 peptide intensity
731:738 440:445 77:77
3:49260646:T:C_T
p = 3.3e−19, beta = 0.407, N = 1248
G3XAK1 NP3
log10 protein intensity
241:241 73:73 11:11
rs4955427
p = 0.11, beta = 0.163, N = 325
CGPLMDCR pc2
log10 peptide intensity
240:241 73:73 11:11
rs4955427
p = 0.05, beta = 0.205, N = 324
WDAQIPHQHR pc3
log10 peptide intensity
238:241 73:73 11:11
rs4955427
p = 0.07, beta = 0.189, N = 322
NPDGSEAPWCFTLRPGMR pc3
log10 peptide intensity
237:241 72:73 11:11
rs4955427
p = 0.078, beta = 0.185, N = 320
QEATTVSCFR pc2
log10 peptide intensity
241:241 73:73 11:11
rs4955427
p = 0.084, beta = 0.181, N = 325
Locus: 92 (Sort: 109)
Status: Not replicated
80% power: Yes
SNP: 3:49260646:T:C
rsID: rs4955427
UniProtID: G3XAK1
HGNC gene: MST1
Type: Cis−pQTL
SCWPAVFTR pc2
rs3197999 ALT
log10 peptide intensity
380:738 155:445 3:77
3:49260646:T:C_T
p = 2.7e−15, model = DOM, N = 538
WPAVFTR pc2
rs3197999 REF
log10 peptide intensity
221:738 177:445 38:77
3:49260646:T:C_T
p = 4.3e−05, model = REC, N = 436

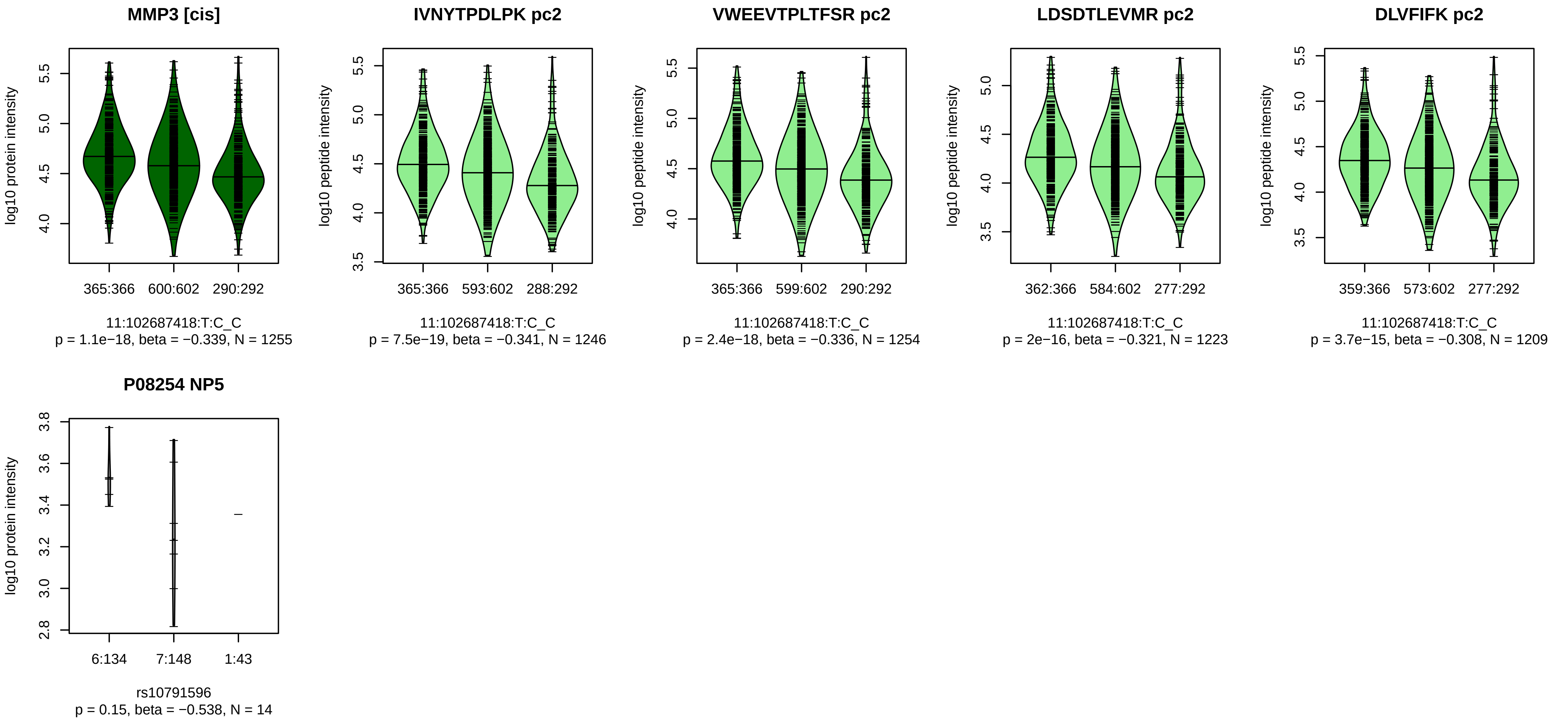


Locus: 93 (Sort: 110)

Status: Not replicated

80% power: No

SNP: 11:102687418:T:C

rsID: rs10791596

UniProtID: P08254

HGNC gene: MMP3

Type: Cis−pQTL

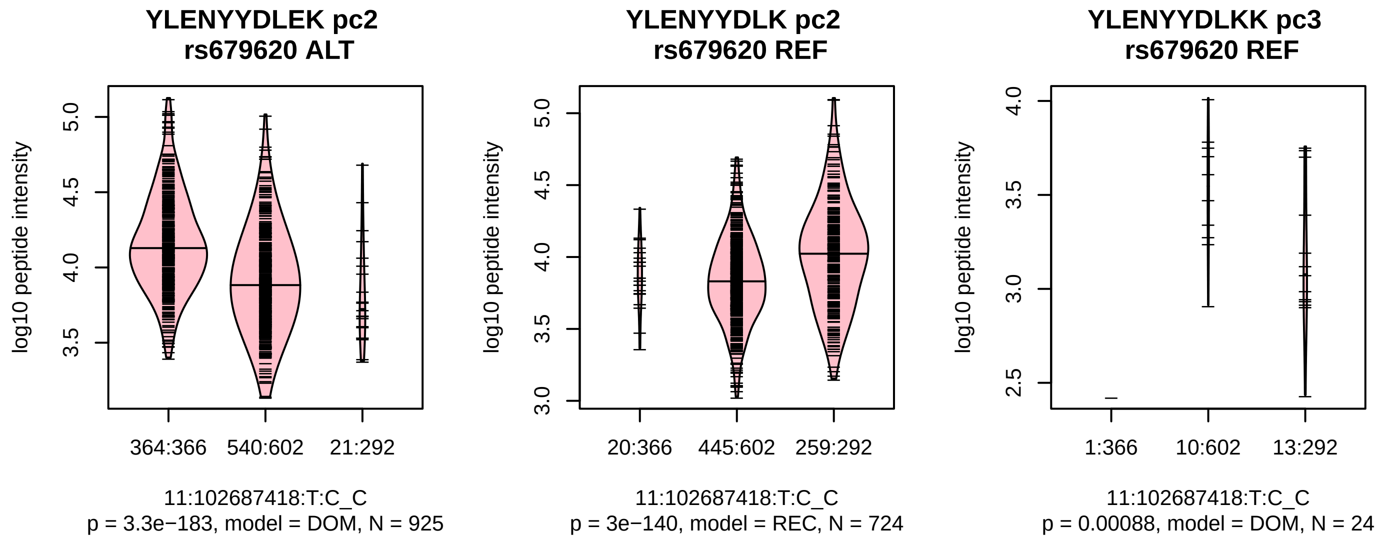

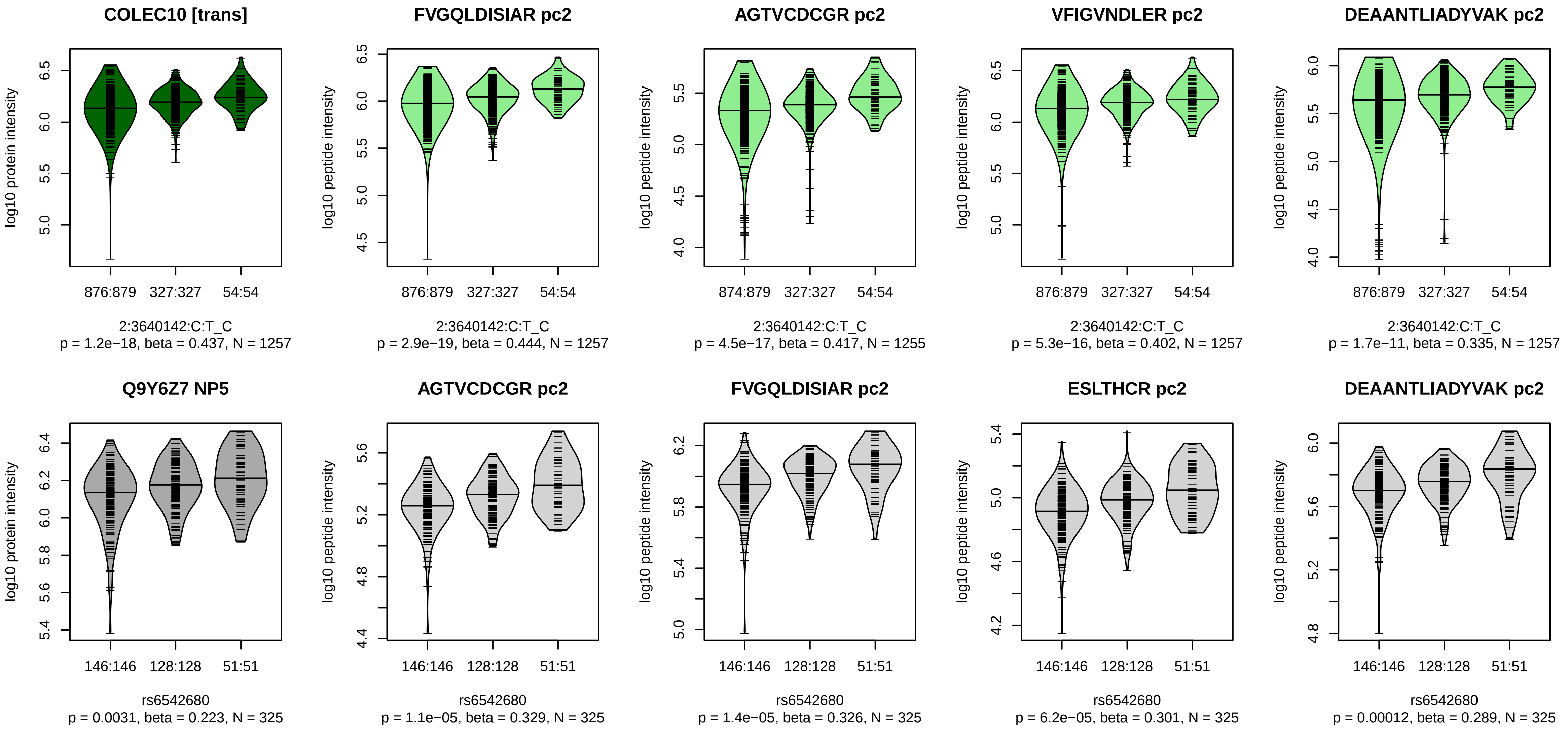


Locus: 94 (Sort: 111)
Status: Not replicated
80% power: No
SNP: 2:3640142:C:T
rsID: rs6542680
UniProtID: Q9Y6Z7
HGNC gene: COLEC10
Type: Trans−pQTL

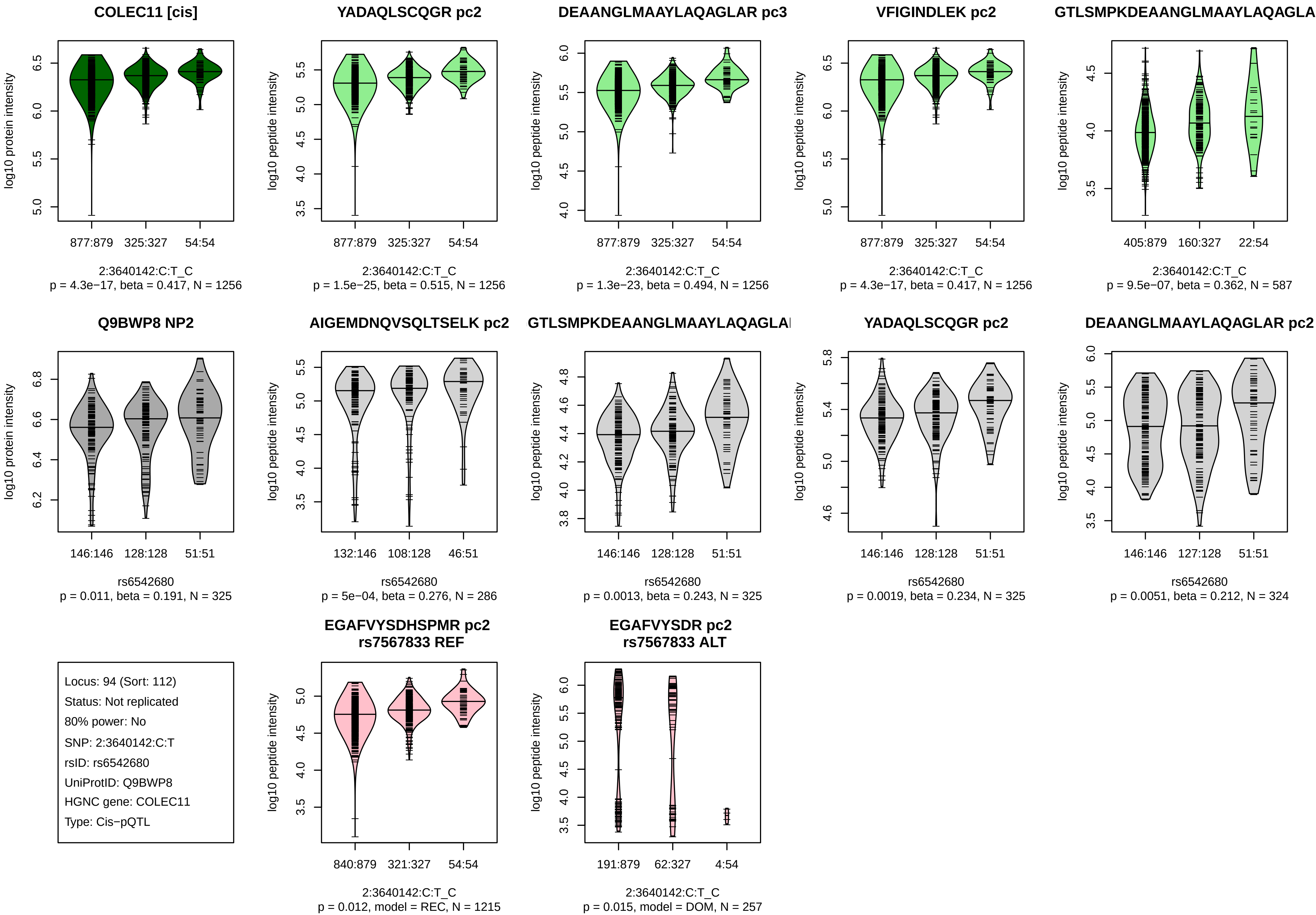
COLEC11 [cis]
log10 protein intensity
877:879
325:327
54:54
2:3640142:C:T_C
p = 4.3e−17, beta = 0.417, N = 1256
YADAQLSCQGR pc2
log10 peptide intensity
877:879
325:327
54:54
2:3640142:C:T_C
p = 1.5e−25, beta = 0.515, N = 1256
DEAANGLMAAYLAQAGLAR pc3
log10 peptide intensity
877:879
325:327
54:54
2:3640142:C:T_C
p = 1.3e−23, beta = 0.494, N = 1256
VFIGINDLEK pc2
log10 peptide intensity
877:879
325:327
54:54
2:3640142:C:T_C
p = 4.3e−17, beta = 0.417, N = 1256
GTLSMPKDEAANGLMAAYLAQAGLA
log10 peptide intensity
405:879
160:327
22:54
2:3640142:C:T_C
p = 9.5e−07, beta = 0.362, N = 587
Q9BWP8 NP2
log10 protein intensity
146:146
128:128
51:51
rs6542680
p = 0.011, beta = 0.191, N = 325
AIGEMDNQVSQLTSELK pc2
log10 peptide intensity
132:146
108:128
46:51
rs6542680
p = 5e−04, beta = 0.276, N = 286
GTLSMPKDEAANGLMAAYLAQAGLA
log10 peptide intensity
146:146
128:128
51:51
rs6542680
p = 0.0013, beta = 0.243, N = 325
YADAQLSCQGR pc2
log10 peptide intensity
146:146
128:128
51:51
rs6542680
p = 0.0019, beta = 0.234, N = 325
DEAANGLMAAYLAQAGLAR pc2
log10 peptide intensity
146:146
127:128
51:51
rs6542680
p = 0.0051, beta = 0.212, N = 324
Locus: 94 (Sort: 112)
Status: Not replicated
80% power: No
SNP: 2:3640142:C:T
rsID: rs6542680
UniProtID: Q9BWP8
HGNC gene: COLEC11
Type: Cis−pQTL
EGAFVYSDHSPMR pc2
rs7567833 REF
log10 peptide intensity
840:879
321:327
54:54
2:3640142:C:T_C
p = 0.012, model = REC, N = 1215
EGAFVYSDR pc2
rs7567833 ALT
log10 peptide intensity
191:879
62:327
4:54
2:3640142:C:T_C
p = 0.015, model = DOM, N = 257

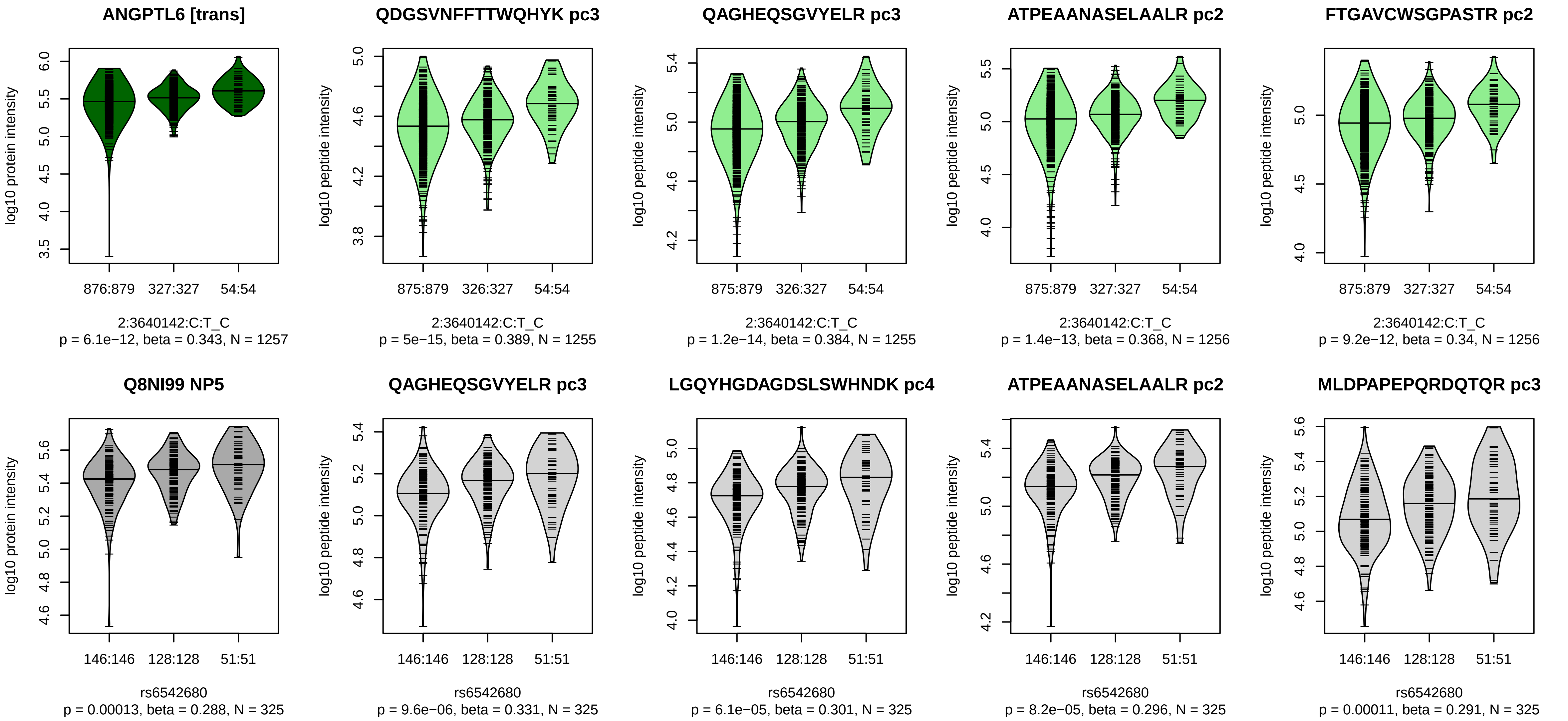


Locus: 94 (Sort: 113)
Status: Replicated
80% power: No
SNP: 2:3640142:C:T
rsID: rs6542680
UniProtID: Q8NI99
HGNC gene: ANGPTL6
Type: Trans−pQTL

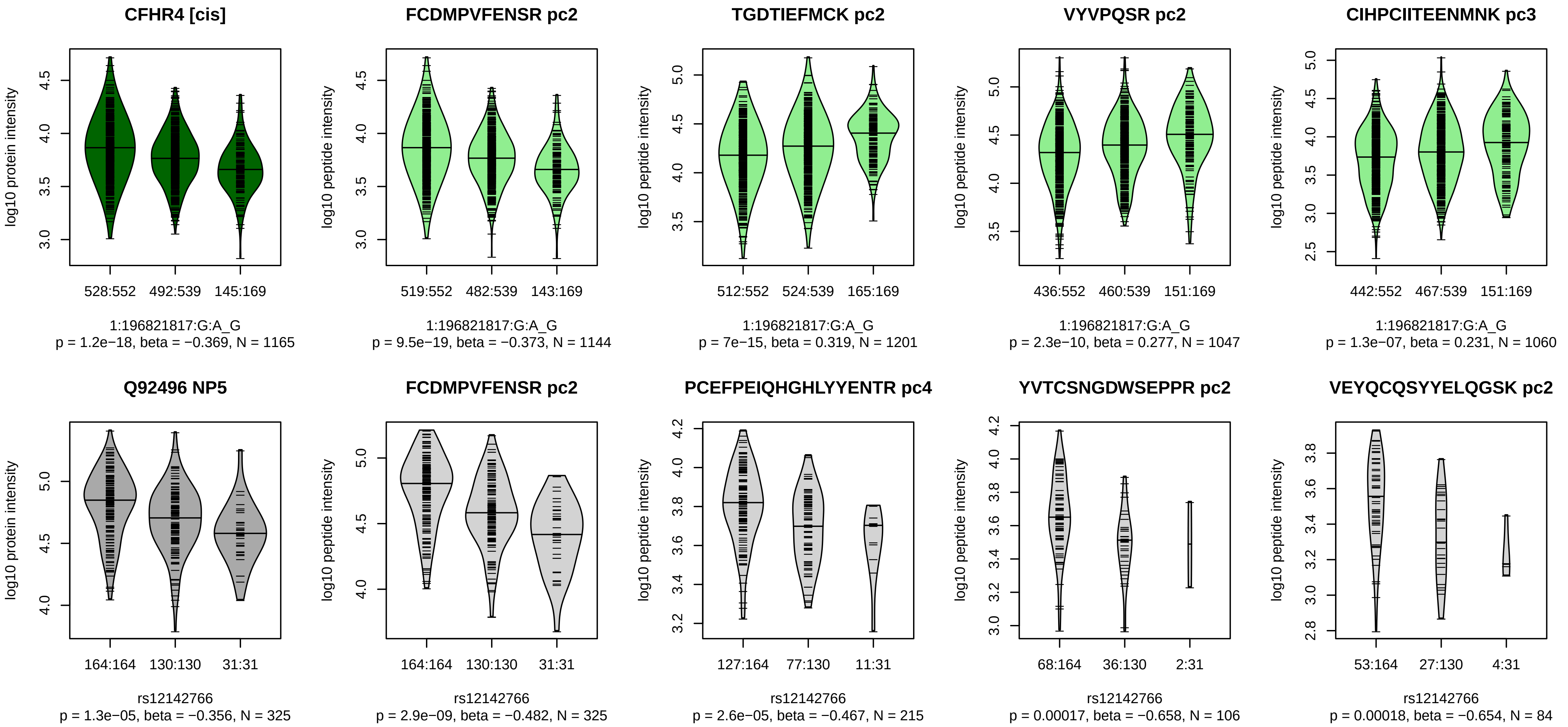


Locus: 95 (Sort: 114)

Status: Replicated

80% power: Yes

SNP: 1:196821817:G:A

rsID: rs12142766

UniProtID: Q92496

HGNC gene: CFHR4

Type: Cis−pQTL

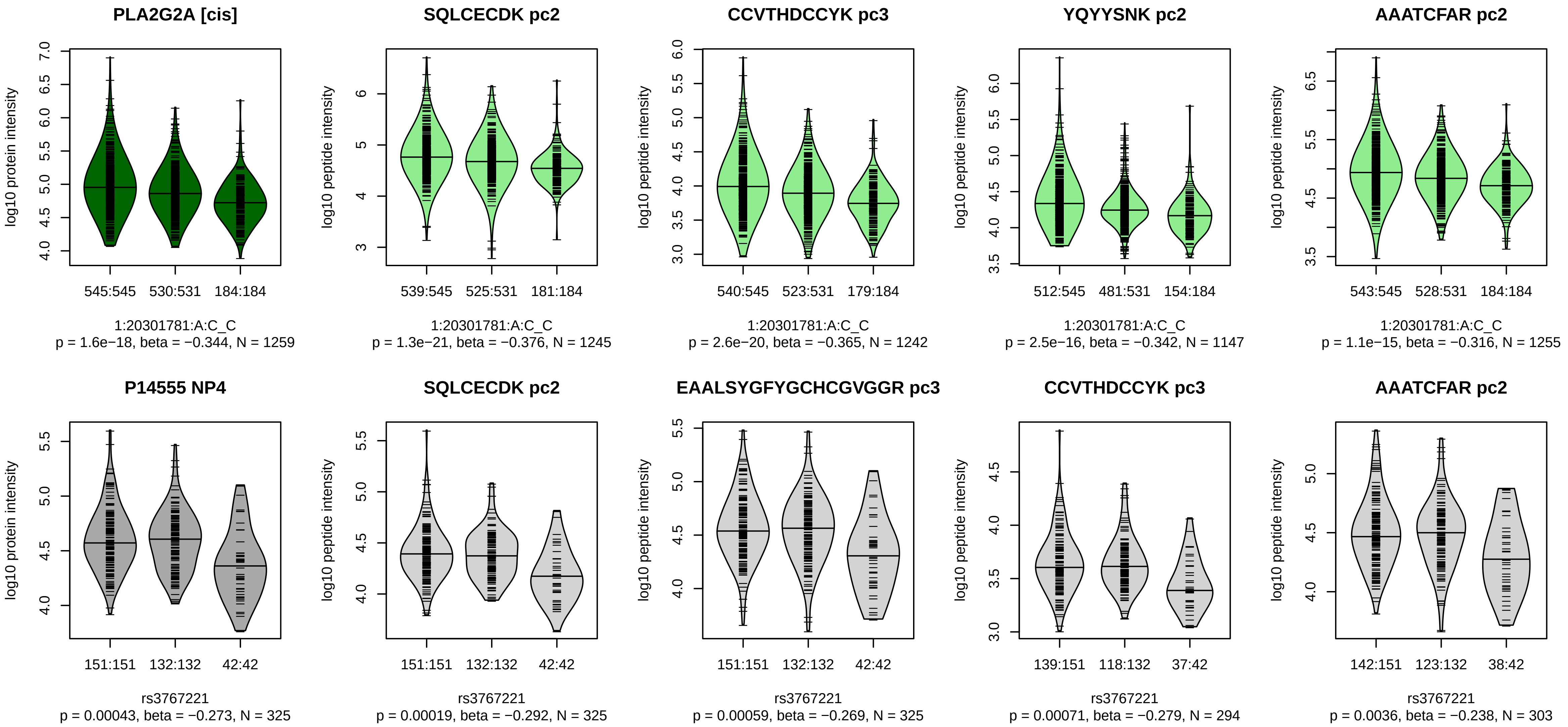


Locus: 96 (Sort: 115)
Status: Not replicated
80% power: No
SNP: 1:20301781:A:C
rsID: rs3767221
UniProtID: P14555
HGNC gene: PLA2G2A
Type: Cis−pQTL

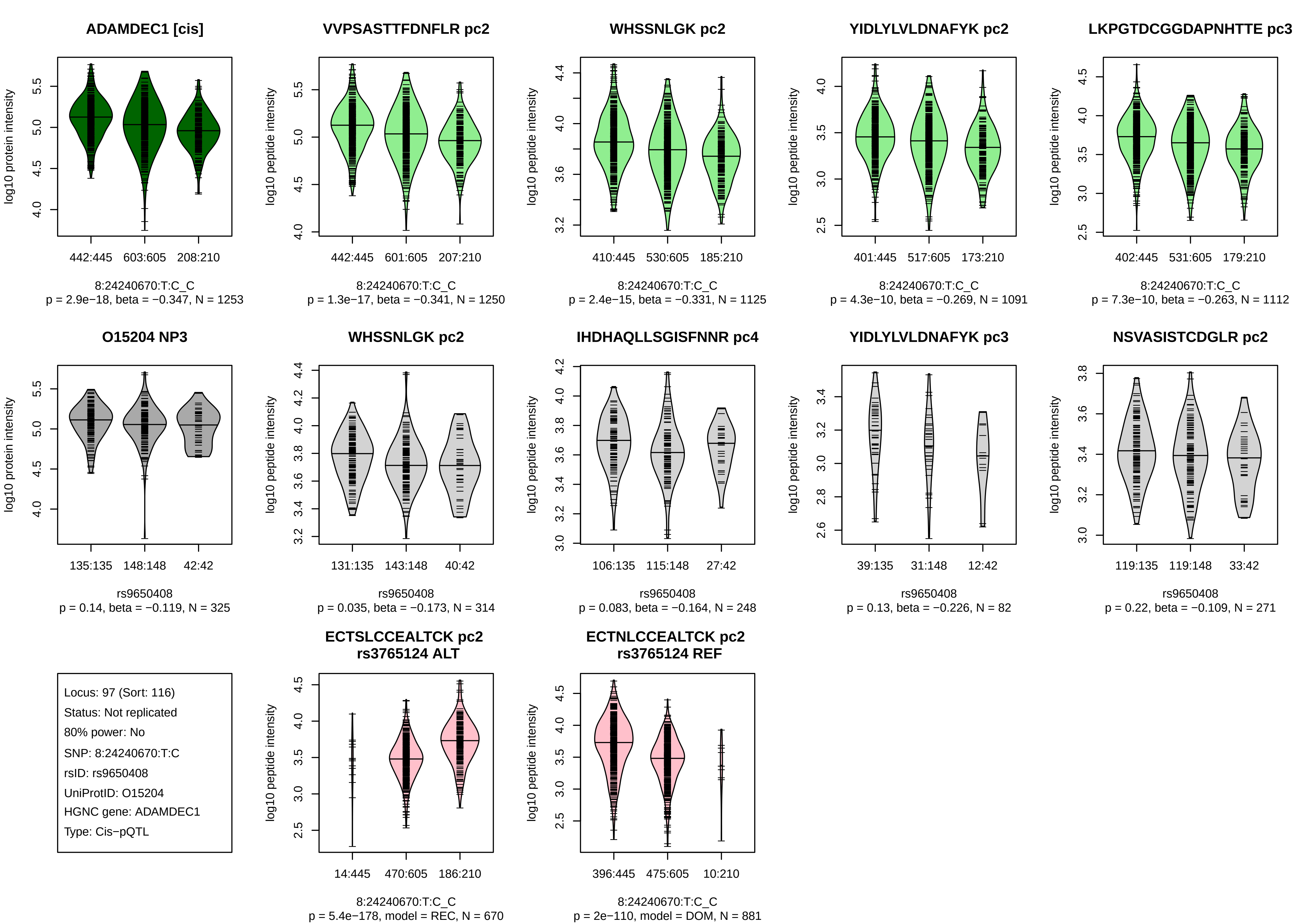
ADAMDEC1 [cis]
log10 protein intensity
442:445
603:605
208:210
8:24240670:T:C_C
p = 2.9e−18, beta = −0.347, N = 1253
VVPSASTTFDNFLR pc2
log10 peptide intensity
442:445
601:605
207:210
8:24240670:T:C_C
p = 1.3e−17, beta = −0.341, N = 1250
WHSSNLGK pc2
log10 peptide intensity
410:445
530:605
185:210
8:24240670:T:C_C
p = 2.4e−15, beta = −0.331, N = 1125
YIDLYLVLDNAFYK pc2
log10 peptide intensity
401:445
517:605
173:210
8:24240670:T:C_C
p = 4.3e−10, beta = −0.269, N = 1091
LKPGTDCGGDAPNHTTE pc3
log10 peptide intensity
402:445
531:605
179:210
8:24240670:T:C_C
p = 7.3e−10, beta = −0.263, N = 1112
O15204 NP3
log10 protein intensity
135:135
148:148
42:42
rs9650408
p = 0.14, beta = −0.119, N = 325
WHSSNLGK pc2
log10 peptide intensity
131:135
143:148
40:42
rs9650408
p = 0.035, beta = −0.173, N = 314
IHDHAQLLSGISFNNR pc4
log10 peptide intensity
106:135
115:148
27:42
rs9650408
p = 0.083, beta = −0.164, N = 248
YIDLYLVLDNAFYK pc3
log10 peptide intensity
39:135
31:148
12:42
rs9650408
p = 0.13, beta = −0.226, N = 82
NSVASISTCDGLR pc2
log10 peptide intensity
119:135
119:148
33:42
rs9650408
p = 0.22, beta = −0.109, N = 271
Locus: 97 (Sort: 116)
Status: Not replicated
80% power: No
SNP: 8:24240670:T:C
rsID: rs9650408
UniProtID: O15204
HGNC gene: ADAMDEC1
Type: Cis−pQTL
ECTSLCCEALTCK pc2
rs3765124 ALT
log10 peptide intensity
14:445
470:605
186:210
8:24240670:T:C_C
p = 5.4e−178, model = REC, N = 670
ECTNLCCEALTCK pc2
rs3765124 REF
log10 peptide intensity
396:445
475:605
10:210
8:24240670:T:C_C
p = 2e−110, model = DOM, N = 881

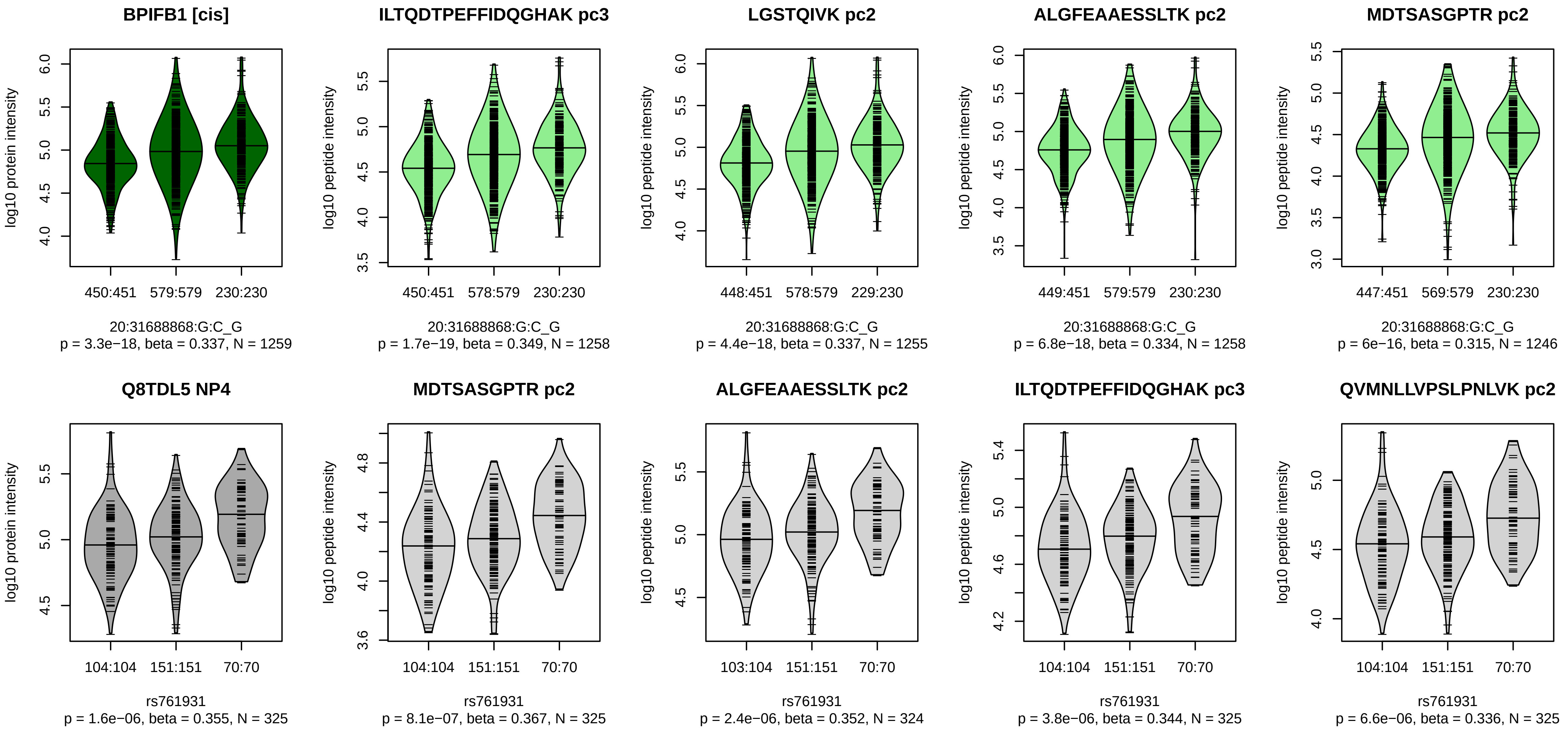


Locus: 98 (Sort: 117)
Status: Replicated
80% power: Yes
SNP: 20:31688868:G:C
rsID: rs761931
UniProtID: Q8TDL5
HGNC gene: BPIFB1
Type: Cis−pQTL

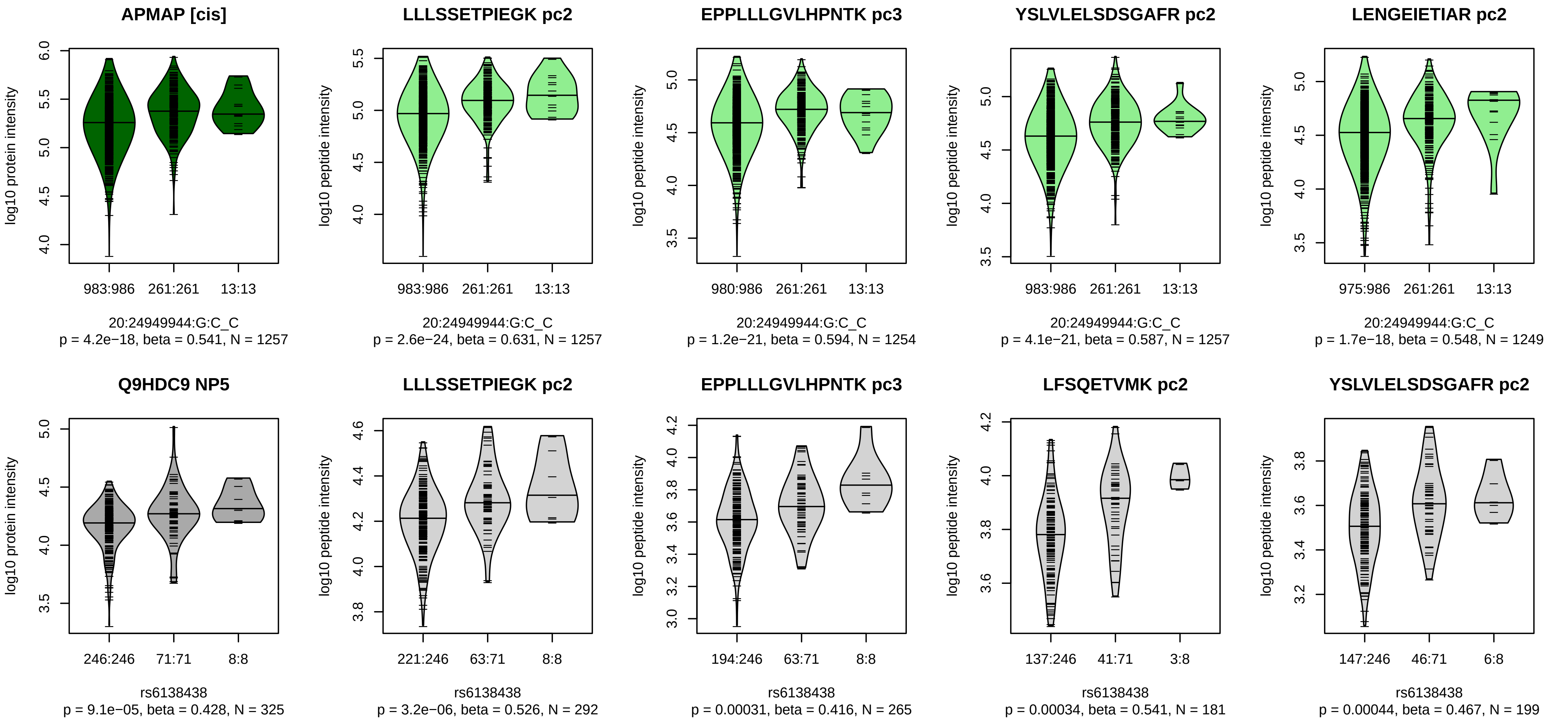


Locus: 99 (Sort: 118)

Status: Replicated

80% power: No

SNP: 20:24949944:G:C

rsID: rs6138438

UniProtID: Q9HDC9

HGNC gene: APMAP

Type: Cis−pQTL

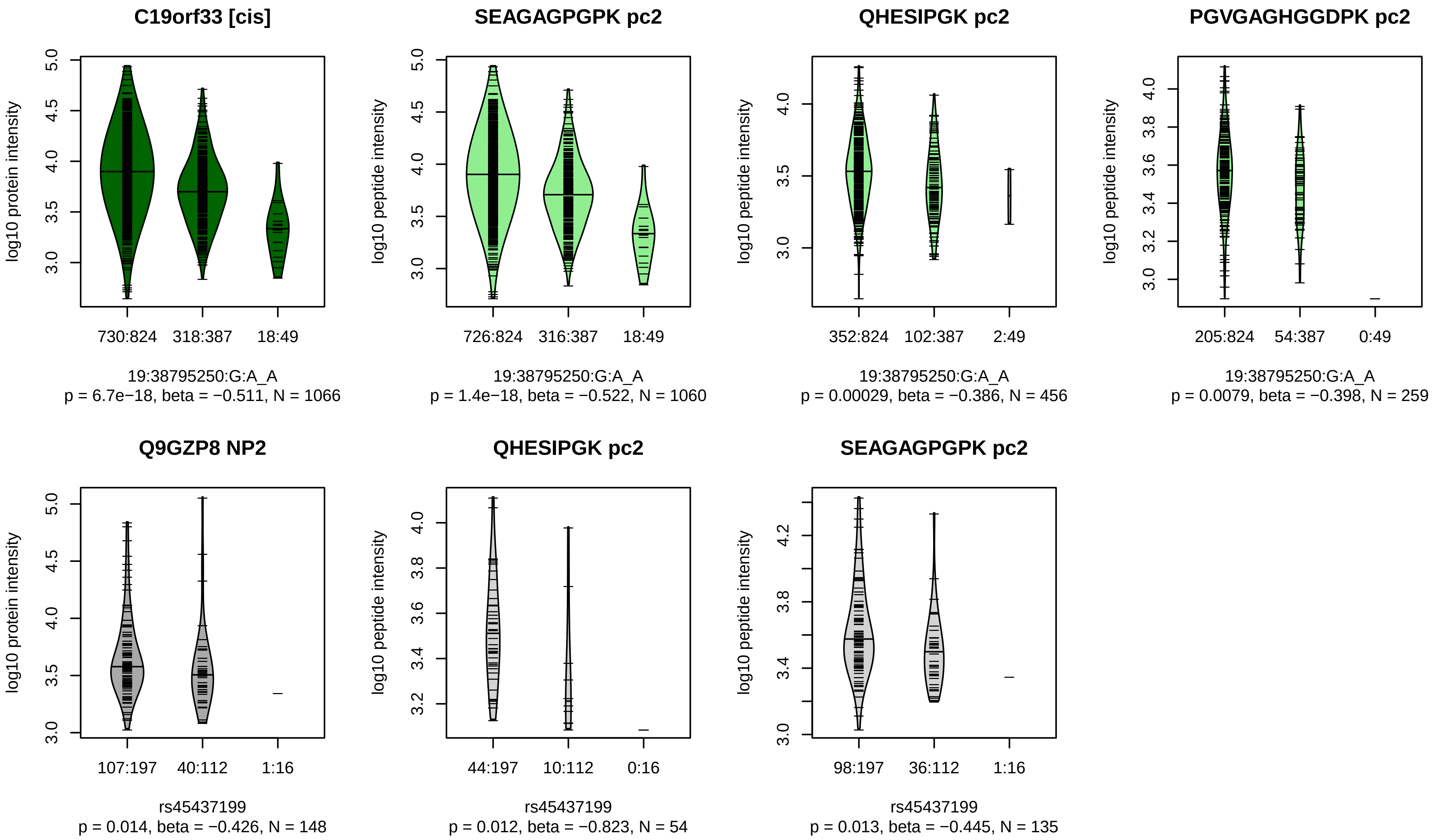


Locus: 100 (Sort: 119)

Status: Not replicated

80% power: No

SNP: 19:38795250:G:A

rsID: rs45437199

UniProtID: Q9GZP8

HGNC gene: C19orf33

Type: Cis−pQTL

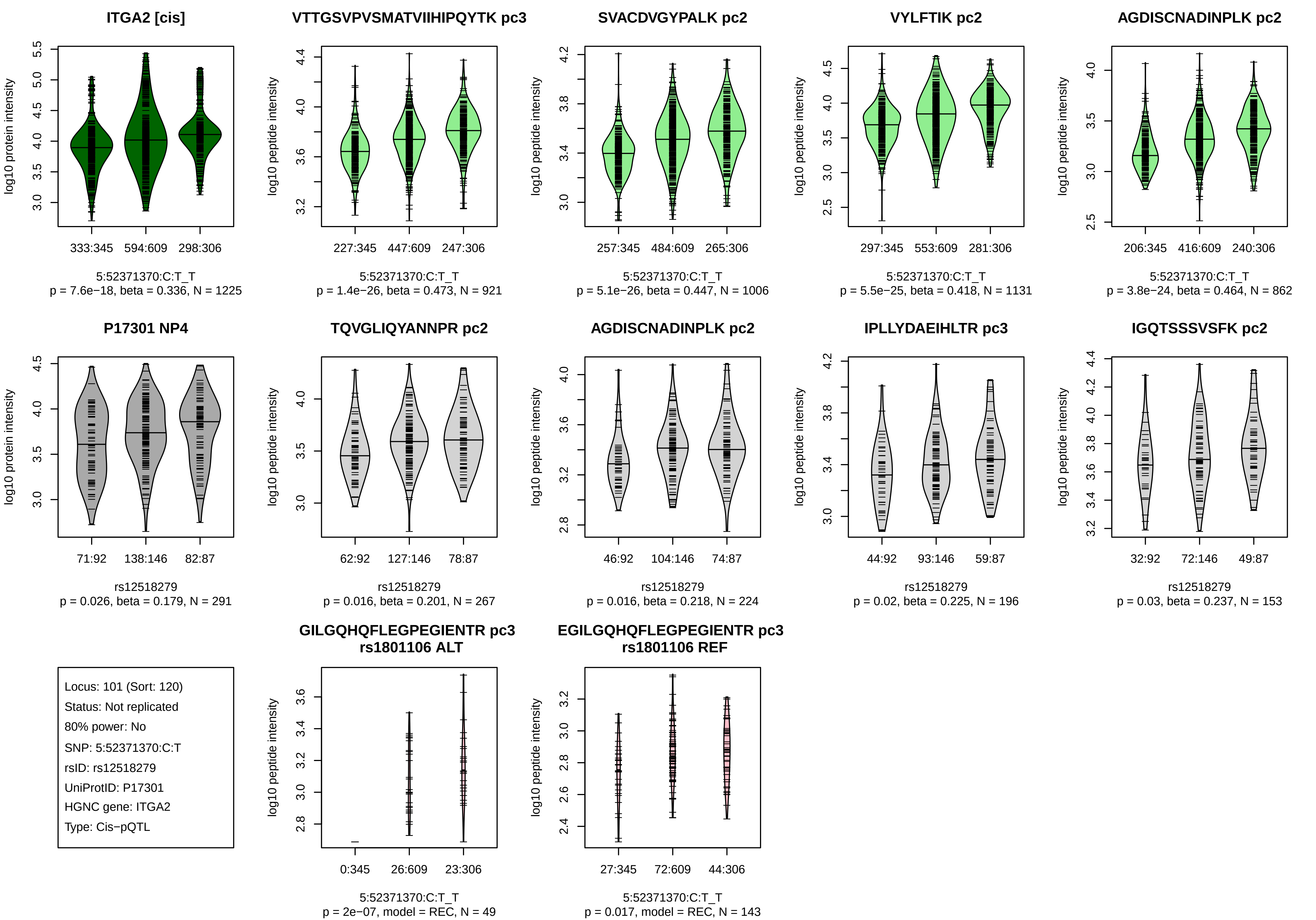
ITGA2 [cis]
log10 protein intensity
333:345
594:609
298:306
5:52371370:C:T_T
p = 7.6e−18, beta = 0.336, N = 1225
VTTGSVPVSMATVIIHIPQYTK pc3
log10 peptide intensity
227:345
447:609
247:306
5:52371370:C:T_T
p = 1.4e−26, beta = 0.473, N = 921
SVACDVGYPALK pc2
log10 peptide intensity
257:345
484:609
265:306
5:52371370:C:T_T
p = 5.1e−26, beta = 0.447, N = 1006
VYLFTIK pc2
log10 peptide intensity
297:345
553:609
281:306
5:52371370:C:T_T
p = 5.5e−25, beta = 0.418, N = 1131
AGDISCNADINPLK pc2
log10 peptide intensity
206:345
416:609
240:306
5:52371370:C:T_T
p = 3.8e−24, beta = 0.464, N = 862
P17301 NP4
log10 protein intensity
71:92
138:146
82:87
rs12518279
p = 0.026, beta = 0.179, N = 291
TQVGLIQYANNPR pc2
log10 peptide intensity
62:92
127:146
78:87
rs12518279
p = 0.016, beta = 0.201, N = 267
AGDISCNADINPLK pc2
log10 peptide intensity
46:92
104:146
74:87
rs12518279
p = 0.016, beta = 0.218, N = 224
IPLLYDAEIHLTR pc3
log10 peptide intensity
44:92
93:146
59:87
rs12518279
p = 0.02, beta = 0.225, N = 196
IGQTSSSVSFK pc2
log10 peptide intensity
32:92
72:146
49:87
rs12518279
p = 0.03, beta = 0.237, N = 153
Locus: 101 (Sort: 120)
Status: Not replicated
80% power: No
SNP: 5:52371370:C:T
rsID: rs12518279
UniProtID: P17301
HGNC gene: ITGA2
Type: Cis−pQTL
GILGQHQFLEGPEGIENTR pc3
rs1801106 ALT
log10 peptide intensity
0:345
26:609
23:306
5:52371370:C:T_T
p = 2e−07, model = REC, N = 49
EGILGQHQFLEGPEGIENTR pc3
rs1801106 REF
log10 peptide intensity
27:345
72:609
44:306
5:52371370:C:T_T
p = 0.017, model = REC, N = 143

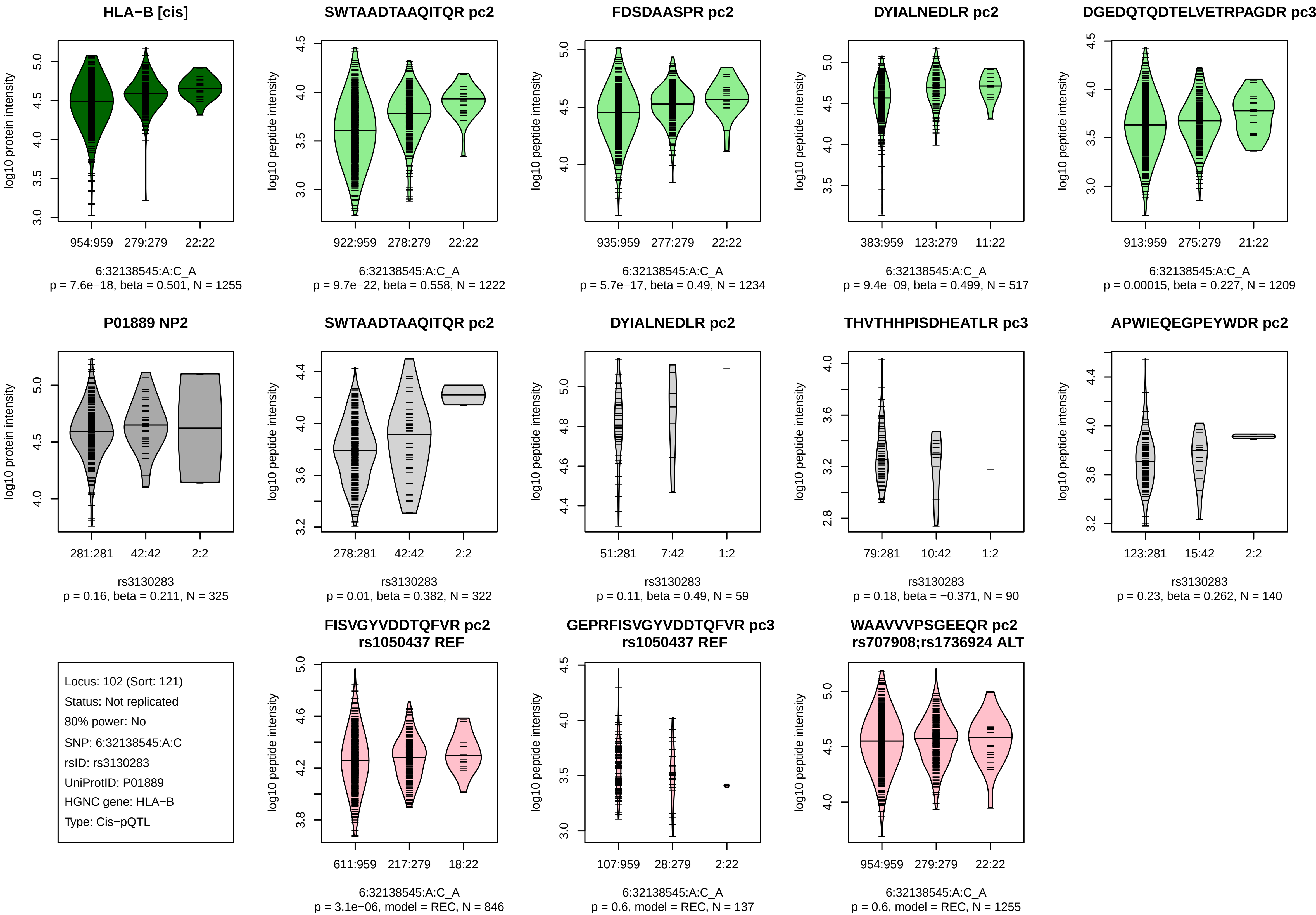
HLA−B [cis]
log10 protein intensity
954:959
279:279
22:22
6:32138545:A:C_A
p = 7.6e−18, beta = 0.501, N = 1255
SWTAADTAAQITQR pc2
log10 peptide intensity
922:959
278:279
22:22
6:32138545:A:C_A
p = 9.7e−22, beta = 0.558, N = 1222
FDSDAASPR pc2
log10 peptide intensity
935:959
277:279
22:22
6:32138545:A:C_A
p = 5.7e−17, beta = 0.49, N = 1234
DYIALNEDLR pc2
log10 peptide intensity
383:959
123:279
11:22
6:32138545:A:C_A
p = 9.4e−09, beta = 0.499, N = 517
DGEDQTQDTELVETRPAGDR pc3
log10 peptide intensity
913:959
275:279
21:22
6:32138545:A:C_A
p = 0.00015, beta = 0.227, N = 1209
P01889 NP2
log10 protein intensity
281:281
42:42
2:2
rs3130283
p = 0.16, beta = 0.211, N = 325
SWTAADTAAQITQR pc2
log10 peptide intensity
278:281
42:42
2:2
rs3130283
p = 0.01, beta = 0.382, N = 322
DYIALNEDLR pc2
log10 peptide intensity
51:281
7:42
1:2
rs3130283
p = 0.11, beta = 0.49, N = 59
THVTHHPISDHEATLR pc3
log10 peptide intensity
79:281
10:42
1:2
rs3130283
p = 0.18, beta = −0.371, N = 90
APWIEQEGPEYWDR pc2
log10 peptide intensity
123:281
15:42
2:2
rs3130283
p = 0.23, beta = 0.262, N = 140
Locus: 102 (Sort: 121)
Status: Not replicated
80% power: No
SNP: 6:32138545:A:C
rsID: rs3130283
UniProtID: P01889
HGNC gene: HLA−B
Type: Cis−pQTL
FISVGYVDDTQFVR pc2
rs1050437 REF
log10 peptide intensity
611:959
217:279
18:22
6:32138545:A:C_A
p = 3.1e−06, model = REC, N = 846
GEPRFISVGYVDDTQFVR pc3
rs1050437 REF
log10 peptide intensity
107:959
28:279
2:22
6:32138545:A:C_A
p = 0.6, model = REC, N = 137
WAAVVVPSGEEQR pc2
rs707908;rs1736924 ALT
log10 peptide intensity
954:959
279:279
22:22
6:32138545:A:C_A
p = 0.6, model = REC, N = 1255

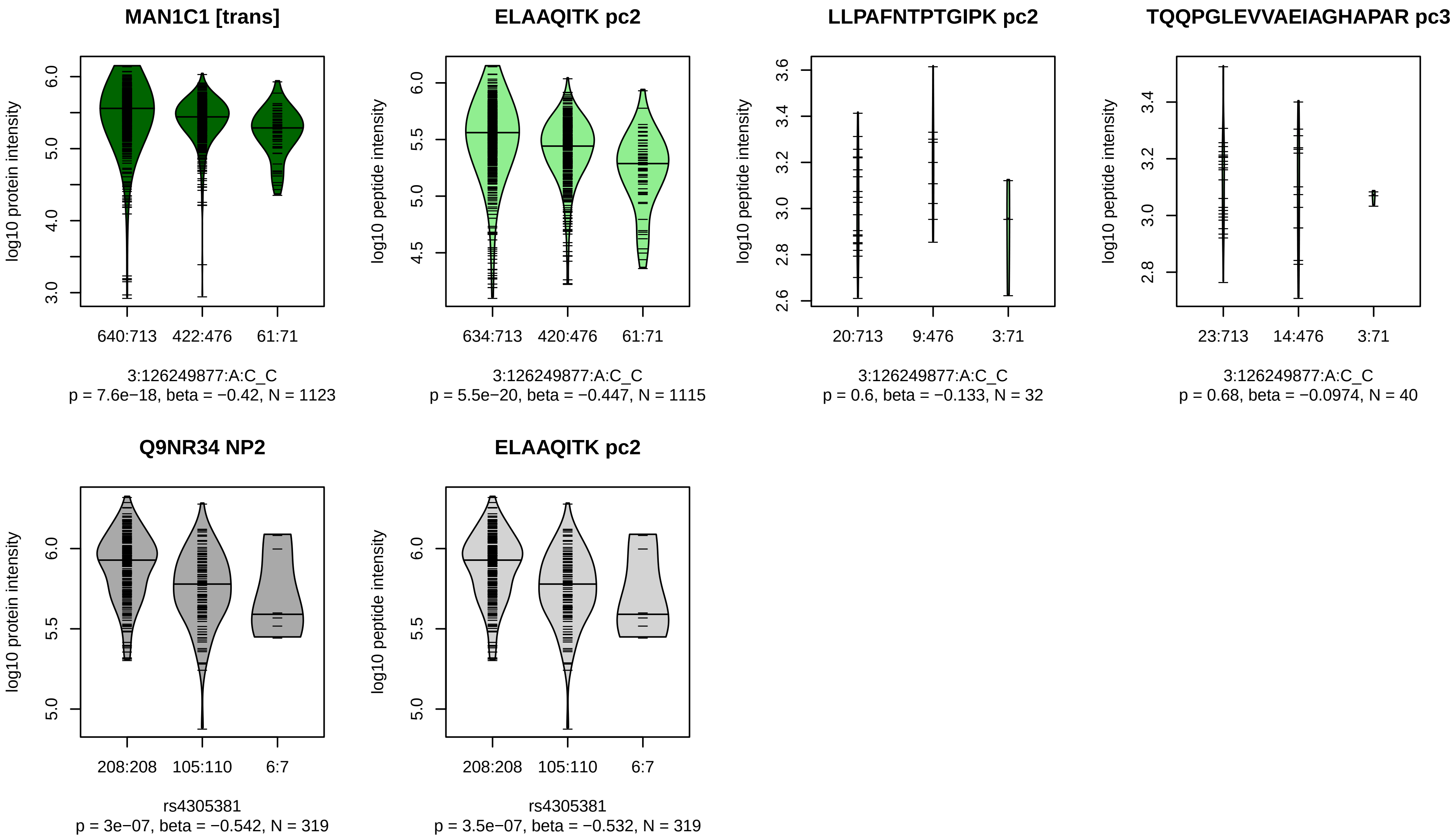


Locus: 103 (Sort: 122)
Status: Replicated
80% power: Yes
SNP: 3:126249877:A:C
rsID: rs4305381
UniProtID: Q9NR34
HGNC gene: MAN1C1
Type: Trans−pQTL

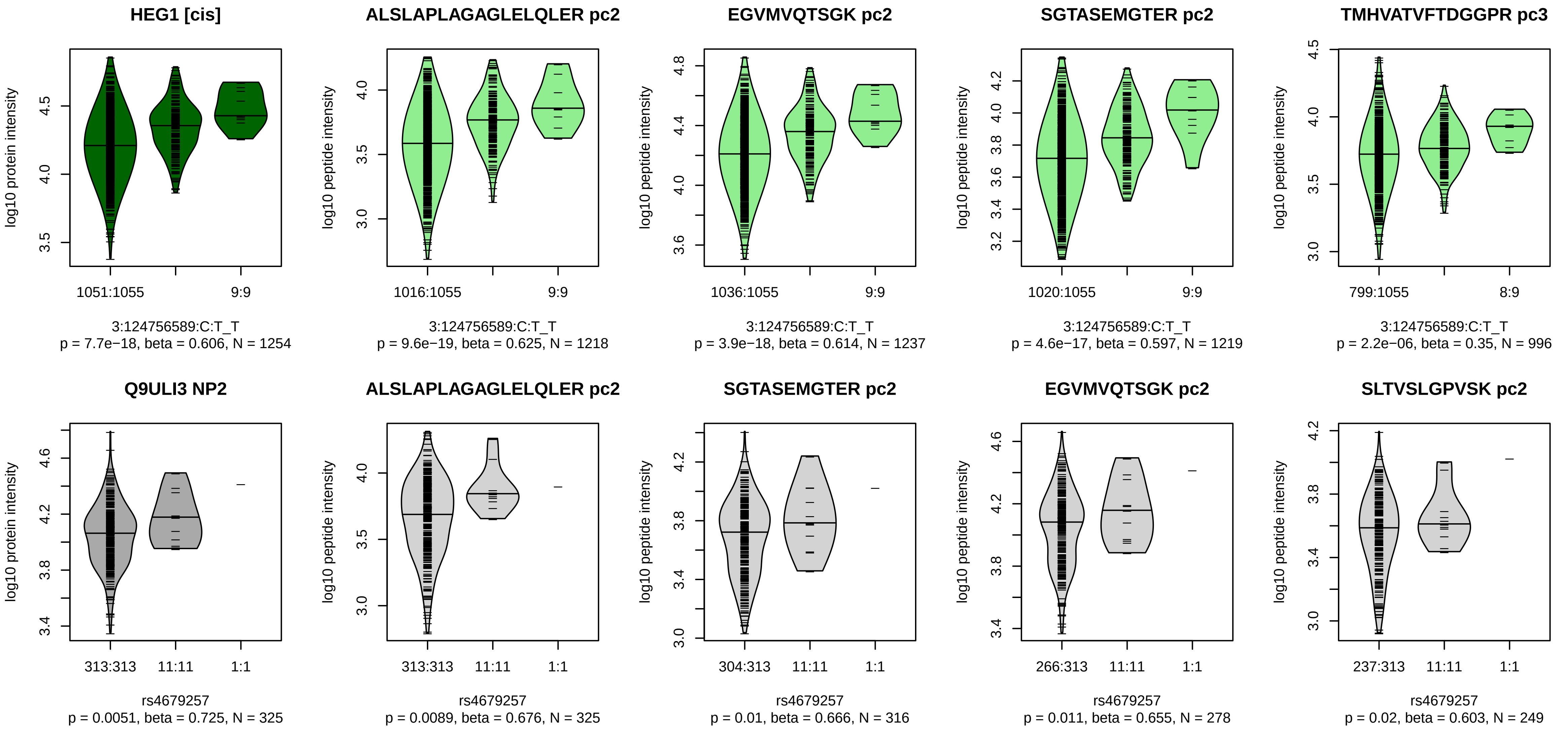


Locus: 104 (Sort: 123)
Status: Not replicated
80% power: No
SNP: 3:124756589:C:T
rsID: rs4679257
UniProtID: Q9ULI3
HGNC gene: HEG1
Type: Cis−pQTL

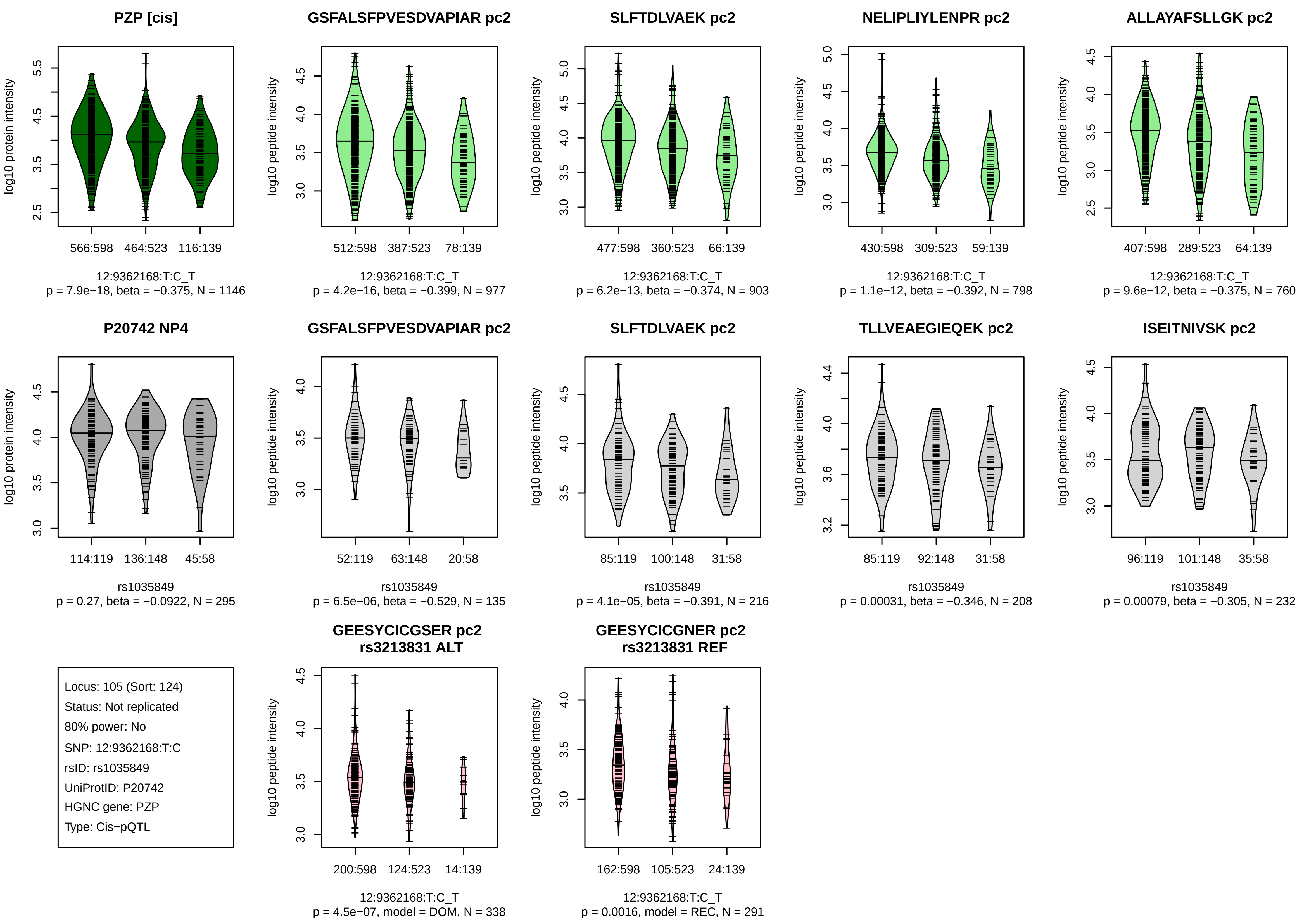

PZP [cis]
log10 protein intensity
566:598
464:523
116:139
12:9362168:T:C_T
p = 7.9e−18, beta = −0.375, N = 1146
GSFALSFPVESDVAPIAR pc2
log10 peptide intensity
512:598
387:523
78:139
12:9362168:T:C_T
p = 4.2e−16, beta = −0.399, N = 977
SLFTDLVAEK pc2
log10 peptide intensity
477:598
360:523
66:139
12:9362168:T:C_T
p = 6.2e−13, beta = −0.374, N = 903
NELIPLIYLENPR pc2
log10 peptide intensity
430:598
309:523
59:139
12:9362168:T:C_T
p = 1.1e−12, beta = −0.392, N = 798
ALLAYAFSLLGK pc2
log10 peptide intensity
407:598
289:523
64:139
12:9362168:T:C_T
p = 9.6e−12, beta = −0.375, N = 760
P20742 NP4
log10 protein intensity
114:119
136:148
45:58
rs1035849
p = 0.27, beta = −0.0922, N = 295
GSFALSFPVESDVAPIAR pc2
log10 peptide intensity
52:119
63:148
20:58
rs1035849
p = 6.5e−06, beta = −0.529, N = 135
SLFTDLVAEK pc2
log10 peptide intensity
85:119
100:148
31:58
rs1035849
p = 4.1e−05, beta = −0.391, N = 216
TLLVEAEGIEQEK pc2
log10 peptide intensity
85:119
92:148
31:58
rs1035849
p = 0.00031, beta = −0.346, N = 208
ISEITNIVSK pc2
log10 peptide intensity
96:119
101:148
35:58
rs1035849
p = 0.00079, beta = −0.305, N = 232
Locus: 105 (Sort: 124)
Status: Not replicated
80% power: No
SNP: 12:9362168:T:C
rsID: rs1035849
UniProtID: P20742
HGNC gene: PZP
Type: Cis−pQTL
GEESYCICGSER pc2
rs3213831 ALT
log10 peptide intensity
200:598
124:523
14:139
12:9362168:T:C_T
p = 4.5e−07, model = DOM, N = 338
GEESYCICGNER pc2
rs3213831 REF
log10 peptide intensity
162:598
105:523
24:139
12:9362168:T:C_T
p = 0.0016, model = REC, N = 291

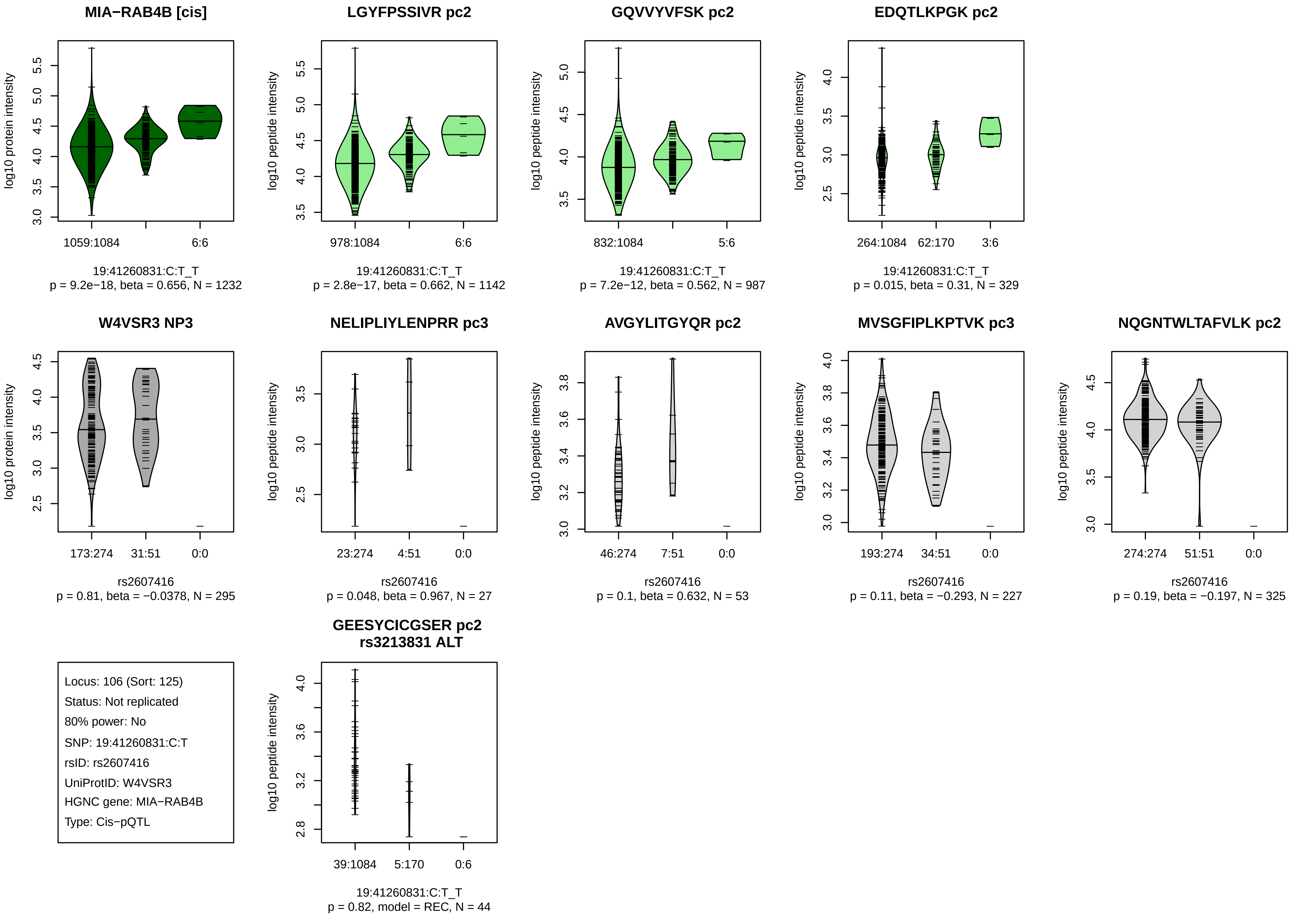
MIA−RAB4B [cis]
log10 protein intensity
1059:1084
6:6
19:41260831:C:T_T
p = 9.2e−18, beta = 0.656, N = 1232
LGYFPSSIVR pc2
log10 peptide intensity
978:1084
6:6
19:41260831:C:T_T
p = 2.8e−17, beta = 0.662, N = 1142
GQVVYVFSK pc2
log10 peptide intensity
832:1084
5:6
19:41260831:C:T_T
p = 7.2e−12, beta = 0.562, N = 987
EDQTLKPGK pc2
log10 peptide intensity
264:1084
62:170
3:6
19:41260831:C:T_T
p = 0.015, beta = 0.31, N = 329
W4VSR3 NP3
log10 protein intensity
173:274
31:51
0:0
rs2607416
p = 0.81, beta = −0.0378, N = 295
NELIPLIYLENPRR pc3
log10 peptide intensity
23:274
4:51
0:0
rs2607416
p = 0.048, beta = 0.967, N = 27
AVGYLITGYQR pc2
log10 peptide intensity
46:274
7:51
0:0
rs2607416
p = 0.1, beta = 0.632, N = 53
MVSGFIPLKPTVK pc3
log10 peptide intensity
193:274
34:51
0:0
rs2607416
p = 0.11, beta = −0.293, N = 227
NQGNTWLTAFVLK pc2
log10 peptide intensity
274:274
51:51
0:0
rs2607416
p = 0.19, beta = −0.197, N = 325
Locus: 106 (Sort: 125)
Status: Not replicated
80% power: No
SNP: 19:41260831:C:T
rsID: rs2607416
UniProtID: W4VSR3
HGNC gene: MIA−RAB4B
Type: Cis−pQTL
GEESYCICGSER pc2
rs3213831 ALT
log10 peptide intensity
39:1084
5:170
0:6
19:41260831:C:T_T
p = 0.82, model = REC, N = 44

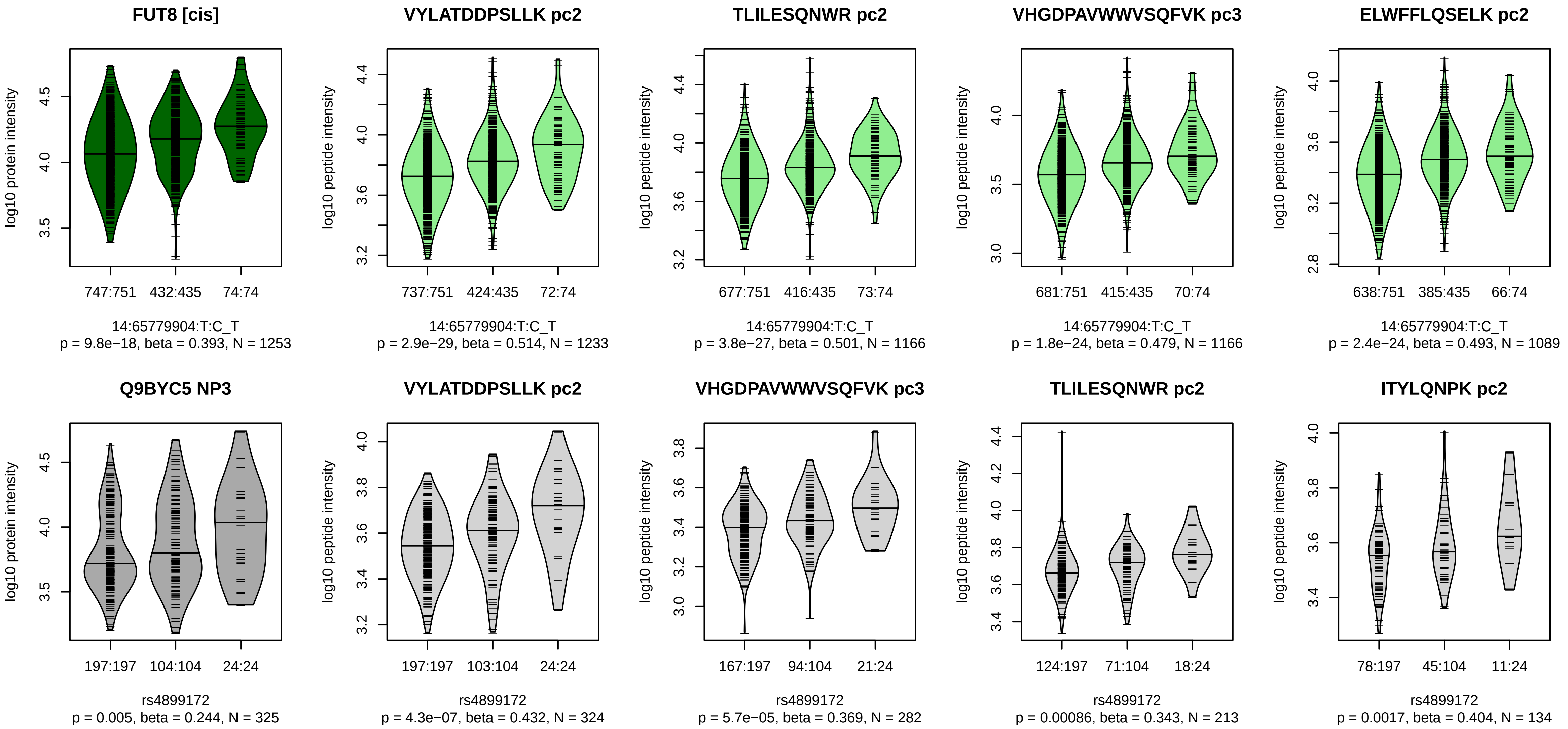


Locus: 107 (Sort: 126)

Status: Not replicated

80% power: No

SNP: 14:65779904:T:C

rsID: rs4899172

UniProtID: Q9BYC5

HGNC gene: FUT8

Type: Cis−pQTL

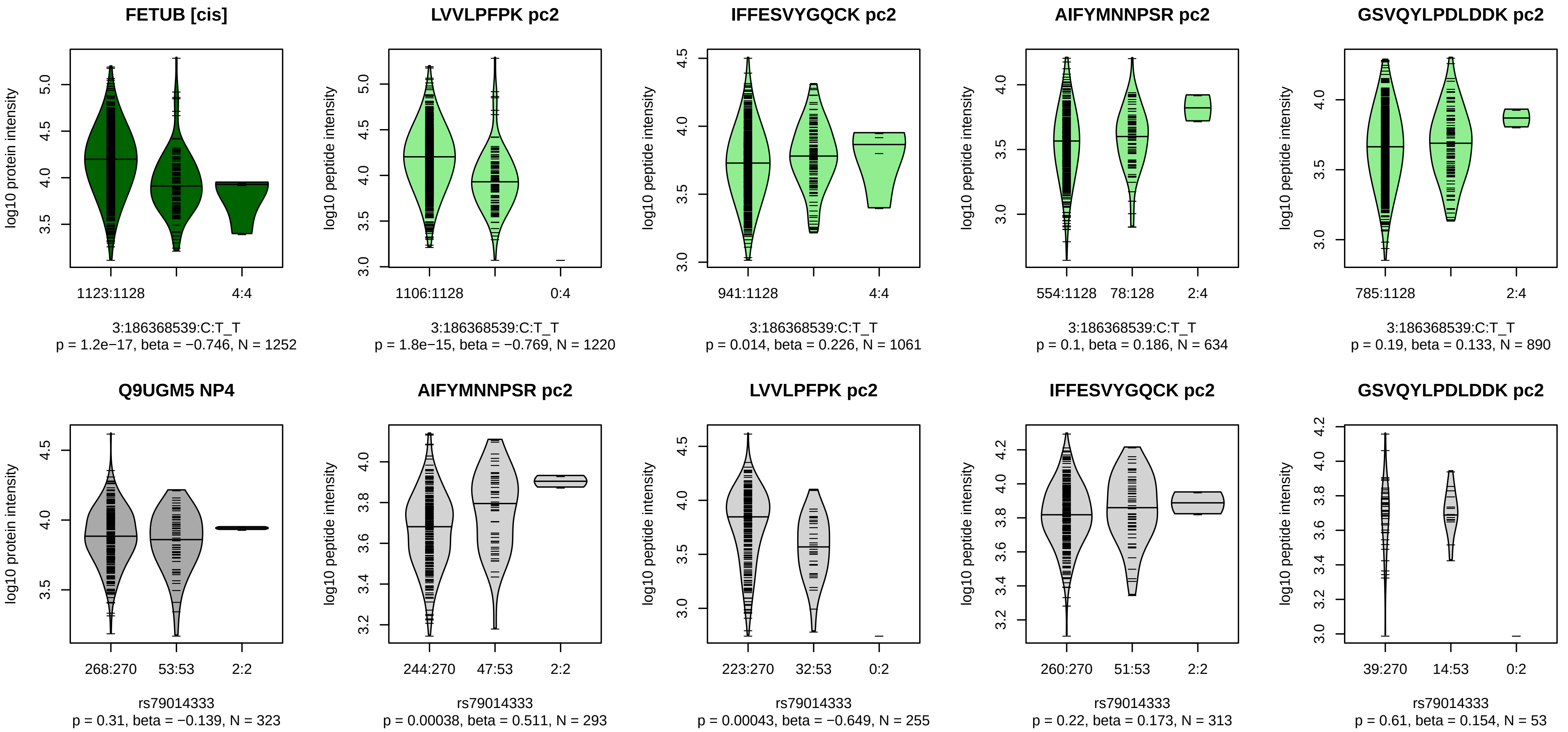


Locus: 108 (Sort: 127)
Status: Not replicated
80% power: No
SNP: 3:186368539:C:T
rsID: rs79014333
UniProtID: Q9UGM5
HGNC gene: FETUB
Type: Cis−pQTL

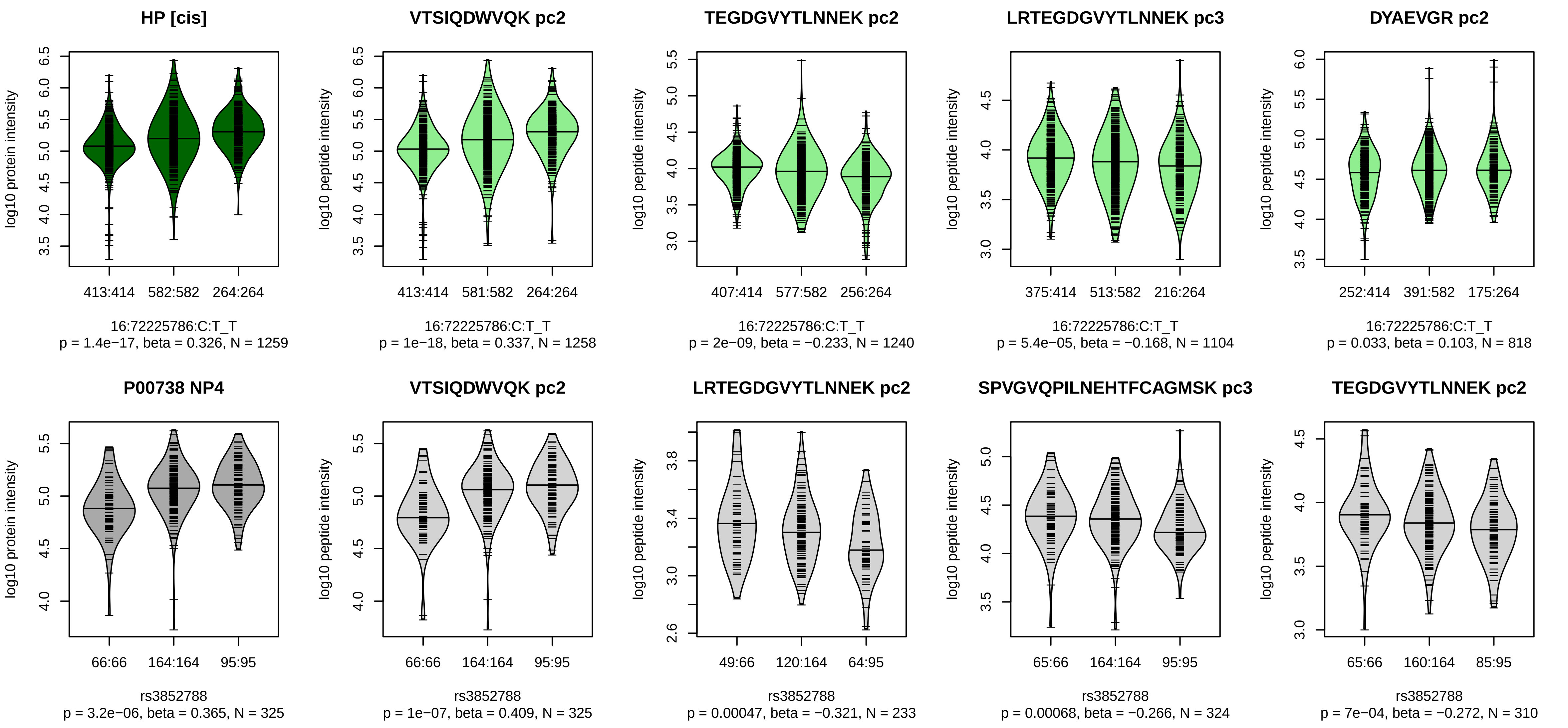


Locus: 109 (Sort: 128)

Status: Replicated

80% power: No

SNP: 16:72225786:C:T

rsID: rs3852788

UniProtID: P00738

HGNC gene: HP

Type: Cis−pQTL

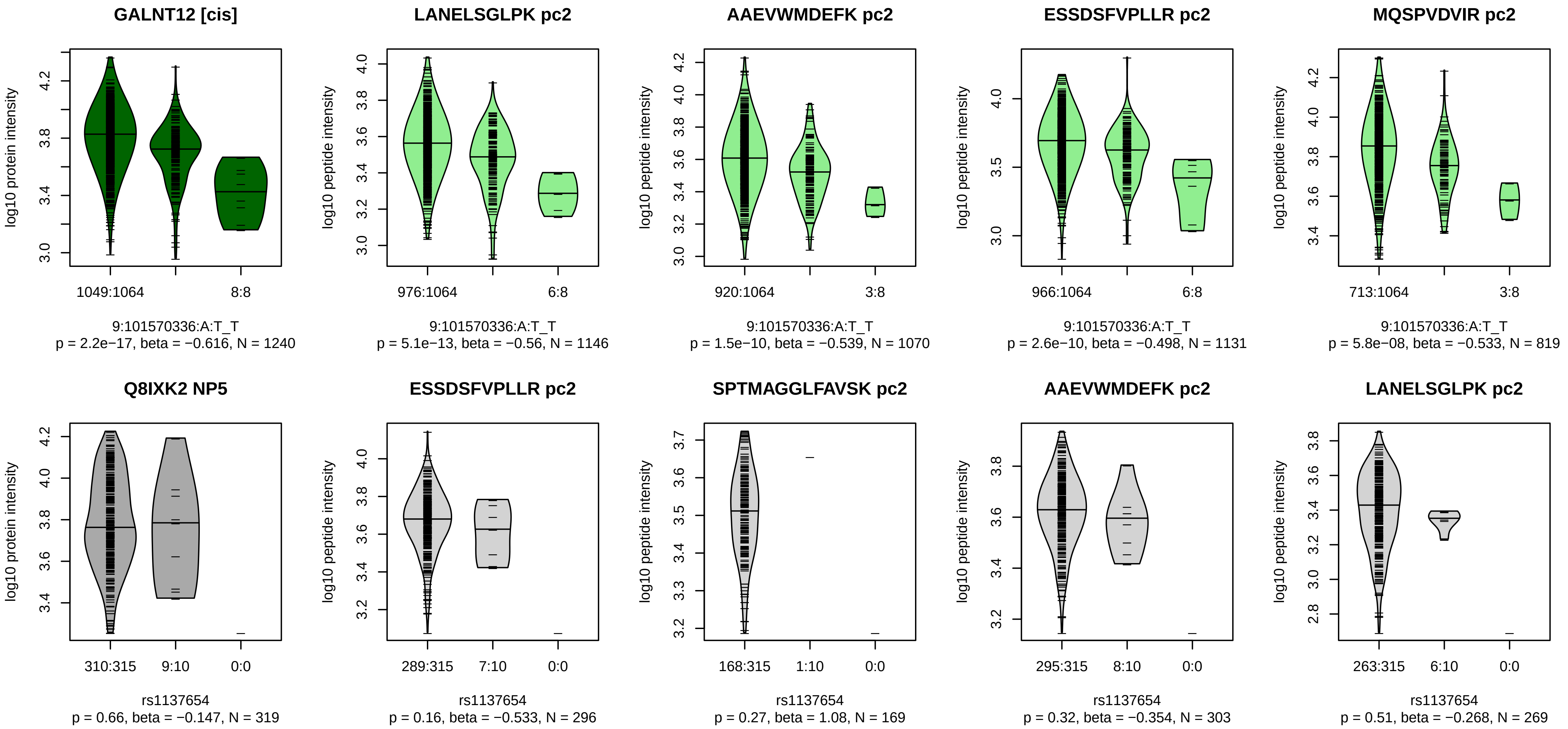


Locus: 110 (Sort: 129)
Status: Not replicated
80% power: No
SNP: 9:101570336:A:T
rsID: rs1137654
UniProtID: Q8IXK2
HGNC gene: GALNT12
Type: Cis−pQTL

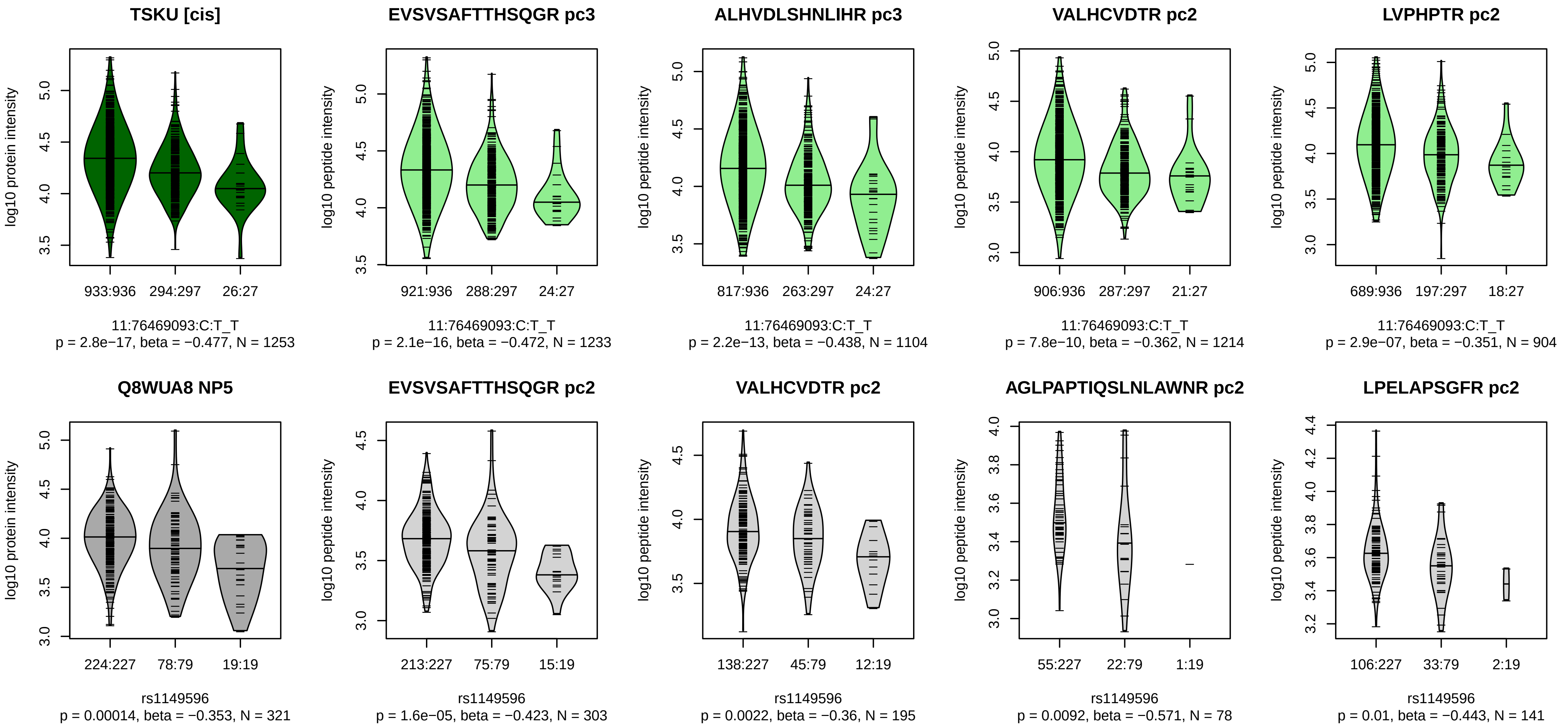


Locus: 111 (Sort: 130)
Status: Replicated
80% power: No
SNP: 11:76469093:C:T
rsID: rs1149596
UniProtID: Q8WUA8
HGNC gene: TSKU
Type: Cis−pQTL

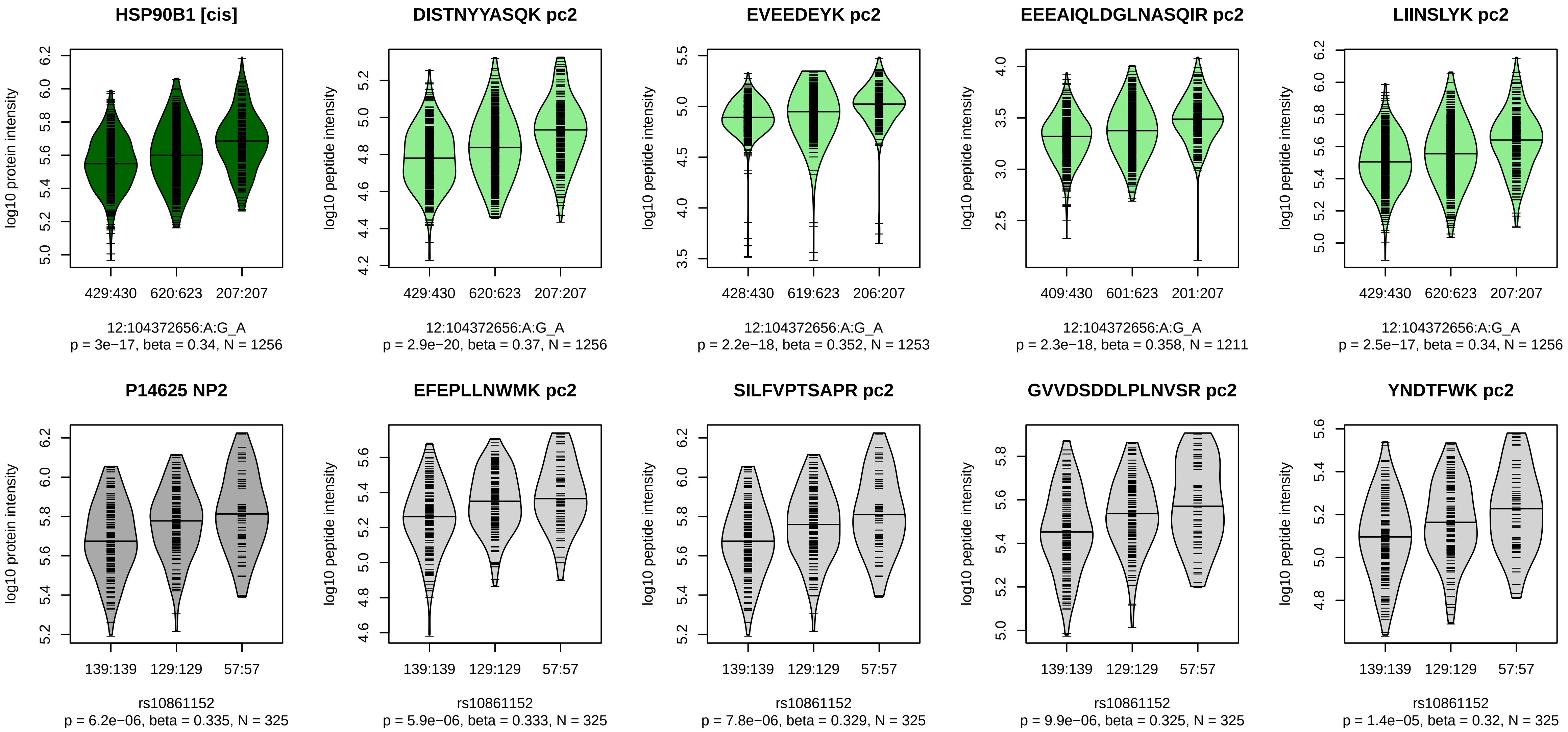


Locus: 112 (Sort: 131)

Status: Replicated

80% power: No

SNP: 12:104372656:A:G

rsID: rs10861152

UniProtID: P14625

HGNC gene: HSP90B1

Type: Cis−pQTL

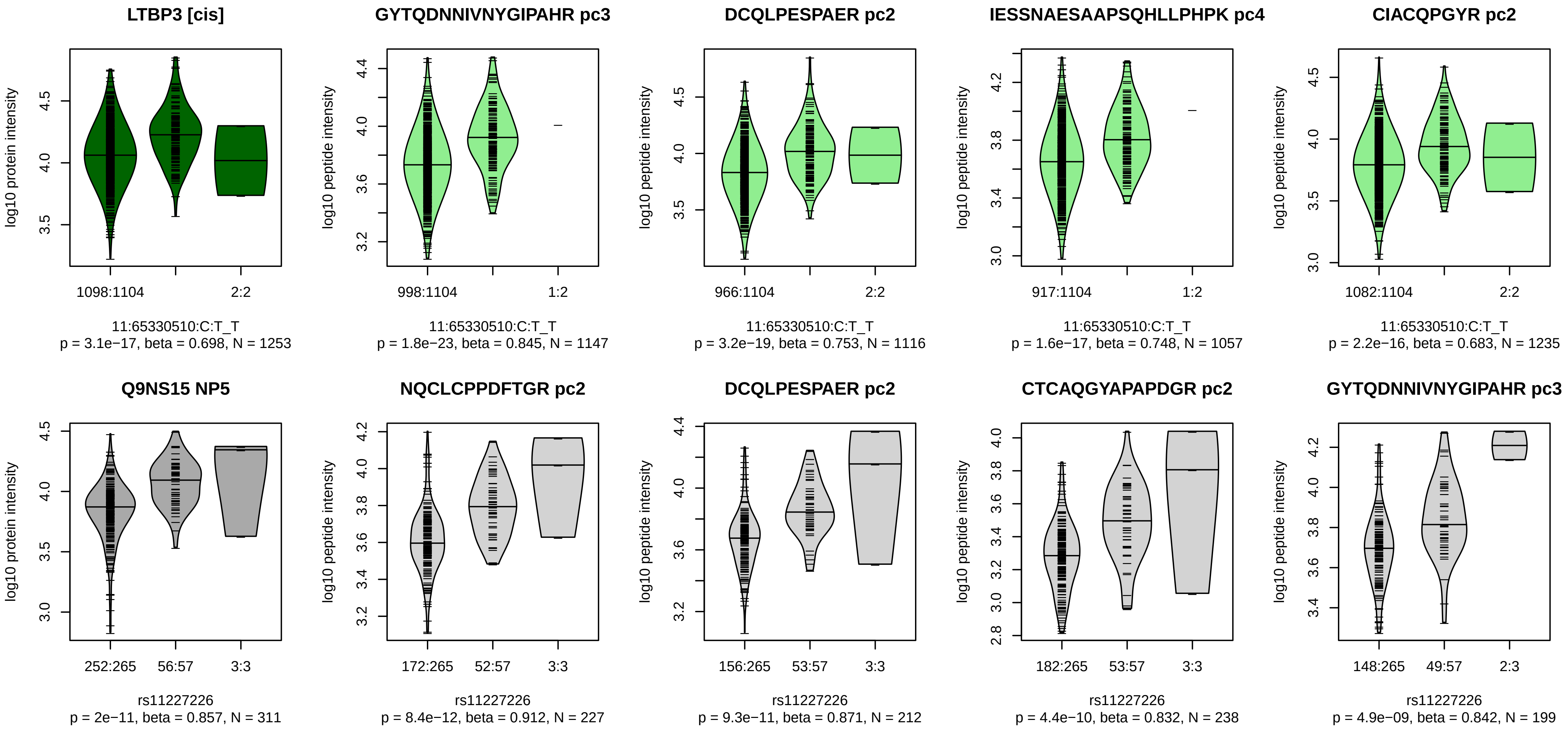


Locus: 113 (Sort: 132)
Status: Replicated
80% power: No
SNP: 11:65330510:C:T
rsID: rs11227226
UniProtID: Q9NS15
HGNC gene: LTBP3
Type: Cis−pQTL

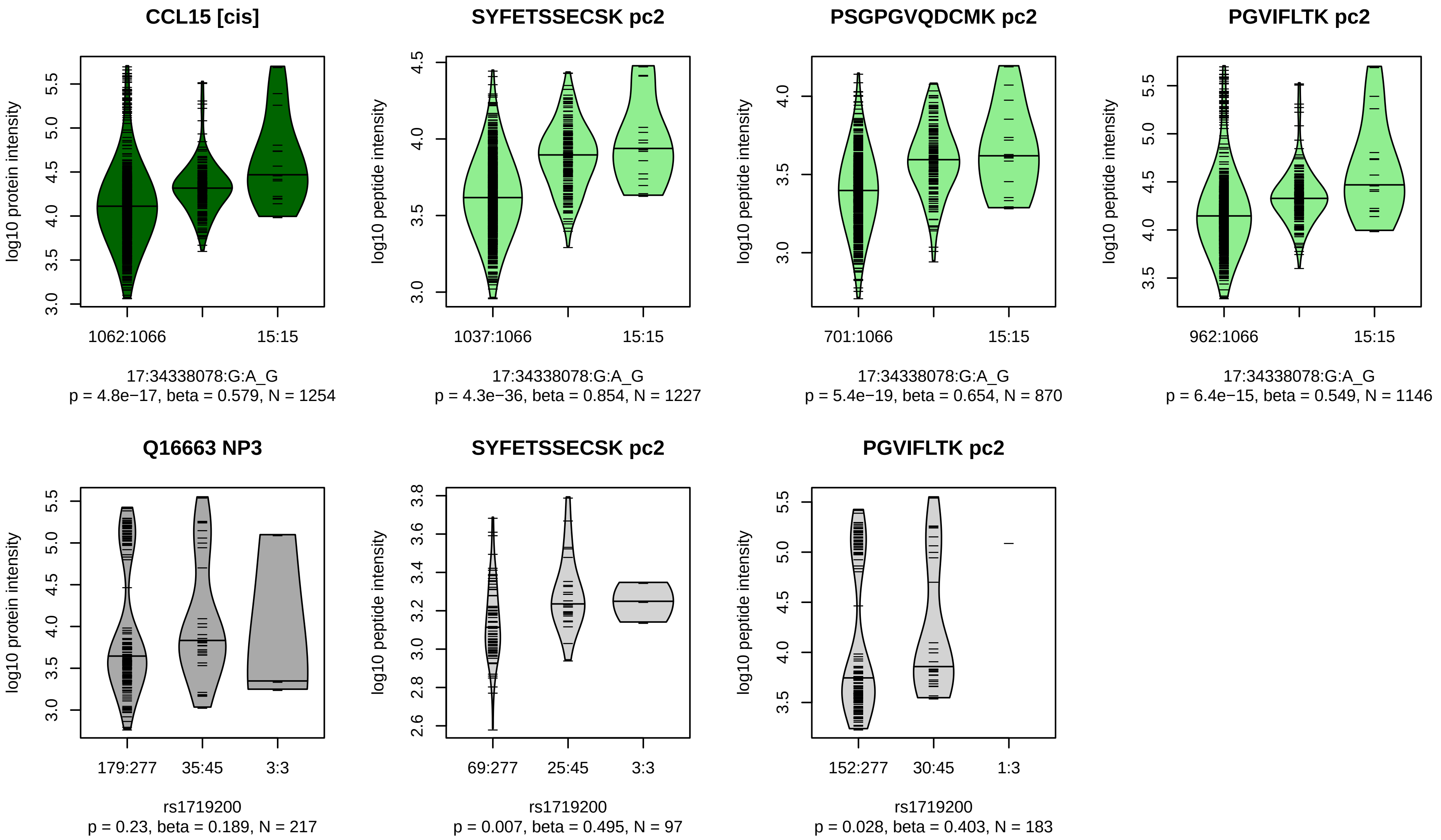


Locus: 114 (Sort: 133)

Status: Not replicated

80% power: No

SNP: 17:34338078:G:A

rsID: rs1719200

UniProtID: Q16663

HGNC gene: CCL15

Type: Cis−pQTL

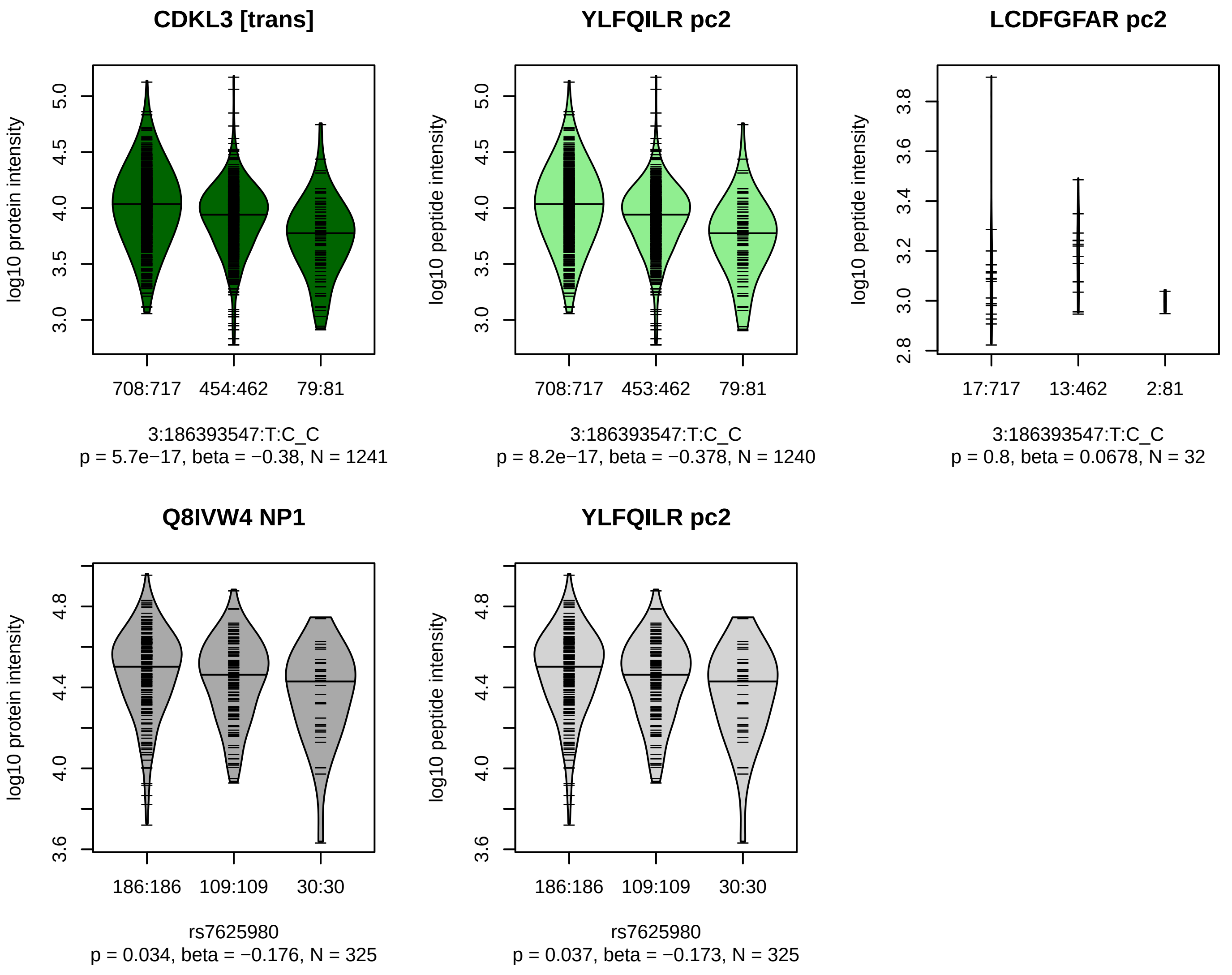


Locus: 115 (Sort: 134)

Status: Not replicated

80% power: No

SNP: 3:186393547:T:C

rsID: rs7625980

UniProtID: Q8IVW4

HGNC gene: CDKL3

Type: Trans−pQTL

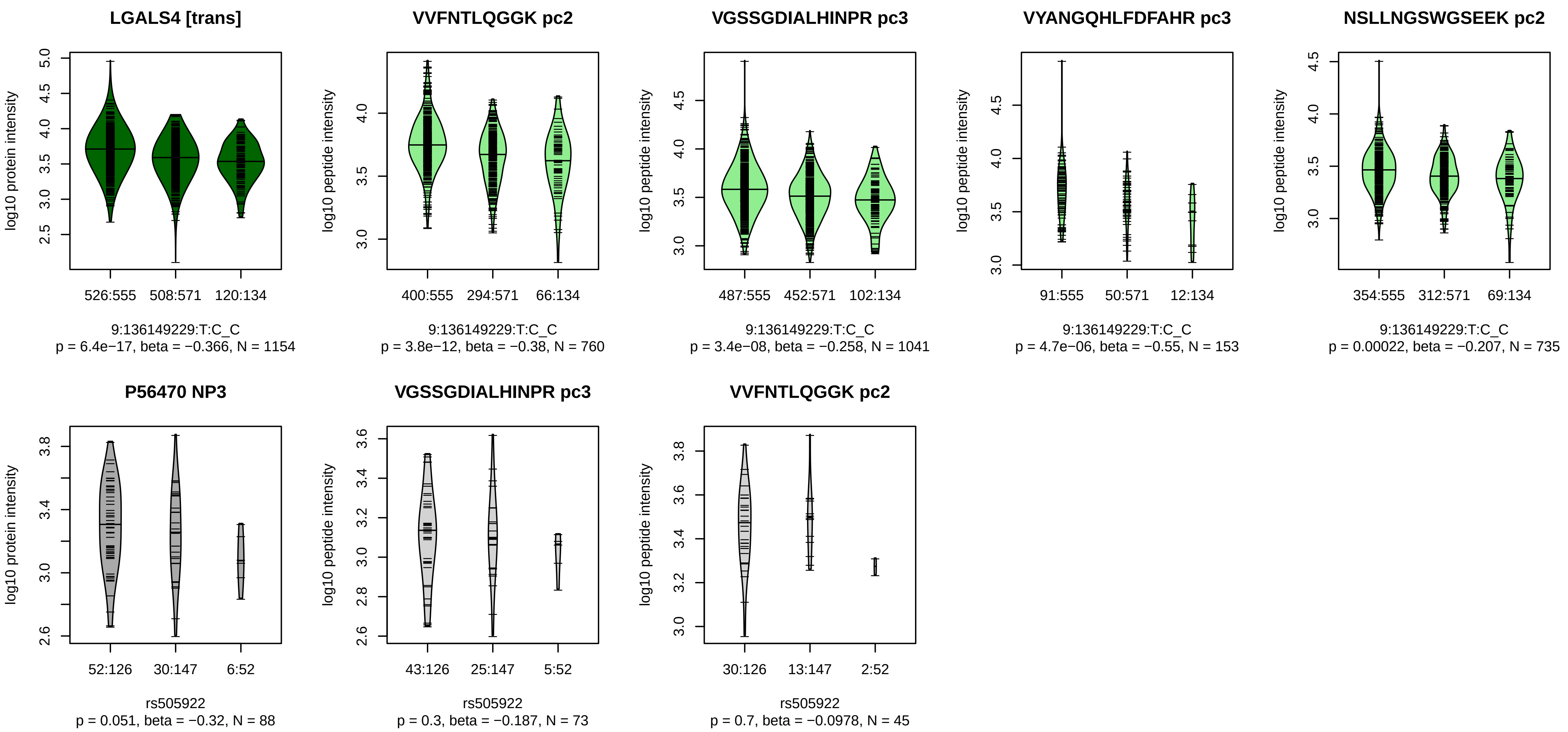


Locus: 116 (Sort: 135)

Status: Not replicated

80% power: No

SNP: 9:136149229:T:C

rsID: rs505922

UniProtID: P56470

HGNC gene: LGALS4

Type: Trans−pQTL

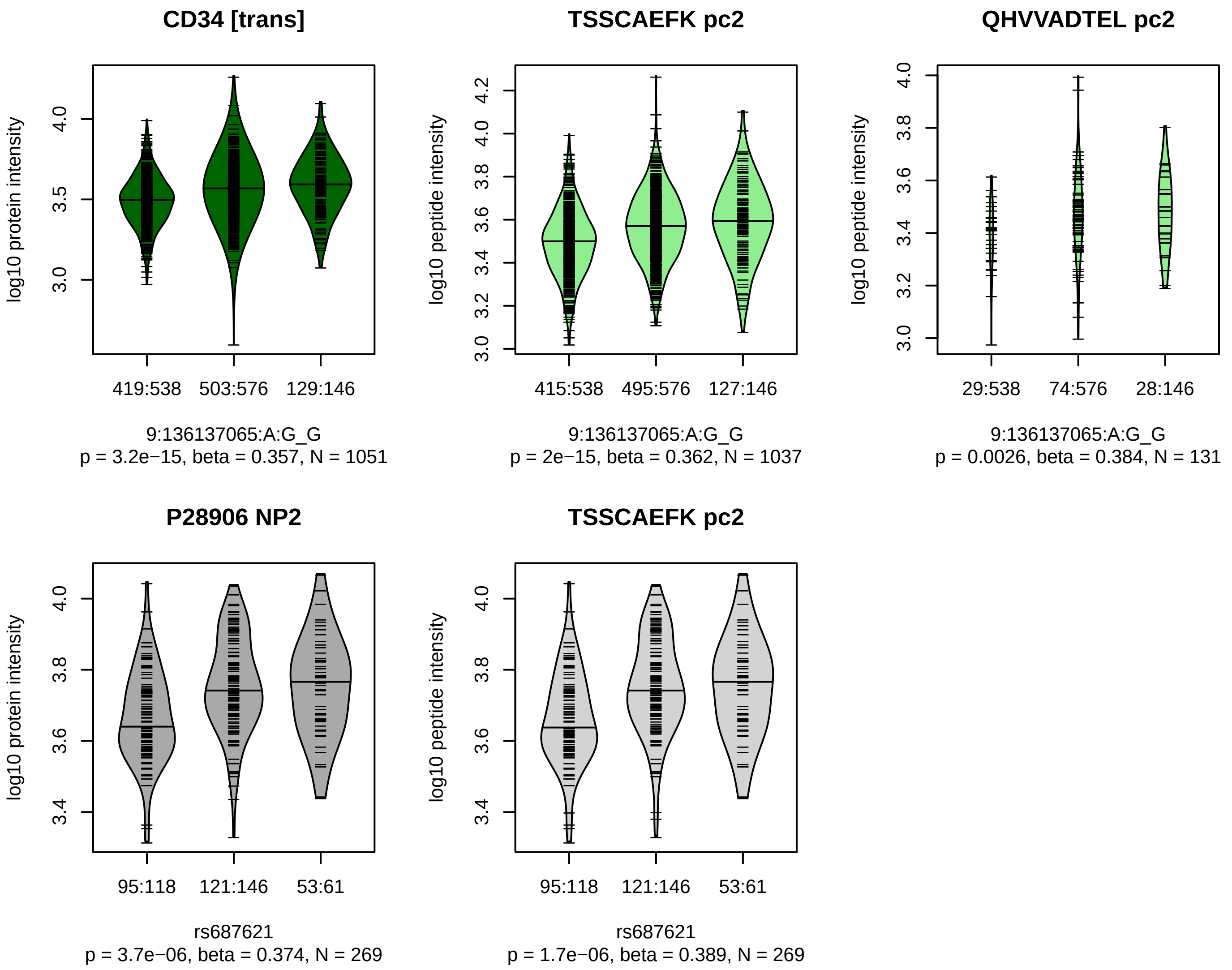


Locus: 116 (Sort: 136)

Status: Replicated

80% power: No

SNP: 9:136137065:A:G

rsID: rs687621

UniProtID: P28906

HGNC gene: CD34

Type: Trans−pQTL

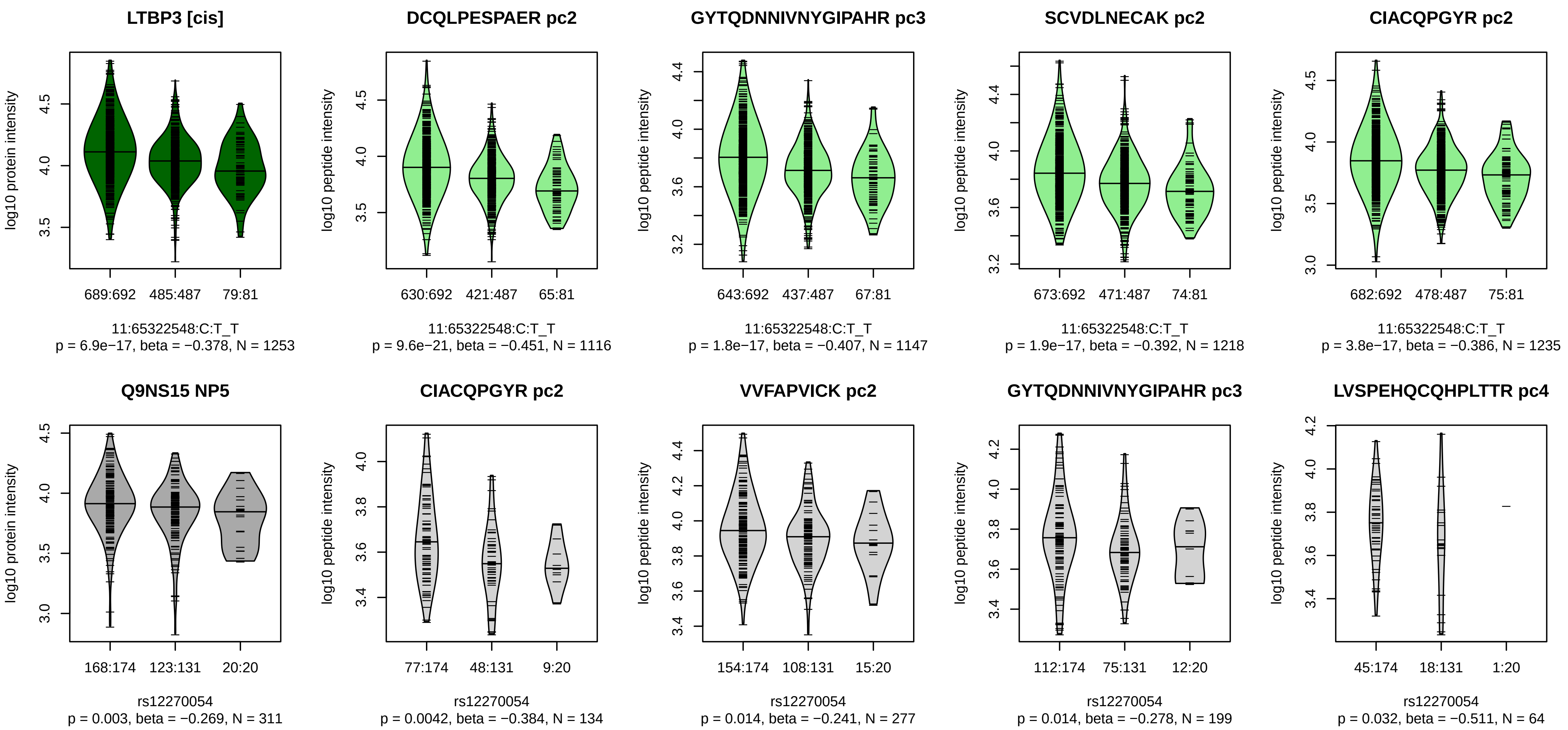


Locus: 117 (Sort: 137)
Status: Not replicated
80% power: No
SNP: 11:65322548:C:T
rsID: rs12270054
UniProtID: Q9NS15
HGNC gene: LTBP3
Type: Cis−pQTL

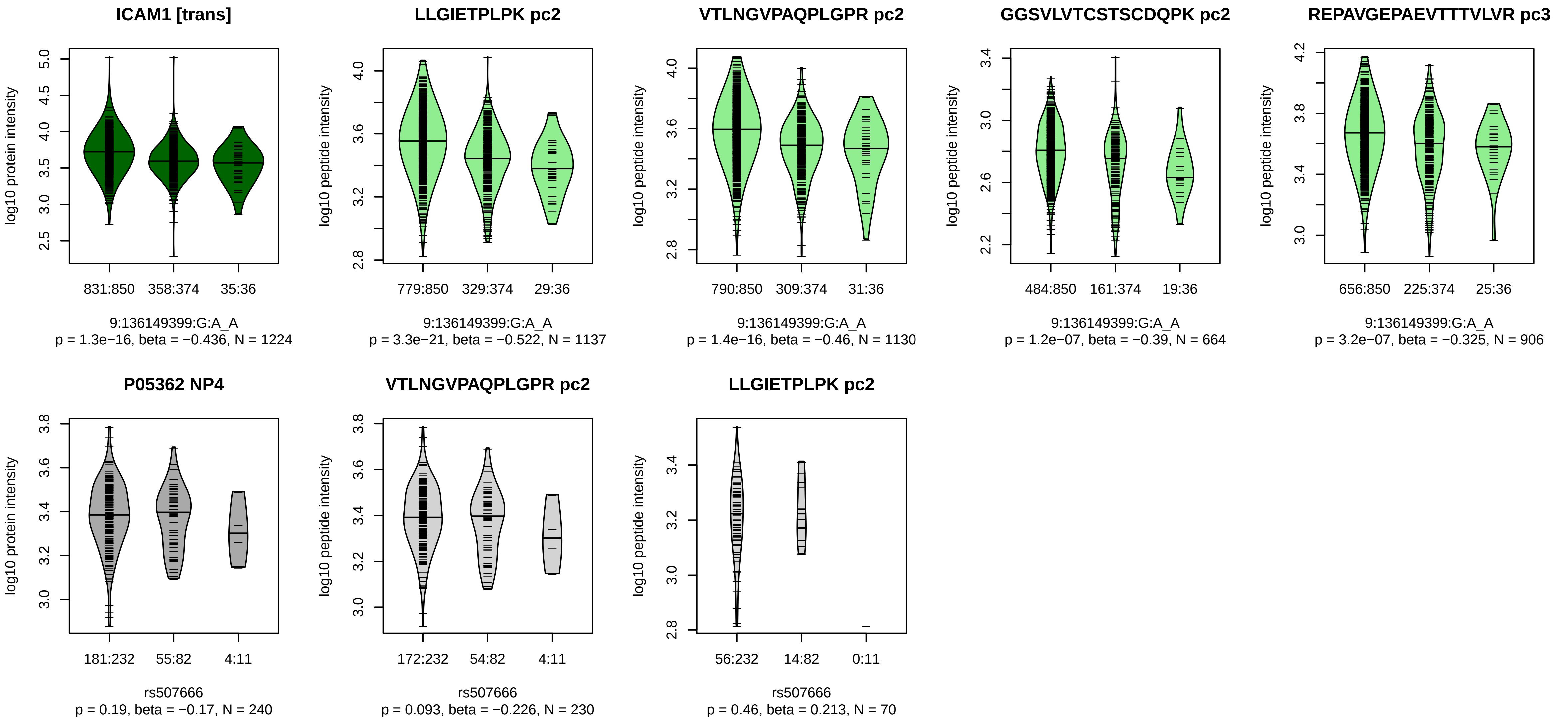


Locus: 118 (Sort: 138)

Status: Not replicated

80% power: No

SNP: 9:136149399:G:A

rsID: rs507666

UniProtID: P05362

HGNC gene: ICAM1

Type: Trans−pQTL

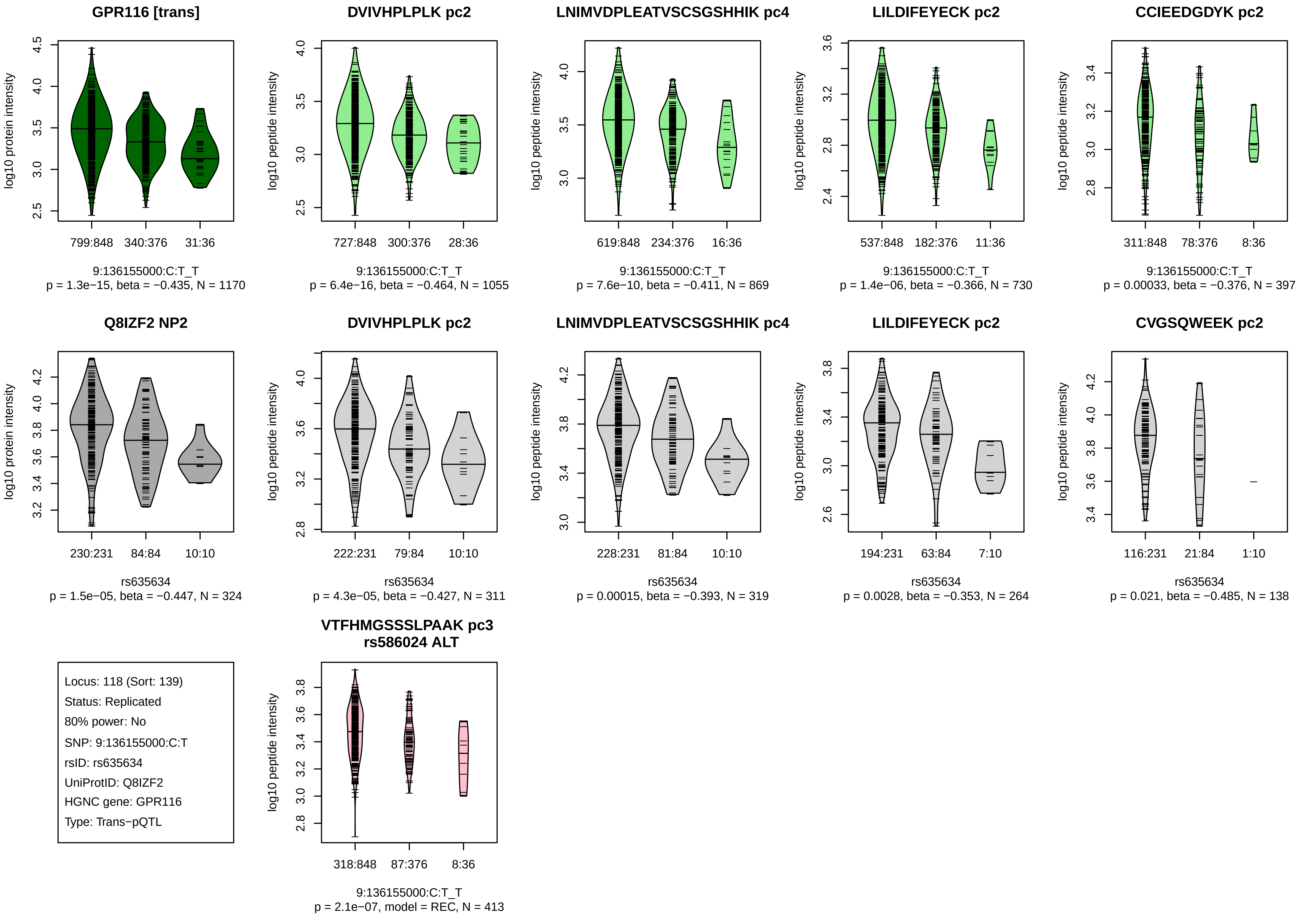
GPR116 [trans]
log10 protein intensity
799:848 340:376 31:36
9:136155000:C:T_T
p = 1.3e−15, beta = −0.435, N = 1170
DVIVHPLPLK pc2
log10 peptide intensity
727:848 300:376 28:36
9:136155000:C:T_T
p = 6.4e−16, beta = −0.464, N = 1055
LNIMVDPLEATVSCSGSHHIK pc4
log10 peptide intensity
619:848 234:376 16:36
9:136155000:C:T_T
p = 7.6e−10, beta = −0.411, N = 869
LILDIFEYECK pc2
log10 peptide intensity
537:848 182:376 11:36
9:136155000:C:T_T
p = 1.4e−06, beta = −0.366, N = 730
CCIEEDGDYK pc2
log10 peptide intensity
311:848 78:376 8:36
9:136155000:C:T_T
p = 0.00033, beta = −0.376, N = 397
Q8IZF2 NP2
log10 protein intensity
230:231 84:84 10:10
rs635634
p = 1.5e−05, beta = −0.447, N = 324
DVIVHPLPLK pc2
log10 peptide intensity
222:231 79:84 10:10
rs635634
p = 4.3e−05, beta = −0.427, N = 311
LNIMVDPLEATVSCSGSHHIK pc4
log10 peptide intensity
228:231 81:84 10:10
rs635634
p = 0.00015, beta = −0.393, N = 319
LILDIFEYECK pc2
log10 peptide intensity
194:231 63:84 7:10
rs635634
p = 0.0028, beta = −0.353, N = 264
CVGSQWEEK pc2
log10 peptide intensity
116:231 21:84 1:10
rs635634
p = 0.021, beta = −0.485, N = 138
Locus: 118 (Sort: 139)
Status: Replicated
80% power: No
SNP: 9:136155000:C:T
rsID: rs635634
UniProtID: Q8IZF2
HGNC gene: GPR116
Type: Trans−pQTL
VTFHMGSSSLPAAK pc3
rs586024 ALT
log10 peptide intensity
318:848 87:376 8:36
9:136155000:C:T_T
p = 2.1e−07, model = REC, N = 413

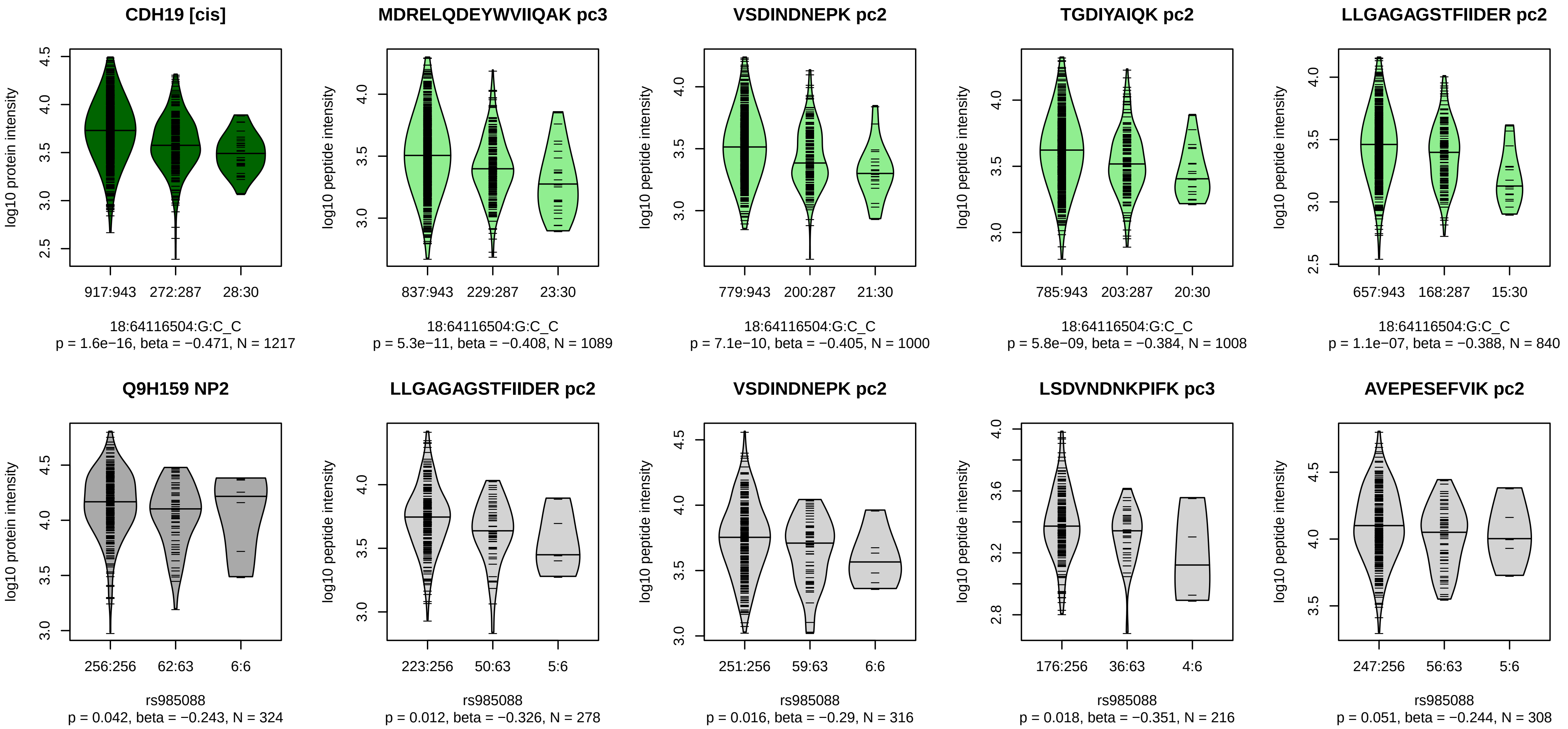


Locus: 119 (Sort: 140)

Status: Not replicated

80% power: No

SNP: 18:64116504:G:C

rsID: rs985088

UniProtID: Q9H159

HGNC gene: CDH19

Type: Cis−pQTL

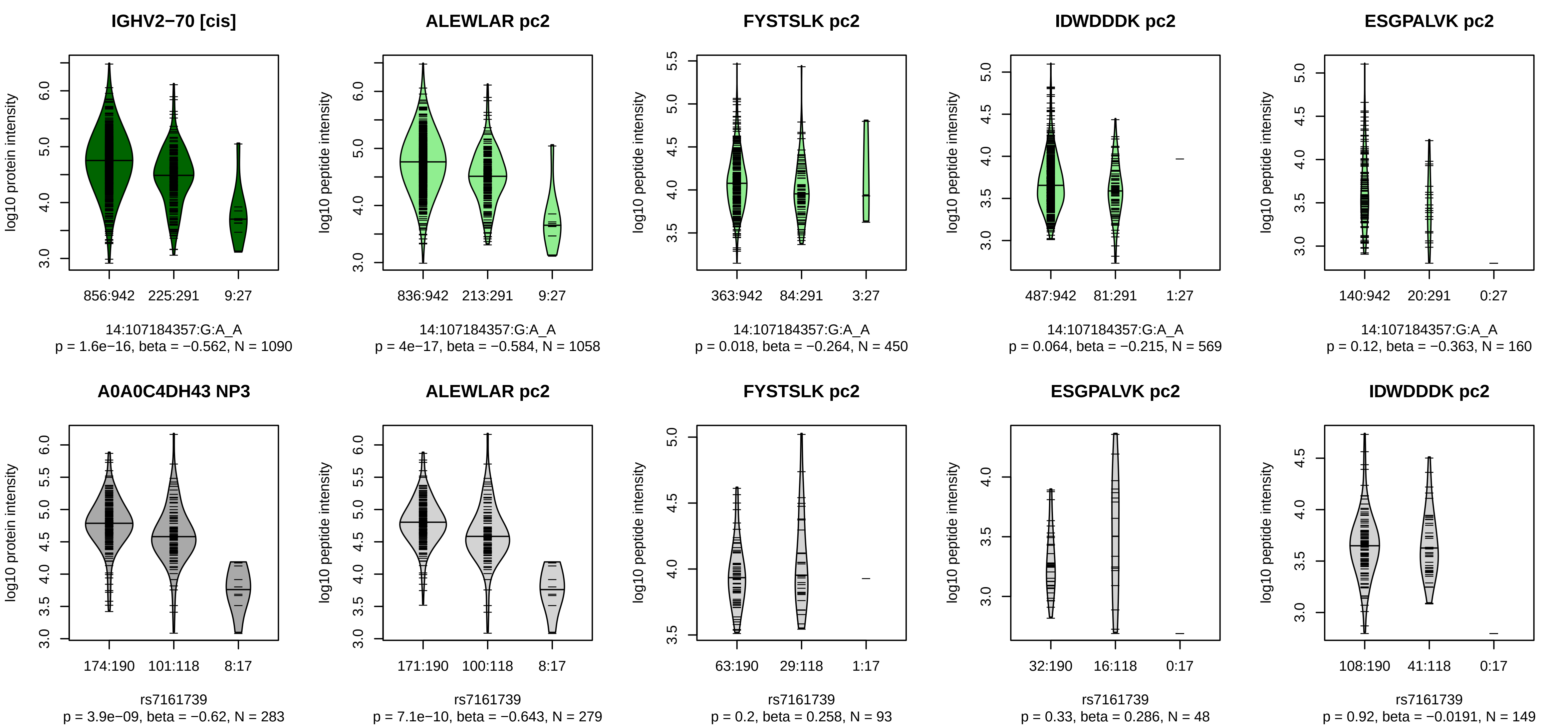


Locus: 120 (Sort: 141)
Status: Replicated
80% power: No
SNP: 14:107184357:G:A
rsID: rs7161739
UniProtID: A0A0C4DH43
HGNC gene: IGHV2−70
Type: Cis−pQTL

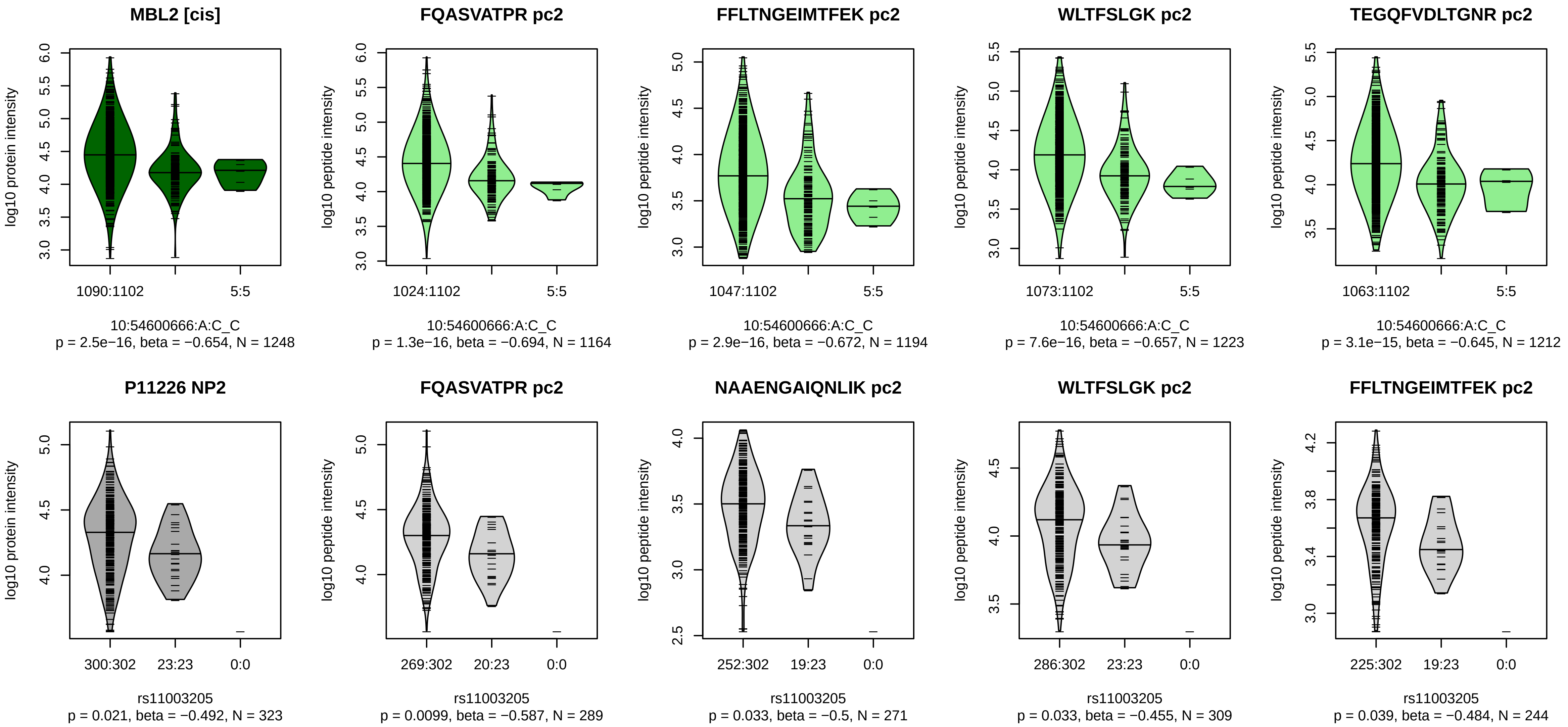


Locus: 121 (Sort: 142)

Status: Not replicated

80% power: No

SNP: 10:54600666:A:C

rsID: rs11003205

UniProtID: P11226

HGNC gene: MBL2

Type: Cis−pQTL

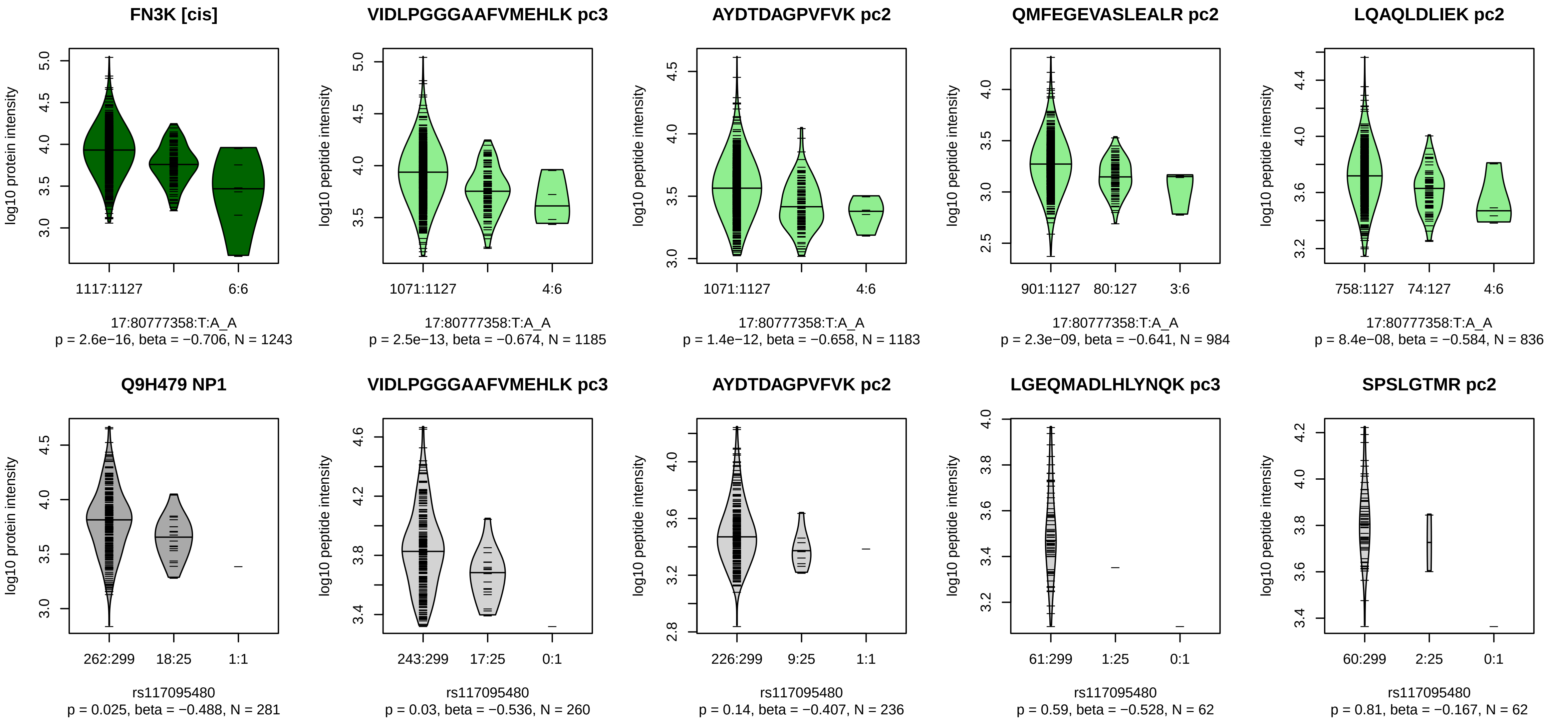


Locus: 122 (Sort: 143)
Status: Not replicated
80% power: No
SNP: 17:80777358:T:A
rsID: rs117095480
UniProtID: Q9H479
HGNC gene: FN3K
Type: Cis−pQTL

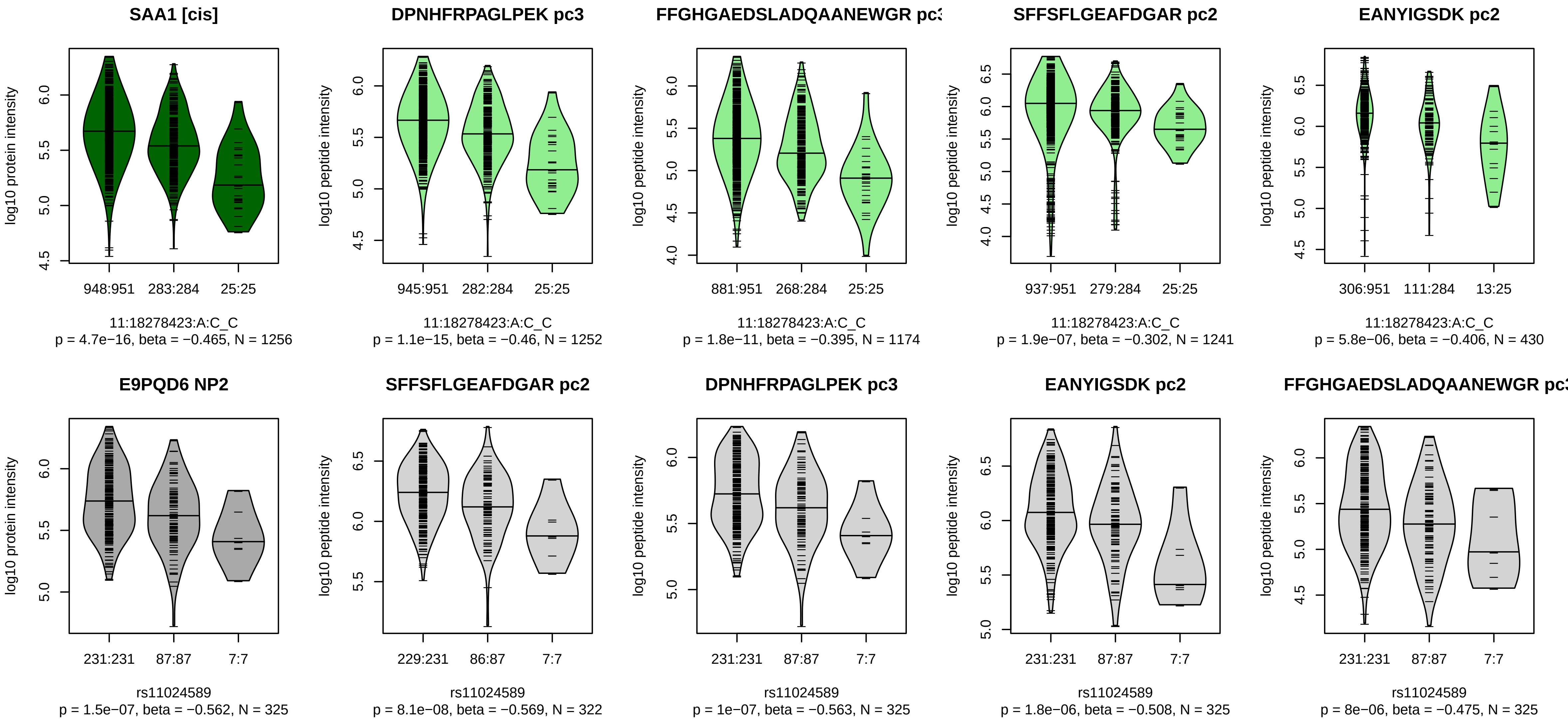


Locus: 123 (Sort: 144)
Status: Replicated
80% power: No
SNP: 11:18278423:A:C
rsID: rs11024589
UniProtID: E9PQD6
HGNC gene: SAA1
Type: Cis−pQTL

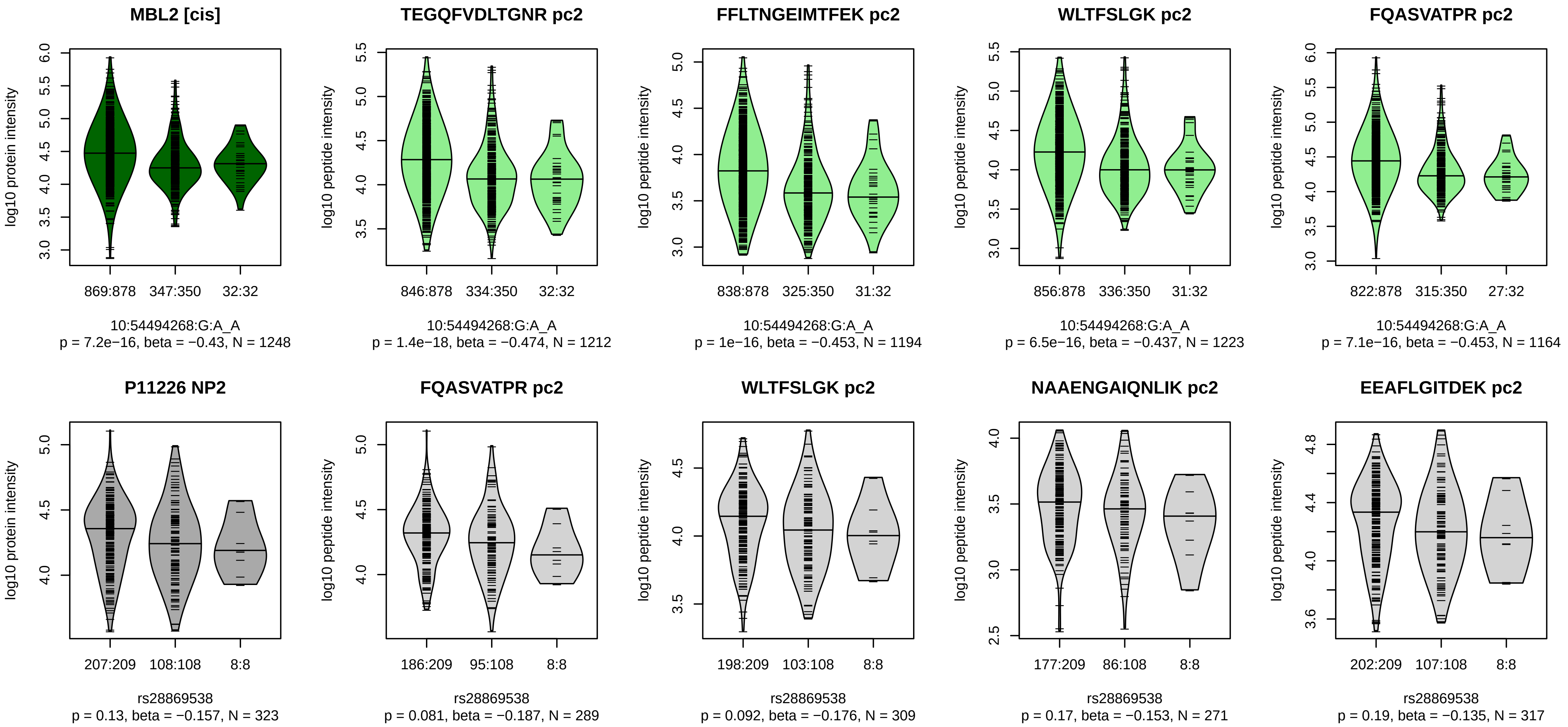


Locus: 124 (Sort: 145)
Status: Not replicated
80% power: No
SNP: 10:54494268:G:A
rsID: rs28869538
UniProtID: P11226
HGNC gene: MBL2
Type: Cis−pQTL

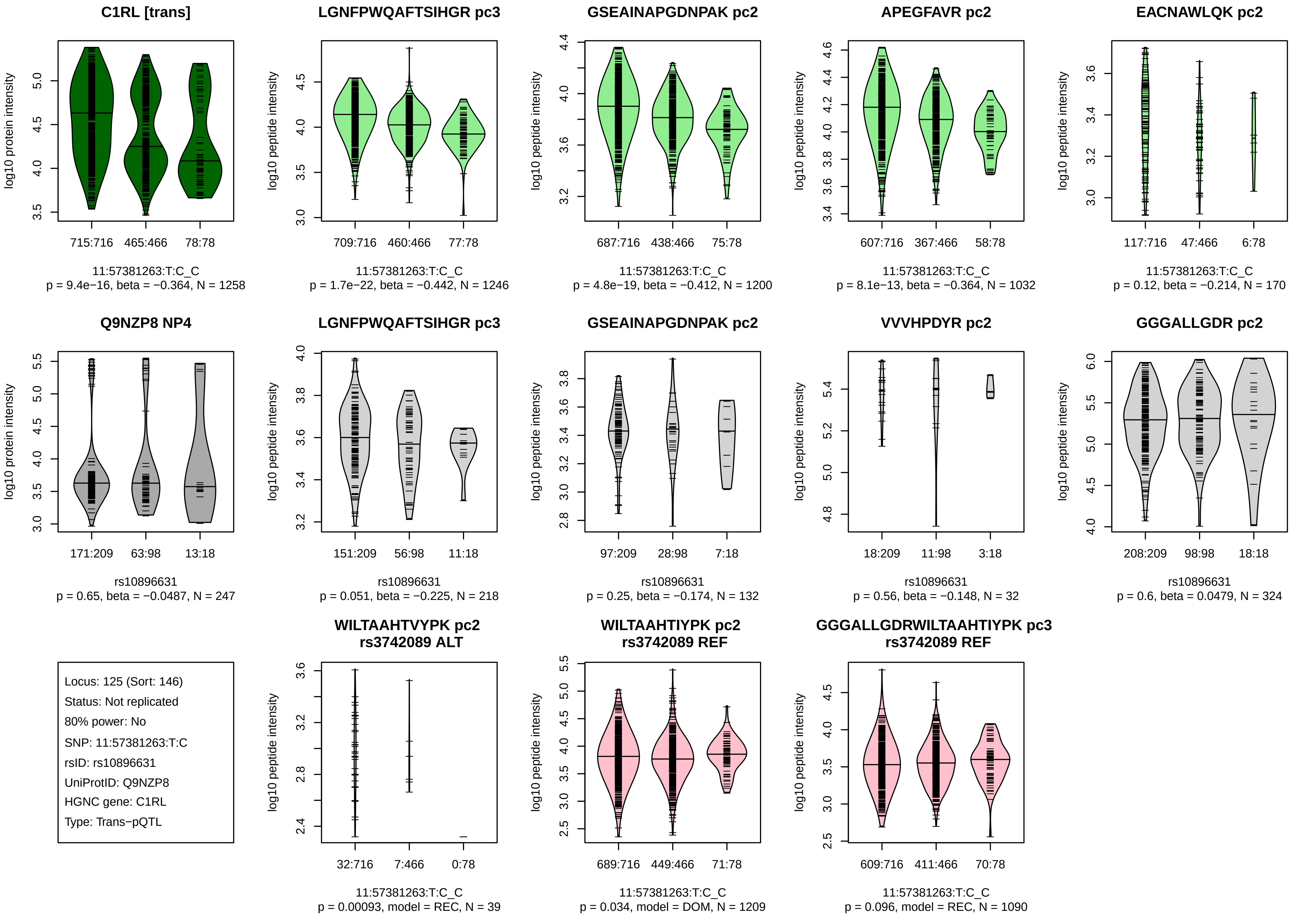
C1RL [trans]
log10 protein intensity
715:716 465:466 78:78
11:57381263:T:C_C
p = 9.4e−16, beta = −0.364, N = 1258
LGNFPWQAFTSIHGR pc3
log10 peptide intensity
709:716 460:466 77:78
11:57381263:T:C_C
p = 1.7e−22, beta = −0.442, N = 1246
GSEAINAPGDNPAK pc2
log10 peptide intensity
687:716 438:466 75:78
11:57381263:T:C_C
p = 4.8e−19, beta = −0.412, N = 1200
APEGFAVR pc2
log10 peptide intensity
607:716 367:466 58:78
11:57381263:T:C_C
p = 8.1e−13, beta = −0.364, N = 1032
EACNAWLQK pc2
log10 peptide intensity
117:716 47:466 6:78
11:57381263:T:C_C
p = 0.12, beta = −0.214, N = 170
Q9NZP8 NP4
log10 protein intensity
171:209 63:98 13:18
rs10896631
p = 0.65, beta = −0.0487, N = 247
LGNFPWQAFTSIHGR pc3
log10 peptide intensity
151:209 56:98 11:18
rs10896631
p = 0.051, beta = −0.225, N = 218
GSEAINAPGDNPAK pc2
log10 peptide intensity
97:209 28:98 7:18
rs10896631
p = 0.25, beta = −0.174, N = 132
VVVHPDYR pc2
log10 peptide intensity
18:209 11:98 3:18
rs10896631
p = 0.56, beta = −0.148, N = 32
GGGALLGDR pc2
log10 peptide intensity
208:209 98:98 18:18
rs10896631
p = 0.6, beta = 0.0479, N = 324
Locus: 125 (Sort: 146)
Status: Not replicated
80% power: No
SNP: 11:57381263:T:C
rsID: rs10896631
UniProtID: Q9NZP8
HGNC gene: C1RL
Type: Trans−pQTL
WILTAAHTVYPK pc2
rs3742089 ALT
log10 peptide intensity
32:716 7:466 0:78
11:57381263:T:C_C
p = 0.00093, model = REC, N = 39
WILTAAHTIYPK pc2
rs3742089 REF
log10 peptide intensity
689:716 449:466 71:78
11:57381263:T:C_C
p = 0.034, model = DOM, N = 1209
GGGALLGDRWILTAAHTIYPK pc3
rs3742089 REF
log10 peptide intensity
609:716 411:466 70:78
11:57381263:T:C_C
p = 0.096, model = REC, N = 1090

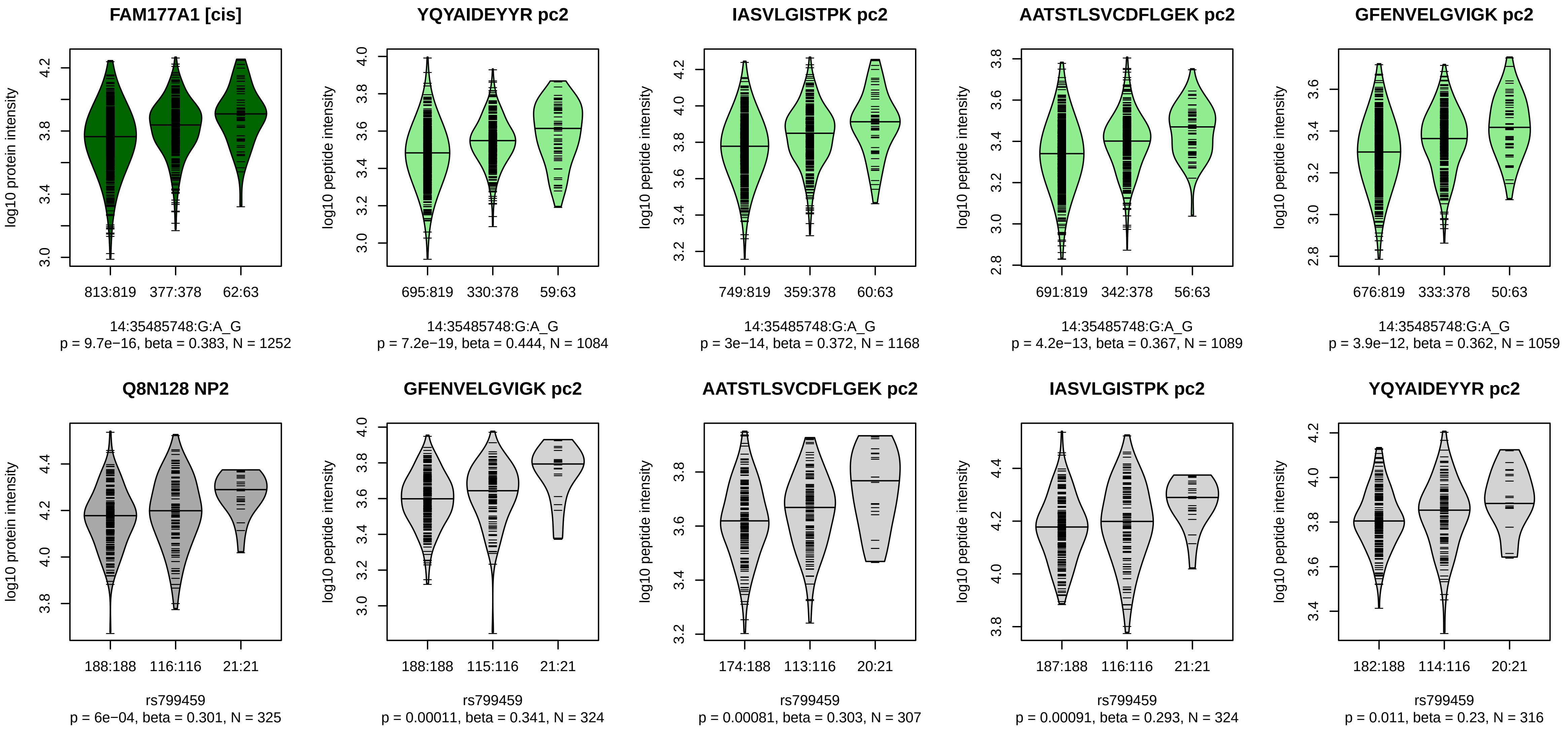


Locus: 126 (Sort: 147)

Status: Not replicated

80% power: No

SNP: 14:35485748:G:A

rsID: rs799459

UniProtID: Q8N128

HGNC gene: FAM177A1

Type: Cis−pQTL

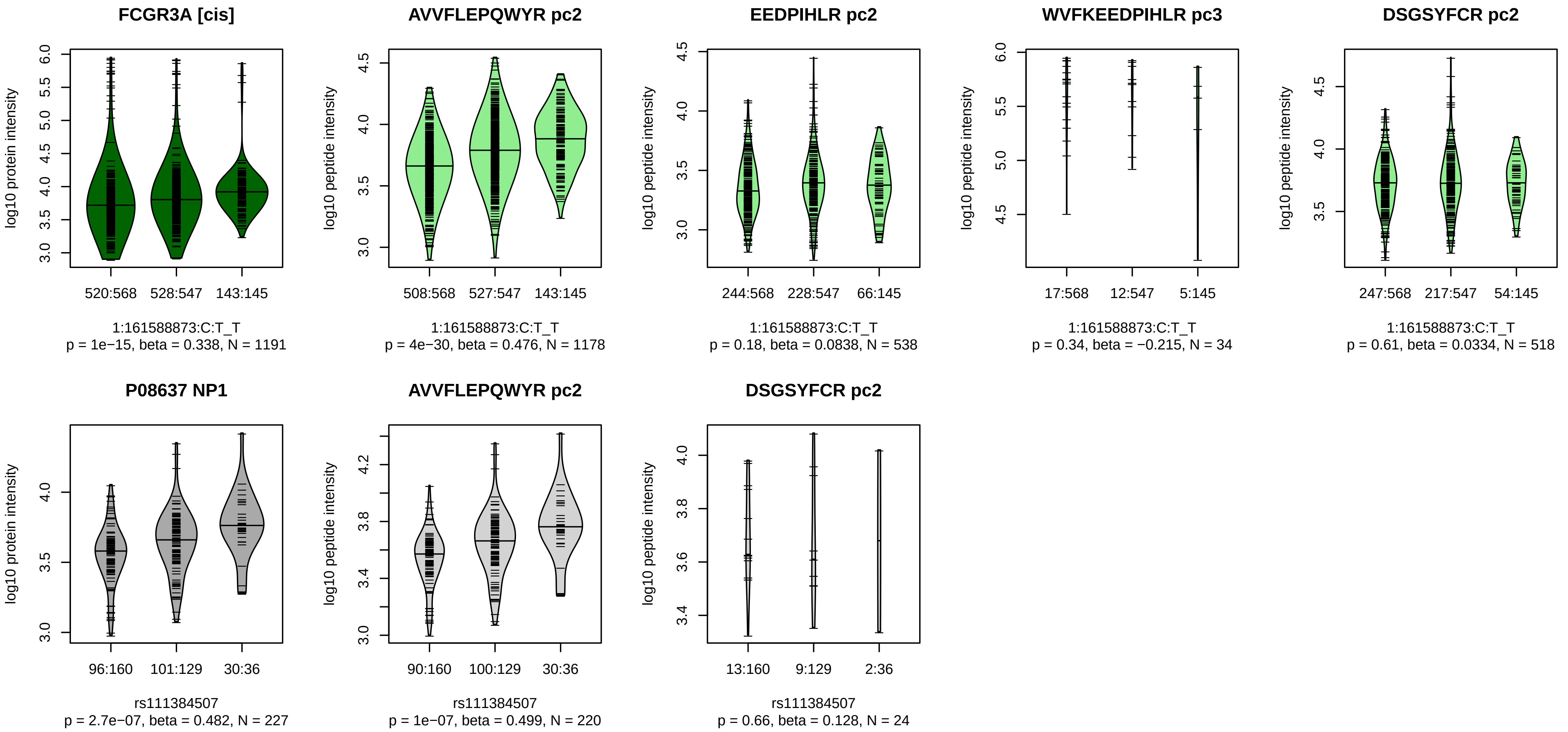


Locus: 127 (Sort: 148)
Status: Replicated
80% power: No
SNP: 1:161588873:C:T
rsID: rs111384507
UniProtID: P08637
HGNC gene: FCGR3A
Type: Cis−pQTL

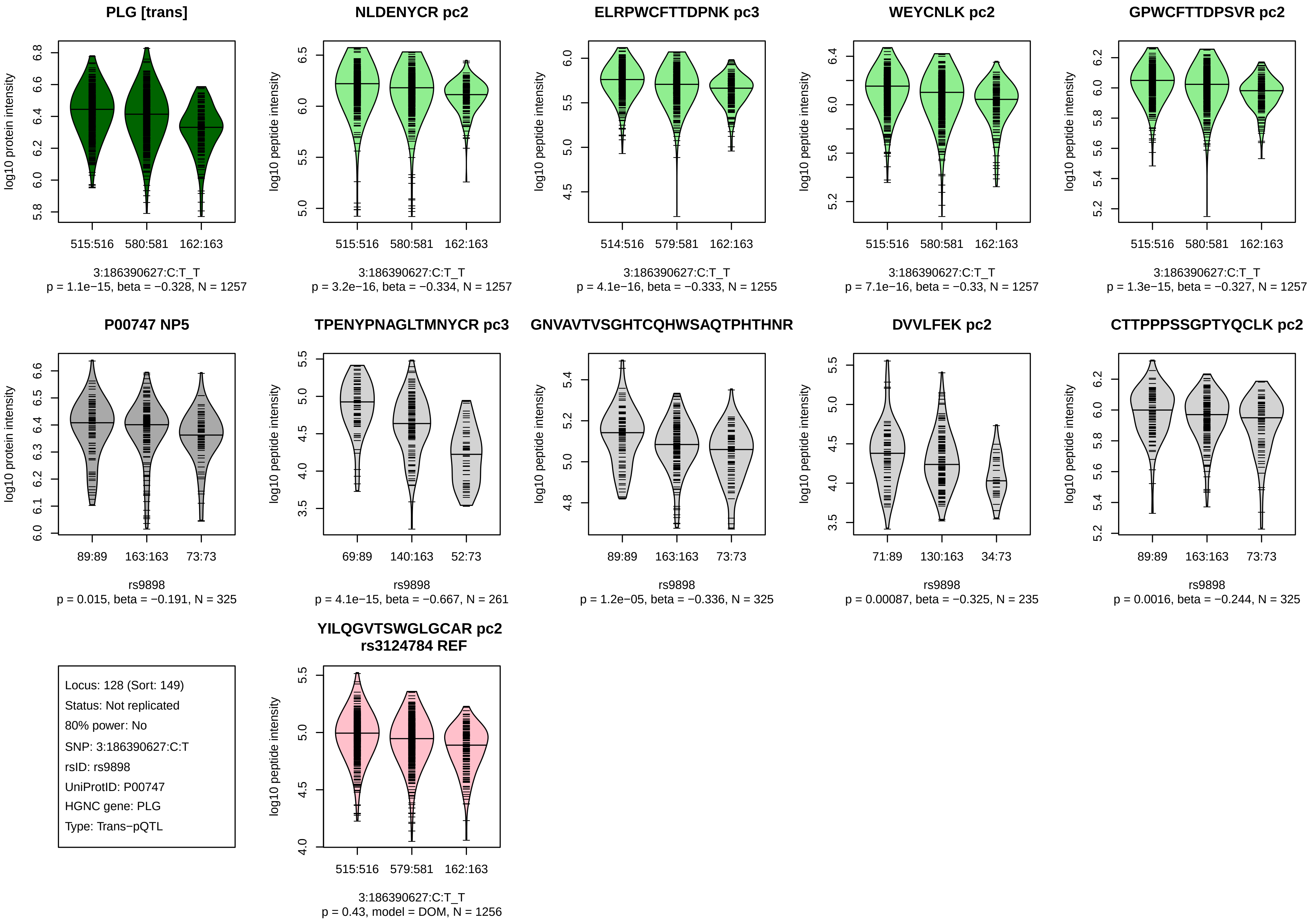

PLG [trans]
log10 protein intensity
515:516
580:581
162:163
3:186390627:C:T_T
p = 1.1e−15, beta = −0.328, N = 1257
NLDENYCR pc2
log10 peptide intensity
515:516
580:581
162:163
3:186390627:C:T_T
p = 3.2e−16, beta = −0.334, N = 1257
ELRPWCFTTDPNK pc3
log10 peptide intensity
514:516
579:581
162:163
3:186390627:C:T_T
p = 4.1e−16, beta = −0.333, N = 1255
WEYCNLK pc2
log10 peptide intensity
515:516
580:581
162:163
3:186390627:C:T_T
p = 7.1e−16, beta = −0.33, N = 1257
GPWCFTTDPSVR pc2
log10 peptide intensity
515:516
580:581
162:163
3:186390627:C:T_T
p = 1.3e−15, beta = −0.327, N = 1257
P00747 NP5
log10 protein intensity
89:89
163:163
73:73
rs9898
p = 0.015, beta = −0.191, N = 325
TPENYPNAGLTMNYCR pc3
log10 peptide intensity
69:89
140:163
52:73
rs9898
p = 4.1e−15, beta = −0.667, N = 261
GNVAVTVSGHTCQHWSAQTPHTHNR
log10 peptide intensity
89:89
163:163
73:73
rs9898
p = 1.2e−05, beta = −0.336, N = 325
DVVLFEK pc2
log10 peptide intensity
71:89
130:163
34:73
rs9898
p = 0.00087, beta = −0.325, N = 235
CTTPPPSSGPTYQCLK pc2
log10 peptide intensity
89:89
163:163
73:73
rs9898
p = 0.0016, beta = −0.244, N = 325
Locus: 128 (Sort: 149)
Status: Not replicated
80% power: No
SNP: 3:186390627:C:T
rsID: rs9898
UniProtID: P00747
HGNC gene: PLG
Type: Trans−pQTL
YILQGVTSWGLGCAR pc2
rs3124784 REF
log10 peptide intensity
515:516
579:581
162:163
3:186390627:C:T_T
p = 0.43, model = DOM, N = 1256

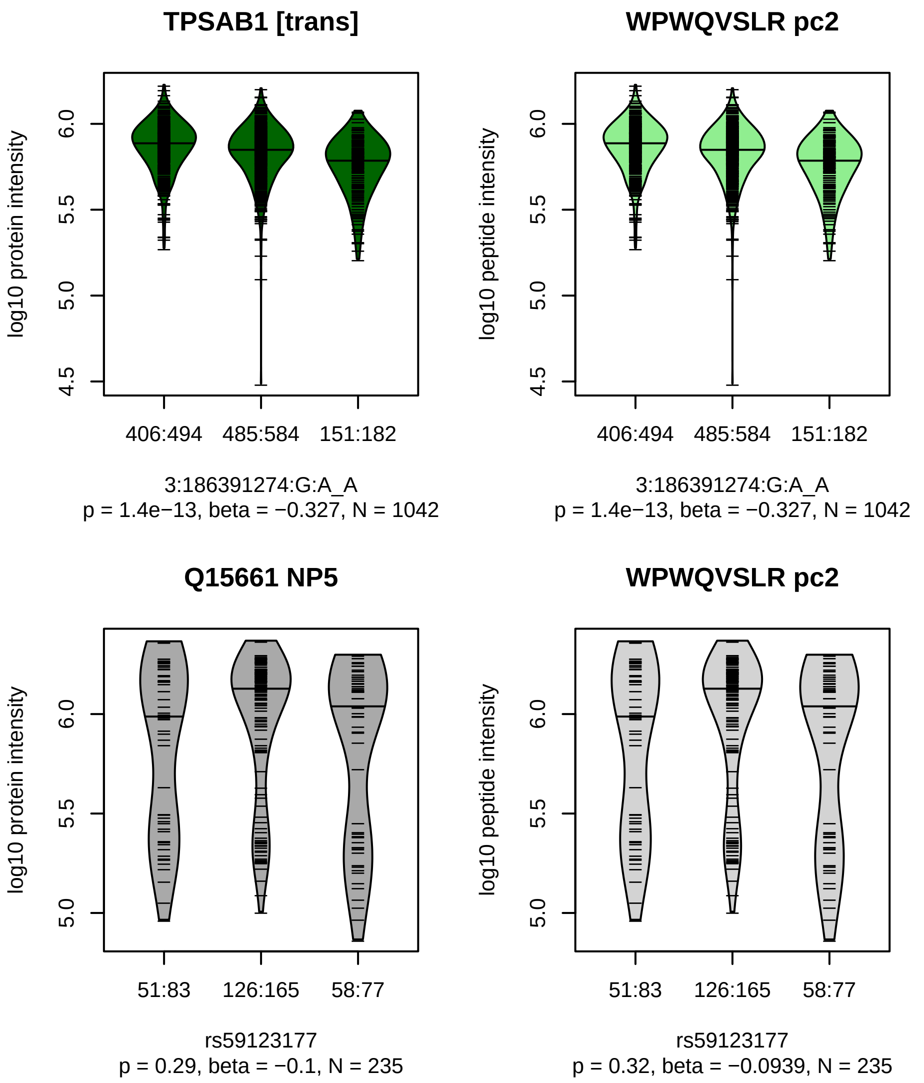


Locus: 128 (Sort: 150)

Status: Not replicated

80% power: No

SNP: 3:186391274:G:A

rsID: rs59123177

UniProtID: Q15661

HGNC gene: TPSAB1

Type: Trans−pQTL

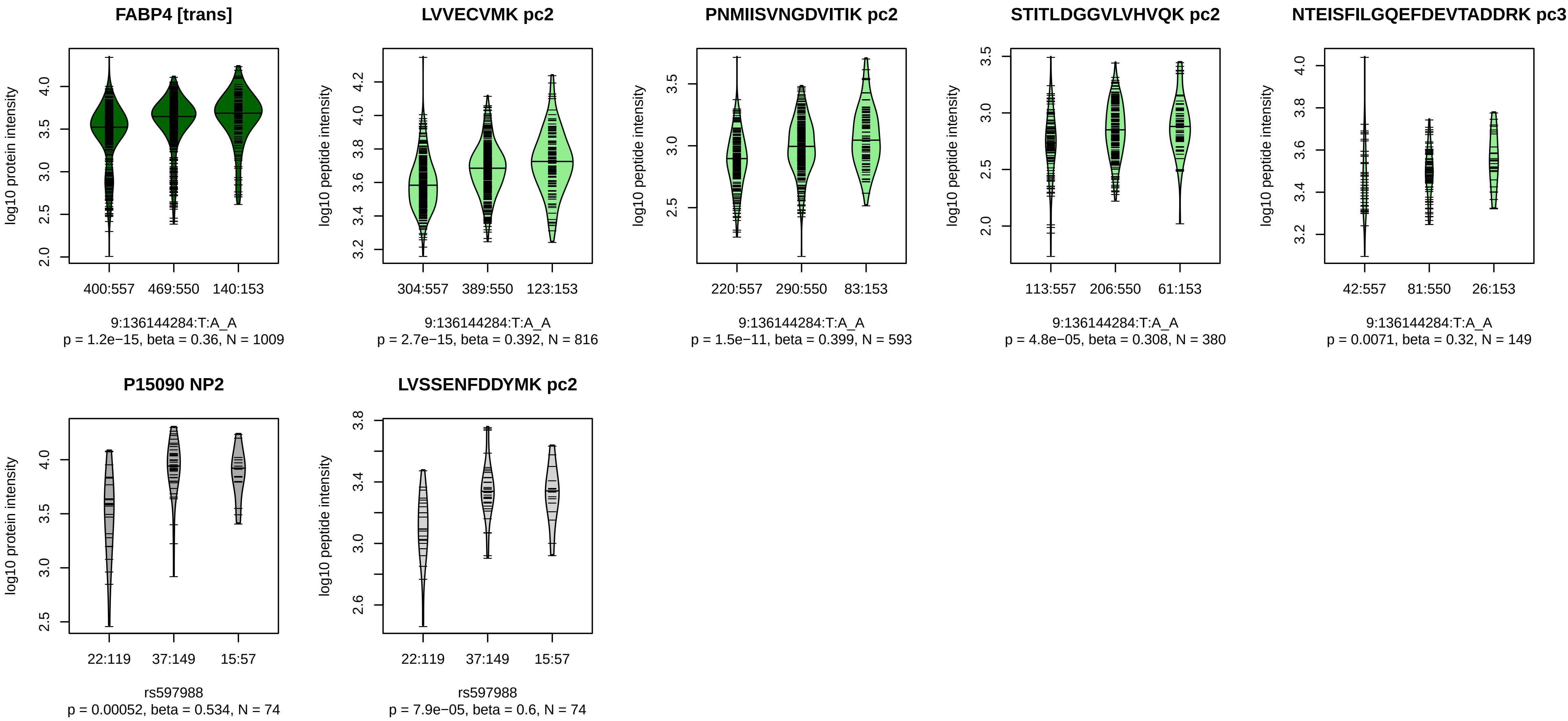


Locus: 129 (Sort: 151)

Status: Not replicated

80% power: No

SNP: 9:136144284:T:A

rsID: rs597988

UniProtID: P15090

HGNC gene: FABP4

Type: Trans−pQTL

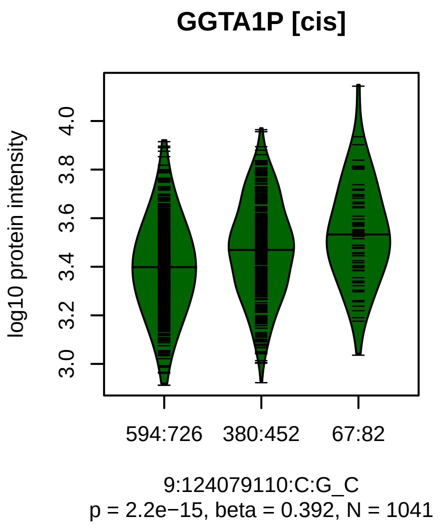


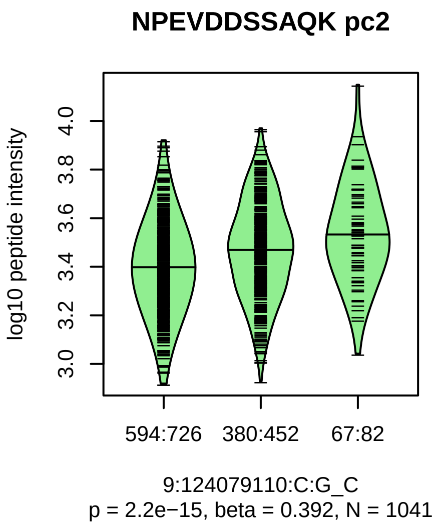


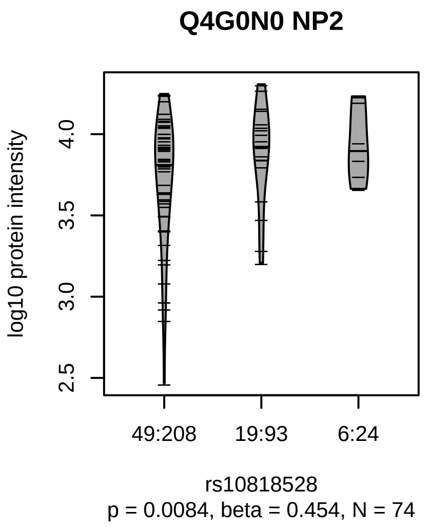


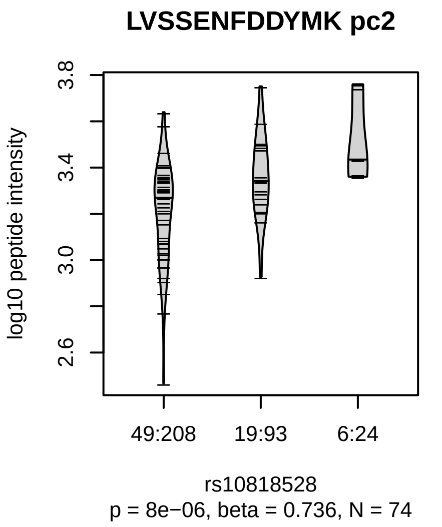


Locus: 130 (Sort: 152)
Status: Not replicated
80% power: No
SNP: 9:124079110:C:G
rsID: rs10818528
UniProtID: Q4G0N0
HGNC gene: GGTA1P
Type: Cis−pQTL

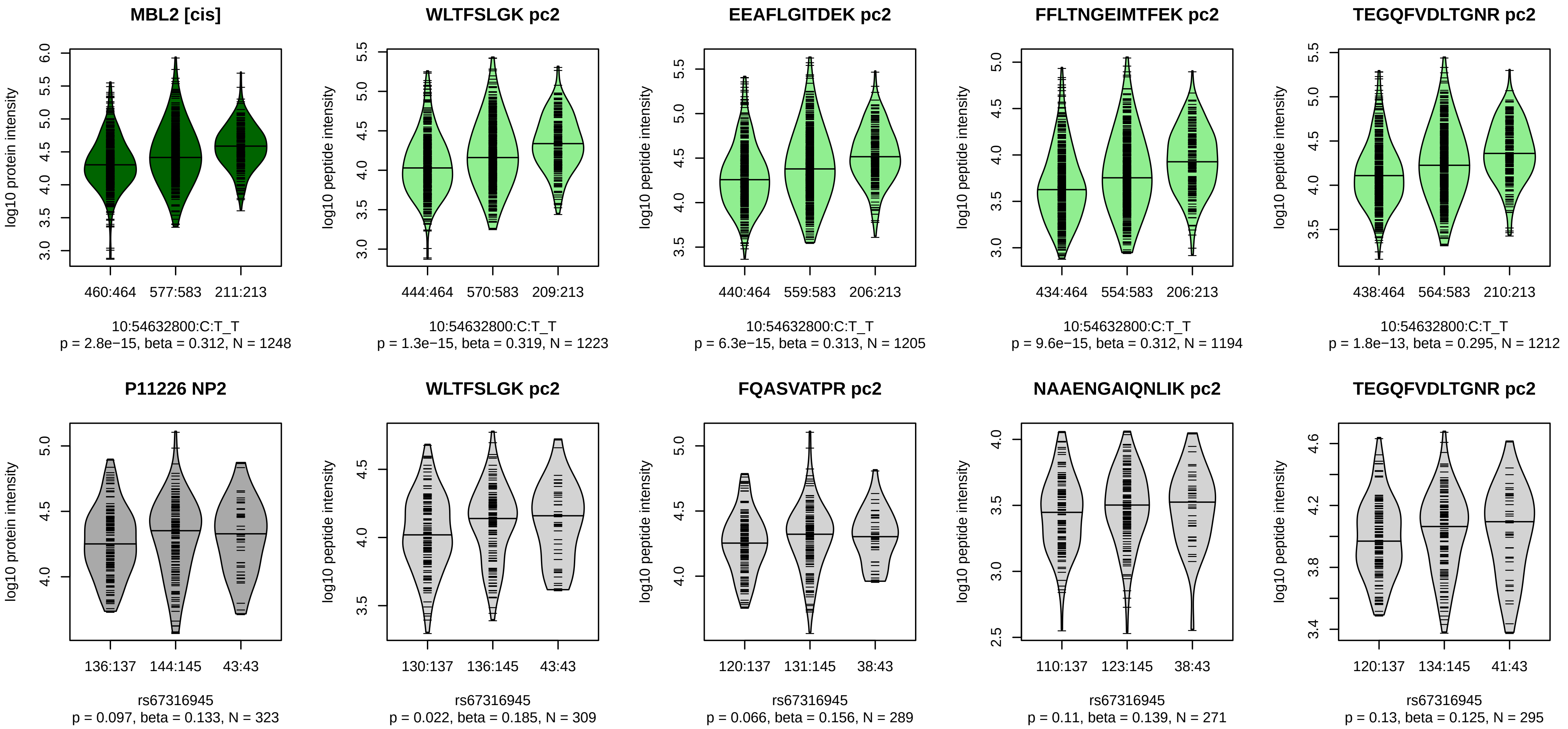


Locus: 131 (Sort: 153)
Status: Not replicated
80% power: No
SNP: 10:54632800:C:T
rsID: rs67316945
UniProtID: P11226
HGNC gene: MBL2
Type: Cis−pQTL

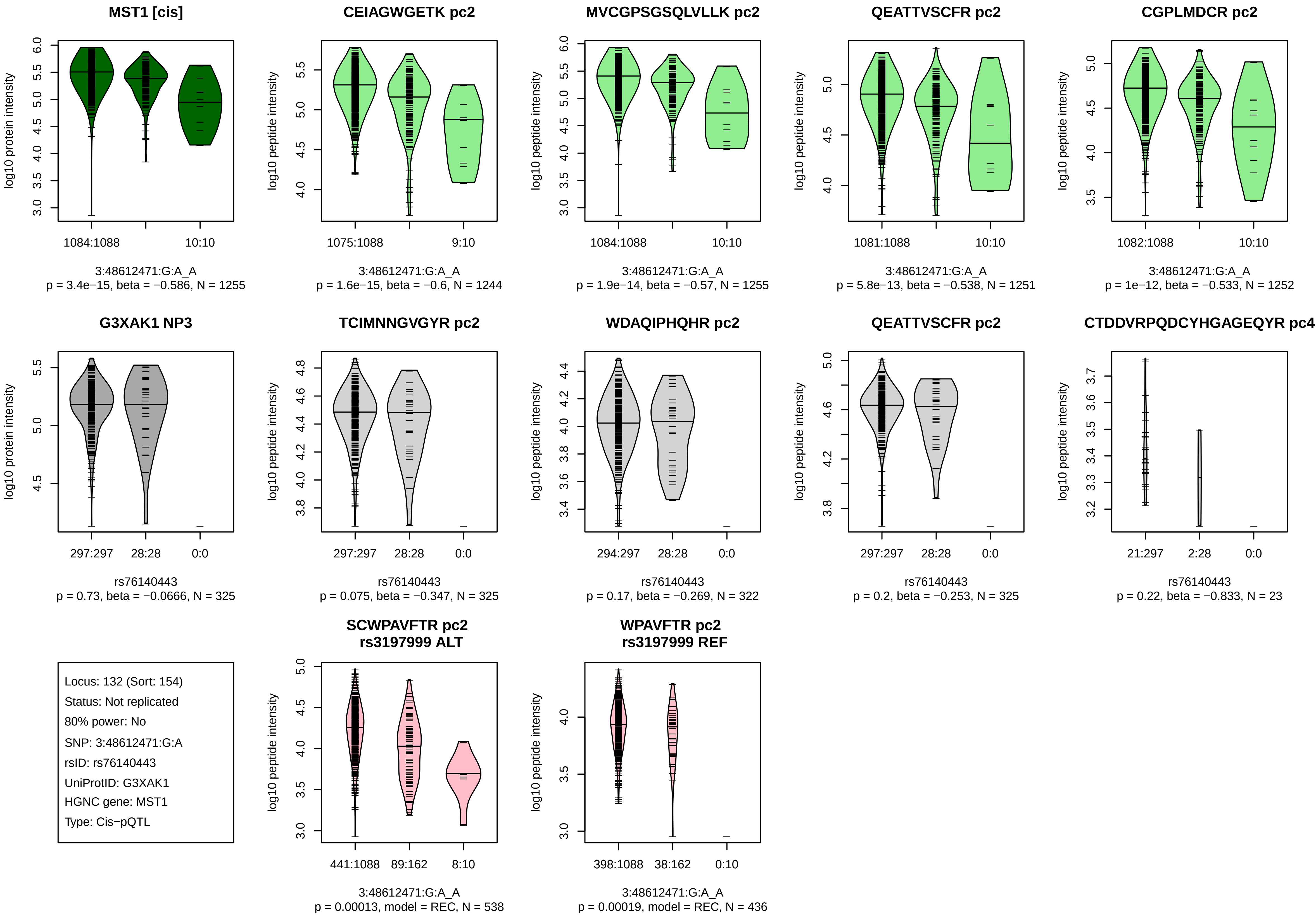
MST1 [cis]
log10 protein intensity
1084:1088
10:10
3:48612471:G:A_A
p = 3.4e−15, beta = −0.586, N = 1255
CEIAGWGETK pc2
log10 peptide intensity
1075:1088
9:10
3:48612471:G:A_A
p = 1.6e−15, beta = −0.6, N = 1244
MVCGPSGSQLVLLK pc2
log10 peptide intensity
1084:1088
10:10
3:48612471:G:A_A
p = 1.9e−14, beta = −0.57, N = 1255
QEATTVSCFR pc2
log10 peptide intensity
1081:1088
10:10
3:48612471:G:A_A
p = 5.8e−13, beta = −0.538, N = 1251
CGPLMDCR pc2
log10 peptide intensity
1082:1088
10:10
3:48612471:G:A_A
p = 1e−12, beta = −0.533, N = 1252
G3XAK1 NP3
log10 protein intensity
297:297
28:28
0:0
rs76140443
p = 0.73, beta = −0.0666, N = 325
TCIMNNGVGYR pc2
log10 peptide intensity
297:297
28:28
0:0
rs76140443
p = 0.075, beta = −0.347, N = 325
WDAQIPHQHR pc2
log10 peptide intensity
294:297
28:28
0:0
rs76140443
p = 0.17, beta = −0.269, N = 322
QEATTVSCFR pc2
log10 peptide intensity
297:297
28:28
0:0
rs76140443
p = 0.2, beta = −0.253, N = 325
CTDDVRPQDCYHGAGEQYR pc4
log10 peptide intensity
21:297
2:28
0:0
rs76140443
p = 0.22, beta = −0.833, N = 23
Locus: 132 (Sort: 154)
Status: Not replicated
80% power: No
SNP: 3:48612471:G:A
rsID: rs76140443
UniProtID: G3XAK1
HGNC gene: MST1
Type: Cis−pQTL
SCWPAVFTR pc2
rs3197999 ALT
log10 peptide intensity
441:1088
89:162
8:10
3:48612471:G:A_A
p = 0.00013, model = REC, N = 538
WPAVFTR pc2
rs3197999 REF
log10 peptide intensity
398:1088
38:162
0:10
3:48612471:G:A_A
p = 0.00019, model = REC, N = 436

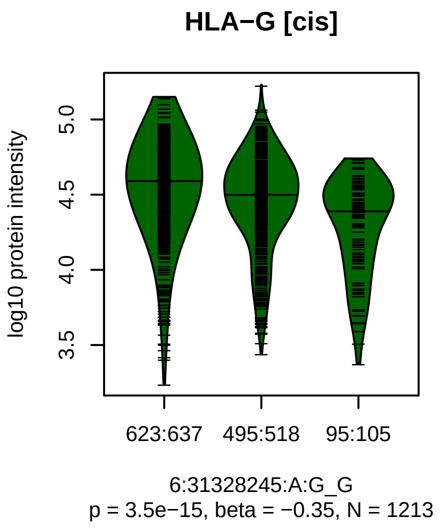


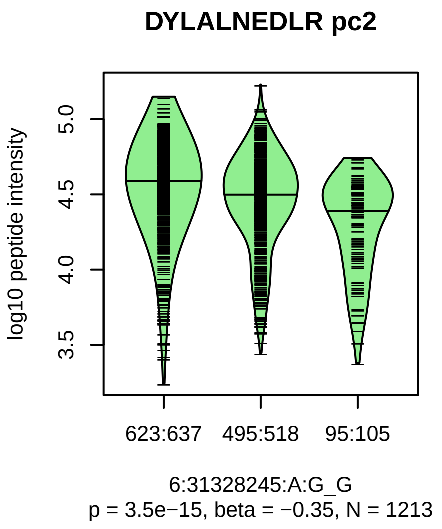


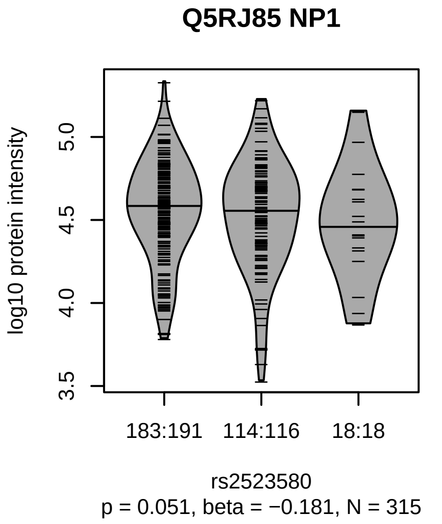


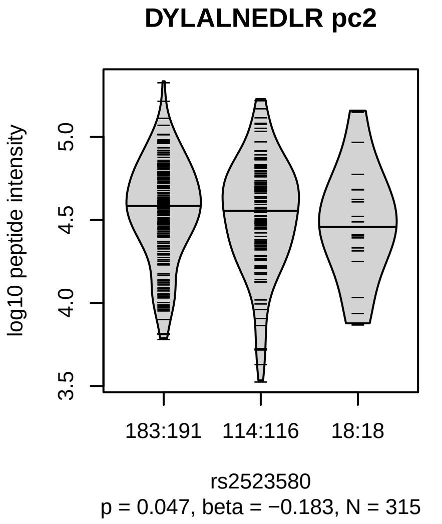


Locus: 133 (Sort: 155)

Status: Not replicated

80% power: No

SNP: 6:31328245:A:G

rsID: rs2523580

UniProtID: Q5RJ85

HGNC gene: HLA−G

Type: Cis−pQTL

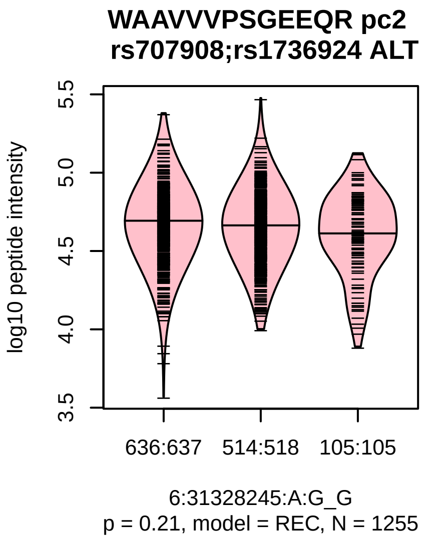

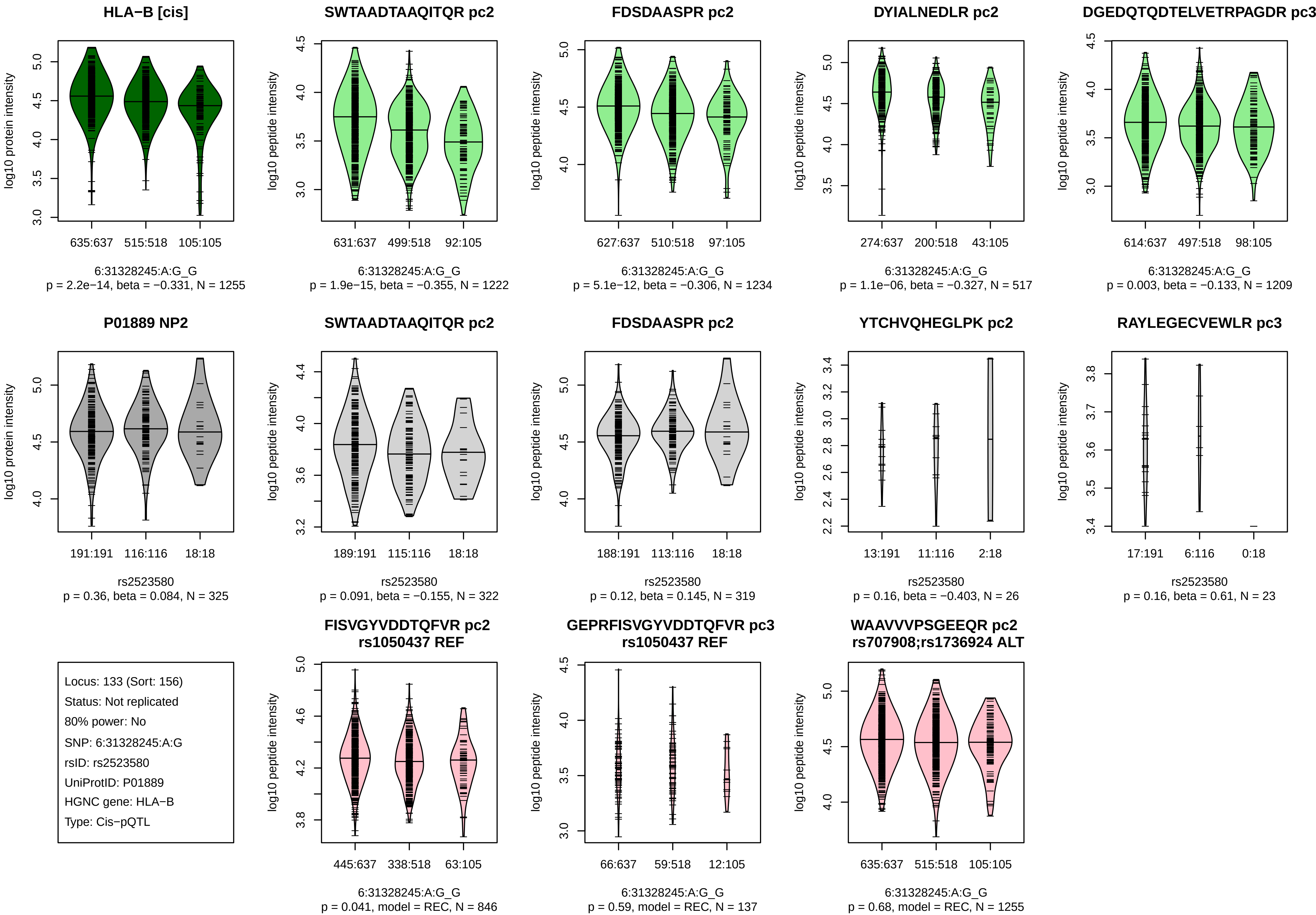
HLA−B [cis]
635:637
515:518
105:105
6:31328245:A:G_G
p = 2.2e−14, beta = −0.331, N = 1255
SWTAADTAAQITQR pc2
631:637
499:518
92:105
6:31328245:A:G_G
p = 1.9e−15, beta = −0.355, N = 1222
FDSDAASPR pc2
627:637
510:518
97:105
6:31328245:A:G_G
p = 5.1e−12, beta = −0.306, N = 1234
DYIALNEDLR pc2
274:637
200:518
43:105
6:31328245:A:G_G
p = 1.1e−06, beta = −0.327, N = 517
DGEDQTQDTELVETRPAGDR pc3
614:637
497:518
98:105
6:31328245:A:G_G
p = 0.003, beta = −0.133, N = 1209
P01889 NP2
191:191
116:116
18:18
rs2523580
p = 0.36, beta = 0.084, N = 325
SWTAADTAAQITQR pc2
189:191
115:116
18:18
rs2523580
p = 0.091, beta = −0.155, N = 322
FDSDAASPR pc2
188:191
113:116
18:18
rs2523580
p = 0.12, beta = 0.145, N = 319
YTCHVQHEGLPK pc2
13:191
11:116
2:18
rs2523580
p = 0.16, beta = −0.403, N = 26
RAYLEGECVEWLR pc3
17:191
6:116
0:18
rs2523580
p = 0.16, beta = 0.61, N = 23
Locus: 133 (Sort: 156)
Status: Not replicated
80% power: No
SNP: 6:31328245:A:G
rsID: rs2523580
UniProtID: P01889
HGNC gene: HLA−B
Type: Cis−pQTL
FISVGYVDDTQFVR pc2
rs1050437 REF
445:637
338:518
63:105
6:31328245:A:G_G
p = 0.041, model = REC, N = 846
GEPRFISVGYVDDTQFVR pc3
rs1050437 REF
66:637
59:518
12:105
6:31328245:A:G_G
p = 0.59, model = REC, N = 137
WAAVVVPSGEEQR pc2
rs707908;rs1736924 ALT
635:637
515:518
105:105
6:31328245:A:G_G
p = 0.68, model = REC, N = 1255
log10 protein intensity
log10 peptide intensity

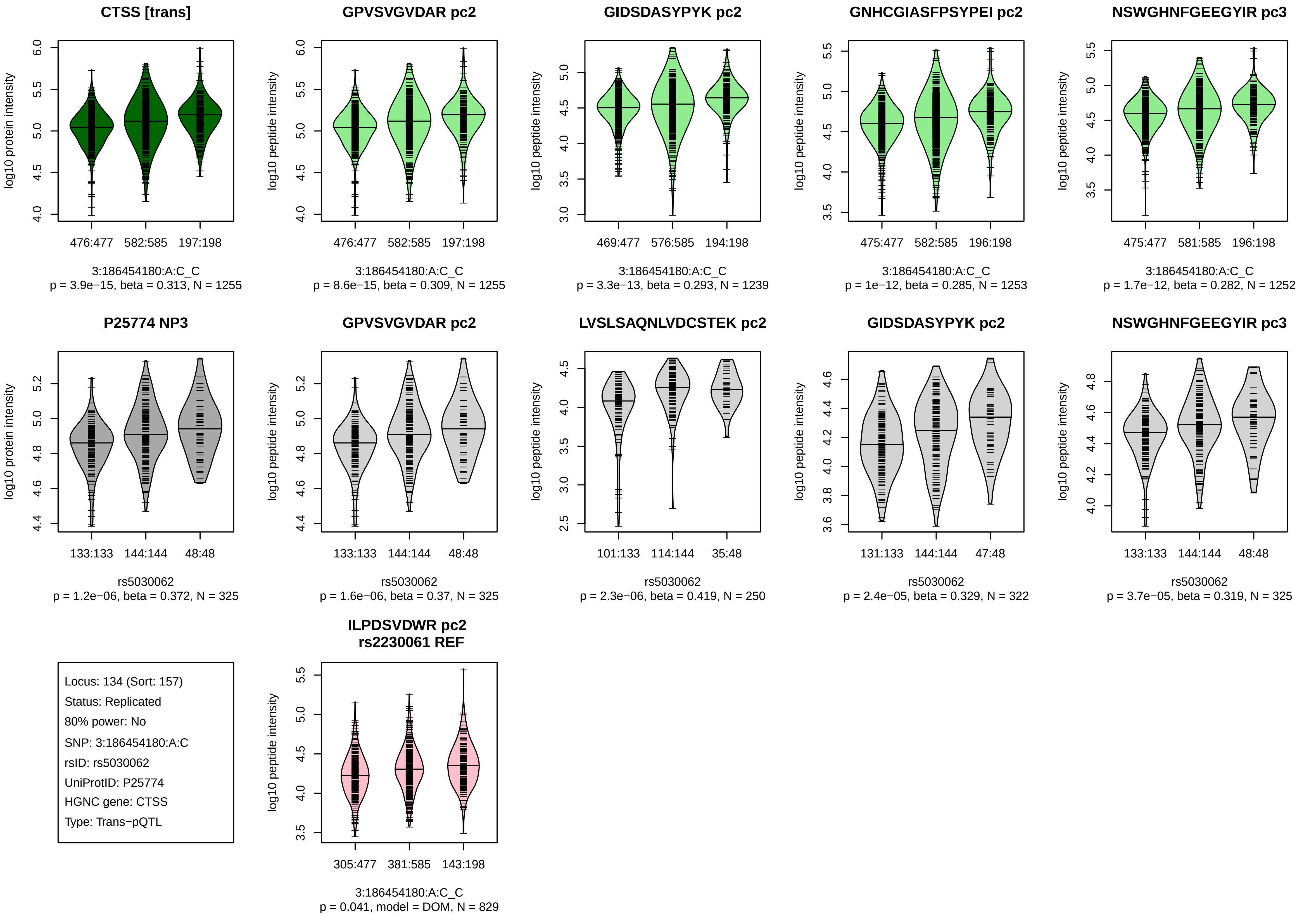
CTSS [trans]
log10 protein intensity
476:477
582:585
197:198
3:186454180:A:C_C
p = 3.9e−15, beta = 0.313, N = 1255
GPVSVGVDAR pc2
log10 peptide intensity
476:477
582:585
197:198
3:186454180:A:C_C
p = 8.6e−15, beta = 0.309, N = 1255
GIDSDASYPYK pc2
log10 peptide intensity
469:477
576:585
194:198
3:186454180:A:C_C
p = 3.3e−13, beta = 0.293, N = 1239
GNHCGIASFPSYPEI pc2
log10 peptide intensity
475:477
582:585
196:198
3:186454180:A:C_C
p = 1e−12, beta = 0.285, N = 1253
NSWGHNFGEEGYIR pc3
log10 peptide intensity
475:477
581:585
196:198
3:186454180:A:C_C
p = 1.7e−12, beta = 0.282, N = 1252
P25774 NP3
log10 protein intensity
133:133
144:144
48:48
rs5030062
p = 1.2e−06, beta = 0.372, N = 325
GPVSVGVDAR pc2
log10 peptide intensity
133:133
144:144
48:48
rs5030062
p = 1.6e−06, beta = 0.37, N = 325
LVSLSAQNLVDCSTEK pc2
log10 peptide intensity
101:133
114:144
35:48
rs5030062
p = 2.3e−06, beta = 0.419, N = 250
GIDSDASYPYK pc2
log10 peptide intensity
131:133
144:144
47:48
rs5030062
p = 2.4e−05, beta = 0.329, N = 322
NSWGHNFGEEGYIR pc3
log10 peptide intensity
133:133
144:144
48:48
rs5030062
p = 3.7e−05, beta = 0.319, N = 325
Locus: 134 (Sort: 157)
Status: Replicated
80% power: No
SNP: 3:186454180:A:C
rsID: rs5030062
UniProtID: P25774
HGNC gene: CTSS
Type: Trans−pQTL
ILPDSVDWR pc2
rs2230061 REF
log10 peptide intensity
305:477
381:585
143:198
3:186454180:A:C_C
p = 0.041, model = DOM, N = 829

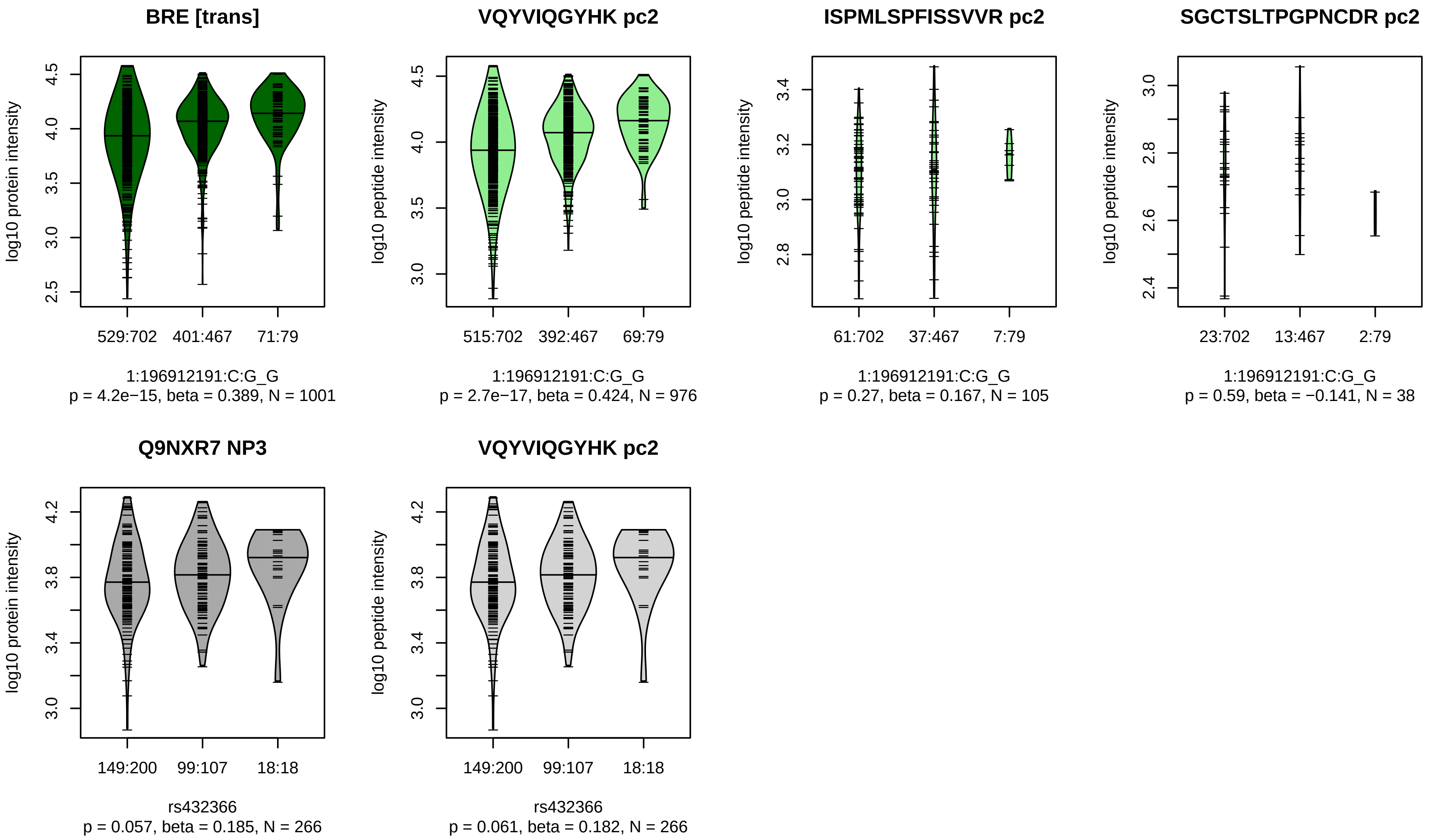


Locus: 135 (Sort: 158)
Status: Not replicated
80% power: No
SNP: 1:196912191:C:G
rsID: rs432366
UniProtID: Q9NXR7
HGNC gene: BRE
Type: Trans−pQTL

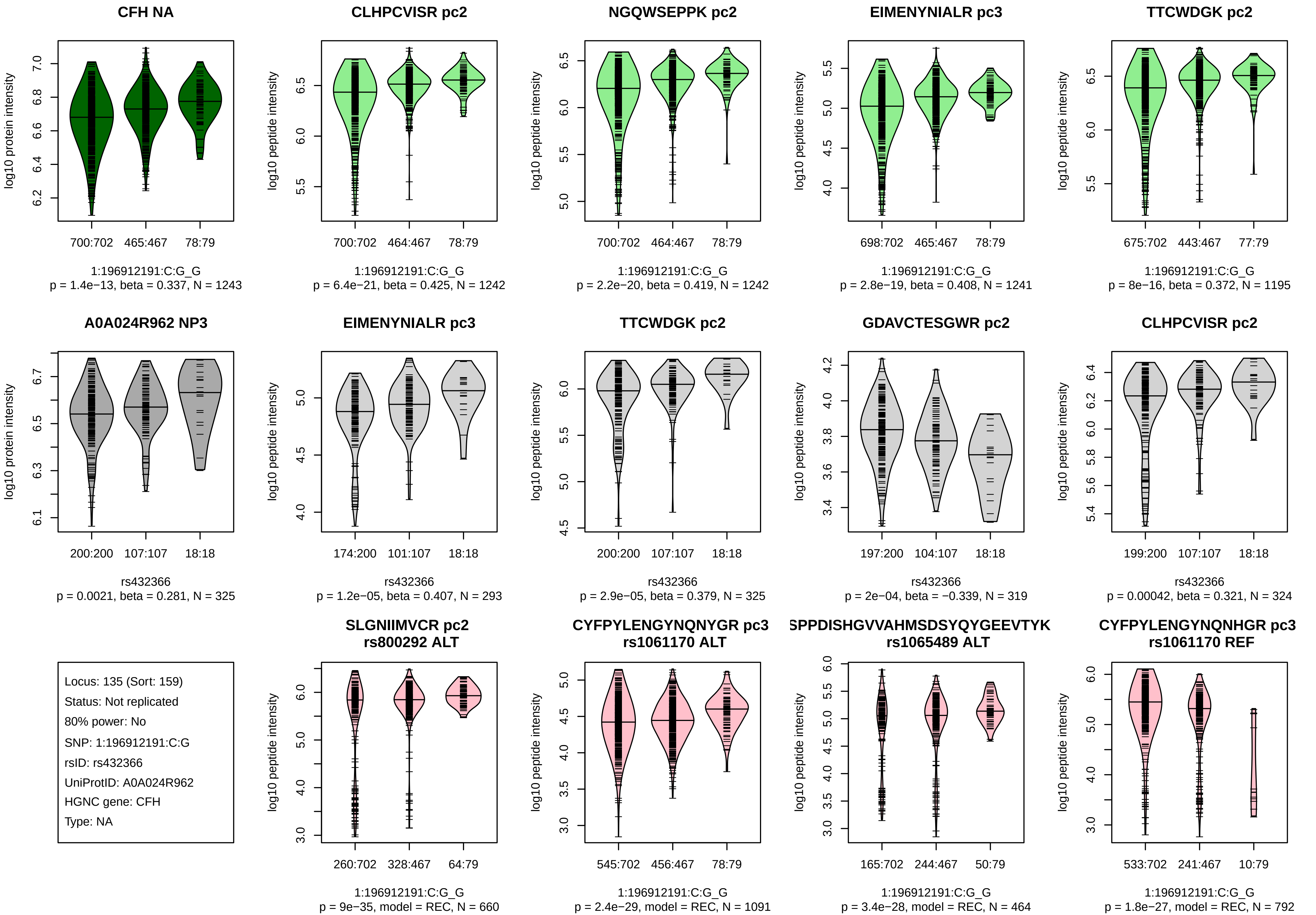
CFH NA
log10 protein intensity
700:702 465:467 78:79
1:196912191:C:G_G
p = 1.4e−13, beta = 0.337, N = 1243
CLHPCVISR pc2
log10 peptide intensity
700:702 464:467 78:79
1:196912191:C:G_G
p = 6.4e−21, beta = 0.425, N = 1242
NGQWSEPPK pc2
log10 peptide intensity
700:702 464:467 78:79
1:196912191:C:G_G
p = 2.2e−20, beta = 0.419, N = 1242
EIMENYNIALR pc3
log10 peptide intensity
698:702 465:467 78:79
1:196912191:C:G_G
p = 2.8e−19, beta = 0.408, N = 1241
TTCWDGK pc2
log10 peptide intensity
675:702 443:467 77:79
1:196912191:C:G_G
p = 8e−16, beta = 0.372, N = 1195
A0A024R962 NP3
log10 protein intensity
200:200 107:107 18:18
rs432366
p = 0.0021, beta = 0.281, N = 325
EIMENYNIALR pc3
log10 peptide intensity
174:200 101:107 18:18
rs432366
p = 1.2e−05, beta = 0.407, N = 293
TTCWDGK pc2
log10 peptide intensity
200:200 107:107 18:18
rs432366
p = 2.9e−05, beta = 0.379, N = 325
GDAVCTESGWR pc2
log10 peptide intensity
197:200 104:107 18:18
rs432366
p = 2e−04, beta = −0.339, N = 319
CLHPCVISR pc2
log10 peptide intensity
199:200 107:107 18:18
rs432366
p = 0.00042, beta = 0.321, N = 324
Locus: 135 (Sort: 159)
Status: Not replicated
80% power: No
SNP: 1:196912191:C:G
rsID: rs432366
UniProtID: A0A024R962
HGNC gene: CFH
Type: NA
SLGNIIMVCR pc2
rs800292 ALT
log10 peptide intensity
260:702 328:467 64:79
1:196912191:C:G_G
p = 9e−35, model = REC, N = 660
CYFPYLENGYNQNYGR pc3
rs1061170 ALT
log10 peptide intensity
545:702 456:467 78:79
1:196912191:C:G_G
p = 2.4e−29, model = REC, N = 1091
SPPDISHGVVAHMSDSYQYGEEVTYK
rs1065489 ALT
log10 peptide intensity
165:702 244:467 50:79
1:196912191:C:G_G
p = 3.4e−28, model = REC, N = 464
CYFPYLENGYNQNHGR pc3
rs1061170 REF
log10 peptide intensity
533:702 241:467 10:79
1:196912191:C:G_G
p = 1.8e−27, model = REC, N = 792

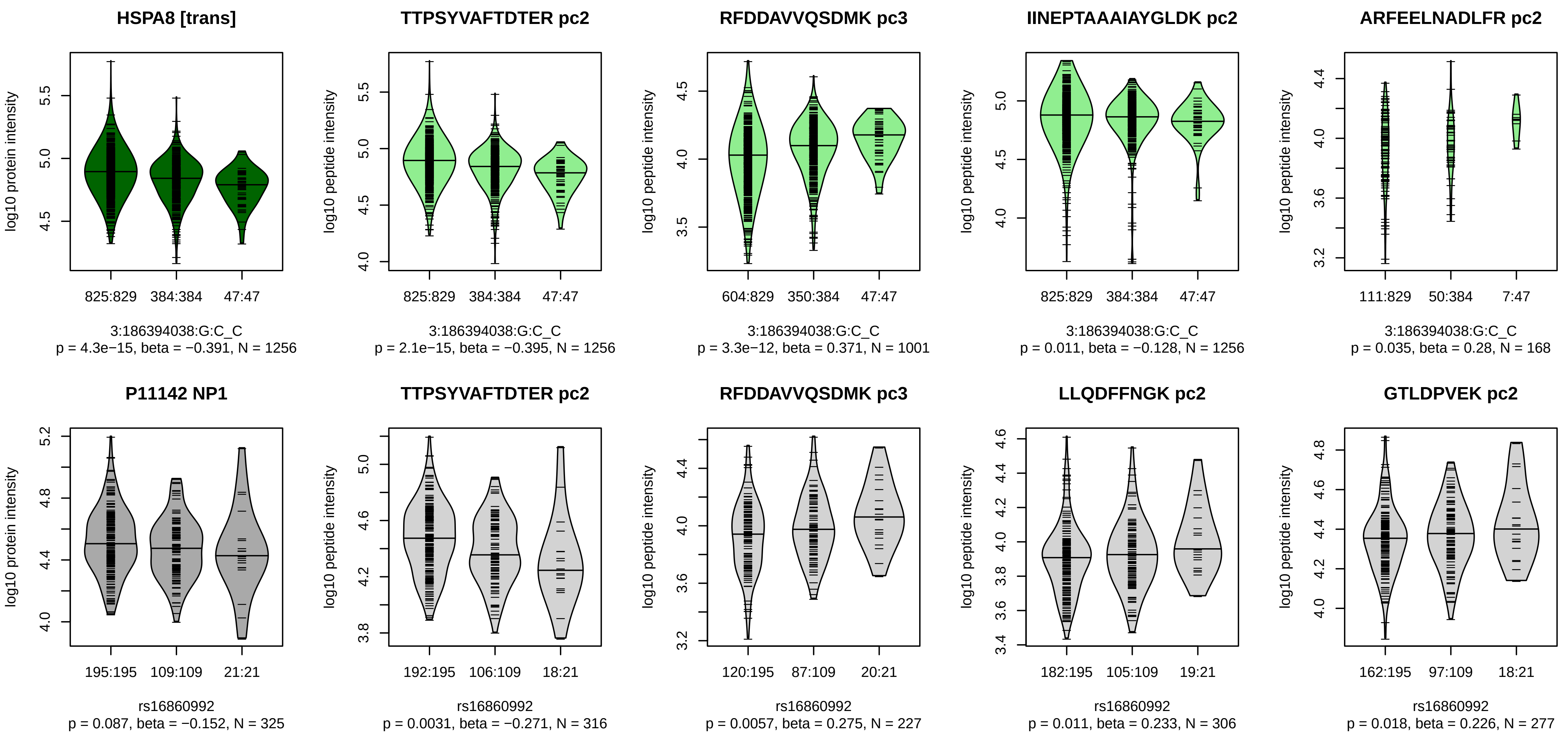


Locus: 136 (Sort: 160)
Status: Not replicated
80% power: No
SNP: 3:186394038:G:C
rsID: rs16860992
UniProtID: P11142
HGNC gene: HSPA8
Type: Trans−pQTL

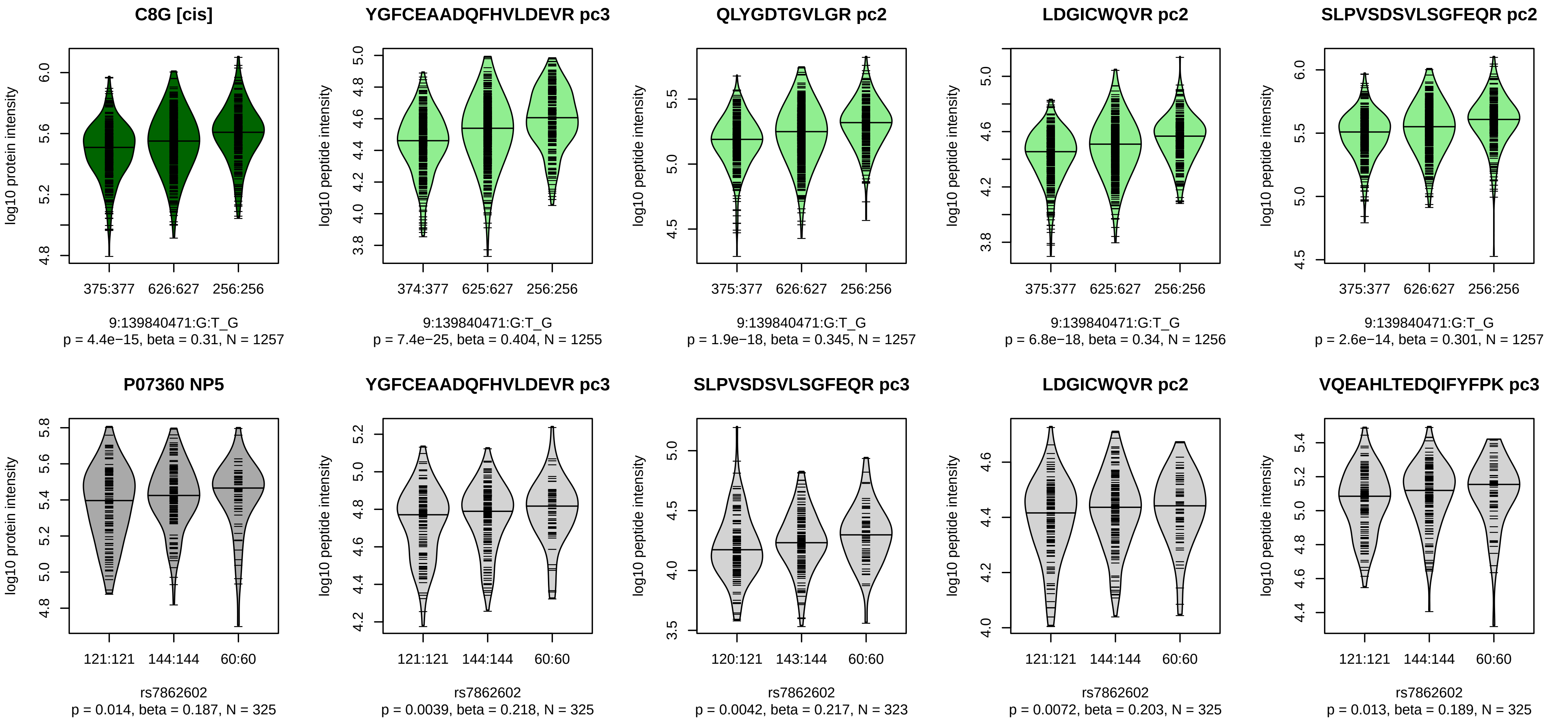


Locus: 137 (Sort: 161)

Status: Not replicated

80% power: No

SNP: 9:139840471:G:T

rsID: rs7862602

UniProtID: P07360

HGNC gene: C8G

Type: Cis−pQTL

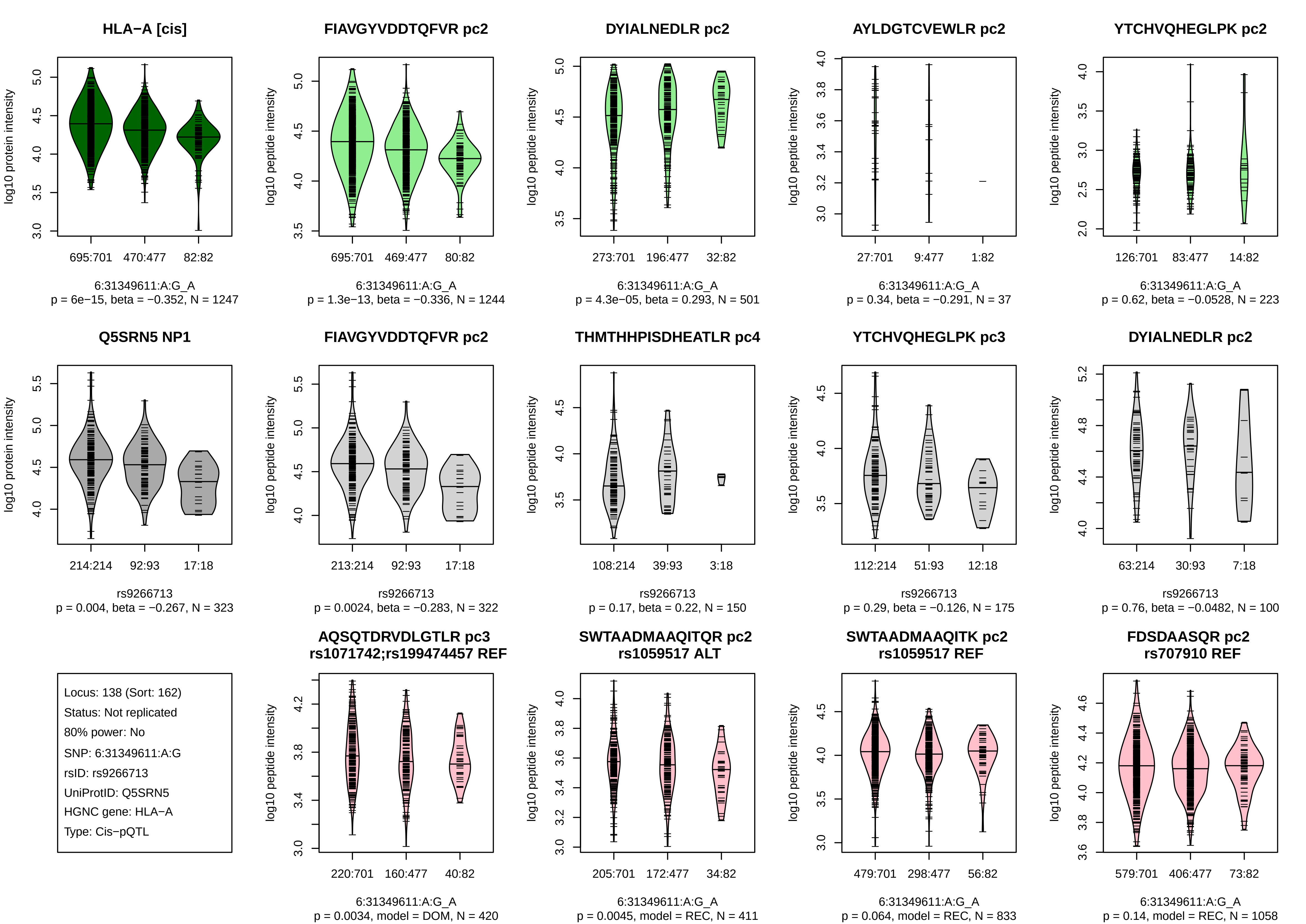

HLA−A [cis]
log10 protein intensity
695:701
470:477
82:82
6:31349611:A:G_A
p = 6e−15, beta = −0.352, N = 1247
FIAVGYVDDTQFVR pc2
log10 peptide intensity
695:701
469:477
80:82
6:31349611:A:G_A
p = 1.3e−13, beta = −0.336, N = 1244
DYIALNEDLR pc2
log10 peptide intensity
273:701
196:477
32:82
6:31349611:A:G_A
p = 4.3e−05, beta = 0.293, N = 501
AYLDGTCVEWLR pc2
log10 peptide intensity
27:701
9:477
1:82
6:31349611:A:G_A
p = 0.34, beta = −0.291, N = 37
YTCHVQHEGLPK pc2
log10 peptide intensity
126:701
83:477
14:82
6:31349611:A:G_A
p = 0.62, beta = −0.0528, N = 223
Q5SRN5 NP1
log10 protein intensity
214:214
92:93
17:18
rs9266713
p = 0.004, beta = −0.267, N = 323
FIAVGYVDDTQFVR pc2
log10 peptide intensity
213:214
92:93
17:18
rs9266713
p = 0.0024, beta = −0.283, N = 322
THMTHHPISDHEATLR pc4
log10 peptide intensity
108:214
39:93
3:18
rs9266713
p = 0.17, beta = 0.22, N = 150
YTCHVQHEGLPK pc3
log10 peptide intensity
112:214
51:93
12:18
rs9266713
p = 0.29, beta = −0.126, N = 175
DYIALNEDLR pc2
log10 peptide intensity
63:214
30:93
7:18
rs9266713
p = 0.76, beta = −0.0482, N = 100
Locus: 138 (Sort: 162)
Status: Not replicated
80% power: No
SNP: 6:31349611:A:G
rsID: rs9266713
UniProtID: Q5SRN5
HGNC gene: HLA−A
Type: Cis−pQTL
AQSQTDRVDLGTLR pc3
rs1071742;rs199474457 REF
log10 peptide intensity
220:701
160:477
40:82
6:31349611:A:G_A
p = 0.0034, model = DOM, N = 420
SWTAADMAAQITQR pc2
rs1059517 ALT
log10 peptide intensity
205:701
172:477
34:82
6:31349611:A:G_A
p = 0.0045, model = REC, N = 411
SWTAADMAAQITK pc2
rs1059517 REF
log10 peptide intensity
479:701
298:477
56:82
6:31349611:A:G_A
p = 0.064, model = REC, N = 833
FDSDAASQR pc2
rs707910 REF
log10 peptide intensity
579:701
406:477
73:82
6:31349611:A:G_A
p = 0.14, model = REC, N = 1058

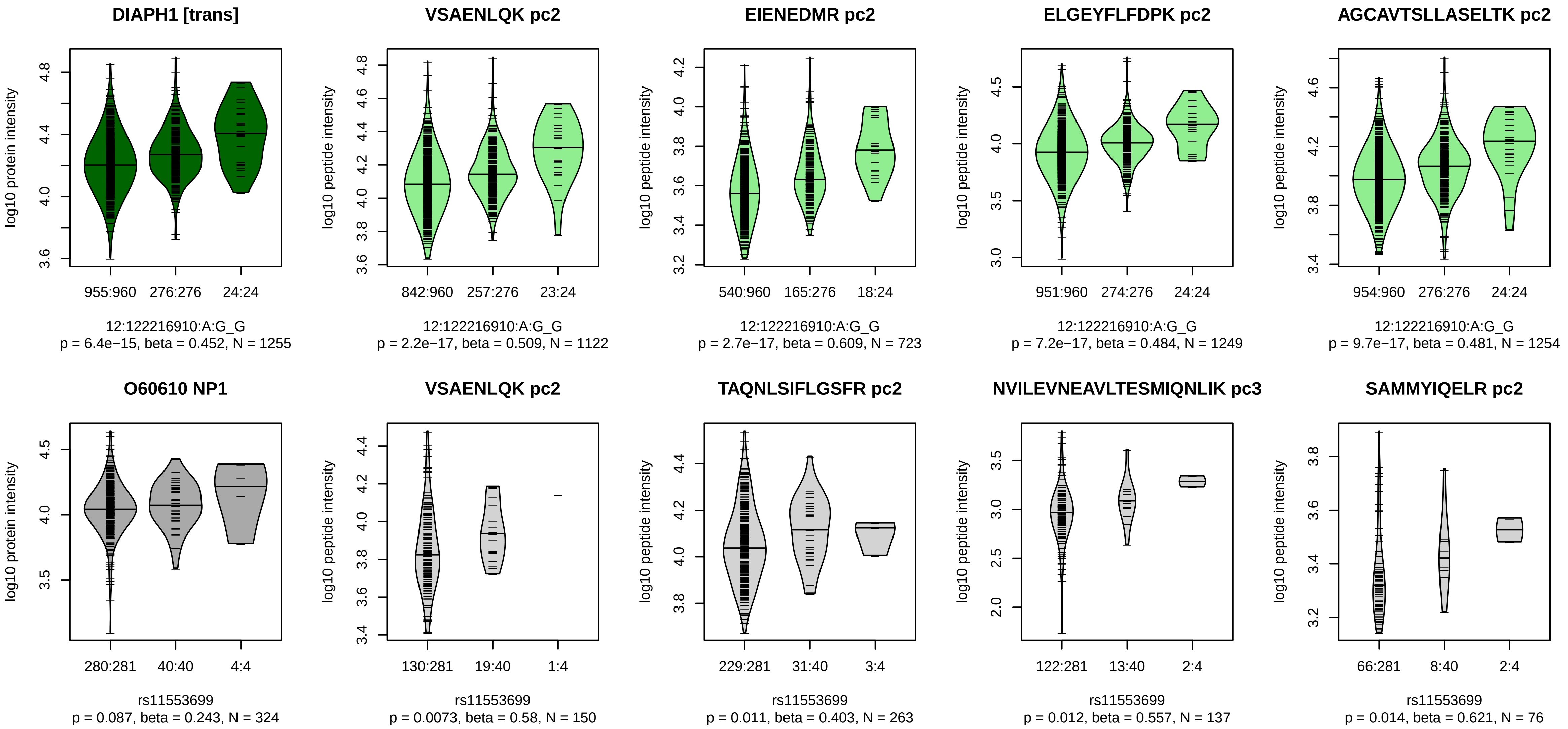


Locus: 139 (Sort: 163)

Status: Not replicated

80% power: No

SNP: 12:122216910:A:G

rsID: rs11553699

UniProtID: O60610

HGNC gene: DIAPH1

Type: Trans−pQTL

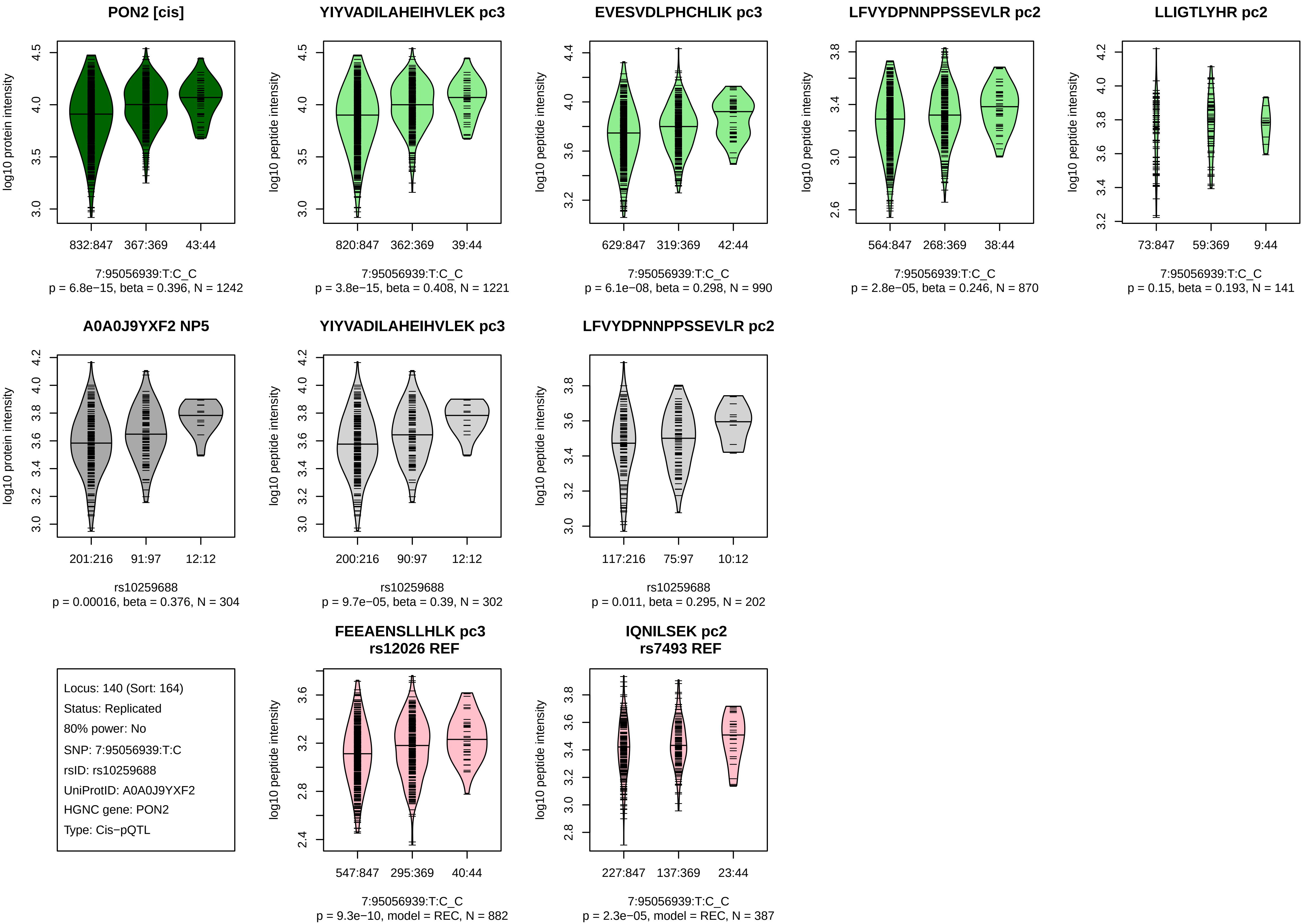
PON2 [cis]
log10 protein intensity
832:847
367:369
43:44
7:95056939:T:C_C
p = 6.8e−15, beta = 0.396, N = 1242
YIYVADILAHEIHVLEK pc3
log10 peptide intensity
820:847
362:369
39:44
7:95056939:T:C_C
p = 3.8e−15, beta = 0.408, N = 1221
EVESVDLPHCHLIK pc3
log10 peptide intensity
629:847
319:369
42:44
7:95056939:T:C_C
p = 6.1e−08, beta = 0.298, N = 990
LFVYDPNNPPSSEVLR pc2
log10 peptide intensity
564:847
268:369
38:44
7:95056939:T:C_C
p = 2.8e−05, beta = 0.246, N = 870
LLIGTLYHR pc2
log10 peptide intensity
73:847
59:369
9:44
7:95056939:T:C_C
p = 0.15, beta = 0.193, N = 141
A0A0J9YXF2 NP5
log10 protein intensity
201:216
91:97
12:12
rs10259688
p = 0.00016, beta = 0.376, N = 304
YIYVADILAHEIHVLEK pc3
log10 peptide intensity
200:216
90:97
12:12
rs10259688
p = 9.7e−05, beta = 0.39, N = 302
LFVYDPNNPPSSEVLR pc2
log10 peptide intensity
117:216
75:97
10:12
rs10259688
p = 0.011, beta = 0.295, N = 202
Locus: 140 (Sort: 164)
Status: Replicated
80% power: No
SNP: 7:95056939:T:C
rsID: rs10259688
UniProtID: A0A0J9YXF2
HGNC gene: PON2
Type: Cis−pQTL
FEEAENSLLHLK pc3
rs12026 REF
log10 peptide intensity
547:847
295:369
40:44
7:95056939:T:C_C
p = 9.3e−10, model = REC, N = 882
IQNILSEK pc2
rs7493 REF
log10 peptide intensity
227:847
137:369
23:44
7:95056939:T:C_C
p = 2.3e−05, model = REC, N = 387

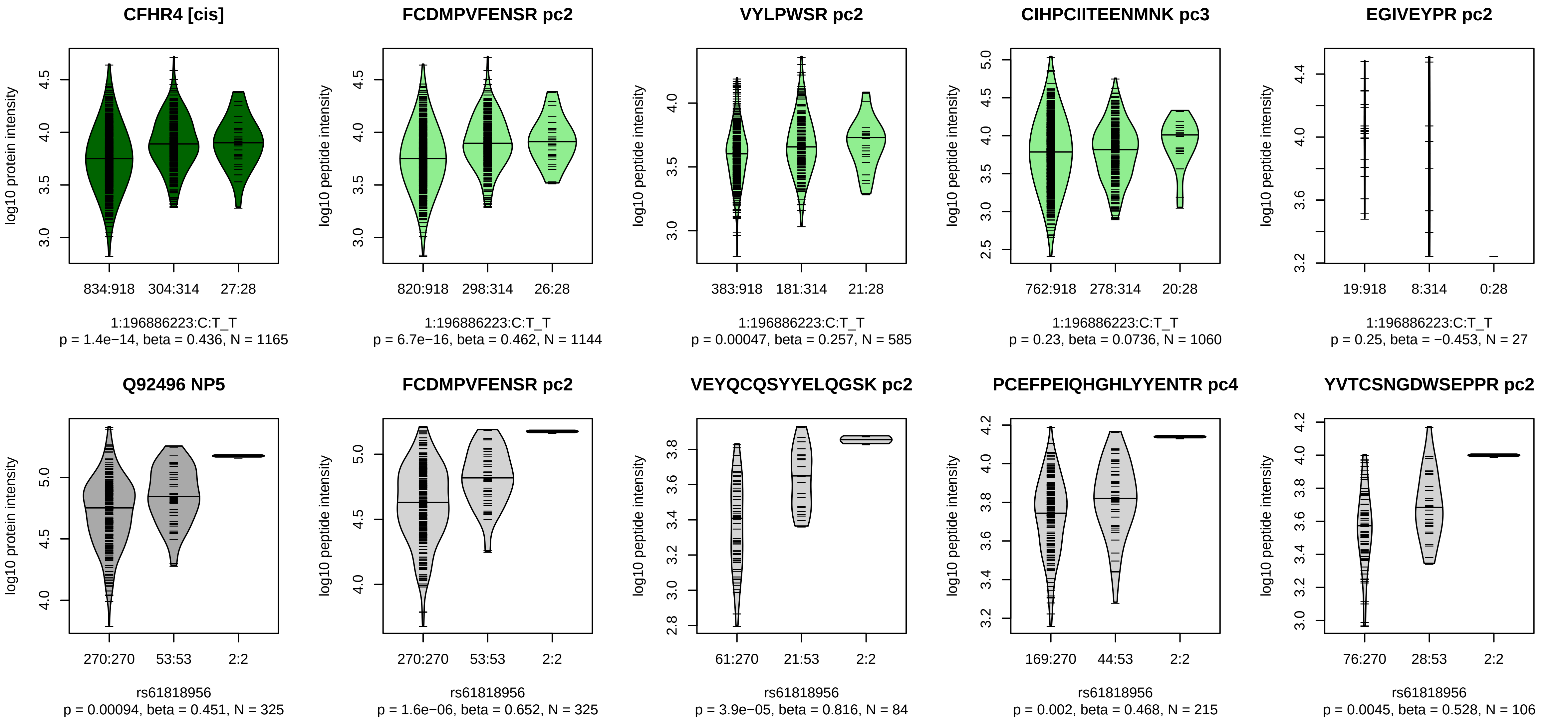


Locus: 141 (Sort: 165)
Status: Not replicated
80% power: No
SNP: 1:196886223:C:T
rsID: rs61818956
UniProtID: Q92496
HGNC gene: CFHR4
Type: Cis−pQTL

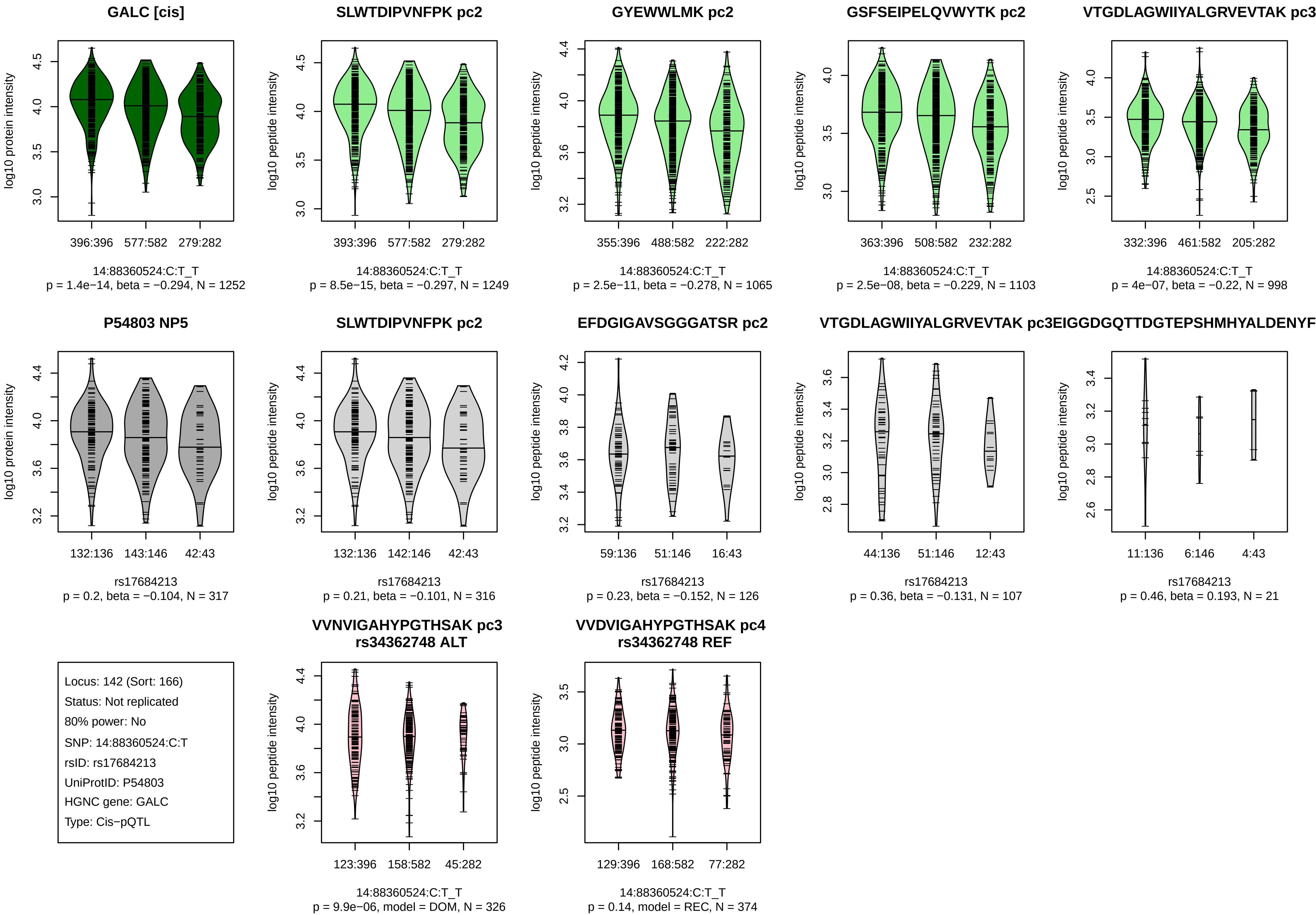
GALC [cis]
log10 protein intensity
396:396
577:582
279:282
14:88360524:C:T_T
p = 1.4e−14, beta = −0.294, N = 1252
SLWTDIPVNFPK pc2
log10 peptide intensity
393:396
577:582
279:282
14:88360524:C:T_T
p = 8.5e−15, beta = −0.297, N = 1249
GYEWWLMK pc2
log10 peptide intensity
355:396
488:582
222:282
14:88360524:C:T_T
p = 2.5e−11, beta = −0.278, N = 1065
GSFSEIPELQVWYTK pc2
log10 peptide intensity
363:396
508:582
232:282
14:88360524:C:T_T
p = 2.5e−08, beta = −0.229, N = 1103
VTGDLAGWIIYALGRVEVTAK pc3
log10 peptide intensity
332:396
461:582
205:282
14:88360524:C:T_T
p = 4e−07, beta = −0.22, N = 998
P54803 NP5
log10 protein intensity
132:136
143:146
42:43
rs17684213
p = 0.2, beta = −0.104, N = 317
SLWTDIPVNFPK pc2
log10 peptide intensity
132:136
142:146
42:43
rs17684213
p = 0.21, beta = −0.101, N = 316
EFDGIGAVSGGGATSR pc2
log10 peptide intensity
59:136
51:146
16:43
rs17684213
p = 0.23, beta = −0.152, N = 126
VTGDLAGWIIYALGRVEVTAK pc3
log10 peptide intensity
44:136
51:146
12:43
rs17684213
p = 0.36, beta = −0.131, N = 107
EIGGDGQTTDGTEPSHMHYALDENYF
log10 peptide intensity
11:136
6:146
4:43
rs17684213
p = 0.46, beta = 0.193, N = 21
Locus: 142 (Sort: 166)
Status: Not replicated
80% power: No
SNP: 14:88360524:C:T
rsID: rs17684213
UniProtID: P54803
HGNC gene: GALC
Type: Cis−pQTL
VVNVIGAHYPGTHSAK pc3
rs34362748 ALT
log10 peptide intensity
123:396
158:582
45:282
14:88360524:C:T_T
p = 9.9e−06, model = DOM, N = 326
VVDVIGAHYPGTHSAK pc4
rs34362748 REF
log10 peptide intensity
129:396
168:582
77:282
14:88360524:C:T_T
p = 0.14, model = REC, N = 374

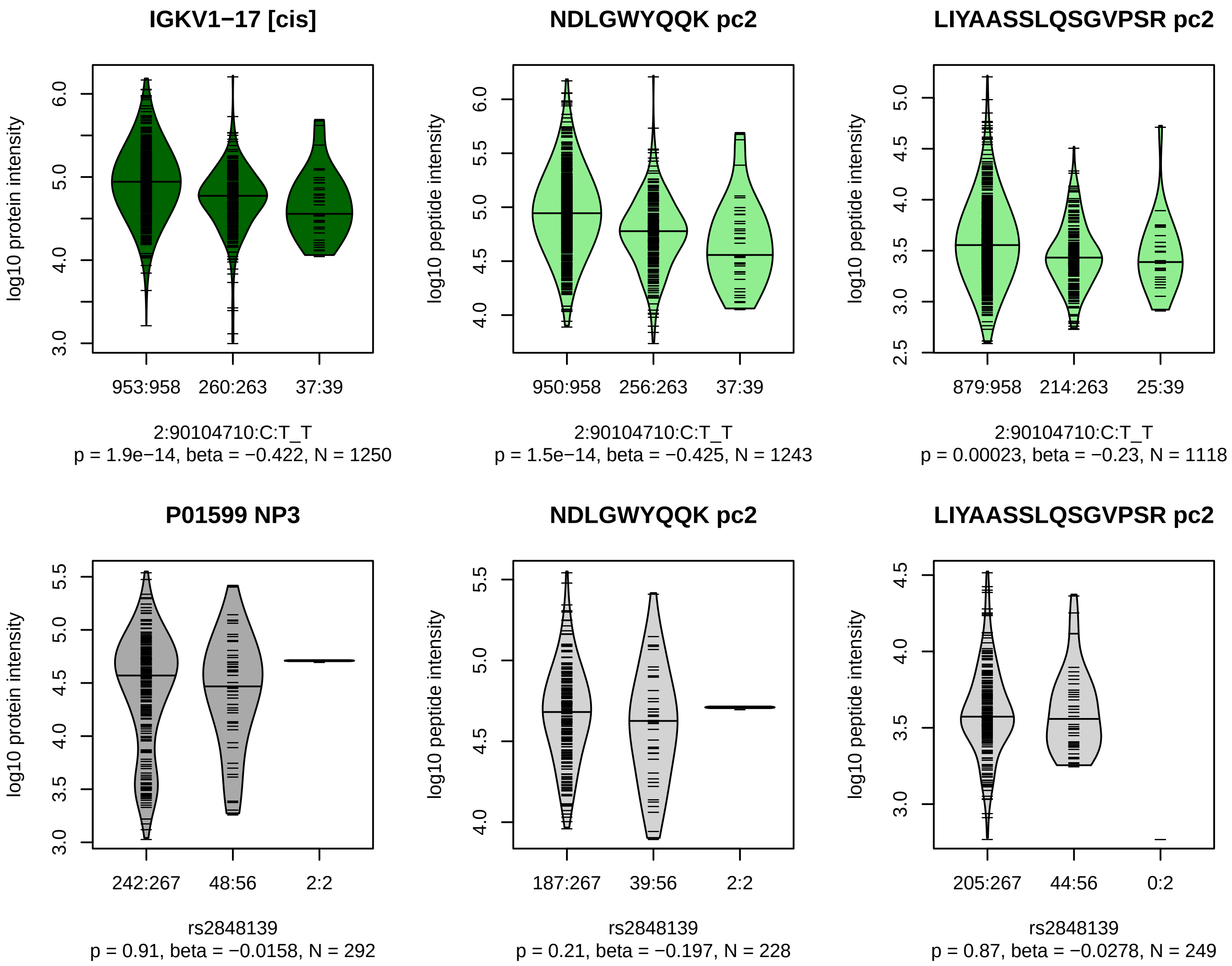


Locus: 143 (Sort: 167)

Status: Not replicated

80% power: No

SNP: 2:90104710:C:T

rsID: rs2848139

UniProtID: P01599

HGNC gene: IGKV1−17

Type: Cis−pQTL

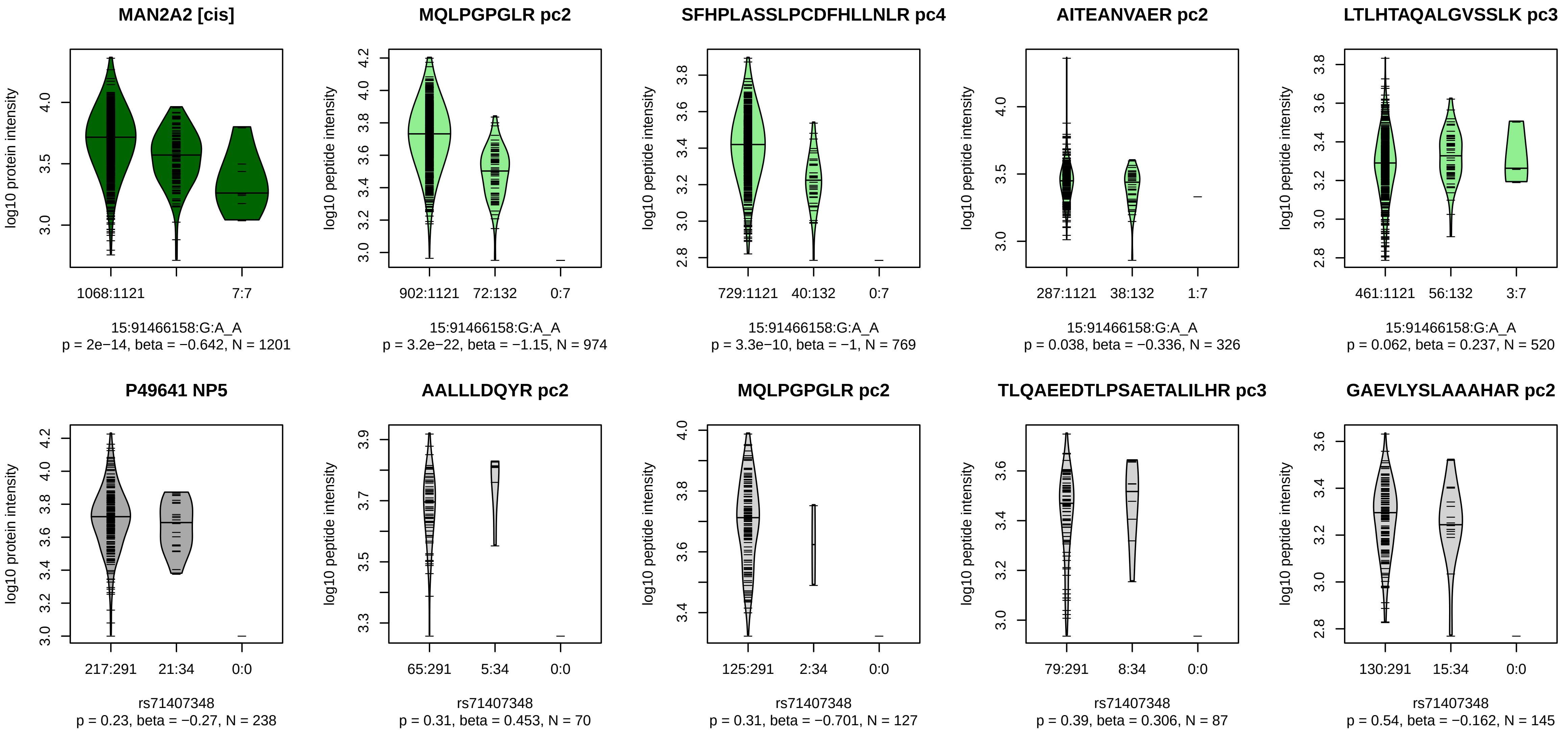


Locus: 144 (Sort: 168)
Status: Not replicated
80% power: No
SNP: 15:91466158:G:A
rsID: rs71407348
UniProtID: P49641
HGNC gene: MAN2A2
Type: Cis−pQTL

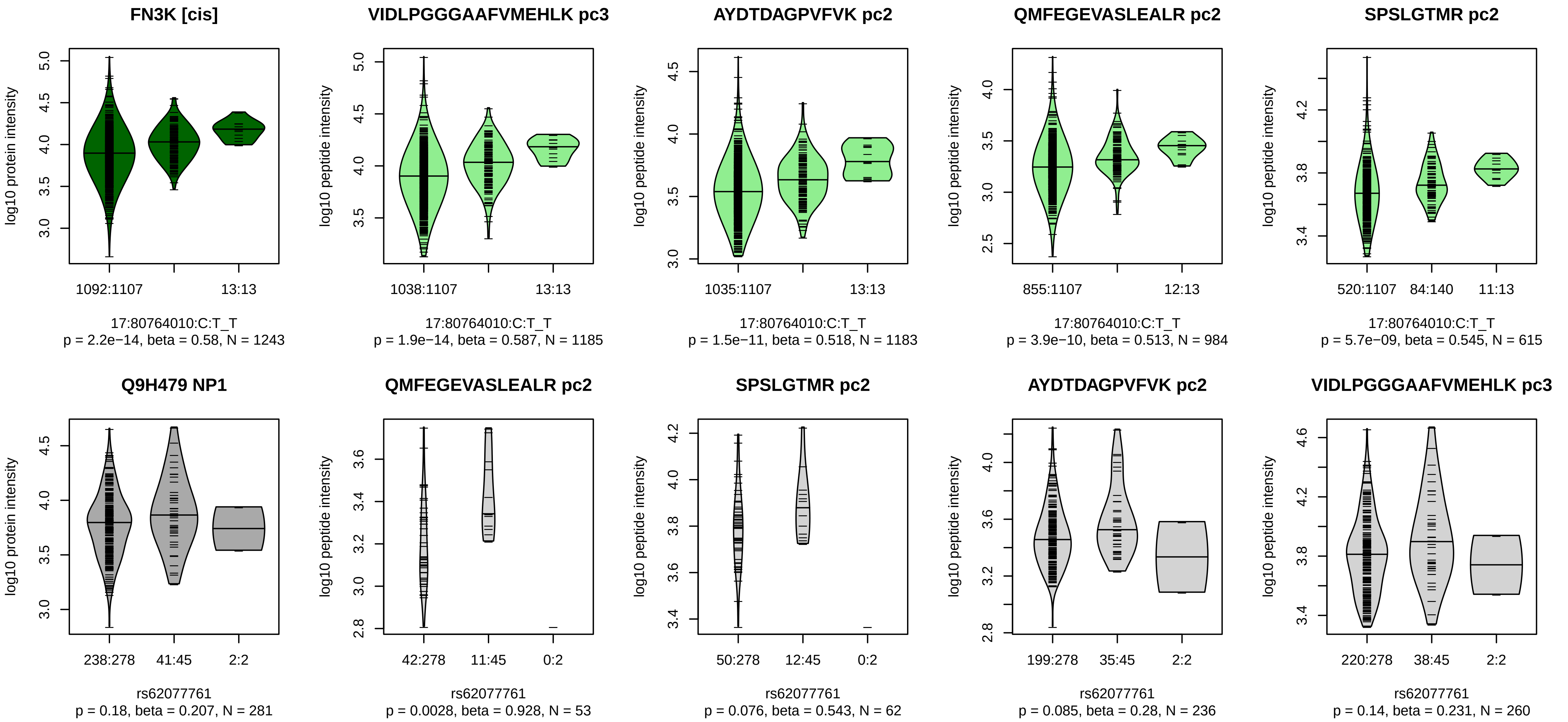


Locus: 145 (Sort: 169)

Status: Not replicated

80% power: No

SNP: 17:80764010:C:T

rsID: rs62077761

UniProtID: Q9H479

HGNC gene: FN3K

Type: Cis−pQTL

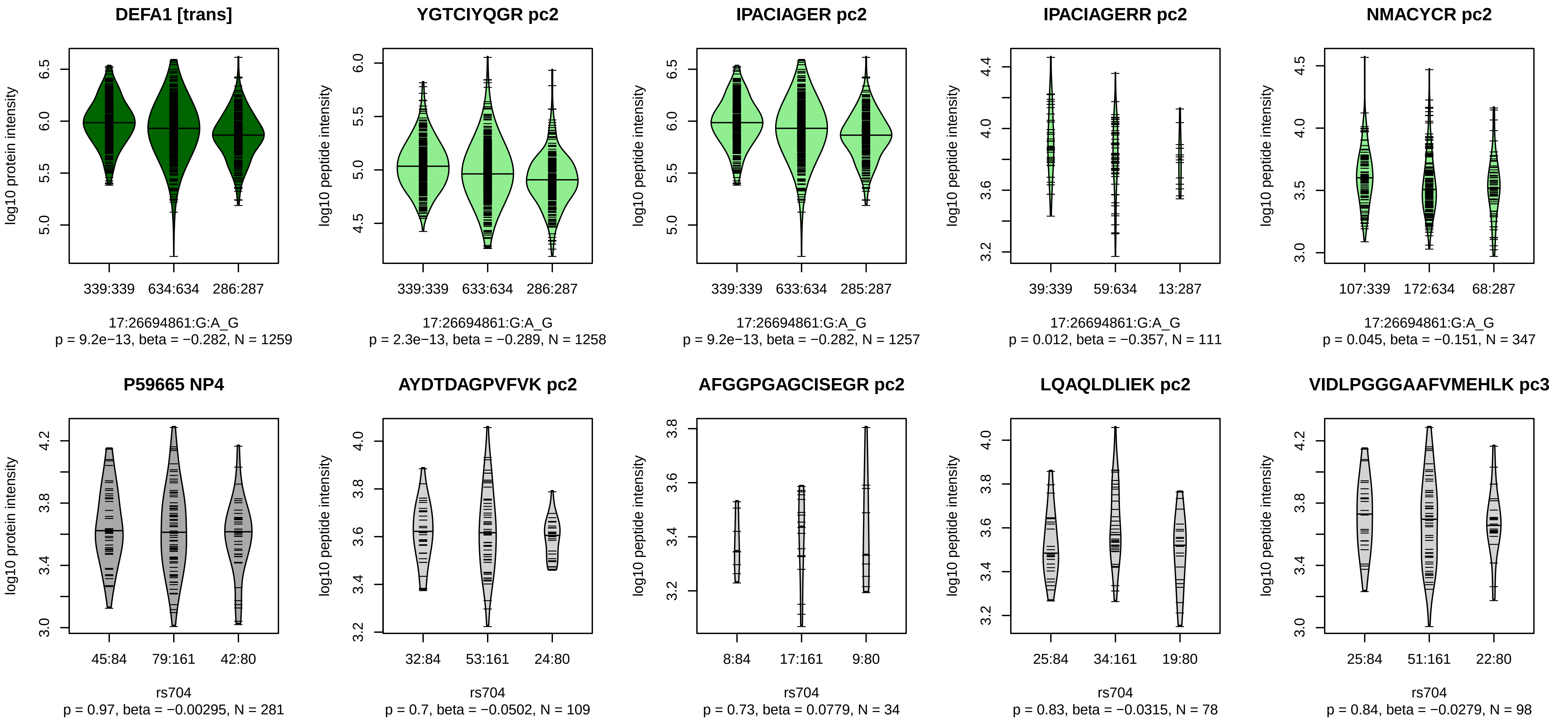


Locus: 146 (Sort: 170)

Status: Not replicated

80% power: No

SNP: 17:26694861:G:A

rsID: rs704

UniProtID: P59665

HGNC gene: DEFA1

Type: Trans−pQTL

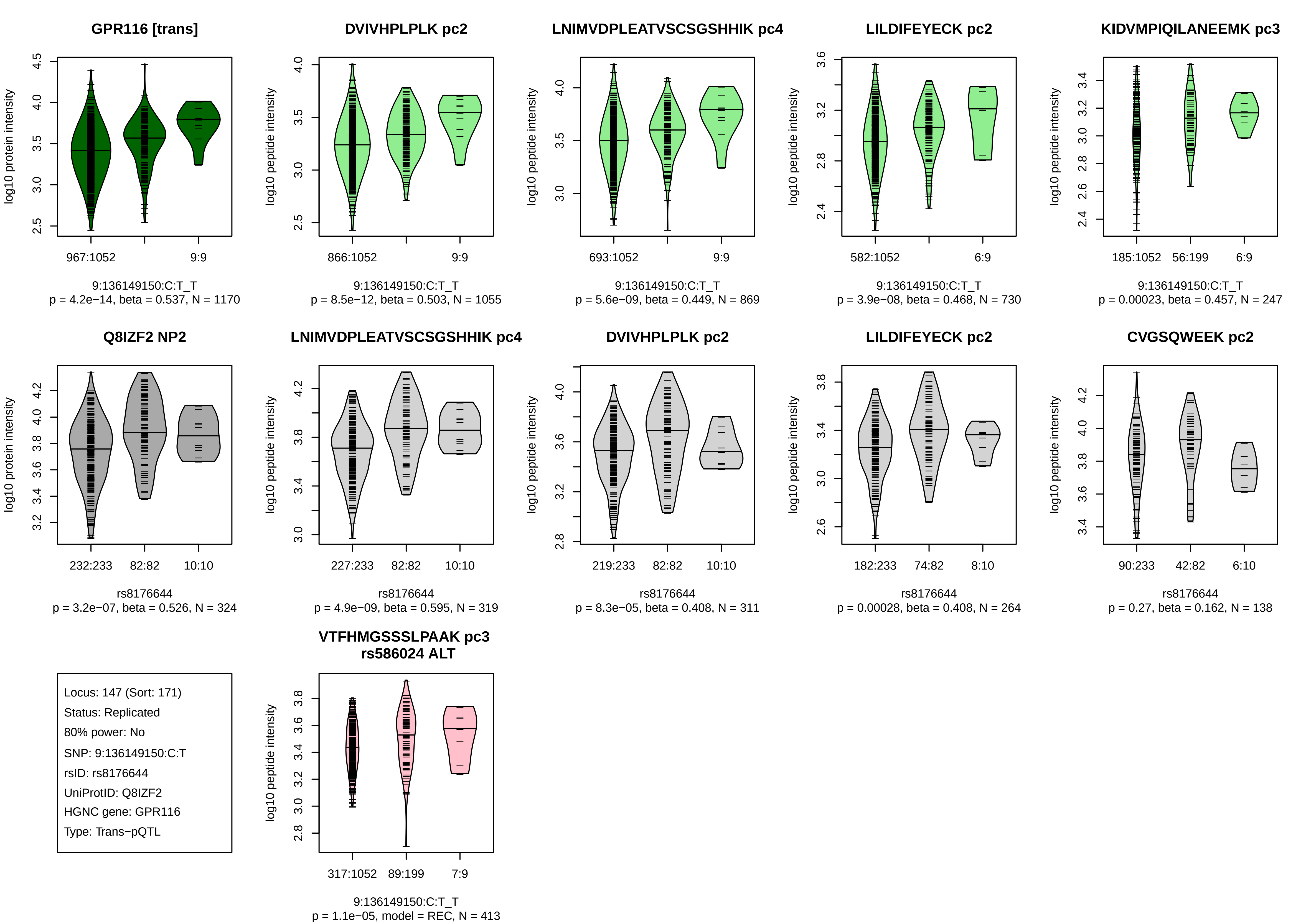
GPR116 [trans]
log10 protein intensity
967:1052
9:9
9:136149150:C:T_T
p = 4.2e−14, beta = 0.537, N = 1170
DVIVHPLPLK pc2
log10 peptide intensity
866:1052
9:9
9:136149150:C:T_T
p = 8.5e−12, beta = 0.503, N = 1055
LNIMVDPLEATVSCSGSHHIK pc4
log10 peptide intensity
693:1052
9:9
9:136149150:C:T_T
p = 5.6e−09, beta = 0.449, N = 869
LILDIFEYECK pc2
log10 peptide intensity
582:1052
6:9
9:136149150:C:T_T
p = 3.9e−08, beta = 0.468, N = 730
KIDVMPIQILANEEMK pc3
log10 peptide intensity
185:1052
56:199
6:9
9:136149150:C:T_T
p = 0.00023, beta = 0.457, N = 247
Q8IZF2 NP2
log10 protein intensity
232:233
82:82
10:10
rs8176644
p = 3.2e−07, beta = 0.526, N = 324
LNIMVDPLEATVSCSGSHHIK pc4
log10 peptide intensity
227:233
82:82
10:10
rs8176644
p = 4.9e−09, beta = 0.595, N = 319
DVIVHPLPLK pc2
log10 peptide intensity
219:233
82:82
10:10
rs8176644
p = 8.3e−05, beta = 0.408, N = 311
LILDIFEYECK pc2
log10 peptide intensity
182:233
74:82
8:10
rs8176644
p = 0.00028, beta = 0.408, N = 264
CVGSQWEEK pc2
log10 peptide intensity
90:233
42:82
6:10
rs8176644
p = 0.27, beta = 0.162, N = 138
Locus: 147 (Sort: 171)
Status: Replicated
80% power: No
SNP: 9:136149150:C:T
rsID: rs8176644
UniProtID: Q8IZF2
HGNC gene: GPR116
Type: Trans−pQTL
VTFHMGSSSLPAAK pc3
rs586024 ALT
log10 peptide intensity
317:1052
89:199
7:9
9:136149150:C:T_T
p = 1.1e−05, model = REC, N = 413

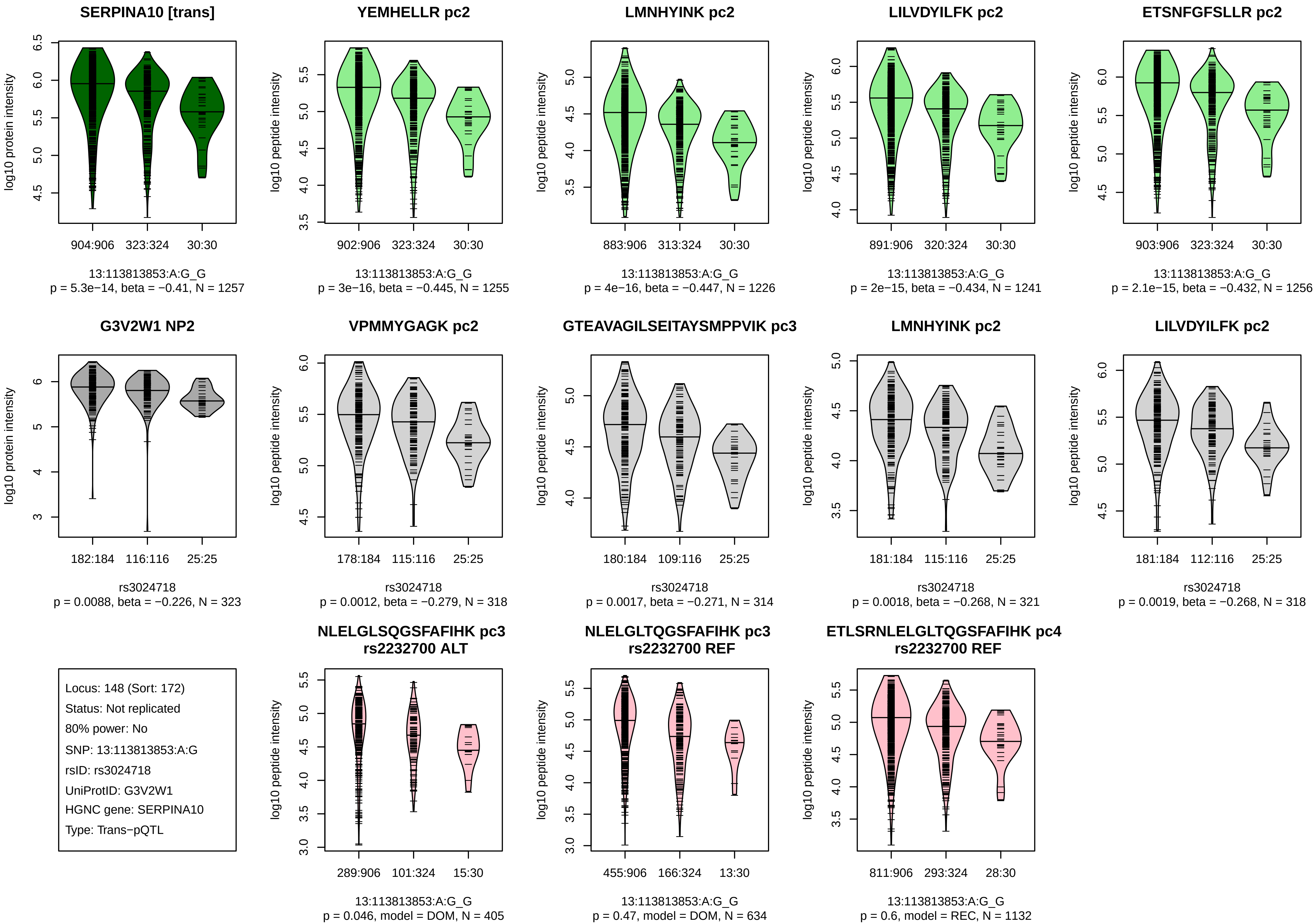
SERPINA10 [trans]
log10 protein intensity
904:906
323:324
30:30
13:113813853:A:G_G
p = 5.3e−14, beta = −0.41, N = 1257
YEMHELLR pc2
log10 peptide intensity
902:906
323:324
30:30
13:113813853:A:G_G
p = 3e−16, beta = −0.445, N = 1255
LMNHYINK pc2
883:906
313:324
30:30
13:113813853:A:G_G
p = 4e−16, beta = −0.447, N = 1226
LILVDYILFK pc2
891:906
320:324
30:30
13:113813853:A:G_G
p = 2e−15, beta = −0.434, N = 1241
ETSNFGFSLLR pc2
903:906
323:324
30:30
13:113813853:A:G_G
p = 2.1e−15, beta = −0.432, N = 1256
G3V2W1 NP2
182:184
116:116
25:25
rs3024718
p = 0.0088, beta = −0.226, N = 323
VPMMYGAGK pc2
178:184
115:116
25:25
rs3024718
p = 0.0012, beta = −0.279, N = 318
GTEAVAGILSEITAYSMPPVIK pc3
180:184
109:116
25:25
rs3024718
p = 0.0017, beta = −0.271, N = 314
LMNHYINK pc2
181:184
115:116
25:25
rs3024718
p = 0.0018, beta = −0.268, N = 321
LILVDYILFK pc2
181:184
112:116
25:25
rs3024718
p = 0.0019, beta = −0.268, N = 318
Locus: 148 (Sort: 172)
Status: Not replicated
80% power: No
SNP: 13:113813853:A:G
rsID: rs3024718
UniProtID: G3V2W1
HGNC gene: SERPINA10
Type: Trans−pQTL
NLELGLSQGSFAFIHK pc3
rs2232700 ALT
289:906
101:324
15:30
13:113813853:A:G_G
p = 0.046, model = DOM, N = 405
NLELGLTQGSFAFIHK pc3
rs2232700 REF
455:906
166:324
13:30
13:113813853:A:G_G
p = 0.47, model = DOM, N = 634
ETLSRNLELGLTQGSFAFIHK pc4
rs2232700 REF
811:906
293:324
28:30
13:113813853:A:G_G
p = 0.6, model = REC, N = 1132

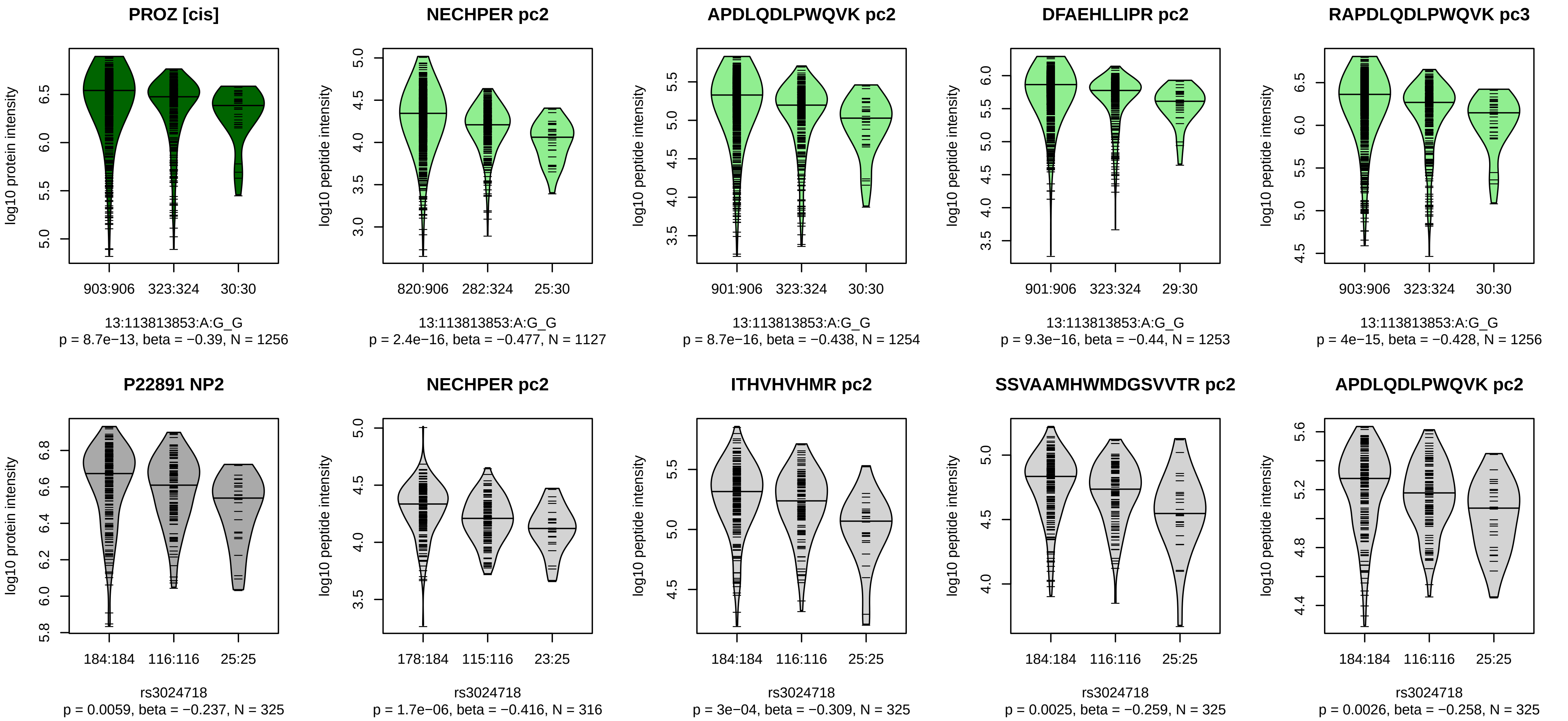


Locus: 148 (Sort: 173)

Status: Not replicated

80% power: No

SNP: 13:113813853:A:G

rsID: rs3024718

UniProtID: P22891

HGNC gene: PROZ

Type: Cis−pQTL

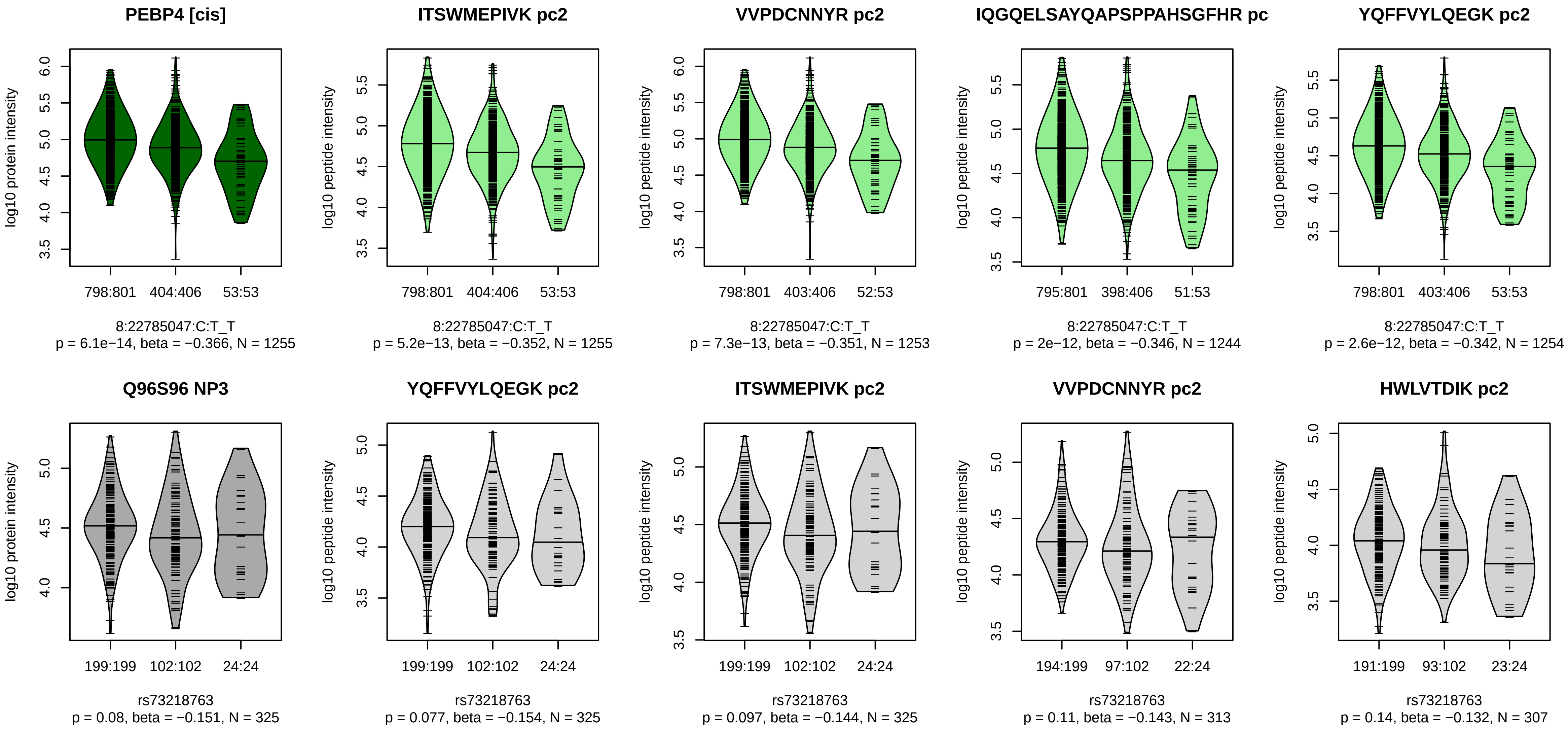


Locus: 149 (Sort: 174)

Status: Not replicated

80% power: No

SNP: 8:22785047:C:T

rsID: rs73218763

UniProtID: Q96S96

HGNC gene: PEBP4

Type: Cis−pQTL

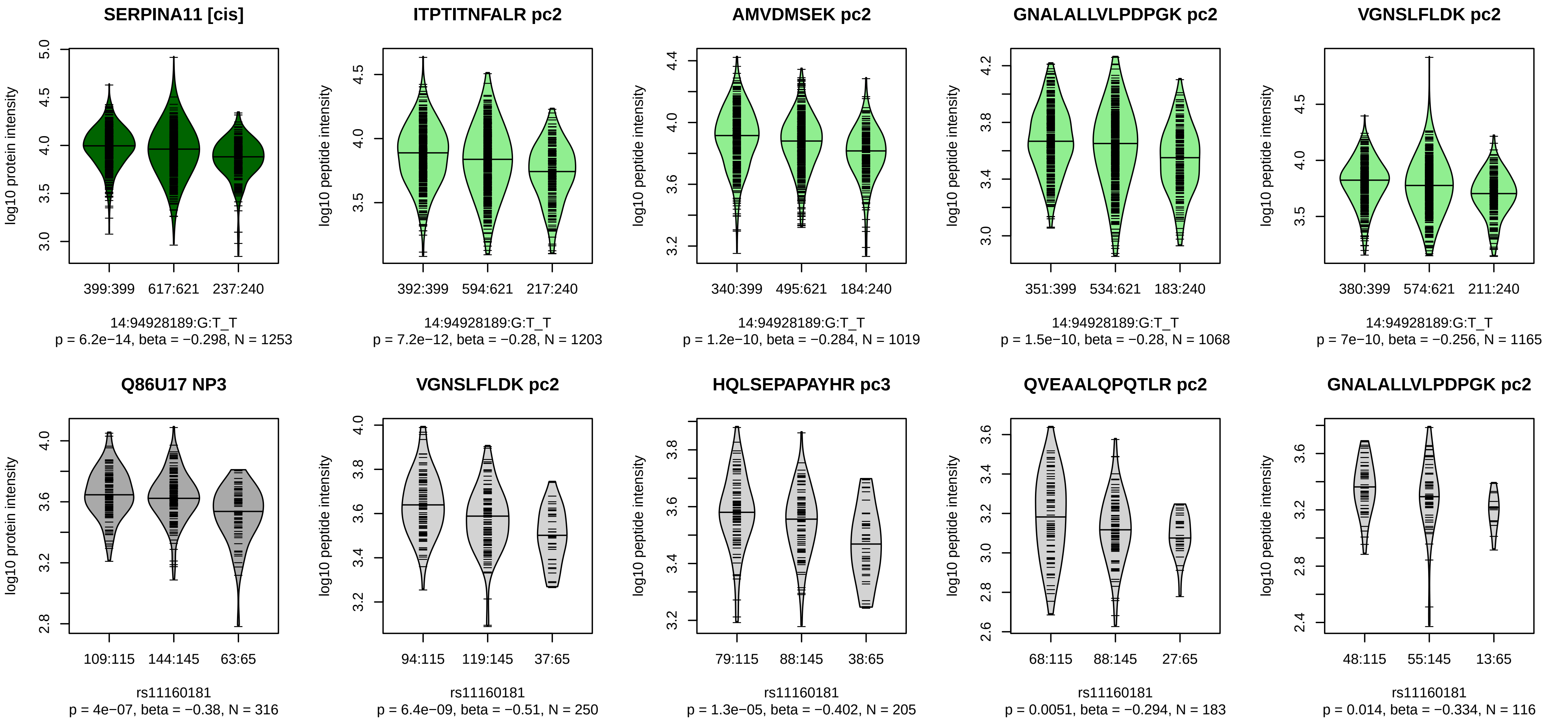


Locus: 150 (Sort: 175)
Status: Replicated
80% power: No
SNP: 14:94928189:G:T
rsID: rs11160181
UniProtID: Q86U17
HGNC gene: SERPINA11
Type: Cis−pQTL

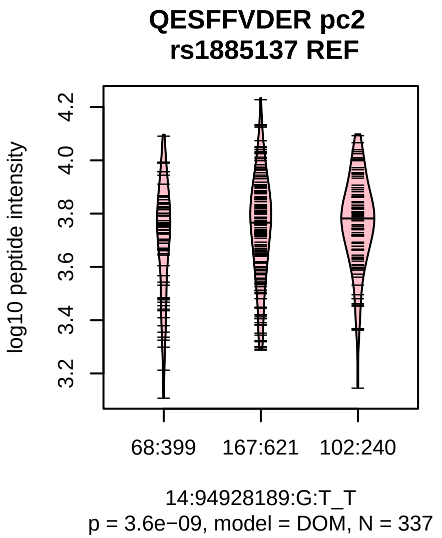

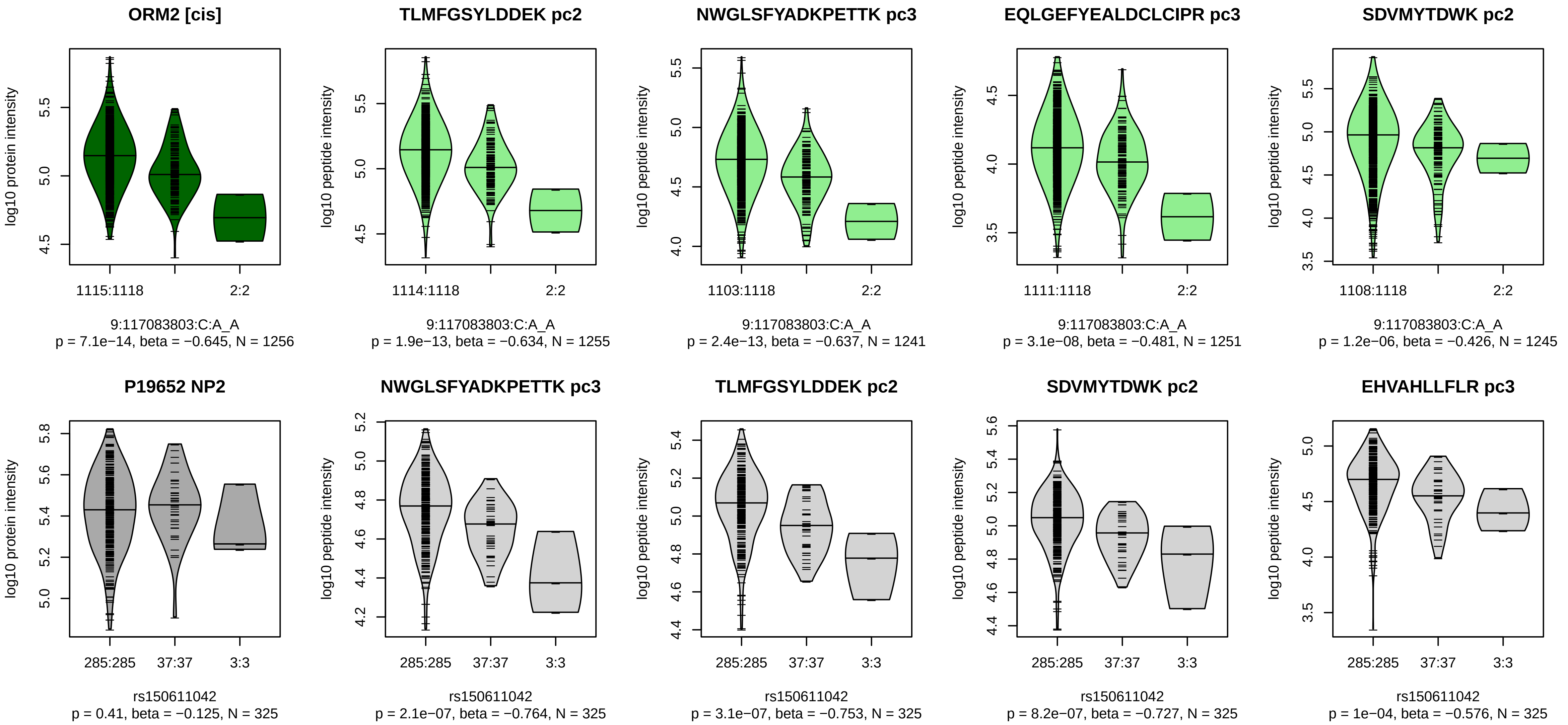


Locus: 151 (Sort: 176)

Status: Not replicated

80% power: No

SNP: 9:117083803:C:A

rsID: rs150611042

UniProtID: P19652

HGNC gene: ORM2

Type: Cis−pQTL

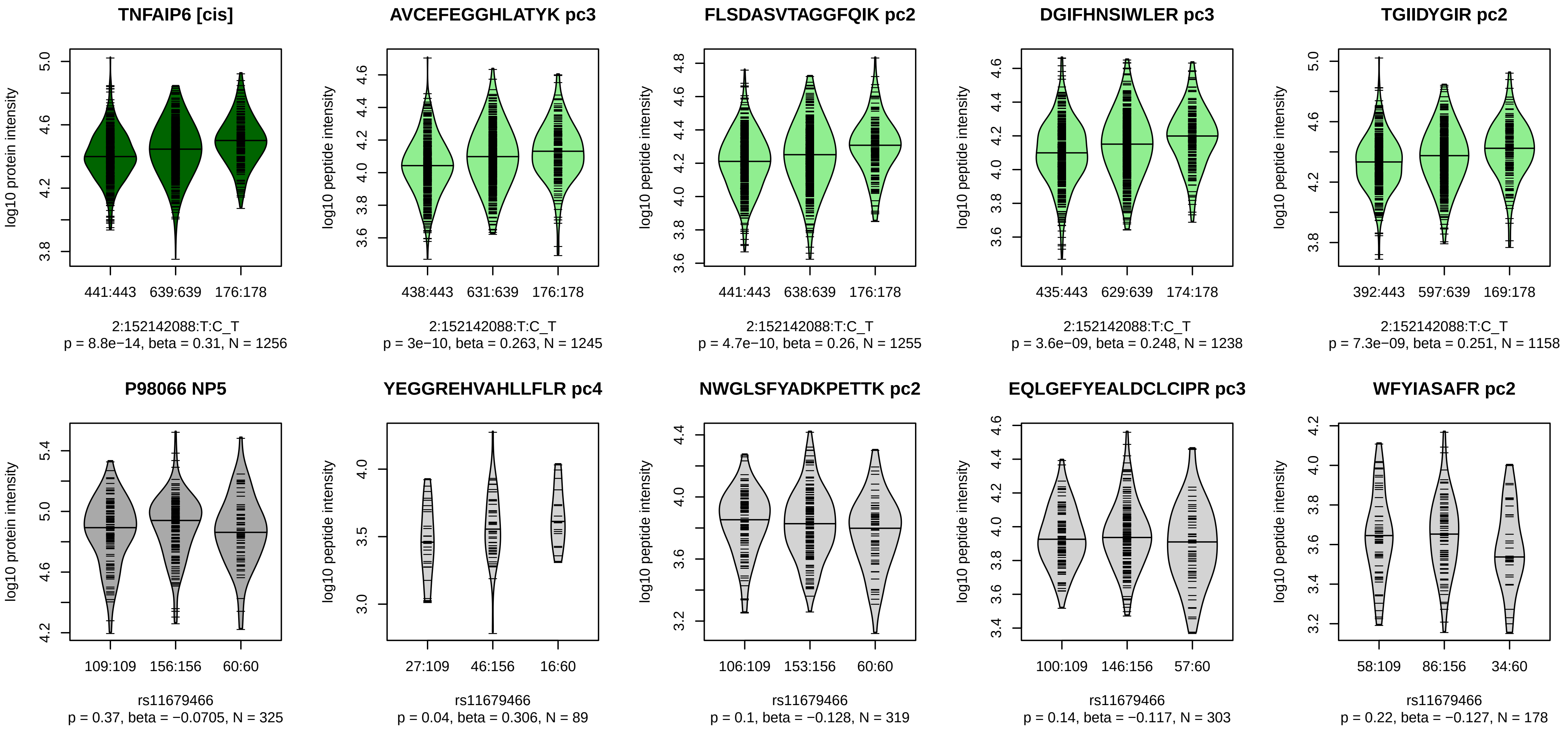


Locus: 152 (Sort: 177)

Status: Not replicated

80% power: No

SNP: 2:152142088:T:C

rsID: rs11679466

UniProtID: P98066

HGNC gene: TNFAIP6

Type: Cis−pQTL

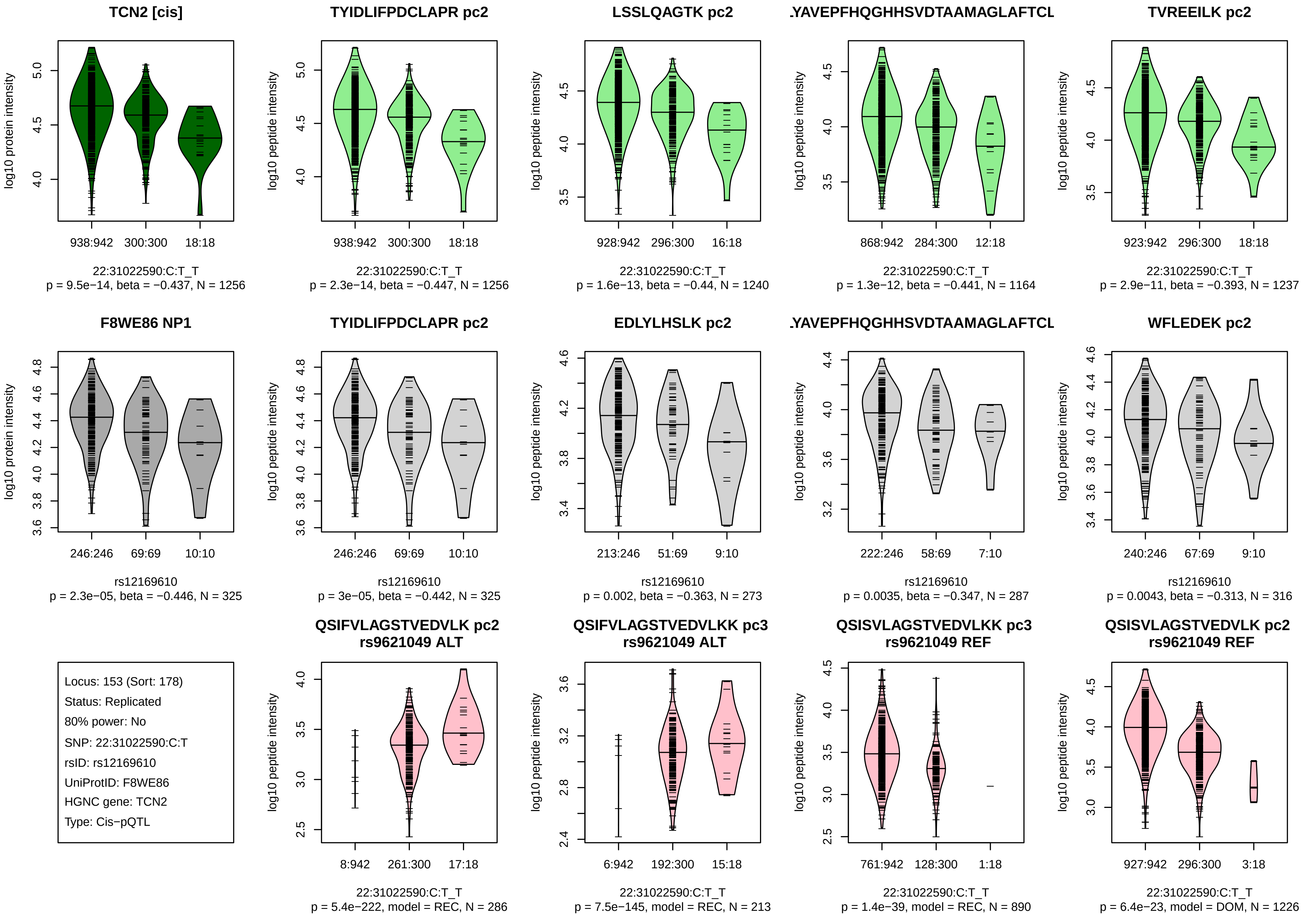
TCN2 [cis]
log10 protein intensity
938:942
300:300
18:18
22:31022590:C:T_T
p = 9.5e−14, beta = −0.437, N = 1256
TYIDLIFPDCLAPR pc2
log10 peptide intensity
938:942
300:300
18:18
22:31022590:C:T_T
p = 2.3e−14, beta = −0.447, N = 1256
LSSLQAGTK pc2
log10 peptide intensity
928:942
296:300
16:18
22:31022590:C:T_T
p = 1.6e−13, beta = −0.44, N = 1240
.YAVEPFHQGHHSVDTAAMAGLAFTCL
log10 peptide intensity
868:942
284:300
12:18
22:31022590:C:T_T
p = 1.3e−12, beta = −0.441, N = 1164
TVREEILK pc2
log10 peptide intensity
923:942
296:300
18:18
22:31022590:C:T_T
p = 2.9e−11, beta = −0.393, N = 1237
F8WE86 NP1
log10 protein intensity
246:246
69:69
10:10
rs12169610
p = 2.3e−05, beta = −0.446, N = 325
TYIDLIFPDCLAPR pc2
log10 peptide intensity
246:246
69:69
10:10
rs12169610
p = 3e−05, beta = −0.442, N = 325
EDLYLHSLK pc2
log10 peptide intensity
213:246
51:69
9:10
rs12169610
p = 0.002, beta = −0.363, N = 273
.YAVEPFHQGHHSVDTAAMAGLAFTCL
log10 peptide intensity
222:246
58:69
7:10
rs12169610
p = 0.0035, beta = −0.347, N = 287
WFLEDEK pc2
log10 peptide intensity
240:246
67:69
9:10
rs12169610
p = 0.0043, beta = −0.313, N = 316
Locus: 153 (Sort: 178)
Status: Replicated
80% power: No
SNP: 22:31022590:C:T
rsID: rs12169610
UniProtID: F8WE86
HGNC gene: TCN2
Type: Cis−pQTL
QSIFVLAGSTVEDVLK pc2
rs9621049 ALT
log10 peptide intensity
8:942
261:300
17:18
22:31022590:C:T_T
p = 5.4e−222, model = REC, N = 286
QSIFVLAGSTVEDVLKK pc3
rs9621049 ALT
log10 peptide intensity
6:942
192:300
15:18
22:31022590:C:T_T
p = 7.5e−145, model = REC, N = 213
QSISVLAGSTVEDVLKK pc3
rs9621049 REF
log10 peptide intensity
761:942
128:300
1:18
22:31022590:C:T_T
p = 1.4e−39, model = REC, N = 890
QSISVLAGSTVEDVLK pc2
rs9621049 REF
log10 peptide intensity
927:942
296:300
3:18
22:31022590:C:T_T
p = 6.4e−23, model = DOM, N = 1226

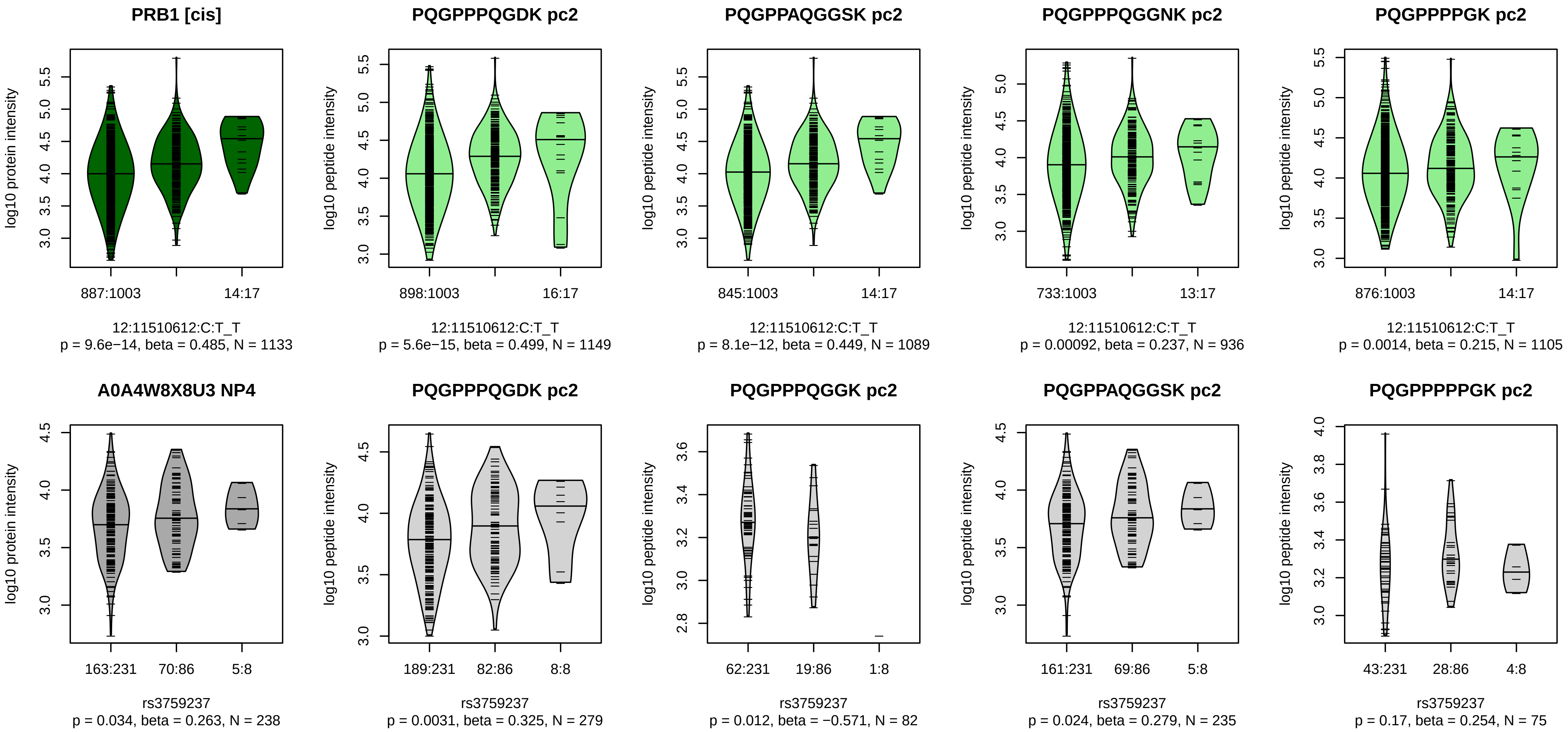


Locus: 154 (Sort: 179)
Status: Not replicated
80% power: No
SNP: 12:11510612:C:T
rsID: rs3759237
UniProtID: A0A4W8X8U3
HGNC gene: PRB1
Type: Cis−pQTL

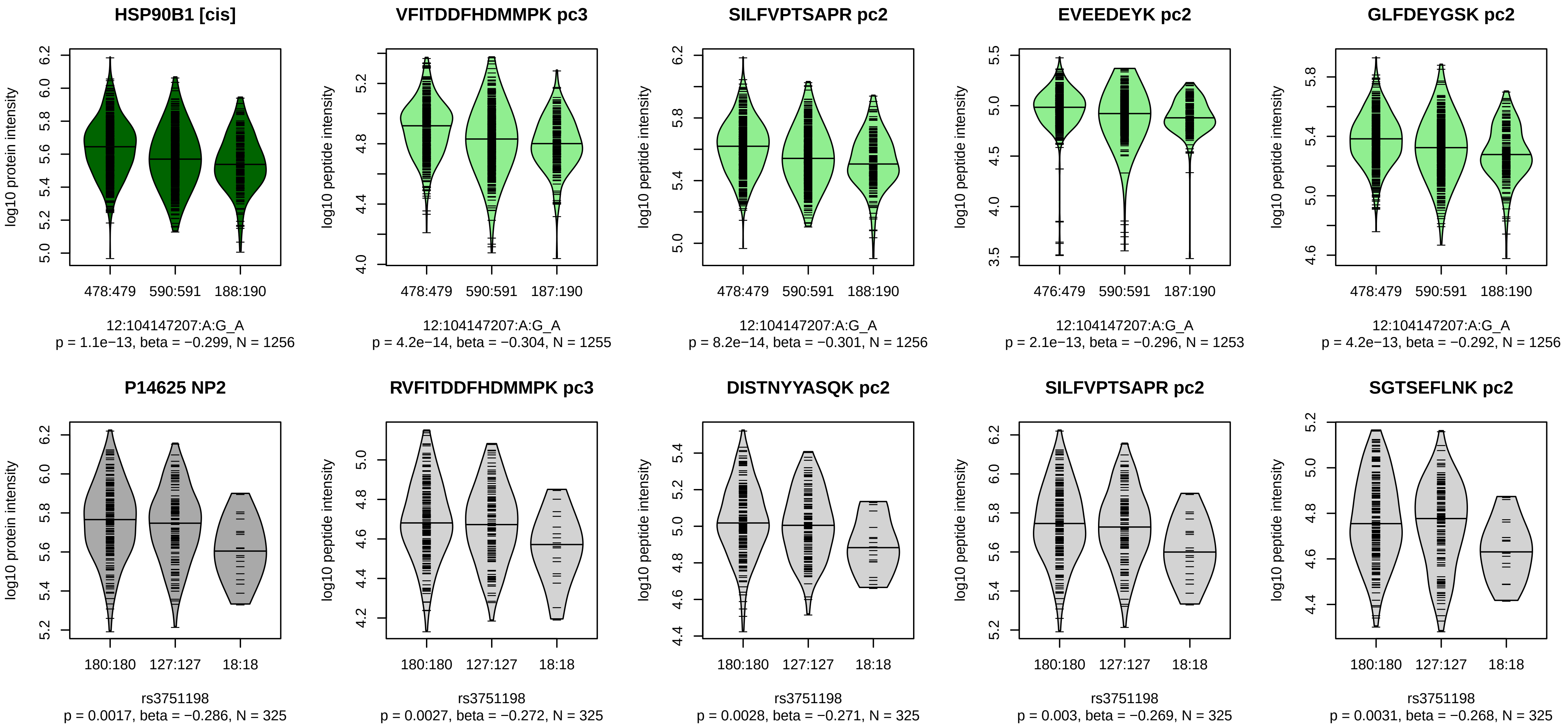


Locus: 155 (Sort: 180)
Status: Not replicated
80% power: No
SNP: 12:104147207:A:G
rsID: rs3751198
UniProtID: P14625
HGNC gene: HSP90B1
Type: Cis−pQTL

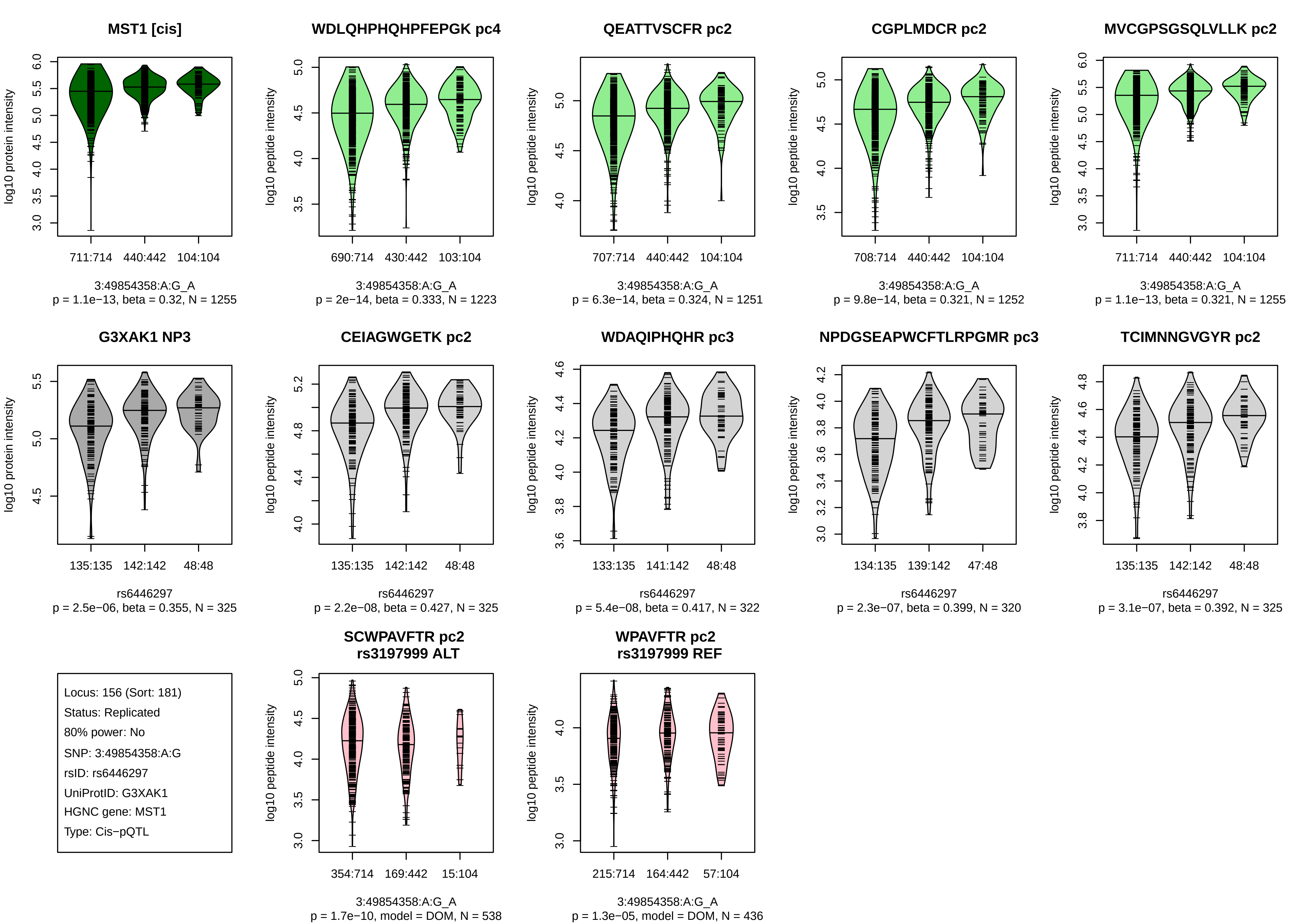

MST1 [cis]
log10 protein intensity
711:714
440:442
104:104
3:49854358:A:G_A
p = 1.1e−13, beta = 0.32, N = 1255
WDLQHPHQHPFEPGK pc4
log10 peptide intensity
690:714
430:442
103:104
3:49854358:A:G_A
p = 2e−14, beta = 0.333, N = 1223
QEATTVSCFR pc2
log10 peptide intensity
707:714
440:442
104:104
3:49854358:A:G_A
p = 6.3e−14, beta = 0.324, N = 1251
CGPLMDCR pc2
log10 peptide intensity
708:714
440:442
104:104
3:49854358:A:G_A
p = 9.8e−14, beta = 0.321, N = 1252
MVCGPSGSQLVLLK pc2
log10 peptide intensity
711:714
440:442
104:104
3:49854358:A:G_A
p = 1.1e−13, beta = 0.321, N = 1255
G3XAK1 NP3
log10 protein intensity
135:135
142:142
48:48
rs6446297
p = 2.5e−06, beta = 0.355, N = 325
CEIAGWGETK pc2
log10 peptide intensity
135:135
142:142
48:48
rs6446297
p = 2.2e−08, beta = 0.427, N = 325
WDAQIPHQHR pc3
log10 peptide intensity
133:135
141:142
48:48
rs6446297
p = 5.4e−08, beta = 0.417, N = 322
NPDGSEAPWCFTLRPGMR pc3
log10 peptide intensity
134:135
139:142
47:48
rs6446297
p = 2.3e−07, beta = 0.399, N = 320
TCIMNNGVGYR pc2
log10 peptide intensity
135:135
142:142
48:48
rs6446297
p = 3.1e−07, beta = 0.392, N = 325
Locus: 156 (Sort: 181)
Status: Replicated
80% power: No
SNP: 3:49854358:A:G
rsID: rs6446297
UniProtID: G3XAK1
HGNC gene: MST1
Type: Cis−pQTL
SCWPAVFTR pc2
rs3197999 ALT
log10 peptide intensity
354:714
169:442
15:104
3:49854358:A:G_A
p = 1.7e−10, model = DOM, N = 538
WPAVFTR pc2
rs3197999 REF
log10 peptide intensity
215:714
164:442
57:104
3:49854358:A:G_A
p = 1.3e−05, model = DOM, N = 436

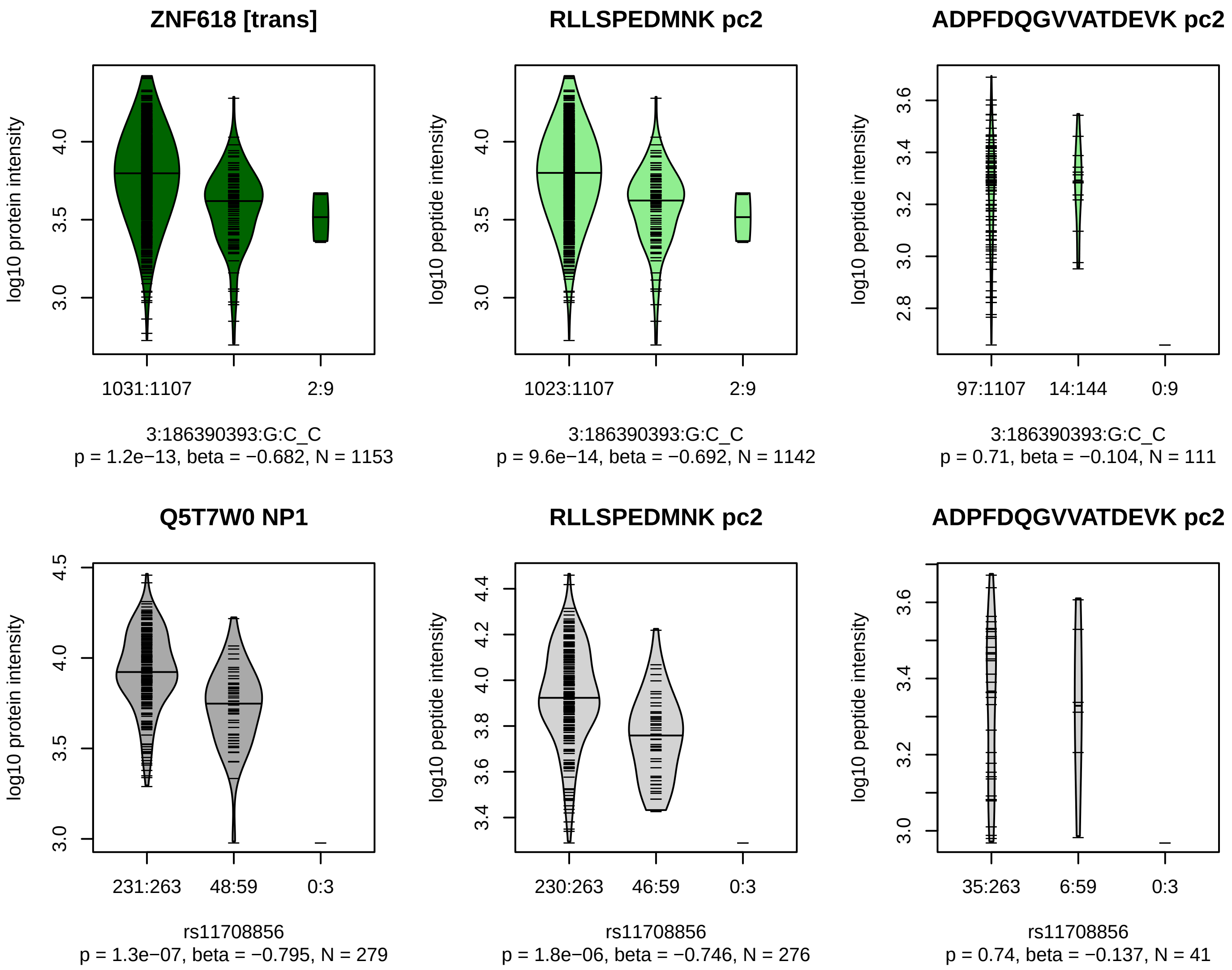


Locus: 157 (Sort: 182)

Status: Replicated

80% power: No

SNP: 3:186390393:G:C

rsID: rs11708856

UniProtID: Q5T7W0

HGNC gene: ZNF618

Type: Trans−pQTL

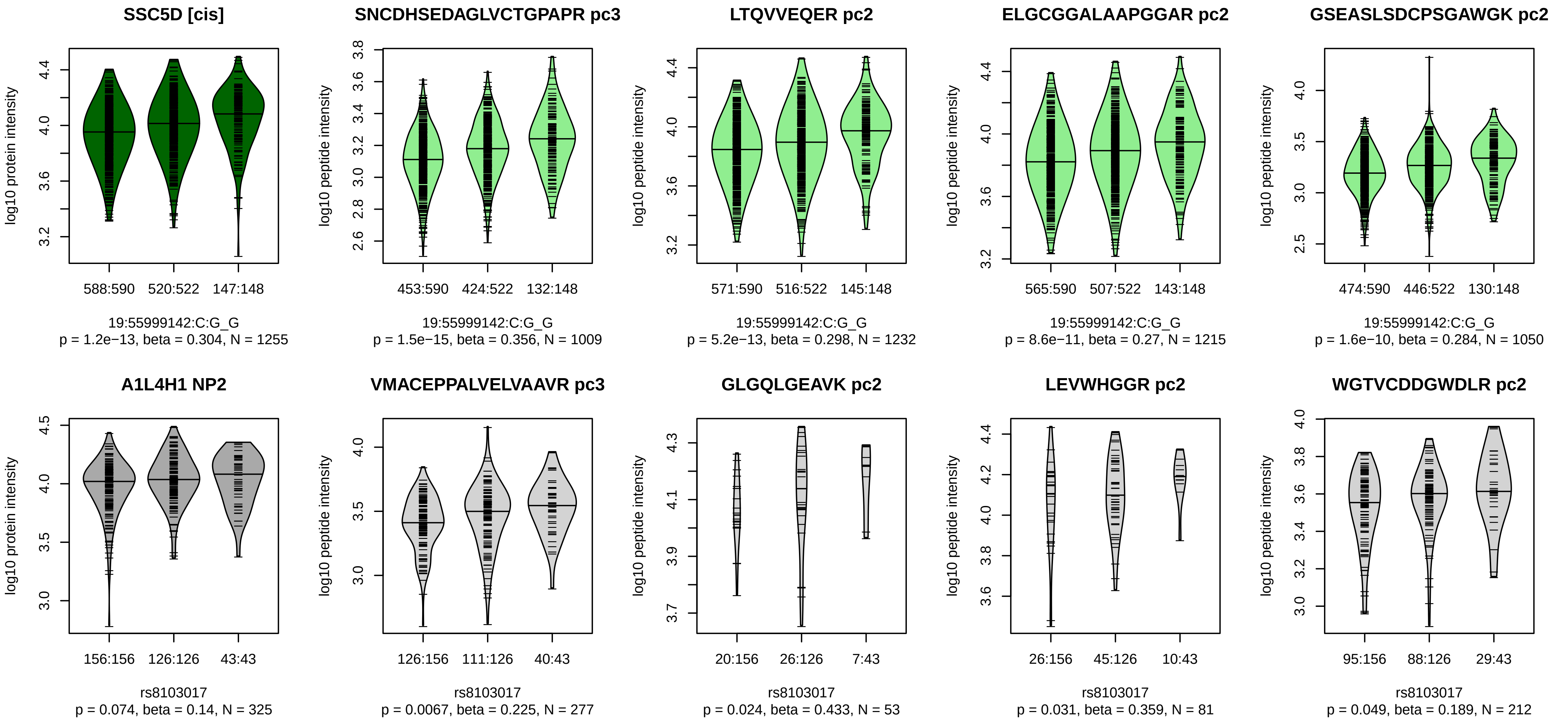


Locus: 158 (Sort: 183)

Status: Not replicated

80% power: No

SNP: 19:55999142:C:G

rsID: rs8103017

UniProtID: A1L4H1

HGNC gene: SSC5D

Type: Cis−pQTL

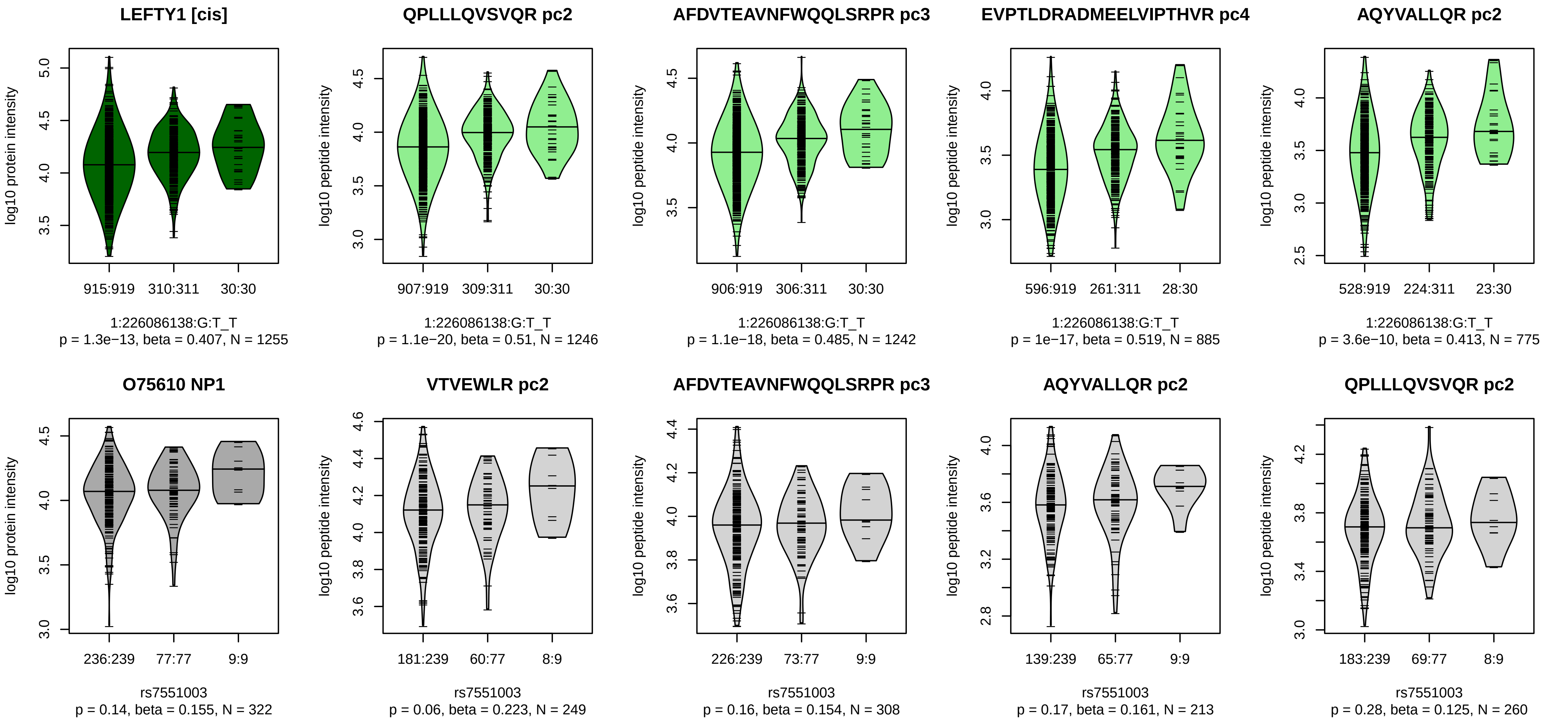


Locus: 159 (Sort: 184)

Status: Not replicated

80% power: No

SNP: 1:226086138:G:T

rsID: rs7551003

UniProtID: O75610

HGNC gene: LEFTY1

Type: Cis−pQTL

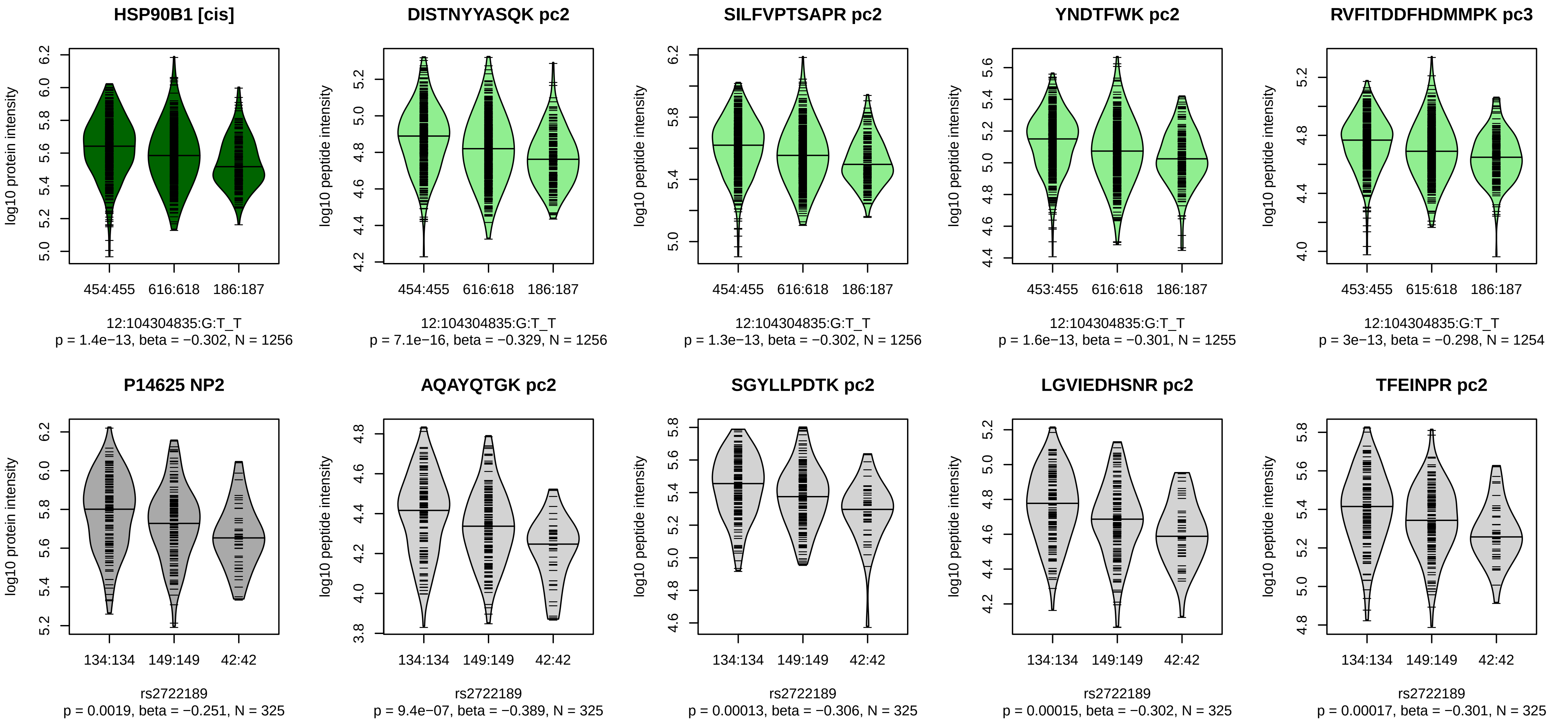


Locus: 160 (Sort: 185)

Status: Not replicated

80% power: No

SNP: 12:104304835:G:T

rsID: rs2722189

UniProtID: P14625

HGNC gene: HSP90B1

Type: Cis−pQTL

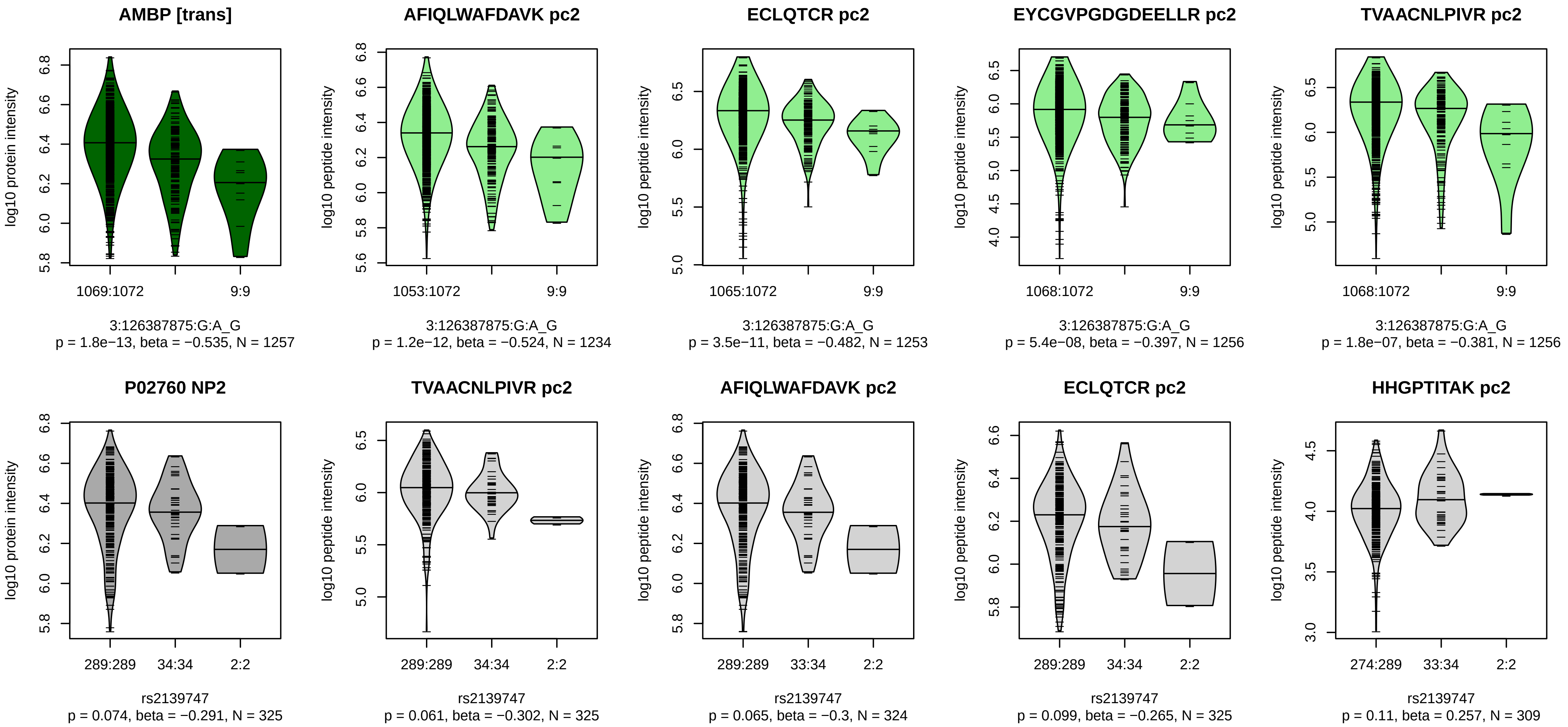


Locus: 161 (Sort: 186)
Status: Not replicated
80% power: No
SNP: 3:126387875:G:A
rsID: rs2139747
UniProtID: P02760
HGNC gene: AMBP
Type: Trans−pQTL

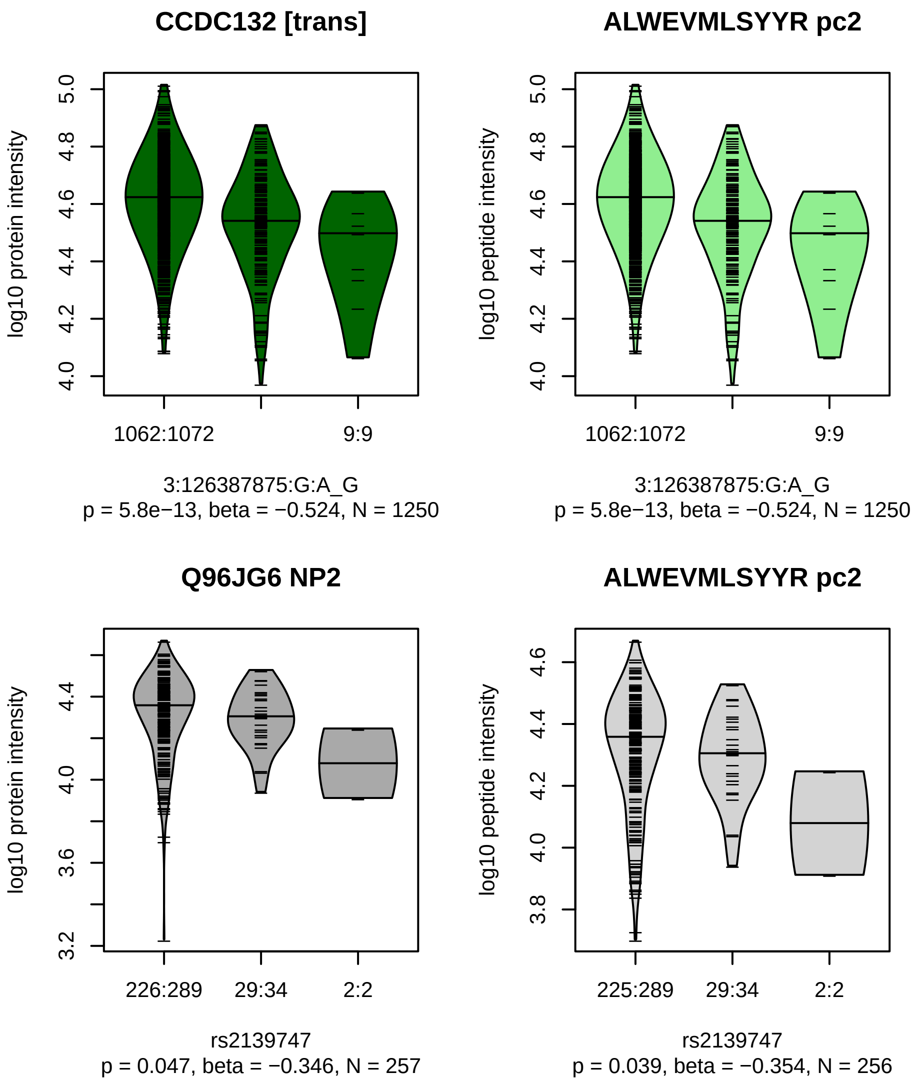


Locus: 161 (Sort: 187)

Status: Not replicated

80% power: No

SNP: 3:126387875:G:A

rsID: rs2139747

UniProtID: Q96JG6

HGNC gene: CCDC132

Type: Trans−pQTL

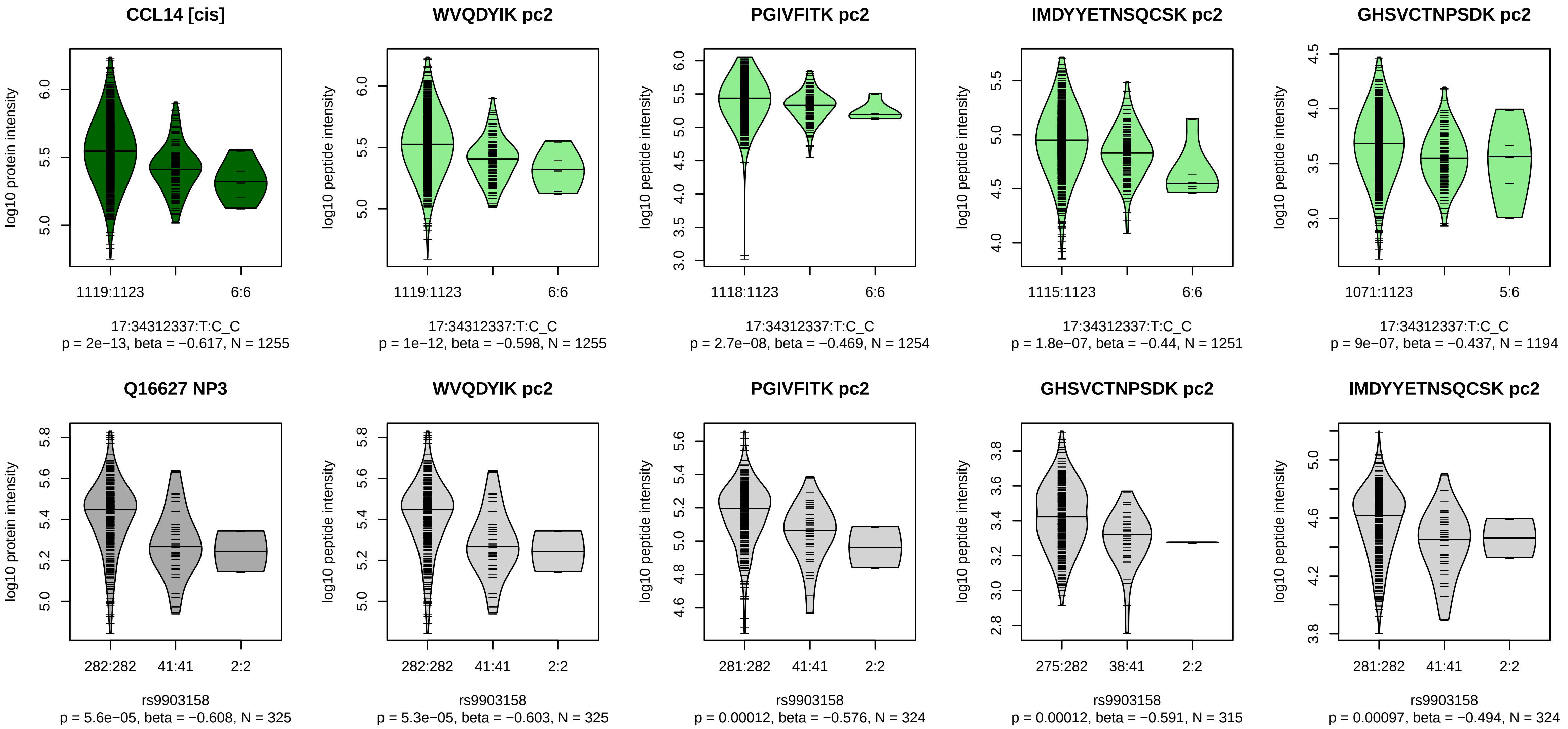


Locus: 162 (Sort: 188)
Status: Replicated
80% power: No
SNP: 17:34312337:T:C
rsID: rs9903158
UniProtID: Q16627
HGNC gene: CCL14
Type: Cis−pQTL

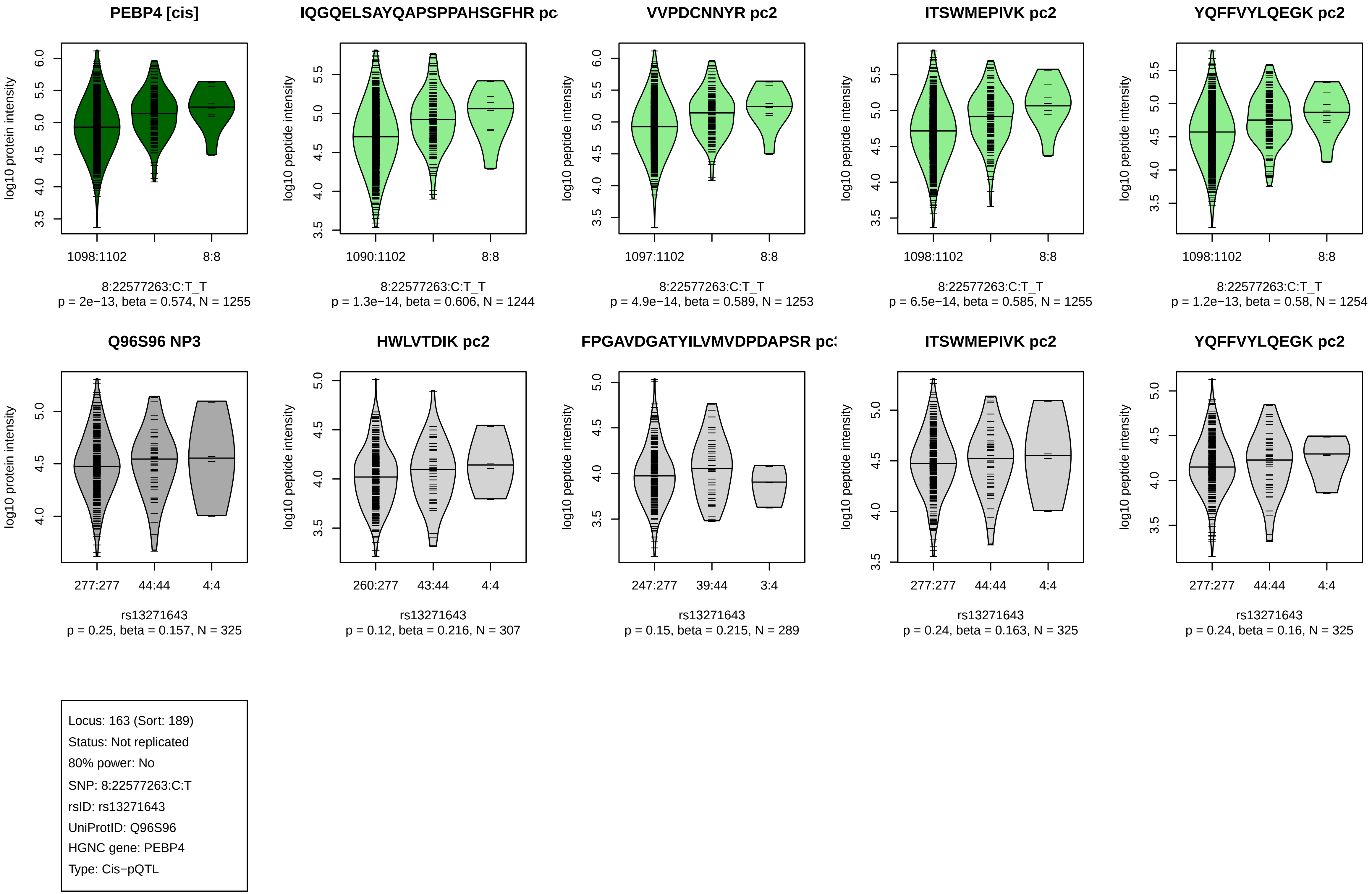
PEBP4 [cis]
log10 protein intensity
1098:1102
8:8
8:22577263:C:T_T
p = 2e−13, beta = 0.574, N = 1255
IQGQELSAYQAPSPPAHSGFHR pc
log10 peptide intensity
1090:1102
8:8
8:22577263:C:T_T
p = 1.3e−14, beta = 0.606, N = 1244
VVPDCNNYR pc2
log10 peptide intensity
1097:1102
8:8
8:22577263:C:T_T
p = 4.9e−14, beta = 0.589, N = 1253
ITSWMEPIVK pc2
log10 peptide intensity
1098:1102
8:8
8:22577263:C:T_T
p = 6.5e−14, beta = 0.585, N = 1255
YQFFVYLQEGK pc2
log10 peptide intensity
1098:1102
8:8
8:22577263:C:T_T
p = 1.2e−13, beta = 0.58, N = 1254
Q96S96 NP3
log10 protein intensity
277:277
44:44
4:4
rs13271643
p = 0.25, beta = 0.157, N = 325
HWLVTDIK pc2
log10 peptide intensity
260:277
43:44
4:4
rs13271643
p = 0.12, beta = 0.216, N = 307
FPGAVDGATYILVMVDPDAPSR pc
log10 peptide intensity
247:277
39:44
3:4
rs13271643
p = 0.15, beta = 0.215, N = 289
ITSWMEPIVK pc2
log10 peptide intensity
277:277
44:44
4:4
rs13271643
p = 0.24, beta = 0.163, N = 325
YQFFVYLQEGK pc2
log10 peptide intensity
277:277
44:44
4:4
rs13271643
p = 0.24, beta = 0.16, N = 325
Locus: 163 (Sort: 189)
Status: Not replicated
80% power: No
SNP: 8:22577263:C:T
rsID: rs13271643
UniProtID: Q96S96
HGNC gene: PEBP4
Type: Cis−pQTL

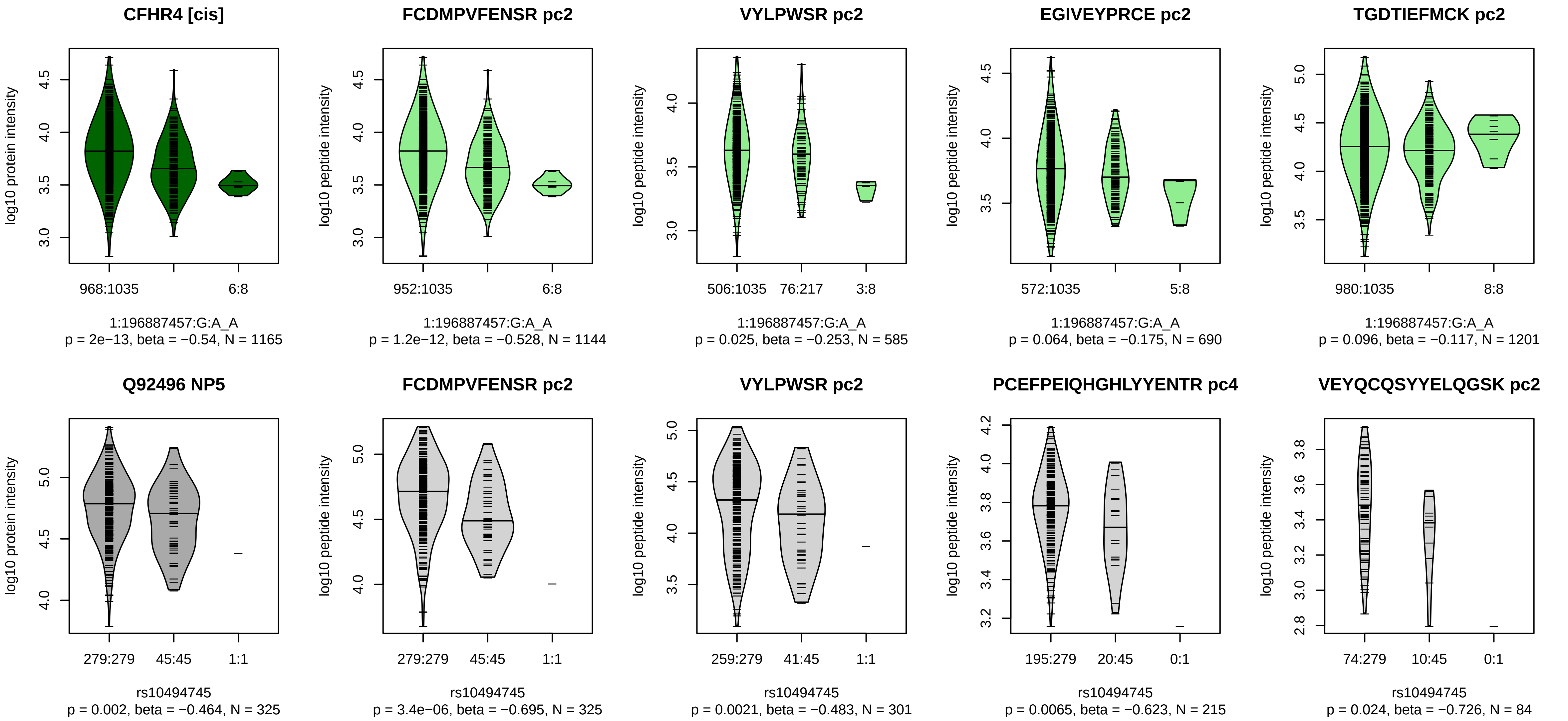


Locus: 164 (Sort: 190)

Status: Not replicated

80% power: No

SNP: 1:196887457:G:A

rsID: rs10494745

UniProtID: Q92496

HGNC gene: CFHR4

Type: Cis−pQTL

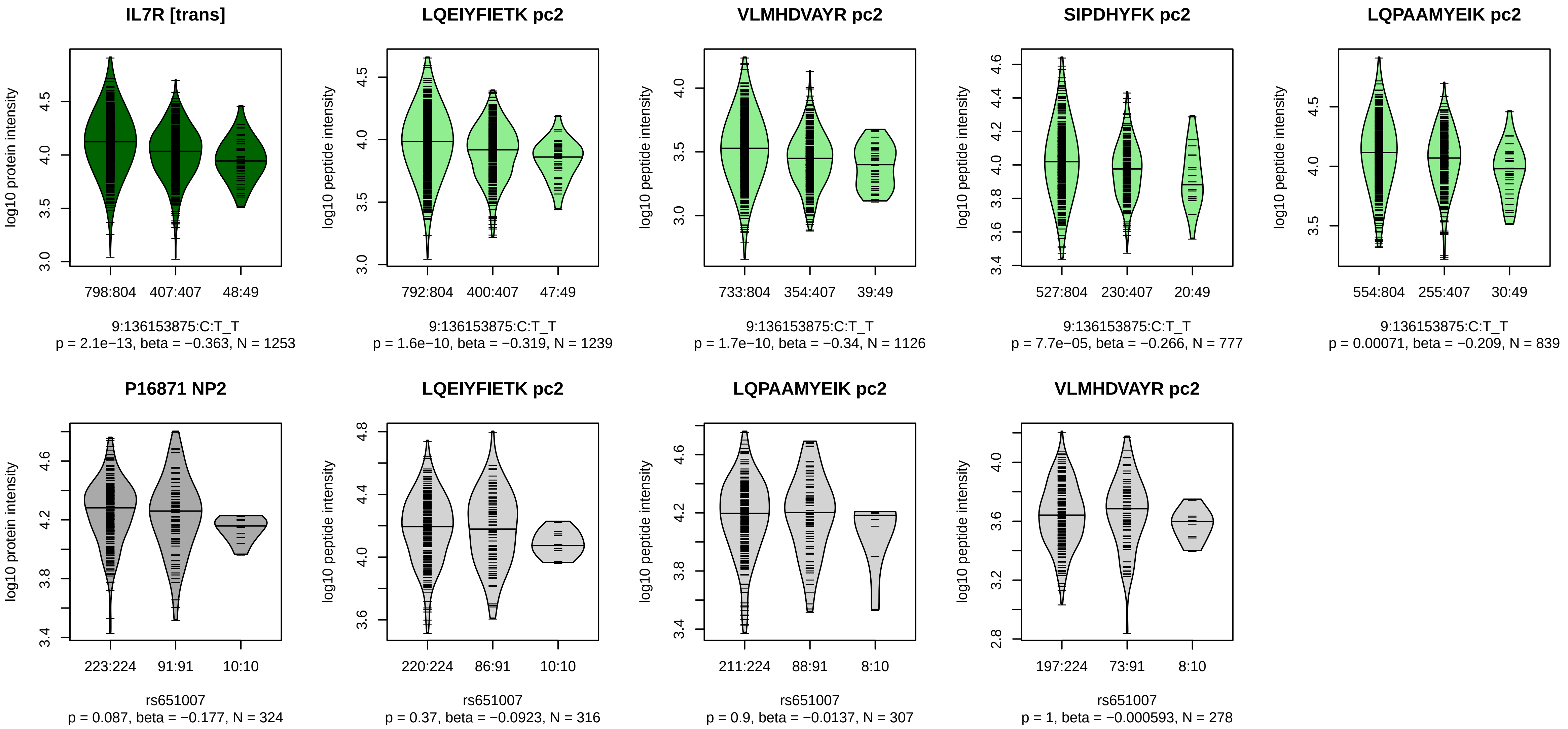


Locus: 165 (Sort: 191)
Status: Not replicated
80% power: No
SNP: 9:136153875:C:T
rsID: rs651007
UniProtID: P16871
HGNC gene: IL7R
Type: Trans−pQTL

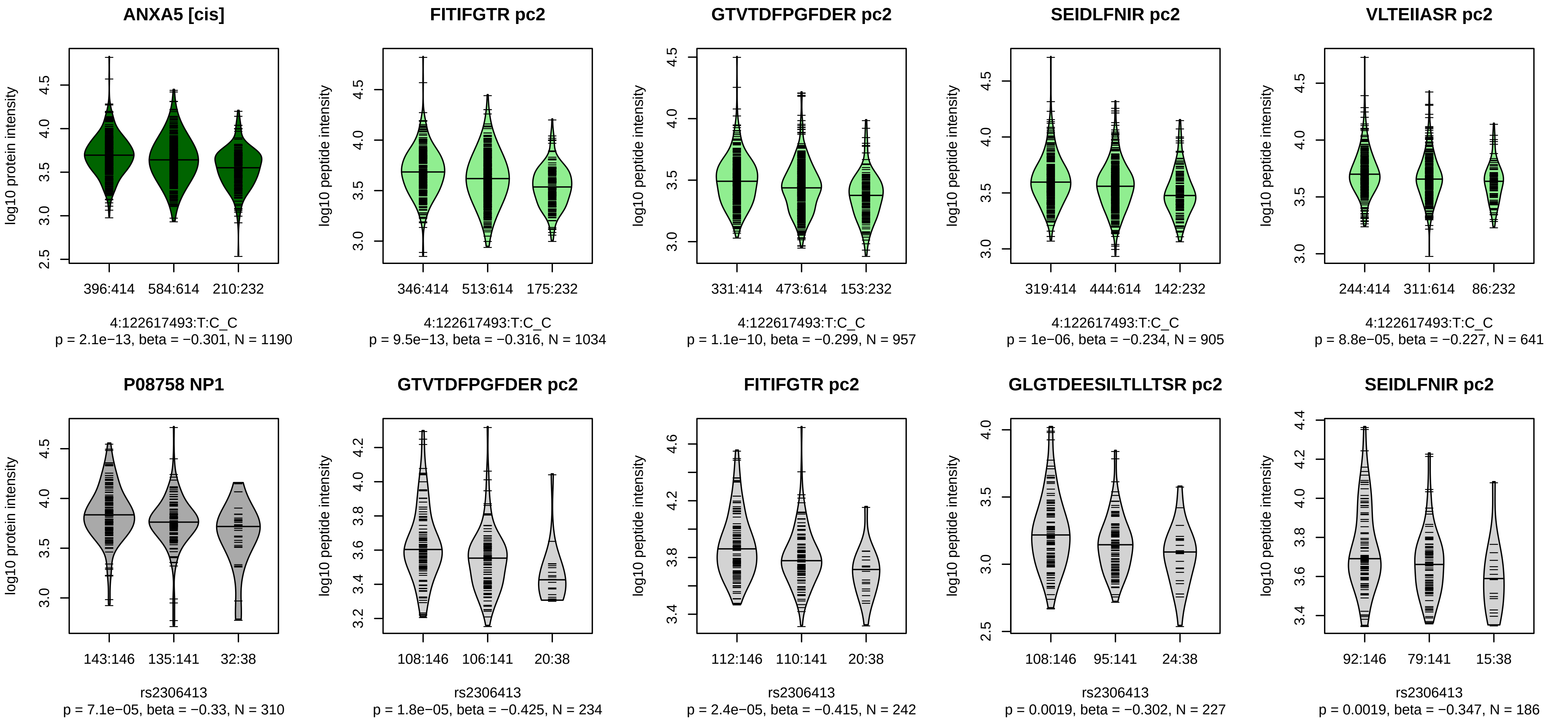


Locus: 166 (Sort: 192)
Status: Replicated
80% power: No
SNP: 4:122617493:T:C
rsID: rs2306413
UniProtID: P08758
HGNC gene: ANXA5
Type: Cis−pQTL

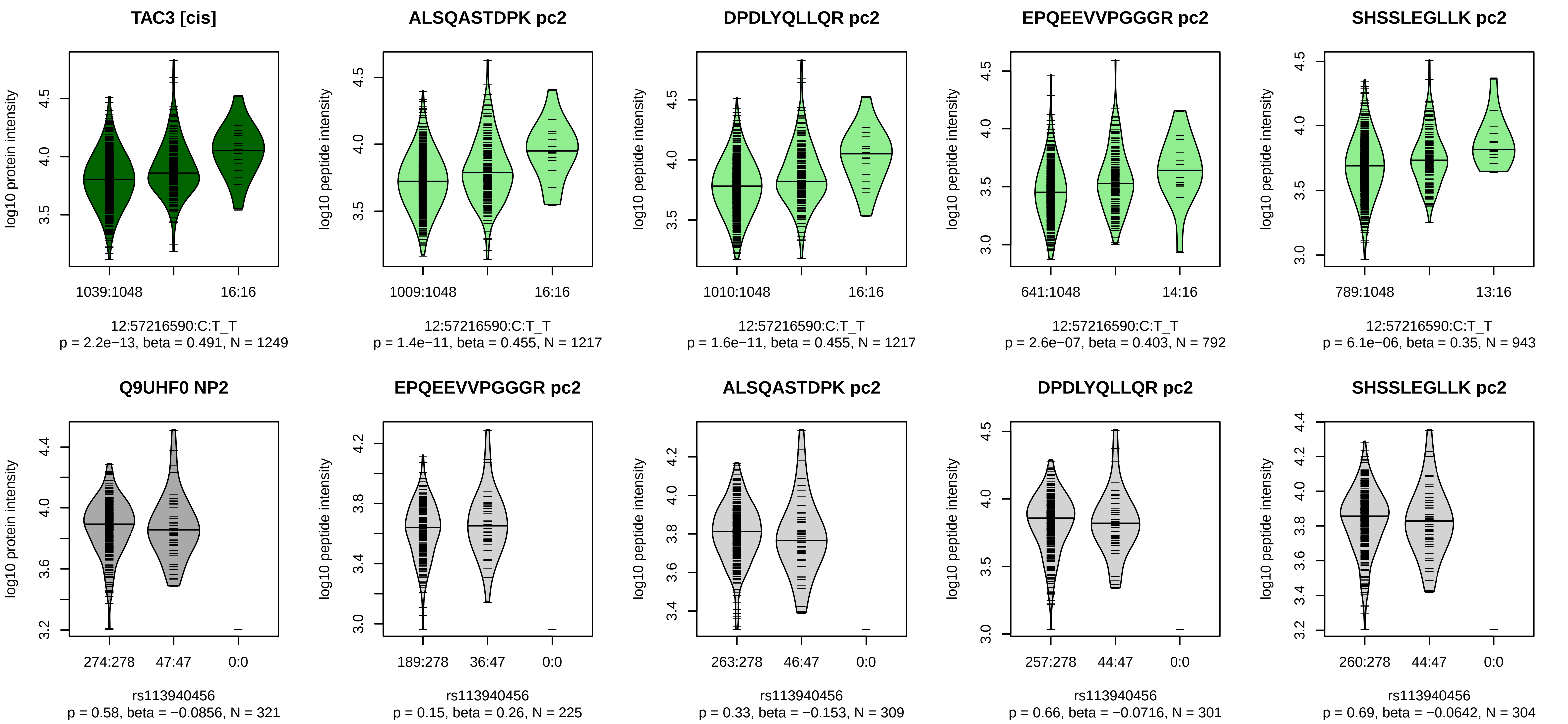


Locus: 167 (Sort: 193)
Status: Not replicated
80% power: No
SNP: 12:57216590:C:T
rsID: rs113940456
UniProtID: Q9UHF0
HGNC gene: TAC3
Type: Cis−pQTL

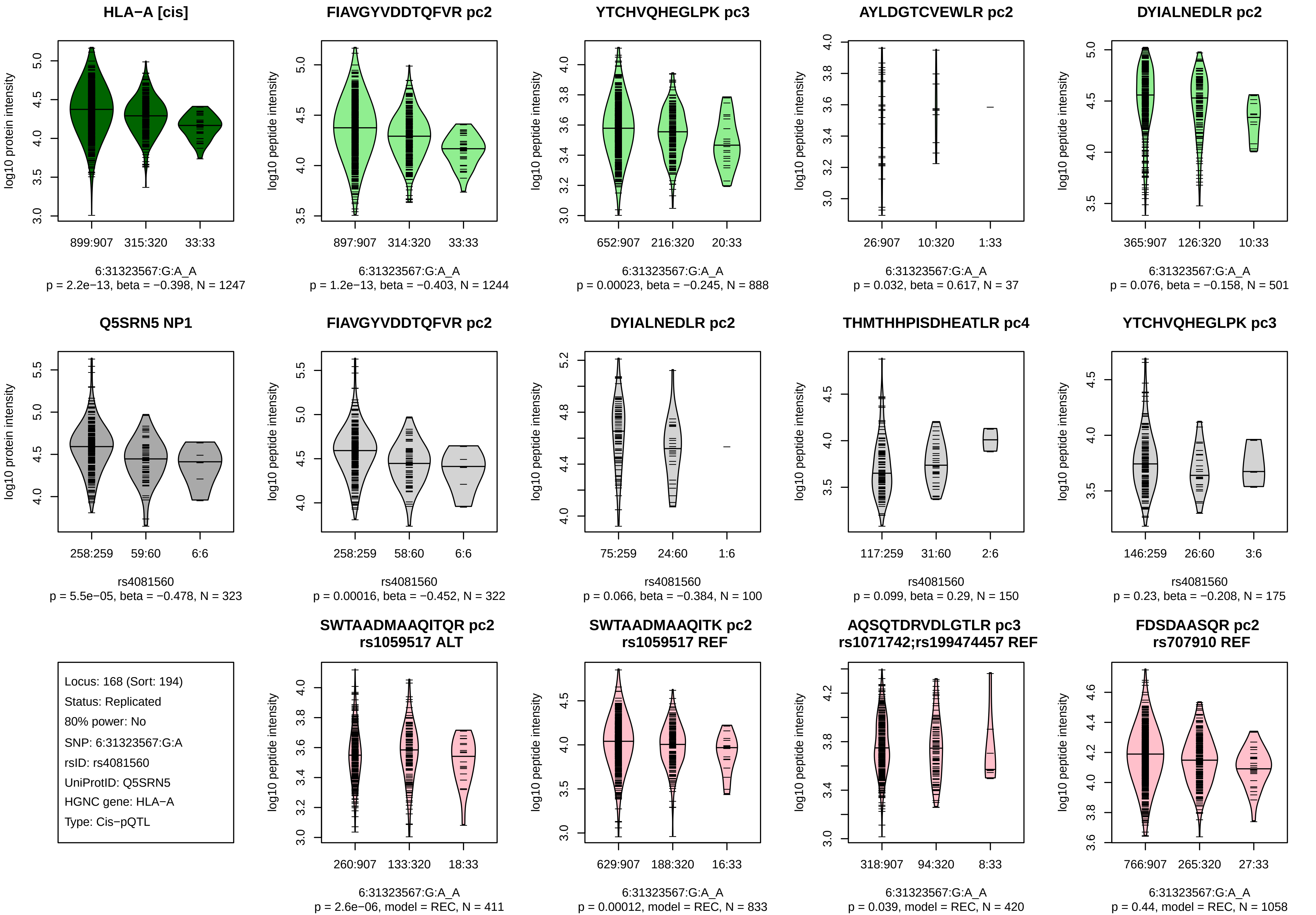

HLA−A [cis]
log10 protein intensity
899:907
315:320
33:33
6:31323567:G:A_A
p = 2.2e−13, beta = −0.398, N = 1247
FIAVGYVDDTQFVR pc2
log10 peptide intensity
897:907
314:320
33:33
6:31323567:G:A_A
p = 1.2e−13, beta = −0.403, N = 1244
YTCHVQHEGLPK pc3
652:907
216:320
20:33
6:31323567:G:A_A
p = 0.00023, beta = −0.245, N = 888
AYLDGTCVEWLR pc2
26:907
10:320
1:33
6:31323567:G:A_A
p = 0.032, beta = 0.617, N = 37
DYIALNEDLR pc2
365:907
126:320
10:33
6:31323567:G:A_A
p = 0.076, beta = −0.158, N = 501
Q5SRN5 NP1
258:259
59:60
6:6
rs4081560
p = 5.5e−05, beta = −0.478, N = 323
FIAVGYVDDTQFVR pc2
258:259
58:60
6:6
rs4081560
p = 0.00016, beta = −0.452, N = 322
DYIALNEDLR pc2
75:259
24:60
1:6
rs4081560
p = 0.066, beta = −0.384, N = 100
THMTHHPISDHEATLR pc4
117:259
31:60
2:6
rs4081560
p = 0.099, beta = 0.29, N = 150
YTCHVQHEGLPK pc3
146:259
26:60
3:6
rs4081560
p = 0.23, beta = −0.208, N = 175
Locus: 168 (Sort: 194)
Status: Replicated
80% power: No
SNP: 6:31323567:G:A
rsID: rs4081560
UniProtID: Q5SRN5
HGNC gene: HLA−A
Type: Cis−pQTL
SWTAADMAAQITQR pc2
rs1059517 ALT
260:907
133:320
18:33
6:31323567:G:A_A
p = 2.6e−06, model = REC, N = 411
SWTAADMAAQITK pc2
rs1059517 REF
629:907
188:320
16:33
6:31323567:G:A_A
p = 0.00012, model = REC, N = 833
AQSQTDRVDLGTLR pc3
rs1071742;rs199474457 REF
318:907
94:320
8:33
6:31323567:G:A_A
p = 0.039, model = REC, N = 420
FDSDAASQR pc2
rs707910 REF
766:907
265:320
27:33
6:31323567:G:A_A
p = 0.44, model = REC, N = 1058

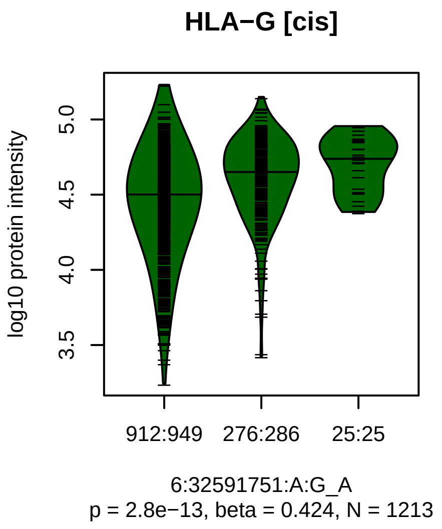


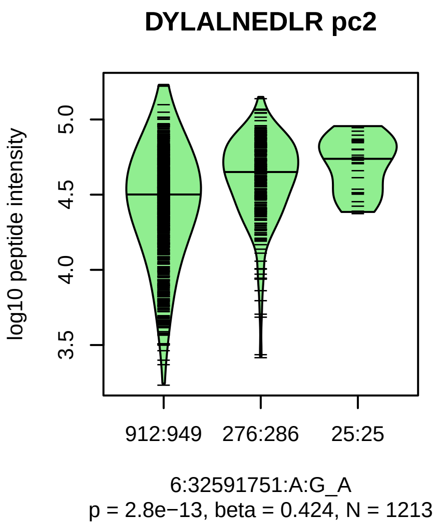


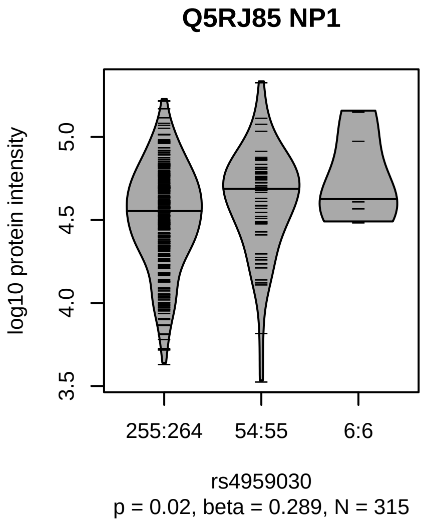


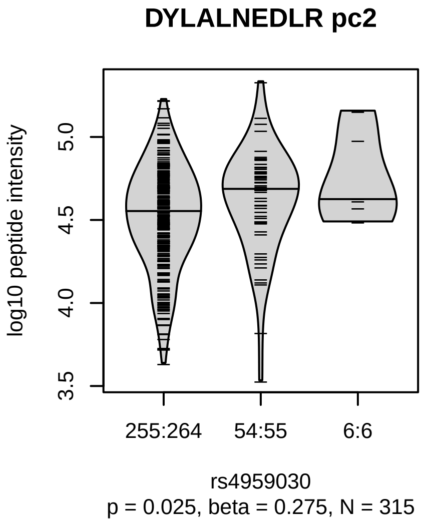


Locus: 169 (Sort: 195)

Status: Not replicated

80% power: No

SNP: 6:32591751:A:G

rsID: rs4959030

UniProtID: Q5RJ85

HGNC gene: HLA−G

Type: Cis−pQTL

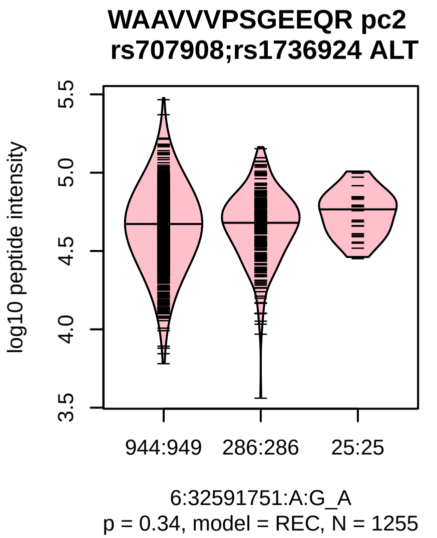

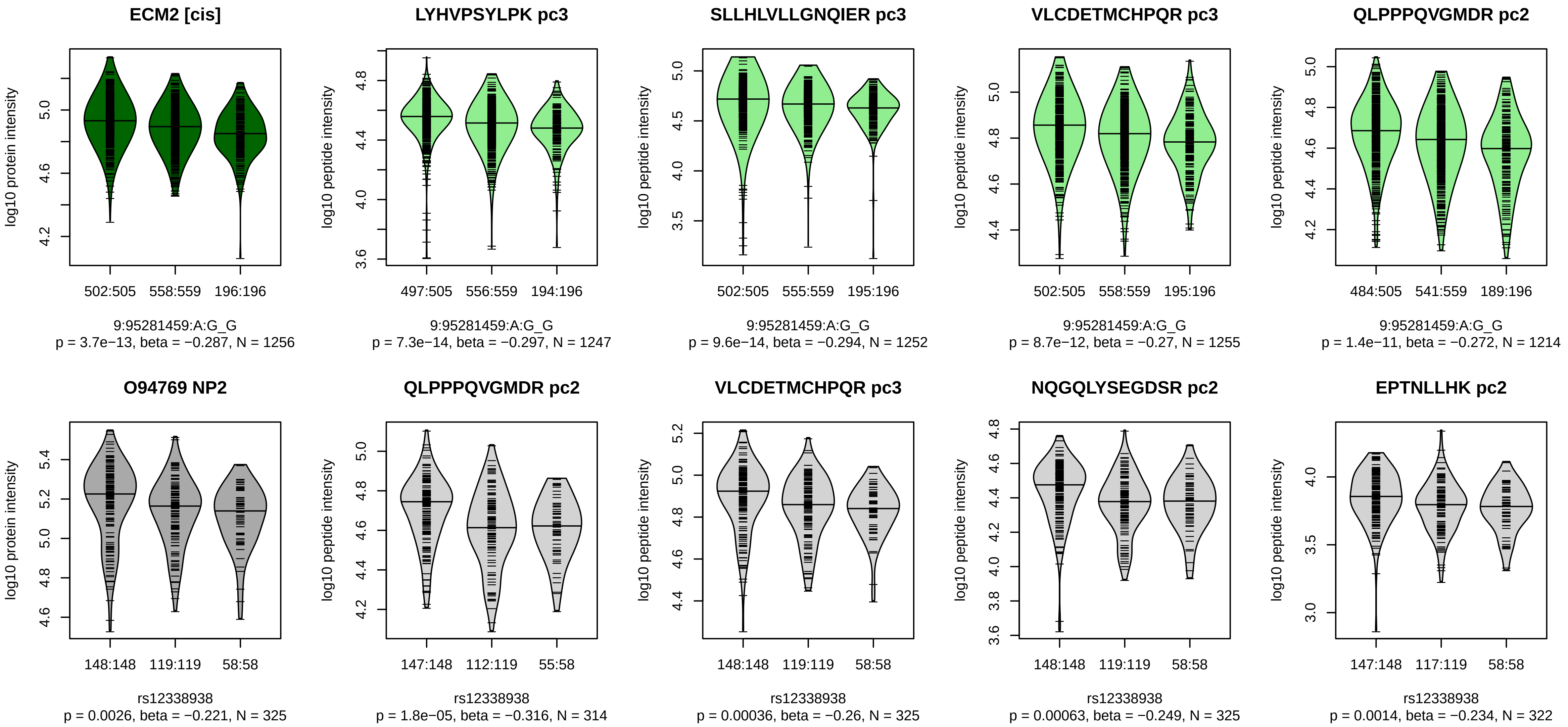


Locus: 170 (Sort: 196)
Status: Not replicated
80% power: No
SNP: 9:95281459:A:G
rsID: rs12338938
UniProtID: O94769
HGNC gene: ECM2
Type: Cis−pQTL

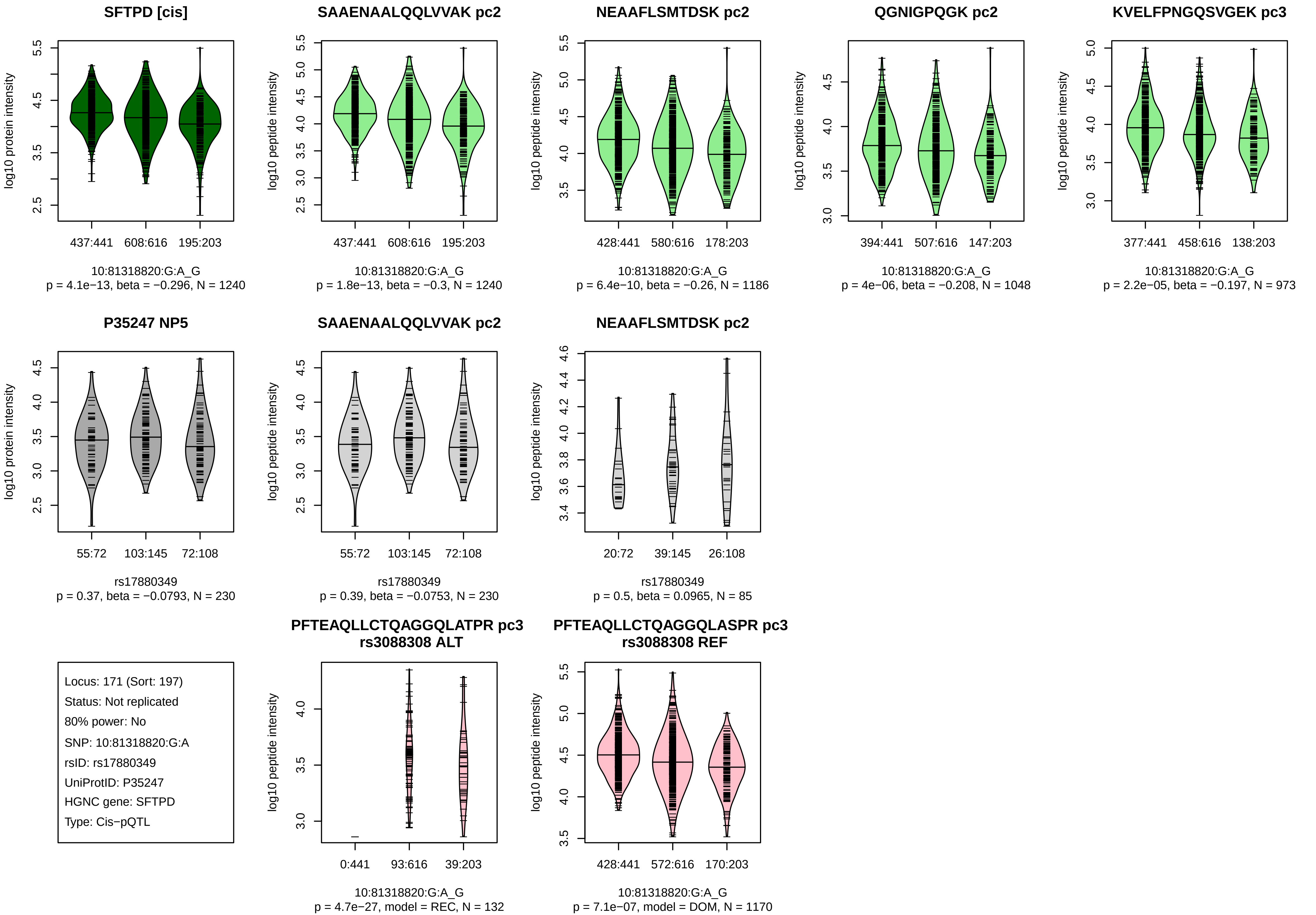

SFTPD [cis]
log10 protein intensity
437:441
608:616
195:203
10:81318820:G:A_G
p = 4.1e−13, beta = −0.296, N = 1240
SAAENAALQQLVVAK pc2
log10 peptide intensity
437:441
608:616
195:203
10:81318820:G:A_G
p = 1.8e−13, beta = −0.3, N = 1240
NEAAFLSMTDSK pc2
log10 peptide intensity
428:441
580:616
178:203
10:81318820:G:A_G
p = 6.4e−10, beta = −0.26, N = 1186
QGNIGPQGK pc2
log10 peptide intensity
394:441
507:616
147:203
10:81318820:G:A_G
p = 4e−06, beta = −0.208, N = 1048
KVELFPNGQSVGEK pc3
log10 peptide intensity
377:441
458:616
138:203
10:81318820:G:A_G
p = 2.2e−05, beta = −0.197, N = 973
P35247 NP5
log10 protein intensity
55:72
103:145
72:108
rs17880349
p = 0.37, beta = −0.0793, N = 230
SAAENAALQQLVVAK pc2
log10 peptide intensity
55:72
103:145
72:108
rs17880349
p = 0.39, beta = −0.0753, N = 230
NEAAFLSMTDSK pc2
log10 peptide intensity
20:72
39:145
26:108
rs17880349
p = 0.5, beta = 0.0965, N = 85
Locus: 171 (Sort: 197)
Status: Not replicated
80% power: No
SNP: 10:81318820:G:A
rsID: rs17880349
UniProtID: P35247
HGNC gene: SFTPD
Type: Cis−pQTL
PFTEAQLLCTQAGGQLATPR pc3
rs3088308 ALT
log10 peptide intensity
0:441
93:616
39:203
10:81318820:G:A_G
p = 4.7e−27, model = REC, N = 132
PFTEAQLLCTQAGGQLASPR pc3
rs3088308 REF
log10 peptide intensity
428:441
572:616
170:203
10:81318820:G:A_G
p = 7.1e−07, model = DOM, N = 1170

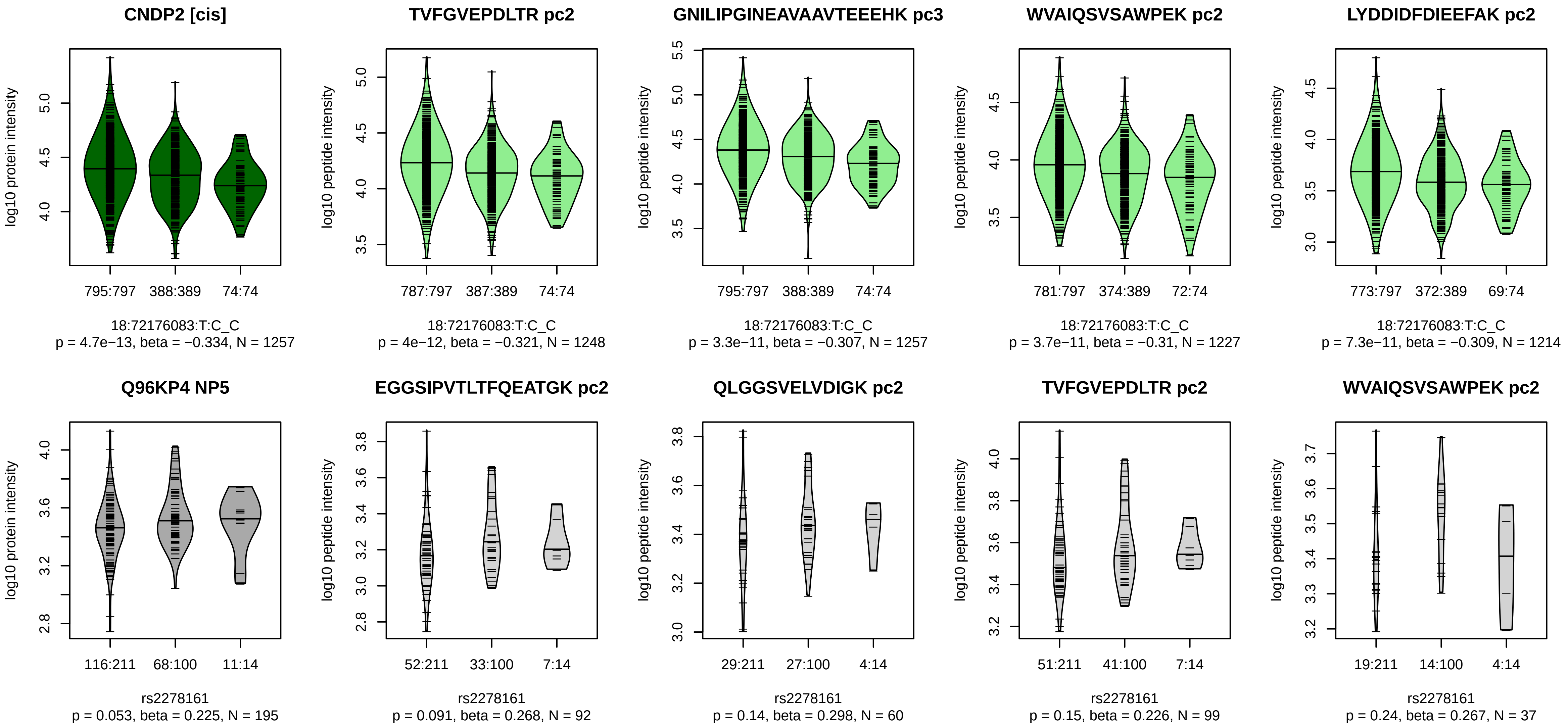


Locus: 172 (Sort: 198)

Status: Not replicated

80% power: No

SNP: 18:72176083:T:C

rsID: rs2278161

UniProtID: Q96KP4

HGNC gene: CNDP2

Type: Cis−pQTL

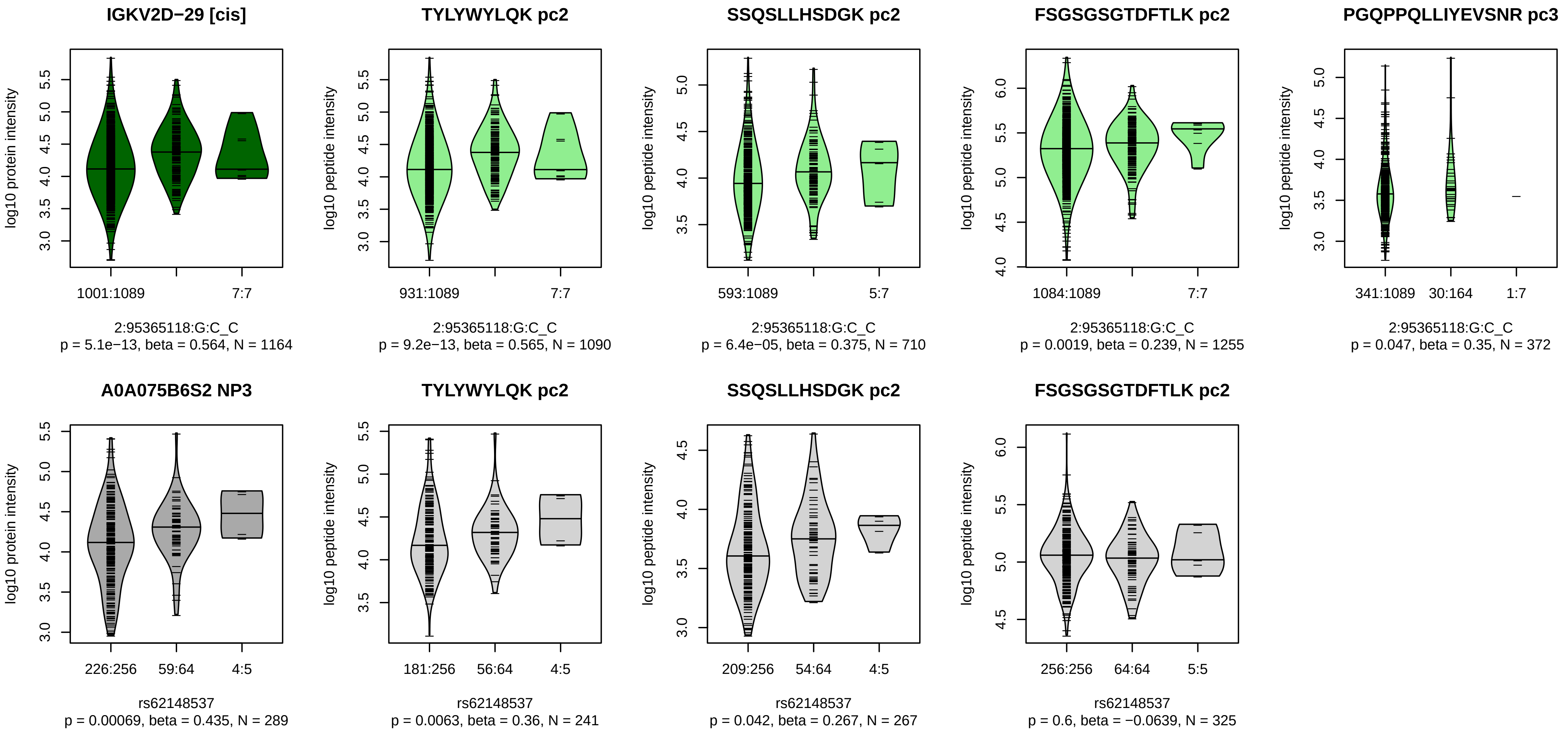


Locus: 173 (Sort: 199)
Status: Not replicated
80% power: No
SNP: 2:95365118:G:C
rsID: rs62148537
UniProtID: A0A075B6S2
HGNC gene: IGKV2D−29
Type: Cis−pQTL

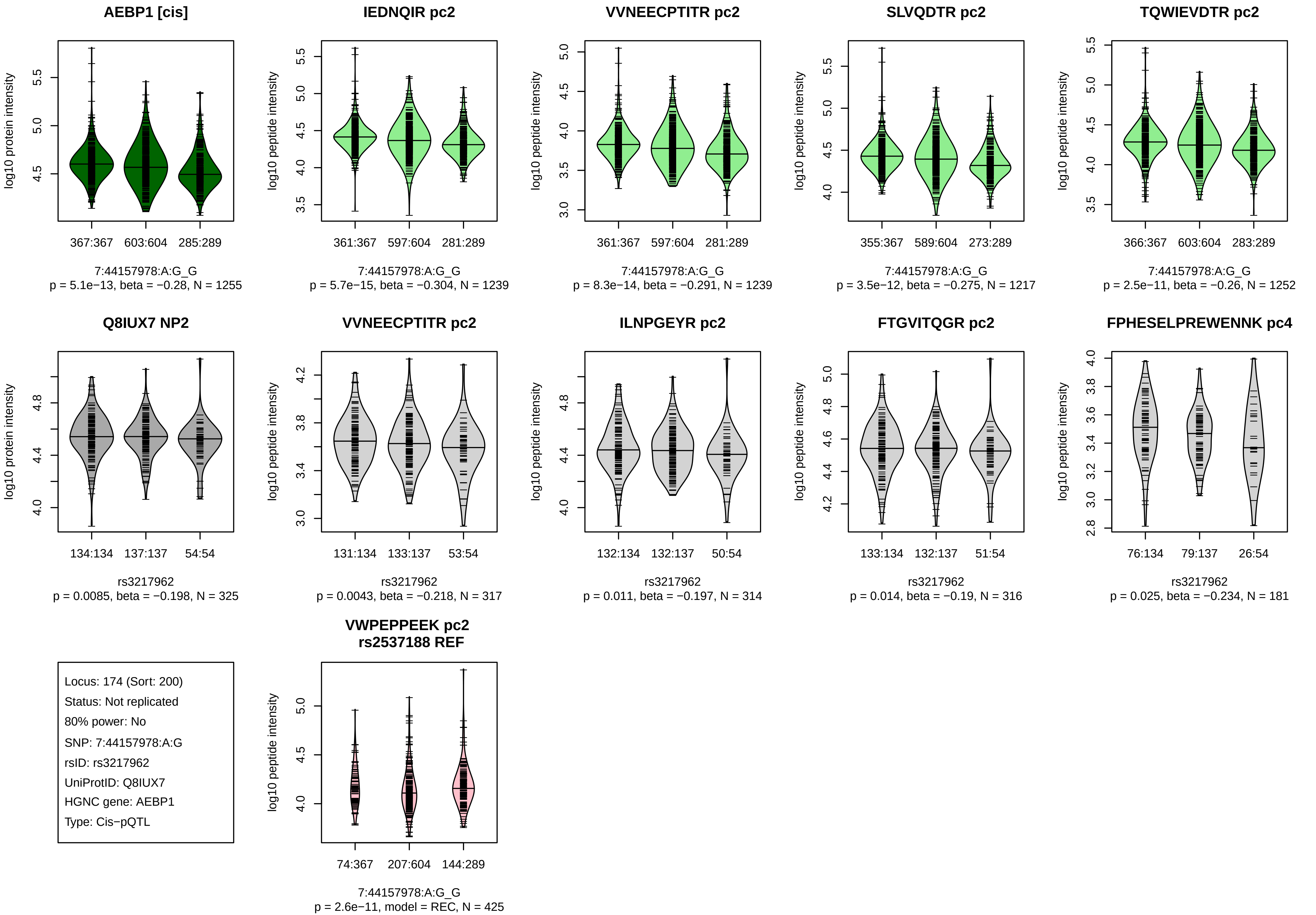
AEBP1 [cis]
log10 protein intensity
367:367 603:604 285:289
7:44157978:A:G_G
p = 5.1e−13, beta = −0.28, N = 1255
IEDNQIR pc2
log10 peptide intensity
361:367 597:604 281:289
7:44157978:A:G_G
p = 5.7e−15, beta = −0.304, N = 1239
VVNEECPTITR pc2
log10 peptide intensity
361:367 597:604 281:289
7:44157978:A:G_G
p = 8.3e−14, beta = −0.291, N = 1239
SLVQDTR pc2
log10 peptide intensity
355:367 589:604 273:289
7:44157978:A:G_G
p = 3.5e−12, beta = −0.275, N = 1217
TQWIEVDTR pc2
log10 peptide intensity
366:367 603:604 283:289
7:44157978:A:G_G
p = 2.5e−11, beta = −0.26, N = 1252
Q8IUX7 NP2
log10 protein intensity
134:134 137:137 54:54
rs3217962
p = 0.0085, beta = −0.198, N = 325
VVNEECPTITR pc2
log10 peptide intensity
131:134 133:137 53:54
rs3217962
p = 0.0043, beta = −0.218, N = 317
ILNPGEYR pc2
log10 peptide intensity
132:134 132:137 50:54
rs3217962
p = 0.011, beta = −0.197, N = 314
FTGVITQGR pc2
log10 peptide intensity
133:134 132:137 51:54
rs3217962
p = 0.014, beta = −0.19, N = 316
FPHESELPREWENNK pc4
log10 peptide intensity
76:134 79:137 26:54
rs3217962
p = 0.025, beta = −0.234, N = 181
Locus: 174 (Sort: 200)
Status: Not replicated
80% power: No
SNP: 7:44157978:A:G
rsID: rs3217962
UniProtID: Q8IUX7
HGNC gene: AEBP1
Type: Cis−pQTL
VWPEPPEEK pc2
rs2537188 REF
log10 peptide intensity
74:367 207:604 144:289
7:44157978:A:G_G
p = 2.6e−11, model = REC, N = 425

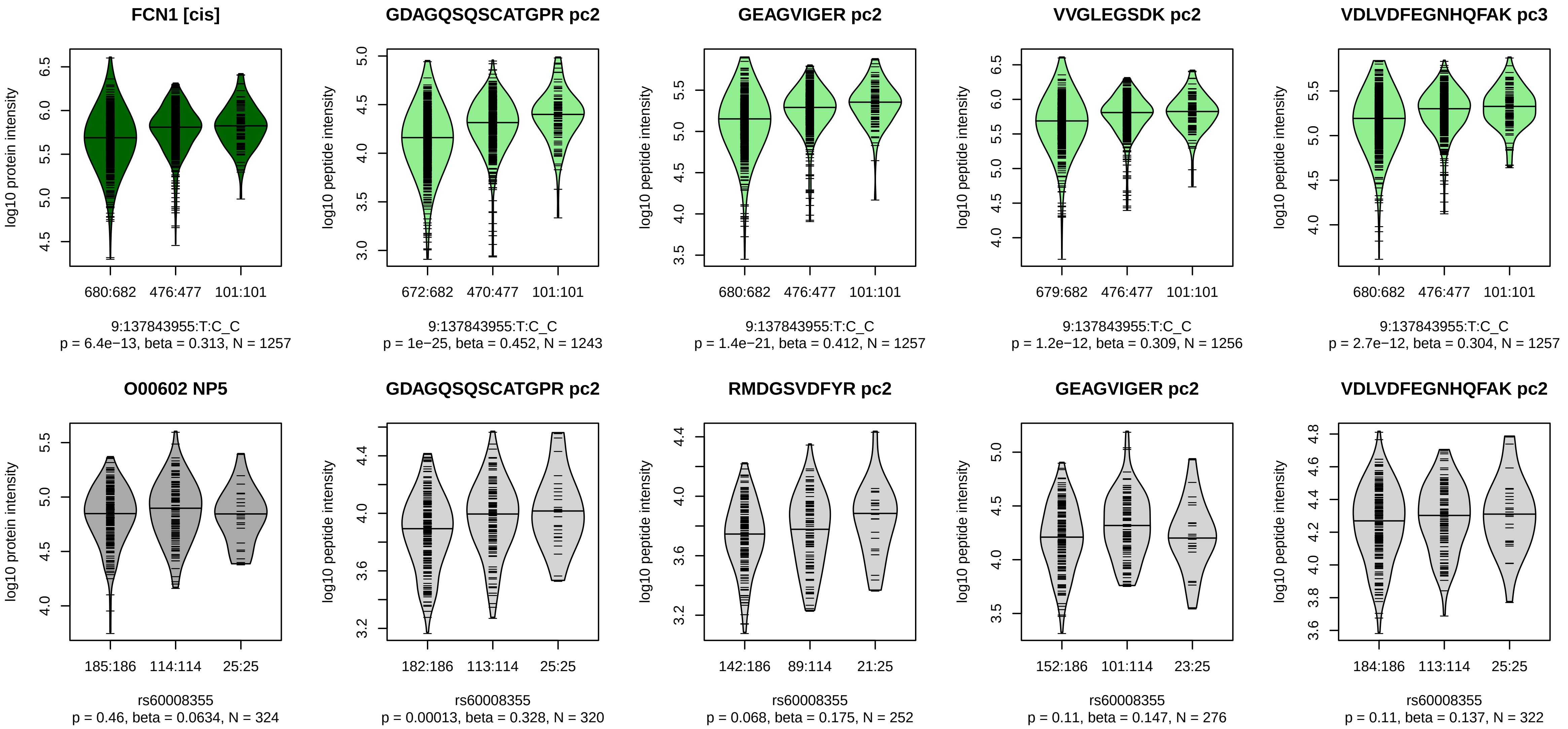


Locus: 175 (Sort: 201)
Status: Not replicated
80% power: No
SNP: 9:137843955:T:C
rsID: rs60008355
UniProtID: O00602
HGNC gene: FCN1
Type: Cis−pQTL

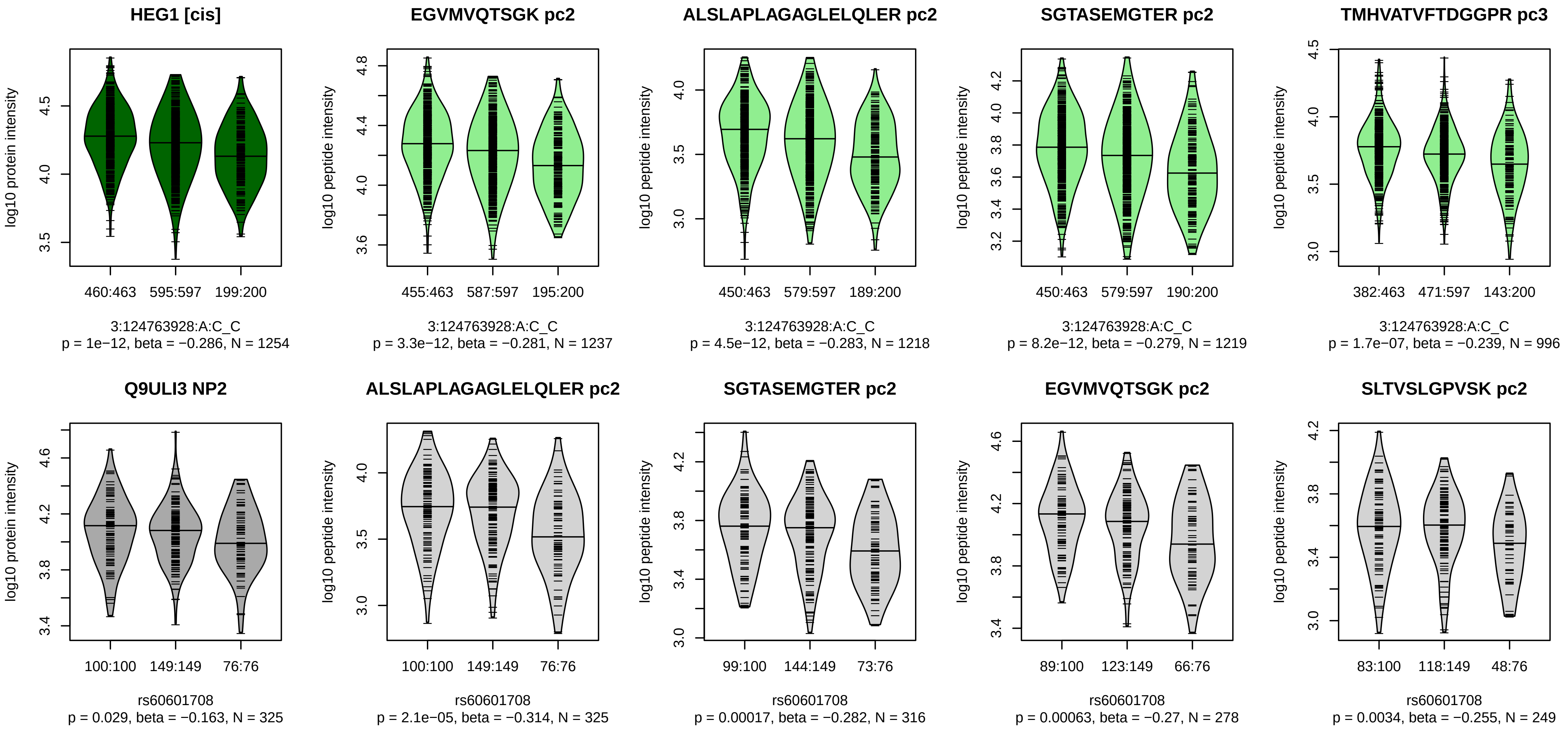


Locus: 176 (Sort: 202)
Status: Not replicated
80% power: No
SNP: 3:124763928:A:C
rsID: rs60601708
UniProtID: Q9ULI3
HGNC gene: HEG1
Type: Cis−pQTL

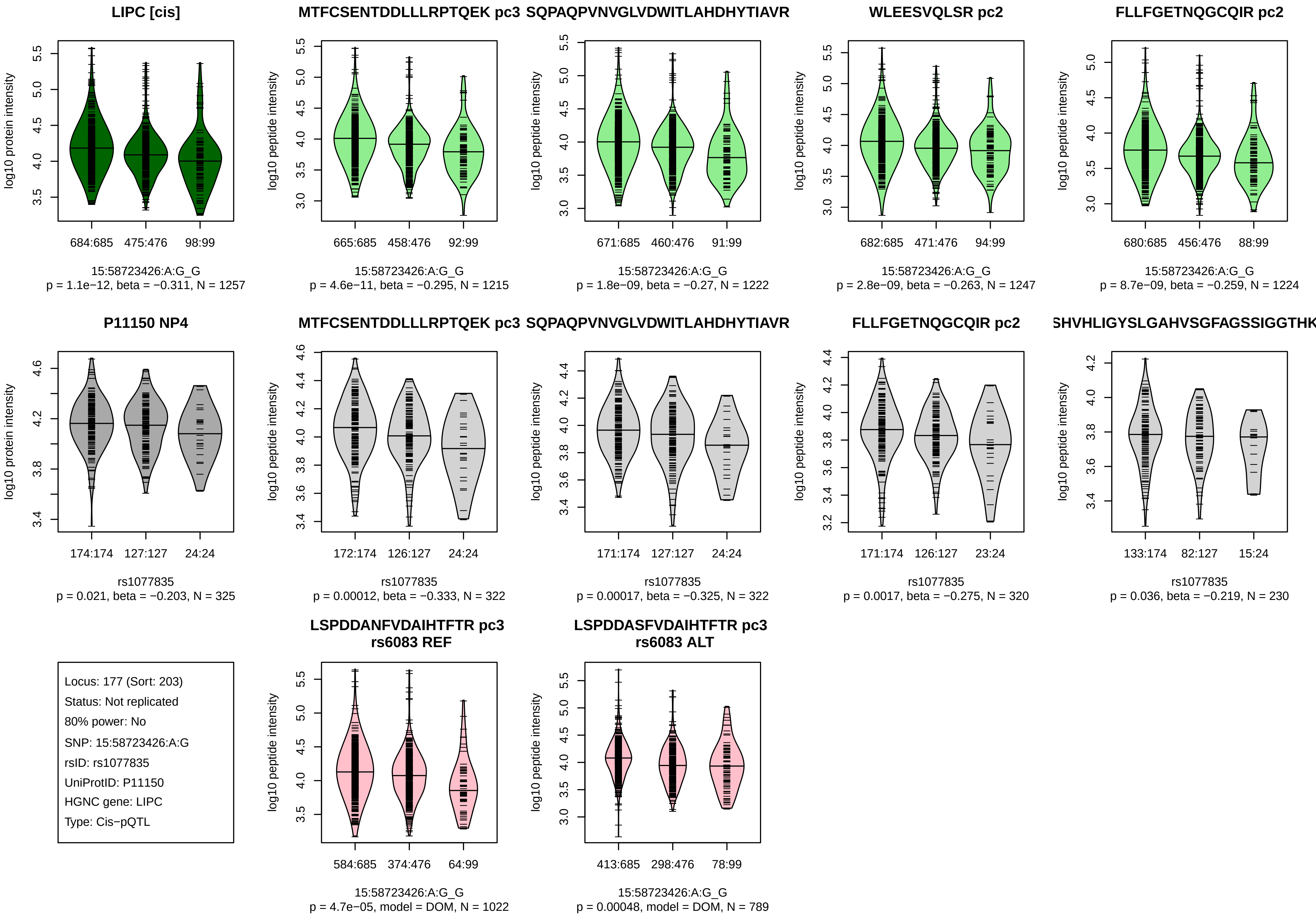
LIPC [cis]
log10 protein intensity
684:685
475:476
98:99
15:58723426:A:G_G
p = 1.1e−12, beta = −0.311, N = 1257
MTFCSENTDDLLLRPTQEK pc3
log10 peptide intensity
665:685
458:476
92:99
15:58723426:A:G_G
p = 4.6e−11, beta = −0.295, N = 1215
SQPAQPVNVGLVDWITLAHDHYTIAVR
log10 peptide intensity
671:685
460:476
91:99
15:58723426:A:G_G
p = 1.8e−09, beta = −0.27, N = 1222
WLEESVQLSR pc2
log10 peptide intensity
682:685
471:476
94:99
15:58723426:A:G_G
p = 2.8e−09, beta = −0.263, N = 1247
FLLFGETNQGCQIR pc2
log10 peptide intensity
680:685
456:476
88:99
15:58723426:A:G_G
p = 8.7e−09, beta = −0.259, N = 1224
P11150 NP4
log10 protein intensity
174:174
127:127
24:24
rs1077835
p = 0.021, beta = −0.203, N = 325
MTFCSENTDDLLLRPTQEK pc3
log10 peptide intensity
172:174
126:127
24:24
rs1077835
p = 0.00012, beta = −0.333, N = 322
SQPAQPVNVGLVDWITLAHDHYTIAVR
log10 peptide intensity
171:174
127:127
24:24
rs1077835
p = 0.00017, beta = −0.325, N = 322
FLLFGETNQGCQIR pc2
log10 peptide intensity
171:174
126:127
23:24
rs1077835
p = 0.0017, beta = −0.275, N = 320
SHVHLIGYSLGAHVSGFAGSSIGGTHK
log10 peptide intensity
133:174
82:127
15:24
rs1077835
p = 0.036, beta = −0.219, N = 230
Locus: 177 (Sort: 203)
Status: Not replicated
80% power: No
SNP: 15:58723426:A:G
rsID: rs1077835
UniProtID: P11150
HGNC gene: LIPC
Type: Cis−pQTL
LSPDDANFVDAIHTFTR pc3
rs6083 REF
log10 peptide intensity
584:685
374:476
64:99
15:58723426:A:G_G
p = 4.7e−05, model = DOM, N = 1022
LSPDDASFVDAIHTFTR pc3
rs6083 ALT
log10 peptide intensity
413:685
298:476
78:99
15:58723426:A:G_G
p = 0.00048, model = DOM, N = 789

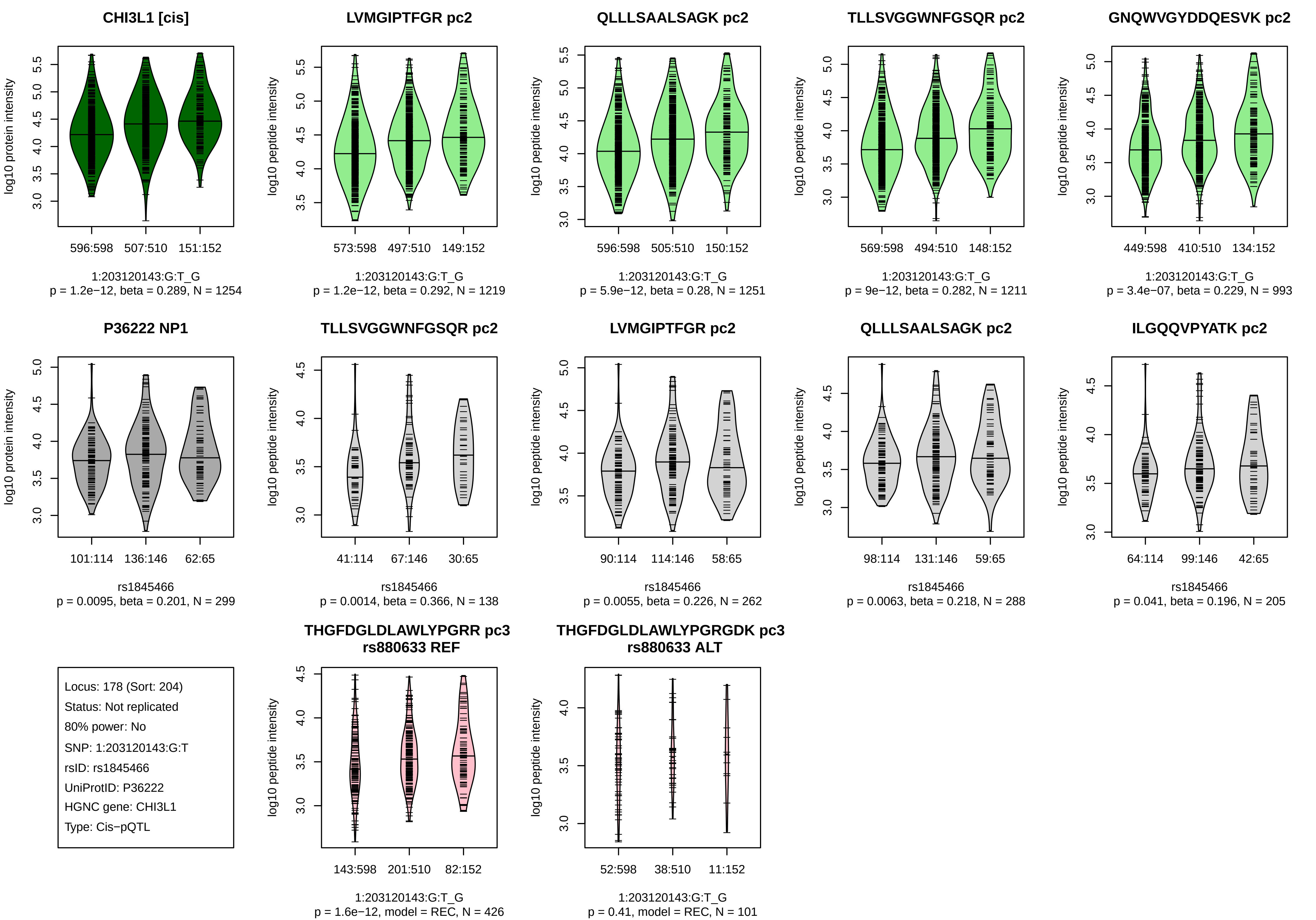
CHI3L1 [cis]
log10 protein intensity
596:598
507:510
151:152
1:203120143:G:T_G
p = 1.2e−12, beta = 0.289, N = 1254
LVMGIPTFGR pc2
log10 peptide intensity
573:598
497:510
149:152
1:203120143:G:T_G
p = 1.2e−12, beta = 0.292, N = 1219
QLLLSAALSAGK pc2
log10 peptide intensity
596:598
505:510
150:152
1:203120143:G:T_G
p = 5.9e−12, beta = 0.28, N = 1251
TLLSVGGWNFGSQR pc2
log10 peptide intensity
569:598
494:510
148:152
1:203120143:G:T_G
p = 9e−12, beta = 0.282, N = 1211
GNQWVGYDDQESVK pc2
log10 peptide intensity
449:598
410:510
134:152
1:203120143:G:T_G
p = 3.4e−07, beta = 0.229, N = 993
P36222 NP1
log10 protein intensity
101:114
136:146
62:65
rs1845466
p = 0.0095, beta = 0.201, N = 299
TLLSVGGWNFGSQR pc2
log10 peptide intensity
41:114
67:146
30:65
rs1845466
p = 0.0014, beta = 0.366, N = 138
LVMGIPTFGR pc2
log10 peptide intensity
90:114
114:146
58:65
rs1845466
p = 0.0055, beta = 0.226, N = 262
QLLLSAALSAGK pc2
log10 peptide intensity
98:114
131:146
59:65
rs1845466
p = 0.0063, beta = 0.218, N = 288
ILGQQVPYATK pc2
log10 peptide intensity
64:114
99:146
42:65
rs1845466
p = 0.041, beta = 0.196, N = 205
Locus: 178 (Sort: 204)
Status: Not replicated
80% power: No
SNP: 1:203120143:G:T
rsID: rs1845466
UniProtID: P36222
HGNC gene: CHI3L1
Type: Cis−pQTL
THGFDGLDLAWLYPGRR pc3
rs880633 REF
log10 peptide intensity
143:598
201:510
82:152
1:203120143:G:T_G
p = 1.6e−12, model = REC, N = 426
THGFDGLDLAWLYPGRGDK pc3
rs880633 ALT
log10 peptide intensity
52:598
38:510
11:152
1:203120143:G:T_G
p = 0.41, model = REC, N = 101

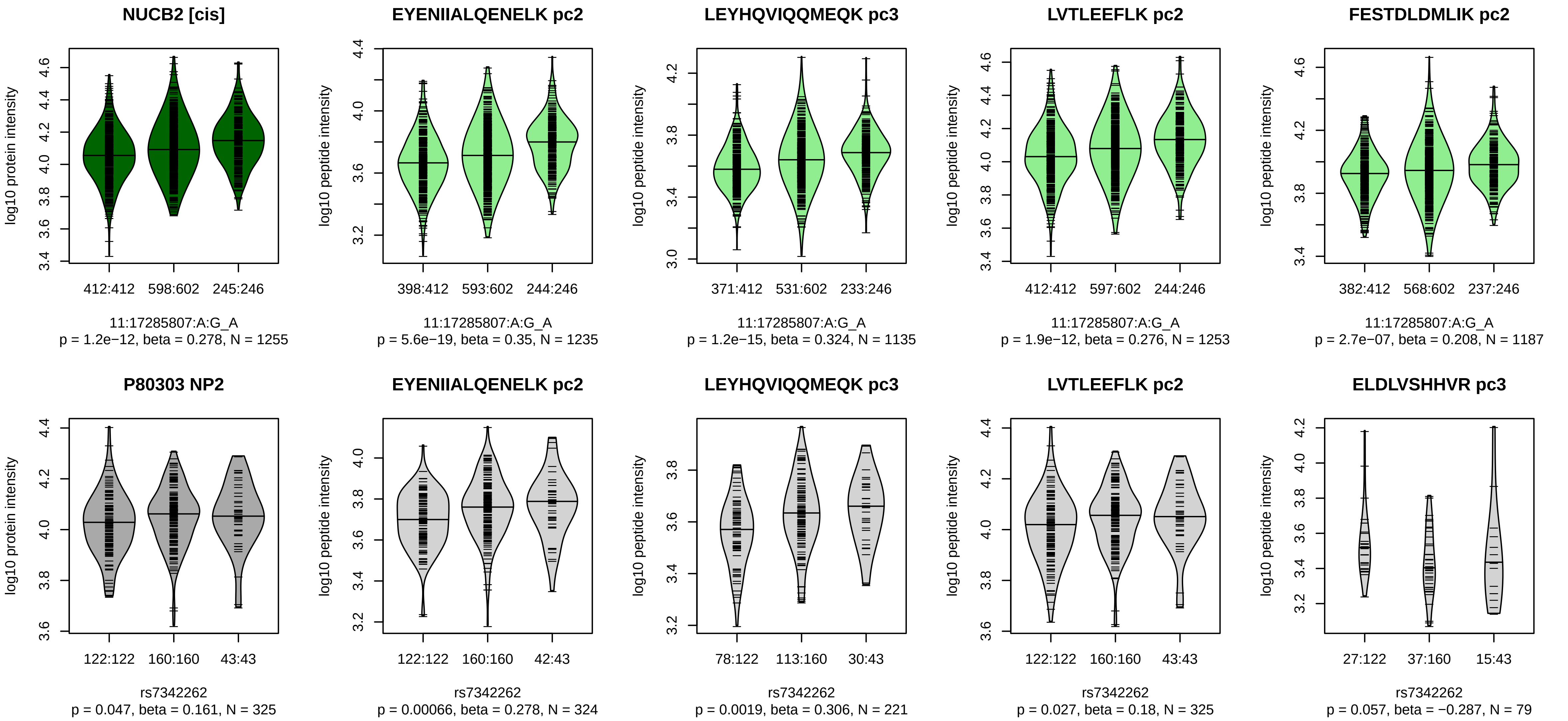


Locus: 179 (Sort: 205)
Status: Not replicated
80% power: No
SNP: 11:17285807:A:G
rsID: rs7342262
UniProtID: P80303
HGNC gene: NUCB2
Type: Cis−pQTL

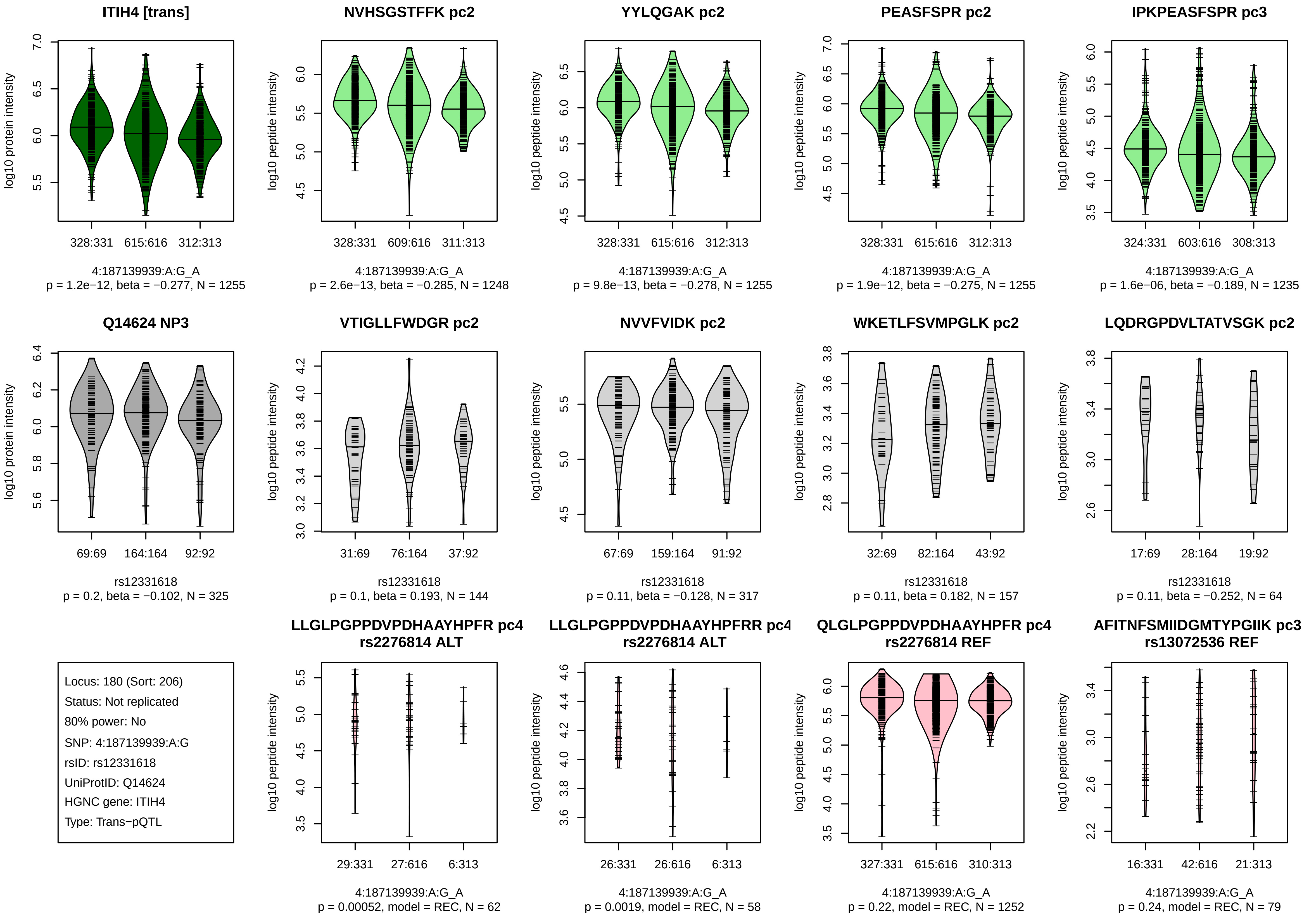

ITIH4 [trans]
log10 protein intensity
328:331
615:616
312:313
4:187139939:A:G_A
p = 1.2e−12, beta = −0.277, N = 1255
NVHSGSTFFK pc2
log10 peptide intensity
328:331
609:616
311:313
4:187139939:A:G_A
p = 2.6e−13, beta = −0.285, N = 1248
YYLQGAK pc2
log10 peptide intensity
328:331
615:616
312:313
4:187139939:A:G_A
p = 9.8e−13, beta = −0.278, N = 1255
PEASFSPR pc2
log10 peptide intensity
328:331
615:616
312:313
4:187139939:A:G_A
p = 1.9e−12, beta = −0.275, N = 1255
IPKPEASFSPR pc3
log10 peptide intensity
324:331
603:616
308:313
4:187139939:A:G_A
p = 1.6e−06, beta = −0.189, N = 1235
Q14624 NP3
log10 protein intensity
69:69
164:164
92:92
rs12331618
p = 0.2, beta = −0.102, N = 325
VTIGLLFWDGR pc2
log10 peptide intensity
31:69
76:164
37:92
rs12331618
p = 0.1, beta = 0.193, N = 144
NVVFVIDK pc2
log10 peptide intensity
67:69
159:164
91:92
rs12331618
p = 0.11, beta = −0.128, N = 317
WKETLFSVMPGLK pc2
log10 peptide intensity
32:69
82:164
43:92
rs12331618
p = 0.11, beta = 0.182, N = 157
LQDRGPDVLTATVSGK pc2
log10 peptide intensity
17:69
28:164
19:92
rs12331618
p = 0.11, beta = −0.252, N = 64
Locus: 180 (Sort: 206)
Status: Not replicated
80% power: No
SNP: 4:187139939:A:G
rsID: rs12331618
UniProtID: Q14624
HGNC gene: ITIH4
Type: Trans−pQTL
LLGLPGPPDVPDHAAYHPFR pc4
rs2276814 ALT
log10 peptide intensity
29:331
27:616
6:313
4:187139939:A:G_A
p = 0.00052, model = REC, N = 62
LLGLPGPPDVPDHAAYHPFRR pc4
rs2276814 ALT
log10 peptide intensity
26:331
26:616
6:313
4:187139939:A:G_A
p = 0.0019, model = REC, N = 58
QLGLPGPPDVPDHAAYHPFR pc4
rs2276814 REF
log10 peptide intensity
327:331
615:616
310:313
4:187139939:A:G_A
p = 0.22, model = REC, N = 1252
AFITNFSMIIDGMTYPGIIK pc3
rs13072536 REF
log10 peptide intensity
16:331
42:616
21:313
4:187139939:A:G_A
p = 0.24, model = REC, N = 79

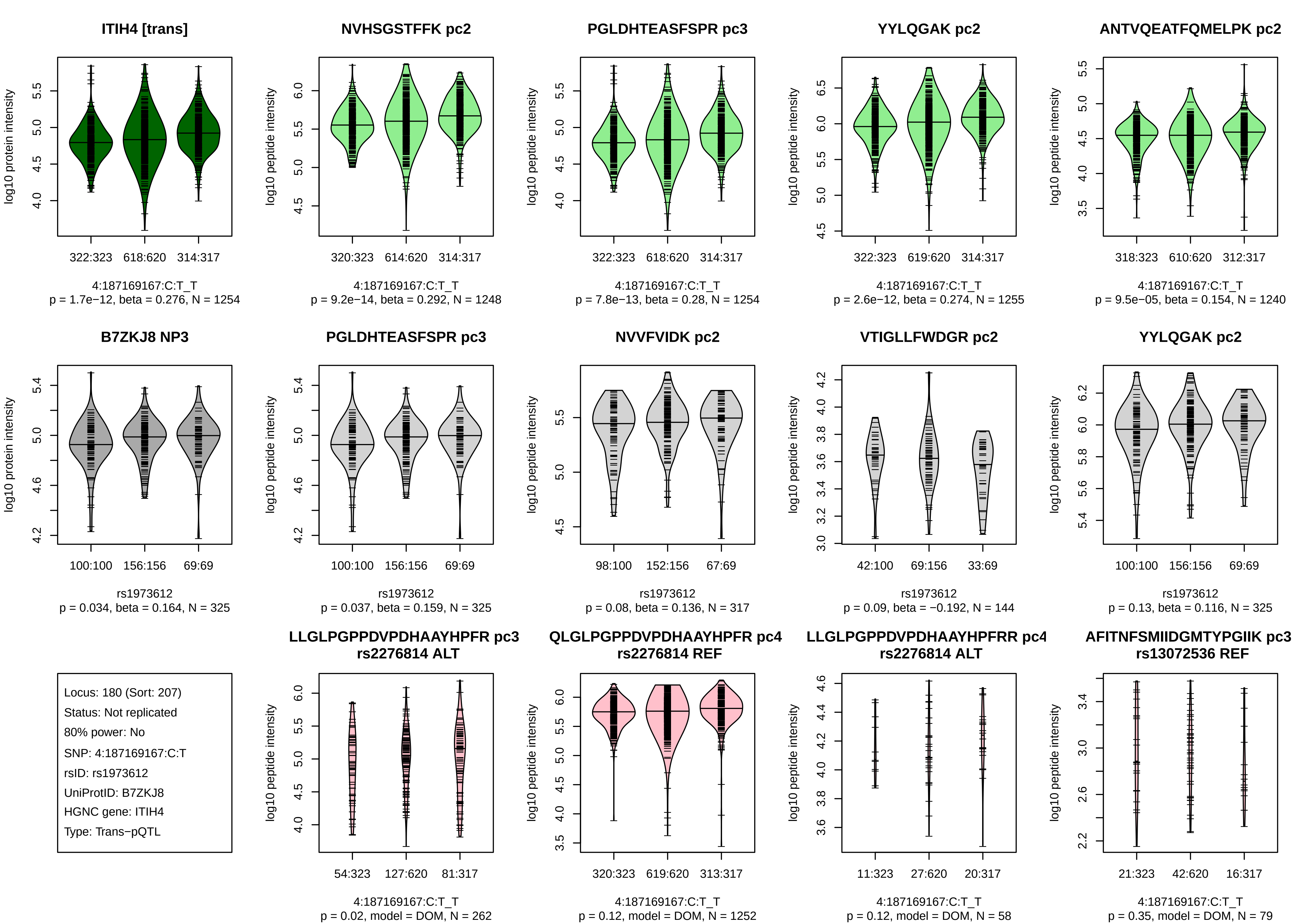

ITIH4 [trans]
log10 protein intensity
322:323
618:620
314:317
4:187169167:C:T_T
p = 1.7e−12, beta = 0.276, N = 1254
NVHSGSTFFK pc2
log10 peptide intensity
320:323
614:620
314:317
4:187169167:C:T_T
p = 9.2e−14, beta = 0.292, N = 1248
PGLDHTEASFSPR pc3
log10 peptide intensity
322:323
618:620
314:317
4:187169167:C:T_T
p = 7.8e−13, beta = 0.28, N = 1254
YYLQGAK pc2
log10 peptide intensity
322:323
619:620
314:317
4:187169167:C:T_T
p = 2.6e−12, beta = 0.274, N = 1255
ANTVQEATFQMELPK pc2
log10 peptide intensity
318:323
610:620
312:317
4:187169167:C:T_T
p = 9.5e−05, beta = 0.154, N = 1240
B7ZKJ8 NP3
log10 protein intensity
100:100
156:156
69:69
rs1973612
p = 0.034, beta = 0.164, N = 325
PGLDHTEASFSPR pc3
log10 peptide intensity
100:100
156:156
69:69
rs1973612
p = 0.037, beta = 0.159, N = 325
NVVFVIDK pc2
log10 peptide intensity
98:100
152:156
67:69
rs1973612
p = 0.08, beta = 0.136, N = 317
VTIGLLFWDGR pc2
log10 peptide intensity
42:100
69:156
33:69
rs1973612
p = 0.09, beta = −0.192, N = 144
YYLQGAK pc2
log10 peptide intensity
100:100
156:156
69:69
rs1973612
p = 0.13, beta = 0.116, N = 325
Locus: 180 (Sort: 207)
Status: Not replicated
80% power: No
SNP: 4:187169167:C:T
rsID: rs1973612
UniProtID: B7ZKJ8
HGNC gene: ITIH4
Type: Trans−pQTL
LLGLPGPPDVPDHAAYHPFR pc3
rs2276814 ALT
log10 peptide intensity
54:323
127:620
81:317
4:187169167:C:T_T
p = 0.02, model = DOM, N = 262
QLGLPGPPDVPDHAAYHPFR pc4
rs2276814 REF
log10 peptide intensity
320:323
619:620
313:317
4:187169167:C:T_T
p = 0.12, model = DOM, N = 1252
LLGLPGPPDVPDHAAYHPFRR pc4
rs2276814 ALT
log10 peptide intensity
11:323
27:620
20:317
4:187169167:C:T_T
p = 0.12, model = DOM, N = 58
AFITNFSMIIDGMTYPGIIK pc3
rs13072536 REF
log10 peptide intensity
21:323
42:620
16:317
4:187169167:C:T_T
p = 0.35, model = DOM, N = 79

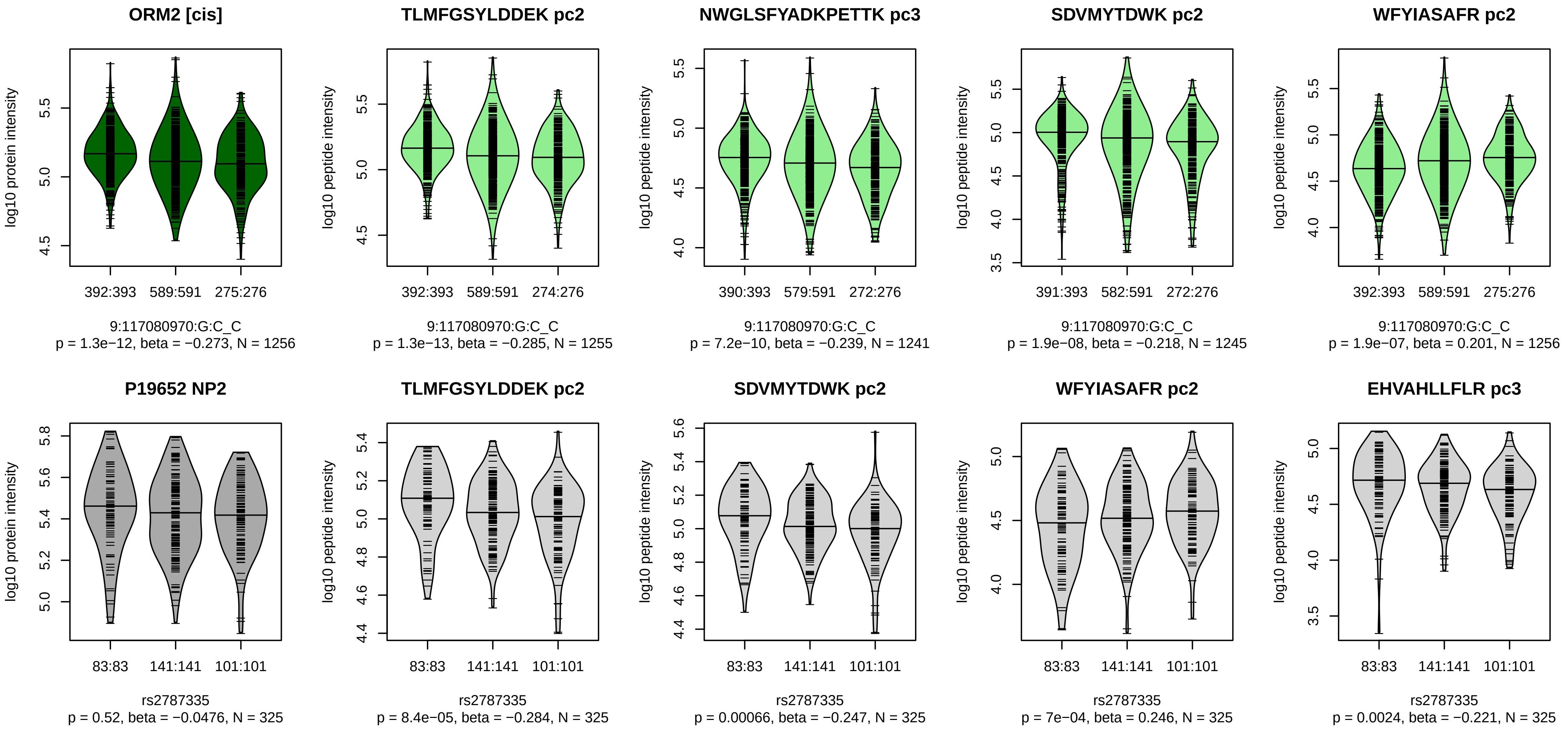


Locus: 181 (Sort: 208)

Status: Not replicated

80% power: No

SNP: 9:117080970:G:C

rsID: rs2787335

UniProtID: P19652

HGNC gene: ORM2

Type: Cis−pQTL

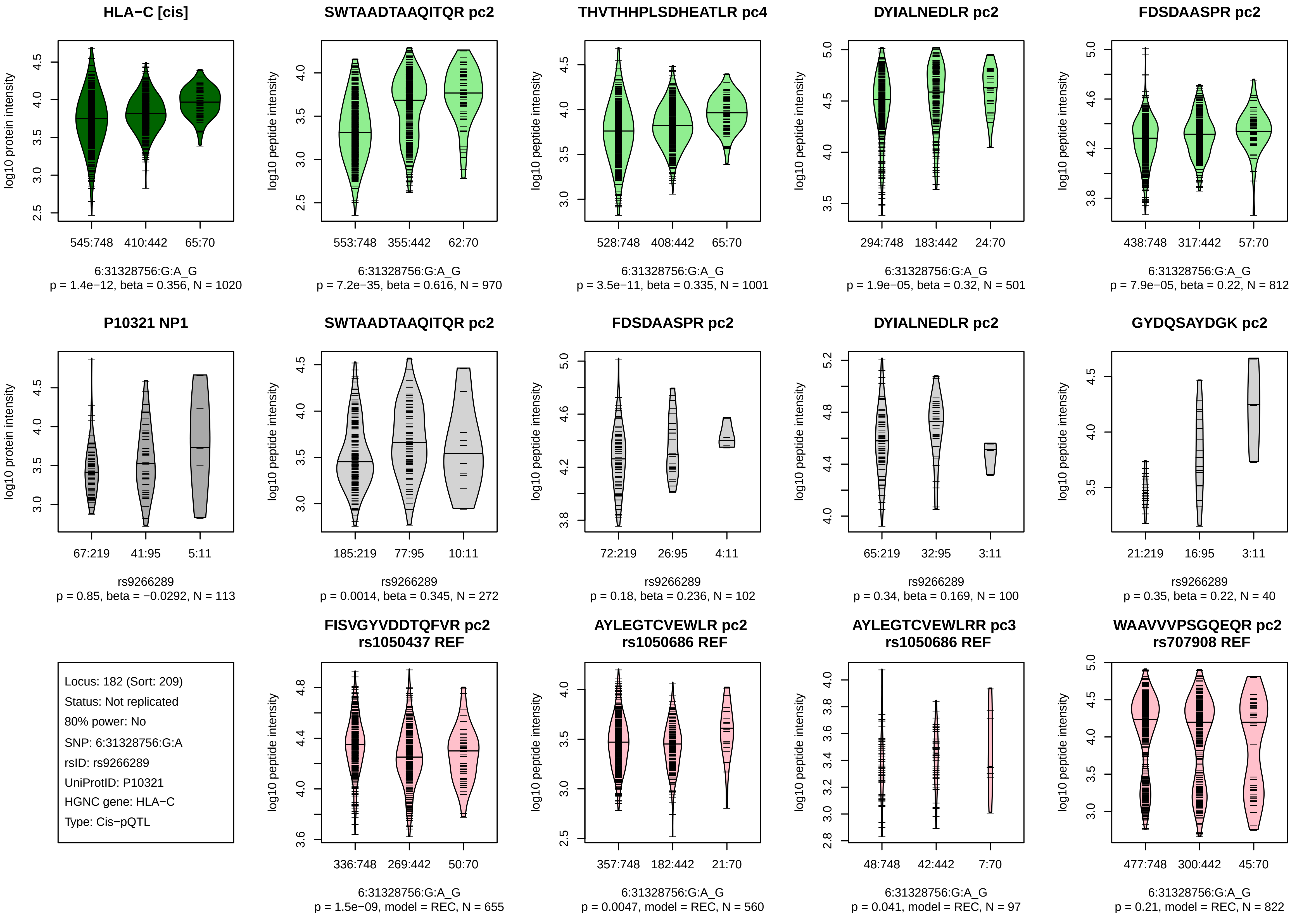
HLA−C [cis]
log10 protein intensity
545:748
410:442
65:70
6:31328756:G:A_G
p = 1.4e−12, beta = 0.356, N = 1020
SWTAADTAAQITQR pc2
log10 peptide intensity
553:748
355:442
62:70
6:31328756:G:A_G
p = 7.2e−35, beta = 0.616, N = 970
THVTHHPLSDHEATLR pc4
log10 peptide intensity
528:748
408:442
65:70
6:31328756:G:A_G
p = 3.5e−11, beta = 0.335, N = 1001
DYIALNEDLR pc2
log10 peptide intensity
294:748
183:442
24:70
6:31328756:G:A_G
p = 1.9e−05, beta = 0.32, N = 501
FDSDAASPR pc2
log10 peptide intensity
438:748
317:442
57:70
6:31328756:G:A_G
p = 7.9e−05, beta = 0.22, N = 812
P10321 NP1
log10 protein intensity
67:219
41:95
5:11
rs9266289
p = 0.85, beta = −0.0292, N = 113
SWTAADTAAQITQR pc2
log10 peptide intensity
185:219
77:95
10:11
rs9266289
p = 0.0014, beta = 0.345, N = 272
FDSDAASPR pc2
log10 peptide intensity
72:219
26:95
4:11
rs9266289
p = 0.18, beta = 0.236, N = 102
DYIALNEDLR pc2
log10 peptide intensity
65:219
32:95
3:11
rs9266289
p = 0.34, beta = 0.169, N = 100
GYDQSAYDGK pc2
log10 peptide intensity
21:219
16:95
3:11
rs9266289
p = 0.35, beta = 0.22, N = 40
Locus: 182 (Sort: 209)
Status: Not replicated
80% power: No
SNP: 6:31328756:G:A
rsID: rs9266289
UniProtID: P10321
HGNC gene: HLA−C
Type: Cis−pQTL
FISVGYVDDTQFVR pc2
rs1050437 REF
log10 peptide intensity
336:748
269:442
50:70
6:31328756:G:A_G
p = 1.5e−09, model = REC, N = 655
AYLEGTCVEWLR pc2
rs1050686 REF
log10 peptide intensity
357:748
182:442
21:70
6:31328756:G:A_G
p = 0.0047, model = REC, N = 560
AYLEGTCVEWLRR pc3
rs1050686 REF
log10 peptide intensity
48:748
42:442
7:70
6:31328756:G:A_G
p = 0.041, model = REC, N = 97
WAAVVVPSGQEQR pc2
rs707908 REF
log10 peptide intensity
477:748
300:442
45:70
6:31328756:G:A_G
p = 0.21, model = REC, N = 822

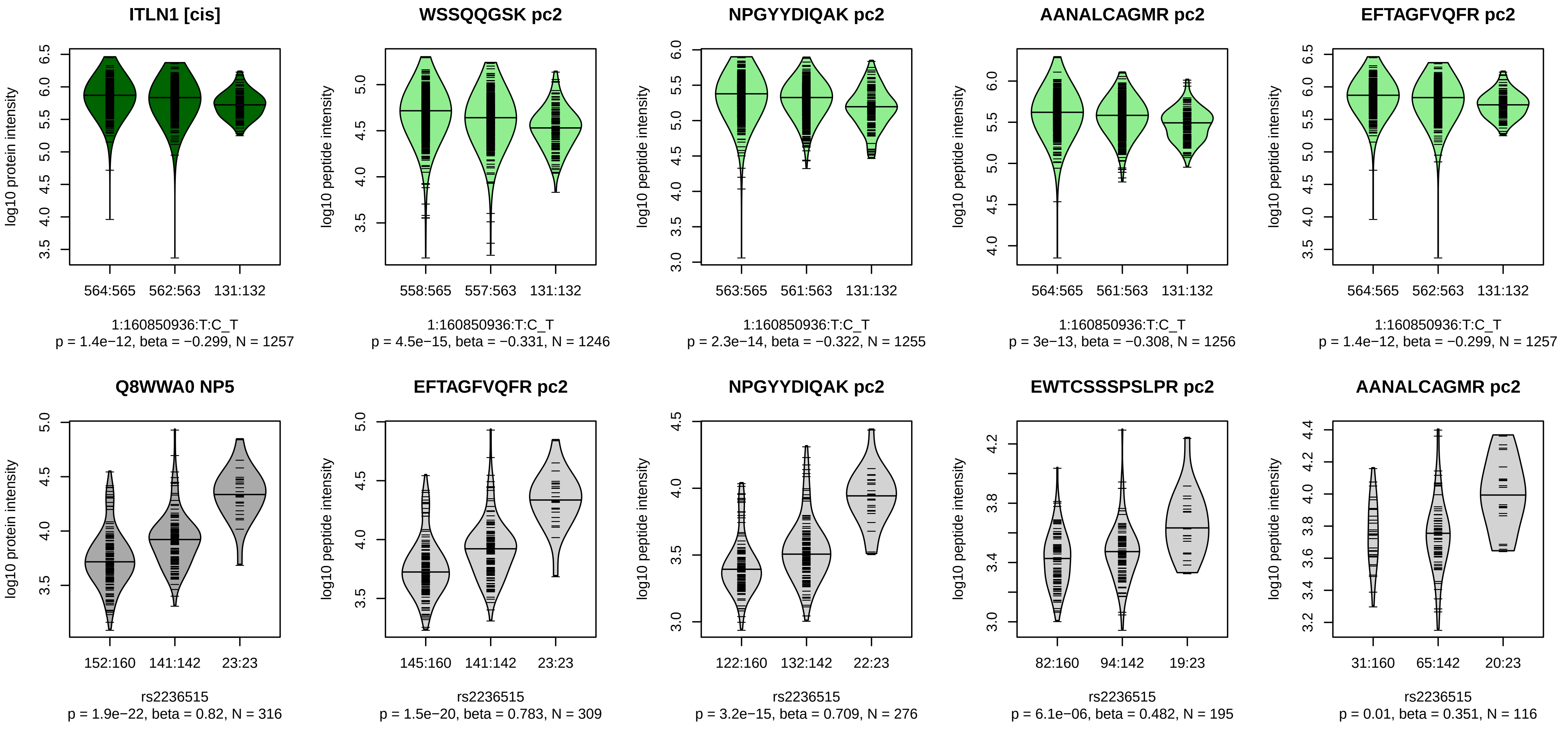


Locus: 183 (Sort: 210)
Status: Not replicated
80% power: No
SNP: 1:160850936:T:C
rsID: rs2236515
UniProtID: Q8WWA0
HGNC gene: ITLN1
Type: Cis−pQTL

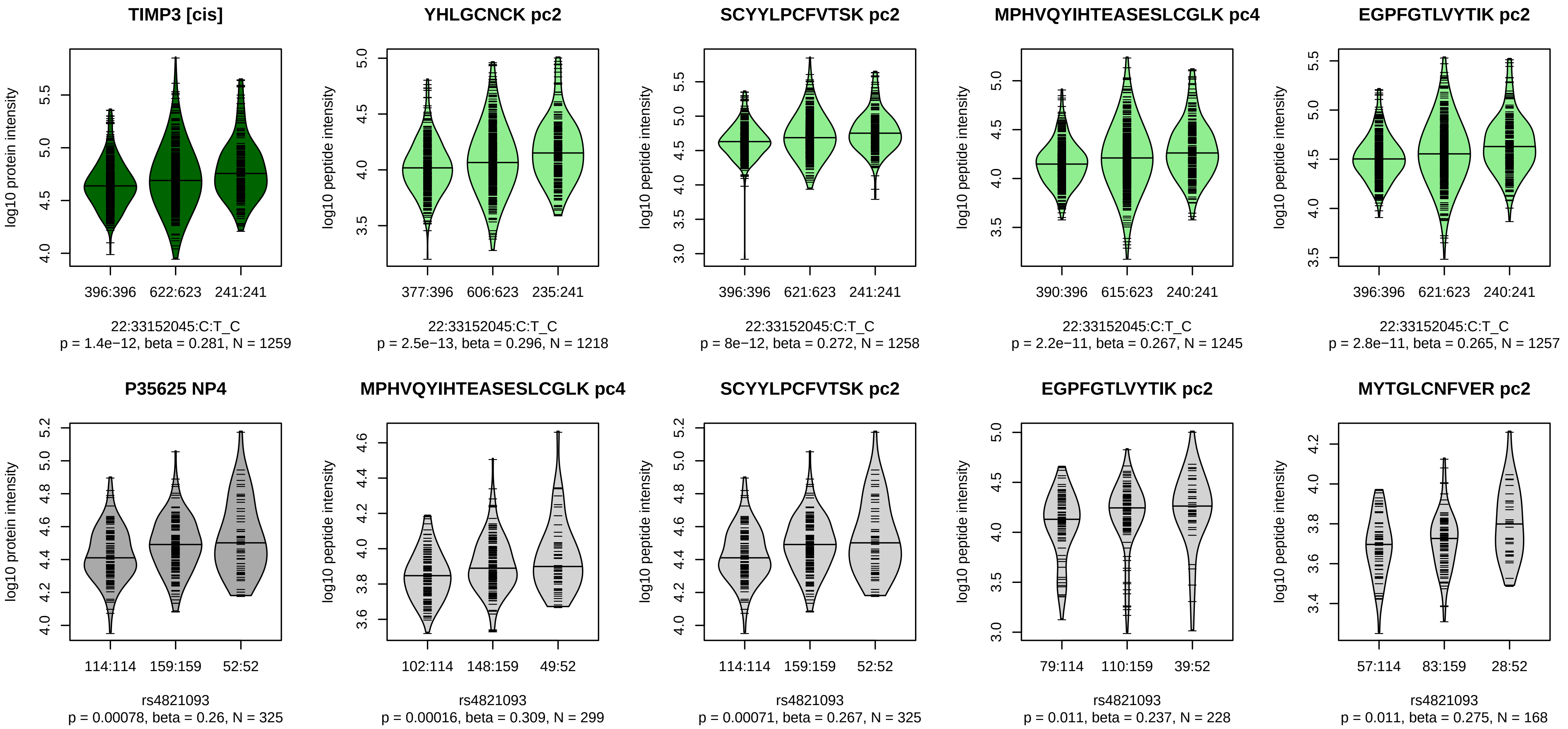


Locus: 184 (Sort: 211)

Status: Not replicated

80% power: No

SNP: 22:33152045:C:T

rsID: rs4821093

UniProtID: P35625

HGNC gene: TIMP3

Type: Cis−pQTL

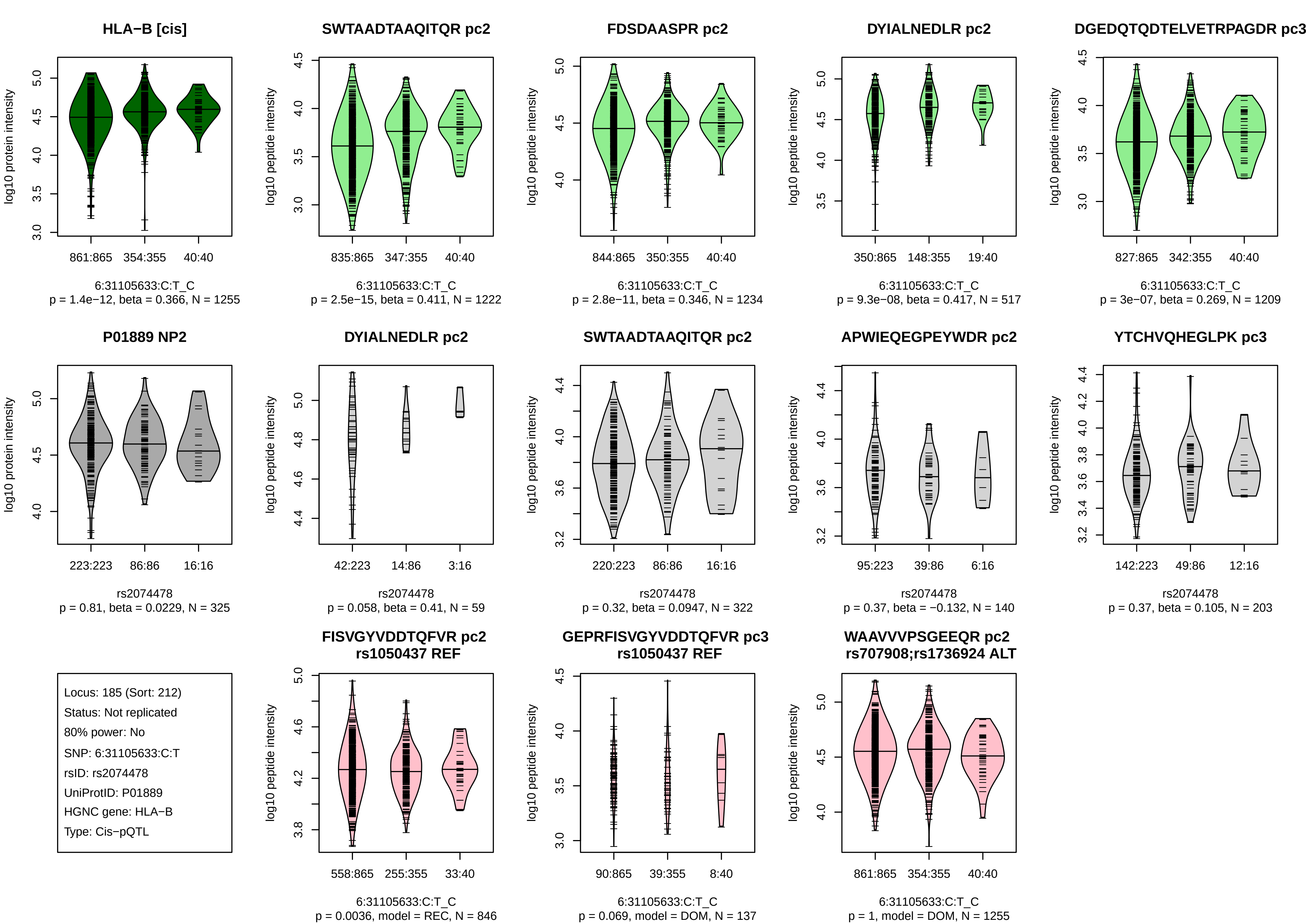

HLA−B [cis]
log10 protein intensity
861:865 354:355 40:40
6:31105633:C:T_C
p = 1.4e−12, beta = 0.366, N = 1255
SWTAADTAAQITQR pc2
log10 peptide intensity
835:865 347:355 40:40
6:31105633:C:T_C
p = 2.5e−15, beta = 0.411, N = 1222
FDSDAASPR pc2
log10 peptide intensity
844:865 350:355 40:40
6:31105633:C:T_C
p = 2.8e−11, beta = 0.346, N = 1234
DYIALNEDLR pc2
log10 peptide intensity
350:865 148:355 19:40
6:31105633:C:T_C
p = 9.3e−08, beta = 0.417, N = 517
DGEDQTQDTELVETRPAGDR pc3
log10 peptide intensity
827:865 342:355 40:40
6:31105633:C:T_C
p = 3e−07, beta = 0.269, N = 1209
P01889 NP2
log10 protein intensity
223:223 86:86 16:16
rs2074478
p = 0.81, beta = 0.0229, N = 325
DYIALNEDLR pc2
log10 peptide intensity
42:223 14:86 3:16
rs2074478
p = 0.058, beta = 0.41, N = 59
SWTAADTAAQITQR pc2
log10 peptide intensity
220:223 86:86 16:16
rs2074478
p = 0.32, beta = 0.0947, N = 322
APWIEQEGPEYWDR pc2
log10 peptide intensity
95:223 39:86 6:16
rs2074478
p = 0.37, beta = −0.132, N = 140
YTCHVQHEGLPK pc3
log10 peptide intensity
142:223 49:86 12:16
rs2074478
p = 0.37, beta = 0.105, N = 203
Locus: 185 (Sort: 212)
Status: Not replicated
80% power: No
SNP: 6:31105633:C:T
rsID: rs2074478
UniProtID: P01889
HGNC gene: HLA−B
Type: Cis−pQTL
FISVGYVDDTQFVR pc2
rs1050437 REF
log10 peptide intensity
558:865 255:355 33:40
6:31105633:C:T_C
p = 0.0036, model = REC, N = 846
GEPRFISVGYVDDTQFVR pc3
rs1050437 REF
log10 peptide intensity
90:865 39:355 8:40
6:31105633:C:T_C
p = 0.069, model = DOM, N = 137
WAAVVVPSGEEQR pc2
rs707908;rs1736924 ALT
log10 peptide intensity
861:865 354:355 40:40
6:31105633:C:T_C
p = 1, model = DOM, N = 1255

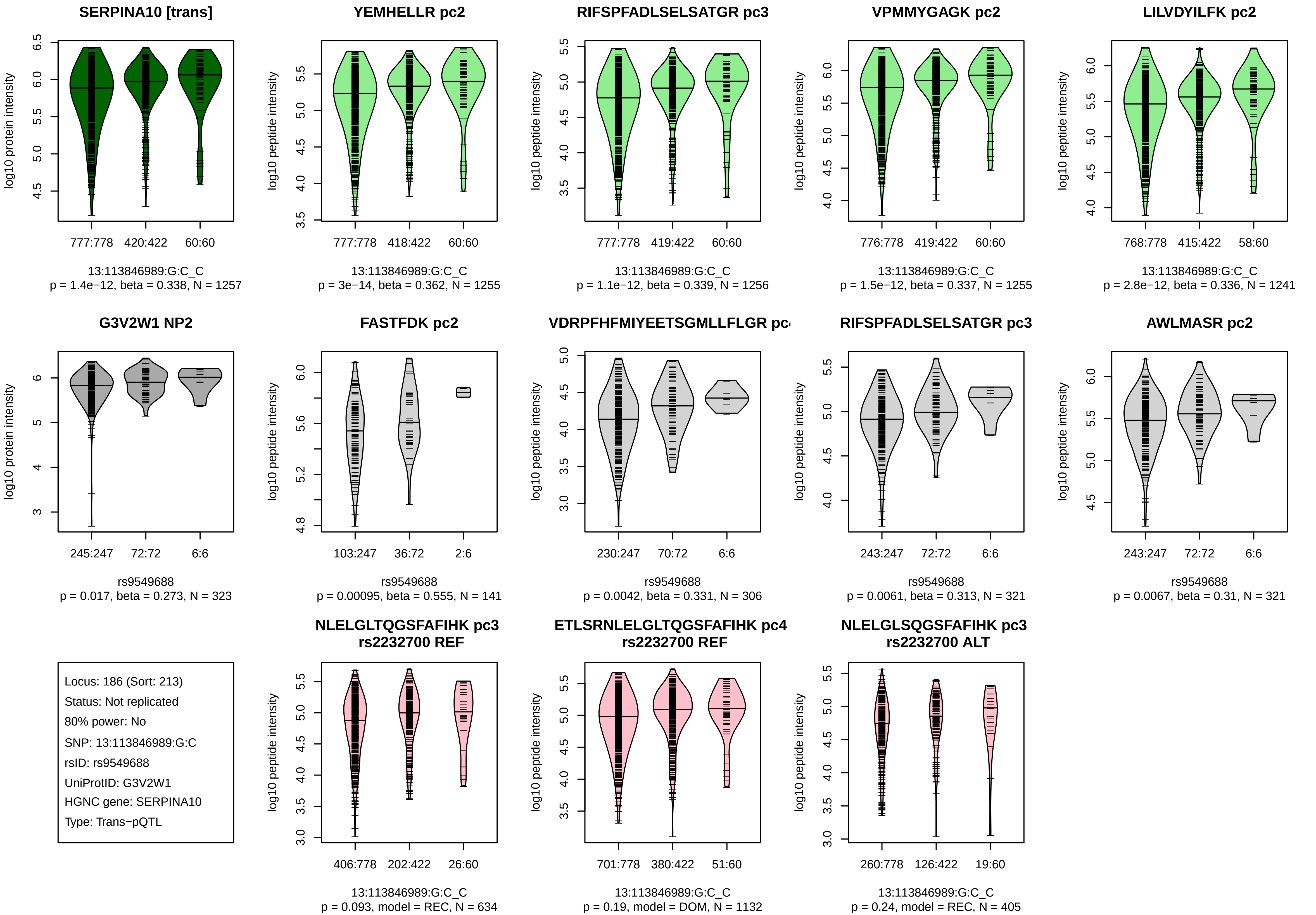
SERPINA10 [trans]
log10 protein intensity
777:778
420:422
60:60
13:113846989:G:C_C
p = 1.4e−12, beta = 0.338, N = 1257
YEMHELLR pc2
log10 peptide intensity
777:778
418:422
60:60
13:113846989:G:C_C
p = 3e−14, beta = 0.362, N = 1255
RIFSPFADLSELSATGR pc3
log10 peptide intensity
777:778
419:422
60:60
13:113846989:G:C_C
p = 1.1e−12, beta = 0.339, N = 1256
VPMMYGAGK pc2
log10 peptide intensity
776:778
419:422
60:60
13:113846989:G:C_C
p = 1.5e−12, beta = 0.337, N = 1255
LILVDYILFK pc2
log10 peptide intensity
768:778
415:422
58:60
13:113846989:G:C_C
p = 2.8e−12, beta = 0.336, N = 1241
G3V2W1 NP2
log10 protein intensity
245:247
72:72
6:6
rs9549688
p = 0.017, beta = 0.273, N = 323
FASTFDK pc2
log10 peptide intensity
103:247
36:72
2:6
rs9549688
p = 0.00095, beta = 0.555, N = 141
VDRPFHFMIYEETSGMLLFLGR pc
log10 peptide intensity
230:247
70:72
6:6
rs9549688
p = 0.0042, beta = 0.331, N = 306
RIFSPFADLSELSATGR pc3
log10 peptide intensity
243:247
72:72
6:6
rs9549688
p = 0.0061, beta = 0.313, N = 321
AWLMASR pc2
log10 peptide intensity
243:247
72:72
6:6
rs9549688
p = 0.0067, beta = 0.31, N = 321
Locus: 186 (Sort: 213)
Status: Not replicated
80% power: No
SNP: 13:113846989:G:C
rsID: rs9549688
UniProtID: G3V2W1
HGNC gene: SERPINA10
Type: Trans−pQTL
NLELGLTQGSFAFIHK pc3
rs2232700 REF
log10 peptide intensity
406:778
202:422
26:60
13:113846989:G:C_C
p = 0.093, model = REC, N = 634
ETLSRNLELGLTQGSFAFIHK pc4
rs2232700 REF
log10 peptide intensity
701:778
380:422
51:60
13:113846989:G:C_C
p = 0.19, model = DOM, N = 1132
NLELGLSQGSFAFIHK pc3
rs2232700 ALT
log10 peptide intensity
260:778
126:422
19:60
13:113846989:G:C_C
p = 0.24, model = REC, N = 405

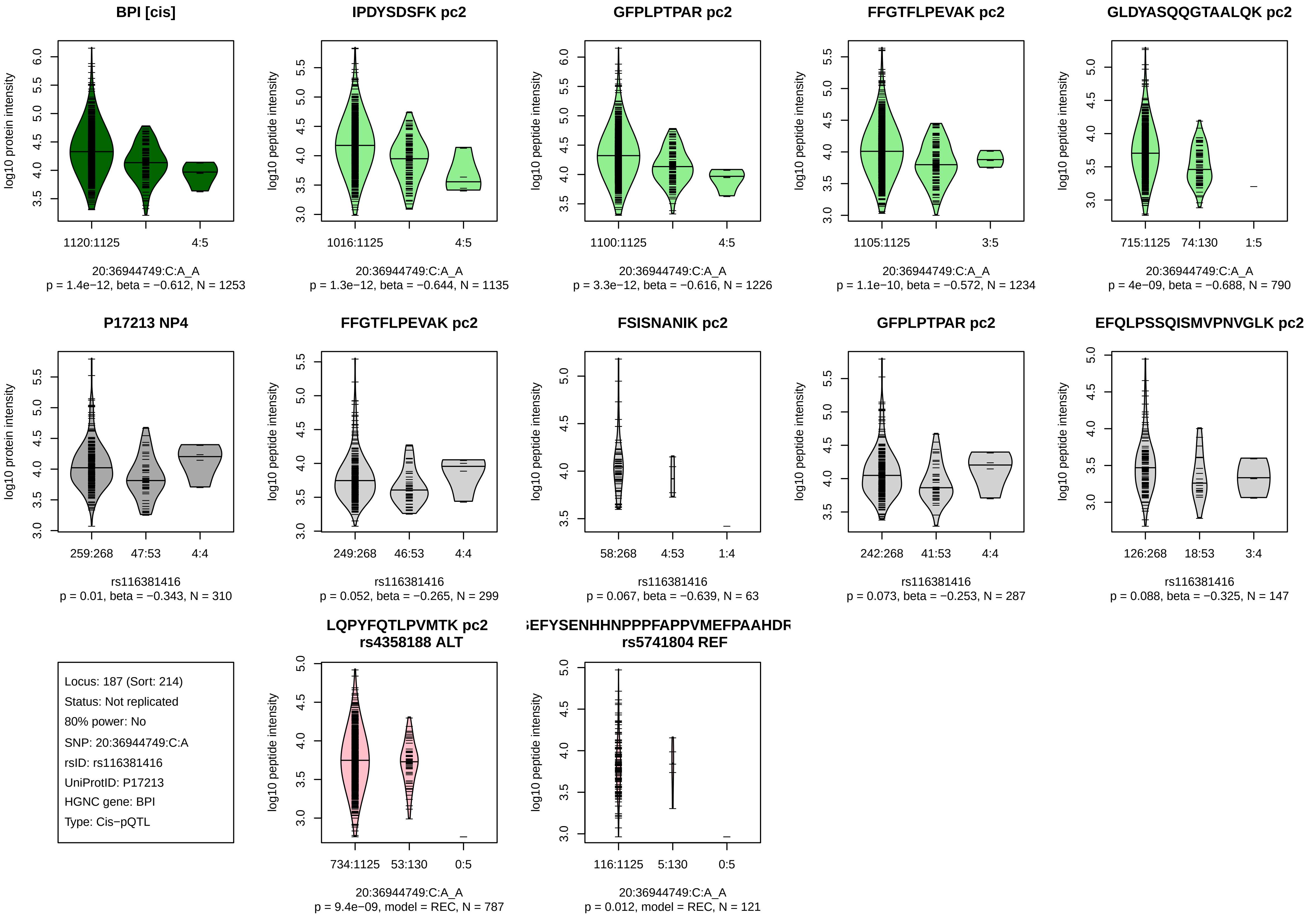
BPI [cis]
log10 protein intensity
1120:1125
4:5
20:36944749:C:A_A
p = 1.4e−12, beta = −0.612, N = 1253
IPDYSDSFK pc2
log10 peptide intensity
1016:1125
4:5
20:36944749:C:A_A
p = 1.3e−12, beta = −0.644, N = 1135
GFPLPTPAR pc2
log10 peptide intensity
1100:1125
4:5
20:36944749:C:A_A
p = 3.3e−12, beta = −0.616, N = 1226
FFGTFLPEVAK pc2
log10 peptide intensity
1105:1125
3:5
20:36944749:C:A_A
p = 1.1e−10, beta = −0.572, N = 1234
GLDYASQQGTAALQK pc2
log10 peptide intensity
715:1125
74:130
1:5
20:36944749:C:A_A
p = 4e−09, beta = −0.688, N = 790
P17213 NP4
log10 protein intensity
259:268
47:53
4:4
rs116381416
p = 0.01, beta = −0.343, N = 310
FFGTFLPEVAK pc2
log10 peptide intensity
249:268
46:53
4:4
rs116381416
p = 0.052, beta = −0.265, N = 299
FSISNANIK pc2
log10 peptide intensity
58:268
4:53
1:4
rs116381416
p = 0.067, beta = −0.639, N = 63
GFPLPTPAR pc2
log10 peptide intensity
242:268
41:53
4:4
rs116381416
p = 0.073, beta = −0.253, N = 287
EFQLPSSQISMVPNVGLK pc2
log10 peptide intensity
126:268
18:53
3:4
rs116381416
p = 0.088, beta = −0.325, N = 147
Locus: 187 (Sort: 214)
Status: Not replicated
80% power: No
SNP: 20:36944749:C:A
rsID: rs116381416
UniProtID: P17213
HGNC gene: BPI
Type: Cis−pQTL
LQPYFQTLPVMTK pc2
rs4358188 ALT
log10 peptide intensity
734:1125
53:130
0:5
20:36944749:C:A_A
p = 9.4e−09, model = REC, N = 787
EFYSENHHNPPPFAPPVMEFPAAHDR
rs5741804 REF
log10 peptide intensity
116:1125
5:130
0:5
20:36944749:C:A_A
p = 0.012, model = REC, N = 121

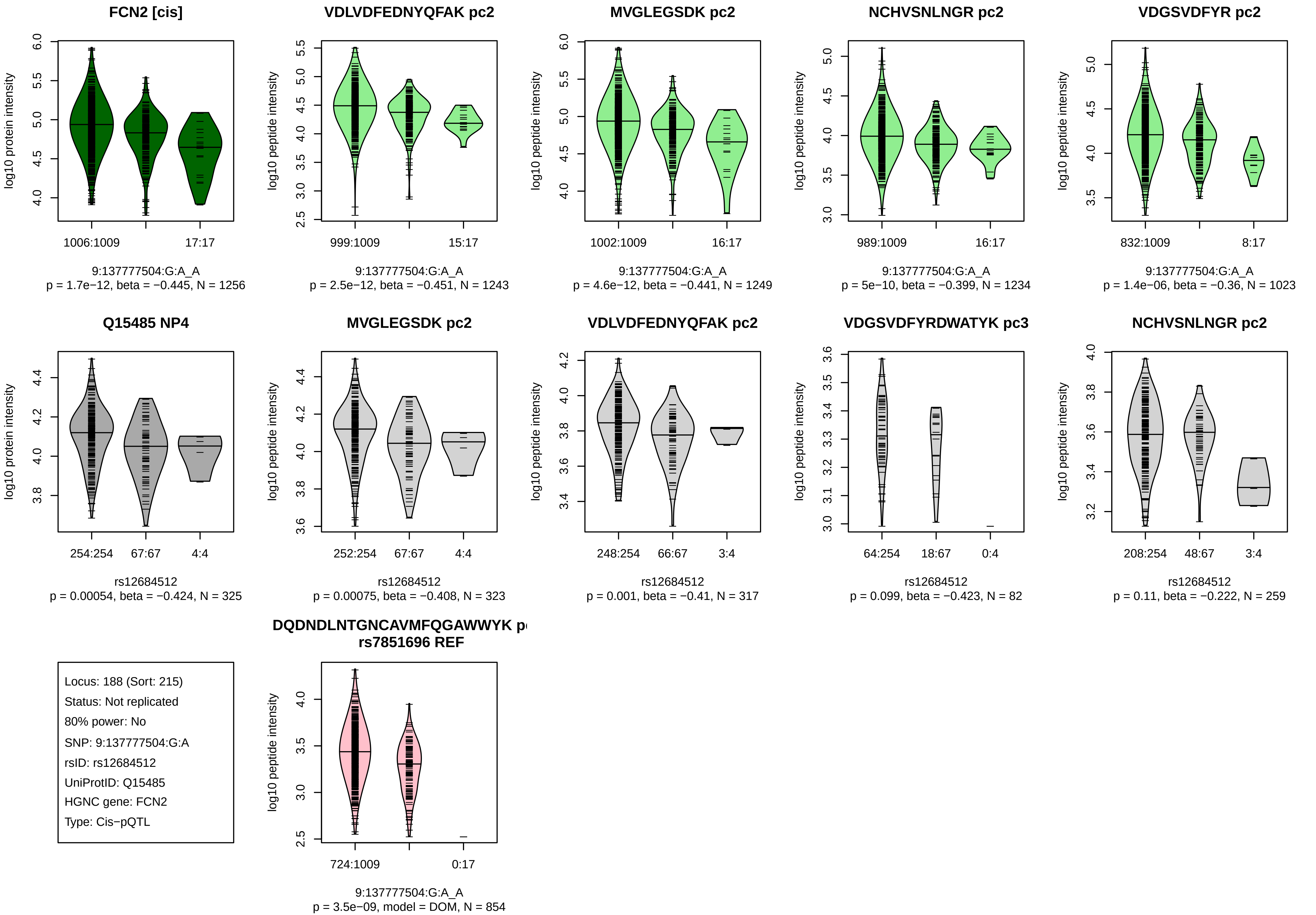
FCN2 [cis]
log10 protein intensity
1006:1009
17:17
9:137777504:G:A_A
p = 1.7e−12, beta = −0.445, N = 1256
VDLVDFEDNYQFAK pc2
log10 peptide intensity
999:1009
15:17
9:137777504:G:A_A
p = 2.5e−12, beta = −0.451, N = 1243
MVGLEGSDK pc2
log10 peptide intensity
1002:1009
16:17
9:137777504:G:A_A
p = 4.6e−12, beta = −0.441, N = 1249
NCHVSNLNGR pc2
log10 peptide intensity
989:1009
16:17
9:137777504:G:A_A
p = 5e−10, beta = −0.399, N = 1234
VDGSVDFYR pc2
log10 peptide intensity
832:1009
8:17
9:137777504:G:A_A
p = 1.4e−06, beta = −0.36, N = 1023
Q15485 NP4
log10 protein intensity
254:254
67:67
4:4
rs12684512
p = 0.00054, beta = −0.424, N = 325
MVGLEGSDK pc2
log10 peptide intensity
252:254
67:67
4:4
rs12684512
p = 0.00075, beta = −0.408, N = 323
VDLVDFEDNYQFAK pc2
log10 peptide intensity
248:254
66:67
3:4
rs12684512
p = 0.001, beta = −0.41, N = 317
VDGSVDFYRDWATYK pc3
log10 peptide intensity
64:254
18:67
0:4
rs12684512
p = 0.099, beta = −0.423, N = 82
NCHVSNLNGR pc2
log10 peptide intensity
208:254
48:67
3:4
rs12684512
p = 0.11, beta = −0.222, N = 259
Locus: 188 (Sort: 215)
Status: Not replicated
80% power: No
SNP: 9:137777504:G:A
rsID: rs12684512
UniProtID: Q15485
HGNC gene: FCN2
Type: Cis−pQTL
DQDNDLNTGNCAVMFQGAWWYK p
rs7851696 REF
log10 peptide intensity
724:1009
0:17
9:137777504:G:A_A
p = 3.5e−09, model = DOM, N = 854

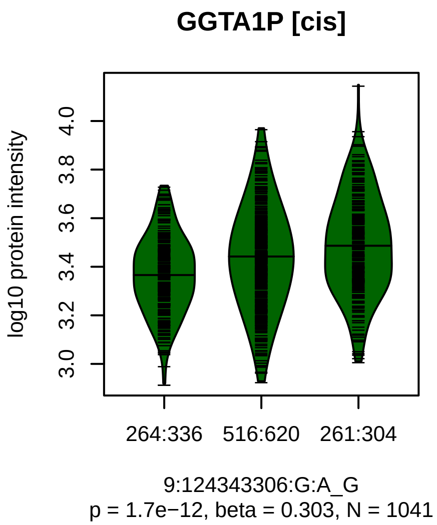


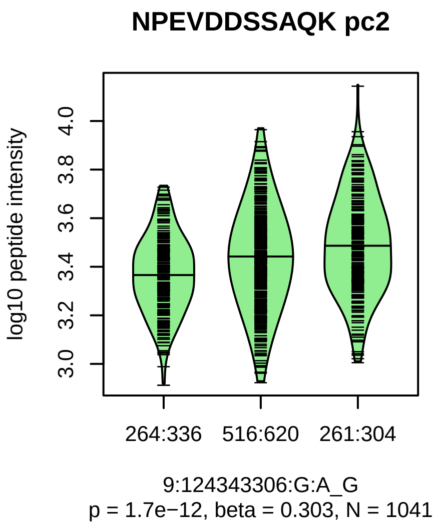


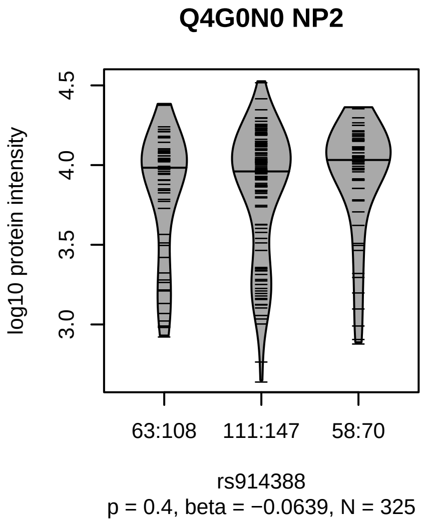


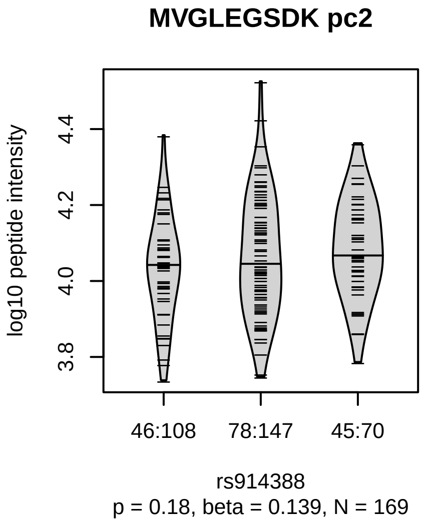


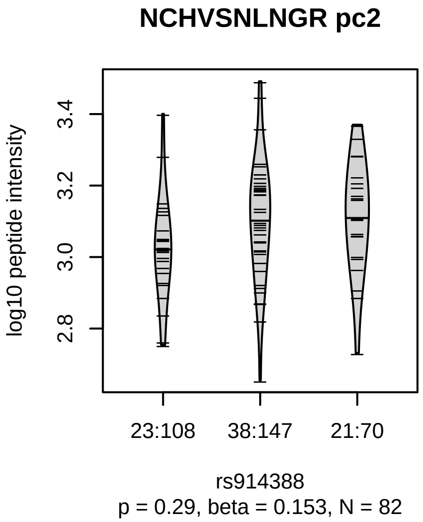


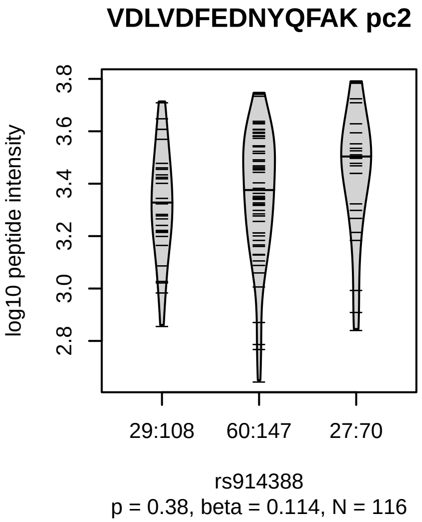


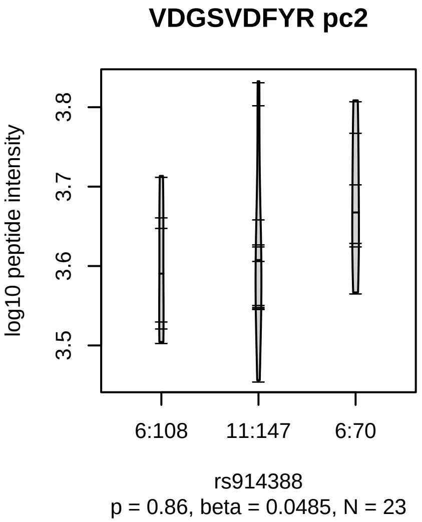


Locus: 189 (Sort: 216)
Status: Not replicated
80% power: No
SNP: 9:124343306:G:A
rsID: rs914388
UniProtID: Q4G0N0
HGNC gene: GGTA1P
Type: Cis−pQTL

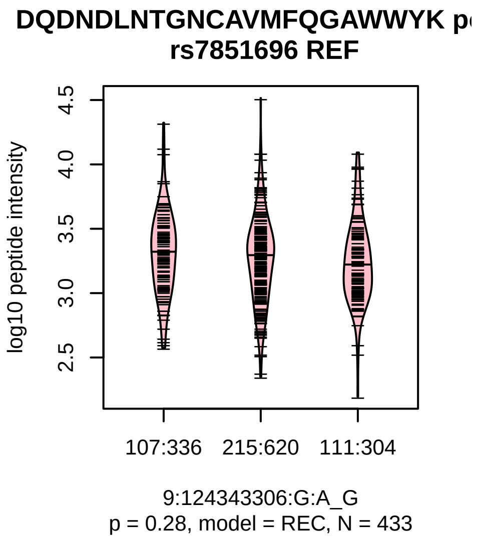

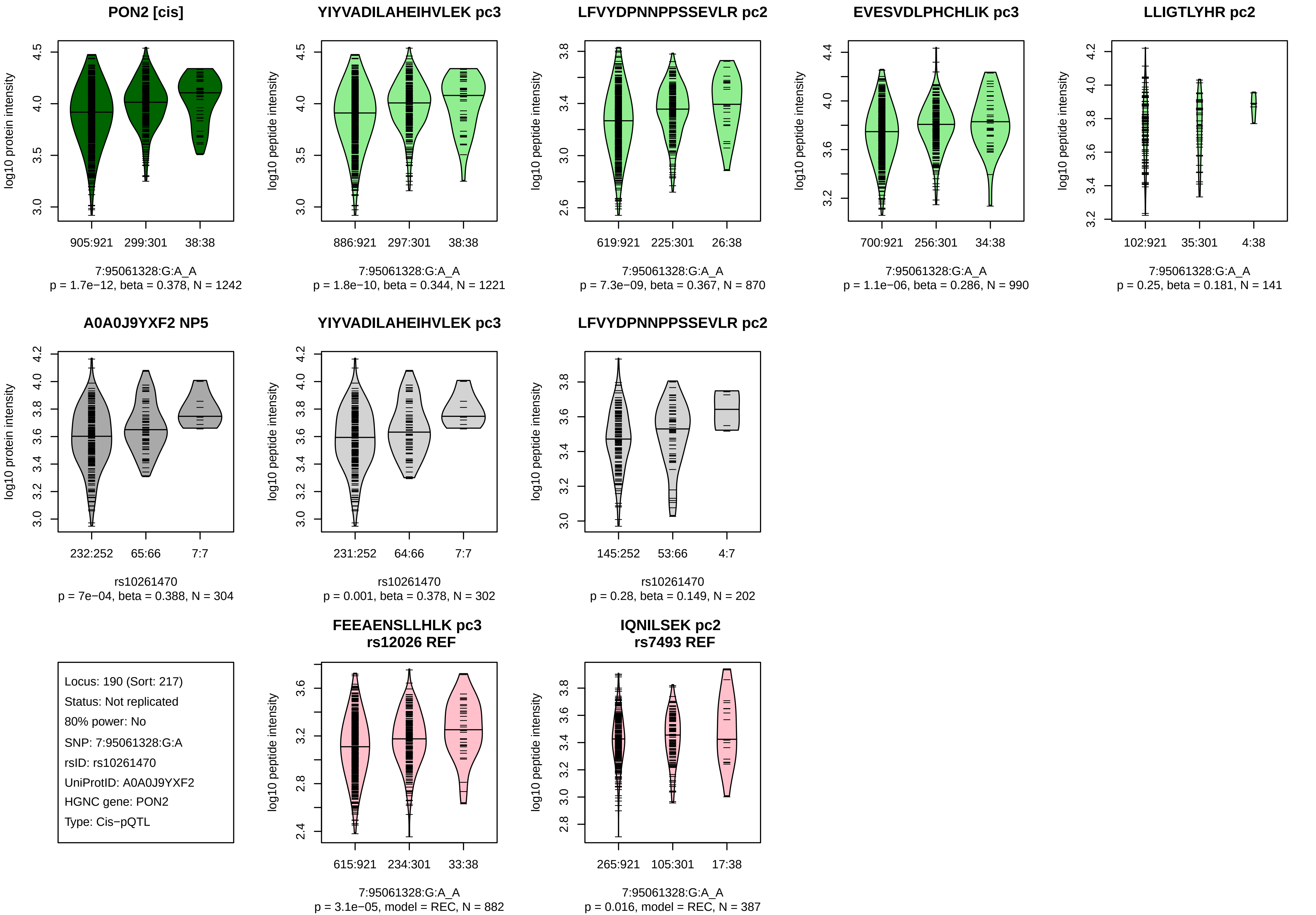
PON2 [cis]
log10 protein intensity
905:921
299:301
38:38
7:95061328:G:A_A
p = 1.7e−12, beta = 0.378, N = 1242
YIYVADILAHEIHVLEK pc3
log10 peptide intensity
886:921
297:301
38:38
7:95061328:G:A_A
p = 1.8e−10, beta = 0.344, N = 1221
LFVYDPNNPPSSEVLR pc2
log10 peptide intensity
619:921
225:301
26:38
7:95061328:G:A_A
p = 7.3e−09, beta = 0.367, N = 870
EVESVDLPHCHLIK pc3
log10 peptide intensity
700:921
256:301
34:38
7:95061328:G:A_A
p = 1.1e−06, beta = 0.286, N = 990
LLIGTLYHR pc2
log10 peptide intensity
102:921
35:301
4:38
7:95061328:G:A_A
p = 0.25, beta = 0.181, N = 141
A0A0J9YXF2 NP5
log10 protein intensity
232:252
65:66
7:7
rs10261470
p = 7e−04, beta = 0.388, N = 304
YIYVADILAHEIHVLEK pc3
log10 peptide intensity
231:252
64:66
7:7
rs10261470
p = 0.001, beta = 0.378, N = 302
LFVYDPNNPPSSEVLR pc2
log10 peptide intensity
145:252
53:66
4:7
rs10261470
p = 0.28, beta = 0.149, N = 202
Locus: 190 (Sort: 217)
Status: Not replicated
80% power: No
SNP: 7:95061328:G:A
rsID: rs10261470
UniProtID: A0A0J9YXF2
HGNC gene: PON2
Type: Cis−pQTL
FEEAENSLLHLK pc3
rs12026 REF
log10 peptide intensity
615:921
234:301
33:38
7:95061328:G:A_A
p = 3.1e−05, model = REC, N = 882
IQNILSEK pc2
rs7493 REF
log10 peptide intensity
265:921
105:301
17:38
7:95061328:G:A_A
p = 0.016, model = REC, N = 387

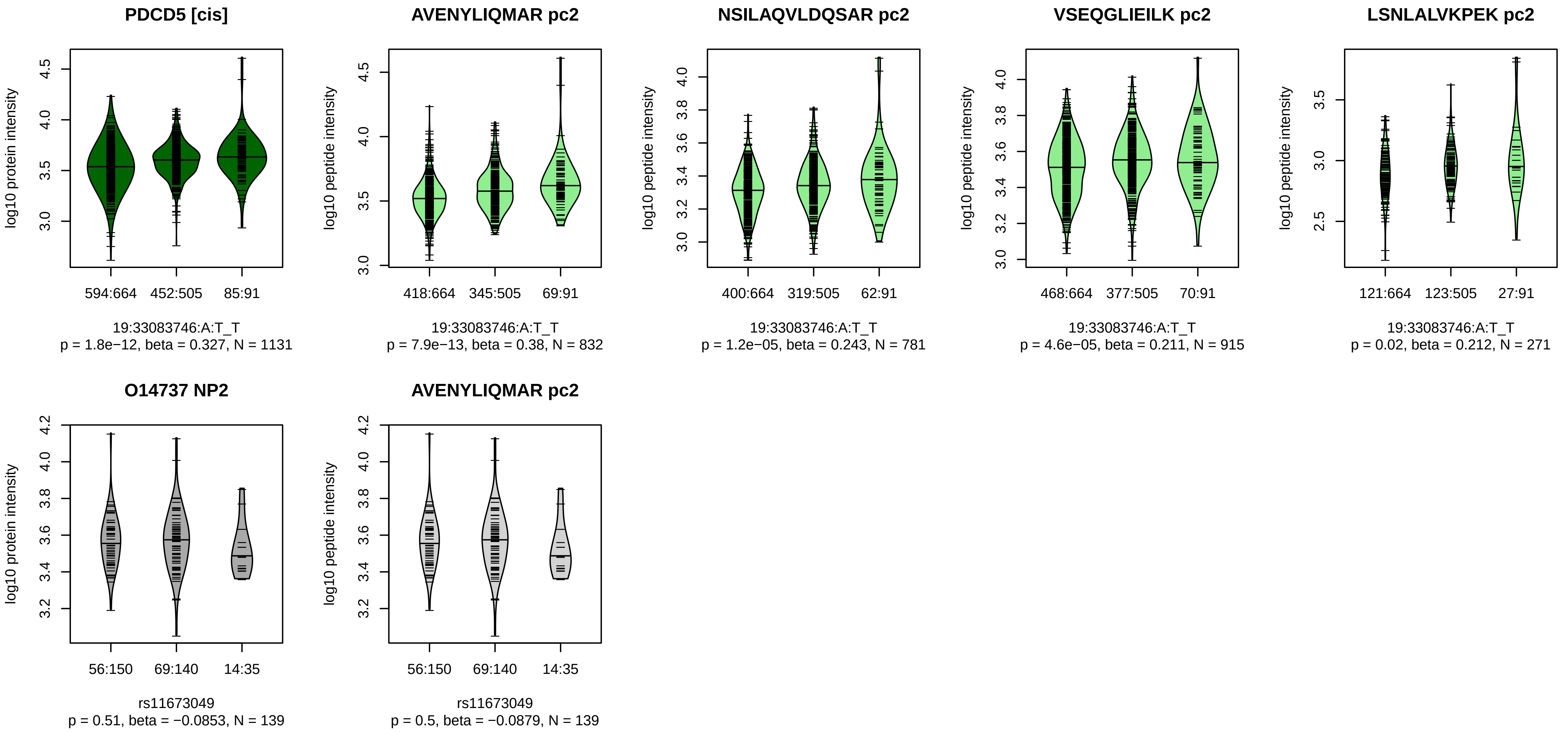


Locus: 191 (Sort: 218)

Status: Not replicated

80% power: No

SNP: 19:33083746:A:T

rsID: rs11673049

UniProtID: O14737

HGNC gene: PDCD5

Type: Cis−pQTL

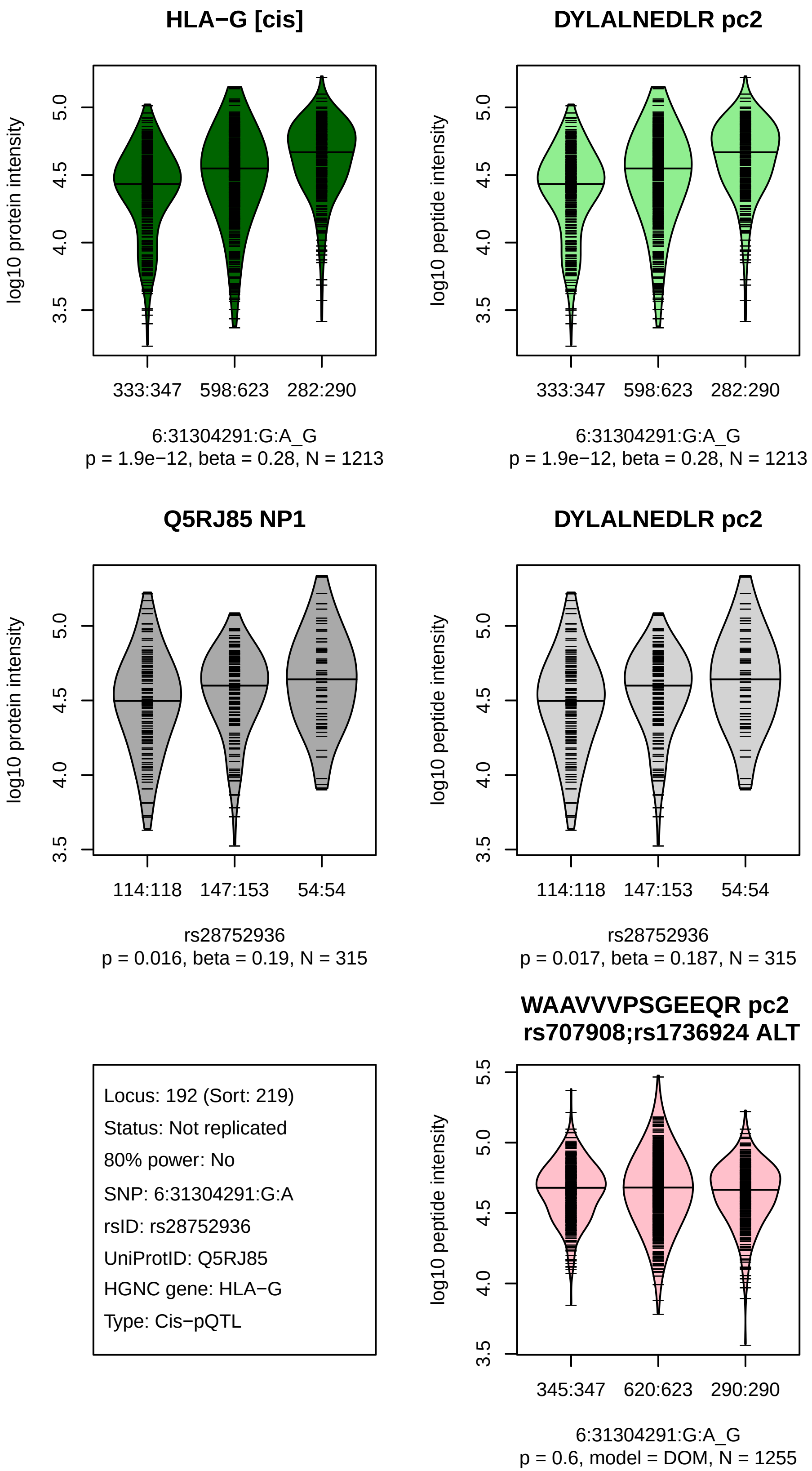
HLA−G [cis]
log10 protein intensity
3.5
4.0
4.5
5.0
333:347
598:623
282:290
6:31304291:G:A_G
p = 1.9e−12, beta = 0.28, N = 1213
DYLALNEDLR pc2
log10 peptide intensity
3.5
4.0
4.5
5.0
333:347
598:623
282:290
6:31304291:G:A_G
p = 1.9e−12, beta = 0.28, N = 1213
Q5RJ85 NP1
log10 protein intensity
3.5
4.0
4.5
5.0
114:118
147:153
54:54
rs28752936
p = 0.016, beta = 0.19, N = 315
DYLALNEDLR pc2
log10 peptide intensity
3.5
4.0
4.5
5.0
114:118
147:153
54:54
rs28752936
p = 0.017, beta = 0.187, N = 315
Locus: 192 (Sort: 219)
Status: Not replicated
80% power: No
SNP: 6:31304291:G:A
rsID: rs28752936
UniProtID: Q5RJ85
HGNC gene: HLA−G
Type: Cis−pQTL
WAAVVVPSGEEQR pc2
rs707908;rs1736924 ALT
log10 peptide intensity
3.5
4.0
4.5
5.0
5.5
345:347
620:623
290:290
6:31304291:G:A_G
p = 0.6, model = DOM, N = 1255

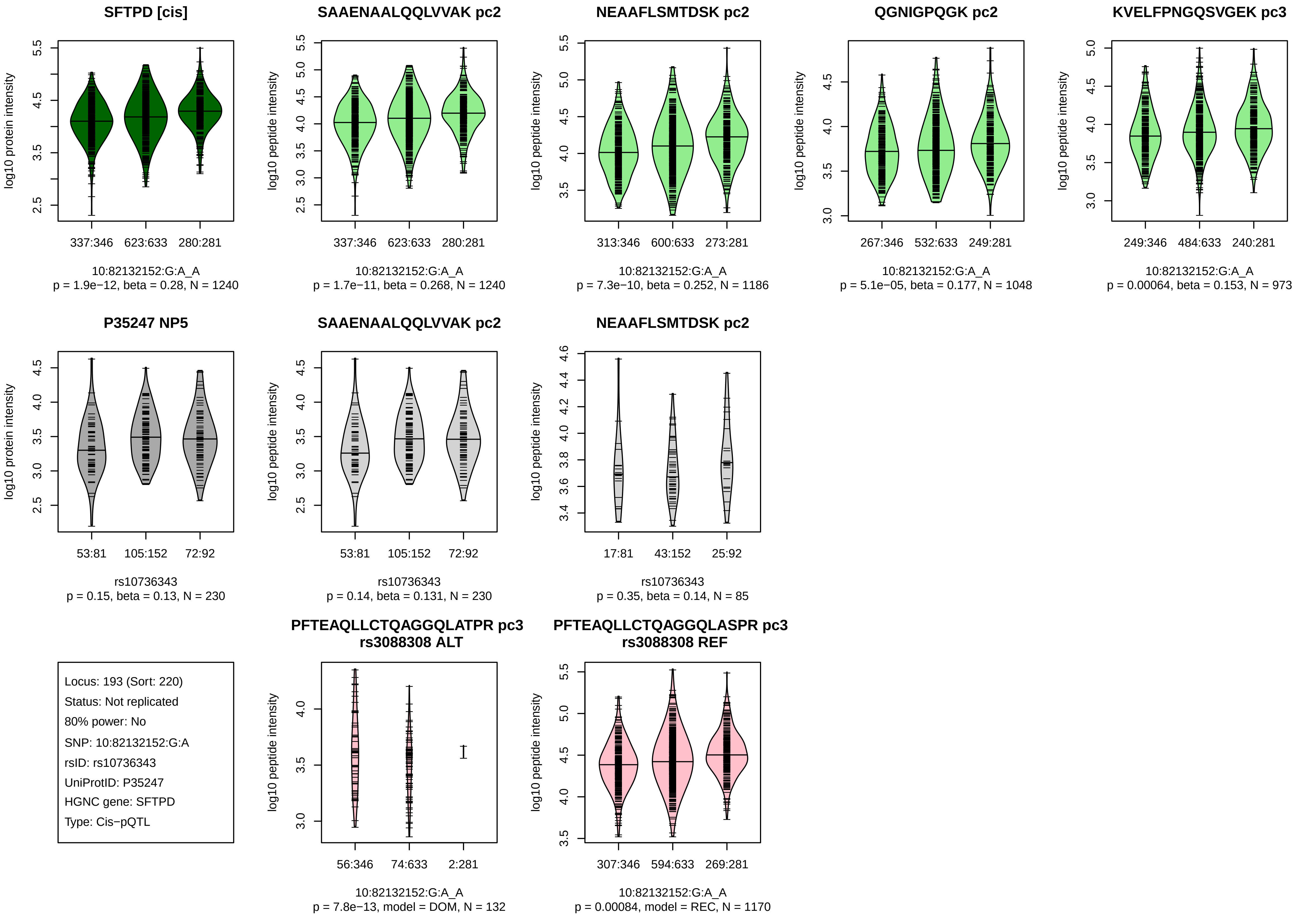
SFTPD [cis]
log10 protein intensity
337:346
623:633
280:281
10:82132152:G:A_A
p = 1.9e−12, beta = 0.28, N = 1240
SAAENAALQQLVVAK pc2
log10 peptide intensity
337:346
623:633
280:281
10:82132152:G:A_A
p = 1.7e−11, beta = 0.268, N = 1240
NEAAFLSMTDSK pc2
log10 peptide intensity
313:346
600:633
273:281
10:82132152:G:A_A
p = 7.3e−10, beta = 0.252, N = 1186
QGNIGPQGK pc2
log10 peptide intensity
267:346
532:633
249:281
10:82132152:G:A_A
p = 5.1e−05, beta = 0.177, N = 1048
KVELFPNGQSVGEK pc3
log10 peptide intensity
249:346
484:633
240:281
10:82132152:G:A_A
p = 0.00064, beta = 0.153, N = 973
P35247 NP5
log10 protein intensity
53:81
105:152
72:92
rs10736343
p = 0.15, beta = 0.13, N = 230
SAAENAALQQLVVAK pc2
log10 peptide intensity
53:81
105:152
72:92
rs10736343
p = 0.14, beta = 0.131, N = 230
NEAAFLSMTDSK pc2
log10 peptide intensity
17:81
43:152
25:92
rs10736343
p = 0.35, beta = 0.14, N = 85
Locus: 193 (Sort: 220)
Status: Not replicated
80% power: No
SNP: 10:82132152:G:A
rsID: rs10736343
UniProtID: P35247
HGNC gene: SFTPD
Type: Cis−pQTL
PFTEAQLLCTQAGGQLATPR pc3
rs3088308 ALT
log10 peptide intensity
56:346
74:633
2:281
10:82132152:G:A_A
p = 7.8e−13, model = DOM, N = 132
PFTEAQLLCTQAGGQLASPR pc3
rs3088308 REF
log10 peptide intensity
307:346
594:633
269:281
10:82132152:G:A_A
p = 0.00084, model = REC, N = 1170

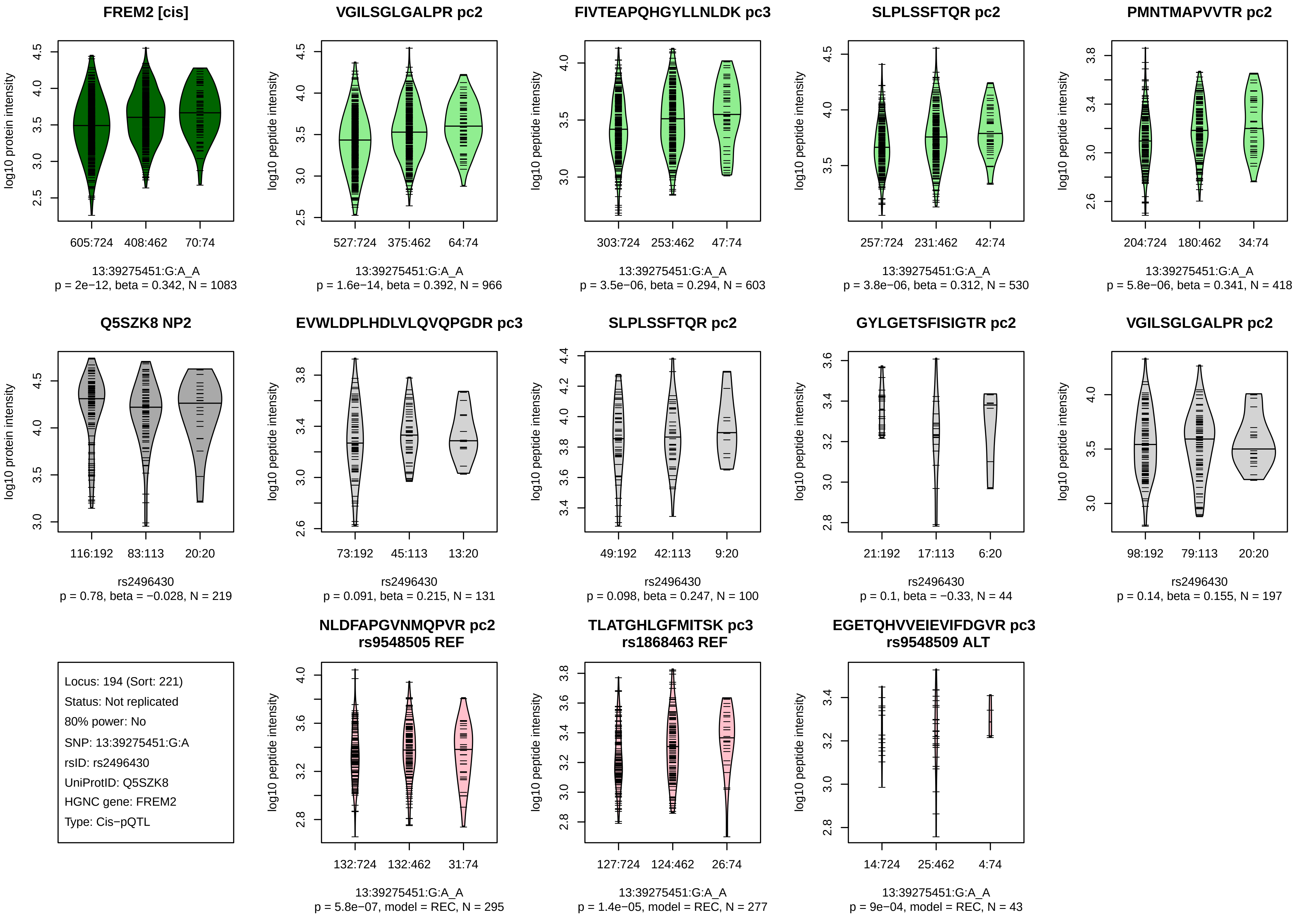
FREM2 [cis]
log10 protein intensity
605:724
408:462
70:74
13:39275451:G:A_A
p = 2e−12, beta = 0.342, N = 1083
VGILSGLGALPR pc2
log10 peptide intensity
527:724
375:462
64:74
13:39275451:G:A_A
p = 1.6e−14, beta = 0.392, N = 966
FIVTEAPQHGYLLNLDK pc3
303:724
253:462
47:74
13:39275451:G:A_A
p = 3.5e−06, beta = 0.294, N = 603
SLPLSSFTQR pc2
257:724
231:462
42:74
13:39275451:G:A_A
p = 3.8e−06, beta = 0.312, N = 530
PMNTMAPVVTR pc2
204:724
180:462
34:74
13:39275451:G:A_A
p = 5.8e−06, beta = 0.341, N = 418
Q5SZK8 NP2
116:192
83:113
20:20
rs2496430
p = 0.78, beta = −0.028, N = 219
EVWLDPLHDLVLQVQPGDR pc3
73:192
45:113
13:20
rs2496430
p = 0.091, beta = 0.215, N = 131
SLPLSSFTQR pc2
49:192
42:113
9:20
rs2496430
p = 0.098, beta = 0.247, N = 100
GYLGETSFISIGTR pc2
21:192
17:113
6:20
rs2496430
p = 0.1, beta = −0.33, N = 44
VGILSGLGALPR pc2
98:192
79:113
20:20
rs2496430
p = 0.14, beta = 0.155, N = 197
Locus: 194 (Sort: 221)
Status: Not replicated
80% power: No
SNP: 13:39275451:G:A
rsID: rs2496430
UniProtID: Q5SZK8
HGNC gene: FREM2
Type: Cis−pQTL
NLDFAPGVNMQPVR pc2
rs9548505 REF
132:724
132:462
31:74
13:39275451:G:A_A
p = 5.8e−07, model = REC, N = 295
TLATGHLGFMITSK pc3
rs1868463 REF
127:724
124:462
26:74
13:39275451:G:A_A
p = 1.4e−05, model = REC, N = 277
EGETQHVVEIEVIFDGVR pc3
rs9548509 ALT
14:724
25:462
4:74
13:39275451:G:A_A
p = 9e−04, model = REC, N = 43

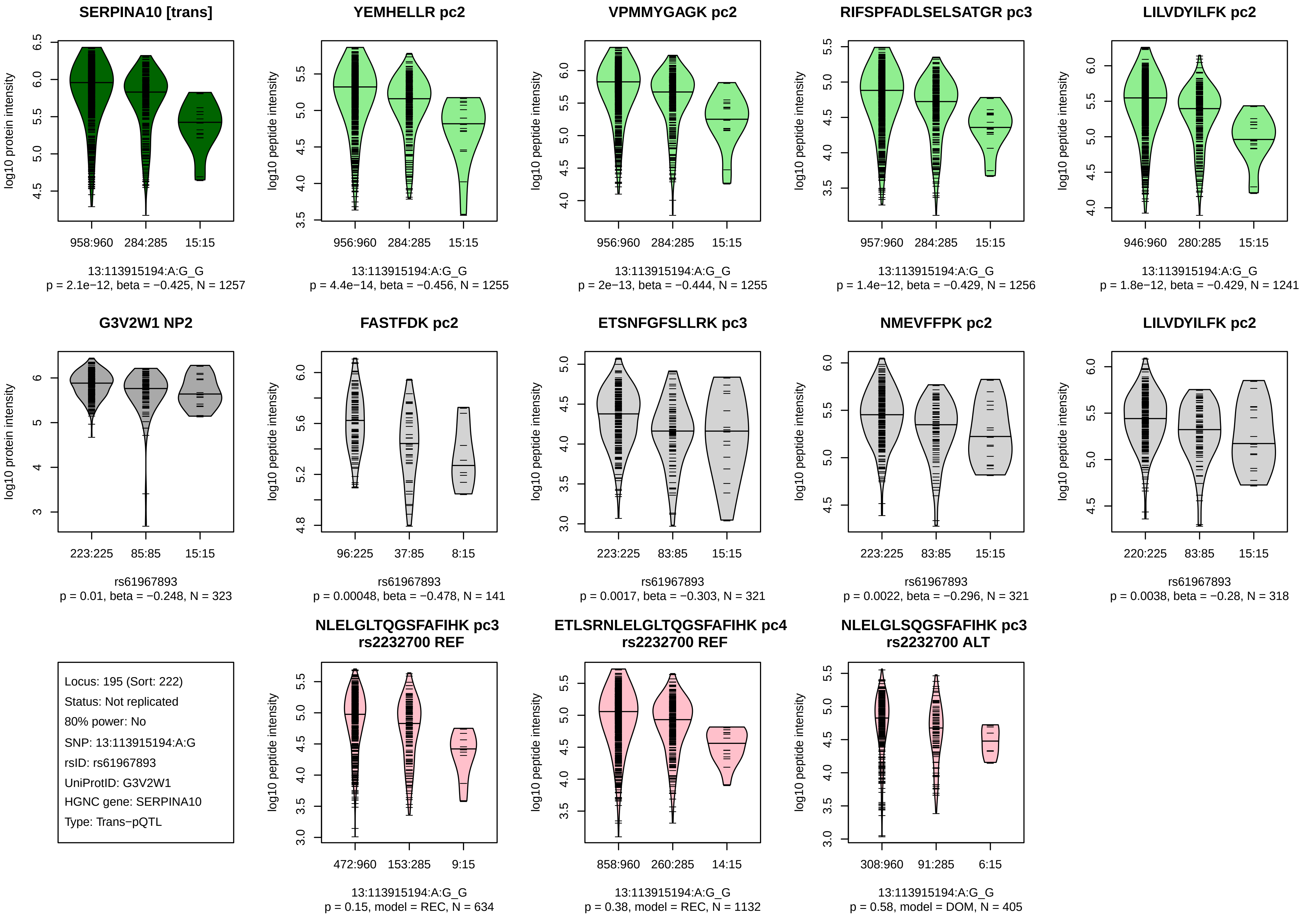
SERPINA10 [trans]
log10 protein intensity
958:960
284:285
15:15
13:113915194:A:G_G
p = 2.1e−12, beta = −0.425, N = 1257
YEMHELLR pc2
log10 peptide intensity
956:960
284:285
15:15
13:113915194:A:G_G
p = 4.4e−14, beta = −0.456, N = 1255
VPMMYGAGK pc2
log10 peptide intensity
956:960
284:285
15:15
13:113915194:A:G_G
p = 2e−13, beta = −0.444, N = 1255
RIFSPFADLSELSATGR pc3
log10 peptide intensity
957:960
284:285
15:15
13:113915194:A:G_G
p = 1.4e−12, beta = −0.429, N = 1256
LILVDYILFK pc2
log10 peptide intensity
946:960
280:285
15:15
13:113915194:A:G_G
p = 1.8e−12, beta = −0.429, N = 1241
G3V2W1 NP2
log10 protein intensity
223:225
85:85
15:15
rs61967893
p = 0.01, beta = −0.248, N = 323
FASTFDK pc2
log10 peptide intensity
96:225
37:85
8:15
rs61967893
p = 0.00048, beta = −0.478, N = 141
ETSNFGFSLLRK pc3
log10 peptide intensity
223:225
83:85
15:15
rs61967893
p = 0.0017, beta = −0.303, N = 321
NMEVFFPK pc2
log10 peptide intensity
223:225
83:85
15:15
rs61967893
p = 0.0022, beta = −0.296, N = 321
LILVDYILFK pc2
log10 peptide intensity
220:225
83:85
15:15
rs61967893
p = 0.0038, beta = −0.28, N = 318
Locus: 195 (Sort: 222)
Status: Not replicated
80% power: No
SNP: 13:113915194:A:G
rsID: rs61967893
UniProtID: G3V2W1
HGNC gene: SERPINA10
Type: Trans−pQTL
NLELGLTQGSFAFIHK pc3
rs2232700 REF
log10 peptide intensity
472:960
153:285
9:15
13:113915194:A:G_G
p = 0.15, model = REC, N = 634
ETLSRNLELGLTQGSFAFIHK pc4
rs2232700 REF
log10 peptide intensity
858:960
260:285
14:15
13:113915194:A:G_G
p = 0.38, model = REC, N = 1132
NLELGLSQGSFAFIHK pc3
rs2232700 ALT
log10 peptide intensity
308:960
91:285
6:15
13:113915194:A:G_G
p = 0.58, model = DOM, N = 405

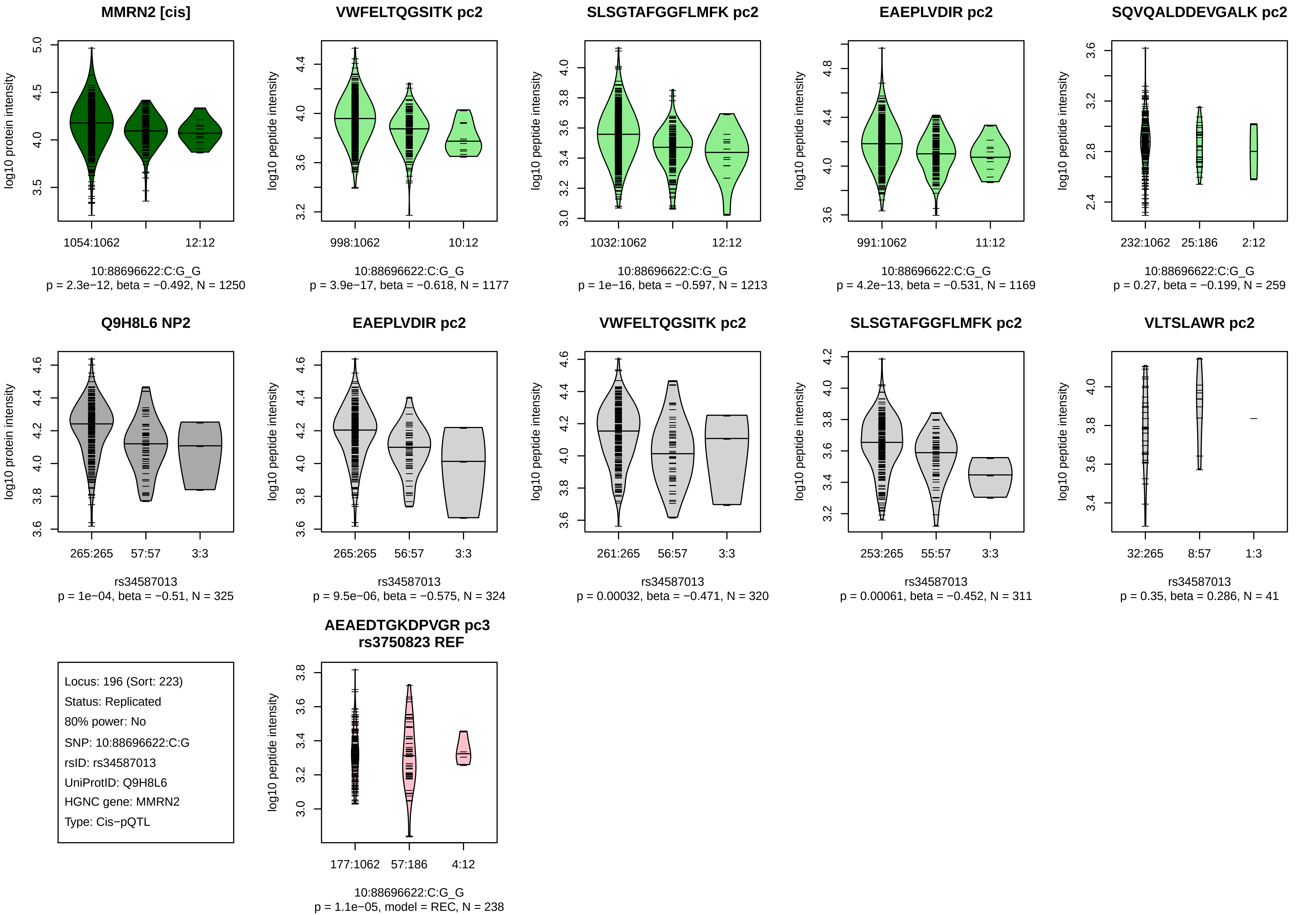
MMRN2 [cis]
log10 protein intensity
1054:1062
12:12
10:88696622:C:G_G
p = 2.3e−12, beta = −0.492, N = 1250
VWFELTQGSITK pc2
log10 peptide intensity
998:1062
10:12
10:88696622:C:G_G
p = 3.9e−17, beta = −0.618, N = 1177
SLSGTAFGGFLMFK pc2
1032:1062
12:12
10:88696622:C:G_G
p = 1e−16, beta = −0.597, N = 1213
EAEPLVDIR pc2
991:1062
11:12
10:88696622:C:G_G
p = 4.2e−13, beta = −0.531, N = 1169
SQVQALDDEVGALK pc2
232:1062
25:186
2:12
10:88696622:C:G_G
p = 0.27, beta = −0.199, N = 259
Q9H8L6 NP2
265:265
57:57
3:3
rs34587013
p = 1e−04, beta = −0.51, N = 325
EAEPLVDIR pc2
265:265
56:57
3:3
rs34587013
p = 9.5e−06, beta = −0.575, N = 324
VWFELTQGSITK pc2
261:265
56:57
3:3
rs34587013
p = 0.00032, beta = −0.471, N = 320
SLSGTAFGGFLMFK pc2
253:265
55:57
3:3
rs34587013
p = 0.00061, beta = −0.452, N = 311
VLTSLAWR pc2
32:265
8:57
1:3
rs34587013
p = 0.35, beta = 0.286, N = 41
Locus: 196 (Sort: 223)
Status: Replicated
80% power: No
SNP: 10:88696622:C:G
rsID: rs34587013
UniProtID: Q9H8L6
HGNC gene: MMRN2
Type: Cis−pQTL
AEAEDTGKDPVGR pc3
rs3750823 REF
177:1062
57:186
4:12
10:88696622:C:G_G
p = 1.1e−05, model = REC, N = 238

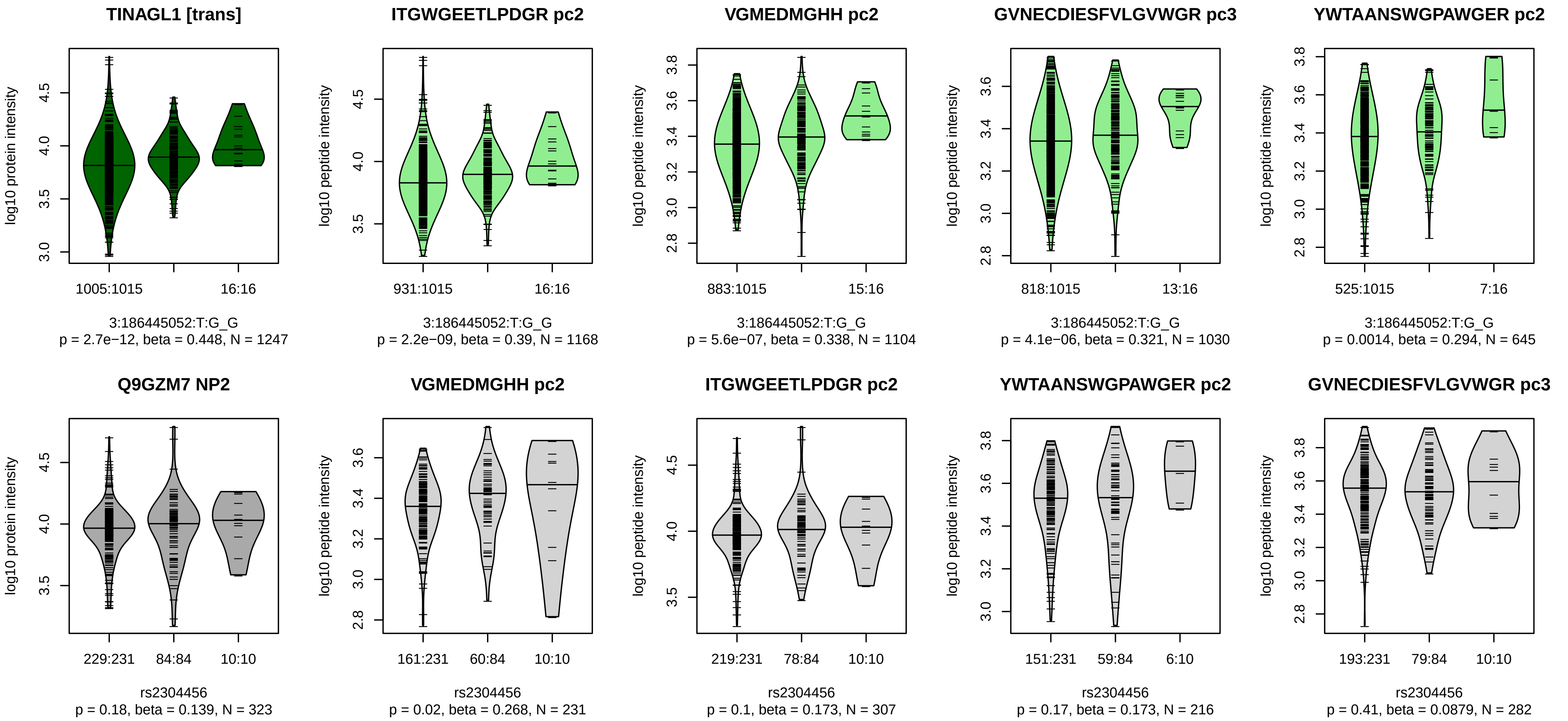


Locus: 197 (Sort: 224)
Status: Not replicated
80% power: No
SNP: 3:186445052:T:G
rsID: rs2304456
UniProtID: Q9GZM7
HGNC gene: TINAGL1
Type: Trans−pQTL

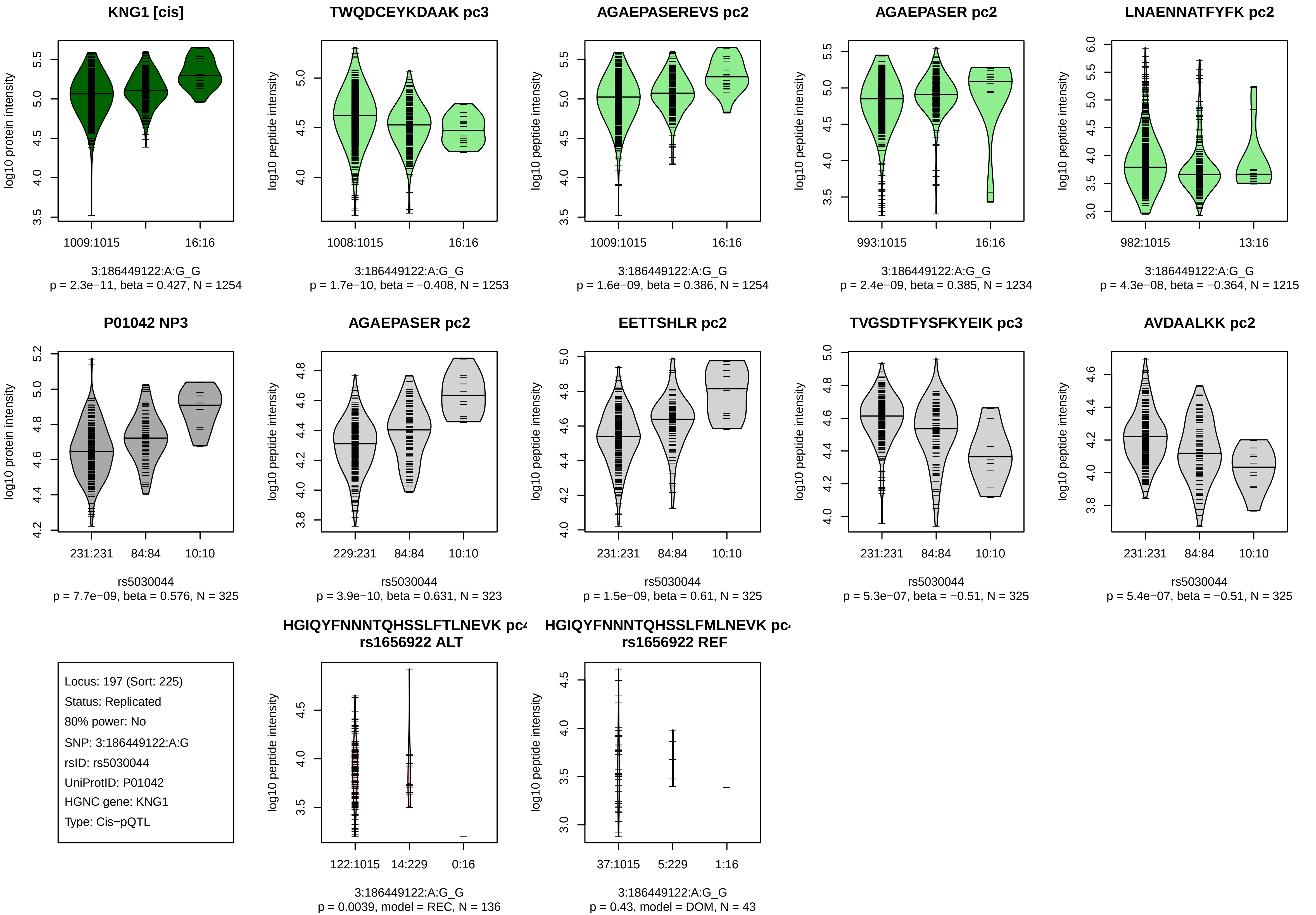
KNG1 [cis]
log10 protein intensity
1009:1015
16:16
3:186449122:A:G_G
p = 2.3e−11, beta = 0.427, N = 1254
TWQDCEYKDAAK pc3
log10 peptide intensity
1008:1015
16:16
3:186449122:A:G_G
p = 1.7e−10, beta = −0.408, N = 1253
AGAEPASEREVS pc2
log10 peptide intensity
1009:1015
16:16
3:186449122:A:G_G
p = 1.6e−09, beta = 0.386, N = 1254
AGAEPASER pc2
log10 peptide intensity
993:1015
16:16
3:186449122:A:G_G
p = 2.4e−09, beta = 0.385, N = 1234
LNAENNATFYFK pc2
log10 peptide intensity
982:1015
13:16
3:186449122:A:G_G
p = 4.3e−08, beta = −0.364, N = 1215
P01042 NP3
log10 protein intensity
231:231
84:84
10:10
rs5030044
p = 7.7e−09, beta = 0.576, N = 325
AGAEPASER pc2
log10 peptide intensity
229:231
84:84
10:10
rs5030044
p = 3.9e−10, beta = 0.631, N = 323
EETTSHLR pc2
log10 peptide intensity
231:231
84:84
10:10
rs5030044
p = 1.5e−09, beta = 0.61, N = 325
TVGSDTFYSFKYEIK pc3
log10 peptide intensity
231:231
84:84
10:10
rs5030044
p = 5.3e−07, beta = −0.51, N = 325
AVDAALKK pc2
log10 peptide intensity
231:231
84:84
10:10
rs5030044
p = 5.4e−07, beta = −0.51, N = 325
Locus: 197 (Sort: 225)
Status: Replicated
80% power: No
SNP: 3:186449122:A:G
rsID: rs5030044
UniProtID: P01042
HGNC gene: KNG1
Type: Cis−pQTL
HGIQYFNNNTQHSSLFTLNEVK pc4
rs1656922 ALT
log10 peptide intensity
122:1015
14:229
0:16
3:186449122:A:G_G
p = 0.0039, model = REC, N = 136
HGIQYFNNNTQHSSLFMLNEVK pc4
rs1656922 REF
log10 peptide intensity
37:1015
5:229
1:16
3:186449122:A:G_G
p = 0.43, model = DOM, N = 43

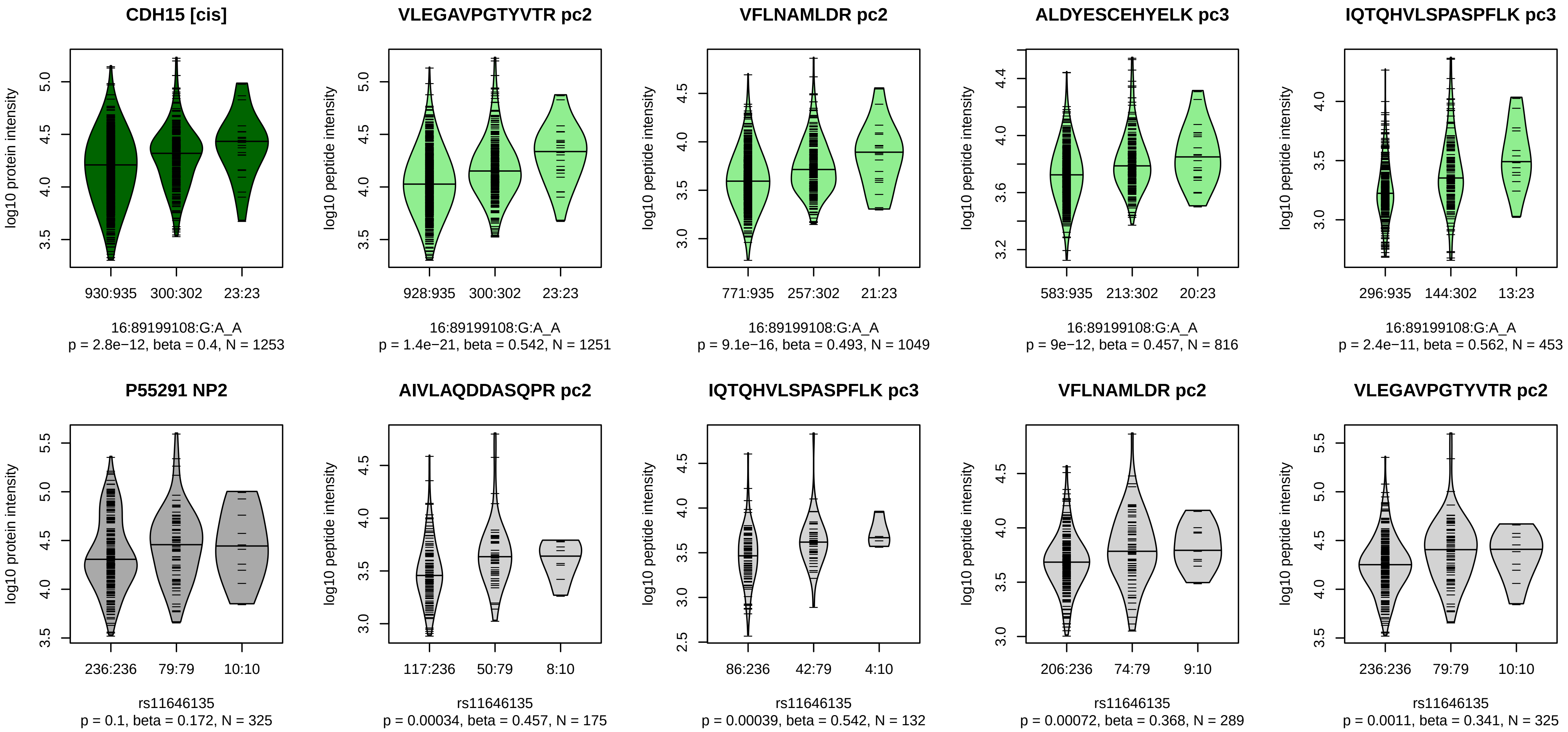


Locus: 198 (Sort: 226)

Status: Not replicated

80% power: No

SNP: 16:89199108:G:A

rsID: rs11646135

UniProtID: P55291

HGNC gene: CDH15

Type: Cis−pQTL

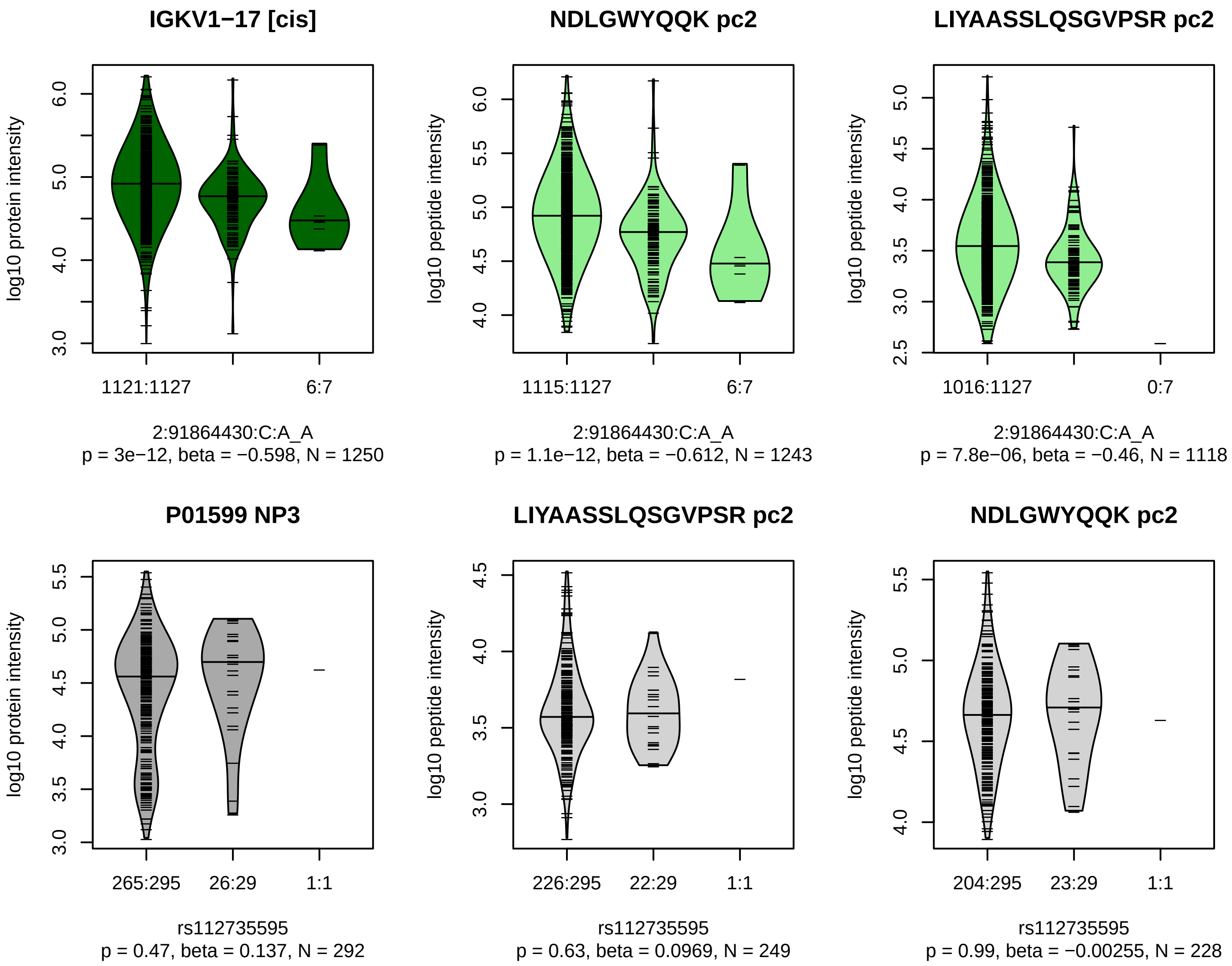


Locus: 199 (Sort: 227)

Status: Not replicated

80% power: No

SNP: 2:91864430:C:A

rsID: rs112735595

UniProtID: P01599

HGNC gene: IGKV1−17

Type: Cis−pQTL

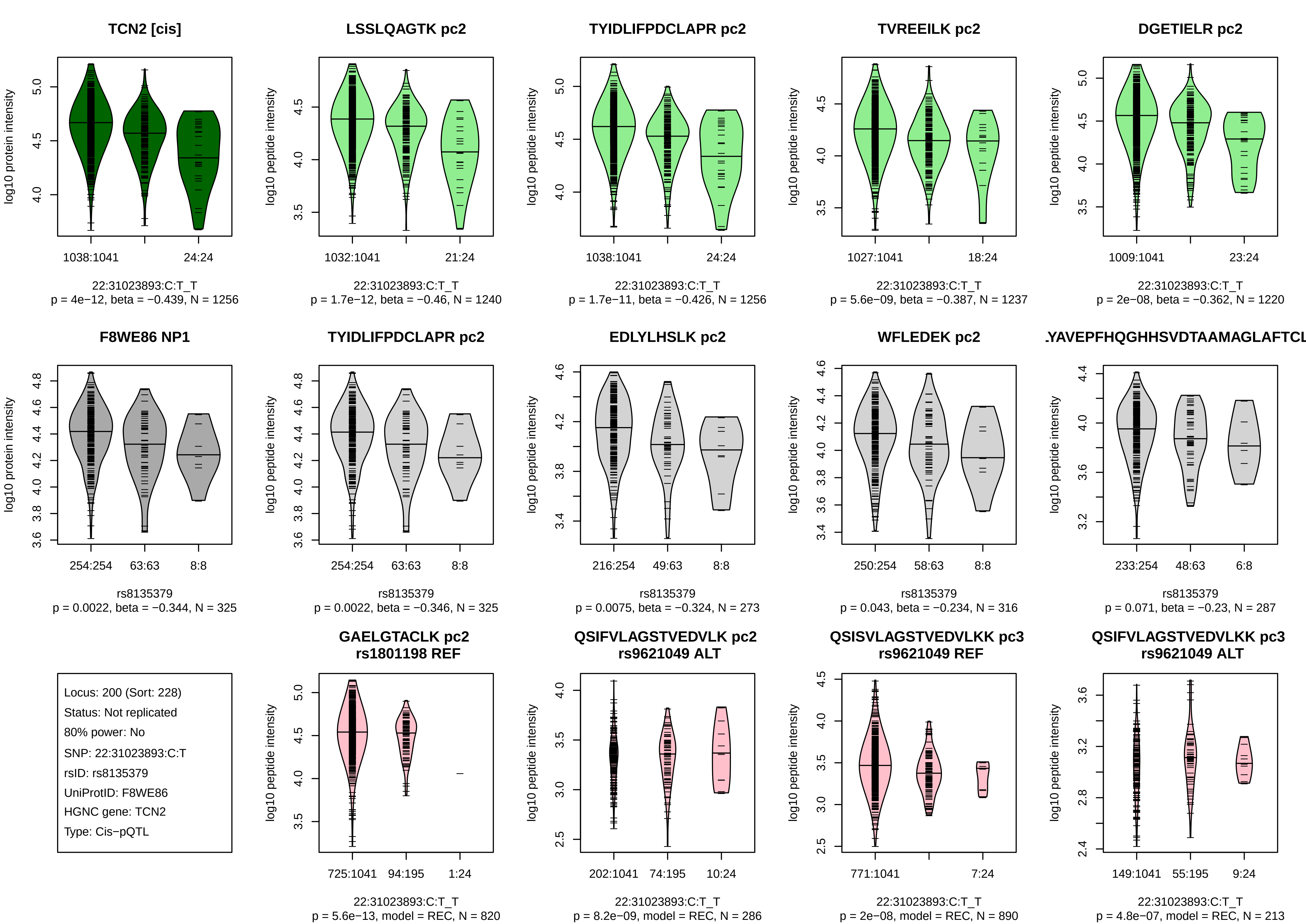
TCN2 [cis]
log10 protein intensity
1038:1041
24:24
22:31023893:C:T_T
p = 4e−12, beta = −0.439, N = 1256
LSSLQAGTK pc2
log10 peptide intensity
1032:1041
21:24
22:31023893:C:T_T
p = 1.7e−12, beta = −0.46, N = 1240
TYIDLIFPDCLAPR pc2
log10 peptide intensity
1038:1041
24:24
22:31023893:C:T_T
p = 1.7e−11, beta = −0.426, N = 1256
TVREEILK pc2
log10 peptide intensity
1027:1041
18:24
22:31023893:C:T_T
p = 5.6e−09, beta = −0.387, N = 1237
DGETIELR pc2
log10 peptide intensity
1009:1041
23:24
22:31023893:C:T_T
p = 2e−08, beta = −0.362, N = 1220
F8WE86 NP1
log10 protein intensity
254:254
63:63
8:8
rs8135379
p = 0.0022, beta = −0.344, N = 325
TYIDLIFPDCLAPR pc2
log10 peptide intensity
254:254
63:63
8:8
rs8135379
p = 0.0022, beta = −0.346, N = 325
EDLYLHSLK pc2
log10 peptide intensity
216:254
49:63
8:8
rs8135379
p = 0.0075, beta = −0.324, N = 273
WFLEDEK pc2
log10 peptide intensity
250:254
58:63
8:8
rs8135379
p = 0.043, beta = −0.234, N = 316
.YAVEPFHQGHHSVDTAAMAGLAFTCL
log10 peptide intensity
233:254
48:63
6:8
rs8135379
p = 0.071, beta = −0.23, N = 287
Locus: 200 (Sort: 228)
Status: Not replicated
80% power: No
SNP: 22:31023893:C:T
rsID: rs8135379
UniProtID: F8WE86
HGNC gene: TCN2
Type: Cis−pQTL
GAELGTACLK pc2
rs1801198 REF
log10 peptide intensity
725:1041
94:195
1:24
22:31023893:C:T_T
p = 5.6e−13, model = REC, N = 820
QSIFVLAGSTVEDVLK pc2
rs9621049 ALT
log10 peptide intensity
202:1041
74:195
10:24
22:31023893:C:T_T
p = 8.2e−09, model = REC, N = 286
QSISVLAGSTVEDVLKK pc3
rs9621049 REF
log10 peptide intensity
771:1041
7:24
22:31023893:C:T_T
p = 2e−08, model = REC, N = 890
QSIFVLAGSTVEDVLKK pc3
rs9621049 ALT
log10 peptide intensity
149:1041
55:195
9:24
22:31023893:C:T_T
p = 4.8e−07, model = REC, N = 213

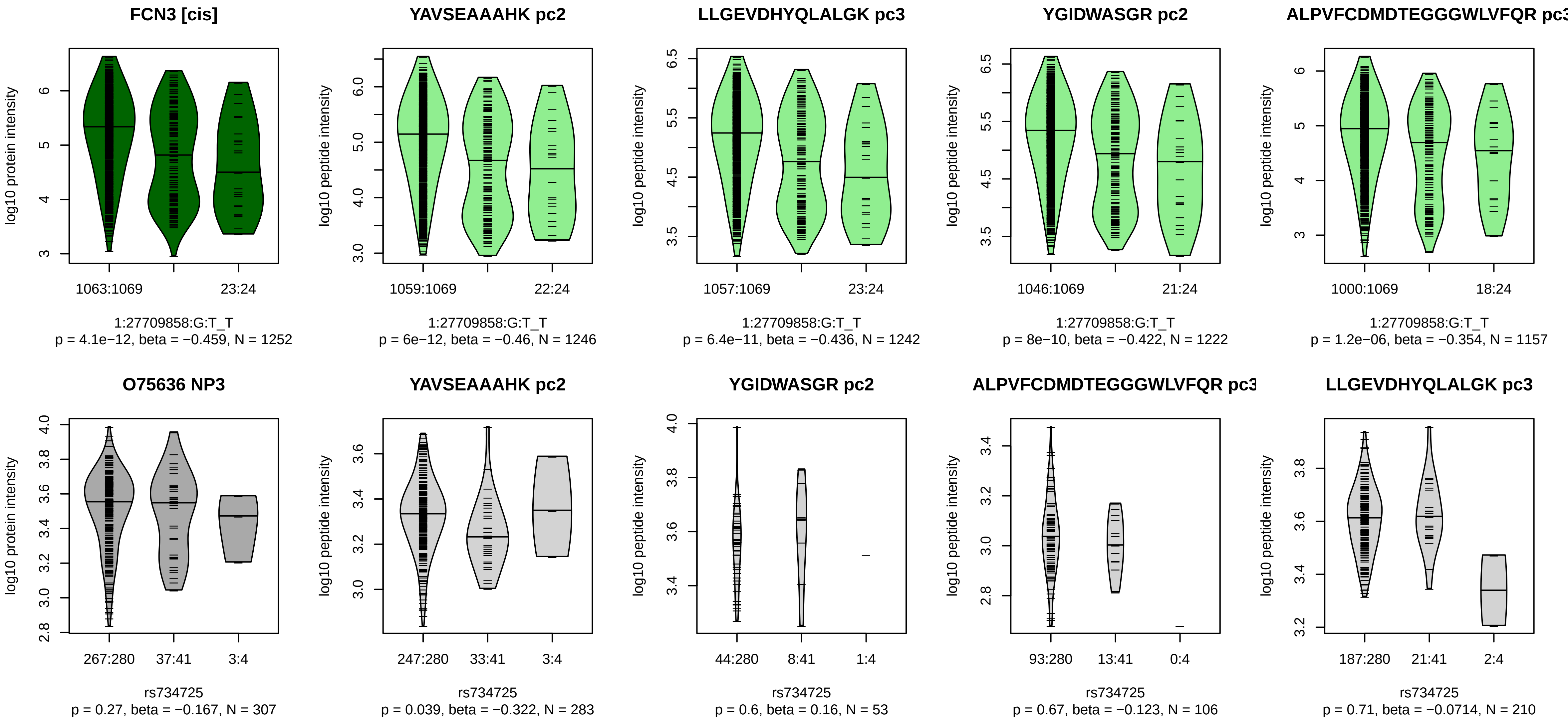


Locus: 201 (Sort: 229)

Status: Not replicated

80% power: No

SNP: 1:27709858:G:T

rsID: rs734725

UniProtID: O75636

HGNC gene: FCN3

Type: Cis−pQTL

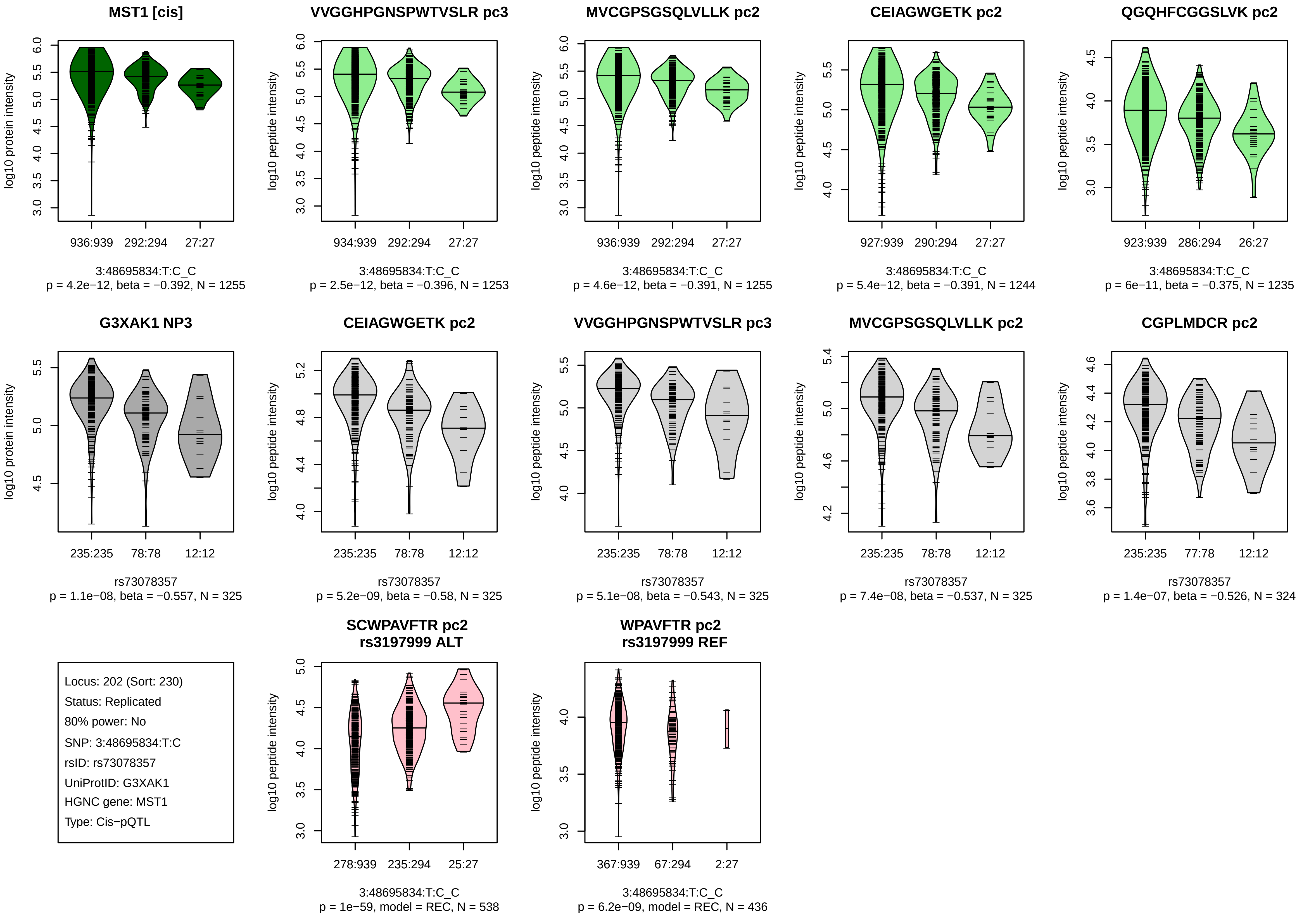
MST1 [cis]
log10 protein intensity
936:939
292:294
27:27
3:48695834:T:C_C
p = 4.2e−12, beta = −0.392, N = 1255
VVGGHPGNSPWTVSLR pc3
log10 peptide intensity
934:939
292:294
27:27
3:48695834:T:C_C
p = 2.5e−12, beta = −0.396, N = 1253
MVCGPSGSQLVLLK pc2
log10 peptide intensity
936:939
292:294
27:27
3:48695834:T:C_C
p = 4.6e−12, beta = −0.391, N = 1255
CEIAGWGETK pc2
log10 peptide intensity
927:939
290:294
27:27
3:48695834:T:C_C
p = 5.4e−12, beta = −0.391, N = 1244
QGQHFCGGSLVK pc2
log10 peptide intensity
923:939
286:294
26:27
3:48695834:T:C_C
p = 6e−11, beta = −0.375, N = 1235
G3XAK1 NP3
log10 protein intensity
235:235
78:78
12:12
rs73078357
p = 1.1e−08, beta = −0.557, N = 325
CEIAGWGETK pc2
log10 peptide intensity
235:235
78:78
12:12
rs73078357
p = 5.2e−09, beta = −0.58, N = 325
VVGGHPGNSPWTVSLR pc3
log10 peptide intensity
235:235
78:78
12:12
rs73078357
p = 5.1e−08, beta = −0.543, N = 325
MVCGPSGSQLVLLK pc2
log10 peptide intensity
235:235
78:78
12:12
rs73078357
p = 7.4e−08, beta = −0.537, N = 325
CGPLMDCR pc2
log10 peptide intensity
235:235
77:78
12:12
rs73078357
p = 1.4e−07, beta = −0.526, N = 324
Locus: 202 (Sort: 230)
Status: Replicated
80% power: No
SNP: 3:48695834:T:C
rsID: rs73078357
UniProtID: G3XAK1
HGNC gene: MST1
Type: Cis−pQTL
SCWPAVFTR pc2
rs3197999 ALT
log10 peptide intensity
278:939
235:294
25:27
3:48695834:T:C_C
p = 1e−59, model = REC, N = 538
WPAVFTR pc2
rs3197999 REF
log10 peptide intensity
367:939
67:294
2:27
3:48695834:T:C_C
p = 6.2e−09, model = REC, N = 436

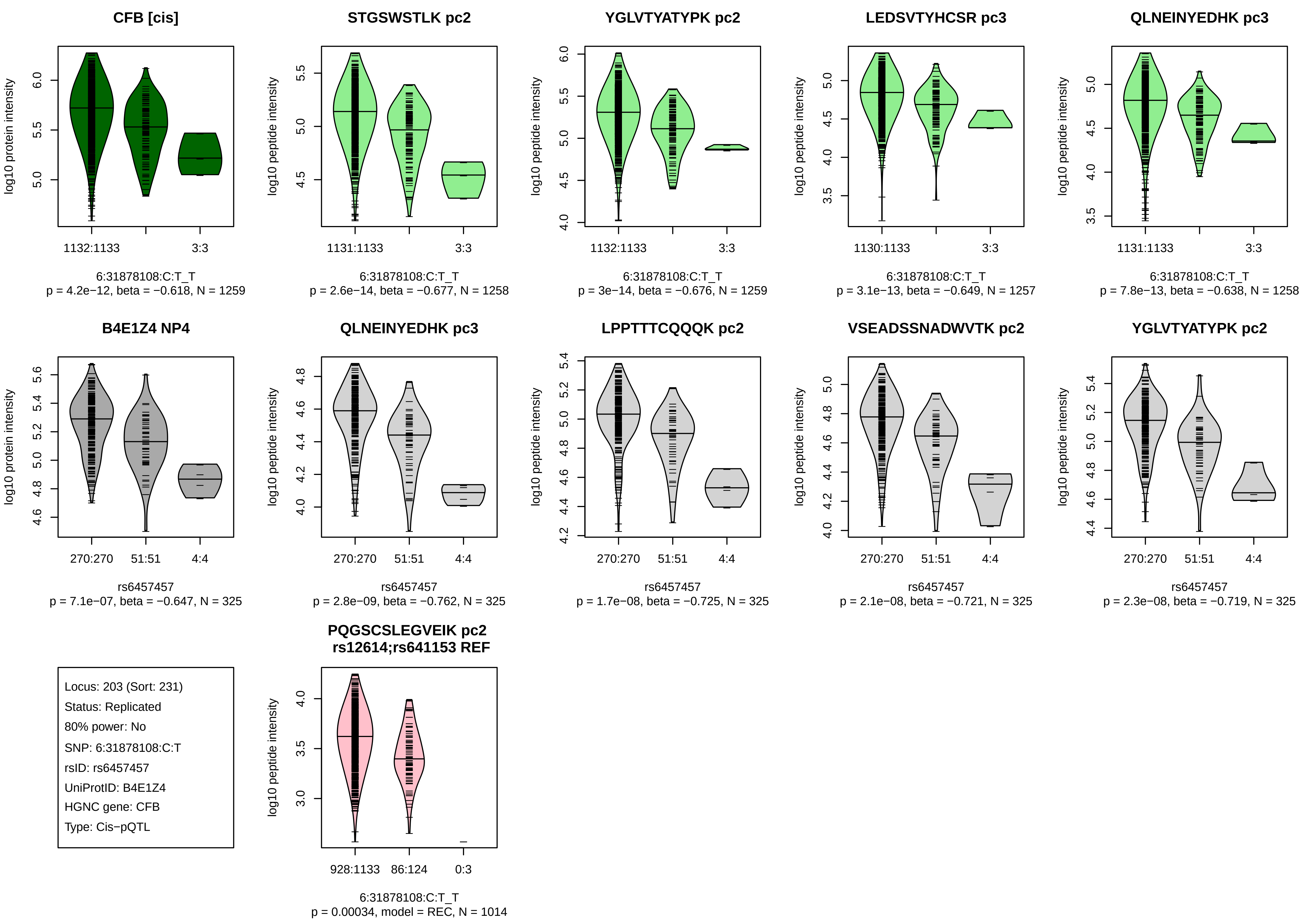
CFB [cis]
log10 protein intensity
1132:1133
3:3
6:31878108:C:T_T
p = 4.2e−12, beta = −0.618, N = 1259
STGSWSTLK pc2
log10 peptide intensity
1131:1133
3:3
6:31878108:C:T_T
p = 2.6e−14, beta = −0.677, N = 1258
YGLVTYATYPK pc2
log10 peptide intensity
1132:1133
3:3
6:31878108:C:T_T
p = 3e−14, beta = −0.676, N = 1259
LEDSVTYHCSR pc3
log10 peptide intensity
1130:1133
3:3
6:31878108:C:T_T
p = 3.1e−13, beta = −0.649, N = 1257
QLNEINYEDHK pc3
log10 peptide intensity
1131:1133
3:3
6:31878108:C:T_T
p = 7.8e−13, beta = −0.638, N = 1258
B4E1Z4 NP4
log10 protein intensity
270:270
51:51
4:4
rs6457457
p = 7.1e−07, beta = −0.647, N = 325
QLNEINYEDHK pc3
log10 peptide intensity
270:270
51:51
4:4
rs6457457
p = 2.8e−09, beta = −0.762, N = 325
LPPTTTCQQQK pc2
log10 peptide intensity
270:270
51:51
4:4
rs6457457
p = 1.7e−08, beta = −0.725, N = 325
VSEADSSNADWVTK pc2
log10 peptide intensity
270:270
51:51
4:4
rs6457457
p = 2.1e−08, beta = −0.721, N = 325
YGLVTYATYPK pc2
log10 peptide intensity
270:270
51:51
4:4
rs6457457
p = 2.3e−08, beta = −0.719, N = 325
Locus: 203 (Sort: 231)
Status: Replicated
80% power: No
SNP: 6:31878108:C:T
rsID: rs6457457
UniProtID: B4E1Z4
HGNC gene: CFB
Type: Cis−pQTL
PQGSCSLEGVEIK pc2
rs12614;rs641153 REF
log10 peptide intensity
928:1133
86:124
0:3
6:31878108:C:T_T
p = 0.00034, model = REC, N = 1014

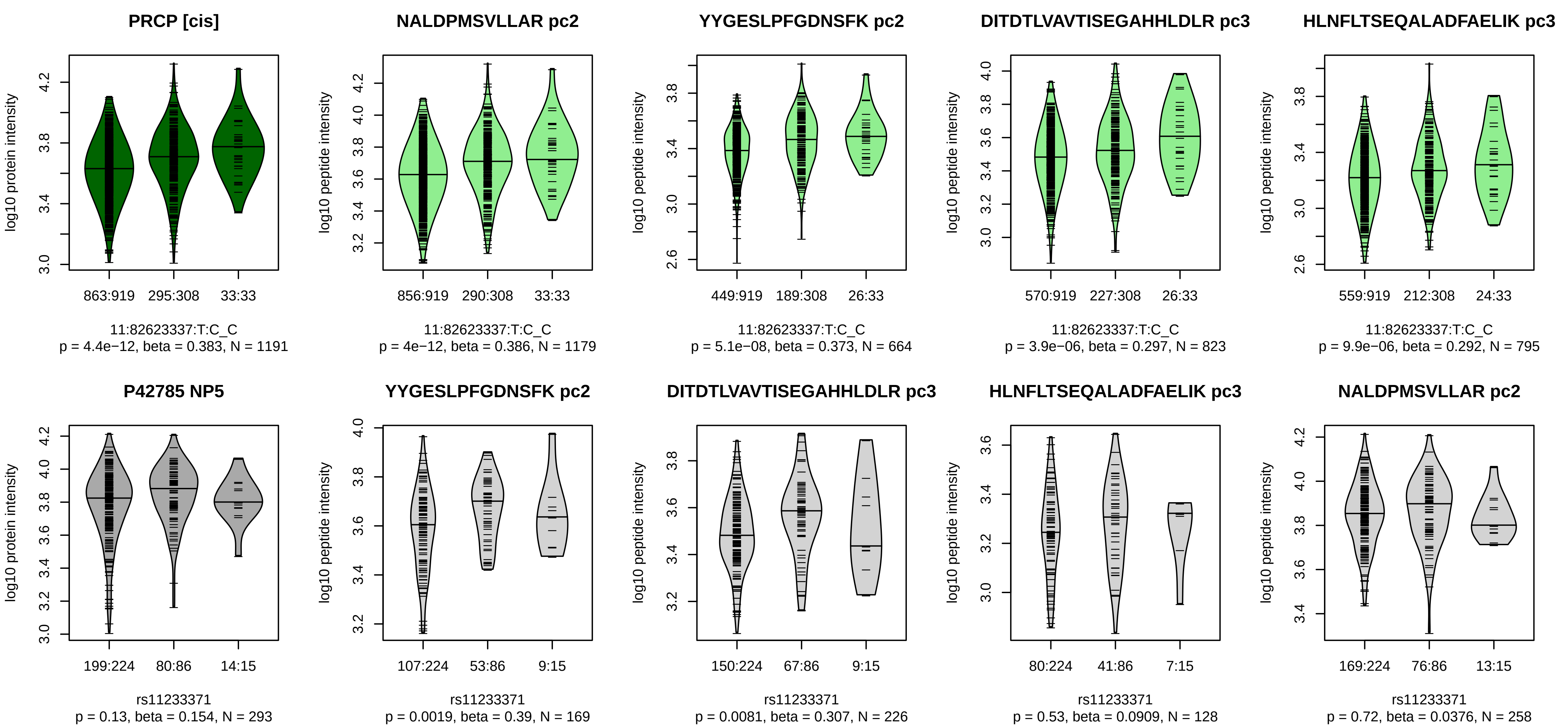


Locus: 204 (Sort: 232)
Status: Not replicated
80% power: No
SNP: 11:82623337:T:C
rsID: rs11233371
UniProtID: P42785
HGNC gene: PRCP
Type: Cis−pQTL

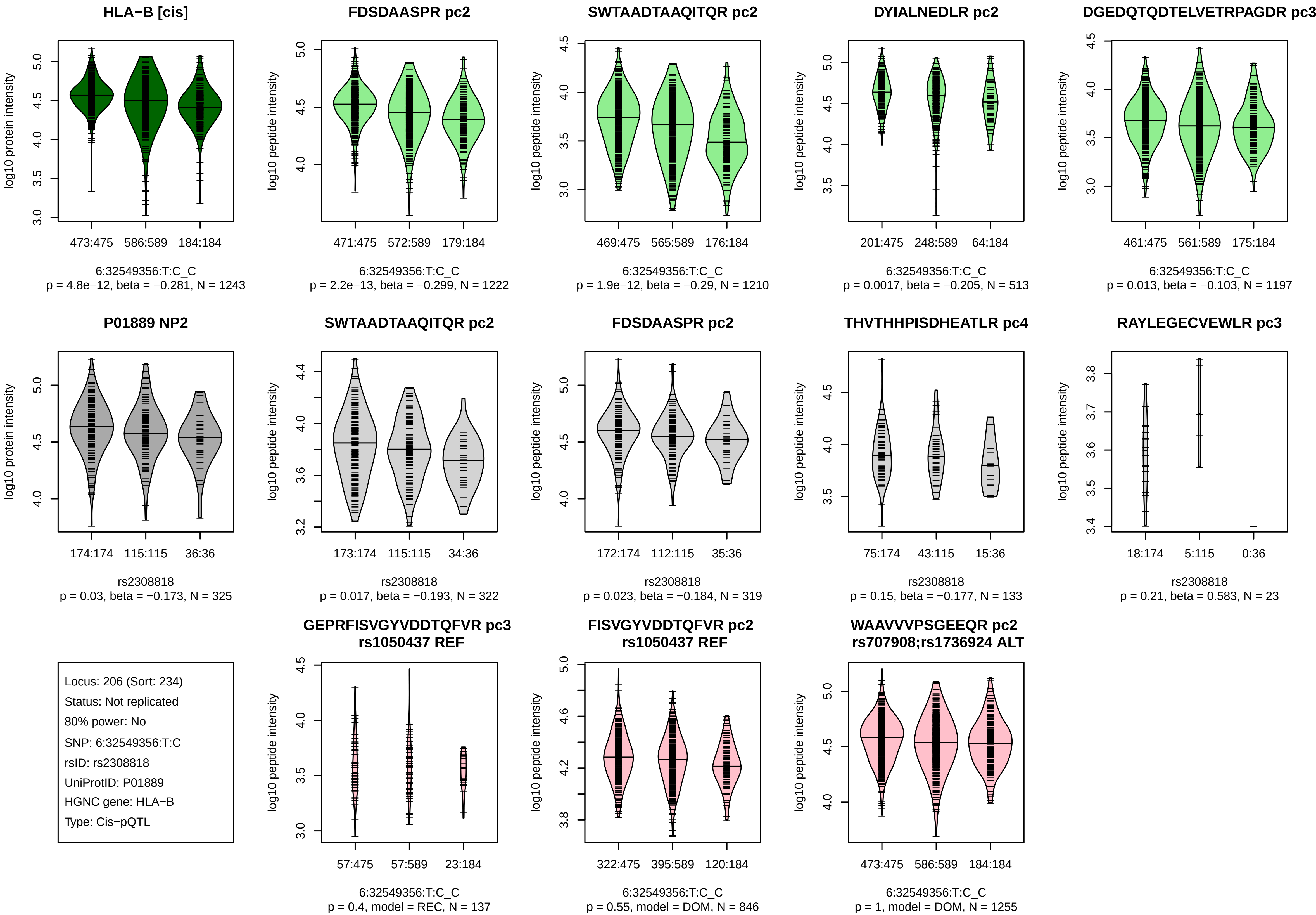
HLA−B [cis]
log10 protein intensity
473:475
586:589
184:184
6:32549356:T:C_C
p = 4.8e−12, beta = −0.281, N = 1243
FDSDAASPR pc2
log10 peptide intensity
471:475
572:589
179:184
6:32549356:T:C_C
p = 2.2e−13, beta = −0.299, N = 1222
SWTAADTAAQITQR pc2
log10 peptide intensity
469:475
565:589
176:184
6:32549356:T:C_C
p = 1.9e−12, beta = −0.29, N = 1210
DYIALNEDLR pc2
log10 peptide intensity
201:475
248:589
64:184
6:32549356:T:C_C
p = 0.0017, beta = −0.205, N = 513
DGEDQTQDTELVETRPAGDR pc3
log10 peptide intensity
461:475
561:589
175:184
6:32549356:T:C_C
p = 0.013, beta = −0.103, N = 1197
P01889 NP2
log10 protein intensity
174:174
115:115
36:36
rs2308818
p = 0.03, beta = −0.173, N = 325
SWTAADTAAQITQR pc2
log10 peptide intensity
173:174
115:115
34:36
rs2308818
p = 0.017, beta = −0.193, N = 322
FDSDAASPR pc2
log10 peptide intensity
172:174
112:115
35:36
rs2308818
p = 0.023, beta = −0.184, N = 319
THVTHHPISDHEATLR pc4
log10 peptide intensity
75:174
43:115
15:36
rs2308818
p = 0.15, beta = −0.177, N = 133
RAYLEGECVEWLR pc3
log10 peptide intensity
18:174
5:115
0:36
rs2308818
p = 0.21, beta = 0.583, N = 23
Locus: 206 (Sort: 234)
Status: Not replicated
80% power: No
SNP: 6:32549356:T:C
rsID: rs2308818
UniProtID: P01889
HGNC gene: HLA−B
Type: Cis−pQTL
GEPRFISVGYVDDTQFVR pc3
rs1050437 REF
log10 peptide intensity
57:475
57:589
23:184
6:32549356:T:C_C
p = 0.4, model = REC, N = 137
FISVGYVDDTQFVR pc2
rs1050437 REF
log10 peptide intensity
322:475
395:589
120:184
6:32549356:T:C_C
p = 0.55, model = DOM, N = 846
WAAVVVPSGEEQR pc2
rs707908;rs1736924 ALT
log10 peptide intensity
473:475
586:589
184:184
6:32549356:T:C_C
p = 1, model = DOM, N = 1255

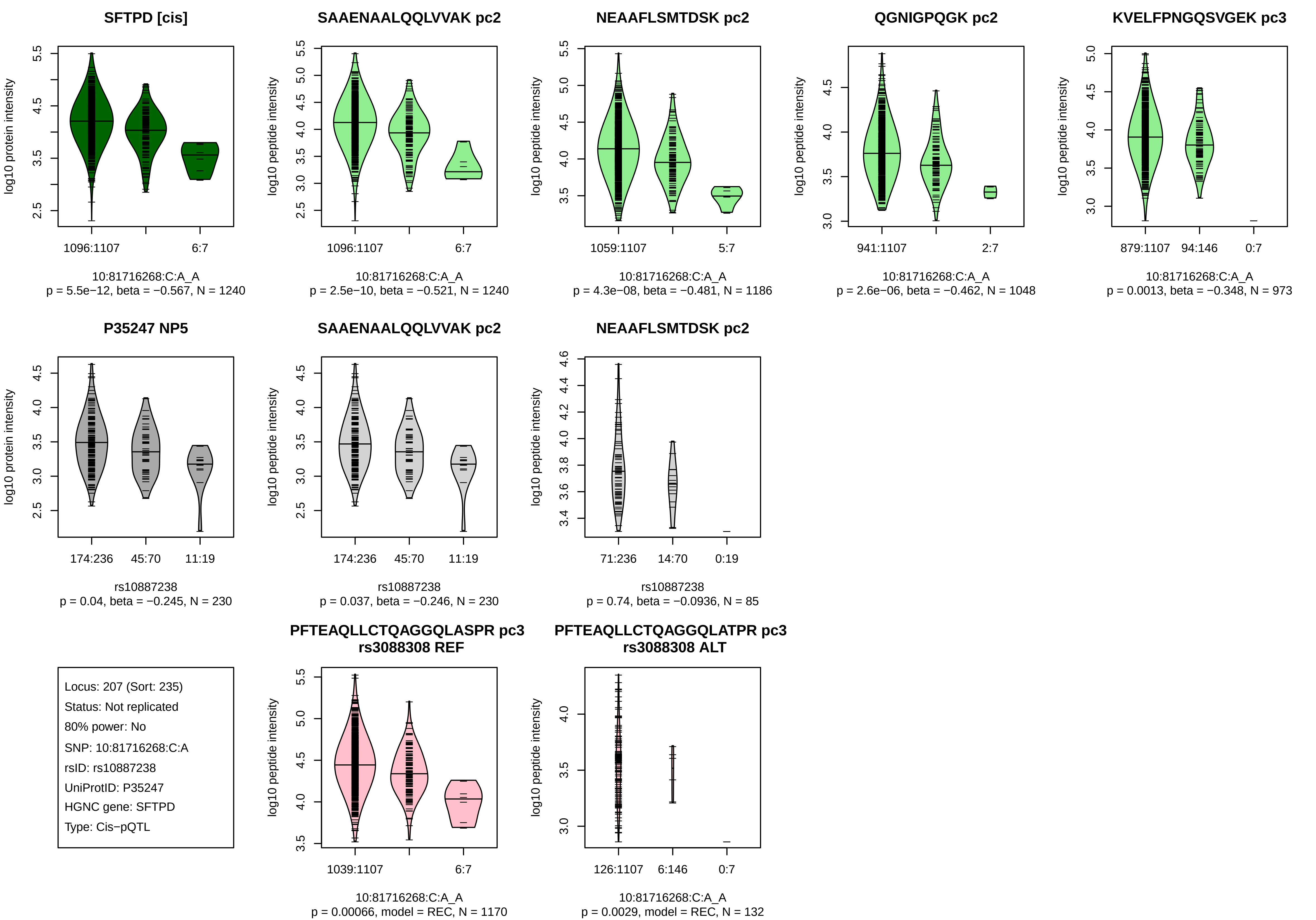
SFTPD [cis]
log10 protein intensity
1096:1107
6:7
10:81716268:C:A_A
p = 5.5e−12, beta = −0.567, N = 1240
SAAENAALQQLVVAK pc2
log10 peptide intensity
1096:1107
6:7
10:81716268:C:A_A
p = 2.5e−10, beta = −0.521, N = 1240
NEAAFLSMTDSK pc2
log10 peptide intensity
1059:1107
5:7
10:81716268:C:A_A
p = 4.3e−08, beta = −0.481, N = 1186
QGNIGPQGK pc2
log10 peptide intensity
941:1107
2:7
10:81716268:C:A_A
p = 2.6e−06, beta = −0.462, N = 1048
KVELFPNGQSVGEK pc3
log10 peptide intensity
879:1107
94:146
0:7
10:81716268:C:A_A
p = 0.0013, beta = −0.348, N = 973
P35247 NP5
log10 protein intensity
174:236
45:70
11:19
rs10887238
p = 0.04, beta = −0.245, N = 230
SAAENAALQQLVVAK pc2
log10 peptide intensity
174:236
45:70
11:19
rs10887238
p = 0.037, beta = −0.246, N = 230
NEAAFLSMTDSK pc2
log10 peptide intensity
71:236
14:70
0:19
rs10887238
p = 0.74, beta = −0.0936, N = 85
Locus: 207 (Sort: 235)
Status: Not replicated
80% power: No
SNP: 10:81716268:C:A
rsID: rs10887238
UniProtID: P35247
HGNC gene: SFTPD
Type: Cis−pQTL
PFTEAQLLCTQAGGQLASPR pc3
rs3088308 REF
log10 peptide intensity
1039:1107
6:7
10:81716268:C:A_A
p = 0.00066, model = REC, N = 1170
PFTEAQLLCTQAGGQLATPR pc3
rs3088308 ALT
log10 peptide intensity
126:1107
6:146
0:7
10:81716268:C:A_A
p = 0.0029, model = REC, N = 132

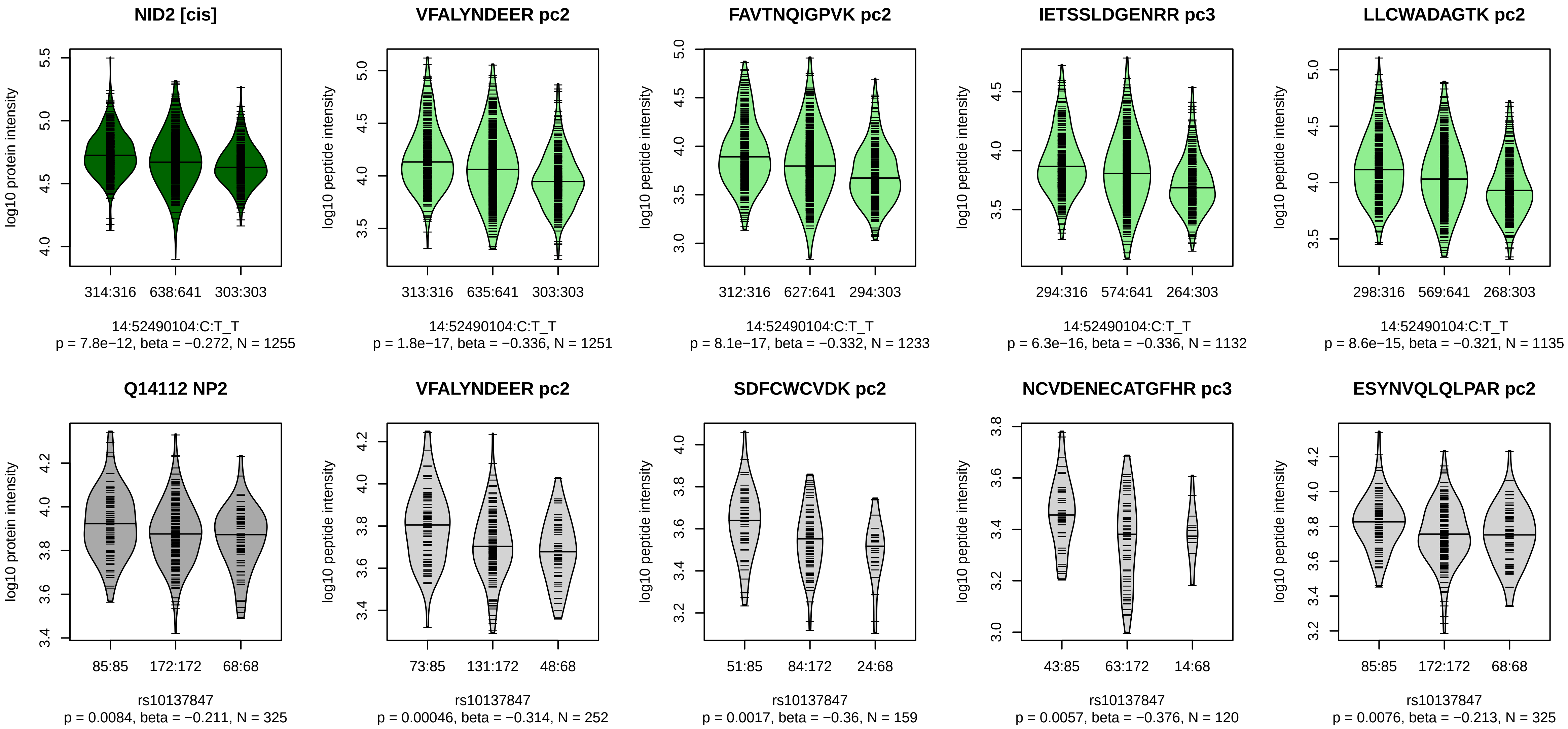


Locus: 208 (Sort: 236)
Status: Not replicated
80% power: No
SNP: 14:52490104:C:T
rsID: rs10137847
UniProtID: Q14112
HGNC gene: NID2
Type: Cis−pQTL

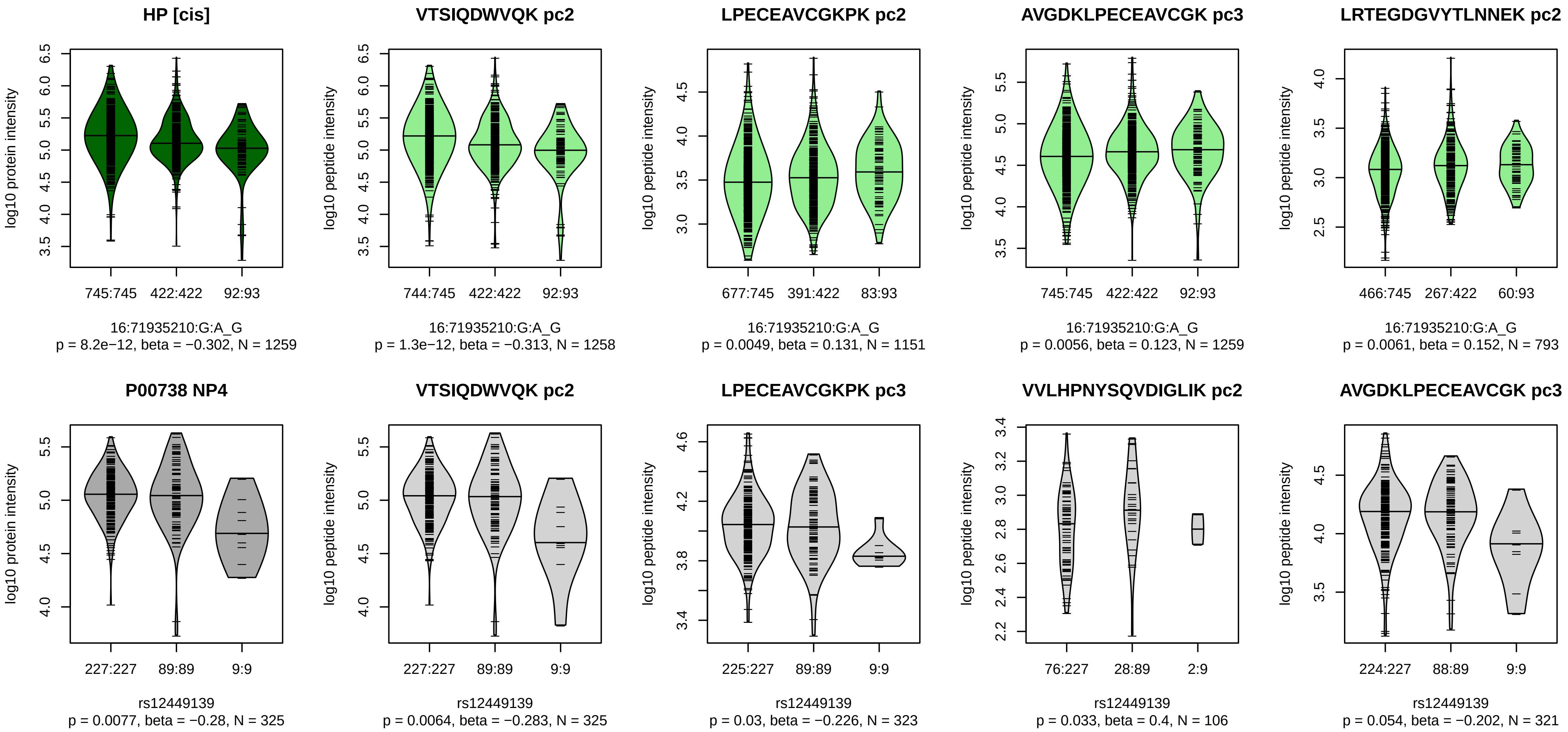


Locus: 209 (Sort: 237)
Status: Not replicated
80% power: No
SNP: 16:71935210:G:A
rsID: rs12449139
UniProtID: P00738
HGNC gene: HP
Type: Cis−pQTL

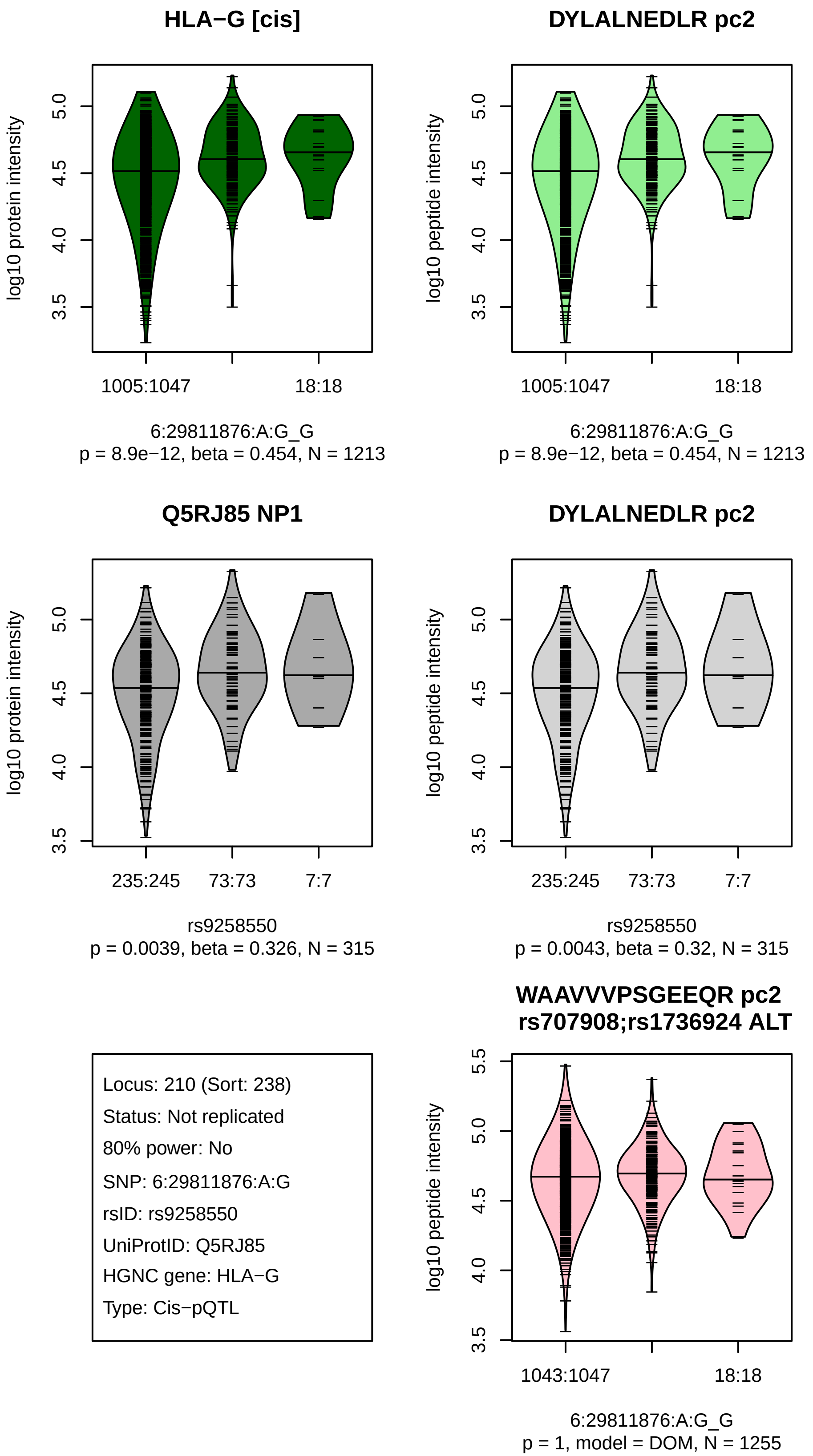
HLA−G [cis]
log10 protein intensity
1005:1047
18:18
6:29811876:A:G_G
p = 8.9e−12, beta = 0.454, N = 1213
DYLALNEDLR pc2
log10 peptide intensity
1005:1047
18:18
6:29811876:A:G_G
p = 8.9e−12, beta = 0.454, N = 1213
Q5RJ85 NP1
log10 protein intensity
235:245
73:73
7:7
rs9258550
p = 0.0039, beta = 0.326, N = 315
DYLALNEDLR pc2
log10 peptide intensity
235:245
73:73
7:7
rs9258550
p = 0.0043, beta = 0.32, N = 315
Locus: 210 (Sort: 238)
Status: Not replicated
80% power: No
SNP: 6:29811876:A:G
rsID: rs9258550
UniProtID: Q5RJ85
HGNC gene: HLA−G
Type: Cis−pQTL
WAAVVVPSGEEQR pc2
rs707908;rs1736924 ALT
log10 peptide intensity
1043:1047
18:18
6:29811876:A:G_G
p = 1, model = DOM, N = 1255

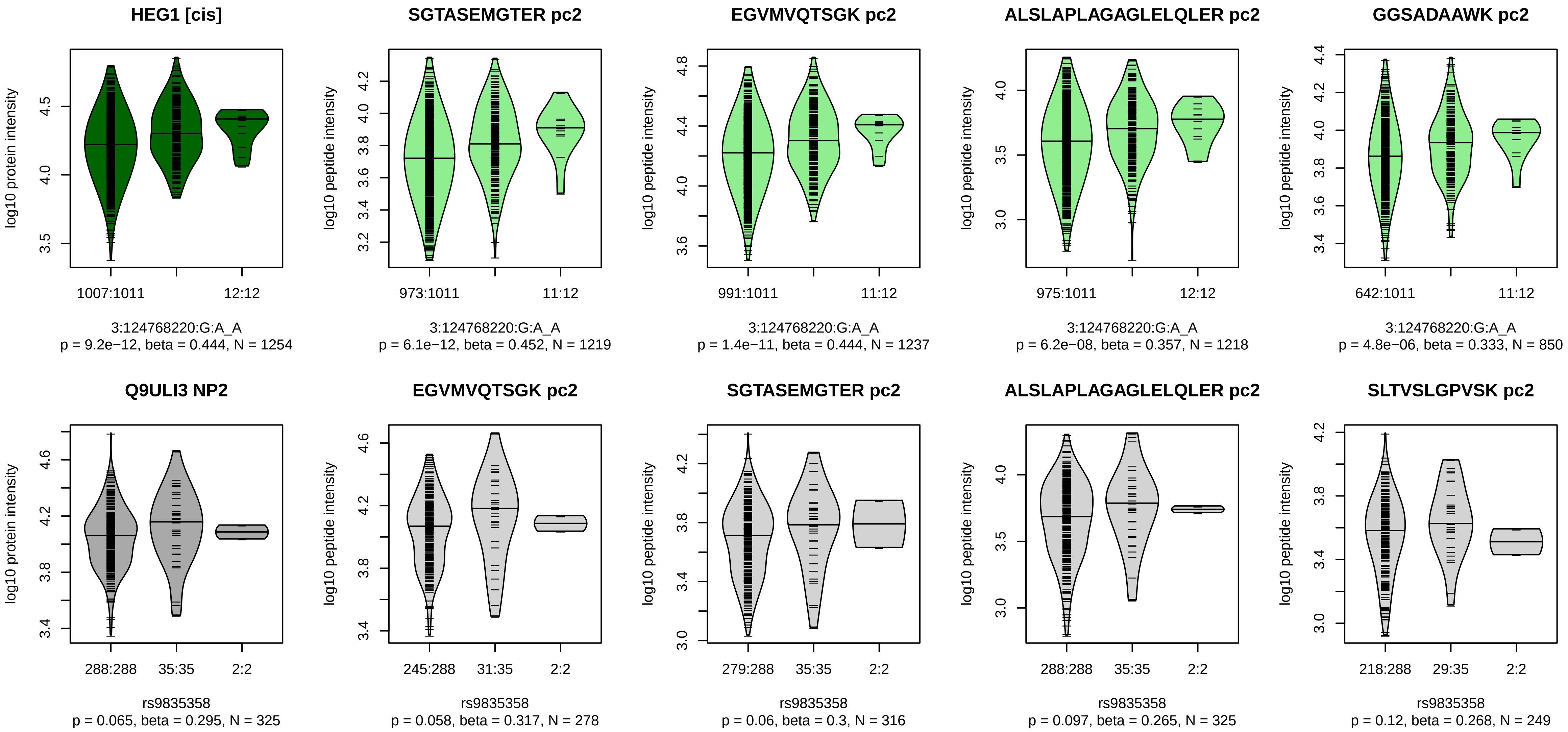


Locus: 211 (Sort: 239)
Status: Not replicated
80% power: No
SNP: 3:124768220:G:A
rsID: rs9835358
UniProtID: Q9ULI3
HGNC gene: HEG1
Type: Cis−pQTL

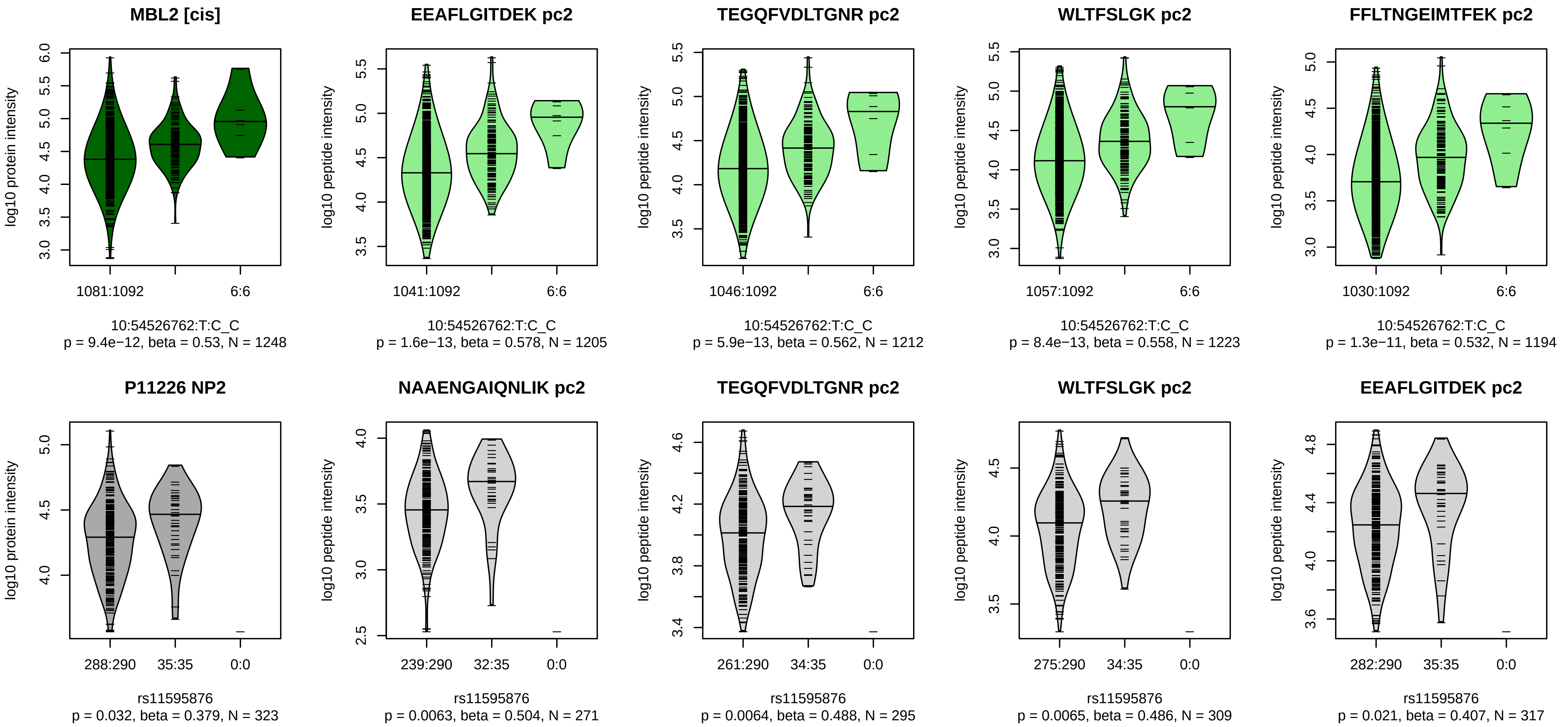


Locus: 212 (Sort: 240)
Status: Not replicated
80% power: No
SNP: 10:54526762:T:C
rsID: rs11595876
UniProtID: P11226
HGNC gene: MBL2
Type: Cis−pQTL

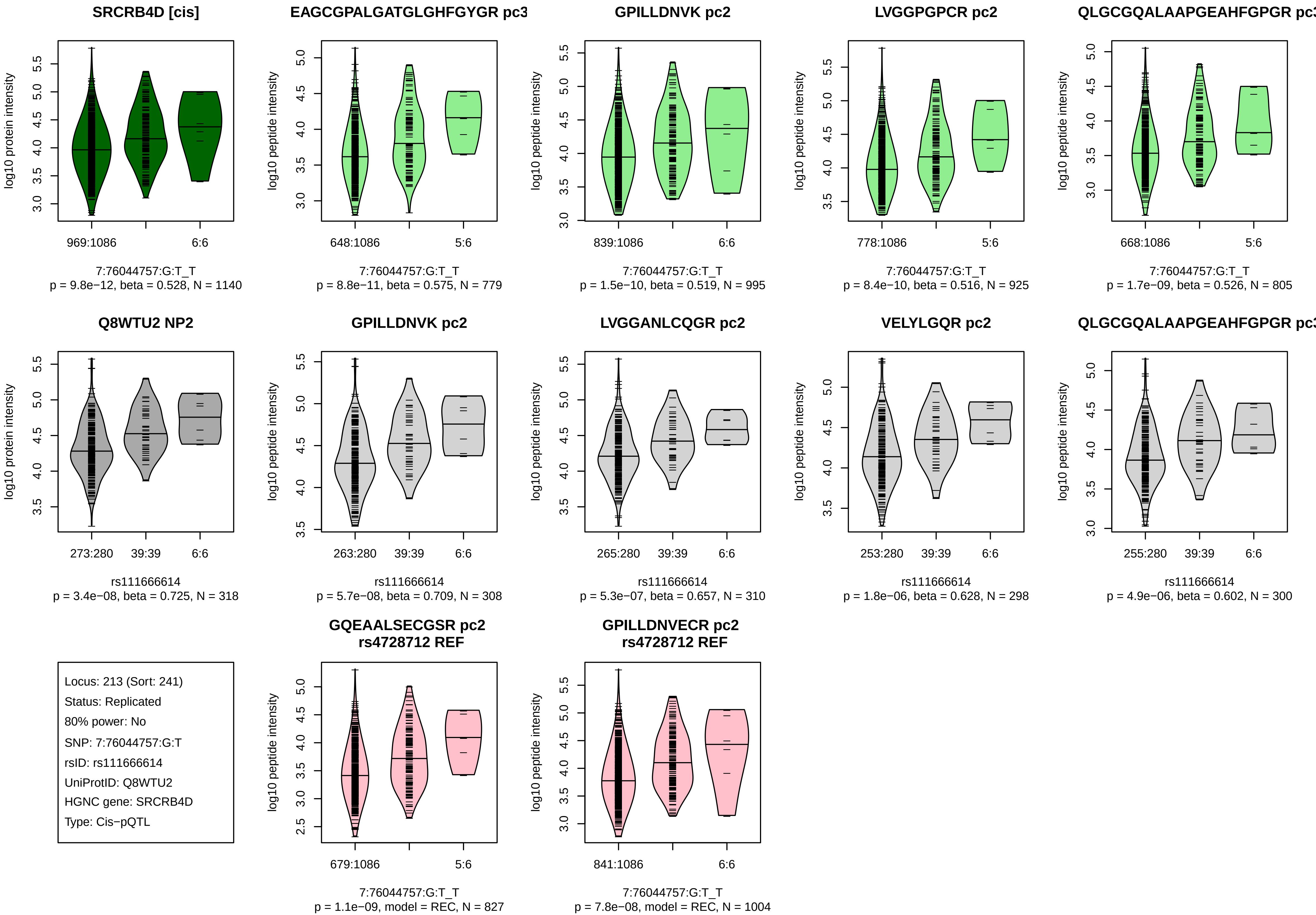

SRCRB4D [cis]
log10 protein intensity
969:1086
6:6
7:76044757:G:T_T
p = 9.8e−12, beta = 0.528, N = 1140
EAGCGPALGATGLGHFGYGR pc3
log10 peptide intensity
648:1086
5:6
7:76044757:G:T_T
p = 8.8e−11, beta = 0.575, N = 779
GPILLDNVK pc2
log10 peptide intensity
839:1086
6:6
7:76044757:G:T_T
p = 1.5e−10, beta = 0.519, N = 995
LVGGPGPCR pc2
log10 peptide intensity
778:1086
5:6
7:76044757:G:T_T
p = 8.4e−10, beta = 0.516, N = 925
QLGCGQALAAPGEAHFGPGR pc3
log10 peptide intensity
668:1086
5:6
7:76044757:G:T_T
p = 1.7e−09, beta = 0.526, N = 805
Q8WTU2 NP2
log10 protein intensity
273:280
39:39
6:6
rs111666614
p = 3.4e−08, beta = 0.725, N = 318
GPILLDNVK pc2
log10 peptide intensity
263:280
39:39
6:6
rs111666614
p = 5.7e−08, beta = 0.709, N = 308
LVGGANLCQGR pc2
log10 peptide intensity
265:280
39:39
6:6
rs111666614
p = 5.3e−07, beta = 0.657, N = 310
VELYLGQR pc2
log10 peptide intensity
253:280
39:39
6:6
rs111666614
p = 1.8e−06, beta = 0.628, N = 298
QLGCGQALAAPGEAHFGPGR pc3
log10 peptide intensity
255:280
39:39
6:6
rs111666614
p = 4.9e−06, beta = 0.602, N = 300
Locus: 213 (Sort: 241)
Status: Replicated
80% power: No
SNP: 7:76044757:G:T
rsID: rs111666614
UniProtID: Q8WTU2
HGNC gene: SRCRB4D
Type: Cis−pQTL
GQEAALSECGSR pc2
rs4728712 REF
log10 peptide intensity
679:1086
5:6
7:76044757:G:T_T
p = 1.1e−09, model = REC, N = 827
GPILLDNVECR pc2
rs4728712 REF
log10 peptide intensity
841:1086
6:6
7:76044757:G:T_T
p = 7.8e−08, model = REC, N = 1004

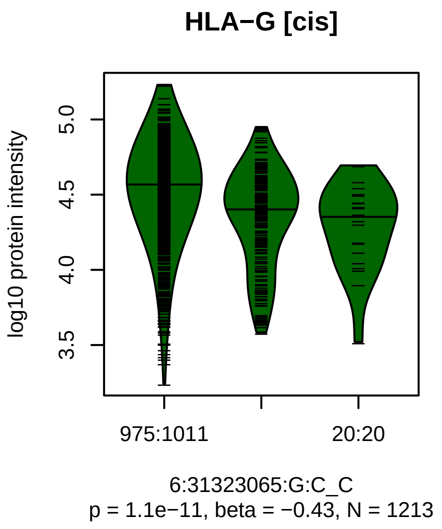


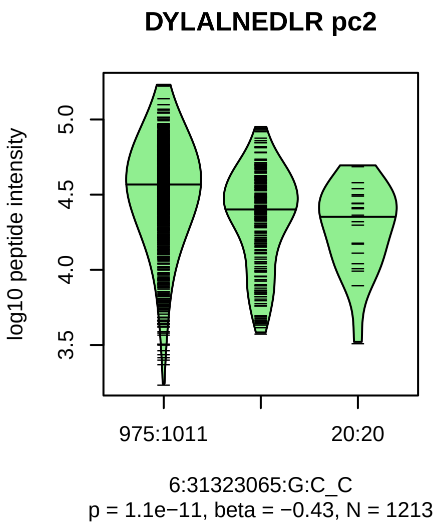


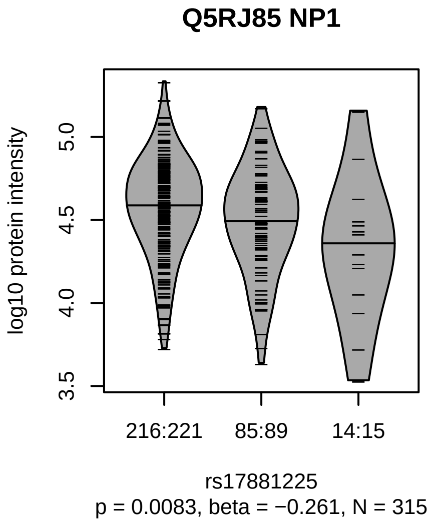


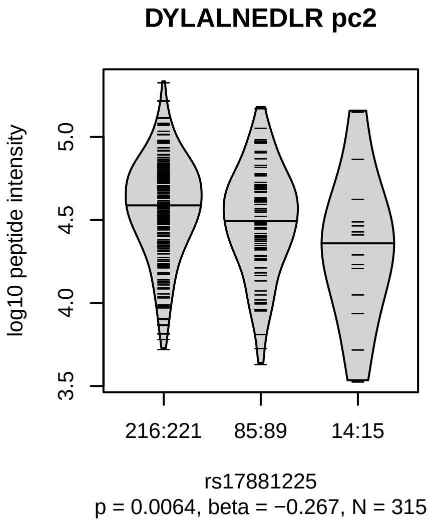


Locus: 214 (Sort: 242)

Status: Not replicated

80% power: No

SNP: 6:31323065:G:C

rsID: rs17881225

UniProtID: Q5RJ85

HGNC gene: HLA−G

Type: Cis−pQTL

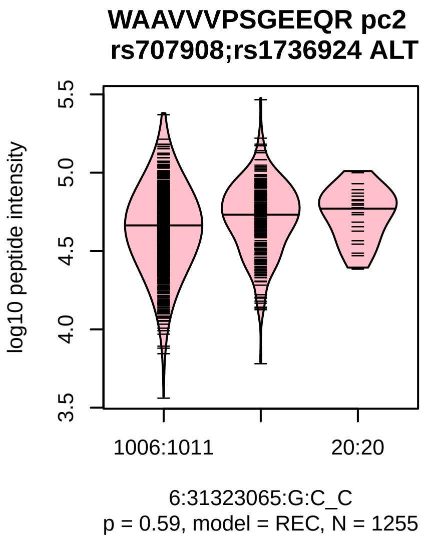

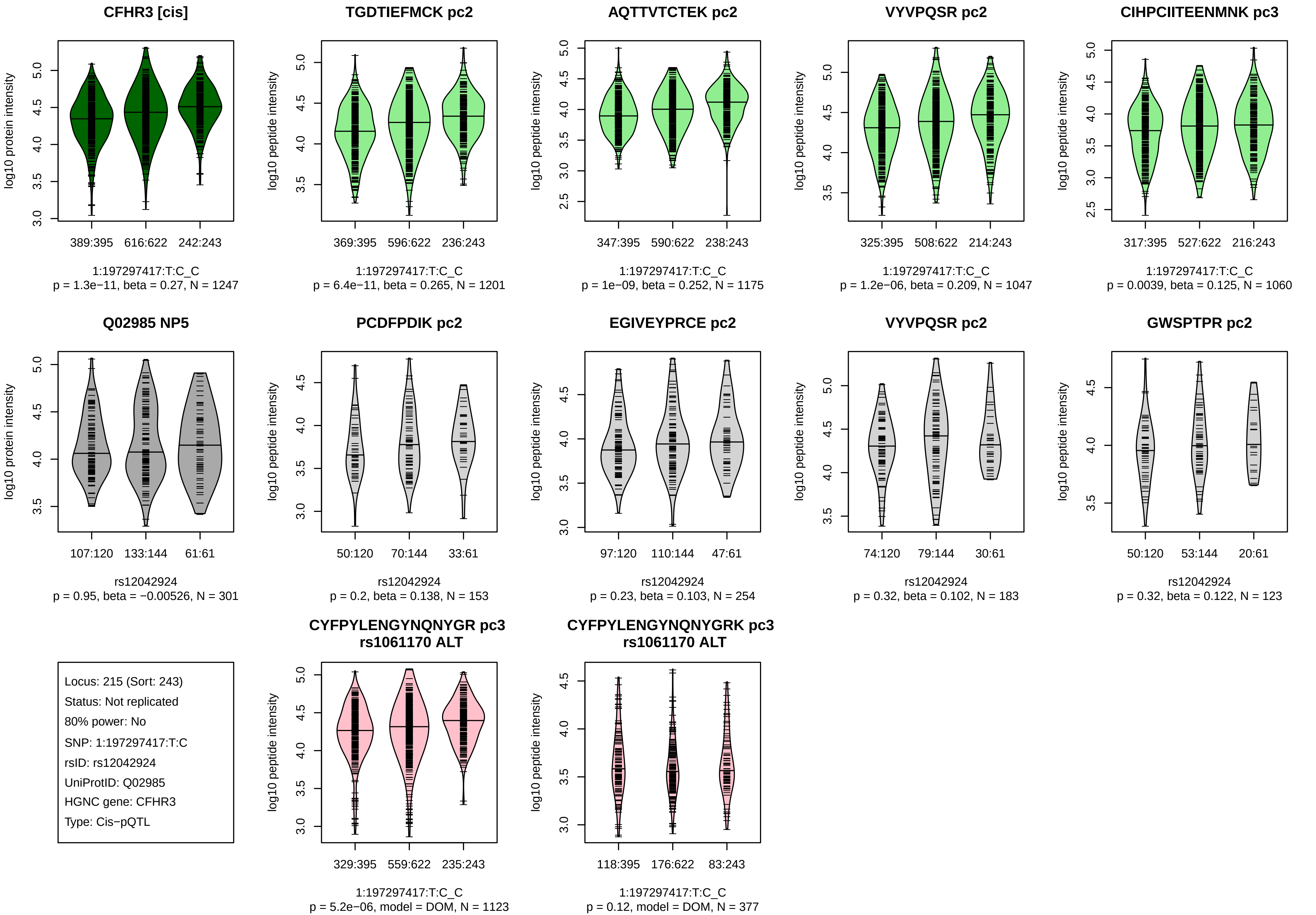
CFHR3 [cis]
log10 protein intensity
389:395
616:622
242:243
1:197297417:T:C_C
p = 1.3e−11, beta = 0.27, N = 1247
TGDTIEFMCK pc2
log10 peptide intensity
369:395
596:622
236:243
1:197297417:T:C_C
p = 6.4e−11, beta = 0.265, N = 1201
AQTTVTCTEK pc2
log10 peptide intensity
347:395
590:622
238:243
1:197297417:T:C_C
p = 1e−09, beta = 0.252, N = 1175
VYVPQSR pc2
log10 peptide intensity
325:395
508:622
214:243
1:197297417:T:C_C
p = 1.2e−06, beta = 0.209, N = 1047
CIHPCIITEENMNK pc3
log10 peptide intensity
317:395
527:622
216:243
1:197297417:T:C_C
p = 0.0039, beta = 0.125, N = 1060
Q02985 NP5
log10 protein intensity
107:120
133:144
61:61
rs12042924
p = 0.95, beta = −0.00526, N = 301
PCDFPDIK pc2
log10 peptide intensity
50:120
70:144
33:61
rs12042924
p = 0.2, beta = 0.138, N = 153
EGIVEYPRCE pc2
log10 peptide intensity
97:120
110:144
47:61
rs12042924
p = 0.23, beta = 0.103, N = 254
VYVPQSR pc2
log10 peptide intensity
74:120
79:144
30:61
rs12042924
p = 0.32, beta = 0.102, N = 183
GWSPTPR pc2
log10 peptide intensity
50:120
53:144
20:61
rs12042924
p = 0.32, beta = 0.122, N = 123
Locus: 215 (Sort: 243)
Status: Not replicated
80% power: No
SNP: 1:197297417:T:C
rsID: rs12042924
UniProtID: Q02985
HGNC gene: CFHR3
Type: Cis−pQTL
CYFPYLENGYNQNYGR pc3
rs1061170 ALT
log10 peptide intensity
329:395
559:622
235:243
1:197297417:T:C_C
p = 5.2e−06, model = DOM, N = 1123
CYFPYLENGYNQNYGRK pc3
rs1061170 ALT
log10 peptide intensity
118:395
176:622
83:243
1:197297417:T:C_C
p = 0.12, model = DOM, N = 377

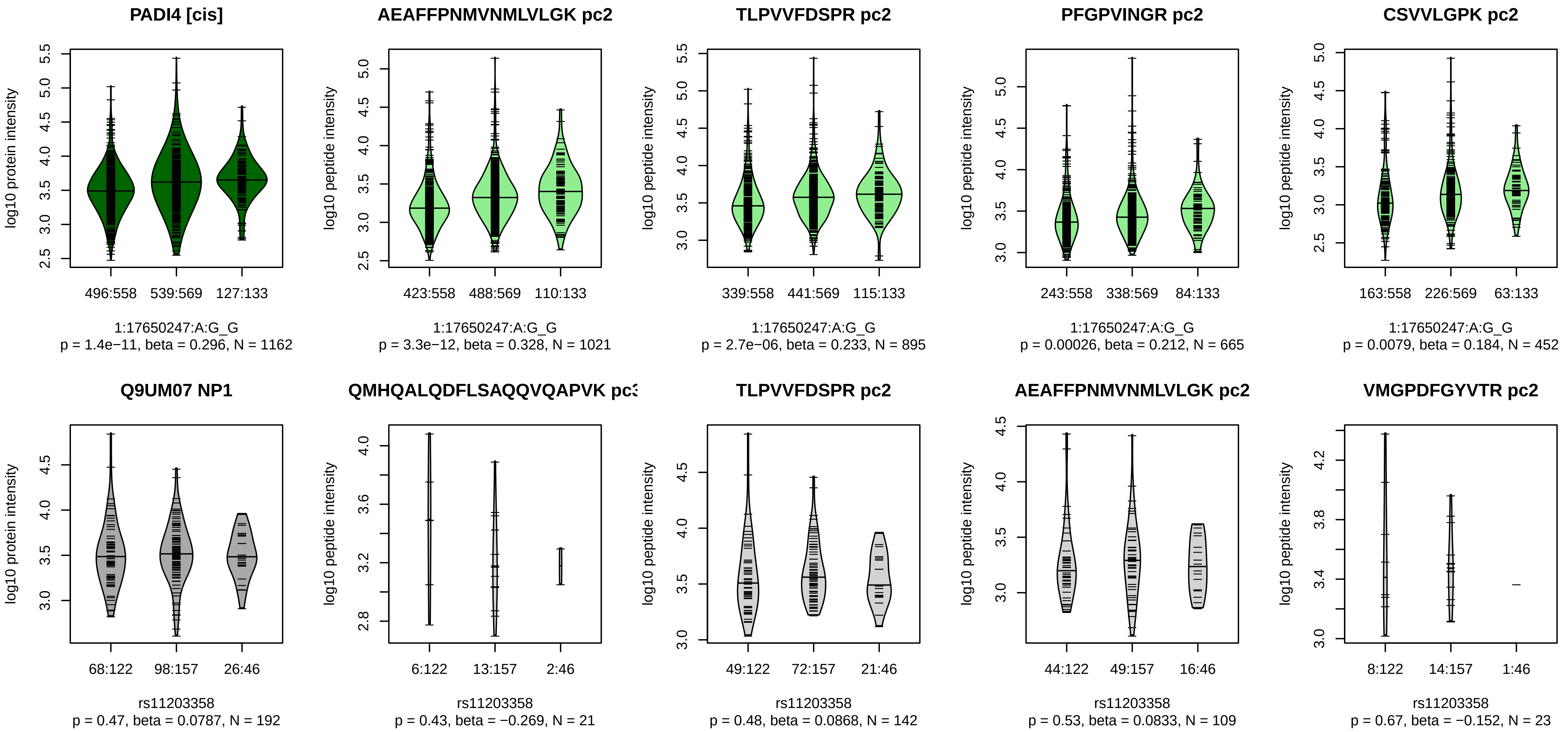


Locus: 216 (Sort: 244)

Status: Not replicated

80% power: No

SNP: 1:17650247:A:G

rsID: rs11203358

UniProtID: Q9UM07

HGNC gene: PADI4

Type: Cis−pQTL

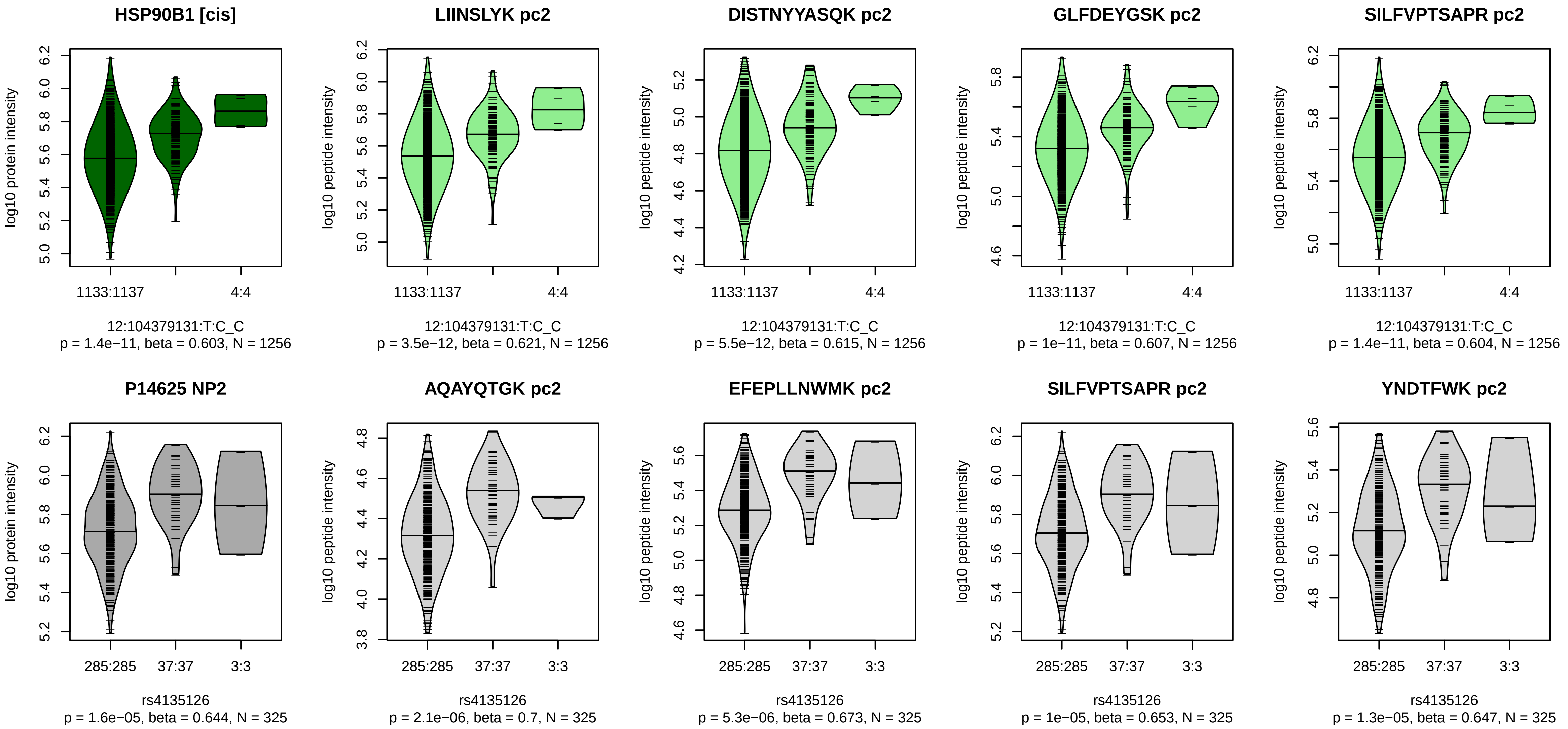


Locus: 217 (Sort: 245)

Status: Replicated

80% power: No

SNP: 12:104379131:T:C

rsID: rs4135126

UniProtID: P14625

HGNC gene: HSP90B1

Type: Cis−pQTL

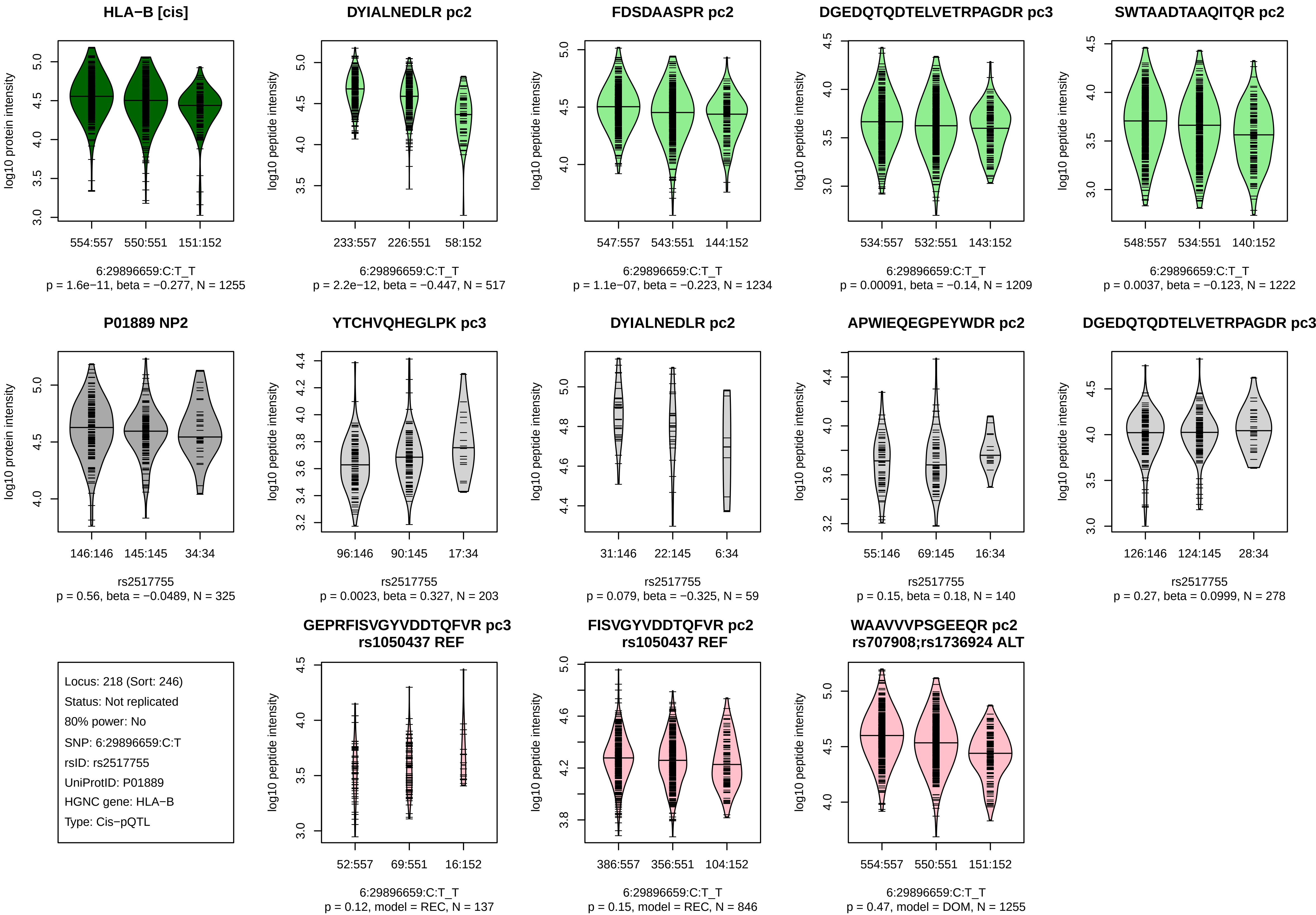

HLA−B [cis]
log10 protein intensity
554:557
550:551
151:152
6:29896659:C:T_T
p = 1.6e−11, beta = −0.277, N = 1255
DYIALNEDLR pc2
log10 peptide intensity
233:557
226:551
58:152
6:29896659:C:T_T
p = 2.2e−12, beta = −0.447, N = 517
FDSDAASPR pc2
547:557
543:551
144:152
6:29896659:C:T_T
p = 1.1e−07, beta = −0.223, N = 1234
DGEDQTQDTELVETRPAGDR pc3
534:557
532:551
143:152
6:29896659:C:T_T
p = 0.00091, beta = −0.14, N = 1209
SWTAADTAAQITQR pc2
548:557
534:551
140:152
6:29896659:C:T_T
p = 0.0037, beta = −0.123, N = 1222
P01889 NP2
146:146
145:145
34:34
rs2517755
p = 0.56, beta = −0.0489, N = 325
YTCHVQHEGLPK pc3
96:146
90:145
17:34
rs2517755
p = 0.0023, beta = 0.327, N = 203
DYIALNEDLR pc2
31:146
22:145
6:34
rs2517755
p = 0.079, beta = −0.325, N = 59
APWIEQEGPEYWDR pc2
55:146
69:145
16:34
rs2517755
p = 0.15, beta = 0.18, N = 140
DGEDQTQDTELVETRPAGDR pc3
126:146
124:145
28:34
rs2517755
p = 0.27, beta = 0.0999, N = 278
Locus: 218 (Sort: 246)
Status: Not replicated
80% power: No
SNP: 6:29896659:C:T
rsID: rs2517755
UniProtID: P01889
HGNC gene: HLA−B
Type: Cis−pQTL
GEPRFISVGYVDDTQFVR pc3
rs1050437 REF
52:557
69:551
16:152
6:29896659:C:T_T
p = 0.12, model = REC, N = 137
FISVGYVDDTQFVR pc2
rs1050437 REF
386:557
356:551
104:152
6:29896659:C:T_T
p = 0.15, model = REC, N = 846
WAAVVVPSGEEQR pc2
rs707908;rs1736924 ALT
554:557
550:551
151:152
6:29896659:C:T_T
p = 0.47, model = DOM, N = 1255

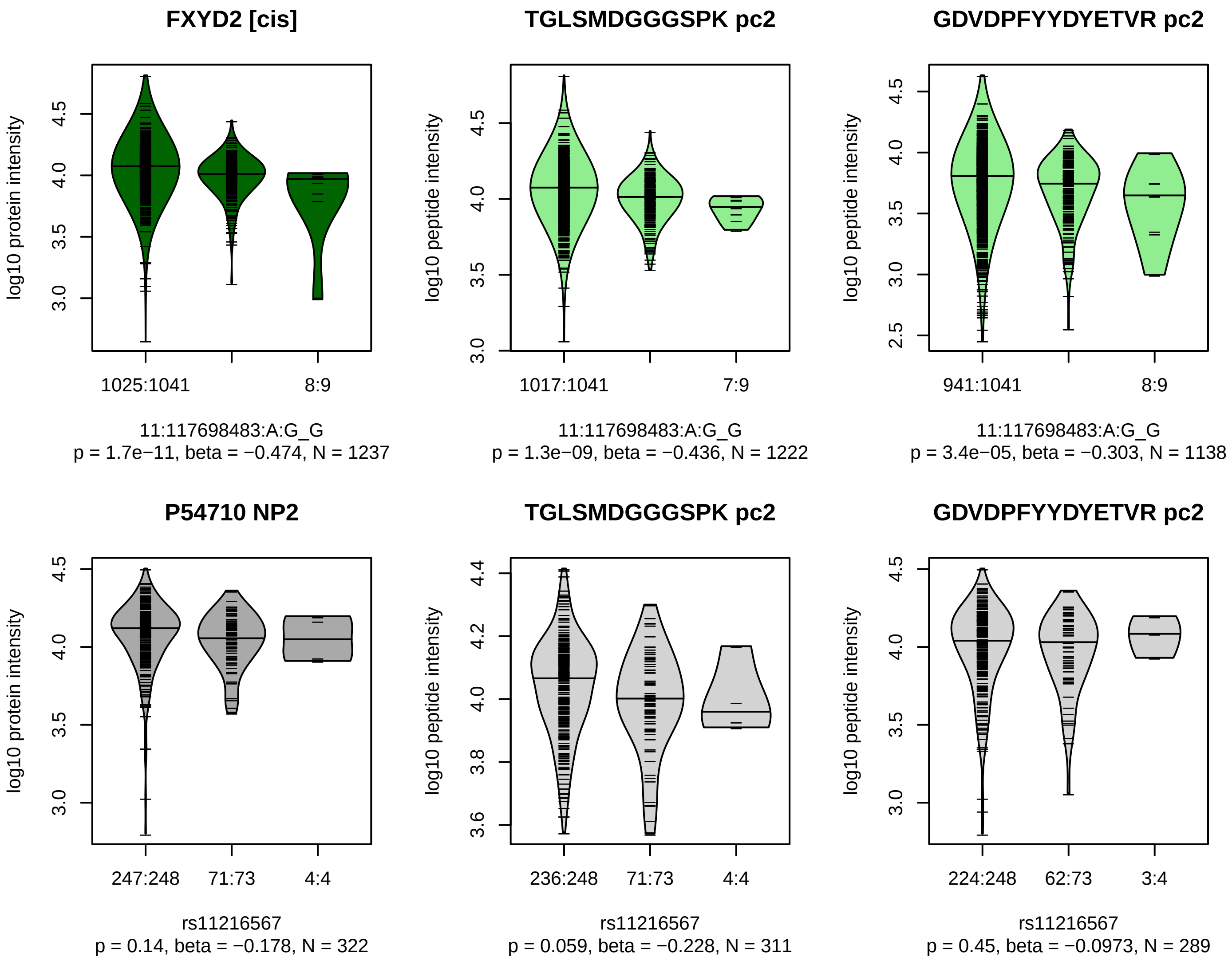


Locus: 219 (Sort: 247)

Status: Not replicated

80% power: No

SNP: 11:117698483:A:G

rsID: rs11216567

UniProtID: P54710

HGNC gene: FXYD2

Type: Cis−pQTL

**HLA−G [cis]**

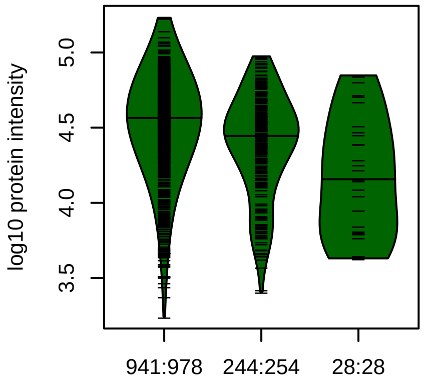


6:29924440:T:C_C
p = 1.8e−11, beta = −0.395, N = 1213

**DYLALNEDLR pc2**

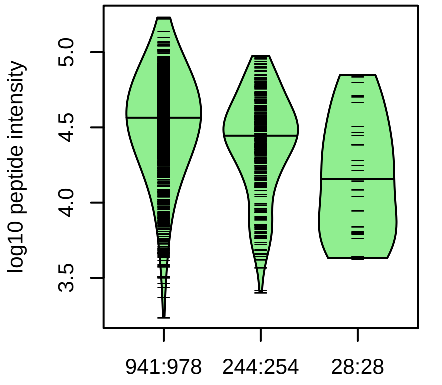


6:29924440:T:C_C
p = 1.8e−11, beta = −0.395, N = 1213

**Q5RJ85 NP1**

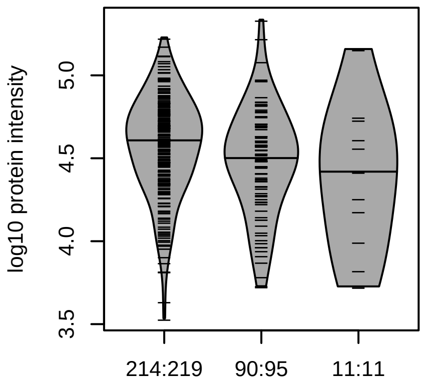


rs17179851
p = 0.0089, beta = −0.267, N = 315

**DYLALNEDLR pc2**

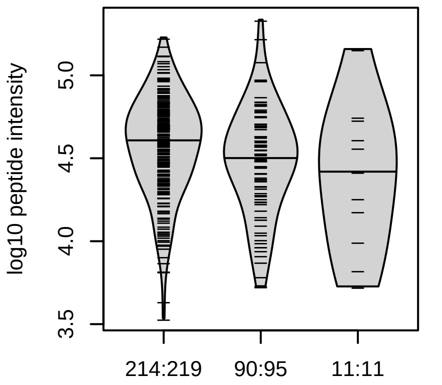


rs17179851
p = 0.0066, beta = −0.275, N = 315

Locus: 220 (Sort: 248)
Status: Not replicated
80% power: No
SNP: 6:29924440:T:C
rsID: rs17179851
UniProtID: Q5RJ85
HGNC gene: HLA−G
Type: Cis−pQTL

**WAAVVVPSGEEQR pc2**
**rs707908;rs1736924 ALT**

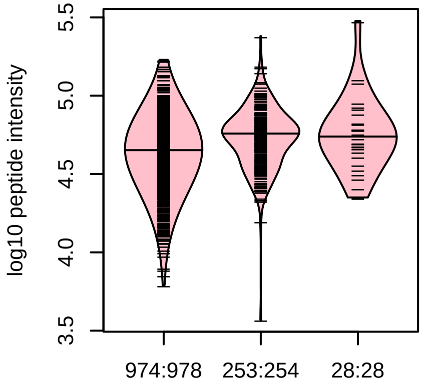


6:29924440:T:C_C
p = 1, model = DOM, N = 1255

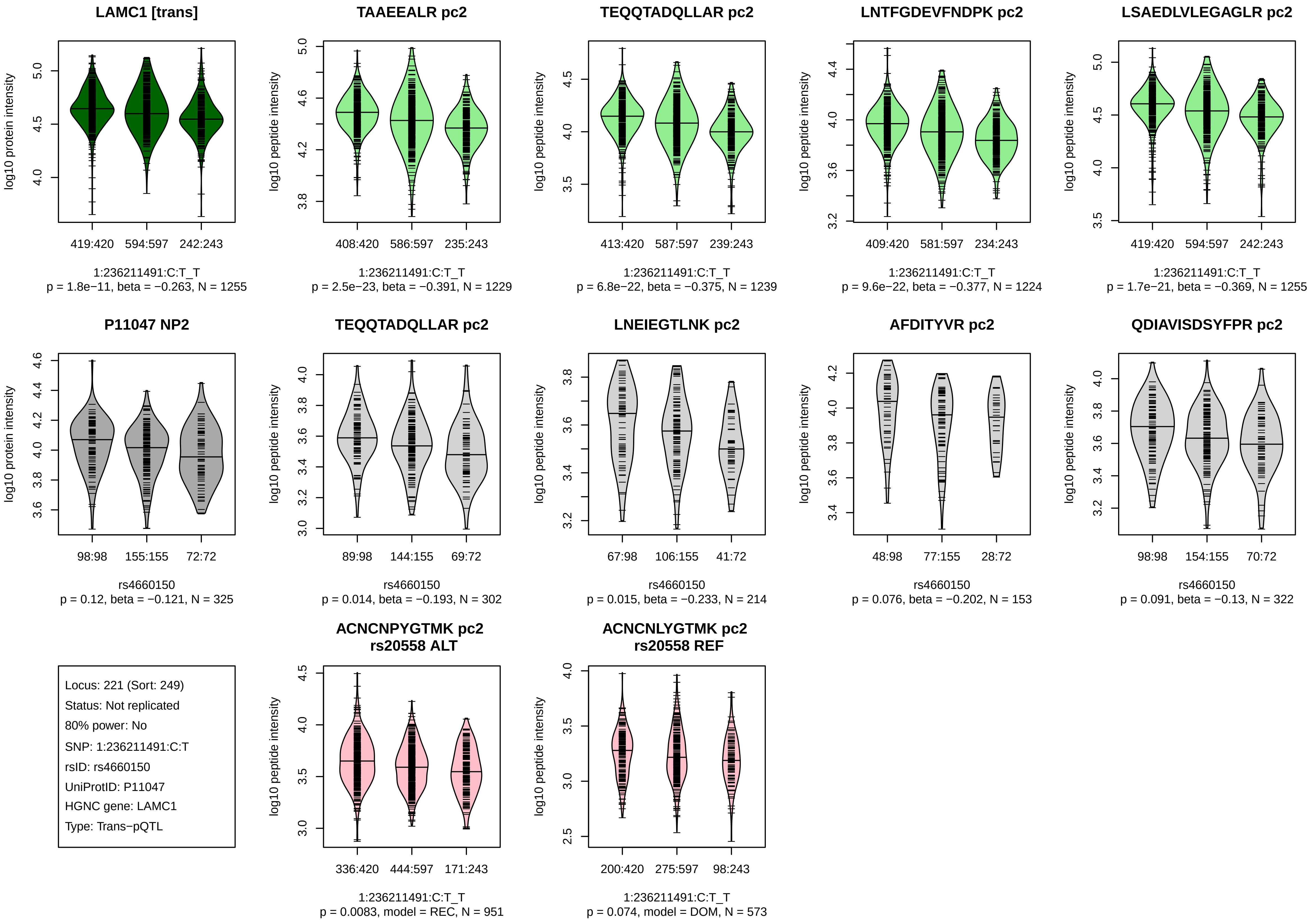

LAMC1 [trans]
log10 protein intensity
419:420
594:597
242:243
1:236211491:C:T_T
p = 1.8e−11, beta = −0.263, N = 1255
TAAEEALR pc2
log10 peptide intensity
408:420
586:597
235:243
1:236211491:C:T_T
p = 2.5e−23, beta = −0.391, N = 1229
TEQQTADQLLAR pc2
log10 peptide intensity
413:420
587:597
239:243
1:236211491:C:T_T
p = 6.8e−22, beta = −0.375, N = 1239
LNTFGDEVFNDPK pc2
log10 peptide intensity
409:420
581:597
234:243
1:236211491:C:T_T
p = 9.6e−22, beta = −0.377, N = 1224
LSAEDLVLEGAGLR pc2
log10 peptide intensity
419:420
594:597
242:243
1:236211491:C:T_T
p = 1.7e−21, beta = −0.369, N = 1255
P11047 NP2
log10 protein intensity
98:98
155:155
72:72
rs4660150
p = 0.12, beta = −0.121, N = 325
TEQQTADQLLAR pc2
log10 peptide intensity
89:98
144:155
69:72
rs4660150
p = 0.014, beta = −0.193, N = 302
LNEIEGTLNK pc2
log10 peptide intensity
67:98
106:155
41:72
rs4660150
p = 0.015, beta = −0.233, N = 214
AFDITYVR pc2
log10 peptide intensity
48:98
77:155
28:72
rs4660150
p = 0.076, beta = −0.202, N = 153
QDIAVISDSYFPR pc2
log10 peptide intensity
98:98
154:155
70:72
rs4660150
p = 0.091, beta = −0.13, N = 322
Locus: 221 (Sort: 249)
Status: Not replicated
80% power: No
SNP: 1:236211491:C:T
rsID: rs4660150
UniProtID: P11047
HGNC gene: LAMC1
Type: Trans−pQTL
ACNCNPYGTMK pc2
rs20558 ALT
log10 peptide intensity
336:420
444:597
171:243
1:236211491:C:T_T
p = 0.0083, model = REC, N = 951
ACNCNLYGTMK pc2
rs20558 REF
log10 peptide intensity
200:420
275:597
98:243
1:236211491:C:T_T
p = 0.074, model = DOM, N = 573

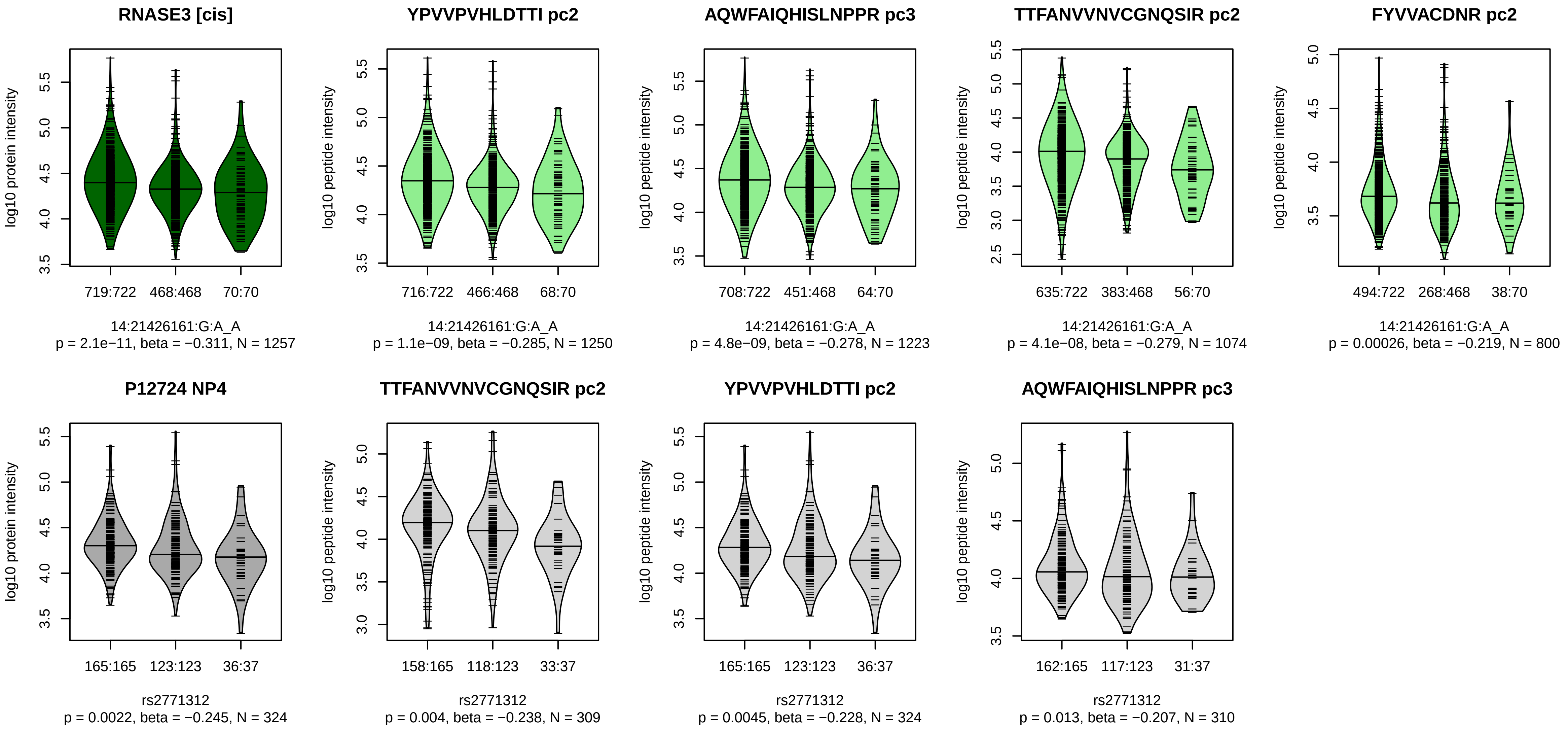


Locus: 222 (Sort: 250)

Status: Not replicated

80% power: No

SNP: 14:21426161:G:A

rsID: rs2771312

UniProtID: P12724

HGNC gene: RNASE3

Type: Cis−pQTL

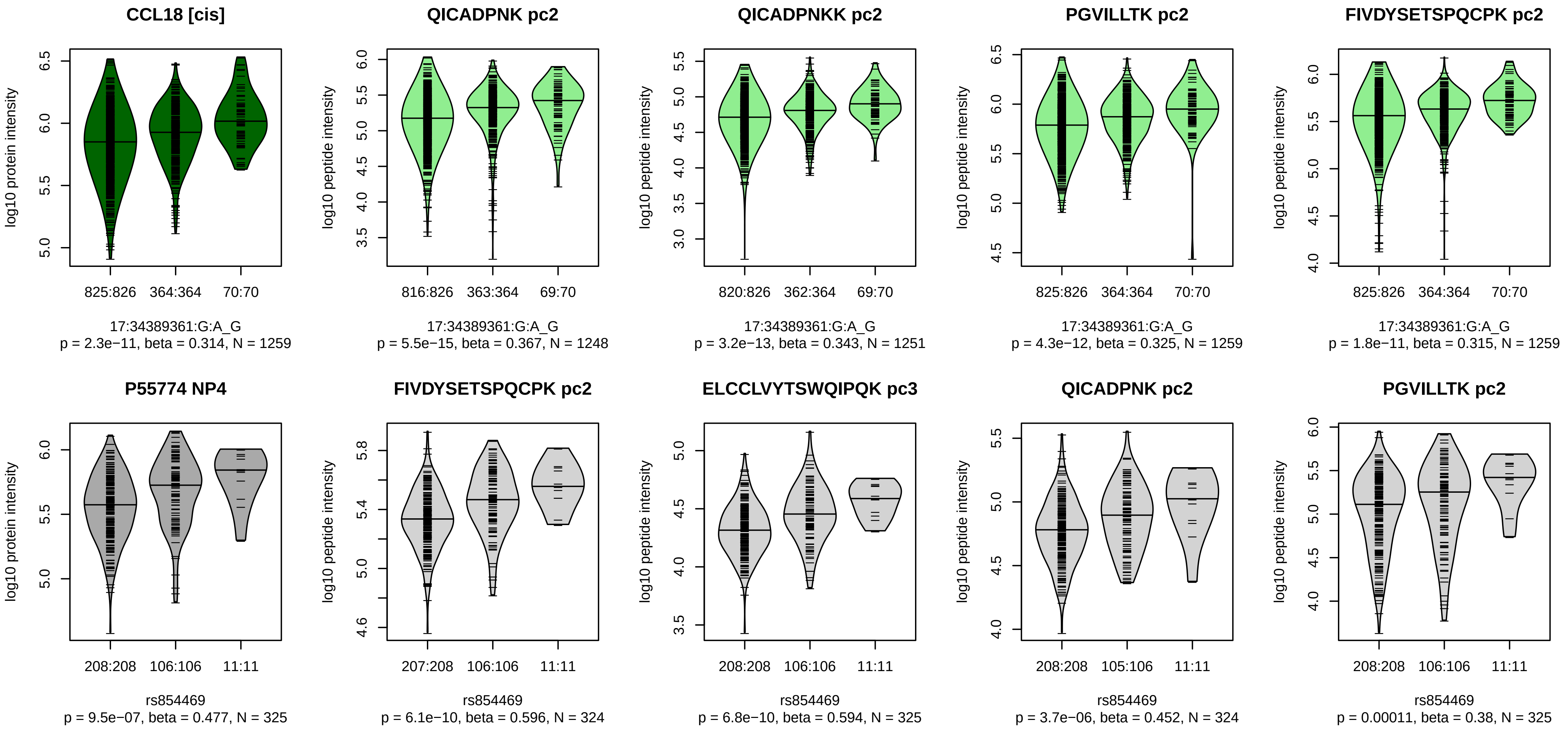


Locus: 223 (Sort: 251)
Status: Replicated
80% power: No
SNP: 17:34389361:G:A
rsID: rs854469
UniProtID: P55774
HGNC gene: CCL18
Type: Cis−pQTL

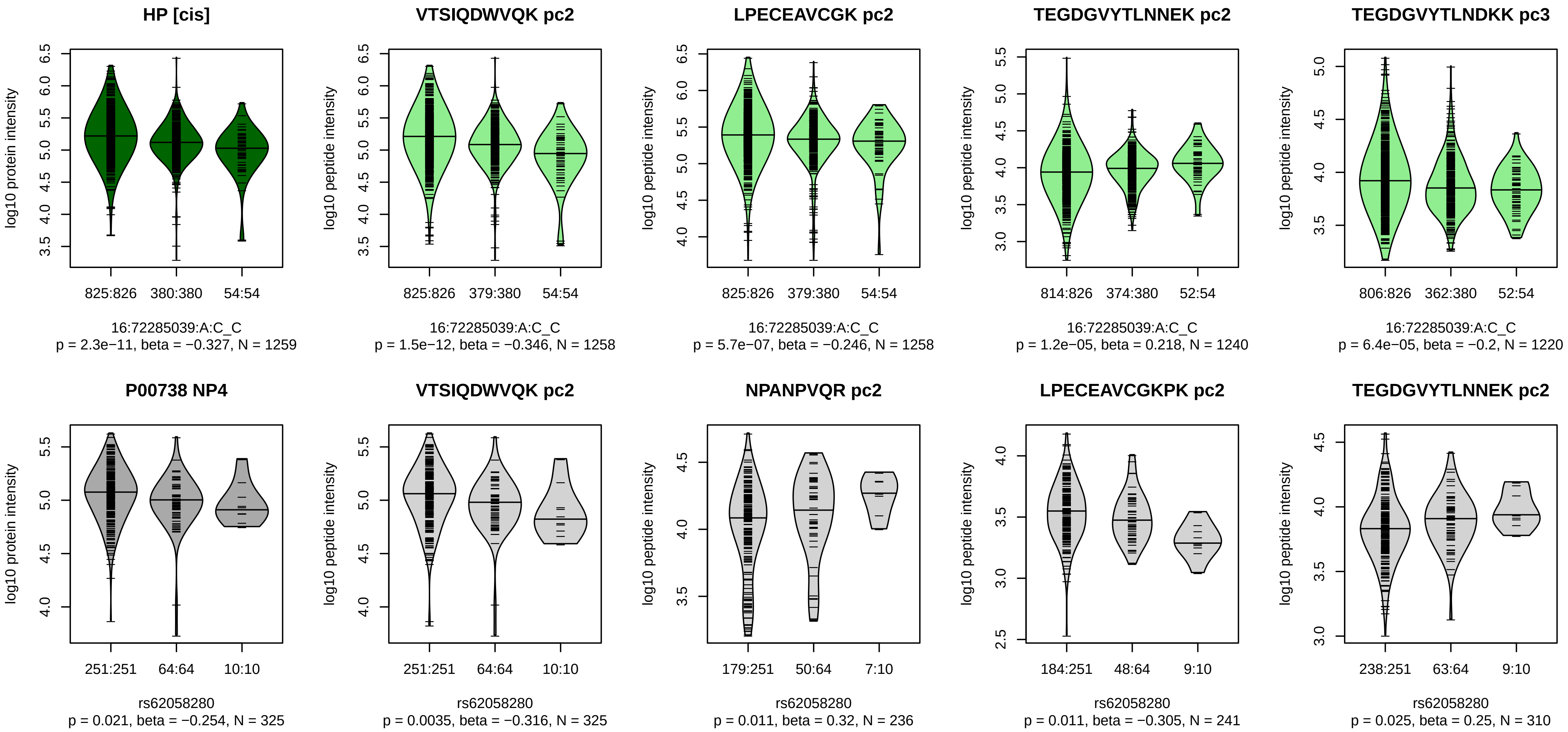


Locus: 224 (Sort: 252)
Status: Not replicated
80% power: No
SNP: 16:72285039:A:C
rsID: rs62058280
UniProtID: P00738
HGNC gene: HP
Type: Cis−pQTL
