## Supplementary material for "A genome-wide association study of mass spectrometry proteomics using the Seer Proteograph platform": Figure S4

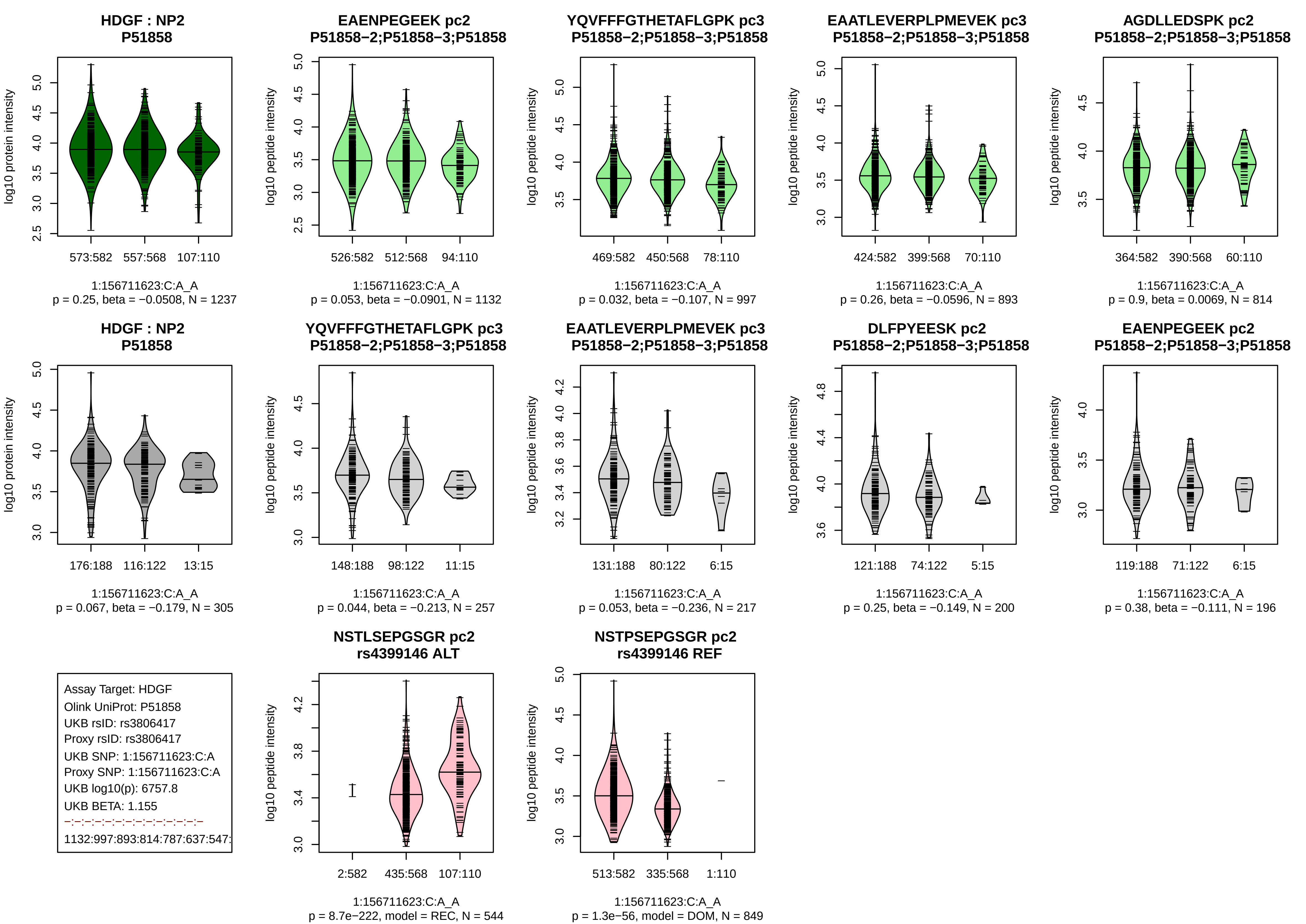
HDGF : NP2
P51858
log10 protein intensity
573:582
557:568
107:110
1:156711623:C:A_A
p = 0.25, beta = −0.0508, N = 1237
EAENPEGEEK pc2
P51858−2;P51858−3;P51858
log10 peptide intensity
526:582
512:568
94:110
1:156711623:C:A_A
p = 0.053, beta = −0.0901, N = 1132
YQVFFFGTHETAFLGPK pc3
P51858−2;P51858−3;P51858
log10 peptide intensity
469:582
450:568
78:110
1:156711623:C:A_A
p = 0.032, beta = −0.107, N = 997
EAATLEVERPLPMEVEK pc3
P51858−2;P51858−3;P51858
log10 peptide intensity
424:582
399:568
70:110
1:156711623:C:A_A
p = 0.26, beta = −0.0596, N = 893
AGDLLEDSPK pc2
P51858−2;P51858−3;P51858
log10 peptide intensity
364:582
390:568
60:110
1:156711623:C:A_A
p = 0.9, beta = 0.0069, N = 814
HDGF : NP2
P51858
log10 protein intensity
176:188
116:122
13:15
1:156711623:C:A_A
p = 0.067, beta = −0.179, N = 305
YQVFFFGTHETAFLGPK pc3
P51858−2;P51858−3;P51858
log10 peptide intensity
148:188
98:122
11:15
1:156711623:C:A_A
p = 0.044, beta = −0.213, N = 257
EAATLEVERPLPMEVEK pc3
P51858−2;P51858−3;P51858
log10 peptide intensity
131:188
80:122
6:15
1:156711623:C:A_A
p = 0.053, beta = −0.236, N = 217
DLFPYEESK pc2
P51858−2;P51858−3;P51858
log10 peptide intensity
121:188
74:122
5:15
1:156711623:C:A_A
p = 0.25, beta = −0.149, N = 200
EAENPEGEEK pc2
P51858−2;P51858−3;P51858
log10 peptide intensity
119:188
71:122
6:15
1:156711623:C:A_A
p = 0.38, beta = −0.111, N = 196
Assay Target: HDGF
Olink UniProt: P51858
UKB rsID: rs3806417
Proxy rsID: rs3806417
UKB SNP: 1:156711623:C:A
Proxy SNP: 1:156711623:C:A
UKB log10(p): 6757.8
UKB BETA: 1.155
−:−:−:−:−:−:−:−:−:−:−:−:−:−
1132:997:893:814:787:637:547:
NSTLSEPGSGR pc2
rs4399146 ALT
log10 peptide intensity
2:582
435:568
107:110
1:156711623:C:A_A
p = 8.7e−222, model = REC, N = 544
NSTPSEPGSGR pc2
rs4399146 REF
log10 peptide intensity
513:582
335:568
1:110
1:156711623:C:A_A
p = 1.3e−56, model = DOM, N = 849

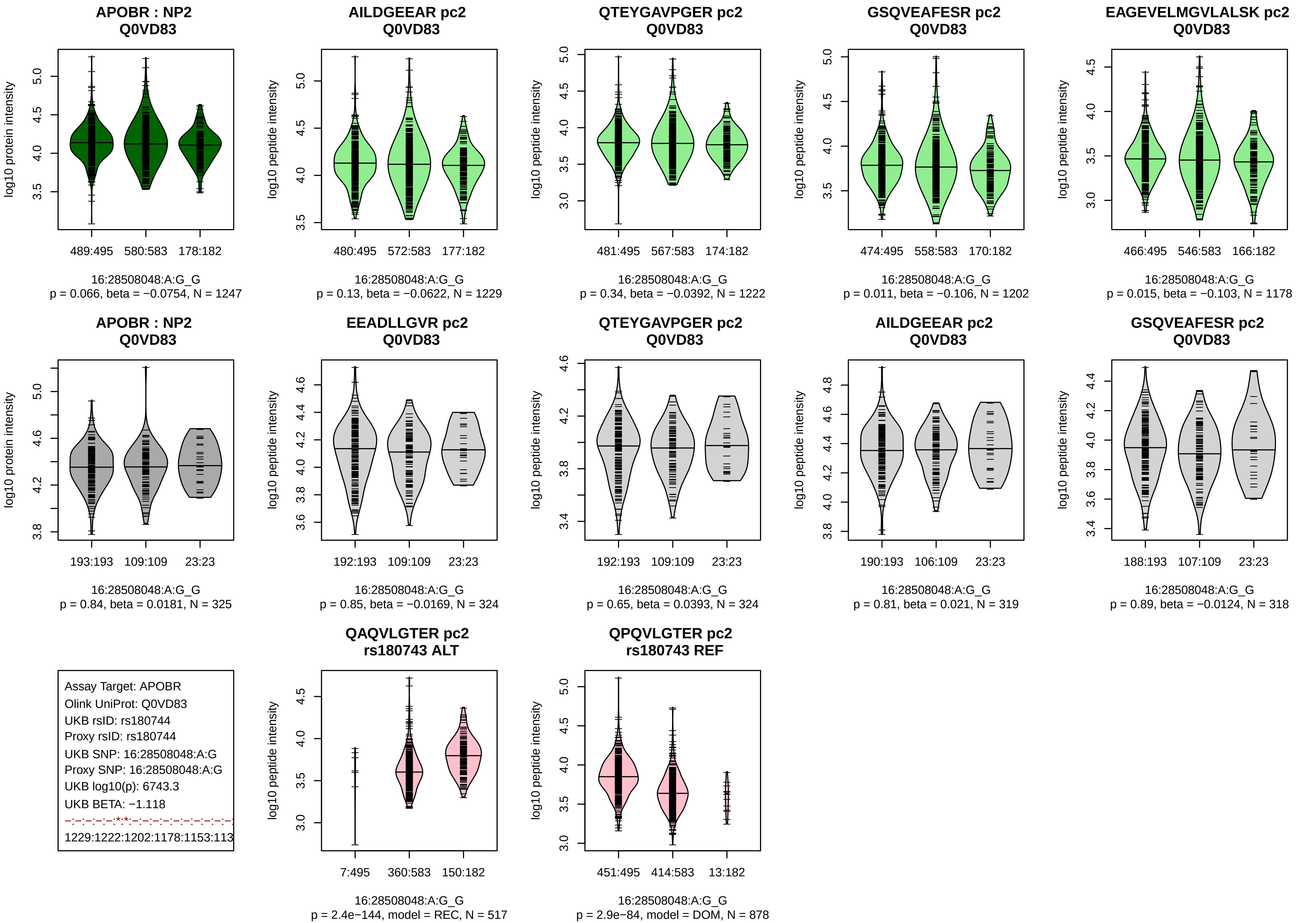

APOBR : NP2
Q0VD83
log10 protein intensity
489:495
580:583
178:182
16:28508048:A:G_G
p = 0.066, beta = −0.0754, N = 1247
AILDGEEAR pc2
Q0VD83
log10 peptide intensity
480:495
572:583
177:182
16:28508048:A:G_G
p = 0.13, beta = −0.0622, N = 1229
QTEYGAVPGER pc2
Q0VD83
481:495
567:583
174:182
16:28508048:A:G_G
p = 0.34, beta = −0.0392, N = 1222
GSQVEAFESR pc2
Q0VD83
474:495
558:583
170:182
16:28508048:A:G_G
p = 0.011, beta = −0.106, N = 1202
EAGEVELMGVLALSK pc2
Q0VD83
466:495
546:583
166:182
16:28508048:A:G_G
p = 0.015, beta = −0.103, N = 1178
APOBR : NP2
Q0VD83
193:193
109:109
23:23
16:28508048:A:G_G
p = 0.84, beta = 0.0181, N = 325
EEADLLGVR pc2
Q0VD83
192:193
109:109
23:23
16:28508048:A:G_G
p = 0.85, beta = −0.0169, N = 324
QTEYGAVPGER pc2
Q0VD83
192:193
109:109
23:23
16:28508048:A:G_G
p = 0.65, beta = 0.0393, N = 324
AILDGEEAR pc2
Q0VD83
190:193
106:109
23:23
16:28508048:A:G_G
p = 0.81, beta = 0.021, N = 319
GSQVEAFESR pc2
Q0VD83
188:193
107:109
23:23
16:28508048:A:G_G
p = 0.89, beta = −0.0124, N = 318
Assay Target: APOBR
Olink UniProt: Q0VD83
UKB rsID: rs180744
Proxy rsID: rs180744
UKB SNP: 16:28508048:A:G
Proxy SNP: 16:28508048:A:G
UKB log10(p): 6743.3
UKB BETA: −1.118
−:−:−:−:−:*:*:−:−:−:−:−:−:−:−:−:−:−
1229:1222:1202:1178:1153:113
QAQVLGTER pc2
rs180743 ALT
7:495
360:583
150:182
16:28508048:A:G_G
p = 2.4e−144, model = REC, N = 517
QPQVLGTER pc2
rs180743 REF
451:495
414:583
13:182
16:28508048:A:G_G
p = 2.9e−84, model = DOM, N = 878

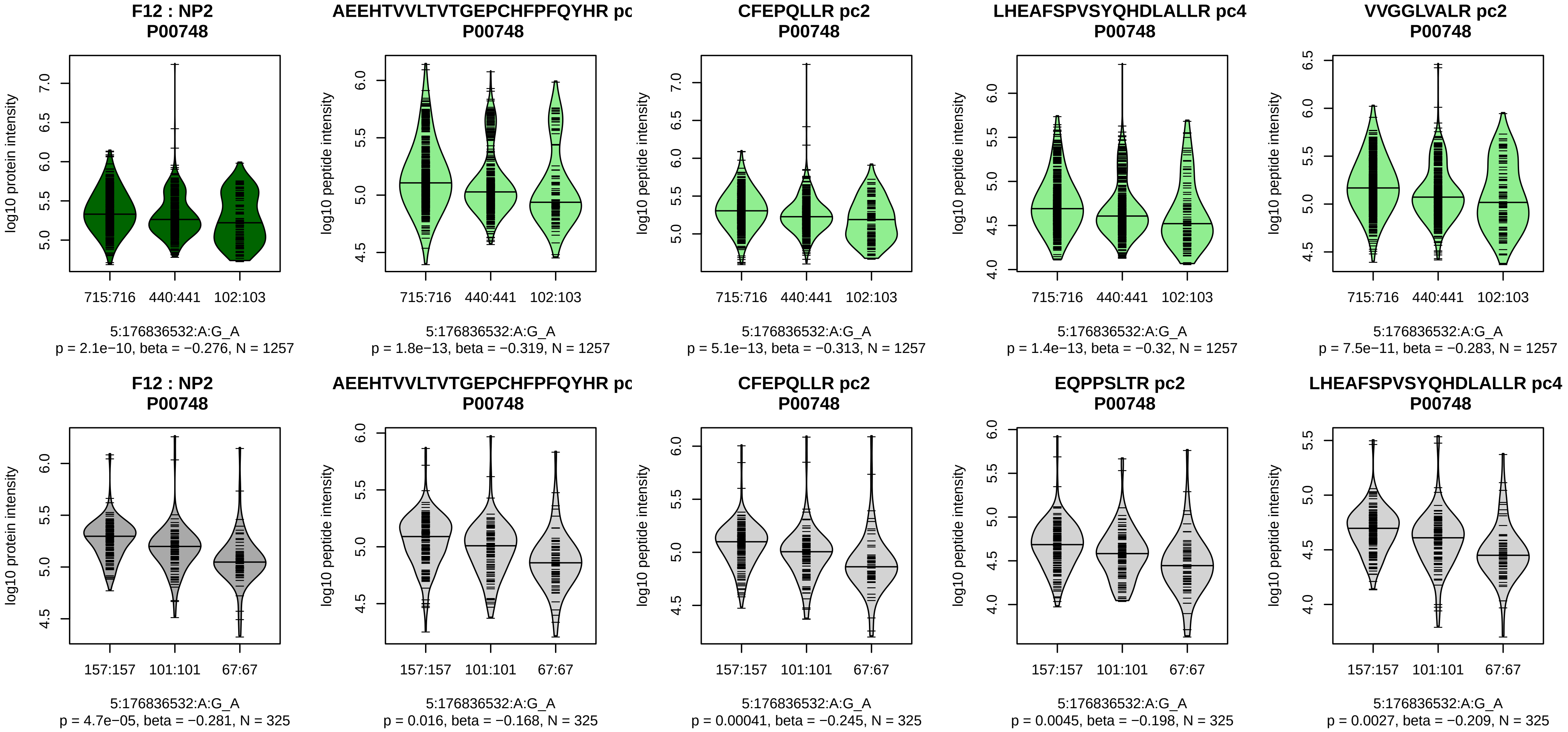


Assay Target: F12
Olink UniProt: P00748
UKB rsID: rs1801020
Proxy rsID: rs1801020
UKB SNP: 5:176836532:A:G
Proxy SNP: 5:176836532:A:G
UKB log10(p): 6066.9
UKB BETA: 1.167
*:*:*:*:*:*:*:*:*:*:*:*:*:*:*:*:*:*:*:*:*:*:–:–:–
1257:1257:1257:1257:1256:125

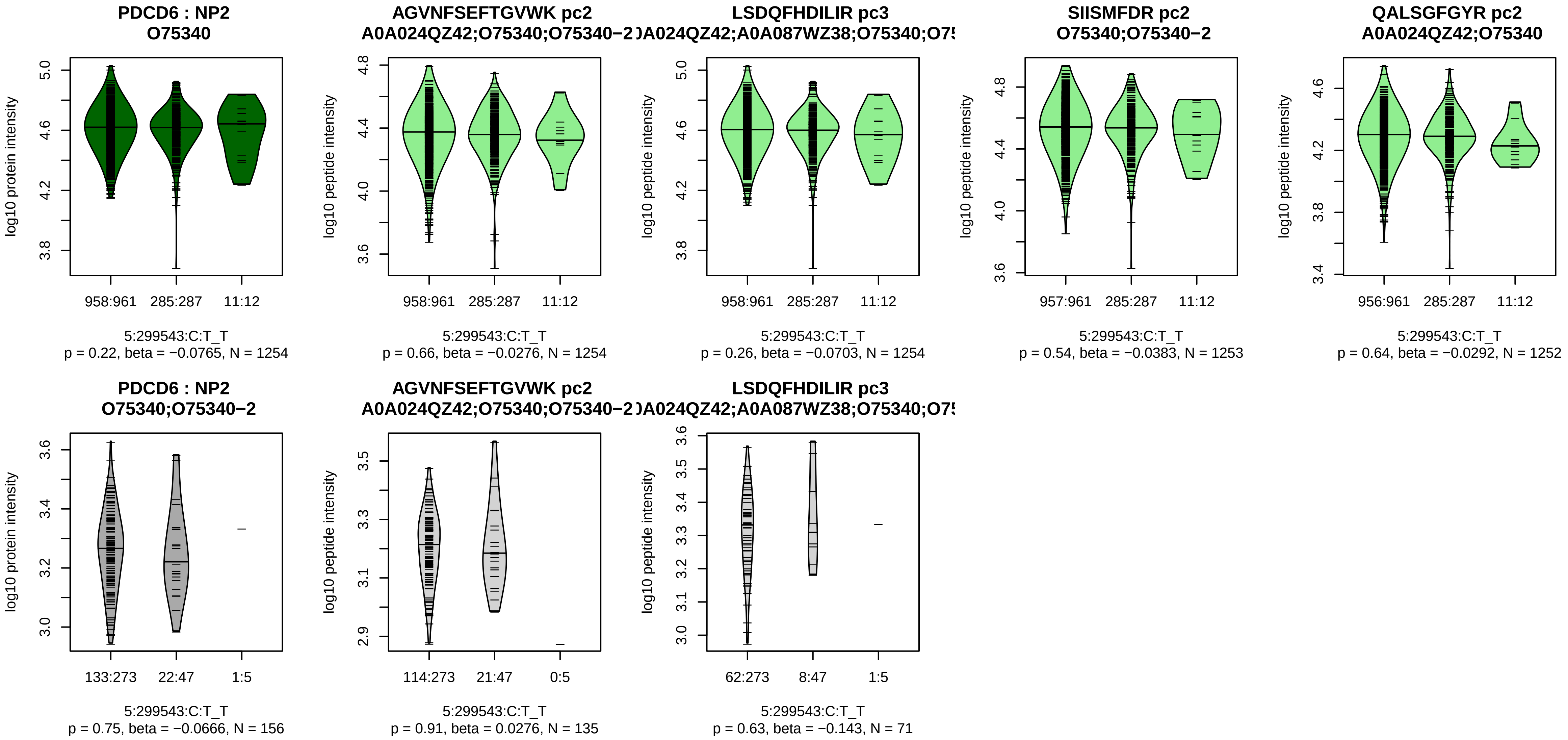


Assay Target: PDCD6
Olink UniProt: O75340
UKB rsID: rs9654451
Proxy rsID: rs9654451
UKB SNP: 5:299543:C:T
Proxy SNP: 5:299543:C:T
UKB log10(p): 5682.7
UKB BETA: −1.506
−:−:−:−:−:−:−:−:−:−:−
1254:1254:1253:1252:1217:121

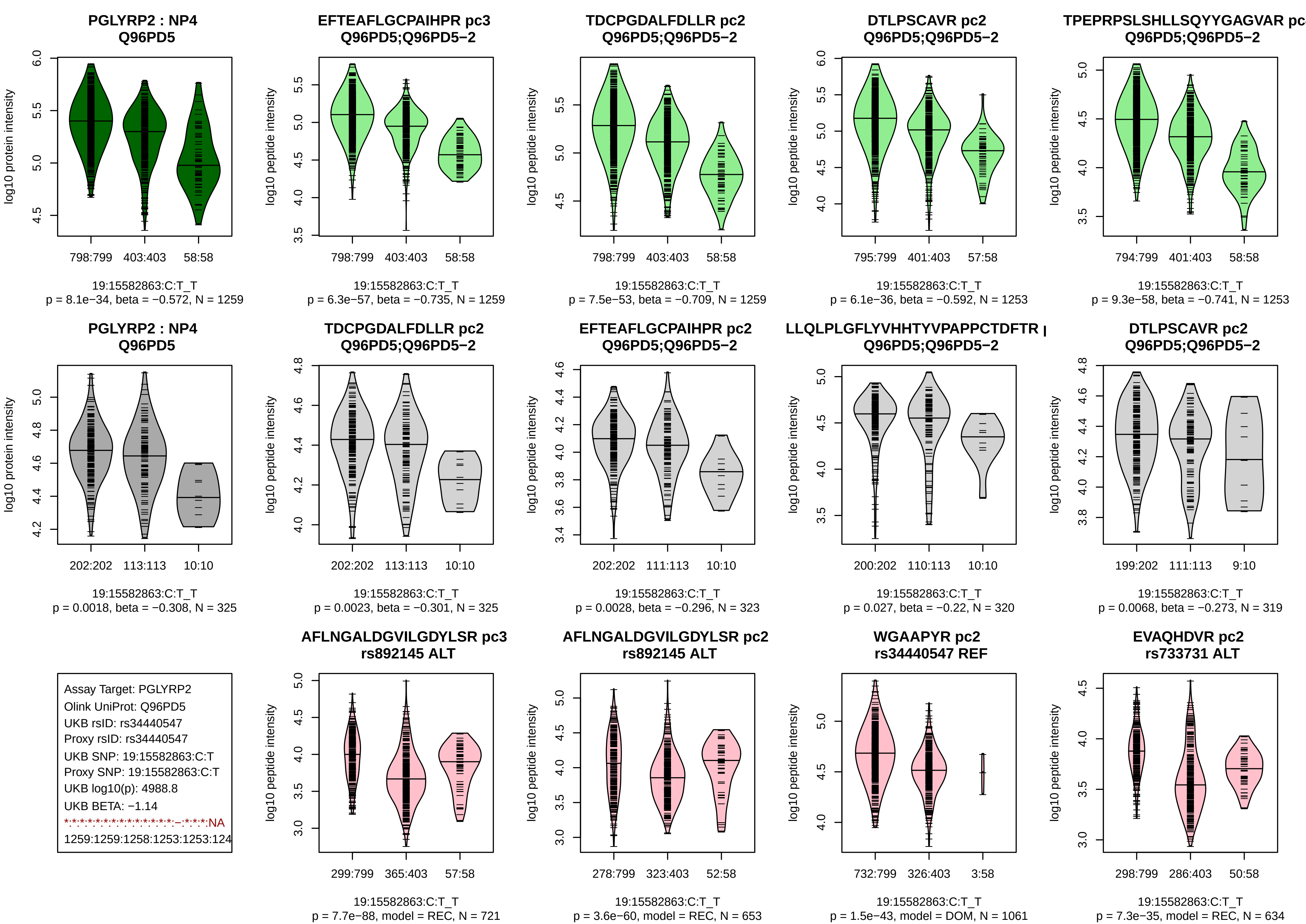
PGLYRP2 : NP4
Q96PD5
log10 protein intensity
798:799
403:403
58:58
19:15582863:C:T_T
p = 8.1e−34, beta = −0.572, N = 1259
EFTEAFLGCPAIHPR pc3
Q96PD5;Q96PD5−2
log10 peptide intensity
798:799
403:403
58:58
19:15582863:C:T_T
p = 6.3e−57, beta = −0.735, N = 1259
TDCPGDALFDLLR pc2
Q96PD5;Q96PD5−2
log10 peptide intensity
798:799
403:403
58:58
19:15582863:C:T_T
p = 7.5e−53, beta = −0.709, N = 1259
DTLPSCAVR pc2
Q96PD5;Q96PD5−2
log10 peptide intensity
795:799
401:403
57:58
19:15582863:C:T_T
p = 6.1e−36, beta = −0.592, N = 1253
TPEPRPSLSHLLSQYYGAGVAR pc
Q96PD5;Q96PD5−2
log10 peptide intensity
794:799
401:403
58:58
19:15582863:C:T_T
p = 9.3e−58, beta = −0.741, N = 1253
PGLYRP2 : NP4
Q96PD5
log10 protein intensity
202:202
113:113
10:10
19:15582863:C:T_T
p = 0.0018, beta = −0.308, N = 325
TDCPGDALFDLLR pc2
Q96PD5;Q96PD5−2
log10 peptide intensity
202:202
113:113
10:10
19:15582863:C:T_T
p = 0.0023, beta = −0.301, N = 325
EFTEAFLGCPAIHPR pc2
Q96PD5;Q96PD5−2
log10 peptide intensity
202:202
111:113
10:10
19:15582863:C:T_T
p = 0.0028, beta = −0.296, N = 323
LLQLPLGFLYVHHTYVPAPPCTDFTR
Q96PD5;Q96PD5−2
log10 peptide intensity
200:202
110:113
10:10
19:15582863:C:T_T
p = 0.027, beta = −0.22, N = 320
DTLPSCAVR pc2
Q96PD5;Q96PD5−2
log10 peptide intensity
199:202
111:113
9:10
19:15582863:C:T_T
p = 0.0068, beta = −0.273, N = 319
Assay Target: PGLYRP2
Olink UniProt: Q96PD5
UKB rsID: rs34440547
Proxy rsID: rs34440547
UKB SNP: 19:15582863:C:T
Proxy SNP: 19:15582863:C:T
UKB log10(p): 4988.8
UKB BETA: −1.14
*:*:*:*:*:*:*:*:*:*:*:*:*:*:–:*:*:*:NA
1259:1259:1258:1253:1253:124
AFLNGALDGVILGDYLSR pc3
rs892145 ALT
log10 peptide intensity
299:799
365:403
57:58
19:15582863:C:T_T
p = 7.7e−88, model = REC, N = 721
AFLNGALDGVILGDYLSR pc2
rs892145 ALT
log10 peptide intensity
278:799
323:403
52:58
19:15582863:C:T_T
p = 3.6e−60, model = REC, N = 653
WGAAPYR pc2
rs34440547 REF
log10 peptide intensity
732:799
326:403
3:58
19:15582863:C:T_T
p = 1.5e−43, model = DOM, N = 1061
EVAQHDVR pc2
rs733731 ALT
log10 peptide intensity
298:799
286:403
50:58
19:15582863:C:T_T
p = 7.3e−35, model = REC, N = 634

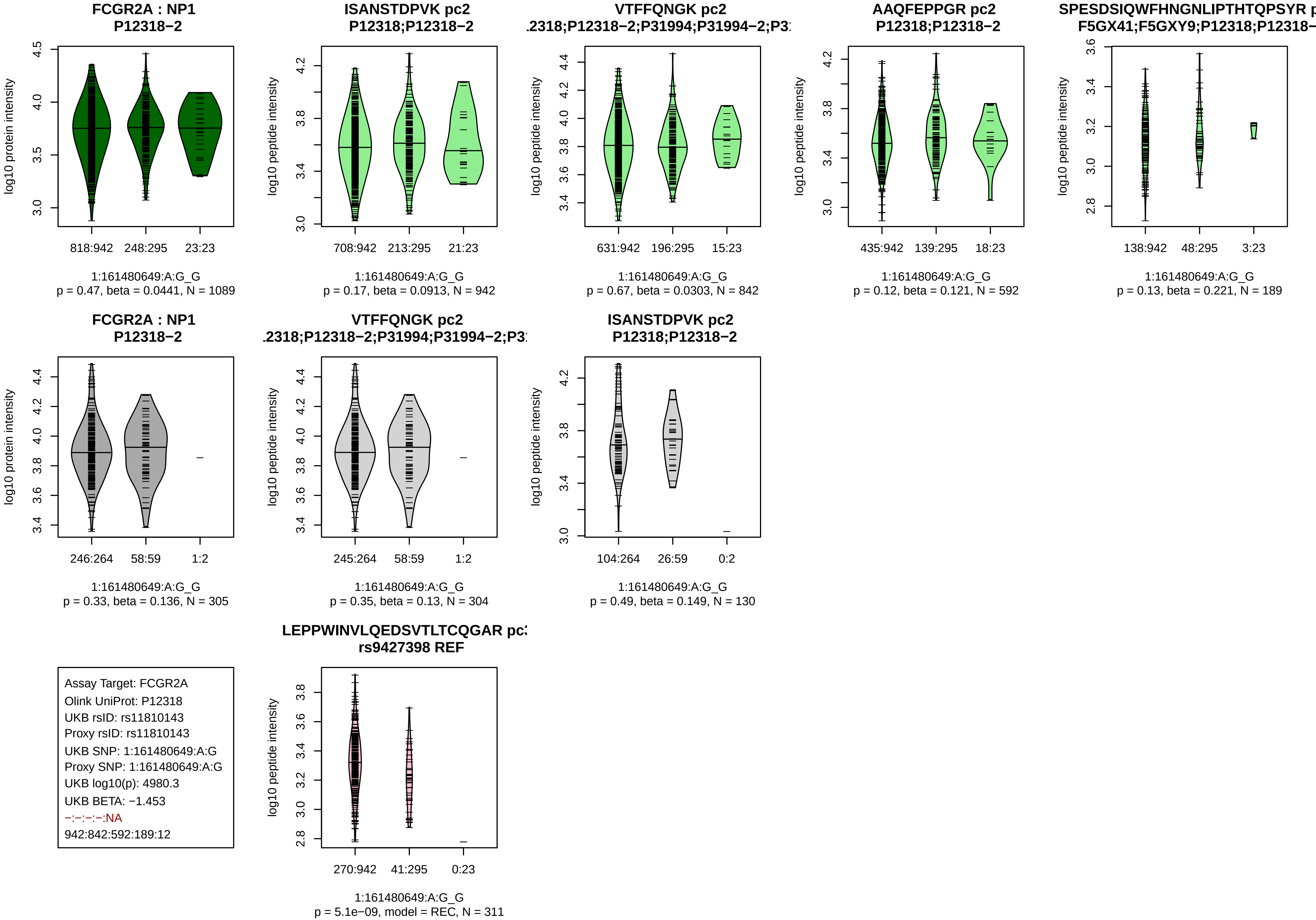

FCGR2A : NP1
P12318−2
log10 protein intensity
818:942
248:295
23:23
1:161480649:A:G_G
p = 0.47, beta = 0.0441, N = 1089
ISANSTDPVK pc2
P12318;P12318−2
log10 peptide intensity
708:942
213:295
21:23
1:161480649:A:G_G
p = 0.17, beta = 0.0913, N = 942
VTFFQNGK pc2
.2318;P12318−2;P31994;P31994−2;P3
log10 peptide intensity
631:942
196:295
15:23
1:161480649:A:G_G
p = 0.67, beta = 0.0303, N = 842
AAQFEPPGR pc2
P12318;P12318−2
log10 peptide intensity
435:942
139:295
18:23
1:161480649:A:G_G
p = 0.12, beta = 0.121, N = 592
SPESDSIQWFHNGNLIPTHTQPSYR p
F5GX41;F5GXY9;P12318;P12318−
log10 peptide intensity
138:942
48:295
3:23
1:161480649:A:G_G
p = 0.13, beta = 0.221, N = 189
FCGR2A : NP1
P12318−2
log10 protein intensity
246:264
58:59
1:2
1:161480649:A:G_G
p = 0.33, beta = 0.136, N = 305
VTFFQNGK pc2
.2318;P12318−2;P31994;P31994−2;P3
log10 peptide intensity
245:264
58:59
1:2
1:161480649:A:G_G
p = 0.35, beta = 0.13, N = 304
ISANSTDPVK pc2
P12318;P12318−2
log10 peptide intensity
104:264
26:59
0:2
1:161480649:A:G_G
p = 0.49, beta = 0.149, N = 130
Assay Target: FCGR2A
Olink UniProt: P12318
UKB rsID: rs11810143
Proxy rsID: rs11810143
UKB SNP: 1:161480649:A:G
Proxy SNP: 1:161480649:A:G
UKB log10(p): 4980.3
UKB BETA: −1.453
−:−:−:−:NA
942:842:592:189:12
LEPPWINVLQEDSVTLTCQGAR pc
rs9427398 REF
log10 peptide intensity
270:942
41:295
0:23
1:161480649:A:G_G
p = 5.1e−09, model = REC, N = 311

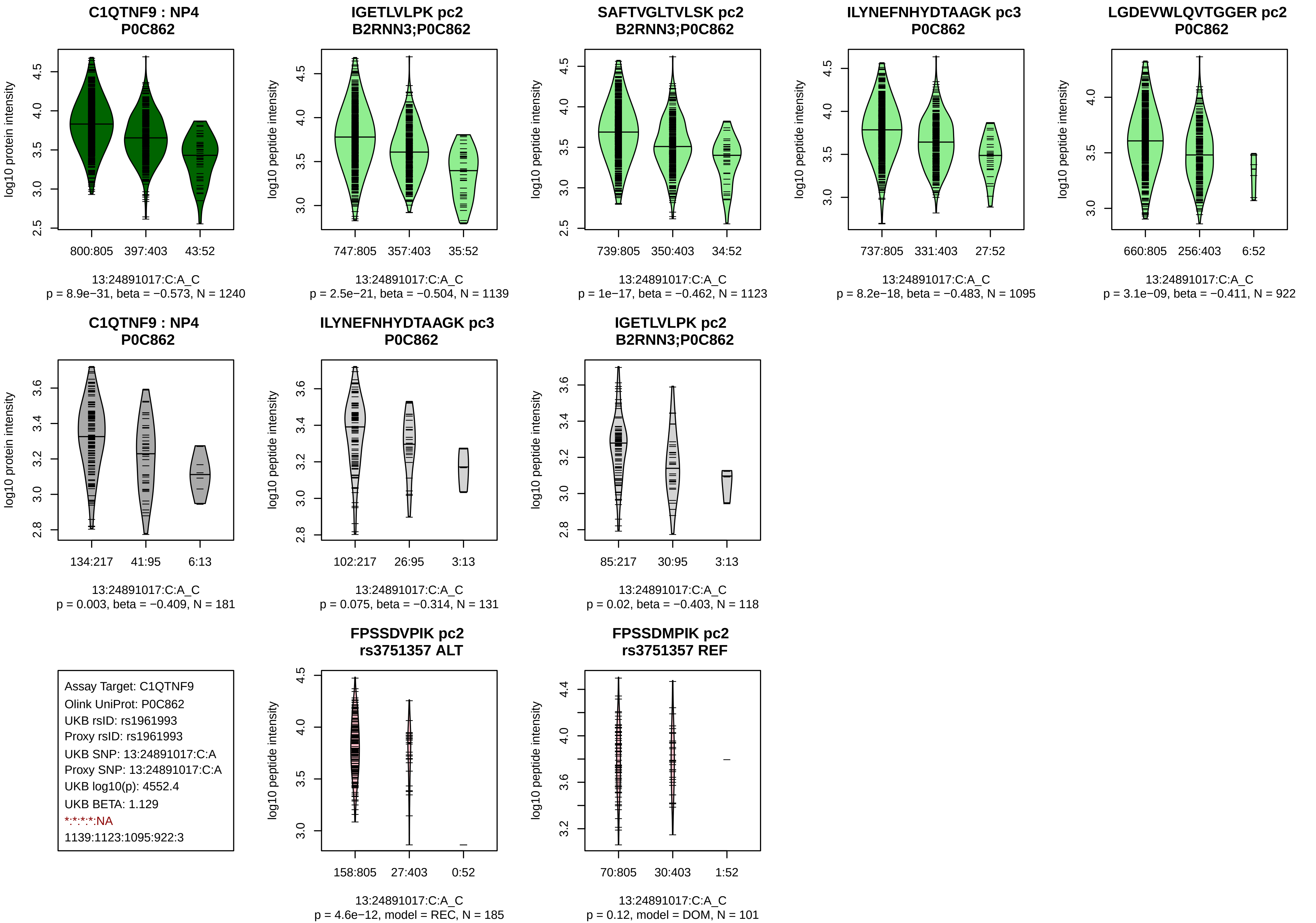
C1QTNF9 : NP4
P0C862
log10 protein intensity
800:805
397:403
43:52
13:24891017:C:A_C
p = 8.9e−31, beta = −0.573, N = 1240
IGETLVLPK pc2
B2RNN3;P0C862
log10 peptide intensity
747:805
357:403
35:52
13:24891017:C:A_C
p = 2.5e−21, beta = −0.504, N = 1139
SAFTVGLTVLSK pc2
B2RNN3;P0C862
log10 peptide intensity
739:805
350:403
34:52
13:24891017:C:A_C
p = 1e−17, beta = −0.462, N = 1123
ILYNEFNHYDTAAGK pc3
P0C862
log10 peptide intensity
737:805
331:403
27:52
13:24891017:C:A_C
p = 8.2e−18, beta = −0.483, N = 1095
LGDEVWLQVTGGER pc2
P0C862
log10 peptide intensity
660:805
256:403
6:52
13:24891017:C:A_C
p = 3.1e−09, beta = −0.411, N = 922
C1QTNF9 : NP4
P0C862
log10 protein intensity
134:217
41:95
6:13
13:24891017:C:A_C
p = 0.003, beta = −0.409, N = 181
ILYNEFNHYDTAAGK pc3
P0C862
log10 peptide intensity
102:217
26:95
3:13
13:24891017:C:A_C
p = 0.075, beta = −0.314, N = 131
IGETLVLPK pc2
B2RNN3;P0C862
log10 peptide intensity
85:217
30:95
3:13
13:24891017:C:A_C
p = 0.02, beta = −0.403, N = 118
Assay Target: C1QTNF9
Olink UniProt: P0C862
UKB rsID: rs1961993
Proxy rsID: rs1961993
UKB SNP: 13:24891017:C:A
Proxy SNP: 13:24891017:C:A
UKB log10(p): 4552.4
UKB BETA: 1.129
*:*:*:*:NA
1139:1123:1095:922:3
FPSSDVPIK pc2
rs3751357 ALT
log10 peptide intensity
158:805
27:403
0:52
13:24891017:C:A_C
p = 4.6e−12, model = REC, N = 185
FPSSDMPIK pc2
rs3751357 REF
log10 peptide intensity
70:805
30:403
1:52
13:24891017:C:A_C
p = 0.12, model = DOM, N = 101

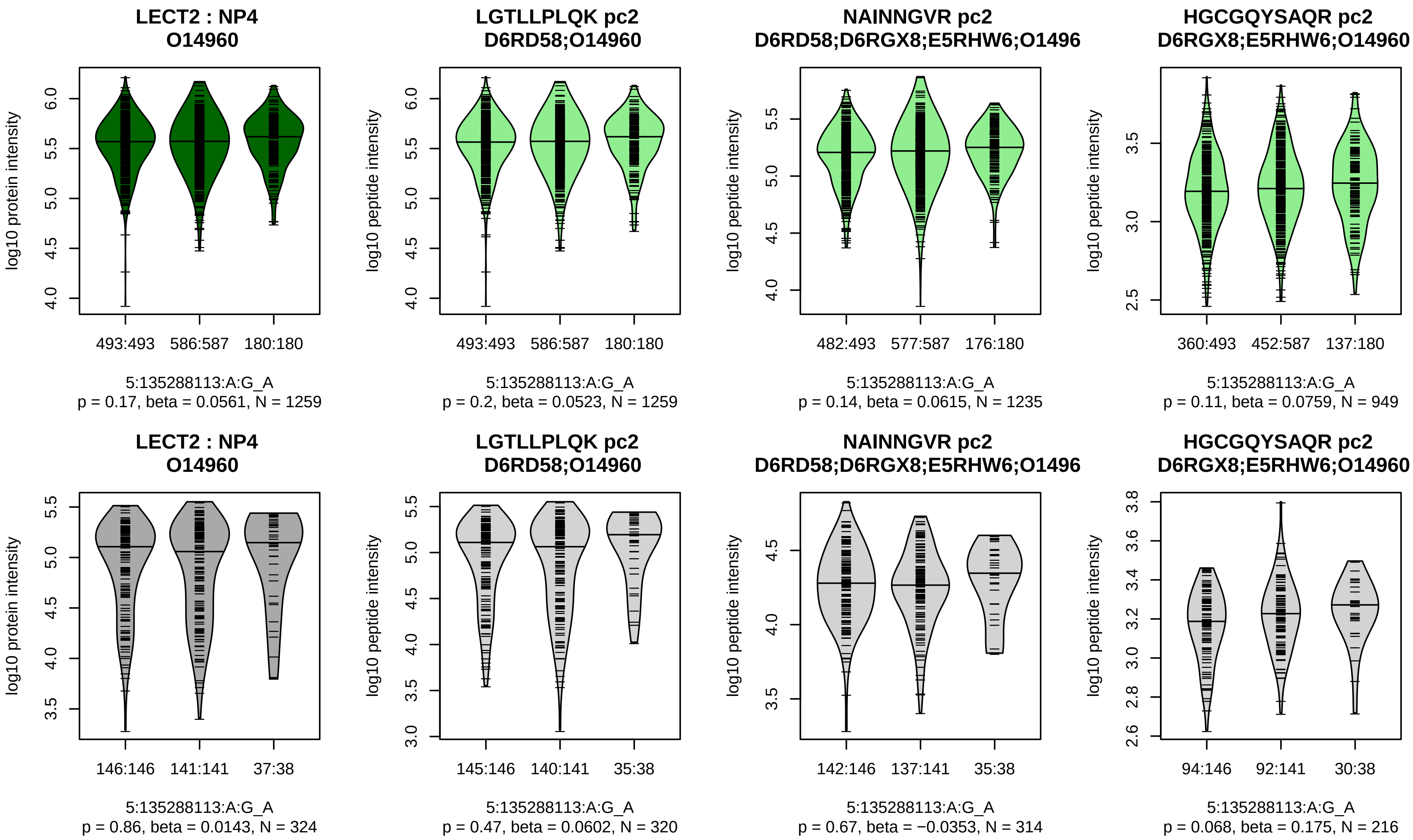


Assay Target: LECT2
Olink UniProt: O14960
UKB rsID: rs31515
Proxy rsID: rs31515
UKB SNP: 5:135288113:A:G
Proxy SNP: 5:135288113:A:G
UKB log10(p): 4434.6
UKB BETA: −0.789
−:−:−
1259:1235:949

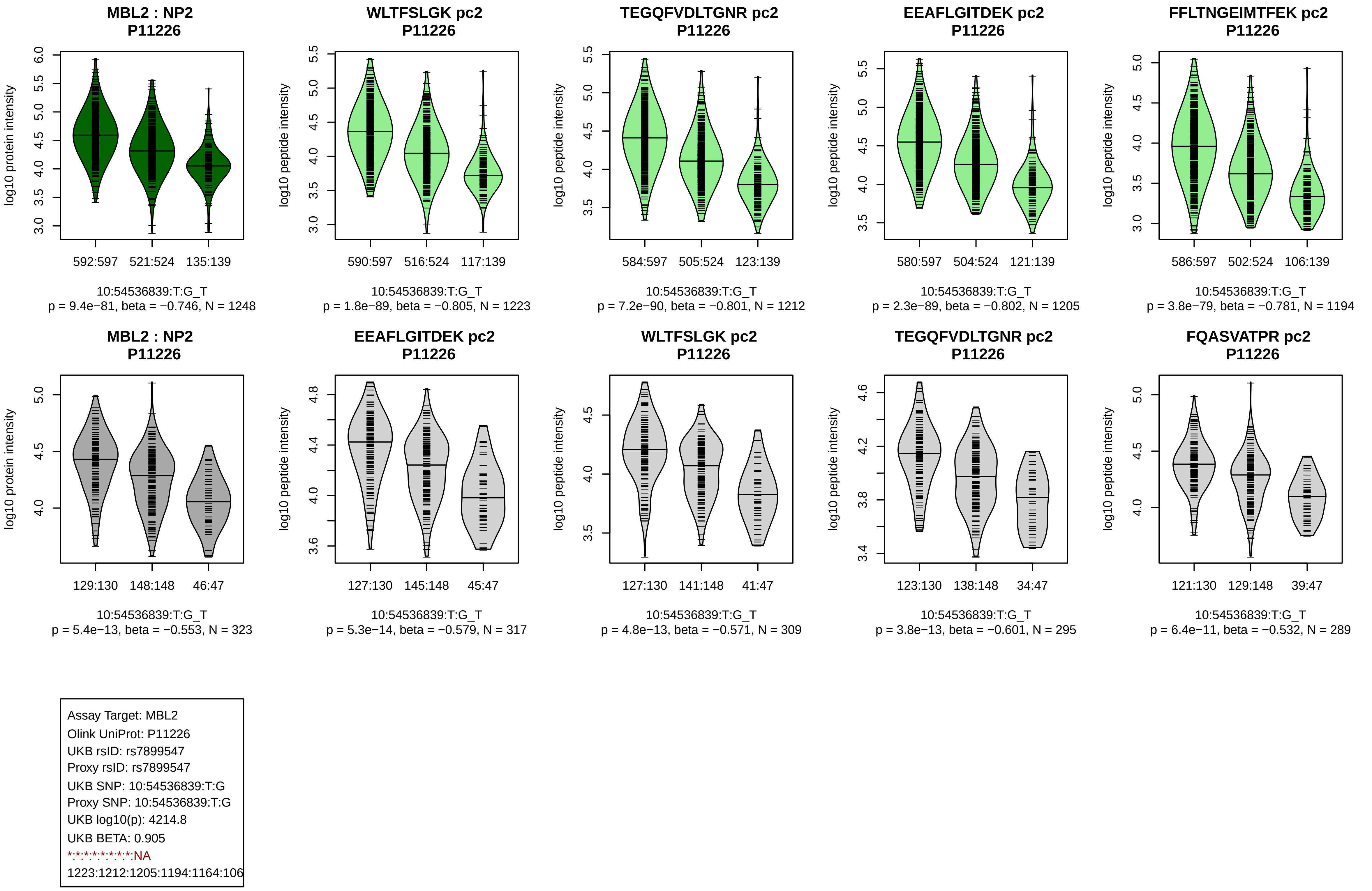
MBL2 : NP2
P11226
log10 protein intensity
592:597
521:524
135:139
10:54536839:T:G_T
p = 9.4e−81, beta = −0.746, N = 1248
WLTFSLGK pc2
P11226
log10 peptide intensity
590:597
516:524
117:139
10:54536839:T:G_T
p = 1.8e−89, beta = −0.805, N = 1223
TEGQFVDLTGNR pc2
P11226
log10 peptide intensity
584:597
505:524
123:139
10:54536839:T:G_T
p = 7.2e−90, beta = −0.801, N = 1212
EEAFLGITDEK pc2
P11226
log10 peptide intensity
580:597
504:524
121:139
10:54536839:T:G_T
p = 2.3e−89, beta = −0.802, N = 1205
FFLTNGEIMTFEK pc2
P11226
log10 peptide intensity
586:597
502:524
106:139
10:54536839:T:G_T
p = 3.8e−79, beta = −0.781, N = 1194
MBL2 : NP2
P11226
log10 protein intensity
129:130
148:148
46:47
10:54536839:T:G_T
p = 5.4e−13, beta = −0.553, N = 323
EEAFLGITDEK pc2
P11226
log10 peptide intensity
127:130
145:148
45:47
10:54536839:T:G_T
p = 5.3e−14, beta = −0.579, N = 317
WLTFSLGK pc2
P11226
log10 peptide intensity
127:130
141:148
41:47
10:54536839:T:G_T
p = 4.8e−13, beta = −0.571, N = 309
TEGQFVDLTGNR pc2
P11226
log10 peptide intensity
123:130
138:148
34:47
10:54536839:T:G_T
p = 3.8e−13, beta = −0.601, N = 295
FQASVATPR pc2
P11226
log10 peptide intensity
121:130
129:148
39:47
10:54536839:T:G_T
p = 6.4e−11, beta = −0.532, N = 289
Assay Target: MBL2
Olink UniProt: P11226
UKB rsID: rs7899547
Proxy rsID: rs7899547
UKB SNP: 10:54536839:T:G
Proxy SNP: 10:54536839:T:G
UKB log10(p): 4214.8
UKB BETA: 0.905
*:*:*:*:*:*:*:NA
1223:1212:1205:1194:1164:106

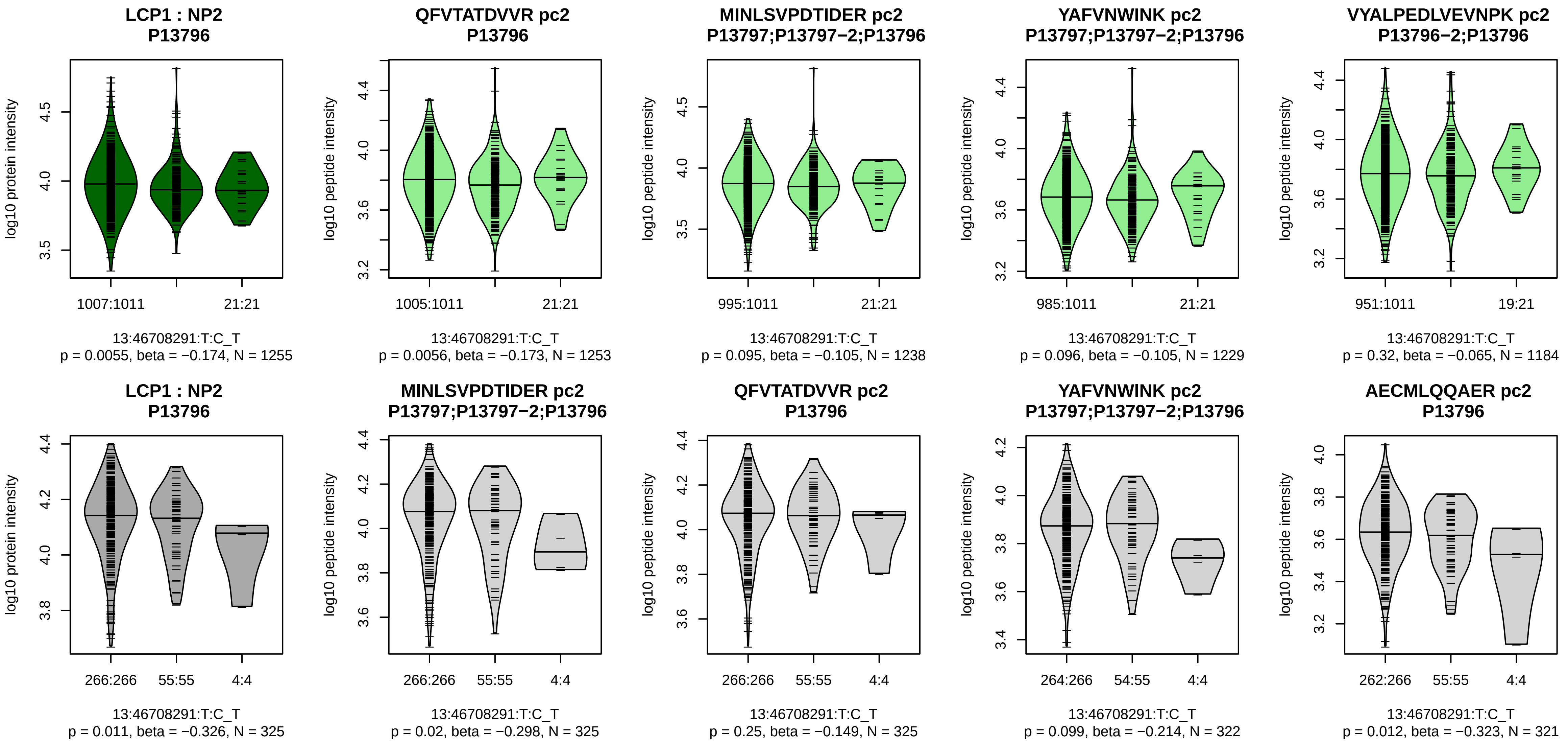


Assay Target: LCP1
Olink UniProt: P13796
UKB rsID: rs4941543
Proxy rsID: rs4941543
UKB SNP: 13:46708291:T:C
Proxy SNP: 13:46708291:T:C
UKB log10(p): 4209.4
UKB BETA: −1.447
*:−:−:−:−:*:−:*:−:−:−:−:−:−:−:
1253:1238:1229:1184:1159:111

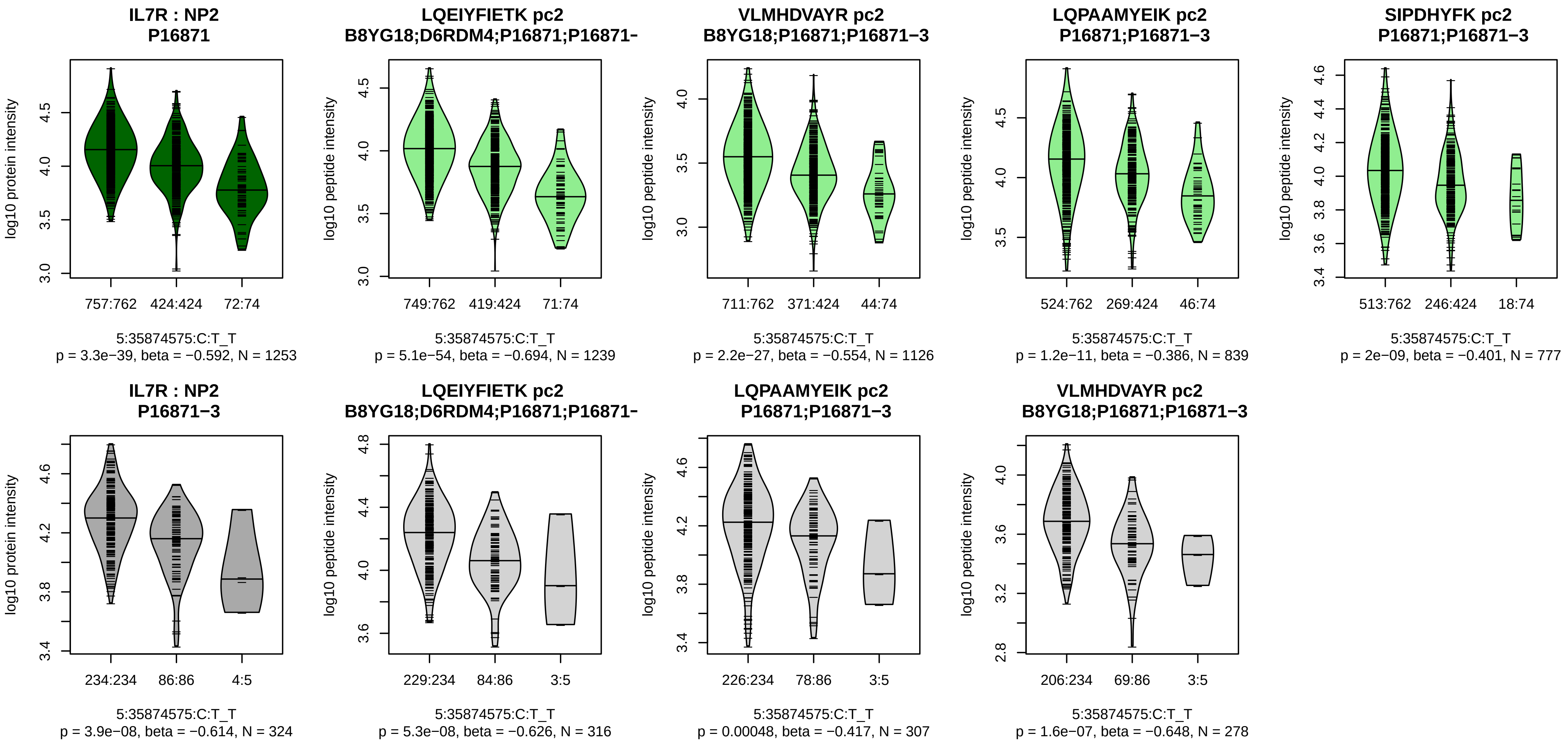


Assay Target: IL7R
Olink UniProt: P16871
UKB rsID: rs6897932
Proxy rsID: rs6897932
UKB SNP: 5:35874575:C:T
Proxy SNP: 5:35874575:C:T
UKB log10(p): 3931.5
UKB BETA: −0.889
*:*:*:*:–:–
1239:1126:839:777:84:38

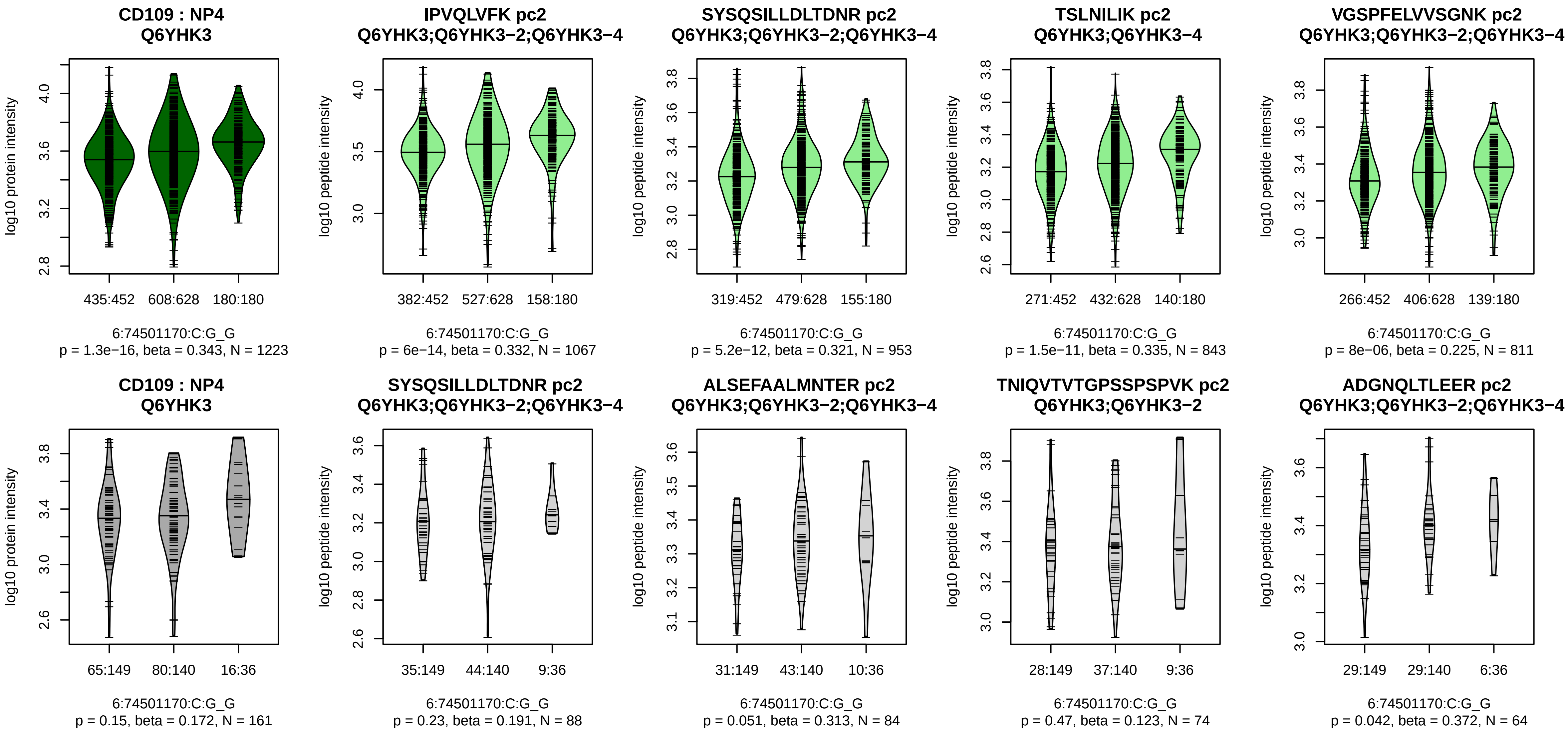


Assay Target: CD109
Olink UniProt: Q6YHK3
UKB rsID: rs9447027
Proxy rsID: rs9447027
UKB SNP: 6:74501170:C:G
Proxy SNP: 6:74501170:C:G
UKB log10(p): 3909.6
UKB BETA: 0.845
*:*:*:*:*:–:–:–:–:*:*:*:–:–:–:–:–:–:–:
1067:953:843:811:609:533:491:

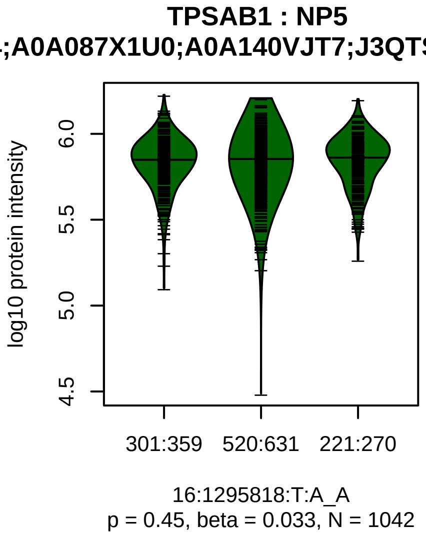


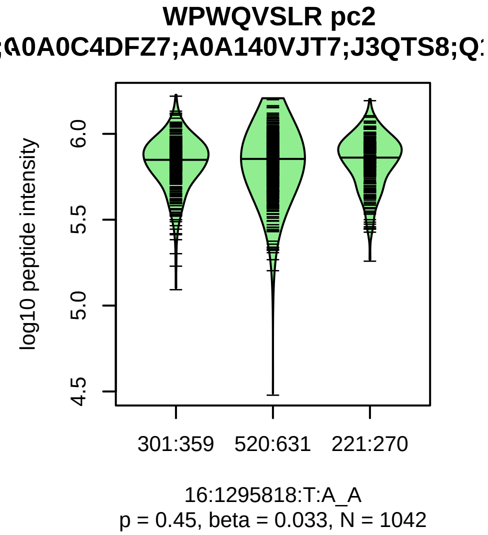


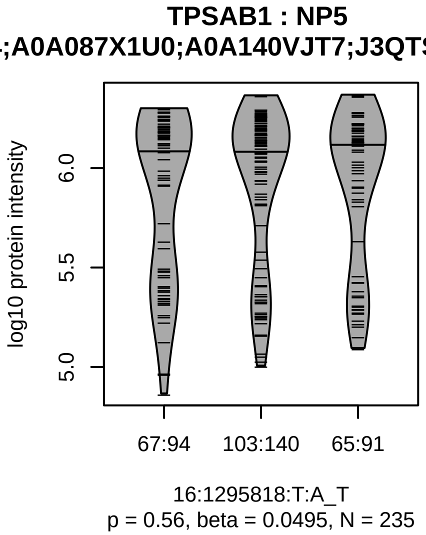


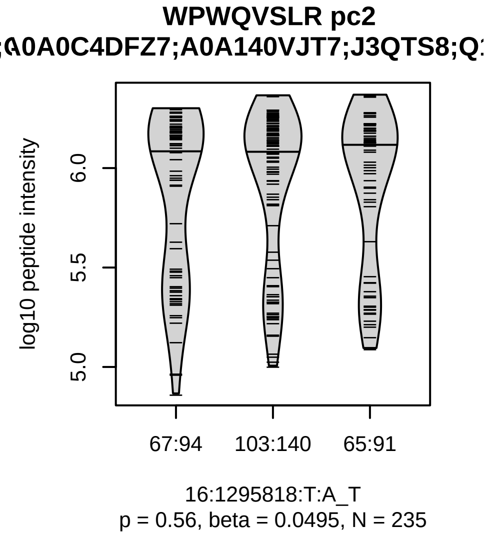


Assay Target: TPSAB1
Olink UniProt: Q15661
UKB rsID: rs8051930
Proxy rsID: rs8051930
UKB SNP: 16:1295818:T:A
Proxy SNP: 16:1295818:T:A
UKB log10(p): 3832.7
UKB BETA: 0.814
–
1042

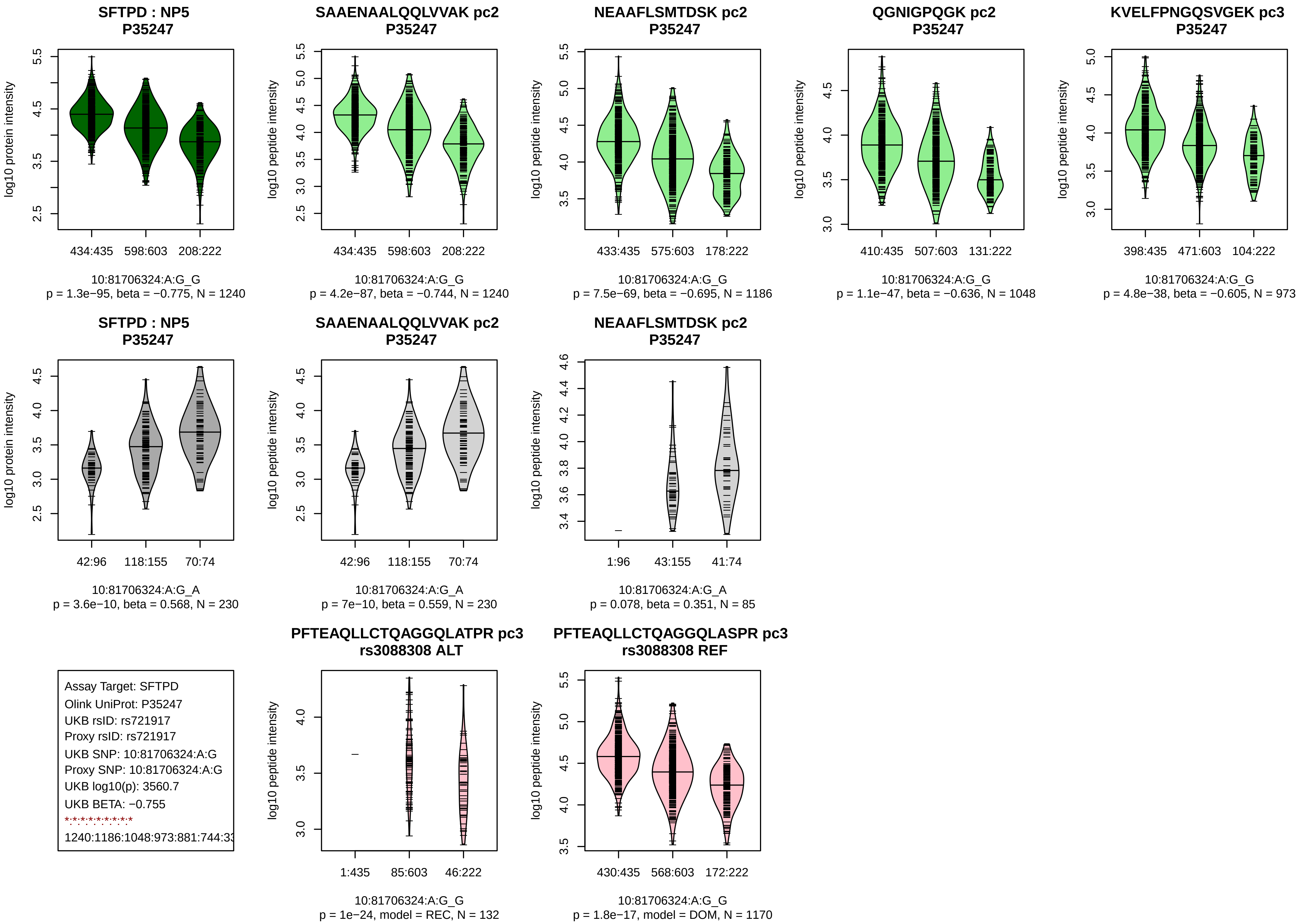

SFTPD : NP5
P35247
log10 protein intensity
434:435
598:603
208:222
10:81706324:A:G_G
p = 1.3e−95, beta = −0.775, N = 1240
SAAENAALQQLVVAK pc2
P35247
log10 peptide intensity
434:435
598:603
208:222
10:81706324:A:G_G
p = 4.2e−87, beta = −0.744, N = 1240
NEAAFLSMTDSK pc2
P35247
log10 peptide intensity
433:435
575:603
178:222
10:81706324:A:G_G
p = 7.5e−69, beta = −0.695, N = 1186
QGNIGPQGK pc2
P35247
log10 peptide intensity
410:435
507:603
131:222
10:81706324:A:G_G
p = 1.1e−47, beta = −0.636, N = 1048
KVELFPNGQSVGEK pc3
P35247
log10 peptide intensity
398:435
471:603
104:222
10:81706324:A:G_G
p = 4.8e−38, beta = −0.605, N = 973
SFTPD : NP5
P35247
log10 protein intensity
42:96
118:155
70:74
10:81706324:A:G_A
p = 3.6e−10, beta = 0.568, N = 230
SAAENAALQQLVVAK pc2
P35247
log10 peptide intensity
42:96
118:155
70:74
10:81706324:A:G_A
p = 7e−10, beta = 0.559, N = 230
NEAAFLSMTDSK pc2
P35247
log10 peptide intensity
1:96
43:155
41:74
10:81706324:A:G_A
p = 0.078, beta = 0.351, N = 85
Assay Target: SFTPD
Olink UniProt: P35247
UKB rsID: rs721917
Proxy rsID: rs721917
UKB SNP: 10:81706324:A:G
Proxy SNP: 10:81706324:A:G
UKB log10(p): 3560.7
UKB BETA: −0.755
1240:1186:1048:973:881:744:33
PFTEAQLLCTQAGGQLATPR pc3
rs3088308 ALT
log10 peptide intensity
1:435
85:603
46:222
10:81706324:A:G_G
p = 1e−24, model = REC, N = 132
PFTEAQLLCTQAGGQLASPR pc3
rs3088308 REF
log10 peptide intensity
430:435
568:603
172:222
10:81706324:A:G_G
p = 1.8e−17, model = DOM, N = 1170

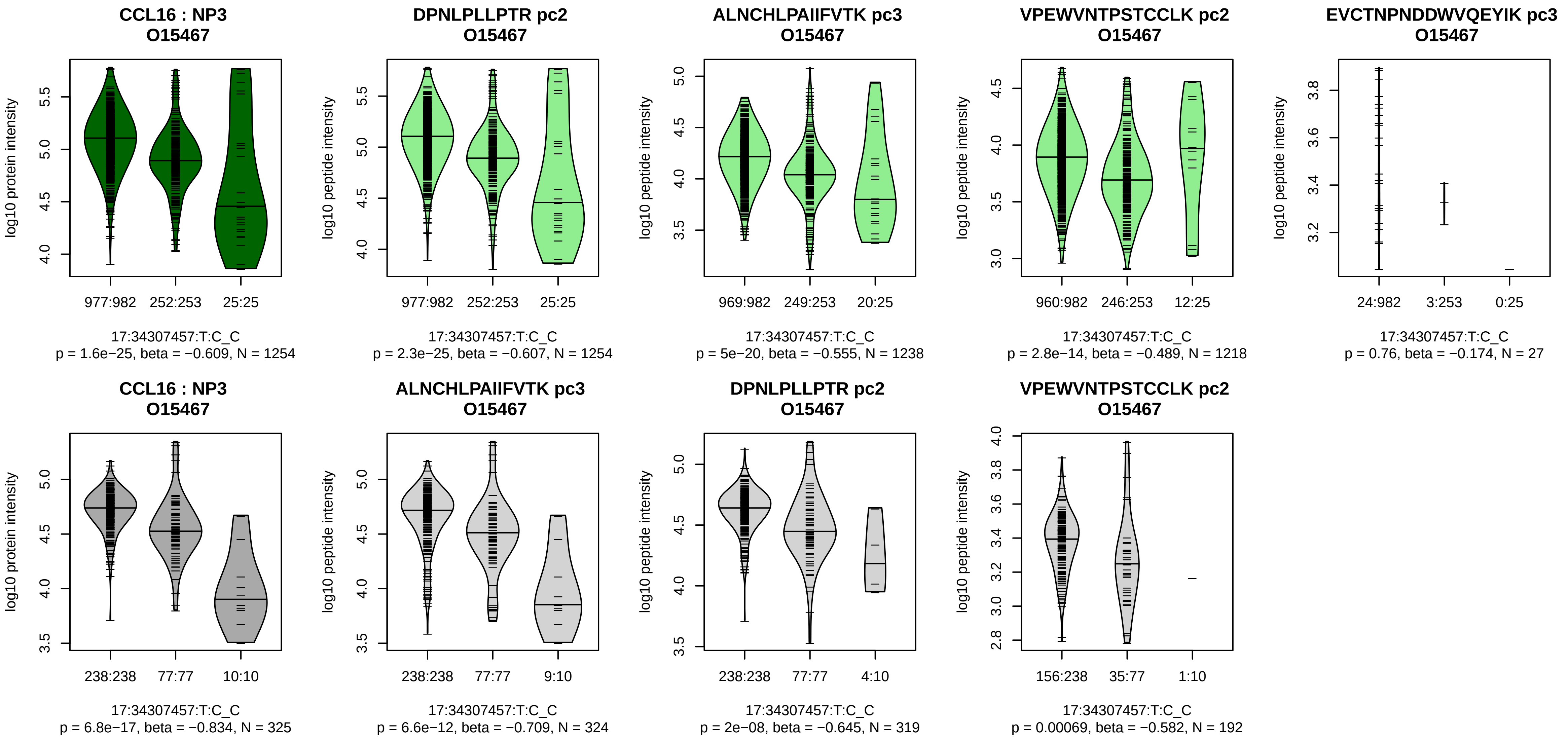


Assay Target: CCL16
Olink UniProt: O15467
UKB rsID: rs112689088
Proxy rsID: rs112689088
UKB SNP: 17:34307457:T:C
Proxy SNP: 17:34307457:T:C
UKB log10(p): 3249.4
UKB BETA: −1.203
*:*:*:*:–
1254:1238:1219:1218:27

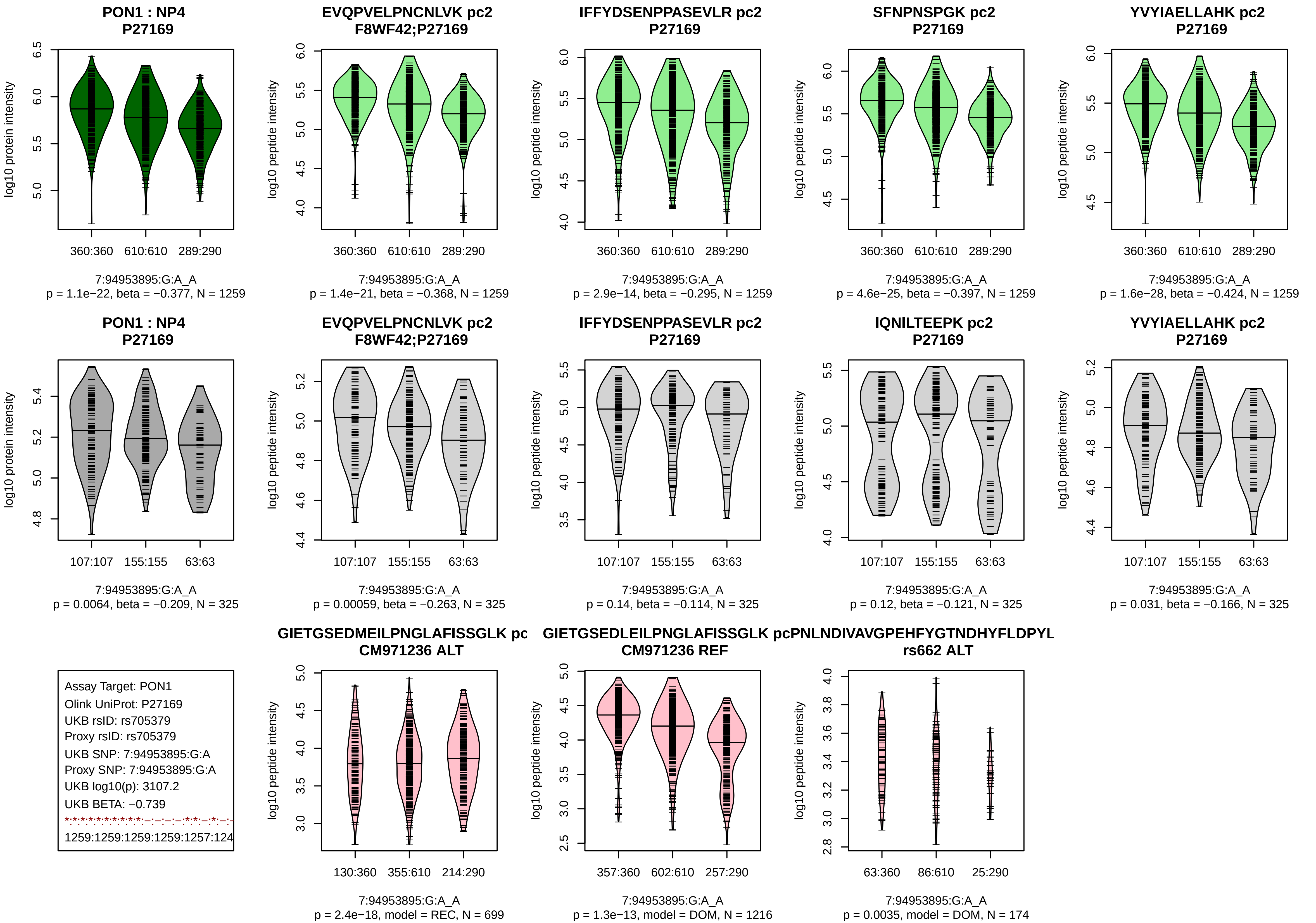

PON1 : NP4
P27169
log10 protein intensity
360:360
610:610
289:290
7:94953895:G:A_A
p = 1.1e−22, beta = −0.377, N = 1259
EVQPVELPNCNLVK pc2
F8WF42;P27169
log10 peptide intensity
360:360
610:610
289:290
7:94953895:G:A_A
p = 1.4e−21, beta = −0.368, N = 1259
IFFYDSENPPASEVLR pc2
P27169
log10 peptide intensity
360:360
610:610
289:290
7:94953895:G:A_A
p = 2.9e−14, beta = −0.295, N = 1259
SFNPNSPGK pc2
P27169
log10 peptide intensity
360:360
610:610
289:290
7:94953895:G:A_A
p = 4.6e−25, beta = −0.397, N = 1259
YVYIAELLAHK pc2
P27169
log10 peptide intensity
360:360
610:610
289:290
7:94953895:G:A_A
p = 1.6e−28, beta = −0.424, N = 1259
PON1 : NP4
P27169
log10 protein intensity
107:107
155:155
63:63
7:94953895:G:A_A
p = 0.0064, beta = −0.209, N = 325
EVQPVELPNCNLVK pc2
F8WF42;P27169
log10 peptide intensity
107:107
155:155
63:63
7:94953895:G:A_A
p = 0.00059, beta = −0.263, N = 325
IFFYDSENPPASEVLR pc2
P27169
log10 peptide intensity
107:107
155:155
63:63
7:94953895:G:A_A
p = 0.14, beta = −0.114, N = 325
IQNILTEEPK pc2
P27169
log10 peptide intensity
107:107
155:155
63:63
7:94953895:G:A_A
p = 0.12, beta = −0.121, N = 325
YVYIAELLAHK pc2
P27169
log10 peptide intensity
107:107
155:155
63:63
7:94953895:G:A_A
p = 0.031, beta = −0.166, N = 325
Assay Target: PON1
Olink UniProt: P27169
UKB rsID: rs705379
Proxy rsID: rs705379
UKB SNP: 7:94953895:G:A
Proxy SNP: 7:94953895:G:A
UKB log10(p): 3107.2
UKB BETA: −0.739
1259:1259:1259:1259:1257:124
GIETGSEDMEILPNGLAFISSGLK pc
CM971236 ALT
log10 peptide intensity
130:360
355:610
214:290
7:94953895:G:A_A
p = 2.4e−18, model = REC, N = 699
GIETGSEDLEILPNGLAFISSGLK pc
CM971236 REF
log10 peptide intensity
357:360
602:610
257:290
7:94953895:G:A_A
p = 1.3e−13, model = DOM, N = 1216
PNLNDIVAVGPEHFYGTNDHYFLDPYL
rs662 ALT
log10 peptide intensity
63:360
86:610
25:290
7:94953895:G:A_A
p = 0.0035, model = DOM, N = 174

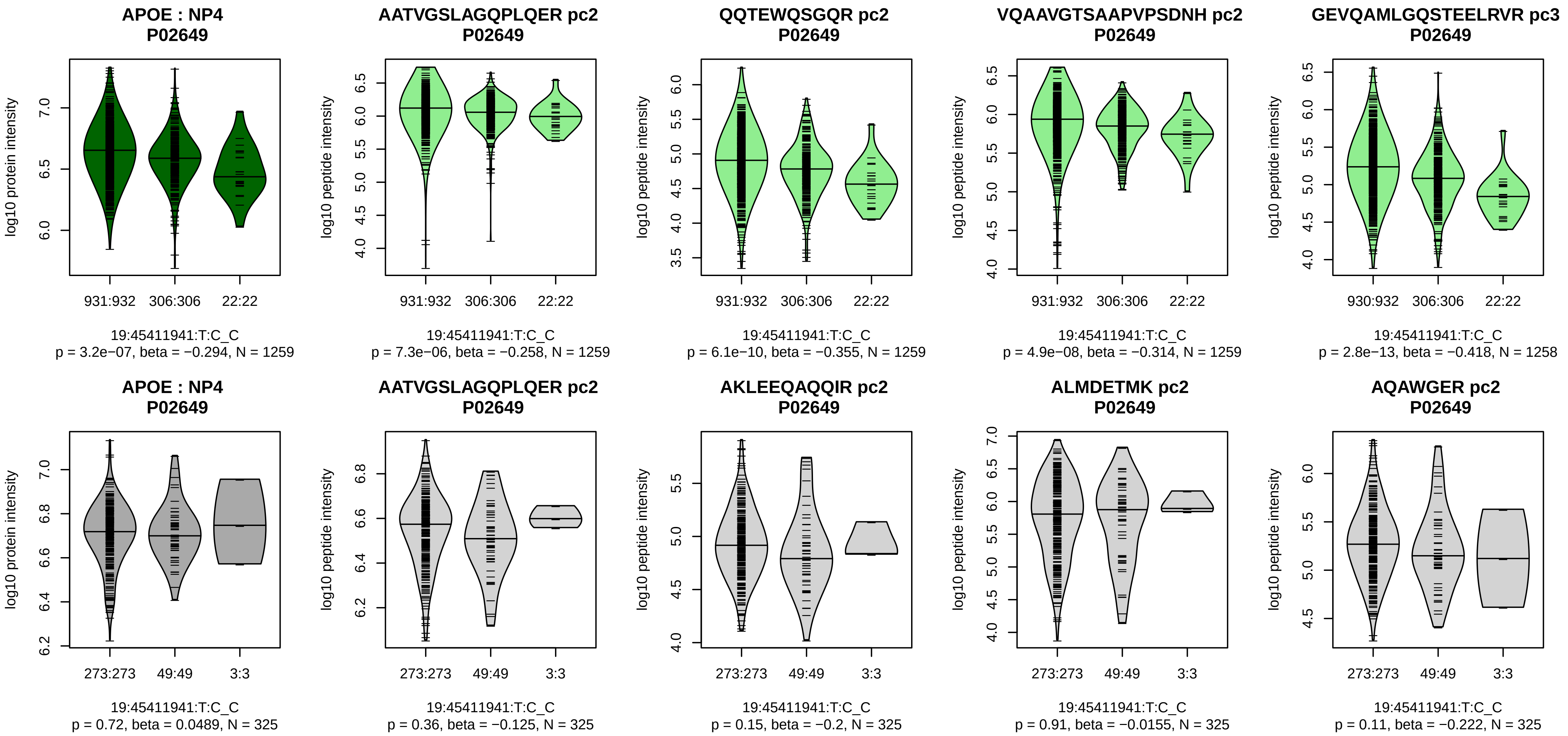


Assay Target: APOE
Olink UniProt: P02649
UKB rsID: rs429358
Proxy rsID: rs429358
UKB SNP: 19:45411941:T:C
Proxy SNP: 19:45411941:T:C
UKB log10(p): 3102.3
UKB BETA: −0.99
*:*:*:*:*:*:*:*:*:*:*:*:*:*:*:*:*:*:*:*:*:*:*:*:*:*:
1259:1259:1259:1258:1258:125

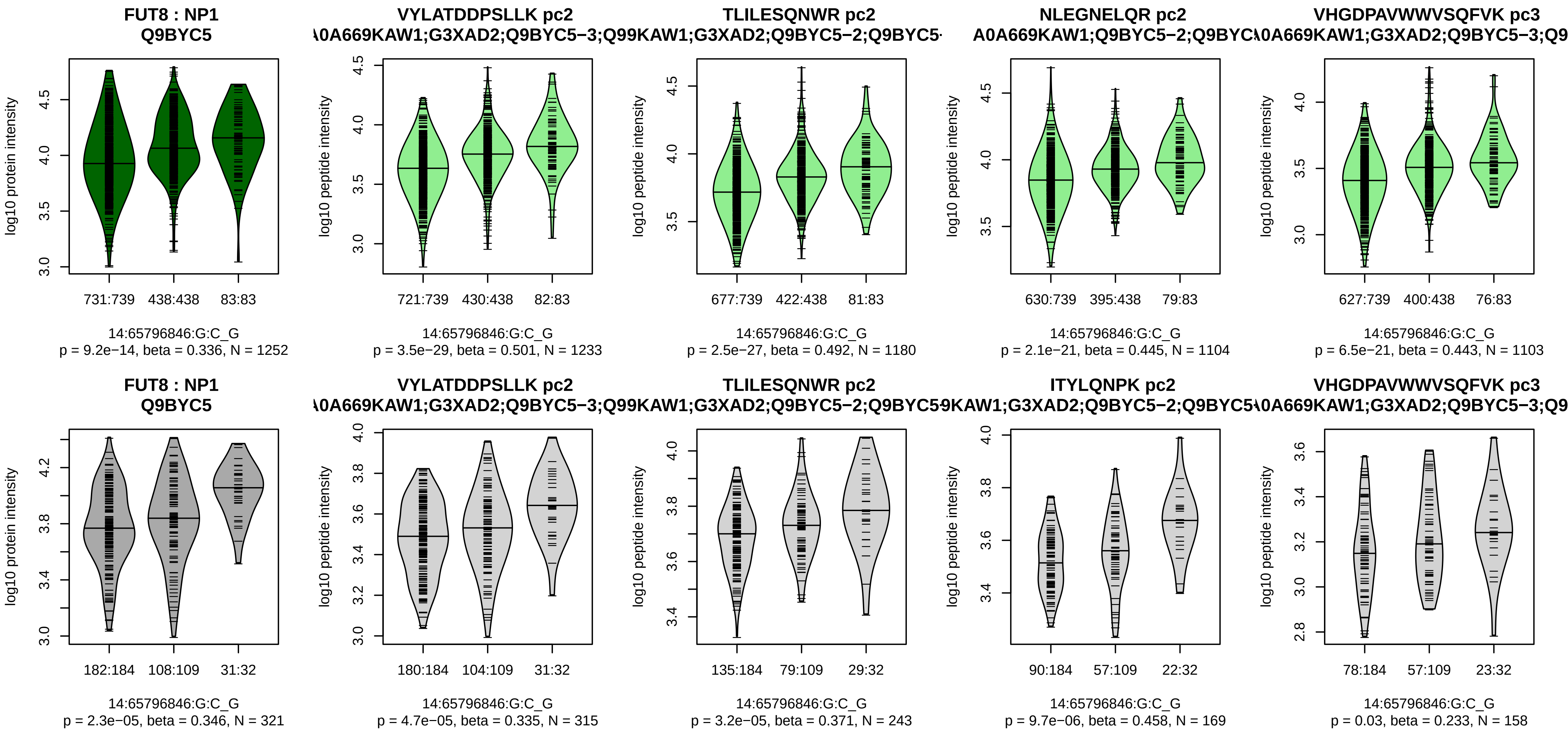


Assay Target: FUT8
Olink UniProt: Q9BYC5
UKB rsID: rs2127870
Proxy rsID: rs2127870
UKB SNP: 14:65796846:G:C
Proxy SNP: 14:65796846:G:C
UKB log10(p): 3097.8
UKB BETA: −0.905
*:*:*:*:*:*:*:*:*:*:–:*:–:*:*:*:*:*:–:–
1233:1180:1104:1103:1050:979

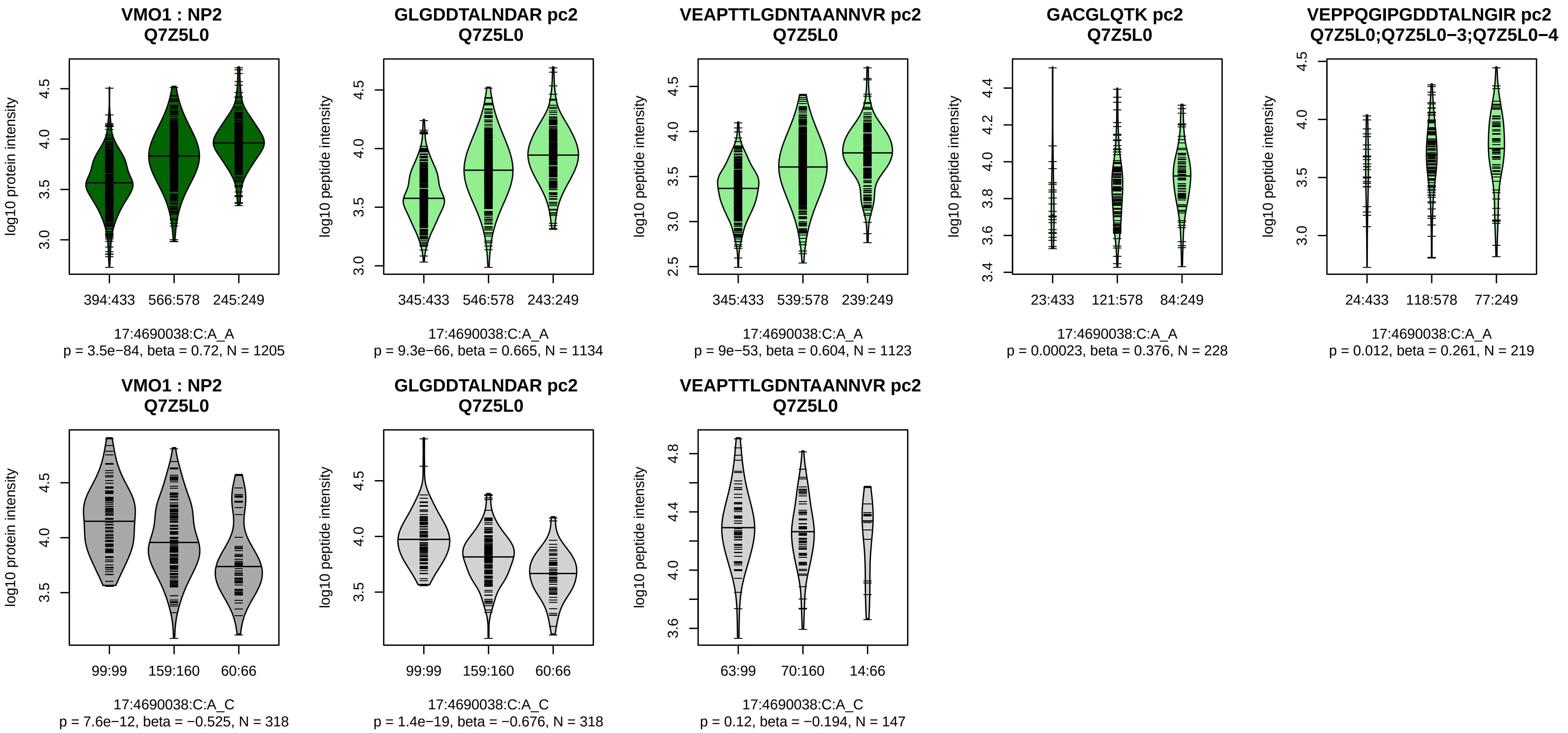


Assay Target: VMO1
Olink UniProt: Q7Z5L0
UKB rsID: rs8078118
Proxy rsID: rs8078118
UKB SNP: 17:4690038:C:A
Proxy SNP: 17:4690038:C:A
UKB log10(p): 3082.1
UKB BETA: 0.732
*:*:*:–:–:*
1134:1123:228:219:144:132

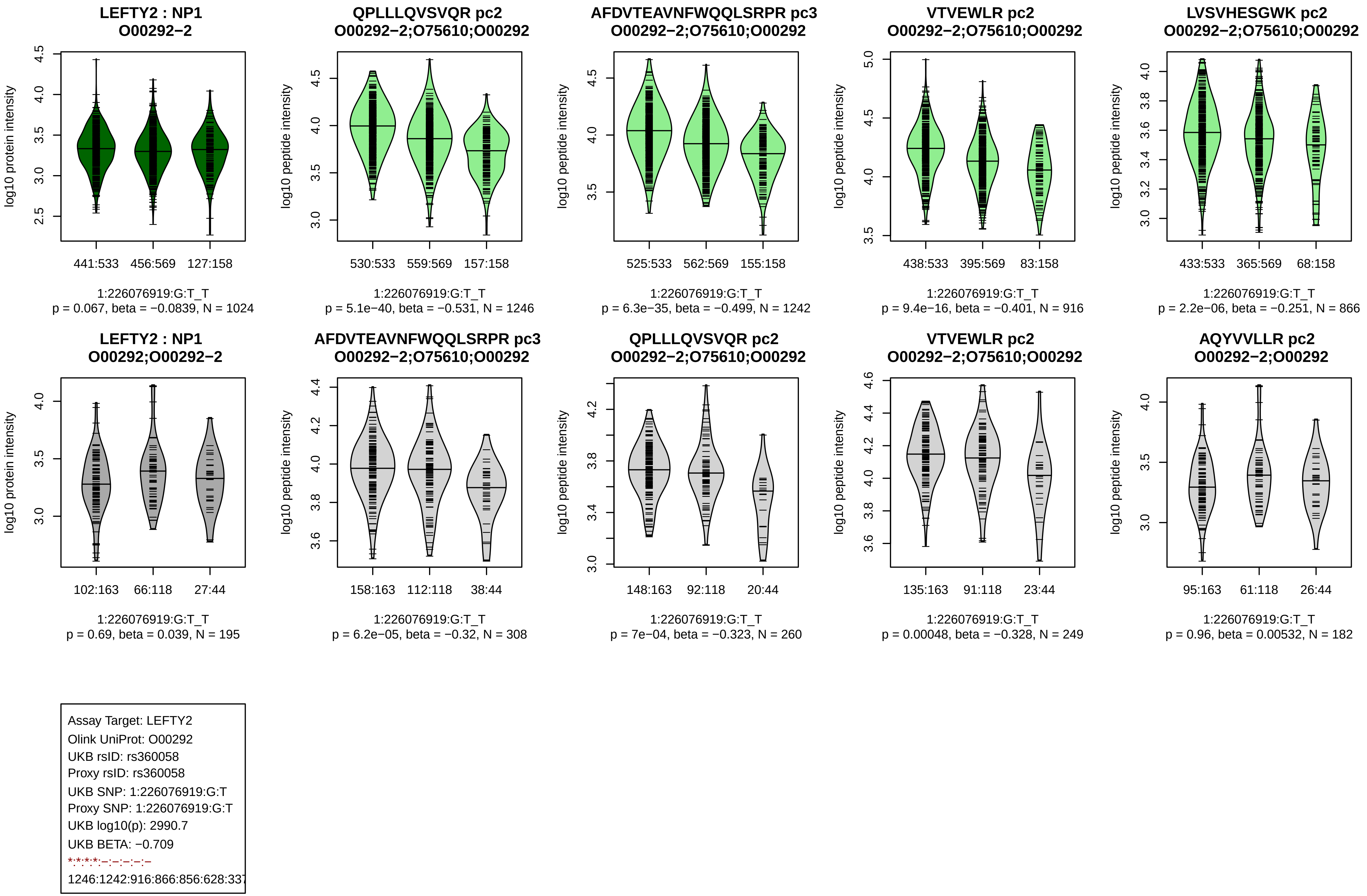
LEFTY2 : NP1
O00292−2
log10 protein intensity
441:533 456:569 127:158
1:226076919:G:T_T
p = 0.067, beta = −0.0839, N = 1024
QPLLLQVSVQR pc2
O00292−2;O75610;O00292
log10 peptide intensity
530:533 559:569 157:158
1:226076919:G:T_T
p = 5.1e−40, beta = −0.531, N = 1246
AFDVTEAVNFWQQLSRPR pc3
O00292−2;O75610;O00292
log10 peptide intensity
525:533 562:569 155:158
1:226076919:G:T_T
p = 6.3e−35, beta = −0.499, N = 1242
VTVEWLR pc2
O00292−2;O75610;O00292
log10 peptide intensity
438:533 395:569 83:158
1:226076919:G:T_T
p = 9.4e−16, beta = −0.401, N = 916
LVSVHESGWK pc2
O00292−2;O75610;O00292
log10 peptide intensity
433:533 365:569 68:158
1:226076919:G:T_T
p = 2.2e−06, beta = −0.251, N = 866
LEFTY2 : NP1
O00292;O00292−2
log10 protein intensity
102:163 66:118 27:44
1:226076919:G:T_T
p = 0.69, beta = 0.039, N = 195
AFDVTEAVNFWQQLSRPR pc3
O00292−2;O75610;O00292
log10 peptide intensity
158:163 112:118 38:44
1:226076919:G:T_T
p = 6.2e−05, beta = −0.32, N = 308
QPLLLQVSVQR pc2
O00292−2;O75610;O00292
log10 peptide intensity
148:163 92:118 20:44
1:226076919:G:T_T
p = 7e−04, beta = −0.323, N = 260
VTVEWLR pc2
O00292−2;O75610;O00292
log10 peptide intensity
135:163 91:118 23:44
1:226076919:G:T_T
p = 0.00048, beta = −0.328, N = 249
AQYVVLLR pc2
O00292−2;O00292
log10 peptide intensity
95:163 61:118 26:44
1:226076919:G:T_T
p = 0.96, beta = 0.00532, N = 182
Assay Target: LEFTY2
Olink UniProt: O00292
UKB rsID: rs360058
Proxy rsID: rs360058
UKB SNP: 1:226076919:G:T
Proxy SNP: 1:226076919:G:T
UKB log10(p): 2990.7
UKB BETA: −0.709
*:*:*:*:–:–:–:–:–
1246:1242:916:866:856:628:337

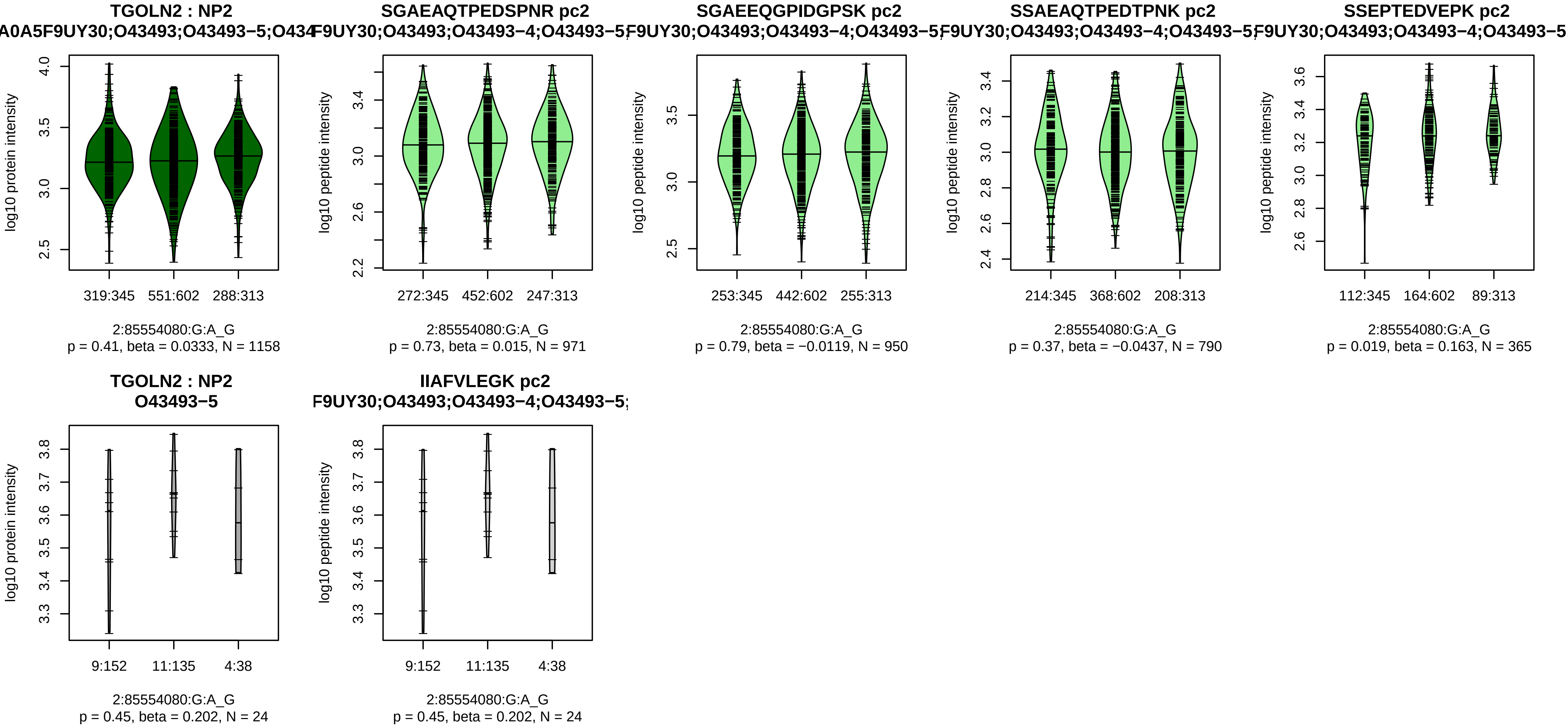


Assay Target: TGOLN2
Olink UniProt: O43493
UKB rsID: rs4247303
Proxy rsID: rs4247303
UKB SNP: 2:85554080:G:A
Proxy SNP: 2:85554080:G:A
UKB log10(p): 2870.9
UKB BETA: −0.688
−:−:−:−:−:NA
971:950:790:365:24:12

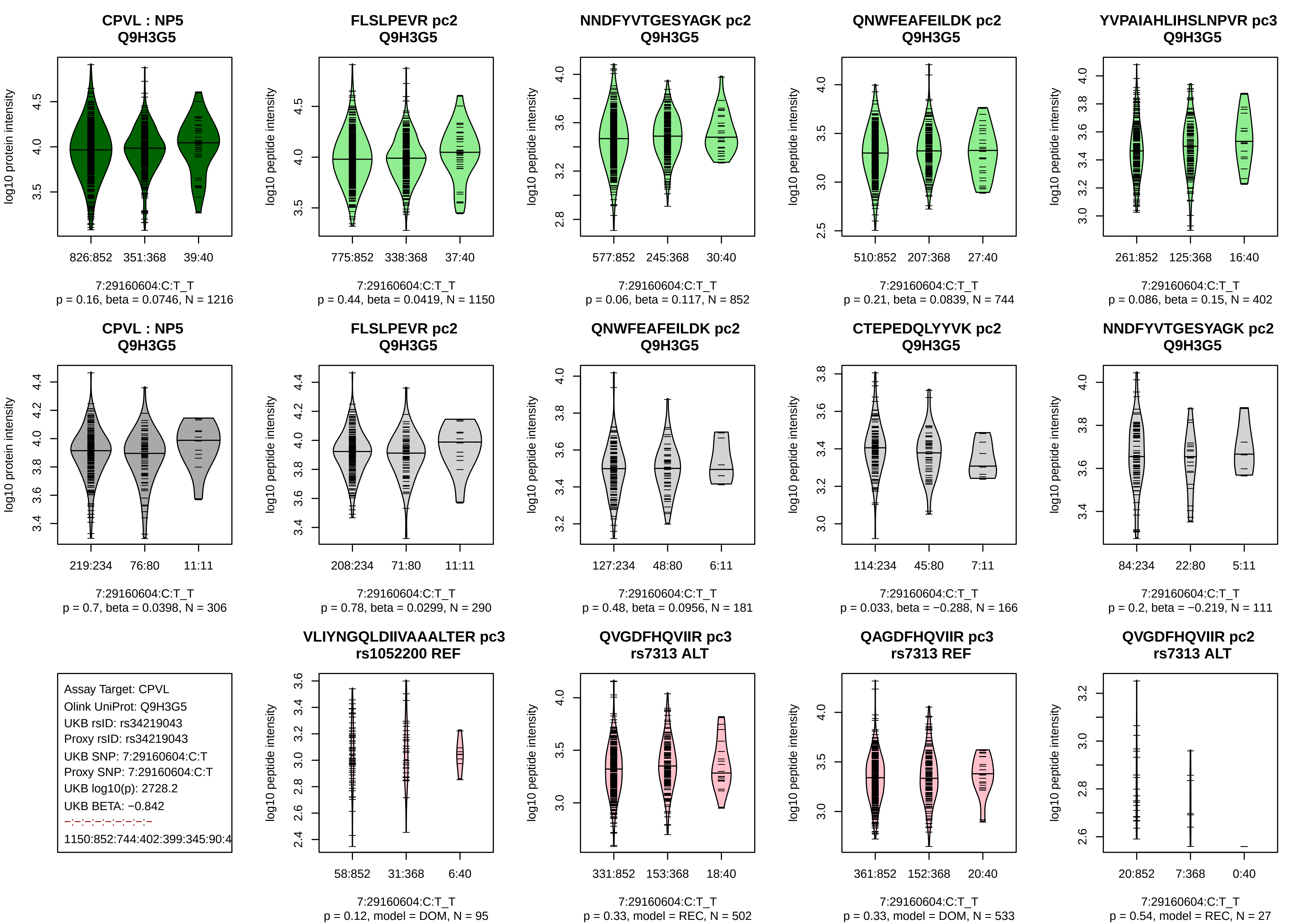
CPVL : NP5
Q9H3G5
log10 protein intensity
826:852 351:368 39:40
7:29160604:C:T_T
p = 0.16, beta = 0.0746, N = 1216
FLSLPEVR pc2
Q9H3G5
log10 peptide intensity
775:852 338:368 37:40
7:29160604:C:T_T
p = 0.44, beta = 0.0419, N = 1150
NNDFYVTGESYAGK pc2
Q9H3G5
log10 peptide intensity
577:852 245:368 30:40
7:29160604:C:T_T
p = 0.06, beta = 0.117, N = 852
QNWFEAFEILDK pc2
Q9H3G5
log10 peptide intensity
510:852 207:368 27:40
7:29160604:C:T_T
p = 0.21, beta = 0.0839, N = 744
YVPAIAHLIHSLNPVR pc3
Q9H3G5
log10 peptide intensity
261:852 125:368 16:40
7:29160604:C:T_T
p = 0.086, beta = 0.15, N = 402
CPVL : NP5
Q9H3G5
log10 protein intensity
219:234 76:80 11:11
7:29160604:C:T_T
p = 0.7, beta = 0.0398, N = 306
FLSLPEVR pc2
Q9H3G5
log10 peptide intensity
208:234 71:80 11:11
7:29160604:C:T_T
p = 0.78, beta = 0.0299, N = 290
QNWFEAFEILDK pc2
Q9H3G5
log10 peptide intensity
127:234 48:80 6:11
7:29160604:C:T_T
p = 0.48, beta = 0.0956, N = 181
CTEPEDQLYYVK pc2
Q9H3G5
log10 peptide intensity
114:234 45:80 7:11
7:29160604:C:T_T
p = 0.033, beta = −0.288, N = 166
NNDFYVTGESYAGK pc2
Q9H3G5
log10 peptide intensity
84:234 22:80 5:11
7:29160604:C:T_T
p = 0.2, beta = −0.219, N = 111
Assay Target: CPVL
Olink UniProt: Q9H3G5
UKB rsID: rs34219043
Proxy rsID: rs34219043
UKB SNP: 7:29160604:C:T
Proxy SNP: 7:29160604:C:T
UKB log10(p): 2728.2
UKB BETA: −0.842
−:−:−:−:−:−:−:−:−
1150:852:744:402:399:345:90:4
VLIYNGQLDIIVAAALTER pc3
rs1052200 REF
log10 peptide intensity
58:852 31:368 6:40
7:29160604:C:T_T
p = 0.12, model = DOM, N = 95
QVGDFHQVIIR pc3
rs7313 ALT
log10 peptide intensity
331:852 153:368 18:40
7:29160604:C:T_T
p = 0.33, model = REC, N = 502
QAGDFHQVIIR pc3
rs7313 REF
log10 peptide intensity
361:852 152:368 20:40
7:29160604:C:T_T
p = 0.33, model = DOM, N = 533
QVGDFHQVIIR pc2
rs7313 ALT
log10 peptide intensity
20:852 7:368 0:40
7:29160604:C:T_T
p = 0.54, model = REC, N = 27

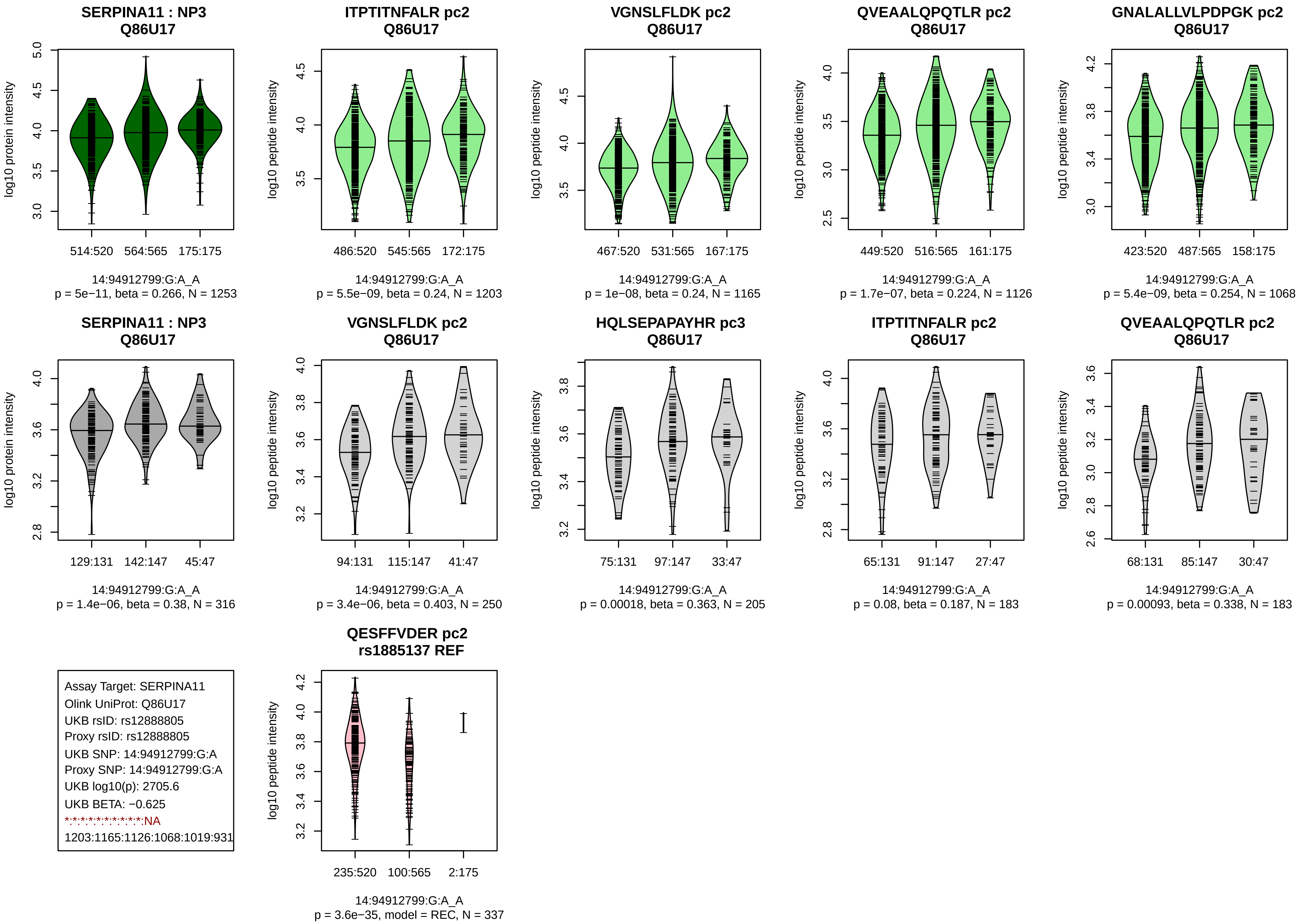
SERPINA11 : NP3
Q86U17
log10 protein intensity
514:520 564:565 175:175
14:94912799:G:A_A
p = 5e−11, beta = 0.266, N = 1253
ITPTITNFALR pc2
Q86U17
log10 peptide intensity
486:520 545:565 172:175
14:94912799:G:A_A
p = 5.5e−09, beta = 0.24, N = 1203
VGNSLFLDK pc2
Q86U17
log10 peptide intensity
467:520 531:565 167:175
14:94912799:G:A_A
p = 1e−08, beta = 0.24, N = 1165
QVEAALQPQTLR pc2
Q86U17
log10 peptide intensity
449:520 516:565 161:175
14:94912799:G:A_A
p = 1.7e−07, beta = 0.224, N = 1126
GNALALLVLPDPGK pc2
Q86U17
log10 peptide intensity
423:520 487:565 158:175
14:94912799:G:A_A
p = 5.4e−09, beta = 0.254, N = 1068
SERPINA11 : NP3
Q86U17
log10 protein intensity
129:131 142:147 45:47
14:94912799:G:A_A
p = 1.4e−06, beta = 0.38, N = 316
VGNSLFLDK pc2
Q86U17
log10 peptide intensity
94:131 115:147 41:47
14:94912799:G:A_A
p = 3.4e−06, beta = 0.403, N = 250
HQLSEPAPAYHR pc3
Q86U17
log10 peptide intensity
75:131 97:147 33:47
14:94912799:G:A_A
p = 0.00018, beta = 0.363, N = 205
ITPTITNFALR pc2
Q86U17
log10 peptide intensity
65:131 91:147 27:47
14:94912799:G:A_A
p = 0.08, beta = 0.187, N = 183
QVEAALQPQTLR pc2
Q86U17
log10 peptide intensity
68:131 85:147 30:47
14:94912799:G:A_A
p = 0.00093, beta = 0.338, N = 183
Assay Target: SERPINA11
Olink UniProt: Q86U17
UKB rsID: rs12888805
Proxy rsID: rs12888805
UKB SNP: 14:94912799:G:A
Proxy SNP: 14:94912799:G:A
UKB log10(p): 2705.6
UKB BETA: −0.625
*:*:*:*:*:*:*:*:*:*:*:NA
1203:1165:1126:1068:1019:931
QESFFVDER pc2
rs1885137 REF
log10 peptide intensity
235:520 100:565 2:175
14:94912799:G:A_A
p = 3.6e−35, model = REC, N = 337

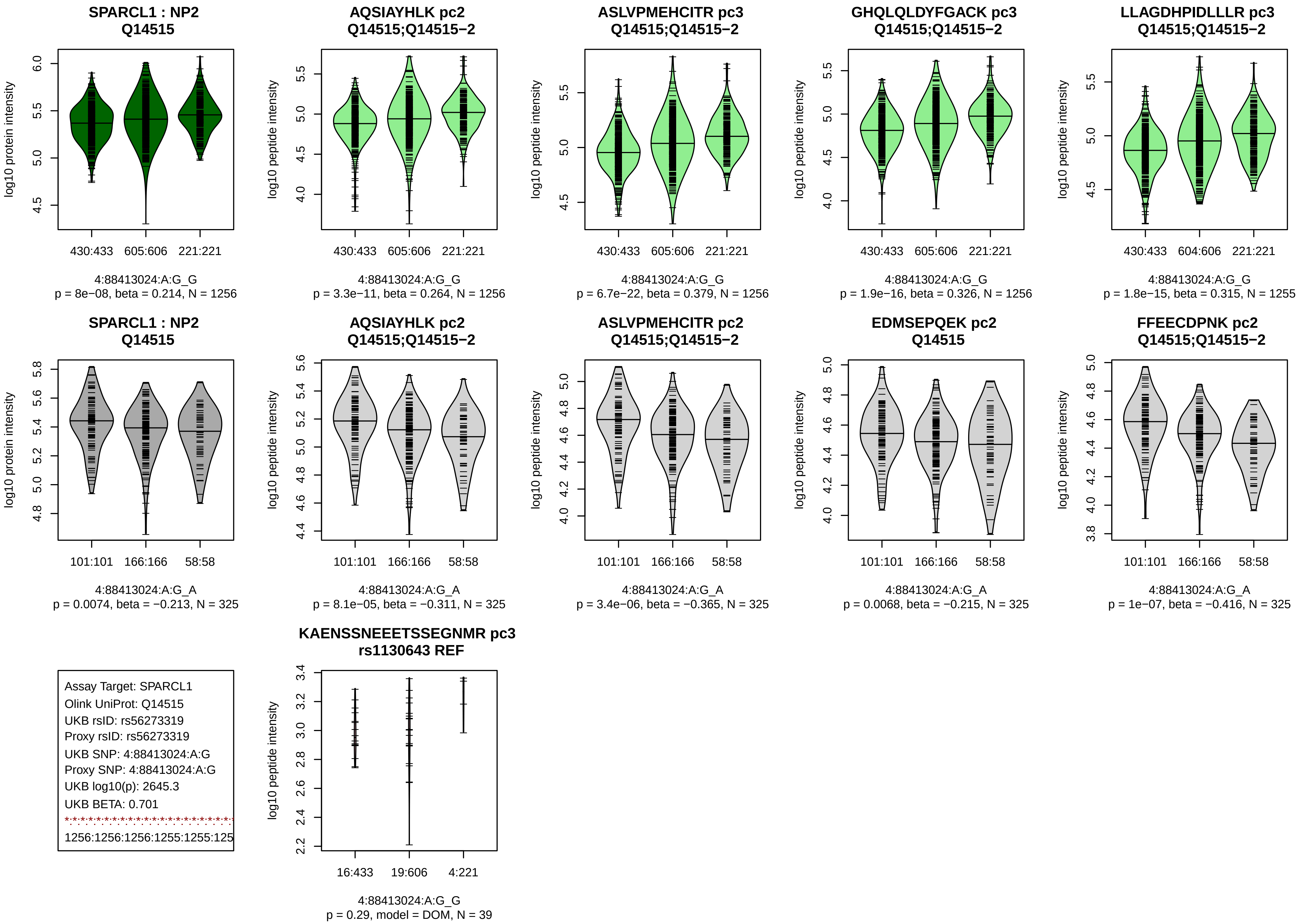
SPARCL1 : NP2
Q14515
log10 protein intensity
430:433 605:606 221:221
4:88413024:A:G_G
p = 8e−08, beta = 0.214, N = 1256
AQSIAYHLK pc2
Q14515;Q14515−2
log10 peptide intensity
430:433 605:606 221:221
4:88413024:A:G_G
p = 3.3e−11, beta = 0.264, N = 1256
ASLVPMEHCITR pc3
Q14515;Q14515−2
log10 peptide intensity
430:433 605:606 221:221
4:88413024:A:G_G
p = 6.7e−22, beta = 0.379, N = 1256
GHQLQLDYFGACK pc3
Q14515;Q14515−2
log10 peptide intensity
430:433 605:606 221:221
4:88413024:A:G_G
p = 1.9e−16, beta = 0.326, N = 1256
LLAGDHPIDLLLR pc3
Q14515;Q14515−2
log10 peptide intensity
430:433 604:606 221:221
4:88413024:A:G_G
p = 1.8e−15, beta = 0.315, N = 1255
SPARCL1 : NP2
Q14515
log10 protein intensity
101:101 166:166 58:58
4:88413024:A:G_A
p = 0.0074, beta = −0.213, N = 325
AQSIAYHLK pc2
Q14515;Q14515−2
log10 peptide intensity
101:101 166:166 58:58
4:88413024:A:G_A
p = 8.1e−05, beta = −0.311, N = 325
ASLVPMEHCITR pc2
Q14515;Q14515−2
log10 peptide intensity
101:101 166:166 58:58
4:88413024:A:G_A
p = 3.4e−06, beta = −0.365, N = 325
EDMSEPQEK pc2
Q14515
log10 peptide intensity
101:101 166:166 58:58
4:88413024:A:G_A
p = 0.0068, beta = −0.215, N = 325
FFEECDPNK pc2
Q14515;Q14515−2
log10 peptide intensity
101:101 166:166 58:58
4:88413024:A:G_A
p = 1e−07, beta = −0.416, N = 325
Assay Target: SPARCL1
Olink UniProt: Q14515
UKB rsID: rs56273319
Proxy rsID: rs56273319
UKB SNP: 4:88413024:A:G
Proxy SNP: 4:88413024:A:G
UKB log10(p): 2645.3
UKB BETA: 0.701
1256:1256:1256:1255:1255:125
KAENSSNEEETSSEGNMR pc3
rs1130643 REF
log10 peptide intensity
16:433 19:606 4:221
4:88413024:A:G_G
p = 0.29, model = DOM, N = 39

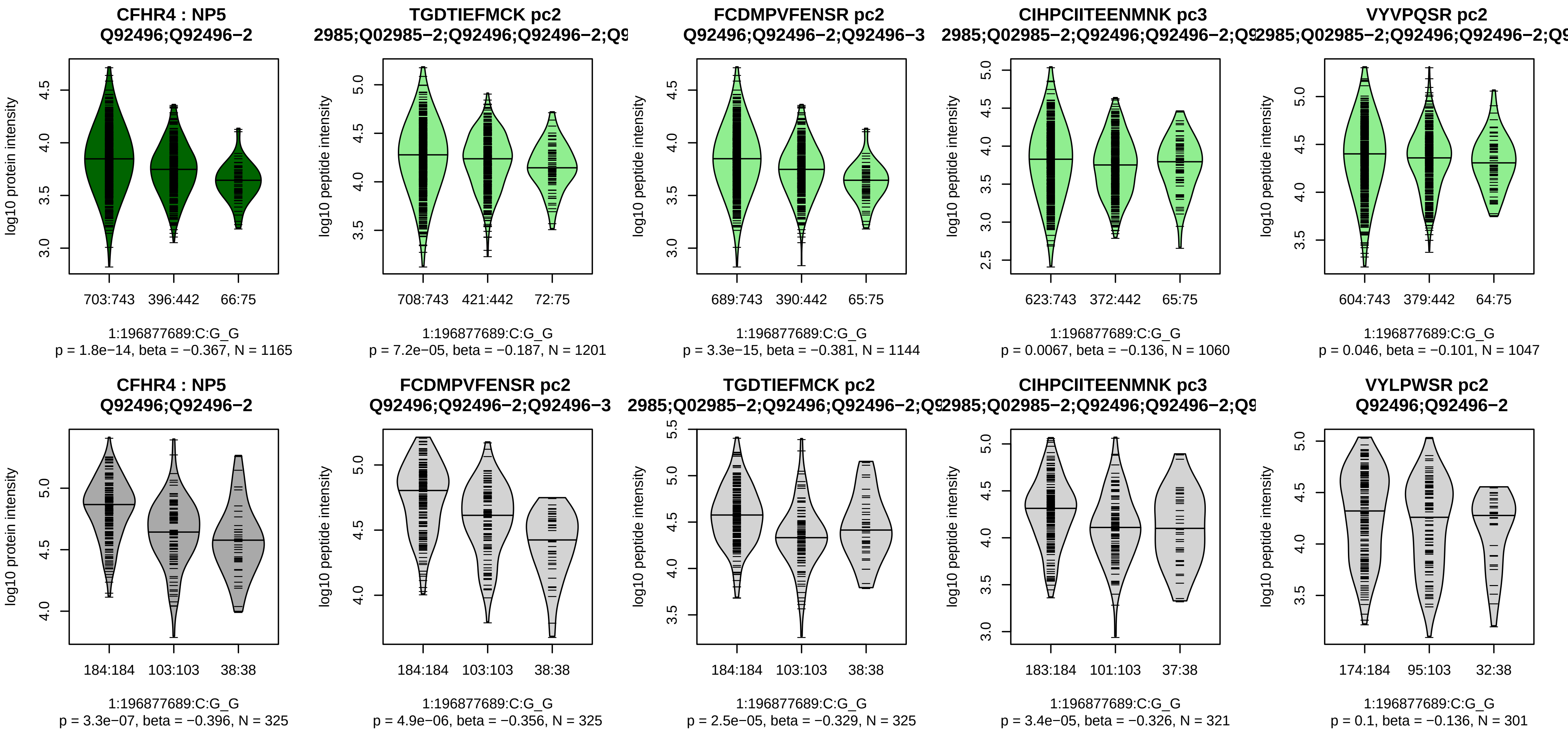


Assay Target: CFHR4
Olink UniProt: Q92496
UKB rsID: rs10922147
Proxy rsID: rs10922147
UKB SNP: 1:196877689:C:G
Proxy SNP: 1:196877689:C:G
UKB log10(p): 2512.3
UKB BETA: −0.785
*:*:*:–:*:*:–:–
1201:1144:1060:1047:690:585:1

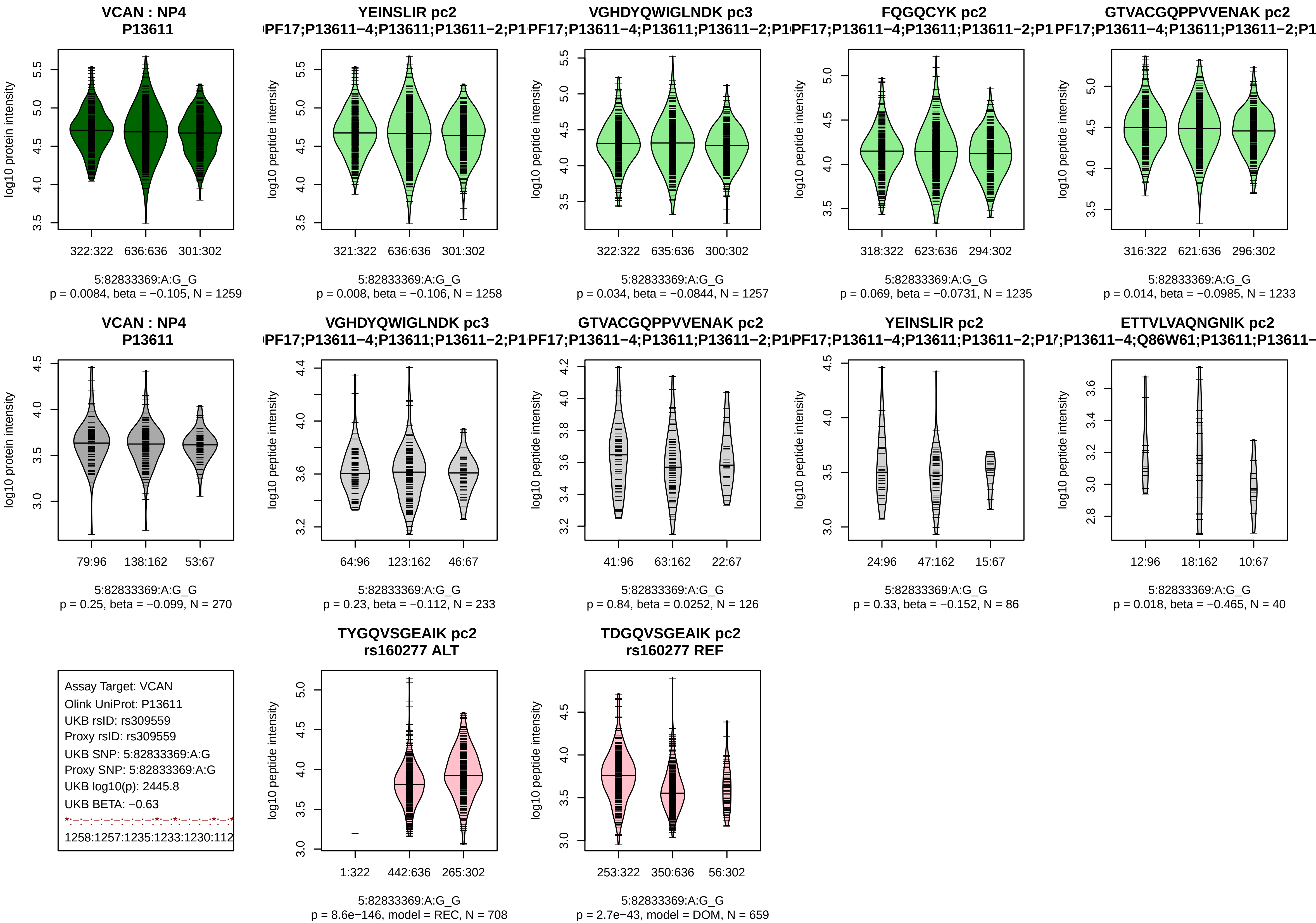
VCAN : NP4
P13611
log10 protein intensity
322:322 636:636 301:302
5:82833369:A:G_G
p = 0.0084, beta = −0.105, N = 1259
YEINSLIR pc2
log10 peptide intensity
321:322 636:636 301:302
5:82833369:A:G_G
p = 0.008, beta = −0.106, N = 1258
VGHDYQWIGLNDK pc3
322:322 635:636 300:302
5:82833369:A:G_G
p = 0.034, beta = −0.0844, N = 1257
FQGQCYK pc2
318:322 623:636 294:302
5:82833369:A:G_G
p = 0.069, beta = −0.0731, N = 1235
GTVACGQPPVVENAK pc2
316:322 621:636 296:302
5:82833369:A:G_G
p = 0.014, beta = −0.0985, N = 1233
VCAN : NP4
P13611
79:96 138:162 53:67
5:82833369:A:G_G
p = 0.25, beta = −0.099, N = 270
VGHDYQWIGLNDK pc3
64:96 123:162 46:67
5:82833369:A:G_G
p = 0.23, beta = −0.112, N = 233
GTVACGQPPVVENAK pc2
41:96 63:162 22:67
5:82833369:A:G_G
p = 0.84, beta = 0.0252, N = 126
YEINSLIR pc2
24:96 47:162 15:67
5:82833369:A:G_G
p = 0.33, beta = −0.152, N = 86
ETTVLVAQNGNIK pc2
12:96 18:162 10:67
5:82833369:A:G_G
p = 0.018, beta = −0.465, N = 40
Assay Target: VCAN
Olink UniProt: P13611
UKB rsID: rs309559
Proxy rsID: rs309559
UKB SNP: 5:82833369:A:G
Proxy SNP: 5:82833369:A:G
UKB log10(p): 2445.8
UKB BETA: −0.63
1258:1257:1235:1233:1230:112
TYGQVSGEAIK pc2
rs160277 ALT
1:322 442:636 265:302
5:82833369:A:G_G
p = 8.6e−146, model = REC, N = 708
TDGQVSGEAIK pc2
rs160277 REF
253:322 350:636 56:302
5:82833369:A:G_G
p = 2.7e−43, model = DOM, N = 659

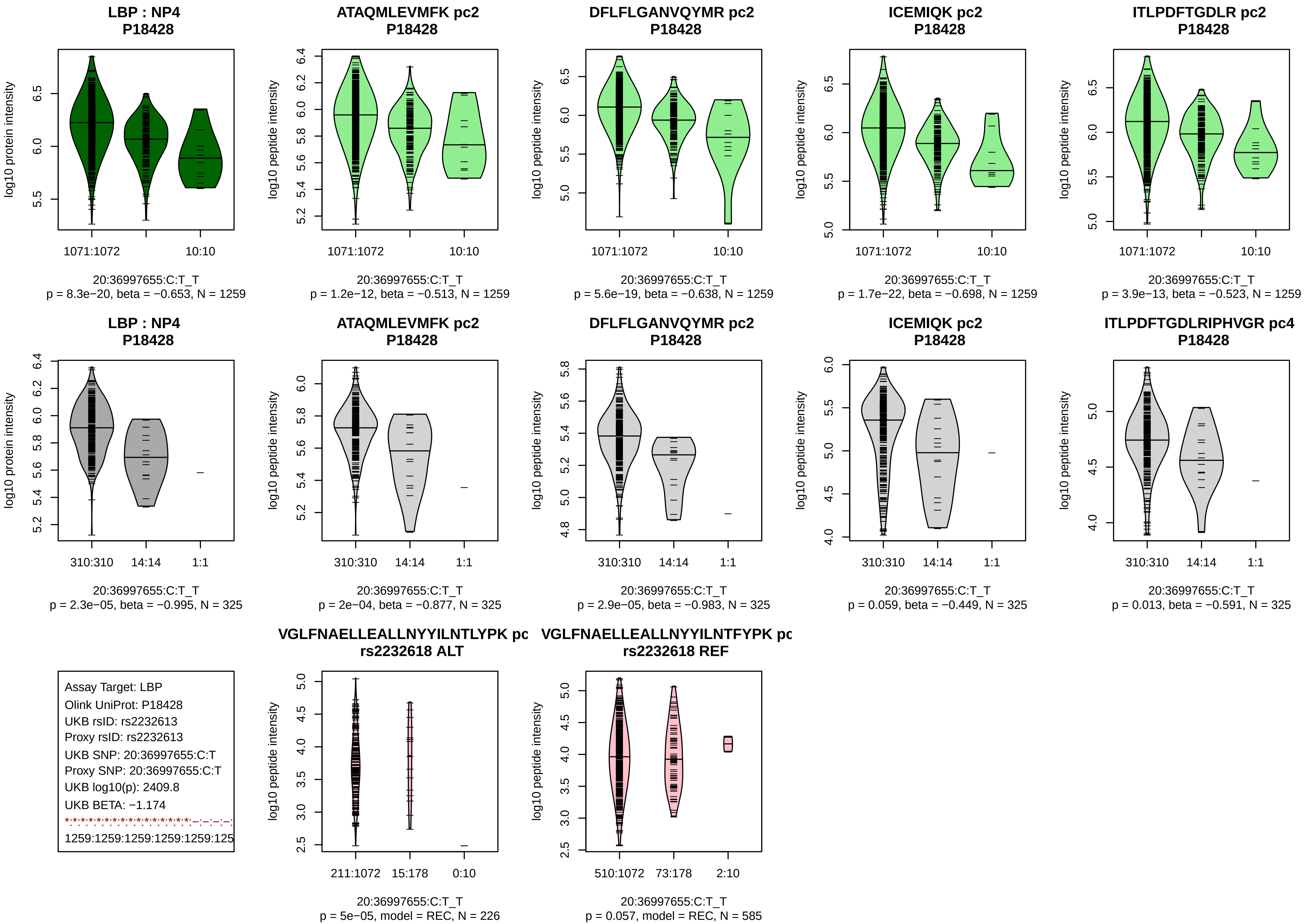
LBP : NP4
P18428
log10 protein intensity
1071:1072
10:10
20:36997655:C:T_T
p = 8.3e−20, beta = −0.653, N = 1259
ATAQMLEVMFK pc2
P18428
log10 peptide intensity
1071:1072
10:10
20:36997655:C:T_T
p = 1.2e−12, beta = −0.513, N = 1259
DFLFLGANVQYMR pc2
P18428
log10 peptide intensity
1071:1072
10:10
20:36997655:C:T_T
p = 5.6e−19, beta = −0.638, N = 1259
ICEMIQK pc2
P18428
log10 peptide intensity
1071:1072
10:10
20:36997655:C:T_T
p = 1.7e−22, beta = −0.698, N = 1259
ITLPDFTGDLR pc2
P18428
log10 peptide intensity
1071:1072
10:10
20:36997655:C:T_T
p = 3.9e−13, beta = −0.523, N = 1259
LBP : NP4
P18428
log10 protein intensity
310:310
14:14
1:1
20:36997655:C:T_T
p = 2.3e−05, beta = −0.995, N = 325
ATAQMLEVMFK pc2
P18428
log10 peptide intensity
310:310
14:14
1:1
20:36997655:C:T_T
p = 2e−04, beta = −0.877, N = 325
DFLFLGANVQYMR pc2
P18428
log10 peptide intensity
310:310
14:14
1:1
20:36997655:C:T_T
p = 2.9e−05, beta = −0.983, N = 325
ICEMIQK pc2
P18428
log10 peptide intensity
310:310
14:14
1:1
20:36997655:C:T_T
p = 0.059, beta = −0.449, N = 325
ITLPDFTGDLRIPHVGR pc4
P18428
log10 peptide intensity
310:310
14:14
1:1
20:36997655:C:T_T
p = 0.013, beta = −0.591, N = 325
Assay Target: LBP
Olink UniProt: P18428
UKB rsID: rs2232613
Proxy rsID: rs2232613
UKB SNP: 20:36997655:C:T
Proxy SNP: 20:36997655:C:T
UKB log10(p): 2409.8
UKB BETA: −1.174
1259:1259:1259:1259:1259:125
VGLFNAELLEALLNYYILNTLYPK pc
rs2232618 ALT
log10 peptide intensity
211:1072
15:178
0:10
20:36997655:C:T_T
p = 5e−05, model = REC, N = 226
VGLFNAELLEALLNYYILNTFYPK pc
rs2232618 REF
log10 peptide intensity
510:1072
73:178
2:10
20:36997655:C:T_T
p = 0.057, model = REC, N = 585

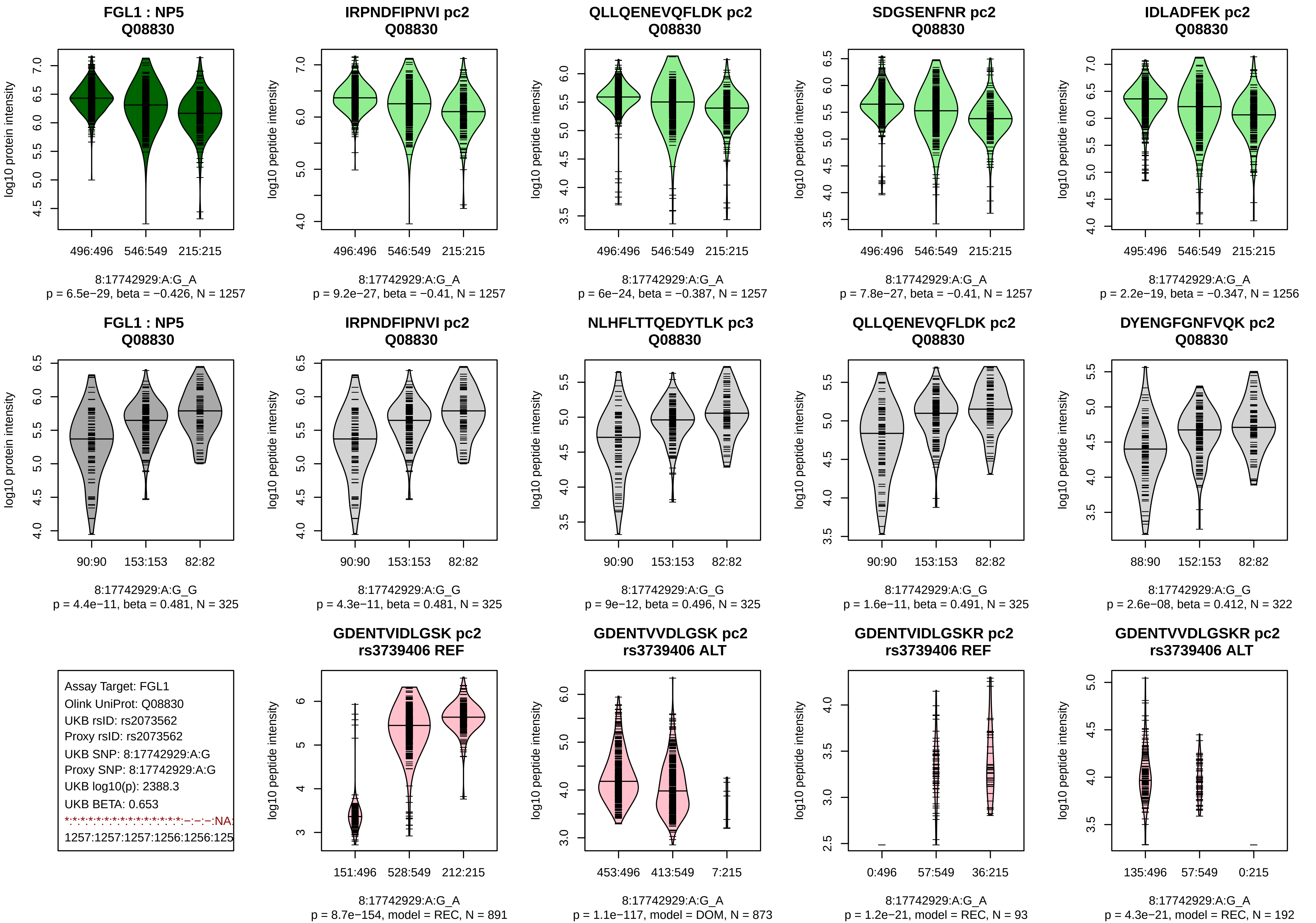

FGL1 : NP5
Q08830
log10 protein intensity
496:496
546:549
215:215
8:17742929:A:G_A
p = 6.5e−29, beta = −0.426, N = 1257
IRPNDFIPNVI pc2
Q08830
log10 peptide intensity
496:496
546:549
215:215
8:17742929:A:G_A
p = 9.2e−27, beta = −0.41, N = 1257
QLLQENEVQFLDK pc2
Q08830
log10 peptide intensity
496:496
546:549
215:215
8:17742929:A:G_A
p = 6e−24, beta = −0.387, N = 1257
SDGSENFNR pc2
Q08830
log10 peptide intensity
496:496
546:549
215:215
8:17742929:A:G_A
p = 7.8e−27, beta = −0.41, N = 1257
IDLADFEK pc2
Q08830
log10 peptide intensity
495:496
546:549
215:215
8:17742929:A:G_A
p = 2.2e−19, beta = −0.347, N = 1256
FGL1 : NP5
Q08830
log10 protein intensity
90:90
153:153
82:82
8:17742929:A:G_G
p = 4.4e−11, beta = 0.481, N = 325
IRPNDFIPNVI pc2
Q08830
log10 peptide intensity
90:90
153:153
82:82
8:17742929:A:G_G
p = 4.3e−11, beta = 0.481, N = 325
NLHFLTTQEDYTLK pc3
Q08830
log10 peptide intensity
90:90
153:153
82:82
8:17742929:A:G_G
p = 9e−12, beta = 0.496, N = 325
QLLQENEVQFLDK pc2
Q08830
log10 peptide intensity
90:90
153:153
82:82
8:17742929:A:G_G
p = 1.6e−11, beta = 0.491, N = 325
DYENGFGNFVQK pc2
Q08830
log10 peptide intensity
88:90
152:153
82:82
8:17742929:A:G_G
p = 2.6e−08, beta = 0.412, N = 322
Assay Target: FGL1
Olink UniProt: Q08830
UKB rsID: rs2073562
Proxy rsID: rs2073562
UKB SNP: 8:17742929:A:G
Proxy SNP: 8:17742929:A:G
UKB log10(p): 2388.3
UKB BETA: 0.653
*:*:*:*:*:*:*:*:*:*:*:*:*:*:*:*:*:*:–:–:–:–:NA:
1257:1257:1257:1256:1256:125
GDENTVIDLGSK pc2
rs3739406 REF
log10 peptide intensity
151:496
528:549
212:215
8:17742929:A:G_A
p = 8.7e−154, model = REC, N = 891
GDENTVVDLGSK pc2
rs3739406 ALT
log10 peptide intensity
453:496
413:549
7:215
8:17742929:A:G_A
p = 1.1e−117, model = DOM, N = 873
GDENTVIDLGSKR pc2
rs3739406 REF
log10 peptide intensity
0:496
57:549
36:215
8:17742929:A:G_A
p = 1.2e−21, model = REC, N = 93
GDENTVVDLGSKR pc2
rs3739406 ALT
log10 peptide intensity
135:496
57:549
0:215
8:17742929:A:G_A
p = 4.3e−21, model = REC, N = 192

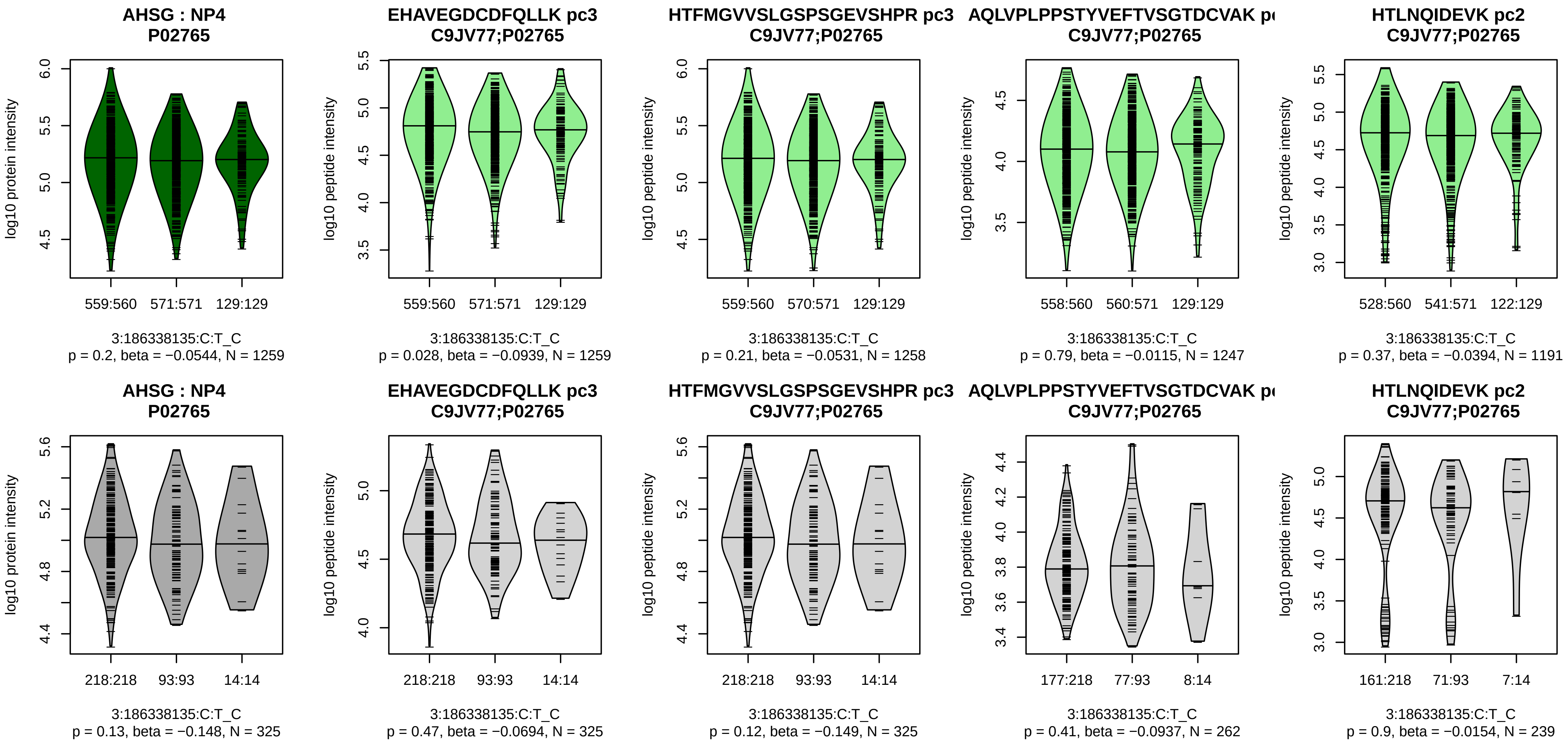


Assay Target: AHSG
Olink UniProt: P02765
UKB rsID: rs1900618
Proxy rsID: rs1900618
UKB SNP: 3:186338135:C:T
Proxy SNP: 3:186338135:C:T
UKB log10(p): 2299.5
UKB BETA: 0.668
−:−:−:−:−:−:−:−:*:*:−:−:NA
1259:1258:1249:1247:1230:119

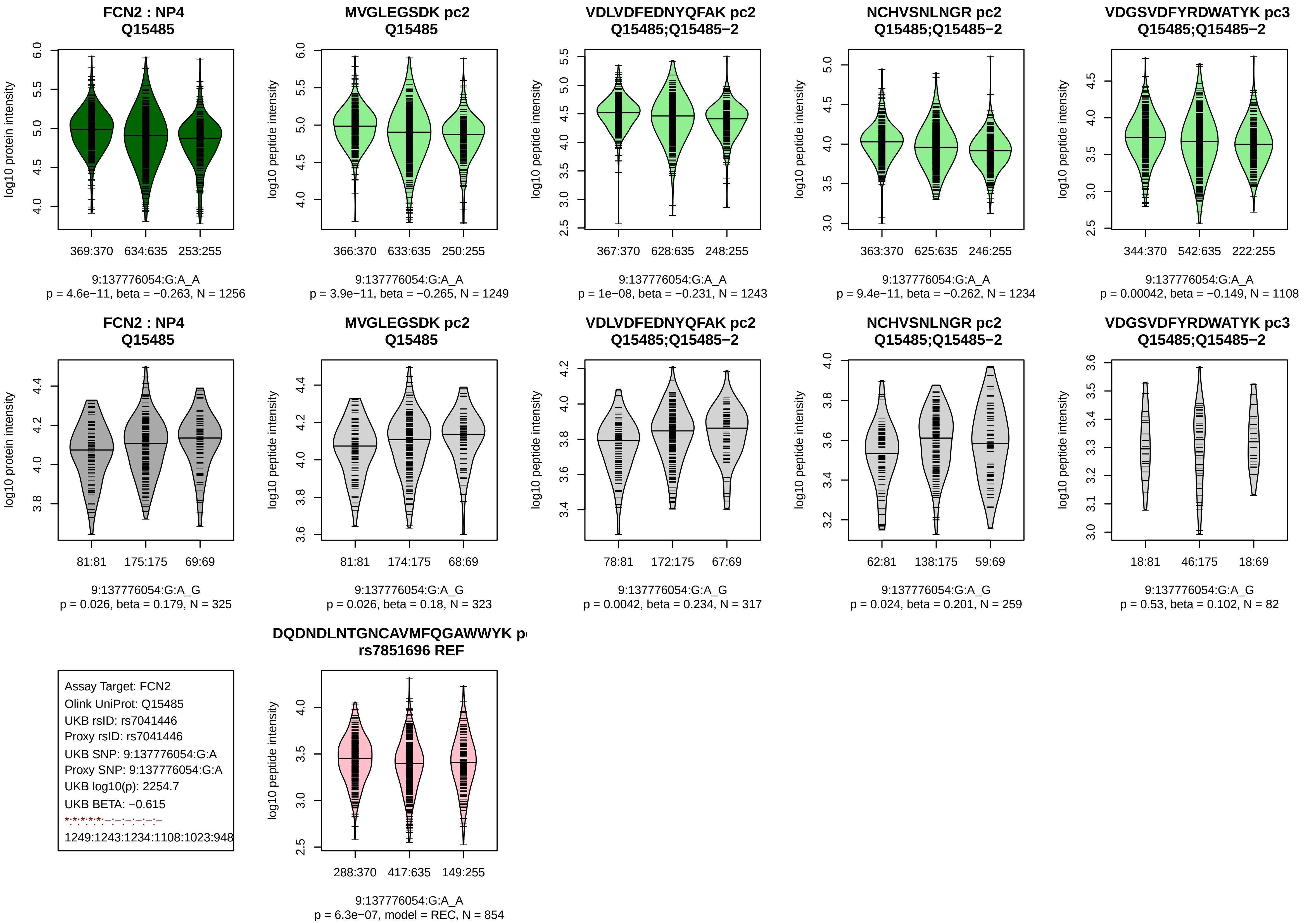

FCN2 : NP4
Q15485
log10 protein intensity
369:370
634:635
253:255
9:137776054:G:A_A
p = 4.6e−11, beta = −0.263, N = 1256
MVGLEGSDK pc2
Q15485
log10 peptide intensity
366:370
633:635
250:255
9:137776054:G:A_A
p = 3.9e−11, beta = −0.265, N = 1249
VDLVDFEDNYQFAK pc2
Q15485;Q15485−2
367:370
628:635
248:255
9:137776054:G:A_A
p = 1e−08, beta = −0.231, N = 1243
NCHVSNLNGR pc2
Q15485;Q15485−2
363:370
625:635
246:255
9:137776054:G:A_A
p = 9.4e−11, beta = −0.262, N = 1234
VDGSVDFYRDWATYK pc3
Q15485;Q15485−2
344:370
542:635
222:255
9:137776054:G:A_A
p = 0.00042, beta = −0.149, N = 1108
FCN2 : NP4
Q15485
81:81
175:175
69:69
9:137776054:G:A_G
p = 0.026, beta = 0.179, N = 325
MVGLEGSDK pc2
Q15485
81:81
174:175
68:69
9:137776054:G:A_G
p = 0.026, beta = 0.18, N = 323
VDLVDFEDNYQFAK pc2
Q15485;Q15485−2
78:81
172:175
67:69
9:137776054:G:A_G
p = 0.0042, beta = 0.234, N = 317
NCHVSNLNGR pc2
Q15485;Q15485−2
62:81
138:175
59:69
9:137776054:G:A_G
p = 0.024, beta = 0.201, N = 259
VDGSVDFYRDWATYK pc3
Q15485;Q15485−2
18:81
46:175
18:69
9:137776054:G:A_G
p = 0.53, beta = 0.102, N = 82
Assay Target: FCN2
Olink UniProt: Q15485
UKB rsID: rs7041446
Proxy rsID: rs7041446
UKB SNP: 9:137776054:G:A
Proxy SNP: 9:137776054:G:A
UKB log10(p): 2254.7
UKB BETA: −0.615
*:*:*:*:*:–:–:–:–:–:–
1249:1243:1234:1108:1023:948
DQDNDLNTGNCAVMFQGAWWYK p
rs7851696 REF
288:370
417:635
149:255
9:137776054:G:A_A
p = 6.3e−07, model = REC, N = 854

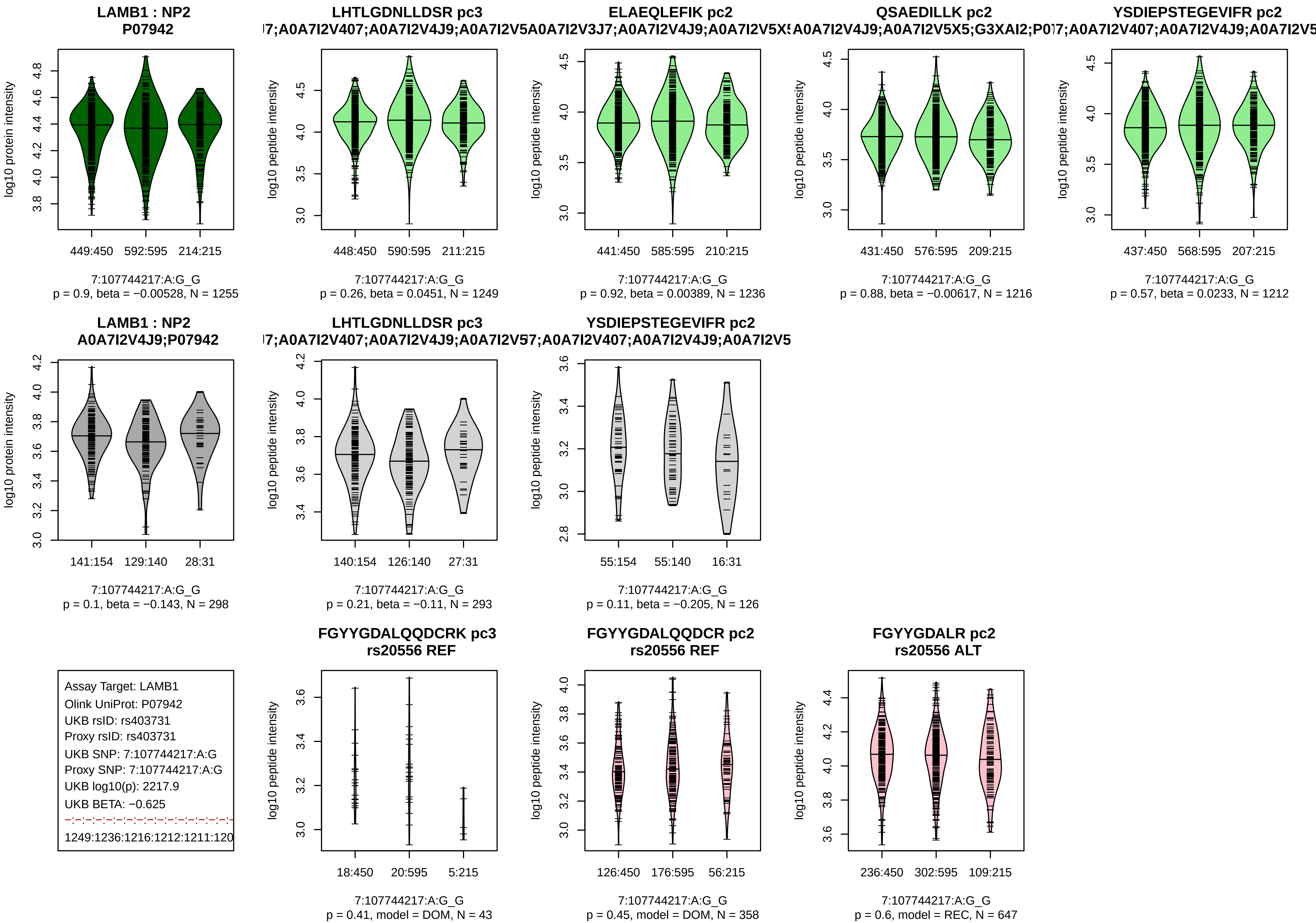

LAMB1 : NP2
P07942
log10 protein intensity
449:450
592:595
214:215
7:107744217:A:G_G
p = 0.9, beta = −0.00528, N = 1255
LHTLGDNLLDSR pc3
l7;A0A7I2V407;A0A7I2V4J9;A0A7I2V5
log10 peptide intensity
448:450
590:595
211:215
7:107744217:A:G_G
p = 0.26, beta = 0.0451, N = 1249
ELAEQLEFIK pc2
A0A7I2V3J7;A0A7I2V4J9;A0A7I2V5X5
log10 peptide intensity
441:450
585:595
210:215
7:107744217:A:G_G
p = 0.92, beta = 0.00389, N = 1236
QSAEDILLK pc2
A0A7I2V4J9;A0A7I2V5X5;G3XAI2;P0
log10 peptide intensity
431:450
576:595
209:215
7:107744217:A:G_G
p = 0.88, beta = −0.00617, N = 1216
YSDIEPSTEGEVIFR pc2
l7;A0A7I2V407;A0A7I2V4J9;A0A7I2V5
log10 peptide intensity
437:450
568:595
207:215
7:107744217:A:G_G
p = 0.57, beta = 0.0233, N = 1212
LAMB1 : NP2
A0A7I2V4J9;P07942
log10 protein intensity
141:154
129:140
28:31
7:107744217:A:G_G
p = 0.1, beta = −0.143, N = 298
LHTLGDNLLDSR pc3
l7;A0A7I2V407;A0A7I2V4J9;A0A7I2V5
log10 peptide intensity
140:154
126:140
27:31
7:107744217:A:G_G
p = 0.21, beta = −0.11, N = 293
YSDIEPSTEGEVIFR pc2
7;A0A7I2V407;A0A7I2V4J9;A0A7I2V5
log10 peptide intensity
55:154
55:140
16:31
7:107744217:A:G_G
p = 0.11, beta = −0.205, N = 126
Assay Target: LAMB1
Olink UniProt: P07942
UKB rsID: rs403731
Proxy rsID: rs403731
UKB SNP: 7:107744217:A:G
Proxy SNP: 7:107744217:A:G
UKB log10(p): 2217.9
UKB BETA: −0.625
1249:1236:1216:1212:1211:120
FGYYGDALQQDCRK pc3
rs20556 REF
log10 peptide intensity
18:450
20:595
5:215
7:107744217:A:G_G
p = 0.41, model = DOM, N = 43
FGYYGDALQQDCR pc2
rs20556 REF
log10 peptide intensity
126:450
176:595
56:215
7:107744217:A:G_G
p = 0.45, model = DOM, N = 358
FGYYGDALR pc2
rs20556 ALT
log10 peptide intensity
236:450
302:595
109:215
7:107744217:A:G_G
p = 0.6, model = REC, N = 647

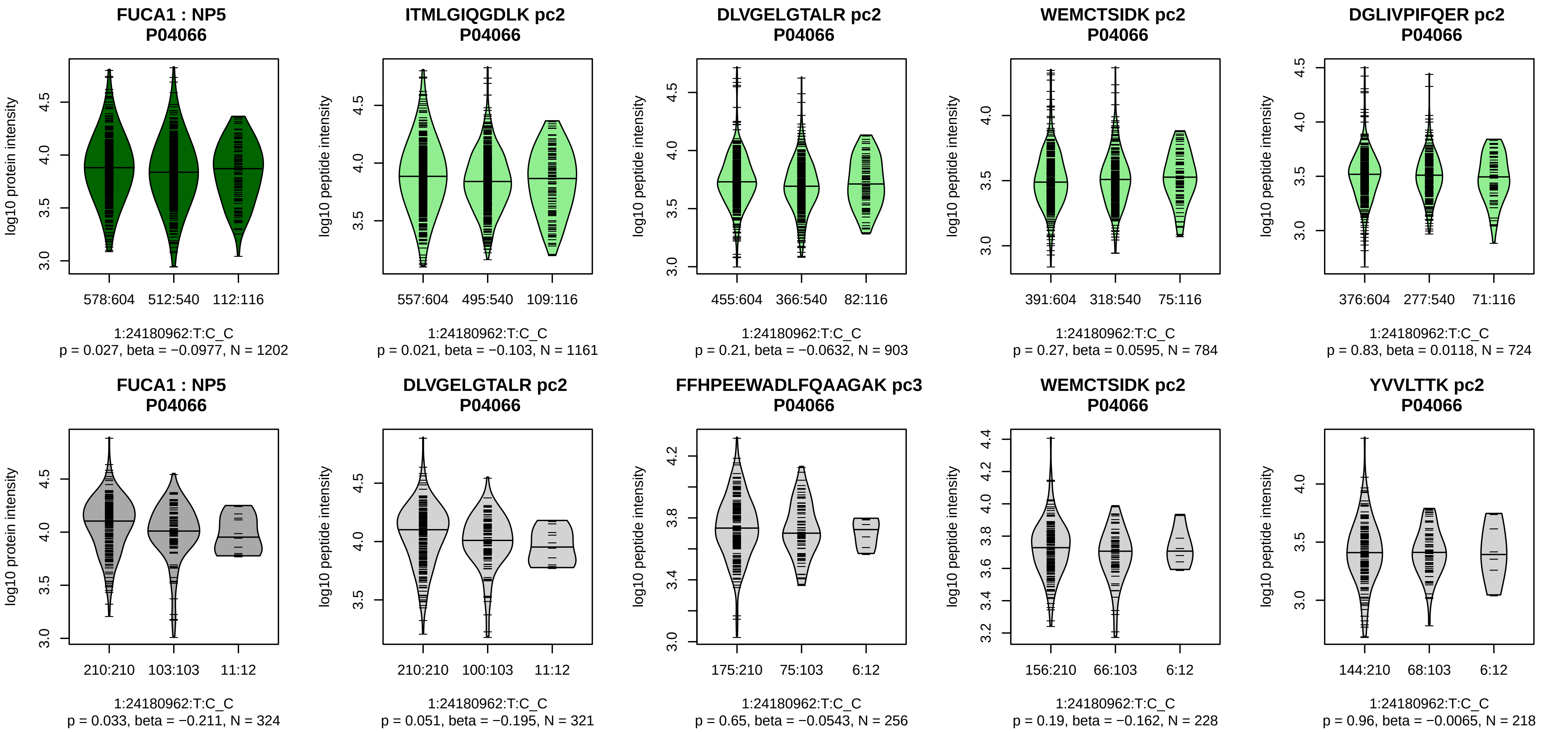


Assay Target: FUCA1
Olink UniProt: P04066
UKB rsID: rs13551
Proxy rsID: rs13551
UKB SNP: 1:24180962:T:C
Proxy SNP: 1:24180962:T:C
UKB log10(p): 2197.8
UKB BETA: −0.38
−:−:−:−:−:−:−:−
1161:903:784:724:682:159:147:

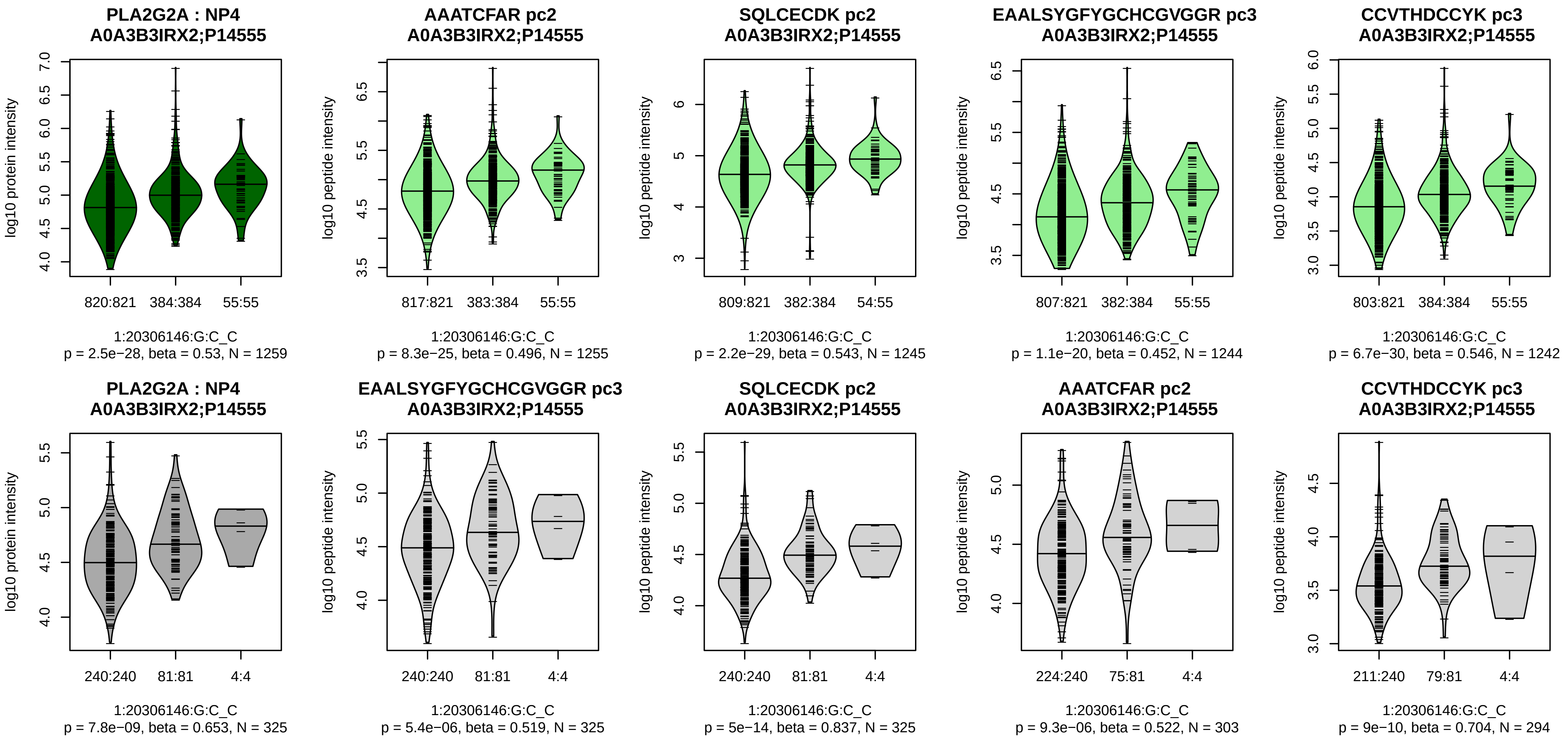


Assay Target: PLA2G2A
Olink UniProt: P14555
UKB rsID: rs11573156
Proxy rsID: rs11573156
UKB SNP: 1:20306146:G:C
Proxy SNP: 1:20306146:G:C
UKB log10(p): 2153.3
UKB BETA: 0.721
*:*:*:*:*:*
1255:1245:1244:1242:1180:114

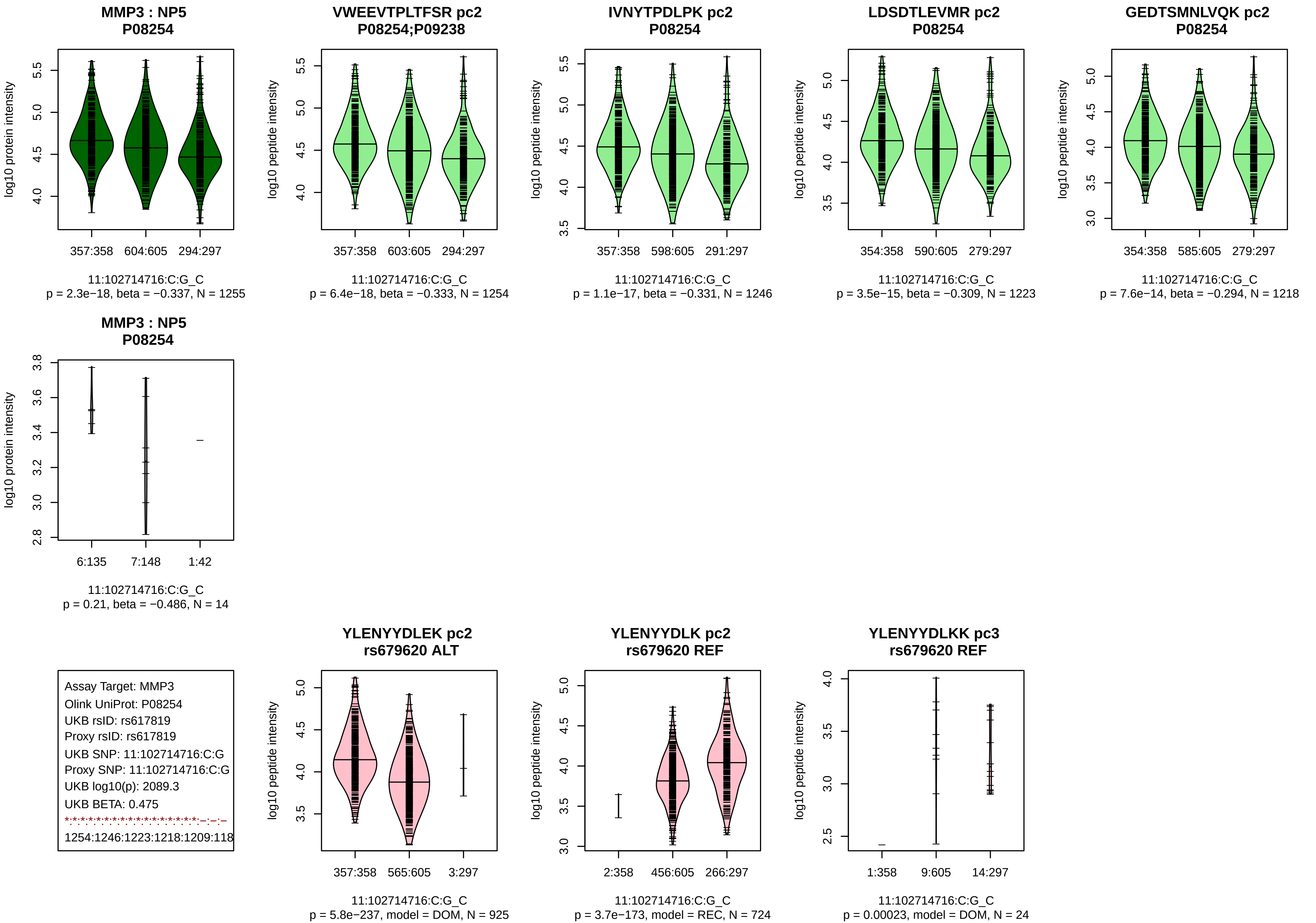
MMP3 : NP5
P08254
log10 protein intensity
357:358
604:605
294:297
11:102714716:C:G_C
p = 2.3e−18, beta = −0.337, N = 1255
VWEEVTPLTFSR pc2
P08254;P09238
log10 peptide intensity
357:358
603:605
294:297
11:102714716:C:G_C
p = 6.4e−18, beta = −0.333, N = 1254
IVNYTPDLPK pc2
P08254
log10 peptide intensity
357:358
598:605
291:297
11:102714716:C:G_C
p = 1.1e−17, beta = −0.331, N = 1246
LDSDTLEVMR pc2
P08254
log10 peptide intensity
354:358
590:605
279:297
11:102714716:C:G_C
p = 3.5e−15, beta = −0.309, N = 1223
GEDTSMNLVQK pc2
P08254
log10 peptide intensity
354:358
585:605
279:297
11:102714716:C:G_C
p = 7.6e−14, beta = −0.294, N = 1218
MMP3 : NP5
P08254
log10 protein intensity
6:135
7:148
1:42
11:102714716:C:G_C
p = 0.21, beta = −0.486, N = 14
Assay Target: MMP3
Olink UniProt: P08254
UKB rsID: rs617819
Proxy rsID: rs617819
UKB SNP: 11:102714716:C:G
Proxy SNP: 11:102714716:C:G
UKB log10(p): 2089.3
UKB BETA: 0.475
1254:1246:1223:1218:1209:118
YLENYYDLEK pc2
rs679620 ALT
log10 peptide intensity
357:358
565:605
3:297
11:102714716:C:G_C
p = 5.8e−237, model = DOM, N = 925
YLENYYDLK pc2
rs679620 REF
log10 peptide intensity
2:358
456:605
266:297
11:102714716:C:G_C
p = 3.7e−173, model = REC, N = 724
YLENYYDLKK pc3
rs679620 REF
log10 peptide intensity
1:358
9:605
14:297
11:102714716:C:G_C
p = 0.00023, model = DOM, N = 24

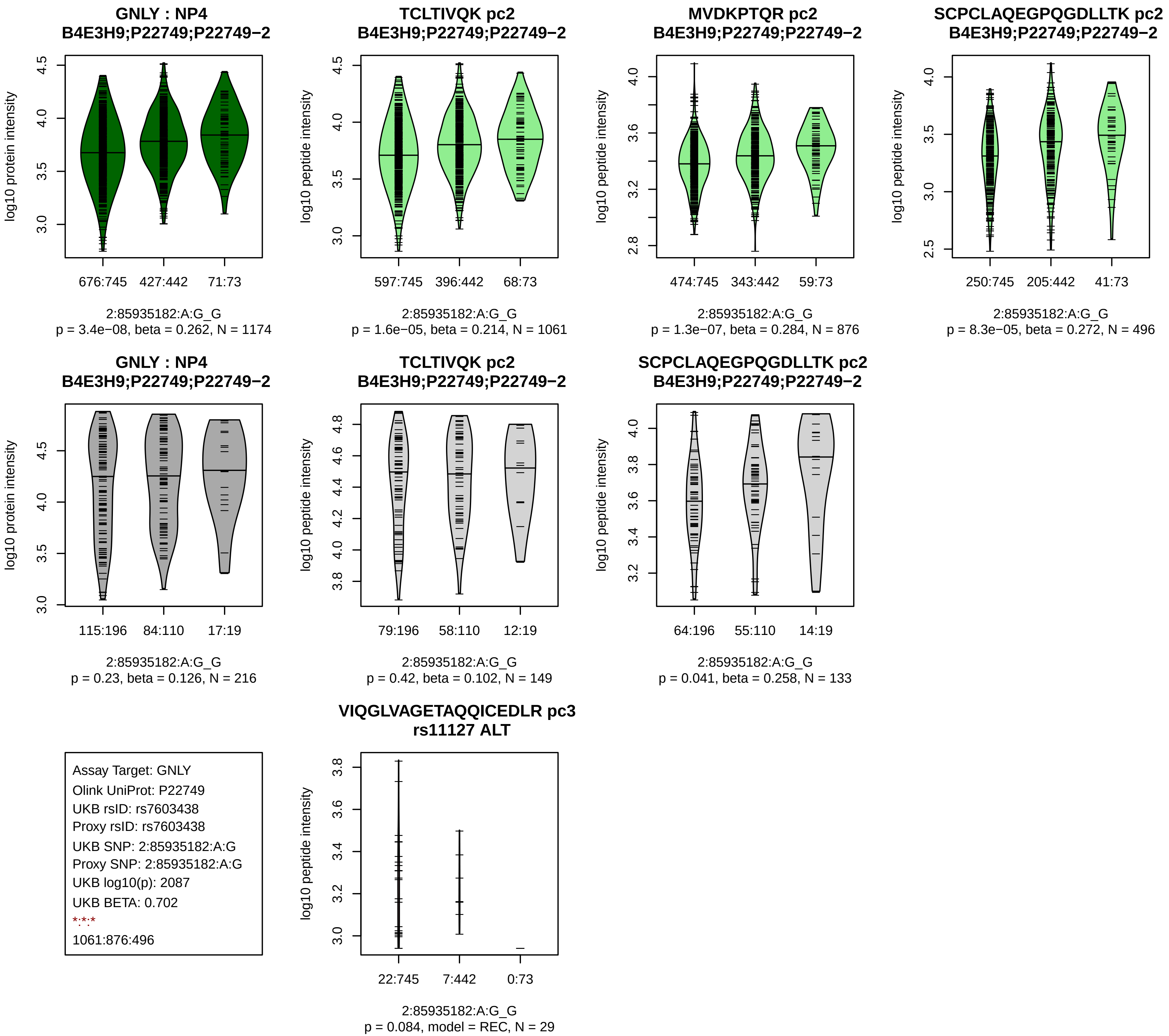
GNLY : NP4
B4E3H9;P22749;P22749−2
log10 protein intensity
676:745
427:442
71:73
2:85935182:A:G_G
p = 3.4e−08, beta = 0.262, N = 1174
TCLTIVQK pc2
B4E3H9;P22749;P22749−2
log10 peptide intensity
597:745
396:442
68:73
2:85935182:A:G_G
p = 1.6e−05, beta = 0.214, N = 1061
MVDKPTQR pc2
B4E3H9;P22749;P22749−2
log10 peptide intensity
474:745
343:442
59:73
2:85935182:A:G_G
p = 1.3e−07, beta = 0.284, N = 876
SCPCLAQEGPQGDLLTK pc2
B4E3H9;P22749;P22749−2
log10 peptide intensity
250:745
205:442
41:73
2:85935182:A:G_G
p = 8.3e−05, beta = 0.272, N = 496
GNLY : NP4
B4E3H9;P22749;P22749−2
log10 protein intensity
115:196
84:110
17:19
2:85935182:A:G_G
p = 0.23, beta = 0.126, N = 216
TCLTIVQK pc2
B4E3H9;P22749;P22749−2
log10 peptide intensity
79:196
58:110
12:19
2:85935182:A:G_G
p = 0.42, beta = 0.102, N = 149
SCPCLAQEGPQGDLLTK pc2
B4E3H9;P22749;P22749−2
log10 peptide intensity
64:196
55:110
14:19
2:85935182:A:G_G
p = 0.041, beta = 0.258, N = 133
Assay Target: GNLY
Olink UniProt: P22749
UKB rsID: rs7603438
Proxy rsID: rs7603438
UKB SNP: 2:85935182:A:G
Proxy SNP: 2:85935182:A:G
UKB log10(p): 2087
UKB BETA: 0.702
*:*:*
1061:876:496
VIQGLVAGETAQQICEDLR pc3
rs11127 ALT
log10 peptide intensity
22:745
7:442
0:73
2:85935182:A:G_G
p = 0.084, model = REC, N = 29

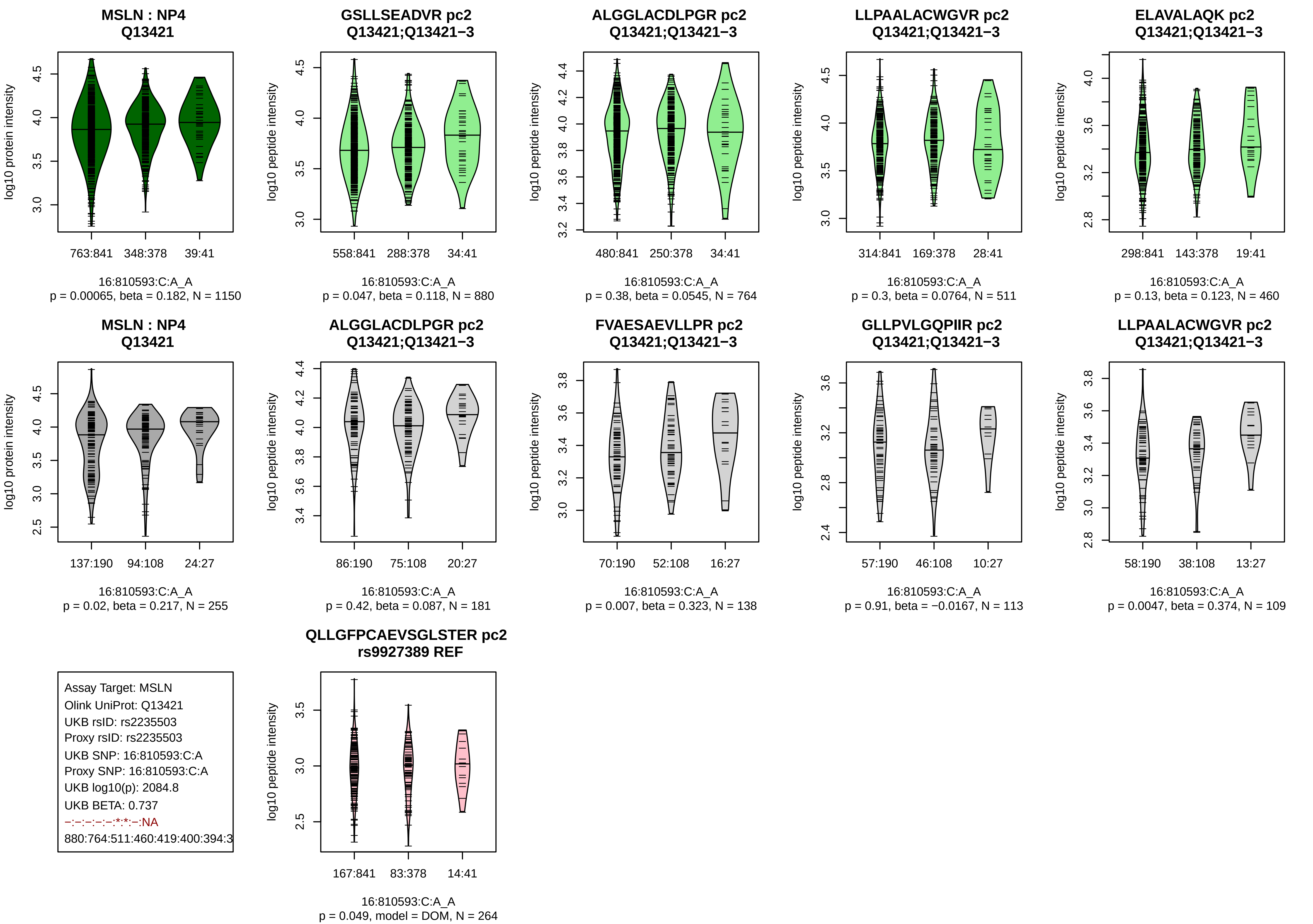
MSLN : NP4
Q13421
log10 protein intensity
763:841
348:378
39:41
16:810593:C:A_A
p = 0.00065, beta = 0.182, N = 1150
GSLLSEADVR pc2
Q13421;Q13421−3
log10 peptide intensity
558:841
288:378
34:41
16:810593:C:A_A
p = 0.047, beta = 0.118, N = 880
ALGGLACDLPGR pc2
Q13421;Q13421−3
log10 peptide intensity
480:841
250:378
34:41
16:810593:C:A_A
p = 0.38, beta = 0.0545, N = 764
LLPAALACWGVR pc2
Q13421;Q13421−3
log10 peptide intensity
314:841
169:378
28:41
16:810593:C:A_A
p = 0.3, beta = 0.0764, N = 511
ELAVALAQK pc2
Q13421;Q13421−3
log10 peptide intensity
298:841
143:378
19:41
16:810593:C:A_A
p = 0.13, beta = 0.123, N = 460
MSLN : NP4
Q13421
log10 protein intensity
137:190
94:108
24:27
16:810593:C:A_A
p = 0.02, beta = 0.217, N = 255
ALGGLACDLPGR pc2
Q13421;Q13421−3
log10 peptide intensity
86:190
75:108
20:27
16:810593:C:A_A
p = 0.42, beta = 0.087, N = 181
FVAESAEVLLPR pc2
Q13421;Q13421−3
log10 peptide intensity
70:190
52:108
16:27
16:810593:C:A_A
p = 0.007, beta = 0.323, N = 138
GLLPVLGQPIIR pc2
Q13421;Q13421−3
log10 peptide intensity
57:190
46:108
10:27
16:810593:C:A_A
p = 0.91, beta = −0.0167, N = 113
LLPAALACWGVR pc2
Q13421;Q13421−3
log10 peptide intensity
58:190
38:108
13:27
16:810593:C:A_A
p = 0.0047, beta = 0.374, N = 109
Assay Target: MSLN
Olink UniProt: Q13421
UKB rsID: rs2235503
Proxy rsID: rs2235503
UKB SNP: 16:810593:C:A
Proxy SNP: 16:810593:C:A
UKB log10(p): 2084.8
UKB BETA: 0.737
−:−:−:−:−:*:*:−:NA
880:764:511:460:419:400:394:3
QLLGFPCAEVSGLSTER pc2
rs9927389 REF
log10 peptide intensity
167:841
83:378
14:41
16:810593:C:A_A
p = 0.049, model = DOM, N = 264

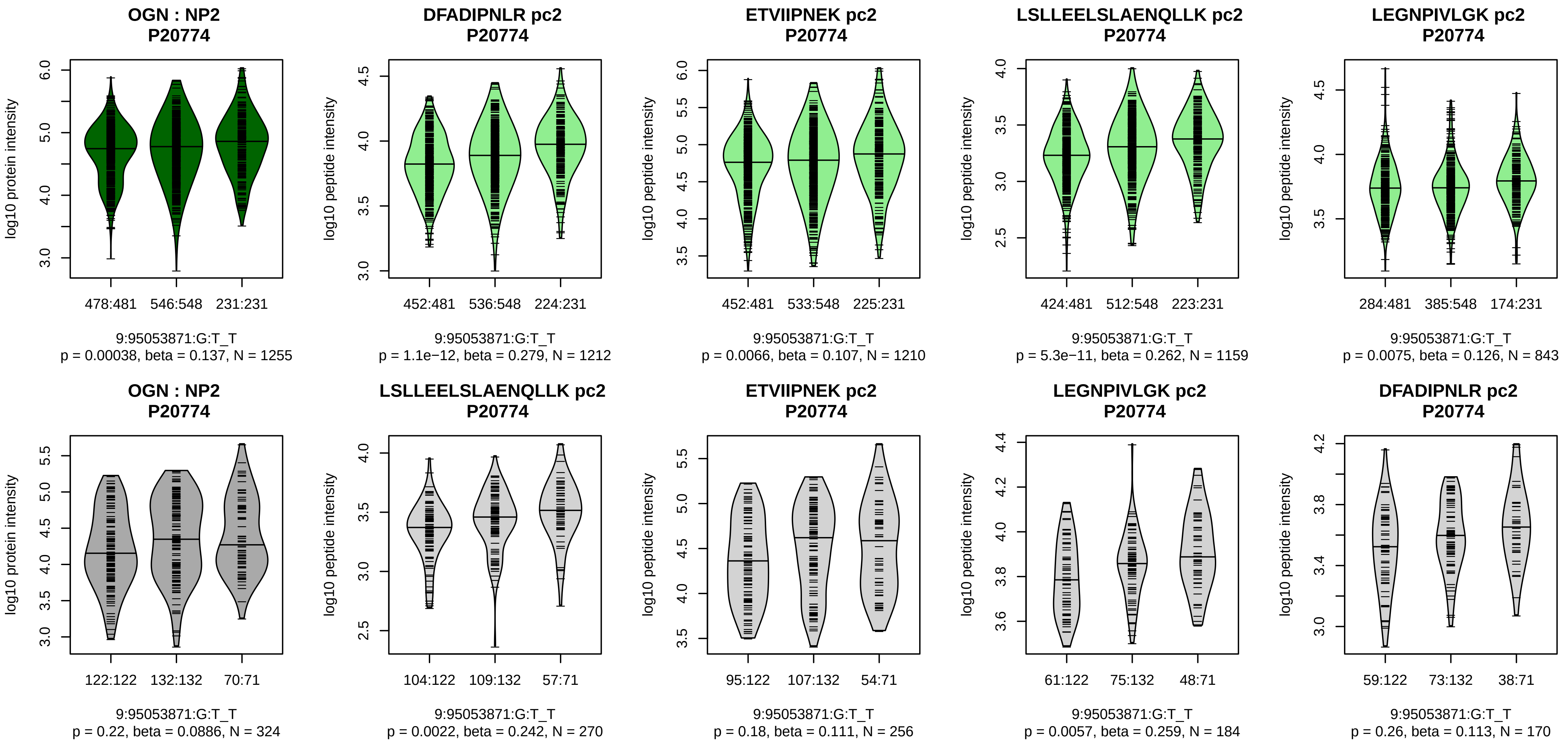


Assay Target: OGN
Olink UniProt: P20774
UKB rsID: rs6479416
Proxy rsID: rs6479416
UKB SNP: 9:95053871:G:T
Proxy SNP: 9:95053871:G:T
UKB log10(p): 2084.4
UKB BETA: 0.579
*:*:*:*:–:–:–:–:–
1212:1210:1159:843:383:235:22

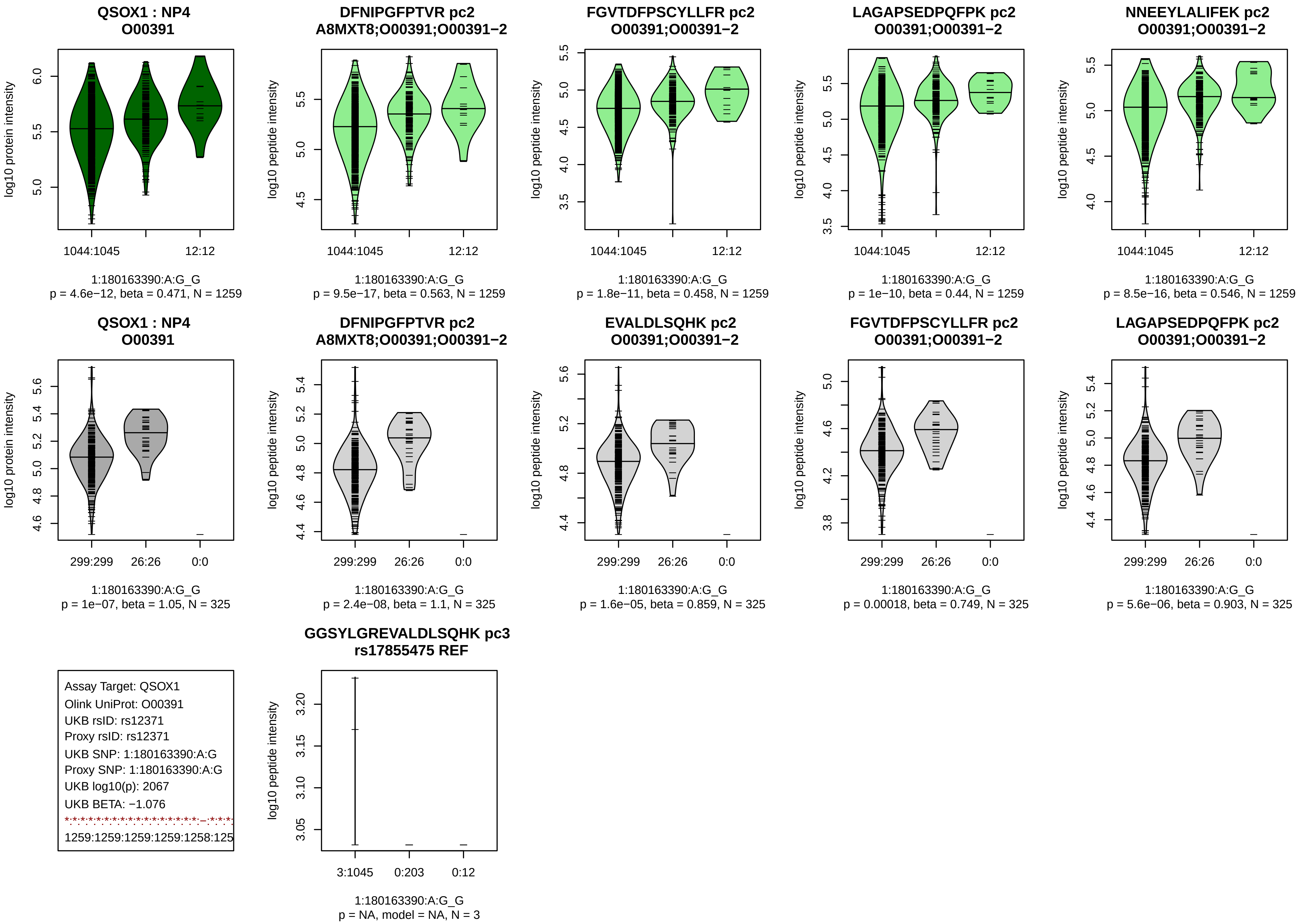
QSOX1 : NP4
O00391
log10 protein intensity
1044:1045
12:12
1:180163390:A:G_G
p = 4.6e−12, beta = 0.471, N = 1259
DFNIPGFPTVR pc2
A8MXT8;O00391;O00391−2
log10 peptide intensity
1044:1045
12:12
1:180163390:A:G_G
p = 9.5e−17, beta = 0.563, N = 1259
FGVTDFPSCYLLFR pc2
O00391;O00391−2
log10 peptide intensity
1044:1045
12:12
1:180163390:A:G_G
p = 1.8e−11, beta = 0.458, N = 1259
LAGAPSEDPQFPK pc2
O00391;O00391−2
log10 peptide intensity
1044:1045
12:12
1:180163390:A:G_G
p = 1e−10, beta = 0.44, N = 1259
NNEEYLALIFEK pc2
O00391;O00391−2
log10 peptide intensity
1044:1045
12:12
1:180163390:A:G_G
p = 8.5e−16, beta = 0.546, N = 1259
QSOX1 : NP4
O00391
log10 protein intensity
299:299
26:26
0:0
1:180163390:A:G_G
p = 1e−07, beta = 1.05, N = 325
DFNIPGFPTVR pc2
A8MXT8;O00391;O00391−2
log10 peptide intensity
299:299
26:26
0:0
1:180163390:A:G_G
p = 2.4e−08, beta = 1.1, N = 325
EVALDLSQHK pc2
O00391;O00391−2
log10 peptide intensity
299:299
26:26
0:0
1:180163390:A:G_G
p = 1.6e−05, beta = 0.859, N = 325
FGVTDFPSCYLLFR pc2
O00391;O00391−2
log10 peptide intensity
299:299
26:26
0:0
1:180163390:A:G_G
p = 0.00018, beta = 0.749, N = 325
LAGAPSEDPQFPK pc2
O00391;O00391−2
log10 peptide intensity
299:299
26:26
0:0
1:180163390:A:G_G
p = 5.6e−06, beta = 0.903, N = 325
Assay Target: QSOX1
Olink UniProt: O00391
UKB rsID: rs12371
Proxy rsID: rs12371
UKB SNP: 1:180163390:A:G
Proxy SNP: 1:180163390:A:G
UKB log10(p): 2067
UKB BETA: −1.076
1259:1259:1259:1259:1258:125
GGSYLGREVALDLSQHK pc3
rs17855475 REF
log10 peptide intensity
3:1045
0:203
0:12
1:180163390:A:G_G
p = NA, model = NA, N = 3

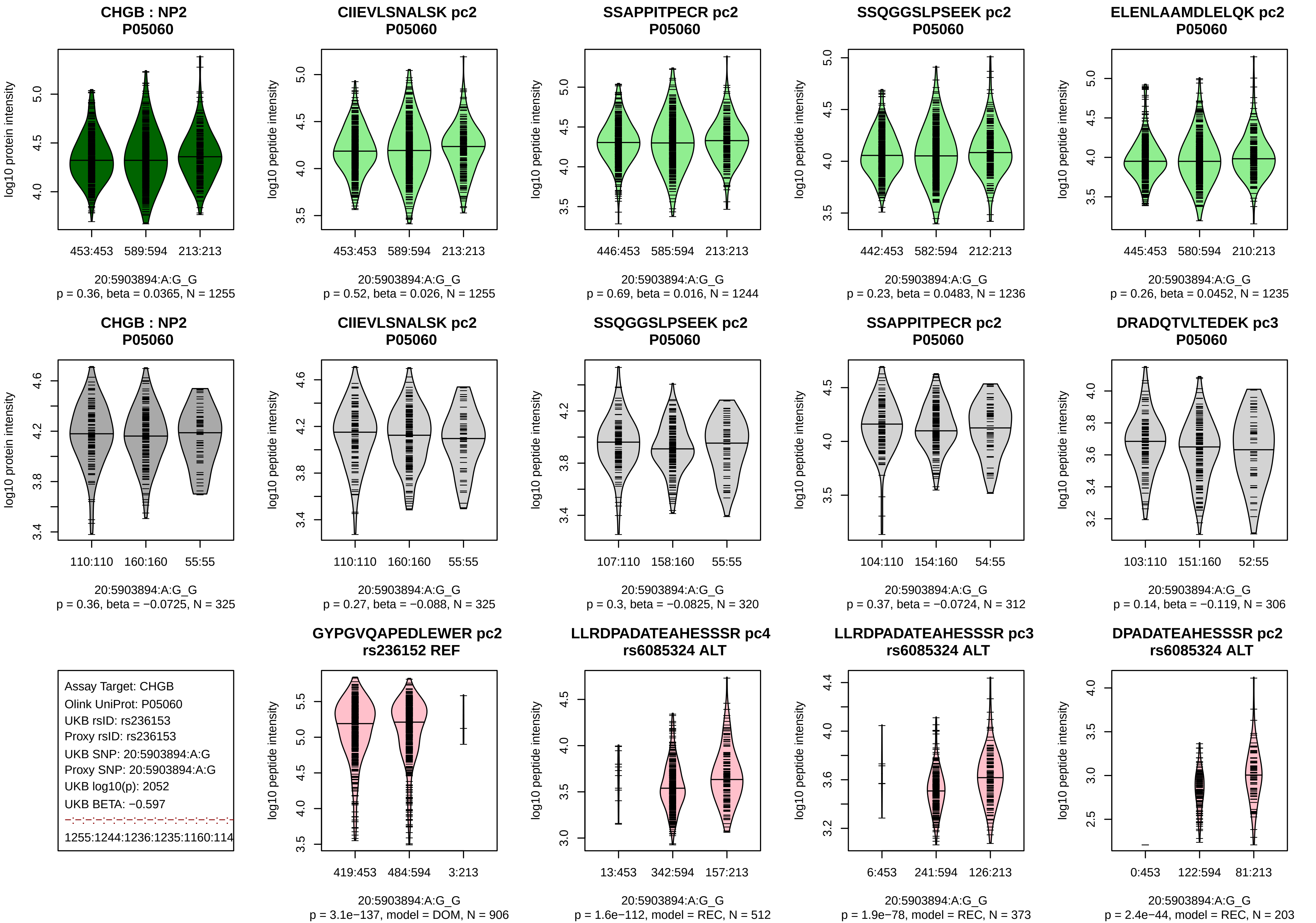
CHGB : NP2
P05060
log10 protein intensity
453:453
589:594
213:213
20:5903894:A:G_G
p = 0.36, beta = 0.0365, N = 1255
CIIEVLSNALSK pc2
P05060
log10 peptide intensity
453:453
589:594
213:213
20:5903894:A:G_G
p = 0.52, beta = 0.026, N = 1255
SSAPPITPECR pc2
P05060
log10 peptide intensity
446:453
585:594
213:213
20:5903894:A:G_G
p = 0.69, beta = 0.016, N = 1244
SSQGGSLPSEEK pc2
P05060
log10 peptide intensity
442:453
582:594
212:213
20:5903894:A:G_G
p = 0.23, beta = 0.0483, N = 1236
ELENLAAMDLELQK pc2
P05060
log10 peptide intensity
445:453
580:594
210:213
20:5903894:A:G_G
p = 0.26, beta = 0.0452, N = 1235
CHGB : NP2
P05060
log10 protein intensity
110:110
160:160
55:55
20:5903894:A:G_G
p = 0.36, beta = −0.0725, N = 325
CIIEVLSNALSK pc2
P05060
log10 peptide intensity
110:110
160:160
55:55
20:5903894:A:G_G
p = 0.27, beta = −0.088, N = 325
SSQGGSLPSEEK pc2
P05060
log10 peptide intensity
107:110
158:160
55:55
20:5903894:A:G_G
p = 0.3, beta = −0.0825, N = 320
SSAPPITPECR pc2
P05060
log10 peptide intensity
104:110
154:160
54:55
20:5903894:A:G_G
p = 0.37, beta = −0.0724, N = 312
DRADQTVLTEDEK pc3
P05060
log10 peptide intensity
103:110
151:160
52:55
20:5903894:A:G_G
p = 0.14, beta = −0.119, N = 306
Assay Target: CHGB
Olink UniProt: P05060
UKB rsID: rs236153
Proxy rsID: rs236153
UKB SNP: 20:5903894:A:G
Proxy SNP: 20:5903894:A:G
UKB log10(p): 2052
UKB BETA: −0.597
1255:1244:1236:1235:1160:114
GYPGVQAPEDLEWER pc2
rs236152 REF
log10 peptide intensity
419:453
484:594
3:213
20:5903894:A:G_G
p = 3.1e−137, model = DOM, N = 906
LLRDPADATEAHESSSR pc4
rs6085324 ALT
log10 peptide intensity
13:453
342:594
157:213
20:5903894:A:G_G
p = 1.6e−112, model = REC, N = 512
LLRDPADATEAHESSSR pc3
rs6085324 ALT
log10 peptide intensity
6:453
241:594
126:213
20:5903894:A:G_G
p = 1.9e−78, model = REC, N = 373
DPADATEAHESSSR pc2
rs6085324 ALT
log10 peptide intensity
0:453
122:594
81:213
20:5903894:A:G_G
p = 2.4e−44, model = REC, N = 203

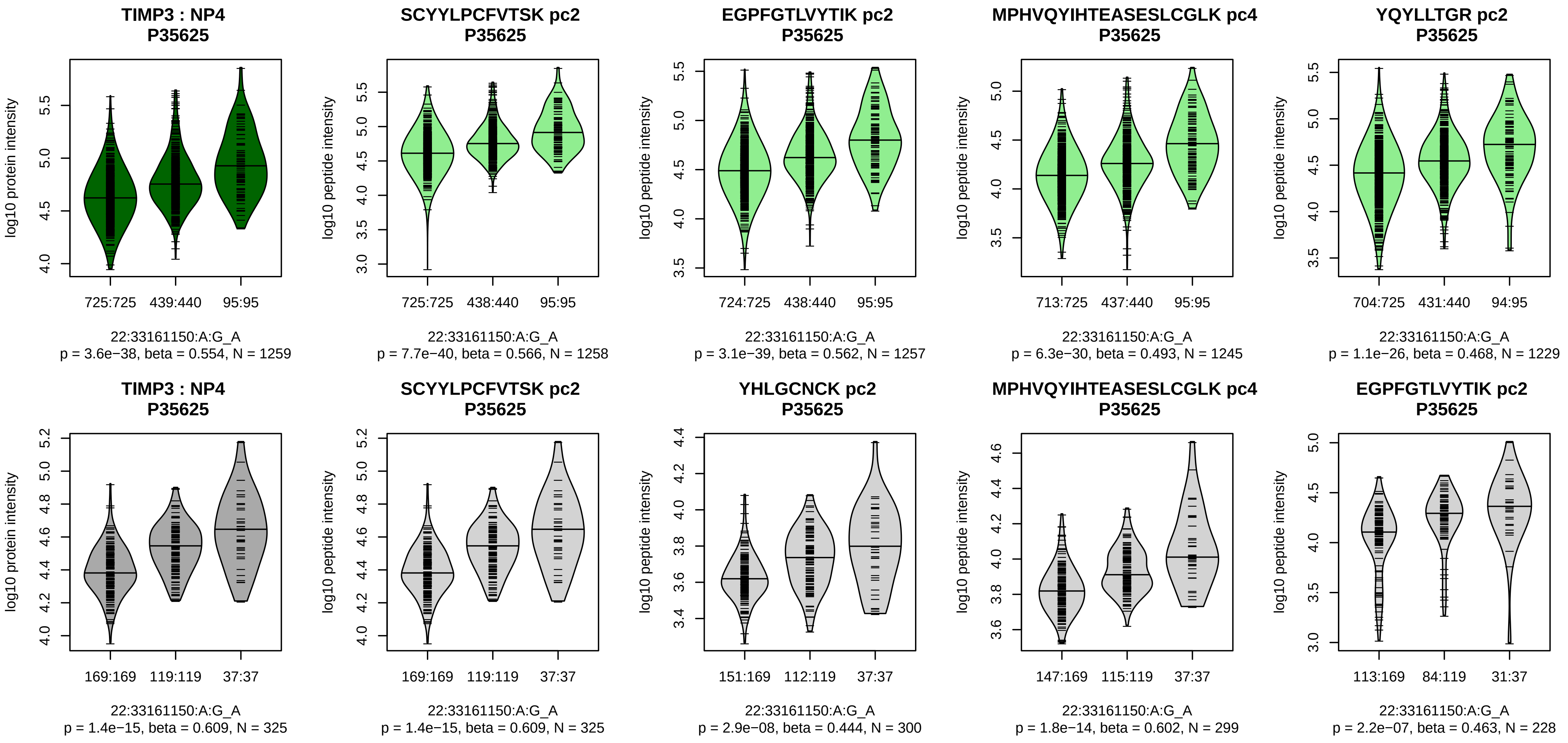


Assay Target: TIMP3
Olink UniProt: P35625
UKB rsID: rs11404116
Proxy rsID: rs929015
UKB SNP: 22:33164633:A:AT
Proxy SNP: 22:33161150:A:G
UKB log10(p): 1997.5
UKB BETA: −0.652
*:*:*:*:*:*:*:*:*
1258:1257:1245:1229:1218:121

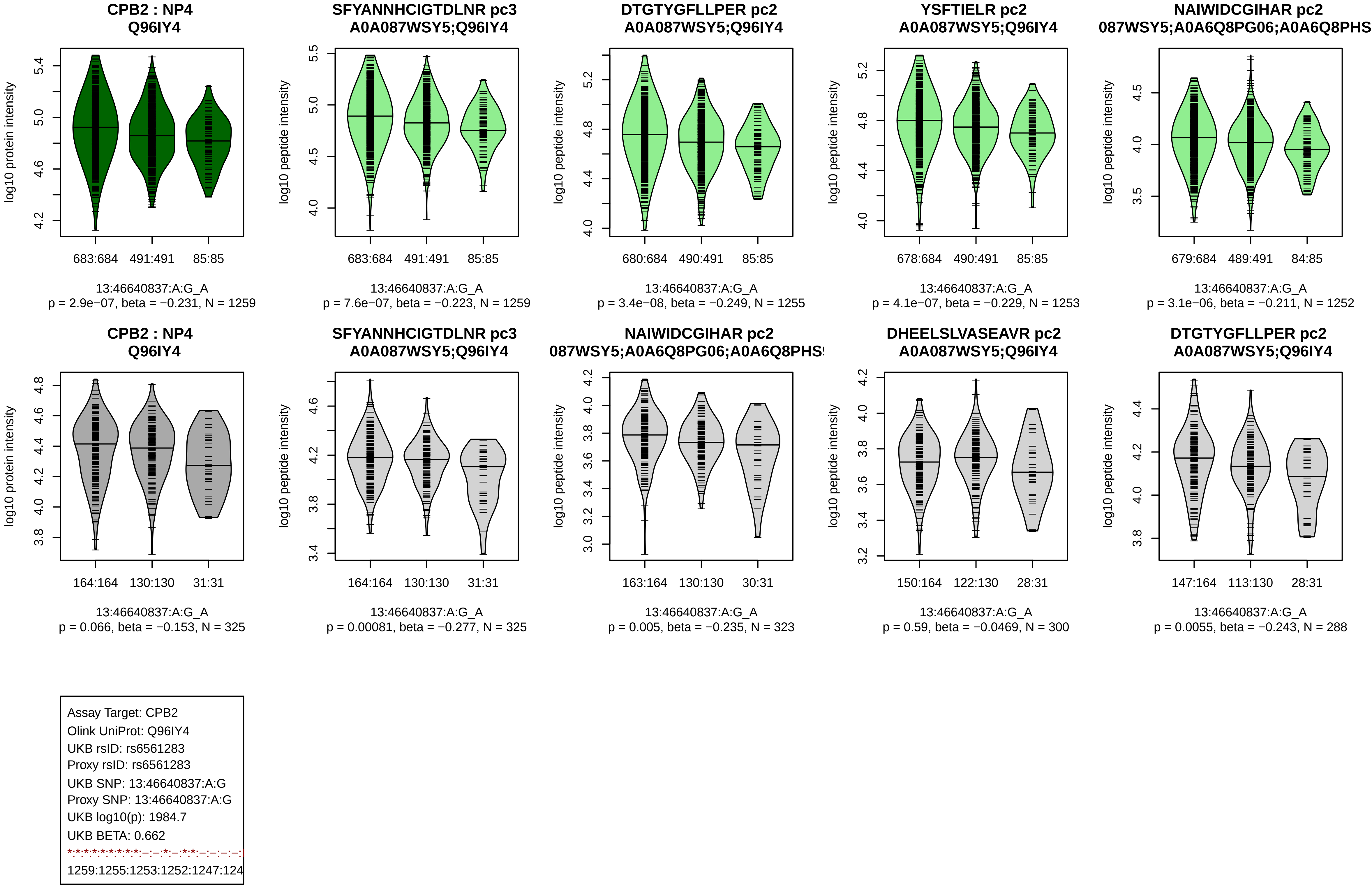
CPB2 : NP4
Q96IY4
log10 protein intensity
683:684 491:491 85:85
13:46640837:A:G_A
p = 2.9e−07, beta = −0.231, N = 1259
SFYANNHCIGTDLNR pc3
A0A087WSY5;Q96IY4
log10 peptide intensity
683:684 491:491 85:85
13:46640837:A:G_A
p = 7.6e−07, beta = −0.223, N = 1259
DTGTYGFLLPER pc2
A0A087WSY5;Q96IY4
log10 peptide intensity
680:684 490:491 85:85
13:46640837:A:G_A
p = 3.4e−08, beta = −0.249, N = 1255
YSFTIELR pc2
A0A087WSY5;Q96IY4
log10 peptide intensity
678:684 490:491 85:85
13:46640837:A:G_A
p = 4.1e−07, beta = −0.229, N = 1253
NAIWIDCGIHAR pc2
087WSY5;A0A6Q8PG06;A0A6Q8PHS
log10 peptide intensity
679:684 489:491 84:85
13:46640837:A:G_A
p = 3.1e−06, beta = −0.211, N = 1252
CPB2 : NP4
Q96IY4
log10 protein intensity
164:164 130:130 31:31
13:46640837:A:G_A
p = 0.066, beta = −0.153, N = 325
SFYANNHCIGTDLNR pc3
A0A087WSY5;Q96IY4
log10 peptide intensity
164:164 130:130 31:31
13:46640837:A:G_A
p = 0.00081, beta = −0.277, N = 325
NAIWIDCGIHAR pc2
087WSY5;A0A6Q8PG06;A0A6Q8PHS
log10 peptide intensity
163:164 130:130 30:31
13:46640837:A:G_A
p = 0.005, beta = −0.235, N = 323
DHEELSLVASEAVR pc2
A0A087WSY5;Q96IY4
log10 peptide intensity
150:164 122:130 28:31
13:46640837:A:G_A
p = 0.59, beta = −0.0469, N = 300
DTGTYGFLLPER pc2
A0A087WSY5;Q96IY4
log10 peptide intensity
147:164 113:130 28:31
13:46640837:A:G_A
p = 0.0055, beta = −0.243, N = 288
Assay Target: CPB2
Olink UniProt: Q96IY4
UKB rsID: rs6561283
Proxy rsID: rs6561283
UKB SNP: 13:46640837:A:G
Proxy SNP: 13:46640837:A:G
UKB log10(p): 1984.7
UKB BETA: 0.662
1259:1255:1253:1252:1247:124

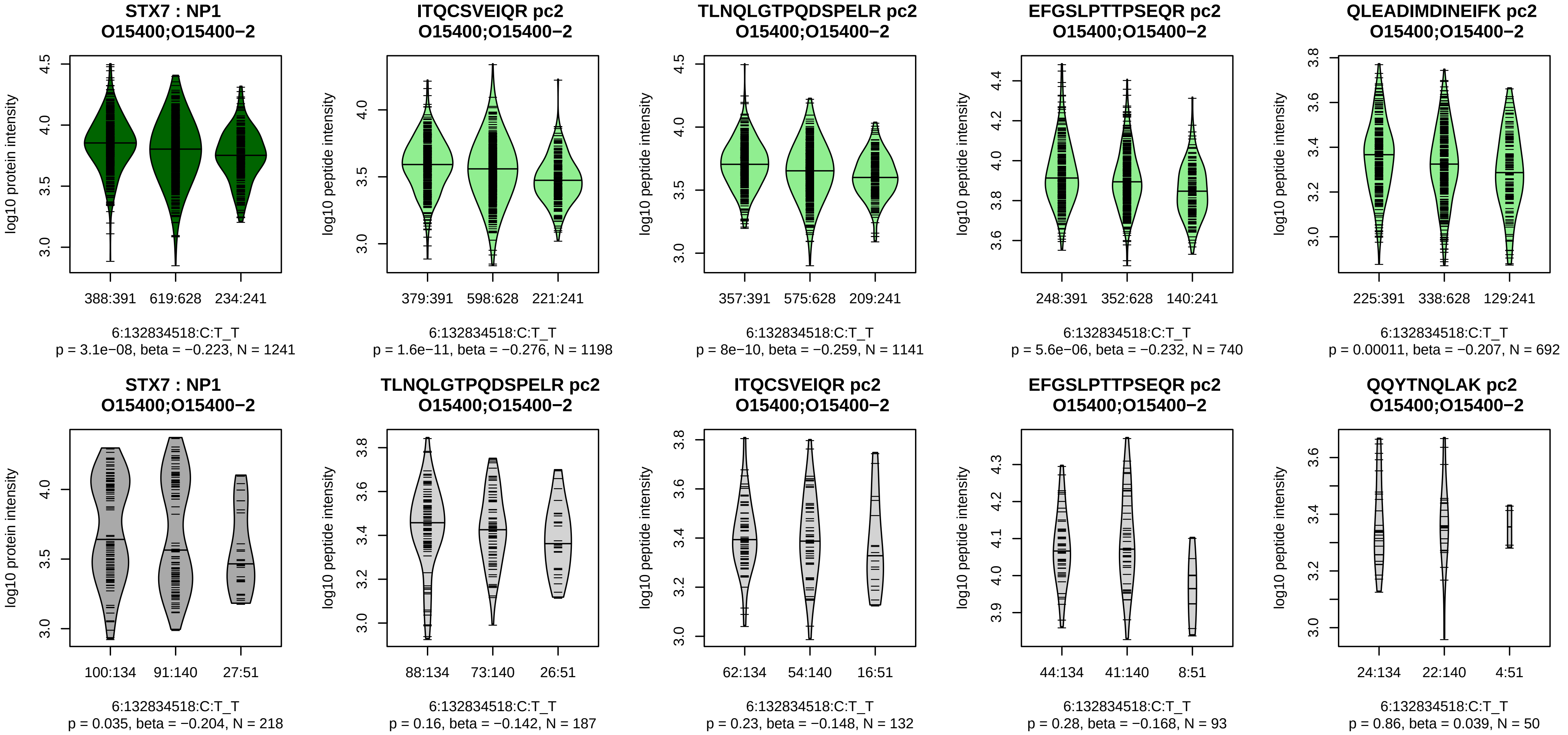


Assay Target: STX7
Olink UniProt: O15400
UKB rsID: rs3813356
Proxy rsID: rs3813356
UKB SNP: 6:132834518:C:T
Proxy SNP: 6:132834518:C:T
UKB log10(p): 1960.4
UKB BETA: −0.582
*:*:*:*:*:–:*:*
1198:1141:740:692:667:431:427

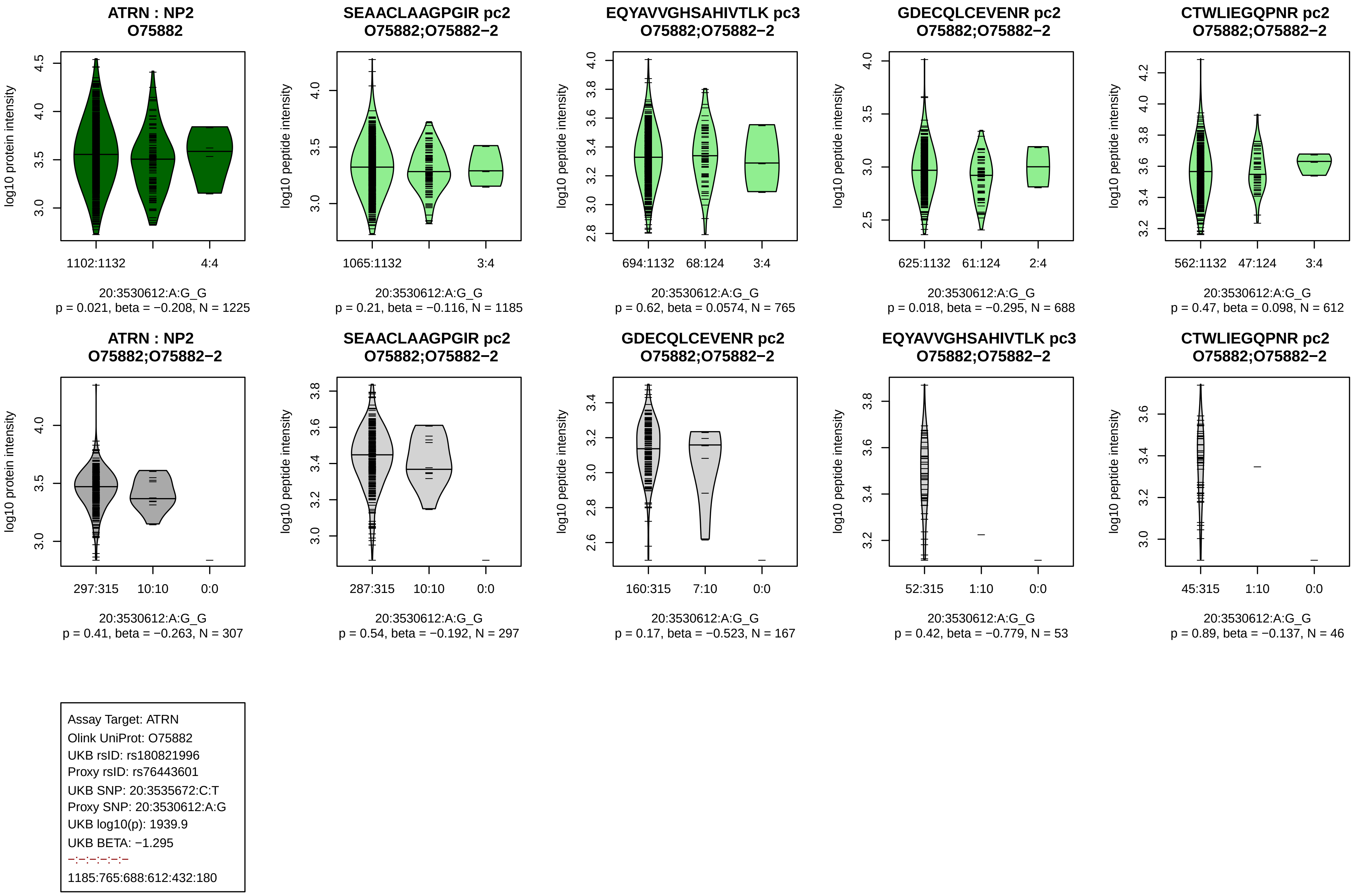
ATRN : NP2
O75882
SEAACLAAGPGIR pc2
O75882;O75882−2
EQYAVVGHSAHIVTLK pc3
O75882;O75882−2
GDECQLCEVENR pc2
O75882;O75882−2
CTWLIEGQPNR pc2
O75882;O75882−2
log10 protein intensity
log10 peptide intensity
1102:1132
4:4
1065:1132
3:4
694:1132
68:124
3:4
625:1132
61:124
2:4
562:1132
47:124
3:4
20:3530612:A:G_G
p = 0.021, beta = −0.208, N = 1225
20:3530612:A:G_G
p = 0.21, beta = −0.116, N = 1185
20:3530612:A:G_G
p = 0.62, beta = 0.0574, N = 765
20:3530612:A:G_G
p = 0.018, beta = −0.295, N = 688
20:3530612:A:G_G
p = 0.47, beta = 0.098, N = 612
ATRN : NP2
O75882;O75882−2
SEAACLAAGPGIR pc2
O75882;O75882−2
GDECQLCEVENR pc2
O75882;O75882−2
EQYAVVGHSAHIVTLK pc3
O75882;O75882−2
CTWLIEGQPNR pc2
O75882;O75882−2
297:315
10:10
0:0
287:315
10:10
0:0
160:315
7:10
0:0
52:315
1:10
0:0
45:315
1:10
0:0
20:3530612:A:G_G
p = 0.41, beta = −0.263, N = 307
20:3530612:A:G_G
p = 0.54, beta = −0.192, N = 297
20:3530612:A:G_G
p = 0.17, beta = −0.523, N = 167
20:3530612:A:G_G
p = 0.42, beta = −0.779, N = 53
20:3530612:A:G_G
p = 0.89, beta = −0.137, N = 46
Assay Target: ATRN
Olink UniProt: O75882
UKB rsID: rs180821996
Proxy rsID: rs76443601
UKB SNP: 20:3535672:C:T
Proxy SNP: 20:3530612:A:G
UKB log10(p): 1939.9
UKB BETA: −1.295
−:−:−:−:−:−
1185:765:688:612:432:180

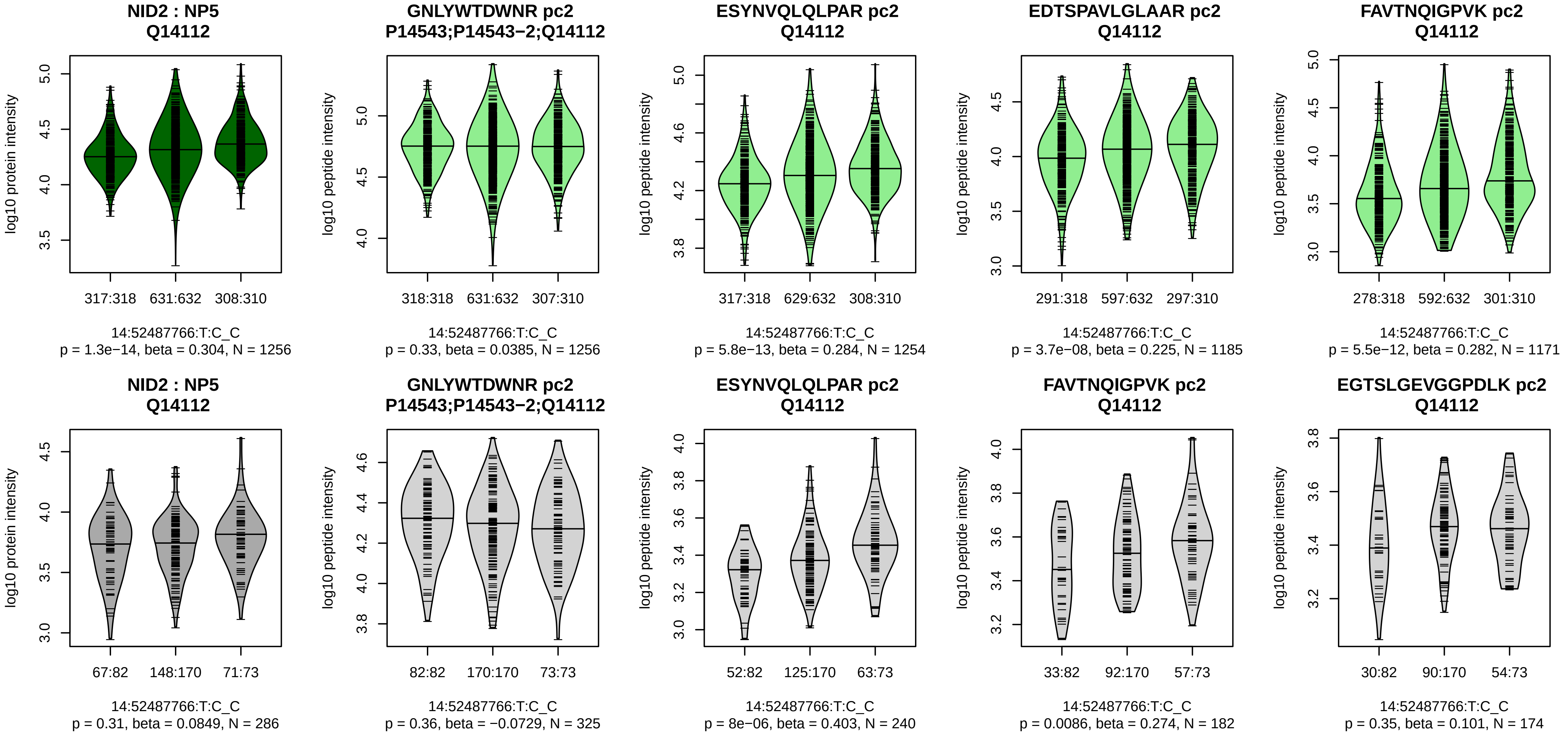


Assay Target: NID2
Olink UniProt: Q14112
UKB rsID: rs2516600
Proxy rsID: rs2516600
UKB SNP: 14:52487766:T:C
Proxy SNP: 14:52487766:T:C
UKB log10(p): 1914.1
UKB BETA: 0.574

1256:1254:1185:1171:1164:107

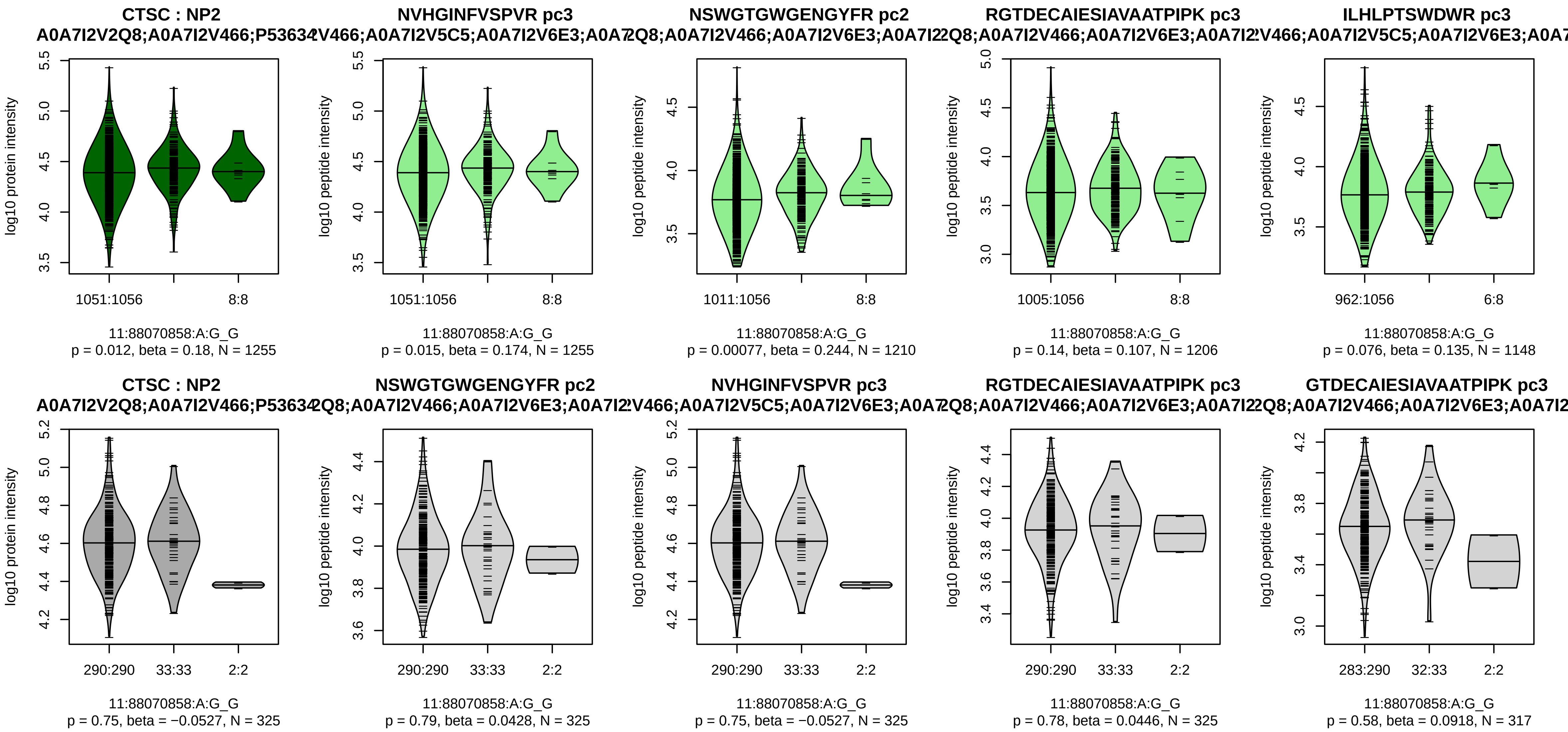


Assay Target: CTSC
Olink UniProt: P53634
UKB rsID: rs11019400
Proxy rsID: rs11019400
UKB SNP: 11:88070858:A:G
Proxy SNP: 11:88070858:A:G
UKB log10(p): 1890.8
UKB BETA: 0.971
−:−:*:−:−:−:−:−:−:−:NA
1255:1249:1210:1206:1148:108

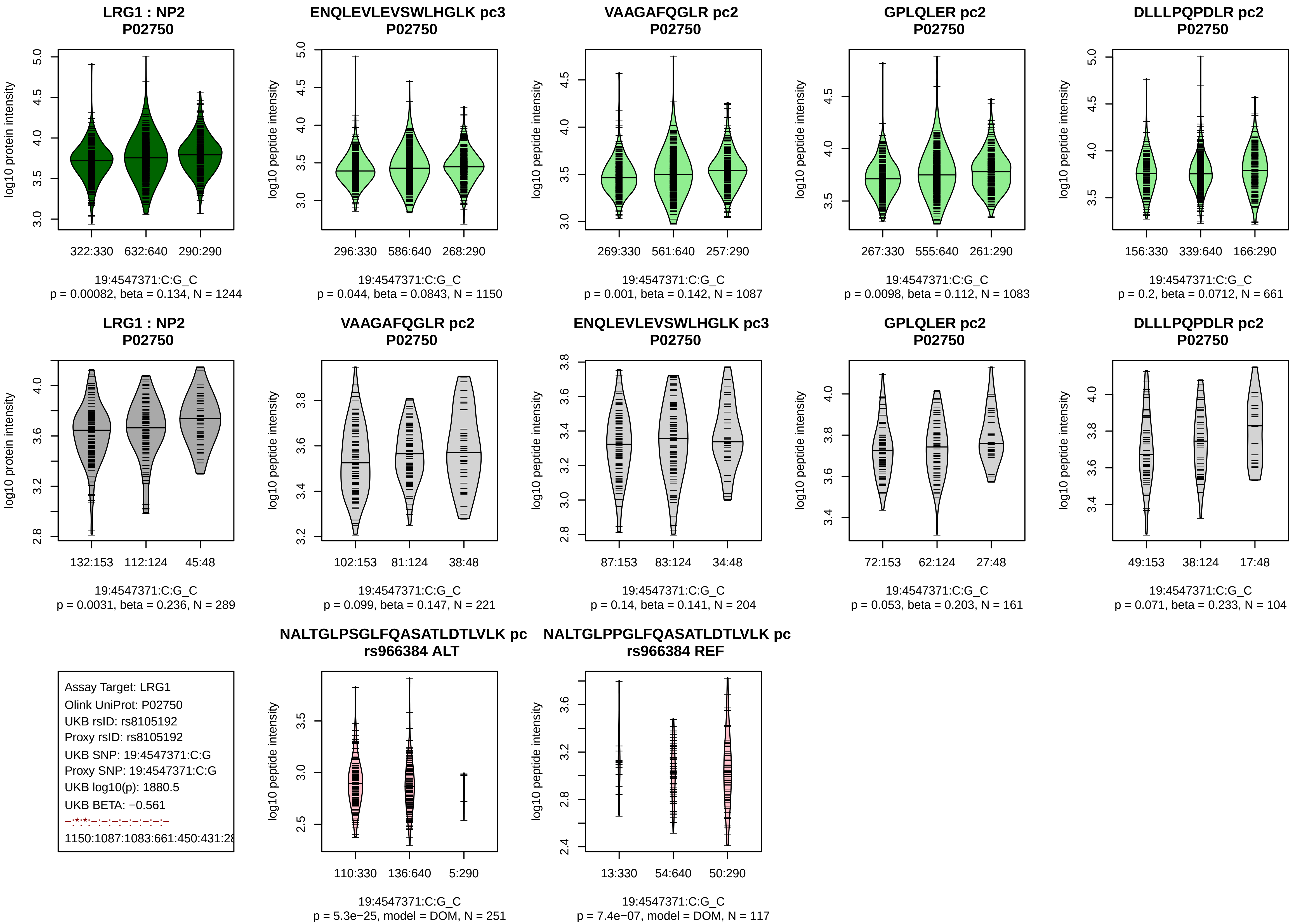
LRG1 : NP2
P02750
log10 protein intensity
322:330 632:640 290:290
19:4547371:C:G_C
p = 0.00082, beta = 0.134, N = 1244
ENQLEVLEVSWLHGLK pc3
P02750
log10 peptide intensity
296:330 586:640 268:290
19:4547371:C:G_C
p = 0.044, beta = 0.0843, N = 1150
VAAGAFQGLR pc2
P02750
log10 peptide intensity
269:330 561:640 257:290
19:4547371:C:G_C
p = 0.001, beta = 0.142, N = 1087
GPLQLER pc2
P02750
log10 peptide intensity
267:330 555:640 261:290
19:4547371:C:G_C
p = 0.0098, beta = 0.112, N = 1083
DLLLPQPDLR pc2
P02750
log10 peptide intensity
156:330 339:640 166:290
19:4547371:C:G_C
p = 0.2, beta = 0.0712, N = 661
LRG1 : NP2
P02750
log10 protein intensity
132:153 112:124 45:48
19:4547371:C:G_C
p = 0.0031, beta = 0.236, N = 289
VAAGAFQGLR pc2
P02750
log10 peptide intensity
102:153 81:124 38:48
19:4547371:C:G_C
p = 0.099, beta = 0.147, N = 221
ENQLEVLEVSWLHGLK pc3
P02750
log10 peptide intensity
87:153 83:124 34:48
19:4547371:C:G_C
p = 0.14, beta = 0.141, N = 204
GPLQLER pc2
P02750
log10 peptide intensity
72:153 62:124 27:48
19:4547371:C:G_C
p = 0.053, beta = 0.203, N = 161
DLLLPQPDLR pc2
P02750
log10 peptide intensity
49:153 38:124 17:48
19:4547371:C:G_C
p = 0.071, beta = 0.233, N = 104
Assay Target: LRG1
Olink UniProt: P02750
UKB rsID: rs8105192
Proxy rsID: rs8105192
UKB SNP: 19:4547371:C:G
Proxy SNP: 19:4547371:C:G
UKB log10(p): 1880.5
UKB BETA: −0.561
−:*:*:−:−:−:−:−:−:−
1150:1087:1083:661:450:431:28
NALTGLPSGLFQASATLDTLVLK pc
rs966384 ALT
log10 peptide intensity
110:330 136:640 5:290
19:4547371:C:G_C
p = 5.3e−25, model = DOM, N = 251
NALTGLPPGLFQASATLDTLVLK pc
rs966384 REF
log10 peptide intensity
13:330 54:640 50:290
19:4547371:C:G_C
p = 7.4e−07, model = DOM, N = 117

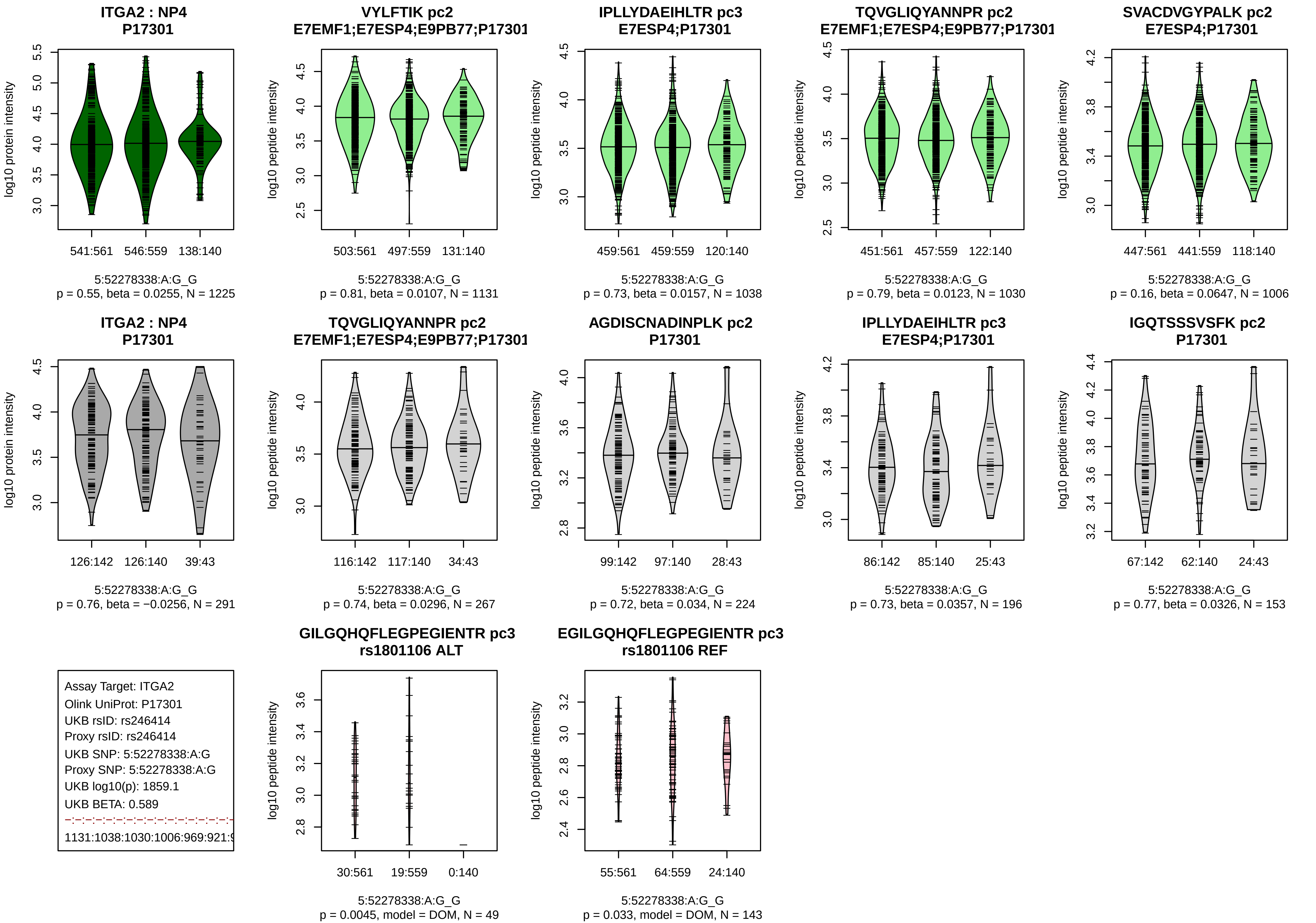
ITGA2 : NP4
P17301
log10 protein intensity
541:561
546:559
138:140
5:52278338:A:G_G
p = 0.55, beta = 0.0255, N = 1225
VYLFTIK pc2
E7EMF1;E7ESP4;E9PB77;P17301
log10 peptide intensity
503:561
497:559
131:140
5:52278338:A:G_G
p = 0.81, beta = 0.0107, N = 1131
IPLLYDAEIHLTR pc3
E7ESP4;P17301
log10 peptide intensity
459:561
459:559
120:140
5:52278338:A:G_G
p = 0.73, beta = 0.0157, N = 1038
TQVGLIQYANNPR pc2
E7EMF1;E7ESP4;E9PB77;P17301
log10 peptide intensity
451:561
457:559
122:140
5:52278338:A:G_G
p = 0.79, beta = 0.0123, N = 1030
SVACDVGYPALK pc2
E7ESP4;P17301
log10 peptide intensity
447:561
441:559
118:140
5:52278338:A:G_G
p = 0.16, beta = 0.0647, N = 1006
ITGA2 : NP4
P17301
log10 protein intensity
126:142
126:140
39:43
5:52278338:A:G_G
p = 0.76, beta = −0.0256, N = 291
TQVGLIQYANNPR pc2
E7EMF1;E7ESP4;E9PB77;P17301
log10 peptide intensity
116:142
117:140
34:43
5:52278338:A:G_G
p = 0.74, beta = 0.0296, N = 267
AGDISCNADINPLK pc2
P17301
log10 peptide intensity
99:142
97:140
28:43
5:52278338:A:G_G
p = 0.72, beta = 0.034, N = 224
IPLLYDAEIHLTR pc3
E7ESP4;P17301
log10 peptide intensity
86:142
85:140
25:43
5:52278338:A:G_G
p = 0.73, beta = 0.0357, N = 196
IGQTSSSVSFK pc2
P17301
log10 peptide intensity
67:142
62:140
24:43
5:52278338:A:G_G
p = 0.77, beta = 0.0326, N = 153
Assay Target: ITGA2
Olink UniProt: P17301
UKB rsID: rs246414
Proxy rsID: rs246414
UKB SNP: 5:52278338:A:G
Proxy SNP: 5:52278338:A:G
UKB log10(p): 1859.1
UKB BETA: 0.589
1131:1038:1030:1006:969:921:
GILGQHQFLEGPEGIENTR pc3
rs1801106 ALT
log10 peptide intensity
30:561
19:559
0:140
5:52278338:A:G_G
p = 0.0045, model = DOM, N = 49
EGILGQHQFLEGPEGIENTR pc3
rs1801106 REF
log10 peptide intensity
55:561
64:559
24:140
5:52278338:A:G_G
p = 0.033, model = DOM, N = 143

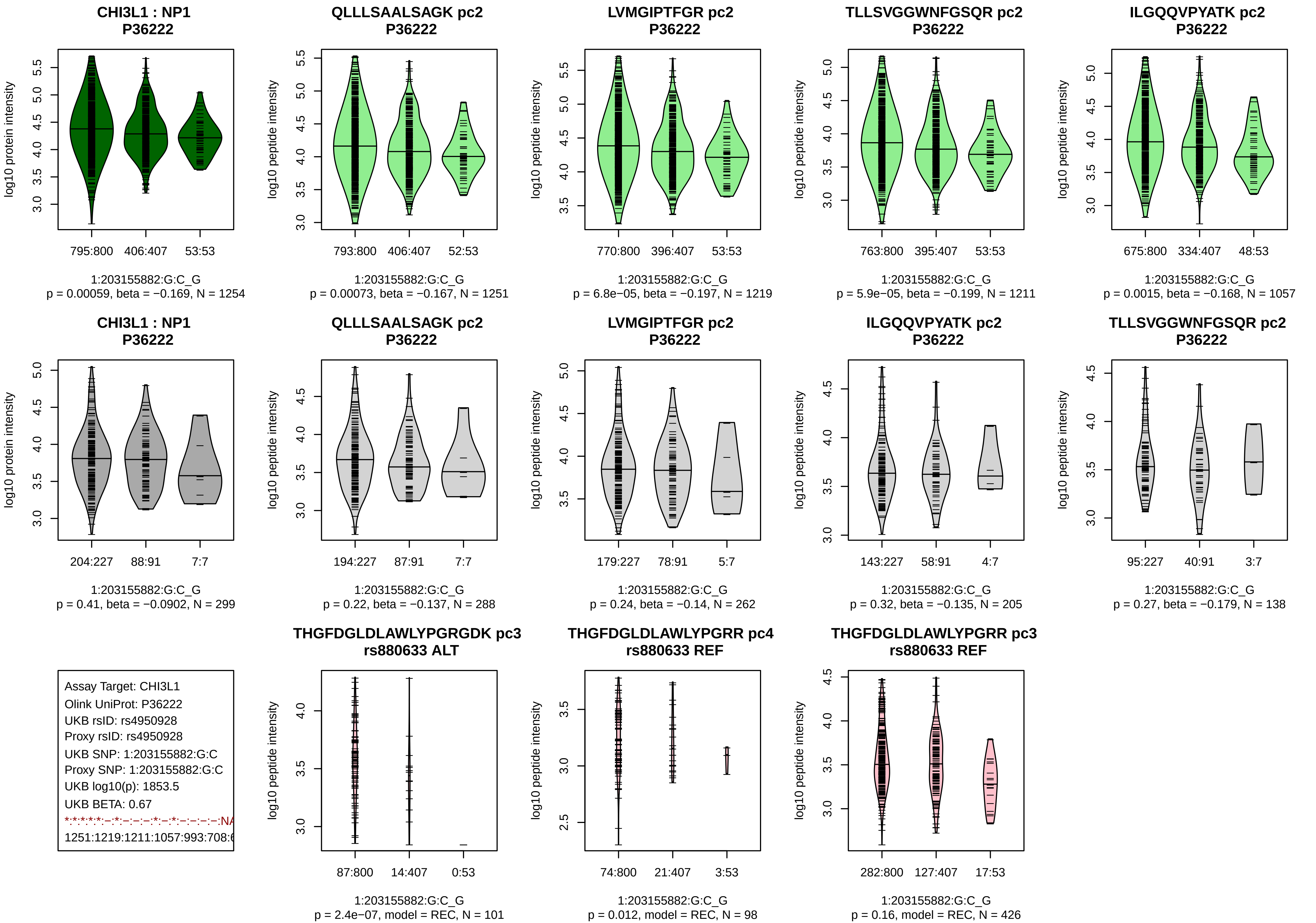
CHI3L1 : NP1
P36222
log10 protein intensity
795:800 406:407 53:53
1:203155882:G:C_G
p = 0.00059, beta = −0.169, N = 1254
QLLLSAALSAGK pc2
P36222
log10 peptide intensity
793:800 406:407 52:53
1:203155882:G:C_G
p = 0.00073, beta = −0.167, N = 1251
LVMGIPTFGR pc2
P36222
log10 peptide intensity
770:800 396:407 53:53
1:203155882:G:C_G
p = 6.8e−05, beta = −0.197, N = 1219
TLLSVGGWNFGSQR pc2
P36222
log10 peptide intensity
763:800 395:407 53:53
1:203155882:G:C_G
p = 5.9e−05, beta = −0.199, N = 1211
ILGQQVPYATK pc2
P36222
log10 peptide intensity
675:800 334:407 48:53
1:203155882:G:C_G
p = 0.0015, beta = −0.168, N = 1057
CHI3L1 : NP1
P36222
log10 protein intensity
204:227 88:91 7:7
1:203155882:G:C_G
p = 0.41, beta = −0.0902, N = 299
QLLLSAALSAGK pc2
P36222
log10 peptide intensity
194:227 87:91 7:7
1:203155882:G:C_G
p = 0.22, beta = −0.137, N = 288
LVMGIPTFGR pc2
P36222
log10 peptide intensity
179:227 78:91 5:7
1:203155882:G:C_G
p = 0.24, beta = −0.14, N = 262
ILGQQVPYATK pc2
P36222
log10 peptide intensity
143:227 58:91 4:7
1:203155882:G:C_G
p = 0.32, beta = −0.135, N = 205
TLLSVGGWNFGSQR pc2
P36222
log10 peptide intensity
95:227 40:91 3:7
1:203155882:G:C_G
p = 0.27, beta = −0.179, N = 138
Assay Target: CHI3L1
Olink UniProt: P36222
UKB rsID: rs4950928
Proxy rsID: rs4950928
UKB SNP: 1:203155882:G:C
Proxy SNP: 1:203155882:G:C
UKB log10(p): 1853.5
UKB BETA: 0.67
*:*:*:*:*:–:*:–:–:–:*:–:*:–:–:–:–:NA
1251:1219:1211:1057:993:708:6
THGFDGLDLAWLYPGRGDK pc3
rs880633 ALT
log10 peptide intensity
87:800 14:407 0:53
1:203155882:G:C_G
p = 2.4e−07, model = REC, N = 101
THGFDGLDLAWLYPGRR pc4
rs880633 REF
log10 peptide intensity
74:800 21:407 3:53
1:203155882:G:C_G
p = 0.012, model = REC, N = 98
THGFDGLDLAWLYPGRR pc3
rs880633 REF
log10 peptide intensity
282:800 127:407 17:53
1:203155882:G:C_G
p = 0.16, model = REC, N = 426

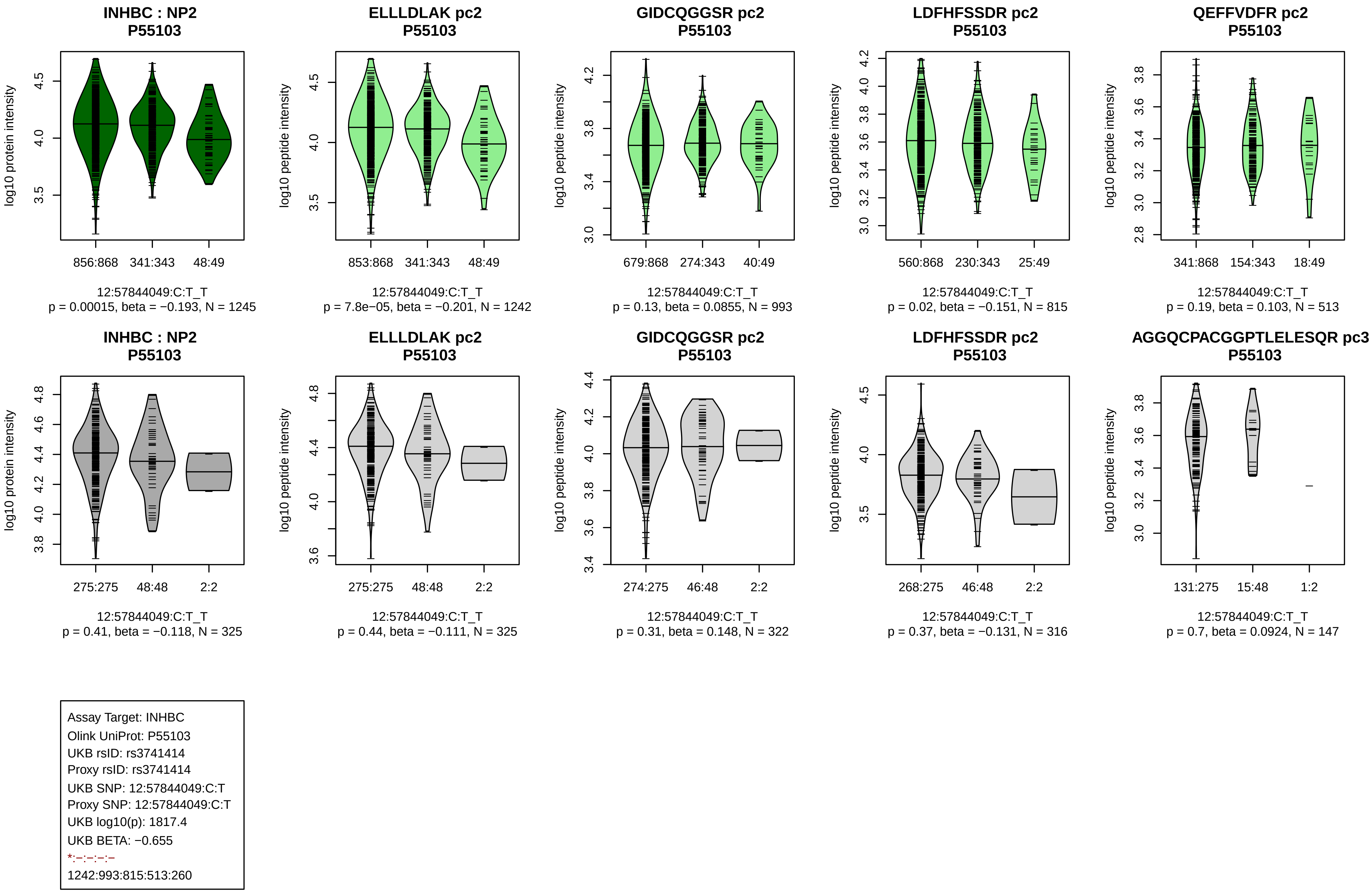
INHBC : NP2
P55103
log10 protein intensity
856:868 341:343 48:49
12:57844049:C:T_T
p = 0.00015, beta = −0.193, N = 1245
ELLLDLAK pc2
P55103
log10 peptide intensity
853:868 341:343 48:49
12:57844049:C:T_T
p = 7.8e−05, beta = −0.201, N = 1242
GIDCQGGSR pc2
P55103
log10 peptide intensity
679:868 274:343 40:49
12:57844049:C:T_T
p = 0.13, beta = 0.0855, N = 993
LDFHFSSDR pc2
P55103
log10 peptide intensity
560:868 230:343 25:49
12:57844049:C:T_T
p = 0.02, beta = −0.151, N = 815
QEFFVDFR pc2
P55103
log10 peptide intensity
341:868 154:343 18:49
12:57844049:C:T_T
p = 0.19, beta = 0.103, N = 513
INHBC : NP2
P55103
log10 protein intensity
275:275 48:48 2:2
12:57844049:C:T_T
p = 0.41, beta = −0.118, N = 325
ELLLDLAK pc2
P55103
log10 peptide intensity
275:275 48:48 2:2
12:57844049:C:T_T
p = 0.44, beta = −0.111, N = 325
GIDCQGGSR pc2
P55103
log10 peptide intensity
274:275 46:48 2:2
12:57844049:C:T_T
p = 0.31, beta = 0.148, N = 322
LDFHFSSDR pc2
P55103
log10 peptide intensity
268:275 46:48 2:2
12:57844049:C:T_T
p = 0.37, beta = −0.131, N = 316
AGGQCPACGGPTLELESQR pc3
P55103
log10 peptide intensity
131:275 15:48 1:2
12:57844049:C:T_T
p = 0.7, beta = 0.0924, N = 147
Assay Target: INHBC
Olink UniProt: P55103
UKB rsID: rs3741414
Proxy rsID: rs3741414
UKB SNP: 12:57844049:C:T
Proxy SNP: 12:57844049:C:T
UKB log10(p): 1817.4
UKB BETA: −0.655
*:−:−:−:−
1242:993:815:513:260

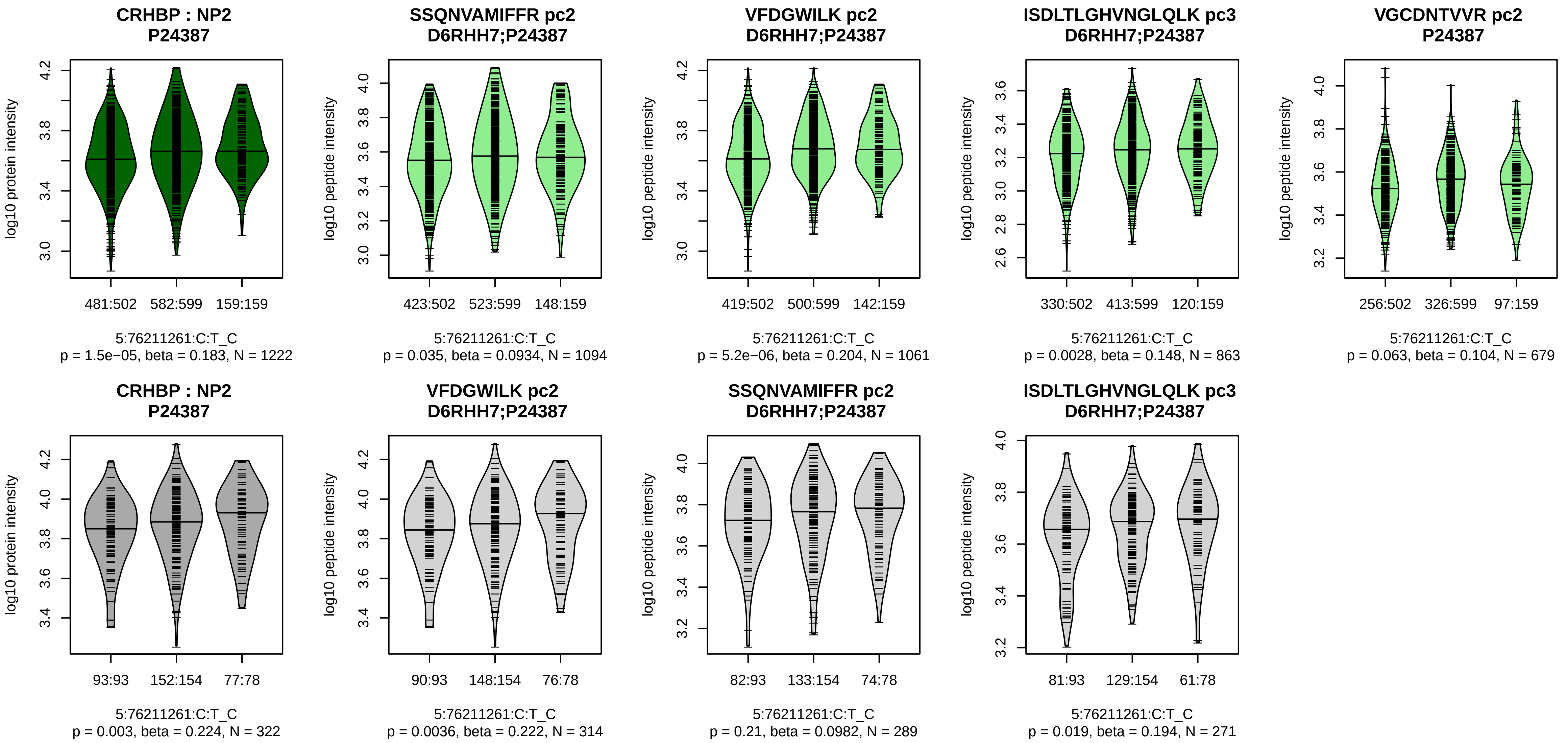


Assay Target: CRHBP
Olink UniProt: P24387
UKB rsID: rs1700657
Proxy rsID: rs1700657
UKB SNP: 5:76211261:C:T
Proxy SNP: 5:76211261:C:T
UKB log10(p): 1809.2
UKB BETA: −0.569
−:*:*:−:−
1094:1061:863:679:181

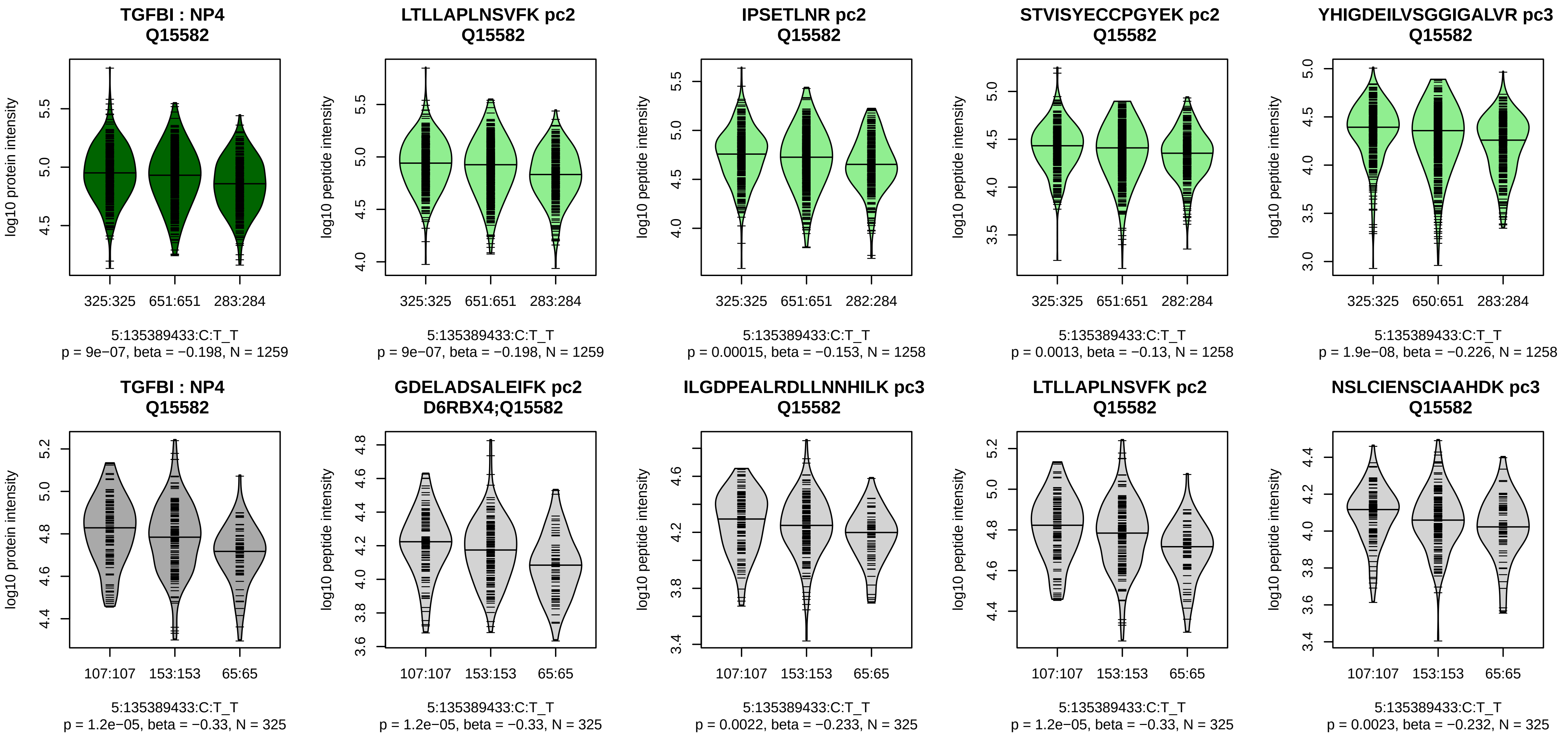


Assay Target: TGFBI
Olink UniProt: Q15582
UKB rsID: rs13159365
Proxy rsID: rs13159365
UKB SNP: 5:135389433:C:T
Proxy SNP: 5:135389433:C:T
UKB log10(p): 1803.9
UKB BETA: −0.547
*:*:*:*:*:*:*:*:*:*:*:*:*:*:*:*:_:*:*:_:*
1259:1258:1258:1258:1258:125

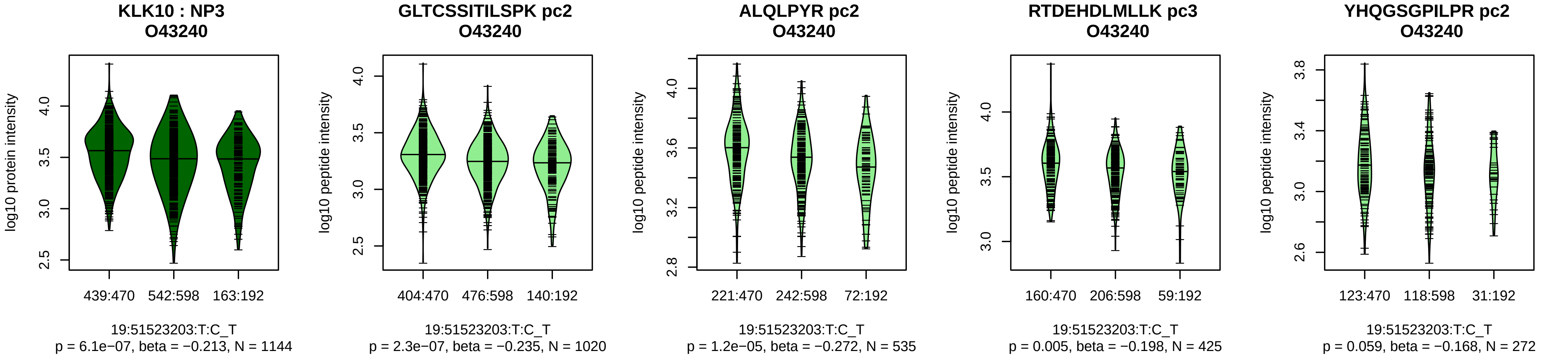


Assay Target: KLK10
Olink UniProt: O43240
UKB rsID: rs2569454
Proxy rsID: rs2569454
UKB SNP: 19:51523203:T:C
Proxy SNP: 19:51523203:T:C
UKB log10(p): 1762.7
UKB BETA: 0.576
*:*:*:−:−:−:−:NA:NA
1020:535:425:272:248:86:76:5:4

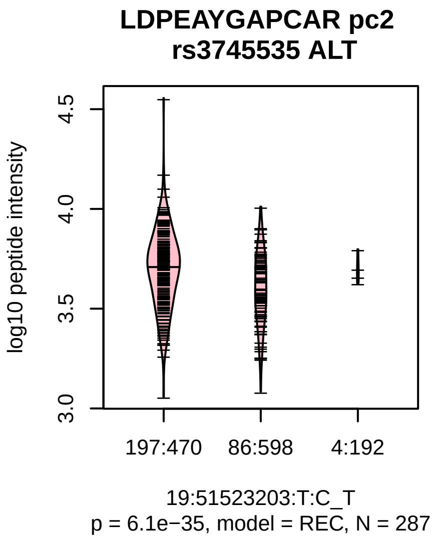

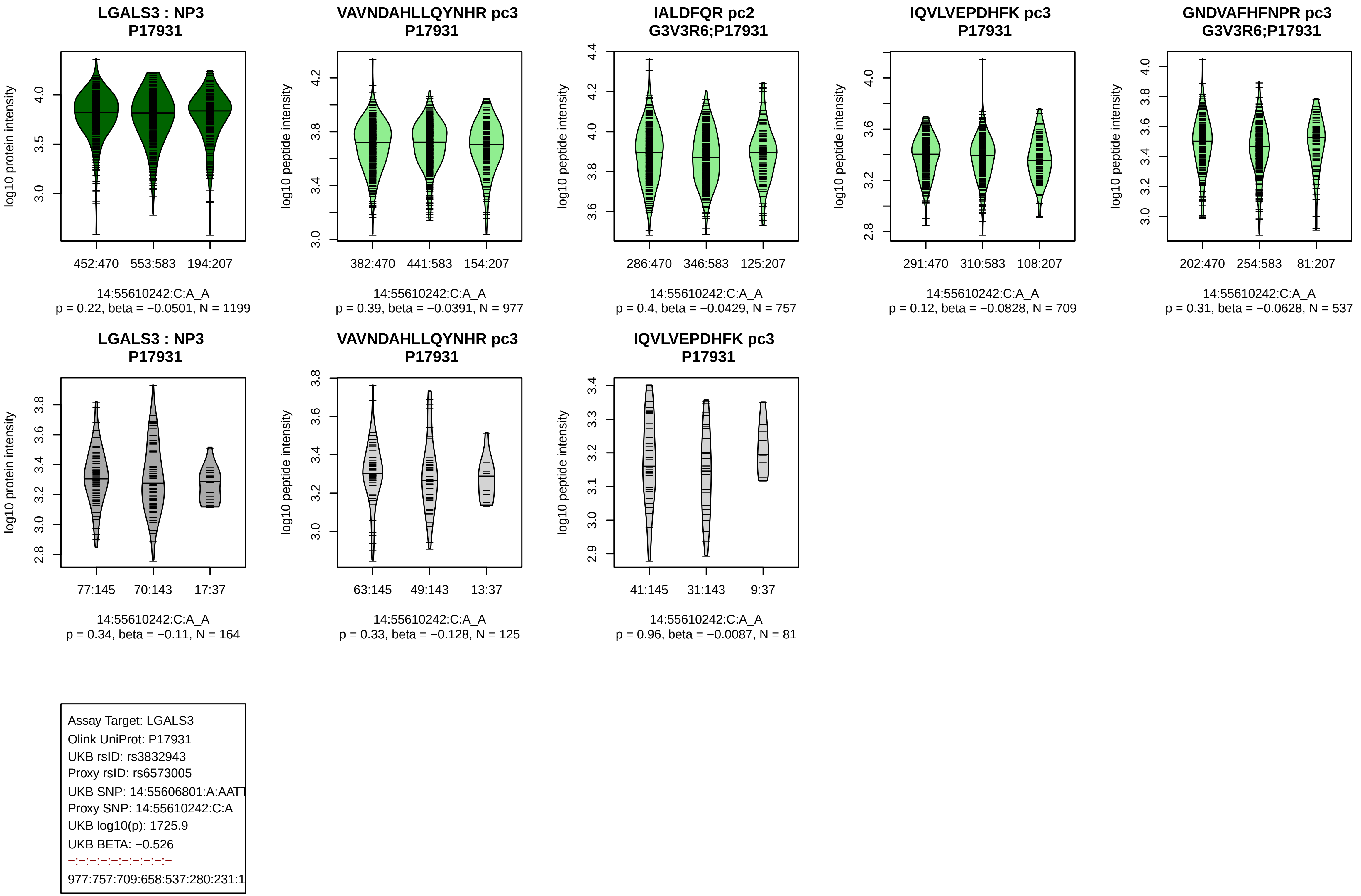

LGALS3 : NP3
P17931
log10 protein intensity
452:470
553:583
194:207
14:55610242:C:A_A
p = 0.22, beta = −0.0501, N = 1199
VAVNDAHLLQYNHR pc3
P17931
log10 peptide intensity
382:470
441:583
154:207
14:55610242:C:A_A
p = 0.39, beta = −0.0391, N = 977
IALDFQR pc2
G3V3R6;P17931
log10 peptide intensity
286:470
346:583
125:207
14:55610242:C:A_A
p = 0.4, beta = −0.0429, N = 757
IQVLVEPDHFK pc3
P17931
log10 peptide intensity
291:470
310:583
108:207
14:55610242:C:A_A
p = 0.12, beta = −0.0828, N = 709
GNDVAFHFNPR pc3
G3V3R6;P17931
log10 peptide intensity
202:470
254:583
81:207
14:55610242:C:A_A
p = 0.31, beta = −0.0628, N = 537
LGALS3 : NP3
P17931
log10 protein intensity
77:145
70:143
17:37
14:55610242:C:A_A
p = 0.34, beta = −0.11, N = 164
VAVNDAHLLQYNHR pc3
P17931
log10 peptide intensity
63:145
49:143
13:37
14:55610242:C:A_A
p = 0.33, beta = −0.128, N = 125
IQVLVEPDHFK pc3
P17931
log10 peptide intensity
41:145
31:143
9:37
14:55610242:C:A_A
p = 0.96, beta = −0.0087, N = 81
Assay Target: LGALS3
Olink UniProt: P17931
UKB rsID: rs3832943
Proxy rsID: rs6573005
UKB SNP: 14:55606801:A:AATT
Proxy SNP: 14:55610242:C:A
UKB log10(p): 1725.9
UKB BETA: −0.526
977:757:709:658:537:280:231:1

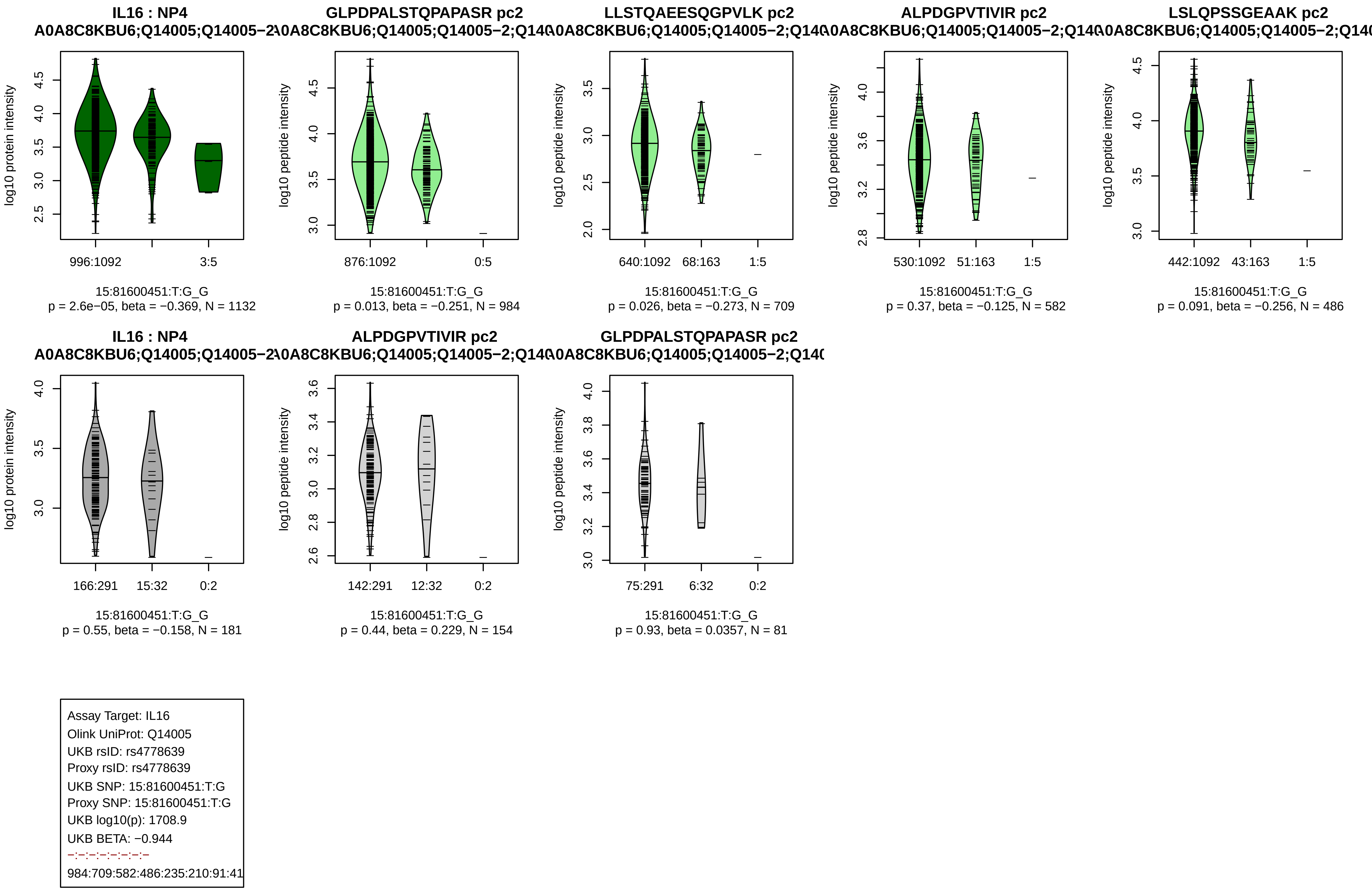

IL16 : NP4
A0A8C8KBU6;Q14005;Q14005−2
log10 protein intensity
996:1092
3:5
15:81600451:T:G_G
p = 2.6e−05, beta = −0.369, N = 1132
GLPDPALSTQPAPASR pc2
A0A8C8KBU6;Q14005;Q14005−2;Q140
log10 peptide intensity
876:1092
0:5
15:81600451:T:G_G
p = 0.013, beta = −0.251, N = 984
LLSTQAEESQGPVLK pc2
A0A8C8KBU6;Q14005;Q14005−2;Q140
log10 peptide intensity
640:1092
68:163
1:5
15:81600451:T:G_G
p = 0.026, beta = −0.273, N = 709
ALPDGPVTIVIR pc2
A0A8C8KBU6;Q14005;Q14005−2;Q140
log10 peptide intensity
530:1092
51:163
1:5
15:81600451:T:G_G
p = 0.37, beta = −0.125, N = 582
LSLQPSSGEAAK pc2
A0A8C8KBU6;Q14005;Q14005−2;Q140
log10 peptide intensity
442:1092
43:163
1:5
15:81600451:T:G_G
p = 0.091, beta = −0.256, N = 486
IL16 : NP4
A0A8C8KBU6;Q14005;Q14005−2
log10 protein intensity
166:291
15:32
0:2
15:81600451:T:G_G
p = 0.55, beta = −0.158, N = 181
ALPDGPVTIVIR pc2
A0A8C8KBU6;Q14005;Q14005−2;Q140
log10 peptide intensity
142:291
12:32
0:2
15:81600451:T:G_G
p = 0.44, beta = 0.229, N = 154
GLPDPALSTQPAPASR pc2
A0A8C8KBU6;Q14005;Q14005−2;Q140
log10 peptide intensity
75:291
6:32
0:2
15:81600451:T:G_G
p = 0.93, beta = 0.0357, N = 81
Assay Target: IL16
Olink UniProt: Q14005
UKB rsID: rs4778639
Proxy rsID: rs4778639
UKB SNP: 15:81600451:T:G
Proxy SNP: 15:81600451:T:G
UKB log10(p): 1708.9
UKB BETA: −0.944
−:−:−:−:−:−:−:−
984:709:582:486:235:210:91:41

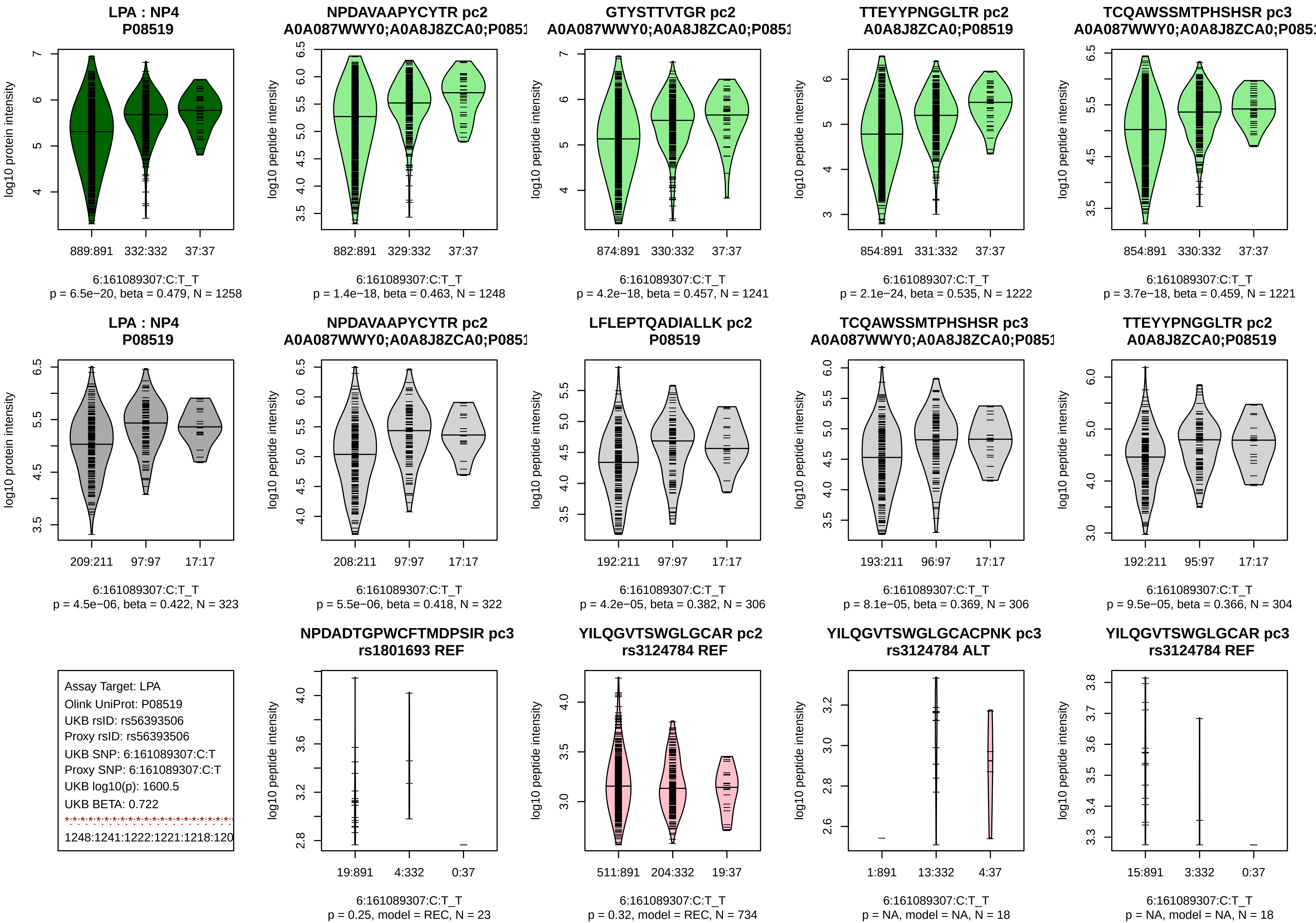
LPA : NP4
P08519
log10 protein intensity
889:891 332:332 37:37
6:161089307:C:T_T
p = 6.5e−20, beta = 0.479, N = 1258
NPDAVAAPYCYTR pc2
A0A087WWY0;A0A8J8ZCA0;P0851
log10 peptide intensity
882:891 329:332 37:37
6:161089307:C:T_T
p = 1.4e−18, beta = 0.463, N = 1248
GTYSTTVTGR pc2
A0A087WWY0;A0A8J8ZCA0;P0851
log10 peptide intensity
874:891 330:332 37:37
6:161089307:C:T_T
p = 4.2e−18, beta = 0.457, N = 1241
TTEYYPNGGLTR pc2
A0A8J8ZCA0;P08519
log10 peptide intensity
854:891 331:332 37:37
6:161089307:C:T_T
p = 2.1e−24, beta = 0.535, N = 1222
TCQAWSSMTPHSHSR pc3
A0A087WWY0;A0A8J8ZCA0;P0851
log10 peptide intensity
854:891 330:332 37:37
6:161089307:C:T_T
p = 3.7e−18, beta = 0.459, N = 1221
LPA : NP4
P08519
log10 protein intensity
209:211 97:97 17:17
6:161089307:C:T_T
p = 4.5e−06, beta = 0.422, N = 323
NPDAVAAPYCYTR pc2
A0A087WWY0;A0A8J8ZCA0;P0851
log10 peptide intensity
208:211 97:97 17:17
6:161089307:C:T_T
p = 5.5e−06, beta = 0.418, N = 322
LFLEPTQADIALLK pc2
P08519
log10 peptide intensity
192:211 97:97 17:17
6:161089307:C:T_T
p = 4.2e−05, beta = 0.382, N = 306
TCQAWSSMTPHSHSR pc3
A0A087WWY0;A0A8J8ZCA0;P0851
log10 peptide intensity
193:211 96:97 17:17
6:161089307:C:T_T
p = 8.1e−05, beta = 0.369, N = 306
TTEYYPNGGLTR pc2
A0A8J8ZCA0;P08519
log10 peptide intensity
192:211 95:97 17:17
6:161089307:C:T_T
p = 9.5e−05, beta = 0.366, N = 304
Assay Target: LPA
Olink UniProt: P08519
UKB rsID: rs56393506
Proxy rsID: rs56393506
UKB SNP: 6:161089307:C:T
Proxy SNP: 6:161089307:C:T
UKB log10(p): 1600.5
UKB BETA: 0.722
1248:1241:1222:1221:1218:120
NPDADTGPWCFTMDPSIR pc3
rs1801693 REF
log10 peptide intensity
19:891 4:332 0:37
6:161089307:C:T_T
p = 0.25, model = REC, N = 23
YILQGVTSWGLGCAR pc2
rs3124784 REF
log10 peptide intensity
511:891 204:332 19:37
6:161089307:C:T_T
p = 0.32, model = REC, N = 734
YILQGVTSWGLGCACPNK pc3
rs3124784 ALT
log10 peptide intensity
1:891 13:332 4:37
6:161089307:C:T_T
p = NA, model = NA, N = 18
YILQGVTSWGLGCAR pc3
rs3124784 REF
log10 peptide intensity
15:891 3:332 0:37
6:161089307:C:T_T
p = NA, model = NA, N = 18

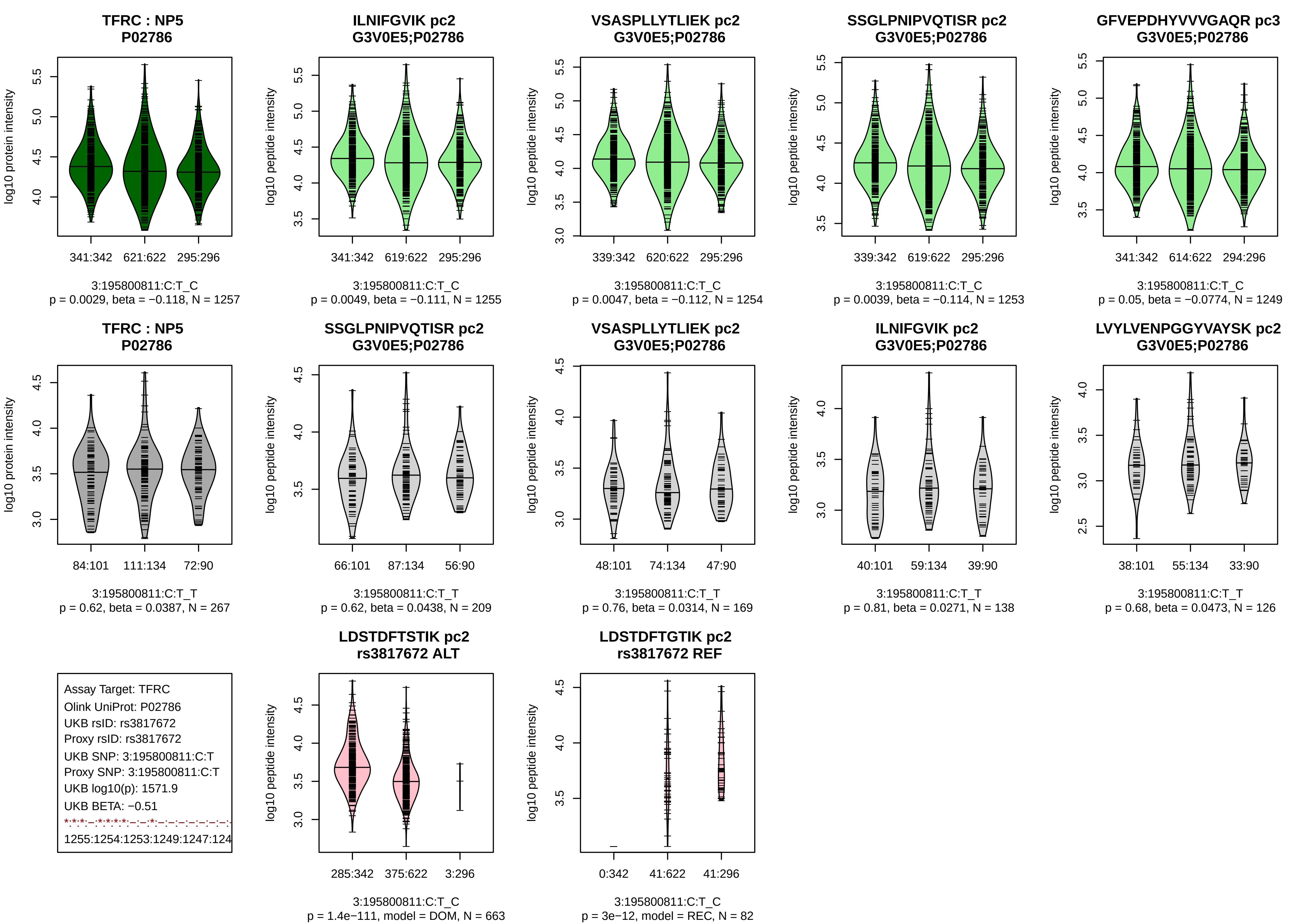
TFRC : NP5
P02786
log10 protein intensity
341:342 621:622 295:296
3:195800811:C:T_C
p = 0.0029, beta = −0.118, N = 1257
ILNIFGVIK pc2
G3V0E5;P02786
log10 peptide intensity
341:342 619:622 295:296
3:195800811:C:T_C
p = 0.0049, beta = −0.111, N = 1255
VSASPLLYTLIEK pc2
G3V0E5;P02786
log10 peptide intensity
339:342 620:622 295:296
3:195800811:C:T_C
p = 0.0047, beta = −0.112, N = 1254
SSGLPNIPVQTISR pc2
G3V0E5;P02786
log10 peptide intensity
339:342 619:622 295:296
3:195800811:C:T_C
p = 0.0039, beta = −0.114, N = 1253
GFVEPDHYVVVGAQR pc3
G3V0E5;P02786
log10 peptide intensity
341:342 614:622 294:296
3:195800811:C:T_C
p = 0.05, beta = −0.0774, N = 1249
TFRC : NP5
P02786
log10 protein intensity
84:101 111:134 72:90
3:195800811:C:T_T
p = 0.62, beta = 0.0387, N = 267
SSGLPNIPVQTISR pc2
G3V0E5;P02786
log10 peptide intensity
66:101 87:134 56:90
3:195800811:C:T_T
p = 0.62, beta = 0.0438, N = 209
VSASPLLYTLIEK pc2
G3V0E5;P02786
log10 peptide intensity
48:101 74:134 47:90
3:195800811:C:T_T
p = 0.76, beta = 0.0314, N = 169
ILNIFGVIK pc2
G3V0E5;P02786
log10 peptide intensity
40:101 59:134 39:90
3:195800811:C:T_T
p = 0.81, beta = 0.0271, N = 138
LVYLVENPGGYVAYSK pc2
G3V0E5;P02786
log10 peptide intensity
38:101 55:134 33:90
3:195800811:C:T_T
p = 0.68, beta = 0.0473, N = 126
Assay Target: TFRC
Olink UniProt: P02786
UKB rsID: rs3817672
Proxy rsID: rs3817672
UKB SNP: 3:195800811:C:T
Proxy SNP: 3:195800811:C:T
UKB log10(p): 1571.9
UKB BETA: −0.51
1255:1254:1253:1249:1247:124
LDSTDFTSTIK pc2
rs3817672 ALT
log10 peptide intensity
285:342 375:622 3:296
3:195800811:C:T_C
p = 1.4e−111, model = DOM, N = 663
LDSTDFTGTIK pc2
rs3817672 REF
log10 peptide intensity
0:342 41:622 41:296
3:195800811:C:T_C
p = 3e−12, model = REC, N = 82

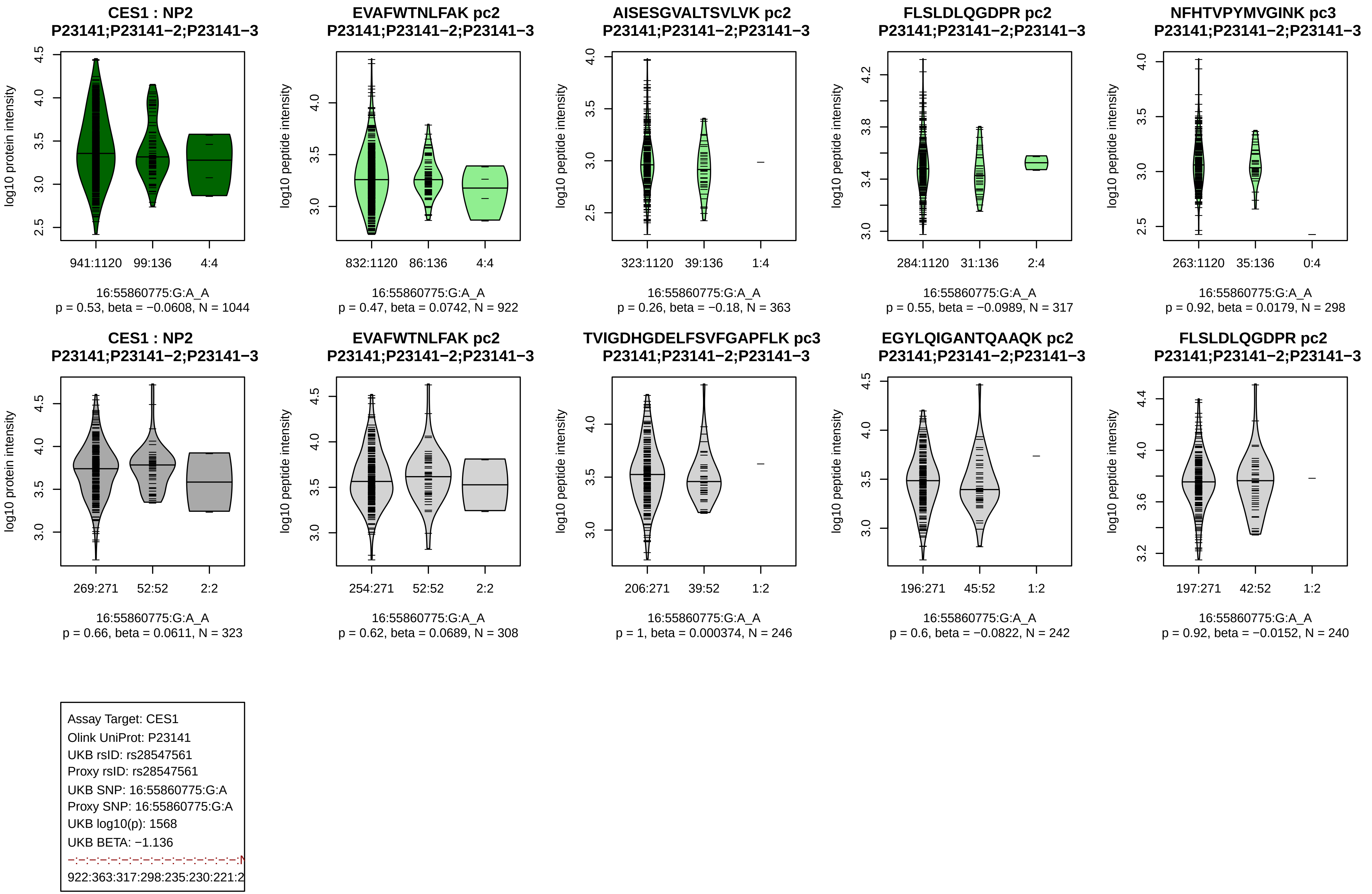
CES1 : NP2
P23141;P23141−2;P23141−3
log10 protein intensity
941:1120
99:136
4:4
16:55860775:G:A_A
p = 0.53, beta = −0.0608, N = 1044
EVAFWTNLFAK pc2
P23141;P23141−2;P23141−3
log10 peptide intensity
832:1120
86:136
4:4
16:55860775:G:A_A
p = 0.47, beta = 0.0742, N = 922
AISESGVALTSVLVK pc2
P23141;P23141−2;P23141−3
log10 peptide intensity
323:1120
39:136
1:4
16:55860775:G:A_A
p = 0.26, beta = −0.18, N = 363
FLSLDLQGDPR pc2
P23141;P23141−2;P23141−3
log10 peptide intensity
284:1120
31:136
2:4
16:55860775:G:A_A
p = 0.55, beta = −0.0989, N = 317
NFHTVPYMVGINK pc3
P23141;P23141−2;P23141−3
log10 peptide intensity
263:1120
35:136
0:4
16:55860775:G:A_A
p = 0.92, beta = 0.0179, N = 298
CES1 : NP2
P23141;P23141−2;P23141−3
log10 protein intensity
269:271
52:52
2:2
16:55860775:G:A_A
p = 0.66, beta = 0.0611, N = 323
EVAFWTNLFAK pc2
P23141;P23141−2;P23141−3
log10 peptide intensity
254:271
52:52
2:2
16:55860775:G:A_A
p = 0.62, beta = 0.0689, N = 308
TVIGDHGDELFSVFGAPFLK pc3
P23141;P23141−2;P23141−3
log10 peptide intensity
206:271
39:52
1:2
16:55860775:G:A_A
p = 1, beta = 0.000374, N = 246
EGYLQIGANTQAAQK pc2
P23141;P23141−2;P23141−3
log10 peptide intensity
196:271
45:52
1:2
16:55860775:G:A_A
p = 0.6, beta = −0.0822, N = 242
FLSLDLQGDPR pc2
P23141;P23141−2;P23141−3
log10 peptide intensity
197:271
42:52
1:2
16:55860775:G:A_A
p = 0.92, beta = −0.0152, N = 240
Assay Target: CES1
Olink UniProt: P23141
UKB rsID: rs28547561
Proxy rsID: rs28547561
UKB SNP: 16:55860775:G:A
Proxy SNP: 16:55860775:G:A
UKB log10(p): 1568
UKB BETA: −1.136
−:−:−:−:−:−:−:−:−:−:−:−:−:−:−:−:N
922:363:317:298:235:230:221:2

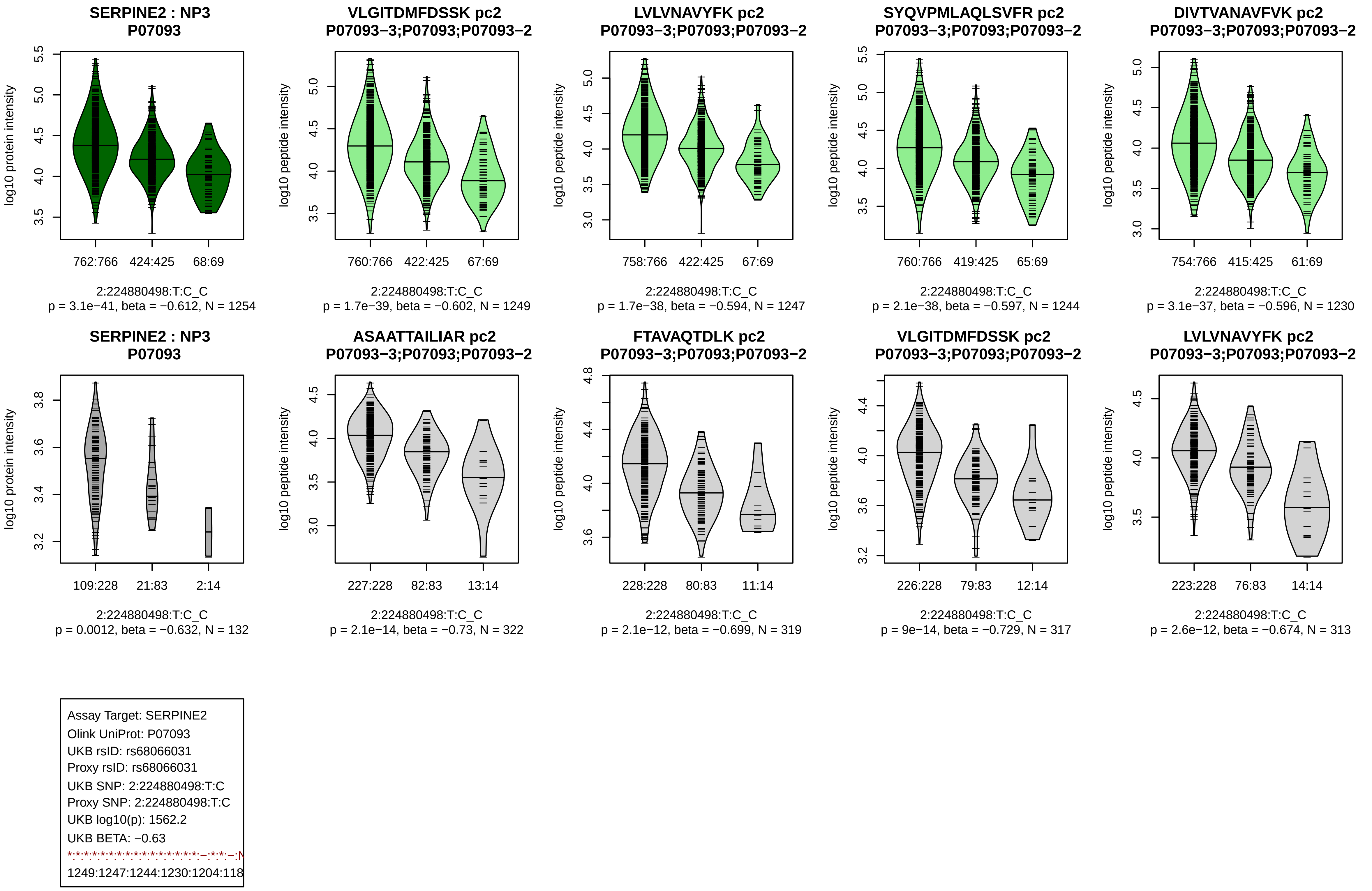
SERPINE2 : NP3
P07093
log10 protein intensity
762:766 424:425 68:69
2:224880498:T:C_C
p = 3.1e−41, beta = −0.612, N = 1254
VLGITDMFDSSK pc2
P07093−3;P07093;P07093−2
log10 peptide intensity
760:766 422:425 67:69
2:224880498:T:C_C
p = 1.7e−39, beta = −0.602, N = 1249
LVLVNAVYFK pc2
P07093−3;P07093;P07093−2
log10 peptide intensity
758:766 422:425 67:69
2:224880498:T:C_C
p = 1.7e−38, beta = −0.594, N = 1247
SYQVPMLAQLSVFR pc2
P07093−3;P07093;P07093−2
log10 peptide intensity
760:766 419:425 65:69
2:224880498:T:C_C
p = 2.1e−38, beta = −0.597, N = 1244
DIVTVANAVFVK pc2
P07093−3;P07093;P07093−2
log10 peptide intensity
754:766 415:425 61:69
2:224880498:T:C_C
p = 3.1e−37, beta = −0.596, N = 1230
SERPINE2 : NP3
P07093
log10 protein intensity
109:228 21:83 2:14
2:224880498:T:C_C
p = 0.0012, beta = −0.632, N = 132
ASAATTAILIAR pc2
P07093−3;P07093;P07093−2
log10 peptide intensity
227:228 82:83 13:14
2:224880498:T:C_C
p = 2.1e−14, beta = −0.73, N = 322
FTAVAQTDLK pc2
P07093−3;P07093;P07093−2
log10 peptide intensity
228:228 80:83 11:14
2:224880498:T:C_C
p = 2.1e−12, beta = −0.699, N = 319
VLGITDMFDSSK pc2
P07093−3;P07093;P07093−2
log10 peptide intensity
226:228 79:83 12:14
2:224880498:T:C_C
p = 9e−14, beta = −0.729, N = 317
LVLVNAVYFK pc2
P07093−3;P07093;P07093−2
log10 peptide intensity
223:228 76:83 14:14
2:224880498:T:C_C
p = 2.6e−12, beta = −0.674, N = 313
Assay Target: SERPINE2
Olink UniProt: P07093
UKB rsID: rs68066031
Proxy rsID: rs68066031
UKB SNP: 2:224880498:T:C
Proxy SNP: 2:224880498:T:C
UKB log10(p): 1562.2
UKB BETA: −0.63
1249:1247:1244:1230:1204:118

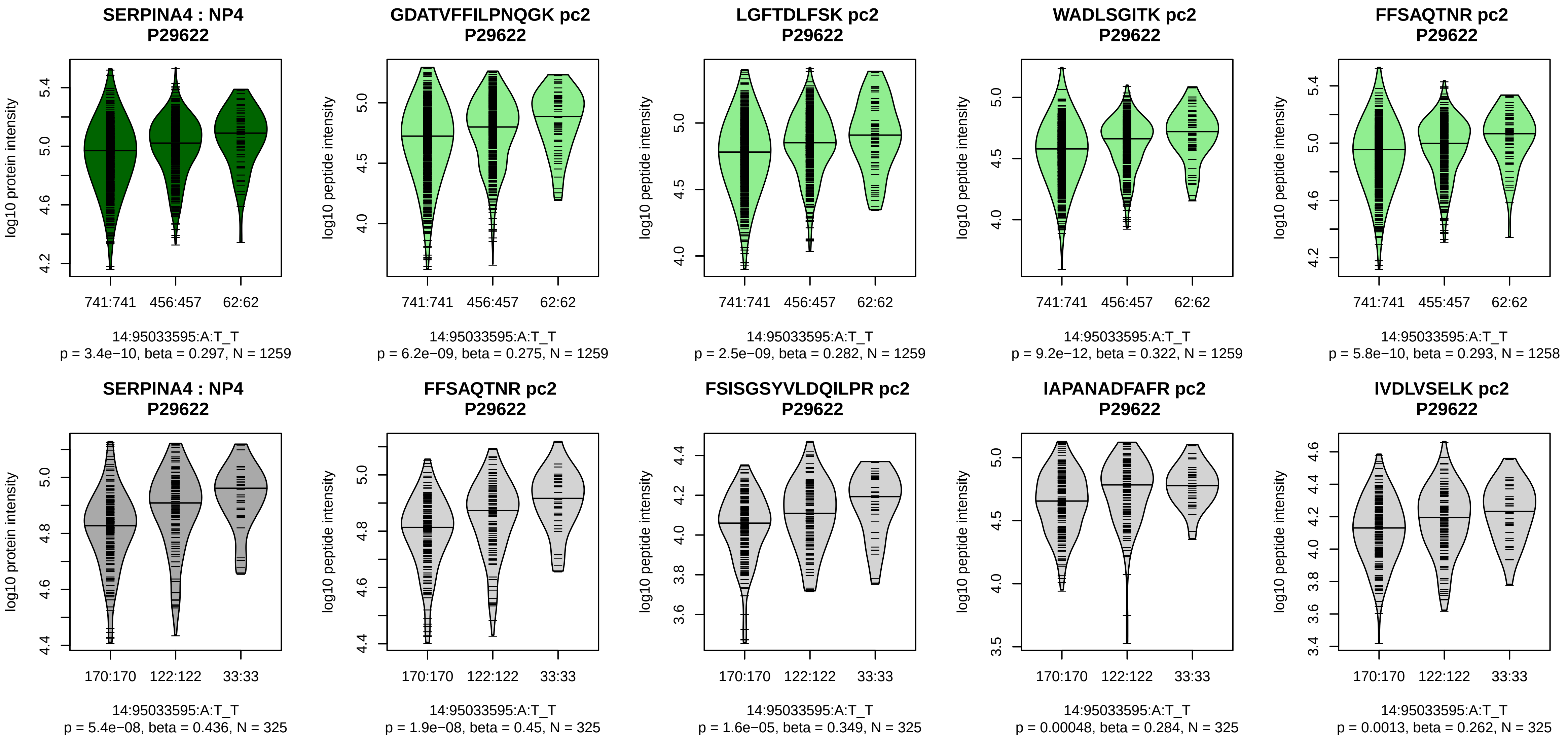


Assay Target: SERPINA4
Olink UniProt: P29622
UKB rsID: rs5511
Proxy rsID: rs5511
UKB SNP: 14:95033595:A:T
Proxy SNP: 14:95033595:A:T
UKB log10(p): 1529.9
UKB BETA: 0.612
*:*:*:*:*:*:*:*:*:*:*:*:*:*:*:*:*:*:*:*:*:–:*:*:
1259:1259:1259:1258:1258:125

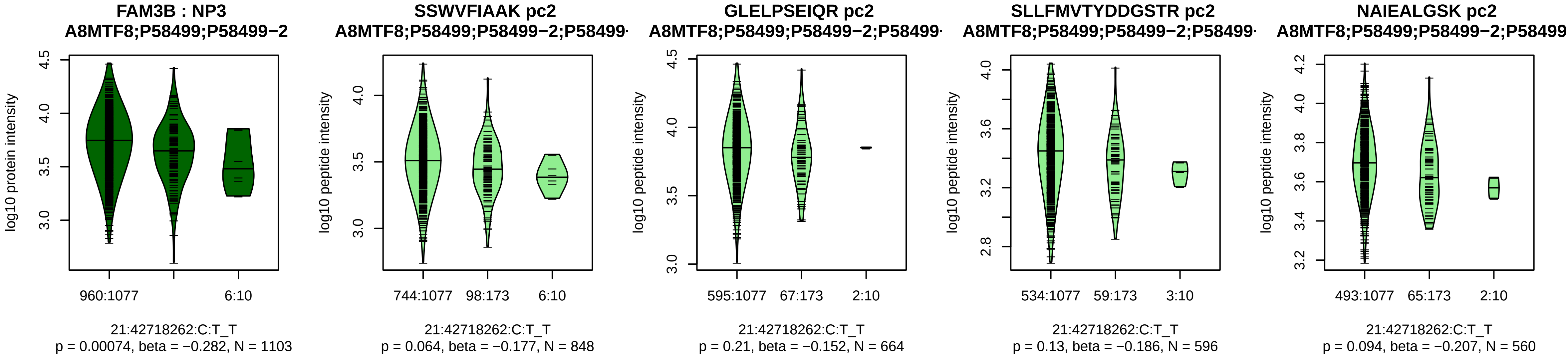


Assay Target: FAM3B
Olink UniProt: P58499
UKB rsID: rs57529409
Proxy rsID: rs57529409
UKB SNP: 21:42718262:C:T
Proxy SNP: 21:42718262:C:T
UKB log10(p): 1516.9
UKB BETA: −1.087
−:−:−:−:*:−
848:664:596:560:520:92

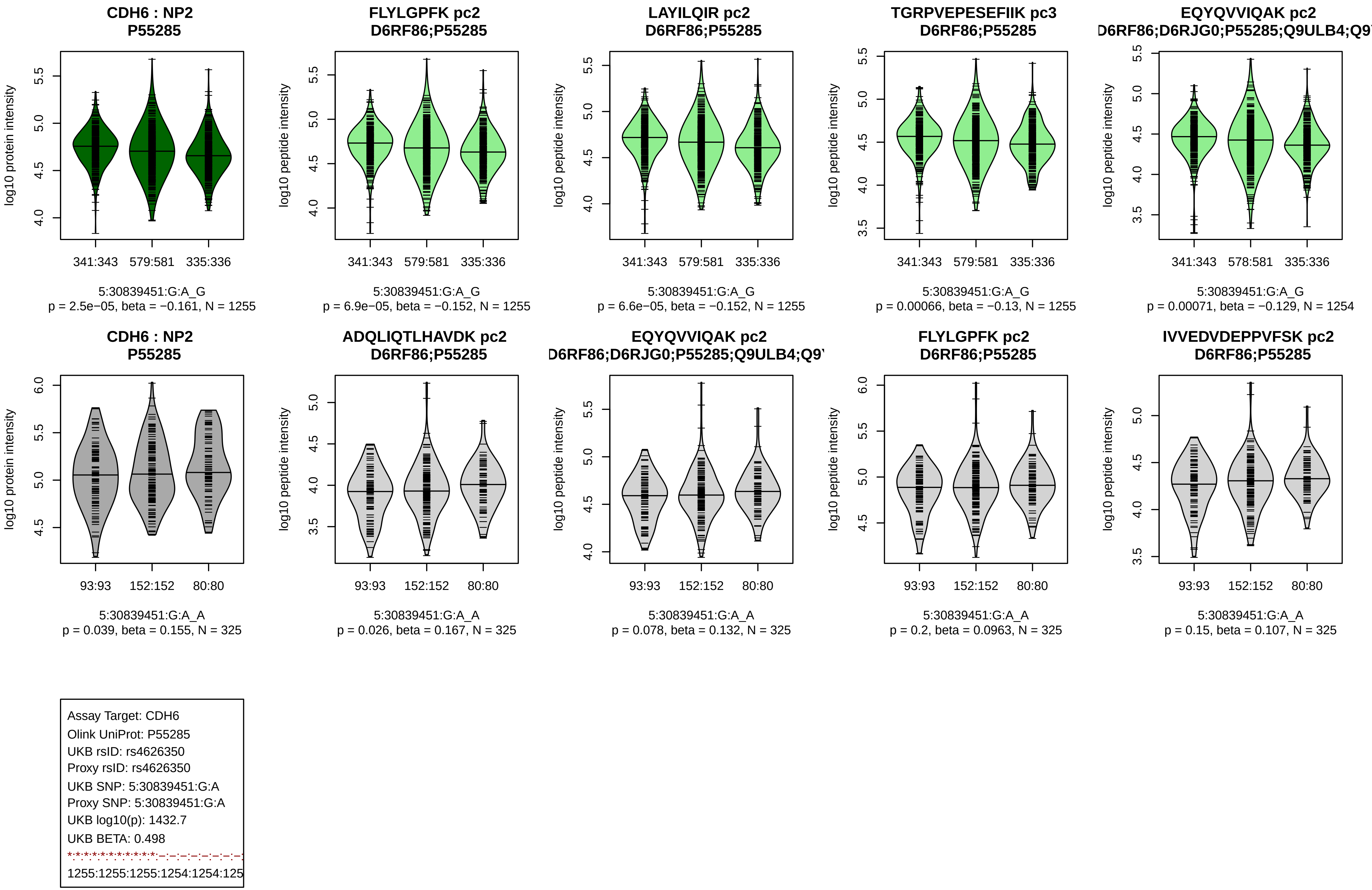

CDH6 : NP2
P55285
log10 protein intensity
341:343
579:581
335:336
5:30839451:G:A_G
p = 2.5e−05, beta = −0.161, N = 1255
FLYLGPFK pc2
D6RF86;P55285
log10 peptide intensity
341:343
579:581
335:336
5:30839451:G:A_G
p = 6.9e−05, beta = −0.152, N = 1255
LAYILQIR pc2
D6RF86;P55285
log10 peptide intensity
341:343
579:581
335:336
5:30839451:G:A_G
p = 6.6e−05, beta = −0.152, N = 1255
TGRPVEPESEFIIK pc3
D6RF86;P55285
log10 peptide intensity
341:343
579:581
335:336
5:30839451:G:A_G
p = 0.00066, beta = −0.13, N = 1255
EQYQVVIQAK pc2
D6RF86;D6RJG0;P55285;Q9ULB4;Q9
log10 peptide intensity
341:343
578:581
335:336
5:30839451:G:A_G
p = 0.00071, beta = −0.129, N = 1254
CDH6 : NP2
P55285
log10 protein intensity
93:93
152:152
80:80
5:30839451:G:A_A
p = 0.039, beta = 0.155, N = 325
ADQLIQTLHAVDK pc2
D6RF86;P55285
log10 peptide intensity
93:93
152:152
80:80
5:30839451:G:A_A
p = 0.026, beta = 0.167, N = 325
EQYQVVIQAK pc2
D6RF86;D6RJG0;P55285;Q9ULB4;Q9
log10 peptide intensity
93:93
152:152
80:80
5:30839451:G:A_A
p = 0.078, beta = 0.132, N = 325
FLYLGPFK pc2
D6RF86;P55285
log10 peptide intensity
93:93
152:152
80:80
5:30839451:G:A_A
p = 0.2, beta = 0.0963, N = 325
IVVEDVDEPPVFSK pc2
D6RF86;P55285
log10 peptide intensity
93:93
152:152
80:80
5:30839451:G:A_A
p = 0.15, beta = 0.107, N = 325
Assay Target: CDH6
Olink UniProt: P55285
UKB rsID: rs4626350
Proxy rsID: rs4626350
UKB SNP: 5:30839451:G:A
Proxy SNP: 5:30839451:G:A
UKB log10(p): 1432.7
UKB BETA: 0.498
1255:1255:1255:1254:1254:125

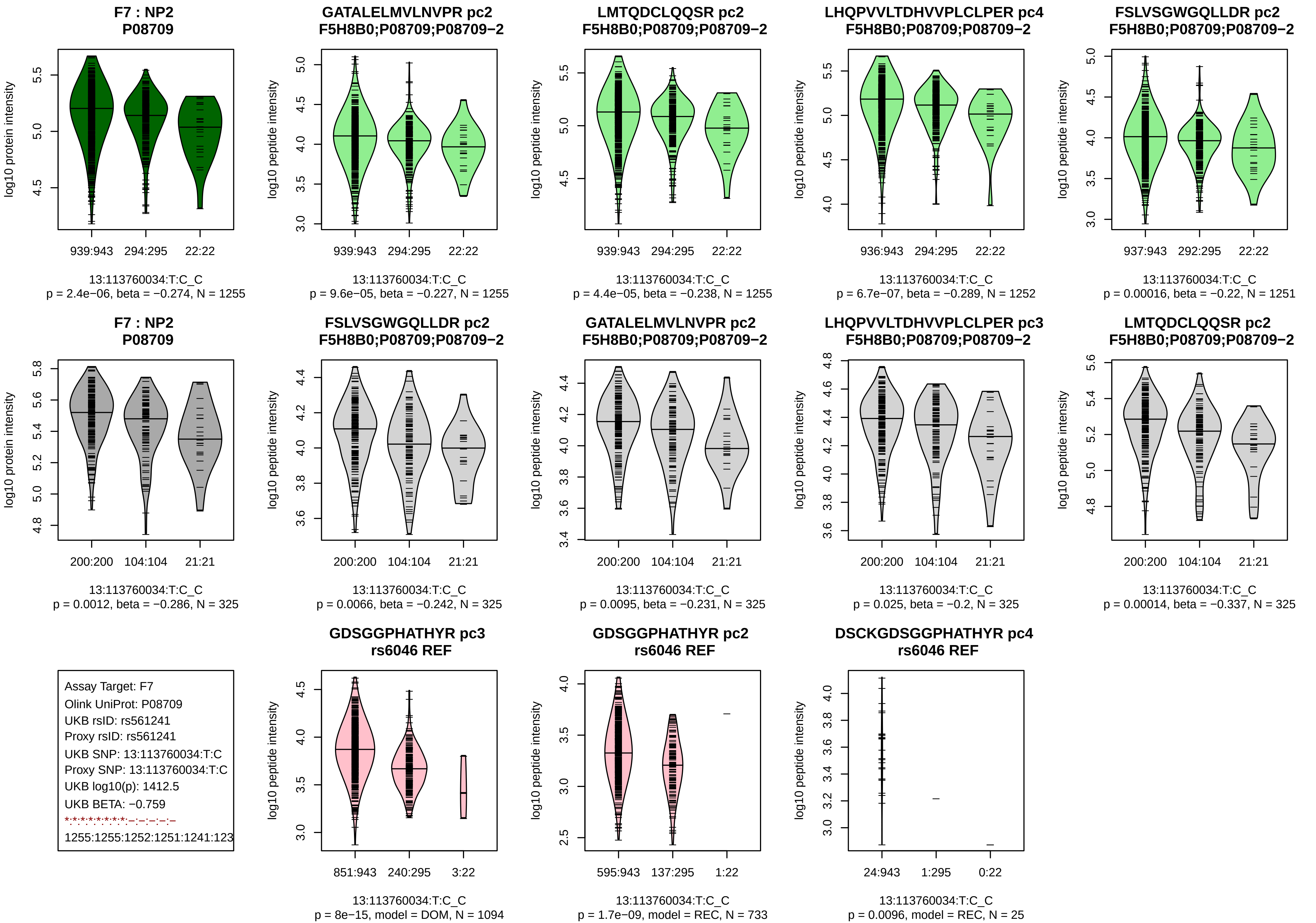
F7 : NP2
P08709
log10 protein intensity
939:943 294:295 22:22
13:113760034:T:C_C
p = 2.4e−06, beta = −0.274, N = 1255
GATALELMVLNVPR pc2
F5H8B0;P08709;P08709−2
log10 peptide intensity
939:943 294:295 22:22
13:113760034:T:C_C
p = 9.6e−05, beta = −0.227, N = 1255
LMTQDCLQQSR pc2
F5H8B0;P08709;P08709−2
939:943 294:295 22:22
13:113760034:T:C_C
p = 4.4e−05, beta = −0.238, N = 1255
LHQPVVLTDHVVPLCLPER pc4
F5H8B0;P08709;P08709−2
936:943 294:295 22:22
13:113760034:T:C_C
p = 6.7e−07, beta = −0.289, N = 1252
FSLVSGWGQLLDR pc2
F5H8B0;P08709;P08709−2
937:943 292:295 22:22
13:113760034:T:C_C
p = 0.00016, beta = −0.22, N = 1251
F7 : NP2
P08709
200:200 104:104 21:21
13:113760034:T:C_C
p = 0.0012, beta = −0.286, N = 325
FSLVSGWGQLLDR pc2
F5H8B0;P08709;P08709−2
200:200 104:104 21:21
13:113760034:T:C_C
p = 0.0066, beta = −0.242, N = 325
GATALELMVLNVPR pc2
F5H8B0;P08709;P08709−2
200:200 104:104 21:21
13:113760034:T:C_C
p = 0.0095, beta = −0.231, N = 325
LHQPVVLTDHVVPLCLPER pc3
F5H8B0;P08709;P08709−2
200:200 104:104 21:21
13:113760034:T:C_C
p = 0.025, beta = −0.2, N = 325
LMTQDCLQQSR pc2
F5H8B0;P08709;P08709−2
200:200 104:104 21:21
13:113760034:T:C_C
p = 0.00014, beta = −0.337, N = 325
Assay Target: F7
Olink UniProt: P08709
UKB rsID: rs561241
Proxy rsID: rs561241
UKB SNP: 13:113760034:T:C
Proxy SNP: 13:113760034:T:C
UKB log10(p): 1412.5
UKB BETA: −0.759
1255:1255:1252:1251:1241:123
GDSGGPHATHYR pc3
rs6046 REF
851:943 240:295 3:22
13:113760034:T:C_C
p = 8e−15, model = DOM, N = 1094
GDSGGPHATHYR pc2
rs6046 REF
595:943 137:295 1:22
13:113760034:T:C_C
p = 1.7e−09, model = REC, N = 733
DSCKGDSGGPHATHYR pc4
rs6046 REF
24:943 1:295 0:22
13:113760034:T:C_C
p = 0.0096, model = REC, N = 25

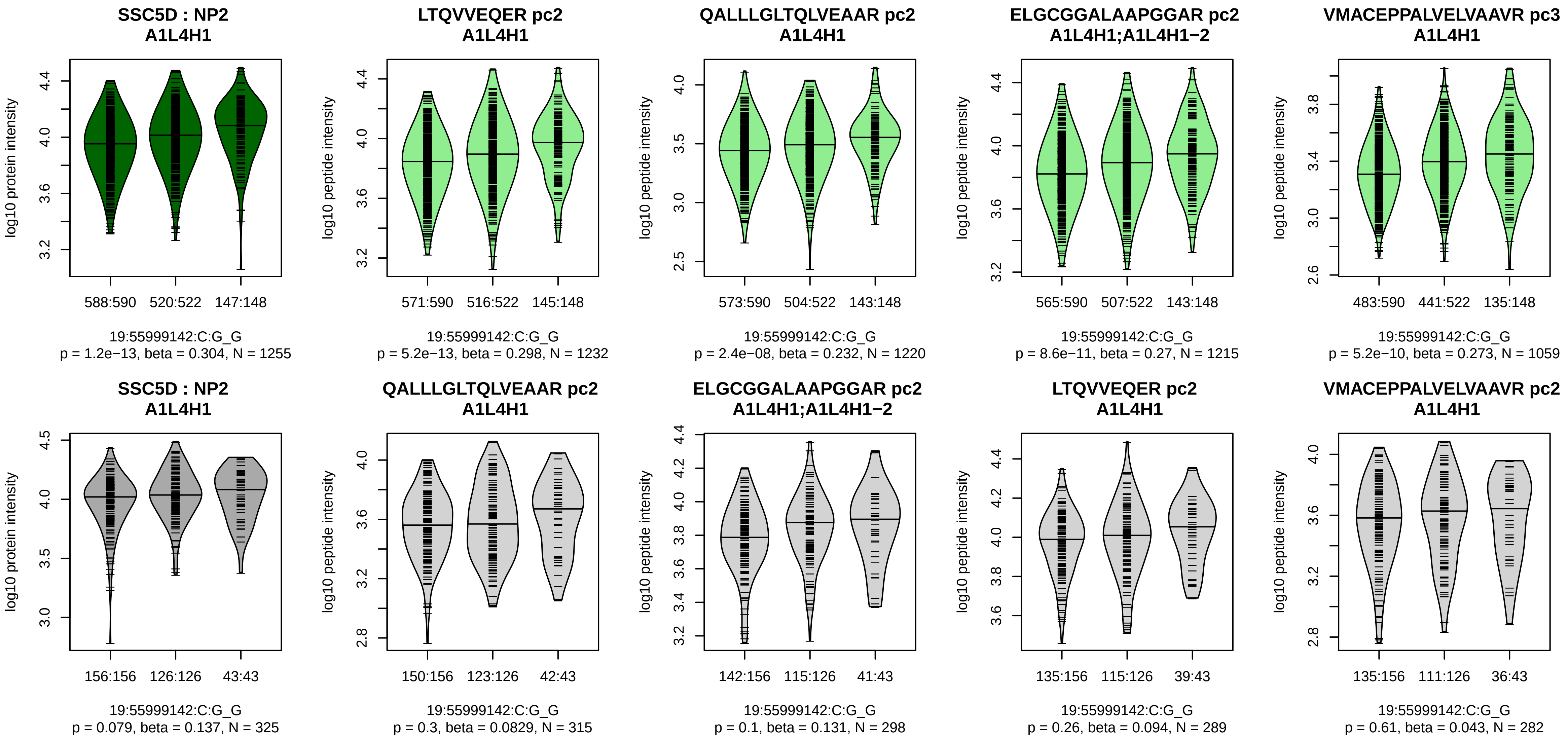


Assay Target: SSC5D
Olink UniProt: A1L4H1
UKB rsID: rs55799523
Proxy rsID: rs8103017
UKB SNP: 19:55999545:C:A
Proxy SNP: 19:55999142:C:G
UKB log10(p): 1347.2
UKB BETA: 0.53
1232:1220:1215:1059:1050:100

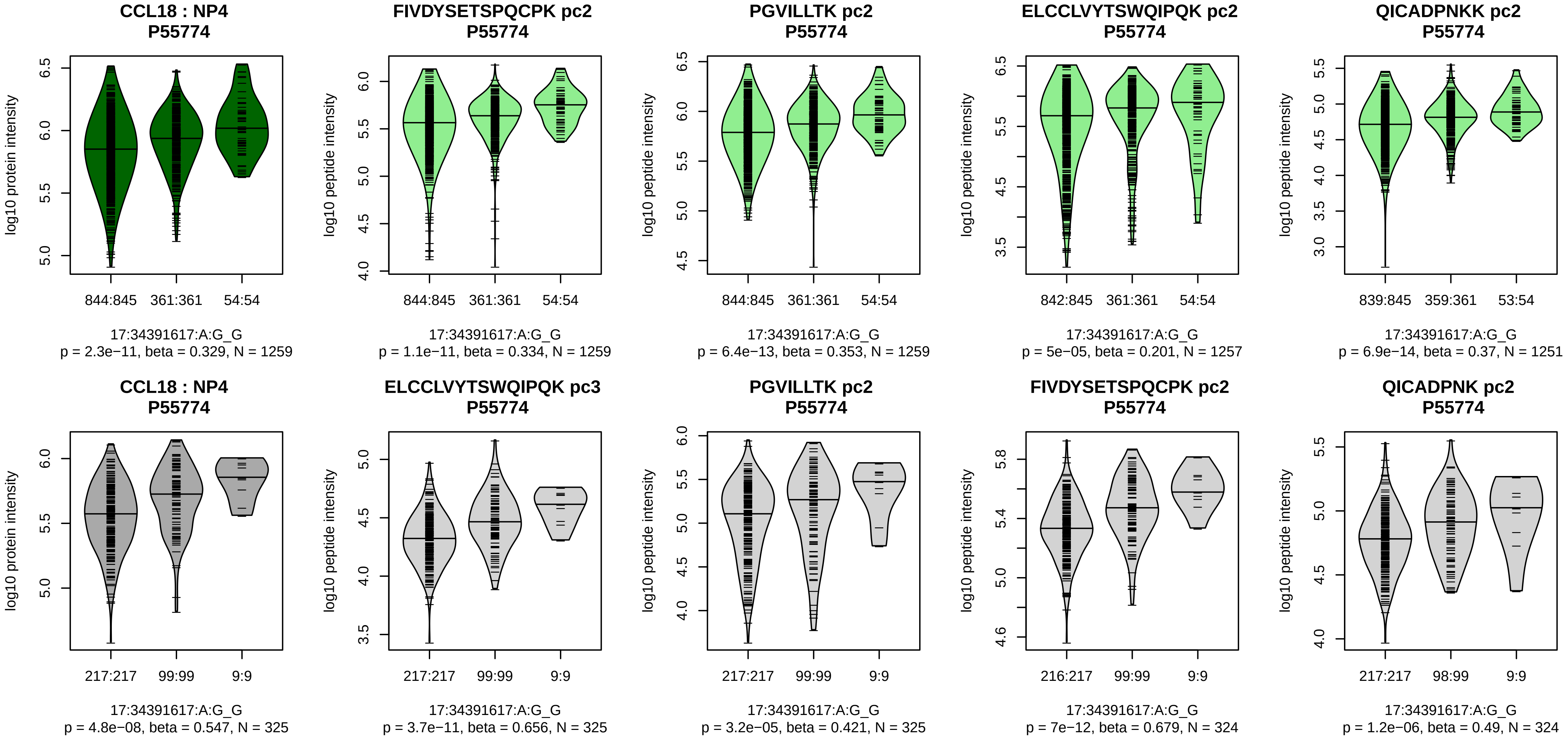


Assay Target: CCL18
Olink UniProt: P55774
UKB rsID: rs2015086
Proxy rsID: rs2015086
UKB SNP: 17:34391617:A:G
Proxy SNP: 17:34391617:A:G
UKB log10(p): 1335.8
UKB BETA: 0.68
*:*:*:*:*:*:–
1259:1259:1257:1257:1251:124

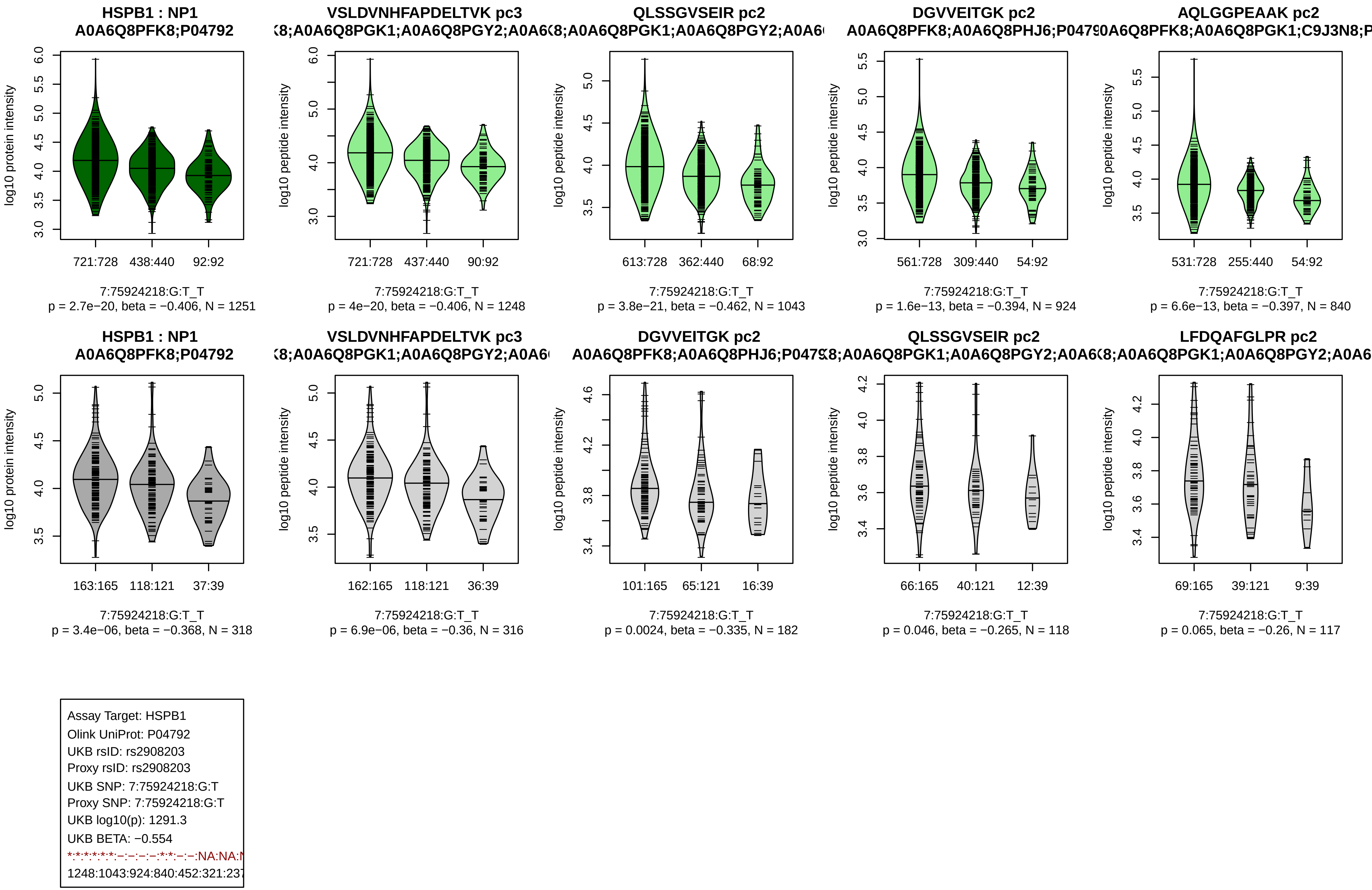
HSPB1 : NP1
A0A6Q8PFK8;P04792
log10 protein intensity
721:728
438:440
92:92
7:75924218:G:T_T
p = 2.7e−20, beta = −0.406, N = 1251
VSLDVNHFAPDELTVK pc3
(8;A0A6Q8PGK1;A0A6Q8PGY2;A0A6(
log10 peptide intensity
721:728
437:440
90:92
7:75924218:G:T_T
p = 4e−20, beta = −0.406, N = 1248
QLSSGVSEIR pc2
8;A0A6Q8PGK1;A0A6Q8PGY2;A0A6(
log10 peptide intensity
613:728
362:440
68:92
7:75924218:G:T_T
p = 3.8e−21, beta = −0.462, N = 1043
DGVVEITGK pc2
A0A6Q8PFK8;A0A6Q8PHJ6;P047(
log10 peptide intensity
561:728
309:440
54:92
7:75924218:G:T_T
p = 1.6e−13, beta = −0.394, N = 924
AQLGGPEAAK pc2
0A6Q8PFK8;A0A6Q8PGK1;C9J3N8;P
log10 peptide intensity
531:728
255:440
54:92
7:75924218:G:T_T
p = 6.6e−13, beta = −0.397, N = 840
HSPB1 : NP1
A0A6Q8PFK8;P04792
log10 protein intensity
163:165
118:121
37:39
7:75924218:G:T_T
p = 3.4e−06, beta = −0.368, N = 318
VSLDVNHFAPDELTVK pc3
(8;A0A6Q8PGK1;A0A6Q8PGY2;A0A6(
log10 peptide intensity
162:165
118:121
36:39
7:75924218:G:T_T
p = 6.9e−06, beta = −0.36, N = 316
DGVVEITGK pc2
A0A6Q8PFK8;A0A6Q8PHJ6;P047(
log10 peptide intensity
101:165
65:121
16:39
7:75924218:G:T_T
p = 0.0024, beta = −0.335, N = 182
QLSSGVSEIR pc2
8;A0A6Q8PGK1;A0A6Q8PGY2;A0A6(
log10 peptide intensity
66:165
40:121
12:39
7:75924218:G:T_T
p = 0.046, beta = −0.265, N = 118
LFDQAFGLPR pc2
8;A0A6Q8PGK1;A0A6Q8PGY2;A0A6
log10 peptide intensity
69:165
39:121
9:39
7:75924218:G:T_T
p = 0.065, beta = −0.26, N = 117
Assay Target: HSPB1
Olink UniProt: P04792
UKB rsID: rs2908203
Proxy rsID: rs2908203
UKB SNP: 7:75924218:G:T
Proxy SNP: 7:75924218:G:T
UKB log10(p): 1291.3
UKB BETA: −0.554
*:*:*:*:*:*:−:−:−:−:*:*:−:−:NA:NA:N
1248:1043:924:840:452:321:237

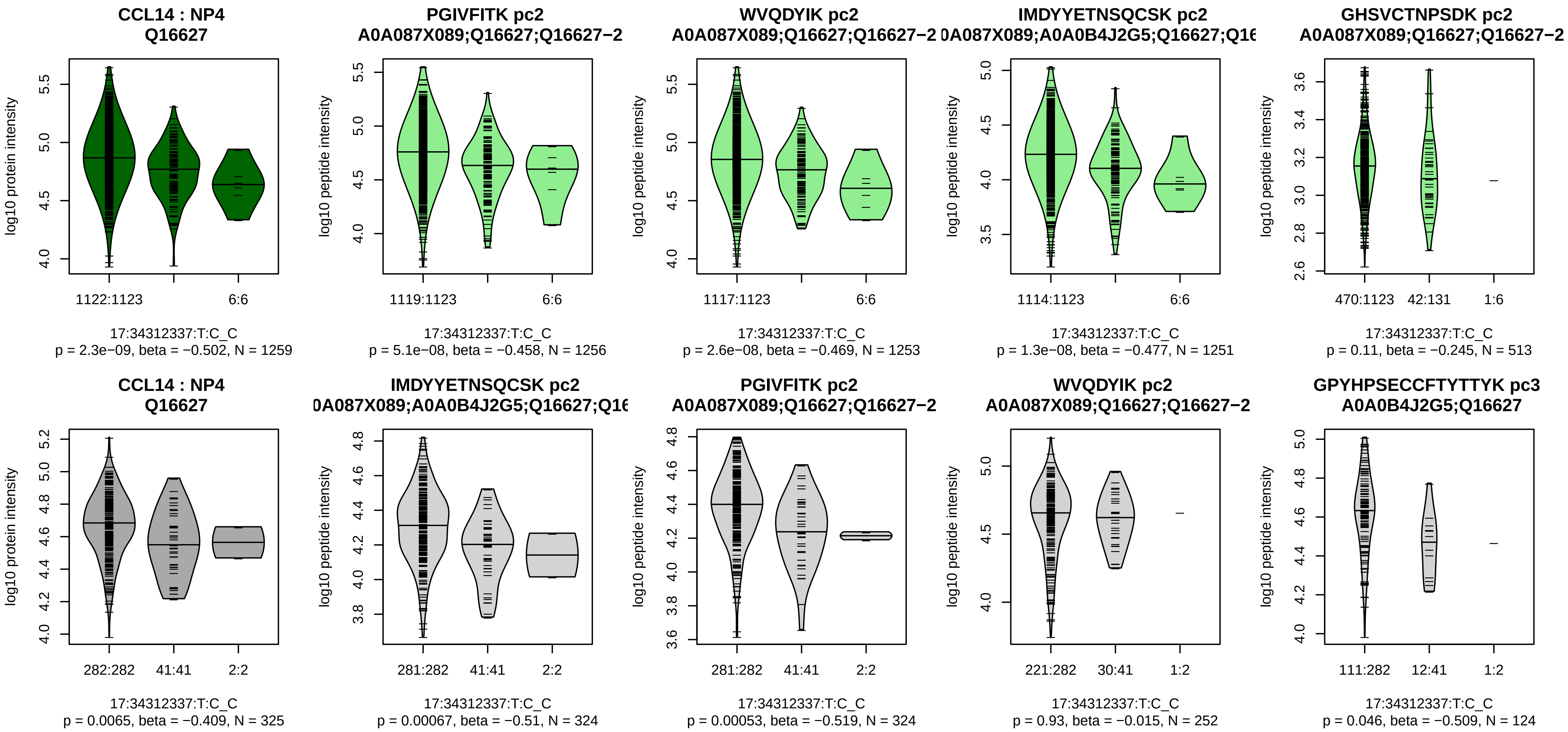


Assay Target: CCL14
Olink UniProt: Q16627
UKB rsID: rs9903158
Proxy rsID: rs9903158
UKB SNP: 17:34312337:T:C
Proxy SNP: 17:34312337:T:C
UKB log10(p): 1253.1
UKB BETA: −0.997
*:*:*:−:−:NA
1256:1253:1251:513:239:32

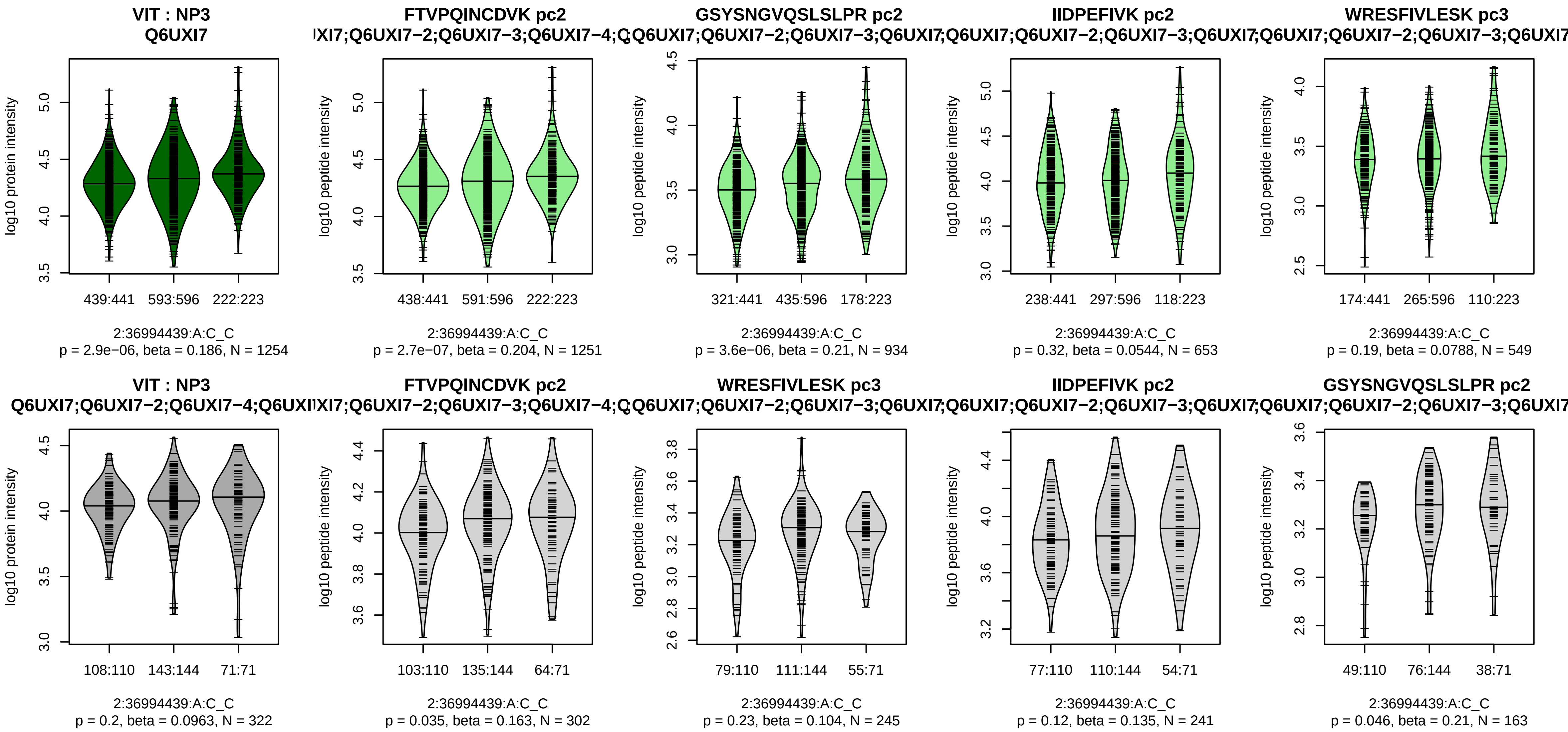


Assay Target: VIT
Olink UniProt: Q6UXI7
UKB rsID: rs11124542
Proxy rsID: rs11124542
UKB SNP: 2:36994439:A:C
Proxy SNP: 2:36994439:A:C
UKB log10(p): 1223.6
UKB BETA: 0.413
*:*:−:−:−:−:−:−:−:−:−:NA:NA:NA
1251:934:653:549:480:465:415:

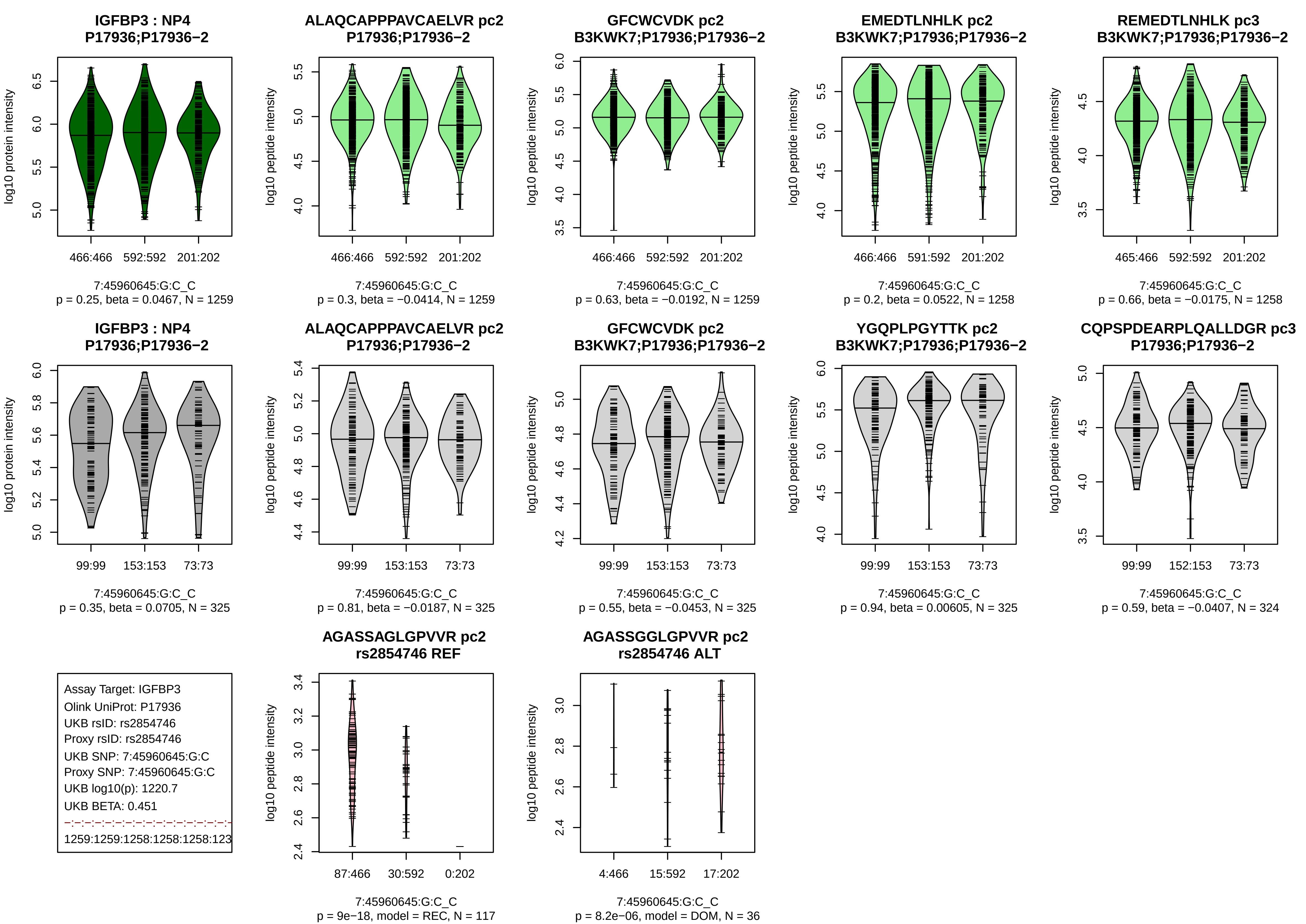
IGFBP3 : NP4
P17936;P17936−2
log10 protein intensity
466:466 592:592 201:202
7:45960645:G:C_C
p = 0.25, beta = 0.0467, N = 1259
ALAQCAPPPAVCAELVR pc2
P17936;P17936−2
log10 peptide intensity
466:466 592:592 201:202
7:45960645:G:C_C
p = 0.3, beta = −0.0414, N = 1259
GFCWCVDK pc2
B3KWK7;P17936;P17936−2
log10 peptide intensity
466:466 592:592 201:202
7:45960645:G:C_C
p = 0.63, beta = −0.0192, N = 1259
EMEDTLNHLK pc2
B3KWK7;P17936;P17936−2
log10 peptide intensity
466:466 591:592 201:202
7:45960645:G:C_C
p = 0.2, beta = 0.0522, N = 1258
REMEDTLNHLK pc3
B3KWK7;P17936;P17936−2
log10 peptide intensity
465:466 592:592 201:202
7:45960645:G:C_C
p = 0.66, beta = −0.0175, N = 1258
IGFBP3 : NP4
P17936;P17936−2
log10 protein intensity
99:99 153:153 73:73
7:45960645:G:C_C
p = 0.35, beta = 0.0705, N = 325
ALAQCAPPPAVCAELVR pc2
P17936;P17936−2
log10 peptide intensity
99:99 153:153 73:73
7:45960645:G:C_C
p = 0.81, beta = −0.0187, N = 325
GFCWCVDK pc2
B3KWK7;P17936;P17936−2
log10 peptide intensity
99:99 153:153 73:73
7:45960645:G:C_C
p = 0.55, beta = −0.0453, N = 325
YGQPLPGYTTK pc2
B3KWK7;P17936;P17936−2
log10 peptide intensity
99:99 153:153 73:73
7:45960645:G:C_C
p = 0.94, beta = 0.00605, N = 325
CQPSPDEARPLQALLDGR pc3
P17936;P17936−2
log10 peptide intensity
99:99 152:153 73:73
7:45960645:G:C_C
p = 0.59, beta = −0.0407, N = 324
Assay Target: IGFBP3
Olink UniProt: P17936
UKB rsID: rs2854746
Proxy rsID: rs2854746
UKB SNP: 7:45960645:G:C
Proxy SNP: 7:45960645:G:C
UKB log10(p): 1220.7
UKB BETA: 0.451
1259:1259:1258:1258:1258:123
AGASSAGLGPVVR pc2
rs2854746 REF
log10 peptide intensity
87:466 30:592 0:202
7:45960645:G:C_C
p = 9e−18, model = REC, N = 117
AGASSGGLGPVVR pc2
rs2854746 ALT
log10 peptide intensity
4:466 15:592 17:202
7:45960645:G:C_C
p = 8.2e−06, model = DOM, N = 36

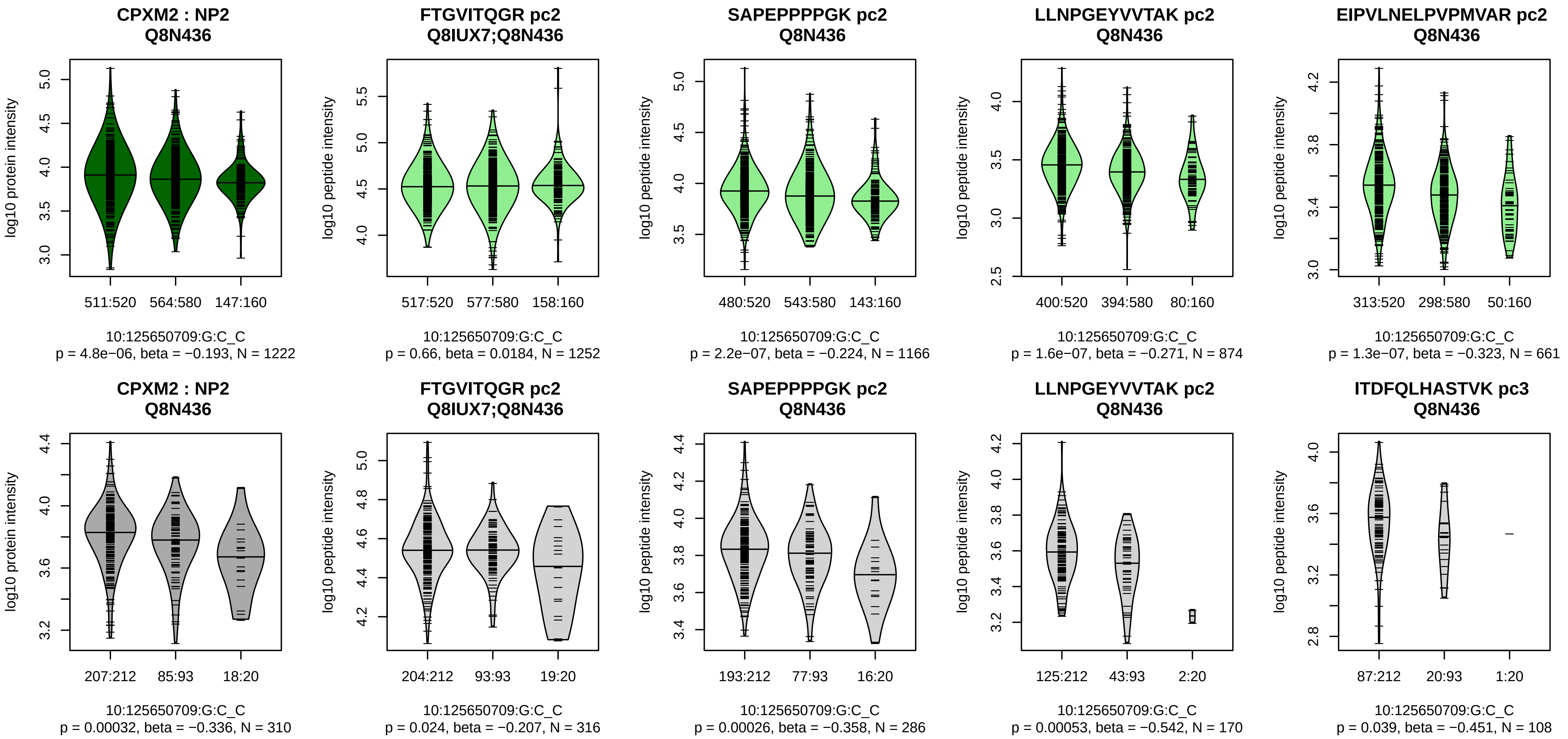


Assay Target: CPXM2
Olink UniProt: Q8N436
UKB rsID: rs28741041
Proxy rsID: rs28741041
UKB SNP: 10:125650709:G:C
Proxy SNP: 10:125650709:G:C
UKB log10(p): 1208
UKB BETA: −0.456
1252:1166:874:661:396:383:366 [illegible]

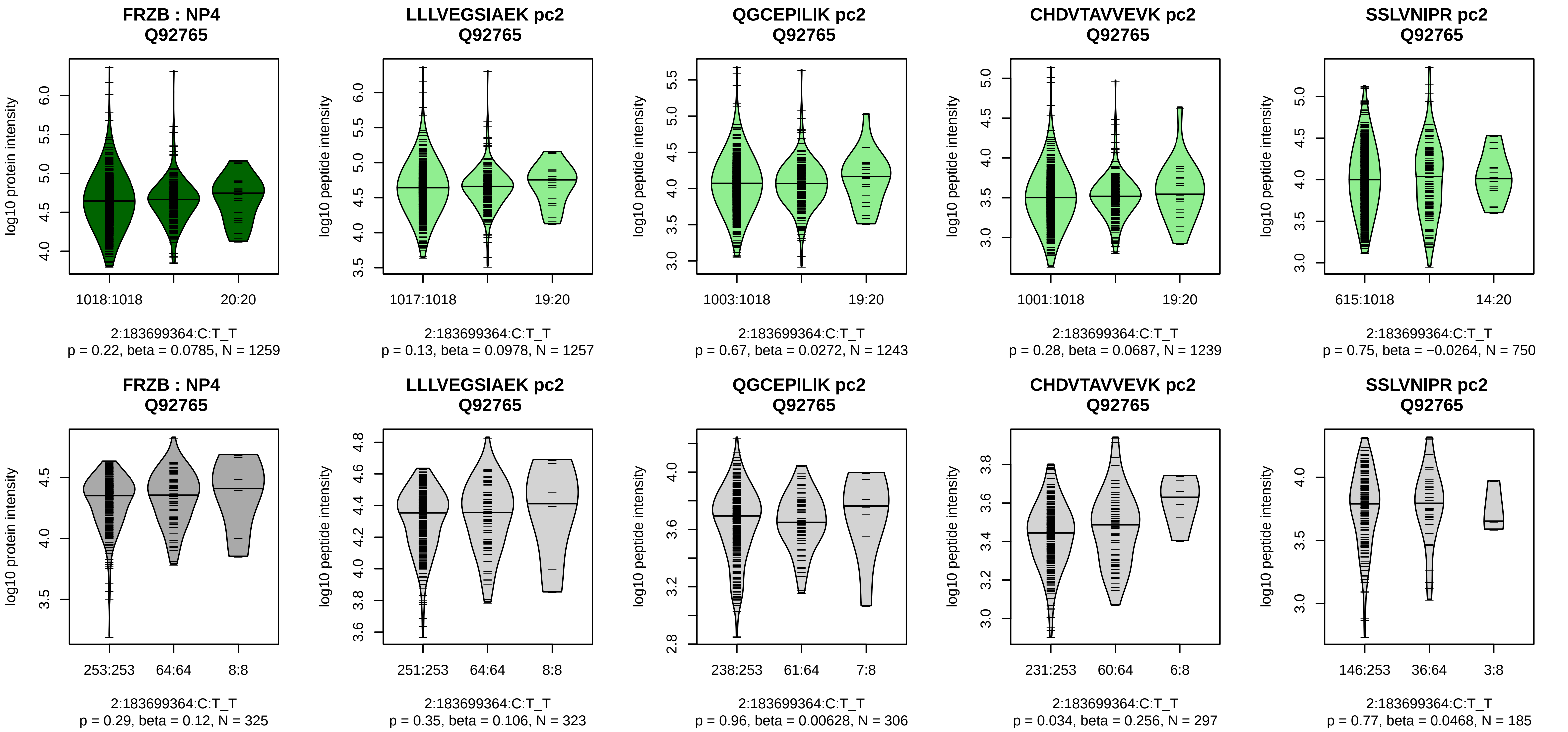


Assay Target: FRZB
Olink UniProt: Q92765
UKB rsID: rs13009
Proxy rsID: rs13009
UKB SNP: 2:183699364:C:T
Proxy SNP: 2:183699364:C:T
UKB log10(p): 1192.8
UKB BETA: −0.783
−:−:−:−:−:−:−:NA
1257:1243:1239:750:522:294:53

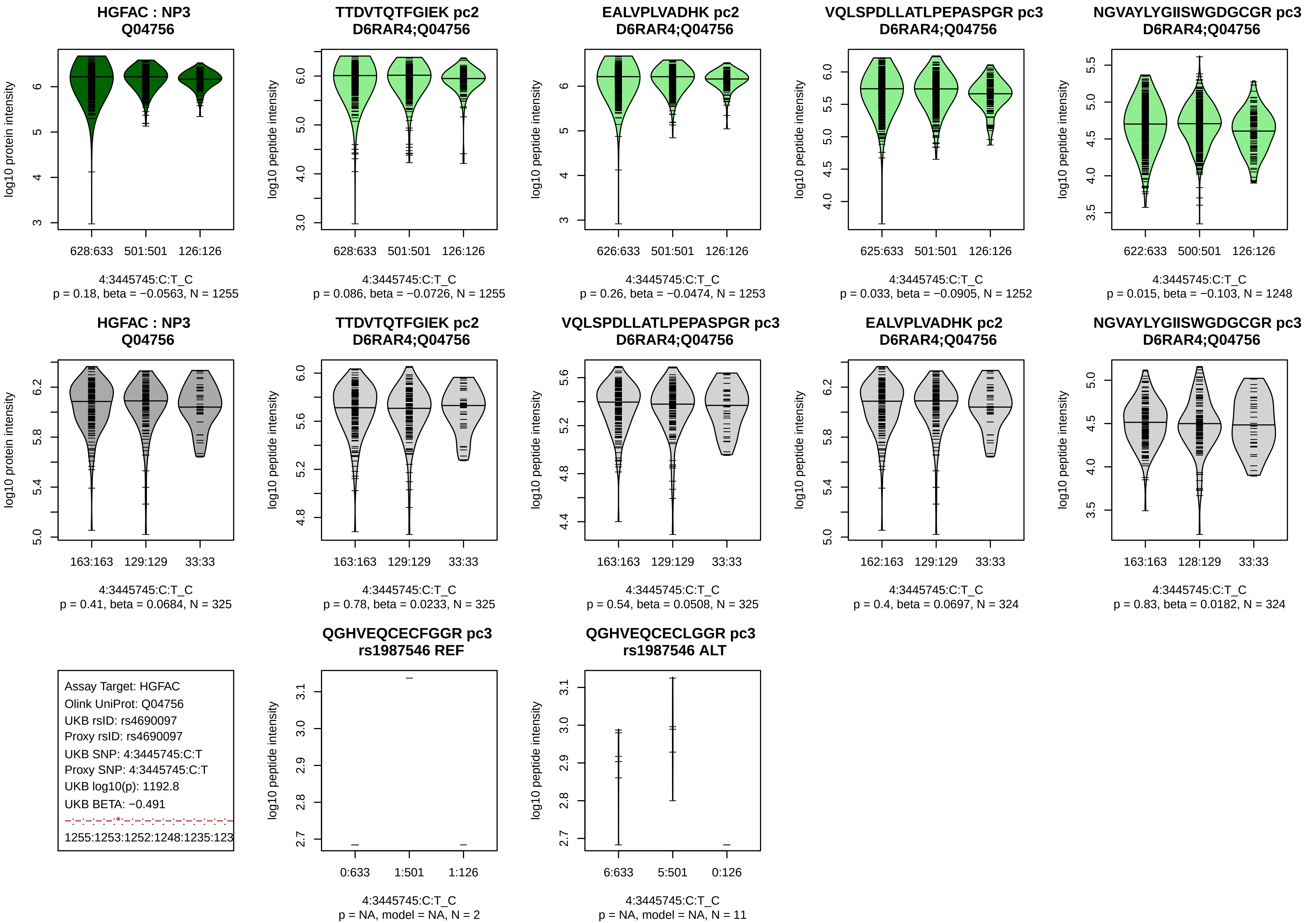
HGFAC : NP3
Q04756
log10 protein intensity
628:633
501:501
126:126
4:3445745:C:T_C
p = 0.18, beta = −0.0563, N = 1255
TTDVTQTFGIEK pc2
D6RAR4;Q04756
log10 peptide intensity
628:633
501:501
126:126
4:3445745:C:T_C
p = 0.086, beta = −0.0726, N = 1255
EALVPLVADHK pc2
D6RAR4;Q04756
log10 peptide intensity
626:633
501:501
126:126
4:3445745:C:T_C
p = 0.26, beta = −0.0474, N = 1253
VQLSPDLLATLPEPASPGR pc3
D6RAR4;Q04756
log10 peptide intensity
625:633
501:501
126:126
4:3445745:C:T_C
p = 0.033, beta = −0.0905, N = 1252
NGVAYLYGIISWGDGCGR pc3
D6RAR4;Q04756
log10 peptide intensity
622:633
500:501
126:126
4:3445745:C:T_C
p = 0.015, beta = −0.103, N = 1248
HGFAC : NP3
Q04756
log10 protein intensity
163:163
129:129
33:33
4:3445745:C:T_C
p = 0.41, beta = 0.0684, N = 325
TTDVTQTFGIEK pc2
D6RAR4;Q04756
log10 peptide intensity
163:163
129:129
33:33
4:3445745:C:T_C
p = 0.78, beta = 0.0233, N = 325
VQLSPDLLATLPEPASPGR pc3
D6RAR4;Q04756
log10 peptide intensity
163:163
129:129
33:33
4:3445745:C:T_C
p = 0.54, beta = 0.0508, N = 325
EALVPLVADHK pc2
D6RAR4;Q04756
log10 peptide intensity
162:163
129:129
33:33
4:3445745:C:T_C
p = 0.4, beta = 0.0697, N = 324
NGVAYLYGIISWGDGCGR pc3
D6RAR4;Q04756
log10 peptide intensity
163:163
128:129
33:33
4:3445745:C:T_C
p = 0.83, beta = 0.0182, N = 324
Assay Target: HGFAC
Olink UniProt: Q04756
UKB rsID: rs4690097
Proxy rsID: rs4690097
UKB SNP: 4:3445745:C:T
Proxy SNP: 4:3445745:C:T
UKB log10(p): 1192.8
UKB BETA: −0.491
−:−:−:−:−:*:−:−:−:−:−:−:−:−:−:−:−
1255:1253:1252:1248:1235:123
QGHVEQCECFGGR pc3
rs1987546 REF
log10 peptide intensity
0:633
1:501
1:126
4:3445745:C:T_C
p = NA, model = NA, N = 2
QGHVEQCECLGGR pc3
rs1987546 ALT
log10 peptide intensity
6:633
5:501
0:126
4:3445745:C:T_C
p = NA, model = NA, N = 11

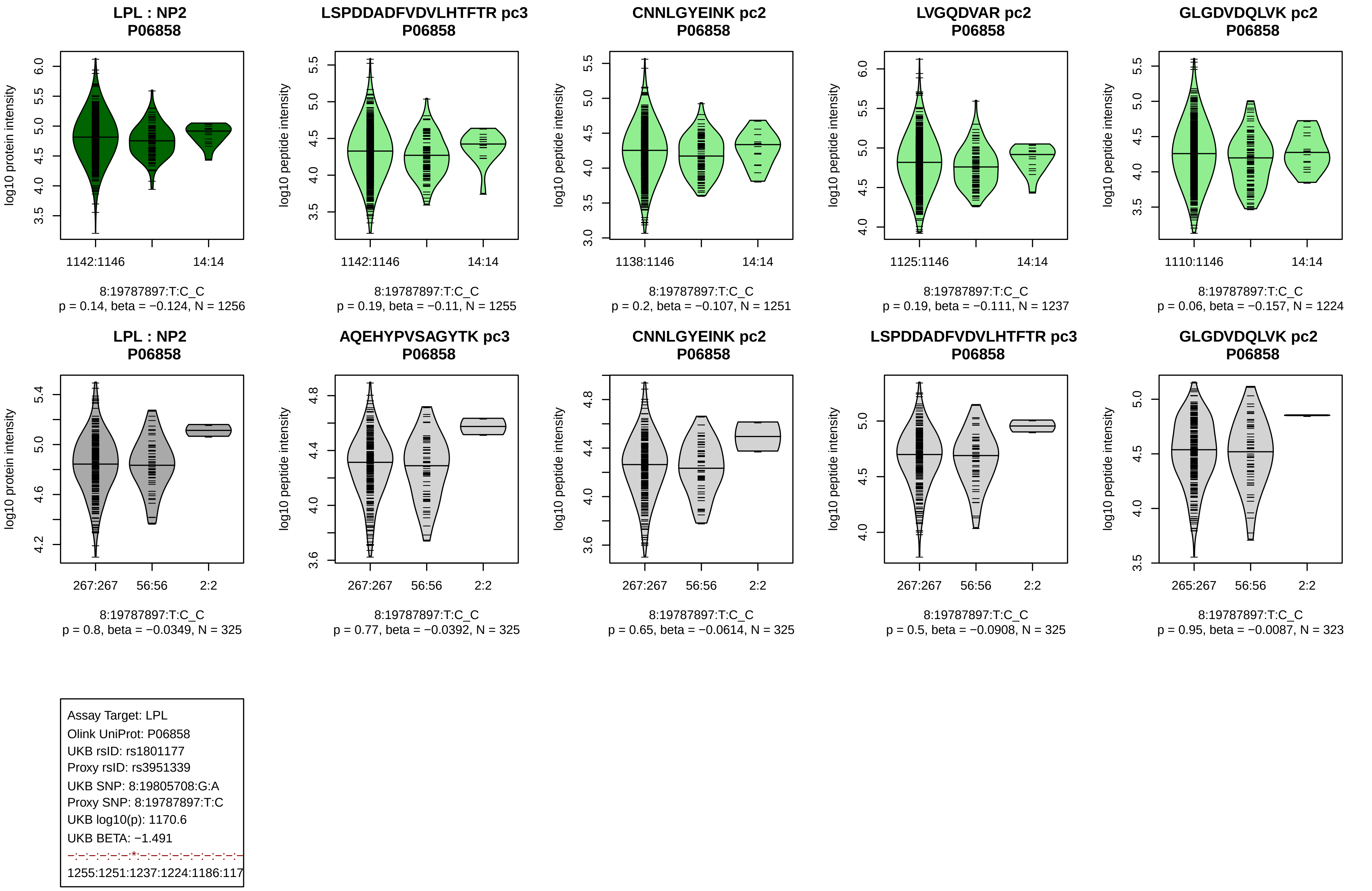
LPL : NP2
P06858
log10 protein intensity
1142:1146
14:14
8:19787897:T:C_C
p = 0.14, beta = −0.124, N = 1256
LSPDDADFVDVLHTFTR pc3
P06858
log10 peptide intensity
1142:1146
14:14
8:19787897:T:C_C
p = 0.19, beta = −0.11, N = 1255
CNNLGYEINK pc2
P06858
log10 peptide intensity
1138:1146
14:14
8:19787897:T:C_C
p = 0.2, beta = −0.107, N = 1251
LVGQDVAR pc2
P06858
log10 peptide intensity
1125:1146
14:14
8:19787897:T:C_C
p = 0.19, beta = −0.111, N = 1237
GLGDVDQLVK pc2
P06858
log10 peptide intensity
1110:1146
14:14
8:19787897:T:C_C
p = 0.06, beta = −0.157, N = 1224
LPL : NP2
P06858
log10 protein intensity
267:267
56:56
2:2
8:19787897:T:C_C
p = 0.8, beta = −0.0349, N = 325
AQEHYPVSAGYTK pc3
P06858
log10 peptide intensity
267:267
56:56
2:2
8:19787897:T:C_C
p = 0.77, beta = −0.0392, N = 325
CNNLGYEINK pc2
P06858
log10 peptide intensity
267:267
56:56
2:2
8:19787897:T:C_C
p = 0.65, beta = −0.0614, N = 325
LSPDDADFVDVLHTFTR pc3
P06858
log10 peptide intensity
267:267
56:56
2:2
8:19787897:T:C_C
p = 0.5, beta = −0.0908, N = 325
GLGDVDQLVK pc2
P06858
log10 peptide intensity
265:267
56:56
2:2
8:19787897:T:C_C
p = 0.95, beta = −0.0087, N = 323
Assay Target: LPL
Olink UniProt: P06858
UKB rsID: rs1801177
Proxy rsID: rs3951339
UKB SNP: 8:19805708:G:A
Proxy SNP: 8:19787897:T:C
UKB log10(p): 1170.6
UKB BETA: −1.491
−:−:−:−:−:−:*:−:−:−:−:−:−:−:−:−:−:−
1255:1251:1237:1224:1186:117

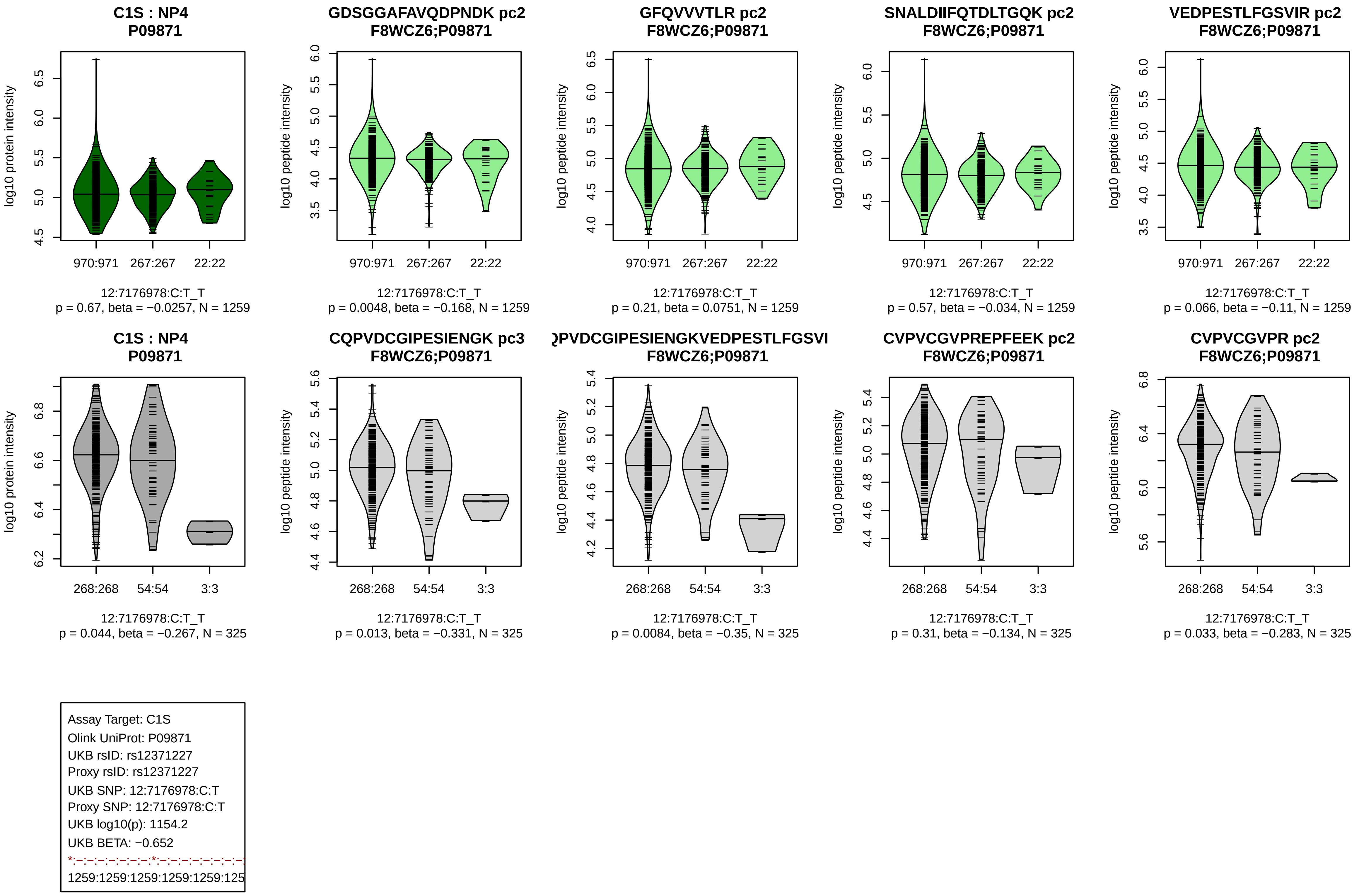

C1S : NP4
P09871
GDSGGAFAVQDPNDK pc2
F8WCZ6;P09871
GFQVVVTLR pc2
F8WCZ6;P09871
SNALDIIFQTDLTGQK pc2
F8WCZ6;P09871
VEDPESTLFGSVIR pc2
F8WCZ6;P09871
log10 protein intensity
log10 peptide intensity
970:971
267:267
22:22
12:7176978:C:T_T
p = 0.67, beta = −0.0257, N = 1259
p = 0.0048, beta = −0.168, N = 1259
p = 0.21, beta = 0.0751, N = 1259
p = 0.57, beta = −0.034, N = 1259
p = 0.066, beta = −0.11, N = 1259
C1S : NP4
P09871
CQPVDCGIPESIENGK pc3
F8WCZ6;P09871
QPVDCGIPESIENGKVEDPESTLFGSVI
F8WCZ6;P09871
CVPVCGVPREPFEEK pc2
F8WCZ6;P09871
CVPVCGVPR pc2
F8WCZ6;P09871
268:268
54:54
3:3
p = 0.044, beta = −0.267, N = 325
p = 0.013, beta = −0.331, N = 325
p = 0.0084, beta = −0.35, N = 325
p = 0.31, beta = −0.134, N = 325
p = 0.033, beta = −0.283, N = 325
Assay Target: C1S
Olink UniProt: P09871
UKB rsID: rs12371227
Proxy rsID: rs12371227
UKB SNP: 12:7176978:C:T
Proxy SNP: 12:7176978:C:T
UKB log10(p): 1154.2
UKB BETA: −0.652
1259:1259:1259:1259:1259:125

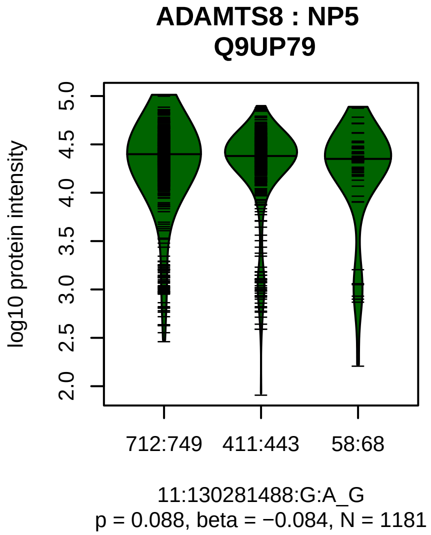


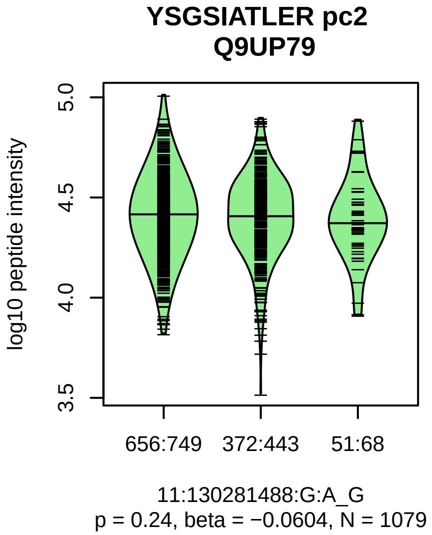


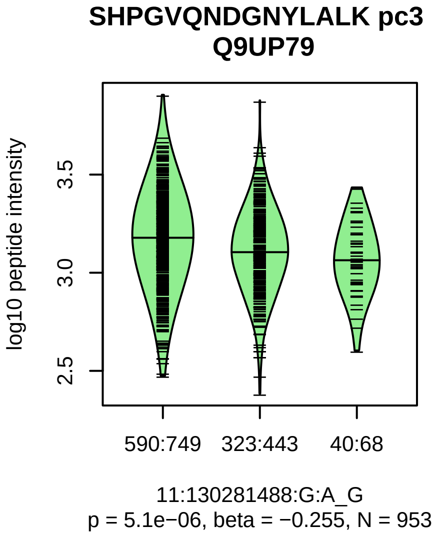


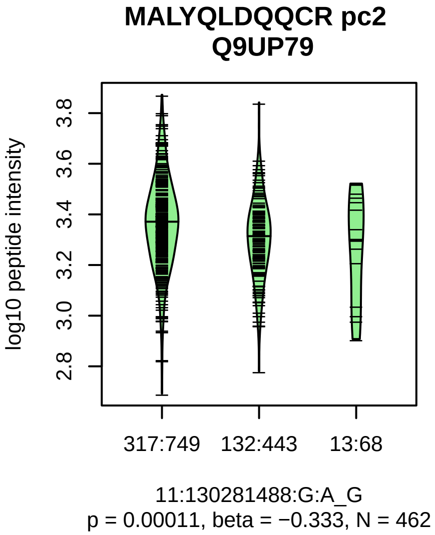


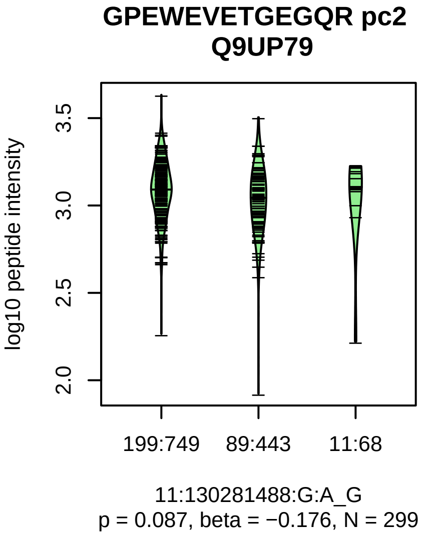


Assay Target: ADAMTS8
Olink UniProt: Q9UP79
UKB rsID: rs2131535
Proxy rsID: rs2131535
UKB SNP: 11:130281488:G:A
Proxy SNP: 11:130281488:G:A
UKB log10(p): 1117.7
UKB BETA: 0.494
–:*:*:–:*
1079:953:462:299:270

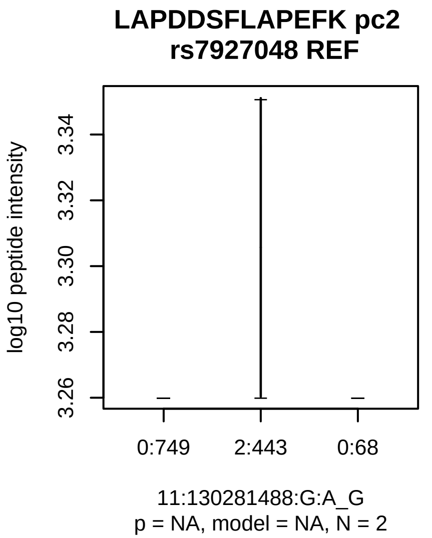

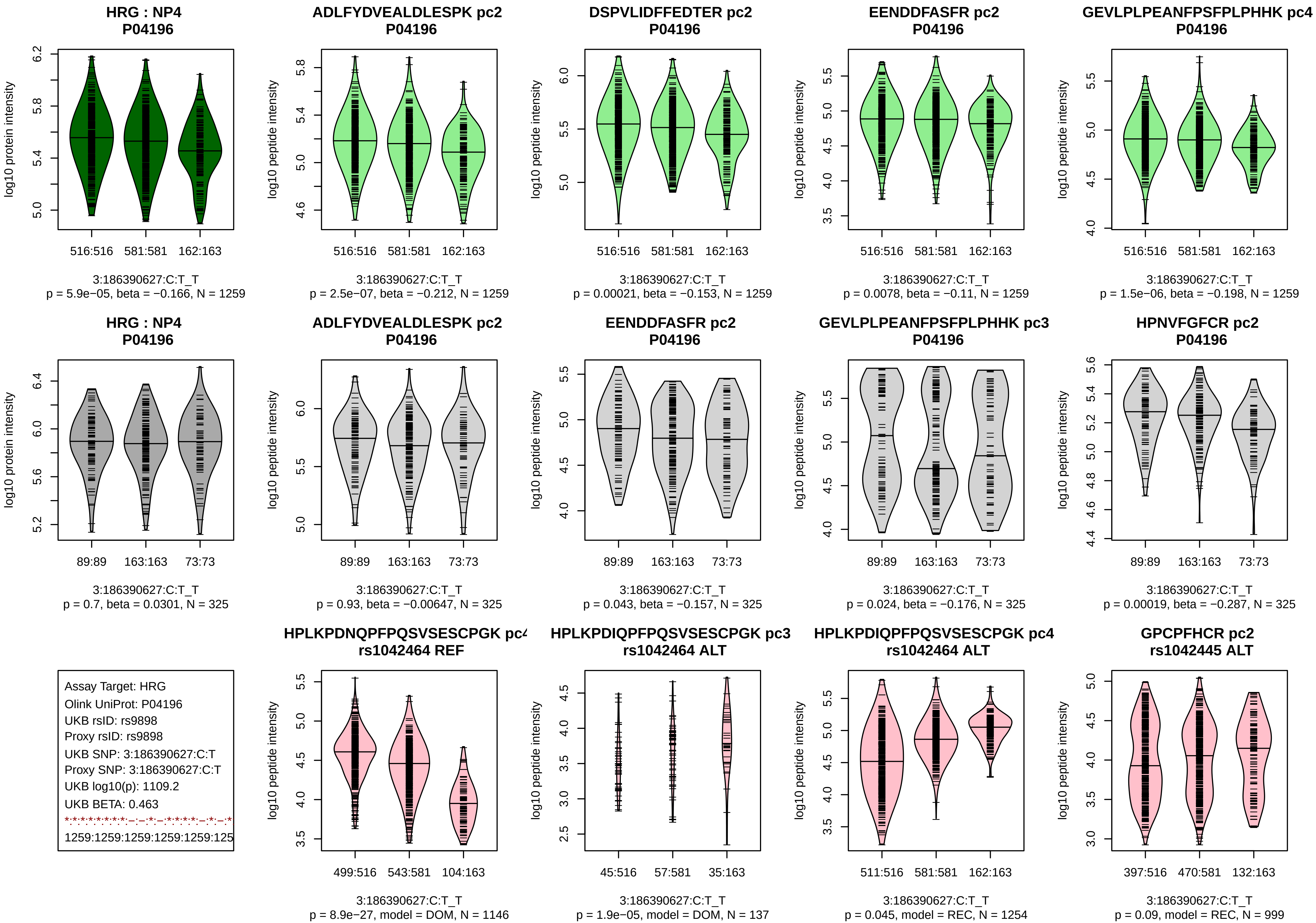
HRG : NP4
P04196
log10 protein intensity
516:516
581:581
162:163
3:186390627:C:T_T
p = 5.9e−05, beta = −0.166, N = 1259
ADLFYDVEALDLESPK pc2
P04196
log10 peptide intensity
516:516
581:581
162:163
3:186390627:C:T_T
p = 2.5e−07, beta = −0.212, N = 1259
DSPVLIDFFEDTER pc2
P04196
log10 peptide intensity
516:516
581:581
162:163
3:186390627:C:T_T
p = 0.00021, beta = −0.153, N = 1259
EENDDFASFR pc2
P04196
log10 peptide intensity
516:516
581:581
162:163
3:186390627:C:T_T
p = 0.0078, beta = −0.11, N = 1259
GEVLPLPEANFPSFPLPHHK pc4
P04196
log10 peptide intensity
516:516
581:581
162:163
3:186390627:C:T_T
p = 1.5e−06, beta = −0.198, N = 1259
HRG : NP4
P04196
log10 protein intensity
89:89
163:163
73:73
3:186390627:C:T_T
p = 0.7, beta = 0.0301, N = 325
ADLFYDVEALDLESPK pc2
P04196
log10 peptide intensity
89:89
163:163
73:73
3:186390627:C:T_T
p = 0.93, beta = −0.00647, N = 325
EENDDFASFR pc2
P04196
log10 peptide intensity
89:89
163:163
73:73
3:186390627:C:T_T
p = 0.043, beta = −0.157, N = 325
GEVLPLPEANFPSFPLPHHK pc3
P04196
log10 peptide intensity
89:89
163:163
73:73
3:186390627:C:T_T
p = 0.024, beta = −0.176, N = 325
HPNVFGFCR pc2
P04196
log10 peptide intensity
89:89
163:163
73:73
3:186390627:C:T_T
p = 0.00019, beta = −0.287, N = 325
Assay Target: HRG
Olink UniProt: P04196
UKB rsID: rs9898
Proxy rsID: rs9898
UKB SNP: 3:186390627:C:T
Proxy SNP: 3:186390627:C:T
UKB log10(p): 1109.2
UKB BETA: 0.463
1259:1259:1259:1259:1259:125
HPLKPDNQPFPQSVSESCPGK pc4
rs1042464 REF
log10 peptide intensity
499:516
543:581
104:163
3:186390627:C:T_T
p = 8.9e−27, model = DOM, N = 1146
HPLKPDIQPFPQSVSESCPGK pc3
rs1042464 ALT
log10 peptide intensity
45:516
57:581
35:163
3:186390627:C:T_T
p = 1.9e−05, model = DOM, N = 137
HPLKPDIQPFPQSVSESCPGK pc4
rs1042464 ALT
log10 peptide intensity
511:516
581:581
162:163
3:186390627:C:T_T
p = 0.045, model = REC, N = 1254
GPCPFHCR pc2
rs1042445 ALT
log10 peptide intensity
397:516
470:581
132:163
3:186390627:C:T_T
p = 0.09, model = REC, N = 999

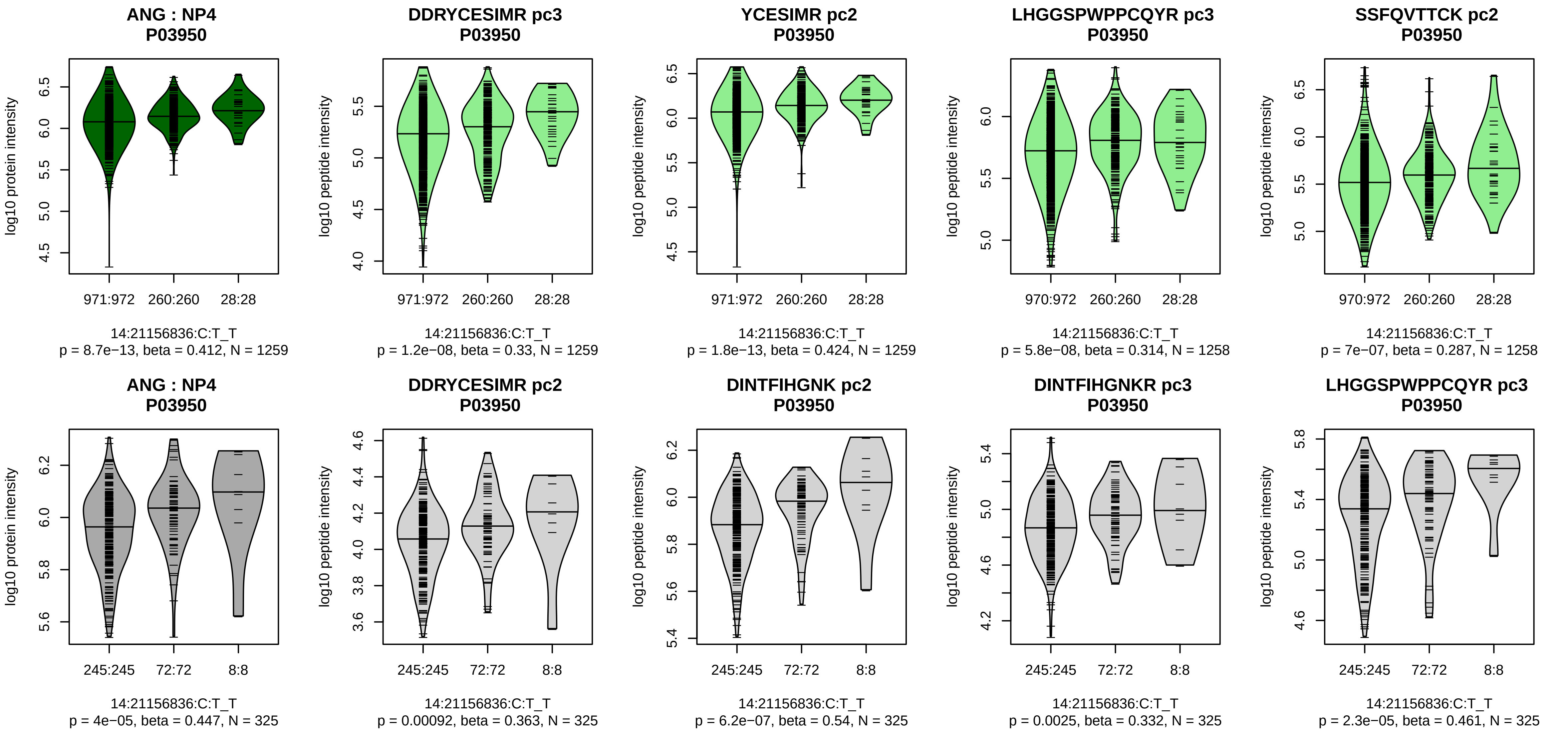


Assay Target: ANG
Olink UniProt: P03950
UKB rsID: rs4982325
Proxy rsID: rs4982325
UKB SNP: 14:21156836:C:T
Proxy SNP: 14:21156836:C:T
UKB log10(p): 1100.5
UKB BETA: 0.652
*:*:*:*:*:*:*:*:–:*:–:*:–:*:–:*:–
1259:1259:1258:1258:1253:124

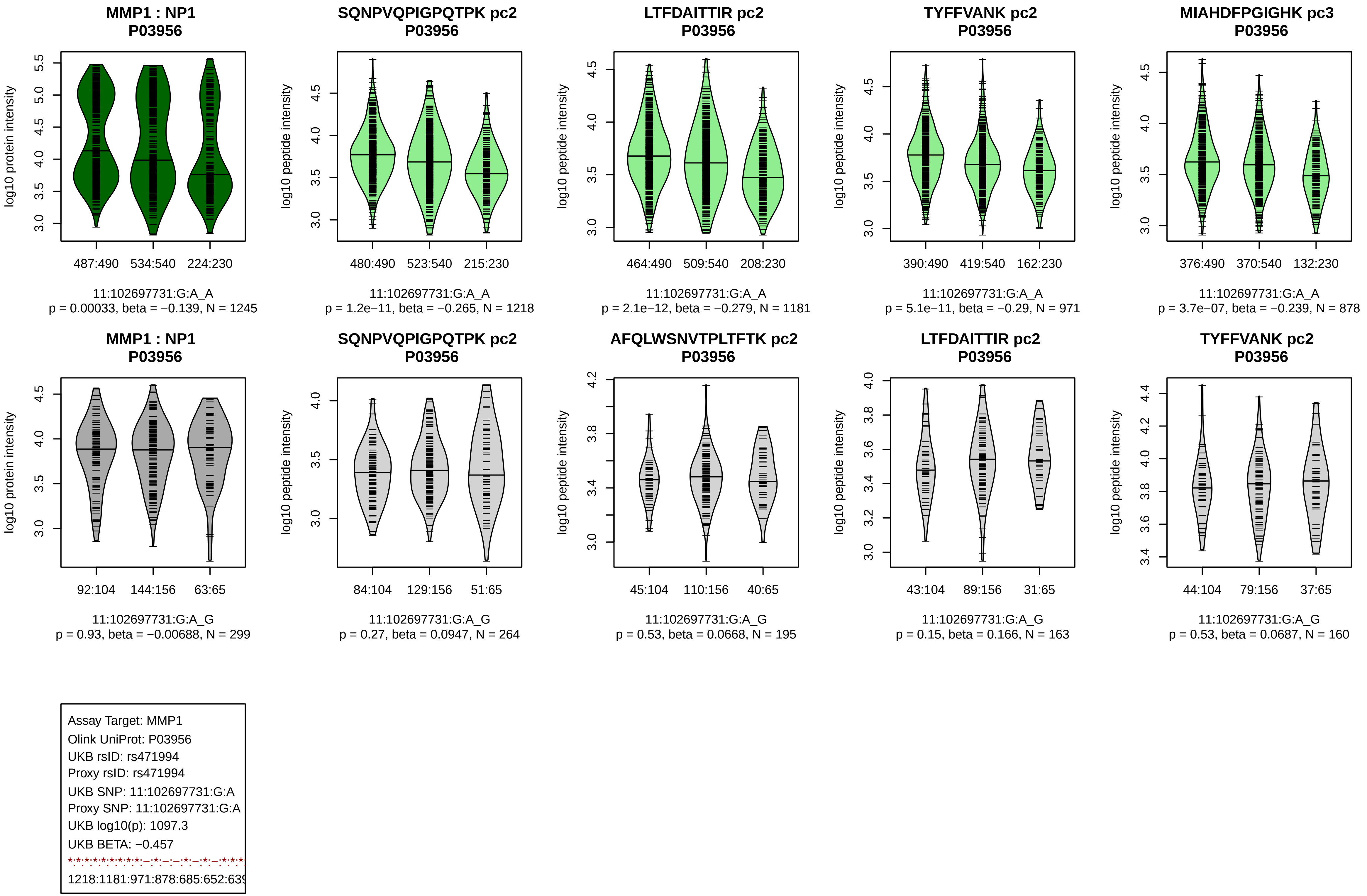

MMP1 : NP1
P03956
log10 protein intensity
487:490
534:540
224:230
11:102697731:G:A_A
p = 0.00033, beta = −0.139, N = 1245
SQNPVQPIGPQTPK pc2
P03956
log10 peptide intensity
480:490
523:540
215:230
11:102697731:G:A_A
p = 1.2e−11, beta = −0.265, N = 1218
LTFDAITTIR pc2
P03956
log10 peptide intensity
464:490
509:540
208:230
11:102697731:G:A_A
p = 2.1e−12, beta = −0.279, N = 1181
TYFFVANK pc2
P03956
log10 peptide intensity
390:490
419:540
162:230
11:102697731:G:A_A
p = 5.1e−11, beta = −0.29, N = 971
MIAHDFPGIGHK pc3
P03956
log10 peptide intensity
376:490
370:540
132:230
11:102697731:G:A_A
p = 3.7e−07, beta = −0.239, N = 878
MMP1 : NP1
P03956
log10 protein intensity
92:104
144:156
63:65
11:102697731:G:A_G
p = 0.93, beta = −0.00688, N = 299
SQNPVQPIGPQTPK pc2
P03956
log10 peptide intensity
84:104
129:156
51:65
11:102697731:G:A_G
p = 0.27, beta = 0.0947, N = 264
AFQLWSNVTPLTFTK pc2
P03956
log10 peptide intensity
45:104
110:156
40:65
11:102697731:G:A_G
p = 0.53, beta = 0.0668, N = 195
LTFDAITTIR pc2
P03956
log10 peptide intensity
43:104
89:156
31:65
11:102697731:G:A_G
p = 0.15, beta = 0.166, N = 163
TYFFVANK pc2
P03956
log10 peptide intensity
44:104
79:156
37:65
11:102697731:G:A_G
p = 0.53, beta = 0.0687, N = 160
Assay Target: MMP1
Olink UniProt: P03956
UKB rsID: rs471994
Proxy rsID: rs471994
UKB SNP: 11:102697731:G:A
Proxy SNP: 11:102697731:G:A
UKB log10(p): 1097.3
UKB BETA: −0.457
1218:1181:971:878:685:652:63

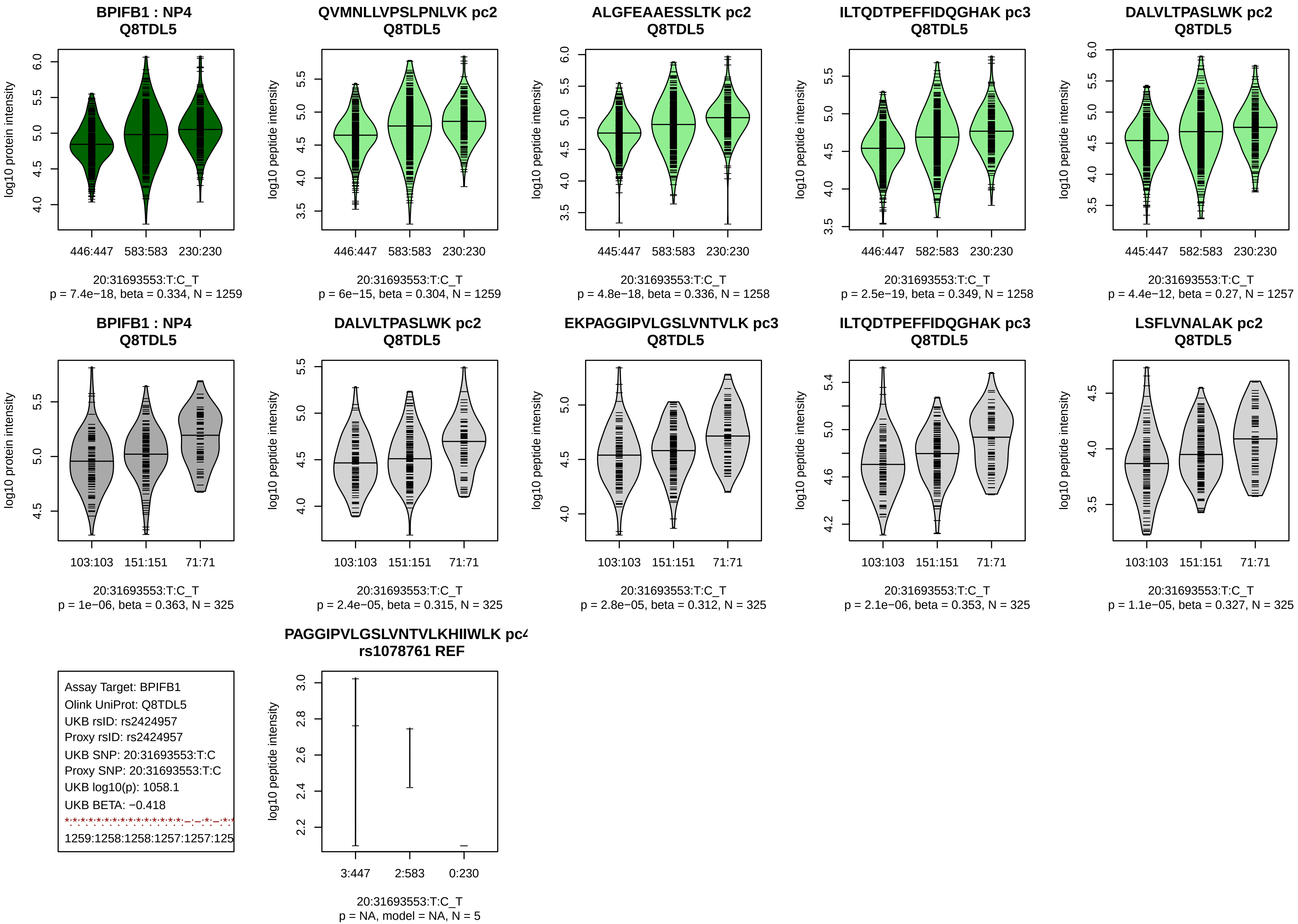

BPIFB1 : NP4
Q8TDL5
log10 protein intensity
446:447
583:583
230:230
20:31693553:T:C_T
p = 7.4e−18, beta = 0.334, N = 1259
QVMNLLVPSLPNLVK pc2
Q8TDL5
log10 peptide intensity
446:447
583:583
230:230
20:31693553:T:C_T
p = 6e−15, beta = 0.304, N = 1259
ALGFEAAESSLTK pc2
Q8TDL5
log10 peptide intensity
445:447
583:583
230:230
20:31693553:T:C_T
p = 4.8e−18, beta = 0.336, N = 1258
ILTQDTPEFFIDQGHAK pc3
Q8TDL5
log10 peptide intensity
446:447
582:583
230:230
20:31693553:T:C_T
p = 2.5e−19, beta = 0.349, N = 1258
DALVLTPASLWK pc2
Q8TDL5
log10 peptide intensity
445:447
582:583
230:230
20:31693553:T:C_T
p = 4.4e−12, beta = 0.27, N = 1257
BPIFB1 : NP4
Q8TDL5
log10 protein intensity
103:103
151:151
71:71
20:31693553:T:C_T
p = 1e−06, beta = 0.363, N = 325
DALVLTPASLWK pc2
Q8TDL5
log10 peptide intensity
103:103
151:151
71:71
20:31693553:T:C_T
p = 2.4e−05, beta = 0.315, N = 325
EKPAGGIPVLGSLVNTVLK pc3
Q8TDL5
log10 peptide intensity
103:103
151:151
71:71
20:31693553:T:C_T
p = 2.8e−05, beta = 0.312, N = 325
ILTQDTPEFFIDQGHAK pc3
Q8TDL5
log10 peptide intensity
103:103
151:151
71:71
20:31693553:T:C_T
p = 2.1e−06, beta = 0.353, N = 325
LSFLVNALAK pc2
Q8TDL5
log10 peptide intensity
103:103
151:151
71:71
20:31693553:T:C_T
p = 1.1e−05, beta = 0.327, N = 325
Assay Target: BPIFB1
Olink UniProt: Q8TDL5
UKB rsID: rs2424957
Proxy rsID: rs2424957
UKB SNP: 20:31693553:T:C
Proxy SNP: 20:31693553:T:C
UKB log10(p): 1058.1
UKB BETA: −0.418
1259:1258:1258:1257:1257:125
PAGGIPVLGSLVNTVLKHIIWLK pc4
rs1078761 REF
log10 peptide intensity
3:447
2:583
0:230
20:31693553:T:C_T
p = NA, model = NA, N = 5

**NOMO1 : NP1**
**Q15155**

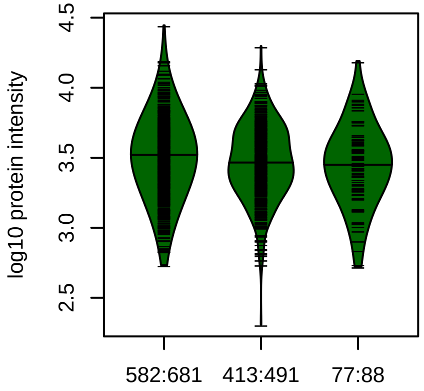


16:16294465:T:C_T
p = 0.11, beta = −0.0785, N = 1072

**GQPLGPAGVQVSLR pc2**
**\0A7I2V4T6;A0A7I2V686;A0A7I2V6B0**

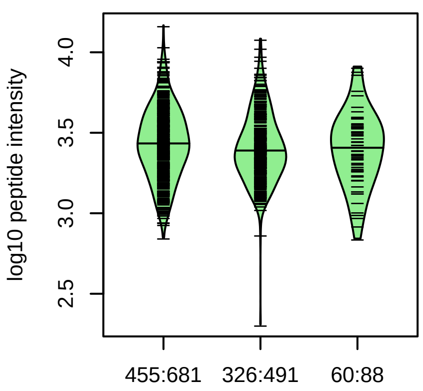


16:16294465:T:C_T
p = 0.28, beta = −0.0586, N = 841

**FLLFSSLVTK pc2**
**2V4T6;A0A7I2V686;A0A7I2V6B0;J3KN**

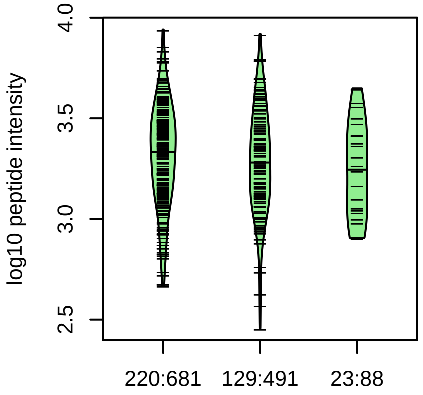


16:16294465:T:C_T
p = 0.13, beta = −0.128, N = 372

**VQVMVPEAETR pc2**
**\A0A7I2V4T6;A0A7I2V6B0;J3KN36;P6**

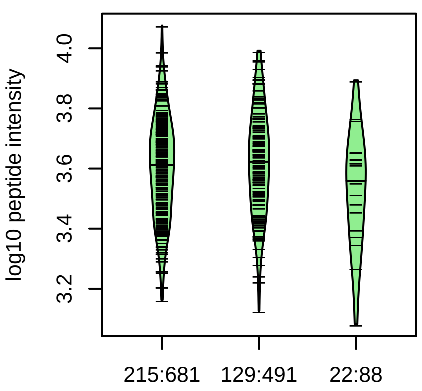


16:16294465:T:C_T
p = 0.66, beta = −0.0369, N = 366

**SQYDYILPQVSFTAVGYHK pc3**
**2V4T6;A0A7I2V6B0;J3KN36;P69849;**

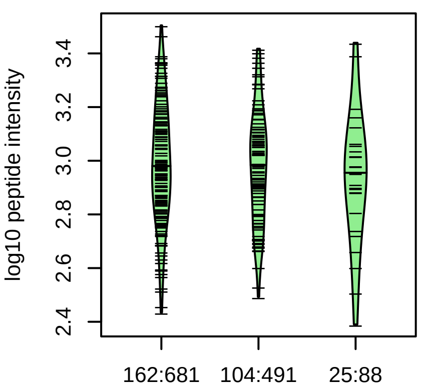


16:16294465:T:C_T
p = 0.61, beta = −0.0452, N = 291

**NOMO1 : NP1**
**Q15155**

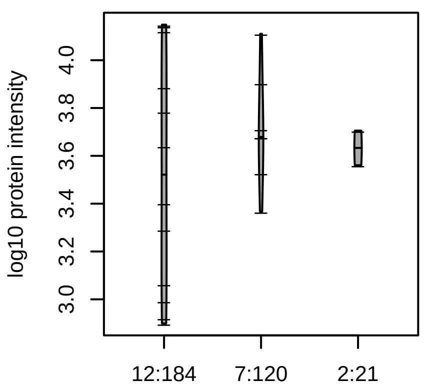


16:16294465:T:C_T
p = 0.87, beta = 0.049, N = 21

Assay Target: NOMO1
Olink UniProt: Q15155
UKB rsID: rs3891245
Proxy rsID: rs7184653
UKB SNP: 16:16348287:T:G
Proxy SNP: 16:16294465:T:C
UKB log10(p): 1051.8
UKB BETA: −0.503
−:−:−:−:−:−:−:−:−:−:−:−:−:−:−
841:372:366:291:252:244:216:1

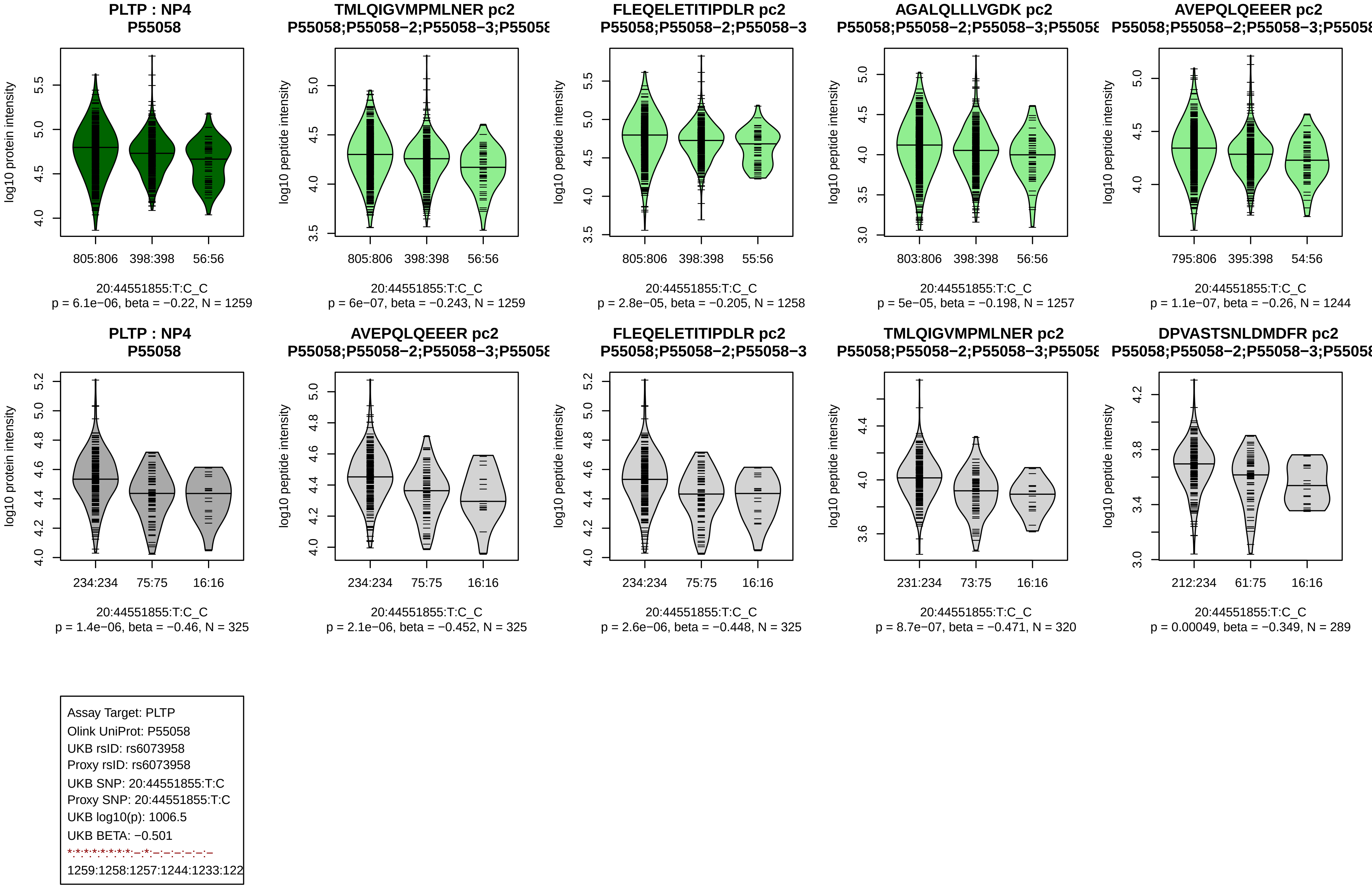
PLTP : NP4
P55058
log10 protein intensity
805:806
398:398
56:56
20:44551855:T:C_C
p = 6.1e−06, beta = −0.22, N = 1259
TMLQIGVMPMLNER pc2
P55058;P55058−2;P55058−3;P5505
log10 peptide intensity
805:806
398:398
56:56
20:44551855:T:C_C
p = 6e−07, beta = −0.243, N = 1259
FLEQELETITIPDLR pc2
P55058;P55058−2;P55058−3
log10 peptide intensity
805:806
398:398
55:56
20:44551855:T:C_C
p = 2.8e−05, beta = −0.205, N = 1258
AGALQLLLVGDK pc2
P55058;P55058−2;P55058−3;P5505
log10 peptide intensity
803:806
398:398
56:56
20:44551855:T:C_C
p = 5e−05, beta = −0.198, N = 1257
AVEPQLQEEER pc2
P55058;P55058−2;P55058−3;P5505
log10 peptide intensity
795:806
395:398
54:56
20:44551855:T:C_C
p = 1.1e−07, beta = −0.26, N = 1244
PLTP : NP4
P55058
log10 protein intensity
234:234
75:75
16:16
20:44551855:T:C_C
p = 1.4e−06, beta = −0.46, N = 325
AVEPQLQEEER pc2
P55058;P55058−2;P55058−3;P5505
log10 peptide intensity
234:234
75:75
16:16
20:44551855:T:C_C
p = 2.1e−06, beta = −0.452, N = 325
FLEQELETITIPDLR pc2
P55058;P55058−2;P55058−3
log10 peptide intensity
234:234
75:75
16:16
20:44551855:T:C_C
p = 2.6e−06, beta = −0.448, N = 325
TMLQIGVMPMLNER pc2
P55058;P55058−2;P55058−3;P5505
log10 peptide intensity
231:234
73:75
16:16
20:44551855:T:C_C
p = 8.7e−07, beta = −0.471, N = 320
DPVASTSNLDMDFR pc2
P55058;P55058−2;P55058−3;P5505
log10 peptide intensity
212:234
61:75
16:16
20:44551855:T:C_C
p = 0.00049, beta = −0.349, N = 289
Assay Target: PLTP
Olink UniProt: P55058
UKB rsID: rs6073958
Proxy rsID: rs6073958
UKB SNP: 20:44551855:T:C
Proxy SNP: 20:44551855:T:C
UKB log10(p): 1006.5
UKB BETA: −0.501
*:*:*:*:*:*:*:*:–:*:–:–:–:–:–:–
1259:1258:1257:1244:1233:122

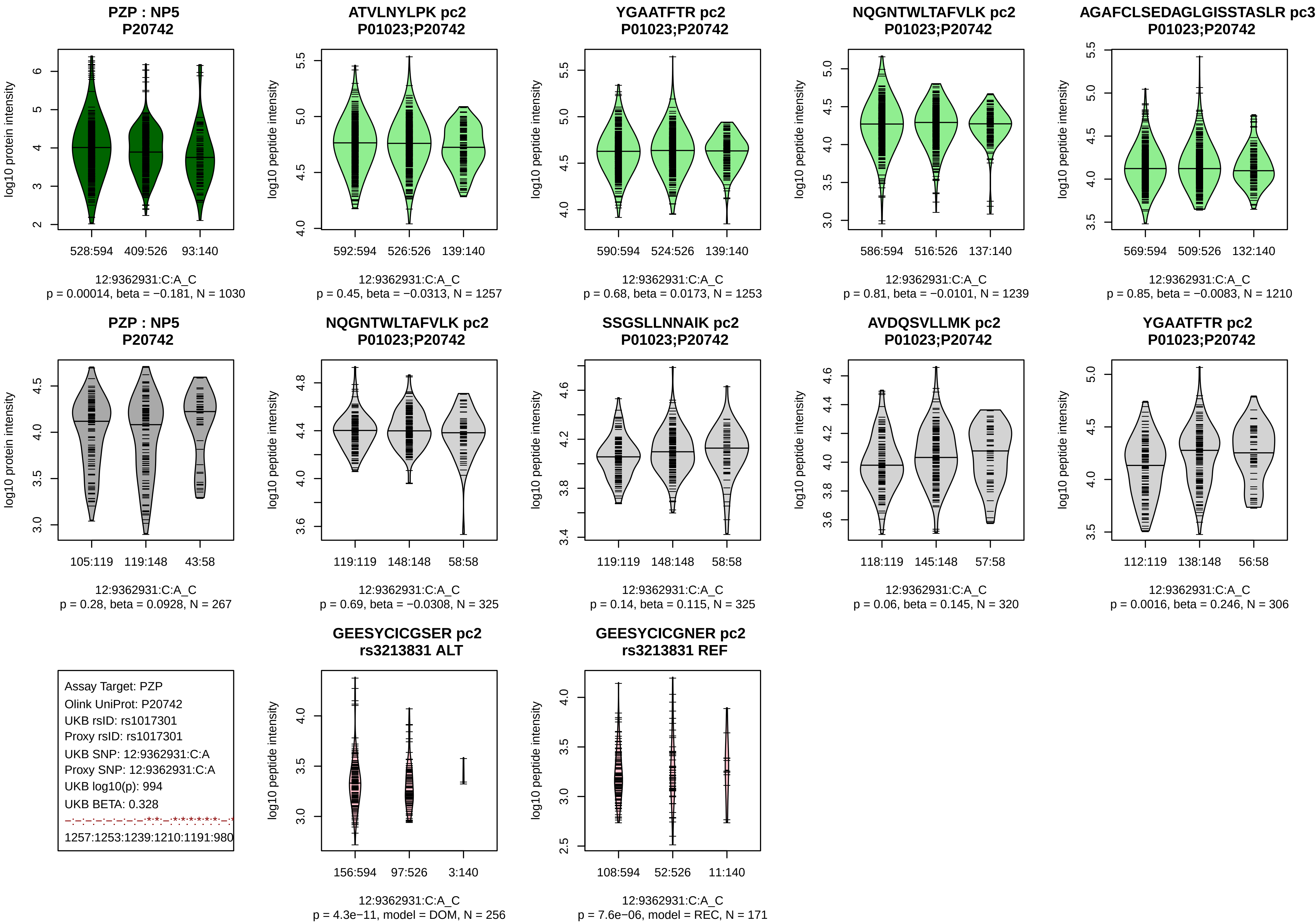

PZP : NP5
P20742
log10 protein intensity
528:594 409:526 93:140
12:9362931:C:A_C
p = 0.00014, beta = −0.181, N = 1030
ATVLNYLPK pc2
P01023;P20742
log10 peptide intensity
592:594 526:526 139:140
12:9362931:C:A_C
p = 0.45, beta = −0.0313, N = 1257
YGAATFTR pc2
P01023;P20742
590:594 524:526 139:140
12:9362931:C:A_C
p = 0.68, beta = 0.0173, N = 1253
NQGNTWLTAFVLK pc2
P01023;P20742
586:594 516:526 137:140
12:9362931:C:A_C
p = 0.81, beta = −0.0101, N = 1239
AGAFCLSEDAGLGISSTASLR pc3
P01023;P20742
569:594 509:526 132:140
12:9362931:C:A_C
p = 0.85, beta = −0.0083, N = 1210
PZP : NP5
P20742
105:119 119:148 43:58
12:9362931:C:A_C
p = 0.28, beta = 0.0928, N = 267
NQGNTWLTAFVLK pc2
P01023;P20742
119:119 148:148 58:58
12:9362931:C:A_C
p = 0.69, beta = −0.0308, N = 325
SSGSLLNNAIK pc2
P01023;P20742
119:119 148:148 58:58
12:9362931:C:A_C
p = 0.14, beta = 0.115, N = 325
AVDQSVLLMK pc2
P01023;P20742
118:119 145:148 57:58
12:9362931:C:A_C
p = 0.06, beta = 0.145, N = 320
YGAATFTR pc2
P01023;P20742
112:119 138:148 56:58
12:9362931:C:A_C
p = 0.0016, beta = 0.246, N = 306
Assay Target: PZP
Olink UniProt: P20742
UKB rsID: rs1017301
Proxy rsID: rs1017301
UKB SNP: 12:9362931:C:A
Proxy SNP: 12:9362931:C:A
UKB log10(p): 994
UKB BETA: 0.328
1257:1253:1239:1210:1191:980
GEESYCICGSER pc2
rs3213831 ALT
156:594 97:526 3:140
12:9362931:C:A_C
p = 4.3e−11, model = DOM, N = 256
GEESYCICGNER pc2
rs3213831 REF
108:594 52:526 11:140
12:9362931:C:A_C
p = 7.6e−06, model = REC, N = 171

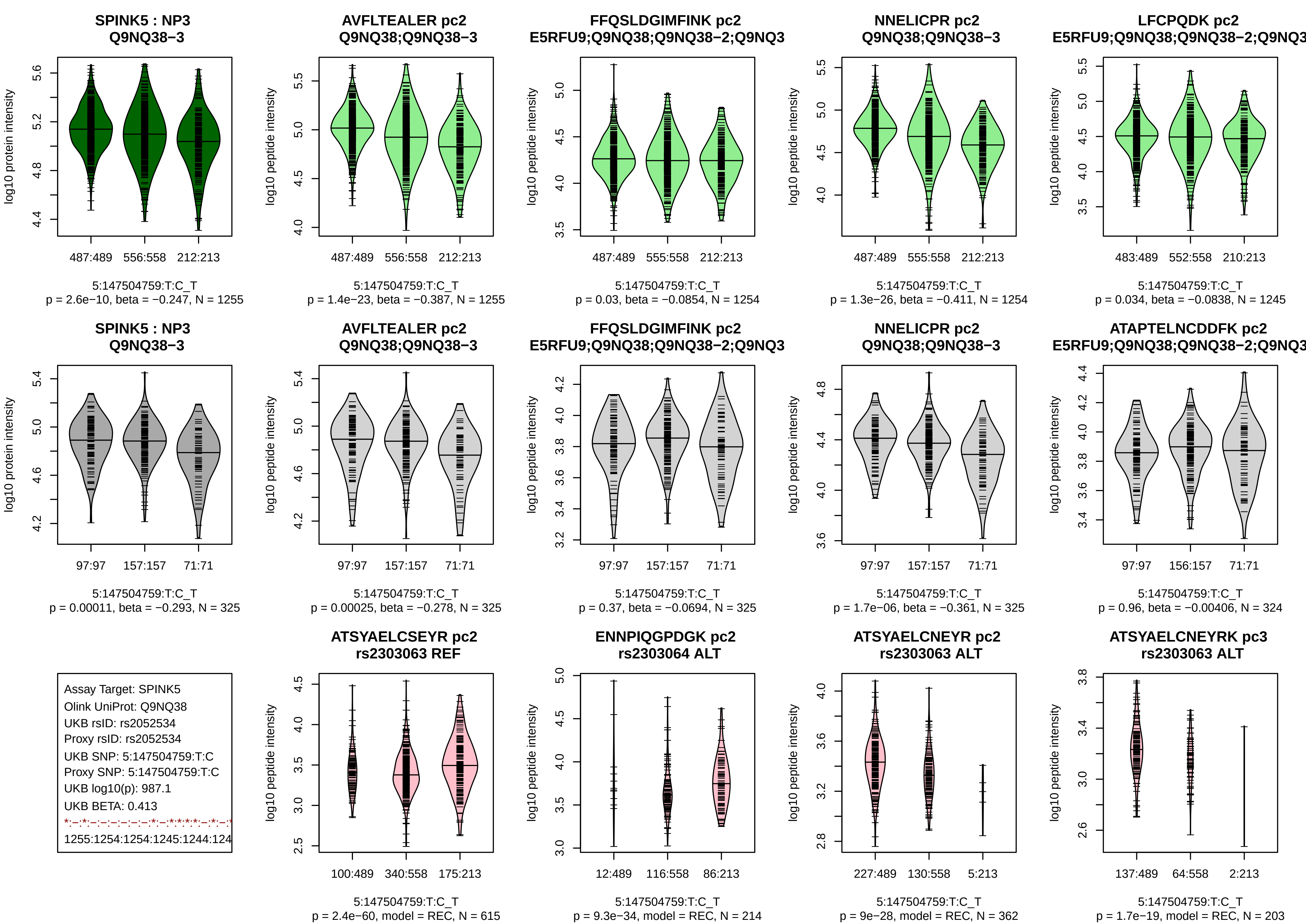

SPINK5 : NP3
Q9NQ38−3
log10 protein intensity
487:489 556:558 212:213
5:147504759:T:C_T
p = 2.6e−10, beta = −0.247, N = 1255
AVFLTEALER pc2
Q9NQ38;Q9NQ38−3
log10 peptide intensity
487:489 556:558 212:213
5:147504759:T:C_T
p = 1.4e−23, beta = −0.387, N = 1255
FFQSLDGIMFINK pc2
E5RFU9;Q9NQ38;Q9NQ38−2;Q9NQ3
log10 peptide intensity
487:489 555:558 212:213
5:147504759:T:C_T
p = 0.03, beta = −0.0854, N = 1254
NNELICPR pc2
Q9NQ38;Q9NQ38−3
log10 peptide intensity
487:489 555:558 212:213
5:147504759:T:C_T
p = 1.3e−26, beta = −0.411, N = 1254
LFCPQDK pc2
E5RFU9;Q9NQ38;Q9NQ38−2;Q9NQ3
log10 peptide intensity
483:489 552:558 210:213
5:147504759:T:C_T
p = 0.034, beta = −0.0838, N = 1245
SPINK5 : NP3
Q9NQ38−3
log10 protein intensity
97:97 157:157 71:71
5:147504759:T:C_T
p = 0.00011, beta = −0.293, N = 325
AVFLTEALER pc2
Q9NQ38;Q9NQ38−3
log10 peptide intensity
97:97 157:157 71:71
5:147504759:T:C_T
p = 0.00025, beta = −0.278, N = 325
FFQSLDGIMFINK pc2
E5RFU9;Q9NQ38;Q9NQ38−2;Q9NQ3
log10 peptide intensity
97:97 157:157 71:71
5:147504759:T:C_T
p = 0.37, beta = −0.0694, N = 325
NNELICPR pc2
Q9NQ38;Q9NQ38−3
log10 peptide intensity
97:97 157:157 71:71
5:147504759:T:C_T
p = 1.7e−06, beta = −0.361, N = 325
ATAPTELNCDDFK pc2
E5RFU9;Q9NQ38;Q9NQ38−2;Q9NQ3
log10 peptide intensity
97:97 156:157 71:71
5:147504759:T:C_T
p = 0.96, beta = −0.00406, N = 324
Assay Target: SPINK5
Olink UniProt: Q9NQ38
UKB rsID: rs2052534
Proxy rsID: rs2052534
UKB SNP: 5:147504759:T:C
Proxy SNP: 5:147504759:T:C
UKB log10(p): 987.1
UKB BETA: 0.413
*:−:*:−:−:−:−:−:*:−:*:*:*:*:−:*:−:*
1255:1254:1254:1245:1244:124
ATSYAELCSEYR pc2
rs2303063 REF
log10 peptide intensity
100:489 340:558 175:213
5:147504759:T:C_T
p = 2.4e−60, model = REC, N = 615
ENNPIQGPDGK pc2
rs2303064 ALT
log10 peptide intensity
12:489 116:558 86:213
5:147504759:T:C_T
p = 9.3e−34, model = REC, N = 214
ATSYAELCNEYR pc2
rs2303063 ALT
log10 peptide intensity
227:489 130:558 5:213
5:147504759:T:C_T
p = 9e−28, model = REC, N = 362
ATSYAELCNEYRK pc3
rs2303063 ALT
log10 peptide intensity
137:489 64:558 2:213
5:147504759:T:C_T
p = 1.7e−19, model = REC, N = 203

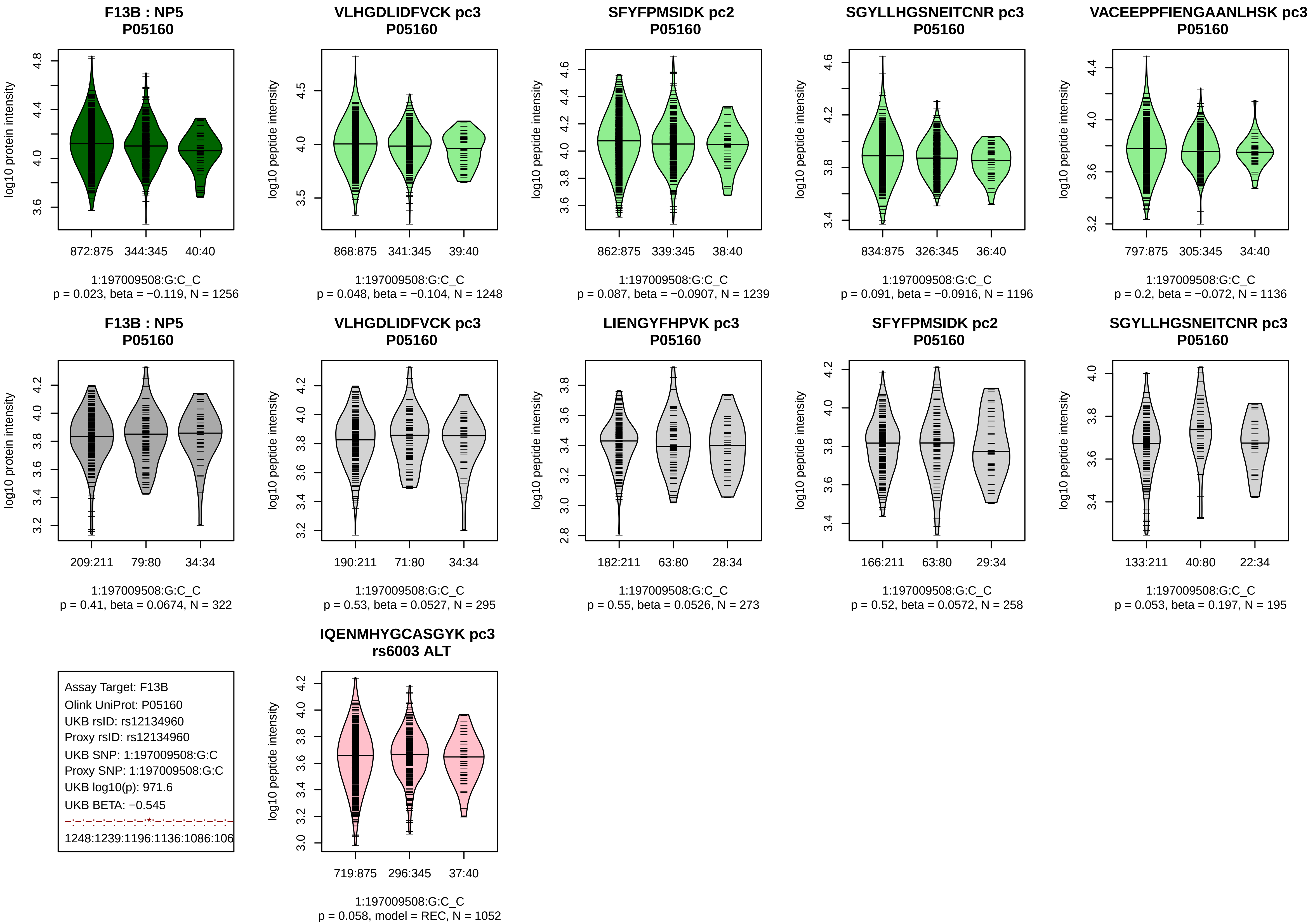

F13B : NP5
P05160
log10 protein intensity
872:875 344:345 40:40
1:197009508:G:C_C
p = 0.023, beta = −0.119, N = 1256
VLHGDLIDFVCK pc3
P05160
log10 peptide intensity
868:875 341:345 39:40
1:197009508:G:C_C
p = 0.048, beta = −0.104, N = 1248
SFYFPMSIDK pc2
P05160
log10 peptide intensity
862:875 339:345 38:40
1:197009508:G:C_C
p = 0.087, beta = −0.0907, N = 1239
SGYLLHGSNEITCNR pc3
P05160
log10 peptide intensity
834:875 326:345 36:40
1:197009508:G:C_C
p = 0.091, beta = −0.0916, N = 1196
VACEEPPFIENGAANLHSK pc3
P05160
log10 peptide intensity
797:875 305:345 34:40
1:197009508:G:C_C
p = 0.2, beta = −0.072, N = 1136
F13B : NP5
P05160
log10 protein intensity
209:211 79:80 34:34
1:197009508:G:C_C
p = 0.41, beta = 0.0674, N = 322
VLHGDLIDFVCK pc3
P05160
log10 peptide intensity
190:211 71:80 34:34
1:197009508:G:C_C
p = 0.53, beta = 0.0527, N = 295
LIENGYFHPVK pc3
P05160
log10 peptide intensity
182:211 63:80 28:34
1:197009508:G:C_C
p = 0.55, beta = 0.0526, N = 273
SFYFPMSIDK pc2
P05160
log10 peptide intensity
166:211 63:80 29:34
1:197009508:G:C_C
p = 0.52, beta = 0.0572, N = 258
SGYLLHGSNEITCNR pc3
P05160
log10 peptide intensity
133:211 40:80 22:34
1:197009508:G:C_C
p = 0.053, beta = 0.197, N = 195
Assay Target: F13B
Olink UniProt: P05160
UKB rsID: rs12134960
Proxy rsID: rs12134960
UKB SNP: 1:197009508:G:C
Proxy SNP: 1:197009508:G:C
UKB log10(p): 971.6
UKB BETA: −0.545
−:−:−:−:−:−:−:*:−:−:−:−:−:−:−:−
1248:1239:1196:1136:1086:106
IQENMHYGCASGYK pc3
rs6003 ALT
log10 peptide intensity
719:875 296:345 37:40
1:197009508:G:C_C
p = 0.058, model = REC, N = 1052

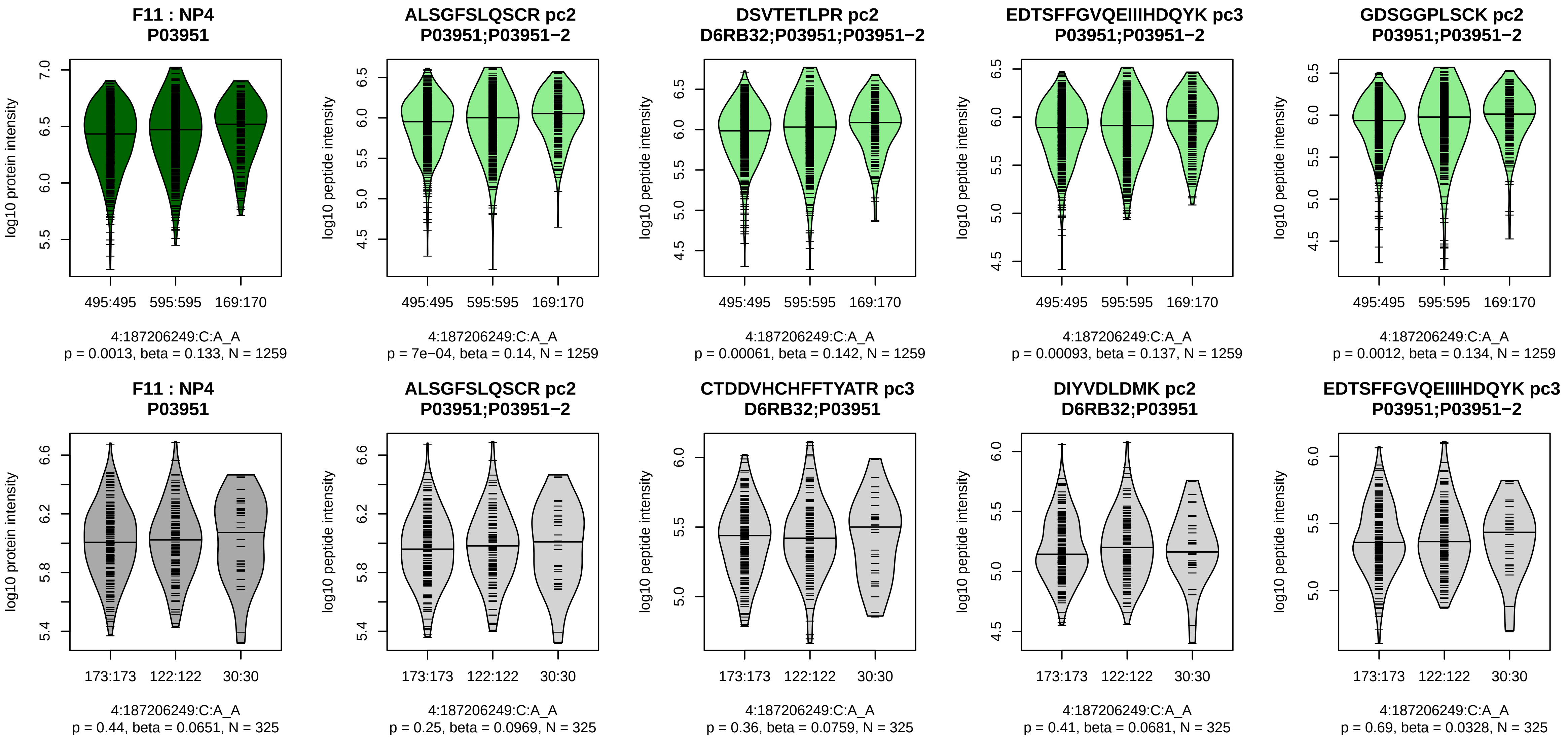


Assay Target: F11
Olink UniProt: P03951
UKB rsID: rs3756011
Proxy rsID: rs3756011
UKB SNP: 4:187206249:C:A
Proxy SNP: 4:187206249:C:A
UKB log10(p): 971.5
UKB BETA: 0.385
*:*:*:*:*:*:*:*:*:*:*:*:*:*:_:*:*:*:_:*:*
1259:1259:1259:1259:1259:125

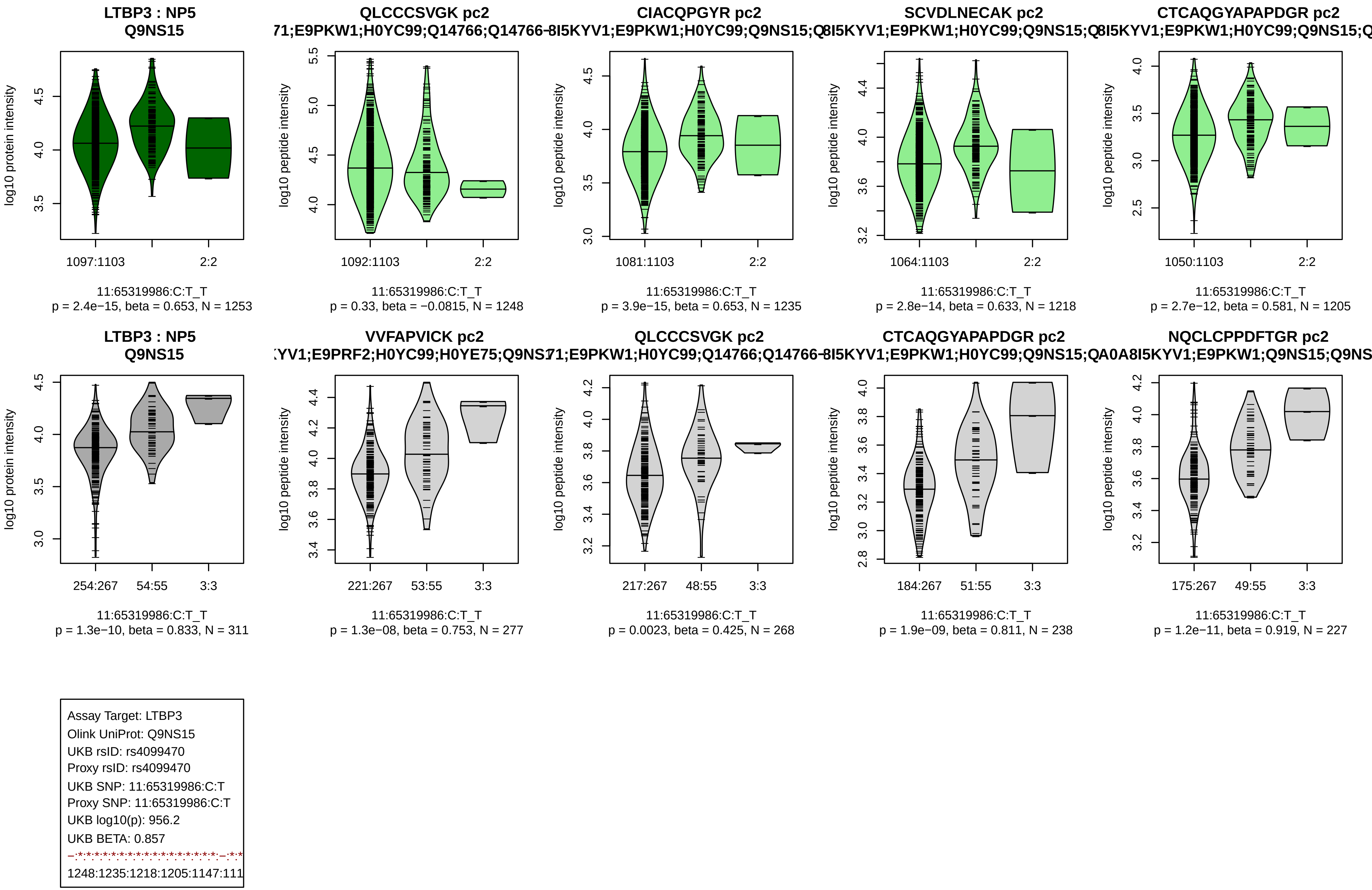

LTBP3 : NP5
Q9NS15
log10 protein intensity
1097:1103
2:2
11:65319986:C:T_T
p = 2.4e−15, beta = 0.653, N = 1253
QLCCCSVGK pc2
71;E9PKW1;H0YC99;Q14766;Q14766-
log10 peptide intensity
1092:1103
2:2
11:65319986:C:T_T
p = 0.33, beta = −0.0815, N = 1248
CIACQPGYR pc2
8I5KYV1;E9PKW1;H0YC99;Q9NS15;Q
log10 peptide intensity
1081:1103
2:2
11:65319986:C:T_T
p = 3.9e−15, beta = 0.653, N = 1235
SCVDLNECAK pc2
8I5KYV1;E9PKW1;H0YC99;Q9NS15;Q
log10 peptide intensity
1064:1103
2:2
11:65319986:C:T_T
p = 2.8e−14, beta = 0.633, N = 1218
CTCAQGYAPAPDGR pc2
8I5KYV1;E9PKW1;H0YC99;Q9NS15;Q
log10 peptide intensity
1050:1103
2:2
11:65319986:C:T_T
p = 2.7e−12, beta = 0.581, N = 1205
LTBP3 : NP5
Q9NS15
log10 protein intensity
254:267
54:55
3:3
11:65319986:C:T_T
p = 1.3e−10, beta = 0.833, N = 311
VVFAPVICK pc2
YV1;E9PRF2;H0YC99;H0YE75;Q9NS1
log10 peptide intensity
221:267
53:55
3:3
11:65319986:C:T_T
p = 1.3e−08, beta = 0.753, N = 277
QLCCCSVGK pc2
71;E9PKW1;H0YC99;Q14766;Q14766-
log10 peptide intensity
217:267
48:55
3:3
11:65319986:C:T_T
p = 0.0023, beta = 0.425, N = 268
CTCAQGYAPAPDGR pc2
8I5KYV1;E9PKW1;H0YC99;Q9NS15;QA0A
log10 peptide intensity
184:267
51:55
3:3
11:65319986:C:T_T
p = 1.9e−09, beta = 0.811, N = 238
NQCLCPPDFTGR pc2
8I5KYV1;E9PKW1;Q9NS15;Q9NS
log10 peptide intensity
175:267
49:55
3:3
11:65319986:C:T_T
p = 1.2e−11, beta = 0.919, N = 227
Assay Target: LTBP3
Olink UniProt: Q9NS15
UKB rsID: rs4099470
Proxy rsID: rs4099470
UKB SNP: 11:65319986:C:T
Proxy SNP: 11:65319986:C:T
UKB log10(p): 956.2
UKB BETA: 0.857
1248:1235:1218:1205:1147:111

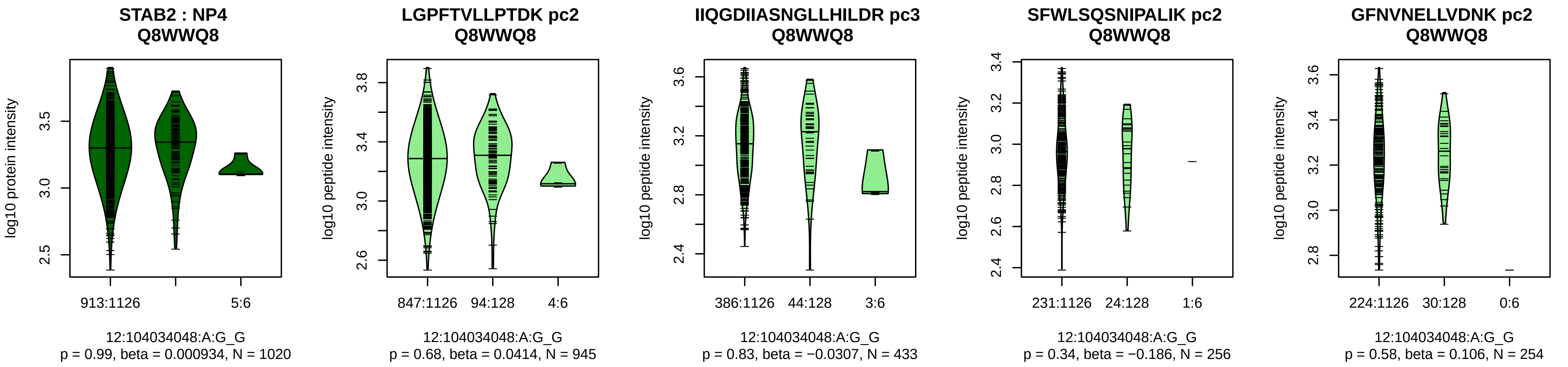


Assay Target: STAB2
Olink UniProt: Q8WWQ8
UKB rsID: rs115840458
Proxy rsID: rs115840458
UKB SNP: 12:104034048:A:G
Proxy SNP: 12:104034048:A:G
UKB log10(p): 952.3
UKB BETA: 0.786
−:−:−:−:−:−:−:−
945:433:256:254:226:219:145:7

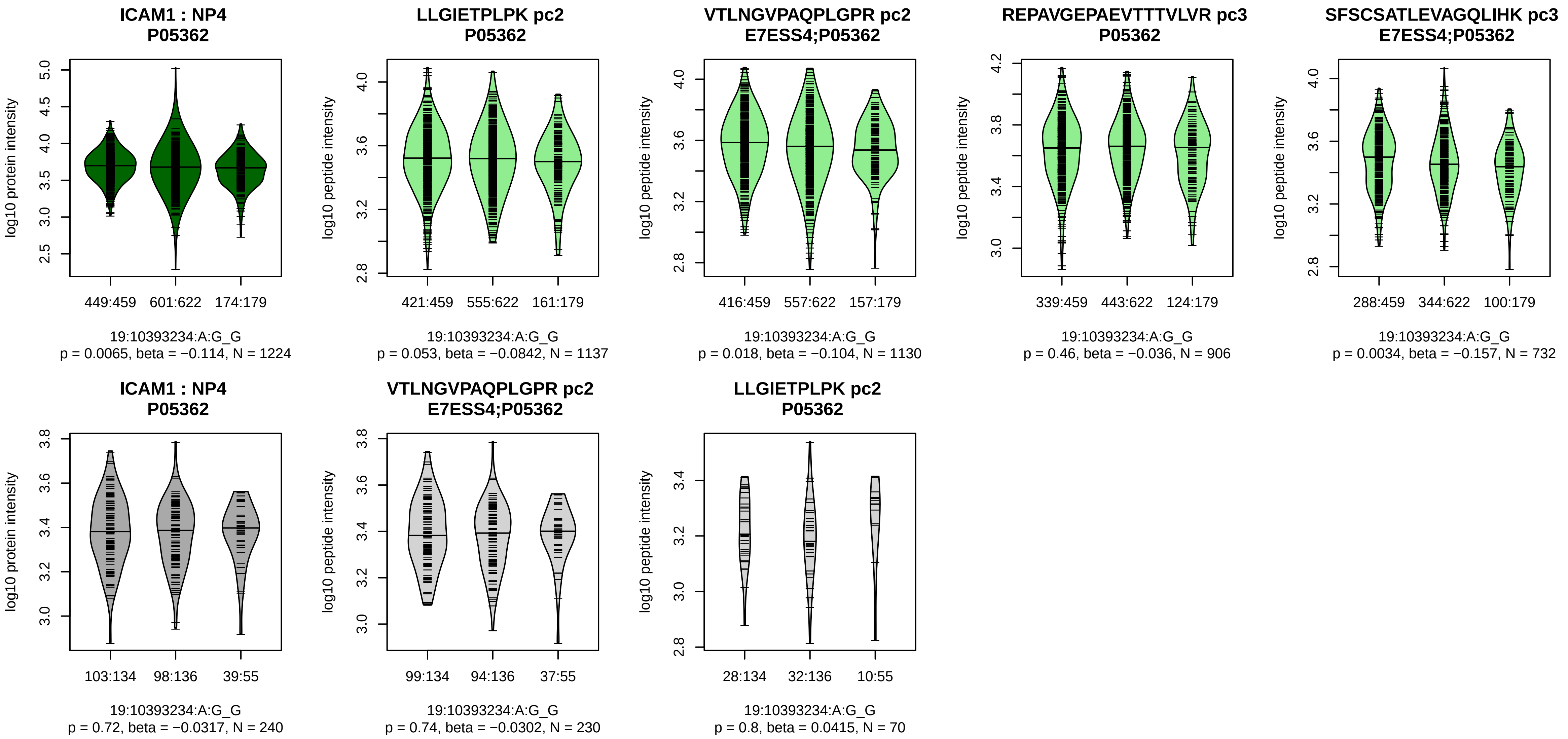


Assay Target: ICAM1
Olink UniProt: P05362
UKB rsID: rs5030379
Proxy rsID: rs5030377
UKB SNP: 19:10393605:C:CTT(
Proxy SNP: 19:10393234:A:G
UKB log10(p): 945.1
UKB BETA: 0.413
−:−:−:*:−:−
1137:1130:906:732:664:564

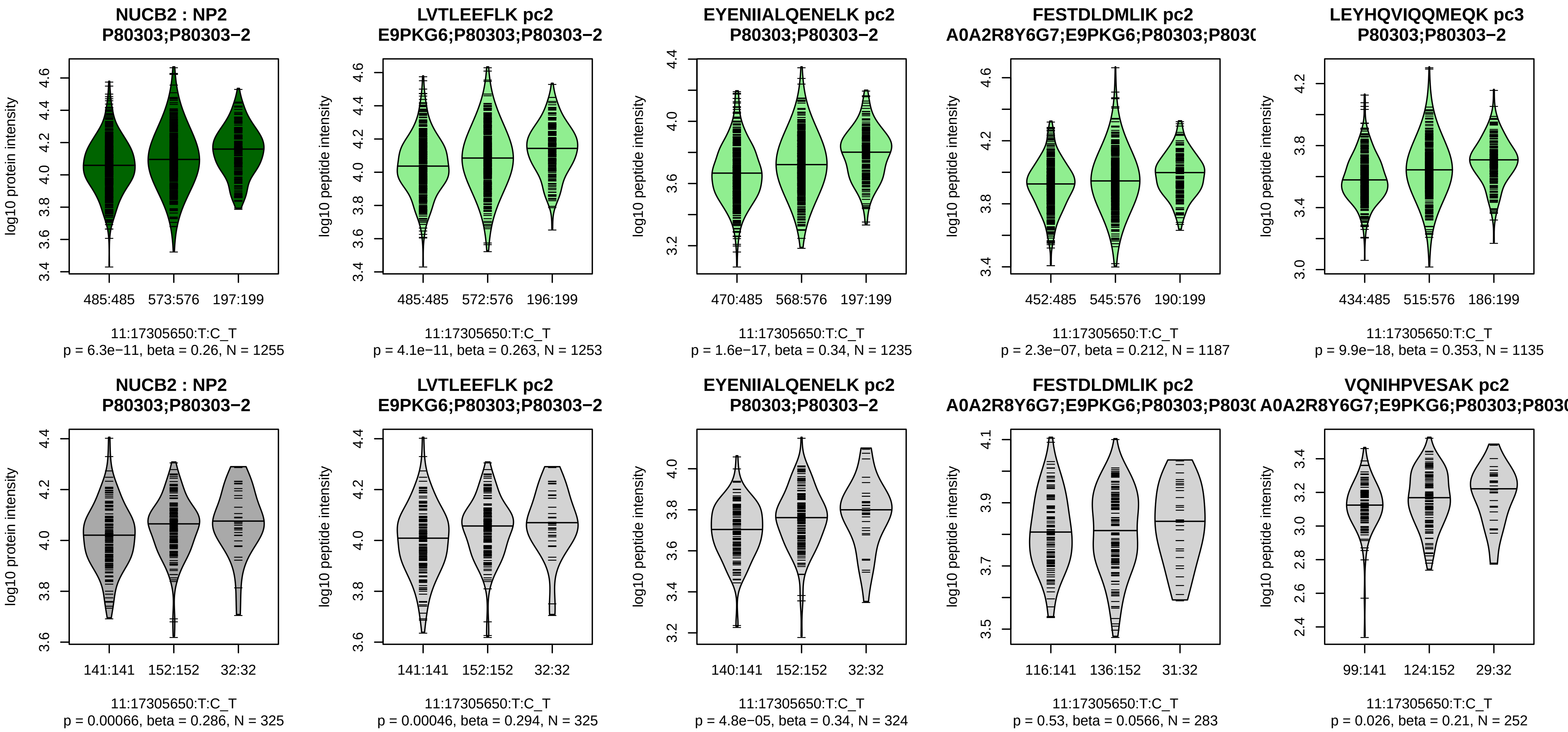


Assay Target: NUCB2
Olink UniProt: P80303
UKB rsID: rs214068
Proxy rsID: rs214068
UKB SNP: 11:17305650:T:C
Proxy SNP: 11:17305650:T:C
UKB log10(p): 922.3
UKB BETA: −0.402
*:*:*:*:*:*:*:–:–:–:–:–:–:–:–:–:
1253:1235:1187:1135:955:899:7

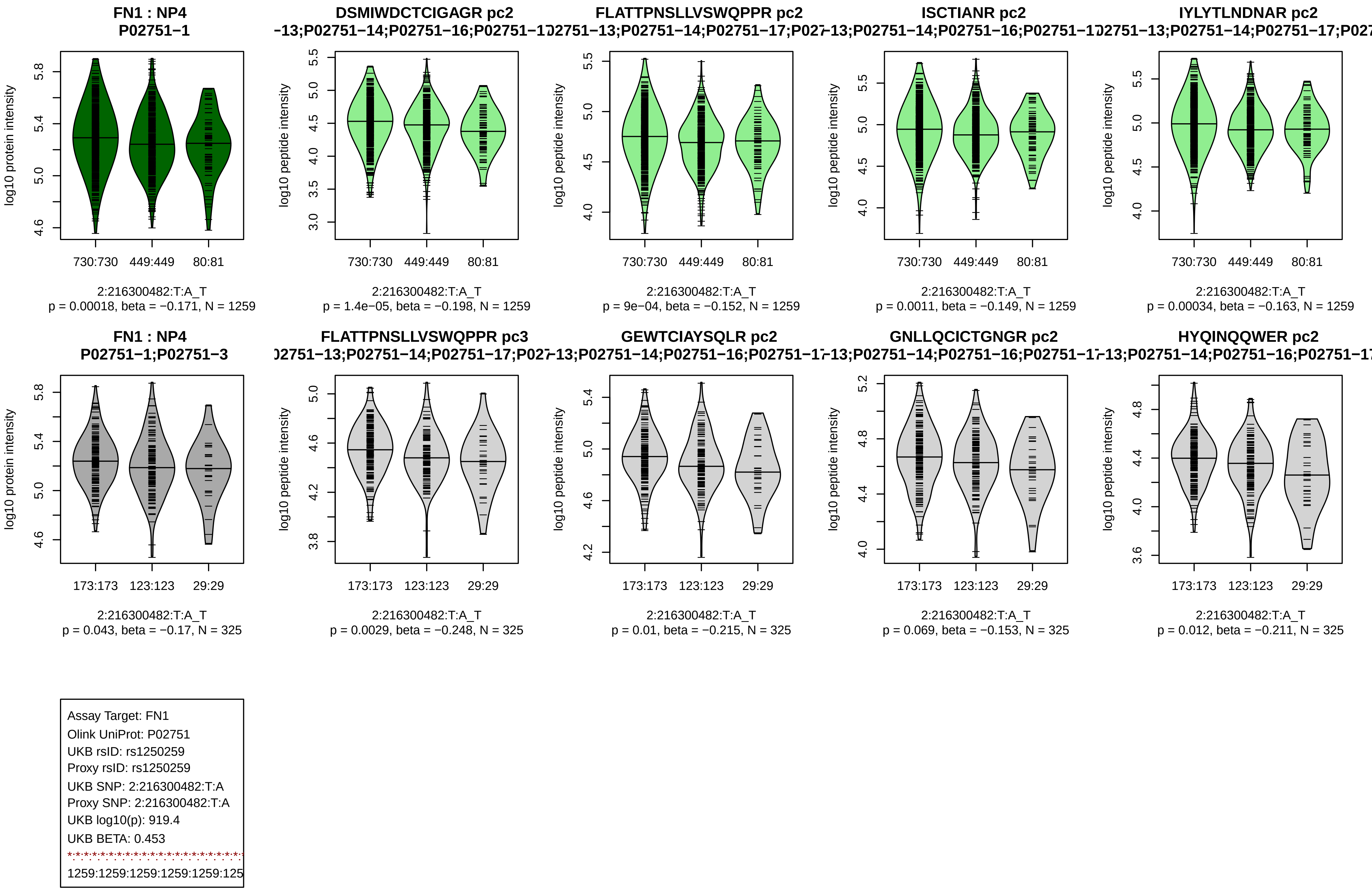
FN1 : NP4
P02751−1
log10 protein intensity
730:730
449:449
80:81
2:216300482:T:A_T
p = 0.00018, beta = −0.171, N = 1259
DSMIWDCTCIGAGR pc2
−13;P02751−14;P02751−16;P02751−1
log10 peptide intensity
2:216300482:T:A_T
p = 1.4e−05, beta = −0.198, N = 1259
FLATTPNSLLVSWQPPR pc2
)2751−13;P02751−14;P02751−17;P027
2:216300482:T:A_T
p = 9e−04, beta = −0.152, N = 1259
ISCTIANR pc2
−13;P02751−14;P02751−16;P02751−1
2:216300482:T:A_T
p = 0.0011, beta = −0.149, N = 1259
IYLYTLNDNAR pc2
)2751−13;P02751−14;P02751−17;P027
2:216300482:T:A_T
p = 0.00034, beta = −0.163, N = 1259
FN1 : NP4
P02751−1;P02751−3
173:173
123:123
29:29
2:216300482:T:A_T
p = 0.043, beta = −0.17, N = 325
FLATTPNSLLVSWQPPR pc3
)2751−13;P02751−14;P02751−17;P027
2:216300482:T:A_T
p = 0.0029, beta = −0.248, N = 325
GEWTCIAYSQLR pc2
−13;P02751−14;P02751−16;P02751−1
2:216300482:T:A_T
p = 0.01, beta = −0.215, N = 325
GNLLQCICTGNGR pc2
−13;P02751−14;P02751−16;P02751−1
2:216300482:T:A_T
p = 0.069, beta = −0.153, N = 325
HYQINQQWER pc2
−13;P02751−14;P02751−16;P02751−1
2:216300482:T:A_T
p = 0.012, beta = −0.211, N = 325
Assay Target: FN1
Olink UniProt: P02751
UKB rsID: rs1250259
Proxy rsID: rs1250259
UKB SNP: 2:216300482:T:A
Proxy SNP: 2:216300482:T:A
UKB log10(p): 919.4
UKB BETA: 0.453
1259:1259:1259:1259:1259:125

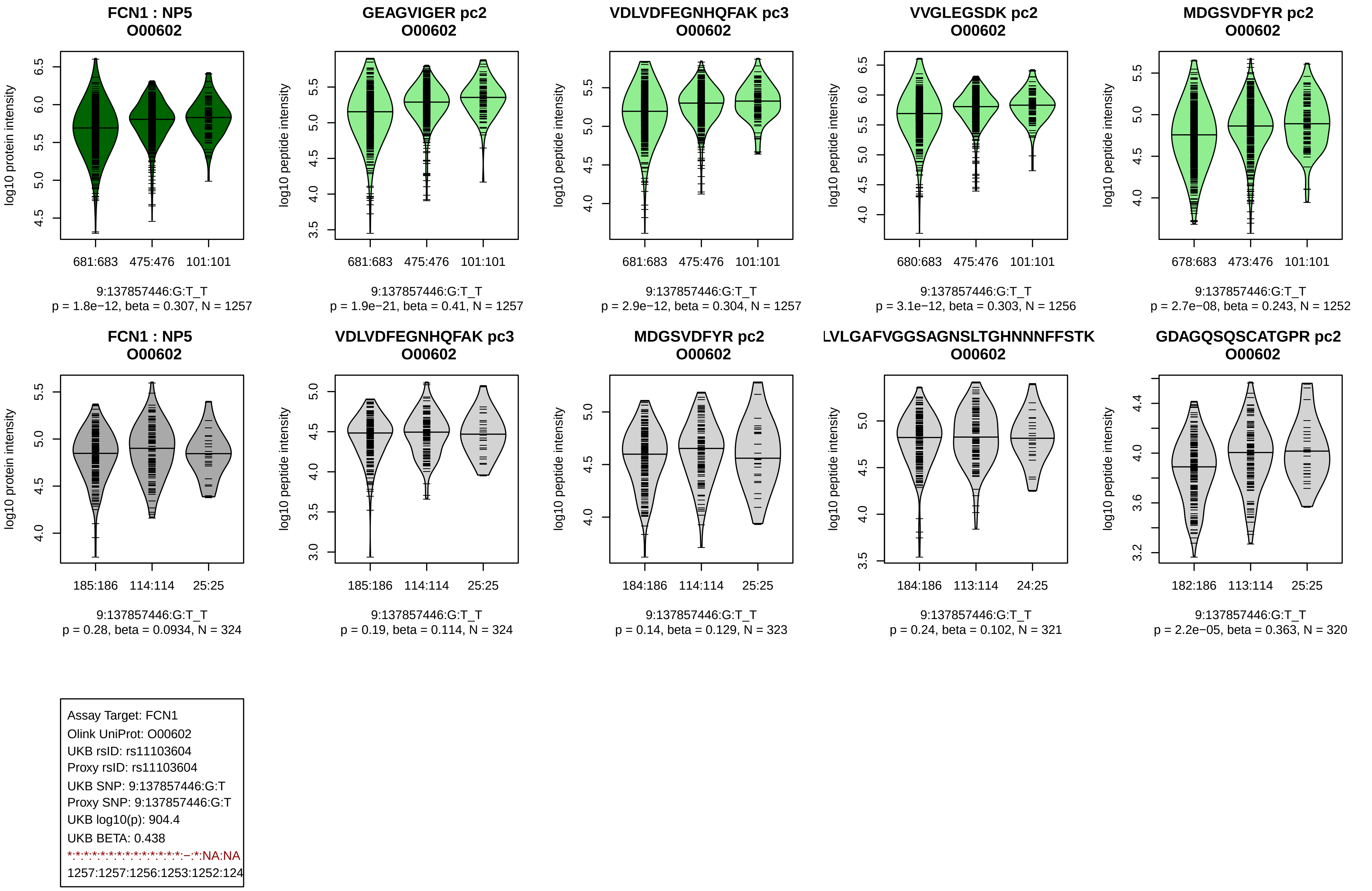

FCN1 : NP5
O00602
log10 protein intensity
681:683 475:476 101:101
9:137857446:G:T_T
p = 1.8e−12, beta = 0.307, N = 1257
GEAGVIGER pc2
O00602
log10 peptide intensity
681:683 475:476 101:101
9:137857446:G:T_T
p = 1.9e−21, beta = 0.41, N = 1257
VDLVDFEGNHQFAK pc3
O00602
log10 peptide intensity
681:683 475:476 101:101
9:137857446:G:T_T
p = 2.9e−12, beta = 0.304, N = 1257
VVGLEGSDK pc2
O00602
log10 peptide intensity
680:683 475:476 101:101
9:137857446:G:T_T
p = 3.1e−12, beta = 0.303, N = 1256
MDGSVDFYR pc2
O00602
log10 peptide intensity
678:683 473:476 101:101
9:137857446:G:T_T
p = 2.7e−08, beta = 0.243, N = 1252
FCN1 : NP5
O00602
log10 protein intensity
185:186 114:114 25:25
9:137857446:G:T_T
p = 0.28, beta = 0.0934, N = 324
VDLVDFEGNHQFAK pc3
O00602
log10 peptide intensity
185:186 114:114 25:25
9:137857446:G:T_T
p = 0.19, beta = 0.114, N = 324
MDGSVDFYR pc2
O00602
log10 peptide intensity
184:186 114:114 25:25
9:137857446:G:T_T
p = 0.14, beta = 0.129, N = 323
LVLGAFVGGSAGNSLTGHNNNFFSTK
O00602
log10 peptide intensity
184:186 113:114 24:25
9:137857446:G:T_T
p = 0.24, beta = 0.102, N = 321
GDAGQSQSCATGPR pc2
O00602
log10 peptide intensity
182:186 113:114 25:25
9:137857446:G:T_T
p = 2.2e−05, beta = 0.363, N = 320
Assay Target: FCN1
Olink UniProt: O00602
UKB rsID: rs11103604
Proxy rsID: rs11103604
UKB SNP: 9:137857446:G:T
Proxy SNP: 9:137857446:G:T
UKB log10(p): 904.4
UKB BETA: 0.438
*:*:*:*:*:*:*:*:*:*:*:*:*:*:*:–:*:NA:NA
1257:1257:1256:1253:1252:124

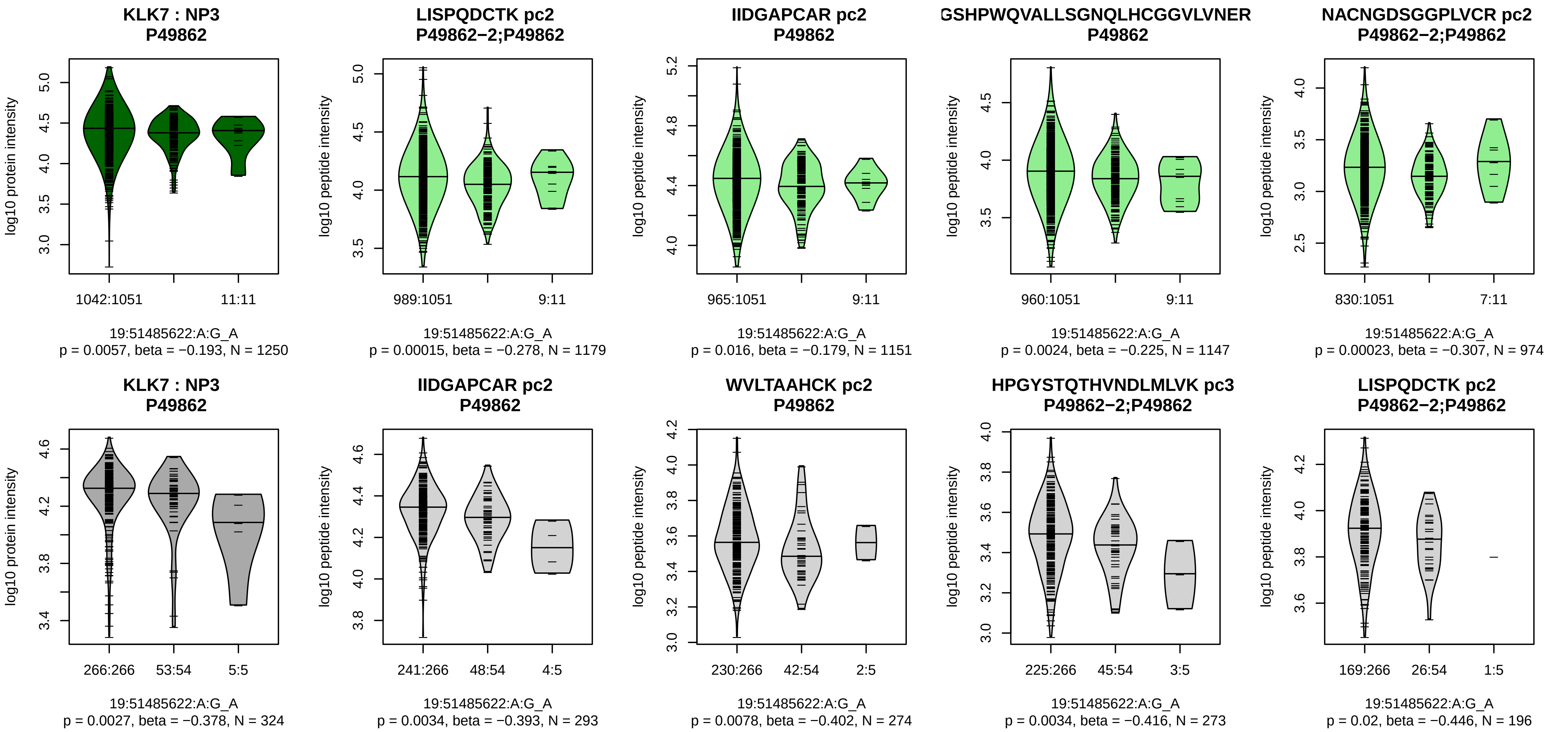


Assay Target: KLK7
Olink UniProt: P49862
UKB rsID: rs2659067
Proxy rsID: rs2659067
UKB SNP: 19:51485622:A:G
Proxy SNP: 19:51485622:A:G
UKB log10(p): 900.3
UKB BETA: 0.681
*:–:*:*:*:*:–:–:–:–:–
1179:1151:1147:974:918:702:32

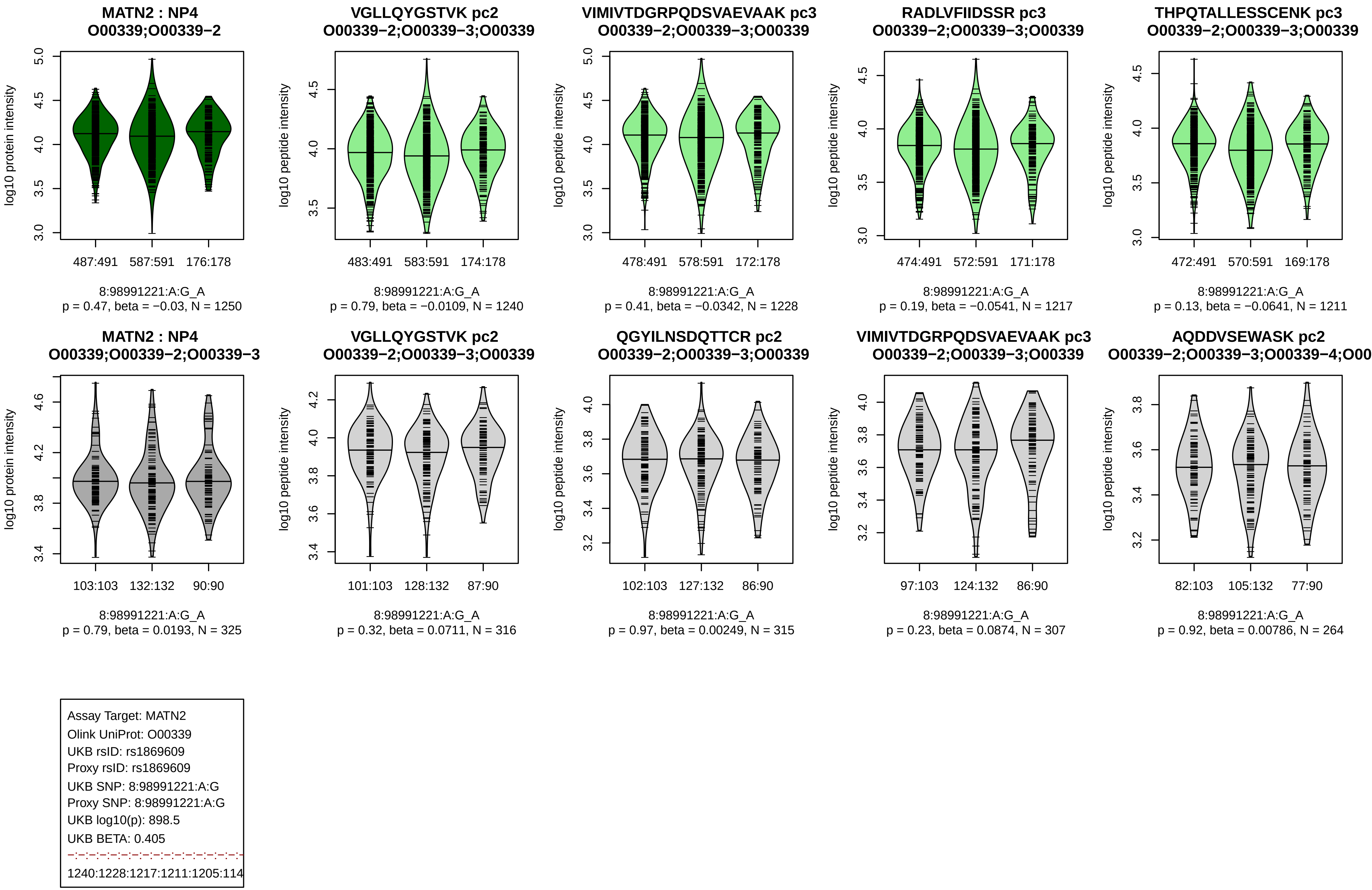
MATN2 : NP4
O00339;O00339−2
log10 protein intensity
487:491 587:591 176:178
8:98991221:A:G_A
p = 0.47, beta = −0.03, N = 1250
VGLLQYGSTVK pc2
O00339−2;O00339−3;O00339
log10 peptide intensity
483:491 583:591 174:178
8:98991221:A:G_A
p = 0.79, beta = −0.0109, N = 1240
VIMIVTDGRPQDSVAEVAAK pc3
O00339−2;O00339−3;O00339
log10 peptide intensity
478:491 578:591 172:178
8:98991221:A:G_A
p = 0.41, beta = −0.0342, N = 1228
RADLVFIIDSSR pc3
O00339−2;O00339−3;O00339
log10 peptide intensity
474:491 572:591 171:178
8:98991221:A:G_A
p = 0.19, beta = −0.0541, N = 1217
THPQTALLESSCENK pc3
O00339−2;O00339−3;O00339
log10 peptide intensity
472:491 570:591 169:178
8:98991221:A:G_A
p = 0.13, beta = −0.0641, N = 1211
MATN2 : NP4
O00339;O00339−2;O00339−3
log10 protein intensity
103:103 132:132 90:90
8:98991221:A:G_A
p = 0.79, beta = 0.0193, N = 325
VGLLQYGSTVK pc2
O00339−2;O00339−3;O00339
log10 peptide intensity
101:103 128:132 87:90
8:98991221:A:G_A
p = 0.32, beta = 0.0711, N = 316
QGYILNSDQTTCR pc2
O00339−2;O00339−3;O00339
log10 peptide intensity
102:103 127:132 86:90
8:98991221:A:G_A
p = 0.97, beta = 0.00249, N = 315
VIMIVTDGRPQDSVAEVAAK pc3
O00339−2;O00339−3;O00339
log10 peptide intensity
97:103 124:132 86:90
8:98991221:A:G_A
p = 0.23, beta = 0.0874, N = 307
AQDDVSEWASK pc2
O00339−2;O00339−3;O00339−4;O00
log10 peptide intensity
82:103 105:132 77:90
8:98991221:A:G_A
p = 0.92, beta = 0.00786, N = 264
Assay Target: MATN2
Olink UniProt: O00339
UKB rsID: rs1869609
Proxy rsID: rs1869609
UKB SNP: 8:98991221:A:G
Proxy SNP: 8:98991221:A:G
UKB log10(p): 898.5
UKB BETA: 0.405
1240:1228:1217:1211:1205:114

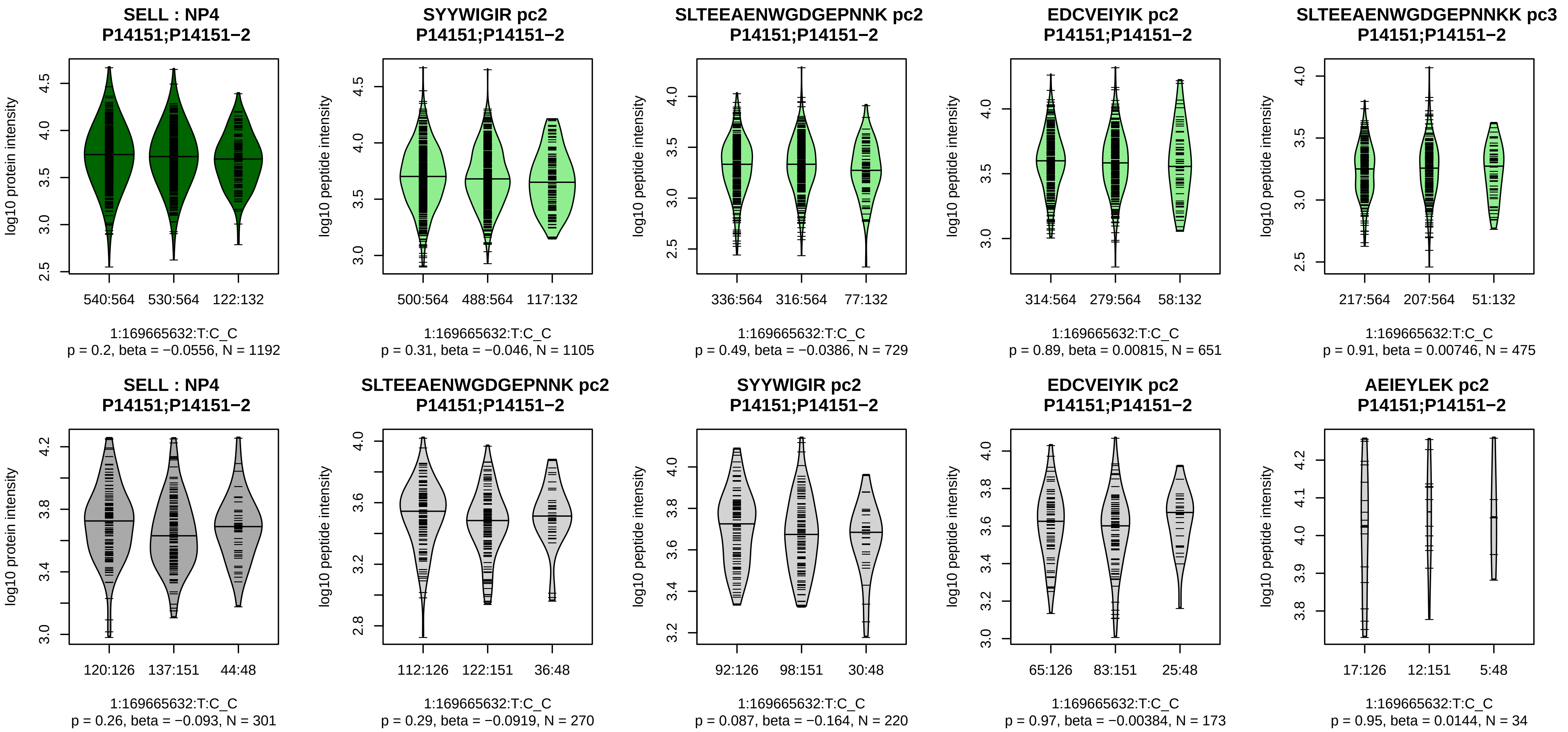


Assay Target: SELL
Olink UniProt: P14151
UKB rsID: rs2223286
Proxy rsID: rs2223286
UKB SNP: 1:169665632:T:C
Proxy SNP: 1:169665632:T:C
UKB log10(p): 888.4
UKB BETA: −0.407
−:−:−:−:−
1105:729:651:475:449

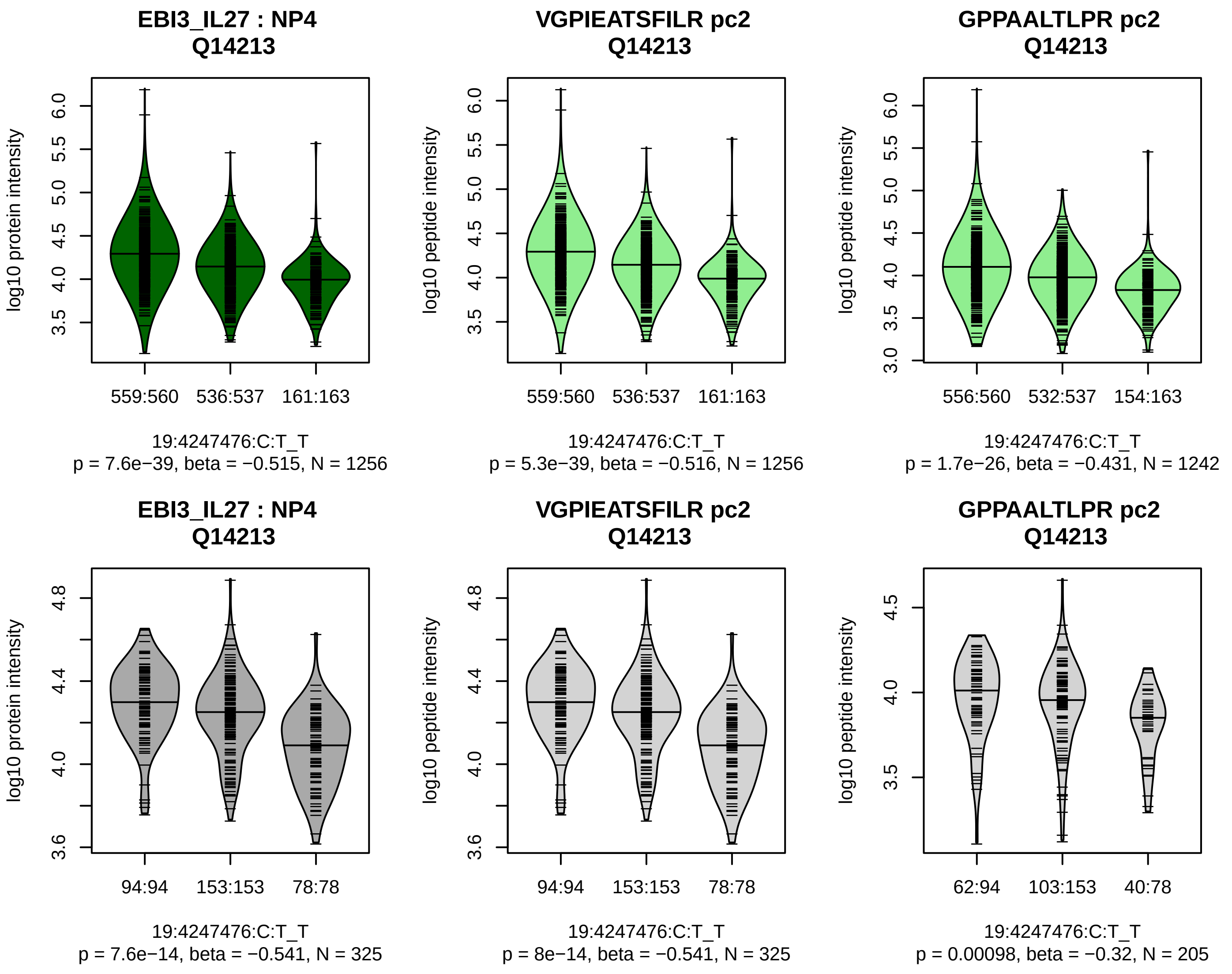


Assay Target: EBI3_IL27
Olink UniProt: Q14213
UKB rsID: rs8109160
Proxy rsID: rs8109160
UKB SNP: 19:4247476:C:T
Proxy SNP: 19:4247476:C:T
UKB log10(p): 876.3
UKB BETA: −0.401
*:*
1256:1242

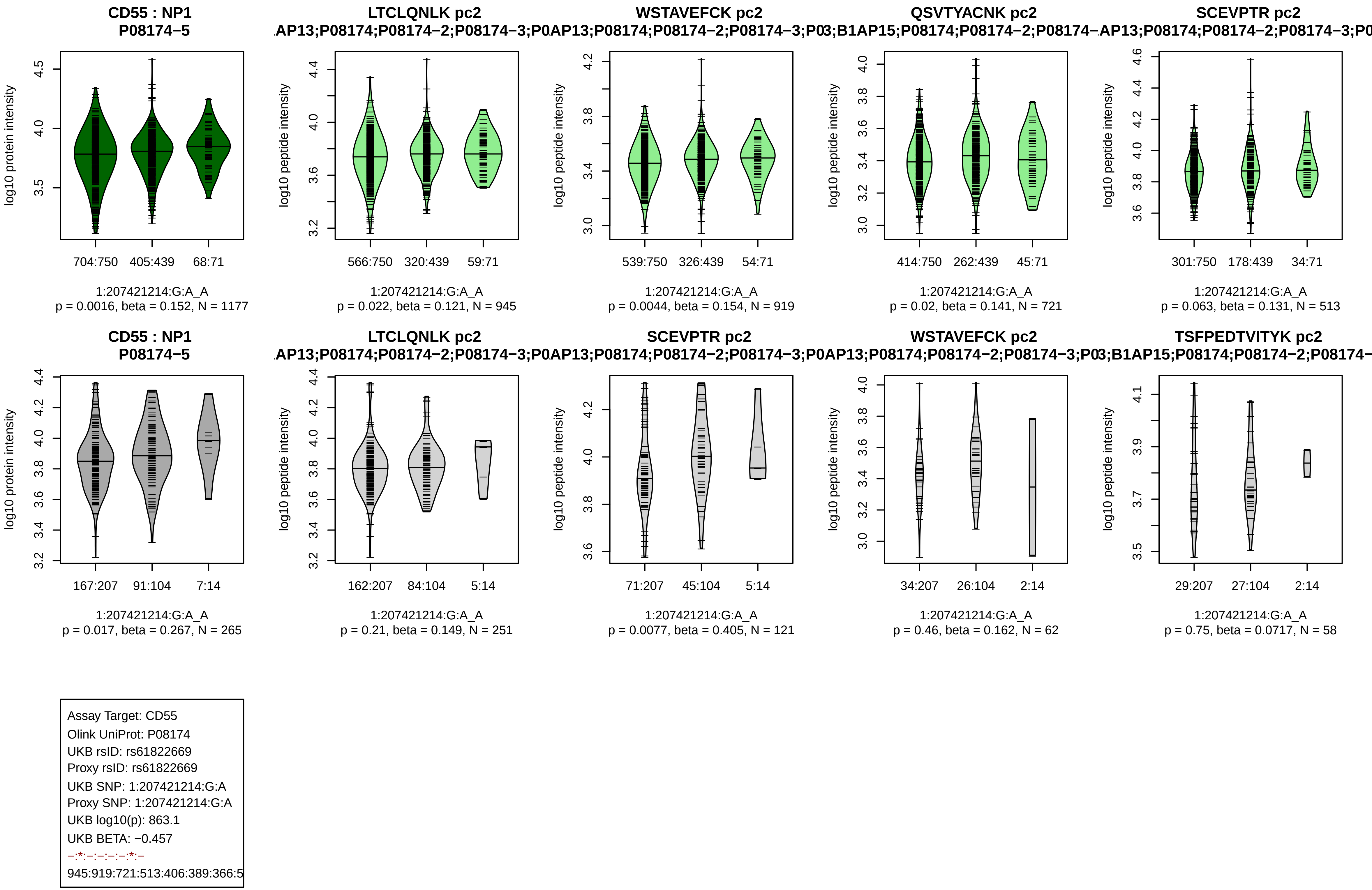
CD55 : NP1
P08174−5
log10 protein intensity
704:750
405:439
68:71
1:207421214:G:A_A
p = 0.0016, beta = 0.152, N = 1177
LTCLQNLK pc2
AP13;P08174;P08174−2;P08174−3;P0
log10 peptide intensity
566:750
320:439
59:71
1:207421214:G:A_A
p = 0.022, beta = 0.121, N = 945
WSTAVEFCK pc2
AP13;P08174;P08174−2;P08174−3;P0
log10 peptide intensity
539:750
326:439
54:71
1:207421214:G:A_A
p = 0.0044, beta = 0.154, N = 919
QSVTYACNK pc2
3;B1AP15;P08174;P08174−2;P08174−
log10 peptide intensity
414:750
262:439
45:71
1:207421214:G:A_A
p = 0.02, beta = 0.141, N = 721
SCEVPTR pc2
AP13;P08174;P08174−2;P08174−3;P0
log10 peptide intensity
301:750
178:439
34:71
1:207421214:G:A_A
p = 0.063, beta = 0.131, N = 513
CD55 : NP1
P08174−5
log10 protein intensity
167:207
91:104
7:14
1:207421214:G:A_A
p = 0.017, beta = 0.267, N = 265
LTCLQNLK pc2
AP13;P08174;P08174−2;P08174−3;P0
log10 peptide intensity
162:207
84:104
5:14
1:207421214:G:A_A
p = 0.21, beta = 0.149, N = 251
SCEVPTR pc2
AP13;P08174;P08174−2;P08174−3;P0
log10 peptide intensity
71:207
45:104
5:14
1:207421214:G:A_A
p = 0.0077, beta = 0.405, N = 121
WSTAVEFCK pc2
AP13;P08174;P08174−2;P08174−3;P0
log10 peptide intensity
34:207
26:104
2:14
1:207421214:G:A_A
p = 0.46, beta = 0.162, N = 62
TSFPEDTVITYK pc2
3;B1AP15;P08174;P08174−2;P08174−
log10 peptide intensity
29:207
27:104
2:14
1:207421214:G:A_A
p = 0.75, beta = 0.0717, N = 58
Assay Target: CD55
Olink UniProt: P08174
UKB rsID: rs61822669
Proxy rsID: rs61822669
UKB SNP: 1:207421214:G:A
Proxy SNP: 1:207421214:G:A
UKB log10(p): 863.1
UKB BETA: −0.457
−:*:−:−:−:−:*:−
945:919:721:513:406:389:366:5

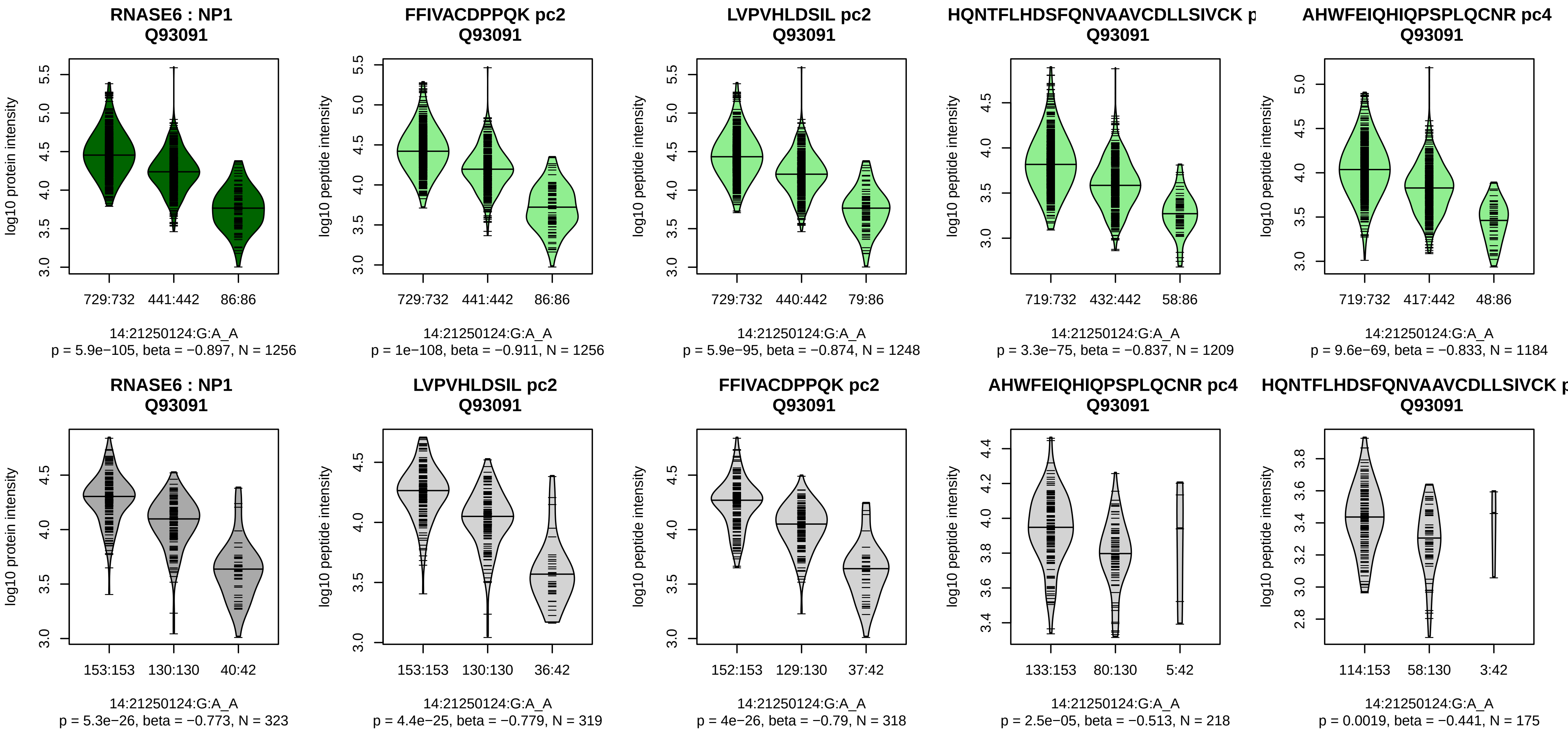


Assay Target: RNASE6
Olink UniProt: Q93091
UKB rsID: rs1045922
Proxy rsID: rs1045922
UKB SNP: 14:21250124:G:A
Proxy SNP: 14:21250124:G:A
UKB log10(p): 853.8
UKB BETA: −0.427
*:*:*:*:*:*:−:NA
1256:1248:1209:1184:1166:493

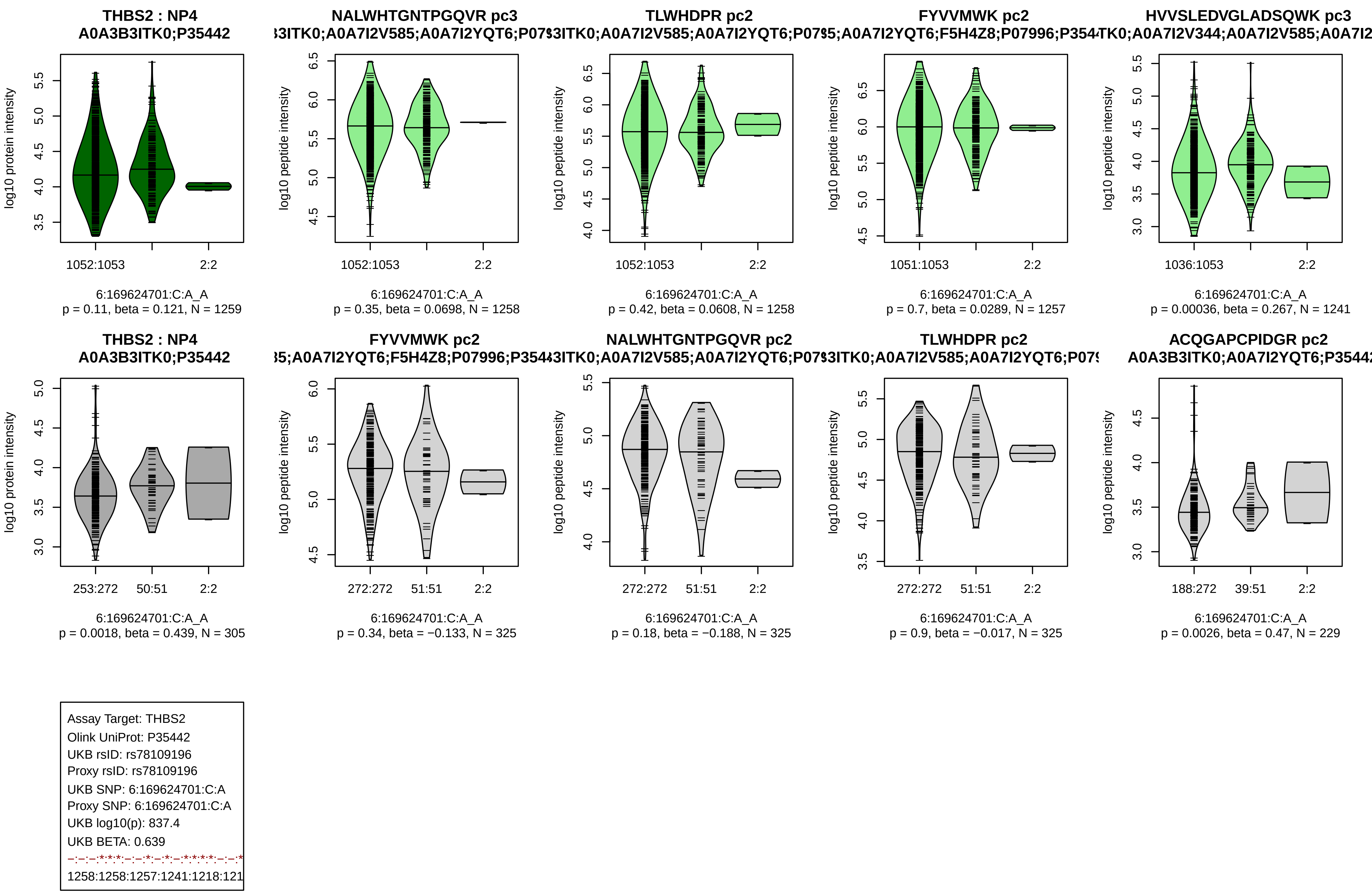
THBS2 : NP4
A0A3B3ITK0;P35442
log10 protein intensity
1052:1053
2:2
6:169624701:C:A_A
p = 0.11, beta = 0.121, N = 1259
NALWHTGNTPGQVR pc3
log10 peptide intensity
1052:1053
2:2
6:169624701:C:A_A
p = 0.35, beta = 0.0698, N = 1258
TLWHDPR pc2
log10 peptide intensity
1052:1053
2:2
6:169624701:C:A_A
p = 0.42, beta = 0.0608, N = 1258
FYVVMWK pc2
log10 peptide intensity
1051:1053
2:2
6:169624701:C:A_A
p = 0.7, beta = 0.0289, N = 1257
HVVSLEDVGLADSQWK pc3
log10 peptide intensity
1036:1053
2:2
6:169624701:C:A_A
p = 0.00036, beta = 0.267, N = 1241
THBS2 : NP4
A0A3B3ITK0;P35442
log10 protein intensity
253:272
50:51
2:2
6:169624701:C:A_A
p = 0.0018, beta = 0.439, N = 305
FYVVMWK pc2
log10 peptide intensity
272:272
51:51
2:2
6:169624701:C:A_A
p = 0.34, beta = −0.133, N = 325
NALWHTGNTPGQVR pc2
log10 peptide intensity
272:272
51:51
2:2
6:169624701:C:A_A
p = 0.18, beta = −0.188, N = 325
TLWHDPR pc2
log10 peptide intensity
272:272
51:51
2:2
6:169624701:C:A_A
p = 0.9, beta = −0.017, N = 325
ACQGAPCPIDGR pc2
A0A3B3ITK0;A0A7I2YQT6;P35442
log10 peptide intensity
188:272
39:51
2:2
6:169624701:C:A_A
p = 0.0026, beta = 0.47, N = 229
Assay Target: THBS2
Olink UniProt: P35442
UKB rsID: rs78109196
Proxy rsID: rs78109196
UKB SNP: 6:169624701:C:A
Proxy SNP: 6:169624701:C:A
UKB log10(p): 837.4
UKB BETA: 0.639
1258:1258:1257:1241:1218:121

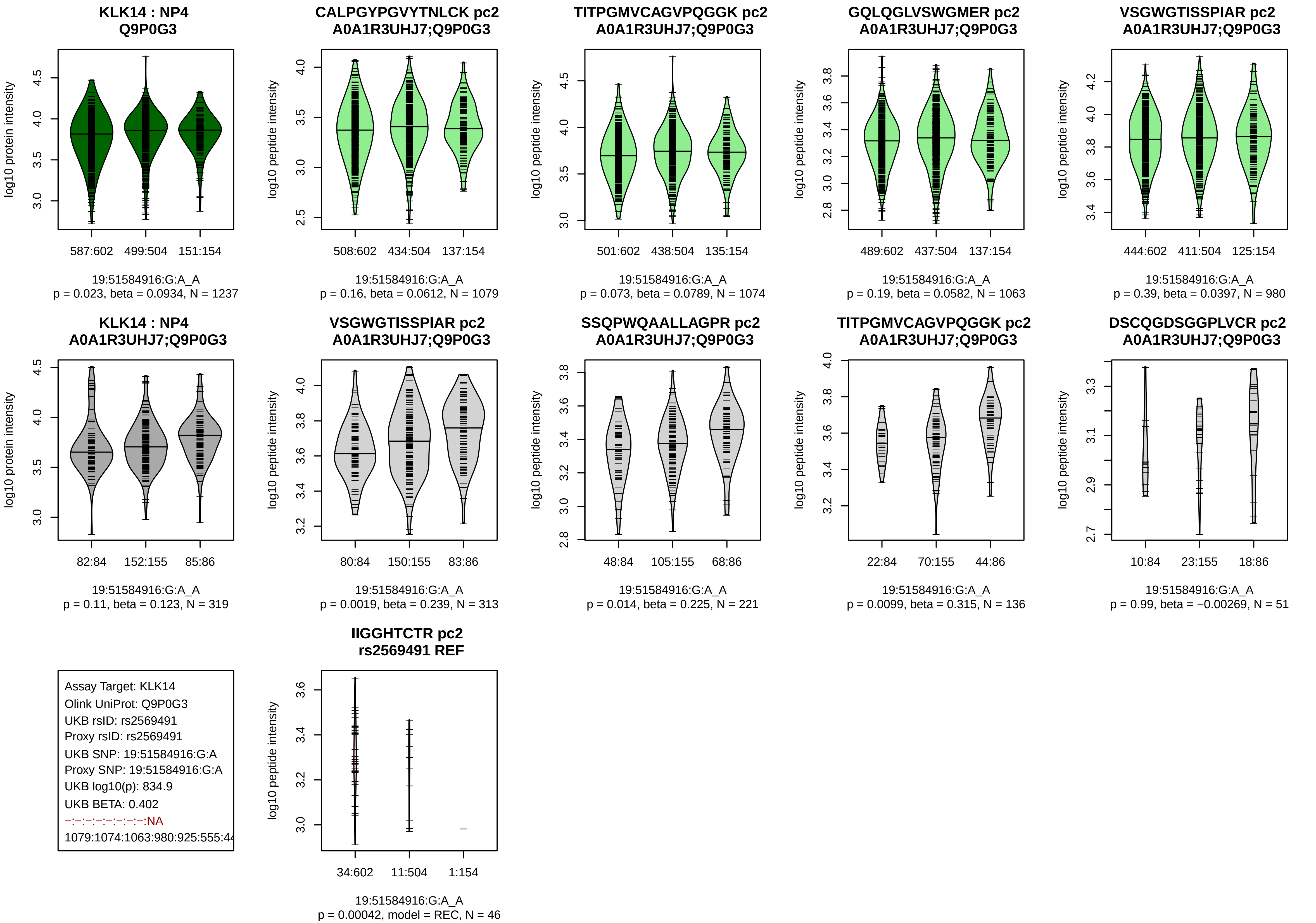

KLK14 : NP4
Q9P0G3
log10 protein intensity
587:602 499:504 151:154
19:51584916:G:A_A
p = 0.023, beta = 0.0934, N = 1237
CALPGYPGVYTNLCK pc2
A0A1R3UHJ7;Q9P0G3
log10 peptide intensity
508:602 434:504 137:154
19:51584916:G:A_A
p = 0.16, beta = 0.0612, N = 1079
TITPGMVCAGVPQGGK pc2
A0A1R3UHJ7;Q9P0G3
log10 peptide intensity
501:602 438:504 135:154
19:51584916:G:A_A
p = 0.073, beta = 0.0789, N = 1074
GQLQGLVSWGMER pc2
A0A1R3UHJ7;Q9P0G3
log10 peptide intensity
489:602 437:504 137:154
19:51584916:G:A_A
p = 0.19, beta = 0.0582, N = 1063
VSGWGTISSPIAR pc2
A0A1R3UHJ7;Q9P0G3
log10 peptide intensity
444:602 411:504 125:154
19:51584916:G:A_A
p = 0.39, beta = 0.0397, N = 980
KLK14 : NP4
A0A1R3UHJ7;Q9P0G3
log10 protein intensity
82:84 152:155 85:86
19:51584916:G:A_A
p = 0.11, beta = 0.123, N = 319
VSGWGTISSPIAR pc2
A0A1R3UHJ7;Q9P0G3
log10 peptide intensity
80:84 150:155 83:86
19:51584916:G:A_A
p = 0.0019, beta = 0.239, N = 313
SSQPWQAALLAGPR pc2
A0A1R3UHJ7;Q9P0G3
log10 peptide intensity
48:84 105:155 68:86
19:51584916:G:A_A
p = 0.014, beta = 0.225, N = 221
TITPGMVCAGVPQGGK pc2
A0A1R3UHJ7;Q9P0G3
log10 peptide intensity
22:84 70:155 44:86
19:51584916:G:A_A
p = 0.0099, beta = 0.315, N = 136
DSCQGDSGGPLVCR pc2
A0A1R3UHJ7;Q9P0G3
log10 peptide intensity
10:84 23:155 18:86
19:51584916:G:A_A
p = 0.99, beta = −0.00269, N = 51
Assay Target: KLK14
Olink UniProt: Q9P0G3
UKB rsID: rs2569491
Proxy rsID: rs2569491
UKB SNP: 19:51584916:G:A
Proxy SNP: 19:51584916:G:A
UKB log10(p): 834.9
UKB BETA: 0.402
−:−:−:−:−:−:−:−:NA
1079:1074:1063:980:925:555:44
IIGGHTCTR pc2
rs2569491 REF
log10 peptide intensity
34:602 11:504 1:154
19:51584916:G:A_A
p = 0.00042, model = REC, N = 46

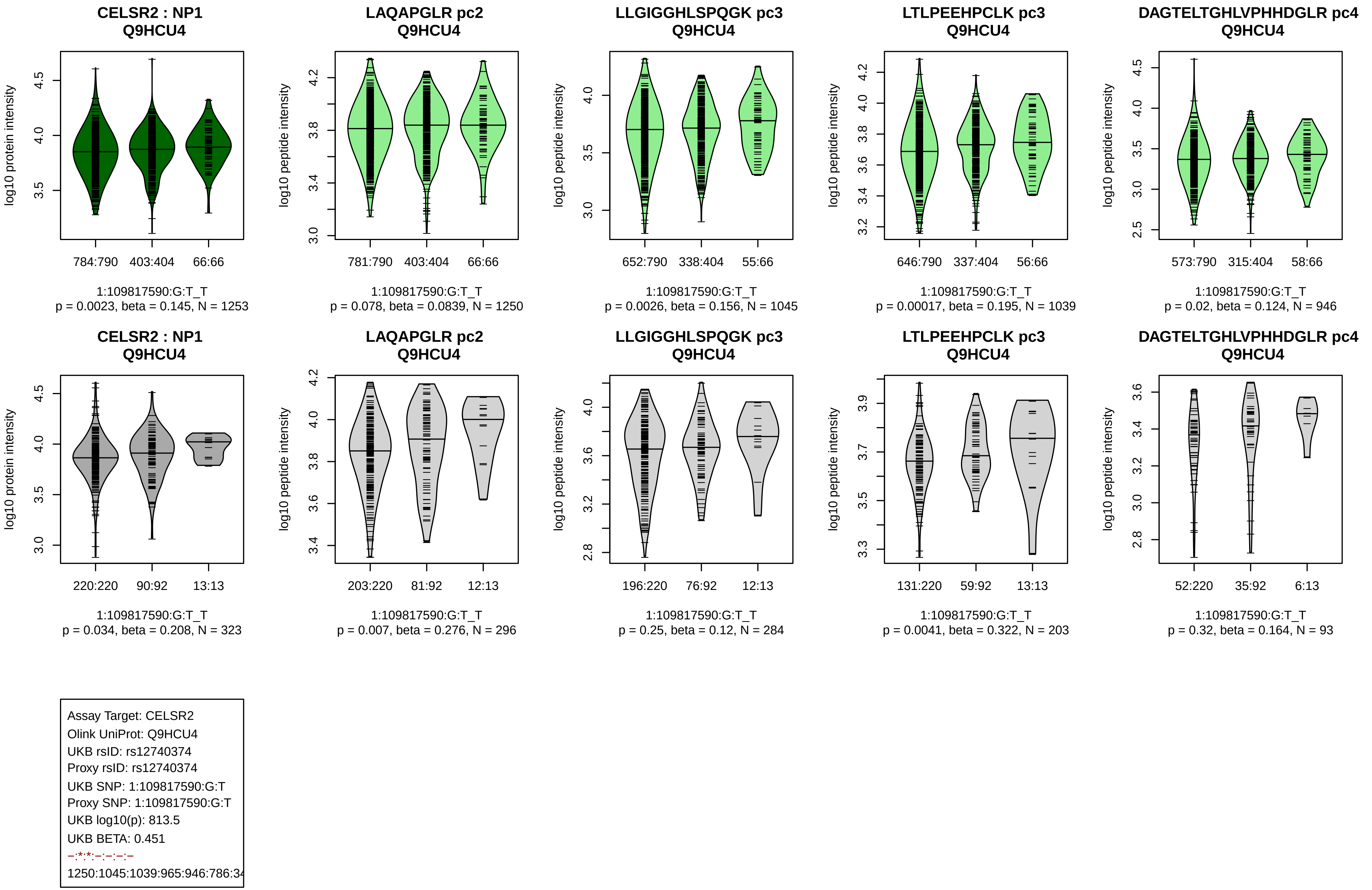
CELSR2 : NP1
Q9HCU4
log10 protein intensity
784:790 403:404 66:66
1:109817590:G:T_T
p = 0.0023, beta = 0.145, N = 1253
LAQAPGLR pc2
Q9HCU4
log10 peptide intensity
781:790 403:404 66:66
1:109817590:G:T_T
p = 0.078, beta = 0.0839, N = 1250
LLGIGGHLSPQGK pc3
Q9HCU4
log10 peptide intensity
652:790 338:404 55:66
1:109817590:G:T_T
p = 0.0026, beta = 0.156, N = 1045
LTLPEEHPCLK pc3
Q9HCU4
log10 peptide intensity
646:790 337:404 56:66
1:109817590:G:T_T
p = 0.00017, beta = 0.195, N = 1039
DAGTELTGHLVPHHDGLR pc4
Q9HCU4
log10 peptide intensity
573:790 315:404 58:66
1:109817590:G:T_T
p = 0.02, beta = 0.124, N = 946
CELSR2 : NP1
Q9HCU4
log10 protein intensity
220:220 90:92 13:13
1:109817590:G:T_T
p = 0.034, beta = 0.208, N = 323
LAQAPGLR pc2
Q9HCU4
log10 peptide intensity
203:220 81:92 12:13
1:109817590:G:T_T
p = 0.007, beta = 0.276, N = 296
LLGIGGHLSPQGK pc3
Q9HCU4
log10 peptide intensity
196:220 76:92 12:13
1:109817590:G:T_T
p = 0.25, beta = 0.12, N = 284
LTLPEEHPCLK pc3
Q9HCU4
log10 peptide intensity
131:220 59:92 13:13
1:109817590:G:T_T
p = 0.0041, beta = 0.322, N = 203
DAGTELTGHLVPHHDGLR pc4
Q9HCU4
log10 peptide intensity
52:220 35:92 6:13
1:109817590:G:T_T
p = 0.32, beta = 0.164, N = 93
Assay Target: CELSR2
Olink UniProt: Q9HCU4
UKB rsID: rs12740374
Proxy rsID: rs12740374
UKB SNP: 1:109817590:G:T
Proxy SNP: 1:109817590:G:T
UKB log10(p): 813.5
UKB BETA: 0.451
–:*:*:–:–:–:–
1250:1045:1039:965:946:786:34

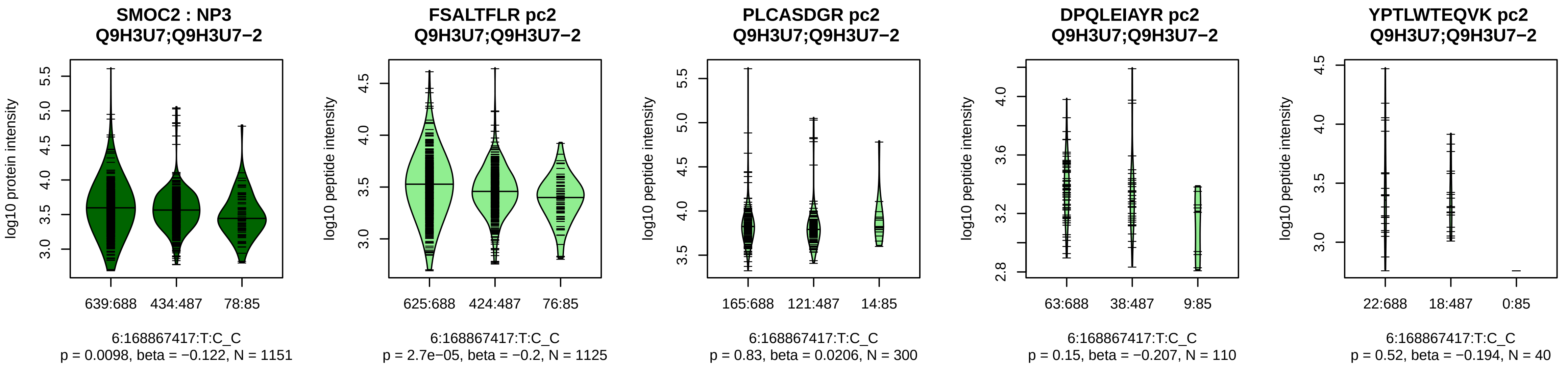


Assay Target: SMOC2
Olink UniProt: Q9H3U7
UKB rsID: rs11759500
Proxy rsID: rs11759500
UKB SNP: 6:168867417:T:C
Proxy SNP: 6:168867417:T:C
UKB log10(p): 784.1
UKB BETA: −0.384
*:−:−:−:−:−:NA:NA:NA:NA:NA:N
1125:300:110:40:38:21:15:11:3:

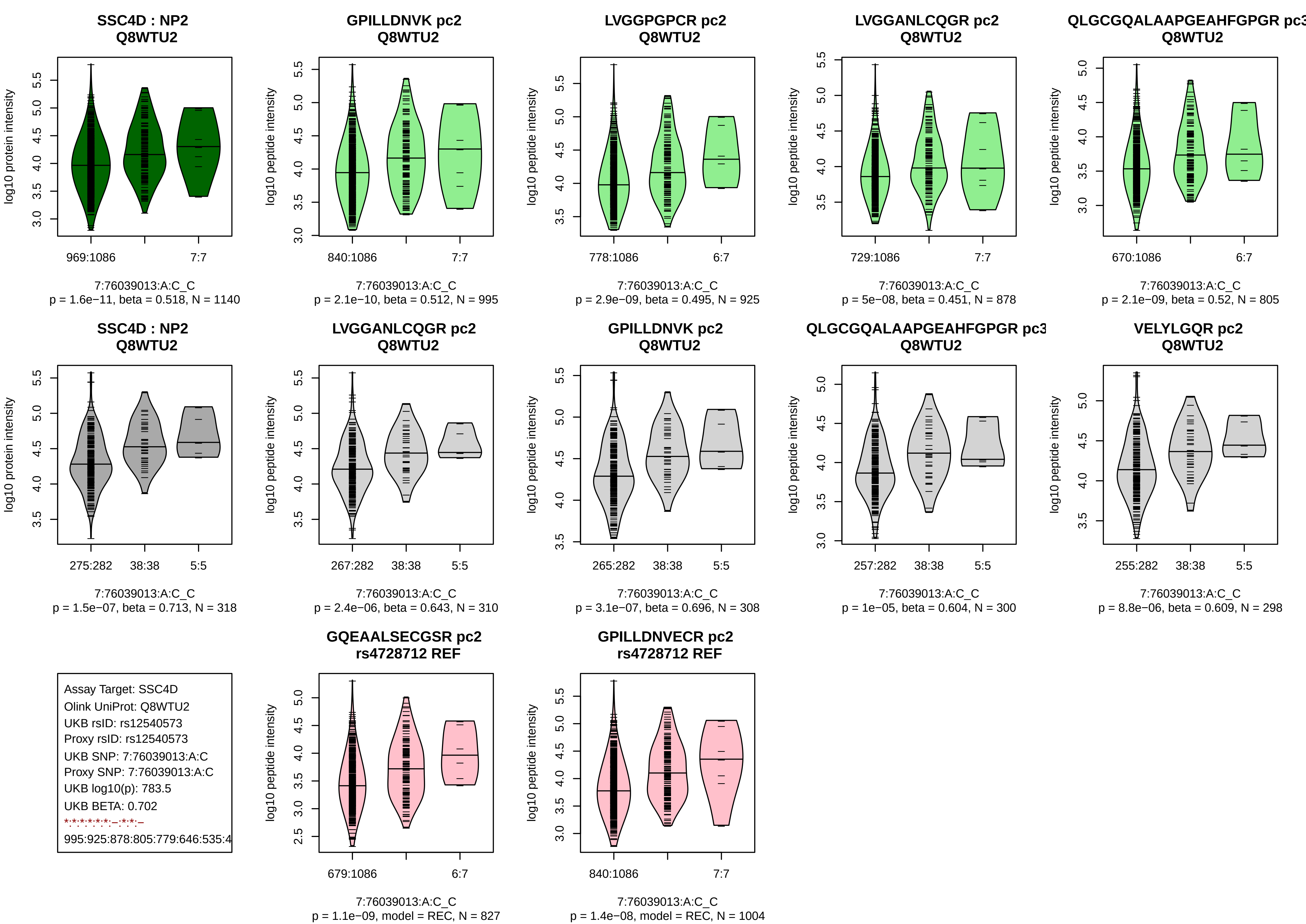
SSC4D : NP2
Q8WTU2
log10 protein intensity
969:1086
7:7
7:76039013:A:C_C
p = 1.6e−11, beta = 0.518, N = 1140
GPILLDNVK pc2
Q8WTU2
log10 peptide intensity
840:1086
7:7
7:76039013:A:C_C
p = 2.1e−10, beta = 0.512, N = 995
LVGGPGPCR pc2
Q8WTU2
log10 peptide intensity
778:1086
6:7
7:76039013:A:C_C
p = 2.9e−09, beta = 0.495, N = 925
LVGGANLCQGR pc2
Q8WTU2
log10 peptide intensity
729:1086
7:7
7:76039013:A:C_C
p = 5e−08, beta = 0.451, N = 878
QLGCGQALAAPGEAHFGPGR pc3
Q8WTU2
log10 peptide intensity
670:1086
6:7
7:76039013:A:C_C
p = 2.1e−09, beta = 0.52, N = 805
SSC4D : NP2
Q8WTU2
log10 protein intensity
275:282
38:38
5:5
7:76039013:A:C_C
p = 1.5e−07, beta = 0.713, N = 318
LVGGANLCQGR pc2
Q8WTU2
log10 peptide intensity
267:282
38:38
5:5
7:76039013:A:C_C
p = 2.4e−06, beta = 0.643, N = 310
GPILLDNVK pc2
Q8WTU2
log10 peptide intensity
265:282
38:38
5:5
7:76039013:A:C_C
p = 3.1e−07, beta = 0.696, N = 308
QLGCGQALAAPGEAHFGPGR pc3
Q8WTU2
log10 peptide intensity
257:282
38:38
5:5
7:76039013:A:C_C
p = 1e−05, beta = 0.604, N = 300
VELYLGQR pc2
Q8WTU2
log10 peptide intensity
255:282
38:38
5:5
7:76039013:A:C_C
p = 8.8e−06, beta = 0.609, N = 298
Assay Target: SSC4D
Olink UniProt: Q8WTU2
UKB rsID: rs12540573
Proxy rsID: rs12540573
UKB SNP: 7:76039013:A:C
Proxy SNP: 7:76039013:A:C
UKB log10(p): 783.5
UKB BETA: 0.702
*:*:*:*:*:*:–:*:*:–
995:925:878:805:779:646:535:4
GQEAALSECGSR pc2
rs4728712 REF
log10 peptide intensity
679:1086
6:7
7:76039013:A:C_C
p = 1.1e−09, model = REC, N = 827
GPILLDNVECR pc2
rs4728712 REF
log10 peptide intensity
840:1086
7:7
7:76039013:A:C_C
p = 1.4e−08, model = REC, N = 1004

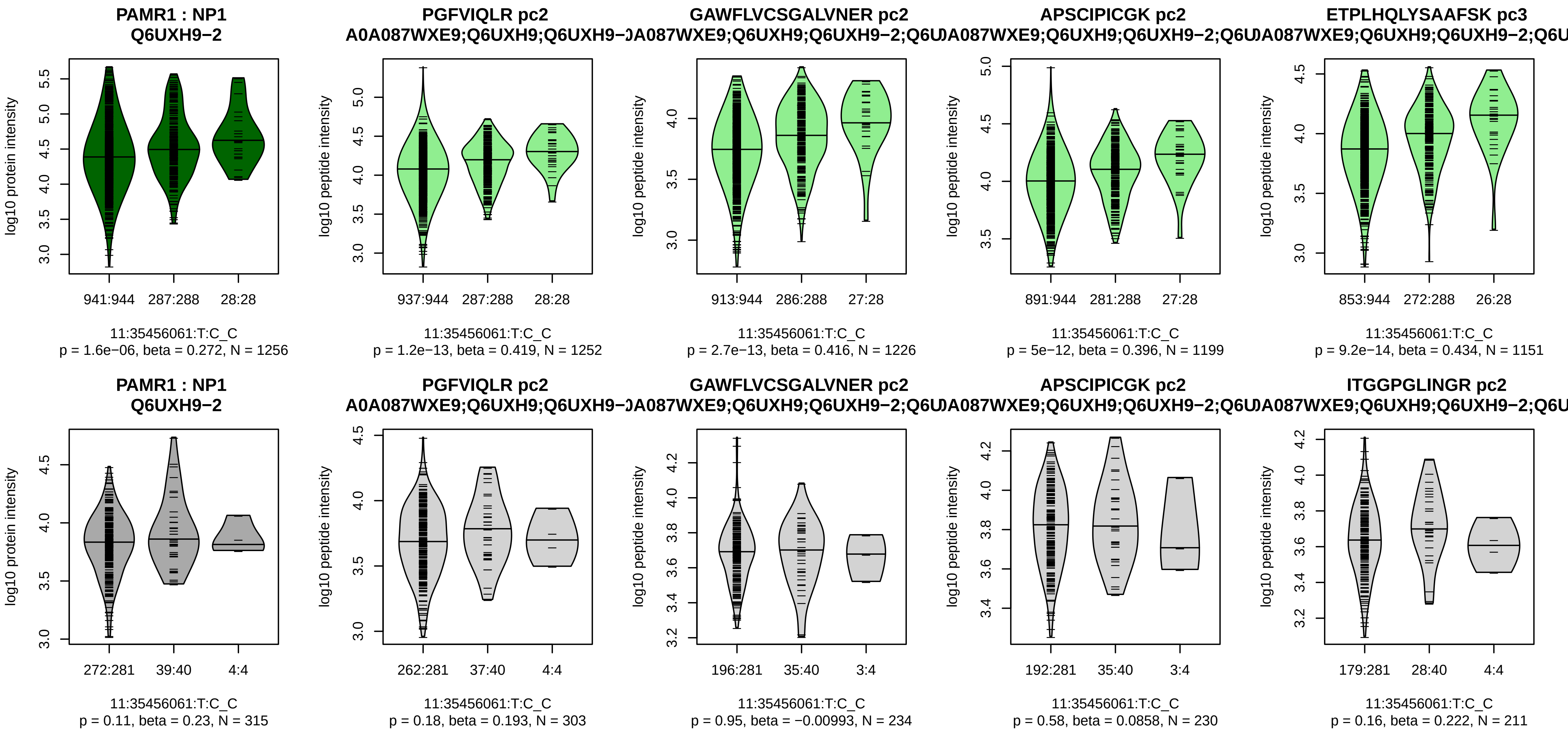


Assay Target: PAMR1
Olink UniProt: Q6UXH9
UKB rsID: rs61736408
Proxy rsID: rs61736408
UKB SNP: 11:35456061:T:C
Proxy SNP: 11:35456061:T:C
UKB log10(p): 772.5
UKB BETA: −0.481
*:*:*:*:*:–:*:*:*:*:–:–:*:*:*:–:–:–:*:
1252:1226:1199:1151:1133:106

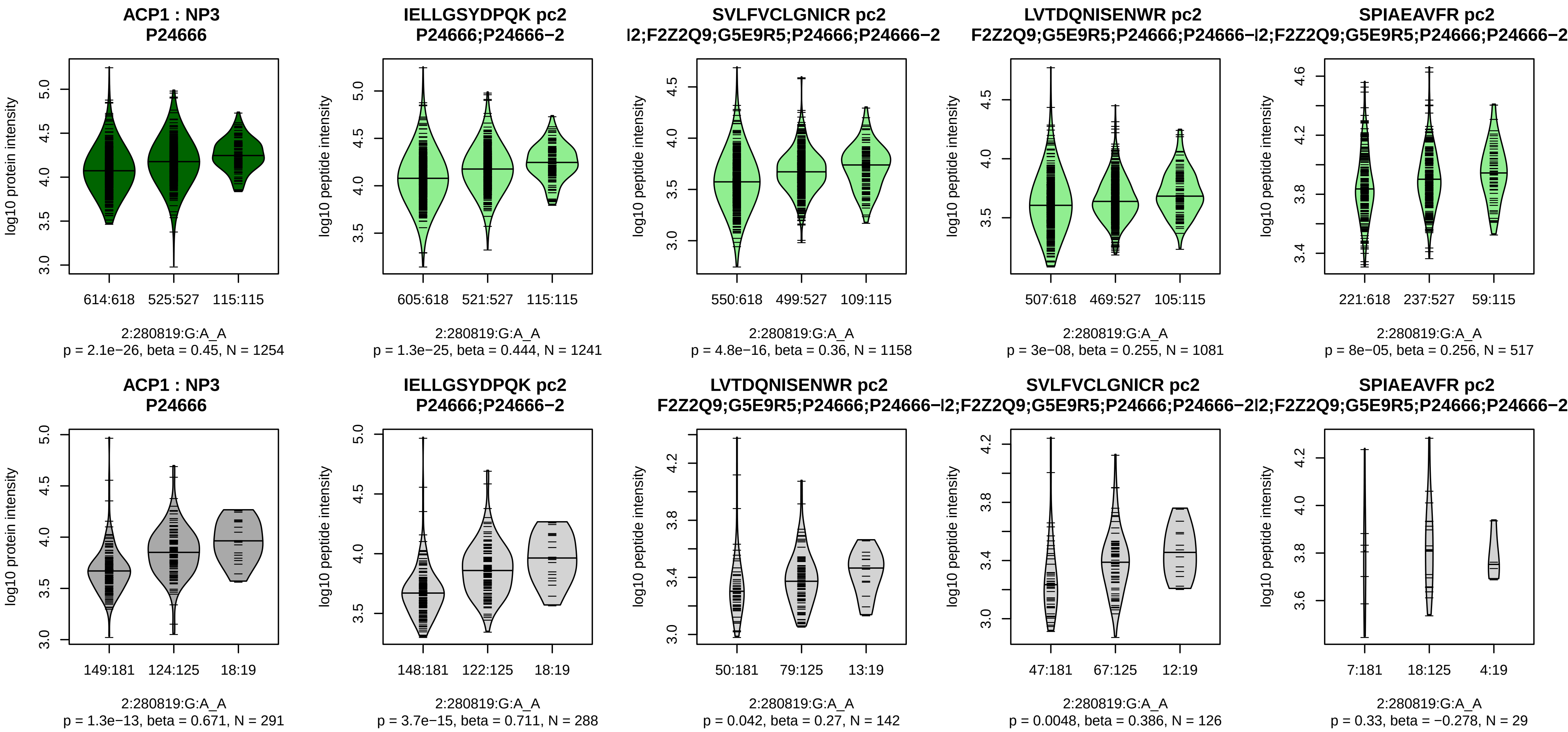


Assay Target: ACP1
Olink UniProt: P24666
UKB rsID: rs7605824
Proxy rsID: rs7605824
UKB SNP: 2:280819:G:A
Proxy SNP: 2:280819:G:A
UKB log10(p): 736.8
UKB BETA: −0.376
*:*:*:*:*:−:−:−:−:NA:NA:NA
1241:1158:1081:517:226:165:11

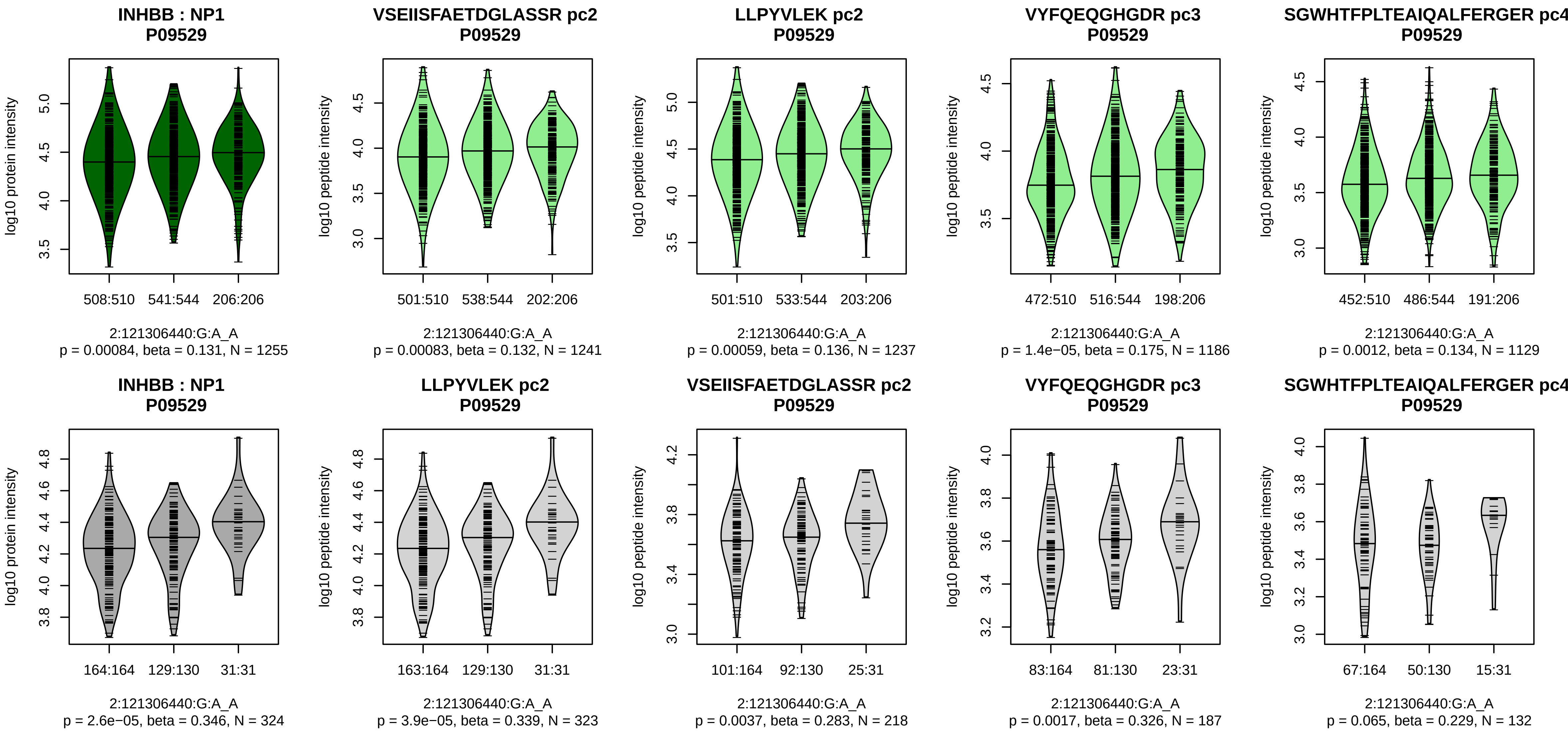


Assay Target: INHBB
Olink UniProt: P09529
UKB rsID: rs17050272
Proxy rsID: rs17050272
UKB SNP: 2:121306440:G:A
Proxy SNP: 2:121306440:G:A
UKB log10(p): 735
UKB BETA: 0.355
*:*:*:*:−:−:−:−:−:−:−:−:−:−:NA
1241:1237:1186:1129:751:500:3

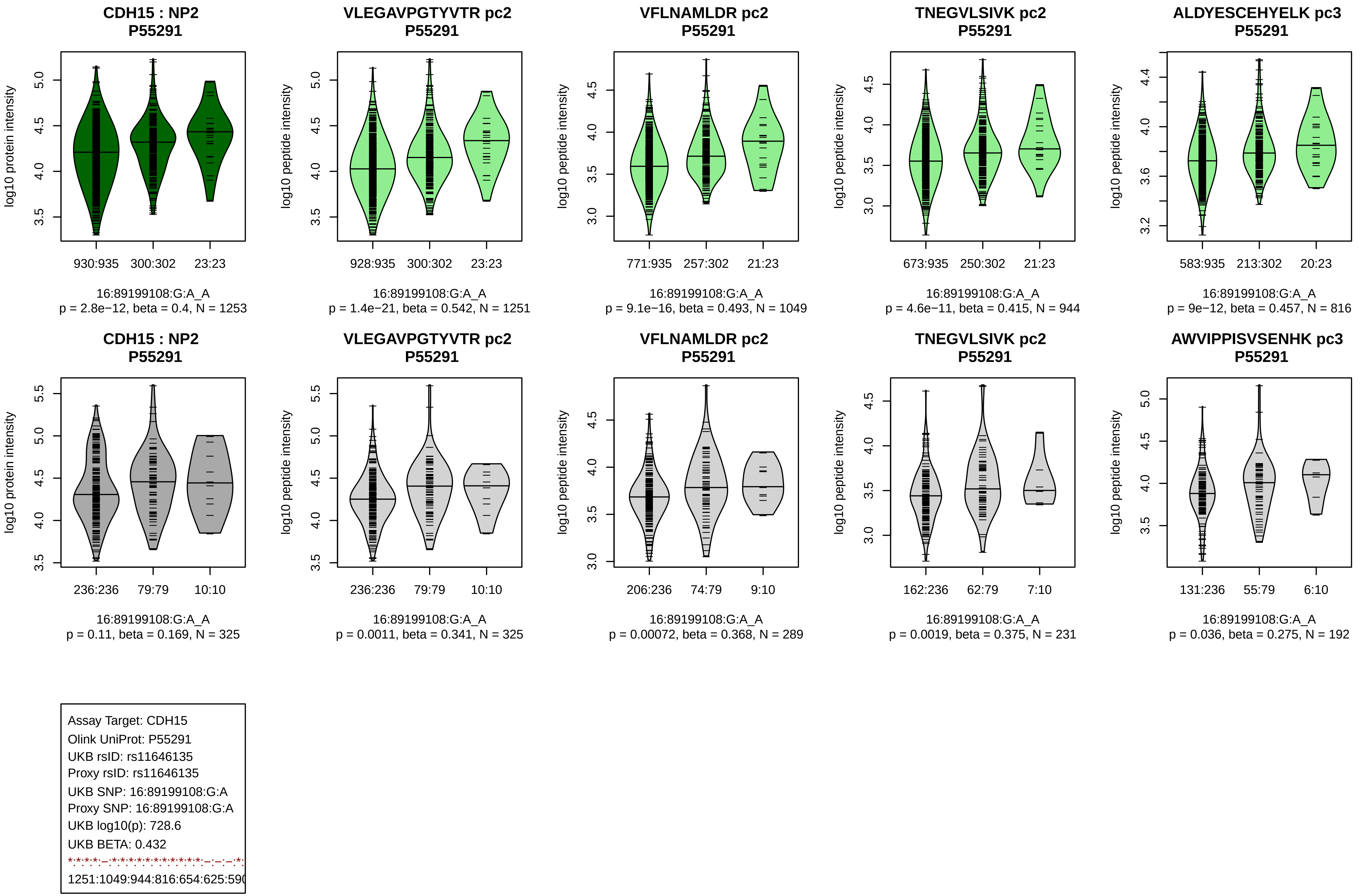
CDH15 : NP2
P55291
log10 protein intensity
930:935
300:302
23:23
16:89199108:G:A_A
p = 2.8e−12, beta = 0.4, N = 1253
VLEGAVPGTYVTR pc2
P55291
log10 peptide intensity
928:935
300:302
23:23
16:89199108:G:A_A
p = 1.4e−21, beta = 0.542, N = 1251
VFLNAMLDR pc2
P55291
log10 peptide intensity
771:935
257:302
21:23
16:89199108:G:A_A
p = 9.1e−16, beta = 0.493, N = 1049
TNEGVLSIVK pc2
P55291
log10 peptide intensity
673:935
250:302
21:23
16:89199108:G:A_A
p = 4.6e−11, beta = 0.415, N = 944
ALDYESCEHYELK pc3
P55291
log10 peptide intensity
583:935
213:302
20:23
16:89199108:G:A_A
p = 9e−12, beta = 0.457, N = 816
CDH15 : NP2
P55291
log10 protein intensity
236:236
79:79
10:10
16:89199108:G:A_A
p = 0.11, beta = 0.169, N = 325
VLEGAVPGTYVTR pc2
P55291
log10 peptide intensity
236:236
79:79
10:10
16:89199108:G:A_A
p = 0.0011, beta = 0.341, N = 325
VFLNAMLDR pc2
P55291
log10 peptide intensity
206:236
74:79
9:10
16:89199108:G:A_A
p = 0.00072, beta = 0.368, N = 289
TNEGVLSIVK pc2
P55291
log10 peptide intensity
162:236
62:79
7:10
16:89199108:G:A_A
p = 0.0019, beta = 0.375, N = 231
AWVIPPISVSENHK pc3
P55291
log10 peptide intensity
131:236
55:79
6:10
16:89199108:G:A_A
p = 0.036, beta = 0.275, N = 192
Assay Target: CDH15
Olink UniProt: P55291
UKB rsID: rs11646135
Proxy rsID: rs11646135
UKB SNP: 16:89199108:G:A
Proxy SNP: 16:89199108:G:A
UKB log10(p): 728.6
UKB BETA: 0.432
1251:1049:944:816:654:625:59

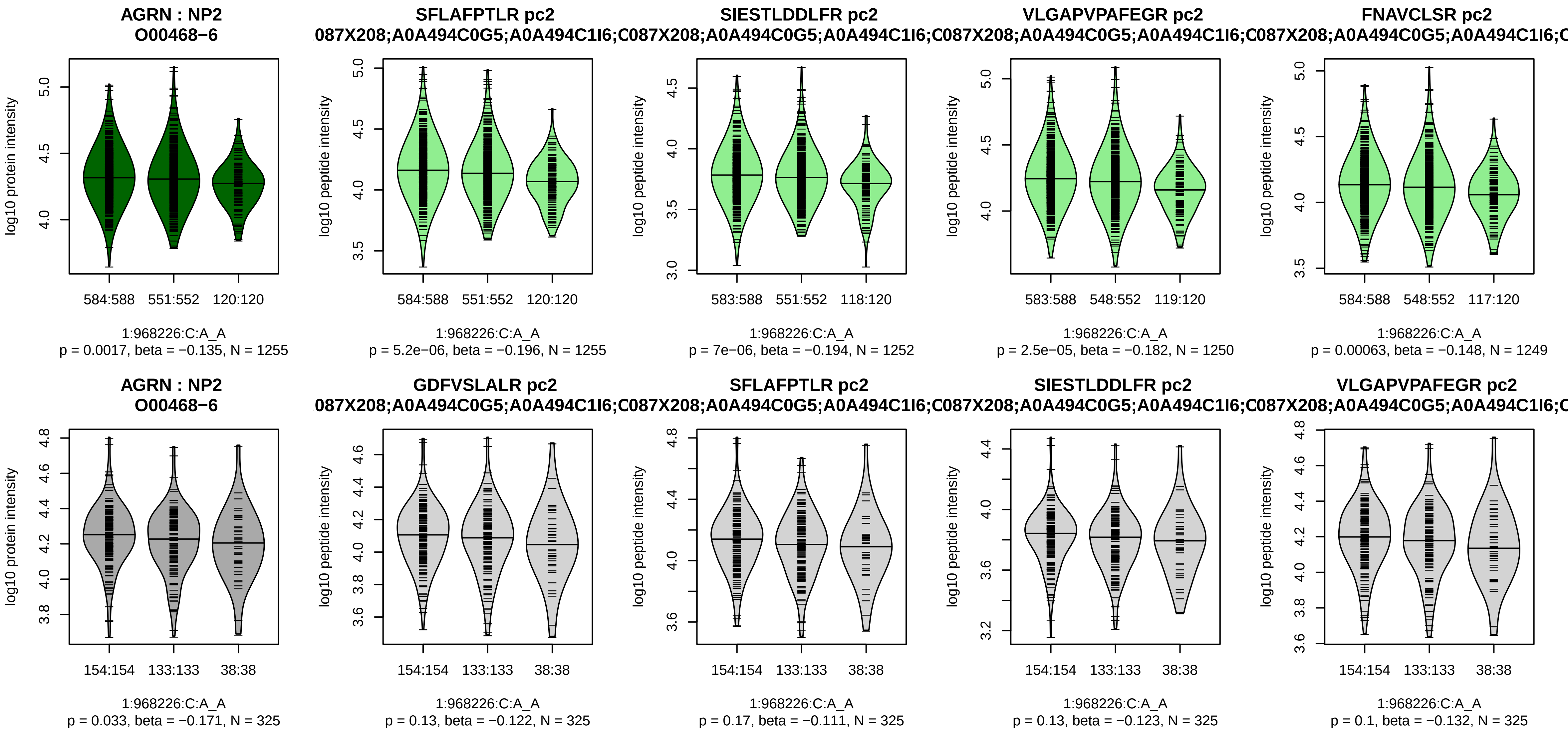


Assay Target: AGRN
Olink UniProt: O00468
UKB rsID: rs3121560
Proxy rsID: rs3121560
UKB SNP: 1:968226:C:A
Proxy SNP: 1:968226:C:A
UKB log10(p): 720.7
UKB BETA: −0.364
*:*:*:*:*:*:*:*:*:*:*:*:*:–:*:–:*:–:*:*:*
1255:1252:1250:1249:1248:124

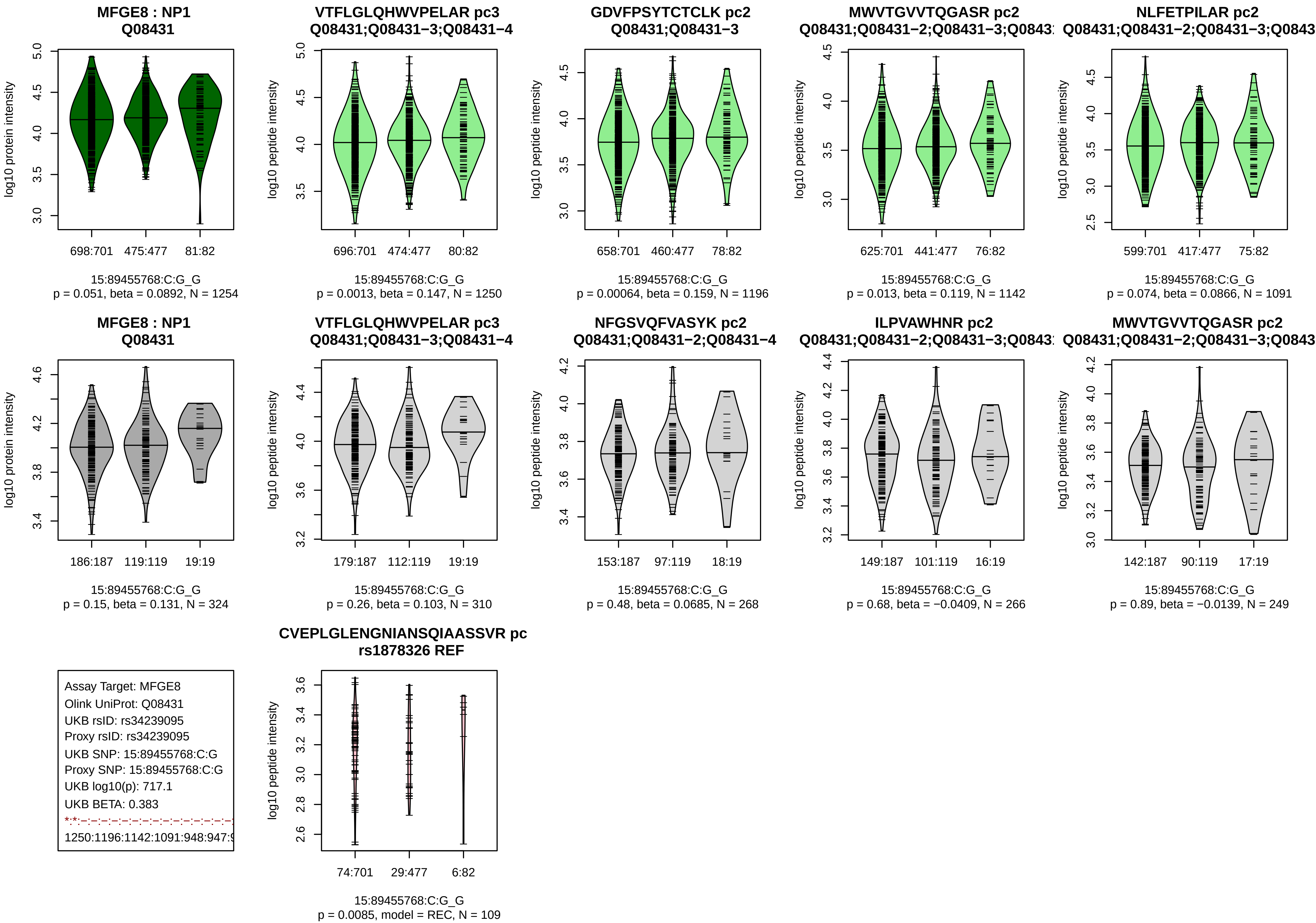
MFGE8 : NP1
Q08431
log10 protein intensity
698:701 475:477 81:82
15:89455768:C:G_G
p = 0.051, beta = 0.0892, N = 1254
VTFLGLQHWVPELAR pc3
Q08431;Q08431−3;Q08431−4
log10 peptide intensity
696:701 474:477 80:82
15:89455768:C:G_G
p = 0.0013, beta = 0.147, N = 1250
GDVFPSYTCTCLK pc2
Q08431;Q08431−3
658:701 460:477 78:82
15:89455768:C:G_G
p = 0.00064, beta = 0.159, N = 1196
MWVTGVVTQGASR pc2
625:701 441:477 76:82
15:89455768:C:G_G
p = 0.013, beta = 0.119, N = 1142
NLFETPILAR pc2
599:701 417:477 75:82
15:89455768:C:G_G
p = 0.074, beta = 0.0866, N = 1091
MFGE8 : NP1
Q08431
186:187 119:119 19:19
15:89455768:C:G_G
p = 0.15, beta = 0.131, N = 324
VTFLGLQHWVPELAR pc3
Q08431;Q08431−3;Q08431−4
179:187 112:119 19:19
15:89455768:C:G_G
p = 0.26, beta = 0.103, N = 310
NFGSVQFVASYK pc2
Q08431;Q08431−2;Q08431−4
153:187 97:119 18:19
15:89455768:C:G_G
p = 0.48, beta = 0.0685, N = 268
ILPVAWHNR pc2
149:187 101:119 16:19
15:89455768:C:G_G
p = 0.68, beta = −0.0409, N = 266
MWVTGVVTQGASR pc2
142:187 90:119 17:19
15:89455768:C:G_G
p = 0.89, beta = −0.0139, N = 249
Assay Target: MFGE8
Olink UniProt: Q08431
UKB rsID: rs34239095
Proxy rsID: rs34239095
UKB SNP: 15:89455768:C:G
Proxy SNP: 15:89455768:C:G
UKB log10(p): 717.1
UKB BETA: 0.383
1250:1196:1142:1091:948:947:
CVEPLGLENGNIANSQIAASSVR pc
rs1878326 REF
74:701 29:477 6:82
15:89455768:C:G_G
p = 0.0085, model = REC, N = 109

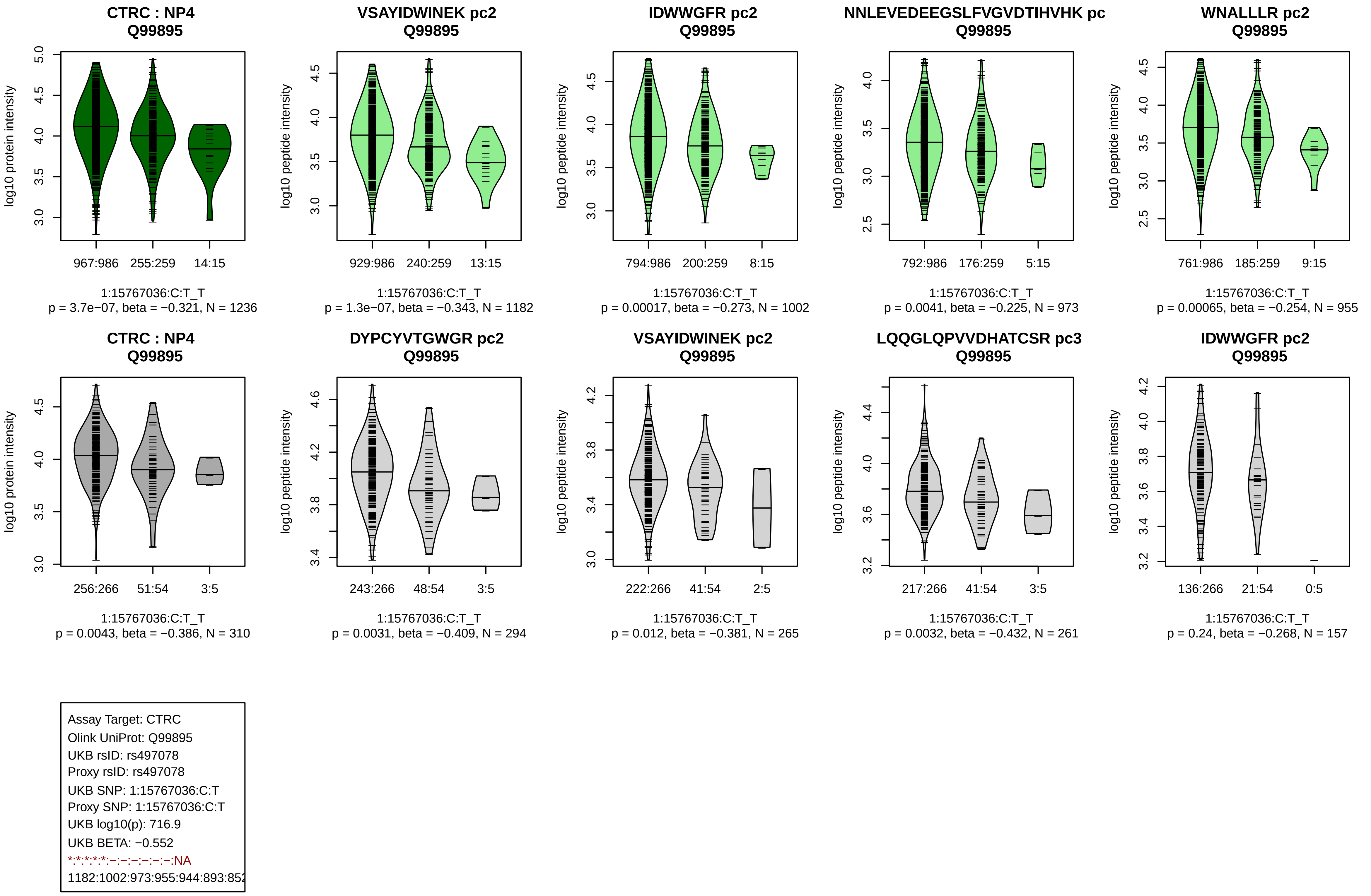
CTRC : NP4
Q99895
log10 protein intensity
967:986
255:259
14:15
1:15767036:C:T_T
p = 3.7e−07, beta = −0.321, N = 1236
VSAYIDWINEK pc2
Q99895
log10 peptide intensity
929:986
240:259
13:15
1:15767036:C:T_T
p = 1.3e−07, beta = −0.343, N = 1182
IDWWGFR pc2
Q99895
log10 peptide intensity
794:986
200:259
8:15
1:15767036:C:T_T
p = 0.00017, beta = −0.273, N = 1002
NNLEVEDEEGSLFVGVDTIHVHK pc
Q99895
log10 peptide intensity
792:986
176:259
5:15
1:15767036:C:T_T
p = 0.0041, beta = −0.225, N = 973
WNALLLR pc2
Q99895
log10 peptide intensity
761:986
185:259
9:15
1:15767036:C:T_T
p = 0.00065, beta = −0.254, N = 955
CTRC : NP4
Q99895
log10 protein intensity
256:266
51:54
3:5
1:15767036:C:T_T
p = 0.0043, beta = −0.386, N = 310
DYPCYVTGWGR pc2
Q99895
log10 peptide intensity
243:266
48:54
3:5
1:15767036:C:T_T
p = 0.0031, beta = −0.409, N = 294
VSAYIDWINEK pc2
Q99895
log10 peptide intensity
222:266
41:54
2:5
1:15767036:C:T_T
p = 0.012, beta = −0.381, N = 265
LQQGLQPVVDHATCSR pc3
Q99895
log10 peptide intensity
217:266
41:54
3:5
1:15767036:C:T_T
p = 0.0032, beta = −0.432, N = 261
IDWWGFR pc2
Q99895
log10 peptide intensity
136:266
21:54
0:5
1:15767036:C:T_T
p = 0.24, beta = −0.268, N = 157
Assay Target: CTRC
Olink UniProt: Q99895
UKB rsID: rs497078
Proxy rsID: rs497078
UKB SNP: 1:15767036:C:T
Proxy SNP: 1:15767036:C:T
UKB log10(p): 716.9
UKB BETA: −0.552
*:*:*:*:*:−:−:−:−:−:−:−:NA
1182:1002:973:955:944:893:85

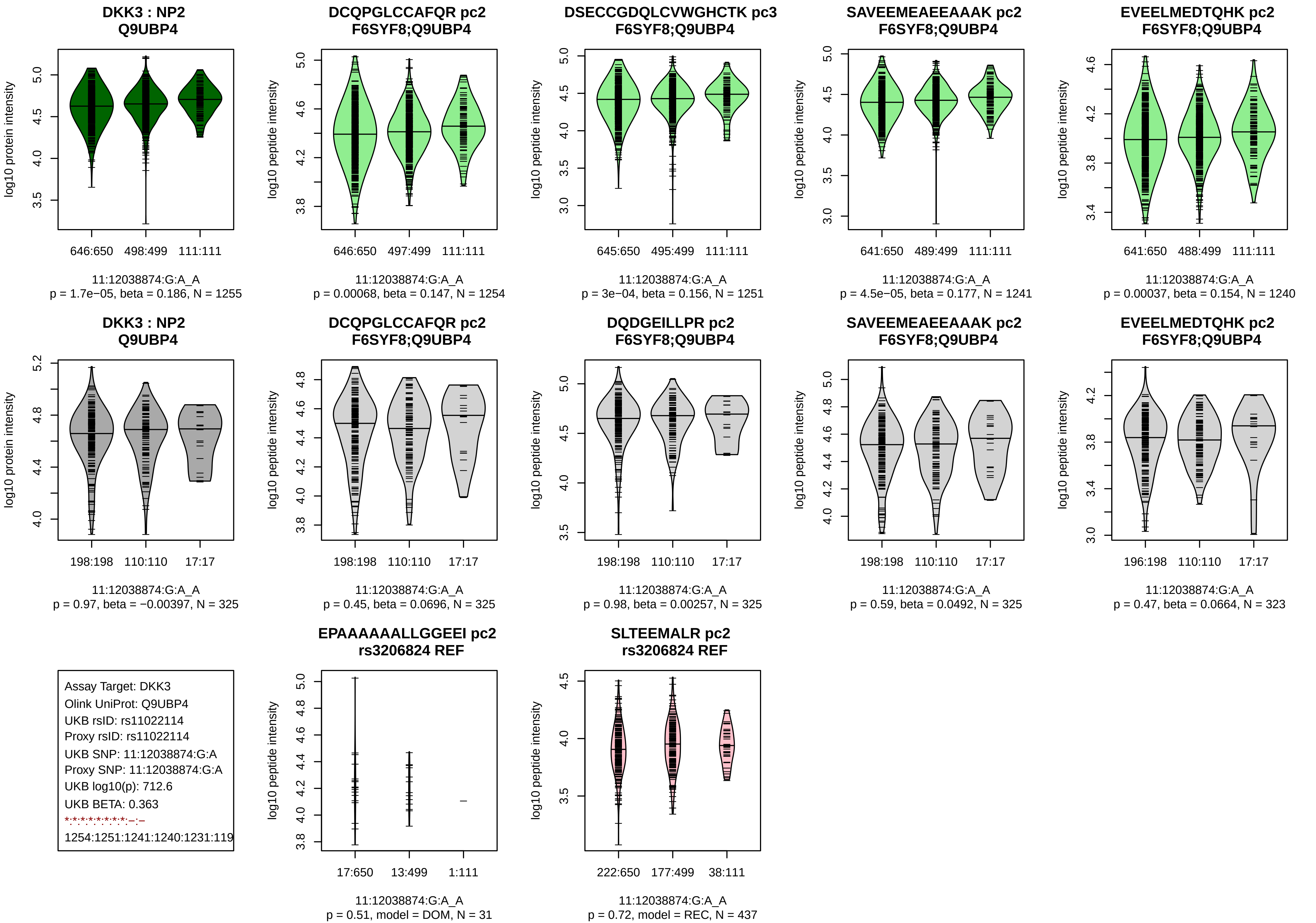
DKK3 : NP2
Q9UBP4
log10 protein intensity
646:650 498:499 111:111
11:12038874:G:A_A
p = 1.7e−05, beta = 0.186, N = 1255
DCQPGLCCAFQR pc2
F6SYF8;Q9UBP4
log10 peptide intensity
646:650 497:499 111:111
11:12038874:G:A_A
p = 0.00068, beta = 0.147, N = 1254
DSECCGDQLCVWGHCTK pc3
F6SYF8;Q9UBP4
645:650 495:499 111:111
11:12038874:G:A_A
p = 3e−04, beta = 0.156, N = 1251
SAVEEMEAEEAAAK pc2
F6SYF8;Q9UBP4
641:650 489:499 111:111
11:12038874:G:A_A
p = 4.5e−05, beta = 0.177, N = 1241
EVEELMEDTQHK pc2
F6SYF8;Q9UBP4
641:650 488:499 111:111
11:12038874:G:A_A
p = 0.00037, beta = 0.154, N = 1240
DKK3 : NP2
Q9UBP4
198:198 110:110 17:17
11:12038874:G:A_A
p = 0.97, beta = −0.00397, N = 325
DCQPGLCCAFQR pc2
F6SYF8;Q9UBP4
198:198 110:110 17:17
11:12038874:G:A_A
p = 0.45, beta = 0.0696, N = 325
DQDGEILLPR pc2
F6SYF8;Q9UBP4
198:198 110:110 17:17
11:12038874:G:A_A
p = 0.98, beta = 0.00257, N = 325
SAVEEMEAEEAAAK pc2
F6SYF8;Q9UBP4
198:198 110:110 17:17
11:12038874:G:A_A
p = 0.59, beta = 0.0492, N = 325
EVEELMEDTQHK pc2
F6SYF8;Q9UBP4
196:198 110:110 17:17
11:12038874:G:A_A
p = 0.47, beta = 0.0664, N = 323
Assay Target: DKK3
Olink UniProt: Q9UBP4
UKB rsID: rs11022114
Proxy rsID: rs11022114
UKB SNP: 11:12038874:G:A
Proxy SNP: 11:12038874:G:A
UKB log10(p): 712.6
UKB BETA: 0.363
1254:1251:1241:1240:1231:119
EPAAAAAALLGGEEI pc2
rs3206824 REF
17:650 13:499 1:111
11:12038874:G:A_A
p = 0.51, model = DOM, N = 31
SLTEEMALR pc2
rs3206824 REF
222:650 177:499 38:111
11:12038874:G:A_A
p = 0.72, model = REC, N = 437

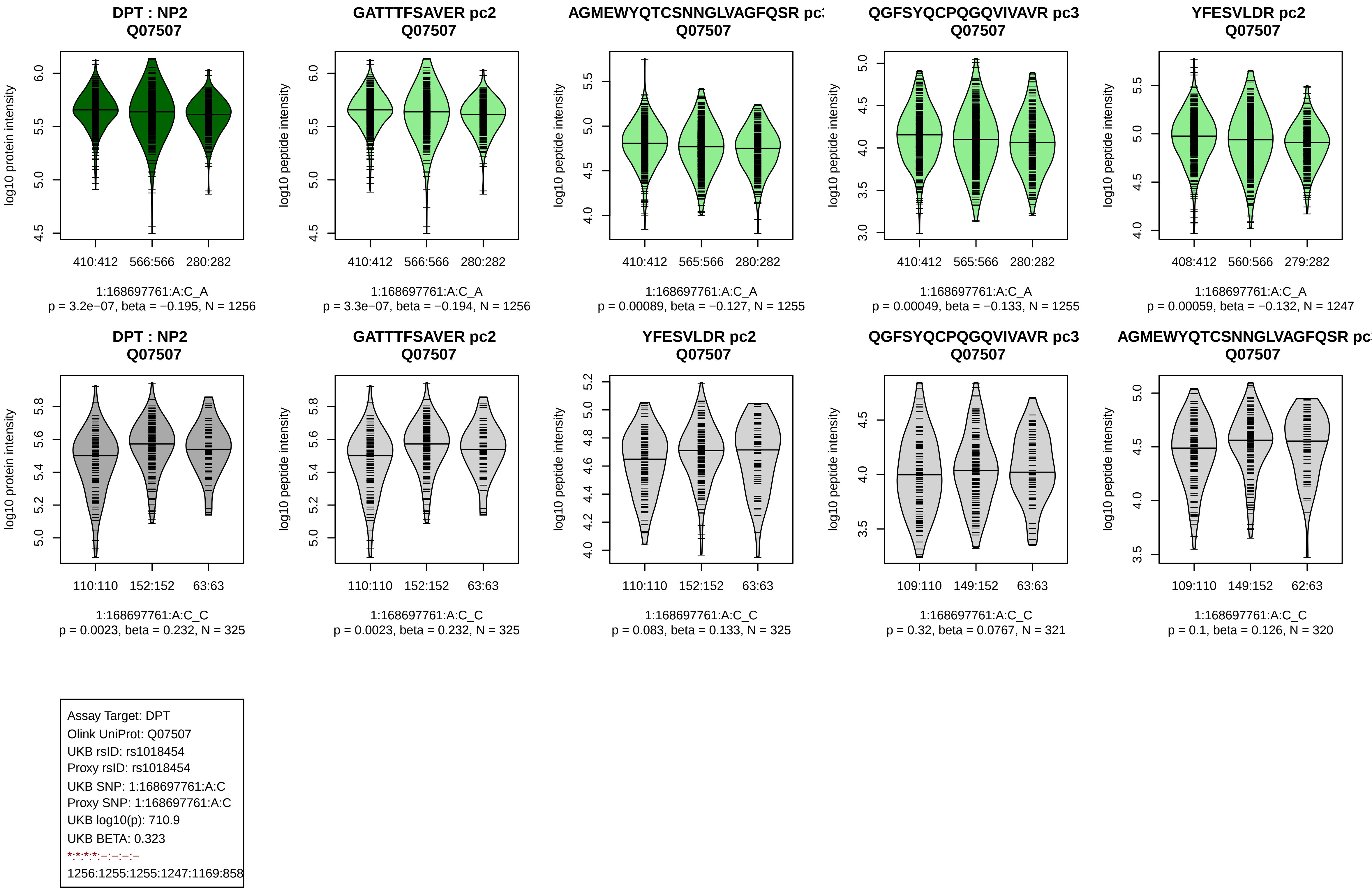
DPT : NP2
Q07507
log10 protein intensity
410:412
566:566
280:282
1:168697761:A:C_A
p = 3.2e−07, beta = −0.195, N = 1256
GATTTFSAVER pc2
Q07507
log10 peptide intensity
410:412
566:566
280:282
1:168697761:A:C_A
p = 3.3e−07, beta = −0.194, N = 1256
AGMEWYQTCSNNGLVAGFQSR pc
Q07507
log10 peptide intensity
410:412
565:566
280:282
1:168697761:A:C_A
p = 0.00089, beta = −0.127, N = 1255
QGFSYQCPQGQVIVAVR pc3
Q07507
log10 peptide intensity
410:412
565:566
280:282
1:168697761:A:C_A
p = 0.00049, beta = −0.133, N = 1255
YFESVLDR pc2
Q07507
log10 peptide intensity
408:412
560:566
279:282
1:168697761:A:C_A
p = 0.00059, beta = −0.132, N = 1247
DPT : NP2
Q07507
log10 protein intensity
110:110
152:152
63:63
1:168697761:A:C_C
p = 0.0023, beta = 0.232, N = 325
GATTTFSAVER pc2
Q07507
log10 peptide intensity
110:110
152:152
63:63
1:168697761:A:C_C
p = 0.0023, beta = 0.232, N = 325
YFESVLDR pc2
Q07507
log10 peptide intensity
110:110
152:152
63:63
1:168697761:A:C_C
p = 0.083, beta = 0.133, N = 325
QGFSYQCPQGQVIVAVR pc3
Q07507
log10 peptide intensity
109:110
149:152
63:63
1:168697761:A:C_C
p = 0.32, beta = 0.0767, N = 321
AGMEWYQTCSNNGLVAGFQSR pc
Q07507
log10 peptide intensity
109:110
149:152
62:63
1:168697761:A:C_C
p = 0.1, beta = 0.126, N = 320
Assay Target: DPT
Olink UniProt: Q07507
UKB rsID: rs1018454
Proxy rsID: rs1018454
UKB SNP: 1:168697761:A:C
Proxy SNP: 1:168697761:A:C
UKB log10(p): 710.9
UKB BETA: 0.323
*:*:*:*:–:–:–:–
1256:1255:1255:1247:1169:858

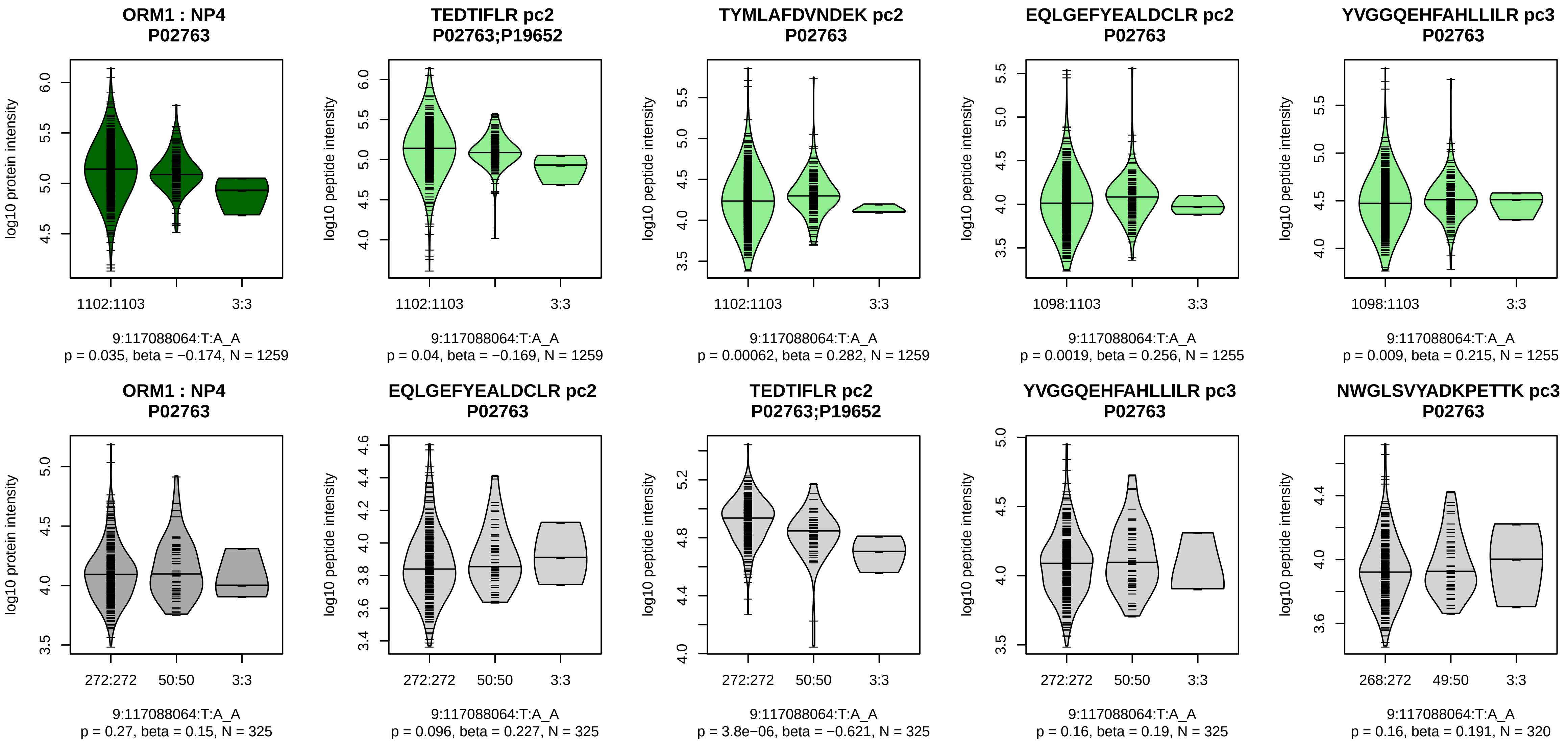


Assay Target: ORM1
Olink UniProt: P02763
UKB rsID: rs10982156
Proxy rsID: rs10982156
UKB SNP: 9:117088064:T:A
Proxy SNP: 9:117088064:T:A
UKB log10(p): 706.3
UKB BETA: 0.721
−:*:*:*:−:−:−:−:−:−:−:−:−:NA:NA:I
1259:1259:1255:1255:1153:110

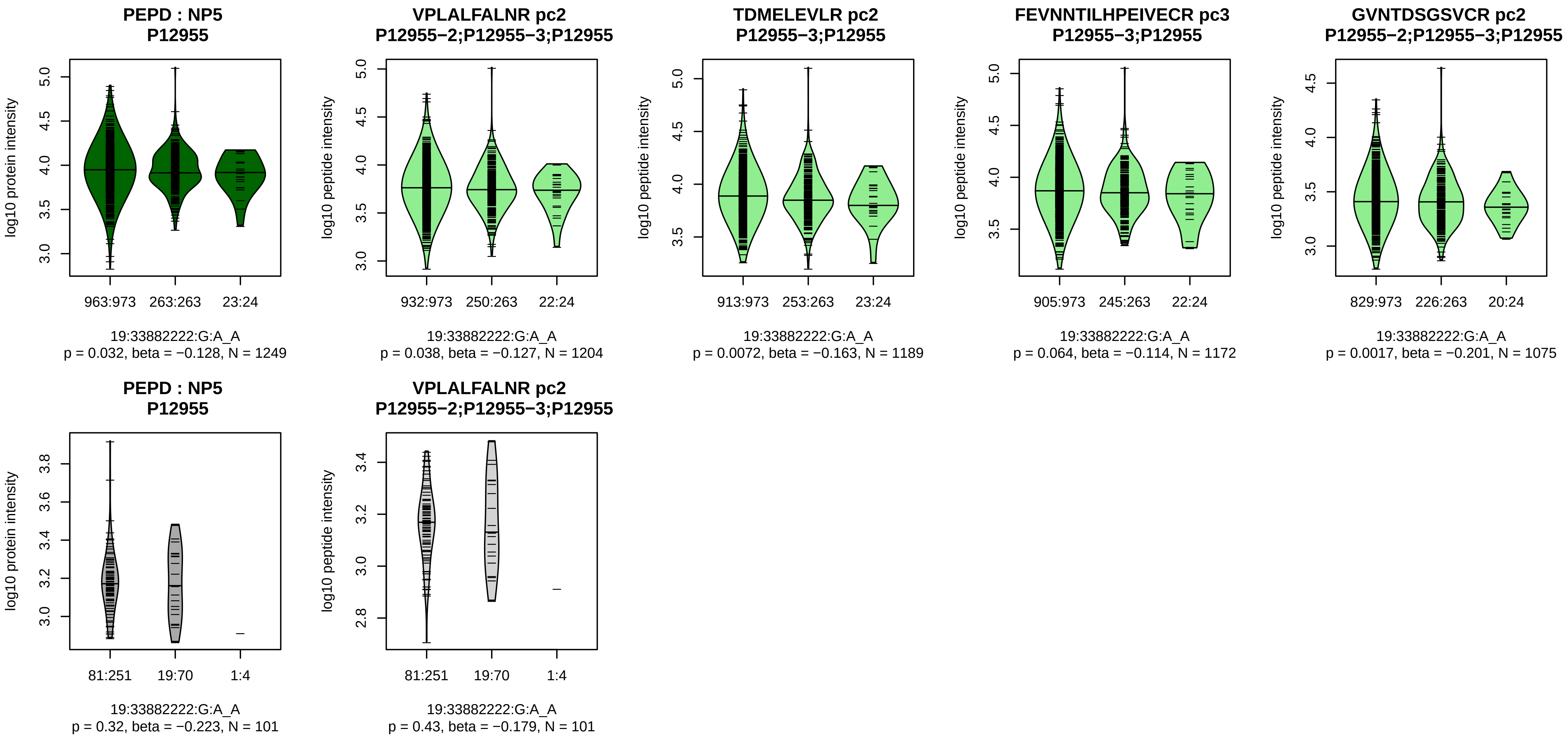


Assay Target: PEPD
Olink UniProt: P12955
UKB rsID: rs17569
Proxy rsID: rs17569
UKB SNP: 19:33882222:G:A
Proxy SNP: 19:33882222:G:A
UKB log10(p): 697.8
UKB BETA: −0.478
−:*:−:*:−:−:−:−:−:−:−:−:−:−:−:−:−
1204:1189:1172:1075:967:937:[illegible]

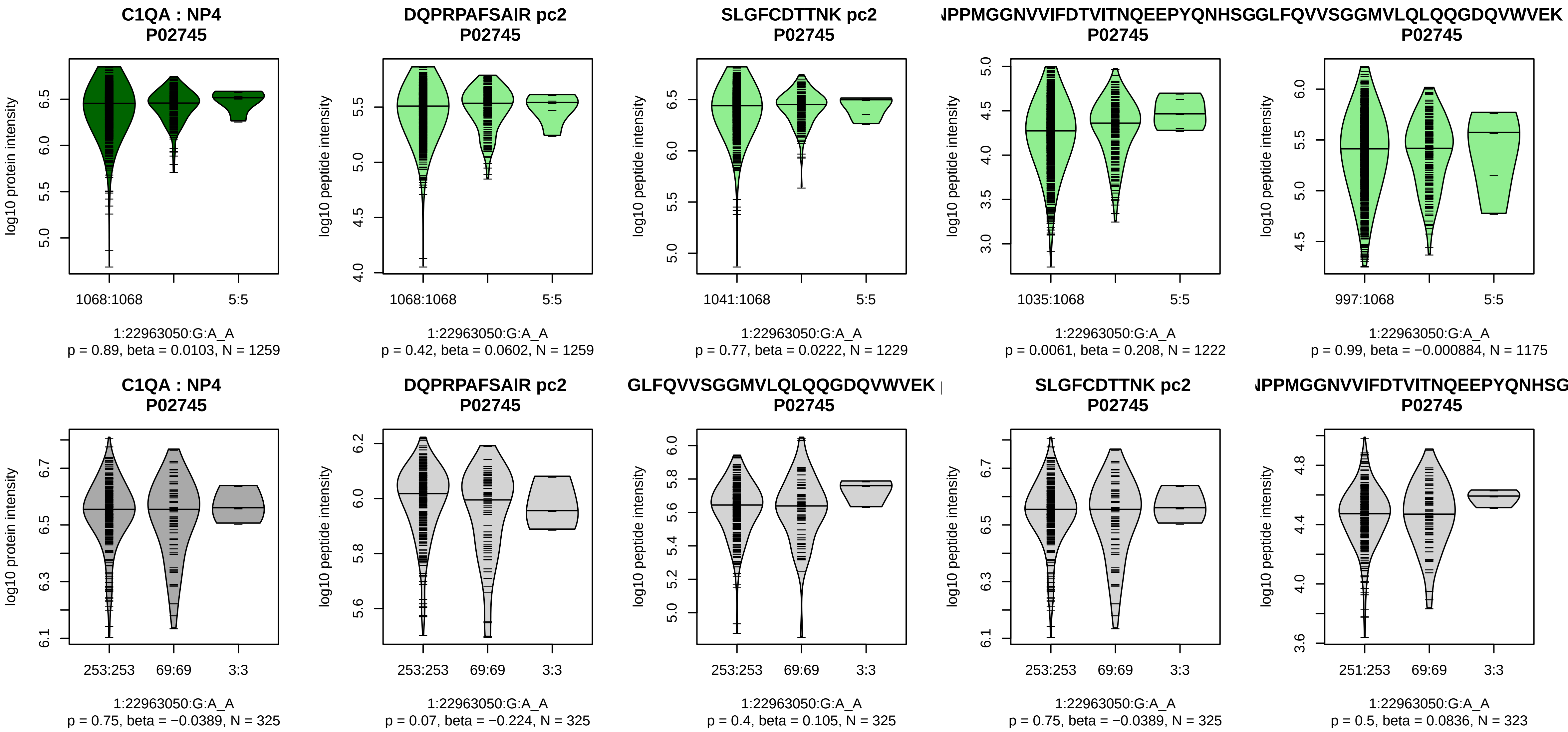


Assay Target: C1QA
Olink UniProt: P02745
UKB rsID: rs12058824
Proxy rsID: rs12058824
UKB SNP: 1:22963050:G:A
Proxy SNP: 1:22963050:G:A
UKB log10(p): 696.2
UKB BETA: −0.621
−:−:*:−:−:−:−:−:−:−:−:−:−:−:−:−:NA
1259:1229:1222:1175:1091:108

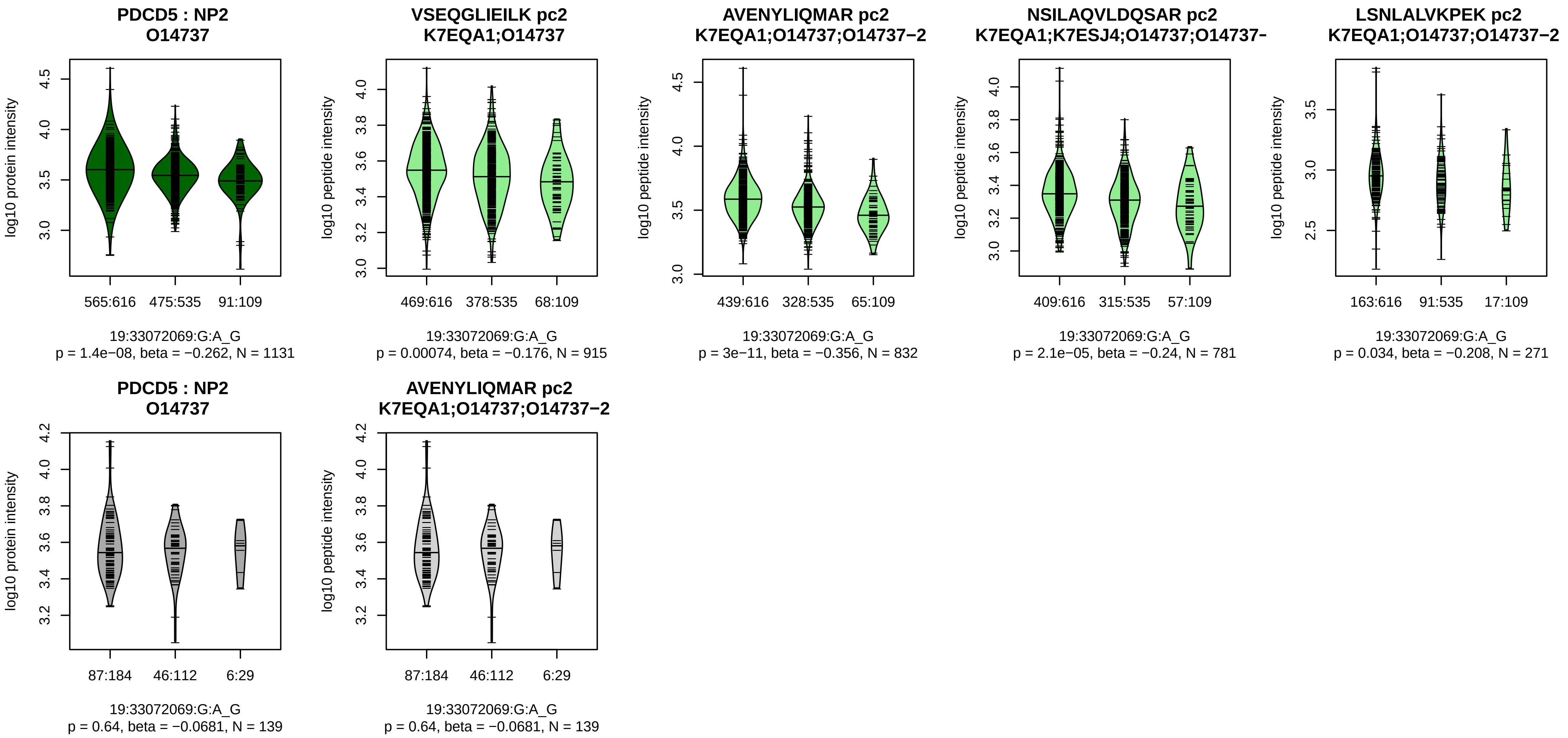


Assay Target: PDCD5
Olink UniProt: O14737
UKB rsID: rs10402931
Proxy rsID: rs10402931
UKB SNP: 19:33072069:G:A
Proxy SNP: 19:33072069:G:A
UKB log10(p): 691.9
UKB BETA: 0.376
*:*:*:−:−:NA:NA
915:832:781:271:121:1:2

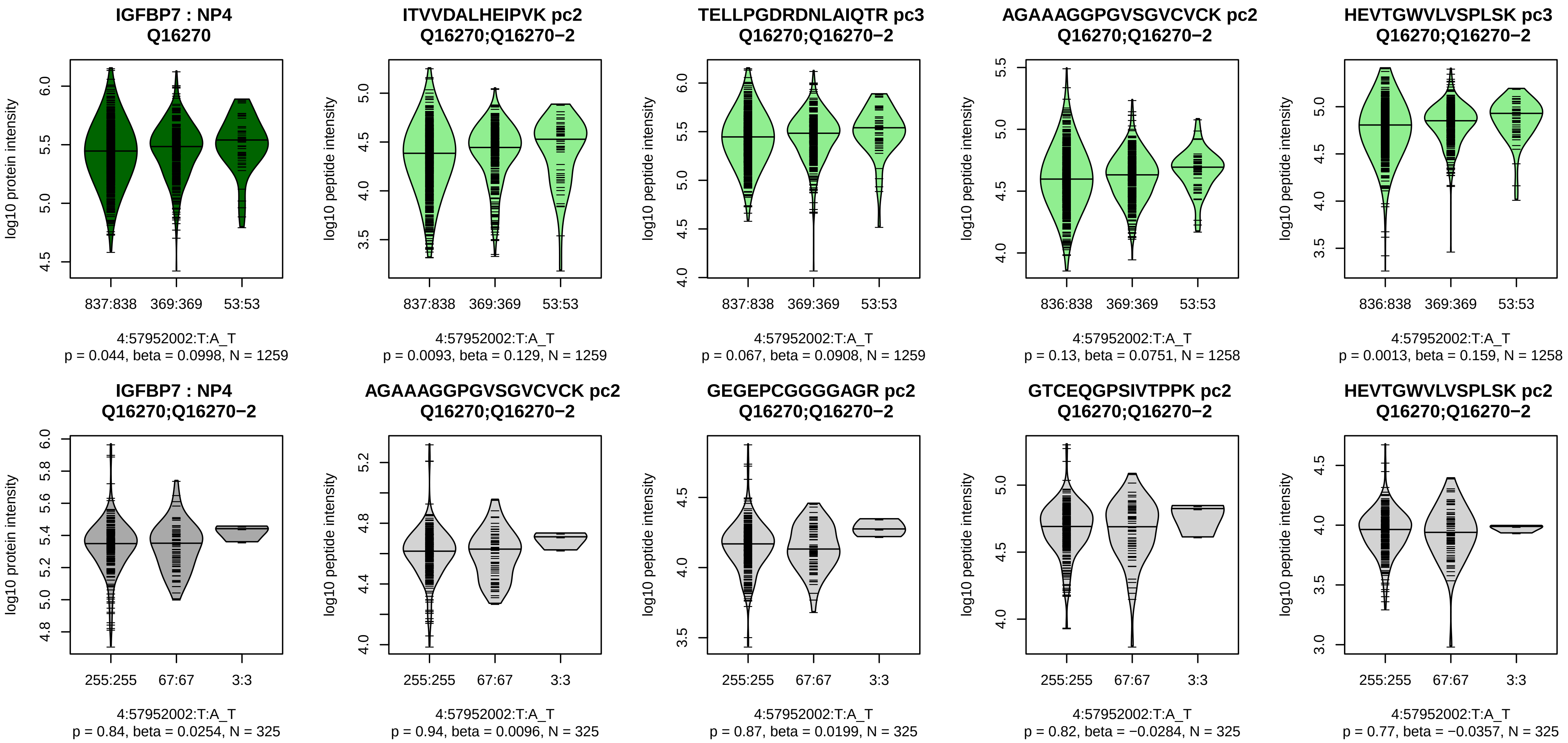


Assay Target: IGFBP7
Olink UniProt: Q16270
UKB rsID: rs6827768
Proxy rsID: rs6827768
UKB SNP: 4:57952002:T:A
Proxy SNP: 4:57952002:T:A
UKB log10(p): 684.7
UKB BETA: −0.393
*:−:−:*:*:−:*:−:*:−:−:−:−:NA
1259:1259:1258:1258:1258:125

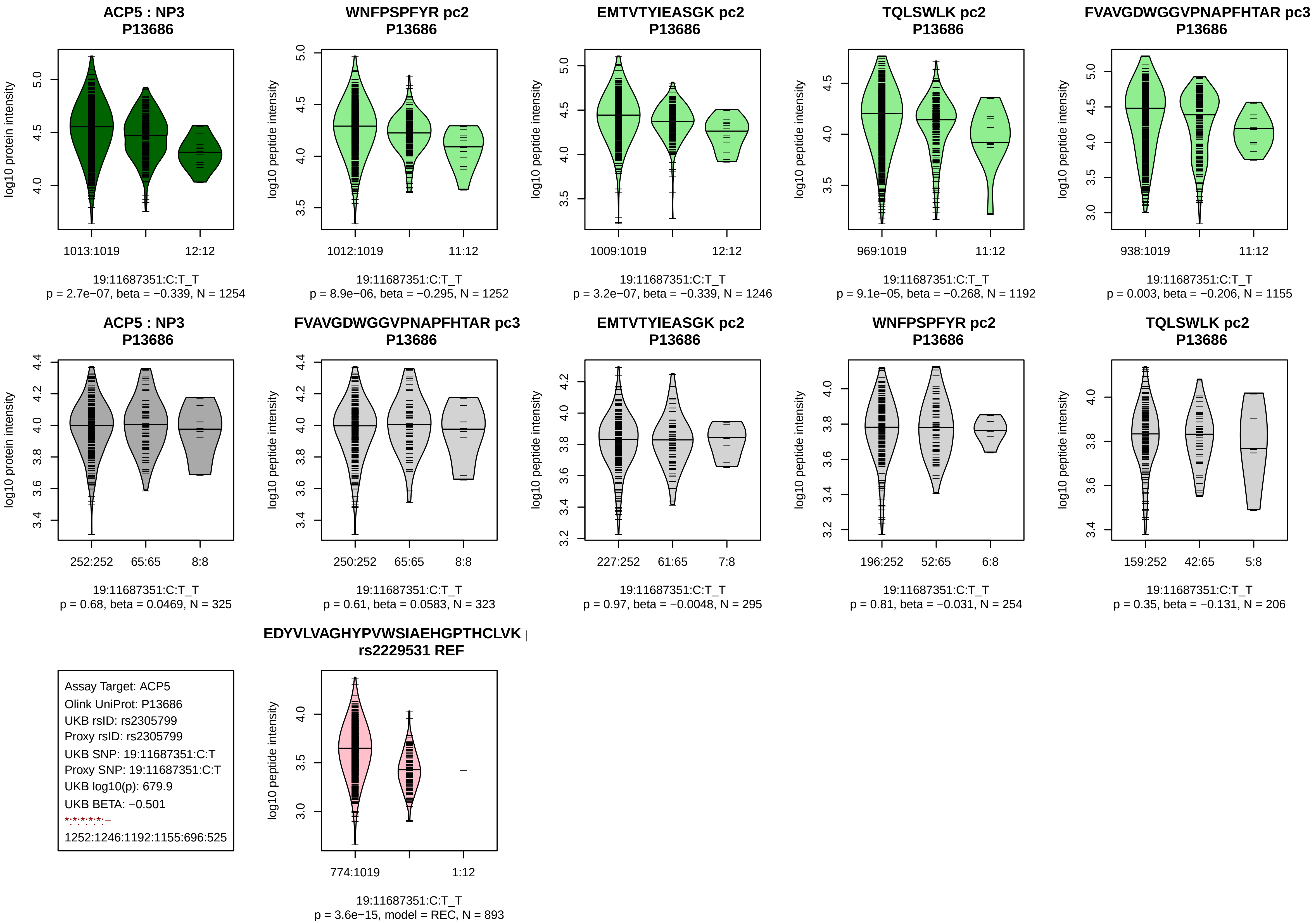
ACP5 : NP3
P13686
log10 protein intensity
1013:1019
12:12
19:11687351:C:T_T
p = 2.7e−07, beta = −0.339, N = 1254
WNFPSPFYR pc2
P13686
log10 peptide intensity
1012:1019
11:12
19:11687351:C:T_T
p = 8.9e−06, beta = −0.295, N = 1252
EMTVTYIEASGK pc2
P13686
log10 peptide intensity
1009:1019
12:12
19:11687351:C:T_T
p = 3.2e−07, beta = −0.339, N = 1246
TQLSWLK pc2
P13686
log10 peptide intensity
969:1019
11:12
19:11687351:C:T_T
p = 9.1e−05, beta = −0.268, N = 1192
FVAVGDWGGVPNAPFHTAR pc3
P13686
log10 peptide intensity
938:1019
11:12
19:11687351:C:T_T
p = 0.003, beta = −0.206, N = 1155
ACP5 : NP3
P13686
log10 protein intensity
252:252
65:65
8:8
19:11687351:C:T_T
p = 0.68, beta = 0.0469, N = 325
FVAVGDWGGVPNAPFHTAR pc3
P13686
log10 peptide intensity
250:252
65:65
8:8
19:11687351:C:T_T
p = 0.61, beta = 0.0583, N = 323
EMTVTYIEASGK pc2
P13686
log10 peptide intensity
227:252
61:65
7:8
19:11687351:C:T_T
p = 0.97, beta = −0.0048, N = 295
WNFPSPFYR pc2
P13686
log10 peptide intensity
196:252
52:65
6:8
19:11687351:C:T_T
p = 0.81, beta = −0.031, N = 254
TQLSWLK pc2
P13686
log10 peptide intensity
159:252
42:65
5:8
19:11687351:C:T_T
p = 0.35, beta = −0.131, N = 206
Assay Target: ACP5
Olink UniProt: P13686
UKB rsID: rs2305799
Proxy rsID: rs2305799
UKB SNP: 19:11687351:C:T
Proxy SNP: 19:11687351:C:T
UKB log10(p): 679.9
UKB BETA: −0.501
*:*:*:*:*:–
1252:1246:1192:1155:696:525
EDYVLVAGHYPVWSIAEHGPTHCLVK
rs2229531 REF
log10 peptide intensity
774:1019
1:12
19:11687351:C:T_T
p = 3.6e−15, model = REC, N = 893

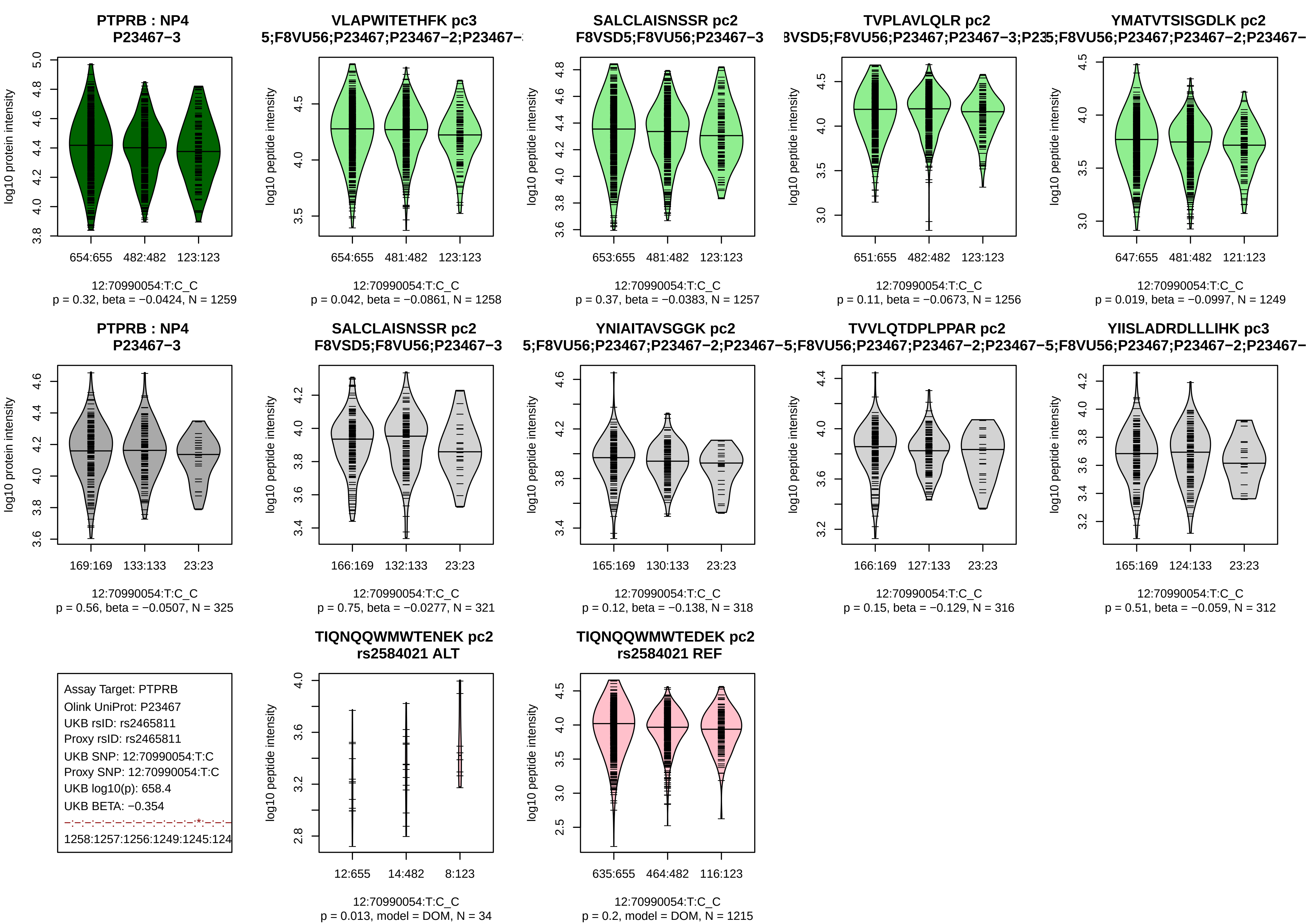

PTPRB : NP4
P23467−3
log10 protein intensity
654:655
482:482
123:123
12:70990054:T:C_C
p = 0.32, beta = −0.0424, N = 1259
VLAPWITETHFK pc3
5;F8VU56;P23467;P23467−2;P23467−
log10 peptide intensity
654:655
481:482
123:123
12:70990054:T:C_C
p = 0.042, beta = −0.0861, N = 1258
SALCLAISNSSR pc2
F8VSD5;F8VU56;P23467−3
653:655
481:482
123:123
12:70990054:T:C_C
p = 0.37, beta = −0.0383, N = 1257
TVPLAVLQLR pc2
3VSD5;F8VU56;P23467;P23467−3;P23
651:655
482:482
123:123
12:70990054:T:C_C
p = 0.11, beta = −0.0673, N = 1256
YMATVTSISGDLK pc2
5;F8VU56;P23467;P23467−2;P23467−
647:655
481:482
121:123
12:70990054:T:C_C
p = 0.019, beta = −0.0997, N = 1249
PTPRB : NP4
P23467−3
169:169
133:133
23:23
12:70990054:T:C_C
p = 0.56, beta = −0.0507, N = 325
SALCLAISNSSR pc2
F8VSD5;F8VU56;P23467−3
166:169
132:133
23:23
12:70990054:T:C_C
p = 0.75, beta = −0.0277, N = 321
YNIAITAVSGGK pc2
5;F8VU56;P23467;P23467−2;P23467−
165:169
130:133
23:23
12:70990054:T:C_C
p = 0.12, beta = −0.138, N = 318
TVVLQTDPLPPAR pc2
5;F8VU56;P23467;P23467−2;P23467−
166:169
127:133
23:23
12:70990054:T:C_C
p = 0.15, beta = −0.129, N = 316
YIISLADRDLLLIHK pc3
5;F8VU56;P23467;P23467−2;P23467−
165:169
124:133
23:23
12:70990054:T:C_C
p = 0.51, beta = −0.059, N = 312
Assay Target: PTPRB
Olink UniProt: P23467
UKB rsID: rs2465811
Proxy rsID: rs2465811
UKB SNP: 12:70990054:T:C
Proxy SNP: 12:70990054:T:C
UKB log10(p): 658.4
UKB BETA: −0.354
−:−:−:−:−:−:−:−:−:−:−:−:*:−:−:−
1258:1257:1256:1249:1245:124
TIQNQQWMWTENEK pc2
rs2584021 ALT
12:655
14:482
8:123
12:70990054:T:C_C
p = 0.013, model = DOM, N = 34
TIQNQQWMWTEDEK pc2
rs2584021 REF
635:655
464:482
116:123
12:70990054:T:C_C
p = 0.2, model = DOM, N = 1215

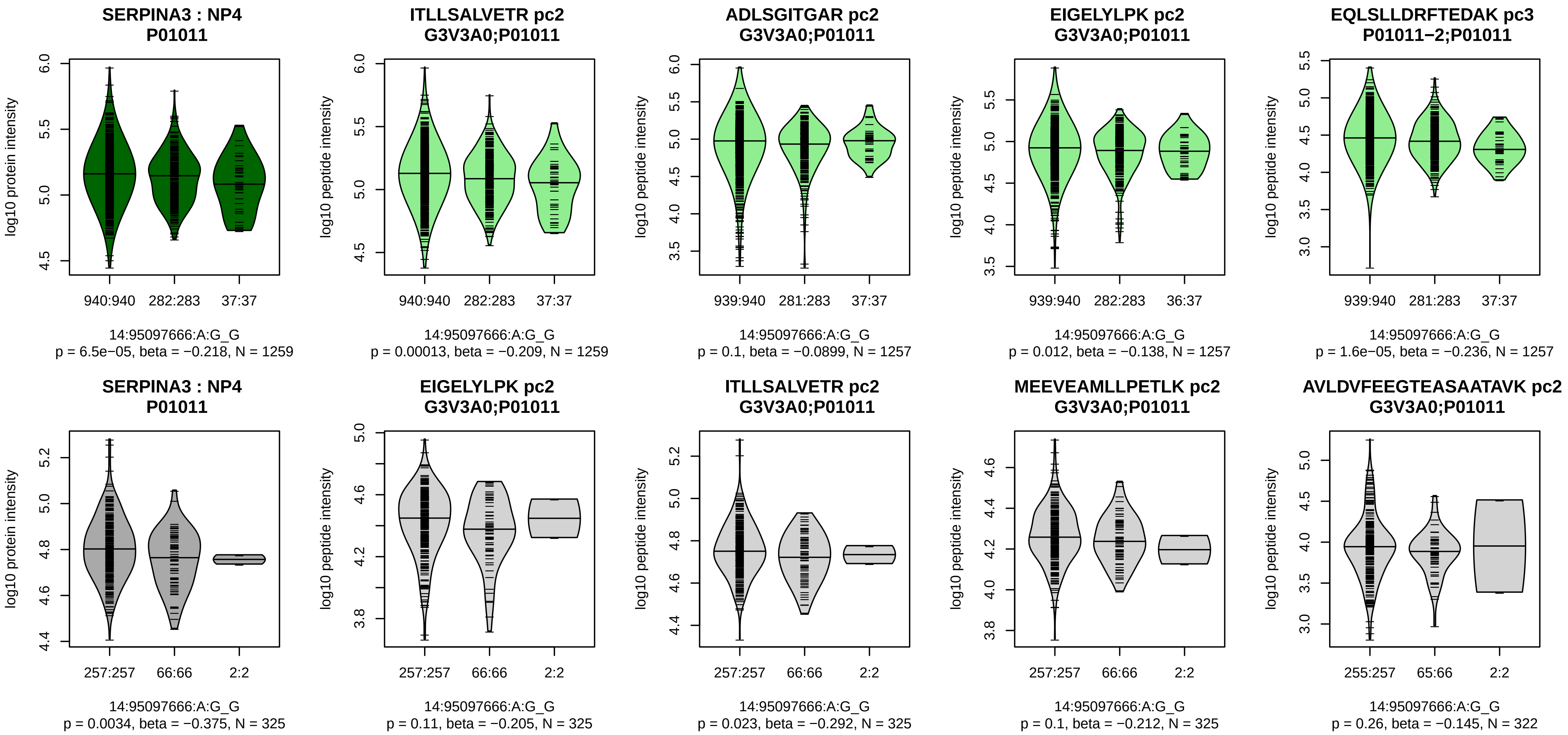


Assay Target: SERPINA3
Olink UniProt: P01011
UKB rsID: rs61976127
Proxy rsID: rs61976127
UKB SNP: 14:95097666:A:G
Proxy SNP: 14:95097666:A:G
UKB log10(p): 645.2
UKB BETA: −0.445
*:–:–:*:*:–:*:*:–:*:–:–:–:–:–:–:–:–
1259:1257:1257:1257:1248:124

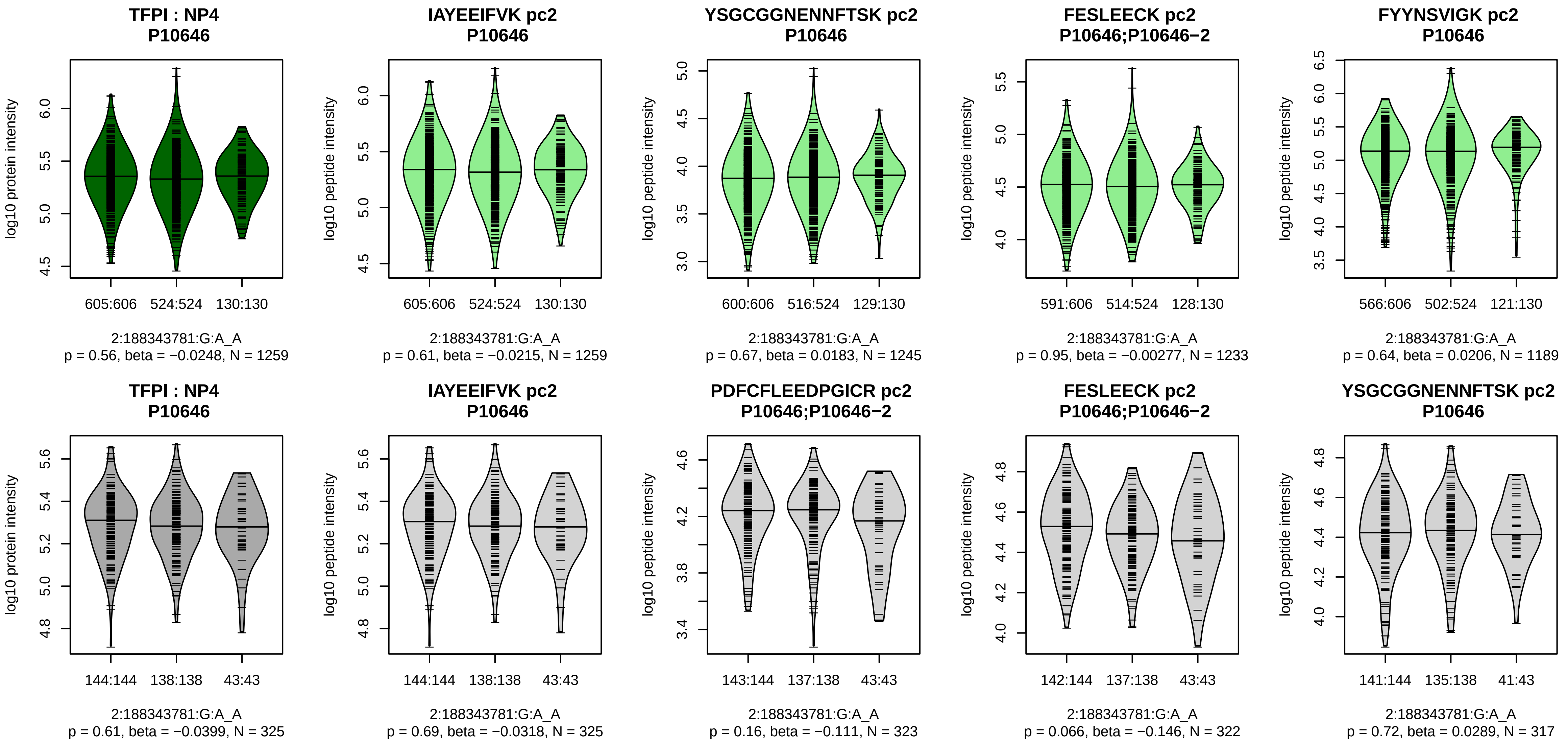


Assay Target: TFPI
Olink UniProt: P10646
UKB rsID: rs7576066
Proxy rsID: rs7576066
UKB SNP: 2:188343781:G:A
Proxy SNP: 2:188343781:G:A
UKB log10(p): 643.7
UKB BETA: 0.35
−:−:−:−:−:−:−:−:−:−:−:−:−:−:NA
1259:1245:1233:1189:1173:107

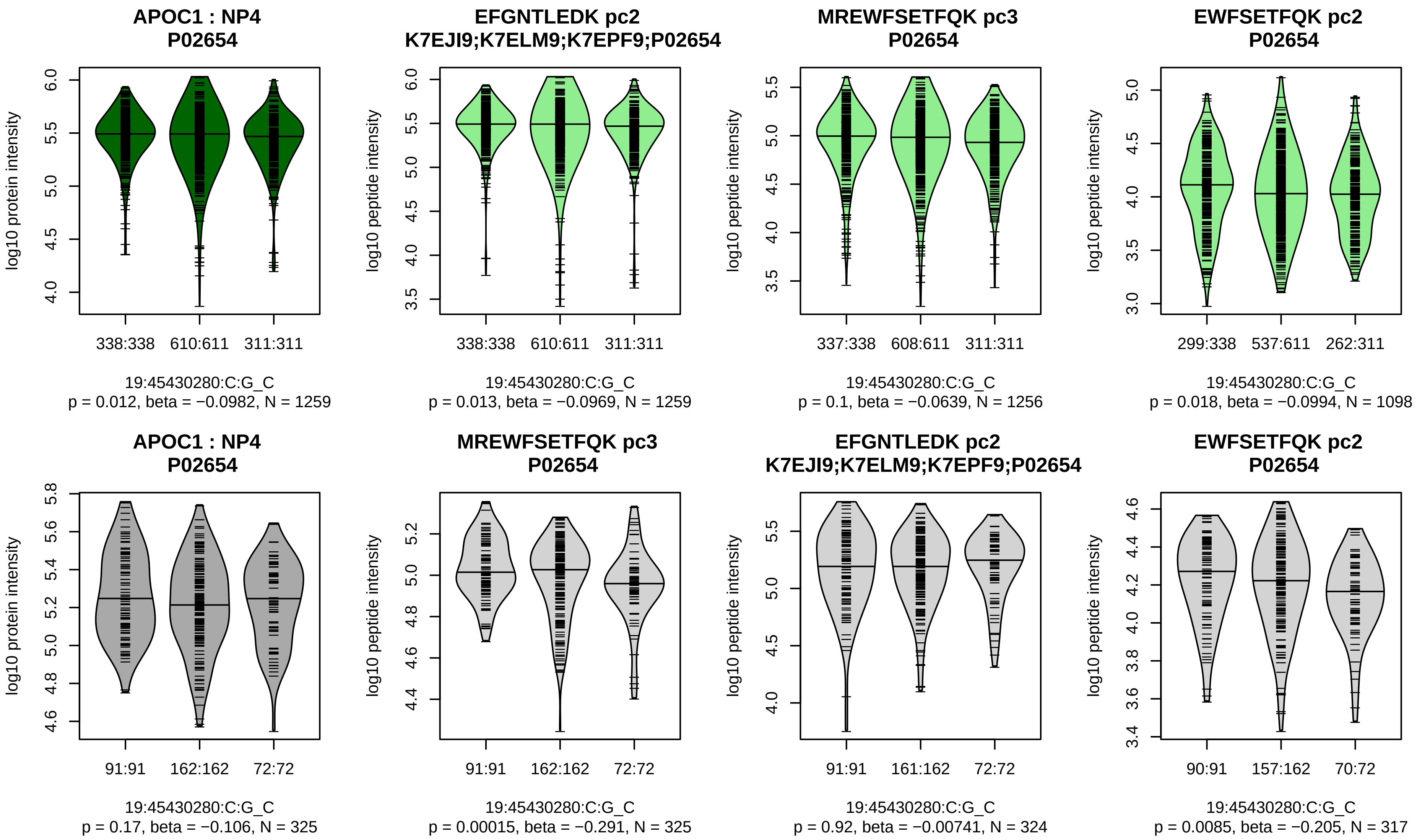


Assay Target: APOC1
Olink UniProt: P02654
UKB rsID: rs5112
Proxy rsID: rs5112
UKB SNP: 19:45430280:C:G
Proxy SNP: 19:45430280:C:G
UKB log10(p): 624.8
UKB BETA: 0.333
−:−:−:−:NA:NA:NA:NA
1259:1256:1143:1098:6:7:4:10

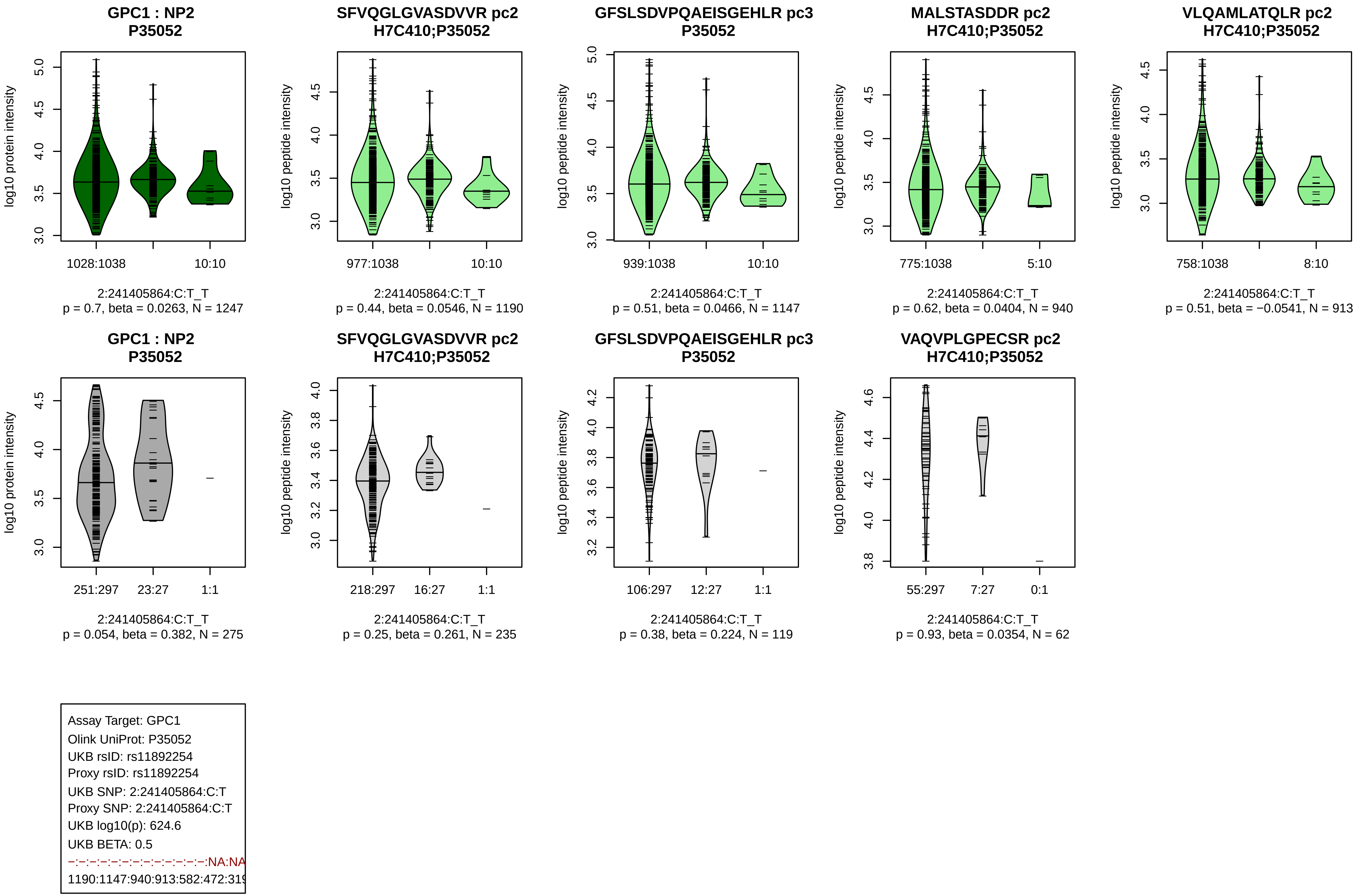
GPC1 : NP2
P35052
log10 protein intensity
1028:1038
10:10
2:241405864:C:T_T
p = 0.7, beta = 0.0263, N = 1247
SFVQGLGVASDVVR pc2
H7C410;P35052
log10 peptide intensity
977:1038
10:10
2:241405864:C:T_T
p = 0.44, beta = 0.0546, N = 1190
GFSLSDVPQAEISGEHLR pc3
P35052
939:1038
10:10
2:241405864:C:T_T
p = 0.51, beta = 0.0466, N = 1147
MALSTASDDR pc2
H7C410;P35052
775:1038
5:10
2:241405864:C:T_T
p = 0.62, beta = 0.0404, N = 940
VLQAMLATQLR pc2
H7C410;P35052
758:1038
8:10
2:241405864:C:T_T
p = 0.51, beta = −0.0541, N = 913
GPC1 : NP2
P35052
251:297
23:27
1:1
2:241405864:C:T_T
p = 0.054, beta = 0.382, N = 275
SFVQGLGVASDVVR pc2
H7C410;P35052
218:297
16:27
1:1
2:241405864:C:T_T
p = 0.25, beta = 0.261, N = 235
GFSLSDVPQAEISGEHLR pc3
P35052
106:297
12:27
1:1
2:241405864:C:T_T
p = 0.38, beta = 0.224, N = 119
VAQVPLGPECSR pc2
H7C410;P35052
55:297
7:27
0:1
2:241405864:C:T_T
p = 0.93, beta = 0.0354, N = 62
Assay Target: GPC1
Olink UniProt: P35052
UKB rsID: rs11892254
Proxy rsID: rs11892254
UKB SNP: 2:241405864:C:T
Proxy SNP: 2:241405864:C:T
UKB log10(p): 624.6
UKB BETA: 0.5
−:−:−:−:−:−:−:−:−:−:−:−:−:−:NA:NA
1190:1147:940:913:582:472:319

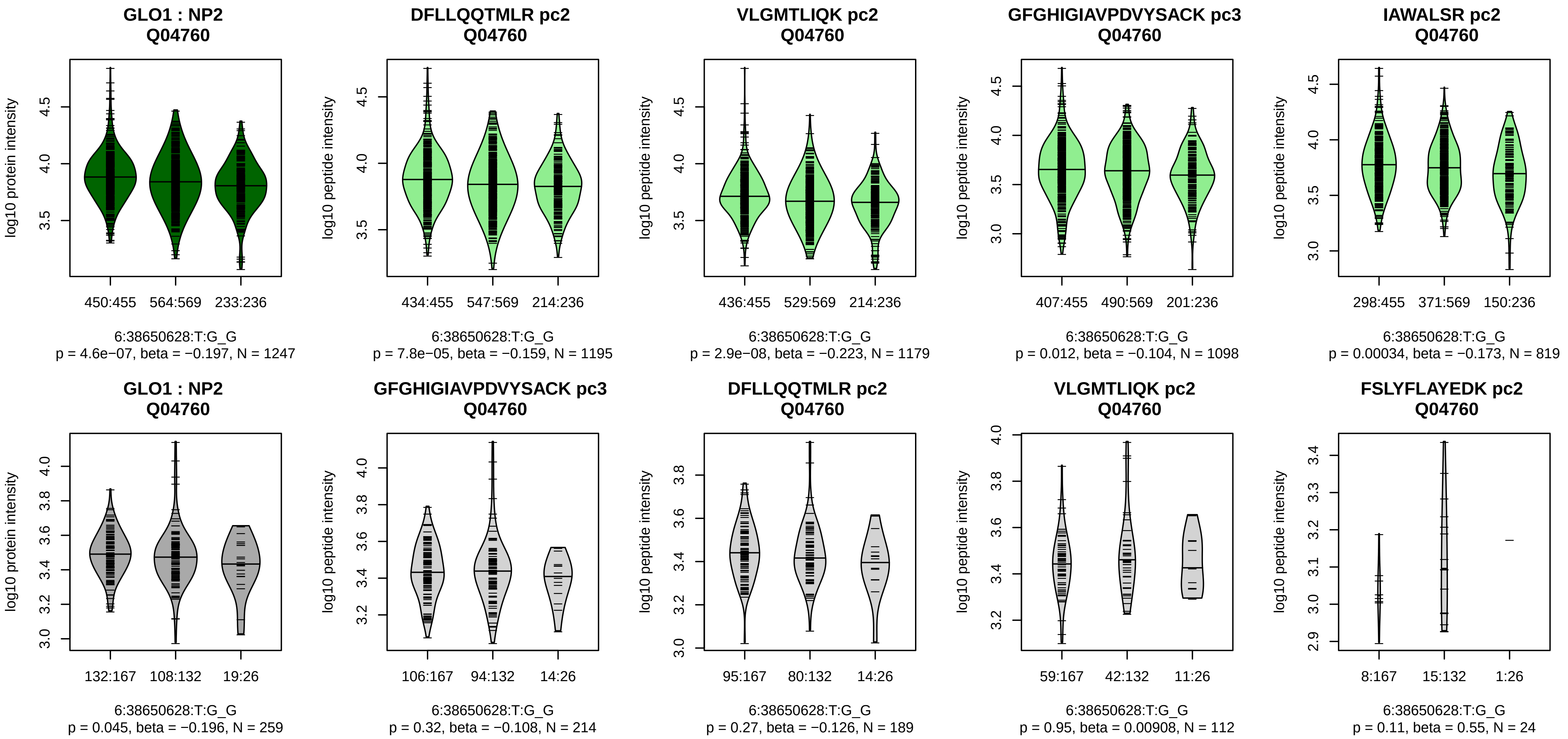


Assay Target: GLO1
Olink UniProt: Q04760
UKB rsID: rs4746
Proxy rsID: rs4746
UKB SNP: 6:38650628:T:G
Proxy SNP: 6:38650628:T:G
UKB log10(p): 616.9
UKB BETA: −0.326
*:*:–:*:–:–:*:–
1195:1179:1098:819:650:199:18

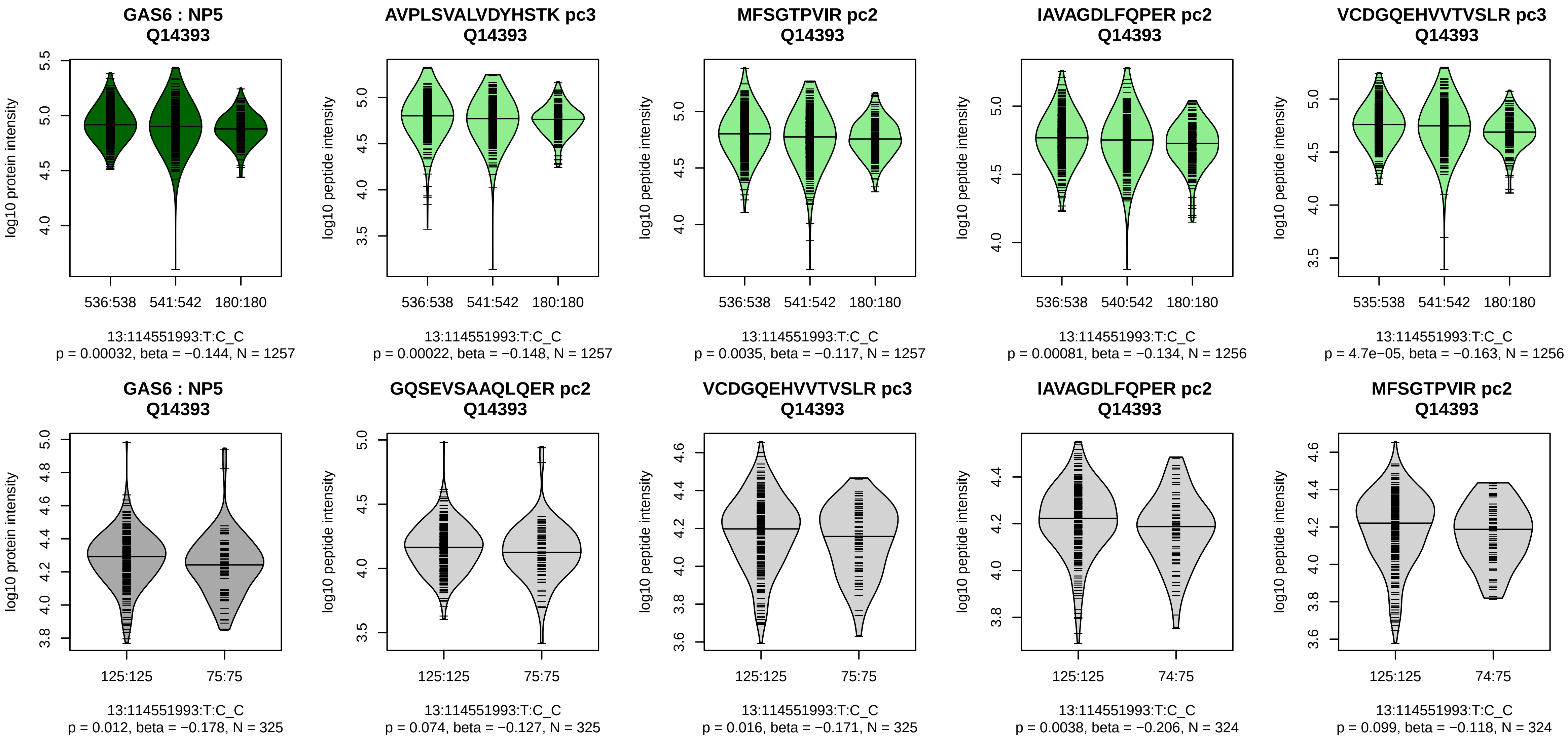


Assay Target: GAS6
Olink UniProt: Q14393
UKB rsID: rs6602909
Proxy rsID: rs6602909
UKB SNP: 13:114551993:T:C
Proxy SNP: 13:114551993:T:C
UKB log10(p): 606.9
UKB BETA: −0.333
*:*:*:*:*:–:*:*:*:*:–:–:–:–:–:–:*:–:–
1257:1257:1256:1256:1256:125

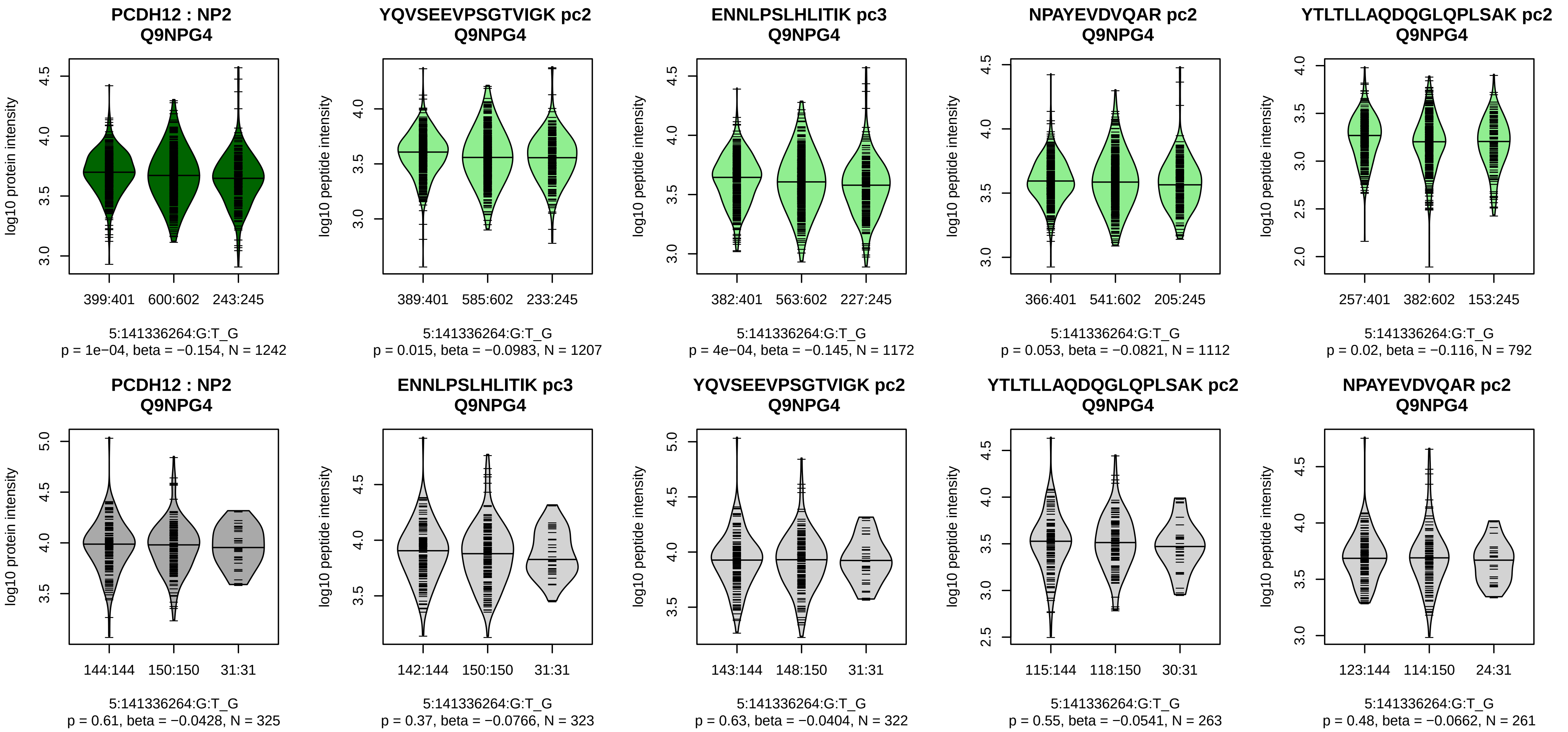


Assay Target: PCDH12
Olink UniProt: Q9NPG4
UKB rsID: rs164075
Proxy rsID: rs164075
UKB SNP: 5:141336264:G:T
Proxy SNP: 5:141336264:G:T
UKB log10(p): 600.4
UKB BETA: 0.325
–:*:–:–:–:–:*:–:–:–:–
1207:1172:1112:792:756:687:68

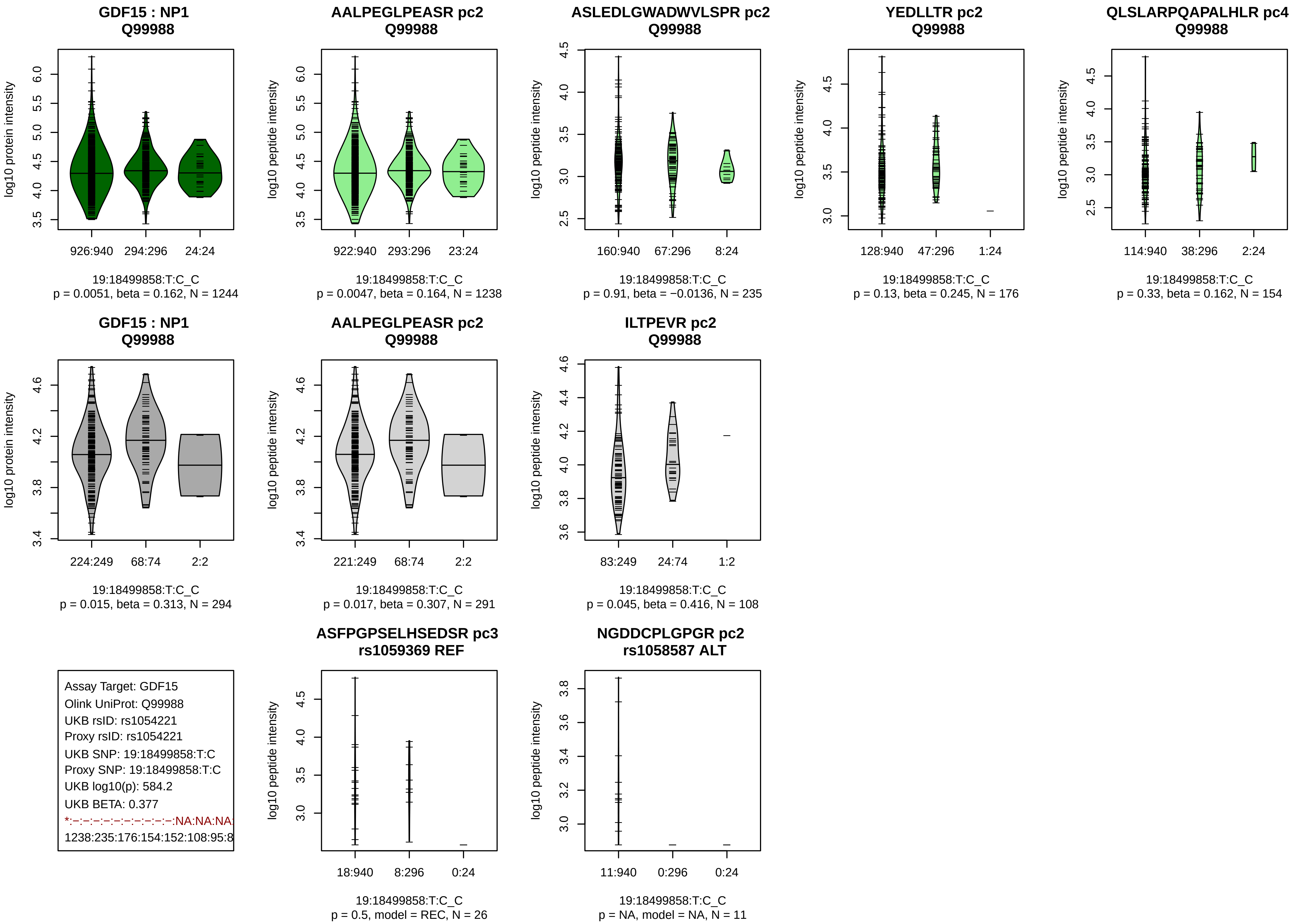
GDF15 : NP1
Q99988
log10 protein intensity
926:940
294:296
24:24
19:18499858:T:C_C
p = 0.0051, beta = 0.162, N = 1244
AALPEGLPEASR pc2
Q99988
log10 peptide intensity
922:940
293:296
23:24
19:18499858:T:C_C
p = 0.0047, beta = 0.164, N = 1238
ASLEDLGWADWVLSPR pc2
Q99988
log10 peptide intensity
160:940
67:296
8:24
19:18499858:T:C_C
p = 0.91, beta = −0.0136, N = 235
YEDLLTR pc2
Q99988
log10 peptide intensity
128:940
47:296
1:24
19:18499858:T:C_C
p = 0.13, beta = 0.245, N = 176
QLSLARPQAPALHLR pc4
Q99988
log10 peptide intensity
114:940
38:296
2:24
19:18499858:T:C_C
p = 0.33, beta = 0.162, N = 154
GDF15 : NP1
Q99988
log10 protein intensity
224:249
68:74
2:2
19:18499858:T:C_C
p = 0.015, beta = 0.313, N = 294
AALPEGLPEASR pc2
Q99988
log10 peptide intensity
221:249
68:74
2:2
19:18499858:T:C_C
p = 0.017, beta = 0.307, N = 291
ILTPEVR pc2
Q99988
log10 peptide intensity
83:249
24:74
1:2
19:18499858:T:C_C
p = 0.045, beta = 0.416, N = 108
Assay Target: GDF15
Olink UniProt: Q99988
UKB rsID: rs1054221
Proxy rsID: rs1054221
UKB SNP: 19:18499858:T:C
Proxy SNP: 19:18499858:T:C
UKB log10(p): 584.2
UKB BETA: 0.377
*:−:−:−:−:−:−:−:−:−:−:NA:NA:NA
1238:235:176:154:152:108:95:8
ASFPGPSELHSEDSR pc3
rs1059369 REF
log10 peptide intensity
18:940
8:296
0:24
19:18499858:T:C_C
p = 0.5, model = REC, N = 26
NGDDCPLGPGR pc2
rs1058587 ALT
log10 peptide intensity
11:940
0:296
0:24
19:18499858:T:C_C
p = NA, model = NA, N = 11

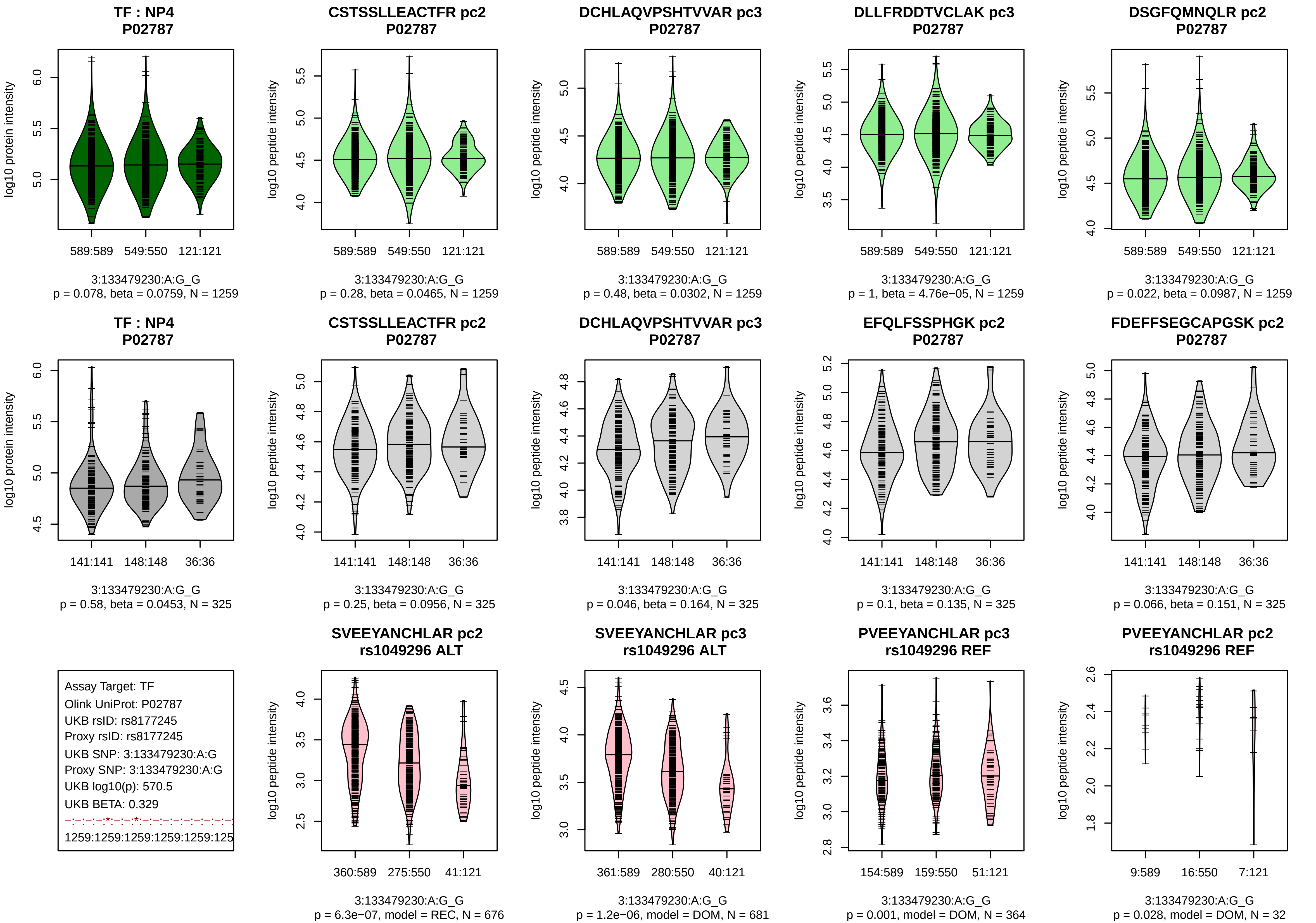
TF : NP4
P02787
log10 protein intensity
589:589 549:550 121:121
3:133479230:A:G_G
p = 0.078, beta = 0.0759, N = 1259
CSTSSLLEACTFR pc2
P02787
log10 peptide intensity
589:589 549:550 121:121
3:133479230:A:G_G
p = 0.28, beta = 0.0465, N = 1259
DCHLAQVPSHTVVAR pc3
P02787
log10 peptide intensity
589:589 549:550 121:121
3:133479230:A:G_G
p = 0.48, beta = 0.0302, N = 1259
DLLFRDDTVCLAK pc3
P02787
log10 peptide intensity
589:589 549:550 121:121
3:133479230:A:G_G
p = 1, beta = 4.76e−05, N = 1259
DSGFQMNQLR pc2
P02787
log10 peptide intensity
589:589 549:550 121:121
3:133479230:A:G_G
p = 0.022, beta = 0.0987, N = 1259
TF : NP4
P02787
log10 protein intensity
141:141 148:148 36:36
3:133479230:A:G_G
p = 0.58, beta = 0.0453, N = 325
CSTSSLLEACTFR pc2
P02787
log10 peptide intensity
141:141 148:148 36:36
3:133479230:A:G_G
p = 0.25, beta = 0.0956, N = 325
DCHLAQVPSHTVVAR pc3
P02787
log10 peptide intensity
141:141 148:148 36:36
3:133479230:A:G_G
p = 0.046, beta = 0.164, N = 325
EFQLFSSPHGK pc2
P02787
log10 peptide intensity
141:141 148:148 36:36
3:133479230:A:G_G
p = 0.1, beta = 0.135, N = 325
FDEFFSEGCAPGSK pc2
P02787
log10 peptide intensity
141:141 148:148 36:36
3:133479230:A:G_G
p = 0.066, beta = 0.151, N = 325
Assay Target: TF
Olink UniProt: P02787
UKB rsID: rs8177245
Proxy rsID: rs8177245
UKB SNP: 3:133479230:A:G
Proxy SNP: 3:133479230:A:G
UKB log10(p): 570.5
UKB BETA: 0.329
−:−:−:−:*:−:−:*:−:−:−:−:−:−:−:−:−:−
1259:1259:1259:1259:1259:125
SVEEYANCHLAR pc2
rs1049296 ALT
log10 peptide intensity
360:589 275:550 41:121
3:133479230:A:G_G
p = 6.3e−07, model = REC, N = 676
SVEEYANCHLAR pc3
rs1049296 ALT
log10 peptide intensity
361:589 280:550 40:121
3:133479230:A:G_G
p = 1.2e−06, model = DOM, N = 681
PVEEYANCHLAR pc3
rs1049296 REF
log10 peptide intensity
154:589 159:550 51:121
3:133479230:A:G_G
p = 0.001, model = DOM, N = 364
PVEEYANCHLAR pc2
rs1049296 REF
log10 peptide intensity
9:589 16:550 7:121
3:133479230:A:G_G
p = 0.028, model = DOM, N = 32

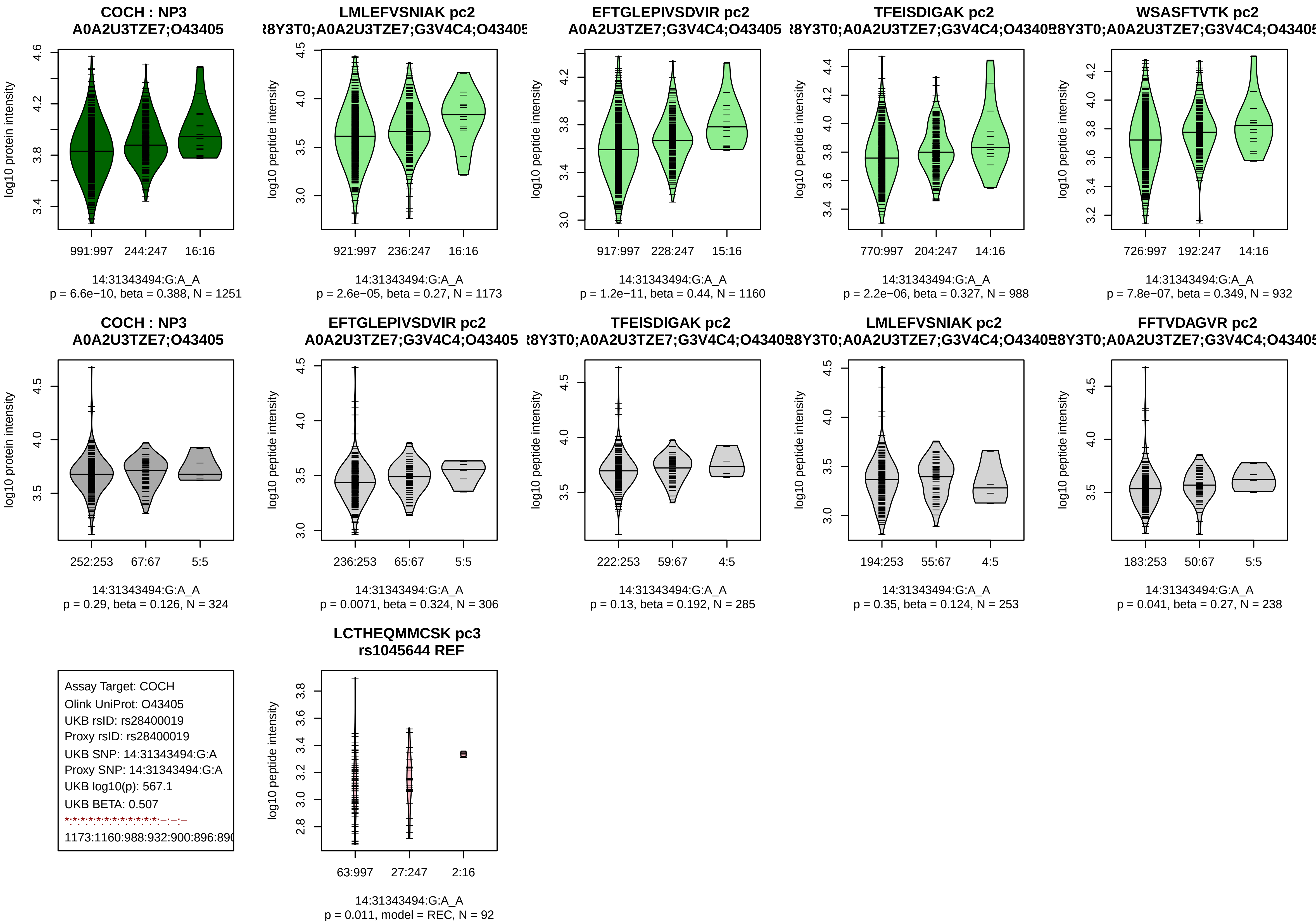

COCH : NP3
A0A2U3TZE7;O43405
log10 protein intensity
991:997 244:247 16:16
14:31343494:G:A_A
p = 6.6e−10, beta = 0.388, N = 1251
LMLEFVSNIAK pc2
log10 peptide intensity
921:997 236:247 16:16
14:31343494:G:A_A
p = 2.6e−05, beta = 0.27, N = 1173
EFTGLEPIVSDVIR pc2
A0A2U3TZE7;G3V4C4;O43405
log10 peptide intensity
917:997 228:247 15:16
14:31343494:G:A_A
p = 1.2e−11, beta = 0.44, N = 1160
TFEISDIGAK pc2
log10 peptide intensity
770:997 204:247 14:16
14:31343494:G:A_A
p = 2.2e−06, beta = 0.327, N = 988
WSASFTVTK pc2
log10 peptide intensity
726:997 192:247 14:16
14:31343494:G:A_A
p = 7.8e−07, beta = 0.349, N = 932
COCH : NP3
A0A2U3TZE7;O43405
log10 protein intensity
252:253 67:67 5:5
14:31343494:G:A_A
p = 0.29, beta = 0.126, N = 324
EFTGLEPIVSDVIR pc2
A0A2U3TZE7;G3V4C4;O43405
log10 peptide intensity
236:253 65:67 5:5
14:31343494:G:A_A
p = 0.0071, beta = 0.324, N = 306
TFEISDIGAK pc2
log10 peptide intensity
222:253 59:67 4:5
14:31343494:G:A_A
p = 0.13, beta = 0.192, N = 285
LMLEFVSNIAK pc2
log10 peptide intensity
194:253 55:67 4:5
14:31343494:G:A_A
p = 0.35, beta = 0.124, N = 253
FFTVDAGVR pc2
log10 peptide intensity
183:253 50:67 5:5
14:31343494:G:A_A
p = 0.041, beta = 0.27, N = 238
Assay Target: COCH
Olink UniProt: O43405
UKB rsID: rs28400019
Proxy rsID: rs28400019
UKB SNP: 14:31343494:G:A
Proxy SNP: 14:31343494:G:A
UKB log10(p): 567.1
UKB BETA: 0.507
LCTHEQMMCSK pc3
rs1045644 REF
log10 peptide intensity
63:997 27:247 2:16
14:31343494:G:A_A
p = 0.011, model = REC, N = 92

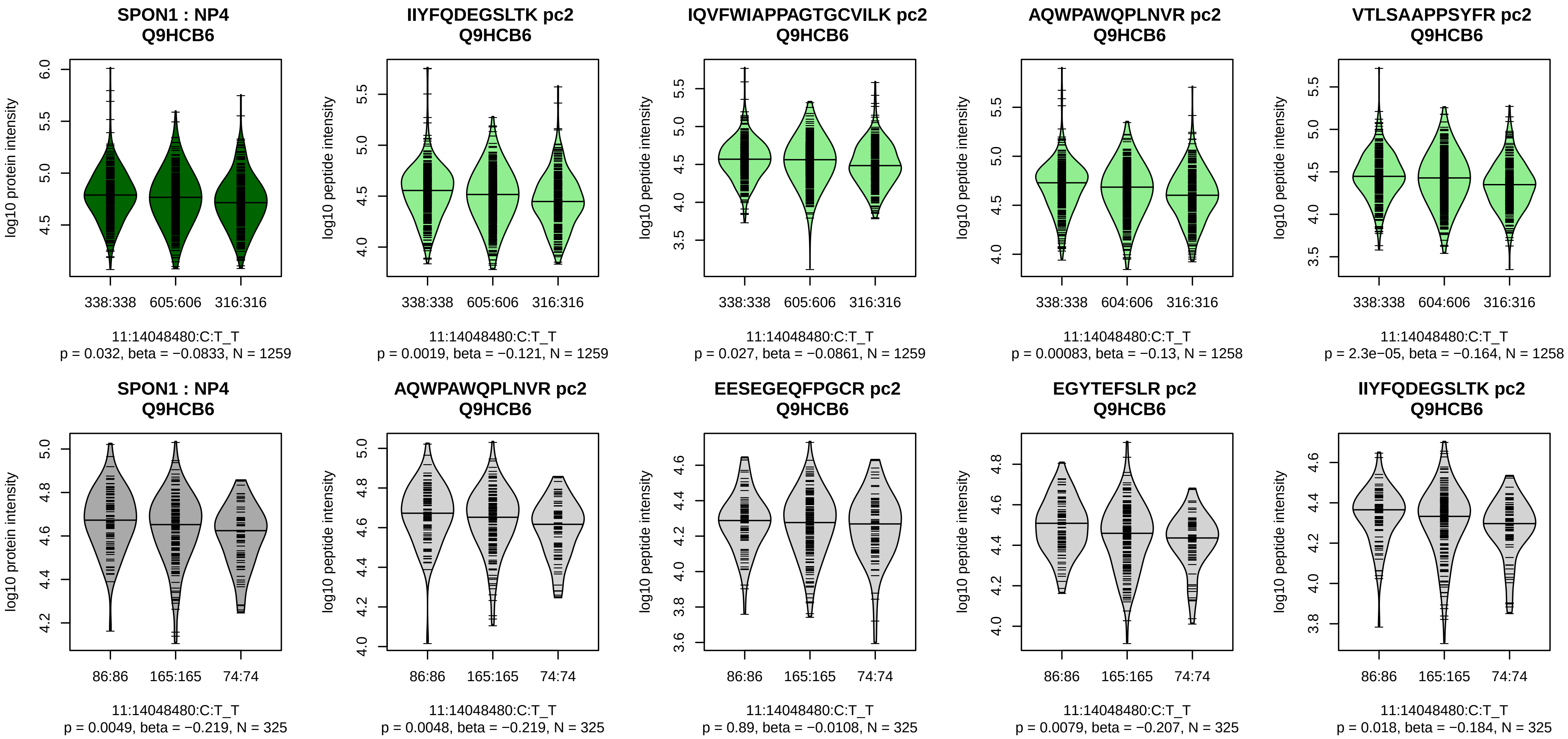


Assay Target: SPON1
Olink UniProt: Q9HCB6
UKB rsID: rs10832164
Proxy rsID: rs10832164
UKB SNP: 11:14048480:C:T
Proxy SNP: 11:14048480:C:T
UKB log10(p): 541.1
UKB BETA: −0.288
*:–:*:*:*:–:–:*:*:–:*:–:*:–:–:–:–:–:
1259:1259:1258:1258:1246:124

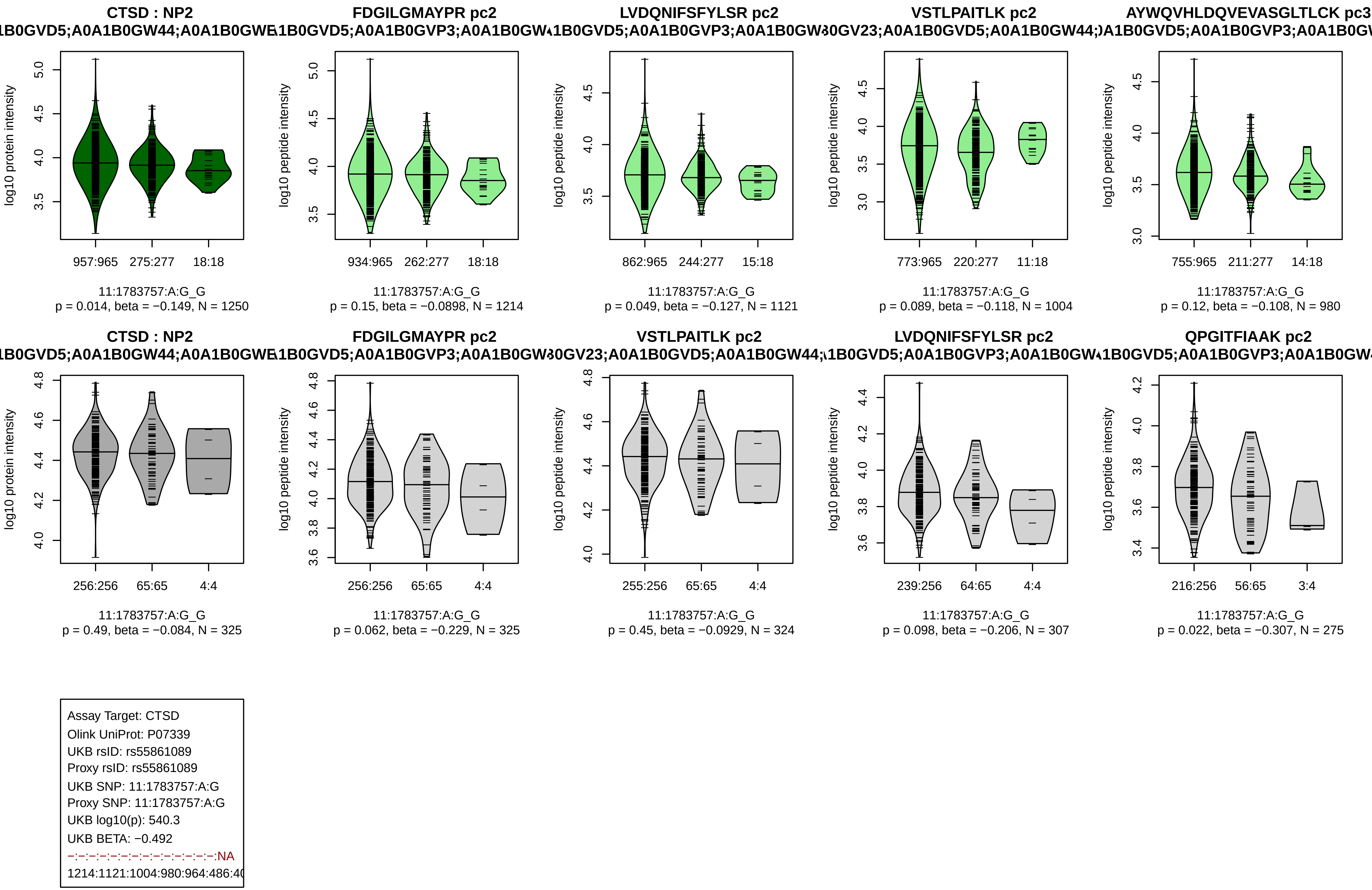
CTSD : NP2
FDGILGMAYPR pc2
LVDQNIFSFYLSR pc2
VSTLPAITLK pc2
AYWQVHLDQVEVASGLTLCK pc3
log10 protein intensity
log10 peptide intensity
957:965 275:277 18:18
934:965 262:277 18:18
862:965 244:277 15:18
773:965 220:277 11:18
755:965 211:277 14:18
11:1783757:A:G_G
p = 0.014, beta = −0.149, N = 1250
p = 0.15, beta = −0.0898, N = 1214
p = 0.049, beta = −0.127, N = 1121
p = 0.089, beta = −0.118, N = 1004
p = 0.12, beta = −0.108, N = 980
QPGITFIAAK pc2
256:256 65:65 4:4
255:256 65:65 4:4
239:256 64:65 4:4
216:256 56:65 3:4
p = 0.49, beta = −0.084, N = 325
p = 0.062, beta = −0.229, N = 325
p = 0.45, beta = −0.0929, N = 324
p = 0.098, beta = −0.206, N = 307
p = 0.022, beta = −0.307, N = 275
Assay Target: CTSD
Olink UniProt: P07339
UKB rsID: rs55861089
Proxy rsID: rs55861089
UKB SNP: 11:1783757:A:G
Proxy SNP: 11:1783757:A:G
UKB log10(p): 540.3
UKB BETA: −0.492
−:−:−:−:−:−:−:−:−:−:−:−:−:−:−:NA
1214:1121:1004:980:964:486:4

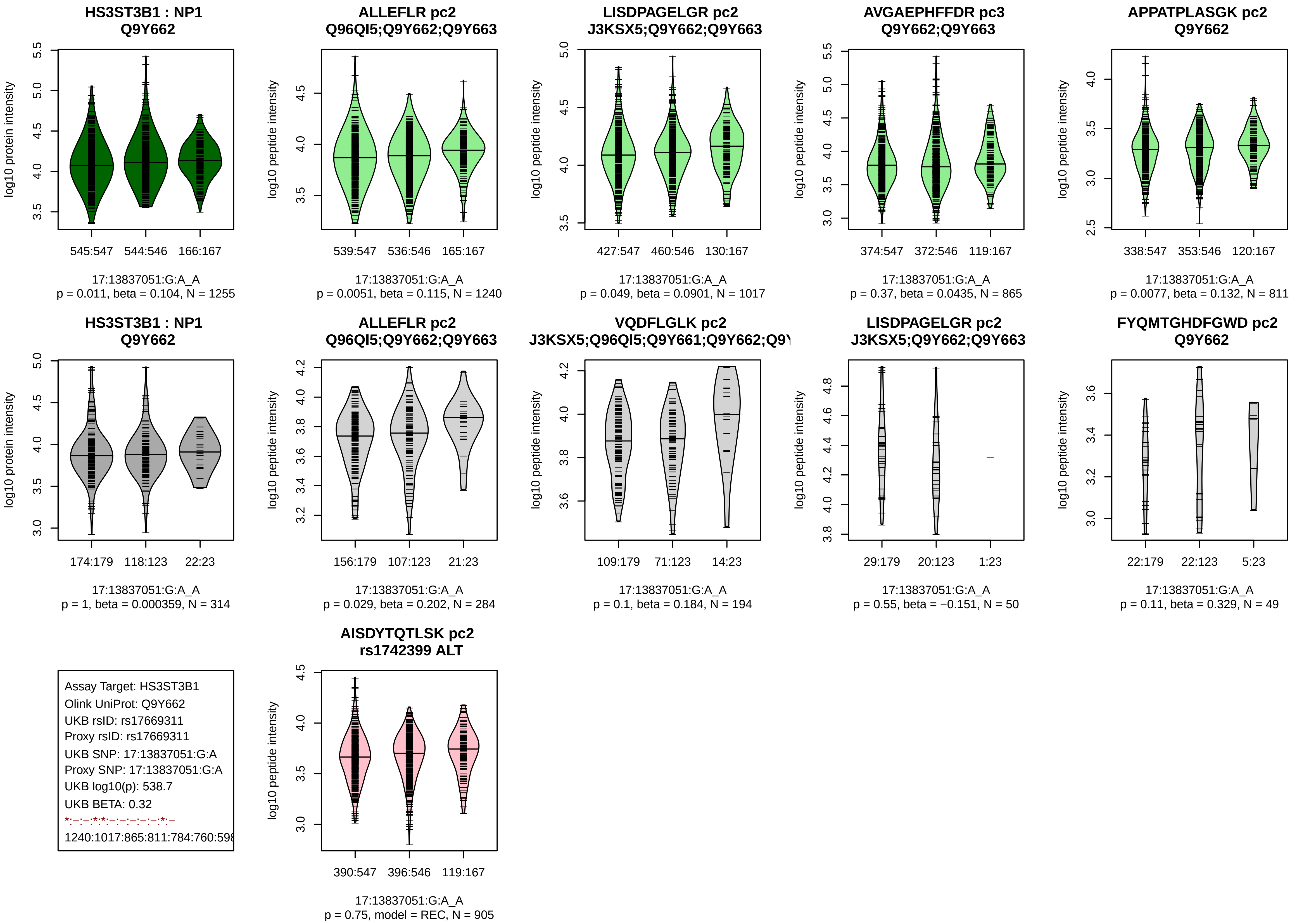

HS3ST3B1 : NP1
Q9Y662
log10 protein intensity
545:547 544:546 166:167
17:13837051:G:A_A
p = 0.011, beta = 0.104, N = 1255
ALLEFLR pc2
Q96QI5;Q9Y662;Q9Y663
log10 peptide intensity
539:547 536:546 165:167
17:13837051:G:A_A
p = 0.0051, beta = 0.115, N = 1240
LISDPAGELGR pc2
J3KSX5;Q9Y662;Q9Y663
log10 peptide intensity
427:547 460:546 130:167
17:13837051:G:A_A
p = 0.049, beta = 0.0901, N = 1017
AVGAEPHFFDR pc3
Q9Y662;Q9Y663
log10 peptide intensity
374:547 372:546 119:167
17:13837051:G:A_A
p = 0.37, beta = 0.0435, N = 865
APPATPLASGK pc2
Q9Y662
log10 peptide intensity
338:547 353:546 120:167
17:13837051:G:A_A
p = 0.0077, beta = 0.132, N = 811
HS3ST3B1 : NP1
Q9Y662
log10 protein intensity
174:179 118:123 22:23
17:13837051:G:A_A
p = 1, beta = 0.000359, N = 314
ALLEFLR pc2
Q96QI5;Q9Y662;Q9Y663
log10 peptide intensity
156:179 107:123 21:23
17:13837051:G:A_A
p = 0.029, beta = 0.202, N = 284
VQDFLGLK pc2
J3KSX5;Q96QI5;Q9Y661;Q9Y662;Q9Y
log10 peptide intensity
109:179 71:123 14:23
17:13837051:G:A_A
p = 0.1, beta = 0.184, N = 194
LISDPAGELGR pc2
J3KSX5;Q9Y662;Q9Y663
log10 peptide intensity
29:179 20:123 1:23
17:13837051:G:A_A
p = 0.55, beta = −0.151, N = 50
FYQMTGHDFGWD pc2
Q9Y662
log10 peptide intensity
22:179 22:123 5:23
17:13837051:G:A_A
p = 0.11, beta = 0.329, N = 49
Assay Target: HS3ST3B1
Olink UniProt: Q9Y662
UKB rsID: rs17669311
Proxy rsID: rs17669311
UKB SNP: 17:13837051:G:A
Proxy SNP: 17:13837051:G:A
UKB log10(p): 538.7
UKB BETA: 0.32
*:–:–:*:*:–:–:–:–:–:*:–
1240:1017:865:811:784:760:598
AISDYTQTLSK pc2
rs1742399 ALT
log10 peptide intensity
390:547 396:546 119:167
17:13837051:G:A_A
p = 0.75, model = REC, N = 905

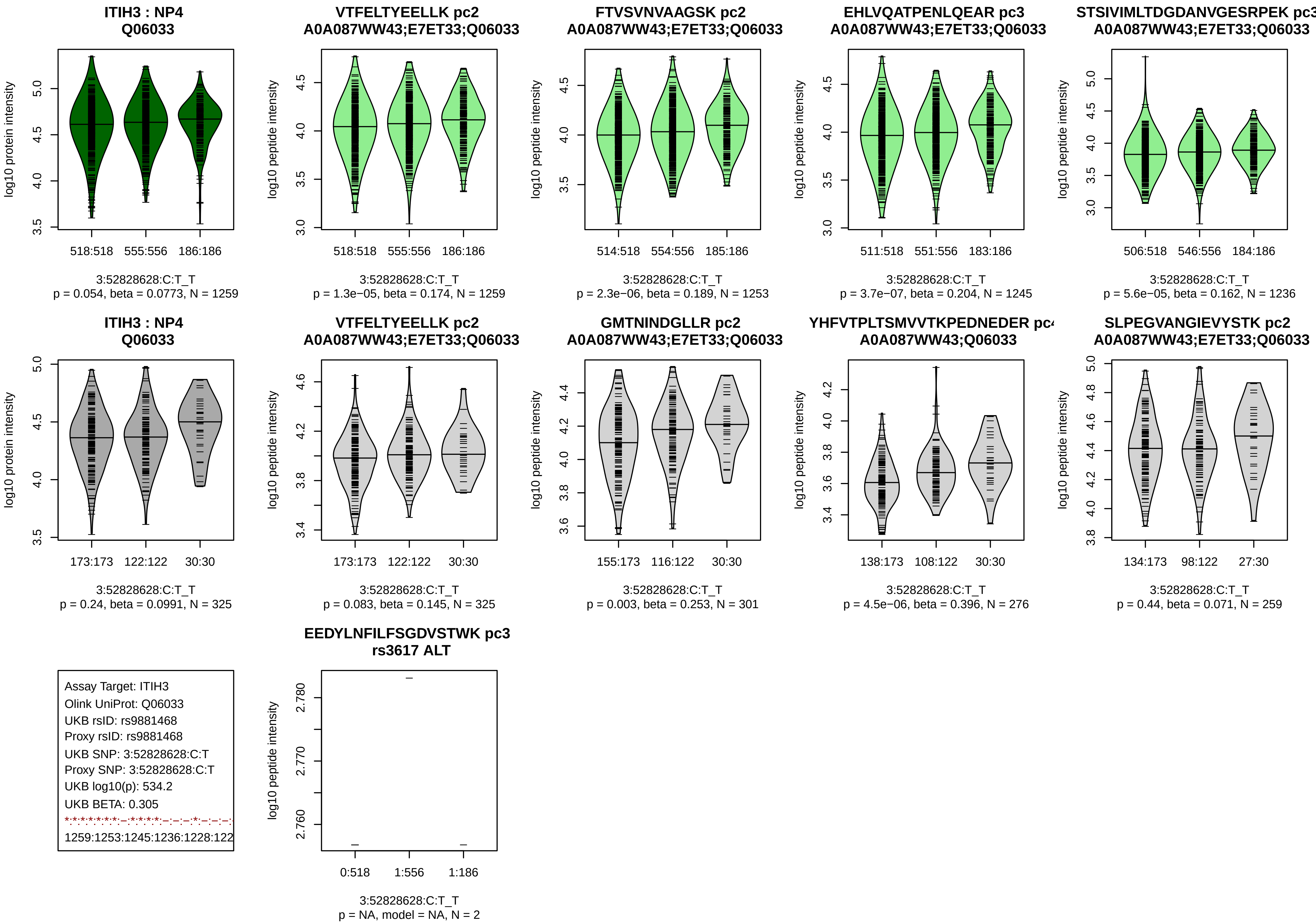
ITIH3 : NP4
Q06033
log10 protein intensity
518:518 555:556 186:186
3:52828628:C:T_T
p = 0.054, beta = 0.0773, N = 1259
VTFELTYEELLK pc2
A0A087WW43;E7ET33;Q06033
log10 peptide intensity
518:518 555:556 186:186
3:52828628:C:T_T
p = 1.3e−05, beta = 0.174, N = 1259
FTVSVNVAAGSK pc2
A0A087WW43;E7ET33;Q06033
514:518 554:556 185:186
3:52828628:C:T_T
p = 2.3e−06, beta = 0.189, N = 1253
EHLVQATPENLQEAR pc3
A0A087WW43;E7ET33;Q06033
511:518 551:556 183:186
3:52828628:C:T_T
p = 3.7e−07, beta = 0.204, N = 1245
STSIVIMLTDGDANVGESRPEK pc3
A0A087WW43;E7ET33;Q06033
506:518 546:556 184:186
3:52828628:C:T_T
p = 5.6e−05, beta = 0.162, N = 1236
ITIH3 : NP4
Q06033
173:173 122:122 30:30
3:52828628:C:T_T
p = 0.24, beta = 0.0991, N = 325
VTFELTYEELLK pc2
A0A087WW43;E7ET33;Q06033
173:173 122:122 30:30
3:52828628:C:T_T
p = 0.083, beta = 0.145, N = 325
GMTNINDGLLR pc2
A0A087WW43;E7ET33;Q06033
155:173 116:122 30:30
3:52828628:C:T_T
p = 0.003, beta = 0.253, N = 301
YHFVTPLTSMVVTKPEDNEDER pc
A0A087WW43;Q06033
138:173 108:122 30:30
3:52828628:C:T_T
p = 4.5e−06, beta = 0.396, N = 276
SLPEGVANGIEVYSTK pc2
A0A087WW43;E7ET33;Q06033
134:173 98:122 27:30
3:52828628:C:T_T
p = 0.44, beta = 0.071, N = 259
Assay Target: ITIH3
Olink UniProt: Q06033
UKB rsID: rs9881468
Proxy rsID: rs9881468
UKB SNP: 3:52828628:C:T
Proxy SNP: 3:52828628:C:T
UKB log10(p): 534.2
UKB BETA: 0.305
1259:1253:1245:1236:1228:122
EEDYLNFILFSGDVSTWK pc3
rs3617 ALT
0:518 1:556 1:186
3:52828628:C:T_T
p = NA, model = NA, N = 2

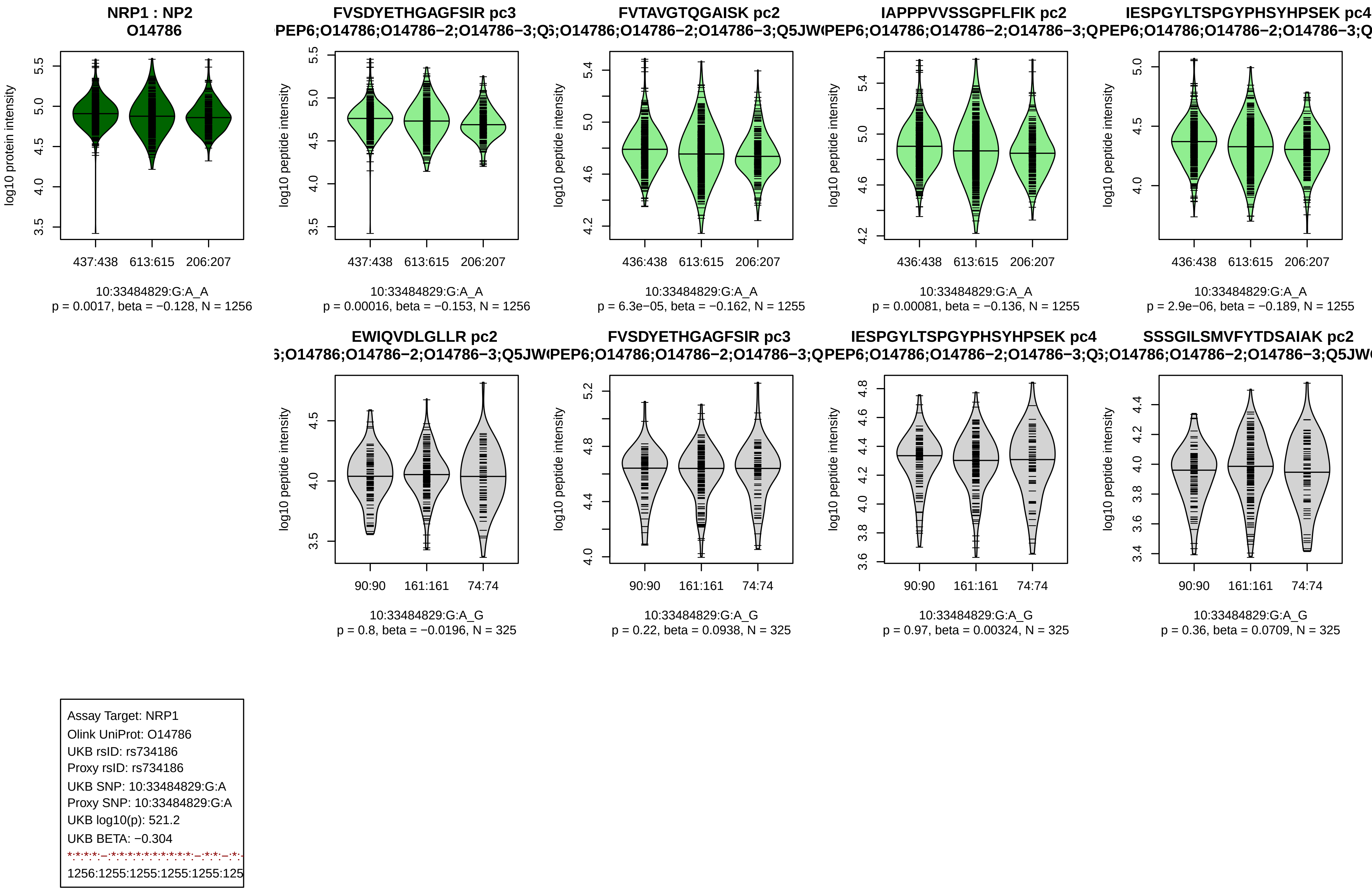

NRP1 : NP2
O14786
log10 protein intensity
437:438
613:615
206:207
10:33484829:G:A_A
p = 0.0017, beta = −0.128, N = 1256
FVSDYETHGAGFSIR pc3
log10 peptide intensity
437:438
613:615
206:207
10:33484829:G:A_A
p = 0.00016, beta = −0.153, N = 1256
FVTAVGTQGAISK pc2
436:438
613:615
206:207
10:33484829:G:A_A
p = 6.3e−05, beta = −0.162, N = 1255
IAPPPVVSSGPFLFIK pc2
436:438
613:615
206:207
10:33484829:G:A_A
p = 0.00081, beta = −0.136, N = 1255
IESPGYLTSPGYPHSYHPSEK pc4
436:438
613:615
206:207
10:33484829:G:A_A
p = 2.9e−06, beta = −0.189, N = 1255
EWIQVDLGLLR pc2
90:90
161:161
74:74
10:33484829:G:A_G
p = 0.8, beta = −0.0196, N = 325
FVSDYETHGAGFSIR pc3
90:90
161:161
74:74
10:33484829:G:A_G
p = 0.22, beta = 0.0938, N = 325
IESPGYLTSPGYPHSYHPSEK pc4
90:90
161:161
74:74
10:33484829:G:A_G
p = 0.97, beta = 0.00324, N = 325
SSSGILSMVFYTDSAIAK pc2
90:90
161:161
74:74
10:33484829:G:A_G
p = 0.36, beta = 0.0709, N = 325
Assay Target: NRP1
Olink UniProt: O14786
UKB rsID: rs734186
Proxy rsID: rs734186
UKB SNP: 10:33484829:G:A
Proxy SNP: 10:33484829:G:A
UKB log10(p): 521.2
UKB BETA: −0.304
1256:1255:1255:1255:1255:125

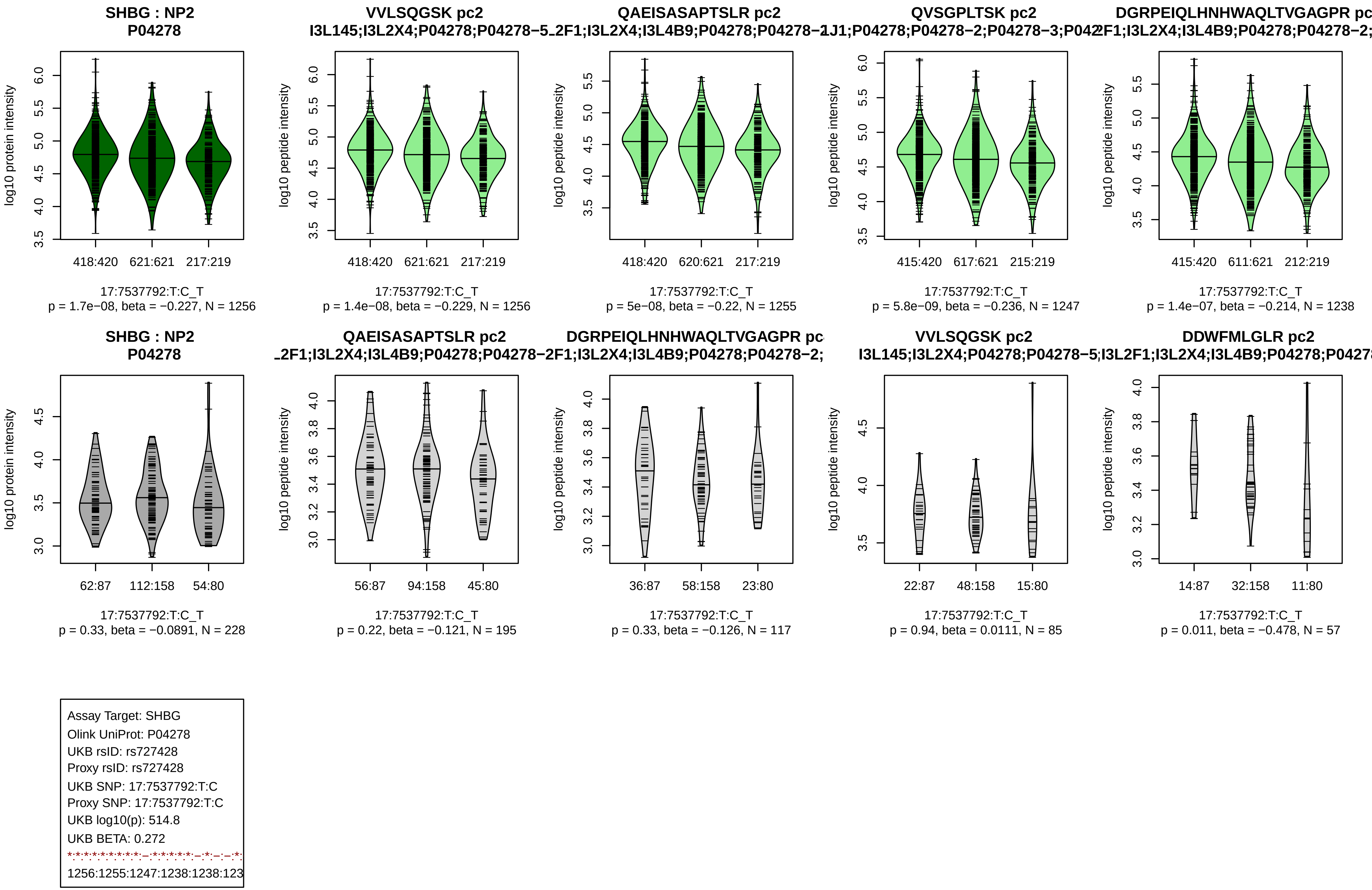
SHBG : NP2
P04278
VVLSQGSK pc2
QAEISASAPTSLR pc2
QVSGPLTSK pc2
log10 protein intensity
log10 peptide intensity
418:420 621:621 217:219
418:420 621:621 217:219
418:420 620:621 217:219
415:420 617:621 215:219
415:420 611:621 212:219
17:7537792:T:C_T
p = 1.7e−08, beta = −0.227, N = 1256
p = 1.4e−08, beta = −0.229, N = 1256
p = 5e−08, beta = −0.22, N = 1255
p = 5.8e−09, beta = −0.236, N = 1247
p = 1.4e−07, beta = −0.214, N = 1238
DDWFMLGLR pc2
62:87 112:158 54:80
56:87 94:158 45:80
36:87 58:158 23:80
22:87 48:158 15:80
14:87 32:158 11:80
p = 0.33, beta = −0.0891, N = 228
p = 0.22, beta = −0.121, N = 195
p = 0.33, beta = −0.126, N = 117
p = 0.94, beta = 0.0111, N = 85
p = 0.011, beta = −0.478, N = 57
Assay Target: SHBG
Olink UniProt: P04278
UKB rsID: rs727428
Proxy rsID: rs727428
UKB SNP: 17:7537792:T:C
Proxy SNP: 17:7537792:T:C
UKB log10(p): 514.8
UKB BETA: 0.272
1256:1255:1247:1238:1238:123

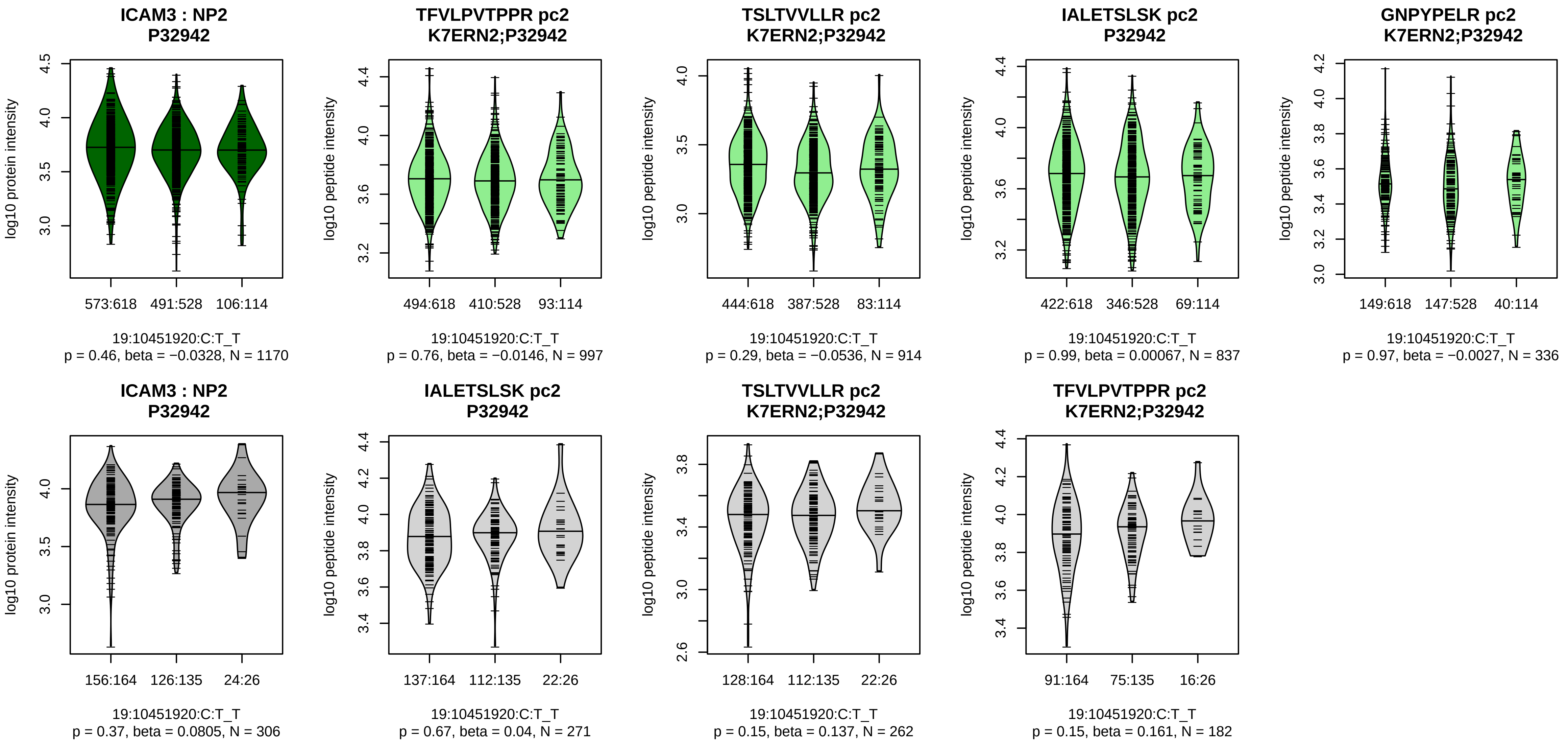


Assay Target: ICAM3
Olink UniProt: P32942
UKB rsID: rs61451658
Proxy rsID: rs61451658
UKB SNP: 19:10451920:C:T
Proxy SNP: 19:10451920:C:T
UKB log10(p): 510
UKB BETA: 0.309
–:–:–:–:–:–:–
997:914:837:336:144:141:71

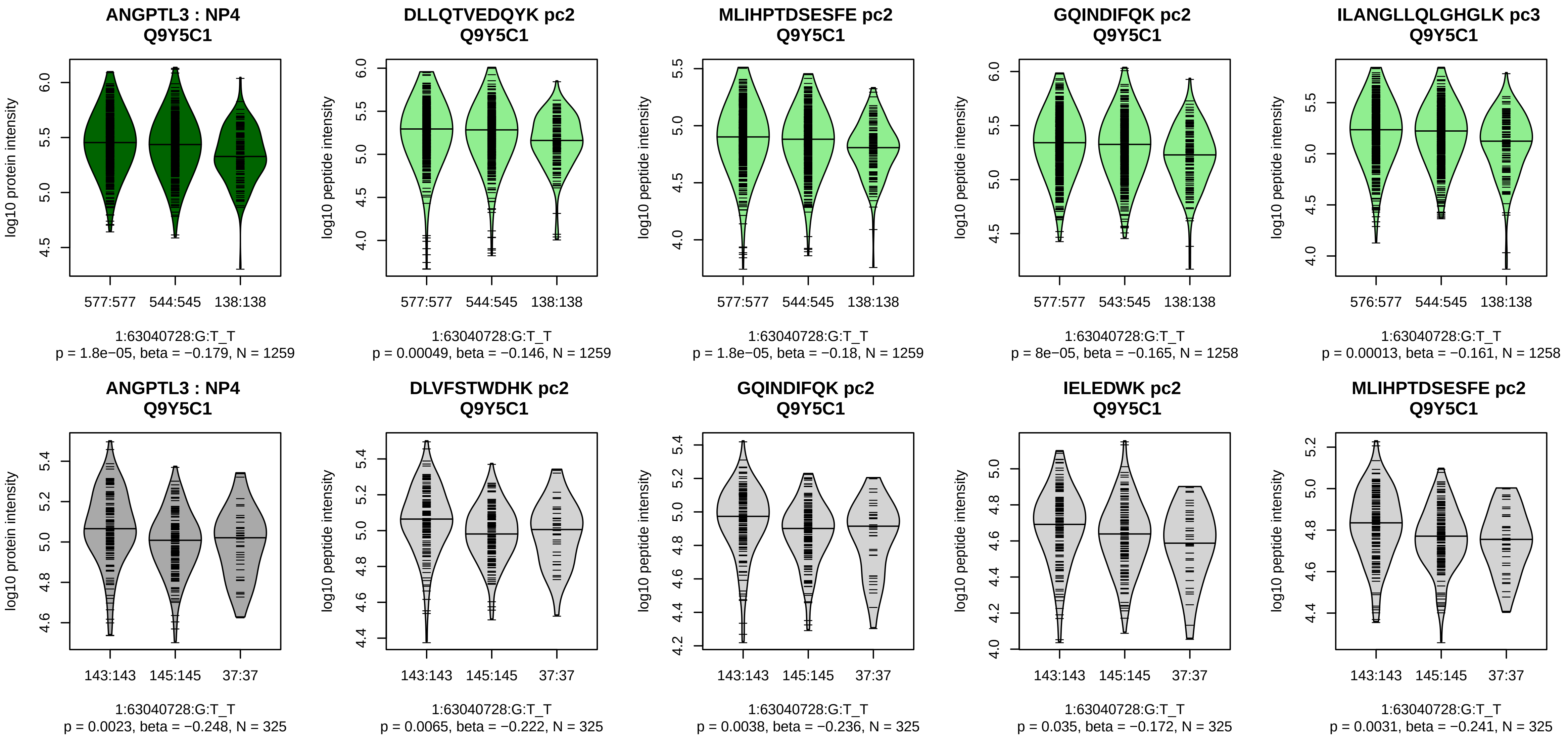


Assay Target: ANGPTL3
Olink UniProt: Q9Y5C1
UKB rsID: rs775677524
Proxy rsID: rs12038768
UKB SNP: 1:63070537:AGTTAA
Proxy SNP: 1:63040728:G:T
UKB log10(p): 500.3
UKB BETA: −0.282
*:*:*:*:*:–:*:*:*:*:*:*:*:*:*:–:–:*:*:*:*:
1259:1259:1258:1258:1257:125

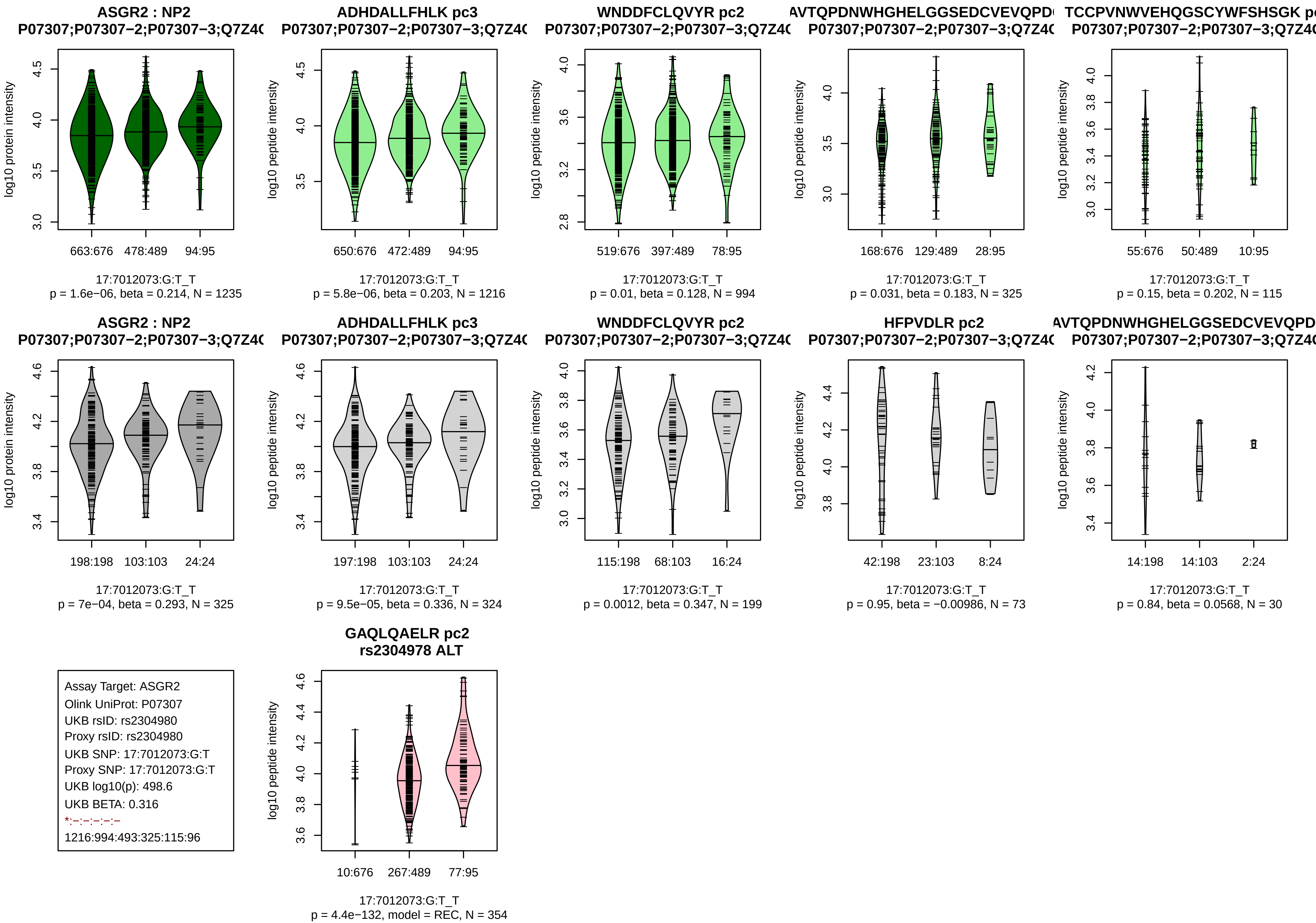

ASGR2 : NP2
P07307;P07307−2;P07307−3;Q7Z4(
log10 protein intensity
663:676 478:489 94:95
17:7012073:G:T_T
p = 1.6e−06, beta = 0.214, N = 1235
ADHDALLFHLK pc3
P07307;P07307−2;P07307−3;Q7Z4(
log10 peptide intensity
650:676 472:489 94:95
17:7012073:G:T_T
p = 5.8e−06, beta = 0.203, N = 1216
WNDDFCLQVYR pc2
P07307;P07307−2;P07307−3;Q7Z4(
log10 peptide intensity
519:676 397:489 78:95
17:7012073:G:T_T
p = 0.01, beta = 0.128, N = 994
AVTQPDNWHGHELGGSEDCVEVQPD(
P07307;P07307−2;P07307−3;Q7Z4(
log10 peptide intensity
168:676 129:489 28:95
17:7012073:G:T_T
p = 0.031, beta = 0.183, N = 325
TCCPVNWVEHQGSCYWFSHSGK p(
P07307;P07307−2;P07307−3;Q7Z4(
log10 peptide intensity
55:676 50:489 10:95
17:7012073:G:T_T
p = 0.15, beta = 0.202, N = 115
ASGR2 : NP2
P07307;P07307−2;P07307−3;Q7Z4(
log10 protein intensity
198:198 103:103 24:24
17:7012073:G:T_T
p = 7e−04, beta = 0.293, N = 325
ADHDALLFHLK pc3
P07307;P07307−2;P07307−3;Q7Z4(
log10 peptide intensity
197:198 103:103 24:24
17:7012073:G:T_T
p = 9.5e−05, beta = 0.336, N = 324
WNDDFCLQVYR pc2
P07307;P07307−2;P07307−3;Q7Z4(
log10 peptide intensity
115:198 68:103 16:24
17:7012073:G:T_T
p = 0.0012, beta = 0.347, N = 199
HFPVDLR pc2
P07307;P07307−2;P07307−3;Q7Z4(
log10 peptide intensity
42:198 23:103 8:24
17:7012073:G:T_T
p = 0.95, beta = −0.00986, N = 73
AVTQPDNWHGHELGGSEDCVEVQPD
P07307;P07307−2;P07307−3;Q7Z4(
log10 peptide intensity
14:198 14:103 2:24
17:7012073:G:T_T
p = 0.84, beta = 0.0568, N = 30
Assay Target: ASGR2
Olink UniProt: P07307
UKB rsID: rs2304980
Proxy rsID: rs2304980
UKB SNP: 17:7012073:G:T
Proxy SNP: 17:7012073:G:T
UKB log10(p): 498.6
UKB BETA: 0.316
*:−:−:−:−:−
1216:994:493:325:115:96
GAQLQAELR pc2
rs2304978 ALT
log10 peptide intensity
10:676 267:489 77:95
17:7012073:G:T_T
p = 4.4e−132, model = REC, N = 354

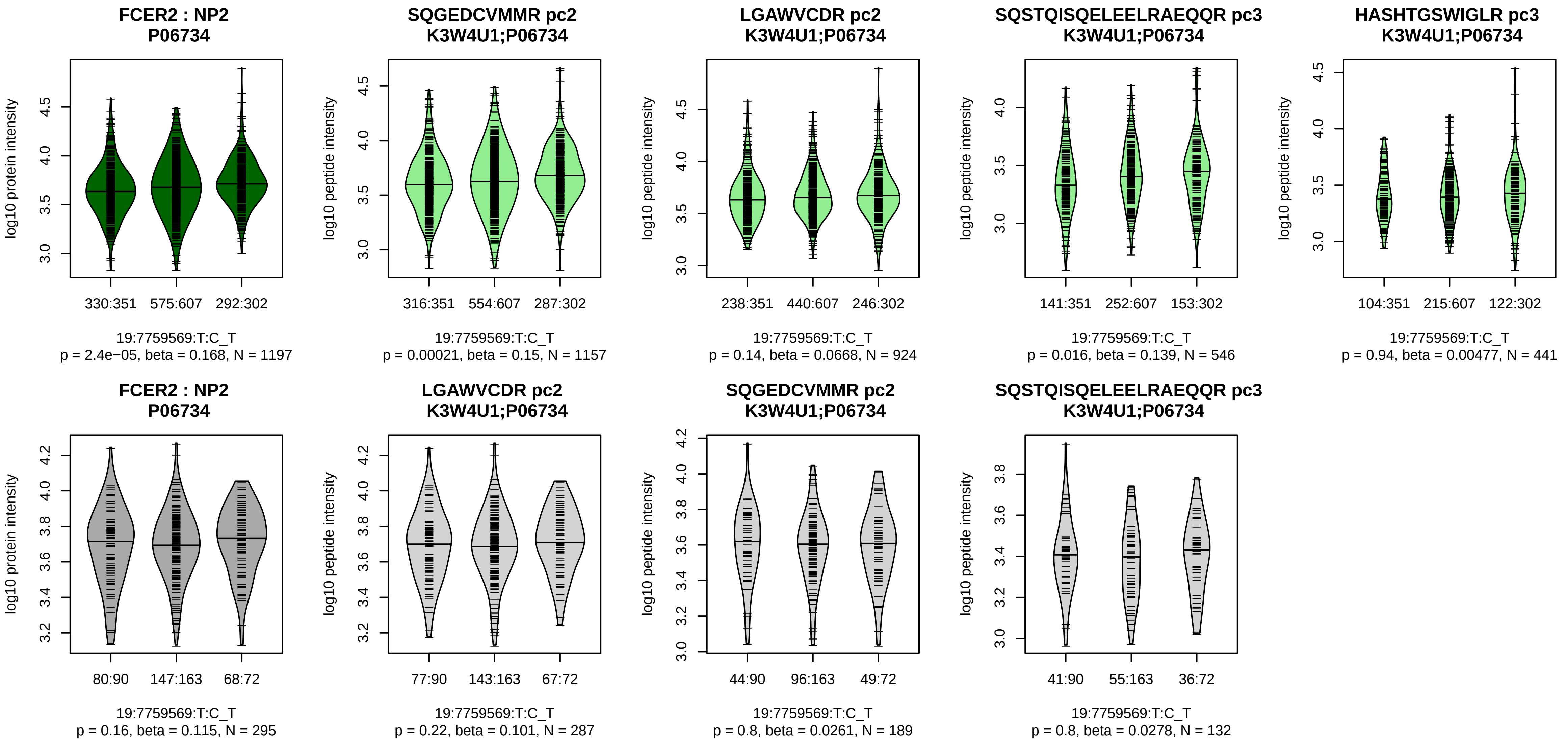


Assay Target: FCER2
Olink UniProt: P06734
UKB rsID: rs11260012
Proxy rsID: rs11260012
UKB SNP: 19:7759569:T:C
Proxy SNP: 19:7759569:T:C
UKB log10(p): 497.2
UKB BETA: −0.289
*:−:−:−:−
1157:924:546:441:205

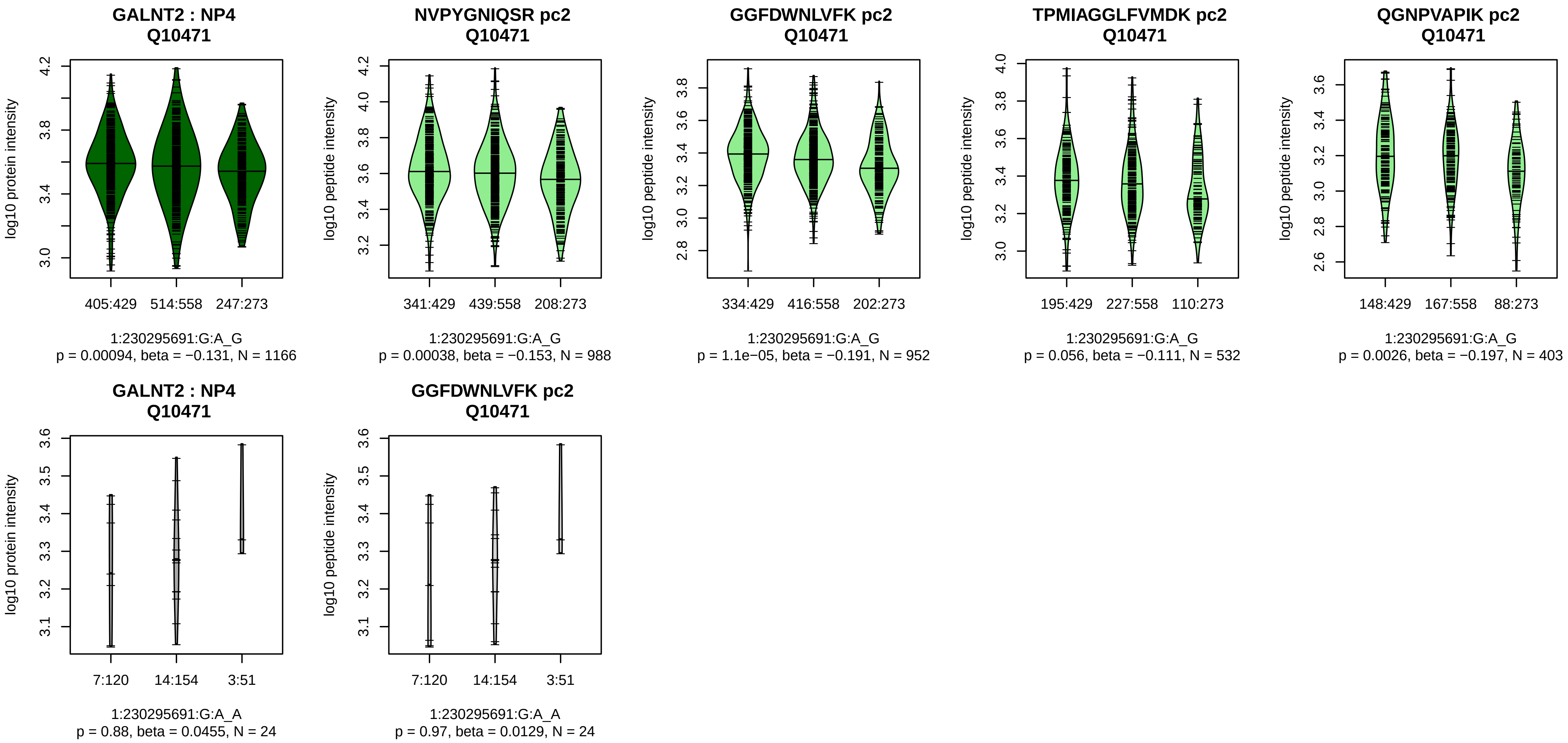


Assay Target: GALNT2
Olink UniProt: Q10471
UKB rsID: rs4846914
Proxy rsID: rs4846914
UKB SNP: 1:230295691:G:A
Proxy SNP: 1:230295691:G:A
UKB log10(p): 490
UKB BETA: 0.287
*:*:−:*:−:−:−:NA
988:952:532:403:337:85:27:14

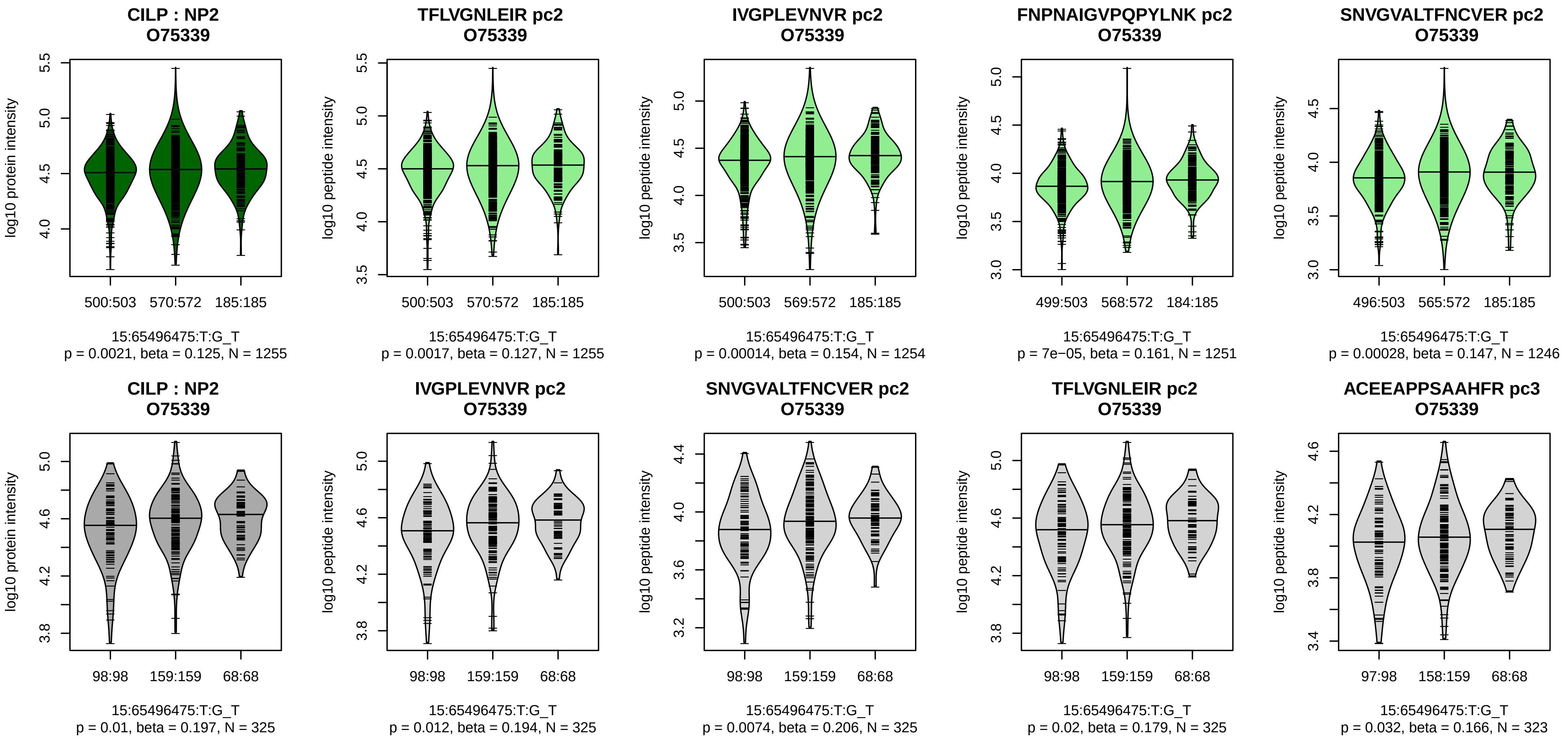


Assay Target: CILP
Olink UniProt: O75339
UKB rsID: rs2585035
Proxy rsID: rs2585035
UKB SNP: 15:65496475:T:G
Proxy SNP: 15:65496475:T:G
UKB log10(p): 484.6
UKB BETA: −0.288
*:*:*:*:*:–:*:–:*:*:–:–:–:–:–:–:–:–:
1255:1254:1251:1246:1239:123

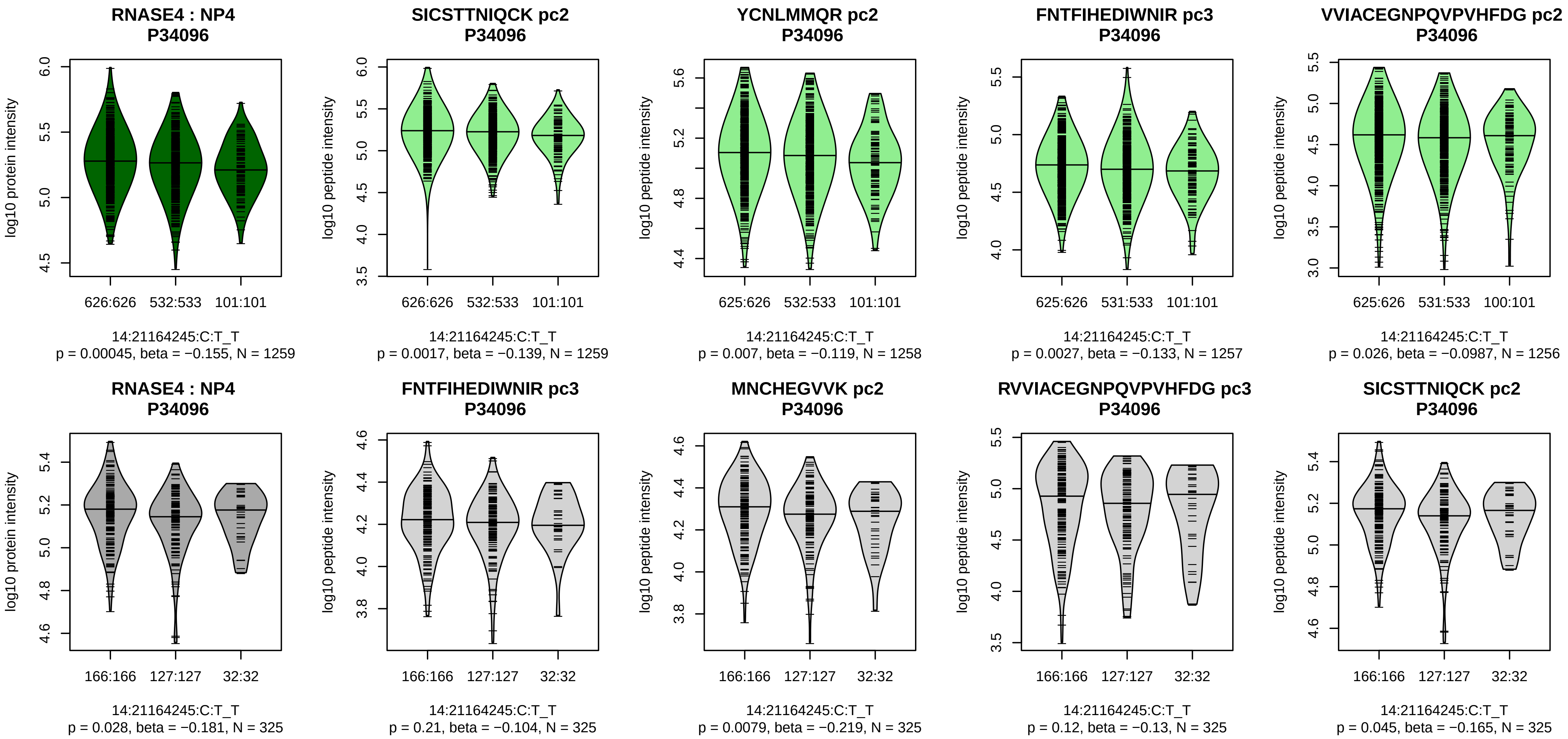


Assay Target: RNASE4
Olink UniProt: P34096
UKB rsID: rs1888560
Proxy rsID: rs1888560
UKB SNP: 14:21164245:C:T
Proxy SNP: 14:21164245:C:T
UKB log10(p): 484.2
UKB BETA: −0.304
*:*:*:−:*:−:−:−:−:−:−:−:−:−:−:NA
1259:1258:1257:1256:1252:123

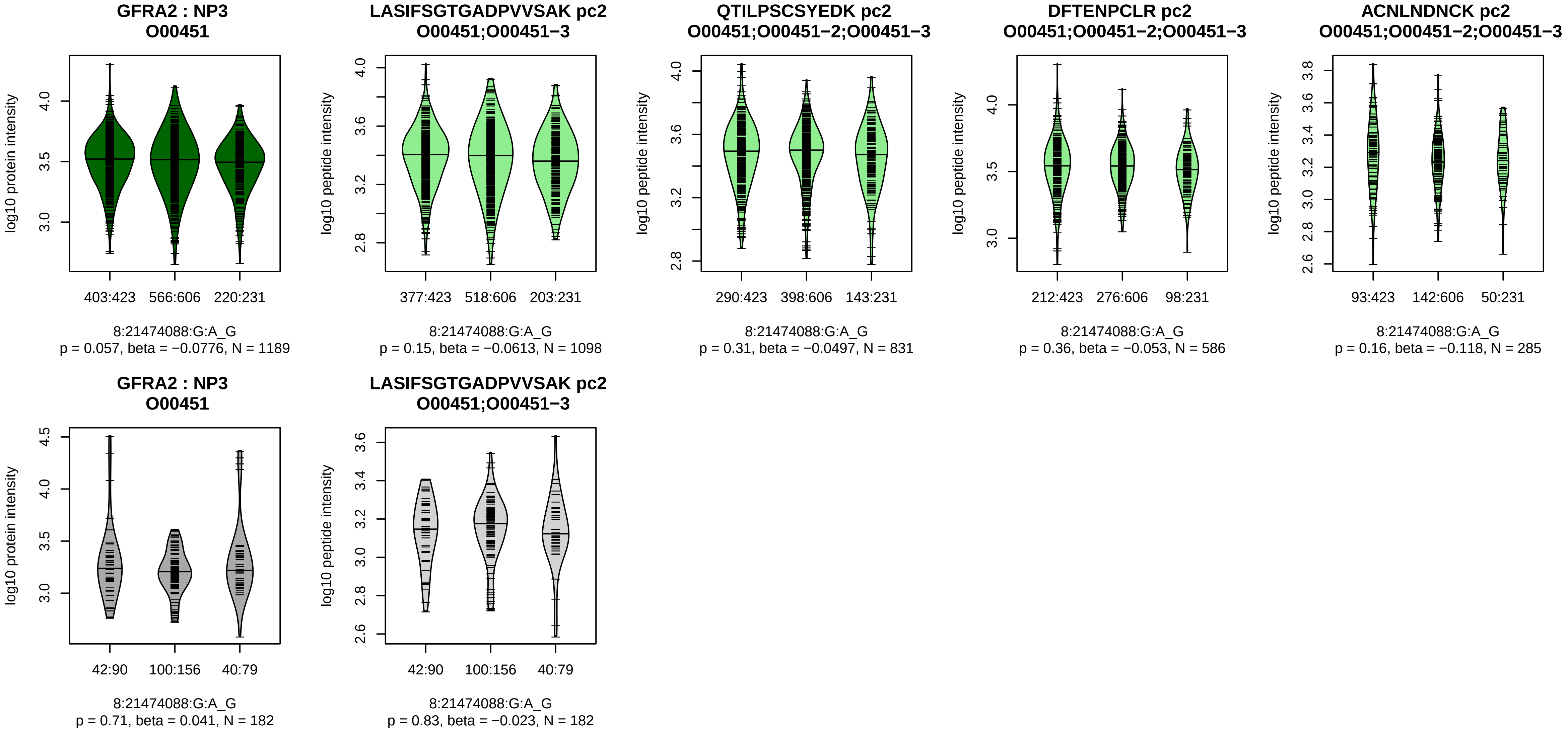


Assay Target: GFRA2
Olink UniProt: O00451
UKB rsID: rs1480999
Proxy rsID: rs1480999
UKB SNP: 8:21474088:G:A
Proxy SNP: 8:21474088:G:A
UKB log10(p): 474.1
UKB BETA: 0.294
−:−:−:−:−:−:−
1098:831:586:285:211:169:139

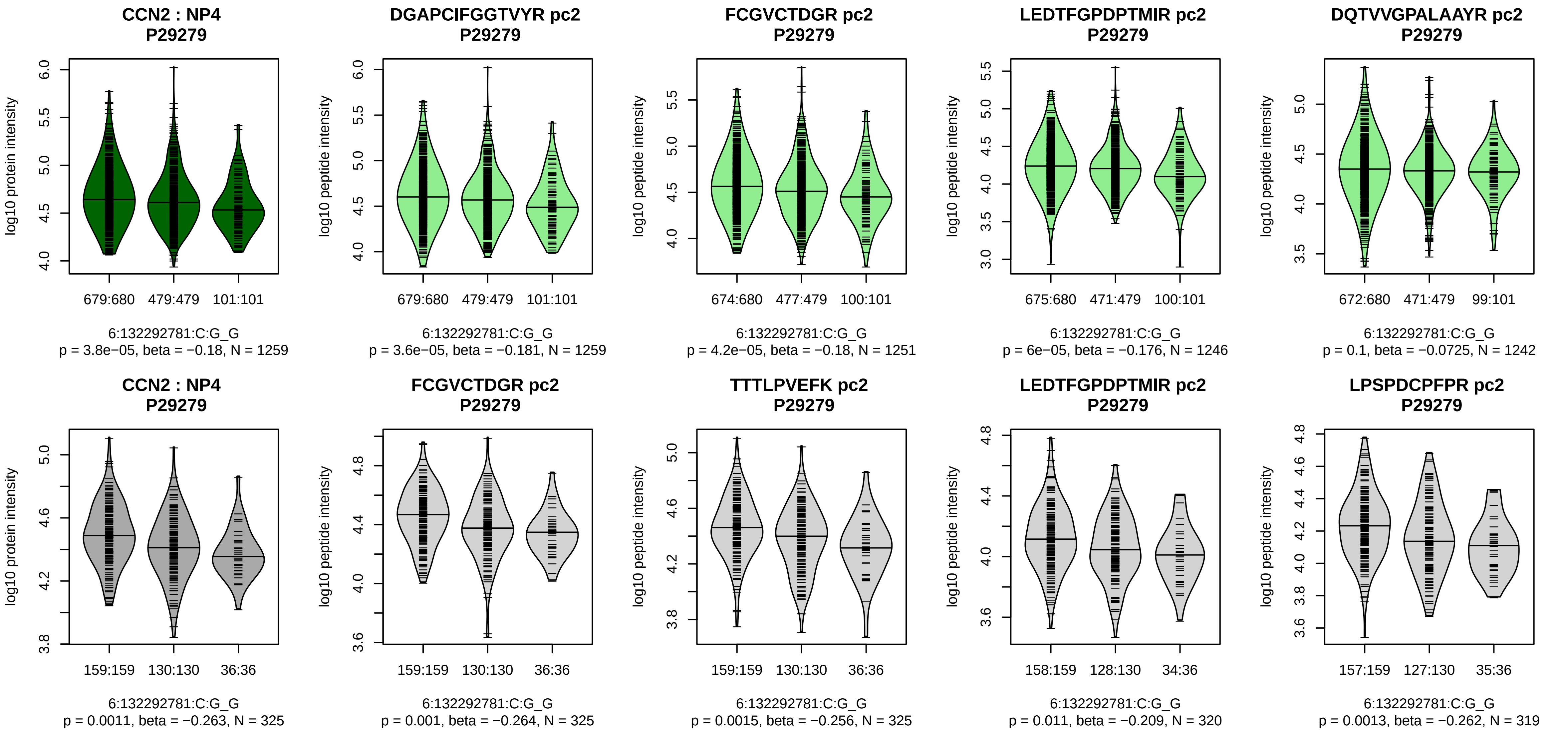


Assay Target: CCN2
Olink UniProt: P29279
UKB rsID: rs7767139
Proxy rsID: rs7767139
UKB SNP: 6:132292781:C:G
Proxy SNP: 6:132292781:C:G
UKB log10(p): 471.2
UKB BETA: −0.343
*:*:*:–:*:*:*:*:*:–:–:–:*:*:–:–:–:–:–
1259:1251:1246:1242:1242:121

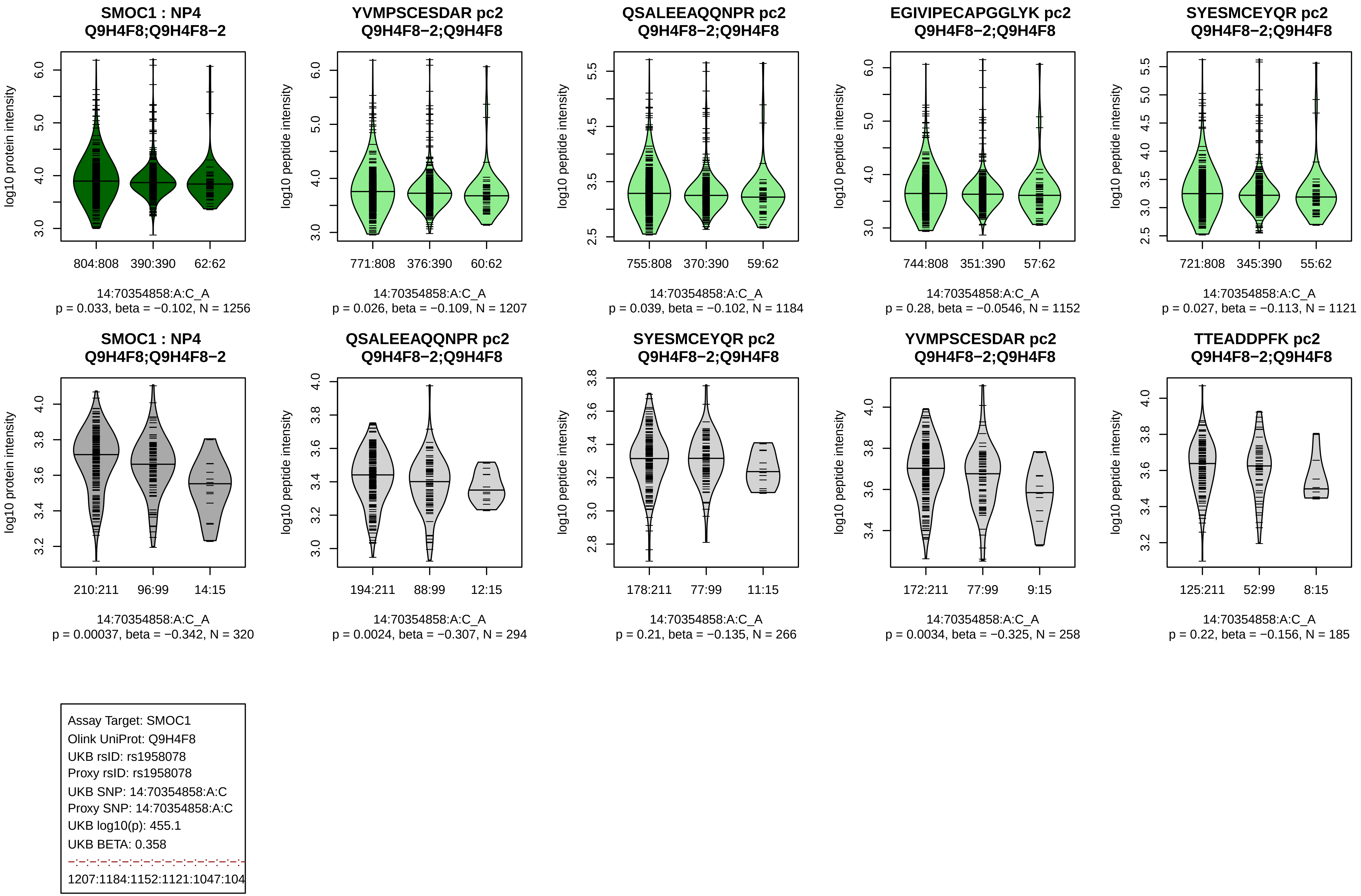

SMOC1 : NP4 Q9H4F8;Q9H4F8−2
log10 protein intensity
804:808 390:390 62:62
14:70354858:A:C_A
p = 0.033, beta = −0.102, N = 1256
YVMPSCESDAR pc2 Q9H4F8−2;Q9H4F8
log10 peptide intensity
771:808 376:390 60:62
14:70354858:A:C_A
p = 0.026, beta = −0.109, N = 1207
QSALEEAQQNPR pc2 Q9H4F8−2;Q9H4F8
log10 peptide intensity
755:808 370:390 59:62
14:70354858:A:C_A
p = 0.039, beta = −0.102, N = 1184
EGIVIPECAPGGLYK pc2 Q9H4F8−2;Q9H4F8
log10 peptide intensity
744:808 351:390 57:62
14:70354858:A:C_A
p = 0.28, beta = −0.0546, N = 1152
SYESMCEYQR pc2 Q9H4F8−2;Q9H4F8
log10 peptide intensity
721:808 345:390 55:62
14:70354858:A:C_A
p = 0.027, beta = −0.113, N = 1121
SMOC1 : NP4 Q9H4F8;Q9H4F8−2
log10 protein intensity
210:211 96:99 14:15
14:70354858:A:C_A
p = 0.00037, beta = −0.342, N = 320
QSALEEAQQNPR pc2 Q9H4F8−2;Q9H4F8
log10 peptide intensity
194:211 88:99 12:15
14:70354858:A:C_A
p = 0.0024, beta = −0.307, N = 294
SYESMCEYQR pc2 Q9H4F8−2;Q9H4F8
log10 peptide intensity
178:211 77:99 11:15
14:70354858:A:C_A
p = 0.21, beta = −0.135, N = 266
YVMPSCESDAR pc2 Q9H4F8−2;Q9H4F8
log10 peptide intensity
172:211 77:99 9:15
14:70354858:A:C_A
p = 0.0034, beta = −0.325, N = 258
TTEADDPFK pc2 Q9H4F8−2;Q9H4F8
log10 peptide intensity
125:211 52:99 8:15
14:70354858:A:C_A
p = 0.22, beta = −0.156, N = 185
Assay Target: SMOC1
Olink UniProt: Q9H4F8
UKB rsID: rs1958078
Proxy rsID: rs1958078
UKB SNP: 14:70354858:A:C
Proxy SNP: 14:70354858:A:C
UKB log10(p): 455.1
UKB BETA: 0.358
1207:1184:1152:1121:1047:104

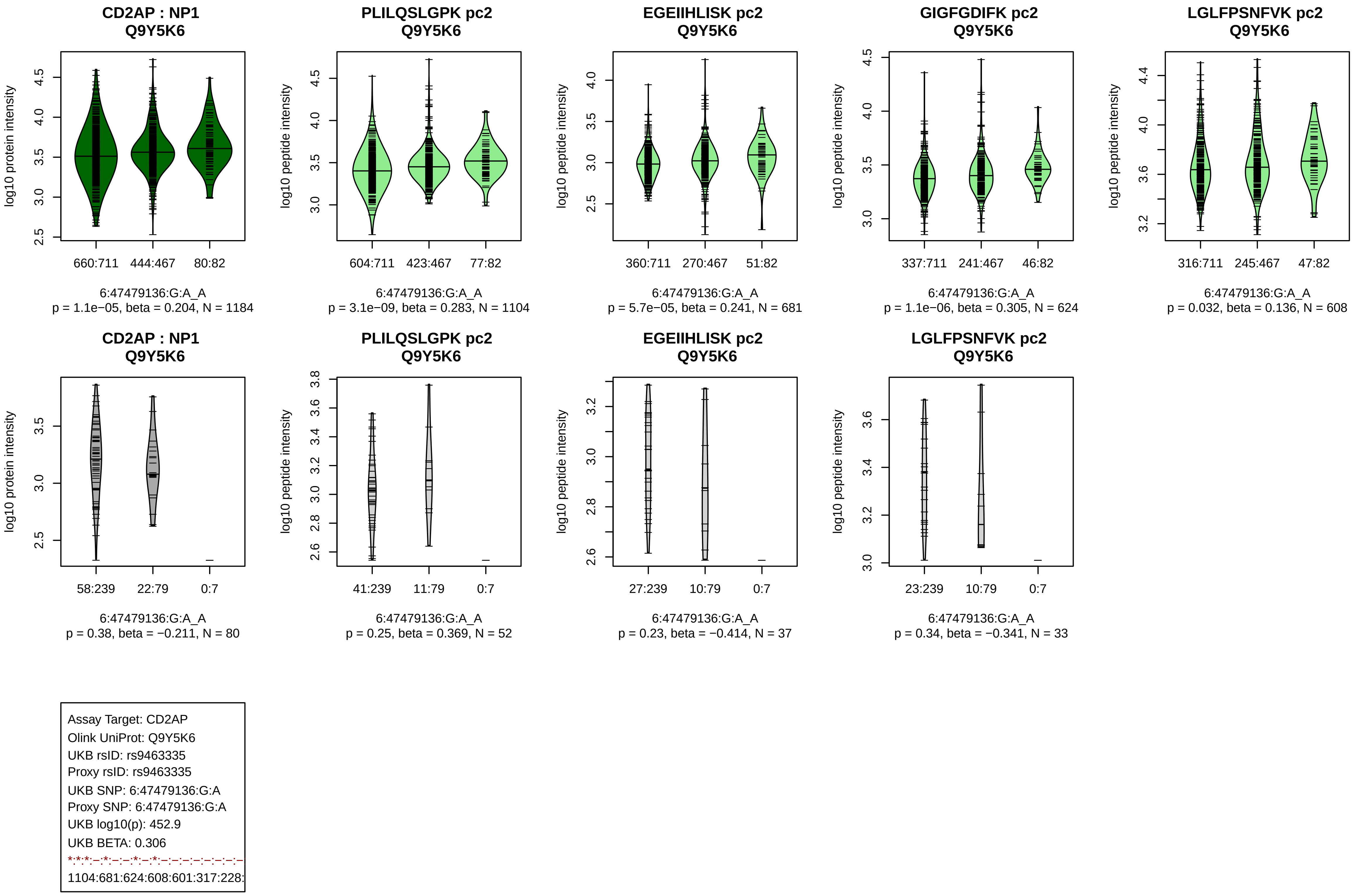
CD2AP : NP1
Q9Y5K6
log10 protein intensity
660:711
444:467
80:82
6:47479136:G:A_A
p = 1.1e−05, beta = 0.204, N = 1184
PLILQSLGPK pc2
Q9Y5K6
log10 peptide intensity
604:711
423:467
77:82
6:47479136:G:A_A
p = 3.1e−09, beta = 0.283, N = 1104
EGEIIHLISK pc2
Q9Y5K6
log10 peptide intensity
360:711
270:467
51:82
6:47479136:G:A_A
p = 5.7e−05, beta = 0.241, N = 681
GIGFGDIFK pc2
Q9Y5K6
log10 peptide intensity
337:711
241:467
46:82
6:47479136:G:A_A
p = 1.1e−06, beta = 0.305, N = 624
LGLFPSNFVK pc2
Q9Y5K6
log10 peptide intensity
316:711
245:467
47:82
6:47479136:G:A_A
p = 0.032, beta = 0.136, N = 608
CD2AP : NP1
Q9Y5K6
log10 protein intensity
58:239
22:79
0:7
6:47479136:G:A_A
p = 0.38, beta = −0.211, N = 80
PLILQSLGPK pc2
Q9Y5K6
log10 peptide intensity
41:239
11:79
0:7
6:47479136:G:A_A
p = 0.25, beta = 0.369, N = 52
EGEIIHLISK pc2
Q9Y5K6
log10 peptide intensity
27:239
10:79
0:7
6:47479136:G:A_A
p = 0.23, beta = −0.414, N = 37
LGLFPSNFVK pc2
Q9Y5K6
log10 peptide intensity
23:239
10:79
0:7
6:47479136:G:A_A
p = 0.34, beta = −0.341, N = 33
Assay Target: CD2AP
Olink UniProt: Q9Y5K6
UKB rsID: rs9463335
Proxy rsID: rs9463335
UKB SNP: 6:47479136:G:A
Proxy SNP: 6:47479136:G:A
UKB log10(p): 452.9
UKB BETA: 0.306
*:*:*:–:*:–:–:*:–:*:–:–:–:–:–:–:–:–:–
1104:681:624:608:601:317:228:

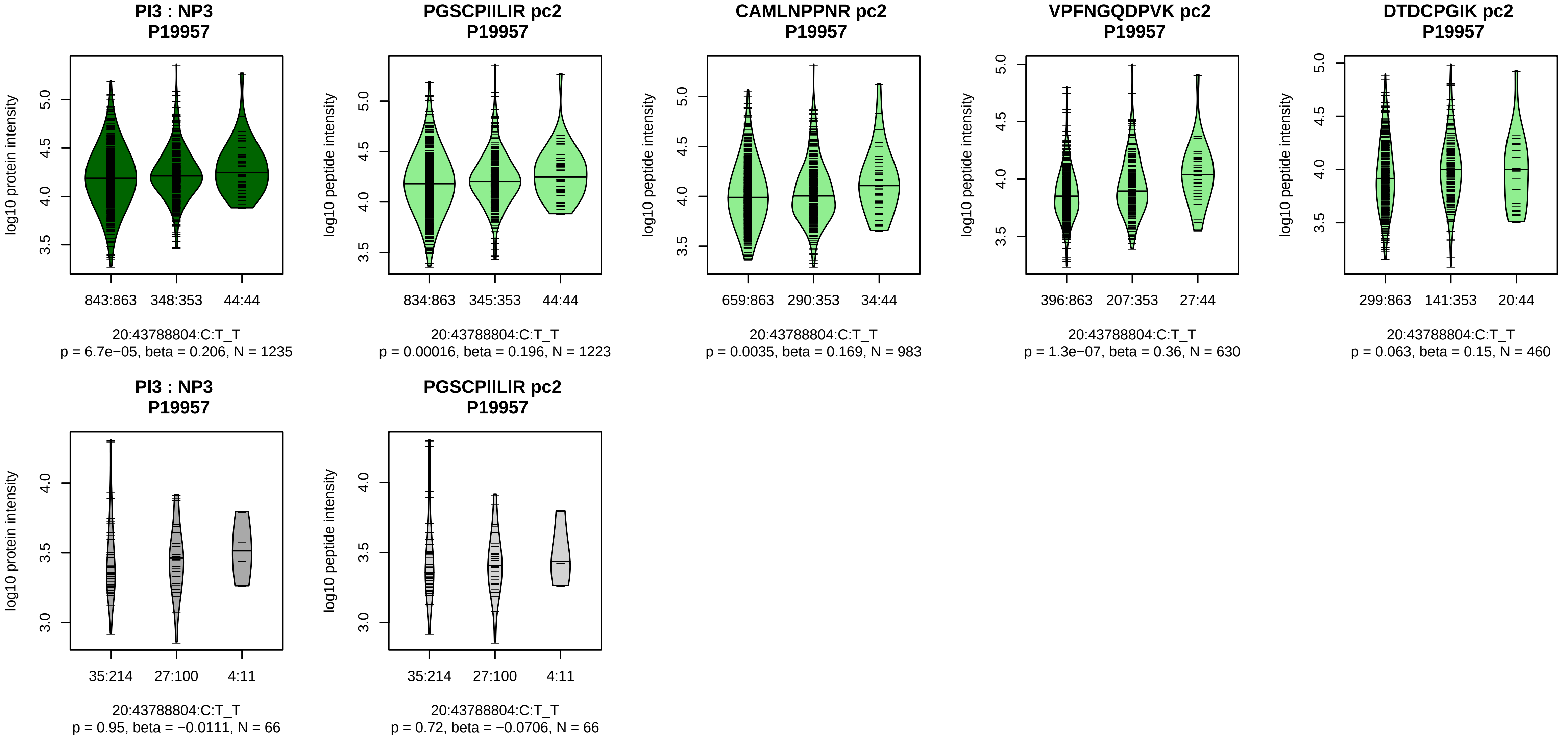


Assay Target: PI3
Olink UniProt: P19957
UKB rsID: rs35770435
Proxy rsID: rs35770435
UKB SNP: 20:43788804:C:T
Proxy SNP: 20:43788804:C:T
UKB log10(p): 450.3
UKB BETA: 0.345
*:*:*:–:–
1223:983:630:460:31

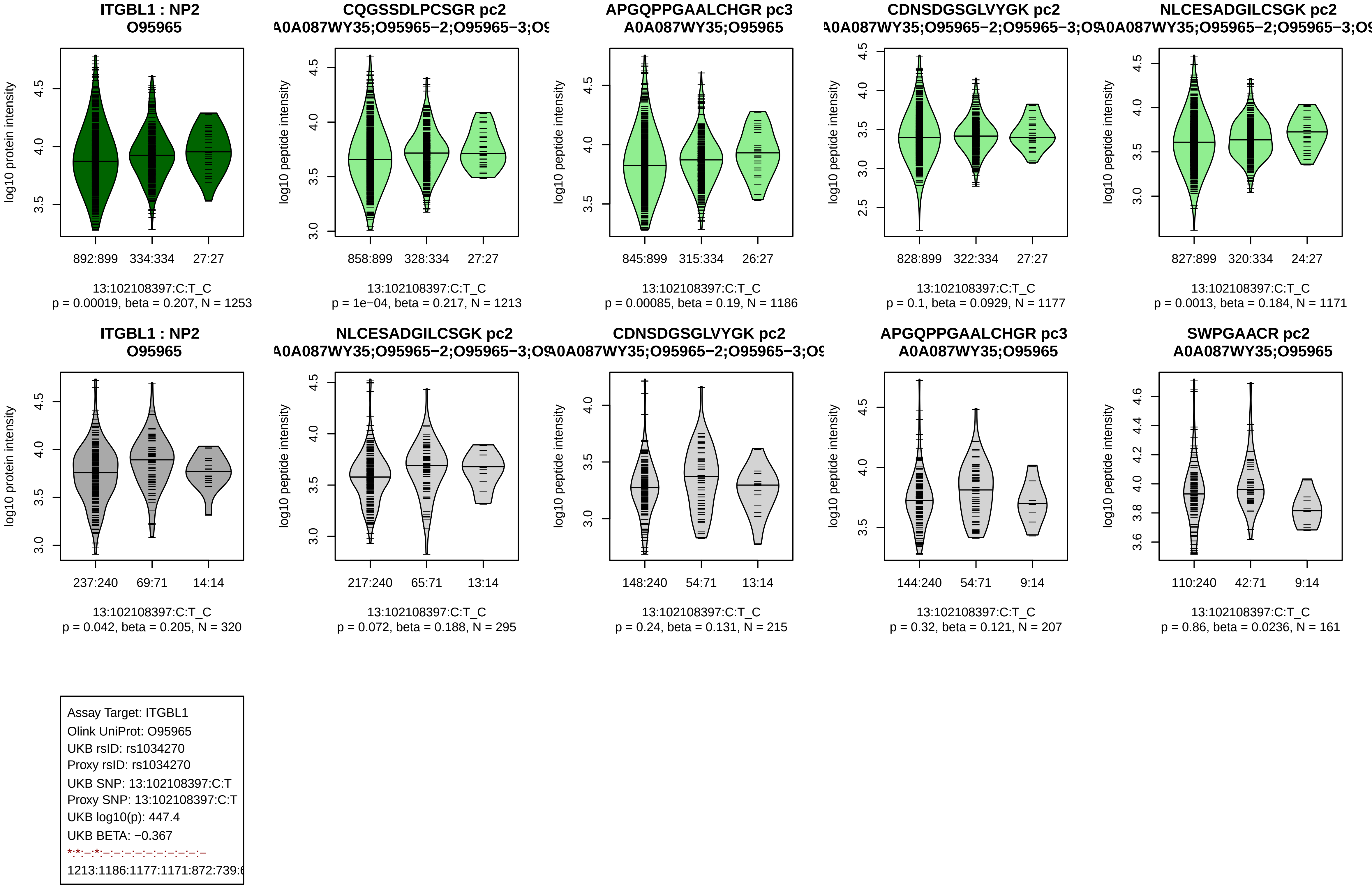
ITGBL1 : NP2
O95965
log10 protein intensity
892:899 334:334 27:27
13:102108397:C:T_C
p = 0.00019, beta = 0.207, N = 1253
CQGSSDLPCSGR pc2
A0A087WY35;O95965−2;O95965−3;O9
log10 peptide intensity
858:899 328:334 27:27
13:102108397:C:T_C
p = 1e−04, beta = 0.217, N = 1213
APGQPPGAALCHGR pc3
A0A087WY35;O95965
log10 peptide intensity
845:899 315:334 26:27
13:102108397:C:T_C
p = 0.00085, beta = 0.19, N = 1186
CDNSDGSGLVYGK pc2
A0A087WY35;O95965−2;O95965−3;O9
log10 peptide intensity
828:899 322:334 27:27
13:102108397:C:T_C
p = 0.1, beta = 0.0929, N = 1177
NLCESADGILCSGK pc2
A0A087WY35;O95965−2;O95965−3;O9
log10 peptide intensity
827:899 320:334 24:27
13:102108397:C:T_C
p = 0.0013, beta = 0.184, N = 1171
ITGBL1 : NP2
O95965
log10 protein intensity
237:240 69:71 14:14
13:102108397:C:T_C
p = 0.042, beta = 0.205, N = 320
NLCESADGILCSGK pc2
A0A087WY35;O95965−2;O95965−3;O9
log10 peptide intensity
217:240 65:71 13:14
13:102108397:C:T_C
p = 0.072, beta = 0.188, N = 295
CDNSDGSGLVYGK pc2
A0A087WY35;O95965−2;O95965−3;O9
log10 peptide intensity
148:240 54:71 13:14
13:102108397:C:T_C
p = 0.24, beta = 0.131, N = 215
APGQPPGAALCHGR pc3
A0A087WY35;O95965
log10 peptide intensity
144:240 54:71 9:14
13:102108397:C:T_C
p = 0.32, beta = 0.121, N = 207
SWPGAACR pc2
A0A087WY35;O95965
log10 peptide intensity
110:240 42:71 9:14
13:102108397:C:T_C
p = 0.86, beta = 0.0236, N = 161
Assay Target: ITGBL1
Olink UniProt: O95965
UKB rsID: rs1034270
Proxy rsID: rs1034270
UKB SNP: 13:102108397:C:T
Proxy SNP: 13:102108397:C:T
UKB log10(p): 447.4
UKB BETA: −0.367
*:*:−:*:−:−:−:−:−:−:−:−:−:−
1213:1186:1177:1171:872:739:6

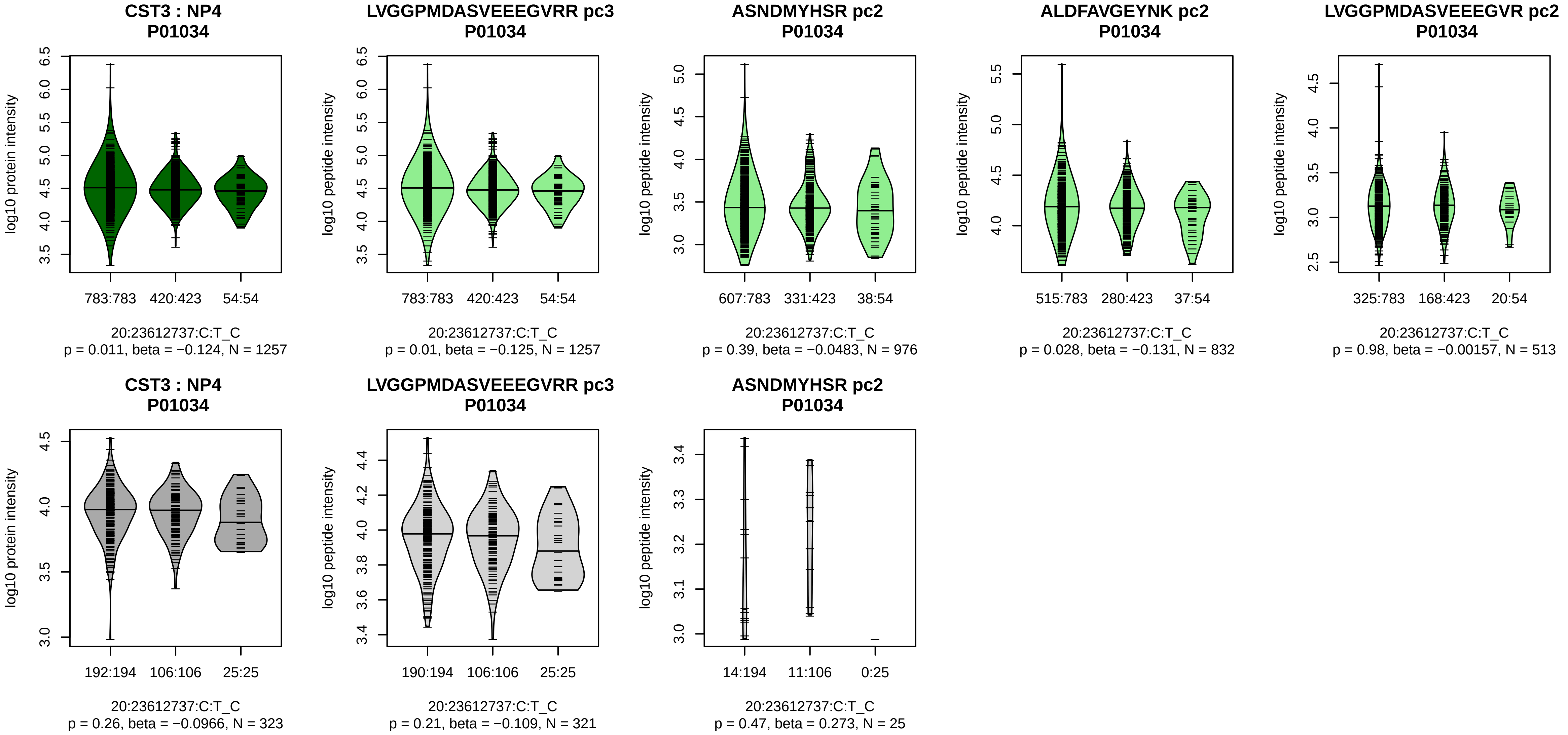


Assay Target: CST3
Olink UniProt: P01034
UKB rsID: rs911119
Proxy rsID: rs911119
UKB SNP: 20:23612737:C:T
Proxy SNP: 20:23612737:C:T
UKB log10(p): 447.4
UKB BETA: 0.307
−:−:−:−:−:−:−:−:−:−:NA:NA:NA
1257:976:832:513:510:184:160:

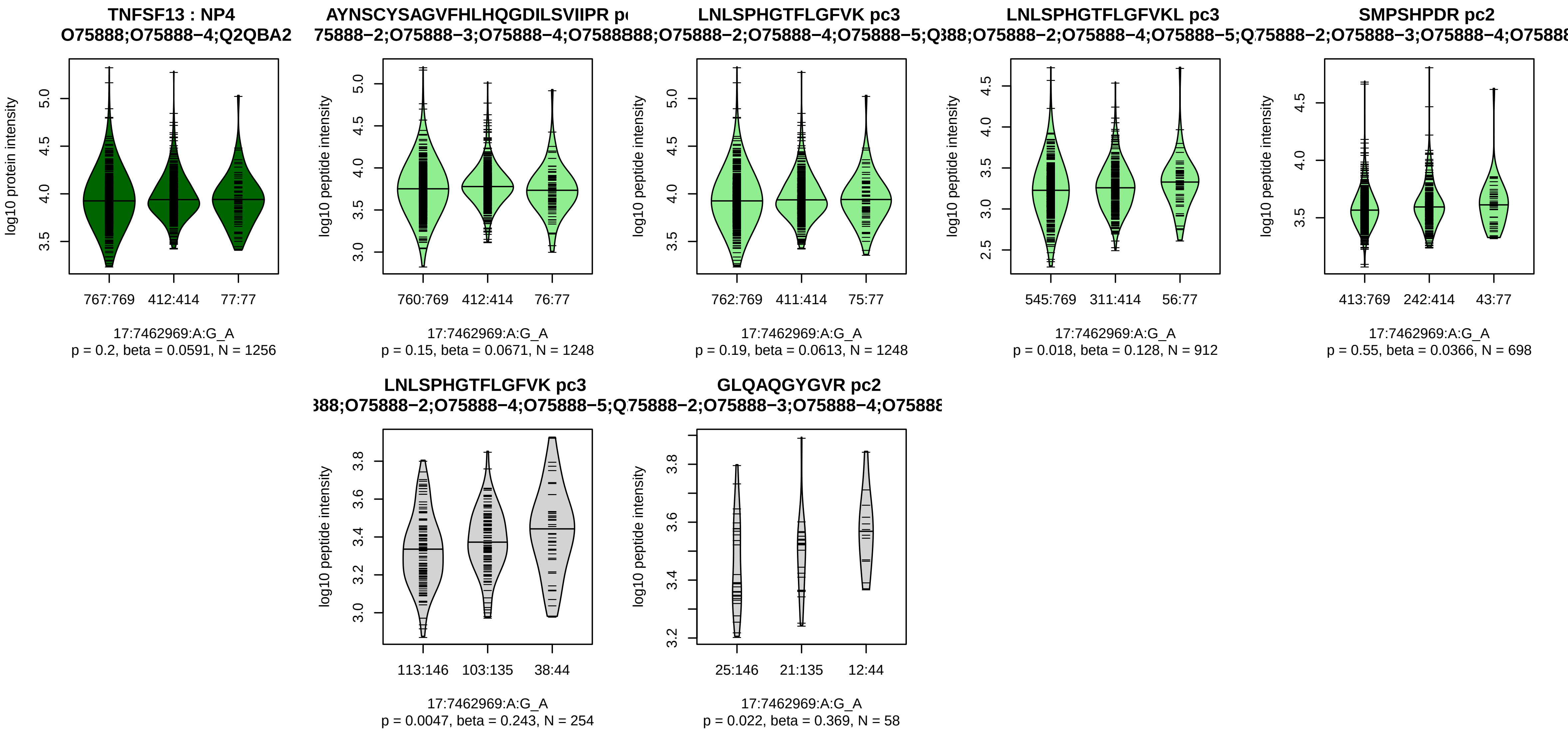


Assay Target: TNFSF13
Olink UniProt: O75888
UKB rsID: rs3803800
Proxy rsID: rs3803800
UKB SNP: 17:7462969:A:G
Proxy SNP: 17:7462969:A:G
UKB log10(p): 444
UKB BETA: −0.313
−:−:−:−:−:−:−:−:NA
1248:1248:912:698:235:114:90:

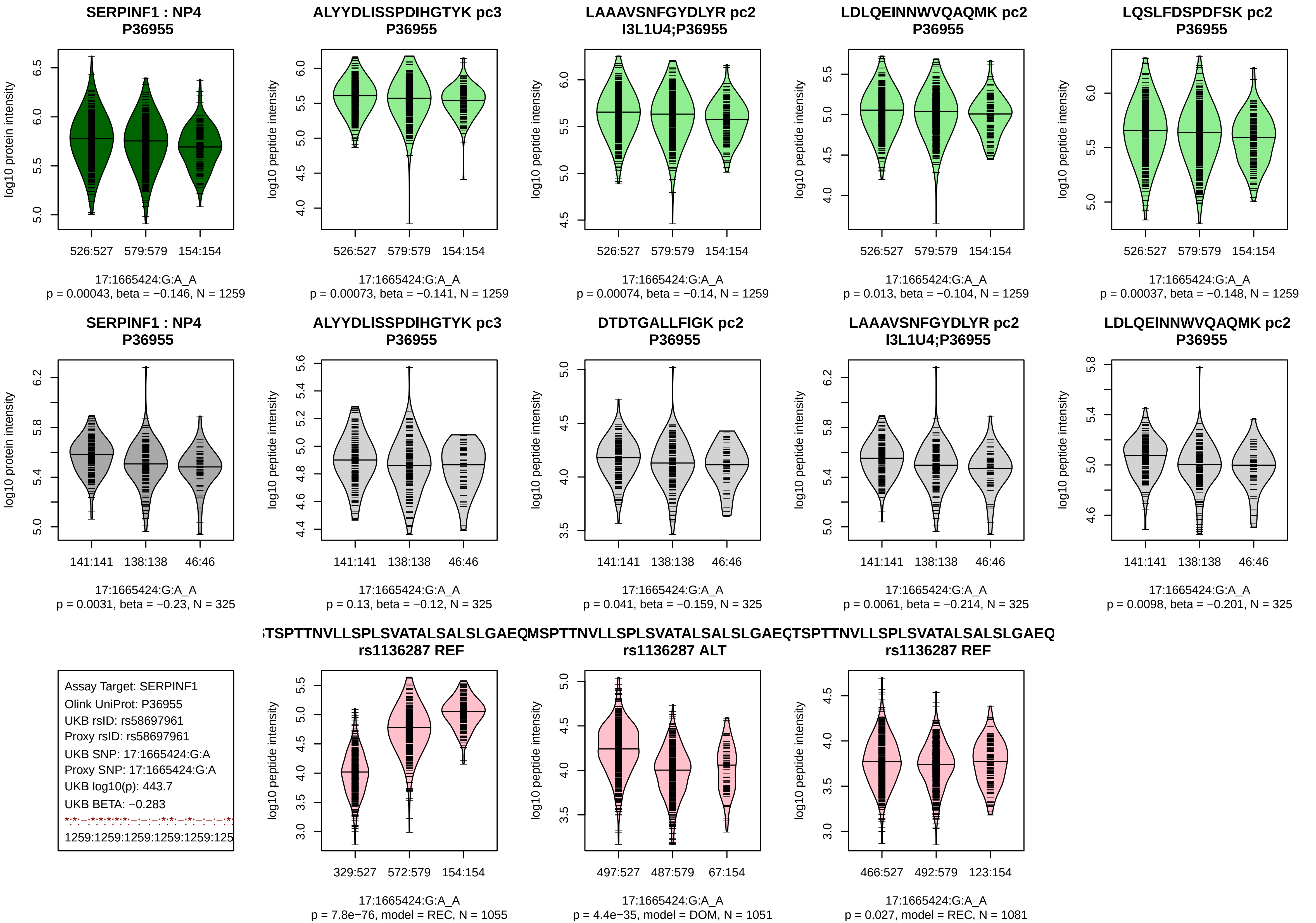

SERPINF1 : NP4
P36955
log10 protein intensity
526:527 579:579 154:154
17:1665424:G:A_A
p = 0.00043, beta = −0.146, N = 1259
ALYYDLISSPDIHGTYK pc3
P36955
log10 peptide intensity
526:527 579:579 154:154
17:1665424:G:A_A
p = 0.00073, beta = −0.141, N = 1259
LAAAVSNFGYDLYR pc2
I3L1U4;P36955
log10 peptide intensity
526:527 579:579 154:154
17:1665424:G:A_A
p = 0.00074, beta = −0.14, N = 1259
LDLQEINNWVQAQMK pc2
P36955
log10 peptide intensity
526:527 579:579 154:154
17:1665424:G:A_A
p = 0.013, beta = −0.104, N = 1259
LQSLFDSPDFSK pc2
P36955
log10 peptide intensity
526:527 579:579 154:154
17:1665424:G:A_A
p = 0.00037, beta = −0.148, N = 1259
SERPINF1 : NP4
P36955
log10 protein intensity
141:141 138:138 46:46
17:1665424:G:A_A
p = 0.0031, beta = −0.23, N = 325
ALYYDLISSPDIHGTYK pc3
P36955
log10 peptide intensity
141:141 138:138 46:46
17:1665424:G:A_A
p = 0.13, beta = −0.12, N = 325
DTDTGALLFIGK pc2
P36955
log10 peptide intensity
141:141 138:138 46:46
17:1665424:G:A_A
p = 0.041, beta = −0.159, N = 325
LAAAVSNFGYDLYR pc2
I3L1U4;P36955
log10 peptide intensity
141:141 138:138 46:46
17:1665424:G:A_A
p = 0.0061, beta = −0.214, N = 325
LDLQEINNWVQAQMK pc2
P36955
log10 peptide intensity
141:141 138:138 46:46
17:1665424:G:A_A
p = 0.0098, beta = −0.201, N = 325
Assay Target: SERPINF1
Olink UniProt: P36955
UKB rsID: rs58697961
Proxy rsID: rs58697961
UKB SNP: 17:1665424:G:A
Proxy SNP: 17:1665424:G:A
UKB log10(p): 443.7
UKB BETA: −0.283
1259:1259:1259:1259:1259:125
TSPTTNVLLSPLSVATALSALSLGAEQ
rs1136287 REF
log10 peptide intensity
329:527 572:579 154:154
17:1665424:G:A_A
p = 7.8e−76, model = REC, N = 1055
MSPTTNVLLSPLSVATALSALSLGAEQ
rs1136287 ALT
log10 peptide intensity
497:527 487:579 67:154
17:1665424:G:A_A
p = 4.4e−35, model = DOM, N = 1051
TSPTTNVLLSPLSVATALSALSLGAEQ
rs1136287 REF
log10 peptide intensity
466:527 492:579 123:154
17:1665424:G:A_A
p = 0.027, model = REC, N = 1081

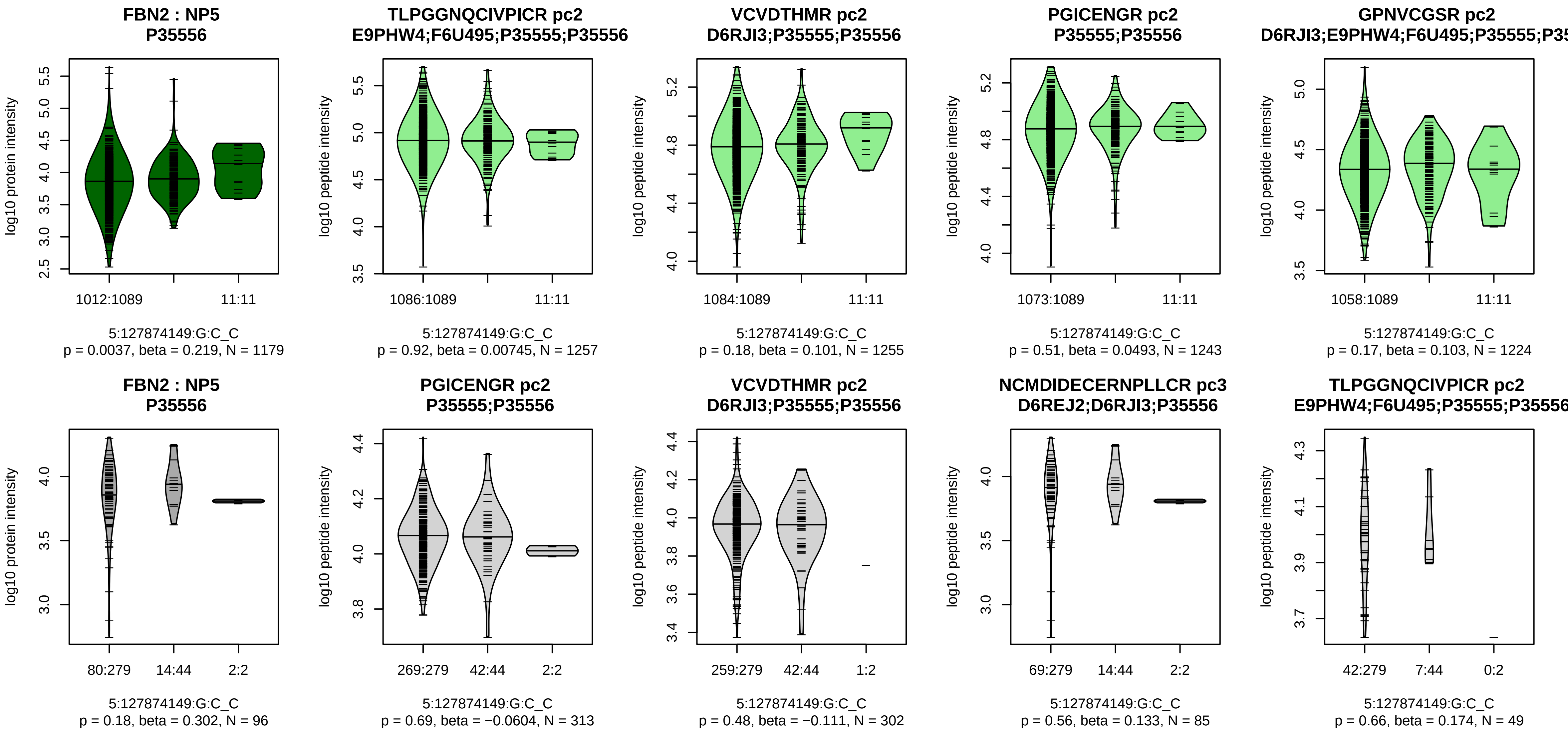


Assay Target: FBN2
Olink UniProt: P35556
UKB rsID: rs78821460
Proxy rsID: rs78821460
UKB SNP: 5:127874149:G:C
Proxy SNP: 5:127874149:G:C
UKB log10(p): 443
UKB BETA: 0.523
–:–:–:–:–:–:*:*:*:*:*:–:–:–:–:–:–
1257:1255:1243:1224:1113:107

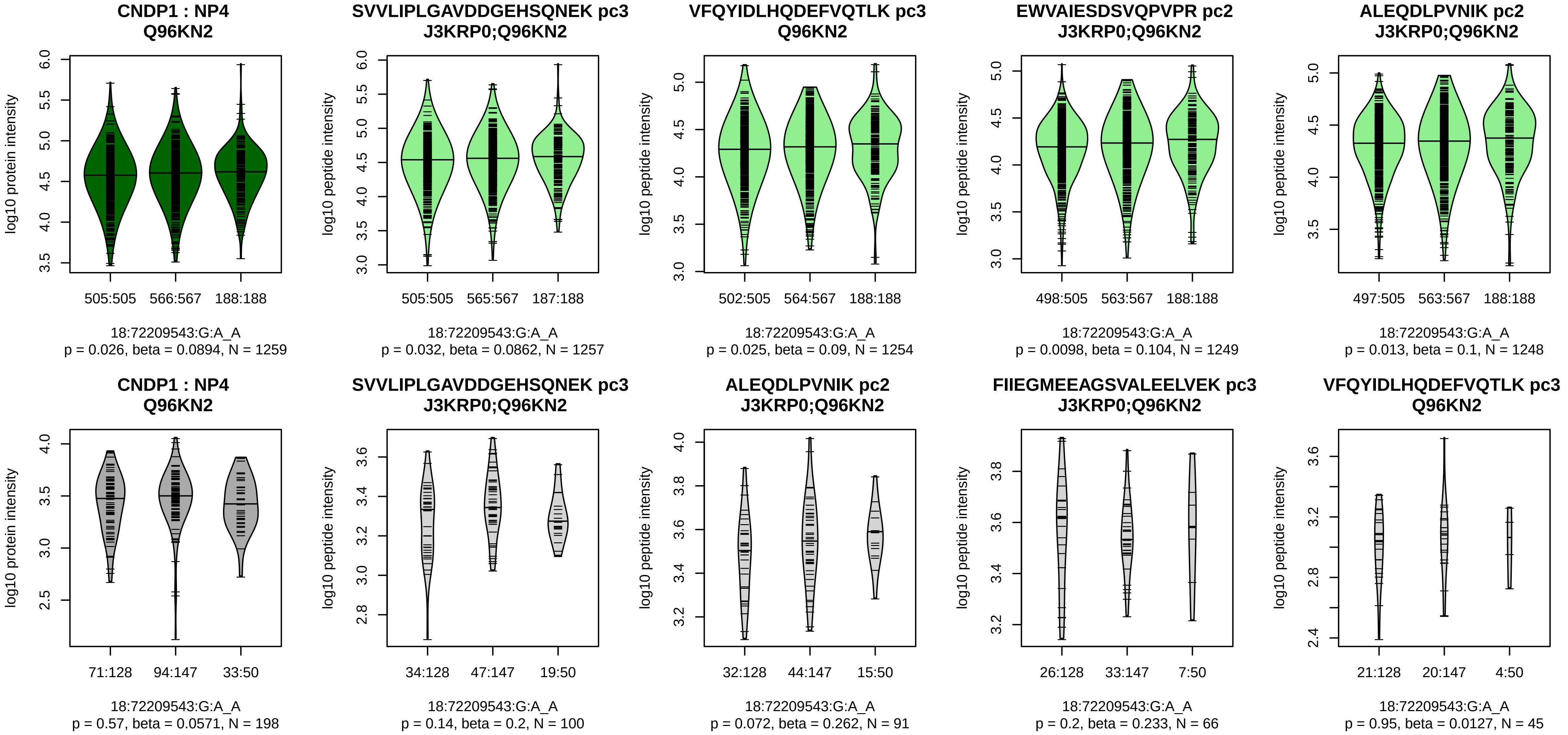


Assay Target: CNDP1
Olink UniProt: Q96KN2
UKB rsID: rs17817077
Proxy rsID: rs17817077
UKB SNP: 18:72209543:G:A
Proxy SNP: 18:72209543:G:A
UKB log10(p): 442.1
UKB BETA: 0.281
1257:1254:1249:1248:1235:123

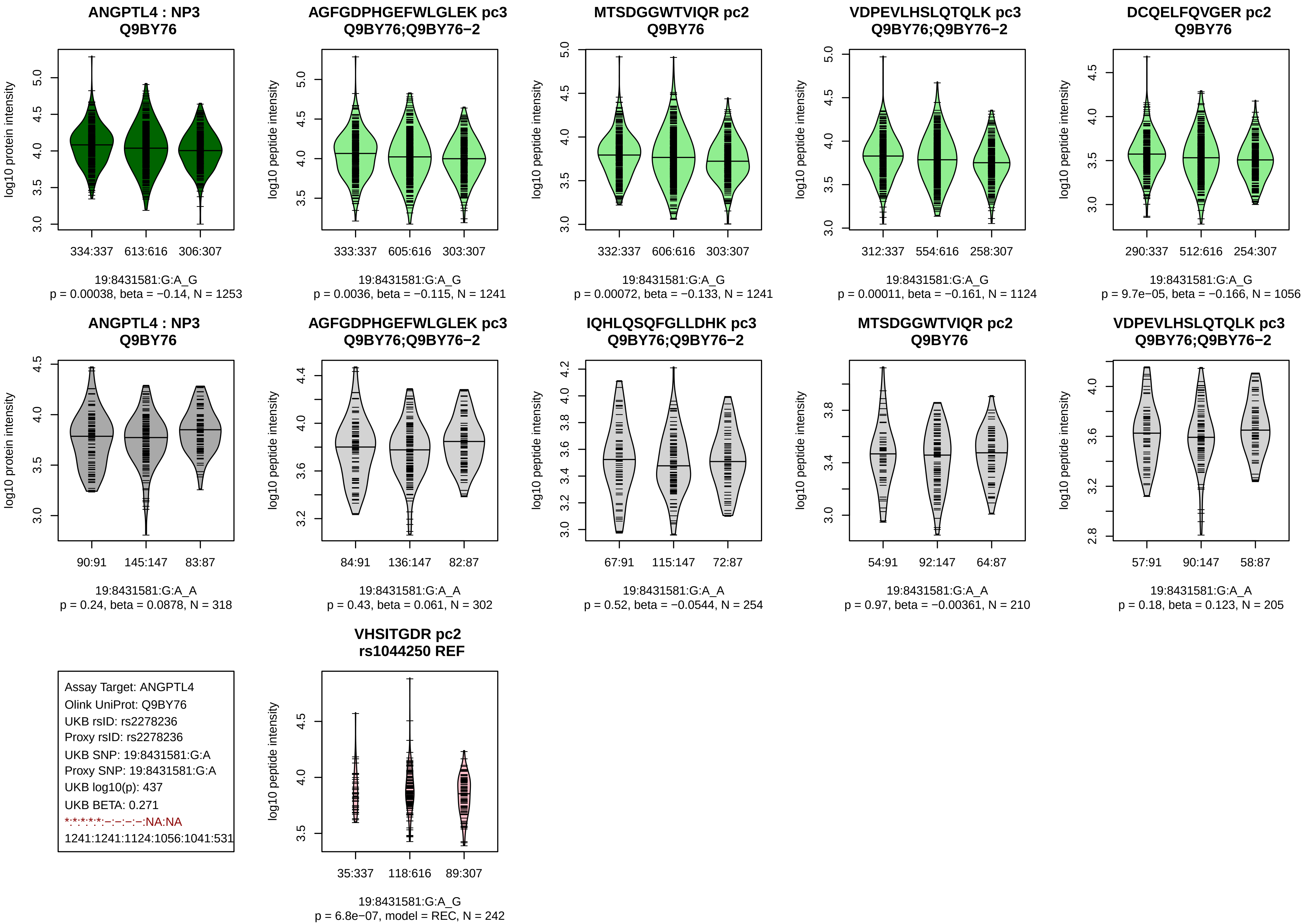

ANGPTL4 : NP3
Q9BY76
log10 protein intensity
334:337
613:616
306:307
19:8431581:G:A_G
p = 0.00038, beta = −0.14, N = 1253
AGFGDPHGEFWLGLEK pc3
Q9BY76;Q9BY76−2
log10 peptide intensity
333:337
605:616
303:307
19:8431581:G:A_G
p = 0.0036, beta = −0.115, N = 1241
MTSDGGWTVIQR pc2
Q9BY76
log10 peptide intensity
332:337
606:616
303:307
19:8431581:G:A_G
p = 0.00072, beta = −0.133, N = 1241
VDPEVLHSLQTQLK pc3
Q9BY76;Q9BY76−2
log10 peptide intensity
312:337
554:616
258:307
19:8431581:G:A_G
p = 0.00011, beta = −0.161, N = 1124
DCQELFQVGER pc2
Q9BY76
log10 peptide intensity
290:337
512:616
254:307
19:8431581:G:A_G
p = 9.7e−05, beta = −0.166, N = 1056
ANGPTL4 : NP3
Q9BY76
log10 protein intensity
90:91
145:147
83:87
19:8431581:G:A_A
p = 0.24, beta = 0.0878, N = 318
AGFGDPHGEFWLGLEK pc3
Q9BY76;Q9BY76−2
log10 peptide intensity
84:91
136:147
82:87
19:8431581:G:A_A
p = 0.43, beta = 0.061, N = 302
IQHLQSQFGLLDHK pc3
Q9BY76;Q9BY76−2
log10 peptide intensity
67:91
115:147
72:87
19:8431581:G:A_A
p = 0.52, beta = −0.0544, N = 254
MTSDGGWTVIQR pc2
Q9BY76
log10 peptide intensity
54:91
92:147
64:87
19:8431581:G:A_A
p = 0.97, beta = −0.00361, N = 210
VDPEVLHSLQTQLK pc3
Q9BY76;Q9BY76−2
log10 peptide intensity
57:91
90:147
58:87
19:8431581:G:A_A
p = 0.18, beta = 0.123, N = 205
Assay Target: ANGPTL4
Olink UniProt: Q9BY76
UKB rsID: rs2278236
Proxy rsID: rs2278236
UKB SNP: 19:8431581:G:A
Proxy SNP: 19:8431581:G:A
UKB log10(p): 437
UKB BETA: 0.271
*:*:*:*:*:−:−:−:−:NA:NA
1241:1241:1124:1056:1041:531
VHSITGDR pc2
rs1044250 REF
log10 peptide intensity
35:337
118:616
89:307
19:8431581:G:A_G
p = 6.8e−07, model = REC, N = 242

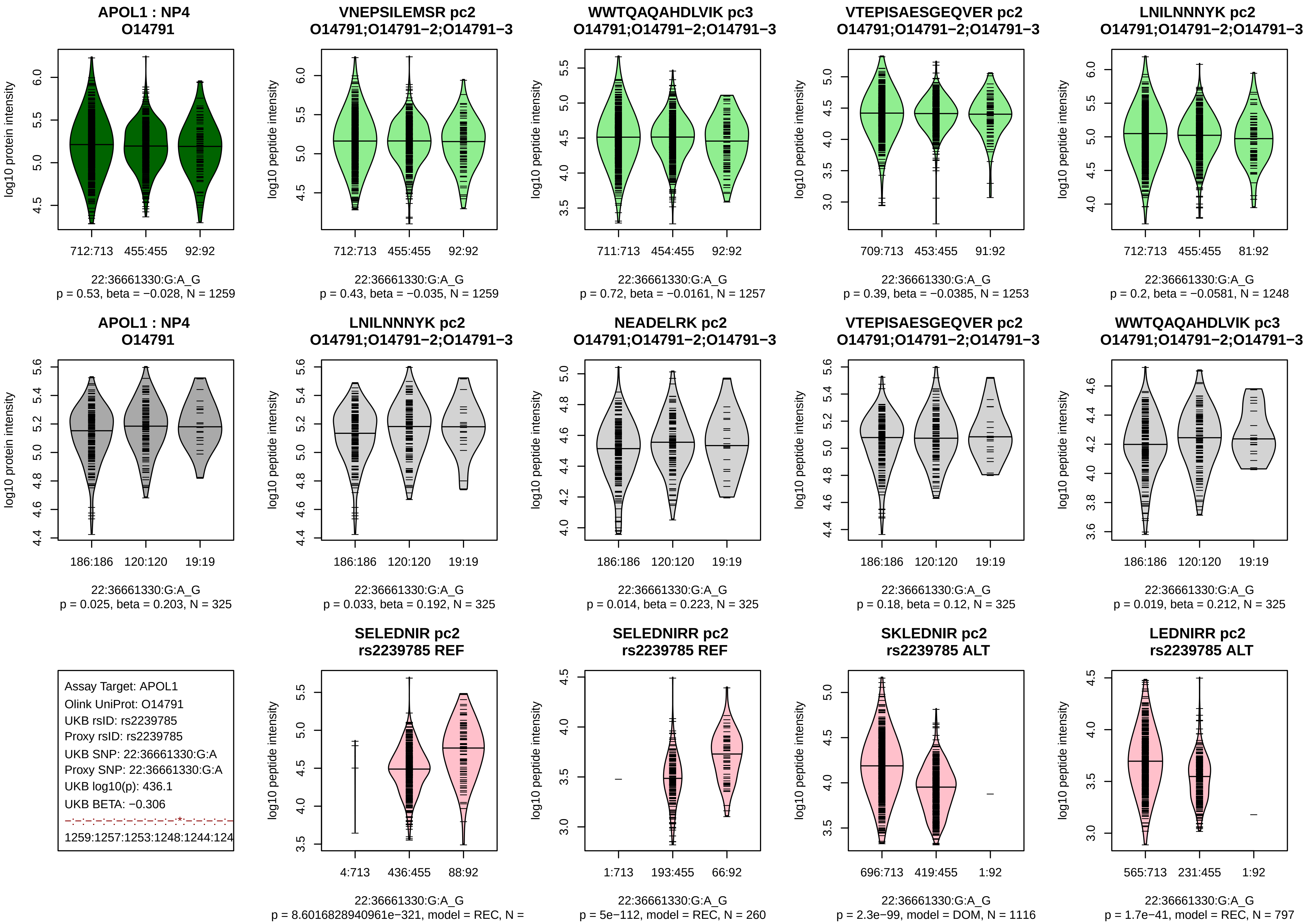
APOL1 : NP4
O14791
log10 protein intensity
712:713
455:455
92:92
22:36661330:G:A_G
p = 0.53, beta = −0.028, N = 1259
VNEPSILEMSR pc2
O14791;O14791−2;O14791−3
log10 peptide intensity
712:713
455:455
92:92
22:36661330:G:A_G
p = 0.43, beta = −0.035, N = 1259
WWTQAQAHDLVIK pc3
O14791;O14791−2;O14791−3
log10 peptide intensity
711:713
454:455
92:92
22:36661330:G:A_G
p = 0.72, beta = −0.0161, N = 1257
VTEPISAESGEQVER pc2
O14791;O14791−2;O14791−3
log10 peptide intensity
709:713
453:455
91:92
22:36661330:G:A_G
p = 0.39, beta = −0.0385, N = 1253
LNILNNNYK pc2
O14791;O14791−2;O14791−3
log10 peptide intensity
712:713
455:455
81:92
22:36661330:G:A_G
p = 0.2, beta = −0.0581, N = 1248
APOL1 : NP4
O14791
log10 protein intensity
186:186
120:120
19:19
22:36661330:G:A_G
p = 0.025, beta = 0.203, N = 325
LNILNNNYK pc2
O14791;O14791−2;O14791−3
log10 peptide intensity
186:186
120:120
19:19
22:36661330:G:A_G
p = 0.033, beta = 0.192, N = 325
NEADELRK pc2
O14791;O14791−2;O14791−3
log10 peptide intensity
186:186
120:120
19:19
22:36661330:G:A_G
p = 0.014, beta = 0.223, N = 325
VTEPISAESGEQVER pc2
O14791;O14791−2;O14791−3
log10 peptide intensity
186:186
120:120
19:19
22:36661330:G:A_G
p = 0.18, beta = 0.12, N = 325
WWTQAQAHDLVIK pc3
O14791;O14791−2;O14791−3
log10 peptide intensity
186:186
120:120
19:19
22:36661330:G:A_G
p = 0.019, beta = 0.212, N = 325
Assay Target: APOL1
Olink UniProt: O14791
UKB rsID: rs2239785
Proxy rsID: rs2239785
UKB SNP: 22:36661330:G:A
Proxy SNP: 22:36661330:G:A
UKB log10(p): 436.1
UKB BETA: −0.306
−:−:−:−:−:−:−:−:−:−:*:−:−:−:−:−
1259:1257:1253:1248:1244:124
SELEDNIR pc2
rs2239785 REF
log10 peptide intensity
4:713
436:455
88:92
22:36661330:G:A_G
p = 8.6016828940961e−321, model = REC, N =
SELEDNIRR pc2
rs2239785 REF
log10 peptide intensity
1:713
193:455
66:92
22:36661330:G:A_G
p = 5e−112, model = REC, N = 260
SKLEDNIR pc2
rs2239785 ALT
log10 peptide intensity
696:713
419:455
1:92
22:36661330:G:A_G
p = 2.3e−99, model = DOM, N = 1116
LEDNIRR pc2
rs2239785 ALT
log10 peptide intensity
565:713
231:455
1:92
22:36661330:G:A_G
p = 1.7e−41, model = REC, N = 797

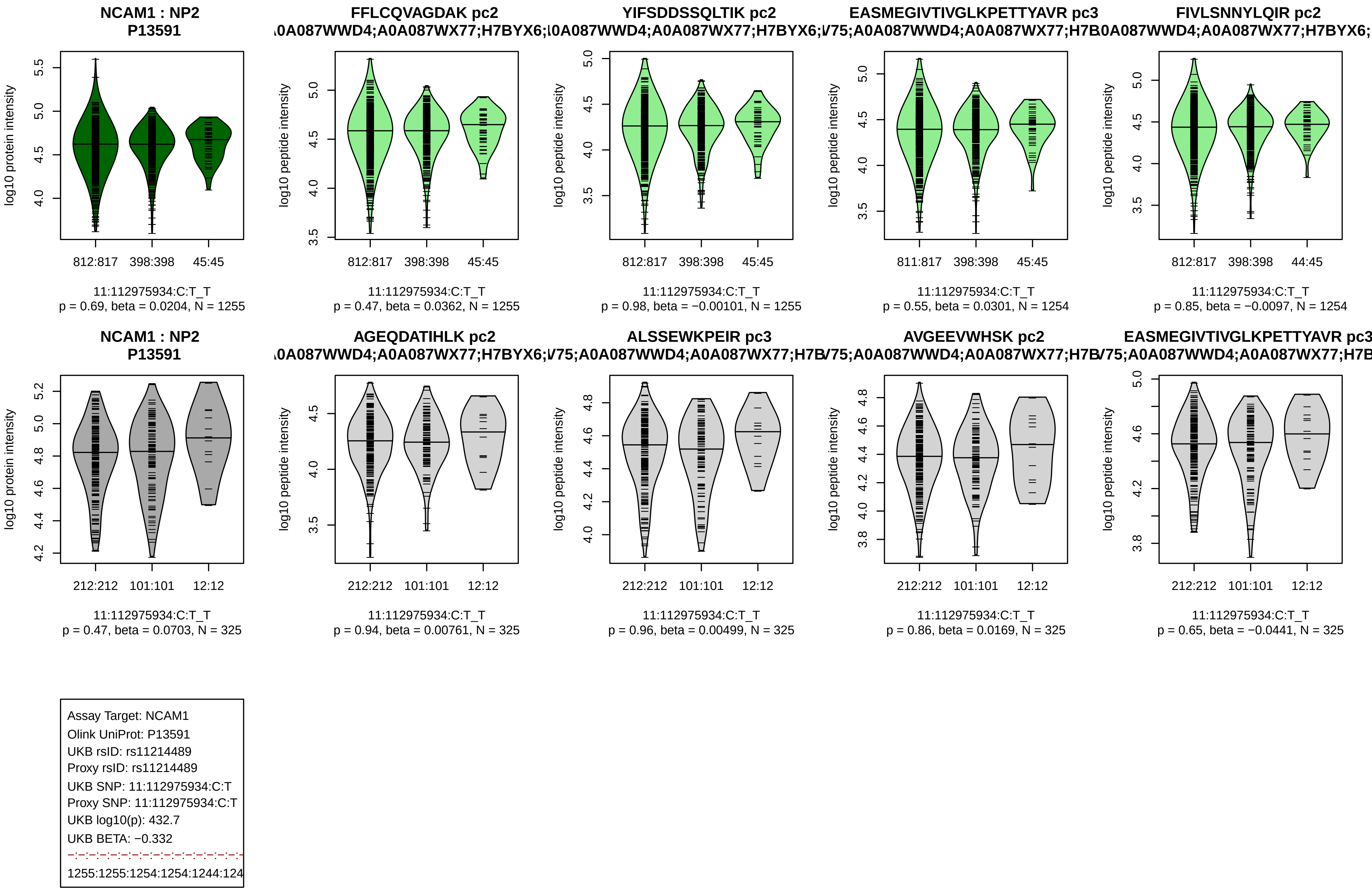
NCAM1 : NP2
P13591
log10 protein intensity
812:817 398:398 45:45
11:112975934:C:T_T
p = 0.69, beta = 0.0204, N = 1255
FFLCQVAGDAK pc2
A0A087WWD4;A0A087WX77;H7BYX6;
log10 peptide intensity
812:817 398:398 45:45
11:112975934:C:T_T
p = 0.47, beta = 0.0362, N = 1255
YIFSDDSSQLTIK pc2
A0A087WWD4;A0A087WX77;H7BYX6;
log10 peptide intensity
812:817 398:398 45:45
11:112975934:C:T_T
p = 0.98, beta = −0.00101, N = 1255
EASMEGIVTIVGLKPETTYAVR pc3
V75;A0A087WWD4;A0A087WX77;H7B
log10 peptide intensity
811:817 398:398 45:45
11:112975934:C:T_T
p = 0.55, beta = 0.0301, N = 1254
FIVLSNNYLQIR pc2
0A087WWD4;A0A087WX77;H7BYX6;
log10 peptide intensity
812:817 398:398 44:45
11:112975934:C:T_T
p = 0.85, beta = −0.0097, N = 1254
NCAM1 : NP2
P13591
log10 protein intensity
212:212 101:101 12:12
11:112975934:C:T_T
p = 0.47, beta = 0.0703, N = 325
AGEQDATIHLK pc2
A0A087WWD4;A0A087WX77;H7BYX6;
log10 peptide intensity
212:212 101:101 12:12
11:112975934:C:T_T
p = 0.94, beta = 0.00761, N = 325
ALSSEWKPEIR pc3
V75;A0A087WWD4;A0A087WX77;H7B
log10 peptide intensity
212:212 101:101 12:12
11:112975934:C:T_T
p = 0.96, beta = 0.00499, N = 325
AVGEEVWHSK pc2
V75;A0A087WWD4;A0A087WX77;H7B
log10 peptide intensity
212:212 101:101 12:12
11:112975934:C:T_T
p = 0.86, beta = 0.0169, N = 325
EASMEGIVTIVGLKPETTYAVR pc3
V75;A0A087WWD4;A0A087WX77;H7B
log10 peptide intensity
212:212 101:101 12:12
11:112975934:C:T_T
p = 0.65, beta = −0.0441, N = 325
Assay Target: NCAM1
Olink UniProt: P13591
UKB rsID: rs11214489
Proxy rsID: rs11214489
UKB SNP: 11:112975934:C:T
Proxy SNP: 11:112975934:C:T
UKB log10(p): 432.7
UKB BETA: −0.332
1255:1255:1254:1254:1244:124

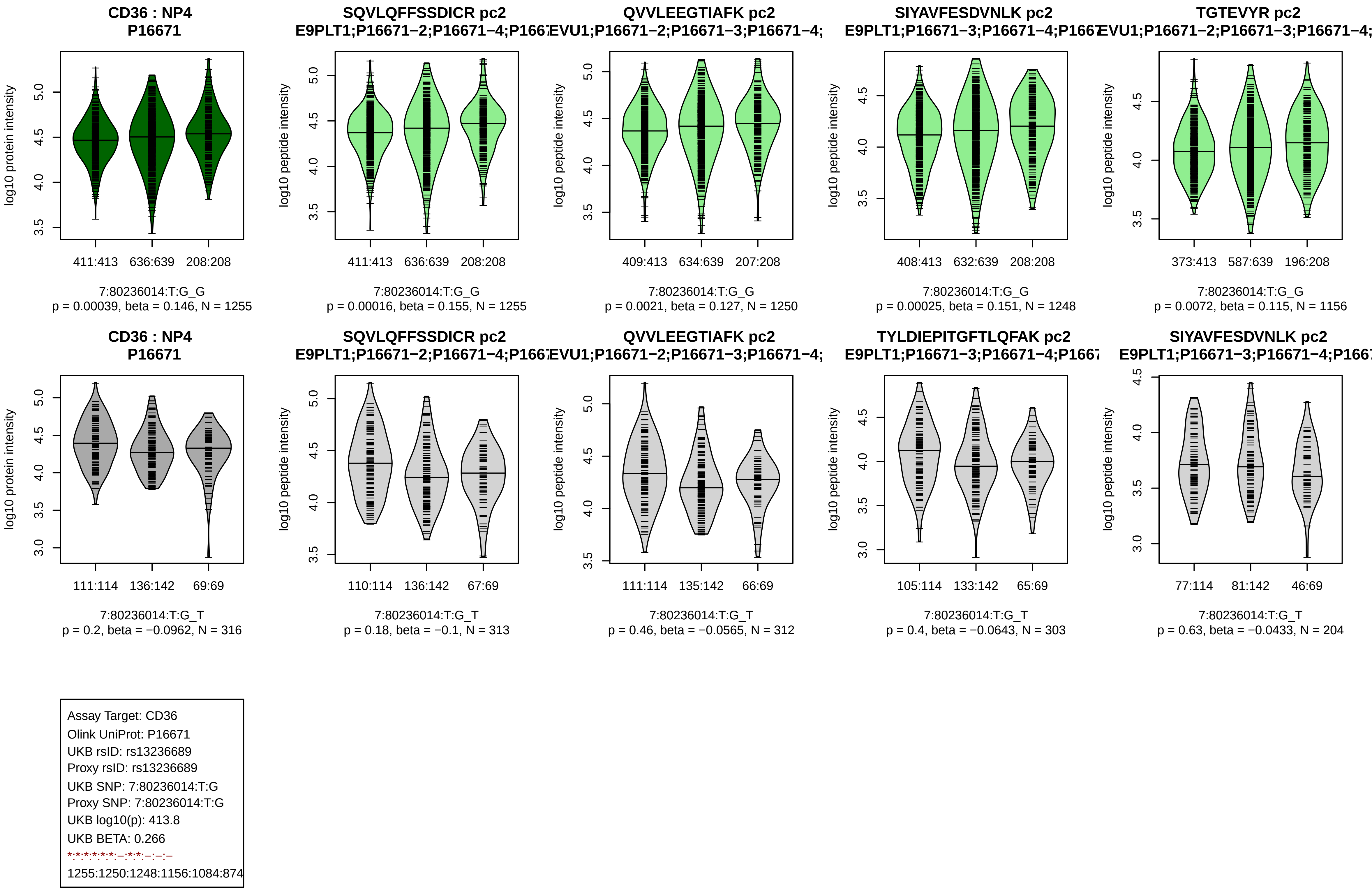
CD36 : NP4
P16671
SQVLQFFSSDICR pc2
E9PLT1;P16671−2;P16671−4;P1667
QVVLEEGTIAFK pc2
EVU1;P16671−2;P16671−3;P16671−4;
SIYAVFESDVNLK pc2
E9PLT1;P16671−3;P16671−4;P1667
TGTEVYR pc2
EVU1;P16671−2;P16671−3;P16671−4;
log10 protein intensity
log10 peptide intensity
411:413 636:639 208:208
411:413 636:639 208:208
409:413 634:639 207:208
408:413 632:639 208:208
373:413 587:639 196:208
7:80236014:T:G_G
p = 0.00039, beta = 0.146, N = 1255
7:80236014:T:G_G
p = 0.00016, beta = 0.155, N = 1255
7:80236014:T:G_G
p = 0.0021, beta = 0.127, N = 1250
7:80236014:T:G_G
p = 0.00025, beta = 0.151, N = 1248
7:80236014:T:G_G
p = 0.0072, beta = 0.115, N = 1156
CD36 : NP4
P16671
SQVLQFFSSDICR pc2
E9PLT1;P16671−2;P16671−4;P1667
QVVLEEGTIAFK pc2
EVU1;P16671−2;P16671−3;P16671−4;
TYLDIEPITGFTLQFAK pc2
E9PLT1;P16671−3;P16671−4;P1667
SIYAVFESDVNLK pc2
E9PLT1;P16671−3;P16671−4;P1667
111:114 136:142 69:69
110:114 136:142 67:69
111:114 135:142 66:69
105:114 133:142 65:69
77:114 81:142 46:69
7:80236014:T:G_T
p = 0.2, beta = −0.0962, N = 316
7:80236014:T:G_T
p = 0.18, beta = −0.1, N = 313
7:80236014:T:G_T
p = 0.46, beta = −0.0565, N = 312
7:80236014:T:G_T
p = 0.4, beta = −0.0643, N = 303
7:80236014:T:G_T
p = 0.63, beta = −0.0433, N = 204
Assay Target: CD36
Olink UniProt: P16671
UKB rsID: rs13236689
Proxy rsID: rs13236689
UKB SNP: 7:80236014:T:G
Proxy SNP: 7:80236014:T:G
UKB log10(p): 413.8
UKB BETA: 0.266
*:*:*:*:*:*:–:*:*:–:–:–
1255:1250:1248:1156:1084:874

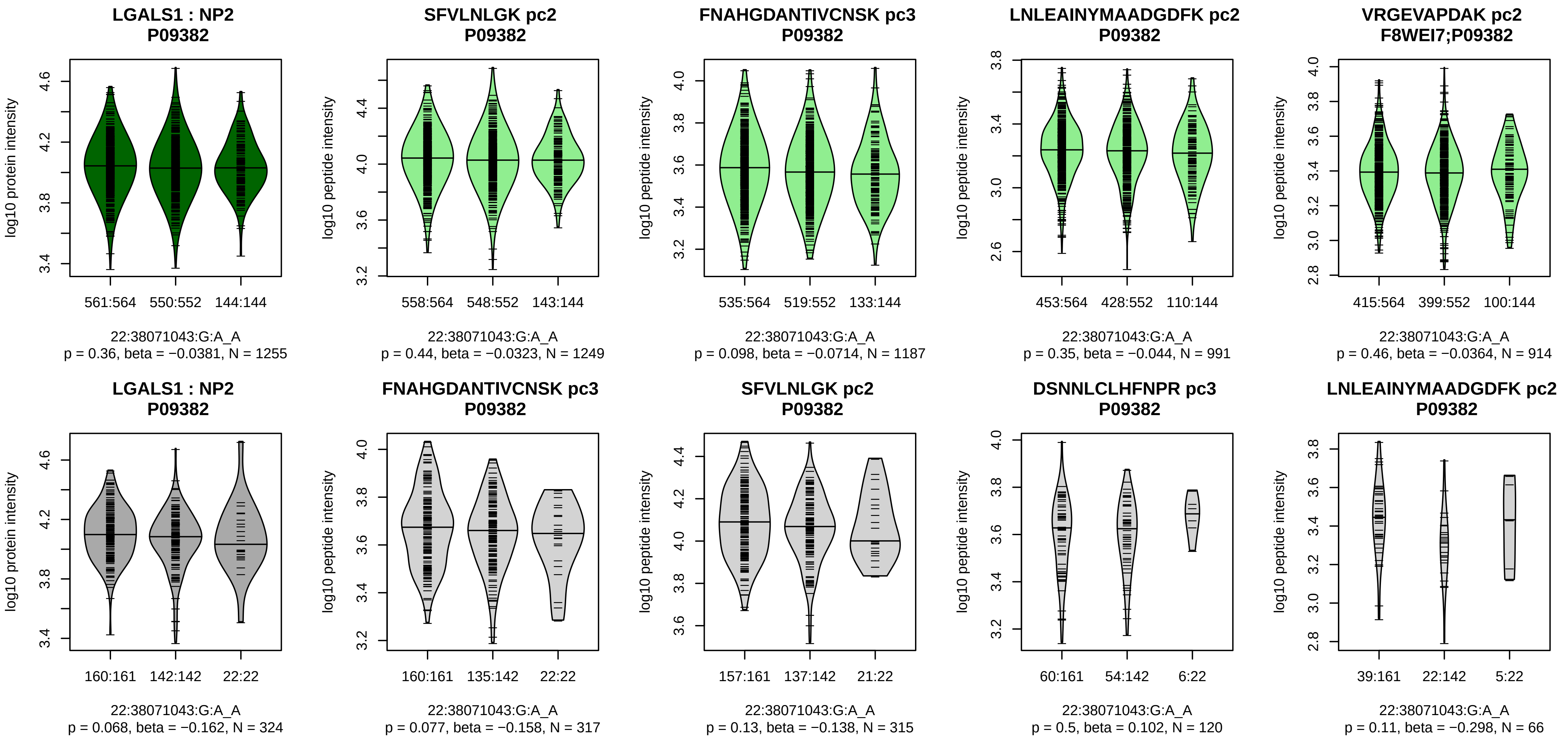


Assay Target: LGALS1
Olink UniProt: P09382
UKB rsID: rs4820294
Proxy rsID: rs4820294
UKB SNP: 22:38071043:G:A
Proxy SNP: 22:38071043:G:A
UKB log10(p): 413
UKB BETA: −0.279
−:−:−:−:*:−:−:−:−
1249:1187:991:914:730:664:123

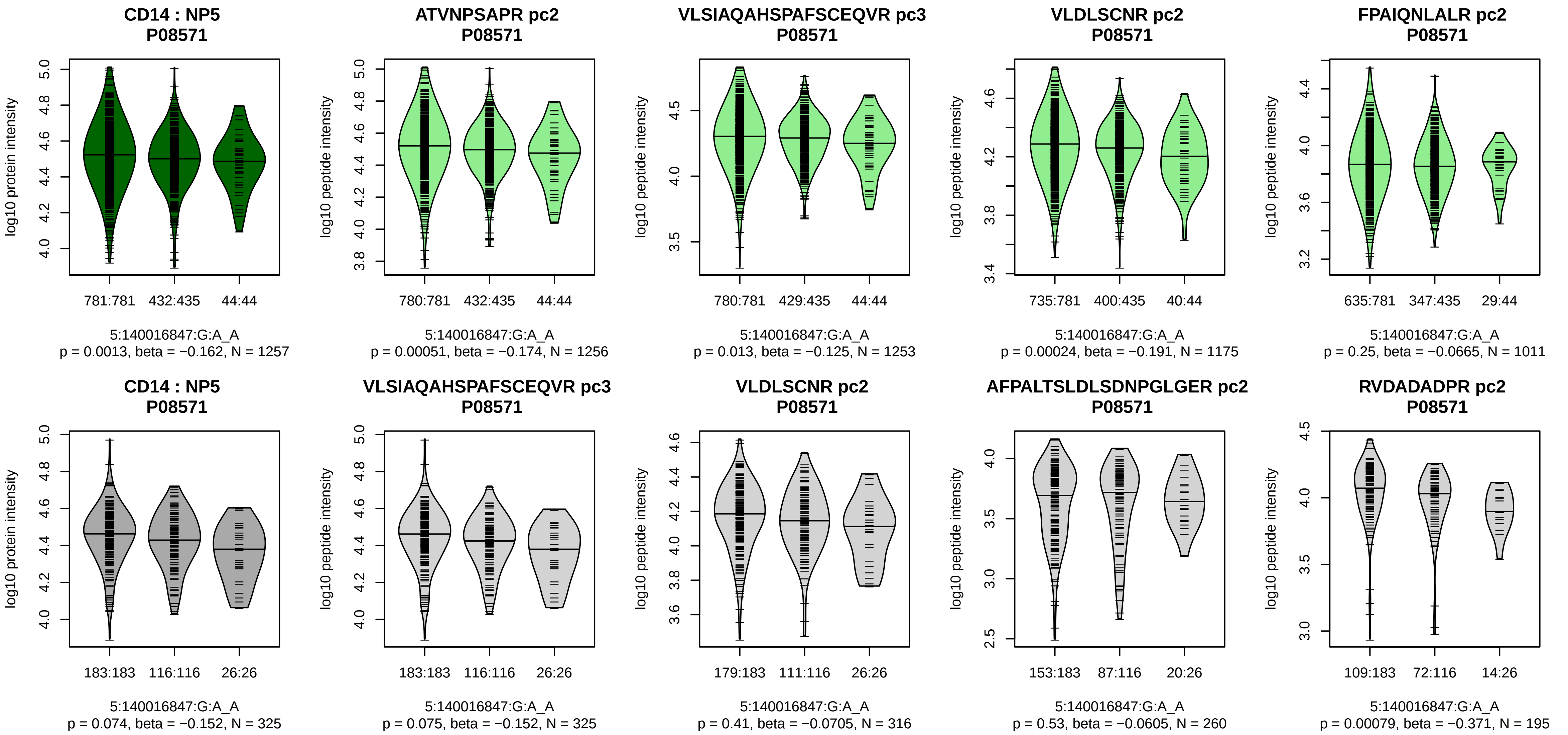


Assay Target: CD14
Olink UniProt: P08571
UKB rsID: rs5744441
Proxy rsID: rs5744441
UKB SNP: 5:140016847:G:A
Proxy SNP: 5:140016847:G:A
UKB log10(p): 411.3
UKB BETA: −0.313
*:–:*:–:*:–:–:–:–:–:–:–:–:–
1256:1253:1175:1011:982:929:8

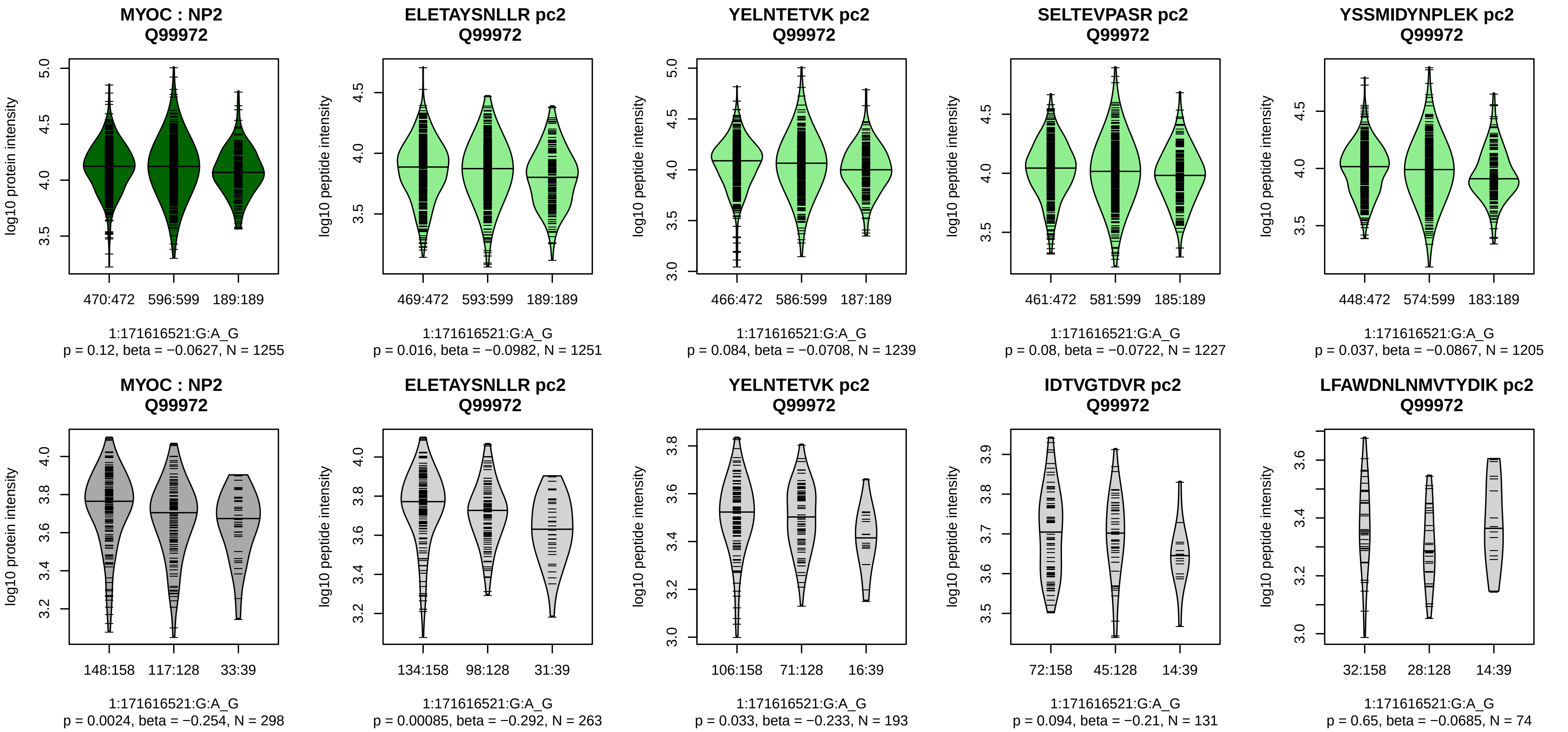


Assay Target: MYOC
Olink UniProt: Q99972
UKB rsID: rs171003
Proxy rsID: rs171003
UKB SNP: 1:171616521:G:A
Proxy SNP: 1:171616521:G:A
UKB log10(p): 404.1
UKB BETA: 0.255
–:–:–:–:–:–:–:–:–:–:–:–:–:–:–:
1251:1239:1227:1205:1191:114

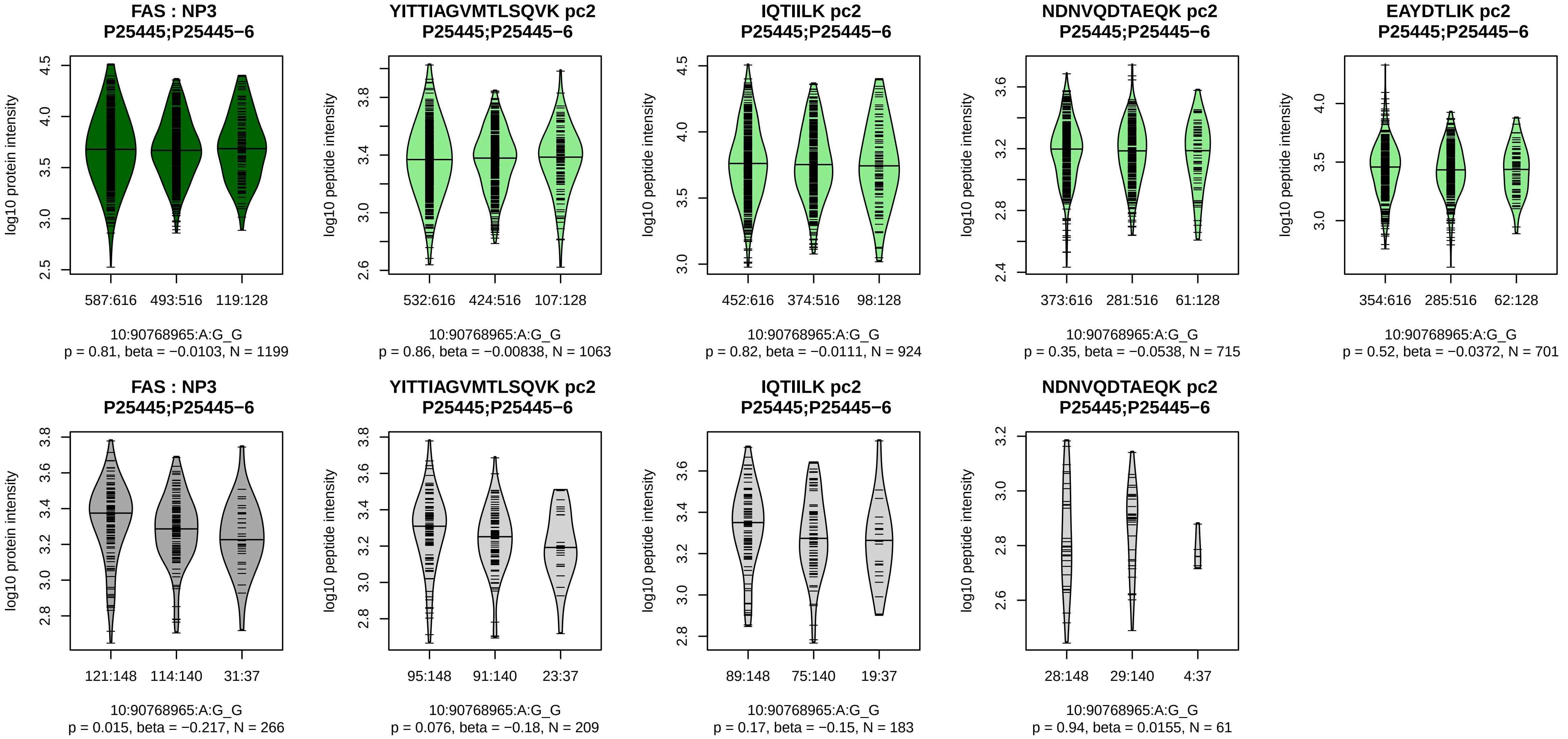


Assay Target: FAS
Olink UniProt: P25445
UKB rsID: rs7911226
Proxy rsID: rs7911226
UKB SNP: 10:90768965:A:G
Proxy SNP: 10:90768965:A:G
UKB log10(p): 399.8
UKB BETA: −0.26
−:−:−:−
1063:924:715:701

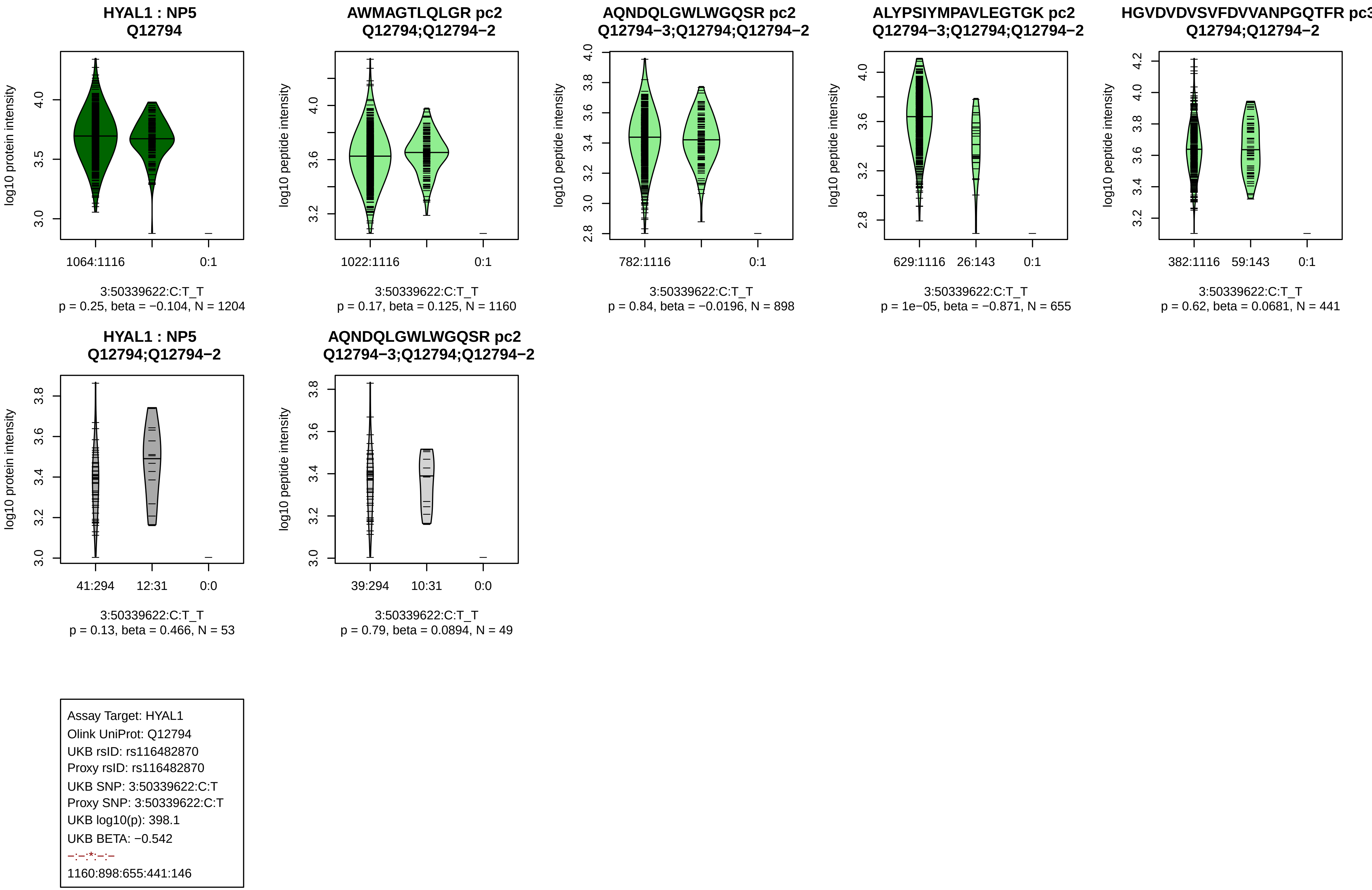
HYAL1 : NP5
Q12794
log10 protein intensity
1064:1116
0:1
3:50339622:C:T_T
p = 0.25, beta = −0.104, N = 1204
AWMAGTLQLGR pc2
Q12794;Q12794−2
log10 peptide intensity
1022:1116
0:1
3:50339622:C:T_T
p = 0.17, beta = 0.125, N = 1160
AQNDQLGWLWGQSR pc2
Q12794−3;Q12794;Q12794−2
log10 peptide intensity
782:1116
0:1
3:50339622:C:T_T
p = 0.84, beta = −0.0196, N = 898
ALYPSIYMPAVLEGTGK pc2
Q12794−3;Q12794;Q12794−2
log10 peptide intensity
629:1116
26:143
0:1
3:50339622:C:T_T
p = 1e−05, beta = −0.871, N = 655
HGVDVDVSVFDVVANPGQTFR pc3
Q12794;Q12794−2
log10 peptide intensity
382:1116
59:143
0:1
3:50339622:C:T_T
p = 0.62, beta = 0.0681, N = 441
HYAL1 : NP5
Q12794;Q12794−2
log10 protein intensity
41:294
12:31
0:0
3:50339622:C:T_T
p = 0.13, beta = 0.466, N = 53
AQNDQLGWLWGQSR pc2
Q12794−3;Q12794;Q12794−2
log10 peptide intensity
39:294
10:31
0:0
3:50339622:C:T_T
p = 0.79, beta = 0.0894, N = 49
Assay Target: HYAL1
Olink UniProt: Q12794
UKB rsID: rs116482870
Proxy rsID: rs116482870
UKB SNP: 3:50339622:C:T
Proxy SNP: 3:50339622:C:T
UKB log10(p): 398.1
UKB BETA: −0.542
−:−:*:−:−
1160:898:655:441:146

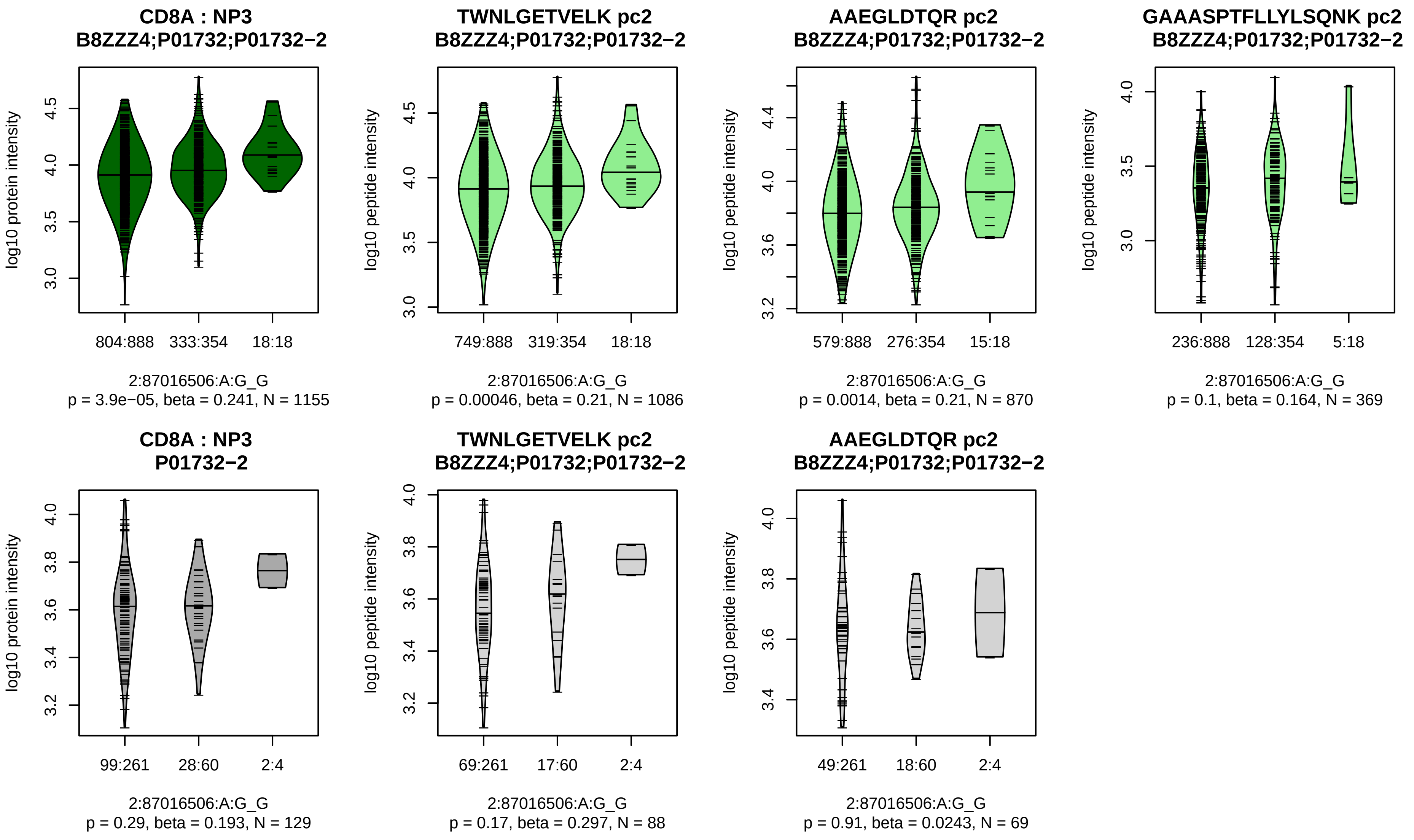


Assay Target: CD8A
Olink UniProt: P01732
UKB rsID: rs3020726
Proxy rsID: rs3020726
UKB SNP: 2:87016506:A:G
Proxy SNP: 2:87016506:A:G
UKB log10(p): 383.5
UKB BETA: 0.345
*:*:–:–
1086:870:369:65

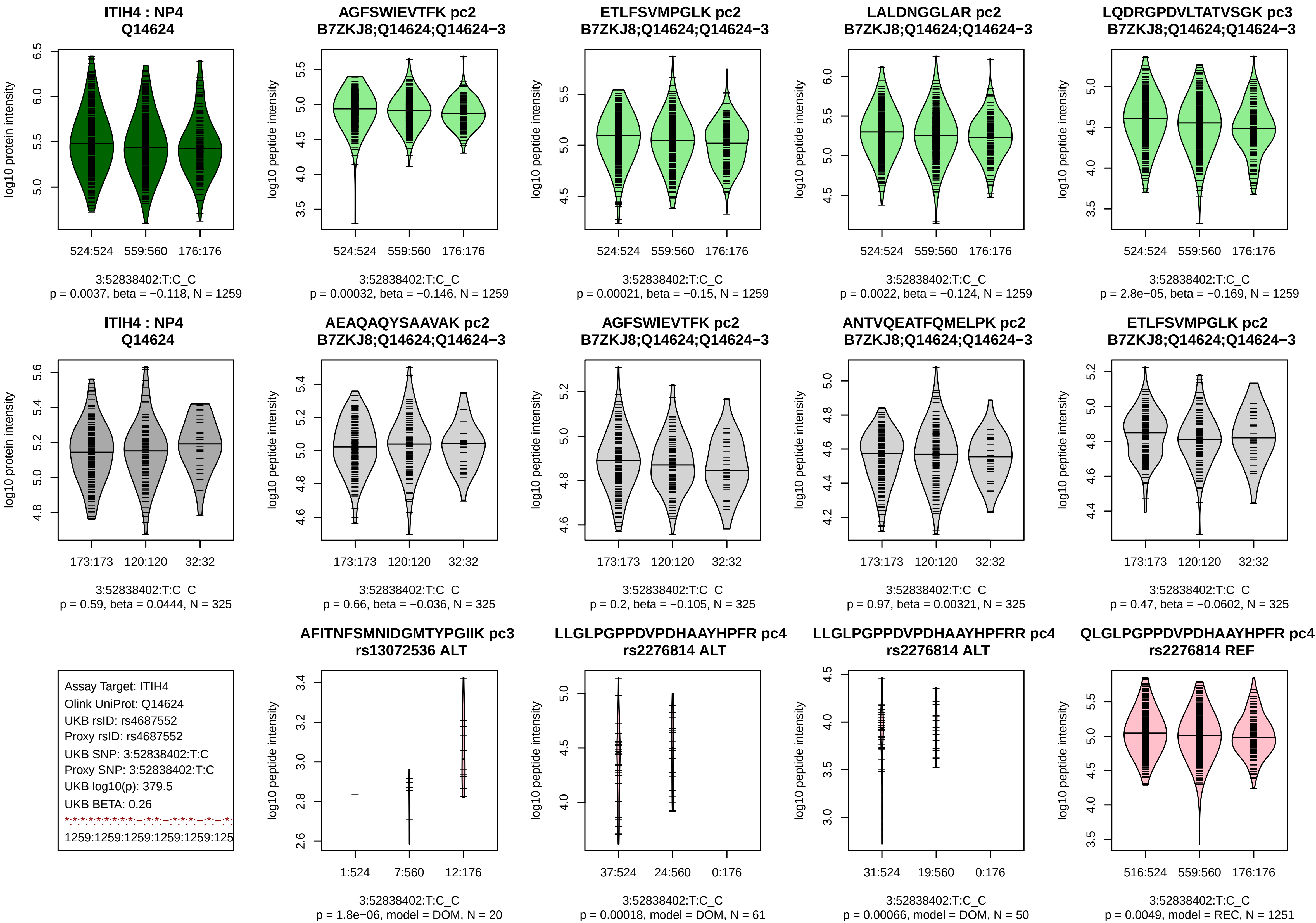

ITIH4 : NP4
Q14624
log10 protein intensity
524:524 559:560 176:176
3:52838402:T:C_C
p = 0.0037, beta = −0.118, N = 1259
AGFSWIEVTFK pc2
B7ZKJ8;Q14624;Q14624−3
log10 peptide intensity
524:524 559:560 176:176
3:52838402:T:C_C
p = 0.00032, beta = −0.146, N = 1259
ETLFSVMPGLK pc2
B7ZKJ8;Q14624;Q14624−3
log10 peptide intensity
524:524 559:560 176:176
3:52838402:T:C_C
p = 0.00021, beta = −0.15, N = 1259
LALDNGGLAR pc2
B7ZKJ8;Q14624;Q14624−3
log10 peptide intensity
524:524 559:560 176:176
3:52838402:T:C_C
p = 0.0022, beta = −0.124, N = 1259
LQDRGPDVLTATVSGK pc3
B7ZKJ8;Q14624;Q14624−3
log10 peptide intensity
524:524 559:560 176:176
3:52838402:T:C_C
p = 2.8e−05, beta = −0.169, N = 1259
ITIH4 : NP4
Q14624
log10 protein intensity
173:173 120:120 32:32
3:52838402:T:C_C
p = 0.59, beta = 0.0444, N = 325
AEAQAQYSAAVAK pc2
B7ZKJ8;Q14624;Q14624−3
log10 peptide intensity
173:173 120:120 32:32
3:52838402:T:C_C
p = 0.66, beta = −0.036, N = 325
AGFSWIEVTFK pc2
B7ZKJ8;Q14624;Q14624−3
log10 peptide intensity
173:173 120:120 32:32
3:52838402:T:C_C
p = 0.2, beta = −0.105, N = 325
ANTVQEATFQMELPK pc2
B7ZKJ8;Q14624;Q14624−3
log10 peptide intensity
173:173 120:120 32:32
3:52838402:T:C_C
p = 0.97, beta = 0.00321, N = 325
ETLFSVMPGLK pc2
B7ZKJ8;Q14624;Q14624−3
log10 peptide intensity
173:173 120:120 32:32
3:52838402:T:C_C
p = 0.47, beta = −0.0602, N = 325
Assay Target: ITIH4
Olink UniProt: Q14624
UKB rsID: rs4687552
Proxy rsID: rs4687552
UKB SNP: 3:52838402:T:C
Proxy SNP: 3:52838402:T:C
UKB log10(p): 379.5
UKB BETA: 0.26
1259:1259:1259:1259:1259:125
AFITNFSMNIDGMTYPGIIK pc3
rs13072536 ALT
log10 peptide intensity
1:524 7:560 12:176
3:52838402:T:C_C
p = 1.8e−06, model = DOM, N = 20
LLGLPGPPDVPDHAAYHPFR pc4
rs2276814 ALT
log10 peptide intensity
37:524 24:560 0:176
3:52838402:T:C_C
p = 0.00018, model = DOM, N = 61
LLGLPGPPDVPDHAAYHPFRR pc4
rs2276814 ALT
log10 peptide intensity
31:524 19:560 0:176
3:52838402:T:C_C
p = 0.00066, model = DOM, N = 50
QLGLPGPPDVPDHAAYHPFR pc4
rs2276814 REF
log10 peptide intensity
516:524 559:560 176:176
3:52838402:T:C_C
p = 0.0049, model = REC, N = 1251

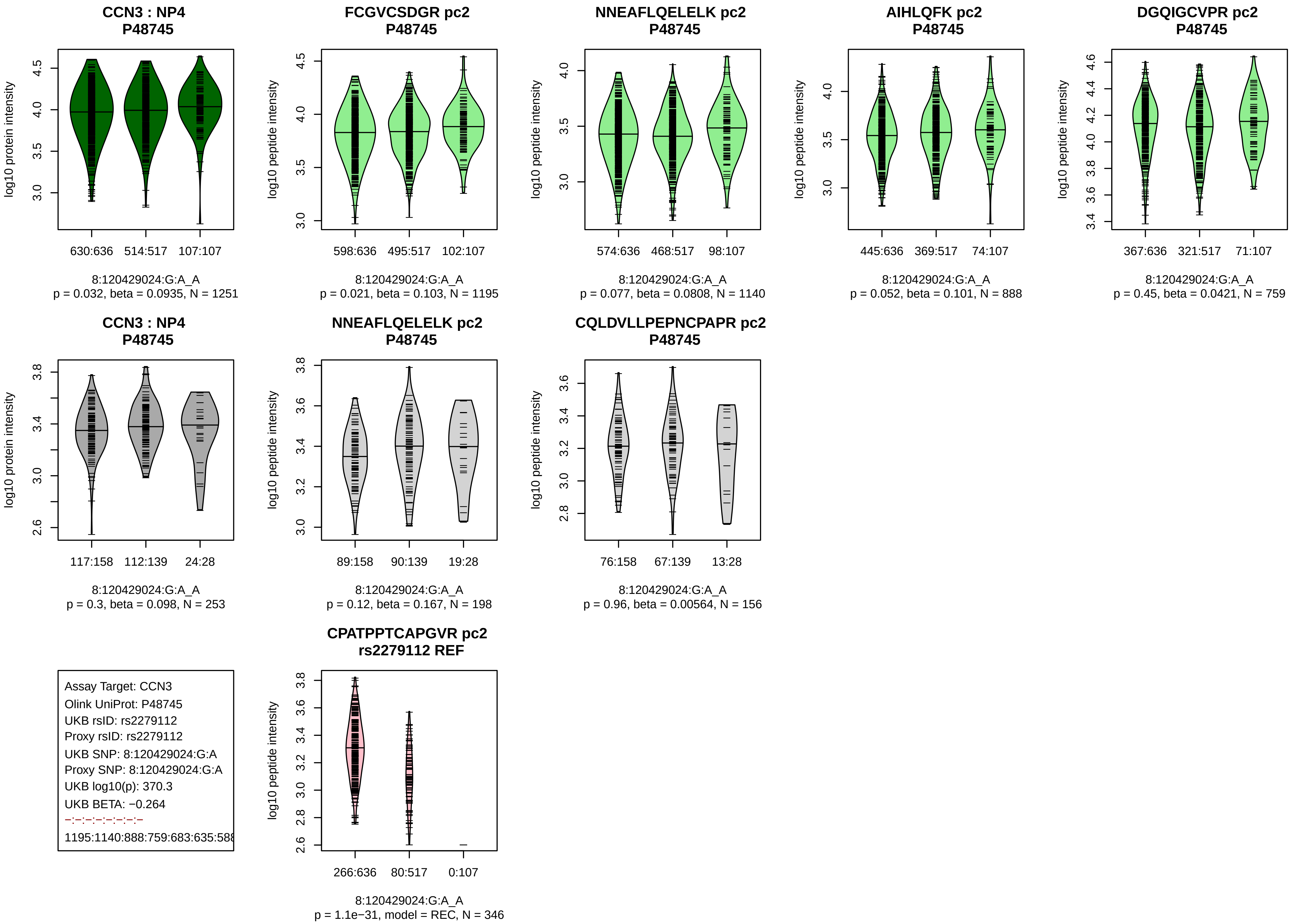
CCN3 : NP4
P48745
log10 protein intensity
630:636
514:517
107:107
8:120429024:G:A_A
p = 0.032, beta = 0.0935, N = 1251
FCGVCSDGR pc2
P48745
log10 peptide intensity
598:636
495:517
102:107
8:120429024:G:A_A
p = 0.021, beta = 0.103, N = 1195
NNEAFLQELELK pc2
P48745
log10 peptide intensity
574:636
468:517
98:107
8:120429024:G:A_A
p = 0.077, beta = 0.0808, N = 1140
AIHLQFK pc2
P48745
log10 peptide intensity
445:636
369:517
74:107
8:120429024:G:A_A
p = 0.052, beta = 0.101, N = 888
DGQIGCVPR pc2
P48745
log10 peptide intensity
367:636
321:517
71:107
8:120429024:G:A_A
p = 0.45, beta = 0.0421, N = 759
CCN3 : NP4
P48745
log10 protein intensity
117:158
112:139
24:28
8:120429024:G:A_A
p = 0.3, beta = 0.098, N = 253
NNEAFLQELELK pc2
P48745
log10 peptide intensity
89:158
90:139
19:28
8:120429024:G:A_A
p = 0.12, beta = 0.167, N = 198
CQLDVLLPEPNCPAPR pc2
P48745
log10 peptide intensity
76:158
67:139
13:28
8:120429024:G:A_A
p = 0.96, beta = 0.00564, N = 156
Assay Target: CCN3
Olink UniProt: P48745
UKB rsID: rs2279112
Proxy rsID: rs2279112
UKB SNP: 8:120429024:G:A
Proxy SNP: 8:120429024:G:A
UKB log10(p): 370.3
UKB BETA: −0.264
−:−:−:−:−:−:−:−
1195:1140:888:759:683:635:58
CPATPPTCAPGVR pc2
rs2279112 REF
log10 peptide intensity
266:636
80:517
0:107
8:120429024:G:A_A
p = 1.1e−31, model = REC, N = 346

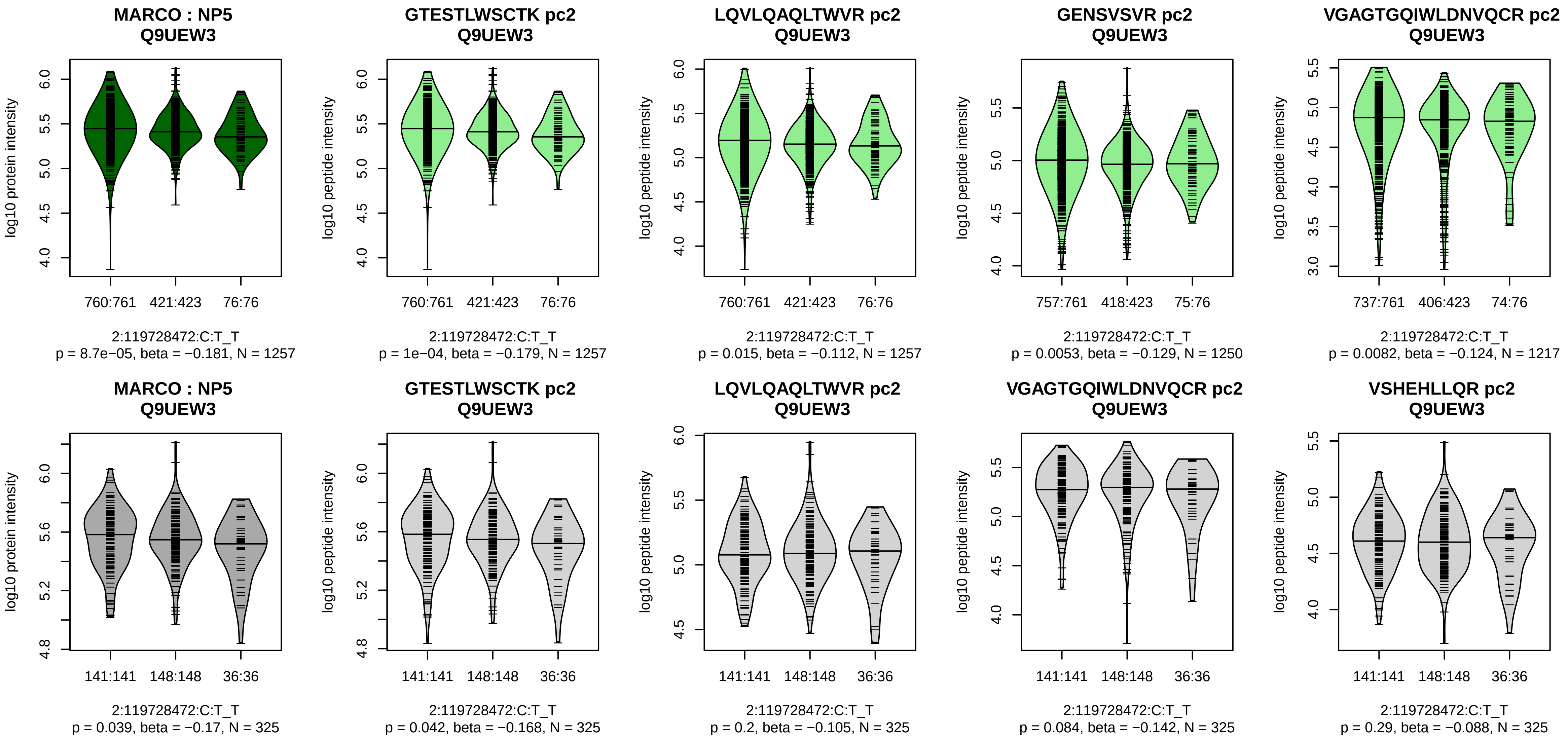


Assay Target: MARCO
Olink UniProt: Q9UEW3
UKB rsID: rs35288880
Proxy rsID: rs35288880
UKB SNP: 2:119728472:C:T
Proxy SNP: 2:119728472:C:T
UKB log10(p): 367.5
UKB BETA: −0.324
*:–:*:*:–:–:–:–
1257:1257:1250:1217:1211:110

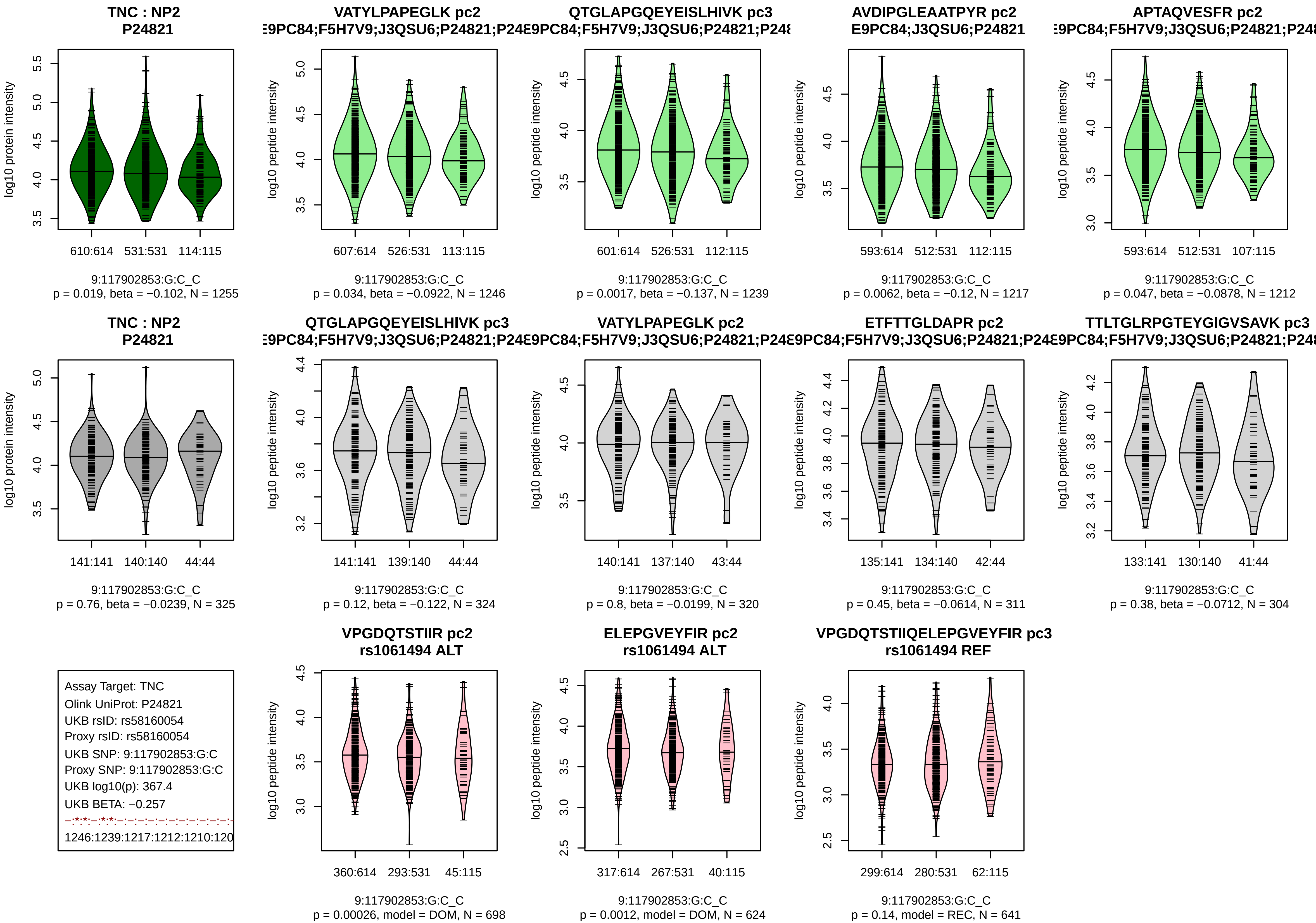

TNC : NP2
P24821
log10 protein intensity
610:614 531:531 114:115
9:117902853:G:C_C
p = 0.019, beta = −0.102, N = 1255
VATYLPAPEGLK pc2
log10 peptide intensity
607:614 526:531 113:115
9:117902853:G:C_C
p = 0.034, beta = −0.0922, N = 1246
QTGLAPGQEYEISLHIVK pc3
601:614 526:531 112:115
9:117902853:G:C_C
p = 0.0017, beta = −0.137, N = 1239
AVDIPGLEAATPYR pc2
E9PC84;J3QSU6;P24821
593:614 512:531 112:115
9:117902853:G:C_C
p = 0.0062, beta = −0.12, N = 1217
APTAQVESFR pc2
593:614 512:531 107:115
9:117902853:G:C_C
p = 0.047, beta = −0.0878, N = 1212
TNC : NP2
P24821
141:141 140:140 44:44
9:117902853:G:C_C
p = 0.76, beta = −0.0239, N = 325
QTGLAPGQEYEISLHIVK pc3
141:141 139:140 44:44
9:117902853:G:C_C
p = 0.12, beta = −0.122, N = 324
VATYLPAPEGLK pc2
140:141 137:140 43:44
9:117902853:G:C_C
p = 0.8, beta = −0.0199, N = 320
ETFTTGLDAPR pc2
135:141 134:140 42:44
9:117902853:G:C_C
p = 0.45, beta = −0.0614, N = 311
TTLTGLRPGTEYGIGVSAVK pc3
133:141 130:140 41:44
9:117902853:G:C_C
p = 0.38, beta = −0.0712, N = 304
Assay Target: TNC
Olink UniProt: P24821
UKB rsID: rs58160054
Proxy rsID: rs58160054
UKB SNP: 9:117902853:G:C
Proxy SNP: 9:117902853:G:C
UKB log10(p): 367.4
UKB BETA: −0.257
1246:1239:1217:1212:1210:120
VPGDQTSTIIR pc2
rs1061494 ALT
360:614 293:531 45:115
9:117902853:G:C_C
p = 0.00026, model = DOM, N = 698
ELEPGVEYFIR pc2
rs1061494 ALT
317:614 267:531 40:115
9:117902853:G:C_C
p = 0.0012, model = DOM, N = 624
VPGDQTSTIIQELEPGVEYFIR pc3
rs1061494 REF
299:614 280:531 62:115
9:117902853:G:C_C
p = 0.14, model = REC, N = 641

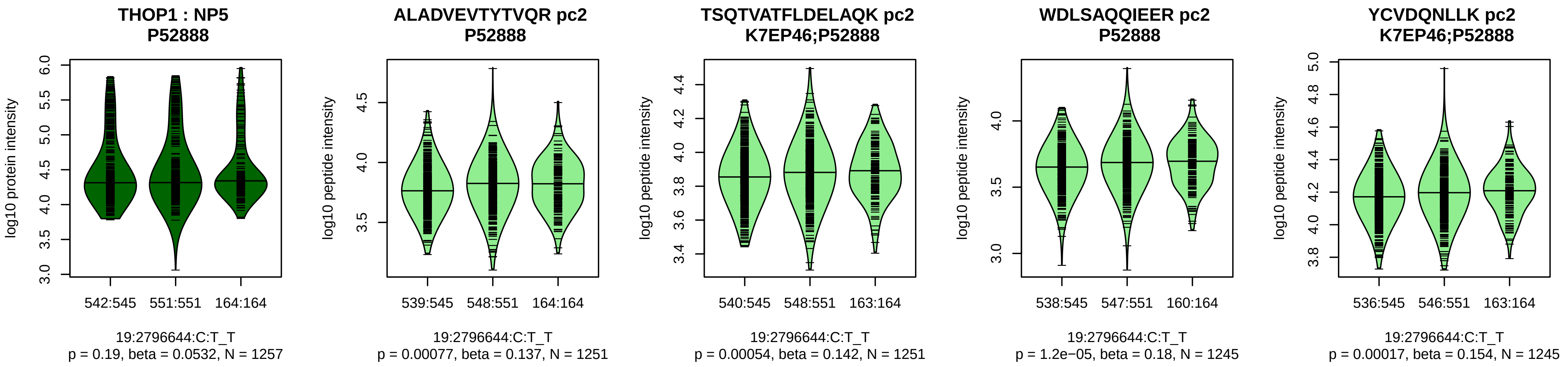


Assay Target: THOP1
Olink UniProt: P52888
UKB rsID: rs550767741
Proxy rsID: rs2741990
UKB SNP: 19:2786763:A:ATTTT
Proxy SNP: 19:2796644:C:T
UKB log10(p): 366.5
UKB BETA: 0.266
*:*:*:*:*:*:*:*:*:*:_:*:*:_:*:*:_:_:*:*:
1251:1251:1245:1245:1236:123

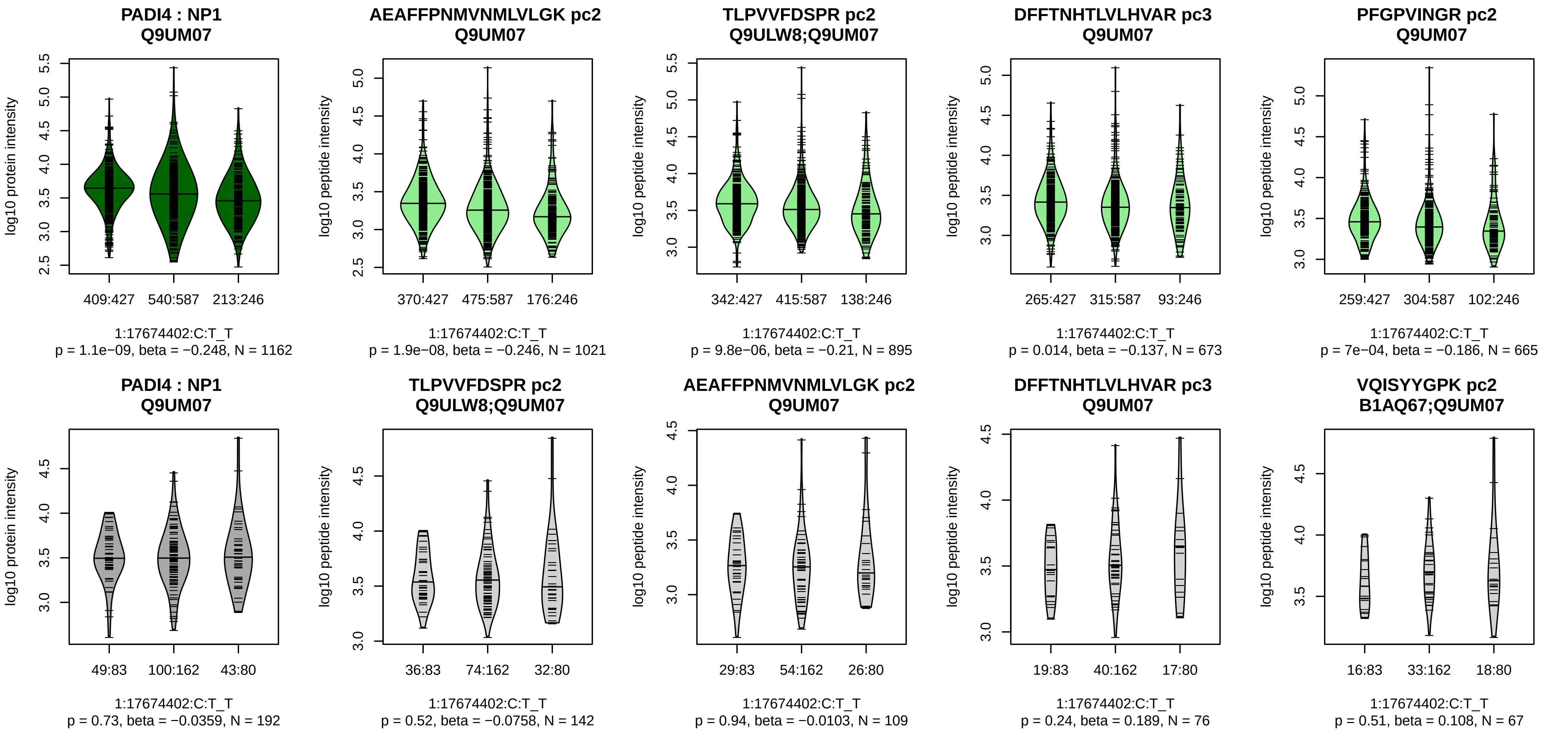


Assay Target: PADI4
Olink UniProt: Q9UM07
UKB rsID: rs2240336
Proxy rsID: rs2240336
UKB SNP: 1:17674402:C:T
Proxy SNP: 1:17674402:C:T
UKB log10(p): 366.3
UKB BETA: −0.253
*:*:–:*:–:–:–:–:–:–:–:–:–:–:–:–:–:–:
1021:895:673:665:604:452:358:

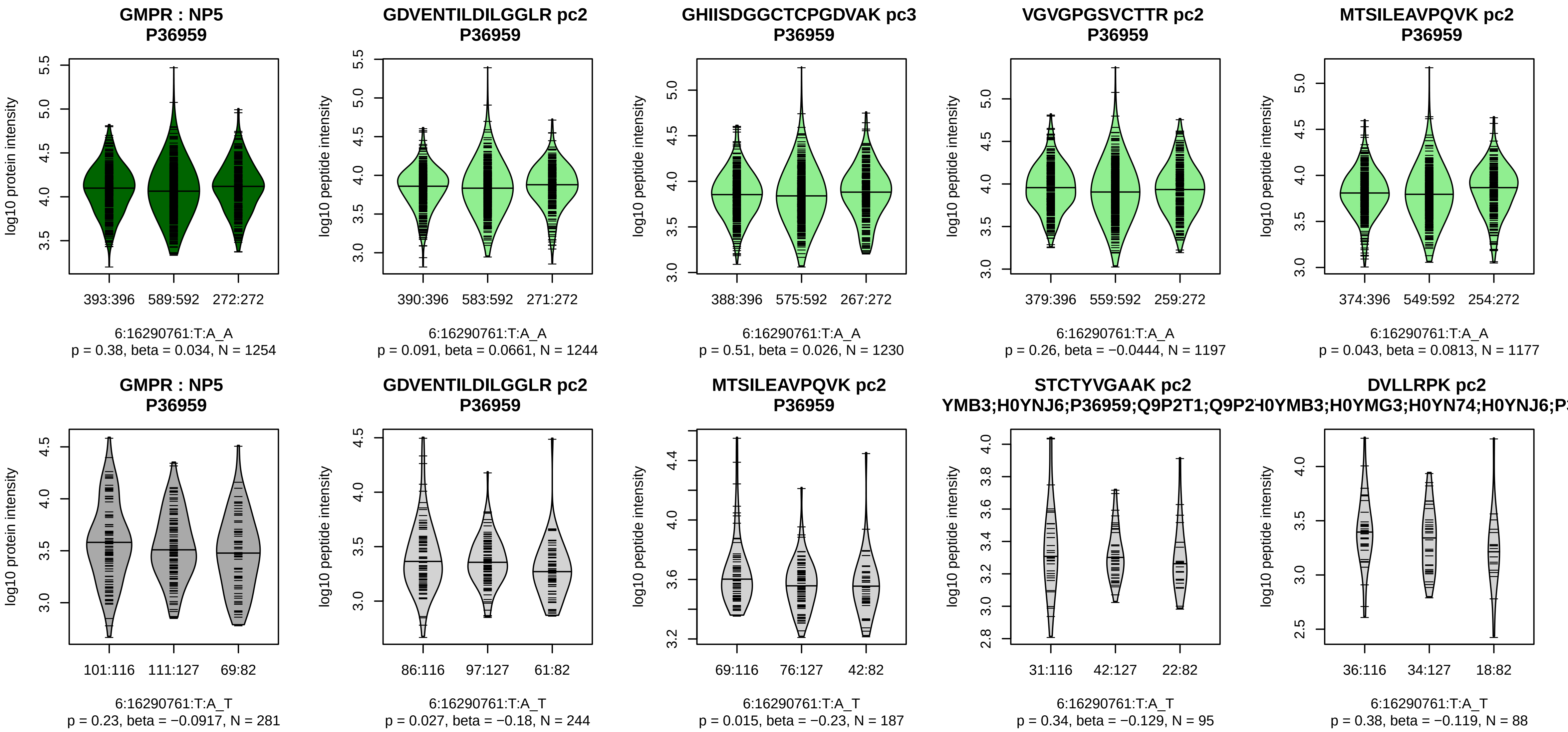


Assay Target: GMPR
Olink UniProt: P36959
UKB rsID: rs1042391
Proxy rsID: rs1042391
UKB SNP: 6:16290761:T:A
Proxy SNP: 6:16290761:T:A
UKB log10(p): 362.5
UKB BETA: 0.252
−:−:−:−:−:−:−:−:−:−:−:−:−:−:−:NA
1244:1230:1197:1177:1174:109

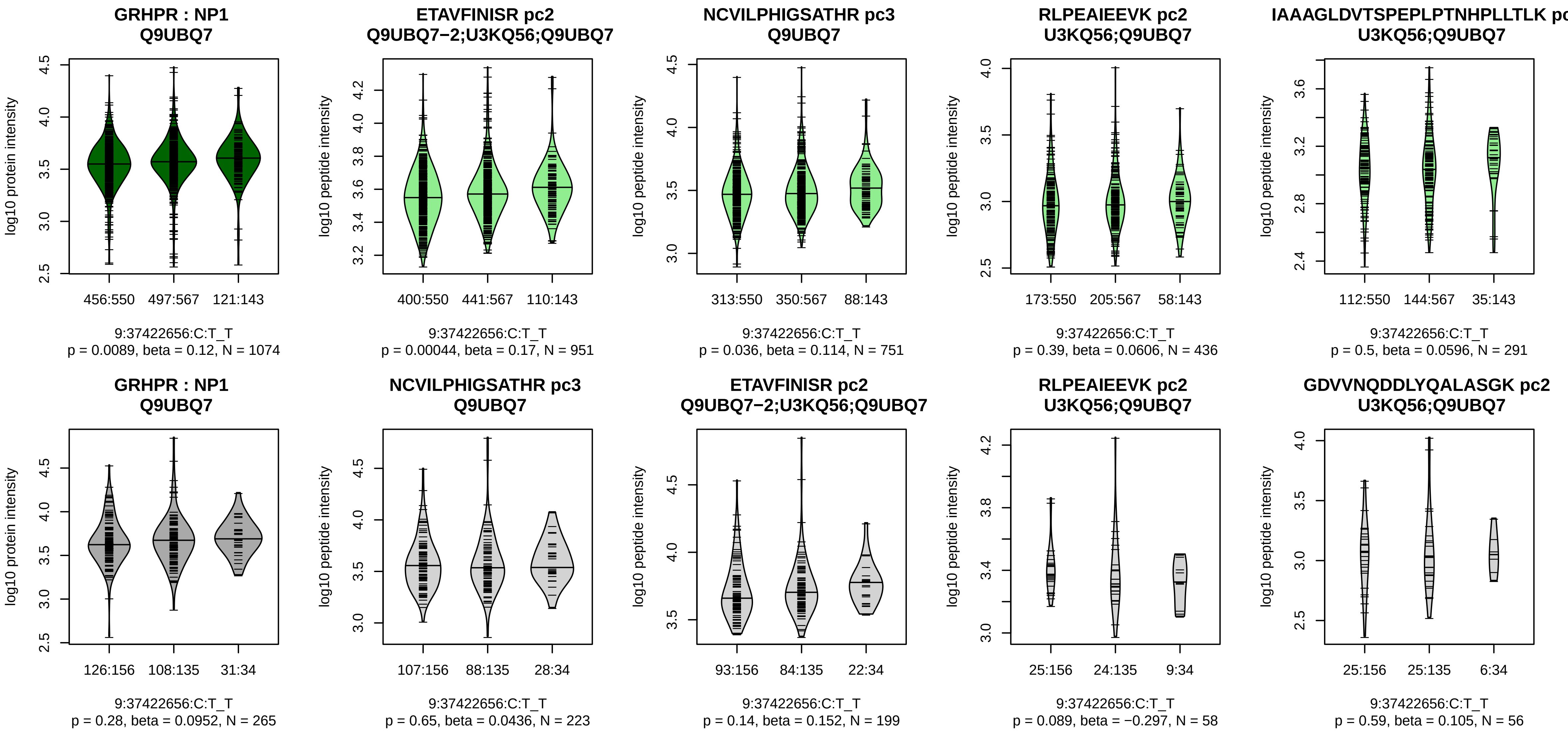


Assay Target: GRHPR
Olink UniProt: Q9UBQ7
UKB rsID: rs4878690
Proxy rsID: rs4878690
UKB SNP: 9:37422656:C:T
Proxy SNP: 9:37422656:C:T
UKB log10(p): 355.1
UKB BETA: 0.264
*:−:−:−:−:−:−:−:−:−:NA:NA
951:751:436:291:225:220:198:1

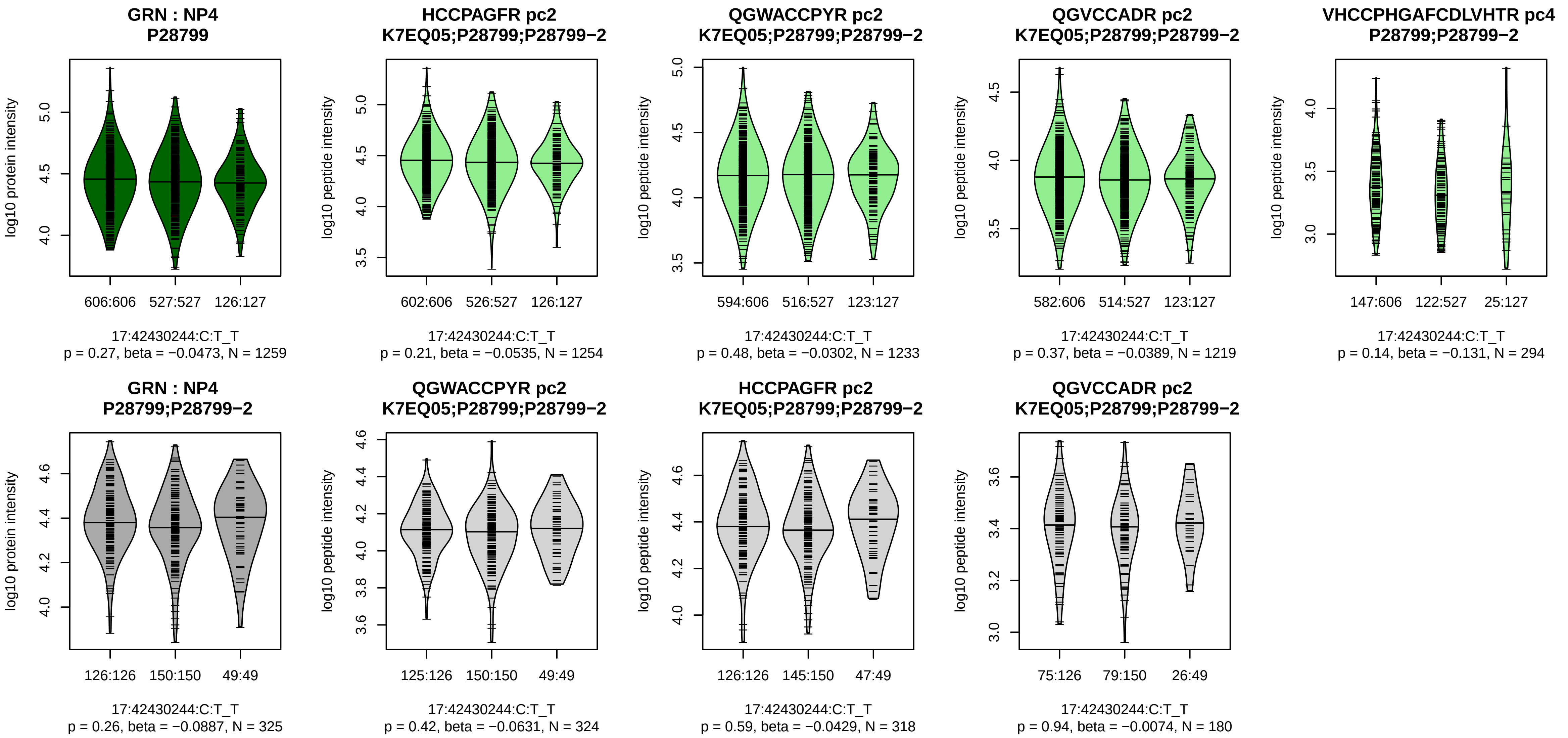


Assay Target: GRN
Olink UniProt: P28799
UKB rsID: rs5848
Proxy rsID: rs5848
UKB SNP: 17:42430244:C:T
Proxy SNP: 17:42430244:C:T
UKB log10(p): 351.4
UKB BETA: −0.25
−:−:−:−:−:−:NA
1254:1233:1219:294:280:58:16

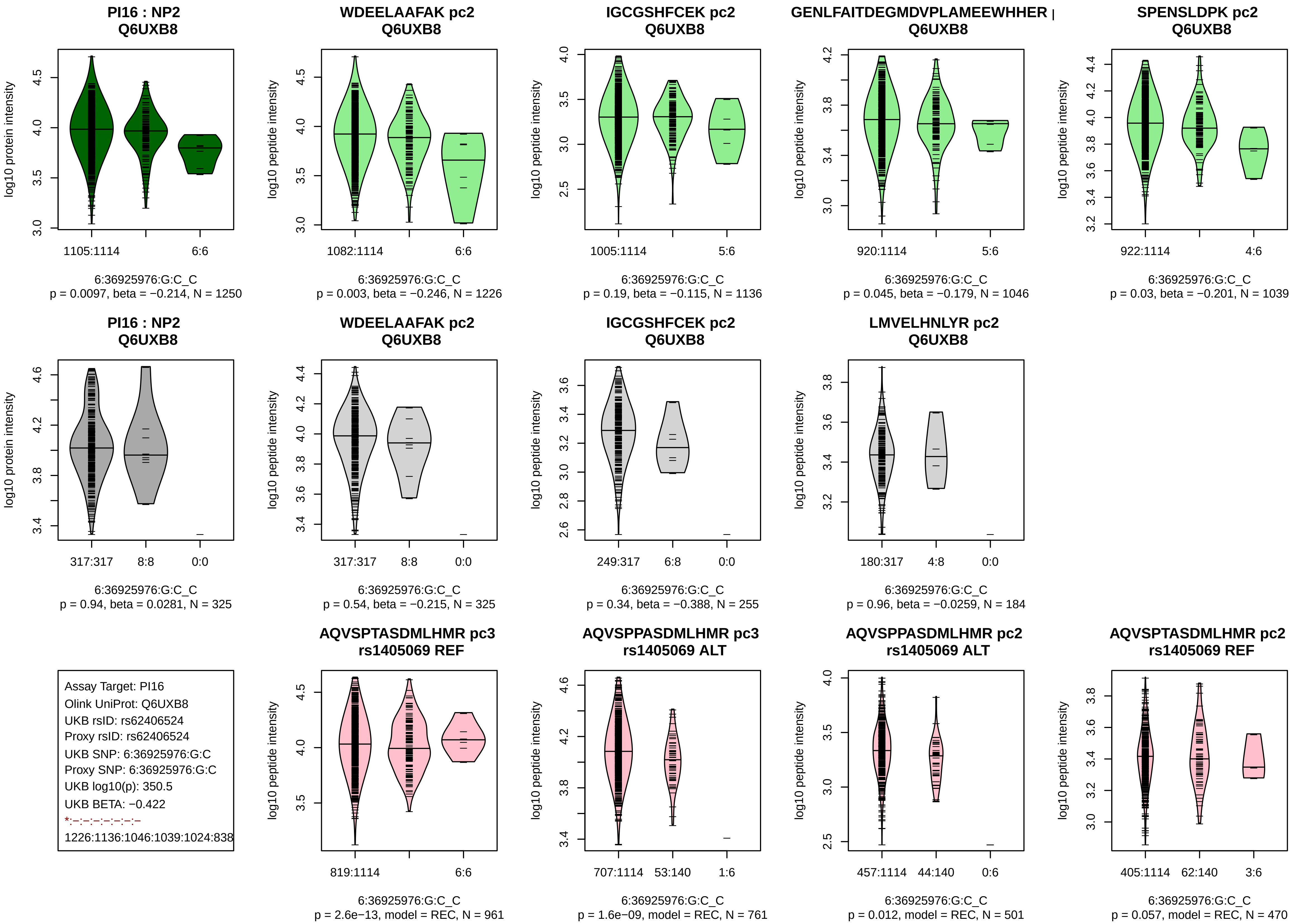
PI16 : NP2
Q6UXB8
log10 protein intensity
1105:1114
6:6
6:36925976:G:C_C
p = 0.0097, beta = −0.214, N = 1250
WDEELAAFAK pc2
Q6UXB8
log10 peptide intensity
1082:1114
6:6
6:36925976:G:C_C
p = 0.003, beta = −0.246, N = 1226
IGCGSHFCEK pc2
Q6UXB8
log10 peptide intensity
1005:1114
5:6
6:36925976:G:C_C
p = 0.19, beta = −0.115, N = 1136
GENLFAITDEGMDVPLAMEEWHHER |
Q6UXB8
log10 peptide intensity
920:1114
5:6
6:36925976:G:C_C
p = 0.045, beta = −0.179, N = 1046
SPENSLDPK pc2
Q6UXB8
log10 peptide intensity
922:1114
4:6
6:36925976:G:C_C
p = 0.03, beta = −0.201, N = 1039
PI16 : NP2
Q6UXB8
log10 protein intensity
317:317
8:8
0:0
6:36925976:G:C_C
p = 0.94, beta = 0.0281, N = 325
WDEELAAFAK pc2
Q6UXB8
log10 peptide intensity
317:317
8:8
0:0
6:36925976:G:C_C
p = 0.54, beta = −0.215, N = 325
IGCGSHFCEK pc2
Q6UXB8
log10 peptide intensity
249:317
6:8
0:0
6:36925976:G:C_C
p = 0.34, beta = −0.388, N = 255
LMVELHNLYR pc2
Q6UXB8
log10 peptide intensity
180:317
4:8
0:0
6:36925976:G:C_C
p = 0.96, beta = −0.0259, N = 184
Assay Target: PI16
Olink UniProt: Q6UXB8
UKB rsID: rs62406524
Proxy rsID: rs62406524
UKB SNP: 6:36925976:G:C
Proxy SNP: 6:36925976:G:C
UKB log10(p): 350.5
UKB BETA: −0.422
*:–:–:–:–:–:–:–
1226:1136:1046:1039:1024:838
AQVSPTASDMLHMR pc3
rs1405069 REF
log10 peptide intensity
819:1114
6:6
6:36925976:G:C_C
p = 2.6e−13, model = REC, N = 961
AQVSPPASDMLHMR pc3
rs1405069 ALT
log10 peptide intensity
707:1114
53:140
1:6
6:36925976:G:C_C
p = 1.6e−09, model = REC, N = 761
AQVSPPASDMLHMR pc2
rs1405069 ALT
log10 peptide intensity
457:1114
44:140
0:6
6:36925976:G:C_C
p = 0.012, model = REC, N = 501
AQVSPTASDMLHMR pc2
rs1405069 REF
log10 peptide intensity
405:1114
62:140
3:6
6:36925976:G:C_C
p = 0.057, model = REC, N = 470

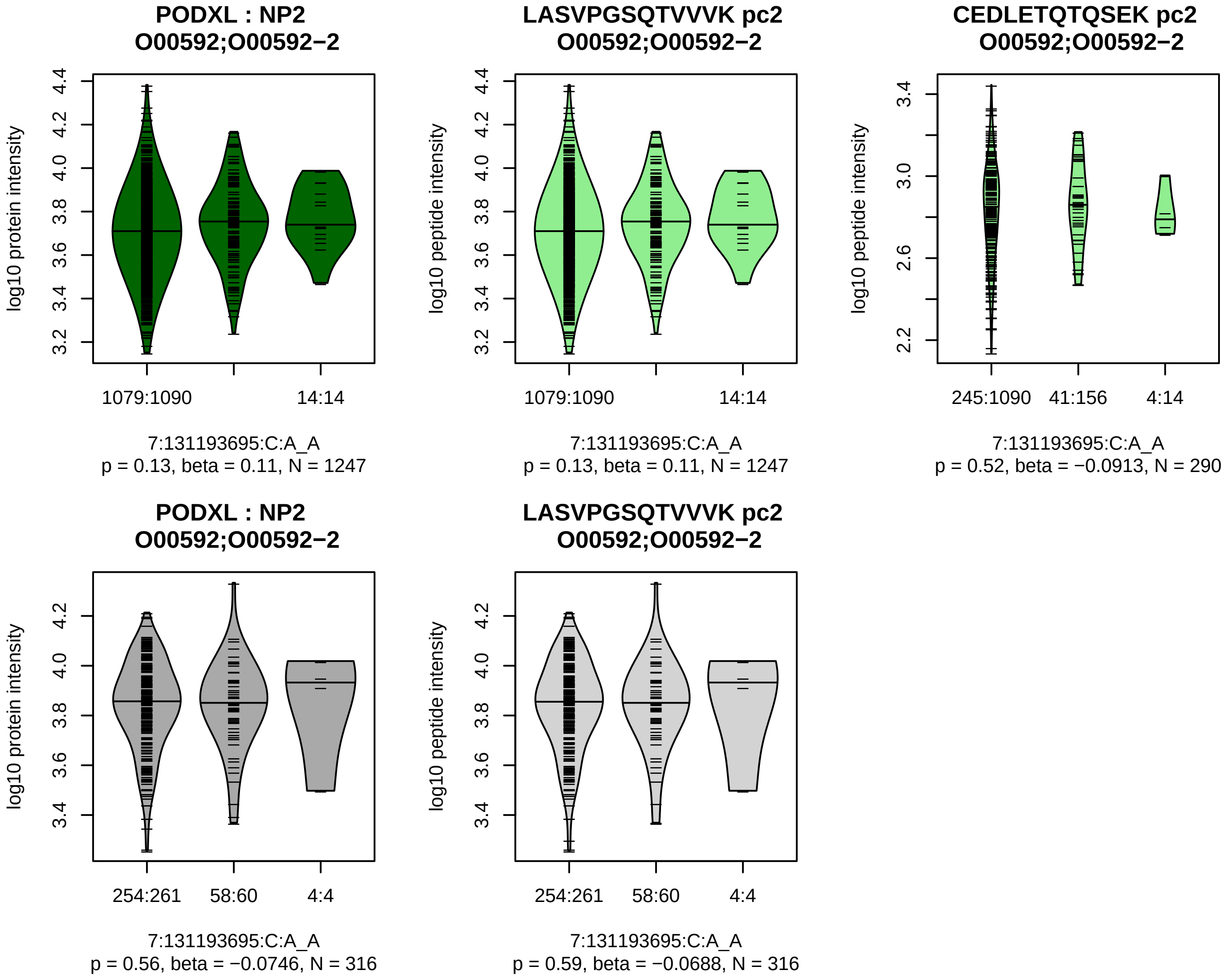


Assay Target: PODXL
Olink UniProt: O00592
UKB rsID: rs3212298
Proxy rsID: rs3212299
UKB SNP: 7:131193739:C:T
Proxy SNP: 7:131193695:C:A
UKB log10(p): 325.3
UKB BETA: −0.555
−:−
1247:290

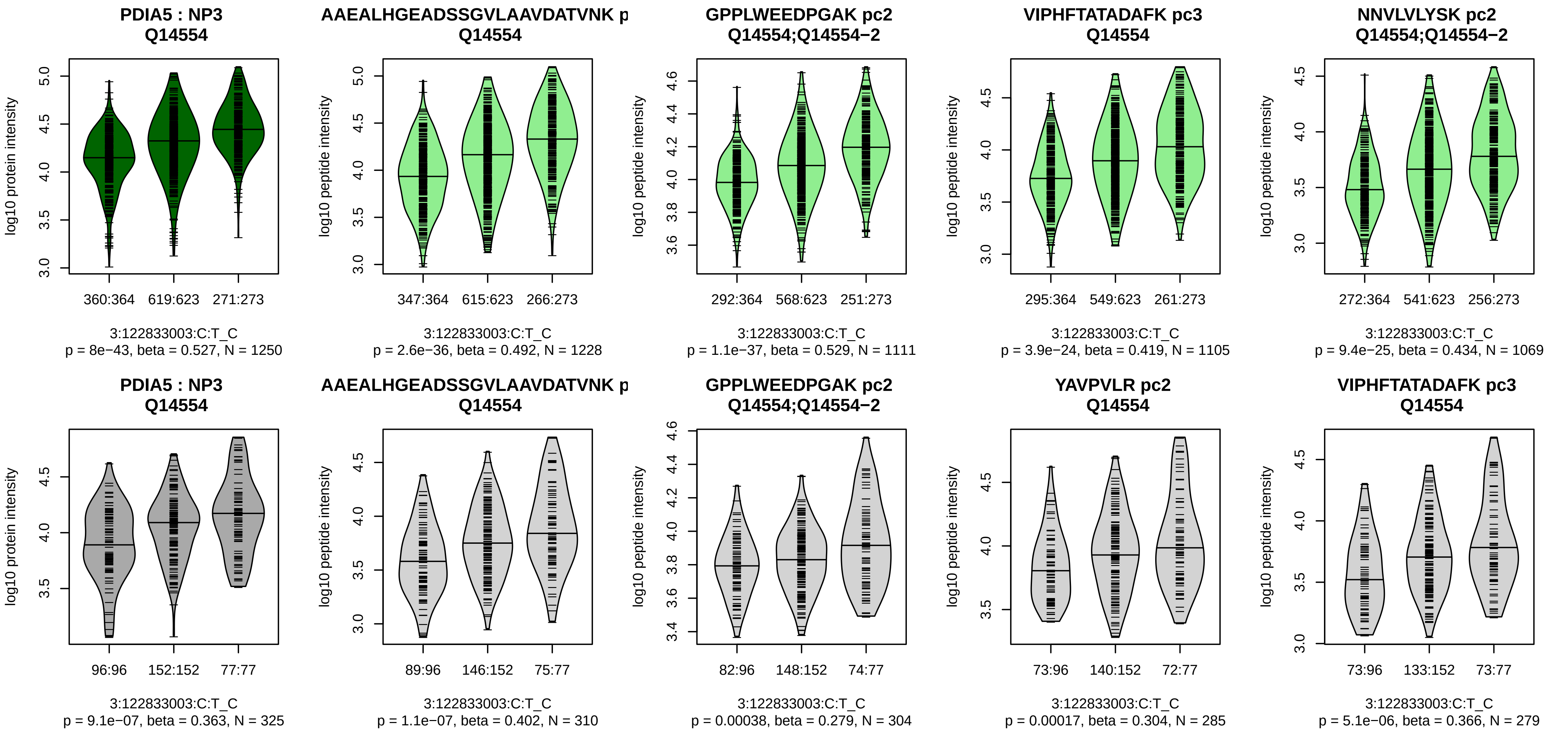


Assay Target: PDIA5
Olink UniProt: Q14554
UKB rsID: rs3804749
Proxy rsID: rs3804749
UKB SNP: 3:122833003:C:T
Proxy SNP: 3:122833003:C:T
UKB log10(p): 321.3
UKB BETA: −0.247
*:*:*:*:*:*:*:*:*:*:*:*:*:*:*:*:*:*:*:*:*:*:*:*:*:−:
1228:1111:1105:1069:1051:100

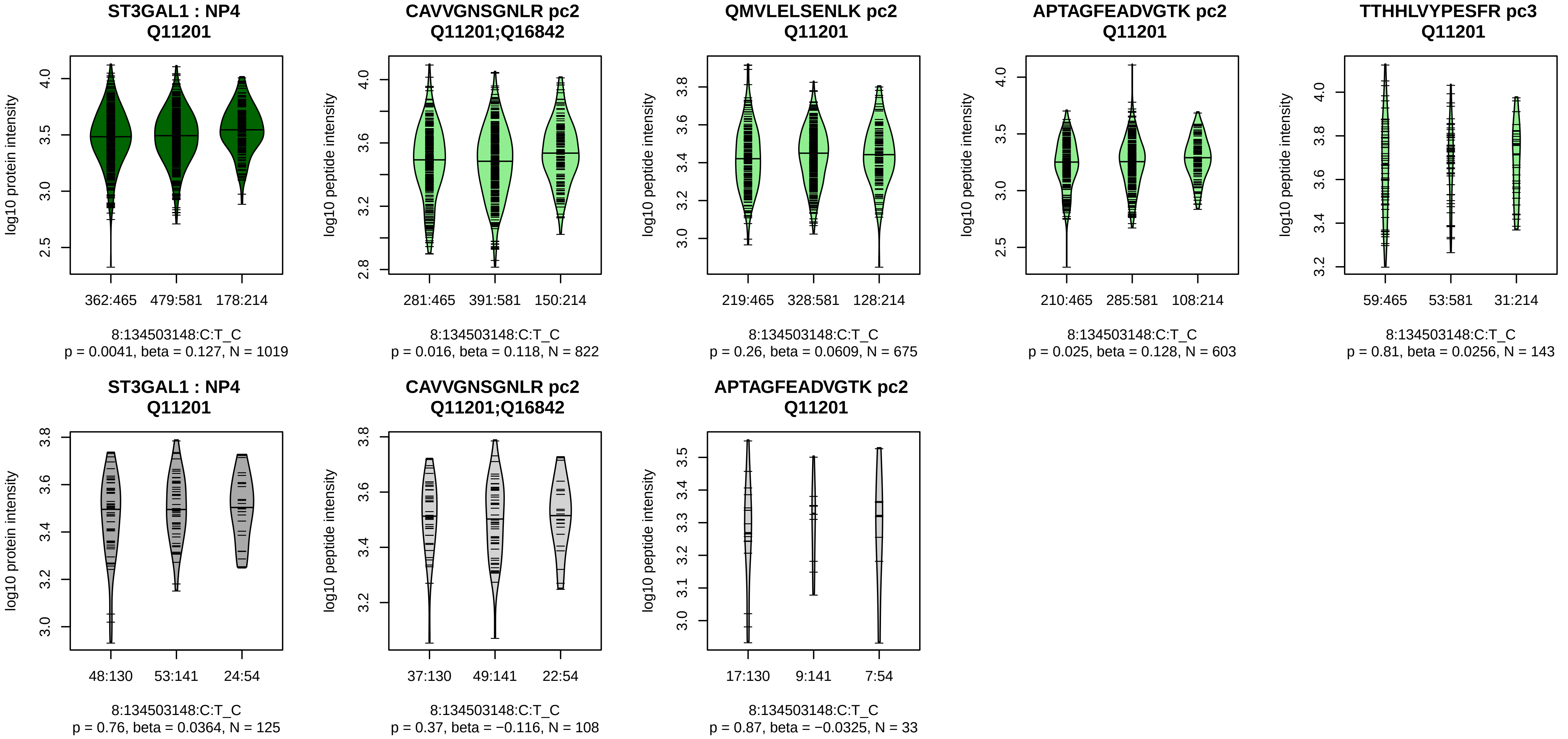


Assay Target: ST3GAL1
Olink UniProt: Q11201
UKB rsID: rs9643300
Proxy rsID: rs9643300
UKB SNP: 8:134503148:C:T
Proxy SNP: 8:134503148:C:T
UKB log10(p): 312.8
UKB BETA: −0.227
−:−:−:−
822:675:603:143

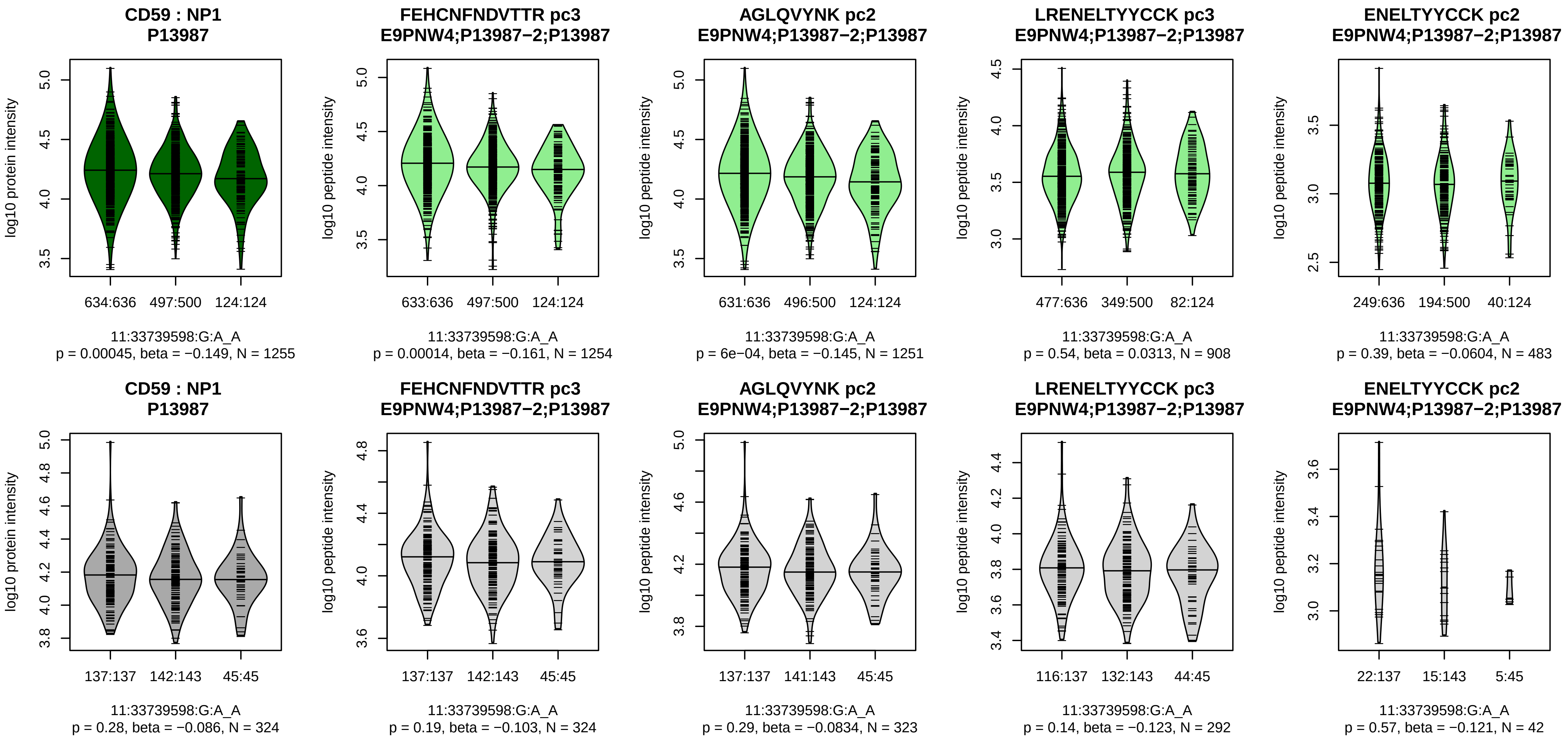


Assay Target: CD59
Olink UniProt: P13987
UKB rsID: rs831636
Proxy rsID: rs831636
UKB SNP: 11:33739598:G:A
Proxy SNP: 11:33739598:G:A
UKB log10(p): 305.8
UKB BETA: −0.229
*:*:−:−:−:−
1254:1251:908:589:483:60

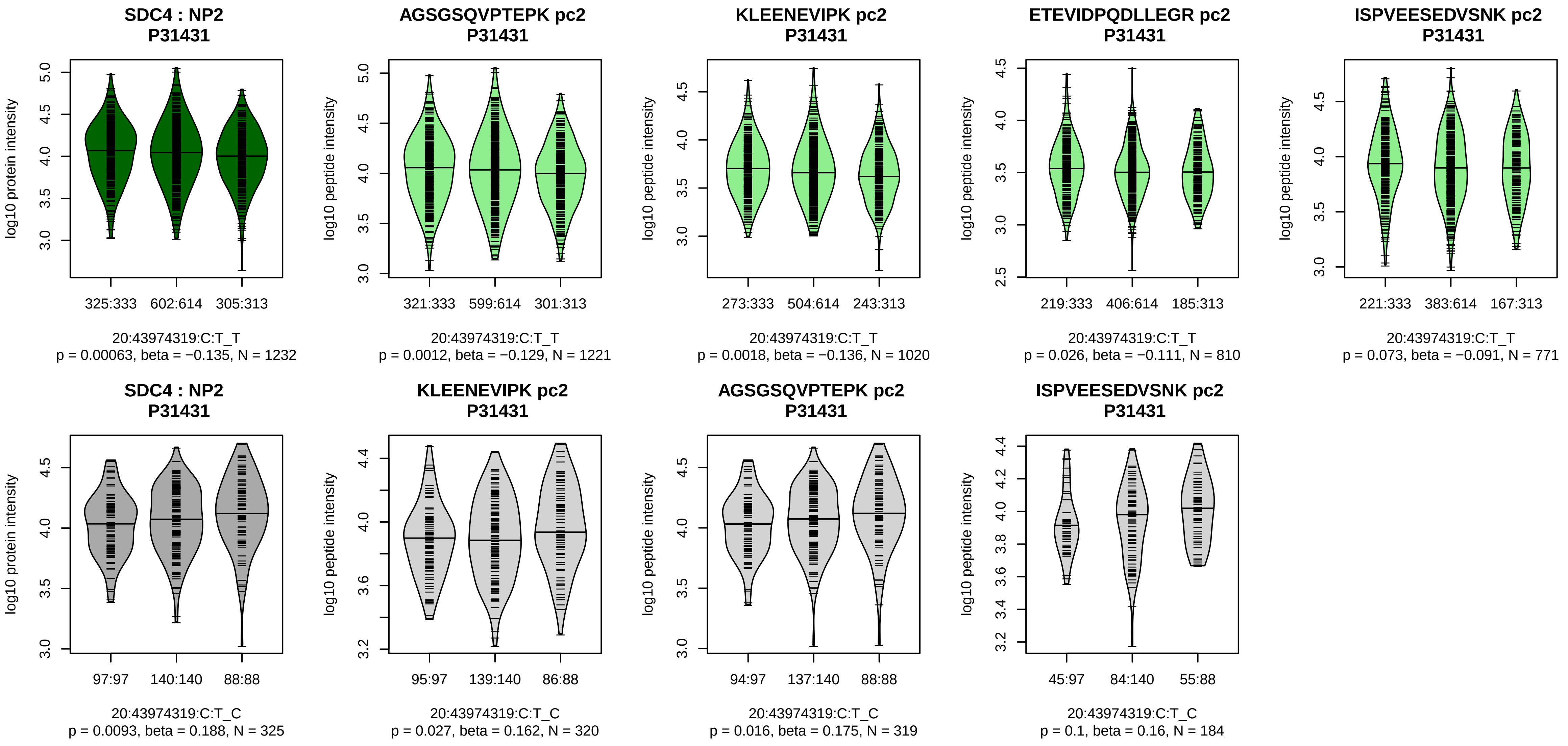


Assay Target: SDC4
Olink UniProt: P31431
UKB rsID: rs2741454
Proxy rsID: rs2741454
UKB SNP: 20:43974319:C:T
Proxy SNP: 20:43974319:C:T
UKB log10(p): 302.4
UKB BETA: −0.226
*:*:–:–:–:–
1221:1020:810:771:544:99

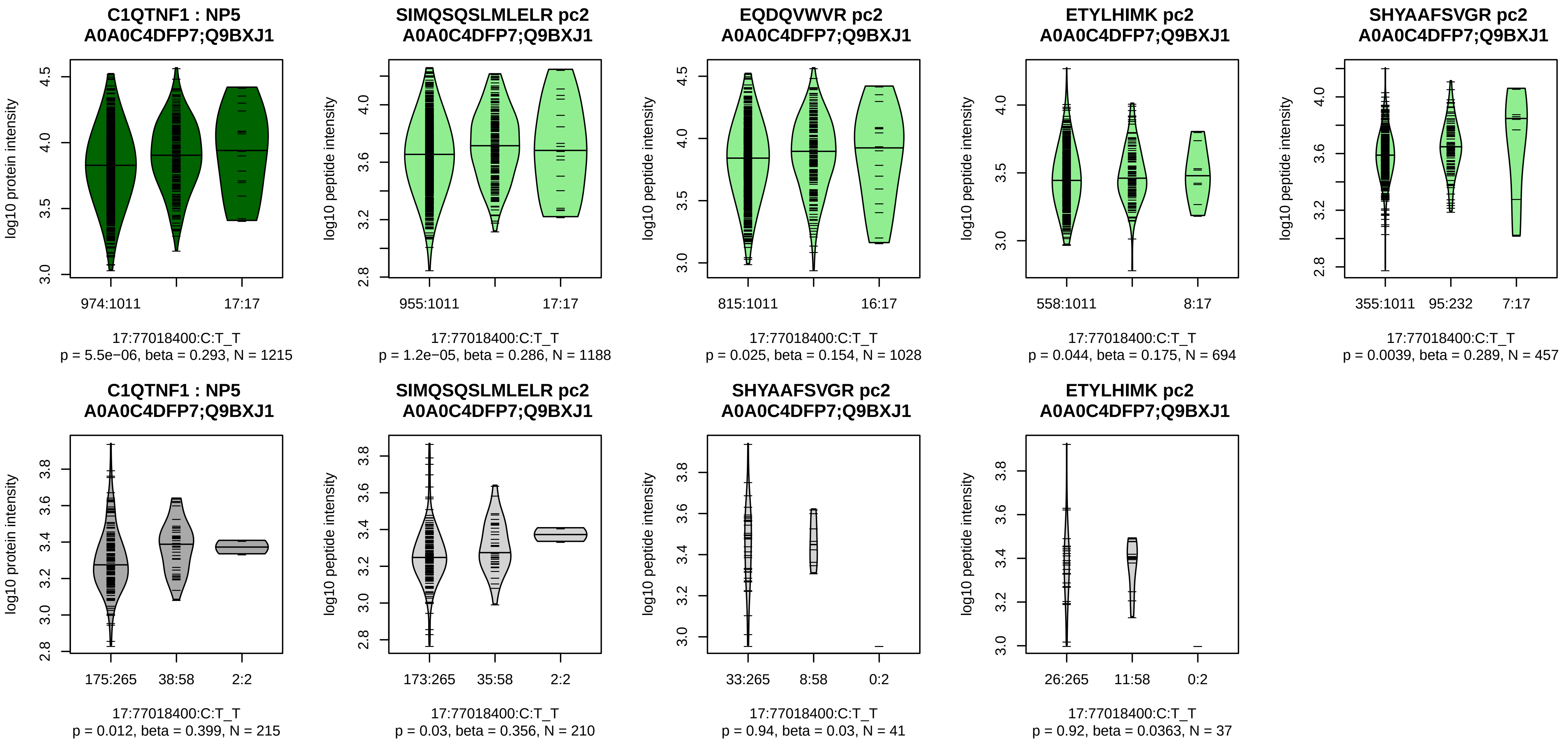


Assay Target: C1QTNF1
Olink UniProt: Q9BXJ1
UKB rsID: rs116539064
Proxy rsID: rs116539064
UKB SNP: 17:77018400:C:T
Proxy SNP: 17:77018400:C:T
UKB log10(p): 297.8
UKB BETA: 0.387
*:−:−:*:−:NA
1188:1028:694:457:250:4

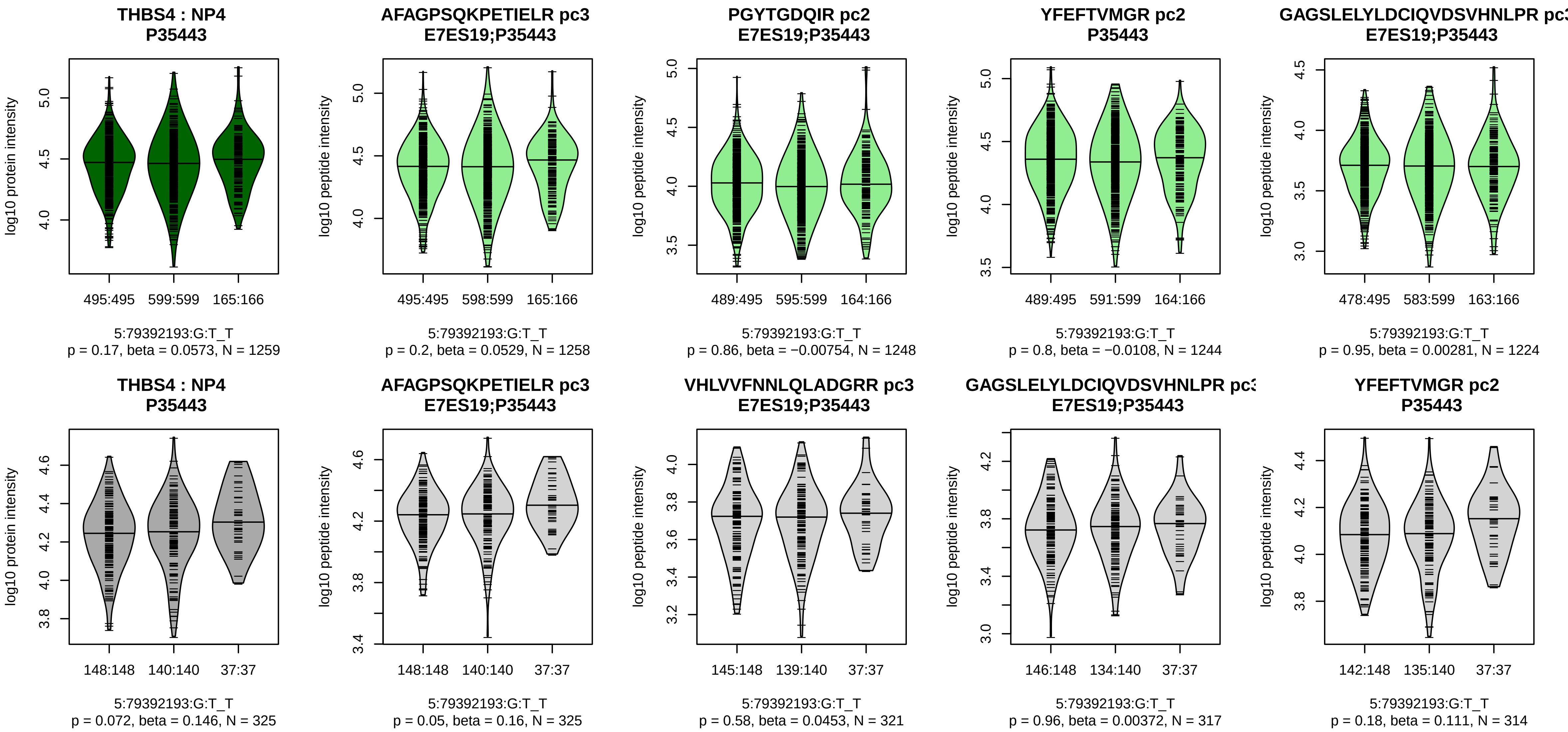


Assay Target: THBS4
Olink UniProt: P35443
UKB rsID: rs2438632
Proxy rsID: rs2438632
UKB SNP: 5:79392193:G:T
Proxy SNP: 5:79392193:G:T
UKB log10(p): 289.3
UKB BETA: 0.225
–:–:–:–:–:–:–:–:–:–:*:–:–:–:–:–:–
1258:1248:1244:1224:1215:120

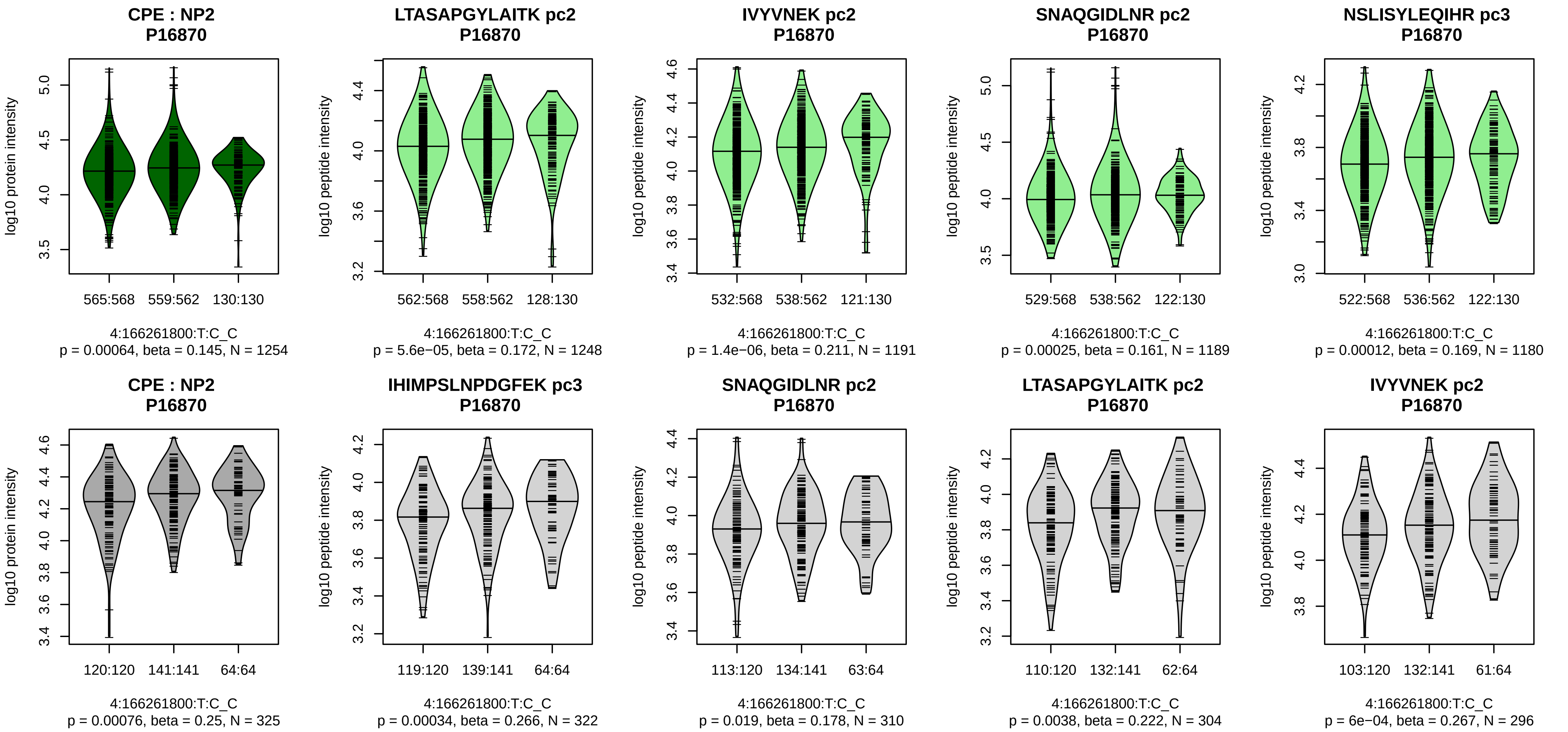


Assay Target: CPE
Olink UniProt: P16870
UKB rsID: rs1550270
Proxy rsID: rs1550270
UKB SNP: 4:166261800:T:C
Proxy SNP: 4:166261800:T:C
UKB log10(p): 287.3
UKB BETA: 0.244
*:*:*:*:*:*:*:*:–:–:–
1248:1191:1189:1180:1167:114

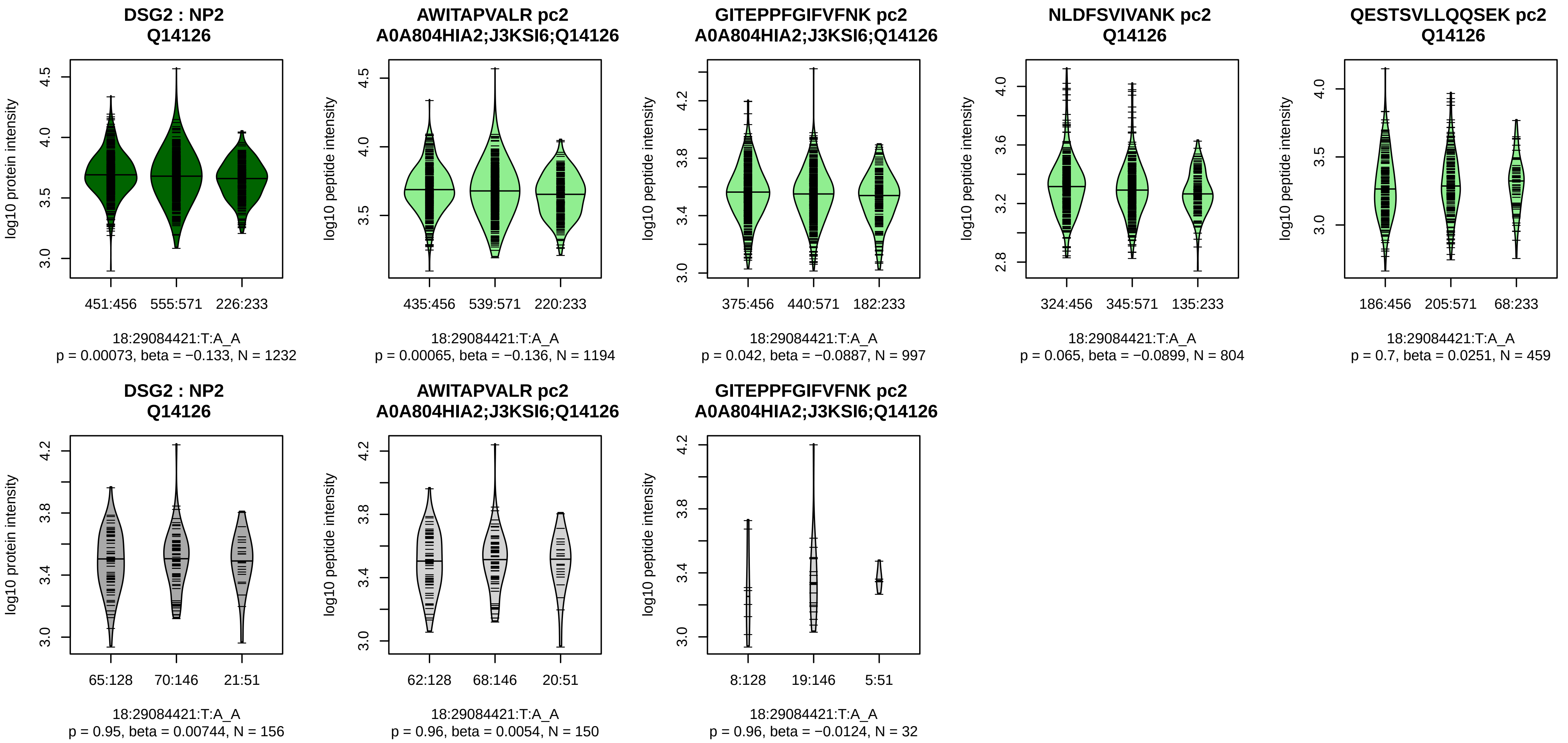


Assay Target: DSG2
Olink UniProt: Q14126
UKB rsID: rs1841396
Proxy rsID: rs1841396
UKB SNP: 18:29084421:T:A
Proxy SNP: 18:29084421:T:A
UKB log10(p): 274.8
UKB BETA: −0.213
*:−:−:−:−:−:−:−
1194:997:804:459:313:275:252:

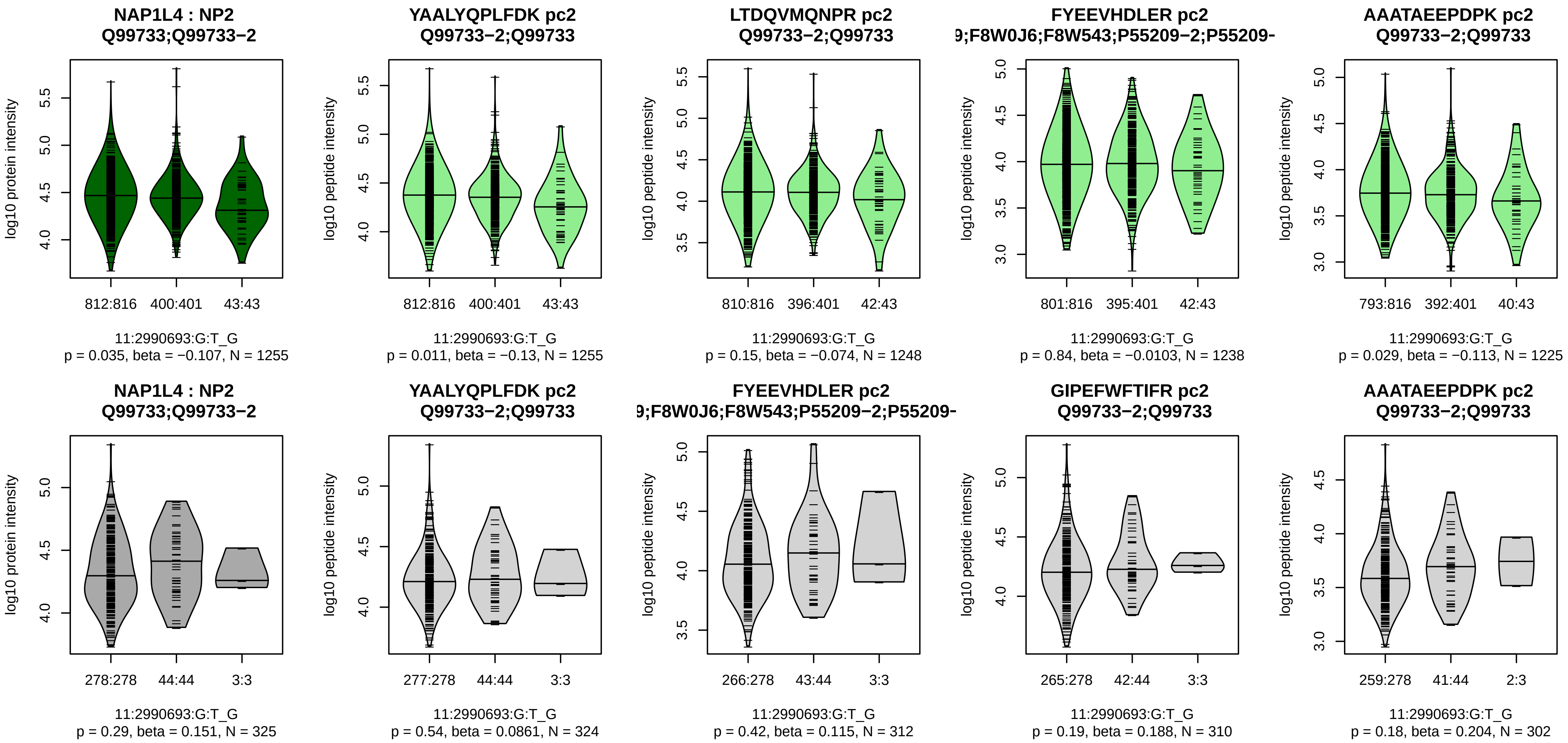


Assay Target: NAP1L4
Olink UniProt: Q99733
UKB rsID: rs57530484
Proxy rsID: rs4758501
UKB SNP: 11:2987643:T:TAACA
Proxy SNP: 11:2990693:G:T
UKB log10(p): 266.3
UKB BETA: 0.251
−:−:−:−:−:−:−:−:−:−:−:−:−:−:NA
1255:1248:1238:1225:1225:118

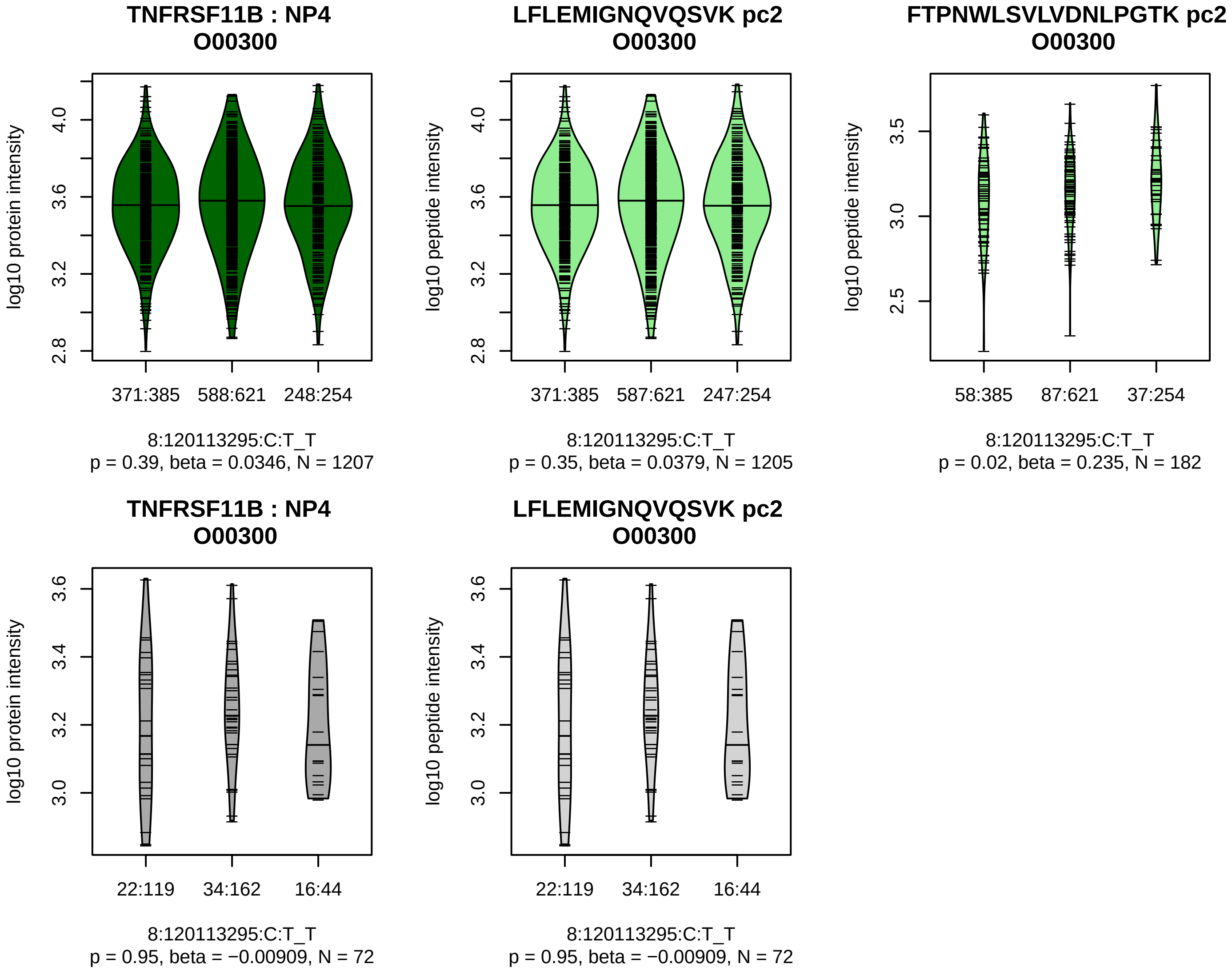


Assay Target: TNFRSF11B
Olink UniProt: O00300
UKB rsID: rs748443516
Proxy rsID: rs2465390
UKB SNP: 8:120101198:CT:C
Proxy SNP: 8:120113295:C:T
UKB log10(p): 264.7
UKB BETA: 0.193
–:–:–
1205:182:126

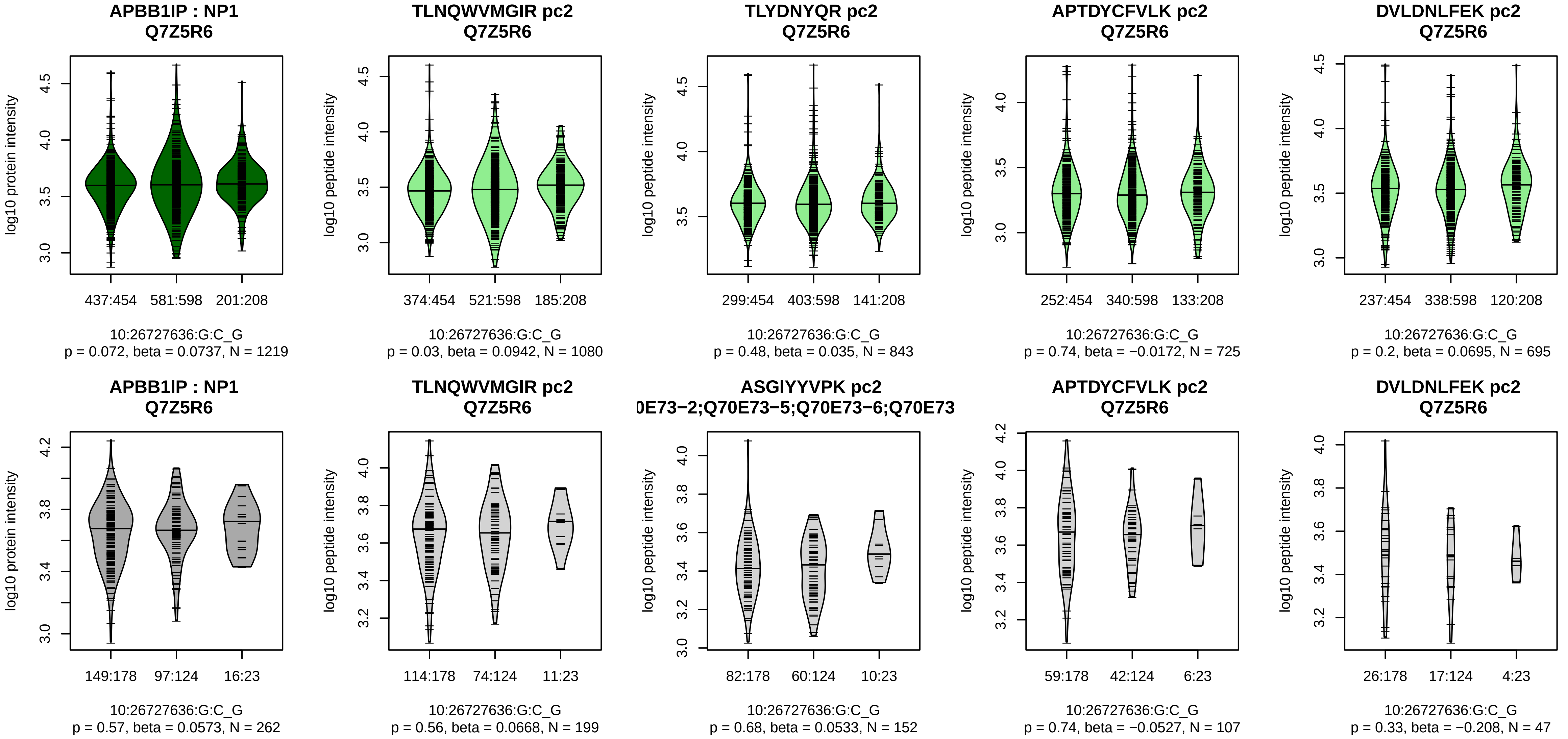


Assay Target: APBB1IP
Olink UniProt: Q7Z5R6
UKB rsID: rs4749125
Proxy rsID: rs4749125
UKB SNP: 10:26727636:G:C
Proxy SNP: 10:26727636:G:C
UKB log10(p): 264.2
UKB BETA: −0.216
−:−:−:−:−:−:−:−:−:−:−:−:−:NA
1080:843:725:695:604:494:487:

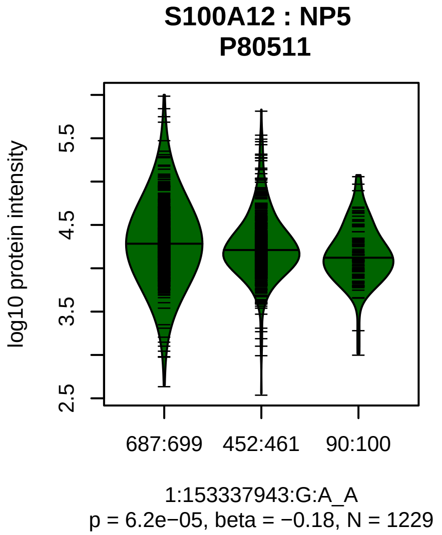


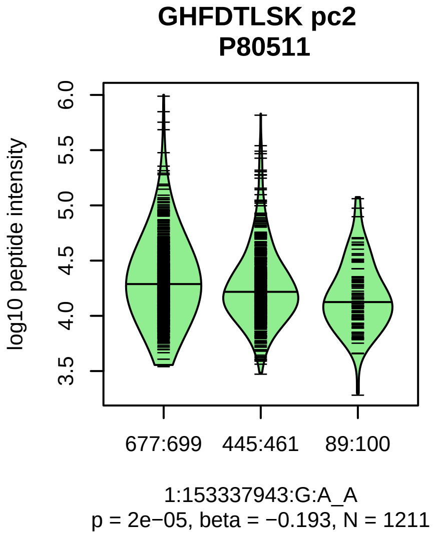


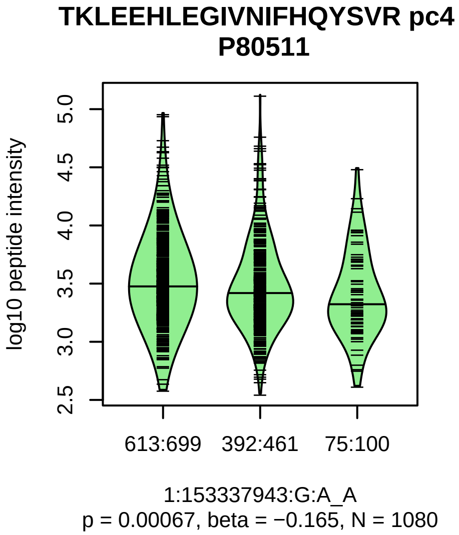


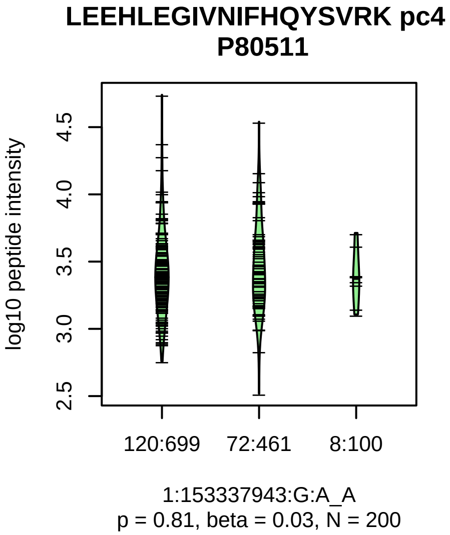


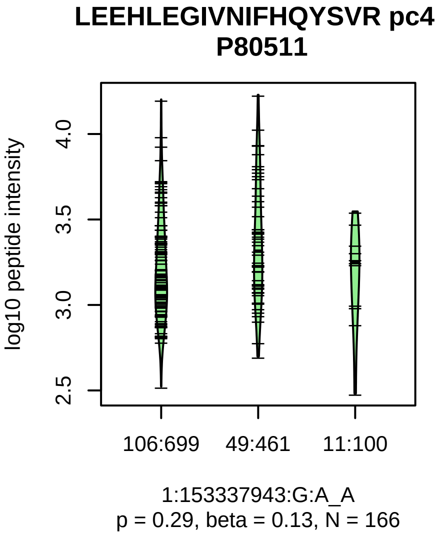


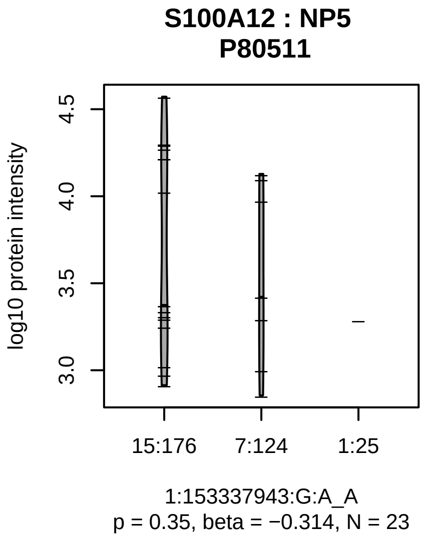


Assay Target: S100A12
Olink UniProt: P80511
UKB rsID: rs3014874
Proxy rsID: rs3014874
UKB SNP: 1:153337943:G:A
Proxy SNP: 1:153337943:G:A
UKB log10(p): 256.4
UKB BETA: −0.235
*:*:−:−:−:−
1211:1080:200:166:55:39

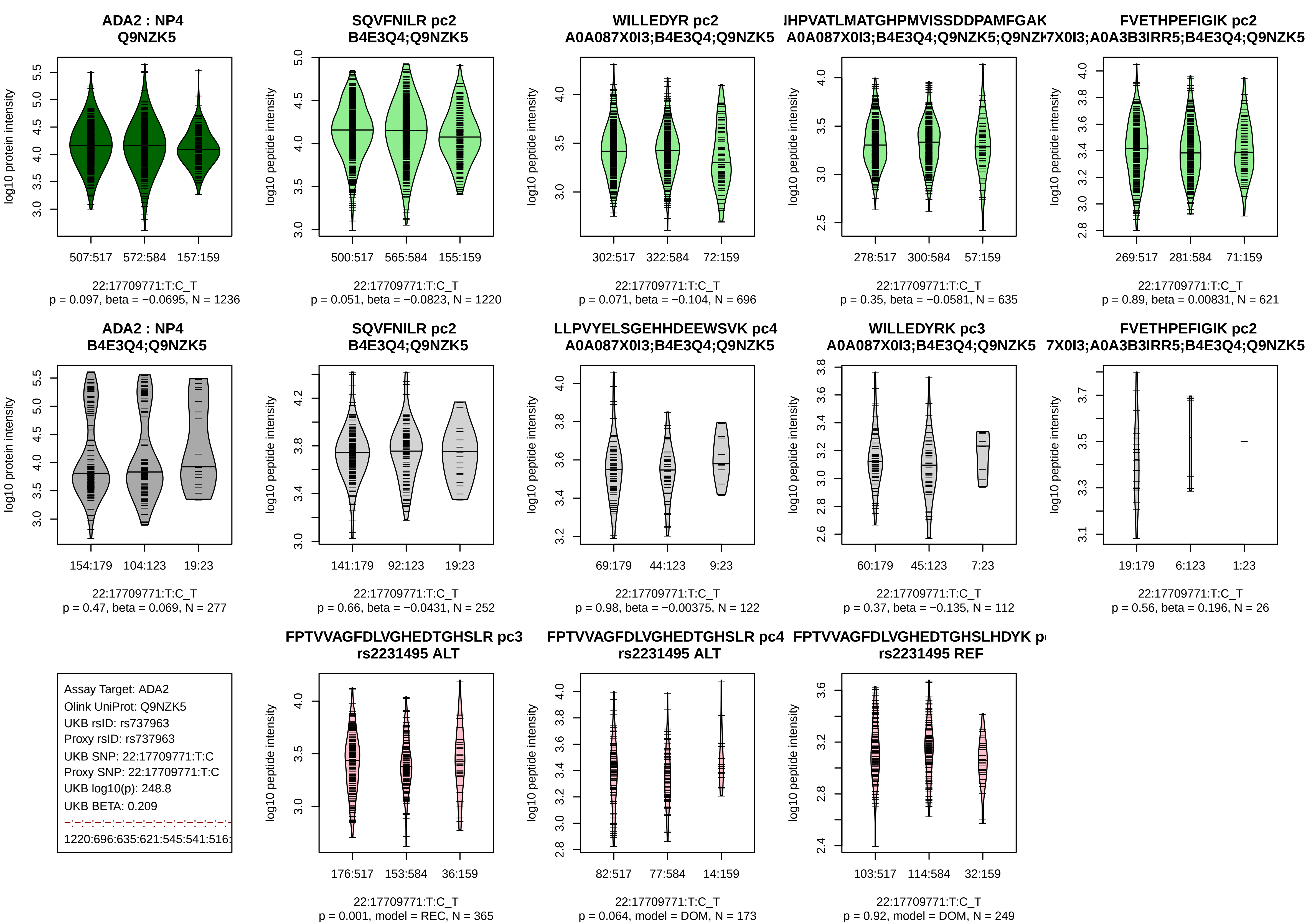
ADA2 : NP4
Q9NZK5
log10 protein intensity
507:517
572:584
157:159
22:17709771:T:C_T
p = 0.097, beta = −0.0695, N = 1236
SQVFNILR pc2
B4E3Q4;Q9NZK5
log10 peptide intensity
500:517
565:584
155:159
22:17709771:T:C_T
p = 0.051, beta = −0.0823, N = 1220
WILLEDYR pc2
A0A087X0I3;B4E3Q4;Q9NZK5
302:517
322:584
72:159
22:17709771:T:C_T
p = 0.071, beta = −0.104, N = 696
IHPVATLMATGHPMVISSDDPAMFGAK
A0A087X0I3;B4E3Q4;Q9NZK5;Q9NZI
278:517
300:584
57:159
22:17709771:T:C_T
p = 0.35, beta = −0.0581, N = 635
FVETHPEFIGIK pc2
7X0I3;A0A3B3IRR5;B4E3Q4;Q9NZK5
269:517
281:584
71:159
22:17709771:T:C_T
p = 0.89, beta = 0.00831, N = 621
ADA2 : NP4
B4E3Q4;Q9NZK5
154:179
104:123
19:23
22:17709771:T:C_T
p = 0.47, beta = 0.069, N = 277
SQVFNILR pc2
B4E3Q4;Q9NZK5
141:179
92:123
19:23
22:17709771:T:C_T
p = 0.66, beta = −0.0431, N = 252
LLPVYELSGEHHDEEWSVK pc4
A0A087X0I3;B4E3Q4;Q9NZK5
69:179
44:123
9:23
22:17709771:T:C_T
p = 0.98, beta = −0.00375, N = 122
WILLEDYRK pc3
A0A087X0I3;B4E3Q4;Q9NZK5
60:179
45:123
7:23
22:17709771:T:C_T
p = 0.37, beta = −0.135, N = 112
FVETHPEFIGIK pc2
7X0I3;A0A3B3IRR5;B4E3Q4;Q9NZK5
19:179
6:123
1:23
22:17709771:T:C_T
p = 0.56, beta = 0.196, N = 26
Assay Target: ADA2
Olink UniProt: Q9NZK5
UKB rsID: rs737963
Proxy rsID: rs737963
UKB SNP: 22:17709771:T:C
Proxy SNP: 22:17709771:T:C
UKB log10(p): 248.8
UKB BETA: 0.209
1220:696:635:621:545:541:516:
FPTVVAGFDLVGHEDTGHSLR pc3
rs2231495 ALT
176:517
153:584
36:159
22:17709771:T:C_T
p = 0.001, model = REC, N = 365
FPTVVAGFDLVGHEDTGHSLR pc4
rs2231495 ALT
82:517
77:584
14:159
22:17709771:T:C_T
p = 0.064, model = DOM, N = 173
FPTVVAGFDLVGHEDTGHSLHDYK p
rs2231495 REF
103:517
114:584
32:159
22:17709771:T:C_T
p = 0.92, model = DOM, N = 249

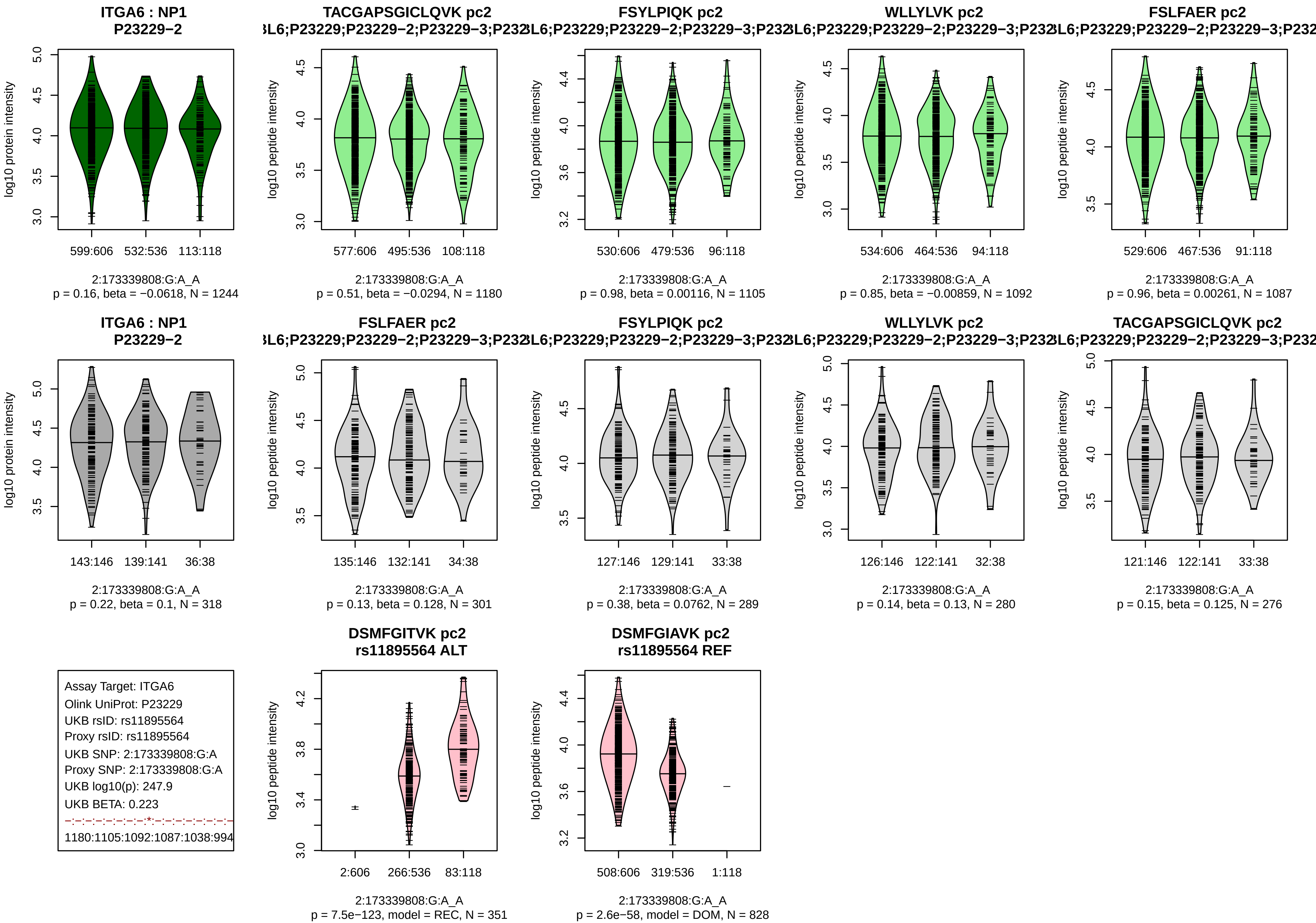

ITGA6 : NP1
P23229−2
log10 protein intensity
599:606 532:536 113:118
2:173339808:G:A_A
p = 0.16, beta = −0.0618, N = 1244
TACGAPSGICLQVK pc2
L6;P23229;P23229−2;P23229−3;P232
log10 peptide intensity
577:606 495:536 108:118
2:173339808:G:A_A
p = 0.51, beta = −0.0294, N = 1180
FSYLPIQK pc2
L6;P23229;P23229−2;P23229−3;P232
log10 peptide intensity
530:606 479:536 96:118
2:173339808:G:A_A
p = 0.98, beta = 0.00116, N = 1105
WLLYLVK pc2
L6;P23229;P23229−2;P23229−3;P232
log10 peptide intensity
534:606 464:536 94:118
2:173339808:G:A_A
p = 0.85, beta = −0.00859, N = 1092
FSLFAER pc2
L6;P23229;P23229−2;P23229−3;P232
log10 peptide intensity
529:606 467:536 91:118
2:173339808:G:A_A
p = 0.96, beta = 0.00261, N = 1087
ITGA6 : NP1
P23229−2
log10 protein intensity
143:146 139:141 36:38
2:173339808:G:A_A
p = 0.22, beta = 0.1, N = 318
FSLFAER pc2
L6;P23229;P23229−2;P23229−3;P232
log10 peptide intensity
135:146 132:141 34:38
2:173339808:G:A_A
p = 0.13, beta = 0.128, N = 301
FSYLPIQK pc2
L6;P23229;P23229−2;P23229−3;P232
log10 peptide intensity
127:146 129:141 33:38
2:173339808:G:A_A
p = 0.38, beta = 0.0762, N = 289
WLLYLVK pc2
L6;P23229;P23229−2;P23229−3;P232
log10 peptide intensity
126:146 122:141 32:38
2:173339808:G:A_A
p = 0.14, beta = 0.13, N = 280
TACGAPSGICLQVK pc2
L6;P23229;P23229−2;P23229−3;P232
log10 peptide intensity
121:146 122:141 33:38
2:173339808:G:A_A
p = 0.15, beta = 0.125, N = 276
Assay Target: ITGA6
Olink UniProt: P23229
UKB rsID: rs11895564
Proxy rsID: rs11895564
UKB SNP: 2:173339808:G:A
Proxy SNP: 2:173339808:G:A
UKB log10(p): 247.9
UKB BETA: 0.223
1180:1105:1092:1087:1038:994
DSMFGITVK pc2
rs11895564 ALT
log10 peptide intensity
2:606 266:536 83:118
2:173339808:G:A_A
p = 7.5e−123, model = REC, N = 351
DSMFGIAVK pc2
rs11895564 REF
log10 peptide intensity
508:606 319:536 1:118
2:173339808:G:A_A
p = 2.6e−58, model = DOM, N = 828

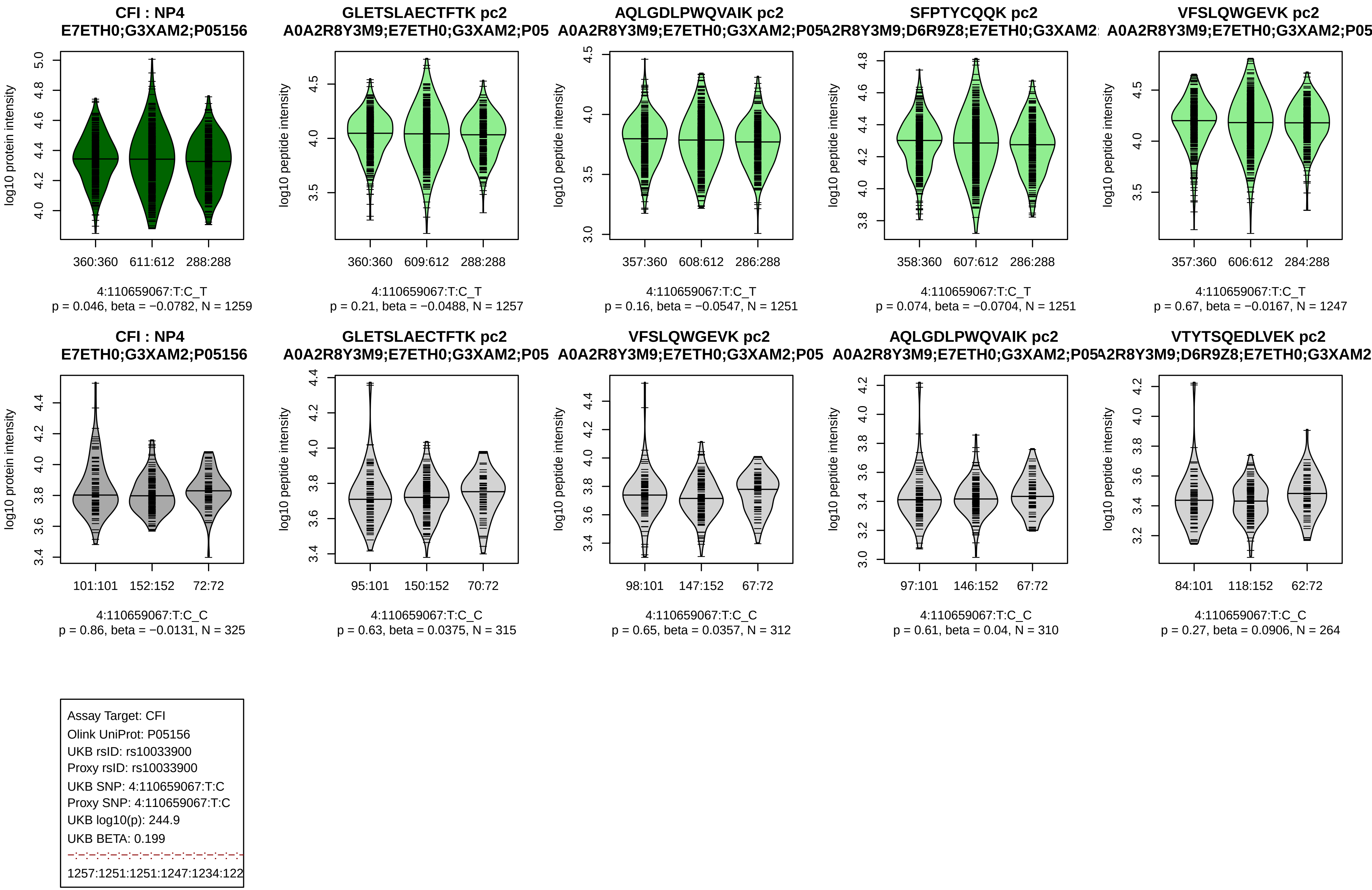

CFI : NP4
E7ETH0;G3XAM2;P05156
log10 protein intensity
360:360 611:612 288:288
4:110659067:T:C_T
p = 0.046, beta = −0.0782, N = 1259
GLETSLAECTFTK pc2
A0A2R8Y3M9;E7ETH0;G3XAM2;P05
log10 peptide intensity
360:360 609:612 288:288
4:110659067:T:C_T
p = 0.21, beta = −0.0488, N = 1257
AQLGDLPWQVAIK pc2
A0A2R8Y3M9;E7ETH0;G3XAM2;P05
log10 peptide intensity
357:360 608:612 286:288
4:110659067:T:C_T
p = 0.16, beta = −0.0547, N = 1251
SFPTYCQQK pc2
A2R8Y3M9;D6R9Z8;E7ETH0;G3XAM2
log10 peptide intensity
358:360 607:612 286:288
4:110659067:T:C_T
p = 0.074, beta = −0.0704, N = 1251
VFSLQWGEVK pc2
A0A2R8Y3M9;E7ETH0;G3XAM2;P05
log10 peptide intensity
357:360 606:612 284:288
4:110659067:T:C_T
p = 0.67, beta = −0.0167, N = 1247
CFI : NP4
E7ETH0;G3XAM2;P05156
log10 protein intensity
101:101 152:152 72:72
4:110659067:T:C_C
p = 0.86, beta = −0.0131, N = 325
GLETSLAECTFTK pc2
A0A2R8Y3M9;E7ETH0;G3XAM2;P05
log10 peptide intensity
95:101 150:152 70:72
4:110659067:T:C_C
p = 0.63, beta = 0.0375, N = 315
VFSLQWGEVK pc2
A0A2R8Y3M9;E7ETH0;G3XAM2;P05
log10 peptide intensity
98:101 147:152 67:72
4:110659067:T:C_C
p = 0.65, beta = 0.0357, N = 312
AQLGDLPWQVAIK pc2
A0A2R8Y3M9;E7ETH0;G3XAM2;P05
log10 peptide intensity
97:101 146:152 67:72
4:110659067:T:C_C
p = 0.61, beta = 0.04, N = 310
VTYTSQEDLVEK pc2
A2R8Y3M9;D6R9Z8;E7ETH0;G3XAM2
log10 peptide intensity
84:101 118:152 62:72
4:110659067:T:C_C
p = 0.27, beta = 0.0906, N = 264
Assay Target: CFI
Olink UniProt: P05156
UKB rsID: rs10033900
Proxy rsID: rs10033900
UKB SNP: 4:110659067:T:C
Proxy SNP: 4:110659067:T:C
UKB log10(p): 244.9
UKB BETA: 0.199
1257:1251:1251:1247:1234:122

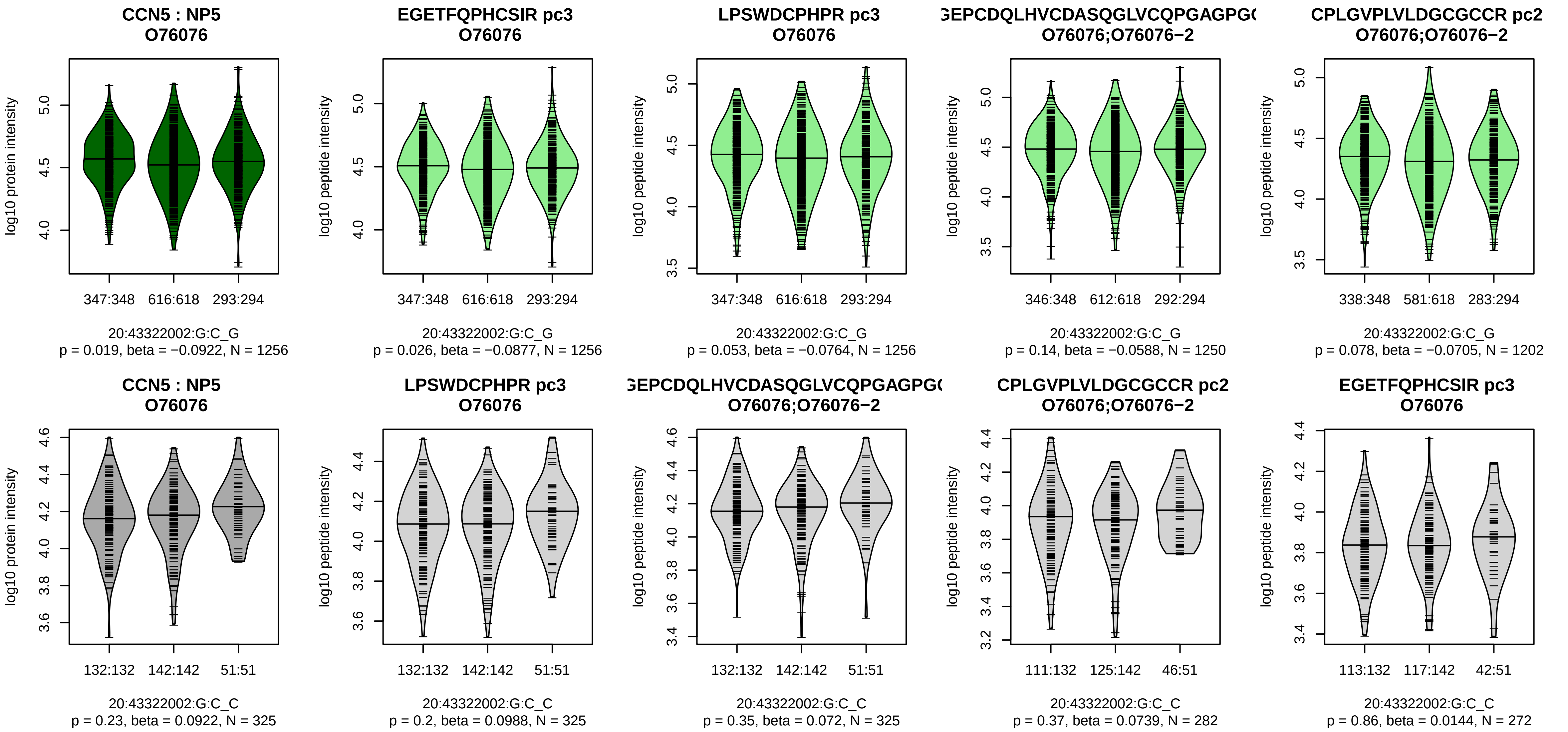


Assay Target: CCN5
Olink UniProt: O76076
UKB rsID: rs6065746
Proxy rsID: rs6065746
UKB SNP: 20:43322002:G:C
Proxy SNP: 20:43322002:G:C
UKB log10(p): 243.7
UKB BETA: 0.189
−:−:−:−:−:−:−:−:−:−:−:−:−:−:NA
1256:1256:1250:1202:1095:964

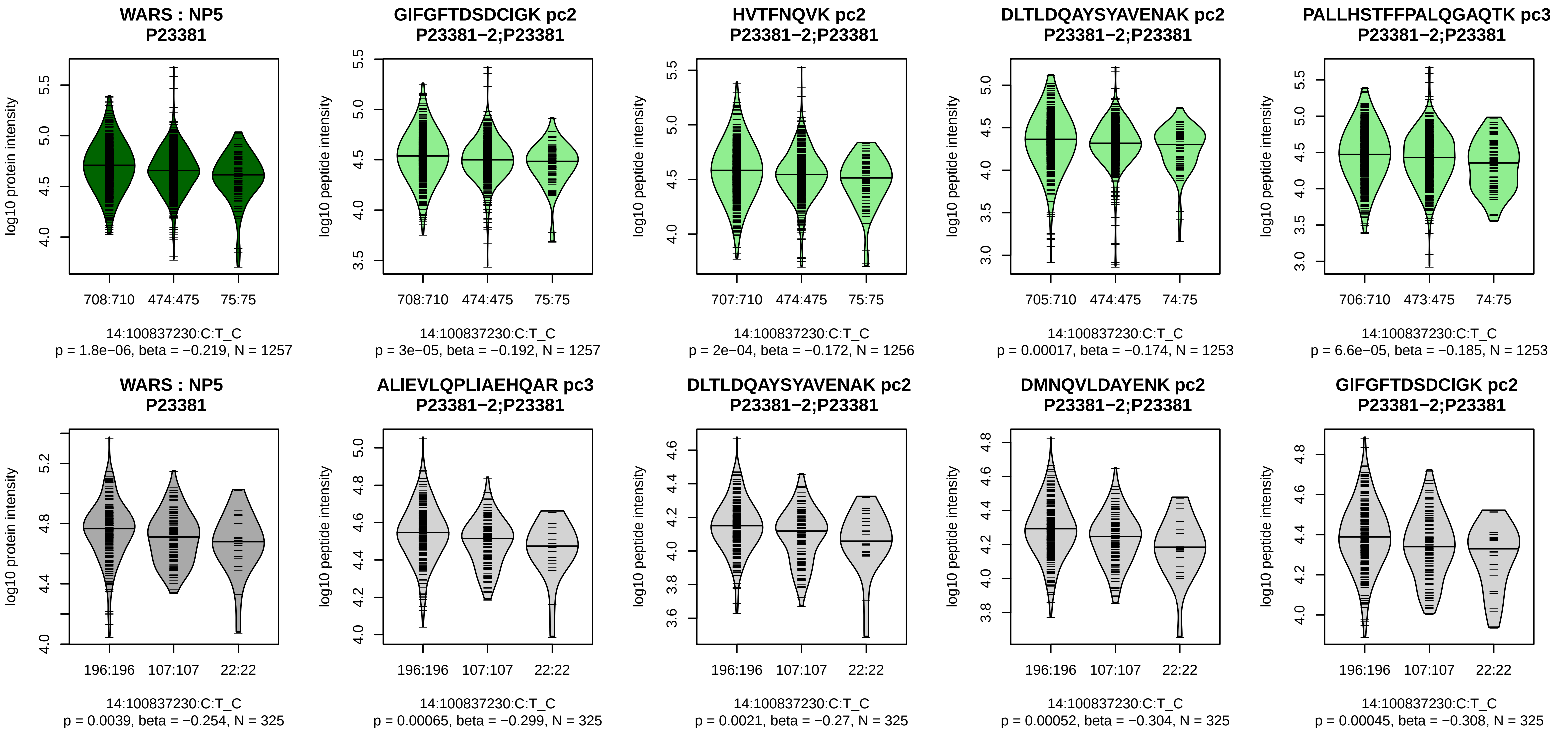


Assay Target: WARS
Olink UniProt: P23381
UKB rsID: rs4905957
Proxy rsID: rs4905957
UKB SNP: 14:100837230:C:T
Proxy SNP: 14:100837230:C:T
UKB log10(p): 240.8
UKB BETA: 0.232
*:*:*:*:*:*:*:*:*:*:*:*:–:*:–:–:*:–:–:–:*
1257:1256:1253:1253:1241:124

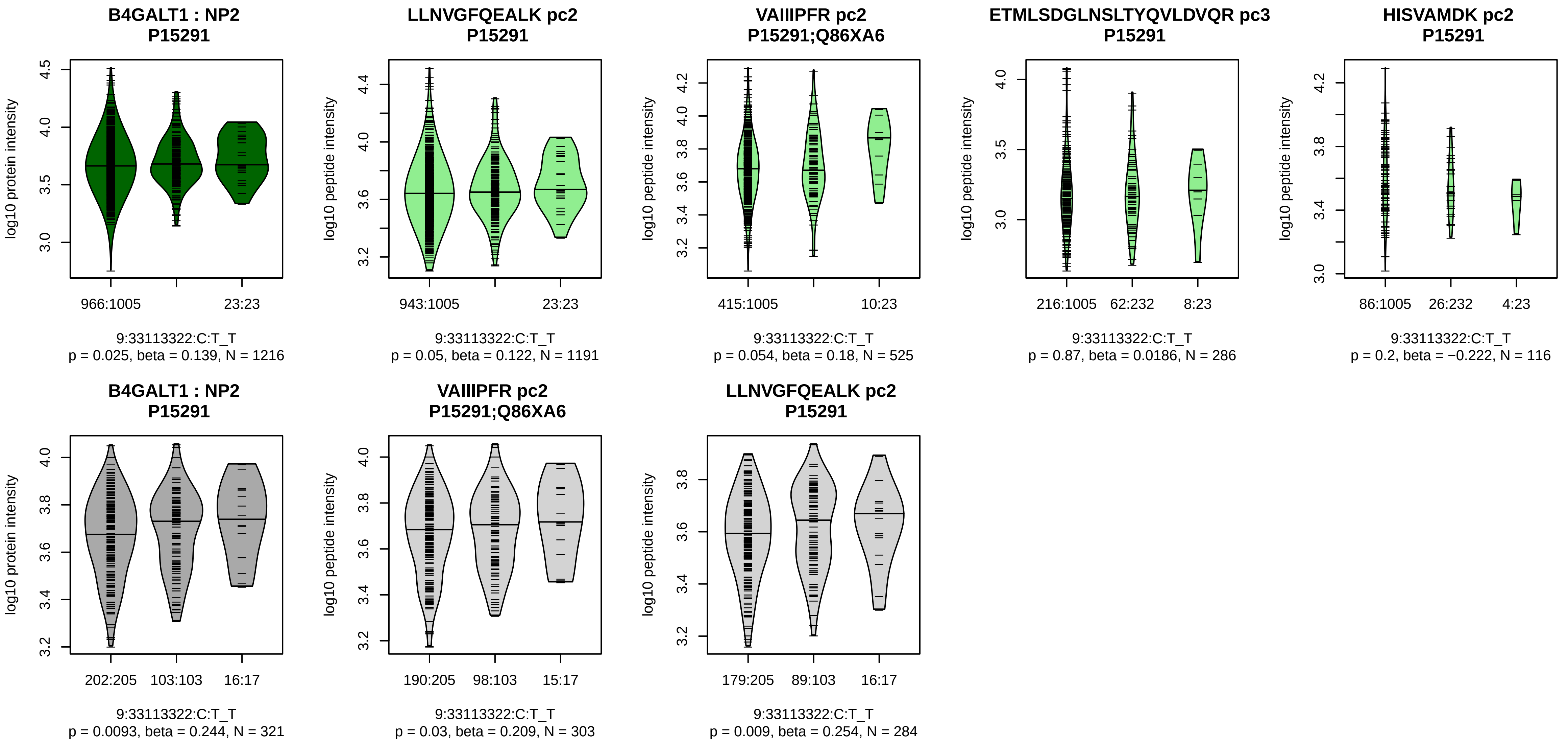


Assay Target: B4GALT1
Olink UniProt: P15291
UKB rsID: rs7019909
Proxy rsID: rs7019909
UKB SNP: 9:33113322:C:T
Proxy SNP: 9:33113322:C:T
UKB log10(p): 240.1
UKB BETA: 0.314
−:−:−:−:−:−:NA
1191:525:286:116:89:74:9

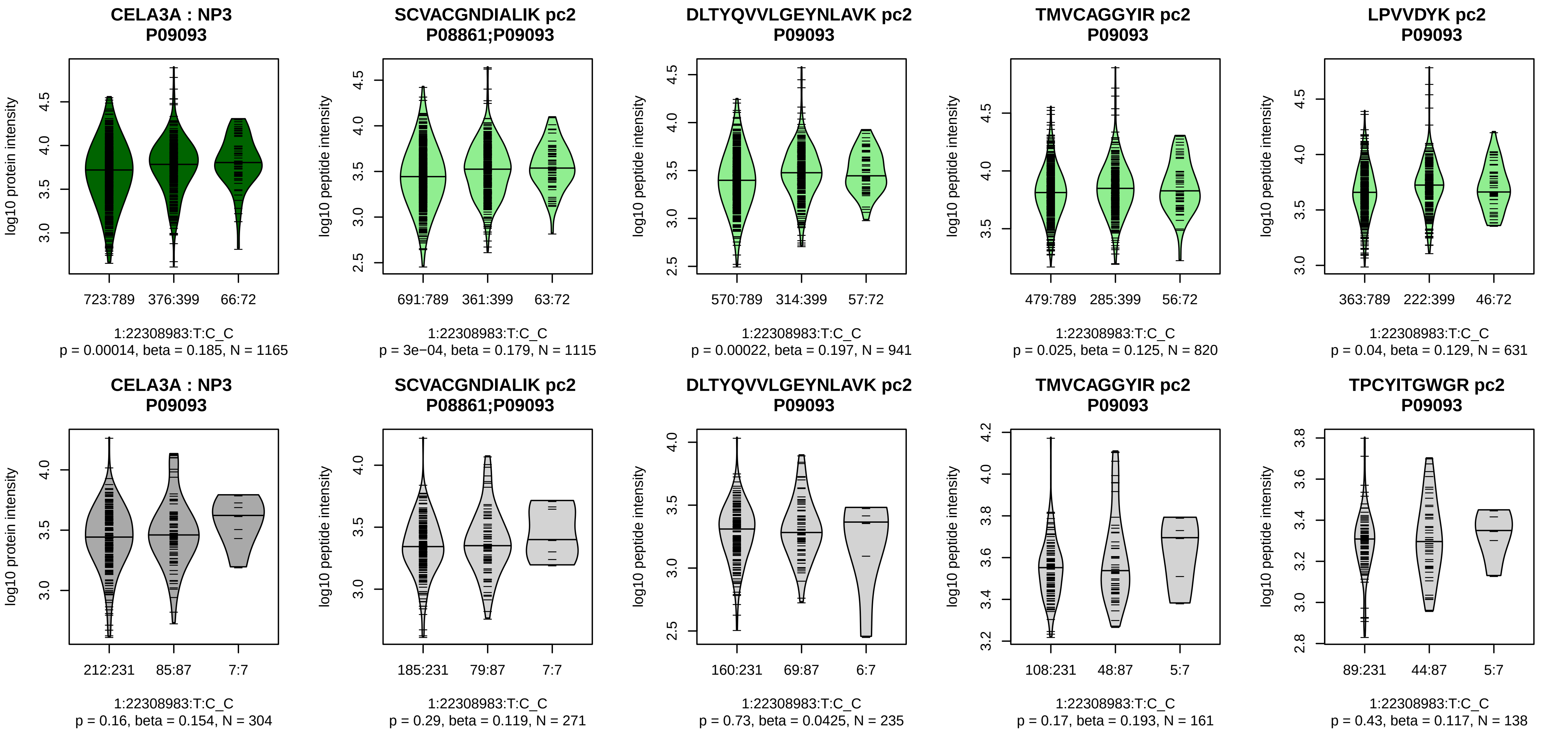


Assay Target: CELA3A
Olink UniProt: P09093
UKB rsID: rs34262568
Proxy rsID: rs34637160
UKB SNP: 1:22313371:G:A
Proxy SNP: 1:22308983:T:C
UKB log10(p): 233.1
UKB BETA: 0.231
*:*:−:−:−:−:−:−:−:−
1115:941:820:631:580:564:515:

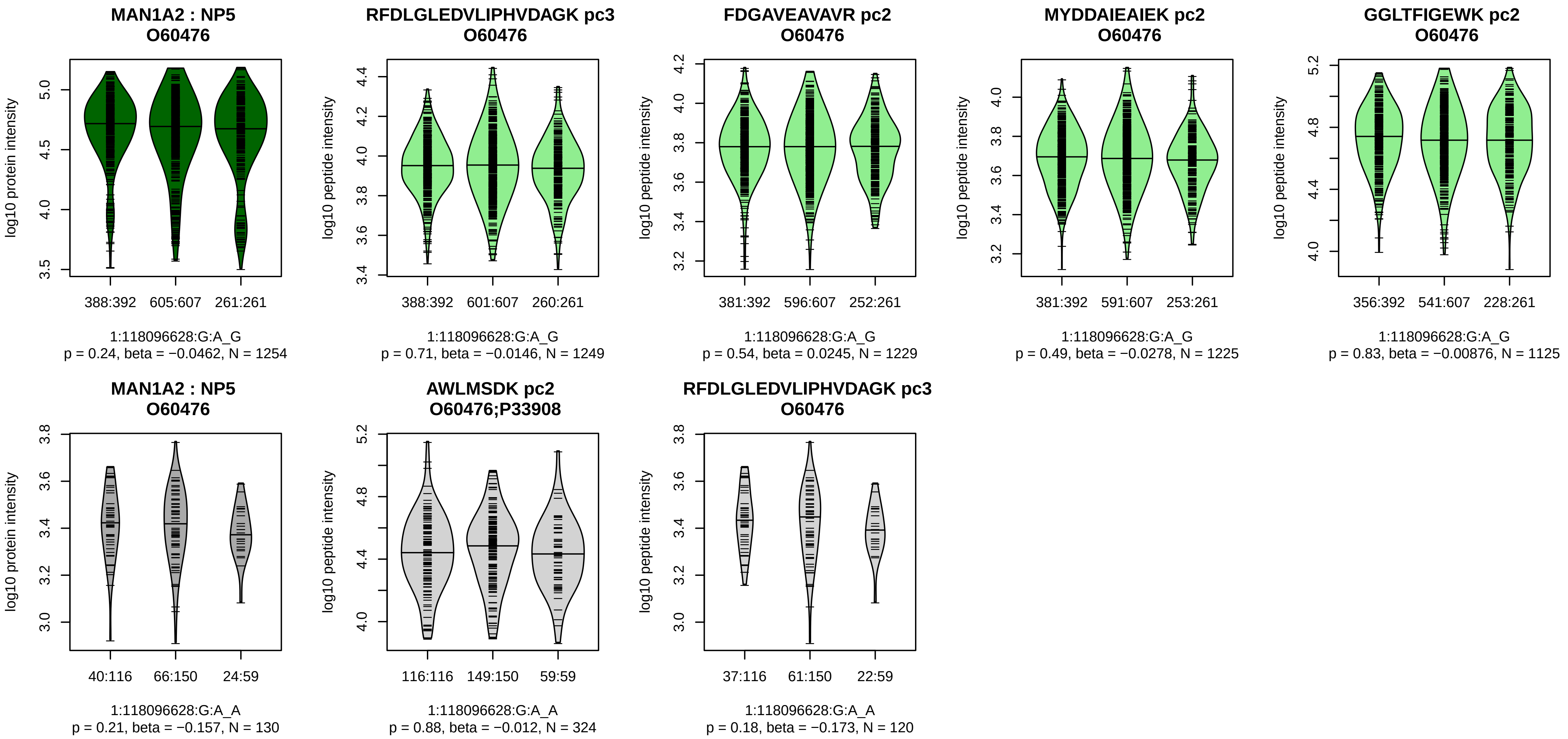


Assay Target: MAN1A2
Olink UniProt: O60476
UKB rsID: rs10923344
Proxy rsID: rs10923344
UKB SNP: 1:118096628:G:A
Proxy SNP: 1:118096628:G:A
UKB log10(p): 233.1
UKB BETA: 0.195
–:–:–:–:–:–:–:–:–:–:–
1249:1229:1225:1125:1124:998

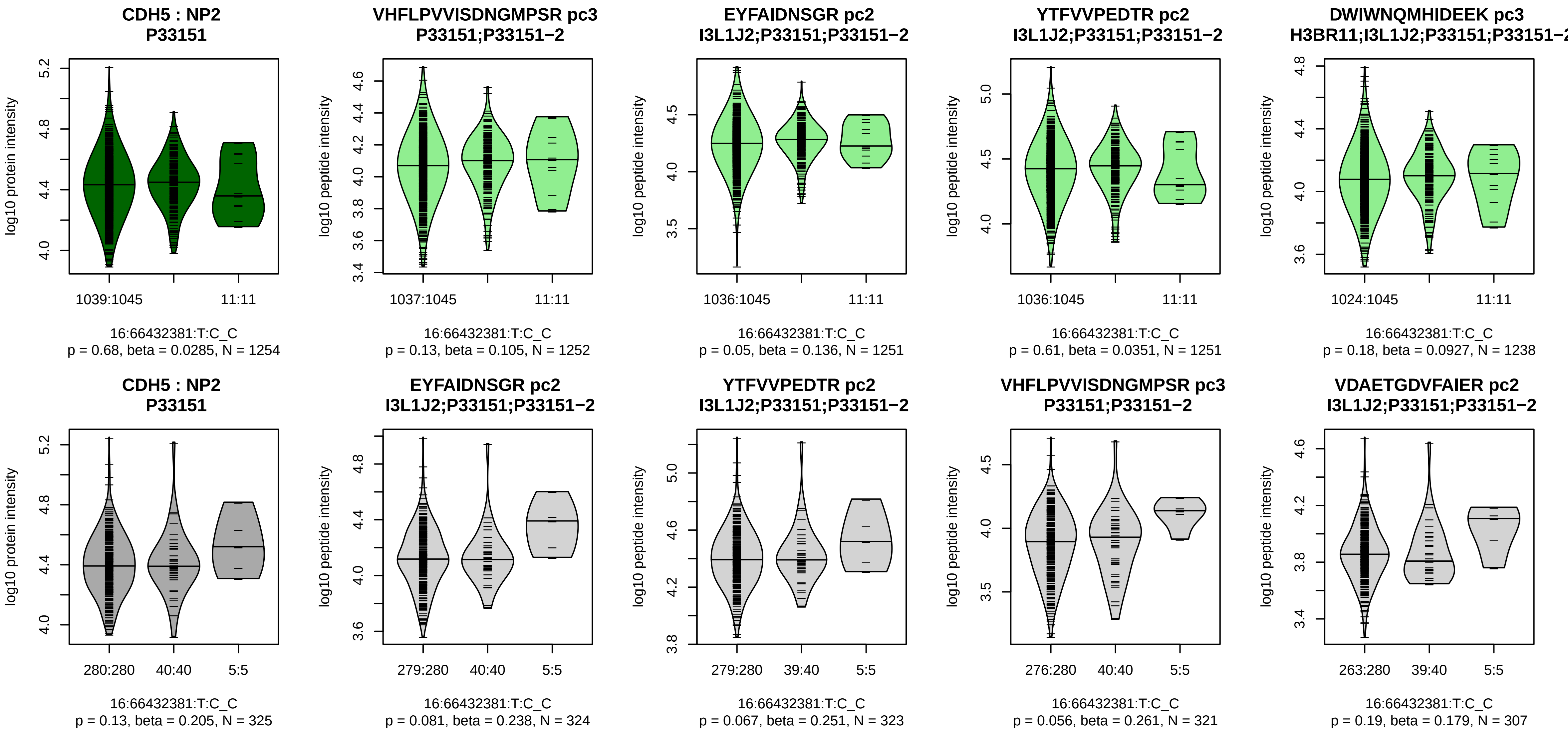


Assay Target: CDH5
Olink UniProt: P33151
UKB rsID: rs16956504
Proxy rsID: rs16956504
UKB SNP: 16:66432381:T:C
Proxy SNP: 16:66432381:T:C
UKB log10(p): 230.2
UKB BETA: 0.296
−:−:−:−:−:−:−:−:−:−:−:−:−:*:−:−:−
1252:1251:1251:1238:1238:122

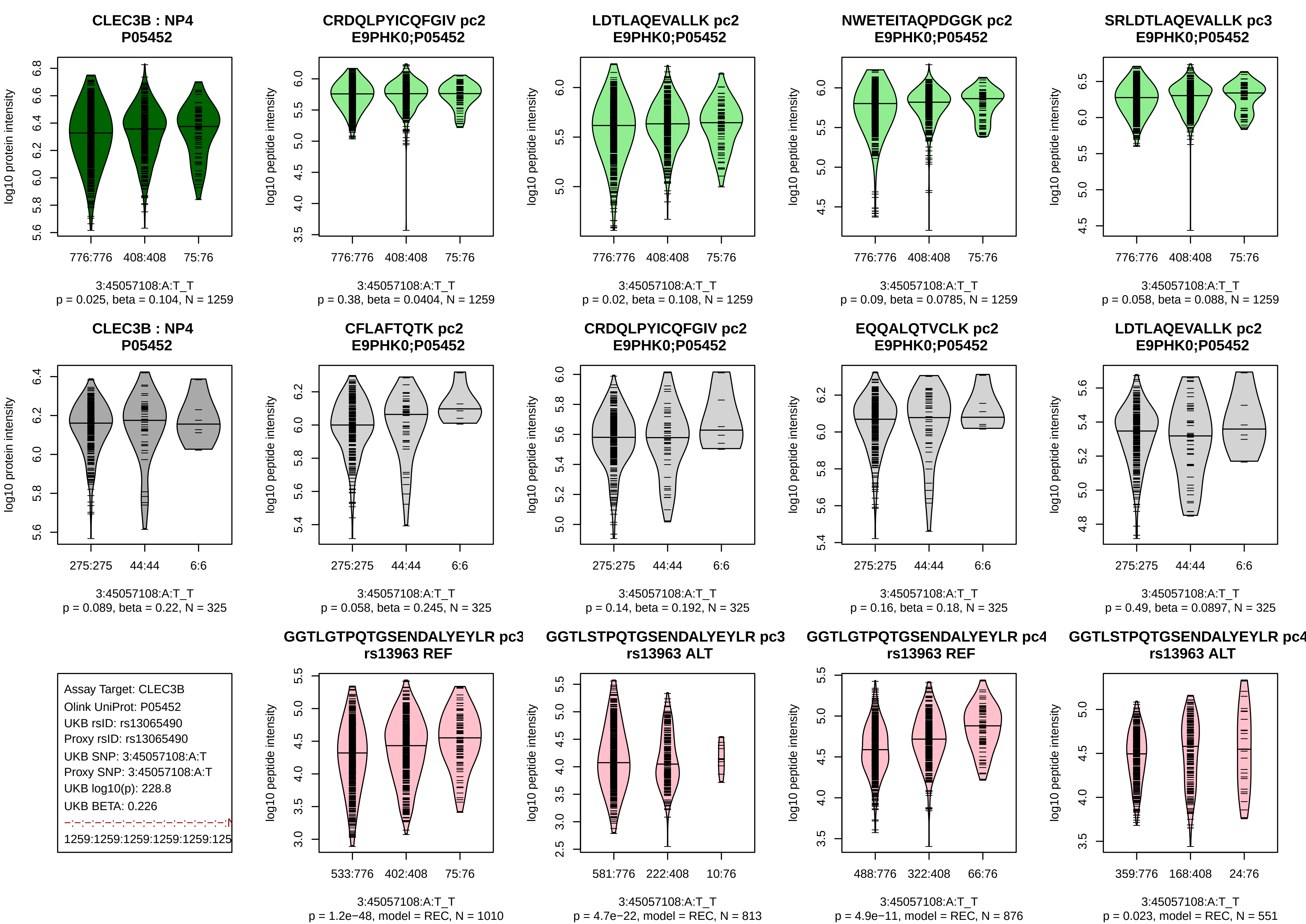
CLEC3B : NP4
P05452
log10 protein intensity
776:776
408:408
75:76
3:45057108:A:T_T
p = 0.025, beta = 0.104, N = 1259
CRDQLPYICQFGIV pc2
E9PHK0;P05452
log10 peptide intensity
776:776
408:408
75:76
3:45057108:A:T_T
p = 0.38, beta = 0.0404, N = 1259
LDTLAQEVALLK pc2
E9PHK0;P05452
log10 peptide intensity
776:776
408:408
75:76
3:45057108:A:T_T
p = 0.02, beta = 0.108, N = 1259
NWETEITAQPDGGK pc2
E9PHK0;P05452
log10 peptide intensity
776:776
408:408
75:76
3:45057108:A:T_T
p = 0.09, beta = 0.0785, N = 1259
SRLDTLAQEVALLK pc3
E9PHK0;P05452
log10 peptide intensity
776:776
408:408
75:76
3:45057108:A:T_T
p = 0.058, beta = 0.088, N = 1259
CLEC3B : NP4
P05452
log10 protein intensity
275:275
44:44
6:6
3:45057108:A:T_T
p = 0.089, beta = 0.22, N = 325
CFLAFTQTK pc2
E9PHK0;P05452
log10 peptide intensity
275:275
44:44
6:6
3:45057108:A:T_T
p = 0.058, beta = 0.245, N = 325
CRDQLPYICQFGIV pc2
E9PHK0;P05452
log10 peptide intensity
275:275
44:44
6:6
3:45057108:A:T_T
p = 0.14, beta = 0.192, N = 325
EQQALQTVCLK pc2
E9PHK0;P05452
log10 peptide intensity
275:275
44:44
6:6
3:45057108:A:T_T
p = 0.16, beta = 0.18, N = 325
LDTLAQEVALLK pc2
E9PHK0;P05452
log10 peptide intensity
275:275
44:44
6:6
3:45057108:A:T_T
p = 0.49, beta = 0.0897, N = 325
Assay Target: CLEC3B
Olink UniProt: P05452
UKB rsID: rs13065490
Proxy rsID: rs13065490
UKB SNP: 3:45057108:A:T
Proxy SNP: 3:45057108:A:T
UKB log10(p): 228.8
UKB BETA: 0.226
1259:1259:1259:1259:1259:125
GGTLGTPQTGSENDALYEYLR pc3
rs13963 REF
log10 peptide intensity
533:776
402:408
75:76
3:45057108:A:T_T
p = 1.2e−48, model = REC, N = 1010
GGTLSTPQTGSENDALYEYLR pc3
rs13963 ALT
log10 peptide intensity
581:776
222:408
10:76
3:45057108:A:T_T
p = 4.7e−22, model = REC, N = 813
GGTLGTPQTGSENDALYEYLR pc4
rs13963 REF
log10 peptide intensity
488:776
322:408
66:76
3:45057108:A:T_T
p = 4.9e−11, model = REC, N = 876
GGTLSTPQTGSENDALYEYLR pc4
rs13963 ALT
log10 peptide intensity
359:776
168:408
24:76
3:45057108:A:T_T
p = 0.023, model = REC, N = 551

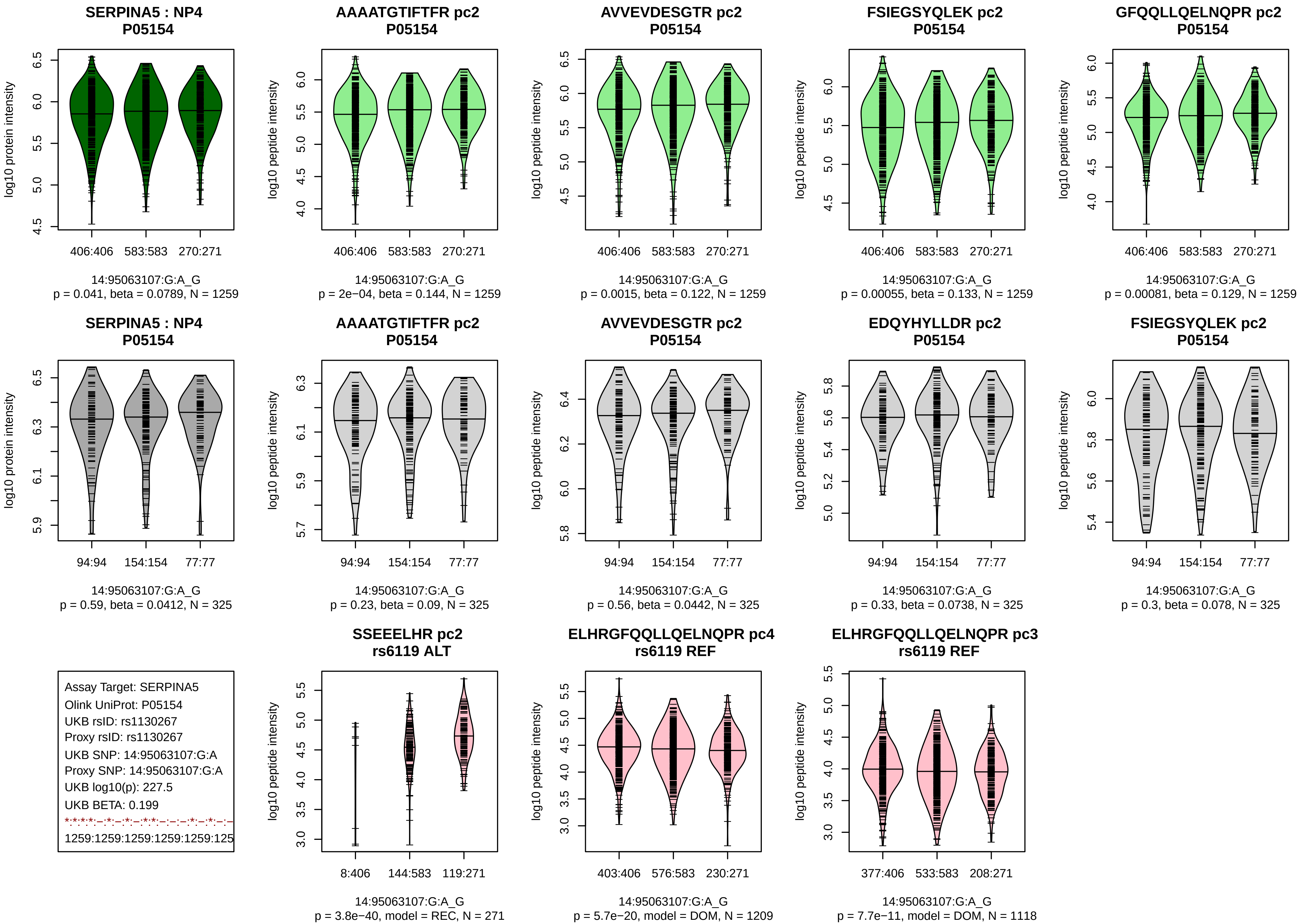

SERPINA5 : NP4
P05154
log10 protein intensity
406:406 583:583 270:271
14:95063107:G:A_G
p = 0.041, beta = 0.0789, N = 1259
AAAATGTIFTFR pc2
P05154
log10 peptide intensity
406:406 583:583 270:271
14:95063107:G:A_G
p = 2e−04, beta = 0.144, N = 1259
AVVEVDESGTR pc2
P05154
log10 peptide intensity
406:406 583:583 270:271
14:95063107:G:A_G
p = 0.0015, beta = 0.122, N = 1259
FSIEGSYQLEK pc2
P05154
log10 peptide intensity
406:406 583:583 270:271
14:95063107:G:A_G
p = 0.00055, beta = 0.133, N = 1259
GFQQLLQELNQPR pc2
P05154
log10 peptide intensity
406:406 583:583 270:271
14:95063107:G:A_G
p = 0.00081, beta = 0.129, N = 1259
SERPINA5 : NP4
P05154
log10 protein intensity
94:94 154:154 77:77
14:95063107:G:A_G
p = 0.59, beta = 0.0412, N = 325
AAAATGTIFTFR pc2
P05154
log10 peptide intensity
94:94 154:154 77:77
14:95063107:G:A_G
p = 0.23, beta = 0.09, N = 325
AVVEVDESGTR pc2
P05154
log10 peptide intensity
94:94 154:154 77:77
14:95063107:G:A_G
p = 0.56, beta = 0.0442, N = 325
EDQYHYLLDR pc2
P05154
log10 peptide intensity
94:94 154:154 77:77
14:95063107:G:A_G
p = 0.33, beta = 0.0738, N = 325
FSIEGSYQLEK pc2
P05154
log10 peptide intensity
94:94 154:154 77:77
14:95063107:G:A_G
p = 0.3, beta = 0.078, N = 325
Assay Target: SERPINA5
Olink UniProt: P05154
UKB rsID: rs1130267
Proxy rsID: rs1130267
UKB SNP: 14:95063107:G:A
Proxy SNP: 14:95063107:G:A
UKB log10(p): 227.5
UKB BETA: 0.199
1259:1259:1259:1259:1259:125
SSEEELHR pc2
rs6119 ALT
log10 peptide intensity
8:406 144:583 119:271
14:95063107:G:A_G
p = 3.8e−40, model = REC, N = 271
ELHRGFQQLLQELNQPR pc4
rs6119 REF
log10 peptide intensity
403:406 576:583 230:271
14:95063107:G:A_G
p = 5.7e−20, model = DOM, N = 1209
ELHRGFQQLLQELNQPR pc3
rs6119 REF
log10 peptide intensity
377:406 533:583 208:271
14:95063107:G:A_G
p = 7.7e−11, model = DOM, N = 1118

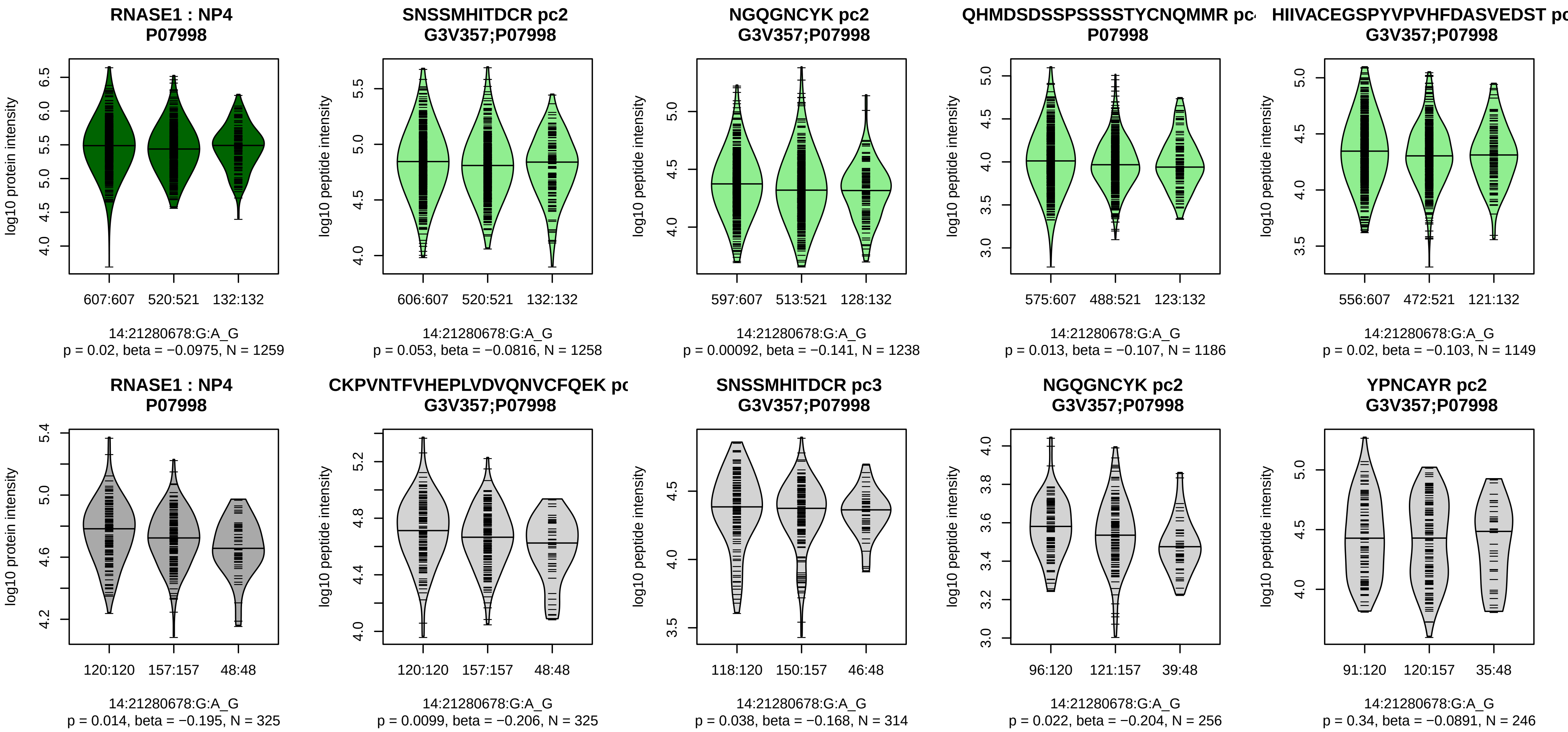


Assay Target: RNASE1
Olink UniProt: P07998
UKB rsID: rs17254387
Proxy rsID: rs17254387
UKB SNP: 14:21280678:G:A
Proxy SNP: 14:21280678:G:A
UKB log10(p): 226.5
UKB BETA: 0.206
−:−:*:−:−:−:−:−
1258:1256:1238:1186:1149:114

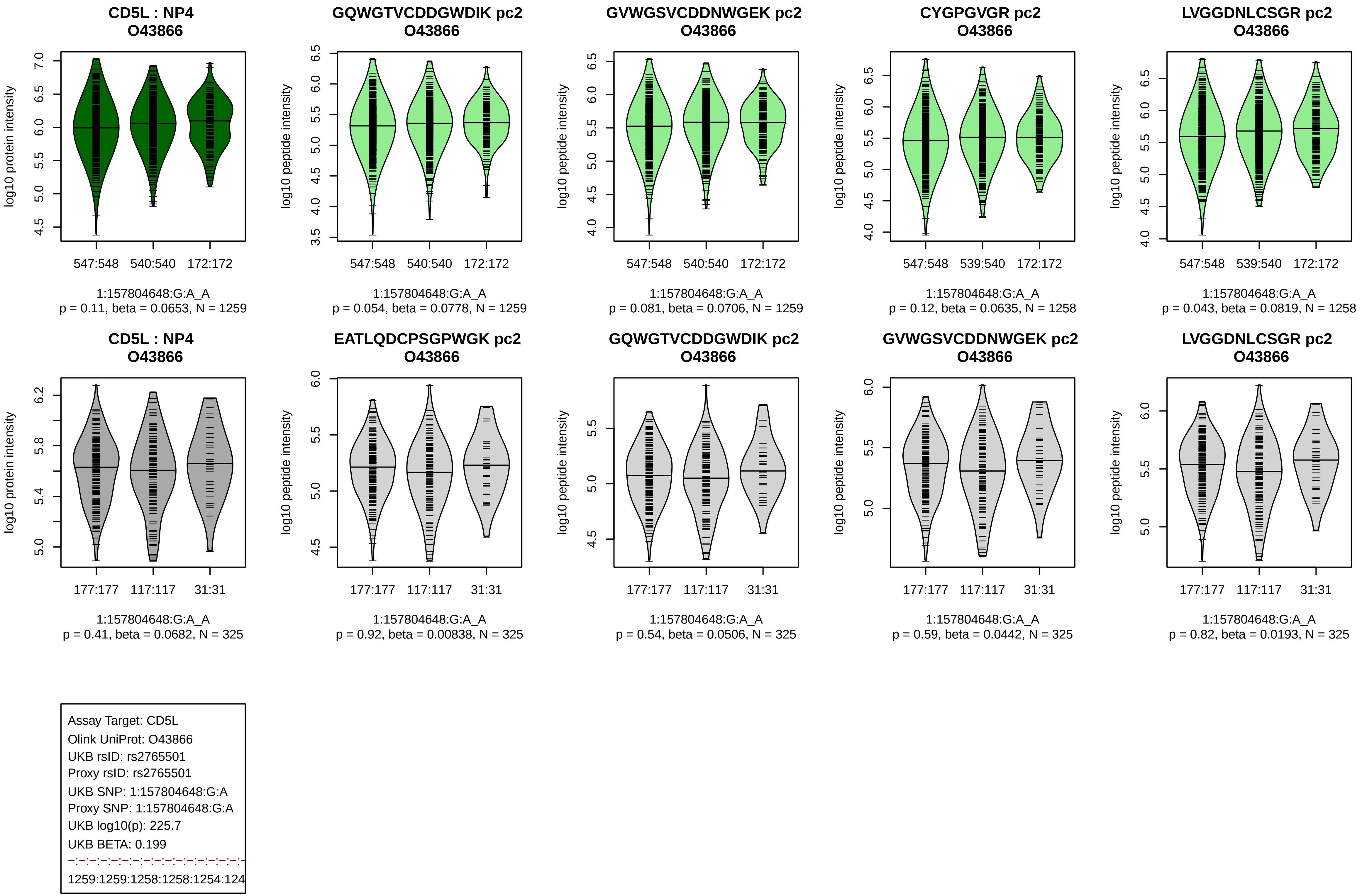
CD5L : NP4
O43866
log10 protein intensity
547:548 540:540 172:172
1:157804648:G:A_A
p = 0.11, beta = 0.0653, N = 1259
GQWGTVCDDGWDIK pc2
O43866
log10 peptide intensity
547:548 540:540 172:172
1:157804648:G:A_A
p = 0.054, beta = 0.0778, N = 1259
GVWGSVCDDNWGEK pc2
O43866
log10 peptide intensity
547:548 540:540 172:172
1:157804648:G:A_A
p = 0.081, beta = 0.0706, N = 1259
CYGPGVGR pc2
O43866
log10 peptide intensity
547:548 539:540 172:172
1:157804648:G:A_A
p = 0.12, beta = 0.0635, N = 1258
LVGGDNLCSGR pc2
O43866
log10 peptide intensity
547:548 539:540 172:172
1:157804648:G:A_A
p = 0.043, beta = 0.0819, N = 1258
CD5L : NP4
O43866
log10 protein intensity
177:177 117:117 31:31
1:157804648:G:A_A
p = 0.41, beta = 0.0682, N = 325
EATLQDCPSGPWGK pc2
O43866
log10 peptide intensity
177:177 117:117 31:31
1:157804648:G:A_A
p = 0.92, beta = 0.00838, N = 325
GQWGTVCDDGWDIK pc2
O43866
log10 peptide intensity
177:177 117:117 31:31
1:157804648:G:A_A
p = 0.54, beta = 0.0506, N = 325
GVWGSVCDDNWGEK pc2
O43866
log10 peptide intensity
177:177 117:117 31:31
1:157804648:G:A_A
p = 0.59, beta = 0.0442, N = 325
LVGGDNLCSGR pc2
O43866
log10 peptide intensity
177:177 117:117 31:31
1:157804648:G:A_A
p = 0.82, beta = 0.0193, N = 325
Assay Target: CD5L
Olink UniProt: O43866
UKB rsID: rs2765501
Proxy rsID: rs2765501
UKB SNP: 1:157804648:G:A
Proxy SNP: 1:157804648:G:A
UKB log10(p): 225.7
UKB BETA: 0.199
1259:1259:1258:1258:1254:124

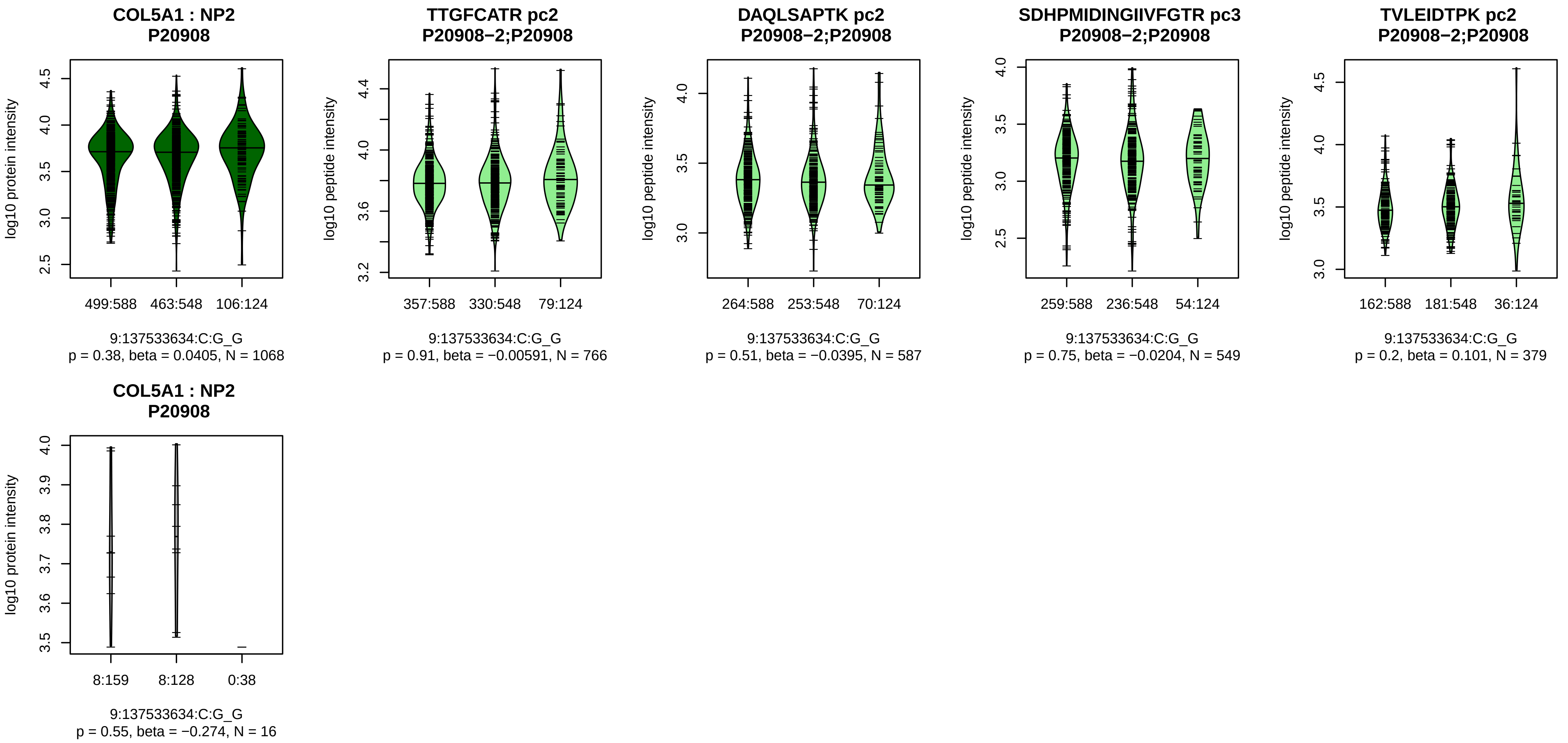


Assay Target: COL5A1
Olink UniProt: P20908
UKB rsID: rs12002679
Proxy rsID: rs12002679
UKB SNP: 9:137533634:C:G
Proxy SNP: 9:137533634:C:G
UKB log10(p): 223.2
UKB BETA: 0.221
−:−:−:−:−:−:−:−
766:587:549:379:237:136:119:4

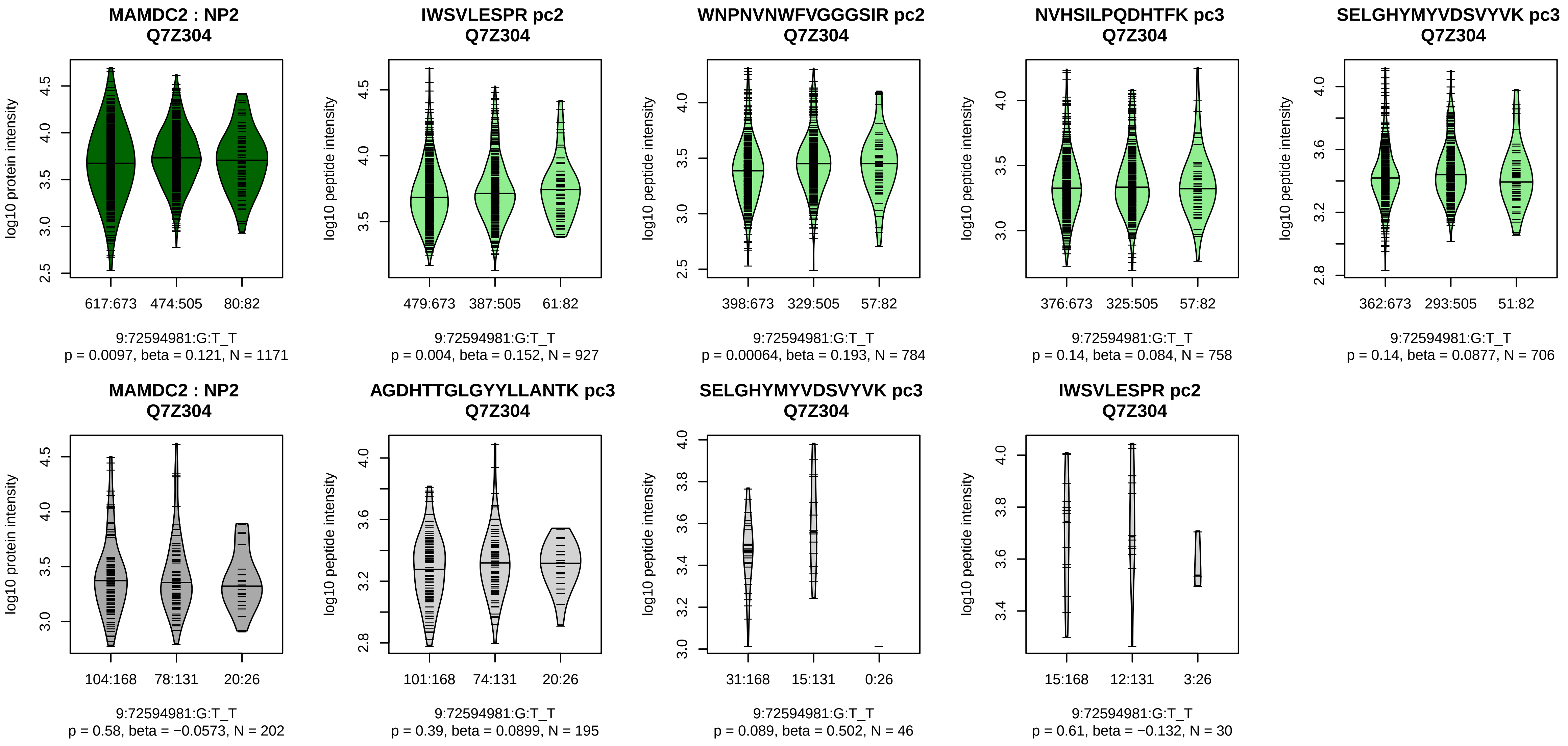


Assay Target: MAMDC2
Olink UniProt: Q7Z304
UKB rsID: rs12683555
Proxy rsID: rs12683555
UKB SNP: 9:72594981:G:T
Proxy SNP: 9:72594981:G:T
UKB log10(p): 222
UKB BETA: 0.213
*:*:–:–:–:–:–:–:–:–:–:–:–:–:–:–:–:
927:784:758:706:692:371:345:2

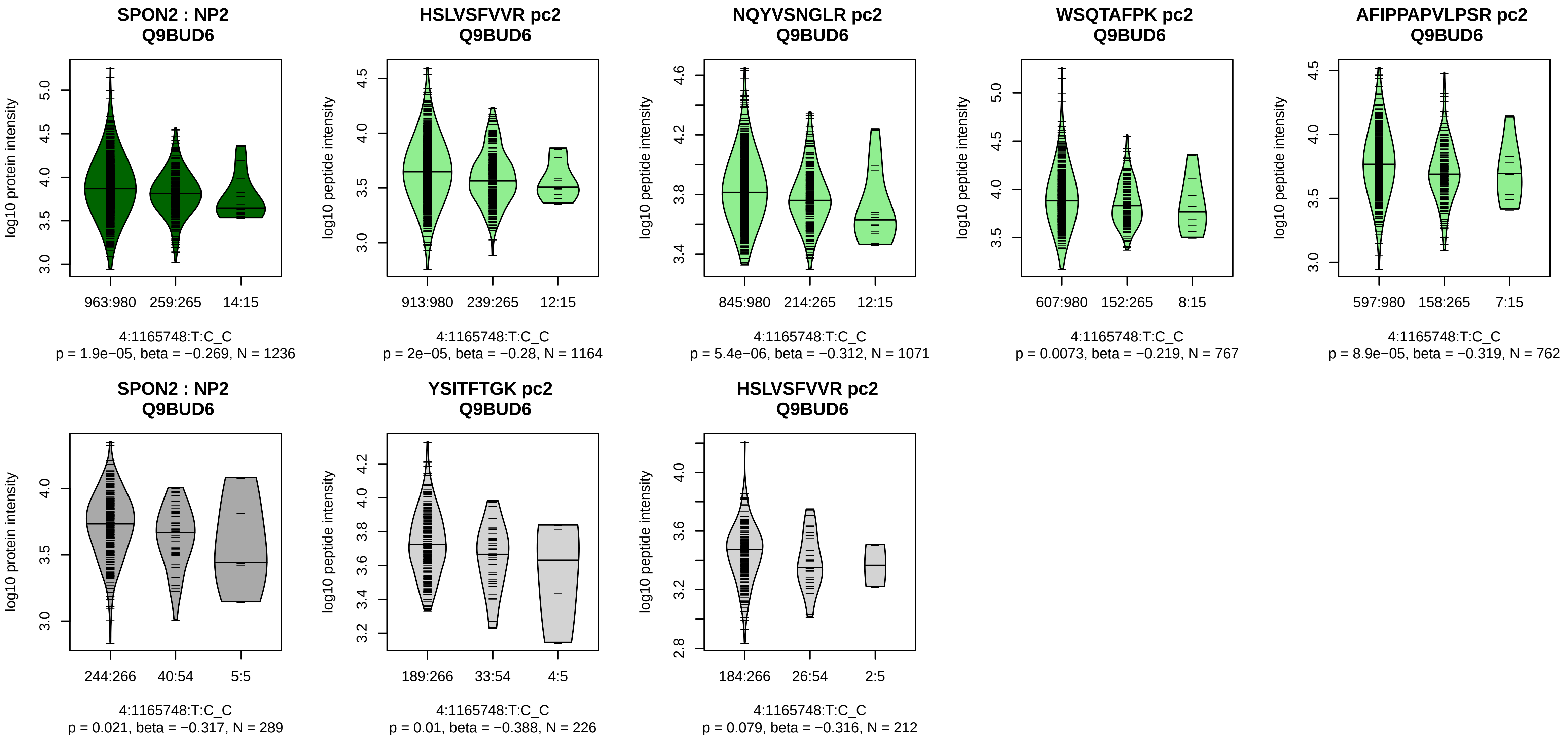


Assay Target: SPON2
Olink UniProt: Q9BUD6
UKB rsID: rs6836335
Proxy rsID: rs6836335
UKB SNP: 4:1165748:T:C
Proxy SNP: 4:1165748:T:C
UKB log10(p): 221.2
UKB BETA: 0.293
*:*:*:*:*:–
1164:1071:767:762:733:478

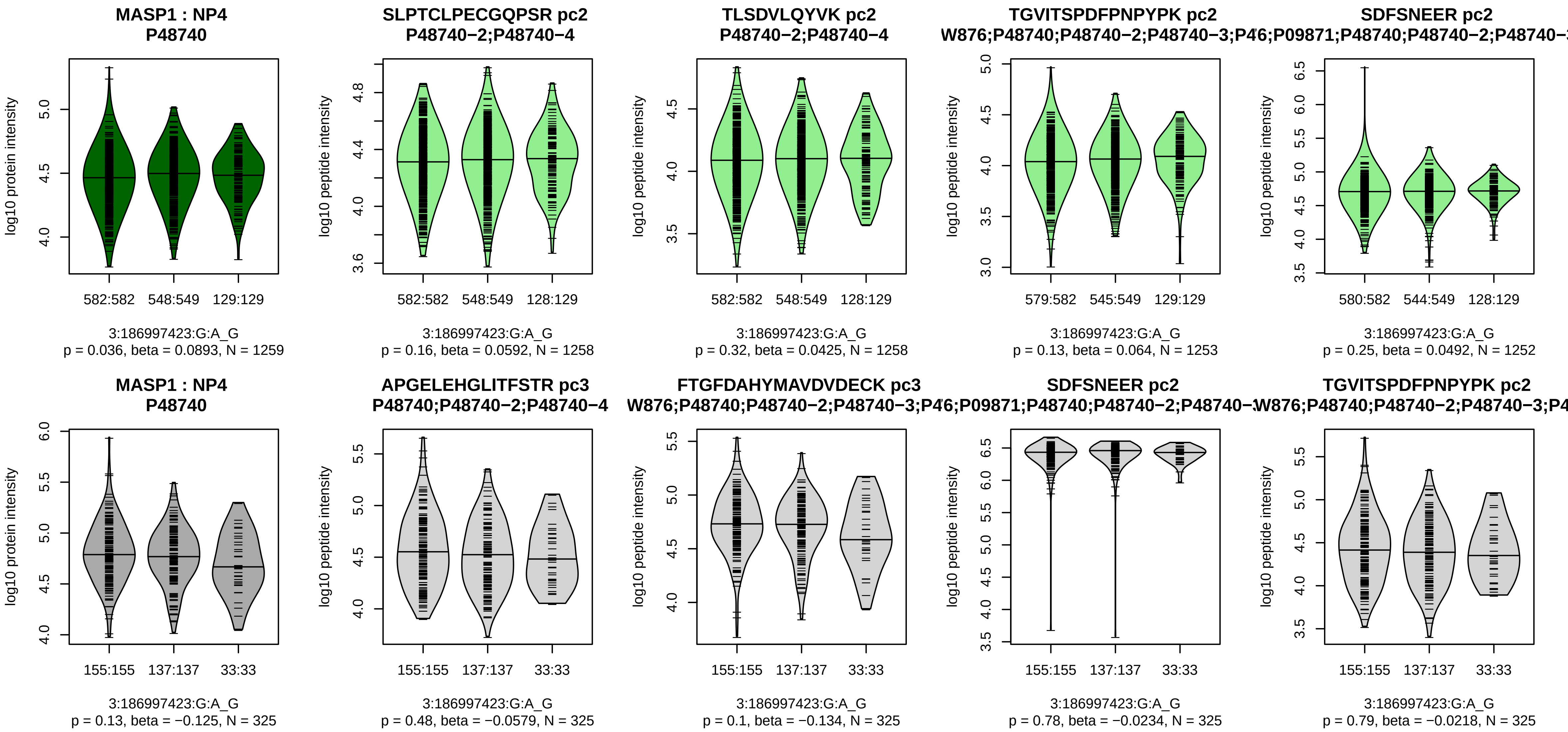


Assay Target: MASP1
Olink UniProt: P48740
UKB rsID: rs34053057
Proxy rsID: rs698085
UKB SNP: 3:186982263:T:TA
Proxy SNP: 3:186997423:G:A
UKB log10(p): 219.9
UKB BETA: −0.205
−:−:−:−:−:−:−:−:*:−:−:*:−:−:−:*:−:
1258:1258:1253:1252:1249:124

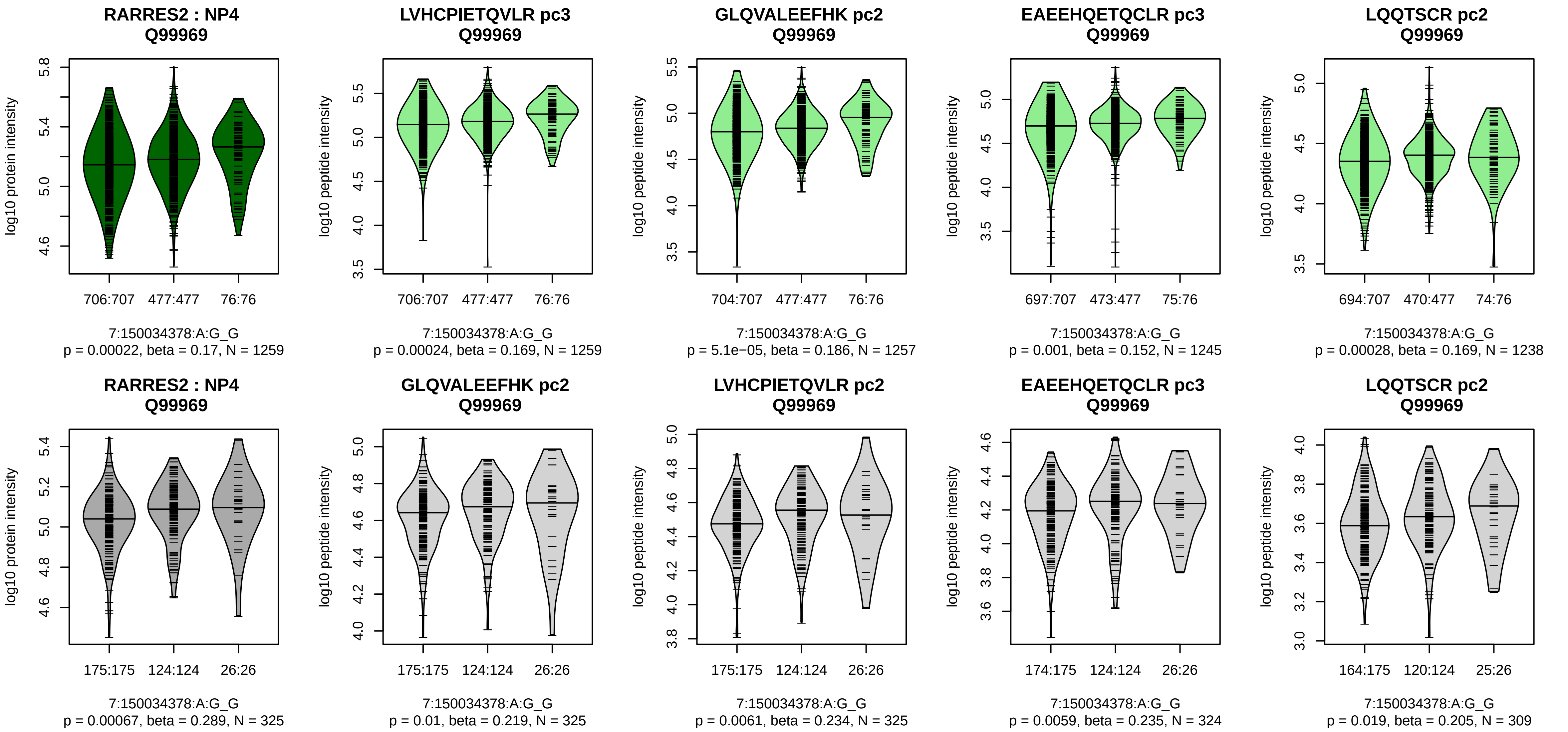


Assay Target: RARRES2
Olink UniProt: Q99969
UKB rsID: rs3735169
Proxy rsID: rs3735169
UKB SNP: 7:150034378:A:G
Proxy SNP: 7:150034378:A:G
UKB log10(p): 217.9
UKB BETA: 0.217
*:*:*:*:*:*:–:–:*:–
1259:1257:1257:1250:1245:123

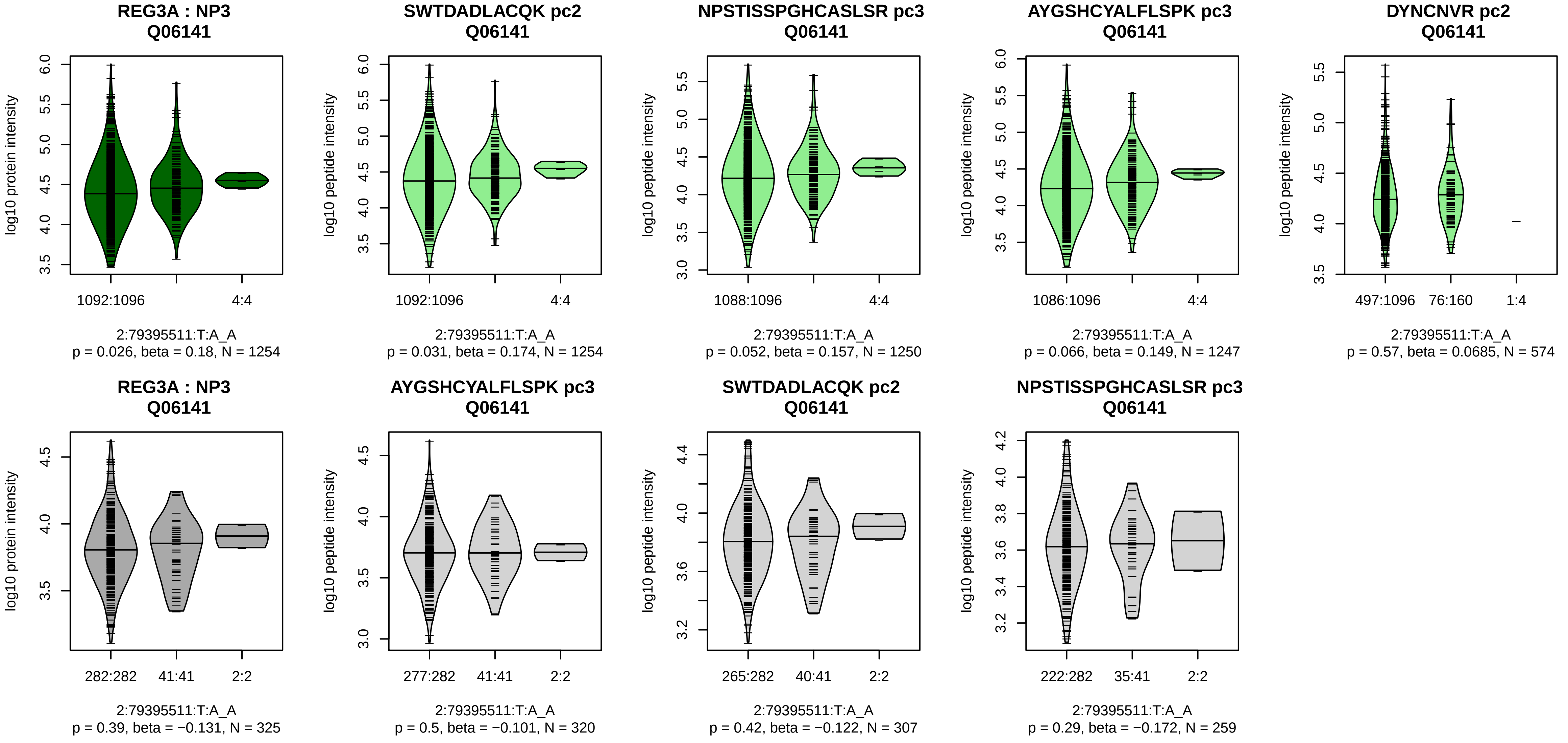


Assay Target: REG3A
Olink UniProt: Q06141
UKB rsID: rs147616162
Proxy rsID: rs147616162
UKB SNP: 2:79395511:T:A
Proxy SNP: 2:79395511:T:A
UKB log10(p): 215.7
UKB BETA: 0.396
–:–:–:–:–:–:–
1254:1250:1247:694:574:324:17

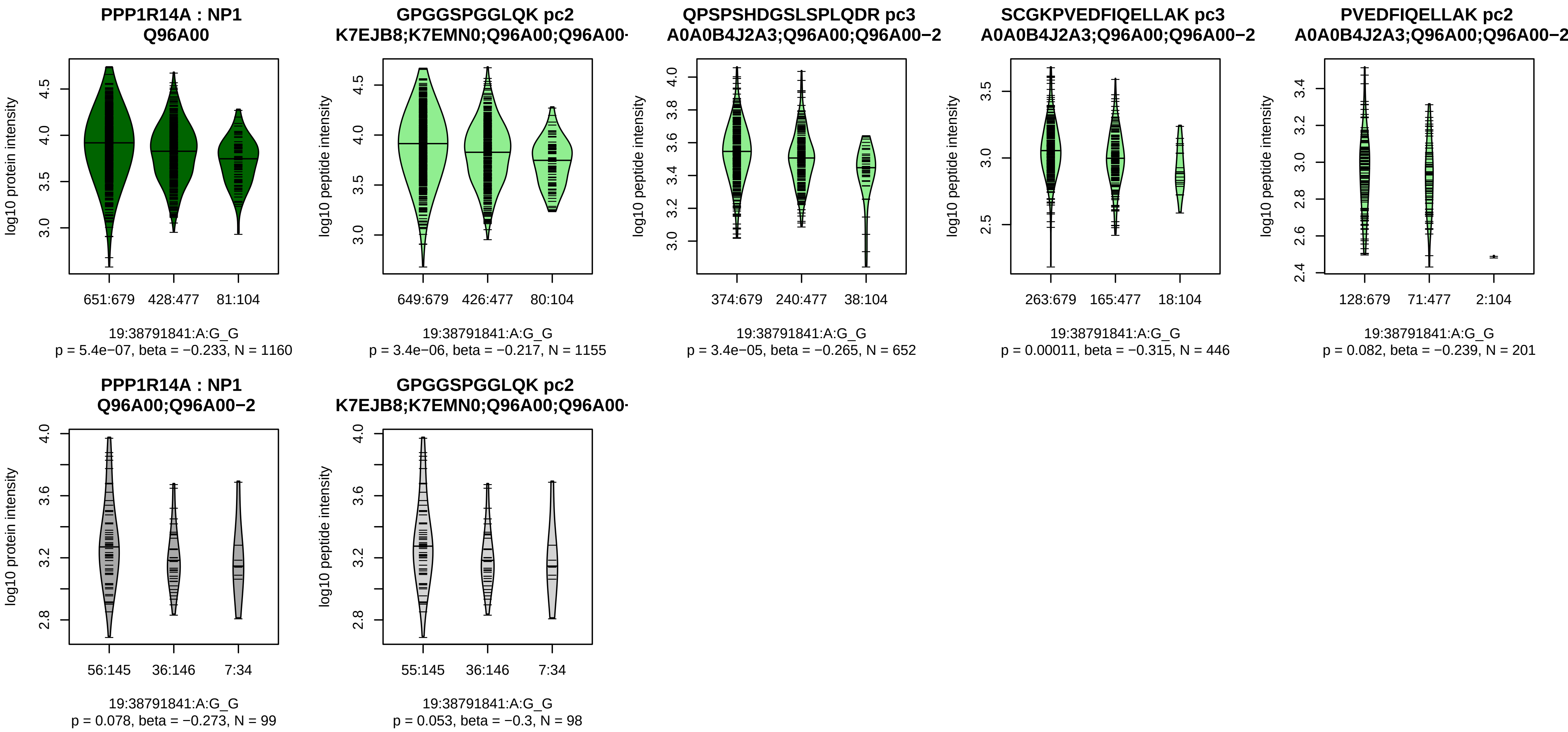


Assay Target: PPP1R14A
Olink UniProt: Q96A00
UKB rsID: rs71354995
Proxy rsID: rs71354995
UKB SNP: 19:38791841:A:G
Proxy SNP: 19:38791841:A:G
UKB log10(p): 215
UKB BETA: −0.221
*:*:*:–:–
1155:652:446:201:90

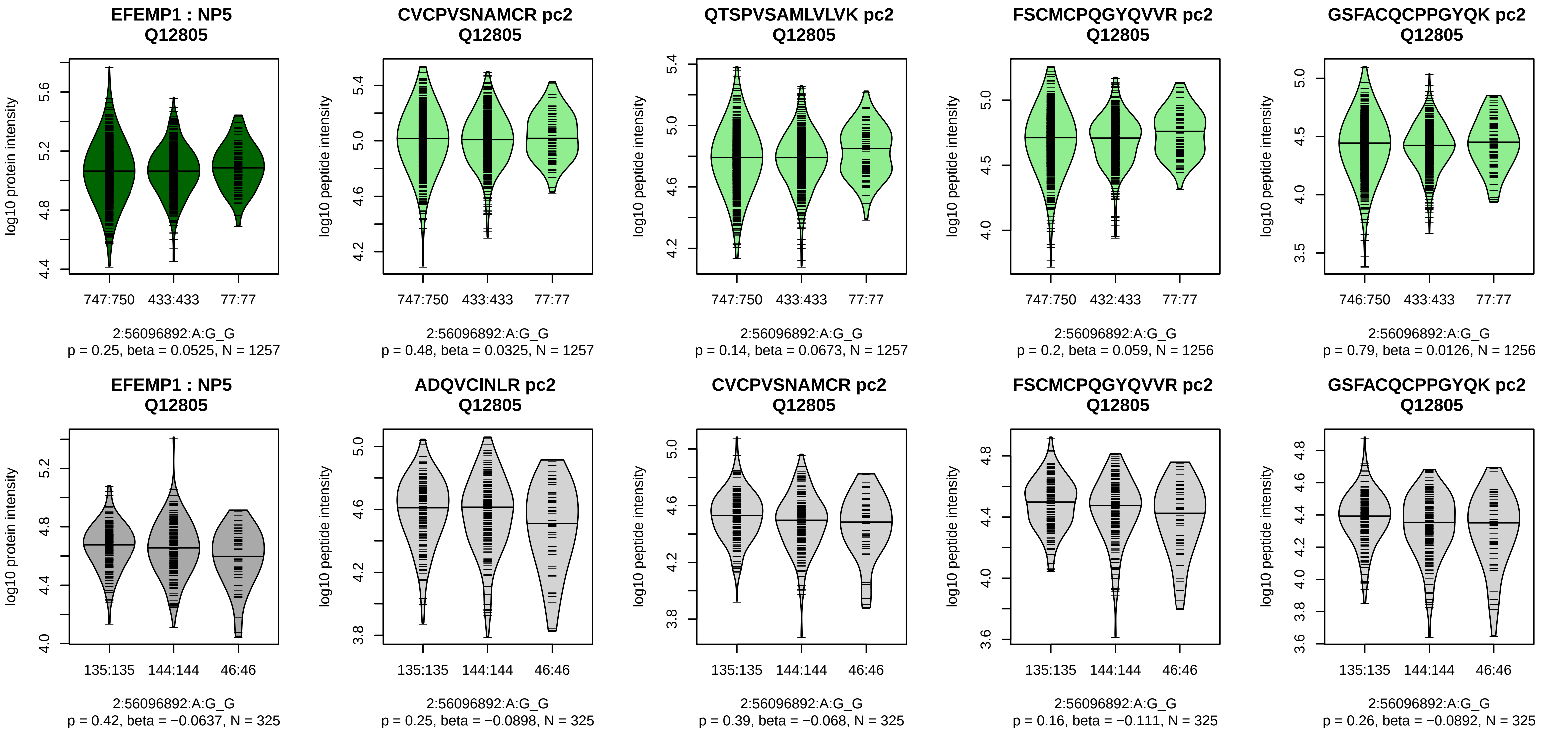


Assay Target: EFEMP1
Olink UniProt: Q12805
UKB rsID: rs3791679
Proxy rsID: rs3791679
UKB SNP: 2:56096892:A:G
Proxy SNP: 2:56096892:A:G
UKB log10(p): 208.1
UKB BETA: 0.205
–:–:–:–:–:–:–:–:–:–:–:–:–:–:–:–:–
1257:1257:1256:1256:1256:125

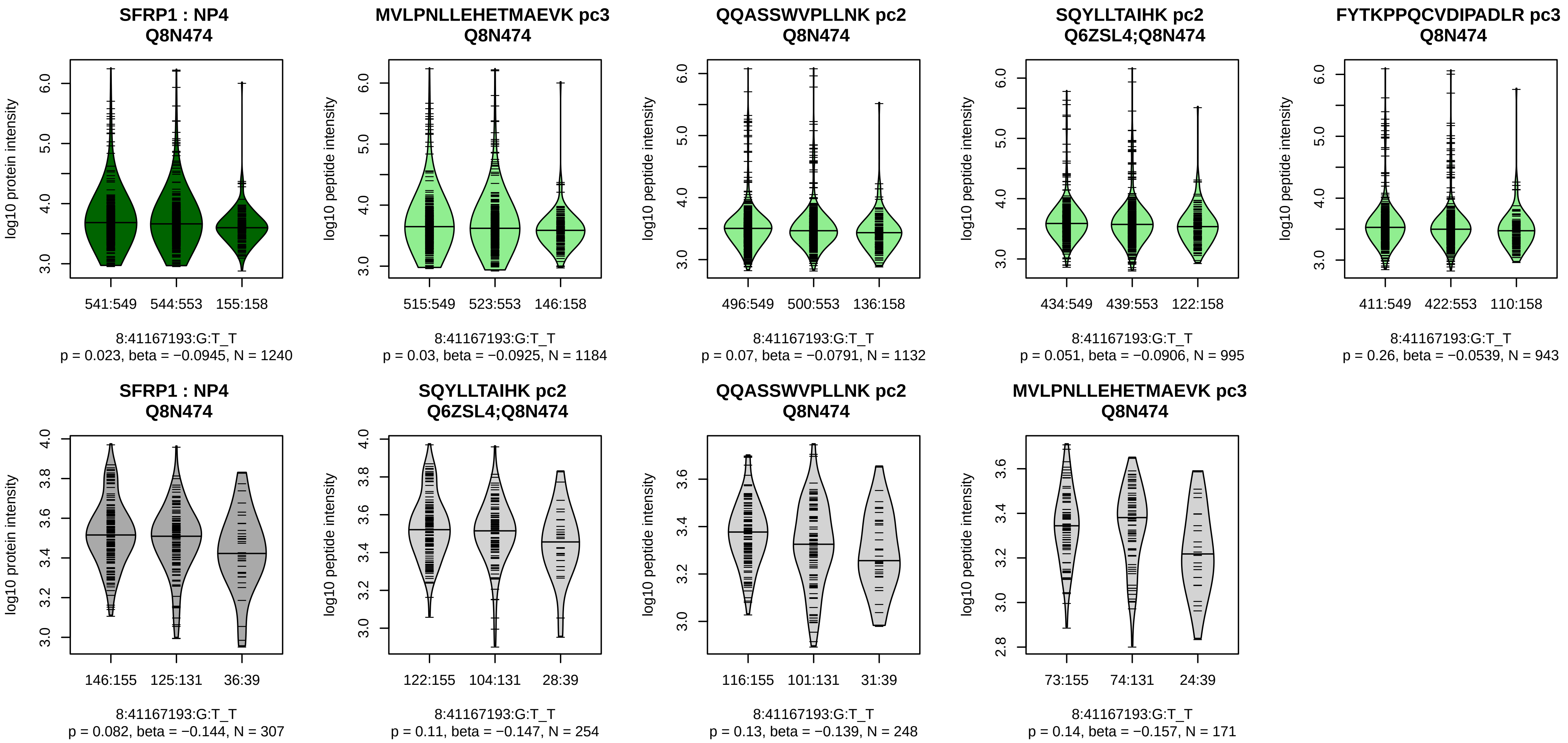


Assay Target: SFRP1
Olink UniProt: Q8N474
UKB rsID: rs72643819
Proxy rsID: rs72643819
UKB SNP: 8:41167193:G:T
Proxy SNP: 8:41167193:G:T
UKB log10(p): 206.1
UKB BETA: −0.188
−:−:−:−:−:−:−:−:−:−:−:−:NA:NA
1184:1132:995:943:637:260:82:

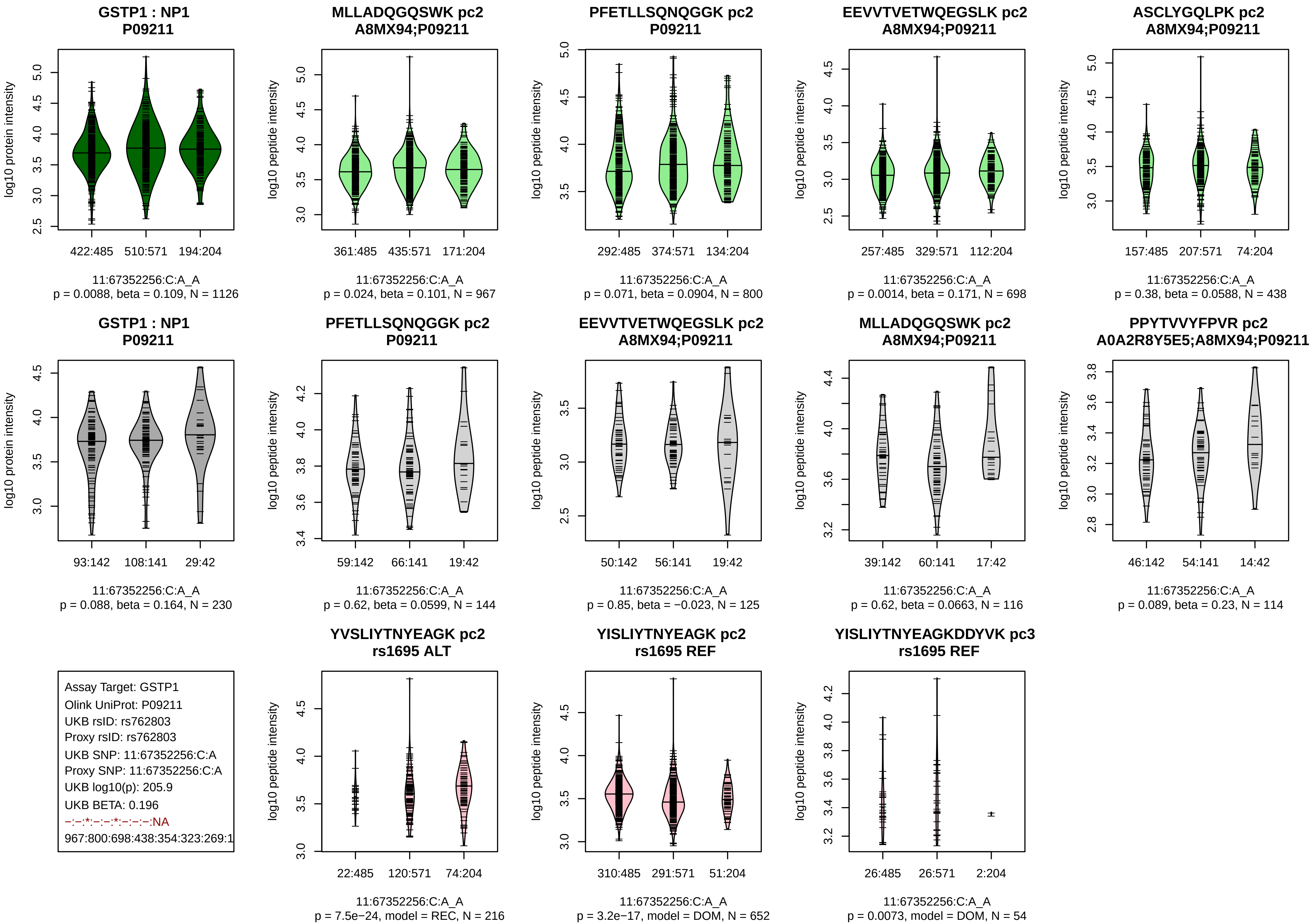
GSTP1 : NP1
P09211
log10 protein intensity
422:485
510:571
194:204
11:67352256:C:A_A
p = 0.0088, beta = 0.109, N = 1126
MLLADQGQSWK pc2
A8MX94;P09211
log10 peptide intensity
361:485
435:571
171:204
11:67352256:C:A_A
p = 0.024, beta = 0.101, N = 967
PFETLLSQNQGGK pc2
P09211
292:485
374:571
134:204
11:67352256:C:A_A
p = 0.071, beta = 0.0904, N = 800
EEVVTVETWQEGSLK pc2
A8MX94;P09211
257:485
329:571
112:204
11:67352256:C:A_A
p = 0.0014, beta = 0.171, N = 698
ASCLYGQLPK pc2
A8MX94;P09211
157:485
207:571
74:204
11:67352256:C:A_A
p = 0.38, beta = 0.0588, N = 438
GSTP1 : NP1
P09211
93:142
108:141
29:42
11:67352256:C:A_A
p = 0.088, beta = 0.164, N = 230
PFETLLSQNQGGK pc2
P09211
59:142
66:141
19:42
11:67352256:C:A_A
p = 0.62, beta = 0.0599, N = 144
EEVVTVETWQEGSLK pc2
A8MX94;P09211
50:142
56:141
19:42
11:67352256:C:A_A
p = 0.85, beta = −0.023, N = 125
MLLADQGQSWK pc2
A8MX94;P09211
39:142
60:141
17:42
11:67352256:C:A_A
p = 0.62, beta = 0.0663, N = 116
PPYTVVYFPVR pc2
A0A2R8Y5E5;A8MX94;P09211
46:142
54:141
14:42
11:67352256:C:A_A
p = 0.089, beta = 0.23, N = 114
Assay Target: GSTP1
Olink UniProt: P09211
UKB rsID: rs762803
Proxy rsID: rs762803
UKB SNP: 11:67352256:C:A
Proxy SNP: 11:67352256:C:A
UKB log10(p): 205.9
UKB BETA: 0.196
−:−:*:−:−:*:−:−:−:NA
967:800:698:438:354:323:269:1
YVSLIYTNYEAGK pc2
rs1695 ALT
22:485
120:571
74:204
11:67352256:C:A_A
p = 7.5e−24, model = REC, N = 216
YISLIYTNYEAGK pc2
rs1695 REF
310:485
291:571
51:204
11:67352256:C:A_A
p = 3.2e−17, model = DOM, N = 652
YISLIYTNYEAGKDDYVK pc3
rs1695 REF
26:485
26:571
2:204
11:67352256:C:A_A
p = 0.0073, model = DOM, N = 54

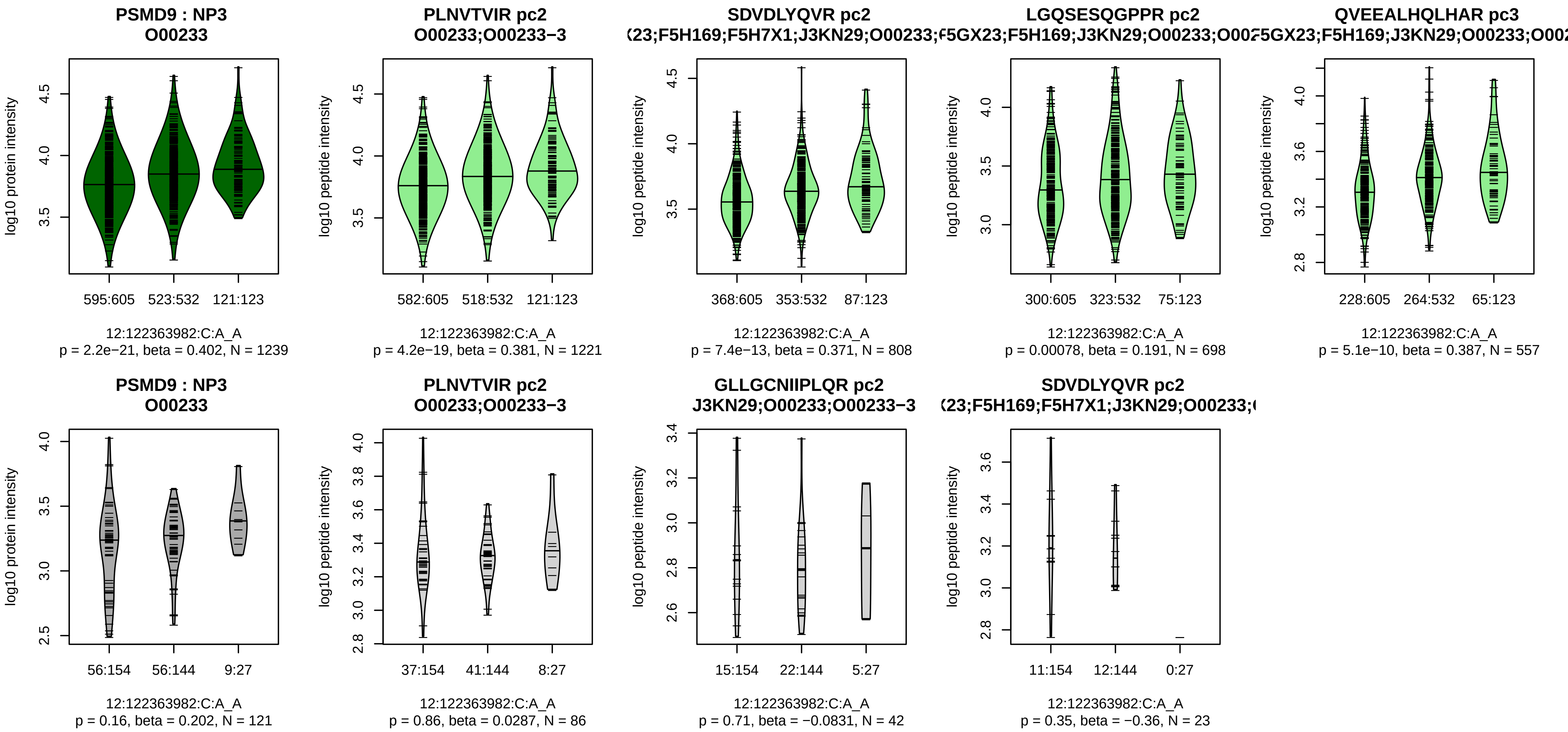


Assay Target: PSMD9
Olink UniProt: O00233
UKB rsID: rs1720066
Proxy rsID: rs1720066
UKB SNP: 12:122363982:C:A
Proxy SNP: 12:122363982:C:A
UKB log10(p): 203.8
UKB BETA: −0.199
*:*:*:*:*:−:*:−:NA:NA:NA
1221:808:698:557:483:419:371:

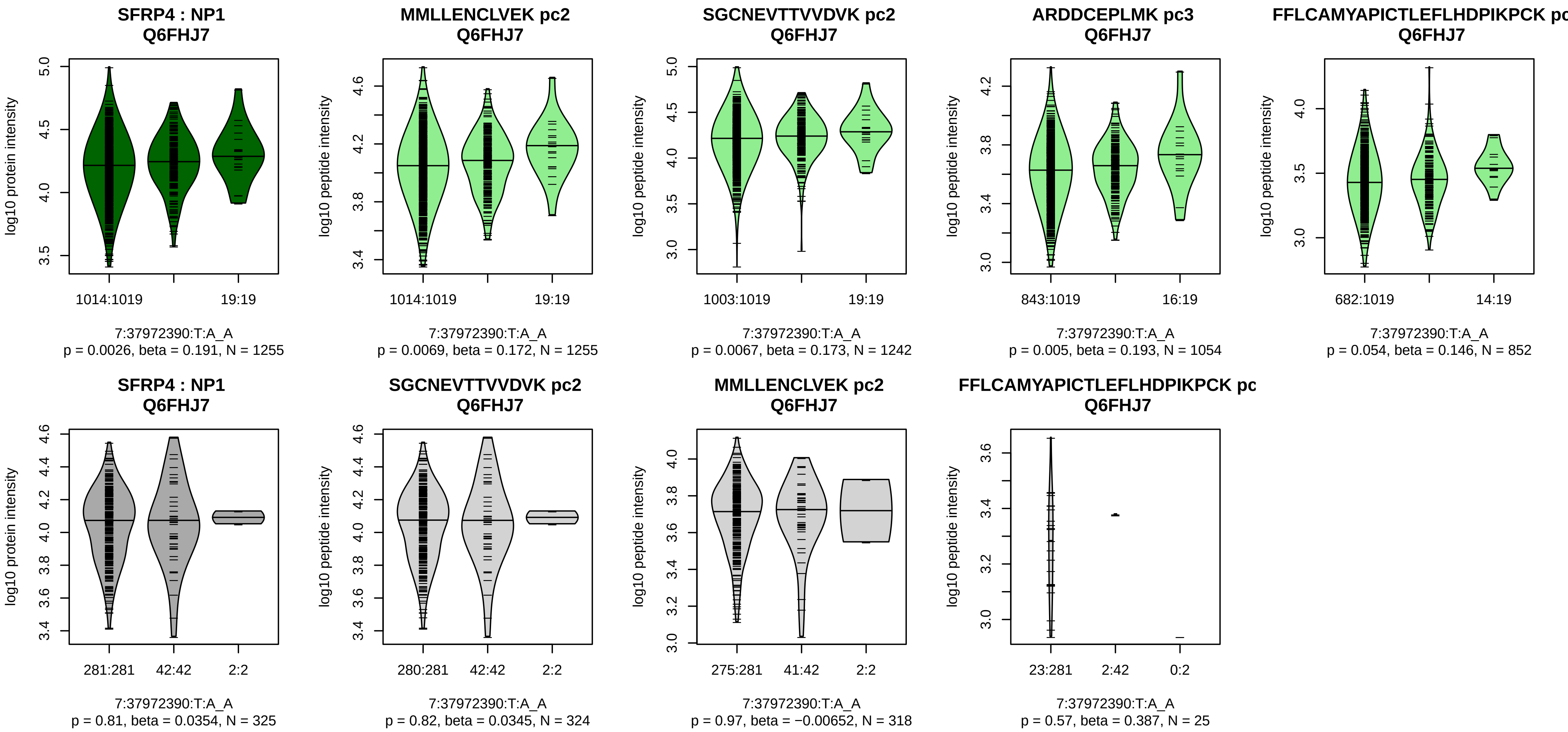


Assay Target: SFRP4
Olink UniProt: Q6FHJ7
UKB rsID: rs75207237
Proxy rsID: rs75207237
UKB SNP: 7:37972390:T:A
Proxy SNP: 7:37972390:T:A
UKB log10(p): 201
UKB BETA: 0.28
*:*:*:−:−:−:NA
1255:1242:1054:852:662:103:1[illegible]

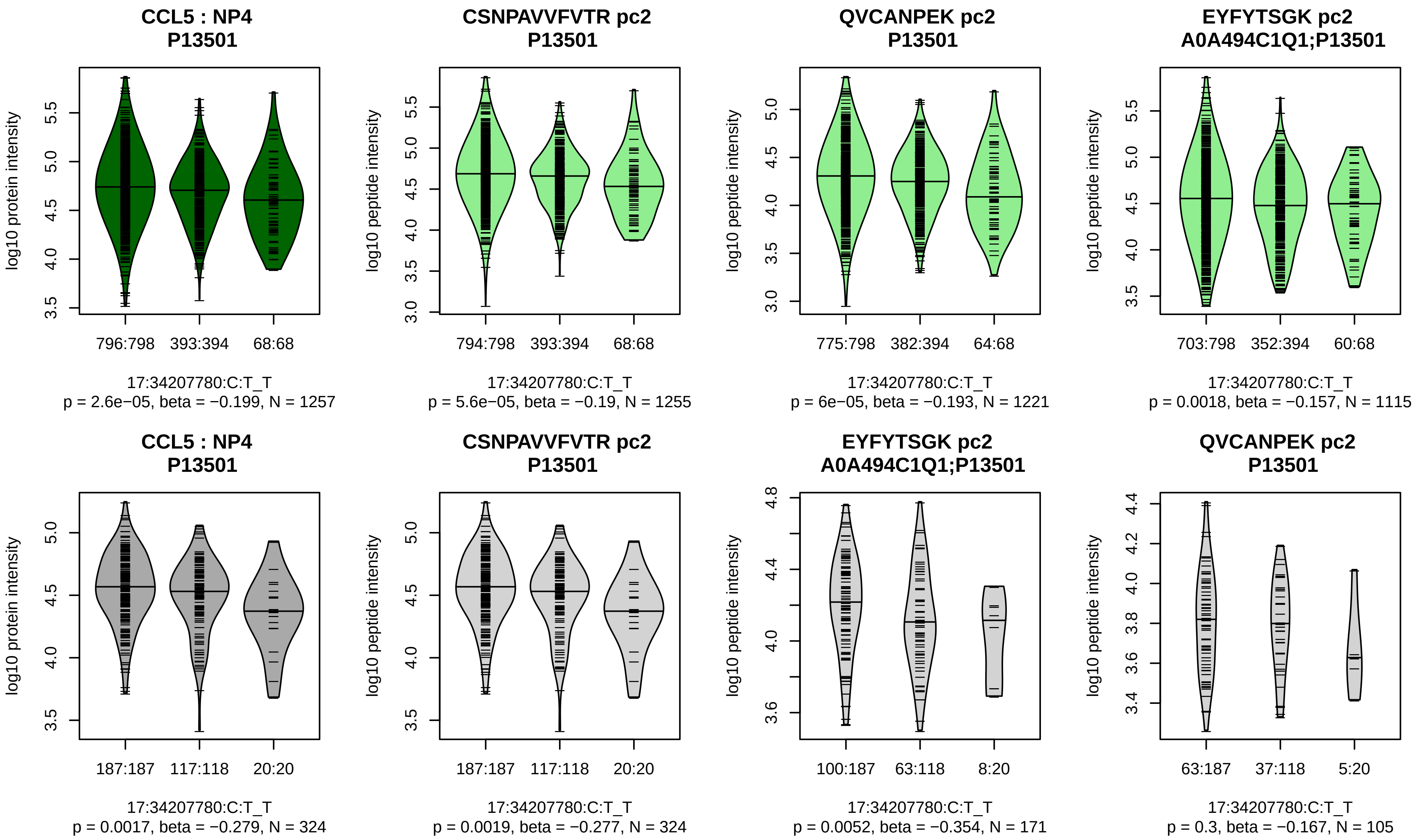


Assay Target: CCL5
Olink UniProt: P13501
UKB rsID: rs2107538
Proxy rsID: rs2107538
UKB SNP: 17:34207780:C:T
Proxy SNP: 17:34207780:C:T
UKB log10(p): 196.7
UKB BETA: −0.231
*:*:*
1255:1221:1115

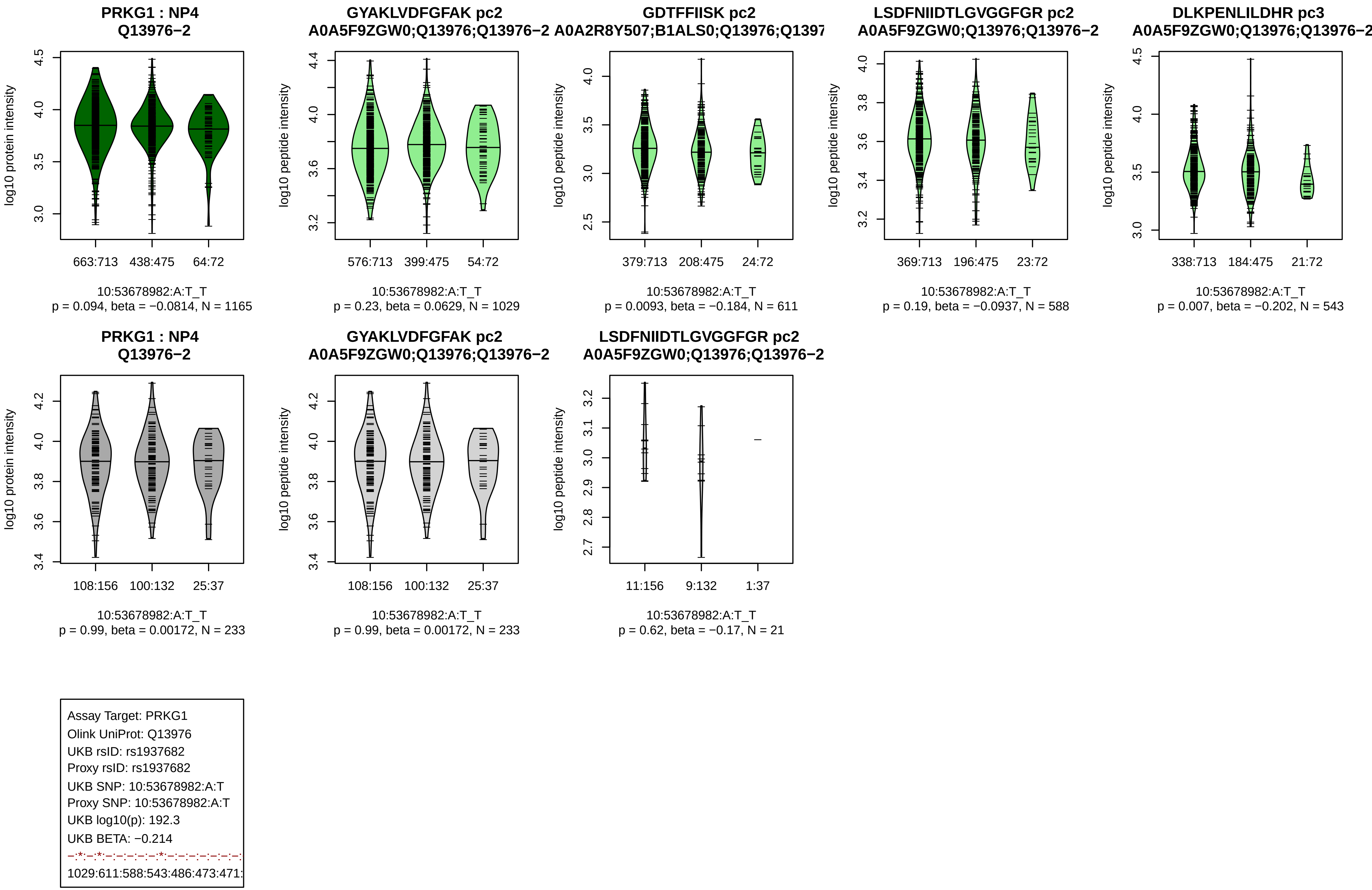

PRKG1 : NP4
Q13976−2
log10 protein intensity
663:713
438:475
64:72
10:53678982:A:T_T
p = 0.094, beta = −0.0814, N = 1165
GYAKLVDFGFAK pc2
A0A5F9ZGW0;Q13976;Q13976−2
log10 peptide intensity
576:713
399:475
54:72
10:53678982:A:T_T
p = 0.23, beta = 0.0629, N = 1029
GDTFFIISK pc2
A0A2R8Y507;B1ALS0;Q13976;Q1397
log10 peptide intensity
379:713
208:475
24:72
10:53678982:A:T_T
p = 0.0093, beta = −0.184, N = 611
LSDFNIIDTLGVGGFGR pc2
A0A5F9ZGW0;Q13976;Q13976−2
log10 peptide intensity
369:713
196:475
23:72
10:53678982:A:T_T
p = 0.19, beta = −0.0937, N = 588
DLKPENLILDHR pc3
A0A5F9ZGW0;Q13976;Q13976−2
log10 peptide intensity
338:713
184:475
21:72
10:53678982:A:T_T
p = 0.007, beta = −0.202, N = 543
PRKG1 : NP4
Q13976−2
log10 protein intensity
108:156
100:132
25:37
10:53678982:A:T_T
p = 0.99, beta = 0.00172, N = 233
GYAKLVDFGFAK pc2
A0A5F9ZGW0;Q13976;Q13976−2
log10 peptide intensity
108:156
100:132
25:37
10:53678982:A:T_T
p = 0.99, beta = 0.00172, N = 233
LSDFNIIDTLGVGGFGR pc2
A0A5F9ZGW0;Q13976;Q13976−2
log10 peptide intensity
11:156
9:132
1:37
10:53678982:A:T_T
p = 0.62, beta = −0.17, N = 21
Assay Target: PRKG1
Olink UniProt: Q13976
UKB rsID: rs1937682
Proxy rsID: rs1937682
UKB SNP: 10:53678982:A:T
Proxy SNP: 10:53678982:A:T
UKB log10(p): 192.3
UKB BETA: −0.214
−:*:−:*:−:−:−:−:*:−:−:−:−:−:−:
1029:611:588:543:486:473:471:

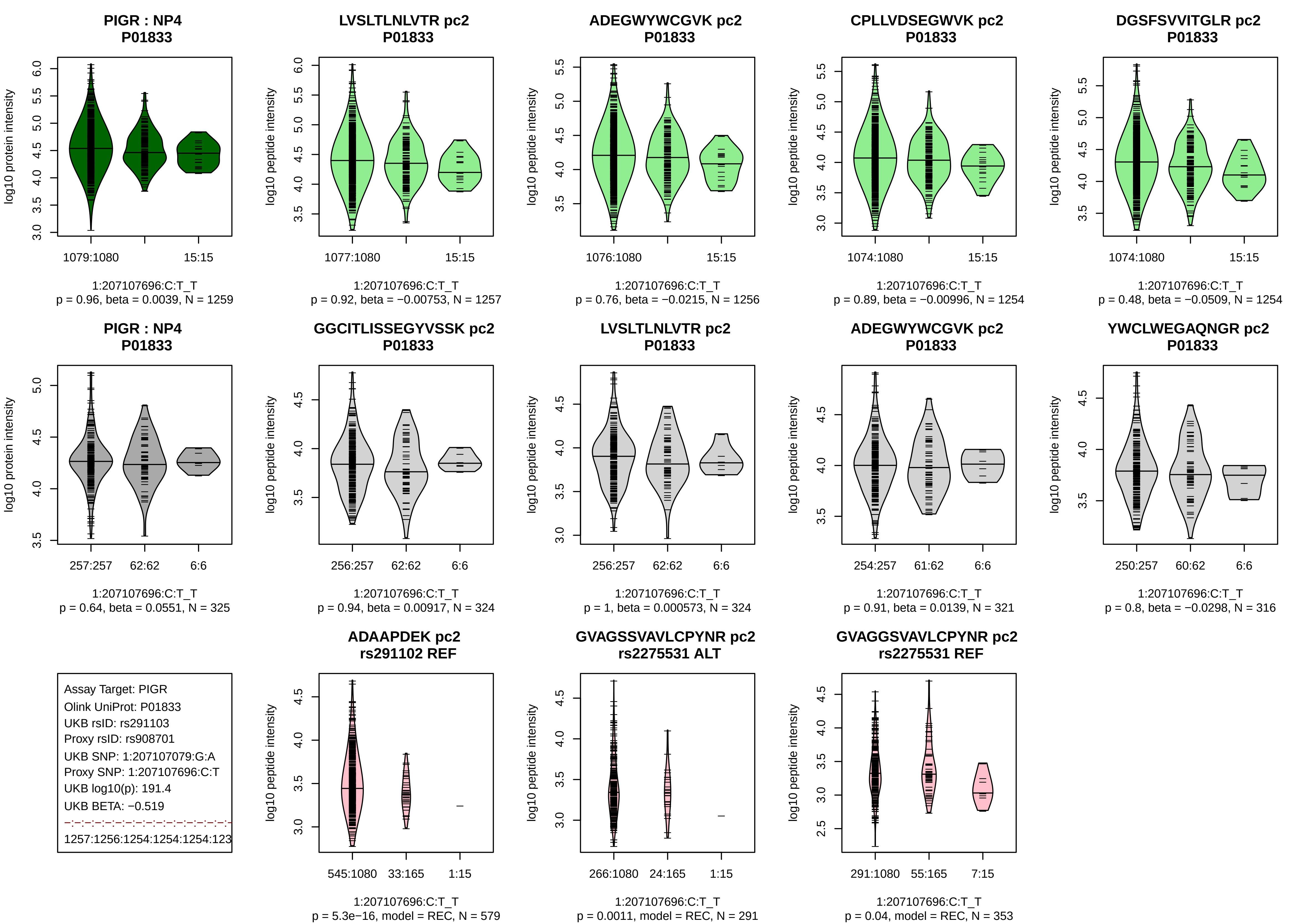
PIGR : NP4
P01833
log10 protein intensity
1079:1080
15:15
1:207107696:C:T_T
p = 0.96, beta = 0.0039, N = 1259
LVSLTLNLVTR pc2
P01833
log10 peptide intensity
1077:1080
15:15
1:207107696:C:T_T
p = 0.92, beta = −0.00753, N = 1257
ADEGWYWCGVK pc2
P01833
1076:1080
15:15
1:207107696:C:T_T
p = 0.76, beta = −0.0215, N = 1256
CPLLVDSEGWVK pc2
P01833
1074:1080
15:15
1:207107696:C:T_T
p = 0.89, beta = −0.00996, N = 1254
DGSFSVVITGLR pc2
P01833
1074:1080
15:15
1:207107696:C:T_T
p = 0.48, beta = −0.0509, N = 1254
PIGR : NP4
P01833
257:257
62:62
6:6
1:207107696:C:T_T
p = 0.64, beta = 0.0551, N = 325
GGCITLISSEGYVSSK pc2
P01833
256:257
62:62
6:6
1:207107696:C:T_T
p = 0.94, beta = 0.00917, N = 324
LVSLTLNLVTR pc2
P01833
256:257
62:62
6:6
1:207107696:C:T_T
p = 1, beta = 0.000573, N = 324
ADEGWYWCGVK pc2
P01833
254:257
61:62
6:6
1:207107696:C:T_T
p = 0.91, beta = 0.0139, N = 321
YWCLWEGAQNGR pc2
P01833
250:257
60:62
6:6
1:207107696:C:T_T
p = 0.8, beta = −0.0298, N = 316
Assay Target: PIGR
Olink UniProt: P01833
UKB rsID: rs291103
Proxy rsID: rs908701
UKB SNP: 1:207107079:G:A
Proxy SNP: 1:207107696:C:T
UKB log10(p): 191.4
UKB BETA: −0.519
1257:1256:1254:1254:1254:123
ADAAPDEK pc2
rs291102 REF
545:1080
33:165
1:15
1:207107696:C:T_T
p = 5.3e−16, model = REC, N = 579
GVAGSSVAVLCPYNR pc2
rs2275531 ALT
266:1080
24:165
1:15
1:207107696:C:T_T
p = 0.0011, model = REC, N = 291
GVAGGSVAVLCPYNR pc2
rs2275531 REF
291:1080
55:165
7:15
1:207107696:C:T_T
p = 0.04, model = REC, N = 353

**AHNAK : NP1**
**Q09666**

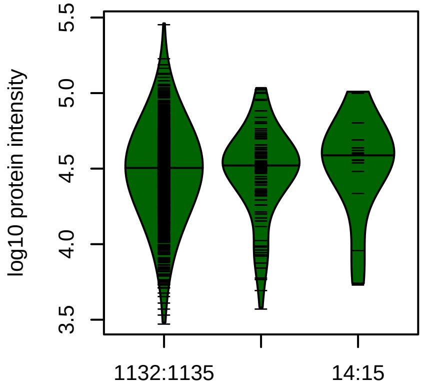


11:62269774:G:A_A
p = 0.47, beta = −0.0589, N = 1255

**FSMPGFK pc2**
**Q09666**

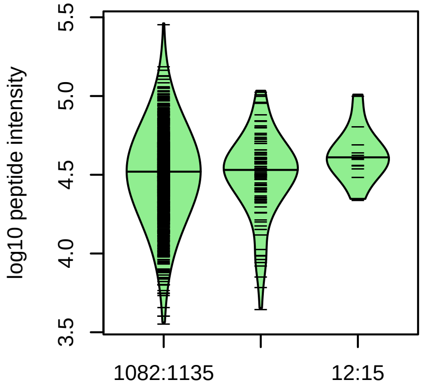


11:62269774:G:A_A
p = 0.84, beta = −0.017, N = 1199

**ISMPDVDLHLK pc2**
**Q09666**

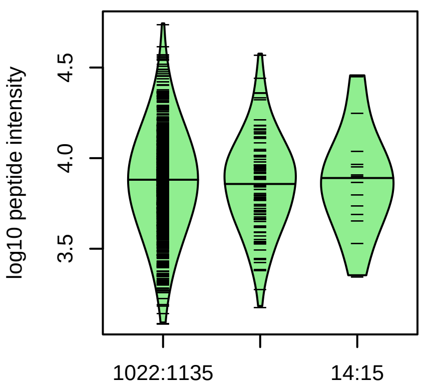


11:62269774:G:A_A
p = 0.66, beta = −0.0363, N = 1137

**ADVVVSGPK pc2**
**Q09666**

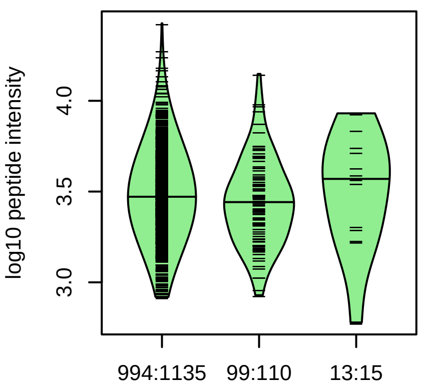


11:62269774:G:A_A
p = 0.37, beta = −0.0764, N = 1106

**GDMDVSVPK pc2**
**Q09666**

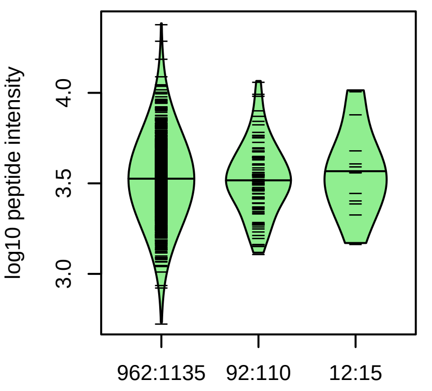


11:62269774:G:A_A
p = 0.53, beta = 0.0551, N = 1066

**AHNAK : NP1**
**Q09666**

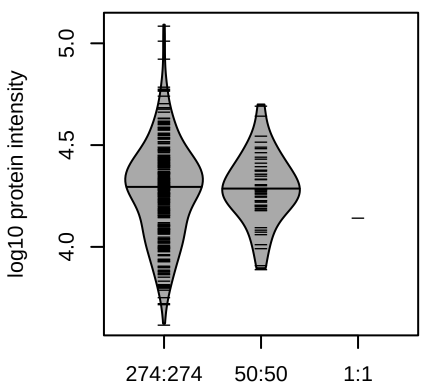


11:62269774:G:A_A
p = 0.93, beta = −0.0131, N = 325

**FSMPGFK pc2**
**Q09666**

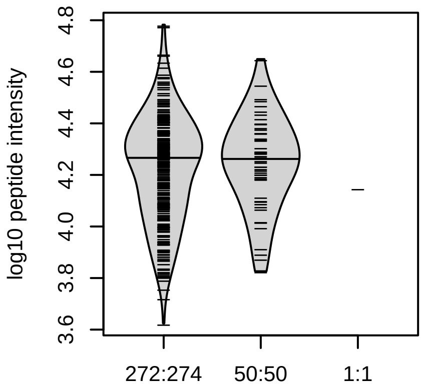


11:62269774:G:A_A
p = 0.98, beta = −0.004, N = 323

**GDVDVSLPK pc2**
**Q09666**

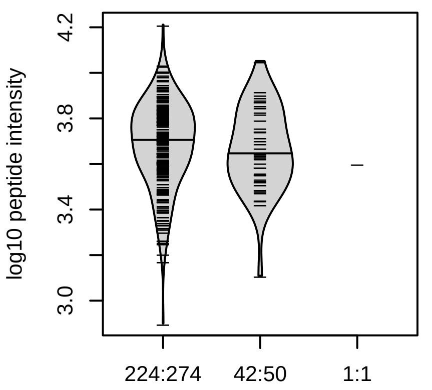


11:62269774:G:A_A
p = 0.24, beta = −0.184, N = 267

**ISMPDVDLHLK pc2**
**Q09666**

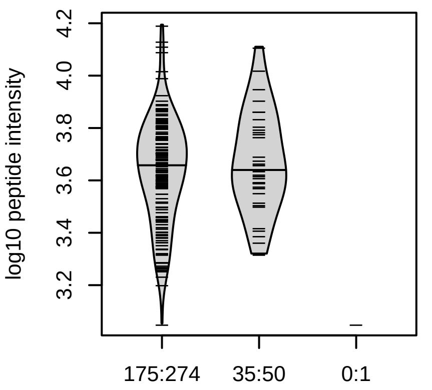


11:62269774:G:A_A
p = 0.49, beta = 0.127, N = 210

**MPEMNIK pc2**
**Q09666**

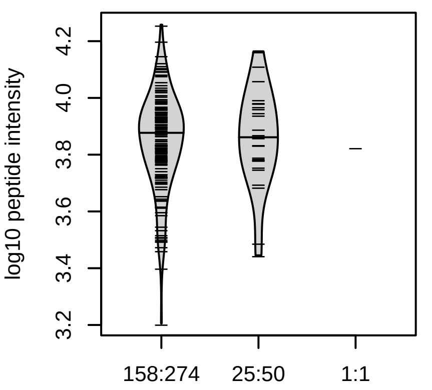


11:62269774:G:A_A
p = 0.76, beta = −0.0589, N = 184

Assay Target: AHNAK
Olink UniProt: Q09666
UKB rsID: rs145590149
Proxy rsID: rs11231121
UKB SNP: 11:62303555:T:TCT[illegible]
Proxy SNP: 11:62269774:G:A
UKB log10(p): 189.7
UKB BETA: −0.496
–:–:–:–:–:–:–:–:–:–:–:–:–:–:–:–
1199:1137:1106:1066:1008:100[illegible]

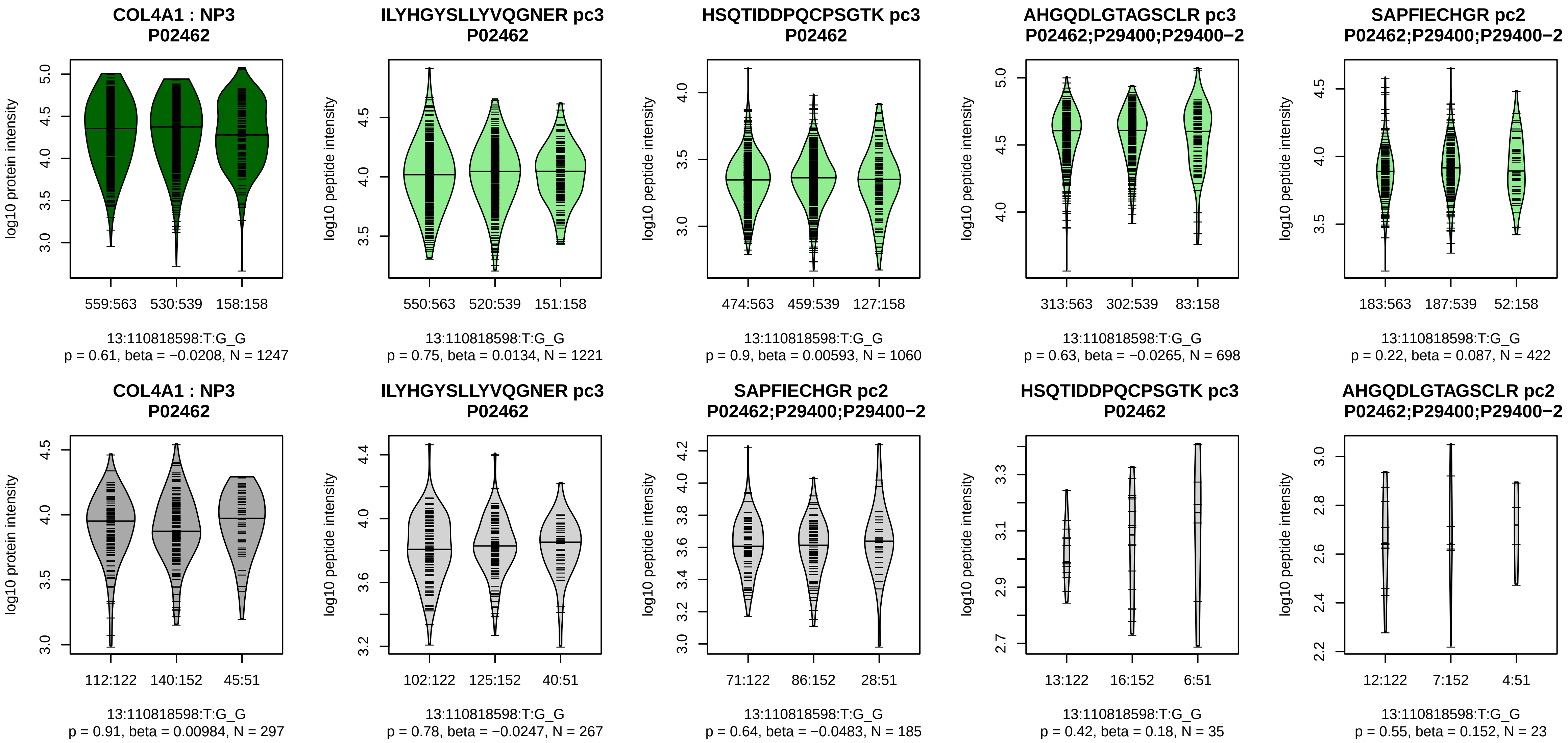


Assay Target: COL4A1
Olink UniProt: P02462
UKB rsID: rs3742207
Proxy rsID: rs3742207
UKB SNP: 13:110818598:T:G
Proxy SNP: 13:110818598:T:G
UKB log10(p): 186.6
UKB BETA: −0.184
−:−:−:−:−:−:NA:NA
1221:1060:698:422:283:130:2:1

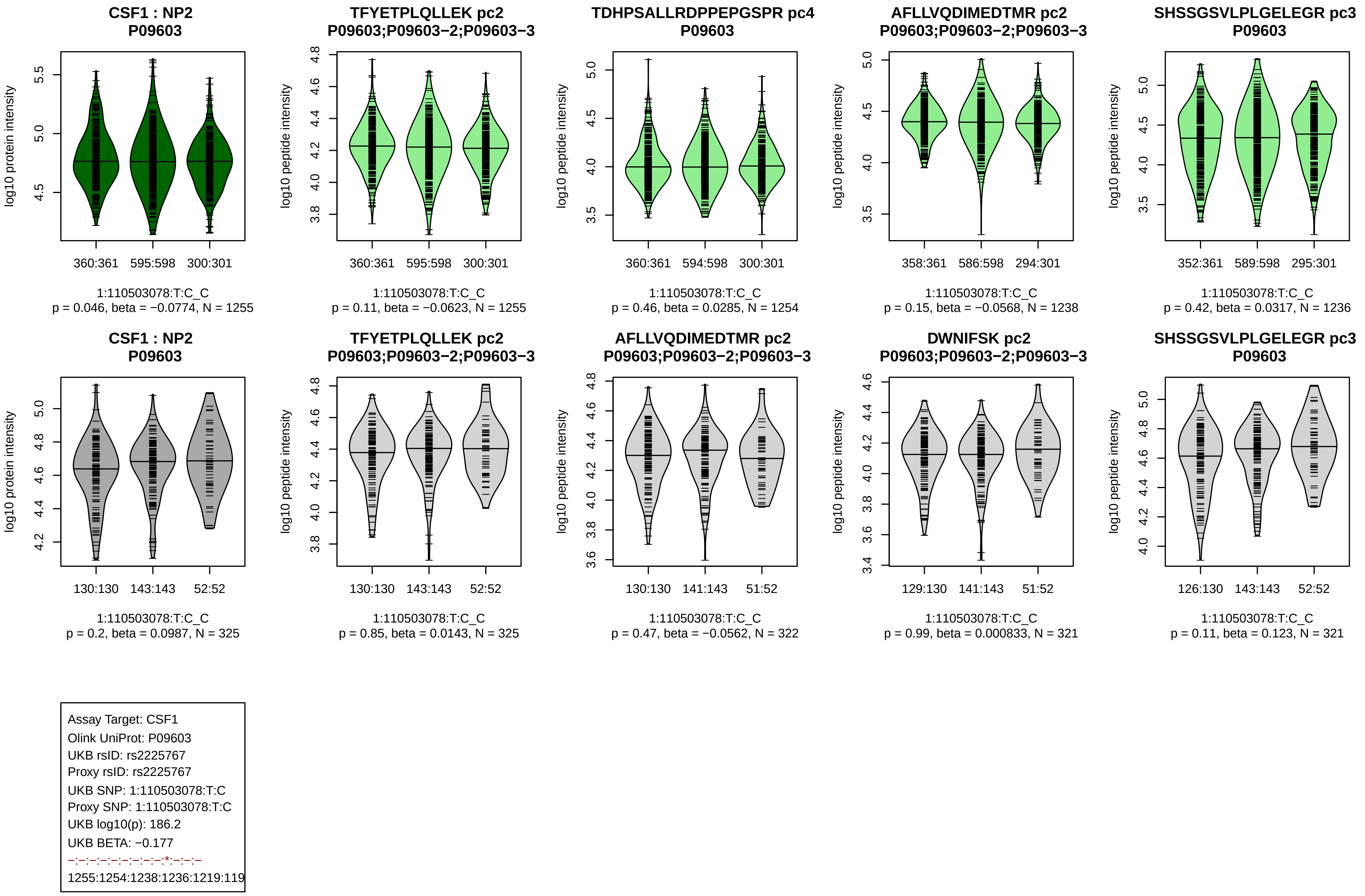
CSF1 : NP2
P09603
TFYETPLQLLEK pc2
P09603;P09603−2;P09603−3
TDHPSALLRDPPEPGSPR pc4
P09603
AFLLVQDIMEDTMR pc2
P09603;P09603−2;P09603−3
SHSSGSVLPLGELEGR pc3
P09603
log10 protein intensity
log10 peptide intensity
360:361 595:598 300:301
360:361 595:598 300:301
360:361 594:598 300:301
358:361 586:598 294:301
352:361 589:598 295:301
1:110503078:T:C_C
p = 0.046, beta = −0.0774, N = 1255
1:110503078:T:C_C
p = 0.11, beta = −0.0623, N = 1255
1:110503078:T:C_C
p = 0.46, beta = 0.0285, N = 1254
1:110503078:T:C_C
p = 0.15, beta = −0.0568, N = 1238
1:110503078:T:C_C
p = 0.42, beta = 0.0317, N = 1236
CSF1 : NP2
P09603
TFYETPLQLLEK pc2
P09603;P09603−2;P09603−3
AFLLVQDIMEDTMR pc2
P09603;P09603−2;P09603−3
DWNIFSK pc2
P09603;P09603−2;P09603−3
SHSSGSVLPLGELEGR pc3
P09603
130:130 143:143 52:52
130:130 143:143 52:52
130:130 141:143 51:52
129:130 141:143 51:52
126:130 143:143 52:52
1:110503078:T:C_C
p = 0.2, beta = 0.0987, N = 325
1:110503078:T:C_C
p = 0.85, beta = 0.0143, N = 325
1:110503078:T:C_C
p = 0.47, beta = −0.0562, N = 322
1:110503078:T:C_C
p = 0.99, beta = 0.000833, N = 321
1:110503078:T:C_C
p = 0.11, beta = 0.123, N = 321
Assay Target: CSF1
Olink UniProt: P09603
UKB rsID: rs2225767
Proxy rsID: rs2225767
UKB SNP: 1:110503078:T:C
Proxy SNP: 1:110503078:T:C
UKB log10(p): 186.2
UKB BETA: −0.177
−:−:−:−:−:−:−:−:−:*:−:−:−
1255:1254:1238:1236:1219:119

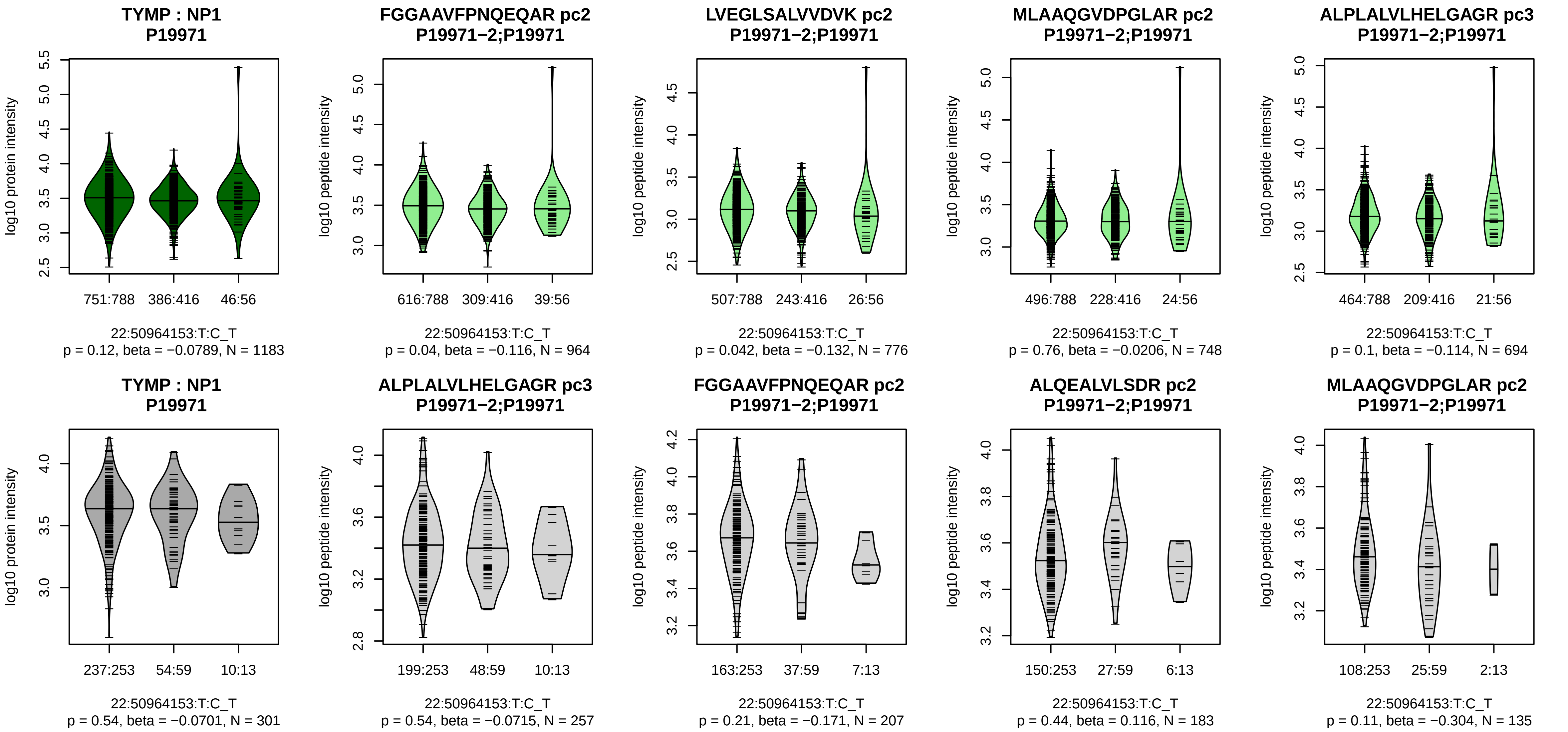


Assay Target: TYMP
Olink UniProt: P19971
UKB rsID: rs131805
Proxy rsID: rs131805
UKB SNP: 22:50964153:T:C
Proxy SNP: 22:50964153:T:C
UKB log10(p): 185.4
UKB BETA: 0.22
−:−:−:−:−:−:−:−:−:−:−:−:−:−:−:NA
964:776:748:694:601:521:495:4

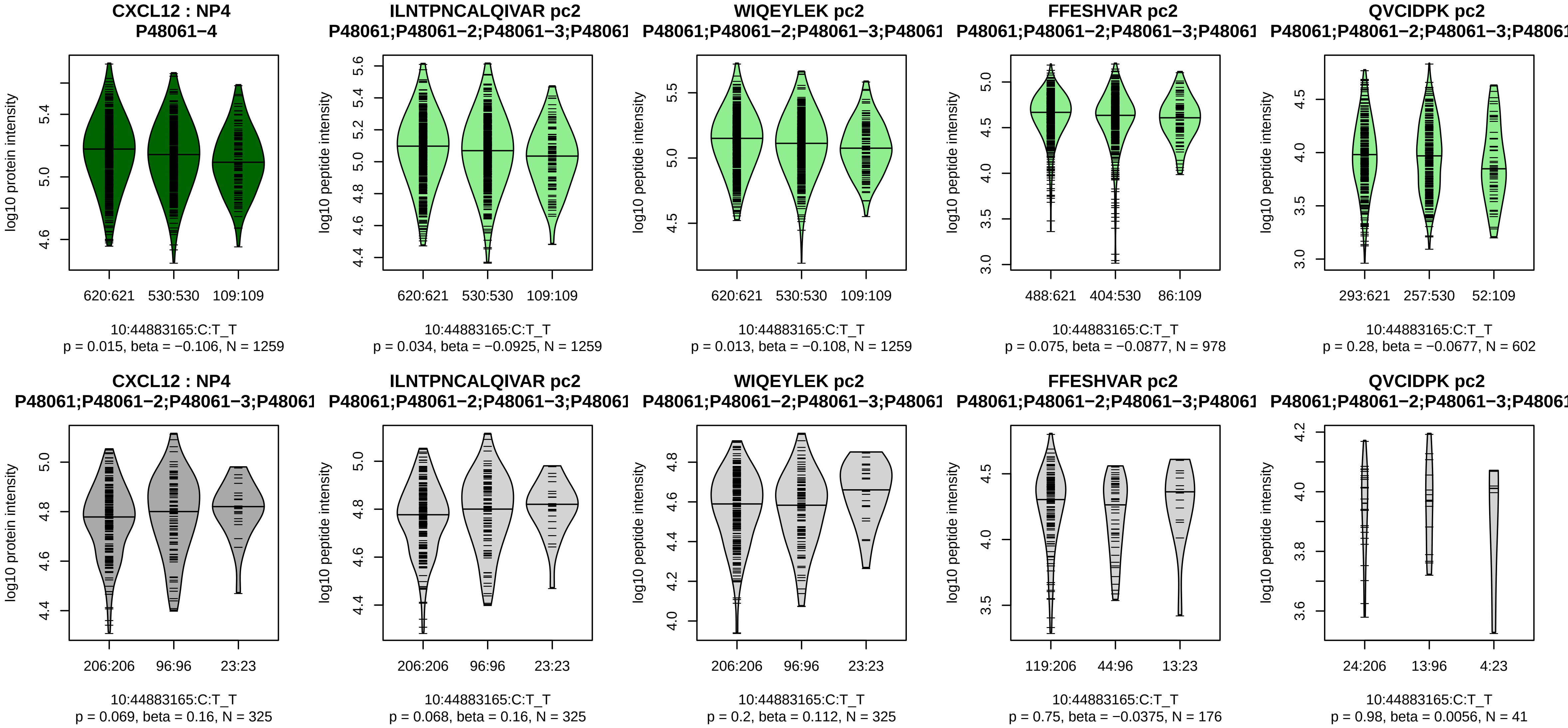


Assay Target: CXCL12
Olink UniProt: P48061
UKB rsID: rs11239028
Proxy rsID: rs11239028
UKB SNP: 10:44883165:C:T
Proxy SNP: 10:44883165:C:T
UKB log10(p): 184.6
UKB BETA: 0.191
−:−:−:−
1259:1259:978:602

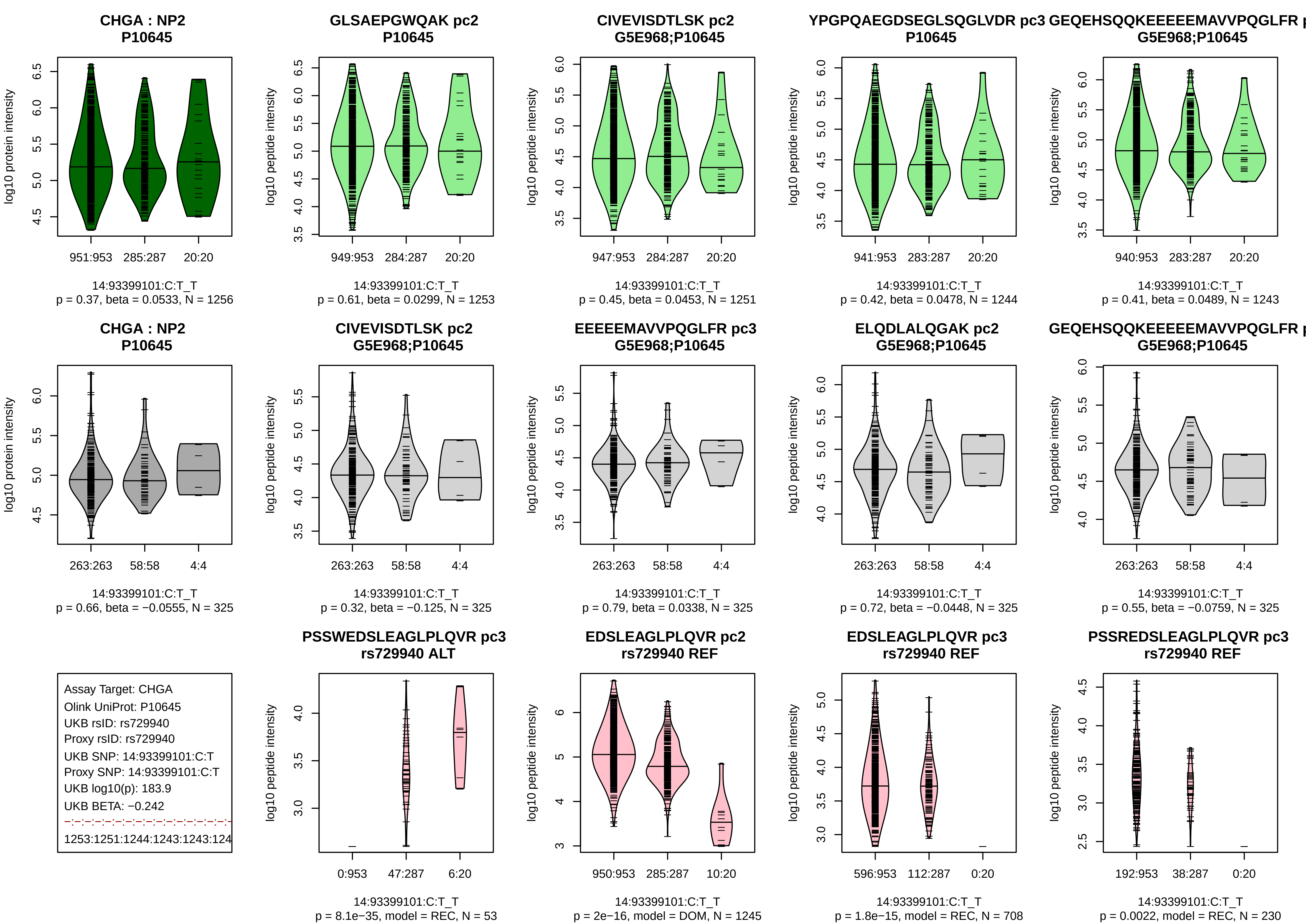
CHGA : NP2
P10645
log10 protein intensity
951:953
285:287
20:20
14:93399101:C:T_T
p = 0.37, beta = 0.0533, N = 1256
GLSAEPGWQAK pc2
P10645
log10 peptide intensity
949:953
284:287
20:20
14:93399101:C:T_T
p = 0.61, beta = 0.0299, N = 1253
CIVEVISDTLSK pc2
G5E968;P10645
947:953
284:287
20:20
14:93399101:C:T_T
p = 0.45, beta = 0.0453, N = 1251
YPGPQAEGDSEGLSQGLVDR pc3
P10645
941:953
283:287
20:20
14:93399101:C:T_T
p = 0.42, beta = 0.0478, N = 1244
GEQEHSQQKEEEEEMAVVPQGLFR p
G5E968;P10645
940:953
283:287
20:20
14:93399101:C:T_T
p = 0.41, beta = 0.0489, N = 1243
CHGA : NP2
P10645
263:263
58:58
4:4
14:93399101:C:T_T
p = 0.66, beta = −0.0555, N = 325
CIVEVISDTLSK pc2
G5E968;P10645
263:263
58:58
4:4
14:93399101:C:T_T
p = 0.32, beta = −0.125, N = 325
EEEEEMAVVPQGLFR pc3
G5E968;P10645
263:263
58:58
4:4
14:93399101:C:T_T
p = 0.79, beta = 0.0338, N = 325
ELQDLALQGAK pc2
G5E968;P10645
263:263
58:58
4:4
14:93399101:C:T_T
p = 0.72, beta = −0.0448, N = 325
GEQEHSQQKEEEEEMAVVPQGLFR p
G5E968;P10645
263:263
58:58
4:4
14:93399101:C:T_T
p = 0.55, beta = −0.0759, N = 325
Assay Target: CHGA
Olink UniProt: P10645
UKB rsID: rs729940
Proxy rsID: rs729940
UKB SNP: 14:93399101:C:T
Proxy SNP: 14:93399101:C:T
UKB log10(p): 183.9
UKB BETA: −0.242
1253:1251:1244:1243:1243:124
PSSWEDSLEAGLPLQVR pc3
rs729940 ALT
0:953
47:287
6:20
14:93399101:C:T_T
p = 8.1e−35, model = REC, N = 53
EDSLEAGLPLQVR pc2
rs729940 REF
950:953
285:287
10:20
14:93399101:C:T_T
p = 2e−16, model = DOM, N = 1245
EDSLEAGLPLQVR pc3
rs729940 REF
596:953
112:287
0:20
14:93399101:C:T_T
p = 1.8e−15, model = REC, N = 708
PSSREDSLEAGLPLQVR pc3
rs729940 REF
192:953
38:287
0:20
14:93399101:C:T_T
p = 0.0022, model = REC, N = 230

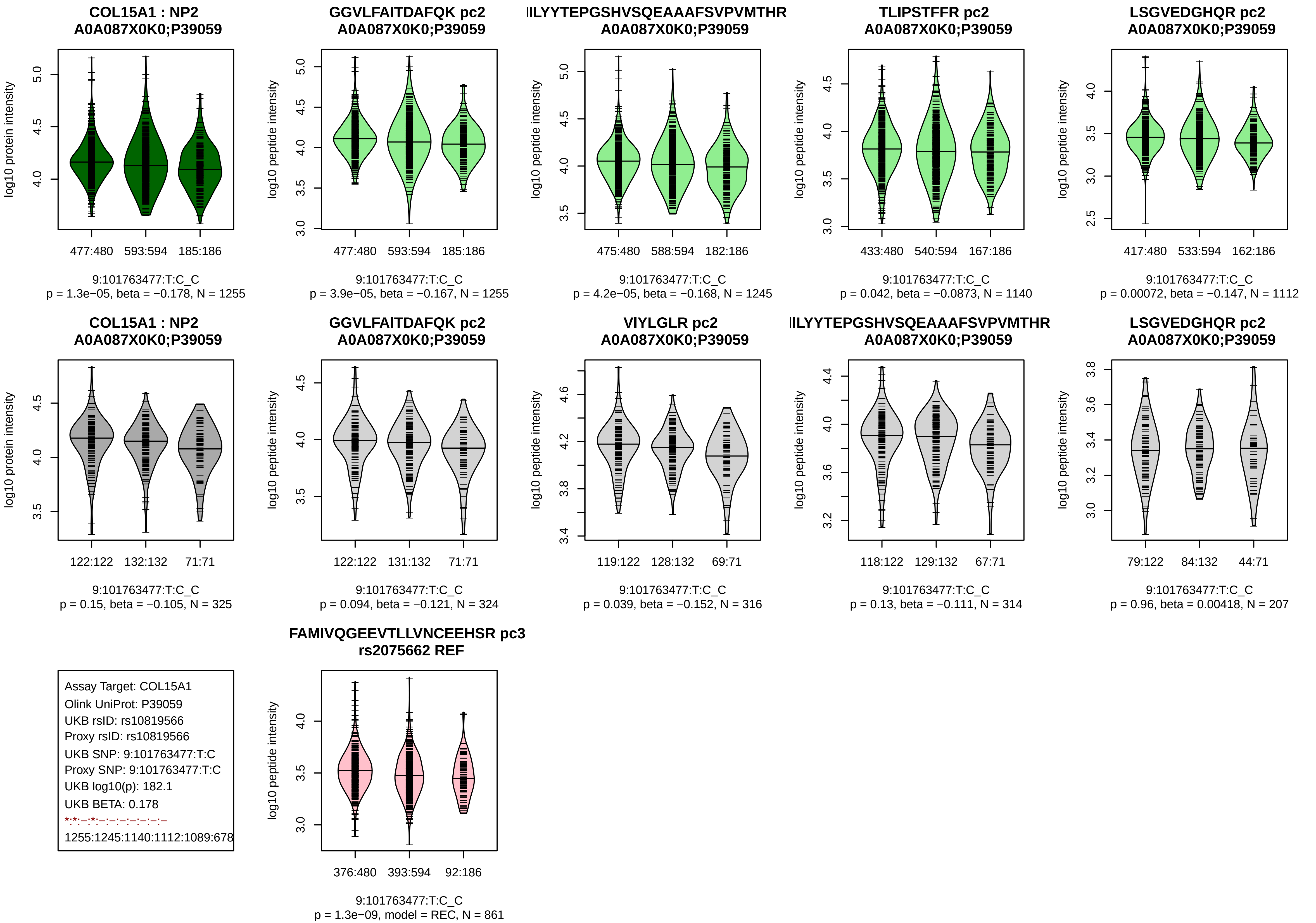

COL15A1 : NP2
A0A087X0K0;P39059
log10 protein intensity
477:480 593:594 185:186
9:101763477:T:C_C
p = 1.3e−05, beta = −0.178, N = 1255
GGVLFAITDAFQK pc2
A0A087X0K0;P39059
log10 peptide intensity
477:480 593:594 185:186
9:101763477:T:C_C
p = 3.9e−05, beta = −0.167, N = 1255
ILYYTEPGSHVSQEAAAFSVPVMTHR
A0A087X0K0;P39059
log10 peptide intensity
475:480 588:594 182:186
9:101763477:T:C_C
p = 4.2e−05, beta = −0.168, N = 1245
TLIPSTFFR pc2
A0A087X0K0;P39059
log10 peptide intensity
433:480 540:594 167:186
9:101763477:T:C_C
p = 0.042, beta = −0.0873, N = 1140
LSGVEDGHQR pc2
A0A087X0K0;P39059
log10 peptide intensity
417:480 533:594 162:186
9:101763477:T:C_C
p = 0.00072, beta = −0.147, N = 1112
COL15A1 : NP2
A0A087X0K0;P39059
log10 protein intensity
122:122 132:132 71:71
9:101763477:T:C_C
p = 0.15, beta = −0.105, N = 325
GGVLFAITDAFQK pc2
A0A087X0K0;P39059
log10 peptide intensity
122:122 131:132 71:71
9:101763477:T:C_C
p = 0.094, beta = −0.121, N = 324
VIYLGLR pc2
A0A087X0K0;P39059
log10 peptide intensity
119:122 128:132 69:71
9:101763477:T:C_C
p = 0.039, beta = −0.152, N = 316
ILYYTEPGSHVSQEAAAFSVPVMTHR
A0A087X0K0;P39059
log10 peptide intensity
118:122 129:132 67:71
9:101763477:T:C_C
p = 0.13, beta = −0.111, N = 314
LSGVEDGHQR pc2
A0A087X0K0;P39059
log10 peptide intensity
79:122 84:132 44:71
9:101763477:T:C_C
p = 0.96, beta = 0.00418, N = 207
Assay Target: COL15A1
Olink UniProt: P39059
UKB rsID: rs10819566
Proxy rsID: rs10819566
UKB SNP: 9:101763477:T:C
Proxy SNP: 9:101763477:T:C
UKB log10(p): 182.1
UKB BETA: 0.178
*:*:–:*:–:–:–:–:–:–:–
1255:1245:1140:1112:1089:678
FAMIVQGEEVTLLVNCEEHSR pc3
rs2075662 REF
log10 peptide intensity
376:480 393:594 92:186
9:101763477:T:C_C
p = 1.3e−09, model = REC, N = 861

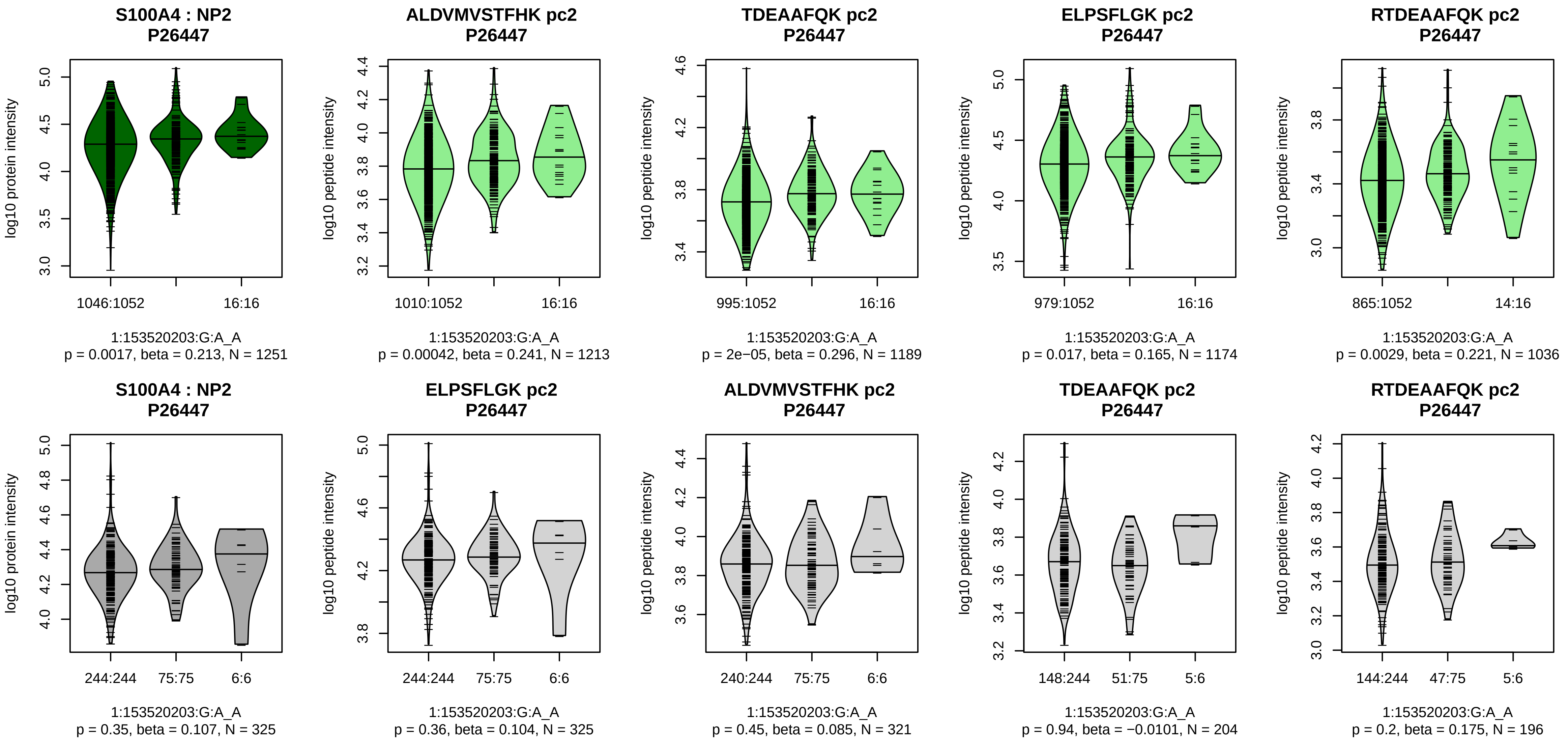


Assay Target: S100A4
Olink UniProt: P26447
UKB rsID: rs41265164
Proxy rsID: rs41265164
UKB SNP: 1:153520203:G:A
Proxy SNP: 1:153520203:G:A
UKB log10(p): 178.9
UKB BETA: 0.373
*:*:−:*:NA
1213:1189:1174:1036:17

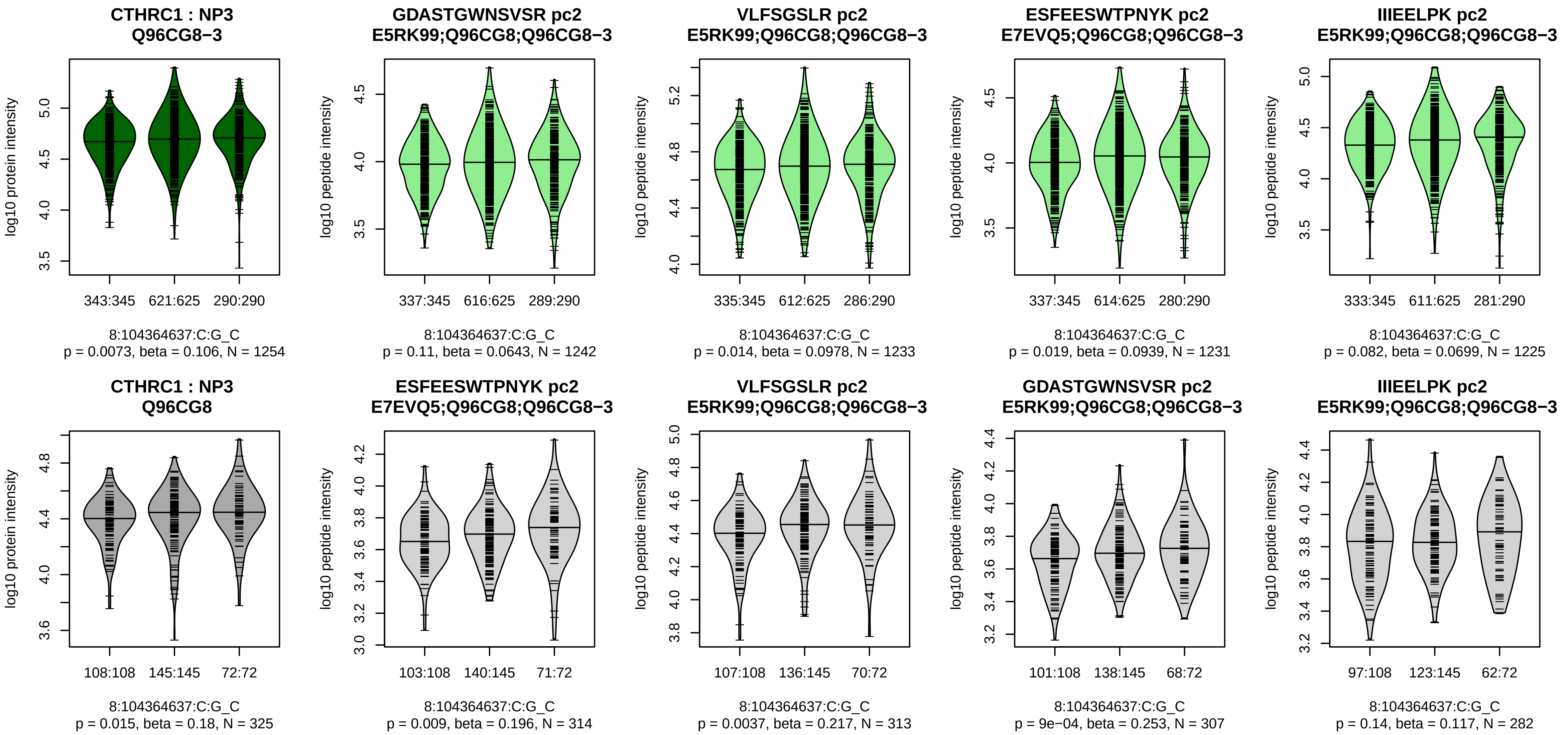


Assay Target: CTHRC1
Olink UniProt: Q96CG8
UKB rsID: rs827592
Proxy rsID: rs827592
UKB SNP: 8:104364637:C:G
Proxy SNP: 8:104364637:C:G
UKB log10(p): 177.8
UKB BETA: −0.157
−:−:−:−:−:−
1242:1233:1231:1225:1143:966

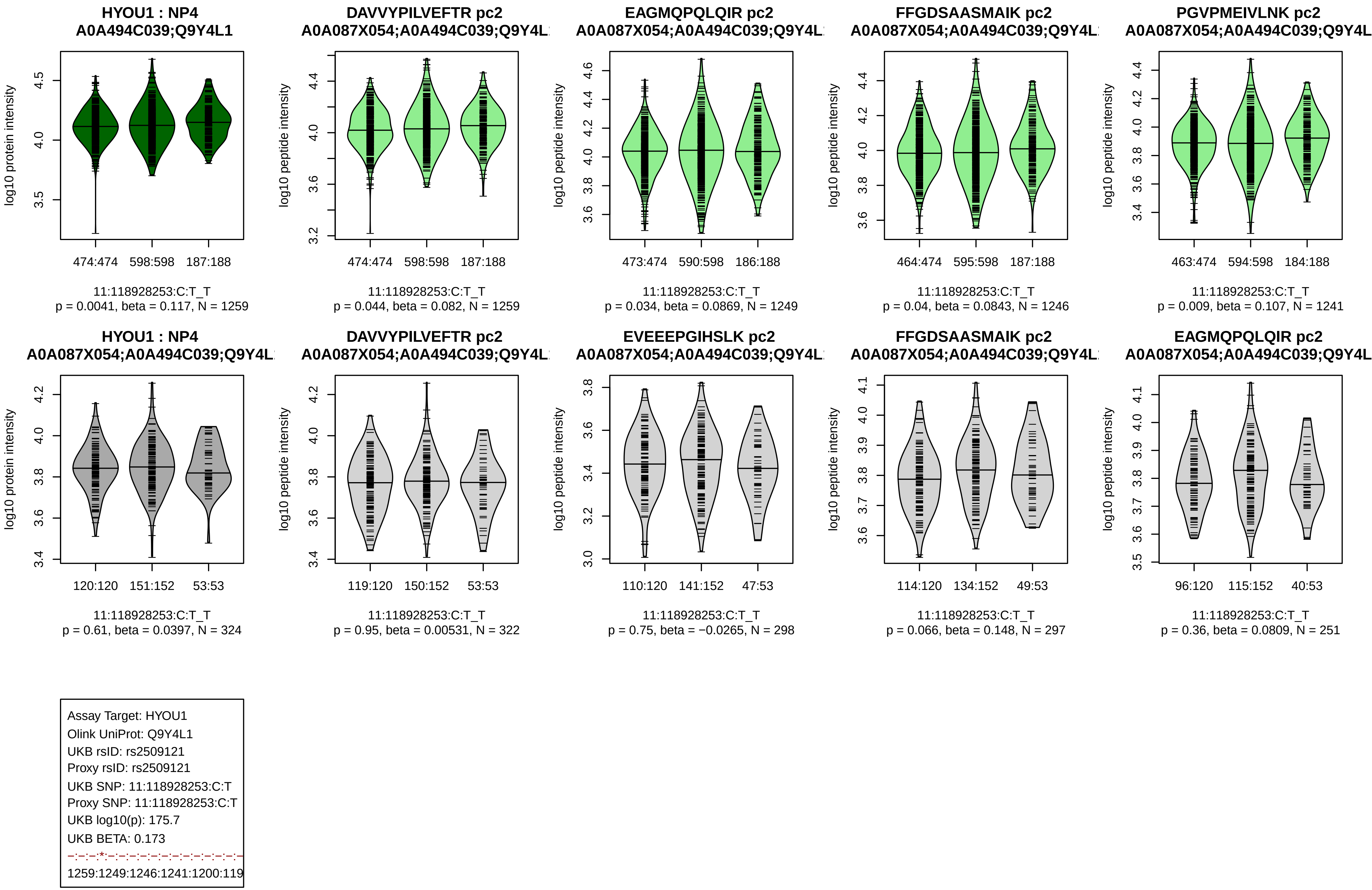
HYOU1 : NP4
A0A494C039;Q9Y4L1
log10 protein intensity
474:474
598:598
187:188
11:118928253:C:T_T
p = 0.0041, beta = 0.117, N = 1259
DAVVYPILVEFTR pc2
A0A087X054;A0A494C039;Q9Y4L
log10 peptide intensity
474:474
598:598
187:188
11:118928253:C:T_T
p = 0.044, beta = 0.082, N = 1259
EAGMQPQLQIR pc2
A0A087X054;A0A494C039;Q9Y4L
log10 peptide intensity
473:474
590:598
186:188
11:118928253:C:T_T
p = 0.034, beta = 0.0869, N = 1249
FFGDSAASMAIK pc2
A0A087X054;A0A494C039;Q9Y4L
log10 peptide intensity
464:474
595:598
187:188
11:118928253:C:T_T
p = 0.04, beta = 0.0843, N = 1246
PGVPMEIVLNK pc2
A0A087X054;A0A494C039;Q9Y4L
log10 peptide intensity
463:474
594:598
184:188
11:118928253:C:T_T
p = 0.009, beta = 0.107, N = 1241
HYOU1 : NP4
A0A087X054;A0A494C039;Q9Y4L
log10 protein intensity
120:120
151:152
53:53
11:118928253:C:T_T
p = 0.61, beta = 0.0397, N = 324
DAVVYPILVEFTR pc2
A0A087X054;A0A494C039;Q9Y4L
log10 peptide intensity
119:120
150:152
53:53
11:118928253:C:T_T
p = 0.95, beta = 0.00531, N = 322
EVEEEPGIHSLK pc2
A0A087X054;A0A494C039;Q9Y4L
log10 peptide intensity
110:120
141:152
47:53
11:118928253:C:T_T
p = 0.75, beta = −0.0265, N = 298
FFGDSAASMAIK pc2
A0A087X054;A0A494C039;Q9Y4L
log10 peptide intensity
114:120
134:152
49:53
11:118928253:C:T_T
p = 0.066, beta = 0.148, N = 297
EAGMQPQLQIR pc2
A0A087X054;A0A494C039;Q9Y4L
log10 peptide intensity
96:120
115:152
40:53
11:118928253:C:T_T
p = 0.36, beta = 0.0809, N = 251
Assay Target: HYOU1
Olink UniProt: Q9Y4L1
UKB rsID: rs2509121
Proxy rsID: rs2509121
UKB SNP: 11:118928253:C:T
Proxy SNP: 11:118928253:C:T
UKB log10(p): 175.7
UKB BETA: 0.173
–:–:–:*:–:–:–:–:–:–:–:–:–:–:–:–:–
1259:1249:1246:1241:1200:119

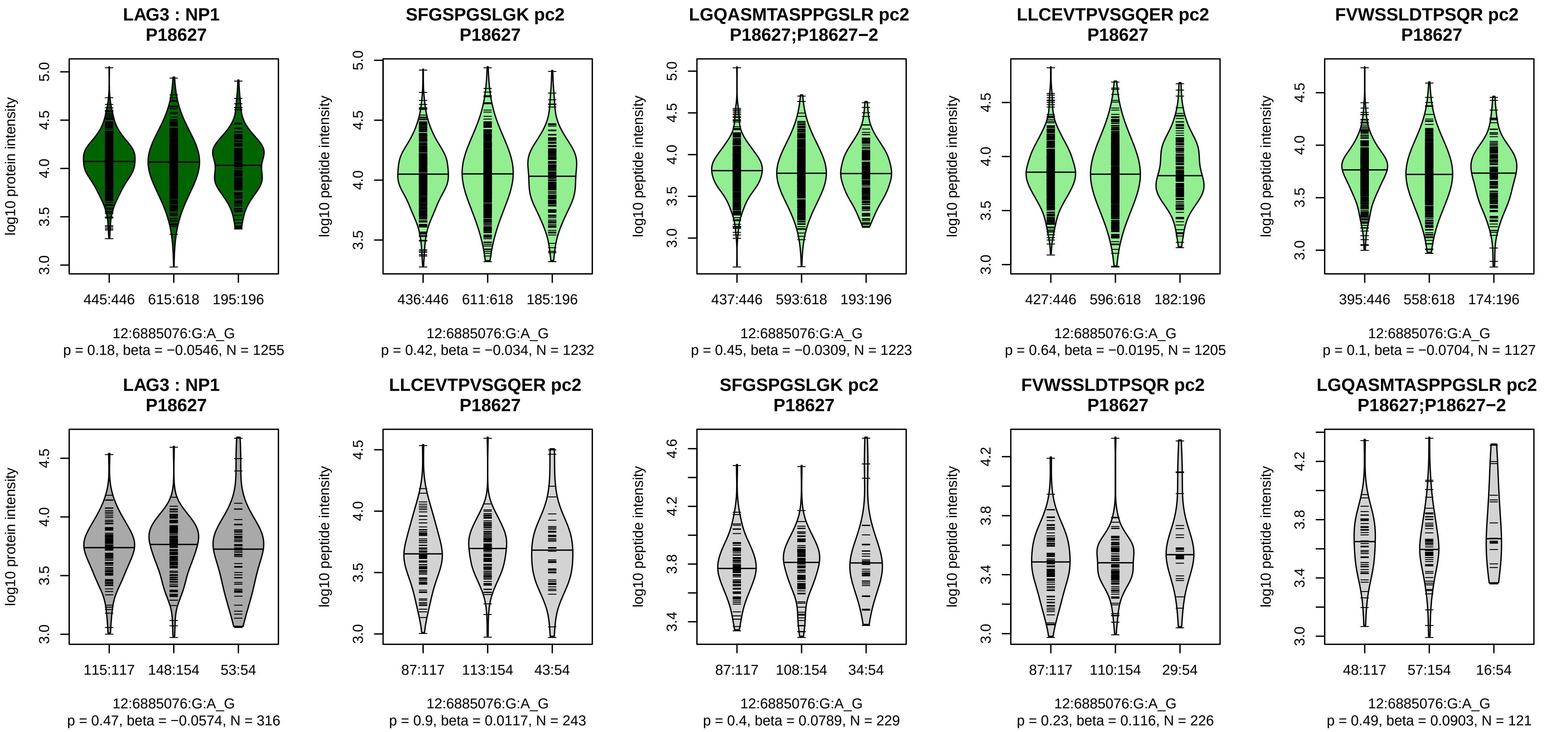


Assay Target: LAG3
Olink UniProt: P18627
UKB rsID: rs3782735
Proxy rsID: rs3782735
UKB SNP: 12:6885076:G:A
Proxy SNP: 12:6885076:G:A
UKB log10(p): 173.6
UKB BETA: 0.174
−:−:−:−:−:−:−
1232:1223:1205:1127:651:92:59

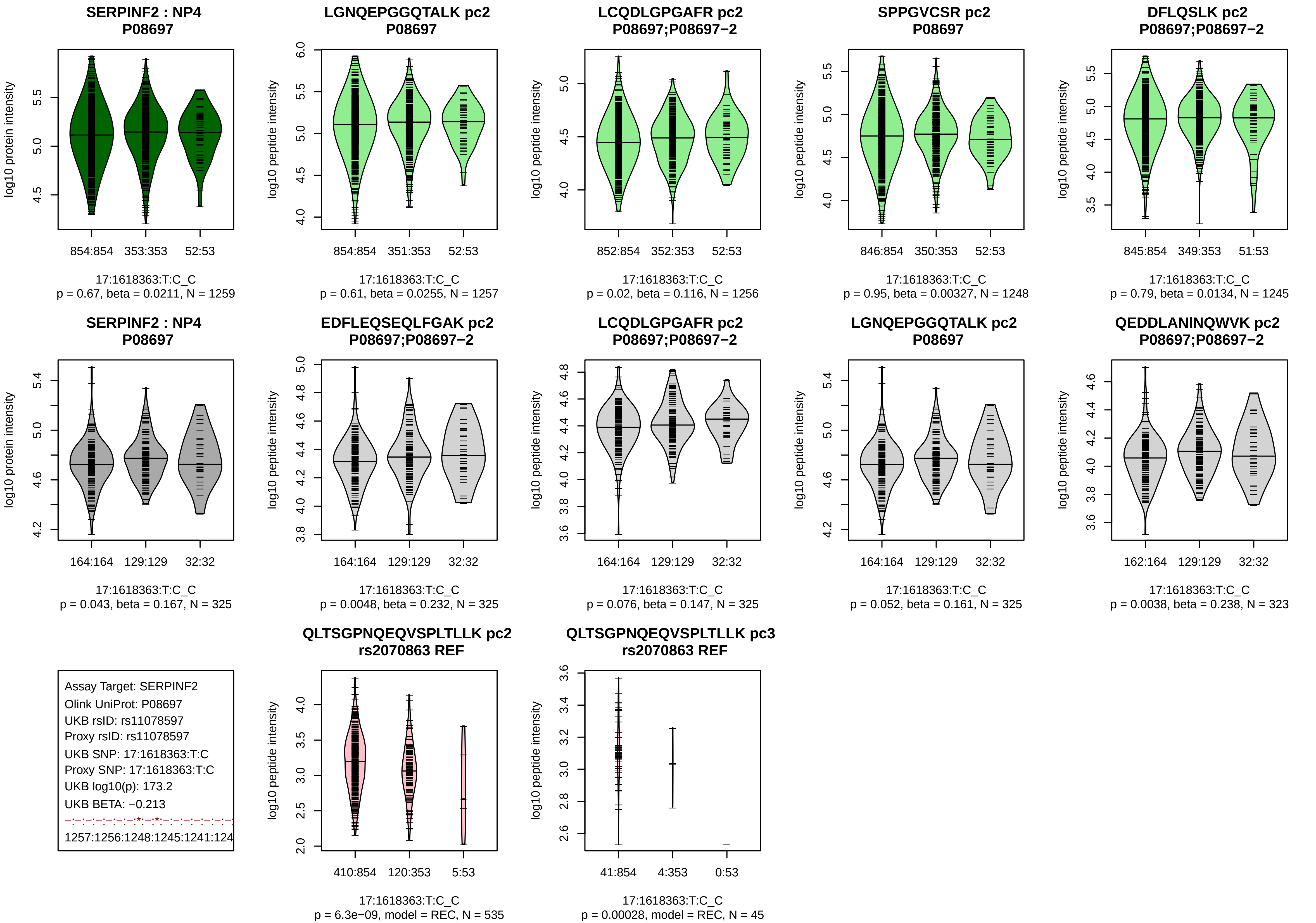
SERPINF2 : NP4
P08697
log10 protein intensity
854:854
353:353
52:53
17:1618363:T:C_C
p = 0.67, beta = 0.0211, N = 1259
LGNQEPGGQTALK pc2
P08697
log10 peptide intensity
854:854
351:353
52:53
17:1618363:T:C_C
p = 0.61, beta = 0.0255, N = 1257
LCQDLGPGAFR pc2
P08697;P08697−2
log10 peptide intensity
852:854
352:353
52:53
17:1618363:T:C_C
p = 0.02, beta = 0.116, N = 1256
SPPGVCSR pc2
P08697
log10 peptide intensity
846:854
350:353
52:53
17:1618363:T:C_C
p = 0.95, beta = 0.00327, N = 1248
DFLQSLK pc2
P08697;P08697−2
log10 peptide intensity
845:854
349:353
51:53
17:1618363:T:C_C
p = 0.79, beta = 0.0134, N = 1245
SERPINF2 : NP4
P08697
log10 protein intensity
164:164
129:129
32:32
17:1618363:T:C_C
p = 0.043, beta = 0.167, N = 325
EDFLEQSEQLFGAK pc2
P08697;P08697−2
log10 peptide intensity
164:164
129:129
32:32
17:1618363:T:C_C
p = 0.0048, beta = 0.232, N = 325
LCQDLGPGAFR pc2
P08697;P08697−2
log10 peptide intensity
164:164
129:129
32:32
17:1618363:T:C_C
p = 0.076, beta = 0.147, N = 325
LGNQEPGGQTALK pc2
P08697
log10 peptide intensity
164:164
129:129
32:32
17:1618363:T:C_C
p = 0.052, beta = 0.161, N = 325
QEDDLANINQWVK pc2
P08697;P08697−2
log10 peptide intensity
162:164
129:129
32:32
17:1618363:T:C_C
p = 0.0038, beta = 0.238, N = 323
Assay Target: SERPINF2
Olink UniProt: P08697
UKB rsID: rs11078597
Proxy rsID: rs11078597
UKB SNP: 17:1618363:T:C
Proxy SNP: 17:1618363:T:C
UKB log10(p): 173.2
UKB BETA: −0.213
−:−:−:−:−:−:*:−:*:−:−:−:−:−:−:−
1257:1256:1248:1245:1241:124
QLTSGPNQEQVSPLTLLK pc2
rs2070863 REF
log10 peptide intensity
410:854
120:353
5:53
17:1618363:T:C_C
p = 6.3e−09, model = REC, N = 535
QLTSGPNQEQVSPLTLLK pc3
rs2070863 REF
log10 peptide intensity
41:854
4:353
0:53
17:1618363:T:C_C
p = 0.00028, model = REC, N = 45

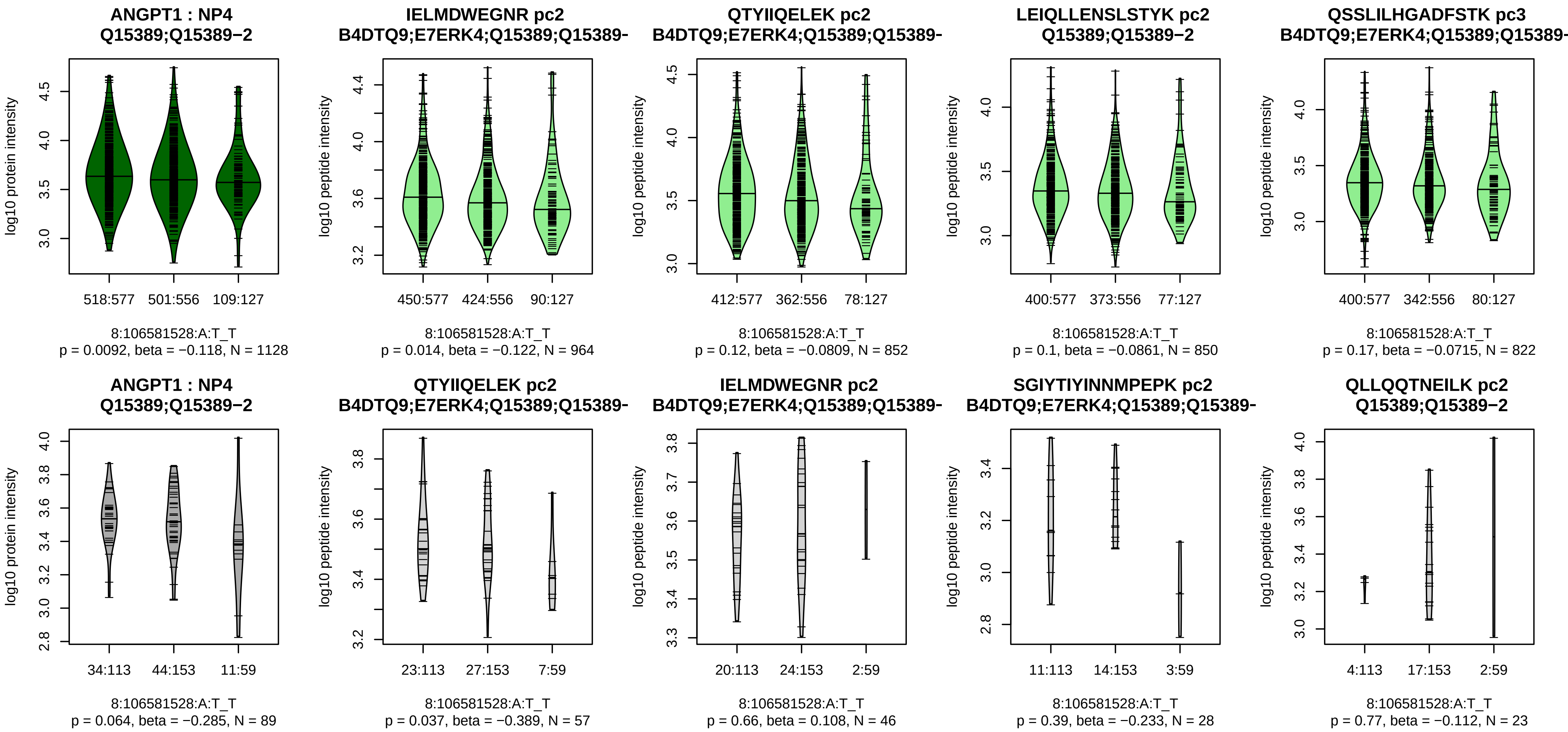


Assay Target: ANGPT1
Olink UniProt: Q15389
UKB rsID: rs6993770
Proxy rsID: rs6993770
UKB SNP: 8:106581528:A:T
Proxy SNP: 8:106581528:A:T
UKB log10(p): 172.5
UKB BETA: −0.187
−:−:−:−:−:−:−:−:−:−:−:−:−:−:−:−:
964:852:850:822:732:624:507:4

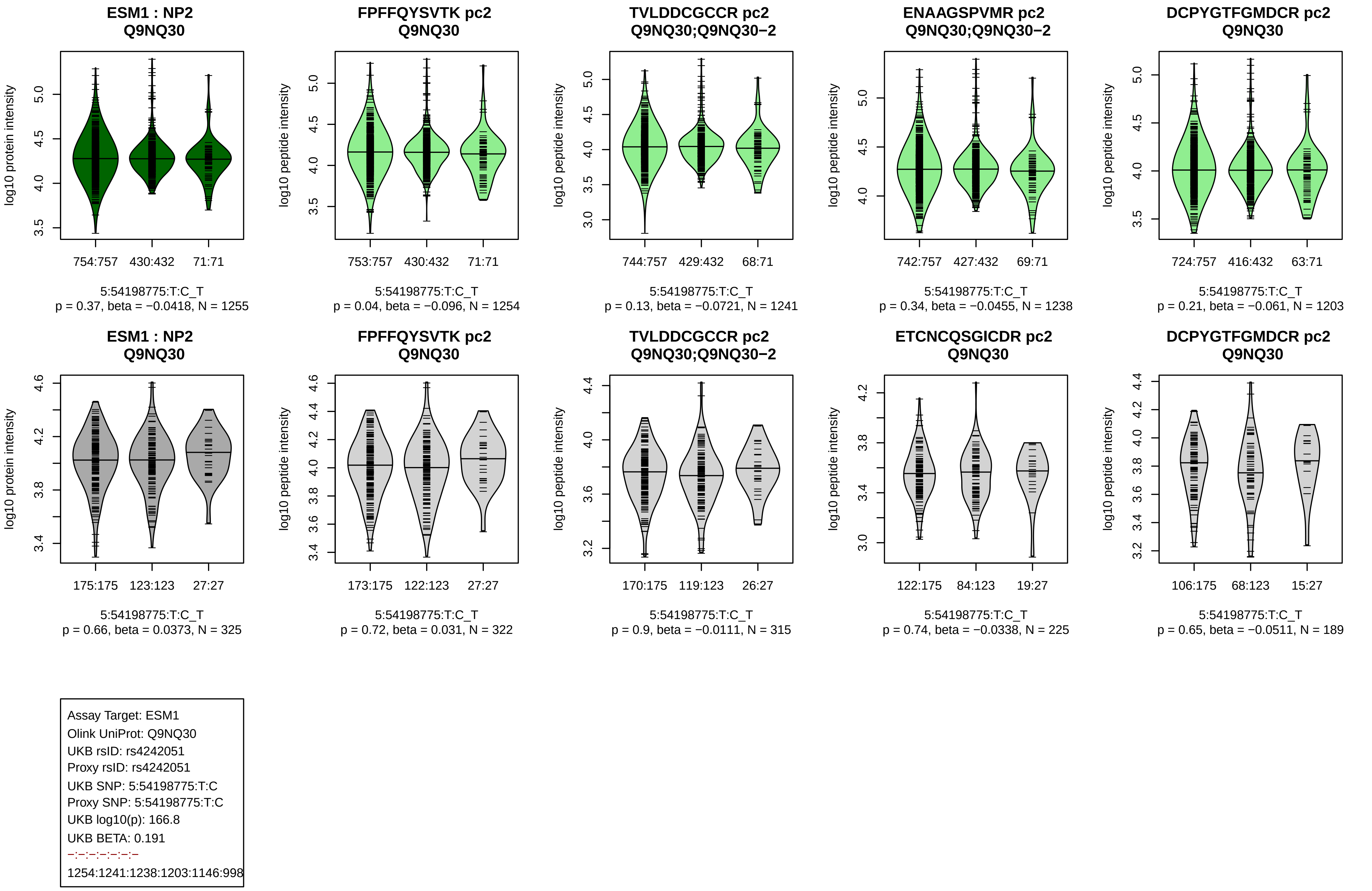
ESM1 : NP2
Q9NQ30
754:757 430:432 71:71
5:54198775:T:C_T
p = 0.37, beta = −0.0418, N = 1255
FPFFQYSVTK pc2
Q9NQ30
753:757 430:432 71:71
5:54198775:T:C_T
p = 0.04, beta = −0.096, N = 1254
TVLDDCGCCR pc2
Q9NQ30;Q9NQ30−2
744:757 429:432 68:71
5:54198775:T:C_T
p = 0.13, beta = −0.0721, N = 1241
ENAAGSPVMR pc2
Q9NQ30;Q9NQ30−2
742:757 427:432 69:71
5:54198775:T:C_T
p = 0.34, beta = −0.0455, N = 1238
DCPYGTFGMDCR pc2
Q9NQ30
724:757 416:432 63:71
5:54198775:T:C_T
p = 0.21, beta = −0.061, N = 1203
ESM1 : NP2
Q9NQ30
175:175 123:123 27:27
5:54198775:T:C_T
p = 0.66, beta = 0.0373, N = 325
FPFFQYSVTK pc2
Q9NQ30
173:175 122:123 27:27
5:54198775:T:C_T
p = 0.72, beta = 0.031, N = 322
TVLDDCGCCR pc2
Q9NQ30;Q9NQ30−2
170:175 119:123 26:27
5:54198775:T:C_T
p = 0.9, beta = −0.0111, N = 315
ETCNCQSGICDR pc2
Q9NQ30
122:175 84:123 19:27
5:54198775:T:C_T
p = 0.74, beta = −0.0338, N = 225
DCPYGTFGMDCR pc2
Q9NQ30
106:175 68:123 15:27
5:54198775:T:C_T
p = 0.65, beta = −0.0511, N = 189
log10 protein intensity
log10 peptide intensity
Assay Target: ESM1
Olink UniProt: Q9NQ30
UKB rsID: rs4242051
Proxy rsID: rs4242051
UKB SNP: 5:54198775:T:C
Proxy SNP: 5:54198775:T:C
UKB log10(p): 166.8
UKB BETA: 0.191
−:−:−:−:−:−:−
1254:1241:1238:1203:1146:998

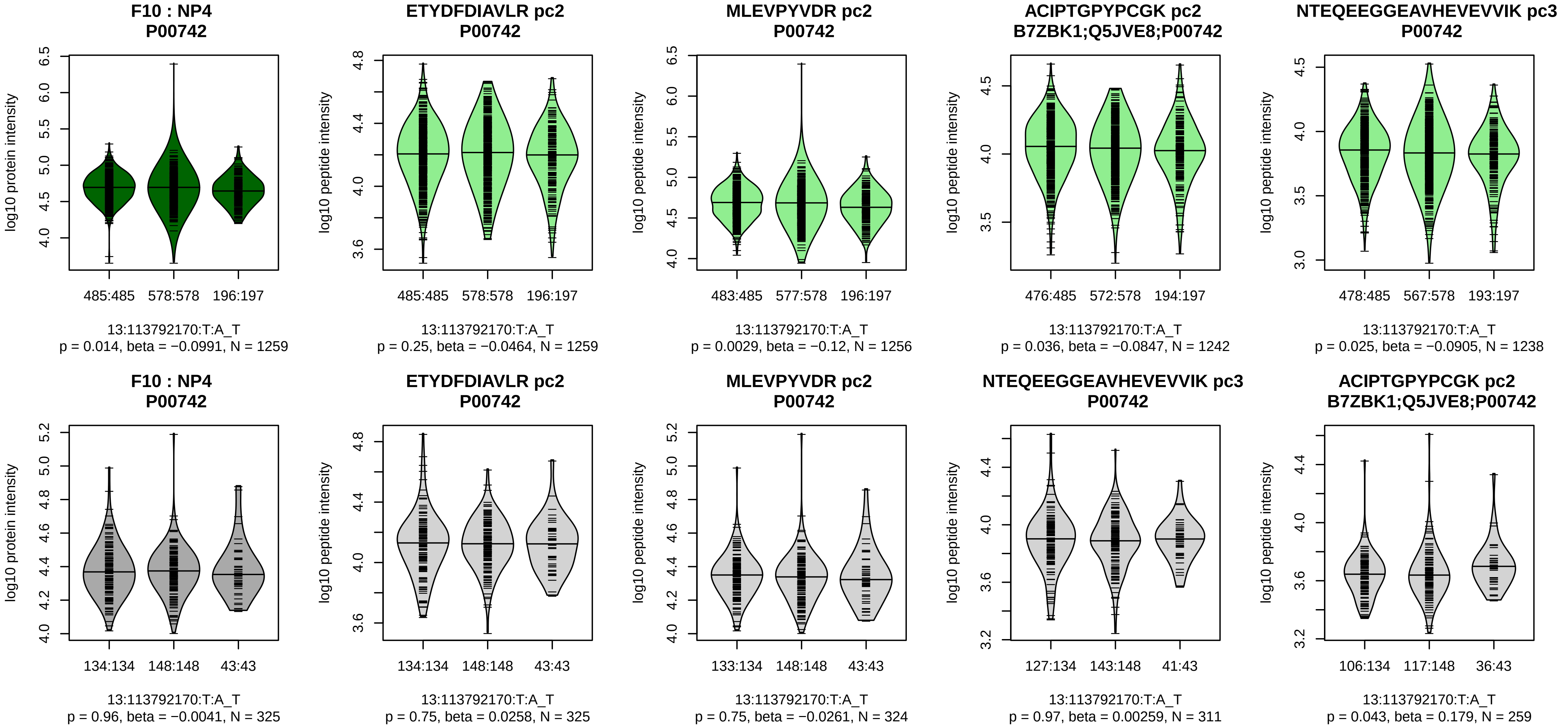


Assay Target: F10
Olink UniProt: P00742
UKB rsID: rs547138
Proxy rsID: rs547138
UKB SNP: 13:113792170:T:A
Proxy SNP: 13:113792170:T:A
UKB log10(p): 159.6
UKB BETA: 0.17
−:*:−:−:−:−:−:−:−:−:−:−:−:−:−:−:NA
1259:1256:1242:1238:1216:106

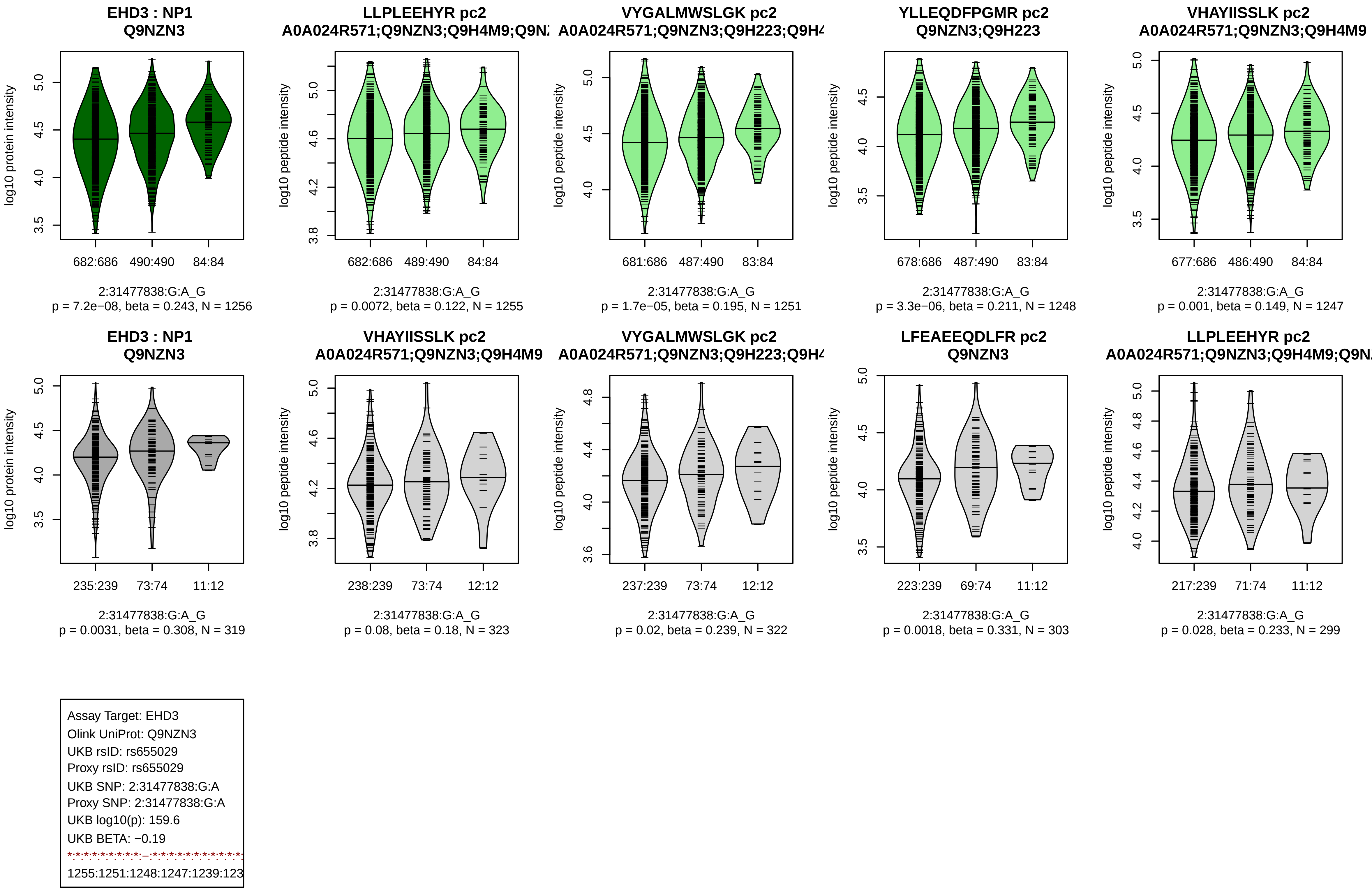

EHD3 : NP1
Q9NZN3
log10 protein intensity
682:686
490:490
84:84
2:31477838:G:A_G
p = 7.2e−08, beta = 0.243, N = 1256
LLPLEEHYR pc2
A0A024R571;Q9NZN3;Q9H4M9;Q9N
log10 peptide intensity
682:686
489:490
84:84
2:31477838:G:A_G
p = 0.0072, beta = 0.122, N = 1255
VYGALMWSLGK pc2
A0A024R571;Q9NZN3;Q9H223;Q9H4
log10 peptide intensity
681:686
487:490
83:84
2:31477838:G:A_G
p = 1.7e−05, beta = 0.195, N = 1251
YLLEQDFPGMR pc2
Q9NZN3;Q9H223
log10 peptide intensity
678:686
487:490
83:84
2:31477838:G:A_G
p = 3.3e−06, beta = 0.211, N = 1248
VHAYIISSLK pc2
A0A024R571;Q9NZN3;Q9H4M9
log10 peptide intensity
677:686
486:490
84:84
2:31477838:G:A_G
p = 0.001, beta = 0.149, N = 1247
EHD3 : NP1
Q9NZN3
log10 protein intensity
235:239
73:74
11:12
2:31477838:G:A_G
p = 0.0031, beta = 0.308, N = 319
VHAYIISSLK pc2
A0A024R571;Q9NZN3;Q9H4M9
log10 peptide intensity
238:239
73:74
12:12
2:31477838:G:A_G
p = 0.08, beta = 0.18, N = 323
VYGALMWSLGK pc2
A0A024R571;Q9NZN3;Q9H223;Q9H4
log10 peptide intensity
237:239
73:74
12:12
2:31477838:G:A_G
p = 0.02, beta = 0.239, N = 322
LFEAEEQDLFR pc2
Q9NZN3
log10 peptide intensity
223:239
69:74
11:12
2:31477838:G:A_G
p = 0.0018, beta = 0.331, N = 303
LLPLEEHYR pc2
A0A024R571;Q9NZN3;Q9H4M9;Q9N
log10 peptide intensity
217:239
71:74
11:12
2:31477838:G:A_G
p = 0.028, beta = 0.233, N = 299
Assay Target: EHD3
Olink UniProt: Q9NZN3
UKB rsID: rs655029
Proxy rsID: rs655029
UKB SNP: 2:31477838:G:A
Proxy SNP: 2:31477838:G:A
UKB log10(p): 159.6
UKB BETA: −0.19
1255:1251:1248:1247:1239:123

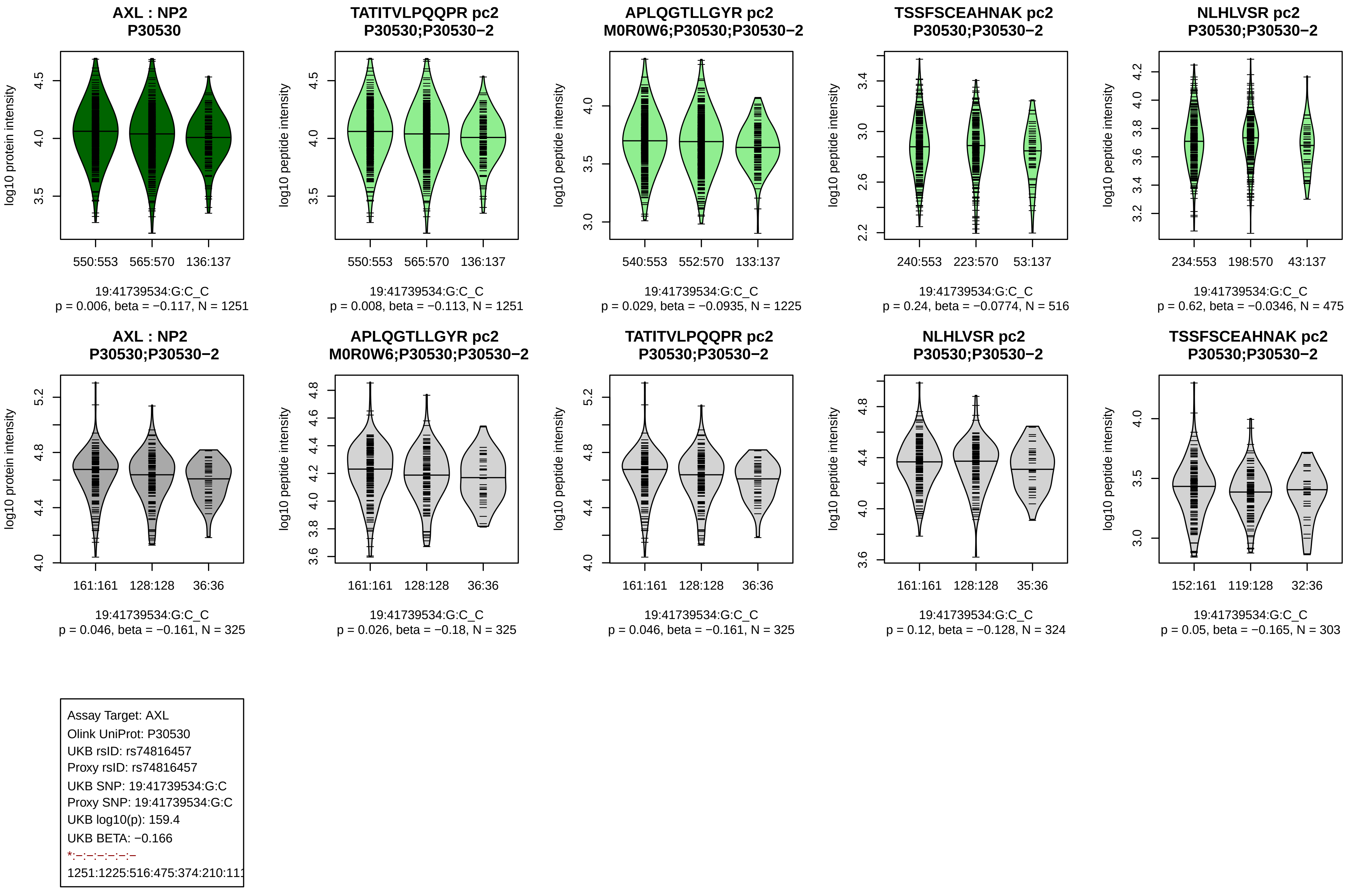
AXL : NP2
P30530
log10 protein intensity
550:553
565:570
136:137
19:41739534:G:C_C
p = 0.006, beta = −0.117, N = 1251
TATITVLPQQPR pc2
P30530;P30530−2
log10 peptide intensity
550:553
565:570
136:137
19:41739534:G:C_C
p = 0.008, beta = −0.113, N = 1251
APLQGTLLGYR pc2
M0R0W6;P30530;P30530−2
log10 peptide intensity
540:553
552:570
133:137
19:41739534:G:C_C
p = 0.029, beta = −0.0935, N = 1225
TSSFSCEAHNAK pc2
P30530;P30530−2
log10 peptide intensity
240:553
223:570
53:137
19:41739534:G:C_C
p = 0.24, beta = −0.0774, N = 516
NLHLVSR pc2
P30530;P30530−2
log10 peptide intensity
234:553
198:570
43:137
19:41739534:G:C_C
p = 0.62, beta = −0.0346, N = 475
AXL : NP2
P30530;P30530−2
log10 protein intensity
161:161
128:128
36:36
19:41739534:G:C_C
p = 0.046, beta = −0.161, N = 325
APLQGTLLGYR pc2
M0R0W6;P30530;P30530−2
log10 peptide intensity
161:161
128:128
36:36
19:41739534:G:C_C
p = 0.026, beta = −0.18, N = 325
TATITVLPQQPR pc2
P30530;P30530−2
log10 peptide intensity
161:161
128:128
36:36
19:41739534:G:C_C
p = 0.046, beta = −0.161, N = 325
NLHLVSR pc2
P30530;P30530−2
log10 peptide intensity
161:161
128:128
35:36
19:41739534:G:C_C
p = 0.12, beta = −0.128, N = 324
TSSFSCEAHNAK pc2
P30530;P30530−2
log10 peptide intensity
152:161
119:128
32:36
19:41739534:G:C_C
p = 0.05, beta = −0.165, N = 303
Assay Target: AXL
Olink UniProt: P30530
UKB rsID: rs74816457
Proxy rsID: rs74816457
UKB SNP: 19:41739534:G:C
Proxy SNP: 19:41739534:G:C
UKB log10(p): 159.4
UKB BETA: −0.166
*:−:−:−:−:−:−
1251:1225:516:475:374:210:111

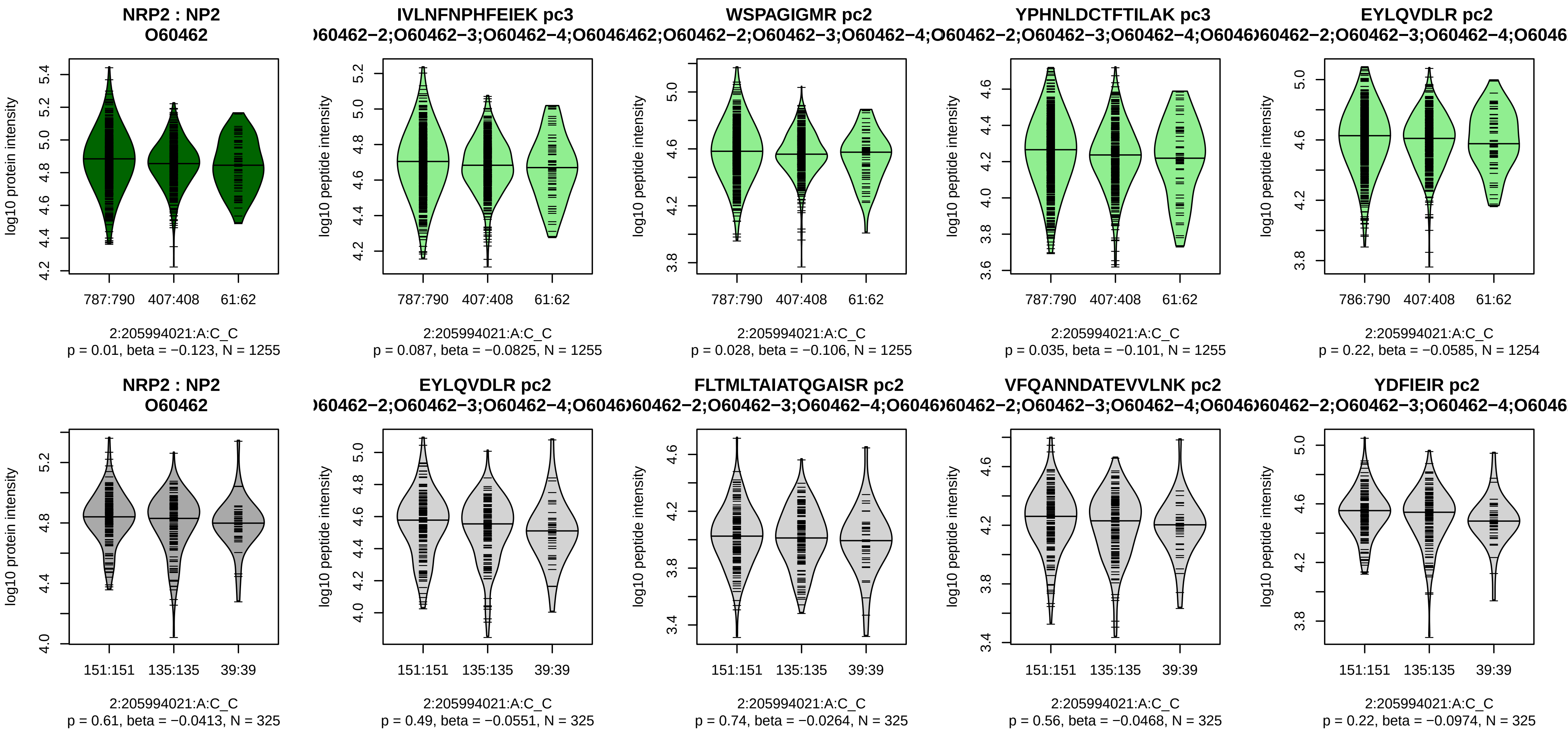


Assay Target: NRP2
Olink UniProt: O60462
UKB rsID: rs761913954
Proxy rsID: rs10932097
UKB SNP: 2:205979199:AGGG
Proxy SNP: 2:205994021:A:C
UKB log10(p): 158.6
UKB BETA: −0.189
−:−:−:−:−:−:−:−:−:−:−:−:−:−:−:
1255:1255:1255:1254:1254:125

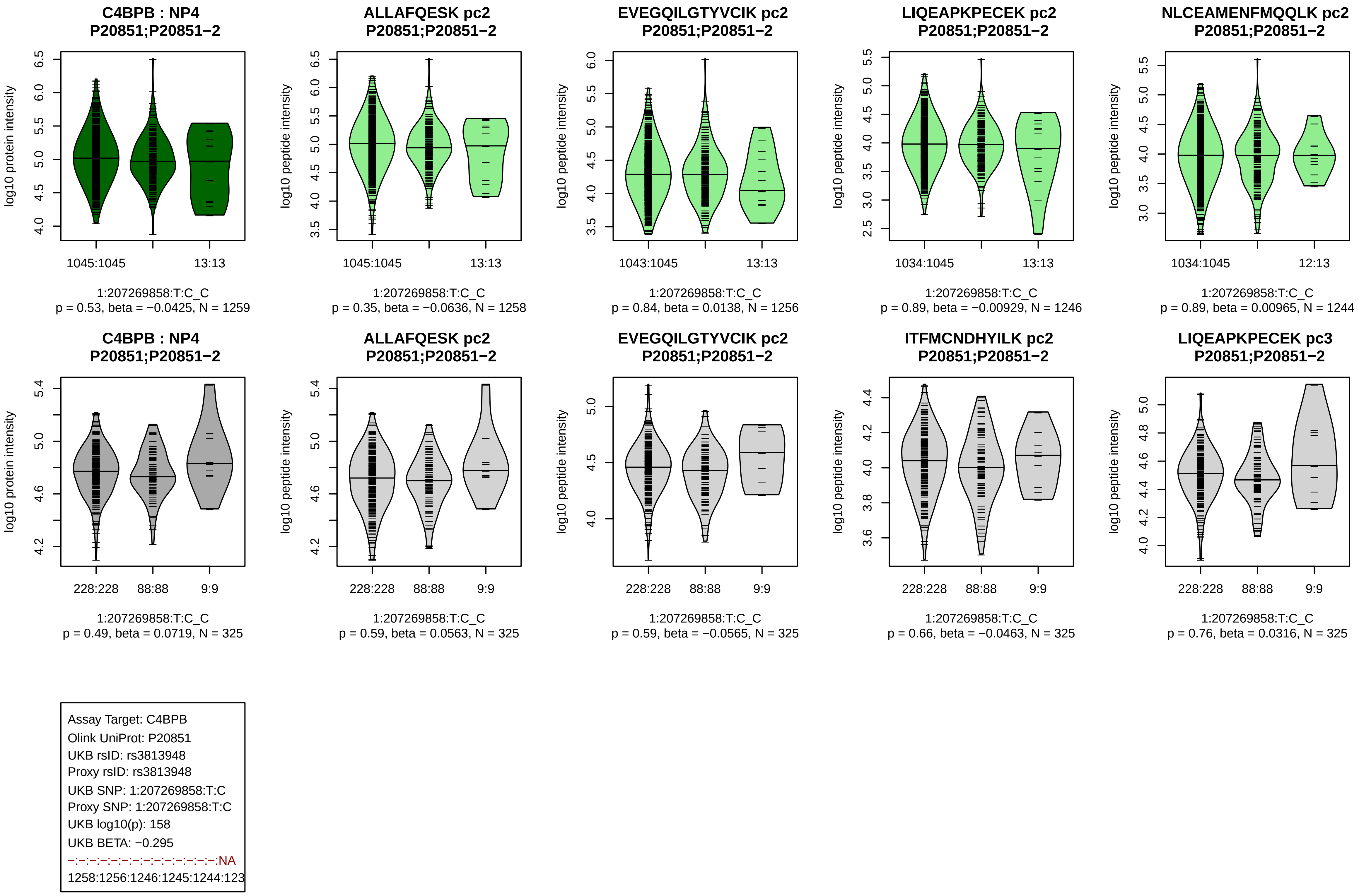

C4BPB : NP4
P20851;P20851−2
log10 protein intensity
1045:1045
13:13
1:207269858:T:C_C
p = 0.53, beta = −0.0425, N = 1259
ALLAFQESK pc2
P20851;P20851−2
log10 peptide intensity
1045:1045
13:13
1:207269858:T:C_C
p = 0.35, beta = −0.0636, N = 1258
EVEGQILGTYVCIK pc2
P20851;P20851−2
log10 peptide intensity
1043:1045
13:13
1:207269858:T:C_C
p = 0.84, beta = 0.0138, N = 1256
LIQEAPKPECEK pc2
P20851;P20851−2
log10 peptide intensity
1034:1045
13:13
1:207269858:T:C_C
p = 0.89, beta = −0.00929, N = 1246
NLCEAMENFMQQLK pc2
P20851;P20851−2
log10 peptide intensity
1034:1045
12:13
1:207269858:T:C_C
p = 0.89, beta = 0.00965, N = 1244
C4BPB : NP4
P20851;P20851−2
log10 protein intensity
228:228
88:88
9:9
1:207269858:T:C_C
p = 0.49, beta = 0.0719, N = 325
ALLAFQESK pc2
P20851;P20851−2
log10 peptide intensity
228:228
88:88
9:9
1:207269858:T:C_C
p = 0.59, beta = 0.0563, N = 325
EVEGQILGTYVCIK pc2
P20851;P20851−2
log10 peptide intensity
228:228
88:88
9:9
1:207269858:T:C_C
p = 0.59, beta = −0.0565, N = 325
ITFMCNDHYILK pc2
P20851;P20851−2
log10 peptide intensity
228:228
88:88
9:9
1:207269858:T:C_C
p = 0.66, beta = −0.0463, N = 325
LIQEAPKPECEK pc3
P20851;P20851−2
log10 peptide intensity
228:228
88:88
9:9
1:207269858:T:C_C
p = 0.76, beta = 0.0316, N = 325
Assay Target: C4BPB
Olink UniProt: P20851
UKB rsID: rs3813948
Proxy rsID: rs3813948
UKB SNP: 1:207269858:T:C
Proxy SNP: 1:207269858:T:C
UKB log10(p): 158
UKB BETA: −0.295
−:−:−:−:−:−:−:−:−:−:−:−:−:−:−:NA
1258:1256:1246:1245:1244:123

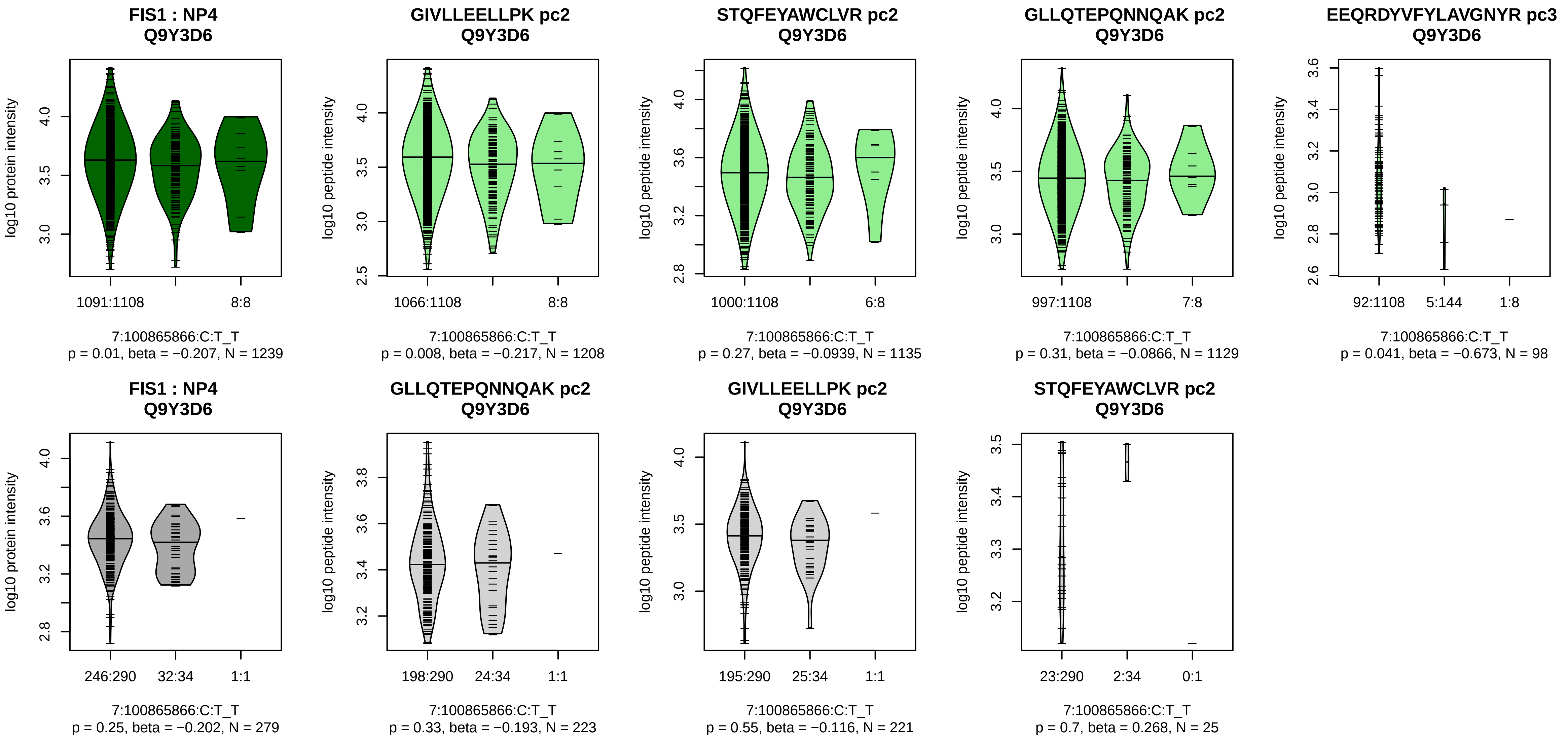


Assay Target: FIS1
Olink UniProt: Q9Y3D6
UKB rsID: rs74662330
Proxy rsID: rs74662330
UKB SNP: 7:100865866:C:T
Proxy SNP: 7:100865866:C:T
UKB log10(p): 155.2
UKB BETA: −0.353
*:−:−:−
1208:1135:1129:98

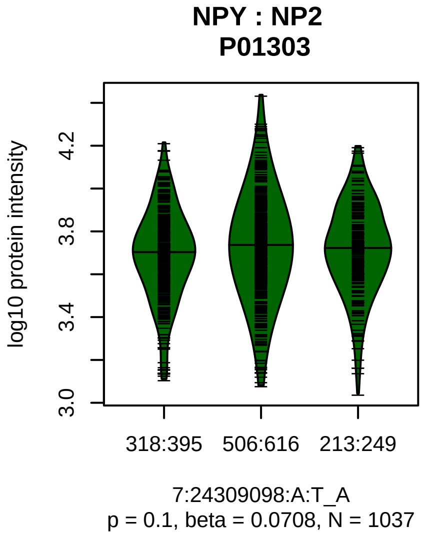


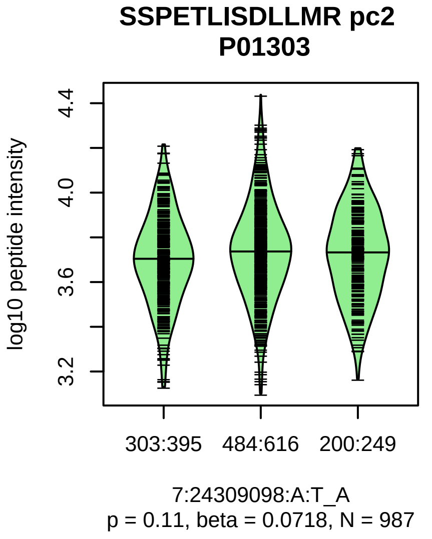


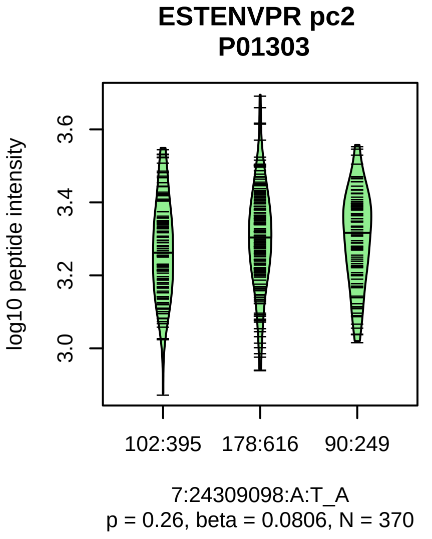


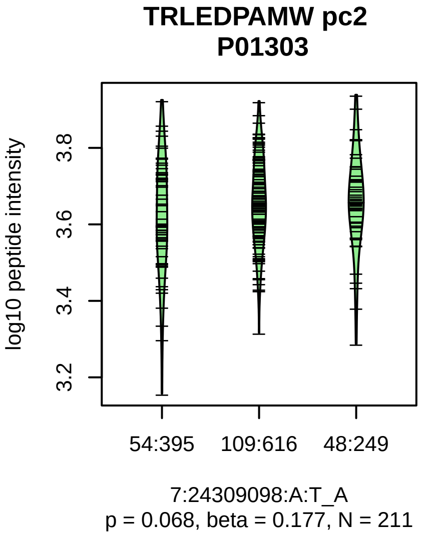


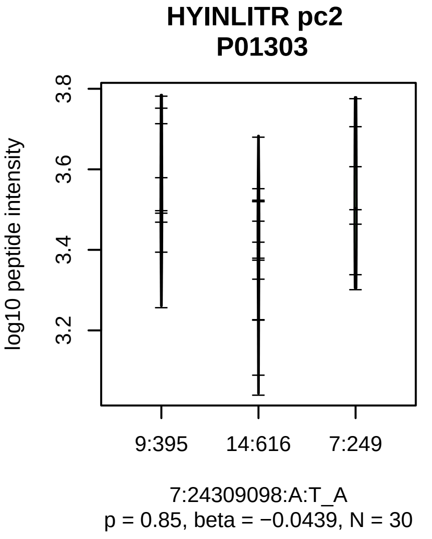


Assay Target: NPY
Olink UniProt: P01303
UKB rsID: rs4722337
Proxy rsID: rs4722337
UKB SNP: 7:24309098:A:T
Proxy SNP: 7:24309098:A:T
UKB log10(p): 152.7
UKB BETA: −0.157
−:−:−:−
987:370:211:30

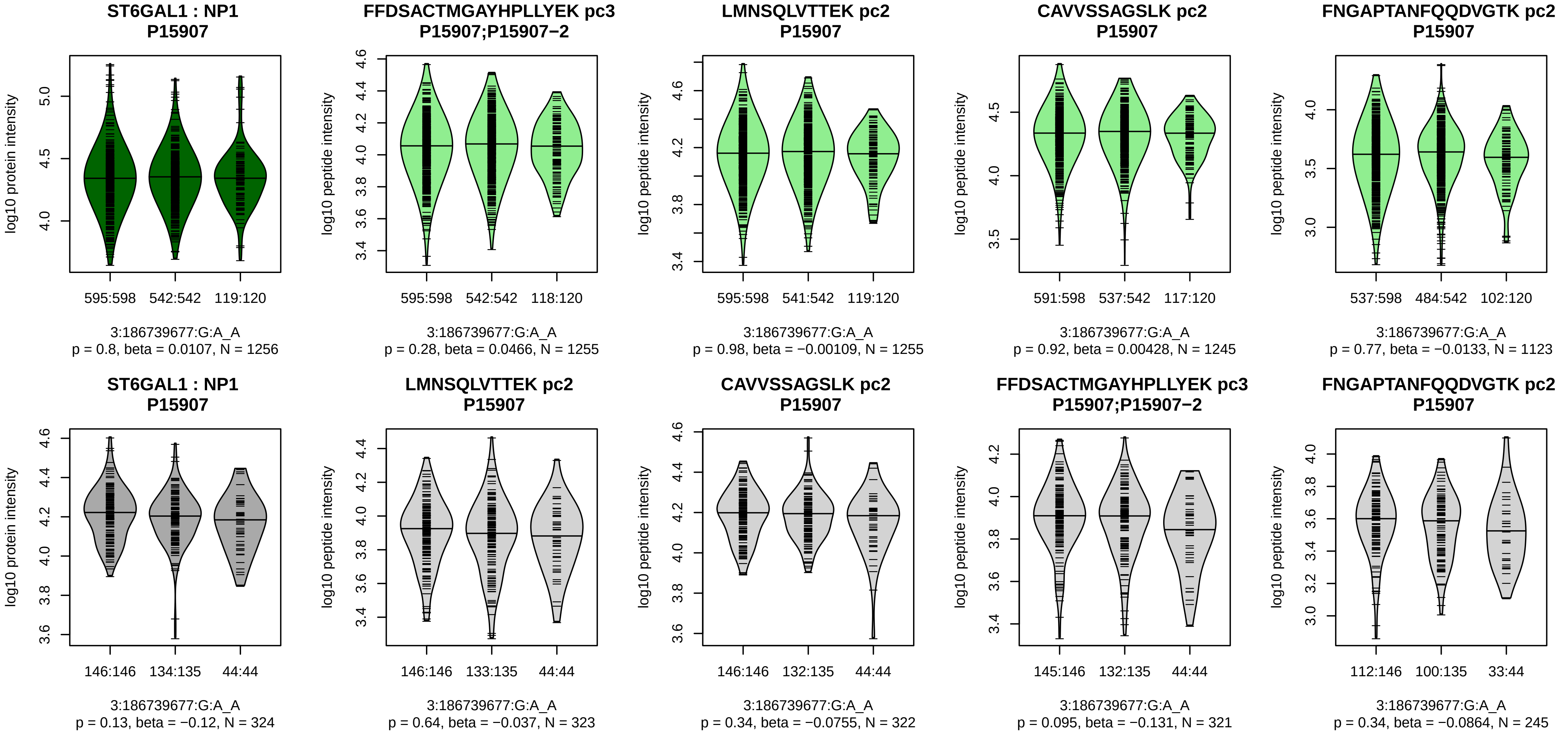


Assay Target: ST6GAL1
Olink UniProt: P15907
UKB rsID: rs4686837
Proxy rsID: rs4686837
UKB SNP: 3:186739677:G:A
Proxy SNP: 3:186739677:G:A
UKB log10(p): 152.6
UKB BETA: −0.17
−:−:−:−:−:−:−:−:−:−:−:−:−:NA:NA
1255:1255:1245:1123:658:500:3

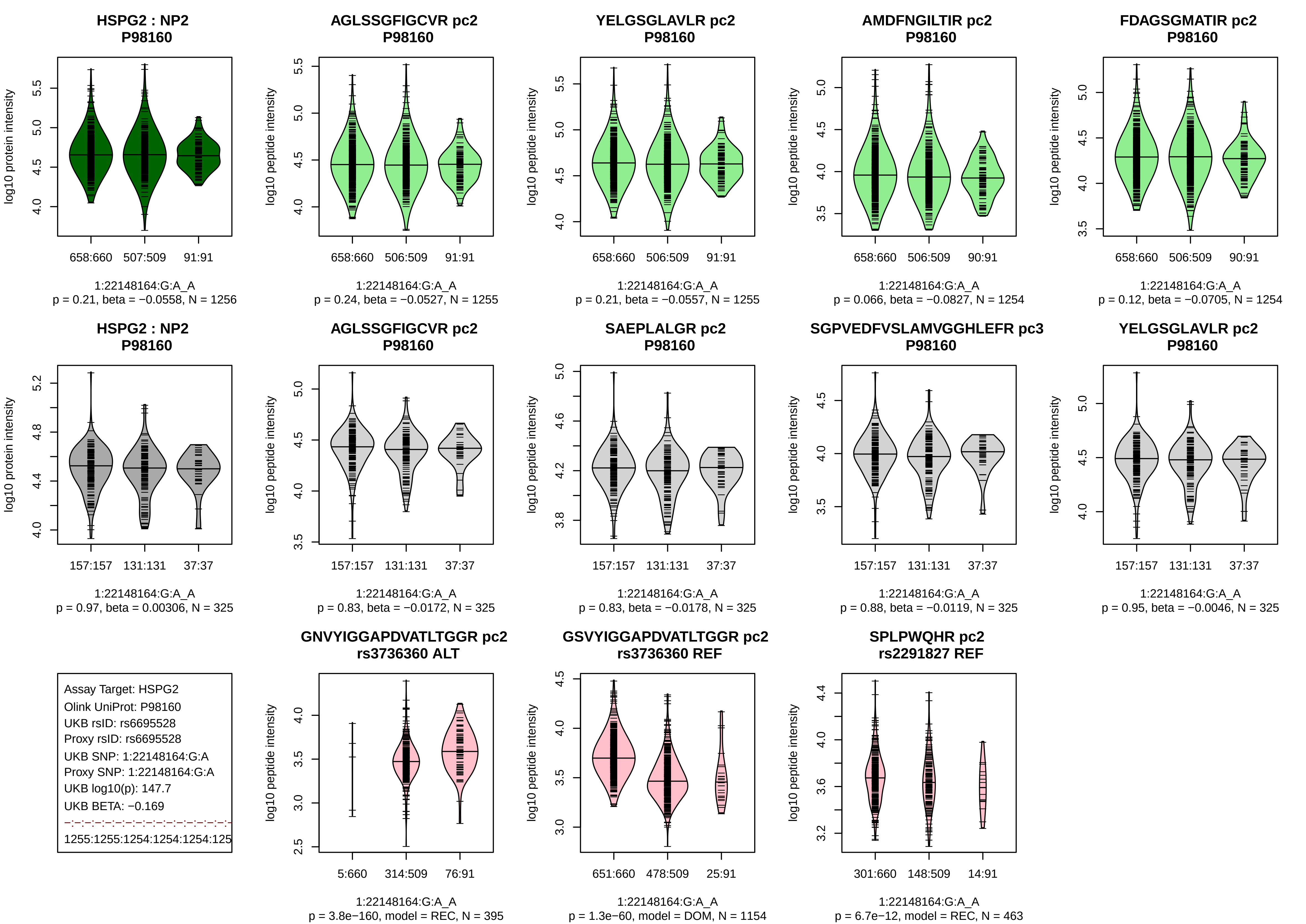
HSPG2 : NP2
P98160
log10 protein intensity
658:660
507:509
91:91
1:22148164:G:A_A
p = 0.21, beta = −0.0558, N = 1256
AGLSSGFIGCVR pc2
P98160
log10 peptide intensity
658:660
506:509
91:91
1:22148164:G:A_A
p = 0.24, beta = −0.0527, N = 1255
YELGSGLAVLR pc2
P98160
log10 peptide intensity
658:660
506:509
91:91
1:22148164:G:A_A
p = 0.21, beta = −0.0557, N = 1255
AMDFNGILTIR pc2
P98160
log10 peptide intensity
658:660
506:509
90:91
1:22148164:G:A_A
p = 0.066, beta = −0.0827, N = 1254
FDAGSGMATIR pc2
P98160
log10 peptide intensity
658:660
506:509
90:91
1:22148164:G:A_A
p = 0.12, beta = −0.0705, N = 1254
HSPG2 : NP2
P98160
log10 protein intensity
157:157
131:131
37:37
1:22148164:G:A_A
p = 0.97, beta = 0.00306, N = 325
AGLSSGFIGCVR pc2
P98160
log10 peptide intensity
157:157
131:131
37:37
1:22148164:G:A_A
p = 0.83, beta = −0.0172, N = 325
SAEPLALGR pc2
P98160
log10 peptide intensity
157:157
131:131
37:37
1:22148164:G:A_A
p = 0.83, beta = −0.0178, N = 325
SGPVEDFVSLAMVGGHLEFR pc3
P98160
log10 peptide intensity
157:157
131:131
37:37
1:22148164:G:A_A
p = 0.88, beta = −0.0119, N = 325
YELGSGLAVLR pc2
P98160
log10 peptide intensity
157:157
131:131
37:37
1:22148164:G:A_A
p = 0.95, beta = −0.0046, N = 325
Assay Target: HSPG2
Olink UniProt: P98160
UKB rsID: rs6695528
Proxy rsID: rs6695528
UKB SNP: 1:22148164:G:A
Proxy SNP: 1:22148164:G:A
UKB log10(p): 147.7
UKB BETA: −0.169
1255:1255:1254:1254:1254:125
GNVYIGGAPDVATLTGGR pc2
rs3736360 ALT
log10 peptide intensity
5:660
314:509
76:91
1:22148164:G:A_A
p = 3.8e−160, model = REC, N = 395
GSVYIGGAPDVATLTGGR pc2
rs3736360 REF
log10 peptide intensity
651:660
478:509
25:91
1:22148164:G:A_A
p = 1.3e−60, model = DOM, N = 1154
SPLPWQHR pc2
rs2291827 REF
log10 peptide intensity
301:660
148:509
14:91
1:22148164:G:A_A
p = 6.7e−12, model = REC, N = 463

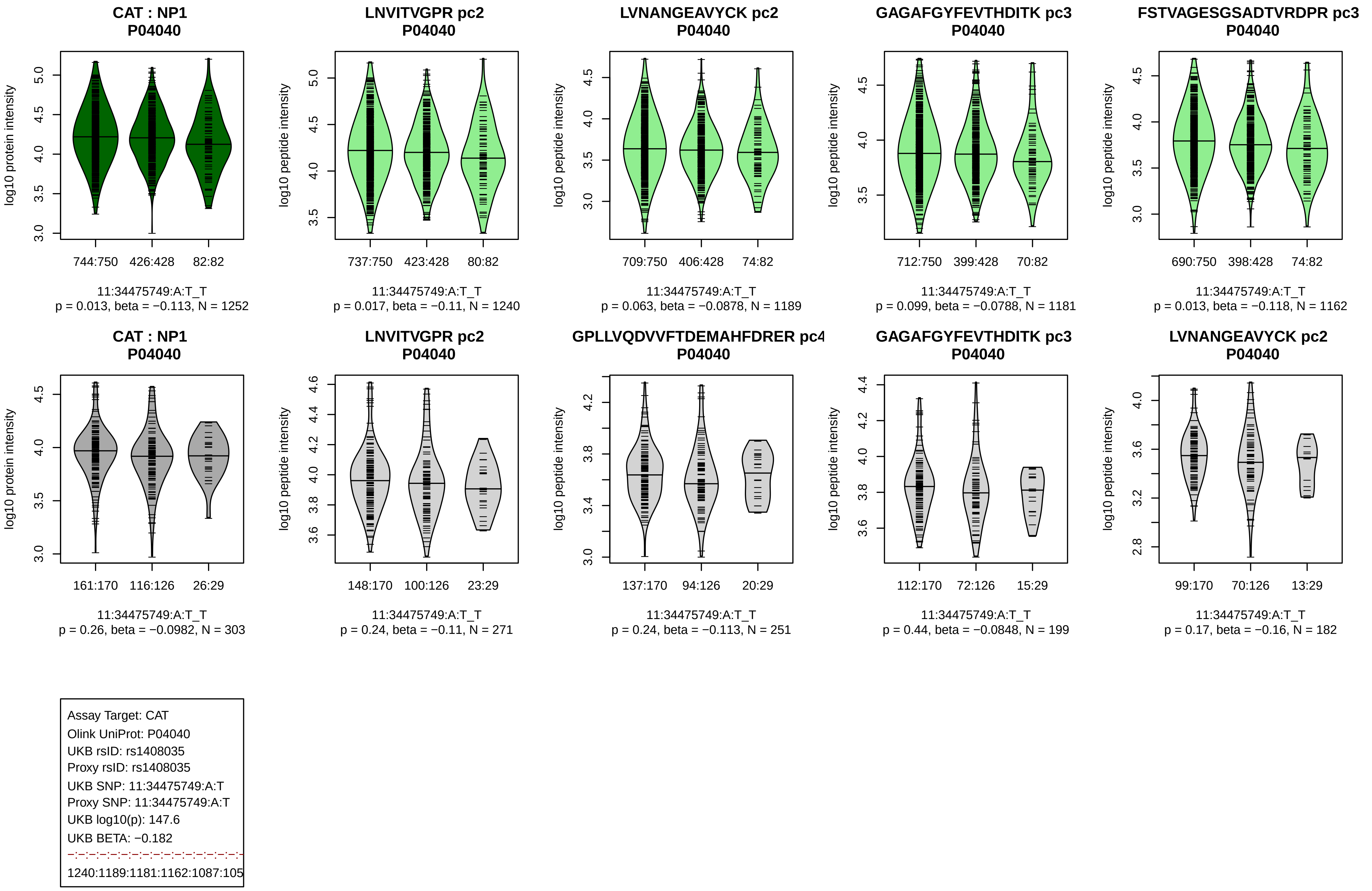
CAT : NP1
P04040
log10 protein intensity
744:750
426:428
82:82
11:34475749:A:T_T
p = 0.013, beta = −0.113, N = 1252
LNVITVGPR pc2
P04040
log10 peptide intensity
737:750
423:428
80:82
11:34475749:A:T_T
p = 0.017, beta = −0.11, N = 1240
LVNANGEAVYCK pc2
P04040
log10 peptide intensity
709:750
406:428
74:82
11:34475749:A:T_T
p = 0.063, beta = −0.0878, N = 1189
GAGAFGYFEVTHDITK pc3
P04040
log10 peptide intensity
712:750
399:428
70:82
11:34475749:A:T_T
p = 0.099, beta = −0.0788, N = 1181
FSTVAGESGSADTVRDPR pc3
P04040
log10 peptide intensity
690:750
398:428
74:82
11:34475749:A:T_T
p = 0.013, beta = −0.118, N = 1162
CAT : NP1
P04040
log10 protein intensity
161:170
116:126
26:29
11:34475749:A:T_T
p = 0.26, beta = −0.0982, N = 303
LNVITVGPR pc2
P04040
log10 peptide intensity
148:170
100:126
23:29
11:34475749:A:T_T
p = 0.24, beta = −0.11, N = 271
GPLLVQDVVFTDEMAHFDRER pc4
P04040
log10 peptide intensity
137:170
94:126
20:29
11:34475749:A:T_T
p = 0.24, beta = −0.113, N = 251
GAGAFGYFEVTHDITK pc3
P04040
log10 peptide intensity
112:170
72:126
15:29
11:34475749:A:T_T
p = 0.44, beta = −0.0848, N = 199
LVNANGEAVYCK pc2
P04040
log10 peptide intensity
99:170
70:126
13:29
11:34475749:A:T_T
p = 0.17, beta = −0.16, N = 182
Assay Target: CAT
Olink UniProt: P04040
UKB rsID: rs1408035
Proxy rsID: rs1408035
UKB SNP: 11:34475749:A:T
Proxy SNP: 11:34475749:A:T
UKB log10(p): 147.6
UKB BETA: −0.182
1240:1189:1181:1162:1087:105

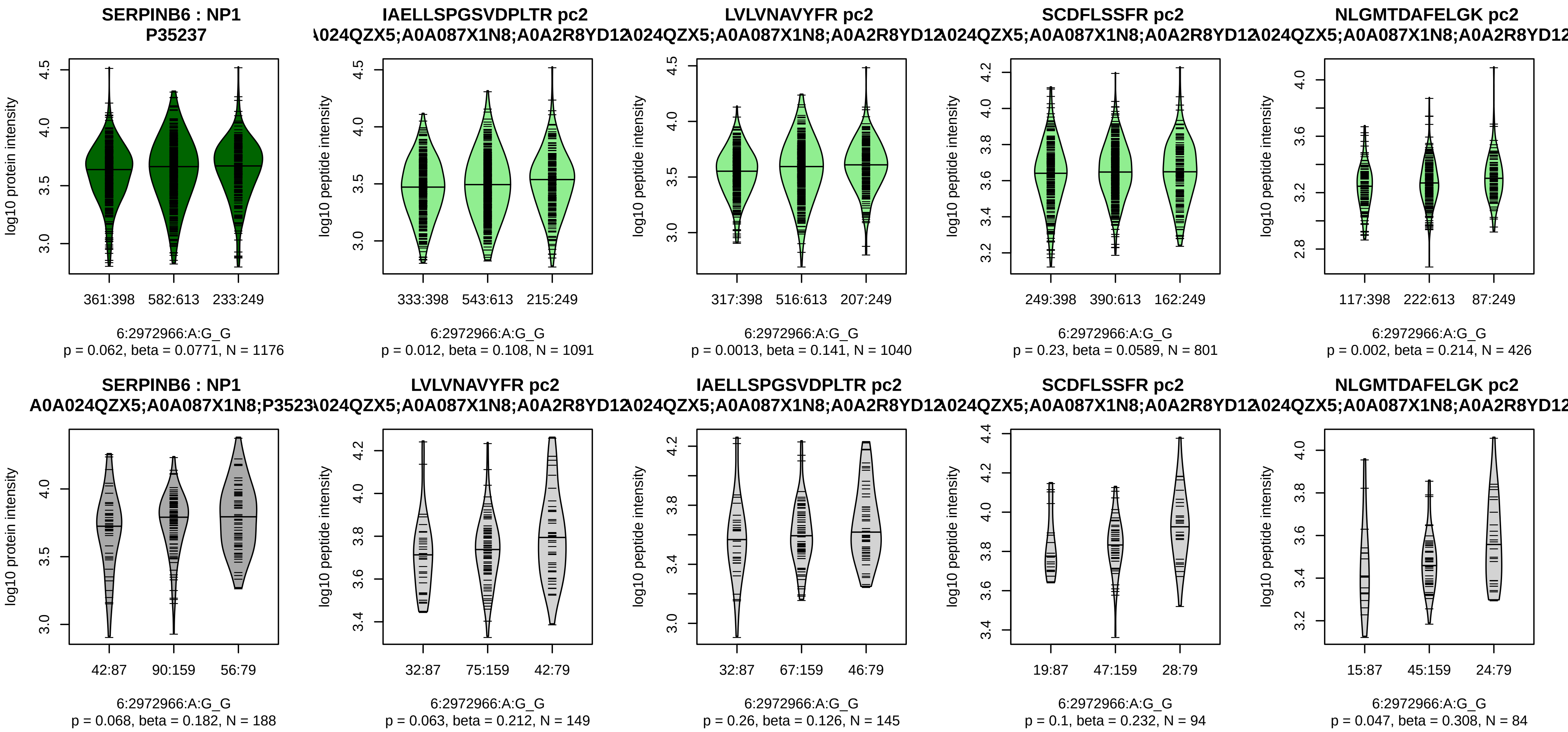


Assay Target: SERPINB6
Olink UniProt: P35237
UKB rsID: rs9405601
Proxy rsID: rs7775515
UKB SNP: 6:2972897:G:A
Proxy SNP: 6:2972966:A:G
UKB log10(p): 147.3
UKB BETA: 0.159
–·*·–·*·–·–·–·–·–·–·–·–·–·–·–·–·–·–·–
1091:1040:801:426:325:255:244

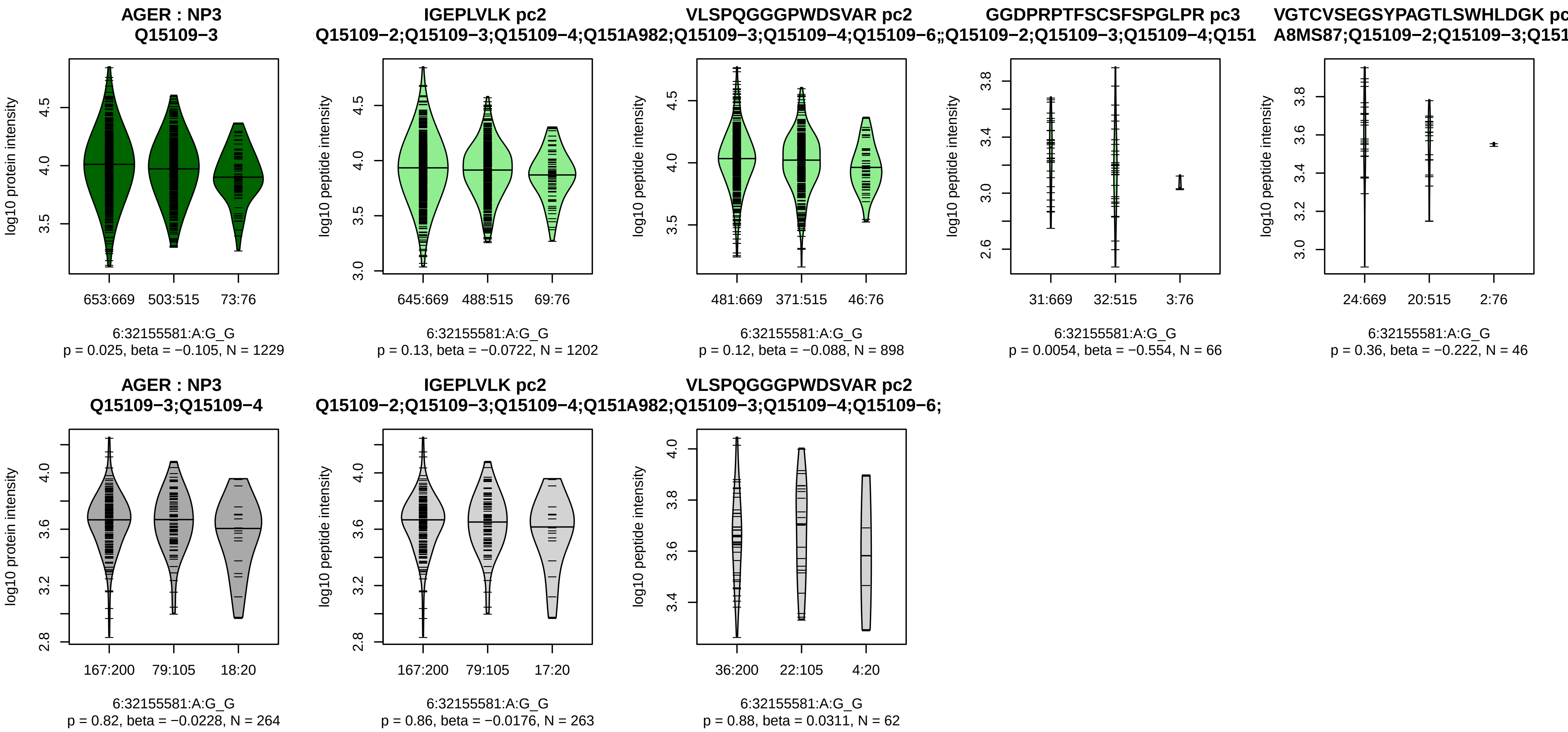


Assay Target: AGER
Olink UniProt: Q15109
UKB rsID: rs204993
Proxy rsID: rs204993
UKB SNP: 6:32155581:A:G
Proxy SNP: 6:32155581:A:G
UKB log10(p): 142.4
UKB BETA: −0.16
−:−:*:−
1202:898:66:46

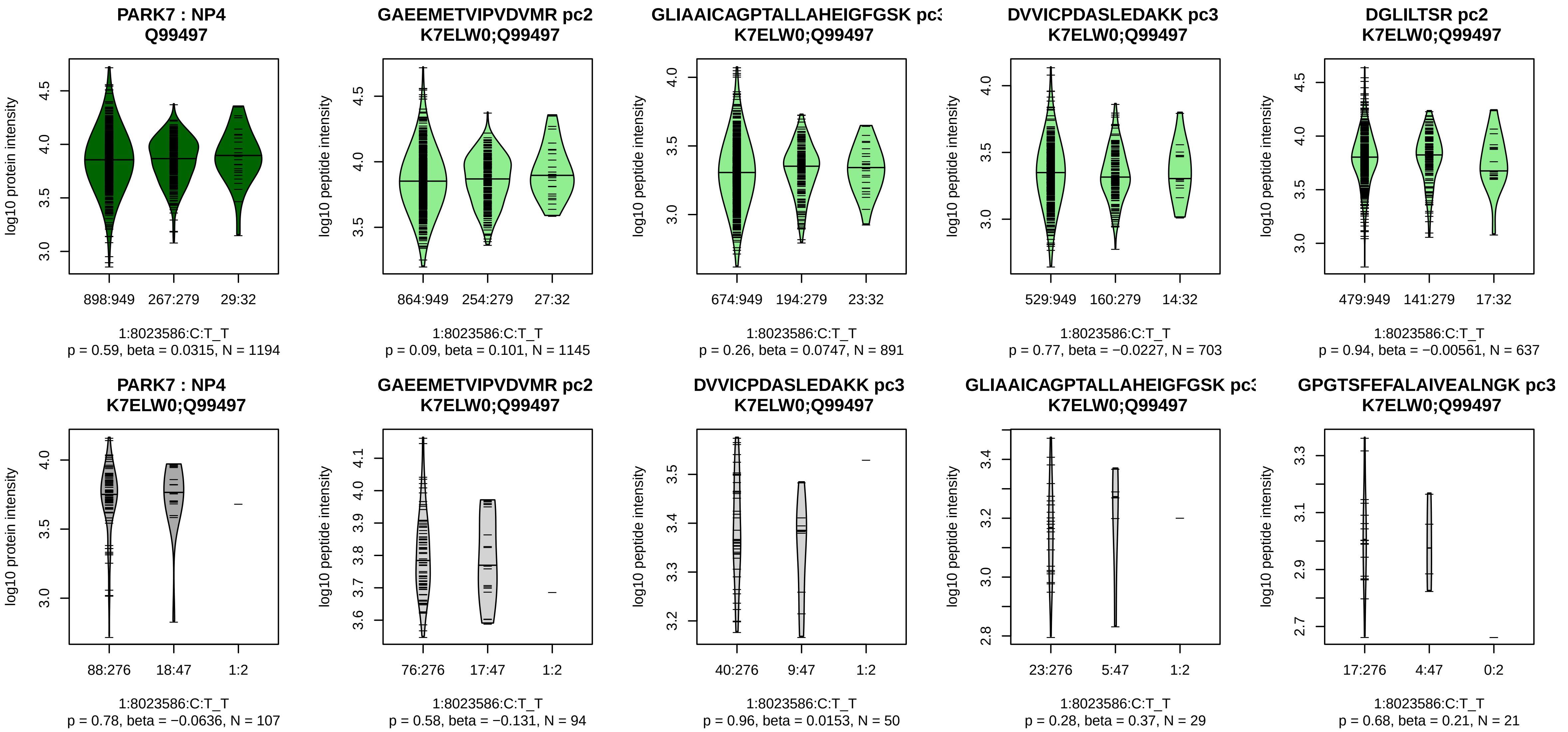


Assay Target: PARK7
Olink UniProt: Q99497
UKB rsID: rs34157438
Proxy rsID: rs34157438
UKB SNP: 1:8023586:C:T
Proxy SNP: 1:8023586:C:T
UKB log10(p): 141.4
UKB BETA: 0.206
−:−:−:−:−:−:−:−:−:−:NA
1145:891:703:637:500:334:244:

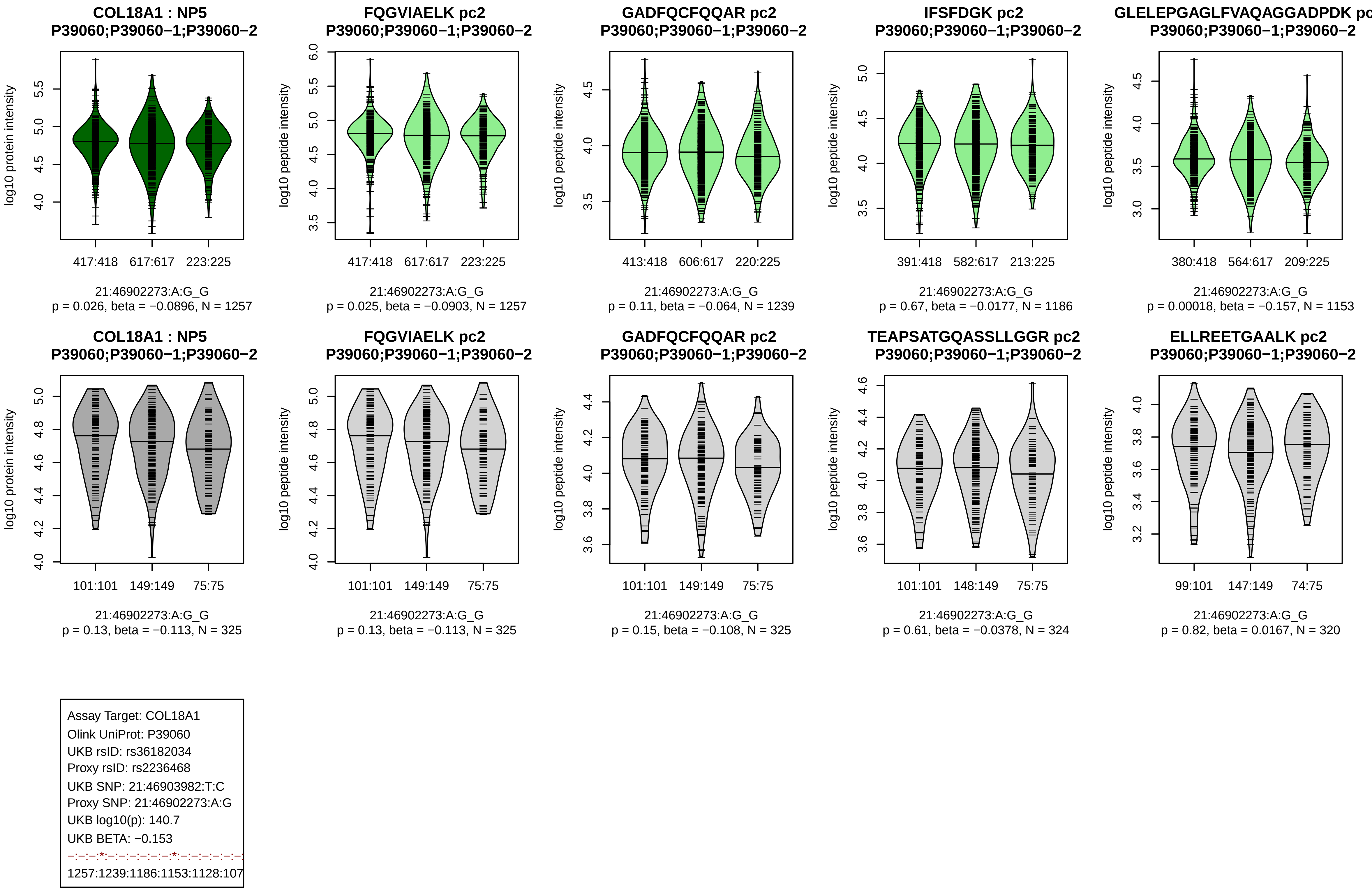
COL18A1 : NP5
P39060;P39060−1;P39060−2
log10 protein intensity
417:418 617:617 223:225
21:46902273:A:G_G
p = 0.026, beta = −0.0896, N = 1257
FQGVIAELK pc2
P39060;P39060−1;P39060−2
log10 peptide intensity
417:418 617:617 223:225
21:46902273:A:G_G
p = 0.025, beta = −0.0903, N = 1257
GADFQCFQQAR pc2
P39060;P39060−1;P39060−2
log10 peptide intensity
413:418 606:617 220:225
21:46902273:A:G_G
p = 0.11, beta = −0.064, N = 1239
IFSFDGK pc2
P39060;P39060−1;P39060−2
log10 peptide intensity
391:418 582:617 213:225
21:46902273:A:G_G
p = 0.67, beta = −0.0177, N = 1186
GLELEPGAGLFVAQAGGADPDK pc
P39060;P39060−1;P39060−2
log10 peptide intensity
380:418 564:617 209:225
21:46902273:A:G_G
p = 0.00018, beta = −0.157, N = 1153
COL18A1 : NP5
P39060;P39060−1;P39060−2
log10 protein intensity
101:101 149:149 75:75
21:46902273:A:G_G
p = 0.13, beta = −0.113, N = 325
FQGVIAELK pc2
P39060;P39060−1;P39060−2
log10 peptide intensity
101:101 149:149 75:75
21:46902273:A:G_G
p = 0.13, beta = −0.113, N = 325
GADFQCFQQAR pc2
P39060;P39060−1;P39060−2
log10 peptide intensity
101:101 149:149 75:75
21:46902273:A:G_G
p = 0.15, beta = −0.108, N = 325
TEAPSATGQASSLLGGR pc2
P39060;P39060−1;P39060−2
log10 peptide intensity
101:101 148:149 75:75
21:46902273:A:G_G
p = 0.61, beta = −0.0378, N = 324
ELLREETGAALK pc2
P39060;P39060−1;P39060−2
log10 peptide intensity
99:101 147:149 74:75
21:46902273:A:G_G
p = 0.82, beta = 0.0167, N = 320
Assay Target: COL18A1
Olink UniProt: P39060
UKB rsID: rs36182034
Proxy rsID: rs2236468
UKB SNP: 21:46903982:T:C
Proxy SNP: 21:46902273:A:G
UKB log10(p): 140.7
UKB BETA: −0.153
−:−:−:*:−:−:−:−:−:−:*:−:−:−:−:−:−:
1257:1239:1186:1153:1128:107

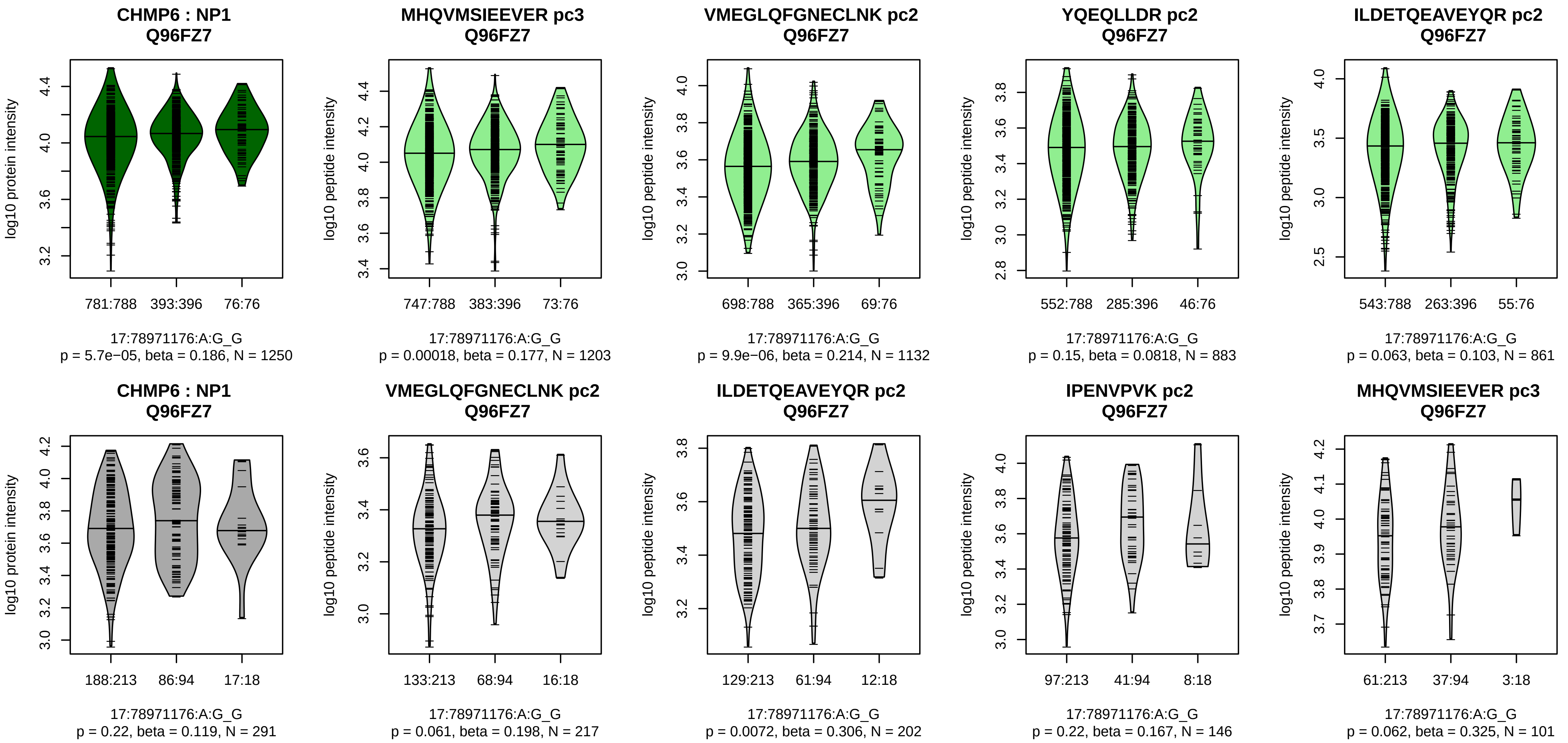


Assay Target: CHMP6
Olink UniProt: Q96FZ7
UKB rsID: rs9989475
Proxy rsID: rs9989475
UKB SNP: 17:78971176:A:G
Proxy SNP: 17:78971176:A:G
UKB log10(p): 139.3
UKB BETA: 0.201
*:*:−:−:−:−
1203:1132:883:861:564:118

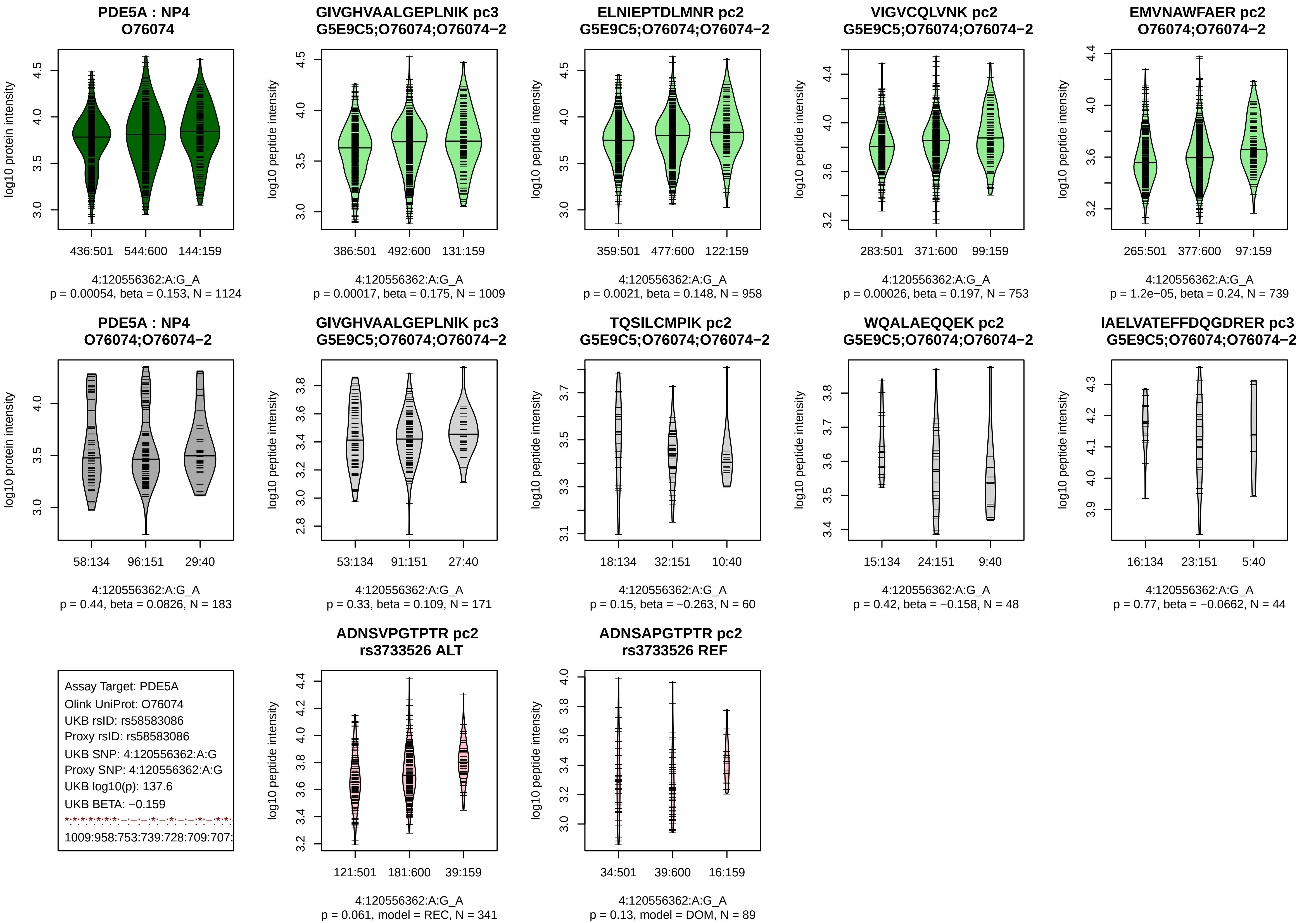

PDE5A : NP4
O76074
log10 protein intensity
436:501
544:600
144:159
4:120556362:A:G_A
p = 0.00054, beta = 0.153, N = 1124
GIVGHVAALGEPLNIK pc3
G5E9C5;O76074;O76074−2
log10 peptide intensity
386:501
492:600
131:159
4:120556362:A:G_A
p = 0.00017, beta = 0.175, N = 1009
ELNIEPTDLMNR pc2
G5E9C5;O76074;O76074−2
log10 peptide intensity
359:501
477:600
122:159
4:120556362:A:G_A
p = 0.0021, beta = 0.148, N = 958
VIGVCQLVNK pc2
G5E9C5;O76074;O76074−2
log10 peptide intensity
283:501
371:600
99:159
4:120556362:A:G_A
p = 0.00026, beta = 0.197, N = 753
EMVNAWFAER pc2
O76074;O76074−2
log10 peptide intensity
265:501
377:600
97:159
4:120556362:A:G_A
p = 1.2e−05, beta = 0.24, N = 739
PDE5A : NP4
O76074;O76074−2
log10 protein intensity
58:134
96:151
29:40
4:120556362:A:G_A
p = 0.44, beta = 0.0826, N = 183
GIVGHVAALGEPLNIK pc3
G5E9C5;O76074;O76074−2
log10 peptide intensity
53:134
91:151
27:40
4:120556362:A:G_A
p = 0.33, beta = 0.109, N = 171
TQSILCMPIK pc2
G5E9C5;O76074;O76074−2
log10 peptide intensity
18:134
32:151
10:40
4:120556362:A:G_A
p = 0.15, beta = −0.263, N = 60
WQALAEQQEK pc2
G5E9C5;O76074;O76074−2
log10 peptide intensity
15:134
24:151
9:40
4:120556362:A:G_A
p = 0.42, beta = −0.158, N = 48
IAELVATEFFDQGDRER pc3
G5E9C5;O76074;O76074−2
log10 peptide intensity
16:134
23:151
5:40
4:120556362:A:G_A
p = 0.77, beta = −0.0662, N = 44
Assay Target: PDE5A
Olink UniProt: O76074
UKB rsID: rs58583086
Proxy rsID: rs58583086
UKB SNP: 4:120556362:A:G
Proxy SNP: 4:120556362:A:G
UKB log10(p): 137.6
UKB BETA: −0.159
*:*:*:*:*:*:–:–:–:*:–:*:–:–:*:–:*:*:
1009:958:753:739:728:709:707:
ADNSVPGTPTR pc2
rs3733526 ALT
log10 peptide intensity
121:501
181:600
39:159
4:120556362:A:G_A
p = 0.061, model = REC, N = 341
ADNSAPGTPTR pc2
rs3733526 REF
log10 peptide intensity
34:501
39:600
16:159
4:120556362:A:G_A
p = 0.13, model = DOM, N = 89

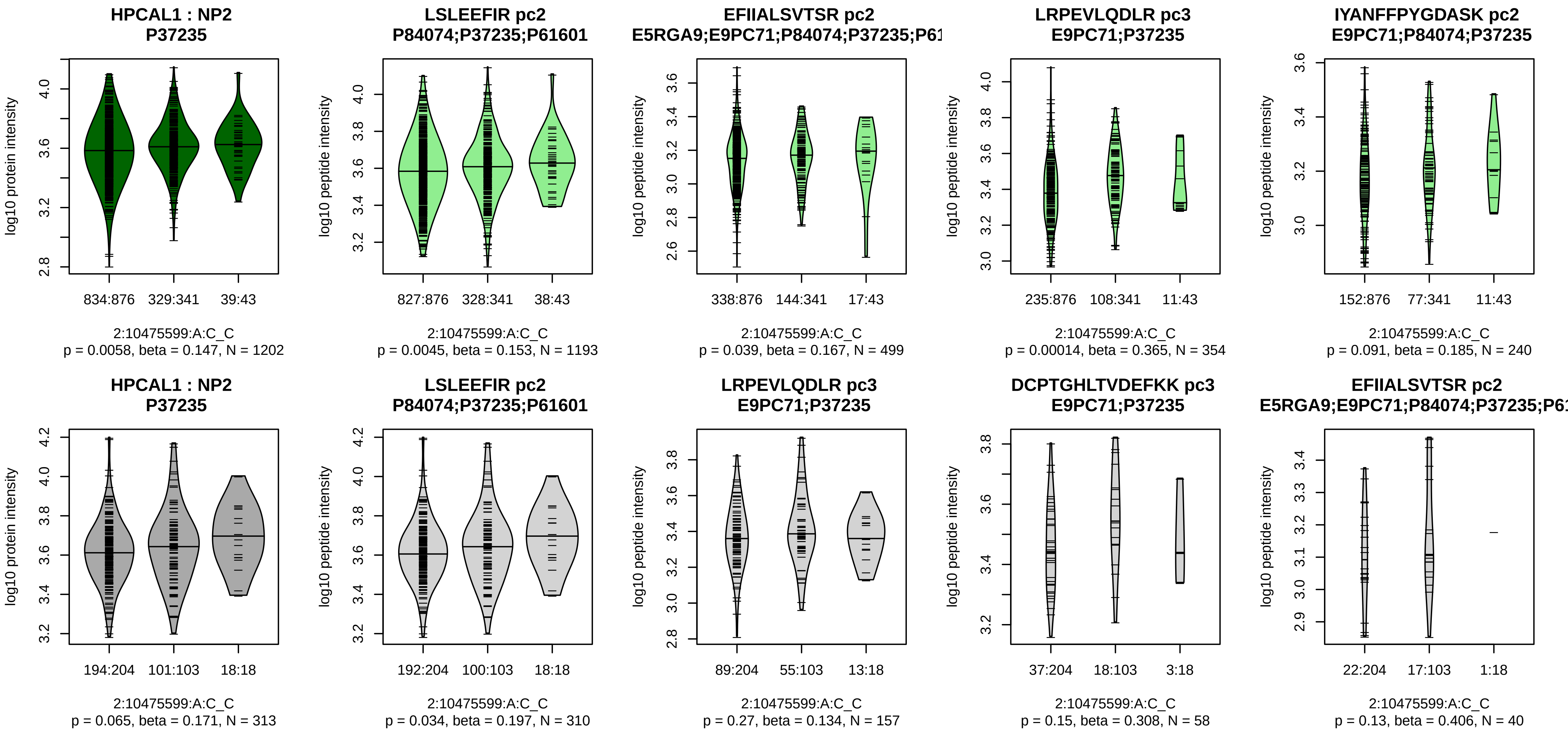


Assay Target: HPCAL1
Olink UniProt: P37235
UKB rsID: rs375285744
Proxy rsID: rs11891604
UKB SNP: 2:10476228:T:C
Proxy SNP: 2:10475599:A:C
UKB log10(p): 137
UKB BETA: 0.213
*:−:*:−:−:−:NA
1193:499:354:240:130:95:2

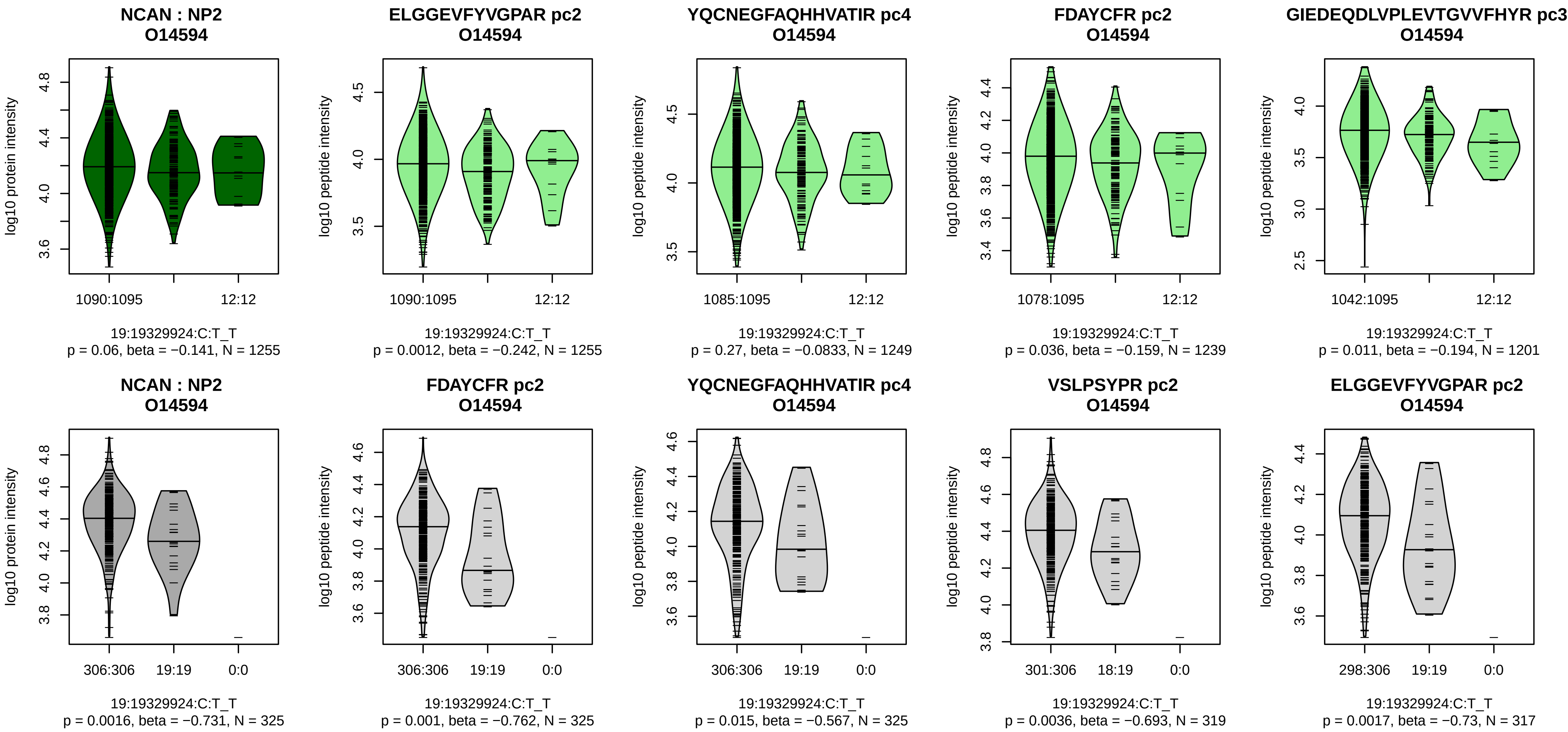


Assay Target: NCAN
Olink UniProt: O14594
UKB rsID: rs2228603
Proxy rsID: rs2228603
UKB SNP: 19:19329924:C:T
Proxy SNP: 19:19329924:C:T
UKB log10(p): 135.3
UKB BETA: −0.272
*:–:–:–:–:–:–:–:–:*:–:–:–:–:*:–:–:
1255:1249:1239:1201:1189:117

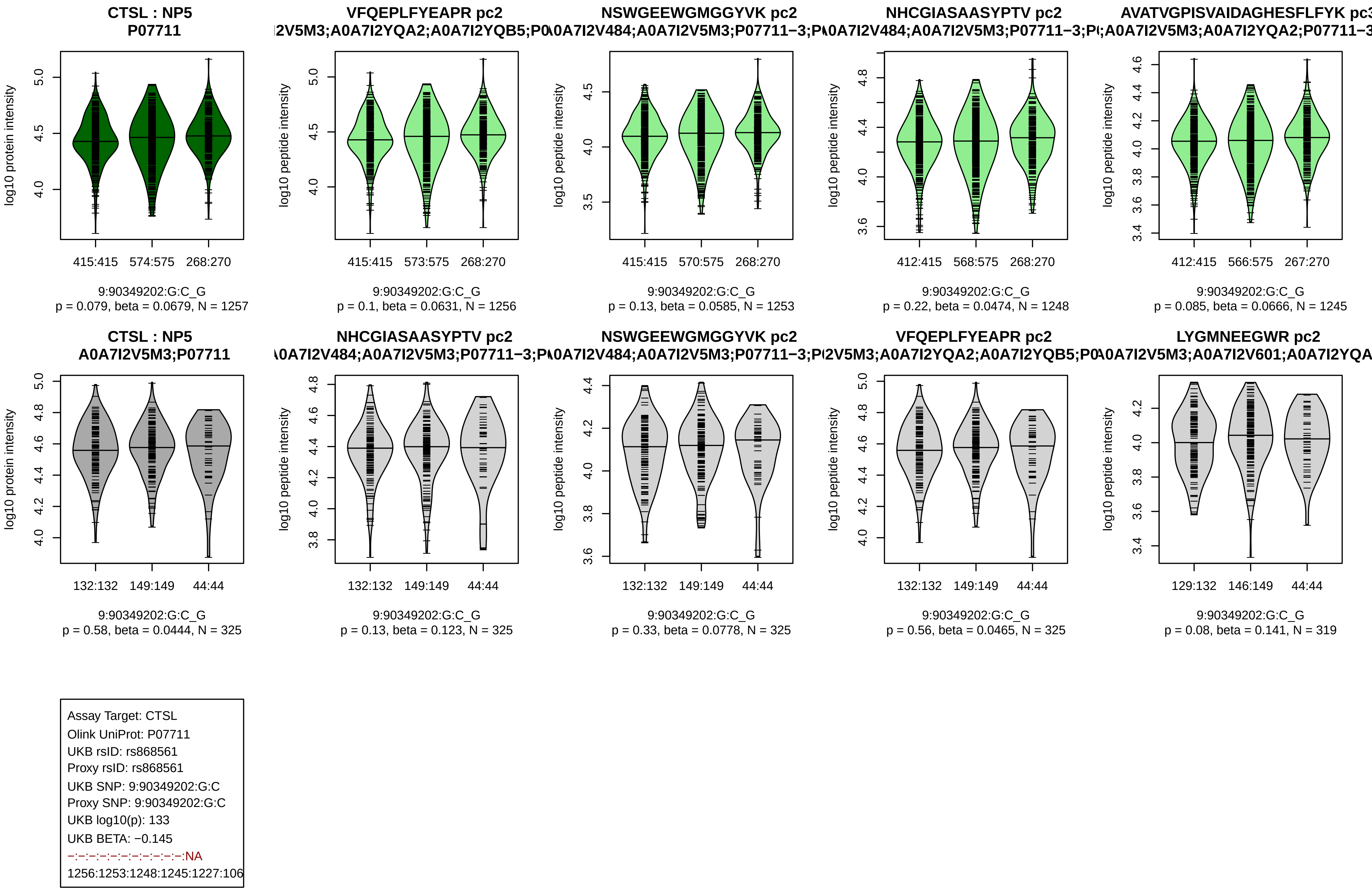

CTSL : NP5
P07711
log10 protein intensity
415:415
574:575
268:270
9:90349202:G:C_G
p = 0.079, beta = 0.0679, N = 1257
VFQEPLFYEAPR pc2
I2V5M3;A0A7I2YQA2;A0A7I2YQB5;P0
log10 peptide intensity
415:415
573:575
268:270
9:90349202:G:C_G
p = 0.1, beta = 0.0631, N = 1256
NSWGEEWGMGGYVK pc2
A0A7I2V484;A0A7I2V5M3;P07711−3;P
log10 peptide intensity
415:415
570:575
268:270
9:90349202:G:C_G
p = 0.13, beta = 0.0585, N = 1253
NHCGIASAASYPTV pc2
A0A7I2V484;A0A7I2V5M3;P07711−3;P
log10 peptide intensity
412:415
568:575
268:270
9:90349202:G:C_G
p = 0.22, beta = 0.0474, N = 1248
AVATVGPISVAIDAGHESFLFYK pc3
A0A7I2V5M3;A0A7I2YQA2;P07711−3
log10 peptide intensity
412:415
566:575
267:270
9:90349202:G:C_G
p = 0.085, beta = 0.0666, N = 1245
CTSL : NP5
A0A7I2V5M3;P07711
log10 protein intensity
132:132
149:149
44:44
9:90349202:G:C_G
p = 0.58, beta = 0.0444, N = 325
NHCGIASAASYPTV pc2
A0A7I2V484;A0A7I2V5M3;P07711−3;P
log10 peptide intensity
132:132
149:149
44:44
9:90349202:G:C_G
p = 0.13, beta = 0.123, N = 325
NSWGEEWGMGGYVK pc2
A0A7I2V484;A0A7I2V5M3;P07711−3;P
log10 peptide intensity
132:132
149:149
44:44
9:90349202:G:C_G
p = 0.33, beta = 0.0778, N = 325
VFQEPLFYEAPR pc2
I2V5M3;A0A7I2YQA2;A0A7I2YQB5;P0
log10 peptide intensity
132:132
149:149
44:44
9:90349202:G:C_G
p = 0.56, beta = 0.0465, N = 325
LYGMNEEGWR pc2
A0A7I2V5M3;A0A7I2V601;A0A7I2YQA
log10 peptide intensity
129:132
146:149
44:44
9:90349202:G:C_G
p = 0.08, beta = 0.141, N = 319
Assay Target: CTSL
Olink UniProt: P07711
UKB rsID: rs868561
Proxy rsID: rs868561
UKB SNP: 9:90349202:G:C
Proxy SNP: 9:90349202:G:C
UKB log10(p): 133
UKB BETA: −0.145
−:−:−:−:−:−:−:−:−:−:−:NA
1256:1253:1248:1245:1227:106

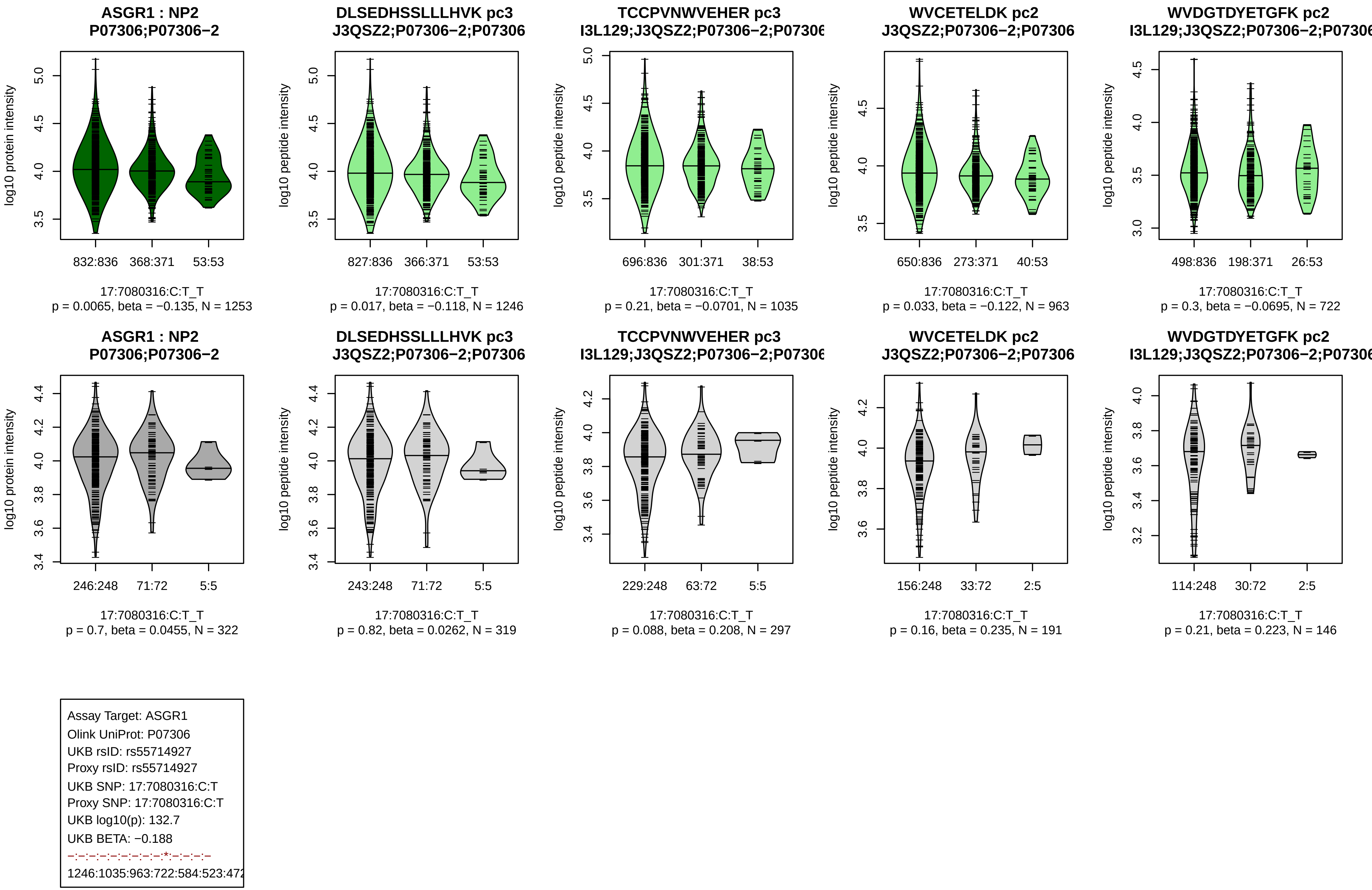
ASGR1 : NP2
P07306;P07306−2
log10 protein intensity
832:836
368:371
53:53
17:7080316:C:T_T
p = 0.0065, beta = −0.135, N = 1253
DLSEDHSSLLLHVK pc3
J3QSZ2;P07306−2;P07306
log10 peptide intensity
827:836
366:371
53:53
17:7080316:C:T_T
p = 0.017, beta = −0.118, N = 1246
TCCPVNWVEHER pc3
I3L129;J3QSZ2;P07306−2;P07306
log10 peptide intensity
696:836
301:371
38:53
17:7080316:C:T_T
p = 0.21, beta = −0.0701, N = 1035
WVCETELDK pc2
J3QSZ2;P07306−2;P07306
log10 peptide intensity
650:836
273:371
40:53
17:7080316:C:T_T
p = 0.033, beta = −0.122, N = 963
WVDGTDYETGFK pc2
I3L129;J3QSZ2;P07306−2;P07306
log10 peptide intensity
498:836
198:371
26:53
17:7080316:C:T_T
p = 0.3, beta = −0.0695, N = 722
ASGR1 : NP2
P07306;P07306−2
log10 protein intensity
246:248
71:72
5:5
17:7080316:C:T_T
p = 0.7, beta = 0.0455, N = 322
DLSEDHSSLLLHVK pc3
J3QSZ2;P07306−2;P07306
log10 peptide intensity
243:248
71:72
5:5
17:7080316:C:T_T
p = 0.82, beta = 0.0262, N = 319
TCCPVNWVEHER pc3
I3L129;J3QSZ2;P07306−2;P07306
log10 peptide intensity
229:248
63:72
5:5
17:7080316:C:T_T
p = 0.088, beta = 0.208, N = 297
WVCETELDK pc2
J3QSZ2;P07306−2;P07306
log10 peptide intensity
156:248
33:72
2:5
17:7080316:C:T_T
p = 0.16, beta = 0.235, N = 191
WVDGTDYETGFK pc2
I3L129;J3QSZ2;P07306−2;P07306
log10 peptide intensity
114:248
30:72
2:5
17:7080316:C:T_T
p = 0.21, beta = 0.223, N = 146
Assay Target: ASGR1
Olink UniProt: P07306
UKB rsID: rs55714927
Proxy rsID: rs55714927
UKB SNP: 17:7080316:C:T
Proxy SNP: 17:7080316:C:T
UKB log10(p): 132.7
UKB BETA: −0.188
−:−:−:−:−:−:−:−:−:*:−:−:−:−
1246:1035:963:722:584:523:472

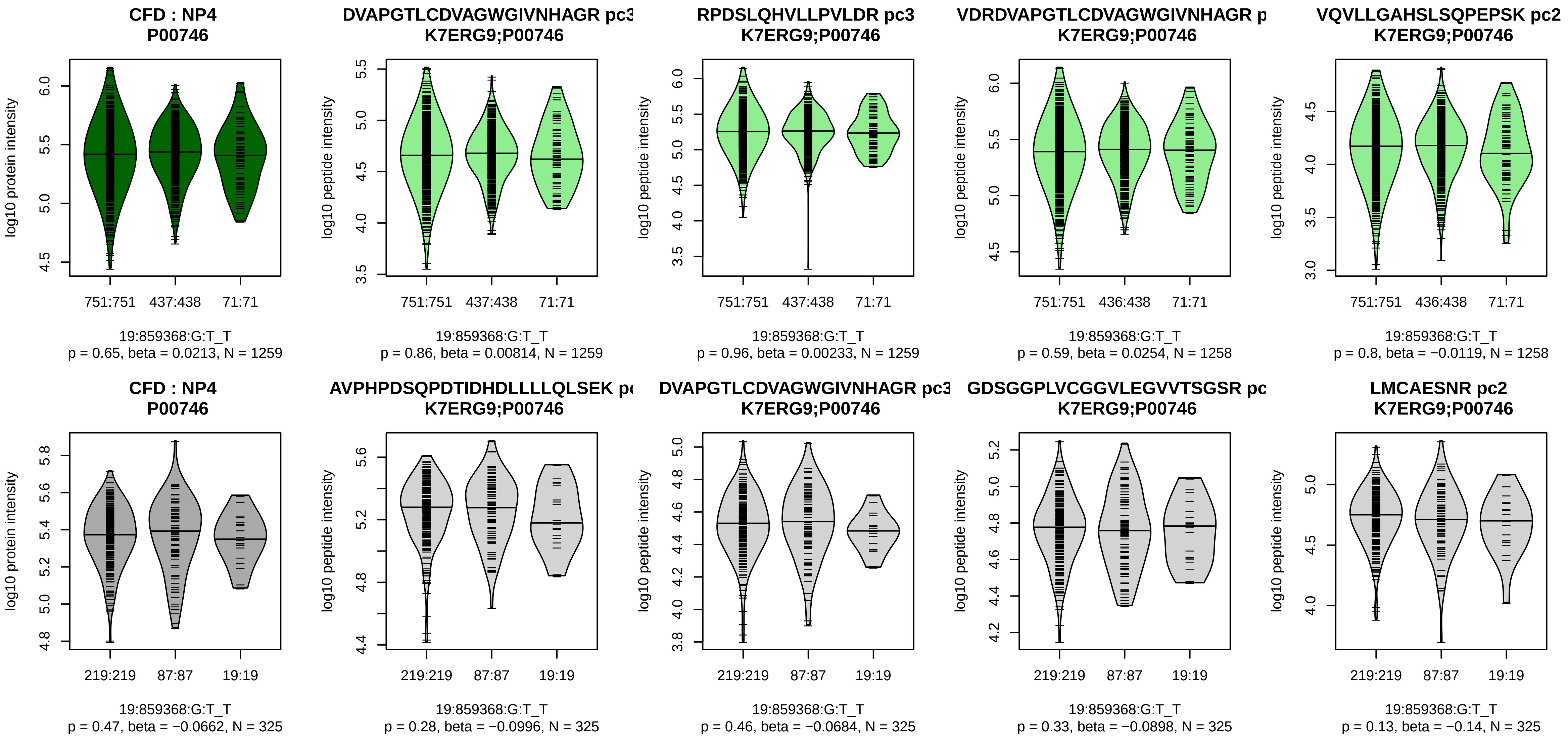


Assay Target: CFD
Olink UniProt: P00746
UKB rsID: rs71335276
Proxy rsID: rs71335276
UKB SNP: 19:859368:G:T
Proxy SNP: 19:859368:G:T
UKB log10(p): 131.4
UKB BETA: −0.181

1259:1259:1258:1258:1258:125

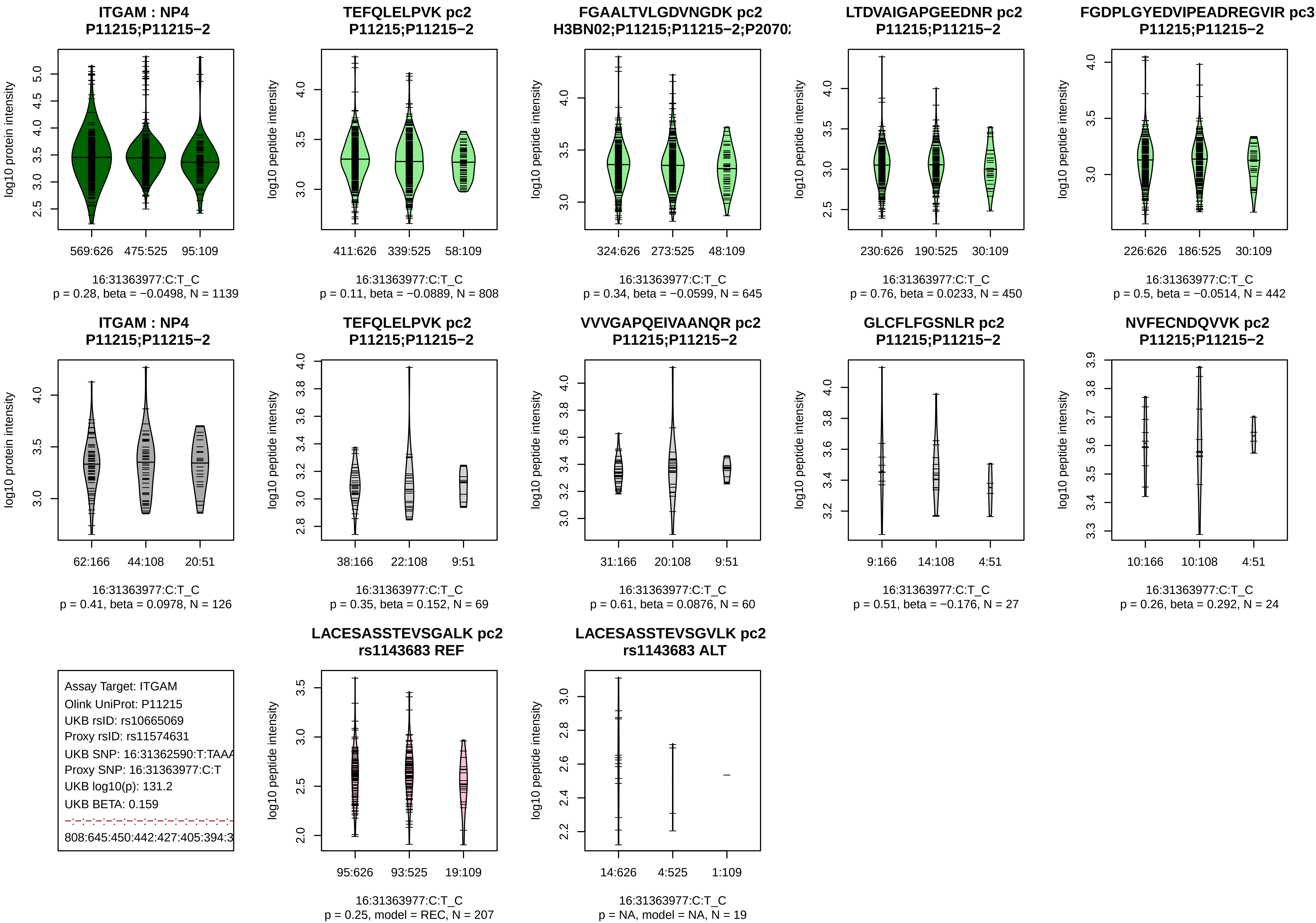

ITGAM : NP4
P11215;P11215−2
log10 protein intensity
569:626
475:525
95:109
16:31363977:C:T_C
p = 0.28, beta = −0.0498, N = 1139
TEFQLELPVK pc2
P11215;P11215−2
log10 peptide intensity
411:626
339:525
58:109
16:31363977:C:T_C
p = 0.11, beta = −0.0889, N = 808
FGAALTVLGDVNGDK pc2
H3BN02;P11215;P11215−2;P2070
log10 peptide intensity
324:626
273:525
48:109
16:31363977:C:T_C
p = 0.34, beta = −0.0599, N = 645
LTDVAIGAPGEEDNR pc2
P11215;P11215−2
log10 peptide intensity
230:626
190:525
30:109
16:31363977:C:T_C
p = 0.76, beta = 0.0233, N = 450
FGDPLGYEDVIPEADREGVIR pc3
P11215;P11215−2
log10 peptide intensity
226:626
186:525
30:109
16:31363977:C:T_C
p = 0.5, beta = −0.0514, N = 442
ITGAM : NP4
P11215;P11215−2
log10 protein intensity
62:166
44:108
20:51
16:31363977:C:T_C
p = 0.41, beta = 0.0978, N = 126
TEFQLELPVK pc2
P11215;P11215−2
log10 peptide intensity
38:166
22:108
9:51
16:31363977:C:T_C
p = 0.35, beta = 0.152, N = 69
VVVGAPQEIVAANQR pc2
P11215;P11215−2
log10 peptide intensity
31:166
20:108
9:51
16:31363977:C:T_C
p = 0.61, beta = 0.0876, N = 60
GLCFLFGSNLR pc2
P11215;P11215−2
log10 peptide intensity
9:166
14:108
4:51
16:31363977:C:T_C
p = 0.51, beta = −0.176, N = 27
NVFECNDQVVK pc2
P11215;P11215−2
log10 peptide intensity
10:166
10:108
4:51
16:31363977:C:T_C
p = 0.26, beta = 0.292, N = 24
Assay Target: ITGAM
Olink UniProt: P11215
UKB rsID: rs10665069
Proxy rsID: rs11574631
UKB SNP: 16:31362590:T:TAAA
Proxy SNP: 16:31363977:C:T
UKB log10(p): 131.2
UKB BETA: 0.159
808:645:450:442:427:405:394:3
LACESASSTEVSGALK pc2
rs1143683 REF
log10 peptide intensity
95:626
93:525
19:109
16:31363977:C:T_C
p = 0.25, model = REC, N = 207
LACESASSTEVSGVLK pc2
rs1143683 ALT
log10 peptide intensity
14:626
4:525
1:109
16:31363977:C:T_C
p = NA, model = NA, N = 19

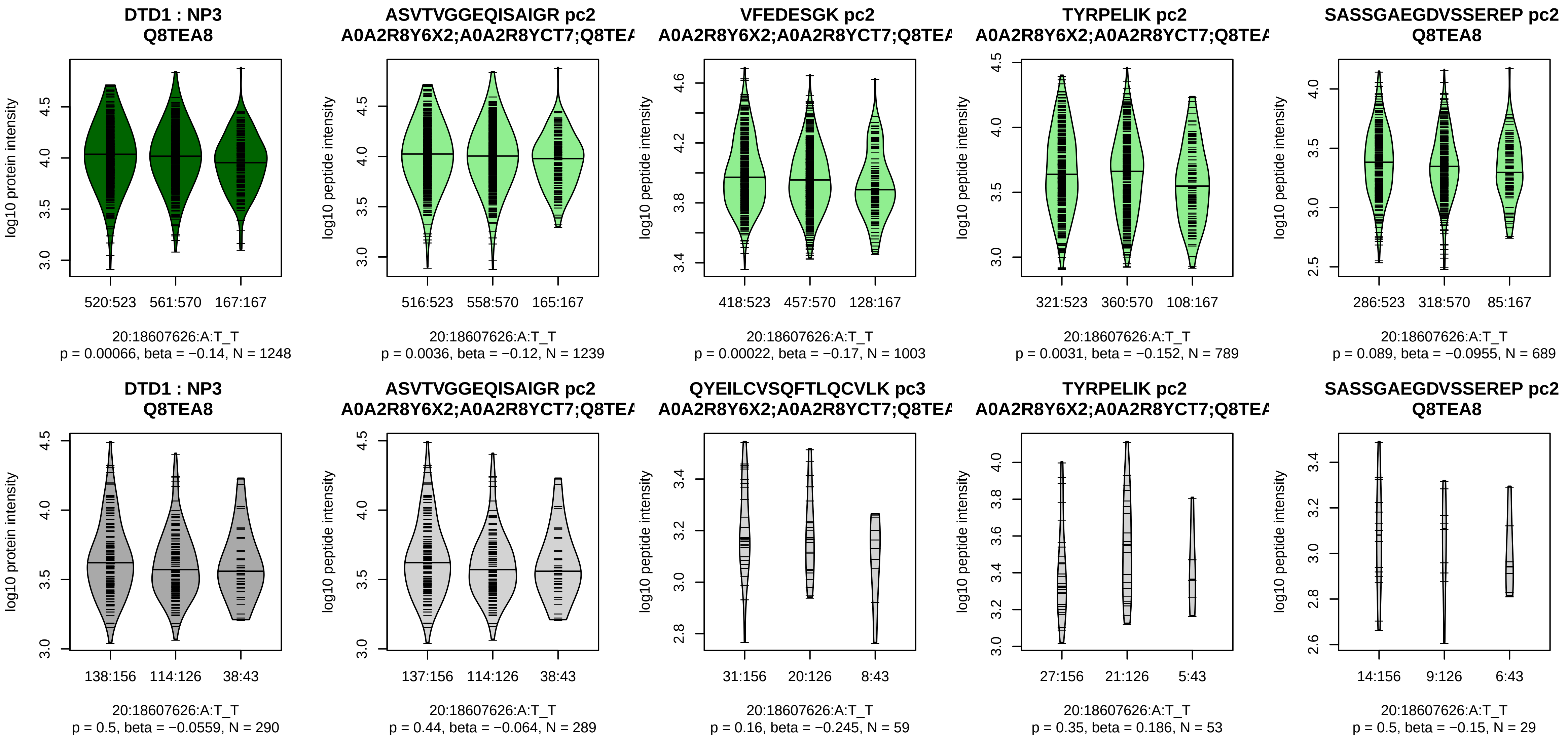


Assay Target: DTD1
Olink UniProt: Q8TEA8
UKB rsID: rs6136442
Proxy rsID: rs6136442
UKB SNP: 20:18607626:A:T
Proxy SNP: 20:18607626:A:T
UKB log10(p): 131
UKB BETA: −0.156
*:*:*:–:*:–:–
1239:1003:789:689:604:456:254

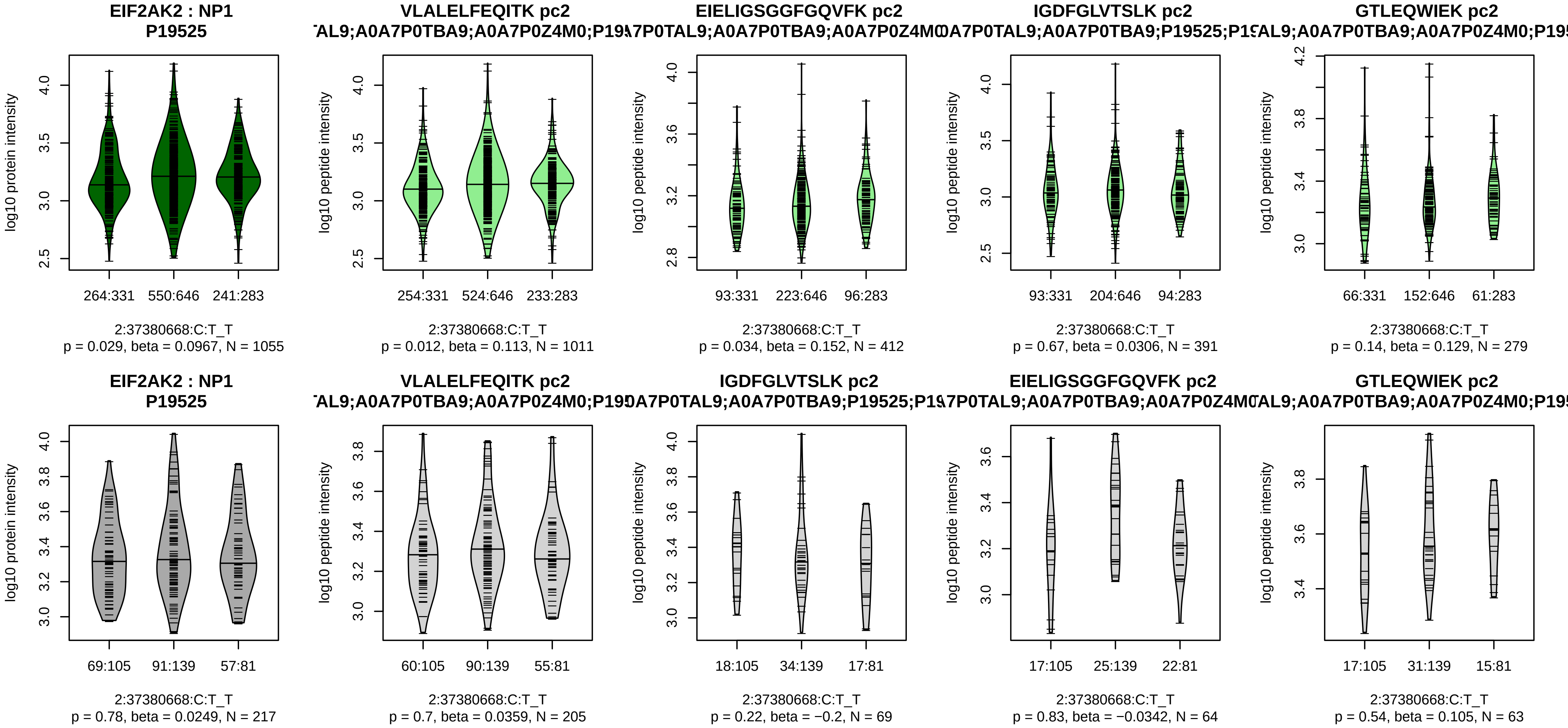


Assay Target: EIF2AK2
Olink UniProt: P19525
UKB rsID: rs11894449
Proxy rsID: rs11894449
UKB SNP: 2:37380668:C:T
Proxy SNP: 2:37380668:C:T
UKB log10(p): 130.4
UKB BETA: 0.148
–:–:–:–:–:–:–:–:–:–:–:–:–:–
1011:412:391:279:260:250:204:

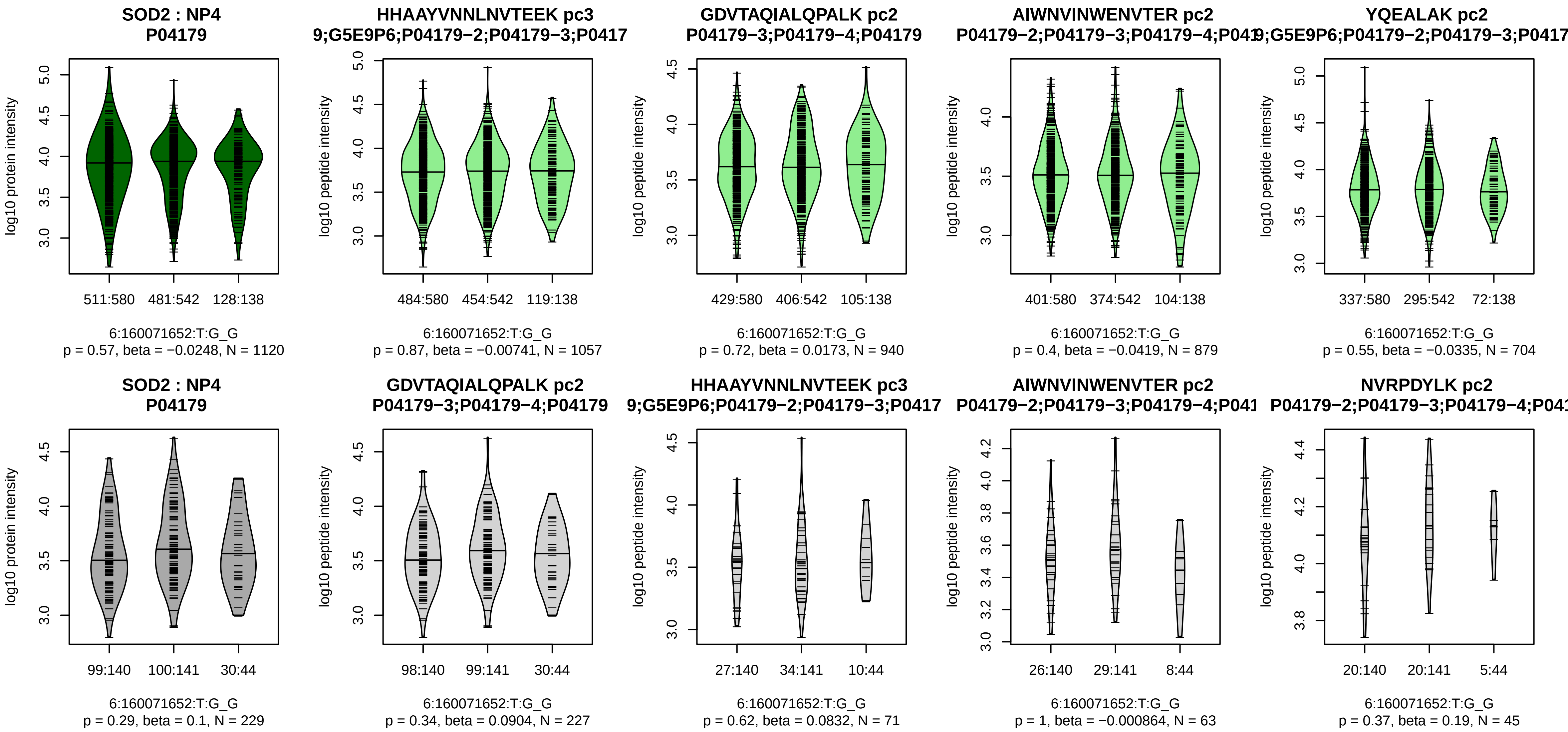


Assay Target: SOD2
Olink UniProt: P04179
UKB rsID: rs9457699
Proxy rsID: rs9457699
UKB SNP: 6:160071652:T:G
Proxy SNP: 6:160071652:T:G
UKB log10(p): 130.2
UKB BETA: 0.157
−:−:−:−:−:−:−:−:−:−:−
1057:940:879:704:669:475:439:

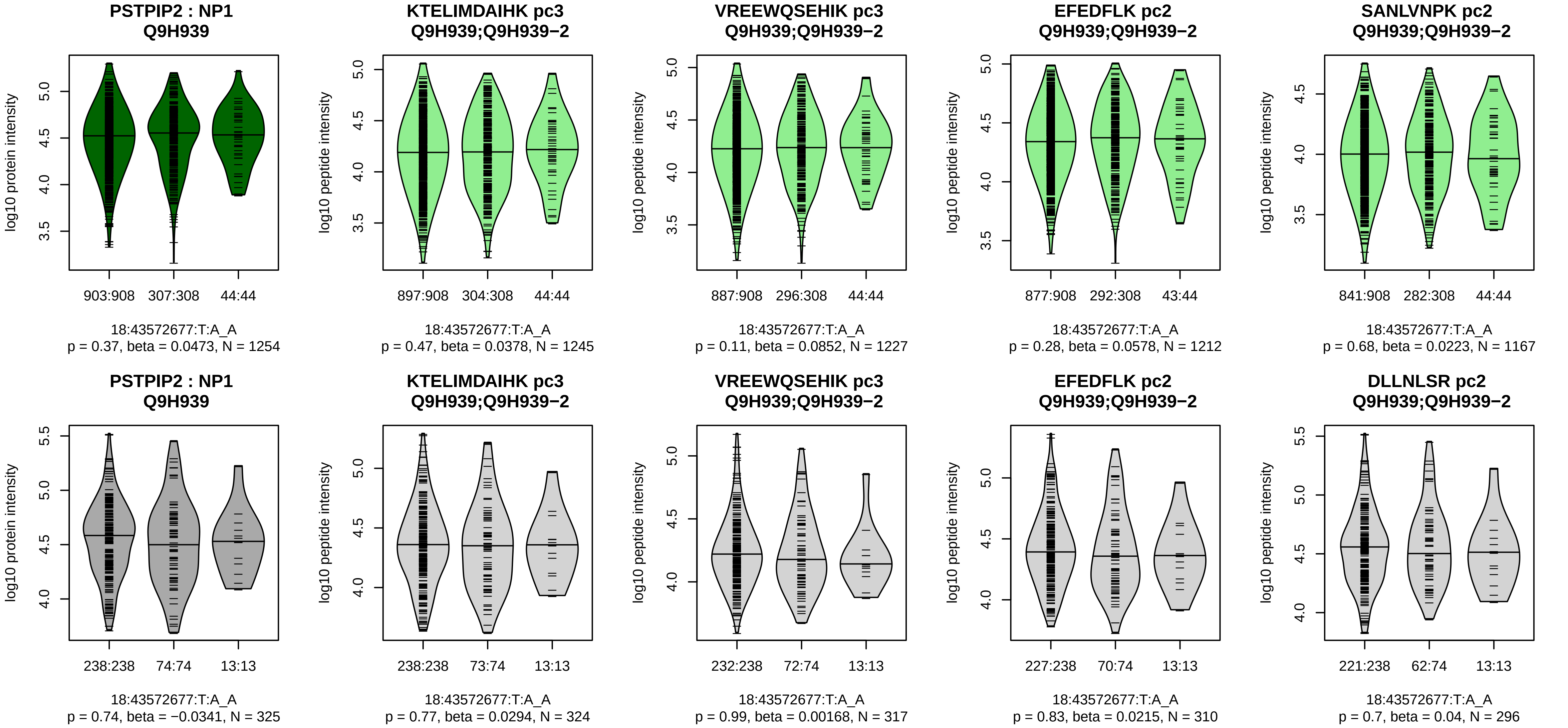


Assay Target: PSTPIP2
Olink UniProt: Q9H939
UKB rsID: rs72642434
Proxy rsID: rs72642434
UKB SNP: 18:43572677:T:A
Proxy SNP: 18:43572677:T:A
UKB log10(p): 130.1
UKB BETA: −0.222
−:−:−:−:−:−:−:−:−:−:−:−:−:−:−:−:
1245:1227:1212:1167:1128:104

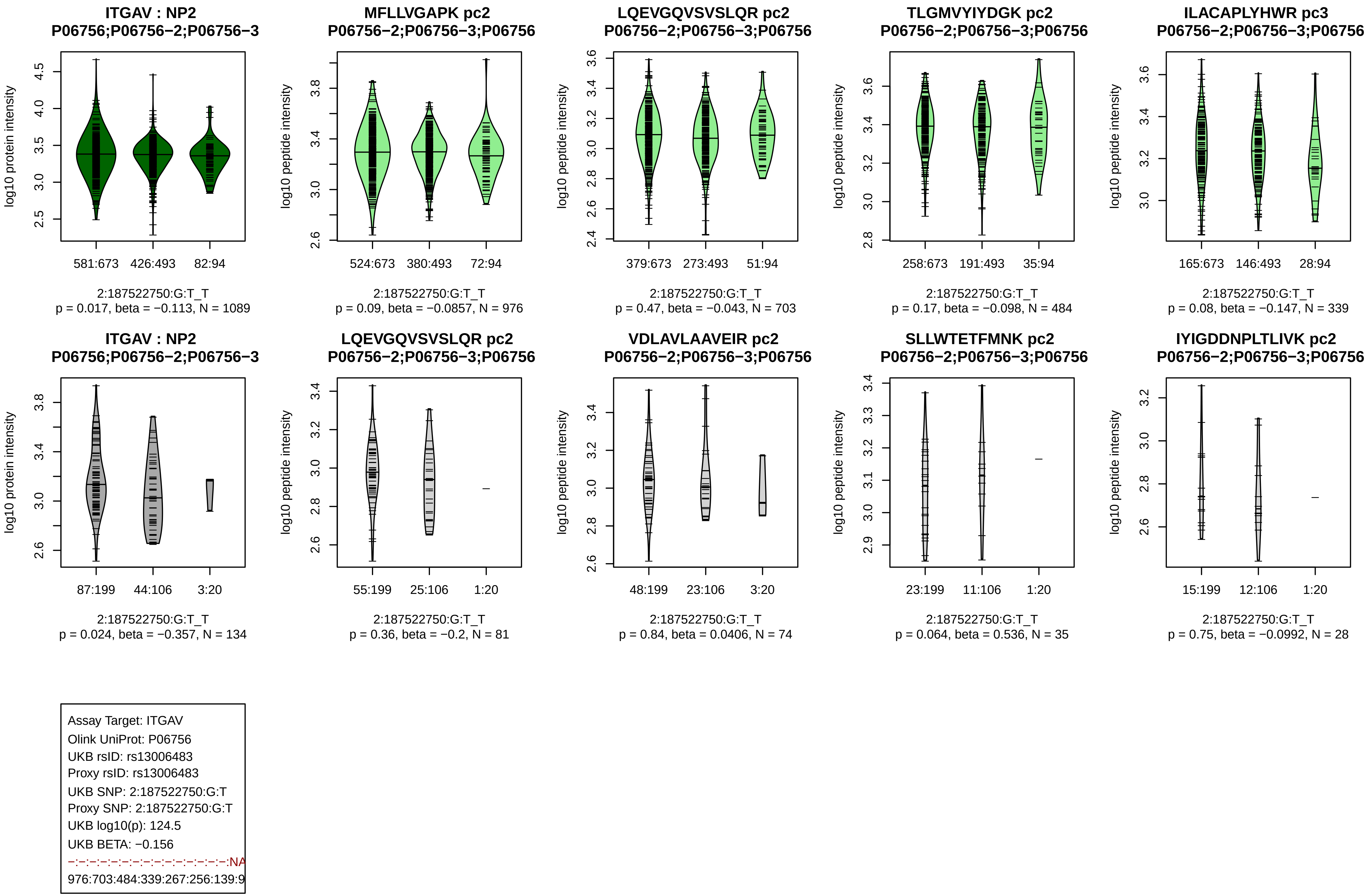
ITGAV : NP2
P06756;P06756−2;P06756−3
log10 protein intensity
581:673
426:493
82:94
2:187522750:G:T_T
p = 0.017, beta = −0.113, N = 1089
MFLLVGAPK pc2
P06756−2;P06756−3;P06756
log10 peptide intensity
524:673
380:493
72:94
2:187522750:G:T_T
p = 0.09, beta = −0.0857, N = 976
LQEVGQVSVSLQR pc2
P06756−2;P06756−3;P06756
log10 peptide intensity
379:673
273:493
51:94
2:187522750:G:T_T
p = 0.47, beta = −0.043, N = 703
TLGMVYIYDGK pc2
P06756−2;P06756−3;P06756
log10 peptide intensity
258:673
191:493
35:94
2:187522750:G:T_T
p = 0.17, beta = −0.098, N = 484
ILACAPLYHWR pc3
P06756−2;P06756−3;P06756
log10 peptide intensity
165:673
146:493
28:94
2:187522750:G:T_T
p = 0.08, beta = −0.147, N = 339
ITGAV : NP2
P06756;P06756−2;P06756−3
log10 protein intensity
87:199
44:106
3:20
2:187522750:G:T_T
p = 0.024, beta = −0.357, N = 134
LQEVGQVSVSLQR pc2
P06756−2;P06756−3;P06756
log10 peptide intensity
55:199
25:106
1:20
2:187522750:G:T_T
p = 0.36, beta = −0.2, N = 81
VDLAVLAAVEIR pc2
P06756−2;P06756−3;P06756
log10 peptide intensity
48:199
23:106
3:20
2:187522750:G:T_T
p = 0.84, beta = 0.0406, N = 74
SLLWTETFMNK pc2
P06756−2;P06756−3;P06756
log10 peptide intensity
23:199
11:106
1:20
2:187522750:G:T_T
p = 0.064, beta = 0.536, N = 35
IYIGDDNPLTLIVK pc2
P06756−2;P06756−3;P06756
log10 peptide intensity
15:199
12:106
1:20
2:187522750:G:T_T
p = 0.75, beta = −0.0992, N = 28
Assay Target: ITGAV
Olink UniProt: P06756
UKB rsID: rs13006483
Proxy rsID: rs13006483
UKB SNP: 2:187522750:G:T
Proxy SNP: 2:187522750:G:T
UKB log10(p): 124.5
UKB BETA: −0.156
−:−:−:−:−:−:−:−:−:−:−:−:−:−:−:−:NA
976:703:484:339:267:256:139:9

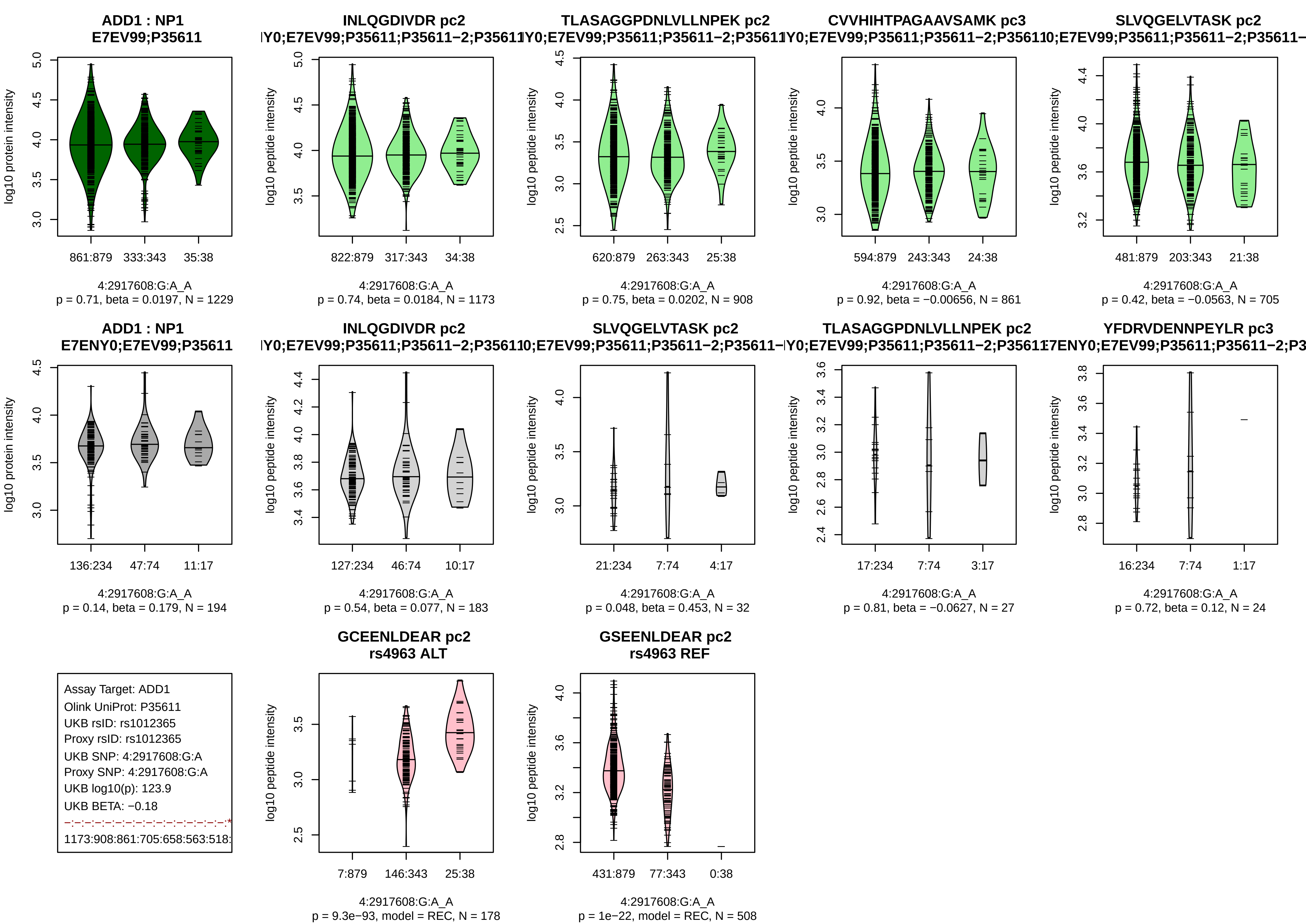
ADD1 : NP1
E7EV99;P35611
log10 protein intensity
861:879
333:343
35:38
4:2917608:G:A_A
p = 0.71, beta = 0.0197, N = 1229
INLQGDIVDR pc2
Y0;E7EV99;P35611;P35611−2;P35611
log10 peptide intensity
822:879
317:343
34:38
4:2917608:G:A_A
p = 0.74, beta = 0.0184, N = 1173
TLASAGGPDNLVLLNPEK pc2
Y0;E7EV99;P35611;P35611−2;P35611
log10 peptide intensity
620:879
263:343
25:38
4:2917608:G:A_A
p = 0.75, beta = 0.0202, N = 908
CVVHIHTPAGAAVSAMK pc3
Y0;E7EV99;P35611;P35611−2;P3561
log10 peptide intensity
594:879
243:343
24:38
4:2917608:G:A_A
p = 0.92, beta = −0.00656, N = 861
SLVQGELVTASK pc2
0;E7EV99;P35611;P35611−2;P35611−
log10 peptide intensity
481:879
203:343
21:38
4:2917608:G:A_A
p = 0.42, beta = −0.0563, N = 705
ADD1 : NP1
E7ENY0;E7EV99;P35611
log10 protein intensity
136:234
47:74
11:17
4:2917608:G:A_A
p = 0.14, beta = 0.179, N = 194
INLQGDIVDR pc2
Y0;E7EV99;P35611;P35611−2;P3561
log10 peptide intensity
127:234
46:74
10:17
4:2917608:G:A_A
p = 0.54, beta = 0.077, N = 183
SLVQGELVTASK pc2
0;E7EV99;P35611;P35611−2;P35611−
log10 peptide intensity
21:234
7:74
4:17
4:2917608:G:A_A
p = 0.048, beta = 0.453, N = 32
TLASAGGPDNLVLLNPEK pc2
Y0;E7EV99;P35611;P35611−2;P3561
log10 peptide intensity
17:234
7:74
3:17
4:2917608:G:A_A
p = 0.81, beta = −0.0627, N = 27
YFDRVDENNPEYLR pc3
7ENY0;E7EV99;P35611;P35611−2;P3
log10 peptide intensity
16:234
7:74
1:17
4:2917608:G:A_A
p = 0.72, beta = 0.12, N = 24
Assay Target: ADD1
Olink UniProt: P35611
UKB rsID: rs1012365
Proxy rsID: rs1012365
UKB SNP: 4:2917608:G:A
Proxy SNP: 4:2917608:G:A
UKB log10(p): 123.9
UKB BETA: −0.18
1173:908:861:705:658:563:518:
GCEENLDEAR pc2
rs4963 ALT
log10 peptide intensity
7:879
146:343
25:38
4:2917608:G:A_A
p = 9.3e−93, model = REC, N = 178
GSEENLDEAR pc2
rs4963 REF
log10 peptide intensity
431:879
77:343
0:38
4:2917608:G:A_A
p = 1e−22, model = REC, N = 508

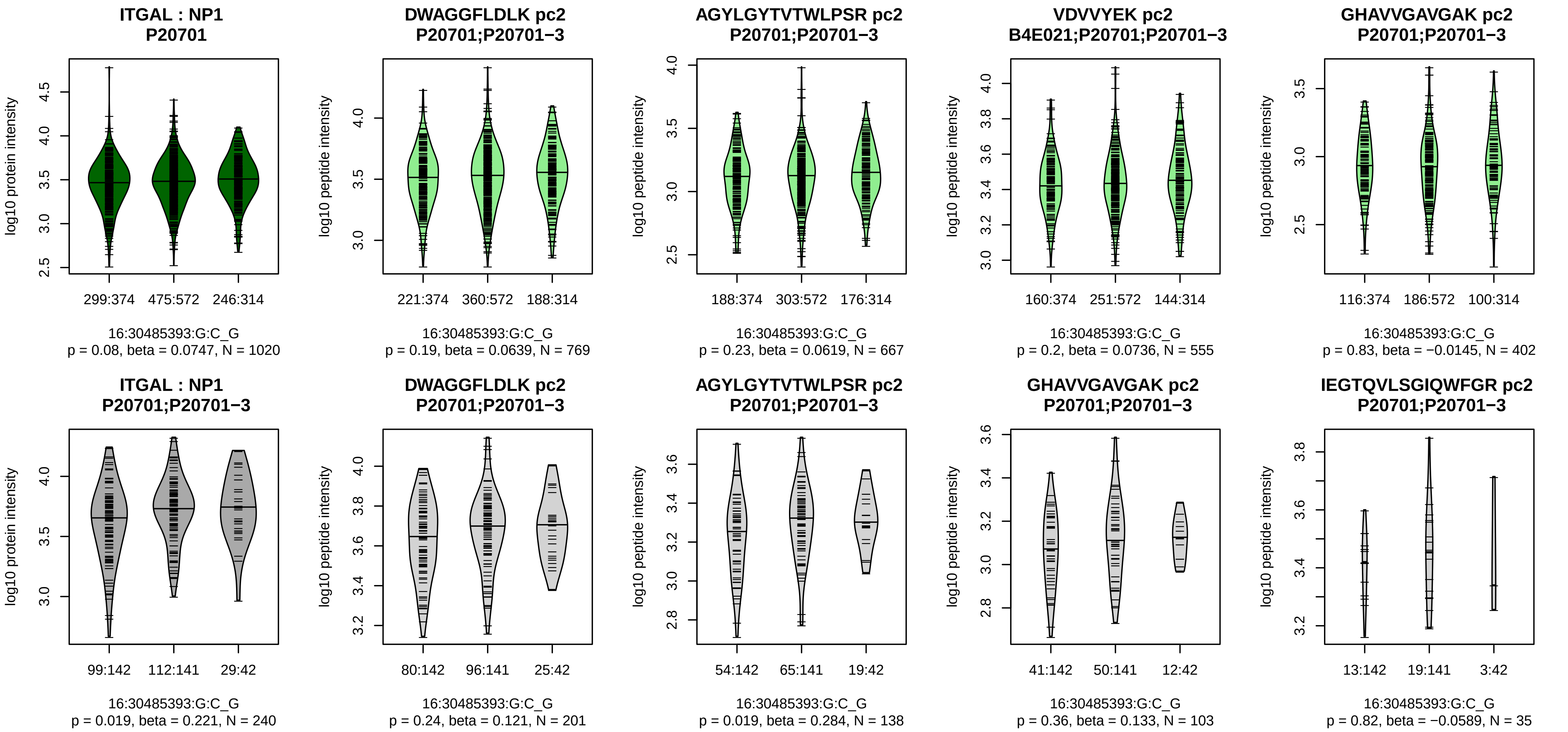


Assay Target: ITGAL
Olink UniProt: P20701
UKB rsID: rs11574938
Proxy rsID: rs11574938
UKB SNP: 16:30485393:G:C
Proxy SNP: 16:30485393:G:C
UKB log10(p): 123.7
UKB BETA: −0.149
−:−:−:−:−:−:−:−:NA
769:667:555:402:337:264:123:9

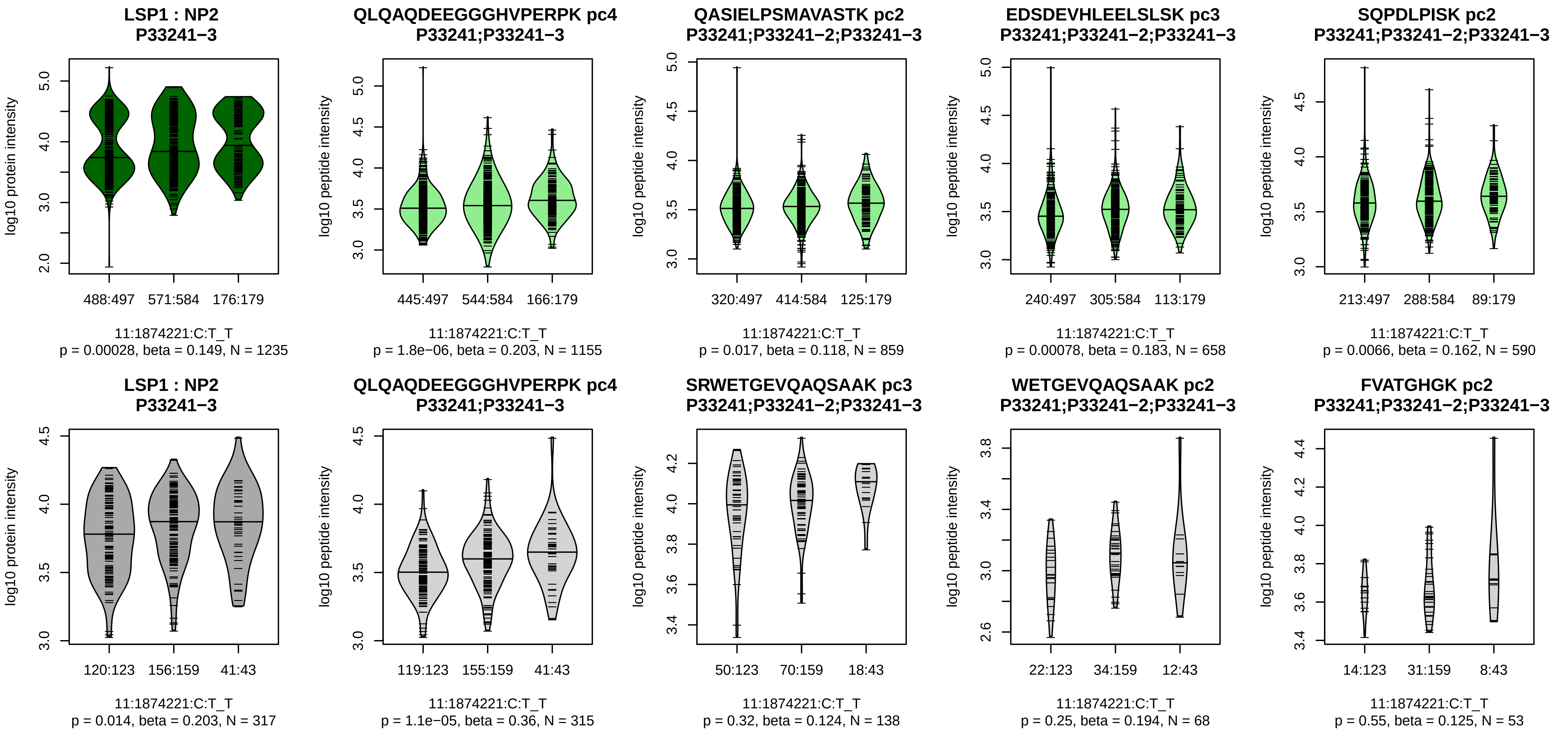


Assay Target: LSP1
Olink UniProt: P33241
UKB rsID: rs907612
Proxy rsID: rs907612
UKB SNP: 11:1874221:C:T
Proxy SNP: 11:1874221:C:T
UKB log10(p): 121.7
UKB BETA: 0.146
*:−:*:*:−:−:−:−:−:−:−:−:NA:NA
1155:859:658:590:572:519:413:

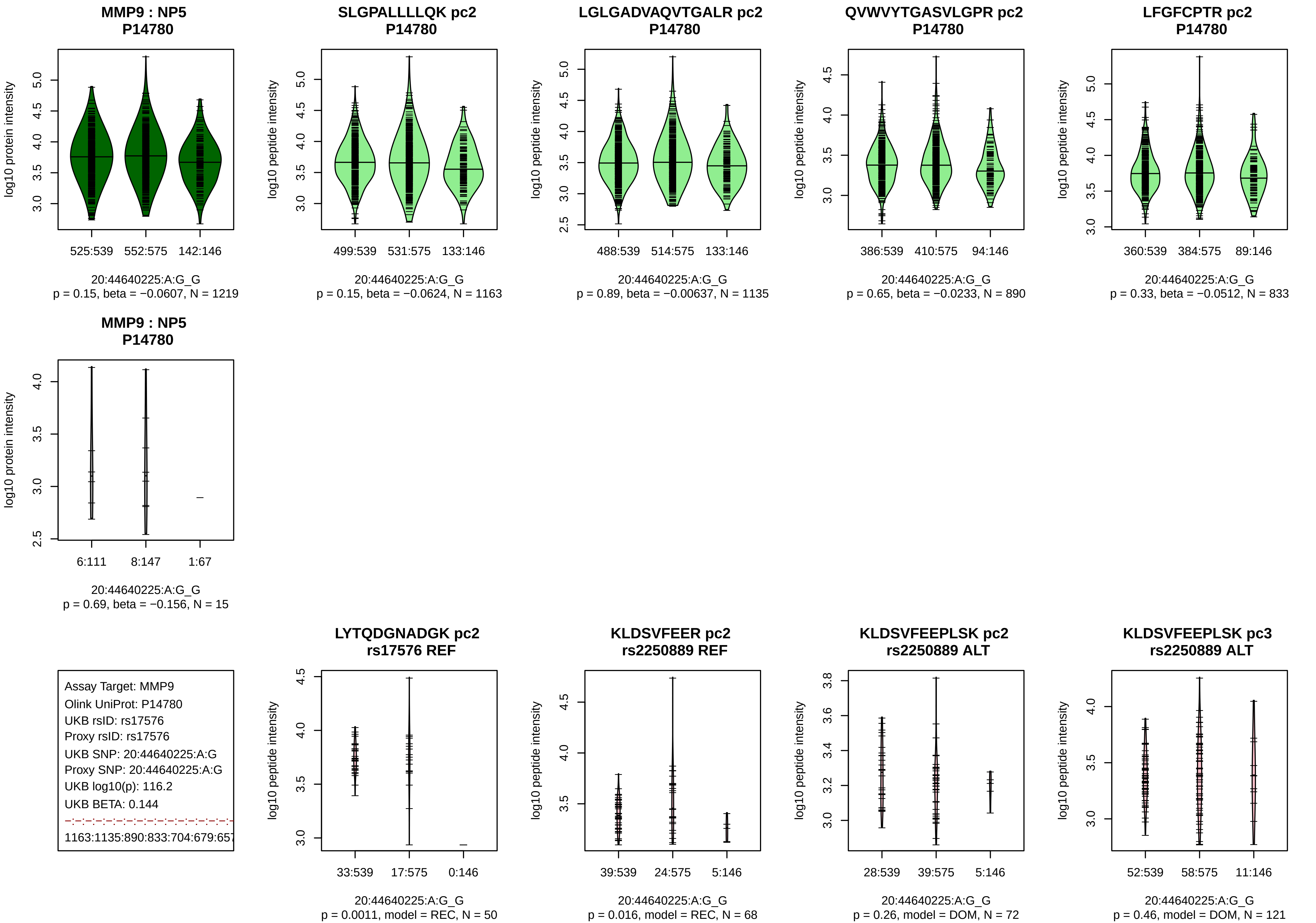
MMP9 : NP5
P14780
log10 protein intensity
525:539
552:575
142:146
20:44640225:A:G_G
p = 0.15, beta = −0.0607, N = 1219
SLGPALLLLQK pc2
P14780
log10 peptide intensity
499:539
531:575
133:146
20:44640225:A:G_G
p = 0.15, beta = −0.0624, N = 1163
LGLGADVAQVTGALR pc2
P14780
log10 peptide intensity
488:539
514:575
133:146
20:44640225:A:G_G
p = 0.89, beta = −0.00637, N = 1135
QVWVYTGASVLGPR pc2
P14780
log10 peptide intensity
386:539
410:575
94:146
20:44640225:A:G_G
p = 0.65, beta = −0.0233, N = 890
LFGFCPTR pc2
P14780
log10 peptide intensity
360:539
384:575
89:146
20:44640225:A:G_G
p = 0.33, beta = −0.0512, N = 833
MMP9 : NP5
P14780
log10 protein intensity
6:111
8:147
1:67
20:44640225:A:G_G
p = 0.69, beta = −0.156, N = 15
Assay Target: MMP9
Olink UniProt: P14780
UKB rsID: rs17576
Proxy rsID: rs17576
UKB SNP: 20:44640225:A:G
Proxy SNP: 20:44640225:A:G
UKB log10(p): 116.2
UKB BETA: 0.144
1163:1135:890:833:704:679:657
LYTQDGNADGK pc2
rs17576 REF
log10 peptide intensity
33:539
17:575
0:146
20:44640225:A:G_G
p = 0.0011, model = REC, N = 50
KLDSVFEER pc2
rs2250889 REF
log10 peptide intensity
39:539
24:575
5:146
20:44640225:A:G_G
p = 0.016, model = REC, N = 68
KLDSVFEEPLSK pc2
rs2250889 ALT
log10 peptide intensity
28:539
39:575
5:146
20:44640225:A:G_G
p = 0.26, model = DOM, N = 72
KLDSVFEEPLSK pc3
rs2250889 ALT
log10 peptide intensity
52:539
58:575
11:146
20:44640225:A:G_G
p = 0.46, model = DOM, N = 121

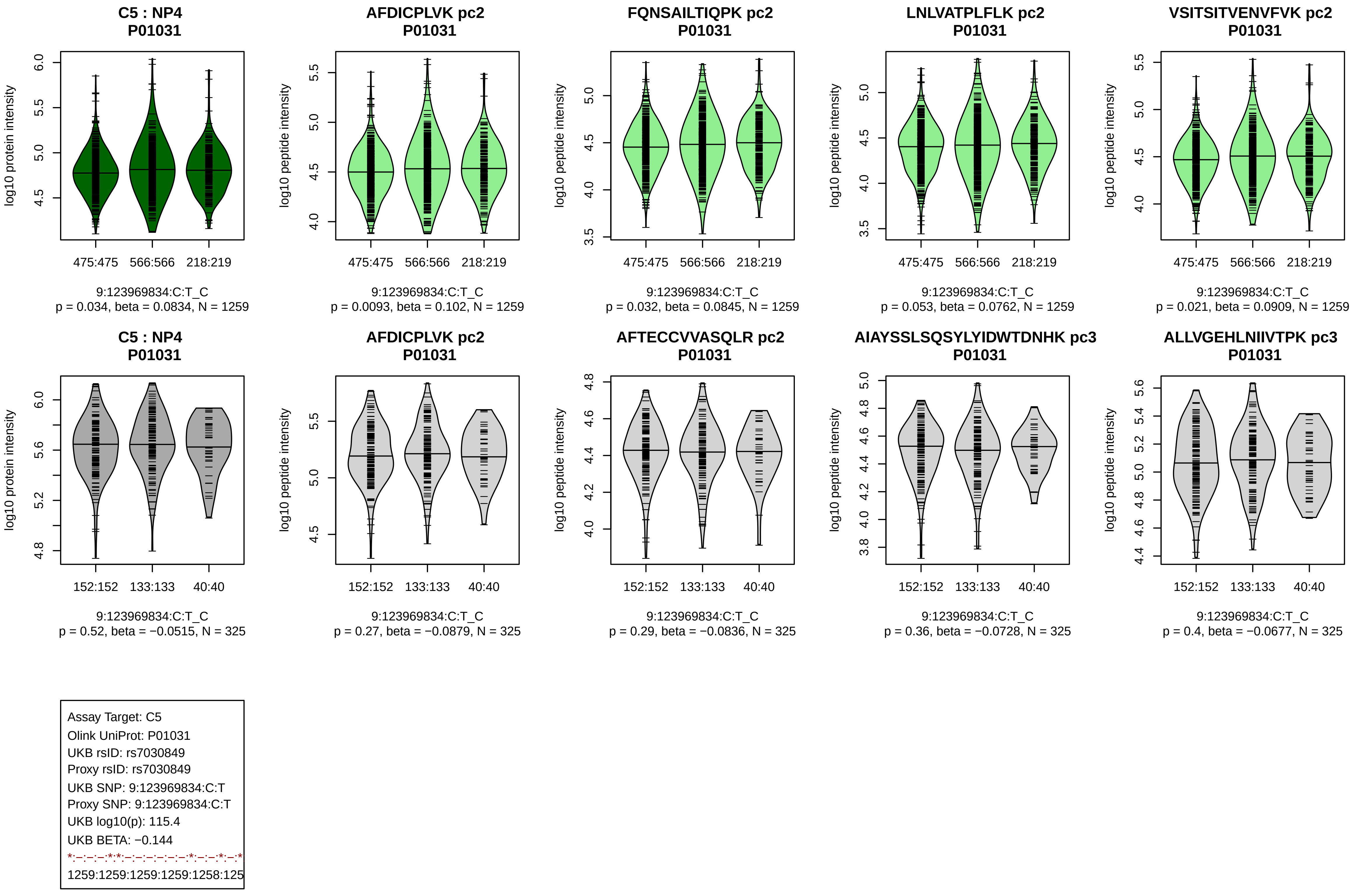
C5 : NP4
P01031
log10 protein intensity
475:475 566:566 218:219
9:123969834:C:T_C
p = 0.034, beta = 0.0834, N = 1259
AFDICPLVK pc2
P01031
log10 peptide intensity
475:475 566:566 218:219
9:123969834:C:T_C
p = 0.0093, beta = 0.102, N = 1259
FQNSAILTIQPK pc2
P01031
log10 peptide intensity
475:475 566:566 218:219
9:123969834:C:T_C
p = 0.032, beta = 0.0845, N = 1259
LNLVATPLFLK pc2
P01031
log10 peptide intensity
475:475 566:566 218:219
9:123969834:C:T_C
p = 0.053, beta = 0.0762, N = 1259
VSITSITVENVFVK pc2
P01031
log10 peptide intensity
475:475 566:566 218:219
9:123969834:C:T_C
p = 0.021, beta = 0.0909, N = 1259
C5 : NP4
P01031
log10 protein intensity
152:152 133:133 40:40
9:123969834:C:T_C
p = 0.52, beta = −0.0515, N = 325
AFDICPLVK pc2
P01031
log10 peptide intensity
152:152 133:133 40:40
9:123969834:C:T_C
p = 0.27, beta = −0.0879, N = 325
AFTECCVVASQLR pc2
P01031
log10 peptide intensity
152:152 133:133 40:40
9:123969834:C:T_C
p = 0.29, beta = −0.0836, N = 325
AIAYSSLSQSYLYIDWTDNHK pc3
P01031
log10 peptide intensity
152:152 133:133 40:40
9:123969834:C:T_C
p = 0.36, beta = −0.0728, N = 325
ALLVGEHLNIIVTPK pc3
P01031
log10 peptide intensity
152:152 133:133 40:40
9:123969834:C:T_C
p = 0.4, beta = −0.0677, N = 325
Assay Target: C5
Olink UniProt: P01031
UKB rsID: rs7030849
Proxy rsID: rs7030849
UKB SNP: 9:123969834:C:T
Proxy SNP: 9:123969834:C:T
UKB log10(p): 115.4
UKB BETA: −0.144
*:–:–:–:*:*:–:–:–:–:–:–:*:–:–:*:–:*
1259:1259:1259:1259:1258:125

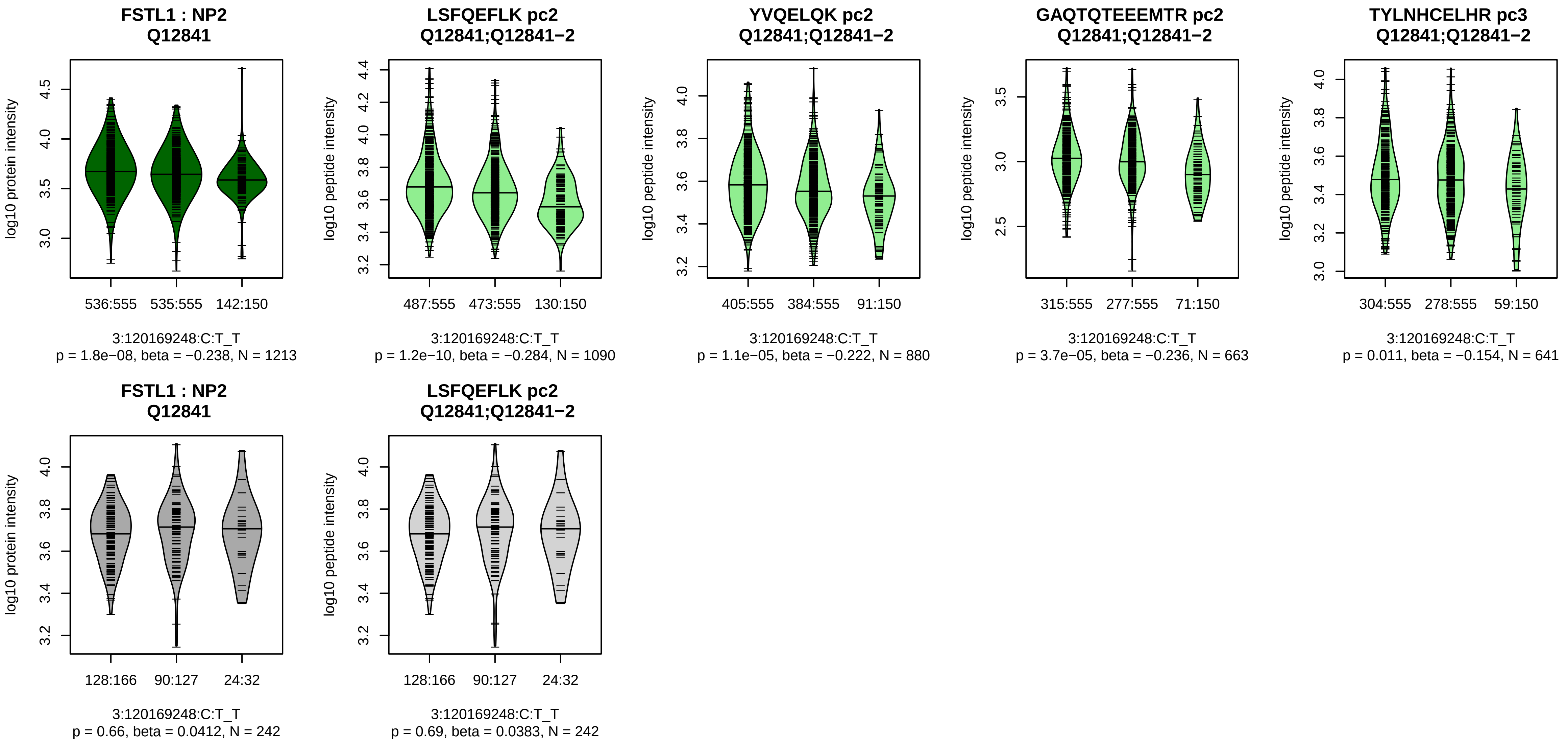


Assay Target: FSTL1
Olink UniProt: Q12841
UKB rsID: rs1147707
Proxy rsID: rs1147707
UKB SNP: 3:120169248:C:T
Proxy SNP: 3:120169248:C:T
UKB log10(p): 114.4
UKB BETA: −0.147
*:*:*:–:*:*:–:–:–
1090:880:663:641:545:538:361:

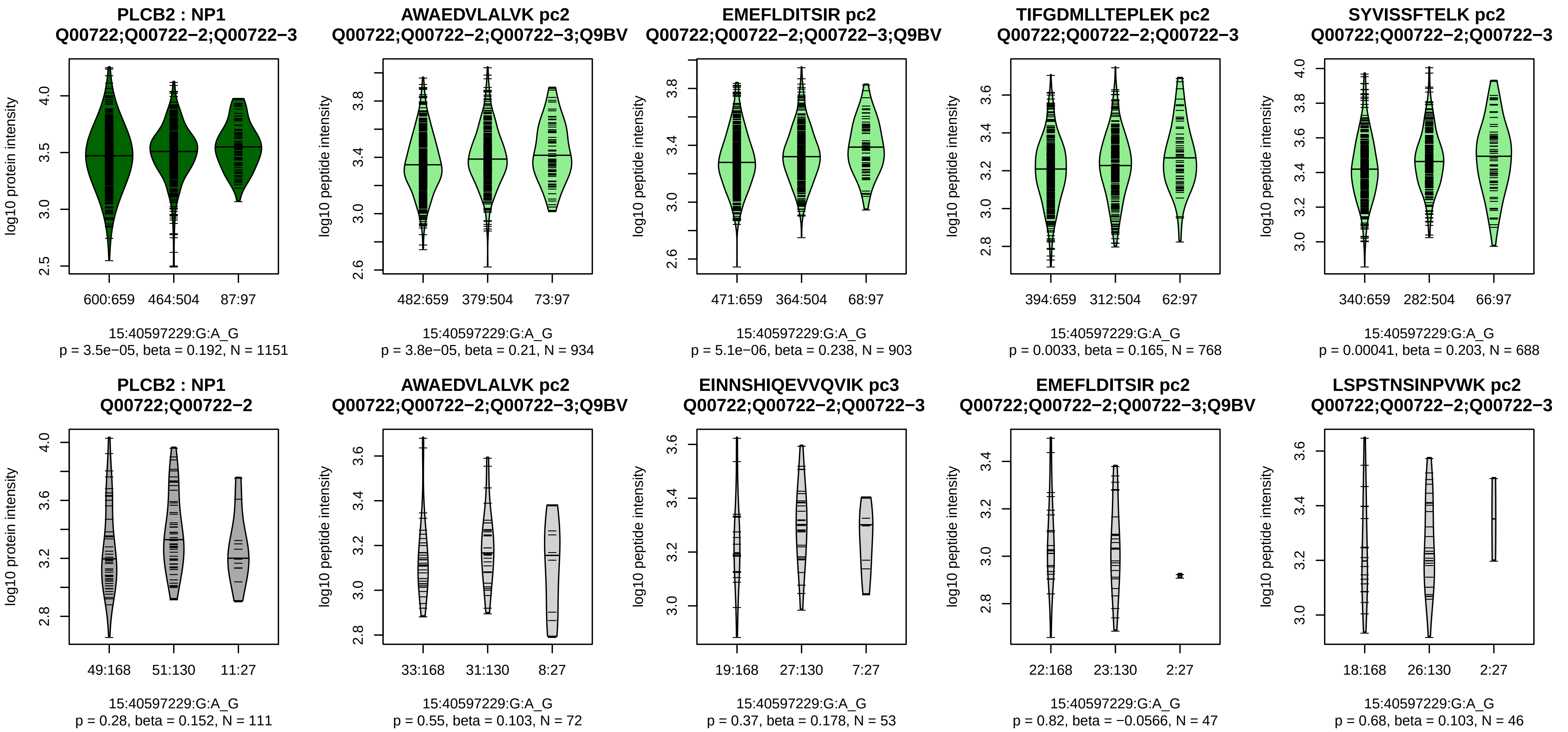


Assay Target: PLCB2
Olink UniProt: Q00722
UKB rsID: rs4924450
Proxy rsID: rs4924450
UKB SNP: 15:40597229:G:A
Proxy SNP: 15:40597229:G:A
UKB log10(p): 112.8
UKB BETA: −0.152
*:*:*:*:*:–:*:–:*:–:–:–:–:–:–:–:–:–:
934:903:768:688:602:582:543:4

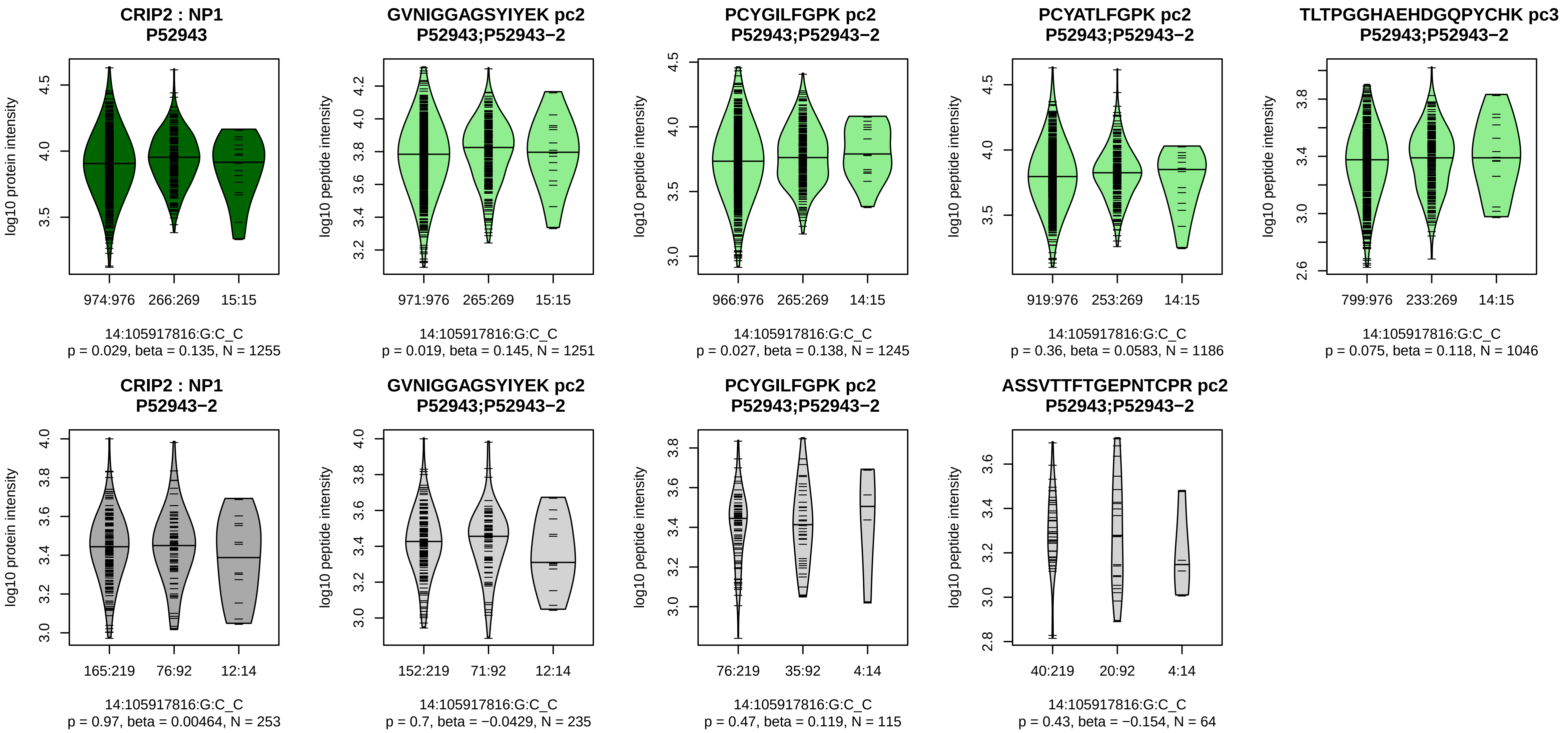


Assay Target: CRIP2
Olink UniProt: P52943
UKB rsID: rs80324969
Proxy rsID: rs80324969
UKB SNP: 14:105917816:G:C
Proxy SNP: 14:105917816:G:C
UKB log10(p): 112.5
UKB BETA: 0.185
−:−:−:−:−:−:−:−:−:−
1251:1245:1186:1046:1013:674

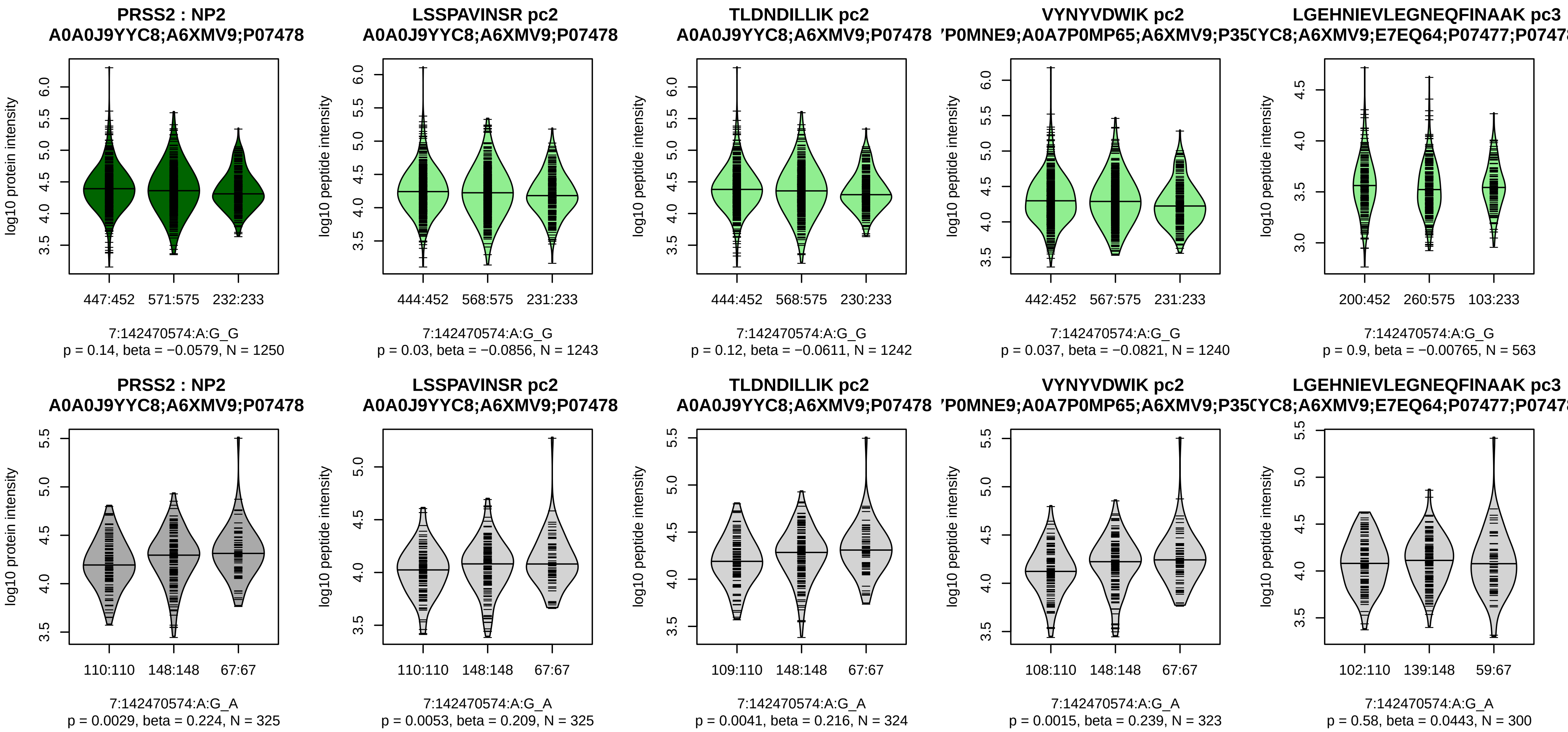


Assay Target: PRSS2
Olink UniProt: P07478
UKB rsID: rs3752404
Proxy rsID: rs3752404
UKB SNP: 7:142470574:A:G
Proxy SNP: 7:142470574:A:G
UKB log10(p): 110
UKB BETA: −0.137
−:−:−:−:−
1243:1242:1240:563:425

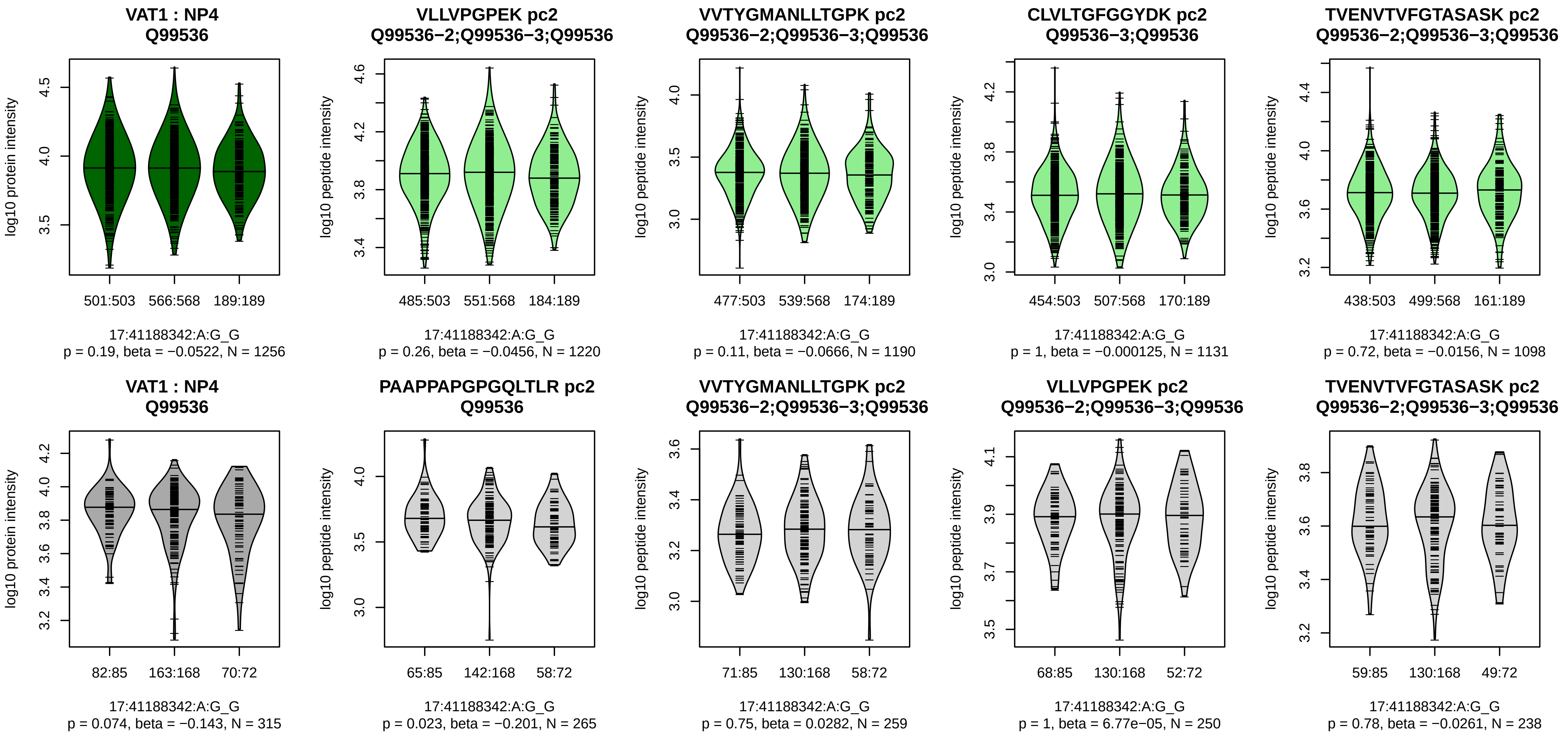


Assay Target: VAT1
Olink UniProt: Q99536
UKB rsID: rs9911630
Proxy rsID: rs9911630
UKB SNP: 17:41188342:A:G
Proxy SNP: 17:41188342:A:G
UKB log10(p): 109.3
UKB BETA: −0.145
−:−:−:−:−:−:−:−:−:−
1220:1190:1131:1098:1028:973

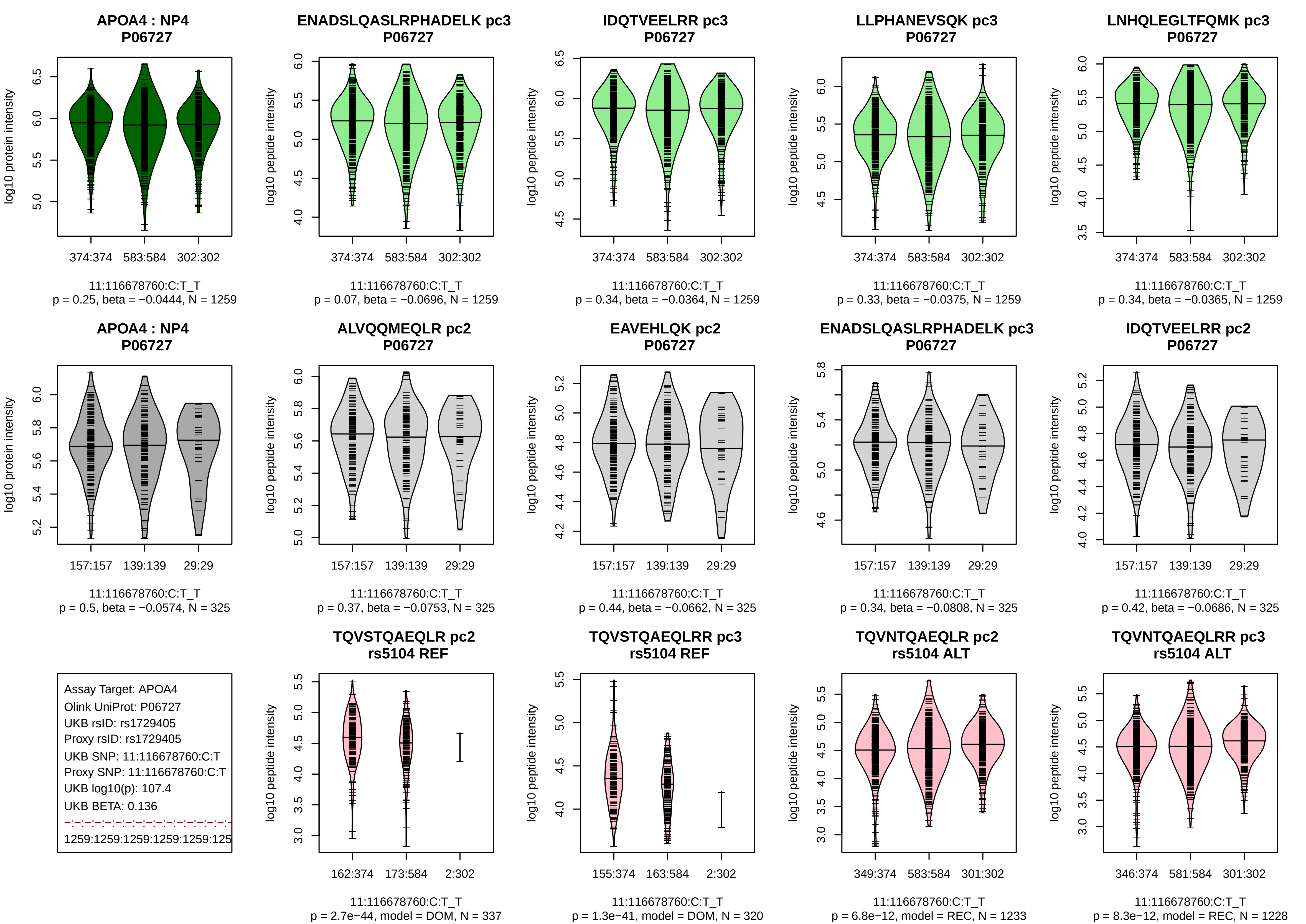
APOA4 : NP4
P06727
log10 protein intensity
374:374
583:584
302:302
11:116678760:C:T_T
p = 0.25, beta = −0.0444, N = 1259
ENADSLQASLRPHADELK pc3
P06727
log10 peptide intensity
374:374
583:584
302:302
11:116678760:C:T_T
p = 0.07, beta = −0.0696, N = 1259
IDQTVEELRR pc3
P06727
log10 peptide intensity
374:374
583:584
302:302
11:116678760:C:T_T
p = 0.34, beta = −0.0364, N = 1259
LLPHANEVSQK pc3
P06727
log10 peptide intensity
374:374
583:584
302:302
11:116678760:C:T_T
p = 0.33, beta = −0.0375, N = 1259
LNHQLEGLTFQMK pc3
P06727
log10 peptide intensity
374:374
583:584
302:302
11:116678760:C:T_T
p = 0.34, beta = −0.0365, N = 1259
APOA4 : NP4
P06727
log10 protein intensity
157:157
139:139
29:29
11:116678760:C:T_T
p = 0.5, beta = −0.0574, N = 325
ALVQQMEQLR pc2
P06727
log10 peptide intensity
157:157
139:139
29:29
11:116678760:C:T_T
p = 0.37, beta = −0.0753, N = 325
EAVEHLQK pc2
P06727
log10 peptide intensity
157:157
139:139
29:29
11:116678760:C:T_T
p = 0.44, beta = −0.0662, N = 325
ENADSLQASLRPHADELK pc3
P06727
log10 peptide intensity
157:157
139:139
29:29
11:116678760:C:T_T
p = 0.34, beta = −0.0808, N = 325
IDQTVEELRR pc2
P06727
log10 peptide intensity
157:157
139:139
29:29
11:116678760:C:T_T
p = 0.42, beta = −0.0686, N = 325
Assay Target: APOA4
Olink UniProt: P06727
UKB rsID: rs1729405
Proxy rsID: rs1729405
UKB SNP: 11:116678760:C:T
Proxy SNP: 11:116678760:C:T
UKB log10(p): 107.4
UKB BETA: 0.136
1259:1259:1259:1259:1259:125
TQVSTQAEQLR pc2
rs5104 REF
log10 peptide intensity
162:374
173:584
2:302
11:116678760:C:T_T
p = 2.7e−44, model = DOM, N = 337
TQVSTQAEQLRR pc3
rs5104 REF
log10 peptide intensity
155:374
163:584
2:302
11:116678760:C:T_T
p = 1.3e−41, model = DOM, N = 320
TQVNTQAEQLR pc2
rs5104 ALT
log10 peptide intensity
349:374
583:584
301:302
11:116678760:C:T_T
p = 6.8e−12, model = REC, N = 1233
TQVNTQAEQLRR pc3
rs5104 ALT
log10 peptide intensity
346:374
581:584
301:302
11:116678760:C:T_T
p = 8.3e−12, model = REC, N = 1228

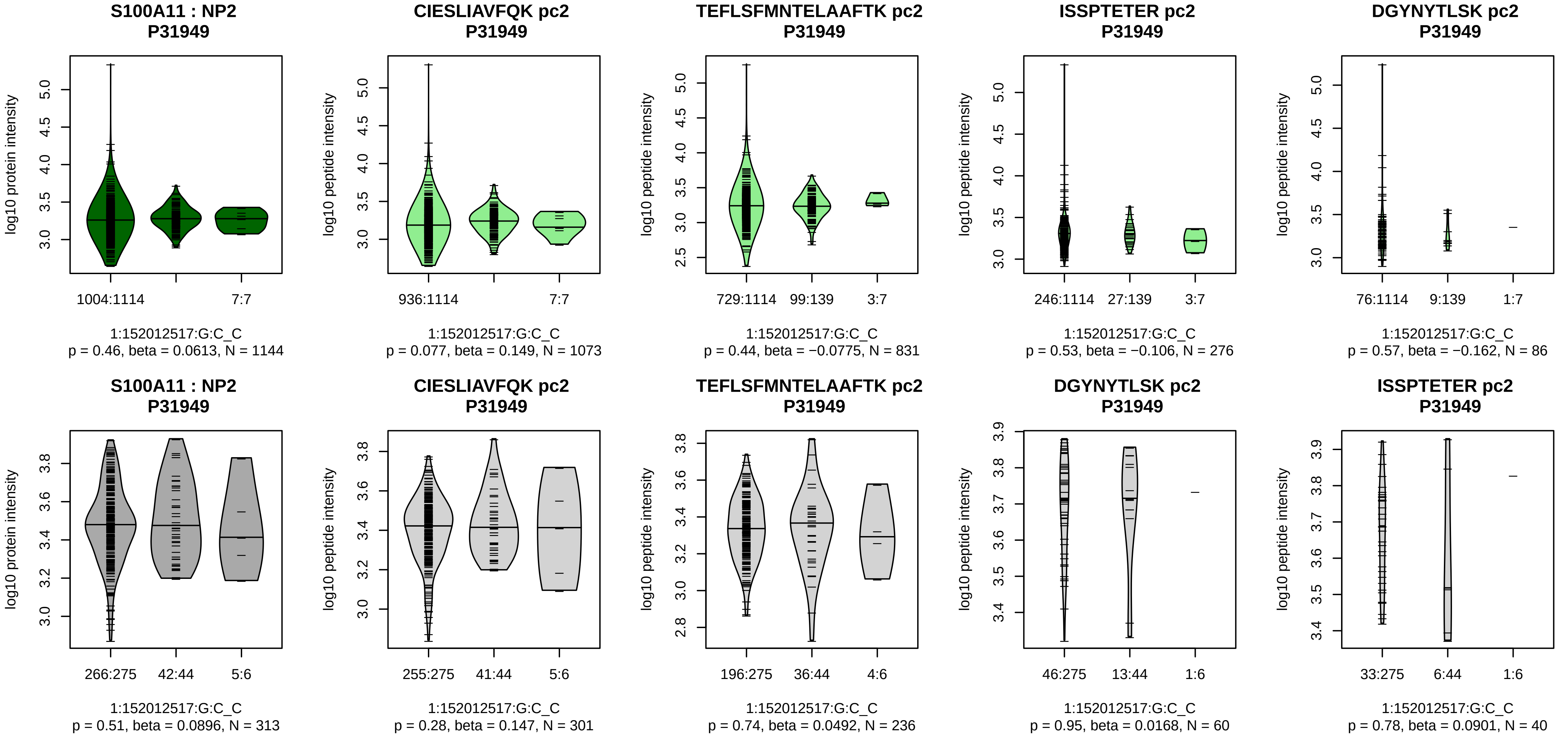


Assay Target: S100A11
Olink UniProt: P31949
UKB rsID: rs3007678
Proxy rsID: rs3007678
UKB SNP: 1:152012517:G:C
Proxy SNP: 1:152012517:G:C
UKB log10(p): 106.8
UKB BETA: 0.31
–:–:–:–
1073:831:276:86

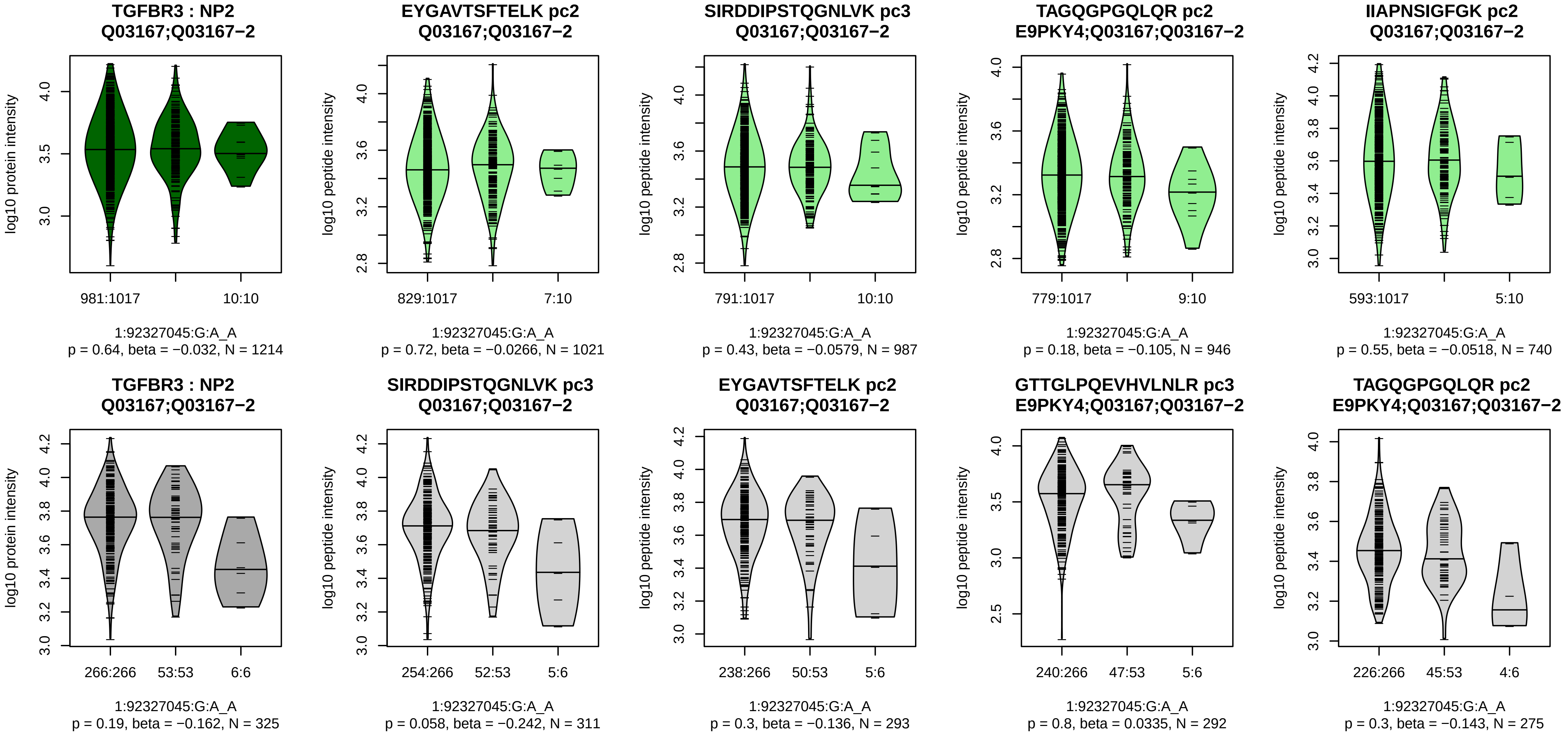


Assay Target: TGFBR3
Olink UniProt: Q03167
UKB rsID: rs1805110
Proxy rsID: rs1805110
UKB SNP: 1:92327045:G:A
Proxy SNP: 1:92327045:G:A
UKB log10(p): 106.6
UKB BETA: −0.234
−:−:−:−:−:−:−
1021:987:946:740:437:392:251

**SDC1 : NP2**
**P18827**

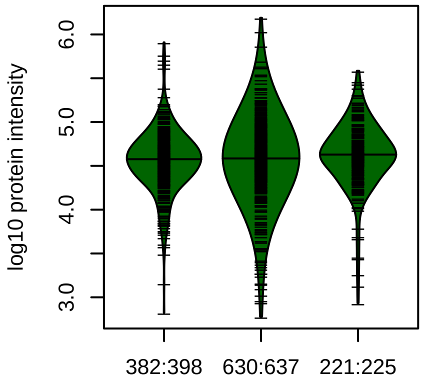


2:20371380:G:A_A
p = 0.037, beta = 0.0859, N = 1233

**EGEAVVLPEVEPGLTAR pc2**
**E9PHH3;P18827**

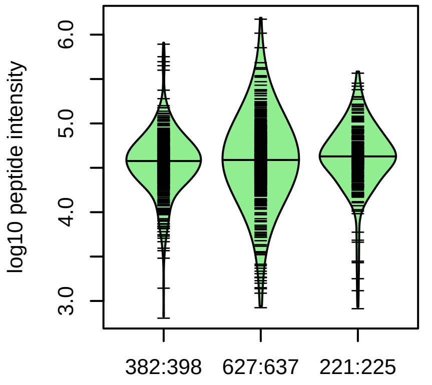


2:20371380:G:A_A
p = 0.032, beta = 0.0882, N = 1230

**NQSPVDQGATGASQGLLDRK pc3**
**P18827**

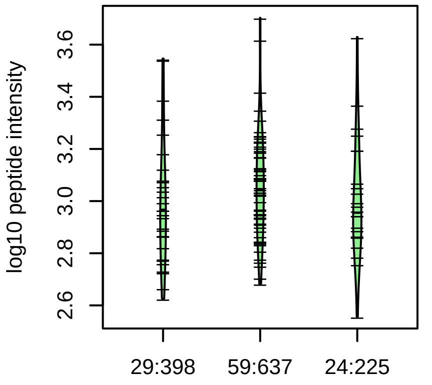


2:20371380:G:A_A
p = 0.58, beta = 0.0738, N = 112

**SDC1 : NP2**
**P18827**

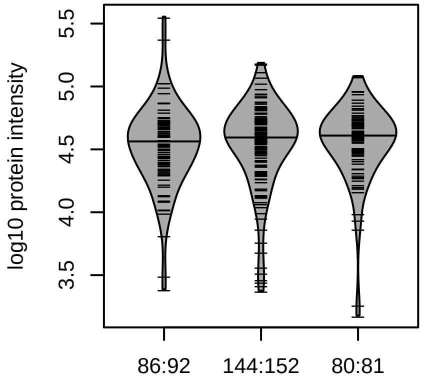


2:20371380:G:A_A
p = 0.51, beta = 0.0509, N = 310

**EGEAVVLPEVEPGLTAR pc2**
**E9PHH3;P18827**

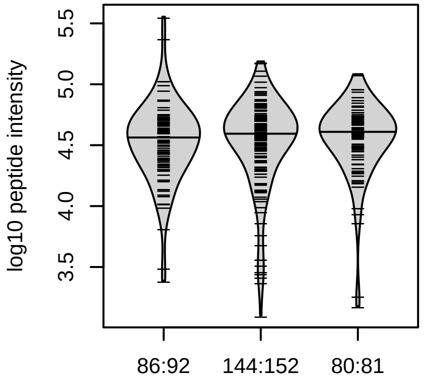


2:20371380:G:A_A
p = 0.52, beta = 0.049, N = 310

Assay Target: SDC1
Olink UniProt: P18827
UKB rsID: rs907866
Proxy rsID: rs907866
UKB SNP: 2:20371380:G:A
Proxy SNP: 2:20371380:G:A
UKB log10(p): 106.1
UKB BETA: 0.131
–:–:–
1230:112:53

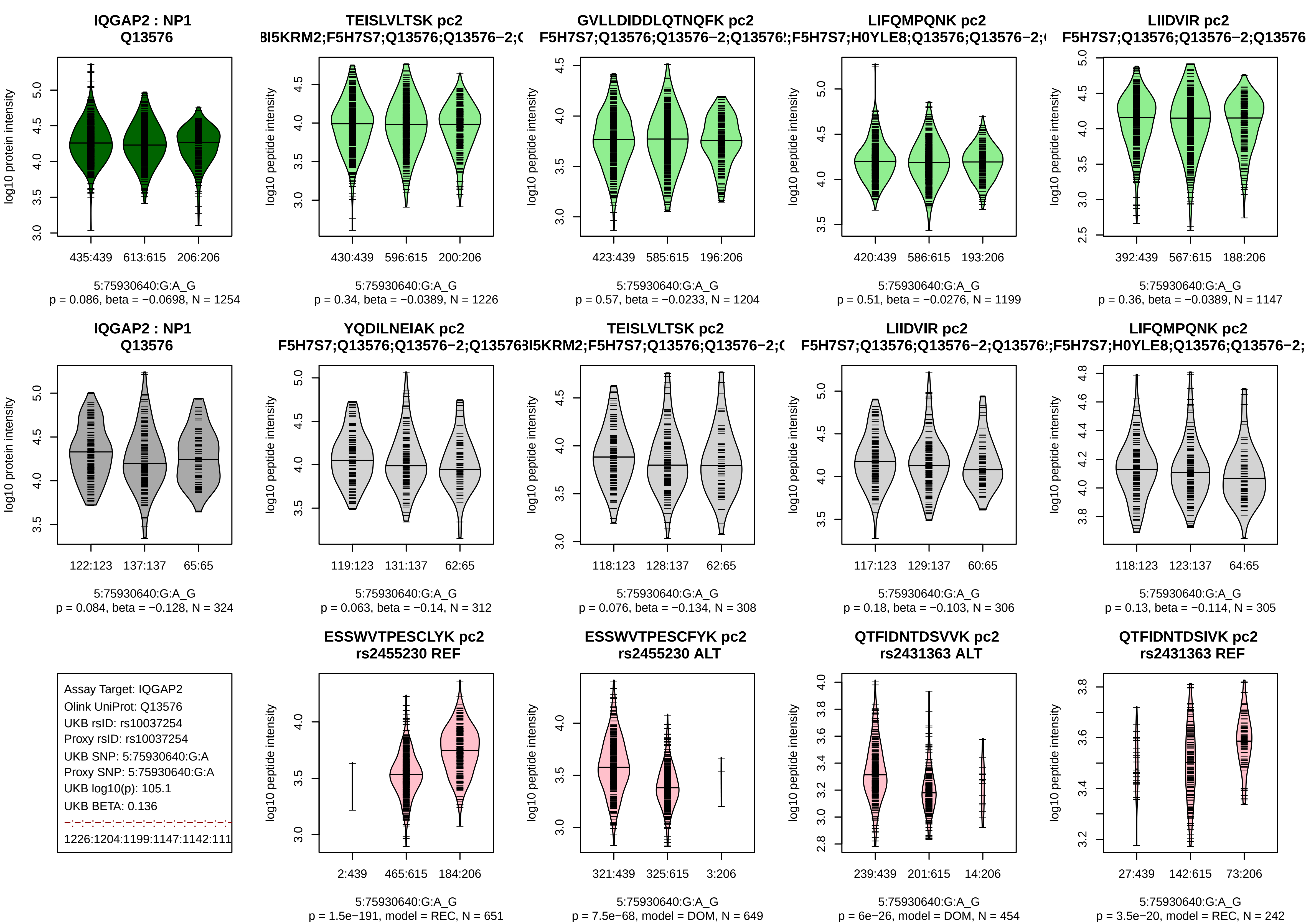
IQGAP2 : NP1
Q13576
log10 protein intensity
435:439 613:615 206:206
5:75930640:G:A_G
p = 0.086, beta = −0.0698, N = 1254
TEISLVLTSK pc2
3I5KRM2;F5H7S7;Q13576;Q13576−2;(
log10 peptide intensity
430:439 596:615 200:206
5:75930640:G:A_G
p = 0.34, beta = −0.0389, N = 1226
GVLLDIDDLQTNQFK pc2
F5H7S7;Q13576;Q13576−2;Q13576
log10 peptide intensity
423:439 585:615 196:206
5:75930640:G:A_G
p = 0.57, beta = −0.0233, N = 1204
LIFQMPQNK pc2
;F5H7S7;H0YLE8;Q13576;Q13576−2;
log10 peptide intensity
420:439 586:615 193:206
5:75930640:G:A_G
p = 0.51, beta = −0.0276, N = 1199
LIIDVIR pc2
F5H7S7;Q13576;Q13576−2;Q13576
log10 peptide intensity
392:439 567:615 188:206
5:75930640:G:A_G
p = 0.36, beta = −0.0389, N = 1147
IQGAP2 : NP1
Q13576
log10 protein intensity
122:123 137:137 65:65
5:75930640:G:A_G
p = 0.084, beta = −0.128, N = 324
YQDILNEIAK pc2
F5H7S7;Q13576;Q13576−2;Q13576
log10 peptide intensity
119:123 131:137 62:65
5:75930640:G:A_G
p = 0.063, beta = −0.14, N = 312
TEISLVLTSK pc2
3I5KRM2;F5H7S7;Q13576;Q13576−2;(
log10 peptide intensity
118:123 128:137 62:65
5:75930640:G:A_G
p = 0.076, beta = −0.134, N = 308
LIIDVIR pc2
F5H7S7;Q13576;Q13576−2;Q13576
log10 peptide intensity
117:123 129:137 60:65
5:75930640:G:A_G
p = 0.18, beta = −0.103, N = 306
LIFQMPQNK pc2
;F5H7S7;H0YLE8;Q13576;Q13576−2;
log10 peptide intensity
118:123 123:137 64:65
5:75930640:G:A_G
p = 0.13, beta = −0.114, N = 305
Assay Target: IQGAP2
Olink UniProt: Q13576
UKB rsID: rs10037254
Proxy rsID: rs10037254
UKB SNP: 5:75930640:G:A
Proxy SNP: 5:75930640:G:A
UKB log10(p): 105.1
UKB BETA: 0.136
1226:1204:1199:1147:1142:111
ESSWVTPESCLYK pc2
rs2455230 REF
log10 peptide intensity
2:439 465:615 184:206
5:75930640:G:A_G
p = 1.5e−191, model = REC, N = 651
ESSWVTPESCFYK pc2
rs2455230 ALT
log10 peptide intensity
321:439 325:615 3:206
5:75930640:G:A_G
p = 7.5e−68, model = DOM, N = 649
QTFIDNTDSVVK pc2
rs2431363 ALT
log10 peptide intensity
239:439 201:615 14:206
5:75930640:G:A_G
p = 6e−26, model = DOM, N = 454
QTFIDNTDSIVK pc2
rs2431363 REF
log10 peptide intensity
27:439 142:615 73:206
5:75930640:G:A_G
p = 3.5e−20, model = REC, N = 242

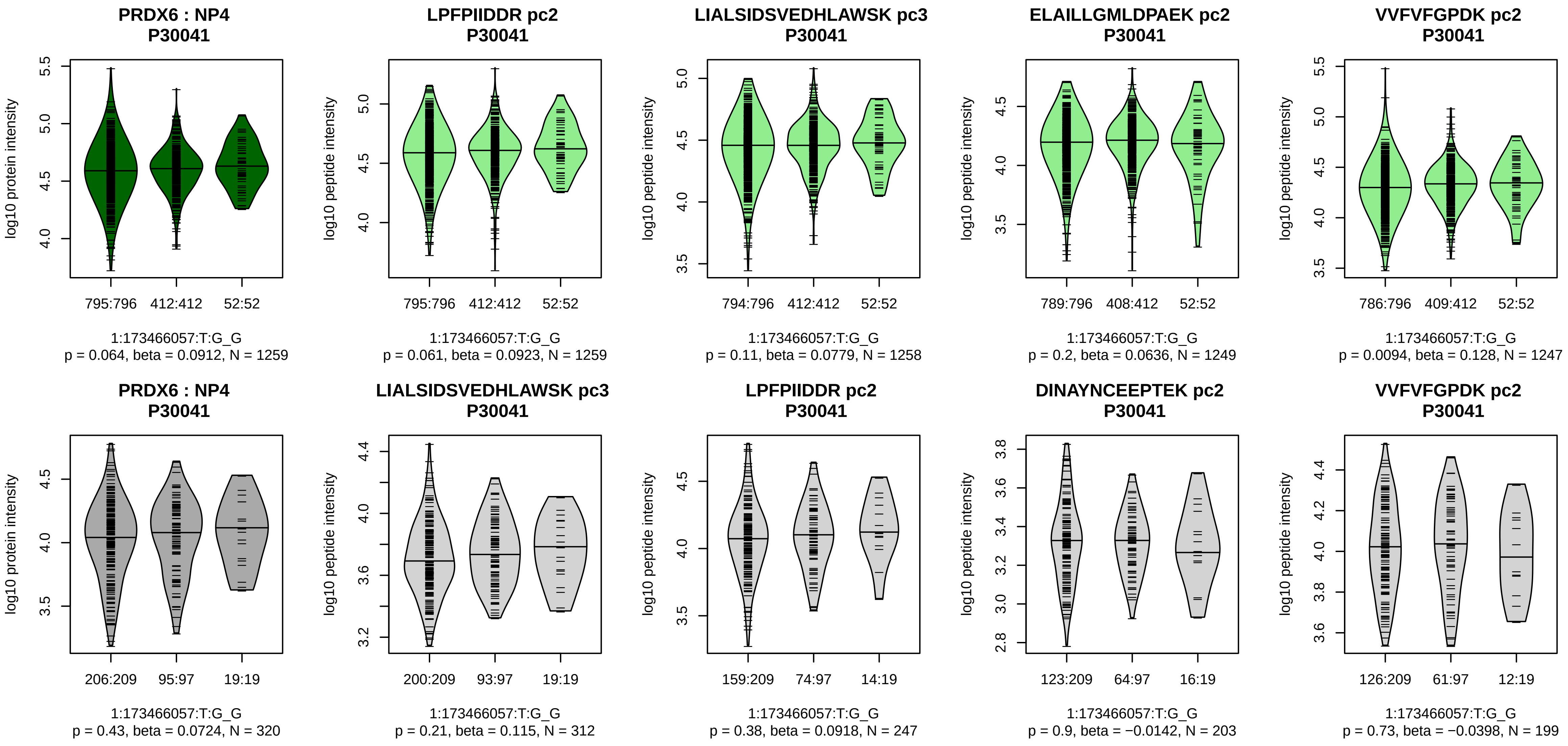


Assay Target: PRDX6
Olink UniProt: P30041
UKB rsID: rs7524592
Proxy rsID: rs7524592
UKB SNP: 1:173466057:T:G
Proxy SNP: 1:173466057:T:G
UKB log10(p): 104.6
UKB BETA: 0.169
–:–:–:*:–:–:–:–:–:–:–:–:–:–:–:–:–:–
1259:1258:1249:1247:1154:112

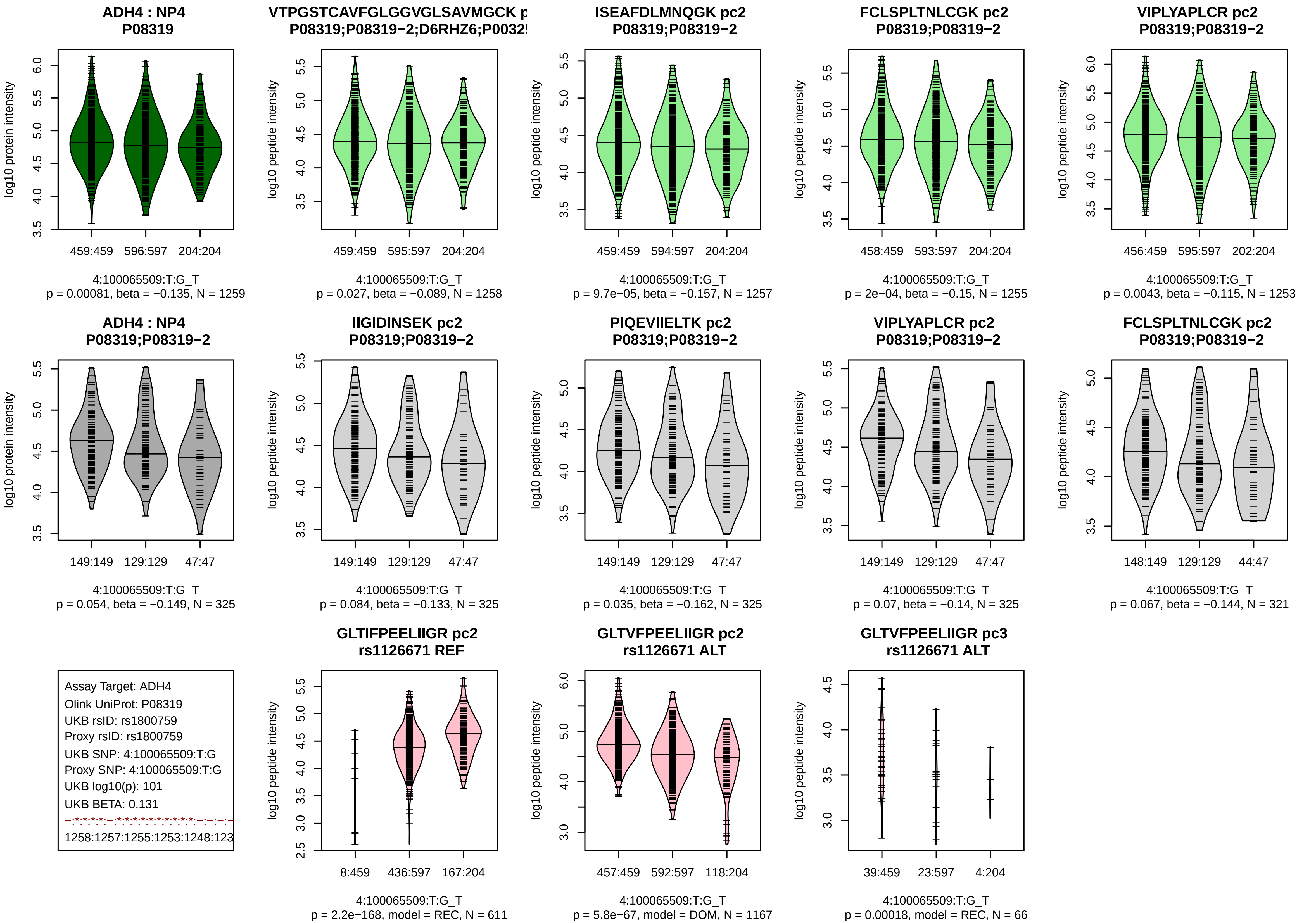

ADH4 : NP4
P08319
log10 protein intensity
459:459
596:597
204:204
4:100065509:T:G_T
p = 0.00081, beta = −0.135, N = 1259
VTPGSTCAVFGLGGVGLSAVMGCK p
P08319;P08319−2;D6RHZ6;P0032
log10 peptide intensity
459:459
595:597
204:204
4:100065509:T:G_T
p = 0.027, beta = −0.089, N = 1258
ISEAFDLMNQGK pc2
P08319;P08319−2
log10 peptide intensity
459:459
594:597
204:204
4:100065509:T:G_T
p = 9.7e−05, beta = −0.157, N = 1257
FCLSPLTNLCGK pc2
P08319;P08319−2
log10 peptide intensity
458:459
593:597
204:204
4:100065509:T:G_T
p = 2e−04, beta = −0.15, N = 1255
VIPLYAPLCR pc2
P08319;P08319−2
log10 peptide intensity
456:459
595:597
202:204
4:100065509:T:G_T
p = 0.0043, beta = −0.115, N = 1253
ADH4 : NP4
P08319;P08319−2
log10 protein intensity
149:149
129:129
47:47
4:100065509:T:G_T
p = 0.054, beta = −0.149, N = 325
IIGIDINSEK pc2
P08319;P08319−2
log10 peptide intensity
149:149
129:129
47:47
4:100065509:T:G_T
p = 0.084, beta = −0.133, N = 325
PIQEVIIELTK pc2
P08319;P08319−2
log10 peptide intensity
149:149
129:129
47:47
4:100065509:T:G_T
p = 0.035, beta = −0.162, N = 325
VIPLYAPLCR pc2
P08319;P08319−2
log10 peptide intensity
149:149
129:129
47:47
4:100065509:T:G_T
p = 0.07, beta = −0.14, N = 325
FCLSPLTNLCGK pc2
P08319;P08319−2
log10 peptide intensity
148:149
129:129
44:47
4:100065509:T:G_T
p = 0.067, beta = −0.144, N = 321
Assay Target: ADH4
Olink UniProt: P08319
UKB rsID: rs1800759
Proxy rsID: rs1800759
UKB SNP: 4:100065509:T:G
Proxy SNP: 4:100065509:T:G
UKB log10(p): 101
UKB BETA: 0.131
1258:1257:1255:1253:1248:123
GLTIFPEELIIGR pc2
rs1126671 REF
log10 peptide intensity
8:459
436:597
167:204
4:100065509:T:G_T
p = 2.2e−168, model = REC, N = 611
GLTVFPEELIIGR pc2
rs1126671 ALT
log10 peptide intensity
457:459
592:597
118:204
4:100065509:T:G_T
p = 5.8e−67, model = DOM, N = 1167
GLTVFPEELIIGR pc3
rs1126671 ALT
log10 peptide intensity
39:459
23:597
4:204
4:100065509:T:G_T
p = 0.00018, model = REC, N = 66

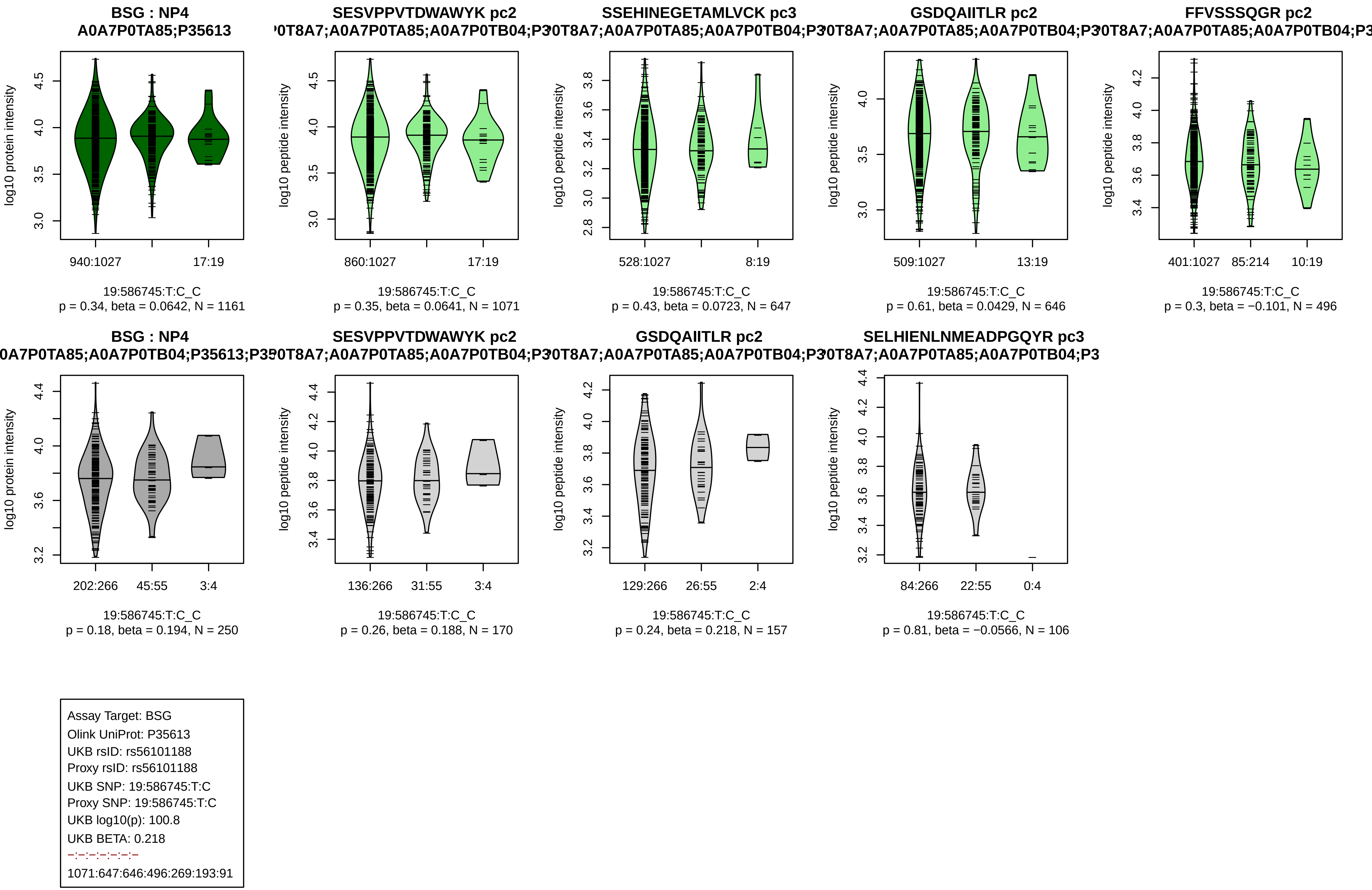

BSG : NP4
A0A7P0TA85;P35613
log10 protein intensity
940:1027
17:19
19:586745:T:C_C
p = 0.34, beta = 0.0642, N = 1161
SESVPPVTDWAWYK pc2
log10 peptide intensity
860:1027
17:19
19:586745:T:C_C
p = 0.35, beta = 0.0641, N = 1071
SSEHINEGETAMLVCK pc3
528:1027
8:19
19:586745:T:C_C
p = 0.43, beta = 0.0723, N = 647
GSDQAIITLR pc2
509:1027
13:19
19:586745:T:C_C
p = 0.61, beta = 0.0429, N = 646
FFVSSSQGR pc2
401:1027
85:214
10:19
19:586745:T:C_C
p = 0.3, beta = −0.101, N = 496
BSG : NP4
202:266
45:55
3:4
19:586745:T:C_C
p = 0.18, beta = 0.194, N = 250
SESVPPVTDWAWYK pc2
136:266
31:55
3:4
19:586745:T:C_C
p = 0.26, beta = 0.188, N = 170
GSDQAIITLR pc2
129:266
26:55
2:4
19:586745:T:C_C
p = 0.24, beta = 0.218, N = 157
SELHIENLNMEADPGQYR pc3
84:266
22:55
0:4
19:586745:T:C_C
p = 0.81, beta = −0.0566, N = 106
Assay Target: BSG
Olink UniProt: P35613
UKB rsID: rs56101188
Proxy rsID: rs56101188
UKB SNP: 19:586745:T:C
Proxy SNP: 19:586745:T:C
UKB log10(p): 100.8
UKB BETA: 0.218
–:–:–:–:–:–:–
1071:647:646:496:269:193:91

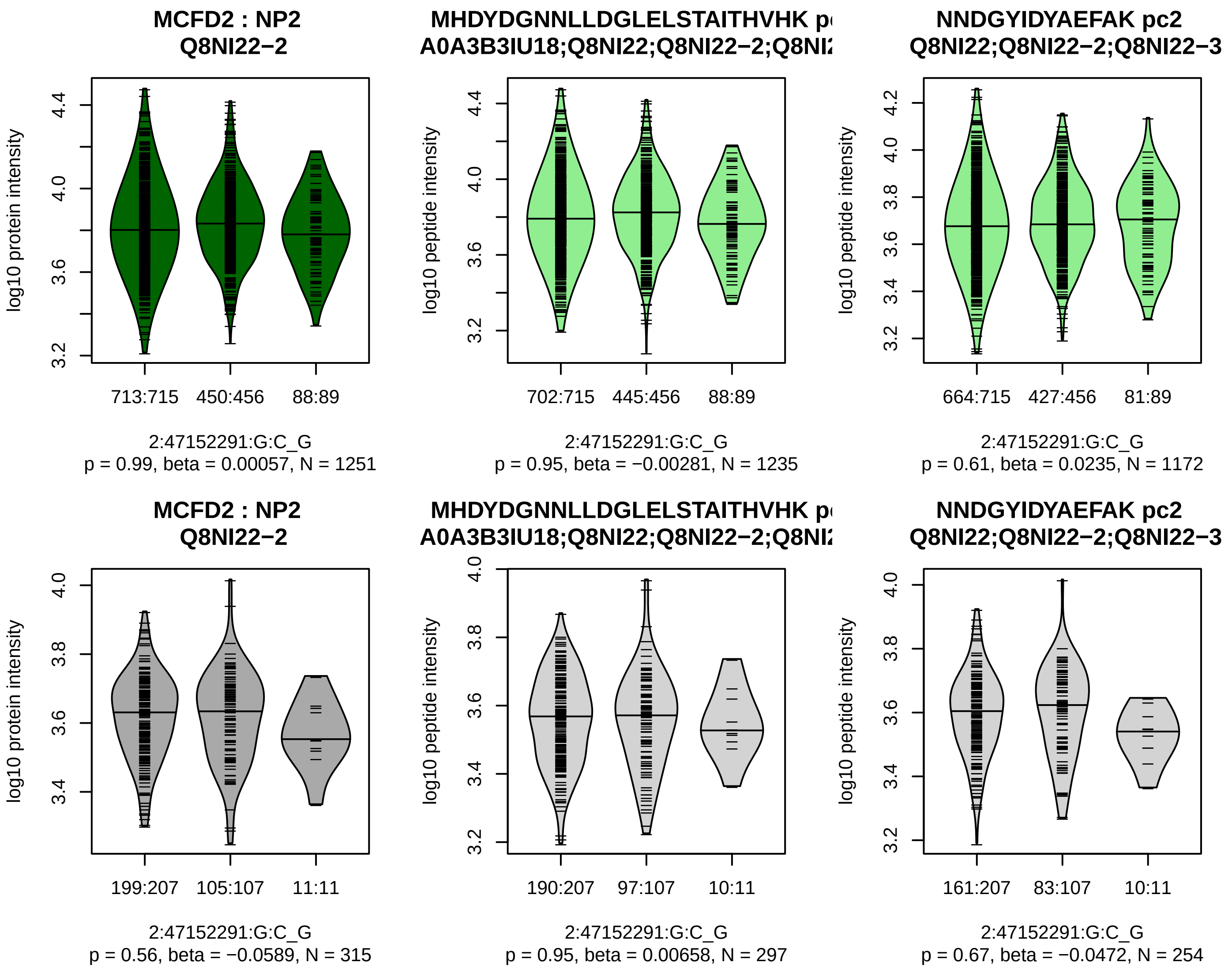


Assay Target: MCFD2
Olink UniProt: Q8NI22
UKB rsID: rs6419615
Proxy rsID: rs6419615
UKB SNP: 2:47152291:G:C
Proxy SNP: 2:47152291:G:C
UKB log10(p): 99.3
UKB BETA: −0.146
−:−
1235:1172

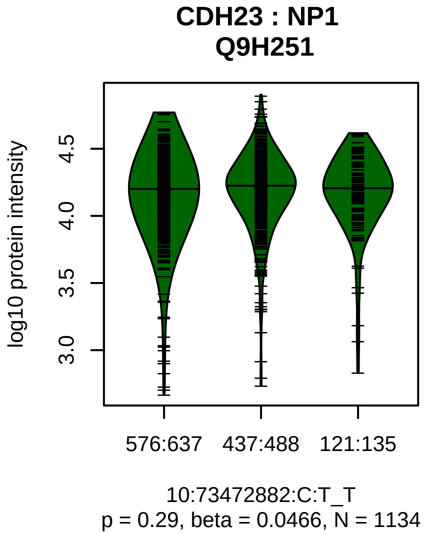


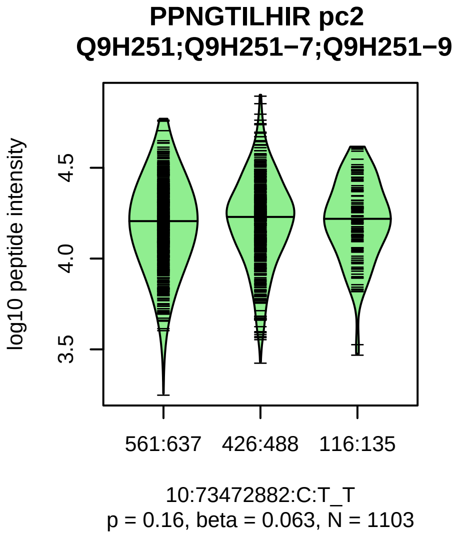


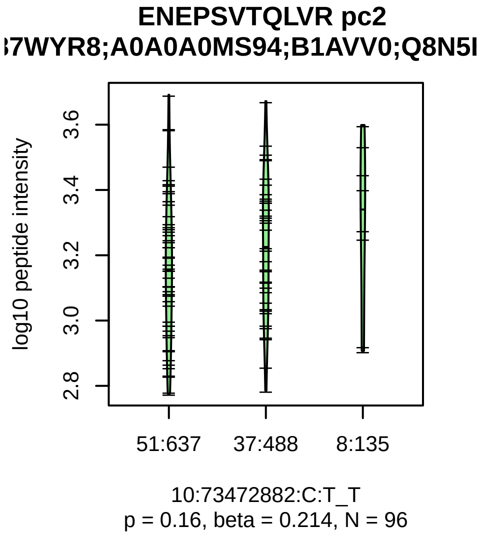


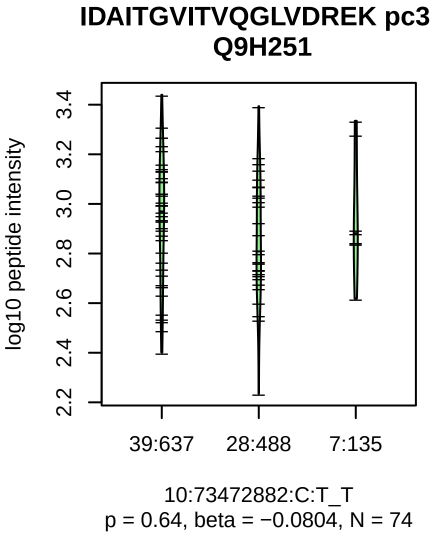


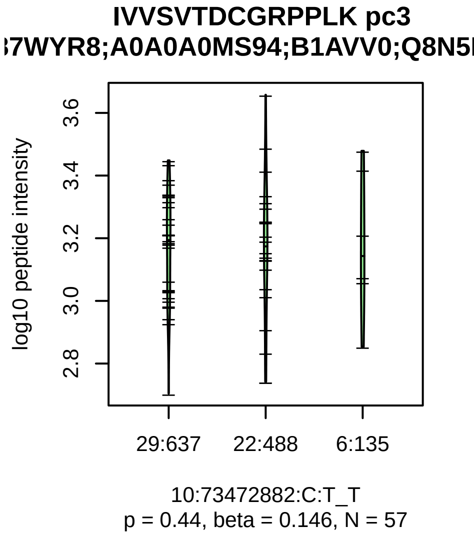


Assay Target: CDH23
Olink UniProt: Q9H251
UKB rsID: rs2166631
Proxy rsID: rs2166631
UKB SNP: 10:73472882:C:T
Proxy SNP: 10:73472882:C:T
UKB log10(p): 96.8
UKB BETA: −0.157
–:–:–:–:–:*:–:–:–:–
1103:96:74:57:55:42:39:38:36:3

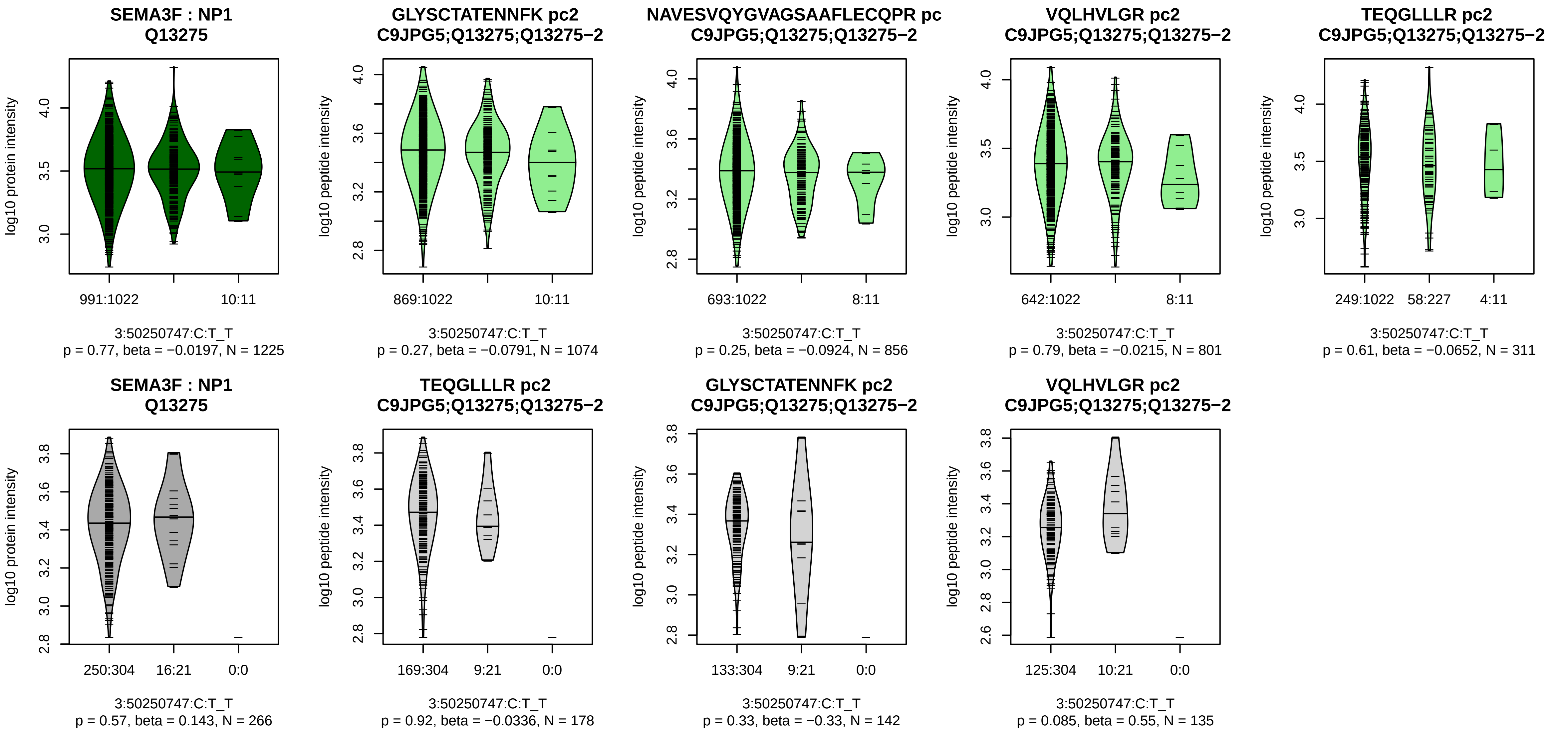


Assay Target: SEMA3F
Olink UniProt: Q13275
UKB rsID: rs74461473
Proxy rsID: rs74461473
UKB SNP: 3:50250747:C:T
Proxy SNP: 3:50250747:C:T
UKB log10(p): 95.6
UKB BETA: −0.201
−:−:−:−:−
1074:856:801:311:24

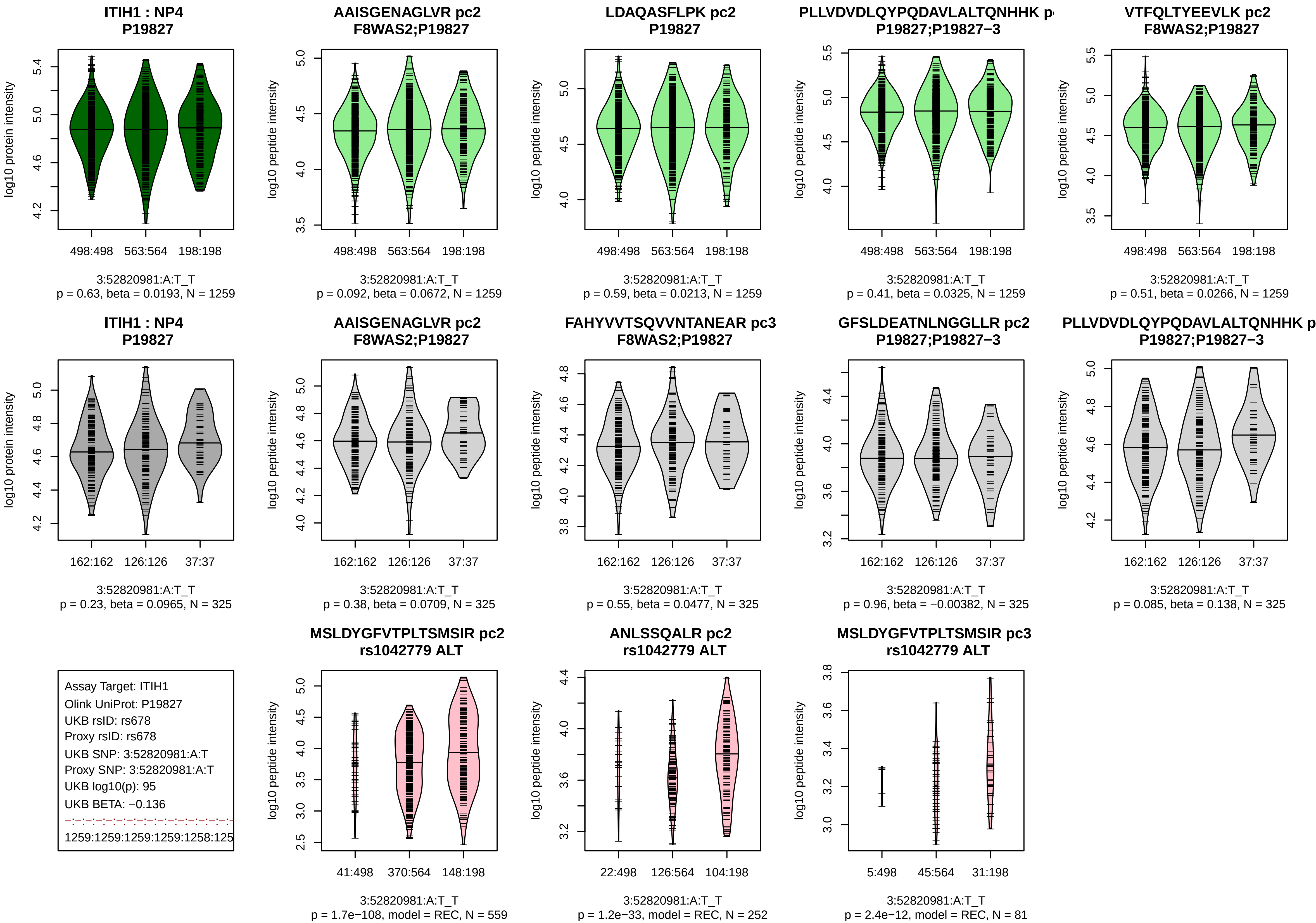

ITIH1 : NP4
P19827
log10 protein intensity
498:498
563:564
198:198
3:52820981:A:T_T
p = 0.63, beta = 0.0193, N = 1259
AAISGENAGLVR pc2
F8WAS2;P19827
log10 peptide intensity
3:52820981:A:T_T
p = 0.092, beta = 0.0672, N = 1259
LDAQASFLPK pc2
P19827
3:52820981:A:T_T
p = 0.59, beta = 0.0213, N = 1259
PLLVDVDLQYPQDAVLALTQNHHK p
P19827;P19827−3
3:52820981:A:T_T
p = 0.41, beta = 0.0325, N = 1259
VTFQLTYEEVLK pc2
F8WAS2;P19827
3:52820981:A:T_T
p = 0.51, beta = 0.0266, N = 1259
ITIH1 : NP4
P19827
162:162
126:126
37:37
3:52820981:A:T_T
p = 0.23, beta = 0.0965, N = 325
AAISGENAGLVR pc2
F8WAS2;P19827
3:52820981:A:T_T
p = 0.38, beta = 0.0709, N = 325
FAHYVVTSQVVNTANEAR pc3
F8WAS2;P19827
3:52820981:A:T_T
p = 0.55, beta = 0.0477, N = 325
GFSLDEATNLNGGLLR pc2
P19827;P19827−3
3:52820981:A:T_T
p = 0.96, beta = −0.00382, N = 325
PLLVDVDLQYPQDAVLALTQNHHK p
P19827;P19827−3
3:52820981:A:T_T
p = 0.085, beta = 0.138, N = 325
Assay Target: ITIH1
Olink UniProt: P19827
UKB rsID: rs678
Proxy rsID: rs678
UKB SNP: 3:52820981:A:T
Proxy SNP: 3:52820981:A:T
UKB log10(p): 95
UKB BETA: −0.136
1259:1259:1259:1259:1258:125
MSLDYGFVTPLTSMSIR pc2
rs1042779 ALT
41:498
370:564
148:198
3:52820981:A:T_T
p = 1.7e−108, model = REC, N = 559
ANLSSQALR pc2
rs1042779 ALT
22:498
126:564
104:198
3:52820981:A:T_T
p = 1.2e−33, model = REC, N = 252
MSLDYGFVTPLTSMSIR pc3
rs1042779 ALT
5:498
45:564
31:198
3:52820981:A:T_T
p = 2.4e−12, model = REC, N = 81

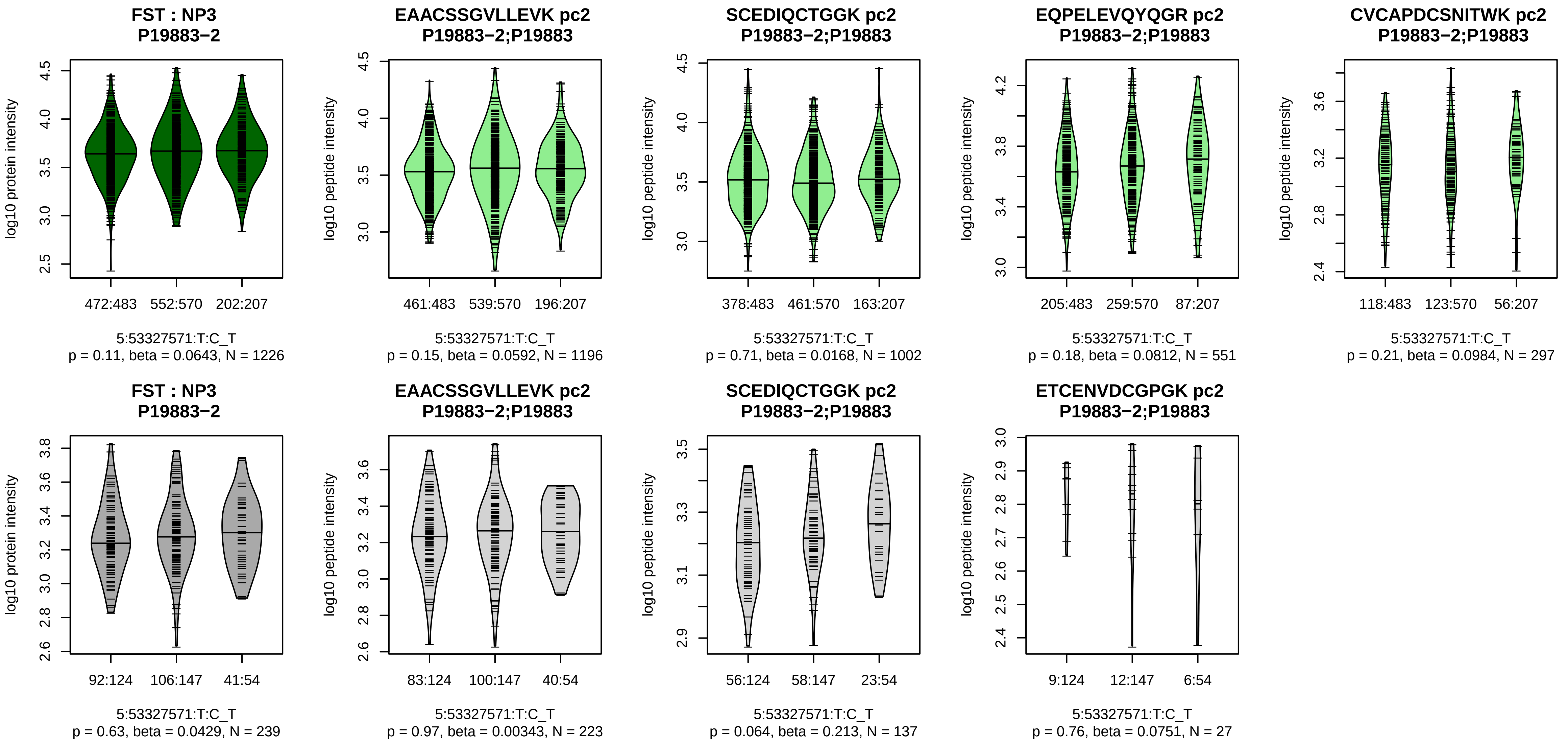


Assay Target: FST
Olink UniProt: P19883
UKB rsID: rs31226
Proxy rsID: rs31226
UKB SNP: 5:53327571:T:C
Proxy SNP: 5:53327571:T:C
UKB log10(p): 94.3
UKB BETA: −0.128
−:−:−:−:−:−:−:−:−:NA
1196:1002:551:297:280:276:172

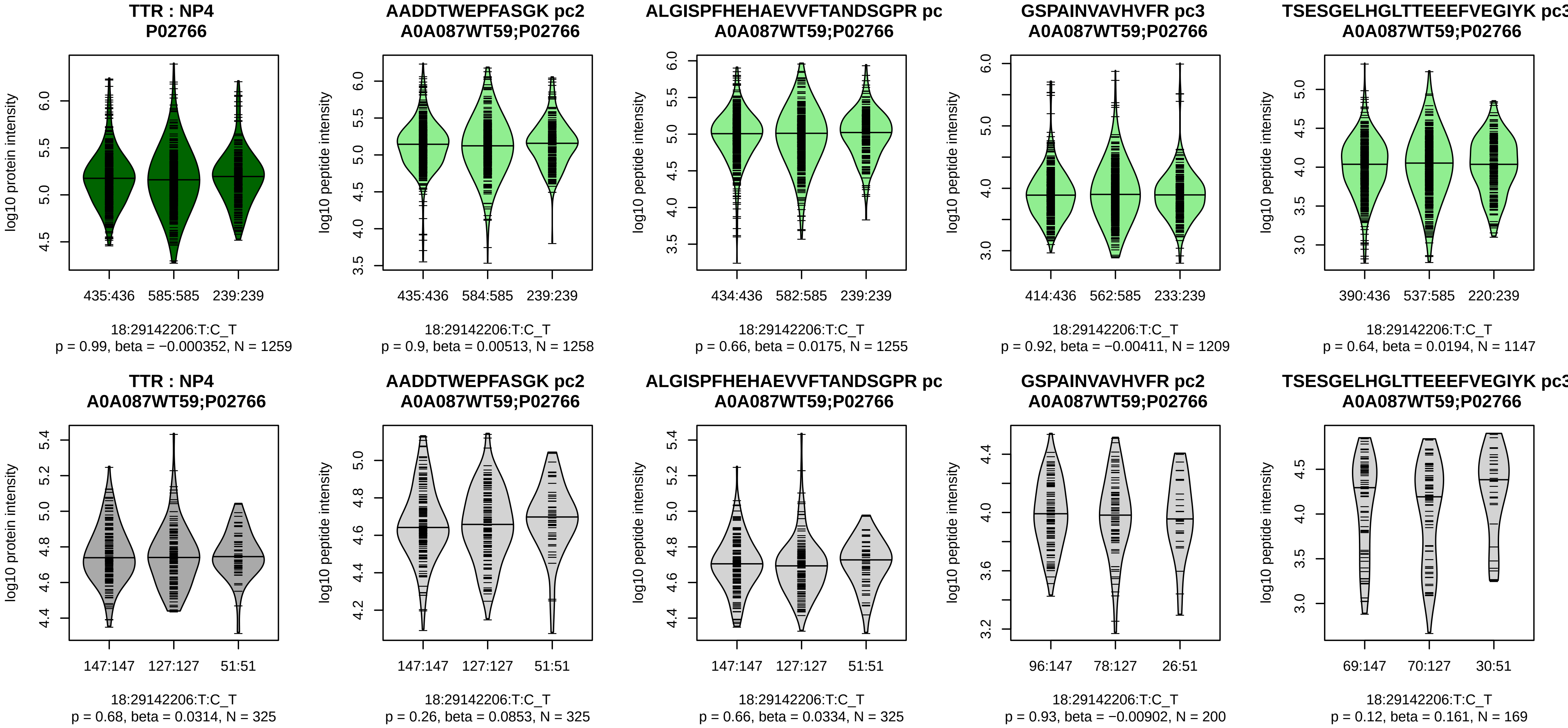


Assay Target: TTR
Olink UniProt: P02766
UKB rsID: rs1667229
Proxy rsID: rs1667229
UKB SNP: 18:29142206:T:C
Proxy SNP: 18:29142206:T:C
UKB log10(p): 94.2
UKB BETA: 0.121
−:−:−:−:−:−:−:−:−:−:−:−:NA:NA:N
1258:1255:1209:1147:1111:978

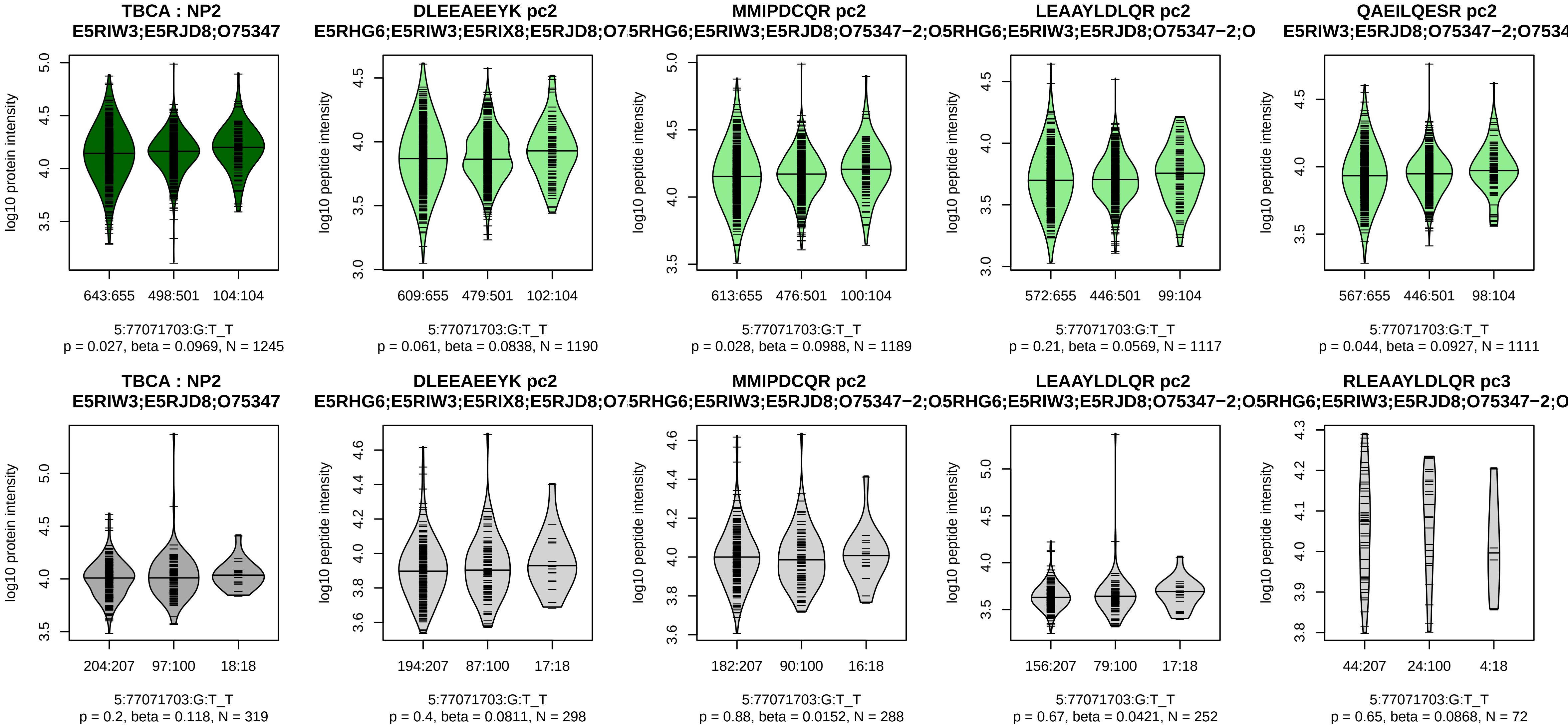


Assay Target: TBCA
Olink UniProt: O75347
UKB rsID: rs254415
Proxy rsID: rs254415
UKB SNP: 5:77071703:G:T
Proxy SNP: 5:77071703:G:T
UKB log10(p): 94.1
UKB BETA: 0.14
−:−:−:−:−:−:−:−:NA
1190:1189:1117:1111:741:601:[illegible]

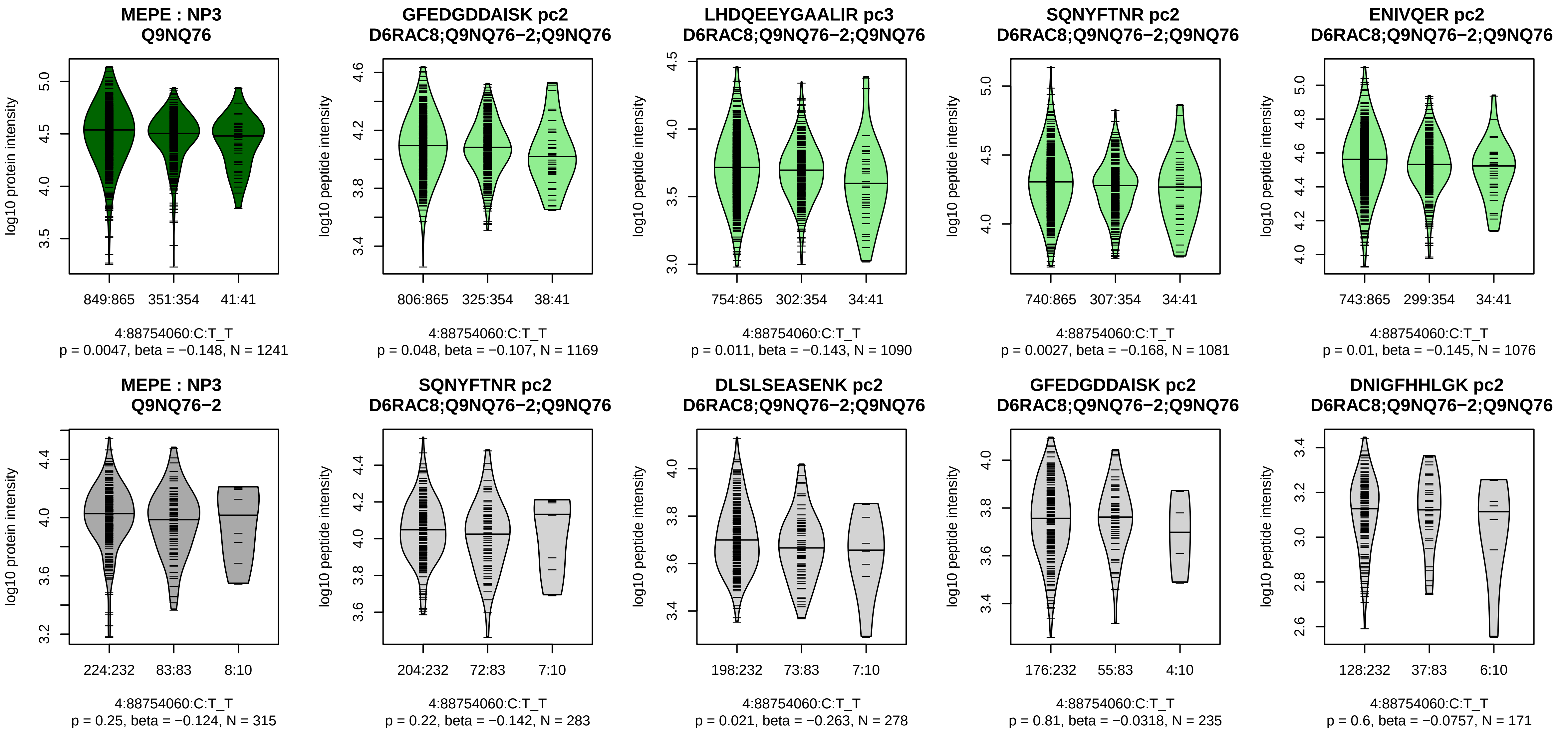


Assay Target: MEPE
Olink UniProt: Q9NQ76
UKB rsID: rs17013212
Proxy rsID: rs17013212
UKB SNP: 4:88754060:C:T
Proxy SNP: 4:88754060:C:T
UKB log10(p): 92.2
UKB BETA: −0.168
−:−:*:*:−:−:−:−:−:−:NA:NA
1169:1090:1081:1076:1038:954

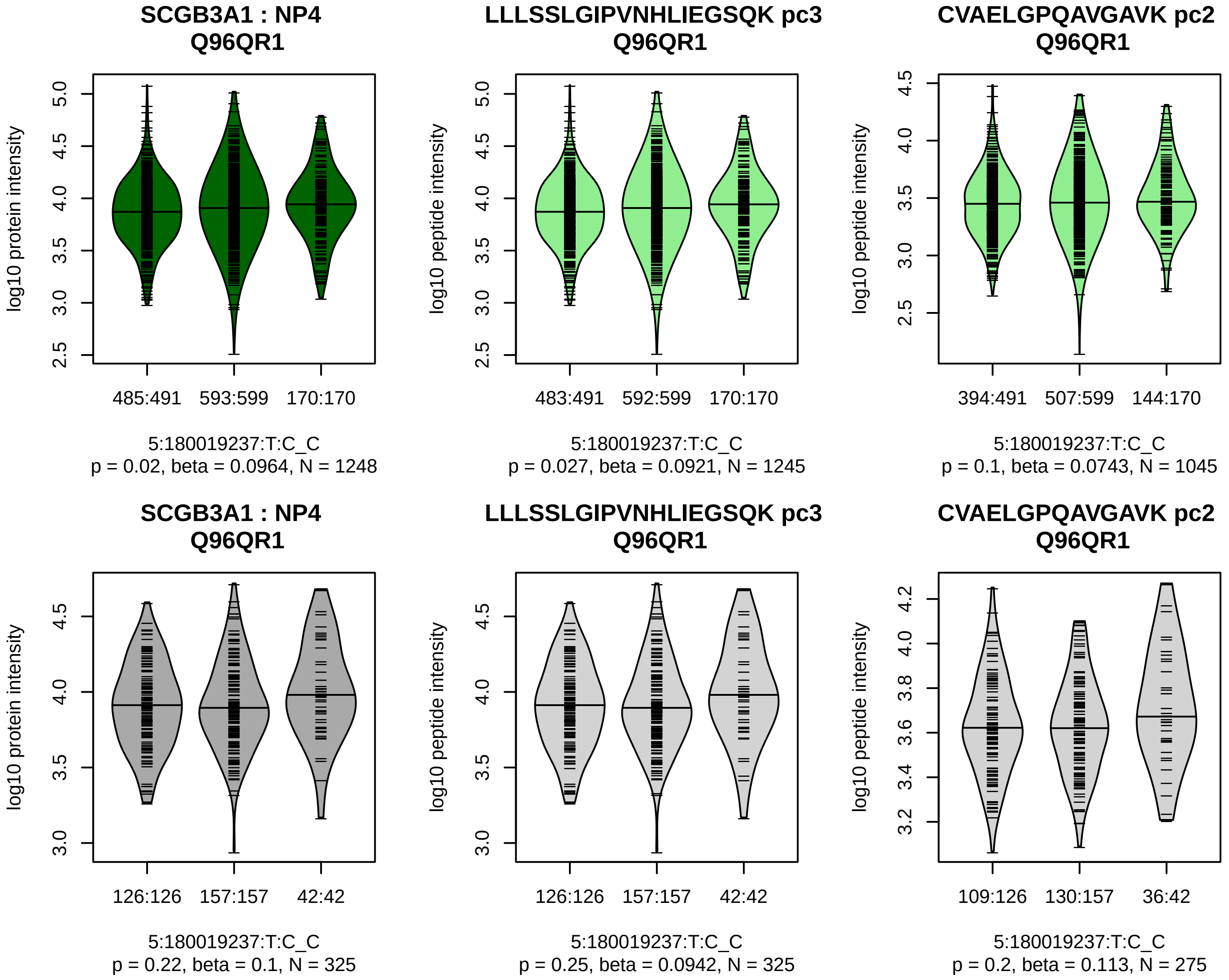


Assay Target: SCGB3A1
Olink UniProt: Q96QR1
UKB rsID: rs307802
Proxy rsID: rs307802
UKB SNP: 5:180019237:T:C
Proxy SNP: 5:180019237:T:C
UKB log10(p): 91.5
UKB BETA: 0.123
–:–
1245:1045

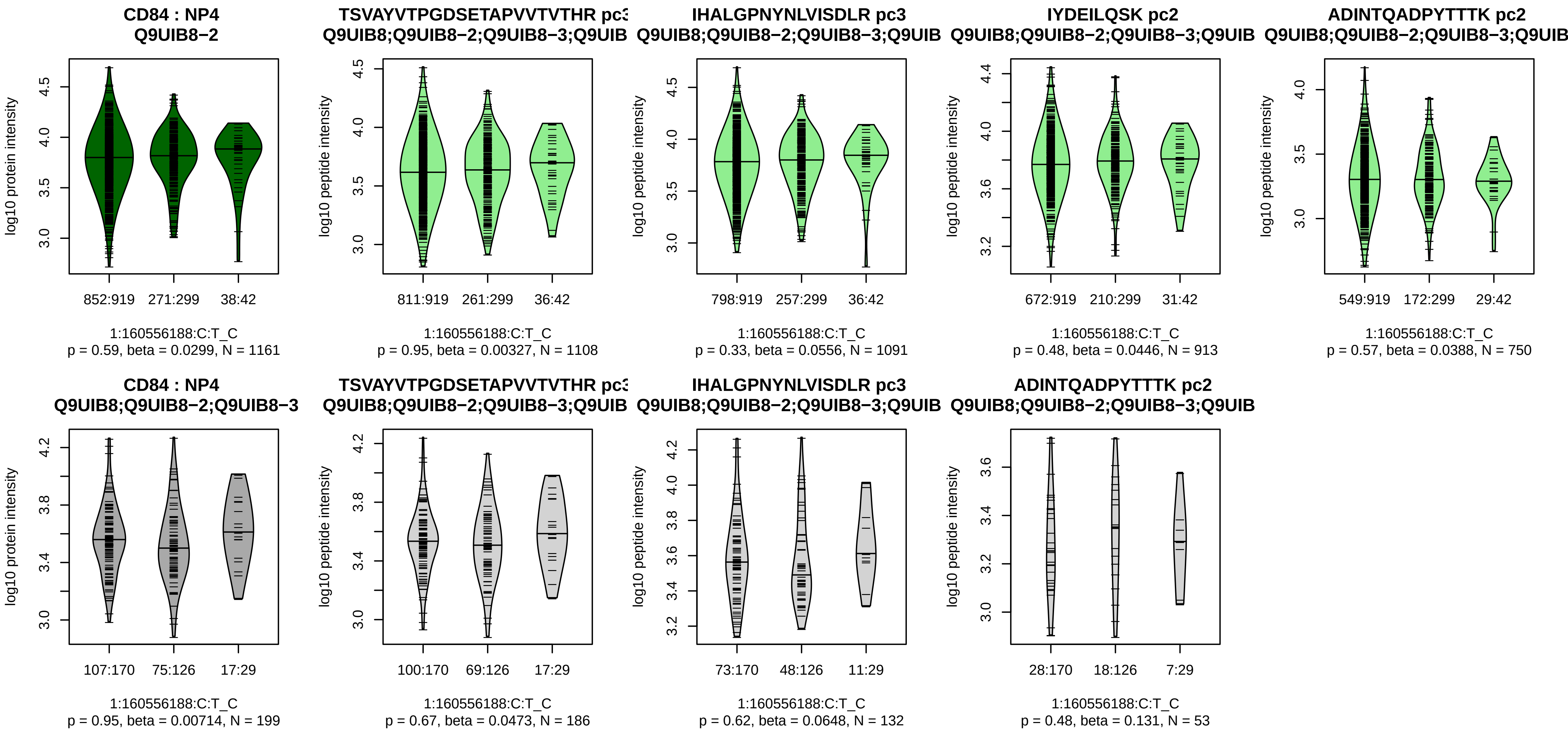


Assay Target: CD84
Olink UniProt: Q9UIB8
UKB rsID: rs200682634
Proxy rsID: rs7539149
UKB SNP: 1:160560109:CAT:C
Proxy SNP: 1:160556188:C:T
UKB log10(p): 88.6
UKB BETA: −0.211
−:−:−:−:−:−:−:−:−
1108:1091:913:750:715:409:161

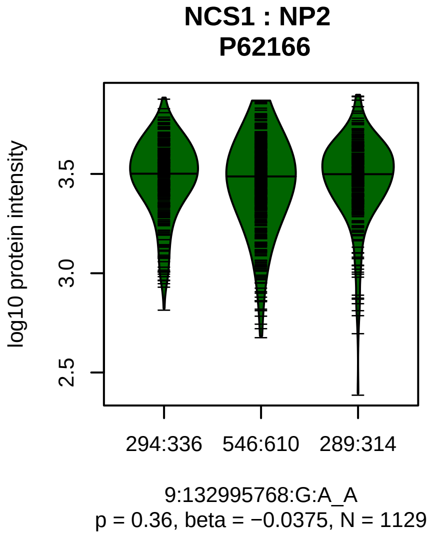


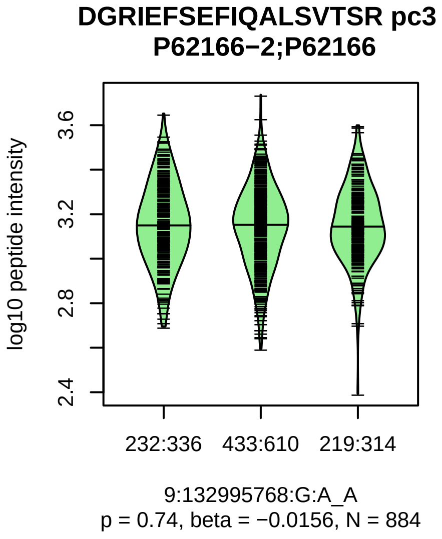


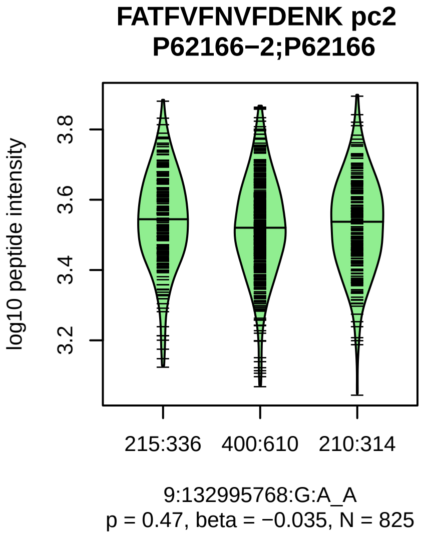


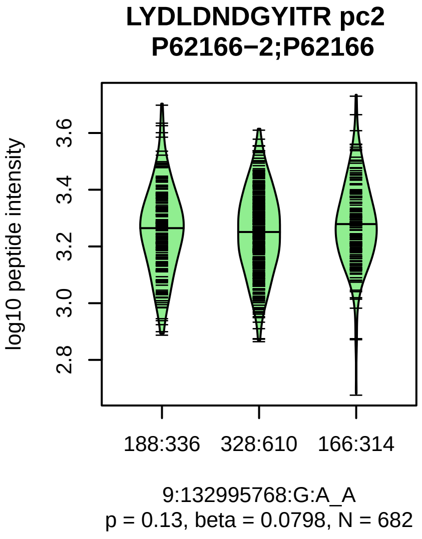


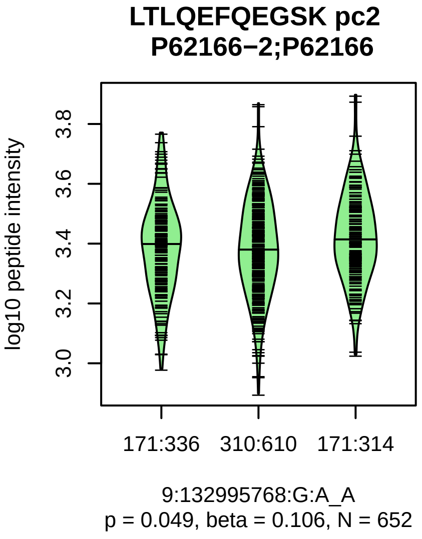


Assay Target: NCS1
Olink UniProt: P62166
UKB rsID: rs1054879
Proxy rsID: rs1054879
UKB SNP: 9:132995768:G:A
Proxy SNP: 9:132995768:G:A
UKB log10(p): 88
UKB BETA: 0.11
−:−:−:−
884:825:682:652

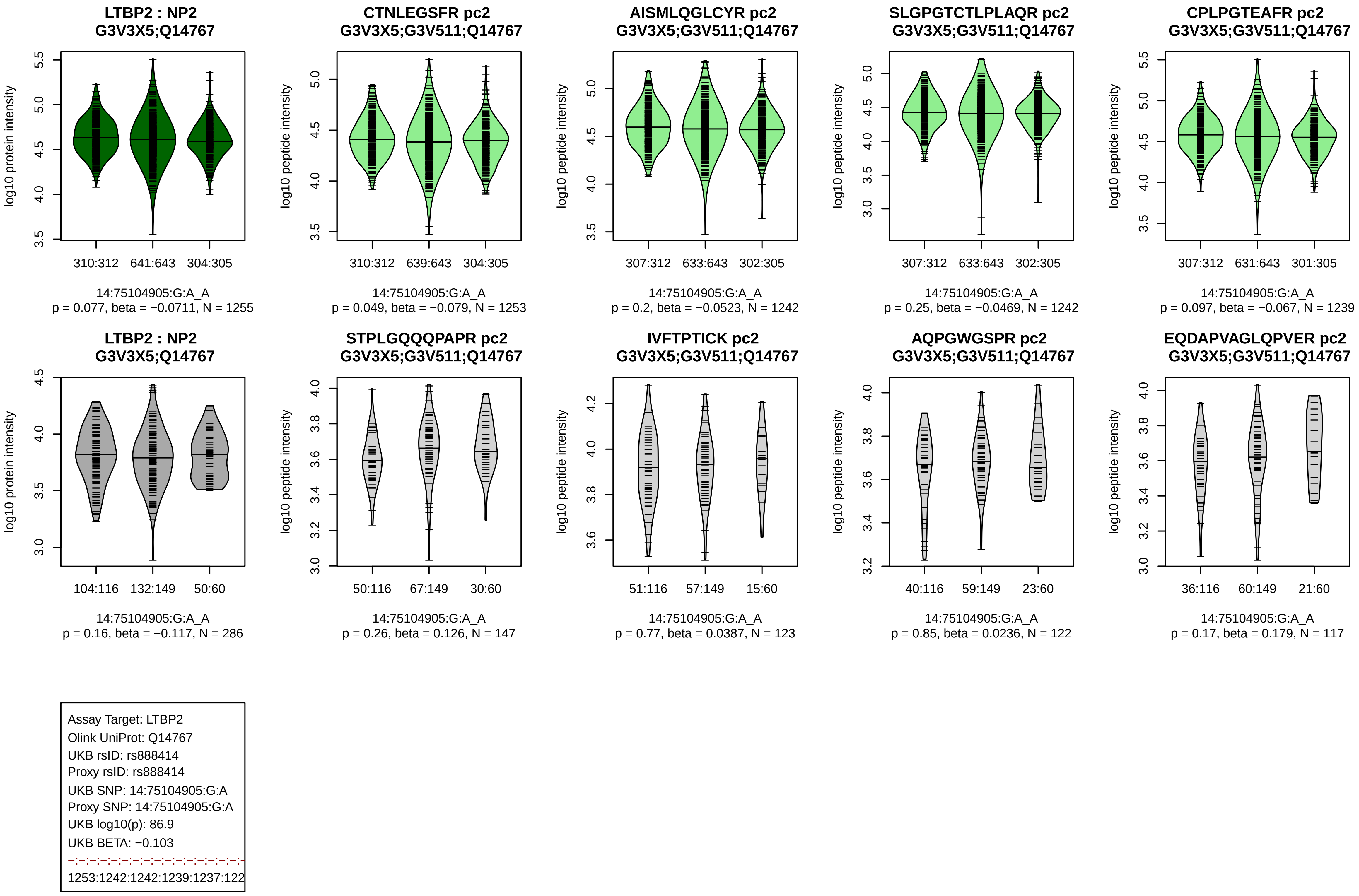

LTBP2 : NP2
G3V3X5;Q14767
log10 protein intensity
310:312
641:643
304:305
14:75104905:G:A_A
p = 0.077, beta = −0.0711, N = 1255
CTNLEGSFR pc2
G3V3X5;G3V511;Q14767
log10 peptide intensity
310:312
639:643
304:305
14:75104905:G:A_A
p = 0.049, beta = −0.079, N = 1253
AISMLQGLCYR pc2
G3V3X5;G3V511;Q14767
log10 peptide intensity
307:312
633:643
302:305
14:75104905:G:A_A
p = 0.2, beta = −0.0523, N = 1242
SLGPGTCTLPLAQR pc2
G3V3X5;G3V511;Q14767
log10 peptide intensity
307:312
633:643
302:305
14:75104905:G:A_A
p = 0.25, beta = −0.0469, N = 1242
CPLPGTEAFR pc2
G3V3X5;G3V511;Q14767
log10 peptide intensity
307:312
631:643
301:305
14:75104905:G:A_A
p = 0.097, beta = −0.067, N = 1239
LTBP2 : NP2
G3V3X5;Q14767
log10 protein intensity
104:116
132:149
50:60
14:75104905:G:A_A
p = 0.16, beta = −0.117, N = 286
STPLGQQQPAPR pc2
G3V3X5;G3V511;Q14767
log10 peptide intensity
50:116
67:149
30:60
14:75104905:G:A_A
p = 0.26, beta = 0.126, N = 147
IVFTPTICK pc2
G3V3X5;G3V511;Q14767
log10 peptide intensity
51:116
57:149
15:60
14:75104905:G:A_A
p = 0.77, beta = 0.0387, N = 123
AQPGWGSPR pc2
G3V3X5;G3V511;Q14767
log10 peptide intensity
40:116
59:149
23:60
14:75104905:G:A_A
p = 0.85, beta = 0.0236, N = 122
EQDAPVAGLQPVER pc2
G3V3X5;G3V511;Q14767
log10 peptide intensity
36:116
60:149
21:60
14:75104905:G:A_A
p = 0.17, beta = 0.179, N = 117
Assay Target: LTBP2
Olink UniProt: Q14767
UKB rsID: rs888414
Proxy rsID: rs888414
UKB SNP: 14:75104905:G:A
Proxy SNP: 14:75104905:G:A
UKB log10(p): 86.9
UKB BETA: −0.103
1253:1242:1242:1239:1237:122

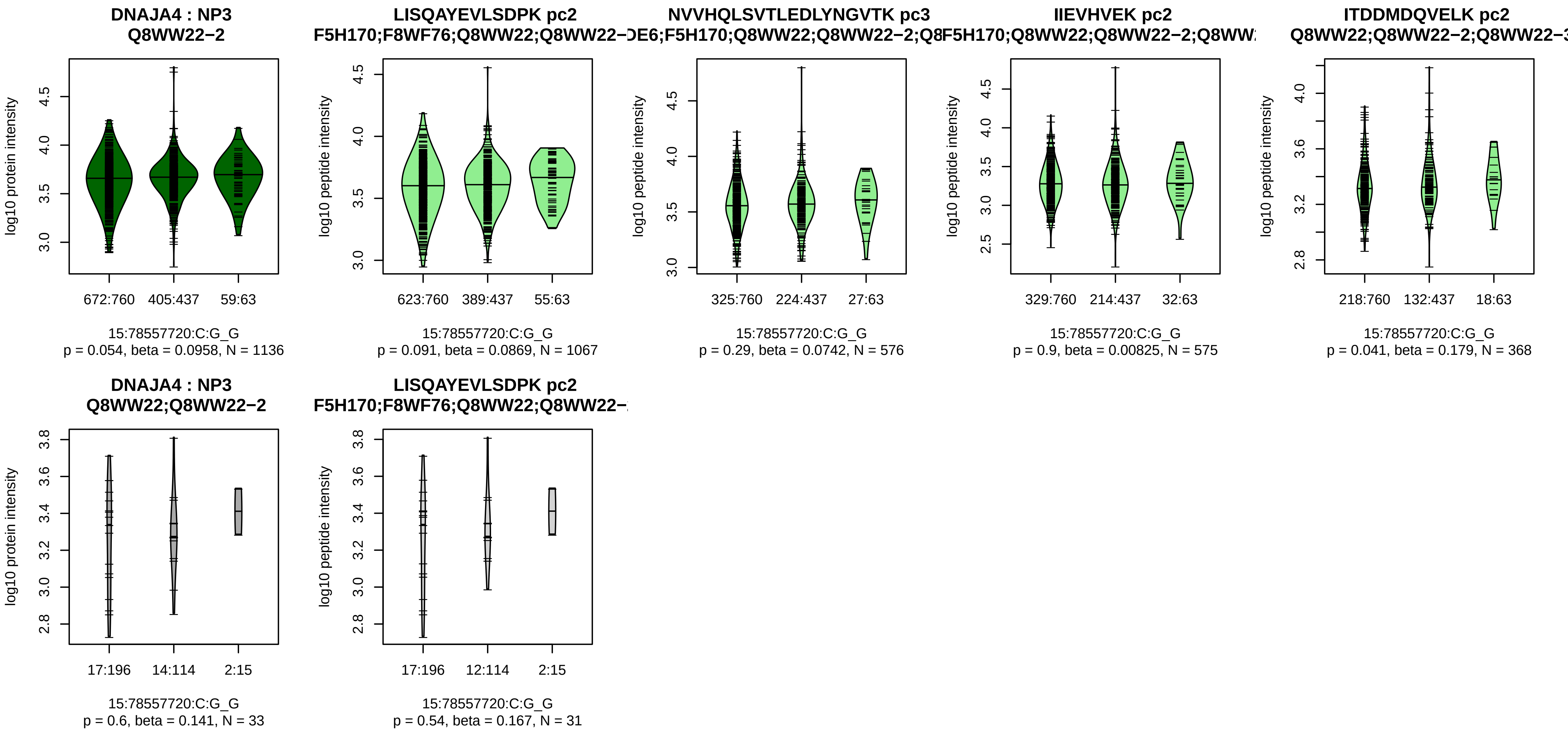


Assay Target: DNAJA4
Olink UniProt: Q8WW22
UKB rsID: rs62007832
Proxy rsID: rs62007832
UKB SNP: 15:78557720:C:G
Proxy SNP: 15:78557720:C:G
UKB log10(p): 86
UKB BETA: 0.143
−:−:−:−:−:−:−:−:−:−:−:−:−:−:NA:N
1067:576:575:368:271:250:205:

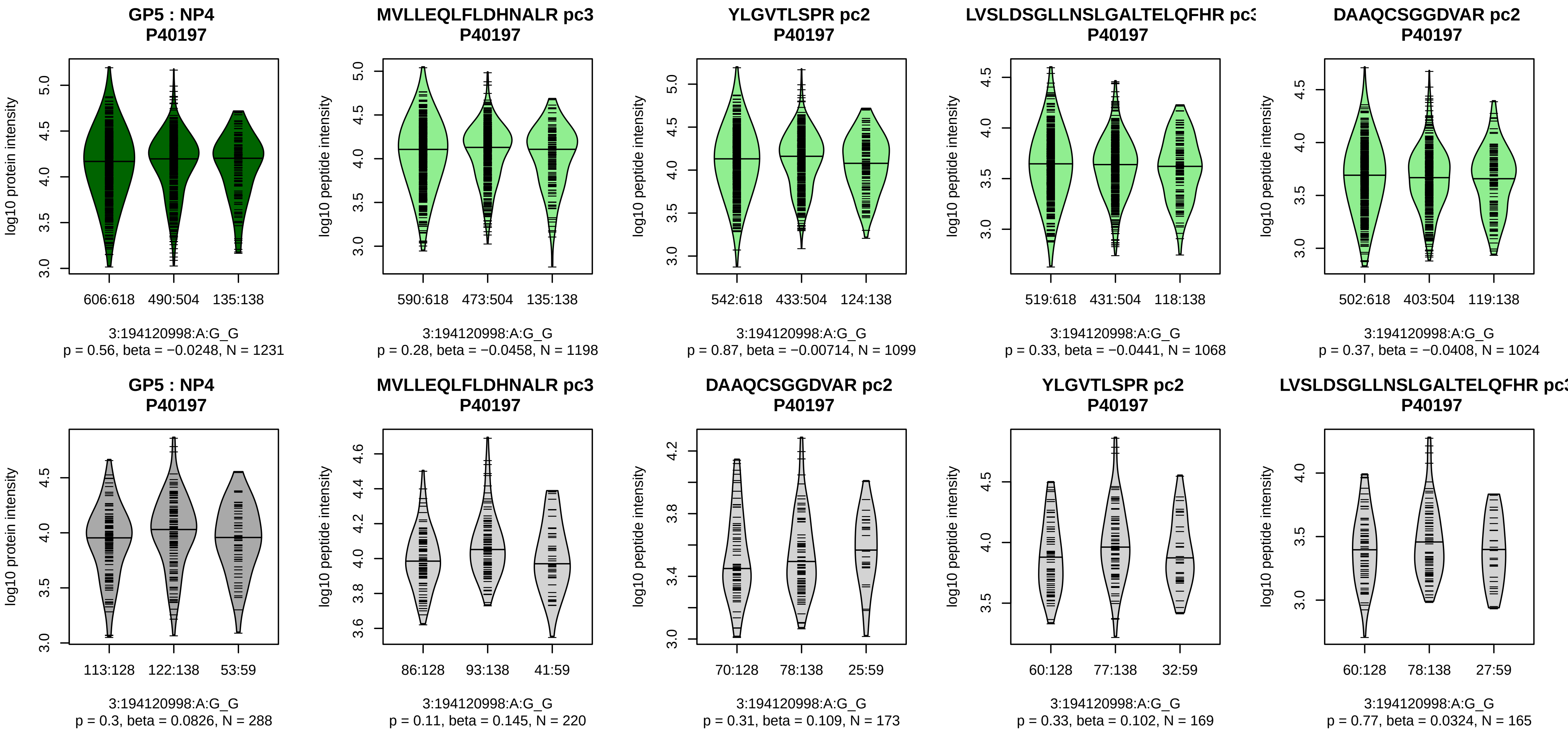


Assay Target: GP5
Olink UniProt: P40197
UKB rsID: rs544985059
Proxy rsID: rs1466733
UKB SNP: 3:194121175:G:T
Proxy SNP: 3:194120998:A:G
UKB log10(p): 85.5
UKB BETA: −0.145
−:−:−:−:−:−:−:−:−:−:−:−:−:−:−:−:
1198:1099:1068:1024:815:785:7

**CDHR2 : NP2**
**Q9BYE9**

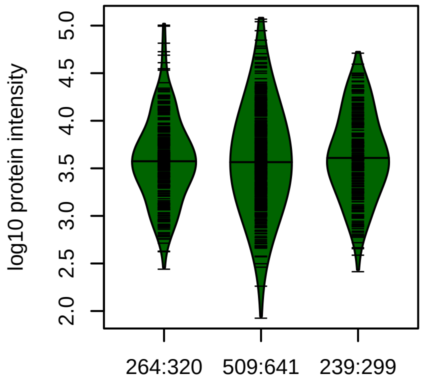


5:175945908:C:G_C
p = 0.072, beta = 0.0799, N = 1012

**TFVIIPELVLPNR pc2**
**Q9BYE9**

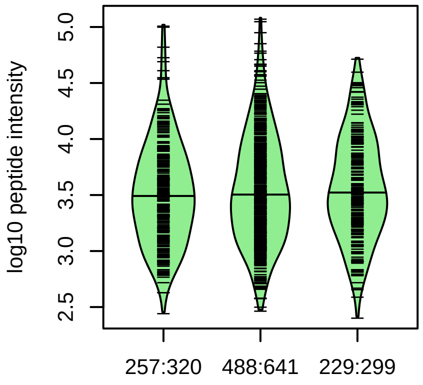


5:175945908:C:G_C
p = 0.1, beta = 0.0736, N = 974

**VTISVSDPYIQVQR pc2**
**Q9BYE9**

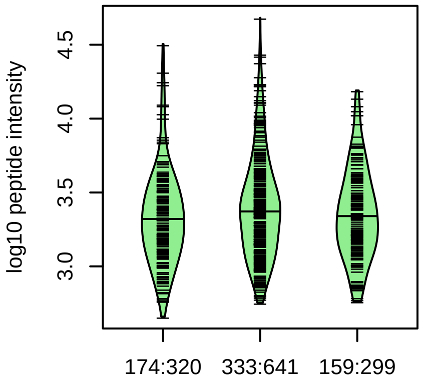


5:175945908:C:G_C
p = 0.1, beta = 0.0889, N = 666

**VSALVDYER pc2**
**Q9BYE9**

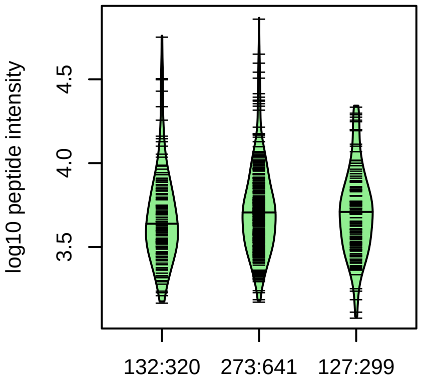


5:175945908:C:G_C
p = 0.099, beta = 0.102, N = 532

**GLVLGAGWAEGYLR pc2**
**Q9BYE9**

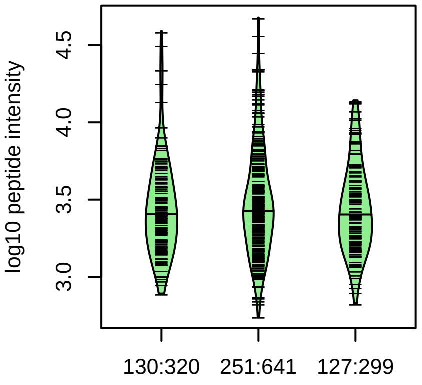


5:175945908:C:G_C
p = 0.83, beta = 0.0133, N = 508

**CDHR2 : NP2**
**Q9BYE9**

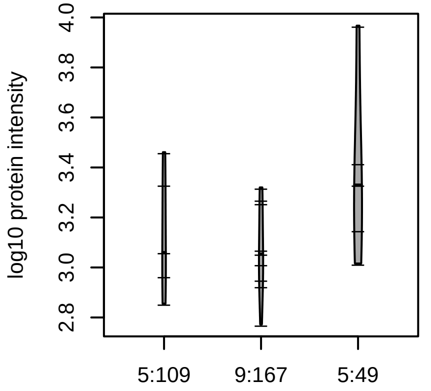


5:175945908:C:G_G
p = 0.13, beta = 0.432, N = 19

Assay Target: CDHR2
Olink UniProt: Q9BYE9
UKB rsID: rs2913909
Proxy rsID: rs2913909
UKB SNP: 5:175945908:C:G
Proxy SNP: 5:175945908:C:G
UKB log10(p): 81.7
UKB BETA: −0.1
−:−:−:−:−:−:−:−:−:−:−:−:−:−:−:−:
974:666:532:508:487:415:408:3

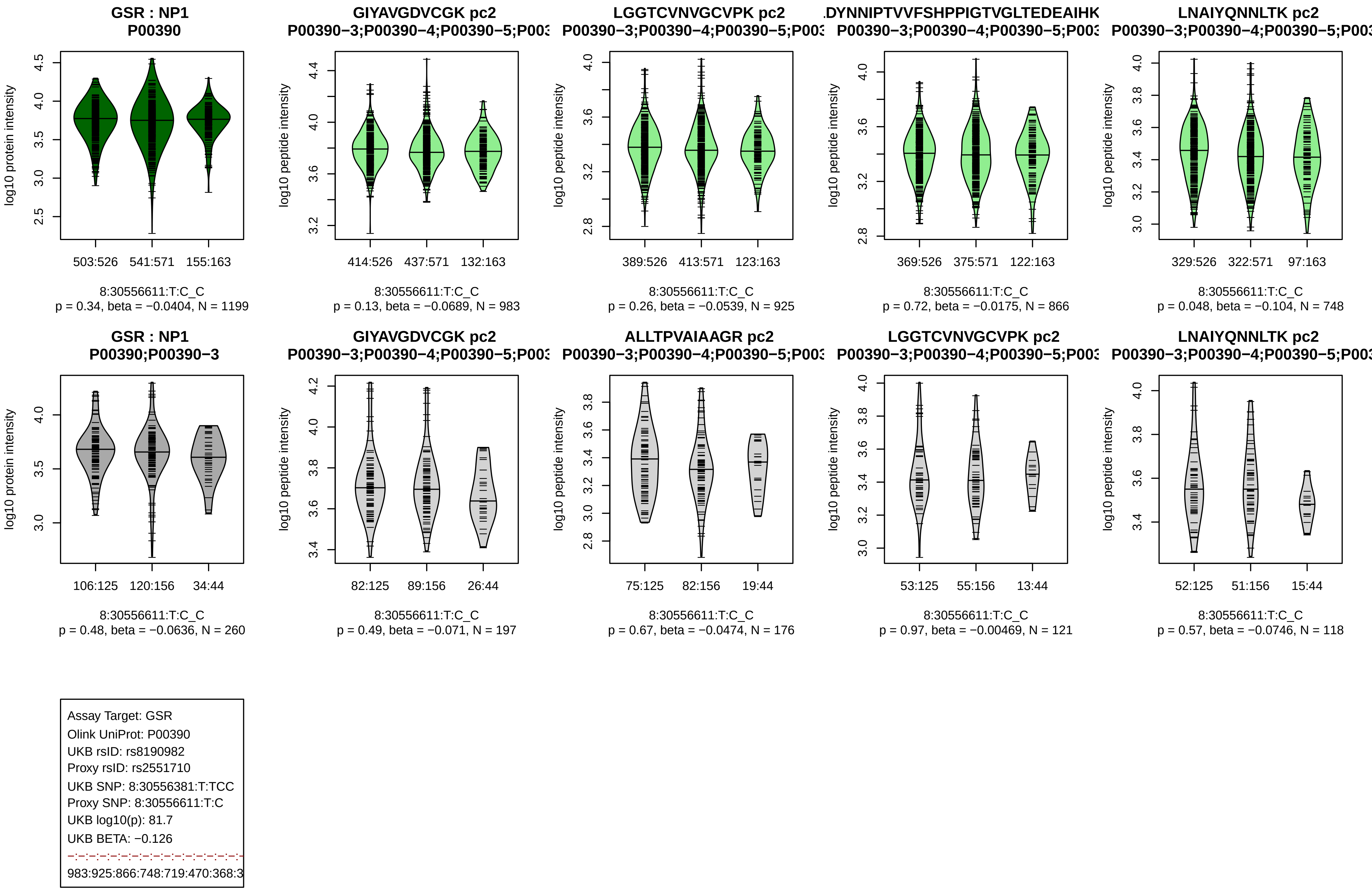
GSR : NP1
P00390
log10 protein intensity
503:526 541:571 155:163
8:30556611:T:C_C
p = 0.34, beta = −0.0404, N = 1199
GIYAVGDVCGK pc2
P00390−3;P00390−4;P00390−5;P003
log10 peptide intensity
414:526 437:571 132:163
8:30556611:T:C_C
p = 0.13, beta = −0.0689, N = 983
LGGTCVNVGCVPK pc2
P00390−3;P00390−4;P00390−5;P003
389:526 413:571 123:163
8:30556611:T:C_C
p = 0.26, beta = −0.0539, N = 925
DYNNIPTVVFSHPPIGTVGLTEDEAIHK
P00390−3;P00390−4;P00390−5;P003
369:526 375:571 122:163
8:30556611:T:C_C
p = 0.72, beta = −0.0175, N = 866
LNAIYQNNLTK pc2
P00390−3;P00390−4;P00390−5;P003
329:526 322:571 97:163
8:30556611:T:C_C
p = 0.048, beta = −0.104, N = 748
GSR : NP1
P00390;P00390−3
106:125 120:156 34:44
8:30556611:T:C_C
p = 0.48, beta = −0.0636, N = 260
GIYAVGDVCGK pc2
P00390−3;P00390−4;P00390−5;P003
82:125 89:156 26:44
8:30556611:T:C_C
p = 0.49, beta = −0.071, N = 197
ALLTPVAIAAGR pc2
P00390−3;P00390−4;P00390−5;P003
75:125 82:156 19:44
8:30556611:T:C_C
p = 0.67, beta = −0.0474, N = 176
LGGTCVNVGCVPK pc2
P00390−3;P00390−4;P00390−5;P003
53:125 55:156 13:44
8:30556611:T:C_C
p = 0.97, beta = −0.00469, N = 121
LNAIYQNNLTK pc2
P00390−3;P00390−4;P00390−5;P003
52:125 51:156 15:44
8:30556611:T:C_C
p = 0.57, beta = −0.0746, N = 118
Assay Target: GSR
Olink UniProt: P00390
UKB rsID: rs8190982
Proxy rsID: rs2551710
UKB SNP: 8:30556381:T:TCC
Proxy SNP: 8:30556611:T:C
UKB log10(p): 81.7
UKB BETA: −0.126
983:925:866:748:719:470:368:3

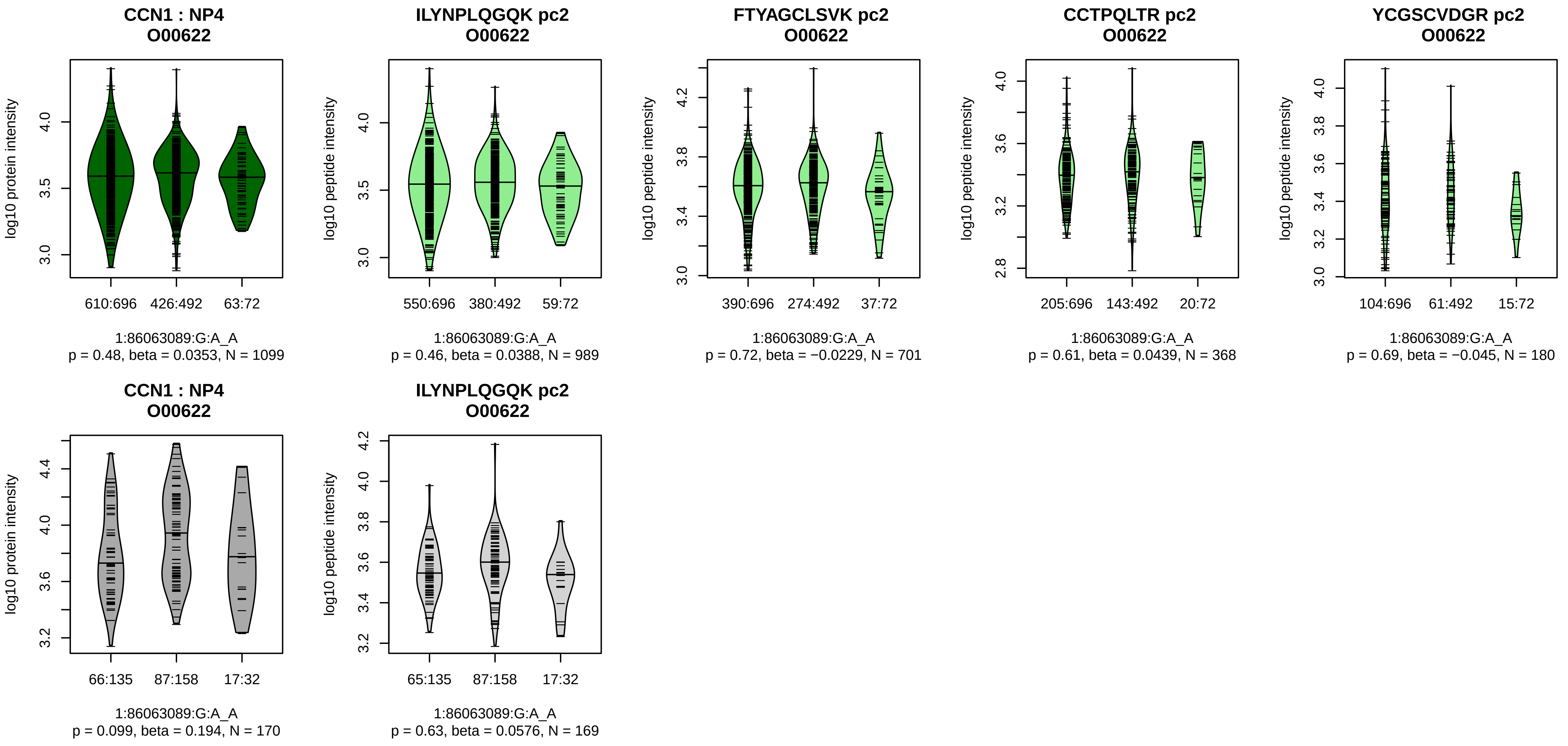


Assay Target: CCN1
Olink UniProt: O00622
UKB rsID: rs4949912
Proxy rsID: rs4949912
UKB SNP: 1:86063089:G:A
Proxy SNP: 1:86063089:G:A
UKB log10(p): 81
UKB BETA: −0.13
−:−:−:−:−:−
989:701:368:180:121:75

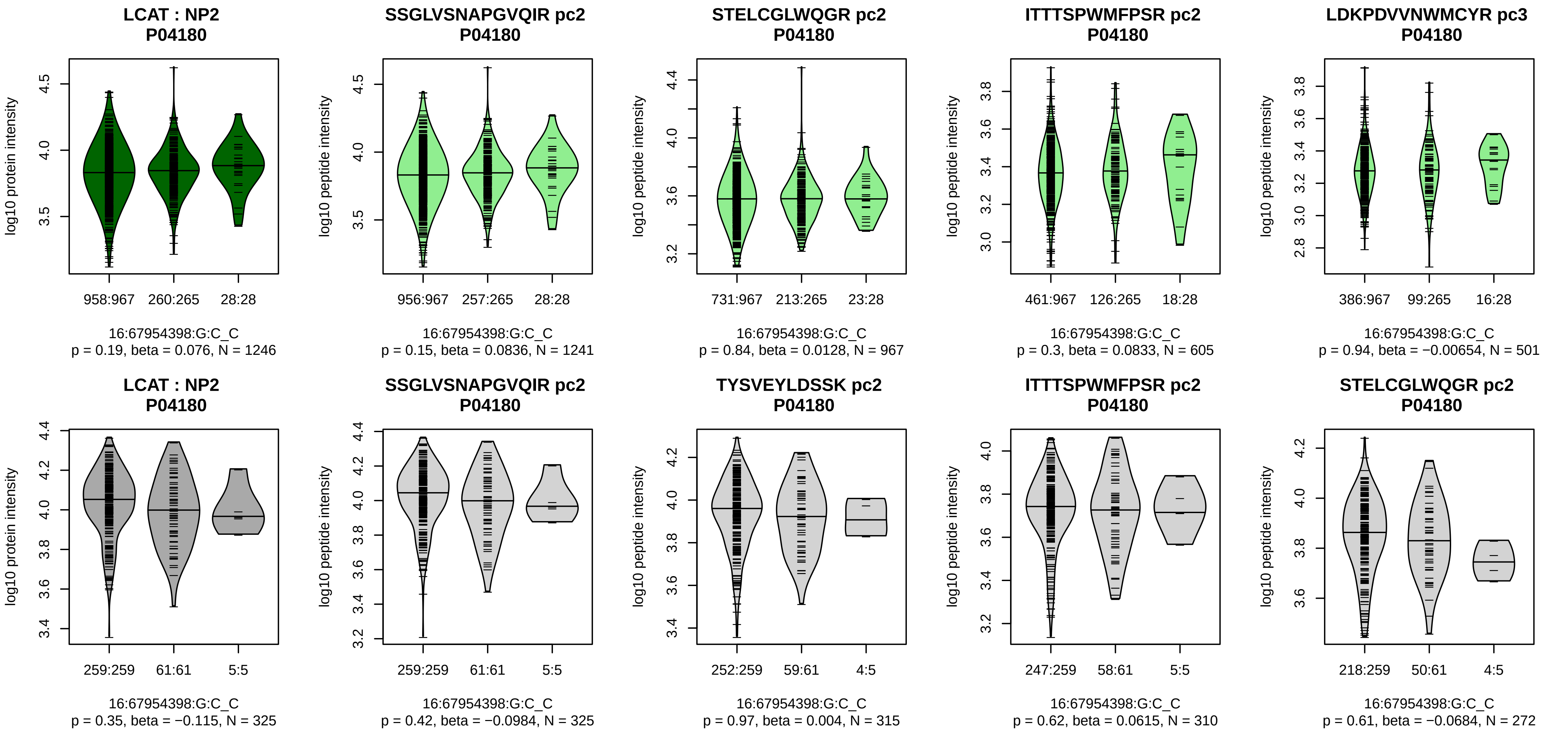


Assay Target: LCAT
Olink UniProt: P04180
UKB rsID: rs112434994
Proxy rsID: rs112434994
UKB SNP: 16:67954398:G:C
Proxy SNP: 16:67954398:G:C
UKB log10(p): 79
UKB BETA: 0.175
–:–:–:–:–:–
1241:967:605:501:354:298

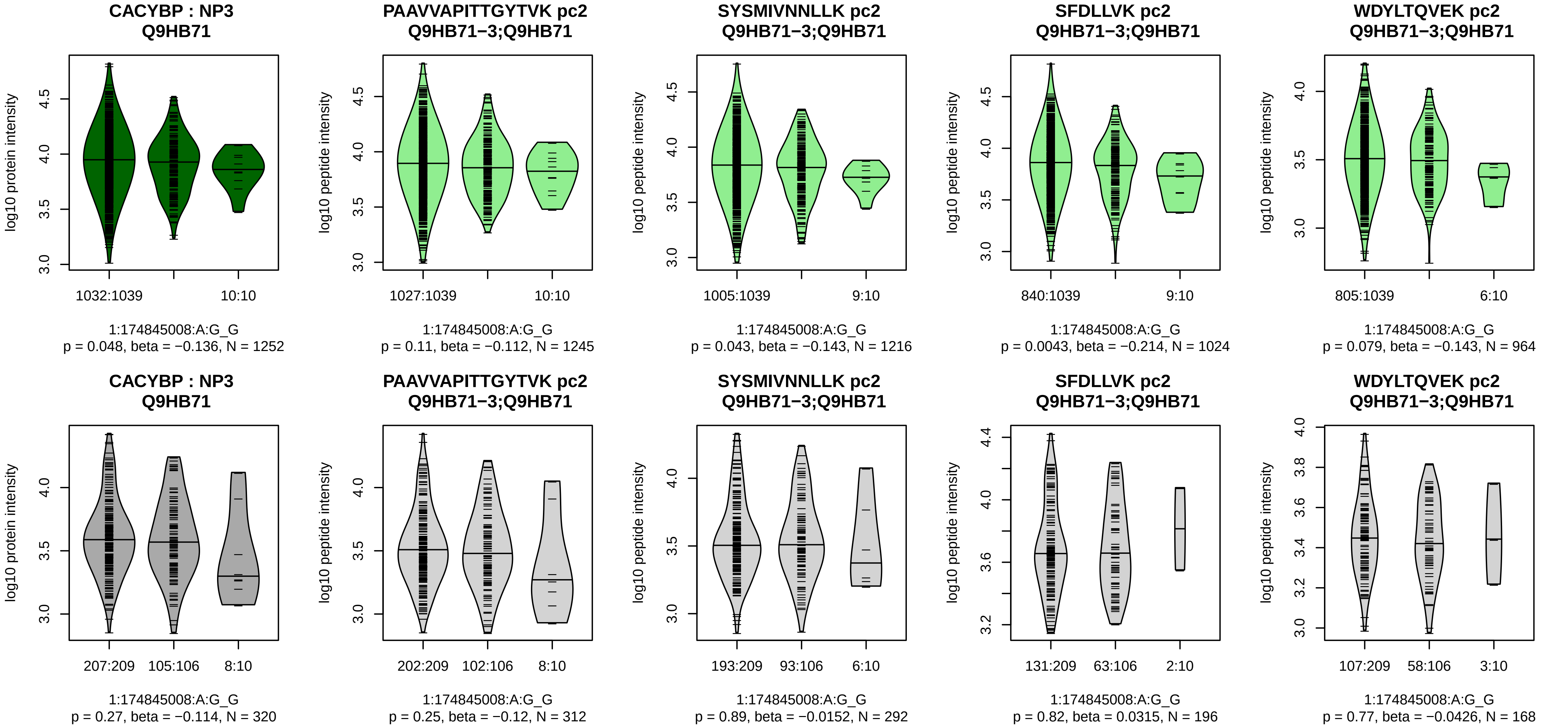


Assay Target: CACYBP
Olink UniProt: Q9HB71
UKB rsID: rs76088187
Proxy rsID: rs76088187
UKB SNP: 1:174845008:A:G
Proxy SNP: 1:174845008:A:G
UKB log10(p): 74.4
UKB BETA: −0.177
−:−:*:−:−:−:−:−:−:−:−:−:NA
1245:1216:1024:964:914:691:47

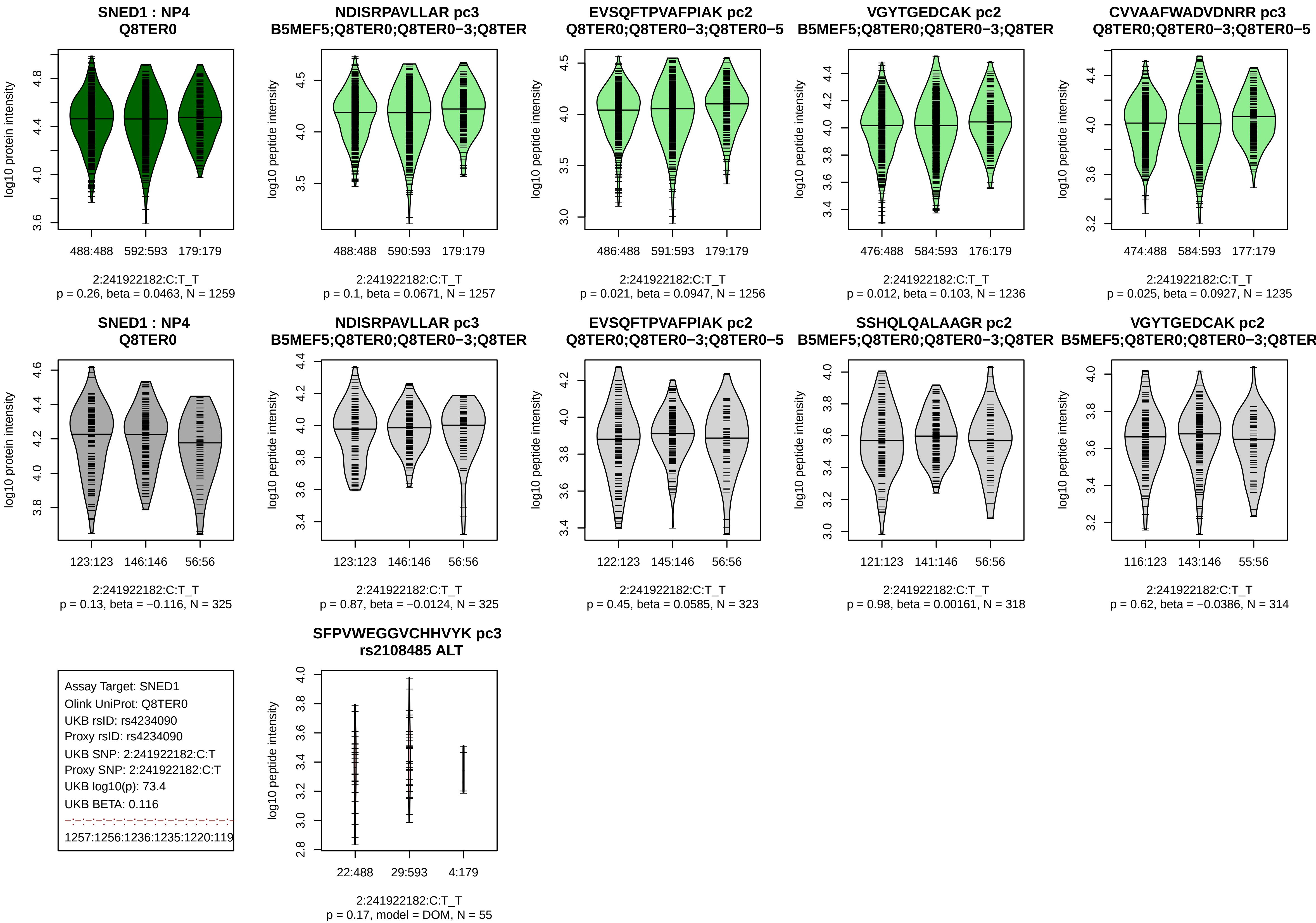
SNED1 : NP4
Q8TER0
log10 protein intensity
488:488 592:593 179:179
2:241922182:C:T_T
p = 0.26, beta = 0.0463, N = 1259
NDISRPAVLLAR pc3
B5MEF5;Q8TER0;Q8TER0−3;Q8TER
log10 peptide intensity
488:488 590:593 179:179
2:241922182:C:T_T
p = 0.1, beta = 0.0671, N = 1257
EVSQFTPVAFPIAK pc2
Q8TER0;Q8TER0−3;Q8TER0−5
log10 peptide intensity
486:488 591:593 179:179
2:241922182:C:T_T
p = 0.021, beta = 0.0947, N = 1256
VGYTGEDCAK pc2
B5MEF5;Q8TER0;Q8TER0−3;Q8TER
log10 peptide intensity
476:488 584:593 176:179
2:241922182:C:T_T
p = 0.012, beta = 0.103, N = 1236
CVVAAFWADVDNRR pc3
Q8TER0;Q8TER0−3;Q8TER0−5
log10 peptide intensity
474:488 584:593 177:179
2:241922182:C:T_T
p = 0.025, beta = 0.0927, N = 1235
SNED1 : NP4
Q8TER0
log10 protein intensity
123:123 146:146 56:56
2:241922182:C:T_T
p = 0.13, beta = −0.116, N = 325
NDISRPAVLLAR pc3
B5MEF5;Q8TER0;Q8TER0−3;Q8TER
log10 peptide intensity
123:123 146:146 56:56
2:241922182:C:T_T
p = 0.87, beta = −0.0124, N = 325
EVSQFTPVAFPIAK pc2
Q8TER0;Q8TER0−3;Q8TER0−5
log10 peptide intensity
122:123 145:146 56:56
2:241922182:C:T_T
p = 0.45, beta = 0.0585, N = 323
SSHQLQALAAGR pc2
B5MEF5;Q8TER0;Q8TER0−3;Q8TER
log10 peptide intensity
121:123 141:146 56:56
2:241922182:C:T_T
p = 0.98, beta = 0.00161, N = 318
VGYTGEDCAK pc2
B5MEF5;Q8TER0;Q8TER0−3;Q8TER
log10 peptide intensity
116:123 143:146 55:56
2:241922182:C:T_T
p = 0.62, beta = −0.0386, N = 314
Assay Target: SNED1
Olink UniProt: Q8TER0
UKB rsID: rs4234090
Proxy rsID: rs4234090
UKB SNP: 2:241922182:C:T
Proxy SNP: 2:241922182:C:T
UKB log10(p): 73.4
UKB BETA: 0.116
1257:1256:1236:1235:1220:119
SFPVWEGGVCHHVYK pc3
rs2108485 ALT
log10 peptide intensity
22:488 29:593 4:179
2:241922182:C:T_T
p = 0.17, model = DOM, N = 55

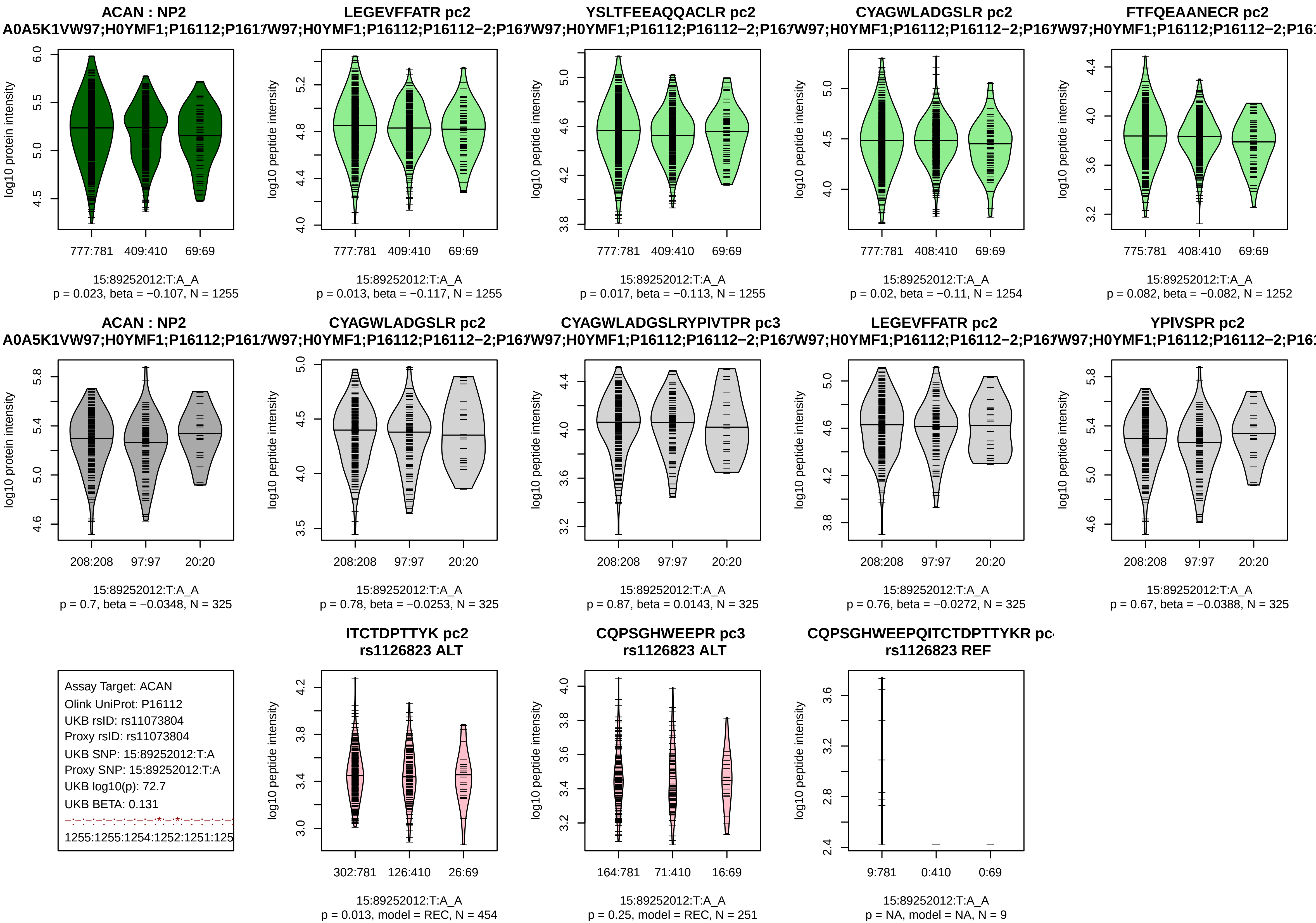

ACAN : NP2
A0A5K1VW97;H0YMF1;P16112;P161
log10 protein intensity
777:781
409:410
69:69
15:89252012:T:A_A
p = 0.023, beta = −0.107, N = 1255
LEGEVFFATR pc2
W97;H0YMF1;P16112;P16112−2;P16
log10 peptide intensity
777:781
409:410
69:69
15:89252012:T:A_A
p = 0.013, beta = −0.117, N = 1255
YSLTFEEAQQACLR pc2
W97;H0YMF1;P16112;P16112−2;P16
777:781
409:410
69:69
15:89252012:T:A_A
p = 0.017, beta = −0.113, N = 1255
CYAGWLADGSLR pc2
W97;H0YMF1;P16112;P16112−2;P16
777:781
408:410
69:69
15:89252012:T:A_A
p = 0.02, beta = −0.11, N = 1254
FTFQEAANECR pc2
W97;H0YMF1;P16112;P16112−2;P16
775:781
408:410
69:69
15:89252012:T:A_A
p = 0.082, beta = −0.082, N = 1252
ACAN : NP2
A0A5K1VW97;H0YMF1;P16112;P161
208:208
97:97
20:20
15:89252012:T:A_A
p = 0.7, beta = −0.0348, N = 325
CYAGWLADGSLR pc2
W97;H0YMF1;P16112;P16112−2;P16
208:208
97:97
20:20
15:89252012:T:A_A
p = 0.78, beta = −0.0253, N = 325
CYAGWLADGSLRYPIVTPR pc3
W97;H0YMF1;P16112;P16112−2;P16
208:208
97:97
20:20
15:89252012:T:A_A
p = 0.87, beta = 0.0143, N = 325
LEGEVFFATR pc2
W97;H0YMF1;P16112;P16112−2;P16
208:208
97:97
20:20
15:89252012:T:A_A
p = 0.76, beta = −0.0272, N = 325
YPIVSPR pc2
W97;H0YMF1;P16112;P16112−2;P16
208:208
97:97
20:20
15:89252012:T:A_A
p = 0.67, beta = −0.0388, N = 325
Assay Target: ACAN
Olink UniProt: P16112
UKB rsID: rs11073804
Proxy rsID: rs11073804
UKB SNP: 15:89252012:T:A
Proxy SNP: 15:89252012:T:A
UKB log10(p): 72.7
UKB BETA: 0.131
−:−:−:−:−:−:−:−:*:−:*:−:−:−:−:−
1255:1255:1254:1252:1251:125
ITCTDPTTYK pc2
rs1126823 ALT
302:781
126:410
26:69
15:89252012:T:A_A
p = 0.013, model = REC, N = 454
CQPSGHWEEPR pc3
rs1126823 ALT
164:781
71:410
16:69
15:89252012:T:A_A
p = 0.25, model = REC, N = 251
CQPSGHWEEPQITCTDPTTYKR pc
rs1126823 REF
9:781
0:410
0:69
15:89252012:T:A_A
p = NA, model = NA, N = 9

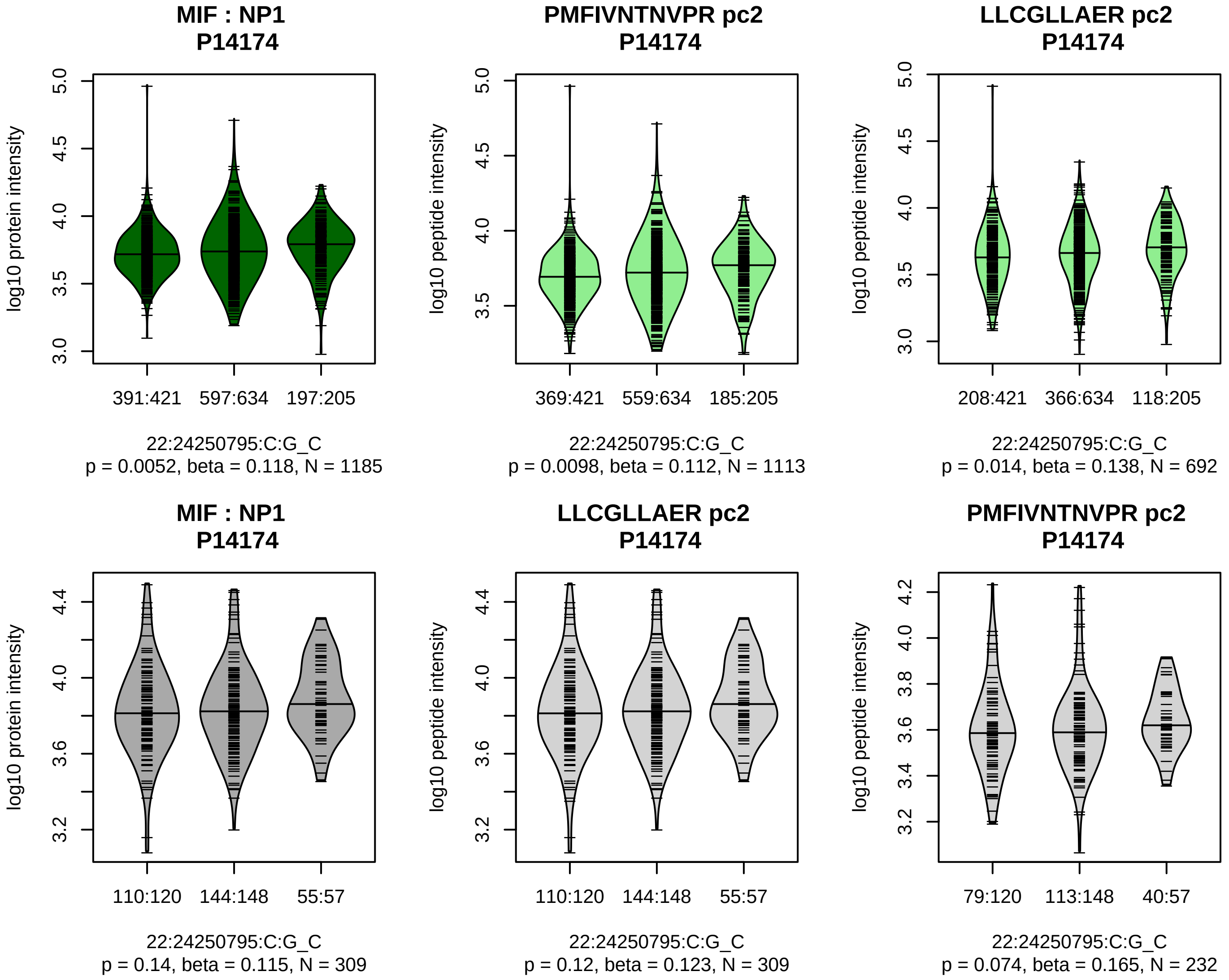


Assay Target: MIF
Olink UniProt: P14174
UKB rsID: rs2330634
Proxy rsID: rs2330634
UKB SNP: 22:24250795:C:G
Proxy SNP: 22:24250795:C:G
UKB log10(p): 71.1
UKB BETA: −0.112
*:–
1113:692

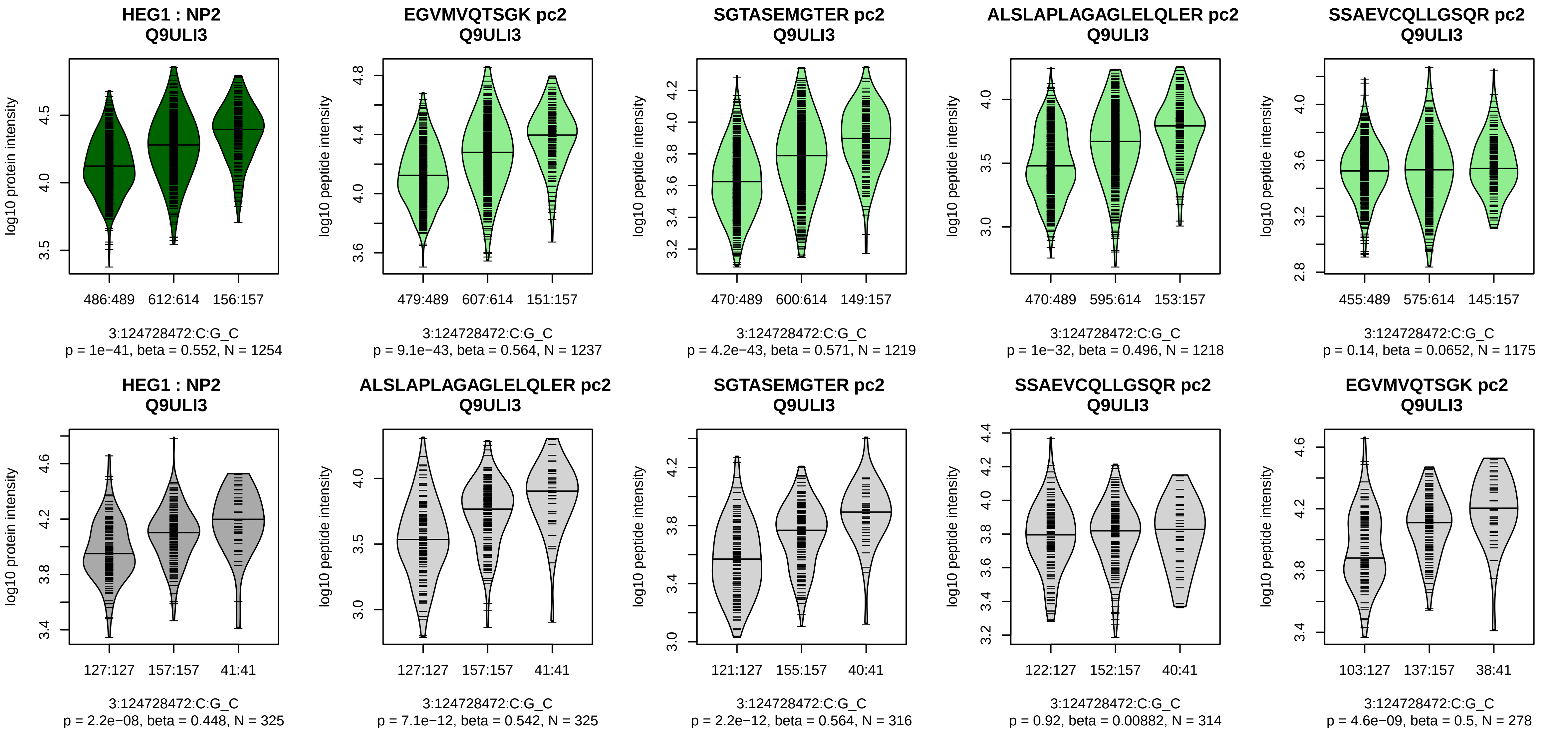


Assay Target: HEG1
Olink UniProt: Q9ULI3
UKB rsID: rs6438868
Proxy rsID: rs6438868
UKB SNP: 3:124728472:C:G
Proxy SNP: 3:124728472:C:G
UKB log10(p): 71.1
UKB BETA: 0.104
*:*:*:−:*:*:−:−:−:*:*:−:−:*:−:−:NA
1237:1219:1218:1175:996:850:8

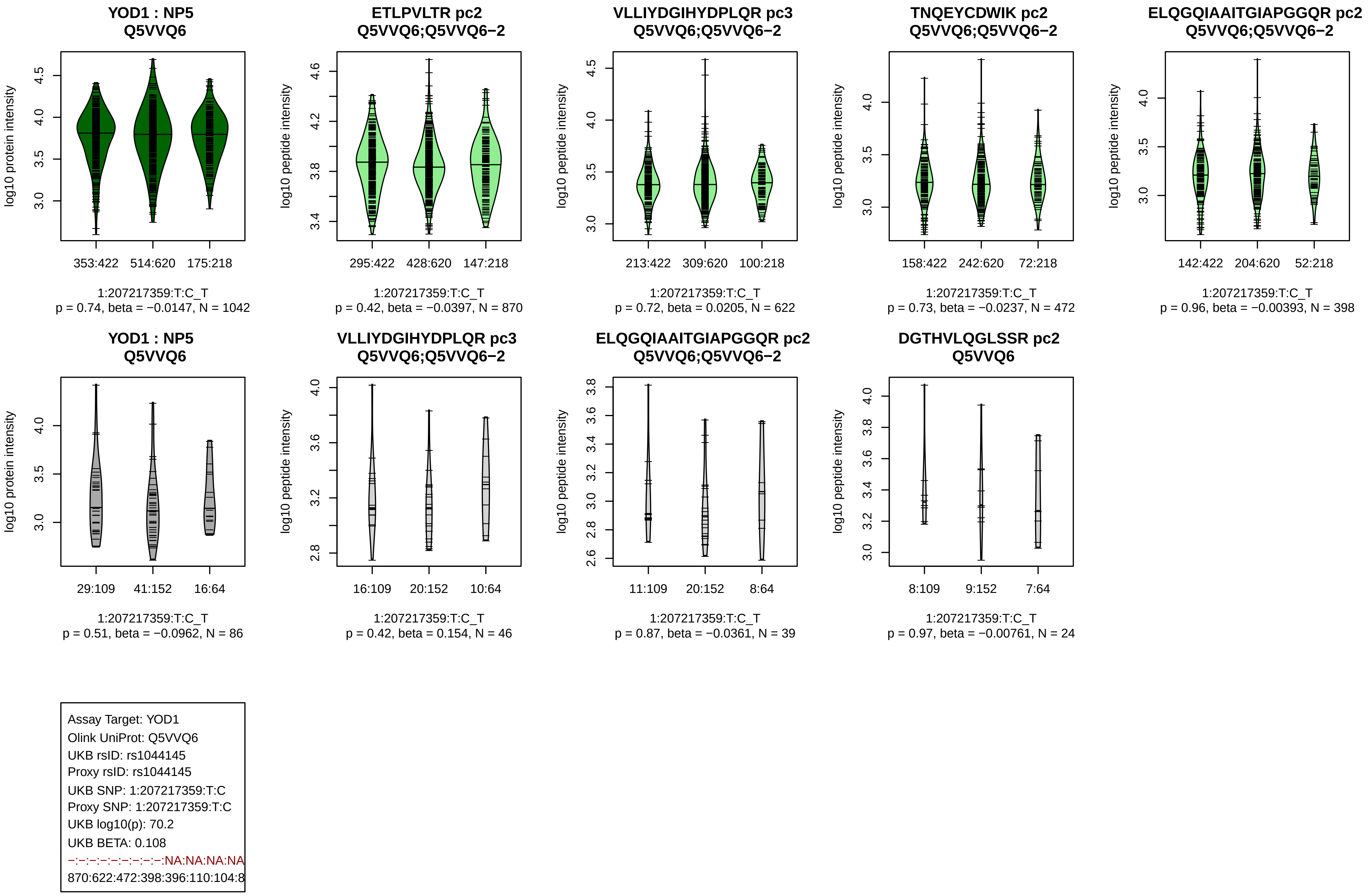
YOD1 : NP5
Q5VVQ6
log10 protein intensity
353:422
514:620
175:218
1:207217359:T:C_T
p = 0.74, beta = −0.0147, N = 1042
ETLPVLTR pc2
Q5VVQ6;Q5VVQ6−2
log10 peptide intensity
295:422
428:620
147:218
1:207217359:T:C_T
p = 0.42, beta = −0.0397, N = 870
VLLIYDGIHYDPLQR pc3
Q5VVQ6;Q5VVQ6−2
log10 peptide intensity
213:422
309:620
100:218
1:207217359:T:C_T
p = 0.72, beta = 0.0205, N = 622
TNQEYCDWIK pc2
Q5VVQ6;Q5VVQ6−2
log10 peptide intensity
158:422
242:620
72:218
1:207217359:T:C_T
p = 0.73, beta = −0.0237, N = 472
ELQGQIAAITGIAPGGQR pc2
Q5VVQ6;Q5VVQ6−2
log10 peptide intensity
142:422
204:620
52:218
1:207217359:T:C_T
p = 0.96, beta = −0.00393, N = 398
YOD1 : NP5
Q5VVQ6
log10 protein intensity
29:109
41:152
16:64
1:207217359:T:C_T
p = 0.51, beta = −0.0962, N = 86
VLLIYDGIHYDPLQR pc3
Q5VVQ6;Q5VVQ6−2
log10 peptide intensity
16:109
20:152
10:64
1:207217359:T:C_T
p = 0.42, beta = 0.154, N = 46
ELQGQIAAITGIAPGGQR pc2
Q5VVQ6;Q5VVQ6−2
log10 peptide intensity
11:109
20:152
8:64
1:207217359:T:C_T
p = 0.87, beta = −0.0361, N = 39
DGTHVLQGLSSR pc2
Q5VVQ6
log10 peptide intensity
8:109
9:152
7:64
1:207217359:T:C_T
p = 0.97, beta = −0.00761, N = 24
Assay Target: YOD1
Olink UniProt: Q5VVQ6
UKB rsID: rs1044145
Proxy rsID: rs1044145
UKB SNP: 1:207217359:T:C
Proxy SNP: 1:207217359:T:C
UKB log10(p): 70.2
UKB BETA: 0.108
−:−:−:−:−:−:−:−:−:NA:NA:NA:NA
870:622:472:398:396:110:104:8

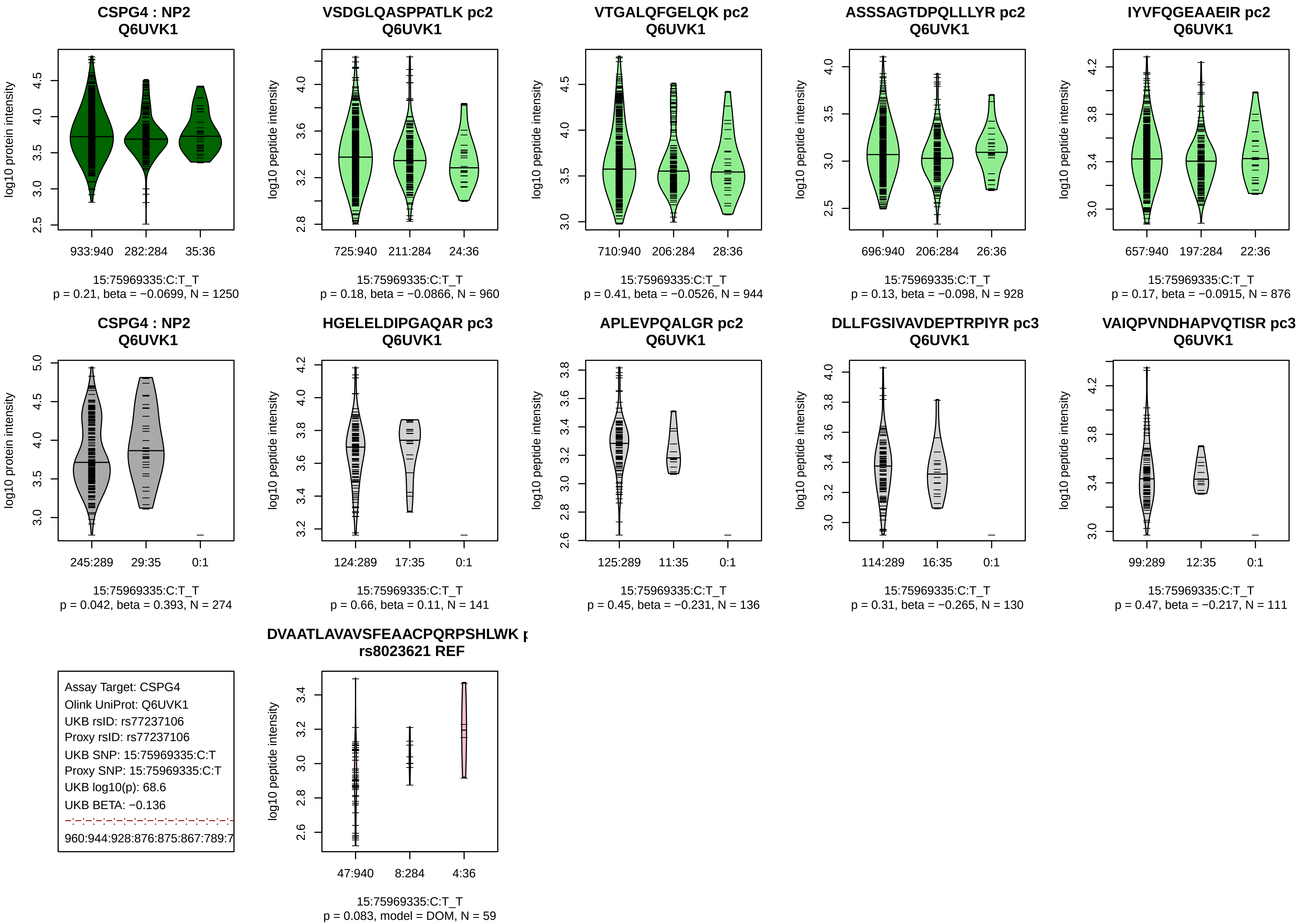

CSPG4 : NP2
Q6UVK1
log10 protein intensity
933:940
282:284
35:36
15:75969335:C:T_T
p = 0.21, beta = −0.0699, N = 1250
VSDGLQASPPATLK pc2
Q6UVK1
log10 peptide intensity
725:940
211:284
24:36
15:75969335:C:T_T
p = 0.18, beta = −0.0866, N = 960
VTGALQFGELQK pc2
Q6UVK1
log10 peptide intensity
710:940
206:284
28:36
15:75969335:C:T_T
p = 0.41, beta = −0.0526, N = 944
ASSSAGTDPQLLLYR pc2
Q6UVK1
log10 peptide intensity
696:940
206:284
26:36
15:75969335:C:T_T
p = 0.13, beta = −0.098, N = 928
IYVFQGEAAEIR pc2
Q6UVK1
log10 peptide intensity
657:940
197:284
22:36
15:75969335:C:T_T
p = 0.17, beta = −0.0915, N = 876
CSPG4 : NP2
Q6UVK1
log10 protein intensity
245:289
29:35
0:1
15:75969335:C:T_T
p = 0.042, beta = 0.393, N = 274
HGELELDIPGAQAR pc3
Q6UVK1
log10 peptide intensity
124:289
17:35
0:1
15:75969335:C:T_T
p = 0.66, beta = 0.11, N = 141
APLEVPQALGR pc2
Q6UVK1
log10 peptide intensity
125:289
11:35
0:1
15:75969335:C:T_T
p = 0.45, beta = −0.231, N = 136
DLLFGSIVAVDEPTRPIYR pc3
Q6UVK1
log10 peptide intensity
114:289
16:35
0:1
15:75969335:C:T_T
p = 0.31, beta = −0.265, N = 130
VAIQPVNDHAPVQTISR pc3
Q6UVK1
log10 peptide intensity
99:289
12:35
0:1
15:75969335:C:T_T
p = 0.47, beta = −0.217, N = 111
Assay Target: CSPG4
Olink UniProt: Q6UVK1
UKB rsID: rs77237106
Proxy rsID: rs77237106
UKB SNP: 15:75969335:C:T
Proxy SNP: 15:75969335:C:T
UKB log10(p): 68.6
UKB BETA: −0.136
960:944:928:876:875:867:789:7
DVAATLAVAVSFEAACPQRPSHLWK
rs8023621 REF
log10 peptide intensity
47:940
8:284
4:36
15:75969335:C:T_T
p = 0.083, model = DOM, N = 59

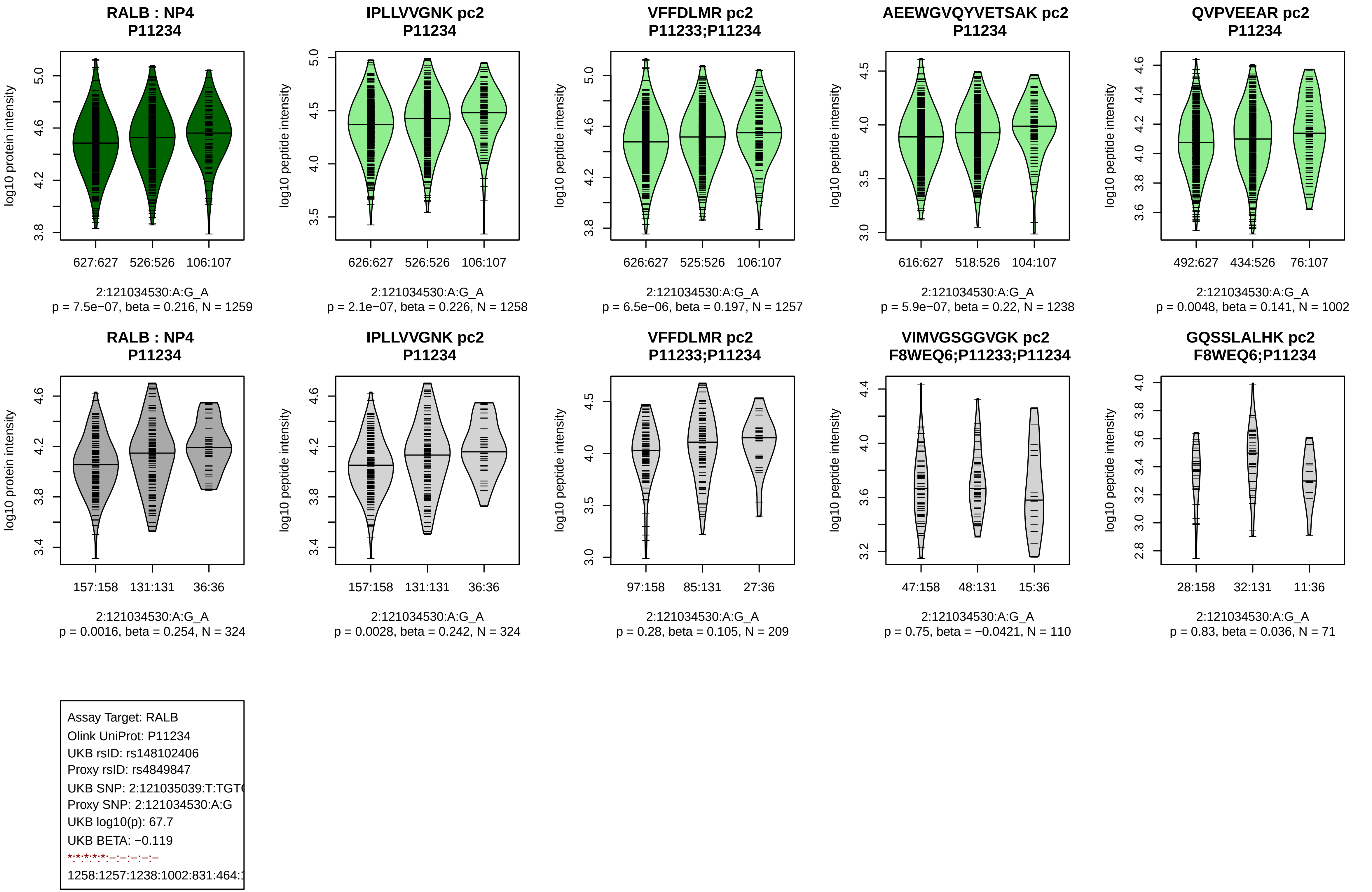

RALB : NP4
P11234
log10 protein intensity
627:627
526:526
106:107
2:121034530:A:G_A
p = 7.5e−07, beta = 0.216, N = 1259
IPLLVVGNK pc2
P11234
log10 peptide intensity
626:627
526:526
106:107
2:121034530:A:G_A
p = 2.1e−07, beta = 0.226, N = 1258
VFFDLMR pc2
P11233;P11234
log10 peptide intensity
626:627
525:526
106:107
2:121034530:A:G_A
p = 6.5e−06, beta = 0.197, N = 1257
AEEWGVQYVETSAK pc2
P11234
log10 peptide intensity
616:627
518:526
104:107
2:121034530:A:G_A
p = 5.9e−07, beta = 0.22, N = 1238
QVPVEEAR pc2
P11234
log10 peptide intensity
492:627
434:526
76:107
2:121034530:A:G_A
p = 0.0048, beta = 0.141, N = 1002
RALB : NP4
P11234
log10 protein intensity
157:158
131:131
36:36
2:121034530:A:G_A
p = 0.0016, beta = 0.254, N = 324
IPLLVVGNK pc2
P11234
log10 peptide intensity
157:158
131:131
36:36
2:121034530:A:G_A
p = 0.0028, beta = 0.242, N = 324
VFFDLMR pc2
P11233;P11234
log10 peptide intensity
97:158
85:131
27:36
2:121034530:A:G_A
p = 0.28, beta = 0.105, N = 209
VIMVGSGGVGK pc2
F8WEQ6;P11233;P11234
log10 peptide intensity
47:158
48:131
15:36
2:121034530:A:G_A
p = 0.75, beta = −0.0421, N = 110
GQSSLALHK pc2
F8WEQ6;P11234
log10 peptide intensity
28:158
32:131
11:36
2:121034530:A:G_A
p = 0.83, beta = 0.036, N = 71
Assay Target: RALB
Olink UniProt: P11234
UKB rsID: rs148102406
Proxy rsID: rs4849847
UKB SNP: 2:121035039:T:TGT
Proxy SNP: 2:121034530:A:G
UKB log10(p): 67.7
UKB BETA: −0.119
*:*:*:*:*:–:–:–:–:–
1258:1257:1238:1002:831:464:

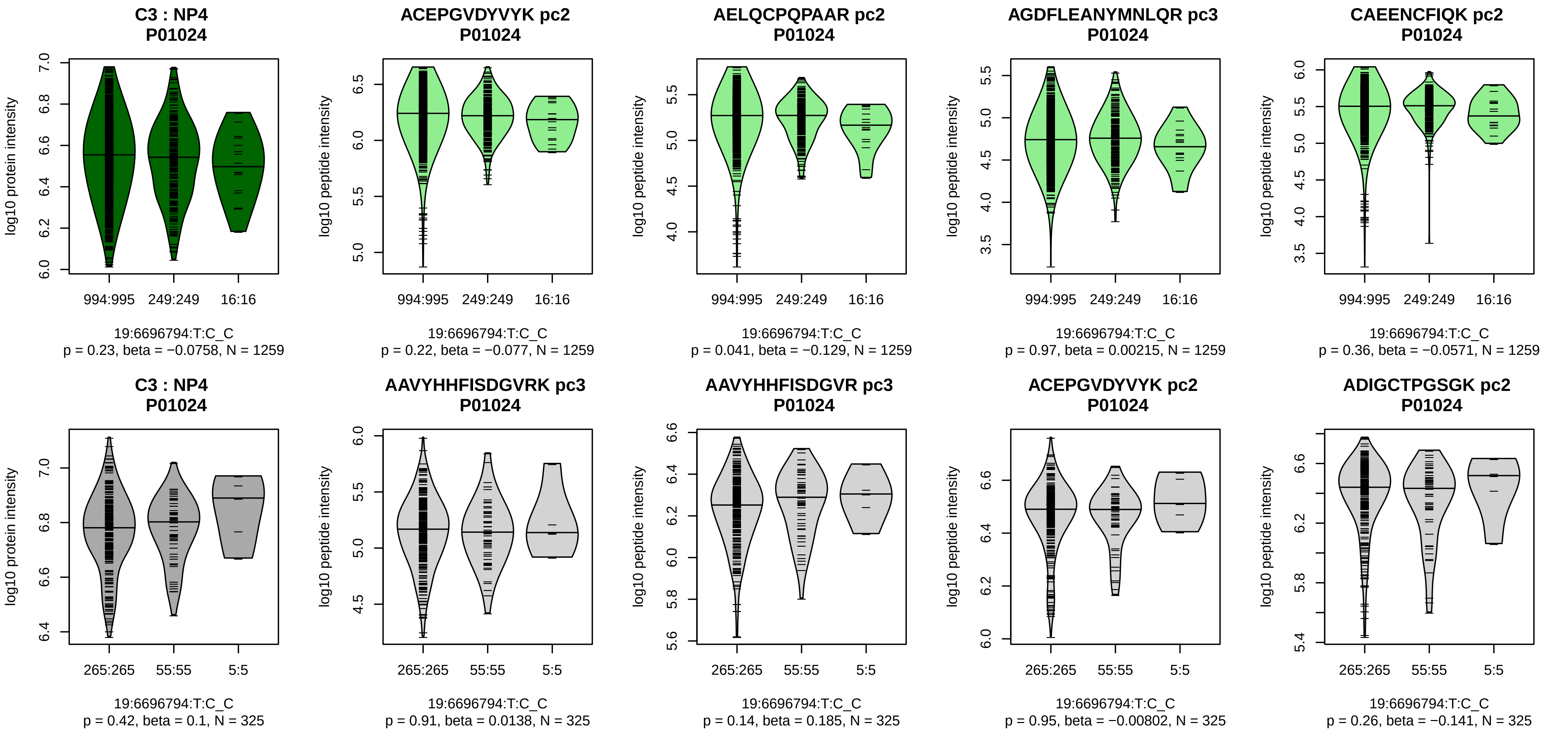


Assay Target: C3
Olink UniProt: P01024
UKB rsID: rs5826945
Proxy rsID: rs11569479
UKB SNP: 19:6697088:A:T
Proxy SNP: 19:6696794:T:C
UKB log10(p): 66.9
UKB BETA: −0.157
−:−:−:−:−:−:−:−:−:−:−:−:−:−:−:*:−
1259:1259:1259:1259:1259:125

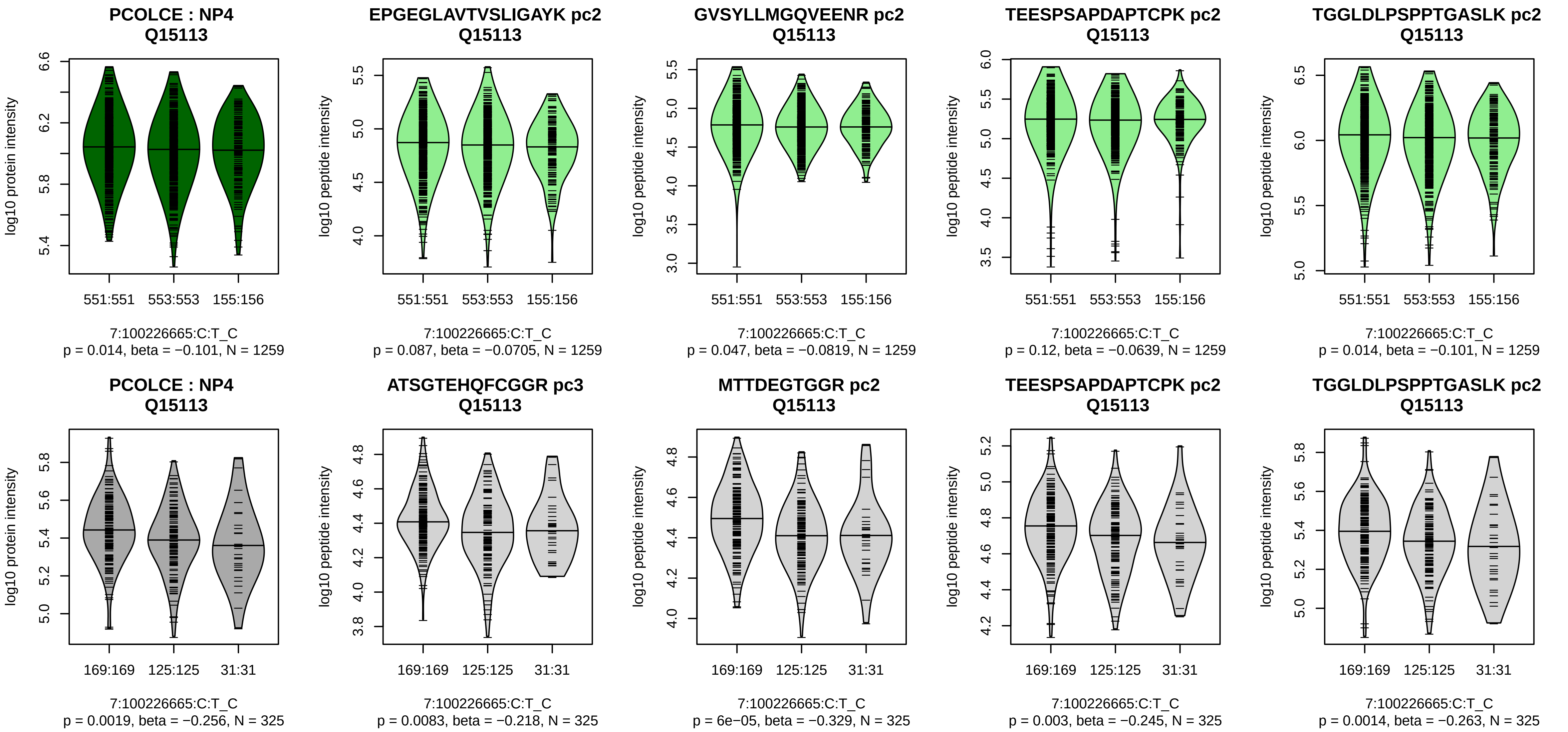


Assay Target: PCOLCE
Olink UniProt: Q15113
UKB rsID: rs4548095
Proxy rsID: rs4548095
UKB SNP: 7:100226665:C:T
Proxy SNP: 7:100226665:C:T
UKB log10(p): 63.7
UKB BETA: 0.169
–:–:–:–:*:–:–:–:–:–:–:–:–:–:–:–:–
1259:1259:1259:1259:1259:125

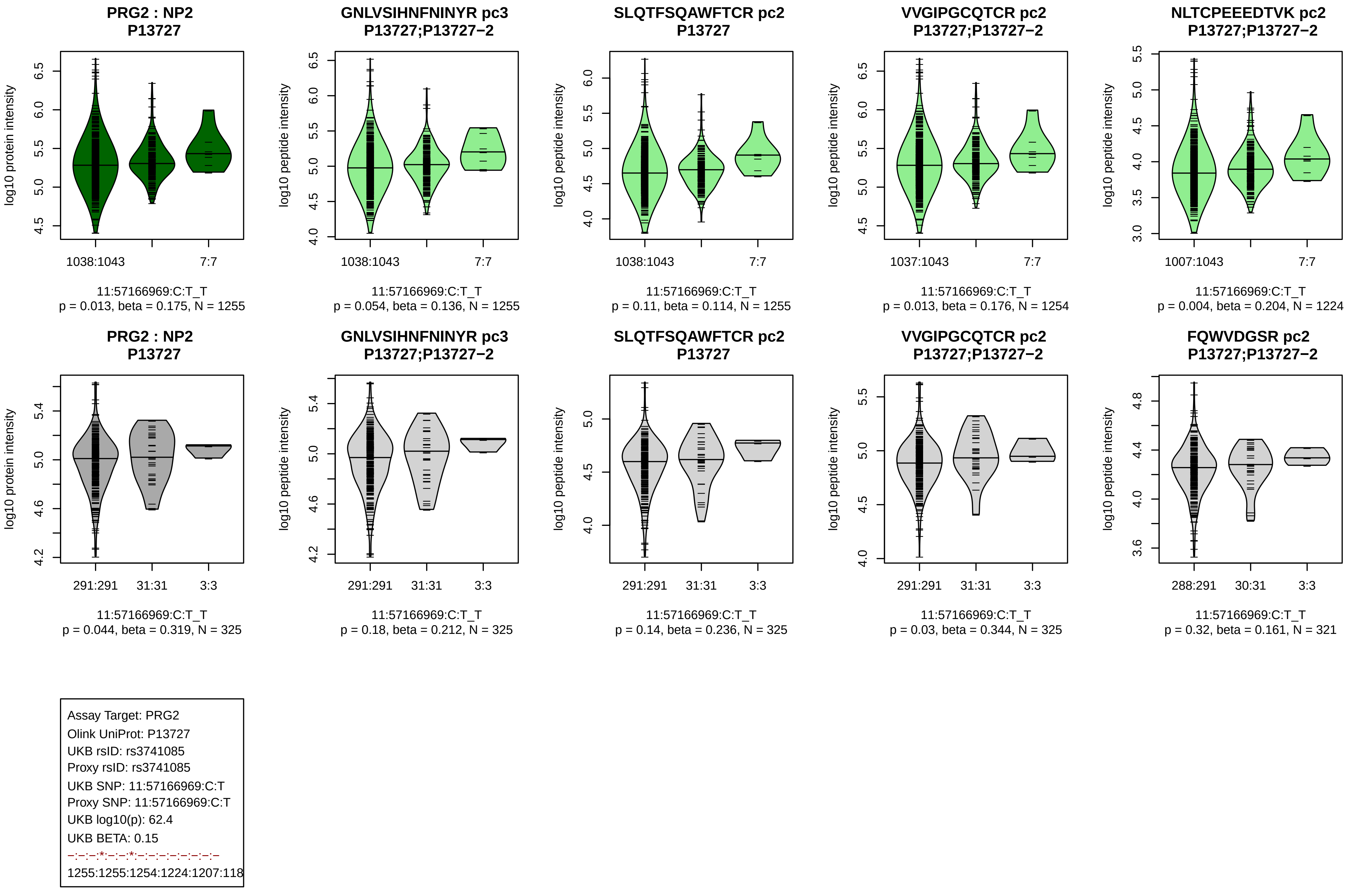

PRG2 : NP2
P13727
log10 protein intensity
1038:1043
7:7
11:57166969:C:T_T
p = 0.013, beta = 0.175, N = 1255
GNLVSIHNFNINYR pc3
P13727;P13727−2
log10 peptide intensity
1038:1043
7:7
11:57166969:C:T_T
p = 0.054, beta = 0.136, N = 1255
SLQTFSQAWFTCR pc2
P13727
log10 peptide intensity
1038:1043
7:7
11:57166969:C:T_T
p = 0.11, beta = 0.114, N = 1255
VVGIPGCQTCR pc2
P13727;P13727−2
log10 peptide intensity
1037:1043
7:7
11:57166969:C:T_T
p = 0.013, beta = 0.176, N = 1254
NLTCPEEEDTVK pc2
P13727;P13727−2
log10 peptide intensity
1007:1043
7:7
11:57166969:C:T_T
p = 0.004, beta = 0.204, N = 1224
PRG2 : NP2
P13727
log10 protein intensity
291:291
31:31
3:3
11:57166969:C:T_T
p = 0.044, beta = 0.319, N = 325
GNLVSIHNFNINYR pc3
P13727;P13727−2
log10 peptide intensity
291:291
31:31
3:3
11:57166969:C:T_T
p = 0.18, beta = 0.212, N = 325
SLQTFSQAWFTCR pc2
P13727
log10 peptide intensity
291:291
31:31
3:3
11:57166969:C:T_T
p = 0.14, beta = 0.236, N = 325
VVGIPGCQTCR pc2
P13727;P13727−2
log10 peptide intensity
291:291
31:31
3:3
11:57166969:C:T_T
p = 0.03, beta = 0.344, N = 325
FQWVDGSR pc2
P13727;P13727−2
log10 peptide intensity
288:291
30:31
3:3
11:57166969:C:T_T
p = 0.32, beta = 0.161, N = 321
Assay Target: PRG2
Olink UniProt: P13727
UKB rsID: rs3741085
Proxy rsID: rs3741085
UKB SNP: 11:57166969:C:T
Proxy SNP: 11:57166969:C:T
UKB log10(p): 62.4
UKB BETA: 0.15
–:–:–:*:–:–:*:–:–:–:–:–:–:–:–
1255:1255:1254:1224:1207:118

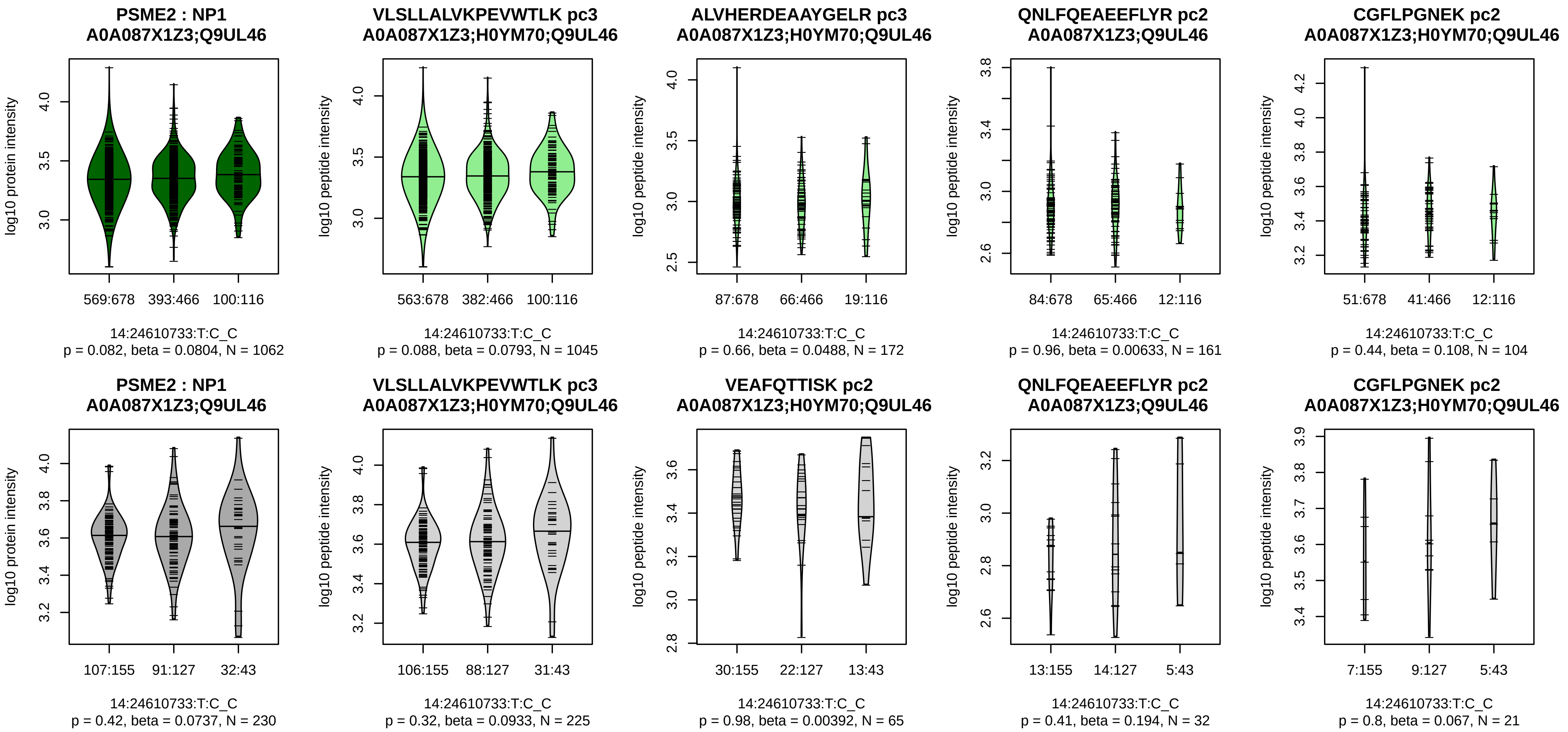


Assay Target: PSME2
Olink UniProt: Q9UL46
UKB rsID: rs2236352
Proxy rsID: rs2236352
UKB SNP: 14:24610733:T:C
Proxy SNP: 14:24610733:T:C
UKB log10(p): 58.8
UKB BETA: 0.111
–:–:–:–:–:–:–:–:–
1045:172:161:104:92:91:54:45:3

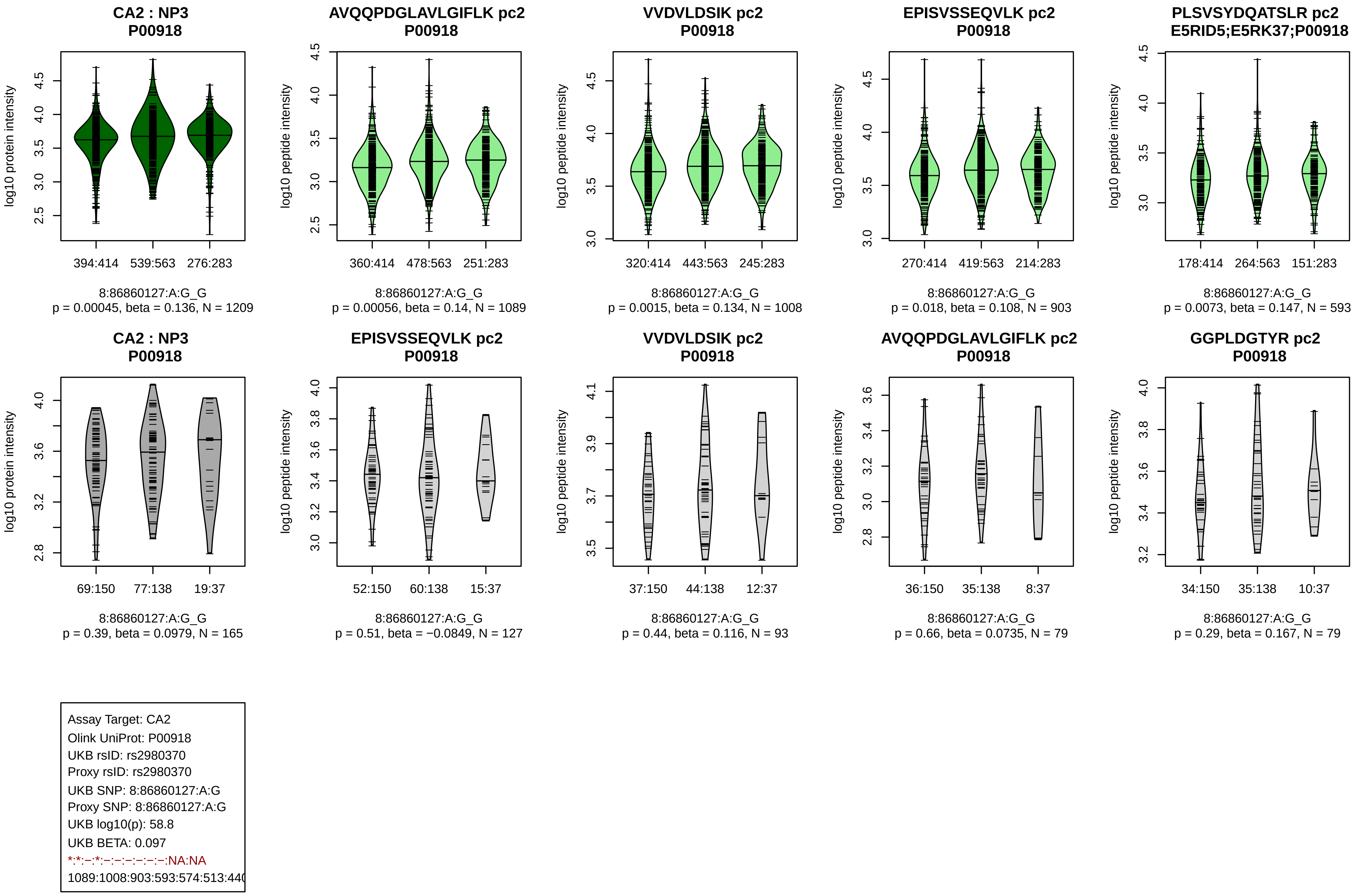
CA2 : NP3
P00918
log10 protein intensity
394:414 539:563 276:283
8:86860127:A:G_G
p = 0.00045, beta = 0.136, N = 1209
AVQQPDGLAVLGIFLK pc2
P00918
log10 peptide intensity
360:414 478:563 251:283
8:86860127:A:G_G
p = 0.00056, beta = 0.14, N = 1089
VVDVLDSIK pc2
P00918
log10 peptide intensity
320:414 443:563 245:283
8:86860127:A:G_G
p = 0.0015, beta = 0.134, N = 1008
EPISVSSEQVLK pc2
P00918
log10 peptide intensity
270:414 419:563 214:283
8:86860127:A:G_G
p = 0.018, beta = 0.108, N = 903
PLSVSYDQATSLR pc2
E5RID5;E5RK37;P00918
log10 peptide intensity
178:414 264:563 151:283
8:86860127:A:G_G
p = 0.0073, beta = 0.147, N = 593
CA2 : NP3
P00918
log10 protein intensity
69:150 77:138 19:37
8:86860127:A:G_G
p = 0.39, beta = 0.0979, N = 165
EPISVSSEQVLK pc2
P00918
log10 peptide intensity
52:150 60:138 15:37
8:86860127:A:G_G
p = 0.51, beta = −0.0849, N = 127
VVDVLDSIK pc2
P00918
log10 peptide intensity
37:150 44:138 12:37
8:86860127:A:G_G
p = 0.44, beta = 0.116, N = 93
AVQQPDGLAVLGIFLK pc2
P00918
log10 peptide intensity
36:150 35:138 8:37
8:86860127:A:G_G
p = 0.66, beta = 0.0735, N = 79
GGPLDGTYR pc2
P00918
log10 peptide intensity
34:150 35:138 10:37
8:86860127:A:G_G
p = 0.29, beta = 0.167, N = 79
Assay Target: CA2
Olink UniProt: P00918
UKB rsID: rs2980370
Proxy rsID: rs2980370
UKB SNP: 8:86860127:A:G
Proxy SNP: 8:86860127:A:G
UKB log10(p): 58.8
UKB BETA: 0.097
*:*:–:*:–:–:–:–:–:–:NA:NA
1089:1008:903:593:574:513:440

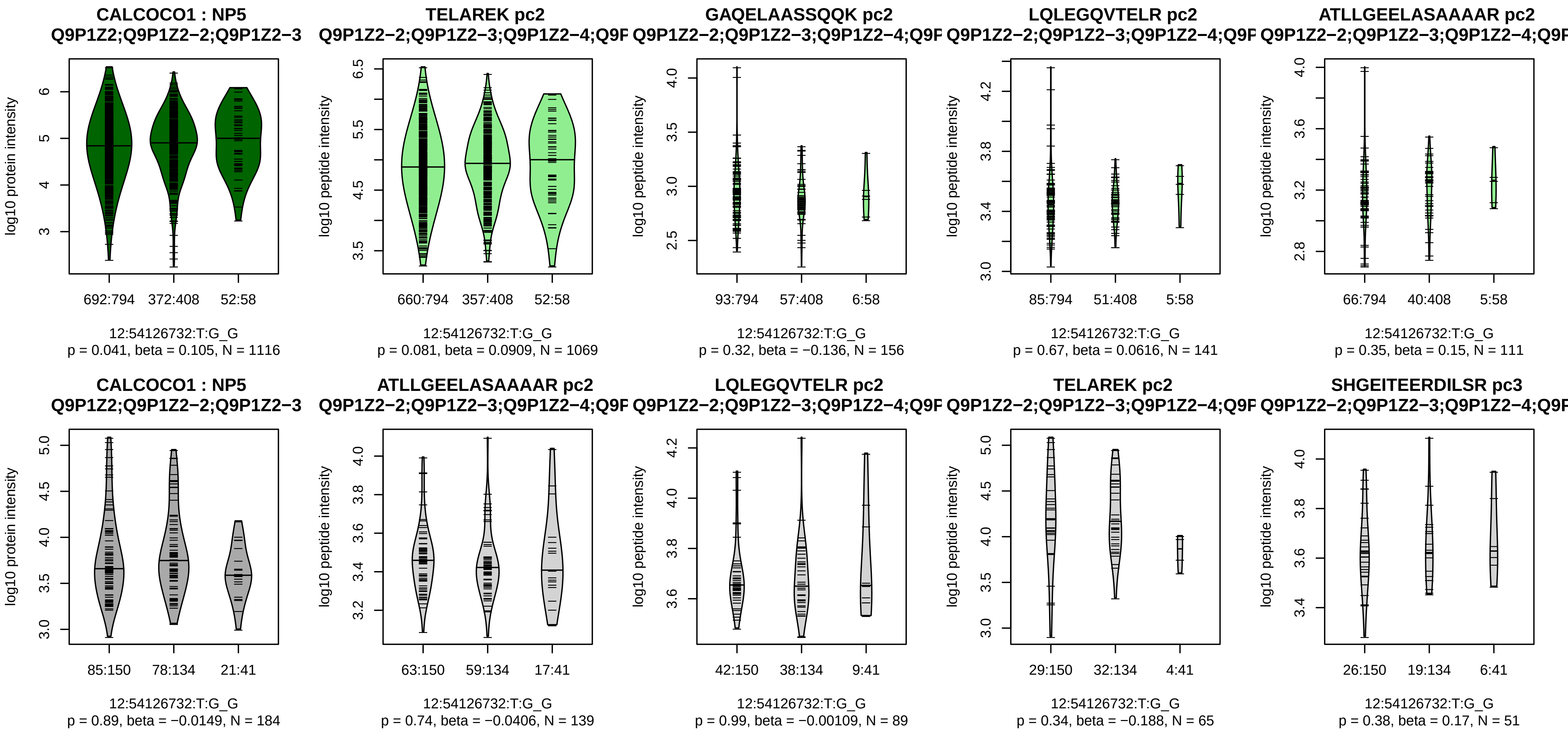


Assay Target: CALCOCO1
Olink UniProt: Q9P1Z2
UKB rsID: rs17102261
Proxy rsID: rs17102261
UKB SNP: 12:54126732:T:G
Proxy SNP: 12:54126732:T:G
UKB log10(p): 58.7
UKB BETA: 0.133
−:−:−:−:−:−:−:−:−:−:−:NA:NA:NA
1069:156:141:111:110:110:98:6

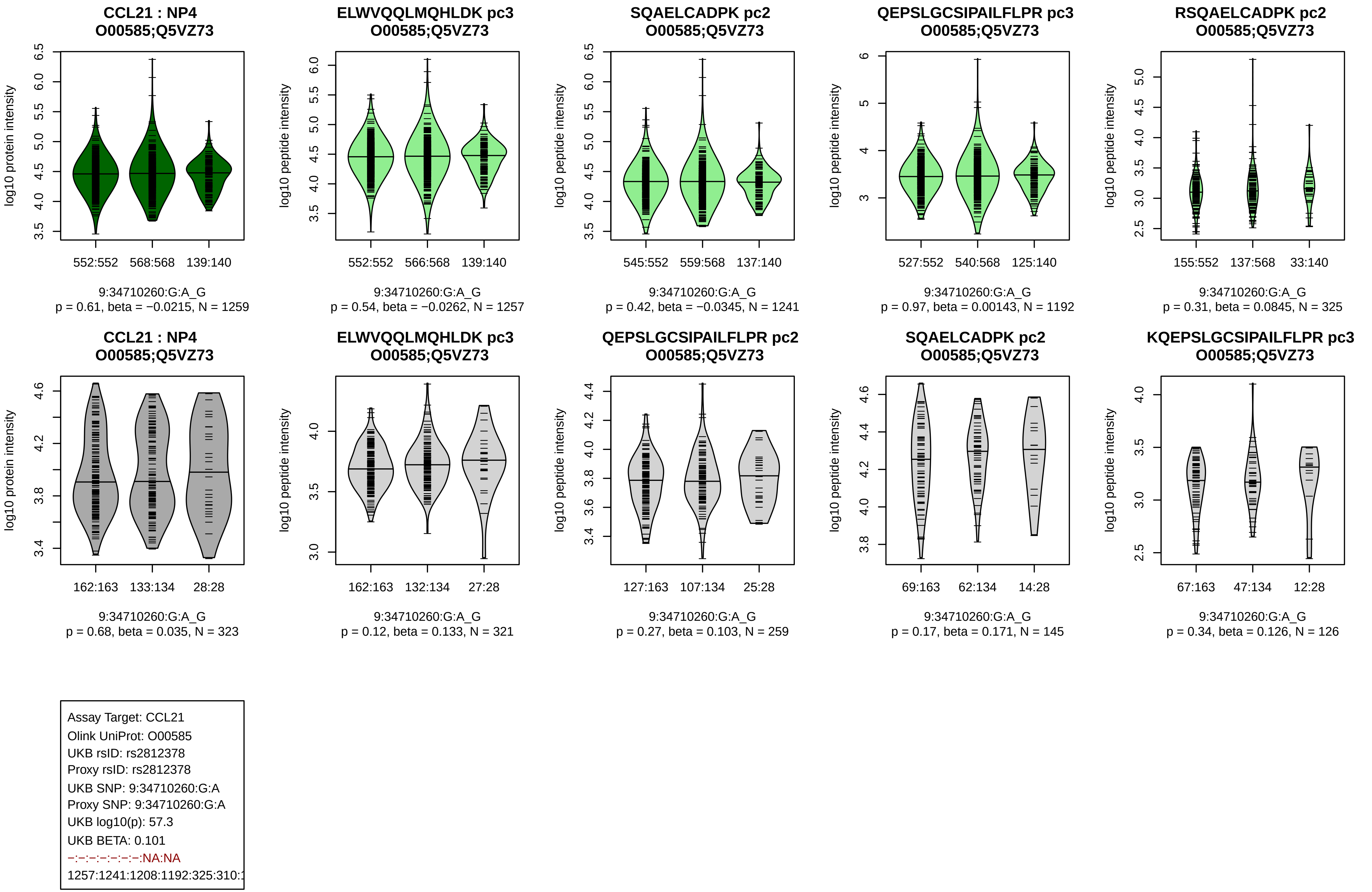
CCL21 : NP4
O00585;Q5VZ73
log10 protein intensity
552:552
568:568
139:140
9:34710260:G:A_G
p = 0.61, beta = −0.0215, N = 1259
ELWVQQLMQHLDK pc3
O00585;Q5VZ73
log10 peptide intensity
552:552
566:568
139:140
9:34710260:G:A_G
p = 0.54, beta = −0.0262, N = 1257
SQAELCADPK pc2
O00585;Q5VZ73
log10 peptide intensity
545:552
559:568
137:140
9:34710260:G:A_G
p = 0.42, beta = −0.0345, N = 1241
QEPSLGCSIPAILFLPR pc3
O00585;Q5VZ73
log10 peptide intensity
527:552
540:568
125:140
9:34710260:G:A_G
p = 0.97, beta = 0.00143, N = 1192
RSQAELCADPK pc2
O00585;Q5VZ73
log10 peptide intensity
155:552
137:568
33:140
9:34710260:G:A_G
p = 0.31, beta = 0.0845, N = 325
CCL21 : NP4
O00585;Q5VZ73
log10 protein intensity
162:163
133:134
28:28
9:34710260:G:A_G
p = 0.68, beta = 0.035, N = 323
ELWVQQLMQHLDK pc3
O00585;Q5VZ73
log10 peptide intensity
162:163
132:134
27:28
9:34710260:G:A_G
p = 0.12, beta = 0.133, N = 321
QEPSLGCSIPAILFLPR pc2
O00585;Q5VZ73
log10 peptide intensity
127:163
107:134
25:28
9:34710260:G:A_G
p = 0.27, beta = 0.103, N = 259
SQAELCADPK pc2
O00585;Q5VZ73
log10 peptide intensity
69:163
62:134
14:28
9:34710260:G:A_G
p = 0.17, beta = 0.171, N = 145
KQEPSLGCSIPAILFLPR pc3
O00585;Q5VZ73
log10 peptide intensity
67:163
47:134
12:28
9:34710260:G:A_G
p = 0.34, beta = 0.126, N = 126
Assay Target: CCL21
Olink UniProt: O00585
UKB rsID: rs2812378
Proxy rsID: rs2812378
UKB SNP: 9:34710260:G:A
Proxy SNP: 9:34710260:G:A
UKB log10(p): 57.3
UKB BETA: 0.101
−:−:−:−:−:−:−:NA:NA
1257:1241:1208:1192:325:310:1

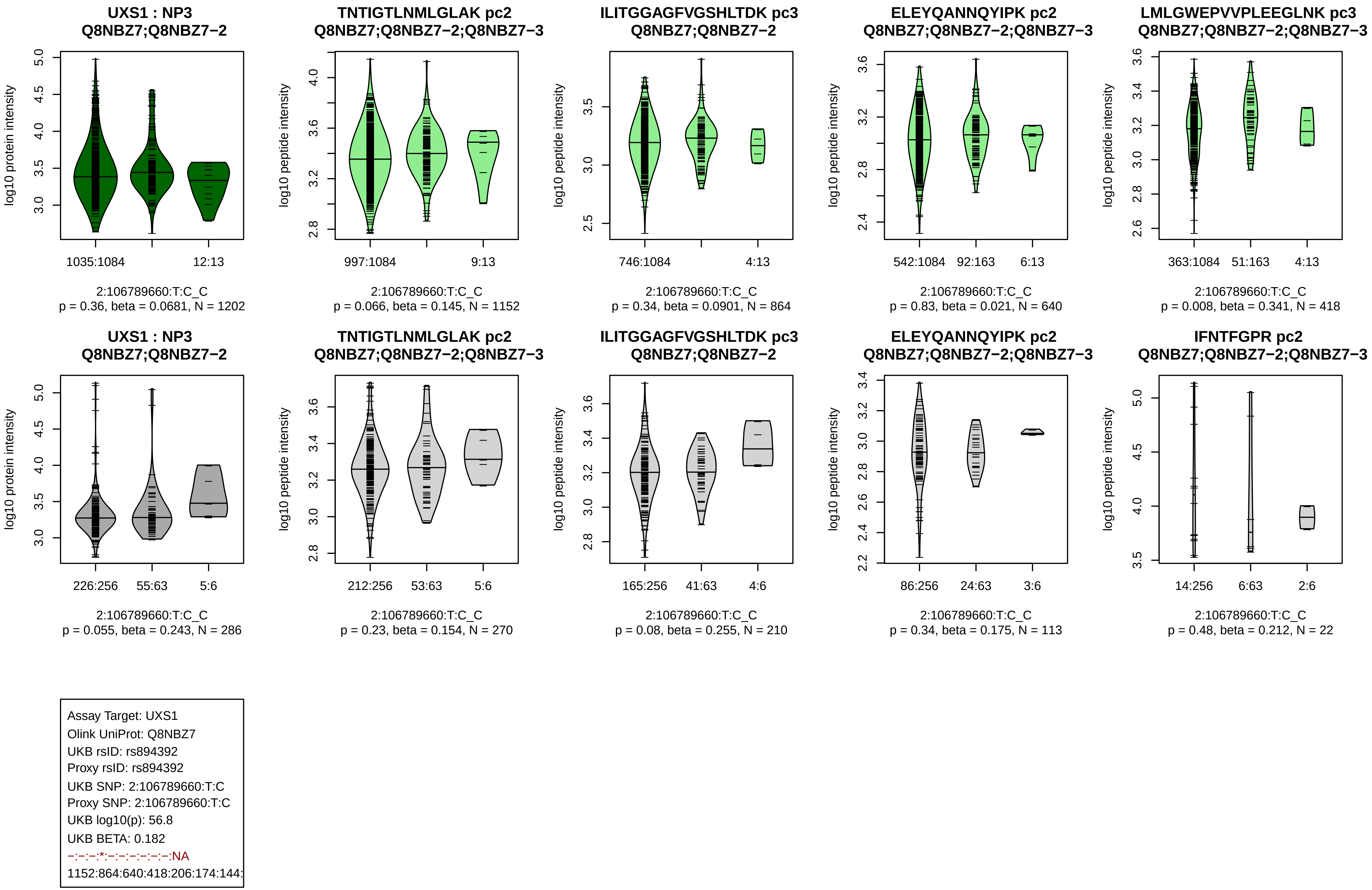
UXS1 : NP3
Q8NBZ7;Q8NBZ7−2
log10 protein intensity
1035:1084
12:13
2:106789660:T:C_C
p = 0.36, beta = 0.0681, N = 1202
TNTIGTLNMLGLAK pc2
Q8NBZ7;Q8NBZ7−2;Q8NBZ7−3
log10 peptide intensity
997:1084
9:13
2:106789660:T:C_C
p = 0.066, beta = 0.145, N = 1152
ILITGGAGFVGSHLTDK pc3
Q8NBZ7;Q8NBZ7−2
log10 peptide intensity
746:1084
4:13
2:106789660:T:C_C
p = 0.34, beta = 0.0901, N = 864
ELEYQANNQYIPK pc2
Q8NBZ7;Q8NBZ7−2;Q8NBZ7−3
log10 peptide intensity
542:1084
92:163
6:13
2:106789660:T:C_C
p = 0.83, beta = 0.021, N = 640
LMLGWEPVVPLEEGLNK pc3
Q8NBZ7;Q8NBZ7−2;Q8NBZ7−3
log10 peptide intensity
363:1084
51:163
4:13
2:106789660:T:C_C
p = 0.008, beta = 0.341, N = 418
UXS1 : NP3
Q8NBZ7;Q8NBZ7−2
log10 protein intensity
226:256
55:63
5:6
2:106789660:T:C_C
p = 0.055, beta = 0.243, N = 286
TNTIGTLNMLGLAK pc2
Q8NBZ7;Q8NBZ7−2;Q8NBZ7−3
log10 peptide intensity
212:256
53:63
5:6
2:106789660:T:C_C
p = 0.23, beta = 0.154, N = 270
ILITGGAGFVGSHLTDK pc3
Q8NBZ7;Q8NBZ7−2
log10 peptide intensity
165:256
41:63
4:6
2:106789660:T:C_C
p = 0.08, beta = 0.255, N = 210
ELEYQANNQYIPK pc2
Q8NBZ7;Q8NBZ7−2;Q8NBZ7−3
log10 peptide intensity
86:256
24:63
3:6
2:106789660:T:C_C
p = 0.34, beta = 0.175, N = 113
IFNTFGPR pc2
Q8NBZ7;Q8NBZ7−2;Q8NBZ7−3
log10 peptide intensity
14:256
6:63
2:6
2:106789660:T:C_C
p = 0.48, beta = 0.212, N = 22
Assay Target: UXS1
Olink UniProt: Q8NBZ7
UKB rsID: rs894392
Proxy rsID: rs894392
UKB SNP: 2:106789660:T:C
Proxy SNP: 2:106789660:T:C
UKB log10(p): 56.8
UKB BETA: 0.182
−:−:−:*:−:−:−:−:−:−:NA
1152:864:640:418:206:174:144:

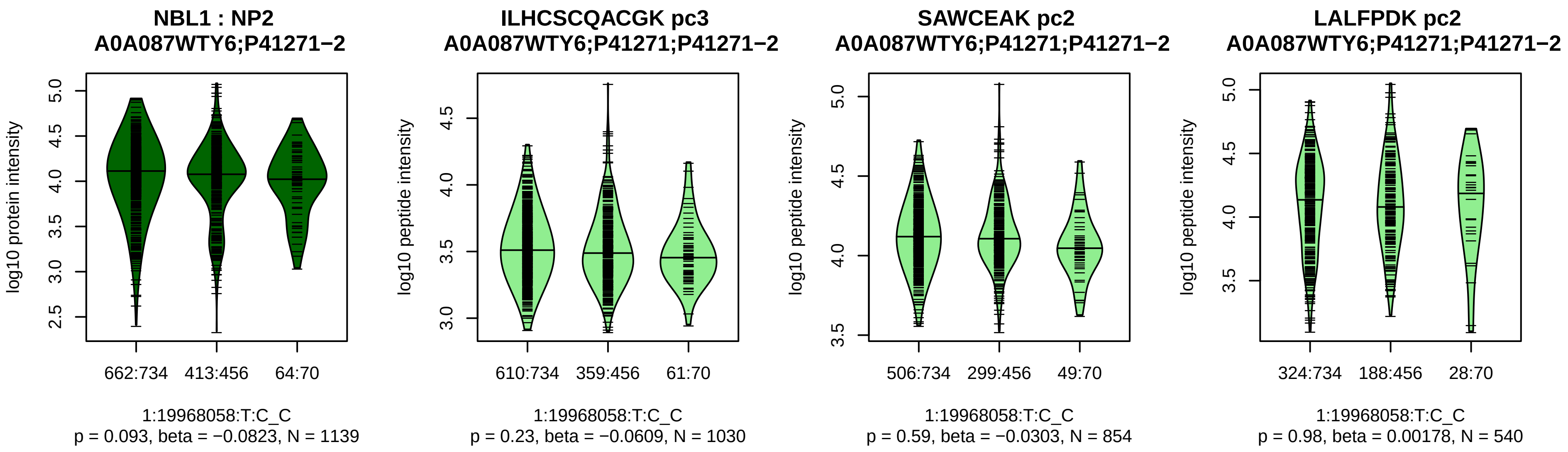


Assay Target: NBL1
Olink UniProt: P41271
UKB rsID: rs12408663
Proxy rsID: rs12408663
UKB SNP: 1:19968058:T:C
Proxy SNP: 1:19968058:T:C
UKB log10(p): 55.9
UKB BETA: −0.114
−:−:−:−
1030:854:540:517

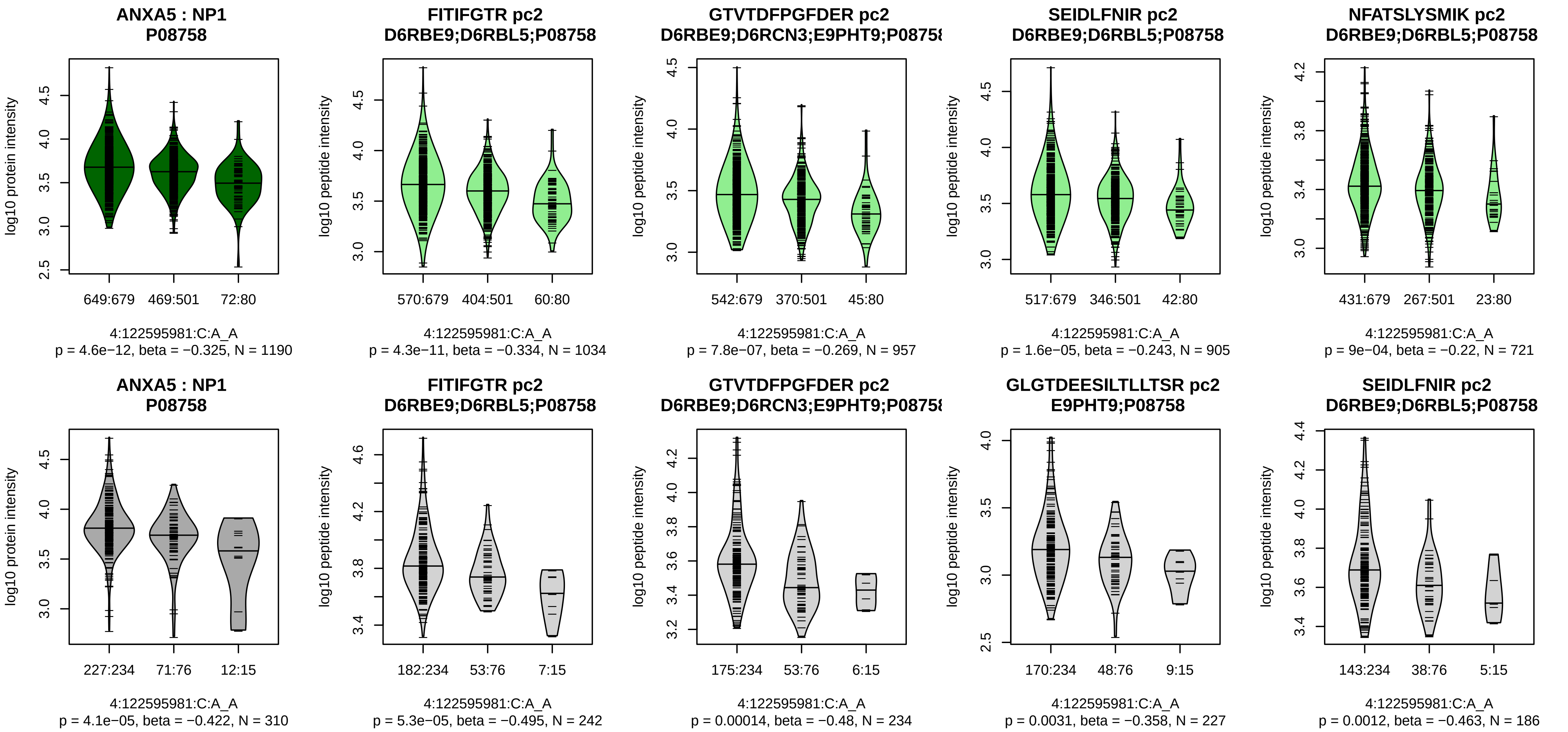


Assay Target: ANXA5
Olink UniProt: P08758
UKB rsID: rs73844249
Proxy rsID: rs73844249
UKB SNP: 4:122595981:C:A
Proxy SNP: 4:122595981:C:A
UKB log10(p): 55.5
UKB BETA: −0.11
*:*:*:*:*:*:−:−:−:−:−:−:NA
1034:957:905:721:641:579:208:

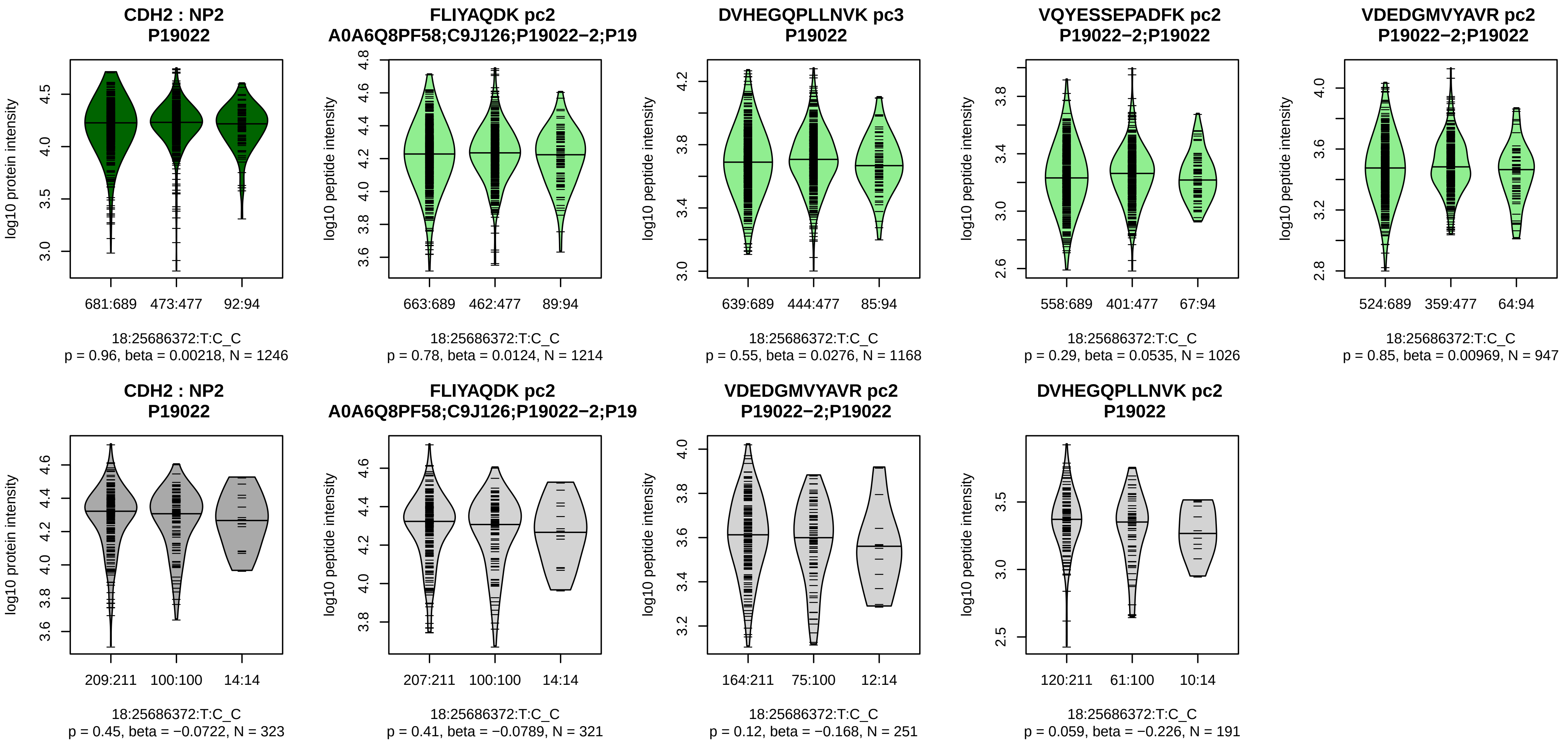


Assay Target: CDH2
Olink UniProt: P19022
UKB rsID: rs12457737
Proxy rsID: rs12457737
UKB SNP: 18:25686372:T:C
Proxy SNP: 18:25686372:T:C
UKB log10(p): 55.1
UKB BETA: −0.096
−:−:−:−:−:−:−:−
1214:1168:1026:947:940:308:2[illegible]

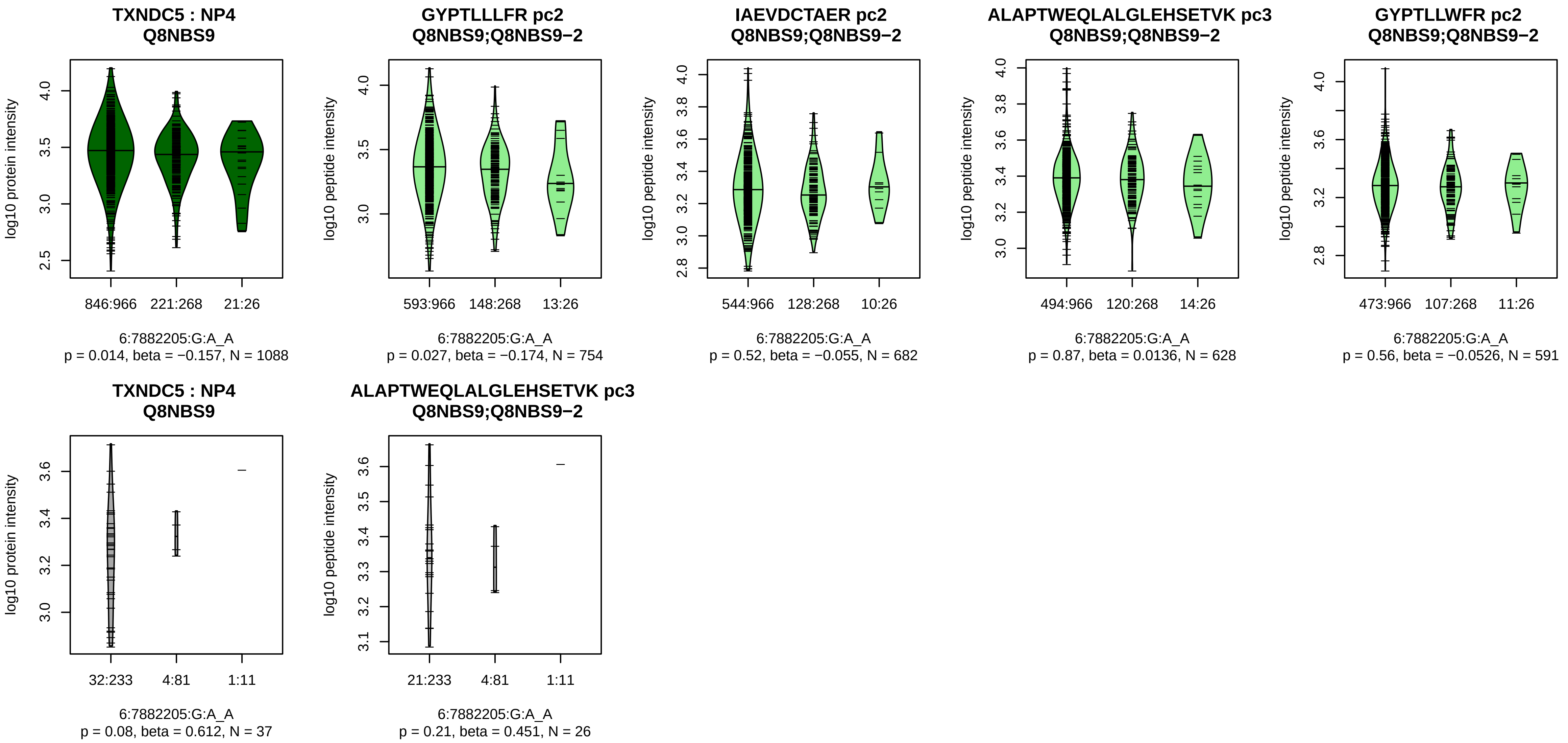


Assay Target: TXNDC5
Olink UniProt: Q8NBS9
UKB rsID: rs7764128
Proxy rsID: rs7764128
UKB SNP: 6:7882205:G:A
Proxy SNP: 6:7882205:G:A
UKB log10(p): 53.8
UKB BETA: −0.148
−:−:−:−:−:−:−:−:−:−:−:−
754:682:628:591:582:500:471:3

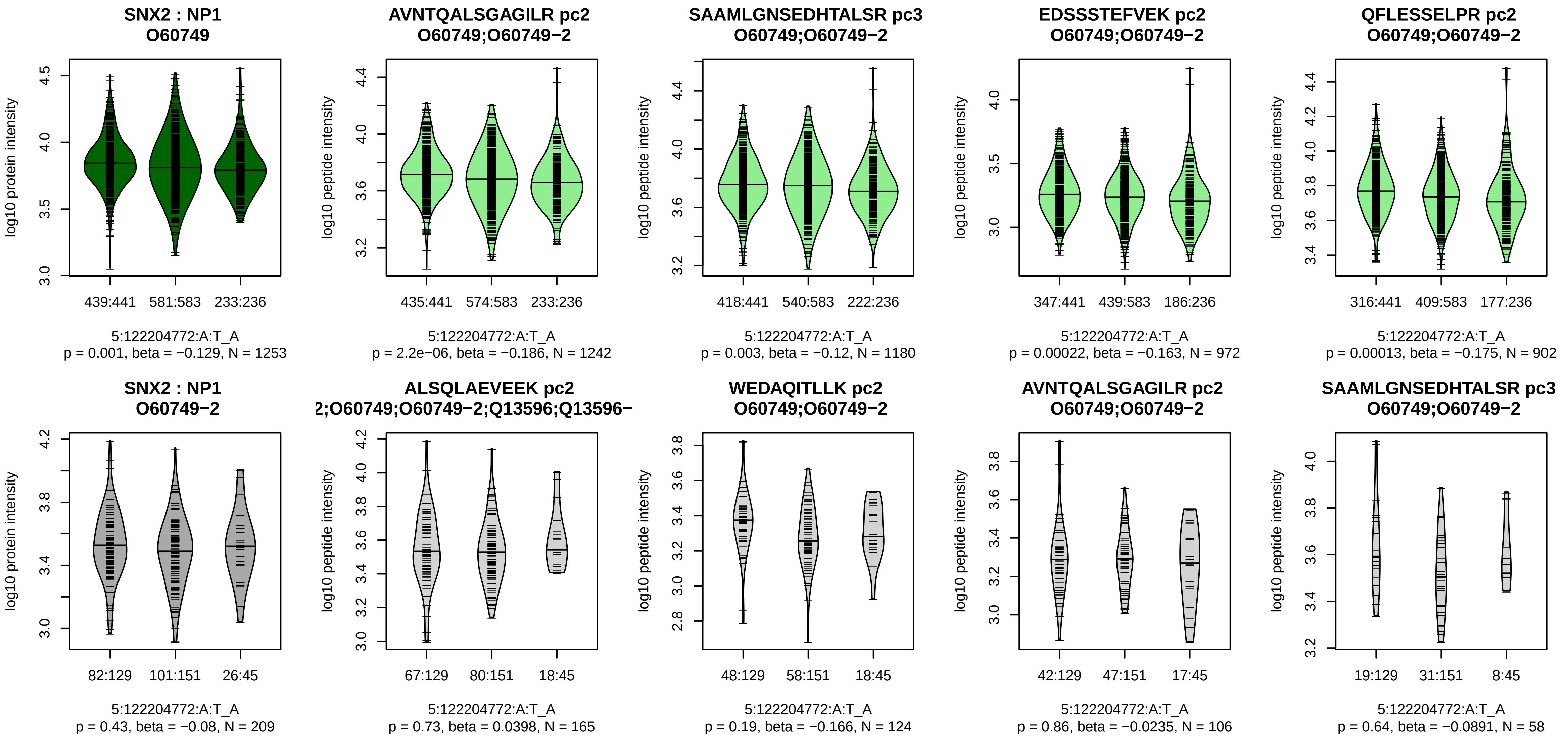


Assay Target: SNX2
Olink UniProt: O60749
UKB rsID: rs27740
Proxy rsID: rs27740
UKB SNP: 5:122204772:A:T
Proxy SNP: 5:122204772:A:T
UKB log10(p): 53.4
UKB BETA: 0.096
*:*:*:*:*:*:–:–:–:*:–:*:–:*:–:–:–:–:–
1242:1180:972:902:887:852:837

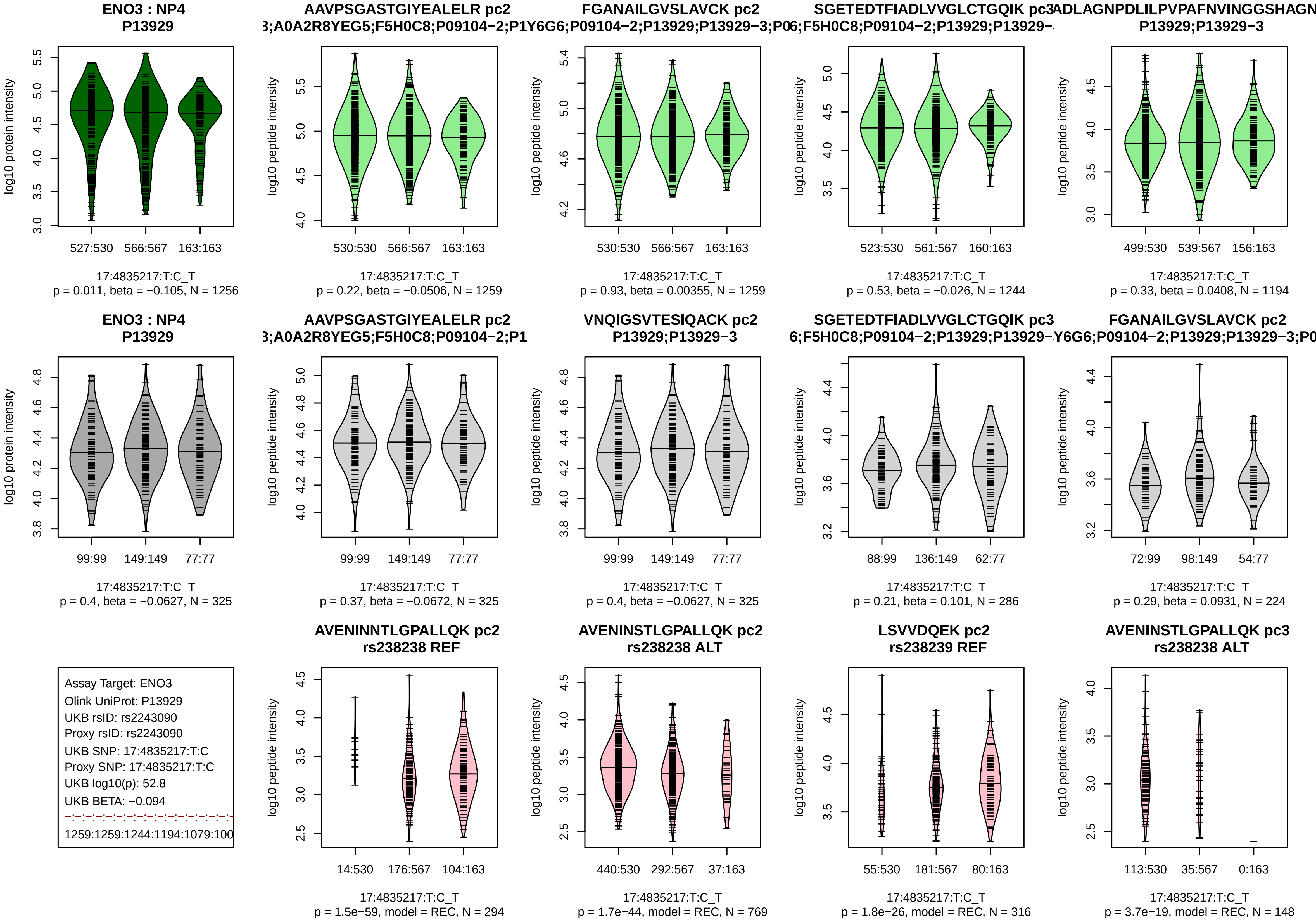
ENO3 : NP4
P13929
log10 protein intensity
527:530
566:567
163:163
17:4835217:T:C_T
p = 0.011, beta = −0.105, N = 1256
AAVPSGASTGIYEALELR pc2
3;A0A2R8YEG5;F5H0C8;P09104−2;P1Y6G6;P09104−2;P13929;P13929−3;P06;F5H0C8;P09104−2;P13929;P13929−
log10 peptide intensity
530:530
566:567
163:163
17:4835217:T:C_T
p = 0.22, beta = −0.0506, N = 1259
FGANAILGVSLAVCK pc2
530:530
566:567
163:163
17:4835217:T:C_T
p = 0.93, beta = 0.00355, N = 1259
SGETEDTFIADLVVGLCTGQIK pc3
523:530
561:567
160:163
17:4835217:T:C_T
p = 0.53, beta = −0.026, N = 1244
ADLAGNPDLILPVPAFNVINGGSHAGN
P13929;P13929−3
499:530
539:567
156:163
17:4835217:T:C_T
p = 0.33, beta = 0.0408, N = 1194
ENO3 : NP4
P13929
99:99
149:149
77:77
17:4835217:T:C_T
p = 0.4, beta = −0.0627, N = 325
AAVPSGASTGIYEALELR pc2
3;A0A2R8YEG5;F5H0C8;P09104−2;P1
99:99
149:149
77:77
17:4835217:T:C_T
p = 0.37, beta = −0.0672, N = 325
VNQIGSVTESIQACK pc2
P13929;P13929−3
99:99
149:149
77:77
17:4835217:T:C_T
p = 0.4, beta = −0.0627, N = 325
SGETEDTFIADLVVGLCTGQIK pc3
6;F5H0C8;P09104−2;P13929;P13929−Y6G6;P09104−2;P13929;P13929−3;P0
88:99
136:149
62:77
17:4835217:T:C_T
p = 0.21, beta = 0.101, N = 286
FGANAILGVSLAVCK pc2
72:99
98:149
54:77
17:4835217:T:C_T
p = 0.29, beta = 0.0931, N = 224
Assay Target: ENO3
Olink UniProt: P13929
UKB rsID: rs2243090
Proxy rsID: rs2243090
UKB SNP: 17:4835217:T:C
Proxy SNP: 17:4835217:T:C
UKB log10(p): 52.8
UKB BETA: −0.094
1259:1259:1244:1194:1079:100
AVENINNTLGPALLQK pc2
rs238238 REF
14:530
176:567
104:163
17:4835217:T:C_T
p = 1.5e−59, model = REC, N = 294
AVENINSTLGPALLQK pc2
rs238238 ALT
440:530
292:567
37:163
17:4835217:T:C_T
p = 1.7e−44, model = REC, N = 769
LSVVDQEK pc2
rs238239 REF
55:530
181:567
80:163
17:4835217:T:C_T
p = 1.8e−26, model = REC, N = 316
AVENINSTLGPALLQK pc3
rs238238 ALT
113:530
35:567
0:163
17:4835217:T:C_T
p = 3.7e−19, model = REC, N = 148

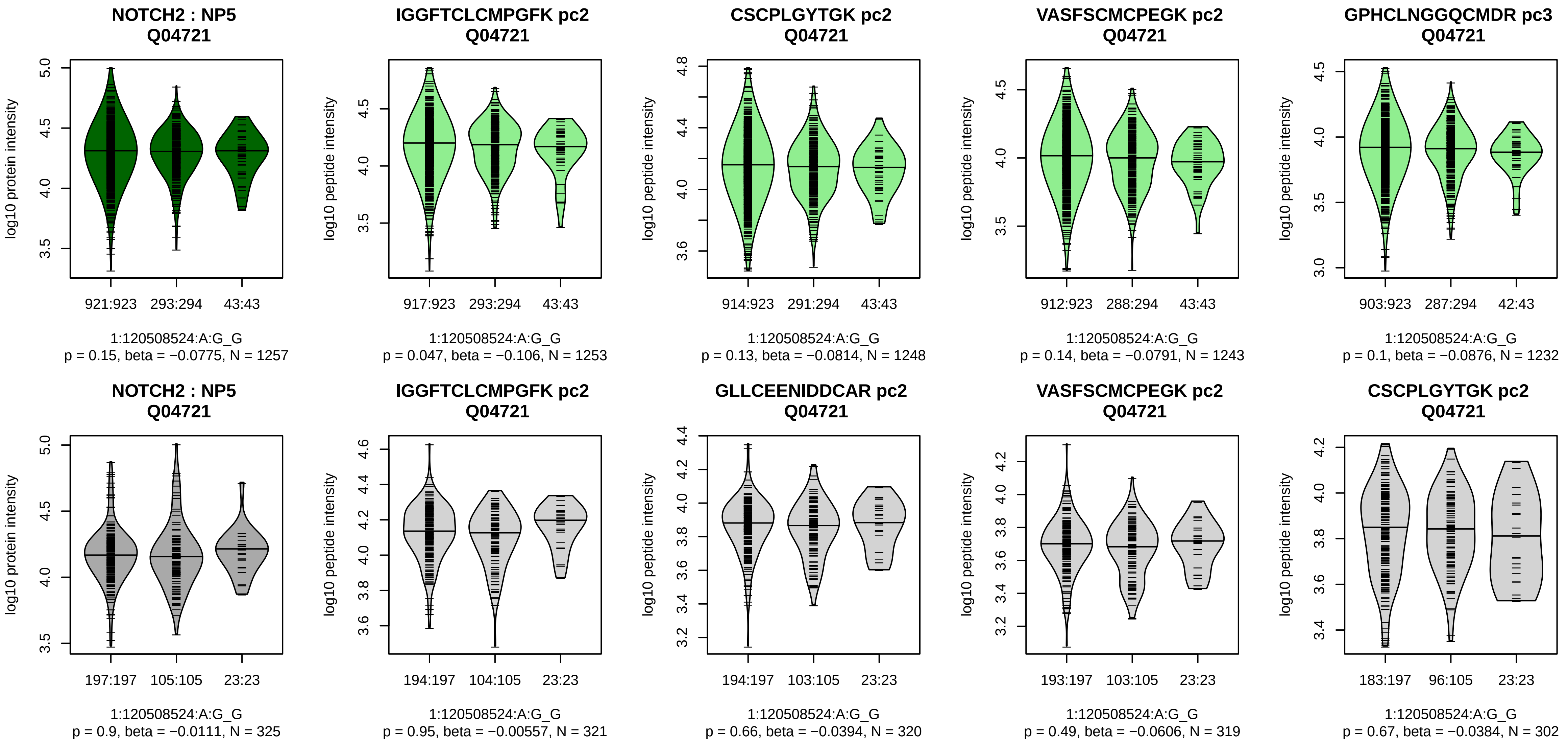


Assay Target: NOTCH2
Olink UniProt: Q04721
UKB rsID: rs2453044
Proxy rsID: rs2453044
UKB SNP: 1:120508524:A:G
Proxy SNP: 1:120508524:A:G
UKB log10(p): 50.4
UKB BETA: −0.131

1253:1248:1243:1232:1198:118

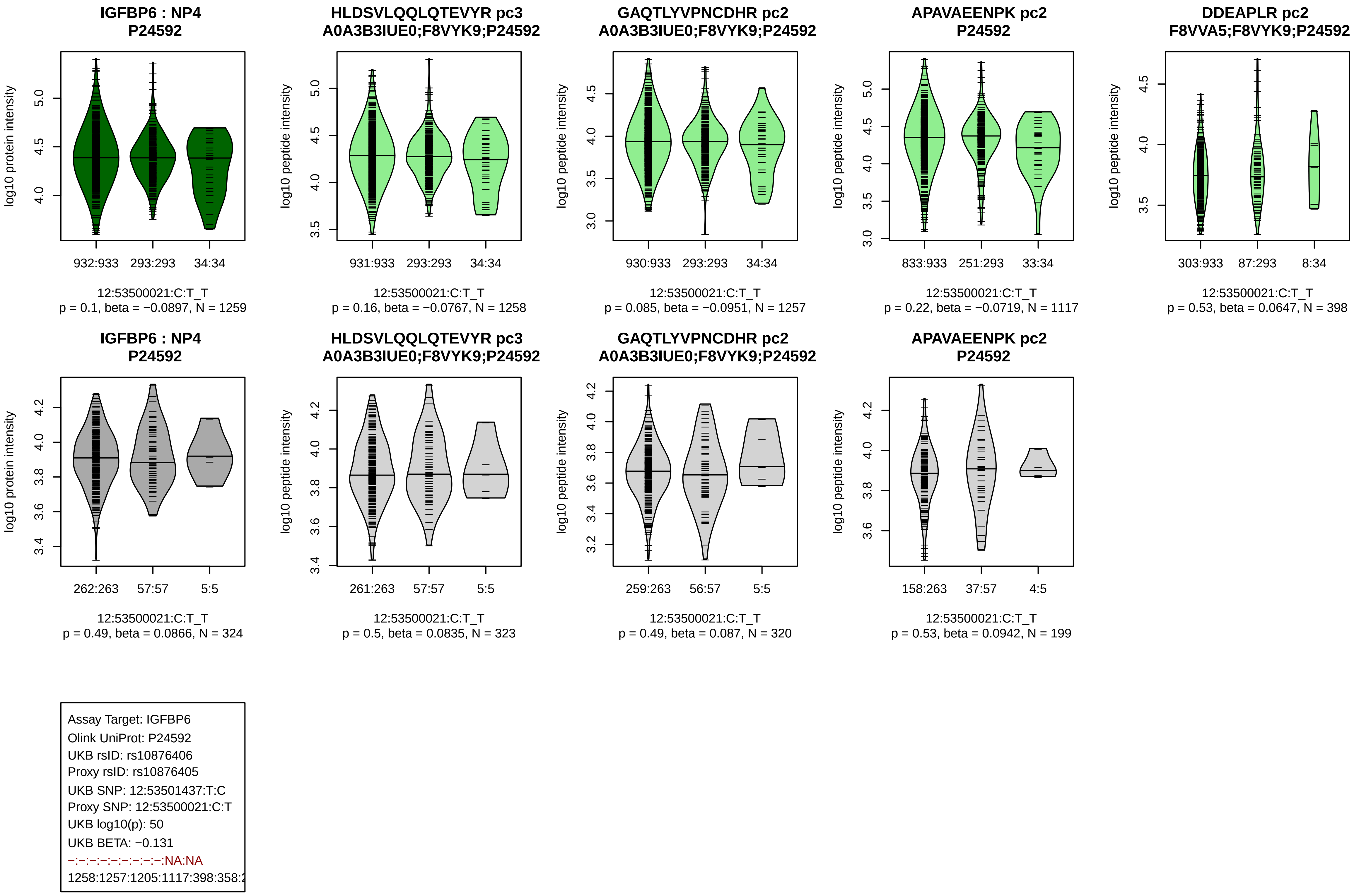
IGFBP6 : NP4
P24592
log10 protein intensity
932:933
293:293
34:34
12:53500021:C:T_T
p = 0.1, beta = −0.0897, N = 1259
HLDSVLQQLQTEVYR pc3
A0A3B3IUE0;F8VYK9;P24592
log10 peptide intensity
931:933
293:293
34:34
12:53500021:C:T_T
p = 0.16, beta = −0.0767, N = 1258
GAQTLYVPNCDHR pc2
A0A3B3IUE0;F8VYK9;P24592
log10 peptide intensity
930:933
293:293
34:34
12:53500021:C:T_T
p = 0.085, beta = −0.0951, N = 1257
APAVAEENPK pc2
P24592
log10 peptide intensity
833:933
251:293
33:34
12:53500021:C:T_T
p = 0.22, beta = −0.0719, N = 1117
DDEAPLR pc2
F8VVA5;F8VYK9;P24592
log10 peptide intensity
303:933
87:293
8:34
12:53500021:C:T_T
p = 0.53, beta = 0.0647, N = 398
IGFBP6 : NP4
P24592
log10 protein intensity
262:263
57:57
5:5
12:53500021:C:T_T
p = 0.49, beta = 0.0866, N = 324
HLDSVLQQLQTEVYR pc3
A0A3B3IUE0;F8VYK9;P24592
log10 peptide intensity
261:263
57:57
5:5
12:53500021:C:T_T
p = 0.5, beta = 0.0835, N = 323
GAQTLYVPNCDHR pc2
A0A3B3IUE0;F8VYK9;P24592
log10 peptide intensity
259:263
56:57
5:5
12:53500021:C:T_T
p = 0.49, beta = 0.087, N = 320
APAVAEENPK pc2
P24592
log10 peptide intensity
158:263
37:57
4:5
12:53500021:C:T_T
p = 0.53, beta = 0.0942, N = 199
Assay Target: IGFBP6
Olink UniProt: P24592
UKB rsID: rs10876406
Proxy rsID: rs10876405
UKB SNP: 12:53501437:T:C
Proxy SNP: 12:53500021:C:T
UKB log10(p): 50
UKB BETA: −0.131
−:−:−:−:−:−:−:−:−:NA:NA
1258:1257:1205:1117:398:358:2

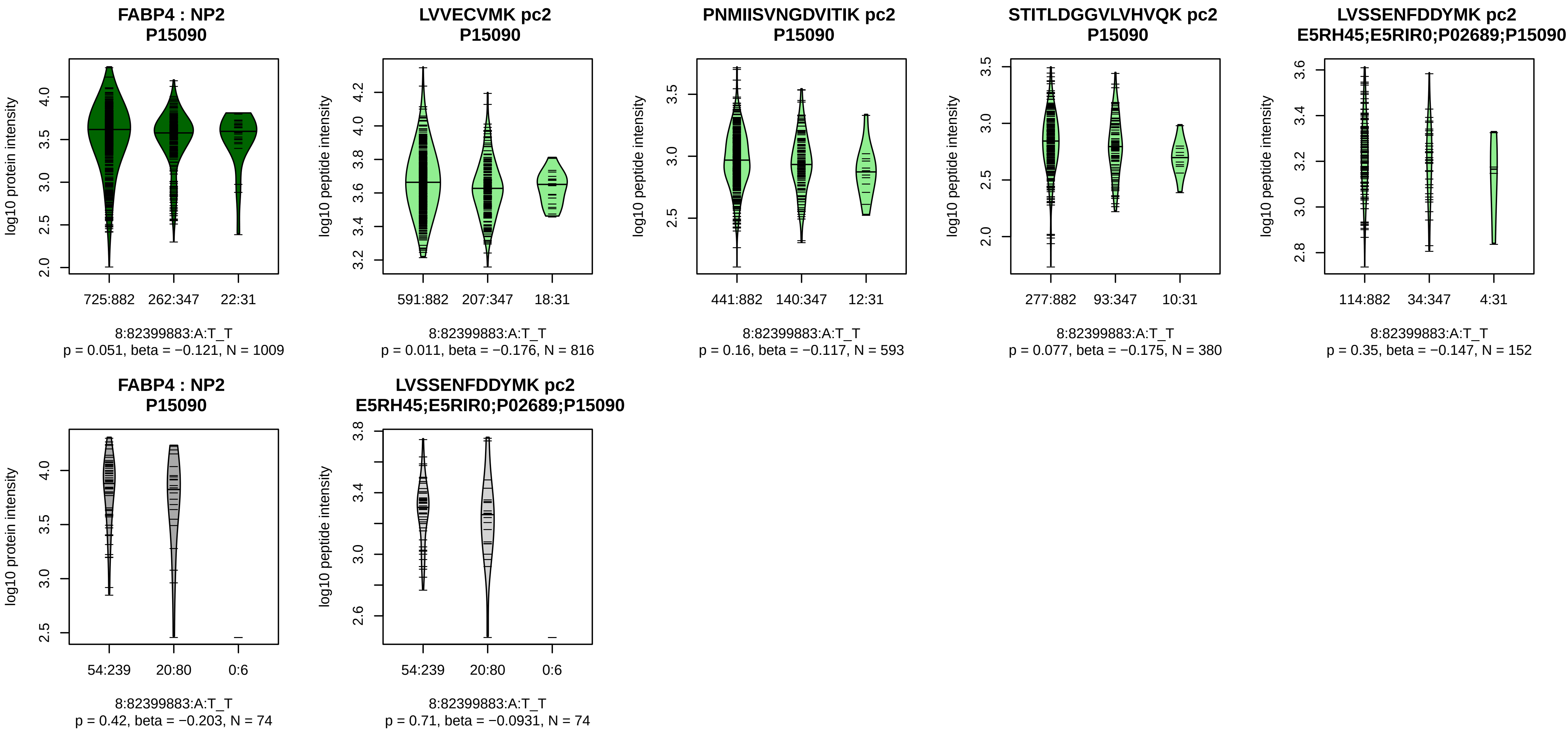


Assay Target: FABP4
Olink UniProt: P15090
UKB rsID: rs72684474
Proxy rsID: rs72684474
UKB SNP: 8:82399883:A:T
Proxy SNP: 8:82399883:A:T
UKB log10(p): 50
UKB BETA: −0.115
−:−:−:−:*
816:593:380:152:149

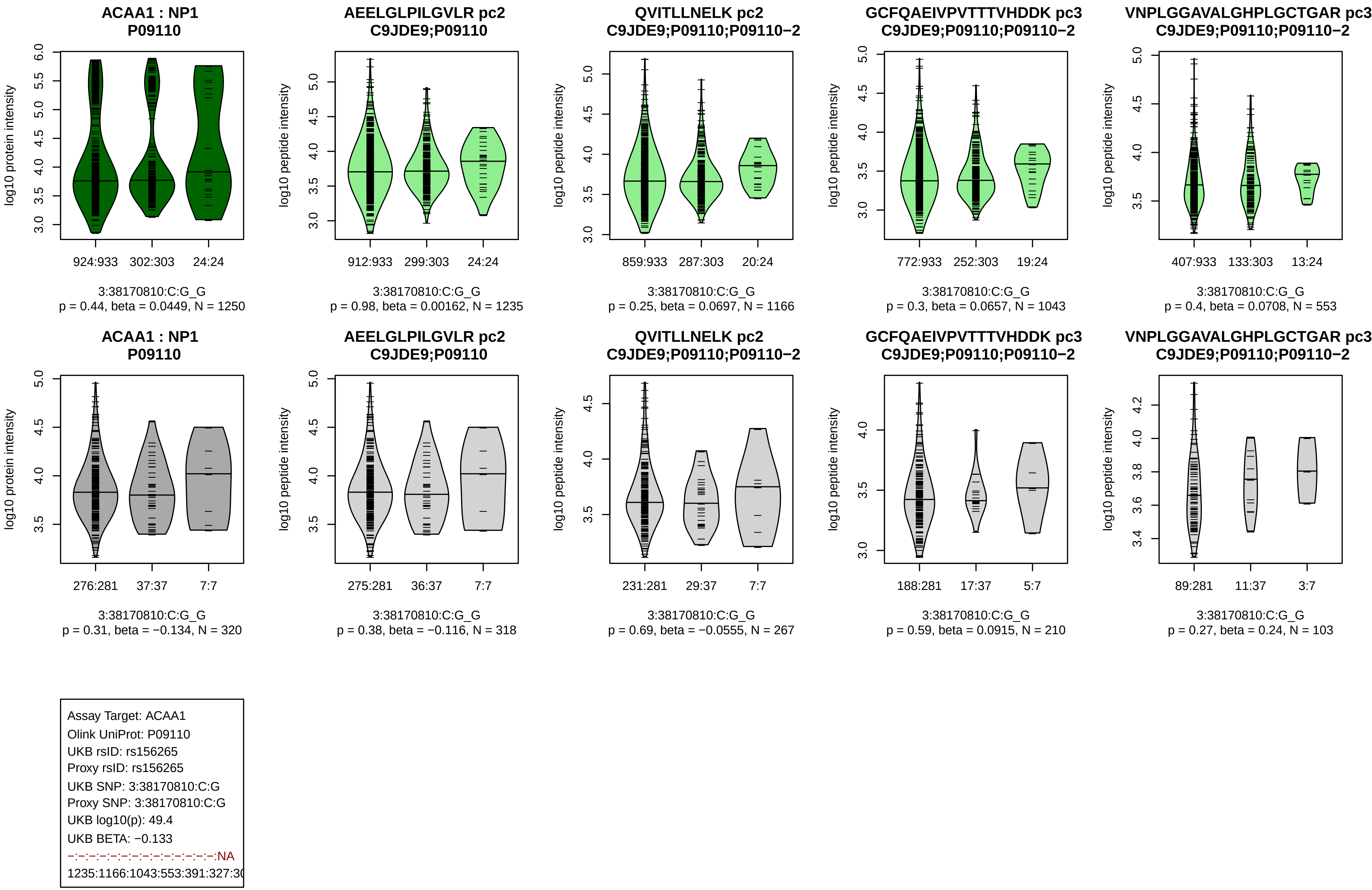
ACAA1 : NP1
P09110
log10 protein intensity
924:933
302:303
24:24
3:38170810:C:G_G
p = 0.44, beta = 0.0449, N = 1250
AEELGLPILGVLR pc2
C9JDE9;P09110
log10 peptide intensity
912:933
299:303
24:24
3:38170810:C:G_G
p = 0.98, beta = 0.00162, N = 1235
QVITLLNELK pc2
C9JDE9;P09110;P09110−2
log10 peptide intensity
859:933
287:303
20:24
3:38170810:C:G_G
p = 0.25, beta = 0.0697, N = 1166
GCFQAEIVPVTTTVHDDK pc3
C9JDE9;P09110;P09110−2
log10 peptide intensity
772:933
252:303
19:24
3:38170810:C:G_G
p = 0.3, beta = 0.0657, N = 1043
VNPLGGAVALGHPLGCTGAR pc3
C9JDE9;P09110;P09110−2
log10 peptide intensity
407:933
133:303
13:24
3:38170810:C:G_G
p = 0.4, beta = 0.0708, N = 553
ACAA1 : NP1
P09110
log10 protein intensity
276:281
37:37
7:7
3:38170810:C:G_G
p = 0.31, beta = −0.134, N = 320
AEELGLPILGVLR pc2
C9JDE9;P09110
log10 peptide intensity
275:281
36:37
7:7
3:38170810:C:G_G
p = 0.38, beta = −0.116, N = 318
QVITLLNELK pc2
C9JDE9;P09110;P09110−2
log10 peptide intensity
231:281
29:37
7:7
3:38170810:C:G_G
p = 0.69, beta = −0.0555, N = 267
GCFQAEIVPVTTTVHDDK pc3
C9JDE9;P09110;P09110−2
log10 peptide intensity
188:281
17:37
5:7
3:38170810:C:G_G
p = 0.59, beta = 0.0915, N = 210
VNPLGGAVALGHPLGCTGAR pc3
C9JDE9;P09110;P09110−2
log10 peptide intensity
89:281
11:37
3:7
3:38170810:C:G_G
p = 0.27, beta = 0.24, N = 103
Assay Target: ACAA1
Olink UniProt: P09110
UKB rsID: rs156265
Proxy rsID: rs156265
UKB SNP: 3:38170810:C:G
Proxy SNP: 3:38170810:C:G
UKB log10(p): 49.4
UKB BETA: −0.133
−:−:−:−:−:−:−:−:−:−:−:−:−:−:NA
1235:1166:1043:553:391:327:3(

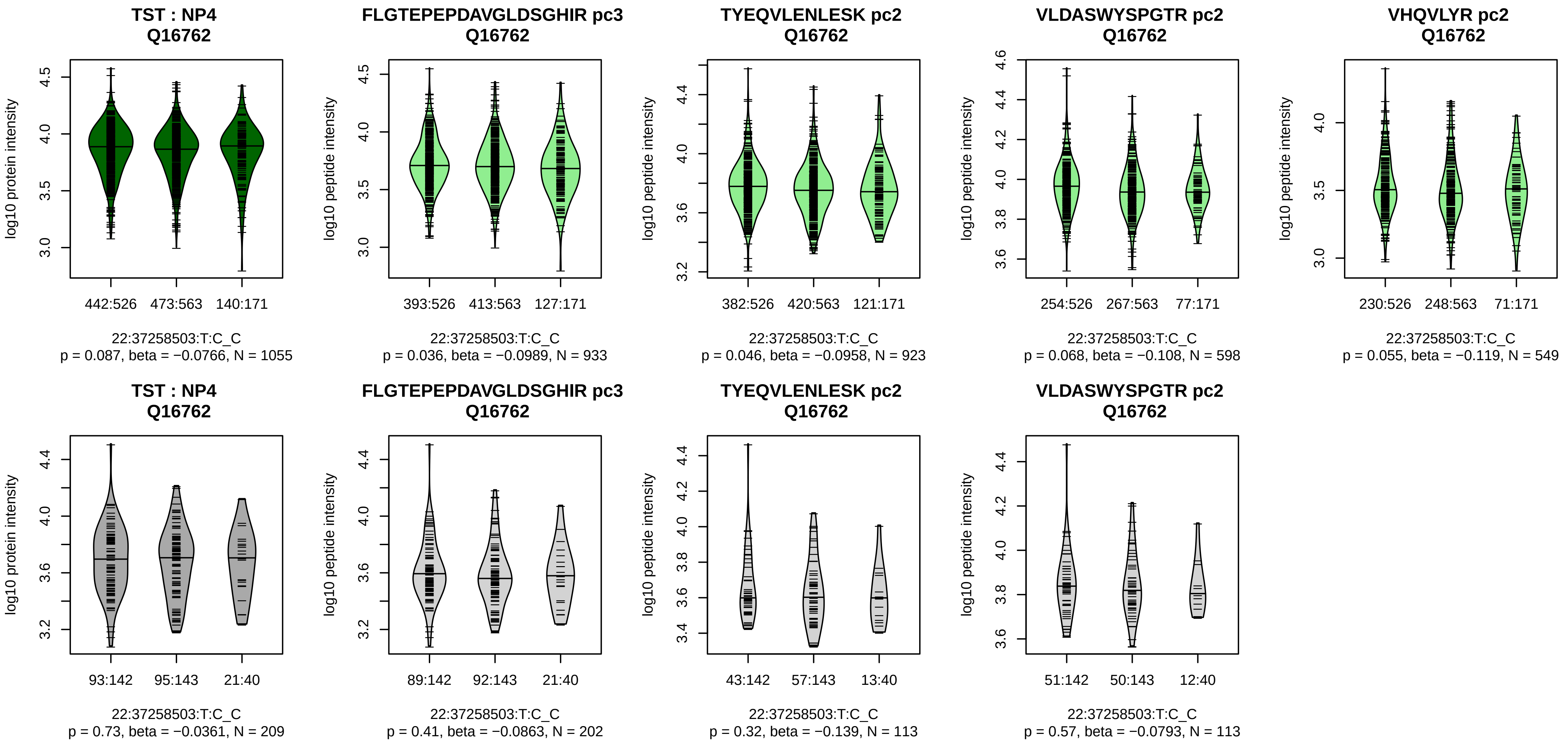


Assay Target: TST
Olink UniProt: Q16762
UKB rsID: rs4821544
Proxy rsID: rs4821544
UKB SNP: 22:37258503:T:C
Proxy SNP: 22:37258503:T:C
UKB log10(p): 49.3
UKB BETA: −0.101
−:−:−:−:−:−:−:−
933:923:598:549:402:133:71:53

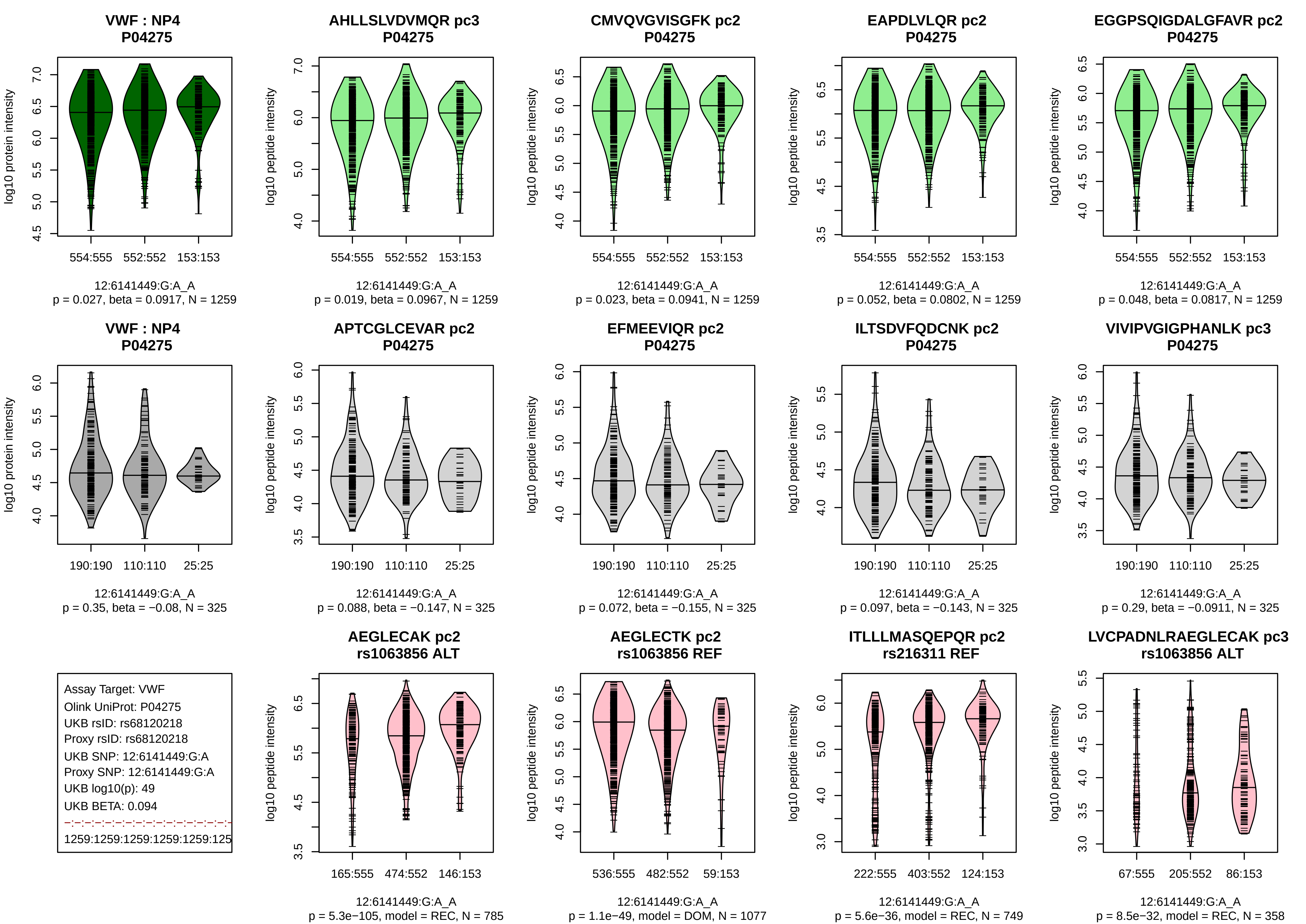
VWF : NP4
P04275
log10 protein intensity
554:555 552:552 153:153
12:6141449:G:A_A
p = 0.027, beta = 0.0917, N = 1259
AHLLSLVDVMQR pc3
P04275
log10 peptide intensity
554:555 552:552 153:153
12:6141449:G:A_A
p = 0.019, beta = 0.0967, N = 1259
CMVQVGVISGFK pc2
P04275
log10 peptide intensity
554:555 552:552 153:153
12:6141449:G:A_A
p = 0.023, beta = 0.0941, N = 1259
EAPDLVLQR pc2
P04275
log10 peptide intensity
554:555 552:552 153:153
12:6141449:G:A_A
p = 0.052, beta = 0.0802, N = 1259
EGGPSQIGDALGFAVR pc2
P04275
log10 peptide intensity
554:555 552:552 153:153
12:6141449:G:A_A
p = 0.048, beta = 0.0817, N = 1259
VWF : NP4
P04275
log10 protein intensity
190:190 110:110 25:25
12:6141449:G:A_A
p = 0.35, beta = −0.08, N = 325
APTCGLCEVAR pc2
P04275
log10 peptide intensity
190:190 110:110 25:25
12:6141449:G:A_A
p = 0.088, beta = −0.147, N = 325
EFMEEVIQR pc2
P04275
log10 peptide intensity
190:190 110:110 25:25
12:6141449:G:A_A
p = 0.072, beta = −0.155, N = 325
ILTSDVFQDCNK pc2
P04275
log10 peptide intensity
190:190 110:110 25:25
12:6141449:G:A_A
p = 0.097, beta = −0.143, N = 325
VIVIPVGIGPHANLK pc3
P04275
log10 peptide intensity
190:190 110:110 25:25
12:6141449:G:A_A
p = 0.29, beta = −0.0911, N = 325
Assay Target: VWF
Olink UniProt: P04275
UKB rsID: rs68120218
Proxy rsID: rs68120218
UKB SNP: 12:6141449:G:A
Proxy SNP: 12:6141449:G:A
UKB log10(p): 49
UKB BETA: 0.094
1259:1259:1259:1259:1259:125
AEGLECAK pc2
rs1063856 ALT
log10 peptide intensity
165:555 474:552 146:153
12:6141449:G:A_A
p = 5.3e−105, model = REC, N = 785
AEGLECTK pc2
rs1063856 REF
log10 peptide intensity
536:555 482:552 59:153
12:6141449:G:A_A
p = 1.1e−49, model = DOM, N = 1077
ITLLLMASQEPQR pc2
rs216311 REF
log10 peptide intensity
222:555 403:552 124:153
12:6141449:G:A_A
p = 5.6e−36, model = REC, N = 749
LVCPADNLRAEGLECAK pc3
rs1063856 ALT
log10 peptide intensity
67:555 205:552 86:153
12:6141449:G:A_A
p = 8.5e−32, model = REC, N = 358

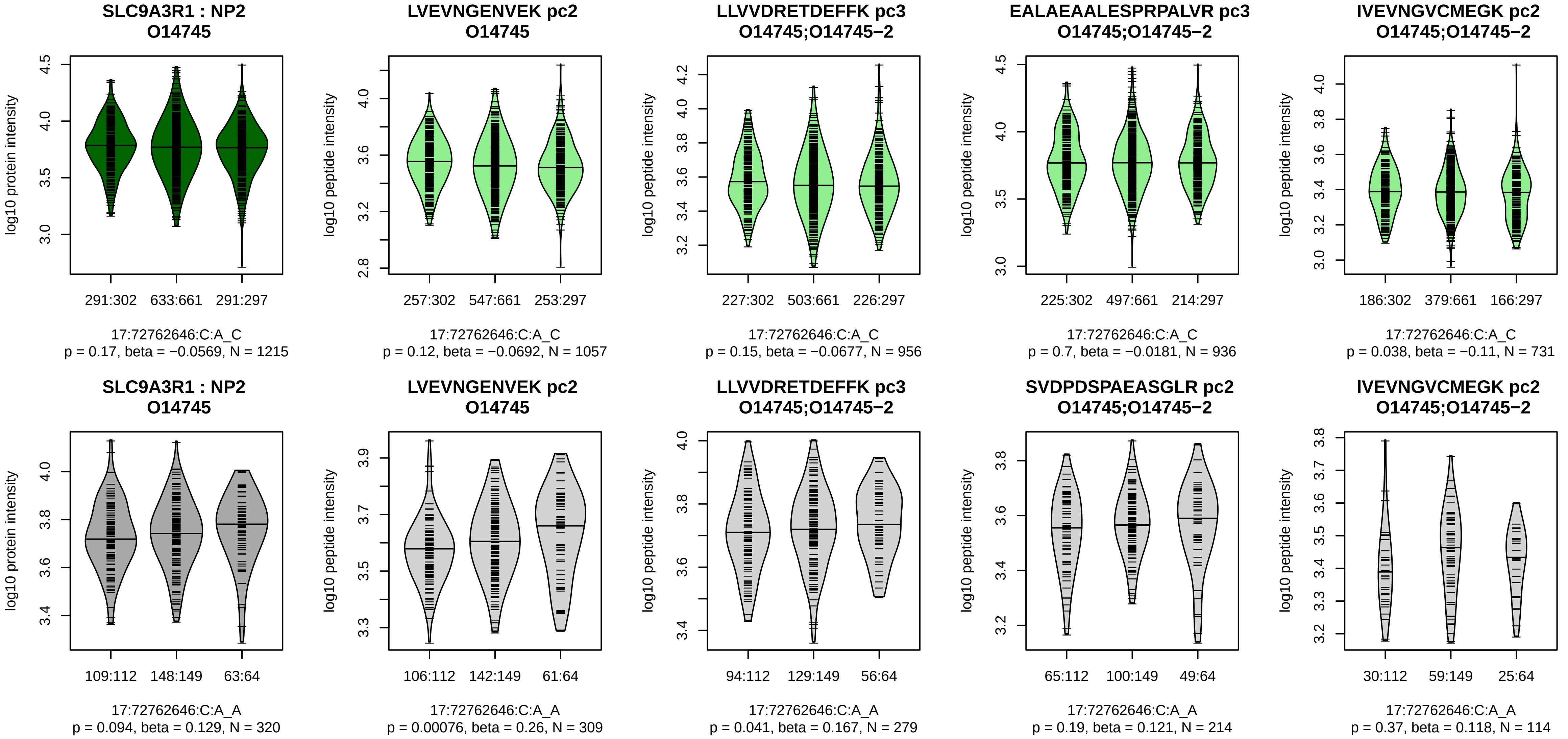


Assay Target: SLC9A3R1
Olink UniProt: O14745
UKB rsID: rs8064299
Proxy rsID: rs8064299
UKB SNP: 17:72762646:C:A
Proxy SNP: 17:72762646:C:A
UKB log10(p): 48.8
UKB BETA: 0.092
−:−:−:−:−:−:−:−
1057:956:936:731:677:603:190:

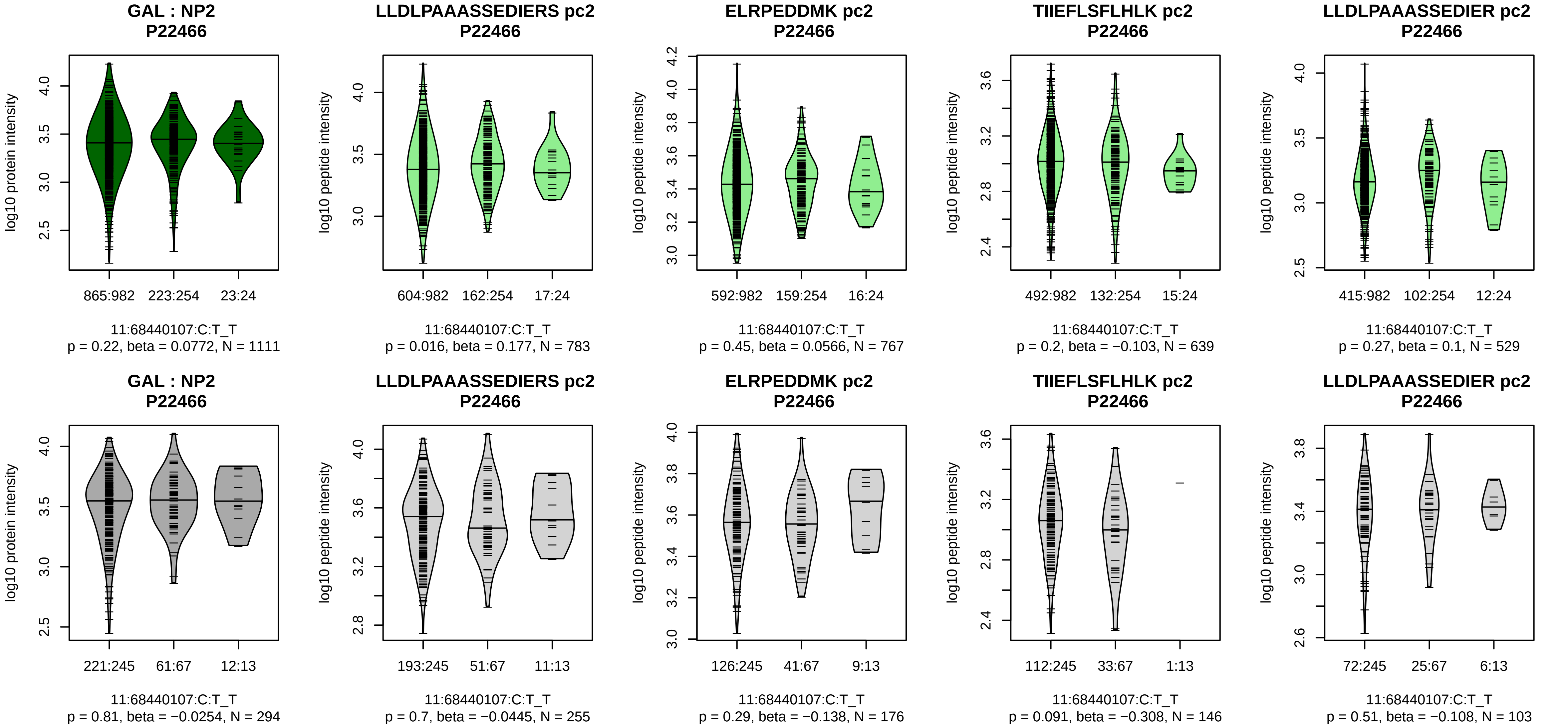


Assay Target: GAL
Olink UniProt: P22466
UKB rsID: rs2012605
Proxy rsID: rs2012605
UKB SNP: 11:68440107:C:T
Proxy SNP: 11:68440107:C:T
UKB log10(p): 47.9
UKB BETA: 0.148
–:–:–:–:–
783:767:639:529:190

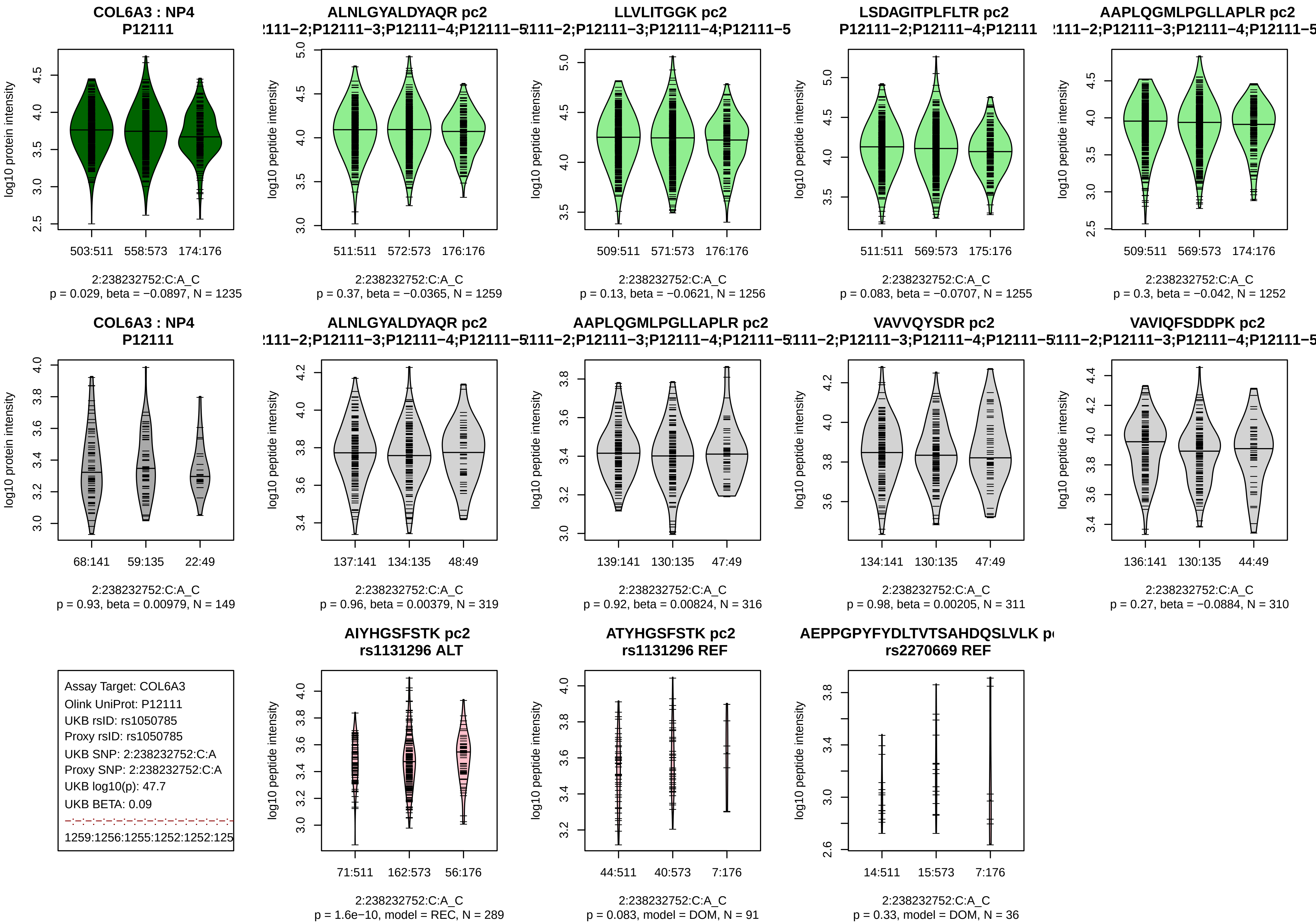
COL6A3 : NP4
P12111
log10 protein intensity
503:511
558:573
174:176
2:238232752:C:A_C
p = 0.029, beta = −0.0897, N = 1235
ALNLGYALDYAQR pc2
log10 peptide intensity
511:511
572:573
176:176
2:238232752:C:A_C
p = 0.37, beta = −0.0365, N = 1259
LLVLITGGK pc2
509:511
571:573
176:176
2:238232752:C:A_C
p = 0.13, beta = −0.0621, N = 1256
LSDAGITPLFLTR pc2
P12111−2;P12111−4;P12111
511:511
569:573
175:176
2:238232752:C:A_C
p = 0.083, beta = −0.0707, N = 1255
AAPLQGMLPGLLAPLR pc2
509:511
569:573
174:176
2:238232752:C:A_C
p = 0.3, beta = −0.042, N = 1252
COL6A3 : NP4
P12111
68:141
59:135
22:49
2:238232752:C:A_C
p = 0.93, beta = 0.00979, N = 149
ALNLGYALDYAQR pc2
137:141
134:135
48:49
2:238232752:C:A_C
p = 0.96, beta = 0.00379, N = 319
AAPLQGMLPGLLAPLR pc2
139:141
130:135
47:49
2:238232752:C:A_C
p = 0.92, beta = 0.00824, N = 316
VAVVQYSDR pc2
134:141
130:135
47:49
2:238232752:C:A_C
p = 0.98, beta = 0.00205, N = 311
VAVIQFSDDPK pc2
136:141
130:135
44:49
2:238232752:C:A_C
p = 0.27, beta = −0.0884, N = 310
Assay Target: COL6A3
Olink UniProt: P12111
UKB rsID: rs1050785
Proxy rsID: rs1050785
UKB SNP: 2:238232752:C:A
Proxy SNP: 2:238232752:C:A
UKB log10(p): 47.7
UKB BETA: 0.09
1259:1256:1255:1252:1252:125
AIYHGSFSTK pc2
rs1131296 ALT
71:511
162:573
56:176
2:238232752:C:A_C
p = 1.6e−10, model = REC, N = 289
ATYHGSFSTK pc2
rs1131296 REF
44:511
40:573
7:176
2:238232752:C:A_C
p = 0.083, model = DOM, N = 91
rs2270669 REF
14:511
15:573
7:176
2:238232752:C:A_C
p = 0.33, model = DOM, N = 36

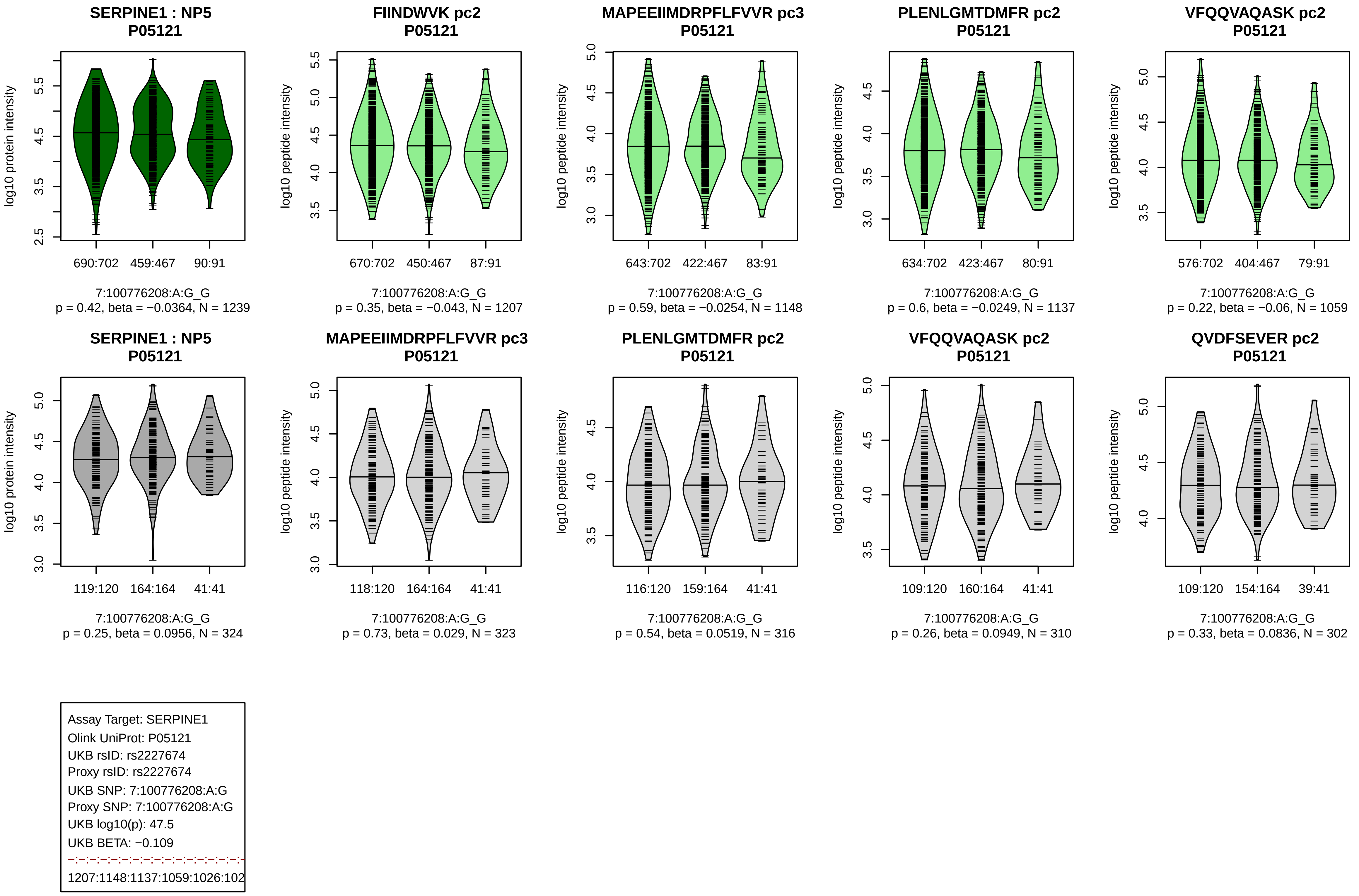

SERPINE1 : NP5
P05121
log10 protein intensity
690:702
459:467
90:91
7:100776208:A:G_G
p = 0.42, beta = −0.0364, N = 1239
FIINDWVK pc2
P05121
log10 peptide intensity
670:702
450:467
87:91
7:100776208:A:G_G
p = 0.35, beta = −0.043, N = 1207
MAPEEIIMDRPFLFVVR pc3
P05121
log10 peptide intensity
643:702
422:467
83:91
7:100776208:A:G_G
p = 0.59, beta = −0.0254, N = 1148
PLENLGMTDMFR pc2
P05121
log10 peptide intensity
634:702
423:467
80:91
7:100776208:A:G_G
p = 0.6, beta = −0.0249, N = 1137
VFQQVAQASK pc2
P05121
log10 peptide intensity
576:702
404:467
79:91
7:100776208:A:G_G
p = 0.22, beta = −0.06, N = 1059
SERPINE1 : NP5
P05121
log10 protein intensity
119:120
164:164
41:41
7:100776208:A:G_G
p = 0.25, beta = 0.0956, N = 324
MAPEEIIMDRPFLFVVR pc3
P05121
log10 peptide intensity
118:120
164:164
41:41
7:100776208:A:G_G
p = 0.73, beta = 0.029, N = 323
PLENLGMTDMFR pc2
P05121
log10 peptide intensity
116:120
159:164
41:41
7:100776208:A:G_G
p = 0.54, beta = 0.0519, N = 316
VFQQVAQASK pc2
P05121
log10 peptide intensity
109:120
160:164
41:41
7:100776208:A:G_G
p = 0.26, beta = 0.0949, N = 310
QVDFSEVER pc2
P05121
log10 peptide intensity
109:120
154:164
39:41
7:100776208:A:G_G
p = 0.33, beta = 0.0836, N = 302
Assay Target: SERPINE1
Olink UniProt: P05121
UKB rsID: rs2227674
Proxy rsID: rs2227674
UKB SNP: 7:100776208:A:G
Proxy SNP: 7:100776208:A:G
UKB log10(p): 47.5
UKB BETA: −0.109
1207:1148:1137:1059:1026:102

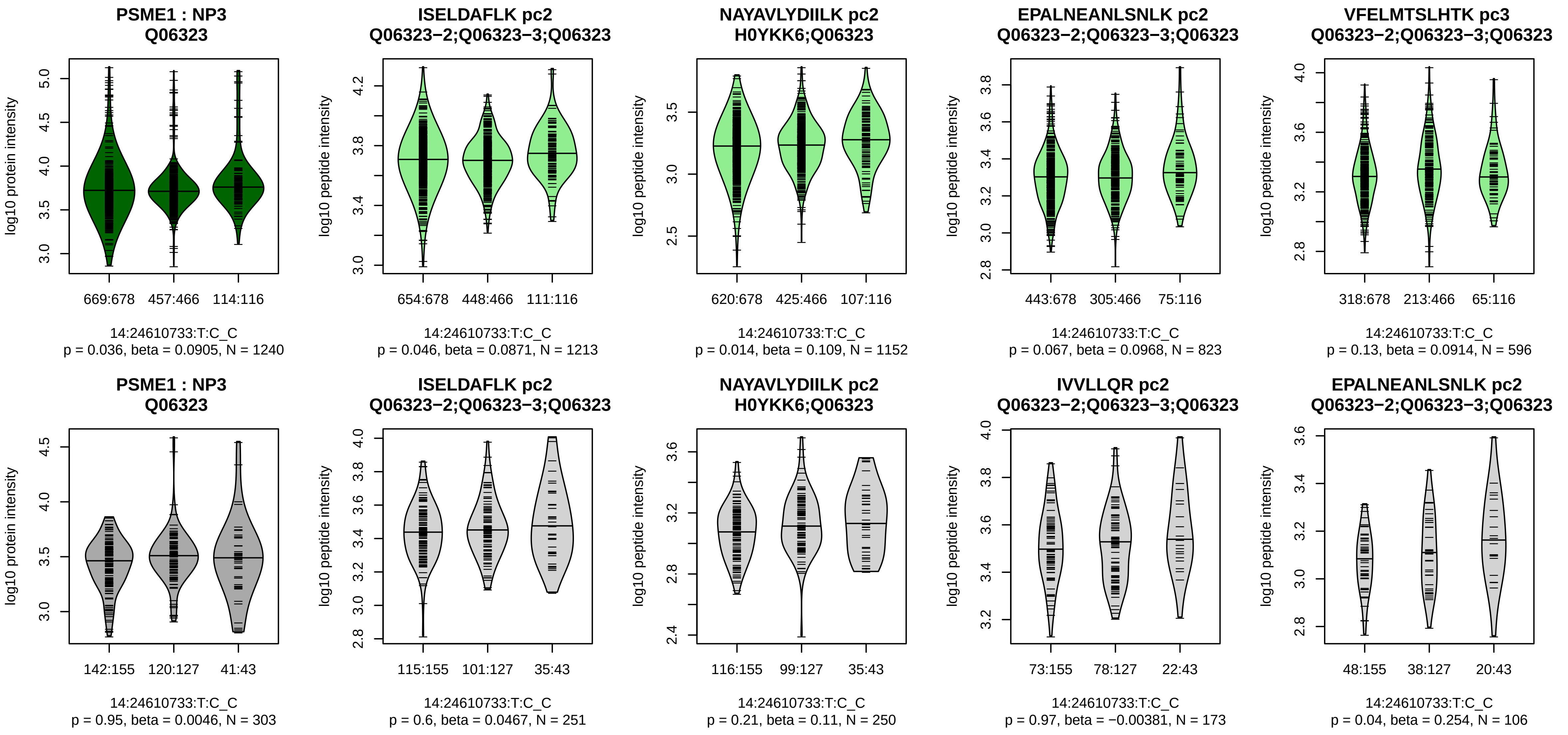


Assay Target: PSME1
Olink UniProt: Q06323
UKB rsID: rs2236352
Proxy rsID: rs2236352
UKB SNP: 14:24610733:T:C
Proxy SNP: 14:24610733:T:C
UKB log10(p): 47.5
UKB BETA: 0.1
−:−:−:−:*:−:−:−:−:−:−:−:−:−:−:NA
1213:1152:823:596:552:510:502

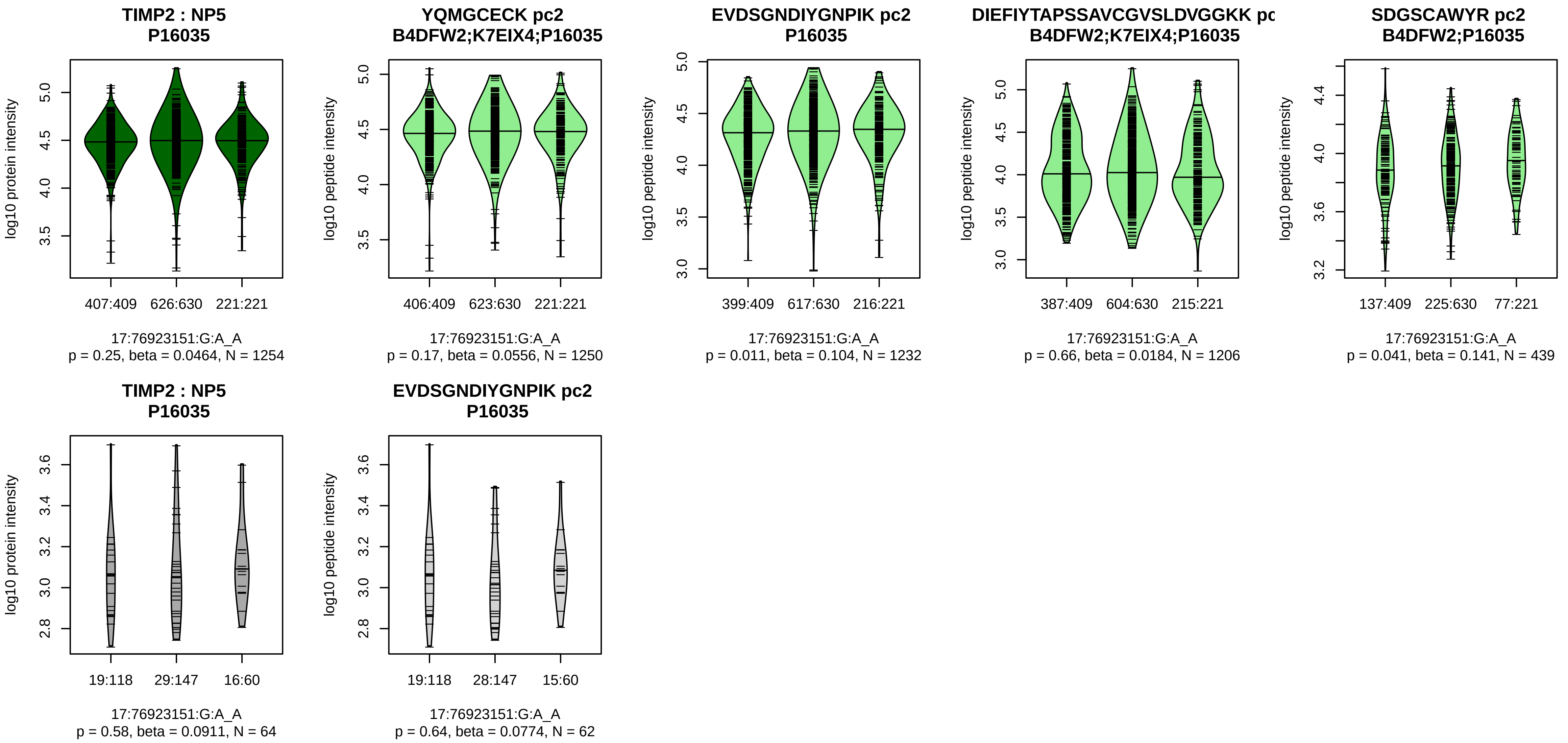


Assay Target: TIMP2
Olink UniProt: P16035
UKB rsID: rs7405712
Proxy rsID: rs7405712
UKB SNP: 17:76923151:G:A
Proxy SNP: 17:76923151:G:A
UKB log10(p): 46.6
UKB BETA: 0.089
−:−:−:−:−:−:−:NA
1250:1232:1206:439:237:34:24:

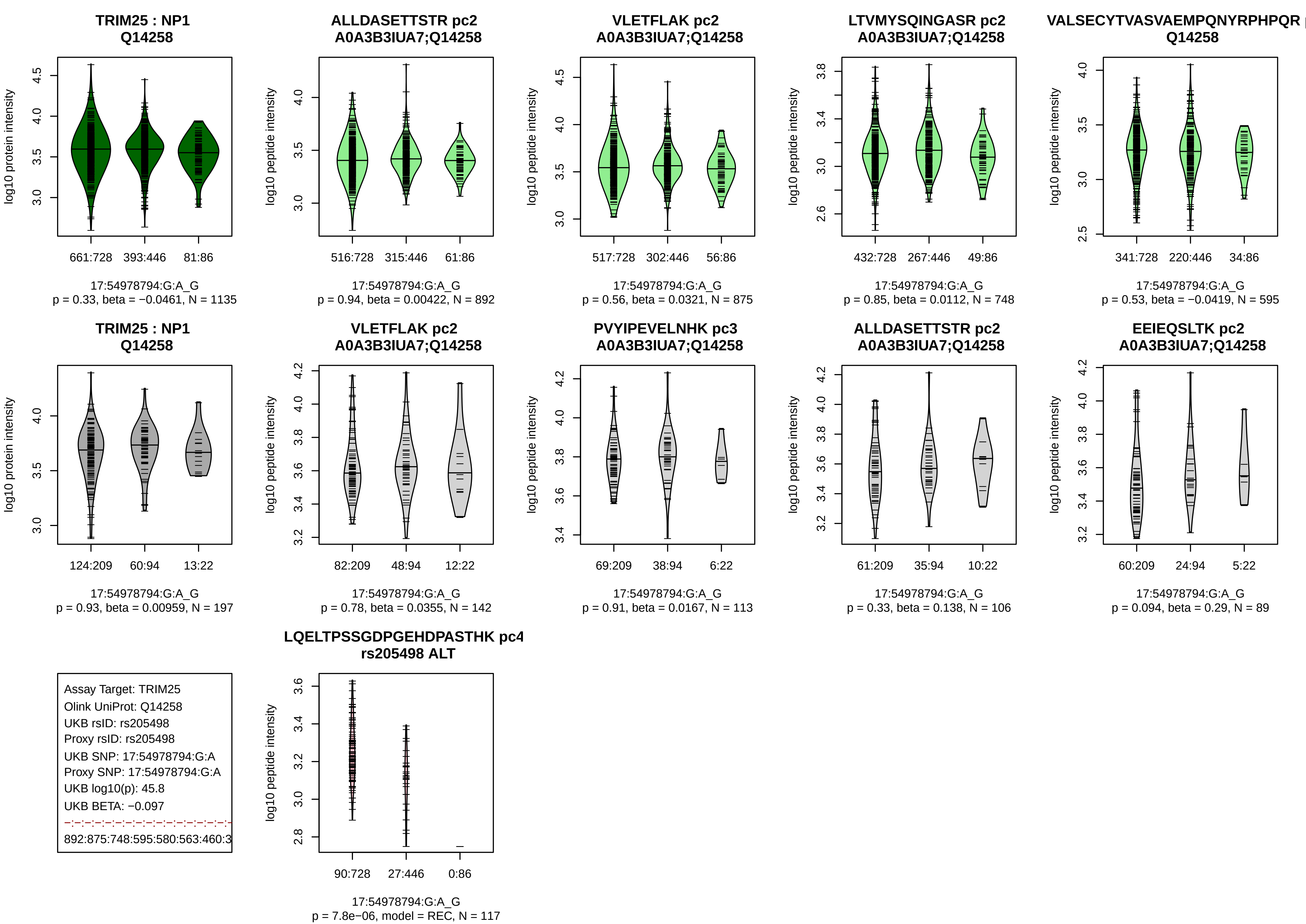
TRIM25 : NP1
Q14258
log10 protein intensity
661:728
393:446
81:86
17:54978794:G:A_G
p = 0.33, beta = −0.0461, N = 1135
ALLDASETTSTR pc2
A0A3B3IUA7;Q14258
log10 peptide intensity
516:728
315:446
61:86
17:54978794:G:A_G
p = 0.94, beta = 0.00422, N = 892
VLETFLAK pc2
A0A3B3IUA7;Q14258
log10 peptide intensity
517:728
302:446
56:86
17:54978794:G:A_G
p = 0.56, beta = 0.0321, N = 875
LTVMYSQINGASR pc2
A0A3B3IUA7;Q14258
log10 peptide intensity
432:728
267:446
49:86
17:54978794:G:A_G
p = 0.85, beta = 0.0112, N = 748
VALSECYTVASVAEMPQNYRPHPQR
Q14258
log10 peptide intensity
341:728
220:446
34:86
17:54978794:G:A_G
p = 0.53, beta = −0.0419, N = 595
TRIM25 : NP1
Q14258
log10 protein intensity
124:209
60:94
13:22
17:54978794:G:A_G
p = 0.93, beta = 0.00959, N = 197
VLETFLAK pc2
A0A3B3IUA7;Q14258
log10 peptide intensity
82:209
48:94
12:22
17:54978794:G:A_G
p = 0.78, beta = 0.0355, N = 142
PVYIPEVELNHK pc3
A0A3B3IUA7;Q14258
log10 peptide intensity
69:209
38:94
6:22
17:54978794:G:A_G
p = 0.91, beta = 0.0167, N = 113
ALLDASETTSTR pc2
A0A3B3IUA7;Q14258
log10 peptide intensity
61:209
35:94
10:22
17:54978794:G:A_G
p = 0.33, beta = 0.138, N = 106
EEIEQSLTK pc2
A0A3B3IUA7;Q14258
log10 peptide intensity
60:209
24:94
5:22
17:54978794:G:A_G
p = 0.094, beta = 0.29, N = 89
Assay Target: TRIM25
Olink UniProt: Q14258
UKB rsID: rs205498
Proxy rsID: rs205498
UKB SNP: 17:54978794:G:A
Proxy SNP: 17:54978794:G:A
UKB log10(p): 45.8
UKB BETA: −0.097
892:875:748:595:580:563:460:3
LQELTPSSGDPGEHDPASTHK pc4
rs205498 ALT
log10 peptide intensity
90:728
27:446
0:86
17:54978794:G:A_G
p = 7.8e−06, model = REC, N = 117

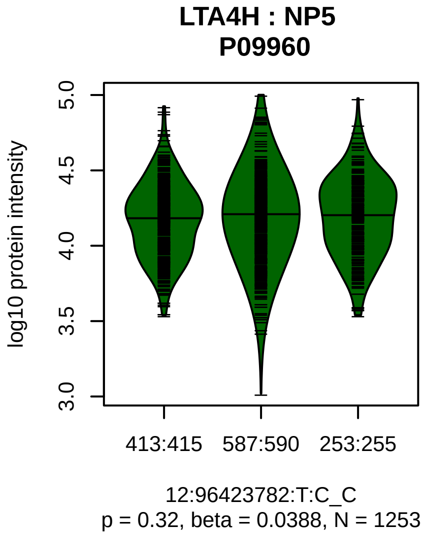


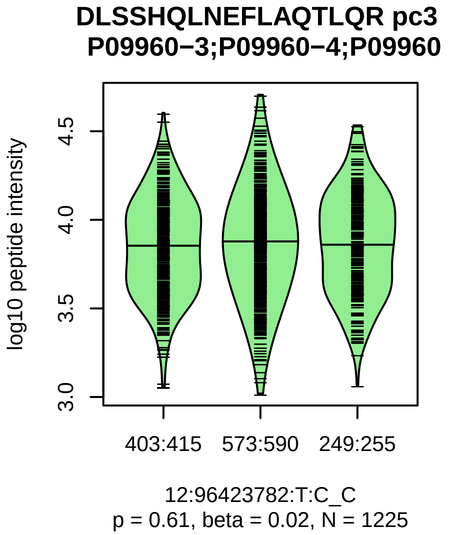


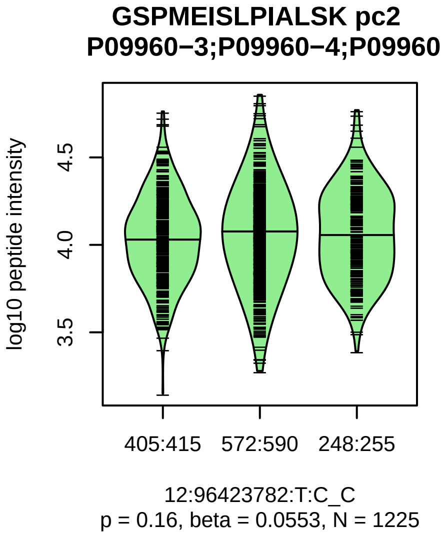


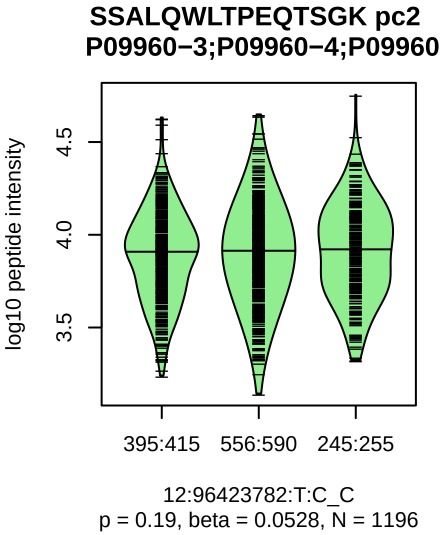


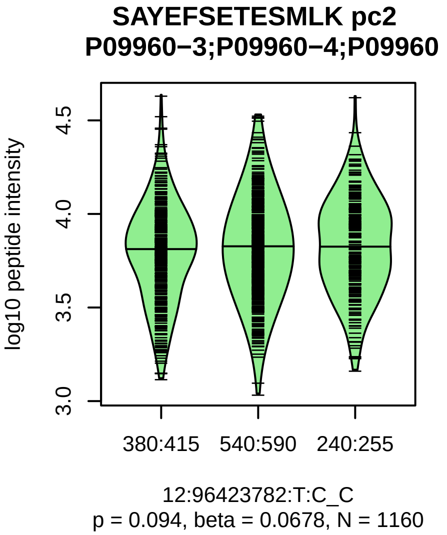


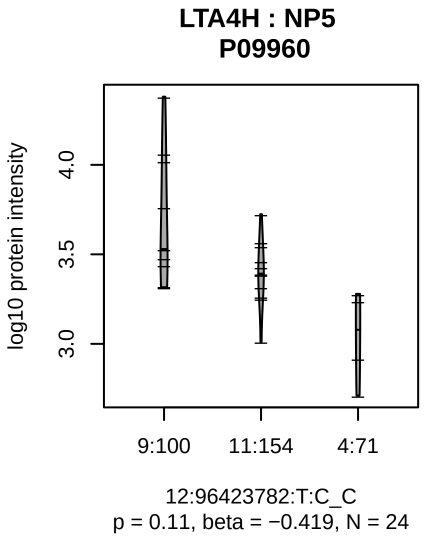


Assay Target: LTA4H
Olink UniProt: P09960
UKB rsID: rs2660896
Proxy rsID: rs2660896
UKB SNP: 12:96423782:T:C
Proxy SNP: 12:96423782:T:C
UKB log10(p): 45.7
UKB BETA: 0.087
−:−:−:−:−:−:−:−:−:−:−:−:−:−:−:−:
1225:1225:1196:1160:1067:104

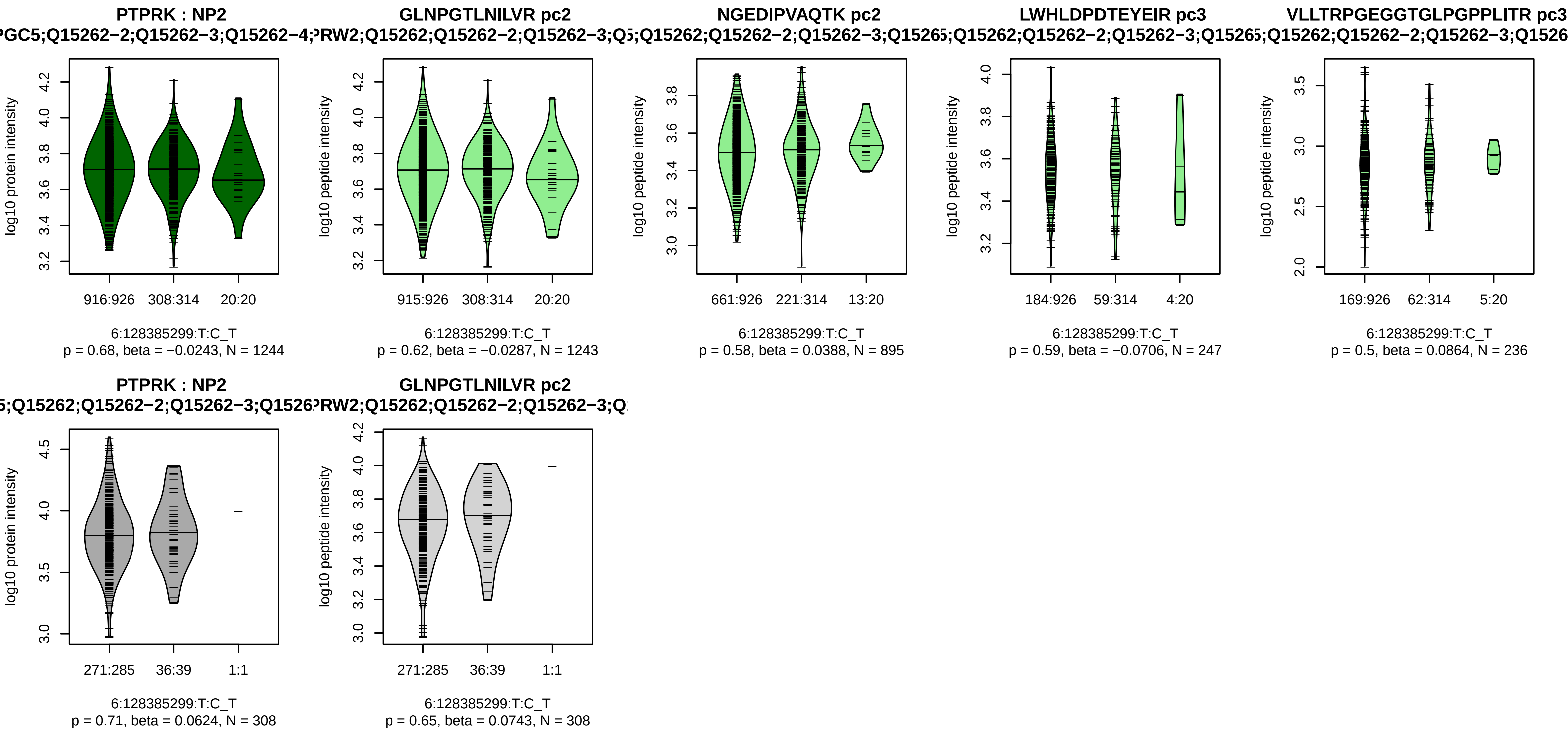


Assay Target: PTPRK
Olink UniProt: Q15262
UKB rsID: rs11422080
Proxy rsID: rs7769691
UKB SNP: 6:128383562:G:GA
Proxy SNP: 6:128385299:T:C
UKB log10(p): 44.7
UKB BETA: −0.12
−:−:−:−
1243:895:247:236

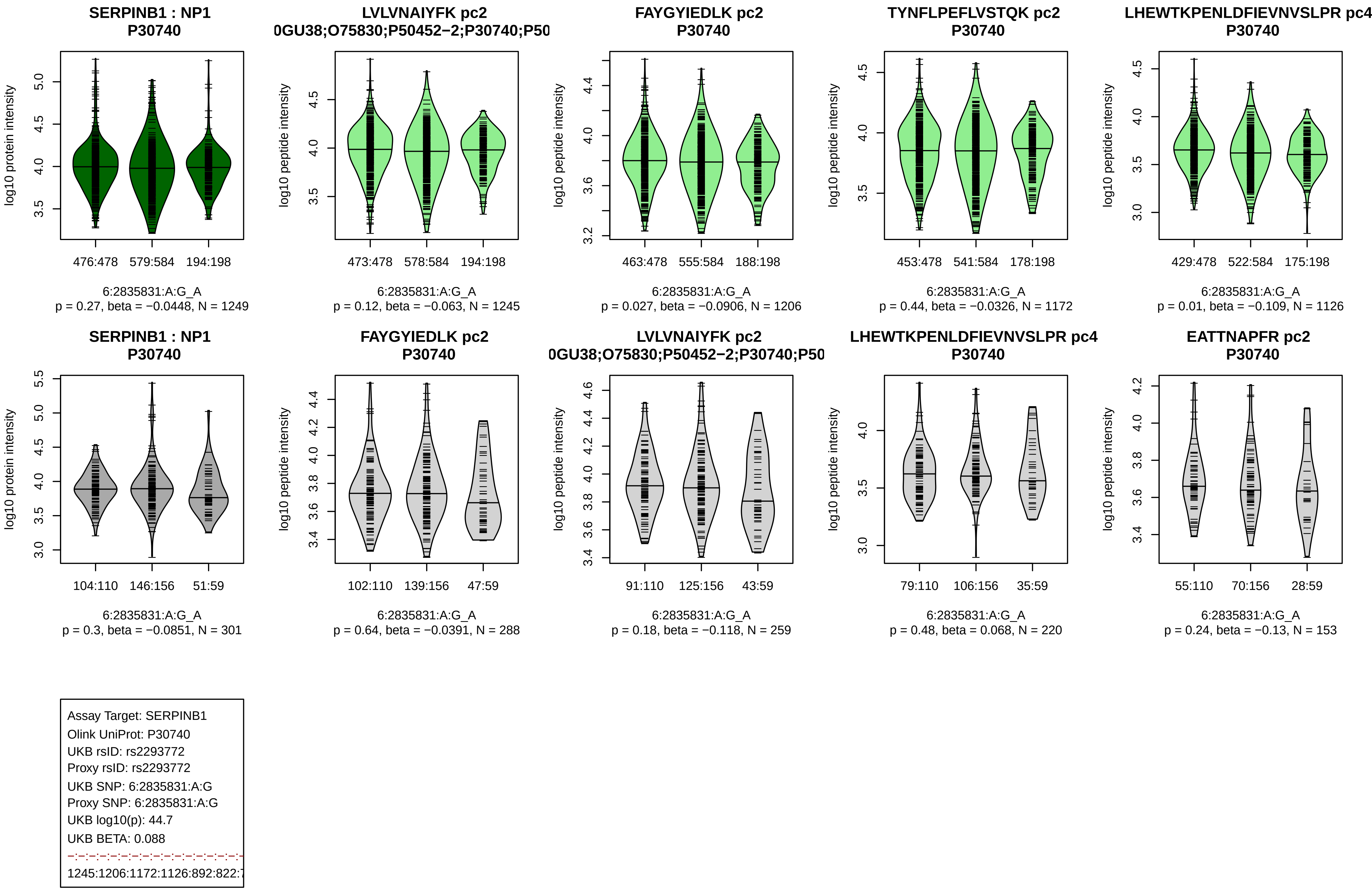
SERPINB1 : NP1
P30740
log10 protein intensity
476:478
579:584
194:198
6:2835831:A:G_A
p = 0.27, beta = −0.0448, N = 1249
LVLVNAIYFK pc2
0GU38;O75830;P50452−2;P30740;P50
log10 peptide intensity
473:478
578:584
194:198
6:2835831:A:G_A
p = 0.12, beta = −0.063, N = 1245
FAYGYIEDLK pc2
P30740
log10 peptide intensity
463:478
555:584
188:198
6:2835831:A:G_A
p = 0.027, beta = −0.0906, N = 1206
TYNFLPEFLVSTQK pc2
P30740
log10 peptide intensity
453:478
541:584
178:198
6:2835831:A:G_A
p = 0.44, beta = −0.0326, N = 1172
LHEWTKPENLDFIEVNVSLPR pc4
P30740
log10 peptide intensity
429:478
522:584
175:198
6:2835831:A:G_A
p = 0.01, beta = −0.109, N = 1126
SERPINB1 : NP1
P30740
log10 protein intensity
104:110
146:156
51:59
6:2835831:A:G_A
p = 0.3, beta = −0.0851, N = 301
FAYGYIEDLK pc2
P30740
log10 peptide intensity
102:110
139:156
47:59
6:2835831:A:G_A
p = 0.64, beta = −0.0391, N = 288
LVLVNAIYFK pc2
0GU38;O75830;P50452−2;P30740;P50
log10 peptide intensity
91:110
125:156
43:59
6:2835831:A:G_A
p = 0.18, beta = −0.118, N = 259
LHEWTKPENLDFIEVNVSLPR pc4
P30740
log10 peptide intensity
79:110
106:156
35:59
6:2835831:A:G_A
p = 0.48, beta = 0.068, N = 220
EATTNAPFR pc2
P30740
log10 peptide intensity
55:110
70:156
28:59
6:2835831:A:G_A
p = 0.24, beta = −0.13, N = 153
Assay Target: SERPINB1
Olink UniProt: P30740
UKB rsID: rs2293772
Proxy rsID: rs2293772
UKB SNP: 6:2835831:A:G
Proxy SNP: 6:2835831:A:G
UKB log10(p): 44.7
UKB BETA: 0.088
1245:1206:1172:1126:892:822:7

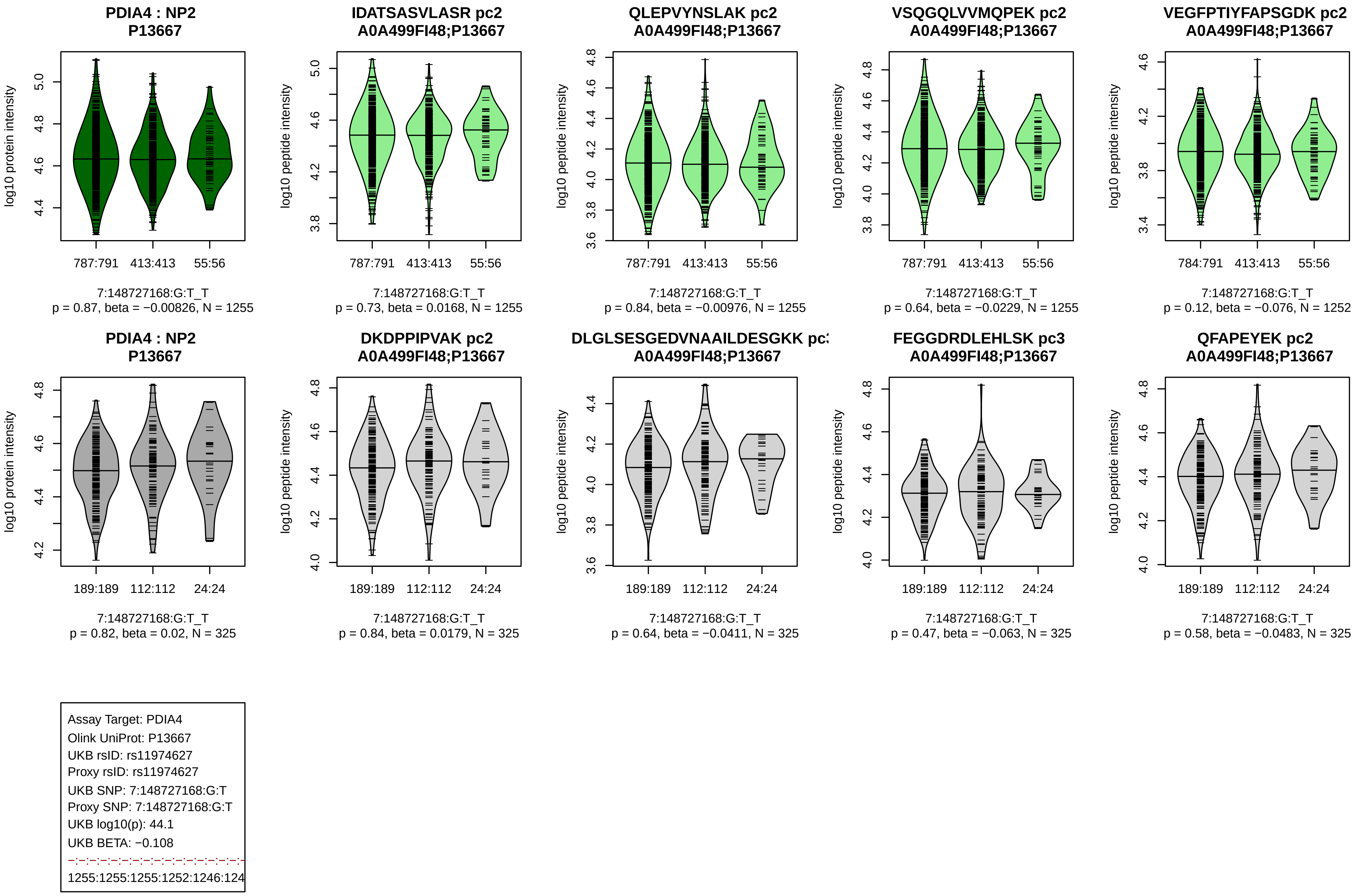
PDIA4 : NP2
P13667
log10 protein intensity
787:791 413:413 55:56
7:148727168:G:T_T
p = 0.87, beta = −0.00826, N = 1255
IDATSASVLASR pc2
A0A499FI48;P13667
log10 peptide intensity
787:791 413:413 55:56
7:148727168:G:T_T
p = 0.73, beta = 0.0168, N = 1255
QLEPVYNSLAK pc2
A0A499FI48;P13667
log10 peptide intensity
787:791 413:413 55:56
7:148727168:G:T_T
p = 0.84, beta = −0.00976, N = 1255
VSQGQLVVMQPEK pc2
A0A499FI48;P13667
log10 peptide intensity
787:791 413:413 55:56
7:148727168:G:T_T
p = 0.64, beta = −0.0229, N = 1255
VEGFPTIYFAPSGDK pc2
A0A499FI48;P13667
log10 peptide intensity
784:791 413:413 55:56
7:148727168:G:T_T
p = 0.12, beta = −0.076, N = 1252
PDIA4 : NP2
P13667
log10 protein intensity
189:189 112:112 24:24
7:148727168:G:T_T
p = 0.82, beta = 0.02, N = 325
DKDPPIPVAK pc2
A0A499FI48;P13667
log10 peptide intensity
189:189 112:112 24:24
7:148727168:G:T_T
p = 0.84, beta = 0.0179, N = 325
DLGLSESGEDVNAAILDESGKK pc
A0A499FI48;P13667
log10 peptide intensity
189:189 112:112 24:24
7:148727168:G:T_T
p = 0.64, beta = −0.0411, N = 325
FEGGDRDLEHLSK pc3
A0A499FI48;P13667
log10 peptide intensity
189:189 112:112 24:24
7:148727168:G:T_T
p = 0.47, beta = −0.063, N = 325
QFAPEYEK pc2
A0A499FI48;P13667
log10 peptide intensity
189:189 112:112 24:24
7:148727168:G:T_T
p = 0.58, beta = −0.0483, N = 325
Assay Target: PDIA4
Olink UniProt: P13667
UKB rsID: rs11974627
Proxy rsID: rs11974627
UKB SNP: 7:148727168:G:T
Proxy SNP: 7:148727168:G:T
UKB log10(p): 44.1
UKB BETA: −0.108
1255:1255:1255:1252:1246:124

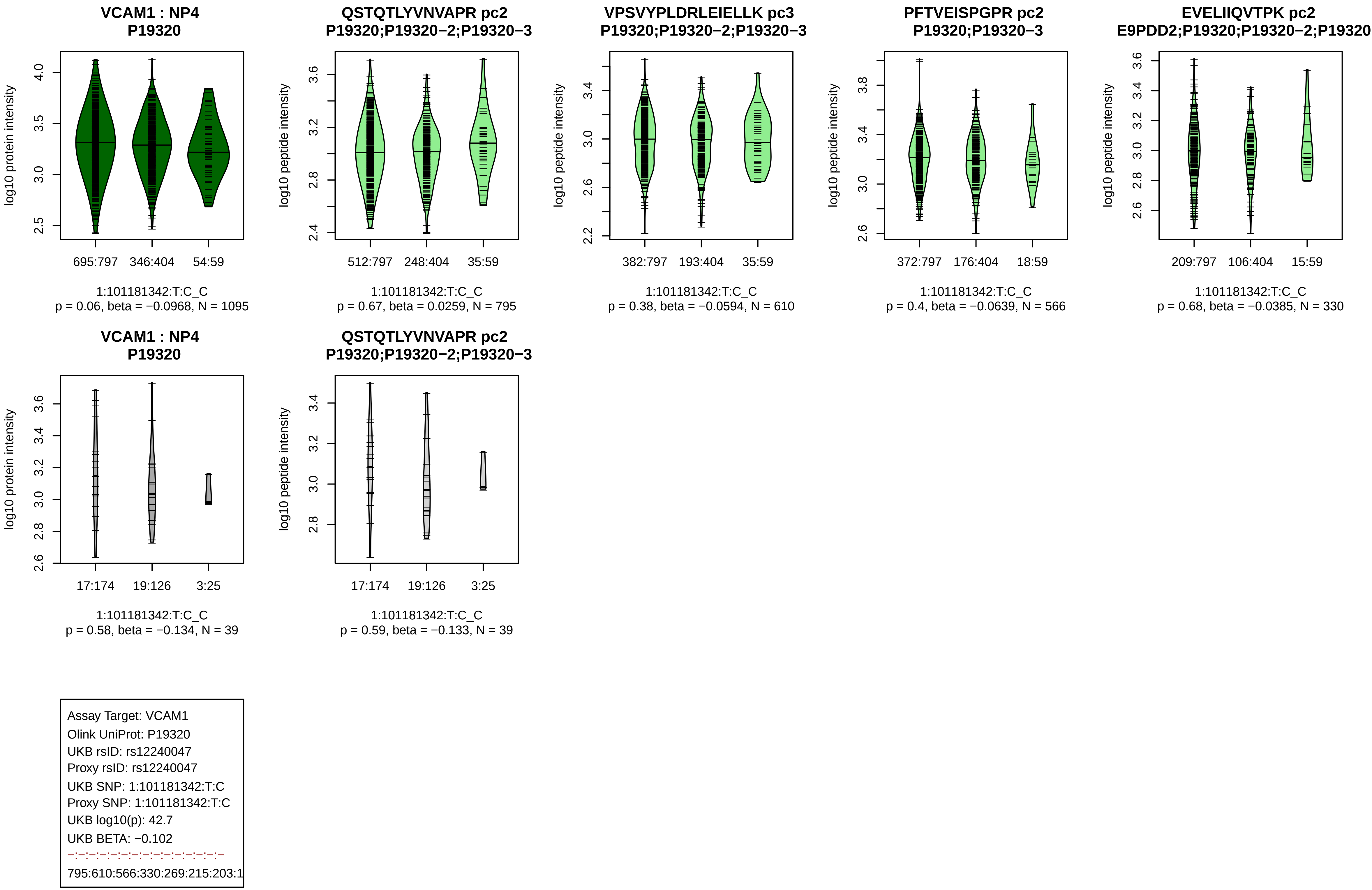

VCAM1 : NP4
P19320
log10 protein intensity
695:797
346:404
54:59
1:101181342:T:C_C
p = 0.06, beta = −0.0968, N = 1095
QSTQTLYVNVAPR pc2
P19320;P19320−2;P19320−3
log10 peptide intensity
512:797
248:404
35:59
1:101181342:T:C_C
p = 0.67, beta = 0.0259, N = 795
VPSVYPLDRLEIELLK pc3
P19320;P19320−2;P19320−3
log10 peptide intensity
382:797
193:404
35:59
1:101181342:T:C_C
p = 0.38, beta = −0.0594, N = 610
PFTVEISPGPR pc2
P19320;P19320−3
log10 peptide intensity
372:797
176:404
18:59
1:101181342:T:C_C
p = 0.4, beta = −0.0639, N = 566
EVELIIQVTPK pc2
E9PDD2;P19320;P19320−2;P19320
log10 peptide intensity
209:797
106:404
15:59
1:101181342:T:C_C
p = 0.68, beta = −0.0385, N = 330
VCAM1 : NP4
P19320
log10 protein intensity
17:174
19:126
3:25
1:101181342:T:C_C
p = 0.58, beta = −0.134, N = 39
QSTQTLYVNVAPR pc2
P19320;P19320−2;P19320−3
log10 peptide intensity
17:174
19:126
3:25
1:101181342:T:C_C
p = 0.59, beta = −0.133, N = 39
Assay Target: VCAM1
Olink UniProt: P19320
UKB rsID: rs12240047
Proxy rsID: rs12240047
UKB SNP: 1:101181342:T:C
Proxy SNP: 1:101181342:T:C
UKB log10(p): 42.7
UKB BETA: −0.102
−:−:−:−:−:−:−:−:−:−:−:−:−:−
795:610:566:330:269:215:203:1

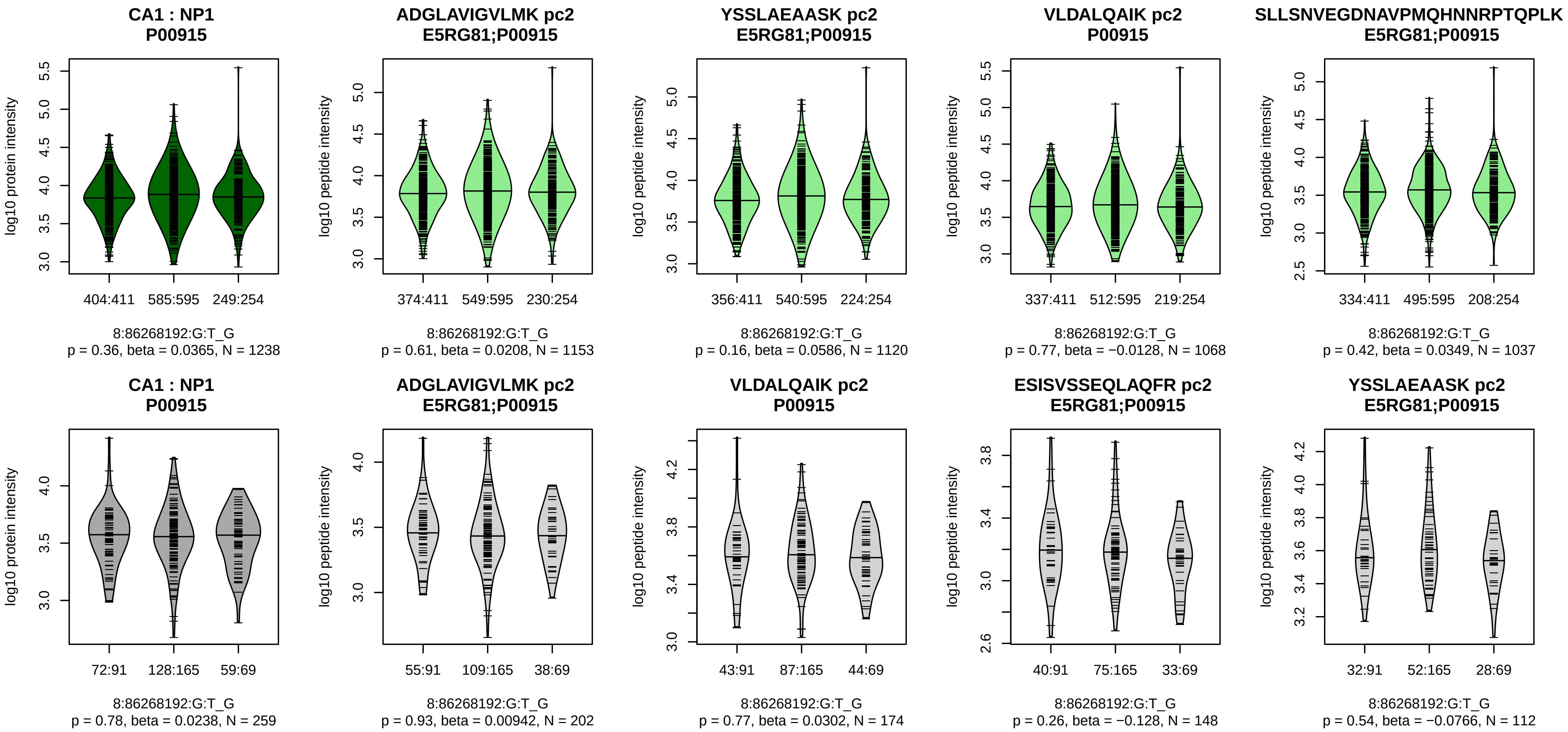


Assay Target: CA1
Olink UniProt: P00915
UKB rsID: rs1532424
Proxy rsID: rs1532424
UKB SNP: 8:86268192:G:T
Proxy SNP: 8:86268192:G:T
UKB log10(p): 41.7
UKB BETA: −0.081
−:−:−:−:−:−:−:−:−:−:−:NA:NA:NA
1153:1120:1068:1037:927:894:8

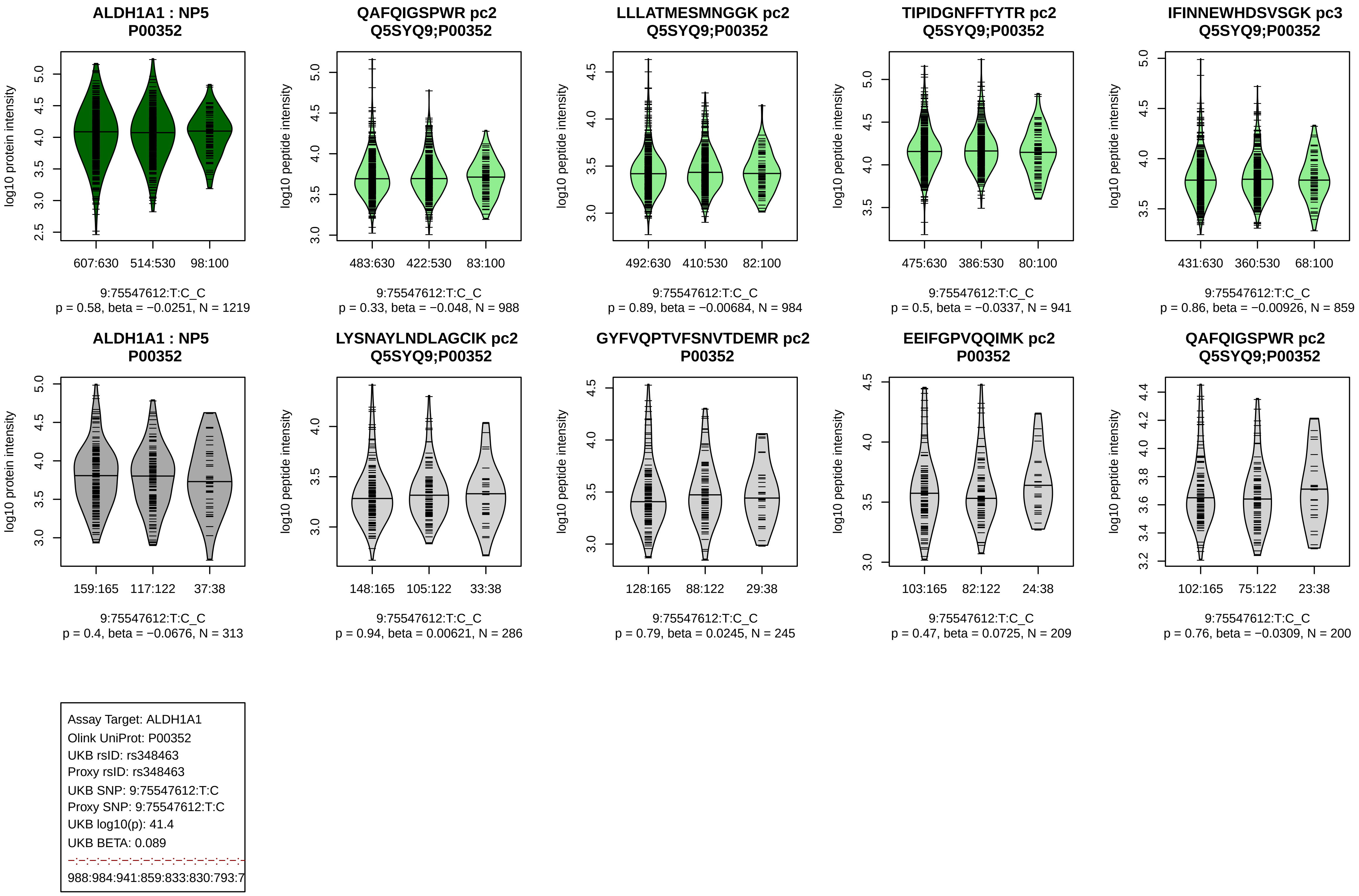
ALDH1A1 : NP5
P00352
log10 protein intensity
607:630
514:530
98:100
9:75547612:T:C_C
p = 0.58, beta = −0.0251, N = 1219
QAFQIGSPWR pc2
Q5SYQ9;P00352
log10 peptide intensity
483:630
422:530
83:100
9:75547612:T:C_C
p = 0.33, beta = −0.048, N = 988
LLLATMESMNGGK pc2
Q5SYQ9;P00352
log10 peptide intensity
492:630
410:530
82:100
9:75547612:T:C_C
p = 0.89, beta = −0.00684, N = 984
TIPIDGNFFTYTR pc2
Q5SYQ9;P00352
log10 peptide intensity
475:630
386:530
80:100
9:75547612:T:C_C
p = 0.5, beta = −0.0337, N = 941
IFINNEWHDSVSGK pc3
Q5SYQ9;P00352
log10 peptide intensity
431:630
360:530
68:100
9:75547612:T:C_C
p = 0.86, beta = −0.00926, N = 859
ALDH1A1 : NP5
P00352
log10 protein intensity
159:165
117:122
37:38
9:75547612:T:C_C
p = 0.4, beta = −0.0676, N = 313
LYSNAYLNDLAGCIK pc2
Q5SYQ9;P00352
log10 peptide intensity
148:165
105:122
33:38
9:75547612:T:C_C
p = 0.94, beta = 0.00621, N = 286
GYFVQPTVFSNVTDEMR pc2
P00352
log10 peptide intensity
128:165
88:122
29:38
9:75547612:T:C_C
p = 0.79, beta = 0.0245, N = 245
EEIFGPVQQIMK pc2
P00352
log10 peptide intensity
103:165
82:122
24:38
9:75547612:T:C_C
p = 0.47, beta = 0.0725, N = 209
QAFQIGSPWR pc2
Q5SYQ9;P00352
log10 peptide intensity
102:165
75:122
23:38
9:75547612:T:C_C
p = 0.76, beta = −0.0309, N = 200
Assay Target: ALDH1A1
Olink UniProt: P00352
UKB rsID: rs348463
Proxy rsID: rs348463
UKB SNP: 9:75547612:T:C
Proxy SNP: 9:75547612:T:C
UKB log10(p): 41.4
UKB BETA: 0.089
988:984:941:859:833:830:793:7

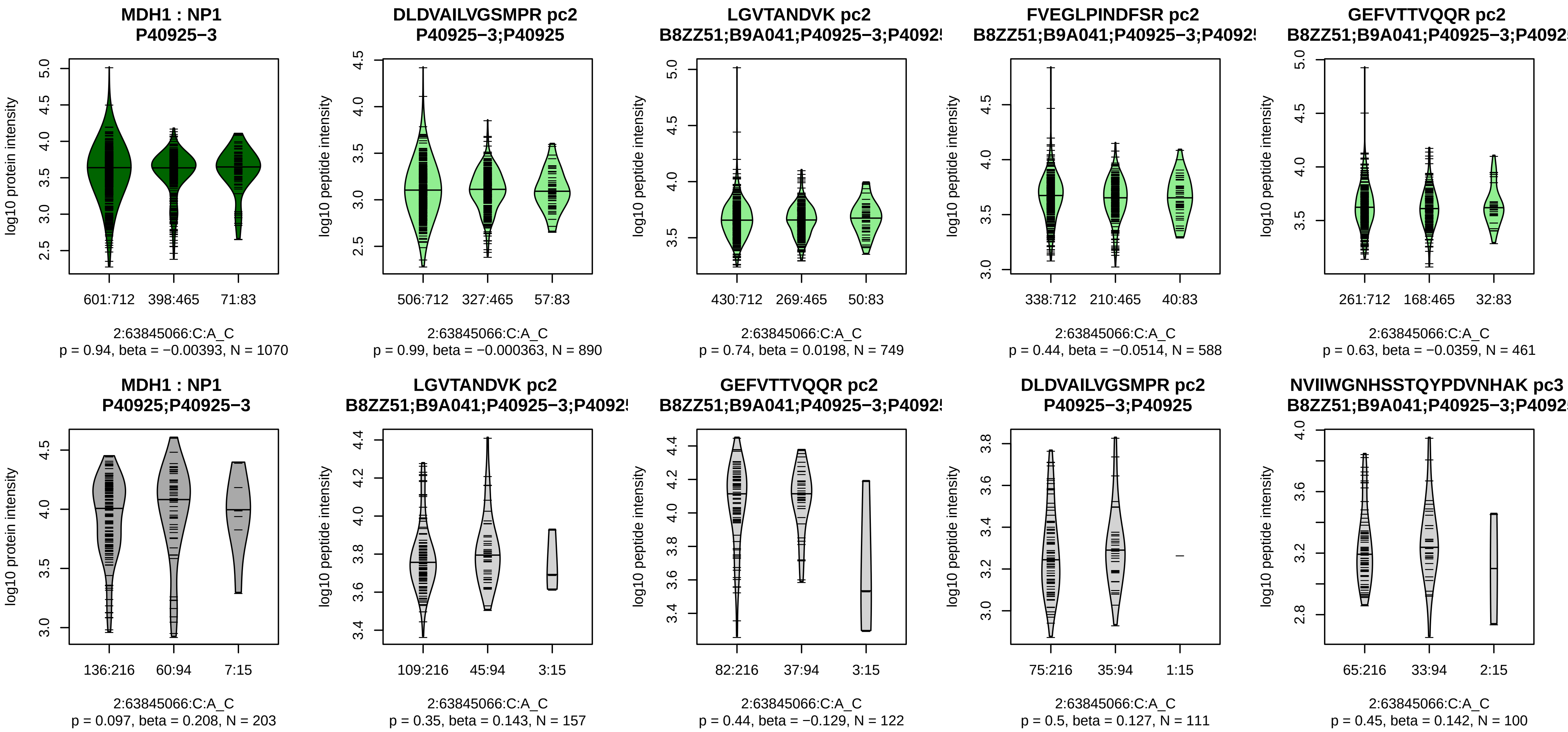


Assay Target: MDH1
Olink UniProt: P40925
UKB rsID: rs262493
Proxy rsID: rs262493
UKB SNP: 2:63845066:C:A
Proxy SNP: 2:63845066:C:A
UKB log10(p): 41.3
UKB BETA: −0.092
−:−:−:−:−:−:−
890:749:588:461:411:152:111

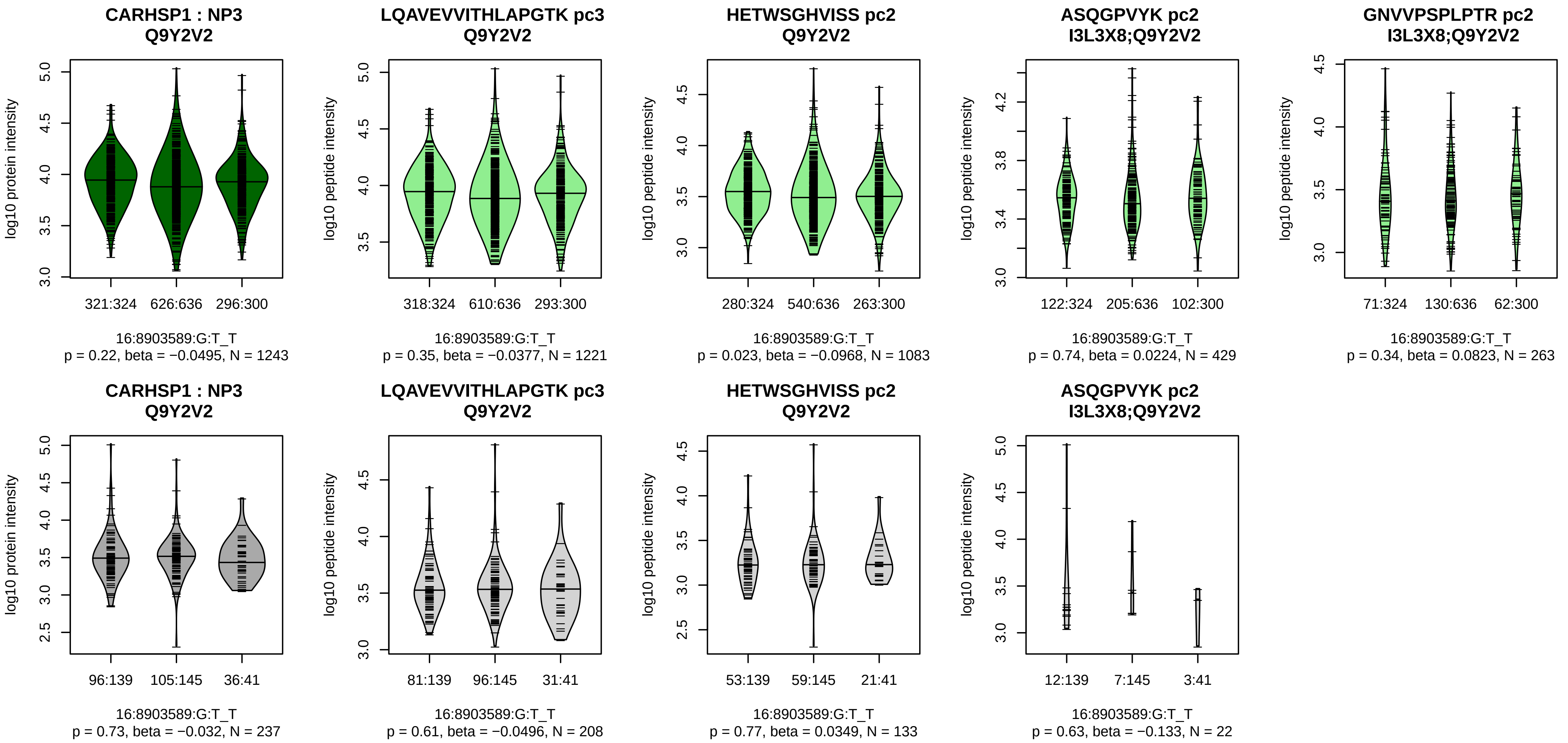


Assay Target: CARHSP1
Olink UniProt: Q9Y2V2
UKB rsID: rs12597479
Proxy rsID: rs12597479
UKB SNP: 16:8903589:G:T
Proxy SNP: 16:8903589:G:T
UKB log10(p): 40.5
UKB BETA: −0.081
−:−:−:−:−
1221:1083:429:338:263

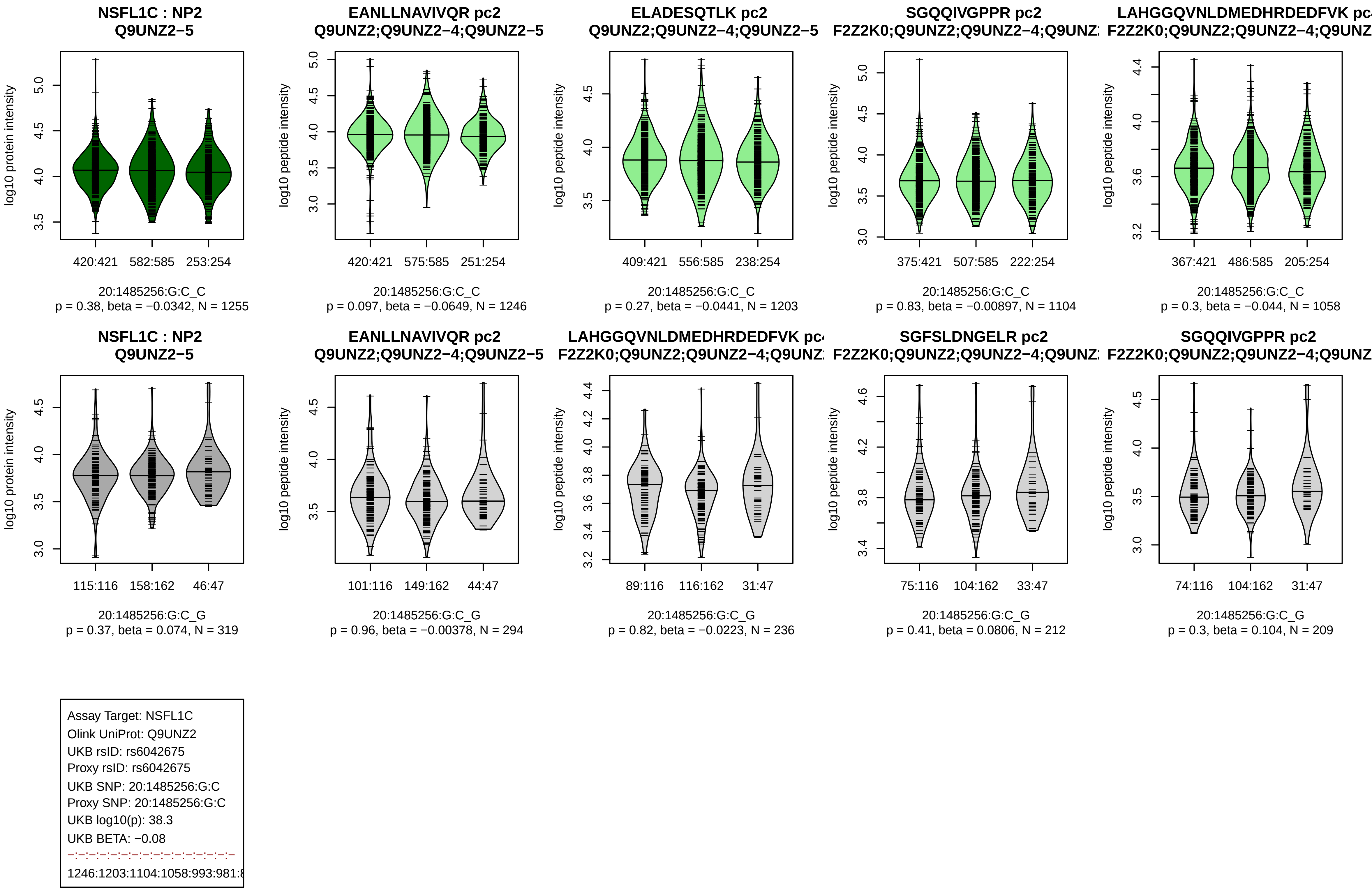
NSFL1C : NP2
Q9UNZ2−5
log10 protein intensity
420:421 582:585 253:254
20:1485256:G:C_C
p = 0.38, beta = −0.0342, N = 1255
EANLLNAVIVQR pc2
Q9UNZ2;Q9UNZ2−4;Q9UNZ2−5
log10 peptide intensity
420:421 575:585 251:254
20:1485256:G:C_C
p = 0.097, beta = −0.0649, N = 1246
ELADESQTLK pc2
Q9UNZ2;Q9UNZ2−4;Q9UNZ2−5
409:421 556:585 238:254
20:1485256:G:C_C
p = 0.27, beta = −0.0441, N = 1203
SGQQIVGPPR pc2
375:421 507:585 222:254
20:1485256:G:C_C
p = 0.83, beta = −0.00897, N = 1104
367:421 486:585 205:254
20:1485256:G:C_C
p = 0.3, beta = −0.044, N = 1058
NSFL1C : NP2
Q9UNZ2−5
115:116 158:162 46:47
20:1485256:G:C_G
p = 0.37, beta = 0.074, N = 319
EANLLNAVIVQR pc2
Q9UNZ2;Q9UNZ2−4;Q9UNZ2−5
101:116 149:162 44:47
20:1485256:G:C_G
p = 0.96, beta = −0.00378, N = 294
89:116 116:162 31:47
20:1485256:G:C_G
p = 0.82, beta = −0.0223, N = 236
SGFSLDNGELR pc2
75:116 104:162 33:47
20:1485256:G:C_G
p = 0.41, beta = 0.0806, N = 212
SGQQIVGPPR pc2
74:116 104:162 31:47
20:1485256:G:C_G
p = 0.3, beta = 0.104, N = 209
Assay Target: NSFL1C
Olink UniProt: Q9UNZ2
UKB rsID: rs6042675
Proxy rsID: rs6042675
UKB SNP: 20:1485256:G:C
Proxy SNP: 20:1485256:G:C
UKB log10(p): 38.3
UKB BETA: −0.08

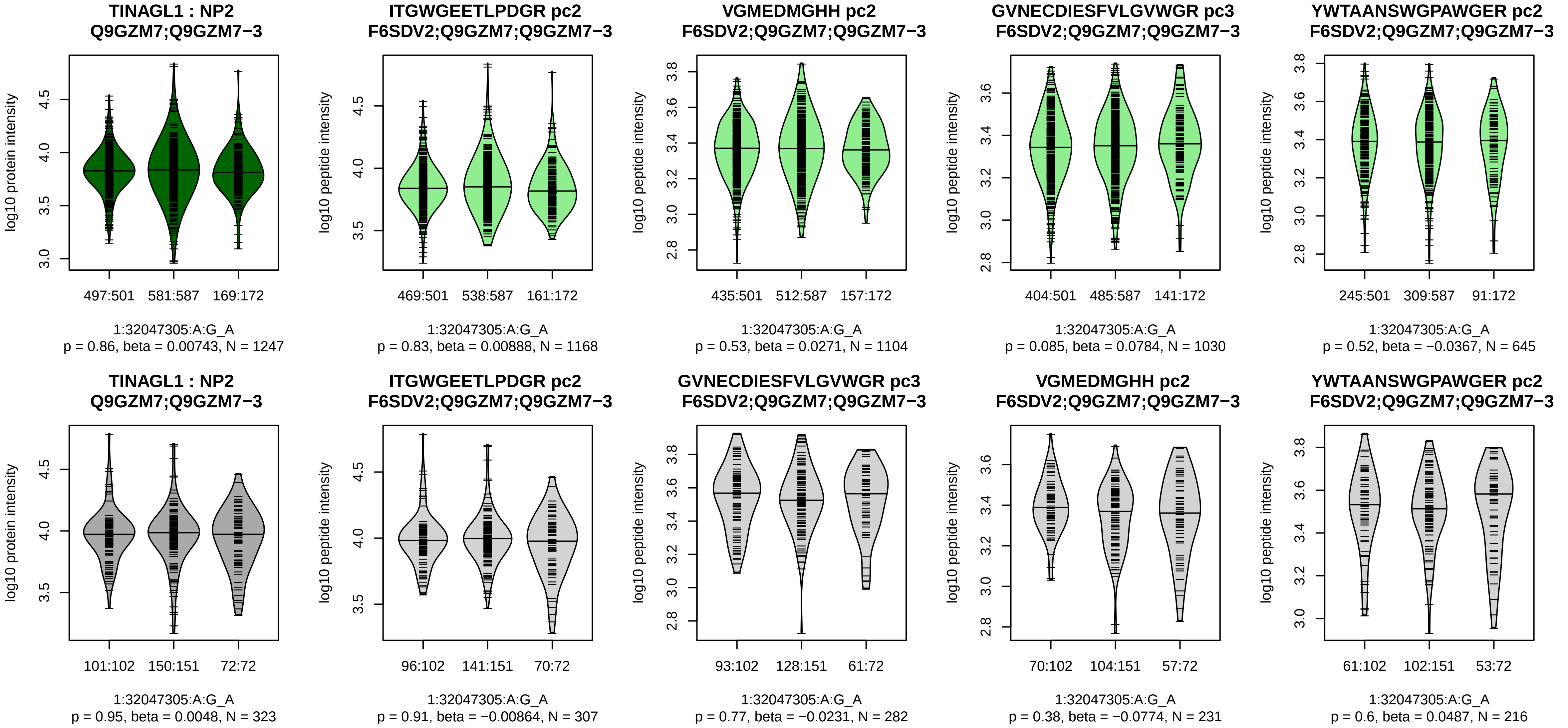


Assay Target: TINAGL1
Olink UniProt: Q9GZM7
UKB rsID: rs945214
Proxy rsID: rs945214
UKB SNP: 1:32047305:A:G
Proxy SNP: 1:32047305:A:G
UKB log10(p): 38
UKB BETA: −0.082
−:−:−:−:−:−:−:−
1168:1104:1030:645:530:427:35

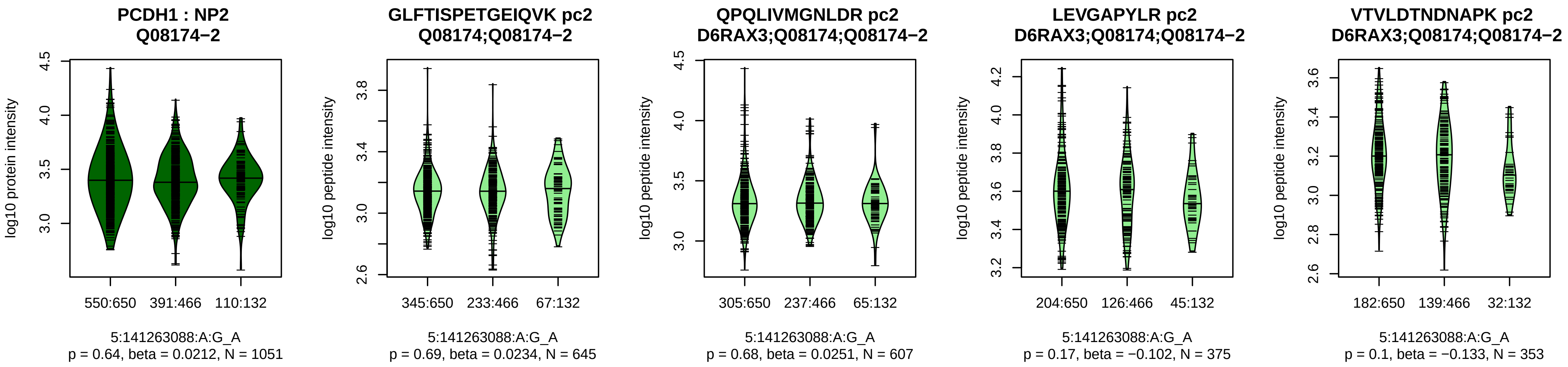


Assay Target: PCDH1
Olink UniProt: Q08174
UKB rsID: rs1124066
Proxy rsID: rs1124066
UKB SNP: 5:141263088:A:G
Proxy SNP: 5:141263088:A:G
UKB log10(p): 37
UKB BETA: 0.078
−:−:−:−:−:−:−:−:−:NA:NA
645:607:375:353:331:162:160:1

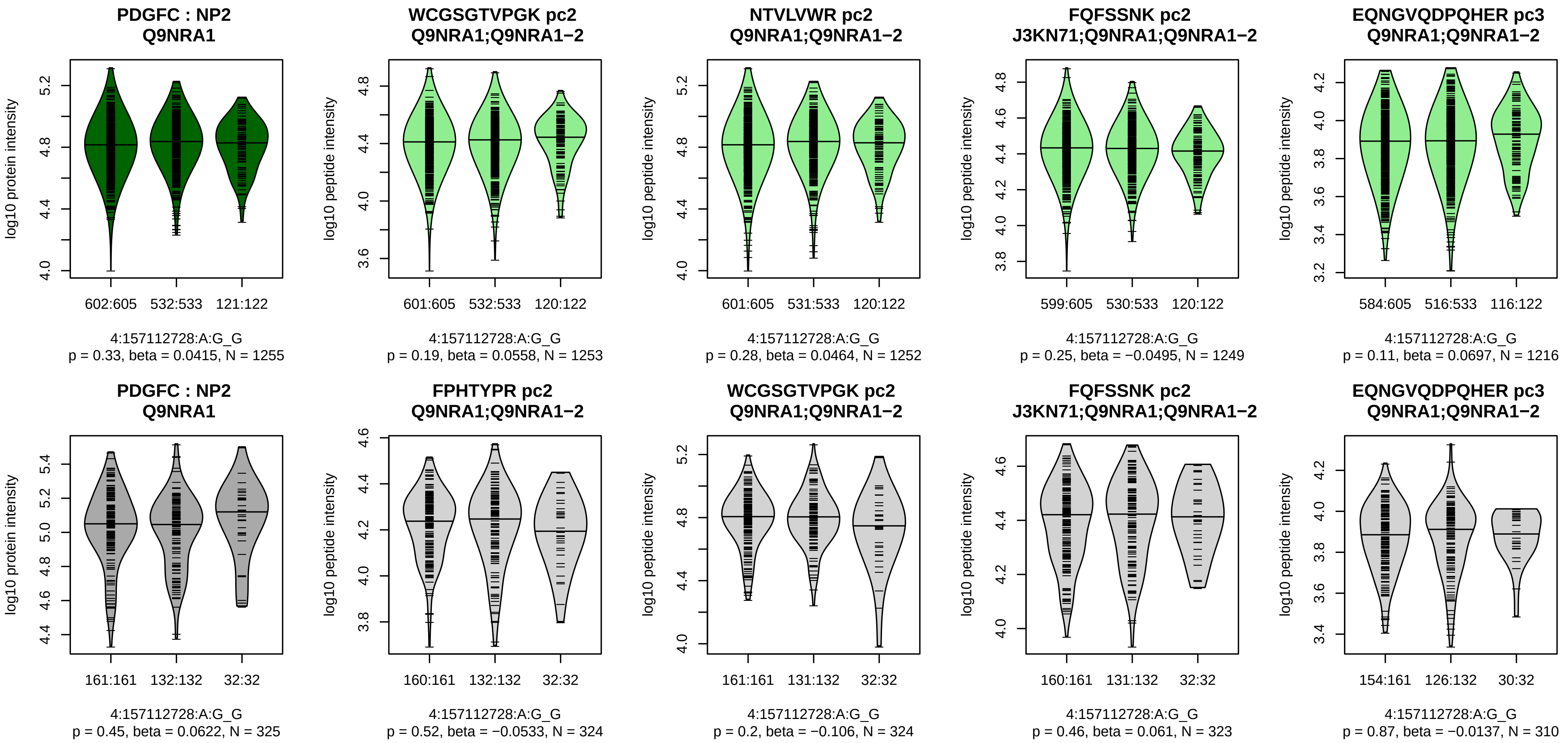


Assay Target: PDGFC
Olink UniProt: Q9NRA1
UKB rsID: rs2341218
Proxy rsID: rs2341218
UKB SNP: 4:157112728:A:G
Proxy SNP: 4:157112728:A:G
UKB log10(p): 36.6
UKB BETA: −0.086
−:−:−:−:−:−:−:−:−:−:−:−:−:NA:NA:N
1253:1252:1249:1216:1181:106

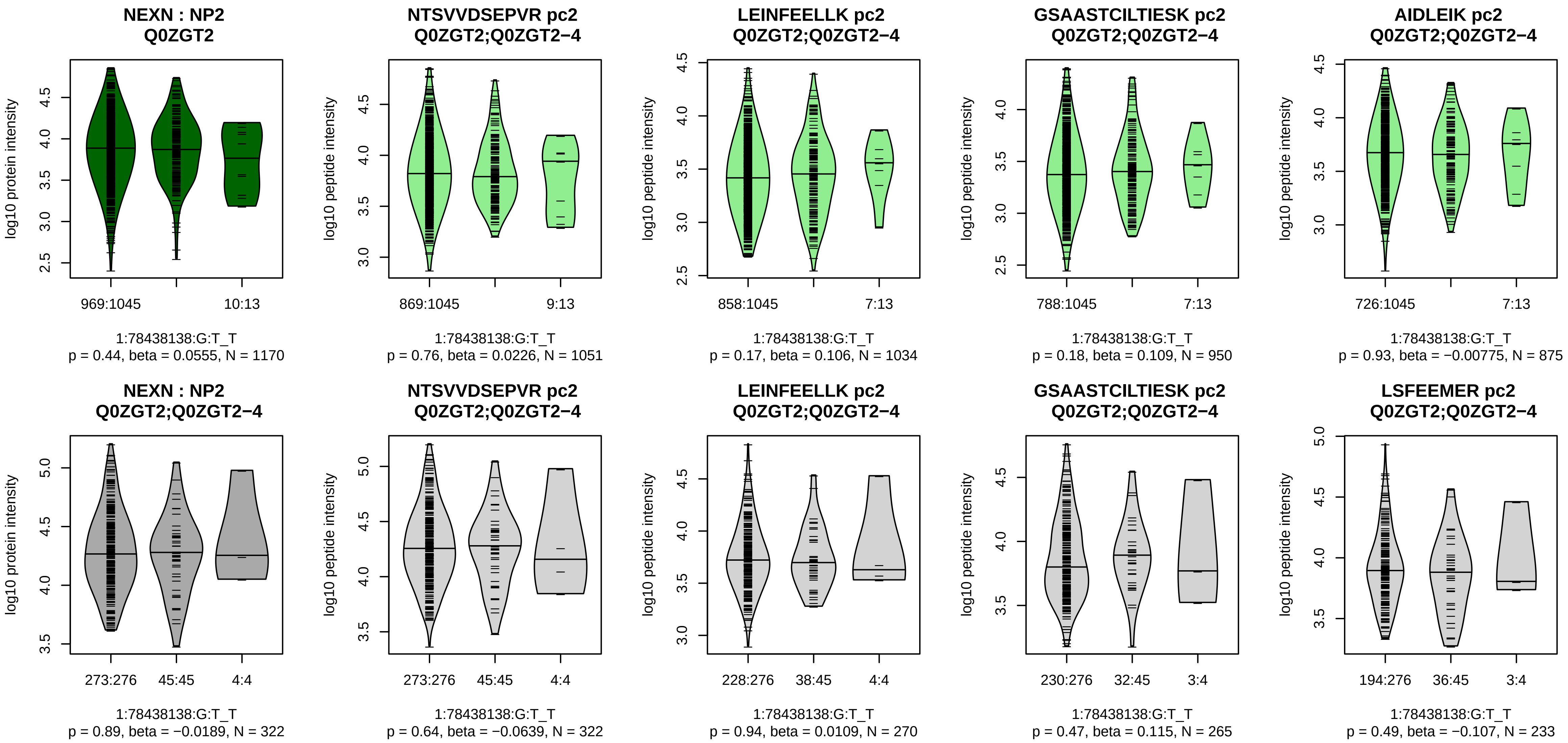


Assay Target: NEXN
Olink UniProt: Q0ZGT2
UKB rsID: rs17387365
Proxy rsID: rs17387365
UKB SNP: 1:78438138:G:T
Proxy SNP: 1:78438138:G:T
UKB log10(p): 35.4
UKB BETA: −0.143
−·:−·:−·:−·:−·:−·:−·:−·:−·:−·:−·:−·:−·:
1051:1034:950:875:837:792:564

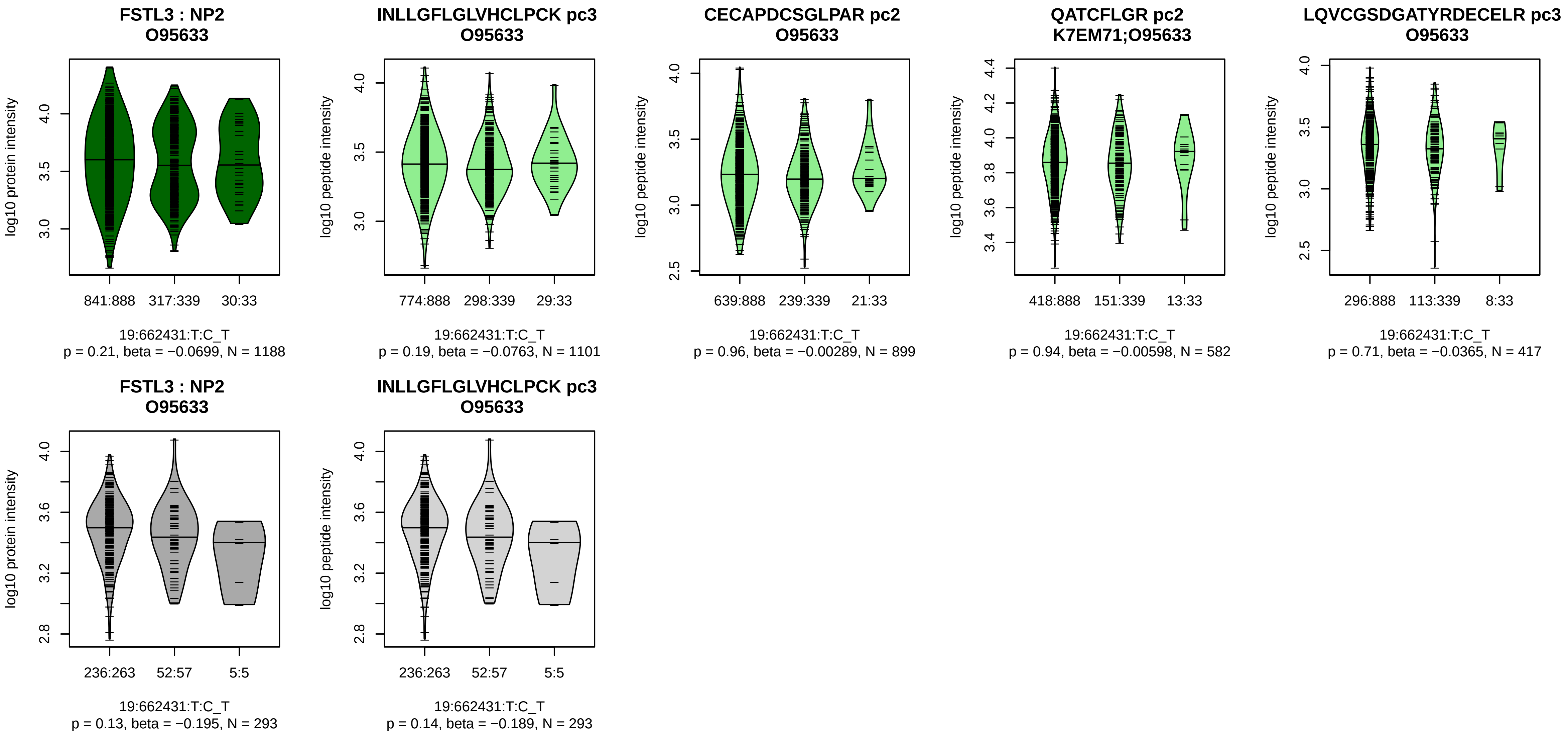


Assay Target: FSTL3
Olink UniProt: O95633
UKB rsID: rs929263
Proxy rsID: rs4919882
UKB SNP: 19:663373:C:T
Proxy SNP: 19:662431:T:C
UKB log10(p): 34.9
UKB BETA: 0.099
–:–:–:–
1101:899:582:417

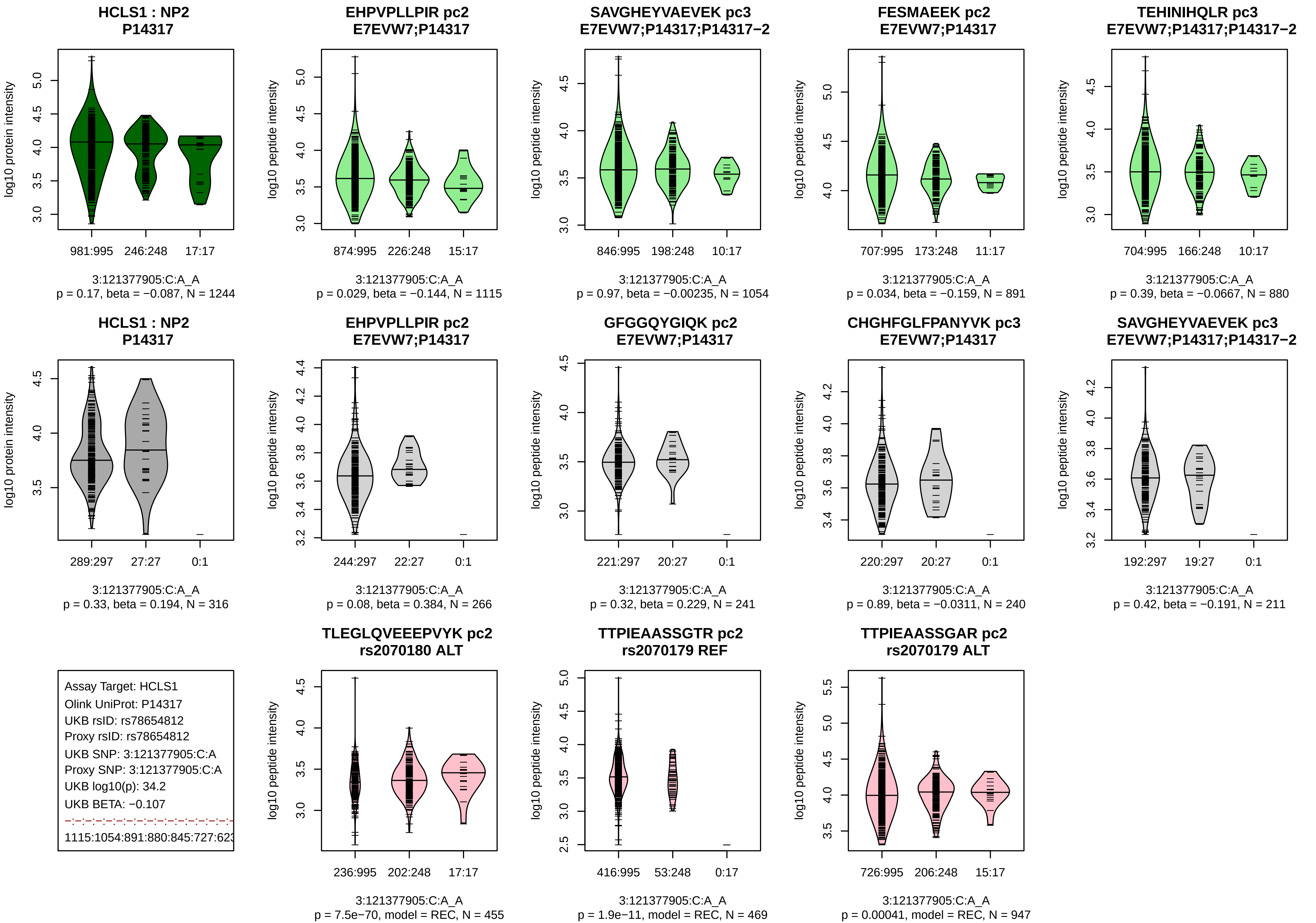
HCLS1 : NP2
P14317
log10 protein intensity
981:995
246:248
17:17
3:121377905:C:A_A
p = 0.17, beta = −0.087, N = 1244
EHPVPLLPIR pc2
E7EVW7;P14317
log10 peptide intensity
874:995
226:248
15:17
3:121377905:C:A_A
p = 0.029, beta = −0.144, N = 1115
SAVGHEYVAEVEK pc3
E7EVW7;P14317;P14317−2
846:995
198:248
10:17
3:121377905:C:A_A
p = 0.97, beta = −0.00235, N = 1054
FESMAEEK pc2
E7EVW7;P14317
707:995
173:248
11:17
3:121377905:C:A_A
p = 0.034, beta = −0.159, N = 891
TEHINIHQLR pc3
E7EVW7;P14317;P14317−2
704:995
166:248
10:17
3:121377905:C:A_A
p = 0.39, beta = −0.0667, N = 880
HCLS1 : NP2
P14317
289:297
27:27
0:1
3:121377905:C:A_A
p = 0.33, beta = 0.194, N = 316
EHPVPLLPIR pc2
E7EVW7;P14317
244:297
22:27
0:1
3:121377905:C:A_A
p = 0.08, beta = 0.384, N = 266
GFGGQYGIQK pc2
E7EVW7;P14317
221:297
20:27
0:1
3:121377905:C:A_A
p = 0.32, beta = 0.229, N = 241
CHGHFGLFPANYVK pc3
E7EVW7;P14317
220:297
20:27
0:1
3:121377905:C:A_A
p = 0.89, beta = −0.0311, N = 240
SAVGHEYVAEVEK pc3
E7EVW7;P14317;P14317−2
192:297
19:27
0:1
3:121377905:C:A_A
p = 0.42, beta = −0.191, N = 211
Assay Target: HCLS1
Olink UniProt: P14317
UKB rsID: rs78654812
Proxy rsID: rs78654812
UKB SNP: 3:121377905:C:A
Proxy SNP: 3:121377905:C:A
UKB log10(p): 34.2
UKB BETA: −0.107
1115:1054:891:880:845:727:623
TLEGLQVEEEPVYK pc2
rs2070180 ALT
236:995
202:248
17:17
3:121377905:C:A_A
p = 7.5e−70, model = REC, N = 455
TTPIEAASSGTR pc2
rs2070179 REF
416:995
53:248
0:17
3:121377905:C:A_A
p = 1.9e−11, model = REC, N = 469
TTPIEAASSGAR pc2
rs2070179 ALT
726:995
206:248
15:17
3:121377905:C:A_A
p = 0.00041, model = REC, N = 947

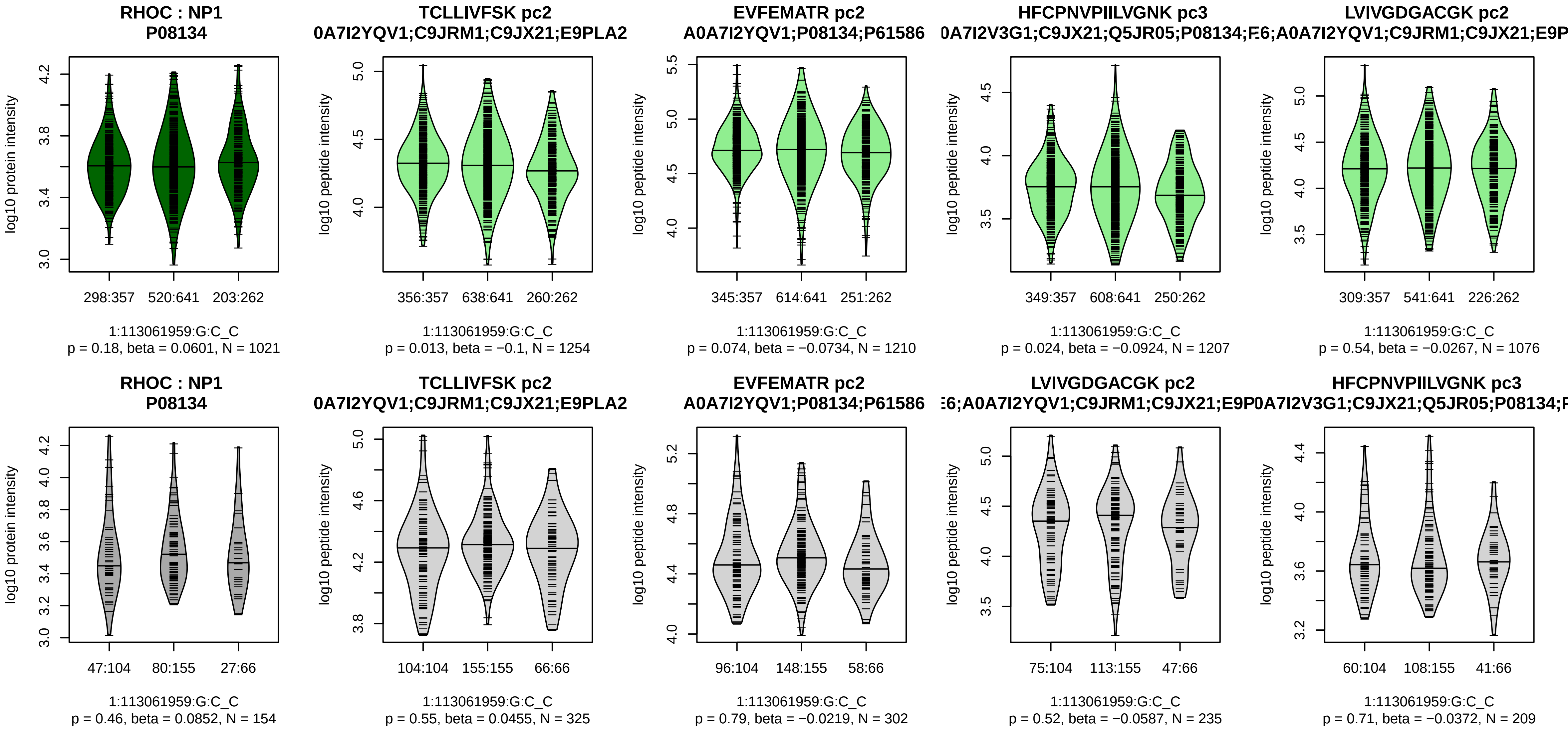


Assay Target: RHOC
Olink UniProt: P08134
UKB rsID: rs1006298
Proxy rsID: rs1006298
UKB SNP: 1:113061959:G:C
Proxy SNP: 1:113061959:G:C
UKB log10(p): 33.9
UKB BETA: 0.076
–:–:–:–:–:–:–:–
1254:1210:1207:1076:1009:842

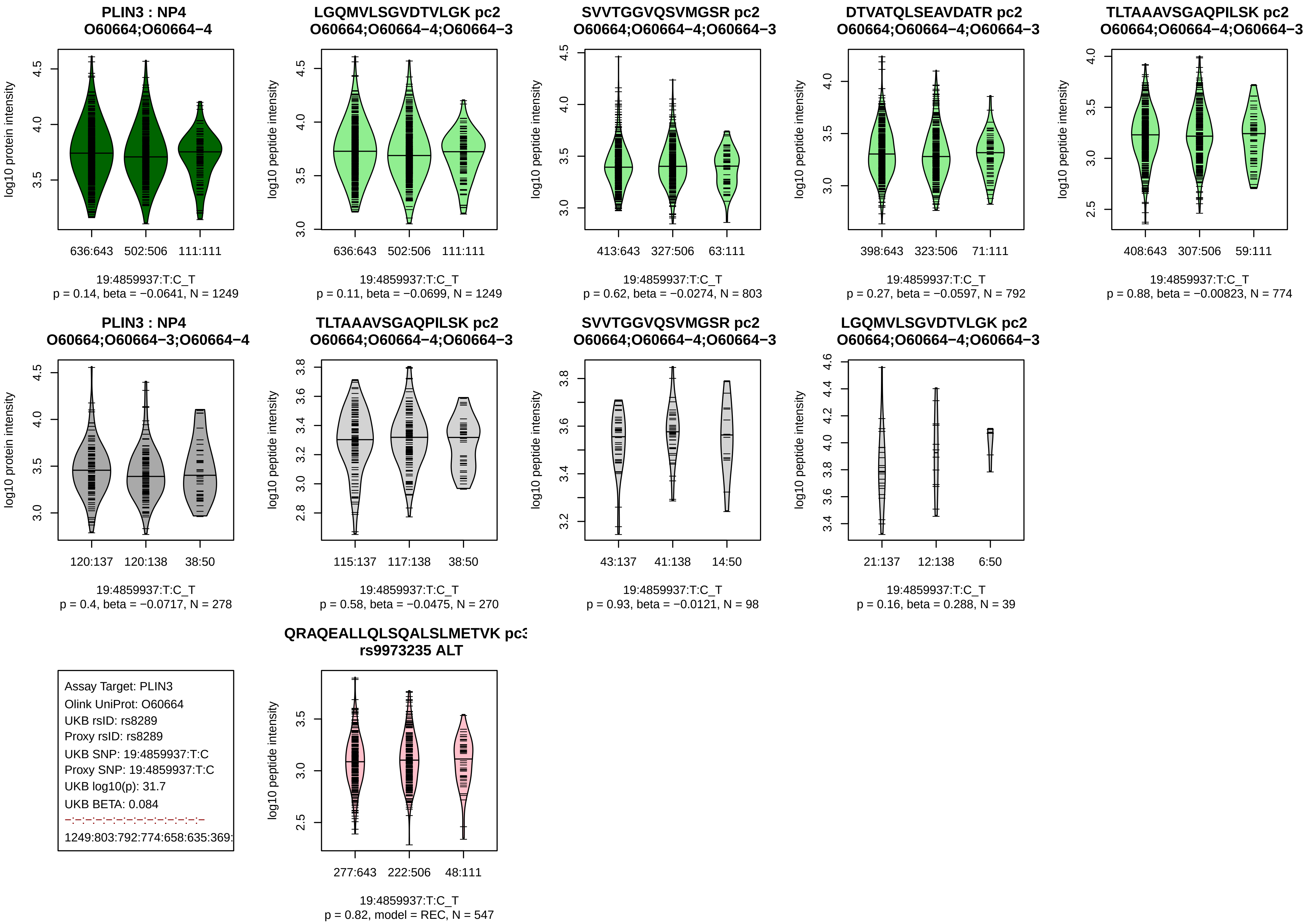

PLIN3 : NP4
O60664;O60664−4
log10 protein intensity
636:643 502:506 111:111
19:4859937:T:C_T
p = 0.14, beta = −0.0641, N = 1249
LGQMVLSGVDTVLGK pc2
O60664;O60664−4;O60664−3
log10 peptide intensity
636:643 502:506 111:111
19:4859937:T:C_T
p = 0.11, beta = −0.0699, N = 1249
SVVTGGVQSVMGSR pc2
O60664;O60664−4;O60664−3
413:643 327:506 63:111
19:4859937:T:C_T
p = 0.62, beta = −0.0274, N = 803
DTVATQLSEAVDATR pc2
O60664;O60664−4;O60664−3
398:643 323:506 71:111
19:4859937:T:C_T
p = 0.27, beta = −0.0597, N = 792
TLTAAAVSGAQPILSK pc2
O60664;O60664−4;O60664−3
408:643 307:506 59:111
19:4859937:T:C_T
p = 0.88, beta = −0.00823, N = 774
PLIN3 : NP4
O60664;O60664−3;O60664−4
120:137 120:138 38:50
19:4859937:T:C_T
p = 0.4, beta = −0.0717, N = 278
TLTAAAVSGAQPILSK pc2
O60664;O60664−4;O60664−3
115:137 117:138 38:50
19:4859937:T:C_T
p = 0.58, beta = −0.0475, N = 270
SVVTGGVQSVMGSR pc2
O60664;O60664−4;O60664−3
43:137 41:138 14:50
19:4859937:T:C_T
p = 0.93, beta = −0.0121, N = 98
LGQMVLSGVDTVLGK pc2
O60664;O60664−4;O60664−3
21:137 12:138 6:50
19:4859937:T:C_T
p = 0.16, beta = 0.288, N = 39
Assay Target: PLIN3
Olink UniProt: O60664
UKB rsID: rs8289
Proxy rsID: rs8289
UKB SNP: 19:4859937:T:C
Proxy SNP: 19:4859937:T:C
UKB log10(p): 31.7
UKB BETA: 0.084
1249:803:792:774:658:635:369:
QRAQEALLQLSQALSLMETVK pc3
rs9973235 ALT
277:643 222:506 48:111
19:4859937:T:C_T
p = 0.82, model = REC, N = 547

**CD34 : NP2**
**P28906**

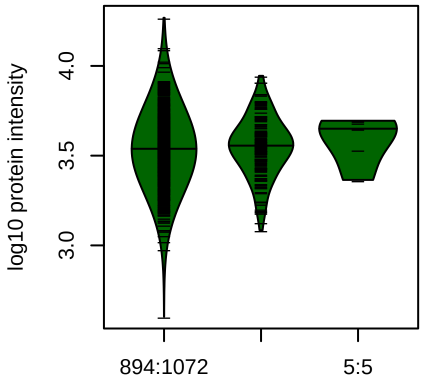


1:208057800:C:T_T
p = 0.35, beta = 0.0772, N = 1051

**TSSCAEFK pc2**
**P28906;P28906−2**

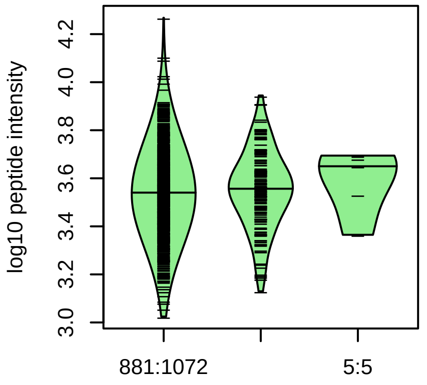


1:208057800:C:T_T
p = 0.32, beta = 0.0824, N = 1037

**QHVVADTEL pc2**
**P28906;Q5JTA5**

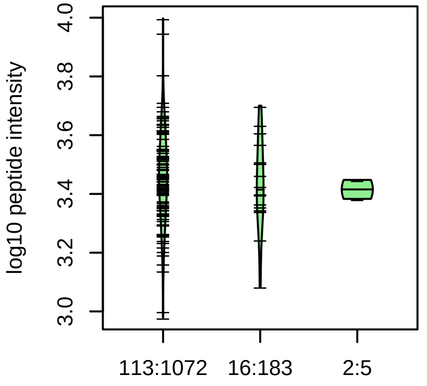


1:208057800:C:T_T
p = 0.41, beta = −0.175, N = 131

**CD34 : NP2**
**P28906**

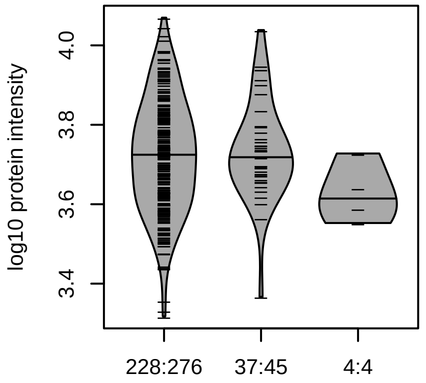


1:208057800:C:T_T
p = 0.87, beta = −0.0249, N = 269

**TSSCAEFK pc2**
**P28906;P28906−2**

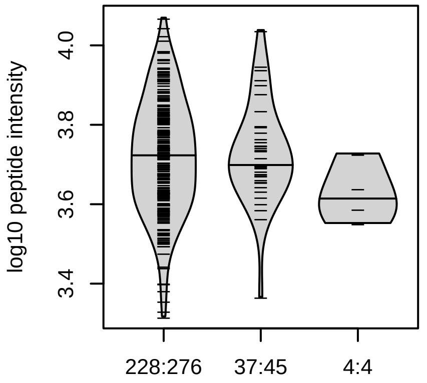


1:208057800:C:T_T
p = 0.77, beta = −0.0428, N = 269

Assay Target: CD34
Olink UniProt: P28906
UKB rsID: rs6671850
Proxy rsID: rs6671850
UKB SNP: 1:208057800:C:T
Proxy SNP: 1:208057800:C:T
UKB log10(p): 30.4
UKB BETA: 0.14
−:−:NA
1037:131:9

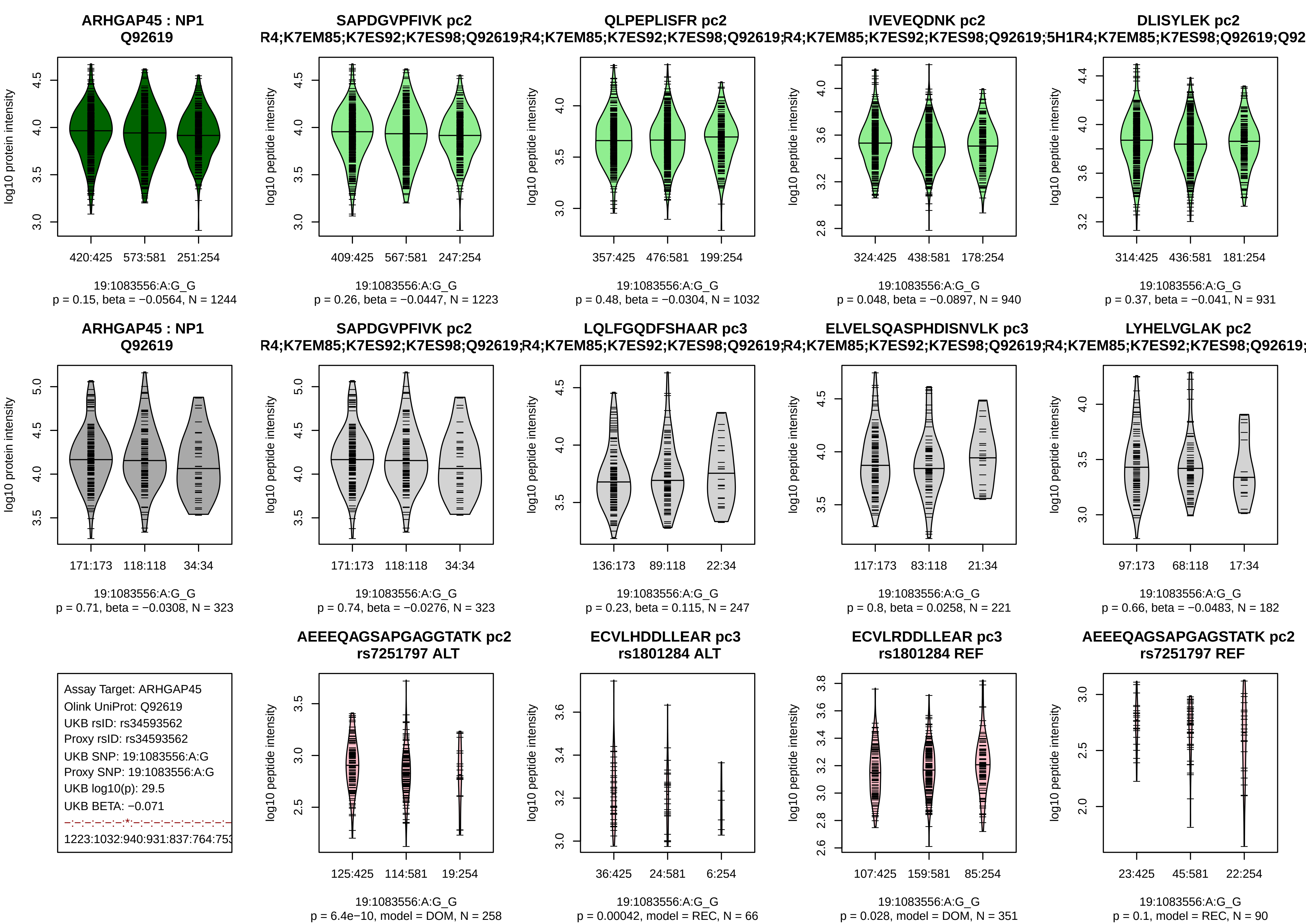
ARHGAP45 : NP1
Q92619
log10 protein intensity
420:425
573:581
251:254
19:1083556:A:G_G
p = 0.15, beta = −0.0564, N = 1244
SAPDGVPFIVK pc2
R4;K7EM85;K7ES92;K7ES98;Q92619;
log10 peptide intensity
409:425
567:581
247:254
19:1083556:A:G_G
p = 0.26, beta = −0.0447, N = 1223
QLPEPLISFR pc2
R4;K7EM85;K7ES92;K7ES98;Q92619;
357:425
476:581
199:254
19:1083556:A:G_G
p = 0.48, beta = −0.0304, N = 1032
IVEVEQDNK pc2
R4;K7EM85;K7ES92;K7ES98;Q92619;5H1
324:425
438:581
178:254
19:1083556:A:G_G
p = 0.048, beta = −0.0897, N = 940
DLISYLEK pc2
R4;K7EM85;K7ES98;Q92619;Q92
314:425
436:581
181:254
19:1083556:A:G_G
p = 0.37, beta = −0.041, N = 931
ARHGAP45 : NP1
Q92619
171:173
118:118
34:34
19:1083556:A:G_G
p = 0.71, beta = −0.0308, N = 323
SAPDGVPFIVK pc2
R4;K7EM85;K7ES92;K7ES98;Q92619;
171:173
118:118
34:34
19:1083556:A:G_G
p = 0.74, beta = −0.0276, N = 323
LQLFGQDFSHAAR pc3
R4;K7EM85;K7ES92;K7ES98;Q92619;
136:173
89:118
22:34
19:1083556:A:G_G
p = 0.23, beta = 0.115, N = 247
ELVELSQASPHDISNVLK pc3
R4;K7EM85;K7ES92;K7ES98;Q92619;
117:173
83:118
21:34
19:1083556:A:G_G
p = 0.8, beta = 0.0258, N = 221
LYHELVGLAK pc2
R4;K7EM85;K7ES92;K7ES98;Q92619;
97:173
68:118
17:34
19:1083556:A:G_G
p = 0.66, beta = −0.0483, N = 182
Assay Target: ARHGAP45
Olink UniProt: Q92619
UKB rsID: rs34593562
Proxy rsID: rs34593562
UKB SNP: 19:1083556:A:G
Proxy SNP: 19:1083556:A:G
UKB log10(p): 29.5
UKB BETA: −0.071
1223:1032:940:931:837:764:753
AEEEQAGSAPGAGGTATK pc2
rs7251797 ALT
125:425
114:581
19:254
19:1083556:A:G_G
p = 6.4e−10, model = DOM, N = 258
ECVLHDDLLEAR pc3
rs1801284 ALT
36:425
24:581
6:254
19:1083556:A:G_G
p = 0.00042, model = REC, N = 66
ECVLRDDLLEAR pc3
rs1801284 REF
107:425
159:581
85:254
19:1083556:A:G_G
p = 0.028, model = DOM, N = 351
AEEEQAGSAPGAGSTATK pc2
rs7251797 REF
23:425
45:581
22:254
19:1083556:A:G_G
p = 0.1, model = REC, N = 90

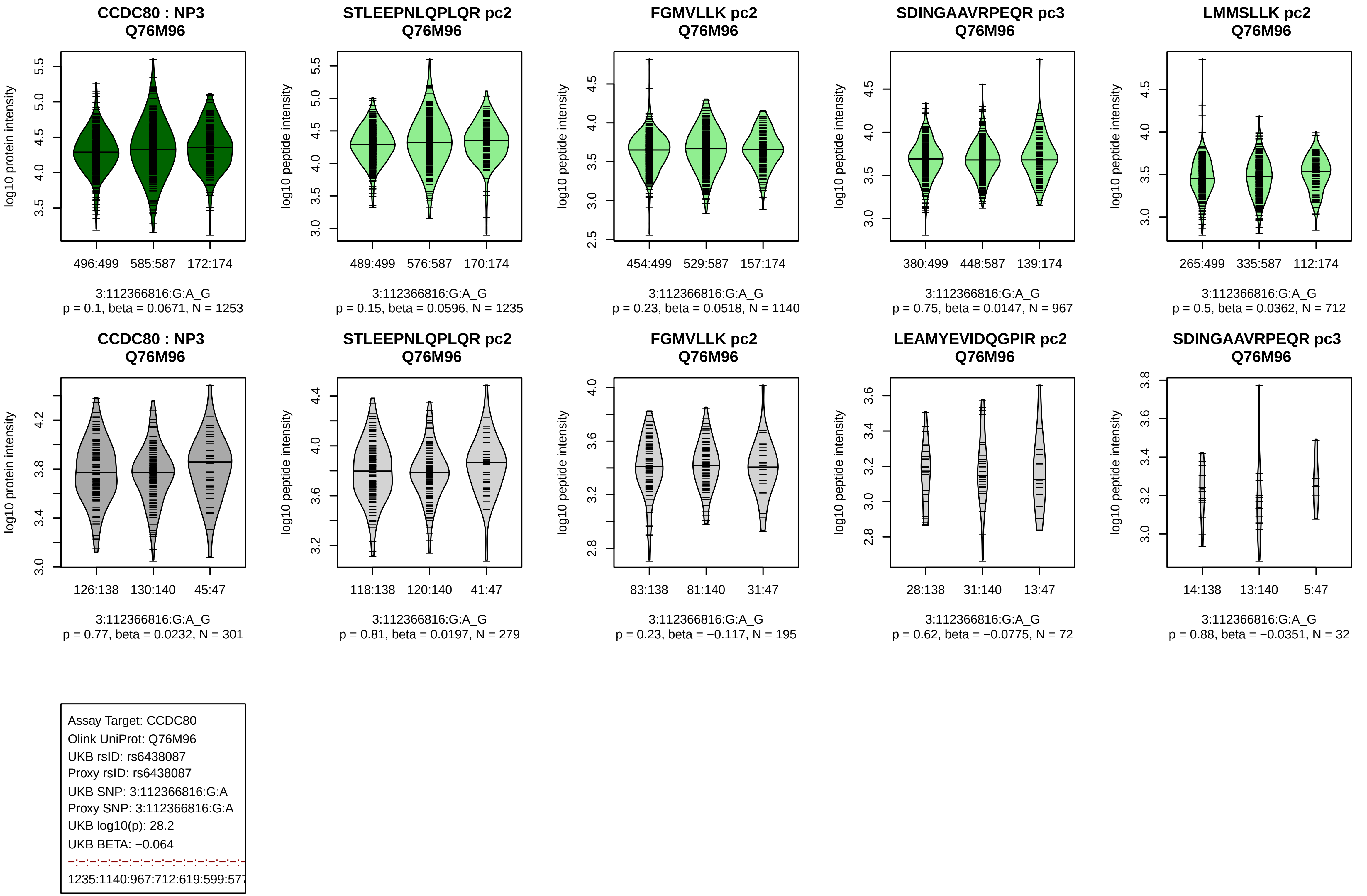
CCDC80 : NP3
Q76M96
log10 protein intensity
496:499 585:587 172:174
3:112366816:G:A_G
p = 0.1, beta = 0.0671, N = 1253
STLEEPNLQPLQR pc2
Q76M96
log10 peptide intensity
489:499 576:587 170:174
3:112366816:G:A_G
p = 0.15, beta = 0.0596, N = 1235
FGMVLLK pc2
Q76M96
log10 peptide intensity
454:499 529:587 157:174
3:112366816:G:A_G
p = 0.23, beta = 0.0518, N = 1140
SDINGAAVRPEQR pc3
Q76M96
log10 peptide intensity
380:499 448:587 139:174
3:112366816:G:A_G
p = 0.75, beta = 0.0147, N = 967
LMMSLLK pc2
Q76M96
log10 peptide intensity
265:499 335:587 112:174
3:112366816:G:A_G
p = 0.5, beta = 0.0362, N = 712
CCDC80 : NP3
Q76M96
log10 protein intensity
126:138 130:140 45:47
3:112366816:G:A_G
p = 0.77, beta = 0.0232, N = 301
STLEEPNLQPLQR pc2
Q76M96
log10 peptide intensity
118:138 120:140 41:47
3:112366816:G:A_G
p = 0.81, beta = 0.0197, N = 279
FGMVLLK pc2
Q76M96
log10 peptide intensity
83:138 81:140 31:47
3:112366816:G:A_G
p = 0.23, beta = −0.117, N = 195
LEAMYEVIDQGPIR pc2
Q76M96
log10 peptide intensity
28:138 31:140 13:47
3:112366816:G:A_G
p = 0.62, beta = −0.0775, N = 72
SDINGAAVRPEQR pc3
Q76M96
log10 peptide intensity
14:138 13:140 5:47
3:112366816:G:A_G
p = 0.88, beta = −0.0351, N = 32
Assay Target: CCDC80
Olink UniProt: Q76M96
UKB rsID: rs6438087
Proxy rsID: rs6438087
UKB SNP: 3:112366816:G:A
Proxy SNP: 3:112366816:G:A
UKB log10(p): 28.2
UKB BETA: −0.064
1235:1140:967:712:619:599:577

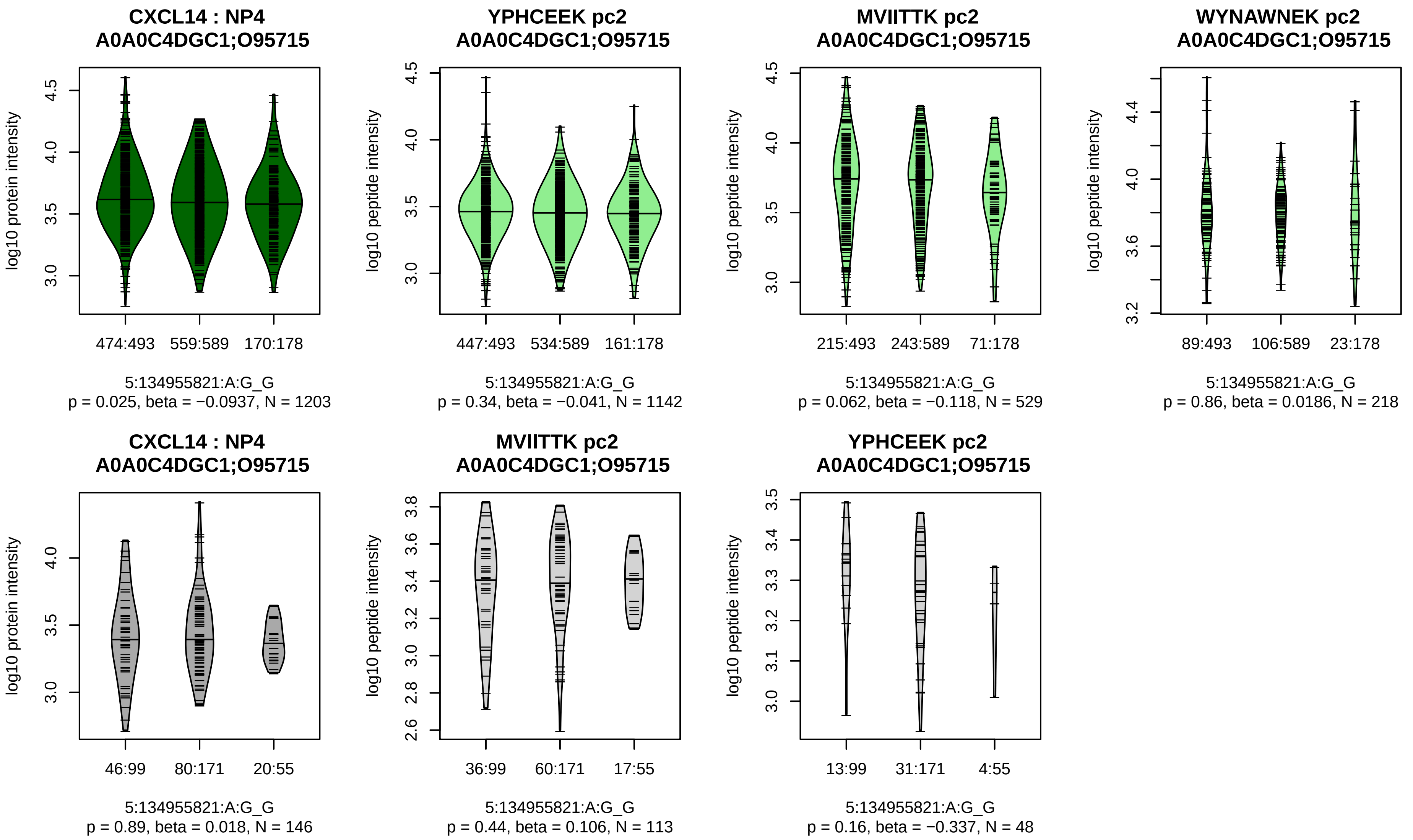


Assay Target: CXCL14
Olink UniProt: O95715
UKB rsID: rs6596251
Proxy rsID: rs6596251
UKB SNP: 5:134955821:A:G
Proxy SNP: 5:134955821:A:G
UKB log10(p): 27.1
UKB BETA: −0.063
−:−:−
1142:529:218

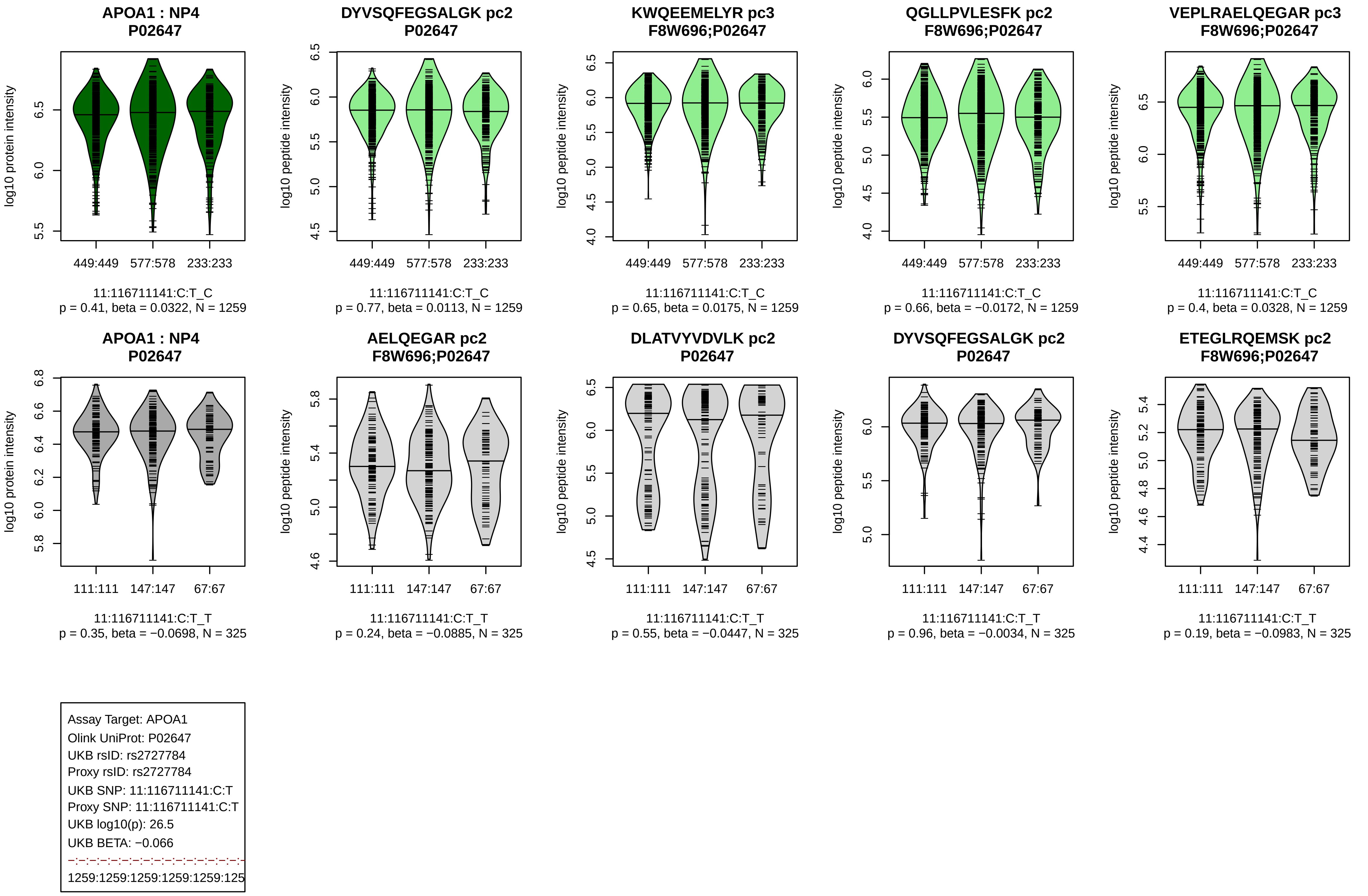
APOA1 : NP4
P02647
log10 protein intensity
449:449 577:578 233:233
11:116711141:C:T_C
p = 0.41, beta = 0.0322, N = 1259
DYVSQFEGSALGK pc2
P02647
log10 peptide intensity
449:449 577:578 233:233
11:116711141:C:T_C
p = 0.77, beta = 0.0113, N = 1259
KWQEEMELYR pc3
F8W696;P02647
log10 peptide intensity
449:449 577:578 233:233
11:116711141:C:T_C
p = 0.65, beta = 0.0175, N = 1259
QGLLPVLESFK pc2
F8W696;P02647
log10 peptide intensity
449:449 577:578 233:233
11:116711141:C:T_C
p = 0.66, beta = −0.0172, N = 1259
VEPLRAELQEGAR pc3
F8W696;P02647
log10 peptide intensity
449:449 577:578 233:233
11:116711141:C:T_C
p = 0.4, beta = 0.0328, N = 1259
APOA1 : NP4
P02647
log10 protein intensity
111:111 147:147 67:67
11:116711141:C:T_T
p = 0.35, beta = −0.0698, N = 325
AELQEGAR pc2
F8W696;P02647
log10 peptide intensity
111:111 147:147 67:67
11:116711141:C:T_T
p = 0.24, beta = −0.0885, N = 325
DLATVYVDVLK pc2
P02647
log10 peptide intensity
111:111 147:147 67:67
11:116711141:C:T_T
p = 0.55, beta = −0.0447, N = 325
DYVSQFEGSALGK pc2
P02647
log10 peptide intensity
111:111 147:147 67:67
11:116711141:C:T_T
p = 0.96, beta = −0.0034, N = 325
ETEGLRQEMSK pc2
F8W696;P02647
log10 peptide intensity
111:111 147:147 67:67
11:116711141:C:T_T
p = 0.19, beta = −0.0983, N = 325
Assay Target: APOA1
Olink UniProt: P02647
UKB rsID: rs2727784
Proxy rsID: rs2727784
UKB SNP: 11:116711141:C:T
Proxy SNP: 11:116711141:C:T
UKB log10(p): 26.5
UKB BETA: −0.066
1259:1259:1259:1259:1259:125

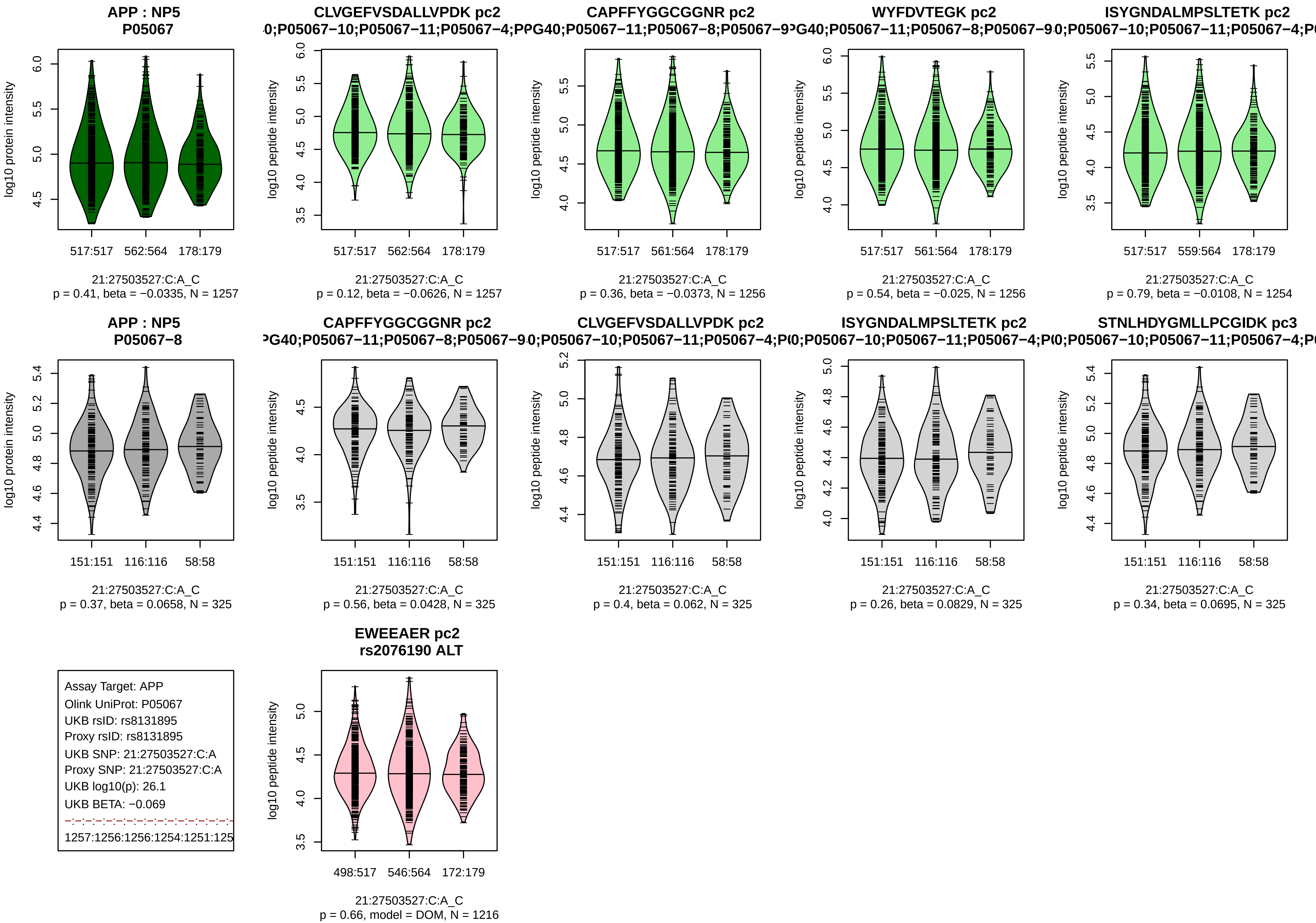

APP : NP5
P05067
log10 protein intensity
517:517
562:564
178:179
21:27503527:C:A_C
p = 0.41, beta = −0.0335, N = 1257
CLVGEFVSDALLVPDK pc2
0;P05067−10;P05067−11;P05067−4;P
log10 peptide intensity
517:517
562:564
178:179
21:27503527:C:A_C
p = 0.12, beta = −0.0626, N = 1257
CAPFFYGGCGGNR pc2
PG40;P05067−11;P05067−8;P05067−9
log10 peptide intensity
517:517
561:564
178:179
21:27503527:C:A_C
p = 0.36, beta = −0.0373, N = 1256
WYFDVTEGK pc2
G40;P05067−11;P05067−8;P05067−9
log10 peptide intensity
517:517
561:564
178:179
21:27503527:C:A_C
p = 0.54, beta = −0.025, N = 1256
ISYGNDALMPSLTETK pc2
0;P05067−10;P05067−11;P05067−4;P
log10 peptide intensity
517:517
559:564
178:179
21:27503527:C:A_C
p = 0.79, beta = −0.0108, N = 1254
APP : NP5
P05067−8
log10 protein intensity
151:151
116:116
58:58
21:27503527:C:A_C
p = 0.37, beta = 0.0658, N = 325
CAPFFYGGCGGNR pc2
PG40;P05067−11;P05067−8;P05067−9
log10 peptide intensity
151:151
116:116
58:58
21:27503527:C:A_C
p = 0.56, beta = 0.0428, N = 325
CLVGEFVSDALLVPDK pc2
0;P05067−10;P05067−11;P05067−4;P
log10 peptide intensity
151:151
116:116
58:58
21:27503527:C:A_C
p = 0.4, beta = 0.062, N = 325
ISYGNDALMPSLTETK pc2
0;P05067−10;P05067−11;P05067−4;P
log10 peptide intensity
151:151
116:116
58:58
21:27503527:C:A_C
p = 0.26, beta = 0.0829, N = 325
STNLHDYGMLLPCGIDK pc3
0;P05067−10;P05067−11;P05067−4;P
log10 peptide intensity
151:151
116:116
58:58
21:27503527:C:A_C
p = 0.34, beta = 0.0695, N = 325
Assay Target: APP
Olink UniProt: P05067
UKB rsID: rs8131895
Proxy rsID: rs8131895
UKB SNP: 21:27503527:C:A
Proxy SNP: 21:27503527:C:A
UKB log10(p): 26.1
UKB BETA: −0.069
1257:1256:1256:1254:1251:125
EWEEAER pc2
rs2076190 ALT
log10 peptide intensity
498:517
546:564
172:179
21:27503527:C:A_C
p = 0.66, model = DOM, N = 1216

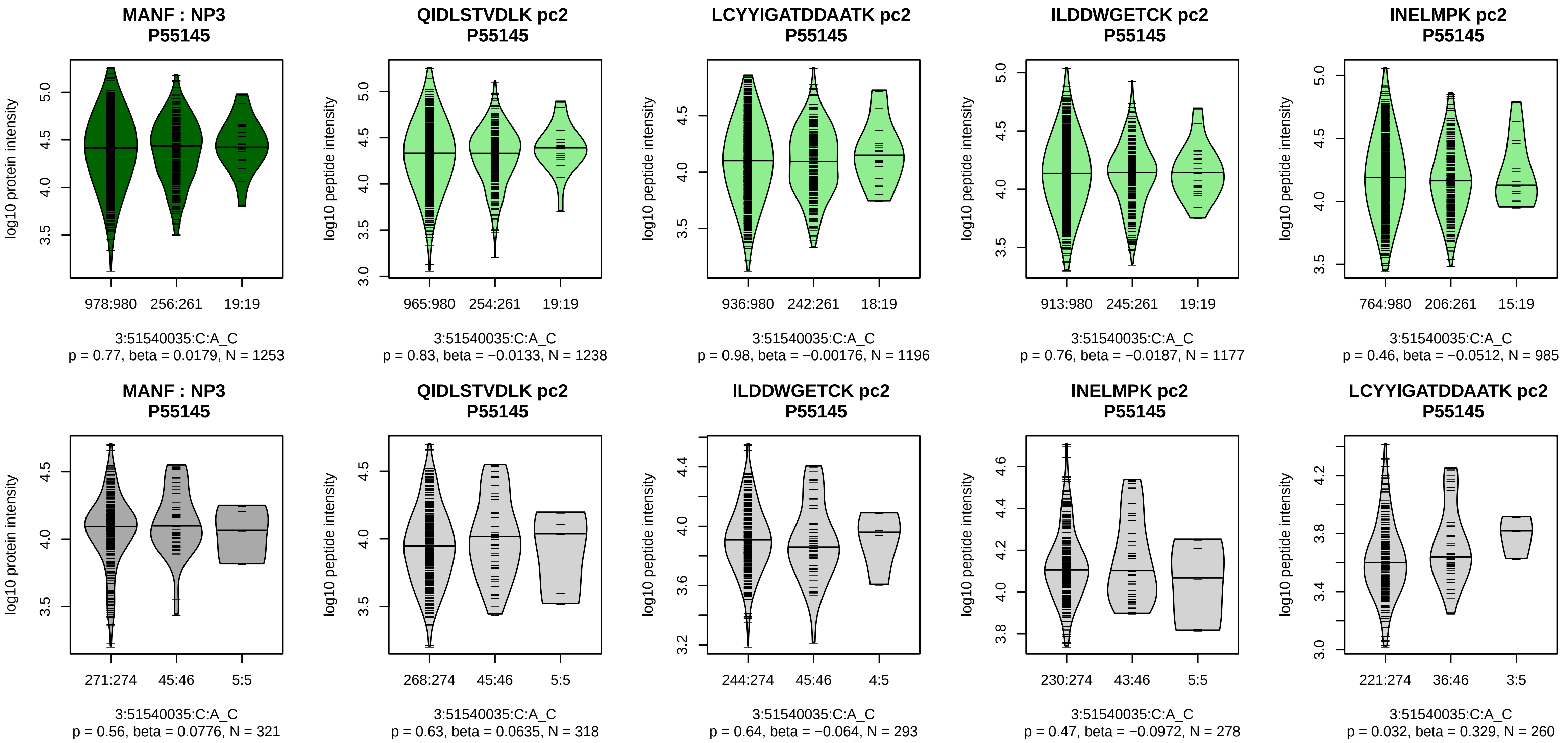


Assay Target: MANF
Olink UniProt: P55145
UKB rsID: rs9311469
Proxy rsID: rs9311469
UKB SNP: 3:51540035:C:A
Proxy SNP: 3:51540035:C:A
UKB log10(p): 26.1
UKB BETA: 0.097
−:−:−:−:−:−:−:−:−:−:−:−:−:NA
1238:1196:1177:985:764:558:54

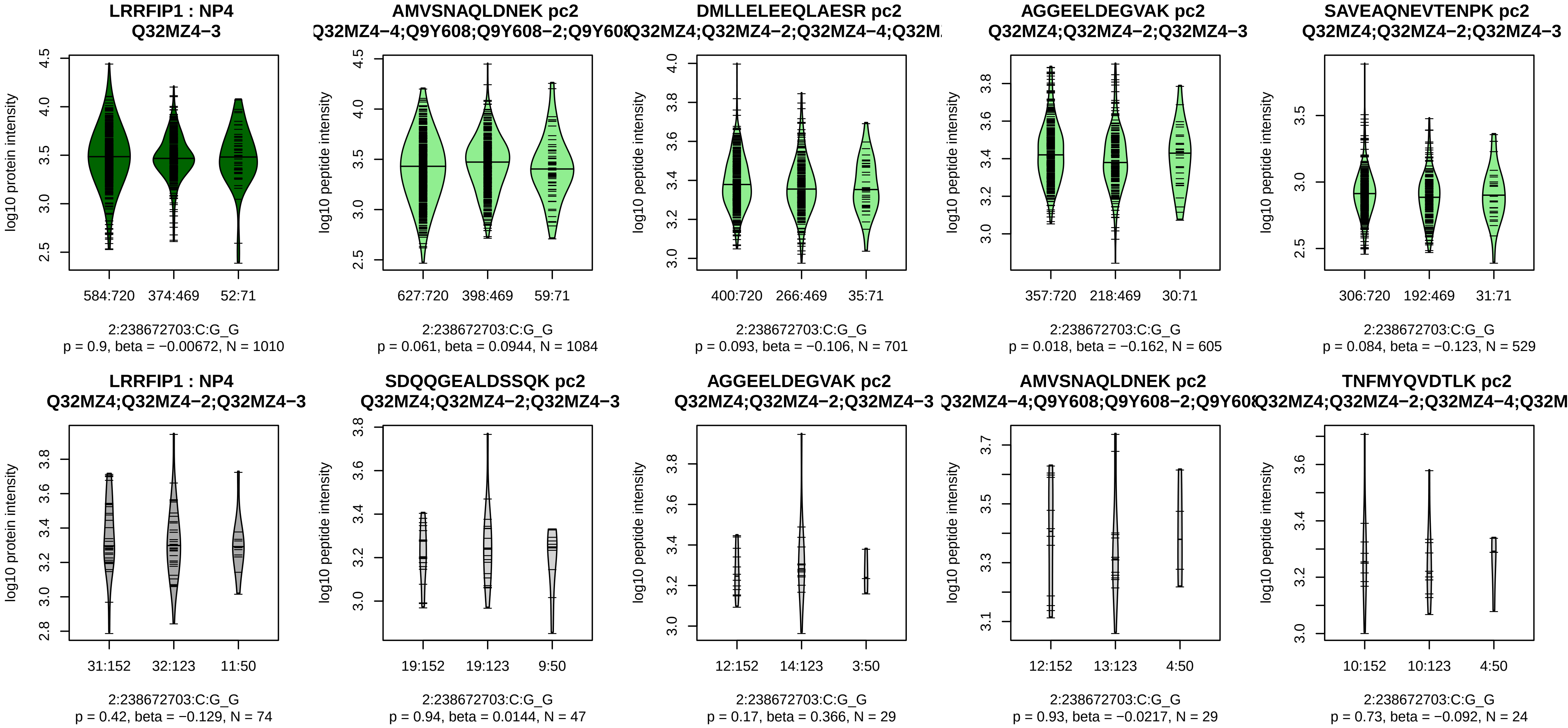


Assay Target: LRRFIP1
Olink UniProt: Q32MZ4
UKB rsID: rs3739038
Proxy rsID: rs3739038
UKB SNP: 2:238672703:C:G
Proxy SNP: 2:238672703:C:G
UKB log10(p): 25.2
UKB BETA: −0.074
−:−:−:−:−:−:−:−:−:−:−:−:−:−:−:−:
1084:701:605:529:452:418:387:

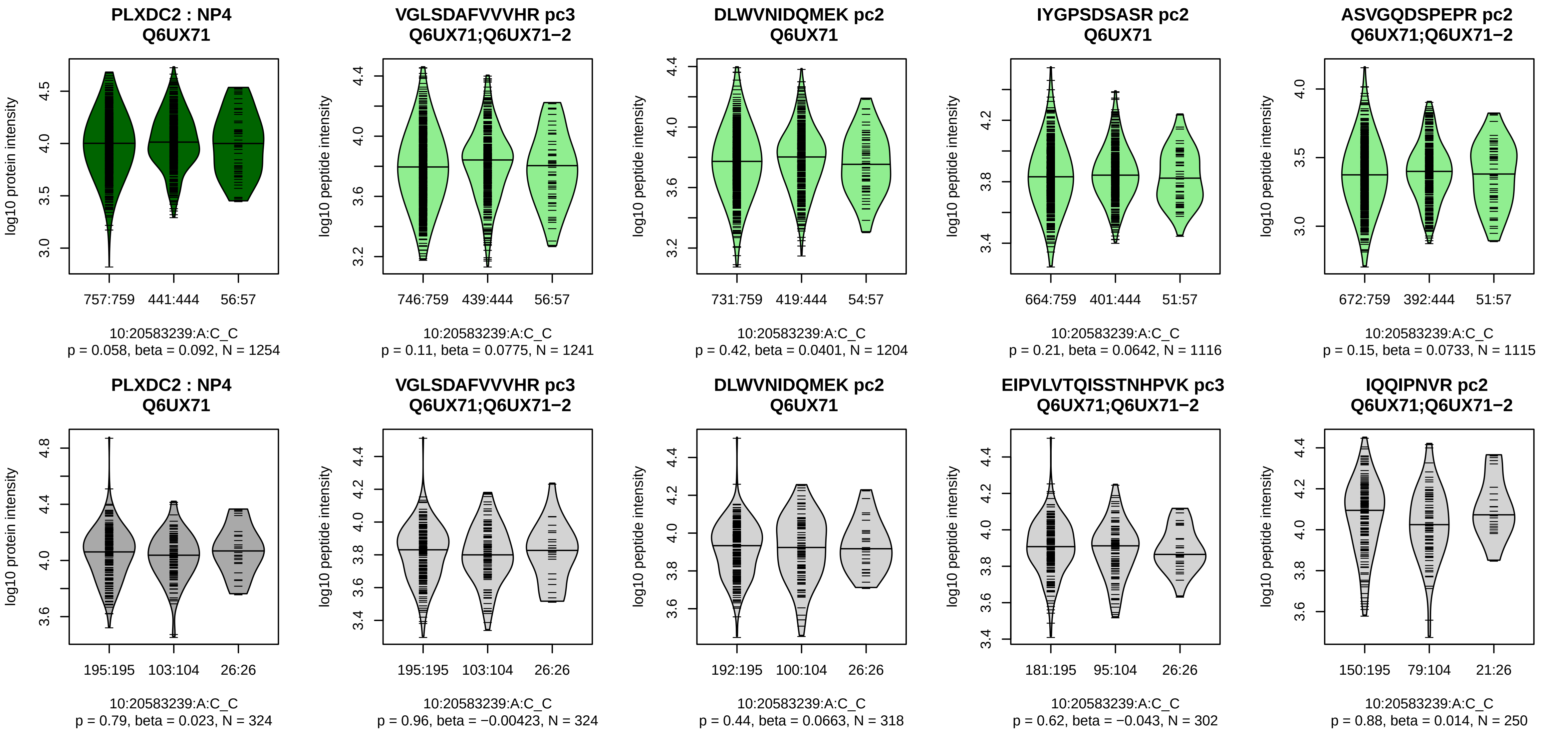


Assay Target: PLXDC2
Olink UniProt: Q6UX71
UKB rsID: rs4748650
Proxy rsID: rs4748650
UKB SNP: 10:20583239:A:C
Proxy SNP: 10:20583239:A:C
UKB log10(p): 25
UKB BETA: 0.078
−:−:−:−:−:*:−:−:−:−:−:−:−
1241:1204:1116:1115:793:720:5

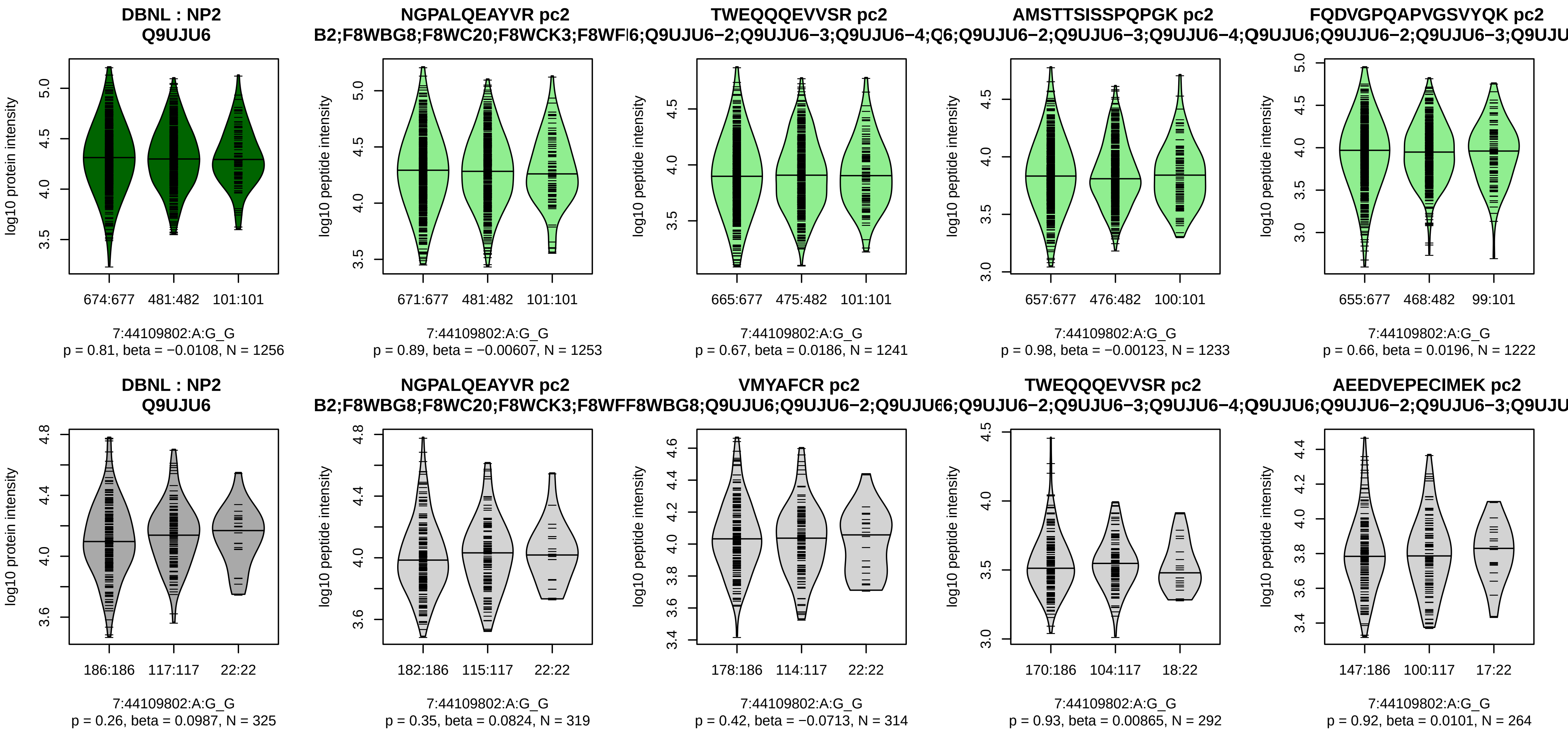


Assay Target: DBNL
Olink UniProt: Q9UJU6
UKB rsID: rs7799945
Proxy rsID: rs7799945
UKB SNP: 7:44109802:A:G
Proxy SNP: 7:44109802:A:G
UKB log10(p): 24.9
UKB BETA: 0.068
−:−:−:−:−:−:−:−:−:−:−:−:−:−:−:
1253:1241:1233:1222:1182:112

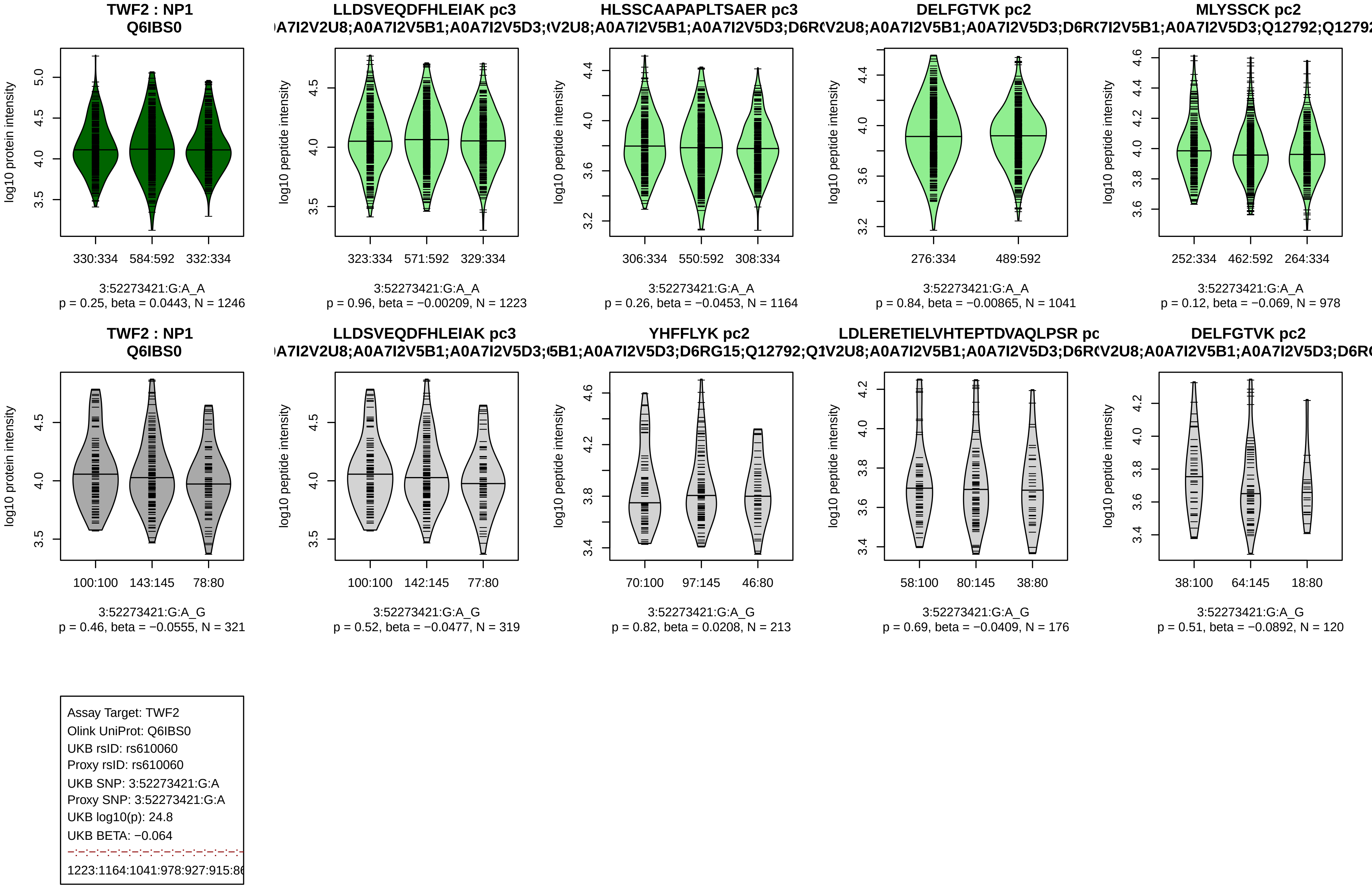

TWF2 : NP1
Q6IBS0
LLDSVEQDFHLEIAK pc3
HLSSCAAPAPLTSAER pc3
DELFGTVK pc2
MLYSSCK pc2
330:334 584:592 332:334
323:334 571:592 329:334
306:334 550:592 308:334
276:334 489:592
252:334 462:592 264:334
3:52273421:G:A_A
p = 0.25, beta = 0.0443, N = 1246
p = 0.96, beta = −0.00209, N = 1223
p = 0.26, beta = −0.0453, N = 1164
p = 0.84, beta = −0.00865, N = 1041
p = 0.12, beta = −0.069, N = 978
YHFFLYK pc2
LDLERETIELVHTEPTDVAQLPSR pc
100:100 143:145 78:80
100:100 142:145 77:80
70:100 97:145 46:80
58:100 80:145 38:80
38:100 64:145 18:80
3:52273421:G:A_G
p = 0.46, beta = −0.0555, N = 321
p = 0.52, beta = −0.0477, N = 319
p = 0.82, beta = 0.0208, N = 213
p = 0.69, beta = −0.0409, N = 176
p = 0.51, beta = −0.0892, N = 120
log10 protein intensity
log10 peptide intensity
Assay Target: TWF2
Olink UniProt: Q6IBS0
UKB rsID: rs610060
Proxy rsID: rs610060
UKB SNP: 3:52273421:G:A
Proxy SNP: 3:52273421:G:A
UKB log10(p): 24.8
UKB BETA: −0.064
1223:1164:1041:978:927:915:86

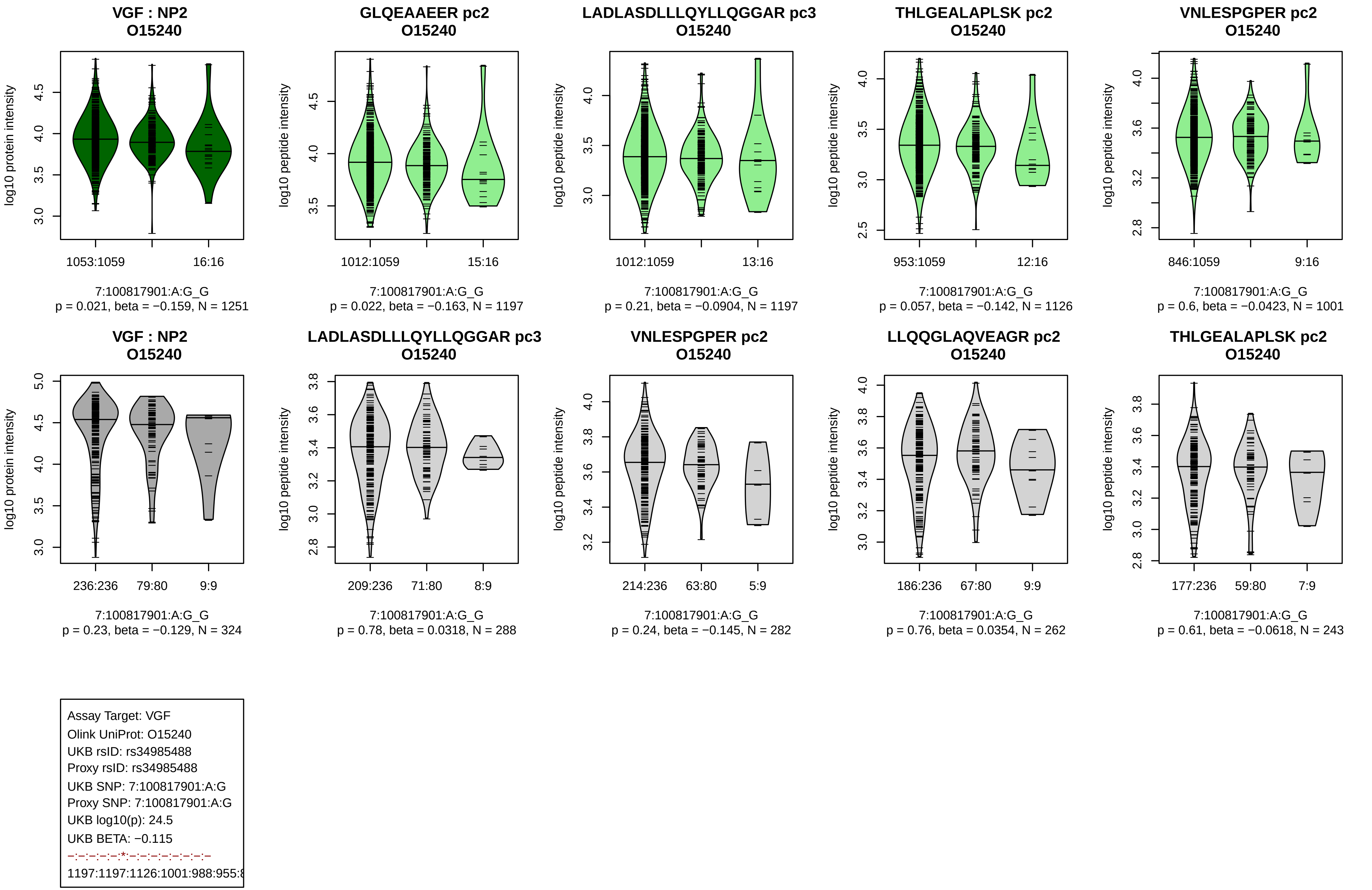
VGF : NP2
O15240
log10 protein intensity
1053:1059
16:16
7:100817901:A:G_G
p = 0.021, beta = −0.159, N = 1251
GLQEAAEER pc2
O15240
log10 peptide intensity
1012:1059
15:16
7:100817901:A:G_G
p = 0.022, beta = −0.163, N = 1197
LADLASDLLLQYLLQGGAR pc3
O15240
log10 peptide intensity
1012:1059
13:16
7:100817901:A:G_G
p = 0.21, beta = −0.0904, N = 1197
THLGEALAPLSK pc2
O15240
log10 peptide intensity
953:1059
12:16
7:100817901:A:G_G
p = 0.057, beta = −0.142, N = 1126
VNLESPGPER pc2
O15240
log10 peptide intensity
846:1059
9:16
7:100817901:A:G_G
p = 0.6, beta = −0.0423, N = 1001
VGF : NP2
O15240
log10 protein intensity
236:236
79:80
9:9
7:100817901:A:G_G
p = 0.23, beta = −0.129, N = 324
LADLASDLLLQYLLQGGAR pc3
O15240
log10 peptide intensity
209:236
71:80
8:9
7:100817901:A:G_G
p = 0.78, beta = 0.0318, N = 288
VNLESPGPER pc2
O15240
log10 peptide intensity
214:236
63:80
5:9
7:100817901:A:G_G
p = 0.24, beta = −0.145, N = 282
LLQQGLAQVEAGR pc2
O15240
log10 peptide intensity
186:236
67:80
9:9
7:100817901:A:G_G
p = 0.76, beta = 0.0354, N = 262
THLGEALAPLSK pc2
O15240
log10 peptide intensity
177:236
59:80
7:9
7:100817901:A:G_G
p = 0.61, beta = −0.0618, N = 243
Assay Target: VGF
Olink UniProt: O15240
UKB rsID: rs34985488
Proxy rsID: rs34985488
UKB SNP: 7:100817901:A:G
Proxy SNP: 7:100817901:A:G
UKB log10(p): 24.5
UKB BETA: −0.115
–:–:–:–:–:*:–:–:–:–:–:–:–:–
1197:1197:1126:1001:988:955:8

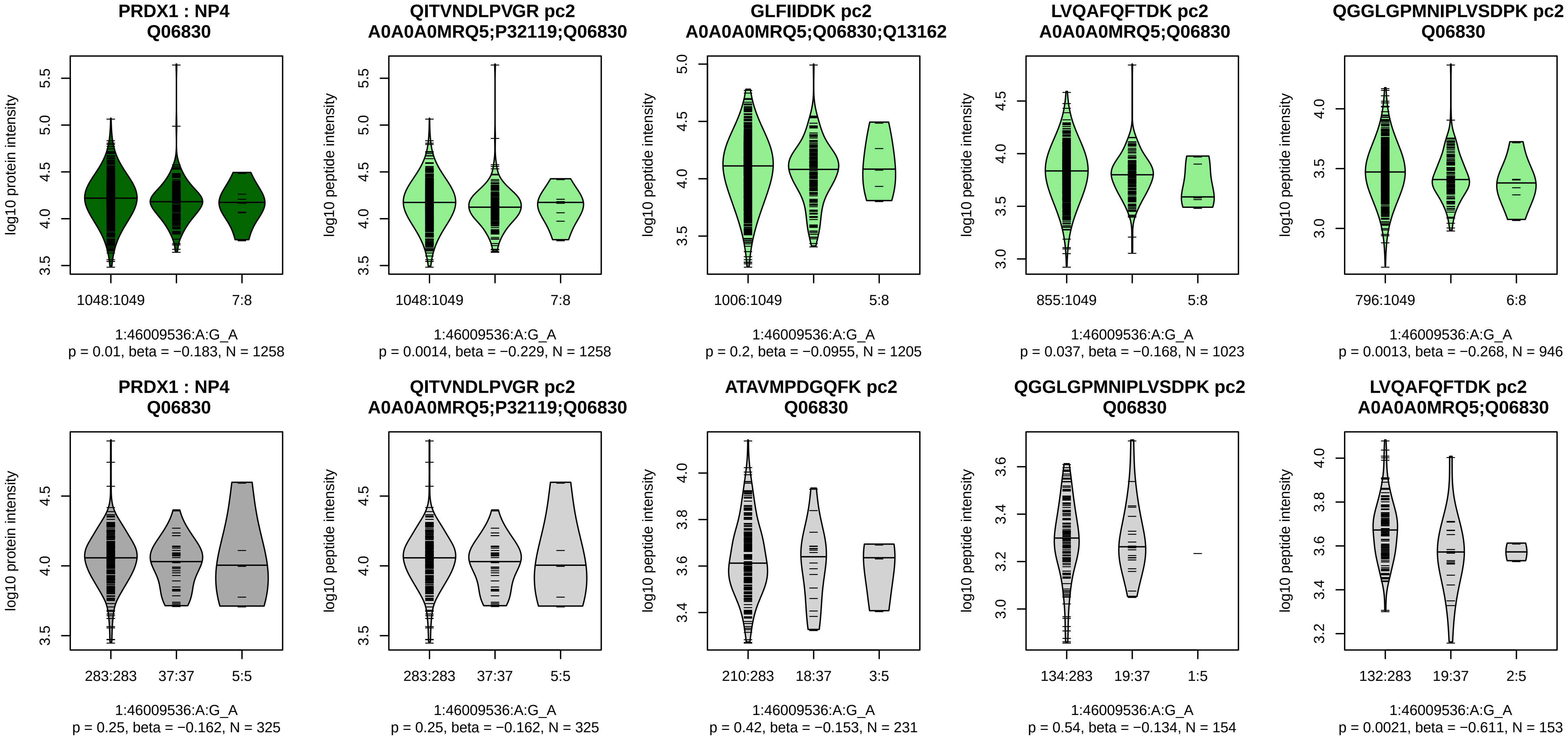


Assay Target: PRDX1
Olink UniProt: Q06830
UKB rsID: rs487174
Proxy rsID: rs487174
UKB SNP: 1:46009536:A:G
Proxy SNP: 1:46009536:A:G
UKB log10(p): 22.1
UKB BETA: 0.102
*:–:–:*:*:*:*:–:–:–:–:–:–
1258:1205:1023:946:934:733:71

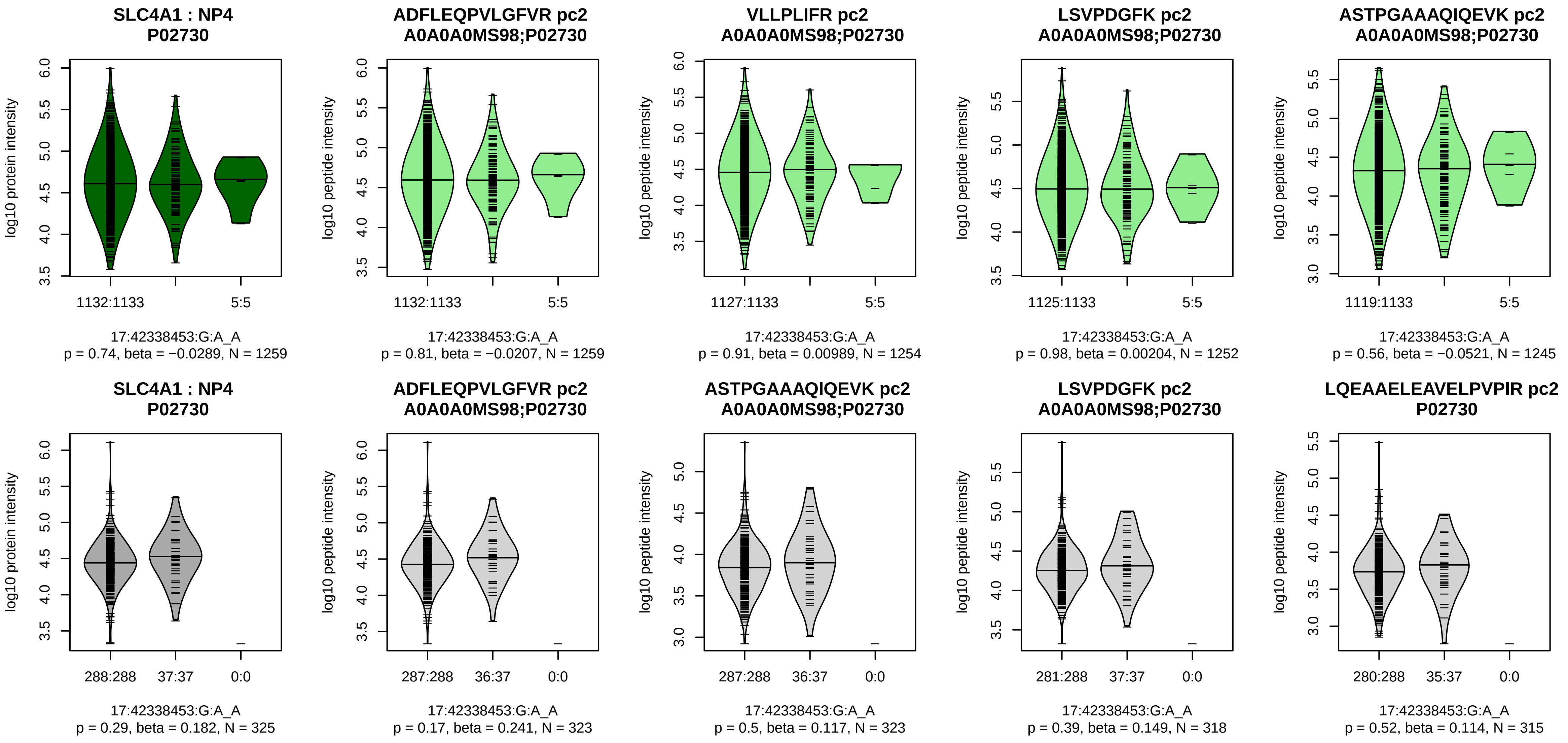


Assay Target: SLC4A1
Olink UniProt: P02730
UKB rsID: rs5035
Proxy rsID: rs45574837
UKB SNP: 17:42338998:T:G
Proxy SNP: 17:42338453:G:A
UKB log10(p): 21.9
UKB BETA: −0.146

1259:1254:1252:1245:1222:121

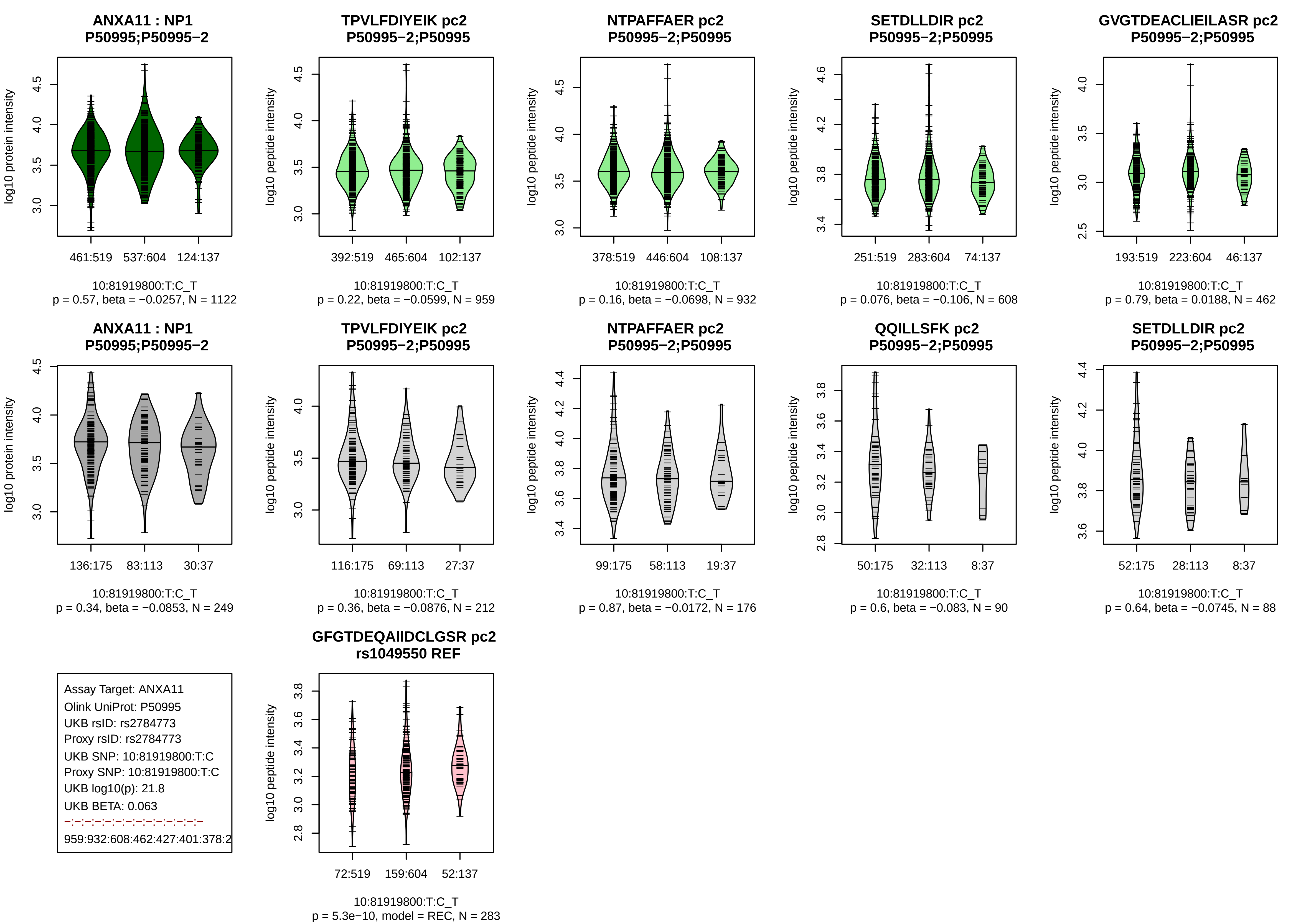
ANXA11 : NP1
P50995;P50995−2
log10 protein intensity
461:519 537:604 124:137
10:81919800:T:C_T
p = 0.57, beta = −0.0257, N = 1122
TPVLFDIYEIK pc2
P50995−2;P50995
log10 peptide intensity
392:519 465:604 102:137
10:81919800:T:C_T
p = 0.22, beta = −0.0599, N = 959
NTPAFFAER pc2
P50995−2;P50995
log10 peptide intensity
378:519 446:604 108:137
10:81919800:T:C_T
p = 0.16, beta = −0.0698, N = 932
SETDLLDIR pc2
P50995−2;P50995
log10 peptide intensity
251:519 283:604 74:137
10:81919800:T:C_T
p = 0.076, beta = −0.106, N = 608
GVGTDEACLIEILASR pc2
P50995−2;P50995
log10 peptide intensity
193:519 223:604 46:137
10:81919800:T:C_T
p = 0.79, beta = 0.0188, N = 462
ANXA11 : NP1
P50995;P50995−2
log10 protein intensity
136:175 83:113 30:37
10:81919800:T:C_T
p = 0.34, beta = −0.0853, N = 249
TPVLFDIYEIK pc2
P50995−2;P50995
log10 peptide intensity
116:175 69:113 27:37
10:81919800:T:C_T
p = 0.36, beta = −0.0876, N = 212
NTPAFFAER pc2
P50995−2;P50995
log10 peptide intensity
99:175 58:113 19:37
10:81919800:T:C_T
p = 0.87, beta = −0.0172, N = 176
QQILLSFK pc2
P50995−2;P50995
log10 peptide intensity
50:175 32:113 8:37
10:81919800:T:C_T
p = 0.6, beta = −0.083, N = 90
SETDLLDIR pc2
P50995−2;P50995
log10 peptide intensity
52:175 28:113 8:37
10:81919800:T:C_T
p = 0.64, beta = −0.0745, N = 88
Assay Target: ANXA11
Olink UniProt: P50995
UKB rsID: rs2784773
Proxy rsID: rs2784773
UKB SNP: 10:81919800:T:C
Proxy SNP: 10:81919800:T:C
UKB log10(p): 21.8
UKB BETA: 0.063
−:−:−:−:−:−:−:−:−:−:−:−:−
959:932:608:462:427:401:378:2
GFGTDEQAIIDCLGSR pc2
rs1049550 REF
log10 peptide intensity
72:519 159:604 52:137
10:81919800:T:C_T
p = 5.3e−10, model = REC, N = 283

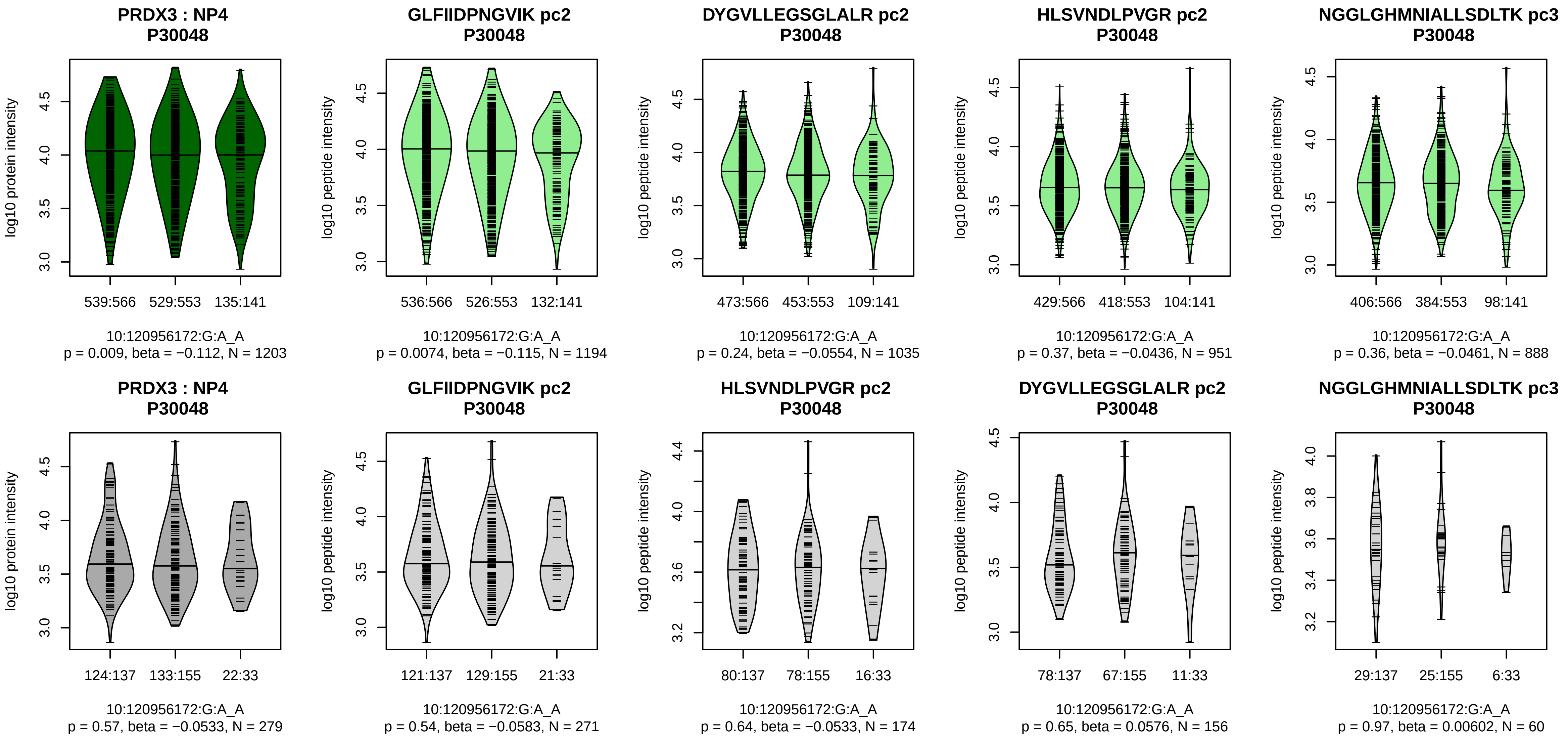


Assay Target: PRDX3
Olink UniProt: P30048
UKB rsID: rs747334131
Proxy rsID: rs7068698
UKB SNP: 10:120953662:TCTG
Proxy SNP: 10:120956172:G:A
UKB log10(p): 21.5
UKB BETA: −0.065
*:−:−:−:−:−:−
1194:1035:951:888:501:467:32

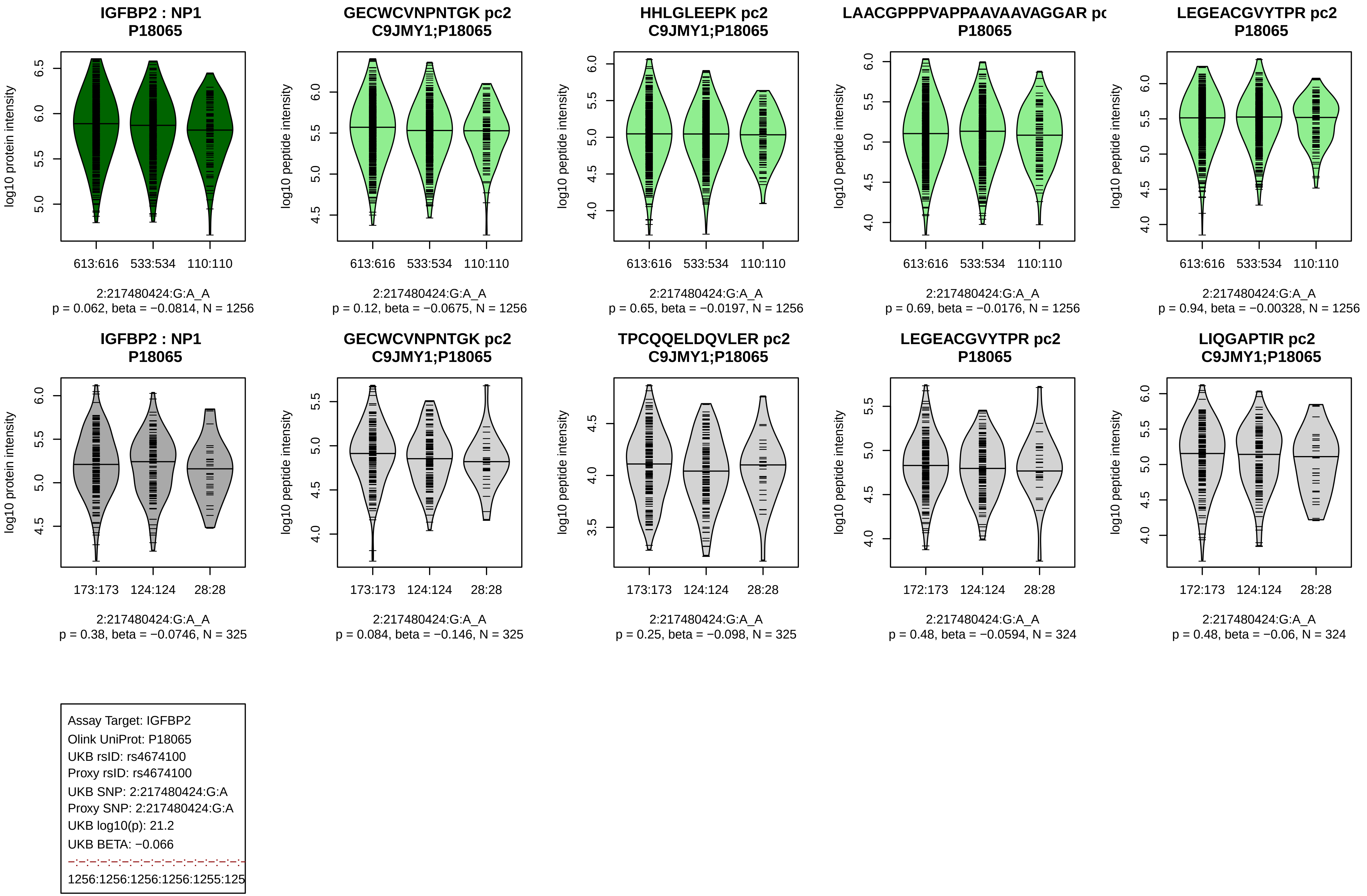

IGFBP2 : NP1
P18065
log10 protein intensity
613:616 533:534 110:110
2:217480424:G:A_A
p = 0.062, beta = −0.0814, N = 1256
GECWCVNPNTGK pc2
C9JMY1;P18065
log10 peptide intensity
613:616 533:534 110:110
2:217480424:G:A_A
p = 0.12, beta = −0.0675, N = 1256
HHLGLEEPK pc2
C9JMY1;P18065
log10 peptide intensity
613:616 533:534 110:110
2:217480424:G:A_A
p = 0.65, beta = −0.0197, N = 1256
LAACGPPPVAPPAAVAAVAGGAR pc
P18065
log10 peptide intensity
613:616 533:534 110:110
2:217480424:G:A_A
p = 0.69, beta = −0.0176, N = 1256
LEGEACGVYTPR pc2
P18065
log10 peptide intensity
613:616 533:534 110:110
2:217480424:G:A_A
p = 0.94, beta = −0.00328, N = 1256
IGFBP2 : NP1
P18065
log10 protein intensity
173:173 124:124 28:28
2:217480424:G:A_A
p = 0.38, beta = −0.0746, N = 325
GECWCVNPNTGK pc2
C9JMY1;P18065
log10 peptide intensity
173:173 124:124 28:28
2:217480424:G:A_A
p = 0.084, beta = −0.146, N = 325
TPCQQELDQVLER pc2
C9JMY1;P18065
log10 peptide intensity
173:173 124:124 28:28
2:217480424:G:A_A
p = 0.25, beta = −0.098, N = 325
LEGEACGVYTPR pc2
P18065
log10 peptide intensity
172:173 124:124 28:28
2:217480424:G:A_A
p = 0.48, beta = −0.0594, N = 324
LIQGAPTIR pc2
C9JMY1;P18065
log10 peptide intensity
172:173 124:124 28:28
2:217480424:G:A_A
p = 0.48, beta = −0.06, N = 324
Assay Target: IGFBP2
Olink UniProt: P18065
UKB rsID: rs4674100
Proxy rsID: rs4674100
UKB SNP: 2:217480424:G:A
Proxy SNP: 2:217480424:G:A
UKB log10(p): 21.2
UKB BETA: −0.066
1256:1256:1256:1256:1255:125

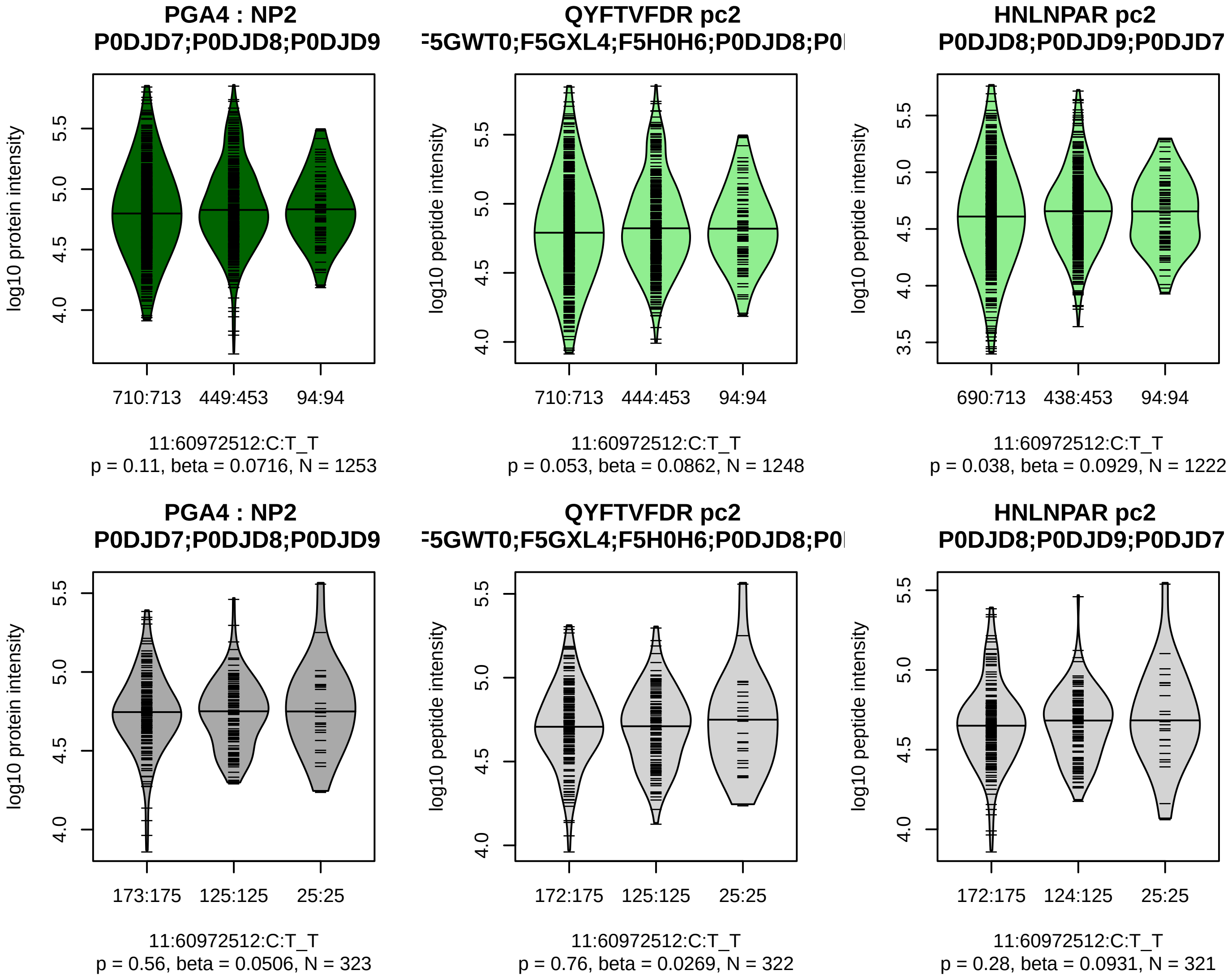


Assay Target: PGA4
Olink UniProt: P0DJD7
UKB rsID: rs4414241
Proxy rsID: rs4414241
UKB SNP: 11:60972512:C:T
Proxy SNP: 11:60972512:C:T
UKB log10(p): 21
UKB BETA: 0.075
–:–
1248:1222

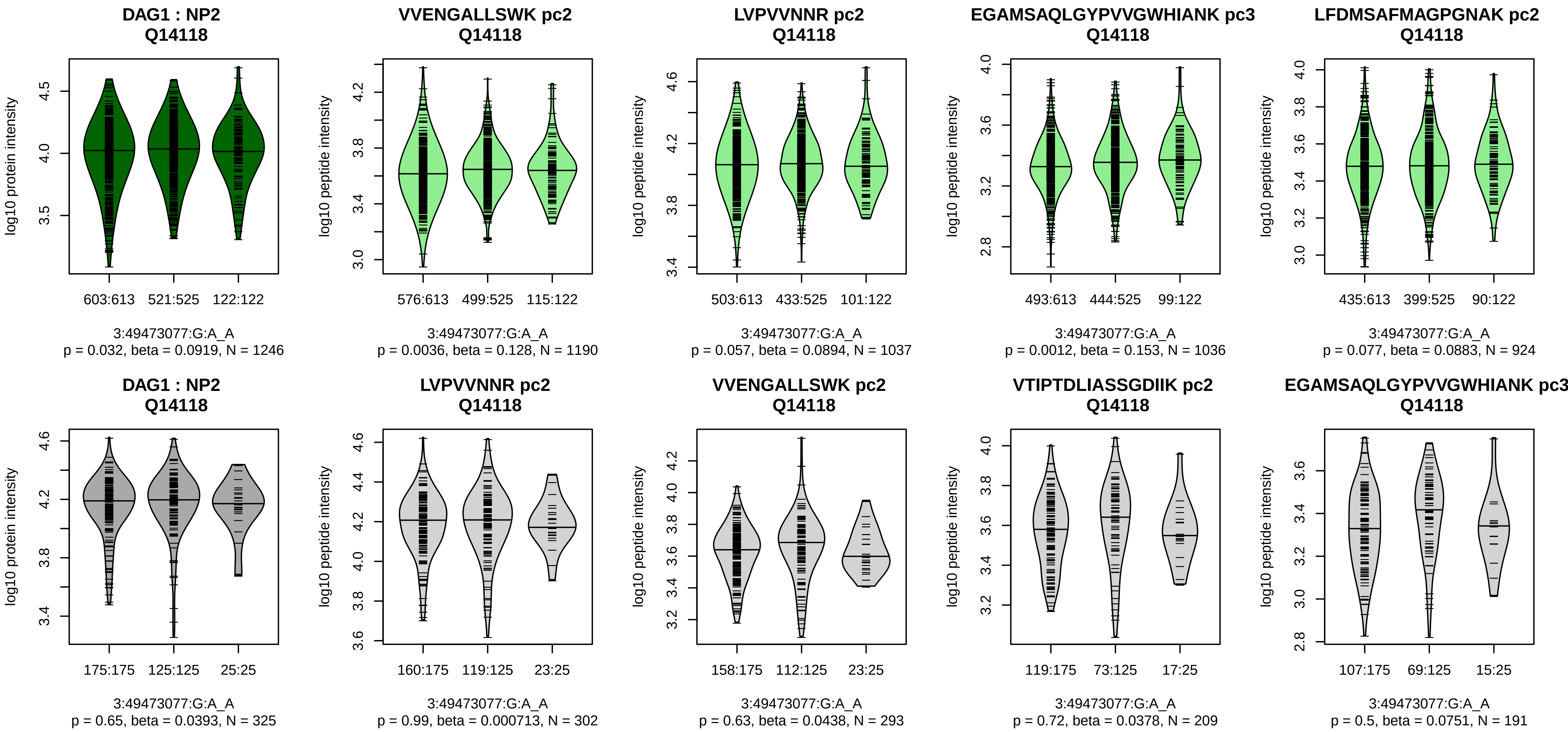


Assay Target: DAG1
Olink UniProt: Q14118
UKB rsID: rs6446277
Proxy rsID: rs6446277
UKB SNP: 3:49473077:G:A
Proxy SNP: 3:49473077:G:A
UKB log10(p): 21
UKB BETA: 0.064
*:–:*:–:–:*:–:–:–:–
1190:1037:1036:924:837:810:6[illegible]

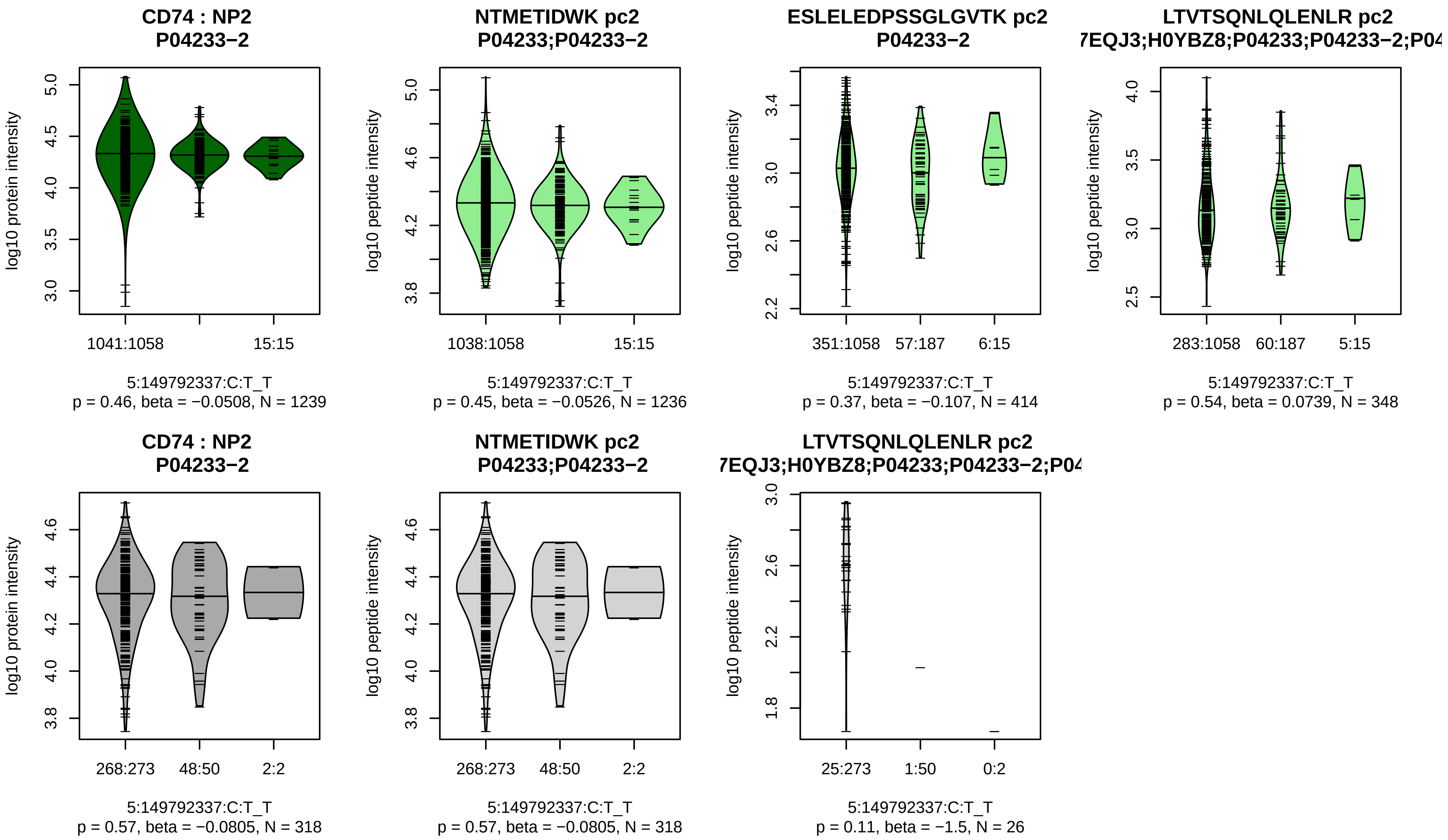


Assay Target: CD74
Olink UniProt: P04233
UKB rsID: rs55796634
Proxy rsID: rs55796634
UKB SNP: 5:149792337:C:T
Proxy SNP: 5:149792337:C:T
UKB log10(p): 20.1
UKB BETA: −0.087
−:−:−
1236:414:348

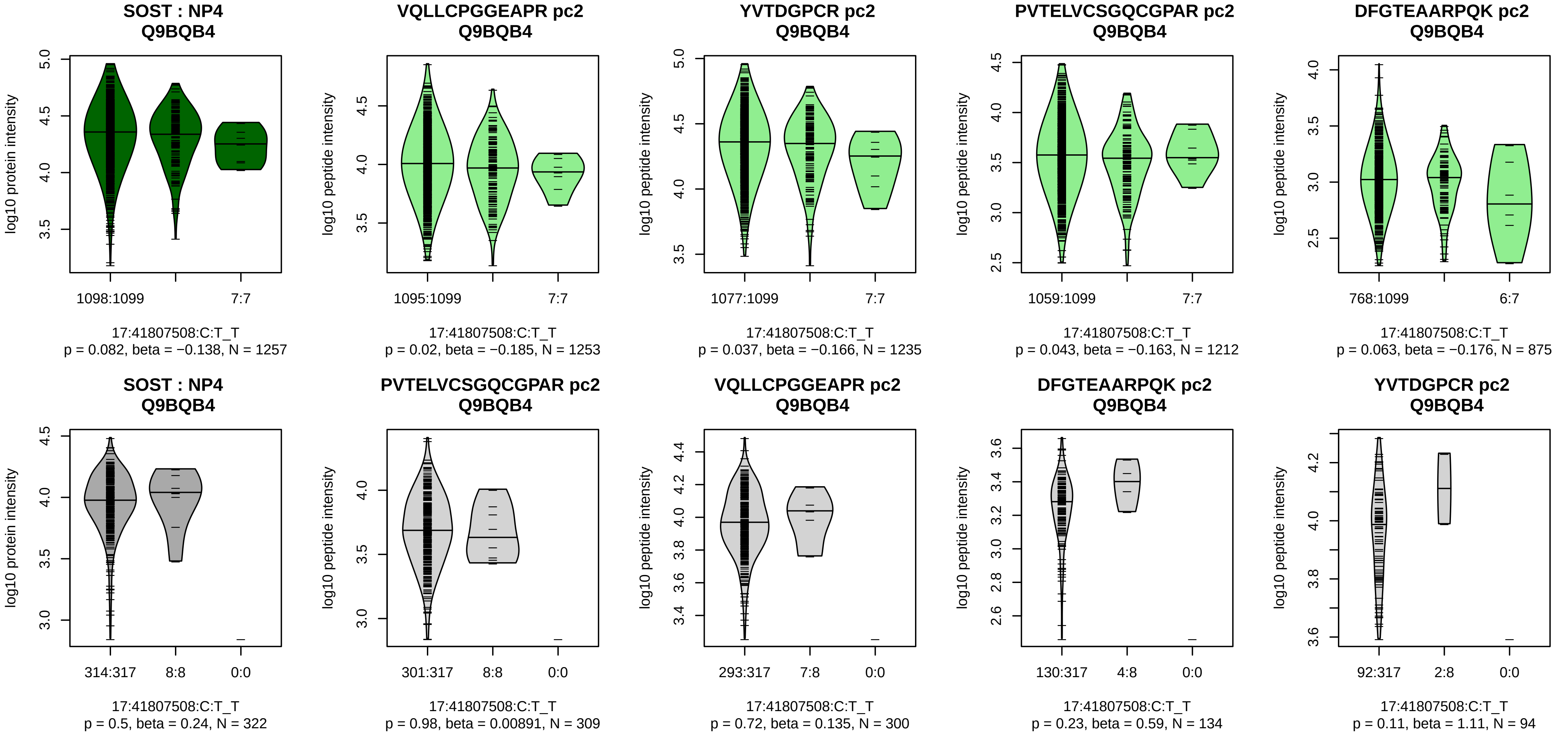


Assay Target: SOST
Olink UniProt: Q9BQB4
UKB rsID: rs76410205
Proxy rsID: rs76410205
UKB SNP: 17:41807508:C:T
Proxy SNP: 17:41807508:C:T
UKB log10(p): 19.7
UKB BETA: −0.097
–:–:–:–:–:–:–
1253:1235:1212:875:871:272:38

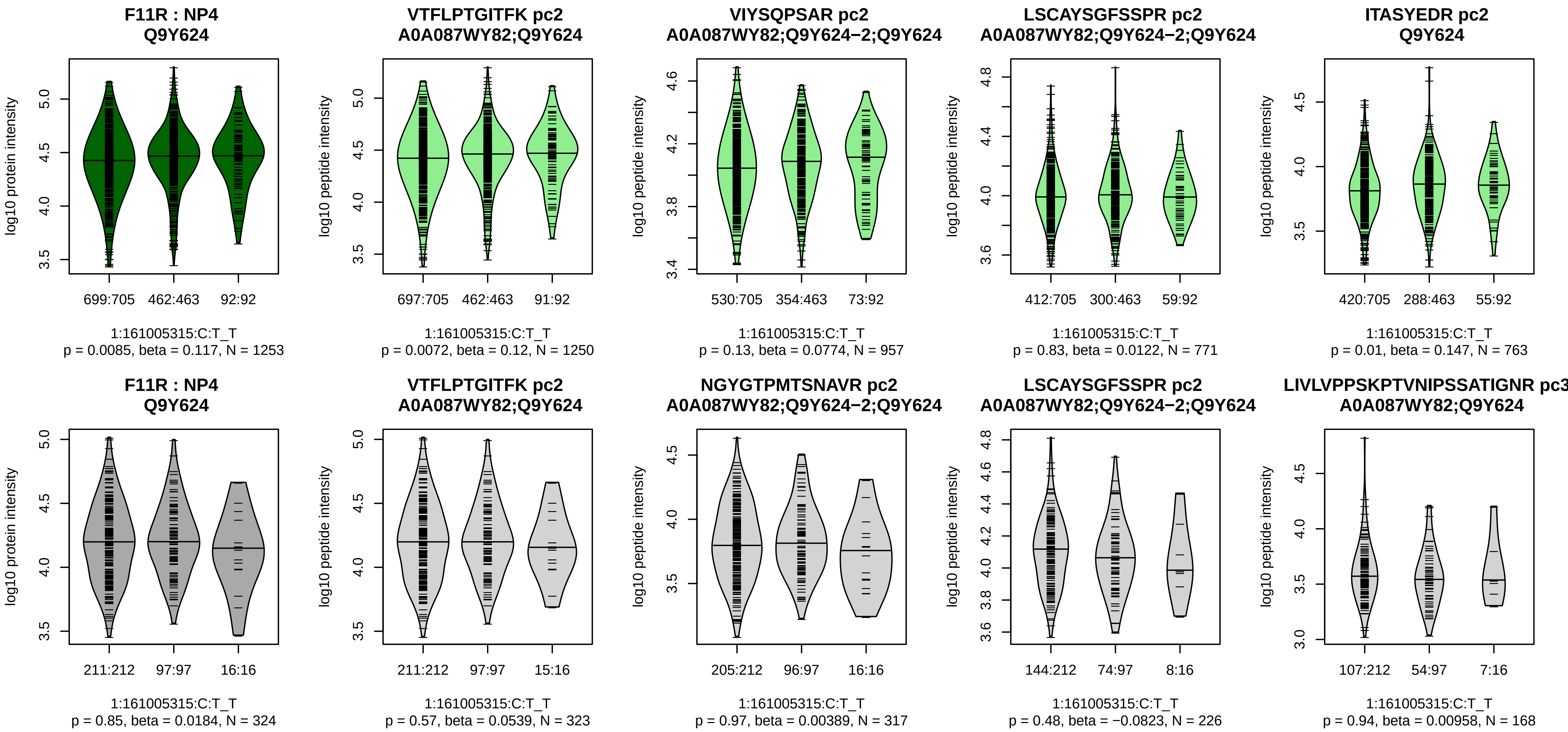


Assay Target: F11R
Olink UniProt: Q9Y624
UKB rsID: rs143551603
Proxy rsID: rs138818128
UKB SNP: 1:161005266:T:TA
Proxy SNP: 1:161005315:C:T
UKB log10(p): 18.7
UKB BETA: 0.067
*:–:–:–:–:–:–:–:–:–
1250:957:771:763:756:667:530:

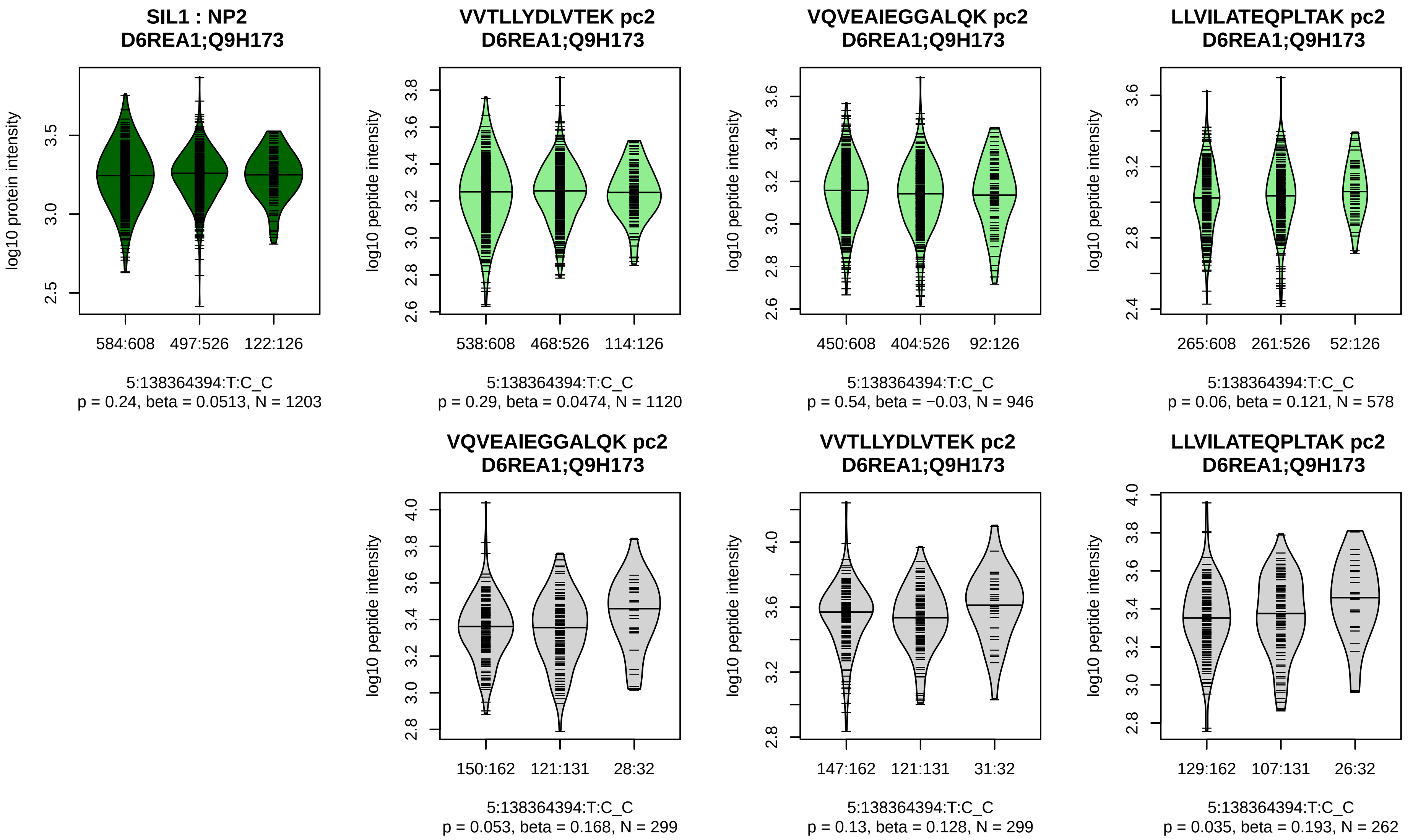


Assay Target: SIL1
Olink UniProt: Q9H173
UKB rsID: rs11957633
Proxy rsID: rs11957633
UKB SNP: 5:138364394:T:C
Proxy SNP: 5:138364394:T:C
UKB log10(p): 18.7
UKB BETA: 0.06
–:–:–
1120:946:578

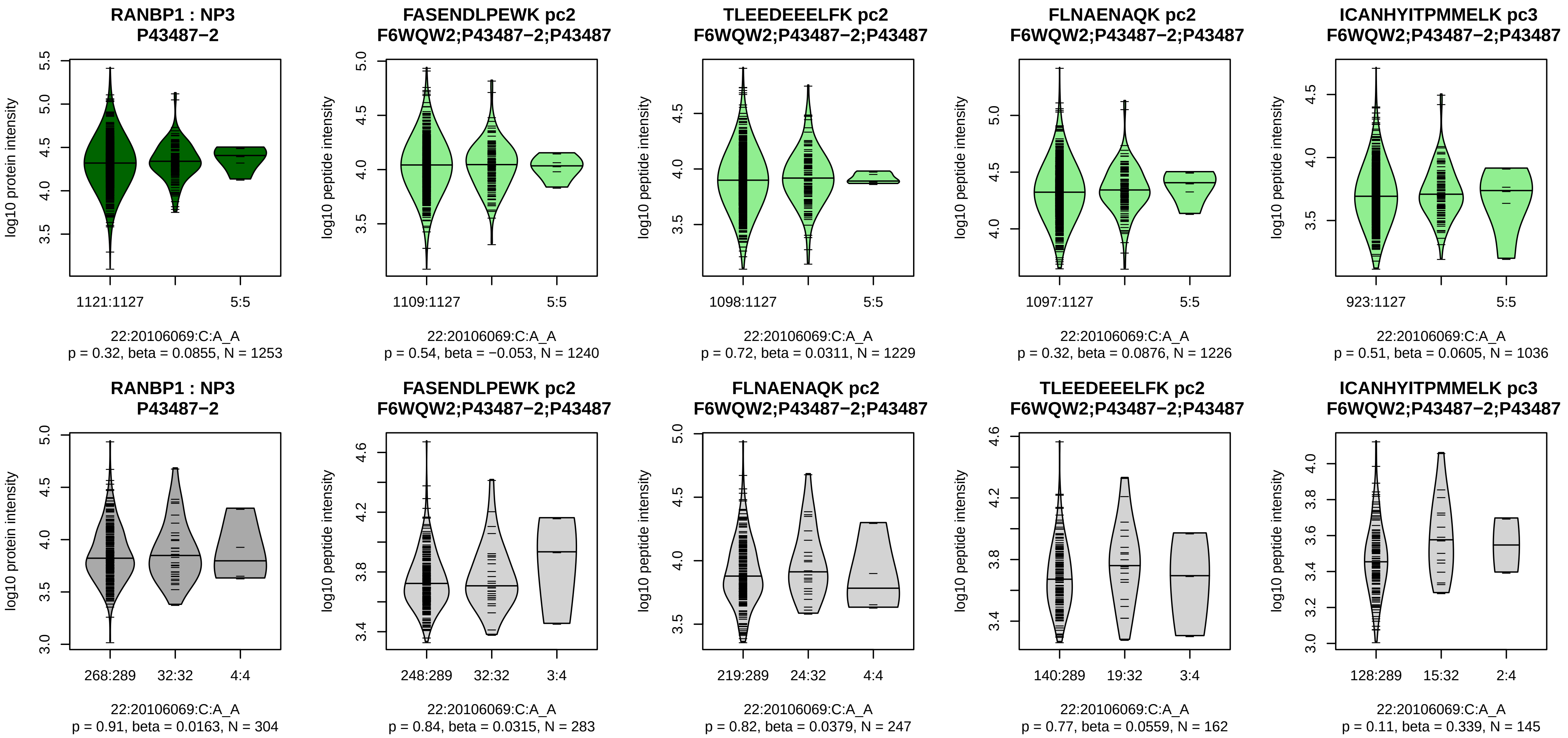


Assay Target: RANBP1
Olink UniProt: P43487
UKB rsID: rs117718692
Proxy rsID: rs4589855
UKB SNP: 22:20117344:G:T
Proxy SNP: 22:20106069:C:A
UKB log10(p): 18.4
UKB BETA: 0.122
−:−:−:−:−:−:NA
1240:1229:1226:1036:950:827:1

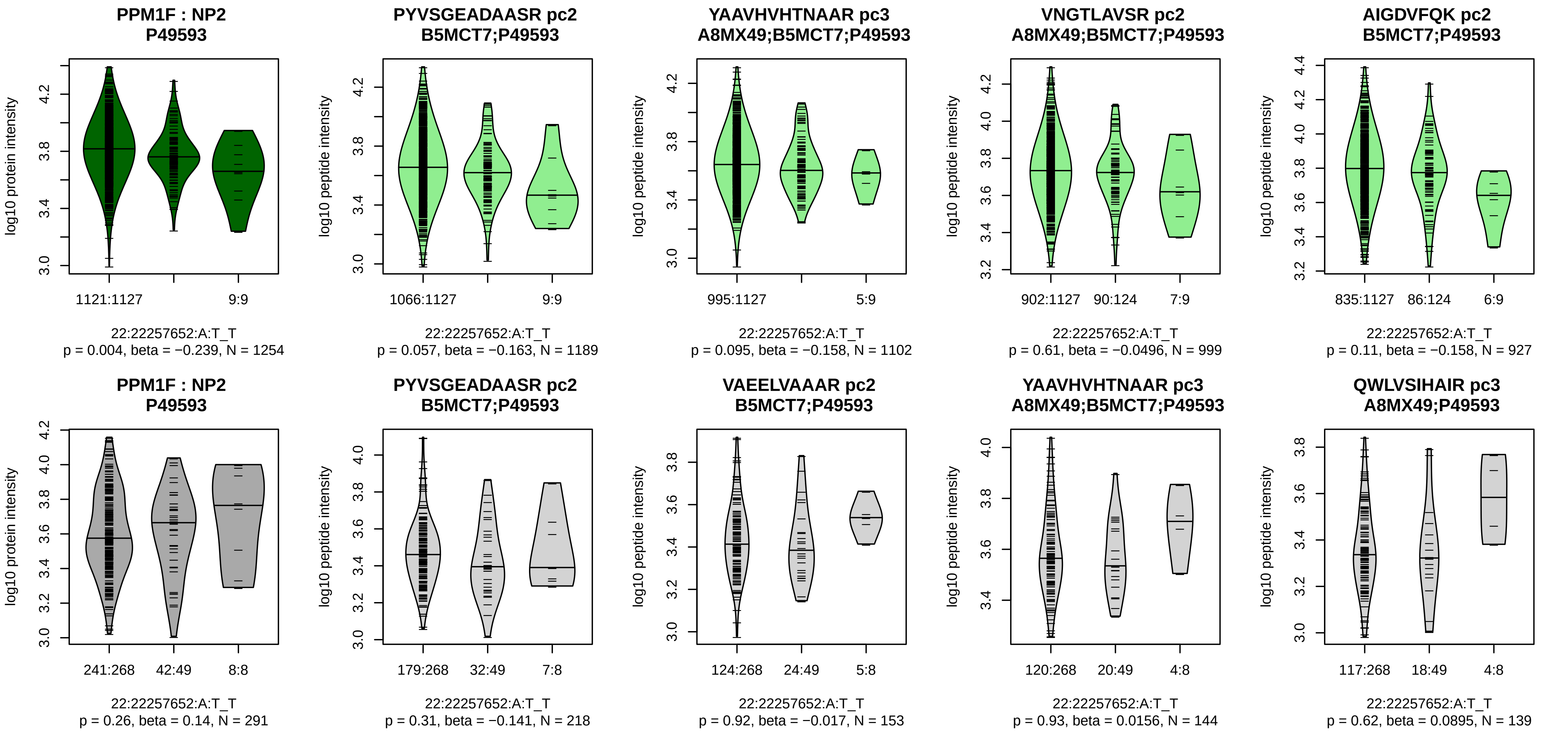


Assay Target: PPM1F
Olink UniProt: P49593
UKB rsID: rs5750281
Proxy rsID: rs5755996
UKB SNP: 22:22327787:C:T
Proxy SNP: 22:22257652:A:T
UKB log10(p): 18.3
UKB BETA: −0.132
−:−:−:−:−:−:−:−:−:−:−:−:−:−:−
1189:1102:999:927:860:842:71[illegible]

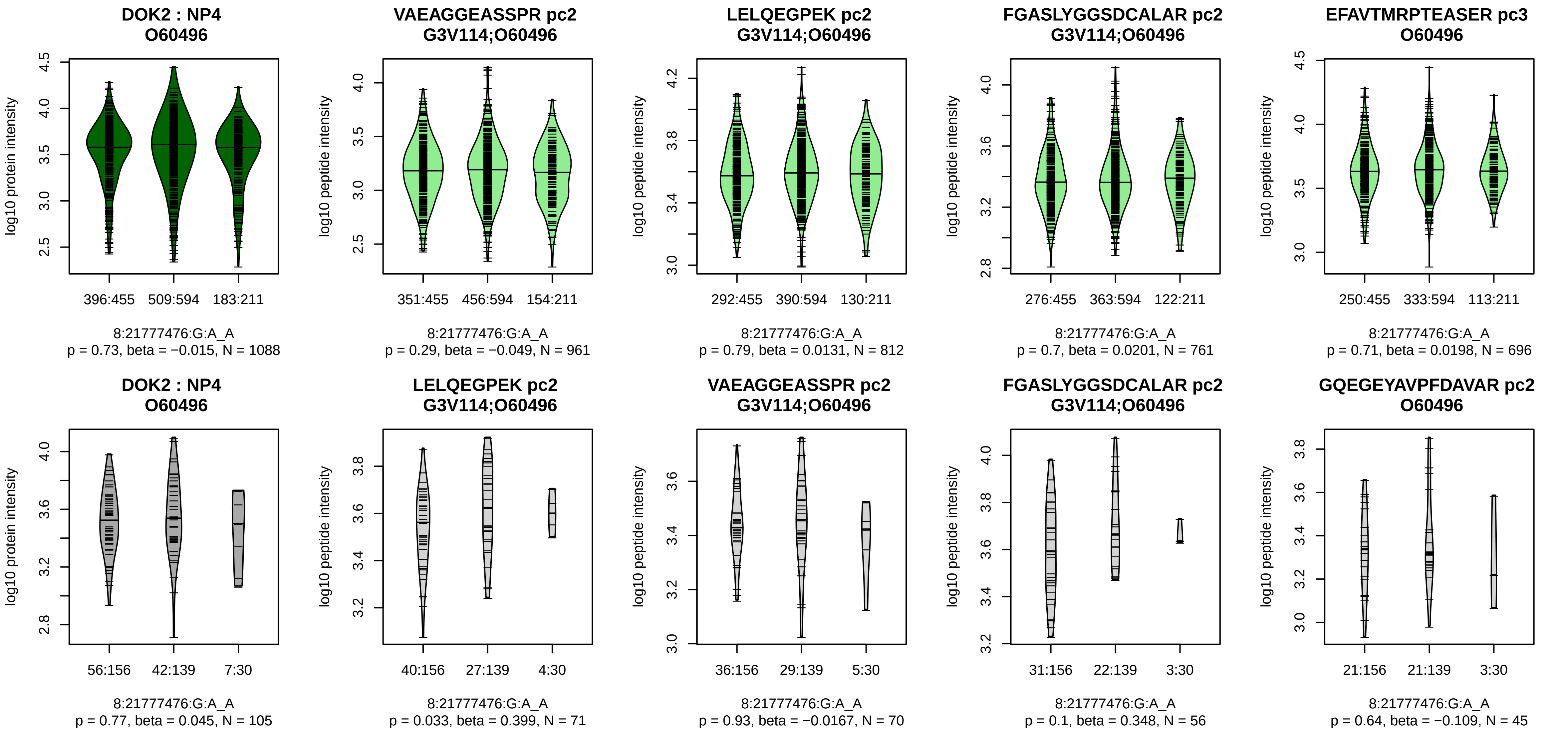


Assay Target: DOK2
Olink UniProt: O60496
UKB rsID: rs4357289
Proxy rsID: rs4357289
UKB SNP: 8:21777476:G:A
Proxy SNP: 8:21777476:G:A
UKB log10(p): 17.8
UKB BETA: 0.054
−:−:−:−:−:−:−:−:−:−:−:−:−:−:−:NA
961:812:761:696:531:498:369:3

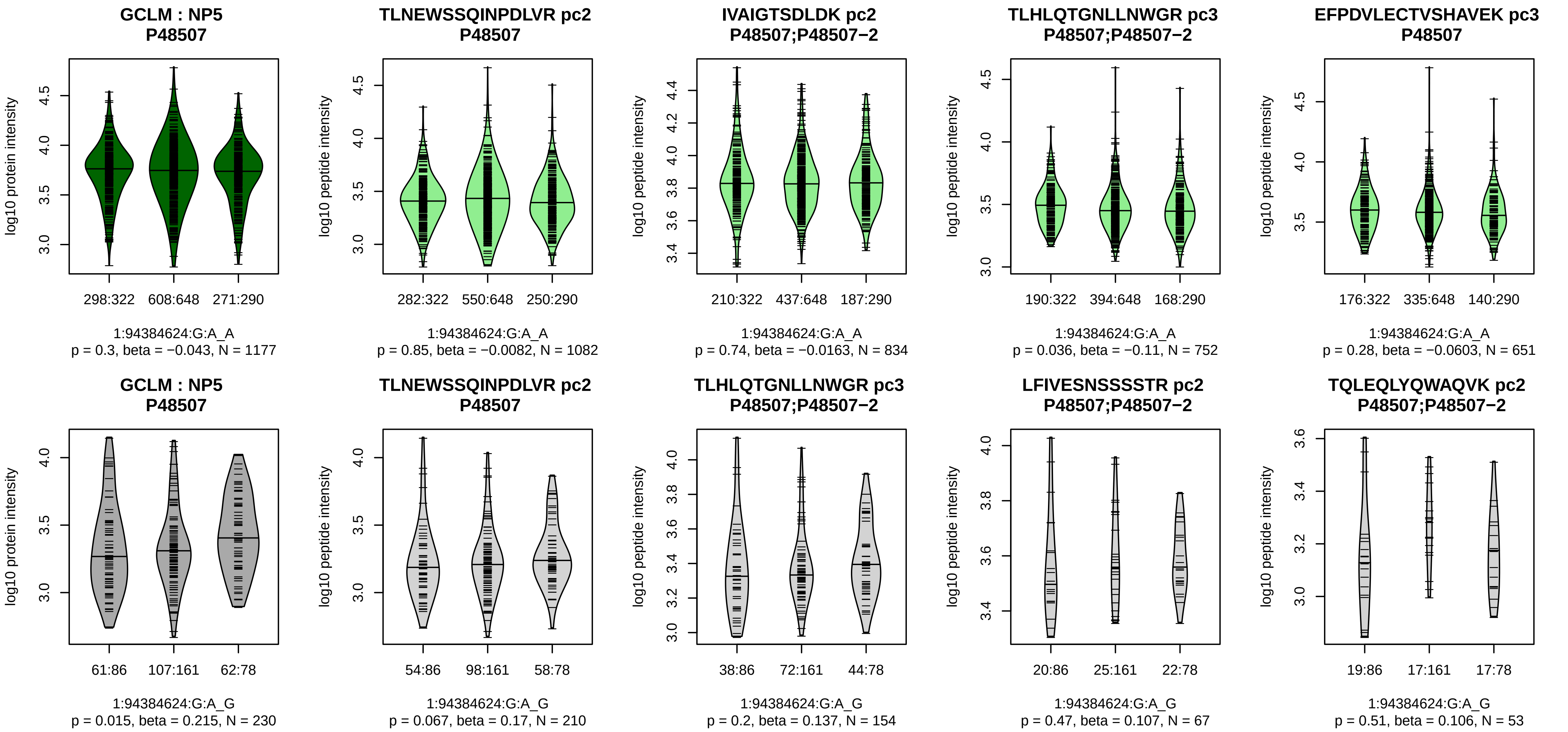


Assay Target: GCLM
Olink UniProt: P48507
UKB rsID: rs774698707
Proxy rsID: rs12563495
UKB SNP: 1:94383448:TTAA:T
Proxy SNP: 1:94384624:G:A
UKB log10(p): 17.6
UKB BETA: −0.055
−:−:−:−:−:−:−:−:−:−:−:−:−:NA:NA
1082:834:752:651:444:227:219:

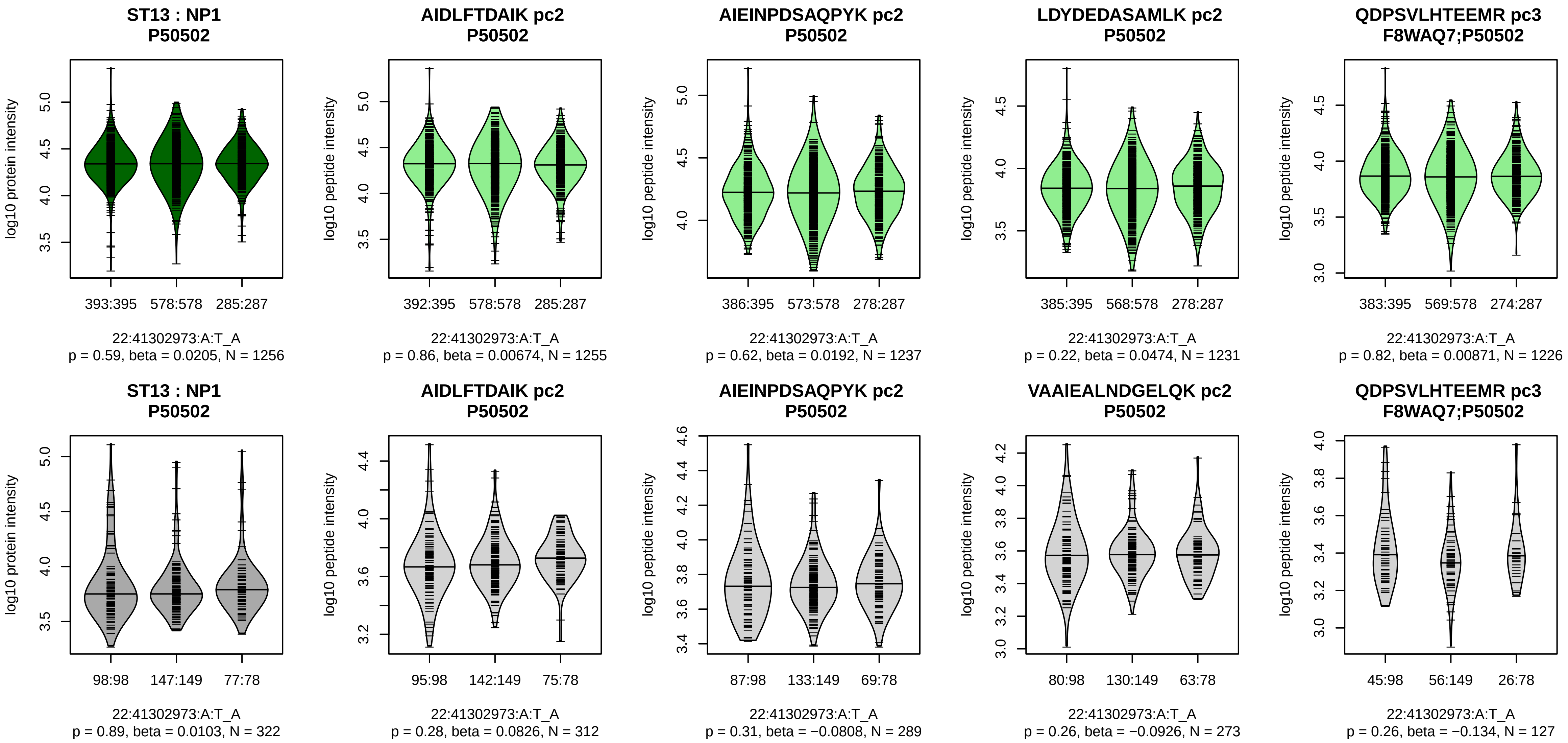


Assay Target: ST13
Olink UniProt: P50502
UKB rsID: rs5995963
Proxy rsID: rs5995963
UKB SNP: 22:41302973:A:T
Proxy SNP: 22:41302973:A:T
UKB log10(p): 17.1
UKB BETA: −0.052
−:−:−:−:−:−:−:−:−:−:−:−:−:−:NA
1255:1237:1231:1226:1202:972

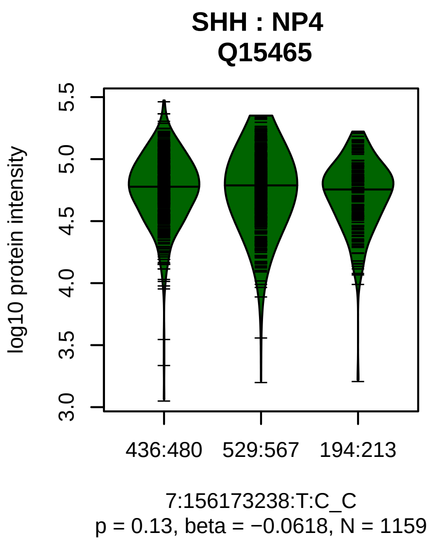


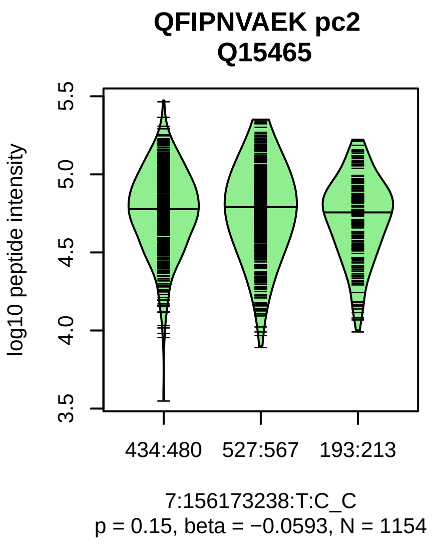


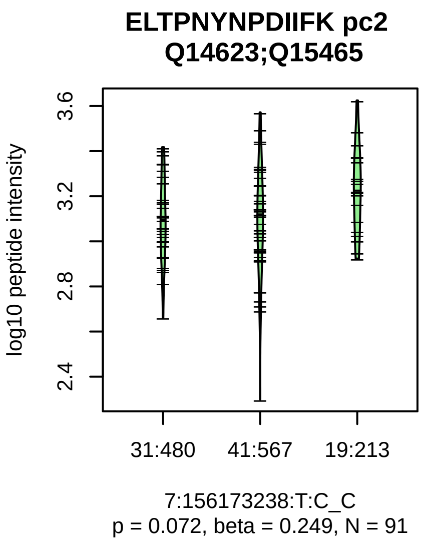


Assay Target: SHH
Olink UniProt: Q15465
UKB rsID: rs10225292
Proxy rsID: rs10225292
UKB SNP: 7:156173238:T:C
Proxy SNP: 7:156173238:T:C
UKB log10(p): 16.6
UKB BETA: −0.055
−:−
1154:91

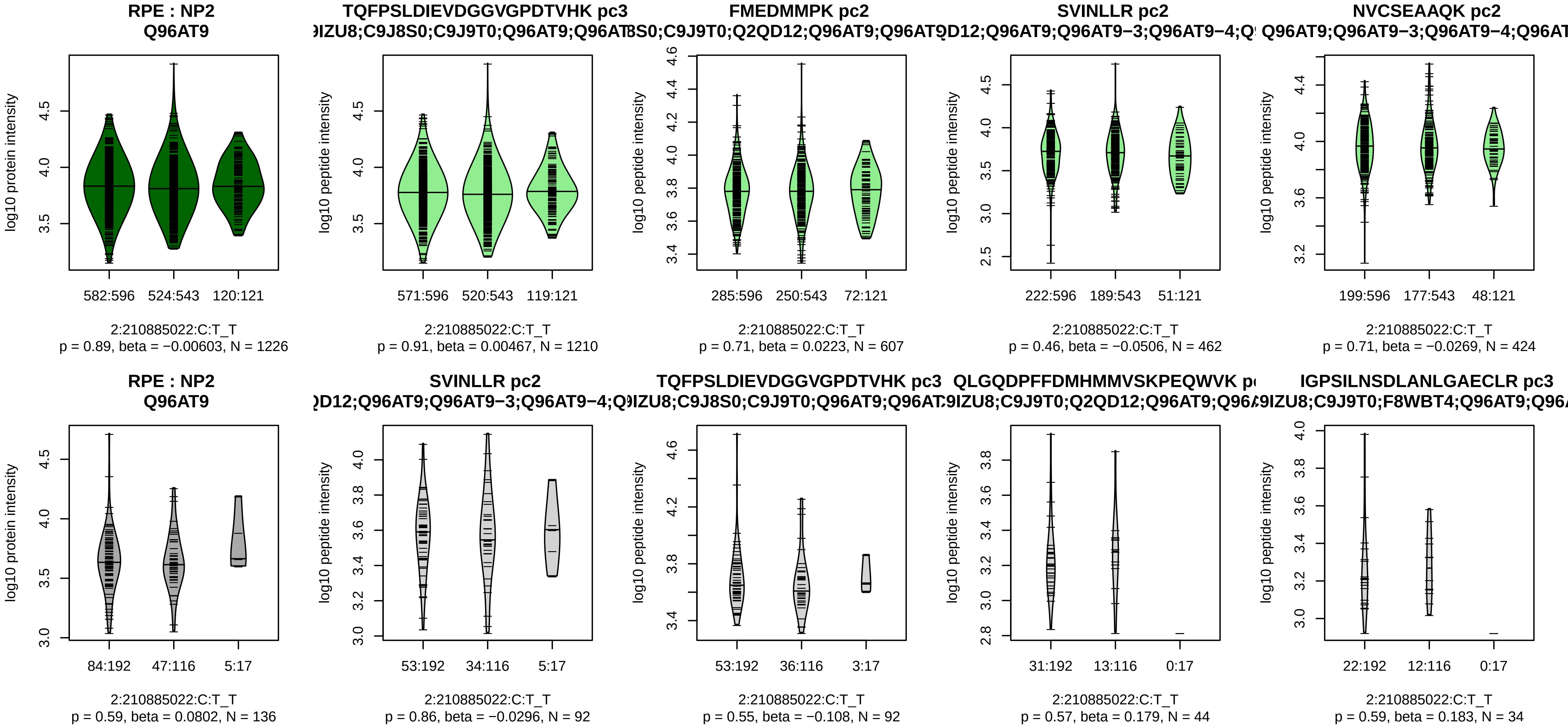


Assay Target: RPE
Olink UniProt: Q96AT9
UKB rsID: rs7585265
Proxy rsID: rs7585265
UKB SNP: 2:210885022:C:T
Proxy SNP: 2:210885022:C:T
UKB log10(p): 16.3
UKB BETA: −0.054
−:−:−:−:−:−:−:NA
1210:607:462:424:353:269:116:

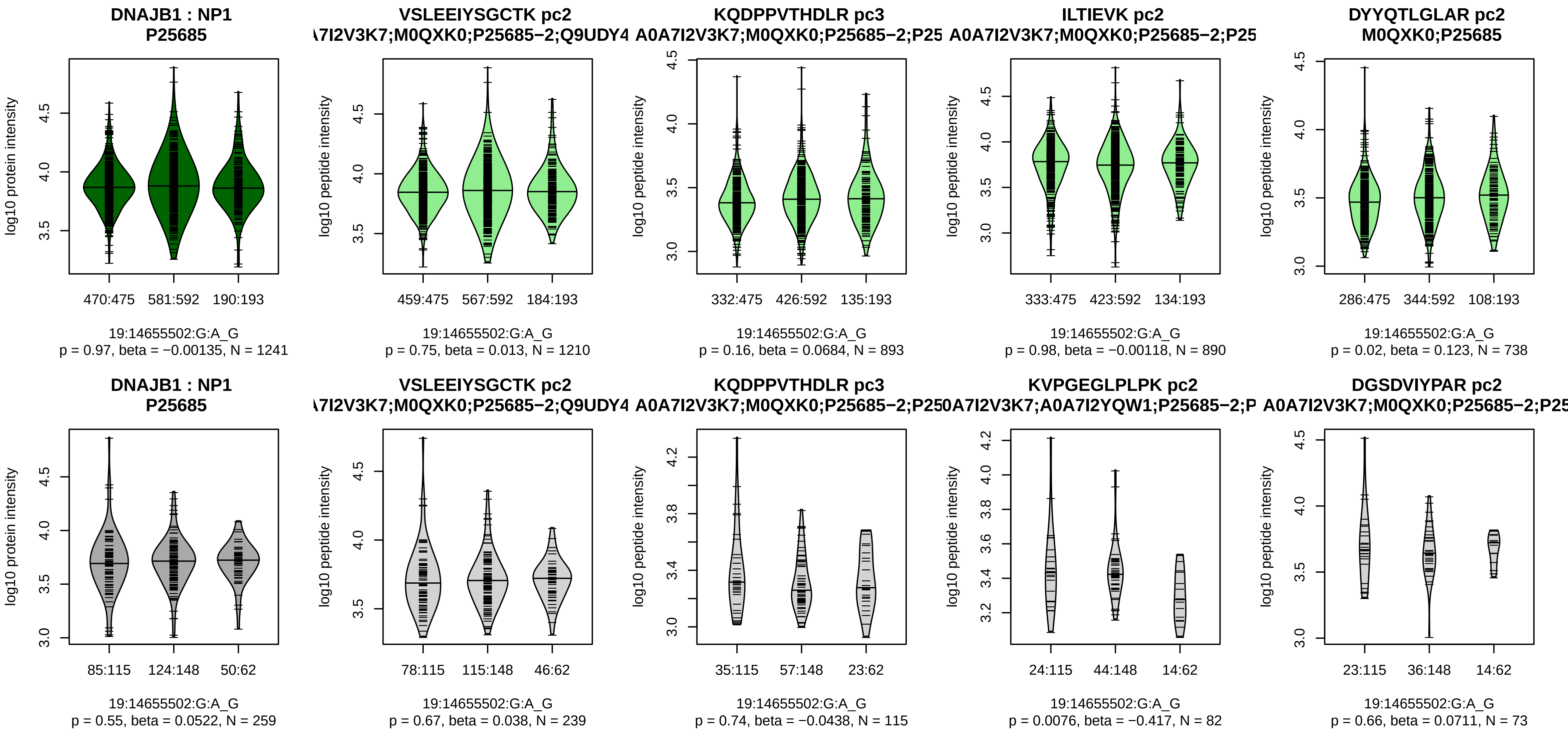


Assay Target: DNAJB1
Olink UniProt: P25685
UKB rsID: rs7248009
Proxy rsID: rs7248009
UKB SNP: 19:14655502:G:A
Proxy SNP: 19:14655502:G:A
UKB log10(p): 15.7
UKB BETA: −0.051
−:−:−:−:−:−:−:−:−:−:−:−:−:−:NA:N
1210:893:890:738:385:348:304:

**SLIT2 : NP4**
**O94813;X6R3P0**

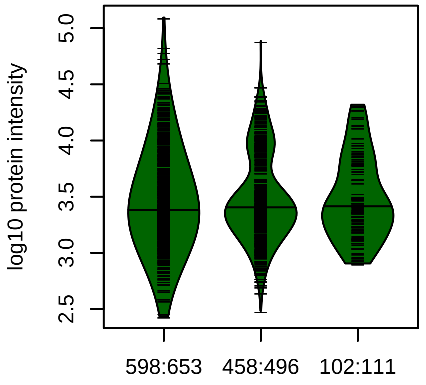


4:20190972:C:T_C
p = 0.7, beta = 0.0175, N = 1158

**VPMQTGILPGCEPCHK pc3**
**87WYV5;O94813;O94813−2;O94813−**

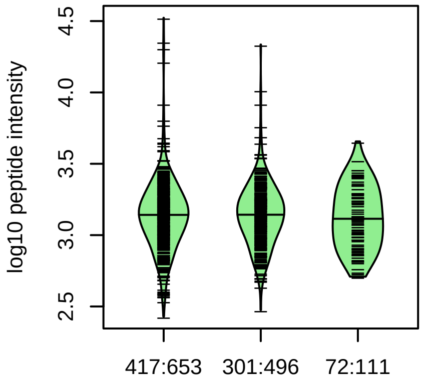


4:20190972:C:T_C
p = 0.64, beta = −0.0256, N = 790

**ESYLQIPSAK pc2**
**87WYV5;O94813;O94813−2;O94813−**

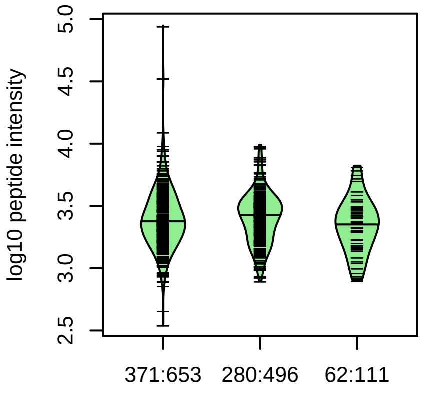


4:20190972:C:T_C
p = 0.15, beta = −0.0819, N = 713

**NLYINSELQDFQK pc2**
**87WYV5;O94813;O94813−2;O94813−**

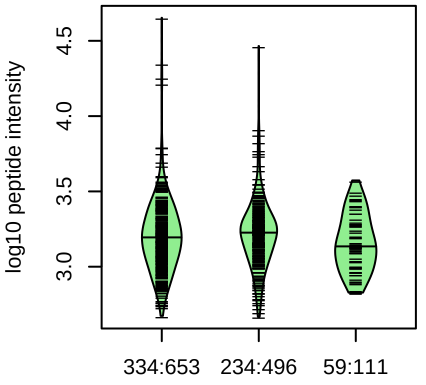


4:20190972:C:T_C
p = 0.47, beta = −0.043, N = 627

**TNDPCLGNK pc2**
**87WYV5;O94813;O94813−2;O94813−**

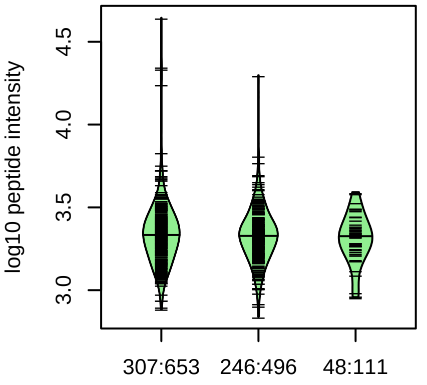


4:20190972:C:T_C
p = 0.15, beta = −0.0914, N = 601

**SLIT2 : NP4**
**87WYV5;O94813;O94813−2;O94813−**

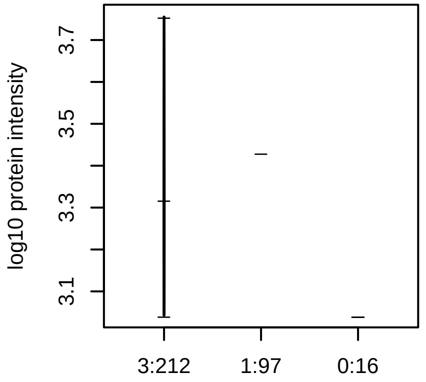


4:20190972:C:T_C
p = NA, beta = NA, N = 4

Assay Target: SLIT2
Olink UniProt: O94813
UKB rsID: rs587668
Proxy rsID: rs587668
UKB SNP: 4:20190972:C:T
Proxy SNP: 4:20190972:C:T
UKB log10(p): 15.6
UKB BETA: −0.056
−:−:−:−:−:−:−:−:−:−:−:*:−:−:−:−:−
790:713:627:601:381:330:292:2
